# Single-cell Transcriptomic Analysis of Salivary Gland Endothelial Cells

**DOI:** 10.1101/2023.06.22.545817

**Authors:** Amber L. Altrieth, Emily Suarez, Deirdre A. Nelson, Sergo Gabunia, Melinda Larsen

## Abstract

Vascular endothelial cells have important functions in fibrosis via direct and indirect methods and in regeneration through secretion of tissue-specific, paracrine-acting angiocrine factors. In the salivary gland, endothelial cells are required for proper development, but their roles within adult glands are largely unknown. The goal of this work was to identify ligand-receptor interactions between endothelial cells and other cell types that are important during homeostasis, fibrosis, and regeneration. To model salivary gland fibrosis and regeneration, we utilized a reversible ductal ligation. To induce injury, a clip was applied to the primary ducts for 14 days, and to induce a regenerative response, the clip was subsequently removed for 5 days. To identify endothelial cell-produced factors, we used single-cell RNA-sequencing of stromal-enriched cells from adult submandibular and sublingual salivary glands. Transcriptional profiles of homeostatic salivary gland endothelial cells were compared to endothelial cells of other organs. Salivary gland endothelial cells were found to express unique genes and displayed the highest overlap in gene expression with other fenestrated endothelial cells from the colon, small intestine, and kidney. Comparison of the 14-day ligated, mock ligated, and 5-day deligated stromal-enriched transcripts and lineage tracing were used to identify evidence for a partial endoMT phenotype, which was observed in a small number of endothelial cell subsets with ligation. CellChat was used to predict changes in ligand-receptor interactions in response to ligation and deligation. CellChat predicted that after ligation, endothelial cells are sources of protein tyrosine phosphatase receptor type m, tumor necrosis factor ligand superfamily member 13, and myelin protein zero signaling and targets for tumor necrosis factor signaling. Following deligation, CellChat predicted that endothelial cells are sources of chemokine (C-X-C motif) and EPH signaling to promote regenerative responses. These studies will inform future endothelial cell-based regenerative therapies.

## 1.1. Introduction

Endothelial cells have many important functions during salivary gland development (Kwon et al. 2017). Endothelial cells also have critical functions in maintaining and promoting homeostasis, and endothelial cell dysfunction can contribute to disease and organ dysfunction. In the salivary glands of a subset of patients with the autoimmune disease, Sjögren’s Disease (SjD), there are vascular defects including: decreased microvessel density, aberrant vessels, and increased atherosclerosis (Tektonidou et al. 1999; Demirci et al. 2016; Valim et al. 2016). Microvascular endothelial cells are known to be sensitive to radiation damage (Wijerathne et al. 2021), and patients who receive radiation therapy as a treatment for head and neck cancer have decreased microvessel density (Lombaert et al. 2017). Thus, vascular-based therapies may provide protective effects for patients suffering from salivary hypofunction.

Studies in other organs have linked endothelial cell dysfunction with tissue damage. In mouse lungs, endothelial cell injury is thought to be an initiator of fibrosis. Following bleomycin treatment, endothelial cells secreted pro-fibrotic factors which recruited inflammatory cells (Leach et al. 2013), and many underwent an endothelial to mesenchymal transition (endoMT) (Hashimoto et al. 2010). In addition, studies in the heart and kidney also found that endothelial cells underwent an endoMT, contributing directly to organ fibrosis (Zeisberg et al. 2007; Zeisberg et al. 2008; Li et al. 2009; Li et al. 2010). Salivary gland fibrosis has been associated with sialadenitis, or swelling, of the salivary glands (Adhikari and Soni 2022), irradiation damage, and in a subset of autoimmune disease patients (J. D. Harrison et al. 1997; Bookman et al. 2011; Llamas-Gutierrez et al. 2014; Straub et al. 2015; Leehan et al. 2018), but the cellular contribution of endothelial cells to salivary gland fibrosis are poorly understood.

Endothelial cells can promote regeneration through the secretion of angiocrine factors. Angiocrine factors maintain stem and progenitor cells in a quiescent state during homeostasis and can promote regeneration following injury in a tissue-specific manner. Angiocrine factors include growth factors, adhesion molecules, and chemokines (Rafii et al. 2016) and are tissue-specific in their expression patterns and in their responses to injury (Nolan et al. 2013; Kalucka et al. 2020); however, salivary gland endothelial cells were not examined.

To design effective therapeutics to restore salivary gland function following injury or disease, it is critical to understand endothelial cell-produced factors and the interactions endothelial cells have with other cell types both in homeostasis and after injury. In this study, we utilized single-cell RNA-sequencing (scRNA-seq) to profile the transcriptomes of salivary gland endothelial cells during homeostasis, fibrotic injury, and the regenerative response following injury to identify pathways that may promote recovery of salivary glands from fibrotic injury.

## 1.2. Materials and Methods

### 1.2.1. Animal husbandry

All animal husbandry, surgical procedures, and tissue collection were performed in accordance with protocols approved by the University at Albany, SUNY IACUC committee. Mice were housed in 12-hour light/dark cycle with access to water and dry food. C57BL/6J (JAX #000664) mice were purchased from The Jackson Laboratory. C57BL/6-Tg(Cdh5-cre/ERT2)1Rha (Taconic #13073) male breeders were purchased from Taconic Biosciences and maintained by crossing to C57BL/6J females. For lineage tracing experiments, C57BL/6-Tg(Cdh5-cre/ERT2)1Rha Cre^+^ males were crossed with B6.Cg-Gt(ROSA)26Sor^tm14(CAG-tdTomato)Hze^/J (JAX #007914) homozygous females to generate female C57BL/6-Tg(Cdh5-cre/ERT2)1Rha; B6.Cg-Gt(ROSA)26Sor^tm14(CAG-tdTomato)Hze^/J (Cdh5-cre/ERT2; R26^tdT^) mice. Mice were assigned a unique identifier at the time of surgery or between postnatal day 7 and 10. Cdh5-cre/ERT2 mice were genotyped with PCR to detect Cre and Cdh5-cre/ERT2; R26^tdT^ mice were genotyped with PCR to detect Cre and tdTomato.

### 1.2.2. Tamoxifen induction

For lineage tracing experiments, Cdh5-cre/ERT2; R26^tdT^ female mice were induced at 11 weeks old for three consecutive days with 100 µL of 10 mg/mL tamoxifen (Sigma cat #T5648) dissolved in corn oil (Sörensen et al. 2009). Surgeries were performed on mice one week after the first induction. As controls for tamoxifen-induced changes in lineage tracing experiments, 1-week induced submandibular salivary glands were used.

### 1.2.3. Salivary gland ductal ligation and deligation

Only female mice were used for surgical manipulations due to their similarities in tissue architecture with human salivary glands (Amano et al. 2012; Maruyama et al. 2019). Adult female mice at 12-weeks-old were used for surgeries. A unique identifier was assigned to each mouse either 7 to 10 days after birth or at the time of surgery. The mice were anesthetized using 100 mg/kg ketamine and 10 mg/kg xylazine by intraperitoneal injection from a stock concentration of 10 mg/mL ketamine and 1 mg/mL xylazine solution in sterile water for injection and were given 100 µL of buprenorphine at a concentration of 0.015mg/mL by subcutaneous injection as an analgesic for post-operative pain management. An incision was made to visualize the main ducts of the submandibular and sublingual glands, Wharton’s and Bartholin’s ducts, respectively. A vascular clamp (Vitalitec/Peter’s Surgical) was applied to these ducts, and the incision was closed using two to four interrupted sutures. Mice were continuously monitored under anesthesia and post-operatively for pain, distress, and changes in weight for a minimum of 48 hours following surgery. The ducts were ligated for 14 days. Successful 14-day ligation was determined if the gland weight was between 30 and 70% of the average historical control gland weight, and samples outside of the range were excluded from downstream analyses. As controls for the C57BL/6J experiments, time point zero (homeostatic) mice were euthanized at 12-weeks old and had no surgical manipulations. Mock surgery control mice received an incision, the ducts were located, but not ligated, and mice were euthanized at 14-days post-surgery.

For deligation surgeries, the mice were anesthetized as described above, and the vascular clamp was removed; the incision was closed using two to four interrupted sutures. Mice were monitored post-operatively, as described above. Mice were harvested five days after deligation. All mice were euthanized at the desired time point under CO2 with secondary cervical dislocation. The submandibular and sublingual glands were immediately weighed upon removal. All subsequent sample processing was performed blind with reference only to the unique identifiers.

### 1.2.4. Single-cell isolation for homeostatic, mock, ligated, and deligated mouse salivary glands

Two submandibular and sublingual glands were harvested, and excess fat and interstitial tissue was removed. The glands were then transferred to a dish containing 1X phosphate buffered saline (1X PBS), liberase TL Research Grade low Thermolysin (Roche), DNase I, and dispase and microdissected for 7 to 9 minutes. The sample was then incubated in a 37°C incubator for 15 minutes and triturated. After trituration, the sample was incubated for an additional 5 minutes in a 37°C incubator and triturated once more. A 15 mL conical tube was placed on ice and the entire sample was transferred to the conical tube and incubated for 10 minutes. The supernatant was isolated and transferred to a fresh conical tube and was centrifuged for 5 minutes at 450xg. The supernatant was removed and discarded, and the cell pellet was resuspended in isolation buffer composed of sterile-filtered Ca^2+^- and Mg^2+^-free 1X PBS, 0.1% bovine serum albumin, and 2mM ethylenediaminetetraacetic acid. 5 μg of EpCAM-A647 (catalog: 118212, BioLegend) and Ter119 (catalog: 50-133-27, eBioscience/ThermoFisher) was added to the cell suspension and incubated at 4°C for 10 minutes for subsequent depletion of epithelial and red blood cells, respectively. The cell suspension was washed using isolation buffer and centrifuged at 450xg for 5 minutes. The supernatant was discarded, and the cell pellet was resuspended in 1 mL of isolation buffer and 25 μL of sheep anti-rat Dynabeads, which bind to the EpCAM and Ter119 labeled cells, (catalog: 11035, Invitrogen) before incubating on ice for 20 minutes on a rocker. The sample was placed on a microcentrifuge magnet for 2 minutes, and the supernatant was transferred to a fresh microcentrifuge tube to remove labeled epithelial and red blood cells. One additional epithelial and red blood cell depletion was performed. The sample was then depleted of dead cells using a dead cell removal kit (catalog: 130-090-101, Miltenyi Biotec) following manufacturer’s instructions. Cells were counted and viability was measured. Only samples with an 80% or higher viability were used for generating sequencing libraries. Viable cells were resuspended to a concentration of 1000 cells/μL. Following the manufacturer’s protocol for the Chromium Next GEM Single Cell 3’ Reagent kits v3.1, scRNA-seq libraries were generated.

### 1.2.5. Single-cell RNA-sequencing analysis using Seurat

Homeostatic, mock, ligated, and 5-day deligated samples were sequenced (the ligated sample was sequenced on the Illumina Nextseq500 and all others were sequenced on the Illumina Nextseq2000), and initial processing of the datasets was performed at the Center for Functional Genomics at the University at Albany. Initial processing steps included generating FASTQ files, aligning to the genome and generating counts files, which were completed using CellRanger version 6.0.1 for all surgically manipulated samples and 7.0.0 for the homeostatic sample. Data files were imported using Seurat v4.1.1 in R v4.2.1 (Stuart et al. 2019; Hao et al. 2021; R: The R Project for Statistical Computing 2021). Data clusters were calculated following the default pipeline (Seurat - Guided Clustering Tutorial 2023). Dead or apoptotic cells were removed if >6% of unique molecular identifiers (UMIs) mapped to mitochondrial genes. Any cells with <200 or >6000 genes were excluded to select for single cells. All samples used 16 dimensions. Samples were processed using clustree to determine resolution and a resolution of 0.8 was used for all samples (Zappia and Oshlack 2018). The processed datasets were saved as RDS files.

The homeostatic sample included two technical replicates and 5-day deligated sample included two biological replicates, which were merged after initial processing. Dead or apoptotic cells were removed if >6% of unique molecular identifiers (UMIs) mapped to mitochondrial genes. Any cells with <200 or >6000 genes were excluded to select for single live cells. The NormalizeData and ScaleData functions in Seurat were used after merging (Satija et al. 2015; Butler et al. 2018; Stuart et al. 2019; Hao et al. 2021). Merged samples used 16 dimensions and were processed using clustree to determine resolution; a resolution of 0.8 was used (Zappia and Oshlack 2018). The merged datasets were saved as RDS files.

To examine differences between sample groups, integration was used. Samples were integrated using the Seurat integration tutorial (Stuart et al. 2019; Introduction to scRNA-seq integration 2023). For mock, 14-day ligated, and 5-day deligated integration, 40 dimensions were used and a resolution of 0.5 as determined by clustree (Zappia and Oshlack 2018). To perform cluster identifications, previously published and annotated scRNA-seq data from salivary glands was used and compared to a list of differentially expressed genes per cluster from the integrated samples (Hauser et al. 2020; Horeth et al. 2021).

### 1.2.6. CellChat Analysis

CellChat analysis was performed to predict ligand-receptor interactions in homeostatic glands, ligated vs mock glands, ligated vs deligated glands, and mock vs deligated. Analysis was performed using the guided tutorial (Jin et al. 2021; Comparison analysis of multiple datasets using CellChat 2022; Inference and analysis of cell-cell communication using CellChat 2022).

### 1.2.7. Preparation of cryosections

Submandibular and sublingual salivary glands were removed together, weighed, and fixed in 4% paraformaldehyde (PFA) in 1X phosphate buffered saline (1X PBS) for 2 hours at 4°C. Glands were washed in three consecutive 1X PBS washes, and incubated in an increasing sucrose gradient progressing from 5%, to 10%, to 15% sucrose for 1 hour each and then transferred to 30% sucrose overnight at 4°C. After overnight incubation, glands were transferred to 15% sucrose, 50% tissue freezing media (Tissue Freezing Medium, Electron Microscopy Sciences) overnight at 4°C. Glands were transferred to tissue freezing medium and then frozen over liquid nitrogen. Ten-micron (10 µm) sections were collected on Superfrost Plus glass slides (Electron Microscopy Sciences) using a Leica CM 1860 cryostat. Slides were collected by taking three consecutive 10 µm sections per slide until the entire tissue was sectioned. Sections were dried for 30 minutes at room temperature and stored at −80°C. Slides were returned to room temperature before staining.

### 1.2.8. Immunohistochemistry (IHC)

Slides were removed from −80°C storage and brought to room temperature and washed in 1X PBS for five minutes. Next, slides were post-fixed in 4% PFA in 1X PBS for 18 minutes at room temperature followed by two five-minute washes in 1X PBS. Slides were permeabilized in 0.5% Triton-X-100 in 1X PBS for 18 minutes, followed by three five-minute washes in 1X PBS. Sections on slides were encircled with a hydrophobic pen and blocked using 10% donkey serum in 3% bovine serum albumin (BSA) in 1X PBS for 60 minutes at room temperature in a humidified chamber. Blocking solution was dabbed off with a Kimtech wipe. Primary antibodies were diluted in 3% BSA in 1X PBS and incubated on sections overnight at 4°C. The following primary antibodies and dilutions were used: anti-goat red fluorescent protein Rockland Biosciences (cat:200-101-379 lot: 37733) 1:1000, and anti-rat CD31 BD Pharmigen (cat:553370 lot: 1005853) 1:200. Three five-minute washes were performed after each round of primary antibody. Secondary antibodies (Jackson ImmunoResearch) were applied at a concentration of 1:500 in 3% BSA in 1X PBS for 60 minutes at room temperature, followed by three five-minute 1X PBS washes. DAPI staining (Life Technologies, 1 ug/ml) was performed for ten minutes followed by two additional five-minute 1X PBS washes. Coverslips were applied with a glycerol-based mounting media containing 4% n-propyl gallate and triethylenediamine (DABCO) as an antifade (Nelson et al. 2013).

### 1.2.9. Image processing and quantification

For quantification of each marker by IHC, sections obtained from similar tissue depths and similar regions of the submandibular glands were compared. Both the submandibular and sublingual glands were stained, but only the submandibular gland was imaged and included in quantifications. For IHC staining, sections at approximately 40% tissue depth were stained, imaged, and quantified. On average, three images were collected, one from the proximal, medial, and distal regions of the submandibular gland from each of three tissue sections on a slide, for a total of 9 images per mouse. IHC sections were imaged on a Zeiss Z1 Cell Observer widefield with an Axio712 mono camera (Carl Zeiss, LLC) using a Plan-Neofluar 20X/0.50 Ph2 M27 objective. All images for one imaging session were captured using one microscope with identical microscope settings and regions containing folds, tears, or unequal staining were avoided.

All image processing and quantification was performed using the freeware, FIJI, an imaging processing package of ImageJ (Schindelin et al. 2012). To estimate the area positive for the expression of a marker, the image was background subtracted using rolling ball background subtraction with a 25-pixel rolling ball radius and disabled smoothing. Images from each day imaged on the same microscope were equally thresholded, and FIJI was used to quantify the tissue area that was positive for a given marker relative to total tissue area. For fluorescent images, the green channel with no background subtraction was used to detect the total issue area by autofluorescence.

### 1.2.10. Statistical analysis

All tabulation of quantifications was performed in Microsoft Excel with graphs generated and statistics run using GraphPad Prism. For comparisons of more than two conditions, an Ordinary one-way analysis of variance (ANOVA) with Tukey’s multiple comparisons test was performed using GraphPad Prism version 9.5.1 for Mac or Windows, GraphPad Software, San Diego, California USA, www.graphpad.com. Statistical significance is indicated with stars over the corresponding bars with p values indicated in the figure legends. Analyses with no statistical significance (p > 0.05) are unmarked.

## 1.3. Results

### 1.3.1. Characterization of homeostatic salivary gland endothelial cells

To profile the transcriptomes of adult salivary gland endothelial cells, we performed a stromal cell enrichment on pooled submandibular and sublingual salivary glands from 12-week-old female C57BL/6J mice followed by scRNA-seq (**Appendix Figure 1A**). We sequenced a total of 23,434 cells, which were grouped into 27 clusters using Seurat, with a majority of the cells sequenced being endothelial (**Figure 1A, Appendix Figure 1B, Appendix Table 1**). To identify endothelial subtypes based on their transcriptional differences, we used a list of the most conserved endothelial cell subtype markers (Kalucka et al. 2020) and generated a heatmap (**Appendix Figure 1C**). A majority of endothelial cells were capillary endothelial cells (85%), 13% were arterial, and 2% were venous, with no lymphatic endothelial cells detected (**Figure 1B, Appendix Figure 1C).**

**Figure 1.**
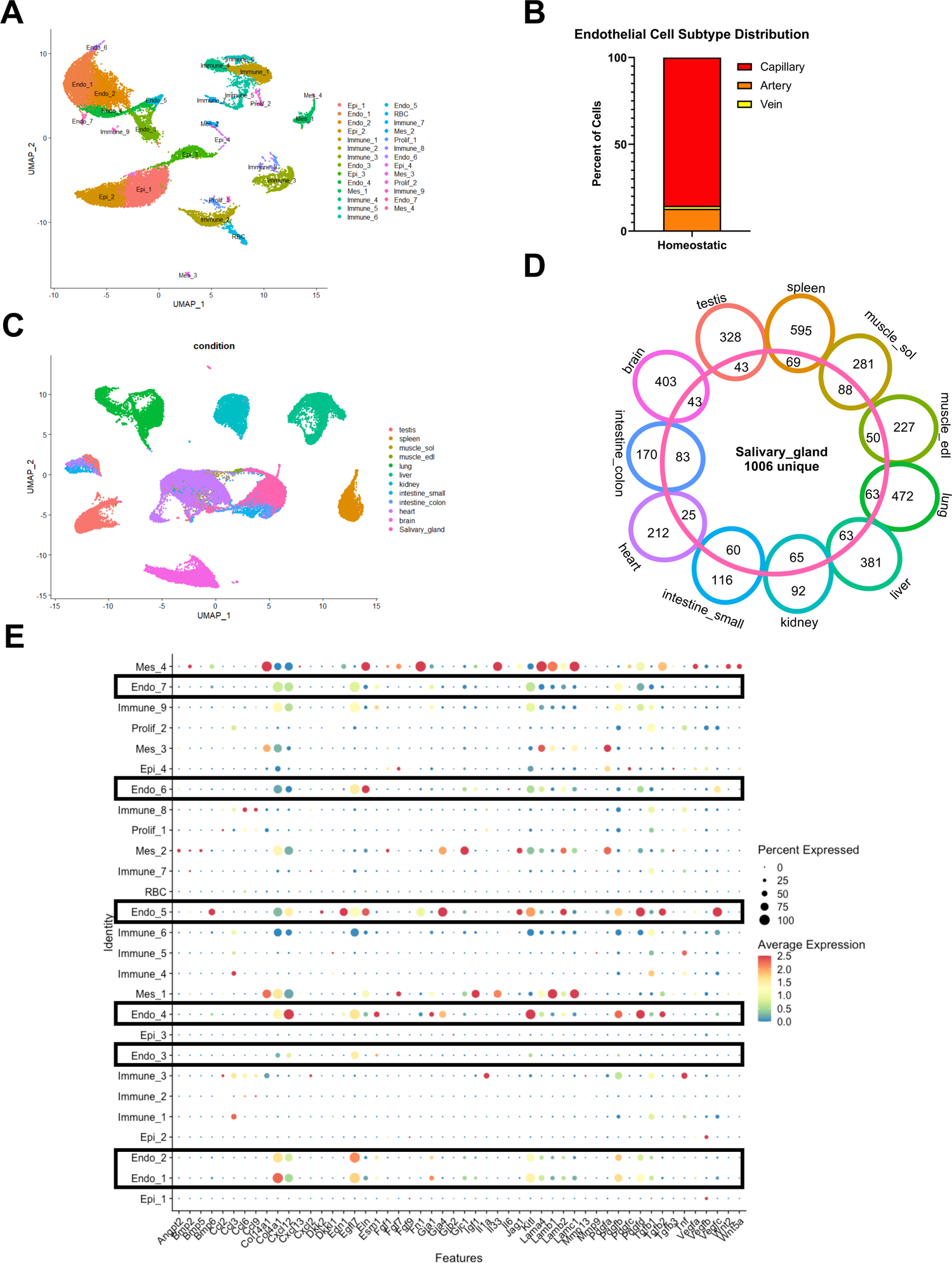
Profiling the homeostatic salivary gland endothelial cell transcriptome. (A) Uniform Manifold Approximation and Projection (UMAP) for two technical replicates from 12-week-old C57BL/6J female mice which were merged, and cell populations were labeled using the following abbreviations: endothelial cells (Endo), epithelial cells (Epi), immune cells (Immune), stromal/mesenchymal cells (Mes), red blood cells (RBC), glial cells (Glial), and dividing cells (Prolif). (B) Graph showing the relative percentages of endothelial cell subtypes out of the total endothelial cells sequenced. (C) Uniform Manifold Approximation and Projection (UMAP) of 11 different organ’s endothelial cells integrated with salivary gland endothelial cells which were subsetted from a stromal-enriched dataset using expression of endothelial cell marker genes and colored based on the organ it was isolated from. (D) Venn diagram showing the number of shared and non-shared genes from each of the organ’s endothelial cells compared to salivary gland endothelial cells, with the total number of unique salivary gland endothelial cell genes in the center of the diagram using the RNA assay on the integrated data. (E) Dot plot of putative angiocrine factors in a homeostatic control submandibular salivary gland with endothelial cell clusters indicated by a black box.

To compare salivary gland endothelial cell transcriptomes to other organs, the endothelial cells were subsetted from the stromal-enriched dataset. The subsetted salivary gland endothelial cells were integrated with published endothelial cell transcriptomes from the brain, heart, colon, small intestine, kidney, liver, lung, extensor digitorum longus muscle, soleus muscle, spleen, and testis (Kalucka et al. 2020) (**Figure 1C**). Salivary gland endothelial cells clustered most closely with colon endothelial cells and a subset of small intestine endothelial cells (**Figure 1C**). The top 10 marker genes were largely unique per organ, with some overlap between salivary glands and colon (**Appendix Figure 2A**). Salivary gland endothelial cells expressed 1006 unique genes that were not shared by any other organ-specific endothelial cell (**Figure 1D, Appendix Table 2**) but shared 83 differentially expressed genes with the colon. In addition, the salivary gland endothelial cells showed significant overlap with endothelial cells of the small intestine (60 differentially expressed genes) and kidney (65 differentially expressed genes) (**Figure 1D**).

**Figure 2.**
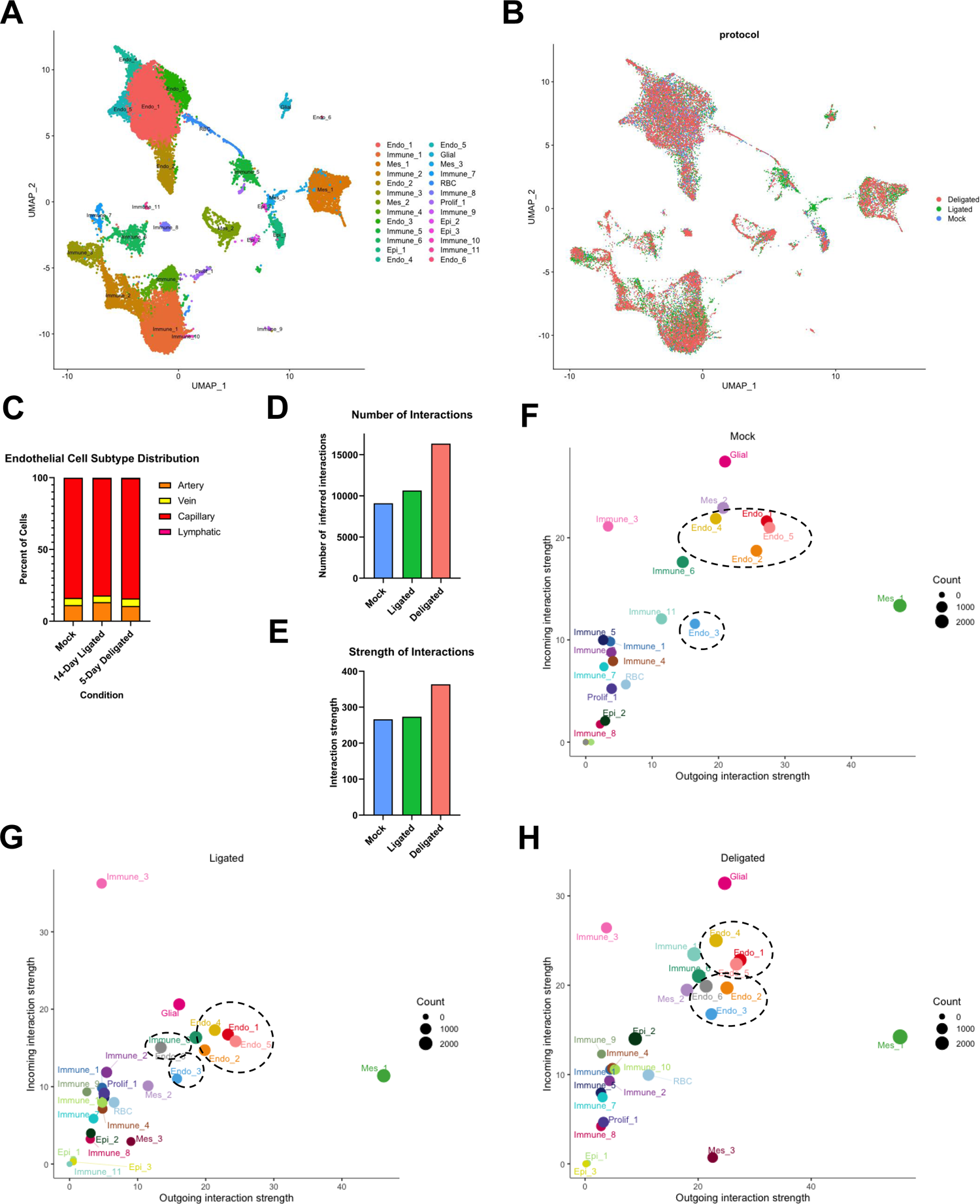
Endothelial cells are important signal senders and receivers in the salivary gland following surgical manipulation. (A) Uniform Manifold Approximation and Projection (UMAP) of stromal-enriched submandibular and sublingual gland mock surgery control, 14-day ligated, and 5-day deligated samples that were integrated and cell populations were labeled using the following abbreviations: endothelial cells (Endo), epithelial cells (Epi), immune cells (Immune), stromal/mesenchymal cells (Mes), red blood cells (RBC), glial cells (Glial), and dividing cells (Prolif). (B) UMAP of the integrated samples in (A) color coded based on the surgical condition. (C) Graph showing the relative percentages of endothelial cell subtypes out of the total endothelial cells sequenced per condition. Bar graph of the (D) number of inferred interactions predicted with CellChat and (E) interaction strength. Graphs of incoming and outgoing interaction strength generated using CellChat grouped by cluster with endothelial cell clusters noted with a black dotted circle in (F) Mock, (G) Ligated, and (H) Deligated.

### 1.3.2. Shifts in cell abundances following surgical manipulation

To identify alterations in the transcriptomes of adult salivary gland endothelial cells following fibrotic injury and regeneration, we used a reversible model of fibrotic injury, in which a ligature is applied to the primary ducts of the submandibular and sublingual salivary glands, to induce injury (Woods et al. 2015; Altrieth et al. 2023) or a regenerative response with ligature removal (Cotroneo et al. 2008; Cotroneo et al. 2010; Aure et al. 2015). We harvested the salivary glands that had been subjected to injury or allowed to repair and compared them to a mock surgery control. Mock surgery (Mock), 14-day ligation surgery (14-Day Ligation), or 14-day ligation followed by 5-day deligation surgery (5-Day Deligation) samples were prepared using stromal-cell enrichment (**Appendix Figure 1A**) and subjected to scRNA-seq. Two biological replicates for the 5-Day Deligation were merged and integrated with the Mock and 14-Day Ligation samples. After integration, we identified 26 cell populations using Seurat (**Figure 2A, B**). Since relative endothelial cell abundance was altered following ligation and deligation (**Appendix Figure 3A**), we compared the abundance of particular endothelial subtypes (Kalucka et al. 2020) with a heatmap (**Appendix Figure 3B**). Endo_1, Endo_3, and Endo_5 were identified as capillary populations, Endo_2 was identified as an artery population, Endo_4 was identified as a vein population, and Endo_6 was identified as a lymphatic population (**Figure 2C, Appendix Figure 3B**). As the overall percent distribution of each endothelial cell subtype was similar across all conditions, transcriptomic changes, rather than cell-type abundance changes, may contribute to fibrosis and regeneration.

**Figure 3.**
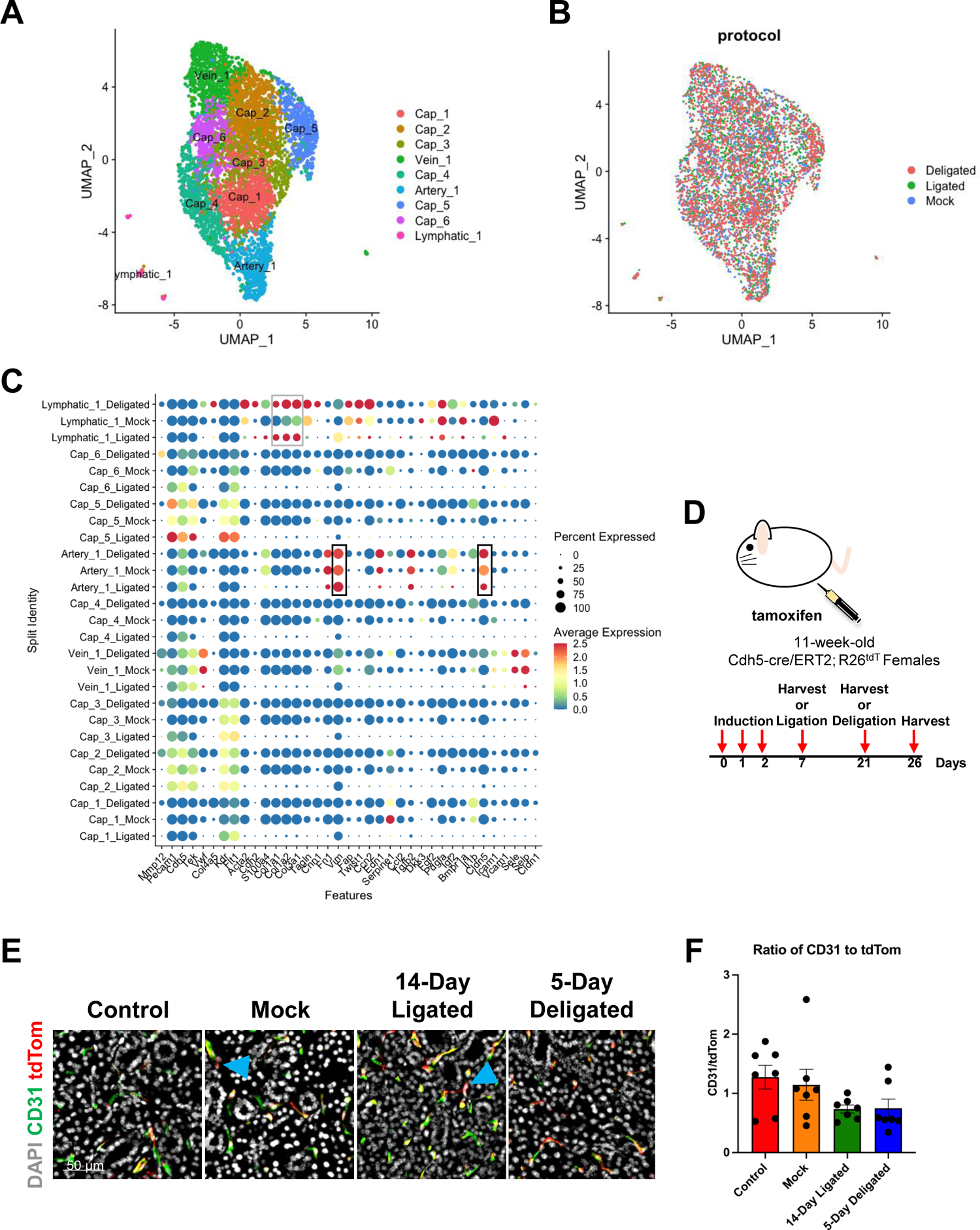
Small subpopulation of endothelial cells show transcriptomic and protein changes consistent with partial or full endothelial to mesenchymal transition. (A) Uniform Manifold Approximation and Projection (UMAP) of stromal-enriched submandibular and sublingual gland mock surgery control, 14-day ligated, and 5-day deligated samples that were integrated and subsetted for endothelial cells using the following gene expression cutoffs for subsetting prior to processing: *Ptprc* < 0.25, *Cdh5*, *Tie1*, and *Kdr* > 1, *Pecam1* > 1.5, *Hba-a1*, *Hba-a2*, and Hbb-bs < 1. (B) UMAP of the integrated samples in (A) color coded based on the surgical condition. (C) Dot plot of genes associated with endothelial to mesenchymal transition in surgically manipulated and endothelial cell subclustered glands, split based on condition with black boxes indicating increased vimentin expression and decreased percent of cells expressing claudin5 in Artery_1 and grey boxes indicating expression of collagen genes in Lymphatic_1 following surgical manipulation. (D) Schematic for induction and surgery time points for Cdh5-cre/ERT2; R26^tdT^ mice with induction occurring at 11-weeks-old for three consecutive days 1 week prior to harvest or surgery. (E) Immunohistochemistry for CD31 (endothelial cells) in green and tdTomato (Cdh5-cre/ERT2; R26^tdT^-derived cells) in red with nuclear staining (DAPI) in blue with blue arrows indicating instances of cells that are Cdh5-cre/ERT2; R26^tdT^-derived, but no longer retain expression of CD31. (F) Graph showing the average ratio of CD31 to tdTom area. N=7 per condition. Error bars: S.E.M. One-way ANOVA followed by Tukey’s multiple comparisons test was performed using GraphPad Prism version 9.5.1.

### 1.3.3. Endothelial cell subsets may undergo a partial endoMT

To look for evidence of a partial endoMT, we subsetted the endothelial cells (**Figure 3A, B**), and generated a dot plot using marker genes that were previously associated with endoMT (Piera-Velazquez and Jimenez 2019) (**Figure 3C**). Artery_1 had increased expression of vimentin (*Vim)* and a decreased percentage of cells expressing claudin 5 (*Cldn5)* (**Figure 3C, black boxes**). In addition, Lymphatic_1 had increased expression of collagen1a1, collagen1a2, and collagen3a2 (*Col1a1, Col1a2*, and *Col3a2)* in both the ligated and deligated samples and had a very distinct transcriptome (**Figure 3C, grey box**). Together, this suggests that following injury, the Atery_1 and Lymphatic_1 cell subsets may undergo a partial endoMT and do not return to homeostatic conditions within 5 days of recovery.

To detect endoMT at the protein level, we utilized an endothelial cell lineage tracing mouse model in which Cre-ERT2 fusion protein is expressed under control of the Cdh5 promoter and is crossed with a tdTomato reporter strain (Cdh5-cre/ERT2; R26^tdT^). Mice were induced with tamoxifen to label Cdh5-expressing cells and their progeny with tdTomato (**Figure 3D**). Cdh5-cre/ERT2; R26^tdT^ glands from 1-week induced (Control), 14-day mock surgery-induced (Mock), 14-Day ligated (14-Day Ligated), and 5-Day deligated (5-Day Deligated) mice were immunostained to detect tdTomato in red, labeling endothelial-derived cells, and CD31 in green, labeling endothelial cells, with nuclear staining (DAPI) in grey (**Figure 3E**). We quantified the ratio of CD31 to tdTom stained area as an indirect readout; 14-Day Ligated and 5-Day Deligated had non-significantly lower ratios (**Figure 3F**). Overall, this data confirms that Cdh5-expressing cells generally retain endothelial cell gene expression following surgical manipulation.

### 1.3.4. Putative salivary gland angiocrine factors

Angiocrine factors play important roles in promoting regeneration and are altered in response to injury (Ding et al. 2010; Nolan et al. 2013; Ding et al. 2014; Rafii et al. 2016). We hypothesized that endothelial cells produce angiocrine factors following deligation to stimulate regeneration. To address this hypothesis, we generated dot plots of reported angiocrine factors (Nolan et al. 2013) in surgically manipulated glands (**Figure 4A**). Similar to homeostatic states, the arterial population expressed the largest number of angiocrine factors (**Figures 1E, 4A**). We searched for cell types that increased expression of and/or showed increased percentages of cells that expressed angiocrine receptors in deligation compared to ligation (**Figure 4B**). The receptor for chemokine C-X-C motif 12 (*Cxcl12)*, atypical chemokine receptor 3 (*Ackr3)*, was expressed by all endothelial clusters. The receptor for endothelin 1 (*Edn1)*, endothelin receptor type a (*Ednra)*, was expressed in Mes_2, the pericyte population. The receptors for platelet derived growth factor alpha (*Pdgfa)*, platelet derived growth factor receptors alpha and beta (*Pdgfra* and *Pdgfrb)*, were both expressed by Mes_1, while *Pdgfrb* was only expressed by the Mes_2 pericyte population. *Notch3*, the receptor for jagged canonical notch ligand 1 (*Jag1)*, was also expressed by the Mes_2 pericytes.

**Figure 4.**
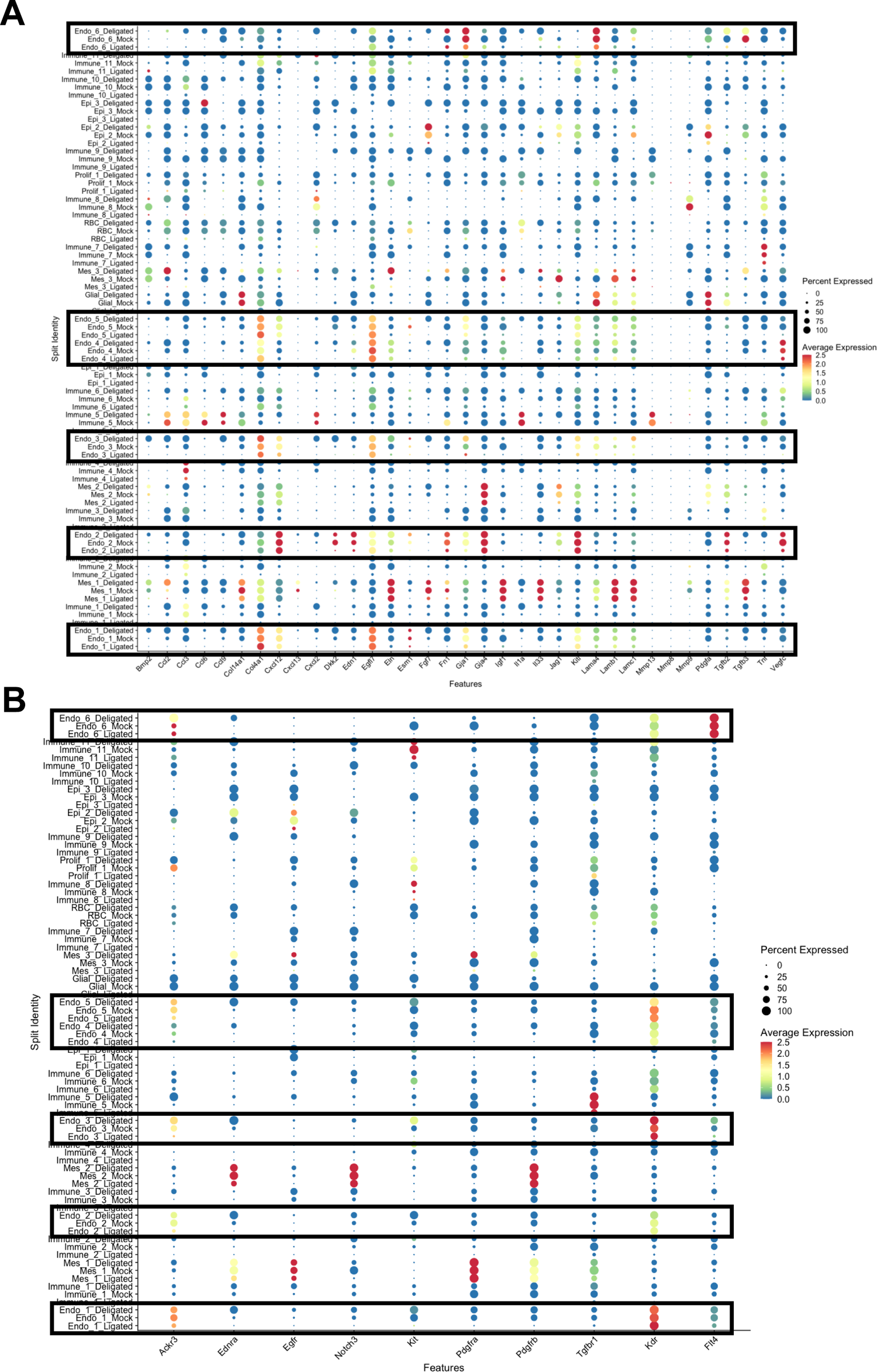
Putative angiocrine factor expression in submandibular salivary glands. (A) Dot plot of putative angiocrine factors in Mock, 14-Day Ligated (Ligated), and 5-Day Deligated (Deligated) integrated stromal-enriched data set with endothelial cell clusters indicated by a black box. (B) Dot plot of receptors for putative angiocrine factors that were expressed in endothelial cell clusters within the Mock, 14-Day Ligated (Ligated), and 5-Day Deligated (Deligated) integrated stromal-enriched data set.

Transforming growth factor beta receptor 1 (*Tgfbr1)*, the receptor for transforming growth factor beta 2 (*Tgfb2)*, was expressed by Immune_5. Kinase insert domain receptor (Kdr) and vascular endothelial growth factor receptor 3 (Flt4) are both receptors for vascular endothelial growth factor c (Vegfc). *Kdr* was expressed by all endothelial subpopulations, while *Flt4* was expressed by Endo_6, the lymphatic population. The receptor for KIT ligand (*Kitl)*, *Kit*, was expressed by Endo_3, Immune_4, Epi_1, and Immune_8. The receptor for C-C motif chemokine ligand (*Ccl2*), C-C motif chemokine receptor 2 (*Ccr2*), is expressed by many immune cell populations. These data suggests that endothelial-pericyte, endothelial-endothelial, and endothelial-immune interactions may play a functional role in regeneration.

### 1.3.5. Subsets of cell signaling pathways are increased in ligation and deligation

As proper cell-cell communication is important for maintaining gland function, and alterations in communication pathways can promote dysfunction, we sought to identify how cell-cell signaling pathways were altered following injury or recovery from injury. We used CellChat to take an unbiased approach to examine the number and strength of interactions and to predict senders and receivers of signaling. Endothelial cells had the highest number of inferred interactions and interaction strength in deligation, followed by ligation, and mock (**Figure 2D, E**). Endothelial cells were found to have both high incoming and outgoing interaction strength in all three conditions compared to all other cell types but overall signaling was lowest with ligation (**Figure 2F-H**). These data suggest that critical endothelial signals may be lost following fibrotic injury and restored with recovery.

While endothelial cell incoming and outgoing signaling is generally decreased in ligation, pathways that promote fibrosis may increase. We identified putative pro-fibrotic pathways in endothelial cells by identifying overall signaling pattern increases in ligated endothelial cells compared to mock. In Endo_1, amyloid beta precursor protein (APP), semaphorin 4 (SEMA4), angiopoietin like (ANGPTL), tumor necrosis factor (TNF), and tumor necrosis factor ligand superfamily member 13 (APRIL) signaling increased in relative strength, with a larger increase in semaphorin 6 (SEMA6), myelin protein zero (MPZ), protein tyrosine phosphatase receptor type m (PTPRM), and neurotrophin (NT) (**Figure 5A, B**). Endo_2 showed increased strength in fas ligand (FALSG), MPZ, vascular endothelial growth factor (VEGF), and semaphorin 7 (SEMA7) (**Figure 5A, B**). Endo_3 had increased endothelial cell adhesion molecule (ESAM) signaling (**Figure 5A, B**). Endo_4 had increased signaling strength for APRIL, insulin growth factor (IGF), PTPRM, TNF, and ANGPTL (**Figure 5A, B**). Endo_5 had increased strength of PTPRM and nicotinamide phosphoribosyltransferase (VISFATIN) (**Figure 5A, B**). PTPRM signaling increased in strength in half of the endothelial cell clusters. MPZ, TNF, and APRIL had increased strength in one third of the endothelial cell clusters, while other pathways were unique to a particular cluster, suggesting that endothelial cell subsets may have some conserved, and some unique responses to injury.

**Figure 5.**
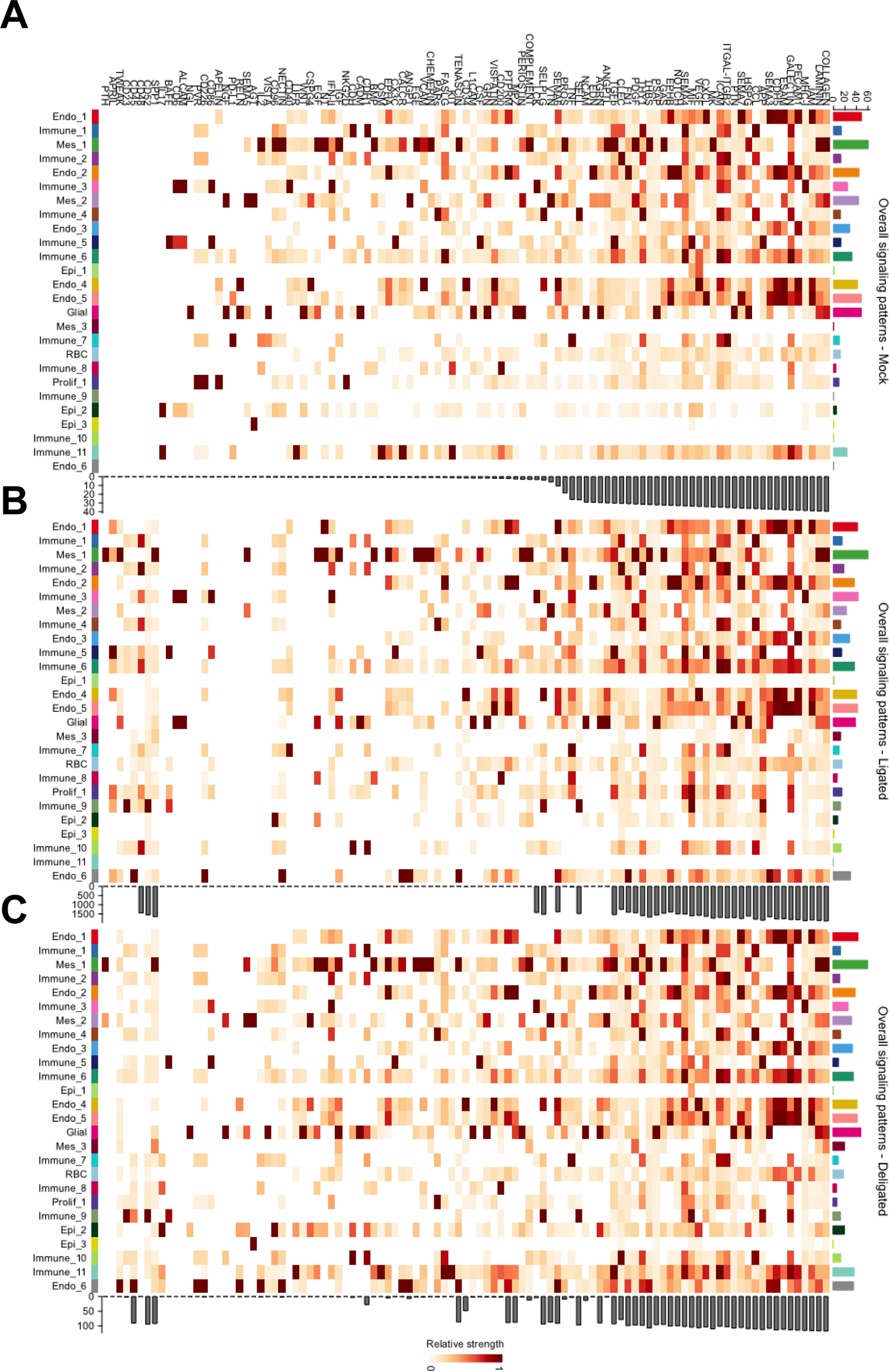
Alterations in cell communication pathways following surgical manipulation Overall signaling pattern heatmap generated using CellChat in (A) Mock, (B) Ligated, and (C) Deligated with pathways listed across the top and cell populations shown on the left, with darker colors indicating a higher relative strength.

Using pathways that were increased in strength with ligation compared to mock (**Figure 5A, B**), we examined ligand-receptor pairs using CellChat to predict sources and targets in multiple endothelial populations. In PTPRM signaling, interactions were predicted to occur with endothelial cells being both the source and target, with both populations expressing *Ptrpm*. TNF signaling interactions were predicted to occur with Immune_2, 3, 5, 7, and 8, and Prolif_1 being sources of *Tnf* with all endothelial cell clusters, Mes_1, and Mes_2 being the predicted target populations expressing the receptor, tumor necrosis factor receptor superfamily member 1A *(Tnfrsf1a)*. APRIL signaling interactions were predicted to occur between Endo_1, 4, and 6, Mes_1, and Immune_6 as sources of tumor necrosis factor ligand superfamily member 13 *(Tnfsf13)* with Immune_5 and 9, and Prolif_1 being target cell populations expressing the receptor, tumor necrosis factor receptor superfamily member 13B (*Tnfrsf13b)*. MPZ interactions were predicted to occur between myelin protein zero like 1 (*Mpzl1)* and *Mpzl1,* both on endothelial cells. Multiple endothelial cell subpopulations are sources of PTPRM, APRIL, and MPZ signaling and are targets of TNF signaling, which may promote fibrosis.

Next, we sought to identify pro-regenerative signaling pathways using CellChat, by identifying pathways which were increased in strength in deligated endothelial cells compared to ligated. In Endo_1, chemokine C-X-C motif ligand (CXCL), EPH receptor A (EPHA), and EPH receptor B (EPHB) were increased with deligation (**Figure 5B, C**). EPHA and fibronectin 1 (FN1) were increased in deligation in Endo_2 (**Figure 5B, C**). Chondroitin sulfate proteoglycan 4 (CSPG4) and VISFATIN were increased in deligation in Endo_4 (**Figure 5B, C**). EPHA and CXCL were increased in deligation in Endo_5. ESAM, thrombospondin 1 (THBS), thy-1 cell surface antigen (THY1), SEMA6, MPZ, interleukin 2 (IL2), reelin (RELN), TNF-related weak inducer of apoptosis (TWEAK), protein tyrosine phosphatase receptor type c (CD45), and PVR cell adhesion molecule (PVR) were increased in deligation in Endo_6 (**Figure 5B, C**). We used CellChat to identify predicted sources and targets for CXCL and EPHA, the pathways conserved in one half to one third of the endothelial cell clusters. CXCL predicted interactions were between *Cxcl12* from endothelial cells, mesenchymal cells, and immune cells and atypical chemokine receptor 3 (*Ackr3)* on Endo_1, Endo_5, and Immune_11. EPHA interactions were predicted to occur with endothelial cells as a source of ephrin a1 (*Efna1)* signaling with Mes_1 a predicted target, expressing the receptor EPH receptor a3 (*Epha3).* Endothelial cells may be acting as sources of both CXCL and EPHA signaling and predicted targets of CXCL signaling to promote regeneration.

## 1.4. Discussion

We used a stromal enrichment strategy combined with scRNA-seq to profile the transcriptome of salivary gland endothelial cells, which were neglected in prior studies (Kalucka et al. 2020). We sequenced endothelial cells, along with other cell types they may be interacting with, including pericytes (**Figure 1A, Appendix Figure 1A, B**). We also compared the transcriptome of salivary gland endothelial cells to 11 other organ’s endothelial cells. Although we identified gland-specific expression patterns, we identified high overlap of the salivary gland endothelial cell gene expression with other fenestrated endothelial organs, like the kidney, small intestine, and colon endothelial cells (**Figure 1D**). This may be due to the shared structure and function of these cells (Abe et al. 1984; Oki et al. 1999; Pi et al. 2018; Gifre-Renom et al. 2022). This provides the first characterization of salivary gland endothelial cells during homeostasis and allows for future studies examining transcriptomic changes following salivary gland manipulation.

Endothelial cells can contribute directly to fibrosis by undergoing endoMT or indirectly through secretion of pro-fibrotic and pro-inflammatory mediators (Sun et al. 2020). Studies in the lung, kidney, heart, and liver have implicated endoMT in the fibrotic phenotype with varying degrees (Zeisberg et al. 2007; Zeisberg et al. 2008; Li et al. 2009; Hashimoto et al. 2010; Piera-Velazquez and Jimenez 2019). With scRNA-seq, we found that Artery_1 showed increased *Vim* expression and decreased percent of cells expressing *Cldn5* following ligation, suggestive of a partial endoMT (**Figure 3B**). There is also a small lymphatic population that expresses collagen genes in both ligated and deligated conditions; however, this is a very small population of less than 30 cells, and further studies to increase the overall abundance of this population would be needed. Previous studies have implicated lymphangiogenesis in kidney and lung fibrosis, suggesting that this population may be of interest for understanding fibrotic mechanisms (El-Chemaly et al. 2009; Zhang et al. 2021). Using an endothelial cell lineage-reporting mouse, we found little evidence for an endoMT, similar to our prior work revealing little evidence for a myofibroblast transition in stromal cells in this same injury model (Altrieth et al. 2023).

Using an unbiased approach to determine indirect contributions to fibrosis via cell-cell communication, we determined that endothelial cells may be acting as sources of PTPRM, APRIL, and MPZ signaling and targets of TNF signaling to promote fibrosis (**Figure 5A, B**). PTPRM is associated with poor prognosis in cervical cancer by promoting epithelial to mesenchymal transition (Liu et al. 2020). Due to high overlap between epithelial to mesenchymal transition and endoMT, PTPRM signaling may also be promoting endoMT. Following spinal cord injury, APRIL was found to increase, and genetic deletion reduced scar formation through decreased expression of TNFα and CCL2, with reduced macrophage and B cell infiltration (Funk et al. 2016). APRIL serum levels in the blood were found to be higher in patients with primary Sjögren’s Syndrome compared to healthy individuals (Jonsson et al. 2005). MPZL1 increased cell proliferation, migration, and invasion of cancer cells, promoting metastasis (Jia et al. 2014; Chen et al. 2019; Liu et al. 2019). TNFα stimulated collagen 1 accumulation and decreased collagen degradation in intestinal fibrosis, and inhibition of TNFα in kidney fibrosis decreased collagen I and III expression (Theiss et al. 2005; Taguchi et al. 2021). These studies suggest that endothelial cells may be contributing both directly and indirectly to salivary gland ductal ligation-induced fibrosis.

Endothelial cells promote regeneration in other organs through secretion of angiocrine factors. We identified a salivary gland angiocrine factor profile during homeostasis and following deligation, which may be important for promoting regeneration (**Figure 4A, B**). We also took an unbiased approach using CellChat. This analysis predicted endothelial-pericyte, endothelial-endothelial, and endothelial-immune interactions as important for regeneration. Specifically, endothelial cells are predicted to be sources of both CXCL and EPHA signaling and targets of CXCL signaling to promote regeneration (**Figure 5B, C**). Chemokines, including CXCL chemokines, have a unique role in inflammation and regeneration, where too much or sustained signaling can be inhibitory for regeneration, and certain levels are necessary to promote regeneration (Van Sweringen et al. 2011; Yamashiro et al. 2020). EPHA signaling is a guidance cue during optic nerve regeneration following optic crush in goldfish (Rodger et al. 2004). These studies suggest that endothelial cells may promote regeneration via signaling to the nerves, which are known to be critical for salivary gland regeneration (Knox et al. 2013; Wang et al. 2021; Li et al. 2022). Endothelial-pericyte interactions are important to endothelial cell homeostasis and new blood vessel formation during regeneration. Pericytes, extend process around the vasculature, especially capillaries, and can regulate blood flow (Attwell et al. 2016).

Pericytes help provide structure to newly forming blood vessels by promoting vascular basement membrane formation and restricting vessel diameter, allowing for increased vessel stability (Stratman et al. 2009). Following surgical manipulation, Endo_1, a capillary subpopulation, had increased *Jag1* expression in mock and deligated glands, which decreased in ligation, suggesting that *Jag1* may be important for regeneration and endothelial cell maintenance. Jag1 and Jag2 are ligands for Notch signaling, which bind to the Notch family of receptors. Previous studies have implicated Notch signaling in vessel stability regulation and proliferation. One study found that NOTCH3 in pericytes leads to expression of DLL4, which activates NOTCH1 in endothelial cells, highlighting that NOTCH signaling is important in both the pericytes and endothelial cells (Tefft et al. 2022). Another study found that *Jag1* expression and Notch signaling was required for endothelial sprouting and pericyte growth *in vitro*, implying that this cell-cell communication event may be important for proper vessel formation (Tattersall et al. 2016). Endothelial Jag2 is also important for promoting hematopoietic stem cell recovery following myelosuppression (Guo et al. 2017). Our data suggests that Notch signaling interactions between pericytes and endothelial cells may be important for salivary gland homeostasis and regeneration.

## 1.7. Author Contributions

A.A., D.N., and M.L. designed experiments, analyzed data, assembled figures, and wrote and revised the manuscript. A.A., E.S., and S.G. performed experiments, analyzed data, and revised the manuscript. All authors gave their final approval and agree to be accountable for all aspects of the work.

## Acknowledgments

The authors are grateful to Douglas Cohn, DVM, and Timothy Quinn for helpful suggestions, insight, and assistance in implementing this surgical model. Research reported in this publication was supported by the National Institute of Dental & Craniofacial Research of the National Institutes of Health under Award Numbers F31DE029688 to A.A., R01DE027953, R01DE030626, and R21DE027571 to M.L. and the RNA Institute for RNA Fellows funding for A.A.. The content is solely the responsibility of the authors and does not necessarily represent the official views of the National Institutes of Health.

## 1.8. Data Availability Statement

All datasets for this study can be found in GEO under SuperSeries GSE226640 and GSE235677. Scripts used to analyze the data can be found on GitHub (https://github.com/MLarsenLab).

## References

Abe K, Takano H, Ito T, Liss R. 1984. Microvasculature of the mouse epididymis, with special reference to fenestrated capillaries localized in the initial segment.

Adhikari R, Soni A. 2022. Submandibular Sialadenitis And Sialadenosis. In: StatPearls. Treasure Island (FL): StatPearls Publishing. [accessed 2022 Jul 19]. http://www.ncbi.nlm.nih.gov/books/NBK562211/.

Altrieth AL, O’Keefe KJ, Gellatly VA, Tavarez JR, Feminella SM, Moskwa NL, Cordi CV, Turrieta JC, Nelson DA, Larsen M. 2023. Identifying fibrogenic cells following salivary gland obstructive injury. Frontiers in Cell and Developmental Biology. 11. [accessed 2023 May 23]. https://www.frontiersin.org/articles/10.3389/fcell.2023.1190386.

Amano O, Mizobe K, Bando Y, Sakiyama K. 2012. Anatomy and Histology of Rodent and Human Major Salivary Glands. Acta Histochem Cytochem. 45(5):241–250. doi:10.1267/ahc.12013.

Attwell D, Mishra A, Hall CN, O’Farrell FM, Dalkara T. 2016. What is a pericyte? J Cereb Blood Flow Metab. 36(2):451–455. doi:10.1177/0271678X15610340.

Aure MH, Konieczny SF, Ovitt CE. 2015. Salivary Gland Homeostasis Is Maintained through Acinar Cell Self-Duplication. Developmental Cell. 33(2):231–237. doi:10.1016/j.devcel.2015.02.013.

Bookman AAM, Shen H, Cook RJ, Bailey D, McComb RJ, Rutka JA, Slomovic AR, Caffery B. 2011. Whole stimulated salivary flow: Correlation with the pathology of inflammation and damage in minor salivary gland biopsy specimens from patients with primary Sjögren’s syndrome but not patients with sicca. Arthritis & Rheumatism. 63(7):2014–2020. doi:10.1002/art.30295.

Butler A, Hoffman P, Smibert P, Papalexi E, Satija R. 2018. Integrating single-cell transcriptomic data across different conditions, technologies, and species. Nat Biotechnol. 36(5):411–420. doi:10.1038/nbt.4096.

Chen D, Cao L, Wang X. 2019. MPZL1 promotes tumor cell proliferation and migration via activation of Src kinase in ovarian cancer. Oncol Rep. 42(2):679–687. doi:10.3892/or.2019.7199.

Comparison analysis of multiple datasets using CellChat. 2022. [accessed 2023 Mar 26]. https://htmlpreview.github.io/?https://github.com/sqjin/CellChat/blob/master/tutorial/Comparison_analysis_of_multiple_datasets.html.

Cotroneo E, Proctor GB, Carpenter GH. 2010. Regeneration of acinar cells following ligation of rat submandibular gland retraces the embryonic-perinatal pathway of cytodifferentiation. Differentiation. 79(2):120–130. doi:10.1016/j.diff.2009.11.005.

Cotroneo E, Proctor GB, Paterson KL, Carpenter GH. 2008. Early Markers of Regeneration Following Ductal Ligation in the Rat Submandibular Gland. Cell Tissue Res. 332(2):227–235. doi:10.1007/s00441-008-0588-6.

Demirci MS, Karabulut G, Gungor O, Celtik A, Ok E, Kabasakal Y. 2016. Is There an Increased Arterial Stiffness in Patients with Primary Sjögren’s Syndrome? Intern Med. 55(5):455–459. doi:10.2169/internalmedicine.55.3472.

Ding B-S, Cao Z, Lis R, Nolan DJ, Guo P, Simons M, Penfold ME, Shido K, Rabbany SY, Rafii S. 2014. Divergent angiocrine signals from vascular niche balance liver regeneration and fibrosis. Nature. 505(7481):97–102. doi:10.1038/nature12681.

Ding B-S, Nolan DJ, Butler JM, James D, Babazadeh AO, Rosenwaks Z, Mittal V, Kobayashi H, Shido K, Lyden D, et al. 2010. Inductive angiocrine signals from sinusoidal endothelium are required for liver regeneration. Nature. 468(7321):310–315. doi:10.1038/nature09493.

El-Chemaly S, Malide D, Zudaire E, Ikeda Y, Weinberg BA, Pacheco-Rodriguez G, Rosas IO, Aparicio M, Ren P, MacDonald SD, et al. 2009. Abnormal lymphangiogenesis in idiopathic pulmonary fibrosis with insights into cellular and molecular mechanisms. Proceedings of the National Academy of Sciences. 106(10):3958–3963. doi:10.1073/pnas.0813368106.

Funk LH, Hackett AR, Bunge MB, Lee JK. 2016. Tumor necrosis factor superfamily member APRIL contributes to fibrotic scar formation after spinal cord injury. Journal of Neuroinflammation. 13(1):87. doi:10.1186/s12974-016-0552-4.

Gifre-Renom L, Daems M, Luttun A, Jones EAV. 2022. Organ-Specific Endothelial Cell Differentiation and Impact of Microenvironmental Cues on Endothelial Heterogeneity. Int J Mol Sci. 23(3):1477. doi:10.3390/ijms23031477.

Guo P, Poulos MG, Palikuqi B, Badwe CR, Lis R, Kunar B, Ding B-S, Rabbany SY, Shido K, Butler JM, et al. 2017. Endothelial jagged-2 sustains hematopoietic stem and progenitor reconstitution after myelosuppression. J Clin Invest. 127(12):4242–4256. doi:10.1172/JCI92309.

Hao Y, Hao S, Andersen-Nissen E, Mauck WM, Zheng S, Butler A, Lee MJ, Wilk AJ, Darby C, Zager M, et al. 2021. Integrated analysis of multimodal single-cell data. Cell. 184(13):3573–3587.e29. doi:10.1016/j.cell.2021.04.048.

Hashimoto N, Phan SH, Imaizumi K, Matsuo M, Nakashima H, Kawabe T, Shimokata K, Hasegawa Y. 2010. Endothelial–Mesenchymal Transition in Bleomycin-Induced Pulmonary Fibrosis. Am J Respir Cell Mol Biol. 43(2):161–172. doi:10.1165/rcmb.2009-0031OC.

Hauser BR, Aure MH, Kelly MC, Hoffman MP, Chibly AM. 2020. Generation of a Single-Cell RNAseq Atlas of Murine Salivary Gland Development. iScience. 23(12):101838. doi:10.1016/j.isci.2020.101838.

Horeth E, Oyelakin A, Song E-AC, Che M, Bard J, Min S, Kiripolsky J, Kramer JM, Sinha S, Romano R-A. 2021. Transcriptomic and Single-Cell Analysis Reveals Regulatory Networks and Cellular Heterogeneity in Mouse Primary Sjögren’s Syndrome Salivary Glands. Front Immunol. 12:729040. doi:10.3389/fimmu.2021.729040.

Inference and analysis of cell-cell communication using CellChat. 2022. [accessed 2023 Mar 26]. https://htmlpreview.github.io/?https://github.com/sqjin/CellChat/blob/master/tutorial/CellChat-vignette.html.

Introduction to scRNA-seq integration. 2023. [accessed 2023 Mar 26]. https://satijalab.org/seurat/articles/integration_introduction.html.

J. D. Harrison, A. Epivatianos, S. N. Bhatia. 1997. Role of microliths in the aetiology of chronic submandibular sialadenitis: a clinicopathological investigation of 154 cases. Histopathology. 31(3):237–251. doi: https://doi.org/10.1046/j.1365-2559.1997.2530856.x.

Jia D, Jing Y, Zhang Z, Liu L, Ding J, Zhao F, Ge C, Wang Q, Chen T, Yao M, et al. 2014. Amplification of MPZL1/PZR promotes tumor cell migration through Src-mediated phosphorylation of cortactin in hepatocellular carcinoma. Cell Res. 24(2):204–217. doi:10.1038/cr.2013.158.

Jin S, Guerrero-Juarez CF, Zhang L, Chang I, Ramos R, Kuan C-H, Myung P, Plikus MV, Nie Q. 2021. Inference and analysis of cell-cell communication using CellChat. Nat Commun. 12(1):1088. doi:10.1038/s41467-021-21246-9.

Jonsson MV, Szodoray P, Jellestad S, Jonsson R, Skarstein K. 2005. Association Between Circulating Levels of the Novel TNF Family Members APRIL and BAFF and Lymphoid Organization in Primary Sjögren’s Syndrome. J Clin Immunol. 25(3):189–201. doi:10.1007/s10875-005-4091-5.

Kalucka J, de Rooij LPMH, Goveia J, Rohlenova K, Dumas SJ, Meta E, Conchinha NV, Taverna F, Teuwen L-A, Veys K, et al. 2020. Single-Cell Transcriptome Atlas of Murine Endothelial Cells. Cell. 180(4):764–779.e20. doi:10.1016/j.cell.2020.01.015.

Knox SM, Lombaert IMA, Haddox CL, Abrams SR, Cotrim A, Wilson AJ, Hoffman MP. 2013. Parasympathetic stimulation improves epithelial organ regeneration. Nat Commun. 4:1494. doi:10.1038/ncomms2493.

Kwon HR, Nelson DA, DeSantis KA, Morrissey JM, Larsen M. 2017. Endothelial cell regulation of salivary gland epithelial patterning. Development. 144(2):211–220. doi:10.1242/dev.142497.

Leach HG, Chrobak I, Han R, Trojanowska M. 2013. Endothelial Cells Recruit Macrophages and Contribute to a Fibrotic Milieu in Bleomycin Lung Injury. Am J Respir Cell Mol Biol. 49(6):1093–1101. doi:10.1165/rcmb.2013-0152OC.

Leehan KM, Pezant NP, Rasmussen A, Grundahl K, Moore JS, Radfar L, Lewis DM, Stone DU, Lessard CJ, Rhodus NL, et al. 2018. Minor salivary gland fibrosis in Sjögren’s syndrome is elevated, associated with focus score and not solely a consequence of aging. Clin Exp Rheumatol. 36(Suppl 112):80–88.

Li J, Qu X, Bertram JF. 2009. Endothelial-Myofibroblast Transition Contributes to the Early Development of Diabetic Renal Interstitial Fibrosis in Streptozotocin-Induced Diabetic Mice. Am J Pathol. 175(4):1380–1388. doi:10.2353/ajpath.2009.090096.

Li J, Qu X, Yao J, Caruana G, Ricardo SD, Yamamoto Y, Yamamoto H, Bertram JF. 2010. Blockade of Endothelial-Mesenchymal Transition by a Smad3 Inhibitor Delays the Early Development of Streptozotocin-Induced Diabetic Nephropathy. Diabetes. 59(10):2612–2624. doi:10.2337/db09-1631.

Li J, Sudiwala S, Berthoin L, Mohabbat S, Gaylord EA, Sinada H, Cruz Pacheco N, Chang JC, Jeon O, Lombaert IMA, et al. 2022. Long-term functional regeneration of radiation-damaged salivary glands through delivery of a neurogenic hydrogel. Sci Adv. 8(51):eadc8753. doi:10.1126/sciadv.adc8753.

Liu P, Zhang C, Liao Y, Liu J, Huang J, Xia M, Chen M, Tan H, He W, Xu M, et al. 2020. High expression of PTPRM predicts poor prognosis and promotes tumor growth and lymph node metastasis in cervical cancer. Cell Death Dis. 11(8):1–17. doi:10.1038/s41419-020-02826-x.

Liu X, Huang J, Liu L, Liu R. 2019. MPZL1 is highly expressed in advanced gallbladder carcinoma and promotes the aggressive behavior of human gallbladder carcinoma GBC-SD cells. Mol Med Rep. 20(3):2725–2733. doi:10.3892/mmr.2019.10506.

Llamas-Gutierrez FJ, Reyes E, Martínez B, Hernández-Molina G. 2014. Histopathological environment besides the focus score in Sjögren’s syndrome. International Journal of Rheumatic Diseases. 17(8):898–903. doi:10.1111/1756-185X.12502.

Lombaert I, Movahednia MM, Adine C, Ferreira JN. 2017. Concise Review: Salivary Gland Regeneration: Therapeutic Approaches from Stem Cells to Tissue Organoids. Stem Cells. 35(1):97–105. doi:10.1002/stem.2455.

Maruyama CL, Monroe M, Hunt J, Buchmann L, Baker OJ. 2019. Comparing Human and Mouse Salivary Glands: A Practice Guide for Salivary Researchers. Oral Dis. 25(2):403–415. doi:10.1111/odi.12840.

Nolan DJ, Ginsberg M, Israely E, Palikuqi B, Poulos MG, James D, Ding B-S, Schachterle W, Liu Y, Rosenwaks Z, et al. 2013. Molecular Signatures of Tissue-Specific Microvascular Endothelial Cell Heterogeneity in Organ Maintenance and Regeneration. Dev Cell. 26(2). doi:10.1016/j.devcel.2013.06.017. [accessed 2018 Jan 1]. https://www.ncbi.nlm.nih.gov/pmc/articles/PMC3873200/.

Oki S, Desaki J, Taguchi Y, Matsuda Y, Shibata T, Okumura H. 1999. Capillary changes with fenestrations in the contralateral soleus muscle of the rat following unilateral limb immobilization. Journal of Orthopaedic Science. 4(1):28–31. doi:10.1007/s007760050070.

Pi X, Xie L, Patterson C. 2018. Emerging roles of vascular endothelium in metabolic homeostasis. Circ Res. 123(4):477–494. doi:10.1161/CIRCRESAHA.118.313237.

Piera-Velazquez S, Jimenez SA. 2019. Endothelial to Mesenchymal Transition: Role in Physiology and in the Pathogenesis of Human Diseases. Physiol Rev. 99(2):1281– 1324. doi:10.1152/physrev.00021.2018.

R: The R Project for Statistical Computing. 2021. [accessed 2023 Jan 10]. https://www.r-project.org/.

Rafii S, Butler JM, Ding B-S. 2016. Angiocrine functions of organ-specific endothelial cells. Nature. 529(7586):316–325. doi:10.1038/nature17040.

Rodger J, Vitale PN, Tee LBG, King CE, Bartlett CA, Fall A, Brennan C, O’Shea JE, Dunlop SA, Beazley LD. 2004. EphA/ephrin-A interactions during optic nerve regeneration: restoration of topography and regulation of ephrin-A2 expression. Molecular and Cellular Neuroscience. 25(1):56–68. doi:10.1016/j.mcn.2003.09.010.

Satija R, Farrell JA, Gennert D, Schier AF, Regev A. 2015. Spatial reconstruction of single-cell gene expression data. Nat Biotechnol. 33(5):495–502. doi:10.1038/nbt.3192.

Schindelin J, Arganda-Carreras I, Frise E, Kaynig V, Longair M, Pietzsch T, Preibisch S, Rueden C, Saalfeld S, Schmid B, et al. 2012. Fiji: an open-source platform for biological-image analysis. Nat Methods. 9(7):676–682. doi:10.1038/nmeth.2019.

Seurat - Guided Clustering Tutorial. 2023. [accessed 2023 Jan 10]. https://satijalab.org/seurat/articles/pbmc3k_tutorial.html.

Sörensen I, Adams RH, Gossler A. 2009. DLL1-mediated Notch activation regulates endothelial identity in mouse fetal arteries. Blood. 113(22):5680–5688. doi:10.1182/blood-2008-08-174508.

Stratman AN, Malotte KM, Mahan RD, Davis MJ, Davis GE. 2009. Pericyte recruitment during vasculogenic tube assembly stimulates endothelial basement membrane matrix formation. Blood. 114(24):5091–5101. doi:10.1182/blood-2009-05-222364.

Straub JM, New J, Hamilton CD, Lominska C, Shnayder Y, Thomas SM. 2015. Radiation-induced fibrosis: mechanisms and implications for therapy. J Cancer Res Clin Oncol. 141(11):1985–1994. doi:10.1007/s00432-015-1974-6.

Stuart T, Butler A, Hoffman P, Hafemeister C, Papalexi E, Mauck WM, Hao Y, Stoeckius M, Smibert P, Satija R. 2019. Comprehensive Integration of Single-Cell Data. Cell. 177(7):1888–1902.e21. doi:10.1016/j.cell.2019.05.031.

Sun X, Nkennor B, Mastikhina O, Soon K, Nunes SS. 2020. Endothelium-mediated contributions to fibrosis. Seminars in Cell & Developmental Biology. 101:78–86. doi:10.1016/j.semcdb.2019.10.015.

Taguchi S, Azushima K, Yamaji T, Urate S, Suzuki T, Abe E, Tanaka S, Tsukamoto S, Kamimura D, Kinguchi S, et al. 2021. Effects of tumor necrosis factor-α inhibition on kidney fibrosis and inflammation in a mouse model of aristolochic acid nephropathy. Sci Rep. 11(1):23587. doi:10.1038/s41598-021-02864-1.

Tattersall IW, Du J, Cong Z, Cho BS, Klein AM, Dieck CL, Chaudhri RA, Cuervo H, Herts JH, Kitajewski J. 2016. In vitro modeling of endothelial interaction with macrophages and pericytes demonstrates Notch signaling function in the vascular microenvironment. Angiogenesis. 19(2):201–215. doi:10.1007/s10456-016-9501-1.

Tefft JB, Bays JL, Lammers A, Kim S, Eyckmans J, Chen CS. 2022. Notch1 and Notch3 coordinate for pericyte-induced stabilization of vasculature. Am J Physiol Cell Physiol. 322(2):C185–C196. doi:10.1152/ajpcell.00320.2021.

Tektonidou M, Kaskani E, Skopouli FN, Moutsopoulos HM. 1999. Microvascular abnormalities in Sjögren’s syndrome: nailfold capillaroscopy. Rheumatology (Oxford). 38(9):826–830. doi:10.1093/rheumatology/38.9.826.

Theiss AL, Simmons JG, Jobin C, Lund PK. 2005. Tumor Necrosis Factor (TNF) α Increases Collagen Accumulation and Proliferation in Intestinal Myofibroblasts via TNF Receptor 2*. Journal of Biological Chemistry. 280(43):36099–36109. doi:10.1074/jbc.M505291200.

Valim V, Gerdts E, Jonsson R, Ferreira G, Brokstad K, Brun J, Midtbo H, Mydel P. 2016. Atherosclerosis in Sjögren’s syndrome: evidence, possible mechanisms and knowledge gaps. Clinical Experimental Rheumatology. 34:133–142.

Van Sweringen HL, Sakai N, Tevar AD, Burns JM, Edwards MJ, Lentsch AB. 2011. CXC chemokine signaling in the liver: Impact on repair and regeneration. Hepatology. 54(4):1445–1453. doi:10.1002/hep.24457.

Wang X, Li Z, Shao Q, Zhang C, Wang J, Han Z, Wang S, Qin L. 2021. The intact parasympathetic nerve promotes submandibular gland regeneration through ductal cell proliferation. Cell Prolif. 54(7):e13078. doi:10.1111/cpr.13078.

Wijerathne H, Langston JC, Yang Q, Sun S, Miyamoto C, Kilpatrick LE, Kiani MF. 2021. Mechanisms of radiation-induced endothelium damage: Emerging models and technologies. Radiother Oncol. 158:21–32. doi:10.1016/j.radonc.2021.02.007.

Woods LT, Camden JM, El-Sayed FG, Khalafalla MG, Petris MJ, Erb L, Weisman GA. 2015. Increased Expression of TGF-β Signaling Components in a Mouse Model of Fibrosis Induced by Submandibular Gland Duct Ligation. PLOS ONE. 10(5):e0123641. doi:10.1371/journal.pone.0123641.

Yamashiro K, Ideguchi H, Aoyagi H, Yoshihara-Hirata C, Hirai A, Suzuki-Kyoshima R, Zhang Y, Wake H, Nishibori M, Yamamoto T, et al. 2020. High Mobility Group Box 1 Expression in Oral Inflammation and Regeneration. Frontiers in Immunology. 11. [accessed 2023 Apr 14]. https://www.frontiersin.org/articles/10.3389/fimmu.2020.01461.

Zappia L, Oshlack A. 2018. Clustering trees: a visualization for evaluating clusterings at multiple resolutions. GigaScience. 7(7):giy083. doi:10.1093/gigascience/giy083.

Zeisberg EM, Potenta SE, Sugimoto H, Zeisberg M, Kalluri R. 2008. Fibroblasts in Kidney Fibrosis Emerge via Endothelial-to-Mesenchymal Transition. J Am Soc Nephrol. 19(12):2282–2287. doi:10.1681/ASN.2008050513.

Zeisberg EM, Tarnavski O, Zeisberg M, Dorfman AL, McMullen JR, Gustafsson E, Chandraker A, Xueli Yuan, Pu WT, Roberts AB, et al. 2007. Endothelial-to-mesenchymal transition contributes to cardiac fibrosis. Nature Medicine. 13(8):952–961. doi:10.1038/nm1613.

Zhang Y, Zhang C, Li L, Liang X, Cheng P, Li Q, Chang X, Wang K, Huang S, Li Y, et al. 2021. Lymphangiogenesis in renal fibrosis arises from macrophages via VEGF-C/VEGFR3-dependent autophagy and polarization. Cell Death Dis. 12(1):1–17. doi:10.1038/s41419-020-03385-x.

