## Supplementary figures and tables for "Single-cell Transcriptomic Analysis of Salivary Gland Endothelial Cells"

**A**

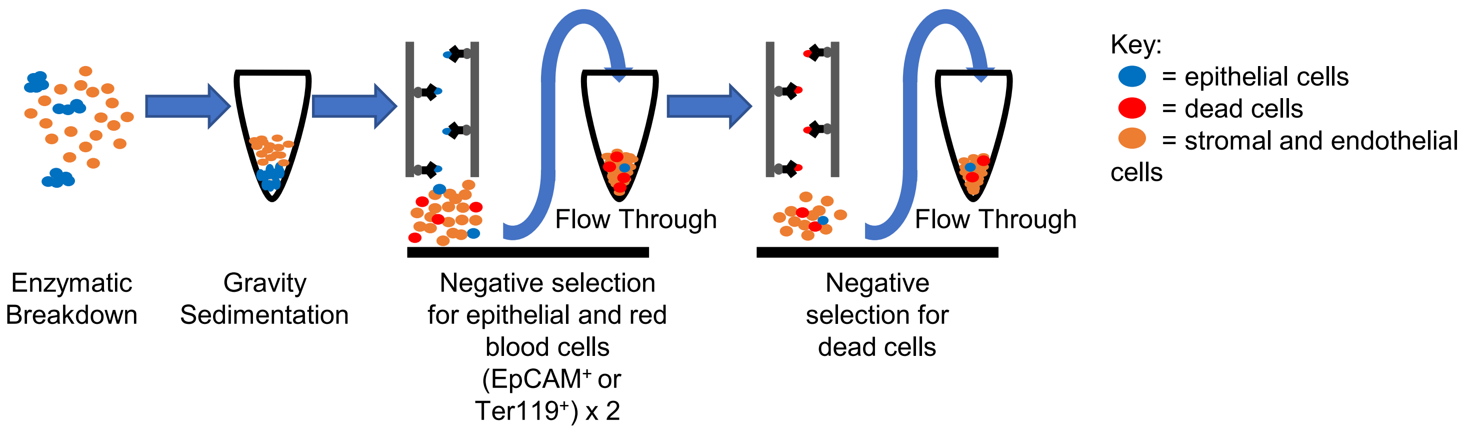
Key:
= epithelial cells
= dead cells
= stromal and endothelial cells
Flow Through
Flow Through
Enzymatic Breakdown
Gravity Sedimentation
Negative selection for epithelial and red blood cells (EpCAM⁺ or Ter119⁺) x 2
Negative selection for dead cells

**B**

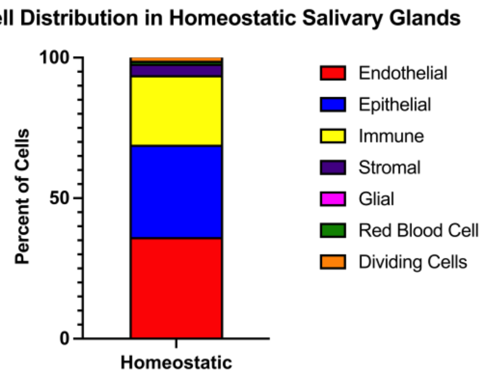
Percent Cell Distribution in Homeostatic Salivary Glands
Percent of Cells
100
50
0
Homeostatic
Endothelial
Epithelial
Immune
Stromal
Glial
Red Blood Cell
Dividing Cells

**C**

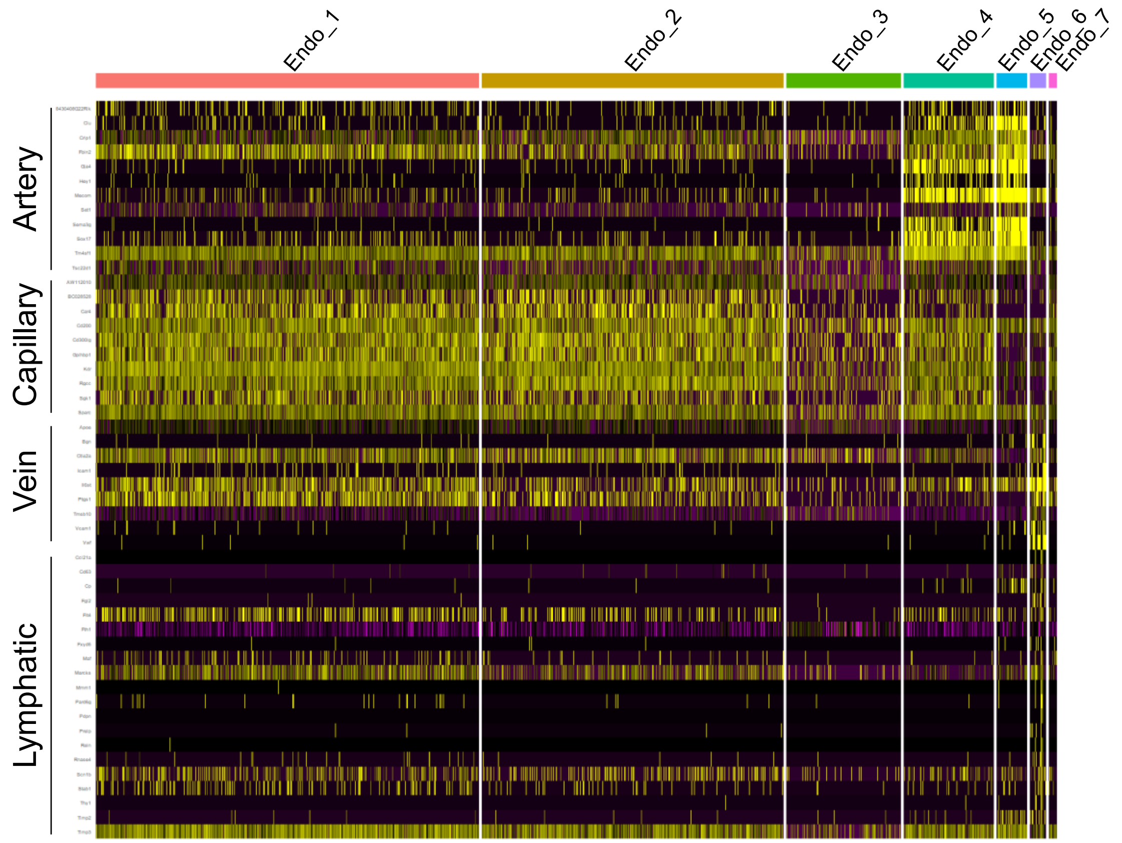
Endo_1
Endo_2
Endo_3
Endo_4
Endo_5
Endo_6
Endo_7
Artery
Capillary
Vein
Lymphatic

**Appendix Figure 1 Stromal-enriched salivary glands contain capillary, artery, and vein subpopulations of endothelial cells with distinct gene ontology terms**

(A) Schematic depicting stromal cell enrichment from two control submandibular and sublingual salivary glands utilizing enzymatic breakdown, gravity sedimentation, and magnetic activated cell sorting to deplete epithelial and red blood cells. The final flow through is counted and the viability is measured and utilized for single-cell RNA-sequencing libraries. (B) Graph showing the relative percentages of each cell type out of the total cells sequenced in homeostatic stromal-enriched salivary glands. (C) Heatmap showing conserved artery, vein, capillary, and lymphatic markers (Kalucka et al. 2020) within the endothelial cell populations of homeostatic stromal-enriched salivary glands.

**A**

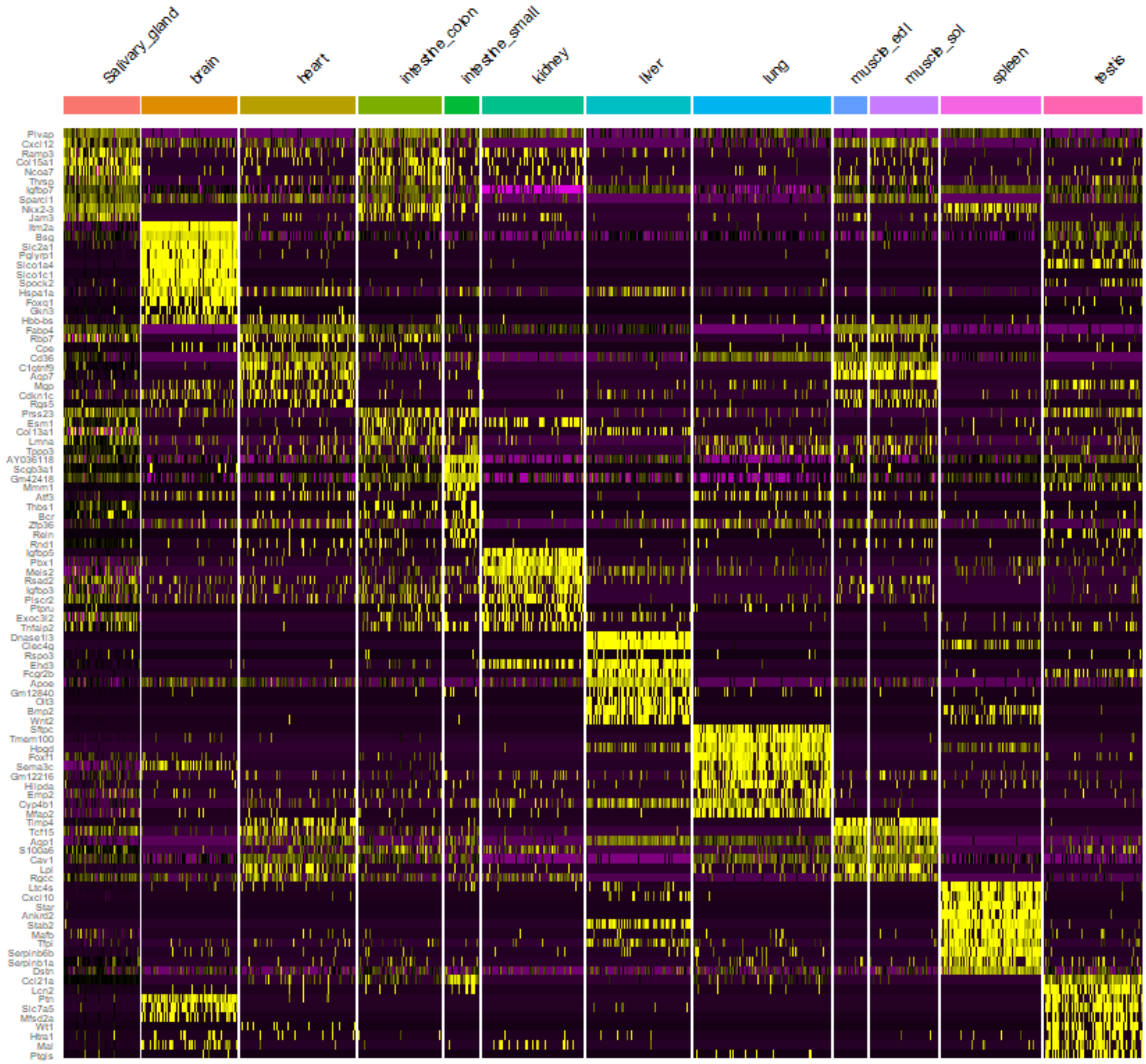
Salivary_gland
brain
heart
intestine_colon
intestine_small
kidney
liver
lung
muscle_edl
muscle_sol
spleen
testis
Plvap
Cxcl12
Ramp3
Col15a1
Ncoa7
Thrsp
Igfbp7
Sparcl1
Nkx2-3
Jam3
Itm2a
Bsg
Slc2a1
Pglyrp1
Slco1a4
Slco1c1
Spock2
Hspa1a
Foxq1
Gkn3
Hbb-bs
Fabp4
Rbp7
Cpe
Cd36
C1qtnf9
Aqp7
Mgp
Cdkn1c
Rgs5
Prss23
Esm1
Col13a1
Lmna
Tppp3
AY036118
Scgb3a1
Gm42418
Mmrn1
Atf3
Thbs1
Bcr
Zfp36
Reln
Rnd1
Igfbp5
Pbx1
Meis2
Rsad2
Igfbp3
Plscr2
Ptpru
Exoc3l2
Tnfaip2
Dnase1l3
Clec4g
Rspo3
Ehd3
Fcgr2b
Apoe
Gm12840
Oit3
Bmp2
Wnt2
Sftpc
Tmem100
Hpgd
Foxf1
Sema3c
Gm12216
Hilpda
Emp2
Cyp4b1
Mfap2
Timp4
Tcf15
Aqp1
S100a6
Cav1
Lpl
Rgcc
Ltc4s
Cxcl10
Star
Ankrd2
Stab2
Mafb
Tfpi
Serpinb6b
Serpinb1a
Dstn
Ccl21a
Lcn2
Ptn
Slc7a5
Mfsd2a
Wt1
Htra1
Mal
Ptgis

**Appendix Figure 2 Salivary gland endothelial cell gene signature**

(A) Heatmap showing the top 10 differentially expressed genes per organ from salivary gland endothelial cells integrated with published endothelial cell transcriptomes from the brain, heart, colon, small intestine, kidney, liver, lung, extensor digitorum longus muscle, soleus muscle, spleen, and testis (Kalucka et al. 2020).

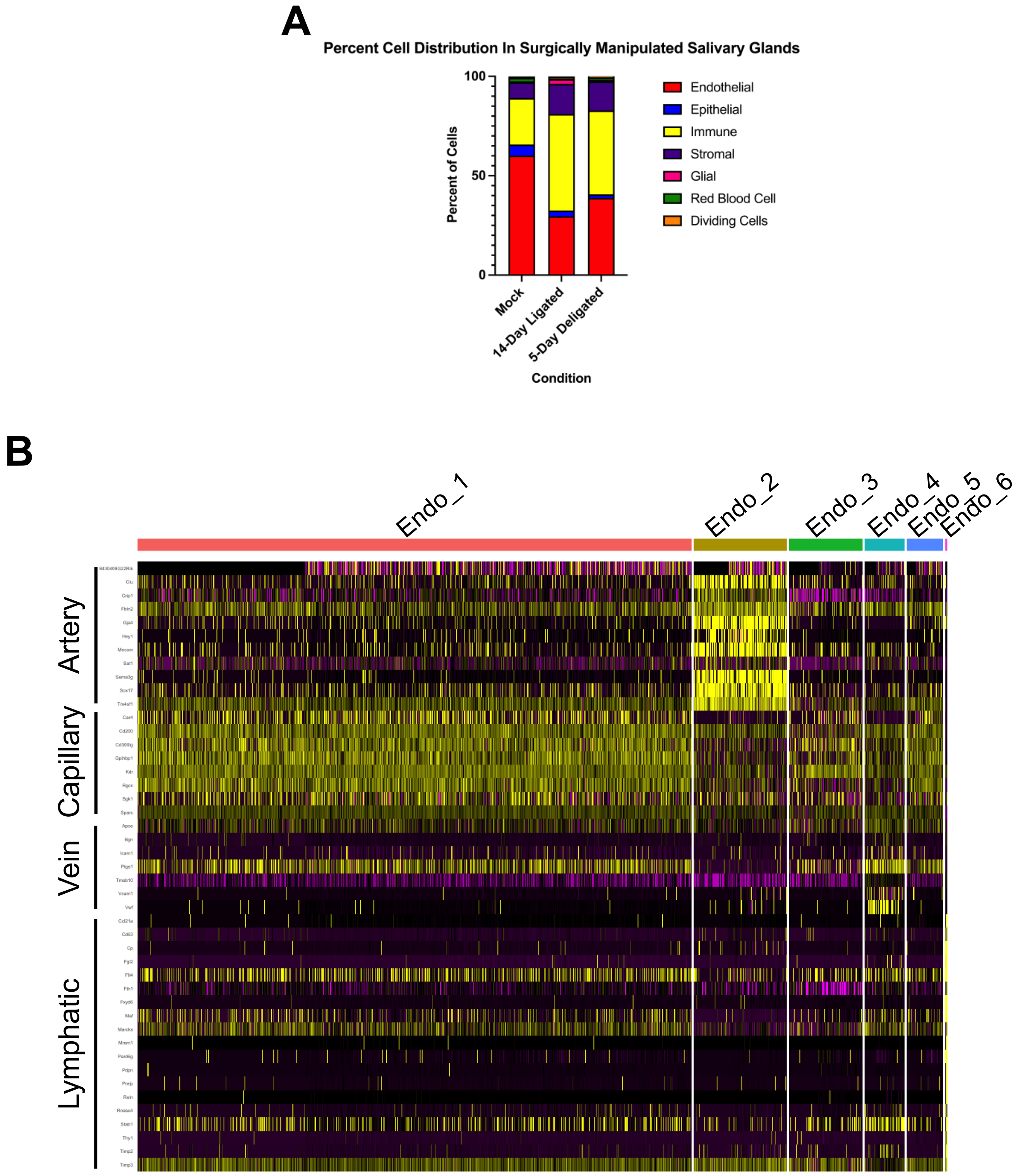
A
Percent Cell Distribution In Surgically Manipulated Salivary Glands
Percent of Cells
100
50
0
Mock
14-Day Ligated
5-Day Deligated
Condition
Endothelial
Epithelial
Immune
Stromal
Glial
Red Blood Cell
Dividing Cells
B
Endo_1
Endo_2
Endo_3
Endo_4
Endo_5
Endo_6
Artery
Capillary
Vein
Lymphatic
8430408G22Rik
Clu
Crip1
Fbln2
Gja4
Hey1
Mecom
Sat1
Sema3g
Sox17
Tm4sf1
Car4
Cd200
Cd300lg
Gpihbp1
Kdr
Rgcc
Sgk1
Sparc
Apoe
Bgn
Icam1
Ptgs1
Tmsb10
Vcam1
Vwf
Ccl21a
Cd63
Cp
Fgl2
Flt4
Fth1
Fxyd6
Maf
Marcks
Mmrn1
Pard6g
Pdpn
Prelp
Reln
Rnase4
Stab1
Thy1
Timp2
Timp3

**Appendix Figure 3 Cell distributions are altered following salivary gland injury and regeneration**

(A) Graph showing the relative percentages of each cell type out of the total cells sequenced in surgically manipulated stromal-enriched salivary glands. (B) Heatmap showing conserved artery, vein, capillary, and lymphatic markers (Kalucka et al. 2020) within the endothelial cell populations of surgically manipulated stromal-enriched salivary glands.

### Appendix Table 1 Positively differentially expressed genes in stromal-enriched salivary gland clusters

The FindAllMarkers function in Seurat was used to identify positively differentially expressed genes per cluster using min.pct = 0.25 and logfc.threshold = 0.25. The clusters in order from 0 to 26 correspond with Epi_1, Endo_1, Endo_2, Epi_2, Immune_1, Immune_2, Immune_3, Endo_3, Epi_3, Endo_4, Mes_1, Immune_4, Immune_5, Immune_6, Endo_5, RBC, Immune_7, Mes_2, Prolif_1, Immune_8, Endo_6, Epi_4, Mes_3, Prolif_2, Immune_9, Endo_7, and Mes_4, respectively.

| | p_val | avg_log2FC | pct.1 | pct.2 | p_val_adj | cluster | gene |
|---|---|---|---|---|---|---|---|
| Fxyd2 | 0 | 2.68016876 | 0.994 | 0.744 | 0 | 0 | Fxyd2 |
| Chchd10 | 0 | 2.323243855 | 0.85 | 0.354 | 0 | 0 | Chchd10 |
| S100a1 | 0 | 2.205721748 | 0.885 | 0.524 | 0 | 0 | S100a1 |
| Mt1 | 0 | 2.161766796 | 0.755 | 0.366 | 0 | 0 | Mt1 |
| Ldhb | 0 | 2.13952224 | 0.551 | 0.208 | 0 | 0 | Ldhb |
| Tesc | 0 | 2.075133785 | 0.759 | 0.315 | 0 | 0 | Tesc |
| Atp5g1 | 0 | 1.930970184 | 0.824 | 0.653 | 0 | 0 | Atp5g1 |
| Mpc2 | 0 | 1.905076298 | 0.666 | 0.498 | 0 | 0 | Mpc2 |
| 2010107E04Rik | 0 | 1.884433827 | 0.697 | 0.509 | 0 | 0 | 2010107E04Rik |
| Tfcp2l1 | 0 | 1.883135065 | 0.535 | 0.224 | 0 | 0 | Tfcp2l1 |
| Uqcr10 | 0 | 1.851856398 | 0.782 | 0.659 | 0 | 0 | Uqcr10 |
| Krt7 | 0 | 1.846299583 | 0.395 | 0.134 | 0 | 0 | Krt7 |
| Mpc1 | 0 | 1.844351153 | 0.672 | 0.5 | 0 | 0 | Mpc1 |
| Atp5g3 | 0 | 1.834355901 | 0.739 | 0.588 | 0 | 0 | Atp5g3 |
| Slc25a5 | 0 | 1.820598728 | 0.792 | 0.65 | 0 | 0 | Slc25a5 |
| Ndufab1 | 0 | 1.799742316 | 0.659 | 0.515 | 0 | 0 | Ndufab1 |
| Sod2 | 0 | 1.795102708 | 0.594 | 0.41 | 0 | 0 | Sod2 |
| Mif | 0 | 1.786226916 | 0.68 | 0.492 | 0 | 0 | Mif |
| Cox7a1 | 0 | 1.784894989 | 0.316 | 0.094 | 0 | 0 | Cox7a1 |
| Cycs | 0 | 1.783883461 | 0.573 | 0.423 | 0 | 0 | Cycs |
| Cox7b | 0 | 1.781877585 | 0.802 | 0.65 | 0 | 0 | Cox7b |
| Ndufc1 | 0 | 1.776300656 | 0.649 | 0.522 | 0 | 0 | Ndufc1 |
| Atp5e | 0 | 1.775253528 | 0.912 | 0.822 | 0 | 0 | Atp5e |
| Atp5k | 0 | 1.767744448 | 0.711 | 0.592 | 0 | 0 | Atp5k |
| Atp5j | 0 | 1.75609426 | 0.901 | 0.82 | 0 | 0 | Atp5j |
| Atp5o.1 | 0 | 1.740742979 | 0.659 | 0.54 | 0 | 0 | Atp5o.1 |
| Cox6c | 0 | 1.728777913 | 0.895 | 0.818 | 0 | 0 | Cox6c |
| Uqcrh | 0 | 1.708952427 | 0.861 | 0.795 | 0 | 0 | Uqcrh |
| Slc25a3 | 0 | 1.708574115 | 0.772 | 0.677 | 0 | 0 | Slc25a3 |
| Uqcrb | 0 | 1.704762507 | 0.798 | 0.719 | 0 | 0 | Uqcrb |
| Atp5b | 0 | 1.70364671 | 0.784 | 0.705 | 0 | 0 | Atp5b |
| Cox7c | 0 | 1.702055437 | 0.913 | 0.831 | 0 | 0 | Cox7c |
| Atp5l | 0 | 1.695153646 | 0.865 | 0.797 | 0 | 0 | Atp5l |
| Atp5h | 0 | 1.693644309 | 0.879 | 0.807 | 0 | 0 | Atp5h |
| Ckmt1 | 0 | 1.669102379 | 0.357 | 0.128 | 0 | 0 | Ckmt1 |
| Prdx5 | 0 | 1.664763443 | 0.638 | 0.512 | 0 | 0 | Prdx5 |
| Krt18 | 0 | 1.662255146 | 0.502 | 0.228 | 0 | 0 | Krt18 |
| Atp5d | 0 | 1.655769727 | 0.735 | 0.67 | 0 | 0 | Atp5d |
| Cox8a | 0 | 1.648250924 | 0.943 | 0.885 | 0 | 0 | Cox8a |
| Atp5c1 | 0 | 1.644634175 | 0.692 | 0.631 | 0 | 0 | Atp5c1 |
| Ndufb10 | 0 | 1.643037498 | 0.692 | 0.62 | 0 | 0 | Ndufb10 |
| Uqcrq | 0 | 1.641564748 | 0.704 | 0.648 | 0 | 0 | Uqcrq |

| | | | | | | | |
|---|---|---|---|---|---|---|---|
| Atp5f1 | 0 | 1.640445361 | 0.736 | 0.678 | 0 | 0 | Atp5f1 |
| Cox5b | 0 | 1.63456496 | 0.773 | 0.698 | 0 | 0 | Cox5b |
| Cox5a | 0 | 1.630384813 | 0.806 | 0.741 | 0 | 0 | Cox5a |
| Ndufb4 | 0 | 1.618447504 | 0.636 | 0.57 | 0 | 0 | Ndufb4 |
| Ndufa4 | 0 | 1.605121868 | 0.85 | 0.812 | 0 | 0 | Ndufa4 |
| Uqcr11 | 0 | 1.594695685 | 0.73 | 0.673 | 0 | 0 | Uqcr11 |
| Aldoa | 0 | 1.580913453 | 0.69 | 0.631 | 0 | 0 | Aldoa |
| Cox4i1 | 0 | 1.546967313 | 0.864 | 0.849 | 0 | 0 | Cox4i1 |
| Atp5j2 | 0 | 1.536745369 | 0.775 | 0.763 | 0 | 0 | Atp5j2 |
| Cox7a2 | 0 | 1.536701616 | 0.791 | 0.767 | 0 | 0 | Cox7a2 |
| Dstn | 0 | 1.523597639 | 0.662 | 0.557 | 0 | 0 | Dstn |
| Cox6b1 | 0 | 1.410944707 | 0.735 | 0.735 | 0 | 0 | Cox6b1 |
| Calm2 | 0 | 1.386246806 | 0.791 | 0.776 | 0 | 0 | Calm2 |
| Klk1b9 | 0 | 1.348514134 | 0.637 | 0.309 | 0 | 0 | Klk1b9 |
| Chchd2 | 0 | 1.335948053 | 0.909 | 0.907 | 0 | 0 | Chchd2 |
| Atp5g2 | 0 | 1.315238715 | 0.712 | 0.735 | 0 | 0 | Atp5g2 |
| H3f3b | 0 | 0.994998445 | 0.88 | 0.913 | 0 | 0 | H3f3b |
| Rplp1 | 0 | 0.989074479 | 0.983 | 0.983 | 0 | 0 | Rplp1 |
| Rplp0 | 0 | 0.91666619 | 0.939 | 0.946 | 0 | 0 | Rplp0 |
| Klk1 | 0 | 0.774749307 | 0.981 | 0.85 | 0 | 0 | Klk1 |
| Rps2 | 0 | 0.75474468 | 0.885 | 0.928 | 0 | 0 | Rps2 |
| Rpl26 | 0 | 0.750162906 | 0.931 | 0.963 | 0 | 0 | Rpl26 |
| Rpl41 | 0 | 0.694869425 | 0.933 | 0.968 | 0 | 0 | Rpl41 |
| Fth1 | 0 | 0.452809834 | 0.962 | 0.971 | 0 | 0 | Fth1 |
| Mucl2 | 0 | 0.325365808 | 0.999 | 0.999 | 0 | 0 | Mucl2 |
| Uqcrfs1 | 1.34E-297 | 1.769891496 | 0.541 | 0.383 | 2.71E-293 | 0 | Uqcrfs1 |
| Gpx1 | 5.80E-294 | 1.454909737 | 0.648 | 0.587 | 1.17E-289 | 0 | Gpx1 |
| Ftl1 | 9.49E-293 | 0.580298663 | 0.864 | 0.922 | 1.92E-288 | 0 | Ftl1 |
| Mdh1 | 1.01E-292 | 1.762341324 | 0.553 | 0.403 | 2.04E-288 | 0 | Mdh1 |
| Ndufa11 | 1.64E-286 | 1.676238679 | 0.6 | 0.511 | 3.32E-282 | 0 | Ndufa11 |
| Ppia | 9.24E-266 | 0.627181266 | 0.898 | 0.947 | 1.87E-261 | 0 | Ppia |
| Nupr1 | 1.01E-264 | 1.192833082 | 0.47 | 0.273 | 2.04E-260 | 0 | Nupr1 |
| Tpt1 | 4.73E-262 | 0.437571089 | 0.984 | 0.991 | 9.58E-258 | 0 | Tpt1 |
| Usmg5 | 6.30E-252 | 1.578600715 | 0.601 | 0.54 | 1.28E-247 | 0 | Usmg5 |
| Atp5a1 | 9.90E-252 | 1.514647237 | 0.628 | 0.627 | 2.00E-247 | 0 | Atp5a1 |
| Phyh | 1.74E-245 | 1.700832956 | 0.377 | 0.19 | 3.52E-241 | 0 | Phyh |
| 2410015M20Rik | 1.24E-243 | 1.572257903 | 0.579 | 0.512 | 2.52E-239 | 0 | 2410015M20Rik |
| Cystm1 | 2.37E-242 | 1.578766403 | 0.339 | 0.152 | 4.79E-238 | 0 | Cystm1 |
| Naca | 1.45E-240 | 0.884139124 | 0.747 | 0.853 | 2.93E-236 | 0 | Naca |
| Aes | 3.22E-238 | 1.331316308 | 0.63 | 0.625 | 6.52E-234 | 0 | Aes |
| Ndufa5 | 6.66E-234 | 1.589765527 | 0.57 | 0.496 | 1.35E-229 | 0 | Ndufa5 |
| Rpl23 | 6.06E-220 | 0.48958324 | 0.953 | 0.975 | 1.23E-215 | 0 | Rpl23 |
| Ndufa12 | 7.46E-215 | 1.621037649 | 0.536 | 0.447 | 1.51E-210 | 0 | Ndufa12 |

| | | | | | | | |
|---|---|---|---|---|---|---|---|
| Rpl10 | 1.76E-214 | 0.664266067 | 0.829 | 0.912 | 3.56E-210 | 0 | Rpl10 |
| Gsta4 | 7.41E-211 | 1.557722454 | 0.288 | 0.123 | 1.50E-206 | 0 | Gsta4 |
| G0s2 | 1.26E-205 | 1.25050093 | 0.459 | 0.274 | 2.54E-201 | 0 | G0s2 |
| Atp6v1f | 5.56E-202 | 1.429749358 | 0.603 | 0.616 | 1.13E-197 | 0 | Atp6v1f |
| Ndufa6 | 1.69E-199 | 1.458136183 | 0.588 | 0.58 | 3.41E-195 | 0 | Ndufa6 |
| Gm42418 | 5.37E-196 | 0.353138963 | 1 | 1 | 1.09E-191 | 0 | Gm42418 |
| Rps8 | 5.09E-194 | 0.500661144 | 0.928 | 0.973 | 1.03E-189 | 0 | Rps8 |
| Ndufb11 | 2.67E-179 | 1.29219646 | 0.609 | 0.654 | 5.41E-175 | 0 | Ndufb11 |
| Oxct1 | 7.72E-179 | 1.742780563 | 0.467 | 0.364 | 1.56E-174 | 0 | Oxct1 |
| Cdk8 | 9.47E-179 | 0.499456538 | 0.919 | 0.981 | 1.92E-174 | 0 | Cdk8 |
| Rps3a1 | 2.84E-176 | 0.495756495 | 0.886 | 0.946 | 5.75E-172 | 0 | Rps3a1 |
| Rpl21 | 5.91E-176 | 0.544732899 | 0.857 | 0.938 | 1.20E-171 | 0 | Rpl21 |
| Suclg1 | 2.41E-175 | 1.545517928 | 0.49 | 0.392 | 4.88E-171 | 0 | Suclg1 |
| Ndufb5 | 1.25E-172 | 1.460882577 | 0.558 | 0.545 | 2.54E-168 | 0 | Ndufb5 |
| Fuom | 1.40E-170 | 1.482526519 | 0.296 | 0.147 | 2.84E-166 | 0 | Fuom |
| Ndufa13 | 3.02E-170 | 1.228506877 | 0.613 | 0.681 | 6.12E-166 | 0 | Ndufa13 |
| Rpl19 | 7.48E-170 | 0.427876023 | 0.934 | 0.97 | 1.51E-165 | 0 | Rpl19 |
| Gapdh | 8.45E-165 | 1.093814984 | 0.634 | 0.726 | 1.71E-160 | 0 | Gapdh |
| S100a11 | 8.46E-162 | 1.188185672 | 0.59 | 0.572 | 1.71E-157 | 0 | S100a11 |
| Slc25a4 | 9.24E-162 | 1.434064766 | 0.559 | 0.563 | 1.87E-157 | 0 | Slc25a4 |
| Klk1b27 | 1.67E-161 | 1.266417174 | 0.284 | 0.132 | 3.38E-157 | 0 | Klk1b27 |
| Prdx2 | 2.71E-161 | 1.462462963 | 0.545 | 0.534 | 5.49E-157 | 0 | Prdx2 |
| Mt2 | 2.48E-155 | 1.406906697 | 0.263 | 0.122 | 5.01E-151 | 0 | Mt2 |
| Rpl24 | 1.23E-154 | 0.582619497 | 0.817 | 0.928 | 2.49E-150 | 0 | Rpl24 |
| Rpl8 | 1.91E-150 | 0.489043375 | 0.86 | 0.94 | 3.88E-146 | 0 | Rpl8 |
| Rpl10a | 5.27E-150 | 0.66649502 | 0.767 | 0.852 | 1.07E-145 | 0 | Rpl10a |
| Lars2 | 1.69E-149 | 0.779886592 | 0.783 | 0.921 | 3.43E-145 | 0 | Lars2 |
| Ndufb9 | 1.96E-148 | 1.238030134 | 0.603 | 0.686 | 3.96E-144 | 0 | Ndufb9 |
| Bri3 | 2.91E-148 | 1.345245537 | 0.575 | 0.619 | 5.88E-144 | 0 | Bri3 |
| Rpl28 | 5.57E-148 | 0.499865086 | 0.861 | 0.946 | 1.13E-143 | 0 | Rpl28 |
| Rpl32 | 2.82E-147 | 0.383483093 | 0.921 | 0.961 | 5.72E-143 | 0 | Rpl32 |
| Cmss1 | 2.14E-146 | 0.388051508 | 0.984 | 0.996 | 4.33E-142 | 0 | Cmss1 |
| Rps26 | 2.56E-144 | 0.640641852 | 0.78 | 0.885 | 5.19E-140 | 0 | Rps26 |
| Ndufb8 | 2.88E-144 | 1.388160719 | 0.554 | 0.564 | 5.84E-140 | 0 | Ndufb8 |
| Rpl11 | 4.41E-144 | 0.47504558 | 0.865 | 0.941 | 8.93E-140 | 0 | Rpl11 |
| Mcee | 1.74E-142 | 1.518521237 | 0.385 | 0.264 | 3.53E-138 | 0 | Mcee |
| Rpl3 | 2.39E-140 | 0.493440302 | 0.847 | 0.924 | 4.85E-136 | 0 | Rpl3 |
| Rpsa | 2.93E-137 | 0.43119759 | 0.852 | 0.914 | 5.93E-133 | 0 | Rpsa |
| H3f3a | 9.66E-135 | 0.627895383 | 0.741 | 0.889 | 1.96E-130 | 0 | H3f3a |
| Vdac1 | 1.78E-134 | 1.530264131 | 0.429 | 0.339 | 3.60E-130 | 0 | Vdac1 |
| Rpl9 | 1.79E-134 | 0.43560547 | 0.896 | 0.955 | 3.63E-130 | 0 | Rpl9 |
| Cyc1 | 2.14E-134 | 1.556669369 | 0.426 | 0.336 | 4.34E-130 | 0 | Cyc1 |
| Ndufs6 | 1.14E-132 | 1.43076029 | 0.508 | 0.484 | 2.30E-128 | 0 | Ndufs6 |

| | | | | | | | |
|---|---|---|---|---|---|---|---|
| Ndufb2 | 1.57E-132 | 1.505219198 | 0.462 | 0.396 | 3.18E-128 | 0 | Ndufb2 |
| Nudt4 | 8.25E-131 | 1.331984944 | 0.479 | 0.398 | 1.67E-126 | 0 | Nudt4 |
| Coq7 | 2.59E-129 | 1.433576549 | 0.268 | 0.142 | 5.24E-125 | 0 | Coq7 |
| Sdhb | 3.86E-128 | 1.511352188 | 0.478 | 0.431 | 7.81E-124 | 0 | Sdhb |
| Eif1 | 7.78E-127 | 0.546607312 | 0.79 | 0.92 | 1.58E-122 | 0 | Eif1 |
| Ndufs5 | 2.16E-120 | 1.32477969 | 0.549 | 0.6 | 4.37E-116 | 0 | Ndufs5 |
| Rpl13 | 2.19E-120 | 0.326537166 | 0.955 | 0.982 | 4.44E-116 | 0 | Rpl13 |
| Paip2 | 8.34E-120 | 0.254862487 | 0.181 | 0.458 | 1.69E-115 | 0 | Paip2 |
| Rps7 | 1.14E-119 | 0.440475126 | 0.852 | 0.931 | 2.30E-115 | 0 | Rps7 |
| Klk1b11 | 6.52E-117 | 0.549548145 | 0.385 | 0.239 | 1.32E-112 | 0 | Klk1b11 |
| Pdcd4 | 2.90E-116 | 1.378187956 | 0.487 | 0.449 | 5.88E-112 | 0 | Pdcd4 |
| Gabarapl1 | 8.01E-115 | 1.506073169 | 0.33 | 0.219 | 1.62E-110 | 0 | Gabarapl1 |
| Rpl36 | 5.84E-113 | 0.494341129 | 0.817 | 0.922 | 1.18E-108 | 0 | Rpl36 |
| Psmd4 | 8.13E-113 | 0.264588857 | 0.144 | 0.381 | 1.65E-108 | 0 | Psmd4 |
| Stxbp6 | 4.55E-111 | 1.201192356 | 0.292 | 0.168 | 9.22E-107 | 0 | Stxbp6 |
| Oaz1 | 1.48E-110 | 0.70202347 | 0.676 | 0.836 | 3.00E-106 | 0 | Oaz1 |
| Atpif1 | 1.72E-109 | 1.264387901 | 0.538 | 0.568 | 3.47E-105 | 0 | Atpif1 |
| Pdcd10 | 2.78E-108 | 0.352009291 | 0.167 | 0.42 | 5.62E-104 | 0 | Pdcd10 |
| Acadl | 4.74E-108 | 1.511816417 | 0.426 | 0.364 | 9.59E-104 | 0 | Acadl |
| Rpl17 | 1.32E-107 | 0.455661161 | 0.84 | 0.934 | 2.67E-103 | 0 | Rpl17 |
| Snx10 | 2.65E-107 | 1.445170148 | 0.286 | 0.175 | 5.36E-103 | 0 | Snx10 |
| Vps29 | 6.12E-106 | 0.260137503 | 0.134 | 0.355 | 1.24E-101 | 0 | Vps29 |
| Srp14 | 1.76E-105 | 0.289787489 | 0.221 | 0.516 | 3.56E-101 | 0 | Srp14 |
| Rp9 | 5.51E-105 | 0.256797793 | 0.142 | 0.367 | 1.12E-100 | 0 | Rp9 |
| Rpl15 | 1.17E-104 | 0.470017341 | 0.812 | 0.921 | 2.36E-100 | 0 | Rpl15 |
| Fdx1 | 1.66E-104 | 1.491432493 | 0.382 | 0.299 | 3.37E-100 | 0 | Fdx1 |
| Atp6v1g1 | 1.82E-104 | 0.280104036 | 0.218 | 0.509 | 3.68E-100 | 0 | Atp6v1g1 |
| 1700020I14Rik | 1.42E-103 | 0.257165345 | 0.152 | 0.384 | 2.88E-99 | 0 | 1700020I14Rik |
| Rpl14 | 3.46E-103 | 0.696190304 | 0.689 | 0.828 | 7.01E-99 | 0 | Rpl14 |
| Ubxn1 | 1.04E-101 | 0.273035068 | 0.207 | 0.484 | 2.10E-97 | 0 | Ubxn1 |
| Arf1 | 1.21E-101 | 0.278102083 | 0.21 | 0.488 | 2.45E-97 | 0 | Arf1 |
| 1110004F10Rik | 2.79E-101 | 0.290567073 | 0.134 | 0.349 | 5.65E-97 | 0 | 1110004F10Rik |
| Rps10 | 5.69E-101 | 0.326462828 | 0.893 | 0.956 | 1.15E-96 | 0 | Rps10 |
| Cggbp1 | 1.46E-100 | 0.275364165 | 0.124 | 0.331 | 2.96E-96 | 0 | Cggbp1 |
| Supt4a | 5.80E-100 | 0.250396853 | 0.143 | 0.363 | 1.17E-95 | 0 | Supt4a |
| Trir | 2.03E-99 | 0.277074446 | 0.199 | 0.467 | 4.11E-95 | 0 | Trir |
| Mpp6 | 3.79E-99 | 1.324096521 | 0.365 | 0.268 | 7.67E-95 | 0 | Mpp6 |
| Abcf1 | 6.30E-99 | 0.286407361 | 0.107 | 0.297 | 1.27E-94 | 0 | Abcf1 |
| Rps4x | 6.72E-98 | 0.356357033 | 0.885 | 0.954 | 1.36E-93 | 0 | Rps4x |
| Mrpl42 | 1.96E-97 | 1.441305539 | 0.409 | 0.344 | 3.96E-93 | 0 | Mrpl42 |
| Eif3e | 2.40E-97 | 0.263098006 | 0.199 | 0.461 | 4.86E-93 | 0 | Eif3e |
| R3hdm2 | 4.44E-97 | 0.32028789 | 0.133 | 0.342 | 9.00E-93 | 0 | R3hdm2 |
| Imp3 | 7.44E-97 | 0.287137961 | 0.14 | 0.355 | 1.51E-92 | 0 | Imp3 |

| | | | | | | | |
|---|---|---|---|---|---|---|---|
| Sptssa | 1.67E-96 | 0.25095524 | 0.135 | 0.343 | 3.37E-92 | 0 | Sptssa |
| Minos1 | 2.64E-96 | 1.346001701 | 0.501 | 0.518 | 5.35E-92 | 0 | Minos1 |
| Rps17 | 6.49E-95 | 0.701492679 | 0.667 | 0.826 | 1.31E-90 | 0 | Rps17 |
| Cab39 | 8.15E-95 | 0.290317217 | 0.121 | 0.318 | 1.65E-90 | 0 | Cab39 |
| Gm26561 | 1.06E-93 | 0.280281949 | 0.14 | 0.349 | 2.14E-89 | 0 | Gm26561 |
| Hadh | 3.25E-93 | 1.45069996 | 0.326 | 0.233 | 6.59E-89 | 0 | Hadh |
| Map1lc3a | 8.04E-93 | 1.171988762 | 0.491 | 0.483 | 1.63E-88 | 0 | Map1lc3a |
| Srp19 | 1.36E-90 | 0.277870536 | 0.142 | 0.349 | 2.75E-86 | 0 | Srp19 |
| Sf3b5 | 1.38E-90 | 0.296569573 | 0.141 | 0.349 | 2.79E-86 | 0 | Sf3b5 |
| Cct8 | 8.42E-90 | 0.263837603 | 0.116 | 0.302 | 1.70E-85 | 0 | Cct8 |
| Psmd8 | 1.53E-89 | 0.272139981 | 0.15 | 0.363 | 3.09E-85 | 0 | Psmd8 |
| Map2k2 | 2.41E-89 | 0.307915964 | 0.13 | 0.327 | 4.87E-85 | 0 | Map2k2 |
| Ptges3 | 3.14E-89 | 0.330324119 | 0.188 | 0.432 | 6.35E-85 | 0 | Ptges3 |
| Ran | 1.18E-88 | 0.280795623 | 0.197 | 0.446 | 2.38E-84 | 0 | Ran |
| Rplp2 | 2.31E-88 | 0.43399418 | 0.795 | 0.912 | 4.68E-84 | 0 | Rplp2 |
| Dnajb6 | 2.67E-88 | 0.346422865 | 0.184 | 0.423 | 5.40E-84 | 0 | Dnajb6 |
| Tcea1 | 2.38E-87 | 0.268659309 | 0.122 | 0.309 | 4.82E-83 | 0 | Tcea1 |
| Sf3b6 | 2.80E-87 | 0.334081603 | 0.173 | 0.403 | 5.68E-83 | 0 | Sf3b6 |
| Pcnp | 4.18E-87 | 0.264436869 | 0.106 | 0.282 | 8.46E-83 | 0 | Pcnp |
| Ndufv2 | 9.00E-87 | 1.411531698 | 0.439 | 0.415 | 1.82E-82 | 0 | Ndufv2 |
| Rps9 | 1.20E-86 | 0.365496261 | 0.851 | 0.936 | 2.43E-82 | 0 | Rps9 |
| Bzw1 | 5.29E-86 | 0.343609119 | 0.191 | 0.433 | 1.07E-81 | 0 | Bzw1 |
| Rheb | 4.24E-85 | 0.360844636 | 0.185 | 0.421 | 8.59E-81 | 0 | Rheb |
| Smim11 | 5.01E-85 | 0.328289292 | 0.143 | 0.344 | 1.02E-80 | 0 | Smim11 |
| Nars | 6.41E-85 | 0.255914436 | 0.115 | 0.294 | 1.30E-80 | 0 | Nars |
| 1810058I24Rik | 6.56E-85 | 0.335374784 | 0.142 | 0.343 | 1.33E-80 | 0 | 1810058I24Rik |
| Nsa2 | 9.81E-85 | 0.270356583 | 0.253 | 0.539 | 1.99E-80 | 0 | Nsa2 |
| Cetn3 | 2.11E-84 | 0.302761514 | 0.16 | 0.373 | 4.27E-80 | 0 | Cetn3 |
| Psma4 | 2.12E-84 | 0.336990194 | 0.173 | 0.399 | 4.30E-80 | 0 | Psma4 |
| Actr10 | 2.13E-84 | 0.266823727 | 0.104 | 0.275 | 4.31E-80 | 0 | Actr10 |
| Rab14 | 3.23E-84 | 0.360378825 | 0.226 | 0.496 | 6.55E-80 | 0 | Rab14 |
| Vamp8 | 4.26E-84 | 1.185416354 | 0.5 | 0.513 | 8.63E-80 | 0 | Vamp8 |
| Cct4 | 5.43E-84 | 0.33187835 | 0.149 | 0.354 | 1.10E-79 | 0 | Cct4 |
| Psmc6 | 1.82E-83 | 0.348484157 | 0.144 | 0.344 | 3.68E-79 | 0 | Psmc6 |
| Zfand5 | 1.92E-83 | 0.261962277 | 0.234 | 0.505 | 3.88E-79 | 0 | Zfand5 |
| Ppp1cb | 3.11E-83 | 0.281246016 | 0.124 | 0.308 | 6.30E-79 | 0 | Ppp1cb |
| Arpp19 | 3.34E-83 | 0.329163145 | 0.223 | 0.49 | 6.77E-79 | 0 | Arpp19 |
| Stx8 | 8.07E-83 | 0.304292356 | 0.12 | 0.302 | 1.63E-78 | 0 | Stx8 |
| Rala | 3.15E-82 | 0.30326403 | 0.137 | 0.329 | 6.39E-78 | 0 | Rala |
| Sub1 | 3.32E-82 | 0.277392478 | 0.27 | 0.571 | 6.71E-78 | 0 | Sub1 |
| Rps28 | 5.63E-82 | 0.393015292 | 0.822 | 0.915 | 1.14E-77 | 0 | Rps28 |
| Rps27a | 9.69E-82 | 0.305518307 | 0.895 | 0.962 | 1.96E-77 | 0 | Rps27a |
| Ifi27l2a | 1.59E-81 | 1.819967404 | 0.422 | 0.358 | 3.21E-77 | 0 | Ifi27l2a |

| | | | | | | | |
|---|---|---|---|---|---|---|---|
| Chmp5 | 4.34E-81 | 0.316803278 | 0.171 | 0.387 | 8.78E-77 | 0 | Chmp5 |
| Trappc4 | 1.15E-80 | 0.287749631 | 0.133 | 0.32 | 2.32E-76 | 0 | Trappc4 |
| Trappc6b | 1.67E-80 | 0.285779916 | 0.141 | 0.334 | 3.38E-76 | 0 | Trappc6b |
| Echs1 | 2.26E-80 | 1.429238277 | 0.329 | 0.25 | 4.58E-76 | 0 | Echs1 |
| Ndufb7 | 3.08E-80 | 1.230112579 | 0.528 | 0.608 | 6.23E-76 | 0 | Ndufb7 |
| Acot13 | 5.11E-80 | 1.371893264 | 0.301 | 0.213 | 1.04E-75 | 0 | Acot13 |
| Chmp4b | 8.52E-80 | 0.359610663 | 0.164 | 0.374 | 1.73E-75 | 0 | Chmp4b |
| Mob4 | 1.06E-79 | 0.360578838 | 0.146 | 0.343 | 2.14E-75 | 0 | Mob4 |
| Arl1 | 2.51E-79 | 0.298381685 | 0.134 | 0.321 | 5.09E-75 | 0 | Arl1 |
| Chmp2a | 3.92E-79 | 0.34573045 | 0.254 | 0.543 | 7.94E-75 | 0 | Chmp2a |
| Snrpf | 9.42E-79 | 0.326890609 | 0.192 | 0.422 | 1.91E-74 | 0 | Snrpf |
| Rpl7 | 1.47E-78 | 0.429036016 | 0.791 | 0.916 | 2.97E-74 | 0 | Rpl7 |
| Dctn3 | 2.02E-78 | 0.368157108 | 0.159 | 0.364 | 4.09E-74 | 0 | Dctn3 |
| Metap2 | 2.09E-78 | 0.373097434 | 0.184 | 0.408 | 4.23E-74 | 0 | Metap2 |
| Aimp1 | 4.12E-78 | 0.359845575 | 0.147 | 0.341 | 8.33E-74 | 0 | Aimp1 |
| Cbx1 | 5.28E-78 | 0.35908021 | 0.155 | 0.356 | 1.07E-73 | 0 | Cbx1 |
| Rpl37 | 5.42E-78 | 0.371173373 | 0.85 | 0.945 | 1.10E-73 | 0 | Rpl37 |
| Eef1a1 | 5.66E-78 | 0.307491006 | 0.915 | 0.97 | 1.15E-73 | 0 | Eef1a1 |
| Hdgf | 1.47E-77 | 0.27753449 | 0.112 | 0.28 | 2.97E-73 | 0 | Hdgf |
| Wdr83os | 1.56E-77 | 0.305190993 | 0.134 | 0.318 | 3.15E-73 | 0 | Wdr83os |
| Psmb4 | 2.12E-77 | 0.35272744 | 0.207 | 0.45 | 4.28E-73 | 0 | Psmb4 |
| Arf5 | 4.34E-77 | 0.265304131 | 0.287 | 0.602 | 8.79E-73 | 0 | Arf5 |
| Polr2j | 5.94E-77 | 0.381512058 | 0.142 | 0.333 | 1.20E-72 | 0 | Polr2j |
| Ptma | 7.24E-77 | 0.35619378 | 0.873 | 0.954 | 1.47E-72 | 0 | Ptma |
| Cisd1 | 8.98E-77 | 1.475646876 | 0.33 | 0.257 | 1.82E-72 | 0 | Cisd1 |
| Esd | 1.97E-76 | 0.38378711 | 0.193 | 0.423 | 3.99E-72 | 0 | Esd |
| Sin3b | 2.74E-76 | 0.297992784 | 0.116 | 0.287 | 5.55E-72 | 0 | Sin3b |
| Ost4 | 4.19E-76 | 0.346723778 | 0.227 | 0.484 | 8.47E-72 | 0 | Ost4 |
| St13 | 9.54E-76 | 0.406617444 | 0.185 | 0.409 | 1.93E-71 | 0 | St13 |
| Psma1 | 1.03E-75 | 0.381643593 | 0.16 | 0.363 | 2.09E-71 | 0 | Psma1 |
| AY036118 | 2.34E-75 | 0.351193297 | 0.834 | 0.946 | 4.74E-71 | 0 | AY036118 |
| Pfdn2 | 3.10E-75 | 0.323536817 | 0.151 | 0.345 | 6.27E-71 | 0 | Pfdn2 |
| Cct2 | 2.60E-74 | 0.382702211 | 0.151 | 0.345 | 5.26E-70 | 0 | Cct2 |
| Rpl27a | 3.86E-74 | 0.272726646 | 0.917 | 0.97 | 7.82E-70 | 0 | Rpl27a |
| Eif4a2 | 4.16E-74 | 0.365598697 | 0.148 | 0.34 | 8.42E-70 | 0 | Eif4a2 |
| D8Ertd738e | 6.80E-74 | 0.358007762 | 0.276 | 0.576 | 1.38E-69 | 0 | D8Ertd738e |
| Nenf | 1.13E-73 | 0.296397525 | 0.144 | 0.331 | 2.30E-69 | 0 | Nenf |
| Cope | 1.22E-73 | 0.330513353 | 0.154 | 0.349 | 2.48E-69 | 0 | Cope |
| Csnk2b | 2.59E-73 | 0.307201327 | 0.12 | 0.289 | 5.24E-69 | 0 | Csnk2b |
| Sra1 | 3.41E-73 | 0.321629557 | 0.136 | 0.316 | 6.90E-69 | 0 | Sra1 |
| Nudc | 5.22E-73 | 0.392039811 | 0.178 | 0.391 | 1.06E-68 | 0 | Nudc |
| Erbin | 1.28E-72 | 0.375924859 | 0.154 | 0.346 | 2.60E-68 | 0 | Erbin |
| Vdac2 | 1.36E-72 | 1.276465431 | 0.495 | 0.554 | 2.74E-68 | 0 | Vdac2 |

| | | | | | | | |
|---|---|---|---|---|---|---|---|
| Ube2n | 4.74E-72 | 0.313841013 | 0.123 | 0.292 | 9.60E-68 | 0 | Ube2n |
| Bola2 | 1.07E-71 | 0.340537005 | 0.124 | 0.294 | 2.17E-67 | 0 | Bola2 |
| Chrac1 | 2.19E-71 | 0.31690866 | 0.115 | 0.277 | 4.44E-67 | 0 | Chrac1 |
| Fam104a | 3.21E-71 | 0.46106964 | 0.149 | 0.337 | 6.50E-67 | 0 | Fam104a |
| Mrpl24 | 5.80E-71 | 0.27680972 | 0.11 | 0.268 | 1.18E-66 | 0 | Mrpl24 |
| Arpc5l | 7.80E-71 | 0.373120571 | 0.152 | 0.341 | 1.58E-66 | 0 | Arpc5l |
| Eif3m | 8.48E-71 | 0.385320275 | 0.189 | 0.408 | 1.72E-66 | 0 | Eif3m |
| Rpl6 | 1.39E-70 | 0.45716548 | 0.772 | 0.906 | 2.81E-66 | 0 | Rpl6 |
| Psmb6 | 5.13E-70 | 0.377137506 | 0.222 | 0.466 | 1.04E-65 | 0 | Psmb6 |
| Tpd52 | 5.97E-70 | 1.181978031 | 0.362 | 0.297 | 1.21E-65 | 0 | Tpd52 |
| Psmd12 | 7.27E-70 | 0.389999906 | 0.135 | 0.311 | 1.47E-65 | 0 | Psmd12 |
| Idh2 | 5.98E-69 | 1.389128383 | 0.279 | 0.2 | 1.21E-64 | 0 | Idh2 |
| Nap1l4 | 7.69E-69 | 0.320428273 | 0.109 | 0.265 | 1.56E-64 | 0 | Nap1l4 |
| Hist1h4d | 2.70E-68 | 0.25878673 | 0.124 | 0.288 | 5.46E-64 | 0 | Hist1h4d |
| Mrpl12 | 4.62E-67 | 1.402037443 | 0.357 | 0.307 | 9.36E-63 | 0 | Mrpl12 |
| Snapc5 | 6.56E-67 | 0.330792131 | 0.114 | 0.269 | 1.33E-62 | 0 | Snapc5 |
| Map1lc3b | 8.08E-67 | 0.36456436 | 0.27 | 0.554 | 1.64E-62 | 0 | Map1lc3b |
| Eloc | 1.23E-66 | 0.405232154 | 0.166 | 0.36 | 2.48E-62 | 0 | Eloc |
| Idh3a | 1.28E-66 | 1.344997907 | 0.251 | 0.172 | 2.60E-62 | 0 | Idh3a |
| Ndufs7 | 7.29E-66 | 1.309804308 | 0.446 | 0.463 | 1.48E-61 | 0 | Ndufs7 |
| Rps13 | 8.76E-66 | 0.284674405 | 0.852 | 0.939 | 1.77E-61 | 0 | Rps13 |
| Psma2 | 9.19E-66 | 0.410341039 | 0.257 | 0.528 | 1.86E-61 | 0 | Psma2 |
| Llph | 2.27E-65 | 0.388052741 | 0.161 | 0.346 | 4.60E-61 | 0 | Llph |
| Rpl35a | 2.81E-65 | 0.290764263 | 0.889 | 0.962 | 5.69E-61 | 0 | Rpl35a |
| Rpl18 | 2.88E-65 | 0.368630273 | 0.798 | 0.923 | 5.82E-61 | 0 | Rpl18 |
| Mat2b | 2.33E-64 | 0.334715856 | 0.109 | 0.259 | 4.72E-60 | 0 | Mat2b |
| Psmc3 | 7.14E-64 | 0.453793083 | 0.185 | 0.389 | 1.45E-59 | 0 | Psmc3 |
| Mrfap1 | 8.90E-64 | 0.371902343 | 0.245 | 0.497 | 1.80E-59 | 0 | Mrfap1 |
| Eif5 | 4.42E-63 | 0.413084143 | 0.24 | 0.49 | 8.94E-59 | 0 | Eif5 |
| Snf8 | 7.41E-63 | 0.378375322 | 0.134 | 0.298 | 1.50E-58 | 0 | Snf8 |
| Hnrnpab | 7.63E-63 | 0.426523305 | 0.195 | 0.407 | 1.55E-58 | 0 | Hnrnpab |
| Ppp1r14b | 8.55E-63 | 1.29188087 | 0.365 | 0.321 | 1.73E-58 | 0 | Ppp1r14b |
| Fkbp3 | 8.28E-62 | 0.421532611 | 0.214 | 0.438 | 1.68E-57 | 0 | Fkbp3 |
| Rpl7a | 7.87E-61 | 0.673009329 | 0.621 | 0.783 | 1.59E-56 | 0 | Rpl7a |
| Pdcd5 | 1.25E-60 | 0.464284573 | 0.22 | 0.447 | 2.53E-56 | 0 | Pdcd5 |
| Mrpl20 | 2.57E-60 | 0.473172949 | 0.181 | 0.378 | 5.20E-56 | 0 | Mrpl20 |
| Ivns1abp | 7.06E-60 | 0.277505618 | 0.186 | 0.378 | 1.43E-55 | 0 | Ivns1abp |
| Gtf2h5 | 1.04E-59 | 0.462857707 | 0.172 | 0.361 | 2.10E-55 | 0 | Gtf2h5 |
| Stub1 | 2.96E-59 | 0.378585655 | 0.128 | 0.284 | 5.99E-55 | 0 | Stub1 |
| Ntan1 | 3.87E-59 | 0.410792974 | 0.169 | 0.353 | 7.84E-55 | 0 | Ntan1 |
| Vti1b | 1.31E-58 | 0.360745893 | 0.128 | 0.281 | 2.64E-54 | 0 | Vti1b |
| Cstb | 1.95E-58 | 0.338569498 | 0.127 | 0.278 | 3.96E-54 | 0 | Cstb |
| Rpl39 | 5.21E-58 | 0.322855439 | 0.808 | 0.921 | 1.05E-53 | 0 | Rpl39 |

| | | | | | | | |
|---|---|---|---|---|---|---|---|
| Rps23 | 7.40E-58 | 0.264422542 | 0.899 | 0.965 | 1.50E-53 | 0 | Rps23 |
| Nucks1 | 1.96E-57 | 0.493179314 | 0.213 | 0.431 | 3.96E-53 | 0 | Nucks1 |
| Nedd8 | 5.36E-57 | 0.372649021 | 0.305 | 0.606 | 1.09E-52 | 0 | Nedd8 |
| Cript | 8.25E-57 | 0.38024751 | 0.131 | 0.285 | 1.67E-52 | 0 | Cript |
| Mrpl27 | 3.37E-56 | 0.342862369 | 0.116 | 0.257 | 6.83E-52 | 0 | Mrpl27 |
| Sumo1 | 5.91E-56 | 0.46510509 | 0.247 | 0.492 | 1.20E-51 | 0 | Sumo1 |
| Serf2 | 1.23E-55 | 0.524451467 | 0.66 | 0.855 | 2.49E-51 | 0 | Serf2 |
| Vapa | 1.35E-55 | 0.511795451 | 0.207 | 0.419 | 2.73E-51 | 0 | Vapa |
| Pttg1 | 4.56E-55 | 0.382566852 | 0.157 | 0.325 | 9.24E-51 | 0 | Pttg1 |
| Magoh | 8.58E-55 | 0.399276234 | 0.133 | 0.286 | 1.74E-50 | 0 | Magoh |
| Vps36 | 1.89E-54 | 0.47205546 | 0.132 | 0.283 | 3.83E-50 | 0 | Vps36 |
| Srsf9 | 3.62E-54 | 0.450054238 | 0.18 | 0.365 | 7.34E-50 | 0 | Srsf9 |
| Tpi1 | 1.03E-53 | 1.333923304 | 0.361 | 0.332 | 2.08E-49 | 0 | Tpi1 |
| Dnaja2 | 4.98E-53 | 0.520250818 | 0.184 | 0.372 | 1.01E-48 | 0 | Dnaja2 |
| Gm15564 | 5.75E-53 | 0.458564411 | 0.225 | 0.442 | 1.16E-48 | 0 | Gm15564 |
| Drap1 | 9.48E-53 | 0.47683672 | 0.216 | 0.429 | 1.92E-48 | 0 | Drap1 |
| Snrnp27 | 2.14E-52 | 0.449363309 | 0.154 | 0.316 | 4.33E-48 | 0 | Snrnp27 |
| Use1 | 2.48E-52 | 0.466776192 | 0.148 | 0.307 | 5.01E-48 | 0 | Use1 |
| Rpl38 | 5.39E-52 | 0.352722342 | 0.786 | 0.917 | 1.09E-47 | 0 | Rpl38 |
| Trappc2l | 6.01E-52 | 0.370368385 | 0.117 | 0.253 | 1.22E-47 | 0 | Trappc2l |
| Sap18 | 1.06E-51 | 0.454238928 | 0.144 | 0.3 | 2.14E-47 | 0 | Sap18 |
| Scand1 | 1.39E-51 | 0.484052637 | 0.261 | 0.512 | 2.81E-47 | 0 | Scand1 |
| Pnkd | 1.52E-51 | 0.462813222 | 0.161 | 0.328 | 3.08E-47 | 0 | Pnkd |
| Ech1 | 2.53E-51 | 0.479890578 | 0.194 | 0.386 | 5.12E-47 | 0 | Ech1 |
| Cacybp | 3.53E-51 | 0.445663547 | 0.129 | 0.273 | 7.14E-47 | 0 | Cacybp |
| Ndufa1 | 4.52E-51 | 1.2542648 | 0.449 | 0.491 | 9.15E-47 | 0 | Ndufa1 |
| Pop5 | 5.42E-51 | 0.476265709 | 0.161 | 0.328 | 1.10E-46 | 0 | Pop5 |
| Hsp90aa1 | 1.00E-50 | 0.443288822 | 0.241 | 0.468 | 2.03E-46 | 0 | Hsp90aa1 |
| Psmd13 | 1.08E-50 | 0.392914922 | 0.123 | 0.262 | 2.18E-46 | 0 | Psmd13 |
| Lamtor1 | 3.41E-50 | 0.466714294 | 0.175 | 0.349 | 6.91E-46 | 0 | Lamtor1 |
| Rpl30 | 4.48E-50 | 0.264428619 | 0.869 | 0.953 | 9.06E-46 | 0 | Rpl30 |
| Smim10l1 | 6.55E-50 | 0.443626354 | 0.175 | 0.348 | 1.33E-45 | 0 | Smim10l1 |
| Zmat2 | 1.02E-49 | 0.474211319 | 0.14 | 0.29 | 2.06E-45 | 0 | Zmat2 |
| Cenpx | 1.49E-49 | 0.498077255 | 0.165 | 0.332 | 3.01E-45 | 0 | Cenpx |
| Txnl1 | 2.20E-49 | 0.490595559 | 0.176 | 0.351 | 4.45E-45 | 0 | Txnl1 |
| Guk1 | 2.95E-49 | 0.407941532 | 0.128 | 0.269 | 5.98E-45 | 0 | Guk1 |
| Rab18 | 3.45E-49 | 0.425931009 | 0.155 | 0.314 | 6.99E-45 | 0 | Rab18 |
| Eif1b | 4.74E-49 | 0.542783773 | 0.186 | 0.37 | 9.59E-45 | 0 | Eif1b |
| Chmp3 | 8.24E-49 | 0.486782367 | 0.187 | 0.368 | 1.67E-44 | 0 | Chmp3 |
| Ogdh | 1.49E-48 | 0.520884022 | 0.141 | 0.291 | 3.01E-44 | 0 | Ogdh |
| Txndc9 | 2.30E-48 | 0.440880397 | 0.119 | 0.252 | 4.65E-44 | 0 | Txndc9 |
| Psmb7 | 2.54E-48 | 0.403476752 | 0.122 | 0.257 | 5.15E-44 | 0 | Psmb7 |
| Eif3h | 5.24E-48 | 0.437148593 | 0.275 | 0.527 | 1.06E-43 | 0 | Eif3h |

| | | | | | | | |
|---|---|---|---|---|---|---|---|
| Psmc5 | 5.59E-48 | 0.468804068 | 0.16 | 0.32 | 1.13E-43 | 0 | Psmc5 |
| Eif3i | 1.45E-47 | 0.487683705 | 0.213 | 0.415 | 2.94E-43 | 0 | Eif3i |
| Nsmce4a | 3.50E-47 | 0.439755466 | 0.145 | 0.294 | 7.09E-43 | 0 | Nsmce4a |
| Pdcd6 | 3.56E-47 | 0.480173502 | 0.15 | 0.302 | 7.22E-43 | 0 | Pdcd6 |
| Fmc1 | 3.71E-47 | 1.327891007 | 0.313 | 0.271 | 7.50E-43 | 0 | Fmc1 |
| Lamtor5 | 6.02E-47 | 0.508227109 | 0.185 | 0.362 | 1.22E-42 | 0 | Lamtor5 |
| Tpm1 | 7.89E-47 | 0.303681968 | 0.165 | 0.323 | 1.60E-42 | 0 | Tpm1 |
| Lsm4 | 1.95E-46 | 0.488713291 | 0.181 | 0.355 | 3.94E-42 | 0 | Lsm4 |
| Ybx3 | 2.13E-46 | 0.505077311 | 0.192 | 0.372 | 4.32E-42 | 0 | Ybx3 |
| Serbp1 | 2.80E-46 | 0.36445821 | 0.34 | 0.647 | 5.68E-42 | 0 | Serbp1 |
| Glrx3 | 5.51E-46 | 0.519017711 | 0.167 | 0.33 | 1.12E-41 | 0 | Glrx3 |
| Aldh2 | 7.90E-46 | 0.271666935 | 0.233 | 0.435 | 1.60E-41 | 0 | Aldh2 |
| Hmox2 | 9.81E-45 | 0.424446456 | 0.127 | 0.259 | 1.99E-40 | 0 | Hmox2 |
| Rpl12 | 3.16E-44 | 0.386527785 | 0.729 | 0.867 | 6.39E-40 | 0 | Rpl12 |
| Rbbp7 | 1.02E-43 | 0.468712055 | 0.137 | 0.274 | 2.07E-39 | 0 | Rbbp7 |
| Ccni | 2.09E-43 | 0.487967043 | 0.128 | 0.26 | 4.22E-39 | 0 | Ccni |
| Rpl35 | 2.25E-43 | 0.481672837 | 0.687 | 0.86 | 4.56E-39 | 0 | Rpl35 |
| Eif2s2 | 3.61E-43 | 0.572406298 | 0.278 | 0.523 | 7.31E-39 | 0 | Eif2s2 |
| Rex1bd | 4.14E-43 | 0.534346866 | 0.183 | 0.352 | 8.39E-39 | 0 | Rex1bd |
| Set | 5.85E-43 | 0.454675942 | 0.276 | 0.519 | 1.19E-38 | 0 | Set |
| Cbx3 | 1.51E-42 | 0.559975691 | 0.217 | 0.413 | 3.07E-38 | 0 | Cbx3 |
| Bcas2 | 1.77E-42 | 0.58549897 | 0.165 | 0.321 | 3.58E-38 | 0 | Bcas2 |
| Eny2 | 7.42E-42 | 0.498996452 | 0.165 | 0.319 | 1.50E-37 | 0 | Eny2 |
| Emc2 | 2.79E-41 | 0.47526941 | 0.146 | 0.285 | 5.66E-37 | 0 | Emc2 |
| Tomm22 | 3.19E-41 | 0.574876378 | 0.198 | 0.377 | 6.47E-37 | 0 | Tomm22 |
| Commd3 | 3.21E-41 | 0.476335647 | 0.144 | 0.282 | 6.49E-37 | 0 | Commd3 |
| Slc25a39 | 3.94E-41 | 1.264660494 | 0.284 | 0.239 | 7.98E-37 | 0 | Slc25a39 |
| Denr | 9.50E-41 | 0.554530659 | 0.159 | 0.307 | 1.92E-36 | 0 | Denr |
| Nop10 | 1.83E-40 | 0.552248319 | 0.218 | 0.409 | 3.71E-36 | 0 | Nop10 |
| Dbi | 2.22E-40 | 0.397758664 | 0.256 | 0.473 | 4.50E-36 | 0 | Dbi |
| Tomm20 | 3.56E-40 | 0.588441875 | 0.256 | 0.481 | 7.21E-36 | 0 | Tomm20 |
| Ndufb3 | 4.35E-40 | 1.21195556 | 0.425 | 0.471 | 8.81E-36 | 0 | Ndufb3 |
| Eif3k | 1.35E-39 | 0.459520167 | 0.294 | 0.549 | 2.74E-35 | 0 | Eif3k |
| Bag1 | 2.29E-39 | 0.493332178 | 0.241 | 0.445 | 4.64E-35 | 0 | Bag1 |
| Tbca | 4.88E-39 | 0.557175077 | 0.279 | 0.519 | 9.88E-35 | 0 | Tbca |
| Mrpl57 | 5.50E-39 | 0.509319834 | 0.159 | 0.303 | 1.11E-34 | 0 | Mrpl57 |
| Rps5 | 8.28E-39 | 0.266590376 | 0.83 | 0.94 | 1.68E-34 | 0 | Rps5 |
| Eif5a | 1.15E-38 | 0.363454769 | 0.336 | 0.617 | 2.33E-34 | 0 | Eif5a |
| Mrps18c | 1.40E-38 | 0.471415278 | 0.143 | 0.274 | 2.83E-34 | 0 | Mrps18c |
| Polr2k | 2.63E-38 | 0.517453978 | 0.149 | 0.285 | 5.33E-34 | 0 | Polr2k |
| Tmem256 | 3.95E-38 | 0.543414799 | 0.215 | 0.399 | 8.00E-34 | 0 | Tmem256 |
| Sdhd | 5.48E-38 | 1.260991883 | 0.302 | 0.27 | 1.11E-33 | 0 | Sdhd |
| Cfl2 | 5.54E-38 | 0.482954451 | 0.13 | 0.254 | 1.12E-33 | 0 | Cfl2 |

| | | | | | | | |
|---|---|---|---|---|---|---|---|
| Coa3 | 1.08E-37 | 0.551987523 | 0.194 | 0.362 | 2.19E-33 | 0 | Coa3 |
| Gpi1 | 4.99E-37 | 0.533620527 | 0.203 | 0.375 | 1.01E-32 | 0 | Gpi1 |
| Psma3 | 1.02E-36 | 0.472256965 | 0.307 | 0.568 | 2.07E-32 | 0 | Psma3 |
| Mrpl43 | 6.03E-36 | 0.573542909 | 0.16 | 0.301 | 1.22E-31 | 0 | Mrpl43 |
| Ifi27 | 1.34E-35 | 1.058892082 | 0.465 | 0.554 | 2.72E-31 | 0 | Ifi27 |
| Acat1 | 6.48E-35 | 1.355791251 | 0.296 | 0.269 | 1.31E-30 | 0 | Acat1 |
| Hspe1 | 2.42E-34 | 1.117130011 | 0.448 | 0.523 | 4.90E-30 | 0 | Hspe1 |
| Snrpd2 | 4.01E-34 | 0.570928427 | 0.246 | 0.447 | 8.11E-30 | 0 | Snrpd2 |
| Snrpd3 | 5.17E-34 | 0.559887815 | 0.187 | 0.341 | 1.05E-29 | 0 | Snrpd3 |
| Ndufs3 | 2.96E-33 | 1.246114015 | 0.349 | 0.35 | 5.99E-29 | 0 | Ndufs3 |
| Pfdn1 | 4.98E-32 | 0.628189456 | 0.191 | 0.345 | 1.01E-27 | 0 | Pfdn1 |
| Sem1 | 5.93E-32 | 0.335920963 | 0.382 | 0.696 | 1.20E-27 | 0 | Sem1 |
| Rps12 | 9.01E-32 | 0.274135968 | 0.79 | 0.922 | 1.83E-27 | 0 | Rps12 |
| Pnrc1 | 2.38E-31 | 0.567165014 | 0.25 | 0.448 | 4.81E-27 | 0 | Pnrc1 |
| Rps6 | 3.00E-31 | 0.469647532 | 0.658 | 0.846 | 6.08E-27 | 0 | Rps6 |
| Txndc17 | 2.81E-30 | 0.567694072 | 0.279 | 0.497 | 5.70E-26 | 0 | Txndc17 |
| Pfdn6 | 3.18E-30 | 0.560138045 | 0.147 | 0.268 | 6.43E-26 | 0 | Pfdn6 |
| Zcrb1 | 4.53E-30 | 0.652992167 | 0.204 | 0.362 | 9.17E-26 | 0 | Zcrb1 |
| Chchd1 | 6.47E-30 | 0.621938776 | 0.19 | 0.34 | 1.31E-25 | 0 | Chchd1 |
| Rpl22 | 6.92E-30 | 0.372593522 | 0.706 | 0.885 | 1.40E-25 | 0 | Rpl22 |
| Pcbd2 | 9.61E-30 | 0.542683077 | 0.151 | 0.272 | 1.95E-25 | 0 | Pcbd2 |
| Hsbp1 | 2.73E-29 | 0.586018345 | 0.224 | 0.393 | 5.54E-25 | 0 | Hsbp1 |
| Eif1ax | 3.63E-29 | 0.600573842 | 0.169 | 0.301 | 7.35E-25 | 0 | Eif1ax |
| Hist1h1e | 2.42E-28 | 0.528840058 | 0.232 | 0.405 | 4.90E-24 | 0 | Hist1h1e |
| Fis1 | 2.77E-28 | 0.58078067 | 0.283 | 0.503 | 5.60E-24 | 0 | Fis1 |
| Mdh2 | 3.61E-28 | 1.260790699 | 0.363 | 0.389 | 7.32E-24 | 0 | Mdh2 |
| Pip | 4.70E-28 | 0.26092498 | 0.36 | 0.337 | 9.51E-24 | 0 | Pip |
| Dtnbp1 | 9.23E-28 | 0.508779169 | 0.14 | 0.25 | 1.87E-23 | 0 | Dtnbp1 |
| Eef1d | 1.82E-27 | 0.443281149 | 0.336 | 0.598 | 3.68E-23 | 0 | Eef1d |
| Aurkaip1 | 2.00E-27 | 0.617034828 | 0.218 | 0.381 | 4.05E-23 | 0 | Aurkaip1 |
| Prdx1 | 2.93E-27 | 0.737896832 | 0.543 | 0.727 | 5.94E-23 | 0 | Prdx1 |
| Psmb5 | 5.10E-27 | 0.614353855 | 0.257 | 0.453 | 1.03E-22 | 0 | Psmb5 |
| Polr1d | 7.10E-27 | 0.563699768 | 0.302 | 0.536 | 1.44E-22 | 0 | Polr1d |
| Phb2 | 1.74E-26 | 0.625113965 | 0.192 | 0.333 | 3.52E-22 | 0 | Phb2 |
| Ubc | 1.81E-26 | 0.28054826 | 0.411 | 0.72 | 3.67E-22 | 0 | Ubc |
| Ndufb6 | 3.48E-26 | 1.213638356 | 0.315 | 0.311 | 7.04E-22 | 0 | Ndufb6 |
| Rack1 | 5.04E-26 | 0.436962985 | 0.641 | 0.826 | 1.02E-21 | 0 | Rack1 |
| Mrpl52 | 2.44E-25 | 0.566548265 | 0.276 | 0.479 | 4.94E-21 | 0 | Mrpl52 |
| Hes1 | 3.18E-25 | 0.366676724 | 0.206 | 0.34 | 6.44E-21 | 0 | Hes1 |
| Bola3 | 5.51E-25 | 1.287563085 | 0.3 | 0.291 | 1.12E-20 | 0 | Bola3 |
| Ndufc2 | 7.74E-25 | 1.040932729 | 0.462 | 0.583 | 1.57E-20 | 0 | Ndufc2 |
| Pfdn5 | 1.21E-24 | 0.548481776 | 0.589 | 0.803 | 2.45E-20 | 0 | Pfdn5 |
| Dnajc19 | 2.68E-24 | 0.583977892 | 0.199 | 0.339 | 5.42E-20 | 0 | Dnajc19 |

| | | | | | | | |
|---|---|---|---|---|---|---|---|
| Lsm6 | 2.86E-24 | 0.628901076 | 0.201 | 0.343 | 5.79E-20 | 0 | Lsm6 |
| Timm8b | 3.14E-24 | 1.240095312 | 0.318 | 0.321 | 6.35E-20 | 0 | Timm8b |
| Ndufa8 | 3.44E-24 | 0.954115613 | 0.469 | 0.592 | 6.96E-20 | 0 | Ndufa8 |
| Ndufa2 | 9.11E-24 | 0.994436608 | 0.465 | 0.592 | 1.85E-19 | 0 | Ndufa2 |
| Hcfc1r1 | 1.22E-23 | 0.593202977 | 0.193 | 0.328 | 2.47E-19 | 0 | Hcfc1r1 |
| Ufc1 | 1.71E-23 | 0.642708395 | 0.206 | 0.352 | 3.47E-19 | 0 | Ufc1 |
| Iscu | 1.74E-23 | 0.60780757 | 0.166 | 0.284 | 3.52E-19 | 0 | Iscu |
| Snrpe | 1.90E-23 | 0.559236252 | 0.299 | 0.519 | 3.85E-19 | 0 | Snrpe |
| Arpc1a | 2.58E-23 | 0.555356278 | 0.204 | 0.344 | 5.22E-19 | 0 | Arpc1a |
| Nol7 | 5.04E-23 | 0.634401145 | 0.276 | 0.474 | 1.02E-18 | 0 | Nol7 |
| Mrpl58 | 5.68E-23 | 0.585068142 | 0.15 | 0.256 | 1.15E-18 | 0 | Mrpl58 |
| Rpl4 | 8.55E-23 | 0.272433275 | 0.416 | 0.713 | 1.73E-18 | 0 | Rpl4 |
| Pcp4l1 | 4.39E-22 | 0.422535381 | 0.207 | 0.339 | 8.89E-18 | 0 | Pcp4l1 |
| Pomp | 8.53E-22 | 0.907117021 | 0.483 | 0.627 | 1.73E-17 | 0 | Pomp |
| Thoc7 | 9.86E-22 | 0.692009945 | 0.194 | 0.324 | 2.00E-17 | 0 | Thoc7 |
| Eef1g | 1.36E-21 | 0.524167658 | 0.322 | 0.554 | 2.76E-17 | 0 | Eef1g |
| Abracl | 2.92E-21 | 1.108736352 | 0.385 | 0.441 | 5.92E-17 | 0 | Abracl |
| 1810022K09Rik | 3.37E-21 | 0.63234815 | 0.178 | 0.298 | 6.82E-17 | 0 | 1810022K09Rik |
| Commd6 | 2.32E-20 | 0.613283928 | 0.15 | 0.251 | 4.70E-16 | 0 | Commd6 |
| Ndufaf8 | 2.99E-20 | 0.638061564 | 0.166 | 0.276 | 6.06E-16 | 0 | Ndufaf8 |
| Edf1 | 7.63E-20 | 0.4208625 | 0.386 | 0.678 | 1.54E-15 | 0 | Edf1 |
| Rtraf | 8.34E-20 | 0.654798788 | 0.271 | 0.456 | 1.69E-15 | 0 | Rtraf |
| Pycard | 1.71E-19 | 0.483255809 | 0.185 | 0.3 | 3.46E-15 | 0 | Pycard |
| Cops9 | 1.87E-19 | 0.634622925 | 0.214 | 0.352 | 3.79E-15 | 0 | Cops9 |
| Mrps33 | 2.53E-19 | 0.682836504 | 0.244 | 0.405 | 5.12E-15 | 0 | Mrps33 |
| Rps25 | 7.59E-19 | 0.362495412 | 0.658 | 0.869 | 1.54E-14 | 0 | Rps25 |
| Ndufa10 | 3.33E-18 | 1.18003188 | 0.304 | 0.314 | 6.74E-14 | 0 | Ndufa10 |
| Etfa | 4.74E-18 | 1.195052858 | 0.267 | 0.26 | 9.59E-14 | 0 | Etfa |
| Commd1 | 1.05E-17 | 0.658594239 | 0.177 | 0.287 | 2.13E-13 | 0 | Commd1 |
| Mrpl23 | 2.28E-17 | 0.672266969 | 0.194 | 0.313 | 4.62E-13 | 0 | Mrpl23 |
| Stx7 | 4.53E-17 | 1.129678165 | 0.307 | 0.319 | 9.17E-13 | 0 | Stx7 |
| Dynlt3 | 1.08E-16 | 0.58878462 | 0.177 | 0.282 | 2.19E-12 | 0 | Dynlt3 |
| Cox20 | 2.64E-16 | 1.176538512 | 0.293 | 0.302 | 5.35E-12 | 0 | Cox20 |
| Eif4g2 | 3.58E-16 | 0.509919322 | 0.363 | 0.612 | 7.24E-12 | 0 | Eif4g2 |
| Rpl27 | 4.29E-16 | 0.402556183 | 0.379 | 0.629 | 8.68E-12 | 0 | Rpl27 |
| Skp1a | 1.63E-15 | 0.713574486 | 0.22 | 0.353 | 3.31E-11 | 0 | Skp1a |
| Pkm | 2.66E-15 | 0.743453598 | 0.249 | 0.403 | 5.39E-11 | 0 | Pkm |
| Hmgn1 | 3.98E-15 | 0.713107733 | 0.222 | 0.355 | 8.07E-11 | 0 | Hmgn1 |
| Etfb | 4.37E-15 | 1.11987903 | 0.321 | 0.347 | 8.85E-11 | 0 | Etfb |
| Fkbp4 | 7.78E-15 | 0.673213342 | 0.161 | 0.255 | 1.58E-10 | 0 | Fkbp4 |
| Ndufs4 | 9.24E-15 | 0.753626487 | 0.257 | 0.418 | 1.87E-10 | 0 | Ndufs4 |
| Adh5 | 1.18E-14 | 0.685342151 | 0.176 | 0.277 | 2.39E-10 | 0 | Adh5 |
| Zfp706 | 1.26E-14 | 0.749885506 | 0.229 | 0.365 | 2.55E-10 | 0 | Zfp706 |

| | | | | | | | |
|---|---|---|---|---|---|---|---|
| Pdap1 | 2.28E-13 | 0.748655941 | 0.261 | 0.415 | 4.61E-09 | 0 | Pdap1 |
| Hadhb | 3.15E-13 | 0.708549563 | 0.163 | 0.252 | 6.38E-09 | 0 | Hadhb |
| Tkt | 5.63E-13 | 0.670261392 | 0.176 | 0.272 | 1.14E-08 | 0 | Tkt |
| Polr2m | 1.16E-12 | 0.699809014 | 0.171 | 0.263 | 2.36E-08 | 0 | Polr2m |
| Ywhae | 1.81E-12 | 0.657188838 | 0.347 | 0.579 | 3.66E-08 | 0 | Ywhae |
| Dnajc15 | 2.33E-12 | 1.098409197 | 0.304 | 0.329 | 4.71E-08 | 0 | Dnajc15 |
| Txn2 | 3.04E-12 | 0.713470686 | 0.195 | 0.301 | 6.15E-08 | 0 | Txn2 |
| Tmem126a | 3.61E-12 | 0.705021708 | 0.165 | 0.254 | 7.31E-08 | 0 | Tmem126a |
| Srp9 | 8.41E-12 | 0.687916526 | 0.308 | 0.496 | 1.70E-07 | 0 | Srp9 |
| Higd1a | 9.69E-12 | 1.108730735 | 0.315 | 0.35 | 1.96E-07 | 0 | Higd1a |
| Rpl22l1 | 1.42E-11 | 0.398481383 | 0.391 | 0.621 | 2.88E-07 | 0 | Rpl22l1 |
| Hmgb1 | 2.21E-11 | 0.266734913 | 0.471 | 0.771 | 4.47E-07 | 0 | Hmgb1 |
| Mrps24 | 5.82E-11 | 0.738482297 | 0.292 | 0.468 | 1.18E-06 | 0 | Mrps24 |
| Ube2m | 3.52E-10 | 0.771038528 | 0.209 | 0.318 | 7.13E-06 | 0 | Ube2m |
| Tsc22d1 | 3.67E-10 | 0.502447964 | 0.311 | 0.478 | 7.42E-06 | 0 | Tsc22d1 |
| Hint1 | 4.24E-10 | 0.65556857 | 0.513 | 0.737 | 8.58E-06 | 0 | Hint1 |
| Emc6 | 4.97E-10 | 0.734122478 | 0.193 | 0.29 | 1.01E-05 | 0 | Emc6 |
| Ucp2 | 8.14E-10 | 0.589923531 | 0.236 | 0.354 | 1.65E-05 | 0 | Ucp2 |
| Rpl36al | 1.56E-09 | 0.89866589 | 0.422 | 0.548 | 3.16E-05 | 0 | Rpl36al |
| Pebp1 | 2.44E-09 | 1.05516487 | 0.386 | 0.494 | 4.94E-05 | 0 | Pebp1 |
| Clint1 | 2.71E-09 | 0.764879025 | 0.262 | 0.401 | 5.48E-05 | 0 | Clint1 |
| Rnf7 | 4.33E-09 | 0.732452059 | 0.271 | 0.417 | 8.76E-05 | 0 | Rnf7 |
| Gabarapl2 | 4.77E-09 | 0.698902025 | 0.332 | 0.535 | 9.65E-05 | 0 | Gabarapl2 |
| Mrpl54 | 9.97E-09 | 0.749052804 | 0.195 | 0.287 | 0.000201942 | 0 | Mrpl54 |
| Glrx5 | 1.28E-08 | 1.104283166 | 0.258 | 0.276 | 0.000260158 | 0 | Glrx5 |
| C1qbp | 1.92E-08 | 1.112218113 | 0.247 | 0.259 | 0.00038855 | 0 | C1qbp |
| Polr2f | 1.96E-08 | 0.795592544 | 0.246 | 0.372 | 0.000396811 | 0 | Polr2f |
| Mrpl51 | 4.86E-08 | 1.108540755 | 0.307 | 0.355 | 0.000983507 | 0 | Mrpl51 |
| Elob | 5.24E-08 | 0.689289175 | 0.505 | 0.739 | 0.001061617 | 0 | Elob |
| Serpinb6a | 1.29E-07 | 0.924891492 | 0.355 | 0.422 | 0.002610458 | 0 | Serpinb6a |
| Fam162a | 1.95E-07 | 0.775913886 | 0.189 | 0.274 | 0.003945396 | 0 | Fam162a |
| Nme1 | 2.15E-07 | 0.729754259 | 0.255 | 0.38 | 0.004360927 | 0 | Nme1 |
| Mien1 | 2.26E-07 | 1.025232491 | 0.315 | 0.365 | 0.004567083 | 0 | Mien1 |
| Ndufv1 | 2.31E-07 | 1.035488903 | 0.252 | 0.269 | 0.004675091 | 0 | Ndufv1 |
| Naa20 | 2.39E-07 | 0.789676829 | 0.207 | 0.302 | 0.004841709 | 0 | Naa20 |
| Btf3 | 3.31E-07 | 0.526711371 | 0.528 | 0.766 | 0.006709154 | 0 | Btf3 |
| Cox17 | 3.99E-07 | 0.809025318 | 0.233 | 0.342 | 0.008070605 | 0 | Cox17 |
| Park7 | 9.30E-07 | 0.89808579 | 0.428 | 0.588 | 0.018820092 | 0 | Park7 |
| Tma7 | 1.23E-06 | 0.650576874 | 0.357 | 0.562 | 0.024871765 | 0 | Tma7 |
| Cyb5a | 1.38E-06 | 0.677605331 | 0.342 | 0.535 | 0.027949567 | 0 | Cyb5a |
| Camk2n1 | 1.85E-06 | 0.716743478 | 0.203 | 0.29 | 0.037376375 | 0 | Camk2n1 |
| Lamtor4 | 2.79E-06 | 0.739086395 | 0.293 | 0.444 | 0.056430678 | 0 | Lamtor4 |
| Gpx4 | 2.84E-06 | 0.551663206 | 0.414 | 0.677 | 0.057409642 | 0 | Gpx4 |

| | | | | | | | |
|---|---|---|---|---|---|---|---|
| Dynlrb1 | 3.67E-06 | 0.789596748 | 0.298 | 0.449 | 0.074365766 | 0 | Dynlrb1 |
| Npm1 | 3.70E-06 | 0.574255425 | 0.508 | 0.718 | 0.074982899 | 0 | Npm1 |
| Atp6v1e1 | 4.96E-06 | 1.157225085 | 0.337 | 0.415 | 0.100391608 | 0 | Atp6v1e1 |
| Higd2a | 5.04E-06 | 0.804456318 | 0.258 | 0.38 | 0.102066057 | 0 | Higd2a |
| Mrps14 | 5.14E-06 | 0.835489406 | 0.283 | 0.428 | 0.103984284 | 0 | Mrps14 |
| Cox6a1 | 6.06E-06 | 0.734572682 | 0.344 | 0.538 | 0.122633934 | 0 | Cox6a1 |
| Eno1 | 7.96E-06 | 0.807882266 | 0.233 | 0.338 | 0.161082784 | 0 | Eno1 |
| Anapc13 | 1.02E-05 | 0.984181171 | 0.273 | 0.306 | 0.205688341 | 0 | Anapc13 |
| Vdac3 | 1.03E-05 | 1.046042063 | 0.341 | 0.425 | 0.207618799 | 0 | Vdac3 |
| Smdt1 | 1.12E-05 | 0.840154868 | 0.448 | 0.638 | 0.226840045 | 0 | Smdt1 |
| Eef1b2 | 1.19E-05 | 0.432923822 | 0.56 | 0.776 | 0.241500052 | 0 | Eef1b2 |
| Pabpc1 | 1.29E-05 | 0.836932083 | 0.434 | 0.596 | 0.261948041 | 0 | Pabpc1 |
| Rpl36a | 1.68E-05 | 0.430623438 | 0.565 | 0.794 | 0.341085142 | 0 | Rpl36a |
| Ybx1 | 3.46E-05 | 0.432796509 | 0.552 | 0.797 | 0.700235562 | 0 | Ybx1 |
| Rpl23a | 3.58E-05 | 0.313671897 | 0.469 | 0.726 | 0.724294215 | 0 | Rpl23a |
| Scp2 | 3.70E-05 | 0.800971345 | 0.327 | 0.504 | 0.748184204 | 0 | Scp2 |
| Jund | 5.60E-05 | 0.283818547 | 0.464 | 0.687 | 1 | 0 | Jund |
| Romo1 | 5.76E-05 | 0.829597742 | 0.296 | 0.442 | 1 | 0 | Romo1 |
| Sec61b | 6.23E-05 | 0.565015479 | 0.402 | 0.633 | 1 | 0 | Sec61b |
| Ndufs2 | 6.45E-05 | 1.065104066 | 0.289 | 0.341 | 1 | 0 | Ndufs2 |
| Ppdpf | 7.24E-05 | 0.845744914 | 0.216 | 0.305 | 1 | 0 | Ppdpf |
| Ndufa7 | 8.91E-05 | 0.843348049 | 0.432 | 0.608 | 1 | 0 | Ndufa7 |
| Cox7a2l | 0.000112137 | 0.82152833 | 0.427 | 0.593 | 1 | 0 | Cox7a2l |
| Uqcrc2 | 0.000158936 | 1.059382534 | 0.292 | 0.346 | 1 | 0 | Uqcrc2 |
| Gm10076 | 0.000227718 | 0.579668798 | 0.379 | 0.584 | 1 | 0 | Gm10076 |
| Gng5 | 0.000239839 | 0.500065999 | 0.548 | 0.813 | 1 | 0 | Gng5 |
| Ptms | 0.000245155 | 0.559686329 | 0.401 | 0.625 | 1 | 0 | Ptms |
| Lamtor2 | 0.000282778 | 0.787189087 | 0.327 | 0.496 | 1 | 0 | Lamtor2 |
| Rpl31 | 0.000613198 | 0.565273715 | 0.498 | 0.726 | 1 | 0 | Rpl31 |
| Cmpk1 | 0.000638432 | 0.767715168 | 0.233 | 0.322 | 1 | 0 | Cmpk1 |
| Rpl5 | 0.000758762 | 0.465863735 | 0.521 | 0.753 | 1 | 0 | Rpl5 |
| Timm17a | 0.000778974 | 1.01729055 | 0.248 | 0.282 | 1 | 0 | Timm17a |
| Hspa9 | 0.00091246 | 0.856178239 | 0.191 | 0.259 | 1 | 0 | Hspa9 |
| Ubl5 | 0.000985969 | 0.704713394 | 0.378 | 0.59 | 1 | 0 | Ubl5 |
| Taldo1 | 0.001027335 | 0.882134073 | 0.399 | 0.543 | 1 | 0 | Taldo1 |
| Uba52 | 0.001066098 | 0.652046669 | 0.367 | 0.558 | 1 | 0 | Uba52 |
| Mrpl18 | 0.001092221 | 0.854615578 | 0.238 | 0.332 | 1 | 0 | Mrpl18 |
| Mrpl30 | 0.001099458 | 1.034675576 | 0.293 | 0.354 | 1 | 0 | Mrpl30 |
| Uqcc2 | 0.001475462 | 0.990503181 | 0.332 | 0.422 | 1 | 0 | Uqcc2 |
| Mrps21 | 0.001559318 | 0.848441666 | 0.286 | 0.414 | 1 | 0 | Mrps21 |
| Hsd17b10 | 0.002139131 | 0.987594518 | 0.231 | 0.259 | 1 | 0 | Hsd17b10 |
| Ube2b | 0.006842991 | 0.847438402 | 0.292 | 0.42 | 1 | 0 | Ube2b |
| Timm13 | 0.00788058 | 0.966214591 | 0.354 | 0.471 | 1 | 0 | Timm13 |

| | | | | | | | |
|---|---|---|---|---|---|---|---|
| Tomm7 | 0.008114086 | 0.68148671 | 0.394 | 0.613 | 1 | 0 | Tomm7 |
| Gabarap | 0.008809557 | 0.706266785 | 0.455 | 0.67 | 1 | 0 | Gabarap |
| Tomm5 | 0.009729986 | 0.934295467 | 0.253 | 0.353 | 1 | 0 | Tomm5 |
| Flt1 | 0 | 2.254289638 | 1 | 0.293 | 0 | 1 | Flt1 |
| Adgrl4 | 0 | 2.215192569 | 0.994 | 0.245 | 0 | 1 | Adgrl4 |
| Cyp26b1 | 0 | 2.171481668 | 0.661 | 0.142 | 0 | 1 | Cyp26b1 |
| Plpp1 | 0 | 2.126561276 | 0.995 | 0.281 | 0 | 1 | Plpp1 |
| Plcb1 | 0 | 2.122599834 | 0.983 | 0.234 | 0 | 1 | Plcb1 |
| Kdr | 0 | 2.112152542 | 0.992 | 0.258 | 0 | 1 | Kdr |
| Plpp3 | 0 | 2.106445063 | 0.997 | 0.306 | 0 | 1 | Plpp3 |
| Plvap | 0 | 2.080877536 | 0.996 | 0.283 | 0 | 1 | Plvap |
| Ptprg | 0 | 2.032206714 | 0.959 | 0.243 | 0 | 1 | Ptprg |
| Emcn | 0 | 2.021360403 | 1 | 0.278 | 0 | 1 | Emcn |
| Cyyr1 | 0 | 2.018013074 | 0.977 | 0.238 | 0 | 1 | Cyyr1 |
| Sptbn1 | 0 | 2.005332465 | 0.999 | 0.412 | 0 | 1 | Sptbn1 |
| Ptprb | 0 | 1.99794832 | 0.994 | 0.264 | 0 | 1 | Ptprb |
| Col4a1 | 0 | 1.958078768 | 0.988 | 0.293 | 0 | 1 | Col4a1 |
| Pecam1 | 0 | 1.952890621 | 0.995 | 0.314 | 0 | 1 | Pecam1 |
| Nrp1 | 0 | 1.944605161 | 0.993 | 0.342 | 0 | 1 | Nrp1 |
| Piezo2 | 0 | 1.941489235 | 0.876 | 0.149 | 0 | 1 | Piezo2 |
| Nfib | 0 | 1.919842922 | 0.997 | 0.353 | 0 | 1 | Nfib |
| Timp3 | 0 | 1.911627712 | 0.978 | 0.287 | 0 | 1 | Timp3 |
| Cavin2 | 0 | 1.902006308 | 0.964 | 0.238 | 0 | 1 | Cavin2 |
| Arhgap31 | 0 | 1.892474438 | 0.977 | 0.272 | 0 | 1 | Arhgap31 |
| Rasal2 | 0 | 1.882282426 | 0.908 | 0.217 | 0 | 1 | Rasal2 |
| Tcf4 | 0 | 1.873922058 | 0.997 | 0.385 | 0 | 1 | Tcf4 |
| Fabp4 | 0 | 1.858912692 | 0.993 | 0.343 | 0 | 1 | Fabp4 |
| Syne1 | 0 | 1.843435244 | 0.935 | 0.263 | 0 | 1 | Syne1 |
| Ldb2 | 0 | 1.842842959 | 0.933 | 0.23 | 0 | 1 | Ldb2 |
| Col4a2 | 0 | 1.834733571 | 0.957 | 0.261 | 0 | 1 | Col4a2 |
| Igfbp7 | 0 | 1.834703545 | 1 | 0.38 | 0 | 1 | Igfbp7 |
| Eng | 0 | 1.831737297 | 0.96 | 0.253 | 0 | 1 | Eng |
| Ebf1 | 0 | 1.814430258 | 0.988 | 0.273 | 0 | 1 | Ebf1 |
| Iigp1 | 0 | 1.788211777 | 0.722 | 0.152 | 0 | 1 | Iigp1 |
| Ptprm | 0 | 1.782645706 | 0.946 | 0.227 | 0 | 1 | Ptprm |
| Arhgap29 | 0 | 1.781515578 | 0.954 | 0.267 | 0 | 1 | Arhgap29 |
| Crim1 | 0 | 1.758387905 | 0.932 | 0.28 | 0 | 1 | Crim1 |
| Podxl | 0 | 1.749747745 | 0.963 | 0.243 | 0 | 1 | Podxl |
| Igfbp3 | 0 | 1.741304931 | 0.845 | 0.213 | 0 | 1 | Igfbp3 |
| Adgrf5 | 0 | 1.738832765 | 0.962 | 0.234 | 0 | 1 | Adgrf5 |
| Dlc1 | 0 | 1.734090125 | 0.952 | 0.264 | 0 | 1 | Dlc1 |
| Etl4 | 0 | 1.730515289 | 0.879 | 0.196 | 0 | 1 | Etl4 |
| Btnl9 | 0 | 1.727003215 | 0.836 | 0.176 | 0 | 1 | Btnl9 |

| | | | | | | | |
|---|---|---|---|---|---|---|---|
| Rsad2 | 0 | 1.720761331 | 0.748 | 0.175 | 0 | 1 | Rsad2 |
| Mef2c | 0 | 1.713917671 | 0.973 | 0.332 | 0 | 1 | Mef2c |
| Snrk | 0 | 1.705942488 | 0.926 | 0.277 | 0 | 1 | Snrk |
| App | 0 | 1.698815098 | 0.968 | 0.329 | 0 | 1 | App |
| Scarb1 | 0 | 1.675375178 | 0.842 | 0.171 | 0 | 1 | Scarb1 |
| Pcdh17 | 0 | 1.65567415 | 0.773 | 0.15 | 0 | 1 | Pcdh17 |
| Cdh5 | 0 | 1.655247616 | 0.952 | 0.227 | 0 | 1 | Cdh5 |
| Ly6c1 | 0 | 1.649936087 | 0.99 | 0.302 | 0 | 1 | Ly6c1 |
| Exoc3l2 | 0 | 1.647427885 | 0.868 | 0.171 | 0 | 1 | Exoc3l2 |
| Apbb2 | 0 | 1.646340326 | 0.926 | 0.274 | 0 | 1 | Apbb2 |
| Egfl7 | 0 | 1.639990551 | 0.995 | 0.294 | 0 | 1 | Egfl7 |
| Mcc | 0 | 1.6246285 | 0.831 | 0.193 | 0 | 1 | Mcc |
| Rgcc | 0 | 1.622910483 | 0.96 | 0.312 | 0 | 1 | Rgcc |
| Slfn5 | 0 | 1.62240169 | 0.905 | 0.266 | 0 | 1 | Slfn5 |
| Fbxl7 | 0 | 1.621591647 | 0.781 | 0.171 | 0 | 1 | Fbxl7 |
| Clec14a | 0 | 1.614250402 | 0.881 | 0.21 | 0 | 1 | Clec14a |
| Rbms1 | 0 | 1.609368184 | 0.965 | 0.407 | 0 | 1 | Rbms1 |
| Dock4 | 0 | 1.604824151 | 0.841 | 0.206 | 0 | 1 | Dock4 |
| Stox2 | 0 | 1.603977101 | 0.776 | 0.159 | 0 | 1 | Stox2 |
| Itga1 | 0 | 1.601907083 | 0.945 | 0.327 | 0 | 1 | Itga1 |
| Jam3 | 0 | 1.598199809 | 0.855 | 0.179 | 0 | 1 | Jam3 |
| Id3 | 0 | 1.596082693 | 0.98 | 0.314 | 0 | 1 | Id3 |
| Fmo2 | 0 | 1.591803335 | 0.857 | 0.194 | 0 | 1 | Fmo2 |
| Dysf | 0 | 1.585551633 | 0.792 | 0.147 | 0 | 1 | Dysf |
| Tcim | 0 | 1.584565425 | 0.876 | 0.208 | 0 | 1 | Tcim |
| Clic4 | 0 | 1.582625648 | 0.923 | 0.341 | 0 | 1 | Clic4 |
| Bmpr2 | 0 | 1.579011819 | 0.885 | 0.261 | 0 | 1 | Bmpr2 |
| Rbp7 | 0 | 1.578775336 | 0.922 | 0.252 | 0 | 1 | Rbp7 |
| Pltp | 0 | 1.577601872 | 0.867 | 0.206 | 0 | 1 | Pltp |
| Adgrl2 | 0 | 1.570279857 | 0.78 | 0.147 | 0 | 1 | Adgrl2 |
| Rgl1 | 0 | 1.5618541 | 0.824 | 0.201 | 0 | 1 | Rgl1 |
| Cd34 | 0 | 1.559688986 | 0.899 | 0.225 | 0 | 1 | Cd34 |
| Gpihbp1 | 0 | 1.559582977 | 0.869 | 0.201 | 0 | 1 | Gpihbp1 |
| Ehd4 | 0 | 1.559007267 | 0.927 | 0.322 | 0 | 1 | Ehd4 |
| Rasgrp3 | 0 | 1.558934093 | 0.836 | 0.19 | 0 | 1 | Rasgrp3 |
| Hmcn1 | 0 | 1.556628079 | 0.718 | 0.127 | 0 | 1 | Hmcn1 |
| Fos | 0 | 1.556455203 | 0.687 | 0.256 | 0 | 1 | Fos |
| Calcrl | 0 | 1.555136636 | 0.784 | 0.177 | 0 | 1 | Calcrl |
| Slc43a3 | 0 | 1.550984409 | 0.786 | 0.184 | 0 | 1 | Slc43a3 |
| Hspg2 | 0 | 1.549276381 | 0.882 | 0.217 | 0 | 1 | Hspg2 |
| Ly6a | 0 | 1.548137645 | 0.987 | 0.372 | 0 | 1 | Ly6a |
| Tek | 0 | 1.546673084 | 0.823 | 0.166 | 0 | 1 | Tek |
| Cd200 | 0 | 1.54525552 | 0.945 | 0.251 | 0 | 1 | Cd200 |

| | | | | | | | |
|---|---|---|---|---|---|---|---|
| Cd93 | 0 | 1.544732524 | 0.814 | 0.163 | 0 | 1 | Cd93 |
| Tshz2 | 0 | 1.540755204 | 0.915 | 0.282 | 0 | 1 | Tshz2 |
| Sparcl1 | 0 | 1.53777063 | 0.939 | 0.268 | 0 | 1 | Sparcl1 |
| Samd12 | 0 | 1.536599913 | 0.755 | 0.142 | 0 | 1 | Samd12 |
| Fbln2 | 0 | 1.536028109 | 0.779 | 0.188 | 0 | 1 | Fbln2 |
| Meis2 | 0 | 1.535586302 | 0.749 | 0.205 | 0 | 1 | Meis2 |
| Slc6a6 | 0 | 1.533861803 | 0.788 | 0.251 | 0 | 1 | Slc6a6 |
| Hecw2 | 0 | 1.531168697 | 0.779 | 0.161 | 0 | 1 | Hecw2 |
| Dnm3 | 0 | 1.530760651 | 0.734 | 0.146 | 0 | 1 | Dnm3 |
| Clic5 | 0 | 1.524862245 | 0.866 | 0.196 | 0 | 1 | Clic5 |
| Col13a1 | 0 | 1.51862329 | 0.779 | 0.153 | 0 | 1 | Col13a1 |
| Jun | 0 | 1.516115009 | 0.908 | 0.45 | 0 | 1 | Jun |
| Rapgef4 | 0 | 1.508522911 | 0.814 | 0.165 | 0 | 1 | Rapgef4 |
| Hspb1 | 0 | 1.504375575 | 0.929 | 0.29 | 0 | 1 | Hspb1 |
| Prex2 | 0 | 1.499840377 | 0.813 | 0.183 | 0 | 1 | Prex2 |
| Tnfsf10 | 0 | 1.495283008 | 0.891 | 0.289 | 0 | 1 | Tnfsf10 |
| Prss23 | 0 | 1.494619598 | 0.788 | 0.197 | 0 | 1 | Prss23 |
| Cd300lg | 0 | 1.494001912 | 0.884 | 0.198 | 0 | 1 | Cd300lg |
| Cav1 | 0 | 1.493745512 | 0.881 | 0.208 | 0 | 1 | Cav1 |
| Klf2 | 0 | 1.488246687 | 0.886 | 0.281 | 0 | 1 | Klf2 |
| Aplp2 | 0 | 1.482180385 | 0.926 | 0.332 | 0 | 1 | Aplp2 |
| Plekhg1 | 0 | 1.481832897 | 0.769 | 0.158 | 0 | 1 | Plekhg1 |
| 9530026P05Rik | 0 | 1.481531779 | 0.703 | 0.165 | 0 | 1 | 9530026P05Rik |
| Car4 | 0 | 1.467317599 | 0.711 | 0.145 | 0 | 1 | Car4 |
| Nostrin | 0 | 1.464017422 | 0.793 | 0.17 | 0 | 1 | Nostrin |
| Id1 | 0 | 1.455454145 | 0.868 | 0.235 | 0 | 1 | Id1 |
| Rdx | 0 | 1.448396757 | 0.979 | 0.481 | 0 | 1 | Rdx |
| Ece1 | 0 | 1.446472304 | 0.848 | 0.216 | 0 | 1 | Ece1 |
| Mgll | 0 | 1.442638589 | 0.973 | 0.363 | 0 | 1 | Mgll |
| Kitl | 0 | 1.44255524 | 0.891 | 0.244 | 0 | 1 | Kitl |
| Ramp3 | 0 | 1.437597624 | 0.838 | 0.172 | 0 | 1 | Ramp3 |
| Kctd12b | 0 | 1.433433284 | 0.733 | 0.169 | 0 | 1 | Kctd12b |
| Itga6 | 0 | 1.426462145 | 0.78 | 0.201 | 0 | 1 | Itga6 |
| Tcf15 | 0 | 1.426144632 | 0.752 | 0.178 | 0 | 1 | Tcf15 |
| Ccdc85a | 0 | 1.423379454 | 0.645 | 0.104 | 0 | 1 | Ccdc85a |
| Myo6 | 0 | 1.421973968 | 0.785 | 0.21 | 0 | 1 | Myo6 |
| Klf4 | 0 | 1.419344487 | 0.745 | 0.191 | 0 | 1 | Klf4 |
| 4931406P16Rik | 0 | 1.417440924 | 0.863 | 0.257 | 0 | 1 | 4931406P16Rik |
| Numb | 0 | 1.41518094 | 0.816 | 0.258 | 0 | 1 | Numb |
| Jam2 | 0 | 1.405047031 | 0.836 | 0.195 | 0 | 1 | Jam2 |
| Sparc | 0 | 1.404187628 | 0.97 | 0.309 | 0 | 1 | Sparc |
| Cald1 | 0 | 1.402779355 | 0.877 | 0.23 | 0 | 1 | Cald1 |
| Elk3 | 0 | 1.397007954 | 0.847 | 0.275 | 0 | 1 | Elk3 |

| | | | | | | | |
|---|---|---|---|---|---|---|---|
| Thbd | 0 | 1.395561246 | 0.766 | 0.17 | 0 | 1 | Thbd |
| Lama3 | 0 | 1.394253935 | 0.722 | 0.138 | 0 | 1 | Lama3 |
| Gng11 | 0 | 1.391935575 | 0.935 | 0.272 | 0 | 1 | Gng11 |
| Pdgfd | 0 | 1.3907249 | 0.768 | 0.168 | 0 | 1 | Pdgfd |
| Plscr2 | 0 | 1.389180274 | 0.891 | 0.242 | 0 | 1 | Plscr2 |
| Pde7b | 0 | 1.385417475 | 0.684 | 0.17 | 0 | 1 | Pde7b |
| Tie1 | 0 | 1.385380186 | 0.798 | 0.163 | 0 | 1 | Tie1 |
| Plxna2 | 0 | 1.385246608 | 0.705 | 0.138 | 0 | 1 | Plxna2 |
| Ednrb | 0 | 1.384041648 | 0.68 | 0.114 | 0 | 1 | Ednrb |
| Fgd5 | 0 | 1.380951004 | 0.736 | 0.145 | 0 | 1 | Fgd5 |
| Klf7 | 0 | 1.375032483 | 0.793 | 0.215 | 0 | 1 | Klf7 |
| Thrsp | 0 | 1.359807994 | 0.849 | 0.201 | 0 | 1 | Thrsp |
| Syne2 | 0 | 1.358879683 | 0.757 | 0.192 | 0 | 1 | Syne2 |
| Sptan1 | 0 | 1.358818625 | 0.824 | 0.264 | 0 | 1 | Sptan1 |
| Gm13889 | 0 | 1.352967225 | 0.696 | 0.14 | 0 | 1 | Gm13889 |
| Arhgef12 | 0 | 1.344434687 | 0.85 | 0.265 | 0 | 1 | Arhgef12 |
| Shank3 | 0 | 1.342474974 | 0.68 | 0.11 | 0 | 1 | Shank3 |
| Sema6a | 0 | 1.340313546 | 0.68 | 0.131 | 0 | 1 | Sema6a |
| Tinagl1 | 0 | 1.338887011 | 0.84 | 0.207 | 0 | 1 | Tinagl1 |
| Rflnb | 0 | 1.337209182 | 0.867 | 0.223 | 0 | 1 | Rflnb |
| Esam | 0 | 1.328183975 | 0.873 | 0.216 | 0 | 1 | Esam |
| Fli1 | 0 | 1.32808576 | 0.85 | 0.328 | 0 | 1 | Fli1 |
| Sox18 | 0 | 1.327456778 | 0.814 | 0.187 | 0 | 1 | Sox18 |
| Rai14 | 0 | 1.322332619 | 0.717 | 0.151 | 0 | 1 | Rai14 |
| Ahr | 0 | 1.321597095 | 0.706 | 0.17 | 0 | 1 | Ahr |
| Prkg1 | 0 | 1.321511276 | 0.607 | 0.129 | 0 | 1 | Prkg1 |
| Qk | 0 | 1.319786291 | 0.883 | 0.332 | 0 | 1 | Qk |
| Zfp366 | 0 | 1.313813678 | 0.632 | 0.108 | 0 | 1 | Zfp366 |
| Hmbox1 | 0 | 1.31333848 | 0.73 | 0.198 | 0 | 1 | Hmbox1 |
| Phlpp1 | 0 | 1.309412352 | 0.7 | 0.191 | 0 | 1 | Phlpp1 |
| Sorbs2 | 0 | 1.305782752 | 0.793 | 0.192 | 0 | 1 | Sorbs2 |
| Mmrn2 | 0 | 1.304280371 | 0.738 | 0.148 | 0 | 1 | Mmrn2 |
| Rapgef2 | 0 | 1.301008181 | 0.797 | 0.263 | 0 | 1 | Rapgef2 |
| Cd24a | 0 | 1.299237788 | 0.813 | 0.265 | 0 | 1 | Cd24a |
| Asap1 | 0 | 1.298686728 | 0.829 | 0.278 | 0 | 1 | Asap1 |
| Ramp2 | 0 | 1.290160526 | 0.879 | 0.241 | 0 | 1 | Ramp2 |
| Car8 | 0 | 1.288876543 | 0.63 | 0.113 | 0 | 1 | Car8 |
| Itpkb | 0 | 1.286295054 | 0.805 | 0.282 | 0 | 1 | Itpkb |
| Rnf220 | 0 | 1.286140383 | 0.651 | 0.174 | 0 | 1 | Rnf220 |
| Tspan9 | 0 | 1.281762582 | 0.74 | 0.187 | 0 | 1 | Tspan9 |
| Slc9a3r2 | 0 | 1.280665582 | 0.919 | 0.279 | 0 | 1 | Slc9a3r2 |
| Ahnak | 0 | 1.275745155 | 0.928 | 0.397 | 0 | 1 | Ahnak |
| Ptgs1 | 0 | 1.274037234 | 0.657 | 0.138 | 0 | 1 | Ptgs1 |

| | | | | | | | |
|---|---|---|---|---|---|---|---|
| Ptk2 | 0 | 1.27177722 | 0.756 | 0.185 | 0 | 1 | Ptk2 |
| Epas1 | 0 | 1.270876085 | 0.905 | 0.269 | 0 | 1 | Epas1 |
| Fam167b | 0 | 1.263676318 | 0.684 | 0.133 | 0 | 1 | Fam167b |
| Gbp7 | 0 | 1.262179189 | 0.668 | 0.204 | 0 | 1 | Gbp7 |
| Ackr3 | 0 | 1.257249127 | 0.534 | 0.089 | 0 | 1 | Ackr3 |
| Tmtc2 | 0 | 1.255435226 | 0.578 | 0.12 | 0 | 1 | Tmtc2 |
| Macf1 | 0 | 1.253777623 | 0.931 | 0.432 | 0 | 1 | Macf1 |
| Rapgef5 | 0 | 1.253374906 | 0.657 | 0.15 | 0 | 1 | Rapgef5 |
| Utrn | 0 | 1.242370521 | 0.872 | 0.329 | 0 | 1 | Utrn |
| Fzd4 | 0 | 1.237714528 | 0.703 | 0.151 | 0 | 1 | Fzd4 |
| Ltbp4 | 0 | 1.232665184 | 0.836 | 0.222 | 0 | 1 | Ltbp4 |
| Ifitm3 | 0 | 1.229548323 | 0.987 | 0.464 | 0 | 1 | Ifitm3 |
| Col15a1 | 0 | 1.229161634 | 0.76 | 0.162 | 0 | 1 | Col15a1 |
| Gas6 | 0 | 1.227468887 | 0.806 | 0.214 | 0 | 1 | Gas6 |
| Pitpnc1 | 0 | 1.224091316 | 0.884 | 0.354 | 0 | 1 | Pitpnc1 |
| Tm4sf1 | 0 | 1.223845187 | 0.965 | 0.28 | 0 | 1 | Tm4sf1 |
| Magi1 | 0 | 1.223228079 | 0.677 | 0.158 | 0 | 1 | Magi1 |
| Csgalnact1 | 0 | 1.217648095 | 0.606 | 0.128 | 0 | 1 | Csgalnact1 |
| Chst15 | 0 | 1.215294597 | 0.615 | 0.102 | 0 | 1 | Chst15 |
| Actn4 | 0 | 1.214992465 | 0.84 | 0.299 | 0 | 1 | Actn4 |
| Oxr1 | 0 | 1.214300572 | 0.753 | 0.267 | 0 | 1 | Oxr1 |
| Nrp2 | 0 | 1.213958894 | 0.601 | 0.115 | 0 | 1 | Nrp2 |
| Bcam | 0 | 1.211226803 | 0.743 | 0.173 | 0 | 1 | Bcam |
| Stard8 | 0 | 1.20957641 | 0.636 | 0.125 | 0 | 1 | Stard8 |
| Prkch | 0 | 1.20698794 | 0.876 | 0.321 | 0 | 1 | Prkch |
| Ctla2a | 0 | 1.205795409 | 0.844 | 0.325 | 0 | 1 | Ctla2a |
| Fut8 | 0 | 1.205594728 | 0.678 | 0.225 | 0 | 1 | Fut8 |
| Abcg2 | 0 | 1.204279406 | 0.736 | 0.194 | 0 | 1 | Abcg2 |
| Cmtm8 | 0 | 1.201093552 | 0.772 | 0.19 | 0 | 1 | Cmtm8 |
| Dock1 | 0 | 1.198917232 | 0.735 | 0.205 | 0 | 1 | Dock1 |
| Nav1 | 0 | 1.195573421 | 0.671 | 0.149 | 0 | 1 | Nav1 |
| Vamp5 | 0 | 1.1941142 | 0.9 | 0.3 | 0 | 1 | Vamp5 |
| Ly6e | 0 | 1.190186486 | 0.988 | 0.559 | 0 | 1 | Ly6e |
| Lrmda | 0 | 1.184895337 | 0.767 | 0.303 | 0 | 1 | Lrmda |
| Ets1 | 0 | 1.18236602 | 0.851 | 0.33 | 0 | 1 | Ets1 |
| Cpd | 0 | 1.18189598 | 0.765 | 0.238 | 0 | 1 | Cpd |
| Cd38 | 0 | 1.177545915 | 0.658 | 0.161 | 0 | 1 | Cd38 |
| Gm12002 | 0 | 1.17712651 | 0.604 | 0.12 | 0 | 1 | Gm12002 |
| S1pr1 | 0 | 1.175497928 | 0.74 | 0.187 | 0 | 1 | S1pr1 |
| Ifitm2 | 0 | 1.171455922 | 0.965 | 0.382 | 0 | 1 | Ifitm2 |
| Mcf2l | 0 | 1.168590691 | 0.648 | 0.142 | 0 | 1 | Mcf2l |
| Sh3bp5 | 0 | 1.167728931 | 0.634 | 0.147 | 0 | 1 | Sh3bp5 |
| Cavin3 | 0 | 1.165360749 | 0.796 | 0.226 | 0 | 1 | Cavin3 |

| | | | | | | | |
|---|---|---|---|---|---|---|---|
| Heg1 | 0 | 1.163983688 | 0.728 | 0.199 | 0 | 1 | Heg1 |
| Ncoa7 | 0 | 1.163175779 | 0.704 | 0.229 | 0 | 1 | Ncoa7 |
| Lrrc8c | 0 | 1.162494356 | 0.736 | 0.241 | 0 | 1 | Lrrc8c |
| Myct1 | 0 | 1.160894293 | 0.685 | 0.146 | 0 | 1 | Myct1 |
| Cd47 | 0 | 1.154189268 | 0.935 | 0.44 | 0 | 1 | Cd47 |
| Ushbp1 | 0 | 1.152416508 | 0.685 | 0.142 | 0 | 1 | Ushbp1 |
| Peak1 | 0 | 1.152226588 | 0.8 | 0.32 | 0 | 1 | Peak1 |
| Arl15 | 0 | 1.150610132 | 0.823 | 0.309 | 0 | 1 | Arl15 |
| Zhx3 | 0 | 1.150213279 | 0.604 | 0.136 | 0 | 1 | Zhx3 |
| Prkce | 0 | 1.138669023 | 0.6 | 0.151 | 0 | 1 | Prkce |
| Tspan12 | 0 | 1.137013738 | 0.692 | 0.156 | 0 | 1 | Tspan12 |
| Tmem88 | 0 | 1.136602946 | 0.666 | 0.135 | 0 | 1 | Tmem88 |
| Srgn | 0 | 1.135794707 | 0.971 | 0.475 | 0 | 1 | Srgn |
| Gnai2 | 0 | 1.134358771 | 0.963 | 0.511 | 0 | 1 | Gnai2 |
| Ndrg1 | 0 | 1.133245037 | 0.674 | 0.16 | 0 | 1 | Ndrg1 |
| Ifi44 | 0 | 1.130492456 | 0.505 | 0.087 | 0 | 1 | Ifi44 |
| Rasgrf2 | 0 | 1.127288914 | 0.582 | 0.11 | 0 | 1 | Rasgrf2 |
| Ntf3 | 0 | 1.127050817 | 0.588 | 0.114 | 0 | 1 | Ntf3 |
| Chst2 | 0 | 1.123160979 | 0.576 | 0.097 | 0 | 1 | Chst2 |
| Grrp1 | 0 | 1.120472591 | 0.669 | 0.145 | 0 | 1 | Grrp1 |
| Rasip1 | 0 | 1.117193352 | 0.628 | 0.123 | 0 | 1 | Rasip1 |
| Kank3 | 0 | 1.115400665 | 0.684 | 0.153 | 0 | 1 | Kank3 |
| Phldb2 | 0 | 1.114574073 | 0.633 | 0.132 | 0 | 1 | Phldb2 |
| Sash1 | 0 | 1.112495494 | 0.692 | 0.185 | 0 | 1 | Sash1 |
| Prkd1 | 0 | 1.112313353 | 0.505 | 0.089 | 0 | 1 | Prkd1 |
| Tmem204 | 0 | 1.111106149 | 0.642 | 0.138 | 0 | 1 | Tmem204 |
| Rhoj | 0 | 1.108970499 | 0.683 | 0.169 | 0 | 1 | Rhoj |
| Msn | 0 | 1.107216426 | 0.858 | 0.375 | 0 | 1 | Msn |
| Gm20532 | 0 | 1.106378492 | 0.43 | 0.057 | 0 | 1 | Gm20532 |
| Tcf7l1 | 0 | 1.101842022 | 0.559 | 0.119 | 0 | 1 | Tcf7l1 |
| Lmcd1 | 0 | 1.10142449 | 0.562 | 0.1 | 0 | 1 | Lmcd1 |
| Rapgef3 | 0 | 1.098548193 | 0.583 | 0.1 | 0 | 1 | Rapgef3 |
| Ephb4 | 0 | 1.098093644 | 0.571 | 0.099 | 0 | 1 | Ephb4 |
| Nr3c1 | 0 | 1.096485644 | 0.804 | 0.314 | 0 | 1 | Nr3c1 |
| Malat1 | 0 | 1.083660742 | 1 | 0.681 | 0 | 1 | Malat1 |
| Xdh | 0 | 1.082815508 | 0.566 | 0.126 | 0 | 1 | Xdh |
| Wwtr1 | 0 | 1.082708105 | 0.674 | 0.169 | 0 | 1 | Wwtr1 |
| Robo4 | 0 | 1.078496396 | 0.597 | 0.107 | 0 | 1 | Robo4 |
| Emp2 | 0 | 1.076729208 | 0.675 | 0.163 | 0 | 1 | Emp2 |
| Meox1 | 0 | 1.073935855 | 0.613 | 0.116 | 0 | 1 | Meox1 |
| Zfp36l1 | 0 | 1.071776349 | 0.83 | 0.368 | 0 | 1 | Zfp36l1 |
| Fermt2 | 0 | 1.070884149 | 0.846 | 0.268 | 0 | 1 | Fermt2 |
| Kirrel3 | 0 | 1.070586305 | 0.284 | 0.035 | 0 | 1 | Kirrel3 |

| | | | | | | | |
|---|---|---|---|---|---|---|---|
| Il6st | 0 | 1.070324858 | 0.644 | 0.164 | 0 | 1 | Il6st |
| Junb | 0 | 1.068544999 | 0.72 | 0.34 | 0 | 1 | Junb |
| Cd81 | 0 | 1.067868553 | 0.982 | 0.45 | 0 | 1 | Cd81 |
| St6galnac3 | 0 | 1.066201924 | 0.635 | 0.147 | 0 | 1 | St6galnac3 |
| Eva1b | 0 | 1.060770866 | 0.715 | 0.215 | 0 | 1 | Eva1b |
| Dusp1 | 0 | 1.058487215 | 0.505 | 0.142 | 0 | 1 | Dusp1 |
| Arap3 | 0 | 1.058285991 | 0.611 | 0.153 | 0 | 1 | Arap3 |
| Dhrs3 | 0 | 1.058284801 | 0.633 | 0.165 | 0 | 1 | Dhrs3 |
| Pdgfb | 0 | 1.058201869 | 0.728 | 0.241 | 0 | 1 | Pdgfb |
| Airn | 0 | 1.056443172 | 0.619 | 0.187 | 0 | 1 | Airn |
| Jup | 0 | 1.052043379 | 0.693 | 0.186 | 0 | 1 | Jup |
| Id4 | 0 | 1.05197572 | 0.459 | 0.07 | 0 | 1 | Id4 |
| Filip1 | 0 | 1.048122125 | 0.59 | 0.129 | 0 | 1 | Filip1 |
| Nbea | 0 | 1.045948145 | 0.569 | 0.135 | 0 | 1 | Nbea |
| Uaca | 0 | 1.045927285 | 0.652 | 0.156 | 0 | 1 | Uaca |
| Sntb2 | 0 | 1.045499145 | 0.655 | 0.202 | 0 | 1 | Sntb2 |
| Fmnl2 | 0 | 1.043394957 | 0.611 | 0.182 | 0 | 1 | Fmnl2 |
| Kctd12 | 0 | 1.043259849 | 0.639 | 0.196 | 0 | 1 | Kctd12 |
| Dip2c | 0 | 1.041723016 | 0.602 | 0.16 | 0 | 1 | Dip2c |
| Mkl2 | 0 | 1.041558056 | 0.652 | 0.19 | 0 | 1 | Mkl2 |
| Zeb1 | 0 | 1.037215404 | 0.706 | 0.259 | 0 | 1 | Zeb1 |
| Afdn | 0 | 1.036063823 | 0.736 | 0.246 | 0 | 1 | Afdn |
| H2-T23 | 0 | 1.035386411 | 0.858 | 0.394 | 0 | 1 | H2-T23 |
| Gimap6 | 0 | 1.032887103 | 0.798 | 0.295 | 0 | 1 | Gimap6 |
| Tjp1 | 0 | 1.032851153 | 0.678 | 0.195 | 0 | 1 | Tjp1 |
| Pdlim5 | 0 | 1.032714167 | 0.663 | 0.224 | 0 | 1 | Pdlim5 |
| Hip1 | 0 | 1.031996518 | 0.731 | 0.269 | 0 | 1 | Hip1 |
| Myo1c | 0 | 1.028113889 | 0.597 | 0.151 | 0 | 1 | Myo1c |
| Mndal | 0 | 1.027319063 | 0.784 | 0.344 | 0 | 1 | Mndal |
| Klf13 | 0 | 1.026383787 | 0.756 | 0.284 | 0 | 1 | Klf13 |
| Arhgap18 | 0 | 1.026335132 | 0.711 | 0.281 | 0 | 1 | Arhgap18 |
| Smco4 | 0 | 1.026159652 | 0.854 | 0.314 | 0 | 1 | Smco4 |
| Ifi203 | 0 | 1.024562372 | 0.748 | 0.305 | 0 | 1 | Ifi203 |
| Plk2 | 0 | 1.023360414 | 0.521 | 0.108 | 0 | 1 | Plk2 |
| Rbpms | 0 | 1.022358714 | 0.791 | 0.277 | 0 | 1 | Rbpms |
| Dab2ip | 0 | 1.020287315 | 0.678 | 0.193 | 0 | 1 | Dab2ip |
| Nxn | 0 | 1.020115399 | 0.667 | 0.203 | 0 | 1 | Nxn |
| Ttc28 | 0 | 1.01954287 | 0.655 | 0.224 | 0 | 1 | Ttc28 |
| Adamts9 | 0 | 1.015999691 | 0.422 | 0.075 | 0 | 1 | Adamts9 |
| Egr1 | 0 | 1.014896746 | 0.437 | 0.133 | 0 | 1 | Egr1 |
| Man2a1 | 0 | 1.013226175 | 0.577 | 0.178 | 0 | 1 | Man2a1 |
| Mfge8 | 0 | 1.012493269 | 0.684 | 0.183 | 0 | 1 | Mfge8 |
| Gnaq | 0 | 1.011933147 | 0.761 | 0.293 | 0 | 1 | Gnaq |

| | | | | | | | |
|---|---|---|---|---|---|---|---|
| Colgalt2 | 0 | 1.009432797 | 0.483 | 0.084 | 0 | 1 | Colgalt2 |
| Cpne8 | 0 | 1.00558976 | 0.638 | 0.161 | 0 | 1 | Cpne8 |
| Itih5 | 0 | 1.005378951 | 0.647 | 0.177 | 0 | 1 | Itih5 |
| Myo18a | 0 | 1.001809433 | 0.661 | 0.191 | 0 | 1 | Myo18a |
| St3gal4 | 0 | 1.001625113 | 0.602 | 0.2 | 0 | 1 | St3gal4 |
| Ecscr | 0 | 1.000822314 | 0.666 | 0.156 | 0 | 1 | Ecscr |
| Ccnd1 | 0 | 1.000497982 | 0.572 | 0.17 | 0 | 1 | Ccnd1 |
| Tgm2 | 0 | 0.998864478 | 0.525 | 0.102 | 0 | 1 | Tgm2 |
| Tmem140 | 0 | 0.998115925 | 0.61 | 0.159 | 0 | 1 | Tmem140 |
| Calm1 | 0 | 0.996987946 | 0.992 | 0.746 | 0 | 1 | Calm1 |
| Clstn1 | 0 | 0.995737853 | 0.641 | 0.17 | 0 | 1 | Clstn1 |
| Sgk1 | 0 | 0.990341119 | 0.658 | 0.206 | 0 | 1 | Sgk1 |
| Reep3 | 0 | 0.984990134 | 0.698 | 0.242 | 0 | 1 | Reep3 |
| Aqp1 | 0 | 0.982527066 | 0.313 | 0.058 | 0 | 1 | Aqp1 |
| Arhgef15 | 0 | 0.98150803 | 0.552 | 0.106 | 0 | 1 | Arhgef15 |
| Sh3bgrl | 0 | 0.981461971 | 0.777 | 0.309 | 0 | 1 | Sh3bgrl |
| 9930111J21Rik2 | 0 | 0.98113111 | 0.605 | 0.2 | 0 | 1 | 9930111J21Rik2 |
| Rhob | 0 | 0.979156675 | 0.603 | 0.213 | 0 | 1 | Rhob |
| Zbtb46 | 0 | 0.978400393 | 0.503 | 0.107 | 0 | 1 | Zbtb46 |
| Grb10 | 0 | 0.97802302 | 0.555 | 0.136 | 0 | 1 | Grb10 |
| Acvrl1 | 0 | 0.977345975 | 0.689 | 0.207 | 0 | 1 | Acvrl1 |
| Gngt2 | 0 | 0.97385692 | 0.795 | 0.288 | 0 | 1 | Gngt2 |
| Spred1 | 0 | 0.973269012 | 0.615 | 0.193 | 0 | 1 | Spred1 |
| Amotl1 | 0 | 0.968423243 | 0.546 | 0.126 | 0 | 1 | Amotl1 |
| H2-Q4 | 0 | 0.966576098 | 0.689 | 0.246 | 0 | 1 | H2-Q4 |
| Lims2 | 0 | 0.965937235 | 0.608 | 0.134 | 0 | 1 | Lims2 |
| Aplnr | 0 | 0.965749257 | 0.38 | 0.047 | 0 | 1 | Aplnr |
| Pls3 | 0 | 0.965717072 | 0.609 | 0.15 | 0 | 1 | Pls3 |
| Ets2 | 0 | 0.964913788 | 0.572 | 0.131 | 0 | 1 | Ets2 |
| Wasf2 | 0 | 0.964575305 | 0.821 | 0.36 | 0 | 1 | Wasf2 |
| Plxnd1 | 0 | 0.962193792 | 0.541 | 0.122 | 0 | 1 | Plxnd1 |
| Ctnnb1 | 0 | 0.95477238 | 0.771 | 0.297 | 0 | 1 | Ctnnb1 |
| Icam2 | 0 | 0.954365568 | 0.66 | 0.159 | 0 | 1 | Icam2 |
| Fry | 0 | 0.951041429 | 0.542 | 0.145 | 0 | 1 | Fry |
| Dab2 | 0 | 0.948859348 | 0.56 | 0.148 | 0 | 1 | Dab2 |
| Dennd5b | 0 | 0.946469755 | 0.47 | 0.094 | 0 | 1 | Dennd5b |
| Ralgapa1 | 0 | 0.942991337 | 0.662 | 0.245 | 0 | 1 | Ralgapa1 |
| Lpcat1 | 0 | 0.939166037 | 0.496 | 0.104 | 0 | 1 | Lpcat1 |
| Myo10 | 0 | 0.938780362 | 0.6 | 0.156 | 0 | 1 | Myo10 |
| Irf1 | 0 | 0.938657415 | 0.648 | 0.242 | 0 | 1 | Irf1 |
| Fez2 | 0 | 0.935116391 | 0.709 | 0.213 | 0 | 1 | Fez2 |
| Fmnl3 | 0 | 0.934172157 | 0.579 | 0.184 | 0 | 1 | Fmnl3 |
| Galnt18 | 0 | 0.933223548 | 0.442 | 0.084 | 0 | 1 | Galnt18 |

| | | | | | | | |
|---|---|---|---|---|---|---|---|
| Luzp1 | 0 | 0.932963656 | 0.758 | 0.291 | 0 | 1 | Luzp1 |
| Yes1 | 0 | 0.93251094 | 0.496 | 0.097 | 0 | 1 | Yes1 |
| Ctnna1 | 0 | 0.930808185 | 0.74 | 0.288 | 0 | 1 | Ctnna1 |
| Piezo1 | 0 | 0.930743742 | 0.506 | 0.119 | 0 | 1 | Piezo1 |
| Rarg | 0 | 0.929863161 | 0.551 | 0.137 | 0 | 1 | Rarg |
| Ripply3 | 0 | 0.929796396 | 0.468 | 0.094 | 0 | 1 | Ripply3 |
| Nedd9 | 0 | 0.929248562 | 0.687 | 0.26 | 0 | 1 | Nedd9 |
| Rock2 | 0 | 0.925579834 | 0.712 | 0.277 | 0 | 1 | Rock2 |
| Ehd2 | 0 | 0.921204646 | 0.56 | 0.133 | 0 | 1 | Ehd2 |
| Fndc3b | 0 | 0.920677671 | 0.58 | 0.19 | 0 | 1 | Fndc3b |
| Dusp3 | 0 | 0.91941955 | 0.545 | 0.134 | 0 | 1 | Dusp3 |
| H2-D1 | 0 | 0.91932317 | 0.994 | 0.616 | 0 | 1 | H2-D1 |
| Kif13a | 0 | 0.919003296 | 0.502 | 0.12 | 0 | 1 | Kif13a |
| Tnfaip1 | 0 | 0.918748577 | 0.586 | 0.14 | 0 | 1 | Tnfaip1 |
| H2-K1 | 0 | 0.918409939 | 0.993 | 0.586 | 0 | 1 | H2-K1 |
| Ppp1r12a | 0 | 0.915306921 | 0.804 | 0.364 | 0 | 1 | Ppp1r12a |
| Tspan18 | 0 | 0.912497757 | 0.474 | 0.082 | 0 | 1 | Tspan18 |
| BC028528 | 0 | 0.911312797 | 0.609 | 0.172 | 0 | 1 | BC028528 |
| Oaz2 | 0 | 0.910470893 | 0.595 | 0.178 | 0 | 1 | Oaz2 |
| Gbp4 | 0 | 0.908992755 | 0.471 | 0.13 | 0 | 1 | Gbp4 |
| Myh9 | 0 | 0.908799019 | 0.803 | 0.377 | 0 | 1 | Myh9 |
| Cav2 | 0 | 0.908583556 | 0.602 | 0.144 | 0 | 1 | Cav2 |
| Trib2 | 0 | 0.90839164 | 0.565 | 0.124 | 0 | 1 | Trib2 |
| Dock9 | 0 | 0.907360003 | 0.554 | 0.149 | 0 | 1 | Dock9 |
| Endod1 | 0 | 0.907357891 | 0.481 | 0.098 | 0 | 1 | Endod1 |
| Tns1 | 0 | 0.907089029 | 0.622 | 0.183 | 0 | 1 | Tns1 |
| Serinc3 | 0 | 0.906609104 | 0.875 | 0.424 | 0 | 1 | Serinc3 |
| Serpinh1 | 0 | 0.90612177 | 0.679 | 0.179 | 0 | 1 | Serpinh1 |
| Cavin1 | 0 | 0.90046559 | 0.596 | 0.15 | 0 | 1 | Cavin1 |
| Tmtc1 | 0 | 0.898572847 | 0.44 | 0.093 | 0 | 1 | Tmtc1 |
| Foxo1 | 0 | 0.896940193 | 0.528 | 0.146 | 0 | 1 | Foxo1 |
| Tmed5 | 0 | 0.895423851 | 0.628 | 0.233 | 0 | 1 | Tmed5 |
| Trp53i11 | 0 | 0.894764592 | 0.47 | 0.11 | 0 | 1 | Trp53i11 |
| Sorbs1 | 0 | 0.893165024 | 0.493 | 0.125 | 0 | 1 | Sorbs1 |
| Smad7 | 0 | 0.891513537 | 0.681 | 0.268 | 0 | 1 | Smad7 |
| Clec1a | 0 | 0.891332393 | 0.505 | 0.113 | 0 | 1 | Clec1a |
| Sema7a | 0 | 0.891046528 | 0.482 | 0.096 | 0 | 1 | Sema7a |
| Hdac7 | 0 | 0.890242362 | 0.549 | 0.146 | 0 | 1 | Hdac7 |
| Rtl8a | 0 | 0.889793892 | 0.515 | 0.121 | 0 | 1 | Rtl8a |
| Insr | 0 | 0.889473543 | 0.328 | 0.097 | 0 | 1 | Insr |
| Zfp503 | 0 | 0.889176392 | 0.461 | 0.095 | 0 | 1 | Zfp503 |
| Dgkh | 0 | 0.888919268 | 0.493 | 0.125 | 0 | 1 | Dgkh |
| Notch1 | 0 | 0.887177946 | 0.563 | 0.16 | 0 | 1 | Notch1 |

| | | | | | | | |
|---|---|---|---|---|---|---|---|
| Sbf2 | 0 | 0.886928689 | 0.521 | 0.156 | 0 | 1 | Sbf2 |
| Tanc1 | 0 | 0.886821255 | 0.567 | 0.167 | 0 | 1 | Tanc1 |
| Xiap | 0 | 0.885495759 | 0.654 | 0.242 | 0 | 1 | Xiap |
| Adam15 | 0 | 0.885250165 | 0.543 | 0.136 | 0 | 1 | Adam15 |
| Erg | 0 | 0.882506268 | 0.455 | 0.077 | 0 | 1 | Erg |
| Selenop | 0 | 0.879447706 | 0.787 | 0.293 | 0 | 1 | Selenop |
| Cdc42bpa | 0 | 0.877933949 | 0.522 | 0.13 | 0 | 1 | Cdc42bpa |
| Parp14 | 0 | 0.876901487 | 0.547 | 0.205 | 0 | 1 | Parp14 |
| Efnb1 | 0 | 0.875874785 | 0.472 | 0.104 | 0 | 1 | Efnb1 |
| Entpd1 | 0 | 0.874898639 | 0.636 | 0.217 | 0 | 1 | Entpd1 |
| Nxpe4 | 0 | 0.874374756 | 0.454 | 0.098 | 0 | 1 | Nxpe4 |
| Klhl5 | 0 | 0.873378596 | 0.59 | 0.165 | 0 | 1 | Klhl5 |
| St8sia4 | 0 | 0.873187357 | 0.567 | 0.21 | 0 | 1 | St8sia4 |
| Rap1b | 0 | 0.870941895 | 0.811 | 0.384 | 0 | 1 | Rap1b |
| Mlec | 0 | 0.870033077 | 0.587 | 0.207 | 0 | 1 | Mlec |
| Lama4 | 0 | 0.869174526 | 0.495 | 0.122 | 0 | 1 | Lama4 |
| Laptm4a | 0 | 0.868515597 | 0.897 | 0.416 | 0 | 1 | Laptm4a |
| B4galt4 | 0 | 0.867585428 | 0.507 | 0.106 | 0 | 1 | B4galt4 |
| Sybu | 0 | 0.867508482 | 0.392 | 0.069 | 0 | 1 | Sybu |
| Gcnt2 | 0 | 0.865880952 | 0.445 | 0.093 | 0 | 1 | Gcnt2 |
| Tpm4 | 0 | 0.865756765 | 0.767 | 0.317 | 0 | 1 | Tpm4 |
| Sypl | 0 | 0.864243686 | 0.6 | 0.181 | 0 | 1 | Sypl |
| Rassf9 | 0 | 0.864006117 | 0.382 | 0.084 | 0 | 1 | Rassf9 |
| Map1b | 0 | 0.860623565 | 0.364 | 0.059 | 0 | 1 | Map1b |
| Nfe2l1 | 0 | 0.859932487 | 0.642 | 0.221 | 0 | 1 | Nfe2l1 |
| Tspan13 | 0 | 0.858771654 | 0.626 | 0.23 | 0 | 1 | Tspan13 |
| Zbtb20 | 0 | 0.857808826 | 0.944 | 0.471 | 0 | 1 | Zbtb20 |
| Arhgef3 | 0 | 0.856841954 | 0.55 | 0.19 | 0 | 1 | Arhgef3 |
| Nedd4 | 0 | 0.856588054 | 0.83 | 0.352 | 0 | 1 | Nedd4 |
| Ptprk | 0 | 0.856444448 | 0.426 | 0.093 | 0 | 1 | Ptprk |
| Cxcl12 | 0 | 0.855111865 | 0.784 | 0.246 | 0 | 1 | Cxcl12 |
| Kcnq1 | 0 | 0.854160546 | 0.362 | 0.053 | 0 | 1 | Kcnq1 |
| Ktn1 | 0 | 0.85352149 | 0.639 | 0.254 | 0 | 1 | Ktn1 |
| Tmem30a | 0 | 0.853155898 | 0.629 | 0.23 | 0 | 1 | Tmem30a |
| Cep85l | 0 | 0.853045992 | 0.458 | 0.117 | 0 | 1 | Cep85l |
| Cped1 | 0 | 0.8520633 | 0.509 | 0.125 | 0 | 1 | Cped1 |
| Ceacam1 | 0 | 0.851956311 | 0.492 | 0.101 | 0 | 1 | Ceacam1 |
| Itm2b | 0 | 0.851419085 | 0.996 | 0.681 | 0 | 1 | Itm2b |
| Tmem47 | 0 | 0.850907756 | 0.478 | 0.104 | 0 | 1 | Tmem47 |
| Ces2e | 0 | 0.849081682 | 0.41 | 0.061 | 0 | 1 | Ces2e |
| Sox4 | 0 | 0.848306604 | 0.528 | 0.15 | 0 | 1 | Sox4 |
| Nbeal1 | 0 | 0.847572907 | 0.594 | 0.208 | 0 | 1 | Nbeal1 |
| Ier5l | 0 | 0.847268286 | 0.498 | 0.126 | 0 | 1 | Ier5l |

| | | | | | | | |
|---|---|---|---|---|---|---|---|
| Adam10 | 0 | 0.845084204 | 0.621 | 0.239 | 0 | 1 | Adam10 |
| Ripor1 | 0 | 0.843535834 | 0.482 | 0.108 | 0 | 1 | Ripor1 |
| Vwa3a | 0 | 0.843532012 | 0.339 | 0.044 | 0 | 1 | Vwa3a |
| Svil | 0 | 0.842389967 | 0.547 | 0.194 | 0 | 1 | Svil |
| Fabp5 | 0 | 0.841902858 | 0.685 | 0.266 | 0 | 1 | Fabp5 |
| She | 0 | 0.841807235 | 0.477 | 0.096 | 0 | 1 | She |
| Slco2b1 | 0 | 0.84125561 | 0.477 | 0.132 | 0 | 1 | Slco2b1 |
| Ppfibp1 | 0 | 0.841196595 | 0.517 | 0.129 | 0 | 1 | Ppfibp1 |
| Prrc2c | 0 | 0.839589507 | 0.765 | 0.364 | 0 | 1 | Prrc2c |
| Diaph2 | 0 | 0.835024906 | 0.699 | 0.324 | 0 | 1 | Diaph2 |
| Pttg1ip | 0 | 0.833969005 | 0.627 | 0.21 | 0 | 1 | Pttg1ip |
| Nhsl2 | 0 | 0.833420204 | 0.435 | 0.085 | 0 | 1 | Nhsl2 |
| Herc6 | 0 | 0.833118986 | 0.429 | 0.144 | 0 | 1 | Herc6 |
| Grasp | 0 | 0.83238143 | 0.48 | 0.115 | 0 | 1 | Grasp |
| Gja1 | 0 | 0.832293615 | 0.391 | 0.085 | 0 | 1 | Gja1 |
| Anxa3 | 0 | 0.832283401 | 0.767 | 0.268 | 0 | 1 | Anxa3 |
| Vcl | 0 | 0.831882866 | 0.467 | 0.112 | 0 | 1 | Vcl |
| Spag9 | 0 | 0.830245987 | 0.698 | 0.304 | 0 | 1 | Spag9 |
| Mprip | 0 | 0.829971788 | 0.656 | 0.234 | 0 | 1 | Mprip |
| Atxn1 | 0 | 0.828936199 | 0.615 | 0.243 | 0 | 1 | Atxn1 |
| Pde8a | 0 | 0.8288277 | 0.514 | 0.129 | 0 | 1 | Pde8a |
| Ifit2 | 0 | 0.828477561 | 0.317 | 0.087 | 0 | 1 | Ifit2 |
| Atrx | 0 | 0.825825552 | 0.76 | 0.364 | 0 | 1 | Atrx |
| Fam198b | 0 | 0.824744475 | 0.507 | 0.131 | 0 | 1 | Fam198b |
| Sos1 | 0 | 0.824223371 | 0.46 | 0.135 | 0 | 1 | Sos1 |
| Lrch1 | 0 | 0.823913239 | 0.622 | 0.261 | 0 | 1 | Lrch1 |
| Nid1 | 0 | 0.823532062 | 0.547 | 0.122 | 0 | 1 | Nid1 |
| Nfat5 | 0 | 0.823312908 | 0.569 | 0.218 | 0 | 1 | Nfat5 |
| Kmt2e | 0 | 0.821575937 | 0.739 | 0.352 | 0 | 1 | Kmt2e |
| 2900026A02Rik | 0 | 0.821008925 | 0.517 | 0.171 | 0 | 1 | 2900026A02Rik |
| Elf1 | 0 | 0.820507601 | 0.707 | 0.323 | 0 | 1 | Elf1 |
| B2m | 0 | 0.82000329 | 0.995 | 0.641 | 0 | 1 | B2m |
| Rock1 | 0 | 0.818785434 | 0.773 | 0.377 | 0 | 1 | Rock1 |
| Kcnb1 | 0 | 0.816338125 | 0.32 | 0.041 | 0 | 1 | Kcnb1 |
| Ppp1r2 | 0 | 0.815587067 | 0.779 | 0.33 | 0 | 1 | Ppp1r2 |
| Baz2b | 0 | 0.815102606 | 0.678 | 0.297 | 0 | 1 | Baz2b |
| Rnf213 | 0 | 0.813574181 | 0.527 | 0.214 | 0 | 1 | Rnf213 |
| Nfia | 0 | 0.811558434 | 0.638 | 0.23 | 0 | 1 | Nfia |
| Nxpe2 | 0 | 0.811414587 | 0.378 | 0.06 | 0 | 1 | Nxpe2 |
| Ppp1r13b | 0 | 0.809270598 | 0.536 | 0.164 | 0 | 1 | Ppp1r13b |
| Sppl2a | 0 | 0.808133351 | 0.6 | 0.247 | 0 | 1 | Sppl2a |
| Bvht | 0 | 0.807427481 | 0.553 | 0.126 | 0 | 1 | Bvht |
| Tdrp | 0 | 0.805914074 | 0.419 | 0.067 | 0 | 1 | Tdrp |

| | | | | | | | |
|---|---|---|---|---|---|---|---|
| Gbp2 | 0 | 0.805585288 | 0.4 | 0.102 | 0 | 1 | Gbp2 |
| Tns2 | 0 | 0.805285718 | 0.45 | 0.095 | 0 | 1 | Tns2 |
| Map4k3 | 0 | 0.803068981 | 0.534 | 0.18 | 0 | 1 | Map4k3 |
| Crip2 | 0 | 0.800859445 | 0.874 | 0.341 | 0 | 1 | Crip2 |
| Tcf12 | 0 | 0.80026775 | 0.708 | 0.322 | 0 | 1 | Tcf12 |
| Map4 | 0 | 0.800206436 | 0.629 | 0.251 | 0 | 1 | Map4 |
| Ctsl | 0 | 0.799792474 | 0.765 | 0.277 | 0 | 1 | Ctsl |
| Nck1 | 0 | 0.795454008 | 0.689 | 0.283 | 0 | 1 | Nck1 |
| Plscr4 | 0 | 0.795045601 | 0.488 | 0.141 | 0 | 1 | Plscr4 |
| Akt3 | 0 | 0.794507641 | 0.546 | 0.204 | 0 | 1 | Akt3 |
| Ier3 | 0 | 0.7932652 | 0.342 | 0.078 | 0 | 1 | Ier3 |
| Son | 0 | 0.792702844 | 0.894 | 0.466 | 0 | 1 | Son |
| Hivep2 | 0 | 0.790551902 | 0.596 | 0.25 | 0 | 1 | Hivep2 |
| Limch1 | 0 | 0.789838185 | 0.422 | 0.093 | 0 | 1 | Limch1 |
| Iqgap1 | 0 | 0.789290222 | 0.759 | 0.356 | 0 | 1 | Iqgap1 |
| Nfkbia | 0 | 0.789006686 | 0.754 | 0.342 | 0 | 1 | Nfkbia |
| St3gal6 | 0 | 0.787264425 | 0.664 | 0.261 | 0 | 1 | St3gal6 |
| Smurf2 | 0 | 0.786768184 | 0.522 | 0.193 | 0 | 1 | Smurf2 |
| Rbfox2 | 0 | 0.786506776 | 0.509 | 0.139 | 0 | 1 | Rbfox2 |
| Cyp2d22 | 0 | 0.783922294 | 0.419 | 0.089 | 0 | 1 | Cyp2d22 |
| Klf3 | 0 | 0.783827258 | 0.564 | 0.183 | 0 | 1 | Klf3 |
| Lipe | 0 | 0.782864806 | 0.426 | 0.09 | 0 | 1 | Lipe |
| Rftn1 | 0 | 0.78218638 | 0.543 | 0.204 | 0 | 1 | Rftn1 |
| Lnx2 | 0 | 0.781314097 | 0.406 | 0.089 | 0 | 1 | Lnx2 |
| Znfx1 | 0 | 0.78006813 | 0.454 | 0.13 | 0 | 1 | Znfx1 |
| Dcbld1 | 0 | 0.779865502 | 0.329 | 0.048 | 0 | 1 | Dcbld1 |
| Cds2 | 0 | 0.779264152 | 0.475 | 0.129 | 0 | 1 | Cds2 |
| Tnfaip2 | 0 | 0.778375747 | 0.435 | 0.106 | 0 | 1 | Tnfaip2 |
| Fam43a | 0 | 0.777613965 | 0.45 | 0.092 | 0 | 1 | Fam43a |
| Atp8a1 | 0 | 0.777231046 | 0.523 | 0.196 | 0 | 1 | Atp8a1 |
| Map4k4 | 0 | 0.7770065 | 0.587 | 0.226 | 0 | 1 | Map4k4 |
| Cd151 | 0 | 0.774629186 | 0.562 | 0.161 | 0 | 1 | Cd151 |
| Ncoa3 | 0 | 0.772100858 | 0.549 | 0.204 | 0 | 1 | Ncoa3 |
| Caskin2 | 0 | 0.771751595 | 0.447 | 0.09 | 0 | 1 | Caskin2 |
| Rhoc | 0 | 0.771373481 | 0.641 | 0.24 | 0 | 1 | Rhoc |
| Ephb1 | 0 | 0.771264916 | 0.322 | 0.042 | 0 | 1 | Ephb1 |
| Tcn2 | 0 | 0.770461802 | 0.474 | 0.132 | 0 | 1 | Tcn2 |
| Sp100 | 0 | 0.770028216 | 0.773 | 0.377 | 0 | 1 | Sp100 |
| Dok4 | 0 | 0.769668534 | 0.406 | 0.072 | 0 | 1 | Dok4 |
| Pitpnm2 | 0 | 0.766878213 | 0.397 | 0.078 | 0 | 1 | Pitpnm2 |
| Palmd | 0 | 0.766861032 | 0.606 | 0.172 | 0 | 1 | Palmd |
| Cd36 | 0 | 0.765534058 | 0.607 | 0.213 | 0 | 1 | Cd36 |
| Osmr | 0 | 0.765139107 | 0.421 | 0.094 | 0 | 1 | Osmr |

| | | | | | | | |
|---|---|---|---|---|---|---|---|
| Crispld1 | 0 | 0.764131604 | 0.419 | 0.099 | 0 | 1 | Crispld1 |
| Prickle1 | 0 | 0.764010654 | 0.421 | 0.102 | 0 | 1 | Prickle1 |
| Elmo1 | 0 | 0.763911896 | 0.719 | 0.309 | 0 | 1 | Elmo1 |
| Flt4 | 0 | 0.762848663 | 0.383 | 0.059 | 0 | 1 | Flt4 |
| Rras | 0 | 0.76279398 | 0.67 | 0.237 | 0 | 1 | Rras |
| Afap1 | 0 | 0.762641196 | 0.432 | 0.104 | 0 | 1 | Afap1 |
| Fam174b | 0 | 0.762227875 | 0.347 | 0.064 | 0 | 1 | Fam174b |
| Pcdh12 | 0 | 0.761917748 | 0.368 | 0.053 | 0 | 1 | Pcdh12 |
| Cnn3 | 0 | 0.761055381 | 0.52 | 0.14 | 0 | 1 | Cnn3 |
| Prickle2 | 0 | 0.760713806 | 0.402 | 0.097 | 0 | 1 | Prickle2 |
| Dock6 | 0 | 0.760402985 | 0.422 | 0.094 | 0 | 1 | Dock6 |
| Ptbp3 | 0 | 0.759426458 | 0.725 | 0.34 | 0 | 1 | Ptbp3 |
| Wwc2 | 0 | 0.757994049 | 0.507 | 0.147 | 0 | 1 | Wwc2 |
| Tnfrsf1a | 0 | 0.757863683 | 0.535 | 0.171 | 0 | 1 | Tnfrsf1a |
| Ptpra | 0 | 0.756965212 | 0.613 | 0.255 | 0 | 1 | Ptpra |
| Bnip2 | 0 | 0.756492577 | 0.609 | 0.231 | 0 | 1 | Bnip2 |
| Zfp462 | 0 | 0.75629823 | 0.381 | 0.078 | 0 | 1 | Zfp462 |
| Arhgap26 | 0 | 0.755729611 | 0.504 | 0.177 | 0 | 1 | Arhgap26 |
| Cdc42ep2 | 0 | 0.753961526 | 0.499 | 0.119 | 0 | 1 | Cdc42ep2 |
| Plekho1 | 0 | 0.753181682 | 0.546 | 0.172 | 0 | 1 | Plekho1 |
| Mfsd6 | 0 | 0.752255731 | 0.42 | 0.099 | 0 | 1 | Mfsd6 |
| Pde10a | 0 | 0.750421743 | 0.296 | 0.05 | 0 | 1 | Pde10a |
| Afap1l1 | 0 | 0.749440065 | 0.369 | 0.067 | 0 | 1 | Afap1l1 |
| Gm38399 | 0 | 0.749182933 | 0.345 | 0.069 | 0 | 1 | Gm38399 |
| Ccdc85b | 0 | 0.748623953 | 0.599 | 0.2 | 0 | 1 | Ccdc85b |
| Gbp5 | 0 | 0.748062646 | 0.353 | 0.092 | 0 | 1 | Gbp5 |
| Shroom4 | 0 | 0.747770037 | 0.357 | 0.064 | 0 | 1 | Shroom4 |
| Vat1 | 0 | 0.747682954 | 0.442 | 0.108 | 0 | 1 | Vat1 |
| Zmiz1 | 0 | 0.746718332 | 0.625 | 0.272 | 0 | 1 | Zmiz1 |
| Unc45b | 0 | 0.745849477 | 0.442 | 0.09 | 0 | 1 | Unc45b |
| Nr4a1 | 0 | 0.744629403 | 0.312 | 0.073 | 0 | 1 | Nr4a1 |
| Birc6 | 0 | 0.743710006 | 0.631 | 0.294 | 0 | 1 | Birc6 |
| Rtn3 | 0 | 0.742754568 | 0.631 | 0.274 | 0 | 1 | Rtn3 |
| Mtdh | 0 | 0.74189611 | 0.696 | 0.337 | 0 | 1 | Mtdh |
| Col4a3bp | 0 | 0.741065578 | 0.545 | 0.212 | 0 | 1 | Col4a3bp |
| F11r | 0 | 0.740246484 | 0.618 | 0.208 | 0 | 1 | F11r |
| Sox7 | 0 | 0.739604797 | 0.345 | 0.054 | 0 | 1 | Sox7 |
| Adamts1 | 0 | 0.739433943 | 0.287 | 0.066 | 0 | 1 | Adamts1 |
| Fam117b | 0 | 0.737192667 | 0.414 | 0.124 | 0 | 1 | Fam117b |
| Nova2 | 0 | 0.735519034 | 0.364 | 0.064 | 0 | 1 | Nova2 |
| Tmem59 | 0 | 0.734809963 | 0.842 | 0.433 | 0 | 1 | Tmem59 |
| Zfp608 | 0 | 0.734598265 | 0.38 | 0.102 | 0 | 1 | Zfp608 |
| 2010111I01Rik | 0 | 0.734142142 | 0.708 | 0.291 | 0 | 1 | 2010111I01Rik |

| | | | | | | | |
|---|---|---|---|---|---|---|---|
| Hook3 | 0 | 0.731967586 | 0.585 | 0.242 | 0 | 1 | Hook3 |
| Adgrg1 | 0 | 0.731785389 | 0.434 | 0.092 | 0 | 1 | Adgrg1 |
| Efna1 | 0 | 0.731285652 | 0.445 | 0.094 | 0 | 1 | Efna1 |
| Gbp9 | 0 | 0.730939428 | 0.391 | 0.101 | 0 | 1 | Gbp9 |
| Mdfic | 0 | 0.730675174 | 0.522 | 0.201 | 0 | 1 | Mdfic |
| Ctnnbip1 | 0 | 0.730108963 | 0.527 | 0.151 | 0 | 1 | Ctnnbip1 |
| Srgap2 | 0 | 0.729943335 | 0.608 | 0.255 | 0 | 1 | Srgap2 |
| Exoc6b | 0 | 0.729938338 | 0.504 | 0.198 | 0 | 1 | Exoc6b |
| Plcb4 | 0 | 0.728928055 | 0.671 | 0.229 | 0 | 1 | Plcb4 |
| Irf2bpl | 0 | 0.728083707 | 0.551 | 0.194 | 0 | 1 | Irf2bpl |
| Stim2 | 0 | 0.726086092 | 0.449 | 0.159 | 0 | 1 | Stim2 |
| Serinc1 | 0 | 0.725871548 | 0.542 | 0.202 | 0 | 1 | Serinc1 |
| Midn | 0 | 0.725820354 | 0.447 | 0.15 | 0 | 1 | Midn |
| Lmo2 | 0 | 0.725650584 | 0.516 | 0.158 | 0 | 1 | Lmo2 |
| Ebf3 | 0 | 0.724451143 | 0.387 | 0.084 | 0 | 1 | Ebf3 |
| Lifr | 0 | 0.723287933 | 0.454 | 0.15 | 0 | 1 | Lifr |
| Flnb | 0 | 0.723101669 | 0.429 | 0.119 | 0 | 1 | Flnb |
| Tapbp | 0 | 0.722027312 | 0.595 | 0.258 | 0 | 1 | Tapbp |
| Irgm2 | 0 | 0.720431215 | 0.336 | 0.093 | 0 | 1 | Irgm2 |
| Marcks | 0 | 0.720261108 | 0.74 | 0.276 | 0 | 1 | Marcks |
| Abi3 | 0 | 0.72010903 | 0.542 | 0.196 | 0 | 1 | Abi3 |
| Sptlc2 | 0 | 0.719526373 | 0.52 | 0.2 | 0 | 1 | Sptlc2 |
| Bst2 | 0 | 0.718907262 | 0.88 | 0.442 | 0 | 1 | Bst2 |
| Fcho2 | 0 | 0.718595292 | 0.603 | 0.262 | 0 | 1 | Fcho2 |
| Fkbp1a | 0 | 0.718577484 | 0.766 | 0.347 | 0 | 1 | Fkbp1a |
| Akap12 | 0 | 0.71600303 | 0.429 | 0.117 | 0 | 1 | Akap12 |
| Herc1 | 0 | 0.715933122 | 0.515 | 0.216 | 0 | 1 | Herc1 |
| Cdc42bpb | 0 | 0.715758776 | 0.501 | 0.135 | 0 | 1 | Cdc42bpb |
| Tap2 | 0 | 0.715747861 | 0.521 | 0.195 | 0 | 1 | Tap2 |
| Atp2b4 | 0 | 0.71503198 | 0.456 | 0.157 | 0 | 1 | Atp2b4 |
| Slc44a1 | 0 | 0.714011056 | 0.419 | 0.113 | 0 | 1 | Slc44a1 |
| Snai2 | 0 | 0.713227274 | 0.321 | 0.054 | 0 | 1 | Snai2 |
| Efnb2 | 0 | 0.713216295 | 0.41 | 0.101 | 0 | 1 | Efnb2 |
| Nav2 | 0 | 0.712945335 | 0.421 | 0.124 | 0 | 1 | Nav2 |
| Ddx3x | 0 | 0.712616533 | 0.62 | 0.257 | 0 | 1 | Ddx3x |
| Dync1h1 | 0 | 0.712094038 | 0.586 | 0.238 | 0 | 1 | Dync1h1 |
| Pcnx | 0 | 0.711845463 | 0.429 | 0.138 | 0 | 1 | Pcnx |
| Nuak1 | 0 | 0.711423082 | 0.35 | 0.08 | 0 | 1 | Nuak1 |
| Enah | 0 | 0.711206579 | 0.436 | 0.103 | 0 | 1 | Enah |
| Parvb | 0 | 0.710098975 | 0.383 | 0.081 | 0 | 1 | Parvb |
| Lpar6 | 0 | 0.708316004 | 0.421 | 0.145 | 0 | 1 | Lpar6 |
| Txnip | 0 | 0.707953059 | 0.879 | 0.491 | 0 | 1 | Txnip |
| Msi2 | 0 | 0.707946125 | 0.584 | 0.246 | 0 | 1 | Msi2 |

| | | | | | | | |
|---|---|---|---|---|---|---|---|
| Foxf1 | 0 | 0.707280241 | 0.321 | 0.045 | 0 | 1 | Foxf1 |
| Tll1 | 0 | 0.706124203 | 0.262 | 0.037 | 0 | 1 | Tll1 |
| Epn2 | 0 | 0.705950031 | 0.44 | 0.108 | 0 | 1 | Epn2 |
| Tsc22d3 | 0 | 0.70565049 | 0.499 | 0.16 | 0 | 1 | Tsc22d3 |
| Prdm1 | 0 | 0.705230792 | 0.33 | 0.074 | 0 | 1 | Prdm1 |
| Cdk17 | 0 | 0.704819986 | 0.515 | 0.201 | 0 | 1 | Cdk17 |
| Stag1 | 0 | 0.70428671 | 0.663 | 0.313 | 0 | 1 | Stag1 |
| Smarca2 | 0 | 0.702559489 | 0.565 | 0.234 | 0 | 1 | Smarca2 |
| Mfhas1 | 0 | 0.701961046 | 0.401 | 0.109 | 0 | 1 | Mfhas1 |
| Mat2a | 0 | 0.700999198 | 0.591 | 0.264 | 0 | 1 | Mat2a |
| Swap70 | 0 | 0.700184749 | 0.524 | 0.202 | 0 | 1 | Swap70 |
| Wars | 0 | 0.699691828 | 0.432 | 0.126 | 0 | 1 | Wars |
| Tspan5 | 0 | 0.697121254 | 0.445 | 0.159 | 0 | 1 | Tspan5 |
| Foxp1 | 0 | 0.695239493 | 0.715 | 0.337 | 0 | 1 | Foxp1 |
| Gab1 | 0 | 0.694142221 | 0.35 | 0.076 | 0 | 1 | Gab1 |
| Maml3 | 0 | 0.694061203 | 0.456 | 0.168 | 0 | 1 | Maml3 |
| Ttll7 | 0 | 0.693804282 | 0.352 | 0.062 | 0 | 1 | Ttll7 |
| Epha4 | 0 | 0.693051717 | 0.35 | 0.079 | 0 | 1 | Epha4 |
| Wdfy3 | 0 | 0.690861911 | 0.446 | 0.139 | 0 | 1 | Wdfy3 |
| Tspan15 | 0 | 0.690482908 | 0.405 | 0.085 | 0 | 1 | Tspan15 |
| Fam219a | 0 | 0.689440779 | 0.367 | 0.093 | 0 | 1 | Fam219a |
| Lysmd2 | 0 | 0.689296681 | 0.486 | 0.144 | 0 | 1 | Lysmd2 |
| Anxa2 | 0 | 0.688883879 | 0.779 | 0.357 | 0 | 1 | Anxa2 |
| Rrbp1 | 0 | 0.688584035 | 0.821 | 0.401 | 0 | 1 | Rrbp1 |
| Fam171a1 | 0 | 0.688101107 | 0.33 | 0.07 | 0 | 1 | Fam171a1 |
| Tpst2 | 0 | 0.687943828 | 0.514 | 0.187 | 0 | 1 | Tpst2 |
| Gpr146 | 0 | 0.686720037 | 0.357 | 0.069 | 0 | 1 | Gpr146 |
| Kmt2c | 0 | 0.68618046 | 0.574 | 0.263 | 0 | 1 | Kmt2c |
| Sema6d | 0 | 0.686110812 | 0.477 | 0.172 | 0 | 1 | Sema6d |
| Cbfa2t3 | 0 | 0.685464122 | 0.383 | 0.099 | 0 | 1 | Cbfa2t3 |
| Ier5 | 0 | 0.685330694 | 0.603 | 0.271 | 0 | 1 | Ier5 |
| Ash1l | 0 | 0.685256456 | 0.587 | 0.253 | 0 | 1 | Ash1l |
| Rassf1 | 0 | 0.685075704 | 0.477 | 0.188 | 0 | 1 | Rassf1 |
| Sema3c | 0 | 0.684959528 | 0.406 | 0.124 | 0 | 1 | Sema3c |
| Etnk1 | 0 | 0.68393229 | 0.488 | 0.183 | 0 | 1 | Etnk1 |
| Dennd3 | 0 | 0.681336591 | 0.393 | 0.106 | 0 | 1 | Dennd3 |
| Sipa1l2 | 0 | 0.680721962 | 0.413 | 0.117 | 0 | 1 | Sipa1l2 |
| Gnb1 | 0 | 0.680191433 | 0.765 | 0.373 | 0 | 1 | Gnb1 |
| Rap1a | 0 | 0.679004369 | 0.869 | 0.483 | 0 | 1 | Rap1a |
| 4930402H24Rik | 0 | 0.678200337 | 0.47 | 0.181 | 0 | 1 | 4930402H24Rik |
| Ptpn14 | 0 | 0.677458701 | 0.351 | 0.067 | 0 | 1 | Ptpn14 |
| Psmb9 | 0 | 0.675875894 | 0.73 | 0.353 | 0 | 1 | Psmb9 |
| Nucb1 | 0 | 0.675368906 | 0.49 | 0.185 | 0 | 1 | Nucb1 |

| | | | | | | | |
|---|---|---|---|---|---|---|---|
| Stard9 | 0 | 0.675036802 | 0.386 | 0.114 | 0 | 1 | Stard9 |
| Plcg1 | 0 | 0.67499163 | 0.376 | 0.081 | 0 | 1 | Plcg1 |
| Proser2 | 0 | 0.673877282 | 0.346 | 0.057 | 0 | 1 | Proser2 |
| Zdhhc17 | 0 | 0.673338327 | 0.384 | 0.112 | 0 | 1 | Zdhhc17 |
| Atrnl1 | 0 | 0.6724809 | 0.398 | 0.128 | 0 | 1 | Atrnl1 |
| C2cd2 | 0 | 0.671884766 | 0.344 | 0.073 | 0 | 1 | C2cd2 |
| Ica1 | 0 | 0.671258281 | 0.477 | 0.127 | 0 | 1 | Ica1 |
| Samd9l | 0 | 0.670969111 | 0.52 | 0.214 | 0 | 1 | Samd9l |
| Ptpn12 | 0 | 0.67068696 | 0.465 | 0.173 | 0 | 1 | Ptpn12 |
| Plscr1 | 0 | 0.670427429 | 0.459 | 0.164 | 0 | 1 | Plscr1 |
| Tgoln1 | 0 | 0.670129304 | 0.522 | 0.202 | 0 | 1 | Tgoln1 |
| Sema3f | 0 | 0.669013439 | 0.335 | 0.055 | 0 | 1 | Sema3f |
| Notch4 | 0 | 0.668727251 | 0.374 | 0.073 | 0 | 1 | Notch4 |
| Abcc4 | 0 | 0.667757142 | 0.361 | 0.1 | 0 | 1 | Abcc4 |
| Ccser2 | 0 | 0.665840951 | 0.529 | 0.183 | 0 | 1 | Ccser2 |
| Kmt2a | 0 | 0.665434182 | 0.525 | 0.226 | 0 | 1 | Kmt2a |
| Inpp5a | 0 | 0.66539889 | 0.369 | 0.091 | 0 | 1 | Inpp5a |
| Synpo | 0 | 0.665248075 | 0.334 | 0.062 | 0 | 1 | Synpo |
| Lrrc3b | 0 | 0.663295393 | 0.279 | 0.064 | 0 | 1 | Lrrc3b |
| Zfyve9 | 0 | 0.663106537 | 0.449 | 0.126 | 0 | 1 | Zfyve9 |
| Fas | 0 | 0.662609317 | 0.362 | 0.078 | 0 | 1 | Fas |
| Slco3a1 | 0 | 0.660485067 | 0.338 | 0.087 | 0 | 1 | Slco3a1 |
| Glg1 | 0 | 0.659986284 | 0.507 | 0.211 | 0 | 1 | Glg1 |
| Gata2 | 0 | 0.659798488 | 0.348 | 0.065 | 0 | 1 | Gata2 |
| Pdzd2 | 0 | 0.657638055 | 0.328 | 0.081 | 0 | 1 | Pdzd2 |
| Zcchc6 | 0 | 0.657627244 | 0.537 | 0.237 | 0 | 1 | Zcchc6 |
| Gm32688 | 0 | 0.656787631 | 0.323 | 0.057 | 0 | 1 | Gm32688 |
| Hsp90b1 | 0 | 0.654721732 | 0.874 | 0.451 | 0 | 1 | Hsp90b1 |
| Mertk | 0 | 0.653935067 | 0.36 | 0.095 | 0 | 1 | Mertk |
| Foxn3 | 0 | 0.653856461 | 0.745 | 0.357 | 0 | 1 | Foxn3 |
| 4930578C19Rik | 0 | 0.653315416 | 0.332 | 0.07 | 0 | 1 | 4930578C19Rik |
| Enpep | 0 | 0.652182476 | 0.301 | 0.049 | 0 | 1 | Enpep |
| Fat4 | 0 | 0.652123165 | 0.336 | 0.069 | 0 | 1 | Fat4 |
| St3gal1 | 0 | 0.650611267 | 0.426 | 0.141 | 0 | 1 | St3gal1 |
| Apold1 | 0 | 0.650486096 | 0.279 | 0.038 | 0 | 1 | Apold1 |
| Unc119b | 0 | 0.649605892 | 0.389 | 0.1 | 0 | 1 | Unc119b |
| Itm2a | 0 | 0.649243204 | 0.457 | 0.135 | 0 | 1 | Itm2a |
| Prrg1 | 0 | 0.646010011 | 0.324 | 0.073 | 0 | 1 | Prrg1 |
| Rassf3 | 0 | 0.645685433 | 0.534 | 0.187 | 0 | 1 | Rassf3 |
| Gulp1 | 0 | 0.645501038 | 0.419 | 0.125 | 0 | 1 | Gulp1 |
| Ankrd28 | 0 | 0.644682139 | 0.346 | 0.099 | 0 | 1 | Ankrd28 |
| Rac1 | 0 | 0.64439881 | 0.757 | 0.377 | 0 | 1 | Rac1 |
| Ifi35 | 0 | 0.642382269 | 0.558 | 0.228 | 0 | 1 | Ifi35 |

| | | | | | | | |
|---|---|---|---|---|---|---|---|
| Pros1 | 0 | 0.640372696 | 0.386 | 0.112 | 0 | 1 | Pros1 |
| Fut4 | 0 | 0.640018851 | 0.299 | 0.047 | 0 | 1 | Fut4 |
| Tgfbr3 | 0 | 0.639072442 | 0.432 | 0.132 | 0 | 1 | Tgfbr3 |
| Gm26883 | 0 | 0.637688686 | 0.273 | 0.05 | 0 | 1 | Gm26883 |
| Adcy4 | 0 | 0.637145002 | 0.348 | 0.068 | 0 | 1 | Adcy4 |
| Trim2 | 0 | 0.634466551 | 0.469 | 0.131 | 0 | 1 | Trim2 |
| Npr1 | 0 | 0.634387273 | 0.319 | 0.055 | 0 | 1 | Npr1 |
| Ppp1r16b | 0 | 0.633993156 | 0.449 | 0.162 | 0 | 1 | Ppp1r16b |
| Eml1 | 0 | 0.633758692 | 0.325 | 0.074 | 0 | 1 | Eml1 |
| Gna11 | 0 | 0.63244729 | 0.557 | 0.194 | 0 | 1 | Gna11 |
| Rell1 | 0 | 0.632414739 | 0.365 | 0.1 | 0 | 1 | Rell1 |
| Ifngr1 | 0 | 0.631808287 | 0.706 | 0.342 | 0 | 1 | Ifngr1 |
| B230219D22Rik | 0 | 0.631642877 | 0.661 | 0.29 | 0 | 1 | B230219D22Rik |
| Nes | 0 | 0.630712617 | 0.361 | 0.097 | 0 | 1 | Nes |
| H2-Q7 | 0 | 0.630454132 | 0.752 | 0.34 | 0 | 1 | H2-Q7 |
| Acvr2a | 0 | 0.629767857 | 0.364 | 0.115 | 0 | 1 | Acvr2a |
| Btbd7 | 0 | 0.629401263 | 0.563 | 0.236 | 0 | 1 | Btbd7 |
| Lmna | 0 | 0.628306341 | 0.461 | 0.155 | 0 | 1 | Lmna |
| Tmod3 | 0 | 0.626336145 | 0.67 | 0.311 | 0 | 1 | Tmod3 |
| Aig1 | 0 | 0.625790403 | 0.472 | 0.172 | 0 | 1 | Aig1 |
| Zmynd11 | 0 | 0.625514443 | 0.567 | 0.244 | 0 | 1 | Zmynd11 |
| Calu | 0 | 0.6236154 | 0.459 | 0.161 | 0 | 1 | Calu |
| Bsg | 0 | 0.622560682 | 0.784 | 0.395 | 0 | 1 | Bsg |
| Stat3 | 0 | 0.622547355 | 0.525 | 0.225 | 0 | 1 | Stat3 |
| Slfn3 | 0 | 0.622110533 | 0.26 | 0.042 | 0 | 1 | Slfn3 |
| Zfpm1 | 0 | 0.620712466 | 0.382 | 0.115 | 0 | 1 | Zfpm1 |
| Mgat1 | 0 | 0.620548436 | 0.402 | 0.13 | 0 | 1 | Mgat1 |
| Pik3c2a | 0 | 0.620360015 | 0.437 | 0.156 | 0 | 1 | Pik3c2a |
| Ccnyl1 | 0 | 0.619770363 | 0.334 | 0.075 | 0 | 1 | Ccnyl1 |
| Mpzl1 | 0 | 0.618210628 | 0.392 | 0.102 | 0 | 1 | Mpzl1 |
| Lman1 | 0 | 0.617791118 | 0.51 | 0.195 | 0 | 1 | Lman1 |
| Erc1 | 0 | 0.61709355 | 0.338 | 0.088 | 0 | 1 | Erc1 |
| Psmb10 | 0 | 0.616909862 | 0.704 | 0.351 | 0 | 1 | Psmb10 |
| Cyp4b1 | 0 | 0.616313532 | 0.25 | 0.039 | 0 | 1 | Cyp4b1 |
| Pld1 | 0 | 0.615901907 | 0.36 | 0.114 | 0 | 1 | Pld1 |
| S100a16 | 0 | 0.615766339 | 0.604 | 0.201 | 0 | 1 | S100a16 |
| Lrrc58 | 0 | 0.615434049 | 0.481 | 0.171 | 0 | 1 | Lrrc58 |
| Gm12185 | 0 | 0.614846484 | 0.335 | 0.1 | 0 | 1 | Gm12185 |
| Itgb1 | 0 | 0.614691095 | 0.844 | 0.419 | 0 | 1 | Itgb1 |
| Adarb1 | 0 | 0.614605439 | 0.321 | 0.076 | 0 | 1 | Adarb1 |
| Gjc1 | 0 | 0.614468062 | 0.345 | 0.077 | 0 | 1 | Gjc1 |
| Elf2 | 0 | 0.613404662 | 0.573 | 0.257 | 0 | 1 | Elf2 |
| Rora | 0 | 0.612791903 | 0.451 | 0.143 | 0 | 1 | Rora |

| | | | | | | | |
|---|---|---|---|---|---|---|---|
| Kalrn | 0 | 0.611149205 | 0.314 | 0.073 | 0 | 1 | Kalrn |
| Nckap1 | 0 | 0.61051371 | 0.527 | 0.177 | 0 | 1 | Nckap1 |
| Osbpl9 | 0 | 0.607543244 | 0.551 | 0.234 | 0 | 1 | Osbpl9 |
| Eln | 0 | 0.607426049 | 0.374 | 0.098 | 0 | 1 | Eln |
| Smc1a | 0 | 0.606675237 | 0.586 | 0.268 | 0 | 1 | Smc1a |
| Rin2 | 0 | 0.606381125 | 0.428 | 0.143 | 0 | 1 | Rin2 |
| Smagp | 0 | 0.605810947 | 0.559 | 0.178 | 0 | 1 | Smagp |
| 4-Sep | 0 | 0.605702885 | 0.438 | 0.12 | 0 | 1 | 4-Sep |
| Jcad | 0 | 0.605539859 | 0.358 | 0.076 | 0 | 1 | Jcad |
| Stom | 0 | 0.6053261 | 0.327 | 0.072 | 0 | 1 | Stom |
| Cdkn1a | 0 | 0.603958359 | 0.338 | 0.093 | 0 | 1 | Cdkn1a |
| Ints9 | 0 | 0.603908582 | 0.344 | 0.103 | 0 | 1 | Ints9 |
| Slc50a1 | 0 | 0.60363633 | 0.583 | 0.253 | 0 | 1 | Slc50a1 |
| Rab12 | 0 | 0.603146269 | 0.426 | 0.137 | 0 | 1 | Rab12 |
| Cyb5r3 | 0 | 0.602314121 | 0.499 | 0.171 | 0 | 1 | Cyb5r3 |
| Klf9 | 0 | 0.602283623 | 0.544 | 0.188 | 0 | 1 | Klf9 |
| Sbds | 0 | 0.601249258 | 0.597 | 0.264 | 0 | 1 | Sbds |
| Csf1 | 0 | 0.601069669 | 0.306 | 0.078 | 0 | 1 | Csf1 |
| Galnt15 | 0 | 0.600426618 | 0.299 | 0.055 | 0 | 1 | Galnt15 |
| Mid2 | 0 | 0.600300269 | 0.3 | 0.058 | 0 | 1 | Mid2 |
| Ghr | 0 | 0.599667212 | 0.382 | 0.114 | 0 | 1 | Ghr |
| Mfap2 | 0 | 0.599663709 | 0.323 | 0.072 | 0 | 1 | Mfap2 |
| Dusp7 | 0 | 0.599247156 | 0.378 | 0.096 | 0 | 1 | Dusp7 |
| Phactr4 | 0 | 0.598822551 | 0.408 | 0.141 | 0 | 1 | Phactr4 |
| Spry4 | 0 | 0.598335184 | 0.304 | 0.063 | 0 | 1 | Spry4 |
| Rgs12 | 0 | 0.597684063 | 0.323 | 0.068 | 0 | 1 | Rgs12 |
| Creb3l2 | 0 | 0.595570198 | 0.331 | 0.09 | 0 | 1 | Creb3l2 |
| Fscn1 | 0 | 0.593718769 | 0.279 | 0.045 | 0 | 1 | Fscn1 |
| Pcbp3 | 0 | 0.592981326 | 0.26 | 0.054 | 0 | 1 | Pcbp3 |
| Spred2 | 0 | 0.592890243 | 0.396 | 0.135 | 0 | 1 | Spred2 |
| Mxd4 | 0 | 0.592848906 | 0.649 | 0.304 | 0 | 1 | Mxd4 |
| Tmed10 | 0 | 0.591382731 | 0.754 | 0.385 | 0 | 1 | Tmed10 |
| Sncg | 0 | 0.590257061 | 0.424 | 0.111 | 0 | 1 | Sncg |
| Rtp3 | 0 | 0.589809331 | 0.288 | 0.048 | 0 | 1 | Rtp3 |
| Slco2a1 | 0 | 0.589240499 | 0.251 | 0.049 | 0 | 1 | Slco2a1 |
| 2510009E07Rik | 0 | 0.588417544 | 0.33 | 0.091 | 0 | 1 | 2510009E07Rik |
| Anp32a | 0 | 0.588400837 | 0.786 | 0.414 | 0 | 1 | Anp32a |
| Rnf19a | 0 | 0.586651057 | 0.357 | 0.104 | 0 | 1 | Rnf19a |
| Lats2 | 0 | 0.585807695 | 0.34 | 0.084 | 0 | 1 | Lats2 |
| Mcam | 0 | 0.585071959 | 0.323 | 0.075 | 0 | 1 | Mcam |
| Arhgef7 | 0 | 0.584128871 | 0.368 | 0.119 | 0 | 1 | Arhgef7 |
| Ppic | 0 | 0.582675551 | 0.348 | 0.084 | 0 | 1 | Ppic |
| Slc28a2 | 0 | 0.582667407 | 0.296 | 0.077 | 0 | 1 | Slc28a2 |

| | | | | | | | |
|---|---|---|---|---|---|---|---|
| Lhx6 | 0 | 0.580225105 | 0.255 | 0.031 | 0 | 1 | Lhx6 |
| Ace | 0 | 0.5789338 | 0.341 | 0.083 | 0 | 1 | Ace |
| Acer2 | 0 | 0.578653836 | 0.258 | 0.04 | 0 | 1 | Acer2 |
| Capn2 | 0 | 0.578499368 | 0.416 | 0.141 | 0 | 1 | Capn2 |
| Stx2 | 0 | 0.577227476 | 0.325 | 0.079 | 0 | 1 | Stx2 |
| Klhl24 | 0 | 0.577107232 | 0.462 | 0.175 | 0 | 1 | Klhl24 |
| Klc1 | 0 | 0.575670446 | 0.368 | 0.114 | 0 | 1 | Klc1 |
| Cx3cl1 | 0 | 0.574904705 | 0.315 | 0.07 | 0 | 1 | Cx3cl1 |
| Mxra7 | 0 | 0.574903959 | 0.378 | 0.104 | 0 | 1 | Mxra7 |
| Tbxa2r | 0 | 0.57389037 | 0.29 | 0.056 | 0 | 1 | Tbxa2r |
| Selenom | 0 | 0.570913375 | 0.841 | 0.381 | 0 | 1 | Selenom |
| Sgms1 | 0 | 0.569281462 | 0.336 | 0.095 | 0 | 1 | Sgms1 |
| Myo1b | 0 | 0.56750373 | 0.374 | 0.099 | 0 | 1 | Myo1b |
| 2700081O15Rik | 0 | 0.56644548 | 0.284 | 0.053 | 0 | 1 | 2700081O15Rik |
| Lamc1 | 0 | 0.566337354 | 0.379 | 0.111 | 0 | 1 | Lamc1 |
| Prkd2 | 0 | 0.566091536 | 0.347 | 0.107 | 0 | 1 | Prkd2 |
| Thsd1 | 0 | 0.563285134 | 0.29 | 0.056 | 0 | 1 | Thsd1 |
| Rnf144a | 0 | 0.563101343 | 0.314 | 0.074 | 0 | 1 | Rnf144a |
| Spns2 | 0 | 0.563055647 | 0.289 | 0.061 | 0 | 1 | Spns2 |
| Sh3glb1 | 0 | 0.561318928 | 0.843 | 0.481 | 0 | 1 | Sh3glb1 |
| Rab13 | 0 | 0.561084716 | 0.388 | 0.097 | 0 | 1 | Rab13 |
| St3gal5 | 0 | 0.56068558 | 0.262 | 0.052 | 0 | 1 | St3gal5 |
| Ltbp1 | 0 | 0.560556748 | 0.297 | 0.073 | 0 | 1 | Ltbp1 |
| Cdc42ep1 | 0 | 0.559891997 | 0.363 | 0.091 | 0 | 1 | Cdc42ep1 |
| Gbp3 | 0 | 0.556605896 | 0.313 | 0.087 | 0 | 1 | Gbp3 |
| Msrb3 | 0 | 0.55563091 | 0.359 | 0.096 | 0 | 1 | Msrb3 |
| Mpdz | 0 | 0.555272859 | 0.291 | 0.067 | 0 | 1 | Mpdz |
| Phf11d | 0 | 0.553792336 | 0.348 | 0.104 | 0 | 1 | Phf11d |
| Mmp15 | 0 | 0.551930075 | 0.319 | 0.066 | 0 | 1 | Mmp15 |
| Fam212a | 0 | 0.551221604 | 0.352 | 0.092 | 0 | 1 | Fam212a |
| Emp1 | 0 | 0.549165765 | 0.328 | 0.089 | 0 | 1 | Emp1 |
| Prkar1a | 0 | 0.548036005 | 0.7 | 0.355 | 0 | 1 | Prkar1a |
| Fcgrt | 0 | 0.546372279 | 0.428 | 0.146 | 0 | 1 | Fcgrt |
| Lrrk2 | 0 | 0.544830122 | 0.255 | 0.049 | 0 | 1 | Lrrk2 |
| Trove2 | 0 | 0.54453501 | 0.343 | 0.098 | 0 | 1 | Trove2 |
| Pqlc1 | 0 | 0.542396383 | 0.333 | 0.093 | 0 | 1 | Pqlc1 |
| Atp8b1 | 0 | 0.542306461 | 0.293 | 0.073 | 0 | 1 | Atp8b1 |
| Hspa12b | 0 | 0.540870763 | 0.309 | 0.069 | 0 | 1 | Hspa12b |
| Rtl8b | 0 | 0.539048506 | 0.322 | 0.078 | 0 | 1 | Rtl8b |
| Efr3b | 0 | 0.537553144 | 0.282 | 0.056 | 0 | 1 | Efr3b |
| Card10 | 0 | 0.531561078 | 0.283 | 0.054 | 0 | 1 | Card10 |
| Impdh1 | 0 | 0.5315533 | 0.305 | 0.081 | 0 | 1 | Impdh1 |
| Prpf40b | 0 | 0.529854983 | 0.25 | 0.049 | 0 | 1 | Prpf40b |

| | | | | | | | |
|---|---|---|---|---|---|---|---|
| Rps6ka2 | 0 | 0.529826672 | 0.264 | 0.052 | 0 | 1 | Rps6ka2 |
| Crk | 0 | 0.528648187 | 0.436 | 0.154 | 0 | 1 | Crk |
| Mapk3 | 0 | 0.525716946 | 0.549 | 0.22 | 0 | 1 | Mapk3 |
| Gnb4 | 0 | 0.525540776 | 0.301 | 0.077 | 0 | 1 | Gnb4 |
| Kctd17 | 0 | 0.525079934 | 0.285 | 0.059 | 0 | 1 | Kctd17 |
| Scarf1 | 0 | 0.523595084 | 0.254 | 0.041 | 0 | 1 | Scarf1 |
| Upp1 | 0 | 0.523591 | 0.299 | 0.067 | 0 | 1 | Upp1 |
| Fam114a1 | 0 | 0.521397754 | 0.34 | 0.098 | 0 | 1 | Fam114a1 |
| Flywch1 | 0 | 0.519693885 | 0.315 | 0.081 | 0 | 1 | Flywch1 |
| C130074G19Rik | 0 | 0.518947269 | 0.507 | 0.149 | 0 | 1 | C130074G19Rik |
| Cebpg | 0 | 0.516374409 | 0.486 | 0.182 | 0 | 1 | Cebpg |
| Ccm2l | 0 | 0.516268569 | 0.288 | 0.061 | 0 | 1 | Ccm2l |
| Lrrc8a | 0 | 0.515321777 | 0.279 | 0.064 | 0 | 1 | Lrrc8a |
| Sox13 | 0 | 0.513296001 | 0.279 | 0.06 | 0 | 1 | Sox13 |
| Ctnnd1 | 0 | 0.511817315 | 0.378 | 0.108 | 0 | 1 | Ctnnd1 |
| Tmem184b | 0 | 0.511548527 | 0.307 | 0.083 | 0 | 1 | Tmem184b |
| Tubb4b | 0 | 0.508186558 | 0.75 | 0.375 | 0 | 1 | Tubb4b |
| Lamb2 | 0 | 0.508139297 | 0.334 | 0.09 | 0 | 1 | Lamb2 |
| Csrp1 | 0 | 0.506414297 | 0.422 | 0.144 | 0 | 1 | Csrp1 |
| Layn | 0 | 0.506222797 | 0.261 | 0.053 | 0 | 1 | Layn |
| Pik3c2b | 0 | 0.505918974 | 0.267 | 0.051 | 0 | 1 | Pik3c2b |
| Tuba1a | 0 | 0.50342733 | 0.536 | 0.211 | 0 | 1 | Tuba1a |
| Tmcc3 | 0 | 0.495988168 | 0.654 | 0.265 | 0 | 1 | Tmcc3 |
| Lamb1 | 0 | 0.494385121 | 0.352 | 0.102 | 0 | 1 | Lamb1 |
| Sod1 | 0 | 0.493055427 | 0.583 | 0.254 | 0 | 1 | Sod1 |
| Zfp979 | 0 | 0.491249048 | 0.258 | 0.05 | 0 | 1 | Zfp979 |
| Smtn | 0 | 0.490472908 | 0.26 | 0.057 | 0 | 1 | Smtn |
| Ankrd33b | 0 | 0.488935812 | 0.261 | 0.055 | 0 | 1 | Ankrd33b |
| Rilpl1 | 0 | 0.48852355 | 0.297 | 0.076 | 0 | 1 | Rilpl1 |
| Pcdh1 | 0 | 0.488289294 | 0.281 | 0.061 | 0 | 1 | Pcdh1 |
| Sec14l1 | 0 | 0.486409042 | 0.573 | 0.229 | 0 | 1 | Sec14l1 |
| Fmo1 | 0 | 0.484774846 | 0.481 | 0.156 | 0 | 1 | Fmo1 |
| Mapk12 | 0 | 0.471019335 | 0.262 | 0.054 | 0 | 1 | Mapk12 |
| Agrn | 0 | 0.468021066 | 0.256 | 0.055 | 0 | 1 | Agrn |
| Pim3 | 0 | 0.460432217 | 0.483 | 0.168 | 0 | 1 | Pim3 |
| Hyal2 | 7.95E-308 | 0.468910327 | 0.258 | 0.059 | 1.61E-303 | 1 | Hyal2 |
| Adipor1 | 1.09E-307 | 0.547039072 | 0.515 | 0.216 | 2.20E-303 | 1 | Adipor1 |
| Arih1 | 1.59E-307 | 0.645761364 | 0.585 | 0.269 | 3.23E-303 | 1 | Arih1 |
| Atp11c | 2.09E-307 | 0.646014784 | 0.361 | 0.118 | 4.23E-303 | 1 | Atp11c |
| Mbnl2 | 2.44E-307 | 0.555667521 | 0.733 | 0.368 | 4.94E-303 | 1 | Mbnl2 |
| Pmp22 | 8.85E-307 | 0.494383033 | 0.363 | 0.109 | 1.79E-302 | 1 | Pmp22 |
| Dstyk | 3.87E-306 | 0.531437211 | 0.364 | 0.115 | 7.84E-302 | 1 | Dstyk |
| Grk5 | 1.77E-305 | 0.665156661 | 0.375 | 0.123 | 3.58E-301 | 1 | Grk5 |

| | | | | | | | |
|---|---|---|---|---|---|---|---|
| Amotl2 | 8.75E-305 | 0.479608027 | 0.256 | 0.059 | 1.77E-300 | 1 | Amotl2 |
| Spata13 | 1.43E-304 | 0.644169037 | 0.443 | 0.168 | 2.90E-300 | 1 | Spata13 |
| Lrp5 | 2.29E-304 | 0.503773852 | 0.268 | 0.066 | 4.63E-300 | 1 | Lrp5 |
| Apc | 4.67E-304 | 0.637488228 | 0.499 | 0.208 | 9.46E-300 | 1 | Apc |
| Lfng | 5.96E-304 | 0.603729191 | 0.337 | 0.104 | 1.21E-299 | 1 | Lfng |
| Ar | 8.35E-304 | 0.560234603 | 0.473 | 0.177 | 1.69E-299 | 1 | Ar |
| Fbxl17 | 1.22E-303 | 0.717026334 | 0.582 | 0.267 | 2.47E-299 | 1 | Fbxl17 |
| Ywhab | 1.60E-303 | 0.536515134 | 0.769 | 0.418 | 3.24E-299 | 1 | Ywhab |
| Il1r1 | 5.75E-302 | 0.494580668 | 0.301 | 0.08 | 1.16E-297 | 1 | Il1r1 |
| Tnrc6c | 2.18E-301 | 0.647809487 | 0.583 | 0.272 | 4.42E-297 | 1 | Tnrc6c |
| Dennd5a | 3.43E-301 | 0.582530504 | 0.417 | 0.15 | 6.94E-297 | 1 | Dennd5a |
| Slc12a7 | 9.55E-301 | 0.521078607 | 0.288 | 0.077 | 1.93E-296 | 1 | Slc12a7 |
| Gnas | 2.05E-300 | 0.465238431 | 0.996 | 0.841 | 4.15E-296 | 1 | Gnas |
| Maml2 | 2.18E-300 | 0.728204444 | 0.627 | 0.297 | 4.42E-296 | 1 | Maml2 |
| Pde4d | 3.93E-300 | 0.555803236 | 0.367 | 0.115 | 7.96E-296 | 1 | Pde4d |
| Nrip1 | 5.14E-300 | 0.77182906 | 0.488 | 0.205 | 1.04E-295 | 1 | Nrip1 |
| Leprot | 9.07E-300 | 0.559575355 | 0.401 | 0.141 | 1.84E-295 | 1 | Leprot |
| Samhd1 | 3.35E-299 | 0.571478863 | 0.512 | 0.213 | 6.79E-295 | 1 | Samhd1 |
| Rpn1 | 3.55E-299 | 0.629093458 | 0.458 | 0.184 | 7.19E-295 | 1 | Rpn1 |
| Triobp | 4.02E-299 | 0.524019286 | 0.341 | 0.105 | 8.15E-295 | 1 | Triobp |
| Lamp1 | 4.42E-299 | 0.488586109 | 0.732 | 0.366 | 8.95E-295 | 1 | Lamp1 |
| Slk | 8.77E-299 | 0.620532106 | 0.485 | 0.202 | 1.77E-294 | 1 | Slk |
| Taok1 | 3.32E-298 | 0.602471959 | 0.5 | 0.211 | 6.72E-294 | 1 | Taok1 |
| Cdh13 | 3.18E-297 | 0.494559827 | 0.278 | 0.069 | 6.44E-293 | 1 | Cdh13 |
| Bag5 | 2.39E-296 | 0.477590864 | 0.443 | 0.163 | 4.83E-292 | 1 | Bag5 |
| Zdhhc18 | 2.74E-295 | 0.595551369 | 0.387 | 0.136 | 5.54E-291 | 1 | Zdhhc18 |
| Pola2 | 2.99E-295 | 0.507021238 | 0.27 | 0.069 | 6.05E-291 | 1 | Pola2 |
| Arhgap23 | 4.10E-295 | 0.489051823 | 0.25 | 0.059 | 8.31E-291 | 1 | Arhgap23 |
| Myo1d | 4.83E-295 | 0.51155963 | 0.291 | 0.077 | 9.78E-291 | 1 | Myo1d |
| Ubl3 | 1.35E-294 | 0.517419082 | 0.487 | 0.198 | 2.73E-290 | 1 | Ubl3 |
| Aldh21 | 2.53E-294 | 0.443008574 | 0.733 | 0.342 | 5.12E-290 | 1 | Aldh2 |
| Strn3 | 3.44E-294 | 0.541560719 | 0.625 | 0.3 | 6.97E-290 | 1 | Strn3 |
| Rhoa | 1.48E-293 | 0.535059897 | 0.897 | 0.544 | 2.99E-289 | 1 | Rhoa |
| Igf1r | 2.32E-292 | 0.593717405 | 0.379 | 0.126 | 4.69E-288 | 1 | Igf1r |
| Prdx4 | 2.69E-292 | 0.513322058 | 0.472 | 0.181 | 5.44E-288 | 1 | Prdx4 |
| Fam13c | 3.33E-292 | 0.493238688 | 0.301 | 0.082 | 6.74E-288 | 1 | Fam13c |
| Palm | 4.26E-292 | 0.452194651 | 0.25 | 0.058 | 8.62E-288 | 1 | Palm |
| Hlx | 6.23E-292 | 0.539457083 | 0.279 | 0.073 | 1.26E-287 | 1 | Hlx |
| Helz2 | 2.11E-291 | 0.481199857 | 0.263 | 0.066 | 4.27E-287 | 1 | Helz2 |
| Trim47 | 2.46E-291 | 0.487695405 | 0.353 | 0.11 | 4.99E-287 | 1 | Trim47 |
| Sh3rf1 | 5.37E-291 | 0.601244701 | 0.326 | 0.099 | 1.09E-286 | 1 | Sh3rf1 |
| Usp34 | 8.85E-291 | 0.605273389 | 0.619 | 0.3 | 1.79E-286 | 1 | Usp34 |
| Itsn2 | 1.04E-289 | 0.644439106 | 0.532 | 0.237 | 2.11E-285 | 1 | Itsn2 |

| | | | | | | | |
|---|---|---|---|---|---|---|---|
| Ifnar2 | 2.93E-289 | 0.616675373 | 0.592 | 0.282 | 5.93E-285 | 1 | Ifnar2 |
| Hmg20b | 3.98E-288 | 0.439158438 | 0.525 | 0.218 | 8.05E-284 | 1 | Hmg20b |
| Kif5b | 4.04E-287 | 0.438711787 | 0.704 | 0.358 | 8.18E-283 | 1 | Kif5b |
| Trim56 | 4.56E-287 | 0.508054785 | 0.316 | 0.095 | 9.23E-283 | 1 | Trim56 |
| Clec2d | 4.95E-287 | 0.568956001 | 0.833 | 0.478 | 1.00E-282 | 1 | Clec2d |
| Abcb1a | 5.21E-287 | 0.492773996 | 0.293 | 0.081 | 1.05E-282 | 1 | Abcb1a |
| Ccdc28b | 1.47E-286 | 0.425700981 | 0.536 | 0.218 | 2.98E-282 | 1 | Ccdc28b |
| Srrm2 | 2.53E-286 | 0.576410737 | 0.781 | 0.419 | 5.12E-282 | 1 | Srrm2 |
| Tpm3 | 3.72E-286 | 0.540079016 | 0.786 | 0.43 | 7.53E-282 | 1 | Tpm3 |
| Gng12 | 3.48E-285 | 0.499181628 | 0.437 | 0.163 | 7.04E-281 | 1 | Gng12 |
| Dnajc3 | 5.59E-285 | 0.59337195 | 0.644 | 0.327 | 1.13E-280 | 1 | Dnajc3 |
| Myl12a | 6.45E-285 | 0.507459081 | 0.767 | 0.422 | 1.31E-280 | 1 | Myl12a |
| Plxna4 | 3.81E-284 | 0.562798309 | 0.307 | 0.089 | 7.72E-280 | 1 | Plxna4 |
| Crebbp | 6.17E-284 | 0.620539876 | 0.562 | 0.264 | 1.25E-279 | 1 | Crebbp |
| Stat2 | 1.74E-282 | 0.59975624 | 0.337 | 0.109 | 3.52E-278 | 1 | Stat2 |
| H2-T22 | 2.90E-282 | 0.557306645 | 0.534 | 0.24 | 5.87E-278 | 1 | H2-T22 |
| Plec | 5.91E-282 | 0.571765827 | 0.39 | 0.14 | 1.20E-277 | 1 | Plec |
| Rab11a | 7.05E-282 | 0.423567595 | 0.585 | 0.264 | 1.43E-277 | 1 | Rab11a |
| Ddx5 | 2.24E-281 | 0.56863068 | 0.95 | 0.606 | 4.54E-277 | 1 | Ddx5 |
| Mcl1 | 7.12E-281 | 0.579459288 | 0.65 | 0.329 | 1.44E-276 | 1 | Mcl1 |
| Nf1 | 1.95E-280 | 0.617984172 | 0.454 | 0.184 | 3.96E-276 | 1 | Nf1 |
| Rab6a | 3.34E-280 | 0.526590501 | 0.449 | 0.178 | 6.75E-276 | 1 | Rab6a |
| Nedd4l | 1.16E-279 | 0.676739195 | 0.446 | 0.181 | 2.35E-275 | 1 | Nedd4l |
| 6430548M08Rik | 2.00E-278 | 0.347744944 | 0.275 | 0.071 | 4.05E-274 | 1 | 6430548M08Rik |
| Fkbp9 | 4.55E-278 | 0.461152098 | 0.293 | 0.082 | 9.22E-274 | 1 | Fkbp9 |
| Acap2 | 9.93E-278 | 0.584623178 | 0.518 | 0.23 | 2.01E-273 | 1 | Acap2 |
| Ivns1abp1 | 1.27E-277 | 0.484350162 | 0.626 | 0.295 | 2.57E-273 | 1 | Ivns1abp |
| Nr3c2 | 1.76E-277 | 0.595755663 | 0.308 | 0.091 | 3.57E-273 | 1 | Nr3c2 |
| Uvrag | 2.74E-276 | 0.632597703 | 0.51 | 0.229 | 5.55E-272 | 1 | Uvrag |
| Dnm2 | 3.42E-276 | 0.572426641 | 0.5 | 0.22 | 6.92E-272 | 1 | Dnm2 |
| Vamp3 | 4.76E-276 | 0.450034292 | 0.395 | 0.141 | 9.63E-272 | 1 | Vamp3 |
| Tbl1x | 1.38E-275 | 0.564642773 | 0.508 | 0.223 | 2.78E-271 | 1 | Tbl1x |
| Psmb8 | 1.80E-274 | 0.552820552 | 0.884 | 0.504 | 3.64E-270 | 1 | Psmb8 |
| Cflar | 2.65E-274 | 0.522244455 | 0.466 | 0.194 | 5.36E-270 | 1 | Cflar |
| Cd274 | 3.78E-274 | 0.610751554 | 0.292 | 0.086 | 7.65E-270 | 1 | Cd274 |
| Tm9sf3 | 3.85E-274 | 0.53932837 | 0.599 | 0.293 | 7.80E-270 | 1 | Tm9sf3 |
| Tmem176a | 1.70E-273 | 0.403231703 | 0.762 | 0.361 | 3.44E-269 | 1 | Tmem176a |
| Exoc4 | 2.08E-273 | 0.649937074 | 0.585 | 0.281 | 4.22E-269 | 1 | Exoc4 |
| Map7d1 | 1.49E-272 | 0.509042316 | 0.377 | 0.134 | 3.01E-268 | 1 | Map7d1 |
| Cntln | 1.60E-272 | 0.58143044 | 0.355 | 0.123 | 3.25E-268 | 1 | Cntln |
| Degs1 | 8.60E-272 | 0.565297216 | 0.4 | 0.15 | 1.74E-267 | 1 | Degs1 |
| Hdgfl3 | 8.98E-272 | 0.508772542 | 0.289 | 0.082 | 1.82E-267 | 1 | Hdgfl3 |
| Irf2 | 9.78E-272 | 0.583986433 | 0.608 | 0.299 | 1.98E-267 | 1 | Irf2 |

| | | | | | | | |
|---|---|---|---|---|---|---|---|
| Klf12 | 3.44E-271 | 0.640728892 | 0.435 | 0.171 | 6.96E-267 | 1 | Klf12 |
| Atg4a | 4.43E-271 | 0.619956887 | 0.321 | 0.102 | 8.97E-267 | 1 | Atg4a |
| Pgrmc1 | 5.21E-271 | 0.461438484 | 0.453 | 0.18 | 1.05E-266 | 1 | Pgrmc1 |
| Ckap4 | 7.68E-271 | 0.466126255 | 0.272 | 0.074 | 1.56E-266 | 1 | Ckap4 |
| H2-Q6 | 1.58E-270 | 0.590906269 | 0.487 | 0.207 | 3.20E-266 | 1 | H2-Q6 |
| Lhfpl2 | 4.52E-270 | 0.526269376 | 0.355 | 0.118 | 9.15E-266 | 1 | Lhfpl2 |
| Emc7 | 4.60E-270 | 0.529495866 | 0.516 | 0.233 | 9.32E-266 | 1 | Emc7 |
| N4bp1 | 7.19E-270 | 0.547463941 | 0.386 | 0.142 | 1.46E-265 | 1 | N4bp1 |
| Fnip2 | 9.65E-270 | 0.529014525 | 0.279 | 0.079 | 1.95E-265 | 1 | Fnip2 |
| Gimap1 | 2.32E-269 | 0.486182473 | 0.589 | 0.257 | 4.70E-265 | 1 | Gimap1 |
| mt-Nd2 | 2.89E-269 | 0.540447255 | 0.989 | 0.639 | 5.84E-265 | 1 | mt-Nd2 |
| Xaf1 | 4.60E-269 | 0.496044039 | 0.457 | 0.183 | 9.32E-265 | 1 | Xaf1 |
| Tmbim6 | 6.25E-269 | 0.542579885 | 0.708 | 0.387 | 1.27E-264 | 1 | Tmbim6 |
| Pbx1 | 1.77E-268 | 0.471282065 | 0.435 | 0.161 | 3.59E-264 | 1 | Pbx1 |
| Polr2a | 2.69E-268 | 0.688497836 | 0.522 | 0.243 | 5.45E-264 | 1 | Polr2a |
| Ppp3ca | 6.45E-268 | 0.590456808 | 0.68 | 0.347 | 1.31E-263 | 1 | Ppp3ca |
| Socs2 | 2.92E-267 | 0.473371282 | 0.407 | 0.147 | 5.91E-263 | 1 | Socs2 |
| Myadm | 3.37E-267 | 0.532238885 | 0.3 | 0.091 | 6.81E-263 | 1 | Myadm |
| Fyn | 8.85E-267 | 0.55671866 | 0.524 | 0.225 | 1.79E-262 | 1 | Fyn |
| Nipbl | 1.09E-266 | 0.607996628 | 0.636 | 0.323 | 2.21E-262 | 1 | Nipbl |
| Herc2 | 2.04E-266 | 0.573051699 | 0.427 | 0.172 | 4.12E-262 | 1 | Herc2 |
| Ankrd12 | 3.03E-266 | 0.640481774 | 0.562 | 0.267 | 6.14E-262 | 1 | Ankrd12 |
| Atp2b1 | 8.47E-266 | 0.617240369 | 0.593 | 0.291 | 1.72E-261 | 1 | Atp2b1 |
| Ralb | 8.88E-266 | 0.444512746 | 0.358 | 0.121 | 1.80E-261 | 1 | Ralb |
| Pkp4 | 5.20E-265 | 0.438355883 | 0.551 | 0.244 | 1.05E-260 | 1 | Pkp4 |
| Crybg3 | 6.44E-265 | 0.516979922 | 0.332 | 0.109 | 1.30E-260 | 1 | Crybg3 |
| Phf20l1 | 1.84E-264 | 0.581602199 | 0.533 | 0.248 | 3.72E-260 | 1 | Phf20l1 |
| Pcm1 | 4.55E-264 | 0.562259492 | 0.487 | 0.213 | 9.21E-260 | 1 | Pcm1 |
| Sec62 | 7.45E-264 | 0.47523306 | 0.675 | 0.349 | 1.51E-259 | 1 | Sec62 |
| Bptf | 9.63E-264 | 0.63096632 | 0.537 | 0.255 | 1.95E-259 | 1 | Bptf |
| Mef2a | 7.42E-263 | 0.466114618 | 0.734 | 0.365 | 1.50E-258 | 1 | Mef2a |
| Mast4 | 1.19E-262 | 0.411020194 | 0.418 | 0.15 | 2.42E-258 | 1 | Mast4 |
| Usp25 | 3.41E-262 | 0.558902685 | 0.525 | 0.242 | 6.91E-258 | 1 | Usp25 |
| Susd6 | 4.06E-262 | 0.626431723 | 0.525 | 0.244 | 8.21E-258 | 1 | Susd6 |
| Trim25 | 4.73E-262 | 0.478445091 | 0.348 | 0.118 | 9.58E-258 | 1 | Trim25 |
| Ipo11 | 7.89E-262 | 0.511687327 | 0.325 | 0.107 | 1.60E-257 | 1 | Ipo11 |
| Igtp | 1.38E-261 | 0.62268026 | 0.279 | 0.081 | 2.79E-257 | 1 | Igtp |
| Smad4 | 1.74E-260 | 0.493725576 | 0.45 | 0.186 | 3.51E-256 | 1 | Smad4 |
| Arhgap17 | 1.64E-259 | 0.591308294 | 0.51 | 0.232 | 3.31E-255 | 1 | Arhgap17 |
| Elovl5 | 2.19E-259 | 0.549662401 | 0.344 | 0.12 | 4.42E-255 | 1 | Elovl5 |
| Aida | 2.29E-259 | 0.502829353 | 0.335 | 0.112 | 4.64E-255 | 1 | Aida |
| Rbm39 | 7.17E-259 | 0.517517787 | 0.899 | 0.521 | 1.45E-254 | 1 | Rbm39 |
| Irgm1 | 1.30E-258 | 0.510902781 | 0.343 | 0.117 | 2.62E-254 | 1 | Irgm1 |

| | | | | | | | |
|---|---|---|---|---|---|---|---|
| Cltc | 1.50E-258 | 0.454845307 | 0.594 | 0.289 | 3.03E-254 | 1 | Cltc |
| Socs5 | 5.87E-258 | 0.453110091 | 0.313 | 0.098 | 1.19E-253 | 1 | Socs5 |
| Pcp4l11 | 1.02E-257 | 0.288532346 | 0.619 | 0.264 | 2.06E-253 | 1 | Pcp4l1 |
| Lamp2 | 4.39E-257 | 0.472107212 | 0.574 | 0.262 | 8.90E-253 | 1 | Lamp2 |
| Egln1 | 4.72E-257 | 0.476113362 | 0.452 | 0.188 | 9.56E-253 | 1 | Egln1 |
| Cr1l | 6.12E-257 | 0.557566665 | 0.46 | 0.197 | 1.24E-252 | 1 | Cr1l |
| Stat1 | 4.64E-256 | 0.626325679 | 0.482 | 0.212 | 9.39E-252 | 1 | Stat1 |
| Pja2 | 2.10E-255 | 0.486801928 | 0.424 | 0.169 | 4.26E-251 | 1 | Pja2 |
| Dst | 4.22E-255 | 0.466034841 | 0.414 | 0.155 | 8.54E-251 | 1 | Dst |
| Vgll4 | 8.18E-255 | 0.547054756 | 0.485 | 0.212 | 1.66E-250 | 1 | Vgll4 |
| Tspan4 | 1.07E-254 | 0.428612617 | 0.277 | 0.079 | 2.16E-250 | 1 | Tspan4 |
| Tm9sf2 | 1.14E-254 | 0.552842546 | 0.488 | 0.218 | 2.31E-250 | 1 | Tm9sf2 |
| Asap2 | 4.13E-254 | 0.44030842 | 0.319 | 0.102 | 8.36E-250 | 1 | Asap2 |
| Cd2ap | 4.43E-254 | 0.365318608 | 0.605 | 0.282 | 8.98E-250 | 1 | Cd2ap |
| Etv6 | 4.88E-254 | 0.539090002 | 0.526 | 0.238 | 9.89E-250 | 1 | Etv6 |
| Vwa1 | 9.34E-254 | 0.408386358 | 0.274 | 0.077 | 1.89E-249 | 1 | Vwa1 |
| Snx9 | 1.29E-253 | 0.447393568 | 0.388 | 0.142 | 2.62E-249 | 1 | Snx9 |
| Btbd3 | 3.08E-253 | 0.421660758 | 0.267 | 0.074 | 6.23E-249 | 1 | Btbd3 |
| Tmco1 | 7.29E-253 | 0.505366221 | 0.561 | 0.273 | 1.48E-248 | 1 | Tmco1 |
| S100a13 | 1.84E-252 | 0.386808562 | 0.843 | 0.487 | 3.72E-248 | 1 | S100a13 |
| Ier2 | 2.31E-252 | 0.654995774 | 0.524 | 0.245 | 4.67E-248 | 1 | Ier2 |
| Vezf1 | 6.81E-252 | 0.442546984 | 0.466 | 0.197 | 1.38E-247 | 1 | Vezf1 |
| Cnot6 | 2.80E-251 | 0.476995313 | 0.354 | 0.128 | 5.67E-247 | 1 | Cnot6 |
| Kif1b | 4.14E-250 | 0.517150703 | 0.472 | 0.207 | 8.38E-246 | 1 | Kif1b |
| Pde4b | 1.07E-249 | 0.633115321 | 0.405 | 0.163 | 2.17E-245 | 1 | Pde4b |
| Csnk1a1 | 1.47E-249 | 0.433469965 | 0.743 | 0.412 | 2.98E-245 | 1 | Csnk1a1 |
| Zfp36 | 2.61E-249 | 0.730399461 | 0.405 | 0.165 | 5.29E-245 | 1 | Zfp36 |
| Selenof | 6.65E-249 | 0.424732986 | 0.778 | 0.445 | 1.35E-244 | 1 | Selenof |
| Ginm1 | 8.53E-249 | 0.461674987 | 0.386 | 0.146 | 1.73E-244 | 1 | Ginm1 |
| Lrrfip1 | 1.21E-248 | 0.519204279 | 0.649 | 0.335 | 2.46E-244 | 1 | Lrrfip1 |
| Slfn2 | 2.57E-248 | 0.637394752 | 0.492 | 0.224 | 5.20E-244 | 1 | Slfn2 |
| Gsk3b | 1.18E-247 | 0.51445788 | 0.585 | 0.29 | 2.40E-243 | 1 | Gsk3b |
| Trim24 | 1.42E-247 | 0.532443636 | 0.373 | 0.141 | 2.87E-243 | 1 | Trim24 |
| Camsap2 | 1.47E-247 | 0.458743029 | 0.274 | 0.082 | 2.98E-243 | 1 | Camsap2 |
| Cmip | 1.10E-246 | 0.573925296 | 0.523 | 0.248 | 2.23E-242 | 1 | Cmip |
| Eepd1 | 1.17E-246 | 0.384685519 | 0.341 | 0.112 | 2.36E-242 | 1 | Eepd1 |
| Gtf2i | 1.39E-245 | 0.441524593 | 0.484 | 0.213 | 2.81E-241 | 1 | Gtf2i |
| Tagln2 | 1.46E-245 | 0.443123883 | 0.724 | 0.381 | 2.95E-241 | 1 | Tagln2 |
| Zdhhc20 | 1.69E-245 | 0.568293807 | 0.449 | 0.195 | 3.42E-241 | 1 | Zdhhc20 |
| Slc23a2 | 1.94E-245 | 0.55264583 | 0.32 | 0.11 | 3.93E-241 | 1 | Slc23a2 |
| Rapgef1 | 8.28E-244 | 0.61067862 | 0.462 | 0.206 | 1.68E-239 | 1 | Rapgef1 |
| Bag3 | 1.12E-243 | 0.455660734 | 0.284 | 0.087 | 2.27E-239 | 1 | Bag3 |
| Zfp609 | 1.14E-243 | 0.507722882 | 0.311 | 0.103 | 2.31E-239 | 1 | Zfp609 |

| | | | | | | | |
|---|---|---|---|---|---|---|---|
| Tcaf1 | 2.30E-243 | 0.421561562 | 0.268 | 0.077 | 4.65E-239 | 1 | Tcaf1 |
| Scn1b | 4.72E-243 | 0.403756972 | 0.415 | 0.161 | 9.56E-239 | 1 | Scn1b |
| Hdac9 | 2.23E-242 | 0.6299443 | 0.355 | 0.127 | 4.52E-238 | 1 | Hdac9 |
| Cast | 2.49E-242 | 0.507080228 | 0.558 | 0.272 | 5.05E-238 | 1 | Cast |
| Grn | 2.70E-242 | 0.475919923 | 0.453 | 0.188 | 5.46E-238 | 1 | Grn |
| Gimap4 | 3.22E-242 | 0.45781691 | 0.512 | 0.216 | 6.53E-238 | 1 | Gimap4 |
| Igf2r | 5.08E-242 | 0.516140148 | 0.313 | 0.106 | 1.03E-237 | 1 | Igf2r |
| Chrm3 | 6.21E-242 | 0.715236842 | 0.26 | 0.074 | 1.26E-237 | 1 | Chrm3 |
| Dhrs7 | 1.02E-241 | 0.505258656 | 0.43 | 0.181 | 2.06E-237 | 1 | Dhrs7 |
| Ankhd1 | 1.45E-241 | 0.592360854 | 0.462 | 0.209 | 2.94E-237 | 1 | Ankhd1 |
| Ext2 | 2.14E-241 | 0.495930182 | 0.326 | 0.114 | 4.34E-237 | 1 | Ext2 |
| Iqsec2 | 2.93E-241 | 0.478474984 | 0.259 | 0.074 | 5.92E-237 | 1 | Iqsec2 |
| Dync1i2 | 9.24E-241 | 0.412886865 | 0.693 | 0.375 | 1.87E-236 | 1 | Dync1i2 |
| Ifih1 | 6.66E-240 | 0.470726636 | 0.306 | 0.101 | 1.35E-235 | 1 | Ifih1 |
| Nras | 1.48E-239 | 0.463044752 | 0.338 | 0.121 | 2.99E-235 | 1 | Nras |
| Ncor1 | 2.42E-239 | 0.497237738 | 0.718 | 0.393 | 4.89E-235 | 1 | Ncor1 |
| Selenok | 6.52E-239 | 0.403485712 | 0.842 | 0.508 | 1.32E-234 | 1 | Selenok |
| Wac | 1.62E-238 | 0.506952573 | 0.569 | 0.283 | 3.28E-234 | 1 | Wac |
| Bicc1 | 4.16E-238 | 0.354822034 | 0.264 | 0.073 | 8.42E-234 | 1 | Bicc1 |
| Fam76a | 5.70E-238 | 0.416857174 | 0.37 | 0.138 | 1.15E-233 | 1 | Fam76a |
| Ccdc47 | 6.12E-238 | 0.503440763 | 0.395 | 0.16 | 1.24E-233 | 1 | Ccdc47 |
| Rsu1 | 8.46E-238 | 0.472290663 | 0.486 | 0.217 | 1.71E-233 | 1 | Rsu1 |
| Xpc | 9.34E-238 | 0.428312954 | 0.265 | 0.078 | 1.89E-233 | 1 | Xpc |
| Agap1 | 1.64E-237 | 0.461815365 | 0.333 | 0.115 | 3.32E-233 | 1 | Agap1 |
| Smpdl3a | 3.34E-237 | 0.419909441 | 0.445 | 0.183 | 6.77E-233 | 1 | Smpdl3a |
| Trim30a | 5.02E-237 | 0.589724178 | 0.432 | 0.185 | 1.02E-232 | 1 | Trim30a |
| Ddx17 | 1.37E-236 | 0.50109477 | 0.504 | 0.236 | 2.78E-232 | 1 | Ddx17 |
| Rab5c | 3.95E-236 | 0.369675105 | 0.533 | 0.252 | 7.99E-232 | 1 | Rab5c |
| Zfp710 | 1.73E-235 | 0.511059291 | 0.346 | 0.125 | 3.50E-231 | 1 | Zfp710 |
| Mrrf | 7.73E-235 | 0.372696508 | 0.278 | 0.084 | 1.56E-230 | 1 | Mrrf |
| Srpr | 1.30E-233 | 0.493342136 | 0.466 | 0.211 | 2.64E-229 | 1 | Srpr |
| Ywhaz | 1.46E-233 | 0.438238531 | 0.78 | 0.446 | 2.95E-229 | 1 | Ywhaz |
| Eif4ebp2 | 7.23E-233 | 0.454004492 | 0.463 | 0.206 | 1.46E-228 | 1 | Eif4ebp2 |
| Ifit1 | 2.05E-232 | 0.69743886 | 0.258 | 0.076 | 4.16E-228 | 1 | Ifit1 |
| Mllt10 | 2.17E-232 | 0.561877269 | 0.501 | 0.239 | 4.40E-228 | 1 | Mllt10 |
| Fus | 7.40E-232 | 0.47759055 | 0.649 | 0.347 | 1.50E-227 | 1 | Fus |
| Dip2b | 1.21E-231 | 0.520882084 | 0.507 | 0.24 | 2.45E-227 | 1 | Dip2b |
| Anxa7 | 1.59E-231 | 0.426077117 | 0.389 | 0.154 | 3.23E-227 | 1 | Anxa7 |
| Pxn | 2.57E-231 | 0.478061117 | 0.355 | 0.135 | 5.20E-227 | 1 | Pxn |
| Ccrl2 | 4.90E-231 | 0.45259835 | 0.272 | 0.083 | 9.92E-227 | 1 | Ccrl2 |
| Gm12216 | 5.94E-231 | 0.444911474 | 0.452 | 0.195 | 1.20E-226 | 1 | Gm12216 |
| Ctdsp2 | 9.85E-231 | 0.447493393 | 0.376 | 0.147 | 1.99E-226 | 1 | Ctdsp2 |
| Dennd6a | 1.20E-230 | 0.472007038 | 0.381 | 0.15 | 2.43E-226 | 1 | Dennd6a |

| | | | | | | | |
|---|---|---|---|---|---|---|---|
| Exoc5 | 2.42E-230 | 0.503667019 | 0.381 | 0.152 | 4.89E-226 | 1 | Exoc5 |
| Ubn1 | 7.76E-230 | 0.505235413 | 0.464 | 0.212 | 1.57E-225 | 1 | Ubn1 |
| Npdc1 | 8.13E-230 | 0.387917235 | 0.314 | 0.105 | 1.65E-225 | 1 | Npdc1 |
| Vav3 | 1.22E-229 | 0.61140147 | 0.481 | 0.22 | 2.47E-225 | 1 | Vav3 |
| Snx3 | 1.44E-229 | 0.389516898 | 0.733 | 0.418 | 2.92E-225 | 1 | Snx3 |
| Siah1a | 1.83E-229 | 0.423986618 | 0.298 | 0.098 | 3.71E-225 | 1 | Siah1a |
| Fam208a | 1.84E-229 | 0.522565316 | 0.403 | 0.168 | 3.72E-225 | 1 | Fam208a |
| Atraid | 5.89E-229 | 0.447275303 | 0.501 | 0.234 | 1.19E-224 | 1 | Atraid |
| Ywhaq | 1.71E-228 | 0.389346616 | 0.765 | 0.443 | 3.46E-224 | 1 | Ywhaq |
| Tor1aip1 | 2.32E-228 | 0.515151544 | 0.51 | 0.245 | 4.70E-224 | 1 | Tor1aip1 |
| Yaf2 | 2.94E-228 | 0.494394461 | 0.424 | 0.182 | 5.96E-224 | 1 | Yaf2 |
| Tmem2 | 3.77E-228 | 0.517197636 | 0.322 | 0.114 | 7.63E-224 | 1 | Tmem2 |
| Fosb | 4.40E-228 | 0.592099574 | 0.389 | 0.158 | 8.91E-224 | 1 | Fosb |
| Pafah1b1 | 5.00E-227 | 0.404325227 | 0.761 | 0.436 | 1.01E-222 | 1 | Pafah1b1 |
| Tmem109 | 1.27E-226 | 0.446118811 | 0.331 | 0.119 | 2.58E-222 | 1 | Tmem109 |
| Dpysl2 | 1.52E-226 | 0.507770152 | 0.367 | 0.144 | 3.07E-222 | 1 | Dpysl2 |
| Pbrm1 | 2.33E-226 | 0.521816636 | 0.516 | 0.251 | 4.73E-222 | 1 | Pbrm1 |
| Lncpint | 3.19E-225 | 0.592991329 | 0.501 | 0.238 | 6.46E-221 | 1 | Lncpint |
| Plod1 | 4.86E-225 | 0.447509262 | 0.289 | 0.094 | 9.83E-221 | 1 | Plod1 |
| Dag1 | 1.06E-224 | 0.460490388 | 0.323 | 0.114 | 2.14E-220 | 1 | Dag1 |
| Pea15a | 1.29E-224 | 0.408694051 | 0.466 | 0.205 | 2.60E-220 | 1 | Pea15a |
| Esm1 | 2.35E-224 | 0.801078512 | 0.262 | 0.078 | 4.76E-220 | 1 | Esm1 |
| Sh2d3c | 3.02E-224 | 0.482187756 | 0.341 | 0.128 | 6.10E-220 | 1 | Sh2d3c |
| Ggta1 | 7.87E-224 | 0.471205855 | 0.318 | 0.112 | 1.59E-219 | 1 | Ggta1 |
| mt-Nd1 | 1.19E-223 | 0.462420973 | 0.927 | 0.545 | 2.41E-219 | 1 | mt-Nd1 |
| Hes11 | 6.55E-223 | 0.44857262 | 0.579 | 0.272 | 1.33E-218 | 1 | Hes1 |
| Itpr1 | 1.16E-222 | 0.486887417 | 0.435 | 0.187 | 2.35E-218 | 1 | Itpr1 |
| Ddx6 | 1.73E-222 | 0.432555107 | 0.586 | 0.302 | 3.51E-218 | 1 | Ddx6 |
| Arglu1 | 2.20E-222 | 0.426821349 | 0.606 | 0.314 | 4.45E-218 | 1 | Arglu1 |
| Nrbp1 | 3.38E-222 | 0.44155779 | 0.417 | 0.178 | 6.84E-218 | 1 | Nrbp1 |
| Cyfip1 | 3.75E-222 | 0.392034396 | 0.519 | 0.24 | 7.60E-218 | 1 | Cyfip1 |
| Lima1 | 4.86E-222 | 0.440234338 | 0.352 | 0.128 | 9.85E-218 | 1 | Lima1 |
| Tsnax | 5.15E-222 | 0.42617343 | 0.356 | 0.135 | 1.04E-217 | 1 | Tsnax |
| Ube2e2 | 6.84E-222 | 0.52607077 | 0.34 | 0.123 | 1.38E-217 | 1 | Ube2e2 |
| Ddx50 | 1.16E-221 | 0.46814578 | 0.524 | 0.256 | 2.34E-217 | 1 | Ddx50 |
| Dram2 | 2.57E-221 | 0.46428991 | 0.338 | 0.125 | 5.21E-217 | 1 | Dram2 |
| Arhgap27 | 3.52E-221 | 0.439254447 | 0.253 | 0.077 | 7.12E-217 | 1 | Arhgap27 |
| Nmi | 6.65E-221 | 0.446069071 | 0.454 | 0.204 | 1.35E-216 | 1 | Nmi |
| Slc29a1 | 6.70E-221 | 0.417194513 | 0.311 | 0.107 | 1.36E-216 | 1 | Slc29a1 |
| Top2b | 9.31E-221 | 0.468097404 | 0.509 | 0.247 | 1.88E-216 | 1 | Top2b |
| Prrc2b | 1.77E-220 | 0.461594897 | 0.442 | 0.196 | 3.59E-216 | 1 | Prrc2b |
| Stx12 | 2.91E-220 | 0.414709625 | 0.348 | 0.132 | 5.90E-216 | 1 | Stx12 |
| Lgals9 | 4.80E-220 | 0.550069658 | 0.46 | 0.214 | 9.73E-216 | 1 | Lgals9 |

| | | | | | | | |
|---|---|---|---|---|---|---|---|
| Klf10 | 4.55E-219 | 0.519547543 | 0.349 | 0.134 | 9.21E-215 | 1 | Klf10 |
| Ubr3 | 1.46E-218 | 0.551343242 | 0.395 | 0.17 | 2.96E-214 | 1 | Ubr3 |
| Kat6a | 1.47E-218 | 0.506720948 | 0.435 | 0.195 | 2.97E-214 | 1 | Kat6a |
| Numa1 | 3.77E-218 | 0.519291517 | 0.389 | 0.164 | 7.64E-214 | 1 | Numa1 |
| mt-Nd3 | 1.70E-217 | 0.454372937 | 0.726 | 0.402 | 3.45E-213 | 1 | mt-Nd3 |
| Ppp1r15a | 6.78E-217 | 0.604896849 | 0.331 | 0.127 | 1.37E-212 | 1 | Ppp1r15a |
| 4930523C07Rik | 9.17E-217 | 0.503277434 | 0.43 | 0.186 | 1.86E-212 | 1 | 4930523C07Rik |
| Me3 | 1.01E-216 | 0.371918172 | 0.256 | 0.076 | 2.05E-212 | 1 | Me3 |
| Tmx3 | 1.43E-216 | 0.521916562 | 0.379 | 0.157 | 2.89E-212 | 1 | Tmx3 |
| Map3k3 | 1.73E-216 | 0.524392926 | 0.366 | 0.148 | 3.51E-212 | 1 | Map3k3 |
| Spen | 3.41E-216 | 0.497005954 | 0.337 | 0.13 | 6.89E-212 | 1 | Spen |
| Trps1 | 3.81E-216 | 0.558710657 | 0.451 | 0.202 | 7.71E-212 | 1 | Trps1 |
| Fer | 3.95E-216 | 0.408664151 | 0.286 | 0.095 | 8.00E-212 | 1 | Fer |
| Lipa | 4.56E-216 | 0.477197528 | 0.372 | 0.148 | 9.23E-212 | 1 | Lipa |
| Sun1 | 6.78E-216 | 0.414592846 | 0.253 | 0.077 | 1.37E-211 | 1 | Sun1 |
| Otud7b | 1.20E-215 | 0.468509396 | 0.29 | 0.098 | 2.43E-211 | 1 | Otud7b |
| Tjp2 | 3.95E-215 | 0.451536702 | 0.315 | 0.113 | 7.99E-211 | 1 | Tjp2 |
| Efcab14 | 4.39E-215 | 0.464770784 | 0.318 | 0.117 | 8.88E-211 | 1 | Efcab14 |
| Mib1 | 5.95E-215 | 0.453921381 | 0.349 | 0.136 | 1.20E-210 | 1 | Mib1 |
| Fbxw2 | 1.64E-214 | 0.402210528 | 0.451 | 0.202 | 3.32E-210 | 1 | Fbxw2 |
| Pam | 1.95E-214 | 0.517768175 | 0.421 | 0.181 | 3.95E-210 | 1 | Pam |
| Ephx1 | 3.20E-214 | 0.428589555 | 0.288 | 0.096 | 6.48E-210 | 1 | Ephx1 |
| Thoc2 | 5.23E-214 | 0.516477587 | 0.49 | 0.24 | 1.06E-209 | 1 | Thoc2 |
| Arhgef10 | 6.19E-214 | 0.298194183 | 0.253 | 0.075 | 1.25E-209 | 1 | Arhgef10 |
| Rtn4 | 1.05E-213 | 0.431748356 | 0.589 | 0.299 | 2.13E-209 | 1 | Rtn4 |
| Ubn2 | 2.79E-213 | 0.549336898 | 0.453 | 0.211 | 5.66E-209 | 1 | Ubn2 |
| Git2 | 3.98E-213 | 0.538797452 | 0.442 | 0.202 | 8.06E-209 | 1 | Git2 |
| Zyx | 6.79E-213 | 0.523895865 | 0.417 | 0.184 | 1.37E-208 | 1 | Zyx |
| Ifit3 | 7.59E-213 | 0.612407936 | 0.289 | 0.097 | 1.54E-208 | 1 | Ifit3 |
| Snx13 | 8.08E-213 | 0.508947129 | 0.324 | 0.123 | 1.64E-208 | 1 | Snx13 |
| Ccnd2 | 1.13E-212 | 0.576507496 | 0.514 | 0.25 | 2.29E-208 | 1 | Ccnd2 |
| Cbx5 | 3.94E-212 | 0.424472516 | 0.384 | 0.157 | 7.97E-208 | 1 | Cbx5 |
| Nisch | 8.36E-212 | 0.419107017 | 0.417 | 0.179 | 1.69E-207 | 1 | Nisch |
| Trim44 | 8.36E-212 | 0.445392943 | 0.433 | 0.195 | 1.69E-207 | 1 | Trim44 |
| Tmem50a | 1.13E-211 | 0.452591488 | 0.631 | 0.343 | 2.29E-207 | 1 | Tmem50a |
| Tmem176b | 2.87E-211 | 0.286517549 | 0.764 | 0.386 | 5.81E-207 | 1 | Tmem176b |
| Nfic | 4.41E-211 | 0.383591176 | 0.488 | 0.225 | 8.92E-207 | 1 | Nfic |
| Pkn2 | 1.03E-210 | 0.456965226 | 0.518 | 0.259 | 2.09E-206 | 1 | Pkn2 |
| Brd2 | 1.08E-210 | 0.461339898 | 0.465 | 0.22 | 2.20E-206 | 1 | Brd2 |
| Eif3a | 2.10E-210 | 0.464836145 | 0.615 | 0.334 | 4.25E-206 | 1 | Eif3a |
| Tnrc18 | 3.55E-210 | 0.485166601 | 0.357 | 0.144 | 7.18E-206 | 1 | Tnrc18 |
| Rc3h1 | 3.93E-210 | 0.49407597 | 0.404 | 0.177 | 7.96E-206 | 1 | Rc3h1 |
| Magi3 | 7.88E-210 | 0.489743713 | 0.256 | 0.081 | 1.59E-205 | 1 | Magi3 |

| | | | | | | | |
|---|---|---|---|---|---|---|---|
| Map4k5 | 7.96E-210 | 0.450118239 | 0.297 | 0.104 | 1.61E-205 | 1 | Map4k5 |
| Cdk11b | 1.10E-209 | 0.460897032 | 0.453 | 0.21 | 2.24E-205 | 1 | Cdk11b |
| Pxk | 1.44E-209 | 0.459282777 | 0.31 | 0.114 | 2.91E-205 | 1 | Pxk |
| Rtp4 | 3.59E-209 | 0.436279175 | 0.346 | 0.132 | 7.26E-205 | 1 | Rtp4 |
| Ccny | 5.55E-209 | 0.481817782 | 0.484 | 0.233 | 1.12E-204 | 1 | Ccny |
| Eif4g1 | 7.20E-209 | 0.469983533 | 0.44 | 0.204 | 1.46E-204 | 1 | Eif4g1 |
| 7-Sep | 1.07E-208 | 0.37983372 | 0.742 | 0.417 | 2.17E-204 | 1 | 7-Sep |
| Babam2 | 1.58E-208 | 0.574605891 | 0.44 | 0.206 | 3.21E-204 | 1 | Babam2 |
| Hspa5 | 1.97E-208 | 0.40618953 | 0.716 | 0.39 | 3.99E-204 | 1 | Hspa5 |
| Fosl2 | 4.61E-208 | 0.429450253 | 0.251 | 0.078 | 9.34E-204 | 1 | Fosl2 |
| Picalm | 9.94E-208 | 0.48406206 | 0.523 | 0.261 | 2.01E-203 | 1 | Picalm |
| Elk4 | 1.04E-207 | 0.476675196 | 0.348 | 0.139 | 2.10E-203 | 1 | Elk4 |
| Ino80d | 1.47E-207 | 0.477370377 | 0.345 | 0.137 | 2.97E-203 | 1 | Ino80d |
| Huwe1 | 3.92E-207 | 0.476954131 | 0.505 | 0.25 | 7.94E-203 | 1 | Huwe1 |
| Taf7 | 8.50E-207 | 0.46401537 | 0.311 | 0.114 | 1.72E-202 | 1 | Taf7 |
| Nbdy | 5.86E-206 | 0.352000611 | 0.396 | 0.165 | 1.19E-201 | 1 | Nbdy |
| Top1 | 8.97E-206 | 0.390810234 | 0.68 | 0.384 | 1.82E-201 | 1 | Top1 |
| Canx | 9.76E-206 | 0.461998606 | 0.532 | 0.271 | 1.98E-201 | 1 | Canx |
| Setd5 | 4.61E-205 | 0.536562814 | 0.474 | 0.23 | 9.33E-201 | 1 | Setd5 |
| Lrch3 | 1.10E-204 | 0.525631993 | 0.433 | 0.201 | 2.23E-200 | 1 | Lrch3 |
| Cmtm3 | 5.88E-204 | 0.381101253 | 0.257 | 0.082 | 1.19E-199 | 1 | Cmtm3 |
| mt-Co2 | 8.61E-204 | 0.484498005 | 1 | 0.684 | 1.74E-199 | 1 | mt-Co2 |
| Pura | 1.98E-203 | 0.409551445 | 0.607 | 0.327 | 4.01E-199 | 1 | Pura |
| Tmpo | 2.09E-203 | 0.444758498 | 0.405 | 0.177 | 4.24E-199 | 1 | Tmpo |
| Lyn | 2.41E-203 | 0.313726155 | 0.508 | 0.235 | 4.88E-199 | 1 | Lyn |
| Cdk12 | 2.87E-203 | 0.50391569 | 0.429 | 0.198 | 5.82E-199 | 1 | Cdk12 |
| Calr | 1.01E-202 | 0.417719702 | 0.676 | 0.365 | 2.05E-198 | 1 | Calr |
| Trim37 | 1.27E-202 | 0.37937682 | 0.251 | 0.079 | 2.57E-198 | 1 | Trim37 |
| Tpr | 4.85E-202 | 0.444497886 | 0.683 | 0.384 | 9.82E-198 | 1 | Tpr |
| Clock | 4.92E-202 | 0.452295124 | 0.399 | 0.173 | 9.97E-198 | 1 | Clock |
| Fubp1 | 5.34E-202 | 0.471319219 | 0.485 | 0.239 | 1.08E-197 | 1 | Fubp1 |
| Ano6 | 7.05E-202 | 0.484855101 | 0.366 | 0.151 | 1.43E-197 | 1 | Ano6 |
| Ago3 | 1.32E-201 | 0.526146659 | 0.325 | 0.128 | 2.66E-197 | 1 | Ago3 |
| Kctd10 | 1.59E-201 | 0.34809919 | 0.274 | 0.092 | 3.21E-197 | 1 | Kctd10 |
| Sh3d19 | 2.06E-201 | 0.431232097 | 0.26 | 0.084 | 4.17E-197 | 1 | Sh3d19 |
| Trim35 | 3.65E-201 | 0.459745979 | 0.307 | 0.114 | 7.39E-197 | 1 | Trim35 |
| Morf4l1 | 3.71E-201 | 0.348984273 | 0.862 | 0.545 | 7.51E-197 | 1 | Morf4l1 |
| Gtdc1 | 1.36E-200 | 0.549536837 | 0.322 | 0.124 | 2.76E-196 | 1 | Gtdc1 |
| Zfand3 | 3.40E-200 | 0.498040522 | 0.524 | 0.266 | 6.88E-196 | 1 | Zfand3 |
| Prpf8 | 3.95E-200 | 0.444548209 | 0.366 | 0.154 | 8.01E-196 | 1 | Prpf8 |
| Sec63 | 6.16E-199 | 0.466498654 | 0.393 | 0.173 | 1.25E-194 | 1 | Sec63 |
| Cfap20 | 7.20E-199 | 0.355559494 | 0.378 | 0.156 | 1.46E-194 | 1 | Cfap20 |
| Tacc1 | 1.05E-198 | 0.485030564 | 0.524 | 0.27 | 2.13E-194 | 1 | Tacc1 |

| | | | | | | | |
|---|---|---|---|---|---|---|---|
| Mesd | 1.73E-198 | 0.450414018 | 0.338 | 0.135 | 3.51E-194 | 1 | Mesd |
| Nsd1 | 1.82E-198 | 0.442807768 | 0.586 | 0.312 | 3.68E-194 | 1 | Nsd1 |
| Hnrnpu | 2.10E-198 | 0.379402864 | 0.679 | 0.382 | 4.25E-194 | 1 | Hnrnpu |
| mt-Cytb | 6.35E-198 | 0.414300374 | 0.994 | 0.652 | 1.29E-193 | 1 | mt-Cytb |
| Smchd1 | 1.03E-197 | 0.483237911 | 0.472 | 0.231 | 2.08E-193 | 1 | Smchd1 |
| Oasl2 | 1.10E-197 | 0.472341392 | 0.295 | 0.105 | 2.23E-193 | 1 | Oasl2 |
| Acyp2 | 3.71E-197 | 0.344918646 | 0.317 | 0.115 | 7.52E-193 | 1 | Acyp2 |
| Atxn7l3b | 6.39E-197 | 0.30985551 | 0.656 | 0.365 | 1.29E-192 | 1 | Atxn7l3b |
| mt-Atp6 | 1.04E-196 | 0.467814914 | 1 | 0.714 | 2.10E-192 | 1 | mt-Atp6 |
| Ep400 | 1.95E-196 | 0.520829062 | 0.394 | 0.178 | 3.95E-192 | 1 | Ep400 |
| Rlim | 2.38E-196 | 0.538446004 | 0.38 | 0.169 | 4.82E-192 | 1 | Rlim |
| Scoc | 3.95E-196 | 0.302365062 | 0.345 | 0.134 | 8.00E-192 | 1 | Scoc |
| Ostf1 | 4.98E-196 | 0.349463864 | 0.786 | 0.462 | 1.01E-191 | 1 | Ostf1 |
| mt-Nd4 | 6.04E-196 | 0.436734004 | 0.988 | 0.624 | 1.22E-191 | 1 | mt-Nd4 |
| Tnrc6b | 7.66E-196 | 0.438792138 | 0.622 | 0.331 | 1.55E-191 | 1 | Tnrc6b |
| Tom1l2 | 9.40E-196 | 0.437981905 | 0.291 | 0.105 | 1.90E-191 | 1 | Tom1l2 |
| Bod1l | 1.39E-195 | 0.508230089 | 0.48 | 0.24 | 2.82E-191 | 1 | Bod1l |
| Tubb5 | 1.75E-195 | 0.317759899 | 0.632 | 0.34 | 3.54E-191 | 1 | Tubb5 |
| Ddx58 | 6.30E-195 | 0.474198081 | 0.332 | 0.133 | 1.27E-190 | 1 | Ddx58 |
| Slc3a2 | 6.99E-195 | 0.507170701 | 0.506 | 0.257 | 1.41E-190 | 1 | Slc3a2 |
| Gimap8 | 1.40E-194 | 0.451361339 | 0.349 | 0.14 | 2.84E-190 | 1 | Gimap8 |
| Tln1 | 5.51E-194 | 0.433191272 | 0.663 | 0.364 | 1.11E-189 | 1 | Tln1 |
| Tnks2 | 6.43E-194 | 0.456577237 | 0.493 | 0.246 | 1.30E-189 | 1 | Tnks2 |
| Kansl1 | 6.46E-194 | 0.471739333 | 0.581 | 0.302 | 1.31E-189 | 1 | Kansl1 |
| Pak2 | 9.82E-194 | 0.376566786 | 0.69 | 0.394 | 1.99E-189 | 1 | Pak2 |
| Zbtb7a | 3.13E-193 | 0.452259157 | 0.424 | 0.198 | 6.34E-189 | 1 | Zbtb7a |
| Trappc9 | 5.66E-193 | 0.467635577 | 0.379 | 0.165 | 1.15E-188 | 1 | Trappc9 |
| Csde1 | 9.91E-193 | 0.315373073 | 0.597 | 0.322 | 2.01E-188 | 1 | Csde1 |
| Jmjd1c | 1.61E-192 | 0.502788613 | 0.526 | 0.273 | 3.26E-188 | 1 | Jmjd1c |
| Large1 | 2.24E-192 | 0.498275228 | 0.322 | 0.125 | 4.53E-188 | 1 | Large1 |
| Stk3 | 5.04E-192 | 0.499772338 | 0.338 | 0.139 | 1.02E-187 | 1 | Stk3 |
| Zfr | 9.71E-192 | 0.489976744 | 0.447 | 0.216 | 1.97E-187 | 1 | Zfr |
| Ctcf | 3.06E-191 | 0.413842346 | 0.487 | 0.244 | 6.20E-187 | 1 | Ctcf |
| Ifi47 | 3.45E-191 | 0.500823274 | 0.356 | 0.151 | 6.98E-187 | 1 | Ifi47 |
| Tmed9 | 3.96E-191 | 0.395820787 | 0.647 | 0.365 | 8.03E-187 | 1 | Tmed9 |
| Prdm2 | 5.47E-191 | 0.46618896 | 0.364 | 0.157 | 1.11E-186 | 1 | Prdm2 |
| Ddrgk1 | 7.69E-191 | 0.362409001 | 0.541 | 0.283 | 1.56E-186 | 1 | Ddrgk1 |
| Pdia3 | 7.71E-191 | 0.395856095 | 0.717 | 0.4 | 1.56E-186 | 1 | Pdia3 |
| Eif4g3 | 8.07E-191 | 0.462006062 | 0.5 | 0.252 | 1.63E-186 | 1 | Eif4g3 |
| Rb1cc1 | 1.41E-190 | 0.497137865 | 0.431 | 0.204 | 2.86E-186 | 1 | Rb1cc1 |
| Thra | 2.47E-190 | 0.315797183 | 0.347 | 0.135 | 4.99E-186 | 1 | Thra |
| Tmem205 | 3.83E-190 | 0.363591971 | 0.334 | 0.131 | 7.76E-186 | 1 | Tmem205 |
| Braf | 1.45E-189 | 0.488128936 | 0.328 | 0.132 | 2.95E-185 | 1 | Braf |

| | | | | | | | |
|---|---|---|---|---|---|---|---|
| Sik3 | 2.63E-189 | 0.512587647 | 0.479 | 0.237 | 5.33E-185 | 1 | Sik3 |
| Slc10a7 | 3.71E-189 | 0.487015679 | 0.311 | 0.122 | 7.50E-185 | 1 | Slc10a7 |
| Tra2a | 5.24E-189 | 0.517080207 | 0.478 | 0.243 | 1.06E-184 | 1 | Tra2a |
| Dnaja1 | 7.68E-189 | 0.39408809 | 0.779 | 0.475 | 1.56E-184 | 1 | Dnaja1 |
| Txlna | 7.83E-189 | 0.381591942 | 0.291 | 0.108 | 1.59E-184 | 1 | Txlna |
| Tax1bp3 | 1.46E-188 | 0.25645119 | 0.363 | 0.145 | 2.95E-184 | 1 | Tax1bp3 |
| Gpatch8 | 1.51E-188 | 0.493761375 | 0.458 | 0.227 | 3.06E-184 | 1 | Gpatch8 |
| Atrn | 2.12E-188 | 0.45927745 | 0.275 | 0.099 | 4.28E-184 | 1 | Atrn |
| Eea1 | 2.64E-188 | 0.446489642 | 0.41 | 0.188 | 5.34E-184 | 1 | Eea1 |
| Slc12a2 | 3.59E-188 | 0.312570154 | 0.269 | 0.091 | 7.27E-184 | 1 | Slc12a2 |
| Arid4a | 3.91E-188 | 0.512430501 | 0.448 | 0.221 | 7.92E-184 | 1 | Arid4a |
| Crip1 | 5.35E-188 | 0.312162889 | 0.822 | 0.49 | 1.08E-183 | 1 | Crip1 |
| Sfpq | 1.37E-187 | 0.441396008 | 0.513 | 0.264 | 2.77E-183 | 1 | Sfpq |
| Xist | 2.57E-187 | 0.340836209 | 0.788 | 0.411 | 5.21E-183 | 1 | Xist |
| Mier1 | 4.36E-187 | 0.358954448 | 0.555 | 0.296 | 8.82E-183 | 1 | Mier1 |
| Ablim1 | 8.08E-187 | 0.389674582 | 0.675 | 0.359 | 1.64E-182 | 1 | Ablim1 |
| AU020206 | 2.65E-186 | 0.481862851 | 0.516 | 0.255 | 5.36E-182 | 1 | AU020206 |
| Mkln1 | 3.07E-186 | 0.439494517 | 0.469 | 0.232 | 6.22E-182 | 1 | Mkln1 |
| Myo9a | 3.71E-186 | 0.485092336 | 0.346 | 0.146 | 7.50E-182 | 1 | Myo9a |
| Zfand51 | 1.30E-185 | 0.356861554 | 0.707 | 0.414 | 2.63E-181 | 1 | Zfand5 |
| Sh3pxd2a | 2.14E-185 | 0.382335619 | 0.307 | 0.116 | 4.34E-181 | 1 | Sh3pxd2a |
| Kat6b | 4.81E-185 | 0.450330494 | 0.316 | 0.126 | 9.74E-181 | 1 | Kat6b |
| Glud1 | 7.34E-185 | 0.376796849 | 0.559 | 0.301 | 1.49E-180 | 1 | Glud1 |
| Cdkn1c | 8.49E-185 | 0.381444504 | 0.286 | 0.101 | 1.72E-180 | 1 | Cdkn1c |
| Purb | 2.74E-184 | 0.315107142 | 0.604 | 0.331 | 5.55E-180 | 1 | Purb |
| Srsf11 | 1.01E-183 | 0.424919009 | 0.579 | 0.315 | 2.04E-179 | 1 | Srsf11 |
| Hipk3 | 2.52E-183 | 0.380755306 | 0.314 | 0.125 | 5.10E-179 | 1 | Hipk3 |
| Gatad2a | 2.94E-183 | 0.41972669 | 0.413 | 0.192 | 5.95E-179 | 1 | Gatad2a |
| Med13l | 3.02E-183 | 0.468090717 | 0.371 | 0.166 | 6.11E-179 | 1 | Med13l |
| Cdipt | 4.17E-183 | 0.398692649 | 0.292 | 0.11 | 8.43E-179 | 1 | Cdipt |
| Lmbrd1 | 2.21E-182 | 0.438991406 | 0.33 | 0.136 | 4.48E-178 | 1 | Lmbrd1 |
| Kbtbd2 | 5.49E-182 | 0.368119549 | 0.251 | 0.086 | 1.11E-177 | 1 | Kbtbd2 |
| Dnajc10 | 8.14E-182 | 0.3942255 | 0.298 | 0.114 | 1.65E-177 | 1 | Dnajc10 |
| Pdia6 | 1.32E-181 | 0.365922018 | 0.57 | 0.3 | 2.67E-177 | 1 | Pdia6 |
| Rabac1 | 2.08E-181 | 0.329748148 | 0.833 | 0.521 | 4.22E-177 | 1 | Rabac1 |
| P4hb | 2.55E-181 | 0.425413308 | 0.45 | 0.221 | 5.16E-177 | 1 | P4hb |
| Mbd5 | 4.00E-181 | 0.51970631 | 0.405 | 0.191 | 8.09E-177 | 1 | Mbd5 |
| Smad1 | 4.51E-181 | 0.383510245 | 0.288 | 0.108 | 9.13E-177 | 1 | Smad1 |
| Dpp8 | 5.60E-181 | 0.389211194 | 0.362 | 0.156 | 1.13E-176 | 1 | Dpp8 |
| Gnb2 | 5.83E-181 | 0.31984148 | 0.789 | 0.49 | 1.18E-176 | 1 | Gnb2 |
| Bbx | 5.88E-181 | 0.450505494 | 0.411 | 0.193 | 1.19E-176 | 1 | Bbx |
| Ankrd11 | 1.10E-180 | 0.41301489 | 0.627 | 0.342 | 2.22E-176 | 1 | Ankrd11 |
| Golim4 | 2.32E-180 | 0.472337124 | 0.355 | 0.153 | 4.69E-176 | 1 | Golim4 |

| | | | | | | | |
|---|---|---|---|---|---|---|---|
| mt-Nd5 | 3.23E-180 | 0.369936151 | 0.797 | 0.448 | 6.54E-176 | 1 | mt-Nd5 |
| Akap11 | 4.57E-180 | 0.433605036 | 0.282 | 0.106 | 9.24E-176 | 1 | Akap11 |
| Rab11b | 7.91E-180 | 0.354018922 | 0.468 | 0.233 | 1.60E-175 | 1 | Rab11b |
| Arid5b | 2.65E-179 | 0.472924666 | 0.486 | 0.245 | 5.36E-175 | 1 | Arid5b |
| Erap1 | 3.85E-179 | 0.454417013 | 0.294 | 0.115 | 7.80E-175 | 1 | Erap1 |
| Arid4b | 3.91E-179 | 0.477490445 | 0.498 | 0.26 | 7.92E-175 | 1 | Arid4b |
| Camk2d | 3.97E-178 | 0.457393403 | 0.475 | 0.237 | 8.05E-174 | 1 | Camk2d |
| Wnk1 | 4.84E-178 | 0.395972024 | 0.604 | 0.325 | 9.79E-174 | 1 | Wnk1 |
| Hnrnph1 | 1.04E-177 | 0.383466317 | 0.447 | 0.219 | 2.11E-173 | 1 | Hnrnph1 |
| Phip | 1.09E-177 | 0.487883582 | 0.451 | 0.225 | 2.22E-173 | 1 | Phip |
| Ndfip2 | 1.53E-177 | 0.398497742 | 0.441 | 0.214 | 3.10E-173 | 1 | Ndfip2 |
| Pum2 | 2.50E-177 | 0.42162129 | 0.467 | 0.238 | 5.06E-173 | 1 | Pum2 |
| Ttc3 | 2.60E-177 | 0.434375829 | 0.451 | 0.221 | 5.27E-173 | 1 | Ttc3 |
| Nfe2l2 | 3.60E-177 | 0.3536935 | 0.52 | 0.266 | 7.28E-173 | 1 | Nfe2l2 |
| Brd4 | 3.62E-177 | 0.433635546 | 0.468 | 0.236 | 7.34E-173 | 1 | Brd4 |
| Cuedc1 | 1.41E-176 | 0.375213594 | 0.292 | 0.112 | 2.86E-172 | 1 | Cuedc1 |
| Myl12b | 2.68E-176 | 0.293206276 | 0.817 | 0.504 | 5.42E-172 | 1 | Myl12b |
| Ccnl2 | 2.77E-175 | 0.42505004 | 0.396 | 0.186 | 5.60E-171 | 1 | Ccnl2 |
| Vcp | 8.68E-175 | 0.32777473 | 0.564 | 0.308 | 1.76E-170 | 1 | Vcp |
| Dnajc8 | 1.21E-174 | 0.306311292 | 0.584 | 0.323 | 2.45E-170 | 1 | Dnajc8 |
| Zbtb4 | 2.28E-173 | 0.368506369 | 0.268 | 0.098 | 4.61E-169 | 1 | Zbtb4 |
| Agtrap | 2.81E-173 | 0.391095817 | 0.276 | 0.104 | 5.69E-169 | 1 | Agtrap |
| Mycbp2 | 7.36E-173 | 0.449907896 | 0.613 | 0.337 | 1.49E-168 | 1 | Mycbp2 |
| Epb41 | 8.60E-173 | 0.440459659 | 0.4 | 0.187 | 1.74E-168 | 1 | Epb41 |
| Ppp2r5a | 2.05E-172 | 0.387819324 | 0.445 | 0.22 | 4.15E-168 | 1 | Ppp2r5a |
| Abhd17b | 2.10E-172 | 0.441372495 | 0.367 | 0.165 | 4.24E-168 | 1 | Abhd17b |
| Mgst1 | 2.23E-172 | 0.331004836 | 0.348 | 0.139 | 4.51E-168 | 1 | Mgst1 |
| 4932438A13Rik | 3.79E-172 | 0.4863051 | 0.398 | 0.191 | 7.68E-168 | 1 | 4932438A13Rik |
| Acin1 | 6.41E-172 | 0.396189629 | 0.528 | 0.283 | 1.30E-167 | 1 | Acin1 |
| Mga | 2.46E-171 | 0.440170713 | 0.337 | 0.145 | 4.99E-167 | 1 | Mga |
| H1f0 | 5.48E-171 | 0.271306442 | 0.382 | 0.164 | 1.11E-166 | 1 | H1f0 |
| Tbc1d2b | 6.76E-171 | 0.416404831 | 0.264 | 0.099 | 1.37E-166 | 1 | Tbc1d2b |
| Kras | 1.20E-170 | 0.326354343 | 0.472 | 0.237 | 2.44E-166 | 1 | Kras |
| Caml | 1.72E-170 | 0.355795751 | 0.285 | 0.109 | 3.47E-166 | 1 | Caml |
| Zfp148 | 2.19E-170 | 0.404829677 | 0.46 | 0.234 | 4.44E-166 | 1 | Zfp148 |
| AW554918 | 3.81E-170 | 0.487292495 | 0.334 | 0.145 | 7.72E-166 | 1 | AW554918 |
| Sp1 | 1.45E-169 | 0.385311566 | 0.296 | 0.118 | 2.94E-165 | 1 | Sp1 |
| Pfkp | 1.59E-169 | 0.317175022 | 0.32 | 0.131 | 3.22E-165 | 1 | Pfkp |
| Plekha1 | 2.66E-169 | 0.373629575 | 0.293 | 0.114 | 5.39E-165 | 1 | Plekha1 |
| Gch1 | 3.59E-169 | 0.403079849 | 0.28 | 0.107 | 7.26E-165 | 1 | Gch1 |
| Zfp638 | 6.51E-169 | 0.455952907 | 0.479 | 0.251 | 1.32E-164 | 1 | Zfp638 |
| Wasl | 1.74E-168 | 0.330431264 | 0.428 | 0.207 | 3.53E-164 | 1 | Wasl |
| Dusp6 | 2.67E-168 | 0.424927503 | 0.29 | 0.115 | 5.40E-164 | 1 | Dusp6 |

| | | | | | | | |
|---|---|---|---|---|---|---|---|
| Gcc2 | 3.06E-168 | 0.466102093 | 0.361 | 0.165 | 6.19E-164 | 1 | Gcc2 |
| Pum1 | 3.09E-168 | 0.435529751 | 0.447 | 0.229 | 6.26E-164 | 1 | Pum1 |
| Ppp4r3b | 5.75E-168 | 0.423253119 | 0.415 | 0.203 | 1.16E-163 | 1 | Ppp4r3b |
| Dph6 | 6.68E-168 | 0.382869436 | 0.28 | 0.108 | 1.35E-163 | 1 | Dph6 |
| Ube2r2 | 1.46E-167 | 0.286881863 | 0.495 | 0.255 | 2.95E-163 | 1 | Ube2r2 |
| Txndc12 | 1.01E-166 | 0.377464245 | 0.255 | 0.093 | 2.05E-162 | 1 | Txndc12 |
| Cep83 | 1.06E-166 | 0.410177057 | 0.252 | 0.092 | 2.14E-162 | 1 | Cep83 |
| Ubxn4 | 2.03E-166 | 0.418359428 | 0.464 | 0.242 | 4.10E-162 | 1 | Ubxn4 |
| Rps6ka3 | 2.86E-166 | 0.419342133 | 0.458 | 0.232 | 5.79E-162 | 1 | Rps6ka3 |
| Zscan26 | 8.11E-166 | 0.40594893 | 0.298 | 0.121 | 1.64E-161 | 1 | Zscan26 |
| Lrrc8d | 9.91E-166 | 0.436777964 | 0.279 | 0.11 | 2.01E-161 | 1 | Lrrc8d |
| Pdzd8 | 1.58E-165 | 0.437992638 | 0.284 | 0.113 | 3.20E-161 | 1 | Pdzd8 |
| Tmem165 | 1.91E-165 | 0.381588138 | 0.348 | 0.153 | 3.87E-161 | 1 | Tmem165 |
| Add3 | 2.00E-165 | 0.359951668 | 0.41 | 0.193 | 4.06E-161 | 1 | Add3 |
| Rb1 | 2.13E-165 | 0.479177735 | 0.392 | 0.186 | 4.32E-161 | 1 | Rb1 |
| Rbm25 | 3.83E-165 | 0.442013503 | 0.544 | 0.301 | 7.76E-161 | 1 | Rbm25 |
| Prpf4b | 5.24E-165 | 0.347675449 | 0.546 | 0.3 | 1.06E-160 | 1 | Prpf4b |
| Slc35b1 | 6.27E-165 | 0.360370106 | 0.326 | 0.138 | 1.27E-160 | 1 | Slc35b1 |
| Dazap2 | 1.34E-164 | 0.307917 | 0.506 | 0.268 | 2.71E-160 | 1 | Dazap2 |
| Smg6 | 1.42E-164 | 0.449619901 | 0.453 | 0.23 | 2.87E-160 | 1 | Smg6 |
| Capns1 | 1.62E-164 | 0.266309966 | 0.57 | 0.313 | 3.28E-160 | 1 | Capns1 |
| Isg15 | 4.05E-164 | 0.520679555 | 0.421 | 0.199 | 8.21E-160 | 1 | Isg15 |
| Phactr2 | 5.14E-164 | 0.369531499 | 0.442 | 0.215 | 1.04E-159 | 1 | Phactr2 |
| Tprgl | 7.89E-164 | 0.302847896 | 0.482 | 0.247 | 1.60E-159 | 1 | Tprgl |
| Ppp6r3 | 2.14E-163 | 0.416430612 | 0.359 | 0.165 | 4.33E-159 | 1 | Ppp6r3 |
| Rab10 | 3.70E-163 | 0.325462439 | 0.572 | 0.315 | 7.48E-159 | 1 | Rab10 |
| Myo1e | 3.68E-162 | 0.414908765 | 0.488 | 0.249 | 7.45E-158 | 1 | Myo1e |
| Fgfr2 | 1.02E-161 | 0.405798775 | 0.306 | 0.129 | 2.07E-157 | 1 | Fgfr2 |
| Cnot4 | 1.96E-161 | 0.441791177 | 0.458 | 0.239 | 3.96E-157 | 1 | Cnot4 |
| Zcchc7 | 7.58E-161 | 0.492963812 | 0.501 | 0.27 | 1.54E-156 | 1 | Zcchc7 |
| Psme2 | 1.53E-160 | 0.279403662 | 0.687 | 0.405 | 3.10E-156 | 1 | Psme2 |
| St5 | 2.00E-160 | 0.306354018 | 0.313 | 0.126 | 4.04E-156 | 1 | St5 |
| Ppig | 5.32E-160 | 0.387138764 | 0.517 | 0.284 | 1.08E-155 | 1 | Ppig |
| Dnajc1 | 6.32E-160 | 0.434929732 | 0.538 | 0.289 | 1.28E-155 | 1 | Dnajc1 |
| Pgrmc2 | 8.48E-160 | 0.329184353 | 0.294 | 0.119 | 1.72E-155 | 1 | Pgrmc2 |
| Gpd2 | 1.05E-159 | 0.383154682 | 0.255 | 0.096 | 2.13E-155 | 1 | Gpd2 |
| 2-Mar | 1.36E-159 | 0.27882795 | 0.618 | 0.358 | 2.75E-155 | 1 | 2-Mar |
| Akap13 | 2.89E-159 | 0.356481586 | 0.637 | 0.355 | 5.84E-155 | 1 | Akap13 |
| Pnisr | 3.19E-159 | 0.440080883 | 0.453 | 0.235 | 6.47E-155 | 1 | Pnisr |
| Chd1 | 3.22E-159 | 0.397026952 | 0.371 | 0.175 | 6.52E-155 | 1 | Chd1 |
| Setx | 4.53E-159 | 0.436701393 | 0.334 | 0.15 | 9.16E-155 | 1 | Setx |
| Mark3 | 1.03E-158 | 0.398639283 | 0.385 | 0.185 | 2.08E-154 | 1 | Mark3 |
| Fto | 1.50E-158 | 0.439674617 | 0.357 | 0.165 | 3.03E-154 | 1 | Fto |

| | | | | | | | |
|---|---|---|---|---|---|---|---|
| Arl6ip1 | 1.81E-158 | 0.383976888 | 0.591 | 0.335 | 3.66E-154 | 1 | Arl6ip1 |
| Rab1a | 1.90E-158 | 0.268395513 | 0.472 | 0.242 | 3.84E-154 | 1 | Rab1a |
| Dcaf6 | 1.90E-158 | 0.411771239 | 0.279 | 0.111 | 3.85E-154 | 1 | Dcaf6 |
| Inpp1 | 4.38E-158 | 0.325509194 | 0.254 | 0.094 | 8.87E-154 | 1 | Inpp1 |
| Hpcal1 | 9.55E-158 | 0.258371045 | 0.515 | 0.27 | 1.93E-153 | 1 | Hpcal1 |
| Luc7l2 | 1.03E-157 | 0.352811645 | 0.643 | 0.367 | 2.09E-153 | 1 | Luc7l2 |
| Evi5 | 1.91E-157 | 0.321270307 | 0.291 | 0.118 | 3.86E-153 | 1 | Evi5 |
| Cttnbp2nl | 2.21E-157 | 0.30934114 | 0.255 | 0.093 | 4.47E-153 | 1 | Cttnbp2nl |
| Yy1 | 3.53E-157 | 0.339115729 | 0.463 | 0.243 | 7.14E-153 | 1 | Yy1 |
| Ppp1r10 | 5.79E-157 | 0.429373672 | 0.353 | 0.165 | 1.17E-152 | 1 | Ppp1r10 |
| Dynll1 | 6.22E-157 | 0.265849057 | 0.863 | 0.564 | 1.26E-152 | 1 | Dynll1 |
| Ccdc186 | 8.00E-157 | 0.345280971 | 0.312 | 0.132 | 1.62E-152 | 1 | Ccdc186 |
| Cnbp | 1.13E-156 | 0.273143167 | 0.729 | 0.448 | 2.28E-152 | 1 | Cnbp |
| Cpeb4 | 1.37E-156 | 0.378509431 | 0.261 | 0.101 | 2.77E-152 | 1 | Cpeb4 |
| Anks1 | 5.30E-156 | 0.398843702 | 0.284 | 0.116 | 1.07E-151 | 1 | Anks1 |
| Hif1a | 9.24E-156 | 0.291889303 | 0.373 | 0.173 | 1.87E-151 | 1 | Hif1a |
| Ap1s2 | 1.07E-155 | 0.366384231 | 0.286 | 0.116 | 2.17E-151 | 1 | Ap1s2 |
| Lrp10 | 1.50E-155 | 0.419626134 | 0.35 | 0.163 | 3.04E-151 | 1 | Lrp10 |
| Pnp | 5.22E-155 | 0.338696949 | 0.444 | 0.224 | 1.06E-150 | 1 | Pnp |
| Trpm7 | 1.22E-154 | 0.446330708 | 0.37 | 0.178 | 2.47E-150 | 1 | Trpm7 |
| Ehmt1 | 2.57E-154 | 0.395344415 | 0.346 | 0.16 | 5.21E-150 | 1 | Ehmt1 |
| Rbbp6 | 4.26E-154 | 0.372947827 | 0.466 | 0.247 | 8.63E-150 | 1 | Rbbp6 |
| Btbd9 | 5.25E-154 | 0.504450879 | 0.413 | 0.21 | 1.06E-149 | 1 | Btbd9 |
| Bclaf1 | 5.30E-154 | 0.428593746 | 0.434 | 0.226 | 1.07E-149 | 1 | Bclaf1 |
| Ubr5 | 5.56E-154 | 0.423732814 | 0.364 | 0.172 | 1.13E-149 | 1 | Ubr5 |
| Ep300 | 1.08E-153 | 0.40249069 | 0.371 | 0.178 | 2.20E-149 | 1 | Ep300 |
| Rere | 1.10E-153 | 0.412016829 | 0.459 | 0.242 | 2.23E-149 | 1 | Rere |
| Matr3 | 1.36E-153 | 0.316849747 | 0.457 | 0.237 | 2.75E-149 | 1 | Matr3 |
| Tspan14 | 1.40E-153 | 0.393995626 | 0.268 | 0.107 | 2.83E-149 | 1 | Tspan14 |
| Arpc3 | 3.00E-153 | 0.308805423 | 0.902 | 0.619 | 6.08E-149 | 1 | Arpc3 |
| Auts2 | 5.05E-153 | 0.300266418 | 0.315 | 0.127 | 1.02E-148 | 1 | Auts2 |
| Rbm6 | 6.49E-153 | 0.431252134 | 0.388 | 0.191 | 1.31E-148 | 1 | Rbm6 |
| Sf3b2 | 9.98E-153 | 0.315932623 | 0.572 | 0.325 | 2.02E-148 | 1 | Sf3b2 |
| Srsf4 | 3.73E-152 | 0.345330274 | 0.313 | 0.136 | 7.56E-148 | 1 | Srsf4 |
| Man1a2 | 9.26E-152 | 0.429238787 | 0.398 | 0.2 | 1.87E-147 | 1 | Man1a2 |
| Il10rb | 1.34E-151 | 0.356936789 | 0.432 | 0.22 | 2.71E-147 | 1 | Il10rb |
| Sipa1l1 | 1.66E-151 | 0.434820874 | 0.354 | 0.164 | 3.35E-147 | 1 | Sipa1l1 |
| Golgb1 | 3.59E-151 | 0.386344918 | 0.445 | 0.234 | 7.28E-147 | 1 | Golgb1 |
| Prkd3 | 3.65E-151 | 0.421868496 | 0.321 | 0.144 | 7.39E-147 | 1 | Prkd3 |
| Keap1 | 4.54E-151 | 0.345315678 | 0.283 | 0.116 | 9.18E-147 | 1 | Keap1 |
| Tmem234 | 8.53E-151 | 0.265243667 | 0.612 | 0.358 | 1.73E-146 | 1 | Tmem234 |
| Hacd4 | 1.44E-150 | 0.371471819 | 0.251 | 0.094 | 2.91E-146 | 1 | Hacd4 |
| Rad21 | 1.77E-150 | 0.330782911 | 0.355 | 0.165 | 3.59E-146 | 1 | Rad21 |

| | | | | | | | |
|---|---|---|---|---|---|---|---|
| Itpripl2 | 3.29E-150 | 0.359248368 | 0.253 | 0.098 | 6.67E-146 | 1 | Itpripl2 |
| Fam3c | 8.55E-150 | 0.354840558 | 0.338 | 0.154 | 1.73E-145 | 1 | Fam3c |
| Ccnt1 | 1.30E-149 | 0.443293863 | 0.318 | 0.144 | 2.63E-145 | 1 | Ccnt1 |
| Zfp292 | 1.32E-149 | 0.41249113 | 0.43 | 0.223 | 2.67E-145 | 1 | Zfp292 |
| Ubr4 | 2.02E-149 | 0.402073643 | 0.288 | 0.123 | 4.10E-145 | 1 | Ubr4 |
| Gm20388 | 2.41E-149 | 0.349652809 | 0.642 | 0.363 | 4.88E-145 | 1 | Gm20388 |
| Jak1 | 5.11E-149 | 0.318174174 | 0.595 | 0.329 | 1.03E-144 | 1 | Jak1 |
| Gbp8 | 5.88E-149 | 0.408250855 | 0.28 | 0.116 | 1.19E-144 | 1 | Gbp8 |
| Pcmtd1 | 7.58E-149 | 0.275094921 | 0.532 | 0.295 | 1.54E-144 | 1 | Pcmtd1 |
| Mgat2 | 9.93E-149 | 0.378344291 | 0.304 | 0.132 | 2.01E-144 | 1 | Mgat2 |
| Epb41l2 | 1.71E-148 | 0.305489254 | 0.433 | 0.211 | 3.47E-144 | 1 | Epb41l2 |
| Psmd7 | 3.31E-148 | 0.274603244 | 0.504 | 0.274 | 6.71E-144 | 1 | Psmd7 |
| Slc38a2 | 3.75E-148 | 0.379116401 | 0.516 | 0.279 | 7.59E-144 | 1 | Slc38a2 |
| Tgfb1 | 6.08E-148 | 0.376987673 | 0.414 | 0.208 | 1.23E-143 | 1 | Tgfb1 |
| Secisbp2l | 6.80E-148 | 0.333801418 | 0.288 | 0.121 | 1.38E-143 | 1 | Secisbp2l |
| Ripor2 | 7.28E-148 | 0.428623059 | 0.331 | 0.148 | 1.47E-143 | 1 | Ripor2 |
| Cmtm6 | 5.31E-147 | 0.335565023 | 0.251 | 0.097 | 1.08E-142 | 1 | Cmtm6 |
| Dusp23 | 5.83E-147 | 0.272839774 | 0.252 | 0.095 | 1.18E-142 | 1 | Dusp23 |
| Arl5a | 1.13E-146 | 0.310001291 | 0.31 | 0.135 | 2.28E-142 | 1 | Arl5a |
| Sdf4 | 1.80E-146 | 0.344751486 | 0.502 | 0.279 | 3.65E-142 | 1 | Sdf4 |
| Ogfrl1 | 2.03E-146 | 0.306470928 | 0.276 | 0.112 | 4.11E-142 | 1 | Ogfrl1 |
| Pds5a | 3.13E-146 | 0.411995253 | 0.366 | 0.179 | 6.33E-142 | 1 | Pds5a |
| Tusc3 | 6.87E-146 | 0.393595169 | 0.293 | 0.126 | 1.39E-141 | 1 | Tusc3 |
| Ppp2r5c | 8.25E-146 | 0.352349995 | 0.425 | 0.22 | 1.67E-141 | 1 | Ppp2r5c |
| Tlk2 | 9.85E-146 | 0.383149277 | 0.308 | 0.137 | 1.99E-141 | 1 | Tlk2 |
| Bdp1 | 1.17E-145 | 0.432903176 | 0.314 | 0.143 | 2.37E-141 | 1 | Bdp1 |
| Trio | 1.44E-145 | 0.446722947 | 0.344 | 0.164 | 2.91E-141 | 1 | Trio |
| Zc3h13 | 1.57E-145 | 0.412828458 | 0.382 | 0.19 | 3.18E-141 | 1 | Zc3h13 |
| Ice1 | 1.81E-145 | 0.339025374 | 0.274 | 0.113 | 3.67E-141 | 1 | Ice1 |
| Map3k5 | 1.89E-145 | 0.417551277 | 0.354 | 0.167 | 3.82E-141 | 1 | Map3k5 |
| Map3k1 | 2.42E-145 | 0.350086634 | 0.361 | 0.174 | 4.90E-141 | 1 | Map3k1 |
| Ranbp2 | 2.86E-145 | 0.412642221 | 0.358 | 0.174 | 5.79E-141 | 1 | Ranbp2 |
| Agfg1 | 3.05E-145 | 0.341174195 | 0.314 | 0.14 | 6.17E-141 | 1 | Agfg1 |
| Brd9 | 8.27E-145 | 0.338290861 | 0.327 | 0.149 | 1.67E-140 | 1 | Brd9 |
| Ccdc82 | 2.88E-144 | 0.379994767 | 0.337 | 0.156 | 5.84E-140 | 1 | Ccdc82 |
| Rnf216 | 3.58E-144 | 0.393094967 | 0.335 | 0.156 | 7.25E-140 | 1 | Rnf216 |
| Tpp1 | 5.19E-144 | 0.367047492 | 0.25 | 0.099 | 1.05E-139 | 1 | Tpp1 |
| Yipf4 | 7.27E-144 | 0.308872437 | 0.394 | 0.195 | 1.47E-139 | 1 | Yipf4 |
| Saraf | 3.34E-143 | 0.365674075 | 0.501 | 0.277 | 6.76E-139 | 1 | Saraf |
| Ppp2r2a | 8.89E-143 | 0.366638654 | 0.378 | 0.187 | 1.80E-138 | 1 | Ppp2r2a |
| Fbxl3 | 1.02E-142 | 0.355119269 | 0.303 | 0.133 | 2.07E-138 | 1 | Fbxl3 |
| mt-Co1 | 2.96E-142 | 0.388753809 | 0.999 | 0.674 | 5.99E-138 | 1 | mt-Co1 |
| Rmdn1 | 3.55E-142 | 0.335584698 | 0.345 | 0.161 | 7.18E-138 | 1 | Rmdn1 |

| | | | | | | | |
|---|---|---|---|---|---|---|---|
| Emc3 | 4.08E-142 | 0.354623297 | 0.324 | 0.149 | 8.27E-138 | 1 | Emc3 |
| Msl1 | 2.58E-141 | 0.334211919 | 0.268 | 0.111 | 5.22E-137 | 1 | Msl1 |
| Unc50 | 3.96E-141 | 0.285108733 | 0.379 | 0.187 | 8.01E-137 | 1 | Unc50 |
| Rasa1 | 6.36E-141 | 0.377928593 | 0.423 | 0.22 | 1.29E-136 | 1 | Rasa1 |
| Zfp644 | 8.51E-141 | 0.360000689 | 0.432 | 0.228 | 1.72E-136 | 1 | Zfp644 |
| Kdm2a | 9.07E-141 | 0.403812292 | 0.409 | 0.214 | 1.84E-136 | 1 | Kdm2a |
| Fbxo11 | 1.77E-140 | 0.407570489 | 0.402 | 0.207 | 3.58E-136 | 1 | Fbxo11 |
| G3bp1 | 4.42E-140 | 0.319960305 | 0.493 | 0.272 | 8.95E-136 | 1 | G3bp1 |
| Rsf1 | 6.04E-140 | 0.404078842 | 0.389 | 0.199 | 1.22E-135 | 1 | Rsf1 |
| Vps13c | 7.32E-140 | 0.37971864 | 0.277 | 0.118 | 1.48E-135 | 1 | Vps13c |
| Sgk3 | 8.44E-140 | 0.325284762 | 0.264 | 0.107 | 1.71E-135 | 1 | Sgk3 |
| mt-Atp8 | 1.23E-139 | 0.307720082 | 0.711 | 0.424 | 2.49E-135 | 1 | mt-Atp8 |
| Ankrd17 | 1.34E-139 | 0.397181125 | 0.437 | 0.234 | 2.70E-135 | 1 | Ankrd17 |
| Mosmo | 2.78E-139 | 0.326349349 | 0.274 | 0.115 | 5.62E-135 | 1 | Mosmo |
| Bhlhe40 | 3.42E-139 | 0.346344743 | 0.307 | 0.135 | 6.92E-135 | 1 | Bhlhe40 |
| Cep350 | 6.67E-139 | 0.427610401 | 0.361 | 0.179 | 1.35E-134 | 1 | Cep350 |
| Wapl | 7.49E-139 | 0.345952094 | 0.423 | 0.224 | 1.52E-134 | 1 | Wapl |
| Psme1 | 1.75E-138 | 0.273723246 | 0.835 | 0.526 | 3.53E-134 | 1 | Psme1 |
| Yme1l1 | 2.47E-138 | 0.38842235 | 0.351 | 0.172 | 5.01E-134 | 1 | Yme1l1 |
| Dctn4 | 6.70E-138 | 0.292164658 | 0.377 | 0.186 | 1.36E-133 | 1 | Dctn4 |
| Hnrnph3 | 8.32E-138 | 0.284434865 | 0.426 | 0.221 | 1.68E-133 | 1 | Hnrnph3 |
| Gimap5 | 9.24E-137 | 0.361741751 | 0.323 | 0.147 | 1.87E-132 | 1 | Gimap5 |
| Gapvd1 | 2.65E-136 | 0.36933927 | 0.346 | 0.17 | 5.37E-132 | 1 | Gapvd1 |
| Aff4 | 4.73E-136 | 0.313689294 | 0.412 | 0.214 | 9.57E-132 | 1 | Aff4 |
| Tnpo1 | 5.96E-136 | 0.382892386 | 0.321 | 0.151 | 1.21E-131 | 1 | Tnpo1 |
| Arl6ip5 | 6.42E-136 | 0.392574234 | 0.385 | 0.196 | 1.30E-131 | 1 | Arl6ip5 |
| Tnfsf12 | 7.64E-136 | 0.307478186 | 0.29 | 0.126 | 1.55E-131 | 1 | Tnfsf12 |
| Xrn1 | 8.70E-136 | 0.427732063 | 0.358 | 0.18 | 1.76E-131 | 1 | Xrn1 |
| Snrnp200 | 1.00E-135 | 0.294409681 | 0.295 | 0.13 | 2.03E-131 | 1 | Snrnp200 |
| Zfp652 | 1.02E-135 | 0.364518028 | 0.42 | 0.22 | 2.06E-131 | 1 | Zfp652 |
| Cpne3 | 1.08E-135 | 0.276019182 | 0.377 | 0.186 | 2.19E-131 | 1 | Cpne3 |
| Clk1 | 1.10E-135 | 0.37576158 | 0.477 | 0.263 | 2.23E-131 | 1 | Clk1 |
| Zfp326 | 1.50E-135 | 0.27256344 | 0.29 | 0.125 | 3.03E-131 | 1 | Zfp326 |
| Ythdc1 | 1.57E-135 | 0.336693612 | 0.401 | 0.209 | 3.18E-131 | 1 | Ythdc1 |
| Setd7 | 2.27E-135 | 0.334627912 | 0.297 | 0.132 | 4.59E-131 | 1 | Setd7 |
| Arhgef1 | 2.35E-135 | 0.360678457 | 0.423 | 0.221 | 4.75E-131 | 1 | Arhgef1 |
| Smc3 | 4.65E-135 | 0.334146446 | 0.426 | 0.226 | 9.42E-131 | 1 | Smc3 |
| Sel1l | 4.82E-135 | 0.389718699 | 0.304 | 0.14 | 9.76E-131 | 1 | Sel1l |
| Ccdc88a | 9.29E-135 | 0.449229951 | 0.363 | 0.18 | 1.88E-130 | 1 | Ccdc88a |
| Lrpap1 | 1.21E-134 | 0.346095234 | 0.352 | 0.173 | 2.44E-130 | 1 | Lrpap1 |
| Crebrf | 2.45E-134 | 0.357706169 | 0.343 | 0.167 | 4.96E-130 | 1 | Crebrf |
| Tmed7 | 2.59E-134 | 0.353195872 | 0.288 | 0.128 | 5.24E-130 | 1 | Tmed7 |
| Sppl3 | 4.85E-134 | 0.409540754 | 0.379 | 0.193 | 9.81E-130 | 1 | Sppl3 |

| | | | | | | | |
|---|---|---|---|---|---|---|---|
| Tap1 | 5.67E-134 | 0.393531151 | 0.27 | 0.117 | 1.15E-129 | 1 | Tap1 |
| Kpna4 | 7.41E-134 | 0.362273534 | 0.518 | 0.293 | 1.50E-129 | 1 | Kpna4 |
| Btg2 | 9.44E-134 | 0.280847681 | 0.568 | 0.309 | 1.91E-129 | 1 | Btg2 |
| Sertad2 | 1.00E-133 | 0.33774012 | 0.317 | 0.146 | 2.03E-129 | 1 | Sertad2 |
| Mindy2 | 2.01E-133 | 0.309965425 | 0.332 | 0.157 | 4.07E-129 | 1 | Mindy2 |
| N4bp2l2 | 2.32E-133 | 0.338901083 | 0.417 | 0.221 | 4.69E-129 | 1 | N4bp2l2 |
| Tubb2a | 3.33E-133 | 0.269031849 | 0.317 | 0.141 | 6.74E-129 | 1 | Tubb2a |
| Trip11 | 4.20E-133 | 0.382642445 | 0.384 | 0.199 | 8.50E-129 | 1 | Trip11 |
| Brd1 | 4.74E-133 | 0.350266191 | 0.407 | 0.215 | 9.59E-129 | 1 | Brd1 |
| Nufip2 | 9.53E-133 | 0.36053776 | 0.396 | 0.207 | 1.93E-128 | 1 | Nufip2 |
| Ube3a | 1.15E-132 | 0.318063821 | 0.479 | 0.267 | 2.33E-128 | 1 | Ube3a |
| Helz | 1.59E-132 | 0.396422068 | 0.313 | 0.147 | 3.22E-128 | 1 | Helz |
| Pnn | 1.60E-132 | 0.32758959 | 0.443 | 0.24 | 3.24E-128 | 1 | Pnn |
| Lpp | 1.62E-132 | 0.353512053 | 0.473 | 0.257 | 3.28E-128 | 1 | Lpp |
| Srsf7 | 2.77E-132 | 0.312953064 | 0.527 | 0.302 | 5.60E-128 | 1 | Srsf7 |
| Cdc14a | 3.39E-132 | 0.371082599 | 0.269 | 0.116 | 6.87E-128 | 1 | Cdc14a |
| Add1 | 3.70E-132 | 0.328564757 | 0.374 | 0.19 | 7.49E-128 | 1 | Add1 |
| Hivep1 | 5.00E-132 | 0.322822271 | 0.268 | 0.114 | 1.01E-127 | 1 | Hivep1 |
| Eif2ak2 | 5.45E-132 | 0.370619473 | 0.308 | 0.142 | 1.10E-127 | 1 | Eif2ak2 |
| Actr1a | 5.79E-132 | 0.338253674 | 0.26 | 0.109 | 1.17E-127 | 1 | Actr1a |
| Adam17 | 1.84E-131 | 0.373461841 | 0.293 | 0.132 | 3.73E-127 | 1 | Adam17 |
| U2surp | 2.69E-131 | 0.292790995 | 0.443 | 0.242 | 5.44E-127 | 1 | U2surp |
| Slmap | 3.85E-131 | 0.351440895 | 0.345 | 0.169 | 7.80E-127 | 1 | Slmap |
| Smg1 | 5.26E-131 | 0.387209372 | 0.398 | 0.209 | 1.06E-126 | 1 | Smg1 |
| Tmed2 | 5.92E-131 | 0.296731644 | 0.563 | 0.335 | 1.20E-126 | 1 | Tmed2 |
| Anapc5 | 6.71E-131 | 0.265694387 | 0.373 | 0.186 | 1.36E-126 | 1 | Anapc5 |
| Celf1 | 7.10E-131 | 0.358557665 | 0.38 | 0.195 | 1.44E-126 | 1 | Celf1 |
| G3bp2 | 2.65E-130 | 0.300944413 | 0.456 | 0.254 | 5.37E-126 | 1 | G3bp2 |
| Gnai3 | 2.13E-129 | 0.276883874 | 0.389 | 0.202 | 4.31E-125 | 1 | Gnai3 |
| Rcn2 | 2.16E-129 | 0.347384525 | 0.277 | 0.122 | 4.37E-125 | 1 | Rcn2 |
| Brd8 | 2.45E-129 | 0.323850941 | 0.306 | 0.142 | 4.96E-125 | 1 | Brd8 |
| Tsc22d2 | 3.41E-129 | 0.392735114 | 0.29 | 0.133 | 6.91E-125 | 1 | Tsc22d2 |
| Prkar2a | 3.71E-129 | 0.257327138 | 0.29 | 0.128 | 7.51E-125 | 1 | Prkar2a |
| Shoc2 | 4.37E-129 | 0.25393906 | 0.386 | 0.196 | 8.85E-125 | 1 | Shoc2 |
| Ifngr2 | 7.32E-129 | 0.292286708 | 0.266 | 0.111 | 1.48E-124 | 1 | Ifngr2 |
| Atp2c1 | 1.06E-128 | 0.39962357 | 0.309 | 0.146 | 2.14E-124 | 1 | Atp2c1 |
| Zmynd8 | 1.68E-128 | 0.36017144 | 0.345 | 0.172 | 3.40E-124 | 1 | Zmynd8 |
| Selenot | 2.04E-128 | 0.315363727 | 0.369 | 0.187 | 4.12E-124 | 1 | Selenot |
| Srrm1 | 2.26E-128 | 0.259111003 | 0.545 | 0.321 | 4.57E-124 | 1 | Srrm1 |
| Chd4 | 3.19E-128 | 0.306606561 | 0.516 | 0.298 | 6.46E-124 | 1 | Chd4 |
| Parp9 | 3.59E-128 | 0.335063198 | 0.263 | 0.113 | 7.26E-124 | 1 | Parp9 |
| Mysm1 | 5.00E-128 | 0.354026393 | 0.32 | 0.153 | 1.01E-123 | 1 | Mysm1 |
| Rbms2 | 1.25E-127 | 0.267761747 | 0.304 | 0.137 | 2.54E-123 | 1 | Rbms2 |

| | | | | | | | |
|---|---|---|---|---|---|---|---|
| Appbp2 | 1.26E-127 | 0.356369765 | 0.271 | 0.12 | 2.55E-123 | 1 | Appbp2 |
| Cyth1 | 1.33E-127 | 0.312526271 | 0.337 | 0.164 | 2.70E-123 | 1 | Cyth1 |
| Papola | 2.13E-127 | 0.275083705 | 0.485 | 0.276 | 4.30E-123 | 1 | Papola |
| 2-Mar | 2.17E-127 | 0.267524277 | 0.352 | 0.173 | 4.39E-123 | 1 | 2-Mar |
| Kat2b | 3.27E-127 | 0.374226171 | 0.297 | 0.137 | 6.63E-123 | 1 | Kat2b |
| Tbc1d20 | 2.32E-126 | 0.262224123 | 0.255 | 0.108 | 4.70E-122 | 1 | Tbc1d20 |
| Dctn1 | 3.57E-126 | 0.261057449 | 0.261 | 0.111 | 7.22E-122 | 1 | Dctn1 |
| Ap2b1 | 7.81E-126 | 0.307377467 | 0.429 | 0.235 | 1.58E-121 | 1 | Ap2b1 |
| Dgkz | 1.29E-125 | 0.336480942 | 0.256 | 0.11 | 2.62E-121 | 1 | Dgkz |
| Klf6 | 1.41E-125 | 0.365547777 | 0.441 | 0.24 | 2.86E-121 | 1 | Klf6 |
| Ggps1 | 1.46E-125 | 0.330114197 | 0.315 | 0.15 | 2.96E-121 | 1 | Ggps1 |
| Ubr1 | 1.49E-125 | 0.368019395 | 0.301 | 0.141 | 3.01E-121 | 1 | Ubr1 |
| Sik1 | 1.52E-125 | 0.378895572 | 0.252 | 0.107 | 3.08E-121 | 1 | Sik1 |
| Arhgef2 | 2.48E-125 | 0.359117935 | 0.304 | 0.143 | 5.02E-121 | 1 | Arhgef2 |
| Sf3b1 | 3.69E-125 | 0.293857544 | 0.601 | 0.361 | 7.47E-121 | 1 | Sf3b1 |
| Nab1 | 5.79E-125 | 0.310493104 | 0.309 | 0.146 | 1.17E-120 | 1 | Nab1 |
| Gls | 8.88E-125 | 0.327624925 | 0.417 | 0.225 | 1.80E-120 | 1 | Gls |
| Arid1b | 1.27E-124 | 0.387996736 | 0.424 | 0.233 | 2.58E-120 | 1 | Arid1b |
| Emc10 | 1.31E-124 | 0.347121761 | 0.349 | 0.177 | 2.65E-120 | 1 | Emc10 |
| Med13 | 1.44E-124 | 0.337300106 | 0.357 | 0.182 | 2.93E-120 | 1 | Med13 |
| Irf9 | 1.64E-124 | 0.276732509 | 0.298 | 0.135 | 3.32E-120 | 1 | Irf9 |
| Mtf2 | 2.05E-124 | 0.350504919 | 0.261 | 0.114 | 4.15E-120 | 1 | Mtf2 |
| Trafd1 | 2.42E-124 | 0.365963649 | 0.288 | 0.133 | 4.91E-120 | 1 | Trafd1 |
| Akap9 | 3.48E-124 | 0.43242113 | 0.392 | 0.214 | 7.04E-120 | 1 | Akap9 |
| Epc1 | 3.72E-124 | 0.336953729 | 0.353 | 0.18 | 7.52E-120 | 1 | Epc1 |
| Atp2a2 | 3.98E-124 | 0.256268814 | 0.356 | 0.176 | 8.05E-120 | 1 | Atp2a2 |
| Rpn2 | 6.98E-124 | 0.33941801 | 0.364 | 0.188 | 1.41E-119 | 1 | Rpn2 |
| Tmem57 | 9.60E-124 | 0.368171819 | 0.262 | 0.116 | 1.94E-119 | 1 | Tmem57 |
| Larp4 | 1.71E-123 | 0.293209854 | 0.315 | 0.151 | 3.47E-119 | 1 | Larp4 |
| 9-Sep | 1.76E-123 | 0.345485067 | 0.312 | 0.149 | 3.57E-119 | 1 | 9-Sep |
| Mphosph8 | 2.01E-123 | 0.293779729 | 0.361 | 0.183 | 4.08E-119 | 1 | Mphosph8 |
| Stag2 | 2.58E-122 | 0.274339882 | 0.463 | 0.261 | 5.22E-118 | 1 | Stag2 |
| Sltm | 4.77E-122 | 0.342978961 | 0.438 | 0.245 | 9.66E-118 | 1 | Sltm |
| Zmym5 | 4.83E-122 | 0.321418475 | 0.28 | 0.128 | 9.78E-118 | 1 | Zmym5 |
| Klhdc10 | 2.47E-121 | 0.359904525 | 0.287 | 0.133 | 5.01E-117 | 1 | Klhdc10 |
| Tecpr1 | 4.01E-121 | 0.335902995 | 0.256 | 0.111 | 8.12E-117 | 1 | Tecpr1 |
| Iws1 | 1.41E-120 | 0.336822309 | 0.288 | 0.134 | 2.85E-116 | 1 | Iws1 |
| AI314180 | 1.43E-120 | 0.335649982 | 0.36 | 0.187 | 2.90E-116 | 1 | AI314180 |
| Tm2d2 | 1.31E-119 | 0.265413209 | 0.365 | 0.187 | 2.66E-115 | 1 | Tm2d2 |
| Ubr2 | 1.37E-119 | 0.362762394 | 0.385 | 0.206 | 2.77E-115 | 1 | Ubr2 |
| Rtf1 | 1.84E-119 | 0.260802853 | 0.448 | 0.254 | 3.73E-115 | 1 | Rtf1 |
| Neat1 | 4.20E-119 | 0.375139253 | 0.358 | 0.186 | 8.51E-115 | 1 | Neat1 |
| Terf1 | 4.74E-119 | 0.350325181 | 0.268 | 0.121 | 9.59E-115 | 1 | Terf1 |

| | | | | | | | |
|---|---|---|---|---|---|---|---|
| 9930021J03Rik | 5.29E-119 | 0.368701995 | 0.282 | 0.131 | 1.07E-114 | 1 | 9930021J03Rik |
| Sos2 | 5.44E-119 | 0.352950007 | 0.295 | 0.141 | 1.10E-114 | 1 | Sos2 |
| Cdc42ep3 | 5.47E-119 | 0.343264772 | 0.317 | 0.153 | 1.11E-114 | 1 | Cdc42ep3 |
| Rnf14 | 6.56E-119 | 0.277763901 | 0.278 | 0.127 | 1.33E-114 | 1 | Rnf14 |
| Vti1a | 1.82E-118 | 0.38421841 | 0.417 | 0.23 | 3.69E-114 | 1 | Vti1a |
| Pds5b | 3.84E-118 | 0.388748175 | 0.319 | 0.159 | 7.77E-114 | 1 | Pds5b |
| Arap2 | 7.05E-118 | 0.381192225 | 0.32 | 0.156 | 1.43E-113 | 1 | Arap2 |
| Fam193a | 1.92E-117 | 0.3885123 | 0.316 | 0.159 | 3.89E-113 | 1 | Fam193a |
| Ywhag | 2.31E-117 | 0.318573459 | 0.321 | 0.16 | 4.67E-113 | 1 | Ywhag |
| Tgfbr2 | 1.22E-116 | 0.347333141 | 0.49 | 0.284 | 2.48E-112 | 1 | Tgfbr2 |
| Stk10 | 1.45E-116 | 0.379162549 | 0.324 | 0.161 | 2.94E-112 | 1 | Stk10 |
| Jarid2 | 3.10E-116 | 0.355431641 | 0.348 | 0.182 | 6.28E-112 | 1 | Jarid2 |
| Pacs1 | 4.55E-116 | 0.366887826 | 0.352 | 0.182 | 9.22E-112 | 1 | Pacs1 |
| Ylpm1 | 6.43E-116 | 0.365616628 | 0.264 | 0.121 | 1.30E-111 | 1 | Ylpm1 |
| Lbr | 6.57E-116 | 0.353511463 | 0.282 | 0.133 | 1.33E-111 | 1 | Lbr |
| Fam120a | 9.39E-116 | 0.315819451 | 0.321 | 0.16 | 1.90E-111 | 1 | Fam120a |
| Rnf169 | 1.07E-115 | 0.407869114 | 0.32 | 0.162 | 2.16E-111 | 1 | Rnf169 |
| Pan3 | 1.22E-115 | 0.407914925 | 0.379 | 0.205 | 2.47E-111 | 1 | Pan3 |
| Copb1 | 1.54E-115 | 0.281135563 | 0.351 | 0.182 | 3.13E-111 | 1 | Copb1 |
| Pip4k2a | 2.53E-115 | 0.292700756 | 0.408 | 0.216 | 5.13E-111 | 1 | Pip4k2a |
| Tox4 | 3.60E-115 | 0.325297732 | 0.323 | 0.162 | 7.30E-111 | 1 | Tox4 |
| Baz1b | 4.09E-115 | 0.323503056 | 0.335 | 0.172 | 8.29E-111 | 1 | Baz1b |
| Phf14 | 7.12E-115 | 0.330499076 | 0.399 | 0.22 | 1.44E-110 | 1 | Phf14 |
| Senp6 | 7.17E-115 | 0.270641584 | 0.474 | 0.271 | 1.45E-110 | 1 | Senp6 |
| Casp8 | 8.50E-115 | 0.301619381 | 0.364 | 0.191 | 1.72E-110 | 1 | Casp8 |
| Epc2 | 9.18E-115 | 0.344855307 | 0.294 | 0.142 | 1.86E-110 | 1 | Epc2 |
| Mef2d | 1.12E-114 | 0.327149606 | 0.257 | 0.116 | 2.26E-110 | 1 | Mef2d |
| Bcap31 | 1.30E-114 | 0.273785461 | 0.452 | 0.259 | 2.63E-110 | 1 | Bcap31 |
| Ccar1 | 2.96E-114 | 0.36563727 | 0.352 | 0.186 | 6.00E-110 | 1 | Ccar1 |
| Trip12 | 3.92E-114 | 0.31425819 | 0.415 | 0.232 | 7.93E-110 | 1 | Trip12 |
| Fam168a | 4.79E-114 | 0.340887115 | 0.268 | 0.123 | 9.69E-110 | 1 | Fam168a |
| Chst11 | 6.64E-114 | 0.397913376 | 0.312 | 0.155 | 1.34E-109 | 1 | Chst11 |
| Gatad2b | 8.35E-114 | 0.337465907 | 0.346 | 0.18 | 1.69E-109 | 1 | Gatad2b |
| Sft2d1 | 9.47E-114 | 0.342520063 | 0.291 | 0.14 | 1.92E-109 | 1 | Sft2d1 |
| Flii | 1.22E-113 | 0.303212266 | 0.288 | 0.138 | 2.47E-109 | 1 | Flii |
| Nlk | 1.80E-113 | 0.369980126 | 0.271 | 0.126 | 3.65E-109 | 1 | Nlk |
| Rsbn1 | 3.17E-113 | 0.333164602 | 0.313 | 0.157 | 6.41E-109 | 1 | Rsbn1 |
| Cask | 5.81E-113 | 0.336896198 | 0.265 | 0.12 | 1.18E-108 | 1 | Cask |
| Rbm5 | 1.00E-112 | 0.341794619 | 0.391 | 0.216 | 2.03E-108 | 1 | Rbm5 |
| 7-Mar | 1.08E-112 | 0.301514387 | 0.322 | 0.162 | 2.18E-108 | 1 | 7-Mar |
| Chd6 | 1.10E-112 | 0.347071119 | 0.364 | 0.195 | 2.24E-108 | 1 | Chd6 |
| Sp3 | 1.41E-112 | 0.32367632 | 0.332 | 0.169 | 2.86E-108 | 1 | Sp3 |
| Supt6 | 2.00E-112 | 0.335519939 | 0.289 | 0.14 | 4.06E-108 | 1 | Supt6 |

| | | | | | | | |
|---|---|---|---|---|---|---|---|
| Zzef1 | 2.97E-112 | 0.373482265 | 0.273 | 0.129 | 6.01E-108 | 1 | Zzef1 |
| Atp1b3 | 3.49E-112 | 0.301809471 | 0.404 | 0.225 | 7.06E-108 | 1 | Atp1b3 |
| Hectd1 | 5.43E-112 | 0.27487443 | 0.478 | 0.273 | 1.10E-107 | 1 | Hectd1 |
| Tcf7l2 | 6.47E-112 | 0.273843834 | 0.254 | 0.11 | 1.31E-107 | 1 | Tcf7l2 |
| Esyt2 | 3.83E-111 | 0.328530817 | 0.401 | 0.218 | 7.76E-107 | 1 | Esyt2 |
| Mapk1 | 4.26E-111 | 0.298816587 | 0.388 | 0.212 | 8.62E-107 | 1 | Mapk1 |
| Nek7 | 2.18E-110 | 0.290938063 | 0.404 | 0.224 | 4.41E-106 | 1 | Nek7 |
| Tmx4 | 4.69E-110 | 0.442234986 | 0.275 | 0.132 | 9.50E-106 | 1 | Tmx4 |
| Vps13d | 5.66E-110 | 0.363726676 | 0.299 | 0.149 | 1.15E-105 | 1 | Vps13d |
| Suz12 | 1.16E-109 | 0.269837233 | 0.324 | 0.163 | 2.35E-105 | 1 | Suz12 |
| Kidins220 | 1.63E-109 | 0.36561773 | 0.28 | 0.136 | 3.30E-105 | 1 | Kidins220 |
| Luc7l3 | 2.07E-109 | 0.264301633 | 0.432 | 0.246 | 4.19E-105 | 1 | Luc7l3 |
| Kmt5b | 3.53E-109 | 0.286220268 | 0.304 | 0.15 | 7.14E-105 | 1 | Kmt5b |
| Ppp4r2 | 4.15E-109 | 0.261798731 | 0.372 | 0.201 | 8.41E-105 | 1 | Ppp4r2 |
| Trim12a | 7.14E-109 | 0.341590888 | 0.317 | 0.161 | 1.45E-104 | 1 | Trim12a |
| Cdk13 | 7.89E-109 | 0.344445761 | 0.373 | 0.205 | 1.60E-104 | 1 | Cdk13 |
| Mecp2 | 9.58E-109 | 0.321744017 | 0.288 | 0.14 | 1.94E-104 | 1 | Mecp2 |
| Ppp2r5e | 1.13E-108 | 0.291117132 | 0.296 | 0.145 | 2.29E-104 | 1 | Ppp2r5e |
| Sec24b | 2.48E-108 | 0.339894685 | 0.258 | 0.12 | 5.01E-104 | 1 | Sec24b |
| Znrf1 | 3.90E-108 | 0.34333323 | 0.328 | 0.168 | 7.90E-104 | 1 | Znrf1 |
| Vps13b | 4.28E-108 | 0.381319934 | 0.365 | 0.197 | 8.66E-104 | 1 | Vps13b |
| Smc6 | 6.48E-108 | 0.286293537 | 0.419 | 0.238 | 1.31E-103 | 1 | Smc6 |
| Thrap3 | 7.80E-108 | 0.250302953 | 0.459 | 0.268 | 1.58E-103 | 1 | Thrap3 |
| Rnf130 | 8.63E-108 | 0.301755833 | 0.447 | 0.259 | 1.75E-103 | 1 | Rnf130 |
| Atf7ip | 1.05E-107 | 0.320892 | 0.349 | 0.185 | 2.13E-103 | 1 | Atf7ip |
| Ndel1 | 1.47E-107 | 0.297943955 | 0.307 | 0.154 | 2.97E-103 | 1 | Ndel1 |
| Vamp4 | 2.26E-107 | 0.261163512 | 0.302 | 0.148 | 4.58E-103 | 1 | Vamp4 |
| Rnf38 | 2.60E-107 | 0.324336661 | 0.251 | 0.115 | 5.27E-103 | 1 | Rnf38 |
| Uhrf2 | 3.14E-107 | 0.334602152 | 0.295 | 0.146 | 6.35E-103 | 1 | Uhrf2 |
| Sfswap | 4.70E-107 | 0.318736899 | 0.253 | 0.117 | 9.52E-103 | 1 | Sfswap |
| Zfp280d | 8.16E-107 | 0.339067192 | 0.317 | 0.163 | 1.65E-102 | 1 | Zfp280d |
| Smg7 | 1.50E-106 | 0.324001986 | 0.281 | 0.137 | 3.03E-102 | 1 | Smg7 |
| Hmg20a | 1.51E-106 | 0.348197242 | 0.262 | 0.124 | 3.05E-102 | 1 | Hmg20a |
| Ifrd1 | 1.79E-106 | 0.27238376 | 0.378 | 0.208 | 3.62E-102 | 1 | Ifrd1 |
| Phc2 | 5.89E-106 | 0.258783457 | 0.303 | 0.149 | 1.19E-101 | 1 | Phc2 |
| Arid2 | 1.40E-105 | 0.333451439 | 0.282 | 0.137 | 2.84E-101 | 1 | Arid2 |
| Ncoa2 | 3.17E-105 | 0.373615764 | 0.386 | 0.216 | 6.43E-101 | 1 | Ncoa2 |
| Ssr3 | 3.58E-105 | 0.265384135 | 0.334 | 0.175 | 7.24E-101 | 1 | Ssr3 |
| Arnt | 3.98E-105 | 0.332221489 | 0.263 | 0.124 | 8.05E-101 | 1 | Arnt |
| Cdkn1b | 4.74E-105 | 0.262037555 | 0.359 | 0.193 | 9.59E-101 | 1 | Cdkn1b |
| 6-Mar | 8.08E-105 | 0.316636598 | 0.263 | 0.125 | 1.64E-100 | 1 | 6-Mar |
| Chd8 | 1.40E-104 | 0.34437874 | 0.296 | 0.148 | 2.84E-100 | 1 | Chd8 |
| Eps15 | 1.66E-104 | 0.306937462 | 0.298 | 0.149 | 3.35E-100 | 1 | Eps15 |

| | | | | | | | |
|---|---|---|---|---|---|---|---|
| Actg1 | 1.69E-104 | 0.313570943 | 0.969 | 0.766 | 3.42E-100 | 1 | Actg1 |
| Smarca4 | 1.74E-104 | 0.278556013 | 0.293 | 0.146 | 3.51E-100 | 1 | Smarca4 |
| Usp1 | 2.55E-104 | 0.266416319 | 0.269 | 0.127 | 5.16E-100 | 1 | Usp1 |
| Zmym4 | 3.11E-104 | 0.326497389 | 0.292 | 0.146 | 6.29E-100 | 1 | Zmym4 |
| Rps6kb1 | 3.19E-104 | 0.301532555 | 0.283 | 0.139 | 6.47E-100 | 1 | Rps6kb1 |
| Scaf11 | 8.82E-104 | 0.250294859 | 0.418 | 0.239 | 1.79E-99 | 1 | Scaf11 |
| Atf2 | 8.82E-104 | 0.281417477 | 0.285 | 0.139 | 1.79E-99 | 1 | Atf2 |
| Ergic2 | 8.94E-104 | 0.274361773 | 0.311 | 0.159 | 1.81E-99 | 1 | Ergic2 |
| Eif5b | 1.13E-103 | 0.253494053 | 0.488 | 0.296 | 2.29E-99 | 1 | Eif5b |
| Kdm7a | 1.17E-103 | 0.320539719 | 0.287 | 0.142 | 2.38E-99 | 1 | Kdm7a |
| Tbc1d5 | 1.47E-103 | 0.326736469 | 0.421 | 0.237 | 2.97E-99 | 1 | Tbc1d5 |
| Cdkal1 | 3.26E-103 | 0.370751037 | 0.309 | 0.161 | 6.59E-99 | 1 | Cdkal1 |
| 5031425E22Rik | 3.43E-103 | 0.305807583 | 0.285 | 0.141 | 6.94E-99 | 1 | 5031425E22Rik |
| Srek1 | 6.07E-103 | 0.337110106 | 0.327 | 0.173 | 1.23E-98 | 1 | Srek1 |
| Ddx42 | 7.19E-103 | 0.257529288 | 0.334 | 0.177 | 1.46E-98 | 1 | Ddx42 |
| Mtmr3 | 1.22E-102 | 0.335284283 | 0.336 | 0.18 | 2.46E-98 | 1 | Mtmr3 |
| Fam76b | 1.26E-102 | 0.267880578 | 0.264 | 0.125 | 2.55E-98 | 1 | Fam76b |
| Coro1c | 2.21E-102 | 0.313733006 | 0.258 | 0.123 | 4.46E-98 | 1 | Coro1c |
| Zfp407 | 3.74E-102 | 0.391427964 | 0.284 | 0.143 | 7.57E-98 | 1 | Zfp407 |
| Tex261 | 4.07E-102 | 0.250756076 | 0.322 | 0.168 | 8.23E-98 | 1 | Tex261 |
| Prpf38b | 9.62E-102 | 0.337621214 | 0.329 | 0.176 | 1.95E-97 | 1 | Prpf38b |
| Taf1 | 2.36E-101 | 0.321634184 | 0.26 | 0.125 | 4.78E-97 | 1 | Taf1 |
| Rabep1 | 4.59E-101 | 0.319662668 | 0.375 | 0.209 | 9.30E-97 | 1 | Rabep1 |
| Wdr33 | 4.68E-101 | 0.305901061 | 0.384 | 0.217 | 9.48E-97 | 1 | Wdr33 |
| Arfgef1 | 5.78E-101 | 0.319357014 | 0.368 | 0.206 | 1.17E-96 | 1 | Arfgef1 |
| Glmp | 1.07E-100 | 0.273522547 | 0.268 | 0.129 | 2.17E-96 | 1 | Glmp |
| Atxn2 | 1.55E-100 | 0.285916303 | 0.258 | 0.121 | 3.15E-96 | 1 | Atxn2 |
| Ddb1 | 1.90E-100 | 0.251596455 | 0.269 | 0.13 | 3.85E-96 | 1 | Ddb1 |
| Bcas3 | 2.06E-100 | 0.384110505 | 0.312 | 0.163 | 4.17E-96 | 1 | Bcas3 |
| Appl1 | 2.38E-100 | 0.282768078 | 0.312 | 0.162 | 4.81E-96 | 1 | Appl1 |
| Tmem208 | 2.40E-100 | 0.250305894 | 0.385 | 0.216 | 4.86E-96 | 1 | Tmem208 |
| Ski | 3.98E-100 | 0.284493524 | 0.356 | 0.193 | 8.07E-96 | 1 | Ski |
| Ankfy1 | 4.06E-100 | 0.32529467 | 0.26 | 0.126 | 8.21E-96 | 1 | Ankfy1 |
| Rbm26 | 5.69E-100 | 0.329775476 | 0.366 | 0.206 | 1.15E-95 | 1 | Rbm26 |
| Atxn7 | 6.62E-100 | 0.349367831 | 0.29 | 0.147 | 1.34E-95 | 1 | Atxn7 |
| Kif2a | 7.52E-100 | 0.300519403 | 0.306 | 0.158 | 1.52E-95 | 1 | Kif2a |
| Tram1 | 7.89E-100 | 0.286041698 | 0.357 | 0.198 | 1.60E-95 | 1 | Tram1 |
| Mia3 | 9.59E-100 | 0.327146332 | 0.296 | 0.151 | 1.94E-95 | 1 | Mia3 |
| Mon2 | 2.24E-99 | 0.309458352 | 0.272 | 0.134 | 4.53E-95 | 1 | Mon2 |
| Ttc14 | 2.53E-99 | 0.353205876 | 0.352 | 0.195 | 5.12E-95 | 1 | Ttc14 |
| Asah1 | 2.70E-99 | 0.253740596 | 0.326 | 0.167 | 5.48E-95 | 1 | Asah1 |
| Ctdspl2 | 4.39E-99 | 0.285012933 | 0.298 | 0.153 | 8.89E-95 | 1 | Ctdspl2 |
| Tnks | 4.43E-99 | 0.312161179 | 0.26 | 0.125 | 8.97E-95 | 1 | Tnks |

| | | | | | | | |
|---|---|---|---|---|---|---|---|
| Phf3 | 8.24E-99 | 0.304684978 | 0.402 | 0.23 | 1.67E-94 | 1 | Phf3 |
| 2810474O19Rik | 1.61E-98 | 0.369072263 | 0.344 | 0.19 | 3.27E-94 | 1 | 2810474O19Rik |
| Ulk2 | 2.07E-98 | 0.296525745 | 0.307 | 0.159 | 4.19E-94 | 1 | Ulk2 |
| Pik3ca | 2.32E-98 | 0.311597995 | 0.293 | 0.149 | 4.70E-94 | 1 | Pik3ca |
| Acer3 | 2.89E-98 | 0.350207138 | 0.258 | 0.124 | 5.85E-94 | 1 | Acer3 |
| Usp7 | 3.94E-98 | 0.250612068 | 0.286 | 0.143 | 7.98E-94 | 1 | Usp7 |
| Rbm27 | 4.16E-98 | 0.250095018 | 0.318 | 0.167 | 8.43E-94 | 1 | Rbm27 |
| Sbno1 | 1.53E-97 | 0.284764614 | 0.344 | 0.189 | 3.10E-93 | 1 | Sbno1 |
| Ext1 | 1.93E-97 | 0.286187575 | 0.297 | 0.147 | 3.91E-93 | 1 | Ext1 |
| Dhx36 | 3.82E-97 | 0.25782992 | 0.376 | 0.214 | 7.74E-93 | 1 | Dhx36 |
| Rictor | 5.79E-97 | 0.342634981 | 0.275 | 0.139 | 1.17E-92 | 1 | Rictor |
| Far1 | 7.19E-97 | 0.305280542 | 0.286 | 0.146 | 1.46E-92 | 1 | Far1 |
| Tra2b | 7.45E-97 | 0.253618444 | 0.459 | 0.272 | 1.51E-92 | 1 | Tra2b |
| Abi1 | 1.68E-96 | 0.296050038 | 0.42 | 0.246 | 3.40E-92 | 1 | Abi1 |
| Scaf8 | 1.77E-96 | 0.34111355 | 0.311 | 0.166 | 3.59E-92 | 1 | Scaf8 |
| Larp4b | 2.00E-96 | 0.324661038 | 0.335 | 0.184 | 4.05E-92 | 1 | Larp4b |
| Usp47 | 2.86E-96 | 0.272717289 | 0.317 | 0.168 | 5.78E-92 | 1 | Usp47 |
| Rgs3 | 4.18E-96 | 0.313187215 | 0.319 | 0.167 | 8.47E-92 | 1 | Rgs3 |
| Ggnbp2 | 8.34E-96 | 0.284727265 | 0.384 | 0.22 | 1.69E-91 | 1 | Ggnbp2 |
| Cdc73 | 8.94E-96 | 0.276585526 | 0.293 | 0.152 | 1.81E-91 | 1 | Cdc73 |
| Wwox | 1.39E-95 | 0.432660494 | 0.301 | 0.157 | 2.81E-91 | 1 | Wwox |
| Taf3 | 4.18E-95 | 0.35648268 | 0.26 | 0.13 | 8.45E-91 | 1 | Taf3 |
| Plgrkt | 5.71E-95 | 0.319612243 | 0.348 | 0.193 | 1.16E-90 | 1 | Plgrkt |
| Nudcd3 | 9.86E-95 | 0.256872093 | 0.39 | 0.224 | 2.00E-90 | 1 | Nudcd3 |
| Rabgap1 | 1.08E-94 | 0.289676308 | 0.285 | 0.145 | 2.18E-90 | 1 | Rabgap1 |
| Clip1 | 1.13E-94 | 0.299047824 | 0.257 | 0.126 | 2.29E-90 | 1 | Clip1 |
| Itch | 1.25E-94 | 0.292132283 | 0.425 | 0.251 | 2.53E-90 | 1 | Itch |
| Tle4 | 2.04E-94 | 0.331953624 | 0.365 | 0.204 | 4.12E-90 | 1 | Tle4 |
| Smarcc2 | 3.69E-94 | 0.291589321 | 0.301 | 0.159 | 7.47E-90 | 1 | Smarcc2 |
| Bach2 | 4.40E-94 | 0.337511406 | 0.282 | 0.143 | 8.91E-90 | 1 | Bach2 |
| Kdm6a | 4.47E-94 | 0.348991395 | 0.346 | 0.194 | 9.04E-90 | 1 | Kdm6a |
| Copa | 1.05E-93 | 0.281654356 | 0.337 | 0.185 | 2.12E-89 | 1 | Copa |
| Swt1 | 1.36E-93 | 0.307162055 | 0.298 | 0.156 | 2.76E-89 | 1 | Swt1 |
| Cop1 | 7.21E-93 | 0.28820157 | 0.303 | 0.16 | 1.46E-88 | 1 | Cop1 |
| Prrc2a | 8.67E-93 | 0.251912838 | 0.266 | 0.133 | 1.75E-88 | 1 | Prrc2a |
| mt-Co3 | 9.57E-93 | 0.270223214 | 0.997 | 0.682 | 1.94E-88 | 1 | mt-Co3 |
| Setd2 | 2.78E-92 | 0.311876343 | 0.318 | 0.174 | 5.63E-88 | 1 | Setd2 |
| Rnf115 | 3.54E-92 | 0.253494269 | 0.316 | 0.17 | 7.16E-88 | 1 | Rnf115 |
| Ascc3 | 4.35E-92 | 0.34688213 | 0.336 | 0.188 | 8.81E-88 | 1 | Ascc3 |
| Enox2 | 2.35E-91 | 0.369723607 | 0.255 | 0.126 | 4.75E-87 | 1 | Enox2 |
| Zmym2 | 9.16E-91 | 0.306708408 | 0.25 | 0.123 | 1.85E-86 | 1 | Zmym2 |
| Ap3b1 | 2.17E-90 | 0.295069289 | 0.349 | 0.198 | 4.39E-86 | 1 | Ap3b1 |
| Lnpep | 3.27E-90 | 0.338463831 | 0.401 | 0.238 | 6.61E-86 | 1 | Lnpep |

| | | | | | | | |
|---|---|---|---|---|---|---|---|
| Kdm4c | 6.71E-90 | 0.316868497 | 0.288 | 0.151 | 1.36E-85 | 1 | Kdm4c |
| Zc3h7a | 1.23E-89 | 0.319744737 | 0.278 | 0.145 | 2.49E-85 | 1 | Zc3h7a |
| Fam172a | 1.24E-89 | 0.3162869 | 0.438 | 0.261 | 2.52E-85 | 1 | Fam172a |
| Dyrk1a | 2.09E-89 | 0.308726781 | 0.3 | 0.162 | 4.23E-85 | 1 | Dyrk1a |
| Pkn1 | 2.13E-89 | 0.273887254 | 0.31 | 0.168 | 4.32E-85 | 1 | Pkn1 |
| Wrn | 2.65E-89 | 0.299130634 | 0.258 | 0.13 | 5.36E-85 | 1 | Wrn |
| Nrd1 | 2.93E-89 | 0.263783184 | 0.336 | 0.188 | 5.93E-85 | 1 | Nrd1 |
| Os9 | 4.07E-89 | 0.288381779 | 0.313 | 0.17 | 8.23E-85 | 1 | Os9 |
| Wsb1 | 8.59E-89 | 0.285314151 | 0.296 | 0.157 | 1.74E-84 | 1 | Wsb1 |
| Magt1 | 9.49E-89 | 0.315765223 | 0.28 | 0.147 | 1.92E-84 | 1 | Magt1 |
| Faf1 | 1.14E-88 | 0.284109815 | 0.328 | 0.182 | 2.32E-84 | 1 | Faf1 |
| Brwd1 | 1.94E-88 | 0.289641839 | 0.252 | 0.125 | 3.92E-84 | 1 | Brwd1 |
| Dennd1a | 3.84E-88 | 0.322363666 | 0.394 | 0.232 | 7.77E-84 | 1 | Dennd1a |
| Cnot2 | 1.56E-87 | 0.281087877 | 0.339 | 0.191 | 3.16E-83 | 1 | Cnot2 |
| Tmcc1 | 1.82E-87 | 0.370840379 | 0.346 | 0.198 | 3.68E-83 | 1 | Tmcc1 |
| Rap1gds1 | 1.26E-86 | 0.306873188 | 0.404 | 0.239 | 2.56E-82 | 1 | Rap1gds1 |
| Rnf13 | 1.43E-86 | 0.294279733 | 0.297 | 0.161 | 2.90E-82 | 1 | Rnf13 |
| Cwf19l2 | 2.91E-86 | 0.261288503 | 0.327 | 0.182 | 5.89E-82 | 1 | Cwf19l2 |
| Asxl1 | 1.39E-85 | 0.312830174 | 0.256 | 0.13 | 2.82E-81 | 1 | Asxl1 |
| Nup98 | 3.33E-85 | 0.290406292 | 0.253 | 0.128 | 6.75E-81 | 1 | Nup98 |
| Pdia4 | 1.74E-84 | 0.291911431 | 0.282 | 0.149 | 3.52E-80 | 1 | Pdia4 |
| Scaper | 5.02E-84 | 0.331312979 | 0.297 | 0.162 | 1.02E-79 | 1 | Scaper |
| Akap8 | 6.87E-84 | 0.26437465 | 0.267 | 0.139 | 1.39E-79 | 1 | Akap8 |
| Mdm4 | 6.69E-83 | 0.262564009 | 0.271 | 0.143 | 1.35E-78 | 1 | Mdm4 |
| Krit1 | 4.61E-82 | 0.320450649 | 0.298 | 0.165 | 9.34E-78 | 1 | Krit1 |
| Scarb2 | 5.42E-82 | 0.262582004 | 0.307 | 0.17 | 1.10E-77 | 1 | Scarb2 |
| Prpf40a | 5.87E-82 | 0.261849328 | 0.403 | 0.246 | 1.19E-77 | 1 | Prpf40a |
| Parp4 | 8.01E-82 | 0.279237258 | 0.261 | 0.136 | 1.62E-77 | 1 | Parp4 |
| Rsbn1l | 2.98E-81 | 0.269236085 | 0.411 | 0.251 | 6.04E-77 | 1 | Rsbn1l |
| Stat5b | 1.55E-80 | 0.291428813 | 0.253 | 0.131 | 3.15E-76 | 1 | Stat5b |
| Nptn | 6.36E-80 | 0.253497517 | 0.383 | 0.229 | 1.29E-75 | 1 | Nptn |
| Stim1 | 7.53E-80 | 0.300750428 | 0.309 | 0.173 | 1.53E-75 | 1 | Stim1 |
| Lrba | 8.42E-80 | 0.309239134 | 0.31 | 0.174 | 1.71E-75 | 1 | Lrba |
| Mgea5 | 1.24E-79 | 0.257092763 | 0.281 | 0.152 | 2.51E-75 | 1 | Mgea5 |
| Arid1a | 7.16E-79 | 0.274953494 | 0.357 | 0.211 | 1.45E-74 | 1 | Arid1a |
| Fam208b | 1.75E-76 | 0.322011996 | 0.253 | 0.135 | 3.53E-72 | 1 | Fam208b |
| Synj1 | 4.56E-76 | 0.285349289 | 0.266 | 0.144 | 9.22E-72 | 1 | Synj1 |
| Trappc8 | 7.22E-76 | 0.290483361 | 0.273 | 0.15 | 1.46E-71 | 1 | Trappc8 |
| Odc1 | 1.91E-75 | 0.294850497 | 0.299 | 0.169 | 3.87E-71 | 1 | Odc1 |
| Ube3c | 2.59E-75 | 0.277699339 | 0.265 | 0.143 | 5.24E-71 | 1 | Ube3c |
| Clasp2 | 3.10E-75 | 0.296603711 | 0.294 | 0.166 | 6.29E-71 | 1 | Clasp2 |
| Gigyf2 | 9.96E-75 | 0.271499895 | 0.276 | 0.152 | 2.02E-70 | 1 | Gigyf2 |
| Atad2b | 3.95E-74 | 0.29434687 | 0.318 | 0.186 | 8.00E-70 | 1 | Atad2b |

| | | | | | | | |
|---|---|---|---|---|---|---|---|
| Stt3b | 5.35E-74 | 0.293518644 | 0.271 | 0.149 | 1.08E-69 | 1 | Stt3b |
| Sf3b3 | 6.71E-73 | 0.256288926 | 0.266 | 0.145 | 1.36E-68 | 1 | Sf3b3 |
| Nr2c2 | 2.25E-72 | 0.269029 | 0.263 | 0.143 | 4.56E-68 | 1 | Nr2c2 |
| Gbf1 | 3.02E-72 | 0.280440696 | 0.29 | 0.166 | 6.12E-68 | 1 | Gbf1 |
| Ambra1 | 4.20E-70 | 0.310323344 | 0.308 | 0.182 | 8.50E-66 | 1 | Ambra1 |
| Osbpl8 | 2.14E-69 | 0.251448892 | 0.312 | 0.183 | 4.34E-65 | 1 | Osbpl8 |
| Lcorl | 1.10E-68 | 0.283795996 | 0.27 | 0.151 | 2.22E-64 | 1 | Lcorl |
| Rnf111 | 2.88E-68 | 0.256782921 | 0.256 | 0.141 | 5.82E-64 | 1 | Rnf111 |
| Cog5 | 4.46E-67 | 0.278142783 | 0.288 | 0.167 | 9.03E-63 | 1 | Cog5 |
| Chd2 | 4.93E-67 | 0.250491536 | 0.379 | 0.235 | 9.98E-63 | 1 | Chd2 |
| B4galt1 | 8.59E-65 | 0.280999594 | 0.279 | 0.162 | 1.74E-60 | 1 | B4galt1 |
| Lcor | 1.57E-63 | 0.265885399 | 0.293 | 0.175 | 3.19E-59 | 1 | Lcor |
| Csnk1g1 | 5.52E-63 | 0.280600385 | 0.253 | 0.143 | 1.12E-58 | 1 | Csnk1g1 |
| Rgcc1 | 0 | 2.009203495 | 0.945 | 0.337 | 0 | 2 | Rgcc |
| Plpp11 | 0 | 1.942823146 | 0.98 | 0.308 | 0 | 2 | Plpp1 |
| Igfbp71 | 0 | 1.929131105 | 0.995 | 0.402 | 0 | 2 | Igfbp7 |
| Emcn1 | 0 | 1.925889406 | 0.989 | 0.305 | 0 | 2 | Emcn |
| Plpp31 | 0 | 1.82829422 | 0.989 | 0.331 | 0 | 2 | Plpp3 |
| Plvap1 | 0 | 1.809892587 | 0.984 | 0.309 | 0 | 2 | Plvap |
| Id31 | 0 | 1.805476339 | 0.952 | 0.34 | 0 | 2 | Id3 |
| Rbp71 | 0 | 1.802717084 | 0.876 | 0.282 | 0 | 2 | Rbp7 |
| Egfl71 | 0 | 1.801143625 | 0.984 | 0.32 | 0 | 2 | Egfl7 |
| Ly6c11 | 0 | 1.752599364 | 0.976 | 0.328 | 0 | 2 | Ly6c1 |
| Gpihbp11 | 0 | 1.717109326 | 0.805 | 0.233 | 0 | 2 | Gpihbp1 |
| Kdr1 | 0 | 1.708252121 | 0.961 | 0.288 | 0 | 2 | Kdr |
| Fabp41 | 0 | 1.689929779 | 0.98 | 0.368 | 0 | 2 | Fabp4 |
| Nrp11 | 0 | 1.681910294 | 0.968 | 0.368 | 0 | 2 | Nrp1 |
| Flt11 | 0 | 1.671742679 | 0.995 | 0.318 | 0 | 2 | Flt1 |
| Car41 | 0 | 1.666341538 | 0.692 | 0.167 | 0 | 2 | Car4 |
| Jam31 | 0 | 1.604969215 | 0.763 | 0.215 | 0 | 2 | Jam3 |
| Adgrl41 | 0 | 1.601068167 | 0.946 | 0.277 | 0 | 2 | Adgrl4 |
| Tcim1 | 0 | 1.597015653 | 0.782 | 0.244 | 0 | 2 | Tcim |
| Thrsp1 | 0 | 1.565087772 | 0.794 | 0.23 | 0 | 2 | Thrsp |
| Ramp31 | 0 | 1.540403919 | 0.75 | 0.206 | 0 | 2 | Ramp3 |
| Ly6a1 | 0 | 1.537888585 | 0.965 | 0.397 | 0 | 2 | Ly6a |
| Hspb11 | 0 | 1.534871442 | 0.885 | 0.318 | 0 | 2 | Hspb1 |
| Id11 | 0 | 1.523341323 | 0.815 | 0.264 | 0 | 2 | Id1 |
| Igfbp31 | 0 | 1.522760744 | 0.756 | 0.247 | 0 | 2 | Igfbp3 |
| Ramp21 | 0 | 1.518050637 | 0.811 | 0.272 | 0 | 2 | Ramp2 |
| Arhgap311 | 0 | 1.516840317 | 0.903 | 0.306 | 0 | 2 | Arhgap31 |
| Cd2001 | 0 | 1.506054105 | 0.878 | 0.284 | 0 | 2 | Cd200 |
| Cd300lg1 | 0 | 1.498176984 | 0.789 | 0.235 | 0 | 2 | Cd300lg |
| Pecam11 | 0 | 1.481999005 | 0.968 | 0.341 | 0 | 2 | Pecam1 |

| | | | | | | | |
|---|---|---|---|---|---|---|---|
| Tcf151 | 0 | 1.481352921 | 0.663 | 0.21 | 0 | 2 | Tcf15 |
| Exoc3l21 | 0 | 1.467065298 | 0.738 | 0.212 | 0 | 2 | Exoc3l2 |
| Cyyr11 | 0 | 1.460832172 | 0.91 | 0.273 | 0 | 2 | Cyyr1 |
| Podxl1 | 0 | 1.454907278 | 0.899 | 0.276 | 0 | 2 | Podxl |
| Plscr21 | 0 | 1.454646718 | 0.812 | 0.275 | 0 | 2 | Plscr2 |
| Esam1 | 0 | 1.446210161 | 0.804 | 0.248 | 0 | 2 | Esam |
| Mef2c1 | 0 | 1.428374297 | 0.912 | 0.363 | 0 | 2 | Mef2c |
| Cyp26b11 | 0 | 1.409684742 | 0.532 | 0.177 | 0 | 2 | Cyp26b1 |
| Pltp1 | 0 | 1.408990935 | 0.751 | 0.244 | 0 | 2 | Pltp |
| Slc9a3r21 | 0 | 1.400816644 | 0.865 | 0.309 | 0 | 2 | Slc9a3r2 |
| Col4a11 | 0 | 1.400560314 | 0.945 | 0.323 | 0 | 2 | Col4a1 |
| Tm4sf11 | 0 | 1.391523788 | 0.93 | 0.309 | 0 | 2 | Tm4sf1 |
| Clec14a1 | 0 | 1.389992086 | 0.749 | 0.25 | 0 | 2 | Clec14a |
| Eng1 | 0 | 1.383222828 | 0.865 | 0.29 | 0 | 2 | Eng |
| Btnl91 | 0 | 1.382450564 | 0.718 | 0.215 | 0 | 2 | Btnl9 |
| Snrk1 | 0 | 1.381847572 | 0.828 | 0.312 | 0 | 2 | Snrk |
| Syne11 | 0 | 1.377492722 | 0.819 | 0.302 | 0 | 2 | Syne1 |
| Gng111 | 0 | 1.373923624 | 0.842 | 0.307 | 0 | 2 | Gng11 |
| Sparcl11 | 0 | 1.369140896 | 0.851 | 0.303 | 0 | 2 | Sparcl1 |
| Cavin21 | 0 | 1.367876543 | 0.852 | 0.278 | 0 | 2 | Cavin2 |
| Arhgap291 | 0 | 1.367471405 | 0.855 | 0.304 | 0 | 2 | Arhgap29 |
| Mcc1 | 0 | 1.365825955 | 0.7 | 0.233 | 0 | 2 | Mcc |
| Timp31 | 0 | 1.365298527 | 0.924 | 0.318 | 0 | 2 | Timp3 |
| Scarb11 | 0 | 1.35773941 | 0.698 | 0.213 | 0 | 2 | Scarb1 |
| Col4a21 | 0 | 1.354866782 | 0.866 | 0.297 | 0 | 2 | Col4a2 |
| Nfib1 | 0 | 1.350860052 | 0.966 | 0.38 | 0 | 2 | Nfib |
| Tcf41 | 0 | 1.350484272 | 0.963 | 0.411 | 0 | 2 | Tcf4 |
| Ifitm31 | 0 | 1.344582746 | 0.965 | 0.485 | 0 | 2 | Ifitm3 |
| Fam167b1 | 0 | 1.344524496 | 0.582 | 0.165 | 0 | 2 | Fam167b |
| Ifitm21 | 0 | 1.329418866 | 0.925 | 0.408 | 0 | 2 | Ifitm2 |
| Rsad21 | 0 | 1.329286164 | 0.636 | 0.21 | 0 | 2 | Rsad2 |
| Ehd41 | 0 | 1.32812665 | 0.829 | 0.356 | 0 | 2 | Ehd4 |
| Tinagl11 | 0 | 1.323757708 | 0.721 | 0.244 | 0 | 2 | Tinagl1 |
| Cdh51 | 0 | 1.318143044 | 0.847 | 0.266 | 0 | 2 | Cdh5 |
| Cd341 | 0 | 1.316348383 | 0.778 | 0.264 | 0 | 2 | Cd34 |
| Ctla2a1 | 0 | 1.314612058 | 0.779 | 0.352 | 0 | 2 | Ctla2a |
| Plcb11 | 0 | 1.31381669 | 0.901 | 0.271 | 0 | 2 | Plcb1 |
| Col13a11 | 0 | 1.30344303 | 0.638 | 0.193 | 0 | 2 | Col13a1 |
| Sox181 | 0 | 1.30287518 | 0.699 | 0.224 | 0 | 2 | Sox18 |
| Ptprm1 | 0 | 1.29922094 | 0.815 | 0.269 | 0 | 2 | Ptprm |
| Cmtm81 | 0 | 1.298627966 | 0.666 | 0.225 | 0 | 2 | Cmtm8 |
| Cd811 | 0 | 1.294080355 | 0.955 | 0.473 | 0 | 2 | Cd81 |
| Vamp51 | 0 | 1.293959349 | 0.823 | 0.331 | 0 | 2 | Vamp5 |

| | | | | | | | |
|---|---|---|---|---|---|---|---|
| Crim11 | 0 | 1.289847567 | 0.776 | 0.323 | 0 | 2 | Crim1 |
| Smco41 | 0 | 1.286551047 | 0.794 | 0.341 | 0 | 2 | Smco4 |
| Ptprg1 | 0 | 1.285754007 | 0.831 | 0.284 | 0 | 2 | Ptprg |
| Sptbn11 | 0 | 1.2842749 | 0.983 | 0.434 | 0 | 2 | Sptbn1 |
| Nostrin1 | 0 | 1.278699636 | 0.644 | 0.211 | 0 | 2 | Nostrin |
| Klf21 | 0 | 1.27787565 | 0.798 | 0.314 | 0 | 2 | Klf2 |
| Ptprb1 | 0 | 1.272506822 | 0.954 | 0.295 | 0 | 2 | Ptprb |
| Clic51 | 0 | 1.271954241 | 0.741 | 0.236 | 0 | 2 | Clic5 |
| Gm138891 | 0 | 1.264266258 | 0.557 | 0.178 | 0 | 2 | Gm13889 |
| Slc43a31 | 0 | 1.263091457 | 0.632 | 0.225 | 0 | 2 | Slc43a3 |
| Rasal21 | 0 | 1.261123433 | 0.735 | 0.263 | 0 | 2 | Rasal2 |
| Mgll1 | 0 | 1.259571985 | 0.921 | 0.391 | 0 | 2 | Mgll |
| Ebf11 | 0 | 1.253083778 | 0.926 | 0.306 | 0 | 2 | Ebf1 |
| Sparc1 | 0 | 1.252482652 | 0.924 | 0.338 | 0 | 2 | Sparc |
| Dlc11 | 0 | 1.241947854 | 0.842 | 0.303 | 0 | 2 | Dlc1 |
| Gm120021 | 0 | 1.237470098 | 0.516 | 0.148 | 0 | 2 | Gm12002 |
| Apbb21 | 0 | 1.232134873 | 0.789 | 0.315 | 0 | 2 | Apbb2 |
| Tnfsf101 | 0 | 1.226253647 | 0.768 | 0.326 | 0 | 2 | Tnfsf10 |
| Adgrf51 | 0 | 1.226249936 | 0.843 | 0.275 | 0 | 2 | Adgrf5 |
| Etl41 | 0 | 1.217172272 | 0.7 | 0.244 | 0 | 2 | Etl4 |
| Ldb21 | 0 | 1.209692525 | 0.784 | 0.274 | 0 | 2 | Ldb2 |
| Piezo21 | 0 | 1.198930583 | 0.626 | 0.207 | 0 | 2 | Piezo2 |
| Hmcn11 | 0 | 1.19235314 | 0.523 | 0.173 | 0 | 2 | Hmcn1 |
| Dysf1 | 0 | 1.191567263 | 0.589 | 0.196 | 0 | 2 | Dysf |
| Hspg21 | 0 | 1.186831522 | 0.744 | 0.258 | 0 | 2 | Hspg2 |
| Rbms11 | 0 | 1.186289611 | 0.867 | 0.439 | 0 | 2 | Rbms1 |
| App1 | 0 | 1.184484663 | 0.879 | 0.363 | 0 | 2 | App |
| Tie11 | 0 | 1.181924741 | 0.624 | 0.208 | 0 | 2 | Tie1 |
| Rdx1 | 0 | 1.181226772 | 0.932 | 0.504 | 0 | 2 | Rdx |
| H2-T231 | 0 | 1.173788493 | 0.763 | 0.423 | 0 | 2 | H2-T23 |
| Kank31 | 0 | 1.166761888 | 0.564 | 0.187 | 0 | 2 | Kank3 |
| Cavin31 | 0 | 1.16338835 | 0.686 | 0.261 | 0 | 2 | Cavin3 |
| Srgn1 | 0 | 1.161277629 | 0.929 | 0.497 | 0 | 2 | Srgn |
| Fmo21 | 0 | 1.157343279 | 0.679 | 0.241 | 0 | 2 | Fmo2 |
| Rapgef41 | 0 | 1.157119298 | 0.615 | 0.214 | 0 | 2 | Rapgef4 |
| Myct11 | 0 | 1.151824265 | 0.543 | 0.183 | 0 | 2 | Myct1 |
| Hecw21 | 0 | 1.145191779 | 0.587 | 0.207 | 0 | 2 | Hecw2 |
| Cxcl121 | 0 | 1.141759498 | 0.714 | 0.274 | 0 | 2 | Cxcl12 |
| Jam21 | 0 | 1.139908431 | 0.676 | 0.238 | 0 | 2 | Jam2 |
| Bcam1 | 0 | 1.139439958 | 0.614 | 0.21 | 0 | 2 | Bcam |
| Grrp11 | 0 | 1.138311202 | 0.558 | 0.178 | 0 | 2 | Grrp1 |
| Kctd12b1 | 0 | 1.138070977 | 0.59 | 0.208 | 0 | 2 | Kctd12b |
| Dock41 | 0 | 1.128423356 | 0.632 | 0.256 | 0 | 2 | Dock4 |

| | | | | | | | |
|---|---|---|---|---|---|---|---|
| Ece11 | 0 | 1.128124448 | 0.668 | 0.262 | 0 | 2 | Ece1 |
| Rgl11 | 0 | 1.124867599 | 0.629 | 0.248 | 0 | 2 | Rgl1 |
| Ahr1 | 0 | 1.119625445 | 0.532 | 0.211 | 0 | 2 | Ahr |
| Fbxl71 | 0 | 1.110554656 | 0.581 | 0.218 | 0 | 2 | Fbxl7 |
| Eva1b1 | 0 | 1.110218204 | 0.602 | 0.247 | 0 | 2 | Eva1b |
| Acvrl11 | 0 | 1.108838648 | 0.588 | 0.237 | 0 | 2 | Acvrl1 |
| Tmem881 | 0 | 1.105707499 | 0.522 | 0.172 | 0 | 2 | Tmem88 |
| 4931406P16Rik1 | 0 | 1.10420437 | 0.708 | 0.298 | 0 | 2 | 4931406P16Rik |
| Ly6e1 | 0 | 1.102886472 | 0.952 | 0.578 | 0 | 2 | Ly6e |
| Kitl1 | 0 | 1.102414486 | 0.752 | 0.285 | 0 | 2 | Kitl |
| Rasgrp31 | 0 | 1.099757201 | 0.659 | 0.236 | 0 | 2 | Rasgrp3 |
| Pcdh171 | 0 | 1.098684337 | 0.572 | 0.198 | 0 | 2 | Pcdh17 |
| Ushbp11 | 0 | 1.097438 | 0.537 | 0.18 | 0 | 2 | Ushbp1 |
| Elk31 | 0 | 1.096799702 | 0.685 | 0.317 | 0 | 2 | Elk3 |
| H2-D11 | 0 | 1.095348861 | 0.983 | 0.63 | 0 | 2 | H2-D1 |
| Myo61 | 0 | 1.092902557 | 0.609 | 0.254 | 0 | 2 | Myo6 |
| Fzd41 | 0 | 1.091763919 | 0.541 | 0.191 | 0 | 2 | Fzd4 |
| Itm2b1 | 0 | 1.090799998 | 0.981 | 0.694 | 0 | 2 | Itm2b |
| 9530026P05Rik1 | 0 | 1.090554137 | 0.541 | 0.205 | 0 | 2 | 9530026P05Rik |
| BC0285281 | 0 | 1.089192049 | 0.52 | 0.199 | 0 | 2 | BC028528 |
| Klf71 | 0 | 1.081445738 | 0.614 | 0.259 | 0 | 2 | Klf7 |
| Cd24a1 | 0 | 1.07986594 | 0.701 | 0.299 | 0 | 2 | Cd24a |
| Calm11 | 0 | 1.079226157 | 0.985 | 0.756 | 0 | 2 | Calm1 |
| Cav11 | 0 | 1.078541655 | 0.713 | 0.254 | 0 | 2 | Cav1 |
| Abcg21 | 0 | 1.076737815 | 0.575 | 0.234 | 0 | 2 | Abcg2 |
| Pdgfb1 | 0 | 1.072076137 | 0.589 | 0.276 | 0 | 2 | Pdgfb |
| Icam21 | 0 | 1.070202536 | 0.546 | 0.191 | 0 | 2 | Icam2 |
| Lama31 | 0 | 1.066130038 | 0.513 | 0.186 | 0 | 2 | Lama3 |
| Thbd1 | 0 | 1.060011671 | 0.574 | 0.216 | 0 | 2 | Thbd |
| Tek1 | 0 | 1.05703285 | 0.614 | 0.217 | 0 | 2 | Tek |
| Slfn51 | 0 | 1.048695468 | 0.739 | 0.31 | 0 | 2 | Slfn5 |
| H2-K11 | 0 | 1.045460231 | 0.975 | 0.602 | 0 | 2 | H2-K1 |
| Ntf31 | 0 | 1.044599693 | 0.445 | 0.149 | 0 | 2 | Ntf3 |
| Ednrb1 | 0 | 1.043872935 | 0.483 | 0.159 | 0 | 2 | Ednrb |
| Tshz21 | 0 | 1.043323002 | 0.765 | 0.324 | 0 | 2 | Tshz2 |
| Cd931 | 0 | 1.040773253 | 0.594 | 0.215 | 0 | 2 | Cd93 |
| Calcrl1 | 0 | 1.040127719 | 0.565 | 0.227 | 0 | 2 | Calcrl |
| Cd471 | 0 | 1.037583952 | 0.832 | 0.471 | 0 | 2 | Cd47 |
| Meox11 | 0 | 1.035971372 | 0.47 | 0.153 | 0 | 2 | Meox1 |
| Mcf2l1 | 0 | 1.033945771 | 0.494 | 0.18 | 0 | 2 | Mcf2l |
| Hmbox11 | 0 | 1.030067092 | 0.551 | 0.24 | 0 | 2 | Hmbox1 |
| Itga11 | 0 | 1.029954449 | 0.789 | 0.369 | 0 | 2 | Itga1 |
| Crip21 | 0 | 1.025674604 | 0.798 | 0.369 | 0 | 2 | Crip2 |

| | | | | | | | |
|---|---|---|---|---|---|---|---|
| Numb1 | 0 | 1.024094628 | 0.617 | 0.304 | 0 | 2 | Numb |
| Epas11 | 0 | 1.023068833 | 0.768 | 0.309 | 0 | 2 | Epas1 |
| Ltbp41 | 0 | 1.02112392 | 0.675 | 0.265 | 0 | 2 | Ltbp4 |
| Bmpr21 | 0 | 1.020887326 | 0.703 | 0.306 | 0 | 2 | Bmpr2 |
| Gas61 | 0 | 1.017175068 | 0.645 | 0.255 | 0 | 2 | Gas6 |
| Plekhg11 | 0 | 1.015238793 | 0.548 | 0.208 | 0 | 2 | Plekhg1 |
| Tmem1401 | 0 | 1.012962186 | 0.487 | 0.191 | 0 | 2 | Tmem140 |
| Gngt21 | 0 | 1.006980138 | 0.674 | 0.321 | 0 | 2 | Gngt2 |
| Chst21 | 0 | 1.00616905 | 0.43 | 0.133 | 0 | 2 | Chst2 |
| Sema6a1 | 0 | 1.002264016 | 0.475 | 0.177 | 0 | 2 | Sema6a |
| Fermt21 | 0 | 1.00037091 | 0.705 | 0.307 | 0 | 2 | Fermt2 |
| Clic41 | 0 | 0.9995786 | 0.776 | 0.38 | 0 | 2 | Clic4 |
| Tmem2041 | 0 | 0.994040052 | 0.477 | 0.177 | 0 | 2 | Tmem204 |
| Rai141 | 0 | 0.993249774 | 0.512 | 0.198 | 0 | 2 | Rai14 |
| Aplp21 | 0 | 0.988748133 | 0.775 | 0.372 | 0 | 2 | Aplp2 |
| Rflnb1 | 0 | 0.98869017 | 0.744 | 0.261 | 0 | 2 | Rflnb |
| Adgrl21 | 0 | 0.988371413 | 0.526 | 0.202 | 0 | 2 | Adgrl2 |
| Ecscr1 | 0 | 0.985940108 | 0.54 | 0.19 | 0 | 2 | Ecscr |
| Mmrn21 | 0 | 0.985608883 | 0.533 | 0.196 | 0 | 2 | Mmrn2 |
| Tspan91 | 0 | 0.984227428 | 0.541 | 0.232 | 0 | 2 | Tspan9 |
| Itga61 | 0 | 0.983823377 | 0.569 | 0.249 | 0 | 2 | Itga6 |
| Prss231 | 0 | 0.98121027 | 0.588 | 0.244 | 0 | 2 | Prss23 |
| Fgd51 | 0 | 0.973406548 | 0.521 | 0.193 | 0 | 2 | Fgd5 |
| Lims21 | 0 | 0.96765027 | 0.466 | 0.17 | 0 | 2 | Lims2 |
| Bst21 | 0 | 0.963927873 | 0.851 | 0.461 | 0 | 2 | Bst2 |
| Fez21 | 0 | 0.952058466 | 0.572 | 0.248 | 0 | 2 | Fez2 |
| Mfge81 | 0 | 0.95044939 | 0.53 | 0.221 | 0 | 2 | Mfge8 |
| Sptan11 | 0 | 0.948310577 | 0.631 | 0.309 | 0 | 2 | Sptan1 |
| S1pr11 | 0 | 0.943367393 | 0.562 | 0.229 | 0 | 2 | S1pr1 |
| Tspan121 | 0 | 0.940381434 | 0.52 | 0.197 | 0 | 2 | Tspan12 |
| Cald11 | 0 | 0.933212224 | 0.673 | 0.279 | 0 | 2 | Cald1 |
| Arhgef121 | 0 | 0.921607996 | 0.67 | 0.309 | 0 | 2 | Arhgef12 |
| Malat11 | 0 | 0.910466137 | 0.996 | 0.692 | 0 | 2 | Malat1 |
| Ets11 | 0 | 0.903834974 | 0.705 | 0.367 | 0 | 2 | Ets1 |
| Ctsl1 | 0 | 0.901735615 | 0.647 | 0.31 | 0 | 2 | Ctsl |
| Sorbs21 | 0 | 0.898841312 | 0.579 | 0.241 | 0 | 2 | Sorbs2 |
| Prex21 | 0 | 0.897975966 | 0.585 | 0.235 | 0 | 2 | Prex2 |
| Laptm4a1 | 0 | 0.8843163 | 0.798 | 0.446 | 0 | 2 | Laptm4a |
| Gnai21 | 0 | 0.878357339 | 0.882 | 0.537 | 0 | 2 | Gnai2 |
| Selenom1 | 0 | 0.827026948 | 0.792 | 0.403 | 0 | 2 | Selenom |
| B2m1 | 0 | 0.765156162 | 0.974 | 0.656 | 0 | 2 | B2m |
| Ccdc85a1 | 1.41E-304 | 0.982009363 | 0.434 | 0.151 | 2.85E-300 | 2 | Ccdc85a |
| Asap11 | 1.61E-304 | 0.981256883 | 0.635 | 0.323 | 3.27E-300 | 2 | Asap1 |

| | | | | | | | |
|---|---|---|---|---|---|---|---|
| Clec1a1 | 4.91E-301 | 0.972147377 | 0.407 | 0.14 | 9.94E-297 | 2 | Clec1a |
| Pdgfd1 | 9.27E-297 | 0.875903453 | 0.545 | 0.218 | 1.88E-292 | 2 | Pdgfd |
| Jup1 | 1.20E-296 | 0.909055034 | 0.529 | 0.225 | 2.44E-292 | 2 | Jup |
| Itpkb1 | 6.69E-296 | 0.948612719 | 0.63 | 0.323 | 1.35E-291 | 2 | Itpkb |
| Syne21 | 4.08E-295 | 0.946465639 | 0.549 | 0.239 | 8.26E-291 | 2 | Syne2 |
| Arl151 | 1.13E-293 | 0.92994272 | 0.668 | 0.347 | 2.28E-289 | 2 | Arl15 |
| Bvht1 | 6.55E-293 | 0.857901085 | 0.431 | 0.157 | 1.33E-288 | 2 | Bvht |
| Ptk21 | 1.55E-292 | 0.911195158 | 0.543 | 0.233 | 3.13E-288 | 2 | Ptk2 |
| Cav21 | 5.59E-292 | 0.888039644 | 0.461 | 0.179 | 1.13E-287 | 2 | Cav2 |
| Anxa31 | 7.99E-292 | 0.848811877 | 0.631 | 0.303 | 1.62E-287 | 2 | Anxa3 |
| Klf131 | 4.93E-288 | 0.916476417 | 0.616 | 0.318 | 9.99E-284 | 2 | Klf13 |
| Qk1 | 3.20E-287 | 0.88159019 | 0.691 | 0.376 | 6.48E-283 | 2 | Qk |
| Plxna21 | 5.76E-287 | 0.948775362 | 0.476 | 0.188 | 1.17E-282 | 2 | Plxna2 |
| Serpinh11 | 1.62E-285 | 0.847690716 | 0.522 | 0.217 | 3.28E-281 | 2 | Serpinh1 |
| Actn41 | 7.04E-284 | 0.871406374 | 0.651 | 0.343 | 1.43E-279 | 2 | Actn4 |
| Robo41 | 1.71E-283 | 0.900856016 | 0.411 | 0.148 | 3.46E-279 | 2 | Robo4 |
| Stox21 | 7.40E-283 | 0.937495861 | 0.521 | 0.214 | 1.50E-278 | 2 | Stox2 |
| Tmtc21 | 9.03E-283 | 1.010905911 | 0.425 | 0.156 | 1.83E-278 | 2 | Tmtc2 |
| Tmem591 | 9.45E-283 | 0.876633945 | 0.739 | 0.461 | 1.91E-278 | 2 | Tmem59 |
| Rbpms1 | 9.96E-282 | 0.849715302 | 0.63 | 0.316 | 2.02E-277 | 2 | Rbpms |
| Col15a11 | 5.50E-281 | 0.7917421 | 0.526 | 0.213 | 1.11E-276 | 2 | Col15a1 |
| Fbln21 | 5.82E-279 | 0.846077329 | 0.568 | 0.236 | 1.18E-274 | 2 | Fbln2 |
| Oaz21 | 6.24E-279 | 0.942442097 | 0.491 | 0.206 | 1.26E-274 | 2 | Oaz2 |
| Chst151 | 1.65E-277 | 0.893662267 | 0.407 | 0.147 | 3.33E-273 | 2 | Chst15 |
| Iigp11 | 5.80E-277 | 1.081761693 | 0.5 | 0.201 | 1.17E-272 | 2 | Iigp1 |
| Cdc42ep21 | 2.87E-274 | 0.842942951 | 0.4 | 0.146 | 5.81E-270 | 2 | Cdc42ep2 |
| Rras1 | 3.64E-274 | 0.870167454 | 0.561 | 0.266 | 7.36E-270 | 2 | Rras |
| Cpne81 | 3.79E-273 | 0.861011172 | 0.482 | 0.198 | 7.67E-269 | 2 | Cpne8 |
| H2-Q41 | 6.83E-269 | 0.94141063 | 0.553 | 0.28 | 1.38E-264 | 2 | H2-Q4 |
| Rasip11 | 4.08E-266 | 0.846141252 | 0.433 | 0.166 | 8.26E-262 | 2 | Rasip1 |
| Stard81 | 1.21E-259 | 0.900498326 | 0.43 | 0.17 | 2.45E-255 | 2 | Stard8 |
| Samd121 | 1.69E-259 | 0.860075487 | 0.49 | 0.198 | 3.43E-255 | 2 | Samd12 |
| Ptgs11 | 2.94E-258 | 0.888936335 | 0.448 | 0.184 | 5.95E-254 | 2 | Ptgs1 |
| Cd381 | 2.67E-254 | 0.922918095 | 0.467 | 0.204 | 5.41E-250 | 2 | Cd38 |
| Zfp3661 | 6.08E-251 | 0.924005689 | 0.407 | 0.155 | 1.23E-246 | 2 | Zfp366 |
| Tubb4b1 | 6.28E-251 | 0.835475434 | 0.681 | 0.397 | 1.27E-246 | 2 | Tubb4b |
| Ccdc85b1 | 4.65E-250 | 0.889964533 | 0.489 | 0.229 | 9.42E-246 | 2 | Ccdc85b |
| Utrn1 | 3.88E-248 | 0.777419975 | 0.682 | 0.373 | 7.86E-244 | 2 | Utrn |
| Emp21 | 7.25E-246 | 0.814576103 | 0.478 | 0.207 | 1.47E-241 | 2 | Emp2 |
| B4galt41 | 1.10E-245 | 0.844232152 | 0.371 | 0.137 | 2.22E-241 | 2 | B4galt4 |
| Gimap61 | 1.20E-245 | 0.824255469 | 0.641 | 0.333 | 2.42E-241 | 2 | Gimap6 |
| Lmcd11 | 2.71E-245 | 0.823952525 | 0.38 | 0.139 | 5.50E-241 | 2 | Lmcd1 |
| Shank31 | 8.79E-244 | 0.925832814 | 0.415 | 0.164 | 1.78E-239 | 2 | Shank3 |

| | | | | | | | |
|---|---|---|---|---|---|---|---|
| Cpd1 | 3.53E-242 | 0.863262874 | 0.556 | 0.283 | 7.15E-238 | 2 | Cpd |
| Bsg1 | 1.44E-240 | 0.814782554 | 0.687 | 0.421 | 2.91E-236 | 2 | Bsg |
| Rapgef21 | 5.58E-240 | 0.903018702 | 0.584 | 0.31 | 1.13E-235 | 2 | Rapgef2 |
| Prkch1 | 6.13E-240 | 0.770852484 | 0.693 | 0.364 | 1.24E-235 | 2 | Prkch |
| Dhrs31 | 1.58E-238 | 0.894858632 | 0.459 | 0.204 | 3.20E-234 | 2 | Dhrs3 |
| Meis21 | 9.12E-238 | 0.925080089 | 0.545 | 0.251 | 1.85E-233 | 2 | Meis2 |
| Rapgef31 | 8.60E-237 | 0.871486722 | 0.377 | 0.144 | 1.74E-232 | 2 | Rapgef3 |
| Tspan131 | 1.08E-236 | 0.930164374 | 0.515 | 0.258 | 2.19E-232 | 2 | Tspan13 |
| Slc50a11 | 1.36E-236 | 0.909580952 | 0.52 | 0.273 | 2.76E-232 | 2 | Slc50a1 |
| Arhgef151 | 6.74E-236 | 0.81648465 | 0.379 | 0.144 | 1.37E-231 | 2 | Arhgef15 |
| Dab2ip1 | 9.78E-236 | 0.854573707 | 0.495 | 0.234 | 1.98E-231 | 2 | Dab2ip |
| Sema7a1 | 1.81E-235 | 0.853634447 | 0.352 | 0.126 | 3.66E-231 | 2 | Sema7a |
| Dnm31 | 5.27E-234 | 0.828647127 | 0.476 | 0.201 | 1.07E-229 | 2 | Dnm3 |
| Tspan151 | 8.40E-233 | 0.782463056 | 0.32 | 0.108 | 1.70E-228 | 2 | Tspan15 |
| Car81 | 8.88E-232 | 0.831815383 | 0.409 | 0.16 | 1.80E-227 | 2 | Car8 |
| Nxn1 | 1.32E-231 | 0.844967287 | 0.508 | 0.24 | 2.66E-227 | 2 | Nxn |
| Itih51 | 1.96E-231 | 0.796472917 | 0.482 | 0.215 | 3.97E-227 | 2 | Itih5 |
| H2-Q71 | 1.67E-230 | 0.817745077 | 0.659 | 0.367 | 3.38E-226 | 2 | H2-Q7 |
| Slc6a61 | 8.89E-230 | 0.85829514 | 0.585 | 0.297 | 1.80E-225 | 2 | Slc6a6 |
| Sgk11 | 1.13E-224 | 0.866931436 | 0.514 | 0.241 | 2.29E-220 | 2 | Sgk1 |
| Luzp11 | 5.15E-223 | 0.779337433 | 0.594 | 0.329 | 1.04E-218 | 2 | Luzp1 |
| Rtl8a1 | 7.88E-222 | 0.817458229 | 0.38 | 0.152 | 1.60E-217 | 2 | Rtl8a |
| Klf41 | 3.28E-221 | 0.823324376 | 0.514 | 0.24 | 6.64E-217 | 2 | Klf4 |
| Pttg1ip1 | 3.32E-219 | 0.833848799 | 0.486 | 0.243 | 6.72E-215 | 2 | Pttg1ip |
| Grb101 | 5.27E-219 | 0.855835231 | 0.406 | 0.17 | 1.07E-214 | 2 | Grb10 |
| Magi11 | 9.51E-219 | 0.819470453 | 0.463 | 0.204 | 1.93E-214 | 2 | Magi1 |
| Clstn11 | 9.51E-219 | 0.809734647 | 0.455 | 0.211 | 1.93E-214 | 2 | Clstn1 |
| Fut81 | 1.53E-217 | 0.910561957 | 0.511 | 0.263 | 3.09E-213 | 2 | Fut8 |
| Selenop1 | 1.73E-216 | 0.726029258 | 0.617 | 0.333 | 3.49E-212 | 2 | Selenop |
| Fli11 | 2.06E-216 | 0.793188459 | 0.65 | 0.373 | 4.16E-212 | 2 | Fli1 |
| AW112010 | 1.34E-215 | 0.421613921 | 0.896 | 0.528 | 2.72E-211 | 2 | AW112010 |
| Psmb81 | 2.26E-214 | 0.707870773 | 0.813 | 0.527 | 4.57E-210 | 2 | Psmb8 |
| Ndrg11 | 5.55E-214 | 0.766036735 | 0.459 | 0.206 | 1.12E-209 | 2 | Ndrg1 |
| Cd1511 | 1.52E-213 | 0.821179864 | 0.425 | 0.193 | 3.07E-209 | 2 | Cd151 |
| S100a161 | 2.40E-213 | 0.737431451 | 0.487 | 0.231 | 4.87E-209 | 2 | S100a16 |
| Nfkbia1 | 6.14E-213 | 0.78369539 | 0.634 | 0.373 | 1.24E-208 | 2 | Nfkbia |
| Tnfaip11 | 8.02E-213 | 0.765067257 | 0.413 | 0.178 | 1.62E-208 | 2 | Tnfaip1 |
| Psmb91 | 1.35E-212 | 0.772453748 | 0.63 | 0.379 | 2.74E-208 | 2 | Psmb9 |
| Ppp1r21 | 3.60E-212 | 0.710564749 | 0.638 | 0.364 | 7.28E-208 | 2 | Ppp1r2 |
| Sash11 | 1.41E-211 | 0.763381263 | 0.486 | 0.23 | 2.85E-207 | 2 | Sash1 |
| Ifi351 | 7.23E-211 | 0.839253459 | 0.485 | 0.249 | 1.46E-206 | 2 | Ifi35 |
| Msn1 | 1.21E-210 | 0.73400957 | 0.683 | 0.415 | 2.44E-206 | 2 | Msn |
| Pitpnc11 | 3.80E-209 | 0.778062517 | 0.686 | 0.399 | 7.69E-205 | 2 | Pitpnc1 |

| | | | | | | | |
|---|---|---|---|---|---|---|---|
| Afdn1 | 8.97E-209 | 0.768461221 | 0.545 | 0.288 | 1.82E-204 | 2 | Afdn |
| Esm11 | 2.73E-205 | 1.040723778 | 0.272 | 0.083 | 5.52E-201 | 2 | Esm1 |
| Phlpp11 | 3.65E-205 | 0.878896125 | 0.48 | 0.238 | 7.39E-201 | 2 | Phlpp1 |
| Plxnd11 | 6.19E-205 | 0.824300611 | 0.378 | 0.158 | 1.25E-200 | 2 | Plxnd1 |
| Lrrc8c1 | 1.02E-201 | 0.858010917 | 0.53 | 0.286 | 2.06E-197 | 2 | Lrrc8c |
| Efna11 | 1.87E-201 | 0.782697854 | 0.324 | 0.122 | 3.78E-197 | 2 | Efna1 |
| Psmb101 | 2.24E-201 | 0.81270478 | 0.608 | 0.376 | 4.54E-197 | 2 | Psmb10 |
| Sh3bp51 | 8.82E-201 | 0.78420468 | 0.428 | 0.192 | 1.79E-196 | 2 | Sh3bp5 |
| Serinc31 | 1.13E-200 | 0.691396739 | 0.73 | 0.458 | 2.30E-196 | 2 | Serinc3 |
| Wwtr11 | 1.41E-200 | 0.739695737 | 0.461 | 0.214 | 2.86E-196 | 2 | Wwtr1 |
| Lpcat11 | 4.98E-199 | 0.79342774 | 0.344 | 0.137 | 1.01E-194 | 2 | Lpcat1 |
| Rnf2201 | 6.01E-199 | 0.910799821 | 0.453 | 0.217 | 1.22E-194 | 2 | Rnf220 |
| Ifi2031 | 1.20E-198 | 0.811815568 | 0.585 | 0.342 | 2.43E-194 | 2 | Ifi203 |
| Nr3c11 | 1.47E-198 | 0.766685628 | 0.608 | 0.357 | 2.98E-194 | 2 | Nr3c1 |
| Csgalnact11 | 1.49E-197 | 0.815135748 | 0.399 | 0.172 | 3.02E-193 | 2 | Csgalnact1 |
| Son1 | 1.69E-197 | 0.711463429 | 0.761 | 0.498 | 3.42E-193 | 2 | Son |
| Phldb21 | 3.24E-196 | 0.744232371 | 0.408 | 0.179 | 6.56E-192 | 2 | Phldb2 |
| Ccdc28b1 | 1.29E-194 | 0.698493201 | 0.48 | 0.236 | 2.61E-190 | 2 | Ccdc28b |
| Macf11 | 2.84E-194 | 0.67243153 | 0.772 | 0.471 | 5.74E-190 | 2 | Macf1 |
| Rabac11 | 5.76E-194 | 0.627877061 | 0.773 | 0.54 | 1.17E-189 | 2 | Rabac1 |
| Gnas1 | 1.07E-193 | 0.470302582 | 0.976 | 0.849 | 2.17E-189 | 2 | Gnas |
| Uaca1 | 1.87E-193 | 0.697160494 | 0.442 | 0.201 | 3.79E-189 | 2 | Uaca |
| Dok41 | 2.11E-193 | 0.709151789 | 0.285 | 0.099 | 4.28E-189 | 2 | Dok4 |
| Filip11 | 2.40E-192 | 0.77664527 | 0.395 | 0.171 | 4.86E-188 | 2 | Filip1 |
| Ahnak1 | 6.35E-191 | 0.597680912 | 0.762 | 0.437 | 1.29E-186 | 2 | Ahnak |
| Wasf21 | 6.92E-191 | 0.708303357 | 0.643 | 0.4 | 1.40E-186 | 2 | Wasf2 |
| Rhoj1 | 8.81E-191 | 0.691505281 | 0.466 | 0.215 | 1.78E-186 | 2 | Rhoj |
| S100a131 | 3.52E-188 | 0.593531556 | 0.753 | 0.511 | 7.13E-184 | 2 | S100a13 |
| Oxr11 | 7.07E-188 | 0.828808094 | 0.549 | 0.311 | 1.43E-183 | 2 | Oxr1 |
| Lysmd21 | 2.85E-187 | 0.764008896 | 0.377 | 0.17 | 5.77E-183 | 2 | Lysmd2 |
| Ackr31 | 4.52E-187 | 0.816946923 | 0.334 | 0.131 | 9.15E-183 | 2 | Ackr3 |
| Crip11 | 1.24E-186 | 0.612744954 | 0.774 | 0.508 | 2.50E-182 | 2 | Crip1 |
| Selenof1 | 2.70E-186 | 0.614425437 | 0.699 | 0.467 | 5.48E-182 | 2 | Selenof |
| Mkl21 | 7.00E-184 | 0.798889839 | 0.46 | 0.231 | 1.42E-179 | 2 | Mkl2 |
| Unc45b1 | 1.57E-183 | 0.661863296 | 0.313 | 0.119 | 3.19E-179 | 2 | Unc45b |
| Prkd11 | 2.06E-183 | 0.840669232 | 0.323 | 0.127 | 4.17E-179 | 2 | Prkd1 |
| Heg11 | 6.96E-183 | 0.686089765 | 0.497 | 0.248 | 1.41E-178 | 2 | Heg1 |
| Rhoc1 | 1.46E-182 | 0.756861937 | 0.5 | 0.273 | 2.96E-178 | 2 | Rhoc |
| Anxa21 | 3.28E-182 | 0.684148336 | 0.644 | 0.39 | 6.63E-178 | 2 | Anxa2 |
| Rab131 | 4.83E-182 | 0.653600738 | 0.307 | 0.118 | 9.79E-178 | 2 | Rab13 |
| Gbp71 | 1.35E-181 | 0.885098112 | 0.471 | 0.246 | 2.73E-177 | 2 | Gbp7 |
| Sncg1 | 2.49E-181 | 0.697150087 | 0.338 | 0.133 | 5.04E-177 | 2 | Sncg |
| Prdx41 | 4.23E-181 | 0.749797078 | 0.415 | 0.199 | 8.55E-177 | 2 | Prdx4 |

| | | | | | | | |
|---|---|---|---|---|---|---|---|
| F11r1 | 4.23E-181 | 0.698124955 | 0.475 | 0.241 | 8.57E-177 | 2 | F11r |
| Grasp1 | 1.41E-179 | 0.748311126 | 0.344 | 0.146 | 2.85E-175 | 2 | Grasp |
| Ctnnb11 | 1.95E-179 | 0.672033596 | 0.579 | 0.338 | 3.94E-175 | 2 | Ctnnb1 |
| Adgrg11 | 2.55E-179 | 0.723713898 | 0.308 | 0.12 | 5.16E-175 | 2 | Adgrg1 |
| Itm2a1 | 4.63E-179 | 0.757178744 | 0.366 | 0.158 | 9.38E-175 | 2 | Itm2a |
| Hip11 | 1.10E-177 | 0.787761132 | 0.542 | 0.31 | 2.22E-173 | 2 | Hip1 |
| Tjp11 | 2.96E-177 | 0.745232079 | 0.468 | 0.24 | 6.00E-173 | 2 | Tjp1 |
| Pde7b1 | 1.12E-176 | 0.806538569 | 0.449 | 0.218 | 2.27E-172 | 2 | Pde7b |
| Sep-41 | 5.10E-176 | 0.716250387 | 0.34 | 0.144 | 1.03E-171 | 2 | 4-Sep |
| Ces2e1 | 5.62E-176 | 0.687207862 | 0.265 | 0.092 | 1.14E-171 | 2 | Ces2e |
| Ctnnbip11 | 6.04E-176 | 0.697799655 | 0.391 | 0.182 | 1.22E-171 | 2 | Ctnnbip1 |
| Ets21 | 6.24E-176 | 0.704837945 | 0.383 | 0.172 | 1.26E-171 | 2 | Ets2 |
| Xdh1 | 1.29E-174 | 0.7321509 | 0.373 | 0.166 | 2.61E-170 | 2 | Xdh |
| 1-Jun | 6.34E-174 | 0.790523792 | 0.749 | 0.486 | 1.28E-169 | 2 | Jun |
| mt-Co21 | 2.61E-173 | 0.53225486 | 0.996 | 0.696 | 5.29E-169 | 2 | mt-Co2 |
| Ncoa71 | 1.36E-172 | 0.786420841 | 0.493 | 0.273 | 2.75E-168 | 2 | Ncoa7 |
| Ripply31 | 1.92E-172 | 0.711903596 | 0.314 | 0.127 | 3.89E-168 | 2 | Ripply3 |
| Reep31 | 3.41E-171 | 0.743330547 | 0.505 | 0.283 | 6.90E-167 | 2 | Reep3 |
| Nav11 | 1.09E-170 | 0.743741439 | 0.423 | 0.199 | 2.21E-166 | 2 | Nav1 |
| Plekho11 | 5.31E-170 | 0.709379729 | 0.413 | 0.202 | 1.07E-165 | 2 | Plekho1 |
| Tmed51 | 1.00E-169 | 0.809753323 | 0.471 | 0.268 | 2.03E-165 | 2 | Tmed5 |
| Rarg1 | 4.03E-169 | 0.743109999 | 0.375 | 0.174 | 8.16E-165 | 2 | Rarg |
| Fam43a1 | 2.18E-168 | 0.673402325 | 0.31 | 0.123 | 4.42E-164 | 2 | Fam43a |
| Palmd1 | 1.14E-167 | 0.64051241 | 0.438 | 0.209 | 2.32E-163 | 2 | Palmd |
| Arap31 | 1.20E-167 | 0.777840529 | 0.399 | 0.197 | 2.43E-163 | 2 | Arap3 |
| Tmem176a1 | 1.34E-166 | 0.584807737 | 0.666 | 0.388 | 2.72E-162 | 2 | Tmem176a |
| Fkbp1a1 | 2.08E-166 | 0.628430768 | 0.611 | 0.382 | 4.21E-162 | 2 | Fkbp1a |
| Tdrp1 | 3.84E-166 | 0.667000767 | 0.269 | 0.098 | 7.77E-162 | 2 | Tdrp |
| Ephb41 | 6.47E-165 | 0.737007111 | 0.338 | 0.146 | 1.31E-160 | 2 | Ephb4 |
| Trp53i111 | 5.13E-164 | 0.779054497 | 0.328 | 0.141 | 1.04E-159 | 2 | Trp53i11 |
| Dock11 | 8.24E-164 | 0.732812371 | 0.48 | 0.257 | 1.67E-159 | 2 | Dock1 |
| Sntb21 | 1.85E-163 | 0.74807254 | 0.452 | 0.244 | 3.74E-159 | 2 | Sntb2 |
| mt-Cytb1 | 1.04E-162 | 0.480595761 | 0.976 | 0.667 | 2.11E-158 | 2 | mt-Cytb |
| Smagp1 | 1.84E-162 | 0.592398113 | 0.424 | 0.209 | 3.72E-158 | 2 | Smagp |
| Fabp51 | 4.77E-162 | 0.711785692 | 0.547 | 0.299 | 9.65E-158 | 2 | Fabp5 |
| Trib21 | 1.02E-161 | 0.6594129 | 0.367 | 0.166 | 2.07E-157 | 2 | Trib2 |
| Mndal1 | 1.86E-161 | 0.706505591 | 0.609 | 0.382 | 3.76E-157 | 2 | Mndal |
| Clec2d1 | 2.19E-161 | 0.68416006 | 0.727 | 0.505 | 4.44E-157 | 2 | Clec2d |
| Fam212a1 | 2.71E-161 | 0.685964914 | 0.28 | 0.11 | 5.48E-157 | 2 | Fam212a |
| Tspan181 | 4.40E-161 | 0.696985182 | 0.298 | 0.119 | 8.91E-157 | 2 | Tspan18 |
| Arhgap181 | 9.44E-161 | 0.752710114 | 0.533 | 0.32 | 1.91E-156 | 2 | Arhgap18 |
| Rap1a1 | 1.63E-160 | 0.571400875 | 0.744 | 0.512 | 3.30E-156 | 2 | Rap1a |
| Dusp31 | 2.24E-160 | 0.718086283 | 0.37 | 0.171 | 4.54E-156 | 2 | Dusp3 |

| | | | | | | | |
|---|---|---|---|---|---|---|---|
| Nrp21 | 2.89E-158 | 0.719664992 | 0.362 | 0.163 | 5.86E-154 | 2 | Nrp2 |
| Zbtb461 | 9.35E-158 | 0.777877163 | 0.327 | 0.143 | 1.89E-153 | 2 | Zbtb46 |
| Rapgef51 | 2.30E-157 | 0.708416897 | 0.411 | 0.2 | 4.66E-153 | 2 | Rapgef5 |
| Plk21 | 4.26E-157 | 0.727440041 | 0.337 | 0.147 | 8.63E-153 | 2 | Plk2 |
| Sypl1 | 4.20E-156 | 0.71789187 | 0.417 | 0.219 | 8.49E-152 | 2 | Sypl |
| mt-Nd21 | 1.47E-155 | 0.494278119 | 0.957 | 0.656 | 2.98E-151 | 2 | mt-Nd2 |
| Zhx31 | 2.46E-155 | 0.755935139 | 0.379 | 0.182 | 4.98E-151 | 2 | Zhx3 |
| Aldh22 | 4.96E-155 | 0.515251326 | 0.629 | 0.369 | 1.00E-150 | 2 | Aldh2 |
| Lama41 | 5.30E-155 | 0.741576087 | 0.345 | 0.155 | 1.07E-150 | 2 | Lama4 |
| Lrmda1 | 5.99E-154 | 0.822822976 | 0.574 | 0.344 | 1.21E-149 | 2 | Lrmda |
| mt-Atp61 | 6.13E-154 | 0.49024005 | 0.993 | 0.725 | 1.24E-149 | 2 | mt-Atp6 |
| Ier5l1 | 2.52E-153 | 0.71842673 | 0.346 | 0.159 | 5.10E-149 | 2 | Ier5l |
| Tpm41 | 3.01E-152 | 0.615354358 | 0.582 | 0.356 | 6.10E-148 | 2 | Tpm4 |
| Tmed101 | 4.52E-152 | 0.596208423 | 0.623 | 0.415 | 9.15E-148 | 2 | Tmed10 |
| Nxpe41 | 1.58E-151 | 0.751692599 | 0.306 | 0.129 | 3.20E-147 | 2 | Nxpe4 |
| Endod11 | 1.63E-151 | 0.706988382 | 0.308 | 0.134 | 3.30E-147 | 2 | Endod1 |
| Myl12a1 | 5.31E-151 | 0.581762899 | 0.659 | 0.449 | 1.07E-146 | 2 | Myl12a |
| Tpst21 | 2.09E-150 | 0.759628335 | 0.398 | 0.213 | 4.24E-146 | 2 | Tpst2 |
| 21-Mar | 1.05E-149 | 0.629755541 | 0.573 | 0.373 | 2.12E-145 | 2 | 2-Mar |
| Txnip1 | 1.09E-149 | 0.548731031 | 0.752 | 0.521 | 2.20E-145 | 2 | Txnip |
| Myo18a1 | 1.33E-149 | 0.676128098 | 0.445 | 0.236 | 2.70E-145 | 2 | Myo18a |
| Slco2b11 | 1.41E-148 | 0.785084875 | 0.344 | 0.161 | 2.86E-144 | 2 | Slco2b1 |
| Peak11 | 5.70E-148 | 0.718663453 | 0.578 | 0.365 | 1.15E-143 | 2 | Peak1 |
| Dock61 | 1.41E-147 | 0.715049356 | 0.291 | 0.123 | 2.85E-143 | 2 | Dock6 |
| Tnfaip21 | 8.20E-147 | 0.732248392 | 0.308 | 0.134 | 1.66E-142 | 2 | Tnfaip2 |
| Sybu1 | 8.20E-147 | 0.69849934 | 0.258 | 0.098 | 1.66E-142 | 2 | Sybu |
| Rhoa1 | 8.72E-147 | 0.528836998 | 0.789 | 0.571 | 1.77E-142 | 2 | Rhoa |
| Selenok1 | 1.62E-146 | 0.512684709 | 0.736 | 0.533 | 3.28E-142 | 2 | Selenok |
| Nck11 | 2.88E-146 | 0.635881928 | 0.521 | 0.319 | 5.83E-142 | 2 | Nck1 |
| Pim31 | 3.68E-146 | 0.594769334 | 0.389 | 0.191 | 7.46E-142 | 2 | Pim3 |
| Zfp36l11 | 5.06E-146 | 0.632639619 | 0.637 | 0.409 | 1.02E-141 | 2 | Zfp36l1 |
| Cd361 | 6.63E-146 | 0.604643714 | 0.474 | 0.244 | 1.34E-141 | 2 | Cd36 |
| Tmem176b1 | 1.79E-145 | 0.475555401 | 0.68 | 0.41 | 3.63E-141 | 2 | Tmem176b |
| Hsp90b11 | 2.03E-145 | 0.53954632 | 0.731 | 0.484 | 4.11E-141 | 2 | Hsp90b1 |
| Nedd41 | 3.83E-145 | 0.54515651 | 0.639 | 0.394 | 7.76E-141 | 2 | Nedd4 |
| Erg1 | 5.69E-145 | 0.654287755 | 0.28 | 0.113 | 1.15E-140 | 2 | Erg |
| Caskin21 | 1.12E-144 | 0.620671962 | 0.292 | 0.122 | 2.26E-140 | 2 | Caskin2 |
| Tns11 | 1.57E-143 | 0.655957392 | 0.427 | 0.224 | 3.18E-139 | 2 | Tns1 |
| Rasgrf21 | 1.95E-143 | 0.680378009 | 0.343 | 0.158 | 3.95E-139 | 2 | Rasgrf2 |
| Hdac71 | 5.95E-143 | 0.765443657 | 0.363 | 0.184 | 1.21E-138 | 2 | Hdac7 |
| Pls31 | 9.28E-143 | 0.637761424 | 0.392 | 0.194 | 1.88E-138 | 2 | Pls3 |
| Prickle11 | 3.74E-142 | 0.697821425 | 0.3 | 0.129 | 7.56E-138 | 2 | Prickle1 |
| Colgalt21 | 4.06E-142 | 0.672344336 | 0.291 | 0.123 | 8.21E-138 | 2 | Colgalt2 |

| | | | | | | | |
|---|---|---|---|---|---|---|---|
| Ctnna11 | 5.19E-142 | 0.609182293 | 0.541 | 0.33 | 1.05E-137 | 2 | Ctnna1 |
| Il6st1 | 1.08E-141 | 0.648413582 | 0.414 | 0.211 | 2.18E-137 | 2 | Il6st |
| Prkg11 | 1.93E-141 | 0.654574779 | 0.377 | 0.175 | 3.91E-137 | 2 | Prkg1 |
| Tns21 | 2.03E-141 | 0.657854542 | 0.295 | 0.127 | 4.11E-137 | 2 | Tns2 |
| Notch11 | 1.19E-140 | 0.710186975 | 0.385 | 0.198 | 2.41E-136 | 2 | Notch1 |
| Id41 | 2.21E-140 | 0.729616534 | 0.271 | 0.108 | 4.47E-136 | 2 | Id4 |
| Tap21 | 4.16E-140 | 0.725212081 | 0.401 | 0.222 | 8.42E-136 | 2 | Tap2 |
| She1 | 6.48E-140 | 0.622507102 | 0.304 | 0.132 | 1.31E-135 | 2 | She |
| Fam198b1 | 6.95E-139 | 0.634900796 | 0.354 | 0.164 | 1.41E-134 | 2 | Fam198b |
| Dynll11 | 2.05E-137 | 0.492402358 | 0.785 | 0.585 | 4.15E-133 | 2 | Dynll1 |
| Myo101 | 2.95E-137 | 0.603253778 | 0.397 | 0.198 | 5.98E-133 | 2 | Myo10 |
| Spred11 | 1.21E-136 | 0.693321377 | 0.423 | 0.233 | 2.44E-132 | 2 | Spred1 |
| C130074G19Rik1 | 4.64E-136 | 0.517453136 | 0.373 | 0.179 | 9.40E-132 | 2 | C130074G19Rik |
| Rbm391 | 1.24E-135 | 0.517696856 | 0.789 | 0.549 | 2.51E-131 | 2 | Rbm39 |
| Tsc22d31 | 8.89E-135 | 0.690762876 | 0.367 | 0.189 | 1.80E-130 | 2 | Tsc22d3 |
| Anp32a1 | 1.30E-134 | 0.535321326 | 0.648 | 0.445 | 2.63E-130 | 2 | Anp32a |
| Lmo21 | 2.27E-134 | 0.642370799 | 0.37 | 0.189 | 4.61E-130 | 2 | Lmo2 |
| Dip2c1 | 5.80E-134 | 0.736450278 | 0.389 | 0.204 | 1.18E-129 | 2 | Dip2c |
| Irf11 | 8.42E-134 | 0.707266477 | 0.465 | 0.28 | 1.70E-129 | 2 | Irf1 |
| St6galnac31 | 1.13E-133 | 0.608794354 | 0.394 | 0.196 | 2.29E-129 | 2 | St6galnac3 |
| Ceacam11 | 1.90E-133 | 0.662361964 | 0.308 | 0.139 | 3.86E-129 | 2 | Ceacam1 |
| Sppl2a1 | 3.68E-132 | 0.700701288 | 0.456 | 0.278 | 7.45E-128 | 2 | Sppl2a |
| Ehd21 | 1.17E-130 | 0.612968928 | 0.356 | 0.174 | 2.37E-126 | 2 | Ehd2 |
| Rap1b1 | 2.87E-130 | 0.577130107 | 0.616 | 0.425 | 5.81E-126 | 2 | Rap1b |
| Pde8a1 | 4.48E-130 | 0.67885672 | 0.336 | 0.166 | 9.06E-126 | 2 | Pde8a |
| Bnip21 | 4.51E-130 | 0.636985732 | 0.448 | 0.265 | 9.13E-126 | 2 | Bnip2 |
| Sh3bgrl1 | 1.00E-129 | 0.603130489 | 0.551 | 0.355 | 2.03E-125 | 2 | Sh3bgrl |
| Dennd5b1 | 5.64E-129 | 0.647070823 | 0.293 | 0.13 | 1.14E-124 | 2 | Dennd5b |
| Ifngr11 | 1.33E-128 | 0.597949044 | 0.568 | 0.373 | 2.70E-124 | 2 | Ifngr1 |
| Adam151 | 3.12E-128 | 0.63850584 | 0.351 | 0.175 | 6.32E-124 | 2 | Adam15 |
| Gna111 | 1.46E-127 | 0.585398528 | 0.411 | 0.226 | 2.97E-123 | 2 | Gna11 |
| Nfe2l11 | 5.11E-127 | 0.618333731 | 0.446 | 0.261 | 1.04E-122 | 2 | Nfe2l1 |
| Plscr41 | 5.93E-127 | 0.688170343 | 0.341 | 0.173 | 1.20E-122 | 2 | Plscr4 |
| Tanc11 | 6.66E-127 | 0.677563389 | 0.385 | 0.205 | 1.35E-122 | 2 | Tanc1 |
| Efnb11 | 2.08E-126 | 0.688481764 | 0.3 | 0.139 | 4.21E-122 | 2 | Efnb1 |
| Yes11 | 2.60E-126 | 0.632431877 | 0.3 | 0.137 | 5.27E-122 | 2 | Yes1 |
| Tapbp1 | 4.66E-125 | 0.684354987 | 0.459 | 0.287 | 9.44E-121 | 2 | Tapbp |
| Tcn21 | 8.73E-125 | 0.659676656 | 0.327 | 0.163 | 1.77E-120 | 2 | Tcn2 |
| Tnfrsf1a1 | 3.43E-124 | 0.683418924 | 0.372 | 0.205 | 6.94E-120 | 2 | Tnfrsf1a |
| Mlec1 | 5.80E-124 | 0.724396384 | 0.414 | 0.243 | 1.17E-119 | 2 | Mlec |
| Arpc31 | 6.22E-124 | 0.438482781 | 0.835 | 0.638 | 1.26E-119 | 2 | Arpc3 |
| Ripor11 | 8.38E-123 | 0.628721511 | 0.303 | 0.144 | 1.70E-118 | 2 | Ripor1 |
| Lipe1 | 1.52E-122 | 0.630648051 | 0.273 | 0.122 | 3.08E-118 | 2 | Lipe |

| | | | | | | | |
|---|---|---|---|---|---|---|---|
| Efnb21 | 3.00E-122 | 0.623806068 | 0.289 | 0.128 | 6.08E-118 | 2 | Efnb2 |
| Zbtb201 | 2.74E-121 | 0.37490882 | 0.797 | 0.506 | 5.55E-117 | 2 | Zbtb20 |
| Psme11 | 3.53E-121 | 0.477063475 | 0.759 | 0.547 | 7.15E-117 | 2 | Psme1 |
| Limch11 | 3.77E-121 | 0.602797888 | 0.28 | 0.123 | 7.63E-117 | 2 | Limch1 |
| Gcnt21 | 6.78E-120 | 0.624444244 | 0.282 | 0.127 | 1.37E-115 | 2 | Gcnt2 |
| B230219D22Rik1 | 7.67E-120 | 0.540456878 | 0.504 | 0.324 | 1.55E-115 | 2 | B230219D22Rik |
| Rhob1 | 8.36E-120 | 0.678364368 | 0.428 | 0.249 | 1.69E-115 | 2 | Rhob |
| Gja11 | 9.17E-120 | 0.665751816 | 0.262 | 0.113 | 1.86E-115 | 2 | Gja1 |
| Tmem471 | 1.03E-119 | 0.627254581 | 0.299 | 0.14 | 2.09E-115 | 2 | Tmem47 |
| Hes12 | 1.08E-119 | 0.580202545 | 0.502 | 0.292 | 2.18E-115 | 2 | Hes1 |
| Tmtc11 | 1.49E-119 | 0.670597521 | 0.28 | 0.126 | 3.02E-115 | 2 | Tmtc1 |
| Ivns1abp2 | 1.90E-119 | 0.562857545 | 0.515 | 0.321 | 3.85E-115 | 2 | Ivns1abp |
| Amotl11 | 7.70E-119 | 0.641667731 | 0.336 | 0.168 | 1.56E-114 | 2 | Amotl1 |
| Fmnl31 | 1.57E-117 | 0.673677431 | 0.388 | 0.223 | 3.18E-113 | 2 | Fmnl3 |
| Cyp2d221 | 3.26E-117 | 0.610018934 | 0.268 | 0.12 | 6.59E-113 | 2 | Cyp2d22 |
| Crispld11 | 5.15E-117 | 0.629338843 | 0.282 | 0.128 | 1.04E-112 | 2 | Crispld1 |
| Tmem50a1 | 2.78E-116 | 0.607875232 | 0.529 | 0.367 | 5.63E-112 | 2 | Tmem50a |
| Vat11 | 5.91E-116 | 0.61175471 | 0.291 | 0.139 | 1.20E-111 | 2 | Vat1 |
| Gnaq1 | 7.74E-116 | 0.598580884 | 0.527 | 0.34 | 1.57E-111 | 2 | Gnaq |
| Tcf7l11 | 8.69E-116 | 0.619788798 | 0.328 | 0.164 | 1.76E-111 | 2 | Tcf7l1 |
| Myl12b1 | 1.72E-115 | 0.435373946 | 0.713 | 0.529 | 3.49E-111 | 2 | Myl12b |
| Elf11 | 3.26E-115 | 0.616529766 | 0.532 | 0.359 | 6.61E-111 | 2 | Elf1 |
| Cavin11 | 3.95E-115 | 0.514261247 | 0.375 | 0.195 | 8.00E-111 | 2 | Cavin1 |
| Nbea1 | 4.71E-115 | 0.670024703 | 0.346 | 0.179 | 9.54E-111 | 2 | Nbea |
| Fry1 | 6.29E-115 | 0.678480203 | 0.349 | 0.184 | 1.27E-110 | 2 | Fry |
| Mprip1 | 1.52E-113 | 0.579298158 | 0.453 | 0.275 | 3.09E-109 | 2 | Mprip |
| Zfp5031 | 4.32E-113 | 0.617488439 | 0.282 | 0.131 | 8.76E-109 | 2 | Zfp503 |
| Sod11 | 8.63E-113 | 0.548183043 | 0.454 | 0.282 | 1.75E-108 | 2 | Sod1 |
| Nbeal11 | 1.70E-112 | 0.675698753 | 0.412 | 0.245 | 3.44E-108 | 2 | Nbeal1 |
| Ywhaq1 | 1.81E-112 | 0.450003091 | 0.651 | 0.469 | 3.66E-108 | 2 | Ywhaq |
| Mpzl11 | 2.42E-112 | 0.58805966 | 0.276 | 0.128 | 4.89E-108 | 2 | Mpzl1 |
| Galnt181 | 5.61E-112 | 0.628379532 | 0.267 | 0.119 | 1.13E-107 | 2 | Galnt18 |
| Fmnl21 | 6.98E-112 | 0.622319238 | 0.401 | 0.224 | 1.41E-107 | 2 | Fmnl2 |
| Airn1 | 1.12E-111 | 0.696508554 | 0.405 | 0.23 | 2.28E-107 | 2 | Airn |
| Sipa1l21 | 2.11E-111 | 0.622135977 | 0.296 | 0.143 | 4.28E-107 | 2 | Sipa1l2 |
| Ttc281 | 2.91E-111 | 0.645174281 | 0.44 | 0.267 | 5.89E-107 | 2 | Ttc28 |
| Adam101 | 9.36E-111 | 0.657632548 | 0.438 | 0.276 | 1.90E-106 | 2 | Adam10 |
| Tgm21 | 2.96E-110 | 0.587351981 | 0.303 | 0.146 | 6.00E-106 | 2 | Tgm2 |
| Marcks1 | 1.29E-109 | 0.379409343 | 0.543 | 0.318 | 2.60E-105 | 2 | Marcks |
| Pcp4l12 | 3.04E-109 | 0.319275535 | 0.512 | 0.29 | 6.16E-105 | 2 | Pcp4l1 |
| Dock91 | 1.78E-108 | 0.636823055 | 0.351 | 0.189 | 3.60E-104 | 2 | Dock9 |
| Prkce1 | 1.83E-108 | 0.748058664 | 0.357 | 0.198 | 3.71E-104 | 2 | Prkce |
| Lamp11 | 2.53E-108 | 0.488468156 | 0.582 | 0.399 | 5.13E-104 | 2 | Lamp1 |

| | | | | | | | |
|---|---|---|---|---|---|---|---|
| Slc44a11 | 3.22E-107 | 0.625674505 | 0.287 | 0.141 | 6.52E-103 | 2 | Slc44a1 |
| Pdlim51 | 4.43E-106 | 0.623785345 | 0.435 | 0.269 | 8.96E-102 | 2 | Pdlim5 |
| Abi31 | 5.75E-105 | 0.654055706 | 0.38 | 0.229 | 1.16E-100 | 2 | Abi3 |
| 2010111I01Rik1 | 5.79E-105 | 0.513958044 | 0.509 | 0.331 | 1.17E-100 | 2 | 2010111I01Rik |
| Gimap11 | 7.96E-105 | 0.53507219 | 0.462 | 0.285 | 1.61E-100 | 2 | Gimap1 |
| Tuba1a1 | 1.53E-104 | 0.533512781 | 0.412 | 0.239 | 3.09E-100 | 2 | Tuba1a |
| Selenow | 4.73E-104 | 0.372122487 | 0.799 | 0.61 | 9.57E-100 | 2 | Selenow |
| Wwc21 | 6.84E-104 | 0.608419154 | 0.335 | 0.182 | 1.38E-99 | 2 | Wwc2 |
| Prickle21 | 7.47E-104 | 0.610846367 | 0.267 | 0.125 | 1.51E-99 | 2 | Prickle2 |
| Dab21 | 6.19E-103 | 0.589550973 | 0.348 | 0.191 | 1.25E-98 | 2 | Dab2 |
| Itgb11 | 7.22E-103 | 0.384311093 | 0.69 | 0.454 | 1.46E-98 | 2 | Itgb1 |
| Rrbp11 | 1.21E-102 | 0.446469417 | 0.642 | 0.439 | 2.46E-98 | 2 | Rrbp1 |
| Smad71 | 1.51E-102 | 0.602490795 | 0.476 | 0.31 | 3.06E-98 | 2 | Smad7 |
| Dgkh1 | 1.52E-102 | 0.669911531 | 0.309 | 0.162 | 3.08E-98 | 2 | Dgkh |
| Gnb21 | 3.38E-102 | 0.412273826 | 0.686 | 0.514 | 6.84E-98 | 2 | Gnb2 |
| Baz2b1 | 1.33E-101 | 0.595502674 | 0.496 | 0.334 | 2.70E-97 | 2 | Baz2b |
| Mxra71 | 1.50E-101 | 0.55180558 | 0.269 | 0.127 | 3.03E-97 | 2 | Mxra7 |
| Ints91 | 1.67E-101 | 0.617252565 | 0.258 | 0.123 | 3.39E-97 | 2 | Ints9 |
| Ralgapa11 | 2.04E-101 | 0.642261826 | 0.446 | 0.288 | 4.14E-97 | 2 | Ralgapa1 |
| Ifi441 | 8.21E-101 | 0.585753513 | 0.277 | 0.131 | 1.66E-96 | 2 | Ifi44 |
| Epn21 | 3.65E-100 | 0.57165049 | 0.284 | 0.14 | 7.39E-96 | 2 | Epn2 |
| Bag51 | 1.21E-99 | 0.51506453 | 0.338 | 0.186 | 2.44E-95 | 2 | Bag5 |
| Myo1c1 | 2.51E-99 | 0.573541972 | 0.352 | 0.198 | 5.08E-95 | 2 | Myo1c |
| Tmbim61 | 3.75E-99 | 0.517350467 | 0.569 | 0.416 | 7.58E-95 | 2 | Tmbim6 |
| Xiap1 | 4.86E-99 | 0.642690736 | 0.437 | 0.285 | 9.84E-95 | 2 | Xiap |
| Tmco11 | 5.29E-99 | 0.575767756 | 0.448 | 0.297 | 1.07E-94 | 2 | Tmco1 |
| Rbfox21 | 8.15E-99 | 0.593429528 | 0.328 | 0.175 | 1.65E-94 | 2 | Rbfox2 |
| Ostf11 | 9.69E-99 | 0.41312355 | 0.663 | 0.489 | 1.96E-94 | 2 | Ostf1 |
| Plscr11 | 1.07E-98 | 0.655547333 | 0.337 | 0.19 | 2.16E-94 | 2 | Plscr1 |
| Tagln21 | 1.30E-98 | 0.488561164 | 0.585 | 0.411 | 2.64E-94 | 2 | Tagln2 |
| Sec621 | 2.13E-98 | 0.524149295 | 0.53 | 0.379 | 4.31E-94 | 2 | Sec62 |
| Klf31 | 5.98E-98 | 0.556058684 | 0.379 | 0.22 | 1.21E-93 | 2 | Klf3 |
| Piezo11 | 1.33E-97 | 0.617967154 | 0.301 | 0.159 | 2.69E-93 | 2 | Piezo1 |
| Lpar61 | 1.63E-97 | 0.68013705 | 0.311 | 0.169 | 3.29E-93 | 2 | Lpar6 |
| Klhl51 | 3.71E-97 | 0.551933791 | 0.364 | 0.209 | 7.52E-93 | 2 | Klhl5 |
| mt-Nd11 | 1.37E-96 | 0.394579967 | 0.83 | 0.571 | 2.77E-92 | 2 | mt-Nd1 |
| Ktn11 | 3.81E-96 | 0.643867173 | 0.442 | 0.293 | 7.71E-92 | 2 | Ktn1 |
| Sh3glb11 | 2.18E-95 | 0.402585346 | 0.686 | 0.514 | 4.42E-91 | 2 | Sh3glb1 |
| Rac11 | 2.72E-95 | 0.483432613 | 0.57 | 0.415 | 5.50E-91 | 2 | Rac1 |
| Mapk31 | 2.81E-95 | 0.509213485 | 0.406 | 0.25 | 5.69E-91 | 2 | Mapk3 |
| Cds21 | 3.19E-95 | 0.582985957 | 0.303 | 0.163 | 6.46E-91 | 2 | Cds2 |
| Psme21 | 2.04E-94 | 0.440422826 | 0.59 | 0.427 | 4.13E-90 | 2 | Psme2 |
| Ppp1r13b1 | 4.48E-94 | 0.618581717 | 0.346 | 0.202 | 9.07E-90 | 2 | Ppp1r13b |

| | | | | | | | |
|---|---|---|---|---|---|---|---|
| Cebpg1 | 4.81E-94 | 0.544088337 | 0.355 | 0.209 | 9.74E-90 | 2 | Cebpg |
| St3gal61 | 1.39E-93 | 0.539051306 | 0.462 | 0.302 | 2.82E-89 | 2 | St3gal6 |
| Irf2bpl1 | 1.79E-93 | 0.52855317 | 0.379 | 0.229 | 3.61E-89 | 2 | Irf2bpl |
| Map4k31 | 1.91E-93 | 0.622778057 | 0.365 | 0.214 | 3.87E-89 | 2 | Map4k3 |
| Ptpra1 | 5.32E-93 | 0.594113621 | 0.434 | 0.29 | 1.08E-88 | 2 | Ptpra |
| Tmem30a1 | 7.07E-93 | 0.592684708 | 0.417 | 0.272 | 1.43E-88 | 2 | Tmem30a |
| Rock21 | 1.04E-92 | 0.549555533 | 0.482 | 0.322 | 2.10E-88 | 2 | Rock2 |
| 1810037I17Rik | 1.48E-92 | 0.338042586 | 0.626 | 0.463 | 3.01E-88 | 2 | 1810037I17Rik |
| Txn1 | 1.66E-92 | 0.320690305 | 0.816 | 0.661 | 3.36E-88 | 2 | Txn1 |
| Hmg20b1 | 1.94E-92 | 0.514446791 | 0.397 | 0.246 | 3.93E-88 | 2 | Hmg20b |
| Kmt2e1 | 9.90E-92 | 0.524324305 | 0.55 | 0.391 | 2.00E-87 | 2 | Kmt2e |
| Emc71 | 1.18E-91 | 0.568685623 | 0.398 | 0.258 | 2.38E-87 | 2 | Emc7 |
| Mtdh1 | 1.29E-91 | 0.569984815 | 0.522 | 0.372 | 2.61E-87 | 2 | Mtdh |
| Ywhab1 | 5.98E-91 | 0.425469083 | 0.614 | 0.451 | 1.21E-86 | 2 | Ywhab |
| Foxo11 | 1.45E-90 | 0.592876748 | 0.326 | 0.186 | 2.93E-86 | 2 | Foxo1 |
| Rab11a1 | 1.46E-90 | 0.447633667 | 0.449 | 0.293 | 2.96E-86 | 2 | Rab11a |
| Cdc42bpa1 | 5.70E-90 | 0.556983582 | 0.315 | 0.17 | 1.15E-85 | 2 | Cdc42bpa |
| Ddx51 | 7.49E-90 | 0.39052968 | 0.848 | 0.631 | 1.52E-85 | 2 | Ddx5 |
| Aig11 | 1.01E-89 | 0.54129079 | 0.345 | 0.199 | 2.05E-85 | 2 | Aig1 |
| Gnb11 | 1.09E-89 | 0.446793533 | 0.568 | 0.412 | 2.21E-85 | 2 | Gnb1 |
| Ica11 | 2.43E-89 | 0.452766248 | 0.309 | 0.162 | 4.91E-85 | 2 | Ica1 |
| Ubc1 | 3.52E-88 | 0.338982752 | 0.815 | 0.646 | 7.13E-84 | 2 | Ubc |
| Tmcc31 | 4.66E-88 | 0.353076503 | 0.48 | 0.301 | 9.43E-84 | 2 | Tmcc3 |
| Nfia1 | 8.57E-88 | 0.493118358 | 0.438 | 0.271 | 1.74E-83 | 2 | Nfia |
| H2-Q61 | 2.24E-87 | 0.618013954 | 0.37 | 0.232 | 4.54E-83 | 2 | H2-Q6 |
| H2-T221 | 3.02E-87 | 0.555747217 | 0.406 | 0.267 | 6.12E-83 | 2 | H2-T22 |
| Gbp41 | 9.45E-87 | 0.640008866 | 0.298 | 0.164 | 1.91E-82 | 2 | Gbp4 |
| Rock11 | 2.33E-86 | 0.480753477 | 0.578 | 0.417 | 4.72E-82 | 2 | Rock1 |
| Atraid1 | 2.64E-86 | 0.516260927 | 0.396 | 0.257 | 5.34E-82 | 2 | Atraid |
| Enah1 | 4.80E-86 | 0.494140778 | 0.27 | 0.136 | 9.72E-82 | 2 | Enah |
| Nid11 | 5.34E-86 | 0.411667987 | 0.316 | 0.167 | 1.08E-81 | 2 | Nid1 |
| Trim471 | 8.61E-86 | 0.550475127 | 0.258 | 0.131 | 1.74E-81 | 2 | Trim47 |
| Degs11 | 1.06E-85 | 0.605539376 | 0.3 | 0.172 | 2.15E-81 | 2 | Degs1 |
| Sbds1 | 1.87E-85 | 0.519250796 | 0.438 | 0.296 | 3.80E-81 | 2 | Sbds |
| Mgst11 | 2.29E-85 | 0.494197911 | 0.294 | 0.153 | 4.64E-81 | 2 | Mgst1 |
| Prrc2c1 | 4.28E-85 | 0.497749369 | 0.559 | 0.405 | 8.68E-81 | 2 | Prrc2c |
| Kctd121 | 1.03E-84 | 0.534995375 | 0.39 | 0.244 | 2.09E-80 | 2 | Kctd12 |
| Dhrs71 | 1.87E-84 | 0.588095837 | 0.333 | 0.202 | 3.79E-80 | 2 | Dhrs7 |
| Ccser21 | 1.95E-84 | 0.486293368 | 0.36 | 0.217 | 3.94E-80 | 2 | Ccser2 |
| Snx31 | 3.85E-84 | 0.368582827 | 0.597 | 0.446 | 7.80E-80 | 2 | Snx3 |
| Ier51 | 2.22E-83 | 0.570330994 | 0.443 | 0.304 | 4.50E-79 | 2 | Ier5 |
| Mgat11 | 2.33E-83 | 0.582422094 | 0.28 | 0.155 | 4.72E-79 | 2 | Mgat1 |
| St3gal41 | 4.29E-83 | 0.605144339 | 0.38 | 0.243 | 8.68E-79 | 2 | St3gal4 |

| | | | | | | | |
|---|---|---|---|---|---|---|---|
| Osmr1 | 7.19E-83 | 0.545059487 | 0.252 | 0.127 | 1.46E-78 | 2 | Osmr |
| Nedd91 | 2.43E-82 | 0.514522916 | 0.464 | 0.304 | 4.92E-78 | 2 | Nedd9 |
| Spag91 | 3.40E-82 | 0.525883156 | 0.487 | 0.345 | 6.89E-78 | 2 | Spag9 |
| Klf91 | 1.61E-81 | 0.475059017 | 0.371 | 0.223 | 3.26E-77 | 2 | Klf9 |
| Rtn31 | 2.39E-81 | 0.55793412 | 0.446 | 0.311 | 4.85E-77 | 2 | Rtn3 |
| Sorbs11 | 3.09E-81 | 0.568930656 | 0.296 | 0.164 | 6.25E-77 | 2 | Sorbs1 |
| Xaf11 | 6.12E-81 | 0.494789417 | 0.345 | 0.207 | 1.24E-76 | 2 | Xaf1 |
| Actg11 | 1.05E-80 | 0.379709401 | 0.934 | 0.778 | 2.13E-76 | 2 | Actg1 |
| Smpdl3a1 | 8.70E-80 | 0.478451535 | 0.342 | 0.206 | 1.76E-75 | 2 | Smpdl3a |
| Socs21 | 1.03E-79 | 0.485953466 | 0.304 | 0.169 | 2.08E-75 | 2 | Socs2 |
| Eepd11 | 1.19E-79 | 0.387166316 | 0.259 | 0.13 | 2.42E-75 | 2 | Eepd1 |
| Nckap11 | 1.24E-79 | 0.489896493 | 0.353 | 0.213 | 2.50E-75 | 2 | Nckap1 |
| mt-Co31 | 1.26E-79 | 0.318547408 | 0.989 | 0.694 | 2.56E-75 | 2 | mt-Co3 |
| St8sia41 | 3.28E-79 | 0.604020313 | 0.382 | 0.246 | 6.63E-75 | 2 | St8sia4 |
| Atrx1 | 3.29E-79 | 0.515282147 | 0.554 | 0.405 | 6.66E-75 | 2 | Atrx |
| Zeb11 | 7.03E-79 | 0.559296901 | 0.458 | 0.307 | 1.42E-74 | 2 | Zeb1 |
| Ppp1r12a1 | 8.82E-79 | 0.455544315 | 0.576 | 0.409 | 1.79E-74 | 2 | Ppp1r12a |
| Leprot1 | 1.34E-78 | 0.537735406 | 0.289 | 0.165 | 2.72E-74 | 2 | Leprot |
| Sox41 | 1.68E-78 | 0.455720452 | 0.336 | 0.189 | 3.40E-74 | 2 | Sox4 |
| Serinc11 | 3.98E-78 | 0.515620417 | 0.369 | 0.236 | 8.05E-74 | 2 | Serinc1 |
| Cdc42bpb1 | 9.87E-78 | 0.473776007 | 0.304 | 0.173 | 2.00E-73 | 2 | Cdc42bpb |
| Man2a11 | 6.35E-77 | 0.585942543 | 0.355 | 0.221 | 1.29E-72 | 2 | Man2a1 |
| Ppib | 7.99E-77 | 0.333155272 | 0.701 | 0.537 | 1.62E-72 | 2 | Ppib |
| Pmp221 | 5.93E-76 | 0.468728967 | 0.255 | 0.132 | 1.20E-71 | 2 | Pmp22 |
| Lman11 | 7.35E-76 | 0.569482471 | 0.351 | 0.226 | 1.49E-71 | 2 | Lman1 |
| Myl6 | 7.83E-76 | 0.280809801 | 0.928 | 0.758 | 1.59E-71 | 2 | Myl6 |
| Tmed91 | 3.75E-75 | 0.441792925 | 0.518 | 0.391 | 7.60E-71 | 2 | Tmed9 |
| Pgrmc11 | 4.08E-75 | 0.506284404 | 0.331 | 0.206 | 8.26E-71 | 2 | Pgrmc1 |
| Kif13a1 | 6.21E-74 | 0.562694145 | 0.284 | 0.162 | 1.26E-69 | 2 | Kif13a |
| Osbpl91 | 6.46E-74 | 0.519037004 | 0.393 | 0.266 | 1.31E-69 | 2 | Osbpl9 |
| Entpd11 | 7.56E-74 | 0.460795959 | 0.407 | 0.262 | 1.53E-69 | 2 | Entpd1 |
| Nav21 | 1.62E-73 | 0.586544385 | 0.275 | 0.153 | 3.27E-69 | 2 | Nav2 |
| Hook31 | 1.77E-73 | 0.547735335 | 0.406 | 0.278 | 3.59E-69 | 2 | Hook3 |
| Scn1b1 | 6.37E-73 | 0.501327609 | 0.308 | 0.184 | 1.29E-68 | 2 | Scn1b |
| Map41 | 7.26E-73 | 0.531043331 | 0.422 | 0.291 | 1.47E-68 | 2 | Map4 |
| Mat2a1 | 1.10E-72 | 0.530672918 | 0.423 | 0.297 | 2.22E-68 | 2 | Mat2a |
| Ifnar21 | 2.35E-72 | 0.531228351 | 0.437 | 0.313 | 4.76E-68 | 2 | Ifnar2 |
| mt-Nd41 | 3.68E-72 | 0.296435603 | 0.939 | 0.643 | 7.46E-68 | 2 | mt-Nd4 |
| Mxd41 | 5.40E-72 | 0.459084702 | 0.475 | 0.339 | 1.09E-67 | 2 | Mxd4 |
| Svil1 | 1.09E-71 | 0.578634712 | 0.355 | 0.231 | 2.20E-67 | 2 | Svil |
| Tax1bp31 | 1.22E-71 | 0.356532199 | 0.29 | 0.163 | 2.48E-67 | 2 | Tax1bp3 |
| Pura1 | 1.24E-71 | 0.441601143 | 0.483 | 0.353 | 2.50E-67 | 2 | Pura |
| Tmod31 | 2.63E-71 | 0.44575583 | 0.482 | 0.349 | 5.33E-67 | 2 | Tmod3 |

| | | | | | | | |
|---|---|---|---|---|---|---|---|
| Cnn31 | 2.85E-71 | 0.438489792 | 0.312 | 0.181 | 5.77E-67 | 2 | Cnn3 |
| Zfyve91 | 3.04E-71 | 0.487165265 | 0.282 | 0.159 | 6.16E-67 | 2 | Zfyve9 |
| Lrch11 | 4.06E-71 | 0.566802658 | 0.425 | 0.3 | 8.22E-67 | 2 | Lrch1 |
| Egln11 | 9.86E-71 | 0.479983055 | 0.335 | 0.212 | 2.00E-66 | 2 | Egln1 |
| Ptbp31 | 1.32E-70 | 0.476201447 | 0.516 | 0.381 | 2.68E-66 | 2 | Ptbp3 |
| Xist1 | 1.35E-70 | 0.311256829 | 0.655 | 0.441 | 2.73E-66 | 2 | Xist |
| Afap11 | 2.05E-70 | 0.541370268 | 0.254 | 0.139 | 4.15E-66 | 2 | Afap1 |
| Dnajc31 | 6.71E-70 | 0.490298268 | 0.483 | 0.359 | 1.36E-65 | 2 | Dnajc3 |
| Wars1 | 7.42E-69 | 0.501500882 | 0.274 | 0.157 | 1.50E-64 | 2 | Wars |
| Cyb5r31 | 1.25E-68 | 0.435580545 | 0.331 | 0.205 | 2.53E-64 | 2 | Cyb5r3 |
| St3gal11 | 1.66E-68 | 0.537238392 | 0.285 | 0.169 | 3.36E-64 | 2 | St3gal1 |
| Plcb41 | 1.86E-68 | 0.353625438 | 0.43 | 0.276 | 3.78E-64 | 2 | Plcb4 |
| Ddx501 | 2.68E-68 | 0.482691842 | 0.399 | 0.281 | 5.42E-64 | 2 | Ddx50 |
| 9930111J21Rik21 | 6.01E-68 | 0.546468486 | 0.37 | 0.245 | 1.22E-63 | 2 | 9930111J21Rik2 |
| Dnaja11 | 8.78E-68 | 0.330591759 | 0.651 | 0.502 | 1.78E-63 | 2 | Dnaja1 |
| Dync1i21 | 6.69E-67 | 0.402973599 | 0.53 | 0.407 | 1.36E-62 | 2 | Dync1i2 |
| Prkar1a1 | 1.23E-66 | 0.383328688 | 0.52 | 0.391 | 2.49E-62 | 2 | Prkar1a |
| Mrfap11 | 2.81E-66 | 0.251098129 | 0.575 | 0.437 | 5.69E-62 | 2 | Mrfap1 |
| Snrpb | 4.81E-66 | 0.323282123 | 0.583 | 0.461 | 9.74E-62 | 2 | Snrpb |
| Pttg11 | 4.29E-65 | 0.311293111 | 0.414 | 0.28 | 8.69E-61 | 2 | Pttg1 |
| Atxn7l3b1 | 6.21E-65 | 0.316052517 | 0.522 | 0.392 | 1.26E-60 | 2 | Atxn7l3b |
| Fndc3b1 | 1.48E-64 | 0.522708667 | 0.354 | 0.233 | 3.00E-60 | 2 | Fndc3b |
| Psmd71 | 1.73E-64 | 0.393813732 | 0.415 | 0.294 | 3.51E-60 | 2 | Psmd7 |
| Tsnax1 | 6.51E-64 | 0.457256845 | 0.265 | 0.155 | 1.32E-59 | 2 | Tsnax |
| Trim21 | 7.29E-64 | 0.404971648 | 0.288 | 0.166 | 1.48E-59 | 2 | Trim2 |
| Lamp21 | 2.29E-63 | 0.448838736 | 0.421 | 0.293 | 4.63E-59 | 2 | Lamp2 |
| Ddrgk11 | 2.54E-62 | 0.44993271 | 0.417 | 0.308 | 5.15E-58 | 2 | Ddrgk1 |
| Srsf71 | 4.77E-62 | 0.470635117 | 0.433 | 0.322 | 9.66E-58 | 2 | Srsf7 |
| Tpm31 | 6.14E-62 | 0.329368182 | 0.61 | 0.465 | 1.24E-57 | 2 | Tpm3 |
| Cnbp1 | 1.83E-61 | 0.297064823 | 0.601 | 0.475 | 3.70E-57 | 2 | Cnbp |
| Ppfibp11 | 2.73E-61 | 0.481124263 | 0.287 | 0.173 | 5.54E-57 | 2 | Ppfibp1 |
| Rftn11 | 3.26E-61 | 0.528150158 | 0.354 | 0.241 | 6.60E-57 | 2 | Rftn1 |
| Cep85l1 | 3.54E-61 | 0.500048114 | 0.263 | 0.154 | 7.17E-57 | 2 | Cep85l |
| Nucb11 | 2.99E-60 | 0.542638597 | 0.323 | 0.218 | 6.05E-56 | 2 | Nucb1 |
| mt-Co11 | 6.77E-60 | 0.279913742 | 0.989 | 0.687 | 1.37E-55 | 2 | mt-Co1 |
| Iqgap11 | 9.80E-60 | 0.428273751 | 0.534 | 0.399 | 1.98E-55 | 2 | Iqgap1 |
| Foxn31 | 2.99E-59 | 0.416744394 | 0.532 | 0.398 | 6.06E-55 | 2 | Foxn3 |
| Col4a3bp1 | 3.10E-59 | 0.482076785 | 0.359 | 0.248 | 6.28E-55 | 2 | Col4a3bp |
| Ncoa31 | 9.97E-59 | 0.536577227 | 0.35 | 0.242 | 2.02E-54 | 2 | Ncoa3 |
| Tgfbr31 | 2.06E-58 | 0.496455474 | 0.273 | 0.164 | 4.17E-54 | 2 | Tgfbr3 |
| Myh91 | 6.06E-58 | 0.385376356 | 0.564 | 0.423 | 1.23E-53 | 2 | Myh9 |
| Slc3a21 | 7.65E-58 | 0.5024311 | 0.388 | 0.281 | 1.55E-53 | 2 | Slc3a2 |
| Ccnd11 | 7.78E-58 | 0.423725926 | 0.341 | 0.215 | 1.57E-53 | 2 | Ccnd1 |

| | | | | | | | |
|---|---|---|---|---|---|---|---|
| Rab121 | 7.83E-58 | 0.454037072 | 0.272 | 0.167 | 1.58E-53 | 2 | Rab12 |
| Birc61 | 1.63E-57 | 0.484998929 | 0.439 | 0.331 | 3.30E-53 | 2 | Birc6 |
| Rin21 | 2.78E-57 | 0.464219 | 0.282 | 0.172 | 5.62E-53 | 2 | Rin2 |
| Tubb51 | 4.09E-57 | 0.307773958 | 0.49 | 0.369 | 8.29E-53 | 2 | Tubb5 |
| Lmna1 | 6.74E-57 | 0.442456765 | 0.299 | 0.187 | 1.36E-52 | 2 | Lmna |
| Cped11 | 8.07E-57 | 0.422891179 | 0.285 | 0.168 | 1.63E-52 | 2 | Cped1 |
| Foxp11 | 9.27E-57 | 0.394796416 | 0.507 | 0.377 | 1.88E-52 | 2 | Foxp1 |
| Calu1 | 9.93E-57 | 0.455554668 | 0.3 | 0.192 | 2.01E-52 | 2 | Calu |
| Fcgrt1 | 1.85E-56 | 0.436499963 | 0.285 | 0.175 | 3.74E-52 | 2 | Fcgrt |
| Fus1 | 2.31E-56 | 0.381354427 | 0.489 | 0.379 | 4.69E-52 | 2 | Fus |
| Csnk1a11 | 3.20E-56 | 0.313742842 | 0.566 | 0.446 | 6.48E-52 | 2 | Csnk1a1 |
| Arhgef31 | 4.62E-56 | 0.57305428 | 0.337 | 0.231 | 9.34E-52 | 2 | Arhgef3 |
| Mbnl21 | 6.42E-56 | 0.329713781 | 0.538 | 0.406 | 1.30E-51 | 2 | Mbnl2 |
| Tprgl1 | 9.87E-56 | 0.390967847 | 0.376 | 0.269 | 2.00E-51 | 2 | Tprgl |
| Vamp31 | 1.86E-55 | 0.401055287 | 0.271 | 0.167 | 3.77E-51 | 2 | Vamp3 |
| Map4k41 | 1.16E-54 | 0.493974355 | 0.373 | 0.267 | 2.35E-50 | 2 | Map4k4 |
| Cr1l1 | 1.16E-54 | 0.517267708 | 0.324 | 0.224 | 2.36E-50 | 2 | Cr1l |
| Ddah2 | 1.47E-54 | 0.252526675 | 0.315 | 0.194 | 2.97E-50 | 2 | Ddah2 |
| Samd9l1 | 1.56E-54 | 0.491007649 | 0.351 | 0.247 | 3.16E-50 | 2 | Samd9l |
| Pdia61 | 5.96E-54 | 0.40551955 | 0.434 | 0.328 | 1.21E-49 | 2 | Pdia6 |
| Dnm21 | 8.22E-54 | 0.520866639 | 0.347 | 0.25 | 1.67E-49 | 2 | Dnm2 |
| Fcho21 | 3.12E-53 | 0.468459499 | 0.404 | 0.3 | 6.33E-49 | 2 | Fcho2 |
| Rsu11 | 3.87E-53 | 0.427074889 | 0.349 | 0.244 | 7.83E-49 | 2 | Rsu1 |
| Camk1 | 5.21E-53 | 0.352597007 | 0.327 | 0.217 | 1.05E-48 | 2 | Camk1 |
| 4930523C07Rik1 | 6.13E-53 | 0.502415783 | 0.313 | 0.21 | 1.24E-48 | 2 | 4930523C07Rik |
| Fmo11 | 7.56E-53 | 0.295412542 | 0.311 | 0.19 | 1.53E-48 | 2 | Fmo1 |
| Akt31 | 1.63E-52 | 0.501471586 | 0.348 | 0.242 | 3.30E-48 | 2 | Akt3 |
| mt-Nd31 | 3.15E-52 | 0.380295419 | 0.554 | 0.435 | 6.38E-48 | 2 | mt-Nd3 |
| Tmem1651 | 7.97E-52 | 0.515380844 | 0.264 | 0.171 | 1.61E-47 | 2 | Tmem165 |
| Cfap201 | 2.59E-51 | 0.3940141 | 0.278 | 0.177 | 5.24E-47 | 2 | Cfap20 |
| Tgoln11 | 2.61E-51 | 0.497944 | 0.334 | 0.238 | 5.28E-47 | 2 | Tgoln1 |
| Sbf21 | 2.78E-51 | 0.459274677 | 0.303 | 0.198 | 5.63E-47 | 2 | Sbf2 |
| Atxn11 | 3.58E-51 | 0.473775738 | 0.398 | 0.285 | 7.25E-47 | 2 | Atxn1 |
| Tm9sf21 | 1.13E-50 | 0.485489649 | 0.341 | 0.247 | 2.30E-46 | 2 | Tm9sf2 |
| Rpn11 | 1.52E-50 | 0.49953333 | 0.308 | 0.213 | 3.08E-46 | 2 | Rpn1 |
| Unc501 | 4.63E-50 | 0.420746906 | 0.3 | 0.204 | 9.38E-46 | 2 | Unc50 |
| Parp141 | 9.15E-50 | 0.506268685 | 0.344 | 0.243 | 1.85E-45 | 2 | Parp14 |
| Csrp11 | 9.45E-50 | 0.356485904 | 0.273 | 0.173 | 1.91E-45 | 2 | Csrp1 |
| Tm9sf31 | 1.68E-49 | 0.433034933 | 0.423 | 0.326 | 3.41E-45 | 2 | Tm9sf3 |
| Capn21 | 1.77E-49 | 0.439572515 | 0.268 | 0.17 | 3.58E-45 | 2 | Capn2 |
| Sptlc21 | 1.89E-49 | 0.492563493 | 0.331 | 0.236 | 3.82E-45 | 2 | Sptlc2 |
| Nbdy1 | 2.11E-49 | 0.389252789 | 0.286 | 0.187 | 4.27E-45 | 2 | Nbdy |
| Dad1 | 2.24E-49 | 0.267097536 | 0.559 | 0.449 | 4.54E-45 | 2 | Dad1 |

| | | | | | | | |
|---|---|---|---|---|---|---|---|
| Msi21 | 8.92E-49 | 0.473121798 | 0.386 | 0.283 | 1.81E-44 | 2 | Msi2 |
| 2900026A02Rik1 | 1.31E-48 | 0.491744819 | 0.308 | 0.211 | 2.65E-44 | 2 | 2900026A02Rik |
| Samhd11 | 1.31E-48 | 0.435401615 | 0.346 | 0.245 | 2.65E-44 | 2 | Samhd1 |
| Smc1a1 | 2.07E-48 | 0.441031933 | 0.401 | 0.303 | 4.19E-44 | 2 | Smc1a |
| Tmem2341 | 3.18E-48 | 0.322810469 | 0.478 | 0.385 | 6.44E-44 | 2 | Tmem234 |
| Spr | 5.54E-48 | 0.33956575 | 0.275 | 0.177 | 1.12E-43 | 2 | Spr |
| Btbd71 | 7.96E-48 | 0.446447759 | 0.37 | 0.272 | 1.61E-43 | 2 | Btbd7 |
| Sec14l11 | 8.01E-48 | 0.284558676 | 0.382 | 0.266 | 1.62E-43 | 2 | Sec14l1 |
| Dusp11 | 1.33E-47 | 0.536701433 | 0.283 | 0.184 | 2.69E-43 | 2 | Dusp1 |
| Tuba1b | 2.07E-47 | 0.279837365 | 0.424 | 0.312 | 4.19E-43 | 2 | Tuba1b |
| Elf21 | 2.30E-47 | 0.38830564 | 0.388 | 0.292 | 4.65E-43 | 2 | Elf2 |
| Ginm11 | 3.70E-47 | 0.430212864 | 0.262 | 0.171 | 7.49E-43 | 2 | Ginm1 |
| Pdia31 | 9.45E-47 | 0.319278984 | 0.549 | 0.433 | 1.91E-42 | 2 | Pdia3 |
| Tcf121 | 1.24E-46 | 0.41372253 | 0.476 | 0.366 | 2.52E-42 | 2 | Tcf12 |
| Irf21 | 1.80E-46 | 0.431044218 | 0.425 | 0.333 | 3.64E-42 | 2 | Irf2 |
| Gng121 | 1.85E-46 | 0.344744791 | 0.293 | 0.191 | 3.74E-42 | 2 | Gng12 |
| Srrm21 | 1.89E-46 | 0.339114995 | 0.581 | 0.458 | 3.82E-42 | 2 | Srrm2 |
| Klhl241 | 2.45E-46 | 0.432306629 | 0.301 | 0.207 | 4.95E-42 | 2 | Klhl24 |
| Gulp11 | 2.75E-46 | 0.416162861 | 0.257 | 0.157 | 5.56E-42 | 2 | Gulp1 |
| Zmiz11 | 4.43E-46 | 0.435277642 | 0.405 | 0.313 | 8.96E-42 | 2 | Zmiz1 |
| Arglu11 | 6.22E-46 | 0.385868815 | 0.438 | 0.346 | 1.26E-41 | 2 | Arglu1 |
| Dync1h11 | 6.23E-46 | 0.458545377 | 0.369 | 0.278 | 1.26E-41 | 2 | Dync1h1 |
| Pafah1b11 | 7.37E-46 | 0.260432218 | 0.581 | 0.47 | 1.49E-41 | 2 | Pafah1b1 |
| N4bp11 | 8.49E-46 | 0.483820514 | 0.256 | 0.168 | 1.72E-41 | 2 | N4bp1 |
| Sos11 | 9.72E-46 | 0.460330091 | 0.265 | 0.172 | 1.97E-41 | 2 | Sos1 |
| Swap701 | 1.61E-45 | 0.458123221 | 0.331 | 0.238 | 3.26E-41 | 2 | Swap70 |
| mt-Nd4l | 8.44E-45 | 0.279909464 | 0.607 | 0.479 | 1.71E-40 | 2 | mt-Nd4l |
| Gm122161 | 1.00E-44 | 0.401002941 | 0.318 | 0.222 | 2.03E-40 | 2 | Gm12216 |
| Phactr41 | 2.64E-44 | 0.454837313 | 0.259 | 0.17 | 5.35E-40 | 2 | Phactr4 |
| Tmx31 | 3.46E-44 | 0.48137553 | 0.265 | 0.18 | 7.00E-40 | 2 | Tmx3 |
| Sep-71 | 4.18E-44 | 0.269156311 | 0.568 | 0.451 | 8.47E-40 | 2 | 7-Sep |
| Abhd17b1 | 6.19E-44 | 0.472466904 | 0.273 | 0.184 | 1.25E-39 | 2 | Abhd17b |
| Cltb | 1.07E-43 | 0.296736957 | 0.308 | 0.21 | 2.16E-39 | 2 | Cltb |
| Elmo11 | 1.92E-43 | 0.331985946 | 0.478 | 0.354 | 3.89E-39 | 2 | Elmo1 |
| Hivep21 | 2.05E-43 | 0.455384878 | 0.384 | 0.29 | 4.15E-39 | 2 | Hivep2 |
| Wdfy31 | 2.25E-43 | 0.458777467 | 0.264 | 0.174 | 4.55E-39 | 2 | Wdfy3 |
| Sp1001 | 2.98E-43 | 0.322437317 | 0.533 | 0.422 | 6.03E-39 | 2 | Sp100 |
| Tmed21 | 3.45E-43 | 0.331369845 | 0.445 | 0.358 | 6.99E-39 | 2 | Tmed2 |
| Stag11 | 3.93E-43 | 0.423957775 | 0.447 | 0.353 | 7.95E-39 | 2 | Stag1 |
| Pnp1 | 8.68E-43 | 0.374397569 | 0.341 | 0.245 | 1.76E-38 | 2 | Pnp |
| Znfx11 | 8.72E-43 | 0.451881529 | 0.255 | 0.167 | 1.76E-38 | 2 | Znfx1 |
| Ddx3x1 | 4.93E-42 | 0.368616605 | 0.392 | 0.3 | 9.98E-38 | 2 | Ddx3x |
| Herc11 | 6.04E-42 | 0.469933416 | 0.336 | 0.249 | 1.22E-37 | 2 | Herc1 |

| | | | | | | | |
|---|---|---|---|---|---|---|---|
| Fbxw21 | 1.13E-41 | 0.371481346 | 0.317 | 0.228 | 2.29E-37 | 2 | Fbxw2 |
| Ar1 | 1.62E-41 | 0.372903991 | 0.309 | 0.209 | 3.28E-37 | 2 | Ar |
| Calr1 | 3.97E-41 | 0.308048676 | 0.507 | 0.398 | 8.05E-37 | 2 | Calr |
| Pcnx1 | 4.20E-41 | 0.476063852 | 0.255 | 0.171 | 8.50E-37 | 2 | Pcnx |
| mt-Atp81 | 6.00E-41 | 0.304702298 | 0.564 | 0.453 | 1.21E-36 | 2 | mt-Atp8 |
| Rab6a1 | 1.11E-40 | 0.372903915 | 0.296 | 0.208 | 2.25E-36 | 2 | Rab6a |
| Tnrc6c1 | 1.12E-40 | 0.447093546 | 0.394 | 0.308 | 2.27E-36 | 2 | Tnrc6c |
| Rassf11 | 1.43E-40 | 0.498547769 | 0.303 | 0.221 | 2.90E-36 | 2 | Rassf1 |
| Cnih4 | 1.74E-40 | 0.289373622 | 0.335 | 0.246 | 3.52E-36 | 2 | Cnih4 |
| Dpm3 | 1.84E-40 | 0.324925009 | 0.425 | 0.342 | 3.73E-36 | 2 | Dpm3 |
| Dnajc81 | 2.12E-40 | 0.288228875 | 0.439 | 0.351 | 4.29E-36 | 2 | Dnajc8 |
| Srgap21 | 2.47E-40 | 0.425267902 | 0.387 | 0.296 | 5.00E-36 | 2 | Srgap2 |
| Etnk11 | 3.09E-40 | 0.444596504 | 0.302 | 0.218 | 6.25E-36 | 2 | Etnk1 |
| Maml21 | 3.31E-40 | 0.388719304 | 0.438 | 0.334 | 6.70E-36 | 2 | Maml2 |
| Lrrfip11 | 5.21E-40 | 0.330812183 | 0.466 | 0.37 | 1.06E-35 | 2 | Lrrfip1 |
| Phf20l11 | 7.34E-40 | 0.448290695 | 0.362 | 0.28 | 1.49E-35 | 2 | Phf20l1 |
| Tsen34 | 7.38E-40 | 0.308521651 | 0.256 | 0.169 | 1.49E-35 | 2 | Tsen34 |
| Arl6ip11 | 2.26E-39 | 0.342232196 | 0.449 | 0.363 | 4.57E-35 | 2 | Arl6ip1 |
| Lrrc581 | 8.55E-39 | 0.364231646 | 0.295 | 0.206 | 1.73E-34 | 2 | Lrrc58 |
| Mdfic1 | 1.55E-38 | 0.429172602 | 0.323 | 0.238 | 3.15E-34 | 2 | Mdfic |
| Atp2b11 | 2.32E-38 | 0.392492286 | 0.411 | 0.326 | 4.70E-34 | 2 | Atp2b1 |
| Rassf31 | 2.61E-38 | 0.319161915 | 0.321 | 0.227 | 5.29E-34 | 2 | Rassf3 |
| Diaph21 | 6.77E-38 | 0.383432319 | 0.468 | 0.368 | 1.37E-33 | 2 | Diaph2 |
| Grn1 | 7.38E-38 | 0.390952891 | 0.303 | 0.216 | 1.49E-33 | 2 | Grn |
| 4930402H24Rik1 | 9.87E-38 | 0.437198637 | 0.297 | 0.214 | 2.00E-33 | 2 | 4930402H24Rik |
| Purb1 | 1.20E-37 | 0.252776802 | 0.449 | 0.361 | 2.43E-33 | 2 | Purb |
| Saraf1 | 1.47E-37 | 0.40973958 | 0.377 | 0.301 | 2.98E-33 | 2 | Saraf |
| Git21 | 1.70E-37 | 0.448997181 | 0.307 | 0.228 | 3.44E-33 | 2 | Git2 |
| Rab5c1 | 1.73E-37 | 0.287978546 | 0.369 | 0.283 | 3.50E-33 | 2 | Rab5c |
| Atp2b41 | 2.04E-37 | 0.453380651 | 0.273 | 0.191 | 4.13E-33 | 2 | Atp2b4 |
| Polr2e | 6.21E-37 | 0.27260358 | 0.398 | 0.317 | 1.26E-32 | 2 | Polr2e |
| Tm2d21 | 1.63E-36 | 0.396916088 | 0.282 | 0.205 | 3.29E-32 | 2 | Tm2d2 |
| Sema6d1 | 1.71E-36 | 0.425275172 | 0.293 | 0.206 | 3.46E-32 | 2 | Sema6d |
| H1f01 | 2.23E-36 | 0.26822134 | 0.277 | 0.186 | 4.51E-32 | 2 | H1f0 |
| Nfe2l21 | 2.33E-36 | 0.312512717 | 0.381 | 0.293 | 4.71E-32 | 2 | Nfe2l2 |
| Dennd5a1 | 2.59E-36 | 0.396366954 | 0.261 | 0.18 | 5.24E-32 | 2 | Dennd5a |
| Bcap311 | 2.65E-36 | 0.399099722 | 0.352 | 0.279 | 5.37E-32 | 2 | Bcap31 |
| Snx91 | 2.69E-36 | 0.36181145 | 0.251 | 0.169 | 5.46E-32 | 2 | Snx9 |
| Ptpn121 | 2.71E-36 | 0.466643739 | 0.284 | 0.206 | 5.48E-32 | 2 | Ptpn12 |
| Abhd17a | 1.13E-35 | 0.289529335 | 0.35 | 0.268 | 2.29E-31 | 2 | Abhd17a |
| Rtn41 | 1.49E-35 | 0.32256621 | 0.417 | 0.331 | 3.01E-31 | 2 | Rtn4 |
| Mcl11 | 1.71E-35 | 0.318175204 | 0.449 | 0.366 | 3.46E-31 | 2 | Mcl1 |
| Ash1l1 | 1.78E-35 | 0.399936581 | 0.373 | 0.293 | 3.61E-31 | 2 | Ash1l |

| | | | | | | | |
|---|---|---|---|---|---|---|---|
| Midn1 | 2.35E-35 | 0.402567524 | 0.262 | 0.184 | 4.77E-31 | 2 | Midn |
| Yipf41 | 2.41E-35 | 0.376135597 | 0.291 | 0.215 | 4.88E-31 | 2 | Yipf4 |
| Nmi1 | 3.33E-35 | 0.338888205 | 0.312 | 0.231 | 6.74E-31 | 2 | Nmi |
| Cfdp1 | 3.77E-35 | 0.280317868 | 0.38 | 0.299 | 7.63E-31 | 2 | Cfdp1 |
| Atp8a11 | 4.80E-35 | 0.468359236 | 0.314 | 0.235 | 9.71E-31 | 2 | Atp8a1 |
| Capns11 | 6.88E-35 | 0.267720981 | 0.422 | 0.342 | 1.39E-30 | 2 | Capns1 |
| Anxa71 | 9.56E-35 | 0.386203258 | 0.256 | 0.179 | 1.94E-30 | 2 | Anxa7 |
| Tspan51 | 1.31E-34 | 0.419257288 | 0.272 | 0.191 | 2.66E-30 | 2 | Tspan5 |
| Tmem2081 | 2.19E-34 | 0.414165464 | 0.304 | 0.232 | 4.43E-30 | 2 | Tmem208 |
| Fam96b | 2.44E-34 | 0.301963429 | 0.315 | 0.236 | 4.94E-30 | 2 | Fam96b |
| Yaf21 | 2.59E-34 | 0.410160579 | 0.284 | 0.209 | 5.25E-30 | 2 | Yaf2 |
| Nfat51 | 2.85E-34 | 0.460079652 | 0.337 | 0.261 | 5.77E-30 | 2 | Nfat5 |
| Arid5b1 | 6.01E-34 | 0.400035548 | 0.353 | 0.271 | 1.22E-29 | 2 | Arid5b |
| Arhgap261 | 9.98E-34 | 0.390629299 | 0.297 | 0.215 | 2.02E-29 | 2 | Arhgap26 |
| Spata131 | 1.03E-33 | 0.449356865 | 0.274 | 0.2 | 2.09E-29 | 2 | Spata13 |
| Gtf2i1 | 1.45E-33 | 0.335135325 | 0.322 | 0.244 | 2.94E-29 | 2 | Gtf2i |
| Vgll41 | 1.54E-33 | 0.400135299 | 0.319 | 0.244 | 3.11E-29 | 2 | Vgll4 |
| Hnrnph31 | 2.59E-33 | 0.330855962 | 0.318 | 0.243 | 5.25E-29 | 2 | Hnrnph3 |
| Srsf111 | 2.72E-33 | 0.380257545 | 0.415 | 0.345 | 5.51E-29 | 2 | Srsf11 |
| Itsn21 | 3.12E-33 | 0.433164205 | 0.343 | 0.272 | 6.32E-29 | 2 | Itsn2 |
| Junb1 | 1.03E-32 | 0.560496577 | 0.476 | 0.385 | 2.08E-28 | 2 | Junb |
| Kif1b1 | 1.17E-32 | 0.407831282 | 0.31 | 0.237 | 2.38E-28 | 2 | Kif1b |
| Il10rb1 | 3.08E-32 | 0.392847701 | 0.315 | 0.243 | 6.24E-28 | 2 | Il10rb |
| Strn31 | 3.37E-32 | 0.329399838 | 0.415 | 0.339 | 6.81E-28 | 2 | Strn3 |
| Ssu72 | 3.40E-32 | 0.256946166 | 0.381 | 0.305 | 6.88E-28 | 2 | Ssu72 |
| Susd61 | 3.53E-32 | 0.434578995 | 0.347 | 0.277 | 7.14E-28 | 2 | Susd6 |
| Usp341 | 3.55E-32 | 0.370111434 | 0.412 | 0.338 | 7.19E-28 | 2 | Usp34 |
| Lrpap11 | 4.96E-32 | 0.409877074 | 0.261 | 0.191 | 1.00E-27 | 2 | Lrpap1 |
| Adipor11 | 5.15E-32 | 0.356531642 | 0.325 | 0.251 | 1.04E-27 | 2 | Adipor1 |
| Ubald2 | 5.86E-32 | 0.260200396 | 0.285 | 0.205 | 1.19E-27 | 2 | Ubald2 |
| Ankrd121 | 7.54E-32 | 0.468787134 | 0.375 | 0.302 | 1.53E-27 | 2 | Ankrd12 |
| Zmynd111 | 8.29E-32 | 0.364806942 | 0.357 | 0.283 | 1.68E-27 | 2 | Zmynd11 |
| Raly | 9.43E-32 | 0.316989466 | 0.338 | 0.265 | 1.91E-27 | 2 | Raly |
| Tmpo1 | 1.06E-31 | 0.343259146 | 0.277 | 0.202 | 2.15E-27 | 2 | Tmpo |
| Stim21 | 1.13E-31 | 0.438910279 | 0.265 | 0.193 | 2.28E-27 | 2 | Stim2 |
| Cltc1 | 1.81E-31 | 0.277999349 | 0.399 | 0.325 | 3.66E-27 | 2 | Cltc |
| Pik3c2a1 | 2.67E-31 | 0.413389273 | 0.262 | 0.189 | 5.41E-27 | 2 | Pik3c2a |
| Polr2a1 | 3.07E-31 | 0.445862299 | 0.345 | 0.276 | 6.22E-27 | 2 | Polr2a |
| Tgfbr21 | 1.26E-30 | 0.422179678 | 0.376 | 0.306 | 2.55E-26 | 2 | Tgfbr2 |
| Dip2b1 | 1.26E-30 | 0.398756329 | 0.337 | 0.271 | 2.56E-26 | 2 | Dip2b |
| Fos1 | 1.82E-30 | 0.513636256 | 0.396 | 0.309 | 3.68E-26 | 2 | Fos |
| Srpr1 | 2.40E-30 | 0.388573068 | 0.307 | 0.24 | 4.86E-26 | 2 | Srpr |
| Tra2a1 | 3.03E-30 | 0.423462617 | 0.335 | 0.27 | 6.14E-26 | 2 | Tra2a |

| | | | | | | | |
|---|---|---|---|---|---|---|---|
| Prpf4b1 | 5.69E-30 | 0.331650654 | 0.396 | 0.328 | 1.15E-25 | 2 | Prpf4b |
| Ier21 | 1.28E-29 | 0.428154606 | 0.351 | 0.278 | 2.60E-25 | 2 | Ier2 |
| Gm203881 | 1.34E-29 | 0.277959901 | 0.481 | 0.393 | 2.71E-25 | 2 | Gm20388 |
| G3bp11 | 1.43E-29 | 0.312629308 | 0.366 | 0.297 | 2.90E-25 | 2 | G3bp1 |
| Rapgef11 | 1.45E-29 | 0.465860847 | 0.301 | 0.236 | 2.93E-25 | 2 | Rapgef1 |
| Eif4ebp21 | 1.58E-29 | 0.344191415 | 0.306 | 0.235 | 3.21E-25 | 2 | Eif4ebp2 |
| Pkn21 | 1.69E-29 | 0.341801758 | 0.356 | 0.289 | 3.42E-25 | 2 | Pkn2 |
| Ubn11 | 1.77E-29 | 0.359727475 | 0.308 | 0.241 | 3.59E-25 | 2 | Ubn1 |
| Isg151 | 2.18E-29 | 0.314722973 | 0.31 | 0.222 | 4.42E-25 | 2 | Isg15 |
| Sdf41 | 4.60E-29 | 0.322387569 | 0.369 | 0.304 | 9.32E-25 | 2 | Sdf4 |
| Ppp1r16b1 | 4.98E-29 | 0.379935838 | 0.267 | 0.196 | 1.01E-24 | 2 | Ppp1r16b |
| Rnf2131 | 5.26E-29 | 0.420851363 | 0.322 | 0.251 | 1.06E-24 | 2 | Rnf213 |
| Tbrg1 | 8.21E-29 | 0.260696464 | 0.369 | 0.301 | 1.66E-24 | 2 | Tbrg1 |
| Taok11 | 8.29E-29 | 0.359613023 | 0.315 | 0.246 | 1.68E-24 | 2 | Taok1 |
| Ndfip21 | 1.30E-28 | 0.354959273 | 0.305 | 0.24 | 2.62E-24 | 2 | Ndfip2 |
| Hprt | 1.55E-28 | 0.292144888 | 0.348 | 0.28 | 3.13E-24 | 2 | Hprt |
| Crk1 | 1.88E-28 | 0.318020379 | 0.257 | 0.187 | 3.81E-24 | 2 | Crk |
| Ddx171 | 3.20E-28 | 0.362228031 | 0.332 | 0.268 | 6.48E-24 | 2 | Ddx17 |
| Acap21 | 5.89E-28 | 0.376649761 | 0.33 | 0.264 | 1.19E-23 | 2 | Acap2 |
| Nisch1 | 6.50E-28 | 0.323847202 | 0.275 | 0.206 | 1.32E-23 | 2 | Nisch |
| Pja21 | 1.04E-27 | 0.360415696 | 0.266 | 0.199 | 2.10E-23 | 2 | Pja2 |
| Uvrag1 | 1.41E-27 | 0.396553409 | 0.325 | 0.263 | 2.86E-23 | 2 | Uvrag |
| Fubp11 | 1.47E-27 | 0.346663972 | 0.331 | 0.268 | 2.98E-23 | 2 | Fubp1 |
| Cdk171 | 1.62E-27 | 0.409719719 | 0.305 | 0.24 | 3.28E-23 | 2 | Cdk17 |
| Wasl1 | 3.29E-27 | 0.285372381 | 0.299 | 0.232 | 6.66E-23 | 2 | Wasl |
| Smurf21 | 4.80E-27 | 0.41747717 | 0.298 | 0.233 | 9.72E-23 | 2 | Smurf2 |
| Nipbl1 | 8.86E-27 | 0.357081767 | 0.428 | 0.361 | 1.79E-22 | 2 | Nipbl |
| Gimap41 | 1.22E-26 | 0.255518782 | 0.333 | 0.249 | 2.48E-22 | 2 | Gimap4 |
| Nrbp11 | 1.66E-26 | 0.347160138 | 0.269 | 0.206 | 3.35E-22 | 2 | Nrbp1 |
| Cyfip11 | 1.78E-26 | 0.280033615 | 0.345 | 0.272 | 3.60E-22 | 2 | Cyfip1 |
| Mier11 | 2.15E-26 | 0.263902722 | 0.39 | 0.327 | 4.34E-22 | 2 | Mier1 |
| Prrc2b1 | 2.54E-26 | 0.366843173 | 0.288 | 0.225 | 5.15E-22 | 2 | Prrc2b |
| Tbl1x1 | 9.53E-26 | 0.336821842 | 0.321 | 0.257 | 1.93E-21 | 2 | Tbl1x |
| Bax | 1.25E-25 | 0.261358965 | 0.371 | 0.309 | 2.54E-21 | 2 | Bax |
| Apc1 | 3.97E-25 | 0.366630253 | 0.304 | 0.244 | 8.03E-21 | 2 | Apc |
| Tln11 | 6.57E-25 | 0.25701697 | 0.472 | 0.399 | 1.33E-20 | 2 | Tln1 |
| Slk1 | 6.88E-25 | 0.357027222 | 0.298 | 0.236 | 1.39E-20 | 2 | Slk |
| Cast1 | 1.28E-24 | 0.283107724 | 0.369 | 0.307 | 2.59E-20 | 2 | Cast |
| Kdelr1 | 1.74E-24 | 0.297533507 | 0.293 | 0.233 | 3.51E-20 | 2 | Kdelr1 |
| Clock1 | 3.13E-24 | 0.362359911 | 0.26 | 0.199 | 6.34E-20 | 2 | Clock |
| Vezf11 | 1.80E-23 | 0.259025502 | 0.293 | 0.229 | 3.64E-19 | 2 | Vezf1 |
| Tmbim4 | 2.56E-23 | 0.253037744 | 0.35 | 0.295 | 5.19E-19 | 2 | Tmbim4 |
| Ubn21 | 5.11E-23 | 0.40370153 | 0.295 | 0.241 | 1.04E-18 | 2 | Ubn2 |

| | | | | | | | |
|---|---|---|---|---|---|---|---|
| Pbrm11 | 5.54E-23 | 0.327569514 | 0.341 | 0.283 | 1.12E-18 | 2 | Pbrm1 |
| Ccnl21 | 6.21E-23 | 0.376155715 | 0.265 | 0.21 | 1.26E-18 | 2 | Ccnl2 |
| Dctn41 | 8.31E-23 | 0.291814797 | 0.266 | 0.207 | 1.68E-18 | 2 | Dctn4 |
| Stat31 | 1.07E-22 | 0.363871643 | 0.317 | 0.263 | 2.17E-18 | 2 | Stat3 |
| Klf121 | 1.65E-22 | 0.372534874 | 0.265 | 0.202 | 3.33E-18 | 2 | Klf12 |
| Zdhhc201 | 5.74E-22 | 0.415675245 | 0.277 | 0.227 | 1.16E-17 | 2 | Zdhhc20 |
| Bptf1 | 7.84E-22 | 0.363439125 | 0.342 | 0.29 | 1.59E-17 | 2 | Bptf |
| Ubl31 | 9.91E-22 | 0.259774904 | 0.293 | 0.234 | 2.01E-17 | 2 | Ubl3 |
| Mydgf | 1.30E-21 | 0.328591153 | 0.279 | 0.228 | 2.62E-17 | 2 | Mydgf |
| Wac1 | 3.73E-21 | 0.28993955 | 0.369 | 0.319 | 7.55E-17 | 2 | Wac |
| Herpud1 | 3.99E-21 | 0.334746682 | 0.262 | 0.21 | 8.09E-17 | 2 | Herpud1 |
| Crebbp1 | 4.01E-21 | 0.351337524 | 0.352 | 0.302 | 8.12E-17 | 2 | Crebbp |
| Glg11 | 4.69E-21 | 0.366493013 | 0.3 | 0.249 | 9.50E-17 | 2 | Glg1 |
| Lrch31 | 7.78E-21 | 0.375605328 | 0.28 | 0.229 | 1.58E-16 | 2 | Lrch3 |
| Tacc11 | 1.08E-20 | 0.299182692 | 0.352 | 0.302 | 2.20E-16 | 2 | Tacc1 |
| Exoc41 | 1.20E-20 | 0.369930071 | 0.374 | 0.319 | 2.43E-16 | 2 | Exoc4 |
| Zcchc61 | 1.31E-20 | 0.320075041 | 0.326 | 0.275 | 2.65E-16 | 2 | Zcchc6 |
| Canx1 | 1.61E-20 | 0.285017675 | 0.355 | 0.303 | 3.25E-16 | 2 | Canx |
| Lgals91 | 1.70E-20 | 0.366691021 | 0.294 | 0.244 | 3.44E-16 | 2 | Lgals9 |
| Puf60 | 3.68E-20 | 0.27054419 | 0.264 | 0.211 | 7.44E-16 | 2 | Puf60 |
| Ctcf1 | 3.96E-20 | 0.281987424 | 0.325 | 0.273 | 8.02E-16 | 2 | Ctcf |
| Slfn21 | 4.52E-20 | 0.326205968 | 0.314 | 0.257 | 9.16E-16 | 2 | Slfn2 |
| Gsk3b1 | 6.30E-20 | 0.263982915 | 0.378 | 0.327 | 1.27E-15 | 2 | Gsk3b |
| Rpn21 | 7.09E-20 | 0.367638247 | 0.256 | 0.209 | 1.43E-15 | 2 | Rpn2 |
| Usp251 | 1.02E-19 | 0.291597068 | 0.33 | 0.278 | 2.06E-15 | 2 | Usp25 |
| Tor1aip11 | 1.30E-19 | 0.303488483 | 0.327 | 0.278 | 2.64E-15 | 2 | Tor1aip1 |
| Rab11b1 | 1.33E-19 | 0.263978397 | 0.313 | 0.262 | 2.69E-15 | 2 | Rab11b |
| Rbbp61 | 1.75E-19 | 0.290509597 | 0.324 | 0.273 | 3.55E-15 | 2 | Rbbp6 |
| Arhgap171 | 1.78E-19 | 0.33117405 | 0.317 | 0.267 | 3.60E-15 | 2 | Arhgap17 |
| AU0202061 | 1.90E-19 | 0.307296018 | 0.346 | 0.287 | 3.84E-15 | 2 | AU020206 |
| Casp81 | 1.90E-19 | 0.32782251 | 0.261 | 0.211 | 3.85E-15 | 2 | Casp8 |
| Fam174a | 1.92E-19 | 0.252586425 | 0.251 | 0.198 | 3.89E-15 | 2 | Fam174a |
| Arih11 | 2.46E-19 | 0.297048895 | 0.361 | 0.309 | 4.98E-15 | 2 | Arih1 |
| Pcm11 | 5.45E-19 | 0.327243201 | 0.296 | 0.248 | 1.10E-14 | 2 | Pcm1 |
| Kmt2a1 | 8.83E-19 | 0.382146976 | 0.31 | 0.264 | 1.79E-14 | 2 | Kmt2a |
| Gpatch81 | 3.02E-18 | 0.376226199 | 0.297 | 0.256 | 6.12E-14 | 2 | Gpatch8 |
| Gatad2a1 | 3.93E-18 | 0.29160222 | 0.267 | 0.219 | 7.95E-14 | 2 | Gatad2a |
| Ppig1 | 4.73E-18 | 0.320748353 | 0.356 | 0.314 | 9.57E-14 | 2 | Ppig |
| Cdk121 | 8.81E-18 | 0.345472401 | 0.272 | 0.227 | 1.78E-13 | 2 | Cdk12 |
| Exoc6b1 | 1.30E-17 | 0.356781675 | 0.285 | 0.238 | 2.63E-13 | 2 | Exoc6b |
| Cmip1 | 1.41E-17 | 0.347820951 | 0.325 | 0.283 | 2.85E-13 | 2 | Cmip |
| Arl6ip51 | 1.73E-17 | 0.333533065 | 0.264 | 0.219 | 3.50E-13 | 2 | Arl6ip5 |
| Smarca21 | 4.31E-17 | 0.297527015 | 0.324 | 0.278 | 8.73E-13 | 2 | Smarca2 |

| | | | | | | | |
|---|---|---|---|---|---|---|---|
| Golgb11 | 9.69E-17 | 0.292051449 | 0.303 | 0.26 | 1.96E-12 | 2 | Golgb1 |
| Nf11 | 1.05E-16 | 0.36124173 | 0.264 | 0.218 | 2.12E-12 | 2 | Nf1 |
| Ttc141 | 1.08E-16 | 0.379047519 | 0.253 | 0.213 | 2.18E-12 | 2 | Ttc14 |
| Eea11 | 1.10E-16 | 0.297621032 | 0.262 | 0.215 | 2.23E-12 | 2 | Eea1 |
| Hnrnph11 | 1.50E-16 | 0.26608023 | 0.292 | 0.247 | 3.03E-12 | 2 | Hnrnph1 |
| Jmjd1c1 | 2.27E-16 | 0.314111163 | 0.348 | 0.305 | 4.60E-12 | 2 | Jmjd1c |
| Thoc21 | 3.09E-16 | 0.344748381 | 0.312 | 0.272 | 6.26E-12 | 2 | Thoc2 |
| Nedd4l1 | 4.31E-16 | 0.342793793 | 0.259 | 0.214 | 8.73E-12 | 2 | Nedd4l |
| Ubxn41 | 4.52E-16 | 0.309050049 | 0.309 | 0.27 | 9.14E-12 | 2 | Ubxn4 |
| Kpna41 | 4.87E-16 | 0.264773721 | 0.362 | 0.321 | 9.86E-12 | 2 | Kpna4 |
| Rb1cc11 | 5.26E-16 | 0.330088595 | 0.276 | 0.232 | 1.06E-11 | 2 | Rb1cc1 |
| Pum11 | 6.05E-16 | 0.30847847 | 0.295 | 0.257 | 1.23E-11 | 2 | Pum1 |
| Brd21 | 7.18E-16 | 0.275333155 | 0.293 | 0.251 | 1.45E-11 | 2 | Brd2 |
| Bod1l1 | 1.47E-15 | 0.327640686 | 0.312 | 0.271 | 2.97E-11 | 2 | Bod1l |
| Fyn1 | 1.53E-15 | 0.254419941 | 0.314 | 0.263 | 3.10E-11 | 2 | Fyn |
| Babam21 | 1.63E-15 | 0.367590511 | 0.275 | 0.236 | 3.31E-11 | 2 | Babam2 |
| Tnks21 | 1.64E-15 | 0.265222049 | 0.318 | 0.278 | 3.33E-11 | 2 | Tnks2 |
| Kmt2c1 | 4.06E-15 | 0.286504929 | 0.342 | 0.304 | 8.22E-11 | 2 | Kmt2c |
| Tgfb11 | 5.62E-15 | 0.303779005 | 0.274 | 0.234 | 1.14E-10 | 2 | Tgfb1 |
| Zfr1 | 1.13E-14 | 0.318712532 | 0.284 | 0.245 | 2.30E-10 | 2 | Zfr |
| Atp1b31 | 1.26E-14 | 0.294238222 | 0.285 | 0.247 | 2.55E-10 | 2 | Atp1b3 |
| Stat11 | 2.62E-14 | 0.257040646 | 0.292 | 0.247 | 5.31E-10 | 2 | Stat1 |
| Brd11 | 3.04E-14 | 0.284379075 | 0.275 | 0.239 | 6.15E-10 | 2 | Brd1 |
| Fbxl171 | 4.29E-14 | 0.307633863 | 0.349 | 0.309 | 8.69E-10 | 2 | Fbxl17 |
| Cflar1 | 4.37E-14 | 0.263066292 | 0.269 | 0.23 | 8.85E-10 | 2 | Cflar |
| Rb11 | 5.78E-14 | 0.332687795 | 0.251 | 0.212 | 1.17E-09 | 2 | Rb1 |
| Nrip11 | 8.28E-14 | 0.347023531 | 0.28 | 0.242 | 1.68E-09 | 2 | Nrip1 |
| Setd51 | 1.94E-13 | 0.33102957 | 0.294 | 0.262 | 3.93E-09 | 2 | Setd5 |
| Mllt101 | 3.43E-13 | 0.310617314 | 0.306 | 0.273 | 6.95E-09 | 2 | Mllt10 |
| Zfp6441 | 3.43E-13 | 0.281756338 | 0.287 | 0.254 | 6.95E-09 | 2 | Zfp644 |
| Cdk11b1 | 4.64E-13 | 0.269137693 | 0.277 | 0.242 | 9.40E-09 | 2 | Cdk11b |
| Luc7l31 | 5.57E-13 | 0.274014406 | 0.303 | 0.27 | 1.13E-08 | 2 | Luc7l3 |
| Arhgef11 | 8.71E-13 | 0.320288639 | 0.279 | 0.247 | 1.76E-08 | 2 | Arhgef1 |
| Rbm251 | 1.35E-12 | 0.261302577 | 0.364 | 0.333 | 2.72E-08 | 2 | Rbm25 |
| Camk2d1 | 1.81E-12 | 0.301954905 | 0.304 | 0.268 | 3.67E-08 | 2 | Camk2d |
| Brd41 | 2.06E-12 | 0.326539529 | 0.295 | 0.266 | 4.17E-08 | 2 | Brd4 |
| Pnisr1 | 2.23E-12 | 0.317717012 | 0.293 | 0.263 | 4.51E-08 | 2 | Pnisr |
| Ankhd11 | 1.17E-11 | 0.327717273 | 0.273 | 0.243 | 2.37E-07 | 2 | Ankhd1 |
| Zbtb7a1 | 2.31E-11 | 0.278464938 | 0.259 | 0.228 | 4.68E-07 | 2 | Zbtb7a |
| Trim30a1 | 9.02E-11 | 0.305732321 | 0.25 | 0.218 | 1.83E-06 | 2 | Trim30a |
| Rnf1301 | 1.20E-10 | 0.254905794 | 0.308 | 0.283 | 2.42E-06 | 2 | Rnf130 |
| Ccny1 | 1.91E-10 | 0.266862653 | 0.293 | 0.267 | 3.86E-06 | 2 | Ccny |
| Smchd11 | 6.15E-10 | 0.286463074 | 0.287 | 0.264 | 1.24E-05 | 2 | Smchd1 |

| | | | | | | | |
|---|---|---|---|---|---|---|---|
| Pnn1 | 1.20E-09 | 0.255817053 | 0.289 | 0.267 | 2.43E-05 | 2 | Pnn |
| Trps11 | 1.62E-09 | 0.264650799 | 0.267 | 0.235 | 3.29E-05 | 2 | Trps1 |
| Arid4b1 | 4.74E-09 | 0.262879552 | 0.314 | 0.293 | 9.60E-05 | 2 | Arid4b |
| Kdm2a1 | 1.11E-08 | 0.259674922 | 0.262 | 0.24 | 0.000225335 | 2 | Kdm2a |
| Sik31 | 1.52E-07 | 0.255402125 | 0.291 | 0.27 | 0.003076645 | 2 | Sik3 |
| Arid4a1 | 1.86E-07 | 0.274356146 | 0.269 | 0.252 | 0.0037564 | 2 | Arid4a |
| Phf31 | 2.56E-07 | 0.259092295 | 0.268 | 0.253 | 0.005185265 | 2 | Phf3 |
| Btbd91 | 3.67E-07 | 0.323440954 | 0.254 | 0.238 | 0.007436773 | 2 | Btbd9 |
| Zfp2921 | 6.24E-07 | 0.250745057 | 0.268 | 0.252 | 0.012634169 | 2 | Zfp292 |
| Akap91 | 8.46E-06 | 0.256303688 | 0.251 | 0.238 | 0.171320502 | 2 | Akap9 |
| Klk1b26 | 0 | 3.987975674 | 0.909 | 0.343 | 0 | 3 | Klk1b26 |
| Klk11 | 0 | 3.711582034 | 0.999 | 0.858 | 0 | 3 | Klk1 |
| Crisp3 | 0 | 3.47208559 | 0.795 | 0.231 | 0 | 3 | Crisp3 |
| Klk1b5 | 0 | 3.2404655 | 0.726 | 0.14 | 0 | 3 | Klk1b5 |
| Klk1b111 | 0 | 3.205807653 | 0.83 | 0.196 | 0 | 3 | Klk1b11 |
| Klk1b22 | 0 | 3.054372632 | 0.492 | 0.068 | 0 | 3 | Klk1b22 |
| Klk1b4 | 0 | 2.808748176 | 0.499 | 0.073 | 0 | 3 | Klk1b4 |
| Klk1b91 | 0 | 2.744625666 | 0.928 | 0.299 | 0 | 3 | Klk1b9 |
| G0s21 | 0 | 2.408060272 | 0.815 | 0.245 | 0 | 3 | G0s2 |
| Klk1b3 | 0 | 2.314953691 | 0.556 | 0.091 | 0 | 3 | Klk1b3 |
| Klk1b21 | 0 | 2.224698614 | 0.347 | 0.04 | 0 | 3 | Klk1b21 |
| Wfdc2 | 0 | 2.148941957 | 0.489 | 0.081 | 0 | 3 | Wfdc2 |
| Klk1b271 | 0 | 2.012746737 | 0.564 | 0.11 | 0 | 3 | Klk1b27 |
| Stxbp61 | 0 | 1.931463581 | 0.554 | 0.146 | 0 | 3 | Stxbp6 |
| Fxyd21 | 0 | 1.880158269 | 1 | 0.762 | 0 | 3 | Fxyd2 |
| S100a12 | 0 | 1.872409931 | 0.938 | 0.545 | 0 | 3 | S100a1 |
| Krt181 | 0 | 1.80766995 | 0.728 | 0.221 | 0 | 3 | Krt18 |
| Tesc1 | 0 | 1.80561489 | 0.867 | 0.336 | 0 | 3 | Tesc |
| Nudt41 | 0 | 1.790456566 | 0.73 | 0.373 | 0 | 3 | Nudt4 |
| Tfcp2l11 | 0 | 1.754183407 | 0.69 | 0.229 | 0 | 3 | Tfcp2l1 |
| Chchd101 | 0 | 1.733327287 | 0.891 | 0.387 | 0 | 3 | Chchd10 |
| Scgb1c1 | 0 | 1.669452278 | 0.271 | 0.03 | 0 | 3 | Scgb1c1 |
| Mpp61 | 0 | 1.6580244 | 0.589 | 0.248 | 0 | 3 | Mpp6 |
| Mif1 | 0 | 1.56659159 | 0.813 | 0.49 | 0 | 3 | Mif |
| Mt11 | 0 | 1.564590746 | 0.823 | 0.388 | 0 | 3 | Mt1 |
| Cox7b1 | 0 | 1.551263632 | 0.88 | 0.652 | 0 | 3 | Cox7b |
| Calm21 | 0 | 1.509677457 | 0.905 | 0.763 | 0 | 3 | Calm2 |
| Cox7c1 | 0 | 1.505655509 | 0.952 | 0.832 | 0 | 3 | Cox7c |
| Mt21 | 0 | 1.497498088 | 0.426 | 0.113 | 0 | 3 | Mt2 |
| Serpinb1a | 0 | 1.488146371 | 0.423 | 0.135 | 0 | 3 | Serpinb1a |
| Dstn1 | 0 | 1.473093405 | 0.829 | 0.545 | 0 | 3 | Dstn |
| Vamp81 | 0 | 1.47187122 | 0.714 | 0.486 | 0 | 3 | Vamp8 |
| Mpc11 | 0 | 1.465540316 | 0.786 | 0.5 | 0 | 3 | Mpc1 |

| | | | | | | | |
|---|---|---|---|---|---|---|---|
| Suclg11 | 0 | 1.462787665 | 0.661 | 0.378 | 0 | 3 | Suclg1 |
| Atp5g11 | 0 | 1.448804603 | 0.883 | 0.659 | 0 | 3 | Atp5g1 |
| Ckmt11 | 0 | 1.436736368 | 0.484 | 0.13 | 0 | 3 | Ckmt1 |
| Apoc1 | 0 | 1.433840963 | 0.261 | 0.054 | 0 | 3 | Apoc1 |
| Atp5j1 | 0 | 1.430725111 | 0.945 | 0.821 | 0 | 3 | Atp5j |
| Sod21 | 0 | 1.430020218 | 0.71 | 0.41 | 0 | 3 | Sod2 |
| Atp5g31 | 0 | 1.406727577 | 0.825 | 0.589 | 0 | 3 | Atp5g3 |
| Cox6c1 | 0 | 1.406635277 | 0.941 | 0.818 | 0 | 3 | Cox6c |
| Cox5b1 | 0 | 1.403955487 | 0.863 | 0.693 | 0 | 3 | Cox5b |
| Tpd521 | 0 | 1.401188201 | 0.574 | 0.276 | 0 | 3 | Tpd52 |
| Ndufab11 | 0 | 1.399837386 | 0.765 | 0.513 | 0 | 3 | Ndufab1 |
| Krt71 | 0 | 1.396878423 | 0.511 | 0.139 | 0 | 3 | Krt7 |
| Slc25a51 | 0 | 1.383529445 | 0.863 | 0.652 | 0 | 3 | Slc25a5 |
| Mdh11 | 0 | 1.379681104 | 0.662 | 0.401 | 0 | 3 | Mdh1 |
| Plet1 | 0 | 1.37902715 | 0.255 | 0.05 | 0 | 3 | Plet1 |
| Atp5k1 | 0 | 1.363079356 | 0.817 | 0.588 | 0 | 3 | Atp5k |
| Mpc21 | 0 | 1.361126937 | 0.754 | 0.5 | 0 | 3 | Mpc2 |
| Mapk13 | 0 | 1.357143071 | 0.305 | 0.054 | 0 | 3 | Mapk13 |
| Ldhb1 | 0 | 1.355441761 | 0.588 | 0.23 | 0 | 3 | Ldhb |
| Atp5b1 | 0 | 1.353390884 | 0.863 | 0.702 | 0 | 3 | Atp5b |
| Cox8a1 | 0 | 1.353356088 | 0.964 | 0.887 | 0 | 3 | Cox8a |
| Cox6b11 | 0 | 1.343783934 | 0.855 | 0.72 | 0 | 3 | Cox6b1 |
| Usmg51 | 0 | 1.343695839 | 0.742 | 0.527 | 0 | 3 | Usmg5 |
| Uqcrb1 | 0 | 1.34294351 | 0.866 | 0.717 | 0 | 3 | Uqcrb |
| Slc25a31 | 0 | 1.333978271 | 0.852 | 0.675 | 0 | 3 | Slc25a3 |
| Cystm11 | 0 | 1.333794251 | 0.477 | 0.149 | 0 | 3 | Cystm1 |
| Cox4i11 | 0 | 1.333366954 | 0.944 | 0.84 | 0 | 3 | Cox4i1 |
| Ndufc11 | 0 | 1.329106928 | 0.741 | 0.521 | 0 | 3 | Ndufc1 |
| Cdo1 | 0 | 1.323712142 | 0.32 | 0.069 | 0 | 3 | Cdo1 |
| 2010107E04Rik1 | 0 | 1.312086612 | 0.761 | 0.516 | 0 | 3 | 2010107E04Rik |
| Stard10 | 0 | 1.310820929 | 0.39 | 0.107 | 0 | 3 | Stard10 |
| Cox7a21 | 0 | 1.307749506 | 0.878 | 0.758 | 0 | 3 | Cox7a2 |
| Smim5 | 0 | 1.306220666 | 0.371 | 0.081 | 0 | 3 | Smim5 |
| Ndufb101 | 0 | 1.303974841 | 0.783 | 0.615 | 0 | 3 | Ndufb10 |
| Atp5l1 | 0 | 1.298082443 | 0.908 | 0.797 | 0 | 3 | Atp5l |
| Atp5h1 | 0 | 1.293285133 | 0.921 | 0.808 | 0 | 3 | Atp5h |
| Cox5a1 | 0 | 1.291540014 | 0.874 | 0.738 | 0 | 3 | Cox5a |
| Atp5o.11 | 0 | 1.291429234 | 0.743 | 0.539 | 0 | 3 | Atp5o.1 |
| Ndufb41 | 0 | 1.286800086 | 0.746 | 0.562 | 0 | 3 | Ndufb4 |
| Uqcr111 | 0 | 1.282994447 | 0.818 | 0.667 | 0 | 3 | Uqcr11 |
| Atp5e1 | 0 | 1.282678118 | 0.942 | 0.825 | 0 | 3 | Atp5e |
| Aldoa1 | 0 | 1.278008587 | 0.788 | 0.623 | 0 | 3 | Aldoa |
| Uqcrq1 | 0 | 1.273094588 | 0.798 | 0.641 | 0 | 3 | Uqcrq |

| | | | | | | | |
|---|---|---|---|---|---|---|---|
| Uqcr101 | 0 | 1.259199521 | 0.826 | 0.663 | 0 | 3 | Uqcr10 |
| Atp5f11 | 0 | 1.247057474 | 0.82 | 0.672 | 0 | 3 | Atp5f1 |
| S100a111 | 0 | 1.244772902 | 0.761 | 0.553 | 0 | 3 | S100a11 |
| Ehf | 0 | 1.240200776 | 0.34 | 0.091 | 0 | 3 | Ehf |
| Arg1 | 0 | 1.226989536 | 0.356 | 0.091 | 0 | 3 | Arg1 |
| Atp5d1 | 0 | 1.220239139 | 0.818 | 0.664 | 0 | 3 | Atp5d |
| Ndufa41 | 0 | 1.20670404 | 0.889 | 0.81 | 0 | 3 | Ndufa4 |
| Atp5c11 | 0 | 1.199904117 | 0.785 | 0.625 | 0 | 3 | Atp5c1 |
| Uqcrh1 | 0 | 1.172438316 | 0.901 | 0.795 | 0 | 3 | Uqcrh |
| Atp5j21 | 0 | 1.127781927 | 0.851 | 0.755 | 0 | 3 | Atp5j2 |
| Chchd21 | 0 | 1.067072332 | 0.945 | 0.902 | 0 | 3 | Chchd2 |
| H3f3b1 | 0 | 0.866732741 | 0.93 | 0.904 | 0 | 3 | H3f3b |
| Rplp01 | 0 | 0.825718828 | 0.963 | 0.943 | 0 | 3 | Rplp0 |
| Rplp11 | 0 | 0.78978057 | 0.983 | 0.983 | 0 | 3 | Rplp1 |
| Atp5g21 | 2.54E-306 | 1.050141295 | 0.806 | 0.722 | 5.14E-302 | 3 | Atp5g2 |
| Serpinb6a1 | 2.51E-299 | 1.52962429 | 0.628 | 0.384 | 5.08E-295 | 3 | Serpinb6a |
| Ndufa111 | 6.58E-299 | 1.239340181 | 0.713 | 0.504 | 1.33E-294 | 3 | Ndufa11 |
| Ndufb81 | 7.73E-299 | 1.283304475 | 0.719 | 0.543 | 1.57E-294 | 3 | Ndufb8 |
| Ndufa51 | 9.46E-299 | 1.280395404 | 0.7 | 0.486 | 1.92E-294 | 3 | Ndufa5 |
| Krt8 | 1.14E-298 | 1.148270428 | 0.328 | 0.087 | 2.31E-294 | 3 | Krt8 |
| Gsta41 | 7.93E-295 | 1.185729606 | 0.39 | 0.123 | 1.61E-290 | 3 | Gsta4 |
| Prdx51 | 2.25E-293 | 1.233426028 | 0.719 | 0.512 | 4.55E-289 | 3 | Prdx5 |
| Phyh1 | 8.86E-285 | 1.239473968 | 0.479 | 0.192 | 1.79E-280 | 3 | Phyh |
| Gpx11 | 1.59E-284 | 1.157093533 | 0.752 | 0.579 | 3.21E-280 | 3 | Gpx1 |
| Rpl101 | 1.96E-284 | 0.803468473 | 0.922 | 0.894 | 3.97E-280 | 3 | Rpl10 |
| Cycs1 | 2.09E-271 | 1.25758373 | 0.657 | 0.424 | 4.24E-267 | 3 | Cycs |
| Ndufb111 | 4.71E-269 | 1.173777225 | 0.739 | 0.634 | 9.54E-265 | 3 | Ndufb11 |
| Mcee1 | 1.15E-263 | 1.301533646 | 0.527 | 0.256 | 2.32E-259 | 3 | Mcee |
| Atp5a11 | 8.85E-262 | 1.145088503 | 0.734 | 0.614 | 1.79E-257 | 3 | Atp5a1 |
| Cox7a11 | 1.91E-257 | 1.08724105 | 0.35 | 0.107 | 3.87E-253 | 3 | Cox7a1 |
| Ndufa61 | 1.30E-255 | 1.143271243 | 0.718 | 0.565 | 2.63E-251 | 3 | Ndufa6 |
| Prr15l | 1.35E-249 | 1.038731038 | 0.271 | 0.069 | 2.73E-245 | 3 | Prr15l |
| Map1lc3a1 | 3.69E-249 | 1.251937947 | 0.663 | 0.462 | 7.46E-245 | 3 | Map1lc3a |
| Atp6v0d2 | 4.10E-249 | 1.050053975 | 0.317 | 0.092 | 8.31E-245 | 3 | Atp6v0d2 |
| Fuom1 | 1.18E-246 | 1.167680799 | 0.397 | 0.146 | 2.38E-242 | 3 | Fuom |
| Ndufb91 | 4.00E-245 | 1.103034163 | 0.745 | 0.662 | 8.10E-241 | 3 | Ndufb9 |
| Atp1b1 | 2.27E-244 | 1.696143703 | 0.367 | 0.139 | 4.60E-240 | 3 | Atp1b1 |
| Ndufa121 | 8.87E-241 | 1.215498183 | 0.645 | 0.44 | 1.80E-236 | 3 | Ndufa12 |
| Uqcrfs11 | 1.06E-240 | 1.223207022 | 0.616 | 0.386 | 2.14E-236 | 3 | Uqcrfs1 |
| Rragd | 1.82E-239 | 1.017047107 | 0.291 | 0.081 | 3.68E-235 | 3 | Rragd |
| Sdhb1 | 5.68E-235 | 1.225364122 | 0.622 | 0.417 | 1.15E-230 | 3 | Sdhb |
| Kctd1 | 1.04E-228 | 1.042998176 | 0.288 | 0.085 | 2.10E-224 | 3 | Kctd1 |
| Csrp2 | 1.81E-227 | 1.315644097 | 0.412 | 0.176 | 3.66E-223 | 3 | Csrp2 |

| | | | | | | | |
|---|---|---|---|---|---|---|---|
| Aes1 | 6.74E-225 | 1.064014157 | 0.725 | 0.614 | 1.36E-220 | 3 | Aes |
| 2410015M20Rik1 | 1.85E-222 | 1.122107771 | 0.672 | 0.506 | 3.76E-218 | 3 | 2410015M20Rik |
| Acot131 | 7.07E-220 | 1.237283575 | 0.442 | 0.203 | 1.43E-215 | 3 | Acot13 |
| Vdac11 | 1.33E-219 | 1.226503518 | 0.559 | 0.33 | 2.70E-215 | 3 | Vdac1 |
| Atp6v1f1 | 3.97E-219 | 1.06333278 | 0.712 | 0.602 | 8.03E-215 | 3 | Atp6v1f |
| Rpl261 | 1.11E-212 | 0.566457664 | 0.949 | 0.958 | 2.24E-208 | 3 | Rpl26 |
| Ndufb21 | 1.36E-212 | 1.209116833 | 0.589 | 0.385 | 2.76E-208 | 3 | Ndufb2 |
| Dhrs4 | 7.27E-212 | 1.223689438 | 0.375 | 0.152 | 1.47E-207 | 3 | Dhrs4 |
| Hadh1 | 7.02E-211 | 1.234906352 | 0.459 | 0.224 | 1.42E-206 | 3 | Hadh |
| Mrpl421 | 1.47E-208 | 1.206590796 | 0.552 | 0.332 | 2.98E-204 | 3 | Mrpl42 |
| Ndufb51 | 7.54E-208 | 1.086736759 | 0.67 | 0.532 | 1.53E-203 | 3 | Ndufb5 |
| Ndufs61 | 4.41E-204 | 1.129434622 | 0.636 | 0.47 | 8.93E-200 | 3 | Ndufs6 |
| Snx101 | 2.52E-202 | 1.126830576 | 0.397 | 0.17 | 5.09E-198 | 3 | Snx10 |
| Apoo | 2.72E-202 | 1.052245881 | 0.299 | 0.1 | 5.50E-198 | 3 | Apoo |
| Rab25 | 1.27E-199 | 0.893764987 | 0.251 | 0.071 | 2.58E-195 | 3 | Rab25 |
| Pdcd41 | 3.48E-192 | 1.114697081 | 0.622 | 0.435 | 7.04E-188 | 3 | Pdcd4 |
| Rpl411 | 3.44E-190 | 0.520619841 | 0.955 | 0.963 | 6.96E-186 | 3 | Rpl41 |
| Echs11 | 2.69E-189 | 1.173443966 | 0.46 | 0.24 | 5.45E-185 | 3 | Echs1 |
| Coq71 | 4.29E-188 | 1.050256575 | 0.354 | 0.141 | 8.69E-184 | 3 | Coq7 |
| Ppia1 | 8.32E-185 | 0.522253576 | 0.935 | 0.939 | 1.68E-180 | 3 | Ppia |
| Acadl1 | 1.98E-184 | 1.213736078 | 0.547 | 0.354 | 4.01E-180 | 3 | Acadl |
| Gabarapl11 | 1.69E-180 | 1.160662383 | 0.431 | 0.215 | 3.43E-176 | 3 | Gabarapl1 |
| Minos11 | 6.70E-178 | 1.067630459 | 0.633 | 0.501 | 1.36E-173 | 3 | Minos1 |
| Rpl10a1 | 2.86E-177 | 0.64603453 | 0.875 | 0.832 | 5.80E-173 | 3 | Rpl10a |
| Ndufa14 | 2.12E-173 | 1.162041944 | 0.607 | 0.468 | 4.28E-169 | 3 | Ndufa1 |
| Naca1 | 1.94E-171 | 0.65178069 | 0.837 | 0.834 | 3.93E-167 | 3 | Naca |
| Gapdh1 | 9.39E-168 | 0.85582436 | 0.74 | 0.706 | 1.90E-163 | 3 | Gapdh |
| Ndufa131 | 1.46E-167 | 0.936537957 | 0.707 | 0.664 | 2.96E-163 | 3 | Ndufa13 |
| Ndufb71 | 1.36E-164 | 1.011834388 | 0.667 | 0.585 | 2.76E-160 | 3 | Ndufb7 |
| Fdx11 | 1.70E-163 | 1.086951937 | 0.491 | 0.292 | 3.44E-159 | 3 | Fdx1 |
| Nupr11 | 3.03E-163 | 0.2756996 | 0.516 | 0.282 | 6.13E-159 | 3 | Nupr1 |
| Gm28285 | 8.89E-162 | 0.989966275 | 0.294 | 0.111 | 1.80E-157 | 3 | Gm28285 |
| Zfp467 | 8.05E-161 | 1.158309396 | 0.348 | 0.157 | 1.63E-156 | 3 | Zfp467 |
| Cyc11 | 2.79E-160 | 1.099538661 | 0.517 | 0.331 | 5.66E-156 | 3 | Cyc1 |
| Ifi271 | 7.60E-160 | 1.080913477 | 0.633 | 0.526 | 1.54E-155 | 3 | Ifi27 |
| Ndufs51 | 5.64E-158 | 0.97329725 | 0.663 | 0.582 | 1.14E-153 | 3 | Ndufs5 |
| Ndufv21 | 4.26E-157 | 1.113040229 | 0.557 | 0.402 | 8.62E-153 | 3 | Ndufv2 |
| Prdx21 | 1.50E-150 | 0.954344177 | 0.631 | 0.524 | 3.04E-146 | 3 | Prdx2 |
| Oaz11 | 3.32E-149 | 0.662763298 | 0.796 | 0.81 | 6.72E-145 | 3 | Oaz1 |
| Rps21 | 4.95E-146 | 0.501401136 | 0.915 | 0.921 | 1.00E-141 | 3 | Rps2 |
| Tpi11 | 6.21E-144 | 1.05741824 | 0.492 | 0.318 | 1.26E-139 | 3 | Tpi1 |
| Pdlim4 | 2.62E-142 | 0.947205595 | 0.289 | 0.117 | 5.30E-138 | 3 | Pdlim4 |
| Atpif11 | 1.51E-139 | 0.906330894 | 0.65 | 0.552 | 3.07E-135 | 3 | Atpif1 |

| | | | | | | | |
|---|---|---|---|---|---|---|---|
| Vdac21 | 5.78E-139 | 0.940051875 | 0.623 | 0.534 | 1.17E-134 | 3 | Vdac2 |
| Mien11 | 3.48E-137 | 1.13587228 | 0.494 | 0.339 | 7.04E-133 | 3 | Mien1 |
| 1110008P14Rik | 1.53E-136 | 1.168560881 | 0.358 | 0.184 | 3.10E-132 | 3 | 1110008P14Rik |
| Pabpc11 | 1.05E-135 | 1.030905566 | 0.631 | 0.56 | 2.13E-131 | 3 | Pabpc1 |
| Ftl11 | 5.42E-132 | 0.273915626 | 0.899 | 0.913 | 1.10E-127 | 3 | Ftl1 |
| Ndufb61 | 9.49E-132 | 1.065320663 | 0.46 | 0.294 | 1.92E-127 | 3 | Ndufb6 |
| Ndufs71 | 1.52E-130 | 1.019775749 | 0.565 | 0.447 | 3.08E-126 | 3 | Ndufs7 |
| Tuba4a | 1.79E-130 | 0.956774051 | 0.267 | 0.109 | 3.62E-126 | 3 | Tuba4a |
| H3f3a1 | 3.31E-129 | 0.52984034 | 0.833 | 0.866 | 6.70E-125 | 3 | H3f3a |
| Bri31 | 2.25E-128 | 0.87099119 | 0.659 | 0.605 | 4.56E-124 | 3 | Bri3 |
| Vegfb | 4.89E-128 | 0.931709772 | 0.293 | 0.128 | 9.91E-124 | 3 | Vegfb |
| Etfb1 | 4.46E-126 | 1.101343075 | 0.477 | 0.326 | 9.03E-122 | 3 | Etfb |
| Mrpl121 | 8.16E-126 | 1.023269539 | 0.462 | 0.298 | 1.65E-121 | 3 | Mrpl12 |
| Ift20 | 6.98E-125 | 1.199265545 | 0.508 | 0.379 | 1.41E-120 | 3 | Ift20 |
| Cmss11 | 2.15E-123 | 0.380533827 | 0.994 | 0.993 | 4.36E-119 | 3 | Cmss1 |
| Sec61b1 | 3.42E-123 | 0.993450371 | 0.633 | 0.587 | 6.92E-119 | 3 | Sec61b |
| Pomp1 | 3.53E-123 | 0.887036073 | 0.649 | 0.596 | 7.14E-119 | 3 | Pomp |
| Rps261 | 6.49E-123 | 0.515345576 | 0.867 | 0.866 | 1.31E-118 | 3 | Rps26 |
| Stx71 | 2.92E-120 | 1.070110346 | 0.455 | 0.3 | 5.91E-116 | 3 | Stx7 |
| Rpl141 | 3.58E-120 | 0.578524959 | 0.804 | 0.804 | 7.26E-116 | 3 | Rpl14 |
| Cisd11 | 5.94E-120 | 1.032930426 | 0.418 | 0.252 | 1.20E-115 | 3 | Cisd1 |
| Ndufa81 | 5.72E-119 | 0.899870975 | 0.627 | 0.564 | 1.16E-114 | 3 | Ndufa8 |
| Idh3a1 | 1.76E-117 | 0.942802381 | 0.332 | 0.168 | 3.56E-113 | 3 | Idh3a |
| Rpl36al1 | 7.79E-117 | 0.944618757 | 0.597 | 0.517 | 1.58E-112 | 3 | Rpl36al |
| Ndufb31 | 5.73E-113 | 0.995570043 | 0.553 | 0.452 | 1.16E-108 | 3 | Ndufb3 |
| Sec61g | 1.20E-112 | 0.932653462 | 0.631 | 0.591 | 2.43E-108 | 3 | Sec61g |
| Fmc11 | 4.28E-112 | 1.007422571 | 0.418 | 0.261 | 8.66E-108 | 3 | Fmc1 |
| Oxct11 | 4.22E-111 | 1.028655074 | 0.503 | 0.367 | 8.53E-107 | 3 | Oxct1 |
| Ndufs31 | 3.68E-106 | 1.006380756 | 0.47 | 0.335 | 7.46E-102 | 3 | Ndufs3 |
| Ppp1r14b1 | 6.66E-106 | 0.916534254 | 0.463 | 0.313 | 1.35E-101 | 3 | Ppp1r14b |
| Ndufa21 | 1.29E-104 | 0.897825104 | 0.609 | 0.565 | 2.61E-100 | 3 | Ndufa2 |
| Abracl1 | 1.35E-104 | 0.974619198 | 0.526 | 0.419 | 2.73E-100 | 3 | Abracl |
| Gng51 | 3.09E-103 | 0.662840637 | 0.735 | 0.77 | 6.26E-99 | 3 | Gng5 |
| 4833439L19Rik | 5.07E-103 | 1.023608037 | 0.336 | 0.186 | 1.03E-98 | 3 | 4833439L19Rik |
| Slc25a391 | 1.24E-102 | 0.971907025 | 0.381 | 0.23 | 2.50E-98 | 3 | Slc25a39 |
| Slc25a41 | 1.90E-101 | 0.838524129 | 0.616 | 0.556 | 3.85E-97 | 3 | Slc25a4 |
| Prdx3 | 1.37E-100 | 0.912414562 | 0.306 | 0.159 | 2.76E-96 | 3 | Prdx3 |
| Dnajc151 | 9.05E-96 | 1.010396801 | 0.441 | 0.311 | 1.83E-91 | 3 | Dnajc15 |
| Serf21 | 1.04E-95 | 0.523397526 | 0.791 | 0.824 | 2.11E-91 | 3 | Serf2 |
| Anapc131 | 1.71E-95 | 1.054349513 | 0.419 | 0.285 | 3.46E-91 | 3 | Anapc13 |
| Idh21 | 2.24E-94 | 0.863868473 | 0.347 | 0.198 | 4.53E-90 | 3 | Idh2 |
| Higd1a1 | 3.26E-91 | 0.938099228 | 0.452 | 0.331 | 6.60E-87 | 3 | Higd1a |
| Sdhd1 | 9.79E-91 | 0.945999175 | 0.398 | 0.26 | 1.98E-86 | 3 | Sdhd |

| | | | | | | | |
|---|---|---|---|---|---|---|---|
| Rpl7a1 | 1.08E-88 | 0.575809937 | 0.743 | 0.756 | 2.20E-84 | 3 | Rpl7a |
| Bola31 | 7.48E-87 | 0.919261735 | 0.409 | 0.278 | 1.51E-82 | 3 | Bola3 |
| Hspe11 | 2.23E-86 | 0.802431139 | 0.568 | 0.502 | 4.52E-82 | 3 | Hspe1 |
| Lgals3 | 1.28E-84 | 0.772286643 | 0.351 | 0.204 | 2.59E-80 | 3 | Lgals3 |
| Cdk81 | 5.17E-83 | 0.337142318 | 0.962 | 0.971 | 1.05E-78 | 3 | Cdk8 |
| Pebp11 | 5.13E-82 | 0.894790218 | 0.528 | 0.468 | 1.04E-77 | 3 | Pebp1 |
| Eif11 | 7.20E-82 | 0.396283089 | 0.866 | 0.901 | 1.46E-77 | 3 | Eif1 |
| Rpl111 | 8.83E-82 | 0.330620672 | 0.91 | 0.93 | 1.79E-77 | 3 | Rpl11 |
| Rpl91 | 1.38E-81 | 0.332139544 | 0.931 | 0.947 | 2.79E-77 | 3 | Rpl9 |
| Rps81 | 3.70E-81 | 0.317300454 | 0.957 | 0.966 | 7.50E-77 | 3 | Rps8 |
| Mdh21 | 2.13E-80 | 0.922131123 | 0.471 | 0.373 | 4.31E-76 | 3 | Mdh2 |
| Etfa1 | 2.87E-79 | 0.957114332 | 0.372 | 0.247 | 5.82E-75 | 3 | Etfa |
| Rps171 | 8.45E-79 | 0.544538587 | 0.754 | 0.803 | 1.71E-74 | 3 | Rps17 |
| Rpl33 | 3.71E-78 | 0.353066733 | 0.901 | 0.911 | 7.51E-74 | 3 | Rpl3 |
| Taldo11 | 4.69E-77 | 0.835081297 | 0.556 | 0.513 | 9.49E-73 | 3 | Taldo1 |
| Ndufa101 | 4.73E-75 | 0.911847558 | 0.411 | 0.3 | 9.57E-71 | 3 | Ndufa10 |
| Cox201 | 1.41E-73 | 0.896871914 | 0.402 | 0.288 | 2.86E-69 | 3 | Cox20 |
| Prdx11 | 1.68E-73 | 0.614542055 | 0.675 | 0.696 | 3.40E-69 | 3 | Prdx1 |
| Timm8b1 | 6.49E-72 | 0.899256354 | 0.417 | 0.309 | 1.31E-67 | 3 | Timm8b |
| Sdhc | 2.66E-70 | 0.899954082 | 0.326 | 0.209 | 5.38E-66 | 3 | Sdhc |
| Acadm | 2.33E-69 | 0.924161779 | 0.334 | 0.218 | 4.71E-65 | 3 | Acadm |
| Papss1 | 1.38E-67 | 0.897803828 | 0.279 | 0.165 | 2.79E-63 | 3 | Papss1 |
| Clint11 | 2.48E-67 | 1.025105586 | 0.444 | 0.368 | 5.03E-63 | 3 | Clint1 |
| Rpl231 | 2.05E-65 | 0.258759968 | 0.958 | 0.973 | 4.16E-61 | 3 | Rpl23 |
| Idh3b | 8.56E-64 | 0.895018152 | 0.339 | 0.232 | 1.73E-59 | 3 | Idh3b |
| Ifi27l2a1 | 3.93E-63 | 0.69438978 | 0.48 | 0.356 | 7.96E-59 | 3 | Ifi27l2a |
| Atp6v1e11 | 4.40E-63 | 0.863335403 | 0.465 | 0.393 | 8.90E-59 | 3 | Atp6v1e1 |
| Rps3a11 | 8.20E-63 | 0.279728246 | 0.926 | 0.937 | 1.66E-58 | 3 | Rps3a1 |
| Uqcrc21 | 1.77E-62 | 0.888059129 | 0.415 | 0.327 | 3.58E-58 | 3 | Uqcrc2 |
| Mocs2 | 2.90E-62 | 0.931603904 | 0.306 | 0.199 | 5.87E-58 | 3 | Mocs2 |
| Timm131 | 1.91E-60 | 0.801394498 | 0.494 | 0.445 | 3.86E-56 | 3 | Timm13 |
| Ndufa71 | 7.58E-59 | 0.765798061 | 0.57 | 0.578 | 1.53E-54 | 3 | Ndufa7 |
| Ndufv11 | 6.16E-58 | 0.875327099 | 0.354 | 0.255 | 1.25E-53 | 3 | Ndufv1 |
| Mrpl14 | 8.89E-58 | 0.870368992 | 0.369 | 0.274 | 1.80E-53 | 3 | Mrpl14 |
| Vdac31 | 3.22E-57 | 0.816017486 | 0.464 | 0.404 | 6.52E-53 | 3 | Vdac3 |
| Rpl241 | 4.72E-57 | 0.320672623 | 0.87 | 0.913 | 9.56E-53 | 3 | Rpl24 |
| Pts | 3.65E-55 | 0.796170063 | 0.276 | 0.173 | 7.39E-51 | 3 | Pts |
| Ptma1 | 1.14E-54 | 0.29025935 | 0.933 | 0.941 | 2.32E-50 | 3 | Ptma |
| Ndufc21 | 9.55E-54 | 0.783275253 | 0.56 | 0.562 | 1.93E-49 | 3 | Ndufc2 |
| C1qbp1 | 1.16E-53 | 0.83499113 | 0.342 | 0.246 | 2.34E-49 | 3 | C1qbp |
| Pdha1 | 1.31E-53 | 0.775057103 | 0.285 | 0.184 | 2.65E-49 | 3 | Pdha1 |
| Lsm8 | 3.55E-53 | 0.862358346 | 0.31 | 0.213 | 7.18E-49 | 3 | Lsm8 |
| Glrx51 | 5.87E-53 | 0.833010248 | 0.354 | 0.263 | 1.19E-48 | 3 | Glrx5 |

| | | | | | | | |
|---|---|---|---|---|---|---|---|
| Ccs | 5.08E-52 | 0.868832599 | 0.265 | 0.169 | 1.03E-47 | 3 | Ccs |
| Prdx6 | 1.70E-51 | 0.814450002 | 0.409 | 0.332 | 3.44E-47 | 3 | Prdx6 |
| Lars21 | 3.75E-51 | 0.426241597 | 0.853 | 0.902 | 7.59E-47 | 3 | Lars2 |
| Rpl61 | 4.05E-51 | 0.313739903 | 0.855 | 0.886 | 8.21E-47 | 3 | Rpl6 |
| Rpl211 | 4.64E-51 | 0.288976295 | 0.888 | 0.928 | 9.39E-47 | 3 | Rpl21 |
| Dynlrb11 | 5.16E-50 | 0.839460356 | 0.463 | 0.418 | 1.04E-45 | 3 | Dynlrb1 |
| Acat11 | 2.61E-49 | 0.784804116 | 0.354 | 0.264 | 5.29E-45 | 3 | Acat1 |
| H2afj | 3.70E-49 | 0.741557045 | 0.495 | 0.451 | 7.50E-45 | 3 | H2afj |
| Idh3g | 3.80E-47 | 0.794555532 | 0.3 | 0.21 | 7.69E-43 | 3 | Idh3g |
| Mrpl301 | 4.50E-47 | 0.818095188 | 0.404 | 0.336 | 9.11E-43 | 3 | Mrpl30 |
| Churc1 | 4.71E-47 | 0.809904314 | 0.36 | 0.278 | 9.53E-43 | 3 | Churc1 |
| Rpl371 | 6.71E-47 | 0.261935438 | 0.902 | 0.931 | 1.36E-42 | 3 | Rpl37 |
| Mrps36 | 2.63E-46 | 0.773758448 | 0.317 | 0.227 | 5.32E-42 | 3 | Mrps36 |
| Cox6a11 | 8.23E-46 | 0.827889479 | 0.507 | 0.504 | 1.67E-41 | 3 | Cox6a1 |
| Hras | 2.88E-45 | 0.780563994 | 0.418 | 0.357 | 5.84E-41 | 3 | Hras |
| Smdt11 | 3.62E-45 | 0.663979156 | 0.572 | 0.609 | 7.32E-41 | 3 | Smdt1 |
| Rplp21 | 4.00E-45 | 0.282371331 | 0.858 | 0.895 | 8.09E-41 | 3 | Rplp2 |
| Rps27l | 2.73E-43 | 0.613132247 | 0.572 | 0.596 | 5.53E-39 | 3 | Rps27l |
| Ndufs8 | 2.54E-42 | 0.778123248 | 0.409 | 0.354 | 5.14E-38 | 3 | Ndufs8 |
| Ndufs21 | 3.71E-42 | 0.817134101 | 0.387 | 0.325 | 7.51E-38 | 3 | Ndufs2 |
| Sumo2 | 7.75E-42 | 0.569617449 | 0.607 | 0.667 | 1.57E-37 | 3 | Sumo2 |
| Hsd17b101 | 1.22E-41 | 0.792365551 | 0.325 | 0.245 | 2.46E-37 | 3 | Hsd17b10 |
| Reep5 | 2.80E-40 | 0.925115018 | 0.484 | 0.484 | 5.67E-36 | 3 | Reep5 |
| Dynll2 | 2.84E-40 | 0.829272312 | 0.297 | 0.217 | 5.75E-36 | 3 | Dynll2 |
| Rpl311 | 5.36E-40 | 0.52492776 | 0.64 | 0.691 | 1.08E-35 | 3 | Rpl31 |
| Rps251 | 4.99E-39 | 0.312113328 | 0.787 | 0.837 | 1.01E-34 | 3 | Rps25 |
| Uqcc21 | 1.71E-37 | 0.740796854 | 0.439 | 0.402 | 3.47E-33 | 3 | Uqcc2 |
| Mrpl511 | 9.97E-37 | 0.741158484 | 0.394 | 0.341 | 2.02E-32 | 3 | Mrpl51 |
| Srp91 | 1.28E-36 | 0.784657883 | 0.468 | 0.462 | 2.60E-32 | 3 | Srp9 |
| Rpl71 | 5.36E-36 | 0.250573524 | 0.857 | 0.898 | 1.09E-31 | 3 | Rpl7 |
| Anxa1 | 7.12E-36 | 0.671762827 | 0.301 | 0.22 | 1.44E-31 | 3 | Anxa1 |
| Mtx2 | 2.19E-35 | 0.758345573 | 0.299 | 0.226 | 4.43E-31 | 3 | Mtx2 |
| Tsc22d11 | 3.78E-35 | 0.663114088 | 0.479 | 0.445 | 7.66E-31 | 3 | Tsc22d1 |
| Rps61 | 4.55E-35 | 0.320150824 | 0.774 | 0.818 | 9.21E-31 | 3 | Rps6 |
| Rpl351 | 6.10E-35 | 0.286351856 | 0.782 | 0.835 | 1.23E-30 | 3 | Rpl35 |
| Swi5 | 1.93E-34 | 0.727519197 | 0.456 | 0.442 | 3.91E-30 | 3 | Swi5 |
| Btf31 | 2.22E-34 | 0.457783263 | 0.658 | 0.732 | 4.50E-30 | 3 | Btf3 |
| Hmgn2 | 4.03E-34 | 0.717414058 | 0.263 | 0.188 | 8.17E-30 | 3 | Hmgn2 |
| Cox7a2l1 | 6.08E-34 | 0.612776933 | 0.544 | 0.566 | 1.23E-29 | 3 | Cox7a2l |
| Scp21 | 2.01E-32 | 0.707090026 | 0.47 | 0.473 | 4.07E-28 | 3 | Scp2 |
| Higd2a1 | 5.33E-32 | 0.784951279 | 0.392 | 0.355 | 1.08E-27 | 3 | Higd2a |
| Pfdn51 | 6.15E-32 | 0.365203213 | 0.701 | 0.773 | 1.24E-27 | 3 | Pfdn5 |
| Gabarap1 | 1.79E-31 | 0.578399566 | 0.575 | 0.639 | 3.61E-27 | 3 | Gabarap |

| | | | | | | | |
|---|---|---|---|---|---|---|---|
| Ube2b1 | 1.92E-31 | 0.780509489 | 0.418 | 0.395 | 3.88E-27 | 3 | Ube2b |
| Lamtor41 | 3.44E-31 | 0.763645083 | 0.433 | 0.415 | 6.97E-27 | 3 | Lamtor4 |
| Pgls | 4.77E-31 | 0.731728686 | 0.422 | 0.399 | 9.65E-27 | 3 | Pgls |
| Ddt | 3.85E-30 | 0.734396101 | 0.311 | 0.25 | 7.79E-26 | 3 | Ddt |
| Ndufv3 | 1.03E-29 | 0.752648664 | 0.425 | 0.411 | 2.09E-25 | 3 | Ndufv3 |
| Elob1 | 3.54E-29 | 0.450919837 | 0.626 | 0.707 | 7.16E-25 | 3 | Elob |
| Hint11 | 4.24E-29 | 0.475340329 | 0.626 | 0.706 | 8.59E-25 | 3 | Hint1 |
| Mrpl34 | 5.02E-29 | 0.765799823 | 0.35 | 0.303 | 1.02E-24 | 3 | Mrpl34 |
| Mrpl181 | 8.80E-29 | 0.739297978 | 0.355 | 0.311 | 1.78E-24 | 3 | Mrpl18 |
| Hint2 | 1.96E-28 | 0.737712467 | 0.311 | 0.254 | 3.96E-24 | 3 | Hint2 |
| Kxd1 | 2.96E-28 | 0.74135289 | 0.305 | 0.247 | 5.98E-24 | 3 | Kxd1 |
| Park71 | 3.99E-28 | 0.587170082 | 0.524 | 0.564 | 8.07E-24 | 3 | Park7 |
| Rps15 | 4.22E-28 | 0.395102691 | 0.683 | 0.759 | 8.54E-24 | 3 | Rps15 |
| Hibadh | 7.11E-28 | 0.726268942 | 0.278 | 0.217 | 1.44E-23 | 3 | Hibadh |
| Tomm71 | 1.03E-27 | 0.591132234 | 0.534 | 0.579 | 2.08E-23 | 3 | Tomm7 |
| Ndufa3 | 9.25E-27 | 0.68707605 | 0.411 | 0.391 | 1.87E-22 | 3 | Ndufa3 |
| Npm11 | 9.33E-27 | 0.414809703 | 0.637 | 0.686 | 1.89E-22 | 3 | Npm1 |
| Cox14 | 1.38E-26 | 0.707825773 | 0.386 | 0.357 | 2.80E-22 | 3 | Cox14 |
| Mrpl41 | 3.12E-26 | 0.726053737 | 0.262 | 0.203 | 6.32E-22 | 3 | Mrpl41 |
| Tma71 | 8.27E-26 | 0.620945084 | 0.505 | 0.529 | 1.67E-21 | 3 | Tma7 |
| Romo11 | 1.36E-25 | 0.691584964 | 0.423 | 0.415 | 2.75E-21 | 3 | Romo1 |
| Ubl51 | 2.60E-25 | 0.599925506 | 0.514 | 0.557 | 5.26E-21 | 3 | Ubl5 |
| Sdha | 7.13E-25 | 0.715675573 | 0.348 | 0.31 | 1.44E-20 | 3 | Sdha |
| Grpel1 | 1.13E-24 | 0.70859849 | 0.359 | 0.325 | 2.29E-20 | 3 | Grpel1 |
| Cracr2b | 1.15E-24 | 0.614350635 | 0.294 | 0.234 | 2.33E-20 | 3 | Cracr2b |
| Ndufa9 | 2.96E-24 | 0.68411062 | 0.262 | 0.206 | 6.00E-20 | 3 | Ndufa9 |
| Rbx1 | 3.30E-24 | 0.588014942 | 0.526 | 0.585 | 6.69E-20 | 3 | Rbx1 |
| Ufm1 | 5.68E-24 | 0.716208772 | 0.266 | 0.211 | 1.15E-19 | 3 | Ufm1 |
| Hsp90ab1 | 1.26E-23 | 0.286069379 | 0.795 | 0.847 | 2.56E-19 | 3 | Hsp90ab1 |
| Rack11 | 2.16E-23 | 0.274774645 | 0.741 | 0.8 | 4.37E-19 | 3 | Rack1 |
| H2afz | 8.12E-23 | 0.423056964 | 0.571 | 0.615 | 1.64E-18 | 3 | H2afz |
| Tceal9 | 9.13E-23 | 0.65548337 | 0.406 | 0.387 | 1.85E-18 | 3 | Tceal9 |
| Emc61 | 2.88E-22 | 0.716623612 | 0.309 | 0.268 | 5.84E-18 | 3 | Emc6 |
| Timm17a1 | 7.80E-22 | 0.684561776 | 0.314 | 0.271 | 1.58E-17 | 3 | Timm17a |
| Slc25a11 | 7.89E-22 | 0.678040935 | 0.264 | 0.214 | 1.60E-17 | 3 | Slc25a11 |
| Cmtm7 | 1.95E-21 | 0.660479033 | 0.262 | 0.209 | 3.96E-17 | 3 | Cmtm7 |
| Cebpzos | 3.15E-21 | 0.657084678 | 0.251 | 0.198 | 6.38E-17 | 3 | Cebpzos |
| Mrps211 | 7.27E-21 | 0.692879767 | 0.397 | 0.39 | 1.47E-16 | 3 | Mrps21 |
| Ghitm | 8.24E-21 | 0.726348326 | 0.346 | 0.319 | 1.67E-16 | 3 | Ghitm |
| Polr2f1 | 4.36E-20 | 0.692184699 | 0.365 | 0.348 | 8.83E-16 | 3 | Polr2f |
| Gabarapl21 | 8.52E-20 | 0.642151948 | 0.467 | 0.503 | 1.72E-15 | 3 | Gabarapl2 |
| Cox171 | 1.06E-19 | 0.685425529 | 0.346 | 0.32 | 2.15E-15 | 3 | Cox17 |
| Hspd1 | 1.47E-19 | 0.675716311 | 0.355 | 0.334 | 2.97E-15 | 3 | Hspd1 |

| | | | | | | | |
|---|---|---|---|---|---|---|---|
| Pkm1 | 8.39E-19 | 0.682826217 | 0.381 | 0.376 | 1.70E-14 | 3 | Pkm |
| 0610012G03Rik | 9.68E-19 | 0.656451162 | 0.392 | 0.387 | 1.96E-14 | 3 | 0610012G03Rik |
| Elof1 | 1.07E-17 | 0.631956841 | 0.271 | 0.231 | 2.16E-13 | 3 | Elof1 |
| Pcp4l13 | 3.17E-17 | 0.581772607 | 0.35 | 0.312 | 6.41E-13 | 3 | Pcp4l1 |
| Lamtor21 | 3.55E-17 | 0.609465044 | 0.442 | 0.469 | 7.20E-13 | 3 | Lamtor2 |
| Rpl51 | 1.09E-16 | 0.329749887 | 0.64 | 0.721 | 2.21E-12 | 3 | Rpl5 |
| Fkbp41 | 1.29E-16 | 0.647370242 | 0.27 | 0.234 | 2.61E-12 | 3 | Fkbp4 |
| Pgk1 | 3.16E-16 | 0.660370367 | 0.251 | 0.213 | 6.40E-12 | 3 | Pgk1 |
| Cops91 | 4.03E-16 | 0.63514091 | 0.342 | 0.326 | 8.17E-12 | 3 | Cops9 |
| Pycard1 | 7.72E-16 | 0.589181947 | 0.311 | 0.276 | 1.56E-11 | 3 | Pycard |
| Rpl36a1 | 1.25E-15 | 0.268255014 | 0.686 | 0.762 | 2.53E-11 | 3 | Rpl36a |
| Dnajc191 | 1.35E-15 | 0.717266227 | 0.327 | 0.313 | 2.73E-11 | 3 | Dnajc19 |
| Mrps16 | 1.65E-15 | 0.625501363 | 0.327 | 0.307 | 3.33E-11 | 3 | Mrps16 |
| Skp1a1 | 1.77E-15 | 0.664627049 | 0.341 | 0.328 | 3.59E-11 | 3 | Skp1a |
| Camk2n11 | 2.24E-15 | 0.627940384 | 0.303 | 0.271 | 4.54E-11 | 3 | Camk2n1 |
| Nol71 | 3.27E-15 | 0.630366482 | 0.419 | 0.442 | 6.62E-11 | 3 | Nol7 |
| Chchd3 | 1.46E-14 | 0.621936843 | 0.353 | 0.344 | 2.96E-10 | 3 | Chchd3 |
| Mrps141 | 2.03E-14 | 0.602193204 | 0.392 | 0.404 | 4.10E-10 | 3 | Mrps14 |
| Immt | 3.75E-14 | 0.624454658 | 0.281 | 0.253 | 7.59E-10 | 3 | Immt |
| Uqcrc1 | 5.58E-14 | 0.631686211 | 0.28 | 0.252 | 1.13E-09 | 3 | Uqcrc1 |
| Tkt1 | 5.76E-14 | 0.703670477 | 0.278 | 0.252 | 1.17E-09 | 3 | Tkt |
| Fundc2 | 3.16E-13 | 0.613037499 | 0.293 | 0.269 | 6.40E-09 | 3 | Fundc2 |
| Ywhae1 | 4.45E-13 | 0.524543864 | 0.483 | 0.545 | 9.00E-09 | 3 | Ywhae |
| Hspa91 | 6.00E-13 | 0.624527337 | 0.27 | 0.244 | 1.21E-08 | 3 | Hspa9 |
| 1110004F10Rik1 | 8.36E-13 | 0.268226978 | 0.208 | 0.323 | 1.69E-08 | 3 | 1110004F10Rik |
| Ufc11 | 8.99E-13 | 0.660848336 | 0.33 | 0.326 | 1.82E-08 | 3 | Ufc1 |
| Ybx11 | 1.08E-12 | 0.277742949 | 0.664 | 0.765 | 2.19E-08 | 3 | Ybx1 |
| Rnf71 | 1.30E-12 | 0.593397223 | 0.382 | 0.393 | 2.63E-08 | 3 | Rnf7 |
| Pdap11 | 1.85E-12 | 0.67674684 | 0.377 | 0.389 | 3.74E-08 | 3 | Pdap1 |
| Smap1 | 2.85E-12 | 0.288295057 | 0.246 | 0.377 | 5.78E-08 | 3 | Smap1 |
| Caprin1 | 3.43E-12 | 0.261185684 | 0.193 | 0.298 | 6.94E-08 | 3 | Caprin1 |
| Mrpl541 | 8.21E-12 | 0.641573284 | 0.287 | 0.269 | 1.66E-07 | 3 | Mrpl54 |
| Zc3h15 | 1.48E-11 | 0.29665264 | 0.198 | 0.305 | 3.00E-07 | 3 | Zc3h15 |
| Ube2m1 | 1.65E-11 | 0.606911601 | 0.308 | 0.297 | 3.35E-07 | 3 | Ube2m |
| Txn11 | 2.22E-11 | 0.334619083 | 0.598 | 0.688 | 4.49E-07 | 3 | Txn1 |
| Mrps241 | 2.67E-11 | 0.561536079 | 0.408 | 0.441 | 5.40E-07 | 3 | Mrps24 |
| Eef1b21 | 4.03E-11 | 0.25731302 | 0.667 | 0.747 | 8.15E-07 | 3 | Eef1b2 |
| Polr1d1 | 4.59E-11 | 0.532503206 | 0.447 | 0.501 | 9.28E-07 | 3 | Polr1d |
| Cmpk11 | 6.37E-11 | 0.525810043 | 0.317 | 0.305 | 1.29E-06 | 3 | Cmpk1 |
| Tcf25 | 1.76E-10 | 0.258550187 | 0.231 | 0.346 | 3.56E-06 | 3 | Tcf25 |
| 22-Mar | 2.20E-10 | 0.308446101 | 0.276 | 0.411 | 4.45E-06 | 3 | 2-Mar |
| Uba521 | 2.50E-10 | 0.435544155 | 0.479 | 0.53 | 5.05E-06 | 3 | Uba52 |
| Ndufs41 | 6.39E-10 | 0.621017225 | 0.368 | 0.392 | 1.29E-05 | 3 | Ndufs4 |

| | | | | | | | |
|---|---|---|---|---|---|---|---|
| Sugt1 | 6.56E-10 | 0.26126997 | 0.228 | 0.338 | 1.33E-05 | 3 | Sugt1 |
| Hdgf1 | 7.51E-10 | 0.252367948 | 0.173 | 0.26 | 1.52E-05 | 3 | Hdgf |
| Txn21 | 8.63E-10 | 0.597295094 | 0.29 | 0.281 | 1.75E-05 | 3 | Txn2 |
| Psmc2 | 1.09E-09 | 0.250194619 | 0.18 | 0.27 | 2.21E-05 | 3 | Psmc2 |
| Nme11 | 1.69E-09 | 0.525940458 | 0.353 | 0.359 | 3.42E-05 | 3 | Nme1 |
| Ccdc12 | 3.20E-09 | 0.663136982 | 0.384 | 0.421 | 6.48E-05 | 3 | Ccdc12 |
| Thoc71 | 3.89E-09 | 0.676223637 | 0.302 | 0.301 | 7.88E-05 | 3 | Thoc7 |
| Cwc15 | 4.52E-09 | 0.289862334 | 0.262 | 0.387 | 9.15E-05 | 3 | Cwc15 |
| Sfr1 | 9.08E-09 | 0.293517619 | 0.272 | 0.398 | 0.000183891 | 3 | Sfr1 |
| Cdc37 | 9.56E-09 | 0.269148099 | 0.23 | 0.336 | 0.000193538 | 3 | Cdc37 |
| Sf3b61 | 1.37E-08 | 0.264904483 | 0.257 | 0.376 | 0.000278111 | 3 | Sf3b6 |
| Zfp7061 | 1.64E-08 | 0.62695328 | 0.33 | 0.343 | 0.000332072 | 3 | Zfp706 |
| Trappc41 | 2.19E-08 | 0.263768793 | 0.205 | 0.297 | 0.000443574 | 3 | Trappc4 |
| Tmem126a1 | 3.15E-08 | 0.569709165 | 0.25 | 0.237 | 0.000637628 | 3 | Tmem126a |
| Cct81 | 3.58E-08 | 0.254315563 | 0.191 | 0.278 | 0.000724751 | 3 | Cct8 |
| Hdlbp | 4.79E-08 | 0.300130978 | 0.213 | 0.31 | 0.000970437 | 3 | Hdlbp |
| Aurkaip11 | 6.65E-08 | 0.599560116 | 0.336 | 0.354 | 0.001346817 | 3 | Aurkaip1 |
| Mrps331 | 8.80E-08 | 0.594425752 | 0.354 | 0.38 | 0.001782545 | 3 | Mrps33 |
| Phb21 | 1.43E-07 | 0.597653047 | 0.303 | 0.309 | 0.00290259 | 3 | Phb2 |
| Arl11 | 2.13E-07 | 0.263248288 | 0.209 | 0.298 | 0.004318445 | 3 | Arl1 |
| Dctn31 | 2.59E-07 | 0.266538906 | 0.238 | 0.339 | 0.00523964 | 3 | Dctn3 |
| Cd9 | 4.09E-07 | 0.907271803 | 0.359 | 0.392 | 0.008289521 | 3 | Cd9 |
| Chrac11 | 4.50E-07 | 0.291024182 | 0.18 | 0.257 | 0.009118064 | 3 | Chrac1 |
| Psmb2 | 5.23E-07 | 0.264534649 | 0.29 | 0.419 | 0.010598093 | 3 | Psmb2 |
| Tpm11 | 6.21E-07 | 0.489232002 | 0.294 | 0.295 | 0.012576468 | 3 | Tpm1 |
| Eif5a1 | 6.51E-07 | 0.402792572 | 0.493 | 0.577 | 0.013189324 | 3 | Eif5a |
| Arpc1a1 | 8.30E-07 | 0.548744991 | 0.311 | 0.321 | 0.01680804 | 3 | Arpc1a |
| Dynlt31 | 9.99E-07 | 0.583560181 | 0.268 | 0.263 | 0.020235694 | 3 | Dynlt3 |
| Cyb5a1 | 1.09E-06 | 0.441792521 | 0.448 | 0.508 | 0.022040802 | 3 | Cyb5a |
| Smim111 | 1.62E-06 | 0.266083975 | 0.227 | 0.319 | 0.032812926 | 3 | Smim11 |
| Tbca1 | 1.68E-06 | 0.543611771 | 0.417 | 0.484 | 0.034083378 | 3 | Tbca |
| Esd1 | 1.69E-06 | 0.261935398 | 0.279 | 0.395 | 0.034301294 | 3 | Esd |
| Gstm1 | 1.86E-06 | 0.401258467 | 0.324 | 0.325 | 0.037730957 | 3 | Gstm1 |
| Psma6 | 2.00E-06 | 0.29526576 | 0.214 | 0.303 | 0.040479688 | 3 | Psma6 |
| Atp1a1 | 2.27E-06 | 0.528637371 | 0.206 | 0.291 | 0.045884285 | 3 | Atp1a1 |
| Hist1h1e1 | 2.59E-06 | 0.527928627 | 0.353 | 0.377 | 0.052462628 | 3 | Hist1h1e |
| Ptges31 | 3.02E-06 | 0.268759108 | 0.284 | 0.402 | 0.061233178 | 3 | Ptges3 |
| Pfdn21 | 3.20E-06 | 0.268447406 | 0.229 | 0.321 | 0.064863364 | 3 | Pfdn2 |
| Cbx11 | 3.91E-06 | 0.305547499 | 0.238 | 0.33 | 0.079258666 | 3 | Cbx1 |
| Eef1g1 | 3.99E-06 | 0.442908833 | 0.452 | 0.521 | 0.080685754 | 3 | Eef1g |
| Psma31 | 4.95E-06 | 0.480901654 | 0.449 | 0.531 | 0.100234367 | 3 | Psma3 |
| Psmb3 | 5.09E-06 | 0.255814605 | 0.31 | 0.436 | 0.103134722 | 3 | Psmb3 |
| Zcrb11 | 5.17E-06 | 0.635437107 | 0.316 | 0.336 | 0.104729291 | 3 | Zcrb1 |

| | | | | | | | |
|---|---|---|---|---|---|---|---|
| Ppdpf1 | 5.47E-06 | 0.517480092 | 0.286 | 0.29 | 0.110797927 | 3 | Ppdpf |
| Rpl22l11 | 5.80E-06 | 0.309319778 | 0.526 | 0.587 | 0.117414459 | 3 | Rpl22l1 |
| Banf1 | 6.09E-06 | 0.268232576 | 0.185 | 0.259 | 0.123372362 | 3 | Banf1 |
| Fam104a1 | 6.16E-06 | 0.285729285 | 0.224 | 0.313 | 0.124615435 | 3 | Fam104a |
| Snf81 | 7.65E-06 | 0.275690042 | 0.201 | 0.278 | 0.154947411 | 3 | Snf8 |
| Map2k21 | 8.12E-06 | 0.286212403 | 0.217 | 0.302 | 0.164380983 | 3 | Map2k2 |
| Psma11 | 8.20E-06 | 0.256703765 | 0.242 | 0.338 | 0.166116618 | 3 | Psma1 |
| Mtch1 | 8.36E-06 | 0.27138898 | 0.239 | 0.328 | 0.169269183 | 3 | Mtch1 |
| Guk11 | 8.92E-06 | 0.262355638 | 0.182 | 0.252 | 0.180624398 | 3 | Guk1 |
| 1810022K09Rik1 | 1.13E-05 | 0.549446202 | 0.273 | 0.278 | 0.228416675 | 3 | 1810022K09Rik |
| Pdcd51 | 1.29E-05 | 0.624755339 | 0.365 | 0.413 | 0.260579002 | 3 | Pdcd5 |
| Ogdh1 | 1.78E-05 | 0.317825121 | 0.197 | 0.273 | 0.360539108 | 3 | Ogdh |
| Tomm51 | 1.92E-05 | 0.509257818 | 0.318 | 0.337 | 0.389398429 | 3 | Tomm5 |
| Gm100761 | 2.06E-05 | 0.394764426 | 0.477 | 0.557 | 0.416652897 | 3 | Gm10076 |
| Nars1 | 3.18E-05 | 0.331364842 | 0.197 | 0.271 | 0.644276076 | 3 | Nars |
| Tcea11 | 4.26E-05 | 0.409374615 | 0.206 | 0.284 | 0.861892908 | 3 | Tcea1 |
| Polr2j1 | 5.12E-05 | 0.347398782 | 0.224 | 0.308 | 1 | 3 | Polr2j |
| Txndc171 | 5.72E-05 | 0.49508686 | 0.404 | 0.466 | 1 | 3 | Txndc17 |
| Csnk2b1 | 6.96E-05 | 0.31847218 | 0.196 | 0.267 | 1 | 3 | Csnk2b |
| Trappc6b1 | 9.95E-05 | 0.369198222 | 0.227 | 0.309 | 1 | 3 | Trappc6b |
| Wdr83os1 | 0.000105871 | 0.286550613 | 0.217 | 0.294 | 1 | 3 | Wdr83os |
| Cetn31 | 0.000113791 | 0.275037143 | 0.253 | 0.345 | 1 | 3 | Cetn3 |
| Psma41 | 0.000169315 | 0.32759271 | 0.269 | 0.37 | 1 | 3 | Psma4 |
| Dnajb61 | 0.000201309 | 0.376585765 | 0.283 | 0.393 | 1 | 3 | Dnajb6 |
| Fis11 | 0.000212658 | 0.474035116 | 0.402 | 0.472 | 1 | 3 | Fis1 |
| Ybx31 | 0.000250837 | 0.529743192 | 0.317 | 0.344 | 1 | 3 | Ybx3 |
| Pgam1 | 0.000332305 | 0.493766872 | 0.276 | 0.287 | 1 | 3 | Pgam1 |
| Rbbp71 | 0.000360311 | 0.586476109 | 0.243 | 0.25 | 1 | 3 | Rbbp7 |
| Magoh1 | 0.000401382 | 0.352158807 | 0.199 | 0.266 | 1 | 3 | Magoh |
| Mrpl231 | 0.000419128 | 0.499844613 | 0.28 | 0.294 | 1 | 3 | Mrpl23 |
| Metap21 | 0.00042512 | 0.383189371 | 0.277 | 0.38 | 1 | 3 | Metap2 |
| Vapa1 | 0.000431557 | 0.551185963 | 0.34 | 0.386 | 1 | 3 | Vapa |
| Bag11 | 0.000483047 | 0.454481467 | 0.365 | 0.415 | 1 | 3 | Bag1 |
| Aimp11 | 0.000567516 | 0.354262073 | 0.237 | 0.316 | 1 | 3 | Aimp1 |
| Zmat21 | 0.000628721 | 0.326230037 | 0.204 | 0.271 | 1 | 3 | Zmat2 |
| Tomm201 | 0.0006564 | 0.527397624 | 0.383 | 0.449 | 1 | 3 | Tomm20 |
| Anp32b | 0.000695699 | 0.574487801 | 0.372 | 0.432 | 1 | 3 | Anp32b |
| Cenpx1 | 0.000737004 | 0.338841972 | 0.233 | 0.311 | 1 | 3 | Cenpx |
| Cct21 | 0.001057074 | 0.34437773 | 0.238 | 0.32 | 1 | 3 | Cct2 |
| Lsm61 | 0.001203895 | 0.493526495 | 0.296 | 0.32 | 1 | 3 | Lsm6 |
| Txnl11 | 0.001476898 | 0.536286951 | 0.294 | 0.323 | 1 | 3 | Txnl1 |
| Fam162a1 | 0.001523283 | 0.4871808 | 0.251 | 0.26 | 1 | 3 | Fam162a |
| Eno11 | 0.002132787 | 0.476506905 | 0.297 | 0.322 | 1 | 3 | Eno1 |

| | | | | | | | |
|---|---|---|---|---|---|---|---|
| Naa201 | 0.002366846 | 0.455892724 | 0.272 | 0.287 | 1 | 3 | Naa20 |
| Rpl23a1 | 0.002452139 | 0.256460985 | 0.587 | 0.692 | 1 | 3 | Rpl23a |
| Ucp21 | 0.003036227 | 0.37618505 | 0.316 | 0.335 | 1 | 3 | Ucp2 |
| Sra11 | 0.003216964 | 0.366736206 | 0.221 | 0.292 | 1 | 3 | Sra1 |
| Chchd11 | 0.003394653 | 0.279550534 | 0.246 | 0.322 | 1 | 3 | Chchd1 |
| Psmd121 | 0.003711405 | 0.35442682 | 0.22 | 0.288 | 1 | 3 | Psmd12 |
| Vti1b1 | 0.004626203 | 0.351932578 | 0.201 | 0.261 | 1 | 3 | Vti1b |
| Emc21 | 0.004778949 | 0.320984418 | 0.207 | 0.267 | 1 | 3 | Emc2 |
| Fkbp2 | 0.005142002 | 0.626447287 | 0.259 | 0.281 | 1 | 3 | Fkbp2 |
| Tmem258 | 0.005577138 | 0.393615532 | 0.242 | 0.32 | 1 | 3 | Tmem258 |
| Eny21 | 0.006056202 | 0.522395185 | 0.271 | 0.294 | 1 | 3 | Eny2 |
| Ivns1abp3 | 0.006270103 | 0.482543161 | 0.32 | 0.347 | 1 | 3 | Ivns1abp |
| Hist1h2bc | 0.006993167 | 0.298598334 | 0.205 | 0.258 | 1 | 3 | Hist1h2bc |
| Ntan11 | 0.007022685 | 0.282630118 | 0.254 | 0.329 | 1 | 3 | Ntan1 |
| Chmp4b1 | 0.007982269 | 0.374735805 | 0.262 | 0.346 | 1 | 3 | Chmp4b |
| Sap181 | 0.00809368 | 0.313871926 | 0.217 | 0.279 | 1 | 3 | Sap18 |
| Serbp11 | 0.00879691 | 0.363110386 | 0.491 | 0.605 | 1 | 3 | Serbp1 |
| Gzmc | 0 | 3.970601927 | 0.811 | 0.037 | 0 | 4 | Gzmc |
| Gzmb | 0 | 3.961463885 | 0.832 | 0.035 | 0 | 4 | Gzmb |
| Ccl5 | 0 | 3.840715242 | 0.958 | 0.102 | 0 | 4 | Ccl5 |
| Cd7 | 0 | 3.29747971 | 0.978 | 0.058 | 0 | 4 | Cd7 |
| Xcl1 | 0 | 3.269984922 | 0.976 | 0.077 | 0 | 4 | Xcl1 |
| Inpp4b | 0 | 3.205943566 | 0.997 | 0.154 | 0 | 4 | Inpp4b |
| Nkg7 | 0 | 2.913514086 | 0.987 | 0.074 | 0 | 4 | Nkg7 |
| Ctsw | 0 | 2.725989809 | 0.99 | 0.074 | 0 | 4 | Ctsw |
| Klrd1 | 0 | 2.722423182 | 0.978 | 0.069 | 0 | 4 | Klrd1 |
| Klra4 | 0 | 2.651604862 | 0.705 | 0.04 | 0 | 4 | Klra4 |
| Klra8 | 0 | 2.640452387 | 0.71 | 0.041 | 0 | 4 | Klra8 |
| Ptprc | 0 | 2.597810035 | 0.992 | 0.15 | 0 | 4 | Ptprc |
| Smyd3 | 0 | 2.59502612 | 0.946 | 0.183 | 0 | 4 | Smyd3 |
| Il2rb | 0 | 2.592963879 | 0.972 | 0.067 | 0 | 4 | Il2rb |
| Tox | 0 | 2.55107633 | 0.922 | 0.072 | 0 | 4 | Tox |
| Bcl2 | 0 | 2.548398653 | 0.934 | 0.162 | 0 | 4 | Bcl2 |
| S100a4 | 0 | 2.542496623 | 0.891 | 0.082 | 0 | 4 | S100a4 |
| Clnk | 0 | 2.53287003 | 0.941 | 0.06 | 0 | 4 | Clnk |
| Chn2 | 0 | 2.505248217 | 0.954 | 0.152 | 0 | 4 | Chn2 |
| Fcer1g | 0 | 2.474751001 | 0.998 | 0.225 | 0 | 4 | Fcer1g |
| Dennd4a | 0 | 2.447755926 | 0.964 | 0.14 | 0 | 4 | Dennd4a |
| Ltb | 0 | 2.436968961 | 0.95 | 0.085 | 0 | 4 | Ltb |
| Klra7 | 0 | 2.42526435 | 0.661 | 0.038 | 0 | 4 | Klra7 |
| Klra9 | 0 | 2.417535552 | 0.745 | 0.037 | 0 | 4 | Klra9 |
| Styk1 | 0 | 2.407707858 | 0.894 | 0.048 | 0 | 4 | Styk1 |
| Skap1 | 0 | 2.344839923 | 0.962 | 0.083 | 0 | 4 | Skap1 |

| | | | | | | | |
|---|---|---|---|---|---|---|---|
| S100a6 | 0 | 2.336059316 | 0.938 | 0.231 | 0 | 4 | S100a6 |
| Tmsb10 | 0 | 2.308517766 | 0.994 | 0.471 | 0 | 4 | Tmsb10 |
| Cd226 | 0 | 2.307239467 | 0.769 | 0.048 | 0 | 4 | Cd226 |
| Cd52 | 0 | 2.294305395 | 1 | 0.238 | 0 | 4 | Cd52 |
| Dock10 | 0 | 2.288372361 | 0.973 | 0.162 | 0 | 4 | Dock10 |
| Rgs1 | 0 | 2.240929999 | 0.876 | 0.095 | 0 | 4 | Rgs1 |
| Tyrobp | 0 | 2.22128315 | 0.998 | 0.217 | 0 | 4 | Tyrobp |
| Ptprcap | 0 | 2.202538288 | 0.959 | 0.081 | 0 | 4 | Ptprcap |
| Ccl4 | 0 | 2.20235368 | 0.725 | 0.093 | 0 | 4 | Ccl4 |
| Itgae | 0 | 2.175124109 | 0.727 | 0.025 | 0 | 4 | Itgae |
| Runx3 | 0 | 2.139943573 | 0.907 | 0.102 | 0 | 4 | Runx3 |
| Ms4a4b | 0 | 2.135182476 | 0.929 | 0.072 | 0 | 4 | Ms4a4b |
| Ikzf3 | 0 | 2.112649077 | 0.899 | 0.072 | 0 | 4 | Ikzf3 |
| Klrb1c | 0 | 2.11159796 | 0.869 | 0.049 | 0 | 4 | Klrb1c |
| Dock2 | 0 | 2.103188691 | 0.962 | 0.138 | 0 | 4 | Dock2 |
| AW1120101 | 0 | 2.085829981 | 0.998 | 0.543 | 0 | 4 | AW112010 |
| Rac2 | 0 | 2.08332886 | 0.958 | 0.13 | 0 | 4 | Rac2 |
| Txk | 0 | 2.079687826 | 0.873 | 0.059 | 0 | 4 | Txk |
| Ncr1 | 0 | 2.0746706 | 0.913 | 0.053 | 0 | 4 | Ncr1 |
| Klre1 | 0 | 2.029669184 | 0.883 | 0.055 | 0 | 4 | Klre1 |
| Mbnl1 | 0 | 2.028316543 | 0.996 | 0.427 | 0 | 4 | Mbnl1 |
| Gem | 0 | 1.96638943 | 0.716 | 0.066 | 0 | 4 | Gem |
| Ipcef1 | 0 | 1.945524509 | 0.82 | 0.074 | 0 | 4 | Ipcef1 |
| Tmem163 | 0 | 1.928351634 | 0.765 | 0.046 | 0 | 4 | Tmem163 |
| Stk17b | 0 | 1.907193976 | 0.903 | 0.117 | 0 | 4 | Stk17b |
| Hcst | 0 | 1.889705259 | 0.894 | 0.099 | 0 | 4 | Hcst |
| Mir142hg | 0 | 1.886256252 | 0.91 | 0.125 | 0 | 4 | Mir142hg |
| Id2 | 0 | 1.878052623 | 0.986 | 0.281 | 0 | 4 | Id2 |
| Klri2 | 0 | 1.876087083 | 0.696 | 0.036 | 0 | 4 | Klri2 |
| Osbpl3 | 0 | 1.87151452 | 0.818 | 0.077 | 0 | 4 | Osbpl3 |
| Cd244 | 0 | 1.842399075 | 0.81 | 0.047 | 0 | 4 | Cd244 |
| Rabgap1l | 0 | 1.824580366 | 0.917 | 0.217 | 0 | 4 | Rabgap1l |
| Chsy1 | 0 | 1.815097529 | 0.811 | 0.085 | 0 | 4 | Chsy1 |
| Atp8b4 | 0 | 1.812294834 | 0.771 | 0.055 | 0 | 4 | Atp8b4 |
| Shisa5 | 0 | 1.810249959 | 0.976 | 0.29 | 0 | 4 | Shisa5 |
| Ppp3cc | 0 | 1.805579617 | 0.855 | 0.098 | 0 | 4 | Ppp3cc |
| Ugcg | 0 | 1.792457633 | 0.887 | 0.128 | 0 | 4 | Ugcg |
| Klrk1 | 0 | 1.789372368 | 0.737 | 0.062 | 0 | 4 | Klrk1 |
| Dock8 | 0 | 1.780307652 | 0.908 | 0.176 | 0 | 4 | Dock8 |
| Ptk2b | 0 | 1.766092281 | 0.854 | 0.107 | 0 | 4 | Ptk2b |
| Dusp2 | 0 | 1.761813418 | 0.753 | 0.062 | 0 | 4 | Dusp2 |
| Pde7a | 0 | 1.750049963 | 0.838 | 0.114 | 0 | 4 | Pde7a |
| Ptpn22 | 0 | 1.72600919 | 0.795 | 0.078 | 0 | 4 | Ptpn22 |

| | | | | | | | |
|---|---|---|---|---|---|---|---|
| Dock5 | 0 | 1.707099338 | 0.787 | 0.073 | 0 | 4 | Dock5 |
| 1-Sep | 0 | 1.704293014 | 0.881 | 0.101 | 0 | 4 | 1-Sep |
| Ankrd44 | 0 | 1.706440762 | 0.929 | 0.256 | 0 | 4 | Ankrd44 |
| Gas7 | 0 | 1.70331069 | 0.834 | 0.143 | 0 | 4 | Gas7 |
| Stat4 | 0 | 1.695861282 | 0.792 | 0.065 | 0 | 4 | Stat4 |
| Epsti1 | 0 | 1.678350982 | 0.89 | 0.131 | 0 | 4 | Epsti1 |
| AC163354.1 | 0 | 1.674313374 | 0.616 | 0.028 | 0 | 4 | AC163354.1 |
| Klra3 | 0 | 1.661910104 | 0.47 | 0.02 | 0 | 4 | Klra3 |
| Coro1a | 0 | 1.661143681 | 0.948 | 0.175 | 0 | 4 | Coro1a |
| Selplg | 0 | 1.656674848 | 0.865 | 0.111 | 0 | 4 | Selplg |
| Itgax | 0 | 1.655688612 | 0.795 | 0.065 | 0 | 4 | Itgax |
| Arhgap45 | 0 | 1.651969226 | 0.863 | 0.115 | 0 | 4 | Arhgap45 |
| Lcp1 | 0 | 1.647560771 | 0.936 | 0.195 | 0 | 4 | Lcp1 |
| Arhgdib | 0 | 1.639009395 | 0.938 | 0.191 | 0 | 4 | Arhgdib |
| Lck | 0 | 1.638582924 | 0.806 | 0.06 | 0 | 4 | Lck |
| Gimap3 | 0 | 1.629924982 | 0.78 | 0.058 | 0 | 4 | Gimap3 |
| Ikzf1 | 0 | 1.62696292 | 0.835 | 0.115 | 0 | 4 | Ikzf1 |
| Btg1 | 0 | 1.617937409 | 0.987 | 0.428 | 0 | 4 | Btg1 |
| Cd69 | 0 | 1.617062983 | 0.686 | 0.066 | 0 | 4 | Cd69 |
| Pde3b | 0 | 1.61124483 | 0.729 | 0.091 | 0 | 4 | Pde3b |
| Sema4d | 0 | 1.585419946 | 0.777 | 0.09 | 0 | 4 | Sema4d |
| Dok6 | 0 | 1.568057548 | 0.513 | 0.014 | 0 | 4 | Dok6 |
| Srgap3 | 0 | 1.560918892 | 0.705 | 0.058 | 0 | 4 | Srgap3 |
| Coro2a | 0 | 1.550841666 | 0.701 | 0.082 | 0 | 4 | Coro2a |
| Mef2a1 | 0 | 1.536229579 | 0.948 | 0.385 | 0 | 4 | Mef2a |
| Samd3 | 0 | 1.533301131 | 0.72 | 0.047 | 0 | 4 | Samd3 |
| Btg21 | 0 | 1.532047265 | 0.913 | 0.31 | 0 | 4 | Btg2 |
| Hivep3 | 0 | 1.525358658 | 0.778 | 0.124 | 0 | 4 | Hivep3 |
| Cdh1 | 0 | 1.517070915 | 0.698 | 0.035 | 0 | 4 | Cdh1 |
| Ccr5 | 0 | 1.491231451 | 0.75 | 0.095 | 0 | 4 | Ccr5 |
| Dapk2 | 0 | 1.474974486 | 0.733 | 0.072 | 0 | 4 | Dapk2 |
| Ndfip1 | 0 | 1.460119852 | 0.908 | 0.286 | 0 | 4 | Ndfip1 |
| Prex1 | 0 | 1.460045559 | 0.787 | 0.136 | 0 | 4 | Prex1 |
| Myo1f | 0 | 1.456460846 | 0.757 | 0.1 | 0 | 4 | Myo1f |
| Rps15a | 0 | 1.454379495 | 0.997 | 0.92 | 0 | 4 | Rps15a |
| Car2 | 0 | 1.445166159 | 0.756 | 0.083 | 0 | 4 | Car2 |
| Rapgef6 | 0 | 1.423301913 | 0.893 | 0.261 | 0 | 4 | Rapgef6 |
| Ctsd | 0 | 1.413333372 | 0.924 | 0.277 | 0 | 4 | Ctsd |
| Evl | 0 | 1.399899612 | 0.829 | 0.184 | 0 | 4 | Evl |
| Arhgap15 | 0 | 1.395936464 | 0.885 | 0.145 | 0 | 4 | Arhgap15 |
| Mctp2 | 0 | 1.394670714 | 0.678 | 0.047 | 0 | 4 | Mctp2 |
| Ssbp2 | 0 | 1.389858468 | 0.679 | 0.107 | 0 | 4 | Ssbp2 |
| Pfn1 | 0 | 1.389281847 | 0.996 | 0.664 | 0 | 4 | Pfn1 |

| | | | | | | | |
|---|---|---|---|---|---|---|---|
| Efhd2 | 0 | 1.384824603 | 0.837 | 0.147 | 0 | 4 | Efhd2 |
| Ptpn18 | 0 | 1.382742524 | 0.954 | 0.277 | 0 | 4 | Ptpn18 |
| Il2rg | 0 | 1.377948539 | 0.737 | 0.079 | 0 | 4 | Il2rg |
| Myo1e1 | 0 | 1.375287414 | 0.842 | 0.247 | 0 | 4 | Myo1e |
| Fndc3a | 0 | 1.373957553 | 0.841 | 0.243 | 0 | 4 | Fndc3a |
| Cers6 | 0 | 1.37171073 | 0.684 | 0.081 | 0 | 4 | Cers6 |
| Padi2 | 0 | 1.371452365 | 0.707 | 0.058 | 0 | 4 | Padi2 |
| Rinl | 0 | 1.371132561 | 0.762 | 0.093 | 0 | 4 | Rinl |
| Tnfrsf9 | 0 | 1.368890016 | 0.604 | 0.034 | 0 | 4 | Tnfrsf9 |
| Fnbp1 | 0 | 1.36824571 | 0.834 | 0.193 | 0 | 4 | Fnbp1 |
| Cd3g | 0 | 1.357786506 | 0.288 | 0.03 | 0 | 4 | Cd3g |
| Sh3bgrl3 | 0 | 1.345749589 | 0.993 | 0.534 | 0 | 4 | Sh3bgrl3 |
| Kif13b | 0 | 1.33230355 | 0.787 | 0.181 | 0 | 4 | Kif13b |
| Gimap42 | 0 | 1.327604224 | 0.864 | 0.22 | 0 | 4 | Gimap4 |
| Slc38a21 | 0 | 1.318231991 | 0.85 | 0.279 | 0 | 4 | Slc38a2 |
| Gprin3 | 0 | 1.316873633 | 0.532 | 0.034 | 0 | 4 | Gprin3 |
| Ccl3 | 0 | 1.312758448 | 0.474 | 0.059 | 0 | 4 | Ccl3 |
| Pik3cd | 0 | 1.308208226 | 0.659 | 0.075 | 0 | 4 | Pik3cd |
| Cotl1 | 0 | 1.301277398 | 0.846 | 0.187 | 0 | 4 | Cotl1 |
| Peli1 | 0 | 1.297862709 | 0.759 | 0.2 | 0 | 4 | Peli1 |
| Kcnq5 | 0 | 1.297823723 | 0.408 | 0.048 | 0 | 4 | Kcnq5 |
| Rhoh | 0 | 1.297752562 | 0.638 | 0.071 | 0 | 4 | Rhoh |
| Rps11 | 0 | 1.289775754 | 0.997 | 0.902 | 0 | 4 | Rps11 |
| Serpina3g | 0 | 1.289594422 | 0.559 | 0.037 | 0 | 4 | Serpina3g |
| Cst7 | 0 | 1.288024448 | 0.639 | 0.039 | 0 | 4 | Cst7 |
| 11-Sep | 0 | 1.285966536 | 0.799 | 0.167 | 0 | 4 | 11-Sep |
| Sulf2 | 0 | 1.273766295 | 0.662 | 0.065 | 0 | 4 | Sulf2 |
| Prf1 | 0 | 1.263443869 | 0.572 | 0.032 | 0 | 4 | Prf1 |
| Gm2682 | 0 | 1.258357635 | 0.537 | 0.053 | 0 | 4 | Gm2682 |
| Cd96 | 0 | 1.256185859 | 0.606 | 0.036 | 0 | 4 | Cd96 |
| Arsb | 0 | 1.255235839 | 0.647 | 0.095 | 0 | 4 | Arsb |
| AC149090.1 | 0 | 1.240614939 | 0.792 | 0.198 | 0 | 4 | AC149090.1 |
| Tnik | 0 | 1.240588771 | 0.636 | 0.081 | 0 | 4 | Tnik |
| Trbc1 | 0 | 1.24003719 | 0.616 | 0.054 | 0 | 4 | Trbc1 |
| Lat2 | 0 | 1.225423717 | 0.734 | 0.095 | 0 | 4 | Lat2 |
| S100a10 | 0 | 1.215212346 | 0.978 | 0.439 | 0 | 4 | S100a10 |
| Limd2 | 0 | 1.212293647 | 0.73 | 0.113 | 0 | 4 | Limd2 |
| Aoah | 0 | 1.210673658 | 0.574 | 0.043 | 0 | 4 | Aoah |
| Gng2 | 0 | 1.210192005 | 0.771 | 0.151 | 0 | 4 | Gng2 |
| Prkch2 | 0 | 1.209638846 | 0.95 | 0.367 | 0 | 4 | Prkch |
| Cdk6 | 0 | 1.199505132 | 0.547 | 0.085 | 0 | 4 | Cdk6 |
| Adgre5 | 0 | 1.198904353 | 0.711 | 0.209 | 0 | 4 | Adgre5 |
| Il21r | 0 | 1.189206267 | 0.604 | 0.062 | 0 | 4 | Il21r |

| | | | | | | | |
|---|---|---|---|---|---|---|---|
| Cd37 | 0 | 1.189000269 | 0.658 | 0.081 | 0 | 4 | Cd37 |
| Cyfip2 | 0 | 1.186717402 | 0.621 | 0.059 | 0 | 4 | Cyfip2 |
| Ccnd3 | 0 | 1.18635407 | 0.788 | 0.207 | 0 | 4 | Ccnd3 |
| Ldlrad4 | 0 | 1.184515475 | 0.77 | 0.219 | 0 | 4 | Ldlrad4 |
| Sorl1 | 0 | 1.181473795 | 0.649 | 0.078 | 0 | 4 | Sorl1 |
| Tbx21 | 0 | 1.178253645 | 0.555 | 0.036 | 0 | 4 | Tbx21 |
| Setbp1 | 0 | 1.167785451 | 0.61 | 0.092 | 0 | 4 | Setbp1 |
| Rasal3 | 0 | 1.167483068 | 0.584 | 0.053 | 0 | 4 | Rasal3 |
| Prkcq | 0 | 1.163011798 | 0.594 | 0.057 | 0 | 4 | Prkcq |
| Fyb | 0 | 1.161012339 | 0.85 | 0.143 | 0 | 4 | Fyb |
| Apbb1ip | 0 | 1.148499356 | 0.751 | 0.139 | 0 | 4 | Apbb1ip |
| Ccr2 | 0 | 1.14675308 | 0.521 | 0.06 | 0 | 4 | Ccr2 |
| Malt1 | 0 | 1.139909455 | 0.645 | 0.12 | 0 | 4 | Malt1 |
| Cxcr6 | 0 | 1.137810692 | 0.48 | 0.038 | 0 | 4 | Cxcr6 |
| Laptm5 | 0 | 1.132906597 | 0.907 | 0.168 | 0 | 4 | Laptm5 |
| Nr4a2 | 0 | 1.130658623 | 0.525 | 0.107 | 0 | 4 | Nr4a2 |
| Gimap12 | 0 | 1.130217538 | 0.876 | 0.268 | 0 | 4 | Gimap1 |
| Fam49b | 0 | 1.128479317 | 0.832 | 0.201 | 0 | 4 | Fam49b |
| Prkcb | 0 | 1.127801889 | 0.693 | 0.124 | 0 | 4 | Prkcb |
| Fryl | 0 | 1.125615502 | 0.789 | 0.237 | 0 | 4 | Fryl |
| Vps37b | 0 | 1.12518249 | 0.639 | 0.106 | 0 | 4 | Vps37b |
| Dnajc11 | 0 | 1.116931046 | 0.832 | 0.292 | 0 | 4 | Dnajc1 |
| Adcy7 | 0 | 1.112003984 | 0.636 | 0.082 | 0 | 4 | Adcy7 |
| Rab8b | 0 | 1.111794445 | 0.817 | 0.252 | 0 | 4 | Rab8b |
| Celf2 | 0 | 1.111094137 | 0.837 | 0.163 | 0 | 4 | Celf2 |
| Klrb1a | 0 | 1.108176801 | 0.428 | 0.02 | 0 | 4 | Klrb1a |
| Camkmt | 0 | 1.10204428 | 0.588 | 0.112 | 0 | 4 | Camkmt |
| Rpsa1 | 0 | 1.09787022 | 0.997 | 0.897 | 0 | 4 | Rpsa |
| Pag1 | 0 | 1.095199817 | 0.54 | 0.096 | 0 | 4 | Pag1 |
| Itgb12 | 0 | 1.094185081 | 0.941 | 0.452 | 0 | 4 | Itgb1 |
| 4933406I18Rik | 0 | 1.093945479 | 0.447 | 0.053 | 0 | 4 | 4933406I18Rik |
| Akap131 | 0 | 1.09345563 | 0.904 | 0.364 | 0 | 4 | Akap13 |
| Ptpn6 | 0 | 1.092951937 | 0.618 | 0.101 | 0 | 4 | Ptpn6 |
| Xist2 | 0 | 1.091309036 | 0.976 | 0.433 | 0 | 4 | Xist |
| Sema6d2 | 0 | 1.089136563 | 0.672 | 0.187 | 0 | 4 | Sema6d |
| Mapre2 | 0 | 1.088337114 | 0.741 | 0.221 | 0 | 4 | Mapre2 |
| Arnt2 | 0 | 1.088223253 | 0.479 | 0.018 | 0 | 4 | Arnt2 |
| Rps3 | 0 | 1.086968144 | 0.999 | 0.887 | 0 | 4 | Rps3 |
| Cd2 | 0 | 1.085636466 | 0.574 | 0.054 | 0 | 4 | Cd2 |
| Tmem131 | 0 | 1.07822358 | 0.714 | 0.178 | 0 | 4 | Tmem131 |
| Cblb | 0 | 1.076003505 | 0.654 | 0.142 | 0 | 4 | Cblb |
| Jak11 | 0 | 1.075577943 | 0.883 | 0.335 | 0 | 4 | Jak1 |
| Atp11b | 0 | 1.074502814 | 0.684 | 0.15 | 0 | 4 | Atp11b |

| | | | | | | | |
|---|---|---|---|---|---|---|---|
| Dgkd | 0 | 1.072661267 | 0.655 | 0.131 | 0 | 4 | Dgkd |
| Itgad | 0 | 1.072572644 | 0.391 | 0.016 | 0 | 4 | Itgad |
| Tbc1d10c | 0 | 1.072088308 | 0.58 | 0.042 | 0 | 4 | Tbc1d10c |
| Prkacb | 0 | 1.062704707 | 0.735 | 0.187 | 0 | 4 | Prkacb |
| Sipa1l11 | 0 | 1.054949245 | 0.65 | 0.162 | 0 | 4 | Sipa1l1 |
| Tab2 | 0 | 1.054684419 | 0.741 | 0.2 | 0 | 4 | Tab2 |
| Grap2 | 0 | 1.052941436 | 0.572 | 0.078 | 0 | 4 | Grap2 |
| Fasl | 0 | 1.051928973 | 0.507 | 0.029 | 0 | 4 | Fasl |
| Gm203882 | 0 | 1.049072825 | 0.927 | 0.369 | 0 | 4 | Gm20388 |
| Ripor21 | 0 | 1.045146076 | 0.648 | 0.144 | 0 | 4 | Ripor2 |
| Pglyrp1 | 0 | 1.045058837 | 0.616 | 0.061 | 0 | 4 | Pglyrp1 |
| Card11 | 0 | 1.040954831 | 0.528 | 0.055 | 0 | 4 | Card11 |
| Abr | 0 | 1.040415955 | 0.644 | 0.129 | 0 | 4 | Abr |
| Tnni1 | 0 | 1.039071511 | 0.409 | 0.009 | 0 | 4 | Tnni1 |
| Mllt3 | 0 | 1.034679772 | 0.621 | 0.123 | 0 | 4 | Mllt3 |
| Oprm1 | 0 | 1.034324179 | 0.502 | 0.033 | 0 | 4 | Oprm1 |
| Rbpj | 0 | 1.030958711 | 0.736 | 0.19 | 0 | 4 | Rbpj |
| Fgl2 | 0 | 1.026799821 | 0.569 | 0.08 | 0 | 4 | Fgl2 |
| Lsp1 | 0 | 1.024321781 | 0.65 | 0.108 | 0 | 4 | Lsp1 |
| Lax1 | 0 | 1.021403072 | 0.517 | 0.038 | 0 | 4 | Lax1 |
| Ubac2 | 0 | 1.021401049 | 0.685 | 0.158 | 0 | 4 | Ubac2 |
| Slc38a1 | 0 | 1.021111052 | 0.552 | 0.06 | 0 | 4 | Slc38a1 |
| Tex14 | 0 | 1.020314831 | 0.549 | 0.126 | 0 | 4 | Tex14 |
| Cd48 | 0 | 1.018208166 | 0.643 | 0.104 | 0 | 4 | Cd48 |
| Ptpre | 0 | 1.016354511 | 0.625 | 0.133 | 0 | 4 | Ptpre |
| Itk | 0 | 1.014919542 | 0.531 | 0.05 | 0 | 4 | Itk |
| Rps27 | 0 | 1.011765779 | 0.997 | 0.892 | 0 | 4 | Rps27 |
| Eml4 | 0 | 1.006345522 | 0.772 | 0.228 | 0 | 4 | Eml4 |
| Lgals31 | 0 | 0.998484952 | 0.748 | 0.186 | 0 | 4 | Lgals3 |
| Fxyd5 | 0 | 0.994897169 | 0.944 | 0.379 | 0 | 4 | Fxyd5 |
| Klrb1f | 0 | 0.994201527 | 0.502 | 0.036 | 0 | 4 | Klrb1f |
| Ptprj | 0 | 0.994112566 | 0.671 | 0.12 | 0 | 4 | Ptprj |
| Rpl22l12 | 0 | 0.993883796 | 0.98 | 0.555 | 0 | 4 | Rpl22l1 |
| Rin3 | 0 | 0.99306033 | 0.648 | 0.152 | 0 | 4 | Rin3 |
| Matk | 0 | 0.989789063 | 0.464 | 0.016 | 0 | 4 | Matk |
| Lrrk1 | 0 | 0.988116735 | 0.653 | 0.163 | 0 | 4 | Lrrk1 |
| Neurl3 | 0 | 0.987883051 | 0.561 | 0.088 | 0 | 4 | Neurl3 |
| Parp8 | 0 | 0.986972088 | 0.592 | 0.123 | 0 | 4 | Parp8 |
| Hectd11 | 0 | 0.986680079 | 0.791 | 0.271 | 0 | 4 | Hectd1 |
| Gimap62 | 0 | 0.983702924 | 0.901 | 0.334 | 0 | 4 | Gimap6 |
| Lncpint1 | 0 | 0.982705309 | 0.73 | 0.247 | 0 | 4 | Lncpint |
| Rgs2 | 0 | 0.978081886 | 0.556 | 0.105 | 0 | 4 | Rgs2 |
| Sema4a | 0 | 0.976554864 | 0.514 | 0.05 | 0 | 4 | Sema4a |

| | | | | | | | |
|---|---|---|---|---|---|---|---|
| Emid1 | 0 | 0.976321305 | 0.597 | 0.079 | 0 | 4 | Emid1 |
| Nsd3 | 0 | 0.976088424 | 0.795 | 0.279 | 0 | 4 | Nsd3 |
| Ighm | 0 | 0.975176714 | 0.649 | 0.095 | 0 | 4 | Ighm |
| Lta4h | 0 | 0.974404494 | 0.632 | 0.119 | 0 | 4 | Lta4h |
| Znrf3 | 0 | 0.972823507 | 0.539 | 0.081 | 0 | 4 | Znrf3 |
| Bin2 | 0 | 0.971496566 | 0.62 | 0.098 | 0 | 4 | Bin2 |
| Mycbp21 | 0 | 0.971051861 | 0.872 | 0.345 | 0 | 4 | Mycbp2 |
| Aak1 | 0 | 0.970289421 | 0.658 | 0.156 | 0 | 4 | Aak1 |
| Mgat5 | 0 | 0.969439264 | 0.616 | 0.144 | 0 | 4 | Mgat5 |
| Sec11c | 0 | 0.96678195 | 0.858 | 0.278 | 0 | 4 | Sec11c |
| 1810011H11Rik | 0 | 0.965627266 | 0.496 | 0.048 | 0 | 4 | 1810011H11Rik |
| Ptpn4 | 0 | 0.964584021 | 0.562 | 0.105 | 0 | 4 | Ptpn4 |
| Tnfrsf18 | 0 | 0.959614993 | 0.523 | 0.046 | 0 | 4 | Tnfrsf18 |
| Rps18 | 0 | 0.958088059 | 0.999 | 0.938 | 0 | 4 | Rps18 |
| Chl1 | 0 | 0.950933995 | 0.413 | 0.031 | 0 | 4 | Chl1 |
| Diaph1 | 0 | 0.949591214 | 0.733 | 0.213 | 0 | 4 | Diaph1 |
| Pycard2 | 0 | 0.948141913 | 0.862 | 0.242 | 0 | 4 | Pycard |
| Sik32 | 0 | 0.946511078 | 0.71 | 0.244 | 0 | 4 | Sik3 |
| Apobec3 | 0 | 0.944890671 | 0.758 | 0.193 | 0 | 4 | Apobec3 |
| Leprotl1 | 0 | 0.944332399 | 0.729 | 0.187 | 0 | 4 | Leprotl1 |
| Dscam | 0 | 0.943440066 | 0.339 | 0.029 | 0 | 4 | Dscam |
| Zfp36l2 | 0 | 0.940504434 | 0.782 | 0.277 | 0 | 4 | Zfp36l2 |
| Sytl2 | 0 | 0.939773472 | 0.452 | 0.032 | 0 | 4 | Sytl2 |
| Dock111 | 0 | 0.936316573 | 0.54 | 0.088 | 0 | 4 | Dock11 |
| Fyn2 | 0 | 0.934792443 | 0.76 | 0.237 | 0 | 4 | Fyn |
| Ccnd21 | 0 | 0.934455174 | 0.768 | 0.257 | 0 | 4 | Ccnd2 |
| Ube2h | 0 | 0.934150184 | 0.742 | 0.237 | 0 | 4 | Ube2h |
| Rpl37a | 0 | 0.933811514 | 0.997 | 0.935 | 0 | 4 | Rpl37a |
| B4galnt1 | 0 | 0.932186663 | 0.565 | 0.067 | 0 | 4 | B4galnt1 |
| Sytl3 | 0 | 0.931897059 | 0.465 | 0.033 | 0 | 4 | Sytl3 |
| Traf3ip3 | 0 | 0.93169855 | 0.521 | 0.052 | 0 | 4 | Traf3ip3 |
| A930015D03Rik | 0 | 0.931503038 | 0.47 | 0.085 | 0 | 4 | A930015D03Rik |
| Chd21 | 0 | 0.931068236 | 0.723 | 0.226 | 0 | 4 | Chd2 |
| Gimap7 | 0 | 0.928815914 | 0.467 | 0.034 | 0 | 4 | Gimap7 |
| Kansl11 | 0 | 0.923276315 | 0.809 | 0.312 | 0 | 4 | Kansl1 |
| Acap1 | 0 | 0.922807123 | 0.505 | 0.044 | 0 | 4 | Acap1 |
| Rps16 | 0 | 0.918370647 | 0.997 | 0.923 | 0 | 4 | Rps16 |
| Apol7e | 0 | 0.915937276 | 0.44 | 0.027 | 0 | 4 | Apol7e |
| Rpl321 | 0 | 0.915008344 | 0.999 | 0.951 | 0 | 4 | Rpl32 |
| Rps3a12 | 0 | 0.914867292 | 0.996 | 0.932 | 0 | 4 | Rps3a1 |
| Ccdc88c | 0 | 0.914125551 | 0.549 | 0.073 | 0 | 4 | Ccdc88c |
| Tle41 | 0 | 0.911627643 | 0.68 | 0.199 | 0 | 4 | Tle4 |
| Slc9a9 | 0 | 0.911446885 | 0.602 | 0.124 | 0 | 4 | Slc9a9 |

| | | | | | | | |
|---|---|---|---|---|---|---|---|
| Rps101 | 0 | 0.909375614 | 0.997 | 0.941 | 0 | 4 | Rps10 |
| Tnfaip3 | 0 | 0.906319501 | 0.413 | 0.071 | 0 | 4 | Tnfaip3 |
| Ncf4 | 0 | 0.90549652 | 0.549 | 0.086 | 0 | 4 | Ncf4 |
| Plek | 0 | 0.903822906 | 0.552 | 0.099 | 0 | 4 | Plek |
| Hsd11b1 | 0 | 0.903794858 | 0.528 | 0.066 | 0 | 4 | Hsd11b1 |
| Il12rb2 | 0 | 0.898822508 | 0.439 | 0.026 | 0 | 4 | Il12rb2 |
| Arhgap30 | 0 | 0.89800659 | 0.559 | 0.083 | 0 | 4 | Arhgap30 |
| Itgal | 0 | 0.896521263 | 0.502 | 0.062 | 0 | 4 | Itgal |
| AU0202062 | 0 | 0.895057973 | 0.767 | 0.263 | 0 | 4 | AU020206 |
| Rpl131 | 0 | 0.894404501 | 0.999 | 0.976 | 0 | 4 | Rpl13 |
| Aim2 | 0 | 0.894063143 | 0.519 | 0.078 | 0 | 4 | Aim2 |
| Gm26827 | 0 | 0.891917272 | 0.43 | 0.028 | 0 | 4 | Gm26827 |
| Jaml | 0 | 0.891694604 | 0.439 | 0.035 | 0 | 4 | Jaml |
| Itgb2 | 0 | 0.891507013 | 0.549 | 0.075 | 0 | 4 | Itgb2 |
| Plcl2 | 0 | 0.890464404 | 0.508 | 0.091 | 0 | 4 | Plcl2 |
| Rps20 | 0 | 0.888390894 | 0.997 | 0.949 | 0 | 4 | Rps20 |
| Rps24 | 0 | 0.888164503 | 0.997 | 0.971 | 0 | 4 | Rps24 |
| Bcl2l11 | 0 | 0.886707091 | 0.533 | 0.092 | 0 | 4 | Bcl2l11 |
| Tra2b1 | 0 | 0.886086285 | 0.784 | 0.267 | 0 | 4 | Tra2b |
| Lims1 | 0 | 0.885596736 | 0.749 | 0.274 | 0 | 4 | Lims1 |
| Mta3 | 0 | 0.885020063 | 0.623 | 0.124 | 0 | 4 | Mta3 |
| Tmsb4x | 0 | 0.883053511 | 1 | 0.93 | 0 | 4 | Tmsb4x |
| Gimap51 | 0 | 0.881635879 | 0.635 | 0.143 | 0 | 4 | Gimap5 |
| Rpl381 | 0 | 0.878404063 | 0.994 | 0.888 | 0 | 4 | Rpl38 |
| Rps71 | 0 | 0.878080014 | 0.996 | 0.912 | 0 | 4 | Rps7 |
| Suco | 0 | 0.877544966 | 0.578 | 0.128 | 0 | 4 | Suco |
| Akna | 0 | 0.875829194 | 0.498 | 0.067 | 0 | 4 | Akna |
| Rfx7 | 0 | 0.874682079 | 0.572 | 0.13 | 0 | 4 | Rfx7 |
| Ppm1h | 0 | 0.874221137 | 0.597 | 0.135 | 0 | 4 | Ppm1h |
| Trim14 | 0 | 0.87337444 | 0.528 | 0.096 | 0 | 4 | Trim14 |
| Stk4 | 0 | 0.872625477 | 0.643 | 0.147 | 0 | 4 | Stk4 |
| Nfkb1 | 0 | 0.87202539 | 0.72 | 0.245 | 0 | 4 | Nfkb1 |
| Rpl18a | 0 | 0.869474174 | 0.999 | 0.927 | 0 | 4 | Rpl18a |
| Rpl34 | 0 | 0.868877062 | 0.997 | 0.913 | 0 | 4 | Rpl34 |
| 6-Sep | 0 | 0.867395731 | 0.532 | 0.073 | 0 | 4 | 6-Sep |
| Vav31 | 0 | 0.865314361 | 0.713 | 0.229 | 0 | 4 | Vav3 |
| Zap70 | 0 | 0.864642203 | 0.463 | 0.032 | 0 | 4 | Zap70 |
| Spry2 | 0 | 0.860201031 | 0.465 | 0.059 | 0 | 4 | Spry2 |
| Rpl361 | 0 | 0.859971855 | 0.996 | 0.898 | 0 | 4 | Rpl36 |
| Sh2d1a | 0 | 0.858954035 | 0.44 | 0.033 | 0 | 4 | Sh2d1a |
| Fam107b | 0 | 0.857924584 | 0.564 | 0.094 | 0 | 4 | Fam107b |
| Ubash3b | 0 | 0.85780945 | 0.57 | 0.144 | 0 | 4 | Ubash3b |
| Gramd1a | 0 | 0.85214963 | 0.512 | 0.076 | 0 | 4 | Gramd1a |

| | | | | | | | |
|---|---|---|---|---|---|---|---|
| Gpsm3 | 0 | 0.849037328 | 0.618 | 0.12 | 0 | 4 | Gpsm3 |
| Eya2 | 0 | 0.847301041 | 0.539 | 0.074 | 0 | 4 | Eya2 |
| Dennd1b | 0 | 0.847012394 | 0.613 | 0.159 | 0 | 4 | Dennd1b |
| Lcp2 | 0 | 0.845901172 | 0.491 | 0.071 | 0 | 4 | Lcp2 |
| Sla2 | 0 | 0.845774057 | 0.431 | 0.025 | 0 | 4 | Sla2 |
| Litaf | 0 | 0.84471702 | 0.593 | 0.132 | 0 | 4 | Litaf |
| Ntan12 | 0 | 0.842078703 | 0.837 | 0.287 | 0 | 4 | Ntan1 |
| Zfp318 | 0 | 0.840462452 | 0.493 | 0.074 | 0 | 4 | Zfp318 |
| Rpl391 | 0 | 0.839717566 | 0.997 | 0.895 | 0 | 4 | Rpl39 |
| Zcchc11 | 0 | 0.838727015 | 0.727 | 0.24 | 0 | 4 | Zcchc11 |
| Esyt1 | 0 | 0.83840459 | 0.556 | 0.111 | 0 | 4 | Esyt1 |
| Plcg2 | 0 | 0.837488198 | 0.545 | 0.095 | 0 | 4 | Plcg2 |
| Trip4 | 0 | 0.836653815 | 0.514 | 0.095 | 0 | 4 | Trip4 |
| Vim | 0 | 0.836073176 | 0.852 | 0.274 | 0 | 4 | Vim |
| Rasa11 | 0 | 0.83584474 | 0.683 | 0.221 | 0 | 4 | Rasa1 |
| Ifitm10 | 0 | 0.829770605 | 0.45 | 0.037 | 0 | 4 | Ifitm10 |
| Sesn1 | 0 | 0.82957764 | 0.621 | 0.188 | 0 | 4 | Sesn1 |
| Ncald | 0 | 0.828894967 | 0.504 | 0.077 | 0 | 4 | Ncald |
| Tec | 0 | 0.828800784 | 0.521 | 0.085 | 0 | 4 | Tec |
| Capzb | 0 | 0.826953395 | 0.86 | 0.34 | 0 | 4 | Capzb |
| Rnf43 | 0 | 0.826406121 | 0.398 | 0.029 | 0 | 4 | Rnf43 |
| Rpl372 | 0 | 0.825330026 | 0.998 | 0.923 | 0 | 4 | Rpl37 |
| Arhgap9 | 0 | 0.824526335 | 0.583 | 0.083 | 0 | 4 | Arhgap9 |
| Tbl1xr1 | 0 | 0.823564623 | 0.612 | 0.138 | 0 | 4 | Tbl1xr1 |
| Rps131 | 0 | 0.822075142 | 0.998 | 0.919 | 0 | 4 | Rps13 |
| Saraf2 | 0 | 0.820313506 | 0.769 | 0.28 | 0 | 4 | Saraf |
| Ttc39b | 0 | 0.820062587 | 0.542 | 0.111 | 0 | 4 | Ttc39b |
| Cyth4 | 0 | 0.819947224 | 0.595 | 0.106 | 0 | 4 | Cyth4 |
| Trbc2 | 0 | 0.819672455 | 0.36 | 0.049 | 0 | 4 | Trbc2 |
| Cytip | 0 | 0.815206956 | 0.51 | 0.066 | 0 | 4 | Cytip |
| Prkca | 0 | 0.812508784 | 0.471 | 0.078 | 0 | 4 | Prkca |
| Fau | 0 | 0.812396157 | 0.998 | 0.951 | 0 | 4 | Fau |
| Rps6ka1 | 0 | 0.812111896 | 0.535 | 0.089 | 0 | 4 | Rps6ka1 |
| B4galt5 | 0 | 0.809510181 | 0.455 | 0.072 | 0 | 4 | B4galt5 |
| Mtmr12 | 0 | 0.809396984 | 0.532 | 0.107 | 0 | 4 | Mtmr12 |
| Sntb1 | 0 | 0.807058285 | 0.395 | 0.045 | 0 | 4 | Sntb1 |
| Tpp2 | 0 | 0.803609467 | 0.673 | 0.202 | 0 | 4 | Tpp2 |
| Nup210 | 0 | 0.802140478 | 0.439 | 0.044 | 0 | 4 | Nup210 |
| Gpr174 | 0 | 0.80061916 | 0.407 | 0.031 | 0 | 4 | Gpr174 |
| Psd4 | 0 | 0.800499539 | 0.428 | 0.043 | 0 | 4 | Psd4 |
| Gimap9 | 0 | 0.800395806 | 0.598 | 0.132 | 0 | 4 | Gimap9 |
| Vps13a | 0 | 0.796633169 | 0.567 | 0.149 | 0 | 4 | Vps13a |
| Nlrc3 | 0 | 0.795537444 | 0.424 | 0.036 | 0 | 4 | Nlrc3 |

| | | | | | | | |
|---|---|---|---|---|---|---|---|
| Rps281 | 0 | 0.793863334 | 0.993 | 0.892 | 0 | 4 | Rps28 |
| Camk2g | 0 | 0.792236037 | 0.542 | 0.11 | 0 | 4 | Camk2g |
| Rps51 | 0 | 0.790725249 | 0.997 | 0.916 | 0 | 4 | Rps5 |
| H2afy | 0 | 0.790082998 | 0.832 | 0.308 | 0 | 4 | H2afy |
| Rpl27a1 | 0 | 0.789684836 | 0.999 | 0.958 | 0 | 4 | Rpl27a |
| Gpr171 | 0 | 0.788532447 | 0.411 | 0.025 | 0 | 4 | Gpr171 |
| Tespa1 | 0 | 0.788180406 | 0.415 | 0.057 | 0 | 4 | Tespa1 |
| Jpt1 | 0 | 0.785866552 | 0.706 | 0.193 | 0 | 4 | Jpt1 |
| Runx1 | 0 | 0.783086697 | 0.52 | 0.108 | 0 | 4 | Runx1 |
| Acot7 | 0 | 0.781188907 | 0.444 | 0.054 | 0 | 4 | Acot7 |
| Pip5k1b | 0 | 0.779070271 | 0.472 | 0.077 | 0 | 4 | Pip5k1b |
| Ptger4 | 0 | 0.778883529 | 0.42 | 0.065 | 0 | 4 | Ptger4 |
| Pkp3 | 0 | 0.778858908 | 0.431 | 0.047 | 0 | 4 | Pkp3 |
| Glmn | 0 | 0.778378317 | 0.463 | 0.075 | 0 | 4 | Glmn |
| Rpl310 | 0 | 0.777811878 | 0.996 | 0.905 | 0 | 4 | Rpl3 |
| Pde1c | 0 | 0.777748245 | 0.392 | 0.049 | 0 | 4 | Pde1c |
| Klrc2 | 0 | 0.77510012 | 0.364 | 0.035 | 0 | 4 | Klrc2 |
| Gfod1 | 0 | 0.773504605 | 0.396 | 0.05 | 0 | 4 | Gfod1 |
| Rpl301 | 0 | 0.771657251 | 0.997 | 0.934 | 0 | 4 | Rpl30 |
| Rsbn1l1 | 0 | 0.770789431 | 0.725 | 0.246 | 0 | 4 | Rsbn1l |
| Cyba | 0 | 0.76672929 | 0.957 | 0.41 | 0 | 4 | Cyba |
| Rpl181 | 0 | 0.766005789 | 0.999 | 0.895 | 0 | 4 | Rpl18 |
| Fermt3 | 0 | 0.763082937 | 0.526 | 0.09 | 0 | 4 | Fermt3 |
| Itpr2 | 0 | 0.761358609 | 0.687 | 0.218 | 0 | 4 | Itpr2 |
| Tiam2 | 0 | 0.759232974 | 0.378 | 0.032 | 0 | 4 | Tiam2 |
| Unc119 | 0 | 0.757389382 | 0.468 | 0.061 | 0 | 4 | Unc119 |
| Inpp5d | 0 | 0.755548855 | 0.644 | 0.178 | 0 | 4 | Inpp5d |
| Pik3ap1 | 0 | 0.754180364 | 0.446 | 0.069 | 0 | 4 | Pik3ap1 |
| Cd53 | 0 | 0.752555856 | 0.524 | 0.092 | 0 | 4 | Cd53 |
| Gramd1b | 0 | 0.748499861 | 0.443 | 0.069 | 0 | 4 | Gramd1b |
| Eya1 | 0 | 0.743315493 | 0.339 | 0.022 | 0 | 4 | Eya1 |
| Ust | 0 | 0.74214358 | 0.401 | 0.067 | 0 | 4 | Ust |
| Vipr2 | 0 | 0.741953062 | 0.328 | 0.014 | 0 | 4 | Vipr2 |
| Jakmip1 | 0 | 0.74159679 | 0.377 | 0.029 | 0 | 4 | Jakmip1 |
| Gsap | 0 | 0.741218467 | 0.436 | 0.065 | 0 | 4 | Gsap |
| Otulin | 0 | 0.740930431 | 0.644 | 0.173 | 0 | 4 | Otulin |
| Mvb12b | 0 | 0.738381239 | 0.578 | 0.143 | 0 | 4 | Mvb12b |
| Rbl2 | 0 | 0.736975985 | 0.504 | 0.113 | 0 | 4 | Rbl2 |
| Ppp1r18 | 0 | 0.735673138 | 0.558 | 0.13 | 0 | 4 | Ppp1r18 |
| Eomes | 0 | 0.734358015 | 0.38 | 0.031 | 0 | 4 | Eomes |
| Napsa | 0 | 0.732531153 | 0.386 | 0.032 | 0 | 4 | Napsa |
| Znrf2 | 0 | 0.730800785 | 0.57 | 0.135 | 0 | 4 | Znrf2 |
| Strbp | 0 | 0.728353862 | 0.633 | 0.165 | 0 | 4 | Strbp |

| | | | | | | | |
|---|---|---|---|---|---|---|---|
| Lpxn | 0 | 0.726457609 | 0.45 | 0.072 | 0 | 4 | Lpxn |
| Gimap81 | 0 | 0.7257859 | 0.557 | 0.146 | 0 | 4 | Gimap8 |
| Sdf2l1 | 0 | 0.724214388 | 0.565 | 0.144 | 0 | 4 | Sdf2l1 |
| Slamf7 | 0 | 0.721330625 | 0.456 | 0.078 | 0 | 4 | Slamf7 |
| Asxl2 | 0 | 0.720243829 | 0.558 | 0.152 | 0 | 4 | Asxl2 |
| Mxd1 | 0 | 0.717530504 | 0.384 | 0.044 | 0 | 4 | Mxd1 |
| Sesn3 | 0 | 0.716813505 | 0.549 | 0.104 | 0 | 4 | Sesn3 |
| Rps231 | 0 | 0.712702125 | 0.999 | 0.95 | 0 | 4 | Rps23 |
| Rras2 | 0 | 0.706469459 | 0.415 | 0.065 | 0 | 4 | Rras2 |
| Elf4 | 0 | 0.703723966 | 0.455 | 0.103 | 0 | 4 | Elf4 |
| Rgs31 | 0 | 0.700466625 | 0.591 | 0.163 | 0 | 4 | Rgs3 |
| Gpr68 | 0 | 0.70032627 | 0.352 | 0.027 | 0 | 4 | Gpr68 |
| Rnf149 | 0 | 0.699758333 | 0.453 | 0.083 | 0 | 4 | Rnf149 |
| Tes | 0 | 0.697642801 | 0.537 | 0.105 | 0 | 4 | Tes |
| Gab3 | 0 | 0.696597511 | 0.391 | 0.077 | 0 | 4 | Gab3 |
| Rasgrp1 | 0 | 0.696201132 | 0.384 | 0.043 | 0 | 4 | Rasgrp1 |
| Cdyl | 0 | 0.691712063 | 0.484 | 0.108 | 0 | 4 | Cdyl |
| Rpl191 | 0 | 0.69009186 | 0.999 | 0.961 | 0 | 4 | Rpl19 |
| Klk8 | 0 | 0.689942572 | 0.406 | 0.063 | 0 | 4 | Klk8 |
| Fam189b | 0 | 0.688030392 | 0.384 | 0.037 | 0 | 4 | Fam189b |
| Mob3a | 0 | 0.686891486 | 0.41 | 0.061 | 0 | 4 | Mob3a |
| Satb1 | 0 | 0.685153117 | 0.368 | 0.045 | 0 | 4 | Satb1 |
| Vasp | 0 | 0.684336898 | 0.671 | 0.212 | 0 | 4 | Vasp |
| Slc14a1 | 0 | 0.68358851 | 0.329 | 0.029 | 0 | 4 | Slc14a1 |
| Btbd11 | 0 | 0.681381589 | 0.333 | 0.039 | 0 | 4 | Btbd11 |
| Cited4 | 0 | 0.678915067 | 0.603 | 0.106 | 0 | 4 | Cited4 |
| Ppm1b | 0 | 0.678746563 | 0.517 | 0.116 | 0 | 4 | Ppm1b |
| Atxn10 | 0 | 0.673859049 | 0.582 | 0.151 | 0 | 4 | Atxn10 |
| Rnf166 | 0 | 0.670481937 | 0.448 | 0.087 | 0 | 4 | Rnf166 |
| Hopx | 0 | 0.669943439 | 0.433 | 0.054 | 0 | 4 | Hopx |
| Pstpip1 | 0 | 0.667094909 | 0.377 | 0.036 | 0 | 4 | Pstpip1 |
| Sh2d1b1 | 0 | 0.666277322 | 0.407 | 0.058 | 0 | 4 | Sh2d1b1 |
| Trmt112 | 0 | 0.665297711 | 0.743 | 0.246 | 0 | 4 | Trmt112 |
| Cnot3 | 0 | 0.650270272 | 0.579 | 0.162 | 0 | 4 | Cnot3 |
| Cd82 | 0 | 0.649425736 | 0.419 | 0.082 | 0 | 4 | Cd82 |
| Cysltr2 | 0 | 0.641427572 | 0.297 | 0.024 | 0 | 4 | Cysltr2 |
| Sash3 | 0 | 0.640874546 | 0.365 | 0.042 | 0 | 4 | Sash3 |
| Hvcn1 | 0 | 0.639392515 | 0.352 | 0.038 | 0 | 4 | Hvcn1 |
| Pglyrp2 | 0 | 0.63875428 | 0.356 | 0.025 | 0 | 4 | Pglyrp2 |
| D16Ertd472e | 0 | 0.63772561 | 0.445 | 0.086 | 0 | 4 | D16Ertd472e |
| Vwa8 | 0 | 0.636846638 | 0.445 | 0.101 | 0 | 4 | Vwa8 |
| Rgs18 | 0 | 0.634124853 | 0.315 | 0.015 | 0 | 4 | Rgs18 |
| Gm16341 | 0 | 0.632104537 | 0.321 | 0.019 | 0 | 4 | Gm16341 |

| | | | | | | | |
|---|---|---|---|---|---|---|---|
| Ptpn2 | 0 | 0.625236581 | 0.511 | 0.122 | 0 | 4 | Ptpn2 |
| Traf1 | 0 | 0.623961017 | 0.352 | 0.045 | 0 | 4 | Traf1 |
| Dtnb | 0 | 0.623182314 | 0.437 | 0.095 | 0 | 4 | Dtnb |
| Slc16a10 | 0 | 0.622690163 | 0.305 | 0.033 | 0 | 4 | Slc16a10 |
| Dennd1c | 0 | 0.622494921 | 0.357 | 0.039 | 0 | 4 | Dennd1c |
| Serp2 | 0 | 0.621705383 | 0.335 | 0.024 | 0 | 4 | Serp2 |
| Sla | 0 | 0.619911404 | 0.362 | 0.056 | 0 | 4 | Sla |
| Magohb | 0 | 0.61641335 | 0.497 | 0.099 | 0 | 4 | Magohb |
| Rassf5 | 0 | 0.615238126 | 0.37 | 0.051 | 0 | 4 | Rassf5 |
| P2ry10 | 0 | 0.614850422 | 0.31 | 0.022 | 0 | 4 | P2ry10 |
| Was | 0 | 0.610612748 | 0.386 | 0.054 | 0 | 4 | Was |
| Pla2g16 | 0 | 0.60889601 | 0.695 | 0.216 | 0 | 4 | Pla2g16 |
| Npm3 | 0 | 0.606370838 | 0.487 | 0.113 | 0 | 4 | Npm3 |
| Sms | 0 | 0.606153135 | 0.549 | 0.127 | 0 | 4 | Sms |
| Fam105a | 0 | 0.605599904 | 0.49 | 0.104 | 0 | 4 | Fam105a |
| Plcxd2 | 0 | 0.603059369 | 0.281 | 0.019 | 0 | 4 | Plcxd2 |
| Fgr | 0 | 0.602819935 | 0.298 | 0.028 | 0 | 4 | Fgr |
| Mfsd10 | 0 | 0.602488196 | 0.386 | 0.078 | 0 | 4 | Mfsd10 |
| Pnpla7 | 0 | 0.597604345 | 0.342 | 0.048 | 0 | 4 | Pnpla7 |
| Syk | 0 | 0.595868844 | 0.345 | 0.051 | 0 | 4 | Syk |
| Eef2k | 0 | 0.594838599 | 0.37 | 0.061 | 0 | 4 | Eef2k |
| Sh2d2a | 0 | 0.592664146 | 0.333 | 0.032 | 0 | 4 | Sh2d2a |
| 1810006J02Rik | 0 | 0.591664091 | 0.301 | 0.026 | 0 | 4 | 1810006J02Rik |
| Rab37 | 0 | 0.590306623 | 0.316 | 0.027 | 0 | 4 | Rab37 |
| Cracr2a | 0 | 0.584554431 | 0.446 | 0.068 | 0 | 4 | Cracr2a |
| Isg20 | 0 | 0.583776541 | 0.465 | 0.095 | 0 | 4 | Isg20 |
| Fmnl1 | 0 | 0.579299013 | 0.386 | 0.075 | 0 | 4 | Fmnl1 |
| Bin1 | 0 | 0.577229509 | 0.58 | 0.152 | 0 | 4 | Bin1 |
| Lrmp | 0 | 0.576390096 | 0.34 | 0.044 | 0 | 4 | Lrmp |
| Ptpn7 | 0 | 0.573036364 | 0.322 | 0.043 | 0 | 4 | Ptpn7 |
| Stk26 | 0 | 0.572874802 | 0.326 | 0.028 | 0 | 4 | Stk26 |
| Vav1 | 0 | 0.567743331 | 0.377 | 0.063 | 0 | 4 | Vav1 |
| Gpr34 | 0 | 0.56406087 | 0.333 | 0.043 | 0 | 4 | Gpr34 |
| Kcnip3 | 0 | 0.557392137 | 0.277 | 0.034 | 0 | 4 | Kcnip3 |
| Spin2c | 0 | 0.555826023 | 0.307 | 0.02 | 0 | 4 | Spin2c |
| Gata3 | 0 | 0.55203274 | 0.321 | 0.036 | 0 | 4 | Gata3 |
| Gtpbp1 | 0 | 0.545748087 | 0.331 | 0.056 | 0 | 4 | Gtpbp1 |
| Slc9a3r1 | 0 | 0.539941701 | 0.448 | 0.09 | 0 | 4 | Slc9a3r1 |
| Herc3 | 0 | 0.538458956 | 0.332 | 0.056 | 0 | 4 | Herc3 |
| Zfp831 | 0 | 0.531979264 | 0.271 | 0.019 | 0 | 4 | Zfp831 |
| Armc7 | 0 | 0.526200151 | 0.301 | 0.034 | 0 | 4 | Armc7 |
| N4bp2l1 | 0 | 0.519104714 | 0.298 | 0.041 | 0 | 4 | N4bp2l1 |
| Rhof | 0 | 0.51633243 | 0.28 | 0.023 | 0 | 4 | Rhof |

| | | | | | | | |
|---|---|---|---|---|---|---|---|
| Rtn4rl1 | 0 | 0.515800861 | 0.277 | 0.036 | 0 | 4 | Rtn4rl1 |
| Adamts14 | 0 | 0.511361663 | 0.252 | 0.023 | 0 | 4 | Adamts14 |
| Gm33104 | 0 | 0.50532984 | 0.25 | 0.016 | 0 | 4 | Gm33104 |
| Prkcz | 0 | 0.500202286 | 0.33 | 0.04 | 0 | 4 | Prkcz |
| Il27ra | 0 | 0.498916383 | 0.277 | 0.028 | 0 | 4 | Il27ra |
| Pde4a | 0 | 0.493427952 | 0.324 | 0.048 | 0 | 4 | Pde4a |
| Lamb3 | 0 | 0.493042899 | 0.314 | 0.04 | 0 | 4 | Lamb3 |
| Kif21b | 0 | 0.485794093 | 0.276 | 0.039 | 0 | 4 | Kif21b |
| Fam78a | 0 | 0.483913155 | 0.263 | 0.027 | 0 | 4 | Fam78a |
| Cdca7l | 0 | 0.481787991 | 0.289 | 0.029 | 0 | 4 | Cdca7l |
| Spn | 0 | 0.479400714 | 0.273 | 0.03 | 0 | 4 | Spn |
| Fhl2 | 0 | 0.47261605 | 0.263 | 0.029 | 0 | 4 | Fhl2 |
| Agap2 | 0 | 0.469923889 | 0.26 | 0.024 | 0 | 4 | Agap2 |
| Sh3bp1 | 0 | 0.468962294 | 0.289 | 0.037 | 0 | 4 | Sh3bp1 |
| Ggct | 0 | 0.461701421 | 0.315 | 0.05 | 0 | 4 | Ggct |
| Clec2i | 0 | 0.456763408 | 0.252 | 0.021 | 0 | 4 | Clec2i |
| Cd247 | 0 | 0.446940767 | 0.259 | 0.034 | 0 | 4 | Cd247 |
| Lat | 0 | 0.441378045 | 0.254 | 0.025 | 0 | 4 | Lat |
| Apobr | 0 | 0.437179428 | 0.28 | 0.036 | 0 | 4 | Apobr |
| Myo1g | 0 | 0.431302468 | 0.257 | 0.035 | 0 | 4 | Myo1g |
| Izumo4 | 0 | 0.425862748 | 0.27 | 0.038 | 0 | 4 | Izumo4 |
| Sh3bp2 | 0 | 0.425022787 | 0.256 | 0.03 | 0 | 4 | Sh3bp2 |
| Stap1 | 0 | 0.399965851 | 0.505 | 0.101 | 0 | 4 | Stap1 |
| Nedd92 | 1.37E-305 | 0.885285259 | 0.795 | 0.291 | 2.77E-301 | 4 | Nedd9 |
| Spint2 | 4.67E-305 | 0.30745783 | 0.434 | 0.094 | 9.46E-301 | 4 | Spint2 |
| Metrnl | 4.67E-305 | 0.554268578 | 0.403 | 0.086 | 9.46E-301 | 4 | Metrnl |
| Zfp512 | 2.86E-304 | 0.587657062 | 0.362 | 0.071 | 5.79E-300 | 4 | Zfp512 |
| Rpl121 | 3.82E-304 | 0.803932398 | 0.994 | 0.833 | 7.73E-300 | 4 | Rpl12 |
| Rpap2 | 1.19E-302 | 0.512876709 | 0.321 | 0.056 | 2.41E-298 | 4 | Rpap2 |
| Zfp6521 | 3.06E-302 | 0.775340187 | 0.674 | 0.222 | 6.20E-298 | 4 | Zfp652 |
| Arap21 | 6.32E-302 | 0.680976512 | 0.564 | 0.155 | 1.28E-297 | 4 | Arap2 |
| Arhgef13 | 2.38E-301 | 0.752552297 | 0.677 | 0.223 | 4.82E-297 | 4 | Arhgef1 |
| Cdc42se2 | 5.05E-301 | 0.779372835 | 0.658 | 0.215 | 1.02E-296 | 4 | Cdc42se2 |
| Smc4 | 1.78E-300 | 0.670453601 | 0.527 | 0.142 | 3.61E-296 | 4 | Smc4 |
| Ppil2 | 1.89E-300 | 0.559311034 | 0.502 | 0.129 | 3.83E-296 | 4 | Ppil2 |
| Eef1a11 | 2.31E-300 | 0.655006026 | 1 | 0.957 | 4.68E-296 | 4 | Eef1a1 |
| Lasp1 | 2.63E-300 | 0.597213143 | 0.616 | 0.184 | 5.32E-296 | 4 | Lasp1 |
| AC151836.1 | 3.44E-299 | 0.418997169 | 0.252 | 0.035 | 6.96E-295 | 4 | AC151836.1 |
| Zyx1 | 7.45E-299 | 0.714808954 | 0.628 | 0.192 | 1.51E-294 | 4 | Zyx |
| Rpl232 | 1.32E-297 | 0.68395942 | 0.998 | 0.969 | 2.67E-293 | 4 | Rpl23 |
| Ttc7 | 2.32E-297 | 0.460253499 | 0.398 | 0.085 | 4.70E-293 | 4 | Ttc7 |
| Cbx4 | 3.75E-297 | 0.579922594 | 0.416 | 0.093 | 7.60E-293 | 4 | Cbx4 |
| Zfp827 | 2.27E-294 | 0.489761024 | 0.297 | 0.049 | 4.59E-290 | 4 | Zfp827 |

| | | | | | | | |
|---|---|---|---|---|---|---|---|
| Zfp654 | 1.48E-292 | 0.659495404 | 0.434 | 0.103 | 2.99E-288 | 4 | Zfp654 |
| Senp7 | 2.59E-292 | 0.645601273 | 0.509 | 0.136 | 5.25E-288 | 4 | Senp7 |
| Prr13 | 5.88E-292 | 0.554903797 | 0.618 | 0.186 | 1.19E-287 | 4 | Prr13 |
| Cd164 | 1.42E-290 | 0.619178377 | 0.617 | 0.19 | 2.88E-286 | 4 | Cd164 |
| mt-Nd51 | 1.64E-290 | 0.886061193 | 0.952 | 0.469 | 3.32E-286 | 4 | mt-Nd5 |
| Ankrd111 | 3.11E-290 | 0.8641895 | 0.85 | 0.353 | 6.30E-286 | 4 | Ankrd11 |
| Ptpn1 | 3.54E-289 | 0.664942761 | 0.653 | 0.211 | 7.17E-285 | 4 | Ptpn1 |
| Tln12 | 4.75E-289 | 0.804048881 | 0.887 | 0.377 | 9.61E-285 | 4 | Tln1 |
| Ogt | 1.74E-288 | 0.705388984 | 0.595 | 0.183 | 3.52E-284 | 4 | Ogt |
| Chst111 | 4.90E-288 | 0.717457541 | 0.543 | 0.154 | 9.93E-284 | 4 | Chst11 |
| Dtnbp11 | 6.27E-288 | 0.456817055 | 0.653 | 0.203 | 1.27E-283 | 4 | Dtnbp1 |
| Cnn2 | 1.23E-287 | 0.663118311 | 0.558 | 0.16 | 2.49E-283 | 4 | Cnn2 |
| Rbm3 | 4.08E-287 | 0.792914411 | 0.931 | 0.45 | 8.27E-283 | 4 | Rbm3 |
| Rpl92 | 1.72E-283 | 0.664487313 | 0.998 | 0.941 | 3.49E-279 | 4 | Rpl9 |
| Cdyl2 | 1.40E-282 | 0.623685088 | 0.362 | 0.076 | 2.83E-278 | 4 | Cdyl2 |
| Chka | 3.69E-281 | 0.695782286 | 0.477 | 0.125 | 7.48E-277 | 4 | Chka |
| Arhgap25 | 2.68E-280 | 0.571929291 | 0.4 | 0.091 | 5.42E-276 | 4 | Arhgap25 |
| Capza1 | 2.72E-279 | 0.658616673 | 0.713 | 0.25 | 5.50E-275 | 4 | Capza1 |
| Rps91 | 3.07E-279 | 0.680000865 | 0.997 | 0.916 | 6.21E-275 | 4 | Rps9 |
| Pik3cg | 3.51E-279 | 0.435949981 | 0.266 | 0.042 | 7.10E-275 | 4 | Pik3cg |
| Npc1 | 7.61E-279 | 0.500524702 | 0.328 | 0.063 | 1.54E-274 | 4 | Npc1 |
| Rpl81 | 9.96E-279 | 0.663679074 | 0.998 | 0.921 | 2.02E-274 | 4 | Rpl8 |
| H2-Q72 | 1.51E-278 | 0.8188389 | 0.884 | 0.369 | 3.06E-274 | 4 | H2-Q7 |
| Rpa1 | 2.10E-277 | 0.492255424 | 0.37 | 0.079 | 4.26E-273 | 4 | Rpa1 |
| AL590144.2 | 2.28E-277 | 0.387592014 | 0.252 | 0.038 | 4.62E-273 | 4 | AL590144.2 |
| Fam214a | 4.14E-277 | 0.577364707 | 0.435 | 0.106 | 8.38E-273 | 4 | Fam214a |
| Cntrl | 2.56E-276 | 0.575282171 | 0.433 | 0.106 | 5.19E-272 | 4 | Cntrl |
| Itch1 | 5.27E-276 | 0.769004821 | 0.695 | 0.249 | 1.07E-271 | 4 | Itch |
| Itgb7 | 4.44E-275 | 0.496333099 | 0.287 | 0.049 | 9.00E-271 | 4 | Itgb7 |
| Rps211 | 9.11E-275 | 0.639625402 | 0.995 | 0.923 | 1.85E-270 | 4 | Rps21 |
| Fam172a1 | 1.32E-274 | 0.822988968 | 0.711 | 0.259 | 2.68E-270 | 4 | Fam172a |
| Rps27a1 | 2.31E-274 | 0.639876896 | 0.999 | 0.947 | 4.68E-270 | 4 | Rps27a |
| Baz1a | 4.47E-274 | 0.653869297 | 0.529 | 0.152 | 9.04E-270 | 4 | Baz1a |
| Hcls1 | 5.59E-274 | 0.465183428 | 0.441 | 0.107 | 1.13E-269 | 4 | Hcls1 |
| Sqstm1 | 9.55E-273 | 0.989361258 | 0.723 | 0.28 | 1.93E-268 | 4 | Sqstm1 |
| Plgrkt1 | 1.31E-272 | 0.658326298 | 0.602 | 0.19 | 2.65E-268 | 4 | Plgrkt |
| Atp2a3 | 2.34E-272 | 0.377408928 | 0.287 | 0.049 | 4.74E-268 | 4 | Atp2a3 |
| Elovl6 | 6.32E-272 | 0.432737919 | 0.252 | 0.038 | 1.28E-267 | 4 | Elovl6 |
| Reep51 | 8.01E-272 | 0.677523397 | 0.93 | 0.454 | 1.62E-267 | 4 | Reep5 |
| Arhgef6 | 8.20E-272 | 0.548338541 | 0.442 | 0.11 | 1.66E-267 | 4 | Arhgef6 |
| Arhgap4 | 9.99E-272 | 0.428604731 | 0.273 | 0.046 | 2.02E-267 | 4 | Arhgap4 |
| Exoc2 | 1.09E-270 | 0.611329413 | 0.477 | 0.128 | 2.21E-266 | 4 | Exoc2 |
| Pitpnc12 | 1.38E-270 | 0.842541557 | 0.911 | 0.401 | 2.80E-266 | 4 | Pitpnc1 |

| | | | | | | | |
|---|---|---|---|---|---|---|---|
| Pacs11 | 1.26E-269 | 0.655237974 | 0.586 | 0.183 | 2.55E-265 | 4 | Pacs1 |
| Wipf1 | 8.57E-268 | 0.744375615 | 0.689 | 0.243 | 1.73E-263 | 4 | Wipf1 |
| Rpl13a | 2.26E-267 | 0.54610636 | 0.723 | 0.259 | 4.57E-263 | 4 | Rpl13a |
| Junb2 | 1.51E-265 | 0.745428929 | 0.877 | 0.364 | 3.06E-261 | 4 | Junb |
| Chd7 | 2.09E-265 | 0.589423585 | 0.485 | 0.132 | 4.23E-261 | 4 | Chd7 |
| Hmgb2 | 1.43E-264 | 0.463493037 | 0.802 | 0.3 | 2.90E-260 | 4 | Hmgb2 |
| Tsc22d4 | 2.20E-264 | 0.528014328 | 0.744 | 0.268 | 4.45E-260 | 4 | Tsc22d4 |
| Esyt21 | 3.17E-264 | 0.752058973 | 0.644 | 0.219 | 6.42E-260 | 4 | Esyt2 |
| Vars | 7.48E-264 | 0.444942904 | 0.333 | 0.067 | 1.51E-259 | 4 | Vars |
| Smg61 | 1.09E-263 | 0.775013436 | 0.664 | 0.236 | 2.22E-259 | 4 | Smg6 |
| Nsmaf | 1.64E-263 | 0.513400441 | 0.367 | 0.082 | 3.31E-259 | 4 | Nsmaf |
| Usp48 | 5.63E-262 | 0.548051339 | 0.435 | 0.111 | 1.14E-257 | 4 | Usp48 |
| Ranbp9 | 1.21E-261 | 0.655929134 | 0.57 | 0.178 | 2.45E-257 | 4 | Ranbp9 |
| Nckap1l | 1.69E-261 | 0.451208289 | 0.349 | 0.073 | 3.43E-257 | 4 | Nckap1l |
| Stk24 | 2.44E-261 | 0.533880988 | 0.57 | 0.174 | 4.94E-257 | 4 | Stk24 |
| mt-Co12 | 3.16E-261 | 0.808215554 | 1 | 0.704 | 6.39E-257 | 4 | mt-Co1 |
| Tnrc6b1 | 5.09E-260 | 0.827803828 | 0.813 | 0.345 | 1.03E-255 | 4 | Tnrc6b |
| Rad54l2 | 9.53E-260 | 0.509925463 | 0.361 | 0.079 | 1.93E-255 | 4 | Rad54l2 |
| Haus3 | 1.04E-259 | 0.432885313 | 0.31 | 0.06 | 2.10E-255 | 4 | Haus3 |
| Ppp1r16b2 | 3.89E-259 | 0.702763627 | 0.579 | 0.18 | 7.88E-255 | 4 | Ppp1r16b |
| Rps121 | 5.54E-259 | 0.668927997 | 0.997 | 0.893 | 1.12E-254 | 4 | Rps12 |
| Lbh | 1.07E-258 | 0.537321983 | 0.442 | 0.114 | 2.17E-254 | 4 | Lbh |
| Me2 | 2.24E-258 | 0.441686997 | 0.313 | 0.061 | 4.54E-254 | 4 | Me2 |
| Ramp1 | 6.80E-258 | 0.53487356 | 0.395 | 0.093 | 1.38E-253 | 4 | Ramp1 |
| Rpl271 | 3.56E-256 | 0.72637267 | 0.973 | 0.56 | 7.21E-252 | 4 | Rpl27 |
| Actr3 | 4.90E-255 | 0.643686797 | 0.869 | 0.363 | 9.92E-251 | 4 | Actr3 |
| Actb | 2.60E-253 | 0.795521268 | 0.998 | 0.834 | 5.27E-249 | 4 | Actb |
| Csk | 3.71E-253 | 0.528156717 | 0.43 | 0.112 | 7.50E-249 | 4 | Csk |
| Ablim11 | 6.85E-253 | 0.698868617 | 0.878 | 0.375 | 1.39E-248 | 4 | Ablim1 |
| Rpl112 | 1.54E-252 | 0.63107898 | 0.998 | 0.923 | 3.11E-248 | 4 | Rpl11 |
| Nfatc3 | 1.91E-252 | 0.620759073 | 0.52 | 0.155 | 3.87E-248 | 4 | Nfatc3 |
| Itga4 | 5.95E-252 | 0.468743539 | 0.315 | 0.062 | 1.20E-247 | 4 | Itga4 |
| Cyld | 9.71E-252 | 0.577141861 | 0.511 | 0.151 | 1.97E-247 | 4 | Cyld |
| Rps6ka5 | 1.23E-251 | 0.57492017 | 0.384 | 0.092 | 2.49E-247 | 4 | Rps6ka5 |
| Dap | 1.64E-251 | 0.449352215 | 0.549 | 0.165 | 3.31E-247 | 4 | Dap |
| Tcf122 | 2.60E-251 | 0.839541592 | 0.822 | 0.35 | 5.26E-247 | 4 | Tcf12 |
| Cog51 | 2.73E-251 | 0.676380619 | 0.528 | 0.163 | 5.54E-247 | 4 | Cog5 |
| Rpl171 | 2.68E-250 | 0.617908369 | 0.994 | 0.912 | 5.43E-246 | 4 | Rpl17 |
| Rab27a | 1.25E-249 | 0.430329172 | 0.334 | 0.07 | 2.53E-245 | 4 | Rab27a |
| Chst12 | 5.40E-249 | 0.439715901 | 0.288 | 0.054 | 1.09E-244 | 4 | Chst12 |
| Ern1 | 1.66E-248 | 0.565030713 | 0.328 | 0.071 | 3.37E-244 | 4 | Ern1 |
| Cdk172 | 5.16E-248 | 0.728989282 | 0.638 | 0.222 | 1.04E-243 | 4 | Cdk17 |
| Pld3 | 5.92E-248 | 0.455832811 | 0.334 | 0.072 | 1.20E-243 | 4 | Pld3 |

| | | | | | | | |
|---|---|---|---|---|---|---|---|
| Mlxip | 6.35E-248 | 0.558870559 | 0.423 | 0.11 | 1.29E-243 | 4 | Mlxip |
| Mmgt2 | 8.65E-248 | 0.366948079 | 0.285 | 0.053 | 1.75E-243 | 4 | Mmgt2 |
| Cbl | 8.66E-248 | 0.631440818 | 0.574 | 0.186 | 1.75E-243 | 4 | Cbl |
| Rnaseh2c | 1.37E-247 | 0.514090241 | 0.715 | 0.26 | 2.76E-243 | 4 | Rnaseh2c |
| Rpl62 | 2.32E-247 | 0.650887261 | 0.996 | 0.875 | 4.70E-243 | 4 | Rpl6 |
| Vps28 | 4.55E-247 | 0.547491541 | 0.728 | 0.276 | 9.20E-243 | 4 | Vps28 |
| Nabp1 | 1.07E-246 | 0.500576208 | 0.324 | 0.068 | 2.17E-242 | 4 | Nabp1 |
| Eif3e1 | 7.59E-246 | 0.57444304 | 0.871 | 0.386 | 1.54E-241 | 4 | Eif3e |
| Tor1aip2 | 5.12E-245 | 0.597904265 | 0.529 | 0.162 | 1.04E-240 | 4 | Tor1aip2 |
| 4930453N24Rik | 1.86E-243 | 0.424173117 | 0.514 | 0.15 | 3.76E-239 | 4 | 4930453N24Rik |
| Capg | 2.20E-243 | 0.635446886 | 0.364 | 0.085 | 4.46E-239 | 4 | Capg |
| Phf6 | 3.22E-243 | 0.54187081 | 0.439 | 0.118 | 6.51E-239 | 4 | Phf6 |
| Clk11 | 1.69E-242 | 0.717992725 | 0.702 | 0.268 | 3.43E-238 | 4 | Clk1 |
| Mapkapk3 | 3.22E-242 | 0.410400596 | 0.322 | 0.068 | 6.52E-238 | 4 | Mapkapk3 |
| Luc7l21 | 4.61E-242 | 0.724783186 | 0.854 | 0.379 | 9.34E-238 | 4 | Luc7l2 |
| Fhit | 4.61E-242 | 0.686684274 | 0.314 | 0.066 | 9.34E-238 | 4 | Fhit |
| Smc5 | 6.57E-242 | 0.528847462 | 0.403 | 0.102 | 1.33E-237 | 4 | Smc5 |
| Cdk2ap2 | 1.01E-241 | 0.402622576 | 0.568 | 0.178 | 2.04E-237 | 4 | Cdk2ap2 |
| Fosb1 | 2.08E-241 | 0.766731272 | 0.535 | 0.169 | 4.21E-237 | 4 | Fosb |
| Taf15 | 8.68E-241 | 0.598085075 | 0.623 | 0.215 | 1.76E-236 | 4 | Taf15 |
| Zcchc71 | 1.02E-240 | 0.702775774 | 0.72 | 0.276 | 2.06E-236 | 4 | Zcchc7 |
| Rpl35a1 | 1.11E-240 | 0.551283467 | 0.997 | 0.946 | 2.25E-236 | 4 | Rpl35a |
| Rpl281 | 2.00E-240 | 0.598028541 | 0.999 | 0.926 | 4.05E-236 | 4 | Rpl28 |
| Grina | 2.87E-240 | 0.518687673 | 0.443 | 0.122 | 5.82E-236 | 4 | Grina |
| Ralgapa2 | 5.80E-240 | 0.538190757 | 0.476 | 0.135 | 1.17E-235 | 4 | Ralgapa2 |
| Mtmr1 | 8.95E-240 | 0.424562215 | 0.308 | 0.063 | 1.81E-235 | 4 | Mtmr1 |
| Arf3 | 9.42E-240 | 0.496009356 | 0.44 | 0.12 | 1.91E-235 | 4 | Arf3 |
| Ppp1r12a2 | 7.60E-239 | 0.74232195 | 0.888 | 0.399 | 1.54E-234 | 4 | Ppp1r12a |
| Snx5 | 1.15E-238 | 0.435091928 | 0.593 | 0.189 | 2.34E-234 | 4 | Snx5 |
| Trp53inp1 | 1.29E-238 | 0.436503435 | 0.276 | 0.052 | 2.62E-234 | 4 | Trp53inp1 |
| Pim1 | 1.80E-238 | 0.647555959 | 0.427 | 0.116 | 3.64E-234 | 4 | Pim1 |
| Mast3 | 2.26E-237 | 0.383113007 | 0.252 | 0.044 | 4.58E-233 | 4 | Mast3 |
| Agpat5 | 2.81E-237 | 0.528825536 | 0.449 | 0.124 | 5.69E-233 | 4 | Agpat5 |
| Vrk2 | 6.26E-237 | 0.542452116 | 0.411 | 0.107 | 1.27E-232 | 4 | Vrk2 |
| Rngtt | 5.82E-236 | 0.555400289 | 0.433 | 0.118 | 1.18E-231 | 4 | Rngtt |
| Rplp22 | 6.16E-235 | 0.589525044 | 0.997 | 0.884 | 1.25E-230 | 4 | Rplp2 |
| Tecr | 7.86E-235 | 0.525102588 | 0.599 | 0.203 | 1.59E-230 | 4 | Tecr |
| Heca | 9.18E-235 | 0.523353148 | 0.448 | 0.125 | 1.86E-230 | 4 | Heca |
| Lnpep1 | 1.38E-234 | 0.684726628 | 0.653 | 0.237 | 2.80E-230 | 4 | Lnpep |
| Qpct | 2.18E-234 | 0.384583656 | 0.314 | 0.066 | 4.41E-230 | 4 | Qpct |
| Cln3 | 1.08E-233 | 0.424844126 | 0.284 | 0.056 | 2.19E-229 | 4 | Cln3 |
| Snrnp70 | 1.44E-233 | 0.580318585 | 0.673 | 0.249 | 2.91E-229 | 4 | Snrnp70 |
| Phactr21 | 3.17E-233 | 0.814478799 | 0.611 | 0.225 | 6.43E-229 | 4 | Phactr2 |

| | | | | | | | |
|---|---|---|---|---|---|---|---|
| Pdcd42 | 3.85E-233 | 0.366829933 | 0.939 | 0.424 | 7.80E-229 | 4 | Pdcd4 |
| Rel | 1.26E-232 | 0.58001129 | 0.438 | 0.121 | 2.55E-228 | 4 | Rel |
| H2afj1 | 1.92E-232 | 0.460299093 | 0.911 | 0.426 | 3.89E-228 | 4 | H2afj |
| Dennd4c | 2.52E-230 | 0.527358949 | 0.411 | 0.11 | 5.09E-226 | 4 | Dennd4c |
| Vps13b1 | 4.92E-230 | 0.690528712 | 0.577 | 0.199 | 9.96E-226 | 4 | Vps13b |
| Eif3f | 8.89E-230 | 0.620068647 | 0.918 | 0.458 | 1.80E-225 | 4 | Eif3f |
| Eef1b22 | 1.18E-229 | 0.662908026 | 0.99 | 0.721 | 2.39E-225 | 4 | Eef1b2 |
| Rpl10a2 | 1.73E-229 | 0.647885644 | 0.995 | 0.827 | 3.51E-225 | 4 | Rpl10a |
| Phf32 | 2.03E-229 | 0.607931246 | 0.639 | 0.23 | 4.11E-225 | 4 | Phf3 |
| Rps4x1 | 1.46E-228 | 0.579352297 | 0.997 | 0.938 | 2.96E-224 | 4 | Rps4x |
| Ago2 | 2.18E-228 | 0.529283035 | 0.49 | 0.149 | 4.42E-224 | 4 | Ago2 |
| Cep192 | 7.08E-227 | 0.4456802 | 0.321 | 0.072 | 1.43E-222 | 4 | Cep192 |
| Hdgfl2 | 2.09E-226 | 0.468695985 | 0.325 | 0.073 | 4.23E-222 | 4 | Hdgfl2 |
| Sipa1 | 2.91E-226 | 0.578131862 | 0.52 | 0.166 | 5.90E-222 | 4 | Sipa1 |
| Rps19 | 1.06E-225 | 0.577071508 | 0.996 | 0.923 | 2.14E-221 | 4 | Rps19 |
| Rps29 | 1.25E-225 | 0.494544064 | 0.998 | 0.97 | 2.52E-221 | 4 | Rps29 |
| S100a112 | 6.87E-225 | 0.539528519 | 0.983 | 0.549 | 1.39E-220 | 4 | S100a11 |
| Rps14 | 1.23E-224 | 0.514939378 | 0.998 | 0.94 | 2.49E-220 | 4 | Rps14 |
| Rnf19b | 1.77E-224 | 0.502255663 | 0.317 | 0.071 | 3.59E-220 | 4 | Rnf19b |
| Fam102a | 1.54E-223 | 0.376448376 | 0.357 | 0.085 | 3.11E-219 | 4 | Fam102a |
| Emp3 | 2.62E-223 | 0.48196961 | 0.667 | 0.235 | 5.31E-219 | 4 | Emp3 |
| Slfn8 | 3.62E-223 | 0.405561845 | 0.298 | 0.063 | 7.33E-219 | 4 | Slfn8 |
| Ezr | 3.68E-223 | 0.31781868 | 0.436 | 0.119 | 7.46E-219 | 4 | Ezr |
| Kpna42 | 5.41E-223 | 0.670247189 | 0.74 | 0.299 | 1.10E-218 | 4 | Kpna4 |
| Tex2 | 1.81E-222 | 0.414438144 | 0.329 | 0.075 | 3.66E-218 | 4 | Tex2 |
| Wwp2 | 3.34E-222 | 0.537676709 | 0.407 | 0.111 | 6.77E-218 | 4 | Wwp2 |
| Jund1 | 3.95E-222 | 0.785392091 | 0.989 | 0.626 | 7.99E-218 | 4 | Jund |
| Nktr | 8.45E-222 | 0.595811443 | 0.642 | 0.235 | 1.71E-217 | 4 | Nktr |
| Lrba1 | 1.39E-221 | 0.618780576 | 0.528 | 0.172 | 2.82E-217 | 4 | Lrba |
| Vmp1 | 2.20E-221 | 0.515470987 | 0.574 | 0.195 | 4.45E-217 | 4 | Vmp1 |
| Sp110 | 1.43E-220 | 0.454771557 | 0.47 | 0.139 | 2.90E-216 | 4 | Sp110 |
| Csnk1g3 | 1.70E-220 | 0.548662851 | 0.511 | 0.164 | 3.45E-216 | 4 | Csnk1g3 |
| Rpl212 | 7.45E-220 | 0.549715345 | 0.994 | 0.919 | 1.51E-215 | 4 | Rpl21 |
| Plekhj1 | 1.30E-219 | 0.443885883 | 0.49 | 0.15 | 2.63E-215 | 4 | Plekhj1 |
| Supt4a1 | 2.09E-219 | 0.461298877 | 0.744 | 0.297 | 4.23E-215 | 4 | Supt4a |
| Pbxip1 | 3.43E-219 | 0.483518229 | 0.386 | 0.102 | 6.94E-215 | 4 | Pbxip1 |
| Usp3 | 4.57E-219 | 0.493328483 | 0.516 | 0.163 | 9.26E-215 | 4 | Usp3 |
| Itgav | 1.53E-218 | 0.637169206 | 0.469 | 0.143 | 3.10E-214 | 4 | Itgav |
| Clic1 | 2.20E-218 | 0.674569763 | 0.978 | 0.579 | 4.46E-214 | 4 | Clic1 |
| Slc2a1 | 7.19E-218 | 0.527936794 | 0.321 | 0.074 | 1.46E-213 | 4 | Slc2a1 |
| Nol4l | 1.15E-217 | 0.479688086 | 0.293 | 0.062 | 2.33E-213 | 4 | Nol4l |
| Sirt7 | 1.27E-217 | 0.444410685 | 0.422 | 0.117 | 2.57E-213 | 4 | Sirt7 |
| Sun2 | 1.83E-217 | 0.445445401 | 0.298 | 0.065 | 3.71E-213 | 4 | Sun2 |

| | | | | | | | |
|---|---|---|---|---|---|---|---|
| Coq10b | 3.22E-217 | 0.583179734 | 0.454 | 0.136 | 6.53E-213 | 4 | Coq10b |
| Ptp4a1 | 6.52E-217 | 0.485317862 | 0.439 | 0.127 | 1.32E-212 | 4 | Ptp4a1 |
| Commd7 | 6.41E-216 | 0.473617129 | 0.543 | 0.181 | 1.30E-211 | 4 | Commd7 |
| Mkl1 | 7.22E-216 | 0.513141375 | 0.402 | 0.11 | 1.46E-211 | 4 | Mkl1 |
| Arid1a1 | 8.89E-216 | 0.598085136 | 0.589 | 0.209 | 1.80E-211 | 4 | Arid1a |
| Itpr3 | 2.03E-215 | 0.422992452 | 0.294 | 0.063 | 4.11E-211 | 4 | Itpr3 |
| Glipr1 | 3.42E-215 | 0.355853535 | 0.346 | 0.082 | 6.93E-211 | 4 | Glipr1 |
| Irak2 | 9.31E-214 | 0.548756064 | 0.433 | 0.125 | 1.88E-209 | 4 | Irak2 |
| Ppp3ca1 | 6.74E-213 | 0.695163592 | 0.823 | 0.368 | 1.36E-208 | 4 | Ppp3ca |
| Samsn1 | 1.74E-211 | 0.384015894 | 0.304 | 0.068 | 3.53E-207 | 4 | Samsn1 |
| Mgat4a | 1.88E-211 | 0.477861169 | 0.357 | 0.091 | 3.81E-207 | 4 | Mgat4a |
| Ddx10 | 2.11E-211 | 0.546597479 | 0.41 | 0.116 | 4.28E-207 | 4 | Ddx10 |
| Gbp81 | 2.15E-211 | 0.509494168 | 0.427 | 0.122 | 4.35E-207 | 4 | Gbp8 |
| Taf10 | 7.77E-211 | 0.481376756 | 0.819 | 0.359 | 1.57E-206 | 4 | Taf10 |
| Rpl151 | 1.22E-210 | 0.574564821 | 0.996 | 0.896 | 2.47E-206 | 4 | Rpl15 |
| Rcor1 | 1.31E-209 | 0.518318584 | 0.505 | 0.165 | 2.65E-205 | 4 | Rcor1 |
| Rpl221 | 3.42E-209 | 0.563593757 | 0.994 | 0.844 | 6.92E-205 | 4 | Rpl22 |
| Arid4b2 | 4.92E-209 | 0.605673422 | 0.683 | 0.27 | 9.96E-205 | 4 | Arid4b |
| Znrf11 | 1.52E-208 | 0.545085931 | 0.519 | 0.17 | 3.08E-204 | 4 | Znrf1 |
| Atad2 | 2.86E-208 | 0.460143362 | 0.33 | 0.079 | 5.79E-204 | 4 | Atad2 |
| Snrpg | 3.07E-208 | 0.433631452 | 0.746 | 0.299 | 6.22E-204 | 4 | Snrpg |
| Srrm22 | 5.04E-208 | 0.674778302 | 0.9 | 0.444 | 1.02E-203 | 4 | Srrm2 |
| Fam173a | 6.47E-208 | 0.367662051 | 0.537 | 0.178 | 1.31E-203 | 4 | Fam173a |
| Card6 | 2.18E-207 | 0.370428096 | 0.255 | 0.05 | 4.42E-203 | 4 | Card6 |
| Lrrfip12 | 4.04E-207 | 0.627714962 | 0.802 | 0.354 | 8.18E-203 | 4 | Lrrfip1 |
| Phf21a | 4.60E-207 | 0.677554642 | 0.614 | 0.232 | 9.32E-203 | 4 | Phf21a |
| 2900026A02Rik2 | 1.91E-206 | 0.54701526 | 0.577 | 0.199 | 3.86E-202 | 4 | 2900026A02Rik |
| Trp53 | 3.60E-206 | 0.455426996 | 0.461 | 0.141 | 7.30E-202 | 4 | Trp53 |
| Runx2 | 4.27E-206 | 0.372264123 | 0.328 | 0.078 | 8.64E-202 | 4 | Runx2 |
| Sfxn1 | 4.35E-206 | 0.346822852 | 0.331 | 0.08 | 8.81E-202 | 4 | Sfxn1 |
| Uba522 | 1.18E-204 | 0.492155963 | 0.944 | 0.497 | 2.39E-200 | 4 | Uba52 |
| Fbxl172 | 2.58E-204 | 0.703347967 | 0.699 | 0.288 | 5.23E-200 | 4 | Fbxl17 |
| Ncoa21 | 9.16E-204 | 0.611207206 | 0.596 | 0.218 | 1.85E-199 | 4 | Ncoa2 |
| Bcl2l13 | 1.15E-203 | 0.40397404 | 0.276 | 0.059 | 2.32E-199 | 4 | Bcl2l13 |
| Rps82 | 2.26E-203 | 0.504361678 | 0.999 | 0.963 | 4.58E-199 | 4 | Rps8 |
| Ube2i | 2.80E-203 | 0.460657385 | 0.743 | 0.306 | 5.67E-199 | 4 | Ube2i |
| Hnrnpl | 7.63E-203 | 0.493574159 | 0.594 | 0.214 | 1.55E-198 | 4 | Hnrnpl |
| Xrn2 | 8.00E-203 | 0.482340164 | 0.639 | 0.24 | 1.62E-198 | 4 | Xrn2 |
| Atp6v0a2 | 1.03E-202 | 0.449560283 | 0.366 | 0.098 | 2.09E-198 | 4 | Atp6v0a2 |
| Upf1 | 1.90E-201 | 0.406570965 | 0.368 | 0.098 | 3.84E-197 | 4 | Upf1 |
| Prdx61 | 8.57E-201 | 0.278551234 | 0.774 | 0.312 | 1.74E-196 | 4 | Prdx6 |
| Foxo3 | 1.39E-200 | 0.49161065 | 0.372 | 0.102 | 2.82E-196 | 4 | Foxo3 |
| Btbd92 | 5.12E-200 | 0.614156141 | 0.59 | 0.217 | 1.04E-195 | 4 | Btbd9 |

| | | | | | | | |
|---|---|---|---|---|---|---|---|
| Kdm6a1 | 5.27E-200 | 0.576530135 | 0.551 | 0.194 | 1.07E-195 | 4 | Kdm6a |
| Crlf3 | 5.97E-200 | 0.469416188 | 0.455 | 0.14 | 1.21E-195 | 4 | Crlf3 |
| F2r | 2.48E-199 | 0.421141496 | 0.324 | 0.078 | 5.01E-195 | 4 | F2r |
| Rcsd1 | 2.92E-199 | 0.489806882 | 0.401 | 0.114 | 5.92E-195 | 4 | Rcsd1 |
| Stk39 | 3.74E-199 | 0.39609465 | 0.451 | 0.137 | 7.56E-195 | 4 | Stk39 |
| Rpl42 | 5.34E-199 | 0.611077763 | 0.981 | 0.639 | 1.08E-194 | 4 | Rpl4 |
| Riok1 | 6.44E-199 | 0.456586302 | 0.372 | 0.101 | 1.30E-194 | 4 | Riok1 |
| Tpk1 | 1.13E-198 | 0.482187607 | 0.348 | 0.091 | 2.29E-194 | 4 | Tpk1 |
| Slbp | 1.76E-198 | 0.409363858 | 0.445 | 0.134 | 3.56E-194 | 4 | Slbp |
| Crlf2 | 4.45E-198 | 0.434371677 | 0.463 | 0.144 | 9.00E-194 | 4 | Crlf2 |
| Rbm252 | 5.62E-198 | 0.604572314 | 0.732 | 0.311 | 1.14E-193 | 4 | Rbm25 |
| Ptbp32 | 6.56E-198 | 0.641693207 | 0.814 | 0.369 | 1.33E-193 | 4 | Ptbp3 |
| Dcaf5 | 1.11E-197 | 0.492047513 | 0.397 | 0.114 | 2.25E-193 | 4 | Dcaf5 |
| Spcs2 | 1.21E-197 | 0.559017813 | 0.82 | 0.377 | 2.46E-193 | 4 | Spcs2 |
| Eif3h1 | 1.22E-197 | 0.50086638 | 0.905 | 0.455 | 2.46E-193 | 4 | Eif3h |
| Msh3 | 1.51E-197 | 0.371866753 | 0.266 | 0.056 | 3.06E-193 | 4 | Msh3 |
| Fuca2 | 1.72E-197 | 0.373281945 | 0.263 | 0.056 | 3.48E-193 | 4 | Fuca2 |
| Ggnbp21 | 2.81E-197 | 0.545936582 | 0.594 | 0.222 | 5.68E-193 | 4 | Ggnbp2 |
| Ccdc107 | 9.03E-197 | 0.443457843 | 0.392 | 0.112 | 1.83E-192 | 4 | Ccdc107 |
| Crot | 5.16E-196 | 0.310676398 | 0.417 | 0.12 | 1.04E-191 | 4 | Crot |
| Ebpl | 5.48E-196 | 0.406794628 | 0.375 | 0.103 | 1.11E-191 | 4 | Ebpl |
| Vti1a1 | 8.62E-196 | 0.62261729 | 0.611 | 0.234 | 1.74E-191 | 4 | Vti1a |
| Sppl31 | 1.39E-195 | 0.597551967 | 0.549 | 0.199 | 2.80E-191 | 4 | Sppl3 |
| Lcor1 | 1.69E-195 | 0.539332018 | 0.509 | 0.172 | 3.43E-191 | 4 | Lcor |
| Cap1 | 1.88E-195 | 0.463246543 | 0.655 | 0.25 | 3.80E-191 | 4 | Cap1 |
| Sf3b11 | 6.38E-195 | 0.580727431 | 0.809 | 0.369 | 1.29E-190 | 4 | Sf3b1 |
| Itpk1 | 9.06E-195 | 0.367714195 | 0.272 | 0.06 | 1.83E-190 | 4 | Itpk1 |
| Rev3l | 1.45E-194 | 0.489772871 | 0.451 | 0.141 | 2.94E-190 | 4 | Rev3l |
| Rfc2 | 1.54E-194 | 0.384818304 | 0.391 | 0.11 | 3.12E-190 | 4 | Rfc2 |
| Rnf1302 | 3.66E-193 | 0.548929072 | 0.66 | 0.262 | 7.40E-189 | 4 | Rnf130 |
| Cdk131 | 3.89E-193 | 0.543060279 | 0.567 | 0.207 | 7.88E-189 | 4 | Cdk13 |
| Zbtb202 | 4.33E-193 | 0.712375919 | 0.959 | 0.513 | 8.78E-189 | 4 | Zbtb20 |
| H2-K12 | 6.66E-193 | 0.712525034 | 1 | 0.622 | 1.35E-188 | 4 | H2-K1 |
| Plekha5 | 1.52E-192 | 0.538055825 | 0.444 | 0.139 | 3.08E-188 | 4 | Plekha5 |
| Csnk1g11 | 3.51E-192 | 0.528311365 | 0.447 | 0.141 | 7.11E-188 | 4 | Csnk1g1 |
| Pgm2l1 | 4.29E-192 | 0.370575915 | 0.25 | 0.051 | 8.68E-188 | 4 | Pgm2l1 |
| Mri1 | 7.76E-192 | 0.361438872 | 0.287 | 0.066 | 1.57E-187 | 4 | Mri1 |
| Gm26887 | 1.11E-191 | 0.55799868 | 0.333 | 0.087 | 2.25E-187 | 4 | Gm26887 |
| Rack12 | 2.50E-191 | 0.577613802 | 0.994 | 0.781 | 5.06E-187 | 4 | Rack1 |
| mt-Nd42 | 3.08E-191 | 0.680143505 | 0.999 | 0.657 | 6.23E-187 | 4 | mt-Nd4 |
| Gne | 8.13E-191 | 0.466960615 | 0.384 | 0.11 | 1.65E-186 | 4 | Gne |
| Zfp2922 | 8.14E-191 | 0.538164967 | 0.609 | 0.231 | 1.65E-186 | 4 | Zfp292 |
| Cetn2 | 9.45E-190 | 0.329159211 | 0.536 | 0.185 | 1.91E-185 | 4 | Cetn2 |

| | | | | | | | |
|---|---|---|---|---|---|---|---|
| Tmem2581 | 1.82E-189 | 0.358382361 | 0.699 | 0.286 | 3.68E-185 | 4 | Tmem258 |
| Gm47283 | 3.58E-189 | 0.702580771 | 0.639 | 0.261 | 7.24E-185 | 4 | Gm47283 |
| Kcnq1ot1 | 4.65E-189 | 0.637407891 | 0.398 | 0.119 | 9.41E-185 | 4 | Kcnq1ot1 |
| Atxn12 | 1.13E-188 | 0.654322003 | 0.671 | 0.274 | 2.29E-184 | 4 | Atxn1 |
| Fam204a | 3.04E-188 | 0.380154168 | 0.472 | 0.152 | 6.16E-184 | 4 | Fam204a |
| Arpc2 | 1.79E-187 | 0.604352652 | 0.948 | 0.521 | 3.63E-183 | 4 | Arpc2 |
| Diaph22 | 2.27E-187 | 0.659713014 | 0.776 | 0.353 | 4.60E-183 | 4 | Diaph2 |
| Nup153 | 7.73E-187 | 0.461485176 | 0.364 | 0.102 | 1.56E-182 | 4 | Nup153 |
| Rps22 | 1.11E-186 | 0.587804813 | 0.997 | 0.915 | 2.24E-182 | 4 | Rps2 |
| Usp38 | 1.15E-186 | 0.364413198 | 0.267 | 0.059 | 2.32E-182 | 4 | Usp38 |
| Supt20 | 2.03E-186 | 0.48435444 | 0.45 | 0.145 | 4.12E-182 | 4 | Supt20 |
| Cxxc5 | 2.61E-186 | 0.345455701 | 0.362 | 0.1 | 5.29E-182 | 4 | Cxxc5 |
| Dmac1 | 2.81E-186 | 0.296675926 | 0.446 | 0.139 | 5.68E-182 | 4 | Dmac1 |
| Eif2ak3 | 5.12E-186 | 0.481317676 | 0.294 | 0.072 | 1.04E-181 | 4 | Eif2ak3 |
| Gm11808 | 7.00E-186 | 0.258779667 | 0.444 | 0.137 | 1.42E-181 | 4 | Gm11808 |
| Ubr21 | 1.57E-185 | 0.544156805 | 0.567 | 0.21 | 3.18E-181 | 4 | Ubr2 |
| Rpl23a2 | 9.32E-185 | 0.621629257 | 0.979 | 0.662 | 1.89E-180 | 4 | Rpl23a |
| Dbnl | 1.69E-184 | 0.399466739 | 0.507 | 0.174 | 3.41E-180 | 4 | Dbnl |
| Rlf | 8.58E-184 | 0.470919874 | 0.403 | 0.122 | 1.74E-179 | 4 | Rlf |
| Rnf157 | 3.90E-183 | 0.351338256 | 0.252 | 0.054 | 7.89E-179 | 4 | Rnf157 |
| Usp15 | 1.61E-182 | 0.495040429 | 0.565 | 0.212 | 3.27E-178 | 4 | Usp15 |
| Stim11 | 2.69E-182 | 0.549592 | 0.497 | 0.173 | 5.45E-178 | 4 | Stim1 |
| Mettl23 | 6.60E-182 | 0.303562657 | 0.529 | 0.184 | 1.34E-177 | 4 | Mettl23 |
| Ankrd171 | 6.70E-182 | 0.546224294 | 0.614 | 0.241 | 1.36E-177 | 4 | Ankrd17 |
| Myh92 | 9.52E-182 | 0.581901424 | 0.867 | 0.412 | 1.93E-177 | 4 | Myh9 |
| Ahdc1 | 8.04E-181 | 0.378565909 | 0.354 | 0.098 | 1.63E-176 | 4 | Ahdc1 |
| Cerk | 1.82E-180 | 0.402473646 | 0.358 | 0.101 | 3.68E-176 | 4 | Cerk |
| Ncl | 2.56E-180 | 0.679694021 | 0.913 | 0.493 | 5.18E-176 | 4 | Ncl |
| Ppp1ca | 4.79E-180 | 0.436812495 | 0.815 | 0.372 | 9.71E-176 | 4 | Ppp1ca |
| Nlrc5 | 5.12E-180 | 0.434954677 | 0.352 | 0.098 | 1.04E-175 | 4 | Nlrc5 |
| mt-Atp82 | 5.90E-180 | 0.557238376 | 0.892 | 0.438 | 1.19E-175 | 4 | mt-Atp8 |
| Tm6sf1 | 6.64E-180 | 0.396139428 | 0.414 | 0.126 | 1.34E-175 | 4 | Tm6sf1 |
| Smap2 | 2.26E-179 | 0.381069108 | 0.541 | 0.193 | 4.57E-175 | 4 | Smap2 |
| Nsa21 | 3.34E-179 | 0.424852046 | 0.907 | 0.462 | 6.76E-175 | 4 | Nsa2 |
| Pacsin2 | 8.88E-179 | 0.358309411 | 0.439 | 0.142 | 1.80E-174 | 4 | Pacsin2 |
| Efr3a | 2.16E-178 | 0.438134055 | 0.512 | 0.179 | 4.38E-174 | 4 | Efr3a |
| Iqgap12 | 2.26E-178 | 0.596179181 | 0.833 | 0.387 | 4.57E-174 | 4 | Iqgap1 |
| St8sia42 | 3.43E-178 | 0.562249515 | 0.618 | 0.239 | 6.94E-174 | 4 | St8sia4 |
| Myo9b | 4.40E-178 | 0.452574451 | 0.363 | 0.106 | 8.90E-174 | 4 | Myo9b |
| Plekha2 | 5.68E-178 | 0.444117221 | 0.393 | 0.119 | 1.15E-173 | 4 | Plekha2 |
| Gls1 | 9.86E-178 | 0.504308099 | 0.598 | 0.231 | 2.00E-173 | 4 | Gls |
| Cnot6l | 1.17E-177 | 0.447080672 | 0.588 | 0.225 | 2.37E-173 | 4 | Cnot6l |
| Nub1 | 3.85E-177 | 0.384960681 | 0.492 | 0.169 | 7.79E-173 | 4 | Nub1 |

| | | | | | | | |
|---|---|---|---|---|---|---|---|
| Arf6 | 2.16E-176 | 0.32814966 | 0.479 | 0.161 | 4.36E-172 | 4 | Arf6 |
| Sub11 | 2.44E-176 | 0.468439301 | 0.927 | 0.492 | 4.94E-172 | 4 | Sub1 |
| Stt3b1 | 3.74E-176 | 0.468276007 | 0.451 | 0.149 | 7.57E-172 | 4 | Stt3b |
| Ccdc88a1 | 4.38E-176 | 0.577534997 | 0.514 | 0.187 | 8.86E-172 | 4 | Ccdc88a |
| Ddx21 | 8.97E-176 | 0.458782819 | 0.53 | 0.195 | 1.82E-171 | 4 | Ddx21 |
| Tnfaip8 | 1.35E-175 | 0.403898427 | 0.498 | 0.169 | 2.74E-171 | 4 | Tnfaip8 |
| Sp4 | 1.73E-175 | 0.410115115 | 0.319 | 0.084 | 3.51E-171 | 4 | Sp4 |
| Snrpf1 | 1.89E-175 | 0.39224657 | 0.782 | 0.356 | 3.83E-171 | 4 | Snrpf |
| Rps262 | 1.98E-175 | 0.483251551 | 0.988 | 0.859 | 4.02E-171 | 4 | Rps26 |
| Nsmce2 | 2.85E-175 | 0.471104796 | 0.522 | 0.188 | 5.76E-171 | 4 | Nsmce2 |
| Esco1 | 4.53E-175 | 0.439833574 | 0.477 | 0.162 | 9.17E-171 | 4 | Esco1 |
| Arfgef11 | 6.19E-175 | 0.49401282 | 0.554 | 0.209 | 1.25E-170 | 4 | Arfgef1 |
| Pdlim2 | 1.70E-174 | 0.32112063 | 0.257 | 0.058 | 3.44E-170 | 4 | Pdlim2 |
| Nipbl2 | 1.80E-174 | 0.56326205 | 0.762 | 0.343 | 3.63E-170 | 4 | Nipbl |
| Glrx | 2.18E-174 | 0.309468685 | 0.261 | 0.06 | 4.41E-170 | 4 | Glrx |
| Pvt1 | 5.98E-174 | 0.376296752 | 0.28 | 0.067 | 1.21E-169 | 4 | Pvt1 |
| Spata5 | 6.04E-174 | 0.470770637 | 0.369 | 0.109 | 1.22E-169 | 4 | Spata5 |
| Exoc42 | 6.45E-174 | 0.609558452 | 0.698 | 0.301 | 1.31E-169 | 4 | Exoc4 |
| Capn1 | 7.95E-174 | 0.336270704 | 0.297 | 0.075 | 1.61E-169 | 4 | Capn1 |
| Rreb1 | 8.25E-174 | 0.47797443 | 0.516 | 0.184 | 1.67E-169 | 4 | Rreb1 |
| Immp2l | 2.91E-173 | 0.615364125 | 0.391 | 0.122 | 5.89E-169 | 4 | Immp2l |
| Eef1d1 | 2.49E-172 | 0.453445917 | 0.938 | 0.527 | 5.04E-168 | 4 | Eef1d |
| Tcf20 | 3.41E-172 | 0.457101007 | 0.402 | 0.126 | 6.91E-168 | 4 | Tcf20 |
| Psph | 5.79E-172 | 0.305352544 | 0.254 | 0.057 | 1.17E-167 | 4 | Psph |
| Ncor11 | 9.69E-172 | 0.524328154 | 0.859 | 0.413 | 1.96E-167 | 4 | Ncor1 |
| Iqsec1 | 1.21E-171 | 0.427078636 | 0.368 | 0.108 | 2.45E-167 | 4 | Iqsec1 |
| Rpl29 | 3.46E-171 | 0.546546308 | 0.987 | 0.756 | 7.00E-167 | 4 | Rpl29 |
| Npm12 | 5.23E-171 | 0.566467216 | 0.978 | 0.662 | 1.06E-166 | 4 | Npm1 |
| Tmem123 | 6.03E-171 | 0.440844203 | 0.425 | 0.138 | 1.22E-166 | 4 | Tmem123 |
| Ccnl1 | 6.07E-171 | 0.495116011 | 0.521 | 0.192 | 1.23E-166 | 4 | Ccnl1 |
| Usp24 | 6.10E-171 | 0.438337053 | 0.393 | 0.122 | 1.24E-166 | 4 | Usp24 |
| H2-Q62 | 1.06E-170 | 0.458214032 | 0.593 | 0.225 | 2.16E-166 | 4 | H2-Q6 |
| Sep-91 | 1.09E-170 | 0.484065707 | 0.456 | 0.154 | 2.20E-166 | 4 | 9-Sep |
| Reps1 | 2.13E-170 | 0.415880268 | 0.322 | 0.089 | 4.31E-166 | 4 | Reps1 |
| Atr | 2.67E-170 | 0.373028305 | 0.257 | 0.06 | 5.41E-166 | 4 | Atr |
| Rp91 | 2.72E-170 | 0.378928111 | 0.709 | 0.302 | 5.52E-166 | 4 | Rp9 |
| Arl6ip52 | 4.92E-170 | 0.46902334 | 0.545 | 0.203 | 9.97E-166 | 4 | Arl6ip5 |
| Parp41 | 7.11E-170 | 0.435502662 | 0.423 | 0.137 | 1.44E-165 | 4 | Parp4 |
| Ambra11 | 1.39E-169 | 0.513001155 | 0.499 | 0.181 | 2.81E-165 | 4 | Ambra1 |
| Adipor2 | 2.28E-169 | 0.443059834 | 0.417 | 0.135 | 4.62E-165 | 4 | Adipor2 |
| Tmcc11 | 2.99E-169 | 0.57145641 | 0.53 | 0.2 | 6.05E-165 | 4 | Tmcc1 |
| 1700025G04Rik | 3.60E-169 | 0.504773481 | 0.486 | 0.172 | 7.30E-165 | 4 | 1700025G04Rik |
| Maml22 | 4.11E-169 | 0.655454577 | 0.714 | 0.322 | 8.32E-165 | 4 | Maml2 |

| | | | | | | | |
|---|---|---|---|---|---|---|---|
| Rasa2 | 6.54E-169 | 0.469577134 | 0.393 | 0.124 | 1.32E-164 | 4 | Rasa2 |
| Zfp6381 | 2.18E-168 | 0.521479778 | 0.634 | 0.261 | 4.41E-164 | 4 | Zfp638 |
| Tpr1 | 4.58E-168 | 0.499078404 | 0.842 | 0.401 | 9.27E-164 | 4 | Tpr |
| Memo1 | 4.83E-168 | 0.391322462 | 0.32 | 0.088 | 9.78E-164 | 4 | Memo1 |
| Smg11 | 4.91E-168 | 0.503657175 | 0.56 | 0.216 | 9.95E-164 | 4 | Smg1 |
| Rpl352 | 1.97E-167 | 0.491141276 | 0.991 | 0.819 | 3.98E-163 | 4 | Rpl35 |
| Huwe11 | 7.36E-167 | 0.485759103 | 0.639 | 0.265 | 1.49E-162 | 4 | Huwe1 |
| Rab8a | 8.61E-167 | 0.312603609 | 0.572 | 0.217 | 1.74E-162 | 4 | Rab8a |
| Nasp | 1.14E-166 | 0.391101901 | 0.46 | 0.156 | 2.31E-162 | 4 | Nasp |
| Cdkn2d | 1.15E-166 | 0.320390773 | 0.307 | 0.08 | 2.32E-162 | 4 | Cdkn2d |
| Slc1a5 | 1.83E-166 | 0.352693955 | 0.278 | 0.069 | 3.71E-162 | 4 | Slc1a5 |
| Rps6ka31 | 1.83E-165 | 0.514456983 | 0.603 | 0.243 | 3.69E-161 | 4 | Rps6ka3 |
| Rmnd5b | 4.71E-165 | 0.312369741 | 0.26 | 0.062 | 9.53E-161 | 4 | Rmnd5b |
| Wdr43 | 9.40E-165 | 0.432735606 | 0.413 | 0.136 | 1.90E-160 | 4 | Wdr43 |
| Cyth3 | 1.76E-164 | 0.434677558 | 0.369 | 0.111 | 3.56E-160 | 4 | Cyth3 |
| Fgfr1op2 | 2.25E-164 | 0.360093251 | 0.677 | 0.282 | 4.55E-160 | 4 | Fgfr1op2 |
| Rps27rt | 2.30E-164 | 0.29903414 | 0.336 | 0.095 | 4.65E-160 | 4 | Rps27rt |
| Zfp6442 | 3.06E-164 | 0.45739283 | 0.593 | 0.236 | 6.19E-160 | 4 | Zfp644 |
| Rbm392 | 1.05E-163 | 0.576937089 | 0.965 | 0.551 | 2.12E-159 | 4 | Rbm39 |
| Ankrd122 | 1.31E-163 | 0.516557925 | 0.673 | 0.287 | 2.66E-159 | 4 | Ankrd12 |
| Bhlhe401 | 1.54E-163 | 0.437904448 | 0.431 | 0.143 | 3.12E-159 | 4 | Bhlhe40 |
| Dnajc2 | 3.30E-163 | 0.308768778 | 0.414 | 0.133 | 6.68E-159 | 4 | Dnajc2 |
| Nme12 | 1.22E-162 | 0.454070955 | 0.726 | 0.334 | 2.48E-158 | 4 | Nme1 |
| Sh2d3c1 | 1.58E-162 | 0.407187967 | 0.431 | 0.141 | 3.20E-158 | 4 | Sh2d3c |
| Vps54 | 1.81E-162 | 0.470412176 | 0.475 | 0.17 | 3.66E-158 | 4 | Vps54 |
| Tmem131l | 3.59E-162 | 0.429144089 | 0.274 | 0.07 | 7.26E-158 | 4 | Tmem131l |
| Zfp182 | 3.65E-162 | 0.386563241 | 0.337 | 0.096 | 7.39E-158 | 4 | Zfp182 |
| Arih12 | 1.24E-161 | 0.552152233 | 0.669 | 0.292 | 2.51E-157 | 4 | Arih1 |
| Fcgr3 | 4.03E-161 | 0.287386689 | 0.295 | 0.076 | 8.16E-157 | 4 | Fcgr3 |
| Dcps | 5.10E-161 | 0.303767348 | 0.352 | 0.103 | 1.03E-156 | 4 | Dcps |
| Ndfip22 | 7.83E-161 | 0.4423269 | 0.574 | 0.226 | 1.59E-156 | 4 | Ndfip2 |
| Brd42 | 1.33E-160 | 0.481960918 | 0.606 | 0.248 | 2.70E-156 | 4 | Brd4 |
| Os91 | 2.58E-160 | 0.555417009 | 0.476 | 0.173 | 5.21E-156 | 4 | Os9 |
| Rpl142 | 3.47E-160 | 0.474031653 | 0.993 | 0.791 | 7.03E-156 | 4 | Rpl14 |
| Zc3h131 | 4.42E-160 | 0.448580839 | 0.524 | 0.198 | 8.94E-156 | 4 | Zc3h13 |
| Ppp6r1 | 5.96E-160 | 0.362425988 | 0.317 | 0.089 | 1.21E-155 | 4 | Ppp6r1 |
| Ankrd10 | 6.88E-160 | 0.341914458 | 0.372 | 0.114 | 1.39E-155 | 4 | Ankrd10 |
| Polr2a2 | 7.27E-160 | 0.524795065 | 0.627 | 0.261 | 1.47E-155 | 4 | Polr2a |
| Rapgef12 | 9.08E-160 | 0.475041708 | 0.565 | 0.222 | 1.84E-155 | 4 | Rapgef1 |
| Ypel3 | 2.26E-159 | 0.401820869 | 0.607 | 0.248 | 4.58E-155 | 4 | Ypel3 |
| Phf141 | 2.90E-159 | 0.470648625 | 0.57 | 0.226 | 5.87E-155 | 4 | Phf14 |
| Notch2 | 3.20E-159 | 0.378791276 | 0.273 | 0.07 | 6.48E-155 | 4 | Notch2 |
| Eif4b | 3.46E-159 | 0.349723217 | 0.56 | 0.218 | 7.00E-155 | 4 | Eif4b |

| | | | | | | | |
|---|---|---|---|---|---|---|---|
| Nt5dc1 | 5.14E-159 | 0.395699271 | 0.355 | 0.107 | 1.04E-154 | 4 | Nt5dc1 |
| Nhp2 | 8.97E-159 | 0.292254025 | 0.461 | 0.161 | 1.82E-154 | 4 | Nhp2 |
| Bcas31 | 2.33E-158 | 0.538034331 | 0.461 | 0.166 | 4.73E-154 | 4 | Bcas3 |
| Micu2 | 2.73E-158 | 0.330149368 | 0.484 | 0.172 | 5.53E-154 | 4 | Micu2 |
| Bola21 | 2.81E-158 | 0.276132169 | 0.605 | 0.242 | 5.69E-154 | 4 | Bola2 |
| Hdac1 | 6.14E-158 | 0.371540719 | 0.442 | 0.152 | 1.24E-153 | 4 | Hdac1 |
| Ppm1g | 6.79E-158 | 0.390892139 | 0.419 | 0.141 | 1.38E-153 | 4 | Ppm1g |
| Hspa51 | 7.33E-158 | 0.608516331 | 0.825 | 0.412 | 1.48E-153 | 4 | Hspa5 |
| mt-Co32 | 9.76E-158 | 0.582121537 | 1 | 0.71 | 1.98E-153 | 4 | mt-Co3 |
| Fbl | 1.10E-157 | 0.346598758 | 0.372 | 0.117 | 2.24E-153 | 4 | Fbl |
| Nr2c21 | 1.24E-157 | 0.441759613 | 0.424 | 0.144 | 2.51E-153 | 4 | Nr2c2 |
| Arid1b1 | 1.39E-157 | 0.524088497 | 0.584 | 0.24 | 2.81E-153 | 4 | Arid1b |
| Nfatc1 | 2.28E-157 | 0.400686435 | 0.345 | 0.103 | 4.61E-153 | 4 | Nfatc1 |
| Pan31 | 3.41E-157 | 0.526115658 | 0.538 | 0.211 | 6.90E-153 | 4 | Pan3 |
| Birc3 | 4.13E-157 | 0.406380114 | 0.309 | 0.087 | 8.37E-153 | 4 | Birc3 |
| Scfd2 | 8.50E-157 | 0.425635464 | 0.282 | 0.075 | 1.72E-152 | 4 | Scfd2 |
| Actr2 | 1.41E-156 | 0.412606777 | 0.723 | 0.321 | 2.86E-152 | 4 | Actr2 |
| Dyrk1a1 | 2.03E-156 | 0.419947663 | 0.462 | 0.164 | 4.11E-152 | 4 | Dyrk1a |
| Nosip | 2.39E-156 | 0.31358662 | 0.402 | 0.129 | 4.84E-152 | 4 | Nosip |
| Phip1 | 8.39E-156 | 0.495200335 | 0.584 | 0.237 | 1.70E-151 | 4 | Phip |
| Rbm51 | 1.19E-155 | 0.434424886 | 0.565 | 0.221 | 2.41E-151 | 4 | Rbm5 |
| Rps62 | 2.31E-155 | 0.485108673 | 0.992 | 0.801 | 4.67E-151 | 4 | Rps6 |
| Pik3cb | 4.40E-155 | 0.405533631 | 0.4 | 0.132 | 8.90E-151 | 4 | Pik3cb |
| Tmem21 | 4.62E-155 | 0.473580597 | 0.396 | 0.128 | 9.35E-151 | 4 | Tmem2 |
| Cd84 | 5.18E-155 | 0.357579585 | 0.294 | 0.079 | 1.05E-150 | 4 | Cd84 |
| Icam1 | 6.65E-155 | 0.404776381 | 0.26 | 0.065 | 1.35E-150 | 4 | Icam1 |
| Cnot1 | 6.74E-155 | 0.40536229 | 0.447 | 0.158 | 1.36E-150 | 4 | Cnot1 |
| Cops4 | 7.73E-155 | 0.27422582 | 0.468 | 0.165 | 1.57E-150 | 4 | Cops4 |
| Cirbp | 5.90E-154 | 0.330659893 | 0.579 | 0.232 | 1.19E-149 | 4 | Cirbp |
| Rlim1 | 1.06E-153 | 0.412423785 | 0.491 | 0.181 | 2.14E-149 | 4 | Rlim |
| Avl9 | 2.77E-153 | 0.410218638 | 0.382 | 0.123 | 5.61E-149 | 4 | Avl9 |
| Birc2 | 4.78E-153 | 0.376998893 | 0.33 | 0.098 | 9.67E-149 | 4 | Birc2 |
| Hdac8 | 6.16E-153 | 0.566433901 | 0.374 | 0.122 | 1.25E-148 | 4 | Hdac8 |
| Larp1 | 1.38E-152 | 0.485779543 | 0.49 | 0.187 | 2.79E-148 | 4 | Larp1 |
| 4932438A13Rik1 | 2.34E-152 | 0.446515938 | 0.526 | 0.202 | 4.73E-148 | 4 | 4932438A13Rik |
| Tiprl | 3.84E-152 | 0.296991128 | 0.515 | 0.192 | 7.78E-148 | 4 | Tiprl |
| Topors | 5.03E-152 | 0.41329631 | 0.342 | 0.105 | 1.02E-147 | 4 | Topors |
| Rpl102 | 5.05E-152 | 0.401889487 | 0.995 | 0.891 | 1.02E-147 | 4 | Rpl10 |
| Arrb2 | 5.27E-152 | 0.303690483 | 0.297 | 0.081 | 1.07E-147 | 4 | Arrb2 |
| Hip1r | 6.18E-152 | 0.326235479 | 0.258 | 0.064 | 1.25E-147 | 4 | Hip1r |
| Thada | 2.58E-151 | 0.436265736 | 0.345 | 0.107 | 5.22E-147 | 4 | Thada |
| Rftn12 | 3.06E-151 | 0.495638808 | 0.576 | 0.233 | 6.19E-147 | 4 | Rftn1 |
| Kdm5a | 4.04E-151 | 0.390162207 | 0.436 | 0.152 | 8.17E-147 | 4 | Kdm5a |

| | | | | | | | |
|---|---|---|---|---|---|---|---|
| Dapp1 | 4.69E-151 | 0.338709552 | 0.271 | 0.071 | 9.49E-147 | 4 | Dapp1 |
| Stag21 | 1.81E-150 | 0.4242122 | 0.628 | 0.268 | 3.67E-146 | 4 | Stag2 |
| Srgn2 | 1.97E-149 | 0.571341976 | 0.986 | 0.519 | 3.98E-145 | 4 | Srgn |
| Ss18 | 2.12E-149 | 0.373675182 | 0.444 | 0.159 | 4.30E-145 | 4 | Ss18 |
| Scaper1 | 2.67E-149 | 0.450698116 | 0.453 | 0.164 | 5.41E-145 | 4 | Scaper |
| Creb1 | 4.94E-149 | 0.518803213 | 0.492 | 0.192 | 1.00E-144 | 4 | Creb1 |
| Chd3 | 5.72E-149 | 0.254862494 | 0.417 | 0.141 | 1.16E-144 | 4 | Chd3 |
| Rchy1 | 6.14E-149 | 0.335711577 | 0.421 | 0.145 | 1.24E-144 | 4 | Rchy1 |
| Grcc10 | 2.03E-148 | 0.250058501 | 0.326 | 0.095 | 4.11E-144 | 4 | Grcc10 |
| Eif3m1 | 2.07E-148 | 0.255636646 | 0.758 | 0.344 | 4.19E-144 | 4 | Eif3m |
| Mxd42 | 2.31E-148 | 0.429427931 | 0.727 | 0.331 | 4.67E-144 | 4 | Mxd4 |
| Nubp1 | 3.10E-148 | 0.25684759 | 0.311 | 0.089 | 6.29E-144 | 4 | Nubp1 |
| Smc61 | 5.13E-148 | 0.391790536 | 0.59 | 0.243 | 1.04E-143 | 4 | Smc6 |
| Farp1 | 5.32E-148 | 0.359574197 | 0.324 | 0.096 | 1.08E-143 | 4 | Farp1 |
| Mndal2 | 5.77E-148 | 0.457902576 | 0.808 | 0.382 | 1.17E-143 | 4 | Mndal |
| Cct41 | 6.62E-148 | 0.263132721 | 0.68 | 0.294 | 1.34E-143 | 4 | Cct4 |
| Rpl72 | 8.70E-148 | 0.411597559 | 0.994 | 0.887 | 1.76E-143 | 4 | Rpl7 |
| Ttc31 | 1.66E-147 | 0.450314397 | 0.576 | 0.233 | 3.36E-143 | 4 | Ttc3 |
| Atad2b1 | 2.46E-147 | 0.450892453 | 0.49 | 0.187 | 4.98E-143 | 4 | Atad2b |
| Dis3l2 | 3.73E-147 | 0.420933283 | 0.379 | 0.126 | 7.56E-143 | 4 | Dis3l2 |
| Rpl36a2 | 3.76E-147 | 0.479540266 | 0.987 | 0.738 | 7.61E-143 | 4 | Rpl36a |
| Ccm2 | 4.11E-147 | 0.39708789 | 0.355 | 0.113 | 8.31E-143 | 4 | Ccm2 |
| Ccny2 | 4.24E-147 | 0.456297635 | 0.595 | 0.249 | 8.58E-143 | 4 | Ccny |
| Arl6ip12 | 1.09E-146 | 0.438240438 | 0.749 | 0.348 | 2.22E-142 | 4 | Arl6ip1 |
| Slc44a2 | 1.47E-146 | 0.339554597 | 0.352 | 0.109 | 2.98E-142 | 4 | Slc44a2 |
| Pspc1 | 2.55E-146 | 0.392260444 | 0.34 | 0.106 | 5.16E-142 | 4 | Pspc1 |
| Itga12 | 2.77E-146 | 0.373634459 | 0.871 | 0.388 | 5.60E-142 | 4 | Itga1 |
| Lman2 | 5.01E-146 | 0.387299526 | 0.495 | 0.19 | 1.02E-141 | 4 | Lman2 |
| Ascc31 | 9.62E-146 | 0.44545288 | 0.497 | 0.191 | 1.95E-141 | 4 | Ascc3 |
| Med14 | 9.72E-146 | 0.343222437 | 0.312 | 0.092 | 1.97E-141 | 4 | Med14 |
| Fam120b | 1.28E-145 | 0.346544789 | 0.285 | 0.079 | 2.58E-141 | 4 | Fam120b |
| Bbx1 | 1.53E-145 | 0.408048773 | 0.528 | 0.206 | 3.10E-141 | 4 | Bbx |
| Kdm4c1 | 1.60E-145 | 0.447746892 | 0.429 | 0.154 | 3.24E-141 | 4 | Kdm4c |
| Sik2 | 1.85E-145 | 0.43250444 | 0.341 | 0.107 | 3.75E-141 | 4 | Sik2 |
| Rpl242 | 2.56E-145 | 0.370423271 | 0.994 | 0.903 | 5.18E-141 | 4 | Rpl24 |
| Mapk11 | 3.46E-145 | 0.384415912 | 0.545 | 0.218 | 7.01E-141 | 4 | Mapk1 |
| B4galt11 | 4.60E-145 | 0.452094322 | 0.443 | 0.162 | 9.32E-141 | 4 | B4galt1 |
| Ifi471 | 1.64E-144 | 0.355467096 | 0.455 | 0.163 | 3.33E-140 | 4 | Ifi47 |
| mt-Nd4l1 | 1.89E-144 | 0.413656043 | 0.9 | 0.467 | 3.82E-140 | 4 | mt-Nd4l |
| Tnfsf102 | 6.64E-144 | 0.28191183 | 0.817 | 0.348 | 1.34E-139 | 4 | Tnfsf10 |
| Gmip | 1.56E-143 | 0.361515919 | 0.267 | 0.072 | 3.17E-139 | 4 | Gmip |
| Acin11 | 2.03E-143 | 0.42301066 | 0.662 | 0.296 | 4.11E-139 | 4 | Acin1 |
| Akr1b3 | 3.10E-143 | 0.299872257 | 0.469 | 0.173 | 6.28E-139 | 4 | Akr1b3 |

| | | | | | | | |
|---|---|---|---|---|---|---|---|
| Wnk11 | 3.11E-143 | 0.470008653 | 0.741 | 0.342 | 6.30E-139 | 4 | Wnk1 |
| Fkbp5 | 4.94E-143 | 0.289679689 | 0.366 | 0.118 | 9.99E-139 | 4 | Fkbp5 |
| C2cd3 | 5.29E-143 | 0.330447102 | 0.261 | 0.069 | 1.07E-138 | 4 | C2cd3 |
| 2310035C23Rik | 8.34E-143 | 0.388776094 | 0.378 | 0.127 | 1.69E-138 | 4 | 2310035C23Rik |
| Hnrnpf | 8.46E-143 | 0.404849599 | 0.891 | 0.472 | 1.71E-138 | 4 | Hnrnpf |
| Ctla2a2 | 1.50E-142 | 0.265527377 | 0.847 | 0.372 | 3.04E-138 | 4 | Ctla2a |
| Prpf4b2 | 3.23E-142 | 0.401873437 | 0.686 | 0.313 | 6.53E-138 | 4 | Prpf4b |
| Ints6 | 3.26E-142 | 0.398512648 | 0.382 | 0.129 | 6.60E-138 | 4 | Ints6 |
| Socs1 | 3.95E-142 | 0.374726757 | 0.3 | 0.087 | 7.99E-138 | 4 | Socs1 |
| Wac2 | 4.61E-142 | 0.423941662 | 0.676 | 0.302 | 9.34E-138 | 4 | Wac |
| Sltm1 | 6.30E-142 | 0.368542754 | 0.6 | 0.252 | 1.27E-137 | 4 | Sltm |
| Pdcd6ip | 6.36E-142 | 0.363006352 | 0.533 | 0.213 | 1.29E-137 | 4 | Pdcd6ip |
| Trappc10 | 1.14E-141 | 0.272471387 | 0.379 | 0.125 | 2.30E-137 | 4 | Trappc10 |
| Rsf11 | 3.95E-141 | 0.4245916 | 0.519 | 0.207 | 7.99E-137 | 4 | Rsf1 |
| Synrg | 4.60E-141 | 0.352444168 | 0.382 | 0.129 | 9.31E-137 | 4 | Synrg |
| Gm100762 | 4.70E-141 | 0.360634219 | 0.911 | 0.525 | 9.51E-137 | 4 | Gm10076 |
| Hip12 | 5.77E-141 | 0.531733157 | 0.686 | 0.314 | 1.17E-136 | 4 | Hip1 |
| Ppp1cc | 8.90E-141 | 0.257098515 | 0.526 | 0.205 | 1.80E-136 | 4 | Ppp1cc |
| Upf3b | 1.36E-140 | 0.332142605 | 0.375 | 0.125 | 2.75E-136 | 4 | Upf3b |
| Rai1 | 2.93E-140 | 0.349320763 | 0.32 | 0.098 | 5.94E-136 | 4 | Rai1 |
| Zfand6 | 3.11E-140 | 0.324659399 | 0.641 | 0.28 | 6.30E-136 | 4 | Zfand6 |
| Jak2 | 6.22E-140 | 0.371002824 | 0.354 | 0.114 | 1.26E-135 | 4 | Jak2 |
| Uimc1 | 9.31E-140 | 0.3520279 | 0.394 | 0.136 | 1.89E-135 | 4 | Uimc1 |
| Colgalt1 | 1.36E-139 | 0.381755524 | 0.382 | 0.131 | 2.76E-135 | 4 | Colgalt1 |
| N4bp2l21 | 1.47E-139 | 0.369838002 | 0.558 | 0.229 | 2.97E-135 | 4 | N4bp2l2 |
| B3gat3 | 3.23E-139 | 0.298464751 | 0.367 | 0.12 | 6.55E-135 | 4 | B3gat3 |
| Tbc1d51 | 5.84E-139 | 0.499853391 | 0.575 | 0.244 | 1.18E-134 | 4 | Tbc1d5 |
| Rif1 | 7.64E-139 | 0.389895156 | 0.403 | 0.141 | 1.55E-134 | 4 | Rif1 |
| Fli12 | 1.05E-138 | 0.398672586 | 0.815 | 0.378 | 2.13E-134 | 4 | Fli1 |
| Tma72 | 6.70E-138 | 0.251442122 | 0.917 | 0.501 | 1.36E-133 | 4 | Tma7 |
| Man1a | 8.02E-138 | 0.465987764 | 0.419 | 0.153 | 1.62E-133 | 4 | Man1a |
| Chfr | 1.04E-137 | 0.352961548 | 0.331 | 0.104 | 2.10E-133 | 4 | Chfr |
| Prmt1 | 1.07E-137 | 0.291731399 | 0.508 | 0.199 | 2.17E-133 | 4 | Prmt1 |
| Cbfb | 1.10E-137 | 0.395590067 | 0.401 | 0.142 | 2.22E-133 | 4 | Cbfb |
| Eif3k1 | 2.29E-137 | 0.298217213 | 0.897 | 0.479 | 4.63E-133 | 4 | Eif3k |
| Rbbp4 | 3.55E-137 | 0.306181084 | 0.584 | 0.244 | 7.18E-133 | 4 | Rbbp4 |
| Babam22 | 1.11E-136 | 0.40412783 | 0.542 | 0.221 | 2.25E-132 | 4 | Babam2 |
| Lrrfip2 | 1.46E-136 | 0.360960239 | 0.416 | 0.15 | 2.96E-132 | 4 | Lrrfip2 |
| Rbm61 | 1.47E-136 | 0.412621589 | 0.503 | 0.201 | 2.97E-132 | 4 | Rbm6 |
| Pabpc12 | 1.73E-136 | 0.2705764 | 0.936 | 0.544 | 3.51E-132 | 4 | Pabpc1 |
| Pdia32 | 1.73E-136 | 0.486322176 | 0.835 | 0.421 | 3.51E-132 | 4 | Pdia3 |
| Ddx39b | 2.53E-136 | 0.310461933 | 0.534 | 0.217 | 5.11E-132 | 4 | Ddx39b |
| Rptor | 1.19E-135 | 0.407091787 | 0.37 | 0.126 | 2.42E-131 | 4 | Rptor |

| | | | | | | | |
|---|---|---|---|---|---|---|---|
| Eef1g2 | 1.75E-135 | 0.337208329 | 0.887 | 0.489 | 3.53E-131 | 4 | Eef1g |
| Fam208b1 | 1.92E-135 | 0.404930231 | 0.391 | 0.137 | 3.88E-131 | 4 | Fam208b |
| Snd1 | 2.12E-135 | 0.403266905 | 0.426 | 0.158 | 4.29E-131 | 4 | Snd1 |
| Fyco1 | 4.34E-135 | 0.384750827 | 0.342 | 0.112 | 8.79E-131 | 4 | Fyco1 |
| Arid4a2 | 5.11E-135 | 0.434713535 | 0.559 | 0.234 | 1.04E-130 | 4 | Arid4a |
| Taok3 | 5.40E-135 | 0.316554852 | 0.42 | 0.149 | 1.09E-130 | 4 | Taok3 |
| Maz | 6.14E-135 | 0.29952497 | 0.284 | 0.082 | 1.24E-130 | 4 | Maz |
| Pip4k2a1 | 6.26E-135 | 0.374501146 | 0.55 | 0.224 | 1.27E-130 | 4 | Pip4k2a |
| Gtf2f2 | 7.64E-135 | 0.305276806 | 0.305 | 0.092 | 1.55E-130 | 4 | Gtf2f2 |
| Chd61 | 1.75E-134 | 0.400440271 | 0.504 | 0.201 | 3.54E-130 | 4 | Chd6 |
| Krit11 | 2.10E-134 | 0.386402023 | 0.45 | 0.167 | 4.25E-130 | 4 | Krit1 |
| Bptf2 | 2.81E-134 | 0.430669861 | 0.627 | 0.275 | 5.68E-130 | 4 | Bptf |
| Kdm2a2 | 4.72E-134 | 0.371843017 | 0.546 | 0.223 | 9.56E-130 | 4 | Kdm2a |
| Clta | 1.10E-133 | 0.403017879 | 0.929 | 0.536 | 2.23E-129 | 4 | Clta |
| Pold4 | 1.14E-133 | 0.276581705 | 0.479 | 0.184 | 2.30E-129 | 4 | Pold4 |
| Retreg1 | 1.52E-133 | 0.30859804 | 0.292 | 0.086 | 3.08E-129 | 4 | Retreg1 |
| Hnrnpa1 | 1.57E-133 | 0.294483286 | 0.669 | 0.305 | 3.17E-129 | 4 | Hnrnpa1 |
| Ccar11 | 1.65E-133 | 0.408335849 | 0.487 | 0.192 | 3.34E-129 | 4 | Ccar1 |
| Mpp7 | 2.15E-133 | 0.281863518 | 0.312 | 0.095 | 4.36E-129 | 4 | Mpp7 |
| Kmt2a2 | 2.19E-133 | 0.434119419 | 0.585 | 0.249 | 4.43E-129 | 4 | Kmt2a |
| Arhgap172 | 2.80E-133 | 0.418150245 | 0.59 | 0.252 | 5.67E-129 | 4 | Arhgap17 |
| Gatad2a2 | 2.87E-133 | 0.364175571 | 0.515 | 0.206 | 5.80E-129 | 4 | Gatad2a |
| Gpatch82 | 4.46E-133 | 0.418467765 | 0.568 | 0.24 | 9.02E-129 | 4 | Gpatch8 |
| Senp61 | 6.35E-133 | 0.366831448 | 0.633 | 0.279 | 1.29E-128 | 4 | Senp6 |
| Zdhhc202 | 8.11E-133 | 0.444449228 | 0.521 | 0.214 | 1.64E-128 | 4 | Zdhhc20 |
| Ep4001 | 1.12E-132 | 0.395331996 | 0.488 | 0.192 | 2.26E-128 | 4 | Ep400 |
| Manf | 1.21E-132 | 0.343445606 | 0.676 | 0.308 | 2.46E-128 | 4 | Manf |
| Gbf11 | 1.83E-132 | 0.419271774 | 0.44 | 0.167 | 3.70E-128 | 4 | Gbf1 |
| Sbno11 | 2.54E-132 | 0.337541269 | 0.491 | 0.193 | 5.13E-128 | 4 | Sbno1 |
| Mcm6 | 2.69E-132 | 0.310420304 | 0.259 | 0.071 | 5.45E-128 | 4 | Mcm6 |
| Abi11 | 4.23E-132 | 0.376324591 | 0.588 | 0.251 | 8.56E-128 | 4 | Abi1 |
| Prpf40a1 | 4.58E-132 | 0.366930871 | 0.581 | 0.249 | 9.28E-128 | 4 | Prpf40a |
| Susd62 | 6.50E-132 | 0.437912273 | 0.605 | 0.264 | 1.32E-127 | 4 | Susd6 |
| Nin | 1.11E-131 | 0.399756939 | 0.367 | 0.126 | 2.24E-127 | 4 | Nin |
| Sh3gl1 | 2.18E-131 | 0.356135754 | 0.342 | 0.113 | 4.42E-127 | 4 | Sh3gl1 |
| Afg3l2 | 5.19E-131 | 0.251731471 | 0.252 | 0.068 | 1.05E-126 | 4 | Afg3l2 |
| Psmb82 | 5.26E-131 | 0.427861902 | 0.959 | 0.534 | 1.06E-126 | 4 | Psmb8 |
| Smarcc1 | 6.53E-131 | 0.280230561 | 0.375 | 0.129 | 1.32E-126 | 4 | Smarcc1 |
| Foxn32 | 7.91E-131 | 0.491889554 | 0.776 | 0.39 | 1.60E-126 | 4 | Foxn3 |
| Rnf138 | 1.39E-130 | 0.296487001 | 0.365 | 0.124 | 2.81E-126 | 4 | Rnf138 |
| Kmt2c2 | 5.14E-130 | 0.416774306 | 0.642 | 0.287 | 1.04E-125 | 4 | Kmt2c |
| Swap702 | 1.12E-129 | 0.417520204 | 0.548 | 0.229 | 2.27E-125 | 4 | Swap70 |
| Sri | 2.09E-129 | 0.299157662 | 0.805 | 0.398 | 4.23E-125 | 4 | Sri |

| | | | | | | | |
|---|---|---|---|---|---|---|---|
| Mthfs | 5.95E-129 | 0.263450956 | 0.266 | 0.075 | 1.21E-124 | 4 | Mthfs |
| Gspt1 | 6.27E-129 | 0.286850185 | 0.54 | 0.224 | 1.27E-124 | 4 | Gspt1 |
| Dennd1a1 | 3.12E-128 | 0.422861491 | 0.558 | 0.236 | 6.32E-124 | 4 | Dennd1a |
| Rpl52 | 2.77E-127 | 0.41202393 | 0.977 | 0.695 | 5.62E-123 | 4 | Rpl5 |
| Fbxw4 | 3.88E-127 | 0.358999662 | 0.321 | 0.104 | 7.86E-123 | 4 | Fbxw4 |
| Gnptab | 6.13E-127 | 0.318111341 | 0.261 | 0.075 | 1.24E-122 | 4 | Gnptab |
| Per1 | 1.17E-126 | 0.322800579 | 0.268 | 0.078 | 2.38E-122 | 4 | Per1 |
| Rad50 | 1.53E-126 | 0.319841171 | 0.37 | 0.13 | 3.10E-122 | 4 | Rad50 |
| Cdc42ep31 | 1.73E-126 | 0.350645377 | 0.432 | 0.16 | 3.51E-122 | 4 | Cdc42ep3 |
| Dcaf7 | 2.15E-126 | 0.328198951 | 0.333 | 0.11 | 4.35E-122 | 4 | Dcaf7 |
| Smap11 | 2.37E-126 | 0.262165008 | 0.72 | 0.34 | 4.80E-122 | 4 | Smap1 |
| Ppig2 | 5.98E-126 | 0.305625681 | 0.657 | 0.297 | 1.21E-121 | 4 | Ppig |
| Nsd11 | 6.54E-126 | 0.386313462 | 0.701 | 0.329 | 1.32E-121 | 4 | Nsd1 |
| Zc3hav1 | 7.28E-126 | 0.35955437 | 0.475 | 0.189 | 1.47E-121 | 4 | Zc3hav1 |
| Gnptg | 7.29E-126 | 0.284714387 | 0.288 | 0.087 | 1.48E-121 | 4 | Gnptg |
| Atf6 | 2.62E-125 | 0.363078278 | 0.405 | 0.148 | 5.31E-121 | 4 | Atf6 |
| Chd81 | 4.50E-125 | 0.360140634 | 0.412 | 0.154 | 9.12E-121 | 4 | Chd8 |
| Cab391 | 5.45E-125 | 0.271879973 | 0.597 | 0.262 | 1.10E-120 | 4 | Cab39 |
| Twf2 | 6.72E-125 | 0.303947677 | 0.419 | 0.157 | 1.36E-120 | 4 | Twf2 |
| Tra2a2 | 9.79E-125 | 0.390412168 | 0.586 | 0.257 | 1.98E-120 | 4 | Tra2a |
| Rab3gap1 | 9.58E-124 | 0.32535013 | 0.404 | 0.151 | 1.94E-119 | 4 | Rab3gap1 |
| Arpc1b | 1.03E-123 | 0.402774843 | 0.945 | 0.571 | 2.08E-119 | 4 | Arpc1b |
| Gm37240 | 1.26E-123 | 0.321903105 | 0.299 | 0.094 | 2.55E-119 | 4 | Gm37240 |
| Fars2 | 1.95E-123 | 0.273489417 | 0.468 | 0.188 | 3.94E-119 | 4 | Fars2 |
| Peak12 | 4.94E-123 | 0.411184647 | 0.756 | 0.366 | 1.00E-118 | 4 | Peak1 |
| Cep57 | 7.09E-123 | 0.271544608 | 0.306 | 0.098 | 1.44E-118 | 4 | Cep57 |
| Ankib1 | 1.18E-122 | 0.346506669 | 0.359 | 0.127 | 2.39E-118 | 4 | Ankib1 |
| Scaf81 | 1.65E-122 | 0.375694438 | 0.438 | 0.171 | 3.33E-118 | 4 | Scaf8 |
| Birc62 | 1.71E-122 | 0.389777106 | 0.686 | 0.321 | 3.47E-118 | 4 | Birc6 |
| Exoc6 | 2.00E-122 | 0.330113756 | 0.417 | 0.157 | 4.05E-118 | 4 | Exoc6 |
| Setd52 | 2.14E-122 | 0.408400274 | 0.562 | 0.247 | 4.33E-118 | 4 | Setd5 |
| Rsrc1 | 3.26E-122 | 0.359231501 | 0.56 | 0.243 | 6.59E-118 | 4 | Rsrc1 |
| Cnppd1 | 5.68E-122 | 0.311788824 | 0.276 | 0.083 | 1.15E-117 | 4 | Cnppd1 |
| Cep120 | 6.98E-122 | 0.373031256 | 0.319 | 0.106 | 1.41E-117 | 4 | Cep120 |
| Impact | 8.53E-122 | 0.372289554 | 0.331 | 0.112 | 1.73E-117 | 4 | Impact |
| Elmsan1 | 1.15E-121 | 0.344100325 | 0.298 | 0.095 | 2.33E-117 | 4 | Elmsan1 |
| Gm26510 | 1.27E-121 | 0.342704505 | 0.31 | 0.099 | 2.57E-117 | 4 | Gm26510 |
| Galnt1 | 1.62E-121 | 0.349161151 | 0.438 | 0.17 | 3.27E-117 | 4 | Galnt1 |
| Psmb31 | 1.94E-121 | 0.251139976 | 0.798 | 0.398 | 3.93E-117 | 4 | Psmb3 |
| Srsf112 | 2.40E-121 | 0.380045524 | 0.695 | 0.331 | 4.86E-117 | 4 | Srsf11 |
| Virma | 2.84E-121 | 0.323745756 | 0.346 | 0.12 | 5.74E-117 | 4 | Virma |
| Zc3h7a1 | 2.93E-121 | 0.352423176 | 0.4 | 0.149 | 5.94E-117 | 4 | Zc3h7a |
| Eef2 | 3.30E-121 | 0.358360641 | 0.914 | 0.523 | 6.68E-117 | 4 | Eef2 |

| | | | | | | | |
|---|---|---|---|---|---|---|---|
| Sec11a | 6.34E-121 | 0.28789609 | 0.604 | 0.267 | 1.28E-116 | 4 | Sec11a |
| Cmip2 | 7.09E-121 | 0.409492007 | 0.601 | 0.268 | 1.44E-116 | 4 | Cmip |
| Herc4 | 7.98E-121 | 0.378100179 | 0.379 | 0.139 | 1.62E-116 | 4 | Herc4 |
| Rfx3 | 9.79E-121 | 0.38515158 | 0.3 | 0.097 | 1.98E-116 | 4 | Rfx3 |
| Usp342 | 3.64E-120 | 0.38021916 | 0.683 | 0.325 | 7.37E-116 | 4 | Usp34 |
| Ap3b11 | 4.93E-120 | 0.38884239 | 0.485 | 0.203 | 9.98E-116 | 4 | Ap3b1 |
| Xpr1 | 5.60E-120 | 0.376200541 | 0.382 | 0.141 | 1.13E-115 | 4 | Xpr1 |
| Pnn2 | 1.06E-119 | 0.362211564 | 0.567 | 0.25 | 2.15E-115 | 4 | Pnn |
| Plekhm3 | 1.11E-119 | 0.358970039 | 0.258 | 0.077 | 2.25E-115 | 4 | Plekhm3 |
| Pi4ka | 1.27E-119 | 0.310505112 | 0.338 | 0.116 | 2.56E-115 | 4 | Pi4ka |
| Eif4g31 | 1.42E-119 | 0.338394258 | 0.604 | 0.268 | 2.88E-115 | 4 | Eif4g3 |
| Prkdc | 2.61E-119 | 0.318192767 | 0.268 | 0.081 | 5.29E-115 | 4 | Prkdc |
| Trip121 | 3.16E-119 | 0.348259629 | 0.553 | 0.24 | 6.40E-115 | 4 | Trip12 |
| Stag12 | 3.44E-119 | 0.421300106 | 0.711 | 0.341 | 6.97E-115 | 4 | Stag1 |
| Faf11 | 7.86E-119 | 0.336429108 | 0.463 | 0.187 | 1.59E-114 | 4 | Faf1 |
| Adamts10 | 8.04E-119 | 0.313185132 | 0.288 | 0.09 | 1.63E-114 | 4 | Adamts10 |
| Sf1 | 1.58E-118 | 0.287697946 | 0.443 | 0.173 | 3.19E-114 | 4 | Sf1 |
| Cnot41 | 2.34E-118 | 0.347421404 | 0.571 | 0.252 | 4.74E-114 | 4 | Cnot4 |
| Thrap31 | 2.54E-118 | 0.305527271 | 0.609 | 0.276 | 5.14E-114 | 4 | Thrap3 |
| Pdxdc1 | 2.62E-118 | 0.294193545 | 0.383 | 0.141 | 5.31E-114 | 4 | Pdxdc1 |
| Rtf11 | 4.05E-118 | 0.273170672 | 0.593 | 0.262 | 8.21E-114 | 4 | Rtf1 |
| Sik11 | 4.06E-118 | 0.358825108 | 0.334 | 0.115 | 8.21E-114 | 4 | Sik1 |
| Tnks22 | 4.91E-118 | 0.349470144 | 0.589 | 0.262 | 9.95E-114 | 4 | Tnks2 |
| Bclaf11 | 5.15E-118 | 0.352545986 | 0.548 | 0.237 | 1.04E-113 | 4 | Bclaf1 |
| Rnaset2b | 6.26E-118 | 0.278122321 | 0.337 | 0.116 | 1.27E-113 | 4 | Rnaset2b |
| Pkn11 | 7.51E-118 | 0.325088342 | 0.44 | 0.172 | 1.52E-113 | 4 | Pkn1 |
| Nlk1 | 8.38E-118 | 0.38200355 | 0.365 | 0.133 | 1.70E-113 | 4 | Nlk |
| Trappc91 | 9.73E-118 | 0.40250806 | 0.449 | 0.18 | 1.97E-113 | 4 | Trappc9 |
| Kpna1 | 1.25E-117 | 0.279294791 | 0.327 | 0.111 | 2.53E-113 | 4 | Kpna1 |
| Gon4l | 1.44E-117 | 0.321491799 | 0.373 | 0.137 | 2.91E-113 | 4 | Gon4l |
| Hdac4 | 1.47E-117 | 0.335975714 | 0.296 | 0.096 | 2.99E-113 | 4 | Hdac4 |
| Crtc3 | 2.08E-117 | 0.382292396 | 0.397 | 0.152 | 4.21E-113 | 4 | Crtc3 |
| Rabep11 | 2.53E-117 | 0.354148465 | 0.509 | 0.215 | 5.12E-113 | 4 | Rabep1 |
| Ssh2 | 3.31E-117 | 0.339730427 | 0.509 | 0.21 | 6.69E-113 | 4 | Ssh2 |
| Bmp2k | 5.00E-117 | 0.267632854 | 0.306 | 0.1 | 1.01E-112 | 4 | Bmp2k |
| Gatad2b1 | 1.09E-116 | 0.331630891 | 0.463 | 0.188 | 2.21E-112 | 4 | Gatad2b |
| Wbp1l | 2.51E-116 | 0.307776685 | 0.275 | 0.086 | 5.09E-112 | 4 | Wbp1l |
| Elf12 | 4.04E-116 | 0.342601403 | 0.737 | 0.356 | 8.17E-112 | 4 | Elf1 |
| Rbm261 | 4.07E-116 | 0.341705466 | 0.5 | 0.211 | 8.24E-112 | 4 | Rbm26 |
| Tmx41 | 4.52E-116 | 0.423884867 | 0.374 | 0.139 | 9.15E-112 | 4 | Tmx4 |
| Cstf3 | 6.35E-116 | 0.324401025 | 0.332 | 0.115 | 1.28E-111 | 4 | Cstf3 |
| Pdpk1 | 6.59E-116 | 0.356731201 | 0.361 | 0.131 | 1.33E-111 | 4 | Pdpk1 |
| Sep-72 | 8.13E-116 | 0.328096602 | 0.842 | 0.44 | 1.65E-111 | 4 | 7-Sep |

| | | | | | | | |
|---|---|---|---|---|---|---|---|
| Kpna3 | 1.18E-115 | 0.290804152 | 0.385 | 0.144 | 2.39E-111 | 4 | Kpna3 |
| Lbr1 | 1.50E-115 | 0.323869525 | 0.381 | 0.14 | 3.04E-111 | 4 | Lbr |
| Kdm1a | 1.75E-115 | 0.252554593 | 0.508 | 0.213 | 3.55E-111 | 4 | Kdm1a |
| Agpat3 | 2.22E-115 | 0.284519251 | 0.336 | 0.118 | 4.50E-111 | 4 | Agpat3 |
| Ssu721 | 2.37E-115 | 0.252523778 | 0.636 | 0.293 | 4.80E-111 | 4 | Ssu72 |
| Ddx39 | 3.74E-115 | 0.257286288 | 0.333 | 0.114 | 7.58E-111 | 4 | Ddx39 |
| Rpl412 | 5.17E-115 | 0.325861225 | 0.999 | 0.96 | 1.05E-110 | 4 | Rpl41 |
| Jarid21 | 7.78E-115 | 0.323755492 | 0.464 | 0.189 | 1.58E-110 | 4 | Jarid2 |
| Pnisr2 | 8.46E-115 | 0.353214388 | 0.561 | 0.248 | 1.71E-110 | 4 | Pnisr |
| Akt32 | 1.17E-114 | 0.418549944 | 0.541 | 0.235 | 2.37E-110 | 4 | Akt3 |
| Atf7ip1 | 1.60E-114 | 0.365519462 | 0.465 | 0.193 | 3.24E-110 | 4 | Atf7ip |
| Akap92 | 3.50E-114 | 0.357405167 | 0.516 | 0.222 | 7.08E-110 | 4 | Akap9 |
| Sf3b21 | 3.72E-114 | 0.283141894 | 0.708 | 0.339 | 7.54E-110 | 4 | Sf3b2 |
| Taf1d | 4.38E-114 | 0.278205163 | 0.364 | 0.132 | 8.86E-110 | 4 | Taf1d |
| Nop58 | 5.55E-114 | 0.314697855 | 0.349 | 0.126 | 1.12E-109 | 4 | Nop58 |
| Trio1 | 6.80E-114 | 0.368235441 | 0.436 | 0.174 | 1.38E-109 | 4 | Trio |
| Crebrf1 | 9.27E-114 | 0.335371925 | 0.442 | 0.177 | 1.88E-109 | 4 | Crebrf |
| Dusp111 | 9.29E-114 | 0.267176269 | 0.59 | 0.264 | 1.88E-109 | 4 | Dusp11 |
| Nsd2 | 9.83E-114 | 0.275056566 | 0.27 | 0.084 | 1.99E-109 | 4 | Nsd2 |
| Wrn1 | 1.00E-113 | 0.329246733 | 0.365 | 0.134 | 2.03E-109 | 4 | Wrn |
| Txndc11 | 1.63E-113 | 0.320485129 | 0.27 | 0.085 | 3.30E-109 | 4 | Txndc11 |
| Sdf2 | 1.78E-113 | 0.306716884 | 0.408 | 0.158 | 3.61E-109 | 4 | Sdf2 |
| Pfdn52 | 1.82E-113 | 0.354019636 | 0.984 | 0.751 | 3.69E-109 | 4 | Pfdn5 |
| Smad72 | 3.10E-113 | 0.385791162 | 0.66 | 0.307 | 6.28E-109 | 4 | Smad7 |
| Mbd1 | 4.76E-113 | 0.27502013 | 0.283 | 0.09 | 9.63E-109 | 4 | Mbd1 |
| Chd41 | 5.76E-113 | 0.308347465 | 0.658 | 0.309 | 1.17E-108 | 4 | Chd4 |
| Map3k8 | 6.79E-113 | 0.276839195 | 0.252 | 0.075 | 1.38E-108 | 4 | Map3k8 |
| Mbd51 | 9.06E-113 | 0.393121229 | 0.485 | 0.205 | 1.83E-108 | 4 | Mbd5 |
| Dnmt3a | 2.00E-112 | 0.294395742 | 0.365 | 0.135 | 4.05E-108 | 4 | Dnmt3a |
| Galnt7 | 1.34E-111 | 0.340266455 | 0.379 | 0.143 | 2.72E-107 | 4 | Galnt7 |
| Lrmda2 | 1.58E-111 | 0.371597958 | 0.711 | 0.349 | 3.20E-107 | 4 | Lrmda |
| Hnrnpa2b1 | 2.35E-111 | 0.369675443 | 0.874 | 0.494 | 4.76E-107 | 4 | Hnrnpa2b1 |
| Dad11 | 3.35E-111 | 0.31474922 | 0.827 | 0.438 | 6.79E-107 | 4 | Dad1 |
| Ttc7b | 4.40E-111 | 0.344638645 | 0.32 | 0.111 | 8.90E-107 | 4 | Ttc7b |
| Lpp1 | 7.04E-111 | 0.376345892 | 0.586 | 0.269 | 1.43E-106 | 4 | Lpp |
| Lcorl1 | 1.37E-110 | 0.379065204 | 0.393 | 0.154 | 2.77E-106 | 4 | Lcorl |
| Fbxo111 | 1.63E-110 | 0.425759663 | 0.495 | 0.218 | 3.30E-106 | 4 | Fbxo11 |
| Setd21 | 2.42E-110 | 0.340672438 | 0.439 | 0.179 | 4.90E-106 | 4 | Setd2 |
| Snrnp48 | 2.69E-110 | 0.283719231 | 0.429 | 0.172 | 5.45E-106 | 4 | Snrnp48 |
| Ctps2 | 6.32E-110 | 0.255627446 | 0.258 | 0.079 | 1.28E-105 | 4 | Ctps2 |
| Rictor1 | 8.86E-110 | 0.357646857 | 0.378 | 0.145 | 1.79E-105 | 4 | Rictor |
| Ssr4 | 1.04E-109 | 0.27132924 | 0.827 | 0.447 | 2.10E-105 | 4 | Ssr4 |
| Rsbn11 | 1.38E-109 | 0.313672198 | 0.415 | 0.164 | 2.79E-105 | 4 | Rsbn1 |

| | | | | | | | |
|---|---|---|---|---|---|---|---|
| Mtmr31 | 2.51E-109 | 0.388875283 | 0.445 | 0.187 | 5.08E-105 | 4 | Mtmr3 |
| Kdm2b | 2.51E-109 | 0.285545396 | 0.258 | 0.08 | 5.08E-105 | 4 | Kdm2b |
| Tmem243 | 8.05E-109 | 0.283658607 | 0.39 | 0.15 | 1.63E-104 | 4 | Tmem243 |
| Vgll42 | 8.20E-109 | 0.381675093 | 0.526 | 0.235 | 1.66E-104 | 4 | Vgll4 |
| Ttc142 | 9.73E-109 | 0.348555899 | 0.474 | 0.201 | 1.97E-104 | 4 | Ttc14 |
| Ifngr12 | 1.20E-108 | 0.402144039 | 0.747 | 0.373 | 2.42E-104 | 4 | Ifngr1 |
| Cobll1 | 1.68E-108 | 0.269563855 | 0.252 | 0.076 | 3.39E-104 | 4 | Cobll1 |
| Rnf1691 | 1.90E-108 | 0.369805922 | 0.421 | 0.17 | 3.85E-104 | 4 | Rnf169 |
| Tgfb12 | 2.84E-108 | 0.386372564 | 0.503 | 0.221 | 5.74E-104 | 4 | Tgfb1 |
| Sdccag8 | 4.90E-108 | 0.292224667 | 0.317 | 0.111 | 9.91E-104 | 4 | Sdccag8 |
| Serbp12 | 7.51E-108 | 0.348532977 | 0.923 | 0.571 | 1.52E-103 | 4 | Serbp1 |
| Prrc2c2 | 1.06E-107 | 0.360924813 | 0.784 | 0.4 | 2.15E-103 | 4 | Prrc2c |
| Stard3nl | 1.09E-107 | 0.253950608 | 0.368 | 0.139 | 2.21E-103 | 4 | Stard3nl |
| Pum21 | 1.90E-107 | 0.296020578 | 0.56 | 0.253 | 3.85E-103 | 4 | Pum2 |
| Aatf | 3.90E-107 | 0.276499445 | 0.287 | 0.095 | 7.91E-103 | 4 | Aatf |
| Rap1gds11 | 6.48E-107 | 0.390275957 | 0.542 | 0.245 | 1.31E-102 | 4 | Rap1gds1 |
| Sfpq1 | 6.52E-107 | 0.329917482 | 0.606 | 0.281 | 1.32E-102 | 4 | Sfpq |
| Ncoa1 | 1.56E-106 | 0.369414735 | 0.463 | 0.196 | 3.15E-102 | 4 | Ncoa1 |
| Ran1 | 2.21E-106 | 0.281833824 | 0.738 | 0.381 | 4.47E-102 | 4 | Ran |
| Cdkn1b1 | 2.46E-106 | 0.252758475 | 0.477 | 0.2 | 4.99E-102 | 4 | Cdkn1b |
| Ubn22 | 7.24E-106 | 0.382408409 | 0.514 | 0.23 | 1.47E-101 | 4 | Ubn2 |
| Pbx3 | 1.34E-105 | 0.359908348 | 0.346 | 0.128 | 2.72E-101 | 4 | Pbx3 |
| Zufsp | 1.52E-105 | 0.322377566 | 0.34 | 0.125 | 3.07E-101 | 4 | Zufsp |
| Stxbp5 | 2.05E-105 | 0.330062008 | 0.328 | 0.119 | 4.15E-101 | 4 | Stxbp5 |
| Ikbkb | 5.85E-105 | 0.291679924 | 0.354 | 0.133 | 1.18E-100 | 4 | Ikbkb |
| Phf20l12 | 1.13E-104 | 0.281461232 | 0.59 | 0.27 | 2.29E-100 | 4 | Phf20l1 |
| Mllt102 | 1.15E-104 | 0.337091374 | 0.563 | 0.259 | 2.34E-100 | 4 | Mllt10 |
| Pak21 | 1.57E-104 | 0.296737561 | 0.798 | 0.414 | 3.17E-100 | 4 | Pak2 |
| mt-Nd12 | 1.59E-104 | 0.41734239 | 0.969 | 0.577 | 3.22E-100 | 4 | mt-Nd1 |
| Rere1 | 1.97E-104 | 0.306631049 | 0.562 | 0.255 | 3.98E-100 | 4 | Rere |
| Scai | 2.15E-104 | 0.324723332 | 0.279 | 0.092 | 4.35E-100 | 4 | Scai |
| Ythdf3 | 3.51E-104 | 0.29474411 | 0.438 | 0.182 | 7.10E-100 | 4 | Ythdf3 |
| Trim12a1 | 7.56E-104 | 0.298190738 | 0.416 | 0.169 | 1.53E-99 | 4 | Trim12a |
| Cdkal11 | 8.84E-104 | 0.346433623 | 0.412 | 0.168 | 1.79E-99 | 4 | Cdkal1 |
| Snx29 | 1.53E-103 | 0.356609212 | 0.294 | 0.103 | 3.09E-99 | 4 | Snx29 |
| Osbpl81 | 2.10E-103 | 0.322318883 | 0.444 | 0.186 | 4.26E-99 | 4 | Osbpl8 |
| Tbc1d9b | 6.72E-103 | 0.298602411 | 0.312 | 0.112 | 1.36E-98 | 4 | Tbc1d9b |
| Trim33 | 9.12E-103 | 0.259077303 | 0.383 | 0.15 | 1.85E-98 | 4 | Trim33 |
| 4930523C07Rik2 | 1.01E-102 | 0.270089284 | 0.486 | 0.205 | 2.05E-98 | 4 | 4930523C07Rik |
| Fam129a | 1.60E-102 | 0.287046036 | 0.355 | 0.134 | 3.24E-98 | 4 | Fam129a |
| Ube2w | 1.78E-102 | 0.262270273 | 0.396 | 0.159 | 3.61E-98 | 4 | Ube2w |
| Mapk14 | 2.16E-102 | 0.260633019 | 0.41 | 0.166 | 4.38E-98 | 4 | Mapk14 |
| Rock12 | 2.47E-102 | 0.321754982 | 0.801 | 0.412 | 4.99E-98 | 4 | Rock1 |

| | | | | | | | |
|---|---|---|---|---|---|---|---|
| Copg2 | 3.24E-102 | 0.273610397 | 0.304 | 0.107 | 6.56E-98 | 4 | Copg2 |
| Hnrnpa0 | 3.29E-102 | 0.27090332 | 0.866 | 0.472 | 6.67E-98 | 4 | Hnrnpa0 |
| Zfp4071 | 5.61E-102 | 0.337866842 | 0.379 | 0.15 | 1.14E-97 | 4 | Zfp407 |
| Traf5 | 7.02E-102 | 0.285390502 | 0.307 | 0.108 | 1.42E-97 | 4 | Traf5 |
| Arih2 | 2.16E-101 | 0.269090396 | 0.289 | 0.099 | 4.37E-97 | 4 | Arih2 |
| Lmtk2 | 2.28E-101 | 0.274349776 | 0.256 | 0.082 | 4.61E-97 | 4 | Lmtk2 |
| Dlg1 | 2.35E-101 | 0.295551484 | 0.46 | 0.198 | 4.76E-97 | 4 | Dlg1 |
| Xpa | 2.75E-101 | 0.252082713 | 0.314 | 0.113 | 5.57E-97 | 4 | Xpa |
| Rplp02 | 3.87E-101 | 0.311057978 | 0.997 | 0.942 | 7.83E-97 | 4 | Rplp0 |
| Kdm3b | 4.89E-101 | 0.317947533 | 0.334 | 0.125 | 9.91E-97 | 4 | Kdm3b |
| Syncrip | 7.49E-101 | 0.275798275 | 0.467 | 0.202 | 1.52E-96 | 4 | Syncrip |
| Cnot21 | 1.30E-100 | 0.310343702 | 0.459 | 0.196 | 2.64E-96 | 4 | Cnot2 |
| Rnf2 | 1.46E-100 | 0.281508613 | 0.281 | 0.096 | 2.96E-96 | 4 | Rnf2 |
| Asb3 | 2.58E-100 | 0.323941954 | 0.289 | 0.1 | 5.23E-96 | 4 | Asb3 |
| Eif4a1 | 4.50E-100 | 0.343514833 | 0.76 | 0.412 | 9.11E-96 | 4 | Eif4a1 |
| Ostf12 | 5.25E-100 | 0.301921557 | 0.872 | 0.486 | 1.06E-95 | 4 | Ostf1 |
| Nop56 | 5.32E-100 | 0.285663981 | 0.327 | 0.121 | 1.08E-95 | 4 | Nop56 |
| Grk2 | 5.44E-100 | 0.271268339 | 0.314 | 0.114 | 1.10E-95 | 4 | Grk2 |
| Itsn22 | 9.12E-100 | 0.346338764 | 0.563 | 0.262 | 1.85E-95 | 4 | Itsn2 |
| Tbc1d1 | 1.45E-99 | 0.277909068 | 0.384 | 0.153 | 2.94E-95 | 4 | Tbc1d1 |
| Pam1 | 3.13E-99 | 0.396268669 | 0.458 | 0.2 | 6.34E-95 | 4 | Pam |
| Brd12 | 4.00E-99 | 0.28756681 | 0.503 | 0.226 | 8.09E-95 | 4 | Brd1 |
| Wdr82 | 4.24E-99 | 0.265011025 | 0.305 | 0.109 | 8.59E-95 | 4 | Wdr82 |
| Pcna | 4.44E-99 | 0.250940991 | 0.31 | 0.112 | 8.98E-95 | 4 | Pcna |
| Pcca | 5.67E-99 | 0.268426018 | 0.307 | 0.111 | 1.15E-94 | 4 | Pcca |
| Wdr331 | 9.91E-99 | 0.275962045 | 0.502 | 0.225 | 2.01E-94 | 4 | Wdr33 |
| Mgea51 | 2.73E-98 | 0.28495871 | 0.389 | 0.157 | 5.53E-94 | 4 | Mgea5 |
| Pds5a1 | 4.06E-98 | 0.286316276 | 0.447 | 0.191 | 8.22E-94 | 4 | Pds5a |
| Nufip21 | 4.26E-98 | 0.298977713 | 0.489 | 0.218 | 8.62E-94 | 4 | Nufip2 |
| Pum12 | 4.66E-98 | 0.263340781 | 0.539 | 0.243 | 9.44E-94 | 4 | Pum1 |
| Ppp4r3b1 | 9.27E-98 | 0.254152414 | 0.496 | 0.217 | 1.88E-93 | 4 | Ppp4r3b |
| Abcb7 | 9.76E-98 | 0.296673414 | 0.29 | 0.102 | 1.98E-93 | 4 | Abcb7 |
| Map2k4 | 1.15E-97 | 0.27595233 | 0.371 | 0.147 | 2.32E-93 | 4 | Map2k4 |
| Kdm5b | 1.21E-97 | 0.283090435 | 0.323 | 0.12 | 2.46E-93 | 4 | Kdm5b |
| Xpo7 | 1.41E-97 | 0.283272928 | 0.362 | 0.142 | 2.85E-93 | 4 | Xpo7 |
| Fnip1 | 3.88E-97 | 0.309035676 | 0.347 | 0.135 | 7.85E-93 | 4 | Fnip1 |
| Helz1 | 4.78E-97 | 0.292263313 | 0.389 | 0.157 | 9.69E-93 | 4 | Helz |
| Gigyf21 | 5.89E-97 | 0.278620297 | 0.385 | 0.156 | 1.19E-92 | 4 | Gigyf2 |
| Lrif1 | 9.53E-97 | 0.259413437 | 0.338 | 0.128 | 1.93E-92 | 4 | Lrif1 |
| Son2 | 1.02E-96 | 0.327804286 | 0.927 | 0.503 | 2.06E-92 | 4 | Son |
| Lgals3bp | 2.50E-96 | 0.30919256 | 0.349 | 0.136 | 5.07E-92 | 4 | Lgals3bp |
| Stx16 | 2.67E-96 | 0.291584253 | 0.374 | 0.149 | 5.40E-92 | 4 | Stx16 |
| Spata132 | 3.44E-96 | 0.302575621 | 0.451 | 0.193 | 6.97E-92 | 4 | Spata13 |

| | | | | | | | |
|---|---|---|---|---|---|---|---|
| Naa15 | 4.02E-96 | 0.305260619 | 0.379 | 0.154 | 8.13E-92 | 4 | Naa15 |
| Ppp2r2a1 | 4.84E-96 | 0.291888176 | 0.457 | 0.2 | 9.79E-92 | 4 | Ppp2r2a |
| Rps252 | 8.06E-96 | 0.292193758 | 0.99 | 0.821 | 1.63E-91 | 4 | Rps25 |
| Fam13b | 3.34E-95 | 0.285524785 | 0.349 | 0.136 | 6.77E-91 | 4 | Fam13b |
| M6pr | 3.88E-95 | 0.287109809 | 0.372 | 0.151 | 7.86E-91 | 4 | M6pr |
| Dnajc32 | 5.92E-95 | 0.311199223 | 0.695 | 0.352 | 1.20E-90 | 4 | Dnajc3 |
| AI3141801 | 1.87E-94 | 0.282895932 | 0.449 | 0.197 | 3.78E-90 | 4 | AI314180 |
| Cfl1 | 7.38E-94 | 0.368795642 | 0.947 | 0.589 | 1.49E-89 | 4 | Cfl1 |
| Klhl6 | 1.10E-93 | 0.292556993 | 0.303 | 0.111 | 2.23E-89 | 4 | Klhl6 |
| mt-Nd32 | 2.53E-93 | 0.286064621 | 0.808 | 0.426 | 5.13E-89 | 4 | mt-Nd3 |
| Akap10 | 3.64E-93 | 0.32331539 | 0.342 | 0.135 | 7.36E-89 | 4 | Akap10 |
| Zfand31 | 5.75E-93 | 0.336650764 | 0.584 | 0.286 | 1.16E-88 | 4 | Zfand3 |
| Ccdc821 | 6.36E-93 | 0.254858678 | 0.406 | 0.168 | 1.29E-88 | 4 | Ccdc82 |
| Ncoa4 | 1.48E-92 | 0.267467198 | 0.266 | 0.093 | 2.99E-88 | 4 | Ncoa4 |
| Ppp1r101 | 2.36E-92 | 0.295634567 | 0.416 | 0.178 | 4.79E-88 | 4 | Ppp1r10 |
| Zzef11 | 3.59E-92 | 0.321637412 | 0.346 | 0.138 | 7.26E-88 | 4 | Zzef1 |
| Pign | 4.69E-92 | 0.284574699 | 0.253 | 0.086 | 9.50E-88 | 4 | Pign |
| Tcerg1 | 5.67E-92 | 0.290645779 | 0.326 | 0.126 | 1.15E-87 | 4 | Tcerg1 |
| Papd5 | 6.45E-92 | 0.257213989 | 0.298 | 0.109 | 1.31E-87 | 4 | Papd5 |
| Neat11 | 1.96E-91 | 0.295942998 | 0.443 | 0.196 | 3.97E-87 | 4 | Neat1 |
| Map3k2 | 4.31E-91 | 0.312648898 | 0.354 | 0.143 | 8.73E-87 | 4 | Map3k2 |
| Pias1 | 4.39E-91 | 0.295549792 | 0.412 | 0.176 | 8.88E-87 | 4 | Pias1 |
| Trip111 | 5.75E-91 | 0.264352731 | 0.468 | 0.21 | 1.16E-86 | 4 | Trip11 |
| Dmxl1 | 7.68E-91 | 0.274436032 | 0.407 | 0.173 | 1.56E-86 | 4 | Dmxl1 |
| Rpl7a2 | 9.08E-91 | 0.28563542 | 0.984 | 0.739 | 1.84E-86 | 4 | Rpl7a |
| Dmtf1 | 1.18E-90 | 0.251753523 | 0.302 | 0.112 | 2.40E-86 | 4 | Dmtf1 |
| Pdss2 | 1.24E-90 | 0.250169519 | 0.263 | 0.091 | 2.51E-86 | 4 | Pdss2 |
| Psen1 | 1.44E-90 | 0.271323755 | 0.263 | 0.092 | 2.92E-86 | 4 | Psen1 |
| Zfr2 | 2.09E-90 | 0.296559745 | 0.504 | 0.233 | 4.23E-86 | 4 | Zfr |
| Sos21 | 3.57E-90 | 0.272351289 | 0.367 | 0.15 | 7.23E-86 | 4 | Sos2 |
| Taf31 | 4.22E-90 | 0.321854774 | 0.341 | 0.136 | 8.54E-86 | 4 | Taf3 |
| Zdhhc181 | 6.13E-90 | 0.266985477 | 0.387 | 0.159 | 1.24E-85 | 4 | Zdhhc18 |
| Lsm14a | 1.11E-89 | 0.251125962 | 0.37 | 0.152 | 2.25E-85 | 4 | Lsm14a |
| Pear1 | 3.17E-89 | 0.26401463 | 0.266 | 0.093 | 6.42E-85 | 4 | Pear1 |
| Tox41 | 4.94E-89 | 0.251264088 | 0.404 | 0.172 | 1.00E-84 | 4 | Tox4 |
| Thoc22 | 6.30E-89 | 0.260047612 | 0.546 | 0.259 | 1.28E-84 | 4 | Thoc2 |
| Psme12 | 6.85E-89 | 0.28727376 | 0.921 | 0.548 | 1.39E-84 | 4 | Psme1 |
| Cdk122 | 1.27E-88 | 0.326390967 | 0.474 | 0.216 | 2.57E-84 | 4 | Cdk12 |
| Zfp398 | 2.55E-88 | 0.260557574 | 0.25 | 0.086 | 5.16E-84 | 4 | Zfp398 |
| 2610035D17Rik | 2.96E-88 | 0.283561556 | 0.287 | 0.105 | 5.99E-84 | 4 | 2610035D17Rik |
| Jmjd1c2 | 3.14E-88 | 0.317241546 | 0.596 | 0.291 | 6.36E-84 | 4 | Jmjd1c |
| Xrn11 | 3.23E-88 | 0.286632526 | 0.433 | 0.191 | 6.53E-84 | 4 | Xrn1 |
| Tnfrsf1b | 4.43E-88 | 0.263622563 | 0.291 | 0.107 | 8.98E-84 | 4 | Tnfrsf1b |

| | | | | | | | |
|---|---|---|---|---|---|---|---|
| Abcg1 | 5.06E-88 | 0.287067807 | 0.301 | 0.114 | 1.02E-83 | 4 | Abcg1 |
| Lrp101 | 5.62E-88 | 0.296035598 | 0.407 | 0.176 | 1.14E-83 | 4 | Lrp10 |
| Crebbp2 | 6.87E-88 | 0.287659042 | 0.593 | 0.289 | 1.39E-83 | 4 | Crebbp |
| Ep3001 | 8.07E-88 | 0.296920988 | 0.432 | 0.192 | 1.63E-83 | 4 | Ep300 |
| Bbs9 | 1.06E-87 | 0.361129215 | 0.262 | 0.093 | 2.15E-83 | 4 | Bbs9 |
| Eif5a2 | 3.60E-87 | 0.334696368 | 0.874 | 0.548 | 7.28E-83 | 4 | Eif5a |
| Atp9b | 3.98E-87 | 0.275147165 | 0.356 | 0.147 | 8.06E-83 | 4 | Atp9b |
| Irf22 | 1.23E-86 | 0.268475609 | 0.638 | 0.325 | 2.49E-82 | 4 | Irf2 |
| Fbxo38 | 2.43E-86 | 0.255750062 | 0.29 | 0.109 | 4.91E-82 | 4 | Fbxo38 |
| Sf3b31 | 6.35E-86 | 0.278921431 | 0.361 | 0.15 | 1.29E-81 | 4 | Sf3b3 |
| Klf122 | 8.99E-86 | 0.313621769 | 0.44 | 0.195 | 1.82E-81 | 4 | Klf12 |
| Slc3a22 | 1.07E-84 | 0.339005204 | 0.553 | 0.276 | 2.16E-80 | 4 | Slc3a2 |
| Baz2a | 3.47E-84 | 0.265306143 | 0.312 | 0.122 | 7.02E-80 | 4 | Baz2a |
| Kidins2201 | 7.03E-83 | 0.276343709 | 0.35 | 0.145 | 1.42E-78 | 4 | Kidins220 |
| Arhgap10 | 1.36E-82 | 0.260671686 | 0.303 | 0.118 | 2.76E-78 | 4 | Arhgap10 |
| Gpd21 | 1.88E-82 | 0.257746863 | 0.288 | 0.109 | 3.81E-78 | 4 | Gpd2 |
| Traf3 | 1.28E-81 | 0.309149544 | 0.268 | 0.101 | 2.60E-77 | 4 | Traf3 |
| mt-Co22 | 1.46E-81 | 0.419630406 | 1 | 0.713 | 2.96E-77 | 4 | mt-Co2 |
| Bach21 | 2.77E-81 | 0.327854762 | 0.354 | 0.151 | 5.61E-77 | 4 | Bach2 |
| Slmap1 | 4.81E-81 | 0.261787107 | 0.407 | 0.181 | 9.73E-77 | 4 | Slmap |
| Ube3c1 | 5.07E-81 | 0.271411194 | 0.353 | 0.148 | 1.03E-76 | 4 | Ube3c |
| Stat32 | 3.13E-80 | 0.276385415 | 0.523 | 0.253 | 6.33E-76 | 4 | Stat3 |
| Rps151 | 4.08E-80 | 0.292294419 | 0.985 | 0.735 | 8.27E-76 | 4 | Rps15 |
| Malat12 | 5.26E-80 | 0.382025509 | 1 | 0.71 | 1.07E-75 | 4 | Malat1 |
| Kansl1l | 5.87E-80 | 0.280270372 | 0.319 | 0.129 | 1.19E-75 | 4 | Kansl1l |
| Ddx52 | 1.07E-79 | 0.372853443 | 0.974 | 0.636 | 2.17E-75 | 4 | Ddx5 |
| Arid5b2 | 1.08E-79 | 0.33232413 | 0.529 | 0.264 | 2.19E-75 | 4 | Arid5b |
| Stk101 | 1.41E-79 | 0.273665634 | 0.391 | 0.172 | 2.85E-75 | 4 | Stk10 |
| Nrf1 | 2.16E-79 | 0.261334701 | 0.312 | 0.126 | 4.37E-75 | 4 | Nrf1 |
| Rnf1111 | 3.41E-79 | 0.253861819 | 0.347 | 0.146 | 6.91E-75 | 4 | Rnf111 |
| mt-Atp62 | 8.66E-79 | 0.404766462 | 1 | 0.74 | 1.75E-74 | 4 | mt-Atp6 |
| Prpf38b1 | 1.16E-78 | 0.262392545 | 0.408 | 0.185 | 2.34E-74 | 4 | Prpf38b |
| Prkd31 | 1.74E-78 | 0.267697393 | 0.363 | 0.157 | 3.52E-74 | 4 | Prkd3 |
| Pibf1 | 2.58E-78 | 0.260394999 | 0.306 | 0.123 | 5.23E-74 | 4 | Pibf1 |
| Wwox1 | 6.92E-78 | 0.396456694 | 0.371 | 0.165 | 1.40E-73 | 4 | Wwox |
| Trpm71 | 4.70E-77 | 0.254252234 | 0.42 | 0.192 | 9.51E-73 | 4 | Trpm7 |
| Map3k14 | 2.44E-76 | 0.265590033 | 0.259 | 0.098 | 4.95E-72 | 4 | Map3k14 |
| Arhgef32 | 7.56E-76 | 0.256930837 | 0.479 | 0.228 | 1.53E-71 | 4 | Arhgef3 |
| Bmt2 | 2.00E-75 | 0.261550215 | 0.282 | 0.112 | 4.06E-71 | 4 | Bmt2 |
| Ppp6r31 | 1.08E-74 | 0.251324783 | 0.396 | 0.18 | 2.19E-70 | 4 | Ppp6r3 |
| mt-Nd22 | 4.05E-73 | 0.343715636 | 0.997 | 0.671 | 8.21E-69 | 4 | mt-Nd2 |
| Ifnar1 | 9.09E-73 | 0.251723321 | 0.319 | 0.135 | 1.84E-68 | 4 | Ifnar1 |
| Ski1 | 4.20E-72 | 0.257200713 | 0.429 | 0.203 | 8.51E-68 | 4 | Ski |

| | | | | | | | |
|---|---|---|---|---|---|---|---|
| Cep3501 | 9.95E-72 | 0.254443029 | 0.412 | 0.192 | 2.01E-67 | 4 | Cep350 |
| H2-D12 | 8.78E-71 | 0.358736847 | 0.999 | 0.65 | 1.78E-66 | 4 | H2-D1 |
| Tsc22d21 | 1.87E-70 | 0.263876255 | 0.331 | 0.144 | 3.78E-66 | 4 | Tsc22d2 |
| Acap22 | 2.09E-67 | 0.277838682 | 0.503 | 0.257 | 4.22E-63 | 4 | Acap2 |
| Fchsd2 | 8.63E-67 | 0.250140878 | 0.335 | 0.151 | 1.75E-62 | 4 | Fchsd2 |
| Arhgap21 | 1.03E-66 | 0.254349879 | 0.256 | 0.103 | 2.08E-62 | 4 | Arhgap21 |
| Tmem164 | 1.08E-66 | 0.266712181 | 0.342 | 0.154 | 2.20E-62 | 4 | Tmem164 |
| Srgap22 | 7.20E-64 | 0.25860467 | 0.554 | 0.29 | 1.46E-59 | 4 | Srgap2 |
| Myl61 | 1.35E-62 | 0.280693361 | 0.994 | 0.764 | 2.74E-58 | 4 | Myl6 |
| Abl2 | 1.61E-60 | 0.254876299 | 0.263 | 0.112 | 3.25E-56 | 4 | Abl2 |
| mt-Cytb2 | 9.54E-56 | 0.27688031 | 1 | 0.683 | 1.93E-51 | 4 | mt-Cytb |
| B2m2 | 3.87E-48 | 0.277438729 | 0.997 | 0.673 | 7.83E-44 | 4 | B2m |
| Cd74 | 0 | 3.597029173 | 0.98 | 0.21 | 0 | 5 | Cd74 |
| H2-Eb1 | 0 | 3.577957793 | 0.969 | 0.151 | 0 | 5 | H2-Eb1 |
| H2-Aa | 0 | 3.506526286 | 0.956 | 0.144 | 0 | 5 | H2-Aa |
| C1qb | 0 | 3.500983798 | 0.925 | 0.111 | 0 | 5 | C1qb |
| H2-Ab1 | 0 | 3.364326196 | 0.909 | 0.162 | 0 | 5 | H2-Ab1 |
| C1qa | 0 | 3.354803534 | 0.906 | 0.104 | 0 | 5 | C1qa |
| Apoe | 0 | 3.300022945 | 0.949 | 0.512 | 0 | 5 | Apoe |
| Ctss | 0 | 3.273796311 | 0.841 | 0.128 | 0 | 5 | Ctss |
| Aif1 | 0 | 3.242983946 | 0.747 | 0.086 | 0 | 5 | Aif1 |
| C1qc | 0 | 3.121824789 | 0.837 | 0.095 | 0 | 5 | C1qc |
| Lyz2 | 0 | 3.034709473 | 0.609 | 0.084 | 0 | 5 | Lyz2 |
| Cst3 | 0 | 2.851157233 | 0.968 | 0.583 | 0 | 5 | Cst3 |
| Tyrobp1 | 0 | 2.676277878 | 0.914 | 0.226 | 0 | 5 | Tyrobp |
| Fth11 | 0 | 2.638502022 | 0.998 | 0.968 | 0 | 5 | Fth1 |
| Ms4a7 | 0 | 2.636820358 | 0.539 | 0.075 | 0 | 5 | Ms4a7 |
| Rgs10 | 0 | 2.566420147 | 0.599 | 0.137 | 0 | 5 | Rgs10 |
| Lst1 | 0 | 2.448622748 | 0.498 | 0.09 | 0 | 5 | Lst1 |
| Wfdc17 | 0 | 2.341897226 | 0.385 | 0.051 | 0 | 5 | Wfdc17 |
| Bcl2a1b | 0 | 2.313837724 | 0.449 | 0.088 | 0 | 5 | Bcl2a1b |
| Fcer1g1 | 0 | 2.299020721 | 0.881 | 0.236 | 0 | 5 | Fcer1g |
| Ftl12 | 0 | 2.215085072 | 0.993 | 0.907 | 0 | 5 | Ftl1 |
| Cd300c2 | 0 | 2.084050104 | 0.407 | 0.068 | 0 | 5 | Cd300c2 |
| H2-DMa | 0 | 2.078218006 | 0.431 | 0.104 | 0 | 5 | H2-DMa |
| Cd521 | 0 | 2.014651085 | 0.883 | 0.249 | 0 | 5 | Cd52 |
| Tmsb4x1 | 0 | 2.000069572 | 1 | 0.931 | 0 | 5 | Tmsb4x |
| Ctsh | 0 | 1.939195632 | 0.47 | 0.119 | 0 | 5 | Ctsh |
| Ly86 | 0 | 1.936303186 | 0.436 | 0.079 | 0 | 5 | Ly86 |
| Rps291 | 0 | 1.390275746 | 0.994 | 0.97 | 0 | 5 | Rps29 |
| Fau1 | 0 | 1.361921258 | 0.986 | 0.951 | 0 | 5 | Fau |
| Rpl35a2 | 6.30E-307 | 1.060916639 | 0.981 | 0.947 | 1.28E-302 | 5 | Rpl35a |
| Rps27a2 | 2.96E-306 | 1.046651576 | 0.983 | 0.949 | 5.99E-302 | 5 | Rps27a |

| | | | | | | | |
|---|---|---|---|---|---|---|---|
| Cd63 | 2.69E-301 | 2.01121864 | 0.535 | 0.17 | 5.44E-297 | 5 | Cd63 |
| Cd68 | 3.81E-298 | 1.809873041 | 0.335 | 0.066 | 7.71E-294 | 5 | Cd68 |
| Rps241 | 1.05E-289 | 0.968171416 | 0.996 | 0.971 | 2.13E-285 | 5 | Rps24 |
| Rps282 | 1.32E-289 | 1.132034035 | 0.965 | 0.894 | 2.66E-285 | 5 | Rps28 |
| Alox5ap | 5.11E-283 | 1.759010679 | 0.314 | 0.061 | 1.03E-278 | 5 | Alox5ap |
| Spi1 | 2.29E-263 | 1.763322341 | 0.328 | 0.07 | 4.63E-259 | 5 | Spi1 |
| Rps92 | 3.82E-261 | 1.064953995 | 0.962 | 0.919 | 7.73E-257 | 5 | Rps9 |
| Npc2 | 7.59E-253 | 2.145521895 | 0.673 | 0.401 | 1.54E-248 | 5 | Npc2 |
| H2-DMb1 | 1.29E-251 | 1.704754157 | 0.343 | 0.079 | 2.62E-247 | 5 | H2-DMb1 |
| Rpl322 | 7.05E-250 | 0.960108314 | 0.979 | 0.952 | 1.43E-245 | 5 | Rpl32 |
| Sh3bgrl31 | 1.27E-249 | 1.610017223 | 0.792 | 0.548 | 2.56E-245 | 5 | Sh3bgrl3 |
| Rps4x2 | 1.13E-243 | 0.961082925 | 0.968 | 0.94 | 2.28E-239 | 5 | Rps4x |
| Rps132 | 1.57E-233 | 0.933241743 | 0.963 | 0.922 | 3.19E-229 | 5 | Rps13 |
| Rpl302 | 2.95E-230 | 0.953956919 | 0.964 | 0.937 | 5.97E-226 | 5 | Rpl30 |
| Rps201 | 4.98E-226 | 0.881720716 | 0.982 | 0.95 | 1.01E-221 | 5 | Rps20 |
| Trem2 | 3.17E-225 | 1.642190008 | 0.275 | 0.056 | 6.41E-221 | 5 | Trem2 |
| Rpl37a1 | 1.88E-216 | 0.88231898 | 0.973 | 0.936 | 3.80E-212 | 5 | Rpl37a |
| Rps122 | 2.07E-215 | 1.009089353 | 0.943 | 0.896 | 4.19E-211 | 5 | Rps12 |
| Cyba1 | 2.24E-211 | 1.867593115 | 0.679 | 0.429 | 4.53E-207 | 5 | Cyba |
| Rpl392 | 8.33E-206 | 1.000997634 | 0.939 | 0.899 | 1.69E-201 | 5 | Rpl39 |
| Rpl113 | 9.16E-203 | 0.907170448 | 0.955 | 0.926 | 1.85E-198 | 5 | Rpl11 |
| Rpl27a2 | 5.24E-199 | 0.806804818 | 0.98 | 0.959 | 1.06E-194 | 5 | Rpl27a |
| Rps263 | 2.31E-198 | 1.046055234 | 0.907 | 0.864 | 4.67E-194 | 5 | Rps26 |
| Rps52 | 1.99E-197 | 0.888212345 | 0.956 | 0.918 | 4.04E-193 | 5 | Rps5 |
| Rps232 | 2.00E-195 | 0.771834136 | 0.98 | 0.951 | 4.04E-191 | 5 | Rps23 |
| Rpl103 | 6.29E-191 | 0.905596598 | 0.927 | 0.896 | 1.27E-186 | 5 | Rpl10 |
| Slamf9 | 1.68E-187 | 1.445976254 | 0.265 | 0.061 | 3.41E-183 | 5 | Slamf9 |
| Basp1 | 3.82E-187 | 1.548894064 | 0.333 | 0.093 | 7.73E-183 | 5 | Basp1 |
| Ckb | 1.77E-184 | 1.894069466 | 0.472 | 0.203 | 3.58E-180 | 5 | Ckb |
| Cd72 | 3.82E-184 | 1.725164557 | 0.325 | 0.094 | 7.73E-180 | 5 | Cd72 |
| Ifi27l2a2 | 4.28E-184 | 1.142817286 | 0.666 | 0.352 | 8.67E-180 | 5 | Ifi27l2a |
| Rps111 | 5.93E-183 | 0.855993597 | 0.949 | 0.905 | 1.20E-178 | 5 | Rps11 |
| Rpl18a1 | 7.46E-183 | 0.818961098 | 0.962 | 0.929 | 1.51E-178 | 5 | Rpl18a |
| Rps161 | 1.99E-178 | 0.797688679 | 0.958 | 0.926 | 4.03E-174 | 5 | Rps16 |
| Rps181 | 2.90E-177 | 0.782079148 | 0.97 | 0.94 | 5.88E-173 | 5 | Rps18 |
| Rpl132 | 2.83E-173 | 0.681851435 | 0.983 | 0.977 | 5.73E-169 | 5 | Rpl13 |
| Rpl233 | 2.46E-172 | 0.718075554 | 0.981 | 0.97 | 4.98E-168 | 5 | Rpl23 |
| Rpl262 | 3.96E-172 | 0.73906376 | 0.973 | 0.956 | 8.02E-168 | 5 | Rpl26 |
| Rps102 | 7.66E-169 | 0.726351864 | 0.972 | 0.943 | 1.55E-164 | 5 | Rps10 |
| Ms4a6c | 7.94E-163 | 1.562539855 | 0.276 | 0.074 | 1.61E-158 | 5 | Ms4a6c |
| Rpl341 | 7.65E-161 | 0.802674305 | 0.934 | 0.917 | 1.55E-156 | 5 | Rpl34 |
| Tpt11 | 2.73E-160 | 0.592189544 | 0.995 | 0.99 | 5.52E-156 | 5 | Tpt1 |
| Cxcl16 | 1.04E-159 | 1.609383532 | 0.323 | 0.102 | 2.11E-155 | 5 | Cxcl16 |

| | | | | | | | |
|---|---|---|---|---|---|---|---|
| Hpgd | 7.14E-159 | 1.470731848 | 0.284 | 0.078 | 1.45E-154 | 5 | Hpgd |
| Hexb | 1.44E-158 | 1.621188112 | 0.686 | 0.51 | 2.92E-154 | 5 | Hexb |
| Rpl353 | 1.31E-155 | 0.949694091 | 0.867 | 0.827 | 2.66E-151 | 5 | Rpl35 |
| Ctsz | 2.47E-152 | 1.878312015 | 0.567 | 0.35 | 5.01E-148 | 5 | Ctsz |
| Rpl192 | 1.01E-151 | 0.663260243 | 0.972 | 0.963 | 2.05E-147 | 5 | Rpl19 |
| Rps15a1 | 8.11E-146 | 0.69211073 | 0.956 | 0.923 | 1.64E-141 | 5 | Rps15a |
| Lgmn | 1.46E-144 | 1.500698843 | 0.317 | 0.104 | 2.97E-140 | 5 | Lgmn |
| Rplp12 | 1.70E-143 | 0.572419708 | 0.981 | 0.983 | 3.44E-139 | 5 | Rplp1 |
| Rpl122 | 2.75E-142 | 0.857078955 | 0.894 | 0.839 | 5.57E-138 | 5 | Rpl12 |
| Rpl373 | 6.58E-139 | 0.730632001 | 0.936 | 0.927 | 1.33E-134 | 5 | Rpl37 |
| Rplp23 | 1.86E-138 | 0.7913001 | 0.916 | 0.889 | 3.77E-134 | 5 | Rplp2 |
| Rpl222 | 4.05E-135 | 0.839784586 | 0.879 | 0.852 | 8.21E-131 | 5 | Rpl22 |
| Ntpcr | 8.25E-134 | 1.532389429 | 0.29 | 0.095 | 1.67E-129 | 5 | Ntpcr |
| Bst22 | 4.86E-133 | 1.366257391 | 0.681 | 0.495 | 9.84E-129 | 5 | Bst2 |
| Ptpn181 | 5.62E-131 | 1.499992675 | 0.534 | 0.305 | 1.14E-126 | 5 | Ptpn18 |
| Rpl382 | 1.49E-130 | 0.749542948 | 0.929 | 0.892 | 3.01E-126 | 5 | Rpl38 |
| Rps212 | 1.74E-128 | 0.680849383 | 0.934 | 0.927 | 3.52E-124 | 5 | Rps21 |
| Rps253 | 2.94E-128 | 0.888542959 | 0.844 | 0.831 | 5.95E-124 | 5 | Rps25 |
| Rpl10a3 | 2.22E-126 | 0.826266791 | 0.873 | 0.835 | 4.49E-122 | 5 | Rpl10a |
| Rps72 | 2.48E-126 | 0.676088549 | 0.942 | 0.916 | 5.02E-122 | 5 | Rps7 |
| Rps141 | 1.62E-121 | 0.627876294 | 0.952 | 0.943 | 3.27E-117 | 5 | Rps14 |
| Coro1a1 | 1.02E-116 | 1.393455562 | 0.436 | 0.209 | 2.07E-112 | 5 | Coro1a |
| Rpl282 | 1.29E-115 | 0.626967939 | 0.934 | 0.931 | 2.61E-111 | 5 | Rpl28 |
| Rps31 | 2.91E-112 | 0.650688636 | 0.919 | 0.892 | 5.88E-108 | 5 | Rps3 |
| Rpl213 | 7.95E-112 | 0.633127549 | 0.938 | 0.923 | 1.61E-107 | 5 | Rpl21 |
| Prol1 | 1.19E-111 | 0.324164011 | 0.877 | 0.841 | 2.40E-107 | 5 | Prol1 |
| Rpl413 | 2.34E-111 | 0.547277944 | 0.962 | 0.962 | 4.73E-107 | 5 | Rpl41 |
| Actb1 | 1.56E-104 | 0.721991023 | 0.94 | 0.838 | 3.17E-100 | 5 | Actb |
| Rps3a13 | 8.53E-103 | 0.567843286 | 0.949 | 0.935 | 1.73E-98 | 5 | Rps3a1 |
| Rps83 | 1.79E-102 | 0.506164075 | 0.96 | 0.965 | 3.62E-98 | 5 | Rps8 |
| Rps271 | 7.26E-102 | 0.650269339 | 0.928 | 0.897 | 1.47E-97 | 5 | Rps27 |
| Ctsc | 2.44E-101 | 1.312129197 | 0.336 | 0.142 | 4.93E-97 | 5 | Ctsc |
| Rpl172 | 7.70E-100 | 0.639562996 | 0.913 | 0.917 | 1.56E-95 | 5 | Rpl17 |
| Rplp03 | 1.23E-98 | 0.53964116 | 0.937 | 0.946 | 2.49E-94 | 5 | Rplp0 |
| Rpl312 | 1.66E-95 | 0.606689897 | 0.911 | 0.91 | 3.37E-91 | 5 | Rpl3 |
| Eef1a12 | 2.20E-95 | 0.540806908 | 0.965 | 0.96 | 4.46E-91 | 5 | Eef1a1 |
| Rps191 | 1.84E-92 | 0.55155675 | 0.932 | 0.927 | 3.72E-88 | 5 | Rps19 |
| Atox1 | 8.51E-92 | 1.140279127 | 0.67 | 0.618 | 1.72E-87 | 5 | Atox1 |
| Rpl93 | 1.91E-88 | 0.501746975 | 0.957 | 0.944 | 3.86E-84 | 5 | Rpl9 |
| Pfn11 | 7.86E-88 | 0.919747705 | 0.744 | 0.68 | 1.59E-83 | 5 | Pfn1 |
| Rpl73 | 1.17E-85 | 0.628416031 | 0.882 | 0.895 | 2.38E-81 | 5 | Rpl7 |
| Rpl182 | 1.69E-84 | 0.588116816 | 0.905 | 0.901 | 3.43E-80 | 5 | Rpl18 |
| Rpsa2 | 8.95E-84 | 0.558155198 | 0.914 | 0.903 | 1.81E-79 | 5 | Rpsa |

| | | | | | | | |
|---|---|---|---|---|---|---|---|
| Serf22 | 6.15E-83 | 0.744015 | 0.793 | 0.822 | 1.25E-78 | 5 | Serf2 |
| Rpl243 | 1.79E-80 | 0.574221039 | 0.888 | 0.909 | 3.62E-76 | 5 | Rpl24 |
| Laptm51 | 3.38E-78 | 1.254801278 | 0.382 | 0.203 | 6.84E-74 | 5 | Laptm5 |
| Rack13 | 3.47E-76 | 0.731779962 | 0.796 | 0.794 | 7.02E-72 | 5 | Rack1 |
| Rpl63 | 4.30E-75 | 0.588039176 | 0.87 | 0.883 | 8.71E-71 | 5 | Rpl6 |
| Rps210 | 1.90E-72 | 0.480571237 | 0.915 | 0.921 | 3.84E-68 | 5 | Rps2 |
| Rpl362 | 2.80E-71 | 0.538061418 | 0.896 | 0.904 | 5.66E-67 | 5 | Rpl36 |
| Tmem176a2 | 2.38E-68 | 1.255457014 | 0.536 | 0.413 | 4.81E-64 | 5 | Tmem176a |
| Ctsb | 2.30E-63 | 1.482987761 | 0.425 | 0.293 | 4.65E-59 | 5 | Ctsb |
| Tmem176b2 | 6.34E-63 | 1.180092158 | 0.543 | 0.436 | 1.28E-58 | 5 | Tmem176b |
| Rpl152 | 1.79E-57 | 0.490181206 | 0.871 | 0.904 | 3.63E-53 | 5 | Rpl15 |
| Ucp22 | 2.57E-57 | 1.144916802 | 0.453 | 0.326 | 5.20E-53 | 5 | Ucp2 |
| Clic11 | 4.91E-54 | 0.938872482 | 0.616 | 0.602 | 9.94E-50 | 5 | Clic1 |
| Rpl291 | 1.80E-49 | 0.625451035 | 0.752 | 0.771 | 3.64E-45 | 5 | Rpl29 |
| Rpl7a3 | 1.01E-46 | 0.626535342 | 0.722 | 0.756 | 2.04E-42 | 5 | Rpl7a |
| Blvrb | 1.51E-46 | 1.285843006 | 0.355 | 0.239 | 3.06E-42 | 5 | Blvrb |
| Rpl36a3 | 2.99E-44 | 0.651532328 | 0.729 | 0.755 | 6.05E-40 | 5 | Rpl36a |
| Rpl82 | 3.87E-41 | 0.339414868 | 0.916 | 0.926 | 7.83E-37 | 5 | Rpl8 |
| Arhgdib1 | 9.72E-40 | 1.092180304 | 0.344 | 0.23 | 1.97E-35 | 5 | Arhgdib |
| B2m3 | 3.70E-39 | 0.596663657 | 0.775 | 0.688 | 7.49E-35 | 5 | B2m |
| Cotl11 | 2.56E-38 | 1.136952139 | 0.326 | 0.221 | 5.18E-34 | 5 | Cotl1 |
| Rps63 | 1.30E-37 | 0.465409631 | 0.776 | 0.815 | 2.63E-33 | 5 | Rps6 |
| Eef1b23 | 2.60E-37 | 0.614801991 | 0.713 | 0.739 | 5.26E-33 | 5 | Eef1b2 |
| Lgals32 | 1.27E-36 | 1.421663911 | 0.327 | 0.214 | 2.58E-32 | 5 | Lgals3 |
| AY0361181 | 7.12E-34 | 0.273162326 | 0.889 | 0.929 | 1.44E-29 | 5 | AY036118 |
| Pmepa1 | 1.33E-30 | 1.04714607 | 0.339 | 0.249 | 2.68E-26 | 5 | Pmepa1 |
| Rps172 | 1.47E-29 | 0.465718153 | 0.73 | 0.802 | 2.98E-25 | 5 | Rps17 |
| Psap | 5.85E-28 | 1.116188233 | 0.442 | 0.399 | 1.18E-23 | 5 | Psap |
| Rps152 | 6.08E-26 | 0.565350537 | 0.686 | 0.754 | 1.23E-21 | 5 | Rps15 |
| Cfl11 | 2.25E-25 | 0.807752173 | 0.573 | 0.613 | 4.55E-21 | 5 | Cfl1 |
| mt-Co33 | 1.06E-23 | 0.472894673 | 0.803 | 0.724 | 2.14E-19 | 5 | mt-Co3 |
| Fxyd51 | 2.89E-23 | 1.039449487 | 0.434 | 0.413 | 5.86E-19 | 5 | Fxyd5 |
| Gpx12 | 8.19E-22 | 0.453300654 | 0.605 | 0.597 | 1.66E-17 | 5 | Gpx1 |
| Rpl313 | 9.03E-21 | 0.555808953 | 0.623 | 0.69 | 1.83E-16 | 5 | Rpl31 |
| H2afz1 | 1.88E-20 | 0.652039933 | 0.577 | 0.612 | 3.80E-16 | 5 | H2afz |
| Sat1 | 2.36E-20 | 0.945277833 | 0.388 | 0.346 | 4.78E-16 | 5 | Sat1 |
| Arpc32 | 6.25E-20 | 0.59787162 | 0.6 | 0.664 | 1.27E-15 | 5 | Arpc3 |
| Atpif12 | 7.93E-20 | 0.670856494 | 0.547 | 0.564 | 1.60E-15 | 5 | Atpif1 |
| Clta1 | 6.67E-19 | 0.795023901 | 0.518 | 0.562 | 1.35E-14 | 5 | Clta |
| Gng52 | 2.74E-18 | 0.471820066 | 0.672 | 0.772 | 5.54E-14 | 5 | Gng5 |
| Gngt22 | 1.83E-17 | 0.833055568 | 0.397 | 0.36 | 3.71E-13 | 5 | Gngt2 |
| Prdx12 | 1.43E-16 | 0.703538263 | 0.612 | 0.699 | 2.89E-12 | 5 | Prdx1 |
| Itm2b2 | 4.27E-16 | 0.332380006 | 0.787 | 0.723 | 8.64E-12 | 5 | Itm2b |

| | | | | | | | |
|---|---|---|---|---|---|---|---|
| Spcs1 | 4.37E-15 | 0.256454743 | 0.161 | 0.298 | 8.85E-11 | 5 | Spcs1 |
| Hexa | 8.20E-15 | 1.075256786 | 0.266 | 0.22 | 1.66E-10 | 5 | Hexa |
| Arpc1b1 | 9.27E-15 | 0.658744096 | 0.535 | 0.597 | 1.88E-10 | 5 | Arpc1b |
| Pfdn53 | 1.50E-14 | 0.411839525 | 0.68 | 0.771 | 3.03E-10 | 5 | Pfdn5 |
| Ostc | 3.26E-14 | 0.253467992 | 0.147 | 0.272 | 6.60E-10 | 5 | Ostc |
| Gdi2 | 1.05E-13 | 0.260560397 | 0.232 | 0.396 | 2.12E-09 | 5 | Gdi2 |
| mt-Atp63 | 2.36E-13 | 0.310734923 | 0.845 | 0.75 | 4.78E-09 | 5 | mt-Atp6 |
| Ier3ip1 | 4.85E-13 | 0.302255533 | 0.164 | 0.289 | 9.81E-09 | 5 | Ier3ip1 |
| Capzb1 | 1.27E-12 | 0.270733665 | 0.224 | 0.38 | 2.57E-08 | 5 | Capzb |
| Ssr41 | 5.33E-12 | 0.835044344 | 0.435 | 0.472 | 1.08E-07 | 5 | Ssr4 |
| Erp29 | 5.65E-12 | 1.007439059 | 0.339 | 0.33 | 1.14E-07 | 5 | Erp29 |
| mt-Co23 | 1.83E-11 | 0.3141129 | 0.804 | 0.726 | 3.71E-07 | 5 | mt-Co2 |
| Rpl272 | 3.02E-11 | 0.548802302 | 0.53 | 0.589 | 6.11E-07 | 5 | Rpl27 |
| Bax1 | 8.79E-11 | 0.26298283 | 0.194 | 0.323 | 1.78E-06 | 5 | Bax |
| Nsa22 | 2.64E-10 | 0.807953247 | 0.451 | 0.491 | 5.35E-06 | 5 | Nsa2 |
| Psmb21 | 5.82E-10 | 0.268320521 | 0.256 | 0.414 | 1.18E-05 | 5 | Psmb2 |
| Prdx52 | 1.21E-09 | 0.336442067 | 0.523 | 0.535 | 2.45E-05 | 5 | Prdx5 |
| Nap1l1 | 1.22E-09 | 0.335043251 | 0.198 | 0.324 | 2.47E-05 | 5 | Nap1l1 |
| Tmem160 | 3.88E-09 | 0.267435735 | 0.23 | 0.367 | 7.85E-05 | 5 | Tmem160 |
| Arpp191 | 4.22E-09 | 0.269527759 | 0.288 | 0.453 | 8.54E-05 | 5 | Arpp19 |
| Tmem14c | 4.72E-09 | 0.273421697 | 0.241 | 0.384 | 9.56E-05 | 5 | Tmem14c |
| Oaz12 | 1.91E-08 | 0.273699348 | 0.686 | 0.815 | 0.000387096 | 5 | Oaz1 |
| Psmb83 | 3.08E-08 | 0.523102926 | 0.517 | 0.562 | 0.000623988 | 5 | Psmb8 |
| Rpl13a1 | 3.96E-08 | 0.379239044 | 0.183 | 0.293 | 0.000802527 | 5 | Rpl13a |
| Esd2 | 1.12E-07 | 0.327224282 | 0.251 | 0.39 | 0.002260696 | 5 | Esd |
| Pitpna | 2.75E-07 | 0.359311622 | 0.216 | 0.337 | 0.005559213 | 5 | Pitpna |
| Eif4a11 | 4.90E-07 | 0.296111087 | 0.291 | 0.441 | 0.00991877 | 5 | Eif4a1 |
| Brk1 | 6.75E-07 | 0.715746988 | 0.402 | 0.447 | 0.013673655 | 5 | Brk1 |
| Snrpe1 | 7.78E-07 | 0.256960576 | 0.324 | 0.489 | 0.015759624 | 5 | Snrpe |
| Atp6v0b | 7.92E-07 | 0.390989641 | 0.244 | 0.377 | 0.016043366 | 5 | Atp6v0b |
| Hspe12 | 1.10E-06 | 0.544269309 | 0.463 | 0.512 | 0.022196967 | 5 | Hspe1 |
| Sec11c1 | 1.14E-06 | 0.275017616 | 0.213 | 0.319 | 0.023052936 | 5 | Sec11c |
| Actr31 | 1.98E-06 | 0.750499882 | 0.373 | 0.395 | 0.040087501 | 5 | Actr3 |
| Snrpf2 | 8.18E-06 | 0.309391782 | 0.262 | 0.389 | 0.165523104 | 5 | Snrpf |
| Tmem2561 | 8.95E-06 | 0.267196011 | 0.254 | 0.373 | 0.181249459 | 5 | Tmem256 |
| Atp6v1g11 | 9.35E-06 | 0.271111462 | 0.314 | 0.466 | 0.189339539 | 5 | Atp6v1g1 |
| Rpl23a3 | 1.11E-05 | 0.391009602 | 0.587 | 0.687 | 0.225039047 | 5 | Rpl23a |
| Calm3 | 1.18E-05 | 0.342302077 | 0.185 | 0.277 | 0.239327329 | 5 | Calm3 |
| Ybx12 | 1.22E-05 | 0.32545174 | 0.643 | 0.76 | 0.246635006 | 5 | Ybx1 |
| mt-Nd4l2 | 1.61E-05 | 0.33608587 | 0.338 | 0.503 | 0.325212026 | 5 | mt-Nd4l |
| Atp6v0e | 2.10E-05 | 0.390786166 | 0.29 | 0.436 | 0.424980406 | 5 | Atp6v0e |
| Cenpx2 | 2.28E-05 | 0.317237852 | 0.21 | 0.308 | 0.461895363 | 5 | Cenpx |
| Ywhah | 3.06E-05 | 0.338369306 | 0.216 | 0.318 | 0.619220875 | 5 | Ywhah |

| | | | | | | | |
|---|---|---|---|---|---|---|---|
| Wdr83os2 | 3.51E-05 | 0.351057427 | 0.199 | 0.291 | 0.710119597 | 5 | Wdr83os |
| Arpc21 | 4.16E-05 | 0.597894811 | 0.468 | 0.552 | 0.84234097 | 5 | Arpc2 |
| Selenop2 | 5.06E-05 | 0.327709494 | 0.263 | 0.372 | 1 | 5 | Selenop |
| Ost41 | 5.06E-05 | 0.282469591 | 0.307 | 0.446 | 1 | 5 | Ost4 |
| Rpl53 | 5.34E-05 | 0.363264766 | 0.601 | 0.719 | 1 | 5 | Rpl5 |
| Nop101 | 6.53E-05 | 0.265168594 | 0.266 | 0.382 | 1 | 5 | Nop10 |
| Srsf91 | 9.92E-05 | 0.311347277 | 0.235 | 0.338 | 1 | 5 | Srsf9 |
| Cnih41 | 0.000132255 | 0.458223771 | 0.179 | 0.261 | 1 | 5 | Cnih4 |
| Hnrnpa11 | 0.000174464 | 0.422640954 | 0.231 | 0.333 | 1 | 5 | Hnrnpa1 |
| Taldo12 | 0.000262505 | 0.491971212 | 0.45 | 0.522 | 1 | 5 | Taldo1 |
| Bola22 | 0.000263216 | 0.290548177 | 0.191 | 0.269 | 1 | 5 | Bola2 |
| Psenen | 0.000291861 | 0.371015907 | 0.271 | 0.393 | 1 | 5 | Psenen |
| Pomp2 | 0.000303985 | 0.405152574 | 0.508 | 0.607 | 1 | 5 | Pomp |
| Aprt | 0.000453659 | 0.401668 | 0.177 | 0.251 | 1 | 5 | Aprt |
| Tmem2582 | 0.000569182 | 0.38423095 | 0.225 | 0.316 | 1 | 5 | Tmem258 |
| Uba523 | 0.000878346 | 0.461660389 | 0.457 | 0.528 | 1 | 5 | Uba52 |
| Cdc42 | 0.000902831 | 0.543480781 | 0.507 | 0.625 | 1 | 5 | Cdc42 |
| Lamp12 | 0.001056873 | 0.466046296 | 0.298 | 0.427 | 1 | 5 | Lamp1 |
| Snrpg1 | 0.00137769 | 0.409057278 | 0.236 | 0.332 | 1 | 5 | Snrpg |
| Timm10b | 0.001654394 | 0.437396047 | 0.183 | 0.254 | 1 | 5 | Timm10b |
| Nmt1 | 0.001835948 | 0.449333619 | 0.213 | 0.298 | 1 | 5 | Nmt1 |
| Gm100763 | 0.001983191 | 0.458217768 | 0.466 | 0.553 | 1 | 5 | Gm10076 |
| Smim14 | 0.002180231 | 0.308827677 | 0.292 | 0.404 | 1 | 5 | Smim14 |
| Mrpl232 | 0.003731443 | 0.376247432 | 0.219 | 0.297 | 1 | 5 | Mrpl23 |
| Ubl52 | 0.00608372 | 0.255315885 | 0.403 | 0.561 | 1 | 5 | Ubl5 |
| Dbi1 | 0.006120193 | 0.280901018 | 0.322 | 0.442 | 1 | 5 | Dbi |
| Cd741 | 0 | 3.73665285 | 0.998 | 0.214 | 0 | 6 | Cd74 |
| H2-Ab11 | 0 | 3.701444274 | 0.996 | 0.162 | 0 | 6 | H2-Ab1 |
| C1qa1 | 0 | 3.584505254 | 0.994 | 0.105 | 0 | 6 | C1qa |
| H2-Eb11 | 0 | 3.560532973 | 0.997 | 0.155 | 0 | 6 | H2-Eb1 |
| H2-Aa1 | 0 | 3.555736047 | 0.997 | 0.147 | 0 | 6 | H2-Aa |
| Cx3cr1 | 0 | 3.552570629 | 0.976 | 0.039 | 0 | 6 | Cx3cr1 |
| Tgfbr1 | 0 | 3.478296176 | 0.978 | 0.167 | 0 | 6 | Tgfbr1 |
| Ctss1 | 0 | 3.462270741 | 0.998 | 0.125 | 0 | 6 | Ctss |
| Apoe1 | 0 | 3.430505355 | 0.997 | 0.512 | 0 | 6 | Apoe |
| C1qb1 | 0 | 3.414667221 | 0.996 | 0.113 | 0 | 6 | C1qb |
| C1qc1 | 0 | 3.392834046 | 0.993 | 0.092 | 0 | 6 | C1qc |
| Apobec1 | 0 | 3.103986335 | 0.986 | 0.059 | 0 | 6 | Apobec1 |
| Cybb | 0 | 2.996167119 | 0.957 | 0.031 | 0 | 6 | Cybb |
| Ctsc1 | 0 | 2.95648676 | 0.982 | 0.109 | 0 | 6 | Ctsc |
| Ly861 | 0 | 2.901474712 | 0.976 | 0.053 | 0 | 6 | Ly86 |
| Lyz21 | 0 | 2.887092951 | 0.91 | 0.072 | 0 | 6 | Lyz2 |
| Ms4a71 | 0 | 2.858330726 | 0.947 | 0.057 | 0 | 6 | Ms4a7 |

| | | | | | | | |
|---|---|---|---|---|---|---|---|
| Tanc2 | 0 | 2.798173808 | 0.94 | 0.121 | 0 | 6 | Tanc2 |
| Zeb2 | 0 | 2.752716563 | 0.947 | 0.103 | 0 | 6 | Zeb2 |
| Frmpd4 | 0 | 2.710868099 | 0.654 | 0.011 | 0 | 6 | Frmpd4 |
| Fyb1 | 0 | 2.693534147 | 0.967 | 0.145 | 0 | 6 | Fyb |
| Basp11 | 0 | 2.630591434 | 0.951 | 0.062 | 0 | 6 | Basp1 |
| Slc8a1 | 0 | 2.584397391 | 0.859 | 0.021 | 0 | 6 | Slc8a1 |
| Csf1r | 0 | 2.572775234 | 0.934 | 0.031 | 0 | 6 | Csf1r |
| Gab2 | 0 | 2.563632906 | 0.935 | 0.169 | 0 | 6 | Gab2 |
| Arhgap151 | 0 | 2.561185024 | 0.961 | 0.15 | 0 | 6 | Arhgap15 |
| Ctsh1 | 0 | 2.538707565 | 0.969 | 0.095 | 0 | 6 | Ctsh |
| Laptm52 | 0 | 2.522578338 | 0.978 | 0.173 | 0 | 6 | Laptm5 |
| Lgmn1 | 0 | 2.513974949 | 0.95 | 0.072 | 0 | 6 | Lgmn |
| Cst31 | 0 | 2.505233751 | 0.997 | 0.584 | 0 | 6 | Cst3 |
| Pmepa11 | 0 | 2.443552701 | 0.946 | 0.218 | 0 | 6 | Pmepa1 |
| Cadm1 | 0 | 2.438059616 | 0.87 | 0.031 | 0 | 6 | Cadm1 |
| Ptprj1 | 0 | 2.366080059 | 0.89 | 0.114 | 0 | 6 | Ptprj |
| Psap1 | 0 | 2.34962563 | 0.986 | 0.37 | 0 | 6 | Psap |
| Ctsb1 | 0 | 2.329282218 | 0.956 | 0.266 | 0 | 6 | Ctsb |
| Hpgd1 | 0 | 2.328617416 | 0.852 | 0.05 | 0 | 6 | Hpgd |
| Ccl41 | 0 | 2.316024156 | 0.74 | 0.099 | 0 | 6 | Ccl4 |
| Mpeg1 | 0 | 2.284732534 | 0.906 | 0.032 | 0 | 6 | Mpeg1 |
| H2-DMb11 | 0 | 2.269892372 | 0.883 | 0.052 | 0 | 6 | H2-DMb1 |
| Tmcc32 | 0 | 2.266109978 | 0.95 | 0.288 | 0 | 6 | Tmcc3 |
| Lair1 | 0 | 2.246101552 | 0.875 | 0.022 | 0 | 6 | Lair1 |
| Adgre1 | 0 | 2.228075071 | 0.845 | 0.017 | 0 | 6 | Adgre1 |
| Cd86 | 0 | 2.189550069 | 0.791 | 0.02 | 0 | 6 | Cd86 |
| H2-DMa1 | 0 | 2.174535468 | 0.927 | 0.08 | 0 | 6 | H2-DMa |
| Aif11 | 0 | 2.164122792 | 0.95 | 0.08 | 0 | 6 | Aif1 |
| Fat3 | 0 | 2.156747464 | 0.731 | 0.009 | 0 | 6 | Fat3 |
| Unc93b1 | 0 | 2.15497567 | 0.93 | 0.118 | 0 | 6 | Unc93b1 |
| Ctsz1 | 0 | 2.148144755 | 0.986 | 0.33 | 0 | 6 | Ctsz |
| Hexb1 | 0 | 2.145939671 | 0.993 | 0.495 | 0 | 6 | Hexb |
| Cd300c21 | 0 | 2.140762691 | 0.911 | 0.044 | 0 | 6 | Cd300c2 |
| Klra2 | 0 | 2.137635599 | 0.814 | 0.015 | 0 | 6 | Klra2 |
| Itgb5 | 0 | 2.137020303 | 0.891 | 0.108 | 0 | 6 | Itgb5 |
| Lyn1 | 0 | 2.126359205 | 0.951 | 0.239 | 0 | 6 | Lyn |
| Zfhx3 | 0 | 2.124990743 | 0.909 | 0.147 | 0 | 6 | Zfhx3 |
| Plxdc2 | 0 | 2.100287058 | 0.809 | 0.057 | 0 | 6 | Plxdc2 |
| Pid1 | 0 | 2.080717767 | 0.773 | 0.034 | 0 | 6 | Pid1 |
| Gatm | 0 | 2.053824939 | 0.831 | 0.035 | 0 | 6 | Gatm |
| Gpr65 | 0 | 2.044149427 | 0.825 | 0.051 | 0 | 6 | Gpr65 |
| Prkcb1 | 0 | 2.029017936 | 0.877 | 0.121 | 0 | 6 | Prkcb |
| Mafb | 0 | 2.006316551 | 0.886 | 0.118 | 0 | 6 | Mafb |

| | | | | | | | |
|---|---|---|---|---|---|---|---|
| Hdac91 | 0 | 1.988944255 | 0.827 | 0.125 | 0 | 6 | Hdac9 |
| Trem21 | 0 | 1.985663895 | 0.774 | 0.031 | 0 | 6 | Trem2 |
| Slamf91 | 0 | 1.977442536 | 0.835 | 0.032 | 0 | 6 | Slamf9 |
| Cd721 | 0 | 1.975826509 | 0.831 | 0.069 | 0 | 6 | Cd72 |
| Fgf13 | 0 | 1.951807383 | 0.605 | 0.053 | 0 | 6 | Fgf13 |
| Spi11 | 0 | 1.944144083 | 0.873 | 0.043 | 0 | 6 | Spi1 |
| Inpp5d1 | 0 | 1.943468729 | 0.881 | 0.171 | 0 | 6 | Inpp5d |
| Cd681 | 0 | 1.931968022 | 0.845 | 0.041 | 0 | 6 | Cd68 |
| Gpr183 | 0 | 1.910207492 | 0.711 | 0.027 | 0 | 6 | Gpr183 |
| Slc9a91 | 0 | 1.906086116 | 0.815 | 0.119 | 0 | 6 | Slc9a9 |
| Cyba2 | 0 | 1.905817808 | 0.981 | 0.415 | 0 | 6 | Cyba |
| Ms4a6c1 | 0 | 1.903772891 | 0.82 | 0.047 | 0 | 6 | Ms4a6c |
| Pip4k2a2 | 0 | 1.898740713 | 0.895 | 0.209 | 0 | 6 | Pip4k2a |
| Creb5 | 0 | 1.884586007 | 0.724 | 0.028 | 0 | 6 | Creb5 |
| Rgs101 | 0 | 1.884091055 | 0.913 | 0.124 | 0 | 6 | Rgs10 |
| Lst11 | 0 | 1.877111726 | 0.886 | 0.073 | 0 | 6 | Lst1 |
| Fam105a1 | 0 | 1.860846178 | 0.842 | 0.09 | 0 | 6 | Fam105a |
| Cxcl161 | 0 | 1.841397365 | 0.86 | 0.076 | 0 | 6 | Cxcl16 |
| Bcl2a1b1 | 0 | 1.841037212 | 0.854 | 0.069 | 0 | 6 | Bcl2a1b |
| Rgs11 | 0 | 1.833180421 | 0.641 | 0.117 | 0 | 6 | Rgs1 |
| Tyrobp2 | 0 | 1.823310265 | 0.991 | 0.226 | 0 | 6 | Tyrobp |
| Hck | 0 | 1.816029805 | 0.774 | 0.021 | 0 | 6 | Hck |
| Lpcat2 | 0 | 1.806466843 | 0.787 | 0.03 | 0 | 6 | Lpcat2 |
| Dock81 | 0 | 1.803013291 | 0.89 | 0.186 | 0 | 6 | Dock8 |
| Myo5a | 0 | 1.802073039 | 0.758 | 0.051 | 0 | 6 | Myo5a |
| Adap2 | 0 | 1.800966522 | 0.739 | 0.023 | 0 | 6 | Adap2 |
| Pld4 | 0 | 1.78462578 | 0.779 | 0.023 | 0 | 6 | Pld4 |
| Apbb1ip1 | 0 | 1.778137217 | 0.854 | 0.14 | 0 | 6 | Apbb1ip |
| Arhgap22 | 0 | 1.775098 | 0.703 | 0.011 | 0 | 6 | Arhgap22 |
| Ldlrad41 | 0 | 1.769014545 | 0.868 | 0.221 | 0 | 6 | Ldlrad4 |
| Gm26740 | 0 | 1.761024229 | 0.723 | 0.027 | 0 | 6 | Gm26740 |
| Fcer1g2 | 0 | 1.758969178 | 0.991 | 0.235 | 0 | 6 | Fcer1g |
| Rgs21 | 0 | 1.758718616 | 0.785 | 0.098 | 0 | 6 | Rgs2 |
| Nlrp1b | 0 | 1.755322185 | 0.671 | 0.008 | 0 | 6 | Nlrp1b |
| Cd631 | 0 | 1.748004401 | 0.911 | 0.152 | 0 | 6 | Cd63 |
| Skap2 | 0 | 1.745266253 | 0.87 | 0.133 | 0 | 6 | Skap2 |
| Cd14 | 0 | 1.741182061 | 0.753 | 0.027 | 0 | 6 | Cd14 |
| Man2b1 | 0 | 1.72333227 | 0.848 | 0.13 | 0 | 6 | Man2b1 |
| Actr32 | 0 | 1.721661237 | 0.96 | 0.364 | 0 | 6 | Actr3 |
| Blnk | 0 | 1.705570443 | 0.768 | 0.026 | 0 | 6 | Blnk |
| Scimp | 0 | 1.703733376 | 0.769 | 0.042 | 0 | 6 | Scimp |
| Runx11 | 0 | 1.702917851 | 0.747 | 0.101 | 0 | 6 | Runx1 |
| Epb41l21 | 0 | 1.702367836 | 0.866 | 0.211 | 0 | 6 | Epb41l2 |

| | | | | | | | |
|---|---|---|---|---|---|---|---|
| Il10ra | 0 | 1.693100274 | 0.77 | 0.048 | 0 | 6 | Il10ra |
| Hpgds | 0 | 1.689542262 | 0.738 | 0.025 | 0 | 6 | Hpgds |
| Plek1 | 0 | 1.689000758 | 0.763 | 0.093 | 0 | 6 | Plek |
| Csf2ra | 0 | 1.672987274 | 0.781 | 0.038 | 0 | 6 | Csf2ra |
| Frmd4b | 0 | 1.671804117 | 0.821 | 0.146 | 0 | 6 | Frmd4b |
| Glipr11 | 0 | 1.667480726 | 0.708 | 0.066 | 0 | 6 | Glipr1 |
| Npc21 | 0 | 1.652878721 | 0.96 | 0.388 | 0 | 6 | Npc2 |
| Fam49b1 | 0 | 1.651357554 | 0.901 | 0.205 | 0 | 6 | Fam49b |
| Cd180 | 0 | 1.646224562 | 0.688 | 0.014 | 0 | 6 | Cd180 |
| Elmo12 | 0 | 1.640104343 | 0.919 | 0.34 | 0 | 6 | Elmo1 |
| Abhd12 | 0 | 1.637324079 | 0.79 | 0.096 | 0 | 6 | Abhd12 |
| Alox5ap1 | 0 | 1.621345139 | 0.776 | 0.038 | 0 | 6 | Alox5ap |
| Cyth41 | 0 | 1.621343735 | 0.814 | 0.1 | 0 | 6 | Cyth4 |
| Plbd1 | 0 | 1.619931586 | 0.748 | 0.025 | 0 | 6 | Plbd1 |
| Tbxas1 | 0 | 1.616344872 | 0.671 | 0.015 | 0 | 6 | Tbxas1 |
| Ifi207 | 0 | 1.615187304 | 0.734 | 0.076 | 0 | 6 | Ifi207 |
| Negr1 | 0 | 1.613271484 | 0.542 | 0.019 | 0 | 6 | Negr1 |
| Pou2f2 | 0 | 1.613270468 | 0.65 | 0.016 | 0 | 6 | Pou2f2 |
| Fcgr1 | 0 | 1.613025222 | 0.704 | 0.019 | 0 | 6 | Fcgr1 |
| Lcp11 | 0 | 1.608349291 | 0.891 | 0.206 | 0 | 6 | Lcp1 |
| Xylt1 | 0 | 1.608292054 | 0.65 | 0.049 | 0 | 6 | Xylt1 |
| Lpl | 0 | 1.60791834 | 0.765 | 0.053 | 0 | 6 | Lpl |
| Ssh21 | 0 | 1.595355957 | 0.801 | 0.198 | 0 | 6 | Ssh2 |
| Axl | 0 | 1.589369334 | 0.749 | 0.045 | 0 | 6 | Axl |
| Mctp1 | 0 | 1.583336079 | 0.655 | 0.045 | 0 | 6 | Mctp1 |
| Cpq | 0 | 1.570154616 | 0.754 | 0.089 | 0 | 6 | Cpq |
| P2ry6 | 0 | 1.56623034 | 0.699 | 0.018 | 0 | 6 | P2ry6 |
| Slc29a3 | 0 | 1.564935552 | 0.706 | 0.037 | 0 | 6 | Slc29a3 |
| Lilra5 | 0 | 1.559804843 | 0.712 | 0.019 | 0 | 6 | Lilra5 |
| Irf8 | 0 | 1.559050208 | 0.763 | 0.077 | 0 | 6 | Irf8 |
| Atf3 | 0 | 1.556629718 | 0.591 | 0.062 | 0 | 6 | Atf3 |
| Ccr51 | 0 | 1.553333131 | 0.738 | 0.103 | 0 | 6 | Ccr5 |
| Hk2 | 0 | 1.549362552 | 0.689 | 0.033 | 0 | 6 | Hk2 |
| Slc15a3 | 0 | 1.547324303 | 0.692 | 0.045 | 0 | 6 | Slc15a3 |
| Dock101 | 0 | 1.547259567 | 0.895 | 0.175 | 0 | 6 | Dock10 |
| 1-Mar | 0 | 1.53288825 | 0.64 | 0.011 | 0 | 6 | 1-Mar |
| Taok31 | 0 | 1.530987907 | 0.785 | 0.133 | 0 | 6 | Taok3 |
| Ophn1 | 0 | 1.530916557 | 0.727 | 0.103 | 0 | 6 | Ophn1 |
| Sat11 | 0 | 1.524485134 | 0.927 | 0.318 | 0 | 6 | Sat1 |
| Tmem119 | 0 | 1.516891268 | 0.694 | 0.028 | 0 | 6 | Tmem119 |
| Fcgr31 | 0 | 1.51397961 | 0.749 | 0.054 | 0 | 6 | Fcgr3 |
| Tmem176b3 | 0 | 1.509739561 | 0.974 | 0.413 | 0 | 6 | Tmem176b |
| Ckb1 | 0 | 1.507653656 | 0.869 | 0.184 | 0 | 6 | Ckb |

| | | | | | | | |
|---|---|---|---|---|---|---|---|
| C3ar1 | 0 | 1.503761705 | 0.635 | 0.01 | 0 | 6 | C3ar1 |
| Cdk61 | 0 | 1.502641691 | 0.669 | 0.084 | 0 | 6 | Cdk6 |
| Rnf180 | 0 | 1.496464751 | 0.622 | 0.01 | 0 | 6 | Rnf180 |
| Myo1f1 | 0 | 1.495580713 | 0.758 | 0.107 | 0 | 6 | Myo1f |
| Ccl12 | 0 | 1.490221685 | 0.449 | 0.02 | 0 | 6 | Ccl12 |
| Sirpa | 0 | 1.489219864 | 0.671 | 0.02 | 0 | 6 | Sirpa |
| Tnf | 0 | 1.474989692 | 0.536 | 0.05 | 0 | 6 | Tnf |
| Mir142hg1 | 0 | 1.468959592 | 0.823 | 0.138 | 0 | 6 | Mir142hg |
| Tbc1d52 | 0 | 1.466127245 | 0.825 | 0.234 | 0 | 6 | Tbc1d5 |
| Tmem176a3 | 0 | 1.465760944 | 0.967 | 0.391 | 0 | 6 | Tmem176a |
| Cd841 | 0 | 1.464891591 | 0.692 | 0.061 | 0 | 6 | Cd84 |
| Gna12 | 0 | 1.460239351 | 0.741 | 0.124 | 0 | 6 | Gna12 |
| Pik3ap11 | 0 | 1.454881135 | 0.656 | 0.062 | 0 | 6 | Pik3ap1 |
| Trps12 | 0 | 1.454143367 | 0.81 | 0.208 | 0 | 6 | Trps1 |
| Acer31 | 0 | 1.451183973 | 0.733 | 0.112 | 0 | 6 | Acer3 |
| Asph | 0 | 1.44264818 | 0.688 | 0.071 | 0 | 6 | Asph |
| Ntpcr1 | 0 | 1.442101631 | 0.774 | 0.071 | 0 | 6 | Ntpcr |
| Rbm47 | 0 | 1.433709493 | 0.701 | 0.044 | 0 | 6 | Rbm47 |
| Itga9 | 0 | 1.417534757 | 0.667 | 0.074 | 0 | 6 | Itga9 |
| Wnk12 | 0 | 1.414776816 | 0.895 | 0.338 | 0 | 6 | Wnk1 |
| Fam129a1 | 0 | 1.398367154 | 0.701 | 0.118 | 0 | 6 | Fam129a |
| Snx51 | 0 | 1.393821198 | 0.831 | 0.181 | 0 | 6 | Snx5 |
| Sh3kbp1 | 0 | 1.388190946 | 0.704 | 0.096 | 0 | 6 | Sh3kbp1 |
| Itgb8 | 0 | 1.385543183 | 0.517 | 0.012 | 0 | 6 | Itgb8 |
| Plau | 0 | 1.380889909 | 0.476 | 0.042 | 0 | 6 | Plau |
| Il1a | 0 | 1.380153887 | 0.49 | 0.006 | 0 | 6 | Il1a |
| Pvt11 | 0 | 1.37432164 | 0.597 | 0.053 | 0 | 6 | Pvt1 |
| Rap2a | 0 | 1.370174952 | 0.692 | 0.089 | 0 | 6 | Rap2a |
| Tnfrsf11a | 0 | 1.366516283 | 0.618 | 0.031 | 0 | 6 | Tnfrsf11a |
| Cacna1d | 0 | 1.360251578 | 0.571 | 0.013 | 0 | 6 | Cacna1d |
| Acsl1 | 0 | 1.359059836 | 0.604 | 0.046 | 0 | 6 | Acsl1 |
| Wfdc171 | 0 | 1.357728375 | 0.628 | 0.041 | 0 | 6 | Wfdc17 |
| Nfkbiz | 0 | 1.350835221 | 0.628 | 0.114 | 0 | 6 | Nfkbiz |
| Tnfsf13b | 0 | 1.350207133 | 0.615 | 0.016 | 0 | 6 | Tnfsf13b |
| Xist3 | 0 | 1.345603911 | 0.959 | 0.44 | 0 | 6 | Xist |
| Parp81 | 0 | 1.344205825 | 0.681 | 0.124 | 0 | 6 | Parp8 |
| Fgd2 | 0 | 1.34148678 | 0.621 | 0.015 | 0 | 6 | Fgd2 |
| Cdk14 | 0 | 1.341131908 | 0.706 | 0.114 | 0 | 6 | Cdk14 |
| Coro1a2 | 0 | 1.340774627 | 0.899 | 0.186 | 0 | 6 | Coro1a |
| St3gal62 | 0 | 1.340513407 | 0.845 | 0.293 | 0 | 6 | St3gal6 |
| Pag11 | 0 | 1.339996487 | 0.667 | 0.095 | 0 | 6 | Pag1 |
| Bmp2k1 | 0 | 1.338793617 | 0.668 | 0.083 | 0 | 6 | Bmp2k |
| Cd83 | 0 | 1.334317582 | 0.554 | 0.014 | 0 | 6 | Cd83 |

| | | | | | | | |
|---|---|---|---|---|---|---|---|
| Nckap1l1 | 0 | 1.328042085 | 0.658 | 0.06 | 0 | 6 | Nckap1l |
| Hexa1 | 0 | 1.324085177 | 0.823 | 0.191 | 0 | 6 | Hexa |
| Snx24 | 0 | 1.317394399 | 0.609 | 0.032 | 0 | 6 | Snx24 |
| Lhfpl21 | 0 | 1.314971262 | 0.692 | 0.124 | 0 | 6 | Lhfpl2 |
| Mgat51 | 0 | 1.309756467 | 0.723 | 0.144 | 0 | 6 | Mgat5 |
| Lacc1 | 0 | 1.305306801 | 0.604 | 0.017 | 0 | 6 | Lacc1 |
| Celf21 | 0 | 1.302948561 | 0.846 | 0.171 | 0 | 6 | Celf2 |
| Cfh | 0 | 1.301701591 | 0.659 | 0.04 | 0 | 6 | Cfh |
| Fam214a1 | 0 | 1.301182223 | 0.655 | 0.098 | 0 | 6 | Fam214a |
| Srgap23 | 0 | 1.28616879 | 0.807 | 0.28 | 0 | 6 | Srgap2 |
| Itgav1 | 0 | 1.279462423 | 0.668 | 0.136 | 0 | 6 | Itgav |
| Atf61 | 0 | 1.275777199 | 0.666 | 0.138 | 0 | 6 | Atf6 |
| Olfml3 | 0 | 1.272898418 | 0.607 | 0.032 | 0 | 6 | Olfml3 |
| Zmynd15 | 0 | 1.269077324 | 0.57 | 0.026 | 0 | 6 | Zmynd15 |
| Ppm1h1 | 0 | 1.265475612 | 0.691 | 0.135 | 0 | 6 | Ppm1h |
| Retreg11 | 0 | 1.265119725 | 0.629 | 0.071 | 0 | 6 | Retreg1 |
| Mbnl22 | 0 | 1.262907225 | 0.906 | 0.396 | 0 | 6 | Mbnl2 |
| Il2ra | 0 | 1.262474944 | 0.476 | 0.022 | 0 | 6 | Il2ra |
| Rtn42 | 0 | 1.262097884 | 0.877 | 0.313 | 0 | 6 | Rtn4 |
| Chd9 | 0 | 1.253826472 | 0.791 | 0.224 | 0 | 6 | Chd9 |
| Erp291 | 0 | 1.253347387 | 0.889 | 0.301 | 0 | 6 | Erp29 |
| Mitf | 0 | 1.253156784 | 0.59 | 0.05 | 0 | 6 | Mitf |
| Dennd1a2 | 0 | 1.253022844 | 0.773 | 0.228 | 0 | 6 | Dennd1a |
| Tnfaip81 | 0 | 1.250682726 | 0.78 | 0.158 | 0 | 6 | Tnfaip8 |
| Ext11 | 0 | 1.249610341 | 0.69 | 0.142 | 0 | 6 | Ext1 |
| Dst1 | 0 | 1.246157786 | 0.74 | 0.164 | 0 | 6 | Dst |
| Dock21 | 0 | 1.241214363 | 0.86 | 0.153 | 0 | 6 | Dock2 |
| Mef2a2 | 0 | 1.240286631 | 0.933 | 0.392 | 0 | 6 | Mef2a |
| Cyfip12 | 0 | 1.239686991 | 0.849 | 0.251 | 0 | 6 | Cyfip1 |
| Slamf71 | 0 | 1.238897854 | 0.637 | 0.073 | 0 | 6 | Slamf7 |
| Ikzf11 | 0 | 1.235325803 | 0.74 | 0.128 | 0 | 6 | Ikzf1 |
| Zfp7101 | 0 | 1.23494239 | 0.688 | 0.129 | 0 | 6 | Zfp710 |
| Gns | 0 | 1.23151101 | 0.675 | 0.1 | 0 | 6 | Gns |
| Plcg21 | 0 | 1.229695408 | 0.683 | 0.093 | 0 | 6 | Plcg2 |
| Ciita | 0 | 1.228896583 | 0.525 | 0.014 | 0 | 6 | Ciita |
| Cd522 | 0 | 1.22885857 | 0.988 | 0.248 | 0 | 6 | Cd52 |
| Fnbp11 | 0 | 1.222175766 | 0.786 | 0.203 | 0 | 6 | Fnbp1 |
| Cotl12 | 0 | 1.221661045 | 0.803 | 0.197 | 0 | 6 | Cotl1 |
| Tbc1d9 | 0 | 1.21714357 | 0.538 | 0.014 | 0 | 6 | Tbc1d9 |
| Mgat4a1 | 0 | 1.215527529 | 0.626 | 0.08 | 0 | 6 | Mgat4a |
| Maml31 | 0 | 1.209696378 | 0.734 | 0.183 | 0 | 6 | Maml3 |
| Ppfia4 | 0 | 1.206580771 | 0.633 | 0.026 | 0 | 6 | Ppfia4 |
| Irak21 | 0 | 1.206446387 | 0.638 | 0.118 | 0 | 6 | Irak2 |

| | | | | | | | |
|---|---|---|---|---|---|---|---|
| Rreb11 | 0 | 1.203816763 | 0.697 | 0.178 | 0 | 6 | Rreb1 |
| AU0202063 | 0 | 1.198787982 | 0.822 | 0.265 | 0 | 6 | AU020206 |
| Specc1 | 0 | 1.196974479 | 0.569 | 0.056 | 0 | 6 | Specc1 |
| Tns3 | 0 | 1.194008683 | 0.6 | 0.046 | 0 | 6 | Tns3 |
| Ms4a6b | 0 | 1.185130412 | 0.637 | 0.066 | 0 | 6 | Ms4a6b |
| Asah11 | 0 | 1.184729023 | 0.753 | 0.161 | 0 | 6 | Asah1 |
| Rrbp12 | 0 | 1.182832893 | 0.948 | 0.437 | 0 | 6 | Rrbp1 |
| Tec1 | 0 | 1.179154662 | 0.609 | 0.086 | 0 | 6 | Tec |
| Etv61 | 0 | 1.175399027 | 0.805 | 0.253 | 0 | 6 | Etv6 |
| Ptprc1 | 0 | 1.17346551 | 0.931 | 0.162 | 0 | 6 | Ptprc |
| Errfi1 | 0 | 1.172388699 | 0.567 | 0.142 | 0 | 6 | Errfi1 |
| Ncf2 | 0 | 1.169677034 | 0.593 | 0.026 | 0 | 6 | Ncf2 |
| Dapp11 | 0 | 1.168674172 | 0.583 | 0.056 | 0 | 6 | Dapp1 |
| Pkib | 0 | 1.167518505 | 0.567 | 0.017 | 0 | 6 | Pkib |
| Trf | 0 | 1.165823304 | 0.611 | 0.041 | 0 | 6 | Trf |
| Rtn1 | 0 | 1.160390826 | 0.5 | 0.012 | 0 | 6 | Rtn1 |
| Tgfbr22 | 0 | 1.152134531 | 0.82 | 0.287 | 0 | 6 | Tgfbr2 |
| Gm26522 | 0 | 1.1520865 | 0.463 | 0.012 | 0 | 6 | Gm26522 |
| Vrk21 | 0 | 1.151501395 | 0.616 | 0.1 | 0 | 6 | Vrk2 |
| Extl3 | 0 | 1.150954802 | 0.622 | 0.117 | 0 | 6 | Extl3 |
| Snx291 | 0 | 1.148296263 | 0.603 | 0.088 | 0 | 6 | Snx29 |
| Arrb21 | 0 | 1.142844154 | 0.618 | 0.067 | 0 | 6 | Arrb2 |
| Ptpro | 0 | 1.140097029 | 0.478 | 0.006 | 0 | 6 | Ptpro |
| Creg1 | 0 | 1.134069522 | 0.723 | 0.163 | 0 | 6 | Creg1 |
| Grn2 | 0 | 1.133784567 | 0.741 | 0.199 | 0 | 6 | Grn |
| Lpxn1 | 0 | 1.129456347 | 0.564 | 0.07 | 0 | 6 | Lpxn |
| Kcnk13 | 0 | 1.127413068 | 0.48 | 0.007 | 0 | 6 | Kcnk13 |
| Phf11b | 0 | 1.127080327 | 0.617 | 0.067 | 0 | 6 | Phf11b |
| Pak1 | 0 | 1.125418076 | 0.56 | 0.016 | 0 | 6 | Pak1 |
| Ctsa | 0 | 1.125166466 | 0.678 | 0.128 | 0 | 6 | Ctsa |
| Irf5 | 0 | 1.123355057 | 0.585 | 0.03 | 0 | 6 | Irf5 |
| Qpct1 | 0 | 1.122896784 | 0.574 | 0.055 | 0 | 6 | Qpct |
| Pla2g4a | 0 | 1.122059994 | 0.481 | 0.021 | 0 | 6 | Pla2g4a |
| Lrrc25 | 0 | 1.121475649 | 0.549 | 0.012 | 0 | 6 | Lrrc25 |
| Cd481 | 0 | 1.12025875 | 0.705 | 0.107 | 0 | 6 | Cd48 |
| Atp8a12 | 0 | 1.116663038 | 0.747 | 0.218 | 0 | 6 | Atp8a1 |
| C5ar1 | 0 | 1.114668056 | 0.437 | 0.005 | 0 | 6 | C5ar1 |
| Atp6v0b1 | 0 | 1.111704823 | 0.896 | 0.341 | 0 | 6 | Atp6v0b |
| Vsir | 0 | 1.111554383 | 0.614 | 0.079 | 0 | 6 | Vsir |
| Epsti11 | 0 | 1.109226361 | 0.756 | 0.147 | 0 | 6 | Epsti1 |
| Abca1 | 0 | 1.106420144 | 0.608 | 0.106 | 0 | 6 | Abca1 |
| Tmem86a | 0 | 1.105310709 | 0.58 | 0.034 | 0 | 6 | Tmem86a |
| Rasgef1b | 0 | 1.10195629 | 0.448 | 0.016 | 0 | 6 | Rasgef1b |

| | | | | | | | |
|---|---|---|---|---|---|---|---|
| Slco2b12 | 0 | 1.099986613 | 0.715 | 0.154 | 0 | 6 | Slco2b1 |
| Cttnbp2nl1 | 0 | 1.099114628 | 0.624 | 0.09 | 0 | 6 | Cttnbp2nl |
| Hcls11 | 0 | 1.097374681 | 0.634 | 0.101 | 0 | 6 | Hcls1 |
| Rap1gds12 | 0 | 1.094315587 | 0.746 | 0.238 | 0 | 6 | Rap1gds1 |
| Fmnl22 | 0 | 1.09229355 | 0.739 | 0.219 | 0 | 6 | Fmnl2 |
| Havcr2 | 0 | 1.086853887 | 0.508 | 0.008 | 0 | 6 | Havcr2 |
| Arhgap173 | 0 | 1.078364263 | 0.766 | 0.247 | 0 | 6 | Arhgap17 |
| Prkcd | 0 | 1.078307757 | 0.613 | 0.062 | 0 | 6 | Prkcd |
| F11r2 | 0 | 1.076889693 | 0.78 | 0.241 | 0 | 6 | F11r |
| Zfp704 | 0 | 1.074759687 | 0.61 | 0.107 | 0 | 6 | Zfp704 |
| Pde3b1 | 0 | 1.072402236 | 0.603 | 0.105 | 0 | 6 | Pde3b |
| Lamp22 | 0 | 1.062354466 | 0.831 | 0.28 | 0 | 6 | Lamp2 |
| Ifngr21 | 0 | 1.061522574 | 0.648 | 0.107 | 0 | 6 | Ifngr2 |
| Psd3 | 0 | 1.061116446 | 0.517 | 0.099 | 0 | 6 | Psd3 |
| Gng21 | 0 | 1.051501925 | 0.708 | 0.162 | 0 | 6 | Gng2 |
| Slc11a1 | 0 | 1.049017849 | 0.504 | 0.011 | 0 | 6 | Slc11a1 |
| Wipf11 | 0 | 1.040695131 | 0.747 | 0.245 | 0 | 6 | Wipf1 |
| Stab1 | 0 | 1.038714429 | 0.533 | 0.062 | 0 | 6 | Stab1 |
| Fermt31 | 0 | 1.037339008 | 0.603 | 0.091 | 0 | 6 | Fermt3 |
| Cd531 | 0 | 1.036348091 | 0.62 | 0.092 | 0 | 6 | Cd53 |
| Aim21 | 0 | 1.03107676 | 0.558 | 0.081 | 0 | 6 | Aim2 |
| Rel1 | 0 | 1.024982423 | 0.611 | 0.116 | 0 | 6 | Rel |
| Arhgap451 | 0 | 1.023284015 | 0.687 | 0.133 | 0 | 6 | Arhgap45 |
| Lat21 | 0 | 1.021845461 | 0.645 | 0.107 | 0 | 6 | Lat2 |
| Rbpj1 | 0 | 1.020141125 | 0.707 | 0.197 | 0 | 6 | Rbpj |
| Marcks2 | 0 | 1.013600404 | 0.913 | 0.314 | 0 | 6 | Marcks |
| Ccl31 | 0 | 1.006071497 | 0.464 | 0.064 | 0 | 6 | Ccl3 |
| Mapk141 | 0 | 1.004885357 | 0.658 | 0.156 | 0 | 6 | Mapk14 |
| Lipa1 | 0 | 1.004180269 | 0.675 | 0.155 | 0 | 6 | Lipa |
| Nfe2l22 | 0 | 1.00103969 | 0.8 | 0.277 | 0 | 6 | Nfe2l2 |
| Ptpn61 | 0 | 1.000750997 | 0.644 | 0.105 | 0 | 6 | Ptpn6 |
| Crlf31 | 0 | 0.999698159 | 0.637 | 0.134 | 0 | 6 | Crlf3 |
| Gusb | 0 | 0.999486547 | 0.589 | 0.075 | 0 | 6 | Gusb |
| Efhd21 | 0 | 0.990116535 | 0.707 | 0.162 | 0 | 6 | Efhd2 |
| Gm35154 | 0 | 0.987047925 | 0.437 | 0.007 | 0 | 6 | Gm35154 |
| Siglech | 0 | 0.986491285 | 0.414 | 0.007 | 0 | 6 | Siglech |
| Maf | 0 | 0.975268349 | 0.536 | 0.098 | 0 | 6 | Maf |
| Rnase6 | 0 | 0.97412584 | 0.478 | 0.025 | 0 | 6 | Rnase6 |
| Tagap | 0 | 0.969880404 | 0.439 | 0.032 | 0 | 6 | Tagap |
| Arhgap301 | 0 | 0.968411238 | 0.58 | 0.087 | 0 | 6 | Arhgap30 |
| Frmd4a | 0 | 0.967085593 | 0.616 | 0.125 | 0 | 6 | Frmd4a |
| Cers61 | 0 | 0.96642478 | 0.53 | 0.096 | 0 | 6 | Cers6 |
| Evi2a | 0 | 0.965551361 | 0.504 | 0.022 | 0 | 6 | Evi2a |

| | | | | | | | |
|---|---|---|---|---|---|---|---|
| Hspa1a | 0 | 0.963791179 | 0.451 | 0.084 | 0 | 6 | Hspa1a |
| P2ry12 | 0 | 0.954100751 | 0.383 | 0.006 | 0 | 6 | P2ry12 |
| Cyp4f18 | 0 | 0.953869862 | 0.474 | 0.014 | 0 | 6 | Cyp4f18 |
| Stx161 | 0 | 0.950427511 | 0.607 | 0.139 | 0 | 6 | Stx16 |
| 4933406I18Rik1 | 0 | 0.950173184 | 0.458 | 0.057 | 0 | 6 | 4933406I18Rik |
| Rassf4 | 0 | 0.94920839 | 0.476 | 0.022 | 0 | 6 | Rassf4 |
| Bin21 | 0 | 0.949124866 | 0.599 | 0.105 | 0 | 6 | Bin2 |
| Plxna41 | 0 | 0.94424268 | 0.552 | 0.098 | 0 | 6 | Plxna4 |
| Ifi204 | 0 | 0.943620307 | 0.493 | 0.075 | 0 | 6 | Ifi204 |
| Ncf1 | 0 | 0.941537838 | 0.474 | 0.023 | 0 | 6 | Ncf1 |
| Ptk2b1 | 0 | 0.939047757 | 0.631 | 0.127 | 0 | 6 | Ptk2b |
| Slc7a8 | 0 | 0.939033521 | 0.459 | 0.019 | 0 | 6 | Slc7a8 |
| Tnfaip8l2 | 0 | 0.935769566 | 0.539 | 0.049 | 0 | 6 | Tnfaip8l2 |
| Snx20 | 0 | 0.931225945 | 0.507 | 0.04 | 0 | 6 | Snx20 |
| Gadd45g | 0 | 0.9267812 | 0.528 | 0.112 | 0 | 6 | Gadd45g |
| Pik3cb1 | 0 | 0.925936537 | 0.587 | 0.125 | 0 | 6 | Pik3cb |
| Agps | 0 | 0.924064736 | 0.61 | 0.147 | 0 | 6 | Agps |
| Dnajb14 | 0 | 0.923252954 | 0.578 | 0.092 | 0 | 6 | Dnajb14 |
| Grap21 | 0 | 0.918896699 | 0.506 | 0.087 | 0 | 6 | Grap2 |
| Synj11 | 0 | 0.91619288 | 0.58 | 0.14 | 0 | 6 | Synj1 |
| Sdf2l11 | 0 | 0.914749384 | 0.609 | 0.147 | 0 | 6 | Sdf2l1 |
| Igsf6 | 0 | 0.914108679 | 0.458 | 0.01 | 0 | 6 | Igsf6 |
| Vav11 | 0 | 0.911889679 | 0.499 | 0.061 | 0 | 6 | Vav1 |
| Ighm1 | 0 | 0.910917581 | 0.63 | 0.102 | 0 | 6 | Ighm |
| Ppp1r211 | 0 | 0.909882266 | 0.534 | 0.092 | 0 | 6 | Ppp1r21 |
| Atp7a | 0 | 0.908656748 | 0.479 | 0.061 | 0 | 6 | Atp7a |
| Clec12a | 0 | 0.908383385 | 0.412 | 0.015 | 0 | 6 | Clec12a |
| Milr1 | 0 | 0.906838308 | 0.482 | 0.026 | 0 | 6 | Milr1 |
| Skil | 0 | 0.906803235 | 0.623 | 0.152 | 0 | 6 | Skil |
| Prex11 | 0 | 0.904171273 | 0.646 | 0.151 | 0 | 6 | Prex1 |
| Grb2 | 0 | 0.898158951 | 0.786 | 0.258 | 0 | 6 | Grb2 |
| Tlr2 | 0 | 0.896354709 | 0.42 | 0.021 | 0 | 6 | Tlr2 |
| Rab3il1 | 0 | 0.895059761 | 0.499 | 0.048 | 0 | 6 | Rab3il1 |
| Ms4a6d | 0 | 0.890851292 | 0.475 | 0.015 | 0 | 6 | Ms4a6d |
| Neurl31 | 0 | 0.888502409 | 0.534 | 0.095 | 0 | 6 | Neurl3 |
| Cask1 | 0 | 0.888388613 | 0.558 | 0.119 | 0 | 6 | Cask |
| Sgk31 | 0 | 0.888365711 | 0.558 | 0.108 | 0 | 6 | Sgk3 |
| Pf4 | 0 | 0.884572694 | 0.352 | 0.027 | 0 | 6 | Pf4 |
| Ramp11 | 0 | 0.883821555 | 0.585 | 0.087 | 0 | 6 | Ramp1 |
| Arl4c | 0 | 0.883701007 | 0.479 | 0.052 | 0 | 6 | Arl4c |
| Dclre1c | 0 | 0.882883828 | 0.437 | 0.039 | 0 | 6 | Dclre1c |
| Abca9 | 0 | 0.878273304 | 0.426 | 0.024 | 0 | 6 | Abca9 |
| Rap1gap2 | 0 | 0.878065514 | 0.55 | 0.059 | 0 | 6 | Rap1gap2 |

| | | | | | | | |
|---|---|---|---|---|---|---|---|
| Plekhm31 | 0 | 0.8769708 | 0.483 | 0.067 | 0 | 6 | Plekhm3 |
| Gm2a | 0 | 0.876399444 | 0.53 | 0.067 | 0 | 6 | Gm2a |
| Bcl2a1a | 0 | 0.874085692 | 0.445 | 0.017 | 0 | 6 | Bcl2a1a |
| Cd362 | 0 | 0.872078198 | 0.791 | 0.243 | 0 | 6 | Cd36 |
| Soat1 | 0 | 0.870537557 | 0.518 | 0.088 | 0 | 6 | Soat1 |
| Agmo | 0 | 0.870522925 | 0.381 | 0.04 | 0 | 6 | Agmo |
| Tgfbi | 0 | 0.867987525 | 0.467 | 0.036 | 0 | 6 | Tgfbi |
| Cyp27a1 | 0 | 0.866839809 | 0.388 | 0.012 | 0 | 6 | Cyp27a1 |
| Fcgr4 | 0 | 0.866011228 | 0.417 | 0.01 | 0 | 6 | Fcgr4 |
| Gm5086 | 0 | 0.865481877 | 0.395 | 0.004 | 0 | 6 | Gm5086 |
| Tm6sf11 | 0 | 0.864878164 | 0.572 | 0.121 | 0 | 6 | Tm6sf1 |
| Arhgap39 | 0 | 0.864150366 | 0.422 | 0.041 | 0 | 6 | Arhgap39 |
| Themis2 | 0 | 0.86362086 | 0.414 | 0.007 | 0 | 6 | Themis2 |
| Abr1 | 0 | 0.862904932 | 0.558 | 0.139 | 0 | 6 | Abr |
| Fam46c | 0 | 0.857429303 | 0.488 | 0.052 | 0 | 6 | Fam46c |
| Il6ra | 0 | 0.846921055 | 0.406 | 0.013 | 0 | 6 | Il6ra |
| Snx18 | 0 | 0.844970974 | 0.476 | 0.072 | 0 | 6 | Snx18 |
| Arhgap251 | 0 | 0.841187454 | 0.476 | 0.091 | 0 | 6 | Arhgap25 |
| Rnf150 | 0 | 0.840164264 | 0.363 | 0.014 | 0 | 6 | Rnf150 |
| Arhgef61 | 0 | 0.834871188 | 0.508 | 0.11 | 0 | 6 | Arhgef6 |
| Timp2 | 0 | 0.833299042 | 0.637 | 0.105 | 0 | 6 | Timp2 |
| Gsap1 | 0 | 0.833013983 | 0.461 | 0.067 | 0 | 6 | Gsap |
| Rnase4 | 0 | 0.829563446 | 0.571 | 0.062 | 0 | 6 | Rnase4 |
| Ptpre1 | 0 | 0.825279901 | 0.566 | 0.142 | 0 | 6 | Ptpre |
| Ptchd1 | 0 | 0.824392219 | 0.34 | 0.005 | 0 | 6 | Ptchd1 |
| Ppt1 | 0 | 0.824307202 | 0.537 | 0.11 | 0 | 6 | Ppt1 |
| Daglb | 0 | 0.819119581 | 0.423 | 0.033 | 0 | 6 | Daglb |
| Myof | 0 | 0.81810333 | 0.464 | 0.04 | 0 | 6 | Myof |
| Sap30 | 0 | 0.818034702 | 0.547 | 0.068 | 0 | 6 | Sap30 |
| Tmem135 | 0 | 0.817166492 | 0.586 | 0.147 | 0 | 6 | Tmem135 |
| Zeb2os | 0 | 0.812538519 | 0.415 | 0.013 | 0 | 6 | Zeb2os |
| Syk1 | 0 | 0.812524038 | 0.426 | 0.05 | 0 | 6 | Syk |
| Crlf21 | 0 | 0.809158617 | 0.615 | 0.14 | 0 | 6 | Crlf2 |
| Scamp2 | 0 | 0.806463896 | 0.569 | 0.121 | 0 | 6 | Scamp2 |
| Clec5a | 0 | 0.802228294 | 0.425 | 0.012 | 0 | 6 | Clec5a |
| Ch25h | 0 | 0.800884204 | 0.303 | 0.011 | 0 | 6 | Ch25h |
| Hipk2 | 0 | 0.800632786 | 0.492 | 0.1 | 0 | 6 | Hipk2 |
| Prcp | 0 | 0.798242809 | 0.467 | 0.068 | 0 | 6 | Prcp |
| Plekho12 | 0 | 0.79723478 | 0.717 | 0.201 | 0 | 6 | Plekho1 |
| Disc1 | 0 | 0.795988093 | 0.415 | 0.04 | 0 | 6 | Disc1 |
| Gpr341 | 0 | 0.793483443 | 0.364 | 0.045 | 0 | 6 | Gpr34 |
| Sfmbt1 | 0 | 0.78906953 | 0.449 | 0.066 | 0 | 6 | Sfmbt1 |
| Dnmt3a1 | 0 | 0.785647756 | 0.542 | 0.129 | 0 | 6 | Dnmt3a |

| | | | | | | | |
|---|---|---|---|---|---|---|---|
| Blvrb1 | 0 | 0.784296161 | 0.721 | 0.22 | 0 | 6 | Blvrb |
| Sh2d1b11 | 0 | 0.777789111 | 0.457 | 0.059 | 0 | 6 | Sh2d1b1 |
| Zfp599 | 0 | 0.776177007 | 0.358 | 0.008 | 0 | 6 | Zfp599 |
| Dusp16 | 0 | 0.773943454 | 0.497 | 0.101 | 0 | 6 | Dusp16 |
| Batf3 | 0 | 0.771827568 | 0.413 | 0.013 | 0 | 6 | Batf3 |
| Col14a1 | 0 | 0.769677986 | 0.451 | 0.038 | 0 | 6 | Col14a1 |
| Cysltr1 | 0 | 0.766738511 | 0.423 | 0.045 | 0 | 6 | Cysltr1 |
| Clec4a2 | 0 | 0.766096649 | 0.4 | 0.012 | 0 | 6 | Clec4a2 |
| Adcy71 | 0 | 0.765731265 | 0.476 | 0.097 | 0 | 6 | Adcy7 |
| Gtf2h2 | 0 | 0.763174621 | 0.455 | 0.054 | 0 | 6 | Gtf2h2 |
| Hmgcl | 0 | 0.762596113 | 0.574 | 0.112 | 0 | 6 | Hmgcl |
| Tank | 0 | 0.75967378 | 0.541 | 0.124 | 0 | 6 | Tank |
| Tuba1c | 0 | 0.757441379 | 0.521 | 0.101 | 0 | 6 | Tuba1c |
| Sema4d1 | 0 | 0.755923421 | 0.529 | 0.111 | 0 | 6 | Sema4d |
| Siglece | 0 | 0.754304379 | 0.38 | 0.008 | 0 | 6 | Siglece |
| Mcub | 0 | 0.751485412 | 0.45 | 0.048 | 0 | 6 | Mcub |
| Tle1 | 0 | 0.751189856 | 0.478 | 0.073 | 0 | 6 | Tle1 |
| Gmip1 | 0 | 0.750012912 | 0.443 | 0.065 | 0 | 6 | Gmip |
| Rtcb | 0 | 0.747581663 | 0.618 | 0.15 | 0 | 6 | Rtcb |
| Shtn1 | 0 | 0.746847806 | 0.497 | 0.047 | 0 | 6 | Shtn1 |
| Gna15 | 0 | 0.746838706 | 0.374 | 0.014 | 0 | 6 | Gna15 |
| Gm46224 | 0 | 0.743169974 | 0.305 | 0.004 | 0 | 6 | Gm46224 |
| Wdr44 | 0 | 0.742690689 | 0.483 | 0.106 | 0 | 6 | Wdr44 |
| Fgd4 | 0 | 0.742001587 | 0.375 | 0.025 | 0 | 6 | Fgd4 |
| Arhgdib2 | 0 | 0.740273379 | 0.786 | 0.207 | 0 | 6 | Arhgdib |
| Nrros | 0 | 0.738711729 | 0.464 | 0.09 | 0 | 6 | Nrros |
| Cmah | 0 | 0.737912276 | 0.388 | 0.052 | 0 | 6 | Cmah |
| Rab7b | 0 | 0.736350719 | 0.383 | 0.013 | 0 | 6 | Rab7b |
| Ang | 0 | 0.734005688 | 0.39 | 0.021 | 0 | 6 | Ang |
| Ccl6 | 0 | 0.732527747 | 0.394 | 0.016 | 0 | 6 | Ccl6 |
| Heatr5a | 0 | 0.731463575 | 0.495 | 0.1 | 0 | 6 | Heatr5a |
| Cd33 | 0 | 0.726778696 | 0.372 | 0.006 | 0 | 6 | Cd33 |
| Myo9b1 | 0 | 0.725669638 | 0.479 | 0.102 | 0 | 6 | Myo9b |
| Nfxl1 | 0 | 0.715927107 | 0.365 | 0.035 | 0 | 6 | Nfxl1 |
| Susd3 | 0 | 0.712520143 | 0.396 | 0.028 | 0 | 6 | Susd3 |
| Gm33370 | 0 | 0.710321707 | 0.319 | 0.014 | 0 | 6 | Gm33370 |
| P2rx7 | 0 | 0.706621064 | 0.362 | 0.036 | 0 | 6 | P2rx7 |
| Dok3 | 0 | 0.706034681 | 0.37 | 0.012 | 0 | 6 | Dok3 |
| Fcgr2b | 0 | 0.705550251 | 0.38 | 0.023 | 0 | 6 | Fcgr2b |
| Syngr2 | 0 | 0.704278648 | 0.544 | 0.123 | 0 | 6 | Syngr2 |
| Slc4a7 | 0 | 0.703900866 | 0.409 | 0.074 | 0 | 6 | Slc4a7 |
| Btk | 0 | 0.70096612 | 0.355 | 0.009 | 0 | 6 | Btk |
| Bcl2a1d | 0 | 0.698796922 | 0.386 | 0.019 | 0 | 6 | Bcl2a1d |

| | | | | | | | |
|---|---|---|---|---|---|---|---|
| Selplg1 | 0 | 0.698764083 | 0.582 | 0.135 | 0 | 6 | Selplg |
| Apba1 | 0 | 0.697521885 | 0.337 | 0.04 | 0 | 6 | Apba1 |
| Atp6v0a1 | 0 | 0.694988749 | 0.426 | 0.074 | 0 | 6 | Atp6v0a1 |
| Mknk1 | 0 | 0.682514464 | 0.374 | 0.042 | 0 | 6 | Mknk1 |
| Hps3 | 0 | 0.676896822 | 0.391 | 0.053 | 0 | 6 | Hps3 |
| Nfam1 | 0 | 0.674961507 | 0.315 | 0.008 | 0 | 6 | Nfam1 |
| Hps4 | 0 | 0.672044022 | 0.396 | 0.051 | 0 | 6 | Hps4 |
| Ddi2 | 0 | 0.67167925 | 0.48 | 0.101 | 0 | 6 | Ddi2 |
| Tifab | 0 | 0.669386407 | 0.339 | 0.008 | 0 | 6 | Tifab |
| Capn3 | 0 | 0.668221233 | 0.346 | 0.017 | 0 | 6 | Capn3 |
| Asb2 | 0 | 0.667919655 | 0.356 | 0.037 | 0 | 6 | Asb2 |
| St3gal3 | 0 | 0.665217893 | 0.364 | 0.06 | 0 | 6 | St3gal3 |
| Tbc1d16 | 0 | 0.664321967 | 0.325 | 0.03 | 0 | 6 | Tbc1d16 |
| Wdfy4 | 0 | 0.657235214 | 0.319 | 0.008 | 0 | 6 | Wdfy4 |
| Il21r1 | 0 | 0.655969141 | 0.42 | 0.078 | 0 | 6 | Il21r |
| Pfkfb4 | 0 | 0.655544893 | 0.329 | 0.017 | 0 | 6 | Pfkfb4 |
| Ms4a4c | 0 | 0.654890148 | 0.295 | 0.019 | 0 | 6 | Ms4a4c |
| Notch21 | 0 | 0.653501322 | 0.403 | 0.065 | 0 | 6 | Notch2 |
| Clec4a3 | 0 | 0.65349609 | 0.289 | 0.008 | 0 | 6 | Clec4a3 |
| Pla2g7 | 0 | 0.653158123 | 0.302 | 0.011 | 0 | 6 | Pla2g7 |
| Bmyc | 0 | 0.651430085 | 0.618 | 0.149 | 0 | 6 | Bmyc |
| Coro7 | 0 | 0.647682086 | 0.359 | 0.05 | 0 | 6 | Coro7 |
| Pmaip1 | 0 | 0.644953646 | 0.286 | 0.01 | 0 | 6 | Pmaip1 |
| Sdc3 | 0 | 0.644886447 | 0.403 | 0.062 | 0 | 6 | Sdc3 |
| Ncf41 | 0 | 0.644203813 | 0.483 | 0.095 | 0 | 6 | Ncf4 |
| Acox3 | 0 | 0.643593163 | 0.323 | 0.042 | 0 | 6 | Acox3 |
| Grk3 | 0 | 0.638492872 | 0.351 | 0.025 | 0 | 6 | Grk3 |
| Fgd3 | 0 | 0.635034984 | 0.351 | 0.035 | 0 | 6 | Fgd3 |
| Slc16a6 | 0 | 0.627649589 | 0.34 | 0.037 | 0 | 6 | Slc16a6 |
| Fam111a | 0 | 0.6274795 | 0.412 | 0.076 | 0 | 6 | Fam111a |
| Amz1 | 0 | 0.625935668 | 0.32 | 0.009 | 0 | 6 | Amz1 |
| Lilrb4a | 0 | 0.623668234 | 0.302 | 0.01 | 0 | 6 | Lilrb4a |
| Tep1 | 0 | 0.622958521 | 0.384 | 0.043 | 0 | 6 | Tep1 |
| Gm20513 | 0 | 0.622573278 | 0.282 | 0.005 | 0 | 6 | Gm20513 |
| Ppcdc | 0 | 0.622527836 | 0.374 | 0.058 | 0 | 6 | Ppcdc |
| Gpr160 | 0 | 0.622524273 | 0.326 | 0.031 | 0 | 6 | Gpr160 |
| Mapkapk31 | 0 | 0.620920998 | 0.409 | 0.066 | 0 | 6 | Mapkapk3 |
| Pik3cg1 | 0 | 0.620093263 | 0.341 | 0.041 | 0 | 6 | Pik3cg |
| Dtx4 | 0 | 0.616100422 | 0.349 | 0.018 | 0 | 6 | Dtx4 |
| Il7r | 0 | 0.616083566 | 0.354 | 0.024 | 0 | 6 | Il7r |
| Pdcd2l | 0 | 0.613049587 | 0.406 | 0.068 | 0 | 6 | Pdcd2l |
| Abcc3 | 0 | 0.612909723 | 0.301 | 0.007 | 0 | 6 | Abcc3 |
| Xlr | 0 | 0.61137525 | 0.348 | 0.032 | 0 | 6 | Xlr |

| | | | | | | | |
|---|---|---|---|---|---|---|---|
| Pnpla71 | 0 | 0.609370206 | 0.361 | 0.05 | 0 | 6 | Pnpla7 |
| Naip2 | 0 | 0.608621843 | 0.357 | 0.043 | 0 | 6 | Naip2 |
| 5031439G07Rik | 0 | 0.608404844 | 0.397 | 0.064 | 0 | 6 | 5031439G07Rik |
| Abcc5 | 0 | 0.605006769 | 0.38 | 0.066 | 0 | 6 | Abcc5 |
| Tlr7 | 0 | 0.599645631 | 0.281 | 0.004 | 0 | 6 | Tlr7 |
| Lyl1 | 0 | 0.597720113 | 0.317 | 0.01 | 0 | 6 | Lyl1 |
| Sla1 | 0 | 0.596866429 | 0.371 | 0.06 | 0 | 6 | Sla |
| Cmtm71 | 0 | 0.596781185 | 0.694 | 0.19 | 0 | 6 | Cmtm7 |
| Smim3 | 0 | 0.593162985 | 0.31 | 0.014 | 0 | 6 | Smim3 |
| Tbc1d8 | 0 | 0.593066236 | 0.35 | 0.046 | 0 | 6 | Tbc1d8 |
| Snx30 | 0 | 0.590745306 | 0.308 | 0.037 | 0 | 6 | Snx30 |
| 1810011H11Rik1 | 0 | 0.588676818 | 0.363 | 0.06 | 0 | 6 | 1810011H11Rik |
| Ifi209 | 0 | 0.584983465 | 0.314 | 0.045 | 0 | 6 | Ifi209 |
| Rassf2 | 0 | 0.584714982 | 0.327 | 0.041 | 0 | 6 | Rassf2 |
| 0610040J01Rik | 0 | 0.584244916 | 0.33 | 0.02 | 0 | 6 | 0610040J01Rik |
| Il18 | 0 | 0.582325271 | 0.289 | 0.01 | 0 | 6 | Il18 |
| Igf2bp3 | 0 | 0.580432387 | 0.263 | 0.018 | 0 | 6 | Igf2bp3 |
| Icosl | 0 | 0.576780146 | 0.304 | 0.019 | 0 | 6 | Icosl |
| Itga8 | 0 | 0.57397081 | 0.271 | 0.023 | 0 | 6 | Itga8 |
| Arhgap41 | 0 | 0.570703999 | 0.323 | 0.046 | 0 | 6 | Arhgap4 |
| Gm15726 | 0 | 0.563483669 | 0.254 | 0.002 | 0 | 6 | Gm15726 |
| Psen2 | 0 | 0.563129522 | 0.34 | 0.049 | 0 | 6 | Psen2 |
| Tnfrsf13b | 0 | 0.561722797 | 0.3 | 0.01 | 0 | 6 | Tnfrsf13b |
| Gmfg | 0 | 0.557692486 | 0.542 | 0.12 | 0 | 6 | Gmfg |
| P2rx4 | 0 | 0.557130093 | 0.378 | 0.05 | 0 | 6 | P2rx4 |
| Arid3b | 0 | 0.557122545 | 0.268 | 0.029 | 0 | 6 | Arid3b |
| Zfp810 | 0 | 0.555379409 | 0.32 | 0.04 | 0 | 6 | Zfp810 |
| Galnt3 | 0 | 0.549311803 | 0.262 | 0.012 | 0 | 6 | Galnt3 |
| Cmklr1 | 0 | 0.544616125 | 0.283 | 0.02 | 0 | 6 | Cmklr1 |
| B3gnt8 | 0 | 0.541599982 | 0.268 | 0.01 | 0 | 6 | B3gnt8 |
| Adap2os | 0 | 0.541270159 | 0.265 | 0.007 | 0 | 6 | Adap2os |
| Dapk1 | 0 | 0.539378026 | 0.355 | 0.05 | 0 | 6 | Dapk1 |
| Slc12a9 | 0 | 0.537538225 | 0.266 | 0.025 | 0 | 6 | Slc12a9 |
| Pla2g15 | 0 | 0.536770322 | 0.299 | 0.018 | 0 | 6 | Pla2g15 |
| Plk3 | 0 | 0.534837146 | 0.294 | 0.029 | 0 | 6 | Plk3 |
| I830077J02Rik | 0 | 0.533398666 | 0.263 | 0.007 | 0 | 6 | I830077J02Rik |
| Pirb | 0 | 0.531856098 | 0.254 | 0.007 | 0 | 6 | Pirb |
| Nlrp3 | 0 | 0.530697188 | 0.257 | 0.003 | 0 | 6 | Nlrp3 |
| Gpr155 | 0 | 0.528001341 | 0.28 | 0.031 | 0 | 6 | Gpr155 |
| B4galt6 | 0 | 0.527078542 | 0.259 | 0.01 | 0 | 6 | B4galt6 |
| Gk | 0 | 0.522188049 | 0.306 | 0.03 | 0 | 6 | Gk |
| Arl111 | 0 | 0.521032942 | 0.296 | 0.011 | 0 | 6 | Arl11 |
| Mmp14 | 0 | 0.520232255 | 0.37 | 0.045 | 0 | 6 | Mmp14 |

| | | | | | | | |
|---|---|---|---|---|---|---|---|
| Rp2 | 0 | 0.519603005 | 0.334 | 0.041 | 0 | 6 | Rp2 |
| Dock7 | 0 | 0.517071812 | 0.307 | 0.035 | 0 | 6 | Dock7 |
| Fam49a | 0 | 0.515405234 | 0.306 | 0.038 | 0 | 6 | Fam49a |
| Slc37a2 | 0 | 0.512781838 | 0.253 | 0.008 | 0 | 6 | Slc37a2 |
| Ston2 | 0 | 0.5121611 | 0.25 | 0.008 | 0 | 6 | Ston2 |
| A630001G21Rik | 0 | 0.508955878 | 0.283 | 0.03 | 0 | 6 | A630001G21Rik |
| Dcakd | 0 | 0.504513579 | 0.368 | 0.058 | 0 | 6 | Dcakd |
| Cebpa | 0 | 0.502611744 | 0.313 | 0.025 | 0 | 6 | Cebpa |
| St6gal1 | 0 | 0.502424471 | 0.266 | 0.031 | 0 | 6 | St6gal1 |
| Tctex1d2 | 0 | 0.496833975 | 0.374 | 0.048 | 0 | 6 | Tctex1d2 |
| Sh3bp21 | 0 | 0.493483821 | 0.277 | 0.032 | 0 | 6 | Sh3bp2 |
| Dck | 0 | 0.489786026 | 0.256 | 0.023 | 0 | 6 | Dck |
| Slc43a2 | 0 | 0.488895353 | 0.279 | 0.023 | 0 | 6 | Slc43a2 |
| Dglucy | 0 | 0.487567416 | 0.26 | 0.028 | 0 | 6 | Dglucy |
| Frrs1 | 0 | 0.484594493 | 0.26 | 0.017 | 0 | 6 | Frrs1 |
| Galc | 0 | 0.480921001 | 0.274 | 0.033 | 0 | 6 | Galc |
| Zfp992 | 0 | 0.480478248 | 0.255 | 0.015 | 0 | 6 | Zfp992 |
| Slc16a7 | 0 | 0.477058561 | 0.262 | 0.021 | 0 | 6 | Slc16a7 |
| Sh3pxd2b | 0 | 0.47607198 | 0.273 | 0.03 | 0 | 6 | Sh3pxd2b |
| Zdhhc13 | 0 | 0.473679908 | 0.258 | 0.027 | 0 | 6 | Zdhhc13 |
| Cd302 | 0 | 0.469040658 | 0.448 | 0.063 | 0 | 6 | Cd302 |
| Galnt12 | 0 | 0.448794579 | 0.256 | 0.03 | 0 | 6 | Galnt12 |
| Zfp385a | 0 | 0.448475276 | 0.262 | 0.012 | 0 | 6 | Zfp385a |
| Parvg | 0 | 0.439683204 | 0.279 | 0.036 | 0 | 6 | Parvg |
| Fmn1 | 0 | 0.411781369 | 0.289 | 0.035 | 0 | 6 | Fmn1 |
| Lrp1 | 0 | 0.282340679 | 0.327 | 0.043 | 0 | 6 | Lrp1 |
| Serinc32 | 4.12E-306 | 1.160506461 | 0.95 | 0.465 | 8.34E-302 | 6 | Serinc3 |
| Ahcyl2 | 6.71E-305 | 0.696807392 | 0.492 | 0.109 | 1.36E-300 | 6 | Ahcyl2 |
| Cep170 | 7.72E-305 | 0.716566177 | 0.498 | 0.112 | 1.56E-300 | 6 | Cep170 |
| Tifa | 2.98E-303 | 0.359500186 | 0.312 | 0.045 | 6.03E-299 | 6 | Tifa |
| Phka2 | 1.46E-302 | 0.481836405 | 0.27 | 0.035 | 2.96E-298 | 6 | Phka2 |
| Atp6ap2 | 5.43E-301 | 0.808490895 | 0.584 | 0.155 | 1.10E-296 | 6 | Atp6ap2 |
| Fth12 | 1.73E-299 | 0.983727445 | 0.997 | 0.968 | 3.51E-295 | 6 | Fth1 |
| Camk11 | 1.87E-299 | 0.711137615 | 0.7 | 0.205 | 3.79E-295 | 6 | Camk1 |
| Fuca1 | 1.32E-298 | 0.806019281 | 0.65 | 0.187 | 2.67E-294 | 6 | Fuca1 |
| Dock112 | 1.63E-298 | 0.717291499 | 0.456 | 0.098 | 3.31E-294 | 6 | Dock11 |
| Stap11 | 2.86E-298 | 0.39003112 | 0.489 | 0.107 | 5.78E-294 | 6 | Stap1 |
| Zdhhc14 | 2.69E-297 | 0.490753253 | 0.268 | 0.035 | 5.44E-293 | 6 | Zdhhc14 |
| Tor3a | 3.60E-297 | 0.649491325 | 0.435 | 0.088 | 7.29E-293 | 6 | Tor3a |
| Clasp21 | 3.91E-296 | 0.867630862 | 0.591 | 0.163 | 7.91E-292 | 6 | Clasp2 |
| Runx31 | 8.06E-296 | 0.538066022 | 0.563 | 0.13 | 1.63E-291 | 6 | Runx3 |
| Atp2b12 | 8.32E-294 | 1.02901211 | 0.82 | 0.31 | 1.68E-289 | 6 | Atp2b1 |
| Rps6ka11 | 9.36E-294 | 0.642681904 | 0.456 | 0.098 | 1.90E-289 | 6 | Rps6ka1 |

| | | | | | | | |
|---|---|---|---|---|---|---|---|
| Fam46a | 3.45E-292 | 0.60989159 | 0.52 | 0.122 | 6.98E-288 | 6 | Fam46a |
| Dse | 1.32E-290 | 0.542111107 | 0.312 | 0.048 | 2.67E-286 | 6 | Dse |
| Wdfy32 | 7.61E-289 | 0.79197321 | 0.603 | 0.162 | 1.54E-284 | 6 | Wdfy3 |
| Zufsp1 | 5.17E-288 | 0.774586435 | 0.498 | 0.119 | 1.05E-283 | 6 | Zufsp |
| Tmem106a | 9.10E-287 | 0.426094934 | 0.283 | 0.04 | 1.84E-282 | 6 | Tmem106a |
| Lcp21 | 1.94E-285 | 0.633394689 | 0.404 | 0.08 | 3.94E-281 | 6 | Lcp2 |
| Rhog | 1.03E-284 | 0.610403439 | 0.486 | 0.112 | 2.08E-280 | 6 | Rhog |
| Casp1 | 1.57E-284 | 0.441370268 | 0.309 | 0.048 | 3.18E-280 | 6 | Casp1 |
| Fam102b | 2.78E-284 | 0.454972813 | 0.287 | 0.041 | 5.62E-280 | 6 | Fam102b |
| Rnf131 | 1.05E-283 | 0.786959506 | 0.584 | 0.159 | 2.13E-279 | 6 | Rnf13 |
| Fbxw11 | 1.01E-279 | 0.760812833 | 0.564 | 0.152 | 2.05E-275 | 6 | Fbxw11 |
| Dennd1b1 | 1.24E-279 | 0.868911725 | 0.588 | 0.165 | 2.50E-275 | 6 | Dennd1b |
| Ucp23 | 3.41E-279 | 0.741204321 | 0.834 | 0.307 | 6.91E-275 | 6 | Ucp2 |
| Tmem37 | 5.34E-279 | 0.466295204 | 0.271 | 0.038 | 1.08E-274 | 6 | Tmem37 |
| 3110043O21Rik | 1.02E-278 | 0.479305303 | 0.276 | 0.039 | 2.06E-274 | 6 | 3110043O21Rik |
| Tlr4 | 3.21E-278 | 0.467343875 | 0.291 | 0.043 | 6.50E-274 | 6 | Tlr4 |
| Ptpn182 | 9.87E-278 | 0.737017525 | 0.849 | 0.29 | 2.00E-273 | 6 | Ptpn18 |
| Itgb21 | 4.41E-277 | 0.608816875 | 0.418 | 0.087 | 8.93E-273 | 6 | Itgb2 |
| Pgd | 6.80E-277 | 0.489388002 | 0.403 | 0.081 | 1.38E-272 | 6 | Pgd |
| Man1c1 | 1.17E-276 | 0.572488995 | 0.34 | 0.059 | 2.37E-272 | 6 | Man1c1 |
| Ifnar11 | 4.59E-276 | 0.730151032 | 0.508 | 0.127 | 9.29E-272 | 6 | Ifnar1 |
| Itm2c | 6.57E-276 | 0.705352809 | 0.62 | 0.179 | 1.33E-271 | 6 | Itm2c |
| Hbb-bt | 7.11E-275 | 3.188916605 | 0.351 | 0.068 | 1.44E-270 | 6 | Hbb-bt |
| Adam171 | 1.61E-274 | 0.760872194 | 0.527 | 0.136 | 3.26E-270 | 6 | Adam17 |
| Hba-a2 | 2.62E-273 | 3.093874757 | 0.413 | 0.097 | 5.31E-269 | 6 | Hba-a2 |
| Fgl21 | 3.86E-273 | 0.637369224 | 0.431 | 0.093 | 7.81E-269 | 6 | Fgl2 |
| Btg22 | 5.64E-273 | 1.168616599 | 0.823 | 0.322 | 1.14E-268 | 6 | Btg2 |
| Ankrd441 | 1.13E-271 | 0.857912613 | 0.797 | 0.27 | 2.28E-267 | 6 | Ankrd44 |
| Tmem173 | 1.14E-271 | 0.508224089 | 0.335 | 0.059 | 2.30E-267 | 6 | Tmem173 |
| Fes | 4.21E-271 | 0.500952524 | 0.365 | 0.068 | 8.53E-267 | 6 | Fes |
| Cap11 | 5.39E-271 | 0.885611325 | 0.721 | 0.251 | 1.09E-266 | 6 | Cap1 |
| Rab20 | 3.12E-270 | 0.461083357 | 0.342 | 0.061 | 6.33E-266 | 6 | Rab20 |
| Lap3 | 7.85E-269 | 0.473835773 | 0.501 | 0.12 | 1.59E-264 | 6 | Lap3 |
| Tpd522 | 2.71E-267 | 0.391671089 | 0.85 | 0.28 | 5.49E-263 | 6 | Tpd52 |
| Rtn4rl11 | 4.90E-267 | 0.487774547 | 0.269 | 0.039 | 9.92E-263 | 6 | Rtn4rl1 |
| Morc3 | 1.05E-264 | 0.71517101 | 0.536 | 0.145 | 2.12E-260 | 6 | Morc3 |
| Tpp21 | 1.80E-263 | 0.85682471 | 0.648 | 0.208 | 3.65E-259 | 6 | Tpp2 |
| Rps6ka51 | 2.07E-263 | 0.689807331 | 0.423 | 0.093 | 4.20E-259 | 6 | Rps6ka5 |
| Pfkfb3 | 4.17E-262 | 0.565489874 | 0.374 | 0.073 | 8.45E-258 | 6 | Pfkfb3 |
| Fnip11 | 4.62E-262 | 0.74623325 | 0.502 | 0.129 | 9.36E-258 | 6 | Fnip1 |
| Rasa4 | 3.36E-261 | 0.429178445 | 0.298 | 0.047 | 6.79E-257 | 6 | Rasa4 |
| Zfp361 | 1.96E-260 | 0.805463762 | 0.603 | 0.179 | 3.97E-256 | 6 | Zfp36 |
| Rhoh1 | 4.56E-260 | 0.571186721 | 0.414 | 0.089 | 9.24E-256 | 6 | Rhoh |

| | | | | | | | |
|---|---|---|---|---|---|---|---|
| Abcb1b | 4.74E-257 | 0.471601507 | 0.282 | 0.044 | 9.59E-253 | 6 | Abcb1b |
| Tmem1641 | 5.86E-257 | 0.750985986 | 0.536 | 0.146 | 1.19E-252 | 6 | Tmem164 |
| Akap101 | 2.53E-256 | 0.747027684 | 0.497 | 0.129 | 5.12E-252 | 6 | Akap10 |
| Picalm1 | 2.62E-255 | 0.900394194 | 0.749 | 0.275 | 5.30E-251 | 6 | Picalm |
| Foxn33 | 3.69E-255 | 1.011399858 | 0.886 | 0.389 | 7.46E-251 | 6 | Foxn3 |
| Camk2d2 | 3.97E-255 | 0.911968201 | 0.713 | 0.249 | 8.05E-251 | 6 | Camk2d |
| Lrch12 | 4.60E-255 | 0.938727842 | 0.775 | 0.29 | 9.32E-251 | 6 | Lrch1 |
| Alpk1 | 1.13E-252 | 0.431347011 | 0.257 | 0.037 | 2.29E-248 | 6 | Alpk1 |
| Tgif1 | 2.80E-252 | 0.355768373 | 0.26 | 0.038 | 5.67E-248 | 6 | Tgif1 |
| Sesn11 | 2.65E-251 | 0.882742975 | 0.617 | 0.193 | 5.38E-247 | 6 | Sesn1 |
| Jmjd1c3 | 4.34E-251 | 0.967813948 | 0.754 | 0.286 | 8.78E-247 | 6 | Jmjd1c |
| Abi12 | 2.65E-250 | 0.838153567 | 0.713 | 0.248 | 5.36E-246 | 6 | Abi1 |
| Tmem50b | 1.44E-249 | 0.44378831 | 0.329 | 0.06 | 2.91E-245 | 6 | Tmem50b |
| Hba-a1 | 1.50E-249 | 3.128824384 | 0.43 | 0.114 | 3.04E-245 | 6 | Hba-a1 |
| Tbc1d23 | 4.76E-249 | 0.67731082 | 0.444 | 0.107 | 9.65E-245 | 6 | Tbc1d23 |
| Hspa1b | 2.32E-248 | 0.647059055 | 0.314 | 0.056 | 4.69E-244 | 6 | Hspa1b |
| Large11 | 3.63E-248 | 0.833338993 | 0.504 | 0.136 | 7.34E-244 | 6 | Large1 |
| Sgpl1 | 1.21E-247 | 0.474745662 | 0.29 | 0.049 | 2.45E-243 | 6 | Sgpl1 |
| Atxn7l1 | 1.44E-246 | 0.708299894 | 0.469 | 0.118 | 2.91E-242 | 6 | Atxn7l1 |
| Ccnl11 | 3.94E-244 | 0.716321957 | 0.614 | 0.191 | 7.97E-240 | 6 | Ccnl1 |
| Map3k141 | 1.91E-242 | 0.619881593 | 0.406 | 0.092 | 3.86E-238 | 6 | Map3k14 |
| Fxyd52 | 2.56E-242 | 0.861072615 | 0.9 | 0.388 | 5.17E-238 | 6 | Fxyd5 |
| Klhl61 | 4.33E-241 | 0.637229272 | 0.439 | 0.106 | 8.76E-237 | 6 | Klhl6 |
| Enox21 | 1.42E-240 | 0.7286646 | 0.485 | 0.127 | 2.87E-236 | 6 | Enox2 |
| Arhgap6 | 1.98E-238 | 0.339637352 | 0.288 | 0.047 | 4.01E-234 | 6 | Arhgap6 |
| Bst23 | 1.05E-237 | 0.904177721 | 0.957 | 0.482 | 2.12E-233 | 6 | Bst2 |
| Tmem219 | 1.56E-237 | 0.533788354 | 0.445 | 0.108 | 3.15E-233 | 6 | Tmem219 |
| Was1 | 3.03E-237 | 0.447628937 | 0.323 | 0.061 | 6.14E-233 | 6 | Was |
| Rab8b1 | 8.44E-237 | 0.878327433 | 0.723 | 0.264 | 1.71E-232 | 6 | Rab8b |
| Il13ra1 | 3.61E-236 | 0.472431904 | 0.331 | 0.063 | 7.31E-232 | 6 | Il13ra1 |
| Mapre21 | 1.04E-235 | 0.991494055 | 0.657 | 0.231 | 2.11E-231 | 6 | Mapre2 |
| Slx4ip | 6.37E-235 | 0.516965528 | 0.31 | 0.057 | 1.29E-230 | 6 | Slx4ip |
| Pea15a1 | 8.81E-233 | 0.65657416 | 0.666 | 0.22 | 1.78E-228 | 6 | Pea15a |
| Zfp516 | 2.00E-231 | 0.383773388 | 0.273 | 0.045 | 4.05E-227 | 6 | Zfp516 |
| Lrmp1 | 1.07E-229 | 0.455871799 | 0.285 | 0.05 | 2.16E-225 | 6 | Lrmp |
| Hacd2 | 1.87E-229 | 0.537920158 | 0.424 | 0.102 | 3.79E-225 | 6 | Hacd2 |
| Hacd41 | 2.41E-229 | 0.526988389 | 0.427 | 0.101 | 4.89E-225 | 6 | Hacd4 |
| Gnaq2 | 4.42E-229 | 0.901413859 | 0.824 | 0.337 | 8.96E-225 | 6 | Gnaq |
| Ust1 | 6.93E-229 | 0.579018525 | 0.35 | 0.074 | 1.40E-224 | 6 | Ust |
| Fmnl11 | 1.20E-228 | 0.503507155 | 0.368 | 0.08 | 2.44E-224 | 6 | Fmnl1 |
| Stard91 | 1.34E-228 | 0.638560687 | 0.504 | 0.135 | 2.71E-224 | 6 | Stard9 |
| Arap1 | 1.97E-227 | 0.364362935 | 0.282 | 0.049 | 3.98E-223 | 6 | Arap1 |
| Commd8 | 8.84E-227 | 0.493853158 | 0.505 | 0.138 | 1.79E-222 | 6 | Commd8 |

| | | | | | | | |
|---|---|---|---|---|---|---|---|
| Slfn81 | 8.60E-226 | 0.484499724 | 0.324 | 0.064 | 1.74E-221 | 6 | Slfn8 |
| Manba | 1.42E-224 | 0.460189786 | 0.294 | 0.054 | 2.87E-220 | 6 | Manba |
| Actb2 | 1.48E-224 | 0.862758539 | 0.994 | 0.836 | 3.01E-220 | 6 | Actb |
| Ipcef11 | 2.64E-224 | 0.547153923 | 0.431 | 0.103 | 5.34E-220 | 6 | Ipcef1 |
| Dgkd1 | 5.77E-224 | 0.638440471 | 0.511 | 0.145 | 1.17E-219 | 6 | Dgkd |
| Rab31 | 6.36E-224 | 0.529995542 | 0.384 | 0.087 | 1.29E-219 | 6 | Rab31 |
| Soga1 | 2.59E-223 | 0.466018838 | 0.279 | 0.049 | 5.24E-219 | 6 | Soga1 |
| Ero1lb | 3.79E-223 | 0.514810796 | 0.288 | 0.053 | 7.67E-219 | 6 | Ero1lb |
| Dock42 | 1.10E-221 | 0.741739433 | 0.771 | 0.274 | 2.23E-217 | 6 | Dock4 |
| Rabep12 | 2.85E-221 | 0.754537273 | 0.626 | 0.212 | 5.78E-217 | 6 | Rabep1 |
| Twf21 | 3.22E-220 | 0.552072437 | 0.529 | 0.154 | 6.51E-216 | 6 | Twf2 |
| Mid1 | 4.67E-220 | 0.457487463 | 0.347 | 0.073 | 9.45E-216 | 6 | Mid1 |
| Cerk1 | 3.25E-219 | 0.582965218 | 0.41 | 0.101 | 6.58E-215 | 6 | Cerk |
| Nt5c2 | 1.25E-218 | 0.486782537 | 0.369 | 0.083 | 2.54E-214 | 6 | Nt5c2 |
| Tfe3 | 3.70E-218 | 0.398798037 | 0.273 | 0.048 | 7.49E-214 | 6 | Tfe3 |
| Map3k51 | 1.05E-216 | 0.755382785 | 0.556 | 0.176 | 2.12E-212 | 6 | Map3k5 |
| Antxr2 | 6.11E-216 | 0.468432076 | 0.358 | 0.078 | 1.24E-211 | 6 | Antxr2 |
| Mylip | 1.75E-214 | 0.669225077 | 0.587 | 0.192 | 3.55E-210 | 6 | Mylip |
| Comt | 2.91E-214 | 0.522842259 | 0.572 | 0.175 | 5.88E-210 | 6 | Comt |
| AC149090.11 | 2.57E-213 | 0.711293448 | 0.637 | 0.213 | 5.21E-209 | 6 | AC149090.1 |
| Mbnl11 | 2.66E-213 | 0.827027606 | 0.938 | 0.436 | 5.38E-209 | 6 | Mbnl1 |
| Mef2c2 | 4.33E-212 | 0.724693825 | 0.961 | 0.398 | 8.77E-208 | 6 | Mef2c |
| Tmsb4x2 | 3.36E-211 | 0.762723828 | 0.998 | 0.931 | 6.80E-207 | 6 | Tmsb4x |
| Gpsm31 | 2.11E-210 | 0.528469615 | 0.477 | 0.133 | 4.28E-206 | 6 | Gpsm3 |
| Tcirg1 | 2.26E-210 | 0.475106217 | 0.316 | 0.065 | 4.58E-206 | 6 | Tcirg1 |
| Tbc1d14 | 2.92E-210 | 0.448629271 | 0.293 | 0.057 | 5.90E-206 | 6 | Tbc1d14 |
| Plekho2 | 5.80E-208 | 0.495272304 | 0.329 | 0.071 | 1.17E-203 | 6 | Plekho2 |
| Wsb11 | 1.80E-207 | 0.63422579 | 0.524 | 0.159 | 3.64E-203 | 6 | Wsb1 |
| Lgals3bp1 | 9.44E-207 | 0.598483068 | 0.469 | 0.132 | 1.91E-202 | 6 | Lgals3bp |
| Sp140 | 2.20E-205 | 0.383185687 | 0.334 | 0.072 | 4.44E-201 | 6 | Sp140 |
| Ifi213 | 3.69E-205 | 0.595824176 | 0.323 | 0.069 | 7.47E-201 | 6 | Ifi213 |
| Trim30d | 9.68E-205 | 0.514591687 | 0.328 | 0.071 | 1.96E-200 | 6 | Trim30d |
| Lrrk11 | 1.39E-204 | 0.657229576 | 0.553 | 0.174 | 2.82E-200 | 6 | Lrrk1 |
| Chd71 | 7.73E-204 | 0.644990259 | 0.474 | 0.136 | 1.57E-199 | 6 | Chd7 |
| Ifnar22 | 8.40E-203 | 0.708258746 | 0.763 | 0.304 | 1.70E-198 | 6 | Ifnar2 |
| Sdccag81 | 8.59E-203 | 0.520308729 | 0.416 | 0.108 | 1.74E-198 | 6 | Sdccag8 |
| Nptn1 | 4.34E-202 | 0.73978758 | 0.641 | 0.231 | 8.78E-198 | 6 | Nptn |
| Ptpn11 | 4.95E-202 | 0.607640929 | 0.628 | 0.218 | 1.00E-197 | 6 | Ptpn1 |
| Zbp1 | 1.05E-201 | 0.423760536 | 0.296 | 0.059 | 2.12E-197 | 6 | Zbp1 |
| Rnf1491 | 1.19E-201 | 0.479720422 | 0.377 | 0.092 | 2.41E-197 | 6 | Rnf149 |
| Ap1s21 | 2.21E-201 | 0.512009337 | 0.456 | 0.124 | 4.48E-197 | 6 | Ap1s2 |
| Pias11 | 3.32E-201 | 0.640230731 | 0.541 | 0.172 | 6.73E-197 | 6 | Pias1 |
| Fblim1 | 6.89E-201 | 0.343335318 | 0.258 | 0.045 | 1.40E-196 | 6 | Fblim1 |

| | | | | | | | |
|---|---|---|---|---|---|---|---|
| Smox | 1.56E-200 | 0.390351744 | 0.311 | 0.064 | 3.16E-196 | 6 | Smox |
| Ftl13 | 2.13E-199 | 0.711034239 | 0.993 | 0.907 | 4.31E-195 | 6 | Ftl1 |
| Nceh1 | 3.14E-199 | 0.517575781 | 0.349 | 0.081 | 6.36E-195 | 6 | Nceh1 |
| Cpeb2 | 5.19E-199 | 0.46733674 | 0.269 | 0.051 | 1.05E-194 | 6 | Cpeb2 |
| H2-M3 | 8.66E-199 | 0.528039237 | 0.466 | 0.132 | 1.75E-194 | 6 | H2-M3 |
| Bcl2l1 | 1.50E-198 | 0.596759661 | 0.466 | 0.132 | 3.03E-194 | 6 | Bcl2l1 |
| Scarb21 | 1.01E-197 | 0.614470085 | 0.54 | 0.171 | 2.04E-193 | 6 | Scarb2 |
| Selenop3 | 5.53E-197 | 0.774738032 | 0.823 | 0.342 | 1.12E-192 | 6 | Selenop |
| Plekhm2 | 4.71E-194 | 0.429401247 | 0.314 | 0.068 | 9.53E-190 | 6 | Plekhm2 |
| Sft2d11 | 1.01E-193 | 0.54395496 | 0.489 | 0.145 | 2.05E-189 | 6 | Sft2d1 |
| Rasa3 | 5.72E-193 | 0.43901601 | 0.281 | 0.056 | 1.16E-188 | 6 | Rasa3 |
| Lcorl2 | 1.17E-192 | 0.614967896 | 0.497 | 0.151 | 2.36E-188 | 6 | Lcorl |
| Gm203883 | 1.38E-192 | 0.794494371 | 0.851 | 0.38 | 2.79E-188 | 6 | Gm20388 |
| Stambpl1 | 1.98E-192 | 0.438875872 | 0.289 | 0.059 | 4.01E-188 | 6 | Stambpl1 |
| Abl21 | 3.42E-191 | 0.554370659 | 0.402 | 0.106 | 6.93E-187 | 6 | Abl2 |
| Mvb12b1 | 1.82E-190 | 0.553761457 | 0.496 | 0.152 | 3.68E-186 | 6 | Mvb12b |
| Rnf19b1 | 2.66E-190 | 0.516002673 | 0.324 | 0.074 | 5.39E-186 | 6 | Rnf19b |
| Mia2 | 5.98E-190 | 0.598490232 | 0.564 | 0.191 | 1.21E-185 | 6 | Mia2 |
| Uvrag2 | 6.59E-190 | 0.706617438 | 0.666 | 0.249 | 1.33E-185 | 6 | Uvrag |
| Mtss1 | 7.18E-190 | 0.464878453 | 0.363 | 0.088 | 1.45E-185 | 6 | Mtss1 |
| Slc23a21 | 1.10E-189 | 0.577032329 | 0.442 | 0.125 | 2.22E-185 | 6 | Slc23a2 |
| Pitpna1 | 1.11E-189 | 0.643987125 | 0.741 | 0.309 | 2.24E-185 | 6 | Pitpna |
| Nf12 | 2.93E-189 | 0.667624972 | 0.588 | 0.204 | 5.94E-185 | 6 | Nf1 |
| Rnpep | 3.11E-189 | 0.428214546 | 0.397 | 0.103 | 6.30E-185 | 6 | Rnpep |
| Bin11 | 6.50E-189 | 0.465823523 | 0.514 | 0.16 | 1.32E-184 | 6 | Bin1 |
| Ccdc88a2 | 2.38E-187 | 0.67037637 | 0.56 | 0.188 | 4.83E-183 | 6 | Ccdc88a |
| Arpc22 | 1.16E-186 | 0.772001741 | 0.935 | 0.527 | 2.34E-182 | 6 | Arpc2 |
| Tacc12 | 8.18E-186 | 0.70327913 | 0.712 | 0.286 | 1.66E-181 | 6 | Tacc1 |
| Pold41 | 2.07E-185 | 0.481915715 | 0.553 | 0.184 | 4.19E-181 | 6 | Pold4 |
| Tmem229b | 4.73E-185 | 0.409644822 | 0.258 | 0.049 | 9.57E-181 | 6 | Tmem229b |
| Hsd17b12 | 1.60E-183 | 0.479049381 | 0.429 | 0.121 | 3.25E-179 | 6 | Hsd17b12 |
| Tmem55b | 5.22E-183 | 0.444313038 | 0.363 | 0.09 | 1.06E-178 | 6 | Tmem55b |
| Tiparp | 1.87E-182 | 0.516902946 | 0.323 | 0.076 | 3.79E-178 | 6 | Tiparp |
| Tet3 | 3.41E-181 | 0.546213481 | 0.416 | 0.117 | 6.91E-177 | 6 | Tet3 |
| Pbx31 | 9.15E-181 | 0.656046234 | 0.433 | 0.126 | 1.85E-176 | 6 | Pbx3 |
| Elf41 | 3.41E-180 | 0.592498272 | 0.398 | 0.11 | 6.90E-176 | 6 | Elf4 |
| Ubash3b1 | 3.89E-180 | 0.597524102 | 0.489 | 0.153 | 7.88E-176 | 6 | Ubash3b |
| Sorl11 | 5.66E-180 | 0.447556969 | 0.381 | 0.099 | 1.15E-175 | 6 | Sorl1 |
| Itga41 | 1.44E-179 | 0.440696373 | 0.3 | 0.066 | 2.92E-175 | 6 | Itga4 |
| Kdm7a1 | 7.00E-179 | 0.497319937 | 0.482 | 0.147 | 1.42E-174 | 6 | Kdm7a |
| Ski2 | 9.85E-179 | 0.630136687 | 0.564 | 0.199 | 2.00E-174 | 6 | Ski |
| Rnf1112 | 1.03E-178 | 0.541047761 | 0.465 | 0.142 | 2.09E-174 | 6 | Rnf111 |
| Mapkapk2 | 6.49E-178 | 0.419068289 | 0.436 | 0.126 | 1.31E-173 | 6 | Mapkapk2 |

| | | | | | | | |
|---|---|---|---|---|---|---|---|
| Csk1 | 6.52E-178 | 0.421154976 | 0.417 | 0.116 | 1.32E-173 | 6 | Csk |
| Galnt71 | 1.86E-177 | 0.610633227 | 0.459 | 0.142 | 3.77E-173 | 6 | Galnt7 |
| Ogt1 | 3.27E-177 | 0.640924306 | 0.552 | 0.19 | 6.62E-173 | 6 | Ogt |
| Klf61 | 3.62E-177 | 0.699553805 | 0.652 | 0.25 | 7.33E-173 | 6 | Klf6 |
| Nmt11 | 1.51E-176 | 0.496692828 | 0.688 | 0.272 | 3.06E-172 | 6 | Nmt1 |
| Dusp61 | 3.23E-176 | 0.528629328 | 0.433 | 0.125 | 6.54E-172 | 6 | Dusp6 |
| Gnptab1 | 4.08E-176 | 0.450442192 | 0.316 | 0.074 | 8.27E-172 | 6 | Gnptab |
| Maml23 | 6.89E-176 | 0.825976982 | 0.752 | 0.324 | 1.39E-171 | 6 | Maml2 |
| Tmbim1 | 9.13E-176 | 0.473527139 | 0.402 | 0.11 | 1.85E-171 | 6 | Tmbim1 |
| Plekha21 | 1.75E-175 | 0.532256451 | 0.42 | 0.121 | 3.53E-171 | 6 | Plekha2 |
| Trim141 | 1.57E-174 | 0.489318933 | 0.394 | 0.108 | 3.19E-170 | 6 | Trim14 |
| Ubac21 | 3.99E-174 | 0.549113344 | 0.523 | 0.173 | 8.07E-170 | 6 | Ubac2 |
| Myo9a1 | 5.43E-174 | 0.577506325 | 0.495 | 0.159 | 1.10E-169 | 6 | Myo9a |
| Rptor1 | 1.12E-173 | 0.548145066 | 0.426 | 0.126 | 2.27E-169 | 6 | Rptor |
| Tbc1d22a | 4.00E-173 | 0.47266457 | 0.4 | 0.111 | 8.10E-169 | 6 | Tbc1d22a |
| Wdfy2 | 5.17E-173 | 0.459199044 | 0.375 | 0.099 | 1.05E-168 | 6 | Wdfy2 |
| Stk41 | 8.01E-173 | 0.49943225 | 0.501 | 0.16 | 1.62E-168 | 6 | Stk4 |
| Acvr1 | 8.37E-173 | 0.386482478 | 0.27 | 0.056 | 1.70E-168 | 6 | Acvr1 |
| Ccnd12 | 3.52E-172 | 0.532131171 | 0.601 | 0.21 | 7.13E-168 | 6 | Ccnd1 |
| Usp8 | 7.08E-172 | 0.465446896 | 0.456 | 0.139 | 1.43E-167 | 6 | Usp8 |
| Btaf1 | 7.77E-172 | 0.487604805 | 0.401 | 0.113 | 1.57E-167 | 6 | Btaf1 |
| Atp6v1b2 | 5.02E-171 | 0.411001691 | 0.372 | 0.099 | 1.02E-166 | 6 | Atp6v1b2 |
| Adam102 | 8.70E-170 | 0.63081176 | 0.692 | 0.274 | 1.76E-165 | 6 | Adam10 |
| Tmx42 | 1.04E-169 | 0.845816269 | 0.439 | 0.138 | 2.10E-165 | 6 | Tmx4 |
| AI504432 | 1.70E-169 | 0.387436783 | 0.277 | 0.06 | 3.45E-165 | 6 | AI504432 |
| Pik3cd1 | 4.00E-169 | 0.471900319 | 0.369 | 0.097 | 8.10E-165 | 6 | Pik3cd |
| Evl1 | 7.23E-169 | 0.51331685 | 0.588 | 0.204 | 1.46E-164 | 6 | Evl |
| Tgs1 | 9.86E-169 | 0.456302129 | 0.447 | 0.135 | 2.00E-164 | 6 | Tgs1 |
| Edem1 | 4.26E-168 | 0.407117893 | 0.32 | 0.078 | 8.63E-164 | 6 | Edem1 |
| Jpt11 | 1.29E-167 | 0.410561404 | 0.59 | 0.206 | 2.61E-163 | 6 | Jpt1 |
| Lamp13 | 3.06E-167 | 0.670112834 | 0.828 | 0.398 | 6.19E-163 | 6 | Lamp1 |
| Tmem131l1 | 3.53E-167 | 0.412972036 | 0.302 | 0.071 | 7.15E-163 | 6 | Tmem131l |
| Tubgcp5 | 1.55E-166 | 0.440379352 | 0.317 | 0.077 | 3.14E-162 | 6 | Tubgcp5 |
| Rnf1303 | 6.77E-166 | 0.571047045 | 0.671 | 0.266 | 1.37E-161 | 6 | Rnf130 |
| Arpc4 | 4.82E-165 | 0.513806892 | 0.677 | 0.273 | 9.76E-161 | 6 | Arpc4 |
| Egr11 | 8.57E-164 | 0.617436419 | 0.488 | 0.161 | 1.73E-159 | 6 | Egr1 |
| Mkl11 | 1.07E-163 | 0.516280874 | 0.392 | 0.114 | 2.17E-159 | 6 | Mkl1 |
| P4ha1 | 2.80E-163 | 0.523874963 | 0.392 | 0.112 | 5.67E-159 | 6 | P4ha1 |
| Dennd4a1 | 3.57E-163 | 0.266472519 | 0.54 | 0.172 | 7.23E-159 | 6 | Dennd4a |
| Trafd11 | 1.58E-162 | 0.46355033 | 0.452 | 0.14 | 3.20E-158 | 6 | Trafd1 |
| Mbp | 5.84E-162 | 0.304557482 | 0.269 | 0.058 | 1.18E-157 | 6 | Mbp |
| Calr2 | 6.48E-162 | 0.803729502 | 0.801 | 0.39 | 1.31E-157 | 6 | Calr |
| Kdm2b1 | 1.89E-161 | 0.416563102 | 0.317 | 0.079 | 3.83E-157 | 6 | Kdm2b |

| | | | | | | | |
|---|---|---|---|---|---|---|---|
| Ly96 | 2.38E-161 | 0.317323702 | 0.281 | 0.062 | 4.81E-157 | 6 | Ly96 |
| Baz1a1 | 3.93E-161 | 0.560560459 | 0.48 | 0.158 | 7.96E-157 | 6 | Baz1a |
| Sh3bgrl32 | 1.09E-160 | 0.61792909 | 0.951 | 0.541 | 2.20E-156 | 6 | Sh3bgrl3 |
| Plin2 | 2.12E-160 | 0.3324613 | 0.342 | 0.088 | 4.30E-156 | 6 | Plin2 |
| Orai1 | 6.28E-160 | 0.499090961 | 0.397 | 0.117 | 1.27E-155 | 6 | Orai1 |
| Itgax1 | 8.14E-160 | 0.422233973 | 0.361 | 0.096 | 1.65E-155 | 6 | Itgax |
| Slc30a5 | 1.09E-159 | 0.421828867 | 0.309 | 0.076 | 2.20E-155 | 6 | Slc30a5 |
| Sdc4 | 9.57E-159 | 0.313914619 | 0.277 | 0.062 | 1.94E-154 | 6 | Sdc4 |
| Map2k1 | 1.72E-158 | 0.400571327 | 0.375 | 0.105 | 3.48E-154 | 6 | Map2k1 |
| Cited2 | 1.39E-157 | 0.536975144 | 0.491 | 0.166 | 2.81E-153 | 6 | Cited2 |
| Pan32 | 1.48E-157 | 0.611645536 | 0.569 | 0.213 | 2.99E-153 | 6 | Pan3 |
| Cbl1 | 3.05E-157 | 0.596594027 | 0.535 | 0.193 | 6.17E-153 | 6 | Cbl |
| Foxo31 | 5.26E-157 | 0.446348276 | 0.37 | 0.105 | 1.06E-152 | 6 | Foxo3 |
| Nes1 | 2.01E-156 | 0.474565755 | 0.413 | 0.121 | 4.07E-152 | 6 | Nes |
| Epb411 | 3.45E-156 | 0.536881769 | 0.553 | 0.201 | 6.99E-152 | 6 | Epb41 |
| Ambra12 | 3.95E-156 | 0.552302198 | 0.519 | 0.183 | 8.00E-152 | 6 | Ambra1 |
| Stau2 | 4.64E-156 | 0.332413525 | 0.258 | 0.056 | 9.40E-152 | 6 | Stau2 |
| Aftph | 3.02E-155 | 0.473448265 | 0.468 | 0.155 | 6.11E-151 | 6 | Aftph |
| Senp2 | 7.41E-155 | 0.437610872 | 0.4 | 0.12 | 1.50E-150 | 6 | Senp2 |
| Gdi21 | 7.54E-155 | 0.491468428 | 0.793 | 0.366 | 1.53E-150 | 6 | Gdi2 |
| Csnk1g31 | 1.42E-154 | 0.508257788 | 0.494 | 0.169 | 2.88E-150 | 6 | Csnk1g3 |
| Pip5k1c | 1.62E-154 | 0.406998537 | 0.332 | 0.088 | 3.27E-150 | 6 | Pip5k1c |
| Hivep22 | 4.71E-154 | 0.723702643 | 0.669 | 0.281 | 9.54E-150 | 6 | Hivep2 |
| Tfeb | 2.05E-153 | 0.305835476 | 0.273 | 0.063 | 4.14E-149 | 6 | Tfeb |
| Iqgap13 | 4.61E-153 | 0.708238379 | 0.824 | 0.393 | 9.33E-149 | 6 | Iqgap1 |
| Nedd4l2 | 7.63E-153 | 0.55927918 | 0.554 | 0.202 | 1.54E-148 | 6 | Nedd4l |
| Sik21 | 1.07E-152 | 0.456644087 | 0.375 | 0.108 | 2.16E-148 | 6 | Sik2 |
| Man1a21 | 2.21E-152 | 0.560856028 | 0.564 | 0.211 | 4.47E-148 | 6 | Man1a2 |
| Pdia33 | 2.83E-152 | 0.721624158 | 0.833 | 0.426 | 5.73E-148 | 6 | Pdia3 |
| Slc25a25 | 2.57E-151 | 0.338429301 | 0.258 | 0.057 | 5.20E-147 | 6 | Slc25a25 |
| Kctd122 | 3.13E-151 | 0.499726991 | 0.632 | 0.241 | 6.34E-147 | 6 | Kctd12 |
| Coro1b | 3.97E-151 | 0.335748062 | 0.385 | 0.112 | 8.03E-147 | 6 | Coro1b |
| Atp13a3 | 7.83E-151 | 0.421382968 | 0.339 | 0.093 | 1.59E-146 | 6 | Atp13a3 |
| Stk102 | 9.51E-151 | 0.525548592 | 0.493 | 0.169 | 1.93E-146 | 6 | Stk10 |
| Btbd93 | 1.29E-150 | 0.564525582 | 0.58 | 0.222 | 2.62E-146 | 6 | Btbd9 |
| Trim30a2 | 3.70E-150 | 0.614926529 | 0.544 | 0.205 | 7.49E-146 | 6 | Trim30a |
| Washc4 | 1.23E-149 | 0.444700336 | 0.391 | 0.117 | 2.50E-145 | 6 | Washc4 |
| Itpripl21 | 1.71E-149 | 0.398729236 | 0.374 | 0.107 | 3.45E-145 | 6 | Itpripl2 |
| Metrnl1 | 2.52E-149 | 0.37596687 | 0.339 | 0.093 | 5.11E-145 | 6 | Metrnl |
| 2810474O19Rik1 | 4.00E-149 | 0.472999187 | 0.542 | 0.195 | 8.09E-145 | 6 | 2810474O19Rik |
| Tbpl1 | 4.04E-149 | 0.334979992 | 0.374 | 0.108 | 8.18E-145 | 6 | Tbpl1 |
| Tbl1xr11 | 1.81E-148 | 0.496055291 | 0.454 | 0.151 | 3.67E-144 | 6 | Tbl1xr1 |
| Ifi206 | 2.92E-148 | 0.398815895 | 0.27 | 0.063 | 5.92E-144 | 6 | Ifi206 |

| | | | | | | | |
|---|---|---|---|---|---|---|---|
| Pcyt1a | 3.10E-148 | 0.374579425 | 0.306 | 0.077 | 6.27E-144 | 6 | Pcyt1a |
| Ncoa32 | 3.37E-148 | 0.548629019 | 0.605 | 0.236 | 6.83E-144 | 6 | Ncoa3 |
| Map3k81 | 5.75E-148 | 0.343093214 | 0.298 | 0.075 | 1.16E-143 | 6 | Map3k8 |
| 1700025G04Rik1 | 1.52E-147 | 0.584100553 | 0.492 | 0.175 | 3.07E-143 | 6 | 1700025G04Rik |
| Gpcpd1 | 4.95E-147 | 0.424272957 | 0.364 | 0.106 | 1.00E-142 | 6 | Gpcpd1 |
| Atad2b2 | 9.10E-147 | 0.544416691 | 0.519 | 0.189 | 1.84E-142 | 6 | Atad2b |
| Plcl21 | 1.13E-146 | 0.498627619 | 0.358 | 0.104 | 2.29E-142 | 6 | Plcl2 |
| Mcl12 | 1.16E-146 | 0.572291924 | 0.775 | 0.355 | 2.35E-142 | 6 | Mcl1 |
| Smap21 | 1.16E-146 | 0.447922271 | 0.536 | 0.197 | 2.36E-142 | 6 | Smap2 |
| Filip1l | 1.10E-145 | 0.442381205 | 0.432 | 0.14 | 2.22E-141 | 6 | Filip1l |
| Traf31 | 1.94E-145 | 0.430255589 | 0.347 | 0.098 | 3.92E-141 | 6 | Traf3 |
| Fmnl32 | 1.24E-144 | 0.501104358 | 0.591 | 0.224 | 2.51E-140 | 6 | Fmnl3 |
| Rilpl2 | 3.45E-144 | 0.416712308 | 0.271 | 0.065 | 6.99E-140 | 6 | Rilpl2 |
| Fos2 | 5.26E-144 | 0.578983807 | 0.693 | 0.3 | 1.07E-139 | 6 | Fos |
| Sh3pxd2a1 | 7.56E-144 | 0.40150421 | 0.417 | 0.129 | 1.53E-139 | 6 | Sh3pxd2a |
| Vwa5a | 8.69E-144 | 0.364074723 | 0.319 | 0.086 | 1.76E-139 | 6 | Vwa5a |
| Il10rb2 | 2.39E-143 | 0.440965929 | 0.6 | 0.233 | 4.84E-139 | 6 | Il10rb |
| Hsd17b11 | 2.73E-143 | 0.378050627 | 0.369 | 0.107 | 5.53E-139 | 6 | Hsd17b11 |
| Dab22 | 4.40E-143 | 0.569460616 | 0.532 | 0.192 | 8.91E-139 | 6 | Dab2 |
| Tnrc181 | 2.33E-142 | 0.471324222 | 0.467 | 0.16 | 4.73E-138 | 6 | Tnrc18 |
| Fuca21 | 8.09E-142 | 0.295298926 | 0.254 | 0.059 | 1.64E-137 | 6 | Fuca2 |
| Tor1aip12 | 8.84E-142 | 0.537977309 | 0.639 | 0.265 | 1.79E-137 | 6 | Tor1aip1 |
| Tubb2a1 | 1.73E-141 | 0.388730811 | 0.457 | 0.151 | 3.49E-137 | 6 | Tubb2a |
| Slc25a12 | 1.80E-141 | 0.33565094 | 0.392 | 0.121 | 3.65E-137 | 6 | Slc25a12 |
| Ube2f | 1.81E-141 | 0.298834973 | 0.392 | 0.12 | 3.67E-137 | 6 | Ube2f |
| Bmt21 | 2.53E-141 | 0.461201093 | 0.366 | 0.11 | 5.12E-137 | 6 | Bmt2 |
| Fkbp15 | 8.79E-141 | 0.40189296 | 0.348 | 0.1 | 1.78E-136 | 6 | Fkbp15 |
| Lsp11 | 9.00E-141 | 0.478884764 | 0.403 | 0.128 | 1.82E-136 | 6 | Lsp1 |
| Samsn11 | 1.38E-140 | 0.331654572 | 0.284 | 0.071 | 2.80E-136 | 6 | Samsn1 |
| Rgs19 | 2.58E-140 | 0.320849571 | 0.25 | 0.058 | 5.21E-136 | 6 | Rgs19 |
| Cept1 | 5.97E-140 | 0.430706161 | 0.398 | 0.125 | 1.21E-135 | 6 | Cept1 |
| Cmip3 | 6.18E-140 | 0.581719259 | 0.644 | 0.269 | 1.25E-135 | 6 | Cmip |
| Ncoa22 | 7.61E-140 | 0.576298867 | 0.567 | 0.224 | 1.54E-135 | 6 | Ncoa2 |
| Fam172a2 | 1.17E-139 | 0.606211059 | 0.643 | 0.268 | 2.37E-135 | 6 | Fam172a |
| Ankfy11 | 4.11E-139 | 0.419395207 | 0.41 | 0.132 | 8.33E-135 | 6 | Ankfy1 |
| Mycbp22 | 3.01E-138 | 0.615301603 | 0.769 | 0.357 | 6.10E-134 | 6 | Mycbp2 |
| 2610507B11Rik | 8.45E-138 | 0.381898694 | 0.435 | 0.145 | 1.71E-133 | 6 | 2610507B11Rik |
| Ptp4a2 | 9.40E-138 | 0.482906162 | 0.792 | 0.376 | 1.90E-133 | 6 | Ptp4a2 |
| M6pr1 | 1.00E-137 | 0.432453546 | 0.442 | 0.15 | 2.03E-133 | 6 | M6pr |
| Capza2 | 1.16E-137 | 0.465807237 | 0.85 | 0.444 | 2.35E-133 | 6 | Capza2 |
| Ppp1r181 | 2.16E-137 | 0.38375496 | 0.43 | 0.142 | 4.37E-133 | 6 | Ppp1r18 |
| Larp4b1 | 2.43E-137 | 0.471736748 | 0.516 | 0.19 | 4.91E-133 | 6 | Larp4b |
| Vps13b2 | 2.46E-137 | 0.554889735 | 0.534 | 0.205 | 4.98E-133 | 6 | Vps13b |

| | | | | | | | |
|---|---|---|---|---|---|---|---|
| Tgfb13 | 3.34E-137 | 0.487053677 | 0.569 | 0.221 | 6.77E-133 | 6 | Tgfb1 |
| Actr21 | 1.38E-136 | 0.489027266 | 0.723 | 0.326 | 2.79E-132 | 6 | Actr2 |
| Lims11 | 1.44E-136 | 0.534850232 | 0.669 | 0.284 | 2.92E-132 | 6 | Lims1 |
| Ube2l6 | 2.15E-136 | 0.312488595 | 0.277 | 0.068 | 4.35E-132 | 6 | Ube2l6 |
| Mtdh2 | 9.82E-136 | 0.577319797 | 0.78 | 0.369 | 1.99E-131 | 6 | Mtdh |
| Atp2c11 | 1.44E-135 | 0.47707898 | 0.451 | 0.155 | 2.92E-131 | 6 | Atp2c1 |
| Exoc61 | 3.06E-135 | 0.452154416 | 0.456 | 0.158 | 6.19E-131 | 6 | Exoc6 |
| Rnf2161 | 1.93E-134 | 0.451111184 | 0.471 | 0.167 | 3.92E-130 | 6 | Rnf216 |
| Ranbp91 | 3.83E-134 | 0.509143523 | 0.501 | 0.187 | 7.76E-130 | 6 | Ranbp9 |
| Nfatc11 | 6.09E-134 | 0.409556255 | 0.352 | 0.106 | 1.23E-129 | 6 | Nfatc1 |
| Kpna43 | 1.05E-133 | 0.591025828 | 0.695 | 0.306 | 2.12E-129 | 6 | Kpna4 |
| Pnp2 | 6.00E-133 | 0.362396632 | 0.604 | 0.238 | 1.22E-128 | 6 | Pnp |
| Colgalt11 | 7.73E-133 | 0.406057852 | 0.405 | 0.132 | 1.57E-128 | 6 | Colgalt1 |
| Ripk2 | 1.83E-132 | 0.363164709 | 0.277 | 0.072 | 3.70E-128 | 6 | Ripk2 |
| Rnaset2a | 2.02E-132 | 0.313143725 | 0.346 | 0.102 | 4.09E-128 | 6 | Rnaset2a |
| Nav22 | 3.75E-132 | 0.625585869 | 0.439 | 0.153 | 7.59E-128 | 6 | Nav2 |
| Npc11 | 4.46E-132 | 0.3741678 | 0.271 | 0.07 | 9.03E-128 | 6 | Npc1 |
| Diaph23 | 1.00E-131 | 0.66182002 | 0.762 | 0.359 | 2.03E-127 | 6 | Diaph2 |
| Kansl12 | 1.23E-131 | 0.575175146 | 0.719 | 0.323 | 2.49E-127 | 6 | Kansl1 |
| Lpar62 | 2.40E-131 | 0.48674509 | 0.477 | 0.17 | 4.86E-127 | 6 | Lpar6 |
| Galnt11 | 2.72E-131 | 0.467368343 | 0.472 | 0.171 | 5.50E-127 | 6 | Galnt1 |
| Snap23 | 4.62E-131 | 0.32656504 | 0.472 | 0.166 | 9.36E-127 | 6 | Snap23 |
| Junb3 | 1.10E-130 | 0.465938012 | 0.806 | 0.374 | 2.22E-126 | 6 | Junb |
| Malat13 | 2.52E-130 | 0.739421429 | 0.998 | 0.713 | 5.11E-126 | 6 | Malat1 |
| Pkig | 3.57E-130 | 0.302334738 | 0.531 | 0.196 | 7.23E-126 | 6 | Pkig |
| Rasa21 | 4.30E-130 | 0.473235078 | 0.39 | 0.127 | 8.70E-126 | 6 | Rasa2 |
| Ptger41 | 4.50E-130 | 0.432699665 | 0.284 | 0.076 | 9.11E-126 | 6 | Ptger4 |
| Myadm1 | 5.12E-130 | 0.360419528 | 0.361 | 0.109 | 1.04E-125 | 6 | Myadm |
| Cebpb | 2.02E-129 | 0.333549162 | 0.429 | 0.145 | 4.09E-125 | 6 | Cebpb |
| Phc21 | 9.14E-129 | 0.428012286 | 0.447 | 0.157 | 1.85E-124 | 6 | Phc2 |
| Dram21 | 1.17E-127 | 0.363896036 | 0.426 | 0.143 | 2.37E-123 | 6 | Dram2 |
| Phf21a1 | 1.38E-127 | 0.527427711 | 0.58 | 0.238 | 2.80E-123 | 6 | Phf21a |
| Lrrc8d1 | 2.17E-127 | 0.400393364 | 0.381 | 0.122 | 4.39E-123 | 6 | Lrrc8d |
| Arid4a3 | 2.77E-127 | 0.502222306 | 0.583 | 0.237 | 5.61E-123 | 6 | Arid4a |
| Lnpep2 | 4.65E-127 | 0.54309159 | 0.587 | 0.245 | 9.41E-123 | 6 | Lnpep |
| Pign1 | 5.28E-127 | 0.457409905 | 0.299 | 0.085 | 1.07E-122 | 6 | Pign |
| Papd4 | 5.53E-127 | 0.411876863 | 0.386 | 0.127 | 1.12E-122 | 6 | Papd4 |
| Selenos | 6.81E-127 | 0.379117874 | 0.554 | 0.217 | 1.38E-122 | 6 | Selenos |
| Ube2j1 | 8.35E-127 | 0.394151047 | 0.431 | 0.15 | 1.69E-122 | 6 | Ube2j1 |
| Usp12 | 8.44E-127 | 0.406338416 | 0.389 | 0.127 | 1.71E-122 | 6 | Usp12 |
| Gna13 | 9.23E-127 | 0.424965229 | 0.447 | 0.159 | 1.87E-122 | 6 | Gna13 |
| Nfkb11 | 4.29E-126 | 0.518140429 | 0.611 | 0.256 | 8.68E-122 | 6 | Nfkb1 |
| Ywhah1 | 2.06E-125 | 0.360118999 | 0.673 | 0.293 | 4.16E-121 | 6 | Ywhah |

| | | | | | | | |
|---|---|---|---|---|---|---|---|
| Oasl21 | 2.95E-125 | 0.415037024 | 0.378 | 0.12 | 5.98E-121 | 6 | Oasl2 |
| Glrx1 | 3.04E-125 | 0.251027958 | 0.253 | 0.063 | 6.15E-121 | 6 | Glrx |
| Slfn22 | 4.19E-125 | 0.389912219 | 0.601 | 0.245 | 8.49E-121 | 6 | Slfn2 |
| Tet2 | 4.62E-125 | 0.31632244 | 0.332 | 0.099 | 9.35E-121 | 6 | Tet2 |
| Fbrsl1 | 1.28E-124 | 0.335063888 | 0.305 | 0.087 | 2.59E-120 | 6 | Fbrsl1 |
| Lifr1 | 1.43E-124 | 0.354701046 | 0.498 | 0.179 | 2.90E-120 | 6 | Lifr |
| Hsp90b12 | 2.49E-124 | 0.62254227 | 0.897 | 0.493 | 5.03E-120 | 6 | Hsp90b1 |
| Tpk11 | 3.61E-124 | 0.44127279 | 0.319 | 0.095 | 7.31E-120 | 6 | Tpk1 |
| Cdc42se21 | 7.04E-124 | 0.537847495 | 0.554 | 0.225 | 1.42E-119 | 6 | Cdc42se2 |
| Dip2b2 | 4.16E-123 | 0.505555119 | 0.61 | 0.261 | 8.43E-119 | 6 | Dip2b |
| Phkb | 7.57E-123 | 0.406576901 | 0.394 | 0.134 | 1.53E-118 | 6 | Phkb |
| Acbd5 | 1.23E-122 | 0.367645704 | 0.495 | 0.187 | 2.50E-118 | 6 | Acbd5 |
| Litaf1 | 1.45E-122 | 0.344277861 | 0.424 | 0.146 | 2.95E-118 | 6 | Litaf |
| Atxn71 | 1.90E-122 | 0.434046655 | 0.432 | 0.154 | 3.85E-118 | 6 | Atxn7 |
| Abi32 | 3.54E-122 | 0.41950842 | 0.571 | 0.229 | 7.17E-118 | 6 | Abi3 |
| Ell2 | 3.73E-122 | 0.293818415 | 0.281 | 0.077 | 7.56E-118 | 6 | Ell2 |
| Peli11 | 4.21E-122 | 0.48165609 | 0.548 | 0.218 | 8.52E-118 | 6 | Peli1 |
| Tex141 | 4.31E-122 | 0.513345939 | 0.403 | 0.138 | 8.72E-118 | 6 | Tex14 |
| Tlk1 | 1.01E-121 | 0.383090902 | 0.531 | 0.209 | 2.05E-117 | 6 | Tlk1 |
| Cd371 | 2.86E-121 | 0.337443635 | 0.341 | 0.104 | 5.78E-117 | 6 | Cd37 |
| Nipa2 | 3.57E-121 | 0.372593533 | 0.369 | 0.12 | 7.23E-117 | 6 | Nipa2 |
| Grk21 | 4.08E-121 | 0.317352114 | 0.359 | 0.114 | 8.25E-117 | 6 | Grk2 |
| Ifngr13 | 1.25E-120 | 0.507047522 | 0.783 | 0.375 | 2.53E-116 | 6 | Ifngr1 |
| Psme4 | 1.72E-120 | 0.367230944 | 0.489 | 0.184 | 3.47E-116 | 6 | Psme4 |
| Zfp395 | 2.49E-120 | 0.311731792 | 0.273 | 0.074 | 5.04E-116 | 6 | Zfp395 |
| Samhd12 | 9.71E-120 | 0.448749981 | 0.58 | 0.239 | 1.97E-115 | 6 | Samhd1 |
| Ankrd112 | 2.62E-119 | 0.557679675 | 0.765 | 0.363 | 5.30E-115 | 6 | Ankrd11 |
| Mbtd1 | 4.36E-119 | 0.442146242 | 0.371 | 0.124 | 8.83E-115 | 6 | Mbtd1 |
| Tpp11 | 4.65E-119 | 0.360069064 | 0.346 | 0.109 | 9.42E-115 | 6 | Tpp1 |
| Fam45a | 1.73E-118 | 0.28629925 | 0.298 | 0.085 | 3.50E-114 | 6 | Fam45a |
| Cnot22 | 2.24E-118 | 0.443107997 | 0.502 | 0.197 | 4.54E-114 | 6 | Cnot2 |
| B4galt51 | 5.43E-118 | 0.34910553 | 0.294 | 0.085 | 1.10E-113 | 6 | B4galt5 |
| Mtus1 | 7.47E-118 | 0.321056183 | 0.424 | 0.147 | 1.51E-113 | 6 | Mtus1 |
| Atp6ap1 | 7.61E-118 | 0.380320873 | 0.402 | 0.138 | 1.54E-113 | 6 | Atp6ap1 |
| Tmcc12 | 2.14E-117 | 0.52433244 | 0.513 | 0.205 | 4.34E-113 | 6 | Tmcc1 |
| Exoc43 | 5.87E-117 | 0.572623635 | 0.667 | 0.307 | 1.19E-112 | 6 | Exoc4 |
| Ywhag1 | 8.85E-117 | 0.385376319 | 0.455 | 0.169 | 1.79E-112 | 6 | Ywhag |
| Chka1 | 5.16E-116 | 0.53463174 | 0.384 | 0.134 | 1.04E-111 | 6 | Chka |
| Arl6ip13 | 6.57E-116 | 0.469963331 | 0.735 | 0.354 | 1.33E-111 | 6 | Arl6ip1 |
| Limd21 | 1.13E-115 | 0.253864426 | 0.407 | 0.137 | 2.28E-111 | 6 | Limd2 |
| Ascc32 | 1.40E-115 | 0.447317496 | 0.497 | 0.194 | 2.84E-111 | 6 | Ascc3 |
| 2310035C23Rik1 | 1.58E-115 | 0.411727228 | 0.379 | 0.13 | 3.19E-111 | 6 | 2310035C23Rik |
| Itpr21 | 2.80E-115 | 0.477710223 | 0.555 | 0.23 | 5.68E-111 | 6 | Itpr2 |

| | | | | | | | |
|---|---|---|---|---|---|---|---|
| Tcf201 | 3.09E-115 | 0.413893397 | 0.38 | 0.131 | 6.26E-111 | 6 | Tcf20 |
| Snx131 | 3.67E-115 | 0.37792078 | 0.402 | 0.139 | 7.43E-111 | 6 | Snx13 |
| Ap3b12 | 4.03E-115 | 0.411967738 | 0.514 | 0.204 | 8.15E-111 | 6 | Ap3b1 |
| Macf12 | 4.35E-115 | 0.565864067 | 0.907 | 0.484 | 8.82E-111 | 6 | Macf1 |
| Arhgef111 | 7.44E-115 | 0.325453936 | 0.307 | 0.091 | 1.51E-110 | 6 | Arhgef11 |
| Lgals92 | 3.31E-114 | 0.44388878 | 0.56 | 0.233 | 6.70E-110 | 6 | Lgals9 |
| Hivep31 | 6.69E-114 | 0.377773959 | 0.421 | 0.15 | 1.35E-109 | 6 | Hivep3 |
| Supt3 | 7.30E-114 | 0.360687317 | 0.354 | 0.117 | 1.48E-109 | 6 | Supt3 |
| Zfand32 | 8.49E-114 | 0.460154983 | 0.644 | 0.286 | 1.72E-109 | 6 | Zfand3 |
| Acsl4 | 1.11E-113 | 0.319314631 | 0.366 | 0.122 | 2.24E-109 | 6 | Acsl4 |
| Mtmr6 | 1.21E-113 | 0.320849634 | 0.337 | 0.108 | 2.45E-109 | 6 | Mtmr6 |
| Nrip12 | 2.85E-113 | 0.379319188 | 0.56 | 0.23 | 5.77E-109 | 6 | Nrip1 |
| Map3k31 | 3.03E-113 | 0.45578329 | 0.443 | 0.166 | 6.13E-109 | 6 | Map3k3 |
| Lgals33 | 3.16E-113 | 0.522239254 | 0.525 | 0.204 | 6.40E-109 | 6 | Lgals3 |
| Snd11 | 3.22E-113 | 0.421645378 | 0.431 | 0.161 | 6.52E-109 | 6 | Snd1 |
| Map3k11 | 7.93E-113 | 0.375689563 | 0.48 | 0.186 | 1.61E-108 | 6 | Map3k1 |
| Gtdc11 | 2.56E-112 | 0.455547471 | 0.397 | 0.141 | 5.19E-108 | 6 | Gtdc1 |
| Ncoa11 | 4.40E-112 | 0.481041915 | 0.494 | 0.197 | 8.91E-108 | 6 | Ncoa1 |
| Ccdc121 | 1.20E-110 | 0.271033413 | 0.794 | 0.397 | 2.42E-106 | 6 | Ccdc12 |
| Setd3 | 1.54E-110 | 0.26782219 | 0.489 | 0.189 | 3.12E-106 | 6 | Setd3 |
| Fam13b1 | 2.19E-110 | 0.405471207 | 0.386 | 0.137 | 4.44E-106 | 6 | Fam13b |
| Aff1 | 3.28E-110 | 0.418768578 | 0.408 | 0.149 | 6.64E-106 | 6 | Aff1 |
| Psen11 | 4.27E-110 | 0.35613192 | 0.299 | 0.092 | 8.65E-106 | 6 | Psen1 |
| Specc1l | 1.47E-109 | 0.347981798 | 0.352 | 0.119 | 2.98E-105 | 6 | Specc1l |
| Tor1a | 1.56E-109 | 0.273615559 | 0.314 | 0.097 | 3.16E-105 | 6 | Tor1a |
| Emp31 | 4.31E-109 | 0.368213551 | 0.585 | 0.244 | 8.72E-105 | 6 | Emp3 |
| Kansl1l1 | 6.44E-109 | 0.397105272 | 0.371 | 0.128 | 1.30E-104 | 6 | Kansl1l |
| Phf142 | 1.16E-108 | 0.429809237 | 0.547 | 0.231 | 2.34E-104 | 6 | Phf14 |
| 9930111J21Rik22 | 2.06E-108 | 0.452941455 | 0.573 | 0.243 | 4.17E-104 | 6 | 9930111J21Rik2 |
| Spred12 | 2.42E-108 | 0.455325834 | 0.564 | 0.238 | 4.90E-104 | 6 | Spred1 |
| Mcm9 | 3.47E-108 | 0.340132851 | 0.277 | 0.081 | 7.03E-104 | 6 | Mcm9 |
| Clic12 | 3.50E-108 | 0.505249217 | 0.945 | 0.585 | 7.09E-104 | 6 | Clic1 |
| Cnih42 | 4.39E-108 | 0.295458078 | 0.57 | 0.24 | 8.89E-104 | 6 | Cnih4 |
| Lmo22 | 4.66E-108 | 0.266656817 | 0.507 | 0.194 | 9.44E-104 | 6 | Lmo2 |
| Kpna31 | 2.74E-107 | 0.327562899 | 0.402 | 0.146 | 5.55E-103 | 6 | Kpna3 |
| Irak4 | 3.27E-107 | 0.332149956 | 0.294 | 0.09 | 6.62E-103 | 6 | Irak4 |
| Zcchc72 | 5.99E-107 | 0.487470687 | 0.633 | 0.286 | 1.21E-102 | 6 | Zcchc7 |
| Immp2l1 | 8.30E-107 | 0.45443383 | 0.363 | 0.126 | 1.68E-102 | 6 | Immp2l |
| Apbb22 | 9.37E-107 | 0.371391972 | 0.777 | 0.348 | 1.90E-102 | 6 | Apbb2 |
| Ep3002 | 2.61E-106 | 0.410918484 | 0.477 | 0.192 | 5.29E-102 | 6 | Ep300 |
| Akap8l | 5.93E-106 | 0.389643068 | 0.369 | 0.13 | 1.20E-101 | 6 | Akap8l |
| Ccdc50 | 1.45E-105 | 0.316794289 | 0.525 | 0.215 | 2.93E-101 | 6 | Ccdc50 |
| Pdgfb2 | 2.98E-105 | 0.321134421 | 0.671 | 0.293 | 6.03E-101 | 6 | Pdgfb |

| | | | | | | | |
|---|---|---|---|---|---|---|---|
| Ergic3 | 4.16E-105 | 0.277206904 | 0.46 | 0.177 | 8.42E-101 | 6 | Ergic3 |
| Sdcbp | 6.39E-105 | 0.386444697 | 0.676 | 0.321 | 1.29E-100 | 6 | Sdcbp |
| Vps541 | 1.01E-104 | 0.375056534 | 0.45 | 0.175 | 2.05E-100 | 6 | Vps54 |
| Slc38a9 | 1.37E-104 | 0.36422453 | 0.258 | 0.074 | 2.78E-100 | 6 | Slc38a9 |
| Larp11 | 3.05E-104 | 0.374309824 | 0.477 | 0.191 | 6.17E-100 | 6 | Larp1 |
| Zfp6081 | 5.02E-104 | 0.365203401 | 0.375 | 0.13 | 1.02E-99 | 6 | Zfp608 |
| Stard3nl1 | 5.11E-104 | 0.260199081 | 0.391 | 0.14 | 1.04E-99 | 6 | Stard3nl |
| Rb12 | 5.17E-104 | 0.399969182 | 0.496 | 0.202 | 1.05E-99 | 6 | Rb1 |
| Thada1 | 7.00E-104 | 0.372166486 | 0.33 | 0.111 | 1.42E-99 | 6 | Thada |
| Vmp11 | 1.10E-103 | 0.366265561 | 0.498 | 0.203 | 2.23E-99 | 6 | Vmp1 |
| Pkn12 | 1.13E-103 | 0.360350967 | 0.45 | 0.175 | 2.30E-99 | 6 | Pkn1 |
| Cnpy3 | 1.26E-103 | 0.296522479 | 0.299 | 0.094 | 2.55E-99 | 6 | Cnpy3 |
| Hltf | 3.34E-103 | 0.346322771 | 0.346 | 0.119 | 6.77E-99 | 6 | Hltf |
| Ccnt2 | 3.42E-103 | 0.353074215 | 0.323 | 0.106 | 6.93E-99 | 6 | Ccnt2 |
| Ptpra2 | 5.69E-103 | 0.489336465 | 0.627 | 0.29 | 1.15E-98 | 6 | Ptpra |
| Tle42 | 6.66E-103 | 0.470118833 | 0.51 | 0.213 | 1.35E-98 | 6 | Tle4 |
| Prpf38b2 | 7.75E-103 | 0.389580309 | 0.463 | 0.184 | 1.57E-98 | 6 | Prpf38b |
| Nsmce21 | 1.98E-102 | 0.367434582 | 0.479 | 0.195 | 4.01E-98 | 6 | Nsmce2 |
| Mthfs1 | 3.59E-102 | 0.263639722 | 0.265 | 0.078 | 7.27E-98 | 6 | Mthfs |
| Bcap29 | 1.03E-101 | 0.310650981 | 0.296 | 0.094 | 2.08E-97 | 6 | Bcap29 |
| Zfp36l21 | 2.89E-101 | 0.42448336 | 0.637 | 0.291 | 5.86E-97 | 6 | Zfp36l2 |
| Rufy3 | 5.55E-101 | 0.290898816 | 0.258 | 0.075 | 1.12E-96 | 6 | Rufy3 |
| Atg10 | 5.95E-101 | 0.306263145 | 0.397 | 0.147 | 1.20E-96 | 6 | Atg10 |
| Tmed3 | 7.66E-101 | 0.320532525 | 0.539 | 0.23 | 1.55E-96 | 6 | Tmed3 |
| Ier52 | 9.25E-101 | 0.401605278 | 0.652 | 0.302 | 1.87E-96 | 6 | Ier5 |
| Hbb-bs | 1.78E-100 | 3.350294376 | 0.565 | 0.319 | 3.61E-96 | 6 | Hbb-bs |
| Clip11 | 2.07E-100 | 0.306594756 | 0.372 | 0.133 | 4.20E-96 | 6 | Clip1 |
| Htt | 2.39E-100 | 0.3150424 | 0.295 | 0.094 | 4.84E-96 | 6 | Htt |
| Fbxl173 | 3.20E-100 | 0.487260724 | 0.632 | 0.297 | 6.48E-96 | 6 | Fbxl17 |
| Akna1 | 7.67E-100 | 0.316640622 | 0.274 | 0.083 | 1.55E-95 | 6 | Akna |
| Entpd12 | 1.18E-99 | 0.324866195 | 0.604 | 0.261 | 2.38E-95 | 6 | Entpd1 |
| Paxbp1 | 6.87E-99 | 0.343062994 | 0.334 | 0.115 | 1.39E-94 | 6 | Paxbp1 |
| Pdia62 | 2.72E-97 | 0.476678625 | 0.662 | 0.323 | 5.51E-93 | 6 | Pdia6 |
| Eps151 | 7.84E-97 | 0.305524639 | 0.414 | 0.158 | 1.59E-92 | 6 | Eps15 |
| Hspa52 | 7.98E-97 | 0.566088875 | 0.775 | 0.42 | 1.62E-92 | 6 | Hspa5 |
| Ppp4r1 | 1.57E-96 | 0.31090173 | 0.32 | 0.108 | 3.17E-92 | 6 | Ppp4r1 |
| Mta31 | 2.25E-96 | 0.289214004 | 0.384 | 0.142 | 4.56E-92 | 6 | Mta3 |
| Zfp809 | 2.71E-96 | 0.252804747 | 0.26 | 0.078 | 5.50E-92 | 6 | Zfp809 |
| Capza11 | 3.71E-96 | 0.362424519 | 0.585 | 0.262 | 7.52E-92 | 6 | Capza1 |
| Abcg11 | 6.32E-96 | 0.373624874 | 0.33 | 0.115 | 1.28E-91 | 6 | Abcg1 |
| Kdm3b1 | 6.76E-96 | 0.356835852 | 0.351 | 0.127 | 1.37E-91 | 6 | Kdm3b |
| Lrch32 | 9.56E-96 | 0.45825604 | 0.506 | 0.22 | 1.94E-91 | 6 | Lrch3 |
| Rnf2132 | 1.00E-95 | 0.581473565 | 0.536 | 0.245 | 2.03E-91 | 6 | Rnf213 |

| | | | | | | | |
|---|---|---|---|---|---|---|---|
| Mkrn1 | 1.08E-95 | 0.581024304 | 0.395 | 0.157 | 2.18E-91 | 6 | Mkrn1 |
| Rnf1692 | 3.08E-95 | 0.379878119 | 0.435 | 0.172 | 6.23E-91 | 6 | Rnf169 |
| Arid1a2 | 5.27E-95 | 0.365618418 | 0.512 | 0.218 | 1.07E-90 | 6 | Arid1a |
| Pik3r1 | 5.71E-95 | 0.305668774 | 0.545 | 0.241 | 1.16E-90 | 6 | Pik3r1 |
| Tsc22d41 | 2.78E-94 | 0.259436002 | 0.609 | 0.28 | 5.64E-90 | 6 | Tsc22d4 |
| Lcor2 | 4.30E-94 | 0.385236222 | 0.44 | 0.179 | 8.70E-90 | 6 | Lcor |
| Clic42 | 5.00E-93 | 0.3686944 | 0.803 | 0.406 | 1.01E-88 | 6 | Clic4 |
| Ppp3ca2 | 8.28E-93 | 0.465724026 | 0.74 | 0.378 | 1.68E-88 | 6 | Ppp3ca |
| Eif2ak21 | 2.07E-92 | 0.341997304 | 0.397 | 0.154 | 4.19E-88 | 6 | Eif2ak2 |
| Etv3 | 6.55E-92 | 0.312251026 | 0.281 | 0.091 | 1.33E-87 | 6 | Etv3 |
| Clta2 | 9.70E-92 | 0.386642142 | 0.886 | 0.543 | 1.96E-87 | 6 | Clta |
| Cd812 | 1.07E-91 | 0.426726394 | 0.957 | 0.506 | 2.17E-87 | 6 | Cd81 |
| Gas71 | 1.43E-91 | 0.306175775 | 0.437 | 0.171 | 2.89E-87 | 6 | Gas7 |
| Sqstm11 | 1.77E-91 | 0.470807433 | 0.607 | 0.291 | 3.59E-87 | 6 | Sqstm1 |
| Arid1b2 | 1.88E-91 | 0.428163918 | 0.55 | 0.246 | 3.81E-87 | 6 | Arid1b |
| Mfap3 | 6.69E-91 | 0.303063421 | 0.275 | 0.089 | 1.36E-86 | 6 | Mfap3 |
| Ssr42 | 9.89E-91 | 0.328506429 | 0.815 | 0.452 | 2.00E-86 | 6 | Ssr4 |
| Fnip21 | 1.15E-90 | 0.360930196 | 0.296 | 0.098 | 2.33E-86 | 6 | Fnip2 |
| Apc2 | 1.44E-90 | 0.37742802 | 0.53 | 0.236 | 2.91E-86 | 6 | Apc |
| Gapvd11 | 3.37E-90 | 0.329804092 | 0.443 | 0.182 | 6.83E-86 | 6 | Gapvd1 |
| Eml41 | 3.62E-90 | 0.353039649 | 0.552 | 0.246 | 7.33E-86 | 6 | Eml4 |
| Nap1l11 | 7.02E-90 | 0.281807447 | 0.627 | 0.301 | 1.42E-85 | 6 | Nap1l1 |
| Git22 | 1.06E-89 | 0.399066318 | 0.508 | 0.223 | 2.15E-85 | 6 | Git2 |
| Ttc17 | 1.52E-89 | 0.355951802 | 0.309 | 0.108 | 3.08E-85 | 6 | Ttc17 |
| Usp151 | 2.65E-89 | 0.327675725 | 0.504 | 0.219 | 5.37E-85 | 6 | Usp15 |
| Parp142 | 4.10E-89 | 0.462435411 | 0.531 | 0.24 | 8.31E-85 | 6 | Parp14 |
| Far11 | 1.29E-88 | 0.309027722 | 0.395 | 0.155 | 2.62E-84 | 6 | Far1 |
| Fbxw41 | 1.53E-88 | 0.311051091 | 0.308 | 0.107 | 3.09E-84 | 6 | Fbxw4 |
| Mtmr32 | 2.01E-88 | 0.359252541 | 0.452 | 0.19 | 4.06E-84 | 6 | Mtmr3 |
| Kdm6a2 | 2.77E-88 | 0.332572577 | 0.475 | 0.202 | 5.61E-84 | 6 | Kdm6a |
| B2m4 | 3.95E-88 | 0.507466809 | 0.992 | 0.677 | 8.00E-84 | 6 | B2m |
| Tcof1 | 4.61E-88 | 0.294154784 | 0.316 | 0.112 | 9.34E-84 | 6 | Tcof1 |
| Wwp21 | 5.43E-88 | 0.313664197 | 0.329 | 0.119 | 1.10E-83 | 6 | Wwp2 |
| Tnfaip31 | 8.11E-88 | 0.25971487 | 0.261 | 0.083 | 1.64E-83 | 6 | Tnfaip3 |
| Rapgef61 | 1.16E-87 | 0.341272581 | 0.618 | 0.283 | 2.34E-83 | 6 | Rapgef6 |
| Capzb2 | 1.19E-87 | 0.309490848 | 0.711 | 0.354 | 2.41E-83 | 6 | Capzb |
| Txndc15 | 2.58E-87 | 0.264229398 | 0.35 | 0.129 | 5.22E-83 | 6 | Txndc15 |
| Smg12 | 2.71E-87 | 0.422448009 | 0.5 | 0.223 | 5.49E-83 | 6 | Smg1 |
| Pld11 | 3.08E-87 | 0.290126623 | 0.367 | 0.138 | 6.24E-83 | 6 | Pld1 |
| Rnf21 | 1.17E-86 | 0.270301883 | 0.289 | 0.098 | 2.37E-82 | 6 | Rnf2 |
| Stk38 | 1.64E-86 | 0.271020601 | 0.494 | 0.215 | 3.33E-82 | 6 | Stk38 |
| Gm472831 | 4.26E-86 | 0.454189202 | 0.571 | 0.269 | 8.63E-82 | 6 | Gm47283 |
| Lrmda3 | 5.28E-86 | 0.413555395 | 0.701 | 0.353 | 1.07E-81 | 6 | Lrmda |

| | | | | | | | |
|---|---|---|---|---|---|---|---|
| Anks11 | 7.79E-86 | 0.350843686 | 0.346 | 0.13 | 1.58E-81 | 6 | Anks1 |
| R3hcc1l | 6.75E-85 | 0.25816689 | 0.304 | 0.106 | 1.37E-80 | 6 | R3hcc1l |
| Itm2b3 | 8.53E-85 | 0.447717078 | 0.996 | 0.713 | 1.73E-80 | 6 | Itm2b |
| Spast | 1.03E-83 | 0.278310984 | 0.298 | 0.105 | 2.08E-79 | 6 | Spast |
| Chd22 | 1.11E-83 | 0.361322149 | 0.529 | 0.242 | 2.25E-79 | 6 | Chd2 |
| Eed | 1.46E-83 | 0.294490827 | 0.364 | 0.142 | 2.95E-79 | 6 | Eed |
| Hook32 | 3.20E-83 | 0.312534812 | 0.596 | 0.276 | 6.47E-79 | 6 | Hook3 |
| Fam168b | 3.71E-83 | 0.283400018 | 0.391 | 0.157 | 7.51E-79 | 6 | Fam168b |
| Trappc81 | 4.47E-83 | 0.292464547 | 0.391 | 0.156 | 9.05E-79 | 6 | Trappc8 |
| 2-Jun | 1.79E-82 | 0.421663073 | 0.869 | 0.498 | 3.63E-78 | 6 | Jun |
| mt-Co13 | 4.19E-82 | 0.518498068 | 0.992 | 0.707 | 8.49E-78 | 6 | mt-Co1 |
| Itch2 | 1.13E-81 | 0.323097818 | 0.562 | 0.261 | 2.29E-77 | 6 | Itch |
| Gak | 2.37E-81 | 0.266084401 | 0.317 | 0.117 | 4.80E-77 | 6 | Gak |
| Map4k32 | 3.99E-81 | 0.253085446 | 0.502 | 0.218 | 8.08E-77 | 6 | Map4k3 |
| Tln13 | 6.03E-81 | 0.354723215 | 0.747 | 0.39 | 1.22E-76 | 6 | Tln1 |
| Ebp | 8.56E-81 | 0.261636019 | 0.415 | 0.172 | 1.73E-76 | 6 | Ebp |
| Eml6 | 9.76E-81 | 0.381161025 | 0.274 | 0.095 | 1.98E-76 | 6 | Eml6 |
| Sf3b12 | 1.75E-80 | 0.323884911 | 0.74 | 0.378 | 3.54E-76 | 6 | Sf3b1 |
| Srsf113 | 2.83E-80 | 0.38191015 | 0.669 | 0.337 | 5.72E-76 | 6 | Srsf11 |
| Ythdf31 | 3.22E-80 | 0.277639211 | 0.434 | 0.186 | 6.53E-76 | 6 | Ythdf3 |
| 2510009E07Rik1 | 3.39E-80 | 0.307847844 | 0.317 | 0.116 | 6.86E-76 | 6 | 2510009E07Rik |
| Cbfb1 | 7.68E-80 | 0.314457431 | 0.367 | 0.146 | 1.55E-75 | 6 | Cbfb |
| Nsd31 | 8.36E-80 | 0.357543479 | 0.611 | 0.295 | 1.69E-75 | 6 | Nsd3 |
| Ddost | 1.40E-79 | 0.276698586 | 0.467 | 0.205 | 2.84E-75 | 6 | Ddost |
| Susd63 | 4.27E-79 | 0.381271709 | 0.567 | 0.27 | 8.64E-75 | 6 | Susd6 |
| Rin31 | 4.45E-79 | 0.312919901 | 0.411 | 0.17 | 9.01E-75 | 6 | Rin3 |
| Ulk21 | 8.00E-79 | 0.30901263 | 0.408 | 0.169 | 1.62E-74 | 6 | Ulk2 |
| Man1a1 | 4.01E-78 | 0.313980764 | 0.386 | 0.158 | 8.11E-74 | 6 | Man1a |
| Nek71 | 6.61E-78 | 0.273060399 | 0.517 | 0.237 | 1.34E-73 | 6 | Nek7 |
| Rbm52 | 1.04E-77 | 0.308374837 | 0.5 | 0.228 | 2.10E-73 | 6 | Rbm5 |
| Sipa11 | 1.39E-77 | 0.282155422 | 0.417 | 0.176 | 2.82E-73 | 6 | Sipa1 |
| Setd22 | 1.63E-77 | 0.318645179 | 0.424 | 0.183 | 3.30E-73 | 6 | Setd2 |
| Fbxo112 | 3.81E-77 | 0.343907353 | 0.487 | 0.222 | 7.71E-73 | 6 | Fbxo11 |
| Wdr26 | 1.71E-76 | 0.278970083 | 0.452 | 0.198 | 3.47E-72 | 6 | Wdr26 |
| Fosb2 | 2.49E-76 | 0.387297506 | 0.414 | 0.18 | 5.04E-72 | 6 | Fosb |
| Foxn2 | 2.98E-76 | 0.283303944 | 0.331 | 0.128 | 6.03E-72 | 6 | Foxn2 |
| Sptlc22 | 3.80E-76 | 0.296662304 | 0.51 | 0.233 | 7.69E-72 | 6 | Sptlc2 |
| Fam69a | 2.02E-75 | 0.30518927 | 0.312 | 0.119 | 4.09E-71 | 6 | Fam69a |
| Zfand52 | 6.06E-75 | 0.401586795 | 0.769 | 0.44 | 1.23E-70 | 6 | Zfand5 |
| Dpysl21 | 7.99E-75 | 0.298665938 | 0.395 | 0.165 | 1.62E-70 | 6 | Dpysl2 |
| Fam220a | 1.05E-74 | 0.35243956 | 0.257 | 0.091 | 2.12E-70 | 6 | Fam220a |
| Zfc3h1 | 2.54E-74 | 0.279143492 | 0.327 | 0.128 | 5.15E-70 | 6 | Zfc3h1 |
| Herc41 | 3.32E-74 | 0.323822843 | 0.352 | 0.143 | 6.72E-70 | 6 | Herc4 |

| | | | | | | | |
|---|---|---|---|---|---|---|---|
| Canx2 | 3.92E-74 | 0.256789686 | 0.603 | 0.294 | 7.93E-70 | 6 | Canx |
| Tspan141 | 5.97E-74 | 0.256832992 | 0.317 | 0.12 | 1.21E-69 | 6 | Tspan14 |
| mt-Co34 | 9.33E-74 | 0.486744664 | 0.998 | 0.714 | 1.89E-69 | 6 | mt-Co3 |
| Eif5b1 | 1.01E-73 | 0.252059146 | 0.618 | 0.308 | 2.05E-69 | 6 | Eif5b |
| Mcmbp | 1.24E-73 | 0.269213492 | 0.329 | 0.128 | 2.50E-69 | 6 | Mcmbp |
| Magt11 | 3.95E-73 | 0.266788241 | 0.373 | 0.155 | 8.00E-69 | 6 | Magt1 |
| Rap1b2 | 4.81E-73 | 0.290213479 | 0.78 | 0.429 | 9.74E-69 | 6 | Rap1b |
| Herpud11 | 6.89E-73 | 0.316158126 | 0.45 | 0.204 | 1.40E-68 | 6 | Herpud1 |
| Dis3l21 | 7.50E-73 | 0.306916808 | 0.329 | 0.131 | 1.52E-68 | 6 | Dis3l2 |
| Dmxl11 | 7.87E-73 | 0.345122628 | 0.402 | 0.176 | 1.59E-68 | 6 | Dmxl1 |
| Phactr22 | 1.09E-72 | 0.33770186 | 0.502 | 0.235 | 2.21E-68 | 6 | Phactr2 |
| Acap23 | 2.05E-72 | 0.322534755 | 0.537 | 0.258 | 4.15E-68 | 6 | Acap2 |
| Dnajc13 | 3.08E-72 | 0.278471983 | 0.319 | 0.124 | 6.24E-68 | 6 | Dnajc13 |
| Tmbim62 | 3.68E-72 | 0.300436413 | 0.773 | 0.416 | 7.45E-68 | 6 | Tmbim6 |
| Myo1e2 | 5.24E-72 | 0.433793538 | 0.549 | 0.27 | 1.06E-67 | 6 | Myo1e |
| Tmem50a2 | 9.63E-72 | 0.253929193 | 0.717 | 0.368 | 1.95E-67 | 6 | Tmem50a |
| Abl1 | 2.13E-71 | 0.265200245 | 0.302 | 0.116 | 4.32E-67 | 6 | Abl1 |
| Rbm253 | 2.18E-71 | 0.330349988 | 0.637 | 0.321 | 4.41E-67 | 6 | Rbm25 |
| Nipbl3 | 2.95E-71 | 0.297183403 | 0.682 | 0.352 | 5.98E-67 | 6 | Nipbl |
| Cep3502 | 5.79E-71 | 0.312146433 | 0.432 | 0.194 | 1.17E-66 | 6 | Cep350 |
| Sppl2a2 | 8.47E-71 | 0.286159164 | 0.579 | 0.284 | 1.71E-66 | 6 | Sppl2a |
| Cog52 | 1.81E-70 | 0.334178415 | 0.399 | 0.174 | 3.67E-66 | 6 | Cog5 |
| Fndc3a1 | 3.24E-70 | 0.289710867 | 0.551 | 0.265 | 6.56E-66 | 6 | Fndc3a |
| Tmem87a | 7.95E-70 | 0.288244443 | 0.281 | 0.105 | 1.61E-65 | 6 | Tmem87a |
| Tnpo3 | 8.16E-70 | 0.25218179 | 0.339 | 0.137 | 1.65E-65 | 6 | Tnpo3 |
| Znrf31 | 9.86E-70 | 0.275080996 | 0.273 | 0.1 | 2.00E-65 | 6 | Znrf3 |
| Rsrc11 | 1.01E-69 | 0.273036712 | 0.518 | 0.249 | 2.05E-65 | 6 | Rsrc1 |
| Pfn12 | 2.80E-69 | 0.305790674 | 0.957 | 0.67 | 5.67E-65 | 6 | Pfn1 |
| Cdc421 | 5.40E-69 | 0.353687891 | 0.931 | 0.601 | 1.09E-64 | 6 | Cdc42 |
| Prpf40a2 | 1.84E-68 | 0.250486784 | 0.531 | 0.255 | 3.73E-64 | 6 | Prpf40a |
| Stk31 | 1.87E-68 | 0.286427546 | 0.37 | 0.157 | 3.79E-64 | 6 | Stk3 |
| Wwox2 | 2.43E-68 | 0.404931296 | 0.38 | 0.167 | 4.93E-64 | 6 | Wwox |
| Lncpint2 | 5.56E-68 | 0.337604471 | 0.538 | 0.263 | 1.13E-63 | 6 | Lncpint |
| Ptbp33 | 2.20E-67 | 0.305451136 | 0.726 | 0.379 | 4.46E-63 | 6 | Ptbp3 |
| B4galt12 | 4.28E-67 | 0.32143564 | 0.384 | 0.168 | 8.66E-63 | 6 | B4galt1 |
| Stat12 | 6.36E-67 | 0.301863549 | 0.496 | 0.239 | 1.29E-62 | 6 | Stat1 |
| Baz2b2 | 2.66E-66 | 0.283044751 | 0.653 | 0.337 | 5.38E-62 | 6 | Baz2b |
| Vti1a2 | 3.74E-66 | 0.298871172 | 0.506 | 0.244 | 7.57E-62 | 6 | Vti1a |
| Fchsd21 | 2.92E-65 | 0.272977497 | 0.355 | 0.152 | 5.91E-61 | 6 | Fchsd2 |
| Qk2 | 1.88E-64 | 0.25657198 | 0.745 | 0.395 | 3.81E-60 | 6 | Qk |
| Ankrd172 | 2.65E-64 | 0.296017246 | 0.509 | 0.251 | 5.36E-60 | 6 | Ankrd17 |
| Mapk6 | 8.40E-64 | 0.25079803 | 0.258 | 0.096 | 1.70E-59 | 6 | Mapk6 |
| Eif4g32 | 7.13E-63 | 0.255120871 | 0.551 | 0.274 | 1.44E-58 | 6 | Eif4g3 |

| | | | | | | | |
|---|---|---|---|---|---|---|---|
| Neat12 | 4.53E-62 | 0.269619488 | 0.426 | 0.2 | 9.16E-58 | 6 | Neat1 |
| mt-Nd13 | 5.84E-62 | 0.375502934 | 0.93 | 0.584 | 1.18E-57 | 6 | mt-Nd1 |
| Rps6ka32 | 1.42E-61 | 0.351218481 | 0.505 | 0.252 | 2.87E-57 | 6 | Rps6ka3 |
| Pde7b2 | 1.53E-61 | 0.391672666 | 0.479 | 0.233 | 3.11E-57 | 6 | Pde7b |
| Nlrc51 | 2.69E-61 | 0.255414355 | 0.27 | 0.105 | 5.44E-57 | 6 | Nlrc5 |
| Dach1 | 3.24E-61 | 0.423865156 | 0.254 | 0.098 | 6.55E-57 | 6 | Dach1 |
| Crtc31 | 9.67E-61 | 0.266817386 | 0.357 | 0.157 | 1.96E-56 | 6 | Crtc3 |
| Spata51 | 1.92E-60 | 0.268975686 | 0.287 | 0.117 | 3.89E-56 | 6 | Spata5 |
| Smchd12 | 3.46E-60 | 0.259697201 | 0.509 | 0.254 | 7.00E-56 | 6 | Smchd1 |
| Dyrk1a2 | 1.09E-59 | 0.275313399 | 0.376 | 0.172 | 2.21E-55 | 6 | Dyrk1a |
| Rassf12 | 2.16E-59 | 0.262281713 | 0.453 | 0.219 | 4.38E-55 | 6 | Rassf1 |
| Ifit21 | 3.43E-59 | 0.449135801 | 0.278 | 0.112 | 6.94E-55 | 6 | Ifit2 |
| Gm372401 | 1.30E-58 | 0.267886637 | 0.254 | 0.098 | 2.63E-54 | 6 | Gm37240 |
| Clk12 | 2.29E-58 | 0.25765341 | 0.549 | 0.281 | 4.63E-54 | 6 | Clk1 |
| Rbm62 | 6.43E-58 | 0.263327898 | 0.433 | 0.208 | 1.30E-53 | 6 | Rbm6 |
| Gm26917 | 6.89E-57 | 0.26073887 | 0.277 | 0.114 | 1.40E-52 | 6 | Gm26917 |
| Kdm2a3 | 8.84E-57 | 0.28428945 | 0.465 | 0.231 | 1.79E-52 | 6 | Kdm2a |
| Zfp4072 | 2.08E-55 | 0.276524472 | 0.34 | 0.154 | 4.22E-51 | 6 | Zfp407 |
| Zfp36l12 | 7.02E-54 | 0.349680735 | 0.727 | 0.42 | 1.42E-49 | 6 | Zfp36l1 |
| Nsd12 | 1.15E-53 | 0.252355364 | 0.62 | 0.338 | 2.33E-49 | 6 | Nsd1 |
| Copg21 | 7.84E-53 | 0.250065356 | 0.266 | 0.111 | 1.59E-48 | 6 | Copg2 |
| Son3 | 1.94E-51 | 0.271213342 | 0.867 | 0.511 | 3.92E-47 | 6 | Son |
| Tra2a3 | 5.76E-51 | 0.286196281 | 0.506 | 0.265 | 1.17E-46 | 6 | Tra2a |
| Stat21 | 3.19E-49 | 0.271617797 | 0.299 | 0.134 | 6.47E-45 | 6 | Stat2 |
| mt-Co24 | 1.75E-39 | 0.335398755 | 0.992 | 0.717 | 3.54E-35 | 6 | mt-Co2 |
| mt-Nd43 | 2.00E-39 | 0.29858785 | 0.97 | 0.662 | 4.04E-35 | 6 | mt-Nd4 |
| Ctsd1 | 1.69E-18 | 0.37400649 | 0.475 | 0.308 | 3.41E-14 | 6 | Ctsd |
| Fabp42 | 0 | 1.836712637 | 0.953 | 0.415 | 0 | 7 | Fabp4 |
| Gm424181 | 2.32E-298 | 0.971373906 | 0.999 | 1 | 4.71E-294 | 7 | Gm42418 |
| Cmss12 | 9.51E-276 | 1.063831806 | 0.99 | 0.994 | 1.93E-271 | 7 | Cmss1 |
| AY0361182 | 2.46E-230 | 1.413614891 | 0.938 | 0.926 | 4.98E-226 | 7 | AY036118 |
| Cdk82 | 3.31E-217 | 1.104267541 | 0.956 | 0.97 | 6.71E-213 | 7 | Cdk8 |
| Igfbp72 | 3.07E-191 | 1.216913434 | 0.914 | 0.45 | 6.21E-187 | 7 | Igfbp7 |
| Rbp72 | 6.25E-187 | 1.85336795 | 0.69 | 0.335 | 1.27E-182 | 7 | Rbp7 |
| Ly6c12 | 7.37E-179 | 1.385159298 | 0.789 | 0.385 | 1.49E-174 | 7 | Ly6c1 |
| Ifitm22 | 1.17E-178 | 1.589931908 | 0.738 | 0.455 | 2.37E-174 | 7 | Ifitm2 |
| Ifitm32 | 2.96E-156 | 1.302845712 | 0.817 | 0.528 | 5.98E-152 | 7 | Ifitm3 |
| Egfl72 | 1.84E-135 | 1.19132887 | 0.726 | 0.381 | 3.72E-131 | 7 | Egfl7 |
| Lars22 | 2.20E-116 | 1.004430463 | 0.865 | 0.898 | 4.46E-112 | 7 | Lars2 |
| Ly6a2 | 5.28E-107 | 1.155555369 | 0.732 | 0.45 | 1.07E-102 | 7 | Ly6a |
| Calm12 | 1.53E-102 | 0.945773819 | 0.855 | 0.779 | 3.10E-98 | 7 | Calm1 |
| Sptbn12 | 8.86E-97 | 1.131508468 | 0.742 | 0.486 | 1.79E-92 | 7 | Sptbn1 |
| Tmsb4x3 | 4.70E-90 | 0.476283289 | 0.986 | 0.932 | 9.51E-86 | 7 | Tmsb4x |

| | | | | | | | |
|---|---|---|---|---|---|---|---|
| Id32 | 1.57E-89 | 1.101726936 | 0.661 | 0.4 | 3.18E-85 | 7 | Id3 |
| Gng112 | 8.00E-89 | 1.485249778 | 0.552 | 0.361 | 1.62E-84 | 7 | Gng11 |
| mt-Atp64 | 6.85E-88 | 0.838894403 | 0.826 | 0.752 | 1.39E-83 | 7 | mt-Atp6 |
| mt-Co25 | 7.28E-78 | 0.842159783 | 0.816 | 0.726 | 1.47E-73 | 7 | mt-Co2 |
| Ptma2 | 7.29E-75 | 0.617993006 | 0.917 | 0.941 | 1.48E-70 | 7 | Ptma |
| Rdx2 | 4.96E-74 | 1.386137861 | 0.654 | 0.549 | 1.00E-69 | 7 | Rdx |
| Itm2b4 | 6.36E-74 | 0.806587201 | 0.843 | 0.722 | 1.29E-69 | 7 | Itm2b |
| Rflnb2 | 1.55E-71 | 1.733193946 | 0.482 | 0.31 | 3.14E-67 | 7 | Rflnb |
| Tm4sf12 | 2.74E-67 | 1.145270971 | 0.58 | 0.371 | 5.55E-63 | 7 | Tm4sf1 |
| Rgcc2 | 1.01E-66 | 1.087384314 | 0.598 | 0.398 | 2.04E-62 | 7 | Rgcc |
| Tpt12 | 1.79E-66 | 0.498626728 | 0.977 | 0.99 | 3.62E-62 | 7 | Tpt1 |
| Plvap2 | 3.70E-62 | 0.791679787 | 0.645 | 0.375 | 7.49E-58 | 7 | Plvap |
| Flt12 | 1.88E-60 | 0.51345853 | 0.727 | 0.381 | 3.82E-56 | 7 | Flt1 |
| Slc9a3r22 | 1.32E-59 | 1.275908989 | 0.515 | 0.367 | 2.66E-55 | 7 | Slc9a3r2 |
| H2-D13 | 4.51E-55 | 0.725345199 | 0.793 | 0.666 | 9.12E-51 | 7 | H2-D1 |
| Sparc2 | 3.12E-44 | 0.943774892 | 0.548 | 0.399 | 6.32E-40 | 7 | Sparc |
| Hspb12 | 5.94E-44 | 0.944524593 | 0.52 | 0.378 | 1.20E-39 | 7 | Hspb1 |
| mt-Co35 | 4.45E-43 | 0.642477243 | 0.784 | 0.725 | 9.01E-39 | 7 | mt-Co3 |
| Cd2002 | 2.95E-42 | 1.106138452 | 0.472 | 0.348 | 5.98E-38 | 7 | Cd200 |
| Camk1d | 4.36E-42 | 0.788365048 | 0.74 | 0.828 | 8.84E-38 | 7 | Camk1d |
| Emcn2 | 1.18E-38 | 0.585966215 | 0.607 | 0.373 | 2.38E-34 | 7 | Emcn |
| Gnas2 | 1.11E-37 | 0.558980075 | 0.806 | 0.866 | 2.25E-33 | 7 | Gnas |
| mt-Nd23 | 1.86E-35 | 0.778684663 | 0.7 | 0.69 | 3.77E-31 | 7 | mt-Nd2 |
| Rps292 | 3.27E-33 | 0.376829371 | 0.95 | 0.973 | 6.63E-29 | 7 | Rps29 |
| Pecam12 | 5.86E-33 | 0.71121004 | 0.559 | 0.407 | 1.19E-28 | 7 | Pecam1 |
| mt-Cytb3 | 9.04E-33 | 0.66925652 | 0.716 | 0.702 | 1.83E-28 | 7 | mt-Cytb |
| Myl62 | 1.61E-32 | 0.685637468 | 0.694 | 0.781 | 3.26E-28 | 7 | Myl6 |
| Mgll2 | 1.81E-31 | 0.881695094 | 0.537 | 0.448 | 3.66E-27 | 7 | Mgll |
| Rps192 | 1.01E-29 | 0.463363779 | 0.867 | 0.931 | 2.05E-25 | 7 | Rps19 |
| Rps142 | 2.26E-29 | 0.445832966 | 0.876 | 0.947 | 4.58E-25 | 7 | Rps14 |
| Rps272 | 2.63E-28 | 0.443912694 | 0.833 | 0.902 | 5.33E-24 | 7 | Rps27 |
| Tmed92 | 1.99E-27 | 0.312457248 | 0.176 | 0.417 | 4.02E-23 | 7 | Tmed9 |
| Dpm31 | 3.50E-27 | 0.268523272 | 0.146 | 0.361 | 7.08E-23 | 7 | Dpm3 |
| Gpihbp12 | 1.57E-26 | 1.037164909 | 0.393 | 0.295 | 3.19E-22 | 7 | Gpihbp1 |
| mt-Co14 | 7.29E-26 | 0.533258874 | 0.767 | 0.72 | 1.48E-21 | 7 | mt-Co1 |
| Rtf12 | 1.05E-25 | 0.280681664 | 0.11 | 0.29 | 2.13E-21 | 7 | Rtf1 |
| Tmod32 | 1.17E-25 | 0.252298968 | 0.158 | 0.373 | 2.38E-21 | 7 | Tmod3 |
| Gphn | 1.62E-25 | 0.712199413 | 0.678 | 0.8 | 3.29E-21 | 7 | Gphn |
| Tmem50a3 | 1.89E-25 | 0.309078936 | 0.169 | 0.395 | 3.83E-21 | 7 | Tmem50a |
| Hnrnpu1 | 7.30E-25 | 0.272227556 | 0.195 | 0.436 | 1.48E-20 | 7 | Hnrnpu |
| Sfr11 | 1.11E-24 | 0.256321762 | 0.175 | 0.394 | 2.26E-20 | 7 | Sfr1 |
| Kdr2 | 3.07E-24 | 0.558891093 | 0.514 | 0.359 | 6.22E-20 | 7 | Kdr |
| Slk2 | 3.13E-24 | 0.250470848 | 0.089 | 0.25 | 6.33E-20 | 7 | Slk |

| | | | | | | | |
|---|---|---|---|---|---|---|---|
| Bcap312 | 3.41E-24 | 0.265966263 | 0.116 | 0.295 | 6.90E-20 | 7 | Bcap31 |
| Sec622 | 4.09E-24 | 0.394966914 | 0.179 | 0.407 | 8.29E-20 | 7 | Sec62 |
| Dync1h12 | 2.01E-23 | 0.276340036 | 0.118 | 0.297 | 4.06E-19 | 7 | Dync1h1 |
| Emc72 | 2.16E-23 | 0.301730864 | 0.11 | 0.282 | 4.36E-19 | 7 | Emc7 |
| Sbds2 | 5.74E-23 | 0.283951038 | 0.134 | 0.321 | 1.16E-18 | 7 | Sbds |
| Ctla2a3 | 6.23E-23 | 1.116436994 | 0.45 | 0.399 | 1.26E-18 | 7 | Ctla2a |
| Atraid2 | 6.64E-23 | 0.279006802 | 0.111 | 0.281 | 1.34E-18 | 7 | Atraid |
| Tapbp2 | 9.00E-23 | 0.339444662 | 0.131 | 0.315 | 1.82E-18 | 7 | Tapbp |
| Nt5c | 1.04E-22 | 0.271441185 | 0.099 | 0.259 | 2.11E-18 | 7 | Nt5c |
| Pdia63 | 1.24E-22 | 0.294587198 | 0.15 | 0.349 | 2.52E-18 | 7 | Pdia6 |
| Tmem2342 | 1.38E-22 | 0.361560187 | 0.183 | 0.405 | 2.80E-18 | 7 | Tmem234 |
| Gnb12 | 1.67E-22 | 0.312025211 | 0.204 | 0.441 | 3.38E-18 | 7 | Gnb1 |
| Serinc12 | 1.93E-22 | 0.279300775 | 0.099 | 0.258 | 3.90E-18 | 7 | Serinc1 |
| Arhgdia | 2.40E-22 | 0.251929053 | 0.191 | 0.415 | 4.85E-18 | 7 | Arhgdia |
| Mapre1 | 3.22E-22 | 0.265661671 | 0.121 | 0.294 | 6.51E-18 | 7 | Mapre1 |
| Nck12 | 3.22E-22 | 0.306375706 | 0.153 | 0.351 | 6.51E-18 | 7 | Nck1 |
| Zfp91 | 3.77E-22 | 0.357673657 | 0.136 | 0.318 | 7.63E-18 | 7 | Zfp91 |
| Thrsp2 | 3.83E-22 | 1.104241777 | 0.362 | 0.293 | 7.76E-18 | 7 | Thrsp |
| Ktn12 | 5.25E-22 | 0.35939337 | 0.135 | 0.318 | 1.06E-17 | 7 | Ktn1 |
| Ywhah2 | 7.31E-22 | 0.299590626 | 0.137 | 0.32 | 1.48E-17 | 7 | Ywhah |
| Cpd2 | 8.06E-22 | 0.267780478 | 0.139 | 0.323 | 1.63E-17 | 7 | Cpd |
| Mier12 | 1.12E-21 | 0.307385087 | 0.151 | 0.343 | 2.27E-17 | 7 | Mier1 |
| Rps242 | 3.76E-21 | 0.269266848 | 0.95 | 0.974 | 7.61E-17 | 7 | Rps24 |
| Cxcl122 | 5.84E-21 | 1.107403102 | 0.401 | 0.321 | 1.18E-16 | 7 | Cxcl12 |
| Map4k42 | 7.82E-21 | 0.351599228 | 0.119 | 0.287 | 1.58E-16 | 7 | Map4k4 |
| Cwc151 | 9.07E-21 | 0.28106421 | 0.179 | 0.382 | 1.84E-16 | 7 | Cwc15 |
| Prkar1a2 | 1.50E-20 | 0.301815354 | 0.195 | 0.415 | 3.05E-16 | 7 | Prkar1a |
| Psma5 | 1.84E-20 | 0.260582915 | 0.142 | 0.32 | 3.72E-16 | 7 | Psma5 |
| Akr1a1 | 6.19E-20 | 0.263906722 | 0.2 | 0.416 | 1.25E-15 | 7 | Akr1a1 |
| Id12 | 7.99E-20 | 0.905955841 | 0.404 | 0.324 | 1.62E-15 | 7 | Id1 |
| Mxd43 | 8.16E-20 | 0.383163258 | 0.168 | 0.363 | 1.65E-15 | 7 | Mxd4 |
| Tmem1601 | 1.03E-19 | 0.316140607 | 0.171 | 0.368 | 2.08E-15 | 7 | Tmem160 |
| Capns12 | 1.30E-19 | 0.308157953 | 0.167 | 0.36 | 2.63E-15 | 7 | Capns1 |
| Crip22 | 1.47E-19 | 1.017886807 | 0.436 | 0.418 | 2.98E-15 | 7 | Crip2 |
| Nap1l12 | 1.58E-19 | 0.437322649 | 0.147 | 0.325 | 3.21E-15 | 7 | Nap1l1 |
| Tmem30a2 | 2.43E-19 | 0.347544877 | 0.129 | 0.296 | 4.92E-15 | 7 | Tmem30a |
| R3hdm21 | 2.55E-19 | 0.267013785 | 0.142 | 0.312 | 5.17E-15 | 7 | R3hdm2 |
| Ldha | 5.35E-19 | 0.32913233 | 0.207 | 0.424 | 1.08E-14 | 7 | Ldha |
| Krtcap2 | 5.95E-19 | 0.358503913 | 0.222 | 0.455 | 1.20E-14 | 7 | Krtcap2 |
| Arl12 | 8.82E-19 | 0.269544321 | 0.133 | 0.295 | 1.79E-14 | 7 | Arl1 |
| Tprgl2 | 1.12E-18 | 0.366166706 | 0.128 | 0.288 | 2.27E-14 | 7 | Tprgl |
| Cnbp2 | 1.42E-18 | 0.25586903 | 0.25 | 0.5 | 2.87E-14 | 7 | Cnbp |
| Nfkbia2 | 1.59E-18 | 0.292377693 | 0.203 | 0.412 | 3.22E-14 | 7 | Nfkbia |

| | | | | | | | |
|---|---|---|---|---|---|---|---|
| Hnrnpa12 | 2.55E-18 | 0.317589542 | 0.157 | 0.335 | 5.16E-14 | 7 | Hnrnpa1 |
| Kif5b1 | 2.80E-18 | 0.291383066 | 0.208 | 0.418 | 5.67E-14 | 7 | Kif5b |
| Cavin22 | 3.07E-18 | 0.792136559 | 0.418 | 0.341 | 6.22E-14 | 7 | Cavin2 |
| Tap22 | 3.21E-18 | 0.304158762 | 0.105 | 0.25 | 6.51E-14 | 7 | Tap2 |
| Plpp12 | 3.78E-18 | 0.451891627 | 0.516 | 0.38 | 7.66E-14 | 7 | Plpp1 |
| Ncoa72 | 3.88E-18 | 0.272045338 | 0.142 | 0.306 | 7.86E-14 | 7 | Ncoa7 |
| Llph1 | 5.24E-18 | 0.374812699 | 0.151 | 0.321 | 1.06E-13 | 7 | Llph |
| Rbm8a | 5.58E-18 | 0.322149605 | 0.149 | 0.318 | 1.13E-13 | 7 | Rbm8a |
| Bax2 | 8.55E-18 | 0.342421784 | 0.151 | 0.324 | 1.73E-13 | 7 | Bax |
| Trappc6b2 | 9.66E-18 | 0.270068924 | 0.144 | 0.307 | 1.96E-13 | 7 | Trappc6b |
| Smco42 | 1.34E-17 | 1.067186712 | 0.415 | 0.392 | 2.72E-13 | 7 | Smco4 |
| Sep-73 | 1.57E-17 | 0.336388025 | 0.24 | 0.475 | 3.17E-13 | 7 | 7-Sep |
| Eif3a1 | 1.69E-17 | 0.408613526 | 0.185 | 0.384 | 3.42E-13 | 7 | Eif3a |
| Eid1 | 1.85E-17 | 0.430098688 | 0.156 | 0.331 | 3.74E-13 | 7 | Eid1 |
| Mrpl17 | 2.48E-17 | 0.275253583 | 0.129 | 0.282 | 5.02E-13 | 7 | Mrpl17 |
| Tspan132 | 2.86E-17 | 0.284809455 | 0.137 | 0.295 | 5.80E-13 | 7 | Tspan13 |
| Map1lc3b1 | 3.31E-17 | 0.289555727 | 0.265 | 0.515 | 6.70E-13 | 7 | Map1lc3b |
| Calm31 | 3.88E-17 | 0.333454602 | 0.126 | 0.278 | 7.85E-13 | 7 | Calm3 |
| Hnrnpa3 | 3.89E-17 | 0.307909051 | 0.231 | 0.457 | 7.87E-13 | 7 | Hnrnpa3 |
| Raly1 | 4.00E-17 | 0.359799326 | 0.127 | 0.28 | 8.11E-13 | 7 | Raly |
| Cnih43 | 4.96E-17 | 0.336275259 | 0.117 | 0.262 | 1.00E-12 | 7 | Cnih4 |
| Ralbp1 | 5.29E-17 | 0.415010571 | 0.165 | 0.341 | 1.07E-12 | 7 | Ralbp1 |
| Ahnak2 | 5.40E-17 | 1.044470697 | 0.473 | 0.475 | 1.09E-12 | 7 | Ahnak |
| Lamp14 | 6.48E-17 | 0.349616051 | 0.215 | 0.429 | 1.31E-12 | 7 | Lamp1 |
| H2-Q63 | 7.09E-17 | 0.352994683 | 0.113 | 0.254 | 1.44E-12 | 7 | H2-Q6 |
| Adipor12 | 9.13E-17 | 0.313579847 | 0.12 | 0.266 | 1.85E-12 | 7 | Adipor1 |
| Rps213 | 9.69E-17 | 0.341439244 | 0.839 | 0.931 | 1.96E-12 | 7 | Rps21 |
| Tubb52 | 1.10E-16 | 0.348687231 | 0.195 | 0.392 | 2.22E-12 | 7 | Tubb5 |
| Smim112 | 1.30E-16 | 0.362264046 | 0.151 | 0.316 | 2.63E-12 | 7 | Smim11 |
| Spop | 2.11E-16 | 0.321202589 | 0.197 | 0.392 | 4.28E-12 | 7 | Spop |
| Podxl2 | 2.21E-16 | 0.77725037 | 0.413 | 0.345 | 4.47E-12 | 7 | Podxl |
| Kras1 | 4.13E-16 | 0.384814544 | 0.129 | 0.278 | 8.37E-12 | 7 | Kras |
| Rps233 | 4.55E-16 | 0.314320541 | 0.874 | 0.957 | 9.21E-12 | 7 | Rps23 |
| Sdf42 | 6.16E-16 | 0.502743523 | 0.152 | 0.319 | 1.25E-11 | 7 | Sdf4 |
| Hmg20b2 | 6.43E-16 | 0.250790169 | 0.126 | 0.27 | 1.30E-11 | 7 | Hmg20b |
| Csde11 | 7.99E-16 | 0.383872037 | 0.185 | 0.371 | 1.62E-11 | 7 | Csde1 |
| Dab2ip2 | 1.18E-15 | 0.267987163 | 0.127 | 0.27 | 2.39E-11 | 7 | Dab2ip |
| Crip12 | 2.00E-15 | 0.927179224 | 0.508 | 0.54 | 4.04E-11 | 7 | Crip1 |
| Cd813 | 2.15E-15 | 0.670191833 | 0.555 | 0.527 | 4.35E-11 | 7 | Cd81 |
| Chmp2a1 | 2.67E-15 | 0.313538841 | 0.266 | 0.502 | 5.41E-11 | 7 | Chmp2a |
| Mprip2 | 4.56E-15 | 0.403606497 | 0.148 | 0.302 | 9.23E-11 | 7 | Mprip |
| Cetn32 | 4.82E-15 | 0.361687919 | 0.174 | 0.343 | 9.77E-11 | 7 | Cetn3 |
| Zcrb12 | 5.11E-15 | 0.293608044 | 0.174 | 0.341 | 1.04E-10 | 7 | Zcrb1 |

| | | | | | | | |
|---|---|---|---|---|---|---|---|
| Cdv3 | 6.60E-15 | 0.416039331 | 0.147 | 0.299 | 1.34E-10 | 7 | Cdv3 |
| Fam96b1 | 6.70E-15 | 0.353642188 | 0.117 | 0.251 | 1.36E-10 | 7 | Fam96b |
| Tcf152 | 1.16E-14 | 1.064507041 | 0.319 | 0.26 | 2.34E-10 | 7 | Tcf15 |
| Tm9sf32 | 1.57E-14 | 0.571711821 | 0.173 | 0.345 | 3.17E-10 | 7 | Tm9sf3 |
| Canx3 | 1.85E-14 | 0.516400867 | 0.156 | 0.316 | 3.74E-10 | 7 | Canx |
| Ccnd22 | 1.86E-14 | 0.435585025 | 0.146 | 0.295 | 3.77E-10 | 7 | Ccnd2 |
| H2afv | 1.89E-14 | 0.379880831 | 0.202 | 0.39 | 3.82E-10 | 7 | H2afv |
| mt-Nd44 | 1.99E-14 | 0.557399186 | 0.642 | 0.679 | 4.03E-10 | 7 | mt-Nd4 |
| Rpl74 | 2.83E-14 | 0.35234821 | 0.776 | 0.899 | 5.73E-10 | 7 | Rpl7 |
| Rnf187 | 2.90E-14 | 0.375189433 | 0.124 | 0.259 | 5.86E-10 | 7 | Rnf187 |
| mt-Nd33 | 4.33E-14 | 0.404889391 | 0.241 | 0.459 | 8.76E-10 | 7 | mt-Nd3 |
| Rap1a2 | 4.37E-14 | 0.271822958 | 0.295 | 0.55 | 8.85E-10 | 7 | Rap1a |
| Polr2m1 | 6.92E-14 | 0.254582017 | 0.124 | 0.253 | 1.40E-09 | 7 | Polr2m |
| Sparcl12 | 1.05E-13 | 0.78025697 | 0.41 | 0.364 | 2.12E-09 | 7 | Sparcl1 |
| Slc50a12 | 1.46E-13 | 0.477303432 | 0.155 | 0.308 | 2.96E-09 | 7 | Slc50a1 |
| Abhd17a1 | 1.58E-13 | 0.443342613 | 0.142 | 0.284 | 3.20E-09 | 7 | Abhd17a |
| Pttg12 | 1.79E-13 | 0.332711672 | 0.156 | 0.302 | 3.63E-09 | 7 | Pttg1 |
| Med28 | 2.05E-13 | 0.383545546 | 0.146 | 0.288 | 4.16E-09 | 7 | Med28 |
| Dnajc82 | 2.40E-13 | 0.534890569 | 0.192 | 0.369 | 4.87E-09 | 7 | Dnajc8 |
| Vim1 | 4.04E-13 | 0.322040172 | 0.168 | 0.316 | 8.17E-09 | 7 | Vim |
| Rps123 | 4.94E-13 | 0.351694845 | 0.775 | 0.905 | 1.00E-08 | 7 | Rps12 |
| D8Ertd738e1 | 5.44E-13 | 0.382019443 | 0.289 | 0.534 | 1.10E-08 | 7 | D8Ertd738e |
| Rpl13a2 | 5.49E-13 | 0.369476604 | 0.15 | 0.293 | 1.11E-08 | 7 | Rpl13a |
| Pttg1ip2 | 5.93E-13 | 0.460346913 | 0.14 | 0.277 | 1.20E-08 | 7 | Pttg1ip |
| Tmed102 | 6.76E-13 | 0.49588608 | 0.239 | 0.448 | 1.37E-08 | 7 | Tmed10 |
| Ifi2032 | 7.79E-13 | 0.393744729 | 0.202 | 0.378 | 1.58E-08 | 7 | Ifi203 |
| Rpl37a2 | 1.40E-12 | 0.256338089 | 0.863 | 0.942 | 2.84E-08 | 7 | Rpl37a |
| Abcg22 | 2.23E-12 | 0.299487183 | 0.145 | 0.279 | 4.52E-08 | 7 | Abcg2 |
| Syf2 | 2.52E-12 | 0.501103562 | 0.197 | 0.368 | 5.10E-08 | 7 | Syf2 |
| Ptprb2 | 2.99E-12 | 0.424912875 | 0.471 | 0.366 | 6.05E-08 | 7 | Ptprb |
| Tns12 | 3.87E-12 | 0.346777437 | 0.128 | 0.253 | 7.84E-08 | 7 | Tns1 |
| Bag12 | 6.30E-12 | 0.428942414 | 0.23 | 0.417 | 1.28E-07 | 7 | Bag1 |
| Mrfap12 | 6.63E-12 | 0.405432287 | 0.255 | 0.462 | 1.34E-07 | 7 | Mrfap1 |
| Chmp31 | 8.02E-12 | 0.357215504 | 0.187 | 0.343 | 1.62E-07 | 7 | Chmp3 |
| Snx32 | 8.07E-12 | 0.423297279 | 0.259 | 0.473 | 1.63E-07 | 7 | Snx3 |
| Sem11 | 1.27E-11 | 0.250028574 | 0.361 | 0.654 | 2.56E-07 | 7 | Sem1 |
| Gbp72 | 1.44E-11 | 0.334973156 | 0.149 | 0.277 | 2.92E-07 | 7 | Gbp7 |
| F11r3 | 2.32E-11 | 0.359953804 | 0.145 | 0.274 | 4.69E-07 | 7 | F11r |
| Arpc1a2 | 3.26E-11 | 0.379506019 | 0.18 | 0.326 | 6.61E-07 | 7 | Arpc1a |
| Ramp22 | 4.31E-11 | 0.949841484 | 0.358 | 0.334 | 8.72E-07 | 7 | Ramp2 |
| AW1120102 | 5.04E-11 | 0.457212615 | 0.577 | 0.571 | 1.02E-06 | 7 | AW112010 |
| Psma7 | 6.81E-11 | 0.326313018 | 0.332 | 0.591 | 1.38E-06 | 7 | Psma7 |
| Gna112 | 8.12E-11 | 0.430643084 | 0.133 | 0.252 | 1.64E-06 | 7 | Gna11 |

| | | | | | | | |
|---|---|---|---|---|---|---|---|
| Rab11a2 | 8.41E-11 | 0.36981955 | 0.175 | 0.317 | 1.70E-06 | 7 | Rab11a |
| mt-Nd52 | 8.47E-11 | 0.31808764 | 0.286 | 0.508 | 1.71E-06 | 7 | mt-Nd5 |
| Ostf13 | 1.06E-10 | 0.397249425 | 0.291 | 0.519 | 2.15E-06 | 7 | Ostf1 |
| Tagln22 | 1.07E-10 | 0.359374015 | 0.248 | 0.44 | 2.17E-06 | 7 | Tagln2 |
| Selenow1 | 1.15E-10 | 0.786030841 | 0.52 | 0.637 | 2.33E-06 | 7 | Selenow |
| Pdcl3 | 1.84E-10 | 0.450098047 | 0.142 | 0.263 | 3.73E-06 | 7 | Pdcl3 |
| Rpl283 | 1.98E-10 | 0.250276597 | 0.828 | 0.935 | 4.01E-06 | 7 | Rpl28 |
| Ly6e2 | 2.44E-10 | 0.552890079 | 0.578 | 0.624 | 4.94E-06 | 7 | Ly6e |
| Rab5c2 | 3.12E-10 | 0.54197916 | 0.164 | 0.299 | 6.31E-06 | 7 | Rab5c |
| Tinagl12 | 3.86E-10 | 0.979720379 | 0.323 | 0.299 | 7.82E-06 | 7 | Tinagl1 |
| Arhgef122 | 4.13E-10 | 0.323941511 | 0.204 | 0.358 | 8.35E-06 | 7 | Arhgef12 |
| Ddrgk12 | 6.49E-10 | 0.564053362 | 0.181 | 0.327 | 1.31E-05 | 7 | Ddrgk1 |
| Clec2d2 | 7.05E-10 | 0.347702857 | 0.317 | 0.54 | 1.43E-05 | 7 | Clec2d |
| Ybx13 | 8.55E-10 | 0.59578045 | 0.591 | 0.761 | 1.73E-05 | 7 | Ybx1 |
| Ywhaq2 | 9.40E-10 | 0.424954321 | 0.285 | 0.5 | 1.90E-05 | 7 | Ywhaq |
| Smim10l11 | 1.44E-09 | 0.425642767 | 0.185 | 0.324 | 2.91E-05 | 7 | Smim10l1 |
| Vamp52 | 1.48E-09 | 0.961613463 | 0.382 | 0.388 | 2.99E-05 | 7 | Vamp5 |
| Igfbp32 | 2.00E-09 | 0.678226515 | 0.349 | 0.303 | 4.04E-05 | 7 | Igfbp3 |
| Gngt23 | 2.17E-09 | 1.097101281 | 0.354 | 0.363 | 4.40E-05 | 7 | Gngt2 |
| Sh3bgrl2 | 2.63E-09 | 0.508692086 | 0.218 | 0.385 | 5.31E-05 | 7 | Sh3bgrl |
| Serpinh12 | 3.55E-09 | 0.337635082 | 0.145 | 0.257 | 7.18E-05 | 7 | Serpinh1 |
| Tmem176b4 | 4.01E-09 | 0.25323533 | 0.273 | 0.449 | 8.12E-05 | 7 | Tmem176b |
| Gnai22 | 4.75E-09 | 0.705816885 | 0.503 | 0.58 | 9.61E-05 | 7 | Gnai2 |
| Gnb22 | 6.15E-09 | 0.438198467 | 0.31 | 0.544 | 0.000124418 | 7 | Gnb2 |
| Heg12 | 7.66E-09 | 0.374008344 | 0.163 | 0.282 | 0.000155113 | 7 | Heg1 |
| Psmb102 | 8.24E-09 | 0.536727102 | 0.234 | 0.41 | 0.000166902 | 7 | Psmb10 |
| Gstm11 | 1.16E-08 | 0.278059097 | 0.199 | 0.331 | 0.000235707 | 7 | Gstm1 |
| Rps32 | 1.42E-08 | 0.257233156 | 0.791 | 0.898 | 0.00028759 | 7 | Rps3 |
| Bsg2 | 1.86E-08 | 0.538656962 | 0.268 | 0.46 | 0.000376261 | 7 | Bsg |
| Selenop4 | 3.51E-08 | 0.355100909 | 0.222 | 0.372 | 0.000709755 | 7 | Selenop |
| Mfge82 | 3.64E-08 | 0.37863856 | 0.152 | 0.261 | 0.000737345 | 7 | Mfge8 |
| Ubxn11 | 3.87E-08 | 0.502270408 | 0.26 | 0.443 | 0.000783325 | 7 | Ubxn1 |
| H2-K13 | 4.47E-08 | 0.395724508 | 0.648 | 0.645 | 0.000904861 | 7 | H2-K1 |
| mt-Atp83 | 5.84E-08 | 0.521352311 | 0.277 | 0.475 | 0.001182632 | 7 | mt-Atp8 |
| B2m5 | 8.09E-08 | 0.410780757 | 0.681 | 0.693 | 0.00163854 | 7 | B2m |
| Arpc1b2 | 9.70E-08 | 0.327800681 | 0.355 | 0.605 | 0.001964724 | 7 | Arpc1b |
| Plscr22 | 1.08E-07 | 0.85454238 | 0.349 | 0.337 | 0.002184531 | 7 | Plscr2 |
| Sptan12 | 1.54E-07 | 0.458479794 | 0.211 | 0.352 | 0.003110394 | 7 | Sptan1 |
| Psmb84 | 1.80E-07 | 0.79554612 | 0.472 | 0.564 | 0.003648996 | 7 | Psmb8 |
| Psme22 | 2.02E-07 | 0.568550953 | 0.267 | 0.454 | 0.004084316 | 7 | Psme2 |
| Cd300lg2 | 2.96E-07 | 0.847604632 | 0.312 | 0.298 | 0.005986009 | 7 | Cd300lg |
| 1700020I14Rik1 | 3.79E-07 | 0.494146635 | 0.211 | 0.35 | 0.007678457 | 7 | 1700020I14Rik |
| Kctd12b2 | 4.51E-07 | 0.313833448 | 0.155 | 0.256 | 0.009132676 | 7 | Kctd12b |

| | | | | | | | |
|---|---|---|---|---|---|---|---|
| Adgrl42 | 5.86E-07 | 0.349952768 | 0.412 | 0.352 | 0.01185408 | 7 | Adgrl4 |
| Atox11 | 6.19E-07 | 0.725175971 | 0.502 | 0.626 | 0.012530169 | 7 | Atox1 |
| Srgn3 | 6.74E-07 | 0.53210161 | 0.516 | 0.549 | 0.013650976 | 7 | Srgn |
| Tmem176a4 | 7.35E-07 | 0.256766136 | 0.271 | 0.427 | 0.014872914 | 7 | Tmem176a |
| H2-T222 | 8.44E-07 | 0.730141537 | 0.172 | 0.288 | 0.017078952 | 7 | H2-T22 |
| Gimap13 | 8.78E-07 | 0.611539479 | 0.186 | 0.311 | 0.017776866 | 7 | Gimap1 |
| Klf22 | 9.01E-07 | 0.704088556 | 0.382 | 0.369 | 0.018242232 | 7 | Klf2 |
| Sod12 | 1.23E-06 | 0.662297427 | 0.185 | 0.307 | 0.024817269 | 7 | Sod1 |
| Tie12 | 1.42E-06 | 0.436862742 | 0.158 | 0.261 | 0.028787821 | 7 | Tie1 |
| Cox7a2l2 | 2.08E-06 | 0.375059095 | 0.351 | 0.574 | 0.042082796 | 7 | Cox7a2l |
| Hsp90b13 | 2.45E-06 | 0.421539717 | 0.32 | 0.522 | 0.049620096 | 7 | Hsp90b1 |
| Fbln22 | 2.48E-06 | 0.282038784 | 0.179 | 0.279 | 0.050225899 | 7 | Fbln2 |
| Chmp51 | 2.73E-06 | 0.538156605 | 0.218 | 0.355 | 0.05520182 | 7 | Chmp5 |
| Adgrf52 | 2.96E-06 | 0.660273155 | 0.352 | 0.34 | 0.059972629 | 7 | Adgrf5 |
| Eva1b2 | 3.19E-06 | 0.555909125 | 0.179 | 0.293 | 0.064636992 | 7 | Eva1b |
| Mapk32 | 3.23E-06 | 0.580844236 | 0.166 | 0.273 | 0.065356267 | 7 | Mapk3 |
| Bst24 | 3.78E-06 | 0.682689833 | 0.454 | 0.508 | 0.076552708 | 7 | Bst2 |
| Tshz22 | 3.97E-06 | 0.287897739 | 0.251 | 0.38 | 0.080329942 | 7 | Tshz2 |
| Jam32 | 3.98E-06 | 0.28567586 | 0.182 | 0.282 | 0.080513915 | 7 | Jam3 |
| Epas12 | 4.08E-06 | 0.863318734 | 0.351 | 0.363 | 0.082508811 | 7 | Epas1 |
| Cavin32 | 4.56E-06 | 0.949861977 | 0.305 | 0.31 | 0.092253841 | 7 | Cavin3 |
| Sgk12 | 5.49E-06 | 0.430360062 | 0.177 | 0.277 | 0.111149805 | 7 | Sgk1 |
| Slc43a32 | 1.21E-05 | 0.355497061 | 0.18 | 0.277 | 0.244231133 | 7 | Slc43a3 |
| Anxa5 | 1.43E-05 | 0.461773468 | 0.313 | 0.505 | 0.290275657 | 7 | Anxa5 |
| Cyp26b12 | 1.95E-05 | 0.553217781 | 0.254 | 0.217 | 0.394356406 | 7 | Cyp26b1 |
| Tubb4b2 | 2.30E-05 | 0.490250417 | 0.278 | 0.437 | 0.465342274 | 7 | Tubb4b |
| Pltp2 | 2.79E-05 | 0.745186062 | 0.309 | 0.303 | 0.565612253 | 7 | Pltp |
| Tmem592 | 3.45E-05 | 0.572333497 | 0.305 | 0.502 | 0.699092883 | 7 | Tmem59 |
| Cfl12 | 3.97E-05 | 0.327496578 | 0.386 | 0.621 | 0.803160313 | 7 | Cfl1 |
| Tuba1a2 | 4.11E-05 | 0.650950301 | 0.167 | 0.263 | 0.831371438 | 7 | Tuba1a |
| Ccdc28b2 | 4.35E-05 | 0.483232798 | 0.173 | 0.268 | 0.879973742 | 7 | Ccdc28b |
| Tpm32 | 5.09E-05 | 0.571963161 | 0.303 | 0.49 | 1 | 7 | Tpm3 |
| Aldh23 | 6.29E-05 | 0.379883555 | 0.266 | 0.405 | 1 | 7 | Aldh2 |
| Gm265611 | 7.13E-05 | 0.633127485 | 0.204 | 0.317 | 1 | 7 | Gm26561 |
| Aplp22 | 8.00E-05 | 0.41473363 | 0.278 | 0.425 | 1 | 7 | Aplp2 |
| Acvrl12 | 0.000101456 | 0.533233014 | 0.183 | 0.282 | 1 | 7 | Acvrl1 |
| H2-T232 | 0.000158281 | 0.529103542 | 0.302 | 0.469 | 1 | 7 | H2-T23 |
| Thbd2 | 0.00017035 | 0.405718124 | 0.175 | 0.261 | 1 | 7 | Thbd |
| Slfn52 | 0.000214073 | 0.789908068 | 0.337 | 0.361 | 1 | 7 | Slfn5 |
| Clic52 | 0.000249453 | 0.861165604 | 0.288 | 0.294 | 1 | 7 | Clic5 |
| Ppp1r22 | 0.000282033 | 0.461722603 | 0.264 | 0.402 | 1 | 7 | Ppp1r2 |
| Ece12 | 0.000367785 | 0.446143027 | 0.212 | 0.313 | 1 | 7 | Ece1 |
| Snrk2 | 0.000467197 | 0.356695289 | 0.261 | 0.377 | 1 | 7 | Snrk |

| | | | | | | | |
|---|---|---|---|---|---|---|---|
| Psmb92 | 0.000477864 | 0.675093092 | 0.265 | 0.415 | 1 | 7 | Psmb9 |
| Prss232 | 0.000664255 | 0.398278528 | 0.199 | 0.287 | 1 | 7 | Prss23 |
| Rhoc2 | 0.000822442 | 0.71981635 | 0.201 | 0.303 | 1 | 7 | Rhoc |
| Eif4g21 | 0.00098544 | 0.489430711 | 0.365 | 0.578 | 1 | 7 | Eif4g2 |
| Sh3glb12 | 0.000996351 | 0.506990897 | 0.343 | 0.543 | 1 | 7 | Sh3glb1 |
| Psme13 | 0.001238273 | 0.447712554 | 0.368 | 0.58 | 1 | 7 | Psme1 |
| Selenom2 | 0.001421654 | 0.725669731 | 0.384 | 0.451 | 1 | 7 | Selenom |
| Fermt22 | 0.00147691 | 0.493749581 | 0.244 | 0.358 | 1 | 7 | Fermt2 |
| 1810037I17Rik1 | 0.002282979 | 0.646729874 | 0.317 | 0.489 | 1 | 7 | 1810037I17Rik |
| Tpm42 | 0.002670302 | 1.026308469 | 0.328 | 0.385 | 1 | 7 | Tpm4 |
| Sox182 | 0.003062245 | 0.751692233 | 0.27 | 0.279 | 1 | 7 | Sox18 |
| Ctsl2 | 0.004789319 | 0.518166105 | 0.247 | 0.353 | 1 | 7 | Ctsl |
| Rpl54 | 0.005079086 | 0.264944615 | 0.476 | 0.723 | 1 | 7 | Rpl5 |
| Ets12 | 0.007241727 | 0.606450332 | 0.28 | 0.412 | 1 | 7 | Ets1 |
| Rpl292 | 0.008064861 | 0.368569041 | 0.59 | 0.778 | 1 | 7 | Rpl29 |
| Rsad22 | 0.00843992 | 0.363150291 | 0.195 | 0.262 | 1 | 7 | Rsad2 |
| Ccdc85b2 | 0.008698927 | 0.706780323 | 0.183 | 0.262 | 1 | 7 | Ccdc85b |
| Selenok2 | 0.008736205 | 0.551958782 | 0.364 | 0.566 | 1 | 7 | Selenok |
| Bcam2 | 0.009548258 | 0.560264653 | 0.184 | 0.26 | 1 | 7 | Bcam |
| Wfdc12 | 0 | 5.305783846 | 0.842 | 0.143 | 0 | 8 | Wfdc12 |
| Scgb1b27 | 0 | 4.55569757 | 0.986 | 0.887 | 0 | 8 | Scgb1b27 |
| Scgb2b27 | 0 | 4.445987593 | 0.952 | 0.726 | 0 | 8 | Scgb2b27 |
| Prol11 | 0 | 4.264717771 | 0.976 | 0.838 | 0 | 8 | Prol1 |
| Car6 | 0 | 4.121824983 | 0.803 | 0.377 | 0 | 8 | Car6 |
| Pip1 | 0 | 4.022050664 | 0.848 | 0.321 | 0 | 8 | Pip |
| Nupr12 | 0 | 3.668190518 | 0.87 | 0.285 | 0 | 8 | Nupr1 |
| Fxyd3 | 0 | 1.857686297 | 0.674 | 0.066 | 0 | 8 | Fxyd3 |
| Snhg18 | 0 | 1.73895917 | 0.313 | 0.031 | 0 | 8 | Snhg18 |
| A630073D07Rik | 0 | 1.454336316 | 0.444 | 0.014 | 0 | 8 | A630073D07Rik |
| Aqp5 | 0 | 1.346133381 | 0.412 | 0.008 | 0 | 8 | Aqp5 |
| Cldn10 | 0 | 1.295497111 | 0.464 | 0.02 | 0 | 8 | Cldn10 |
| Crlf1 | 0 | 0.993008786 | 0.299 | 0.025 | 0 | 8 | Crlf1 |
| Fkbp11 | 0 | 0.90805354 | 0.307 | 0.021 | 0 | 8 | Fkbp11 |
| Pigr | 0 | 0.899297137 | 0.259 | 0.012 | 0 | 8 | Pigr |
| Mucl21 | 1.68E-296 | 4.320870379 | 1 | 0.999 | 3.39E-292 | 8 | Mucl2 |
| Scgb2b26 | 4.96E-295 | 4.407672273 | 0.827 | 0.46 | 1.00E-290 | 8 | Scgb2b26 |
| A2ml1 | 1.24E-288 | 3.383218793 | 0.281 | 0.034 | 2.52E-284 | 8 | A2ml1 |
| Tpd52l1 | 7.79E-280 | 1.031917136 | 0.359 | 0.053 | 1.58E-275 | 8 | Tpd52l1 |
| Wfdc18 | 2.04E-244 | 3.707704744 | 0.316 | 0.051 | 4.13E-240 | 8 | Wfdc18 |
| Bglap3 | 5.27E-172 | 0.826967914 | 0.292 | 0.055 | 1.07E-167 | 8 | Bglap3 |
| Asns | 9.63E-169 | 0.689074733 | 0.257 | 0.045 | 1.95E-164 | 8 | Asns |
| Chpt1 | 9.65E-157 | 0.827273509 | 0.502 | 0.155 | 1.95E-152 | 8 | Chpt1 |
| Smgc | 1.74E-134 | 3.331584454 | 0.256 | 0.057 | 3.51E-130 | 8 | Smgc |

| | | | | | | | |
|---|---|---|---|---|---|---|---|
| Gfpt1 | 7.63E-130 | 1.717555427 | 0.388 | 0.121 | 1.54E-125 | 8 | Gfpt1 |
| Epcam | 1.72E-124 | 0.399376218 | 0.299 | 0.072 | 3.48E-120 | 8 | Epcam |
| Plet11 | 9.44E-119 | 0.420576841 | 0.277 | 0.064 | 1.91E-114 | 8 | Plet1 |
| Dcpp1 | 9.09E-115 | 5.698826641 | 0.259 | 0.066 | 1.84E-110 | 8 | Dcpp1 |
| Pla2g12a | 3.78E-109 | 0.589060174 | 0.271 | 0.068 | 7.65E-105 | 8 | Pla2g12a |
| Serp1 | 6.34E-103 | 0.894027758 | 0.509 | 0.208 | 1.28E-98 | 8 | Serp1 |
| Copz2 | 2.59E-101 | 1.047369003 | 0.356 | 0.116 | 5.25E-97 | 8 | Copz2 |
| Spint21 | 8.46E-89 | 0.362275249 | 0.334 | 0.106 | 1.71E-84 | 8 | Spint2 |
| Krt182 | 7.90E-74 | 0.517263953 | 0.594 | 0.263 | 1.60E-69 | 8 | Krt18 |
| Eif4ebp1 | 2.37E-72 | 0.741417595 | 0.348 | 0.133 | 4.81E-68 | 8 | Eif4ebp1 |
| Nucb2 | 1.18E-66 | 0.474690408 | 0.315 | 0.115 | 2.39E-62 | 8 | Nucb2 |
| Phlda1 | 1.20E-65 | 0.357798761 | 0.25 | 0.081 | 2.43E-61 | 8 | Phlda1 |
| Slc12a21 | 1.63E-65 | 0.614953457 | 0.298 | 0.11 | 3.29E-61 | 8 | Slc12a2 |
| Stard101 | 3.23E-58 | 0.608913576 | 0.329 | 0.13 | 6.53E-54 | 8 | Stard10 |
| Ss18l2 | 1.67E-56 | 0.292436809 | 0.333 | 0.132 | 3.39E-52 | 8 | Ss18l2 |
| Sec11c2 | 1.23E-52 | 0.637067895 | 0.561 | 0.303 | 2.49E-48 | 8 | Sec11c |
| Nans | 9.64E-51 | 0.73891869 | 0.303 | 0.129 | 1.95E-46 | 8 | Nans |
| Dcpp2 | 1.93E-50 | 6.076100519 | 0.327 | 0.16 | 3.90E-46 | 8 | Dcpp2 |
| Ncald1 | 3.52E-50 | 0.418742115 | 0.255 | 0.097 | 7.13E-46 | 8 | Ncald |
| Dcpp3 | 6.52E-50 | 6.200663543 | 0.392 | 0.208 | 1.32E-45 | 8 | Dcpp3 |
| Ehf1 | 2.93E-42 | 0.320074899 | 0.274 | 0.112 | 5.93E-38 | 8 | Ehf |
| Gde1 | 3.79E-40 | 0.650530067 | 0.348 | 0.17 | 7.67E-36 | 8 | Gde1 |
| Oat | 1.59E-27 | 0.463364248 | 0.265 | 0.133 | 3.21E-23 | 8 | Oat |
| Smim141 | 8.77E-25 | 0.944359655 | 0.566 | 0.391 | 1.78E-20 | 8 | Smim14 |
| Sec61b2 | 1.46E-24 | 0.707144764 | 0.777 | 0.585 | 2.95E-20 | 8 | Sec61b |
| Tceal91 | 8.68E-21 | 0.805347284 | 0.545 | 0.383 | 1.76E-16 | 8 | Tceal9 |
| Hist1h2bc1 | 5.38E-16 | 0.487504177 | 0.379 | 0.247 | 1.09E-11 | 8 | Hist1h2bc |
| H2afj2 | 1.99E-15 | 0.397331123 | 0.629 | 0.449 | 4.02E-11 | 8 | H2afj |
| St3gal12 | 1.11E-14 | 0.466606341 | 0.284 | 0.178 | 2.25E-10 | 8 | St3gal1 |
| Tmed31 | 6.42E-13 | 0.41609397 | 0.357 | 0.241 | 1.30E-08 | 8 | Tmed3 |
| Cmpk12 | 7.95E-12 | 1.076384829 | 0.396 | 0.303 | 1.61E-07 | 8 | Cmpk1 |
| Hdlbp1 | 3.92E-11 | 0.412664876 | 0.408 | 0.296 | 7.94E-07 | 8 | Hdlbp |
| Ufm11 | 9.92E-11 | 0.335785156 | 0.309 | 0.214 | 2.01E-06 | 8 | Ufm1 |
| Rexo2 | 8.30E-08 | 0.394135056 | 0.393 | 0.296 | 0.001681294 | 8 | Rexo2 |
| Xbp1 | 2.58E-06 | 0.341770088 | 0.267 | 0.196 | 0.052320321 | 8 | Xbp1 |
| Cxcl123 | 0 | 2.973642572 | 0.977 | 0.301 | 0 | 9 | Cxcl12 |
| Mecom | 0 | 2.582487759 | 0.79 | 0.074 | 0 | 9 | Mecom |
| Tm4sf13 | 0 | 2.270875551 | 0.986 | 0.359 | 0 | 9 | Tm4sf1 |
| Sox17 | 0 | 2.21377398 | 0.703 | 0.069 | 0 | 9 | Sox17 |
| Arl152 | 0 | 2.12341889 | 0.888 | 0.366 | 0 | 9 | Arl15 |
| Cd363 | 0 | 2.104877488 | 0.857 | 0.249 | 0 | 9 | Cd36 |
| Glul | 0 | 2.06877158 | 0.788 | 0.178 | 0 | 9 | Glul |
| Plcb12 | 0 | 2.028788164 | 0.962 | 0.321 | 0 | 9 | Plcb1 |

| | | | | | | | |
|---|---|---|---|---|---|---|---|
| Gja4 | 0 | 1.980709389 | 0.637 | 0.043 | 0 | 9 | Gja4 |
| Kitl2 | 0 | 1.977972987 | 0.938 | 0.317 | 0 | 9 | Kitl |
| Pdgfd2 | 0 | 1.866845822 | 0.819 | 0.236 | 0 | 9 | Pdgfd |
| Klf23 | 0 | 1.783152834 | 0.941 | 0.349 | 0 | 9 | Klf2 |
| Id13 | 0 | 1.745014894 | 0.925 | 0.307 | 0 | 9 | Id1 |
| S1pr12 | 0 | 1.709273584 | 0.836 | 0.248 | 0 | 9 | S1pr1 |
| Ptprb3 | 0 | 1.67774103 | 0.988 | 0.349 | 0 | 9 | Ptprb |
| Tgfb2 | 0 | 1.674740739 | 0.522 | 0.073 | 0 | 9 | Tgfb2 |
| Ptprm2 | 0 | 1.653250278 | 0.914 | 0.311 | 0 | 9 | Ptprm |
| Alpl | 0 | 1.629756728 | 0.583 | 0.057 | 0 | 9 | Alpl |
| Ltbp42 | 0 | 1.602560832 | 0.864 | 0.292 | 0 | 9 | Ltbp4 |
| Mast41 | 0 | 1.578638851 | 0.697 | 0.171 | 0 | 9 | Mast4 |
| Cldn5 | 0 | 1.569527579 | 0.34 | 0.023 | 0 | 9 | Cldn5 |
| Epas13 | 0 | 1.565573839 | 0.922 | 0.342 | 0 | 9 | Epas1 |
| Sema3g | 0 | 1.547117556 | 0.46 | 0.019 | 0 | 9 | Sema3g |
| Efnb22 | 0 | 1.534788599 | 0.661 | 0.128 | 0 | 9 | Efnb2 |
| Adgrf53 | 0 | 1.533071001 | 0.933 | 0.319 | 0 | 9 | Adgrf5 |
| Lrrc3b1 | 0 | 1.531183102 | 0.545 | 0.079 | 0 | 9 | Lrrc3b |
| Cdh52 | 0 | 1.525650075 | 0.926 | 0.312 | 0 | 9 | Cdh5 |
| Ptprr | 0 | 1.457677631 | 0.501 | 0.051 | 0 | 9 | Ptprr |
| Alox12 | 0 | 1.431391841 | 0.565 | 0.082 | 0 | 9 | Alox12 |
| Clu | 0 | 1.398582532 | 0.448 | 0.053 | 0 | 9 | Clu |
| Cdh131 | 0 | 1.364323572 | 0.502 | 0.085 | 0 | 9 | Cdh13 |
| Col8a1 | 0 | 1.361624135 | 0.366 | 0.018 | 0 | 9 | Col8a1 |
| Stmn2 | 0 | 1.323931193 | 0.292 | 0.021 | 0 | 9 | Stmn2 |
| Slc26a10 | 0 | 1.234463837 | 0.444 | 0.054 | 0 | 9 | Slc26a10 |
| Fn1 | 0 | 1.0477657 | 0.312 | 0.025 | 0 | 9 | Fn1 |
| Hey1 | 0 | 0.859947702 | 0.268 | 0.02 | 0 | 9 | Hey1 |
| Car7 | 0 | 0.844875509 | 0.277 | 0.026 | 0 | 9 | Car7 |
| Efhd1 | 0 | 0.822821829 | 0.256 | 0.015 | 0 | 9 | Efhd1 |
| Myo102 | 2.59E-307 | 1.427298392 | 0.721 | 0.203 | 5.24E-303 | 9 | Myo10 |
| Slc9a3r23 | 2.50E-305 | 1.528444998 | 0.93 | 0.353 | 5.05E-301 | 9 | Slc9a3r2 |
| Cyyr12 | 1.73E-303 | 1.479304243 | 0.942 | 0.325 | 3.51E-299 | 9 | Cyyr1 |
| Fbln5 | 9.22E-298 | 1.197254581 | 0.292 | 0.032 | 1.87E-293 | 9 | Fbln5 |
| Ebf12 | 3.86E-295 | 1.542697092 | 0.951 | 0.357 | 7.82E-291 | 9 | Ebf1 |
| Rflnb3 | 2.74E-292 | 1.676413915 | 0.855 | 0.298 | 5.55E-288 | 9 | Rflnb |
| Esm12 | 9.51E-289 | 1.471808421 | 0.49 | 0.091 | 1.93E-284 | 9 | Esm1 |
| Ly6c13 | 3.00E-285 | 1.456077663 | 0.983 | 0.382 | 6.06E-281 | 9 | Ly6c1 |
| Cav12 | 5.42E-284 | 1.445043346 | 0.856 | 0.287 | 1.10E-279 | 9 | Cav1 |
| Epb41l4a | 1.59E-281 | 1.115231957 | 0.42 | 0.071 | 3.23E-277 | 9 | Epb41l4a |
| Ly6a3 | 1.62E-280 | 1.481044293 | 0.983 | 0.444 | 3.28E-276 | 9 | Ly6a |
| Fam13c1 | 4.02E-279 | 1.224228438 | 0.492 | 0.1 | 8.14E-275 | 9 | Fam13c |
| Podxl3 | 7.03E-274 | 1.360696791 | 0.922 | 0.328 | 1.42E-269 | 9 | Podxl |

| | | | | | | | |
|---|---|---|---|---|---|---|---|
| Heg13 | 5.87E-272 | 1.493964358 | 0.767 | 0.259 | 1.19E-267 | 9 | Heg1 |
| Adarb11 | 2.35E-266 | 1.098094621 | 0.485 | 0.098 | 4.75E-262 | 9 | Adarb1 |
| Fbxl72 | 7.07E-264 | 1.663234786 | 0.748 | 0.242 | 1.43E-259 | 9 | Fbxl7 |
| Etl42 | 1.65E-263 | 1.395710999 | 0.822 | 0.278 | 3.34E-259 | 9 | Etl4 |
| Ecscr2 | 1.04E-262 | 1.294926849 | 0.71 | 0.213 | 2.11E-258 | 9 | Ecscr |
| Clic53 | 1.37E-261 | 1.294508703 | 0.822 | 0.275 | 2.78E-257 | 9 | Clic5 |
| Entpd13 | 2.89E-261 | 1.519622806 | 0.747 | 0.262 | 5.86E-257 | 9 | Entpd1 |
| Sox183 | 6.23E-259 | 1.378741829 | 0.782 | 0.261 | 1.26E-254 | 9 | Sox18 |
| Slc6a62 | 6.88E-256 | 1.716131538 | 0.801 | 0.313 | 1.39E-251 | 9 | Slc6a6 |
| Clec1a2 | 1.15E-254 | 1.272987608 | 0.598 | 0.155 | 2.33E-250 | 9 | Clec1a |
| Sparc3 | 5.11E-253 | 1.359653929 | 0.942 | 0.386 | 1.04E-248 | 9 | Sparc |
| Dnm32 | 2.52E-252 | 1.38126933 | 0.707 | 0.216 | 5.09E-248 | 9 | Dnm3 |
| Id33 | 4.46E-250 | 1.341662645 | 0.964 | 0.391 | 9.04E-246 | 9 | Id3 |
| Plscr23 | 1.01E-249 | 1.389807822 | 0.857 | 0.319 | 2.05E-245 | 9 | Plscr2 |
| Ldb22 | 4.87E-245 | 1.467034425 | 0.841 | 0.315 | 9.87E-241 | 9 | Ldb2 |
| Ptprg2 | 2.08E-243 | 1.275243276 | 0.889 | 0.328 | 4.21E-239 | 9 | Ptprg |
| Crip13 | 2.53E-243 | 1.390188431 | 0.951 | 0.524 | 5.12E-239 | 9 | Crip1 |
| Jag2 | 3.44E-242 | 0.991280538 | 0.398 | 0.073 | 6.97E-238 | 9 | Jag2 |
| Mcc2 | 5.29E-240 | 1.261383772 | 0.799 | 0.269 | 1.07E-235 | 9 | Mcc |
| Egfl73 | 4.48E-238 | 1.241742499 | 0.97 | 0.376 | 9.07E-234 | 9 | Egfl7 |
| Arhgap312 | 4.17E-237 | 1.280687084 | 0.906 | 0.356 | 8.44E-233 | 9 | Arhgap31 |
| Btnl92 | 4.83E-235 | 1.408325318 | 0.764 | 0.255 | 9.78E-231 | 9 | Btnl9 |
| Fbln23 | 1.24E-230 | 1.298726888 | 0.766 | 0.257 | 2.51E-226 | 9 | Fbln2 |
| Ssu2 | 4.26E-230 | 0.710293014 | 0.279 | 0.037 | 8.62E-226 | 9 | Ssu2 |
| Esam2 | 1.29E-229 | 1.173410118 | 0.826 | 0.294 | 2.62E-225 | 9 | Esam |
| Flt13 | 8.68E-229 | 1.272833289 | 0.977 | 0.376 | 1.76E-224 | 9 | Flt1 |
| Hspb13 | 3.47E-227 | 1.279771832 | 0.906 | 0.365 | 7.03E-223 | 9 | Hspb1 |
| Filip12 | 5.99E-227 | 1.244954273 | 0.615 | 0.182 | 1.21E-222 | 9 | Filip1 |
| Uaca2 | 2.27E-225 | 1.178609331 | 0.672 | 0.213 | 4.59E-221 | 9 | Uaca |
| Cd2003 | 8.13E-225 | 1.187575153 | 0.89 | 0.334 | 1.65E-220 | 9 | Cd200 |
| Notch12 | 2.28E-224 | 1.226603482 | 0.637 | 0.204 | 4.62E-220 | 9 | Notch1 |
| Epha41 | 2.47E-224 | 1.08854789 | 0.465 | 0.106 | 4.99E-220 | 9 | Epha4 |
| Tcf42 | 1.43E-222 | 1.265849599 | 0.965 | 0.457 | 2.89E-218 | 9 | Tcf4 |
| Mcf2l2 | 2.55E-222 | 1.256902443 | 0.647 | 0.201 | 5.15E-218 | 9 | Mcf2l |
| Rasgrp32 | 3.21E-222 | 1.191780572 | 0.768 | 0.267 | 6.50E-218 | 9 | Rasgrp3 |
| Rras3 | 5.62E-222 | 1.199512226 | 0.757 | 0.284 | 1.14E-217 | 9 | Rras |
| Dll4 | 8.90E-222 | 0.961552024 | 0.371 | 0.068 | 1.80E-217 | 9 | Dll4 |
| Tcf153 | 1.12E-221 | 1.25706788 | 0.744 | 0.245 | 2.27E-217 | 9 | Tcf15 |
| Ceacam12 | 2.98E-221 | 1.120403881 | 0.545 | 0.144 | 6.04E-217 | 9 | Ceacam1 |
| Ramp23 | 1.03E-220 | 1.206449496 | 0.837 | 0.317 | 2.09E-216 | 9 | Ramp2 |
| Utrn2 | 3.82E-220 | 1.358717558 | 0.846 | 0.393 | 7.73E-216 | 9 | Utrn |
| Jam22 | 8.78E-220 | 1.130036209 | 0.779 | 0.271 | 1.78E-215 | 9 | Jam2 |
| Pls32 | 2.52E-218 | 1.221295185 | 0.641 | 0.202 | 5.10E-214 | 9 | Pls3 |

| | | | | | | | |
|---|---|---|---|---|---|---|---|
| Ly6e3 | 2.65E-217 | 1.189552874 | 0.984 | 0.609 | 5.36E-213 | 9 | Ly6e |
| Dock92 | 1.11E-214 | 1.185357569 | 0.618 | 0.194 | 2.24E-210 | 9 | Dock9 |
| Igfbp33 | 4.82E-213 | 1.519404469 | 0.778 | 0.289 | 9.76E-209 | 9 | Igfbp3 |
| Fabp43 | 5.60E-213 | 1.169995917 | 0.989 | 0.419 | 1.13E-208 | 9 | Fabp4 |
| Nfib2 | 9.12E-213 | 1.210899096 | 0.959 | 0.429 | 1.85E-208 | 9 | Nfib |
| Map4k33 | 2.03E-212 | 1.234014652 | 0.649 | 0.217 | 4.11E-208 | 9 | Map4k3 |
| Cd932 | 3.98E-212 | 1.144994465 | 0.723 | 0.242 | 8.06E-208 | 9 | Cd93 |
| Klf42 | 3.00E-211 | 1.352050052 | 0.715 | 0.256 | 6.07E-207 | 9 | Klf4 |
| Sncaip | 1.11E-210 | 0.905472707 | 0.374 | 0.072 | 2.24E-206 | 9 | Sncaip |
| Magi12 | 6.82E-210 | 1.204767388 | 0.672 | 0.218 | 1.38E-205 | 9 | Magi1 |
| Kctd12b3 | 2.84E-209 | 1.132025097 | 0.711 | 0.236 | 5.74E-205 | 9 | Kctd12b |
| Sparcl13 | 1.47E-207 | 1.175886147 | 0.88 | 0.348 | 2.97E-203 | 9 | Sparcl1 |
| Plk22 | 1.77E-207 | 1.147822688 | 0.554 | 0.155 | 3.57E-203 | 9 | Plk2 |
| Sorbs22 | 3.32E-207 | 1.25610186 | 0.735 | 0.264 | 6.72E-203 | 9 | Sorbs2 |
| Clec2d3 | 4.77E-206 | 1.265566762 | 0.919 | 0.516 | 9.66E-202 | 9 | Clec2d |
| Mgll3 | 1.15E-205 | 1.236740926 | 0.926 | 0.435 | 2.32E-201 | 9 | Mgll |
| Timp32 | 5.17E-205 | 1.137320553 | 0.926 | 0.369 | 1.05E-200 | 9 | Timp3 |
| Klf123 | 5.04E-204 | 1.355208238 | 0.6 | 0.196 | 1.02E-199 | 9 | Klf12 |
| Pecam13 | 1.66E-203 | 1.104007925 | 0.958 | 0.394 | 3.36E-199 | 9 | Pecam1 |
| Icam22 | 3.81E-203 | 1.103083381 | 0.665 | 0.217 | 7.72E-199 | 9 | Icam2 |
| Rassf91 | 6.27E-203 | 1.089970371 | 0.473 | 0.116 | 1.27E-198 | 9 | Rassf9 |
| Sipa1l22 | 2.10E-200 | 1.069697833 | 0.528 | 0.147 | 4.26E-196 | 9 | Sipa1l2 |
| Prex22 | 3.61E-200 | 1.192576278 | 0.723 | 0.26 | 7.30E-196 | 9 | Prex2 |
| Mpzl12 | 9.14E-200 | 1.001332751 | 0.501 | 0.132 | 1.85E-195 | 9 | Mpzl1 |
| Actn42 | 5.07E-198 | 1.217810761 | 0.803 | 0.363 | 1.03E-193 | 9 | Actn4 |
| Sptbn13 | 3.12E-197 | 1.175462244 | 0.969 | 0.481 | 6.31E-193 | 9 | Sptbn1 |
| Gulp12 | 5.20E-197 | 1.114706198 | 0.547 | 0.155 | 1.05E-192 | 9 | Gulp1 |
| Kdr3 | 2.24E-195 | 1.110215845 | 0.914 | 0.346 | 4.53E-191 | 9 | Kdr |
| Cd300lg3 | 2.63E-195 | 1.07829441 | 0.773 | 0.282 | 5.33E-191 | 9 | Cd300lg |
| Syne12 | 3.76E-195 | 1.206721094 | 0.829 | 0.345 | 7.61E-191 | 9 | Syne1 |
| Ushbp12 | 4.18E-195 | 1.09675902 | 0.636 | 0.206 | 8.46E-191 | 9 | Ushbp1 |
| Ece13 | 1.54E-194 | 1.134377056 | 0.763 | 0.293 | 3.12E-190 | 9 | Ece1 |
| Tspan133 | 4.07E-194 | 1.210590102 | 0.698 | 0.274 | 8.24E-190 | 9 | Tspan13 |
| Chrm31 | 9.57E-194 | 1.2008254 | 0.406 | 0.09 | 1.94E-189 | 9 | Chrm3 |
| Mmrn22 | 1.30E-192 | 1.078377597 | 0.667 | 0.219 | 2.64E-188 | 9 | Mmrn2 |
| Sema7a2 | 4.64E-191 | 0.969814326 | 0.515 | 0.139 | 9.40E-187 | 9 | Sema7a |
| Eng2 | 4.95E-190 | 1.063710133 | 0.862 | 0.339 | 1.00E-185 | 9 | Eng |
| Myct12 | 8.37E-190 | 1.004252187 | 0.646 | 0.21 | 1.69E-185 | 9 | Myct1 |
| Akap121 | 1.01E-188 | 1.063295255 | 0.527 | 0.15 | 2.05E-184 | 9 | Akap12 |
| Lims22 | 2.55E-188 | 1.04436124 | 0.598 | 0.19 | 5.15E-184 | 9 | Lims2 |
| Nxpe42 | 4.39E-188 | 1.005253374 | 0.504 | 0.137 | 8.89E-184 | 9 | Nxpe4 |
| Rgcc3 | 1.28E-186 | 1.11749445 | 0.9 | 0.389 | 2.60E-182 | 9 | Rgcc |
| Col4a12 | 1.73E-186 | 1.044547839 | 0.925 | 0.376 | 3.51E-182 | 9 | Col4a1 |

| | | | | | | | |
|---|---|---|---|---|---|---|---|
| 4931406P16Rik2 | 7.88E-186 | 1.086980151 | 0.79 | 0.33 | 1.60E-181 | 9 | 4931406P16Rik |
| App2 | 1.72E-185 | 1.062103862 | 0.914 | 0.405 | 3.48E-181 | 9 | App |
| Sncg2 | 3.67E-185 | 1.006262937 | 0.519 | 0.144 | 7.43E-181 | 9 | Sncg |
| Tanc12 | 4.25E-185 | 1.098894675 | 0.617 | 0.211 | 8.61E-181 | 9 | Tanc1 |
| Ehd22 | 2.77E-184 | 1.071147516 | 0.573 | 0.182 | 5.62E-180 | 9 | Ehd2 |
| Ablim12 | 4.28E-184 | 1.23740634 | 0.821 | 0.391 | 8.66E-180 | 9 | Ablim1 |
| Rapgef22 | 1.50E-182 | 1.181842879 | 0.756 | 0.327 | 3.03E-178 | 9 | Rapgef2 |
| Mkl22 | 5.48E-182 | 1.159056293 | 0.66 | 0.243 | 1.11E-177 | 9 | Mkl2 |
| Crip23 | 1.30E-181 | 1.080092514 | 0.872 | 0.403 | 2.63E-177 | 9 | Crip2 |
| Limch12 | 1.65E-181 | 1.046746173 | 0.478 | 0.129 | 3.34E-177 | 9 | Limch1 |
| Itga13 | 1.87E-180 | 1.155203664 | 0.867 | 0.402 | 3.78E-176 | 9 | Itga1 |
| S100a162 | 5.55E-180 | 0.992229309 | 0.681 | 0.245 | 1.12E-175 | 9 | S100a16 |
| Rasip12 | 1.93E-178 | 0.9556539 | 0.584 | 0.183 | 3.91E-174 | 9 | Rasip1 |
| Itm2b5 | 2.82E-178 | 1.020745763 | 0.996 | 0.717 | 5.71E-174 | 9 | Itm2b |
| 9530026P05Rik2 | 6.28E-178 | 1.197300883 | 0.656 | 0.229 | 1.27E-173 | 9 | 9530026P05Rik |
| Rbp73 | 2.61E-177 | 1.128040928 | 0.832 | 0.333 | 5.28E-173 | 9 | Rbp7 |
| Pdgfb3 | 4.75E-177 | 1.133778027 | 0.721 | 0.297 | 9.62E-173 | 9 | Pdgfb |
| Pcdh172 | 2.28E-176 | 1.046140873 | 0.666 | 0.226 | 4.63E-172 | 9 | Pcdh17 |
| Gng113 | 1.11E-175 | 0.947531372 | 0.867 | 0.351 | 2.25E-171 | 9 | Gng11 |
| Bmpr22 | 2.79E-175 | 1.136920069 | 0.783 | 0.337 | 5.65E-171 | 9 | Bmpr2 |
| Crim12 | 1.10E-174 | 1.181030249 | 0.811 | 0.36 | 2.23E-170 | 9 | Crim1 |
| Stox22 | 6.89E-174 | 1.138046495 | 0.66 | 0.235 | 1.40E-169 | 9 | Stox2 |
| Syne22 | 1.61E-173 | 1.070685186 | 0.692 | 0.26 | 3.26E-169 | 9 | Syne2 |
| Thrsp3 | 5.37E-173 | 0.957012324 | 0.755 | 0.279 | 1.09E-168 | 9 | Thrsp |
| Bcam3 | 3.14E-172 | 0.964038264 | 0.681 | 0.241 | 6.35E-168 | 9 | Bcam |
| Mef2c3 | 1.00E-171 | 1.154191683 | 0.885 | 0.41 | 2.03E-167 | 9 | Mef2c |
| Nck13 | 1.48E-171 | 1.086626467 | 0.74 | 0.328 | 2.99E-167 | 9 | Nck1 |
| Ndrg12 | 3.24E-171 | 1.044067709 | 0.628 | 0.221 | 6.57E-167 | 9 | Ndrg1 |
| Sptan13 | 1.12E-170 | 1.058597356 | 0.764 | 0.331 | 2.26E-166 | 9 | Sptan1 |
| C130074G19Rik2 | 3.10E-170 | 0.874929507 | 0.584 | 0.188 | 6.28E-166 | 9 | C130074G19Rik |
| Rasgrf22 | 2.18E-169 | 0.981148146 | 0.541 | 0.166 | 4.42E-165 | 9 | Rasgrf2 |
| Eogt | 3.81E-169 | 0.84287126 | 0.343 | 0.074 | 7.72E-165 | 9 | Eogt |
| Fbn1 | 3.34E-168 | 0.690817498 | 0.303 | 0.057 | 6.76E-164 | 9 | Fbn1 |
| Rnf152 | 8.01E-168 | 0.896973303 | 0.343 | 0.074 | 1.62E-163 | 9 | Rnf152 |
| Ctnnb12 | 1.07E-167 | 1.029230034 | 0.773 | 0.352 | 2.18E-163 | 9 | Ctnnb1 |
| Csgalnact12 | 2.02E-167 | 1.088822739 | 0.565 | 0.185 | 4.10E-163 | 9 | Csgalnact1 |
| Cavin23 | 6.06E-167 | 0.904988768 | 0.856 | 0.326 | 1.23E-162 | 9 | Cavin2 |
| Col4a22 | 3.48E-166 | 1.005100298 | 0.841 | 0.346 | 7.04E-162 | 9 | Col4a2 |
| Thbd3 | 2.30E-164 | 1.078194088 | 0.66 | 0.243 | 4.66E-160 | 9 | Thbd |
| Lama32 | 3.07E-164 | 1.087350436 | 0.608 | 0.21 | 6.21E-160 | 9 | Lama3 |
| Samd122 | 5.79E-163 | 1.119894583 | 0.621 | 0.218 | 1.17E-158 | 9 | Samd12 |
| Dusp32 | 1.48E-162 | 0.97919652 | 0.551 | 0.181 | 3.00E-158 | 9 | Dusp3 |
| Gm120022 | 2.30E-162 | 1.066257562 | 0.552 | 0.178 | 4.66E-158 | 9 | Gm12002 |

| | | | | | | | |
|---|---|---|---|---|---|---|---|
| Clic43 | 4.80E-162 | 1.091085852 | 0.83 | 0.412 | 9.72E-158 | 9 | Clic4 |
| Enpp3 | 5.00E-162 | 0.676147817 | 0.254 | 0.044 | 1.01E-157 | 9 | Enpp3 |
| Prdx42 | 6.41E-162 | 1.039855474 | 0.588 | 0.211 | 1.30E-157 | 9 | Prdx4 |
| Ifitm33 | 8.36E-162 | 0.999356819 | 0.985 | 0.525 | 1.69E-157 | 9 | Ifitm3 |
| Tinagl13 | 1.36E-161 | 0.92172577 | 0.747 | 0.284 | 2.75E-157 | 9 | Tinagl1 |
| Foxp12 | 8.67E-161 | 1.118744499 | 0.778 | 0.378 | 1.75E-156 | 9 | Foxp1 |
| Hspa12b1 | 5.67E-160 | 0.846867339 | 0.383 | 0.094 | 1.15E-155 | 9 | Hspa12b |
| She2 | 1.04E-159 | 0.896345592 | 0.481 | 0.14 | 2.11E-155 | 9 | She |
| Plpp32 | 2.74E-159 | 0.999496184 | 0.919 | 0.389 | 5.55E-155 | 9 | Plpp3 |
| Ifitm23 | 9.45E-159 | 0.951889292 | 0.938 | 0.451 | 1.91E-154 | 9 | Ifitm2 |
| Mgst12 | 1.74E-158 | 0.933492736 | 0.512 | 0.157 | 3.52E-154 | 9 | Mgst1 |
| Fzd6 | 1.74E-158 | 0.71979063 | 0.279 | 0.053 | 3.53E-154 | 9 | Fzd6 |
| Nrp12 | 3.49E-158 | 0.957170069 | 0.917 | 0.42 | 7.07E-154 | 9 | Nrp1 |
| Crispld12 | 6.61E-158 | 0.95832592 | 0.466 | 0.134 | 1.34E-153 | 9 | Crispld1 |
| Tns13 | 3.57E-157 | 0.944653643 | 0.63 | 0.234 | 7.22E-153 | 9 | Tns1 |
| Dach11 | 3.88E-157 | 1.140587297 | 0.382 | 0.096 | 7.86E-153 | 9 | Dach1 |
| Palmd2 | 2.00E-156 | 0.926930327 | 0.624 | 0.222 | 4.06E-152 | 9 | Palmd |
| Aplp23 | 4.39E-156 | 0.988577288 | 0.841 | 0.404 | 8.88E-152 | 9 | Aplp2 |
| 4930578C19Rik1 | 5.28E-156 | 0.959224378 | 0.388 | 0.098 | 1.07E-151 | 9 | 4930578C19Rik |
| Emcn3 | 2.99E-155 | 0.877382126 | 0.938 | 0.364 | 6.05E-151 | 9 | Emcn |
| Adamts11 | 4.56E-155 | 0.984963339 | 0.367 | 0.089 | 9.22E-151 | 9 | Adamts1 |
| Itpkb2 | 6.76E-155 | 1.055794061 | 0.744 | 0.345 | 1.37E-150 | 9 | Itpkb |
| Ablim3 | 2.05E-153 | 0.783565903 | 0.314 | 0.067 | 4.15E-149 | 9 | Ablim3 |
| Smco43 | 8.87E-153 | 0.921537107 | 0.83 | 0.377 | 1.80E-148 | 9 | Smco4 |
| Gpihbp13 | 1.24E-152 | 0.930774095 | 0.744 | 0.283 | 2.51E-148 | 9 | Gpihbp1 |
| Tmem44 | 3.80E-151 | 0.648669059 | 0.276 | 0.054 | 7.70E-147 | 9 | Tmem44 |
| Klf72 | 4.72E-151 | 1.025593604 | 0.695 | 0.286 | 9.56E-147 | 9 | Klf7 |
| Lima11 | 6.92E-151 | 0.931053839 | 0.482 | 0.149 | 1.40E-146 | 9 | Lima1 |
| Lmo7 | 3.93E-150 | 0.751812211 | 0.309 | 0.066 | 7.95E-146 | 9 | Lmo7 |
| Cavin33 | 3.24E-149 | 0.909366781 | 0.729 | 0.295 | 6.57E-145 | 9 | Cavin3 |
| Rsad23 | 4.13E-149 | 1.089659956 | 0.652 | 0.245 | 8.36E-145 | 9 | Rsad2 |
| Syt15 | 1.33E-148 | 0.735272656 | 0.284 | 0.057 | 2.70E-144 | 9 | Syt15 |
| Fat41 | 1.58E-148 | 0.790778767 | 0.382 | 0.098 | 3.20E-144 | 9 | Fat4 |
| Tmem882 | 4.22E-148 | 0.933867342 | 0.575 | 0.2 | 8.54E-144 | 9 | Tmem88 |
| Mcam1 | 1.45E-147 | 0.803186503 | 0.392 | 0.102 | 2.93E-143 | 9 | Mcam |
| Tspan7 | 2.21E-147 | 0.871381442 | 0.313 | 0.07 | 4.47E-143 | 9 | Tspan7 |
| Fam198b2 | 2.25E-147 | 1.015665566 | 0.525 | 0.174 | 4.55E-143 | 9 | Fam198b |
| Ralgapa12 | 3.33E-147 | 1.087873143 | 0.662 | 0.293 | 6.75E-143 | 9 | Ralgapa1 |
| Ptk22 | 2.47E-146 | 0.931517799 | 0.655 | 0.255 | 5.00E-142 | 9 | Ptk2 |
| Cd814 | 4.55E-145 | 0.937157538 | 0.961 | 0.513 | 9.21E-141 | 9 | Cd81 |
| Wwtr12 | 7.09E-145 | 0.876653923 | 0.619 | 0.229 | 1.44E-140 | 9 | Wwtr1 |
| Efna12 | 2.72E-144 | 0.885285296 | 0.45 | 0.135 | 5.50E-140 | 9 | Efna1 |
| Gas62 | 3.21E-144 | 0.878979574 | 0.708 | 0.286 | 6.50E-140 | 9 | Gas6 |

| | | | | | | | |
|---|---|---|---|---|---|---|---|
| Jcad1 | 3.88E-144 | 0.889705956 | 0.395 | 0.107 | 7.86E-140 | 9 | Jcad |
| Kank32 | 4.10E-144 | 0.910908459 | 0.599 | 0.217 | 8.30E-140 | 9 | Kank3 |
| Igfbp73 | 4.93E-143 | 0.894783554 | 0.983 | 0.452 | 9.98E-139 | 9 | Igfbp7 |
| Slfn53 | 4.24E-142 | 0.906111151 | 0.784 | 0.344 | 8.59E-138 | 9 | Slfn5 |
| Diaph24 | 5.07E-142 | 1.108137945 | 0.742 | 0.366 | 1.03E-137 | 9 | Diaph2 |
| Arid5b3 | 7.38E-141 | 1.125043159 | 0.625 | 0.268 | 1.49E-136 | 9 | Arid5b |
| Bst25 | 1.72E-140 | 1.002010217 | 0.887 | 0.492 | 3.48E-136 | 9 | Bst2 |
| Tpm43 | 6.94E-140 | 0.941409092 | 0.764 | 0.369 | 1.40E-135 | 9 | Tpm4 |
| Adam152 | 8.31E-140 | 0.889072425 | 0.531 | 0.184 | 1.68E-135 | 9 | Adam15 |
| Cst32 | 8.66E-140 | 0.455993128 | 0.968 | 0.592 | 1.75E-135 | 9 | Cst3 |
| Cald12 | 1.49E-139 | 0.846128149 | 0.747 | 0.31 | 3.02E-135 | 9 | Cald1 |
| Oaz22 | 2.11E-139 | 0.951809792 | 0.597 | 0.226 | 4.28E-135 | 9 | Oaz2 |
| Abcb1a1 | 1.98E-138 | 0.846094417 | 0.376 | 0.102 | 4.02E-134 | 9 | Abcb1a |
| Fryl1 | 2.01E-138 | 1.047952679 | 0.62 | 0.258 | 4.08E-134 | 9 | Fryl |
| Malat14 | 3.39E-138 | 0.951411977 | 0.984 | 0.718 | 6.87E-134 | 9 | Malat1 |
| Efr3b1 | 5.54E-138 | 0.775765559 | 0.332 | 0.08 | 1.12E-133 | 9 | Efr3b |
| Notch41 | 6.80E-138 | 0.809559928 | 0.39 | 0.108 | 1.38E-133 | 9 | Notch4 |
| Arhgef28 | 5.33E-137 | 0.702339252 | 0.317 | 0.074 | 1.08E-132 | 9 | Arhgef28 |
| Arhgef152 | 5.51E-137 | 0.827422224 | 0.492 | 0.159 | 1.12E-132 | 9 | Arhgef15 |
| Fgd52 | 6.26E-137 | 0.876825015 | 0.594 | 0.218 | 1.27E-132 | 9 | Fgd5 |
| Stc1 | 6.49E-137 | 0.752855852 | 0.274 | 0.057 | 1.31E-132 | 9 | Stc1 |
| Clec14a2 | 7.88E-137 | 0.854379233 | 0.73 | 0.293 | 1.60E-132 | 9 | Clec14a |
| Tie13 | 9.48E-137 | 0.828667867 | 0.645 | 0.242 | 1.92E-132 | 9 | Tie1 |
| Enah2 | 7.20E-136 | 0.834253441 | 0.455 | 0.141 | 1.46E-131 | 9 | Enah |
| Grb102 | 9.11E-136 | 0.888492382 | 0.534 | 0.185 | 1.84E-131 | 9 | Grb10 |
| Lrrc8c2 | 1.03E-135 | 0.961494675 | 0.686 | 0.3 | 2.09E-131 | 9 | Lrrc8c |
| Pltp3 | 1.05E-135 | 0.848381715 | 0.724 | 0.288 | 2.12E-131 | 9 | Pltp |
| Tgfbr23 | 1.79E-135 | 1.079794845 | 0.66 | 0.301 | 3.62E-131 | 9 | Tgfbr2 |
| Ets13 | 2.27E-135 | 0.952056749 | 0.787 | 0.393 | 4.60E-131 | 9 | Ets1 |
| Plxna22 | 4.32E-135 | 0.905288016 | 0.572 | 0.209 | 8.75E-131 | 9 | Plxna2 |
| Klf132 | 4.98E-135 | 0.920487271 | 0.723 | 0.339 | 1.01E-130 | 9 | Klf13 |
| Mapk33 | 1.27E-133 | 0.866704701 | 0.617 | 0.256 | 2.58E-129 | 9 | Mapk3 |
| Afdn2 | 1.52E-133 | 0.935002052 | 0.678 | 0.305 | 3.08E-129 | 9 | Afdn |
| Rdx3 | 2.15E-132 | 0.875294068 | 0.933 | 0.54 | 4.35E-128 | 9 | Rdx |
| Fut82 | 2.59E-132 | 1.044565877 | 0.641 | 0.279 | 5.25E-128 | 9 | Fut8 |
| B4galt42 | 3.86E-132 | 0.812769732 | 0.472 | 0.153 | 7.81E-128 | 9 | B4galt4 |
| Ahnak3 | 4.23E-132 | 0.953356207 | 0.867 | 0.46 | 8.55E-128 | 9 | Ahnak |
| Cavin12 | 7.77E-132 | 0.909605003 | 0.56 | 0.203 | 1.57E-127 | 9 | Cavin1 |
| Gnai23 | 7.80E-132 | 0.882154976 | 0.921 | 0.565 | 1.58E-127 | 9 | Gnai2 |
| Pcp4l14 | 1.41E-131 | 0.776620197 | 0.718 | 0.302 | 2.86E-127 | 9 | Pcp4l1 |
| Hecw22 | 1.35E-130 | 0.893980886 | 0.617 | 0.238 | 2.74E-126 | 9 | Hecw2 |
| Rapgef42 | 1.74E-130 | 0.939101923 | 0.629 | 0.247 | 3.53E-126 | 9 | Rapgef4 |
| Cav22 | 2.57E-130 | 0.815695949 | 0.55 | 0.199 | 5.20E-126 | 9 | Cav2 |

| | | | | | | | |
|---|---|---|---|---|---|---|---|
| Sgk13 | 3.56E-130 | 1.0043681 | 0.633 | 0.259 | 7.20E-126 | 9 | Sgk1 |
| Efnb12 | 1.50E-129 | 0.914090806 | 0.448 | 0.147 | 3.04E-125 | 9 | Efnb1 |
| Nbea2 | 1.51E-129 | 0.922853568 | 0.519 | 0.187 | 3.05E-125 | 9 | Nbea |
| Luzp12 | 1.65E-129 | 0.901961436 | 0.721 | 0.347 | 3.35E-125 | 9 | Luzp1 |
| Calm13 | 2.12E-129 | 0.77692837 | 0.99 | 0.775 | 4.30E-125 | 9 | Calm1 |
| Apbb23 | 3.51E-129 | 0.879354265 | 0.774 | 0.355 | 7.11E-125 | 9 | Apbb2 |
| Eml11 | 5.23E-129 | 0.758652318 | 0.37 | 0.102 | 1.06E-124 | 9 | Eml1 |
| St6galnac32 | 1.02E-128 | 0.968928306 | 0.555 | 0.206 | 2.06E-124 | 9 | St6galnac3 |
| Fzd42 | 1.66E-128 | 0.890958307 | 0.578 | 0.219 | 3.36E-124 | 9 | Fzd4 |
| Ephx11 | 5.70E-128 | 0.815431322 | 0.39 | 0.114 | 1.15E-123 | 9 | Ephx1 |
| Msrb31 | 2.58E-127 | 0.801473337 | 0.411 | 0.125 | 5.23E-123 | 9 | Msrb3 |
| Prickle22 | 5.22E-127 | 0.832562635 | 0.424 | 0.131 | 1.06E-122 | 9 | Prickle2 |
| Mertk1 | 1.17E-126 | 0.797777165 | 0.408 | 0.124 | 2.36E-122 | 9 | Mertk |
| Rnf2202 | 2.61E-126 | 1.070321278 | 0.577 | 0.232 | 5.29E-122 | 9 | Rnf220 |
| Elk32 | 3.96E-126 | 0.877157804 | 0.729 | 0.346 | 8.02E-122 | 9 | Elk3 |
| Rgs32 | 8.34E-126 | 0.918471967 | 0.486 | 0.179 | 1.69E-121 | 9 | Rgs3 |
| Il6st2 | 1.07E-125 | 0.837741303 | 0.578 | 0.222 | 2.16E-121 | 9 | Il6st |
| Arhgap292 | 1.15E-125 | 0.798091722 | 0.795 | 0.353 | 2.32E-121 | 9 | Arhgap29 |
| Snrk3 | 1.99E-125 | 0.787947898 | 0.781 | 0.357 | 4.03E-121 | 9 | Snrk |
| Tspan92 | 1.09E-123 | 0.894781416 | 0.612 | 0.255 | 2.21E-119 | 9 | Tspan9 |
| Gja12 | 2.88E-123 | 0.885122303 | 0.397 | 0.12 | 5.83E-119 | 9 | Gja1 |
| Tek2 | 3.18E-122 | 0.812310899 | 0.634 | 0.249 | 6.43E-118 | 9 | Tek |
| Abcc41 | 3.45E-122 | 0.805070145 | 0.408 | 0.128 | 6.98E-118 | 9 | Abcc4 |
| Abcg23 | 5.18E-122 | 0.910130309 | 0.619 | 0.261 | 1.05E-117 | 9 | Abcg2 |
| Dab2ip3 | 7.32E-122 | 0.817962474 | 0.607 | 0.251 | 1.48E-117 | 9 | Dab2ip |
| Slc43a33 | 1.13E-121 | 0.789388473 | 0.642 | 0.259 | 2.29E-117 | 9 | Slc43a3 |
| Socs22 | 1.54E-121 | 0.796543131 | 0.492 | 0.174 | 3.12E-117 | 9 | Socs2 |
| Map42 | 2.30E-121 | 0.996510517 | 0.629 | 0.295 | 4.67E-117 | 9 | Map4 |
| Cdc42ep11 | 3.19E-121 | 0.687851682 | 0.402 | 0.121 | 6.45E-117 | 9 | Cdc42ep1 |
| Dock12 | 8.82E-121 | 0.874281985 | 0.633 | 0.27 | 1.79E-116 | 9 | Dock1 |
| Cd1512 | 9.77E-121 | 0.825963282 | 0.543 | 0.208 | 1.98E-116 | 9 | Cd151 |
| Bsg3 | 5.30E-120 | 0.926178489 | 0.795 | 0.439 | 1.07E-115 | 9 | Bsg |
| Acvrl13 | 1.23E-119 | 0.806488662 | 0.626 | 0.265 | 2.48E-115 | 9 | Acvrl1 |
| Hlx1 | 8.82E-118 | 0.724216975 | 0.342 | 0.095 | 1.78E-113 | 9 | Hlx |
| Gcnt22 | 1.31E-117 | 0.736069214 | 0.425 | 0.135 | 2.64E-113 | 9 | Gcnt2 |
| Mfge83 | 3.54E-117 | 0.816529714 | 0.594 | 0.244 | 7.16E-113 | 9 | Mfge8 |
| Plvap3 | 4.99E-117 | 0.720599578 | 0.883 | 0.369 | 1.01E-112 | 9 | Plvap |
| Tspan52 | 8.64E-117 | 0.921857341 | 0.497 | 0.19 | 1.75E-112 | 9 | Tspan5 |
| Wwc22 | 1.62E-116 | 0.827560315 | 0.502 | 0.189 | 3.28E-112 | 9 | Wwc2 |
| Tmed103 | 1.65E-116 | 0.851299938 | 0.769 | 0.427 | 3.34E-112 | 9 | Tmed10 |
| Robo42 | 3.11E-116 | 0.823542294 | 0.48 | 0.168 | 6.30E-112 | 9 | Robo4 |
| Zfp3662 | 1.27E-115 | 0.828404379 | 0.492 | 0.174 | 2.57E-111 | 9 | Zfp366 |
| Anxa32 | 7.05E-115 | 0.750766567 | 0.716 | 0.328 | 1.43E-110 | 9 | Anxa3 |

| | | | | | | | |
|---|---|---|---|---|---|---|---|
| Fermt23 | 1.06E-114 | 0.72626735 | 0.746 | 0.339 | 2.15E-110 | 9 | Fermt2 |
| Anp32a2 | 3.90E-114 | 0.82723752 | 0.798 | 0.456 | 7.89E-110 | 9 | Anp32a |
| Arhgef123 | 5.00E-114 | 0.855626898 | 0.716 | 0.338 | 1.01E-109 | 9 | Arhgef12 |
| Adgrg12 | 7.30E-114 | 0.709007173 | 0.414 | 0.132 | 1.48E-109 | 9 | Adgrg1 |
| Wasf22 | 9.99E-114 | 0.880893127 | 0.751 | 0.416 | 2.02E-109 | 9 | Wasf2 |
| Ctnna12 | 1.18E-112 | 0.79818498 | 0.697 | 0.342 | 2.39E-108 | 9 | Ctnna1 |
| Mfhas11 | 2.05E-112 | 0.780074646 | 0.419 | 0.142 | 4.15E-108 | 9 | Mfhas1 |
| Klf32 | 3.87E-112 | 0.783884425 | 0.556 | 0.227 | 7.84E-108 | 9 | Klf3 |
| Cdc42bpa2 | 5.97E-112 | 0.820900497 | 0.483 | 0.177 | 1.21E-107 | 9 | Cdc42bpa |
| Rbpms2 | 7.06E-112 | 0.730489812 | 0.715 | 0.339 | 1.43E-107 | 9 | Rbpms |
| Tubb2a2 | 8.13E-112 | 0.775275635 | 0.443 | 0.157 | 1.65E-107 | 9 | Tubb2a |
| Gata21 | 1.48E-111 | 0.687331408 | 0.344 | 0.098 | 2.99E-107 | 9 | Gata2 |
| Rbfox22 | 9.63E-110 | 0.795982944 | 0.488 | 0.182 | 1.95E-105 | 9 | Rbfox2 |
| Mmp151 | 6.67E-109 | 0.659456986 | 0.333 | 0.095 | 1.35E-104 | 9 | Mmp15 |
| Tjp12 | 6.68E-109 | 0.847878349 | 0.587 | 0.255 | 1.35E-104 | 9 | Tjp1 |
| Plscr12 | 6.75E-109 | 0.891645125 | 0.496 | 0.197 | 1.37E-104 | 9 | Plscr1 |
| Ccm2l1 | 7.32E-109 | 0.667309983 | 0.316 | 0.086 | 1.48E-104 | 9 | Ccm2l |
| Hip13 | 9.79E-109 | 0.843226917 | 0.676 | 0.325 | 1.98E-104 | 9 | Hip1 |
| Osmr2 | 2.65E-108 | 0.761299131 | 0.403 | 0.132 | 5.36E-104 | 9 | Osmr |
| Clstn12 | 3.47E-108 | 0.760429443 | 0.556 | 0.228 | 7.02E-104 | 9 | Clstn1 |
| Msn2 | 4.66E-108 | 0.832614526 | 0.776 | 0.434 | 9.44E-104 | 9 | Msn |
| Adgrl43 | 7.16E-108 | 0.475697534 | 0.856 | 0.337 | 1.45E-103 | 9 | Adgrl4 |
| Tril | 1.14E-107 | 0.651855379 | 0.302 | 0.081 | 2.31E-103 | 9 | Tril |
| Cdkn1c1 | 2.11E-107 | 0.763429334 | 0.377 | 0.119 | 4.28E-103 | 9 | Cdkn1c |
| Phactr1 | 8.57E-107 | 0.671270295 | 0.256 | 0.062 | 1.74E-102 | 9 | Phactr1 |
| Pcdh11 | 2.45E-106 | 0.613340467 | 0.311 | 0.085 | 4.96E-102 | 9 | Pcdh1 |
| Crybg31 | 5.58E-106 | 0.819441653 | 0.393 | 0.133 | 1.13E-101 | 9 | Crybg3 |
| Fas1 | 4.74E-105 | 0.690874751 | 0.36 | 0.111 | 9.59E-101 | 9 | Fas |
| Aig12 | 5.32E-105 | 0.735217687 | 0.514 | 0.205 | 1.08E-100 | 9 | Aig1 |
| Flnb1 | 5.55E-105 | 0.77326684 | 0.432 | 0.155 | 1.12E-100 | 9 | Flnb |
| Tmtc12 | 5.71E-105 | 0.844397896 | 0.401 | 0.135 | 1.16E-100 | 9 | Tmtc1 |
| Hspg22 | 1.04E-104 | 0.736097144 | 0.689 | 0.301 | 2.10E-100 | 9 | Hspg2 |
| Epn22 | 1.85E-104 | 0.721095029 | 0.424 | 0.147 | 3.75E-100 | 9 | Epn2 |
| Stom1 | 9.46E-104 | 0.661994696 | 0.34 | 0.101 | 1.91E-99 | 9 | Stom |
| Cnn32 | 7.10E-103 | 0.729358188 | 0.487 | 0.185 | 1.44E-98 | 9 | Cnn3 |
| Dysf2 | 1.67E-102 | 0.675277303 | 0.58 | 0.23 | 3.39E-98 | 9 | Dysf |
| Agap11 | 2.22E-102 | 0.717529416 | 0.401 | 0.138 | 4.49E-98 | 9 | Agap1 |
| Smad73 | 1.34E-101 | 0.926600891 | 0.635 | 0.318 | 2.71E-97 | 9 | Smad7 |
| Plekhg12 | 1.94E-101 | 0.806175416 | 0.566 | 0.236 | 3.92E-97 | 9 | Plekhg1 |
| Nfe2l12 | 6.64E-101 | 0.819121293 | 0.588 | 0.272 | 1.34E-96 | 9 | Nfe2l1 |
| Cpd3 | 2.10E-100 | 0.777883369 | 0.639 | 0.303 | 4.26E-96 | 9 | Cpd |
| Bvht2 | 4.53E-100 | 0.705605077 | 0.475 | 0.178 | 9.17E-96 | 9 | Bvht |
| Tcim2 | 4.86E-100 | 0.732331827 | 0.674 | 0.293 | 9.85E-96 | 9 | Tcim |

| | | | | | | | |
|---|---|---|---|---|---|---|---|
| Unc45b2 | 5.39E-100 | 0.654584399 | 0.397 | 0.132 | 1.09E-95 | 9 | Unc45b |
| Lifr2 | 1.06E-99 | 0.835545086 | 0.473 | 0.185 | 2.15E-95 | 9 | Lifr |
| H2-D14 | 1.53E-99 | 0.710406159 | 0.993 | 0.66 | 3.11E-95 | 9 | H2-D1 |
| Ppfibp12 | 1.93E-99 | 0.764819129 | 0.456 | 0.176 | 3.90E-95 | 9 | Ppfibp1 |
| F11r4 | 2.31E-99 | 0.70127966 | 0.587 | 0.257 | 4.69E-95 | 9 | F11r |
| 9930111J21Rik23 | 3.17E-99 | 0.853962377 | 0.556 | 0.249 | 6.41E-95 | 9 | 9930111J21Rik2 |
| Hmbox12 | 4.08E-99 | 0.758631861 | 0.589 | 0.265 | 8.27E-95 | 9 | Hmbox1 |
| Galnt151 | 1.34E-98 | 0.589764461 | 0.3 | 0.084 | 2.71E-94 | 9 | Galnt15 |
| Sox131 | 1.39E-98 | 0.660029817 | 0.3 | 0.085 | 2.81E-94 | 9 | Sox13 |
| Ifi2033 | 1.94E-98 | 0.839243052 | 0.679 | 0.359 | 3.94E-94 | 9 | Ifi203 |
| Trim472 | 3.54E-98 | 0.739487204 | 0.392 | 0.137 | 7.18E-94 | 9 | Trim47 |
| Dip2c2 | 4.25E-98 | 0.845480334 | 0.519 | 0.215 | 8.60E-94 | 9 | Dip2c |
| Grrp12 | 2.54E-97 | 0.68445782 | 0.53 | 0.211 | 5.14E-93 | 9 | Grrp1 |
| Pcnx2 | 2.78E-97 | 0.823827159 | 0.439 | 0.172 | 5.63E-93 | 9 | Pcnx |
| Slc50a13 | 3.26E-97 | 0.827301093 | 0.597 | 0.291 | 6.60E-93 | 9 | Slc50a1 |
| Nedd42 | 3.47E-97 | 0.731456453 | 0.762 | 0.41 | 7.02E-93 | 9 | Nedd4 |
| Nxn2 | 8.00E-97 | 0.804768308 | 0.581 | 0.26 | 1.62E-92 | 9 | Nxn |
| Amotl21 | 9.43E-97 | 0.615182039 | 0.29 | 0.081 | 1.91E-92 | 9 | Amotl2 |
| Rbms12 | 1.13E-96 | 0.716265261 | 0.845 | 0.476 | 2.28E-92 | 9 | Rbms1 |
| Cmtm82 | 1.96E-96 | 0.673029038 | 0.613 | 0.264 | 3.97E-92 | 9 | Cmtm8 |
| Cep85l2 | 2.06E-96 | 0.803104886 | 0.423 | 0.157 | 4.17E-92 | 9 | Cep85l |
| Btbd72 | 2.72E-96 | 0.786114205 | 0.572 | 0.273 | 5.50E-92 | 9 | Btbd7 |
| Ppp1r13b2 | 1.86E-95 | 0.755947291 | 0.497 | 0.208 | 3.76E-91 | 9 | Ppp1r13b |
| Laptm4a2 | 2.87E-95 | 0.725163308 | 0.822 | 0.475 | 5.81E-91 | 9 | Laptm4a |
| Ace1 | 3.92E-95 | 0.740657227 | 0.35 | 0.113 | 7.94E-91 | 9 | Ace |
| Myo1d1 | 1.51E-94 | 0.643258534 | 0.327 | 0.101 | 3.06E-90 | 9 | Myo1d |
| Zhx32 | 2.02E-94 | 0.741484376 | 0.487 | 0.194 | 4.08E-90 | 9 | Zhx3 |
| Asap12 | 2.24E-94 | 0.741361658 | 0.684 | 0.347 | 4.54E-90 | 9 | Asap1 |
| Ccdc85a2 | 2.48E-94 | 0.78426735 | 0.462 | 0.174 | 5.01E-90 | 9 | Ccdc85a |
| Galnt182 | 2.96E-94 | 0.70284008 | 0.38 | 0.128 | 6.00E-90 | 9 | Galnt18 |
| Igf1r1 | 3.97E-94 | 0.744198887 | 0.418 | 0.154 | 8.04E-90 | 9 | Igf1r |
| Dync1h13 | 4.72E-94 | 0.794085502 | 0.575 | 0.279 | 9.55E-90 | 9 | Dync1h1 |
| Macf13 | 4.86E-94 | 0.774180743 | 0.834 | 0.494 | 9.85E-90 | 9 | Macf1 |
| Klf62 | 6.52E-94 | 0.884688886 | 0.55 | 0.26 | 1.32E-89 | 9 | Klf6 |
| Tmem255b | 6.76E-94 | 0.552837401 | 0.26 | 0.068 | 1.37E-89 | 9 | Tmem255b |
| Xdh2 | 2.01E-93 | 0.726785733 | 0.466 | 0.18 | 4.07E-89 | 9 | Xdh |
| Ankrd33b1 | 2.13E-93 | 0.651771434 | 0.28 | 0.078 | 4.32E-89 | 9 | Ankrd33b |
| Tnfsf103 | 2.53E-93 | 0.708606383 | 0.727 | 0.364 | 5.12E-89 | 9 | Tnfsf10 |
| Tmod33 | 6.68E-93 | 0.780734932 | 0.666 | 0.353 | 1.35E-88 | 9 | Tmod3 |
| Abca11 | 8.38E-93 | 0.796922744 | 0.359 | 0.123 | 1.70E-88 | 9 | Abca1 |
| Col13a12 | 1.01E-92 | 0.765561755 | 0.559 | 0.233 | 2.05E-88 | 9 | Col13a1 |
| Gna113 | 1.28E-92 | 0.673592101 | 0.541 | 0.237 | 2.59E-88 | 9 | Gna11 |
| Slc3a23 | 1.74E-92 | 0.9010913 | 0.577 | 0.283 | 3.53E-88 | 9 | Slc3a2 |

| | | | | | | | |
|---|---|---|---|---|---|---|---|
| Rb13 | 1.83E-92 | 0.828019164 | 0.483 | 0.207 | 3.71E-88 | 9 | Rb1 |
| Cx3cl11 | 2.22E-92 | 0.724499313 | 0.317 | 0.098 | 4.50E-88 | 9 | Cx3cl1 |
| Prickle12 | 1.24E-91 | 0.783716855 | 0.391 | 0.14 | 2.51E-87 | 9 | Prickle1 |
| Parvb1 | 1.92E-91 | 0.640721159 | 0.354 | 0.117 | 3.89E-87 | 9 | Parvb |
| Stmn1 | 1.92E-91 | 0.507160975 | 0.371 | 0.124 | 3.89E-87 | 9 | Stmn1 |
| Ccdc28b3 | 7.83E-91 | 0.599723081 | 0.576 | 0.253 | 1.58E-86 | 9 | Ccdc28b |
| Nova21 | 1.05E-90 | 0.621602932 | 0.322 | 0.1 | 2.12E-86 | 9 | Nova2 |
| Sema6a2 | 1.07E-90 | 0.689409238 | 0.504 | 0.201 | 2.16E-86 | 9 | Sema6a |
| Lama42 | 1.80E-90 | 0.730575918 | 0.44 | 0.168 | 3.64E-86 | 9 | Lama4 |
| Ppp1r23 | 1.26E-89 | 0.726641649 | 0.715 | 0.384 | 2.56E-85 | 9 | Ppp1r2 |
| S100a132 | 1.29E-89 | 0.669586547 | 0.838 | 0.528 | 2.61E-85 | 9 | S100a13 |
| Tmem2042 | 3.89E-89 | 0.680187967 | 0.496 | 0.201 | 7.88E-85 | 9 | Tmem204 |
| Dlc12 | 4.37E-89 | 0.553014896 | 0.762 | 0.351 | 8.85E-85 | 9 | Dlc1 |
| Myl12b2 | 5.30E-89 | 0.689700948 | 0.832 | 0.54 | 1.07E-84 | 9 | Myl12b |
| Hes13 | 6.90E-89 | 0.804352128 | 0.63 | 0.305 | 1.40E-84 | 9 | Hes1 |
| Cd2ap1 | 7.87E-89 | 0.680668463 | 0.635 | 0.318 | 1.59E-84 | 9 | Cd2ap |
| Kalrn1 | 1.25E-88 | 0.626582657 | 0.319 | 0.1 | 2.52E-84 | 9 | Kalrn |
| Plcg11 | 1.82E-88 | 0.610743935 | 0.348 | 0.116 | 3.69E-84 | 9 | Plcg1 |
| Mndal3 | 2.11E-88 | 0.785291056 | 0.72 | 0.397 | 4.27E-84 | 9 | Mndal |
| Ifit22 | 1.05E-87 | 0.760968592 | 0.339 | 0.112 | 2.13E-83 | 9 | Ifit2 |
| Shank32 | 1.86E-87 | 0.741102643 | 0.466 | 0.183 | 3.76E-83 | 9 | Shank3 |
| Ktn13 | 1.89E-87 | 0.795523889 | 0.593 | 0.3 | 3.82E-83 | 9 | Ktn1 |
| Qk3 | 2.23E-87 | 0.678515033 | 0.739 | 0.401 | 4.52E-83 | 9 | Qk |
| Rassf32 | 8.94E-87 | 0.731108841 | 0.514 | 0.228 | 1.81E-82 | 9 | Rassf3 |
| Rhob2 | 3.21E-86 | 0.835837693 | 0.554 | 0.26 | 6.49E-82 | 9 | Rhob |
| Rhoj2 | 4.59E-86 | 0.669030524 | 0.546 | 0.233 | 9.28E-82 | 9 | Rhoj |
| Bnip22 | 1.36E-85 | 0.69842022 | 0.568 | 0.276 | 2.75E-81 | 9 | Bnip2 |
| Sorbs12 | 1.85E-85 | 0.681197531 | 0.434 | 0.17 | 3.74E-81 | 9 | Sorbs1 |
| Eln1 | 2.19E-85 | 0.65286827 | 0.371 | 0.13 | 4.43E-81 | 9 | Eln |
| Dennd5b2 | 2.86E-85 | 0.669429983 | 0.387 | 0.14 | 5.79E-81 | 9 | Dennd5b |
| Fam129b | 6.22E-85 | 0.57262798 | 0.274 | 0.08 | 1.26E-80 | 9 | Fam129b |
| Kctd101 | 9.07E-85 | 0.604237526 | 0.329 | 0.111 | 1.84E-80 | 9 | Kctd10 |
| Cd472 | 1.04E-84 | 0.710670229 | 0.826 | 0.501 | 2.12E-80 | 9 | Cd47 |
| Srgn4 | 2.51E-84 | 0.678690063 | 0.912 | 0.534 | 5.08E-80 | 9 | Srgn |
| Nav12 | 3.27E-84 | 0.704134769 | 0.504 | 0.215 | 6.62E-80 | 9 | Nav1 |
| Oxr12 | 5.12E-84 | 0.783316893 | 0.634 | 0.328 | 1.04E-79 | 9 | Oxr1 |
| Nfia2 | 2.96E-83 | 0.771839473 | 0.583 | 0.28 | 6.00E-79 | 9 | Nfia |
| Ctsl3 | 7.88E-83 | 0.525849614 | 0.691 | 0.336 | 1.60E-78 | 9 | Ctsl |
| Maml24 | 9.42E-83 | 0.882014153 | 0.634 | 0.335 | 1.91E-78 | 9 | Maml2 |
| Rai142 | 1.16E-82 | 0.741020389 | 0.513 | 0.224 | 2.35E-78 | 9 | Rai14 |
| Atp13a31 | 2.20E-82 | 0.828754257 | 0.296 | 0.098 | 4.45E-78 | 9 | Atp13a3 |
| Sos12 | 2.32E-82 | 0.716937015 | 0.427 | 0.174 | 4.69E-78 | 9 | Sos1 |
| Rapgef52 | 3.52E-82 | 0.694071988 | 0.499 | 0.215 | 7.12E-78 | 9 | Rapgef5 |

| | | | | | | | |
|---|---|---|---|---|---|---|---|
| Gbp73 | 4.01E-82 | 0.775195483 | 0.549 | 0.262 | 8.12E-78 | 9 | Gbp7 |
| Ccrl21 | 5.05E-82 | 0.627217247 | 0.314 | 0.104 | 1.02E-77 | 9 | Ccrl2 |
| Ccnyl11 | 9.75E-82 | 0.606320246 | 0.318 | 0.105 | 1.97E-77 | 9 | Ccnyl1 |
| Cpne82 | 1.65E-81 | 0.637595276 | 0.512 | 0.221 | 3.34E-77 | 9 | Cpne8 |
| Slc30a1 | 6.98E-81 | 0.589059295 | 0.256 | 0.075 | 1.41E-76 | 9 | Slc30a1 |
| Cd382 | 9.62E-81 | 0.658788371 | 0.509 | 0.224 | 1.95E-76 | 9 | Cd38 |
| Atrx2 | 1.31E-80 | 0.762657737 | 0.708 | 0.412 | 2.66E-76 | 9 | Atrx |
| Phlpp12 | 1.74E-80 | 0.71559601 | 0.547 | 0.256 | 3.53E-76 | 9 | Phlpp1 |
| Sh3rf11 | 1.81E-80 | 0.688635398 | 0.349 | 0.125 | 3.67E-76 | 9 | Sh3rf1 |
| Tmcc33 | 1.81E-80 | 0.584101171 | 0.63 | 0.311 | 3.67E-76 | 9 | Tmcc3 |
| Plscr42 | 1.84E-80 | 0.657925523 | 0.445 | 0.183 | 3.73E-76 | 9 | Plscr4 |
| Tapbp3 | 1.85E-80 | 0.742005106 | 0.572 | 0.298 | 3.75E-76 | 9 | Tapbp |
| Adgrl22 | 2.46E-80 | 0.608224658 | 0.534 | 0.229 | 4.98E-76 | 9 | Adgrl2 |
| H2-K14 | 2.74E-80 | 0.654215118 | 0.974 | 0.634 | 5.54E-76 | 9 | H2-K1 |
| Arap32 | 9.14E-80 | 0.632305778 | 0.485 | 0.211 | 1.85E-75 | 9 | Arap3 |
| Serpinh13 | 1.58E-79 | 0.609135158 | 0.544 | 0.242 | 3.19E-75 | 9 | Serpinh1 |
| Wars2 | 2.91E-79 | 0.657592769 | 0.406 | 0.162 | 5.88E-75 | 9 | Wars |
| Fam171a11 | 2.92E-79 | 0.558122604 | 0.308 | 0.101 | 5.91E-75 | 9 | Fam171a1 |
| Afap12 | 3.20E-79 | 0.610276542 | 0.382 | 0.144 | 6.49E-75 | 9 | Afap1 |
| Tnfaip12 | 3.90E-79 | 0.572849946 | 0.471 | 0.196 | 7.90E-75 | 9 | Tnfaip1 |
| Aida1 | 9.73E-79 | 0.642061158 | 0.364 | 0.137 | 1.97E-74 | 9 | Aida |
| Gnas3 | 1.13E-78 | 0.493870672 | 0.977 | 0.86 | 2.29E-74 | 9 | Gnas |
| Sh3glb13 | 1.85E-78 | 0.631039358 | 0.814 | 0.524 | 3.74E-74 | 9 | Sh3glb1 |
| Baz2b3 | 2.21E-78 | 0.723543008 | 0.634 | 0.343 | 4.47E-74 | 9 | Baz2b |
| Phldb22 | 4.75E-78 | 0.670007055 | 0.467 | 0.196 | 9.61E-74 | 9 | Phldb2 |
| Dennd5a2 | 9.15E-78 | 0.698536559 | 0.424 | 0.181 | 1.85E-73 | 9 | Dennd5a |
| Ehd42 | 9.94E-78 | 0.676848114 | 0.74 | 0.399 | 2.01E-73 | 9 | Ehd4 |
| Fry2 | 2.42E-77 | 0.719660196 | 0.454 | 0.194 | 4.90E-73 | 9 | Fry |
| Rasal22 | 3.72E-77 | 0.546640479 | 0.651 | 0.306 | 7.52E-73 | 9 | Rasal2 |
| Agrn1 | 5.00E-77 | 0.545456388 | 0.261 | 0.078 | 1.01E-72 | 9 | Agrn |
| Arhgap5 | 8.38E-77 | 0.683087818 | 0.564 | 0.282 | 1.70E-72 | 9 | Arhgap5 |
| Isg152 | 1.43E-76 | 0.803322754 | 0.493 | 0.222 | 2.90E-72 | 9 | Isg15 |
| Lamb21 | 5.19E-76 | 0.689369665 | 0.332 | 0.118 | 1.05E-71 | 9 | Lamb2 |
| Rapgef13 | 5.25E-76 | 0.756656812 | 0.49 | 0.234 | 1.06E-71 | 9 | Rapgef1 |
| Osbpl92 | 5.69E-76 | 0.67730308 | 0.543 | 0.271 | 1.15E-71 | 9 | Osbpl9 |
| Atp2b42 | 7.98E-76 | 0.699495577 | 0.44 | 0.192 | 1.62E-71 | 9 | Atp2b4 |
| Iigp12 | 1.08E-75 | 0.652449245 | 0.52 | 0.225 | 2.18E-71 | 9 | Iigp1 |
| Adcy41 | 1.89E-75 | 0.570587526 | 0.305 | 0.102 | 3.83E-71 | 9 | Adcy4 |
| Rgs121 | 2.01E-75 | 0.593018355 | 0.297 | 0.098 | 4.07E-71 | 9 | Rgs12 |
| Gnb13 | 2.02E-75 | 0.621186721 | 0.707 | 0.42 | 4.09E-71 | 9 | Gnb1 |
| Slain2 | 3.10E-75 | 0.684143156 | 0.408 | 0.172 | 6.28E-71 | 9 | Slain2 |
| Dgkh2 | 3.64E-75 | 0.682717302 | 0.412 | 0.171 | 7.36E-71 | 9 | Dgkh |
| Ccser22 | 5.61E-75 | 0.639231354 | 0.49 | 0.224 | 1.14E-70 | 9 | Ccser2 |

| | | | | | | | |
|---|---|---|---|---|---|---|---|
| Fez22 | 6.33E-75 | 0.574265908 | 0.571 | 0.275 | 1.28E-70 | 9 | Fez2 |
| Cbfa2t31 | 8.89E-75 | 0.675084901 | 0.354 | 0.133 | 1.80E-70 | 9 | Cbfa2t3 |
| Ephb42 | 1.39E-74 | 0.532510535 | 0.411 | 0.16 | 2.82E-70 | 9 | Ephb4 |
| Akt33 | 2.73E-74 | 0.790660213 | 0.51 | 0.245 | 5.52E-70 | 9 | Akt3 |
| Sash12 | 3.32E-74 | 0.637947097 | 0.538 | 0.249 | 6.73E-70 | 9 | Sash1 |
| Tuba1a3 | 4.43E-74 | 0.673268461 | 0.524 | 0.249 | 8.98E-70 | 9 | Tuba1a |
| Hist1h2bc2 | 4.97E-74 | 0.652691863 | 0.51 | 0.243 | 1.01E-69 | 9 | Hist1h2bc |
| Hdac72 | 9.89E-74 | 0.669096467 | 0.446 | 0.196 | 2.00E-69 | 9 | Hdac7 |
| Cd342 | 1.81E-73 | 0.531249017 | 0.661 | 0.312 | 3.67E-69 | 9 | Cd34 |
| Caskin22 | 7.69E-73 | 0.569930871 | 0.359 | 0.134 | 1.56E-68 | 9 | Caskin2 |
| Lgals93 | 8.93E-73 | 0.680430968 | 0.496 | 0.241 | 1.81E-68 | 9 | Lgals9 |
| Fam219a1 | 9.37E-73 | 0.655977825 | 0.338 | 0.126 | 1.90E-68 | 9 | Fam219a |
| Pitpnc13 | 1.21E-72 | 0.651649781 | 0.746 | 0.421 | 2.44E-68 | 9 | Pitpnc1 |
| Foxn34 | 1.21E-72 | 0.745422391 | 0.683 | 0.404 | 2.45E-68 | 9 | Foxn3 |
| Calcrl2 | 1.33E-72 | 0.654039229 | 0.547 | 0.256 | 2.69E-68 | 9 | Calcrl |
| Numb2 | 2.03E-72 | 0.611453227 | 0.631 | 0.329 | 4.11E-68 | 9 | Numb |
| Sybu2 | 5.99E-72 | 0.631542097 | 0.314 | 0.109 | 1.21E-67 | 9 | Sybu |
| Ptpn141 | 6.97E-72 | 0.576745816 | 0.298 | 0.102 | 1.41E-67 | 9 | Ptpn14 |
| Ak3 | 1.25E-71 | 0.438352302 | 0.381 | 0.148 | 2.53E-67 | 9 | Ak3 |
| Atxn13 | 1.32E-71 | 0.764465077 | 0.56 | 0.288 | 2.67E-67 | 9 | Atxn1 |
| Nes2 | 1.68E-71 | 0.658282337 | 0.342 | 0.128 | 3.39E-67 | 9 | Nes |
| Cltb1 | 1.83E-71 | 0.548424301 | 0.467 | 0.212 | 3.70E-67 | 9 | Cltb |
| Fli13 | 2.62E-71 | 0.679664466 | 0.695 | 0.394 | 5.30E-67 | 9 | Fli1 |
| 23-Mar | 3.67E-71 | 0.575909775 | 0.668 | 0.387 | 7.43E-67 | 9 | 2-Mar |
| Ttll71 | 4.40E-71 | 0.554304427 | 0.292 | 0.098 | 8.91E-67 | 9 | Ttll7 |
| Kmt2e2 | 7.33E-71 | 0.646475369 | 0.683 | 0.399 | 1.48E-66 | 9 | Kmt2e |
| Cds22 | 1.54E-70 | 0.615119234 | 0.406 | 0.171 | 3.13E-66 | 9 | Cds2 |
| Ttc282 | 1.99E-70 | 0.663970992 | 0.556 | 0.278 | 4.04E-66 | 9 | Ttc28 |
| St3gal13 | 2.01E-70 | 0.621856153 | 0.411 | 0.174 | 4.07E-66 | 9 | St3gal1 |
| Plxnd12 | 2.30E-70 | 0.603974401 | 0.418 | 0.175 | 4.66E-66 | 9 | Plxnd1 |
| Il10rb3 | 5.66E-70 | 0.633399888 | 0.497 | 0.242 | 1.15E-65 | 9 | Il10rb |
| Iqgap14 | 6.16E-70 | 0.677048351 | 0.684 | 0.405 | 1.25E-65 | 9 | Iqgap1 |
| Klhl52 | 6.90E-70 | 0.588534038 | 0.478 | 0.218 | 1.40E-65 | 9 | Klhl5 |
| Lats21 | 1.10E-69 | 0.552417266 | 0.316 | 0.115 | 2.23E-65 | 9 | Lats2 |
| Ccdc85b3 | 1.49E-69 | 0.56945735 | 0.518 | 0.249 | 3.01E-65 | 9 | Ccdc85b |
| Zfyve92 | 2.89E-69 | 0.583980283 | 0.401 | 0.165 | 5.85E-65 | 9 | Zfyve9 |
| Nr3c12 | 3.87E-69 | 0.671560287 | 0.656 | 0.376 | 7.84E-65 | 9 | Nr3c1 |
| Sgms11 | 4.70E-69 | 0.604601851 | 0.33 | 0.124 | 9.51E-65 | 9 | Sgms1 |
| Sypl2 | 9.58E-69 | 0.603733212 | 0.491 | 0.233 | 1.94E-64 | 9 | Sypl |
| Tagln23 | 1.52E-68 | 0.642798301 | 0.71 | 0.422 | 3.08E-64 | 9 | Tagln2 |
| Gnaq3 | 2.89E-68 | 0.62908308 | 0.64 | 0.352 | 5.84E-64 | 9 | Gnaq |
| Cyb5r32 | 6.10E-68 | 0.647959728 | 0.455 | 0.211 | 1.24E-63 | 9 | Cyb5r3 |
| Grasp2 | 6.46E-68 | 0.56859807 | 0.391 | 0.161 | 1.31E-63 | 9 | Grasp |

| | | | | | | | |
|---|---|---|---|---|---|---|---|
| Gbp21 | 9.52E-68 | 0.631180211 | 0.35 | 0.138 | 1.93E-63 | 9 | Gbp2 |
| Sbf22 | 1.02E-67 | 0.623267641 | 0.449 | 0.201 | 2.08E-63 | 9 | Sbf2 |
| Pdlim52 | 1.06E-67 | 0.696920839 | 0.538 | 0.28 | 2.15E-63 | 9 | Pdlim5 |
| Rac12 | 1.89E-67 | 0.629163372 | 0.695 | 0.423 | 3.83E-63 | 9 | Rac1 |
| Kras2 | 1.92E-67 | 0.572646515 | 0.518 | 0.263 | 3.89E-63 | 9 | Kras |
| Tm6sf12 | 2.35E-67 | 0.642986928 | 0.342 | 0.136 | 4.75E-63 | 9 | Tm6sf1 |
| Ssfa2 | 5.42E-67 | 0.64004236 | 0.27 | 0.093 | 1.10E-62 | 9 | Ssfa2 |
| Hdgfl31 | 6.75E-67 | 0.542746731 | 0.297 | 0.106 | 1.37E-62 | 9 | Hdgfl3 |
| Rhoa2 | 1.23E-66 | 0.555165319 | 0.851 | 0.587 | 2.48E-62 | 9 | Rhoa |
| Ccny3 | 1.37E-66 | 0.726967674 | 0.504 | 0.261 | 2.78E-62 | 9 | Ccny |
| Jup2 | 3.58E-66 | 0.575349319 | 0.523 | 0.251 | 7.26E-62 | 9 | Jup |
| Ebf31 | 3.68E-66 | 0.573862826 | 0.324 | 0.121 | 7.46E-62 | 9 | Ebf3 |
| Capn22 | 4.90E-66 | 0.606371127 | 0.398 | 0.173 | 9.91E-62 | 9 | Capn2 |
| Ctnnbip12 | 6.99E-66 | 0.574209491 | 0.44 | 0.198 | 1.42E-61 | 9 | Ctnnbip1 |
| Nbeal12 | 1.43E-65 | 0.700842692 | 0.503 | 0.256 | 2.90E-61 | 9 | Nbeal1 |
| Anxa22 | 1.58E-65 | 0.580728552 | 0.711 | 0.409 | 3.20E-61 | 9 | Anxa2 |
| Git23 | 1.89E-65 | 0.665353291 | 0.464 | 0.229 | 3.82E-61 | 9 | Git2 |
| Nav23 | 3.06E-65 | 0.636976223 | 0.382 | 0.16 | 6.19E-61 | 9 | Nav2 |
| Tmem1402 | 4.29E-65 | 0.633981503 | 0.465 | 0.217 | 8.69E-61 | 9 | Tmem140 |
| Spns21 | 8.87E-65 | 0.464447498 | 0.265 | 0.088 | 1.80E-60 | 9 | Spns2 |
| Ano61 | 9.72E-65 | 0.613174795 | 0.397 | 0.175 | 1.97E-60 | 9 | Ano6 |
| Vps13d1 | 1.23E-64 | 0.607994359 | 0.379 | 0.164 | 2.50E-60 | 9 | Vps13d |
| Tspan41 | 1.27E-64 | 0.542089595 | 0.286 | 0.101 | 2.57E-60 | 9 | Tspan4 |
| Parp143 | 1.34E-64 | 0.675739003 | 0.491 | 0.247 | 2.71E-60 | 9 | Parp14 |
| Arglu12 | 1.40E-64 | 0.612476068 | 0.603 | 0.348 | 2.83E-60 | 9 | Arglu1 |
| Atp8b11 | 1.49E-64 | 0.555413384 | 0.281 | 0.099 | 3.02E-60 | 9 | Atp8b1 |
| Spata133 | 1.55E-64 | 0.706333101 | 0.427 | 0.201 | 3.13E-60 | 9 | Spata13 |
| Nfkbia3 | 7.39E-64 | 0.650755105 | 0.673 | 0.393 | 1.50E-59 | 9 | Nfkbia |
| Dram22 | 1.16E-63 | 0.566561164 | 0.359 | 0.149 | 2.35E-59 | 9 | Dram2 |
| Hacd42 | 3.60E-63 | 0.566631269 | 0.298 | 0.111 | 7.29E-59 | 9 | Hacd4 |
| Csf11 | 4.47E-63 | 0.594166651 | 0.29 | 0.105 | 9.06E-59 | 9 | Csf1 |
| Reep32 | 9.22E-63 | 0.631121188 | 0.56 | 0.299 | 1.87E-58 | 9 | Reep3 |
| Afap1l11 | 1.05E-62 | 0.479011167 | 0.292 | 0.104 | 2.13E-58 | 9 | Afap1l1 |
| Plcb42 | 1.45E-62 | 0.47119649 | 0.568 | 0.284 | 2.94E-58 | 9 | Plcb4 |
| Peak13 | 2.44E-62 | 0.659378189 | 0.642 | 0.381 | 4.94E-58 | 9 | Peak1 |
| Socs51 | 6.34E-62 | 0.523628218 | 0.317 | 0.123 | 1.28E-57 | 9 | Socs5 |
| Dstyk1 | 6.85E-62 | 0.546217767 | 0.35 | 0.145 | 1.39E-57 | 9 | Dstyk |
| Ston1 | 7.93E-62 | 0.498678575 | 0.255 | 0.086 | 1.61E-57 | 9 | Ston1 |
| Pik3c2a2 | 1.36E-61 | 0.640989157 | 0.411 | 0.19 | 2.75E-57 | 9 | Pik3c2a |
| Zbtb462 | 1.44E-61 | 0.615156582 | 0.372 | 0.157 | 2.93E-57 | 9 | Zbtb46 |
| Tmpo2 | 1.47E-61 | 0.544934157 | 0.43 | 0.202 | 2.97E-57 | 9 | Tmpo |
| Ptpra3 | 1.54E-61 | 0.616872641 | 0.547 | 0.298 | 3.11E-57 | 9 | Ptpra |
| Rock13 | 2.96E-61 | 0.59910998 | 0.702 | 0.426 | 5.99E-57 | 9 | Rock1 |

| | | | | | | | |
|---|---|---|---|---|---|---|---|
| Tgm22 | 4.34E-61 | 0.531685677 | 0.379 | 0.156 | 8.79E-57 | 9 | Tgm2 |
| Clock2 | 5.37E-61 | 0.573727709 | 0.423 | 0.198 | 1.09E-56 | 9 | Clock |
| Prss233 | 5.91E-61 | 0.426729399 | 0.578 | 0.273 | 1.20E-56 | 9 | Prss23 |
| Lysmd22 | 8.07E-61 | 0.543082991 | 0.412 | 0.186 | 1.63E-56 | 9 | Lysmd2 |
| Tmem472 | 9.49E-61 | 0.553172589 | 0.366 | 0.151 | 1.92E-56 | 9 | Tmem47 |
| Nckap12 | 1.46E-60 | 0.544012733 | 0.461 | 0.22 | 2.95E-56 | 9 | Nckap1 |
| Shroom41 | 1.59E-60 | 0.538007238 | 0.28 | 0.101 | 3.21E-56 | 9 | Shroom4 |
| Ier5l2 | 2.14E-60 | 0.563413625 | 0.396 | 0.173 | 4.34E-56 | 9 | Ier5l |
| Dock62 | 2.49E-60 | 0.570858994 | 0.338 | 0.135 | 5.04E-56 | 9 | Dock6 |
| Anxa51 | 6.06E-60 | 0.489968904 | 0.773 | 0.486 | 1.23E-55 | 9 | Anxa5 |
| Rock22 | 8.45E-60 | 0.615493153 | 0.589 | 0.331 | 1.71E-55 | 9 | Rock2 |
| Myl12a2 | 1.39E-59 | 0.557552506 | 0.729 | 0.464 | 2.82E-55 | 9 | Myl12a |
| Itga62 | 1.63E-59 | 0.587234395 | 0.541 | 0.277 | 3.30E-55 | 9 | Itga6 |
| Inpp11 | 1.79E-59 | 0.512638913 | 0.293 | 0.111 | 3.62E-55 | 9 | Inpp1 |
| Plec1 | 1.92E-59 | 0.60635432 | 0.379 | 0.169 | 3.89E-55 | 9 | Plec |
| Emp22 | 1.95E-59 | 0.541359122 | 0.482 | 0.229 | 3.95E-55 | 9 | Emp2 |
| Cdc42ep22 | 2.26E-59 | 0.497554867 | 0.39 | 0.167 | 4.57E-55 | 9 | Cdc42ep2 |
| Klf92 | 3.11E-59 | 0.573393036 | 0.477 | 0.231 | 6.30E-55 | 9 | Klf9 |
| Ggta11 | 3.20E-59 | 0.555692658 | 0.332 | 0.136 | 6.49E-55 | 9 | Ggta1 |
| Zbtb203 | 3.47E-59 | 0.482677983 | 0.862 | 0.528 | 7.03E-55 | 9 | Zbtb20 |
| Nhsl21 | 9.42E-59 | 0.551837614 | 0.327 | 0.13 | 1.91E-54 | 9 | Nhsl2 |
| Strn32 | 1.05E-58 | 0.582968989 | 0.593 | 0.339 | 2.12E-54 | 9 | Strn3 |
| Lrrc582 | 1.05E-58 | 0.540680533 | 0.439 | 0.209 | 2.12E-54 | 9 | Lrrc58 |
| Exoc3l22 | 1.07E-58 | 0.460039741 | 0.556 | 0.263 | 2.17E-54 | 9 | Exoc3l2 |
| Spag92 | 1.19E-58 | 0.615042156 | 0.6 | 0.353 | 2.40E-54 | 9 | Spag9 |
| Gbp91 | 1.36E-58 | 0.503443458 | 0.335 | 0.137 | 2.76E-54 | 9 | Gbp9 |
| Eva1b3 | 2.36E-58 | 0.525596403 | 0.543 | 0.279 | 4.79E-54 | 9 | Eva1b |
| Map3k111 | 2.99E-58 | 0.458565687 | 0.252 | 0.088 | 6.06E-54 | 9 | Map3k11 |
| Swap703 | 3.55E-58 | 0.653467923 | 0.465 | 0.241 | 7.18E-54 | 9 | Swap70 |
| Sox71 | 4.64E-58 | 0.558566993 | 0.258 | 0.091 | 9.39E-54 | 9 | Sox7 |
| Usp46 | 5.69E-58 | 0.42520092 | 0.321 | 0.128 | 1.15E-53 | 9 | Usp46 |
| Rtn32 | 8.57E-58 | 0.646765779 | 0.557 | 0.318 | 1.73E-53 | 9 | Rtn3 |
| Azin1 | 1.87E-57 | 0.520852084 | 0.348 | 0.149 | 3.79E-53 | 9 | Azin1 |
| Mbnl23 | 3.39E-57 | 0.575064396 | 0.678 | 0.412 | 6.86E-53 | 9 | Mbnl2 |
| Gbp51 | 5.85E-57 | 0.54775968 | 0.312 | 0.124 | 1.19E-52 | 9 | Gbp5 |
| Map7d11 | 6.92E-57 | 0.574414202 | 0.363 | 0.163 | 1.40E-52 | 9 | Map7d1 |
| Lpar63 | 1.13E-56 | 0.642020191 | 0.386 | 0.179 | 2.28E-52 | 9 | Lpar6 |
| Mprip3 | 1.29E-56 | 0.583451301 | 0.533 | 0.287 | 2.61E-52 | 9 | Mprip |
| Phf11d1 | 2.00E-56 | 0.47033516 | 0.327 | 0.133 | 4.04E-52 | 9 | Phf11d |
| Cmip4 | 2.80E-56 | 0.60295868 | 0.512 | 0.28 | 5.66E-52 | 9 | Cmip |
| Serinc33 | 3.39E-56 | 0.531237105 | 0.763 | 0.48 | 6.87E-52 | 9 | Serinc3 |
| Phactr42 | 3.78E-56 | 0.560067585 | 0.38 | 0.173 | 7.66E-52 | 9 | Phactr4 |
| Itm2c1 | 4.99E-56 | 0.722272102 | 0.397 | 0.194 | 1.01E-51 | 9 | Itm2c |

| | | | | | | | |
|---|---|---|---|---|---|---|---|
| Arhgef21 | 6.72E-56 | 0.588287308 | 0.356 | 0.16 | 1.36E-51 | 9 | Arhgef2 |
| Ints92 | 6.81E-56 | 0.574819508 | 0.318 | 0.132 | 1.38E-51 | 9 | Ints9 |
| Zfp36l13 | 7.99E-56 | 0.587146196 | 0.693 | 0.426 | 1.62E-51 | 9 | Zfp36l1 |
| Psmb103 | 9.60E-56 | 0.545174072 | 0.655 | 0.393 | 1.94E-51 | 9 | Psmb10 |
| Smpdl3a2 | 1.02E-55 | 0.548401606 | 0.439 | 0.214 | 2.06E-51 | 9 | Smpdl3a |
| Prrg11 | 2.32E-55 | 0.578562037 | 0.276 | 0.104 | 4.69E-51 | 9 | Prrg1 |
| Psmb93 | 2.40E-55 | 0.509340697 | 0.661 | 0.399 | 4.86E-51 | 9 | Psmb9 |
| Fcho22 | 2.84E-55 | 0.574436059 | 0.544 | 0.304 | 5.75E-51 | 9 | Fcho2 |
| Hsp90b14 | 3.06E-55 | 0.531958 | 0.785 | 0.503 | 6.20E-51 | 9 | Hsp90b1 |
| Fstl1 | 4.55E-55 | 0.435593295 | 0.273 | 0.099 | 9.20E-51 | 9 | Fstl1 |
| Pde8a2 | 6.85E-55 | 0.550926082 | 0.391 | 0.178 | 1.39E-50 | 9 | Pde8a |
| Elf13 | 7.39E-55 | 0.616999575 | 0.608 | 0.371 | 1.50E-50 | 9 | Elf1 |
| Tnrc6c2 | 1.26E-54 | 0.621831329 | 0.544 | 0.31 | 2.54E-50 | 9 | Tnrc6c |
| Atox12 | 1.28E-54 | 0.450391783 | 0.867 | 0.612 | 2.59E-50 | 9 | Atox1 |
| Cr1l2 | 1.36E-54 | 0.638274577 | 0.441 | 0.228 | 2.76E-50 | 9 | Cr1l |
| Stard92 | 1.45E-54 | 0.593075042 | 0.342 | 0.147 | 2.94E-50 | 9 | Stard9 |
| Itgb13 | 3.04E-54 | 0.488321451 | 0.769 | 0.471 | 6.16E-50 | 9 | Itgb1 |
| Txnip2 | 4.09E-54 | 0.496866281 | 0.804 | 0.538 | 8.27E-50 | 9 | Txnip |
| Psmb85 | 3.01E-53 | 0.488820219 | 0.845 | 0.55 | 6.10E-49 | 9 | Psmb8 |
| Bcl2l12 | 3.24E-53 | 0.564789519 | 0.329 | 0.143 | 6.56E-49 | 9 | Bcl2l1 |
| Piezo12 | 3.58E-53 | 0.535708969 | 0.372 | 0.169 | 7.26E-49 | 9 | Piezo1 |
| Dusp161 | 8.02E-53 | 0.535367696 | 0.286 | 0.115 | 1.62E-48 | 9 | Dusp16 |
| Ptbp34 | 1.35E-52 | 0.556494582 | 0.64 | 0.388 | 2.74E-48 | 9 | Ptbp3 |
| Tap23 | 2.25E-52 | 0.618108898 | 0.45 | 0.236 | 4.55E-48 | 9 | Tap2 |
| Ghr1 | 2.44E-52 | 0.545009662 | 0.342 | 0.147 | 4.95E-48 | 9 | Ghr |
| Lipa2 | 3.92E-52 | 0.542978067 | 0.374 | 0.174 | 7.93E-48 | 9 | Lipa |
| Col4a3bp2 | 3.95E-52 | 0.558210381 | 0.477 | 0.253 | 8.01E-48 | 9 | Col4a3bp |
| Cebpg2 | 3.96E-52 | 0.446956169 | 0.44 | 0.218 | 8.02E-48 | 9 | Cebpg |
| Ripor12 | 4.14E-52 | 0.524073225 | 0.351 | 0.156 | 8.38E-48 | 9 | Ripor1 |
| Rab122 | 8.61E-52 | 0.547412224 | 0.371 | 0.172 | 1.74E-47 | 9 | Rab12 |
| Ddx53 | 1.16E-51 | 0.48695403 | 0.908 | 0.647 | 2.35E-47 | 9 | Ddx5 |
| Eif3a2 | 1.42E-51 | 0.639390718 | 0.594 | 0.367 | 2.87E-47 | 9 | Eif3a |
| Ets22 | 1.73E-51 | 0.522102723 | 0.404 | 0.189 | 3.50E-47 | 9 | Ets2 |
| Cracr2b1 | 1.90E-51 | 0.294553894 | 0.478 | 0.232 | 3.85E-47 | 9 | Cracr2b |
| Tshz23 | 3.75E-51 | 0.370485667 | 0.682 | 0.364 | 7.58E-47 | 9 | Tshz2 |
| Fbxo7 | 5.01E-51 | 0.479688639 | 0.26 | 0.101 | 1.02E-46 | 9 | Fbxo7 |
| Tubb4b3 | 5.22E-51 | 0.475502413 | 0.684 | 0.421 | 1.06E-46 | 9 | Tubb4b |
| Stag13 | 5.29E-51 | 0.586299477 | 0.591 | 0.356 | 1.07E-46 | 9 | Stag1 |
| Cdc42bpb2 | 5.86E-51 | 0.431433359 | 0.391 | 0.181 | 1.19E-46 | 9 | Cdc42bpb |
| Tmem184b1 | 6.29E-51 | 0.486189496 | 0.277 | 0.11 | 1.27E-46 | 9 | Tmem184b |
| Selenof2 | 6.71E-51 | 0.457422267 | 0.741 | 0.485 | 1.36E-46 | 9 | Selenof |
| Yes12 | 9.32E-51 | 0.500811516 | 0.342 | 0.149 | 1.89E-46 | 9 | Yes1 |
| Mfsd61 | 1.14E-50 | 0.516199503 | 0.324 | 0.14 | 2.31E-46 | 9 | Mfsd6 |

| | | | | | | | |
|---|---|---|---|---|---|---|---|
| Kif5b2 | 1.34E-50 | 0.473072131 | 0.641 | 0.4 | 2.71E-46 | 9 | Kif5b |
| Son4 | 1.87E-50 | 0.500729192 | 0.799 | 0.519 | 3.78E-46 | 9 | Son |
| Eepd12 | 3.55E-50 | 0.431182425 | 0.328 | 0.139 | 7.19E-46 | 9 | Eepd1 |
| Rapgef32 | 4.14E-50 | 0.475953615 | 0.367 | 0.164 | 8.39E-46 | 9 | Rapgef3 |
| Ptp4a3 | 4.97E-50 | 0.542997745 | 0.307 | 0.133 | 1.01E-45 | 9 | Ptp4a3 |
| Taok12 | 5.89E-50 | 0.531161415 | 0.461 | 0.246 | 1.19E-45 | 9 | Taok1 |
| Tjp21 | 6.24E-50 | 0.512449794 | 0.316 | 0.136 | 1.26E-45 | 9 | Tjp2 |
| Eif4ebp22 | 6.26E-50 | 0.508896267 | 0.45 | 0.236 | 1.27E-45 | 9 | Eif4ebp2 |
| Cyp2d222 | 6.79E-50 | 0.471690668 | 0.314 | 0.131 | 1.38E-45 | 9 | Cyp2d22 |
| Erc11 | 1.06E-49 | 0.509711344 | 0.289 | 0.119 | 2.15E-45 | 9 | Erc1 |
| Sec623 | 1.57E-49 | 0.535654407 | 0.625 | 0.389 | 3.18E-45 | 9 | Sec62 |
| Gbp42 | 2.04E-49 | 0.570595467 | 0.367 | 0.173 | 4.14E-45 | 9 | Gbp4 |
| Scarb12 | 3.55E-49 | 0.448019095 | 0.524 | 0.26 | 7.18E-45 | 9 | Scarb1 |
| Tnks23 | 4.14E-49 | 0.55524686 | 0.49 | 0.275 | 8.39E-45 | 9 | Tnks2 |
| Gm383991 | 5.02E-49 | 0.48980957 | 0.265 | 0.104 | 1.02E-44 | 9 | Gm38399 |
| Itm2a2 | 5.07E-49 | 0.559420091 | 0.38 | 0.175 | 1.03E-44 | 9 | Itm2a |
| Tmco12 | 7.13E-49 | 0.518261107 | 0.53 | 0.307 | 1.44E-44 | 9 | Tmco1 |
| Rrbp13 | 7.86E-49 | 0.467476811 | 0.714 | 0.453 | 1.59E-44 | 9 | Rrbp1 |
| Irf7 | 1.66E-48 | 0.561684961 | 0.365 | 0.172 | 3.36E-44 | 9 | Irf7 |
| Tspan122 | 1.79E-48 | 0.41880368 | 0.464 | 0.226 | 3.63E-44 | 9 | Tspan12 |
| Sod13 | 1.94E-48 | 0.438224676 | 0.524 | 0.294 | 3.92E-44 | 9 | Sod1 |
| Atrnl11 | 2.04E-48 | 0.608957231 | 0.35 | 0.161 | 4.13E-44 | 9 | Atrnl1 |
| Picalm2 | 2.23E-48 | 0.542133794 | 0.51 | 0.291 | 4.52E-44 | 9 | Picalm |
| Ric1 | 2.43E-48 | 0.508115847 | 0.276 | 0.114 | 4.93E-44 | 9 | Ric1 |
| Gbp31 | 2.47E-48 | 0.551474723 | 0.28 | 0.114 | 5.01E-44 | 9 | Gbp3 |
| Amotl12 | 2.63E-48 | 0.484500983 | 0.388 | 0.181 | 5.32E-44 | 9 | Amotl1 |
| Tmem30a3 | 2.78E-48 | 0.542550548 | 0.499 | 0.281 | 5.63E-44 | 9 | Tmem30a |
| Map4k43 | 3.21E-48 | 0.553322546 | 0.482 | 0.272 | 6.50E-44 | 9 | Map4k4 |
| Grk51 | 5.15E-48 | 0.533610051 | 0.34 | 0.154 | 1.04E-43 | 9 | Grk5 |
| Prrc2c3 | 1.61E-47 | 0.537730788 | 0.651 | 0.415 | 3.26E-43 | 9 | Prrc2c |
| B230219D22Rik2 | 1.69E-47 | 0.475539666 | 0.567 | 0.337 | 3.42E-43 | 9 | B230219D22Rik |
| Tspan182 | 1.95E-47 | 0.450966504 | 0.314 | 0.133 | 3.95E-43 | 9 | Tspan18 |
| Ncoa73 | 2.32E-47 | 0.544864138 | 0.523 | 0.291 | 4.69E-43 | 9 | Ncoa7 |
| Ptpn122 | 2.33E-47 | 0.578081132 | 0.407 | 0.209 | 4.72E-43 | 9 | Ptpn12 |
| Pcmtd11 | 2.53E-47 | 0.462637718 | 0.546 | 0.322 | 5.13E-43 | 9 | Pcmtd1 |
| Rftn13 | 3.02E-47 | 0.556931355 | 0.457 | 0.247 | 6.10E-43 | 9 | Rftn1 |
| Camk2d3 | 3.41E-47 | 0.543712546 | 0.481 | 0.265 | 6.91E-43 | 9 | Camk2d |
| Foxo12 | 3.73E-47 | 0.508034507 | 0.397 | 0.195 | 7.56E-43 | 9 | Foxo1 |
| Myh93 | 3.89E-47 | 0.548046781 | 0.668 | 0.431 | 7.89E-43 | 9 | Myh9 |
| Irf12 | 4.00E-47 | 0.536334545 | 0.517 | 0.294 | 8.09E-43 | 9 | Irf1 |
| Usp343 | 6.37E-47 | 0.551528892 | 0.565 | 0.339 | 1.29E-42 | 9 | Usp34 |
| Smchd13 | 7.03E-47 | 0.56446784 | 0.465 | 0.259 | 1.42E-42 | 9 | Smchd1 |
| Ifih11 | 1.83E-46 | 0.503849893 | 0.291 | 0.125 | 3.71E-42 | 9 | Ifih1 |

| | | | | | | | |
|---|---|---|---|---|---|---|---|
| Xiap2 | 2.03E-46 | 0.529672093 | 0.512 | 0.295 | 4.10E-42 | 9 | Xiap |
| Smagp2 | 3.36E-46 | 0.380160838 | 0.448 | 0.226 | 6.79E-42 | 9 | Smagp |
| Gimap14 | 3.42E-46 | 0.487140348 | 0.536 | 0.297 | 6.93E-42 | 9 | Gimap1 |
| Tmbim63 | 3.44E-46 | 0.498919429 | 0.66 | 0.426 | 6.97E-42 | 9 | Tmbim6 |
| Dip2b3 | 4.62E-46 | 0.549833528 | 0.476 | 0.272 | 9.36E-42 | 9 | Dip2b |
| Zmynd112 | 5.58E-46 | 0.478573159 | 0.499 | 0.284 | 1.13E-41 | 9 | Zmynd11 |
| Lipe2 | 5.60E-46 | 0.440599806 | 0.309 | 0.133 | 1.13E-41 | 9 | Lipe |
| Prkd21 | 6.76E-46 | 0.543192684 | 0.306 | 0.136 | 1.37E-41 | 9 | Prkd2 |
| Klhl242 | 7.77E-46 | 0.489097514 | 0.411 | 0.211 | 1.57E-41 | 9 | Klhl24 |
| Tcf7l12 | 1.03E-45 | 0.448451804 | 0.376 | 0.176 | 2.09E-41 | 9 | Tcf7l1 |
| Dock43 | 1.18E-45 | 0.434301068 | 0.545 | 0.29 | 2.38E-41 | 9 | Dock4 |
| Cdipt1 | 1.86E-45 | 0.472201129 | 0.297 | 0.131 | 3.76E-41 | 9 | Cdipt |
| Prkch3 | 2.63E-45 | 0.439038737 | 0.671 | 0.393 | 5.33E-41 | 9 | Prkch |
| 4930523C07Rik3 | 2.77E-45 | 0.59976718 | 0.409 | 0.215 | 5.61E-41 | 9 | 4930523C07Rik |
| Rsu12 | 2.90E-45 | 0.45081502 | 0.46 | 0.249 | 5.88E-41 | 9 | Rsu1 |
| Lrch13 | 2.94E-45 | 0.575690213 | 0.525 | 0.307 | 5.95E-41 | 9 | Lrch1 |
| Pam2 | 5.39E-45 | 0.583189962 | 0.406 | 0.209 | 1.09E-40 | 9 | Pam |
| Rabac12 | 6.76E-45 | 0.384901286 | 0.815 | 0.558 | 1.37E-40 | 9 | Rabac1 |
| Atp1b32 | 7.51E-45 | 0.540967161 | 0.444 | 0.244 | 1.52E-40 | 9 | Atp1b3 |
| Nr3c21 | 1.28E-44 | 0.499073134 | 0.28 | 0.117 | 2.59E-40 | 9 | Nr3c2 |
| Fbxw22 | 1.71E-44 | 0.445969726 | 0.433 | 0.231 | 3.45E-40 | 9 | Fbxw2 |
| Mxd44 | 1.97E-44 | 0.464422071 | 0.575 | 0.347 | 3.99E-40 | 9 | Mxd4 |
| Vim2 | 2.17E-44 | 0.839111351 | 0.517 | 0.302 | 4.39E-40 | 9 | Vim |
| Dpp81 | 2.94E-44 | 0.462081459 | 0.365 | 0.18 | 5.94E-40 | 9 | Dpp8 |
| Cdk173 | 3.41E-44 | 0.541067667 | 0.443 | 0.24 | 6.90E-40 | 9 | Cdk17 |
| Dusp71 | 3.95E-44 | 0.443270633 | 0.3 | 0.132 | 8.00E-40 | 9 | Dusp7 |
| Arhgef33 | 6.65E-44 | 0.576418524 | 0.436 | 0.236 | 1.35E-39 | 9 | Arhgef3 |
| Gstm12 | 7.24E-44 | 0.28431788 | 0.565 | 0.316 | 1.47E-39 | 9 | Gstm1 |
| Dab23 | 7.55E-44 | 0.468931549 | 0.402 | 0.202 | 1.53E-39 | 9 | Dab2 |
| Klc11 | 9.74E-44 | 0.439317667 | 0.317 | 0.146 | 1.97E-39 | 9 | Klc1 |
| Zfp6091 | 1.30E-43 | 0.5190911 | 0.29 | 0.128 | 2.62E-39 | 9 | Zfp609 |
| Pitpnm21 | 1.38E-43 | 0.454115835 | 0.282 | 0.119 | 2.79E-39 | 9 | Pitpnm2 |
| Dcaf61 | 2.53E-43 | 0.493503281 | 0.292 | 0.13 | 5.13E-39 | 9 | Dcaf6 |
| Tpm33 | 3.86E-43 | 0.437775053 | 0.713 | 0.474 | 7.81E-39 | 9 | Tpm3 |
| Selenow2 | 5.27E-43 | 0.369273373 | 0.843 | 0.624 | 1.07E-38 | 9 | Selenow |
| Vamp53 | 7.39E-43 | 0.298394082 | 0.671 | 0.378 | 1.50E-38 | 9 | Vamp5 |
| Rarg2 | 7.90E-43 | 0.410003241 | 0.39 | 0.191 | 1.60E-38 | 9 | Rarg |
| Ddah21 | 7.90E-43 | 0.316067005 | 0.407 | 0.201 | 1.60E-38 | 9 | Ddah2 |
| Gnpda2 | 1.07E-42 | 0.42568416 | 0.271 | 0.116 | 2.17E-38 | 9 | Gnpda2 |
| Sep-74 | 1.09E-42 | 0.404975656 | 0.695 | 0.456 | 2.21E-38 | 9 | 7-Sep |
| Gsk3b2 | 1.10E-42 | 0.504732738 | 0.529 | 0.326 | 2.22E-38 | 9 | Gsk3b |
| Prkd12 | 1.19E-42 | 0.523185898 | 0.319 | 0.144 | 2.40E-38 | 9 | Prkd1 |
| Sec14l12 | 1.27E-42 | 0.364156965 | 0.496 | 0.272 | 2.56E-38 | 9 | Sec14l1 |

| | | | | | | | |
|---|---|---|---|---|---|---|---|
| Adipor21 | 1.40E-42 | 0.513127382 | 0.309 | 0.147 | 2.84E-38 | 9 | Adipor2 |
| Slc39a10 | 1.48E-42 | 0.49445106 | 0.258 | 0.108 | 3.00E-38 | 9 | Slc39a10 |
| Znfx12 | 3.93E-42 | 0.477447889 | 0.351 | 0.171 | 7.96E-38 | 9 | Znfx1 |
| Ppp1r16b3 | 3.98E-42 | 0.496573375 | 0.386 | 0.198 | 8.06E-38 | 9 | Ppp1r16b |
| Selenom3 | 4.81E-42 | 0.289951153 | 0.739 | 0.438 | 9.74E-38 | 9 | Selenom |
| Hmcn12 | 5.00E-42 | 0.569147767 | 0.408 | 0.206 | 1.01E-37 | 9 | Hmcn1 |
| Tmem1652 | 5.71E-42 | 0.490529053 | 0.349 | 0.176 | 1.16E-37 | 9 | Tmem165 |
| Dag11 | 6.21E-42 | 0.46400144 | 0.305 | 0.139 | 1.26E-37 | 9 | Dag1 |
| Smc1a2 | 6.86E-42 | 0.5180599 | 0.512 | 0.308 | 1.39E-37 | 9 | Smc1a |
| Xaf12 | 7.14E-42 | 0.453228468 | 0.409 | 0.217 | 1.45E-37 | 9 | Xaf1 |
| Cyth31 | 7.37E-42 | 0.515499782 | 0.275 | 0.122 | 1.49E-37 | 9 | Cyth3 |
| Bptf3 | 8.86E-42 | 0.51513393 | 0.488 | 0.289 | 1.79E-37 | 9 | Bptf |
| Anxa72 | 9.34E-42 | 0.405640694 | 0.365 | 0.182 | 1.89E-37 | 9 | Anxa7 |
| Trak1 | 9.49E-42 | 0.457767994 | 0.276 | 0.123 | 1.92E-37 | 9 | Trak1 |
| Rnf2133 | 9.87E-42 | 0.546646324 | 0.448 | 0.253 | 2.00E-37 | 9 | Rnf213 |
| Prkar1a3 | 9.98E-42 | 0.393708845 | 0.625 | 0.398 | 2.02E-37 | 9 | Prkar1a |
| Smarca22 | 1.06E-41 | 0.556298435 | 0.476 | 0.276 | 2.15E-37 | 9 | Smarca2 |
| Ankrd123 | 1.11E-41 | 0.531147417 | 0.51 | 0.303 | 2.26E-37 | 9 | Ankrd12 |
| Trim241 | 1.37E-41 | 0.56024499 | 0.339 | 0.169 | 2.77E-37 | 9 | Trim24 |
| Erg2 | 1.77E-41 | 0.388924285 | 0.293 | 0.127 | 3.58E-37 | 9 | Erg |
| Pttg1ip3 | 1.83E-41 | 0.446770373 | 0.471 | 0.264 | 3.70E-37 | 9 | Pttg1ip |
| H2-Q42 | 1.88E-41 | 0.450567405 | 0.522 | 0.304 | 3.80E-37 | 9 | H2-Q4 |
| Tmem1091 | 2.59E-41 | 0.433005686 | 0.309 | 0.144 | 5.23E-37 | 9 | Tmem109 |
| Pde4d1 | 2.77E-41 | 0.464104742 | 0.319 | 0.146 | 5.61E-37 | 9 | Pde4d |
| Cdk19 | 4.99E-41 | 0.626131594 | 0.301 | 0.144 | 1.01E-36 | 9 | Cdk19 |
| Itsn23 | 6.08E-41 | 0.524862829 | 0.469 | 0.274 | 1.23E-36 | 9 | Itsn2 |
| Lamp15 | 7.35E-41 | 0.429810286 | 0.64 | 0.412 | 1.49E-36 | 9 | Lamp1 |
| Ar2 | 7.60E-41 | 0.419729329 | 0.413 | 0.214 | 1.54E-36 | 9 | Ar |
| Rbm393 | 9.21E-41 | 0.434224736 | 0.826 | 0.568 | 1.87E-36 | 9 | Rbm39 |
| Samhd13 | 1.52E-40 | 0.490387572 | 0.445 | 0.25 | 3.08E-36 | 9 | Samhd1 |
| Sp1002 | 1.78E-40 | 0.467875617 | 0.652 | 0.427 | 3.60E-36 | 9 | Sp100 |
| Plekho13 | 1.82E-40 | 0.442476063 | 0.418 | 0.22 | 3.68E-36 | 9 | Plekho1 |
| Tmed52 | 2.18E-40 | 0.490739031 | 0.483 | 0.284 | 4.42E-36 | 9 | Tmed5 |
| Dnaja12 | 2.39E-40 | 0.411321095 | 0.747 | 0.511 | 4.83E-36 | 9 | Dnaja1 |
| Elovl51 | 2.40E-40 | 0.477281134 | 0.309 | 0.147 | 4.85E-36 | 9 | Elovl5 |
| Nostrin2 | 2.47E-40 | 0.330310069 | 0.49 | 0.253 | 4.99E-36 | 9 | Nostrin |
| BC0285282 | 2.51E-40 | 0.395355829 | 0.44 | 0.229 | 5.07E-36 | 9 | BC028528 |
| mt-Co26 | 2.60E-40 | 0.424702904 | 0.994 | 0.721 | 5.26E-36 | 9 | mt-Co2 |
| Bnip3l | 2.62E-40 | 0.402734905 | 0.536 | 0.327 | 5.30E-36 | 9 | Bnip3l |
| Serinc13 | 3.39E-40 | 0.466546667 | 0.436 | 0.245 | 6.86E-36 | 9 | Serinc1 |
| Tsc22d12 | 4.41E-40 | 0.258449308 | 0.709 | 0.439 | 8.93E-36 | 9 | Tsc22d1 |
| Tmem593 | 5.73E-40 | 0.399864595 | 0.719 | 0.485 | 1.16E-35 | 9 | Tmem59 |
| Birc63 | 6.26E-40 | 0.53555504 | 0.54 | 0.337 | 1.27E-35 | 9 | Birc6 |

| | | | | | | | |
|---|---|---|---|---|---|---|---|
| Tcf123 | 7.88E-40 | 0.462031694 | 0.602 | 0.371 | 1.60E-35 | 9 | Tcf12 |
| Sertad21 | 9.12E-40 | 0.488820064 | 0.332 | 0.165 | 1.85E-35 | 9 | Sertad2 |
| Herc61 | 9.96E-40 | 0.589204062 | 0.353 | 0.18 | 2.02E-35 | 9 | Herc6 |
| Fam212a2 | 1.00E-39 | 0.410661183 | 0.281 | 0.125 | 2.03E-35 | 9 | Fam212a |
| Chst112 | 1.04E-39 | 0.526011017 | 0.342 | 0.172 | 2.11E-35 | 9 | Chst11 |
| Dynll12 | 1.83E-39 | 0.372934061 | 0.835 | 0.6 | 3.70E-35 | 9 | Dynll1 |
| Bag52 | 2.01E-39 | 0.373362103 | 0.382 | 0.197 | 4.08E-35 | 9 | Bag5 |
| Fnbp1l | 2.73E-39 | 0.32063874 | 0.344 | 0.166 | 5.52E-35 | 9 | Fnbp1l |
| Cdc42ep32 | 2.73E-39 | 0.494609958 | 0.342 | 0.171 | 5.52E-35 | 9 | Cdc42ep3 |
| Sep-42 | 2.91E-39 | 0.419017805 | 0.334 | 0.161 | 5.90E-35 | 9 | 4-Sep |
| Tubb53 | 3.31E-39 | 0.40894093 | 0.587 | 0.376 | 6.71E-35 | 9 | Tubb5 |
| Fkbp31 | 4.72E-39 | 0.391621945 | 0.603 | 0.392 | 9.56E-35 | 9 | Fkbp3 |
| Impdh11 | 5.93E-39 | 0.411447253 | 0.255 | 0.109 | 1.20E-34 | 9 | Impdh1 |
| Nuak11 | 7.32E-39 | 0.510479178 | 0.263 | 0.114 | 1.48E-34 | 9 | Nuak1 |
| Tbl1x2 | 9.40E-39 | 0.473204322 | 0.449 | 0.258 | 1.90E-34 | 9 | Tbl1x |
| Gm121851 | 9.62E-39 | 0.449207533 | 0.284 | 0.129 | 1.95E-34 | 9 | Gm12185 |
| Ifit31 | 1.32E-38 | 0.501436802 | 0.271 | 0.12 | 2.67E-34 | 9 | Ifit3 |
| Dst2 | 1.34E-38 | 0.415322832 | 0.37 | 0.187 | 2.71E-34 | 9 | Dst |
| 2900026A02Rik3 | 1.66E-38 | 0.49285684 | 0.398 | 0.215 | 3.36E-34 | 9 | 2900026A02Rik |
| Rhoc3 | 1.89E-38 | 0.470166007 | 0.498 | 0.292 | 3.82E-34 | 9 | Rhoc |
| Zmiz12 | 2.59E-38 | 0.47723984 | 0.514 | 0.317 | 5.23E-34 | 9 | Zmiz1 |
| Fmnl33 | 2.59E-38 | 0.495671112 | 0.424 | 0.236 | 5.25E-34 | 9 | Fmnl3 |
| Gtf2i2 | 3.14E-38 | 0.428269772 | 0.435 | 0.246 | 6.36E-34 | 9 | Gtf2i |
| Rexo21 | 3.21E-38 | 0.405572612 | 0.492 | 0.293 | 6.51E-34 | 9 | Rexo2 |
| Tprgl3 | 3.23E-38 | 0.411714262 | 0.467 | 0.274 | 6.55E-34 | 9 | Tprgl |
| Esyt22 | 3.41E-38 | 0.519542882 | 0.424 | 0.238 | 6.91E-34 | 9 | Esyt2 |
| Kctd123 | 4.60E-38 | 0.435225177 | 0.455 | 0.254 | 9.32E-34 | 9 | Kctd12 |
| Tm9sf22 | 4.60E-38 | 0.507943485 | 0.433 | 0.252 | 9.32E-34 | 9 | Tm9sf2 |
| Elmo13 | 4.86E-38 | 0.491146069 | 0.586 | 0.361 | 9.84E-34 | 9 | Elmo1 |
| Gpr1461 | 5.24E-38 | 0.396805573 | 0.25 | 0.106 | 1.06E-33 | 9 | Gpr146 |
| Myo62 | 5.86E-38 | 0.400487255 | 0.507 | 0.287 | 1.19E-33 | 9 | Myo6 |
| Adam103 | 6.10E-38 | 0.48617517 | 0.482 | 0.288 | 1.24E-33 | 9 | Adam10 |
| Trim251 | 6.62E-38 | 0.407737189 | 0.306 | 0.146 | 1.34E-33 | 9 | Trim25 |
| Samd9l2 | 7.51E-38 | 0.531129458 | 0.436 | 0.253 | 1.52E-33 | 9 | Samd9l |
| Dennd31 | 1.03E-37 | 0.477784438 | 0.3 | 0.143 | 2.10E-33 | 9 | Dennd3 |
| Kmt2a3 | 1.65E-37 | 0.520900225 | 0.448 | 0.263 | 3.35E-33 | 9 | Kmt2a |
| Arhgap182 | 2.35E-37 | 0.462285379 | 0.545 | 0.337 | 4.75E-33 | 9 | Arhgap18 |
| Stard82 | 2.40E-37 | 0.410020224 | 0.382 | 0.193 | 4.87E-33 | 9 | Stard8 |
| Trib22 | 2.41E-37 | 0.352930298 | 0.369 | 0.182 | 4.87E-33 | 9 | Trib2 |
| Ctla2a4 | 2.59E-37 | 0.350213703 | 0.66 | 0.392 | 5.25E-33 | 9 | Ctla2a |
| Ifnar23 | 2.83E-37 | 0.489621623 | 0.514 | 0.321 | 5.73E-33 | 9 | Ifnar2 |
| Tmem50a4 | 3.90E-37 | 0.44612389 | 0.58 | 0.378 | 7.89E-33 | 9 | Tmem50a |
| H2-T233 | 5.26E-37 | 0.383673418 | 0.682 | 0.454 | 1.07E-32 | 9 | H2-T23 |

| | | | | | | | |
|---|---|---|---|---|---|---|---|
| Ier53 | 6.17E-37 | 0.510110445 | 0.508 | 0.313 | 1.25E-32 | 9 | Ier5 |
| Apc3 | 6.63E-37 | 0.573670337 | 0.422 | 0.245 | 1.34E-32 | 9 | Apc |
| Tns22 | 6.89E-37 | 0.389179779 | 0.301 | 0.141 | 1.39E-32 | 9 | Tns2 |
| Jam33 | 7.68E-37 | 0.289956954 | 0.51 | 0.27 | 1.55E-32 | 9 | Jam3 |
| Hook33 | 9.37E-37 | 0.494327737 | 0.476 | 0.286 | 1.90E-32 | 9 | Hook3 |
| Ext12 | 1.26E-36 | 0.496245133 | 0.327 | 0.163 | 2.55E-32 | 9 | Ext1 |
| Ctnnd11 | 1.45E-36 | 0.320574725 | 0.302 | 0.142 | 2.93E-32 | 9 | Ctnnd1 |
| Ywhaq3 | 2.01E-36 | 0.356873997 | 0.699 | 0.483 | 4.06E-32 | 9 | Ywhaq |
| 4930402H24Rik2 | 2.67E-36 | 0.442645825 | 0.396 | 0.217 | 5.40E-32 | 9 | 4930402H24Rik |
| Itpr11 | 2.81E-36 | 0.443634384 | 0.397 | 0.218 | 5.69E-32 | 9 | Itpr1 |
| Meox12 | 3.19E-36 | 0.367586748 | 0.365 | 0.183 | 6.46E-32 | 9 | Meox1 |
| Tsnax2 | 3.75E-36 | 0.357453828 | 0.327 | 0.162 | 7.59E-32 | 9 | Tsnax |
| N4bp12 | 3.86E-36 | 0.454921134 | 0.333 | 0.172 | 7.81E-32 | 9 | N4bp1 |
| Myo18a2 | 4.95E-36 | 0.37067768 | 0.454 | 0.253 | 1.00E-31 | 9 | Myo18a |
| Nrp22 | 4.97E-36 | 0.445176749 | 0.356 | 0.18 | 1.01E-31 | 9 | Nrp2 |
| Fmo22 | 5.51E-36 | 0.328950828 | 0.522 | 0.283 | 1.12E-31 | 9 | Fmo2 |
| Trim30a3 | 7.88E-36 | 0.549320331 | 0.385 | 0.216 | 1.60E-31 | 9 | Trim30a |
| Vps53 | 9.49E-36 | 0.390650724 | 0.264 | 0.121 | 1.92E-31 | 9 | Vps53 |
| Sntb22 | 1.15E-35 | 0.427549129 | 0.456 | 0.262 | 2.33E-31 | 9 | Sntb2 |
| Zfp664 | 1.37E-35 | 0.326727422 | 0.349 | 0.179 | 2.77E-31 | 9 | Zfp664 |
| Npdc11 | 1.64E-35 | 0.379978027 | 0.28 | 0.13 | 3.33E-31 | 9 | Npdc1 |
| Pbrm12 | 2.23E-35 | 0.491311291 | 0.467 | 0.284 | 4.51E-31 | 9 | Pbrm1 |
| Tom1l21 | 2.70E-35 | 0.488845127 | 0.27 | 0.128 | 5.47E-31 | 9 | Tom1l2 |
| Tmx32 | 3.07E-35 | 0.464193402 | 0.345 | 0.184 | 6.22E-31 | 9 | Tmx3 |
| Sema3c1 | 3.55E-35 | 0.529770112 | 0.321 | 0.16 | 7.20E-31 | 9 | Sema3c |
| Ramp32 | 3.90E-35 | 0.270430267 | 0.494 | 0.261 | 7.90E-31 | 9 | Ramp3 |
| Ash1l2 | 3.95E-35 | 0.486581866 | 0.477 | 0.296 | 7.99E-31 | 9 | Ash1l |
| Map4k51 | 4.42E-35 | 0.440029773 | 0.271 | 0.128 | 8.95E-31 | 9 | Map4k5 |
| Lamb11 | 5.45E-35 | 0.306311034 | 0.291 | 0.133 | 1.10E-30 | 9 | Lamb1 |
| Ncor12 | 5.68E-35 | 0.384022297 | 0.646 | 0.433 | 1.15E-30 | 9 | Ncor1 |
| Exoc6b2 | 6.54E-35 | 0.530611304 | 0.411 | 0.237 | 1.32E-30 | 9 | Exoc6b |
| Rab11a3 | 7.90E-35 | 0.3304028 | 0.503 | 0.304 | 1.60E-30 | 9 | Rab11a |
| Wdfy33 | 8.20E-35 | 0.442673193 | 0.343 | 0.179 | 1.66E-30 | 9 | Wdfy3 |
| Plekha11 | 9.24E-35 | 0.418168022 | 0.282 | 0.135 | 1.87E-30 | 9 | Plekha1 |
| Snx33 | 9.95E-35 | 0.305414065 | 0.671 | 0.456 | 2.02E-30 | 9 | Snx3 |
| Fbxo22 | 1.03E-34 | 0.36883095 | 0.29 | 0.14 | 2.09E-30 | 9 | Fbxo22 |
| St3gal42 | 1.42E-34 | 0.492972169 | 0.432 | 0.252 | 2.87E-30 | 9 | St3gal4 |
| Ier22 | 1.44E-34 | 0.457560302 | 0.464 | 0.28 | 2.91E-30 | 9 | Ier2 |
| Nipbl4 | 1.46E-34 | 0.483183423 | 0.556 | 0.362 | 2.96E-30 | 9 | Nipbl |
| B2m6 | 1.61E-34 | 0.382241395 | 0.964 | 0.683 | 3.26E-30 | 9 | B2m |
| Ptprk1 | 2.37E-34 | 0.359809073 | 0.292 | 0.137 | 4.79E-30 | 9 | Ptprk |
| Prkce2 | 2.80E-34 | 0.447912796 | 0.386 | 0.21 | 5.67E-30 | 9 | Prkce |
| Rgl12 | 2.82E-34 | 0.341286692 | 0.497 | 0.285 | 5.71E-30 | 9 | Rgl1 |

| | | | | | | | |
|---|---|---|---|---|---|---|---|
| Zbtb41 | 3.22E-34 | 0.429252405 | 0.255 | 0.118 | 6.52E-30 | 9 | Zbtb4 |
| Rb1cc12 | 3.26E-34 | 0.464878658 | 0.402 | 0.231 | 6.61E-30 | 9 | Rb1cc1 |
| 21-Mar | 3.73E-34 | 0.393748262 | 0.356 | 0.193 | 7.56E-30 | 9 | 2-Mar |
| Ywhab2 | 4.28E-34 | 0.380428884 | 0.666 | 0.463 | 8.66E-30 | 9 | Ywhab |
| Pea15a2 | 6.25E-34 | 0.38299533 | 0.418 | 0.237 | 1.27E-29 | 9 | Pea15a |
| Cast2 | 6.41E-34 | 0.406214289 | 0.494 | 0.308 | 1.30E-29 | 9 | Cast |
| Mier13 | 8.64E-34 | 0.409441157 | 0.518 | 0.328 | 1.75E-29 | 9 | Mier1 |
| Myo1e3 | 9.07E-34 | 0.41621729 | 0.472 | 0.277 | 1.84E-29 | 9 | Myo1e |
| Sppl2a3 | 9.93E-34 | 0.449429801 | 0.476 | 0.293 | 2.01E-29 | 9 | Sppl2a |
| Gimap63 | 1.43E-33 | 0.352478822 | 0.6 | 0.36 | 2.90E-29 | 9 | Gimap6 |
| Msi22 | 1.52E-33 | 0.411319943 | 0.477 | 0.289 | 3.08E-29 | 9 | Msi2 |
| Pak22 | 1.60E-33 | 0.377118657 | 0.635 | 0.431 | 3.24E-29 | 9 | Pak2 |
| Nae1 | 1.81E-33 | 0.370055957 | 0.303 | 0.153 | 3.67E-29 | 9 | Nae1 |
| Ralbp11 | 2.08E-33 | 0.337872652 | 0.518 | 0.327 | 4.20E-29 | 9 | Ralbp1 |
| Ppp2r2d | 2.08E-33 | 0.40787181 | 0.296 | 0.148 | 4.21E-29 | 9 | Ppp2r2d |
| Nisch2 | 2.12E-33 | 0.429721504 | 0.372 | 0.208 | 4.28E-29 | 9 | Nisch |
| Ddx61 | 2.44E-33 | 0.381435179 | 0.524 | 0.337 | 4.95E-29 | 9 | Ddx6 |
| Lrmda4 | 2.49E-33 | 0.56211351 | 0.572 | 0.364 | 5.04E-29 | 9 | Lrmda |
| Oasl22 | 2.66E-33 | 0.401311928 | 0.273 | 0.128 | 5.38E-29 | 9 | Oasl2 |
| Fkbp1a2 | 4.83E-33 | 0.275483108 | 0.633 | 0.401 | 9.78E-29 | 9 | Fkbp1a |
| Nedd93 | 5.45E-33 | 0.429116842 | 0.525 | 0.315 | 1.10E-28 | 9 | Nedd9 |
| Ctdsp21 | 6.84E-33 | 0.367314952 | 0.334 | 0.175 | 1.39E-28 | 9 | Ctdsp2 |
| Srgap24 | 7.59E-33 | 0.469875797 | 0.486 | 0.3 | 1.54E-28 | 9 | Srgap2 |
| Tnfrsf1a2 | 7.75E-33 | 0.459291019 | 0.39 | 0.218 | 1.57E-28 | 9 | Tnfrsf1a |
| Zeb12 | 7.78E-33 | 0.483036371 | 0.515 | 0.318 | 1.58E-28 | 9 | Zeb1 |
| Rtn43 | 1.04E-32 | 0.396724869 | 0.53 | 0.335 | 2.11E-28 | 9 | Rtn4 |
| Cenpb | 1.08E-32 | 0.305553614 | 0.364 | 0.196 | 2.18E-28 | 9 | Cenpb |
| Atg4a1 | 1.09E-32 | 0.452813731 | 0.27 | 0.129 | 2.21E-28 | 9 | Atg4a |
| Herc21 | 1.20E-32 | 0.47675729 | 0.363 | 0.204 | 2.43E-28 | 9 | Herc2 |
| Frmd4b1 | 1.32E-32 | 0.429328426 | 0.334 | 0.174 | 2.68E-28 | 9 | Frmd4b |
| Dhrs32 | 2.07E-32 | 0.37373025 | 0.411 | 0.227 | 4.18E-28 | 9 | Dhrs3 |
| Usp252 | 2.20E-32 | 0.416961357 | 0.454 | 0.278 | 4.46E-28 | 9 | Usp25 |
| Dusp12 | 2.44E-32 | 0.558612959 | 0.351 | 0.19 | 4.94E-28 | 9 | Dusp1 |
| Ppp3ca3 | 2.75E-32 | 0.425534306 | 0.587 | 0.389 | 5.57E-28 | 9 | Ppp3ca |
| Mxra72 | 3.83E-32 | 0.397860368 | 0.286 | 0.139 | 7.76E-28 | 9 | Mxra7 |
| Psme23 | 3.92E-32 | 0.363637858 | 0.636 | 0.439 | 7.93E-28 | 9 | Psme2 |
| Rtp41 | 4.24E-32 | 0.379403143 | 0.311 | 0.158 | 8.59E-28 | 9 | Rtp4 |
| Fam114a11 | 5.46E-32 | 0.380900807 | 0.269 | 0.129 | 1.11E-27 | 9 | Fam114a1 |
| Psmd72 | 5.76E-32 | 0.322793715 | 0.487 | 0.301 | 1.17E-27 | 9 | Psmd7 |
| Sh3bgrl4 | 6.37E-32 | 0.382991498 | 0.571 | 0.371 | 1.29E-27 | 9 | Sh3bgrl |
| Lrch33 | 9.79E-32 | 0.485059077 | 0.39 | 0.229 | 1.98E-27 | 9 | Lrch3 |
| Pgrmc12 | 1.02E-31 | 0.382169634 | 0.379 | 0.214 | 2.08E-27 | 9 | Pgrmc1 |
| Pqlc11 | 1.16E-31 | 0.346993596 | 0.261 | 0.123 | 2.35E-27 | 9 | Pqlc1 |

| | | | | | | | |
|---|---|---|---|---|---|---|---|
| Ostf14 | 1.39E-31 | 0.308989127 | 0.714 | 0.502 | 2.82E-27 | 9 | Ostf1 |
| Xpr11 | 1.54E-31 | 0.472341325 | 0.291 | 0.151 | 3.13E-27 | 9 | Xpr1 |
| Akap93 | 2.00E-31 | 0.501275462 | 0.393 | 0.234 | 4.05E-27 | 9 | Akap9 |
| Glg12 | 2.57E-31 | 0.462706195 | 0.414 | 0.249 | 5.20E-27 | 9 | Glg1 |
| Rab6a2 | 3.17E-31 | 0.356693729 | 0.379 | 0.212 | 6.41E-27 | 9 | Rab6a |
| Vcl1 | 3.66E-31 | 0.37680476 | 0.311 | 0.158 | 7.42E-27 | 9 | Vcl |
| Zcchc62 | 5.00E-31 | 0.451296563 | 0.441 | 0.275 | 1.01E-26 | 9 | Zcchc6 |
| Stat22 | 5.26E-31 | 0.398880335 | 0.276 | 0.137 | 1.07E-26 | 9 | Stat2 |
| Capns13 | 5.45E-31 | 0.320319582 | 0.53 | 0.345 | 1.10E-26 | 9 | Capns1 |
| Csrp12 | 9.44E-31 | 0.291117378 | 0.339 | 0.179 | 1.91E-26 | 9 | Csrp1 |
| Cyfip13 | 9.96E-31 | 0.33827619 | 0.457 | 0.274 | 2.02E-26 | 9 | Cyfip1 |
| Stim22 | 1.09E-30 | 0.486407444 | 0.349 | 0.196 | 2.21E-26 | 9 | Stim2 |
| Anks12 | 1.14E-30 | 0.423022599 | 0.273 | 0.136 | 2.31E-26 | 9 | Anks1 |
| Sptlc23 | 1.28E-30 | 0.416704156 | 0.408 | 0.241 | 2.60E-26 | 9 | Sptlc2 |
| Mcl13 | 1.42E-30 | 0.416902198 | 0.55 | 0.369 | 2.87E-26 | 9 | Mcl1 |
| Rnf19a1 | 2.04E-30 | 0.39658013 | 0.274 | 0.137 | 4.13E-26 | 9 | Rnf19a |
| Ifi472 | 2.09E-30 | 0.423701338 | 0.326 | 0.175 | 4.23E-26 | 9 | Ifi47 |
| Lamc11 | 2.75E-30 | 0.331422618 | 0.293 | 0.145 | 5.56E-26 | 9 | Lamc1 |
| Rps6ka33 | 3.04E-30 | 0.437401689 | 0.428 | 0.259 | 6.15E-26 | 9 | Rps6ka3 |
| Hnrnpu2 | 3.11E-30 | 0.342890971 | 0.604 | 0.419 | 6.30E-26 | 9 | Hnrnpu |
| Elk41 | 3.77E-30 | 0.376059629 | 0.309 | 0.164 | 7.62E-26 | 9 | Elk4 |
| Tgoln12 | 4.64E-30 | 0.440592769 | 0.407 | 0.243 | 9.39E-26 | 9 | Tgoln1 |
| Rassf13 | 4.73E-30 | 0.440150209 | 0.385 | 0.225 | 9.57E-26 | 9 | Rassf1 |
| Chst152 | 4.96E-30 | 0.301841093 | 0.337 | 0.172 | 1.00E-25 | 9 | Chst15 |
| Spr1 | 6.40E-30 | 0.308169632 | 0.337 | 0.183 | 1.30E-25 | 9 | Spr |
| Ppic1 | 7.13E-30 | 0.370681566 | 0.25 | 0.118 | 1.44E-25 | 9 | Ppic |
| Pgrmc21 | 7.19E-30 | 0.428571048 | 0.275 | 0.14 | 1.46E-25 | 9 | Pgrmc2 |
| Myo1c2 | 1.05E-29 | 0.358894652 | 0.377 | 0.21 | 2.13E-25 | 9 | Myo1c |
| Tspan152 | 1.21E-29 | 0.29348242 | 0.269 | 0.127 | 2.45E-25 | 9 | Tspan15 |
| Zbtb7a2 | 1.37E-29 | 0.49243663 | 0.38 | 0.226 | 2.77E-25 | 9 | Zbtb7a |
| Midn2 | 1.46E-29 | 0.403721904 | 0.339 | 0.188 | 2.95E-25 | 9 | Midn |
| Fubp12 | 1.71E-29 | 0.37474833 | 0.438 | 0.269 | 3.46E-25 | 9 | Fubp1 |
| Tsc22d32 | 2.22E-29 | 0.416931053 | 0.363 | 0.204 | 4.50E-25 | 9 | Tsc22d3 |
| Ywhah3 | 2.37E-29 | 0.305602282 | 0.485 | 0.306 | 4.79E-25 | 9 | Ywhah |
| Hspa53 | 2.44E-29 | 0.370471957 | 0.641 | 0.43 | 4.95E-25 | 9 | Hspa5 |
| Pkn22 | 2.66E-29 | 0.385703238 | 0.459 | 0.291 | 5.39E-25 | 9 | Pkn2 |
| Tmem22 | 3.03E-29 | 0.414788762 | 0.275 | 0.14 | 6.14E-25 | 9 | Tmem2 |
| Tpr2 | 4.06E-29 | 0.370477426 | 0.612 | 0.422 | 8.21E-25 | 9 | Tpr |
| Fus2 | 4.77E-29 | 0.348613482 | 0.576 | 0.385 | 9.66E-25 | 9 | Fus |
| Adamts91 | 4.91E-29 | 0.460410658 | 0.252 | 0.121 | 9.95E-25 | 9 | Adamts9 |
| Ppp1r12a3 | 5.46E-29 | 0.366354142 | 0.629 | 0.421 | 1.11E-24 | 9 | Ppp1r12a |
| Cep3503 | 5.59E-29 | 0.417197958 | 0.35 | 0.201 | 1.13E-24 | 9 | Cep350 |
| Ubxn42 | 5.86E-29 | 0.447994924 | 0.428 | 0.269 | 1.19E-24 | 9 | Ubxn4 |

| | | | | | | | |
|---|---|---|---|---|---|---|---|
| Psme14 | 5.86E-29 | 0.297360498 | 0.789 | 0.563 | 1.19E-24 | 9 | Psme1 |
| Ypel31 | 6.39E-29 | 0.384718292 | 0.432 | 0.264 | 1.29E-24 | 9 | Ypel3 |
| Ywhaz1 | 6.59E-29 | 0.281313806 | 0.702 | 0.488 | 1.33E-24 | 9 | Ywhaz |
| Akap111 | 7.65E-29 | 0.400190561 | 0.255 | 0.127 | 1.55E-24 | 9 | Akap11 |
| Tuba1b1 | 9.32E-29 | 0.310299349 | 0.506 | 0.318 | 1.89E-24 | 9 | Tuba1b |
| Spen1 | 9.65E-29 | 0.410347172 | 0.293 | 0.156 | 1.95E-24 | 9 | Spen |
| Csde12 | 1.13E-28 | 0.303615562 | 0.536 | 0.356 | 2.30E-24 | 9 | Csde1 |
| Golgb12 | 1.25E-28 | 0.419655282 | 0.417 | 0.259 | 2.54E-24 | 9 | Golgb1 |
| 3-Jun | 1.35E-28 | 0.404066188 | 0.718 | 0.51 | 2.74E-24 | 9 | Jun |
| Top2b1 | 1.40E-28 | 0.413298525 | 0.44 | 0.28 | 2.84E-24 | 9 | Top2b |
| St8sia43 | 1.90E-28 | 0.468638258 | 0.422 | 0.256 | 3.84E-24 | 9 | St8sia4 |
| Rap1a3 | 2.42E-28 | 0.335051432 | 0.734 | 0.532 | 4.91E-24 | 9 | Rap1a |
| Mkln11 | 2.55E-28 | 0.409468434 | 0.42 | 0.261 | 5.16E-24 | 9 | Mkln1 |
| Dhrs72 | 2.72E-28 | 0.409417734 | 0.361 | 0.212 | 5.50E-24 | 9 | Dhrs7 |
| Pdia34 | 2.72E-28 | 0.356130943 | 0.646 | 0.439 | 5.51E-24 | 9 | Pdia3 |
| Adgre51 | 2.79E-28 | 0.341053165 | 0.4 | 0.234 | 5.65E-24 | 9 | Adgre5 |
| Rin22 | 2.86E-28 | 0.279159611 | 0.335 | 0.179 | 5.79E-24 | 9 | Rin2 |
| Hmg20b3 | 2.90E-28 | 0.258834001 | 0.43 | 0.257 | 5.87E-24 | 9 | Hmg20b |
| Tmem2343 | 2.97E-28 | 0.270166351 | 0.577 | 0.389 | 6.01E-24 | 9 | Tmem234 |
| Tor1aip13 | 2.99E-28 | 0.372483192 | 0.44 | 0.278 | 6.06E-24 | 9 | Tor1aip1 |
| Kat6a1 | 4.49E-28 | 0.461160013 | 0.374 | 0.225 | 9.10E-24 | 9 | Kat6a |
| Atp11c1 | 4.84E-28 | 0.457404582 | 0.281 | 0.149 | 9.80E-24 | 9 | Atp11c |
| Smurf22 | 6.53E-28 | 0.431078041 | 0.39 | 0.236 | 1.32E-23 | 9 | Smurf2 |
| Casp82 | 7.15E-28 | 0.368266763 | 0.36 | 0.212 | 1.45E-23 | 9 | Casp8 |
| Irf23 | 8.02E-28 | 0.350385956 | 0.513 | 0.338 | 1.62E-23 | 9 | Irf2 |
| Dnajc83 | 8.91E-28 | 0.258122454 | 0.533 | 0.355 | 1.80E-23 | 9 | Dnajc8 |
| H1f02 | 1.27E-27 | 0.338283418 | 0.344 | 0.191 | 2.57E-23 | 9 | H1f0 |
| Fam3c1 | 1.60E-27 | 0.339951293 | 0.319 | 0.176 | 3.23E-23 | 9 | Fam3c |
| Calu2 | 1.73E-27 | 0.378634996 | 0.349 | 0.199 | 3.50E-23 | 9 | Calu |
| Rbms21 | 1.82E-27 | 0.327998719 | 0.296 | 0.157 | 3.69E-23 | 9 | Rbms2 |
| Ccnl22 | 2.24E-27 | 0.423123536 | 0.358 | 0.211 | 4.54E-23 | 9 | Ccnl2 |
| Arap22 | 2.57E-27 | 0.431804924 | 0.317 | 0.175 | 5.20E-23 | 9 | Arap2 |
| Slk3 | 2.67E-27 | 0.393930155 | 0.391 | 0.238 | 5.40E-23 | 9 | Slk |
| Cped12 | 2.87E-27 | 0.289643372 | 0.333 | 0.176 | 5.80E-23 | 9 | Cped1 |
| Hnrnph12 | 3.02E-27 | 0.382297752 | 0.403 | 0.247 | 6.11E-23 | 9 | Hnrnph1 |
| Ankrd13a | 3.33E-27 | 0.302327061 | 0.253 | 0.127 | 6.74E-23 | 9 | Ankrd13a |
| Fndc3b2 | 4.49E-27 | 0.371514443 | 0.404 | 0.241 | 9.08E-23 | 9 | Fndc3b |
| Trove21 | 4.52E-27 | 0.352894508 | 0.258 | 0.129 | 9.16E-23 | 9 | Trove2 |
| Bax3 | 5.37E-27 | 0.300171843 | 0.483 | 0.31 | 1.09E-22 | 9 | Bax |
| Fam96b2 | 7.52E-27 | 0.305938775 | 0.397 | 0.24 | 1.52E-22 | 9 | Fam96b |
| Vat12 | 7.60E-27 | 0.291863098 | 0.292 | 0.152 | 1.54E-22 | 9 | Vat1 |
| Mga1 | 7.72E-27 | 0.399096911 | 0.305 | 0.169 | 1.56E-22 | 9 | Mga |
| Ginm12 | 8.54E-27 | 0.351066836 | 0.319 | 0.177 | 1.73E-22 | 9 | Ginm1 |

| | | | | | | | |
|---|---|---|---|---|---|---|---|
| Abhd17a2 | 8.70E-27 | 0.325717011 | 0.432 | 0.272 | 1.76E-22 | 9 | Abhd17a |
| Luc7l22 | 9.66E-27 | 0.358269186 | 0.592 | 0.401 | 1.95E-22 | 9 | Luc7l2 |
| Bod1l2 | 1.30E-26 | 0.442206492 | 0.422 | 0.27 | 2.64E-22 | 9 | Bod1l |
| Cltc2 | 1.35E-26 | 0.28215371 | 0.502 | 0.327 | 2.73E-22 | 9 | Cltc |
| Phf21a2 | 1.37E-26 | 0.456662757 | 0.402 | 0.25 | 2.77E-22 | 9 | Phf21a |
| Ackr32 | 1.40E-26 | 0.326068537 | 0.292 | 0.149 | 2.84E-22 | 9 | Ackr3 |
| Efcab141 | 1.46E-26 | 0.366228124 | 0.271 | 0.142 | 2.95E-22 | 9 | Efcab14 |
| Spred13 | 1.53E-26 | 0.385752865 | 0.413 | 0.249 | 3.10E-22 | 9 | Spred1 |
| Ddx3x2 | 1.70E-26 | 0.342385578 | 0.47 | 0.305 | 3.45E-22 | 9 | Ddx3x |
| Crk2 | 1.83E-26 | 0.364087761 | 0.332 | 0.19 | 3.70E-22 | 9 | Crk |
| Myl63 | 2.43E-26 | 0.297673483 | 0.953 | 0.771 | 4.92E-22 | 9 | Myl6 |
| Brd43 | 2.45E-26 | 0.422260732 | 0.418 | 0.264 | 4.97E-22 | 9 | Brd4 |
| Ankrd113 | 2.50E-26 | 0.351982735 | 0.559 | 0.377 | 5.06E-22 | 9 | Ankrd11 |
| Csnk1e | 2.76E-26 | 0.275302057 | 0.258 | 0.132 | 5.60E-22 | 9 | Csnk1e |
| Stx121 | 2.93E-26 | 0.327975022 | 0.296 | 0.159 | 5.93E-22 | 9 | Stx12 |
| Pmp222 | 3.34E-26 | 0.271902314 | 0.277 | 0.141 | 6.76E-22 | 9 | Pmp22 |
| Fam208a1 | 3.92E-26 | 0.426982104 | 0.335 | 0.197 | 7.94E-22 | 9 | Fam208a |
| Ifnar12 | 3.97E-26 | 0.399675808 | 0.268 | 0.142 | 8.04E-22 | 9 | Ifnar1 |
| Cyp26b13 | 4.01E-26 | 0.540785544 | 0.371 | 0.213 | 8.12E-22 | 9 | Cyp26b1 |
| Gab21 | 4.93E-26 | 0.276843594 | 0.359 | 0.202 | 9.97E-22 | 9 | Gab2 |
| Kif1b2 | 6.16E-26 | 0.369498849 | 0.39 | 0.24 | 1.25E-21 | 9 | Kif1b |
| mt-Cytb4 | 6.51E-26 | 0.284002691 | 0.979 | 0.693 | 1.32E-21 | 9 | mt-Cytb |
| Ubn12 | 7.46E-26 | 0.364119296 | 0.396 | 0.243 | 1.51E-21 | 9 | Ubn1 |
| Nmi2 | 8.55E-26 | 0.367481348 | 0.385 | 0.235 | 1.73E-21 | 9 | Nmi |
| Lmna2 | 1.04E-25 | 0.378393487 | 0.342 | 0.195 | 2.10E-21 | 9 | Lmna |
| Vcp1 | 1.12E-25 | 0.313410814 | 0.507 | 0.34 | 2.26E-21 | 9 | Vcp |
| Prkar2a1 | 1.18E-25 | 0.289904318 | 0.279 | 0.148 | 2.39E-21 | 9 | Prkar2a |
| Ctbp1 | 1.21E-25 | 0.309692212 | 0.332 | 0.191 | 2.44E-21 | 9 | Ctbp1 |
| St3gal63 | 1.35E-25 | 0.383378491 | 0.486 | 0.314 | 2.74E-21 | 9 | St3gal6 |
| Pcm12 | 1.86E-25 | 0.413072604 | 0.397 | 0.248 | 3.77E-21 | 9 | Pcm1 |
| Rcn21 | 2.31E-25 | 0.331914959 | 0.265 | 0.14 | 4.67E-21 | 9 | Rcn2 |
| Phf20l13 | 2.37E-25 | 0.434525323 | 0.436 | 0.284 | 4.79E-21 | 9 | Phf20l1 |
| Cntln1 | 3.15E-25 | 0.425041686 | 0.279 | 0.152 | 6.38E-21 | 9 | Cntln |
| St51 | 3.22E-25 | 0.296823446 | 0.282 | 0.149 | 6.52E-21 | 9 | St5 |
| Vamp32 | 3.23E-25 | 0.275760559 | 0.314 | 0.174 | 6.54E-21 | 9 | Vamp3 |
| Arl6ip53 | 3.35E-25 | 0.352284129 | 0.365 | 0.219 | 6.79E-21 | 9 | Arl6ip5 |
| Wac3 | 3.38E-25 | 0.394197219 | 0.477 | 0.319 | 6.85E-21 | 9 | Wac |
| Nsd13 | 4.07E-25 | 0.414176711 | 0.506 | 0.347 | 8.25E-21 | 9 | Nsd1 |
| Tnfsf121 | 4.79E-25 | 0.326528104 | 0.273 | 0.145 | 9.71E-21 | 9 | Tnfsf12 |
| Dnajc33 | 5.44E-25 | 0.310765774 | 0.54 | 0.367 | 1.10E-20 | 9 | Dnajc3 |
| Nfe2l23 | 7.52E-25 | 0.300351492 | 0.465 | 0.298 | 1.52E-20 | 9 | Nfe2l2 |
| Cpne31 | 1.06E-24 | 0.301681743 | 0.353 | 0.209 | 2.15E-20 | 9 | Cpne3 |
| Eif4g11 | 1.13E-24 | 0.341602205 | 0.377 | 0.233 | 2.30E-20 | 9 | Eif4g1 |

| | | | | | | | |
|---|---|---|---|---|---|---|---|
| Nek72 | 1.60E-24 | 0.356321588 | 0.39 | 0.246 | 3.23E-20 | 9 | Nek7 |
| Srrm23 | 1.61E-24 | 0.3190407 | 0.655 | 0.465 | 3.25E-20 | 9 | Srrm2 |
| Asap21 | 1.93E-24 | 0.300314633 | 0.254 | 0.129 | 3.91E-20 | 9 | Asap2 |
| Dync1i22 | 2.05E-24 | 0.265141587 | 0.591 | 0.415 | 4.15E-20 | 9 | Dync1i2 |
| Fip1l1 | 2.49E-24 | 0.331189809 | 0.334 | 0.197 | 5.04E-20 | 9 | Fip1l1 |
| Srpr2 | 2.59E-24 | 0.369131847 | 0.387 | 0.243 | 5.25E-20 | 9 | Srpr |
| Ubr51 | 2.95E-24 | 0.390978039 | 0.328 | 0.196 | 5.97E-20 | 9 | Ubr5 |
| Rad211 | 3.31E-24 | 0.30956756 | 0.324 | 0.188 | 6.71E-20 | 9 | Rad21 |
| Ssu722 | 4.82E-24 | 0.290969399 | 0.466 | 0.309 | 9.76E-20 | 9 | Ssu72 |
| Uvrag3 | 5.15E-24 | 0.390996265 | 0.412 | 0.265 | 1.04E-19 | 9 | Uvrag |
| Cnih44 | 5.47E-24 | 0.255425248 | 0.401 | 0.251 | 1.11E-19 | 9 | Cnih4 |
| Fxyd53 | 5.92E-24 | 0.287903365 | 0.608 | 0.407 | 1.20E-19 | 9 | Fxyd5 |
| Rtl8a2 | 6.85E-24 | 0.307456957 | 0.313 | 0.174 | 1.39E-19 | 9 | Rtl8a |
| Helz3 | 7.51E-24 | 0.40066733 | 0.291 | 0.167 | 1.52E-19 | 9 | Helz |
| Prrc2b2 | 8.06E-24 | 0.353492607 | 0.366 | 0.227 | 1.63E-19 | 9 | Prrc2b |
| G3bp21 | 8.33E-24 | 0.356550762 | 0.427 | 0.278 | 1.69E-19 | 9 | G3bp2 |
| Slc10a71 | 8.36E-24 | 0.36297332 | 0.266 | 0.145 | 1.69E-19 | 9 | Slc10a7 |
| Ubr31 | 1.04E-23 | 0.373914552 | 0.334 | 0.198 | 2.11E-19 | 9 | Ubr3 |
| mt-Nd14 | 1.44E-23 | 0.294296906 | 0.845 | 0.593 | 2.91E-19 | 9 | mt-Nd1 |
| Stk32 | 1.72E-23 | 0.416170152 | 0.29 | 0.164 | 3.48E-19 | 9 | Stk3 |
| Ifngr14 | 2.04E-23 | 0.294985401 | 0.577 | 0.389 | 4.14E-19 | 9 | Ifngr1 |
| Psmd1 | 2.20E-23 | 0.292214737 | 0.344 | 0.207 | 4.46E-19 | 9 | Psmd1 |
| Ppp4r3b2 | 2.21E-23 | 0.340455925 | 0.367 | 0.23 | 4.47E-19 | 9 | Ppp4r3b |
| Sfpq2 | 2.44E-23 | 0.357400252 | 0.441 | 0.296 | 4.95E-19 | 9 | Sfpq |
| Man1a22 | 2.44E-23 | 0.359753265 | 0.364 | 0.224 | 4.95E-19 | 9 | Man1a2 |
| Znrf12 | 2.83E-23 | 0.382605193 | 0.316 | 0.187 | 5.73E-19 | 9 | Znrf1 |
| Zfand33 | 2.84E-23 | 0.386551365 | 0.449 | 0.299 | 5.75E-19 | 9 | Zfand3 |
| Irf2bpl2 | 2.90E-23 | 0.303618956 | 0.391 | 0.241 | 5.88E-19 | 9 | Irf2bpl |
| Yipf42 | 2.94E-23 | 0.310127997 | 0.358 | 0.219 | 5.96E-19 | 9 | Yipf4 |
| Golim41 | 3.06E-23 | 0.359608794 | 0.307 | 0.178 | 6.19E-19 | 9 | Golim4 |
| Numa11 | 3.38E-23 | 0.351804707 | 0.324 | 0.193 | 6.85E-19 | 9 | Numa1 |
| Usp11 | 3.71E-23 | 0.35344253 | 0.263 | 0.144 | 7.51E-19 | 9 | Usp1 |
| Mllt103 | 4.83E-23 | 0.359797606 | 0.419 | 0.272 | 9.78E-19 | 9 | Mllt10 |
| Arid21 | 5.45E-23 | 0.401619675 | 0.273 | 0.154 | 1.10E-18 | 9 | Arid2 |
| Ica12 | 6.16E-23 | 0.264485286 | 0.313 | 0.174 | 1.25E-18 | 9 | Ica1 |
| Kmt2c3 | 6.47E-23 | 0.38174594 | 0.448 | 0.304 | 1.31E-18 | 9 | Kmt2c |
| Gatad2a3 | 7.38E-23 | 0.371812878 | 0.358 | 0.22 | 1.49E-18 | 9 | Gatad2a |
| Srsf114 | 8.08E-23 | 0.321718575 | 0.507 | 0.348 | 1.64E-18 | 9 | Srsf11 |
| Akap132 | 9.33E-23 | 0.351328612 | 0.564 | 0.391 | 1.89E-18 | 9 | Akap13 |
| Nras1 | 9.46E-23 | 0.333617026 | 0.268 | 0.148 | 1.92E-18 | 9 | Nras |
| Med13l1 | 1.11E-22 | 0.485363974 | 0.317 | 0.192 | 2.25E-18 | 9 | Med13l |
| Spred21 | 1.16E-22 | 0.380656333 | 0.296 | 0.169 | 2.35E-18 | 9 | Spred2 |
| Kif13a2 | 1.29E-22 | 0.308232556 | 0.303 | 0.171 | 2.62E-18 | 9 | Kif13a |

| | | | | | | | |
|---|---|---|---|---|---|---|---|
| Wbp4 | 1.32E-22 | 0.276747441 | 0.366 | 0.227 | 2.66E-18 | 9 | Wbp4 |
| Wnk13 | 1.37E-22 | 0.305431708 | 0.531 | 0.36 | 2.78E-18 | 9 | Wnk1 |
| Baz1b1 | 1.37E-22 | 0.318879617 | 0.322 | 0.191 | 2.78E-18 | 9 | Baz1b |
| Gkap1 | 1.39E-22 | 0.315144454 | 0.259 | 0.14 | 2.82E-18 | 9 | Gkap1 |
| 2010111I01Rik2 | 1.43E-22 | 0.280115484 | 0.518 | 0.346 | 2.89E-18 | 9 | 2010111I01Rik |
| Fbxl31 | 1.79E-22 | 0.347259443 | 0.274 | 0.154 | 3.63E-18 | 9 | Fbxl3 |
| Exoc62 | 2.01E-22 | 0.396096819 | 0.29 | 0.169 | 4.07E-18 | 9 | Exoc6 |
| Pros11 | 2.02E-22 | 0.374724747 | 0.269 | 0.148 | 4.08E-18 | 9 | Pros1 |
| Sft2d12 | 2.09E-22 | 0.347058629 | 0.277 | 0.158 | 4.24E-18 | 9 | Sft2d1 |
| Ddx502 | 2.42E-22 | 0.279220827 | 0.439 | 0.29 | 4.89E-18 | 9 | Ddx50 |
| Atxn21 | 2.57E-22 | 0.308282342 | 0.253 | 0.137 | 5.21E-18 | 9 | Atxn2 |
| Ntf32 | 3.21E-22 | 0.37008418 | 0.316 | 0.179 | 6.51E-18 | 9 | Ntf3 |
| Snrnp2001 | 3.89E-22 | 0.351349348 | 0.266 | 0.15 | 7.88E-18 | 9 | Snrnp200 |
| Ifi352 | 4.34E-22 | 0.320220887 | 0.417 | 0.271 | 8.80E-18 | 9 | Ifi35 |
| Huwe12 | 4.70E-22 | 0.335529844 | 0.43 | 0.282 | 9.51E-18 | 9 | Huwe1 |
| Crebbp3 | 5.30E-22 | 0.357995088 | 0.445 | 0.303 | 1.07E-17 | 9 | Crebbp |
| Herc12 | 5.30E-22 | 0.375214861 | 0.39 | 0.255 | 1.07E-17 | 9 | Herc1 |
| Selenot1 | 5.36E-22 | 0.345124716 | 0.339 | 0.21 | 1.09E-17 | 9 | Selenot |
| mt-Atp65 | 5.48E-22 | 0.290198797 | 0.995 | 0.747 | 1.11E-17 | 9 | mt-Atp6 |
| Ccdc471 | 5.52E-22 | 0.335577919 | 0.317 | 0.19 | 1.12E-17 | 9 | Ccdc47 |
| Tm9sf33 | 5.78E-22 | 0.340399659 | 0.481 | 0.332 | 1.17E-17 | 9 | Tm9sf3 |
| Degs12 | 5.91E-22 | 0.34967559 | 0.309 | 0.182 | 1.20E-17 | 9 | Degs1 |
| Anapc51 | 6.35E-22 | 0.265513275 | 0.346 | 0.209 | 1.28E-17 | 9 | Anapc5 |
| G3bp12 | 6.54E-22 | 0.301144582 | 0.449 | 0.3 | 1.32E-17 | 9 | G3bp1 |
| Dpysl22 | 6.77E-22 | 0.359049141 | 0.296 | 0.172 | 1.37E-17 | 9 | Dpysl2 |
| Acap24 | 6.86E-22 | 0.383281693 | 0.412 | 0.267 | 1.39E-17 | 9 | Acap2 |
| Lmo23 | 7.36E-22 | 0.266428355 | 0.349 | 0.205 | 1.49E-17 | 9 | Lmo2 |
| Wasl2 | 8.00E-22 | 0.279151967 | 0.372 | 0.235 | 1.62E-17 | 9 | Wasl |
| Brd22 | 8.55E-22 | 0.33683621 | 0.387 | 0.251 | 1.73E-17 | 9 | Brd2 |
| Etnk12 | 8.79E-22 | 0.35020815 | 0.355 | 0.223 | 1.78E-17 | 9 | Etnk1 |
| Svil2 | 8.82E-22 | 0.355747632 | 0.379 | 0.241 | 1.78E-17 | 9 | Svil |
| Tmem2051 | 9.43E-22 | 0.279587894 | 0.28 | 0.157 | 1.91E-17 | 9 | Tmem205 |
| Pum13 | 1.04E-21 | 0.336027725 | 0.395 | 0.256 | 2.11E-17 | 9 | Pum1 |
| Thoc23 | 1.10E-21 | 0.394766381 | 0.407 | 0.272 | 2.23E-17 | 9 | Thoc2 |
| Kpna44 | 1.32E-21 | 0.340274782 | 0.47 | 0.321 | 2.67E-17 | 9 | Kpna4 |
| Elf22 | 1.53E-21 | 0.324469529 | 0.444 | 0.298 | 3.09E-17 | 9 | Elf2 |
| Tmed93 | 1.57E-21 | 0.27261858 | 0.562 | 0.4 | 3.18E-17 | 9 | Tmed9 |
| Mydgf1 | 1.59E-21 | 0.35653286 | 0.359 | 0.23 | 3.21E-17 | 9 | Mydgf |
| Setx1 | 1.79E-21 | 0.374621579 | 0.291 | 0.172 | 3.62E-17 | 9 | Setx |
| Exoc44 | 1.98E-21 | 0.384016365 | 0.472 | 0.32 | 4.02E-17 | 9 | Exoc4 |
| Prpf4b3 | 1.99E-21 | 0.267158763 | 0.485 | 0.33 | 4.03E-17 | 9 | Prpf4b |
| Airn2 | 2.19E-21 | 0.380224385 | 0.392 | 0.246 | 4.42E-17 | 9 | Airn |
| Yif1b | 2.38E-21 | 0.283046059 | 0.263 | 0.146 | 4.82E-17 | 9 | Yif1b |

| | | | | | | | |
|---|---|---|---|---|---|---|---|
| Prdm21 | 2.47E-21 | 0.437765211 | 0.3 | 0.183 | 5.01E-17 | 9 | Prdm2 |
| Cdk123 | 2.50E-21 | 0.368297991 | 0.361 | 0.228 | 5.07E-17 | 9 | Cdk12 |
| Stat33 | 2.52E-21 | 0.35224268 | 0.404 | 0.264 | 5.10E-17 | 9 | Stat3 |
| Gimap82 | 2.95E-21 | 0.332970027 | 0.29 | 0.167 | 5.98E-17 | 9 | Gimap8 |
| Brd91 | 2.96E-21 | 0.343861193 | 0.292 | 0.171 | 6.00E-17 | 9 | Brd9 |
| Ubn23 | 3.40E-21 | 0.404074727 | 0.375 | 0.242 | 6.88E-17 | 9 | Ubn2 |
| Setd53 | 3.51E-21 | 0.349794564 | 0.396 | 0.262 | 7.12E-17 | 9 | Setd5 |
| Zdhhc171 | 3.58E-21 | 0.366634577 | 0.264 | 0.148 | 7.26E-17 | 9 | Zdhhc17 |
| Ppp4r21 | 3.67E-21 | 0.299940611 | 0.355 | 0.222 | 7.43E-17 | 9 | Ppp4r2 |
| Fos3 | 4.29E-21 | 0.452389845 | 0.469 | 0.314 | 8.69E-17 | 9 | Fos |
| Calr3 | 5.64E-21 | 0.293589951 | 0.571 | 0.405 | 1.14E-16 | 9 | Calr |
| Fam120a1 | 6.51E-21 | 0.375001822 | 0.3 | 0.18 | 1.32E-16 | 9 | Fam120a |
| Chd62 | 6.71E-21 | 0.400717299 | 0.339 | 0.215 | 1.36E-16 | 9 | Chd6 |
| Thra1 | 6.74E-21 | 0.282956251 | 0.287 | 0.162 | 1.36E-16 | 9 | Thra |
| Map3k12 | 7.08E-21 | 0.309093735 | 0.322 | 0.197 | 1.43E-16 | 9 | Map3k1 |
| Atp2b13 | 7.47E-21 | 0.340943899 | 0.482 | 0.33 | 1.51E-16 | 9 | Atp2b1 |
| Pik3cb2 | 8.12E-21 | 0.331691614 | 0.258 | 0.144 | 1.64E-16 | 9 | Pik3cb |
| Ppig3 | 8.17E-21 | 0.305281748 | 0.457 | 0.314 | 1.65E-16 | 9 | Ppig |
| Mphosph81 | 9.08E-21 | 0.355289243 | 0.33 | 0.205 | 1.84E-16 | 9 | Mphosph8 |
| Map3k52 | 1.02E-20 | 0.359928269 | 0.313 | 0.19 | 2.06E-16 | 9 | Map3k5 |
| Ubl32 | 1.03E-20 | 0.297629663 | 0.372 | 0.236 | 2.09E-16 | 9 | Ubl3 |
| Tra2a4 | 1.21E-20 | 0.363627928 | 0.407 | 0.272 | 2.46E-16 | 9 | Tra2a |
| Unc119b1 | 1.23E-20 | 0.25079576 | 0.255 | 0.138 | 2.49E-16 | 9 | Unc119b |
| Emc73 | 1.45E-20 | 0.303911678 | 0.407 | 0.27 | 2.94E-16 | 9 | Emc7 |
| Tmtc22 | 1.52E-20 | 0.355006888 | 0.313 | 0.183 | 3.08E-16 | 9 | Tmtc2 |
| Rc3h11 | 1.56E-20 | 0.321010297 | 0.33 | 0.206 | 3.16E-16 | 9 | Rc3h1 |
| Xrn12 | 1.63E-20 | 0.355568918 | 0.324 | 0.202 | 3.30E-16 | 9 | Xrn1 |
| Hectd12 | 1.75E-20 | 0.302003621 | 0.444 | 0.298 | 3.55E-16 | 9 | Hectd1 |
| Sh2d3c2 | 1.92E-20 | 0.316802503 | 0.271 | 0.155 | 3.88E-16 | 9 | Sh2d3c |
| Cnot61 | 2.26E-20 | 0.275329125 | 0.274 | 0.157 | 4.57E-16 | 9 | Cnot6 |
| Tln14 | 2.42E-20 | 0.296117746 | 0.57 | 0.402 | 4.90E-16 | 9 | Tln1 |
| Bbx2 | 2.57E-20 | 0.401512803 | 0.346 | 0.221 | 5.20E-16 | 9 | Bbx |
| Irf91 | 2.91E-20 | 0.253902264 | 0.274 | 0.155 | 5.90E-16 | 9 | Irf9 |
| Cflar2 | 2.97E-20 | 0.326151103 | 0.358 | 0.23 | 6.01E-16 | 9 | Cflar |
| Tulp4 | 3.10E-20 | 0.276011197 | 0.308 | 0.186 | 6.27E-16 | 9 | Tulp4 |
| Ttc143 | 3.10E-20 | 0.401646196 | 0.334 | 0.214 | 6.28E-16 | 9 | Ttc14 |
| Blmh | 3.14E-20 | 0.321175272 | 0.261 | 0.15 | 6.37E-16 | 9 | Blmh |
| Trpm72 | 3.53E-20 | 0.329061038 | 0.327 | 0.202 | 7.14E-16 | 9 | Trpm7 |
| Lrp102 | 3.63E-20 | 0.354994569 | 0.305 | 0.186 | 7.35E-16 | 9 | Lrp10 |
| Dennd6a1 | 4.68E-20 | 0.317888432 | 0.301 | 0.18 | 9.48E-16 | 9 | Dennd6a |
| Arhgef14 | 7.46E-20 | 0.359203813 | 0.374 | 0.247 | 1.51E-15 | 9 | Arhgef1 |
| Gpatch83 | 7.59E-20 | 0.409471272 | 0.38 | 0.256 | 1.54E-15 | 9 | Gpatch8 |
| Fam13b2 | 7.98E-20 | 0.379419181 | 0.252 | 0.145 | 1.61E-15 | 9 | Fam13b |

| | | | | | | | |
|---|---|---|---|---|---|---|---|
| Rab11b2 | 1.08E-19 | 0.283427393 | 0.397 | 0.263 | 2.19E-15 | 9 | Rab11b |
| Sec631 | 1.12E-19 | 0.323251413 | 0.323 | 0.201 | 2.26E-15 | 9 | Sec63 |
| Myo9a2 | 1.26E-19 | 0.355873539 | 0.286 | 0.172 | 2.55E-15 | 9 | Myo9a |
| Papola1 | 1.56E-19 | 0.255705985 | 0.439 | 0.302 | 3.17E-15 | 9 | Papola |
| Arl5a1 | 1.87E-19 | 0.256740121 | 0.269 | 0.156 | 3.79E-15 | 9 | Arl5a |
| Prpf81 | 2.37E-19 | 0.316949398 | 0.297 | 0.181 | 4.79E-15 | 9 | Prpf8 |
| Ripply32 | 2.64E-19 | 0.295081222 | 0.259 | 0.145 | 5.35E-15 | 9 | Ripply3 |
| Slc44a12 | 3.02E-19 | 0.288602031 | 0.27 | 0.154 | 6.12E-15 | 9 | Slc44a1 |
| Zscan261 | 3.07E-19 | 0.263128596 | 0.254 | 0.143 | 6.21E-15 | 9 | Zscan26 |
| Lrrfip13 | 3.09E-19 | 0.284922694 | 0.527 | 0.376 | 6.25E-15 | 9 | Lrrfip1 |
| Canx4 | 3.14E-19 | 0.318596392 | 0.441 | 0.304 | 6.35E-15 | 9 | Canx |
| Iws11 | 3.29E-19 | 0.343003147 | 0.261 | 0.153 | 6.67E-15 | 9 | Iws1 |
| H2-T223 | 3.41E-19 | 0.268319172 | 0.413 | 0.279 | 6.91E-15 | 9 | H2-T22 |
| Lman12 | 3.69E-19 | 0.301972404 | 0.366 | 0.236 | 7.47E-15 | 9 | Lman1 |
| Cdk11b2 | 3.87E-19 | 0.311979466 | 0.367 | 0.242 | 7.84E-15 | 9 | Cdk11b |
| Eif2ak22 | 4.07E-19 | 0.321494533 | 0.275 | 0.162 | 8.24E-15 | 9 | Eif2ak2 |
| Zdhhc182 | 4.25E-19 | 0.337060006 | 0.281 | 0.169 | 8.60E-15 | 9 | Zdhhc18 |
| Jmjd1c4 | 4.32E-19 | 0.323836531 | 0.445 | 0.305 | 8.74E-15 | 9 | Jmjd1c |
| Atp2a21 | 4.70E-19 | 0.269910036 | 0.323 | 0.198 | 9.52E-15 | 9 | Atp2a2 |
| Dnajc12 | 5.89E-19 | 0.395161025 | 0.464 | 0.32 | 1.19E-14 | 9 | Dnajc1 |
| Sltm2 | 6.27E-19 | 0.298629676 | 0.401 | 0.269 | 1.27E-14 | 9 | Sltm |
| Rap1b3 | 7.00E-19 | 0.261245805 | 0.599 | 0.441 | 1.42E-14 | 9 | Rap1b |
| Pdia64 | 7.70E-19 | 0.299988909 | 0.478 | 0.335 | 1.56E-14 | 9 | Pdia6 |
| Dnm22 | 8.89E-19 | 0.295917808 | 0.386 | 0.257 | 1.80E-14 | 9 | Dnm2 |
| Nfat52 | 1.12E-18 | 0.344787699 | 0.401 | 0.265 | 2.27E-14 | 9 | Nfat5 |
| Tlk11 | 1.55E-18 | 0.300556005 | 0.339 | 0.221 | 3.13E-14 | 9 | Tlk1 |
| Acvr2a1 | 1.60E-18 | 0.334422206 | 0.255 | 0.148 | 3.24E-14 | 9 | Acvr2a |
| Nptn2 | 1.64E-18 | 0.338062487 | 0.371 | 0.247 | 3.32E-14 | 9 | Nptn |
| Lpp2 | 1.68E-18 | 0.332756413 | 0.422 | 0.284 | 3.39E-14 | 9 | Lpp |
| Sdf43 | 2.06E-18 | 0.285296171 | 0.44 | 0.307 | 4.16E-14 | 9 | Sdf4 |
| Phip2 | 2.58E-18 | 0.368400588 | 0.377 | 0.254 | 5.23E-14 | 9 | Phip |
| Ctcf2 | 3.38E-18 | 0.255335441 | 0.406 | 0.275 | 6.84E-14 | 9 | Ctcf |
| Slc35b11 | 3.50E-18 | 0.291083554 | 0.271 | 0.162 | 7.08E-14 | 9 | Slc35b1 |
| Mia31 | 4.03E-18 | 0.332427774 | 0.276 | 0.168 | 8.17E-14 | 9 | Mia3 |
| Pnisr3 | 4.15E-18 | 0.342967494 | 0.387 | 0.262 | 8.41E-14 | 9 | Pnisr |
| Rbm53 | 4.37E-18 | 0.349149801 | 0.354 | 0.238 | 8.84E-14 | 9 | Rbm5 |
| Ppp6r32 | 4.50E-18 | 0.349227669 | 0.302 | 0.189 | 9.11E-14 | 9 | Ppp6r3 |
| Acin12 | 4.59E-18 | 0.27395214 | 0.45 | 0.314 | 9.29E-14 | 9 | Acin1 |
| Zfp2923 | 5.27E-18 | 0.312640793 | 0.374 | 0.249 | 1.07E-13 | 9 | Zfp292 |
| Mdfic2 | 5.38E-18 | 0.287399311 | 0.374 | 0.243 | 1.09E-13 | 9 | Mdfic |
| Ccnt11 | 5.63E-18 | 0.317057063 | 0.274 | 0.166 | 1.14E-13 | 9 | Ccnt1 |
| Lbr2 | 6.02E-18 | 0.360028292 | 0.254 | 0.151 | 1.22E-13 | 9 | Lbr |
| Ddx172 | 8.36E-18 | 0.320603608 | 0.397 | 0.271 | 1.69E-13 | 9 | Ddx17 |

| | | | | | | | |
|---|---|---|---|---|---|---|---|
| Zdhhc203 | 9.03E-18 | 0.370094378 | 0.348 | 0.228 | 1.83E-13 | 9 | Zdhhc20 |
| Ankhd12 | 9.52E-18 | 0.342920919 | 0.364 | 0.242 | 1.93E-13 | 9 | Ankhd1 |
| Hivep23 | 1.05E-17 | 0.334758958 | 0.43 | 0.296 | 2.13E-13 | 9 | Hivep2 |
| Pitpna2 | 1.13E-17 | 0.255350838 | 0.461 | 0.326 | 2.28E-13 | 9 | Pitpna |
| Pnn3 | 1.13E-17 | 0.275670443 | 0.392 | 0.265 | 2.29E-13 | 9 | Pnn |
| Trim441 | 1.17E-17 | 0.262629136 | 0.348 | 0.225 | 2.37E-13 | 9 | Trim44 |
| Kat2b1 | 1.36E-17 | 0.295736921 | 0.263 | 0.157 | 2.75E-13 | 9 | Kat2b |
| Mbd52 | 1.47E-17 | 0.353332593 | 0.337 | 0.218 | 2.99E-13 | 9 | Mbd5 |
| Shoc21 | 1.65E-17 | 0.273155393 | 0.338 | 0.22 | 3.34E-13 | 9 | Shoc2 |
| Ddx581 | 1.66E-17 | 0.328551572 | 0.264 | 0.158 | 3.36E-13 | 9 | Ddx58 |
| Rbm254 | 1.78E-17 | 0.323479582 | 0.461 | 0.332 | 3.60E-13 | 9 | Rbm25 |
| Thap3 | 1.95E-17 | 0.296970346 | 0.266 | 0.16 | 3.95E-13 | 9 | Thap3 |
| Irgm11 | 2.02E-17 | 0.254771723 | 0.253 | 0.146 | 4.10E-13 | 9 | Irgm1 |
| Atxn72 | 2.59E-17 | 0.300050514 | 0.271 | 0.164 | 5.25E-13 | 9 | Atxn7 |
| Polr2a3 | 2.60E-17 | 0.341304151 | 0.402 | 0.279 | 5.26E-13 | 9 | Polr2a |
| Pja22 | 3.10E-17 | 0.250291098 | 0.321 | 0.203 | 6.28E-13 | 9 | Pja2 |
| Ndel11 | 3.61E-17 | 0.323464265 | 0.279 | 0.173 | 7.31E-13 | 9 | Ndel1 |
| Trappc82 | 4.78E-17 | 0.307749947 | 0.268 | 0.165 | 9.67E-13 | 9 | Trappc8 |
| Rasa12 | 4.90E-17 | 0.33469817 | 0.365 | 0.245 | 9.92E-13 | 9 | Rasa1 |
| Kansl13 | 4.94E-17 | 0.28927326 | 0.476 | 0.338 | 9.99E-13 | 9 | Kansl1 |
| Add11 | 5.02E-17 | 0.29269079 | 0.326 | 0.213 | 1.02E-12 | 9 | Add1 |
| Lnpep3 | 6.56E-17 | 0.341004211 | 0.379 | 0.258 | 1.33E-12 | 9 | Lnpep |
| Kidins2202 | 7.07E-17 | 0.315288264 | 0.254 | 0.154 | 1.43E-12 | 9 | Kidins220 |
| Arid4b3 | 8.58E-17 | 0.334293534 | 0.416 | 0.291 | 1.74E-12 | 9 | Arid4b |
| Krit12 | 1.08E-16 | 0.354748133 | 0.284 | 0.181 | 2.18E-12 | 9 | Krit1 |
| Braf1 | 1.10E-16 | 0.369720087 | 0.26 | 0.157 | 2.23E-12 | 9 | Braf |
| Zcchc73 | 1.18E-16 | 0.368692375 | 0.418 | 0.299 | 2.38E-12 | 9 | Zcchc7 |
| Emc31 | 1.30E-16 | 0.361074737 | 0.275 | 0.172 | 2.63E-12 | 9 | Emc3 |
| mt-Nd34 | 1.42E-16 | 0.25821834 | 0.6 | 0.444 | 2.88E-12 | 9 | mt-Nd3 |
| Lmbrd11 | 1.69E-16 | 0.299175784 | 0.263 | 0.161 | 3.42E-12 | 9 | Lmbrd1 |
| Tmem2082 | 1.82E-16 | 0.291694201 | 0.351 | 0.237 | 3.69E-12 | 9 | Tmem208 |
| Hnrnpr | 1.93E-16 | 0.260646006 | 0.351 | 0.234 | 3.91E-12 | 9 | Hnrnpr |
| Ep4002 | 1.93E-16 | 0.348418869 | 0.312 | 0.206 | 3.91E-12 | 9 | Ep400 |
| Yme1l11 | 2.18E-16 | 0.290658709 | 0.302 | 0.194 | 4.41E-12 | 9 | Yme1l1 |
| Unc502 | 2.29E-16 | 0.258205662 | 0.324 | 0.211 | 4.63E-12 | 9 | Unc50 |
| Tacc13 | 2.43E-16 | 0.26479574 | 0.433 | 0.303 | 4.92E-12 | 9 | Tacc1 |
| Ap2b11 | 2.44E-16 | 0.254315986 | 0.379 | 0.259 | 4.95E-12 | 9 | Ap2b1 |
| Map3k32 | 2.50E-16 | 0.334660702 | 0.281 | 0.177 | 5.05E-12 | 9 | Map3k3 |
| Ifi442 | 3.18E-16 | 0.255006113 | 0.25 | 0.144 | 6.43E-12 | 9 | Ifi44 |
| Fchsd22 | 3.76E-16 | 0.354044859 | 0.259 | 0.158 | 7.61E-12 | 9 | Fchsd2 |
| U2surp1 | 4.40E-16 | 0.289049652 | 0.383 | 0.267 | 8.92E-12 | 9 | U2surp |
| Zfp6382 | 4.93E-16 | 0.303032752 | 0.397 | 0.28 | 9.99E-12 | 9 | Zfp638 |
| Saraf3 | 5.16E-16 | 0.297231497 | 0.428 | 0.306 | 1.05E-11 | 9 | Saraf |

| | | | | | | | |
|---|---|---|---|---|---|---|---|
| Trps13 | 5.26E-16 | 0.31792732 | 0.355 | 0.234 | 1.07E-11 | 9 | Trps1 |
| Ski3 | 7.31E-16 | 0.308474505 | 0.322 | 0.213 | 1.48E-11 | 9 | Ski |
| Esco11 | 7.96E-16 | 0.290073607 | 0.281 | 0.178 | 1.61E-11 | 9 | Esco1 |
| Snx92 | 8.71E-16 | 0.262126467 | 0.281 | 0.174 | 1.76E-11 | 9 | Snx9 |
| Flii1 | 9.25E-16 | 0.302166177 | 0.254 | 0.157 | 1.87E-11 | 9 | Flii |
| Mecp21 | 1.12E-15 | 0.28045955 | 0.258 | 0.159 | 2.27E-11 | 9 | Mecp2 |
| Ccdc822 | 1.15E-15 | 0.317632003 | 0.281 | 0.179 | 2.33E-11 | 9 | Ccdc82 |
| Zfp6443 | 1.17E-15 | 0.25856315 | 0.37 | 0.254 | 2.36E-11 | 9 | Zfp644 |
| Sp31 | 1.29E-15 | 0.310942318 | 0.292 | 0.19 | 2.60E-11 | 9 | Sp3 |
| Trio2 | 1.48E-15 | 0.325991168 | 0.289 | 0.187 | 2.99E-11 | 9 | Trio |
| Gapvd12 | 1.59E-15 | 0.300497327 | 0.297 | 0.192 | 3.23E-11 | 9 | Gapvd1 |
| Rere2 | 1.61E-15 | 0.282451998 | 0.388 | 0.27 | 3.25E-11 | 9 | Rere |
| Fam193a1 | 1.69E-15 | 0.298209278 | 0.28 | 0.178 | 3.43E-11 | 9 | Fam193a |
| Exoc51 | 1.85E-15 | 0.25420064 | 0.286 | 0.182 | 3.76E-11 | 9 | Exoc5 |
| Kdm2a4 | 2.36E-15 | 0.291833385 | 0.35 | 0.239 | 4.78E-11 | 9 | Kdm2a |
| Trip122 | 2.45E-15 | 0.28304585 | 0.367 | 0.255 | 4.95E-11 | 9 | Trip12 |
| Tnrc182 | 2.75E-15 | 0.310178886 | 0.271 | 0.172 | 5.56E-11 | 9 | Tnrc18 |
| Mycbp23 | 3.12E-15 | 0.280161103 | 0.512 | 0.372 | 6.31E-11 | 9 | Mycbp2 |
| Smad41 | 3.58E-15 | 0.266759342 | 0.33 | 0.221 | 7.25E-11 | 9 | Smad4 |
| Cdc731 | 3.93E-15 | 0.26658097 | 0.269 | 0.169 | 7.96E-11 | 9 | Cdc73 |
| Zfp1481 | 3.96E-15 | 0.28918491 | 0.376 | 0.263 | 8.02E-11 | 9 | Zfp148 |
| Ggps11 | 4.09E-15 | 0.288083304 | 0.271 | 0.17 | 8.29E-11 | 9 | Ggps1 |
| Sel1l1 | 4.21E-15 | 0.326852884 | 0.258 | 0.161 | 8.53E-11 | 9 | Sel1l |
| AU0202064 | 4.34E-15 | 0.337913815 | 0.414 | 0.289 | 8.80E-11 | 9 | AU020206 |
| Zfr3 | 4.92E-15 | 0.292712066 | 0.358 | 0.246 | 9.96E-11 | 9 | Zfr |
| Eif5b2 | 5.36E-15 | 0.304442678 | 0.439 | 0.32 | 1.09E-10 | 9 | Eif5b |
| P4hb1 | 5.63E-15 | 0.260892523 | 0.363 | 0.251 | 1.14E-10 | 9 | P4hb |
| Arid1a3 | 6.33E-15 | 0.312540312 | 0.333 | 0.229 | 1.28E-10 | 9 | Arid1a |
| Ankrd173 | 6.51E-15 | 0.304524162 | 0.372 | 0.26 | 1.32E-10 | 9 | Ankrd17 |
| Psme41 | 7.02E-15 | 0.256413317 | 0.3 | 0.196 | 1.42E-10 | 9 | Psme4 |
| Eea12 | 8.35E-15 | 0.264314918 | 0.323 | 0.217 | 1.69E-10 | 9 | Eea1 |
| Ranbp21 | 9.51E-15 | 0.269377423 | 0.3 | 0.197 | 1.93E-10 | 9 | Ranbp2 |
| Ncoa23 | 9.66E-15 | 0.319417498 | 0.345 | 0.237 | 1.96E-10 | 9 | Ncoa2 |
| Rbbp62 | 1.09E-14 | 0.306193681 | 0.386 | 0.275 | 2.20E-10 | 9 | Rbbp6 |
| Ndfip23 | 1.33E-14 | 0.273084237 | 0.353 | 0.244 | 2.70E-10 | 9 | Ndfip2 |
| Luc7l32 | 1.81E-14 | 0.272842727 | 0.376 | 0.27 | 3.67E-10 | 9 | Luc7l3 |
| Ergic21 | 1.90E-14 | 0.268258248 | 0.276 | 0.178 | 3.84E-10 | 9 | Ergic2 |
| Gls2 | 1.93E-14 | 0.265139982 | 0.359 | 0.25 | 3.91E-10 | 9 | Gls |
| Dhx361 | 2.23E-14 | 0.255473072 | 0.34 | 0.234 | 4.51E-10 | 9 | Dhx36 |
| Smc31 | 2.71E-14 | 0.26648269 | 0.36 | 0.251 | 5.48E-10 | 9 | Smc3 |
| Abi13 | 2.91E-14 | 0.284951587 | 0.379 | 0.268 | 5.90E-10 | 9 | Abi1 |
| Usp71 | 2.95E-14 | 0.255307013 | 0.254 | 0.161 | 5.98E-10 | 9 | Usp7 |
| Fbxl174 | 3.31E-14 | 0.285938092 | 0.433 | 0.309 | 6.70E-10 | 9 | Fbxl17 |

| | | | | | | | |
|---|---|---|---|---|---|---|---|
| Phactr23 | 3.51E-14 | 0.251636196 | 0.358 | 0.244 | 7.11E-10 | 9 | Phactr2 |
| Chd82 | 4.01E-14 | 0.315799237 | 0.259 | 0.167 | 8.11E-10 | 9 | Chd8 |
| Ino80d1 | 4.09E-14 | 0.319691118 | 0.256 | 0.164 | 8.29E-10 | 9 | Ino80d |
| Ehmt11 | 4.17E-14 | 0.287071237 | 0.281 | 0.184 | 8.44E-10 | 9 | Ehmt1 |
| Arhgap262 | 5.28E-14 | 0.315078602 | 0.328 | 0.221 | 1.07E-09 | 9 | Arhgap26 |
| Kmt5b1 | 5.83E-14 | 0.284859777 | 0.263 | 0.169 | 1.18E-09 | 9 | Kmt5b |
| Etv62 | 7.52E-14 | 0.258589652 | 0.396 | 0.276 | 1.52E-09 | 9 | Etv6 |
| Galnt13 | 7.97E-14 | 0.268723852 | 0.28 | 0.183 | 1.61E-09 | 9 | Galnt1 |
| Yy11 | 1.52E-13 | 0.283869697 | 0.376 | 0.272 | 3.07E-09 | 9 | Yy1 |
| Ppp2r2a2 | 1.68E-13 | 0.287395862 | 0.311 | 0.212 | 3.40E-09 | 9 | Ppp2r2a |
| Nucb12 | 1.87E-13 | 0.273800737 | 0.329 | 0.227 | 3.79E-09 | 9 | Nucb1 |
| Rap1gds13 | 1.90E-13 | 0.306287579 | 0.367 | 0.26 | 3.85E-09 | 9 | Rap1gds1 |
| Nudcd31 | 1.95E-13 | 0.284157888 | 0.345 | 0.245 | 3.95E-09 | 9 | Nudcd3 |
| Ttc32 | 2.09E-13 | 0.262317411 | 0.361 | 0.25 | 4.23E-09 | 9 | Ttc3 |
| Prkd32 | 2.67E-13 | 0.259495253 | 0.258 | 0.167 | 5.40E-09 | 9 | Prkd3 |
| Setd23 | 2.85E-13 | 0.295710254 | 0.284 | 0.192 | 5.78E-09 | 9 | Setd2 |
| Bdp11 | 2.86E-13 | 0.321380393 | 0.252 | 0.165 | 5.80E-09 | 9 | Bdp1 |
| Smg13 | 2.93E-13 | 0.308690262 | 0.332 | 0.233 | 5.94E-09 | 9 | Smg1 |
| Arid4a4 | 2.96E-13 | 0.281319495 | 0.354 | 0.25 | 6.00E-09 | 9 | Arid4a |
| AW5549181 | 3.09E-13 | 0.347777443 | 0.259 | 0.17 | 6.25E-09 | 9 | AW554918 |
| Dhx15 | 3.85E-13 | 0.251547614 | 0.296 | 0.201 | 7.80E-09 | 9 | Dhx15 |
| Atf7ip2 | 6.18E-13 | 0.32037385 | 0.301 | 0.206 | 1.25E-08 | 9 | Atf7ip |
| Ppp2r5e1 | 6.47E-13 | 0.288032831 | 0.252 | 0.164 | 1.31E-08 | 9 | Ppp2r5e |
| Chd11 | 7.85E-13 | 0.257577711 | 0.295 | 0.2 | 1.59E-08 | 9 | Chd1 |
| Atp2c12 | 9.43E-13 | 0.270329092 | 0.255 | 0.167 | 1.91E-08 | 9 | Atp2c1 |
| Abi33 | 9.50E-13 | 0.260129434 | 0.348 | 0.243 | 1.92E-08 | 9 | Abi3 |
| Man2a12 | 1.28E-12 | 0.253251435 | 0.339 | 0.233 | 2.59E-08 | 9 | Man2a1 |
| Nfkb12 | 1.75E-12 | 0.263427366 | 0.379 | 0.27 | 3.55E-08 | 9 | Nfkb1 |
| Ccar12 | 1.98E-12 | 0.267497065 | 0.3 | 0.207 | 4.02E-08 | 9 | Ccar1 |
| Brd13 | 2.29E-12 | 0.250437847 | 0.335 | 0.239 | 4.63E-08 | 9 | Brd1 |
| Pkn13 | 2.58E-12 | 0.268084067 | 0.274 | 0.186 | 5.22E-08 | 9 | Pkn1 |
| Emc101 | 5.40E-12 | 0.260610972 | 0.289 | 0.199 | 1.09E-07 | 9 | Emc10 |
| Cdk132 | 6.25E-12 | 0.274350918 | 0.319 | 0.226 | 1.27E-07 | 9 | Cdk13 |
| Fyn3 | 7.68E-12 | 0.254119647 | 0.372 | 0.265 | 1.56E-07 | 9 | Fyn |
| Scaf82 | 1.17E-11 | 0.351023768 | 0.266 | 0.184 | 2.36E-07 | 9 | Scaf8 |
| Prpf38b3 | 1.58E-11 | 0.26739773 | 0.282 | 0.195 | 3.20E-07 | 9 | Prpf38b |
| Nf13 | 1.87E-11 | 0.277497425 | 0.312 | 0.22 | 3.78E-07 | 9 | Nf1 |
| Pan33 | 2.52E-11 | 0.288494348 | 0.318 | 0.228 | 5.11E-07 | 9 | Pan3 |
| Phf143 | 3.45E-11 | 0.273060588 | 0.337 | 0.243 | 6.98E-07 | 9 | Phf14 |
| Trappc92 | 3.79E-11 | 0.256848777 | 0.279 | 0.194 | 7.67E-07 | 9 | Trappc9 |
| Ccdc88a3 | 3.92E-11 | 0.283426888 | 0.292 | 0.204 | 7.94E-07 | 9 | Ccdc88a |
| Ep3003 | 5.73E-11 | 0.256766786 | 0.29 | 0.203 | 1.16E-06 | 9 | Ep300 |
| Mtmr33 | 7.13E-11 | 0.26681139 | 0.284 | 0.2 | 1.44E-06 | 9 | Mtmr3 |

| | | | | | | | |
|---|---|---|---|---|---|---|---|
| Bcas32 | 7.97E-11 | 0.291319897 | 0.264 | 0.182 | 1.61E-06 | 9 | Bcas3 |
| Cnn21 | 8.76E-11 | 0.254199182 | 0.265 | 0.182 | 1.77E-06 | 9 | Cnn2 |
| Trip112 | 1.00E-10 | 0.297624546 | 0.305 | 0.223 | 2.03E-06 | 9 | Trip11 |
| Btbd94 | 2.20E-10 | 0.262169453 | 0.324 | 0.237 | 4.45E-06 | 9 | Btbd9 |
| Trim12a2 | 2.29E-10 | 0.295372669 | 0.259 | 0.181 | 4.63E-06 | 9 | Trim12a |
| Abhd17b2 | 2.61E-10 | 0.279201686 | 0.273 | 0.192 | 5.29E-06 | 9 | Abhd17b |
| Sppl32 | 2.82E-10 | 0.295469816 | 0.301 | 0.218 | 5.71E-06 | 9 | Sppl3 |
| Sept111 | 4.14E-10 | 0.260560543 | 0.29 | 0.203 | 8.38E-06 | 9 | 11-Sep |
| Tmcc13 | 7.24E-10 | 0.313308789 | 0.3 | 0.217 | 1.47E-05 | 9 | Tmcc1 |
| Ubr22 | 2.81E-09 | 0.262030508 | 0.308 | 0.229 | 5.69E-05 | 9 | Ubr2 |
| Kdm6a3 | 2.86E-09 | 0.26792458 | 0.291 | 0.213 | 5.79E-05 | 9 | Kdm6a |
| Clasp22 | 1.12E-08 | 0.25471845 | 0.253 | 0.182 | 0.000227368 | 9 | Clasp2 |
| Rnf1693 | 4.14E-08 | 0.262567354 | 0.252 | 0.183 | 0.000838332 | 9 | Rnf169 |
| Rbm63 | 1.35E-07 | 0.270715908 | 0.282 | 0.217 | 0.002725713 | 9 | Rbm6 |
| 1700025G04Rik2 | 3.07E-07 | 0.261737159 | 0.253 | 0.189 | 0.00621658 | 9 | 1700025G04Rik |
| Dcn | 0 | 6.996344206 | 1 | 0.022 | 0 | 10 | Dcn |
| Gsn | 0 | 6.718310047 | 0.998 | 0.085 | 0 | 10 | Gsn |
| Cxcl14 | 0 | 5.49506869 | 0.944 | 0.03 | 0 | 10 | Cxcl14 |
| Apod | 0 | 5.181737421 | 0.988 | 0.011 | 0 | 10 | Apod |
| Mgp | 0 | 4.891198268 | 0.975 | 0.035 | 0 | 10 | Mgp |
| Col1a2 | 0 | 4.685443132 | 0.992 | 0.014 | 0 | 10 | Col1a2 |
| Col3a1 | 0 | 4.52051358 | 0.989 | 0.013 | 0 | 10 | Col3a1 |
| Lama2 | 0 | 4.460453516 | 0.975 | 0.012 | 0 | 10 | Lama2 |
| Lum | 0 | 4.425875907 | 0.98 | 0.005 | 0 | 10 | Lum |
| Lsamp | 0 | 4.34629254 | 0.953 | 0.014 | 0 | 10 | Lsamp |
| Col1a1 | 0 | 3.824472121 | 0.972 | 0.008 | 0 | 10 | Col1a1 |
| Rbms3 | 0 | 3.755969463 | 0.959 | 0.029 | 0 | 10 | Rbms3 |
| Adamts12 | 0 | 3.730772789 | 0.934 | 0.016 | 0 | 10 | Adamts12 |
| Pdgfra | 0 | 3.71912493 | 0.966 | 0.005 | 0 | 10 | Pdgfra |
| Serpinf1 | 0 | 3.657466525 | 0.973 | 0.025 | 0 | 10 | Serpinf1 |
| Lpl1 | 0 | 3.638332443 | 0.947 | 0.064 | 0 | 10 | Lpl |
| Bgn | 0 | 3.60878057 | 0.969 | 0.02 | 0 | 10 | Bgn |
| Clec3b | 0 | 3.469944171 | 0.93 | 0.007 | 0 | 10 | Clec3b |
| Lrp11 | 0 | 3.395295079 | 0.973 | 0.032 | 0 | 10 | Lrp1 |
| Igfbp4 | 0 | 3.355092917 | 0.961 | 0.075 | 0 | 10 | Igfbp4 |
| Enpp2 | 0 | 3.34900163 | 0.952 | 0.082 | 0 | 10 | Enpp2 |
| Plxdc21 | 0 | 3.297215225 | 0.959 | 0.071 | 0 | 10 | Plxdc2 |
| Ccl11 | 0 | 3.282550993 | 0.909 | 0.008 | 0 | 10 | Ccl11 |
| Arhgap61 | 0 | 3.199542705 | 0.916 | 0.035 | 0 | 10 | Arhgap6 |
| Abca8a | 0 | 3.191836218 | 0.941 | 0.008 | 0 | 10 | Abca8a |
| Fst | 0 | 3.169920169 | 0.794 | 0.006 | 0 | 10 | Fst |
| Serping1 | 0 | 3.164617846 | 0.939 | 0.011 | 0 | 10 | Serping1 |
| Col6a3 | 0 | 3.158771291 | 0.948 | 0.005 | 0 | 10 | Col6a3 |

| | | | | | | | |
|---|---|---|---|---|---|---|---|
| Col6a1 | 0 | 3.115964409 | 0.956 | 0.008 | 0 | 10 | Col6a1 |
| Col5a2 | 0 | 3.112002914 | 0.945 | 0.018 | 0 | 10 | Col5a2 |
| Smoc2 | 0 | 3.109255111 | 0.767 | 0.002 | 0 | 10 | Smoc2 |
| Colec12 | 0 | 3.070987143 | 0.944 | 0.026 | 0 | 10 | Colec12 |
| Cfh1 | 0 | 3.05471321 | 0.952 | 0.046 | 0 | 10 | Cfh |
| Dpep1 | 0 | 3.035935703 | 0.912 | 0.005 | 0 | 10 | Dpep1 |
| C3 | 0 | 3.002412878 | 0.636 | 0.005 | 0 | 10 | C3 |
| C1s1 | 0 | 2.98234043 | 0.928 | 0.01 | 0 | 10 | C1s1 |
| Cdh11 | 0 | 2.970132186 | 0.9 | 0.004 | 0 | 10 | Cdh11 |
| Gas1 | 0 | 2.950305639 | 0.891 | 0.008 | 0 | 10 | Gas1 |
| Ogn | 0 | 2.938336336 | 0.82 | 0.005 | 0 | 10 | Ogn |
| Rora1 | 0 | 2.920008584 | 0.955 | 0.166 | 0 | 10 | Rora |
| Igfbp5 | 0 | 2.881248425 | 0.791 | 0.016 | 0 | 10 | Igfbp5 |
| Crispld2 | 0 | 2.877233697 | 0.889 | 0.003 | 0 | 10 | Crispld2 |
| Itih52 | 0 | 2.852041059 | 0.966 | 0.225 | 0 | 10 | Itih5 |
| Pcolce | 0 | 2.832117915 | 0.928 | 0.022 | 0 | 10 | Pcolce |
| Col15a12 | 0 | 2.827029871 | 0.969 | 0.229 | 0 | 10 | Col15a1 |
| Rnase41 | 0 | 2.823574208 | 0.938 | 0.064 | 0 | 10 | Rnase4 |
| Cd3021 | 0 | 2.791702693 | 0.933 | 0.059 | 0 | 10 | Cd302 |
| Celf22 | 0 | 2.777489693 | 0.95 | 0.184 | 0 | 10 | Celf2 |
| Adamts5 | 0 | 2.762426744 | 0.825 | 0.004 | 0 | 10 | Adamts5 |
| Htra1 | 0 | 2.754369973 | 0.895 | 0.024 | 0 | 10 | Htra1 |
| Bicc11 | 0 | 2.739648661 | 0.934 | 0.078 | 0 | 10 | Bicc1 |
| Fgf10 | 0 | 2.716038501 | 0.839 | 0.01 | 0 | 10 | Fgf10 |
| S100a61 | 0 | 2.680001313 | 0.988 | 0.254 | 0 | 10 | S100a6 |
| Srpx | 0 | 2.675442353 | 0.878 | 0.002 | 0 | 10 | Srpx |
| Cygb | 0 | 2.675177016 | 0.917 | 0.012 | 0 | 10 | Cygb |
| Gpm6b | 0 | 2.600736987 | 0.903 | 0.01 | 0 | 10 | Gpm6b |
| Col6a2 | 0 | 2.597765334 | 0.883 | 0.006 | 0 | 10 | Col6a2 |
| Htra3 | 0 | 2.58650877 | 0.869 | 0.011 | 0 | 10 | Htra3 |
| Mmp2 | 0 | 2.583178667 | 0.858 | 0.005 | 0 | 10 | Mmp2 |
| Cp | 0 | 2.569932816 | 0.884 | 0.024 | 0 | 10 | Cp |
| Crabp1 | 0 | 2.534131379 | 0.645 | 0.001 | 0 | 10 | Crabp1 |
| Timp21 | 0 | 2.533935218 | 0.922 | 0.11 | 0 | 10 | Timp2 |
| Gpc6 | 0 | 2.505231242 | 0.791 | 0.012 | 0 | 10 | Gpc6 |
| Eda | 0 | 2.502945603 | 0.753 | 0.012 | 0 | 10 | Eda |
| Col5a3 | 0 | 2.500841046 | 0.897 | 0.008 | 0 | 10 | Col5a3 |
| Col14a11 | 0 | 2.497164043 | 0.819 | 0.037 | 0 | 10 | Col14a1 |
| Nrxn1 | 0 | 2.490198415 | 0.5 | 0.004 | 0 | 10 | Nrxn1 |
| Rspo3 | 0 | 2.479203885 | 0.797 | 0.024 | 0 | 10 | Rspo3 |
| Pappa | 0 | 2.467213617 | 0.756 | 0.003 | 0 | 10 | Pappa |
| Rarres2 | 0 | 2.436703043 | 0.847 | 0.016 | 0 | 10 | Rarres2 |
| Nid12 | 0 | 2.418803926 | 0.942 | 0.163 | 0 | 10 | Nid1 |

| | | | | | | | |
|---|---|---|---|---|---|---|---|
| Mmp3 | 0 | 2.404963771 | 0.472 | 0.001 | 0 | 10 | Mmp3 |
| Abi3bp | 0 | 2.394225058 | 0.869 | 0.011 | 0 | 10 | Abi3bp |
| Cd632 | 0 | 2.393725555 | 0.972 | 0.168 | 0 | 10 | Cd63 |
| Auts21 | 0 | 2.390608281 | 0.925 | 0.133 | 0 | 10 | Auts2 |
| Lgals1 | 0 | 2.378086294 | 0.942 | 0.114 | 0 | 10 | Lgals1 |
| Marcks3 | 0 | 2.370302499 | 0.972 | 0.326 | 0 | 10 | Marcks |
| Mmp141 | 0 | 2.366571862 | 0.873 | 0.038 | 0 | 10 | Mmp14 |
| Ctsl4 | 0 | 2.363275575 | 0.967 | 0.331 | 0 | 10 | Ctsl |
| Lamb12 | 0 | 2.362881918 | 0.897 | 0.117 | 0 | 10 | Lamb1 |
| Efemp1 | 0 | 2.360748425 | 0.766 | 0.006 | 0 | 10 | Efemp1 |
| Il11ra1 | 0 | 2.360406884 | 0.892 | 0.077 | 0 | 10 | Il11ra1 |
| Prrx1 | 0 | 2.352259384 | 0.875 | 0.01 | 0 | 10 | Prrx1 |
| Islr | 0 | 2.331785464 | 0.844 | 0.005 | 0 | 10 | Islr |
| Cpxm1 | 0 | 2.325055896 | 0.83 | 0.002 | 0 | 10 | Cpxm1 |
| Robo1 | 0 | 2.308943657 | 0.797 | 0.004 | 0 | 10 | Robo1 |
| Cped13 | 0 | 2.30426591 | 0.931 | 0.16 | 0 | 10 | Cped1 |
| Entpd2 | 0 | 2.296253791 | 0.852 | 0.008 | 0 | 10 | Entpd2 |
| Magi2 | 0 | 2.281081897 | 0.709 | 0.02 | 0 | 10 | Magi2 |
| C1ra | 0 | 2.245390938 | 0.863 | 0.01 | 0 | 10 | C1ra |
| Adam19 | 0 | 2.204689121 | 0.798 | 0.062 | 0 | 10 | Adam19 |
| Lhfp | 0 | 2.204645814 | 0.833 | 0.018 | 0 | 10 | Lhfp |
| Antxr21 | 0 | 2.197501077 | 0.873 | 0.07 | 0 | 10 | Antxr2 |
| Astn2 | 0 | 2.185937601 | 0.756 | 0.006 | 0 | 10 | Astn2 |
| Slit3 | 0 | 2.183710039 | 0.825 | 0.063 | 0 | 10 | Slit3 |
| Aebp1 | 0 | 2.181528835 | 0.856 | 0.062 | 0 | 10 | Aebp1 |
| Scara5 | 0 | 2.17425807 | 0.808 | 0.007 | 0 | 10 | Scara5 |
| Adamtsl1 | 0 | 2.159616889 | 0.761 | 0.03 | 0 | 10 | Adamtsl1 |
| Zbtb204 | 0 | 2.157485794 | 0.978 | 0.528 | 0 | 10 | Zbtb20 |
| Dapk11 | 0 | 2.155067573 | 0.823 | 0.044 | 0 | 10 | Dapk1 |
| Fap | 0 | 2.139122792 | 0.798 | 0.016 | 0 | 10 | Fap |
| Ms4a4d | 0 | 2.138917414 | 0.823 | 0.004 | 0 | 10 | Ms4a4d |
| Sned1 | 0 | 2.13116638 | 0.703 | 0.048 | 0 | 10 | Sned1 |
| Egfr | 0 | 2.128967434 | 0.797 | 0.007 | 0 | 10 | Egfr |
| Ank2 | 0 | 2.127741475 | 0.798 | 0.023 | 0 | 10 | Ank2 |
| Spon1 | 0 | 2.111511363 | 0.661 | 0.005 | 0 | 10 | Spon1 |
| Nav3 | 0 | 2.086832507 | 0.839 | 0.052 | 0 | 10 | Nav3 |
| Tgfbi1 | 0 | 2.07439726 | 0.747 | 0.039 | 0 | 10 | Tgfbi |
| Itm2a3 | 0 | 2.064117869 | 0.883 | 0.162 | 0 | 10 | Itm2a |
| Ccdc80 | 0 | 2.05472149 | 0.786 | 0.011 | 0 | 10 | Ccdc80 |
| Pcdh7 | 0 | 2.050901829 | 0.691 | 0.013 | 0 | 10 | Pcdh7 |
| Ank3 | 0 | 2.050046892 | 0.728 | 0.032 | 0 | 10 | Ank3 |
| Rbp1 | 0 | 2.040367671 | 0.814 | 0.083 | 0 | 10 | Rbp1 |
| Scn7a | 0 | 2.039387246 | 0.82 | 0.007 | 0 | 10 | Scn7a |

| | | | | | | | |
|---|---|---|---|---|---|---|---|
| Steap4 | 0 | 2.017888198 | 0.759 | 0.009 | 0 | 10 | Steap4 |
| Pdzrn3 | 0 | 2.016401079 | 0.728 | 0.004 | 0 | 10 | Pdzrn3 |
| Emp11 | 0 | 2.014941629 | 0.869 | 0.103 | 0 | 10 | Emp1 |
| Naaladl2 | 0 | 2.009286529 | 0.711 | 0.023 | 0 | 10 | Naaladl2 |
| Sdc2 | 0 | 2.001080097 | 0.794 | 0.011 | 0 | 10 | Sdc2 |
| Inmt | 0 | 1.99514324 | 0.695 | 0.001 | 0 | 10 | Inmt |
| Serpinh14 | 0 | 1.985116136 | 0.931 | 0.233 | 0 | 10 | Serpinh1 |
| Gpx3 | 0 | 1.985038739 | 0.62 | 0.003 | 0 | 10 | Gpx3 |
| Svep1 | 0 | 1.982419773 | 0.755 | 0.001 | 0 | 10 | Svep1 |
| Igf1 | 0 | 1.968350669 | 0.752 | 0.066 | 0 | 10 | Igf1 |
| Il33 | 0 | 1.944068221 | 0.716 | 0.004 | 0 | 10 | Il33 |
| Mxra8 | 0 | 1.943771855 | 0.795 | 0.022 | 0 | 10 | Mxra8 |
| Nr2f1 | 0 | 1.934517974 | 0.728 | 0.007 | 0 | 10 | Nr2f1 |
| Fstl11 | 0 | 1.917874781 | 0.828 | 0.084 | 0 | 10 | Fstl1 |
| Hsd11b11 | 0 | 1.917819258 | 0.625 | 0.079 | 0 | 10 | Hsd11b1 |
| Col6a6 | 0 | 1.914575676 | 0.634 | 0.002 | 0 | 10 | Col6a6 |
| Fbn11 | 0 | 1.911202536 | 0.812 | 0.045 | 0 | 10 | Fbn1 |
| Mdk | 0 | 1.906631695 | 0.767 | 0.008 | 0 | 10 | Mdk |
| Mafb1 | 0 | 1.902107308 | 0.842 | 0.137 | 0 | 10 | Mafb |
| Fam198b3 | 0 | 1.878312046 | 0.873 | 0.167 | 0 | 10 | Fam198b |
| Rhoj3 | 0 | 1.87178423 | 0.925 | 0.225 | 0 | 10 | Rhoj |
| Robo2 | 0 | 1.869869245 | 0.614 | 0.005 | 0 | 10 | Robo2 |
| Maf1 | 0 | 1.845471684 | 0.805 | 0.101 | 0 | 10 | Maf |
| Meg3 | 0 | 1.842808267 | 0.67 | 0.009 | 0 | 10 | Meg3 |
| Tenm3 | 0 | 1.831953686 | 0.616 | 0.006 | 0 | 10 | Tenm3 |
| Fgfr1 | 0 | 1.82793016 | 0.767 | 0.03 | 0 | 10 | Fgfr1 |
| Igsf10 | 0 | 1.820056169 | 0.73 | 0.002 | 0 | 10 | Igsf10 |
| Nr2f2 | 0 | 1.81026776 | 0.838 | 0.08 | 0 | 10 | Nr2f2 |
| Lamc12 | 0 | 1.804418886 | 0.859 | 0.13 | 0 | 10 | Lamc1 |
| Ror1 | 0 | 1.798157411 | 0.65 | 0.02 | 0 | 10 | Ror1 |
| Fbln7 | 0 | 1.797127755 | 0.7 | 0.001 | 0 | 10 | Fbln7 |
| Eif4g33 | 0 | 1.796964919 | 0.887 | 0.271 | 0 | 10 | Eif4g3 |
| Sema3a | 0 | 1.795384919 | 0.653 | 0.002 | 0 | 10 | Sema3a |
| Barx1 | 0 | 1.787068615 | 0.722 | 0.006 | 0 | 10 | Barx1 |
| B3galt1 | 0 | 1.77308627 | 0.613 | 0.006 | 0 | 10 | B3galt1 |
| Prelp | 0 | 1.772118393 | 0.738 | 0.006 | 0 | 10 | Prelp |
| Zfhx4 | 0 | 1.765182785 | 0.698 | 0.004 | 0 | 10 | Zfhx4 |
| Mrc2 | 0 | 1.763610263 | 0.756 | 0.008 | 0 | 10 | Mrc2 |
| Plac8 | 0 | 1.747253295 | 0.552 | 0.01 | 0 | 10 | Plac8 |
| Rcn3 | 0 | 1.724626211 | 0.764 | 0.035 | 0 | 10 | Rcn3 |
| Plekha6 | 0 | 1.72452017 | 0.752 | 0.023 | 0 | 10 | Plekha6 |
| Col5a1 | 0 | 1.705764933 | 0.703 | 0.005 | 0 | 10 | Col5a1 |
| Arhgap24 | 0 | 1.700276332 | 0.65 | 0.032 | 0 | 10 | Arhgap24 |

| | | | | | | | |
|---|---|---|---|---|---|---|---|
| Pard3b | 0 | 1.692447627 | 0.769 | 0.084 | 0 | 10 | Pard3b |
| Tuba1a4 | 0 | 1.689854222 | 0.878 | 0.241 | 0 | 10 | Tuba1a |
| Tgfbr32 | 0 | 1.688239298 | 0.861 | 0.157 | 0 | 10 | Tgfbr3 |
| Spats2l | 0 | 1.665636159 | 0.778 | 0.063 | 0 | 10 | Spats2l |
| Dchs2 | 0 | 1.660185483 | 0.63 | 0.001 | 0 | 10 | Dchs2 |
| C1qtnf2 | 0 | 1.657879184 | 0.731 | 0.027 | 0 | 10 | C1qtnf2 |
| Aox3 | 0 | 1.653189329 | 0.545 | 0.003 | 0 | 10 | Aox3 |
| Frmd6 | 0 | 1.648677601 | 0.708 | 0.029 | 0 | 10 | Frmd6 |
| Fxyd1 | 0 | 1.642008884 | 0.744 | 0.015 | 0 | 10 | Fxyd1 |
| Pdgfrb | 0 | 1.639442863 | 0.702 | 0.01 | 0 | 10 | Pdgfrb |
| Tcf7l21 | 0 | 1.638870056 | 0.794 | 0.113 | 0 | 10 | Tcf7l2 |
| Atp2a22 | 0 | 1.613897672 | 0.861 | 0.184 | 0 | 10 | Atp2a2 |
| Loxl1 | 0 | 1.613291421 | 0.634 | 0.003 | 0 | 10 | Loxl1 |
| Mast42 | 0 | 1.609190327 | 0.792 | 0.172 | 0 | 10 | Mast4 |
| F3 | 0 | 1.605092621 | 0.611 | 0.005 | 0 | 10 | F3 |
| Pbx11 | 0 | 1.597423492 | 0.85 | 0.183 | 0 | 10 | Pbx1 |
| Nid2 | 0 | 1.587297637 | 0.708 | 0.061 | 0 | 10 | Nid2 |
| Epha3 | 0 | 1.579935344 | 0.577 | 0.003 | 0 | 10 | Epha3 |
| Eps8 | 0 | 1.577448932 | 0.777 | 0.069 | 0 | 10 | Eps8 |
| Mgst13 | 0 | 1.577353996 | 0.845 | 0.15 | 0 | 10 | Mgst1 |
| Ddr2 | 0 | 1.570866789 | 0.7 | 0.009 | 0 | 10 | Ddr2 |
| Atf5 | 0 | 1.568522606 | 0.641 | 0.031 | 0 | 10 | Atf5 |
| Psd31 | 0 | 1.557913855 | 0.742 | 0.102 | 0 | 10 | Psd3 |
| Gstt1 | 0 | 1.555345611 | 0.677 | 0.019 | 0 | 10 | Gstt1 |
| Ctsk | 0 | 1.553989466 | 0.65 | 0.004 | 0 | 10 | Ctsk |
| Smad3 | 0 | 1.53965312 | 0.694 | 0.081 | 0 | 10 | Smad3 |
| Npas2 | 0 | 1.525756699 | 0.613 | 0.012 | 0 | 10 | Npas2 |
| Crlf11 | 0 | 1.51906735 | 0.581 | 0.021 | 0 | 10 | Crlf1 |
| Pmp223 | 0 | 1.517397801 | 0.817 | 0.127 | 0 | 10 | Pmp22 |
| Emp32 | 0 | 1.512658246 | 0.887 | 0.244 | 0 | 10 | Emp3 |
| Scd2 | 0 | 1.512166447 | 0.728 | 0.063 | 0 | 10 | Scd2 |
| Ggt5 | 0 | 1.505836915 | 0.659 | 0.011 | 0 | 10 | Ggt5 |
| Plau1 | 0 | 1.502451777 | 0.677 | 0.046 | 0 | 10 | Plau |
| Nav13 | 0 | 1.50090863 | 0.841 | 0.208 | 0 | 10 | Nav1 |
| Nfix | 0 | 1.496937159 | 0.895 | 0.199 | 0 | 10 | Nfix |
| Oaf | 0 | 1.489067483 | 0.652 | 0.009 | 0 | 10 | Oaf |
| Dpysl3 | 0 | 1.485394027 | 0.72 | 0.05 | 0 | 10 | Dpysl3 |
| Mfap5 | 0 | 1.485031061 | 0.509 | 0.003 | 0 | 10 | Mfap5 |
| Nrxn2 | 0 | 1.470700777 | 0.594 | 0.001 | 0 | 10 | Nrxn2 |
| Rhobtb3 | 0 | 1.458843581 | 0.684 | 0.034 | 0 | 10 | Rhobtb3 |
| Fbln1 | 0 | 1.450295018 | 0.566 | 0.001 | 0 | 10 | Fbln1 |
| Slc29a11 | 0 | 1.449364946 | 0.742 | 0.12 | 0 | 10 | Slc29a1 |
| Bnc2 | 0 | 1.447763089 | 0.577 | 0.003 | 0 | 10 | Bnc2 |

| | | | | | | | |
|---|---|---|---|---|---|---|---|
| Sobp | 0 | 1.446631108 | 0.603 | 0.006 | 0 | 10 | Sobp |
| Lamc3 | 0 | 1.443728032 | 0.595 | 0.003 | 0 | 10 | Lamc3 |
| Zeb21 | 0 | 1.442244977 | 0.8 | 0.127 | 0 | 10 | Zeb2 |
| Ar3 | 0 | 1.440856452 | 0.836 | 0.204 | 0 | 10 | Ar |
| Cdk141 | 0 | 1.429320186 | 0.734 | 0.127 | 0 | 10 | Cdk14 |
| Gli3 | 0 | 1.425180943 | 0.603 | 0.008 | 0 | 10 | Gli3 |
| Igfbp6 | 0 | 1.424987354 | 0.372 | 0.007 | 0 | 10 | Igfbp6 |
| Ntrk2 | 0 | 1.424121958 | 0.564 | 0.005 | 0 | 10 | Ntrk2 |
| Itgbl1 | 0 | 1.415844515 | 0.398 | 0.001 | 0 | 10 | Itgbl1 |
| Naalad2 | 0 | 1.406570248 | 0.613 | 0.005 | 0 | 10 | Naalad2 |
| Setbp11 | 0 | 1.404757226 | 0.697 | 0.108 | 0 | 10 | Setbp1 |
| Akap122 | 0 | 1.404532752 | 0.767 | 0.146 | 0 | 10 | Akap12 |
| Tshz3 | 0 | 1.403170071 | 0.686 | 0.078 | 0 | 10 | Tshz3 |
| Ltbp11 | 0 | 1.393074499 | 0.689 | 0.089 | 0 | 10 | Ltbp1 |
| Cacna1a | 0 | 1.391484247 | 0.602 | 0.022 | 0 | 10 | Cacna1a |
| Fcgrt2 | 0 | 1.389593012 | 0.806 | 0.17 | 0 | 10 | Fcgrt |
| Spry1 | 0 | 1.388297512 | 0.634 | 0.061 | 0 | 10 | Spry1 |
| Cyth32 | 0 | 1.385182494 | 0.72 | 0.11 | 0 | 10 | Cyth3 |
| Gulp13 | 0 | 1.381758958 | 0.773 | 0.151 | 0 | 10 | Gulp1 |
| Lpar1 | 0 | 1.38011582 | 0.63 | 0.008 | 0 | 10 | Lpar1 |
| Mitf1 | 0 | 1.379514467 | 0.637 | 0.061 | 0 | 10 | Mitf |
| Ext13 | 0 | 1.373379001 | 0.745 | 0.153 | 0 | 10 | Ext1 |
| Ssbp21 | 0 | 1.371801178 | 0.766 | 0.125 | 0 | 10 | Ssbp2 |
| Lima12 | 0 | 1.366120506 | 0.772 | 0.143 | 0 | 10 | Lima1 |
| Sgce | 0 | 1.35144878 | 0.658 | 0.028 | 0 | 10 | Sgce |
| Dennd2a | 0 | 1.351204891 | 0.588 | 0.008 | 0 | 10 | Dennd2a |
| Col27a1 | 0 | 1.346392122 | 0.586 | 0.015 | 0 | 10 | Col27a1 |
| Itga81 | 0 | 1.345513578 | 0.566 | 0.021 | 0 | 10 | Itga8 |
| Abca8b | 0 | 1.33462063 | 0.53 | 0.006 | 0 | 10 | Abca8b |
| Bmpr1a | 0 | 1.333751974 | 0.658 | 0.022 | 0 | 10 | Bmpr1a |
| Neo1 | 0 | 1.332315059 | 0.631 | 0.05 | 0 | 10 | Neo1 |
| Egflam | 0 | 1.321756956 | 0.53 | 0.006 | 0 | 10 | Egflam |
| Ccbe1 | 0 | 1.321547443 | 0.481 | 0.002 | 0 | 10 | Ccbe1 |
| Podn | 0 | 1.316239392 | 0.548 | 0.001 | 0 | 10 | Podn |
| Tspan3 | 0 | 1.315457296 | 0.723 | 0.089 | 0 | 10 | Tspan3 |
| Cpq1 | 0 | 1.31529692 | 0.72 | 0.106 | 0 | 10 | Cpq |
| Ptprd | 0 | 1.313878563 | 0.498 | 0.01 | 0 | 10 | Ptprd |
| Adamtsl3 | 0 | 1.309797173 | 0.442 | 0.005 | 0 | 10 | Adamtsl3 |
| Ptges | 0 | 1.299082705 | 0.561 | 0.003 | 0 | 10 | Ptges |
| Twist1 | 0 | 1.297890797 | 0.591 | 0.004 | 0 | 10 | Twist1 |
| Ghr2 | 0 | 1.291624798 | 0.705 | 0.138 | 0 | 10 | Ghr |
| Abca91 | 0 | 1.279825594 | 0.584 | 0.029 | 0 | 10 | Abca9 |
| Col4a5 | 0 | 1.278034463 | 0.491 | 0.004 | 0 | 10 | Col4a5 |

| | | | | | | | |
|---|---|---|---|---|---|---|---|
| Creb3l1 | 0 | 1.275572195 | 0.625 | 0.022 | 0 | 10 | Creb3l1 |
| Steap3 | 0 | 1.274821254 | 0.562 | 0.006 | 0 | 10 | Steap3 |
| Cacna2d1 | 0 | 1.268513065 | 0.566 | 0.01 | 0 | 10 | Cacna2d1 |
| Dpt | 0 | 1.265502229 | 0.339 | 0.003 | 0 | 10 | Dpt |
| Anpep | 0 | 1.265181373 | 0.573 | 0.01 | 0 | 10 | Anpep |
| Bmp4 | 0 | 1.261543141 | 0.483 | 0.007 | 0 | 10 | Bmp4 |
| Ndn | 0 | 1.261136744 | 0.569 | 0.004 | 0 | 10 | Ndn |
| Il1r11 | 0 | 1.260908636 | 0.681 | 0.096 | 0 | 10 | Il1r1 |
| Arhgap32 | 0 | 1.258726411 | 0.588 | 0.04 | 0 | 10 | Arhgap32 |
| Fign | 0 | 1.257896559 | 0.522 | 0.011 | 0 | 10 | Fign |
| Aldh1a1 | 0 | 1.25648532 | 0.603 | 0.019 | 0 | 10 | Aldh1a1 |
| Kcnt2 | 0 | 1.25602821 | 0.488 | 0.003 | 0 | 10 | Kcnt2 |
| Slc24a3 | 0 | 1.253072812 | 0.492 | 0.032 | 0 | 10 | Slc24a3 |
| P3h3 | 0 | 1.251750705 | 0.584 | 0.011 | 0 | 10 | P3h3 |
| Tnxb | 0 | 1.248653621 | 0.503 | 0.008 | 0 | 10 | Tnxb |
| Mxra73 | 0 | 1.245274967 | 0.717 | 0.128 | 0 | 10 | Mxra7 |
| Pid11 | 0 | 1.244789656 | 0.561 | 0.057 | 0 | 10 | Pid1 |
| Tspan11 | 0 | 1.241887965 | 0.562 | 0.001 | 0 | 10 | Tspan11 |
| Rarb | 0 | 1.236820198 | 0.503 | 0.046 | 0 | 10 | Rarb |
| Pard3 | 0 | 1.234651196 | 0.602 | 0.031 | 0 | 10 | Pard3 |
| Peg3 | 0 | 1.233629248 | 0.375 | 0.008 | 0 | 10 | Peg3 |
| P2ry1 | 0 | 1.232048457 | 0.488 | 0.002 | 0 | 10 | P2ry1 |
| Emilin1 | 0 | 1.230325338 | 0.586 | 0.026 | 0 | 10 | Emilin1 |
| Mylk | 0 | 1.221357183 | 0.575 | 0.035 | 0 | 10 | Mylk |
| Thsd7a | 0 | 1.219674954 | 0.458 | 0.026 | 0 | 10 | Thsd7a |
| Prrx2 | 0 | 1.204201579 | 0.542 | 0.001 | 0 | 10 | Prrx2 |
| Axl1 | 0 | 1.20041871 | 0.666 | 0.063 | 0 | 10 | Axl |
| Sec24d | 0 | 1.189984205 | 0.597 | 0.057 | 0 | 10 | Sec24d |
| Spon2 | 0 | 1.1897027 | 0.436 | 0.003 | 0 | 10 | Spon2 |
| S1pr3 | 0 | 1.189104528 | 0.516 | 0.016 | 0 | 10 | S1pr3 |
| Adam12 | 0 | 1.185054783 | 0.417 | 0.027 | 0 | 10 | Adam12 |
| Clmp | 0 | 1.184522738 | 0.522 | 0.011 | 0 | 10 | Clmp |
| Casc4 | 0 | 1.183651143 | 0.555 | 0.021 | 0 | 10 | Casc4 |
| Ifi205 | 0 | 1.182066643 | 0.445 | 0.003 | 0 | 10 | Ifi205 |
| Plekha51 | 0 | 1.177473584 | 0.681 | 0.143 | 0 | 10 | Plekha5 |
| Mmp19 | 0 | 1.176458879 | 0.516 | 0.002 | 0 | 10 | Mmp19 |
| Sgcd | 0 | 1.17603463 | 0.484 | 0.008 | 0 | 10 | Sgcd |
| Gm16263 | 0 | 1.170127244 | 0.467 | 0.002 | 0 | 10 | Gm16263 |
| Kcnk2 | 0 | 1.167495525 | 0.428 | 0.005 | 0 | 10 | Kcnk2 |
| Agmo1 | 0 | 1.166997987 | 0.483 | 0.045 | 0 | 10 | Agmo |
| Ctsh2 | 0 | 1.16013918 | 0.808 | 0.12 | 0 | 10 | Ctsh |
| Antxr1 | 0 | 1.159261903 | 0.539 | 0.009 | 0 | 10 | Antxr1 |
| Tril1 | 0 | 1.15674232 | 0.616 | 0.074 | 0 | 10 | Tril |

| | | | | | | | |
|---|---|---|---|---|---|---|---|
| Zcchc24 | 0 | 1.153390732 | 0.62 | 0.054 | 0 | 10 | Zcchc24 |
| Atrnl12 | 0 | 1.152500389 | 0.728 | 0.152 | 0 | 10 | Atrnl1 |
| Dpyd | 0 | 1.149550848 | 0.527 | 0.036 | 0 | 10 | Dpyd |
| Fkbp7 | 0 | 1.145451196 | 0.605 | 0.068 | 0 | 10 | Fkbp7 |
| Dock71 | 0 | 1.143260237 | 0.567 | 0.034 | 0 | 10 | Dock7 |
| Wdr66 | 0 | 1.142165025 | 0.481 | 0.005 | 0 | 10 | Wdr66 |
| Maged1 | 0 | 1.140379117 | 0.639 | 0.075 | 0 | 10 | Maged1 |
| Pla2r1 | 0 | 1.139119474 | 0.489 | 0.006 | 0 | 10 | Pla2r1 |
| Tpst1 | 0 | 1.134485567 | 0.58 | 0.039 | 0 | 10 | Tpst1 |
| Fcgr2b1 | 0 | 1.127666921 | 0.459 | 0.03 | 0 | 10 | Fcgr2b |
| Rgma | 0 | 1.127444803 | 0.491 | 0.005 | 0 | 10 | Rgma |
| Twsg1 | 0 | 1.118111238 | 0.562 | 0.033 | 0 | 10 | Twsg1 |
| Pkd2 | 0 | 1.117100855 | 0.637 | 0.094 | 0 | 10 | Pkd2 |
| Gfpt2 | 0 | 1.100144029 | 0.517 | 0.031 | 0 | 10 | Gfpt2 |
| Arhgap20 | 0 | 1.099105509 | 0.434 | 0.004 | 0 | 10 | Arhgap20 |
| Lrig1 | 0 | 1.096006257 | 0.516 | 0.013 | 0 | 10 | Lrig1 |
| Atp1a2 | 0 | 1.0890871 | 0.492 | 0.006 | 0 | 10 | Atp1a2 |
| Hunk | 0 | 1.083529982 | 0.445 | 0.003 | 0 | 10 | Hunk |
| Pcdh19 | 0 | 1.07006423 | 0.442 | 0.016 | 0 | 10 | Pcdh19 |
| Nbl1 | 0 | 1.057221053 | 0.409 | 0.013 | 0 | 10 | Nbl1 |
| Plod11 | 0 | 1.054309789 | 0.647 | 0.108 | 0 | 10 | Plod1 |
| Sh3d191 | 0 | 1.046711782 | 0.633 | 0.095 | 0 | 10 | Sh3d19 |
| Flrt2 | 0 | 1.045610802 | 0.456 | 0.034 | 0 | 10 | Flrt2 |
| Chpf | 0 | 1.043666321 | 0.494 | 0.017 | 0 | 10 | Chpf |
| Clcn5 | 0 | 1.041126583 | 0.483 | 0.04 | 0 | 10 | Clcn5 |
| Gpr153 | 0 | 1.040558428 | 0.495 | 0.002 | 0 | 10 | Gpr153 |
| Ptpn13 | 0 | 1.038930517 | 0.509 | 0.019 | 0 | 10 | Ptpn13 |
| Slc1a51 | 0 | 1.034864201 | 0.569 | 0.068 | 0 | 10 | Slc1a5 |
| Zfp385b | 0 | 1.034438538 | 0.414 | 0.01 | 0 | 10 | Zfp385b |
| Snhg181 | 0 | 1.033091637 | 0.558 | 0.027 | 0 | 10 | Snhg18 |
| C2 | 0 | 1.030704939 | 0.445 | 0.004 | 0 | 10 | C2 |
| Acss3 | 0 | 1.024218933 | 0.428 | 0.004 | 0 | 10 | Acss3 |
| Gaa | 0 | 1.021673559 | 0.553 | 0.05 | 0 | 10 | Gaa |
| Maged2 | 0 | 1.020172895 | 0.542 | 0.06 | 0 | 10 | Maged2 |
| P4ha11 | 0 | 1.019893231 | 0.63 | 0.112 | 0 | 10 | P4ha1 |
| Sdk1 | 0 | 1.01973415 | 0.33 | 0.016 | 0 | 10 | Sdk1 |
| Medag | 0 | 1.015386994 | 0.425 | 0.002 | 0 | 10 | Medag |
| Negr11 | 0 | 1.012689861 | 0.403 | 0.035 | 0 | 10 | Negr1 |
| Cd248 | 0 | 1.012018235 | 0.444 | 0.007 | 0 | 10 | Cd248 |
| Dse1 | 0 | 1.009393364 | 0.508 | 0.049 | 0 | 10 | Dse |
| Cmklr11 | 0 | 1.002165512 | 0.48 | 0.021 | 0 | 10 | Cmklr1 |
| Runx1t1 | 0 | 0.998173927 | 0.467 | 0.032 | 0 | 10 | Runx1t1 |
| Thbs1 | 0 | 0.996120622 | 0.377 | 0.02 | 0 | 10 | Thbs1 |

| | | | | | | | |
|---|---|---|---|---|---|---|---|
| Plat | 0 | 0.995899185 | 0.491 | 0.047 | 0 | 10 | Plat |
| Oat1 | 0 | 0.995103717 | 0.692 | 0.123 | 0 | 10 | Oat |
| Col23a1 | 0 | 0.994122789 | 0.411 | 0.005 | 0 | 10 | Col23a1 |
| Fam49a1 | 0 | 0.993058935 | 0.489 | 0.039 | 0 | 10 | Fam49a |
| Fibin | 0 | 0.98298621 | 0.384 | 0.001 | 0 | 10 | Fibin |
| Vcam1 | 0 | 0.98204679 | 0.445 | 0.017 | 0 | 10 | Vcam1 |
| Cdh19 | 0 | 0.979534402 | 0.422 | 0.006 | 0 | 10 | Cdh19 |
| Serpine2 | 0 | 0.978213982 | 0.419 | 0.023 | 0 | 10 | Serpine2 |
| Cacnb2 | 0 | 0.977733567 | 0.308 | 0.012 | 0 | 10 | Cacnb2 |
| Sulf1 | 0 | 0.976323573 | 0.509 | 0.042 | 0 | 10 | Sulf1 |
| Reck | 0 | 0.975285669 | 0.458 | 0.024 | 0 | 10 | Reck |
| Scd1 | 0 | 0.974757921 | 0.447 | 0.014 | 0 | 10 | Scd1 |
| Gpx8 | 0 | 0.968891191 | 0.52 | 0.038 | 0 | 10 | Gpx8 |
| A830082K12Rik | 0 | 0.965728413 | 0.398 | 0.005 | 0 | 10 | A830082K12Rik |
| Loxl2 | 0 | 0.960003586 | 0.447 | 0.011 | 0 | 10 | Loxl2 |
| Slc16a2 | 0 | 0.959815362 | 0.509 | 0.057 | 0 | 10 | Slc16a2 |
| Sh3pxd2b1 | 0 | 0.958933731 | 0.475 | 0.03 | 0 | 10 | Sh3pxd2b |
| Penk | 0 | 0.957118699 | 0.378 | 0.005 | 0 | 10 | Penk |
| Ndufa4l2 | 0 | 0.955279675 | 0.462 | 0.01 | 0 | 10 | Ndufa4l2 |
| Ntrk3 | 0 | 0.952430435 | 0.305 | 0.007 | 0 | 10 | Ntrk3 |
| P2ry14 | 0 | 0.951193944 | 0.42 | 0.02 | 0 | 10 | P2ry14 |
| Ikbip | 0 | 0.950656897 | 0.55 | 0.066 | 0 | 10 | Ikbip |
| Olfml31 | 0 | 0.945878643 | 0.491 | 0.049 | 0 | 10 | Olfml3 |
| Gucy1a2 | 0 | 0.944455191 | 0.347 | 0.008 | 0 | 10 | Gucy1a2 |
| Kirrel | 0 | 0.944213292 | 0.472 | 0.01 | 0 | 10 | Kirrel |
| Fgf7 | 0 | 0.9428411 | 0.414 | 0.003 | 0 | 10 | Fgf7 |
| Adk | 0 | 0.942409337 | 0.672 | 0.111 | 0 | 10 | Adk |
| Glis3 | 0 | 0.940963524 | 0.375 | 0.007 | 0 | 10 | Glis3 |
| Cyp1b1 | 0 | 0.939803052 | 0.38 | 0 | 0 | 10 | Cyp1b1 |
| Gria3 | 0 | 0.93938967 | 0.411 | 0.007 | 0 | 10 | Gria3 |
| Ckap41 | 0 | 0.936676337 | 0.564 | 0.09 | 0 | 10 | Ckap4 |
| Lvrn | 0 | 0.935534477 | 0.35 | 0.001 | 0 | 10 | Lvrn |
| Rgmb | 0 | 0.935234406 | 0.502 | 0.05 | 0 | 10 | Rgmb |
| Txndc5 | 0 | 0.93352772 | 0.517 | 0.071 | 0 | 10 | Txndc5 |
| Vcan | 0 | 0.932402661 | 0.381 | 0.006 | 0 | 10 | Vcan |
| Olfml2a | 0 | 0.930094752 | 0.412 | 0.007 | 0 | 10 | Olfml2a |
| Bmp1 | 0 | 0.925070374 | 0.489 | 0.034 | 0 | 10 | Bmp1 |
| Adamts2 | 0 | 0.917539041 | 0.427 | 0.009 | 0 | 10 | Adamts2 |
| Meis1 | 0 | 0.916267397 | 0.416 | 0.028 | 0 | 10 | Meis1 |
| Angptl2 | 0 | 0.91131849 | 0.458 | 0.01 | 0 | 10 | Angptl2 |
| Adamts15 | 0 | 0.910689919 | 0.395 | 0.002 | 0 | 10 | Adamts15 |
| Srpx2 | 0 | 0.90922056 | 0.402 | 0.001 | 0 | 10 | Srpx2 |
| Pla2g4a1 | 0 | 0.894987086 | 0.447 | 0.033 | 0 | 10 | Pla2g4a |

| | | | | | | | |
|---|---|---|---|---|---|---|---|
| Scarf2 | 0 | 0.891246124 | 0.448 | 0.015 | 0 | 10 | Scarf2 |
| Kdelr3 | 0 | 0.887957702 | 0.405 | 0.007 | 0 | 10 | Kdelr3 |
| CT025619.1 | 0 | 0.883789994 | 0.406 | 0.007 | 0 | 10 | CT025619.1 |
| Adcyap1r1 | 0 | 0.880332148 | 0.416 | 0.003 | 0 | 10 | Adcyap1r1 |
| Lin7a | 0 | 0.879903636 | 0.342 | 0.009 | 0 | 10 | Lin7a |
| Gm29683 | 0 | 0.875414128 | 0.323 | 0.002 | 0 | 10 | Gm29683 |
| Dnm1 | 0 | 0.87488863 | 0.398 | 0.002 | 0 | 10 | Dnm1 |
| Ecm2 | 0 | 0.870595233 | 0.453 | 0.039 | 0 | 10 | Ecm2 |
| Ryk | 0 | 0.865056296 | 0.605 | 0.098 | 0 | 10 | Ryk |
| Asph1 | 0 | 0.860832422 | 0.566 | 0.089 | 0 | 10 | Asph |
| Col18a1 | 0 | 0.854178998 | 0.394 | 0.013 | 0 | 10 | Col18a1 |
| Ptgir | 0 | 0.853436629 | 0.394 | 0.005 | 0 | 10 | Ptgir |
| Boc | 0 | 0.848612612 | 0.425 | 0.017 | 0 | 10 | Boc |
| Lrrtm3 | 0 | 0.846600436 | 0.291 | 0.001 | 0 | 10 | Lrrtm3 |
| Ecm1 | 0 | 0.84549929 | 0.378 | 0.026 | 0 | 10 | Ecm1 |
| Ppp1r9a | 0 | 0.841612615 | 0.427 | 0.025 | 0 | 10 | Ppp1r9a |
| Ltbp3 | 0 | 0.8340962 | 0.428 | 0.016 | 0 | 10 | Ltbp3 |
| Hic1 | 0 | 0.831088227 | 0.491 | 0.059 | 0 | 10 | Hic1 |
| Scpep1 | 0 | 0.830781606 | 0.459 | 0.043 | 0 | 10 | Scpep1 |
| Fbln51 | 0 | 0.829540121 | 0.472 | 0.029 | 0 | 10 | Fbln5 |
| Ednra | 0 | 0.828050986 | 0.372 | 0.008 | 0 | 10 | Ednra |
| Tnfaip6 | 0 | 0.827054343 | 0.317 | 0.002 | 0 | 10 | Tnfaip6 |
| Adcy5 | 0 | 0.826987794 | 0.362 | 0.002 | 0 | 10 | Adcy5 |
| C1qtnf7 | 0 | 0.825885748 | 0.264 | 0.003 | 0 | 10 | C1qtnf7 |
| Epb41l3 | 0 | 0.825655618 | 0.381 | 0.005 | 0 | 10 | Epb41l3 |
| Il17d | 0 | 0.823138413 | 0.38 | 0.005 | 0 | 10 | Il17d |
| Scara3 | 0 | 0.822380899 | 0.367 | 0.006 | 0 | 10 | Scara3 |
| Fads1 | 0 | 0.821949611 | 0.445 | 0.045 | 0 | 10 | Fads1 |
| Pcdh18 | 0 | 0.812905429 | 0.377 | 0.004 | 0 | 10 | Pcdh18 |
| Cspg4 | 0 | 0.810934659 | 0.361 | 0.006 | 0 | 10 | Cspg4 |
| Srebf1 | 0 | 0.806651204 | 0.452 | 0.046 | 0 | 10 | Srebf1 |
| Rspo1 | 0 | 0.79583857 | 0.347 | 0.001 | 0 | 10 | Rspo1 |
| Ptgfrn | 0 | 0.792075935 | 0.403 | 0.017 | 0 | 10 | Ptgfrn |
| Ces2g | 0 | 0.790842018 | 0.348 | 0.033 | 0 | 10 | Ces2g |
| Sdc41 | 0 | 0.784296897 | 0.458 | 0.062 | 0 | 10 | Sdc4 |
| Reps2 | 0 | 0.782172114 | 0.352 | 0.014 | 0 | 10 | Reps2 |
| Basp12 | 0 | 0.774633041 | 0.592 | 0.093 | 0 | 10 | Basp1 |
| Mrvi1 | 0 | 0.77427233 | 0.345 | 0.009 | 0 | 10 | Mrvi1 |
| Tbx15 | 0 | 0.772725793 | 0.336 | 0.001 | 0 | 10 | Tbx15 |
| Sox5 | 0 | 0.772484467 | 0.316 | 0.014 | 0 | 10 | Sox5 |
| Tmem158 | 0 | 0.767604627 | 0.323 | 0.019 | 0 | 10 | Tmem158 |
| Mme | 0 | 0.76554198 | 0.38 | 0.014 | 0 | 10 | Mme |
| Vstm4 | 0 | 0.764140625 | 0.378 | 0.006 | 0 | 10 | Vstm4 |

| | | | | | | | |
|---|---|---|---|---|---|---|---|
| Spock2 | 0 | 0.763761125 | 0.353 | 0.012 | 0 | 10 | Spock2 |
| Efemp2 | 0 | 0.761347296 | 0.388 | 0.01 | 0 | 10 | Efemp2 |
| Sfrp1 | 0 | 0.760438359 | 0.331 | 0.006 | 0 | 10 | Sfrp1 |
| Cdon | 0 | 0.758284931 | 0.333 | 0.013 | 0 | 10 | Cdon |
| Tmem1191 | 0 | 0.75736078 | 0.458 | 0.05 | 0 | 10 | Tmem119 |
| Rab34 | 0 | 0.755022984 | 0.392 | 0.012 | 0 | 10 | Rab34 |
| Nnmt | 0 | 0.754036319 | 0.358 | 0.003 | 0 | 10 | Nnmt |
| Adgra2 | 0 | 0.750318385 | 0.373 | 0.01 | 0 | 10 | Adgra2 |
| Cpe | 0 | 0.746969634 | 0.338 | 0.013 | 0 | 10 | Cpe |
| Lgi2 | 0 | 0.745911012 | 0.312 | 0.002 | 0 | 10 | Lgi2 |
| Nr1h3 | 0 | 0.745732725 | 0.369 | 0.011 | 0 | 10 | Nr1h3 |
| Tnfrsf21 | 0 | 0.742976337 | 0.317 | 0.017 | 0 | 10 | Tnfrsf21 |
| H6pd | 0 | 0.739520521 | 0.35 | 0.024 | 0 | 10 | H6pd |
| Aox1 | 0 | 0.739056717 | 0.348 | 0.01 | 0 | 10 | Aox1 |
| Snai1 | 0 | 0.731350967 | 0.403 | 0.037 | 0 | 10 | Snai1 |
| Arap11 | 0 | 0.728257231 | 0.411 | 0.05 | 0 | 10 | Arap1 |
| Gk5 | 0 | 0.726231232 | 0.342 | 0.022 | 0 | 10 | Gk5 |
| Aoc3 | 0 | 0.725008985 | 0.345 | 0.008 | 0 | 10 | Aoc3 |
| Rassf21 | 0 | 0.72445065 | 0.4 | 0.046 | 0 | 10 | Rassf2 |
| Dsel | 0 | 0.724193773 | 0.397 | 0.035 | 0 | 10 | Dsel |
| Tmed1 | 0 | 0.724150033 | 0.43 | 0.051 | 0 | 10 | Tmed1 |
| Plekhh2 | 0 | 0.717746703 | 0.342 | 0.016 | 0 | 10 | Plekhh2 |
| Pknox2 | 0 | 0.712889726 | 0.322 | 0.006 | 0 | 10 | Pknox2 |
| Arhgap28 | 0 | 0.710497458 | 0.336 | 0.022 | 0 | 10 | Arhgap28 |
| Ptger3 | 0 | 0.701136251 | 0.314 | 0.012 | 0 | 10 | Ptger3 |
| Smo | 0 | 0.697198135 | 0.398 | 0.035 | 0 | 10 | Smo |
| Fgd41 | 0 | 0.694652793 | 0.347 | 0.034 | 0 | 10 | Fgd4 |
| Adam23 | 0 | 0.692821328 | 0.341 | 0.008 | 0 | 10 | Adam23 |
| Pdpn | 0 | 0.682043843 | 0.323 | 0.003 | 0 | 10 | Pdpn |
| Prdm5 | 0 | 0.675100656 | 0.327 | 0.018 | 0 | 10 | Prdm5 |
| Col4a4 | 0 | 0.674237117 | 0.314 | 0.018 | 0 | 10 | Col4a4 |
| Thbs2 | 0 | 0.67245413 | 0.275 | 0.007 | 0 | 10 | Thbs2 |
| Aldh7a1 | 0 | 0.670705084 | 0.462 | 0.052 | 0 | 10 | Aldh7a1 |
| Igsf3 | 0 | 0.667257055 | 0.328 | 0.007 | 0 | 10 | Igsf3 |
| Gxylt2 | 0 | 0.666861825 | 0.295 | 0.006 | 0 | 10 | Gxylt2 |
| Fndc5 | 0 | 0.66298813 | 0.403 | 0.039 | 0 | 10 | Fndc5 |
| Tbx2 | 0 | 0.660356566 | 0.333 | 0.015 | 0 | 10 | Tbx2 |
| Shox2 | 0 | 0.659700852 | 0.291 | 0.002 | 0 | 10 | Shox2 |
| Tenm4 | 0 | 0.659310378 | 0.252 | 0.002 | 0 | 10 | Tenm4 |
| Glis2 | 0 | 0.657316555 | 0.391 | 0.041 | 0 | 10 | Glis2 |
| C1qtnf1 | 0 | 0.65692756 | 0.302 | 0.009 | 0 | 10 | C1qtnf1 |
| Vasn | 0 | 0.649469417 | 0.306 | 0.005 | 0 | 10 | Vasn |
| B3gnt9 | 0 | 0.649426916 | 0.309 | 0.003 | 0 | 10 | B3gnt9 |

| | | | | | | | |
|---|---|---|---|---|---|---|---|
| Enpp1 | 0 | 0.648590886 | 0.281 | 0.022 | 0 | 10 | Enpp1 |
| Sspn | 0 | 0.647132536 | 0.331 | 0.022 | 0 | 10 | Sspn |
| Col16a1 | 0 | 0.646374873 | 0.309 | 0.005 | 0 | 10 | Col16a1 |
| Adh1 | 0 | 0.641912743 | 0.308 | 0.017 | 0 | 10 | Adh1 |
| Pde1a | 0 | 0.641591863 | 0.284 | 0.008 | 0 | 10 | Pde1a |
| Capn6 | 0 | 0.63909615 | 0.291 | 0.002 | 0 | 10 | Capn6 |
| Fzd2 | 0 | 0.637259096 | 0.3 | 0.002 | 0 | 10 | Fzd2 |
| Parp3 | 0 | 0.636483862 | 0.327 | 0.021 | 0 | 10 | Parp3 |
| Arhgef25 | 0 | 0.630632619 | 0.328 | 0.012 | 0 | 10 | Arhgef25 |
| Adcy3 | 0 | 0.626807095 | 0.308 | 0.019 | 0 | 10 | Adcy3 |
| Ror2 | 0 | 0.625839299 | 0.262 | 0.003 | 0 | 10 | Ror2 |
| Epb41l1 | 0 | 0.625831097 | 0.339 | 0.016 | 0 | 10 | Epb41l1 |
| Arhgap42 | 0 | 0.623216508 | 0.345 | 0.018 | 0 | 10 | Arhgap42 |
| Olfml1 | 0 | 0.621578177 | 0.292 | 0.001 | 0 | 10 | Olfml1 |
| Fzd1 | 0 | 0.621120067 | 0.28 | 0.009 | 0 | 10 | Fzd1 |
| Abca6 | 0 | 0.611631808 | 0.264 | 0.003 | 0 | 10 | Abca6 |
| Phlda3 | 0 | 0.601036578 | 0.403 | 0.046 | 0 | 10 | Phlda3 |
| Abat | 0 | 0.600891589 | 0.305 | 0.014 | 0 | 10 | Abat |
| Emilin2 | 0 | 0.591025419 | 0.28 | 0.005 | 0 | 10 | Emilin2 |
| Cacna1g | 0 | 0.587277062 | 0.259 | 0.001 | 0 | 10 | Cacna1g |
| Adm | 0 | 0.586001327 | 0.294 | 0.017 | 0 | 10 | Adm |
| Dact3 | 0 | 0.581197941 | 0.289 | 0.004 | 0 | 10 | Dact3 |
| Arhgef17 | 0 | 0.581054585 | 0.312 | 0.017 | 0 | 10 | Arhgef17 |
| Myh10 | 0 | 0.578424844 | 0.308 | 0.02 | 0 | 10 | Myh10 |
| Zfp618 | 0 | 0.577008603 | 0.266 | 0.005 | 0 | 10 | Zfp618 |
| Ift81 | 0 | 0.576837253 | 0.309 | 0.02 | 0 | 10 | Ift81 |
| Ptprs | 0 | 0.575573528 | 0.328 | 0.031 | 0 | 10 | Ptprs |
| Mmp23 | 0 | 0.560625014 | 0.25 | 0.002 | 0 | 10 | Mmp23 |
| Asap3 | 0 | 0.55825705 | 0.258 | 0.015 | 0 | 10 | Asap3 |
| Gsc | 0 | 0.55140155 | 0.258 | 0.001 | 0 | 10 | Gsc |
| Pkdcc | 0 | 0.551049731 | 0.258 | 0.01 | 0 | 10 | Pkdcc |
| Fam171b | 0 | 0.546154088 | 0.252 | 0.003 | 0 | 10 | Fam171b |
| Fkbp14 | 0 | 0.541713554 | 0.262 | 0.011 | 0 | 10 | Fkbp14 |
| Evc2 | 0 | 0.538496484 | 0.258 | 0.005 | 0 | 10 | Evc2 |
| Fkbp10 | 0 | 0.529728237 | 0.258 | 0.011 | 0 | 10 | Fkbp10 |
| S1pr2 | 0 | 0.529308663 | 0.286 | 0.017 | 0 | 10 | S1pr2 |
| P3h4 | 0 | 0.515132431 | 0.277 | 0.018 | 0 | 10 | P3h4 |
| Evc | 0 | 0.506593827 | 0.255 | 0.004 | 0 | 10 | Evc |
| P3h1 | 0 | 0.493898087 | 0.252 | 0.014 | 0 | 10 | P3h1 |
| Armcx2 | 0 | 0.487582564 | 0.283 | 0.021 | 0 | 10 | Armcx2 |
| Rhbdf1 | 0 | 0.481181415 | 0.267 | 0.017 | 0 | 10 | Rhbdf1 |
| Nfia3 | 5.23E-308 | 1.578566915 | 0.889 | 0.273 | 1.06E-303 | 10 | Nfia |
| Calu3 | 6.43E-308 | 1.18191071 | 0.784 | 0.188 | 1.30E-303 | 10 | Calu |

| | | | | | | | |
|---|---|---|---|---|---|---|---|
| Eln2 | 8.37E-308 | 1.265939548 | 0.65 | 0.124 | 1.69E-303 | 10 | Eln |
| Rrbp14 | 1.34E-307 | 1.737486147 | 0.969 | 0.448 | 2.72E-303 | 10 | Rrbp1 |
| Ifi2071 | 2.73E-306 | 0.919581417 | 0.575 | 0.096 | 5.52E-302 | 10 | Ifi207 |
| Hspg23 | 6.74E-304 | 1.545952221 | 0.944 | 0.297 | 1.36E-299 | 10 | Hspg2 |
| Klhl13 | 5.53E-301 | 0.593786898 | 0.327 | 0.032 | 1.12E-296 | 10 | Klhl13 |
| Cbr3 | 3.71E-300 | 0.738491195 | 0.461 | 0.064 | 7.50E-296 | 10 | Cbr3 |
| Plxnc1 | 3.04E-299 | 0.506040179 | 0.259 | 0.02 | 6.16E-295 | 10 | Plxnc1 |
| Ldlr | 4.81E-297 | 0.771921737 | 0.377 | 0.044 | 9.75E-293 | 10 | Ldlr |
| Idh1 | 6.97E-297 | 0.661876905 | 0.427 | 0.056 | 1.41E-292 | 10 | Idh1 |
| Lrp6 | 2.80E-296 | 1.038242288 | 0.652 | 0.132 | 5.66E-292 | 10 | Lrp6 |
| Fkbp91 | 1.48E-295 | 0.889653811 | 0.577 | 0.1 | 3.00E-291 | 10 | Fkbp9 |
| Plod2 | 1.53E-294 | 0.653587107 | 0.375 | 0.044 | 3.09E-290 | 10 | Plod2 |
| Itgb51 | 1.61E-293 | 0.967579907 | 0.672 | 0.132 | 3.27E-289 | 10 | Itgb5 |
| Vwa11 | 1.58E-291 | 0.80356592 | 0.558 | 0.093 | 3.21E-287 | 10 | Vwa1 |
| Lamp23 | 2.33E-291 | 1.354289927 | 0.903 | 0.291 | 4.71E-287 | 10 | Lamp2 |
| Man2a2 | 7.26E-287 | 0.635370798 | 0.333 | 0.036 | 1.47E-282 | 10 | Man2a2 |
| Fam102b1 | 3.70E-285 | 0.664927099 | 0.37 | 0.045 | 7.50E-281 | 10 | Fam102b |
| Fhit1 | 1.91E-284 | 1.200173483 | 0.461 | 0.071 | 3.87E-280 | 10 | Fhit |
| Selenop5 | 7.57E-284 | 1.55133641 | 0.941 | 0.35 | 1.53E-279 | 10 | Selenop |
| Sept112 | 2.65E-281 | 1.158039612 | 0.764 | 0.19 | 5.36E-277 | 10 | 11-Sep |
| Cdkn1c2 | 2.62E-280 | 0.995439515 | 0.603 | 0.115 | 5.30E-276 | 10 | Cdkn1c |
| Glt8d1 | 1.54E-279 | 0.714360746 | 0.436 | 0.062 | 3.11E-275 | 10 | Glt8d1 |
| Vim3 | 2.13E-279 | 1.497768039 | 0.908 | 0.292 | 4.30E-275 | 10 | Vim |
| Col4a3 | 6.95E-279 | 0.721284839 | 0.372 | 0.045 | 1.41E-274 | 10 | Col4a3 |
| Wls | 1.34E-278 | 0.935665804 | 0.597 | 0.116 | 2.71E-274 | 10 | Wls |
| Chd91 | 1.85E-275 | 1.279020291 | 0.823 | 0.237 | 3.74E-271 | 10 | Chd9 |
| Sh3pxd2a2 | 1.09E-273 | 1.010627792 | 0.633 | 0.13 | 2.20E-269 | 10 | Sh3pxd2a |
| Pdlim21 | 1.47E-273 | 0.742880657 | 0.422 | 0.061 | 2.98E-269 | 10 | Pdlim2 |
| Cc2d2a | 1.37E-272 | 0.490167262 | 0.297 | 0.029 | 2.78E-268 | 10 | Cc2d2a |
| Plxnb2 | 3.42E-272 | 0.565792758 | 0.352 | 0.042 | 6.93E-268 | 10 | Plxnb2 |
| Ssh22 | 1.67E-270 | 1.280308791 | 0.783 | 0.213 | 3.38E-266 | 10 | Ssh2 |
| Ndrg2 | 1.86E-269 | 0.474395969 | 0.441 | 0.061 | 3.77E-265 | 10 | Ndrg2 |
| Golim42 | 2.03E-268 | 1.062113634 | 0.703 | 0.168 | 4.11E-264 | 10 | Golim4 |
| Dapk21 | 6.47E-268 | 0.877448689 | 0.55 | 0.1 | 1.31E-263 | 10 | Dapk2 |
| Dlc13 | 2.05E-267 | 1.668663635 | 0.952 | 0.349 | 4.15E-263 | 10 | Dlc1 |
| Mid11 | 1.60E-265 | 1.077364391 | 0.462 | 0.076 | 3.24E-261 | 10 | Mid1 |
| Tulp3 | 2.40E-262 | 0.525964702 | 0.302 | 0.031 | 4.86E-258 | 10 | Tulp3 |
| Epb41l22 | 4.87E-262 | 1.199743507 | 0.806 | 0.228 | 9.86E-258 | 10 | Epb41l2 |
| Laptm4a3 | 7.76E-262 | 1.508007409 | 0.969 | 0.473 | 1.57E-257 | 10 | Laptm4a |
| P2rx41 | 2.55E-261 | 0.628028433 | 0.405 | 0.057 | 5.16E-257 | 10 | P2rx4 |
| Cyp7b1 | 3.61E-259 | 0.582144272 | 0.295 | 0.03 | 7.31E-255 | 10 | Cyp7b1 |
| Nfic1 | 8.39E-258 | 1.161498485 | 0.827 | 0.248 | 1.70E-253 | 10 | Nfic |
| Tnfsf122 | 5.28E-256 | 0.899337998 | 0.634 | 0.136 | 1.07E-251 | 10 | Tnfsf12 |

| | | | | | | | |
|---|---|---|---|---|---|---|---|
| Nedd43 | 5.84E-253 | 1.43749073 | 0.952 | 0.407 | 1.18E-248 | 10 | Nedd4 |
| Hspa13 | 8.73E-252 | 0.617623851 | 0.388 | 0.054 | 1.77E-247 | 10 | Hspa13 |
| Gab22 | 2.36E-251 | 1.188149146 | 0.745 | 0.192 | 4.77E-247 | 10 | Gab2 |
| Arhgef40 | 5.63E-251 | 0.683446197 | 0.409 | 0.06 | 1.14E-246 | 10 | Arhgef40 |
| Lama43 | 1.43E-250 | 1.030077729 | 0.702 | 0.162 | 2.91E-246 | 10 | Lama4 |
| 8-Sep | 1.00E-248 | 0.570901962 | 0.327 | 0.039 | 2.02E-244 | 10 | 8-Sep |
| Add31 | 1.11E-248 | 1.101698403 | 0.766 | 0.21 | 2.24E-244 | 10 | Add3 |
| Fndc3b3 | 2.49E-246 | 1.153059714 | 0.792 | 0.231 | 5.05E-242 | 10 | Fndc3b |
| Ube2e21 | 2.87E-244 | 0.990583103 | 0.639 | 0.141 | 5.80E-240 | 10 | Ube2e2 |
| Sgcb | 2.25E-242 | 0.562564063 | 0.359 | 0.048 | 4.56E-238 | 10 | Sgcb |
| 2210408I21Rik | 4.31E-242 | 0.730921053 | 0.377 | 0.053 | 8.73E-238 | 10 | 2210408I21Rik |
| Thra2 | 6.90E-242 | 0.858623974 | 0.664 | 0.152 | 1.40E-237 | 10 | Thra |
| Cep112 | 8.30E-242 | 0.703257133 | 0.372 | 0.051 | 1.68E-237 | 10 | Cep112 |
| Ehbp1 | 6.46E-241 | 0.852839931 | 0.525 | 0.101 | 1.31E-236 | 10 | Ehbp1 |
| Olfm1 | 1.38E-240 | 0.505401007 | 0.309 | 0.036 | 2.79E-236 | 10 | Olfm1 |
| Wipi1 | 3.59E-240 | 0.592246698 | 0.362 | 0.049 | 7.26E-236 | 10 | Wipi1 |
| Zfyve21 | 5.39E-240 | 0.509948712 | 0.386 | 0.055 | 1.09E-235 | 10 | Zfyve21 |
| Park2 | 1.91E-238 | 0.849008258 | 0.419 | 0.067 | 3.86E-234 | 10 | Park2 |
| Klf93 | 1.88E-237 | 1.151476478 | 0.784 | 0.224 | 3.81E-233 | 10 | Klf9 |
| Hspa54 | 3.58E-237 | 1.559434993 | 0.941 | 0.423 | 7.25E-233 | 10 | Hspa5 |
| Fam114a12 | 3.33E-236 | 0.840284585 | 0.58 | 0.121 | 6.74E-232 | 10 | Fam114a1 |
| Igf1r2 | 3.86E-236 | 1.031827386 | 0.644 | 0.15 | 7.81E-232 | 10 | Igf1r |
| Itsn1 | 5.96E-235 | 0.713868939 | 0.417 | 0.067 | 1.21E-230 | 10 | Itsn1 |
| Bace1 | 2.50E-234 | 0.488363964 | 0.3 | 0.035 | 5.07E-230 | 10 | Bace1 |
| Cdc42ep5 | 6.56E-234 | 0.516734913 | 0.412 | 0.063 | 1.33E-229 | 10 | Cdc42ep5 |
| Errfi11 | 5.26E-233 | 1.227828544 | 0.623 | 0.15 | 1.06E-228 | 10 | Errfi1 |
| B3glct | 8.25E-233 | 0.538364331 | 0.284 | 0.032 | 1.67E-228 | 10 | B3glct |
| Dcbld2 | 8.46E-233 | 0.570268734 | 0.297 | 0.034 | 1.71E-228 | 10 | Dcbld2 |
| Rbfox23 | 2.58E-232 | 0.931661056 | 0.716 | 0.178 | 5.23E-228 | 10 | Rbfox2 |
| Lifr3 | 1.01E-231 | 1.143511359 | 0.694 | 0.181 | 2.05E-227 | 10 | Lifr |
| Fmo12 | 2.06E-231 | 1.017583694 | 0.73 | 0.189 | 4.18E-227 | 10 | Fmo1 |
| Rcn1 | 8.47E-230 | 0.824924831 | 0.506 | 0.099 | 1.71E-225 | 10 | Rcn1 |
| Cnrip1 | 3.04E-229 | 0.576470606 | 0.327 | 0.042 | 6.16E-225 | 10 | Cnrip1 |
| Tmem263 | 3.27E-229 | 0.64627695 | 0.441 | 0.074 | 6.63E-225 | 10 | Tmem263 |
| Sec31a | 2.72E-228 | 0.85953532 | 0.653 | 0.16 | 5.51E-224 | 10 | Sec31a |
| Malat15 | 4.36E-228 | 1.553818214 | 0.991 | 0.72 | 8.83E-224 | 10 | Malat1 |
| Airn3 | 3.18E-227 | 1.388423557 | 0.78 | 0.236 | 6.43E-223 | 10 | Airn |
| 2900097C17Rik | 7.09E-227 | 0.806949865 | 0.613 | 0.141 | 1.44E-222 | 10 | 2900097C17Rik |
| Ophn11 | 1.70E-226 | 0.912343656 | 0.572 | 0.122 | 3.43E-222 | 10 | Ophn1 |
| Selenos1 | 3.27E-226 | 1.049641892 | 0.745 | 0.219 | 6.61E-222 | 10 | Selenos |
| Lpp3 | 4.27E-224 | 1.191792888 | 0.822 | 0.273 | 8.64E-220 | 10 | Lpp |
| Smim1 | 4.73E-224 | 0.333369996 | 0.258 | 0.026 | 9.58E-220 | 10 | Smim1 |
| Nenf1 | 5.46E-224 | 0.998530359 | 0.855 | 0.282 | 1.10E-219 | 10 | Nenf |

| | | | | | | | |
|---|---|---|---|---|---|---|---|
| Dync2h1 | 6.40E-224 | 0.567294419 | 0.325 | 0.043 | 1.30E-219 | 10 | Dync2h1 |
| Plekhh3 | 1.64E-223 | 0.449144064 | 0.258 | 0.027 | 3.31E-219 | 10 | Plekhh3 |
| Erlec1 | 4.58E-223 | 0.819756937 | 0.605 | 0.14 | 9.28E-219 | 10 | Erlec1 |
| Tubb6 | 2.15E-222 | 0.583098405 | 0.328 | 0.044 | 4.35E-218 | 10 | Tubb6 |
| Ncoa12 | 3.62E-220 | 1.068543728 | 0.709 | 0.198 | 7.34E-216 | 10 | Ncoa1 |
| Pros12 | 4.78E-220 | 0.870421242 | 0.605 | 0.14 | 9.67E-216 | 10 | Pros1 |
| Itgav2 | 1.39E-218 | 0.843944274 | 0.625 | 0.15 | 2.82E-214 | 10 | Itgav |
| Prickle23 | 1.40E-218 | 0.935266422 | 0.583 | 0.129 | 2.84E-214 | 10 | Prickle2 |
| Kdelr2 | 4.71E-218 | 0.900301504 | 0.63 | 0.159 | 9.54E-214 | 10 | Kdelr2 |
| Dag12 | 5.61E-218 | 0.779371482 | 0.591 | 0.132 | 1.14E-213 | 10 | Dag1 |
| Stard13 | 2.55E-216 | 0.496543604 | 0.291 | 0.035 | 5.16E-212 | 10 | Stard13 |
| Dtx41 | 2.22E-215 | 0.47681946 | 0.256 | 0.028 | 4.49E-211 | 10 | Dtx4 |
| Grina1 | 1.66E-214 | 0.783471266 | 0.569 | 0.13 | 3.37E-210 | 10 | Grina |
| Man2a13 | 2.79E-214 | 1.149069982 | 0.745 | 0.223 | 5.65E-210 | 10 | Man2a1 |
| Calr4 | 5.91E-213 | 1.303101342 | 0.92 | 0.396 | 1.20E-208 | 10 | Calr |
| Stt3a | 2.41E-212 | 0.786485169 | 0.55 | 0.123 | 4.88E-208 | 10 | Stt3a |
| Aff3 | 8.53E-211 | 0.528293912 | 0.261 | 0.029 | 1.73E-206 | 10 | Aff3 |
| Slc39a7 | 8.66E-211 | 0.619059705 | 0.458 | 0.085 | 1.75E-206 | 10 | Slc39a7 |
| Il15 | 1.57E-210 | 0.752786963 | 0.477 | 0.093 | 3.18E-206 | 10 | Il15 |
| Ubtd2 | 2.94E-209 | 0.470844767 | 0.281 | 0.034 | 5.95E-205 | 10 | Ubtd2 |
| Ctsb2 | 6.96E-208 | 0.850772306 | 0.856 | 0.285 | 1.41E-203 | 10 | Ctsb |
| Snx93 | 1.10E-204 | 0.852694739 | 0.641 | 0.165 | 2.23E-200 | 10 | Snx9 |
| Lpar4 | 3.64E-204 | 0.590574439 | 0.342 | 0.05 | 7.36E-200 | 10 | Lpar4 |
| Armcx3 | 1.07E-203 | 0.52506644 | 0.386 | 0.064 | 2.16E-199 | 10 | Armcx3 |
| Cald13 | 1.69E-203 | 1.043529058 | 0.903 | 0.309 | 3.42E-199 | 10 | Cald1 |
| Ccdc501 | 3.32E-203 | 0.860490814 | 0.734 | 0.217 | 6.71E-199 | 10 | Ccdc50 |
| Sep-43 | 4.33E-202 | 0.79909597 | 0.631 | 0.154 | 8.76E-198 | 10 | 4-Sep |
| Zfpm2 | 6.07E-202 | 0.953435649 | 0.403 | 0.071 | 1.23E-197 | 10 | Zfpm2 |
| Ntn4 | 1.11E-201 | 0.43374334 | 0.258 | 0.029 | 2.25E-197 | 10 | Ntn4 |
| Por | 1.91E-201 | 0.769339238 | 0.48 | 0.099 | 3.87E-197 | 10 | Por |
| Kdelc2 | 2.93E-201 | 0.526393961 | 0.328 | 0.048 | 5.94E-197 | 10 | Kdelc2 |
| Creg11 | 4.33E-201 | 0.844725613 | 0.658 | 0.178 | 8.77E-197 | 10 | Creg1 |
| Mill2 | 6.36E-201 | 0.654659132 | 0.377 | 0.062 | 1.29E-196 | 10 | Mill2 |
| Scarb22 | 6.90E-201 | 0.887315142 | 0.655 | 0.177 | 1.40E-196 | 10 | Scarb2 |
| Prickle13 | 2.29E-200 | 0.910796438 | 0.58 | 0.136 | 4.63E-196 | 10 | Prickle1 |
| Mettl1 | 2.47E-200 | 0.5152366 | 0.342 | 0.052 | 5.01E-196 | 10 | Mettl1 |
| P4hb2 | 2.94E-200 | 0.943251881 | 0.762 | 0.241 | 5.94E-196 | 10 | P4hb |
| Synm | 4.26E-200 | 0.505603362 | 0.275 | 0.034 | 8.62E-196 | 10 | Synm |
| Itpr12 | 1.31E-199 | 1.062174841 | 0.706 | 0.21 | 2.64E-195 | 10 | Itpr1 |
| Hdgfl32 | 1.37E-199 | 0.678242754 | 0.5 | 0.102 | 2.77E-195 | 10 | Hdgfl3 |
| Cyp20a1 | 2.21E-199 | 0.639632058 | 0.422 | 0.077 | 4.47E-195 | 10 | Cyp20a1 |
| Arhgap211 | 3.66E-199 | 0.771158142 | 0.483 | 0.102 | 7.41E-195 | 10 | Arhgap21 |
| Trf1 | 1.37E-198 | 0.903980049 | 0.369 | 0.062 | 2.77E-194 | 10 | Trf |

| | | | | | | | |
|---|---|---|---|---|---|---|---|
| Dnajc25 | 3.15E-198 | 0.505850276 | 0.327 | 0.048 | 6.38E-194 | 10 | Dnajc25 |
| Hsp90b15 | 5.45E-198 | 1.294575542 | 0.967 | 0.5 | 1.10E-193 | 10 | Hsp90b1 |
| Crtap | 8.18E-197 | 0.618108 | 0.397 | 0.069 | 1.66E-192 | 10 | Crtap |
| Tlr41 | 1.38E-196 | 0.55300988 | 0.325 | 0.048 | 2.79E-192 | 10 | Tlr4 |
| Daam1 | 1.09E-195 | 0.708124844 | 0.481 | 0.1 | 2.22E-191 | 10 | Daam1 |
| Glg13 | 1.45E-195 | 1.003340794 | 0.753 | 0.241 | 2.94E-191 | 10 | Glg1 |
| Ebf32 | 2.81E-195 | 0.741165119 | 0.534 | 0.117 | 5.68E-191 | 10 | Ebf3 |
| Tmed32 | 2.38E-194 | 0.963025643 | 0.734 | 0.232 | 4.82E-190 | 10 | Tmed3 |
| Prss234 | 7.65E-194 | 1.057120969 | 0.831 | 0.268 | 1.55E-189 | 10 | Prss23 |
| Serinc14 | 1.59E-193 | 0.904564409 | 0.761 | 0.237 | 3.23E-189 | 10 | Serinc1 |
| Amotl13 | 9.13E-193 | 0.830226024 | 0.666 | 0.174 | 1.85E-188 | 10 | Amotl1 |
| Mageh1 | 5.02E-192 | 0.489033131 | 0.359 | 0.058 | 1.02E-187 | 10 | Mageh1 |
| Itpripl22 | 5.46E-192 | 0.752540742 | 0.5 | 0.11 | 1.11E-187 | 10 | Itpripl2 |
| Vkorc1 | 2.42E-190 | 0.749798651 | 0.578 | 0.143 | 4.89E-186 | 10 | Vkorc1 |
| Nhs | 2.56E-190 | 0.648759439 | 0.292 | 0.04 | 5.19E-186 | 10 | Nhs |
| Slc39a1 | 4.65E-190 | 0.541665214 | 0.378 | 0.065 | 9.41E-186 | 10 | Slc39a1 |
| mt-Cytb5 | 6.50E-190 | 1.13575847 | 0.988 | 0.694 | 1.32E-185 | 10 | mt-Cytb |
| Mpg | 3.48E-189 | 0.469750809 | 0.32 | 0.048 | 7.04E-185 | 10 | Mpg |
| Arnt1 | 9.07E-189 | 0.753928653 | 0.553 | 0.133 | 1.84E-184 | 10 | Arnt |
| Mettl7a1 | 2.33E-187 | 0.473337279 | 0.306 | 0.044 | 4.71E-183 | 10 | Mettl7a1 |
| Grb103 | 6.67E-186 | 0.825588177 | 0.678 | 0.184 | 1.35E-181 | 10 | Grb10 |
| Spry21 | 2.32E-185 | 0.630808869 | 0.4 | 0.075 | 4.70E-181 | 10 | Spry2 |
| Zeb13 | 2.59E-185 | 1.294142921 | 0.814 | 0.311 | 5.23E-181 | 10 | Zeb1 |
| Sparc4 | 4.18E-185 | 1.212234007 | 0.973 | 0.39 | 8.46E-181 | 10 | Sparc |
| Sash13 | 5.05E-185 | 0.913212815 | 0.781 | 0.245 | 1.02E-180 | 10 | Sash1 |
| Lix1l | 8.20E-185 | 0.591841532 | 0.377 | 0.066 | 1.66E-180 | 10 | Lix1l |
| Cks1b | 5.35E-184 | 0.448284502 | 0.275 | 0.037 | 1.08E-179 | 10 | Cks1b |
| Tgfb21 | 8.98E-184 | 0.63346521 | 0.422 | 0.08 | 1.82E-179 | 10 | Tgfb2 |
| Gnb41 | 4.52E-183 | 0.661448402 | 0.475 | 0.1 | 9.15E-179 | 10 | Gnb4 |
| 9530068E07Rik | 9.02E-183 | 0.591955995 | 0.439 | 0.089 | 1.83E-178 | 10 | 9530068E07Rik |
| Pds5b1 | 1.38E-182 | 0.851244018 | 0.617 | 0.17 | 2.80E-178 | 10 | Pds5b |
| Eri3 | 6.21E-182 | 0.621464022 | 0.452 | 0.095 | 1.26E-177 | 10 | Eri3 |
| Copa1 | 1.58E-181 | 0.812852071 | 0.664 | 0.194 | 3.20E-177 | 10 | Copa |
| Pik3r11 | 4.65E-181 | 0.896648379 | 0.744 | 0.242 | 9.42E-177 | 10 | Pik3r1 |
| Cachd1 | 5.88E-181 | 0.473736645 | 0.256 | 0.033 | 1.19E-176 | 10 | Cachd1 |
| Adamts13 | 1.55E-180 | 0.71475094 | 0.444 | 0.089 | 3.15E-176 | 10 | Adamts1 |
| Camk2n12 | 4.57E-180 | 0.83300444 | 0.789 | 0.26 | 9.26E-176 | 10 | Camk2n1 |
| Ssr31 | 2.06E-178 | 0.843229023 | 0.637 | 0.186 | 4.17E-174 | 10 | Ssr3 |
| Col4a13 | 3.41E-178 | 1.133277548 | 0.969 | 0.379 | 6.91E-174 | 10 | Col4a1 |
| Acvr11 | 1.14E-177 | 0.537535247 | 0.347 | 0.059 | 2.30E-173 | 10 | Acvr1 |
| Xbp11 | 6.60E-177 | 0.734566858 | 0.65 | 0.186 | 1.34E-172 | 10 | Xbp1 |
| Gadd45a | 1.01E-176 | 0.439917218 | 0.358 | 0.061 | 2.04E-172 | 10 | Gadd45a |
| Ccdc71l | 1.46E-176 | 0.620567639 | 0.392 | 0.075 | 2.95E-172 | 10 | Ccdc71l |

| | | | | | | | |
|---|---|---|---|---|---|---|---|
| Arhgap35 | 5.60E-176 | 0.575928285 | 0.411 | 0.081 | 1.13E-171 | 10 | Arhgap35 |
| Nr3c13 | 1.55E-175 | 1.045065191 | 0.889 | 0.372 | 3.15E-171 | 10 | Nr3c1 |
| Tcaf11 | 2.74E-175 | 0.649606997 | 0.455 | 0.095 | 5.56E-171 | 10 | Tcaf1 |
| Mfap21 | 3.19E-174 | 0.638718386 | 0.466 | 0.099 | 6.46E-170 | 10 | Mfap2 |
| Mbtps1 | 3.89E-174 | 0.624573589 | 0.445 | 0.095 | 7.88E-170 | 10 | Mbtps1 |
| Yipf5 | 1.19E-173 | 0.715242558 | 0.511 | 0.123 | 2.41E-169 | 10 | Yipf5 |
| Ccser1 | 1.21E-173 | 0.639575011 | 0.352 | 0.061 | 2.46E-169 | 10 | Ccser1 |
| Pkd1 | 3.13E-173 | 0.58483199 | 0.433 | 0.089 | 6.33E-169 | 10 | Pkd1 |
| Ppic2 | 4.25E-173 | 0.688216635 | 0.495 | 0.112 | 8.61E-169 | 10 | Ppic |
| Samd4 | 4.30E-173 | 0.526857033 | 0.312 | 0.049 | 8.70E-169 | 10 | Samd4 |
| Vav32 | 2.27E-172 | 0.986512114 | 0.741 | 0.245 | 4.59E-168 | 10 | Vav3 |
| Prrc1 | 2.72E-172 | 0.582904974 | 0.417 | 0.085 | 5.50E-168 | 10 | Prrc1 |
| Cbx6 | 9.04E-172 | 0.461601378 | 0.345 | 0.059 | 1.83E-167 | 10 | Cbx6 |
| Tm7sf3 | 1.15E-171 | 0.498361555 | 0.358 | 0.064 | 2.32E-167 | 10 | Tm7sf3 |
| Srebf2 | 1.71E-171 | 0.629770558 | 0.425 | 0.09 | 3.47E-167 | 10 | Srebf2 |
| Pten | 2.03E-171 | 0.935388505 | 0.812 | 0.311 | 4.11E-167 | 10 | Pten |
| Tnik1 | 8.44E-171 | 0.683494198 | 0.473 | 0.105 | 1.71E-166 | 10 | Tnik |
| Abca12 | 1.05E-169 | 0.763431976 | 0.503 | 0.121 | 2.12E-165 | 10 | Abca1 |
| Parva | 1.22E-169 | 0.58419671 | 0.473 | 0.104 | 2.46E-165 | 10 | Parva |
| Tead1 | 1.93E-169 | 0.564847445 | 0.397 | 0.077 | 3.92E-165 | 10 | Tead1 |
| Ptpn9 | 6.01E-169 | 0.669016332 | 0.533 | 0.133 | 1.22E-164 | 10 | Ptpn9 |
| Kdm5b1 | 1.64E-168 | 0.617708857 | 0.508 | 0.122 | 3.31E-164 | 10 | Kdm5b |
| Sec632 | 5.62E-168 | 0.829844134 | 0.648 | 0.193 | 1.14E-163 | 10 | Sec63 |
| Tmem39a | 3.48E-167 | 0.485905827 | 0.291 | 0.045 | 7.05E-163 | 10 | Tmem39a |
| Purg | 7.07E-167 | 0.415434398 | 0.264 | 0.037 | 1.43E-162 | 10 | Purg |
| Rnf217 | 8.28E-166 | 0.430603743 | 0.25 | 0.034 | 1.68E-161 | 10 | Rnf217 |
| Zfp521 | 9.42E-166 | 0.596601061 | 0.367 | 0.068 | 1.91E-161 | 10 | Zfp521 |
| Slc39a13 | 2.10E-165 | 0.494596804 | 0.312 | 0.052 | 4.24E-161 | 10 | Slc39a13 |
| Dnajc101 | 2.96E-165 | 0.700258339 | 0.522 | 0.131 | 5.99E-161 | 10 | Dnajc10 |
| Igf2r1 | 4.61E-165 | 0.749095646 | 0.506 | 0.126 | 9.34E-161 | 10 | Igf2r |
| Sp32 | 7.69E-165 | 0.775568585 | 0.622 | 0.181 | 1.56E-160 | 10 | Sp3 |
| Hgsnat | 7.74E-165 | 0.546534497 | 0.386 | 0.077 | 1.57E-160 | 10 | Hgsnat |
| Kcnq1ot11 | 8.63E-165 | 0.975728822 | 0.498 | 0.126 | 1.75E-160 | 10 | Kcnq1ot1 |
| Tacc14 | 5.04E-163 | 0.889281125 | 0.805 | 0.293 | 1.02E-158 | 10 | Tacc1 |
| Rap2b | 5.57E-163 | 0.556353511 | 0.359 | 0.068 | 1.13E-158 | 10 | Rap2b |
| Ust2 | 2.34E-162 | 0.697811403 | 0.388 | 0.079 | 4.73E-158 | 10 | Ust |
| Clip2 | 3.05E-162 | 0.489138308 | 0.292 | 0.047 | 6.17E-158 | 10 | Clip2 |
| Ifi2041 | 3.10E-161 | 0.65449547 | 0.412 | 0.087 | 6.28E-157 | 10 | Ifi204 |
| Lrrc583 | 7.65E-161 | 0.784831094 | 0.662 | 0.204 | 1.55E-156 | 10 | Lrrc58 |
| Pofut2 | 1.71E-160 | 0.481511178 | 0.375 | 0.073 | 3.47E-156 | 10 | Pofut2 |
| Acp2 | 2.06E-160 | 0.416732548 | 0.258 | 0.037 | 4.16E-156 | 10 | Acp2 |
| B4galt13 | 1.04E-159 | 0.781945748 | 0.588 | 0.167 | 2.10E-155 | 10 | B4galt1 |
| Notch22 | 2.21E-159 | 0.568828806 | 0.372 | 0.074 | 4.47E-155 | 10 | Notch2 |

| | | | | | | | |
|---|---|---|---|---|---|---|---|
| Ano62 | 2.64E-159 | 0.759765555 | 0.598 | 0.171 | 5.34E-155 | 10 | Ano6 |
| mt-Nd45 | 3.16E-159 | 1.014665983 | 0.984 | 0.669 | 6.40E-155 | 10 | mt-Nd4 |
| Tcf7l13 | 3.58E-159 | 0.709505489 | 0.62 | 0.171 | 7.24E-155 | 10 | Tcf7l1 |
| Adgra3 | 3.93E-159 | 0.384776136 | 0.256 | 0.036 | 7.95E-155 | 10 | Adgra3 |
| Ftx | 6.88E-159 | 0.560817274 | 0.277 | 0.043 | 1.39E-154 | 10 | Ftx |
| Maml32 | 1.40E-158 | 0.987758915 | 0.641 | 0.198 | 2.83E-154 | 10 | Maml3 |
| Hikeshi | 6.60E-158 | 0.594562469 | 0.516 | 0.131 | 1.34E-153 | 10 | Hikeshi |
| Ifngr15 | 2.81E-157 | 0.999284503 | 0.881 | 0.382 | 5.68E-153 | 10 | Ifngr1 |
| G0s22 | 1.88E-156 | 0.488793691 | 0.867 | 0.291 | 3.80E-152 | 10 | G0s2 |
| Klhl22 | 5.08E-156 | 0.419466685 | 0.259 | 0.038 | 1.03E-151 | 10 | Klhl22 |
| Rnd3 | 5.51E-156 | 0.527163755 | 0.422 | 0.089 | 1.12E-151 | 10 | Rnd3 |
| Rab3il11 | 1.09E-155 | 0.440683939 | 0.342 | 0.063 | 2.21E-151 | 10 | Rab3il1 |
| Snx241 | 2.58E-155 | 0.480445098 | 0.312 | 0.054 | 5.22E-151 | 10 | Snx24 |
| Xist4 | 4.03E-155 | 1.1178609 | 0.922 | 0.453 | 8.16E-151 | 10 | Xist |
| Tmem106b | 5.36E-155 | 0.574354298 | 0.486 | 0.119 | 1.09E-150 | 10 | Tmem106b |
| Smarca23 | 3.96E-154 | 0.861045354 | 0.756 | 0.27 | 8.01E-150 | 10 | Smarca2 |
| Msmo1 | 4.21E-154 | 0.631685399 | 0.3 | 0.052 | 8.53E-150 | 10 | Msmo1 |
| Manf1 | 8.02E-154 | 1.046579792 | 0.787 | 0.318 | 1.62E-149 | 10 | Manf |
| Slc12a4 | 1.33E-153 | 0.481014584 | 0.297 | 0.049 | 2.69E-149 | 10 | Slc12a4 |
| Scml4 | 1.79E-153 | 0.48758331 | 0.253 | 0.037 | 3.63E-149 | 10 | Scml4 |
| Mpdz1 | 2.09E-153 | 0.569551224 | 0.419 | 0.091 | 4.23E-149 | 10 | Mpdz |
| Mia32 | 1.08E-152 | 0.766279323 | 0.564 | 0.161 | 2.19E-148 | 10 | Mia3 |
| Tram11 | 2.51E-152 | 0.738886094 | 0.652 | 0.209 | 5.08E-148 | 10 | Tram1 |
| Cblb1 | 2.55E-152 | 0.800767816 | 0.564 | 0.162 | 5.16E-148 | 10 | Cblb |
| Lrmda5 | 2.85E-152 | 1.241211302 | 0.825 | 0.358 | 5.78E-148 | 10 | Lrmda |
| Wipf12 | 4.11E-152 | 0.874697172 | 0.734 | 0.257 | 8.32E-148 | 10 | Wipf1 |
| Cyp51 | 4.25E-152 | 0.529601201 | 0.28 | 0.046 | 8.61E-148 | 10 | Cyp51 |
| Adamts101 | 7.97E-152 | 0.621928792 | 0.416 | 0.093 | 1.61E-147 | 10 | Adamts10 |
| Csf12 | 1.69E-151 | 0.555667068 | 0.448 | 0.102 | 3.43E-147 | 10 | Csf1 |
| Nisch3 | 4.39E-151 | 0.685754522 | 0.647 | 0.201 | 8.89E-147 | 10 | Nisch |
| Psap2 | 6.26E-151 | 0.762719262 | 0.903 | 0.387 | 1.27E-146 | 10 | Psap |
| Ebf13 | 3.78E-150 | 1.036824443 | 0.936 | 0.362 | 7.65E-146 | 10 | Ebf1 |
| Sec624 | 8.24E-150 | 0.891260482 | 0.872 | 0.384 | 1.67E-145 | 10 | Sec62 |
| Cd91 | 1.11E-149 | 0.878542486 | 0.87 | 0.375 | 2.24E-145 | 10 | Cd9 |
| Acaca | 1.29E-149 | 0.572842939 | 0.369 | 0.077 | 2.61E-145 | 10 | Acaca |
| Pdia41 | 1.31E-149 | 0.700464282 | 0.553 | 0.158 | 2.66E-145 | 10 | Pdia4 |
| Tmem176b5 | 3.20E-149 | 0.97153793 | 0.923 | 0.428 | 6.49E-145 | 10 | Tmem176b |
| Aldh24 | 4.95E-149 | 0.863385249 | 0.9 | 0.385 | 1.00E-144 | 10 | Aldh2 |
| Ttc33 | 4.98E-149 | 0.779989588 | 0.705 | 0.242 | 1.01E-144 | 10 | Ttc3 |
| Sesn31 | 5.29E-149 | 0.5697556 | 0.483 | 0.121 | 1.07E-144 | 10 | Sesn3 |
| Spred14 | 3.34E-148 | 0.805404084 | 0.719 | 0.242 | 6.76E-144 | 10 | Spred1 |
| Sec11a1 | 3.47E-148 | 0.732798781 | 0.758 | 0.274 | 7.03E-144 | 10 | Sec11a |
| Prkca1 | 5.03E-148 | 0.587955467 | 0.414 | 0.093 | 1.02E-143 | 10 | Prkca |

| | | | | | | | |
|---|---|---|---|---|---|---|---|
| Pdia65 | 5.13E-148 | 0.935381791 | 0.805 | 0.327 | 1.04E-143 | 10 | Pdia6 |
| Phldb1 | 7.78E-148 | 0.454380094 | 0.327 | 0.06 | 1.57E-143 | 10 | Phldb1 |
| Sar1a | 8.95E-148 | 0.642714187 | 0.622 | 0.189 | 1.81E-143 | 10 | Sar1a |
| Surf4 | 2.23E-147 | 0.651969029 | 0.505 | 0.136 | 4.51E-143 | 10 | Surf4 |
| Tnfrsf1a3 | 2.28E-147 | 0.762448569 | 0.664 | 0.212 | 4.61E-143 | 10 | Tnfrsf1a |
| Palld | 5.17E-147 | 0.286219829 | 0.291 | 0.048 | 1.05E-142 | 10 | Palld |
| Ifitm24 | 7.86E-147 | 1.044407092 | 0.958 | 0.454 | 1.59E-142 | 10 | Ifitm2 |
| Atxn101 | 2.08E-146 | 0.62642755 | 0.573 | 0.166 | 4.22E-142 | 10 | Atxn10 |
| Ahnak4 | 7.04E-146 | 0.973883355 | 0.955 | 0.461 | 1.43E-141 | 10 | Ahnak |
| Rsrc12 | 7.48E-146 | 0.866311971 | 0.706 | 0.25 | 1.51E-141 | 10 | Rsrc1 |
| Rev3l1 | 7.66E-146 | 0.714335951 | 0.533 | 0.149 | 1.55E-141 | 10 | Rev3l |
| Cnpy2 | 2.19E-145 | 0.694074961 | 0.719 | 0.253 | 4.43E-141 | 10 | Cnpy2 |
| Pmepa12 | 2.91E-145 | 0.69738817 | 0.719 | 0.241 | 5.89E-141 | 10 | Pmepa1 |
| Hdlbp2 | 2.97E-145 | 0.673346888 | 0.77 | 0.287 | 6.02E-141 | 10 | Hdlbp |
| Srpr3 | 4.10E-145 | 0.698844728 | 0.698 | 0.236 | 8.30E-141 | 10 | Srpr |
| Creb3l21 | 4.27E-145 | 0.613180974 | 0.47 | 0.116 | 8.64E-141 | 10 | Creb3l2 |
| Acsl3 | 2.38E-144 | 0.456207596 | 0.256 | 0.04 | 4.81E-140 | 10 | Acsl3 |
| Grn3 | 3.05E-144 | 0.658248422 | 0.675 | 0.214 | 6.18E-140 | 10 | Grn |
| Ifi211 | 3.47E-144 | 0.431438962 | 0.272 | 0.045 | 7.02E-140 | 10 | Ifi211 |
| Lgals3bp2 | 9.63E-144 | 0.707453062 | 0.505 | 0.139 | 1.95E-139 | 10 | Lgals3bp |
| Stat5b1 | 1.32E-143 | 0.661626846 | 0.509 | 0.138 | 2.67E-139 | 10 | Stat5b |
| Tmem159 | 1.32E-143 | 0.562609888 | 0.434 | 0.103 | 2.68E-139 | 10 | Tmem159 |
| Cxcl124 | 1.76E-142 | 0.294321159 | 0.886 | 0.309 | 3.56E-138 | 10 | Cxcl12 |
| Pdia35 | 6.84E-142 | 0.932191897 | 0.903 | 0.433 | 1.38E-137 | 10 | Pdia3 |
| mt-Co36 | 7.03E-142 | 0.889305083 | 0.989 | 0.721 | 1.42E-137 | 10 | mt-Co3 |
| Sbf23 | 9.79E-142 | 0.753508545 | 0.631 | 0.198 | 1.98E-137 | 10 | Sbf2 |
| Rab11fip2 | 1.04E-141 | 0.557721559 | 0.423 | 0.1 | 2.12E-137 | 10 | Rab11fip2 |
| Morf4l2 | 1.51E-141 | 0.618341103 | 0.637 | 0.204 | 3.05E-137 | 10 | Morf4l2 |
| Tspan53 | 1.81E-141 | 0.862334563 | 0.606 | 0.189 | 3.66E-137 | 10 | Tspan5 |
| Gstm5 | 5.41E-141 | 0.25536509 | 0.317 | 0.058 | 1.09E-136 | 10 | Gstm5 |
| Hlf | 1.43E-140 | 0.498521783 | 0.297 | 0.053 | 2.89E-136 | 10 | Hlf |
| Thy1 | 1.59E-140 | 0.437667554 | 0.303 | 0.055 | 3.22E-136 | 10 | Thy1 |
| Phf21a3 | 3.61E-140 | 0.846809862 | 0.692 | 0.243 | 7.32E-136 | 10 | Phf21a |
| Kdelr11 | 6.14E-140 | 0.67550264 | 0.672 | 0.227 | 1.24E-135 | 10 | Kdelr1 |
| Cebpb1 | 3.18E-138 | 0.653714999 | 0.523 | 0.149 | 6.43E-134 | 10 | Cebpb |
| Tm9sf34 | 7.78E-138 | 0.813221648 | 0.795 | 0.325 | 1.58E-133 | 10 | Tm9sf3 |
| Cep1701 | 8.49E-138 | 0.639516404 | 0.466 | 0.122 | 1.72E-133 | 10 | Cep170 |
| Dnajc34 | 1.15E-137 | 0.88411711 | 0.836 | 0.36 | 2.33E-133 | 10 | Dnajc3 |
| Itm2c2 | 1.20E-137 | 0.739056605 | 0.598 | 0.19 | 2.43E-133 | 10 | Itm2c |
| Prkg12 | 2.79E-137 | 0.84382839 | 0.613 | 0.187 | 5.65E-133 | 10 | Prkg1 |
| Ubr11 | 8.16E-137 | 0.671588448 | 0.536 | 0.154 | 1.65E-132 | 10 | Ubr1 |
| Selenon | 1.38E-136 | 0.408279295 | 0.283 | 0.05 | 2.79E-132 | 10 | Selenon |
| Abcc51 | 5.52E-136 | 0.475036297 | 0.353 | 0.074 | 1.12E-131 | 10 | Abcc5 |

| | | | | | | | |
|---|---|---|---|---|---|---|---|
| Tpcn1 | 6.21E-136 | 0.43153888 | 0.303 | 0.057 | 1.26E-131 | 10 | Tpcn1 |
| Ephx12 | 9.54E-136 | 0.527585994 | 0.461 | 0.114 | 1.93E-131 | 10 | Ephx1 |
| Frmd4a1 | 1.18E-135 | 0.660484183 | 0.5 | 0.14 | 2.38E-131 | 10 | Frmd4a |
| Snx181 | 1.24E-135 | 0.52124028 | 0.375 | 0.084 | 2.50E-131 | 10 | Snx18 |
| Ttc283 | 1.91E-135 | 0.803537289 | 0.745 | 0.275 | 3.87E-131 | 10 | Ttc28 |
| Zfp5032 | 2.06E-135 | 0.60788451 | 0.516 | 0.139 | 4.17E-131 | 10 | Zfp503 |
| Rpn22 | 7.26E-135 | 0.699511393 | 0.619 | 0.203 | 1.47E-130 | 10 | Rpn2 |
| Pam3 | 8.05E-135 | 0.742918128 | 0.631 | 0.205 | 1.63E-130 | 10 | Pam |
| Rhobtb1 | 2.49E-134 | 0.451097763 | 0.303 | 0.057 | 5.03E-130 | 10 | Rhobtb1 |
| Lrpap12 | 2.61E-134 | 0.687294314 | 0.597 | 0.188 | 5.28E-130 | 10 | Lrpap1 |
| Acap25 | 6.64E-134 | 0.760743159 | 0.713 | 0.26 | 1.34E-129 | 10 | Acap2 |
| Tmem176a5 | 9.83E-134 | 0.895480931 | 0.887 | 0.407 | 1.99E-129 | 10 | Tmem176a |
| Tmem110 | 1.30E-133 | 0.46693095 | 0.312 | 0.061 | 2.63E-129 | 10 | Tmem110 |
| Dpy19l1 | 5.20E-133 | 0.529157968 | 0.344 | 0.074 | 1.05E-128 | 10 | Dpy19l1 |
| Ltbr | 5.40E-133 | 0.505581773 | 0.369 | 0.081 | 1.09E-128 | 10 | Ltbr |
| Lman13 | 1.54E-132 | 0.683395373 | 0.664 | 0.229 | 3.12E-128 | 10 | Lman1 |
| Nktr1 | 2.77E-132 | 0.764996924 | 0.686 | 0.248 | 5.61E-128 | 10 | Nktr |
| Fam69a1 | 4.10E-132 | 0.622356759 | 0.455 | 0.12 | 8.29E-128 | 10 | Fam69a |
| Dtx3 | 7.43E-132 | 0.473513219 | 0.3 | 0.057 | 1.50E-127 | 10 | Dtx3 |
| Layn1 | 8.62E-132 | 0.483761906 | 0.356 | 0.076 | 1.75E-127 | 10 | Layn |
| Cdc42bpa3 | 1.49E-131 | 0.622364804 | 0.591 | 0.176 | 3.01E-127 | 10 | Cdc42bpa |
| Dnajc14 | 1.55E-131 | 0.866451475 | 0.78 | 0.313 | 3.13E-127 | 10 | Dnajc1 |
| Rere3 | 2.99E-131 | 0.727190565 | 0.714 | 0.262 | 6.05E-127 | 10 | Rere |
| Shc1 | 8.06E-131 | 0.440268 | 0.297 | 0.057 | 1.63E-126 | 10 | Shc1 |
| Fam129a2 | 1.15E-130 | 0.678158565 | 0.492 | 0.138 | 2.32E-126 | 10 | Fam129a |
| Rnf145 | 1.74E-130 | 0.484489217 | 0.344 | 0.074 | 3.52E-126 | 10 | Rnf145 |
| Rnf144a1 | 8.73E-130 | 0.583580371 | 0.414 | 0.101 | 1.77E-125 | 10 | Rnf144a |
| Zfp260 | 1.81E-129 | 0.444049015 | 0.419 | 0.102 | 3.65E-125 | 10 | Zfp260 |
| Nfat53 | 2.00E-129 | 0.738261083 | 0.709 | 0.257 | 4.05E-125 | 10 | Nfat5 |
| Tcf43 | 3.04E-129 | 0.913440061 | 0.969 | 0.461 | 6.15E-125 | 10 | Tcf4 |
| Npdc12 | 4.18E-129 | 0.507254147 | 0.478 | 0.126 | 8.46E-125 | 10 | Npdc1 |
| Stk33 | 7.77E-129 | 0.58647297 | 0.536 | 0.158 | 1.57E-124 | 10 | Stk3 |
| Fer1 | 8.93E-129 | 0.488207012 | 0.447 | 0.114 | 1.81E-124 | 10 | Fer |
| Psip1 | 9.01E-129 | 0.603046409 | 0.588 | 0.186 | 1.82E-124 | 10 | Psip1 |
| Scmh1 | 1.06E-128 | 0.627495573 | 0.473 | 0.131 | 2.14E-124 | 10 | Scmh1 |
| Zc3hav11 | 1.46E-128 | 1.155201149 | 0.573 | 0.196 | 2.95E-124 | 10 | Zc3hav1 |
| Trp53i13 | 2.36E-128 | 0.417234133 | 0.284 | 0.053 | 4.78E-124 | 10 | Trp53i13 |
| Tm9sf4 | 3.00E-128 | 0.571601059 | 0.461 | 0.124 | 6.07E-124 | 10 | Tm9sf4 |
| Plcl1 | 4.60E-128 | 0.556843481 | 0.267 | 0.048 | 9.31E-124 | 10 | Plcl1 |
| Sec24a | 5.03E-128 | 0.566961094 | 0.353 | 0.08 | 1.02E-123 | 10 | Sec24a |
| Cd815 | 5.91E-128 | 0.951703865 | 0.972 | 0.516 | 1.20E-123 | 10 | Cd81 |
| mt-Nd24 | 7.55E-128 | 0.8839216 | 0.986 | 0.682 | 1.53E-123 | 10 | mt-Nd2 |
| Lmna3 | 8.12E-128 | 0.669965193 | 0.595 | 0.189 | 1.64E-123 | 10 | Lmna |

| | | | | | | | |
|---|---|---|---|---|---|---|---|
| mt-Nd4l3 | 1.65E-127 | 0.848191388 | 0.938 | 0.481 | 3.34E-123 | 10 | mt-Nd4l |
| Arhgap101 | 2.13E-127 | 0.571372119 | 0.452 | 0.12 | 4.32E-123 | 10 | Arhgap10 |
| Eva1b4 | 2.33E-127 | 0.668755886 | 0.756 | 0.275 | 4.72E-123 | 10 | Eva1b |
| Lamp16 | 7.94E-127 | 0.781024021 | 0.873 | 0.407 | 1.61E-122 | 10 | Lamp1 |
| Canx5 | 9.08E-127 | 0.797861164 | 0.731 | 0.297 | 1.84E-122 | 10 | Canx |
| Cul1 | 9.77E-127 | 0.657251892 | 0.72 | 0.267 | 1.98E-122 | 10 | Cul1 |
| Rin23 | 2.78E-126 | 0.644243943 | 0.566 | 0.174 | 5.63E-122 | 10 | Rin2 |
| Spry41 | 3.28E-126 | 0.517368859 | 0.388 | 0.09 | 6.63E-122 | 10 | Spry4 |
| Col4a23 | 5.87E-126 | 0.671463009 | 0.92 | 0.347 | 1.19E-121 | 10 | Col4a2 |
| Ptov1 | 9.70E-126 | 0.387969776 | 0.439 | 0.111 | 1.96E-121 | 10 | Ptov1 |
| Gstm2 | 2.12E-125 | 0.334609031 | 0.438 | 0.108 | 4.30E-121 | 10 | Gstm2 |
| Sec23a | 4.06E-125 | 0.470076551 | 0.312 | 0.064 | 8.22E-121 | 10 | Sec23a |
| Actn1 | 4.87E-125 | 0.497646391 | 0.348 | 0.078 | 9.86E-121 | 10 | Actn1 |
| Hmgcs1 | 1.00E-124 | 0.594700813 | 0.358 | 0.082 | 2.03E-120 | 10 | Hmgcs1 |
| Copb11 | 1.22E-124 | 0.607489815 | 0.591 | 0.196 | 2.48E-120 | 10 | Copb1 |
| Ergic1 | 1.39E-124 | 0.40942089 | 0.341 | 0.075 | 2.82E-120 | 10 | Ergic1 |
| Fto1 | 1.58E-124 | 0.65793786 | 0.57 | 0.182 | 3.19E-120 | 10 | Fto |
| Fbxw8 | 2.09E-124 | 0.43489123 | 0.339 | 0.074 | 4.24E-120 | 10 | Fbxw8 |
| Tigd2 | 3.37E-124 | 0.494468467 | 0.427 | 0.111 | 6.81E-120 | 10 | Tigd2 |
| Ssr1 | 6.93E-124 | 0.584532105 | 0.562 | 0.178 | 1.40E-119 | 10 | Ssr1 |
| Gpr180 | 1.54E-123 | 0.427902002 | 0.302 | 0.061 | 3.12E-119 | 10 | Gpr180 |
| Lsp12 | 2.38E-123 | 0.571284616 | 0.475 | 0.132 | 4.81E-119 | 10 | Lsp1 |
| Pxdn | 3.21E-123 | 0.448048377 | 0.325 | 0.068 | 6.50E-119 | 10 | Pxdn |
| Sdf21 | 6.03E-123 | 0.610345704 | 0.53 | 0.163 | 1.22E-118 | 10 | Sdf2 |
| Rcn22 | 9.54E-123 | 0.580480125 | 0.481 | 0.135 | 1.93E-118 | 10 | Rcn2 |
| Ssbp3 | 5.17E-122 | 0.54528671 | 0.531 | 0.162 | 1.05E-117 | 10 | Ssbp3 |
| Crtc32 | 6.08E-121 | 0.663083932 | 0.512 | 0.157 | 1.23E-116 | 10 | Crtc3 |
| Hook34 | 1.44E-120 | 0.687295074 | 0.728 | 0.28 | 2.91E-116 | 10 | Hook3 |
| mt-Nd35 | 1.97E-120 | 0.816948747 | 0.892 | 0.437 | 3.98E-116 | 10 | mt-Nd3 |
| Dip2c3 | 2.37E-120 | 0.65247777 | 0.636 | 0.214 | 4.79E-116 | 10 | Dip2c |
| Ppp2r3a | 3.22E-120 | 0.542019065 | 0.362 | 0.085 | 6.51E-116 | 10 | Ppp2r3a |
| Nrip13 | 3.24E-120 | 0.71339628 | 0.652 | 0.235 | 6.55E-116 | 10 | Nrip1 |
| Pde10a1 | 6.74E-120 | 0.686159336 | 0.347 | 0.079 | 1.37E-115 | 10 | Pde10a |
| Selenom4 | 1.34E-119 | 0.786297071 | 0.927 | 0.435 | 2.71E-115 | 10 | Selenom |
| Gm2a1 | 1.45E-119 | 0.408317219 | 0.359 | 0.083 | 2.94E-115 | 10 | Gm2a |
| Eps152 | 1.60E-119 | 0.641680787 | 0.52 | 0.161 | 3.25E-115 | 10 | Eps15 |
| Wwtr13 | 4.74E-119 | 0.618482919 | 0.68 | 0.23 | 9.61E-115 | 10 | Wwtr1 |
| St3gal2 | 5.17E-119 | 0.404221843 | 0.275 | 0.053 | 1.05E-114 | 10 | St3gal2 |
| Eif2ak31 | 6.00E-119 | 0.472186374 | 0.342 | 0.078 | 1.22E-114 | 10 | Eif2ak3 |
| Abi2 | 7.77E-119 | 0.501466256 | 0.362 | 0.086 | 1.57E-114 | 10 | Abi2 |
| Comt1 | 2.54E-118 | 0.537013513 | 0.57 | 0.185 | 5.14E-114 | 10 | Comt |
| Tfpi | 2.63E-118 | 0.414586701 | 0.28 | 0.054 | 5.33E-114 | 10 | Tfpi |
| Anxa52 | 3.62E-118 | 0.784683897 | 0.922 | 0.484 | 7.33E-114 | 10 | Anxa5 |

| | | | | | | | |
|---|---|---|---|---|---|---|---|
| Kank2 | 5.74E-118 | 0.397055122 | 0.262 | 0.048 | 1.16E-113 | 10 | Kank2 |
| Tmem5 | 7.10E-118 | 0.50629687 | 0.464 | 0.131 | 1.44E-113 | 10 | Tmem5 |
| Tbc1d2b1 | 1.40E-117 | 0.523277074 | 0.427 | 0.114 | 2.84E-113 | 10 | Tbc1d2b |
| Mid1ip1 | 1.41E-117 | 0.352533385 | 0.322 | 0.069 | 2.85E-113 | 10 | Mid1ip1 |
| 1110051M20Rik | 1.63E-117 | 0.432634243 | 0.253 | 0.046 | 3.30E-113 | 10 | 1110051M20Rik |
| Lamb22 | 3.29E-117 | 0.547418892 | 0.439 | 0.117 | 6.65E-113 | 10 | Lamb2 |
| Rpn12 | 5.77E-117 | 0.629762562 | 0.617 | 0.213 | 1.17E-112 | 10 | Rpn1 |
| Desi2 | 5.81E-117 | 0.517227424 | 0.527 | 0.163 | 1.18E-112 | 10 | Desi2 |
| mt-Nd53 | 5.92E-117 | 0.84202515 | 0.936 | 0.486 | 1.20E-112 | 10 | mt-Nd5 |
| Gng122 | 8.16E-117 | 0.540269816 | 0.591 | 0.192 | 1.65E-112 | 10 | Gng12 |
| Fnbp12 | 1.78E-116 | 0.585998415 | 0.648 | 0.221 | 3.60E-112 | 10 | Fnbp1 |
| Tspan31 | 2.13E-116 | 0.476590672 | 0.433 | 0.117 | 4.31E-112 | 10 | Tspan31 |
| Sgpl11 | 3.73E-116 | 0.424800461 | 0.275 | 0.055 | 7.56E-112 | 10 | Sgpl1 |
| Arl5a2 | 6.35E-116 | 0.511448758 | 0.502 | 0.151 | 1.29E-111 | 10 | Arl5a |
| Tcf124 | 3.89E-115 | 0.788866383 | 0.828 | 0.366 | 7.87E-111 | 10 | Tcf12 |
| Ikzf2 | 1.46E-114 | 0.389166205 | 0.256 | 0.048 | 2.95E-110 | 10 | Ikzf2 |
| Mtdh3 | 3.30E-114 | 0.761823484 | 0.838 | 0.377 | 6.69E-110 | 10 | Mtdh |
| Rtn44 | 9.91E-114 | 0.716936623 | 0.778 | 0.329 | 2.01E-109 | 10 | Rtn4 |
| Plbd2 | 1.01E-113 | 0.421406534 | 0.316 | 0.07 | 2.05E-109 | 10 | Plbd2 |
| Pttg1ip4 | 1.21E-113 | 0.596754642 | 0.703 | 0.259 | 2.46E-109 | 10 | Pttg1ip |
| Rhoq | 2.12E-113 | 0.442150593 | 0.392 | 0.102 | 4.30E-109 | 10 | Rhoq |
| Arntl | 2.98E-113 | 0.467317766 | 0.419 | 0.112 | 6.03E-109 | 10 | Arntl |
| Svil3 | 3.64E-113 | 0.752556803 | 0.637 | 0.234 | 7.36E-109 | 10 | Svil |
| Arhgef112 | 3.90E-113 | 0.521115532 | 0.377 | 0.094 | 7.90E-109 | 10 | Arhgef11 |
| Ocrl | 1.01E-112 | 0.400645846 | 0.27 | 0.054 | 2.04E-108 | 10 | Ocrl |
| H13 | 1.34E-112 | 0.535354341 | 0.489 | 0.15 | 2.72E-108 | 10 | H13 |
| Cers2 | 2.66E-112 | 0.452611863 | 0.42 | 0.114 | 5.39E-108 | 10 | Cers2 |
| Peli2 | 2.82E-112 | 0.449435831 | 0.294 | 0.062 | 5.72E-108 | 10 | Peli2 |
| Magi31 | 3.67E-112 | 0.52601668 | 0.386 | 0.099 | 7.43E-108 | 10 | Magi3 |
| Clcn4 | 4.05E-112 | 0.518482378 | 0.416 | 0.115 | 8.19E-108 | 10 | Clcn4 |
| Tnrc6c3 | 4.19E-112 | 0.685930468 | 0.752 | 0.306 | 8.49E-108 | 10 | Tnrc6c |
| Anxa6 | 4.74E-112 | 0.515261548 | 0.492 | 0.149 | 9.60E-108 | 10 | Anxa6 |
| Gns1 | 1.33E-111 | 0.527300132 | 0.431 | 0.12 | 2.70E-107 | 10 | Gns |
| Pde7b3 | 1.44E-111 | 0.672655062 | 0.661 | 0.233 | 2.92E-107 | 10 | Pde7b |
| Copg1 | 1.66E-111 | 0.413655621 | 0.369 | 0.092 | 3.36E-107 | 10 | Copg1 |
| Ugdh | 2.35E-111 | 0.319435854 | 0.262 | 0.051 | 4.76E-107 | 10 | Ugdh |
| Ifitm34 | 7.47E-111 | 0.831091675 | 0.984 | 0.528 | 1.51E-106 | 10 | Ifitm3 |
| Gjc11 | 1.12E-110 | 0.493319862 | 0.411 | 0.108 | 2.27E-106 | 10 | Gjc1 |
| Smad5 | 1.96E-110 | 0.426132728 | 0.369 | 0.092 | 3.96E-106 | 10 | Smad5 |
| Ahdc11 | 2.54E-110 | 0.488456712 | 0.397 | 0.106 | 5.15E-106 | 10 | Ahdc1 |
| Yipf3 | 4.01E-110 | 0.512003803 | 0.462 | 0.138 | 8.11E-106 | 10 | Yipf3 |
| Pear11 | 4.71E-110 | 0.458638336 | 0.377 | 0.096 | 9.54E-106 | 10 | Pear1 |
| Map3k53 | 5.09E-110 | 0.622253685 | 0.55 | 0.185 | 1.03E-105 | 10 | Map3k5 |

| | | | | | | | |
|---|---|---|---|---|---|---|---|
| Ddost1 | 7.33E-110 | 0.629946581 | 0.584 | 0.208 | 1.48E-105 | 10 | Ddost |
| Atraid3 | 9.13E-110 | 0.644569302 | 0.683 | 0.262 | 1.85E-105 | 10 | Atraid |
| Camkk2 | 1.76E-109 | 0.399674369 | 0.297 | 0.064 | 3.56E-105 | 10 | Camkk2 |
| Nckap13 | 3.00E-109 | 0.479708378 | 0.641 | 0.217 | 6.06E-105 | 10 | Nckap1 |
| Osmr3 | 5.04E-109 | 0.573301499 | 0.462 | 0.133 | 1.02E-104 | 10 | Osmr |
| Ift27 | 5.51E-109 | 0.372644989 | 0.347 | 0.084 | 1.12E-104 | 10 | Ift27 |
| Tex264 | 6.18E-109 | 0.464789262 | 0.369 | 0.093 | 1.25E-104 | 10 | Tex264 |
| Cxxc51 | 7.95E-109 | 0.36481002 | 0.402 | 0.108 | 1.61E-104 | 10 | Cxxc5 |
| Asap13 | 1.50E-108 | 0.676677918 | 0.811 | 0.346 | 3.05E-104 | 10 | Asap1 |
| Tusc31 | 1.83E-108 | 0.503952673 | 0.477 | 0.142 | 3.71E-104 | 10 | Tusc3 |
| Zfp8091 | 2.00E-108 | 0.440406386 | 0.334 | 0.08 | 4.05E-104 | 10 | Zfp809 |
| Ssr2 | 2.64E-108 | 0.566306264 | 0.567 | 0.194 | 5.34E-104 | 10 | Ssr2 |
| Lpgat1 | 6.30E-108 | 0.460484243 | 0.336 | 0.081 | 1.27E-103 | 10 | Lpgat1 |
| Tceal8 | 8.15E-108 | 0.304063549 | 0.452 | 0.126 | 1.65E-103 | 10 | Tceal8 |
| mt-Nd15 | 1.13E-107 | 0.81076668 | 0.953 | 0.591 | 2.28E-103 | 10 | mt-Nd1 |
| Golga3 | 1.54E-107 | 0.379029787 | 0.352 | 0.086 | 3.11E-103 | 10 | Golga3 |
| Sc5d | 1.55E-107 | 0.383288208 | 0.266 | 0.054 | 3.13E-103 | 10 | Sc5d |
| Ctsz2 | 1.60E-107 | 0.411637388 | 0.83 | 0.35 | 3.24E-103 | 10 | Ctsz |
| Vangl1 | 2.65E-107 | 0.390133986 | 0.288 | 0.061 | 5.37E-103 | 10 | Vangl1 |
| Pnisr4 | 2.78E-107 | 0.656106549 | 0.666 | 0.256 | 5.63E-103 | 10 | Pnisr |
| Esyt23 | 4.13E-107 | 0.666828877 | 0.634 | 0.234 | 8.35E-103 | 10 | Esyt2 |
| Cebpd | 4.91E-107 | 0.512930808 | 0.53 | 0.167 | 9.93E-103 | 10 | Cebpd |
| Ginm13 | 5.90E-107 | 0.523561828 | 0.534 | 0.172 | 1.20E-102 | 10 | Ginm1 |
| Myo1b1 | 6.11E-107 | 0.436965697 | 0.458 | 0.131 | 1.24E-102 | 10 | Myo1b |
| Ddb11 | 8.85E-107 | 0.467893545 | 0.47 | 0.142 | 1.79E-102 | 10 | Ddb1 |
| mt-Atp66 | 1.23E-106 | 0.742925285 | 0.992 | 0.749 | 2.49E-102 | 10 | mt-Atp6 |
| Srek11 | 1.85E-106 | 0.574999617 | 0.552 | 0.186 | 3.74E-102 | 10 | Srek1 |
| Ncor2 | 2.65E-106 | 0.485700328 | 0.411 | 0.115 | 5.38E-102 | 10 | Ncor2 |
| Dpysl23 | 3.42E-106 | 0.573343375 | 0.514 | 0.167 | 6.92E-102 | 10 | Dpysl2 |
| Tm9sf23 | 4.89E-106 | 0.666266817 | 0.648 | 0.247 | 9.91E-102 | 10 | Tm9sf2 |
| Rnft1 | 5.29E-106 | 0.427085727 | 0.306 | 0.07 | 1.07E-101 | 10 | Rnft1 |
| Gbf12 | 5.36E-106 | 0.553626566 | 0.528 | 0.174 | 1.09E-101 | 10 | Gbf1 |
| Ostc1 | 9.54E-106 | 0.588970665 | 0.661 | 0.254 | 1.93E-101 | 10 | Ostc |
| Ppib1 | 9.58E-106 | 0.770287728 | 0.925 | 0.546 | 1.94E-101 | 10 | Ppib |
| Sel1l2 | 2.10E-105 | 0.508888673 | 0.495 | 0.155 | 4.26E-101 | 10 | Sel1l |
| Tlcd2 | 2.19E-105 | 0.313332234 | 0.25 | 0.049 | 4.44E-101 | 10 | Tlcd2 |
| Gprasp1 | 2.40E-105 | 0.513781669 | 0.372 | 0.098 | 4.86E-101 | 10 | Gprasp1 |
| Fnip12 | 2.62E-105 | 0.560629256 | 0.459 | 0.139 | 5.30E-101 | 10 | Fnip1 |
| Flna | 2.78E-105 | 0.499947929 | 0.43 | 0.125 | 5.62E-101 | 10 | Flna |
| Il6st3 | 5.66E-105 | 0.419448567 | 0.658 | 0.223 | 1.15E-100 | 10 | Il6st |
| Fkbp21 | 8.30E-105 | 0.437269894 | 0.695 | 0.267 | 1.68E-100 | 10 | Fkbp2 |
| Slc16a9 | 1.21E-104 | 0.386343876 | 0.266 | 0.054 | 2.46E-100 | 10 | Slc16a9 |
| Ston11 | 2.63E-104 | 0.40302766 | 0.347 | 0.085 | 5.32E-100 | 10 | Ston1 |

| | | | | | | | |
|---|---|---|---|---|---|---|---|
| Tmsb101 | 2.72E-104 | 0.46671493 | 0.955 | 0.49 | 5.52E-100 | 10 | Tmsb10 |
| Impad1 | 3.05E-104 | 0.457028834 | 0.367 | 0.095 | 6.17E-100 | 10 | Impad1 |
| Akt2 | 3.33E-104 | 0.328378303 | 0.278 | 0.06 | 6.73E-100 | 10 | Akt2 |
| Usp9x | 3.76E-104 | 0.527719642 | 0.62 | 0.226 | 7.60E-100 | 10 | Usp9x |
| Atxn22 | 3.76E-104 | 0.47332171 | 0.447 | 0.133 | 7.61E-100 | 10 | Atxn2 |
| Tmed71 | 9.77E-104 | 0.507250968 | 0.466 | 0.142 | 1.98E-99 | 10 | Tmed7 |
| Sep-75 | 1.02E-103 | 0.651139754 | 0.897 | 0.452 | 2.07E-99 | 10 | 7-Sep |
| Ddah22 | 1.08E-103 | 0.316746506 | 0.602 | 0.197 | 2.18E-99 | 10 | Ddah2 |
| Arcn1 | 1.17E-103 | 0.438499114 | 0.553 | 0.187 | 2.37E-99 | 10 | Arcn1 |
| Nbea3 | 3.30E-103 | 0.54673274 | 0.569 | 0.188 | 6.69E-99 | 10 | Nbea |
| Pdlim7 | 1.01E-102 | 0.367821105 | 0.302 | 0.069 | 2.05E-98 | 10 | Pdlim7 |
| Adap21 | 1.21E-102 | 0.386027794 | 0.261 | 0.053 | 2.44E-98 | 10 | Adap2 |
| Stam | 1.28E-102 | 0.362218254 | 0.289 | 0.064 | 2.59E-98 | 10 | Stam |
| Degs13 | 1.93E-102 | 0.519667405 | 0.533 | 0.177 | 3.91E-98 | 10 | Degs1 |
| Sgpp1 | 2.73E-102 | 0.378900047 | 0.305 | 0.071 | 5.53E-98 | 10 | Sgpp1 |
| Dennd1a3 | 4.57E-102 | 0.652549884 | 0.639 | 0.245 | 9.26E-98 | 10 | Dennd1a |
| Ski4 | 1.48E-101 | 0.590080517 | 0.575 | 0.207 | 2.99E-97 | 10 | Ski |
| Snd12 | 2.06E-101 | 0.509458467 | 0.506 | 0.165 | 4.18E-97 | 10 | Snd1 |
| Chic2 | 2.65E-101 | 0.543723999 | 0.639 | 0.244 | 5.37E-97 | 10 | Chic2 |
| Adamts92 | 3.61E-101 | 0.560652105 | 0.417 | 0.118 | 7.30E-97 | 10 | Adamts9 |
| Cdc16 | 8.40E-101 | 0.356618249 | 0.298 | 0.069 | 1.70E-96 | 10 | Cdc16 |
| Atp6ap11 | 1.22E-100 | 0.472022526 | 0.461 | 0.143 | 2.46E-96 | 10 | Atp6ap1 |
| Mtch11 | 1.44E-100 | 0.546382899 | 0.733 | 0.307 | 2.91E-96 | 10 | Mtch1 |
| Arid1b3 | 3.04E-100 | 0.652460524 | 0.642 | 0.25 | 6.15E-96 | 10 | Arid1b |
| Pgrmc13 | 3.25E-100 | 0.495202426 | 0.595 | 0.21 | 6.58E-96 | 10 | Pgrmc1 |
| Stambpl11 | 3.96E-100 | 0.447956965 | 0.284 | 0.065 | 8.01E-96 | 10 | Stambpl1 |
| Magt12 | 4.10E-100 | 0.580375994 | 0.48 | 0.157 | 8.29E-96 | 10 | Magt1 |
| Myo9a3 | 4.31E-100 | 0.579008932 | 0.503 | 0.166 | 8.73E-96 | 10 | Myo9a |
| mt-Co15 | 7.38E-100 | 0.747599348 | 0.986 | 0.714 | 1.50E-95 | 10 | mt-Co1 |
| Ppp3ca4 | 1.01E-99 | 0.737057011 | 0.833 | 0.384 | 2.05E-95 | 10 | Ppp3ca |
| Map4k52 | 1.06E-99 | 0.485156125 | 0.427 | 0.124 | 2.14E-95 | 10 | Map4k5 |
| Sep-92 | 1.30E-99 | 0.564835654 | 0.497 | 0.164 | 2.63E-95 | 10 | 9-Sep |
| Btrc | 1.53E-99 | 0.427574145 | 0.384 | 0.105 | 3.09E-95 | 10 | Btrc |
| Add12 | 1.83E-99 | 0.489768185 | 0.581 | 0.206 | 3.70E-95 | 10 | Add1 |
| St52 | 2.08E-99 | 0.448044767 | 0.472 | 0.145 | 4.20E-95 | 10 | St5 |
| Fscn11 | 4.26E-99 | 0.374759749 | 0.311 | 0.073 | 8.63E-95 | 10 | Fscn1 |
| Sec61a1 | 7.44E-99 | 0.393291719 | 0.497 | 0.162 | 1.51E-94 | 10 | Sec61a1 |
| Prrc2b3 | 8.21E-99 | 0.564440791 | 0.602 | 0.222 | 1.66E-94 | 10 | Prrc2b |
| Copz21 | 9.22E-99 | 0.340234418 | 0.412 | 0.117 | 1.87E-94 | 10 | Copz2 |
| Cask2 | 1.00E-98 | 0.548316657 | 0.439 | 0.133 | 2.03E-94 | 10 | Cask |
| Prkcb2 | 1.08E-98 | 0.483890884 | 0.483 | 0.15 | 2.19E-94 | 10 | Prkcb |
| Rhob3 | 1.18E-98 | 0.616289292 | 0.67 | 0.259 | 2.38E-94 | 10 | Rhob |
| L3mbtl3 | 2.04E-98 | 0.408082019 | 0.273 | 0.06 | 4.12E-94 | 10 | L3mbtl3 |

| | | | | | | | |
|---|---|---|---|---|---|---|---|
| 2410089E03Rik | 3.11E-98 | 0.439845497 | 0.345 | 0.089 | 6.29E-94 | 10 | 2410089E03Rik |
| Arl13 | 3.15E-98 | 0.459520567 | 0.689 | 0.277 | 6.38E-94 | 10 | Arl1 |
| Fam172a3 | 3.42E-98 | 0.648620094 | 0.683 | 0.276 | 6.92E-94 | 10 | Fam172a |
| Capg1 | 3.61E-98 | 0.408635109 | 0.361 | 0.095 | 7.30E-94 | 10 | Capg |
| Npepps | 5.15E-98 | 0.461873702 | 0.409 | 0.12 | 1.04E-93 | 10 | Npepps |
| Spaca6 | 5.30E-98 | 0.422324673 | 0.286 | 0.065 | 1.07E-93 | 10 | Spaca6 |
| Kifap3 | 5.40E-98 | 0.386961465 | 0.375 | 0.101 | 1.09E-93 | 10 | Kifap3 |
| Csgalnact2 | 2.27E-97 | 0.378337464 | 0.259 | 0.056 | 4.60E-93 | 10 | Csgalnact2 |
| Colgalt12 | 4.03E-97 | 0.503553015 | 0.442 | 0.138 | 8.16E-93 | 10 | Colgalt1 |
| Dnmt3a2 | 2.70E-96 | 0.519252484 | 0.445 | 0.141 | 5.46E-92 | 10 | Dnmt3a |
| Trim351 | 2.76E-96 | 0.483684324 | 0.439 | 0.134 | 5.58E-92 | 10 | Trim35 |
| App3 | 2.90E-96 | 0.608688016 | 0.93 | 0.408 | 5.86E-92 | 10 | App |
| Tgfbr24 | 4.55E-96 | 0.592145448 | 0.727 | 0.302 | 9.21E-92 | 10 | Tgfbr2 |
| Ttc39b1 | 4.72E-96 | 0.467223379 | 0.428 | 0.129 | 9.55E-92 | 10 | Ttc39b |
| Gja13 | 5.33E-96 | 0.424581545 | 0.425 | 0.122 | 1.08E-91 | 10 | Gja1 |
| Tmem87b | 5.99E-96 | 0.413166578 | 0.309 | 0.076 | 1.21E-91 | 10 | Tmem87b |
| Mesd1 | 6.06E-96 | 0.485656196 | 0.481 | 0.156 | 1.23E-91 | 10 | Mesd |
| Pnp3 | 1.13E-95 | 0.580692196 | 0.639 | 0.245 | 2.29E-91 | 10 | Pnp |
| Luc7l23 | 1.92E-95 | 0.655885171 | 0.842 | 0.395 | 3.88E-91 | 10 | Luc7l2 |
| Gorasp2 | 2.42E-95 | 0.446953352 | 0.403 | 0.12 | 4.90E-91 | 10 | Gorasp2 |
| Slfn54 | 2.68E-95 | 0.52331139 | 0.838 | 0.346 | 5.42E-91 | 10 | Slfn5 |
| Gkap11 | 2.92E-95 | 0.381880486 | 0.442 | 0.136 | 5.91E-91 | 10 | Gkap1 |
| Cyb5a2 | 3.03E-95 | 0.52567344 | 0.928 | 0.489 | 6.12E-91 | 10 | Cyb5a |
| Slc9a1 | 3.66E-95 | 0.406109249 | 0.286 | 0.067 | 7.41E-91 | 10 | Slc9a1 |
| Rufy31 | 4.03E-95 | 0.437347401 | 0.311 | 0.078 | 8.15E-91 | 10 | Rufy3 |
| Dennd4c1 | 1.09E-94 | 0.53386647 | 0.402 | 0.12 | 2.22E-90 | 10 | Dennd4c |
| Grk52 | 1.32E-94 | 0.577290535 | 0.469 | 0.151 | 2.67E-90 | 10 | Grk5 |
| Setd54 | 1.50E-94 | 0.604450513 | 0.642 | 0.256 | 3.04E-90 | 10 | Setd5 |
| Trip123 | 2.00E-94 | 0.54665128 | 0.636 | 0.248 | 4.04E-90 | 10 | Trip12 |
| Arl3 | 2.90E-94 | 0.343090493 | 0.502 | 0.162 | 5.88E-90 | 10 | Arl3 |
| Rnf132 | 2.93E-94 | 0.471555284 | 0.506 | 0.172 | 5.92E-90 | 10 | Rnf13 |
| Cabin1 | 2.96E-94 | 0.436136602 | 0.35 | 0.095 | 6.00E-90 | 10 | Cabin1 |
| Ogt2 | 3.10E-94 | 0.56247432 | 0.553 | 0.199 | 6.27E-90 | 10 | Ogt |
| Cntln2 | 5.13E-94 | 0.492513498 | 0.462 | 0.148 | 1.04E-89 | 10 | Cntln |
| Ctdsp22 | 1.29E-93 | 0.477772731 | 0.506 | 0.171 | 2.61E-89 | 10 | Ctdsp2 |
| Vwa5a1 | 1.63E-93 | 0.415720356 | 0.339 | 0.091 | 3.30E-89 | 10 | Vwa5a |
| Herpud12 | 1.98E-93 | 0.510586582 | 0.564 | 0.206 | 4.01E-89 | 10 | Herpud1 |
| Rbms13 | 2.90E-93 | 0.638849565 | 0.936 | 0.476 | 5.88E-89 | 10 | Rbms1 |
| Sec22b | 5.83E-93 | 0.369759354 | 0.403 | 0.12 | 1.18E-88 | 10 | Sec22b |
| Stt3b2 | 1.02E-92 | 0.522158489 | 0.475 | 0.159 | 2.07E-88 | 10 | Stt3b |
| Nt5c21 | 1.54E-92 | 0.424756038 | 0.336 | 0.09 | 3.12E-88 | 10 | Nt5c2 |
| Hsd17b111 | 1.91E-92 | 0.45066112 | 0.389 | 0.113 | 3.87E-88 | 10 | Hsd17b11 |
| Hmgn3 | 3.02E-92 | 0.267413028 | 0.511 | 0.164 | 6.12E-88 | 10 | Hmgn3 |

| | | | | | | | |
|---|---|---|---|---|---|---|---|
| E130102H24Rik | 3.22E-92 | 0.39225154 | 0.27 | 0.062 | 6.53E-88 | 10 | E130102H24Rik |
| Tuba1b2 | 4.36E-92 | 0.474171213 | 0.745 | 0.313 | 8.82E-88 | 10 | Tuba1b |
| Phip3 | 4.58E-92 | 0.621227157 | 0.628 | 0.248 | 9.27E-88 | 10 | Phip |
| Adprh | 6.55E-92 | 0.393550371 | 0.433 | 0.134 | 1.33E-87 | 10 | Adprh |
| Plscr43 | 6.68E-92 | 0.547132027 | 0.53 | 0.183 | 1.35E-87 | 10 | Plscr4 |
| Copb2 | 7.34E-92 | 0.435103414 | 0.558 | 0.203 | 1.49E-87 | 10 | Copb2 |
| Irf2bp2 | 9.28E-92 | 0.468619052 | 0.589 | 0.221 | 1.88E-87 | 10 | Irf2bp2 |
| mt-Co27 | 9.77E-92 | 0.686048049 | 0.988 | 0.723 | 1.98E-87 | 10 | mt-Co2 |
| Hexa2 | 1.21E-91 | 0.401576923 | 0.589 | 0.213 | 2.46E-87 | 10 | Hexa |
| Stxbp1 | 1.66E-91 | 0.331831286 | 0.266 | 0.06 | 3.37E-87 | 10 | Stxbp1 |
| Nucb13 | 2.13E-91 | 0.488229491 | 0.591 | 0.22 | 4.31E-87 | 10 | Nucb1 |
| Gstm13 | 2.21E-91 | 0.403959079 | 0.748 | 0.313 | 4.47E-87 | 10 | Gstm1 |
| Mydgf2 | 6.24E-91 | 0.48664533 | 0.589 | 0.224 | 1.26E-86 | 10 | Mydgf |
| Spcs11 | 7.55E-91 | 0.532332209 | 0.68 | 0.279 | 1.53E-86 | 10 | Spcs1 |
| Tm2d22 | 1.20E-90 | 0.464850603 | 0.559 | 0.204 | 2.43E-86 | 10 | Tm2d2 |
| Senp62 | 1.40E-90 | 0.508730006 | 0.698 | 0.289 | 2.83E-86 | 10 | Senp6 |
| Hmgn11 | 3.20E-90 | 0.404342996 | 0.744 | 0.32 | 6.49E-86 | 10 | Hmgn1 |
| Ubr52 | 3.43E-90 | 0.497630289 | 0.533 | 0.191 | 6.95E-86 | 10 | Ubr5 |
| Camk2d4 | 4.11E-90 | 0.567541061 | 0.655 | 0.261 | 8.32E-86 | 10 | Camk2d |
| Sqor | 5.39E-90 | 0.327246395 | 0.283 | 0.067 | 1.09E-85 | 10 | Sqor |
| Tgoln13 | 6.58E-90 | 0.487740061 | 0.62 | 0.239 | 1.33E-85 | 10 | Tgoln1 |
| Gmds | 7.10E-90 | 0.509270978 | 0.495 | 0.172 | 1.44E-85 | 10 | Gmds |
| Mon21 | 7.51E-90 | 0.514130626 | 0.445 | 0.146 | 1.52E-85 | 10 | Mon2 |
| Exoc6b3 | 9.24E-90 | 0.517641616 | 0.609 | 0.233 | 1.87E-85 | 10 | Exoc6b |
| Sec13 | 1.32E-89 | 0.296813736 | 0.428 | 0.131 | 2.68E-85 | 10 | Sec13 |
| Nptn3 | 1.42E-89 | 0.537958179 | 0.616 | 0.241 | 2.87E-85 | 10 | Nptn |
| Slc38a10 | 1.54E-89 | 0.467048644 | 0.403 | 0.124 | 3.11E-85 | 10 | Slc38a10 |
| Sec16a | 2.10E-89 | 0.381517726 | 0.302 | 0.076 | 4.25E-85 | 10 | Sec16a |
| Aff11 | 2.15E-89 | 0.559728702 | 0.456 | 0.154 | 4.36E-85 | 10 | Aff1 |
| Ergic31 | 2.24E-89 | 0.448681049 | 0.514 | 0.182 | 4.53E-85 | 10 | Ergic3 |
| Eprs | 2.38E-89 | 0.482923517 | 0.533 | 0.194 | 4.82E-85 | 10 | Eprs |
| Tmed22 | 3.32E-89 | 0.591091248 | 0.767 | 0.357 | 6.73E-85 | 10 | Tmed2 |
| Map3k20 | 4.24E-89 | 0.302151685 | 0.286 | 0.068 | 8.58E-85 | 10 | Map3k20 |
| Fmnl23 | 6.37E-89 | 0.575058468 | 0.611 | 0.235 | 1.29E-84 | 10 | Fmnl2 |
| Man1c11 | 6.55E-89 | 0.397418919 | 0.28 | 0.068 | 1.33E-84 | 10 | Man1c1 |
| Atp8b12 | 8.27E-89 | 0.412030379 | 0.356 | 0.098 | 1.67E-84 | 10 | Atp8b1 |
| Clptm1l | 9.96E-89 | 0.389493443 | 0.353 | 0.1 | 2.02E-84 | 10 | Clptm1l |
| Tmem167 | 1.97E-88 | 0.435441611 | 0.662 | 0.274 | 3.99E-84 | 10 | Tmem167 |
| Smc62 | 2.21E-88 | 0.533566523 | 0.634 | 0.254 | 4.48E-84 | 10 | Smc6 |
| Trps14 | 6.65E-88 | 0.577242639 | 0.595 | 0.228 | 1.35E-83 | 10 | Trps1 |
| Jade1 | 7.80E-88 | 0.379560422 | 0.375 | 0.109 | 1.58E-83 | 10 | Jade1 |
| Raph1 | 8.74E-88 | 0.460433881 | 0.37 | 0.109 | 1.77E-83 | 10 | Raph1 |
| Clasp23 | 1.01E-87 | 0.513535263 | 0.502 | 0.176 | 2.04E-83 | 10 | Clasp2 |

| | | | | | | | |
|---|---|---|---|---|---|---|---|
| Pikfyve | 1.19E-87 | 0.390019196 | 0.283 | 0.069 | 2.41E-83 | 10 | Pikfyve |
| Tnrc183 | 1.32E-87 | 0.46927184 | 0.486 | 0.166 | 2.67E-83 | 10 | Tnrc18 |
| Cnn33 | 2.64E-87 | 0.434442886 | 0.539 | 0.186 | 5.35E-83 | 10 | Cnn3 |
| Iah1 | 2.69E-87 | 0.341566579 | 0.419 | 0.131 | 5.45E-83 | 10 | Iah1 |
| Bcl9l | 3.19E-87 | 0.393233074 | 0.28 | 0.068 | 6.46E-83 | 10 | Bcl9l |
| Ctsd2 | 6.38E-87 | 0.508245914 | 0.72 | 0.305 | 1.29E-82 | 10 | Ctsd |
| Spag93 | 6.78E-87 | 0.587194672 | 0.759 | 0.35 | 1.37E-82 | 10 | Spag9 |
| Nucb21 | 8.74E-87 | 0.363745884 | 0.386 | 0.115 | 1.77E-82 | 10 | Nucb2 |
| Zcchc14 | 1.07E-86 | 0.385253657 | 0.298 | 0.075 | 2.16E-82 | 10 | Zcchc14 |
| Amfr | 3.18E-86 | 0.437142343 | 0.388 | 0.118 | 6.44E-82 | 10 | Amfr |
| mt-Atp84 | 3.86E-86 | 0.619746726 | 0.877 | 0.454 | 7.82E-82 | 10 | mt-Atp8 |
| Tmed104 | 5.32E-86 | 0.605027766 | 0.841 | 0.427 | 1.08E-81 | 10 | Tmed10 |
| Tshz24 | 6.14E-86 | 0.489244494 | 0.848 | 0.361 | 1.24E-81 | 10 | Tshz2 |
| Lrrc32 | 1.36E-85 | 0.318115004 | 0.264 | 0.061 | 2.75E-81 | 10 | Lrrc32 |
| Phka1 | 1.68E-85 | 0.360385732 | 0.258 | 0.06 | 3.40E-81 | 10 | Phka1 |
| Scfd1 | 1.91E-85 | 0.357923498 | 0.484 | 0.166 | 3.86E-81 | 10 | Scfd1 |
| Map3k21 | 2.01E-85 | 0.450603209 | 0.444 | 0.147 | 4.08E-81 | 10 | Map3k2 |
| Fam92a | 2.04E-85 | 0.301932282 | 0.394 | 0.118 | 4.12E-81 | 10 | Fam92a |
| Bfar | 2.34E-85 | 0.455819862 | 0.375 | 0.114 | 4.75E-81 | 10 | Bfar |
| Gpr108 | 2.51E-85 | 0.373241441 | 0.288 | 0.073 | 5.09E-81 | 10 | Gpr108 |
| Plin21 | 2.67E-85 | 0.42580923 | 0.334 | 0.094 | 5.41E-81 | 10 | Plin2 |
| Eif4g12 | 1.15E-84 | 0.520658718 | 0.583 | 0.229 | 2.33E-80 | 10 | Eif4g1 |
| Zfp445 | 1.15E-84 | 0.456017116 | 0.389 | 0.121 | 2.33E-80 | 10 | Zfp445 |
| Ap2a2 | 1.64E-84 | 0.445621794 | 0.438 | 0.144 | 3.32E-80 | 10 | Ap2a2 |
| Chd42 | 2.25E-84 | 0.54228953 | 0.72 | 0.319 | 4.55E-80 | 10 | Chd4 |
| Mbd53 | 2.30E-84 | 0.577066058 | 0.552 | 0.213 | 4.65E-80 | 10 | Mbd5 |
| Ltbp43 | 3.44E-84 | 0.446019449 | 0.762 | 0.299 | 6.96E-80 | 10 | Ltbp4 |
| Insig1 | 4.87E-84 | 0.403892654 | 0.295 | 0.077 | 9.86E-80 | 10 | Insig1 |
| Pnpla8 | 6.59E-84 | 0.386737158 | 0.498 | 0.176 | 1.33E-79 | 10 | Pnpla8 |
| Tor1aip14 | 8.16E-84 | 0.469194559 | 0.664 | 0.273 | 1.65E-79 | 10 | Tor1aip1 |
| Zfp592 | 8.95E-84 | 0.396419824 | 0.364 | 0.108 | 1.81E-79 | 10 | Zfp592 |
| Fuca11 | 1.13E-83 | 0.469495626 | 0.536 | 0.201 | 2.28E-79 | 10 | Fuca1 |
| St7 | 1.50E-83 | 0.438383024 | 0.261 | 0.064 | 3.04E-79 | 10 | St7 |
| Ash1l3 | 1.71E-83 | 0.527158348 | 0.678 | 0.291 | 3.47E-79 | 10 | Ash1l |
| Son5 | 2.50E-83 | 0.647219014 | 0.928 | 0.517 | 5.06E-79 | 10 | Son |
| Ppp1r12b | 4.66E-83 | 0.463864583 | 0.347 | 0.101 | 9.43E-79 | 10 | Ppp1r12b |
| Mast2 | 8.31E-83 | 0.379480356 | 0.272 | 0.068 | 1.68E-78 | 10 | Mast2 |
| Selenof3 | 9.69E-83 | 0.591197885 | 0.877 | 0.483 | 1.96E-78 | 10 | Selenof |
| Pja1 | 9.80E-83 | 0.300391267 | 0.269 | 0.066 | 1.98E-78 | 10 | Pja1 |
| Ahcyl1 | 1.04E-82 | 0.297842739 | 0.486 | 0.168 | 2.11E-78 | 10 | Ahcyl1 |
| Ldb23 | 1.43E-82 | 0.293616781 | 0.814 | 0.32 | 2.89E-78 | 10 | Ldb2 |
| Trip113 | 1.58E-82 | 0.477570476 | 0.559 | 0.217 | 3.19E-78 | 10 | Trip11 |
| Eid11 | 2.02E-82 | 0.481835826 | 0.709 | 0.313 | 4.09E-78 | 10 | Eid1 |

| | | | | | | | |
|---|---|---|---|---|---|---|---|
| Tmed94 | 2.21E-82 | 0.582447557 | 0.789 | 0.395 | 4.48E-78 | 10 | Tmed9 |
| Lpcat3 | 2.69E-82 | 0.443644415 | 0.32 | 0.089 | 5.44E-78 | 10 | Lpcat3 |
| Zfp950 | 4.37E-82 | 0.407235111 | 0.359 | 0.106 | 8.85E-78 | 10 | Zfp950 |
| Dst3 | 7.11E-82 | 0.424608651 | 0.519 | 0.184 | 1.44E-77 | 10 | Dst |
| Abcd3 | 8.94E-82 | 0.287473536 | 0.323 | 0.089 | 1.81E-77 | 10 | Abcd3 |
| Map4k44 | 1.27E-81 | 0.556719012 | 0.642 | 0.269 | 2.57E-77 | 10 | Map4k4 |
| Cbx51 | 2.23E-81 | 0.451578137 | 0.505 | 0.182 | 4.51E-77 | 10 | Cbx5 |
| Ube3a1 | 2.50E-81 | 0.495372656 | 0.67 | 0.287 | 5.07E-77 | 10 | Ube3a |
| Stag14 | 3.94E-81 | 0.589936094 | 0.758 | 0.353 | 7.97E-77 | 10 | Stag1 |
| Scn1b2 | 4.55E-81 | 0.434259867 | 0.52 | 0.189 | 9.22E-77 | 10 | Scn1b |
| Fip1l11 | 6.54E-81 | 0.498860671 | 0.514 | 0.193 | 1.32E-76 | 10 | Fip1l1 |
| Med13l2 | 1.40E-80 | 0.458066292 | 0.512 | 0.187 | 2.83E-76 | 10 | Med13l |
| Tmem1642 | 1.48E-80 | 0.514496658 | 0.452 | 0.158 | 2.99E-76 | 10 | Tmem164 |
| Mycbp24 | 1.83E-80 | 0.668272935 | 0.761 | 0.366 | 3.71E-76 | 10 | Mycbp2 |
| Scamp21 | 2.00E-80 | 0.400164714 | 0.416 | 0.136 | 4.05E-76 | 10 | Scamp2 |
| Zfand34 | 2.94E-80 | 0.540007144 | 0.681 | 0.293 | 5.94E-76 | 10 | Zfand3 |
| Afap13 | 4.15E-80 | 0.36028289 | 0.444 | 0.144 | 8.40E-76 | 10 | Afap1 |
| Nf14 | 6.95E-80 | 0.510366817 | 0.555 | 0.214 | 1.41E-75 | 10 | Nf1 |
| Cyp2d223 | 8.64E-80 | 0.391914053 | 0.412 | 0.129 | 1.75E-75 | 10 | Cyp2d22 |
| Uba5 | 9.54E-80 | 0.334392974 | 0.431 | 0.144 | 1.93E-75 | 10 | Uba5 |
| Fam193a2 | 1.18E-79 | 0.479841393 | 0.481 | 0.173 | 2.38E-75 | 10 | Fam193a |
| Yap1 | 1.29E-79 | 0.265729811 | 0.273 | 0.068 | 2.60E-75 | 10 | Yap1 |
| Mib11 | 3.01E-79 | 0.393242628 | 0.459 | 0.159 | 6.09E-75 | 10 | Mib1 |
| Sigmar1 | 4.53E-79 | 0.315727254 | 0.302 | 0.081 | 9.18E-75 | 10 | Sigmar1 |
| Ttc19 | 4.92E-79 | 0.401572859 | 0.35 | 0.105 | 9.96E-75 | 10 | Ttc19 |
| Cd343 | 1.37E-78 | 0.543796653 | 0.747 | 0.312 | 2.77E-74 | 10 | Cd34 |
| Zhx33 | 1.42E-78 | 0.394848976 | 0.542 | 0.195 | 2.88E-74 | 10 | Zhx3 |
| 2510009E07Rik2 | 1.55E-78 | 0.416150656 | 0.384 | 0.119 | 3.13E-74 | 10 | 2510009E07Rik |
| Bcap313 | 1.57E-78 | 0.497871702 | 0.652 | 0.277 | 3.18E-74 | 10 | Bcap31 |
| Cdip1 | 1.78E-78 | 0.331277111 | 0.258 | 0.064 | 3.61E-74 | 10 | Cdip1 |
| Epc21 | 2.76E-78 | 0.399500793 | 0.452 | 0.157 | 5.58E-74 | 10 | Epc2 |
| Pura2 | 3.32E-78 | 0.46311337 | 0.77 | 0.357 | 6.73E-74 | 10 | Pura |
| Asah12 | 6.34E-78 | 0.424177186 | 0.498 | 0.182 | 1.28E-73 | 10 | Asah1 |
| Aff41 | 8.38E-78 | 0.436553054 | 0.588 | 0.233 | 1.70E-73 | 10 | Aff4 |
| Fam120a2 | 8.43E-78 | 0.395711448 | 0.484 | 0.176 | 1.71E-73 | 10 | Fam120a |
| Prpf19 | 1.18E-77 | 0.320028401 | 0.314 | 0.089 | 2.40E-73 | 10 | Prpf19 |
| Plod3 | 1.43E-77 | 0.305071276 | 0.261 | 0.065 | 2.89E-73 | 10 | Plod3 |
| Reep33 | 1.61E-77 | 0.427186738 | 0.703 | 0.297 | 3.26E-73 | 10 | Reep3 |
| Tpd52l2 | 1.78E-77 | 0.28518987 | 0.469 | 0.164 | 3.60E-73 | 10 | Tpd52l2 |
| Aaed1 | 1.88E-77 | 0.284460928 | 0.42 | 0.137 | 3.82E-73 | 10 | Aaed1 |
| Zfp266 | 2.21E-77 | 0.370781537 | 0.339 | 0.1 | 4.47E-73 | 10 | Zfp266 |
| Zfp3951 | 2.35E-77 | 0.37882679 | 0.289 | 0.079 | 4.76E-73 | 10 | Zfp395 |
| Crebbp4 | 2.35E-77 | 0.502109779 | 0.678 | 0.297 | 4.76E-73 | 10 | Crebbp |

| | | | | | | | |
|---|---|---|---|---|---|---|---|
| Nsd32 | 3.45E-77 | 0.474456313 | 0.694 | 0.3 | 6.99E-73 | 10 | Nsd3 |
| Acbd3 | 3.59E-77 | 0.414737654 | 0.527 | 0.2 | 7.27E-73 | 10 | Acbd3 |
| Yif1a | 3.59E-77 | 0.258370095 | 0.347 | 0.103 | 7.27E-73 | 10 | Yif1a |
| Golgb13 | 5.87E-77 | 0.436800719 | 0.622 | 0.255 | 1.19E-72 | 10 | Golgb1 |
| Ddx173 | 7.11E-77 | 0.512933653 | 0.625 | 0.266 | 1.44E-72 | 10 | Ddx17 |
| Derl1 | 7.68E-77 | 0.42990581 | 0.434 | 0.152 | 1.56E-72 | 10 | Derl1 |
| Gm37494 | 1.20E-76 | 0.294590822 | 0.294 | 0.079 | 2.43E-72 | 10 | Gm37494 |
| Ick | 1.63E-76 | 0.32299143 | 0.295 | 0.08 | 3.30E-72 | 10 | Ick |
| Tmcc14 | 1.80E-76 | 0.61970854 | 0.533 | 0.211 | 3.65E-72 | 10 | Tmcc1 |
| Evi51 | 1.99E-76 | 0.361271891 | 0.409 | 0.135 | 4.03E-72 | 10 | Evi5 |
| Tnpo11 | 3.19E-76 | 0.448173987 | 0.464 | 0.168 | 6.46E-72 | 10 | Tnpo1 |
| Abhd16a | 3.22E-76 | 0.380788722 | 0.364 | 0.114 | 6.53E-72 | 10 | Abhd16a |
| Mlec2 | 3.30E-76 | 0.481191016 | 0.616 | 0.253 | 6.69E-72 | 10 | Mlec |
| Lmbrd12 | 4.25E-76 | 0.444135553 | 0.445 | 0.157 | 8.61E-72 | 10 | Lmbrd1 |
| Psme42 | 5.93E-76 | 0.403060741 | 0.506 | 0.191 | 1.20E-71 | 10 | Psme4 |
| Dlg11 | 6.34E-76 | 0.453675901 | 0.528 | 0.206 | 1.28E-71 | 10 | Dlg1 |
| Snx4 | 6.58E-76 | 0.324540709 | 0.388 | 0.124 | 1.33E-71 | 10 | Snx4 |
| Ext21 | 7.11E-76 | 0.440036415 | 0.409 | 0.138 | 1.44E-71 | 10 | Ext2 |
| AI3141802 | 8.63E-76 | 0.437174654 | 0.525 | 0.204 | 1.75E-71 | 10 | AI314180 |
| Arf4 | 9.57E-76 | 0.486170883 | 0.844 | 0.447 | 1.94E-71 | 10 | Arf4 |
| Cd473 | 1.80E-75 | 0.590000173 | 0.912 | 0.501 | 3.64E-71 | 10 | Cd47 |
| Ppp1r24 | 1.96E-75 | 0.49882093 | 0.802 | 0.384 | 3.96E-71 | 10 | Ppp1r2 |
| Erp44 | 2.77E-75 | 0.385184815 | 0.458 | 0.164 | 5.61E-71 | 10 | Erp44 |
| Phf2 | 3.50E-75 | 0.309732934 | 0.292 | 0.081 | 7.08E-71 | 10 | Phf2 |
| Rbm262 | 4.14E-75 | 0.445276213 | 0.553 | 0.22 | 8.38E-71 | 10 | Rbm26 |
| Lnpep4 | 4.15E-75 | 0.480924803 | 0.608 | 0.252 | 8.41E-71 | 10 | Lnpep |
| Lyst | 4.50E-75 | 0.434625866 | 0.342 | 0.105 | 9.11E-71 | 10 | Lyst |
| Abl11 | 4.94E-75 | 0.403982024 | 0.37 | 0.118 | 1.00E-70 | 10 | Abl1 |
| Kat2b2 | 7.54E-75 | 0.41686506 | 0.438 | 0.153 | 1.53E-70 | 10 | Kat2b |
| Tmem65 | 8.65E-75 | 0.314653082 | 0.317 | 0.091 | 1.75E-70 | 10 | Tmem65 |
| Baz1b2 | 2.19E-74 | 0.428325307 | 0.494 | 0.187 | 4.43E-70 | 10 | Baz1b |
| S100a163 | 2.22E-74 | 0.352049904 | 0.636 | 0.25 | 4.50E-70 | 10 | S100a16 |
| Brwd11 | 2.77E-74 | 0.442119451 | 0.402 | 0.137 | 5.61E-70 | 10 | Brwd1 |
| Sdf44 | 2.98E-74 | 0.483027334 | 0.672 | 0.301 | 6.03E-70 | 10 | Sdf4 |
| Myadm2 | 3.01E-74 | 0.380933392 | 0.366 | 0.115 | 6.10E-70 | 10 | Myadm |
| Cul5 | 4.19E-74 | 0.351581325 | 0.406 | 0.136 | 8.47E-70 | 10 | Cul5 |
| Itgb14 | 4.35E-74 | 0.504779555 | 0.919 | 0.469 | 8.81E-70 | 10 | Itgb1 |
| Snx6 | 5.45E-74 | 0.375692677 | 0.628 | 0.267 | 1.10E-69 | 10 | Snx6 |
| Tug1 | 5.60E-74 | 0.351230228 | 0.291 | 0.081 | 1.13E-69 | 10 | Tug1 |
| Gpatch84 | 8.34E-74 | 0.455585694 | 0.608 | 0.251 | 1.69E-69 | 10 | Gpatch8 |
| Ctdspl | 8.68E-74 | 0.319038928 | 0.309 | 0.087 | 1.76E-69 | 10 | Ctdspl |
| 3-Mar | 9.92E-74 | 0.447602389 | 0.308 | 0.089 | 2.01E-69 | 10 | 3-Mar |
| Stat6 | 1.34E-73 | 0.382927814 | 0.373 | 0.122 | 2.71E-69 | 10 | Stat6 |

| | | | | | | | |
|---|---|---|---|---|---|---|---|
| Ttc144 | 1.62E-73 | 0.461296895 | 0.531 | 0.209 | 3.29E-69 | 10 | Ttc14 |
| Hivep24 | 1.74E-73 | 0.536896022 | 0.666 | 0.29 | 3.52E-69 | 10 | Hivep2 |
| Zmym41 | 2.02E-73 | 0.405015234 | 0.447 | 0.159 | 4.10E-69 | 10 | Zmym4 |
| Susd64 | 2.78E-73 | 0.510085835 | 0.634 | 0.275 | 5.62E-69 | 10 | Susd6 |
| Prkcsh | 2.78E-73 | 0.361471389 | 0.297 | 0.085 | 5.63E-69 | 10 | Prkcsh |
| Mpp1 | 7.01E-73 | 0.323277978 | 0.288 | 0.079 | 1.42E-68 | 10 | Mpp1 |
| Sntb23 | 1.11E-72 | 0.457608027 | 0.623 | 0.258 | 2.24E-68 | 10 | Sntb2 |
| AW549877 | 1.61E-72 | 0.260182456 | 0.309 | 0.088 | 3.25E-68 | 10 | AW549877 |
| Anxa23 | 2.62E-72 | 0.53521193 | 0.819 | 0.408 | 5.30E-68 | 10 | Anxa2 |
| Parp82 | 2.81E-72 | 0.430039964 | 0.417 | 0.144 | 5.68E-68 | 10 | Parp8 |
| Emc74 | 2.85E-72 | 0.429181665 | 0.627 | 0.264 | 5.77E-68 | 10 | Emc7 |
| Klhl9 | 2.87E-72 | 0.270531858 | 0.316 | 0.092 | 5.81E-68 | 10 | Klhl9 |
| Clock3 | 3.36E-72 | 0.423404629 | 0.512 | 0.197 | 6.80E-68 | 10 | Clock |
| Tgfb1i1 | 4.67E-72 | 0.26822468 | 0.302 | 0.084 | 9.45E-68 | 10 | Tgfb1i1 |
| Thap12 | 4.74E-72 | 0.290985397 | 0.383 | 0.126 | 9.59E-68 | 10 | Thap12 |
| Large12 | 4.81E-72 | 0.417787601 | 0.423 | 0.147 | 9.73E-68 | 10 | Large1 |
| Thoc24 | 4.90E-72 | 0.442897654 | 0.623 | 0.267 | 9.92E-68 | 10 | Thoc2 |
| R3hdm4 | 5.95E-72 | 0.315249731 | 0.338 | 0.104 | 1.20E-67 | 10 | R3hdm4 |
| Sgms12 | 7.85E-72 | 0.386388415 | 0.384 | 0.124 | 1.59E-67 | 10 | Sgms1 |
| Mertk2 | 1.11E-71 | 0.387893533 | 0.388 | 0.126 | 2.25E-67 | 10 | Mertk |
| Runx21 | 1.30E-71 | 0.343982792 | 0.302 | 0.088 | 2.64E-67 | 10 | Runx2 |
| Tmem63a | 2.08E-71 | 0.285356241 | 0.261 | 0.069 | 4.22E-67 | 10 | Tmem63a |
| Atxn7l3b2 | 2.59E-71 | 0.394138323 | 0.809 | 0.396 | 5.24E-67 | 10 | Atxn7l3b |
| Soat11 | 3.02E-71 | 0.381047192 | 0.331 | 0.103 | 6.11E-67 | 10 | Soat1 |
| Os92 | 3.14E-71 | 0.50719472 | 0.475 | 0.183 | 6.35E-67 | 10 | Os9 |
| Mafg | 3.69E-71 | 0.357857124 | 0.309 | 0.091 | 7.47E-67 | 10 | Mafg |
| Fus3 | 5.36E-71 | 0.497342102 | 0.777 | 0.38 | 1.08E-66 | 10 | Fus |
| Ulk22 | 7.08E-71 | 0.4327664 | 0.464 | 0.173 | 1.43E-66 | 10 | Ulk2 |
| Anapc52 | 8.00E-71 | 0.30091316 | 0.531 | 0.205 | 1.62E-66 | 10 | Anapc5 |
| Fam168a1 | 1.11E-70 | 0.401479763 | 0.4 | 0.137 | 2.25E-66 | 10 | Fam168a |
| Kif1bp | 1.30E-70 | 0.278463988 | 0.261 | 0.069 | 2.64E-66 | 10 | Kif1bp |
| Pink1 | 1.30E-70 | 0.266398155 | 0.384 | 0.126 | 2.64E-66 | 10 | Pink1 |
| Gpr107 | 1.53E-70 | 0.335494674 | 0.289 | 0.082 | 3.11E-66 | 10 | Gpr107 |
| Wdfy21 | 2.17E-70 | 0.3307898 | 0.342 | 0.107 | 4.38E-66 | 10 | Wdfy2 |
| Ap3d1 | 2.82E-70 | 0.335485163 | 0.319 | 0.096 | 5.70E-66 | 10 | Ap3d1 |
| Rspry1 | 3.47E-70 | 0.369262735 | 0.344 | 0.11 | 7.03E-66 | 10 | Rspry1 |
| Rb1cc13 | 3.51E-70 | 0.435009337 | 0.555 | 0.228 | 7.10E-66 | 10 | Rb1cc1 |
| Plcl22 | 5.47E-70 | 0.50488516 | 0.341 | 0.11 | 1.11E-65 | 10 | Plcl2 |
| Uso1 | 5.93E-70 | 0.377715491 | 0.388 | 0.134 | 1.20E-65 | 10 | Uso1 |
| Tmem2052 | 6.31E-70 | 0.337623383 | 0.434 | 0.153 | 1.28E-65 | 10 | Tmem205 |
| Sfr12 | 6.42E-70 | 0.366721002 | 0.769 | 0.374 | 1.30E-65 | 10 | Sfr1 |
| Cx3cl12 | 9.74E-70 | 0.380904353 | 0.33 | 0.1 | 1.97E-65 | 10 | Cx3cl1 |
| Rer1 | 1.10E-69 | 0.3917807 | 0.689 | 0.318 | 2.23E-65 | 10 | Rer1 |

| | | | | | | | |
|---|---|---|---|---|---|---|---|
| Yme1l12 | 1.59E-69 | 0.428533085 | 0.494 | 0.19 | 3.22E-65 | 10 | Yme1l1 |
| Tnrc6b2 | 1.61E-69 | 0.488291994 | 0.755 | 0.363 | 3.26E-65 | 10 | Tnrc6b |
| Aga | 2.22E-69 | 0.299245947 | 0.275 | 0.076 | 4.50E-65 | 10 | Aga |
| Fbxl175 | 2.35E-69 | 0.54133504 | 0.666 | 0.303 | 4.76E-65 | 10 | Fbxl17 |
| Med131 | 2.39E-69 | 0.41319248 | 0.508 | 0.199 | 4.83E-65 | 10 | Med13 |
| Ankrd174 | 2.50E-69 | 0.484755783 | 0.597 | 0.254 | 5.06E-65 | 10 | Ankrd17 |
| Dmxl12 | 3.06E-69 | 0.413232786 | 0.47 | 0.179 | 6.19E-65 | 10 | Dmxl1 |
| Supt31 | 3.12E-69 | 0.362597557 | 0.367 | 0.122 | 6.32E-65 | 10 | Supt3 |
| Smtn1 | 3.23E-69 | 0.297899618 | 0.289 | 0.081 | 6.55E-65 | 10 | Smtn |
| Washc41 | 3.46E-69 | 0.40808258 | 0.37 | 0.124 | 7.01E-65 | 10 | Washc4 |
| Rbms22 | 3.58E-69 | 0.327130908 | 0.434 | 0.154 | 7.26E-65 | 10 | Rbms2 |
| Hexim1 | 3.73E-69 | 0.326037682 | 0.32 | 0.098 | 7.55E-65 | 10 | Hexim1 |
| Jkamp | 4.65E-69 | 0.340532368 | 0.331 | 0.104 | 9.42E-65 | 10 | Jkamp |
| Cltc3 | 5.01E-69 | 0.408637149 | 0.698 | 0.323 | 1.01E-64 | 10 | Cltc |
| Dhrs73 | 7.24E-69 | 0.413153166 | 0.525 | 0.209 | 1.47E-64 | 10 | Dhrs7 |
| Akap94 | 8.41E-69 | 0.476588448 | 0.55 | 0.231 | 1.70E-64 | 10 | Akap9 |
| Xdh3 | 8.76E-69 | 0.397976299 | 0.495 | 0.182 | 1.77E-64 | 10 | Xdh |
| Efr3a1 | 1.08E-68 | 0.399463714 | 0.491 | 0.192 | 2.18E-64 | 10 | Efr3a |
| Invs | 1.13E-68 | 0.38740935 | 0.258 | 0.07 | 2.29E-64 | 10 | Invs |
| Smg62 | 1.21E-68 | 0.510556368 | 0.586 | 0.253 | 2.45E-64 | 10 | Smg6 |
| 4932438A13Rik2 | 1.22E-68 | 0.459280837 | 0.527 | 0.213 | 2.46E-64 | 10 | 4932438A13Rik |
| Dpy19l4 | 1.50E-68 | 0.351117613 | 0.272 | 0.076 | 3.04E-64 | 10 | Dpy19l4 |
| Ormdl3 | 1.87E-68 | 0.271618588 | 0.289 | 0.083 | 3.78E-64 | 10 | Ormdl3 |
| Itfg1 | 1.98E-68 | 0.431048998 | 0.388 | 0.135 | 4.00E-64 | 10 | Itfg1 |
| Dnajb141 | 2.12E-68 | 0.289110015 | 0.345 | 0.11 | 4.28E-64 | 10 | Dnajb14 |
| Rtn33 | 2.16E-68 | 0.456760379 | 0.689 | 0.316 | 4.38E-64 | 10 | Rtn3 |
| Gnptg1 | 2.27E-68 | 0.352460374 | 0.308 | 0.094 | 4.59E-64 | 10 | Gnptg |
| Kansl1l2 | 2.44E-68 | 0.44894093 | 0.386 | 0.134 | 4.93E-64 | 10 | Kansl1l |
| Top2b2 | 4.44E-68 | 0.414617053 | 0.628 | 0.276 | 8.99E-64 | 10 | Top2b |
| Bet1l | 4.78E-68 | 0.254751423 | 0.362 | 0.118 | 9.68E-64 | 10 | Bet1l |
| Taok13 | 4.89E-68 | 0.376237447 | 0.584 | 0.244 | 9.91E-64 | 10 | Taok1 |
| Rnf6 | 8.01E-68 | 0.292316599 | 0.37 | 0.124 | 1.62E-63 | 10 | Rnf6 |
| Zfr4 | 8.91E-68 | 0.43966374 | 0.572 | 0.241 | 1.80E-63 | 10 | Zfr |
| Ganab | 1.26E-67 | 0.359423364 | 0.342 | 0.111 | 2.55E-63 | 10 | Ganab |
| Wwc23 | 1.34E-67 | 0.374040468 | 0.505 | 0.191 | 2.71E-63 | 10 | Wwc2 |
| Wdfy34 | 1.40E-67 | 0.379617852 | 0.473 | 0.176 | 2.83E-63 | 10 | Wdfy3 |
| Zdhhc5 | 1.64E-67 | 0.297427308 | 0.283 | 0.081 | 3.33E-63 | 10 | Zdhhc5 |
| Tmem55a | 1.75E-67 | 0.27733891 | 0.277 | 0.078 | 3.55E-63 | 10 | Tmem55a |
| Strn33 | 1.85E-67 | 0.394090569 | 0.728 | 0.337 | 3.75E-63 | 10 | Strn3 |
| Rabac13 | 2.02E-67 | 0.523284944 | 0.927 | 0.557 | 4.08E-63 | 10 | Rabac1 |
| Mprip4 | 2.02E-67 | 0.385146633 | 0.661 | 0.285 | 4.09E-63 | 10 | Mprip |
| Man1a3 | 2.04E-67 | 0.579166968 | 0.428 | 0.162 | 4.13E-63 | 10 | Man1a |
| Fbxl32 | 2.06E-67 | 0.376824885 | 0.419 | 0.151 | 4.16E-63 | 10 | Fbxl3 |

| | | | | | | | |
|---|---|---|---|---|---|---|---|
| Ccdc127 | 2.15E-67 | 0.344760955 | 0.347 | 0.112 | 4.36E-63 | 10 | Ccdc127 |
| Pcm13 | 2.70E-67 | 0.451342395 | 0.572 | 0.244 | 5.47E-63 | 10 | Pcm1 |
| Atp9b1 | 2.75E-67 | 0.371167384 | 0.42 | 0.152 | 5.57E-63 | 10 | Atp9b |
| Rps6kb11 | 2.88E-67 | 0.364220242 | 0.422 | 0.153 | 5.83E-63 | 10 | Rps6kb1 |
| Brd3 | 2.92E-67 | 0.285291486 | 0.311 | 0.094 | 5.91E-63 | 10 | Brd3 |
| Bsg4 | 3.79E-67 | 0.458406968 | 0.831 | 0.441 | 7.67E-63 | 10 | Bsg |
| Mga2 | 4.77E-67 | 0.401520726 | 0.445 | 0.166 | 9.66E-63 | 10 | Mga |
| Pon2 | 5.53E-67 | 0.369841438 | 0.445 | 0.166 | 1.12E-62 | 10 | Pon2 |
| Appl11 | 1.76E-66 | 0.433369372 | 0.458 | 0.176 | 3.56E-62 | 10 | Appl1 |
| Ankle2 | 4.54E-66 | 0.316459897 | 0.283 | 0.083 | 9.19E-62 | 10 | Ankle2 |
| Smarcd2 | 5.30E-66 | 0.314775894 | 0.295 | 0.088 | 1.07E-61 | 10 | Smarcd2 |
| Pde4dip | 5.85E-66 | 0.275436063 | 0.259 | 0.072 | 1.18E-61 | 10 | Pde4dip |
| Ogfrl11 | 6.05E-66 | 0.322146497 | 0.378 | 0.129 | 1.22E-61 | 10 | Ogfrl1 |
| Lrp51 | 6.35E-66 | 0.337997923 | 0.3 | 0.089 | 1.29E-61 | 10 | Lrp5 |
| Flot2 | 7.53E-66 | 0.288339897 | 0.275 | 0.079 | 1.52E-61 | 10 | Flot2 |
| Mpv17 | 9.37E-66 | 0.265859149 | 0.253 | 0.069 | 1.90E-61 | 10 | Mpv17 |
| Eml12 | 1.31E-65 | 0.312240105 | 0.336 | 0.105 | 2.66E-61 | 10 | Eml1 |
| Wwox3 | 1.79E-65 | 0.653816726 | 0.439 | 0.171 | 3.63E-61 | 10 | Wwox |
| Krtcap21 | 1.92E-65 | 0.453663888 | 0.825 | 0.434 | 3.89E-61 | 10 | Krtcap2 |
| Emc32 | 2.40E-65 | 0.370539052 | 0.447 | 0.167 | 4.86E-61 | 10 | Emc3 |
| Krit13 | 2.88E-65 | 0.455273161 | 0.455 | 0.177 | 5.83E-61 | 10 | Krit1 |
| Stim12 | 3.86E-65 | 0.465525458 | 0.467 | 0.185 | 7.82E-61 | 10 | Stim1 |
| Dopey1 | 4.36E-65 | 0.384772408 | 0.323 | 0.103 | 8.83E-61 | 10 | Dopey1 |
| Thbd4 | 6.47E-65 | 0.505028151 | 0.595 | 0.248 | 1.31E-60 | 10 | Thbd |
| Foxo32 | 8.17E-65 | 0.36326929 | 0.338 | 0.112 | 1.65E-60 | 10 | Foxo3 |
| Lrrk12 | 1.28E-64 | 0.503078725 | 0.47 | 0.185 | 2.58E-60 | 10 | Lrrk1 |
| Ubr32 | 1.77E-64 | 0.4050095 | 0.489 | 0.195 | 3.58E-60 | 10 | Ubr3 |
| Araf | 1.88E-64 | 0.330103908 | 0.322 | 0.103 | 3.80E-60 | 10 | Araf |
| Smarcc11 | 2.09E-64 | 0.349427858 | 0.384 | 0.137 | 4.22E-60 | 10 | Smarcc1 |
| Tbc1d12 | 3.04E-64 | 0.369116846 | 0.327 | 0.105 | 6.15E-60 | 10 | Tbc1d12 |
| Slc25a24 | 4.02E-64 | 0.278261654 | 0.275 | 0.08 | 8.15E-60 | 10 | Slc25a24 |
| Ppp6r2 | 4.99E-64 | 0.269606824 | 0.25 | 0.069 | 1.01E-59 | 10 | Ppp6r2 |
| Arhgef124 | 6.09E-64 | 0.353254257 | 0.755 | 0.34 | 1.23E-59 | 10 | Arhgef12 |
| Xrn13 | 8.10E-64 | 0.437395393 | 0.495 | 0.198 | 1.64E-59 | 10 | Xrn1 |
| Tax1bp1 | 9.48E-64 | 0.355228582 | 0.802 | 0.409 | 1.92E-59 | 10 | Tax1bp1 |
| Cers5 | 9.82E-64 | 0.376200495 | 0.37 | 0.129 | 1.99E-59 | 10 | Cers5 |
| Cisd2 | 1.16E-63 | 0.28959699 | 0.447 | 0.169 | 2.35E-59 | 10 | Cisd2 |
| Selenok3 | 1.43E-63 | 0.459429831 | 0.916 | 0.547 | 2.89E-59 | 10 | Selenok |
| Fkbp51 | 2.05E-63 | 0.366507078 | 0.362 | 0.127 | 4.16E-59 | 10 | Fkbp5 |
| Cbfa2t2 | 2.10E-63 | 0.344217586 | 0.341 | 0.113 | 4.26E-59 | 10 | Cbfa2t2 |
| Itm2b6 | 2.19E-63 | 0.549631421 | 0.991 | 0.72 | 4.44E-59 | 10 | Itm2b |
| Usp471 | 3.57E-63 | 0.3603795 | 0.467 | 0.182 | 7.23E-59 | 10 | Usp47 |
| Kif13b1 | 4.21E-63 | 0.400301826 | 0.516 | 0.21 | 8.53E-59 | 10 | Kif13b |

| | | | | | | | |
|---|---|---|---|---|---|---|---|
| Mia21 | 4.31E-63 | 0.430235239 | 0.492 | 0.202 | 8.72E-59 | 10 | Mia2 |
| Smarcc21 | 7.66E-63 | 0.336510322 | 0.45 | 0.172 | 1.55E-58 | 10 | Smarcc2 |
| Hdac2 | 8.38E-63 | 0.260179343 | 0.311 | 0.098 | 1.70E-58 | 10 | Hdac2 |
| Vcp2 | 1.23E-62 | 0.397891551 | 0.703 | 0.336 | 2.50E-58 | 10 | Vcp |
| Mex3c | 1.57E-62 | 0.269185566 | 0.286 | 0.086 | 3.18E-58 | 10 | Mex3c |
| Sfpq3 | 1.66E-62 | 0.414747282 | 0.636 | 0.291 | 3.35E-58 | 10 | Sfpq |
| Acsl41 | 1.79E-62 | 0.343326001 | 0.366 | 0.128 | 3.63E-58 | 10 | Acsl4 |
| Specc1l1 | 1.89E-62 | 0.359560785 | 0.358 | 0.124 | 3.83E-58 | 10 | Specc1l |
| Cmtm31 | 2.17E-62 | 0.317127286 | 0.319 | 0.102 | 4.38E-58 | 10 | Cmtm3 |
| Ppp3cb | 3.01E-62 | 0.328355502 | 0.434 | 0.164 | 6.09E-58 | 10 | Ppp3cb |
| Smarcb1 | 3.05E-62 | 0.27616508 | 0.384 | 0.137 | 6.17E-58 | 10 | Smarcb1 |
| Tbce | 3.15E-62 | 0.348609505 | 0.266 | 0.078 | 6.38E-58 | 10 | Tbce |
| Mbnl24 | 3.72E-62 | 0.455975867 | 0.797 | 0.411 | 7.53E-58 | 10 | Mbnl2 |
| Tsc22d13 | 3.79E-62 | 0.422814334 | 0.833 | 0.438 | 7.68E-58 | 10 | Tsc22d1 |
| Myef2 | 4.19E-62 | 0.312680775 | 0.259 | 0.075 | 8.49E-58 | 10 | Myef2 |
| Rab6a3 | 4.40E-62 | 0.301861903 | 0.519 | 0.209 | 8.90E-58 | 10 | Rab6a |
| Slc35a1 | 6.92E-62 | 0.273492507 | 0.25 | 0.07 | 1.40E-57 | 10 | Slc35a1 |
| Crispld13 | 7.30E-62 | 0.612623714 | 0.383 | 0.139 | 1.48E-57 | 10 | Crispld1 |
| Agtrap1 | 7.46E-62 | 0.318210769 | 0.359 | 0.123 | 1.51E-57 | 10 | Agtrap |
| Cd1641 | 1.72E-61 | 0.313300767 | 0.509 | 0.208 | 3.48E-57 | 10 | Cd164 |
| Faf2 | 2.38E-61 | 0.387981299 | 0.348 | 0.121 | 4.82E-57 | 10 | Faf2 |
| Zadh2 | 2.87E-61 | 0.277645401 | 0.309 | 0.099 | 5.81E-57 | 10 | Zadh2 |
| Rabgap11 | 3.22E-61 | 0.344407894 | 0.42 | 0.158 | 6.52E-57 | 10 | Rabgap1 |
| Pdxdc11 | 4.06E-61 | 0.273031979 | 0.405 | 0.149 | 8.21E-57 | 10 | Pdxdc1 |
| Atf4 | 4.44E-61 | 0.315607487 | 0.484 | 0.195 | 8.99E-57 | 10 | Atf4 |
| Socs23 | 4.55E-61 | 0.348696186 | 0.459 | 0.177 | 9.21E-57 | 10 | Socs2 |
| Calm32 | 4.92E-61 | 0.337540269 | 0.586 | 0.263 | 9.95E-57 | 10 | Calm3 |
| Chd63 | 4.96E-61 | 0.41468348 | 0.506 | 0.212 | 1.00E-56 | 10 | Chd6 |
| Lrig2 | 5.13E-61 | 0.362974902 | 0.273 | 0.082 | 1.04E-56 | 10 | Lrig2 |
| Cuedc11 | 6.16E-61 | 0.316016459 | 0.373 | 0.131 | 1.25E-56 | 10 | Cuedc1 |
| Trip10 | 8.82E-61 | 0.295417266 | 0.297 | 0.092 | 1.78E-56 | 10 | Trip10 |
| Tshz1 | 9.99E-61 | 0.331563882 | 0.336 | 0.114 | 2.02E-56 | 10 | Tshz1 |
| Rassf8 | 1.12E-60 | 0.298337308 | 0.267 | 0.078 | 2.27E-56 | 10 | Rassf8 |
| Pi4ka1 | 1.48E-60 | 0.325900842 | 0.355 | 0.123 | 3.00E-56 | 10 | Pi4ka |
| Tecr1 | 1.63E-60 | 0.33045615 | 0.522 | 0.219 | 3.30E-56 | 10 | Tecr |
| Eea13 | 2.37E-60 | 0.338776628 | 0.512 | 0.212 | 4.80E-56 | 10 | Eea1 |
| Braf2 | 3.98E-60 | 0.330070708 | 0.412 | 0.154 | 8.06E-56 | 10 | Braf |
| Scaf83 | 8.39E-60 | 0.386386602 | 0.453 | 0.179 | 1.70E-55 | 10 | Scaf8 |
| Ccdc472 | 1.29E-59 | 0.362027545 | 0.466 | 0.187 | 2.61E-55 | 10 | Ccdc47 |
| Tmco13 | 1.56E-59 | 0.344288059 | 0.658 | 0.305 | 3.16E-55 | 10 | Tmco1 |
| Zfx | 1.66E-59 | 0.328780476 | 0.394 | 0.146 | 3.36E-55 | 10 | Zfx |
| Atp2c13 | 3.32E-59 | 0.379175773 | 0.422 | 0.163 | 6.72E-55 | 10 | Atp2c1 |
| Slc23a22 | 5.13E-59 | 0.343678155 | 0.375 | 0.134 | 1.04E-54 | 10 | Slc23a2 |

| | | | | | | | |
|---|---|---|---|---|---|---|---|
| 4930402H24Rik3 | 5.20E-59 | 0.371416937 | 0.512 | 0.216 | 1.05E-54 | 10 | 4930402H24Rik |
| Cnih1 | 6.62E-59 | 0.265691585 | 0.348 | 0.121 | 1.34E-54 | 10 | Cnih1 |
| Osbpl93 | 6.72E-59 | 0.343319893 | 0.614 | 0.271 | 1.36E-54 | 10 | Osbpl9 |
| Sqstm12 | 7.29E-59 | 0.378306876 | 0.645 | 0.297 | 1.48E-54 | 10 | Sqstm1 |
| Cldnd1 | 8.56E-59 | 0.283349055 | 0.338 | 0.114 | 1.73E-54 | 10 | Cldnd1 |
| Egr12 | 1.04E-58 | 0.852500892 | 0.412 | 0.171 | 2.10E-54 | 10 | Egr1 |
| Scaf111 | 1.37E-58 | 0.394913991 | 0.57 | 0.257 | 2.77E-54 | 10 | Scaf11 |
| Vps13c1 | 1.91E-58 | 0.335030165 | 0.373 | 0.135 | 3.87E-54 | 10 | Vps13c |
| Ubtf | 1.92E-58 | 0.274482974 | 0.456 | 0.181 | 3.88E-54 | 10 | Ubtf |
| Rbm64 | 1.93E-58 | 0.401116826 | 0.503 | 0.212 | 3.92E-54 | 10 | Rbm6 |
| Tmem33 | 2.36E-58 | 0.257203639 | 0.27 | 0.082 | 4.78E-54 | 10 | Tmem33 |
| Smc1a3 | 2.43E-58 | 0.396004599 | 0.647 | 0.305 | 4.93E-54 | 10 | Smc1a |
| Mgat21 | 2.84E-58 | 0.326119288 | 0.403 | 0.15 | 5.74E-54 | 10 | Mgat2 |
| Dis3l22 | 3.18E-58 | 0.396029828 | 0.367 | 0.135 | 6.44E-54 | 10 | Dis3l2 |
| Sil1 | 3.19E-58 | 0.257143894 | 0.261 | 0.078 | 6.45E-54 | 10 | Sil1 |
| Fam45a1 | 5.84E-58 | 0.261056709 | 0.288 | 0.091 | 1.18E-53 | 10 | Fam45a |
| Pign2 | 8.61E-58 | 0.381310717 | 0.281 | 0.091 | 1.74E-53 | 10 | Pign |
| Atl3 | 1.53E-57 | 0.349988548 | 0.35 | 0.124 | 3.11E-53 | 10 | Atl3 |
| Togaram1 | 1.61E-57 | 0.301595403 | 0.275 | 0.086 | 3.27E-53 | 10 | Togaram1 |
| Tnks1 | 2.03E-57 | 0.359576449 | 0.373 | 0.138 | 4.11E-53 | 10 | Tnks |
| Prrc2a1 | 2.03E-57 | 0.290332787 | 0.391 | 0.146 | 4.11E-53 | 10 | Prrc2a |
| Atad1 | 2.24E-57 | 0.292703526 | 0.303 | 0.1 | 4.54E-53 | 10 | Atad1 |
| Tbc1d9b1 | 2.26E-57 | 0.2964243 | 0.339 | 0.118 | 4.58E-53 | 10 | Tbc1d9b |
| Farp11 | 2.46E-57 | 0.313563097 | 0.312 | 0.104 | 4.99E-53 | 10 | Farp1 |
| Ptpra4 | 2.55E-57 | 0.371189231 | 0.639 | 0.298 | 5.17E-53 | 10 | Ptpra |
| Dhx30 | 2.62E-57 | 0.285674682 | 0.295 | 0.096 | 5.30E-53 | 10 | Dhx30 |
| Nipbl5 | 3.11E-57 | 0.403524052 | 0.723 | 0.359 | 6.30E-53 | 10 | Nipbl |
| Tmem50a5 | 3.82E-57 | 0.406378896 | 0.739 | 0.375 | 7.74E-53 | 10 | Tmem50a |
| Jak12 | 4.20E-57 | 0.357573883 | 0.744 | 0.358 | 8.51E-53 | 10 | Jak1 |
| Kmt5b2 | 4.78E-57 | 0.34097018 | 0.422 | 0.166 | 9.68E-53 | 10 | Kmt5b |
| Ptbp2 | 5.11E-57 | 0.354212415 | 0.35 | 0.125 | 1.04E-52 | 10 | Ptbp2 |
| Eif1a | 5.61E-57 | 0.25265059 | 0.395 | 0.148 | 1.14E-52 | 10 | Eif1a |
| Nbeal13 | 5.67E-57 | 0.343449839 | 0.584 | 0.256 | 1.15E-52 | 10 | Nbeal1 |
| Cbfb2 | 5.76E-57 | 0.351576975 | 0.395 | 0.151 | 1.17E-52 | 10 | Cbfb |
| Tmem594 | 5.96E-57 | 0.419096092 | 0.861 | 0.483 | 1.21E-52 | 10 | Tmem59 |
| St131 | 6.54E-57 | 0.266111633 | 0.723 | 0.359 | 1.32E-52 | 10 | St13 |
| Tor2a | 7.87E-57 | 0.317313977 | 0.261 | 0.081 | 1.59E-52 | 10 | Tor2a |
| Akt34 | 8.93E-57 | 0.395433856 | 0.558 | 0.245 | 1.81E-52 | 10 | Akt3 |
| Mtmr34 | 1.24E-56 | 0.368748682 | 0.472 | 0.195 | 2.51E-52 | 10 | Mtmr3 |
| Zfp6082 | 1.50E-56 | 0.354066622 | 0.373 | 0.136 | 3.04E-52 | 10 | Zfp608 |
| Tnks1bp1 | 1.67E-56 | 0.328486318 | 0.277 | 0.086 | 3.38E-52 | 10 | Tnks1bp1 |
| Hnrnpul2 | 2.42E-56 | 0.298239047 | 0.489 | 0.206 | 4.90E-52 | 10 | Hnrnpul2 |
| Agpat31 | 3.59E-56 | 0.257281579 | 0.35 | 0.125 | 7.27E-52 | 10 | Agpat3 |

| | | | | | | | |
|---|---|---|---|---|---|---|---|
| Usp45 | 3.82E-56 | 0.286989459 | 0.259 | 0.079 | 7.73E-52 | 10 | Usp45 |
| Pbxip11 | 4.06E-56 | 0.365605234 | 0.322 | 0.113 | 8.22E-52 | 10 | Pbxip1 |
| Nbas | 4.34E-56 | 0.299555188 | 0.272 | 0.086 | 8.78E-52 | 10 | Nbas |
| Lims12 | 4.46E-56 | 0.407922294 | 0.625 | 0.294 | 9.03E-52 | 10 | Lims1 |
| Rab1a1 | 4.91E-56 | 0.312713843 | 0.586 | 0.267 | 9.95E-52 | 10 | Rab1a |
| Dync1h14 | 6.36E-56 | 0.36638757 | 0.608 | 0.28 | 1.29E-51 | 10 | Dync1h1 |
| Pxk1 | 6.60E-56 | 0.341667222 | 0.369 | 0.136 | 1.34E-51 | 10 | Pxk |
| Dcaf8 | 6.62E-56 | 0.263358862 | 0.47 | 0.192 | 1.34E-51 | 10 | Dcaf8 |
| Dpm32 | 7.52E-56 | 0.40495243 | 0.684 | 0.343 | 1.52E-51 | 10 | Dpm3 |
| Kat6b1 | 8.22E-56 | 0.349425578 | 0.388 | 0.148 | 1.66E-51 | 10 | Kat6b |
| Tmem473 | 1.08E-55 | 0.288931004 | 0.411 | 0.151 | 2.18E-51 | 10 | Tmem47 |
| Acvr2a2 | 1.19E-55 | 0.348746593 | 0.386 | 0.145 | 2.41E-51 | 10 | Acvr2a |
| Sp11 | 1.44E-55 | 0.288228645 | 0.373 | 0.138 | 2.92E-51 | 10 | Sp1 |
| Arhgap12 | 1.51E-55 | 0.270124712 | 0.336 | 0.118 | 3.07E-51 | 10 | Arhgap12 |
| Rsu13 | 1.61E-55 | 0.332700925 | 0.561 | 0.248 | 3.27E-51 | 10 | Rsu1 |
| Pea15a3 | 1.90E-55 | 0.286730405 | 0.548 | 0.234 | 3.84E-51 | 10 | Pea15a |
| Stat34 | 2.41E-55 | 0.383395971 | 0.569 | 0.261 | 4.88E-51 | 10 | Stat3 |
| Ino80 | 2.43E-55 | 0.31071599 | 0.367 | 0.135 | 4.91E-51 | 10 | Ino80 |
| Kif1b3 | 2.81E-55 | 0.359806995 | 0.537 | 0.237 | 5.68E-51 | 10 | Kif1b |
| Pkn23 | 4.07E-55 | 0.357624628 | 0.614 | 0.288 | 8.25E-51 | 10 | Pkn2 |
| Hivep11 | 4.55E-55 | 0.264476875 | 0.361 | 0.131 | 9.21E-51 | 10 | Hivep1 |
| Jmjd1c5 | 4.61E-55 | 0.410521628 | 0.636 | 0.301 | 9.32E-51 | 10 | Jmjd1c |
| Prpsap1 | 5.96E-55 | 0.315553091 | 0.312 | 0.107 | 1.21E-50 | 10 | Prpsap1 |
| Sparcl14 | 6.81E-55 | 0.340073657 | 0.769 | 0.355 | 1.38E-50 | 10 | Sparcl1 |
| Osbpl82 | 7.04E-55 | 0.383512856 | 0.461 | 0.194 | 1.43E-50 | 10 | Osbpl8 |
| Tdrd3 | 7.59E-55 | 0.33741479 | 0.275 | 0.088 | 1.54E-50 | 10 | Tdrd3 |
| Capn23 | 8.14E-55 | 0.284311784 | 0.444 | 0.174 | 1.65E-50 | 10 | Capn2 |
| Birc64 | 8.88E-55 | 0.369605843 | 0.697 | 0.334 | 1.80E-50 | 10 | Birc6 |
| Bclaf12 | 1.05E-54 | 0.347752385 | 0.553 | 0.248 | 2.13E-50 | 10 | Bclaf1 |
| Brip1os | 1.14E-54 | 0.296811678 | 0.273 | 0.087 | 2.32E-50 | 10 | Brip1os |
| Spred22 | 1.17E-54 | 0.332763732 | 0.422 | 0.166 | 2.38E-50 | 10 | Spred2 |
| Ccdc88a4 | 1.23E-54 | 0.429355957 | 0.473 | 0.199 | 2.49E-50 | 10 | Ccdc88a |
| Bcas33 | 1.76E-54 | 0.378195304 | 0.438 | 0.177 | 3.56E-50 | 10 | Bcas3 |
| Zfp397 | 1.85E-54 | 0.273905726 | 0.266 | 0.083 | 3.74E-50 | 10 | Zfp397 |
| Akap112 | 2.25E-54 | 0.333760651 | 0.347 | 0.126 | 4.55E-50 | 10 | Akap11 |
| Atp6v0d1 | 2.36E-54 | 0.277255332 | 0.4 | 0.154 | 4.77E-50 | 10 | Atp6v0d1 |
| Trove22 | 2.51E-54 | 0.290727492 | 0.355 | 0.127 | 5.08E-50 | 10 | Trove2 |
| Gab11 | 2.65E-54 | 0.305225402 | 0.323 | 0.11 | 5.37E-50 | 10 | Gab1 |
| Aak11 | 2.65E-54 | 0.356351345 | 0.445 | 0.18 | 5.37E-50 | 10 | Aak1 |
| Tmem1092 | 3.29E-54 | 0.304072206 | 0.383 | 0.143 | 6.66E-50 | 10 | Tmem109 |
| Cnep1r1 | 3.70E-54 | 0.300445537 | 0.309 | 0.105 | 7.49E-50 | 10 | Cnep1r1 |
| Hnrnph13 | 4.41E-54 | 0.321259359 | 0.545 | 0.244 | 8.92E-50 | 10 | Hnrnph1 |
| Trpm73 | 4.53E-54 | 0.376117835 | 0.469 | 0.199 | 9.17E-50 | 10 | Trpm7 |

| | | | | | | | |
|---|---|---|---|---|---|---|---|
| Erg28 | 4.87E-54 | 0.313584605 | 0.322 | 0.114 | 9.85E-50 | 10 | Erg28 |
| Cpne32 | 5.27E-54 | 0.294689262 | 0.489 | 0.206 | 1.07E-49 | 10 | Cpne3 |
| 9930021J03Rik1 | 5.31E-54 | 0.379539272 | 0.381 | 0.147 | 1.07E-49 | 10 | 9930021J03Rik |
| Zcchc74 | 7.34E-54 | 0.43056338 | 0.62 | 0.295 | 1.49E-49 | 10 | Zcchc7 |
| Nsd14 | 8.52E-54 | 0.379851862 | 0.692 | 0.343 | 1.72E-49 | 10 | Nsd1 |
| Dab24 | 1.05E-53 | 0.370415543 | 0.488 | 0.201 | 2.12E-49 | 10 | Dab2 |
| Pja23 | 1.10E-53 | 0.335799622 | 0.472 | 0.199 | 2.22E-49 | 10 | Pja2 |
| Ankhd13 | 1.38E-53 | 0.406527024 | 0.533 | 0.238 | 2.79E-49 | 10 | Ankhd1 |
| Far12 | 2.01E-53 | 0.293132852 | 0.408 | 0.16 | 4.07E-49 | 10 | Far1 |
| Syncrip1 | 2.36E-53 | 0.324731342 | 0.489 | 0.21 | 4.78E-49 | 10 | Syncrip |
| Arl13b | 2.38E-53 | 0.257221004 | 0.291 | 0.096 | 4.82E-49 | 10 | Arl13b |
| Ccdc6 | 2.69E-53 | 0.267958334 | 0.261 | 0.082 | 5.45E-49 | 10 | Ccdc6 |
| Herpud2 | 2.85E-53 | 0.2748116 | 0.286 | 0.095 | 5.77E-49 | 10 | Herpud2 |
| Slc30a51 | 2.91E-53 | 0.293779077 | 0.261 | 0.083 | 5.88E-49 | 10 | Slc30a5 |
| Tmbim11 | 3.53E-53 | 0.275562228 | 0.334 | 0.119 | 7.14E-49 | 10 | Tmbim1 |
| Arhgef62 | 4.03E-53 | 0.335678549 | 0.339 | 0.125 | 8.16E-49 | 10 | Arhgef6 |
| Tbl1x3 | 4.20E-53 | 0.32582434 | 0.564 | 0.256 | 8.50E-49 | 10 | Tbl1x |
| Pgrmc22 | 4.41E-53 | 0.256751558 | 0.37 | 0.138 | 8.93E-49 | 10 | Pgrmc2 |
| Ambra13 | 5.78E-53 | 0.368208721 | 0.459 | 0.193 | 1.17E-48 | 10 | Ambra1 |
| Snapc3 | 9.68E-53 | 0.274309559 | 0.259 | 0.081 | 1.96E-48 | 10 | Snapc3 |
| Nap1l13 | 1.13E-52 | 0.285975149 | 0.648 | 0.308 | 2.29E-48 | 10 | Nap1l1 |
| Picalm3 | 1.28E-52 | 0.398523563 | 0.606 | 0.29 | 2.60E-48 | 10 | Picalm |
| Fbln24 | 1.82E-52 | 0.263243872 | 0.613 | 0.265 | 3.69E-48 | 10 | Fbln2 |
| 2700081O15Rik1 | 1.85E-52 | 0.251310952 | 0.264 | 0.082 | 3.75E-48 | 10 | 2700081O15Rik |
| Tnks24 | 2.81E-52 | 0.345007608 | 0.591 | 0.274 | 5.69E-48 | 10 | Tnks2 |
| Imp31 | 3.55E-52 | 0.302291833 | 0.634 | 0.309 | 7.18E-48 | 10 | Imp3 |
| Dhrs33 | 3.62E-52 | 0.337830271 | 0.522 | 0.226 | 7.32E-48 | 10 | Dhrs3 |
| Gatad2b2 | 4.33E-52 | 0.33400058 | 0.467 | 0.197 | 8.77E-48 | 10 | Gatad2b |
| Mpzl13 | 5.59E-52 | 0.269951955 | 0.375 | 0.138 | 1.13E-47 | 10 | Mpzl1 |
| Hnrnpr1 | 6.17E-52 | 0.340183662 | 0.512 | 0.23 | 1.25E-47 | 10 | Hnrnpr |
| Hacd21 | 6.31E-52 | 0.298833674 | 0.316 | 0.113 | 1.28E-47 | 10 | Hacd2 |
| Cnpy31 | 6.56E-52 | 0.289276564 | 0.289 | 0.099 | 1.33E-47 | 10 | Cnpy3 |
| Zbtb42 | 6.57E-52 | 0.256921633 | 0.33 | 0.117 | 1.33E-47 | 10 | Zbtb4 |
| Tmem1653 | 6.83E-52 | 0.29595395 | 0.431 | 0.175 | 1.38E-47 | 10 | Tmem165 |
| Epha42 | 7.90E-52 | 0.266675319 | 0.325 | 0.112 | 1.60E-47 | 10 | Epha4 |
| Dnajb11 | 9.12E-52 | 0.3088229 | 0.481 | 0.209 | 1.85E-47 | 10 | Dnajb11 |
| Ppp1cc1 | 9.91E-52 | 0.287286177 | 0.497 | 0.217 | 2.01E-47 | 10 | Ppp1cc |
| Kmt2c4 | 1.45E-51 | 0.363009889 | 0.627 | 0.3 | 2.94E-47 | 10 | Kmt2c |
| Csgalnact13 | 1.49E-51 | 0.325924127 | 0.467 | 0.19 | 3.02E-47 | 10 | Csgalnact1 |
| Txndc151 | 1.65E-51 | 0.301246573 | 0.356 | 0.134 | 3.34E-47 | 10 | Txndc15 |
| Mvp | 1.84E-51 | 0.264173332 | 0.267 | 0.087 | 3.72E-47 | 10 | Mvp |
| Phc3 | 1.84E-51 | 0.360874229 | 0.345 | 0.131 | 3.73E-47 | 10 | Phc3 |
| Herc22 | 2.71E-51 | 0.371324491 | 0.469 | 0.202 | 5.49E-47 | 10 | Herc2 |

| | | | | | | | |
|---|---|---|---|---|---|---|---|
| Hnrnpdl | 2.75E-51 | 0.297483762 | 0.536 | 0.243 | 5.56E-47 | 10 | Hnrnpdl |
| Hs2st1 | 3.07E-51 | 0.304558341 | 0.266 | 0.087 | 6.21E-47 | 10 | Hs2st1 |
| Maml25 | 3.09E-51 | 0.455275924 | 0.673 | 0.336 | 6.25E-47 | 10 | Maml2 |
| Nsmaf1 | 3.58E-51 | 0.308404007 | 0.278 | 0.094 | 7.26E-47 | 10 | Nsmaf |
| Tmem30a4 | 3.61E-51 | 0.332563464 | 0.603 | 0.28 | 7.32E-47 | 10 | Tmem30a |
| Nr3c22 | 3.63E-51 | 0.350519199 | 0.327 | 0.117 | 7.34E-47 | 10 | Nr3c2 |
| Fbxo113 | 4.94E-51 | 0.340044951 | 0.514 | 0.227 | 1.00E-46 | 10 | Fbxo11 |
| Clcc1 | 5.12E-51 | 0.282065393 | 0.272 | 0.09 | 1.04E-46 | 10 | Clcc1 |
| Rc3h2 | 5.72E-51 | 0.266786175 | 0.317 | 0.113 | 1.16E-46 | 10 | Rc3h2 |
| Spcs3 | 7.21E-51 | 0.282946906 | 0.319 | 0.115 | 1.46E-46 | 10 | Spcs3 |
| Rab123 | 8.09E-51 | 0.256992599 | 0.43 | 0.172 | 1.64E-46 | 10 | Rab12 |
| Ankrd124 | 8.86E-51 | 0.363080078 | 0.623 | 0.302 | 1.79E-46 | 10 | Ankrd12 |
| Clk4 | 1.06E-50 | 0.296444097 | 0.359 | 0.137 | 2.15E-46 | 10 | Clk4 |
| Matr31 | 1.61E-50 | 0.297144972 | 0.561 | 0.261 | 3.25E-46 | 10 | Matr3 |
| Ap3b13 | 1.69E-50 | 0.35504401 | 0.484 | 0.212 | 3.42E-46 | 10 | Ap3b1 |
| Gsk3b3 | 1.78E-50 | 0.324160157 | 0.655 | 0.324 | 3.61E-46 | 10 | Gsk3b |
| Wdr261 | 2.00E-50 | 0.312385642 | 0.47 | 0.204 | 4.06E-46 | 10 | Wdr26 |
| Efcab142 | 2.13E-50 | 0.316014989 | 0.367 | 0.141 | 4.32E-46 | 10 | Efcab14 |
| Usp344 | 2.47E-50 | 0.37523772 | 0.672 | 0.338 | 4.99E-46 | 10 | Usp34 |
| Fam13b3 | 2.65E-50 | 0.286668426 | 0.369 | 0.143 | 5.37E-46 | 10 | Fam13b |
| Ago31 | 2.90E-50 | 0.362966081 | 0.378 | 0.151 | 5.87E-46 | 10 | Ago3 |
| Wdr332 | 3.72E-50 | 0.295118858 | 0.522 | 0.234 | 7.53E-46 | 10 | Wdr33 |
| Zfp451 | 3.86E-50 | 0.309338696 | 0.289 | 0.1 | 7.81E-46 | 10 | Zfp451 |
| St7l | 4.49E-50 | 0.25780021 | 0.264 | 0.086 | 9.09E-46 | 10 | St7l |
| Tnrc6a | 5.82E-50 | 0.288026879 | 0.459 | 0.194 | 1.18E-45 | 10 | Tnrc6a |
| Mbtd11 | 6.18E-50 | 0.339400897 | 0.342 | 0.131 | 1.25E-45 | 10 | Mbtd1 |
| Tiparp1 | 6.31E-50 | 0.275213952 | 0.256 | 0.083 | 1.28E-45 | 10 | Tiparp |
| Kansl14 | 9.63E-50 | 0.422819584 | 0.667 | 0.334 | 1.95E-45 | 10 | Kansl1 |
| Rsrp1 | 1.19E-49 | 0.382774385 | 0.902 | 0.564 | 2.41E-45 | 10 | Rsrp1 |
| Pigk | 1.27E-49 | 0.309939641 | 0.317 | 0.116 | 2.57E-45 | 10 | Pigk |
| Dennd1b2 | 1.28E-49 | 0.326847562 | 0.431 | 0.18 | 2.58E-45 | 10 | Dennd1b |
| Ctso | 1.34E-49 | 0.285488041 | 0.311 | 0.111 | 2.71E-45 | 10 | Ctso |
| Rhot1 | 1.83E-49 | 0.314925046 | 0.314 | 0.114 | 3.71E-45 | 10 | Rhot1 |
| Derl2 | 1.95E-49 | 0.312814051 | 0.328 | 0.123 | 3.94E-45 | 10 | Derl2 |
| Txndc111 | 2.15E-49 | 0.309558475 | 0.27 | 0.091 | 4.34E-45 | 10 | Txndc11 |
| Herc42 | 2.28E-49 | 0.365819218 | 0.369 | 0.147 | 4.63E-45 | 10 | Herc4 |
| Gm472832 | 2.35E-49 | 0.396476606 | 0.575 | 0.276 | 4.75E-45 | 10 | Gm47283 |
| Cdkal12 | 3.17E-49 | 0.3731261 | 0.42 | 0.176 | 6.41E-45 | 10 | Cdkal1 |
| Kmt2e3 | 3.17E-49 | 0.378058268 | 0.759 | 0.399 | 6.42E-45 | 10 | Kmt2e |
| Ddx3x3 | 3.70E-49 | 0.313860698 | 0.623 | 0.302 | 7.48E-45 | 10 | Ddx3x |
| Zfp1482 | 4.68E-49 | 0.345834245 | 0.548 | 0.259 | 9.47E-45 | 10 | Zfp148 |
| Abcc42 | 4.77E-49 | 0.308848808 | 0.352 | 0.132 | 9.65E-45 | 10 | Abcc4 |
| Macf14 | 4.97E-49 | 0.437758114 | 0.869 | 0.495 | 1.01E-44 | 10 | Macf1 |

| | | | | | | | |
|---|---|---|---|---|---|---|---|
| Atf7 | 6.43E-49 | 0.255041679 | 0.289 | 0.099 | 1.30E-44 | 10 | Atf7 |
| Ylpm11 | 1.17E-48 | 0.311830695 | 0.35 | 0.136 | 2.37E-44 | 10 | Ylpm1 |
| Zcchc111 | 1.73E-48 | 0.289809241 | 0.567 | 0.262 | 3.50E-44 | 10 | Zcchc11 |
| Mark31 | 1.80E-48 | 0.334886345 | 0.467 | 0.207 | 3.64E-44 | 10 | Mark3 |
| Tmem259 | 2.05E-48 | 0.253661832 | 0.267 | 0.089 | 4.16E-44 | 10 | Tmem259 |
| Extl31 | 2.09E-48 | 0.269237699 | 0.356 | 0.136 | 4.24E-44 | 10 | Extl3 |
| Slk4 | 2.53E-48 | 0.32092374 | 0.52 | 0.236 | 5.13E-44 | 10 | Slk |
| Ncstn | 2.85E-48 | 0.288508676 | 0.305 | 0.11 | 5.77E-44 | 10 | Ncstn |
| Dad12 | 3.74E-48 | 0.310807771 | 0.817 | 0.451 | 7.57E-44 | 10 | Dad1 |
| Phkb1 | 4.14E-48 | 0.268763252 | 0.359 | 0.141 | 8.39E-44 | 10 | Phkb |
| Yipf43 | 4.69E-48 | 0.270151851 | 0.491 | 0.216 | 9.50E-44 | 10 | Yipf4 |
| Cpsf6 | 5.79E-48 | 0.27843274 | 0.37 | 0.147 | 1.17E-43 | 10 | Cpsf6 |
| Dcaf62 | 7.37E-48 | 0.34190735 | 0.339 | 0.13 | 1.49E-43 | 10 | Dcaf6 |
| Ptprk2 | 8.31E-48 | 0.340762555 | 0.355 | 0.136 | 1.68E-43 | 10 | Ptprk |
| Smc32 | 8.43E-48 | 0.306143454 | 0.533 | 0.247 | 1.71E-43 | 10 | Smc3 |
| Atp6ap21 | 8.70E-48 | 0.295200177 | 0.411 | 0.17 | 1.76E-43 | 10 | Atp6ap2 |
| Stk40 | 1.70E-47 | 0.261174413 | 0.261 | 0.088 | 3.43E-43 | 10 | Stk40 |
| Tbrg11 | 1.91E-47 | 0.270907268 | 0.62 | 0.3 | 3.88E-43 | 10 | Tbrg1 |
| Emc102 | 2.57E-47 | 0.272825312 | 0.452 | 0.195 | 5.21E-43 | 10 | Emc10 |
| Cop11 | 2.65E-47 | 0.286276819 | 0.414 | 0.175 | 5.37E-43 | 10 | Cop1 |
| Phactr24 | 2.79E-47 | 0.344201934 | 0.528 | 0.241 | 5.65E-43 | 10 | Phactr2 |
| Pik3ca1 | 3.54E-47 | 0.285453948 | 0.4 | 0.164 | 7.17E-43 | 10 | Pik3ca |
| Nbr1 | 3.98E-47 | 0.258759695 | 0.352 | 0.137 | 8.06E-43 | 10 | Nbr1 |
| Ggnbp22 | 4.11E-47 | 0.290931344 | 0.52 | 0.237 | 8.33E-43 | 10 | Ggnbp2 |
| Ggps12 | 4.30E-47 | 0.27284772 | 0.406 | 0.167 | 8.70E-43 | 10 | Ggps1 |
| Arglu13 | 8.55E-47 | 0.302434312 | 0.691 | 0.347 | 1.73E-42 | 10 | Arglu1 |
| Dnajb9 | 8.62E-47 | 0.30100752 | 0.353 | 0.139 | 1.75E-42 | 10 | Dnajb9 |
| Uvrag4 | 9.31E-47 | 0.299508973 | 0.566 | 0.262 | 1.88E-42 | 10 | Uvrag |
| Rabl6 | 9.56E-47 | 0.267417325 | 0.305 | 0.111 | 1.94E-42 | 10 | Rabl6 |
| Ralgapb | 1.04E-46 | 0.297948158 | 0.306 | 0.114 | 2.11E-42 | 10 | Ralgapb |
| Khdrbs1 | 1.09E-46 | 0.259298739 | 0.556 | 0.26 | 2.22E-42 | 10 | Khdrbs1 |
| St3gal43 | 1.32E-46 | 0.316059984 | 0.541 | 0.251 | 2.68E-42 | 10 | St3gal4 |
| Ube4b | 1.40E-46 | 0.322135395 | 0.289 | 0.105 | 2.84E-42 | 10 | Ube4b |
| Hjurp | 1.49E-46 | 0.262557091 | 0.261 | 0.088 | 3.02E-42 | 10 | Hjurp |
| Thada2 | 1.76E-46 | 0.303124092 | 0.311 | 0.116 | 3.57E-42 | 10 | Thada |
| Phf20 | 1.84E-46 | 0.264252868 | 0.356 | 0.141 | 3.73E-42 | 10 | Phf20 |
| Ttc171 | 2.17E-46 | 0.301770833 | 0.306 | 0.113 | 4.39E-42 | 10 | Ttc17 |
| Setd24 | 2.31E-46 | 0.315119829 | 0.438 | 0.188 | 4.68E-42 | 10 | Setd2 |
| Bptf4 | 2.49E-46 | 0.332439801 | 0.592 | 0.288 | 5.03E-42 | 10 | Bptf |
| Hdac81 | 5.95E-46 | 0.409964884 | 0.334 | 0.132 | 1.20E-41 | 10 | Hdac8 |
| Clptm1 | 6.60E-46 | 0.274183501 | 0.32 | 0.121 | 1.34E-41 | 10 | Clptm1 |
| Kdm2a5 | 6.79E-46 | 0.304925335 | 0.509 | 0.235 | 1.37E-41 | 10 | Kdm2a |
| Zfp6383 | 7.42E-46 | 0.33568643 | 0.573 | 0.276 | 1.50E-41 | 10 | Zfp638 |

| | | | | | | | |
|---|---|---|---|---|---|---|---|
| Zmym21 | 8.24E-46 | 0.300261514 | 0.344 | 0.136 | 1.67E-41 | 10 | Zmym2 |
| Eif3a3 | 8.31E-46 | 0.294941912 | 0.711 | 0.365 | 1.68E-41 | 10 | Eif3a |
| Thap31 | 1.23E-45 | 0.269504107 | 0.386 | 0.158 | 2.48E-41 | 10 | Thap3 |
| Tial1 | 1.26E-45 | 0.275259366 | 0.37 | 0.15 | 2.56E-41 | 10 | Tial1 |
| Ufl1 | 1.29E-45 | 0.290620717 | 0.32 | 0.123 | 2.62E-41 | 10 | Ufl1 |
| Ranbp22 | 1.54E-45 | 0.289272668 | 0.444 | 0.194 | 3.12E-41 | 10 | Ranbp2 |
| Mar-61 | 1.58E-45 | 0.284297107 | 0.352 | 0.139 | 3.20E-41 | 10 | 6-Mar |
| Ncor13 | 1.82E-45 | 0.315569643 | 0.794 | 0.431 | 3.69E-41 | 10 | Ncor1 |
| Ythdf32 | 2.64E-45 | 0.262802239 | 0.439 | 0.191 | 5.35E-41 | 10 | Ythdf3 |
| Prpf40a3 | 3.22E-45 | 0.286183962 | 0.552 | 0.261 | 6.52E-41 | 10 | Prpf40a |
| Sh3gl11 | 3.84E-45 | 0.306408622 | 0.317 | 0.121 | 7.77E-41 | 10 | Sh3gl1 |
| Atf21 | 4.13E-45 | 0.286047206 | 0.377 | 0.155 | 8.36E-41 | 10 | Atf2 |
| Fmr1 | 4.51E-45 | 0.250812143 | 0.328 | 0.127 | 9.13E-41 | 10 | Fmr1 |
| Spcs21 | 7.90E-45 | 0.330210791 | 0.722 | 0.395 | 1.60E-40 | 10 | Spcs2 |
| Srsf115 | 9.18E-45 | 0.328315369 | 0.672 | 0.345 | 1.86E-40 | 10 | Srsf11 |
| Setd71 | 1.02E-44 | 0.298325193 | 0.367 | 0.15 | 2.06E-40 | 10 | Setd7 |
| Srgap25 | 1.09E-44 | 0.374708192 | 0.6 | 0.298 | 2.20E-40 | 10 | Srgap2 |
| Otud7b1 | 1.49E-44 | 0.255434378 | 0.32 | 0.121 | 3.01E-40 | 10 | Otud7b |
| Ncoa6 | 2.01E-44 | 0.312829357 | 0.33 | 0.13 | 4.07E-40 | 10 | Ncoa6 |
| Ddhd2 | 2.26E-44 | 0.26401485 | 0.286 | 0.104 | 4.58E-40 | 10 | Ddhd2 |
| Ckap5 | 4.96E-44 | 0.28717589 | 0.345 | 0.138 | 1.01E-39 | 10 | Ckap5 |
| Ankrd114 | 5.98E-44 | 0.314468988 | 0.72 | 0.374 | 1.21E-39 | 10 | Ankrd11 |
| Prrc2c4 | 6.99E-44 | 0.296698329 | 0.781 | 0.413 | 1.41E-39 | 10 | Prrc2c |
| Gnaq4 | 7.05E-44 | 0.266109597 | 0.702 | 0.352 | 1.43E-39 | 10 | Gnaq |
| Tnpo31 | 7.20E-44 | 0.251604397 | 0.352 | 0.142 | 1.46E-39 | 10 | Tnpo3 |
| Mtss11 | 1.13E-43 | 0.251321544 | 0.272 | 0.097 | 2.29E-39 | 10 | Mtss1 |
| Ube3c2 | 1.20E-43 | 0.304535558 | 0.37 | 0.155 | 2.42E-39 | 10 | Ube3c |
| Baz2b4 | 1.41E-43 | 0.325551171 | 0.667 | 0.344 | 2.86E-39 | 10 | Baz2b |
| Lncpint3 | 1.63E-43 | 0.348439072 | 0.556 | 0.269 | 3.29E-39 | 10 | Lncpint |
| Anapc1 | 1.69E-43 | 0.270333196 | 0.314 | 0.12 | 3.42E-39 | 10 | Anapc1 |
| Galnt14 | 1.80E-43 | 0.293630623 | 0.412 | 0.18 | 3.65E-39 | 10 | Galnt1 |
| Exoc45 | 2.11E-43 | 0.360389805 | 0.63 | 0.317 | 4.28E-39 | 10 | Exoc4 |
| Diaph25 | 2.66E-43 | 0.318735653 | 0.714 | 0.37 | 5.39E-39 | 10 | Diaph2 |
| Psen12 | 3.06E-43 | 0.257004491 | 0.27 | 0.097 | 6.20E-39 | 10 | Psen1 |
| Zmym51 | 3.45E-43 | 0.258264652 | 0.355 | 0.144 | 6.98E-39 | 10 | Zmym5 |
| Pum14 | 3.74E-43 | 0.275882397 | 0.533 | 0.254 | 7.57E-39 | 10 | Pum1 |
| Prpf82 | 4.00E-43 | 0.276583538 | 0.411 | 0.179 | 8.09E-39 | 10 | Prpf8 |
| Sf3b32 | 5.94E-43 | 0.287598727 | 0.375 | 0.157 | 1.20E-38 | 10 | Sf3b3 |
| Numa12 | 9.09E-43 | 0.309323099 | 0.431 | 0.191 | 1.84E-38 | 10 | Numa1 |
| Cast3 | 1.24E-42 | 0.312394424 | 0.603 | 0.306 | 2.51E-38 | 10 | Cast |
| Ppp2r5e2 | 1.40E-42 | 0.27915315 | 0.38 | 0.161 | 2.84E-38 | 10 | Ppp2r5e |
| Itch3 | 1.43E-42 | 0.297962079 | 0.559 | 0.268 | 2.89E-38 | 10 | Itch |
| Herc13 | 2.16E-42 | 0.284963114 | 0.533 | 0.252 | 4.37E-38 | 10 | Herc1 |

| | | | | | | | |
|---|---|---|---|---|---|---|---|
| Hmg20a1 | 2.38E-42 | 0.263438443 | 0.344 | 0.139 | 4.82E-38 | 10 | Hmg20a |
| Nr2c22 | 3.50E-42 | 0.353878744 | 0.367 | 0.155 | 7.09E-38 | 10 | Nr2c2 |
| Trio3 | 5.87E-42 | 0.330356228 | 0.416 | 0.184 | 1.19E-37 | 10 | Trio |
| Atrn1 | 6.07E-42 | 0.299587371 | 0.308 | 0.12 | 1.23E-37 | 10 | Atrn |
| Ankrd281 | 6.72E-42 | 0.280416301 | 0.33 | 0.129 | 1.36E-37 | 10 | Ankrd28 |
| Appbp21 | 9.10E-42 | 0.269990223 | 0.338 | 0.136 | 1.84E-37 | 10 | Appbp2 |
| Tia1 | 1.81E-41 | 0.25877002 | 0.291 | 0.11 | 3.66E-37 | 10 | Tia1 |
| Wnk14 | 2.02E-41 | 0.326469473 | 0.686 | 0.357 | 4.10E-37 | 10 | Wnk1 |
| Frs2 | 2.21E-41 | 0.25688691 | 0.341 | 0.139 | 4.48E-37 | 10 | Frs2 |
| Man1a23 | 2.42E-41 | 0.3194508 | 0.472 | 0.222 | 4.90E-37 | 10 | Man1a2 |
| Cwc27 | 3.32E-41 | 0.255725341 | 0.308 | 0.12 | 6.73E-37 | 10 | Cwc27 |
| Zfp280d1 | 4.76E-41 | 0.261366502 | 0.408 | 0.18 | 9.63E-37 | 10 | Zfp280d |
| Bbx3 | 6.29E-41 | 0.318583169 | 0.466 | 0.218 | 1.27E-36 | 10 | Bbx |
| Btbd73 | 6.52E-41 | 0.257859528 | 0.569 | 0.276 | 1.32E-36 | 10 | Btbd7 |
| Fyco11 | 1.01E-40 | 0.25860113 | 0.306 | 0.121 | 2.05E-36 | 10 | Fyco1 |
| Rap1gds14 | 1.98E-40 | 0.273813821 | 0.531 | 0.256 | 4.02E-36 | 10 | Rap1gds1 |
| Zmynd113 | 2.41E-40 | 0.269574241 | 0.567 | 0.284 | 4.89E-36 | 10 | Zmynd11 |
| Smurf23 | 2.62E-40 | 0.266456819 | 0.495 | 0.234 | 5.30E-36 | 10 | Smurf2 |
| Rbm54 | 2.71E-40 | 0.313785692 | 0.492 | 0.235 | 5.49E-36 | 10 | Rbm5 |
| Ehmt12 | 3.96E-40 | 0.261209145 | 0.411 | 0.181 | 8.03E-36 | 10 | Ehmt1 |
| Nfkb13 | 5.23E-40 | 0.254305996 | 0.552 | 0.266 | 1.06E-35 | 10 | Nfkb1 |
| Ahctf1 | 5.34E-40 | 0.251646537 | 0.306 | 0.12 | 1.08E-35 | 10 | Ahctf1 |
| Sdccag82 | 6.50E-40 | 0.256433206 | 0.302 | 0.119 | 1.32E-35 | 10 | Sdccag8 |
| Cep3504 | 7.17E-40 | 0.267369872 | 0.436 | 0.199 | 1.45E-35 | 10 | Cep350 |
| Rlf1 | 7.84E-40 | 0.323975349 | 0.323 | 0.134 | 1.59E-35 | 10 | Rlf |
| Cnot42 | 8.84E-40 | 0.302018525 | 0.533 | 0.264 | 1.79E-35 | 10 | Cnot4 |
| Golga4 | 9.72E-40 | 0.259031303 | 0.362 | 0.154 | 1.97E-35 | 10 | Golga4 |
| Tmem1311 | 2.66E-39 | 0.316697082 | 0.445 | 0.204 | 5.38E-35 | 10 | Tmem131 |
| Ankib11 | 2.78E-39 | 0.270825993 | 0.328 | 0.135 | 5.64E-35 | 10 | Ankib1 |
| Larp12 | 3.00E-39 | 0.276564625 | 0.434 | 0.199 | 6.07E-35 | 10 | Larp1 |
| Tmx33 | 3.60E-39 | 0.274256471 | 0.411 | 0.183 | 7.29E-35 | 10 | Tmx3 |
| Atp11c2 | 3.94E-39 | 0.339786251 | 0.347 | 0.148 | 7.98E-35 | 10 | Atp11c |
| Map7d12 | 6.60E-39 | 0.285222797 | 0.375 | 0.164 | 1.34E-34 | 10 | Map7d1 |
| 2310035C23Rik2 | 6.64E-39 | 0.268299493 | 0.331 | 0.137 | 1.35E-34 | 10 | 2310035C23Rik |
| Usp241 | 2.32E-38 | 0.260484513 | 0.323 | 0.134 | 4.69E-34 | 10 | Usp24 |
| Trim442 | 5.34E-38 | 0.251540509 | 0.469 | 0.223 | 1.08E-33 | 10 | Trim44 |
| Ccdc1071 | 6.71E-38 | 0.271854521 | 0.306 | 0.125 | 1.36E-33 | 10 | Ccdc107 |
| Celf11 | 9.58E-38 | 0.259092203 | 0.455 | 0.216 | 1.94E-33 | 10 | Celf1 |
| Huwe13 | 1.05E-37 | 0.262579615 | 0.556 | 0.28 | 2.13E-33 | 10 | Huwe1 |
| Coro1c1 | 1.10E-37 | 0.298132074 | 0.325 | 0.137 | 2.23E-33 | 10 | Coro1c |
| Phf33 | 1.27E-37 | 0.25598797 | 0.509 | 0.248 | 2.58E-33 | 10 | Phf3 |
| Colgalt22 | 1.61E-37 | 0.285306754 | 0.334 | 0.137 | 3.25E-33 | 10 | Colgalt2 |
| Smg14 | 2.56E-37 | 0.308475549 | 0.47 | 0.23 | 5.18E-33 | 10 | Smg1 |

| | | | | | | | |
|---|---|---|---|---|---|---|---|
| Gigyf22 | 3.52E-37 | 0.276724364 | 0.369 | 0.164 | 7.12E-33 | 10 | Gigyf2 |
| Akap133 | 4.60E-37 | 0.275485701 | 0.717 | 0.388 | 9.30E-33 | 10 | Akap13 |
| Sik22 | 4.96E-37 | 0.271130518 | 0.291 | 0.117 | 1.00E-32 | 10 | Sik2 |
| Pan34 | 4.97E-37 | 0.286246896 | 0.467 | 0.224 | 1.01E-32 | 10 | Pan3 |
| Rnf2162 | 5.97E-37 | 0.299613336 | 0.389 | 0.177 | 1.21E-32 | 10 | Rnf216 |
| Fnbp4 | 7.83E-37 | 0.252597122 | 0.306 | 0.126 | 1.59E-32 | 10 | Fnbp4 |
| Ccnd23 | 1.05E-36 | 0.305749666 | 0.548 | 0.281 | 2.13E-32 | 10 | Ccnd2 |
| Morf4l11 | 1.22E-36 | 0.276595818 | 0.9 | 0.583 | 2.47E-32 | 10 | Morf4l1 |
| Gpatch2 | 1.26E-36 | 0.293052422 | 0.281 | 0.112 | 2.54E-32 | 10 | Gpatch2 |
| Fgl22 | 2.82E-36 | 0.256055377 | 0.27 | 0.105 | 5.71E-32 | 10 | Fgl2 |
| Scaper2 | 3.22E-36 | 0.292879101 | 0.384 | 0.176 | 6.53E-32 | 10 | Scaper |
| Senp71 | 3.31E-36 | 0.252148852 | 0.35 | 0.153 | 6.70E-32 | 10 | Senp7 |
| Ric11 | 3.39E-36 | 0.271852116 | 0.286 | 0.115 | 6.87E-32 | 10 | Ric1 |
| Dmtf11 | 3.83E-36 | 0.280234443 | 0.289 | 0.119 | 7.75E-32 | 10 | Dmtf1 |
| Lrch34 | 4.01E-36 | 0.258669819 | 0.472 | 0.228 | 8.11E-32 | 10 | Lrch3 |
| Bbs91 | 4.15E-36 | 0.270969298 | 0.258 | 0.099 | 8.40E-32 | 10 | Bbs9 |
| Ascc33 | 4.25E-36 | 0.290105834 | 0.423 | 0.204 | 8.60E-32 | 10 | Ascc3 |
| Ifih12 | 5.21E-36 | 0.254474443 | 0.306 | 0.126 | 1.05E-31 | 10 | Ifih1 |
| Emsy | 8.12E-36 | 0.304655451 | 0.297 | 0.124 | 1.64E-31 | 10 | Emsy |
| Cdk133 | 9.30E-36 | 0.265438086 | 0.455 | 0.223 | 1.88E-31 | 10 | Cdk13 |
| Brd44 | 1.08E-35 | 0.262174726 | 0.523 | 0.263 | 2.18E-31 | 10 | Brd4 |
| Peak14 | 1.67E-35 | 0.317504889 | 0.689 | 0.381 | 3.39E-31 | 10 | Peak1 |
| Mkl12 | 4.42E-35 | 0.267193934 | 0.295 | 0.123 | 8.94E-31 | 10 | Mkl1 |
| Tbc1d53 | 1.15E-34 | 0.268651614 | 0.509 | 0.257 | 2.32E-30 | 10 | Tbc1d5 |
| Csnk1g12 | 1.34E-34 | 0.261421717 | 0.347 | 0.154 | 2.72E-30 | 10 | Csnk1g1 |
| Sulf21 | 1.39E-34 | 0.281283282 | 0.252 | 0.097 | 2.82E-30 | 10 | Sulf2 |
| Ints61 | 1.57E-34 | 0.264249634 | 0.32 | 0.139 | 3.18E-30 | 10 | Ints6 |
| Sh3rf12 | 3.09E-34 | 0.269814554 | 0.305 | 0.128 | 6.25E-30 | 10 | Sh3rf1 |
| Sppl33 | 4.59E-34 | 0.252330067 | 0.447 | 0.214 | 9.30E-30 | 10 | Sppl3 |
| Adcy72 | 4.94E-34 | 0.303364356 | 0.272 | 0.112 | 1.00E-29 | 10 | Adcy7 |
| Rbm33 | 7.53E-34 | 0.254557738 | 0.319 | 0.139 | 1.52E-29 | 10 | Rbm33 |
| Zc3h7a2 | 8.00E-34 | 0.27535717 | 0.35 | 0.159 | 1.62E-29 | 10 | Zc3h7a |
| Kmt2a4 | 2.02E-33 | 0.257444833 | 0.516 | 0.263 | 4.09E-29 | 10 | Kmt2a |
| Ncoa24 | 2.17E-33 | 0.279455901 | 0.473 | 0.234 | 4.39E-29 | 10 | Ncoa2 |
| Neat13 | 3.04E-33 | 0.282888176 | 0.422 | 0.205 | 6.16E-29 | 10 | Neat1 |
| Stat23 | 8.29E-33 | 0.29024262 | 0.317 | 0.137 | 1.68E-28 | 10 | Stat2 |
| Ubn24 | 2.12E-32 | 0.286133512 | 0.472 | 0.241 | 4.29E-28 | 10 | Ubn2 |
| Nrf11 | 6.67E-32 | 0.259061619 | 0.305 | 0.133 | 1.35E-27 | 10 | Nrf1 |
| Abl22 | 1.98E-31 | 0.313712633 | 0.277 | 0.117 | 4.01E-27 | 10 | Abl2 |
| Fgfr21 | 2.73E-31 | 0.250123523 | 0.328 | 0.15 | 5.52E-27 | 10 | Fgfr2 |
| Tcf202 | 4.33E-31 | 0.285803575 | 0.306 | 0.139 | 8.77E-27 | 10 | Tcf20 |
| Cspp1 | 7.88E-31 | 0.258949087 | 0.269 | 0.114 | 1.59E-26 | 10 | Cspp1 |
| Frmd4b2 | 5.71E-30 | 0.378553973 | 0.358 | 0.175 | 1.16E-25 | 10 | Frmd4b |

| | | | | | | | |
|---|---|---|---|---|---|---|---|
| 5031425E22Rik1 | 1.57E-29 | 0.280487295 | 0.334 | 0.157 | 3.18E-25 | 10 | 5031425E22Rik |
| Kdm4c2 | 2.34E-29 | 0.250780886 | 0.348 | 0.166 | 4.74E-25 | 10 | Kdm4c |
| Chd23 | 1.99E-28 | 0.257455665 | 0.472 | 0.25 | 4.02E-24 | 10 | Chd2 |
| Ifi2034 | 7.99E-27 | 0.275588405 | 0.634 | 0.363 | 1.62E-22 | 10 | Ifi203 |
| Iigp13 | 3.03E-19 | 0.3100764 | 0.414 | 0.23 | 6.13E-15 | 10 | Iigp1 |
| Gbp22 | 5.61E-18 | 0.266771733 | 0.277 | 0.142 | 1.14E-13 | 10 | Gbp2 |
| Klra41 | 0 | 3.272451669 | 0.813 | 0.063 | 0 | 11 | Klra4 |
| Klra81 | 0 | 2.942523935 | 0.801 | 0.064 | 0 | 11 | Klra8 |
| Gzma | 0 | 2.917047778 | 0.447 | 0.012 | 0 | 11 | Gzma |
| Ms4a4b1 | 0 | 2.91487483 | 0.978 | 0.104 | 0 | 11 | Ms4a4b |
| Xcl11 | 0 | 2.897198501 | 0.966 | 0.112 | 0 | 11 | Xcl1 |
| Klrk11 | 0 | 2.730521473 | 0.971 | 0.082 | 0 | 11 | Klrk1 |
| Nkg71 | 0 | 2.726419233 | 0.971 | 0.109 | 0 | 11 | Nkg7 |
| Ptprc2 | 0 | 2.703832803 | 0.995 | 0.182 | 0 | 11 | Ptprc |
| Klre11 | 0 | 2.589209955 | 0.975 | 0.084 | 0 | 11 | Klre1 |
| Lgals11 | 0 | 2.579031555 | 0.851 | 0.12 | 0 | 11 | Lgals1 |
| Cd523 | 0 | 2.481611718 | 0.998 | 0.267 | 0 | 11 | Cd52 |
| Skap11 | 0 | 2.387258523 | 0.955 | 0.116 | 0 | 11 | Skap1 |
| Tmsb102 | 0 | 2.384004775 | 0.995 | 0.491 | 0 | 11 | Tmsb10 |
| Klrd11 | 0 | 2.310599221 | 0.964 | 0.104 | 0 | 11 | Klrd1 |
| Dock22 | 0 | 2.301808236 | 0.968 | 0.17 | 0 | 11 | Dock2 |
| Gm26821 | 0 | 2.190215921 | 0.79 | 0.065 | 0 | 11 | Gm2682 |
| Ctsw1 | 0 | 2.173583145 | 0.973 | 0.11 | 0 | 11 | Ctsw |
| Ltb1 | 0 | 2.15214522 | 0.908 | 0.119 | 0 | 11 | Ltb |
| Ncr11 | 0 | 2.134210458 | 0.858 | 0.088 | 0 | 11 | Ncr1 |
| Dennd4a2 | 0 | 2.120609664 | 0.944 | 0.172 | 0 | 11 | Dennd4a |
| Ptprcap1 | 0 | 2.076910506 | 0.932 | 0.115 | 0 | 11 | Ptprcap |
| Ccr21 | 0 | 2.072386376 | 0.824 | 0.07 | 0 | 11 | Ccr2 |
| Clnk1 | 0 | 2.065543317 | 0.855 | 0.096 | 0 | 11 | Clnk |
| Ccl51 | 0 | 2.062260112 | 0.885 | 0.136 | 0 | 11 | Ccl5 |
| Klra71 | 0 | 2.052831487 | 0.621 | 0.063 | 0 | 11 | Klra7 |
| Styk11 | 0 | 2.042950057 | 0.801 | 0.082 | 0 | 11 | Styk1 |
| Ugcg1 | 0 | 2.01792238 | 0.894 | 0.156 | 0 | 11 | Ugcg |
| Tox1 | 0 | 2.009644298 | 0.819 | 0.107 | 0 | 11 | Tox |
| Hcst1 | 0 | 1.995149414 | 0.925 | 0.129 | 0 | 11 | Hcst |
| Stat41 | 0 | 1.976587422 | 0.792 | 0.093 | 0 | 11 | Stat4 |
| Txk1 | 0 | 1.975781617 | 0.831 | 0.091 | 0 | 11 | Txk |
| Samd31 | 0 | 1.96824777 | 0.817 | 0.071 | 0 | 11 | Samd3 |
| Rac21 | 0 | 1.964123725 | 0.944 | 0.162 | 0 | 11 | Rac2 |
| Klri21 | 0 | 1.943795079 | 0.706 | 0.061 | 0 | 11 | Klri2 |
| Il2rb1 | 0 | 1.92830234 | 0.919 | 0.103 | 0 | 11 | Il2rb |
| Ikzf31 | 0 | 1.925786956 | 0.876 | 0.104 | 0 | 11 | Ikzf3 |
| Tyrobp3 | 0 | 1.911731317 | 0.998 | 0.247 | 0 | 11 | Tyrobp |

| | | | | | | | |
|---|---|---|---|---|---|---|---|
| Kcnq51 | 0 | 1.90893839 | 0.528 | 0.058 | 0 | 11 | Kcnq5 |
| Runx32 | 0 | 1.901540434 | 0.846 | 0.134 | 0 | 11 | Runx3 |
| Klra91 | 0 | 1.889367799 | 0.605 | 0.067 | 0 | 11 | Klra9 |
| Dusp21 | 0 | 1.871931059 | 0.754 | 0.088 | 0 | 11 | Dusp2 |
| Fcer1g3 | 0 | 1.865479388 | 1 | 0.255 | 0 | 11 | Fcer1g |
| Bcl21 | 0 | 1.858399367 | 0.867 | 0.194 | 0 | 11 | Bcl2 |
| Mir142hg2 | 0 | 1.851330894 | 0.88 | 0.155 | 0 | 11 | Mir142hg |
| Shisa51 | 0 | 1.830697507 | 0.982 | 0.316 | 0 | 11 | Shisa5 |
| Arhgdib3 | 0 | 1.825366888 | 0.953 | 0.219 | 0 | 11 | Arhgdib |
| Eomes1 | 0 | 1.799864023 | 0.752 | 0.035 | 0 | 11 | Eomes |
| 12-Sep | 0 | 1.788046289 | 0.869 | 0.131 | 0 | 11 | 1-Sep |
| Klrb1c1 | 0 | 1.768938251 | 0.763 | 0.083 | 0 | 11 | Klrb1c |
| Arhgap152 | 0 | 1.720441458 | 0.901 | 0.173 | 0 | 11 | Arhgap15 |
| Adgre52 | 0 | 1.717985744 | 0.851 | 0.225 | 0 | 11 | Adgre5 |
| Epsti12 | 0 | 1.713089307 | 0.858 | 0.161 | 0 | 11 | Epsti1 |
| Dscam1 | 0 | 1.711987901 | 0.433 | 0.039 | 0 | 11 | Dscam |
| Cd21 | 0 | 1.709407289 | 0.747 | 0.07 | 0 | 11 | Cd2 |
| Setbp12 | 0 | 1.700691764 | 0.711 | 0.109 | 0 | 11 | Setbp1 |
| Ikzf12 | 0 | 1.692113657 | 0.81 | 0.143 | 0 | 11 | Ikzf1 |
| Gem1 | 0 | 1.644323386 | 0.688 | 0.091 | 0 | 11 | Gem |
| Ccr52 | 0 | 1.640323867 | 0.749 | 0.12 | 0 | 11 | Ccr5 |
| Coro1a3 | 0 | 1.638132582 | 0.93 | 0.205 | 0 | 11 | Coro1a |
| Arsb1 | 0 | 1.632128598 | 0.74 | 0.114 | 0 | 11 | Arsb |
| Ptpn221 | 0 | 1.63070525 | 0.776 | 0.105 | 0 | 11 | Ptpn22 |
| Dock102 | 0 | 1.630399815 | 0.899 | 0.194 | 0 | 11 | Dock10 |
| Arhgap452 | 0 | 1.608815281 | 0.819 | 0.145 | 0 | 11 | Arhgap45 |
| Chsy11 | 0 | 1.604721159 | 0.781 | 0.114 | 0 | 11 | Chsy1 |
| Fam49b2 | 0 | 1.554738274 | 0.887 | 0.224 | 0 | 11 | Fam49b |
| Prf11 | 0 | 1.548291708 | 0.621 | 0.051 | 0 | 11 | Prf1 |
| Selplg2 | 0 | 1.536923925 | 0.826 | 0.141 | 0 | 11 | Selplg |
| Atp8b41 | 0 | 1.536105801 | 0.684 | 0.085 | 0 | 11 | Atp8b4 |
| Sh2d1a1 | 0 | 1.525938719 | 0.645 | 0.044 | 0 | 11 | Sh2d1a |
| Slc9a92 | 0 | 1.524644741 | 0.754 | 0.139 | 0 | 11 | Slc9a9 |
| Gas72 | 0 | 1.512580811 | 0.783 | 0.17 | 0 | 11 | Gas7 |
| Prkcq1 | 0 | 1.479503056 | 0.632 | 0.076 | 0 | 11 | Prkcq |
| Aff31 | 0 | 1.461437213 | 0.53 | 0.023 | 0 | 11 | Aff3 |
| Cd160 | 0 | 1.458481827 | 0.625 | 0.031 | 0 | 11 | Cd160 |
| Tnik2 | 0 | 1.456623919 | 0.706 | 0.101 | 0 | 11 | Tnik |
| Trbc11 | 0 | 1.446907118 | 0.695 | 0.073 | 0 | 11 | Trbc1 |
| Klrb1f1 | 0 | 1.436503643 | 0.648 | 0.05 | 0 | 11 | Klrb1f |
| Klrc21 | 0 | 1.41052924 | 0.625 | 0.041 | 0 | 11 | Klrc2 |
| Lck1 | 0 | 1.410126837 | 0.725 | 0.09 | 0 | 11 | Lck |
| Stk17b1 | 0 | 1.407998909 | 0.777 | 0.15 | 0 | 11 | Stk17b |

| | | | | | | | |
|---|---|---|---|---|---|---|---|
| Osbpl31 | 0 | 1.376720896 | 0.713 | 0.107 | 0 | 11 | Osbpl3 |
| Aoah1 | 0 | 1.37077039 | 0.607 | 0.062 | 0 | 11 | Aoah |
| Rinl1 | 0 | 1.347280861 | 0.725 | 0.12 | 0 | 11 | Rinl |
| Mctp21 | 0 | 1.345462952 | 0.627 | 0.073 | 0 | 11 | Mctp2 |
| Limd22 | 0 | 1.32137755 | 0.718 | 0.137 | 0 | 11 | Limd2 |
| Prkcb3 | 0 | 1.305832558 | 0.732 | 0.145 | 0 | 11 | Prkcb |
| Ipcef12 | 0 | 1.271695594 | 0.632 | 0.107 | 0 | 11 | Ipcef1 |
| Satb11 | 0 | 1.269390534 | 0.549 | 0.053 | 0 | 11 | Satb1 |
| Znrf32 | 0 | 1.265392571 | 0.618 | 0.097 | 0 | 11 | Znrf3 |
| Itgal1 | 0 | 1.260684955 | 0.618 | 0.076 | 0 | 11 | Itgal |
| Cd372 | 0 | 1.253395398 | 0.671 | 0.103 | 0 | 11 | Cd37 |
| Car21 | 0 | 1.228527748 | 0.661 | 0.111 | 0 | 11 | Car2 |
| Itga2 | 0 | 1.182953027 | 0.454 | 0.029 | 0 | 11 | Itga2 |
| Cst71 | 0 | 1.173816396 | 0.564 | 0.064 | 0 | 11 | Cst7 |
| Il2rg1 | 0 | 1.167956209 | 0.625 | 0.107 | 0 | 11 | Il2rg |
| Il18rap | 0 | 1.1672148 | 0.438 | 0.015 | 0 | 11 | Il18rap |
| Cyfip21 | 0 | 1.164219213 | 0.589 | 0.081 | 0 | 11 | Cyfip2 |
| Hsd11b12 | 0 | 1.160238824 | 0.607 | 0.081 | 0 | 11 | Hsd11b1 |
| Itga42 | 0 | 1.158342339 | 0.497 | 0.068 | 0 | 11 | Itga4 |
| Kcnip31 | 0 | 1.141473835 | 0.513 | 0.037 | 0 | 11 | Kcnip3 |
| Il18r1 | 0 | 1.135513432 | 0.379 | 0.018 | 0 | 11 | Il18r1 |
| Adamts141 | 0 | 1.130388706 | 0.483 | 0.026 | 0 | 11 | Adamts14 |
| Gpr1741 | 0 | 1.128363681 | 0.492 | 0.043 | 0 | 11 | Gpr174 |
| Serpina3g1 | 0 | 1.126642424 | 0.472 | 0.059 | 0 | 11 | Serpina3g |
| Cd71 | 0 | 1.12639223 | 0.693 | 0.1 | 0 | 11 | Cd7 |
| Cd2441 | 0 | 1.104439341 | 0.582 | 0.082 | 0 | 11 | Cd244 |
| Slc38a11 | 0 | 1.093904987 | 0.555 | 0.079 | 0 | 11 | Slc38a1 |
| Pglyrp11 | 0 | 1.088182819 | 0.605 | 0.082 | 0 | 11 | Pglyrp1 |
| Sh2d2a1 | 0 | 1.087785148 | 0.487 | 0.04 | 0 | 11 | Sh2d2a |
| Card111 | 0 | 1.076249228 | 0.53 | 0.073 | 0 | 11 | Card11 |
| Dok2 | 0 | 1.075751563 | 0.472 | 0.033 | 0 | 11 | Dok2 |
| Sep-61 | 0 | 1.075375767 | 0.594 | 0.089 | 0 | 11 | 6-Sep |
| Cd2471 | 0 | 1.068211655 | 0.472 | 0.038 | 0 | 11 | Cd247 |
| Thy11 | 0 | 1.063748797 | 0.46 | 0.052 | 0 | 11 | Thy1 |
| Sell | 0 | 1.05143974 | 0.366 | 0.004 | 0 | 11 | Sell |
| Crtam | 0 | 1.033562676 | 0.325 | 0.018 | 0 | 11 | Crtam |
| Lax11 | 0 | 0.993064741 | 0.49 | 0.057 | 0 | 11 | Lax1 |
| Rnf431 | 0 | 0.980469066 | 0.406 | 0.043 | 0 | 11 | Rnf43 |
| Gm43647 | 0 | 0.973829381 | 0.354 | 0.014 | 0 | 11 | Gm43647 |
| Tbx211 | 0 | 0.964900495 | 0.469 | 0.058 | 0 | 11 | Tbx21 |
| Tnfrsf181 | 0 | 0.946119983 | 0.49 | 0.065 | 0 | 11 | Tnfrsf18 |
| Tcf7 | 0 | 0.945769426 | 0.384 | 0.014 | 0 | 11 | Tcf7 |
| Fasl1 | 0 | 0.913870077 | 0.436 | 0.049 | 0 | 11 | Fasl |

| | | | | | | | |
|---|---|---|---|---|---|---|---|
| Eya11 | 0 | 0.876466383 | 0.375 | 0.034 | 0 | 11 | Eya1 |
| Glipr2 | 0 | 0.83350455 | 0.39 | 0.036 | 0 | 11 | Glipr2 |
| Cd27 | 0 | 0.83114671 | 0.37 | 0.033 | 0 | 11 | Cd27 |
| D5Ertd615e | 0 | 0.829325253 | 0.291 | 0.016 | 0 | 11 | D5Ertd615e |
| Cers62 | 2.20E-302 | 1.235246032 | 0.61 | 0.105 | 4.46E-298 | 11 | Cers6 |
| Ptpn183 | 1.27E-301 | 1.634734368 | 0.962 | 0.302 | 2.58E-297 | 11 | Ptpn18 |
| Coro2a1 | 2.57E-300 | 1.216493963 | 0.618 | 0.108 | 5.19E-296 | 11 | Coro2a |
| Tespa11 | 1.22E-293 | 0.980639299 | 0.487 | 0.069 | 2.47E-289 | 11 | Tespa1 |
| Tbc1d10c1 | 2.60E-293 | 0.913173914 | 0.474 | 0.065 | 5.27E-289 | 11 | Tbc1d10c |
| Sytl31 | 9.39E-293 | 0.883784574 | 0.417 | 0.05 | 1.90E-288 | 11 | Sytl3 |
| Bin22 | 1.07E-292 | 1.091339872 | 0.634 | 0.117 | 2.16E-288 | 11 | Bin2 |
| Chl11 | 2.73E-292 | 1.079585645 | 0.399 | 0.046 | 5.52E-288 | 11 | Chl1 |
| Ptk2b2 | 1.28E-291 | 1.227490473 | 0.706 | 0.139 | 2.58E-287 | 11 | Ptk2b |
| Ccdc88c1 | 5.09E-289 | 1.014788983 | 0.557 | 0.091 | 1.03E-284 | 11 | Ccdc88c |
| Ncf42 | 3.24E-287 | 1.080423879 | 0.587 | 0.103 | 6.57E-283 | 11 | Ncf4 |
| Celf23 | 5.08E-287 | 1.22789546 | 0.826 | 0.189 | 1.03E-282 | 11 | Celf2 |
| Ppp3cc1 | 1.71E-283 | 1.221444034 | 0.671 | 0.131 | 3.45E-279 | 11 | Ppp3cc |
| Fyb2 | 4.11E-276 | 1.051712088 | 0.799 | 0.172 | 8.31E-272 | 11 | Fyb |
| Fam107b1 | 2.53E-275 | 1.137120737 | 0.598 | 0.111 | 5.13E-271 | 11 | Fam107b |
| Cd691 | 4.45E-274 | 1.228631844 | 0.557 | 0.093 | 9.01E-270 | 11 | Cd69 |
| Jak13 | 5.08E-274 | 1.66836228 | 0.959 | 0.354 | 1.03E-269 | 11 | Jak1 |
| Gm163411 | 1.91E-273 | 0.750542726 | 0.316 | 0.031 | 3.87E-269 | 11 | Gm16341 |
| Lcp12 | 4.98E-271 | 1.432423028 | 0.86 | 0.226 | 1.01E-266 | 11 | Lcp1 |
| Itgb22 | 2.40E-270 | 0.988446379 | 0.542 | 0.093 | 4.85E-266 | 11 | Itgb2 |
| Laptm53 | 3.56E-270 | 1.03145577 | 0.865 | 0.198 | 7.21E-266 | 11 | Laptm5 |
| Cotl13 | 8.83E-270 | 1.393927996 | 0.826 | 0.212 | 1.79E-265 | 11 | Cotl1 |
| AW1120103 | 2.87E-269 | 1.938961321 | 0.998 | 0.56 | 5.81E-265 | 11 | AW112010 |
| Nup2101 | 8.20E-269 | 0.855430532 | 0.431 | 0.059 | 1.66E-264 | 11 | Nup210 |
| Rasgrp11 | 1.25E-268 | 0.849426196 | 0.417 | 0.055 | 2.54E-264 | 11 | Rasgrp1 |
| Tmem1631 | 9.38E-267 | 1.071673923 | 0.512 | 0.079 | 1.90E-262 | 11 | Tmem163 |
| Dock82 | 1.97E-263 | 1.340668404 | 0.82 | 0.206 | 3.98E-259 | 11 | Dock8 |
| Nr4a21 | 1.39E-262 | 1.384066476 | 0.614 | 0.121 | 2.80E-258 | 11 | Nr4a2 |
| Efhd22 | 6.87E-262 | 1.197432729 | 0.749 | 0.176 | 1.39E-257 | 11 | Efhd2 |
| Dock113 | 4.79E-261 | 1.037759244 | 0.571 | 0.105 | 9.70E-257 | 11 | Dock11 |
| Cd55 | 1.32E-260 | 0.593134322 | 0.266 | 0.022 | 2.67E-256 | 11 | Cd55 |
| Ptpn62 | 1.27E-259 | 1.148081205 | 0.614 | 0.121 | 2.57E-255 | 11 | Ptpn6 |
| Lsp13 | 1.85E-257 | 1.003973686 | 0.639 | 0.129 | 3.74E-253 | 11 | Lsp1 |
| Cblb2 | 3.99E-257 | 1.343827207 | 0.688 | 0.16 | 8.08E-253 | 11 | Cblb |
| Arhgap91 | 1.43E-255 | 0.874970614 | 0.566 | 0.102 | 2.89E-251 | 11 | Arhgap9 |
| Plek2 | 5.33E-255 | 1.167019145 | 0.596 | 0.115 | 1.08E-250 | 11 | Plek |
| Malt11 | 8.51E-255 | 1.2489182 | 0.646 | 0.14 | 1.72E-250 | 11 | Malt1 |
| Srgap31 | 3.53E-254 | 1.027936172 | 0.519 | 0.088 | 7.15E-250 | 11 | Srgap3 |
| Clec2i1 | 1.20E-252 | 0.639005475 | 0.294 | 0.029 | 2.43E-248 | 11 | Clec2i |

| | | | | | | | |
|---|---|---|---|---|---|---|---|
| Tiam21 | 2.72E-252 | 0.837317708 | 0.37 | 0.046 | 5.51E-248 | 11 | Tiam2 |
| Sema4d2 | 5.13E-252 | 1.026886064 | 0.614 | 0.121 | 1.04E-247 | 11 | Sema4d |
| Cd842 | 8.14E-252 | 0.950804878 | 0.499 | 0.082 | 1.65E-247 | 11 | Cd84 |
| Itgax2 | 2.21E-251 | 0.957870533 | 0.553 | 0.099 | 4.46E-247 | 11 | Itgax |
| Traf11 | 6.40E-250 | 0.929485989 | 0.406 | 0.056 | 1.29E-245 | 11 | Traf1 |
| Itk1 | 2.03E-248 | 0.894974303 | 0.461 | 0.07 | 4.11E-244 | 11 | Itk |
| Gm268271 | 7.67E-247 | 0.82508545 | 0.363 | 0.045 | 1.55E-242 | 11 | Gm26827 |
| Gpr132 | 7.52E-246 | 0.59908198 | 0.26 | 0.023 | 1.52E-241 | 11 | Gpr132 |
| Vps37b1 | 5.35E-244 | 1.077612048 | 0.618 | 0.127 | 1.08E-239 | 11 | Vps37b |
| Tnfrsf91 | 5.95E-244 | 1.010271668 | 0.418 | 0.06 | 1.21E-239 | 11 | Tnfrsf9 |
| B4galnt11 | 6.83E-244 | 0.899071437 | 0.504 | 0.087 | 1.38E-239 | 11 | B4galnt1 |
| Sh3bgrl33 | 8.36E-244 | 1.649763932 | 0.995 | 0.551 | 1.69E-239 | 11 | Sh3bgrl3 |
| Prex12 | 1.60E-243 | 1.239967269 | 0.695 | 0.163 | 3.25E-239 | 11 | Prex1 |
| Unc1191 | 9.04E-243 | 0.801663966 | 0.47 | 0.076 | 1.83E-238 | 11 | Unc119 |
| Cxcr3 | 1.09E-240 | 0.658223628 | 0.305 | 0.032 | 2.20E-236 | 11 | Cxcr3 |
| Gimap31 | 4.63E-239 | 0.864130189 | 0.53 | 0.091 | 9.37E-235 | 11 | Gimap3 |
| Mbnl12 | 6.83E-239 | 1.5944344 | 0.982 | 0.449 | 1.38E-234 | 11 | Mbnl1 |
| Lcp22 | 1.71E-238 | 0.937585106 | 0.496 | 0.087 | 3.46E-234 | 11 | Lcp2 |
| Runx22 | 4.24E-237 | 0.850936691 | 0.494 | 0.084 | 8.59E-233 | 11 | Runx2 |
| Rabgap1l1 | 8.16E-237 | 1.390359962 | 0.838 | 0.245 | 1.65E-232 | 11 | Rabgap1l |
| Rnf1571 | 3.49E-236 | 0.881145381 | 0.402 | 0.058 | 7.07E-232 | 11 | Rnf157 |
| Acap11 | 3.72E-236 | 0.769002049 | 0.425 | 0.064 | 7.53E-232 | 11 | Acap1 |
| Emb | 5.21E-236 | 0.818573323 | 0.321 | 0.036 | 1.05E-231 | 11 | Emb |
| H2-Q73 | 1.36E-234 | 1.567286747 | 0.957 | 0.387 | 2.76E-230 | 11 | H2-Q7 |
| Mgat52 | 5.57E-234 | 1.258585655 | 0.673 | 0.16 | 1.13E-229 | 11 | Mgat5 |
| Rps15a2 | 1.38E-233 | 1.375304403 | 0.996 | 0.923 | 2.78E-229 | 11 | Rps15a |
| Sulf22 | 9.63E-233 | 0.914896228 | 0.506 | 0.091 | 1.95E-228 | 11 | Sulf2 |
| Fnbp13 | 1.74E-232 | 1.199157883 | 0.801 | 0.219 | 3.53E-228 | 11 | Fnbp1 |
| Ankrd442 | 2.47E-232 | 1.411405321 | 0.876 | 0.283 | 5.00E-228 | 11 | Ankrd44 |
| Gimap71 | 9.68E-232 | 0.83983047 | 0.382 | 0.053 | 1.96E-227 | 11 | Gimap7 |
| Dusp5 | 1.18E-230 | 0.707379946 | 0.318 | 0.037 | 2.40E-226 | 11 | Dusp5 |
| Atp11b1 | 1.28E-230 | 1.192016786 | 0.686 | 0.171 | 2.60E-226 | 11 | Atp11b |
| Id21 | 6.56E-229 | 1.333896456 | 0.932 | 0.31 | 1.33E-224 | 11 | Id2 |
| Nlrc31 | 4.57E-228 | 0.769336887 | 0.373 | 0.052 | 9.26E-224 | 11 | Nlrc3 |
| Klra1 | 8.67E-227 | 1.445873669 | 0.262 | 0.025 | 1.75E-222 | 11 | Klra1 |
| Ptpn71 | 5.64E-226 | 0.732924288 | 0.373 | 0.052 | 1.14E-221 | 11 | Ptpn7 |
| Pfn13 | 7.50E-226 | 1.546472525 | 0.996 | 0.676 | 1.52E-221 | 11 | Pfn1 |
| Myo1f2 | 1.24E-225 | 0.990358206 | 0.61 | 0.128 | 2.50E-221 | 11 | Myo1f |
| Padi21 | 3.42E-224 | 0.91324841 | 0.492 | 0.088 | 6.92E-220 | 11 | Padi2 |
| Cd532 | 6.11E-224 | 0.930433214 | 0.546 | 0.108 | 1.24E-219 | 11 | Cd53 |
| Iqgap2 | 1.56E-222 | 0.820974778 | 0.524 | 0.097 | 3.16E-218 | 11 | Iqgap2 |
| Apbb1ip2 | 4.13E-222 | 1.001607663 | 0.693 | 0.163 | 8.36E-218 | 11 | Apbb1ip |
| Pik3r5 | 1.77E-221 | 0.613465191 | 0.262 | 0.026 | 3.58E-217 | 11 | Pik3r5 |

| | | | | | | | |
|---|---|---|---|---|---|---|---|
| Itgb15 | 6.29E-219 | 1.544590799 | 0.982 | 0.469 | 1.27E-214 | 11 | Itgb1 |
| Zap701 | 1.68E-218 | 0.734166522 | 0.364 | 0.051 | 3.40E-214 | 11 | Zap70 |
| Gpr1711 | 2.87E-218 | 0.686840816 | 0.33 | 0.042 | 5.81E-214 | 11 | Gpr171 |
| Btg11 | 2.49E-217 | 1.533394966 | 0.98 | 0.449 | 5.04E-213 | 11 | Btg1 |
| Fam105a2 | 2.02E-215 | 0.897000086 | 0.569 | 0.117 | 4.09E-211 | 11 | Fam105a |
| Lrrk13 | 5.04E-215 | 1.18254157 | 0.684 | 0.181 | 1.02E-210 | 11 | Lrrk1 |
| Rasal31 | 7.10E-214 | 0.903036742 | 0.445 | 0.077 | 1.44E-209 | 11 | Rasal3 |
| Nfatc12 | 1.97E-213 | 0.939006202 | 0.53 | 0.108 | 4.00E-209 | 11 | Nfatc1 |
| Sytl21 | 2.01E-213 | 0.703613446 | 0.359 | 0.05 | 4.07E-209 | 11 | Sytl2 |
| Neurl32 | 3.61E-212 | 0.910489539 | 0.531 | 0.107 | 7.32E-208 | 11 | Neurl3 |
| Spry22 | 7.28E-212 | 0.87374057 | 0.438 | 0.075 | 1.47E-207 | 11 | Spry2 |
| Cyth42 | 8.83E-211 | 0.934861036 | 0.58 | 0.125 | 1.79E-206 | 11 | Cyth4 |
| Aim22 | 1.21E-210 | 0.940016561 | 0.496 | 0.095 | 2.46E-206 | 11 | Aim2 |
| Sema4a1 | 2.75E-210 | 0.815556081 | 0.42 | 0.07 | 5.56E-206 | 11 | Sema4a |
| Prr5l | 9.08E-210 | 0.702703353 | 0.309 | 0.038 | 1.84E-205 | 11 | Prr5l |
| Rps112 | 6.01E-209 | 1.22781797 | 0.998 | 0.905 | 1.22E-204 | 11 | Rps11 |
| Traf3ip31 | 8.15E-209 | 0.760382924 | 0.427 | 0.072 | 1.65E-204 | 11 | Traf3ip3 |
| Ttc39b2 | 7.64E-208 | 1.008064666 | 0.567 | 0.127 | 1.55E-203 | 11 | Ttc39b |
| Fryl2 | 1.89E-207 | 1.450649925 | 0.797 | 0.258 | 3.83E-203 | 11 | Fryl |
| Chn21 | 8.52E-207 | 1.152999047 | 0.725 | 0.188 | 1.72E-202 | 11 | Chn2 |
| Il10ra1 | 1.01E-206 | 0.828972206 | 0.44 | 0.076 | 2.04E-202 | 11 | Il10ra |
| Cd482 | 1.43E-206 | 0.88863743 | 0.58 | 0.126 | 2.89E-202 | 11 | Cd48 |
| Pde7a1 | 1.56E-206 | 1.017350004 | 0.63 | 0.147 | 3.16E-202 | 11 | Pde7a |
| 1810006J02Rik1 | 9.73E-206 | 0.65634952 | 0.3 | 0.036 | 1.97E-201 | 11 | 1810006J02Rik |
| Rgs13 | 1.03E-204 | 0.969332336 | 0.61 | 0.132 | 2.08E-200 | 11 | Rgs1 |
| Cytip1 | 1.15E-203 | 0.792803414 | 0.463 | 0.085 | 2.33E-199 | 11 | Cytip |
| Lat22 | 1.62E-203 | 0.868074693 | 0.571 | 0.123 | 3.29E-199 | 11 | Lat2 |
| Pde3b2 | 1.09E-202 | 1.02844302 | 0.557 | 0.119 | 2.21E-198 | 11 | Pde3b |
| Fhl21 | 1.12E-202 | 0.634846749 | 0.3 | 0.037 | 2.27E-198 | 11 | Fhl2 |
| Gimap43 | 1.61E-202 | 1.148545221 | 0.808 | 0.246 | 3.25E-198 | 11 | Gimap4 |
| Slc38a22 | 3.98E-202 | 1.506797107 | 0.844 | 0.301 | 8.06E-198 | 11 | Slc38a2 |
| Oprm11 | 2.20E-201 | 0.782084144 | 0.363 | 0.054 | 4.44E-197 | 11 | Oprm1 |
| Ccnd31 | 2.42E-201 | 1.202232523 | 0.765 | 0.23 | 4.89E-197 | 11 | Ccnd3 |
| Scml41 | 2.63E-199 | 0.651492801 | 0.298 | 0.037 | 5.32E-195 | 11 | Scml4 |
| Ccl42 | 3.67E-199 | 1.863875368 | 0.558 | 0.121 | 7.44E-195 | 11 | Ccl4 |
| AB124611 | 4.74E-197 | 0.518326452 | 0.25 | 0.026 | 9.60E-193 | 11 | AB124611 |
| Il12rb21 | 2.42E-195 | 0.677949515 | 0.323 | 0.044 | 4.89E-191 | 11 | Il12rb2 |
| Ripor22 | 3.21E-195 | 1.141457304 | 0.639 | 0.163 | 6.50E-191 | 11 | Ripor2 |
| Gpsm32 | 3.85E-194 | 0.868603559 | 0.591 | 0.139 | 7.80E-190 | 11 | Gpsm3 |
| Jpt12 | 1.48E-193 | 0.927892154 | 0.741 | 0.212 | 2.99E-189 | 11 | Jpt1 |
| Pik3cd2 | 1.53E-193 | 0.910523085 | 0.497 | 0.101 | 3.10E-189 | 11 | Pik3cd |
| Stk42 | 2.32E-193 | 1.025182693 | 0.637 | 0.166 | 4.70E-189 | 11 | Stk4 |
| B4galt52 | 2.66E-193 | 0.904058809 | 0.452 | 0.087 | 5.38E-189 | 11 | B4galt5 |

| | | | | | | | |
|---|---|---|---|---|---|---|---|
| Rps182 | 2.64E-192 | 1.064018189 | 0.998 | 0.941 | 5.35E-188 | 11 | Rps18 |
| Rps33 | 1.49E-190 | 1.088477279 | 1 | 0.891 | 3.02E-186 | 11 | Rps3 |
| Rhof1 | 3.57E-190 | 0.54944973 | 0.275 | 0.033 | 7.22E-186 | 11 | Rhof |
| Fyn4 | 3.77E-190 | 1.188392025 | 0.79 | 0.256 | 7.63E-186 | 11 | Fyn |
| Sh3kbp11 | 4.83E-190 | 0.991044375 | 0.53 | 0.117 | 9.77E-186 | 11 | Sh3kbp1 |
| Rab371 | 9.85E-190 | 0.606616722 | 0.298 | 0.039 | 1.99E-185 | 11 | Rab37 |
| Trdc | 2.30E-189 | 0.483417229 | 0.413 | 0.069 | 4.66E-185 | 11 | Trdc |
| Rps243 | 7.14E-189 | 0.993500577 | 0.998 | 0.972 | 1.45E-184 | 11 | Rps24 |
| Gata31 | 8.98E-189 | 0.656952425 | 0.327 | 0.046 | 1.82E-184 | 11 | Gata3 |
| Rps162 | 1.21E-188 | 1.013262199 | 1 | 0.926 | 2.46E-184 | 11 | Rps16 |
| Rpl133 | 2.58E-188 | 0.98391145 | 0.998 | 0.977 | 5.23E-184 | 11 | Rpl13 |
| Cyba3 | 3.66E-188 | 1.212009147 | 0.962 | 0.431 | 7.40E-184 | 11 | Cyba |
| Rpsa3 | 8.76E-188 | 1.164404894 | 0.998 | 0.901 | 1.77E-183 | 11 | Rpsa |
| Jakmip11 | 1.04E-185 | 0.691705336 | 0.314 | 0.044 | 2.10E-181 | 11 | Jakmip1 |
| Jaml1 | 2.95E-185 | 0.762880326 | 0.346 | 0.053 | 5.98E-181 | 11 | Jaml |
| Vim4 | 2.69E-184 | 1.116998047 | 0.856 | 0.296 | 5.45E-180 | 11 | Vim |
| Kif13b2 | 6.11E-183 | 1.086229464 | 0.706 | 0.206 | 1.24E-178 | 11 | Kif13b |
| Lrmp2 | 9.81E-183 | 0.677812823 | 0.348 | 0.055 | 1.99E-178 | 11 | Lrmp |
| Ctla2a5 | 4.92E-181 | 1.258305826 | 0.964 | 0.387 | 9.97E-177 | 11 | Ctla2a |
| Rpl342 | 5.08E-181 | 0.999804513 | 0.993 | 0.916 | 1.03E-176 | 11 | Rpl34 |
| Nabp11 | 7.32E-181 | 0.811442246 | 0.413 | 0.075 | 1.48E-176 | 11 | Nabp1 |
| Ndfip11 | 1.18E-180 | 1.164891335 | 0.851 | 0.311 | 2.38E-176 | 11 | Ndfip1 |
| Smyd31 | 9.16E-180 | 1.248989515 | 0.724 | 0.217 | 1.85E-175 | 11 | Smyd3 |
| Rps103 | 1.80E-178 | 0.9735178 | 1 | 0.943 | 3.64E-174 | 11 | Rps10 |
| Rapgef62 | 1.65E-177 | 1.130821746 | 0.828 | 0.287 | 3.34E-173 | 11 | Rapgef6 |
| S100a101 | 3.61E-176 | 1.247843079 | 0.975 | 0.46 | 7.31E-172 | 11 | S100a10 |
| Slc9a7 | 3.67E-176 | 0.606505009 | 0.28 | 0.037 | 7.42E-172 | 11 | Slc9a7 |
| Rpl27a3 | 6.82E-175 | 0.913641693 | 1 | 0.959 | 1.38E-170 | 11 | Rpl27a |
| Evl2 | 2.89E-174 | 1.038396106 | 0.707 | 0.211 | 5.84E-170 | 11 | Evl |
| Cd821 | 2.90E-174 | 0.94945617 | 0.451 | 0.094 | 5.87E-170 | 11 | Cd82 |
| Ighm2 | 4.19E-174 | 0.732385343 | 0.524 | 0.119 | 8.48E-170 | 11 | Ighm |
| Arhgap302 | 4.84E-174 | 0.841495308 | 0.478 | 0.103 | 9.80E-170 | 11 | Arhgap30 |
| Peli12 | 8.90E-174 | 1.254039687 | 0.707 | 0.223 | 1.80E-169 | 11 | Peli1 |
| Slc14a11 | 4.65E-173 | 0.65322622 | 0.296 | 0.042 | 9.42E-169 | 11 | Slc14a1 |
| Pik3ap12 | 4.99E-173 | 0.772293076 | 0.431 | 0.084 | 1.01E-168 | 11 | Pik3ap1 |
| Rpl323 | 7.44E-173 | 0.991679328 | 1 | 0.953 | 1.51E-168 | 11 | Rpl32 |
| Btg23 | 1.47E-172 | 1.248413018 | 0.882 | 0.334 | 2.97E-168 | 11 | Btg2 |
| Ust3 | 1.54E-171 | 0.93356284 | 0.409 | 0.08 | 3.12E-167 | 11 | Ust |
| Sash31 | 1.61E-171 | 0.675002444 | 0.338 | 0.055 | 3.27E-167 | 11 | Sash3 |
| Acot71 | 5.60E-171 | 0.703997355 | 0.388 | 0.071 | 1.13E-166 | 11 | Acot7 |
| Otulin1 | 7.94E-171 | 0.915151562 | 0.661 | 0.191 | 1.61E-166 | 11 | Otulin |
| Dock51 | 4.06E-170 | 0.899256074 | 0.49 | 0.108 | 8.22E-166 | 11 | Dock5 |
| Runx12 | 1.09E-169 | 0.931487429 | 0.531 | 0.124 | 2.20E-165 | 11 | Runx1 |

| | | | | | | | |
|---|---|---|---|---|---|---|---|
| Rps273 | 3.12E-169 | 0.999111151 | 0.996 | 0.896 | 6.31E-165 | 11 | Rps27 |
| Esyt11 | 1.03E-168 | 0.866310428 | 0.528 | 0.129 | 2.08E-164 | 11 | Esyt1 |
| Klra31 | 1.58E-168 | 1.174250126 | 0.293 | 0.042 | 3.21E-164 | 11 | Klra3 |
| Ctsd3 | 5.99E-168 | 1.025784193 | 0.837 | 0.304 | 1.21E-163 | 11 | Ctsd |
| Xist5 | 8.64E-168 | 1.185569908 | 0.964 | 0.454 | 1.75E-163 | 11 | Xist |
| Ssbp22 | 1.36E-167 | 0.928868738 | 0.548 | 0.132 | 2.75E-163 | 11 | Ssbp2 |
| Gprin31 | 4.87E-166 | 0.730748469 | 0.345 | 0.057 | 9.85E-162 | 11 | Gprin3 |
| Rpl18a2 | 1.63E-165 | 0.921502144 | 1 | 0.93 | 3.30E-161 | 11 | Rpl18a |
| Rpl383 | 8.47E-164 | 0.990316889 | 0.996 | 0.892 | 1.72E-159 | 11 | Rpl38 |
| Leprotl11 | 5.21E-163 | 0.908086817 | 0.684 | 0.208 | 1.05E-158 | 11 | Leprotl1 |
| Pim2 | 7.83E-163 | 0.522328265 | 0.25 | 0.031 | 1.58E-158 | 11 | Pim2 |
| Akna2 | 1.17E-162 | 0.719484147 | 0.418 | 0.085 | 2.37E-158 | 11 | Akna |
| Gimap91 | 2.61E-162 | 0.829272836 | 0.582 | 0.151 | 5.29E-158 | 11 | Gimap9 |
| Bcl6 | 5.81E-162 | 0.625301194 | 0.318 | 0.051 | 1.18E-157 | 11 | Bcl6 |
| Pstpip11 | 2.26E-161 | 0.59512603 | 0.318 | 0.051 | 4.58E-157 | 11 | Pstpip1 |
| Rpl37a3 | 5.99E-161 | 0.914605158 | 1 | 0.937 | 1.21E-156 | 11 | Rpl37a |
| Pycard3 | 6.83E-161 | 0.907706586 | 0.799 | 0.267 | 1.38E-156 | 11 | Pycard |
| Ezr1 | 7.34E-161 | 0.720791178 | 0.53 | 0.129 | 1.49E-156 | 11 | Ezr |
| Rin32 | 8.78E-160 | 0.899375485 | 0.61 | 0.171 | 1.78E-155 | 11 | Rin3 |
| Fxyd54 | 1.28E-159 | 1.134655011 | 0.916 | 0.402 | 2.58E-155 | 11 | Fxyd5 |
| Rpl183 | 1.99E-159 | 0.933433649 | 0.998 | 0.899 | 4.02E-155 | 11 | Rpl18 |
| Rps73 | 2.57E-159 | 0.953229259 | 0.998 | 0.915 | 5.19E-155 | 11 | Rps7 |
| Dennd1c1 | 1.24E-158 | 0.59372944 | 0.32 | 0.053 | 2.52E-154 | 11 | Dennd1c |
| Rps53 | 1.22E-156 | 0.911992516 | 1 | 0.919 | 2.47E-152 | 11 | Rps5 |
| Zfp5121 | 1.27E-156 | 0.773875036 | 0.402 | 0.081 | 2.58E-152 | 11 | Zfp512 |
| Herc31 | 2.28E-155 | 0.735648541 | 0.359 | 0.066 | 4.62E-151 | 11 | Herc3 |
| Irf81 | 3.71E-155 | 0.837814616 | 0.461 | 0.103 | 7.52E-151 | 11 | Irf8 |
| Rps3a14 | 1.69E-154 | 0.909608119 | 0.998 | 0.934 | 3.42E-150 | 11 | Rps3a1 |
| Sh2d1b12 | 2.53E-154 | 0.641288756 | 0.375 | 0.072 | 5.12E-150 | 11 | Sh2d1b1 |
| Rbl21 | 2.66E-154 | 0.793616187 | 0.513 | 0.128 | 5.39E-150 | 11 | Rbl2 |
| Ubac22 | 5.27E-154 | 0.895518168 | 0.618 | 0.18 | 1.07E-149 | 11 | Ubac2 |
| Psd41 | 6.08E-154 | 0.613012151 | 0.339 | 0.06 | 1.23E-149 | 11 | Psd4 |
| Mllt31 | 9.33E-153 | 0.926967203 | 0.549 | 0.144 | 1.89E-148 | 11 | Mllt3 |
| Ifitm101 | 3.65E-152 | 0.60093953 | 0.325 | 0.055 | 7.38E-148 | 11 | Ifitm10 |
| Samsn12 | 6.35E-152 | 0.695077793 | 0.381 | 0.075 | 1.29E-147 | 11 | Samsn1 |
| Trbc21 | 1.23E-151 | 0.951384465 | 0.345 | 0.061 | 2.48E-147 | 11 | Trbc2 |
| Nfkb14 | 1.84E-151 | 1.138574916 | 0.731 | 0.263 | 3.72E-147 | 11 | Nfkb1 |
| Rps133 | 1.23E-150 | 0.867316156 | 0.998 | 0.922 | 2.49E-146 | 11 | Rps13 |
| Tnfaip32 | 2.05E-150 | 0.797392955 | 0.404 | 0.084 | 4.15E-146 | 11 | Tnfaip3 |
| Atp10a | 2.75E-150 | 0.659024719 | 0.282 | 0.043 | 5.56E-146 | 11 | Atp10a |
| Cnn22 | 4.45E-150 | 0.851507171 | 0.605 | 0.174 | 9.00E-146 | 11 | Cnn2 |
| Inpp4b1 | 5.50E-150 | 1.022576097 | 0.664 | 0.194 | 1.11E-145 | 11 | Inpp4b |
| Gfod11 | 2.14E-149 | 0.625141067 | 0.348 | 0.064 | 4.33E-145 | 11 | Gfod1 |

| | | | | | | | |
|---|---|---|---|---|---|---|---|
| Inpp5d2 | 7.86E-149 | 0.878048479 | 0.652 | 0.195 | 1.59E-144 | 11 | Inpp5d |
| AC149090.12 | 7.93E-149 | 1.055088492 | 0.689 | 0.223 | 1.60E-144 | 11 | AC149090.1 |
| Mob3a1 | 9.38E-149 | 0.650279447 | 0.377 | 0.075 | 1.90E-144 | 11 | Mob3a |
| Rps202 | 1.23E-148 | 0.880038917 | 1 | 0.95 | 2.48E-144 | 11 | Rps20 |
| Ccl32 | 4.14E-148 | 1.460444995 | 0.381 | 0.077 | 8.38E-144 | 11 | Ccl3 |
| Adcy73 | 5.23E-148 | 0.794323053 | 0.458 | 0.108 | 1.06E-143 | 11 | Adcy7 |
| Tnf1 | 1.24E-146 | 0.82064262 | 0.355 | 0.067 | 2.51E-142 | 11 | Tnf |
| Gramd1b1 | 4.90E-146 | 0.787146866 | 0.397 | 0.084 | 9.92E-142 | 11 | Gramd1b |
| Cd961 | 9.95E-146 | 0.698230374 | 0.345 | 0.064 | 2.02E-141 | 11 | Cd96 |
| Magohb1 | 1.31E-145 | 0.697740125 | 0.476 | 0.115 | 2.66E-141 | 11 | Magohb |
| Rpl393 | 2.17E-145 | 0.915857496 | 0.993 | 0.899 | 4.39E-141 | 11 | Rpl39 |
| Cobll11 | 3.05E-145 | 0.688973499 | 0.391 | 0.08 | 6.17E-141 | 11 | Cobll1 |
| Zcchc112 | 5.04E-145 | 1.003603904 | 0.731 | 0.259 | 1.02E-140 | 11 | Zcchc11 |
| Gm203884 | 8.59E-145 | 1.043116745 | 0.908 | 0.391 | 1.74E-140 | 11 | Gm20388 |
| Fermt32 | 1.56E-144 | 0.707417312 | 0.456 | 0.108 | 3.16E-140 | 11 | Fermt3 |
| Rpl303 | 7.81E-144 | 0.810726635 | 0.996 | 0.937 | 1.58E-139 | 11 | Rpl30 |
| Bcl2l111 | 1.51E-143 | 0.978992133 | 0.454 | 0.111 | 3.06E-139 | 11 | Bcl2l11 |
| Bach22 | 2.25E-143 | 1.018610001 | 0.548 | 0.154 | 4.56E-139 | 11 | Bach2 |
| Rpl363 | 5.16E-143 | 0.866205067 | 0.995 | 0.902 | 1.05E-138 | 11 | Rpl36 |
| Dennd1b3 | 3.49E-142 | 0.884223539 | 0.592 | 0.177 | 7.07E-138 | 11 | Dennd1b |
| Gsap2 | 6.36E-141 | 0.687008685 | 0.382 | 0.08 | 1.29E-136 | 11 | Gsap |
| Grap22 | 1.48E-140 | 0.771503353 | 0.434 | 0.1 | 3.00E-136 | 11 | Grap2 |
| Fau2 | 9.10E-140 | 0.778721513 | 1 | 0.952 | 1.84E-135 | 11 | Fau |
| Tmem131l2 | 9.28E-140 | 0.672591709 | 0.368 | 0.075 | 1.88E-135 | 11 | Tmem131l |
| Sla21 | 4.03E-139 | 0.531221273 | 0.278 | 0.045 | 8.16E-135 | 11 | Sla2 |
| Crlf32 | 1.56E-138 | 0.771411724 | 0.54 | 0.15 | 3.15E-134 | 11 | Crlf3 |
| Sept113 | 1.57E-138 | 0.83723007 | 0.636 | 0.195 | 3.17E-134 | 11 | 11-Sep |
| Anxa61 | 5.33E-138 | 0.732711666 | 0.54 | 0.15 | 1.08E-133 | 11 | Anxa6 |
| Rpl193 | 7.27E-138 | 0.743686363 | 1 | 0.963 | 1.47E-133 | 11 | Rpl19 |
| Rpl83 | 1.34E-137 | 0.802376746 | 1 | 0.924 | 2.72E-133 | 11 | Rpl8 |
| H2-Q64 | 1.62E-137 | 0.866226158 | 0.706 | 0.237 | 3.29E-133 | 11 | H2-Q6 |
| Rpl234 | 4.23E-137 | 0.7635716 | 1 | 0.97 | 8.56E-133 | 11 | Rpl23 |
| Tle43 | 3.85E-136 | 0.886249578 | 0.657 | 0.217 | 7.80E-132 | 11 | Tle4 |
| Rpl123 | 2.86E-135 | 0.902795683 | 0.993 | 0.839 | 5.79E-131 | 11 | Rpl12 |
| Slc9a3r11 | 5.21E-134 | 0.604126632 | 0.436 | 0.104 | 1.05E-129 | 11 | Slc9a3r1 |
| H2-K15 | 8.19E-134 | 1.013942361 | 1 | 0.637 | 1.66E-129 | 11 | H2-K1 |
| Sub12 | 5.80E-133 | 0.927032742 | 0.953 | 0.508 | 1.17E-128 | 11 | Sub1 |
| Cd44 | 9.07E-133 | 0.703811885 | 0.427 | 0.102 | 1.84E-128 | 11 | Cd44 |
| Plcl23 | 2.06E-132 | 0.780826628 | 0.44 | 0.108 | 4.18E-128 | 11 | Plcl2 |
| Rpl94 | 2.78E-132 | 0.751371954 | 0.998 | 0.944 | 5.62E-128 | 11 | Rpl9 |
| Stk261 | 6.11E-132 | 0.47284165 | 0.26 | 0.041 | 1.24E-127 | 11 | Stk26 |
| Actr33 | 1.43E-131 | 0.820367099 | 0.905 | 0.381 | 2.90E-127 | 11 | Actr3 |
| Gimap52 | 1.89E-131 | 0.810853421 | 0.564 | 0.164 | 3.83E-127 | 11 | Gimap5 |

| | | | | | | | |
|---|---|---|---|---|---|---|---|
| Dapk22 | 2.49E-131 | 0.6956669 | 0.438 | 0.104 | 5.04E-127 | 11 | Dapk2 |
| Btbd111 | 3.86E-131 | 0.643414456 | 0.291 | 0.052 | 7.82E-127 | 11 | Btbd11 |
| Fgf131 | 8.36E-131 | 0.917065683 | 0.357 | 0.074 | 1.69E-126 | 11 | Fgf13 |
| Rab8b2 | 1.34E-130 | 0.952482612 | 0.743 | 0.276 | 2.72E-126 | 11 | Rab8b |
| Actb3 | 1.69E-130 | 0.960074954 | 1 | 0.84 | 3.42E-126 | 11 | Actb |
| 1700025G04Rik3 | 2.70E-130 | 0.870389249 | 0.582 | 0.181 | 5.46E-126 | 11 | 1700025G04Rik |
| Fam189b1 | 3.55E-130 | 0.567432707 | 0.294 | 0.053 | 7.18E-126 | 11 | Fam189b |
| Ldlrad42 | 7.87E-129 | 1.082833795 | 0.688 | 0.243 | 1.59E-124 | 11 | Ldlrad4 |
| Elf42 | 8.54E-129 | 0.76073223 | 0.452 | 0.116 | 1.73E-124 | 11 | Elf4 |
| Sms1 | 1.93E-128 | 0.649866621 | 0.519 | 0.144 | 3.91E-124 | 11 | Sms |
| Sla3 | 4.22E-128 | 0.671794215 | 0.336 | 0.069 | 8.55E-124 | 11 | Sla |
| Zeb22 | 6.58E-128 | 0.500212135 | 0.526 | 0.136 | 1.33E-123 | 11 | Zeb2 |
| Mtmr121 | 1.48E-127 | 0.754821546 | 0.469 | 0.124 | 2.99E-123 | 11 | Mtmr12 |
| Diaph11 | 2.75E-127 | 0.821858533 | 0.677 | 0.234 | 5.57E-123 | 11 | Diaph1 |
| Rassf51 | 1.17E-126 | 0.534101597 | 0.323 | 0.064 | 2.37E-122 | 11 | Rassf5 |
| Rpl314 | 2.64E-126 | 0.790607097 | 0.998 | 0.908 | 5.34E-122 | 11 | Rpl3 |
| Pglyrp21 | 2.82E-126 | 0.528852234 | 0.251 | 0.04 | 5.71E-122 | 11 | Pglyrp2 |
| Gng22 | 3.75E-126 | 0.772605102 | 0.591 | 0.179 | 7.59E-122 | 11 | Gng2 |
| A930015D03Rik1 | 2.50E-125 | 0.932741483 | 0.411 | 0.101 | 5.05E-121 | 11 | A930015D03Rik |
| Znrf21 | 3.15E-125 | 0.74816908 | 0.524 | 0.152 | 6.38E-121 | 11 | Znrf2 |
| Prkca2 | 6.23E-125 | 0.735746858 | 0.4 | 0.095 | 1.26E-120 | 11 | Prkca |
| Mxd11 | 2.24E-124 | 0.612056336 | 0.307 | 0.059 | 4.53E-120 | 11 | Mxd1 |
| Ern11 | 1.41E-123 | 0.756418084 | 0.359 | 0.08 | 2.85E-119 | 11 | Ern1 |
| Lta4h1 | 3.71E-123 | 0.733407705 | 0.503 | 0.142 | 7.51E-119 | 11 | Lta4h |
| Sorl12 | 4.45E-123 | 0.670761741 | 0.427 | 0.106 | 9.01E-119 | 11 | Sorl1 |
| Vps13a1 | 9.23E-123 | 0.773378297 | 0.544 | 0.166 | 1.87E-118 | 11 | Vps13a |
| Wipf13 | 1.17E-122 | 0.925541415 | 0.707 | 0.26 | 2.36E-118 | 11 | Wipf1 |
| Emid11 | 1.49E-122 | 0.634247064 | 0.422 | 0.103 | 3.02E-118 | 11 | Emid1 |
| Atp1b11 | 2.53E-122 | 0.271449123 | 0.555 | 0.154 | 5.13E-118 | 11 | Atp1b1 |
| Rpl223 | 7.67E-122 | 0.756416724 | 0.993 | 0.85 | 1.55E-117 | 11 | Rpl22 |
| Hgsnat1 | 1.48E-121 | 1.09029091 | 0.352 | 0.079 | 3.00E-117 | 11 | Hgsnat |
| Nfatc31 | 5.14E-121 | 0.797871484 | 0.546 | 0.168 | 1.04E-116 | 11 | Nfatc3 |
| Was2 | 6.47E-121 | 0.526341979 | 0.329 | 0.068 | 1.31E-116 | 11 | Was |
| Syk2 | 7.73E-121 | 0.583858655 | 0.314 | 0.063 | 1.56E-116 | 11 | Syk |
| Zfp36l22 | 1.84E-120 | 0.96366398 | 0.745 | 0.297 | 3.72E-116 | 11 | Zfp36l2 |
| Rpl114 | 2.19E-120 | 0.733903504 | 0.998 | 0.926 | 4.44E-116 | 11 | Rpl11 |
| Gimap15 | 4.86E-120 | 0.865633647 | 0.772 | 0.294 | 9.84E-116 | 11 | Gimap1 |
| Apol7e1 | 4.87E-120 | 0.511077406 | 0.268 | 0.047 | 9.85E-116 | 11 | Apol7e |
| Rpl374 | 1.14E-119 | 0.743463934 | 0.998 | 0.926 | 2.30E-115 | 11 | Rpl37 |
| Baz1a2 | 2.15E-119 | 0.802507364 | 0.539 | 0.166 | 4.36E-115 | 11 | Baz1a |
| Capzb3 | 2.13E-118 | 0.836037532 | 0.826 | 0.36 | 4.32E-114 | 11 | Capzb |
| Pim11 | 2.50E-118 | 0.832246749 | 0.461 | 0.127 | 5.06E-114 | 11 | Pim1 |
| Tes1 | 6.47E-118 | 0.591896342 | 0.461 | 0.124 | 1.31E-113 | 11 | Tes |

| | | | | | | | |
|---|---|---|---|---|---|---|---|
| Il21r2 | 2.50E-117 | 0.690945332 | 0.375 | 0.089 | 5.07E-113 | 11 | Il21r |
| Cbl2 | 2.45E-116 | 0.782362533 | 0.591 | 0.2 | 4.96E-112 | 11 | Cbl |
| Ppm1b1 | 3.75E-116 | 0.666464669 | 0.472 | 0.132 | 7.60E-112 | 11 | Ppm1b |
| Nsd33 | 4.79E-116 | 0.846688507 | 0.749 | 0.3 | 9.70E-112 | 11 | Nsd3 |
| Npm31 | 1.50E-115 | 0.545990923 | 0.467 | 0.127 | 3.04E-111 | 11 | Npm3 |
| Rps234 | 1.65E-115 | 0.645827863 | 0.998 | 0.952 | 3.33E-111 | 11 | Rps23 |
| Tex142 | 2.49E-115 | 0.898478091 | 0.488 | 0.143 | 5.04E-111 | 11 | Tex14 |
| Gmfg1 | 3.04E-115 | 0.609743848 | 0.481 | 0.133 | 6.15E-111 | 11 | Gmfg |
| Me21 | 3.77E-115 | 0.544898928 | 0.329 | 0.07 | 7.62E-111 | 11 | Me2 |
| Kdm2b2 | 1.15E-114 | 0.754757973 | 0.359 | 0.084 | 2.33E-110 | 11 | Kdm2b |
| Tec2 | 2.32E-114 | 0.656569269 | 0.411 | 0.105 | 4.70E-110 | 11 | Tec |
| Napsa1 | 3.17E-114 | 0.477162317 | 0.266 | 0.048 | 6.43E-110 | 11 | Napsa |
| Nlrc52 | 3.69E-114 | 0.680753562 | 0.413 | 0.106 | 7.47E-110 | 11 | Nlrc5 |
| mt-Co16 | 1.67E-113 | 0.838803877 | 1 | 0.715 | 3.37E-109 | 11 | mt-Co1 |
| Sesn32 | 2.37E-113 | 0.591947043 | 0.452 | 0.123 | 4.80E-109 | 11 | Sesn3 |
| Sec11c3 | 5.87E-113 | 0.685833372 | 0.768 | 0.302 | 1.19E-108 | 11 | Sec11c |
| Junb4 | 1.73E-112 | 0.773676883 | 0.898 | 0.383 | 3.50E-108 | 11 | Junb |
| Itgb71 | 4.05E-112 | 0.543081672 | 0.293 | 0.058 | 8.20E-108 | 11 | Itgb7 |
| Elovl61 | 4.96E-112 | 0.493002277 | 0.259 | 0.046 | 1.00E-107 | 11 | Elovl6 |
| Suco1 | 2.22E-111 | 0.732267746 | 0.496 | 0.147 | 4.50E-107 | 11 | Suco |
| Ppm1h2 | 2.97E-110 | 0.747857315 | 0.51 | 0.155 | 6.02E-106 | 11 | Ppm1h |
| Gimap64 | 8.58E-110 | 0.89013946 | 0.844 | 0.357 | 1.74E-105 | 11 | Gimap6 |
| Ifi2091 | 9.66E-110 | 0.578690363 | 0.275 | 0.053 | 1.96E-105 | 11 | Ifi209 |
| Rpl13a3 | 9.06E-109 | 0.65497113 | 0.716 | 0.277 | 1.83E-104 | 11 | Rpl13a |
| Rps27a3 | 1.03E-108 | 0.650450497 | 0.998 | 0.949 | 2.09E-104 | 11 | Rps27a |
| Pkp31 | 2.79E-108 | 0.554932447 | 0.307 | 0.065 | 5.65E-104 | 11 | Pkp3 |
| Clcn3 | 5.24E-108 | 0.696372147 | 0.451 | 0.128 | 1.06E-103 | 11 | Clcn3 |
| Tra2b2 | 7.80E-108 | 0.765823433 | 0.725 | 0.289 | 1.58E-103 | 11 | Tra2b |
| Rpl10a4 | 1.45E-106 | 0.742500313 | 0.995 | 0.833 | 2.94E-102 | 11 | Rpl10a |
| Eef1a13 | 2.17E-106 | 0.605710066 | 1 | 0.959 | 4.38E-102 | 11 | Eef1a1 |
| Ttc71 | 3.19E-106 | 0.485796585 | 0.386 | 0.098 | 6.45E-102 | 11 | Ttc7 |
| Pla2g161 | 7.07E-106 | 0.571420109 | 0.654 | 0.236 | 1.43E-101 | 11 | Pla2g16 |
| Chd24 | 1.05E-105 | 0.843788192 | 0.652 | 0.247 | 2.13E-101 | 11 | Chd2 |
| Rps93 | 2.18E-105 | 0.669404738 | 0.998 | 0.919 | 4.41E-101 | 11 | Rps9 |
| Rps283 | 2.72E-105 | 0.730358226 | 0.995 | 0.896 | 5.50E-101 | 11 | Rps28 |
| Zyx2 | 1.03E-104 | 0.730555404 | 0.6 | 0.209 | 2.08E-100 | 11 | Zyx |
| Anxa24 | 1.11E-104 | 0.83115487 | 0.876 | 0.408 | 2.25E-100 | 11 | Anxa2 |
| Pik3cg2 | 3.91E-104 | 0.537992435 | 0.26 | 0.051 | 7.92E-100 | 11 | Pik3cg |
| Sec11a2 | 8.20E-104 | 0.687128059 | 0.702 | 0.277 | 1.66E-99 | 11 | Sec11a |
| Fmnl12 | 2.28E-103 | 0.5575477 | 0.355 | 0.088 | 4.61E-99 | 11 | Fmnl1 |
| Hopx1 | 4.64E-103 | 0.537857467 | 0.318 | 0.071 | 9.39E-99 | 11 | Hopx |
| Chst113 | 5.94E-103 | 0.743353162 | 0.521 | 0.17 | 1.20E-98 | 11 | Chst11 |
| Hivep32 | 7.17E-103 | 0.72829275 | 0.496 | 0.156 | 1.45E-98 | 11 | Hivep3 |

| | | | | | | | |
|---|---|---|---|---|---|---|---|
| Il16 | 1.11E-102 | 0.452398856 | 0.251 | 0.048 | 2.25E-98 | 11 | Il16 |
| Pvt12 | 1.20E-102 | 0.627883452 | 0.323 | 0.074 | 2.43E-98 | 11 | Pvt1 |
| Bbs92 | 1.04E-101 | 0.698242673 | 0.375 | 0.097 | 2.10E-97 | 11 | Bbs9 |
| Arap23 | 2.42E-101 | 0.710290461 | 0.531 | 0.172 | 4.89E-97 | 11 | Arap2 |
| Rpl22l13 | 3.93E-101 | 0.770245521 | 0.971 | 0.571 | 7.96E-97 | 11 | Rpl22l1 |
| Ptprj2 | 6.07E-101 | 0.592571655 | 0.488 | 0.145 | 1.23E-96 | 11 | Ptprj |
| Rasa13 | 6.22E-101 | 0.798569994 | 0.636 | 0.24 | 1.26E-96 | 11 | Rasa1 |
| Rpl273 | 7.34E-101 | 0.772399915 | 0.953 | 0.576 | 1.49E-96 | 11 | Rpl27 |
| Nsmaf2 | 8.68E-101 | 0.561718328 | 0.364 | 0.092 | 1.76E-96 | 11 | Nsmaf |
| Rps214 | 1.13E-100 | 0.624377805 | 0.995 | 0.926 | 2.28E-96 | 11 | Rps21 |
| Chka2 | 1.21E-100 | 0.768590106 | 0.456 | 0.139 | 2.44E-96 | 11 | Chka |
| Akap134 | 3.79E-100 | 0.79050034 | 0.84 | 0.386 | 7.68E-96 | 11 | Akap13 |
| Slamf72 | 4.30E-100 | 0.593210658 | 0.37 | 0.094 | 8.70E-96 | 11 | Slamf7 |
| Mgat4a2 | 8.40E-100 | 0.597888088 | 0.379 | 0.101 | 1.70E-95 | 11 | Mgat4a |
| Mkl13 | 1.81E-99 | 0.640810913 | 0.42 | 0.121 | 3.67E-95 | 11 | Mkl1 |
| Mapk61 | 2.94E-99 | 0.636447078 | 0.372 | 0.098 | 5.95E-95 | 11 | Mapk6 |
| Rplp24 | 4.39E-99 | 0.639929613 | 0.996 | 0.888 | 8.90E-95 | 11 | Rplp2 |
| Elmo14 | 8.60E-99 | 0.823884273 | 0.808 | 0.358 | 1.74E-94 | 11 | Elmo1 |
| Apobec31 | 1.23E-98 | 0.710177544 | 0.603 | 0.219 | 2.48E-94 | 11 | Apobec3 |
| Ankrd115 | 1.28E-98 | 0.849155076 | 0.824 | 0.373 | 2.60E-94 | 11 | Ankrd11 |
| Ppp1r182 | 1.30E-98 | 0.631608666 | 0.474 | 0.148 | 2.63E-94 | 11 | Ppp1r18 |
| Smc41 | 1.39E-98 | 0.725334899 | 0.492 | 0.157 | 2.81E-94 | 11 | Smc4 |
| Ube2h1 | 2.19E-98 | 0.758121429 | 0.661 | 0.258 | 4.44E-94 | 11 | Ube2h |
| Rack14 | 2.55E-98 | 0.712556991 | 0.995 | 0.789 | 5.16E-94 | 11 | Rack1 |
| Chd72 | 2.63E-98 | 0.649345296 | 0.47 | 0.146 | 5.33E-94 | 11 | Chd7 |
| Rpl153 | 1.08E-97 | 0.652801045 | 0.996 | 0.9 | 2.18E-93 | 11 | Rpl15 |
| Itpk11 | 1.86E-97 | 0.482424361 | 0.298 | 0.067 | 3.76E-93 | 11 | Itpk1 |
| Capza12 | 2.01E-97 | 0.689162923 | 0.673 | 0.268 | 4.06E-93 | 11 | Capza1 |
| Lpp4 | 3.29E-97 | 0.868704006 | 0.679 | 0.279 | 6.66E-93 | 11 | Lpp |
| Rps193 | 3.56E-97 | 0.630615884 | 0.996 | 0.926 | 7.21E-93 | 11 | Rps19 |
| Ssh23 | 2.16E-96 | 0.789303106 | 0.598 | 0.219 | 4.37E-92 | 11 | Ssh2 |
| Cbx41 | 1.40E-95 | 0.567312222 | 0.386 | 0.106 | 2.83E-91 | 11 | Cbx4 |
| Rhoh2 | 1.95E-95 | 0.607671532 | 0.372 | 0.099 | 3.94E-91 | 11 | Rhoh |
| Rps143 | 4.31E-95 | 0.56423163 | 0.996 | 0.942 | 8.73E-91 | 11 | Rps14 |
| Rpl35a3 | 5.28E-95 | 0.567459692 | 0.998 | 0.948 | 1.07E-90 | 11 | Rpl35a |
| Pip5k1b1 | 2.55E-94 | 0.587059664 | 0.361 | 0.095 | 5.17E-90 | 11 | Pip5k1b |
| Rps4x3 | 2.85E-94 | 0.608725034 | 0.998 | 0.94 | 5.76E-90 | 11 | Rps4x |
| Rcsd11 | 3.13E-94 | 0.649952039 | 0.422 | 0.124 | 6.35E-90 | 11 | Rcsd1 |
| Prkch4 | 2.34E-93 | 0.786294468 | 0.858 | 0.391 | 4.74E-89 | 11 | Prkch |
| Rps124 | 2.45E-93 | 0.624033908 | 0.991 | 0.897 | 4.95E-89 | 11 | Rps12 |
| Agpat32 | 3.22E-93 | 0.613952454 | 0.415 | 0.124 | 6.53E-89 | 11 | Agpat3 |
| Gramd1a1 | 3.39E-93 | 0.554149345 | 0.361 | 0.096 | 6.86E-89 | 11 | Gramd1a |
| Rbm31 | 3.84E-93 | 0.72914307 | 0.887 | 0.47 | 7.78E-89 | 11 | Rbm3 |

| | | | | | | | |
|---|---|---|---|---|---|---|---|
| Prkacb1 | 4.81E-93 | 0.670178334 | 0.585 | 0.212 | 9.73E-89 | 11 | Prkacb |
| Rras21 | 5.59E-93 | 0.562273251 | 0.325 | 0.08 | 1.13E-88 | 11 | Rras2 |
| Ggct1 | 2.27E-92 | 0.464214083 | 0.276 | 0.061 | 4.60E-88 | 11 | Ggct |
| Rpl173 | 4.95E-92 | 0.58910019 | 0.995 | 0.915 | 1.00E-87 | 11 | Rpl17 |
| Dnajc16 | 9.01E-92 | 0.786944394 | 0.729 | 0.316 | 1.83E-87 | 11 | Dnajc1 |
| Plekha22 | 9.44E-92 | 0.651751927 | 0.424 | 0.129 | 1.91E-87 | 11 | Plekha2 |
| Eef1b24 | 1.48E-91 | 0.678730937 | 0.984 | 0.732 | 2.99E-87 | 11 | Eef1b2 |
| Slc7a1 | 2.15E-91 | 0.553002907 | 0.266 | 0.058 | 4.36E-87 | 11 | Slc7a1 |
| Sp1101 | 5.57E-91 | 0.603235162 | 0.467 | 0.152 | 1.13E-86 | 11 | Sp110 |
| Csk2 | 6.12E-91 | 0.538425116 | 0.417 | 0.124 | 1.24E-86 | 11 | Csk |
| Cracr2a1 | 8.39E-91 | 0.393591356 | 0.338 | 0.085 | 1.70E-86 | 11 | Cracr2a |
| Camk2g1 | 8.52E-91 | 0.679612463 | 0.424 | 0.13 | 1.73E-86 | 11 | Camk2g |
| Hcls12 | 4.32E-90 | 0.504719322 | 0.411 | 0.12 | 8.74E-86 | 11 | Hcls1 |
| Tab21 | 6.79E-90 | 0.700014782 | 0.589 | 0.225 | 1.37E-85 | 11 | Tab2 |
| Eif3f1 | 8.23E-90 | 0.707656724 | 0.894 | 0.476 | 1.67E-85 | 11 | Eif3f |
| Rgs22 | 9.53E-90 | 0.551612675 | 0.425 | 0.125 | 1.93E-85 | 11 | Rgs2 |
| mt-Nd54 | 2.37E-89 | 0.77152419 | 0.919 | 0.488 | 4.79E-85 | 11 | mt-Nd5 |
| Rpl64 | 6.62E-89 | 0.612946529 | 1 | 0.879 | 1.34E-84 | 11 | Rpl6 |
| Cdk2ap21 | 1.67E-88 | 0.491816054 | 0.548 | 0.193 | 3.39E-84 | 11 | Cdk2ap2 |
| Hmgb21 | 1.69E-88 | 0.483651762 | 0.752 | 0.321 | 3.42E-84 | 11 | Hmgb2 |
| Fam172a4 | 1.70E-88 | 0.805115566 | 0.666 | 0.277 | 3.44E-84 | 11 | Fam172a |
| Trmt1121 | 4.21E-88 | 0.571178519 | 0.659 | 0.267 | 8.52E-84 | 11 | Trmt112 |
| Eya21 | 4.97E-88 | 0.499599603 | 0.357 | 0.096 | 1.01E-83 | 11 | Eya2 |
| Eml42 | 8.54E-88 | 0.787345223 | 0.623 | 0.253 | 1.73E-83 | 11 | Eml4 |
| Vasp1 | 2.69E-87 | 0.64087518 | 0.594 | 0.231 | 5.45E-83 | 11 | Vasp |
| Gm44686 | 6.07E-87 | 0.499561017 | 0.26 | 0.057 | 1.23E-82 | 11 | Gm44686 |
| S100a62 | 6.76E-87 | 0.744197314 | 0.679 | 0.264 | 1.37E-82 | 11 | S100a6 |
| Xrn21 | 7.50E-87 | 0.648924628 | 0.627 | 0.255 | 1.52E-82 | 11 | Xrn2 |
| Rpl75 | 1.22E-86 | 0.564067976 | 0.996 | 0.891 | 2.48E-82 | 11 | Rpl7 |
| Rpl214 | 3.43E-86 | 0.566926151 | 1 | 0.922 | 6.95E-82 | 11 | Rpl21 |
| Vps542 | 4.07E-86 | 0.70509816 | 0.506 | 0.181 | 8.23E-82 | 11 | Vps54 |
| Tmsb4x4 | 5.51E-86 | 0.598126284 | 1 | 0.933 | 1.12E-81 | 11 | Tmsb4x |
| Vav12 | 5.86E-86 | 0.481236122 | 0.307 | 0.077 | 1.19E-81 | 11 | Vav1 |
| Prr131 | 6.10E-86 | 0.544687865 | 0.557 | 0.204 | 1.23E-81 | 11 | Prr13 |
| Strbp1 | 6.74E-86 | 0.503874815 | 0.533 | 0.186 | 1.37E-81 | 11 | Strbp |
| Rfx71 | 1.21E-85 | 0.655646581 | 0.452 | 0.15 | 2.44E-81 | 11 | Rfx7 |
| Phf61 | 1.29E-85 | 0.576843559 | 0.42 | 0.13 | 2.60E-81 | 11 | Phf6 |
| Tmem1731 | 5.77E-85 | 0.479623209 | 0.282 | 0.068 | 1.17E-80 | 11 | Tmem173 |
| Rps293 | 7.26E-85 | 0.490223075 | 1 | 0.971 | 1.47E-80 | 11 | Rps29 |
| Klhl62 | 1.08E-84 | 0.65556923 | 0.388 | 0.116 | 2.18E-80 | 11 | Klhl6 |
| Rsbn1l2 | 1.90E-84 | 0.647359823 | 0.65 | 0.266 | 3.84E-80 | 11 | Rsbn1l |
| Clic13 | 2.67E-84 | 0.704125421 | 0.959 | 0.595 | 5.40E-80 | 11 | Clic1 |
| Ago21 | 3.68E-84 | 0.563837774 | 0.478 | 0.162 | 7.46E-80 | 11 | Ago2 |

| | | | | | | | |
|---|---|---|---|---|---|---|---|
| Retreg12 | 4.14E-84 | 0.483195581 | 0.343 | 0.093 | 8.38E-80 | 11 | Retreg1 |
| Eif3e2 | 5.63E-84 | 0.564619556 | 0.838 | 0.405 | 1.14E-79 | 11 | Eif3e |
| Cmc1 | 7.28E-84 | 0.393652177 | 0.53 | 0.185 | 1.47E-79 | 11 | Cmc1 |
| Ptpn21 | 7.94E-84 | 0.563087175 | 0.431 | 0.138 | 1.61E-79 | 11 | Ptpn2 |
| Ppp1r16b4 | 1.21E-83 | 0.736705855 | 0.535 | 0.196 | 2.46E-79 | 11 | Ppp1r16b |
| Ptpn15 | 1.92E-83 | 0.576052237 | 0.594 | 0.23 | 3.88E-79 | 11 | Ptpn1 |
| Rpl293 | 2.18E-83 | 0.653672776 | 0.986 | 0.765 | 4.40E-79 | 11 | Rpl29 |
| Pld31 | 2.65E-83 | 0.504672251 | 0.314 | 0.082 | 5.36E-79 | 11 | Pld3 |
| Ralgapa21 | 5.36E-83 | 0.626871844 | 0.449 | 0.149 | 1.09E-78 | 11 | Ralgapa2 |
| Rps215 | 8.03E-83 | 0.64329949 | 0.998 | 0.918 | 1.63E-78 | 11 | Rps2 |
| Cdkn2d1 | 8.98E-83 | 0.493254243 | 0.329 | 0.088 | 1.82E-78 | 11 | Cdkn2d |
| Zfp8271 | 1.22E-82 | 0.471271349 | 0.262 | 0.06 | 2.47E-78 | 11 | Zfp827 |
| Rps27rt1 | 3.39E-82 | 0.381428656 | 0.364 | 0.103 | 6.87E-78 | 11 | Rps27rt |
| Birc31 | 1.42E-81 | 0.557505261 | 0.339 | 0.095 | 2.88E-77 | 11 | Birc3 |
| Rps264 | 2.50E-81 | 0.584639755 | 0.995 | 0.863 | 5.05E-77 | 11 | Rps26 |
| Rcor11 | 3.64E-81 | 0.635307262 | 0.492 | 0.178 | 7.37E-77 | 11 | Rcor1 |
| Arpc23 | 6.53E-81 | 0.68824655 | 0.926 | 0.538 | 1.32E-76 | 11 | Arpc2 |
| Atp2a31 | 6.89E-81 | 0.335771786 | 0.259 | 0.059 | 1.40E-76 | 11 | Atp2a3 |
| Gtpbp11 | 1.16E-80 | 0.45225303 | 0.278 | 0.068 | 2.36E-76 | 11 | Gtpbp1 |
| Lrrfip14 | 1.60E-80 | 0.627790034 | 0.788 | 0.371 | 3.23E-76 | 11 | Lrrfip1 |
| mt-Co37 | 1.61E-80 | 0.673252784 | 1 | 0.721 | 3.25E-76 | 11 | mt-Co3 |
| Supt4a2 | 5.93E-80 | 0.50146441 | 0.713 | 0.315 | 1.20E-75 | 11 | Supt4a |
| Arhgap252 | 1.81E-79 | 0.568897419 | 0.354 | 0.104 | 3.67E-75 | 11 | Arhgap25 |
| H2afy1 | 3.43E-79 | 0.549659059 | 0.74 | 0.331 | 6.95E-75 | 11 | H2afy |
| Myl64 | 9.08E-79 | 0.684698758 | 0.991 | 0.772 | 1.84E-74 | 11 | Myl6 |
| Sqstm13 | 8.15E-78 | 0.837555449 | 0.657 | 0.298 | 1.65E-73 | 11 | Sqstm1 |
| Tln15 | 1.12E-77 | 0.688613225 | 0.799 | 0.398 | 2.27E-73 | 11 | Tln1 |
| Gm372402 | 1.74E-77 | 0.52619738 | 0.346 | 0.1 | 3.53E-73 | 11 | Gm37240 |
| Rbpj2 | 2.08E-77 | 0.73314403 | 0.546 | 0.215 | 4.21E-73 | 11 | Rbpj |
| Aak12 | 2.59E-77 | 0.597532379 | 0.497 | 0.18 | 5.24E-73 | 11 | Aak1 |
| Bhlhe402 | 3.83E-77 | 0.618227967 | 0.447 | 0.153 | 7.75E-73 | 11 | Bhlhe40 |
| Iqgap15 | 5.19E-77 | 0.655721737 | 0.837 | 0.404 | 1.05E-72 | 11 | Iqgap1 |
| Vwa81 | 6.37E-77 | 0.621738476 | 0.373 | 0.116 | 1.29E-72 | 11 | Vwa8 |
| Dgkd2 | 7.03E-77 | 0.587268342 | 0.451 | 0.156 | 1.42E-72 | 11 | Dgkd |
| Arhgef16 | 1.01E-76 | 0.646836596 | 0.591 | 0.243 | 2.05E-72 | 11 | Arhgef1 |
| Ptger42 | 2.39E-76 | 0.57505029 | 0.302 | 0.081 | 4.83E-72 | 11 | Ptger4 |
| D16Ertd472e1 | 2.74E-76 | 0.545738674 | 0.346 | 0.102 | 5.54E-72 | 11 | D16Ertd472e |
| Tcf125 | 3.39E-76 | 0.701637995 | 0.779 | 0.369 | 6.87E-72 | 11 | Tcf12 |
| Uba524 | 4.46E-76 | 0.472211509 | 0.944 | 0.514 | 9.03E-72 | 11 | Uba52 |
| mt-Atp85 | 8.21E-76 | 0.624233786 | 0.878 | 0.456 | 1.66E-71 | 11 | mt-Atp8 |
| Ppp1r12a4 | 1.13E-75 | 0.671634644 | 0.844 | 0.418 | 2.29E-71 | 11 | Ppp1r12a |
| Mta32 | 1.20E-75 | 0.479399561 | 0.438 | 0.147 | 2.42E-71 | 11 | Mta3 |
| Mycbp25 | 3.21E-75 | 0.677259469 | 0.779 | 0.367 | 6.49E-71 | 11 | Mycbp2 |

| | | | | | | | |
|---|---|---|---|---|---|---|---|
| Kpna45 | 3.53E-75 | 0.630169828 | 0.7 | 0.317 | 7.14E-71 | 11 | Kpna4 |
| Pitpnc14 | 8.19E-75 | 0.724063045 | 0.838 | 0.422 | 1.66E-70 | 11 | Pitpnc1 |
| Rpl284 | 2.09E-74 | 0.507870226 | 0.998 | 0.929 | 4.22E-70 | 11 | Rpl28 |
| Rpl43 | 2.96E-74 | 0.595604998 | 0.971 | 0.653 | 5.99E-70 | 11 | Rpl4 |
| Lnpep5 | 3.53E-74 | 0.629327932 | 0.61 | 0.254 | 7.15E-70 | 11 | Lnpep |
| Nktr2 | 4.53E-74 | 0.619877181 | 0.603 | 0.252 | 9.17E-70 | 11 | Nktr |
| Fndc3a2 | 6.89E-74 | 0.699280771 | 0.623 | 0.271 | 1.39E-69 | 11 | Fndc3a |
| Taf101 | 6.94E-74 | 0.505887918 | 0.785 | 0.377 | 1.41E-69 | 11 | Taf10 |
| Npm13 | 1.54E-73 | 0.65463971 | 0.98 | 0.674 | 3.12E-69 | 11 | Npm1 |
| Pde4b1 | 2.59E-73 | 0.628189509 | 0.506 | 0.191 | 5.24E-69 | 11 | Pde4b |
| Rpl244 | 3.12E-73 | 0.486959431 | 1 | 0.906 | 6.32E-69 | 11 | Rpl24 |
| Ddx211 | 3.91E-73 | 0.564025364 | 0.531 | 0.208 | 7.93E-69 | 11 | Ddx21 |
| Tpp22 | 6.94E-73 | 0.609361847 | 0.557 | 0.223 | 1.41E-68 | 11 | Tpp2 |
| Rps84 | 1.02E-72 | 0.488445241 | 1 | 0.964 | 2.06E-68 | 11 | Rps8 |
| Ranbp92 | 1.25E-72 | 0.623980009 | 0.503 | 0.195 | 2.52E-68 | 11 | Ranbp9 |
| Rev3l2 | 1.30E-72 | 0.566990825 | 0.436 | 0.153 | 2.63E-68 | 11 | Rev3l |
| Ogt3 | 1.60E-72 | 0.555107084 | 0.522 | 0.201 | 3.25E-68 | 11 | Ogt |
| Camkmt1 | 2.18E-72 | 0.716522604 | 0.4 | 0.135 | 4.41E-68 | 11 | Camkmt |
| Nckap1l2 | 2.24E-72 | 0.389138289 | 0.309 | 0.085 | 4.54E-68 | 11 | Nckap1l |
| Ube2i1 | 2.40E-72 | 0.482937934 | 0.706 | 0.324 | 4.87E-68 | 11 | Ube2i |
| H2afj3 | 5.94E-72 | 0.407551282 | 0.867 | 0.446 | 1.20E-67 | 11 | H2afj |
| Crip14 | 6.18E-72 | 0.541887708 | 0.944 | 0.529 | 1.25E-67 | 11 | Crip1 |
| Rnf1492 | 6.28E-72 | 0.507250062 | 0.334 | 0.1 | 1.27E-67 | 11 | Rnf149 |
| Mbp1 | 1.01E-71 | 0.390931362 | 0.259 | 0.064 | 2.04E-67 | 11 | Mbp |
| Gab31 | 1.18E-71 | 0.613733117 | 0.312 | 0.091 | 2.39E-67 | 11 | Gab3 |
| Plekhj11 | 2.05E-71 | 0.490530863 | 0.456 | 0.164 | 4.15E-67 | 11 | Plekhj1 |
| Mapre22 | 2.29E-71 | 0.659849467 | 0.583 | 0.244 | 4.64E-67 | 11 | Mapre2 |
| Rreb12 | 2.31E-71 | 0.602031295 | 0.508 | 0.197 | 4.67E-67 | 11 | Rreb1 |
| Chd31 | 2.33E-71 | 0.453643248 | 0.434 | 0.151 | 4.73E-67 | 11 | Chd3 |
| Rnf1661 | 2.46E-71 | 0.458789576 | 0.341 | 0.103 | 4.98E-67 | 11 | Rnf166 |
| Fam173a1 | 5.87E-71 | 0.374123827 | 0.512 | 0.192 | 1.19E-66 | 11 | Fam173a |
| Cap12 | 6.04E-71 | 0.52360133 | 0.619 | 0.266 | 1.22E-66 | 11 | Cap1 |
| Cntrl1 | 1.05E-70 | 0.517089119 | 0.372 | 0.12 | 2.12E-66 | 11 | Cntrl |
| Ptbp35 | 1.18E-70 | 0.62311866 | 0.794 | 0.387 | 2.40E-66 | 11 | Ptbp3 |
| Tmcc15 | 1.19E-70 | 0.676872689 | 0.528 | 0.213 | 2.41E-66 | 11 | Tmcc1 |
| Ddx101 | 1.50E-70 | 0.526505154 | 0.386 | 0.127 | 3.04E-66 | 11 | Ddx10 |
| Tmem2431 | 1.90E-70 | 0.499736418 | 0.443 | 0.158 | 3.84E-66 | 11 | Tmem243 |
| Crtc33 | 2.27E-70 | 0.57824223 | 0.442 | 0.16 | 4.60E-66 | 11 | Crtc3 |
| Trim12a3 | 2.47E-70 | 0.517446598 | 0.478 | 0.177 | 5.01E-66 | 11 | Trim12a |
| Rpl354 | 3.18E-70 | 0.522497611 | 0.995 | 0.825 | 6.44E-66 | 11 | Rpl35 |
| Plekha52 | 3.34E-70 | 0.639205404 | 0.425 | 0.151 | 6.77E-66 | 11 | Plekha5 |
| Snrpf3 | 7.10E-70 | 0.452389311 | 0.772 | 0.372 | 1.44E-65 | 11 | Snrpf |
| Usp242 | 7.24E-70 | 0.545869097 | 0.393 | 0.133 | 1.47E-65 | 11 | Usp24 |

| | | | | | | | |
|---|---|---|---|---|---|---|---|
| Rasa31 | 1.07E-69 | 0.413653227 | 0.251 | 0.063 | 2.16E-65 | 11 | Rasa3 |
| Pdcd43 | 1.87E-69 | 0.31691201 | 0.889 | 0.445 | 3.79E-65 | 11 | Pdcd4 |
| Nme13 | 2.11E-69 | 0.581229599 | 0.72 | 0.349 | 4.27E-65 | 11 | Nme1 |
| Gmip2 | 2.51E-69 | 0.455178324 | 0.285 | 0.079 | 5.08E-65 | 11 | Gmip |
| Trim142 | 2.97E-69 | 0.557750815 | 0.363 | 0.117 | 6.00E-65 | 11 | Trim14 |
| Nsa23 | 3.69E-69 | 0.451307748 | 0.89 | 0.479 | 7.46E-65 | 11 | Nsa2 |
| Rpl36a4 | 3.72E-69 | 0.576655745 | 0.989 | 0.748 | 7.54E-65 | 11 | Rpl36a |
| Exoc21 | 5.08E-69 | 0.547161979 | 0.409 | 0.143 | 1.03E-64 | 11 | Exoc2 |
| Stk103 | 7.64E-69 | 0.550660897 | 0.474 | 0.178 | 1.55E-64 | 11 | Stk10 |
| Exoc46 | 8.01E-69 | 0.610247787 | 0.691 | 0.316 | 1.62E-64 | 11 | Exoc4 |
| Rps6ka12 | 1.45E-68 | 0.468049402 | 0.352 | 0.111 | 2.94E-64 | 11 | Rps6ka1 |
| Arl6ip54 | 2.45E-68 | 0.579274032 | 0.537 | 0.216 | 4.97E-64 | 11 | Arl6ip5 |
| Zbtb205 | 4.30E-68 | 0.654961233 | 0.941 | 0.53 | 8.72E-64 | 11 | Zbtb20 |
| Inpp5b | 4.94E-68 | 0.432646365 | 0.268 | 0.071 | 1.00E-63 | 11 | Inpp5b |
| Gm472833 | 8.74E-68 | 0.716593937 | 0.618 | 0.276 | 1.77E-63 | 11 | Gm47283 |
| Sf3b13 | 8.76E-68 | 0.580859531 | 0.779 | 0.387 | 1.77E-63 | 11 | Sf3b1 |
| Zcchc75 | 2.11E-67 | 0.621944992 | 0.655 | 0.295 | 4.28E-63 | 11 | Zcchc7 |
| Snrpg2 | 2.23E-67 | 0.440982606 | 0.691 | 0.318 | 4.51E-63 | 11 | Snrpg |
| Sipa12 | 2.43E-67 | 0.536191133 | 0.474 | 0.181 | 4.91E-63 | 11 | Sipa1 |
| Zdhhc204 | 2.49E-67 | 0.58372176 | 0.549 | 0.225 | 5.05E-63 | 11 | Zdhhc20 |
| Ccm21 | 2.55E-67 | 0.475017841 | 0.372 | 0.122 | 5.16E-63 | 11 | Ccm2 |
| Eif3h2 | 4.81E-67 | 0.462369409 | 0.871 | 0.473 | 9.74E-63 | 11 | Eif3h |
| Cln31 | 5.43E-67 | 0.419667157 | 0.253 | 0.066 | 1.10E-62 | 11 | Cln3 |
| Bin12 | 1.22E-66 | 0.409321613 | 0.461 | 0.171 | 2.47E-62 | 11 | Bin1 |
| Tpm34 | 1.39E-66 | 0.552125448 | 0.876 | 0.472 | 2.81E-62 | 11 | Tpm3 |
| Ppp1ca1 | 2.23E-66 | 0.461014583 | 0.781 | 0.39 | 4.51E-62 | 11 | Ppp1ca |
| Cnot6l1 | 2.56E-66 | 0.522149635 | 0.571 | 0.239 | 5.19E-62 | 11 | Cnot6l |
| Nt5c1 | 2.86E-66 | 0.444022707 | 0.578 | 0.244 | 5.80E-62 | 11 | Nt5c |
| Rplp04 | 4.50E-66 | 0.50976517 | 0.998 | 0.944 | 9.12E-62 | 11 | Rplp0 |
| Smad74 | 9.97E-66 | 0.675903872 | 0.67 | 0.32 | 2.02E-61 | 11 | Smad7 |
| H2-D15 | 1.00E-65 | 0.614470997 | 1 | 0.663 | 2.03E-61 | 11 | H2-D1 |
| Vmp12 | 1.05E-65 | 0.510275255 | 0.519 | 0.211 | 2.13E-61 | 11 | Vmp1 |
| Nhp21 | 1.11E-65 | 0.416159075 | 0.463 | 0.172 | 2.25E-61 | 11 | Nhp2 |
| Rnaseh2c1 | 5.97E-65 | 0.381986666 | 0.639 | 0.28 | 1.21E-60 | 11 | Rnaseh2c |
| Slc2a11 | 8.72E-65 | 0.512634292 | 0.291 | 0.084 | 1.77E-60 | 11 | Slc2a1 |
| Tnrc6b3 | 1.05E-64 | 0.563777489 | 0.747 | 0.365 | 2.12E-60 | 11 | Tnrc6b |
| Mef2a3 | 1.42E-64 | 0.616687431 | 0.811 | 0.41 | 2.87E-60 | 11 | Mef2a |
| Zfp6541 | 1.49E-64 | 0.592194545 | 0.352 | 0.117 | 3.02E-60 | 11 | Zfp654 |
| Sdc42 | 1.56E-64 | 0.335648986 | 0.259 | 0.068 | 3.15E-60 | 11 | Sdc4 |
| Sik23 | 2.21E-64 | 0.522970973 | 0.352 | 0.116 | 4.48E-60 | 11 | Sik2 |
| Tecr2 | 2.82E-64 | 0.468556471 | 0.531 | 0.22 | 5.72E-60 | 11 | Tecr |
| mt-Nd46 | 3.17E-64 | 0.610886729 | 0.996 | 0.67 | 6.41E-60 | 11 | mt-Nd4 |
| Ss181 | 6.83E-64 | 0.477446266 | 0.451 | 0.169 | 1.38E-59 | 11 | Ss18 |

| | | | | | | | |
|---|---|---|---|---|---|---|---|
| Vti1a3 | 6.87E-64 | 0.614692595 | 0.569 | 0.25 | 1.39E-59 | 11 | Vti1a |
| Rngtt1 | 8.52E-64 | 0.482701539 | 0.384 | 0.131 | 1.73E-59 | 11 | Rngtt |
| Tsc22d42 | 2.60E-63 | 0.387854917 | 0.648 | 0.288 | 5.27E-59 | 11 | Tsc22d4 |
| Stk241 | 2.67E-63 | 0.475934559 | 0.485 | 0.191 | 5.40E-59 | 11 | Stk24 |
| Rel2 | 3.15E-63 | 0.574749505 | 0.382 | 0.135 | 6.38E-59 | 11 | Rel |
| Tmem23 | 3.66E-63 | 0.536155012 | 0.395 | 0.139 | 7.40E-59 | 11 | Tmem2 |
| Coq10b1 | 4.20E-63 | 0.586003342 | 0.408 | 0.15 | 8.50E-59 | 11 | Coq10b |
| Fhit2 | 4.66E-63 | 0.658910728 | 0.271 | 0.077 | 9.43E-59 | 11 | Fhit |
| Tgfb14 | 7.47E-63 | 0.672938227 | 0.537 | 0.231 | 1.51E-58 | 11 | Tgfb1 |
| Ran2 | 1.12E-62 | 0.468992347 | 0.767 | 0.394 | 2.26E-58 | 11 | Ran |
| Vars1 | 1.36E-62 | 0.403020649 | 0.276 | 0.079 | 2.76E-58 | 11 | Vars |
| Cdk174 | 1.46E-62 | 0.564499835 | 0.567 | 0.24 | 2.96E-58 | 11 | Cdk17 |
| Creb11 | 1.97E-62 | 0.610813241 | 0.496 | 0.203 | 3.98E-58 | 11 | Creb1 |
| Sppl34 | 3.55E-62 | 0.59628987 | 0.51 | 0.213 | 7.20E-58 | 11 | Sppl3 |
| Arhgef63 | 6.94E-62 | 0.478127769 | 0.364 | 0.125 | 1.41E-57 | 11 | Arhgef6 |
| Snrpe2 | 8.37E-62 | 0.348251772 | 0.874 | 0.47 | 1.69E-57 | 11 | Snrpe |
| Pacs12 | 1.49E-61 | 0.528399332 | 0.497 | 0.2 | 3.01E-57 | 11 | Pacs1 |
| Cog53 | 1.73E-61 | 0.559163538 | 0.456 | 0.179 | 3.51E-57 | 11 | Cog5 |
| Plcg22 | 1.75E-61 | 0.489982211 | 0.35 | 0.117 | 3.54E-57 | 11 | Plcg2 |
| Rad54l21 | 1.89E-61 | 0.445030394 | 0.3 | 0.092 | 3.82E-57 | 11 | Rad54l2 |
| Rnf22 | 1.92E-61 | 0.438911151 | 0.321 | 0.102 | 3.88E-57 | 11 | Rnf2 |
| Ppm1g1 | 2.61E-61 | 0.47720973 | 0.411 | 0.152 | 5.29E-57 | 11 | Ppm1g |
| Rpl23a4 | 2.76E-61 | 0.561988854 | 0.971 | 0.674 | 5.58E-57 | 11 | Rpl23a |
| Sik33 | 3.37E-61 | 0.585721131 | 0.592 | 0.265 | 6.82E-57 | 11 | Sik3 |
| Tor1aip21 | 4.73E-61 | 0.533534879 | 0.454 | 0.178 | 9.57E-57 | 11 | Tor1aip2 |
| Sesn12 | 8.59E-61 | 0.592080175 | 0.503 | 0.208 | 1.74E-56 | 11 | Sesn1 |
| Eef1g3 | 9.31E-61 | 0.428577075 | 0.88 | 0.505 | 1.89E-56 | 11 | Eef1g |
| Vrk22 | 1.05E-60 | 0.459894865 | 0.355 | 0.12 | 2.12E-56 | 11 | Vrk2 |
| Ptpre2 | 1.39E-60 | 0.522385989 | 0.42 | 0.157 | 2.81E-56 | 11 | Ptpre |
| Hectd13 | 1.53E-60 | 0.561781039 | 0.634 | 0.295 | 3.10E-56 | 11 | Hectd1 |
| Itch4 | 1.61E-60 | 0.548601684 | 0.592 | 0.268 | 3.25E-56 | 11 | Itch |
| Spata134 | 2.45E-60 | 0.508969987 | 0.499 | 0.201 | 4.95E-56 | 11 | Spata13 |
| Phf21a4 | 3.33E-60 | 0.649866015 | 0.558 | 0.248 | 6.74E-56 | 11 | Phf21a |
| Fchsd23 | 4.42E-60 | 0.536945281 | 0.413 | 0.156 | 8.94E-56 | 11 | Fchsd2 |
| Psmb86 | 8.17E-60 | 0.522059184 | 0.95 | 0.55 | 1.65E-55 | 11 | Psmb8 |
| Cyld1 | 8.67E-60 | 0.42367196 | 0.438 | 0.166 | 1.75E-55 | 11 | Cyld |
| mt-Atp67 | 8.88E-60 | 0.557902848 | 1 | 0.75 | 1.80E-55 | 11 | mt-Atp6 |
| Esyt24 | 9.32E-60 | 0.603663574 | 0.549 | 0.237 | 1.89E-55 | 11 | Esyt2 |
| Gm100764 | 9.33E-60 | 0.437533228 | 0.914 | 0.539 | 1.89E-55 | 11 | Gm10076 |
| Cdc42se22 | 1.13E-59 | 0.579512576 | 0.537 | 0.234 | 2.28E-55 | 11 | Cdc42se2 |
| Grap | 1.15E-59 | 0.421649476 | 0.341 | 0.112 | 2.33E-55 | 11 | Grap |
| B4galt14 | 2.11E-59 | 0.55106292 | 0.44 | 0.173 | 4.26E-55 | 11 | B4galt1 |
| Taf151 | 2.37E-59 | 0.506717114 | 0.539 | 0.233 | 4.80E-55 | 11 | Taf15 |

| | | | | | | | |
|---|---|---|---|---|---|---|---|
| Arid1b4 | 3.09E-59 | 0.578206957 | 0.569 | 0.253 | 6.26E-55 | 11 | Arid1b |
| Tmem2583 | 4.67E-59 | 0.285908338 | 0.664 | 0.303 | 9.46E-55 | 11 | Tmem258 |
| Rpl143 | 5.92E-59 | 0.481068299 | 0.987 | 0.799 | 1.20E-54 | 11 | Rpl14 |
| Memo11 | 6.26E-59 | 0.444581615 | 0.305 | 0.097 | 1.27E-54 | 11 | Memo1 |
| Kansl15 | 7.12E-59 | 0.605761641 | 0.684 | 0.334 | 1.44E-54 | 11 | Kansl1 |
| Npc12 | 1.25E-58 | 0.417865685 | 0.26 | 0.075 | 2.54E-54 | 11 | Npc1 |
| Rpl414 | 1.43E-58 | 0.404345476 | 0.998 | 0.961 | 2.90E-54 | 11 | Rpl41 |
| Itpr22 | 1.63E-58 | 0.527069571 | 0.551 | 0.239 | 3.31E-54 | 11 | Itpr2 |
| Ncoa25 | 1.67E-58 | 0.587357947 | 0.537 | 0.234 | 3.39E-54 | 11 | Ncoa2 |
| Ifngr16 | 1.76E-58 | 0.615239838 | 0.749 | 0.387 | 3.56E-54 | 11 | Ifngr1 |
| Nedd94 | 2.24E-58 | 0.489005771 | 0.691 | 0.313 | 4.54E-54 | 11 | Nedd9 |
| Fam208b2 | 2.40E-58 | 0.491842409 | 0.397 | 0.146 | 4.86E-54 | 11 | Fam208b |
| Plgrkt2 | 2.45E-58 | 0.492721449 | 0.499 | 0.209 | 4.97E-54 | 11 | Plgrkt |
| Arrb22 | 2.50E-58 | 0.363291254 | 0.293 | 0.09 | 5.06E-54 | 11 | Arrb2 |
| Ubash3b2 | 2.65E-58 | 0.590976041 | 0.424 | 0.164 | 5.37E-54 | 11 | Ubash3b |
| Babam23 | 3.00E-58 | 0.499807184 | 0.546 | 0.233 | 6.07E-54 | 11 | Babam2 |
| Ms4a6b1 | 3.82E-58 | 0.350467667 | 0.294 | 0.09 | 7.74E-54 | 11 | Ms4a6b |
| Fbxl176 | 4.26E-58 | 0.580257536 | 0.645 | 0.305 | 8.63E-54 | 11 | Fbxl17 |
| Sun21 | 4.64E-58 | 0.42713111 | 0.259 | 0.075 | 9.40E-54 | 11 | Sun2 |
| Commd71 | 1.24E-57 | 0.397202094 | 0.479 | 0.196 | 2.51E-53 | 11 | Commd7 |
| Tpt13 | 1.55E-57 | 0.335887243 | 0.995 | 0.99 | 3.13E-53 | 11 | Tpt1 |
| Asxl21 | 1.71E-57 | 0.518423104 | 0.434 | 0.171 | 3.47E-53 | 11 | Asxl2 |
| Saraf4 | 2.68E-57 | 0.50422549 | 0.637 | 0.302 | 5.43E-53 | 11 | Saraf |
| Arl6ip14 | 3.07E-57 | 0.479322182 | 0.734 | 0.364 | 6.22E-53 | 11 | Arl6ip1 |
| Sp1003 | 4.07E-57 | 0.465002584 | 0.815 | 0.426 | 8.24E-53 | 11 | Sp100 |
| Actr22 | 4.56E-57 | 0.447563344 | 0.684 | 0.338 | 9.23E-53 | 11 | Actr2 |
| Cetn21 | 6.17E-57 | 0.317415073 | 0.492 | 0.199 | 1.25E-52 | 11 | Cetn2 |
| Polr2a4 | 6.72E-57 | 0.565367971 | 0.591 | 0.276 | 1.36E-52 | 11 | Polr2a |
| Nudt21 | 6.77E-57 | 0.338350737 | 0.411 | 0.153 | 1.37E-52 | 11 | Nudt21 |
| Cfl13 | 7.56E-57 | 0.573982217 | 0.93 | 0.603 | 1.53E-52 | 11 | Cfl1 |
| Myh94 | 8.16E-57 | 0.549751991 | 0.82 | 0.43 | 1.65E-52 | 11 | Myh9 |
| Nfkbiz1 | 8.71E-57 | 0.451427642 | 0.377 | 0.134 | 1.76E-52 | 11 | Nfkbiz |
| Tpk12 | 1.28E-56 | 0.458626388 | 0.312 | 0.101 | 2.58E-52 | 11 | Tpk1 |
| Exoc63 | 1.39E-56 | 0.454538994 | 0.429 | 0.167 | 2.82E-52 | 11 | Exoc6 |
| Mlxip1 | 1.40E-56 | 0.498242779 | 0.352 | 0.124 | 2.84E-52 | 11 | Mlxip |
| Srrm24 | 2.42E-56 | 0.510832281 | 0.86 | 0.463 | 4.91E-52 | 11 | Srrm2 |
| Ubr23 | 2.64E-56 | 0.51224504 | 0.517 | 0.225 | 5.34E-52 | 11 | Ubr2 |
| Slfn82 | 2.75E-56 | 0.467701964 | 0.251 | 0.073 | 5.56E-52 | 11 | Slfn8 |
| Dad13 | 3.23E-56 | 0.461884274 | 0.826 | 0.453 | 6.54E-52 | 11 | Dad1 |
| Larp13 | 4.56E-56 | 0.484150045 | 0.478 | 0.199 | 9.23E-52 | 11 | Larp1 |
| 4933406I18Rik2 | 4.62E-56 | 0.562220576 | 0.251 | 0.073 | 9.36E-52 | 11 | 4933406I18Rik |
| Nin1 | 5.20E-56 | 0.48273282 | 0.37 | 0.135 | 1.05E-51 | 11 | Nin |
| Lsm41 | 7.18E-56 | 0.295757757 | 0.67 | 0.316 | 1.45E-51 | 11 | Lsm4 |

| | | | | | | | |
|---|---|---|---|---|---|---|---|
| Stk391 | 1.28E-55 | 0.368728245 | 0.402 | 0.15 | 2.58E-51 | 11 | Stk39 |
| Cd474 | 1.49E-55 | 0.508847209 | 0.901 | 0.503 | 3.02E-51 | 11 | Cd47 |
| mt-Nd4l4 | 1.89E-55 | 0.440542114 | 0.889 | 0.484 | 3.83E-51 | 11 | mt-Nd4l |
| Vps281 | 1.90E-55 | 0.366490224 | 0.63 | 0.295 | 3.84E-51 | 11 | Vps28 |
| Def6 | 2.01E-55 | 0.255358713 | 0.291 | 0.09 | 4.08E-51 | 11 | Def6 |
| Arpc5l1 | 2.40E-55 | 0.33715526 | 0.632 | 0.299 | 4.85E-51 | 11 | Arpc5l |
| Ccnd24 | 2.79E-55 | 0.633665407 | 0.6 | 0.281 | 5.64E-51 | 11 | Ccnd2 |
| Arhgap263 | 2.93E-55 | 0.545186512 | 0.508 | 0.218 | 5.93E-51 | 11 | Arhgap26 |
| Emp33 | 5.93E-55 | 0.480790907 | 0.567 | 0.254 | 1.20E-50 | 11 | Emp3 |
| Cdyl1 | 7.54E-55 | 0.452548152 | 0.35 | 0.125 | 1.53E-50 | 11 | Cdyl |
| Elmsan11 | 7.71E-55 | 0.485790099 | 0.307 | 0.103 | 1.56E-50 | 11 | Elmsan1 |
| Atf7ip3 | 1.28E-54 | 0.441826754 | 0.485 | 0.203 | 2.59E-50 | 11 | Atf7ip |
| Clk13 | 1.33E-54 | 0.528182404 | 0.607 | 0.287 | 2.69E-50 | 11 | Clk1 |
| Vps13b3 | 1.99E-54 | 0.50026719 | 0.503 | 0.215 | 4.03E-50 | 11 | Vps13b |
| Sri1 | 3.22E-54 | 0.3795222 | 0.799 | 0.414 | 6.52E-50 | 11 | Sri |
| Btbd95 | 3.58E-54 | 0.520563811 | 0.524 | 0.233 | 7.26E-50 | 11 | Btbd9 |
| Rasa22 | 3.84E-54 | 0.460521222 | 0.366 | 0.135 | 7.78E-50 | 11 | Rasa2 |
| Atf62 | 9.47E-54 | 0.44094389 | 0.408 | 0.158 | 1.92E-49 | 11 | Atf6 |
| Ncl1 | 1.06E-53 | 0.616772763 | 0.871 | 0.511 | 2.15E-49 | 11 | Ncl |
| Lsm5 | 1.16E-53 | 0.296032501 | 0.508 | 0.216 | 2.35E-49 | 11 | Lsm5 |
| Grcc101 | 1.38E-53 | 0.31514004 | 0.314 | 0.104 | 2.79E-49 | 11 | Grcc10 |
| Lcor3 | 1.60E-53 | 0.477638586 | 0.452 | 0.186 | 3.25E-49 | 11 | Lcor |
| Nipbl6 | 1.74E-53 | 0.484704378 | 0.727 | 0.36 | 3.53E-49 | 11 | Nipbl |
| Snrpd21 | 1.88E-53 | 0.268222179 | 0.779 | 0.403 | 3.81E-49 | 11 | Snrpd2 |
| Hivep25 | 2.17E-53 | 0.602174862 | 0.61 | 0.293 | 4.40E-49 | 11 | Hivep2 |
| Cmtm72 | 2.69E-53 | 0.283547433 | 0.501 | 0.208 | 5.44E-49 | 11 | Cmtm7 |
| Atp2b14 | 2.75E-53 | 0.523325079 | 0.662 | 0.327 | 5.56E-49 | 11 | Atp2b1 |
| Sdf2l12 | 4.41E-53 | 0.403787515 | 0.418 | 0.164 | 8.93E-49 | 11 | Sdf2l1 |
| Man1a4 | 4.45E-53 | 0.5007924 | 0.411 | 0.164 | 9.02E-49 | 11 | Man1a |
| Nasp1 | 5.21E-53 | 0.463016502 | 0.422 | 0.168 | 1.06E-48 | 11 | Nasp |
| Tmem1312 | 6.31E-53 | 0.460378503 | 0.481 | 0.204 | 1.28E-48 | 11 | Tmem131 |
| Herc43 | 8.11E-53 | 0.49436948 | 0.384 | 0.148 | 1.64E-48 | 11 | Herc4 |
| Eif3k2 | 1.01E-52 | 0.335895436 | 0.883 | 0.495 | 2.05E-48 | 11 | Eif3k |
| Osbpl83 | 1.11E-52 | 0.501695396 | 0.461 | 0.195 | 2.25E-48 | 11 | Osbpl8 |
| Exosc3 | 1.39E-52 | 0.303147066 | 0.375 | 0.139 | 2.82E-48 | 11 | Exosc3 |
| Ablim13 | 1.53E-52 | 0.43606535 | 0.792 | 0.396 | 3.11E-48 | 11 | Ablim1 |
| Borcs8 | 1.59E-52 | 0.331834652 | 0.311 | 0.104 | 3.22E-48 | 11 | Borcs8 |
| Rbm394 | 1.62E-52 | 0.521854822 | 0.939 | 0.568 | 3.28E-48 | 11 | Rbm39 |
| Lncpint4 | 3.97E-52 | 0.525828417 | 0.58 | 0.269 | 8.04E-48 | 11 | Lncpint |
| Usp31 | 4.23E-52 | 0.445042396 | 0.436 | 0.179 | 8.56E-48 | 11 | Usp3 |
| Rgs33 | 4.77E-52 | 0.43200513 | 0.449 | 0.183 | 9.66E-48 | 11 | Rgs3 |
| Eif3i1 | 5.10E-52 | 0.312910321 | 0.718 | 0.371 | 1.03E-47 | 11 | Eif3i |
| Grina2 | 5.29E-52 | 0.426283707 | 0.363 | 0.136 | 1.07E-47 | 11 | Grina |

| | | | | | | | |
|---|---|---|---|---|---|---|---|
| Arhgef34 | 6.11E-52 | 0.538525745 | 0.528 | 0.236 | 1.24E-47 | 11 | Arhgef3 |
| Smg15 | 6.36E-52 | 0.509432001 | 0.515 | 0.23 | 1.29E-47 | 11 | Smg1 |
| Mettl231 | 9.52E-52 | 0.271319415 | 0.479 | 0.198 | 1.93E-47 | 11 | Mettl23 |
| Wdr89 | 1.02E-51 | 0.316135434 | 0.307 | 0.103 | 2.07E-47 | 11 | Wdr89 |
| Rnf1381 | 1.17E-51 | 0.378720667 | 0.361 | 0.133 | 2.36E-47 | 11 | Rnf138 |
| Mndal4 | 1.32E-51 | 0.607659345 | 0.752 | 0.4 | 2.68E-47 | 11 | Mndal |
| Rpl7a4 | 1.37E-51 | 0.41373579 | 0.98 | 0.749 | 2.77E-47 | 11 | Rpl7a |
| Psmd14 | 1.83E-51 | 0.369947454 | 0.49 | 0.213 | 3.70E-47 | 11 | Psmd14 |
| Cab392 | 1.90E-51 | 0.383013787 | 0.585 | 0.275 | 3.84E-47 | 11 | Cab39 |
| Arfgef12 | 1.93E-51 | 0.413911524 | 0.51 | 0.223 | 3.91E-47 | 11 | Arfgef1 |
| mt-Co28 | 1.95E-51 | 0.52559978 | 1 | 0.724 | 3.94E-47 | 11 | mt-Co2 |
| Lyst1 | 2.12E-51 | 0.469021286 | 0.309 | 0.107 | 4.30E-47 | 11 | Lyst |
| Zfp593 | 2.13E-51 | 0.266824929 | 0.251 | 0.075 | 4.32E-47 | 11 | Zfp593 |
| Akr1b31 | 2.15E-51 | 0.334854769 | 0.449 | 0.185 | 4.36E-47 | 11 | Akr1b3 |
| Ncor14 | 4.01E-51 | 0.434510002 | 0.808 | 0.432 | 8.12E-47 | 11 | Ncor1 |
| Rfx31 | 4.32E-51 | 0.469440302 | 0.305 | 0.104 | 8.74E-47 | 11 | Rfx3 |
| Rps64 | 4.73E-51 | 0.432207431 | 0.991 | 0.809 | 9.58E-47 | 11 | Rps6 |
| Cerk2 | 6.95E-51 | 0.540166834 | 0.311 | 0.112 | 1.41E-46 | 11 | Cerk |
| Rpl55 | 7.19E-51 | 0.428456891 | 0.973 | 0.706 | 1.46E-46 | 11 | Rpl5 |
| Cct82 | 7.65E-51 | 0.289116082 | 0.573 | 0.262 | 1.55E-46 | 11 | Cct8 |
| Tbc1d11 | 7.73E-51 | 0.392666662 | 0.408 | 0.162 | 1.56E-46 | 11 | Tbc1d1 |
| Dbnl1 | 8.50E-51 | 0.330927774 | 0.458 | 0.188 | 1.72E-46 | 11 | Dbnl |
| Gimap83 | 9.56E-51 | 0.504663519 | 0.413 | 0.165 | 1.94E-46 | 11 | Gimap8 |
| Srgn5 | 1.21E-50 | 0.537363888 | 0.969 | 0.537 | 2.46E-46 | 11 | Srgn |
| Sltm3 | 1.23E-50 | 0.456800369 | 0.564 | 0.266 | 2.49E-46 | 11 | Sltm |
| Rapgef14 | 1.23E-50 | 0.428533797 | 0.539 | 0.236 | 2.50E-46 | 11 | Rapgef1 |
| Rbm255 | 1.49E-50 | 0.450769301 | 0.664 | 0.328 | 3.02E-46 | 11 | Rbm25 |
| Ascc34 | 1.51E-50 | 0.512078209 | 0.469 | 0.203 | 3.06E-46 | 11 | Ascc3 |
| Pik3r12 | 2.09E-50 | 0.388434412 | 0.544 | 0.249 | 4.23E-46 | 11 | Pik3r1 |
| Hnrnpa13 | 2.14E-50 | 0.299423218 | 0.657 | 0.319 | 4.33E-46 | 11 | Hnrnpa1 |
| Nsmce22 | 2.56E-50 | 0.404148847 | 0.472 | 0.202 | 5.19E-46 | 11 | Nsmce2 |
| Irak22 | 2.57E-50 | 0.456175787 | 0.366 | 0.139 | 5.21E-46 | 11 | Irak2 |
| Tma73 | 3.64E-50 | 0.252638916 | 0.898 | 0.517 | 7.38E-46 | 11 | Tma7 |
| Dyrk1a3 | 3.95E-50 | 0.460214907 | 0.427 | 0.176 | 8.00E-46 | 11 | Dyrk1a |
| Cdyl21 | 4.09E-50 | 0.430499543 | 0.275 | 0.089 | 8.29E-46 | 11 | Cdyl2 |
| Stag15 | 4.97E-50 | 0.504361449 | 0.695 | 0.356 | 1.01E-45 | 11 | Stag1 |
| Ptpn41 | 7.99E-50 | 0.44499022 | 0.346 | 0.128 | 1.62E-45 | 11 | Ptpn4 |
| Iqsec11 | 8.49E-50 | 0.441185092 | 0.329 | 0.119 | 1.72E-45 | 11 | Iqsec1 |
| Gls3 | 1.26E-49 | 0.507465106 | 0.531 | 0.247 | 2.55E-45 | 11 | Gls |
| Zhx2 | 2.06E-49 | 0.402475355 | 0.253 | 0.079 | 4.17E-45 | 11 | Zhx2 |
| Luc7l24 | 2.92E-49 | 0.463937086 | 0.758 | 0.399 | 5.90E-45 | 11 | Luc7l2 |
| Rab27a1 | 2.93E-49 | 0.307770383 | 0.262 | 0.082 | 5.94E-45 | 11 | Rab27a |
| Gne1 | 6.08E-49 | 0.409379449 | 0.332 | 0.121 | 1.23E-44 | 11 | Gne |

| | | | | | | | |
|---|---|---|---|---|---|---|---|
| Foxp13 | 6.94E-49 | 0.473531673 | 0.738 | 0.384 | 1.41E-44 | 11 | Foxp1 |
| Arid1a4 | 7.78E-49 | 0.463873717 | 0.499 | 0.226 | 1.57E-44 | 11 | Arid1a |
| Arid4b4 | 8.74E-49 | 0.463328475 | 0.594 | 0.288 | 1.77E-44 | 11 | Arid4b |
| Celf12 | 9.02E-49 | 0.419965975 | 0.492 | 0.216 | 1.83E-44 | 11 | Celf1 |
| Riok11 | 1.02E-48 | 0.381300275 | 0.316 | 0.113 | 2.06E-44 | 11 | Riok1 |
| Smc51 | 2.90E-48 | 0.421708082 | 0.32 | 0.116 | 5.87E-44 | 11 | Smc5 |
| Rlf2 | 3.34E-48 | 0.421831376 | 0.35 | 0.134 | 6.75E-44 | 11 | Rlf |
| Grk22 | 3.48E-48 | 0.376505672 | 0.33 | 0.122 | 7.04E-44 | 11 | Grk2 |
| Snd13 | 6.04E-48 | 0.415737725 | 0.411 | 0.169 | 1.22E-43 | 11 | Snd1 |
| Fbl1 | 7.31E-48 | 0.343294273 | 0.341 | 0.127 | 1.48E-43 | 11 | Fbl |
| Nek73 | 8.00E-48 | 0.431698619 | 0.524 | 0.244 | 1.62E-43 | 11 | Nek7 |
| Arih13 | 8.26E-48 | 0.484019955 | 0.618 | 0.308 | 1.67E-43 | 11 | Arih1 |
| Tex21 | 8.82E-48 | 0.39054458 | 0.264 | 0.086 | 1.79E-43 | 11 | Tex2 |
| Eef1d2 | 9.65E-48 | 0.340096654 | 0.901 | 0.544 | 1.95E-43 | 11 | Eef1d |
| Sirt71 | 1.69E-47 | 0.395581254 | 0.343 | 0.131 | 3.43E-43 | 11 | Sirt7 |
| Wwp22 | 1.71E-47 | 0.452694136 | 0.33 | 0.124 | 3.45E-43 | 11 | Wwp2 |
| Tiprl1 | 2.38E-47 | 0.321990442 | 0.472 | 0.206 | 4.82E-43 | 11 | Tiprl |
| Ddx39b1 | 3.16E-47 | 0.31740831 | 0.512 | 0.229 | 6.41E-43 | 11 | Ddx39b |
| mt-Nd16 | 4.06E-47 | 0.509118747 | 0.948 | 0.593 | 8.22E-43 | 11 | mt-Nd1 |
| Reps11 | 9.00E-47 | 0.3609214 | 0.285 | 0.098 | 1.82E-42 | 11 | Reps1 |
| Arpc5 | 9.35E-47 | 0.296114526 | 0.795 | 0.422 | 1.89E-42 | 11 | Arpc5 |
| Ggnbp23 | 1.23E-46 | 0.453383389 | 0.515 | 0.238 | 2.49E-42 | 11 | Ggnbp2 |
| Tbl1xr12 | 1.55E-46 | 0.428789847 | 0.393 | 0.161 | 3.13E-42 | 11 | Tbl1xr1 |
| Klf124 | 2.28E-46 | 0.538240386 | 0.461 | 0.204 | 4.61E-42 | 11 | Klf12 |
| Glipr12 | 3.24E-46 | 0.279139538 | 0.28 | 0.093 | 6.56E-42 | 11 | Glipr1 |
| Tm6sf13 | 3.34E-46 | 0.371202337 | 0.357 | 0.138 | 6.77E-42 | 11 | Tm6sf1 |
| Acin13 | 3.37E-46 | 0.387618324 | 0.627 | 0.311 | 6.83E-42 | 11 | Acin1 |
| Dusp112 | 5.34E-46 | 0.345414333 | 0.58 | 0.277 | 1.08E-41 | 11 | Dusp11 |
| Fosb3 | 5.60E-46 | 0.517702955 | 0.431 | 0.186 | 1.13E-41 | 11 | Fosb |
| Phf34 | 7.82E-46 | 0.422761641 | 0.53 | 0.249 | 1.58E-41 | 11 | Phf3 |
| Eif4a12 | 9.38E-46 | 0.448912566 | 0.752 | 0.425 | 1.90E-41 | 11 | Eif4a1 |
| Sipa1l12 | 9.38E-46 | 0.389442481 | 0.442 | 0.186 | 1.90E-41 | 11 | Sipa1l1 |
| Atp1b33 | 1.05E-45 | 0.406931615 | 0.524 | 0.245 | 2.13E-41 | 11 | Atp1b3 |
| Cxxc52 | 1.06E-45 | 0.278074525 | 0.309 | 0.112 | 2.14E-41 | 11 | Cxxc5 |
| Nab11 | 1.07E-45 | 0.345362671 | 0.402 | 0.164 | 2.17E-41 | 11 | Nab1 |
| Lbh1 | 1.15E-45 | 0.369866853 | 0.339 | 0.129 | 2.32E-41 | 11 | Lbh |
| Wdr431 | 1.26E-45 | 0.407742887 | 0.366 | 0.147 | 2.54E-41 | 11 | Wdr43 |
| Srpk1 | 1.52E-45 | 0.303939574 | 0.391 | 0.158 | 3.08E-41 | 11 | Srpk1 |
| Pbxip12 | 2.04E-45 | 0.415202989 | 0.309 | 0.114 | 4.13E-41 | 11 | Pbxip1 |
| Atxn7l11 | 2.47E-45 | 0.428628508 | 0.338 | 0.13 | 5.00E-41 | 11 | Atxn7l1 |
| Heca1 | 2.57E-45 | 0.402592973 | 0.355 | 0.14 | 5.20E-41 | 11 | Heca |
| Skil1 | 2.81E-45 | 0.389540514 | 0.406 | 0.17 | 5.70E-41 | 11 | Skil |
| Anp32b1 | 3.00E-45 | 0.300294777 | 0.758 | 0.417 | 6.08E-41 | 11 | Anp32b |

| | | | | | | | |
|---|---|---|---|---|---|---|---|
| Kcnq1ot12 | 3.90E-45 | 0.564962513 | 0.338 | 0.131 | 7.89E-41 | 11 | Kcnq1ot1 |
| Taok32 | 4.95E-45 | 0.36195785 | 0.391 | 0.16 | 1.00E-40 | 11 | Taok3 |
| Ttc7b1 | 6.35E-45 | 0.415958715 | 0.32 | 0.119 | 1.29E-40 | 11 | Ttc7b |
| Cct5 | 9.12E-45 | 0.297786803 | 0.587 | 0.29 | 1.85E-40 | 11 | Cct5 |
| Ifnar13 | 9.67E-45 | 0.40582764 | 0.357 | 0.141 | 1.96E-40 | 11 | Ifnar1 |
| Zc3hav12 | 1.13E-44 | 0.406371669 | 0.451 | 0.2 | 2.29E-40 | 11 | Zc3hav1 |
| Rplp13 | 1.33E-44 | 0.278971188 | 1 | 0.982 | 2.68E-40 | 11 | Rplp1 |
| Zfp2924 | 1.55E-44 | 0.461690049 | 0.521 | 0.247 | 3.14E-40 | 11 | Zfp292 |
| Kmt2a5 | 1.63E-44 | 0.429769829 | 0.551 | 0.263 | 3.29E-40 | 11 | Kmt2a |
| Thrap32 | 2.02E-44 | 0.388486139 | 0.585 | 0.289 | 4.08E-40 | 11 | Thrap3 |
| 4930453N24Rik1 | 2.15E-44 | 0.280128804 | 0.404 | 0.167 | 4.35E-40 | 11 | 4930453N24Rik |
| Ppil21 | 4.75E-44 | 0.386877652 | 0.361 | 0.147 | 9.61E-40 | 11 | Ppil2 |
| Smg63 | 6.51E-44 | 0.44035277 | 0.535 | 0.256 | 1.32E-39 | 11 | Smg6 |
| Parp83 | 6.86E-44 | 0.482865726 | 0.364 | 0.146 | 1.39E-39 | 11 | Parp8 |
| Jund2 | 7.53E-44 | 0.500893248 | 0.968 | 0.64 | 1.53E-39 | 11 | Jund |
| Set1 | 8.65E-44 | 0.318849064 | 0.824 | 0.468 | 1.75E-39 | 11 | Set |
| Smap22 | 9.36E-44 | 0.305929086 | 0.467 | 0.208 | 1.90E-39 | 11 | Smap2 |
| Hnrnpf1 | 1.46E-43 | 0.365539382 | 0.84 | 0.49 | 2.95E-39 | 11 | Hnrnpf |
| Ddx391 | 1.74E-43 | 0.286830112 | 0.327 | 0.123 | 3.52E-39 | 11 | Ddx39 |
| Esco12 | 1.83E-43 | 0.375737373 | 0.413 | 0.176 | 3.71E-39 | 11 | Esco1 |
| Eif5a3 | 2.24E-43 | 0.460211059 | 0.887 | 0.56 | 4.54E-39 | 11 | Eif5a |
| Cd1642 | 3.10E-43 | 0.333823778 | 0.467 | 0.21 | 6.28E-39 | 11 | Cd164 |
| Ptp4a11 | 3.56E-43 | 0.396130002 | 0.35 | 0.141 | 7.22E-39 | 11 | Ptp4a1 |
| Dcaf51 | 5.19E-43 | 0.371613683 | 0.33 | 0.127 | 1.05E-38 | 11 | Dcaf5 |
| Litaf2 | 6.19E-43 | 0.469103442 | 0.368 | 0.155 | 1.25E-38 | 11 | Litaf |
| Dtnb1 | 9.07E-43 | 0.334814253 | 0.303 | 0.112 | 1.84E-38 | 11 | Dtnb |
| Jarid22 | 1.42E-42 | 0.361506069 | 0.451 | 0.2 | 2.88E-38 | 11 | Jarid2 |
| Phf144 | 1.70E-42 | 0.404278953 | 0.508 | 0.24 | 3.44E-38 | 11 | Phf14 |
| Rfc21 | 1.85E-42 | 0.308892269 | 0.321 | 0.122 | 3.74E-38 | 11 | Rfc2 |
| Ifi473 | 2.01E-42 | 0.316032857 | 0.417 | 0.175 | 4.07E-38 | 11 | Ifi47 |
| Trp531 | 2.40E-42 | 0.376108229 | 0.373 | 0.155 | 4.85E-38 | 11 | Trp53 |
| Uimc11 | 2.46E-42 | 0.362919336 | 0.359 | 0.146 | 4.98E-38 | 11 | Uimc1 |
| Atxn14 | 8.46E-42 | 0.536442198 | 0.578 | 0.291 | 1.71E-37 | 11 | Atxn1 |
| Zfp3181 | 1.06E-41 | 0.33239995 | 0.271 | 0.095 | 2.14E-37 | 11 | Zfp318 |
| Nup1531 | 1.19E-41 | 0.368737093 | 0.302 | 0.114 | 2.41E-37 | 11 | Nup153 |
| Spata52 | 1.31E-41 | 0.40980003 | 0.312 | 0.121 | 2.65E-37 | 11 | Spata5 |
| Prrc2c5 | 2.06E-41 | 0.361645166 | 0.776 | 0.415 | 4.18E-37 | 11 | Prrc2c |
| Snrnp701 | 2.91E-41 | 0.345201117 | 0.546 | 0.268 | 5.88E-37 | 11 | Snrnp70 |
| Usp481 | 3.05E-41 | 0.363034773 | 0.321 | 0.126 | 6.17E-37 | 11 | Usp48 |
| mt-Nd25 | 3.71E-41 | 0.445963974 | 0.996 | 0.683 | 7.52E-37 | 11 | mt-Nd2 |
| Chd43 | 3.94E-41 | 0.375260959 | 0.621 | 0.323 | 7.98E-37 | 11 | Chd4 |
| Snx52 | 5.02E-41 | 0.277505384 | 0.461 | 0.207 | 1.02E-36 | 11 | Snx5 |
| Arf61 | 6.52E-41 | 0.280300601 | 0.406 | 0.175 | 1.32E-36 | 11 | Arf6 |

| | | | | | | | |
|---|---|---|---|---|---|---|---|
| Ets14 | 7.47E-41 | 0.359383616 | 0.763 | 0.398 | 1.51E-36 | 11 | Ets1 |
| Cnot31 | 8.43E-41 | 0.346344929 | 0.415 | 0.182 | 1.71E-36 | 11 | Cnot3 |
| Zmat5 | 8.80E-41 | 0.250373115 | 0.373 | 0.154 | 1.78E-36 | 11 | Zmat5 |
| Arhgap174 | 9.32E-41 | 0.440753317 | 0.537 | 0.266 | 1.89E-36 | 11 | Arhgap17 |
| Fam102a1 | 1.38E-40 | 0.281142389 | 0.273 | 0.098 | 2.79E-36 | 11 | Fam102a |
| St8sia44 | 1.97E-40 | 0.439668103 | 0.526 | 0.255 | 3.98E-36 | 11 | St8sia4 |
| Pan35 | 3.02E-40 | 0.478489979 | 0.476 | 0.225 | 6.12E-36 | 11 | Pan3 |
| Hnrnpl1 | 5.46E-40 | 0.34968771 | 0.487 | 0.231 | 1.10E-35 | 11 | Hnrnpl |
| Cct42 | 6.26E-40 | 0.272505196 | 0.603 | 0.311 | 1.27E-35 | 11 | Cct4 |
| Rpl104 | 6.75E-40 | 0.2748554 | 0.991 | 0.895 | 1.37E-35 | 11 | Rpl10 |
| Sbno12 | 7.19E-40 | 0.391655956 | 0.445 | 0.206 | 1.46E-35 | 11 | Sbno1 |
| Trip124 | 8.41E-40 | 0.378189103 | 0.515 | 0.253 | 1.70E-35 | 11 | Trip12 |
| Nop581 | 1.04E-39 | 0.377665596 | 0.332 | 0.135 | 2.12E-35 | 11 | Nop58 |
| Rcc2 | 1.25E-39 | 0.289237357 | 0.289 | 0.11 | 2.53E-35 | 11 | Rcc2 |
| Rab8a1 | 1.56E-39 | 0.260086135 | 0.492 | 0.233 | 3.15E-35 | 11 | Rab8a |
| Usp152 | 1.78E-39 | 0.38439162 | 0.478 | 0.227 | 3.61E-35 | 11 | Usp15 |
| Rnf1304 | 1.80E-39 | 0.345578736 | 0.56 | 0.28 | 3.65E-35 | 11 | Rnf130 |
| Tpr3 | 1.96E-39 | 0.385156683 | 0.741 | 0.421 | 3.97E-35 | 11 | Tpr |
| Smad31 | 2.00E-39 | 0.363884336 | 0.26 | 0.094 | 4.05E-35 | 11 | Smad3 |
| Actg12 | 2.07E-39 | 0.475385804 | 0.986 | 0.791 | 4.20E-35 | 11 | Actg1 |
| Arhgap102 | 2.16E-39 | 0.349484358 | 0.316 | 0.125 | 4.37E-35 | 11 | Arhgap10 |
| Tra2a5 | 2.37E-39 | 0.340554337 | 0.551 | 0.27 | 4.80E-35 | 11 | Tra2a |
| Nt5dc11 | 3.10E-39 | 0.351966435 | 0.303 | 0.118 | 6.27E-35 | 11 | Nt5dc1 |
| Tcf203 | 4.30E-39 | 0.388130291 | 0.336 | 0.139 | 8.71E-35 | 11 | Tcf20 |
| Senp72 | 6.09E-39 | 0.386255807 | 0.361 | 0.154 | 1.23E-34 | 11 | Senp7 |
| Lman21 | 7.21E-39 | 0.305465258 | 0.442 | 0.203 | 1.46E-34 | 11 | Lman2 |
| Tbc1d54 | 7.73E-39 | 0.457907132 | 0.515 | 0.258 | 1.56E-34 | 11 | Tbc1d5 |
| Srpk2 | 8.67E-39 | 0.342776005 | 0.42 | 0.19 | 1.76E-34 | 11 | Srpk2 |
| Ankib12 | 9.06E-39 | 0.39470874 | 0.33 | 0.136 | 1.83E-34 | 11 | Ankib1 |
| Cdc42ep33 | 9.16E-39 | 0.389862917 | 0.391 | 0.172 | 1.85E-34 | 11 | Cdc42ep3 |
| Smc63 | 9.18E-39 | 0.307978857 | 0.531 | 0.258 | 1.86E-34 | 11 | Smc6 |
| Map3k22 | 1.06E-38 | 0.403166802 | 0.354 | 0.151 | 2.14E-34 | 11 | Map3k2 |
| Stat35 | 1.10E-38 | 0.403894299 | 0.53 | 0.263 | 2.22E-34 | 11 | Stat3 |
| Prpf4b4 | 1.43E-38 | 0.329210882 | 0.623 | 0.329 | 2.89E-34 | 11 | Prpf4b |
| Rsrc13 | 1.69E-38 | 0.357176207 | 0.521 | 0.256 | 3.41E-34 | 11 | Rsrc1 |
| Tmod34 | 2.03E-38 | 0.327982145 | 0.671 | 0.356 | 4.11E-34 | 11 | Tmod3 |
| Thada3 | 2.07E-38 | 0.386109882 | 0.298 | 0.117 | 4.18E-34 | 11 | Thada |
| Lasp11 | 2.24E-38 | 0.287837816 | 0.443 | 0.205 | 4.53E-34 | 11 | Lasp1 |
| Nptn4 | 2.46E-38 | 0.376447621 | 0.503 | 0.245 | 4.99E-34 | 11 | Nptn |
| Sep-76 | 3.34E-38 | 0.325717375 | 0.801 | 0.456 | 6.75E-34 | 11 | 7-Sep |
| Pdcd6ip1 | 4.00E-38 | 0.339373647 | 0.47 | 0.226 | 8.09E-34 | 11 | Pdcd6ip |
| Tapbpl | 4.18E-38 | 0.327618499 | 0.316 | 0.126 | 8.46E-34 | 11 | Tapbpl |
| Psmb94 | 4.24E-38 | 0.294498762 | 0.741 | 0.4 | 8.58E-34 | 11 | Psmb9 |

| | | | | | | | |
|---|---|---|---|---|---|---|---|
| Tmem128 | 4.32E-38 | 0.292883755 | 0.379 | 0.162 | 8.74E-34 | 11 | Tmem128 |
| Stim13 | 6.25E-38 | 0.432824485 | 0.413 | 0.188 | 1.27E-33 | 11 | Stim1 |
| Ccnl12 | 6.64E-38 | 0.380902798 | 0.438 | 0.207 | 1.35E-33 | 11 | Ccnl1 |
| Psmb1 | 8.20E-38 | 0.269114629 | 0.847 | 0.505 | 1.66E-33 | 11 | Psmb1 |
| Cdk62 | 1.09E-37 | 0.360113016 | 0.287 | 0.109 | 2.20E-33 | 11 | Cdk6 |
| Dennd4c2 | 1.12E-37 | 0.366318817 | 0.307 | 0.124 | 2.26E-33 | 11 | Dennd4c |
| Micu21 | 1.68E-37 | 0.298078529 | 0.411 | 0.186 | 3.39E-33 | 11 | Micu2 |
| Immp2l2 | 1.87E-37 | 0.521989678 | 0.321 | 0.134 | 3.79E-33 | 11 | Immp2l |
| Rap1gds15 | 2.30E-37 | 0.396038678 | 0.521 | 0.257 | 4.67E-33 | 11 | Rap1gds1 |
| Birc65 | 2.64E-37 | 0.358202102 | 0.636 | 0.336 | 5.35E-33 | 11 | Birc6 |
| Vgll43 | 2.77E-37 | 0.37926609 | 0.497 | 0.246 | 5.60E-33 | 11 | Vgll4 |
| Ankrd175 | 2.94E-37 | 0.428315021 | 0.512 | 0.257 | 5.96E-33 | 11 | Ankrd17 |
| Ankrd125 | 3.02E-37 | 0.400623298 | 0.582 | 0.304 | 6.11E-33 | 11 | Ankrd12 |
| Max | 3.28E-37 | 0.301335047 | 0.35 | 0.149 | 6.65E-33 | 11 | Max |
| Ambra14 | 3.73E-37 | 0.419777899 | 0.42 | 0.195 | 7.56E-33 | 11 | Ambra1 |
| Brd45 | 4.21E-37 | 0.409893529 | 0.522 | 0.264 | 8.53E-33 | 11 | Brd4 |
| Sft2d2 | 4.35E-37 | 0.310045401 | 0.269 | 0.101 | 8.80E-33 | 11 | Sft2d2 |
| Nr2c23 | 6.39E-37 | 0.401516417 | 0.357 | 0.156 | 1.29E-32 | 11 | Nr2c2 |
| Cdc422 | 6.70E-37 | 0.355088755 | 0.937 | 0.61 | 1.36E-32 | 11 | Cdc42 |
| Ints62 | 6.84E-37 | 0.353811896 | 0.334 | 0.139 | 1.39E-32 | 11 | Ints6 |
| Vav33 | 6.88E-37 | 0.448637201 | 0.506 | 0.252 | 1.39E-32 | 11 | Vav3 |
| Fam204a1 | 7.11E-37 | 0.25190922 | 0.381 | 0.167 | 1.44E-32 | 11 | Fam204a |
| Kdm4c3 | 7.81E-37 | 0.353818437 | 0.377 | 0.166 | 1.58E-32 | 11 | Kdm4c |
| Traf51 | 8.08E-37 | 0.342984442 | 0.294 | 0.116 | 1.64E-32 | 11 | Traf5 |
| Pnn4 | 9.94E-37 | 0.359189157 | 0.524 | 0.263 | 2.01E-32 | 11 | Pnn |
| Ube2j11 | 1.02E-36 | 0.395226207 | 0.357 | 0.159 | 2.07E-32 | 11 | Ube2j1 |
| Krtcap22 | 1.38E-36 | 0.295784606 | 0.743 | 0.437 | 2.79E-32 | 11 | Krtcap2 |
| Prpf40a4 | 2.18E-36 | 0.345490708 | 0.522 | 0.263 | 4.42E-32 | 11 | Prpf40a |
| Itgav3 | 3.10E-36 | 0.452069599 | 0.357 | 0.158 | 6.27E-32 | 11 | Itgav |
| Birc21 | 3.91E-36 | 0.302524584 | 0.28 | 0.108 | 7.92E-32 | 11 | Birc2 |
| Wnk15 | 3.96E-36 | 0.33947504 | 0.666 | 0.359 | 8.03E-32 | 11 | Wnk1 |
| Faf12 | 3.98E-36 | 0.358563138 | 0.422 | 0.198 | 8.05E-32 | 11 | Faf1 |
| Gon4l1 | 5.26E-36 | 0.324398926 | 0.343 | 0.146 | 1.06E-31 | 11 | Gon4l |
| Atad21 | 7.08E-36 | 0.301459123 | 0.25 | 0.091 | 1.43E-31 | 11 | Atad2 |
| Senp63 | 9.56E-36 | 0.381903101 | 0.555 | 0.294 | 1.94E-31 | 11 | Senp6 |
| Gm4070 | 1.00E-35 | 0.335465644 | 0.259 | 0.097 | 2.03E-31 | 11 | Gm4070 |
| Mpp71 | 1.07E-35 | 0.284415101 | 0.273 | 0.105 | 2.17E-31 | 11 | Mpp7 |
| Crlf22 | 1.33E-35 | 0.264387495 | 0.364 | 0.159 | 2.70E-31 | 11 | Crlf2 |
| Trappc83 | 1.65E-35 | 0.382583643 | 0.366 | 0.163 | 3.35E-31 | 11 | Trappc8 |
| Metrnl2 | 2.25E-35 | 0.31084363 | 0.264 | 0.101 | 4.55E-31 | 11 | Metrnl |
| Psip11 | 2.31E-35 | 0.300873663 | 0.413 | 0.191 | 4.68E-31 | 11 | Psip1 |
| Efr3a2 | 2.39E-35 | 0.319538012 | 0.415 | 0.195 | 4.85E-31 | 11 | Efr3a |
| Serbp13 | 2.42E-35 | 0.324798923 | 0.894 | 0.585 | 4.91E-31 | 11 | Serbp1 |

| | | | | | | | |
|---|---|---|---|---|---|---|---|
| Chfr1 | 3.08E-35 | 0.32700684 | 0.287 | 0.114 | 6.24E-31 | 11 | Chfr |
| Taf1d1 | 3.21E-35 | 0.302478659 | 0.334 | 0.142 | 6.50E-31 | 11 | Taf1d |
| Jmjd1c6 | 3.41E-35 | 0.391100598 | 0.573 | 0.303 | 6.90E-31 | 11 | Jmjd1c |
| Psme15 | 4.07E-35 | 0.347392387 | 0.898 | 0.563 | 8.24E-31 | 11 | Psme1 |
| Srsf116 | 4.08E-35 | 0.329178747 | 0.639 | 0.347 | 8.26E-31 | 11 | Srsf11 |
| Lims13 | 4.62E-35 | 0.387900079 | 0.56 | 0.297 | 9.36E-31 | 11 | Lims1 |
| Traf32 | 5.27E-35 | 0.408040255 | 0.271 | 0.107 | 1.07E-30 | 11 | Traf3 |
| 4932438A13Rik3 | 6.27E-35 | 0.387059085 | 0.445 | 0.216 | 1.27E-30 | 11 | 4932438A13Rik |
| Mllt104 | 6.30E-35 | 0.36866999 | 0.528 | 0.271 | 1.28E-30 | 11 | Mllt10 |
| Sh2d3c3 | 7.55E-35 | 0.35424475 | 0.352 | 0.154 | 1.53E-30 | 11 | Sh2d3c |
| Bcor | 7.59E-35 | 0.298388449 | 0.296 | 0.118 | 1.54E-30 | 11 | Bcor |
| Ppp6r11 | 8.49E-35 | 0.27993611 | 0.26 | 0.099 | 1.72E-30 | 11 | Ppp6r1 |
| B2m7 | 1.12E-34 | 0.410612692 | 0.996 | 0.685 | 2.26E-30 | 11 | B2m |
| Cdk134 | 1.13E-34 | 0.393237858 | 0.452 | 0.224 | 2.29E-30 | 11 | Cdk13 |
| Gpatch85 | 1.67E-34 | 0.358923618 | 0.504 | 0.255 | 3.37E-30 | 11 | Gpatch8 |
| Nipa21 | 1.78E-34 | 0.262100923 | 0.311 | 0.128 | 3.60E-30 | 11 | Nipa2 |
| Dennd1a4 | 1.78E-34 | 0.403207232 | 0.496 | 0.25 | 3.61E-30 | 11 | Dennd1a |
| Cdkal13 | 2.12E-34 | 0.333922697 | 0.39 | 0.178 | 4.28E-30 | 11 | Cdkal1 |
| Thoc1 | 2.54E-34 | 0.294327018 | 0.289 | 0.115 | 5.14E-30 | 11 | Thoc1 |
| Hspa55 | 3.47E-34 | 0.378430593 | 0.736 | 0.43 | 7.02E-30 | 11 | Hspa5 |
| Plp2 | 3.65E-34 | 0.267007173 | 0.262 | 0.1 | 7.38E-30 | 11 | Plp2 |
| Map3k142 | 4.52E-34 | 0.382226927 | 0.264 | 0.104 | 9.16E-30 | 11 | Map3k14 |
| Zfp800 | 5.48E-34 | 0.355059969 | 0.287 | 0.116 | 1.11E-29 | 11 | Zfp800 |
| H2afz2 | 5.65E-34 | 0.266740265 | 0.923 | 0.603 | 1.14E-29 | 11 | H2afz |
| mt-Nd36 | 7.22E-34 | 0.314561802 | 0.774 | 0.441 | 1.46E-29 | 11 | mt-Nd3 |
| Scaper3 | 8.27E-34 | 0.407196572 | 0.381 | 0.177 | 1.68E-29 | 11 | Scaper |
| N4bp2l22 | 8.50E-34 | 0.306804894 | 0.488 | 0.244 | 1.72E-29 | 11 | N4bp2l2 |
| Tnfrsf1b1 | 9.34E-34 | 0.361727593 | 0.282 | 0.114 | 1.89E-29 | 11 | Tnfrsf1b |
| Slbp1 | 9.98E-34 | 0.268374885 | 0.343 | 0.149 | 2.02E-29 | 11 | Slbp |
| Gnl3 | 1.06E-33 | 0.29799 | 0.273 | 0.108 | 2.15E-29 | 11 | Gnl3 |
| Gm268871 | 1.08E-33 | 0.420337782 | 0.255 | 0.098 | 2.18E-29 | 11 | Gm26887 |
| Zfp6444 | 1.38E-33 | 0.307522235 | 0.497 | 0.252 | 2.79E-29 | 11 | Zfp644 |
| Atxn102 | 1.44E-33 | 0.267453281 | 0.379 | 0.173 | 2.92E-29 | 11 | Atxn10 |
| Plscr13 | 1.67E-33 | 0.292160954 | 0.434 | 0.202 | 3.38E-29 | 11 | Plscr1 |
| Lrmda6 | 1.94E-33 | 0.331526455 | 0.661 | 0.364 | 3.93E-29 | 11 | Lrmda |
| Btbd1 | 2.07E-33 | 0.253680868 | 0.479 | 0.237 | 4.19E-29 | 11 | Btbd1 |
| Cnot43 | 2.22E-33 | 0.284485718 | 0.528 | 0.265 | 4.49E-29 | 11 | Cnot4 |
| Pear12 | 3.20E-33 | 0.289422456 | 0.26 | 0.1 | 6.48E-29 | 11 | Pear1 |
| Kpna32 | 3.33E-33 | 0.271261121 | 0.348 | 0.154 | 6.75E-29 | 11 | Kpna3 |
| Jak21 | 4.73E-33 | 0.349228016 | 0.298 | 0.125 | 9.58E-29 | 11 | Jak2 |
| 5031425E22Rik2 | 5.00E-33 | 0.311323134 | 0.354 | 0.157 | 1.01E-28 | 11 | 5031425E22Rik |
| Pcca1 | 5.40E-33 | 0.301629791 | 0.289 | 0.119 | 1.09E-28 | 11 | Pcca |
| Sik12 | 6.14E-33 | 0.345498717 | 0.298 | 0.124 | 1.24E-28 | 11 | Sik1 |

| | | | | | | | |
|---|---|---|---|---|---|---|---|
| Abr2 | 7.31E-33 | 0.329385202 | 0.35 | 0.156 | 1.48E-28 | 11 | Abr |
| Rps6ka34 | 9.70E-33 | 0.41978305 | 0.492 | 0.26 | 1.96E-28 | 11 | Rps6ka3 |
| Arpc1b3 | 1.02E-32 | 0.291114657 | 0.912 | 0.586 | 2.06E-28 | 11 | Arpc1b |
| Rhoq1 | 1.02E-32 | 0.278656991 | 0.268 | 0.106 | 2.07E-28 | 11 | Rhoq |
| Atp9b2 | 1.16E-32 | 0.328360324 | 0.346 | 0.155 | 2.34E-28 | 11 | Atp9b |
| Lrrfip21 | 1.25E-32 | 0.306516642 | 0.357 | 0.162 | 2.52E-28 | 11 | Lrrfip2 |
| Rps6ka52 | 1.53E-32 | 0.376685327 | 0.266 | 0.106 | 3.09E-28 | 11 | Rps6ka5 |
| Lrba2 | 1.55E-32 | 0.372833988 | 0.397 | 0.189 | 3.13E-28 | 11 | Lrba |
| Myo9b2 | 1.57E-32 | 0.360406381 | 0.282 | 0.117 | 3.17E-28 | 11 | Myo9b |
| Map4k45 | 1.63E-32 | 0.308442269 | 0.528 | 0.273 | 3.29E-28 | 11 | Map4k4 |
| Zfand35 | 2.16E-32 | 0.383669875 | 0.551 | 0.298 | 4.37E-28 | 11 | Zfand3 |
| Flna1 | 2.86E-32 | 0.2758488 | 0.305 | 0.129 | 5.80E-28 | 11 | Flna |
| Csnk1g32 | 3.05E-32 | 0.28418695 | 0.388 | 0.18 | 6.18E-28 | 11 | Csnk1g3 |
| Pold42 | 3.45E-32 | 0.287088171 | 0.406 | 0.197 | 6.98E-28 | 11 | Pold4 |
| Gatad2a4 | 3.56E-32 | 0.296992262 | 0.447 | 0.219 | 7.21E-28 | 11 | Gatad2a |
| Cd2741 | 4.04E-32 | 0.276413862 | 0.28 | 0.112 | 8.17E-28 | 11 | Cd274 |
| Atad2b3 | 6.44E-32 | 0.371317326 | 0.411 | 0.2 | 1.30E-27 | 11 | Atad2b |
| Supt201 | 7.38E-32 | 0.331342721 | 0.35 | 0.159 | 1.49E-27 | 11 | Supt20 |
| Sdf22 | 7.82E-32 | 0.298032503 | 0.364 | 0.168 | 1.58E-27 | 11 | Sdf2 |
| Cyth11 | 8.70E-32 | 0.276188595 | 0.393 | 0.184 | 1.76E-27 | 11 | Cyth1 |
| Pspc11 | 9.31E-32 | 0.29104434 | 0.282 | 0.116 | 1.88E-27 | 11 | Pspc1 |
| Itsn24 | 1.04E-31 | 0.35960448 | 0.521 | 0.274 | 2.11E-27 | 11 | Itsn2 |
| Bcap314 | 1.10E-31 | 0.283014097 | 0.526 | 0.282 | 2.23E-27 | 11 | Bcap31 |
| Arhgap212 | 1.17E-31 | 0.284936988 | 0.271 | 0.108 | 2.36E-27 | 11 | Arhgap21 |
| Wac4 | 1.26E-31 | 0.32840994 | 0.583 | 0.319 | 2.54E-27 | 11 | Wac |
| Trim12c | 1.26E-31 | 0.275836515 | 0.316 | 0.136 | 2.54E-27 | 11 | Trim12c |
| Srsf72 | 1.57E-31 | 0.29848769 | 0.598 | 0.329 | 3.17E-27 | 11 | Srsf7 |
| Abi14 | 1.90E-31 | 0.330004048 | 0.508 | 0.266 | 3.85E-27 | 11 | Abi1 |
| Kif2a1 | 2.25E-31 | 0.291005978 | 0.375 | 0.175 | 4.56E-27 | 11 | Kif2a |
| Fli14 | 2.40E-31 | 0.272913364 | 0.729 | 0.397 | 4.87E-27 | 11 | Fli1 |
| Nop561 | 2.41E-31 | 0.280132846 | 0.303 | 0.129 | 4.89E-27 | 11 | Nop56 |
| Zc3h7a3 | 2.88E-31 | 0.346966121 | 0.348 | 0.16 | 5.84E-27 | 11 | Zc3h7a |
| Synrg1 | 3.70E-31 | 0.264354405 | 0.32 | 0.141 | 7.50E-27 | 11 | Synrg |
| Kdm6a4 | 4.03E-31 | 0.35656953 | 0.422 | 0.211 | 8.15E-27 | 11 | Kdm6a |
| Setd25 | 4.98E-31 | 0.316745097 | 0.395 | 0.19 | 1.01E-26 | 11 | Setd2 |
| Mapk15 | 6.59E-31 | 0.308330051 | 0.458 | 0.233 | 1.33E-26 | 11 | Mapk1 |
| Nudcd32 | 7.46E-31 | 0.275402844 | 0.472 | 0.243 | 1.51E-26 | 11 | Nudcd3 |
| Wdr333 | 8.80E-31 | 0.293174918 | 0.463 | 0.236 | 1.78E-26 | 11 | Wdr33 |
| Ccar13 | 1.15E-30 | 0.290931271 | 0.417 | 0.205 | 2.34E-26 | 11 | Ccar1 |
| Rere4 | 1.29E-30 | 0.263913718 | 0.515 | 0.268 | 2.60E-26 | 11 | Rere |
| Pak23 | 1.45E-30 | 0.253128575 | 0.736 | 0.43 | 2.93E-26 | 11 | Pak2 |
| Kdm5a1 | 1.54E-30 | 0.25807203 | 0.359 | 0.165 | 3.12E-26 | 11 | Kdm5a |
| Gbf13 | 1.74E-30 | 0.334949199 | 0.373 | 0.179 | 3.52E-26 | 11 | Gbf1 |

| | | | | | | | |
|---|---|---|---|---|---|---|---|
| Pi4ka2 | 2.29E-30 | 0.295704666 | 0.293 | 0.126 | 4.63E-26 | 11 | Pi4ka |
| Rock14 | 2.84E-30 | 0.323911797 | 0.74 | 0.428 | 5.75E-26 | 11 | Rock1 |
| Tmbim64 | 2.88E-30 | 0.276301899 | 0.736 | 0.426 | 5.84E-26 | 11 | Tmbim6 |
| Myo1e4 | 2.94E-30 | 0.339634413 | 0.528 | 0.278 | 5.96E-26 | 11 | Myo1e |
| Tecpr11 | 3.71E-30 | 0.283828623 | 0.3 | 0.128 | 7.51E-26 | 11 | Tecpr1 |
| Sos22 | 4.28E-30 | 0.321311401 | 0.343 | 0.159 | 8.66E-26 | 11 | Sos2 |
| Pip4k2a3 | 4.28E-30 | 0.288641324 | 0.47 | 0.238 | 8.67E-26 | 11 | Pip4k2a |
| Usp345 | 4.51E-30 | 0.311210554 | 0.612 | 0.34 | 9.13E-26 | 11 | Usp34 |
| Abcb71 | 4.55E-30 | 0.32060764 | 0.266 | 0.11 | 9.22E-26 | 11 | Abcb7 |
| Upf11 | 5.09E-30 | 0.265023874 | 0.268 | 0.111 | 1.03E-25 | 11 | Upf1 |
| Macf15 | 5.94E-30 | 0.303516682 | 0.851 | 0.497 | 1.20E-25 | 11 | Macf1 |
| Ifi2035 | 7.44E-30 | 0.300964644 | 0.654 | 0.363 | 1.51E-25 | 11 | Ifi203 |
| Fbxw7 | 7.53E-30 | 0.334164196 | 0.296 | 0.13 | 1.53E-25 | 11 | Fbxw7 |
| Lrrc8c3 | 8.25E-30 | 0.283136561 | 0.585 | 0.307 | 1.67E-25 | 11 | Lrrc8c |
| Hdac82 | 1.10E-29 | 0.403174465 | 0.302 | 0.133 | 2.23E-25 | 11 | Hdac8 |
| Csnk1g13 | 1.13E-29 | 0.342811503 | 0.338 | 0.155 | 2.28E-25 | 11 | Csnk1g1 |
| Bcas34 | 1.18E-29 | 0.348272373 | 0.373 | 0.18 | 2.38E-25 | 11 | Bcas3 |
| Scamp3 | 1.18E-29 | 0.256008318 | 0.257 | 0.105 | 2.39E-25 | 11 | Scamp3 |
| Son6 | 1.19E-29 | 0.26371497 | 0.887 | 0.52 | 2.40E-25 | 11 | Son |
| Kdm3b2 | 1.32E-29 | 0.298968406 | 0.305 | 0.134 | 2.68E-25 | 11 | Kdm3b |
| Klf101 | 1.36E-29 | 0.417161777 | 0.339 | 0.161 | 2.76E-25 | 11 | Klf10 |
| Slfn23 | 1.75E-29 | 0.296583883 | 0.492 | 0.258 | 3.53E-25 | 11 | Slfn2 |
| Smg71 | 2.03E-29 | 0.293266117 | 0.334 | 0.154 | 4.11E-25 | 11 | Smg7 |
| Ep4003 | 2.16E-29 | 0.325826349 | 0.409 | 0.205 | 4.37E-25 | 11 | Ep400 |
| Wdr262 | 2.46E-29 | 0.264184257 | 0.413 | 0.206 | 4.98E-25 | 11 | Wdr26 |
| Adipor22 | 2.46E-29 | 0.295595059 | 0.325 | 0.148 | 4.98E-25 | 11 | Adipor2 |
| Isg201 | 2.65E-29 | 0.273953088 | 0.269 | 0.114 | 5.36E-25 | 11 | Isg20 |
| Fubp13 | 3.02E-29 | 0.275251947 | 0.506 | 0.269 | 6.11E-25 | 11 | Fubp1 |
| Neat14 | 3.37E-29 | 0.346810766 | 0.411 | 0.206 | 6.81E-25 | 11 | Neat1 |
| Sep-93 | 4.46E-29 | 0.307322327 | 0.359 | 0.168 | 9.03E-25 | 11 | 9-Sep |
| Mcl14 | 4.52E-29 | 0.262747472 | 0.657 | 0.369 | 9.14E-25 | 11 | Mcl1 |
| Susd65 | 4.75E-29 | 0.330371103 | 0.519 | 0.279 | 9.61E-25 | 11 | Susd6 |
| Huwe14 | 1.08E-28 | 0.269174431 | 0.524 | 0.282 | 2.19E-24 | 11 | Huwe1 |
| Chd83 | 1.75E-28 | 0.32358461 | 0.346 | 0.166 | 3.55E-24 | 11 | Chd8 |
| Krit14 | 1.91E-28 | 0.349846374 | 0.37 | 0.18 | 3.87E-24 | 11 | Krit1 |
| Stag22 | 2.63E-28 | 0.272753144 | 0.524 | 0.285 | 5.33E-24 | 11 | Stag2 |
| Yaf22 | 4.89E-28 | 0.261769788 | 0.424 | 0.212 | 9.89E-24 | 11 | Yaf2 |
| Bmt22 | 6.27E-28 | 0.300374035 | 0.275 | 0.119 | 1.27E-23 | 11 | Bmt2 |
| Parp42 | 6.41E-28 | 0.276452287 | 0.325 | 0.15 | 1.30E-23 | 11 | Parp4 |
| Hdac41 | 6.90E-28 | 0.275576326 | 0.251 | 0.104 | 1.40E-23 | 11 | Hdac4 |
| Bptf5 | 6.99E-28 | 0.262715412 | 0.539 | 0.29 | 1.42E-23 | 11 | Bptf |
| Slc3a24 | 7.37E-28 | 0.328214698 | 0.517 | 0.288 | 1.49E-23 | 11 | Slc3a2 |
| Rbm55 | 9.47E-28 | 0.286100747 | 0.452 | 0.237 | 1.92E-23 | 11 | Rbm5 |

| | | | | | | | |
|---|---|---|---|---|---|---|---|
| Akap102 | 9.85E-28 | 0.319092896 | 0.311 | 0.144 | 2.00E-23 | 11 | Akap10 |
| Papd51 | 1.01E-27 | 0.306777414 | 0.269 | 0.117 | 2.04E-23 | 11 | Papd5 |
| Zfp6384 | 1.04E-27 | 0.354638355 | 0.508 | 0.279 | 2.11E-23 | 11 | Zfp638 |
| Ppp3ca5 | 1.28E-27 | 0.311793438 | 0.673 | 0.389 | 2.60E-23 | 11 | Ppp3ca |
| Mbd54 | 1.37E-27 | 0.335946342 | 0.42 | 0.217 | 2.78E-23 | 11 | Mbd5 |
| Wrn2 | 1.40E-27 | 0.28986953 | 0.312 | 0.145 | 2.84E-23 | 11 | Wrn |
| Prpf38b4 | 1.42E-27 | 0.278828158 | 0.391 | 0.194 | 2.88E-23 | 11 | Prpf38b |
| Rai11 | 2.03E-27 | 0.282520402 | 0.255 | 0.108 | 4.11E-23 | 11 | Rai1 |
| Airn4 | 2.24E-27 | 0.290121268 | 0.47 | 0.245 | 4.54E-23 | 11 | Airn |
| Virma1 | 2.26E-27 | 0.314876684 | 0.289 | 0.13 | 4.58E-23 | 11 | Virma |
| Zfp1483 | 2.43E-27 | 0.268522724 | 0.487 | 0.262 | 4.93E-23 | 11 | Zfp148 |
| Lcorl3 | 2.47E-27 | 0.309411427 | 0.345 | 0.164 | 5.01E-23 | 11 | Lcorl |
| Peak15 | 2.69E-27 | 0.26398721 | 0.671 | 0.383 | 5.46E-23 | 11 | Peak1 |
| Ythdf33 | 2.73E-27 | 0.279085157 | 0.386 | 0.193 | 5.53E-23 | 11 | Ythdf3 |
| Dmtf12 | 2.91E-27 | 0.266474204 | 0.275 | 0.12 | 5.88E-23 | 11 | Dmtf1 |
| Hnrnpa2b11 | 3.01E-27 | 0.254534298 | 0.808 | 0.511 | 6.09E-23 | 11 | Hnrnpa2b1 |
| Raph11 | 3.06E-27 | 0.268544123 | 0.264 | 0.112 | 6.20E-23 | 11 | Raph1 |
| Klf63 | 3.58E-27 | 0.345728414 | 0.488 | 0.264 | 7.25E-23 | 11 | Klf6 |
| Naa151 | 4.82E-27 | 0.282124042 | 0.341 | 0.164 | 9.76E-23 | 11 | Naa15 |
| Phf20l14 | 5.13E-27 | 0.260945913 | 0.517 | 0.284 | 1.04E-22 | 11 | Phf20l1 |
| Rictor2 | 5.28E-27 | 0.292800258 | 0.327 | 0.155 | 1.07E-22 | 11 | Rictor |
| Ube3a2 | 5.55E-27 | 0.251931467 | 0.531 | 0.292 | 1.12E-22 | 11 | Ube3a |
| Ttc145 | 7.78E-27 | 0.274164432 | 0.415 | 0.213 | 1.58E-22 | 11 | Ttc14 |
| Eed1 | 9.70E-27 | 0.298975739 | 0.316 | 0.149 | 1.96E-22 | 11 | Eed |
| Rbm263 | 1.12E-26 | 0.275304382 | 0.431 | 0.224 | 2.26E-22 | 11 | Rbm26 |
| Pced1b | 1.24E-26 | 0.265585602 | 0.269 | 0.116 | 2.50E-22 | 11 | Pced1b |
| Abhd17b3 | 1.50E-26 | 0.303450039 | 0.379 | 0.19 | 3.04E-22 | 11 | Abhd17b |
| Rsbn12 | 1.70E-26 | 0.269464775 | 0.359 | 0.175 | 3.44E-22 | 11 | Rsbn1 |
| Trip41 | 2.10E-26 | 0.299096716 | 0.268 | 0.117 | 4.25E-22 | 11 | Trip4 |
| Ier54 | 2.75E-26 | 0.264416428 | 0.558 | 0.314 | 5.57E-22 | 11 | Ier5 |
| Igf2r2 | 2.96E-26 | 0.279178068 | 0.294 | 0.132 | 5.99E-22 | 11 | Igf2r |
| Galnt16 | 3.43E-26 | 0.277648094 | 0.366 | 0.182 | 6.95E-22 | 11 | Galnt1 |
| Zfp4073 | 3.50E-26 | 0.34517445 | 0.33 | 0.16 | 7.08E-22 | 11 | Zfp407 |
| Dip2b4 | 4.01E-26 | 0.258568849 | 0.503 | 0.274 | 8.12E-22 | 11 | Dip2b |
| Ncoa13 | 4.73E-26 | 0.287253877 | 0.404 | 0.208 | 9.57E-22 | 11 | Ncoa1 |
| Arl5b | 5.07E-26 | 0.280498102 | 0.264 | 0.115 | 1.03E-21 | 11 | Arl5b |
| Pdpk11 | 5.28E-26 | 0.255295243 | 0.305 | 0.142 | 1.07E-21 | 11 | Pdpk1 |
| Rabep13 | 5.57E-26 | 0.252419874 | 0.436 | 0.228 | 1.13E-21 | 11 | Rabep1 |
| Cdc732 | 6.20E-26 | 0.26565206 | 0.345 | 0.168 | 1.26E-21 | 11 | Cdc73 |
| Nedd4l3 | 8.44E-26 | 0.300136617 | 0.417 | 0.215 | 1.71E-21 | 11 | Nedd4l |
| Lrp103 | 8.80E-26 | 0.284133388 | 0.372 | 0.186 | 1.78E-21 | 11 | Lrp10 |
| Cbfb3 | 9.26E-26 | 0.308707031 | 0.32 | 0.154 | 1.87E-21 | 11 | Cbfb |
| Scaf84 | 1.00E-25 | 0.253127807 | 0.368 | 0.182 | 2.02E-21 | 11 | Scaf8 |

| | | | | | | | |
|---|---|---|---|---|---|---|---|
| Mgea52 | 1.11E-25 | 0.274018819 | 0.341 | 0.167 | 2.25E-21 | 11 | Mgea5 |
| Akap95 | 1.15E-25 | 0.272959484 | 0.445 | 0.235 | 2.32E-21 | 11 | Akap9 |
| Irf24 | 1.33E-25 | 0.255094285 | 0.592 | 0.338 | 2.69E-21 | 11 | Irf2 |
| Ifrd11 | 1.59E-25 | 0.253475165 | 0.429 | 0.228 | 3.22E-21 | 11 | Ifrd1 |
| Topors1 | 1.71E-25 | 0.301428084 | 0.26 | 0.116 | 3.46E-21 | 11 | Topors |
| Trim26 | 1.87E-25 | 0.267019616 | 0.251 | 0.109 | 3.79E-21 | 11 | Trim26 |
| 2310035C23Rik3 | 2.26E-25 | 0.259077218 | 0.298 | 0.138 | 4.58E-21 | 11 | 2310035C23Rik |
| Dmxl13 | 3.02E-25 | 0.289981056 | 0.364 | 0.183 | 6.11E-21 | 11 | Dmxl1 |
| 4930523C07Rik4 | 3.30E-25 | 0.310613324 | 0.413 | 0.217 | 6.68E-21 | 11 | 4930523C07Rik |
| Phip4 | 3.81E-25 | 0.291088319 | 0.47 | 0.253 | 7.72E-21 | 11 | Phip |
| Rftn14 | 3.92E-25 | 0.292540014 | 0.467 | 0.249 | 7.95E-21 | 11 | Rftn1 |
| Sf3b33 | 5.12E-25 | 0.252361252 | 0.329 | 0.159 | 1.04E-20 | 11 | Sf3b3 |
| Zdhhc183 | 5.71E-25 | 0.251517566 | 0.348 | 0.168 | 1.16E-20 | 11 | Zdhhc18 |
| Rptor2 | 7.03E-25 | 0.308286052 | 0.291 | 0.137 | 1.42E-20 | 11 | Rptor |
| Dis3l23 | 7.58E-25 | 0.296219621 | 0.294 | 0.137 | 1.53E-20 | 11 | Dis3l2 |
| Setd55 | 9.14E-25 | 0.250124677 | 0.481 | 0.261 | 1.85E-20 | 11 | Setd5 |
| Ccdc88a5 | 1.27E-24 | 0.335714724 | 0.39 | 0.202 | 2.58E-20 | 11 | Ccdc88a |
| Xpo71 | 3.08E-24 | 0.267151345 | 0.318 | 0.152 | 6.23E-20 | 11 | Xpo7 |
| Stxbp51 | 4.24E-24 | 0.257018336 | 0.278 | 0.128 | 8.59E-20 | 11 | Stxbp5 |
| Wdr441 | 5.02E-24 | 0.275543084 | 0.266 | 0.121 | 1.02E-19 | 11 | Wdr44 |
| Agpat51 | 5.24E-24 | 0.263512203 | 0.296 | 0.141 | 1.06E-19 | 11 | Agpat5 |
| Gtdc12 | 5.59E-24 | 0.393004808 | 0.307 | 0.15 | 1.13E-19 | 11 | Gtdc1 |
| Grk53 | 5.77E-24 | 0.250302695 | 0.327 | 0.156 | 1.17E-19 | 11 | Grk5 |
| Mdm41 | 6.81E-24 | 0.261823695 | 0.321 | 0.158 | 1.38E-19 | 11 | Mdm4 |
| Kmt2c5 | 1.21E-23 | 0.278836589 | 0.531 | 0.303 | 2.46E-19 | 11 | Kmt2c |
| Cep3505 | 1.41E-23 | 0.27196854 | 0.386 | 0.201 | 2.85E-19 | 11 | Cep350 |
| Os93 | 1.96E-23 | 0.293399009 | 0.364 | 0.187 | 3.97E-19 | 11 | Os9 |
| Lbr3 | 2.27E-23 | 0.305181953 | 0.309 | 0.151 | 4.59E-19 | 11 | Lbr |
| Cnot11 | 2.30E-23 | 0.274075269 | 0.339 | 0.171 | 4.66E-19 | 11 | Cnot1 |
| Galnt72 | 2.38E-23 | 0.276722715 | 0.312 | 0.154 | 4.82E-19 | 11 | Galnt7 |
| Pam4 | 2.68E-23 | 0.299623017 | 0.397 | 0.212 | 5.43E-19 | 11 | Pam |
| Pds5b2 | 6.22E-23 | 0.307918949 | 0.346 | 0.178 | 1.26E-18 | 11 | Pds5b |
| Rnf1694 | 6.64E-23 | 0.281313739 | 0.352 | 0.181 | 1.34E-18 | 11 | Rnf169 |
| Elp4 | 6.91E-23 | 0.316810572 | 0.269 | 0.126 | 1.40E-18 | 11 | Elp4 |
| Epc11 | 7.08E-23 | 0.279221926 | 0.379 | 0.201 | 1.43E-18 | 11 | Epc1 |
| Arid5b4 | 1.16E-22 | 0.31524389 | 0.485 | 0.275 | 2.35E-18 | 11 | Arid5b |
| Malat16 | 1.63E-22 | 0.252898982 | 1 | 0.721 | 3.30E-18 | 11 | Malat1 |
| Zzef12 | 2.50E-22 | 0.292790809 | 0.296 | 0.147 | 5.07E-18 | 11 | Zzef1 |
| Mtmr35 | 3.26E-22 | 0.263793954 | 0.375 | 0.199 | 6.59E-18 | 11 | Mtmr3 |
| Gigyf23 | 4.06E-22 | 0.264986508 | 0.325 | 0.166 | 8.23E-18 | 11 | Gigyf2 |
| Znrf13 | 9.68E-22 | 0.269404558 | 0.357 | 0.187 | 1.96E-17 | 11 | Znrf1 |
| Avl91 | 1.27E-21 | 0.25227322 | 0.28 | 0.136 | 2.57E-17 | 11 | Avl9 |
| Dgkz1 | 3.16E-21 | 0.276297685 | 0.268 | 0.128 | 6.40E-17 | 11 | Dgkz |

| | | | | | | | |
|---|---|---|---|---|---|---|---|
| AW5549182 | 3.32E-21 | 0.263265361 | 0.33 | 0.169 | 6.73E-17 | 11 | AW554918 |
| Arid4a5 | 5.09E-21 | 0.266683457 | 0.438 | 0.25 | 1.03E-16 | 11 | Arid4a |
| Nrf12 | 2.54E-20 | 0.274039339 | 0.271 | 0.134 | 5.15E-16 | 11 | Nrf1 |
| Pou2f1 | 9.76E-20 | 0.301715092 | 0.257 | 0.125 | 1.98E-15 | 11 | Pou2f1 |
| Zufsp2 | 1.41E-19 | 0.261265972 | 0.271 | 0.135 | 2.84E-15 | 11 | Zufsp |
| Fnbp41 | 1.69E-19 | 0.263757312 | 0.259 | 0.127 | 3.42E-15 | 11 | Fnbp4 |
| Nfat54 | 2.50E-19 | 0.299095361 | 0.454 | 0.265 | 5.06E-15 | 11 | Nfat5 |
| Tnpo12 | 5.27E-19 | 0.258470279 | 0.321 | 0.172 | 1.07E-14 | 11 | Tnpo1 |
| Ddx582 | 1.69E-18 | 0.266141575 | 0.302 | 0.159 | 3.41E-14 | 11 | Ddx58 |
| Trim30a4 | 1.70E-18 | 0.251235432 | 0.39 | 0.217 | 3.43E-14 | 11 | Trim30a |
| Wwox4 | 1.07E-17 | 0.302685225 | 0.32 | 0.174 | 2.17E-13 | 11 | Wwox |
| Fbxo114 | 1.15E-17 | 0.282084358 | 0.395 | 0.231 | 2.34E-13 | 11 | Fbxo11 |
| Pbx32 | 1.74E-15 | 0.315442207 | 0.259 | 0.139 | 3.52E-11 | 11 | Pbx3 |
| Cd3g1 | 0 | 3.393594905 | 0.742 | 0.03 | 0 | 12 | Cd3g |
| Ccl52 | 0 | 2.93569949 | 0.804 | 0.139 | 0 | 12 | Ccl5 |
| Trbc22 | 0 | 2.904379561 | 0.871 | 0.049 | 0 | 12 | Trbc2 |
| S100a41 | 0 | 2.876648935 | 0.92 | 0.113 | 0 | 12 | S100a4 |
| Cd3d | 0 | 2.830090353 | 0.686 | 0.006 | 0 | 12 | Cd3d |
| Ltb2 | 0 | 2.585502914 | 0.953 | 0.119 | 0 | 12 | Ltb |
| Gm26822 | 0 | 2.527449203 | 0.774 | 0.066 | 0 | 12 | Gm2682 |
| Skap12 | 0 | 2.471290925 | 0.964 | 0.117 | 0 | 12 | Skap1 |
| Tmsb103 | 0 | 2.417976252 | 0.998 | 0.491 | 0 | 12 | Tmsb10 |
| Ms4a4b2 | 0 | 2.407218923 | 0.89 | 0.107 | 0 | 12 | Ms4a4b |
| Cd2261 | 0 | 2.373613388 | 0.864 | 0.074 | 0 | 12 | Cd226 |
| Inpp4b2 | 0 | 2.241297063 | 0.929 | 0.189 | 0 | 12 | Inpp4b |
| Dock23 | 0 | 2.239176524 | 0.963 | 0.17 | 0 | 12 | Dock2 |
| Xcl12 | 0 | 2.238860163 | 0.757 | 0.117 | 0 | 12 | Xcl1 |
| Cd524 | 0 | 2.211137306 | 0.994 | 0.268 | 0 | 12 | Cd52 |
| Hcst2 | 0 | 2.204020249 | 0.923 | 0.13 | 0 | 12 | Hcst |
| Cxcr61 | 0 | 2.19928442 | 0.798 | 0.048 | 0 | 12 | Cxcr6 |
| Cd3e | 0 | 2.182997436 | 0.628 | 0.002 | 0 | 12 | Cd3e |
| Arhgap153 | 0 | 2.142204068 | 0.942 | 0.173 | 0 | 12 | Arhgap15 |
| Ptprc3 | 0 | 2.096976822 | 0.976 | 0.183 | 0 | 12 | Ptprc |
| Nkg72 | 0 | 2.085244437 | 0.843 | 0.113 | 0 | 12 | Nkg7 |
| Rac22 | 0 | 2.079984807 | 0.948 | 0.163 | 0 | 12 | Rac2 |
| Dock103 | 0 | 2.050180679 | 0.951 | 0.194 | 0 | 12 | Dock10 |
| Ctsw2 | 0 | 1.990248755 | 0.826 | 0.114 | 0 | 12 | Ctsw |
| Ptprcap2 | 0 | 1.94604426 | 0.933 | 0.116 | 0 | 12 | Ptprcap |
| Gimap32 | 0 | 1.943071183 | 0.811 | 0.085 | 0 | 12 | Gimap3 |
| Trbc12 | 0 | 1.932618514 | 0.596 | 0.076 | 0 | 12 | Trbc1 |
| Dennd4a3 | 0 | 1.885565052 | 0.807 | 0.176 | 0 | 12 | Dennd4a |
| Arhgdib4 | 0 | 1.862981165 | 0.957 | 0.219 | 0 | 12 | Arhgdib |
| Lgals12 | 0 | 1.837872859 | 0.688 | 0.124 | 0 | 12 | Lgals1 |

| | | | | | | | |
|---|---|---|---|---|---|---|---|
| Coro1a4 | 0 | 1.837812929 | 0.942 | 0.205 | 0 | 12 | Coro1a |
| Ptpn222 | 0 | 1.818500189 | 0.798 | 0.106 | 0 | 12 | Ptpn22 |
| Ly6c2 | 0 | 1.813642743 | 0.308 | 0.004 | 0 | 12 | Ly6c2 |
| Lck2 | 0 | 1.802230037 | 0.804 | 0.089 | 0 | 12 | Lck |
| Stk17b2 | 0 | 1.78600679 | 0.884 | 0.148 | 0 | 12 | Stk17b |
| Stat42 | 0 | 1.779514794 | 0.776 | 0.094 | 0 | 12 | Stat4 |
| Mir142hg3 | 0 | 1.778247179 | 0.869 | 0.156 | 0 | 12 | Mir142hg |
| Epsti13 | 0 | 1.77791187 | 0.809 | 0.163 | 0 | 12 | Epsti1 |
| Klrd12 | 0 | 1.76054386 | 0.68 | 0.112 | 0 | 12 | Klrd1 |
| Itk2 | 0 | 1.760488934 | 0.746 | 0.064 | 0 | 12 | Itk |
| Ikzf13 | 0 | 1.71010214 | 0.828 | 0.143 | 0 | 12 | Ikzf1 |
| Pde7a2 | 0 | 1.709014687 | 0.804 | 0.143 | 0 | 12 | Pde7a |
| Tox2 | 0 | 1.690063294 | 0.731 | 0.109 | 0 | 12 | Tox |
| Arhgap453 | 0 | 1.667273883 | 0.862 | 0.144 | 0 | 12 | Arhgap45 |
| Grap23 | 0 | 1.662832873 | 0.69 | 0.094 | 0 | 12 | Grap2 |
| 13-Sep | 0 | 1.65709525 | 0.847 | 0.132 | 0 | 12 | 1-Sep |
| Klrk12 | 0 | 1.642248169 | 0.634 | 0.091 | 0 | 12 | Klrk1 |
| Trac | 0 | 1.63609198 | 0.46 | 0.005 | 0 | 12 | Trac |
| Themis | 0 | 1.631552395 | 0.385 | 0.002 | 0 | 12 | Themis |
| Bcl11b | 0 | 1.630451955 | 0.555 | 0.017 | 0 | 12 | Bcl11b |
| Slc9a93 | 0 | 1.629447916 | 0.776 | 0.139 | 0 | 12 | Slc9a9 |
| Il2rb2 | 0 | 1.60758489 | 0.764 | 0.107 | 0 | 12 | Il2rb |
| Cd692 | 0 | 1.587507276 | 0.675 | 0.091 | 0 | 12 | Cd69 |
| Prkcq2 | 0 | 1.579297699 | 0.682 | 0.076 | 0 | 12 | Prkcq |
| Thy12 | 0 | 1.57838556 | 0.621 | 0.049 | 0 | 12 | Thy1 |
| Il18r11 | 0 | 1.578019215 | 0.389 | 0.018 | 0 | 12 | Il18r1 |
| Ikzf32 | 0 | 1.57623141 | 0.748 | 0.108 | 0 | 12 | Ikzf3 |
| Cd22 | 0 | 1.572768557 | 0.727 | 0.071 | 0 | 12 | Cd2 |
| Atp8b42 | 0 | 1.532846503 | 0.656 | 0.086 | 0 | 12 | Atp8b4 |
| Runx33 | 0 | 1.52099156 | 0.761 | 0.137 | 0 | 12 | Runx3 |
| Il7r1 | 0 | 1.513283114 | 0.387 | 0.033 | 0 | 12 | Il7r |
| Lsp14 | 0 | 1.495909719 | 0.807 | 0.126 | 0 | 12 | Lsp1 |
| Tnik3 | 0 | 1.473653896 | 0.69 | 0.102 | 0 | 12 | Tnik |
| Selplg3 | 0 | 1.458875483 | 0.785 | 0.143 | 0 | 12 | Selplg |
| Txk2 | 0 | 1.439907705 | 0.634 | 0.097 | 0 | 12 | Txk |
| Rinl2 | 0 | 1.435391274 | 0.763 | 0.119 | 0 | 12 | Rinl |
| Ikzf21 | 0 | 1.434699915 | 0.471 | 0.044 | 0 | 12 | Ikzf2 |
| Itgb72 | 0 | 1.387522506 | 0.622 | 0.05 | 0 | 12 | Itgb7 |
| Ms4a6b2 | 0 | 1.386774241 | 0.579 | 0.083 | 0 | 12 | Ms4a6b |
| Fyb3 | 0 | 1.343770497 | 0.824 | 0.172 | 0 | 12 | Fyb |
| Pag12 | 0 | 1.338646367 | 0.652 | 0.111 | 0 | 12 | Pag1 |
| Cd2472 | 0 | 1.337477977 | 0.529 | 0.037 | 0 | 12 | Cd247 |
| Apbb1ip3 | 0 | 1.303322082 | 0.779 | 0.162 | 0 | 12 | Apbb1ip |

| | | | | | | | |
|---|---|---|---|---|---|---|---|
| Rhoh3 | 0 | 1.263255414 | 0.647 | 0.093 | 0 | 12 | Rhoh |
| Gimap72 | 0 | 1.254527995 | 0.505 | 0.05 | 0 | 12 | Gimap7 |
| Cd373 | 0 | 1.235626284 | 0.679 | 0.103 | 0 | 12 | Cd37 |
| Adam191 | 0 | 1.233024936 | 0.523 | 0.072 | 0 | 12 | Adam19 |
| Cyfip22 | 0 | 1.224940085 | 0.622 | 0.081 | 0 | 12 | Cyfip2 |
| Pik3cd3 | 0 | 1.22290568 | 0.613 | 0.099 | 0 | 12 | Pik3cd |
| Il2rg2 | 0 | 1.222158122 | 0.665 | 0.106 | 0 | 12 | Il2rg |
| Sorl13 | 0 | 1.22006712 | 0.63 | 0.101 | 0 | 12 | Sorl1 |
| Sntb11 | 0 | 1.219649682 | 0.507 | 0.056 | 0 | 12 | Sntb1 |
| Cd822 | 0 | 1.205553888 | 0.581 | 0.091 | 0 | 12 | Cd82 |
| Jaml2 | 0 | 1.204617488 | 0.514 | 0.049 | 0 | 12 | Jaml |
| Cd483 | 0 | 1.202418611 | 0.703 | 0.124 | 0 | 12 | Cd48 |
| Emb1 | 0 | 1.165374497 | 0.402 | 0.034 | 0 | 12 | Emb |
| Itgal2 | 0 | 1.150645806 | 0.609 | 0.076 | 0 | 12 | Itgal |
| Tbx212 | 0 | 1.10999199 | 0.499 | 0.058 | 0 | 12 | Tbx21 |
| Slc38a12 | 0 | 1.107776043 | 0.548 | 0.079 | 0 | 12 | Slc38a1 |
| Tespa12 | 0 | 1.097863471 | 0.533 | 0.068 | 0 | 12 | Tespa1 |
| Cxcr31 | 0 | 1.092810095 | 0.458 | 0.029 | 0 | 12 | Cxcr3 |
| Lat1 | 0 | 1.089764814 | 0.464 | 0.029 | 0 | 12 | Lat |
| Tnfrsf182 | 0 | 1.074918017 | 0.52 | 0.065 | 0 | 12 | Tnfrsf18 |
| Hsd11b13 | 0 | 1.057149114 | 0.564 | 0.083 | 0 | 12 | Hsd11b1 |
| Cd1601 | 0 | 1.045747434 | 0.393 | 0.037 | 0 | 12 | Cd160 |
| Cd962 | 0 | 1.0420258 | 0.508 | 0.06 | 0 | 12 | Cd96 |
| Lef1 | 0 | 1.009735366 | 0.286 | 0.009 | 0 | 12 | Lef1 |
| Tbc1d10c2 | 0 | 0.988506288 | 0.514 | 0.065 | 0 | 12 | Tbc1d10c |
| Hopx2 | 0 | 0.9744299 | 0.516 | 0.067 | 0 | 12 | Hopx |
| Traf12 | 0 | 0.97223971 | 0.452 | 0.055 | 0 | 12 | Traf1 |
| Lax12 | 0 | 0.966634615 | 0.479 | 0.058 | 0 | 12 | Lax1 |
| Klrc1 | 0 | 0.948435732 | 0.297 | 0.022 | 0 | 12 | Klrc1 |
| Camk4 | 0 | 0.931547761 | 0.325 | 0.014 | 0 | 12 | Camk4 |
| Cd200r1 | 0 | 0.922750239 | 0.336 | 0.021 | 0 | 12 | Cd200r1 |
| Cd28 | 0 | 0.909910721 | 0.305 | 0.013 | 0 | 12 | Cd28 |
| Ifng | 0 | 0.905181585 | 0.316 | 0.02 | 0 | 12 | Ifng |
| Gpr681 | 0 | 0.859893995 | 0.383 | 0.039 | 0 | 12 | Gpr68 |
| Apol7e2 | 0 | 0.853537224 | 0.406 | 0.044 | 0 | 12 | Apol7e |
| Tcrg-C4 | 0 | 0.832956585 | 0.293 | 0.007 | 0 | 12 | Tcrg-C4 |
| Il27ra1 | 0 | 0.799653594 | 0.376 | 0.036 | 0 | 12 | Il27ra |
| Acp5 | 0 | 0.61369435 | 0.252 | 0.015 | 0 | 12 | Acp5 |
| Lcp23 | 8.14E-307 | 1.106971737 | 0.559 | 0.086 | 1.65E-302 | 12 | Lcp2 |
| Rasal32 | 8.50E-306 | 1.075460984 | 0.527 | 0.075 | 1.72E-301 | 12 | Rasal3 |
| Itgb23 | 1.04E-304 | 1.06284207 | 0.581 | 0.092 | 2.11E-300 | 12 | Itgb2 |
| Lpxn2 | 5.34E-304 | 1.104195833 | 0.553 | 0.084 | 1.08E-299 | 12 | Lpxn |
| Sema4d3 | 1.02E-302 | 1.271510635 | 0.66 | 0.12 | 2.07E-298 | 12 | Sema4d |

| | | | | | | | |
|---|---|---|---|---|---|---|---|
| Nt5e | 1.41E-302 | 0.718683703 | 0.288 | 0.022 | 2.85E-298 | 12 | Nt5e |
| Rgs14 | 2.07E-302 | 1.582892274 | 0.699 | 0.13 | 4.19E-298 | 12 | Rgs1 |
| Sh3kbp12 | 7.70E-302 | 1.313318717 | 0.643 | 0.115 | 1.56E-297 | 12 | Sh3kbp1 |
| S100a63 | 3.93E-301 | 2.107509313 | 0.92 | 0.259 | 7.96E-297 | 12 | S100a6 |
| Jakmip12 | 3.69E-299 | 0.805326554 | 0.394 | 0.043 | 7.47E-295 | 12 | Jakmip1 |
| Gata32 | 1.03E-295 | 0.80936482 | 0.404 | 0.045 | 2.09E-291 | 12 | Gata3 |
| Gpr1321 | 5.82E-294 | 0.635046866 | 0.288 | 0.022 | 1.18E-289 | 12 | Gpr132 |
| Dusp22 | 7.52E-291 | 1.230254179 | 0.579 | 0.093 | 1.52E-286 | 12 | Dusp2 |
| Ppp3cc2 | 1.67E-290 | 1.292683845 | 0.682 | 0.131 | 3.37E-286 | 12 | Ppp3cc |
| Fgl23 | 2.68E-290 | 1.314280338 | 0.583 | 0.099 | 5.42E-286 | 12 | Fgl2 |
| Lcp13 | 3.64E-290 | 1.515869524 | 0.88 | 0.226 | 7.36E-286 | 12 | Lcp1 |
| Ptpn184 | 5.98E-290 | 1.675501184 | 0.959 | 0.303 | 1.21E-285 | 12 | Ptpn18 |
| Tcf71 | 2.42E-289 | 0.646125353 | 0.252 | 0.017 | 4.89E-285 | 12 | Tcf7 |
| Ankrd443 | 5.53E-285 | 1.659266657 | 0.92 | 0.282 | 1.12E-280 | 12 | Ankrd44 |
| Clnk2 | 1.14E-284 | 1.429579292 | 0.611 | 0.102 | 2.30E-280 | 12 | Clnk |
| Laptm54 | 3.41E-284 | 1.147826077 | 0.884 | 0.198 | 6.90E-280 | 12 | Laptm5 |
| Celf24 | 1.40E-283 | 1.212881775 | 0.839 | 0.19 | 2.83E-279 | 12 | Celf2 |
| Limd23 | 9.48E-283 | 1.282720295 | 0.686 | 0.138 | 1.92E-278 | 12 | Limd2 |
| Cotl14 | 1.99E-282 | 1.456836382 | 0.849 | 0.212 | 4.02E-278 | 12 | Cotl1 |
| Fam107b2 | 4.88E-282 | 1.083786855 | 0.619 | 0.111 | 9.89E-278 | 12 | Fam107b |
| Prex13 | 7.05E-281 | 1.335218211 | 0.738 | 0.163 | 1.43E-276 | 12 | Prex1 |
| Bcl22 | 3.03E-280 | 1.746201506 | 0.8 | 0.196 | 6.14E-276 | 12 | Bcl2 |
| Slc14a12 | 2.90E-279 | 0.812028237 | 0.37 | 0.04 | 5.87E-275 | 12 | Slc14a1 |
| Dock114 | 1.16E-277 | 1.111149099 | 0.596 | 0.105 | 2.35E-273 | 12 | Dock11 |
| Mbp2 | 4.03E-277 | 0.848146391 | 0.45 | 0.06 | 8.17E-273 | 12 | Mbp |
| Rasgrp12 | 2.17E-276 | 0.843155544 | 0.432 | 0.055 | 4.39E-272 | 12 | Rasgrp1 |
| Serpina3g2 | 3.12E-276 | 1.078312502 | 0.45 | 0.06 | 6.33E-272 | 12 | Serpina3g |
| Bin23 | 3.80E-276 | 1.047211204 | 0.636 | 0.118 | 7.70E-272 | 12 | Bin2 |
| Adcy74 | 1.88E-275 | 1.035535179 | 0.594 | 0.105 | 3.80E-271 | 12 | Adcy7 |
| Sep-62 | 4.74E-267 | 0.929354409 | 0.546 | 0.09 | 9.59E-263 | 12 | 6-Sep |
| Pkp32 | 1.04E-266 | 0.88057472 | 0.452 | 0.062 | 2.10E-262 | 12 | Pkp3 |
| Il18rap1 | 2.75E-265 | 0.620102015 | 0.258 | 0.02 | 5.57E-261 | 12 | Il18rap |
| Itgae1 | 1.20E-264 | 1.357209955 | 0.441 | 0.059 | 2.43E-260 | 12 | Itgae |
| Pde3b3 | 2.61E-260 | 1.231638374 | 0.619 | 0.118 | 5.29E-256 | 12 | Pde3b |
| Gramd1b2 | 1.17E-258 | 1.028296382 | 0.51 | 0.082 | 2.37E-254 | 12 | Gramd1b |
| Card112 | 4.37E-258 | 0.949076816 | 0.488 | 0.075 | 8.84E-254 | 12 | Card11 |
| Cd73 | 9.72E-251 | 1.57999064 | 0.581 | 0.103 | 1.97E-246 | 12 | Cd7 |
| Ccnd32 | 1.01E-249 | 1.368492638 | 0.828 | 0.229 | 2.04E-245 | 12 | Ccnd3 |
| Gng23 | 2.44E-249 | 1.245546274 | 0.736 | 0.176 | 4.94E-245 | 12 | Gng2 |
| Mllt32 | 5.21E-248 | 1.167977301 | 0.664 | 0.141 | 1.06E-243 | 12 | Mllt3 |
| Zap702 | 1.80E-247 | 0.754656483 | 0.393 | 0.05 | 3.65E-243 | 12 | Zap70 |
| Smyd32 | 1.90E-245 | 1.634078121 | 0.806 | 0.216 | 3.85E-241 | 12 | Smyd3 |
| Rab372 | 2.44E-245 | 0.627175027 | 0.34 | 0.038 | 4.94E-241 | 12 | Rab37 |

| | | | | | | | |
|---|---|---|---|---|---|---|---|
| Ubash3a | 2.76E-245 | 0.58713501 | 0.25 | 0.02 | 5.59E-241 | 12 | Ubash3a |
| Ipcef13 | 2.77E-245 | 1.149091611 | 0.583 | 0.109 | 5.62E-241 | 12 | Ipcef1 |
| Shisa52 | 6.73E-245 | 1.481133649 | 0.921 | 0.318 | 1.36E-240 | 12 | Shisa5 |
| Sytl22 | 4.99E-244 | 0.807201768 | 0.387 | 0.05 | 1.01E-239 | 12 | Sytl2 |
| Cd533 | 2.30E-242 | 0.973274949 | 0.574 | 0.108 | 4.66E-238 | 12 | Cd53 |
| Nup2102 | 3.55E-241 | 0.798650379 | 0.422 | 0.06 | 7.19E-237 | 12 | Nup210 |
| Pstpip12 | 1.03E-240 | 0.721554778 | 0.383 | 0.05 | 2.09E-236 | 12 | Pstpip1 |
| Itga43 | 5.70E-239 | 1.368877727 | 0.447 | 0.069 | 1.15E-234 | 12 | Itga4 |
| Traf3ip32 | 8.81E-239 | 0.850622773 | 0.462 | 0.072 | 1.78E-234 | 12 | Traf3ip3 |
| Fnbp14 | 1.13E-238 | 1.32471059 | 0.8 | 0.219 | 2.29E-234 | 12 | Fnbp1 |
| Pglyrp22 | 6.04E-237 | 0.632922551 | 0.336 | 0.039 | 1.22E-232 | 12 | Pglyrp2 |
| Dscam2 | 9.85E-237 | 1.800946669 | 0.342 | 0.041 | 1.99E-232 | 12 | Dscam |
| Tnfaip82 | 2.89E-236 | 1.14809523 | 0.733 | 0.176 | 5.86E-232 | 12 | Tnfaip8 |
| Cd271 | 1.64E-235 | 0.652174077 | 0.316 | 0.034 | 3.33E-231 | 12 | Cd27 |
| Coro2a2 | 2.84E-235 | 1.13889515 | 0.57 | 0.109 | 5.75E-231 | 12 | Coro2a |
| Nlrc32 | 6.87E-235 | 0.75079883 | 0.387 | 0.052 | 1.39E-230 | 12 | Nlrc3 |
| Fam189b2 | 9.47E-235 | 0.70287399 | 0.385 | 0.051 | 1.92E-230 | 12 | Fam189b |
| Klk81 | 1.21E-233 | 0.87257124 | 0.471 | 0.075 | 2.45E-229 | 12 | Klk8 |
| Gimap44 | 1.81E-231 | 1.543505503 | 0.813 | 0.246 | 3.66E-227 | 12 | Gimap4 |
| Parp84 | 3.19E-231 | 1.283141318 | 0.634 | 0.14 | 6.46E-227 | 12 | Parp8 |
| Osbpl32 | 3.91E-229 | 1.05770569 | 0.576 | 0.111 | 7.92E-225 | 12 | Osbpl3 |
| Rapgef63 | 8.42E-227 | 1.436371808 | 0.854 | 0.287 | 1.71E-222 | 12 | Rapgef6 |
| Adgre53 | 8.30E-226 | 1.433437301 | 0.793 | 0.227 | 1.68E-221 | 12 | Adgre5 |
| Tnf2 | 5.02E-225 | 1.169247262 | 0.428 | 0.066 | 1.02E-220 | 12 | Tnf |
| Mbnl13 | 5.23E-224 | 1.645476803 | 0.981 | 0.449 | 1.06E-219 | 12 | Mbnl1 |
| B4galnt12 | 1.22E-223 | 0.876688431 | 0.497 | 0.088 | 2.48E-219 | 12 | B4galnt1 |
| Glipr21 | 5.65E-222 | 0.701135255 | 0.321 | 0.038 | 1.14E-217 | 12 | Glipr2 |
| Ptk2b3 | 3.18E-220 | 1.070338377 | 0.645 | 0.141 | 6.43E-216 | 12 | Ptk2b |
| Arhgap303 | 3.50E-220 | 0.983823481 | 0.531 | 0.102 | 7.08E-216 | 12 | Arhgap30 |
| A430078G23Rik | 1.59E-219 | 0.558517453 | 0.254 | 0.024 | 3.21E-215 | 12 | A430078G23Rik |
| Iqgap21 | 4.25E-217 | 0.915519873 | 0.525 | 0.097 | 8.60E-213 | 12 | Iqgap2 |
| Gramd3 | 1.12E-216 | 0.946437972 | 0.288 | 0.031 | 2.26E-212 | 12 | Gramd3 |
| Ccdc88c2 | 4.89E-215 | 0.85532595 | 0.507 | 0.092 | 9.91E-211 | 12 | Ccdc88c |
| Gm267401 | 3.44E-213 | 0.953198203 | 0.381 | 0.054 | 6.97E-209 | 12 | Gm26740 |
| Tnfaip33 | 8.81E-213 | 1.045371339 | 0.467 | 0.083 | 1.78E-208 | 12 | Tnfaip3 |
| Arhgap92 | 7.08E-212 | 0.76227289 | 0.536 | 0.103 | 1.43E-207 | 12 | Arhgap9 |
| Sh2d2a2 | 9.28E-212 | 0.64972793 | 0.34 | 0.044 | 1.88E-207 | 12 | Sh2d2a |
| Cd2442 | 1.63E-211 | 1.019889688 | 0.48 | 0.085 | 3.29E-207 | 12 | Cd244 |
| Gmfg2 | 1.75E-211 | 1.022529577 | 0.602 | 0.131 | 3.54E-207 | 12 | Gmfg |
| Runx13 | 3.00E-211 | 1.124199415 | 0.583 | 0.123 | 6.07E-207 | 12 | Runx1 |
| Cytip2 | 6.21E-211 | 0.86575298 | 0.477 | 0.085 | 1.26E-206 | 12 | Cytip |
| Acap12 | 1.78E-210 | 0.798224301 | 0.413 | 0.065 | 3.60E-206 | 12 | Acap1 |
| Myo1f3 | 9.24E-210 | 1.045079956 | 0.598 | 0.129 | 1.87E-205 | 12 | Myo1f |

| | | | | | | | |
|---|---|---|---|---|---|---|---|
| Pglyrp12 | 7.39E-209 | 0.880877831 | 0.482 | 0.086 | 1.50E-204 | 12 | Pglyrp1 |
| Fasl2 | 8.64E-209 | 0.727568648 | 0.366 | 0.051 | 1.75E-204 | 12 | Fasl |
| Rabgap1l2 | 3.34E-208 | 1.374954032 | 0.802 | 0.247 | 6.77E-204 | 12 | Rabgap1l |
| Rps15a3 | 1.35E-207 | 1.307935977 | 0.996 | 0.923 | 2.73E-203 | 12 | Rps15a |
| Ctse | 1.82E-207 | 0.564014659 | 0.262 | 0.027 | 3.68E-203 | 12 | Ctse |
| H2-Q74 | 2.65E-207 | 1.470797819 | 0.95 | 0.388 | 5.37E-203 | 12 | H2-Q7 |
| Ppm1h3 | 4.84E-207 | 1.09776344 | 0.637 | 0.152 | 9.81E-203 | 12 | Ppm1h |
| Capg2 | 1.38E-206 | 1.375813867 | 0.486 | 0.094 | 2.80E-202 | 12 | Capg |
| Nlrc53 | 2.80E-206 | 0.956853199 | 0.523 | 0.104 | 5.67E-202 | 12 | Nlrc5 |
| Zfp8311 | 6.35E-205 | 0.535455339 | 0.271 | 0.029 | 1.29E-200 | 12 | Zfp831 |
| Ptpn72 | 2.33E-204 | 0.717977281 | 0.366 | 0.053 | 4.71E-200 | 12 | Ptpn7 |
| Acot72 | 3.39E-204 | 0.747589756 | 0.426 | 0.07 | 6.86E-200 | 12 | Acot7 |
| Dok21 | 3.66E-204 | 0.645369797 | 0.307 | 0.037 | 7.42E-200 | 12 | Dok2 |
| AC163354.11 | 4.47E-204 | 1.62353044 | 0.374 | 0.056 | 9.05E-200 | 12 | AC163354.1 |
| Klrc22 | 2.65E-202 | 0.800308114 | 0.348 | 0.048 | 5.37E-198 | 12 | Klrc2 |
| Aak13 | 4.27E-200 | 1.145380574 | 0.671 | 0.176 | 8.64E-196 | 12 | Aak1 |
| Stk43 | 9.47E-199 | 0.974046402 | 0.658 | 0.166 | 1.92E-194 | 12 | Stk4 |
| Evl3 | 1.31E-198 | 1.167382829 | 0.738 | 0.211 | 2.65E-194 | 12 | Evl |
| Asb21 | 8.10E-198 | 0.834045822 | 0.338 | 0.047 | 1.64E-193 | 12 | Asb2 |
| Sema4a2 | 3.14E-197 | 0.773426861 | 0.421 | 0.07 | 6.36E-193 | 12 | Sema4a |
| Il21r3 | 1.23E-194 | 0.914002682 | 0.464 | 0.087 | 2.49E-190 | 12 | Il21r |
| Gpr1742 | 8.34E-194 | 0.689229333 | 0.34 | 0.047 | 1.69E-189 | 12 | Gpr174 |
| Scml42 | 1.51E-193 | 0.721931934 | 0.299 | 0.037 | 3.06E-189 | 12 | Scml4 |
| Spn1 | 6.37E-193 | 0.613838424 | 0.307 | 0.039 | 1.29E-188 | 12 | Spn |
| Gpsm33 | 9.48E-193 | 0.889742081 | 0.6 | 0.139 | 1.92E-188 | 12 | Gpsm3 |
| Btg12 | 3.20E-192 | 1.463545376 | 0.974 | 0.45 | 6.48E-188 | 12 | Btg1 |
| Leprotl12 | 8.53E-192 | 1.056827382 | 0.725 | 0.208 | 1.73E-187 | 12 | Leprotl1 |
| Psd42 | 2.24E-191 | 0.631260149 | 0.379 | 0.059 | 4.53E-187 | 12 | Psd4 |
| Ndfip12 | 1.77E-190 | 1.297948576 | 0.862 | 0.311 | 3.59E-186 | 12 | Ndfip1 |
| Sh3bgrl34 | 3.25E-189 | 1.436978501 | 0.987 | 0.552 | 6.57E-185 | 12 | Sh3bgrl3 |
| Vim5 | 8.16E-188 | 1.256198733 | 0.867 | 0.296 | 1.65E-183 | 12 | Vim |
| Cdh12 | 2.37E-187 | 0.855200522 | 0.406 | 0.068 | 4.80E-183 | 12 | Cdh1 |
| Rpsa4 | 1.96E-186 | 1.23333835 | 0.998 | 0.901 | 3.97E-182 | 12 | Rpsa |
| Nebl | 4.37E-186 | 0.709489832 | 0.323 | 0.044 | 8.84E-182 | 12 | Nebl |
| Znrf33 | 5.60E-186 | 0.966782928 | 0.49 | 0.1 | 1.13E-181 | 12 | Znrf3 |
| Samsn13 | 7.47E-186 | 0.801687531 | 0.421 | 0.074 | 1.51E-181 | 12 | Samsn1 |
| Arl4c1 | 6.32E-185 | 0.827012684 | 0.393 | 0.066 | 1.28E-180 | 12 | Arl4c |
| Sla4 | 3.18E-184 | 0.756907381 | 0.398 | 0.068 | 6.45E-180 | 12 | Sla |
| Chsy12 | 1.28E-183 | 1.006277029 | 0.544 | 0.12 | 2.59E-179 | 12 | Chsy1 |
| Il12rb22 | 1.52E-181 | 0.660717497 | 0.32 | 0.045 | 3.07E-177 | 12 | Il12rb2 |
| Dennd2d | 3.75E-181 | 0.55862701 | 0.273 | 0.033 | 7.59E-177 | 12 | Dennd2d |
| S100a102 | 7.72E-181 | 1.382752476 | 0.974 | 0.46 | 1.56E-176 | 12 | S100a10 |
| Dock83 | 2.62E-180 | 1.023946233 | 0.742 | 0.209 | 5.30E-176 | 12 | Dock8 |

| | | | | | | | |
|---|---|---|---|---|---|---|---|
| Rps163 | 2.93E-180 | 1.049733652 | 0.996 | 0.926 | 5.93E-176 | 12 | Rps16 |
| Prkca3 | 4.47E-180 | 0.937862179 | 0.467 | 0.094 | 9.05E-176 | 12 | Prkca |
| Gprin32 | 1.03E-179 | 0.757878241 | 0.364 | 0.057 | 2.09E-175 | 12 | Gprin3 |
| Cd441 | 1.26E-178 | 0.828537795 | 0.488 | 0.101 | 2.55E-174 | 12 | Cd44 |
| Plcxd21 | 5.50E-178 | 0.535384219 | 0.256 | 0.029 | 1.11E-173 | 12 | Plcxd2 |
| Ramp12 | 5.41E-177 | 1.308033382 | 0.482 | 0.103 | 1.10E-172 | 12 | Ramp1 |
| Rps244 | 2.39E-176 | 1.002712234 | 0.998 | 0.972 | 4.84E-172 | 12 | Rps24 |
| Elf43 | 8.81E-176 | 0.870932738 | 0.521 | 0.115 | 1.78E-171 | 12 | Elf4 |
| Mctp22 | 1.64E-175 | 0.816829756 | 0.424 | 0.078 | 3.33E-171 | 12 | Mctp2 |
| Cblb3 | 1.63E-173 | 1.037968469 | 0.622 | 0.162 | 3.29E-169 | 12 | Cblb |
| Rassf52 | 4.54E-173 | 0.690164054 | 0.372 | 0.063 | 9.19E-169 | 12 | Rassf5 |
| Cst72 | 1.69E-172 | 0.713787248 | 0.394 | 0.068 | 3.43E-168 | 12 | Cst7 |
| Sesn33 | 3.88E-172 | 0.74358463 | 0.536 | 0.121 | 7.86E-168 | 12 | Sesn3 |
| Rps274 | 4.93E-172 | 1.068453722 | 1 | 0.896 | 9.99E-168 | 12 | Rps27 |
| Sla22 | 6.81E-172 | 0.56001532 | 0.31 | 0.044 | 1.38E-167 | 12 | Sla2 |
| Gimap16 | 5.33E-170 | 1.088532925 | 0.85 | 0.293 | 1.08E-165 | 12 | Gimap1 |
| Sash32 | 1.22E-169 | 0.59685174 | 0.346 | 0.055 | 2.47E-165 | 12 | Sash3 |
| Fxyd55 | 1.44E-168 | 1.172999496 | 0.931 | 0.402 | 2.91E-164 | 12 | Fxyd5 |
| Padi22 | 2.52E-168 | 0.797972648 | 0.449 | 0.09 | 5.10E-164 | 12 | Padi2 |
| Aim23 | 8.81E-167 | 0.777043961 | 0.464 | 0.096 | 1.78E-162 | 12 | Aim2 |
| Slamf73 | 1.57E-166 | 0.809809679 | 0.454 | 0.093 | 3.17E-162 | 12 | Slamf7 |
| Emp34 | 1.86E-166 | 0.999337278 | 0.781 | 0.249 | 3.77E-162 | 12 | Emp3 |
| Kif21b1 | 2.93E-166 | 0.581994168 | 0.316 | 0.047 | 5.94E-162 | 12 | Kif21b |
| Cyld2 | 3.87E-166 | 0.847837736 | 0.615 | 0.162 | 7.83E-162 | 12 | Cyld |
| Jak14 | 1.60E-164 | 1.143159593 | 0.888 | 0.356 | 3.24E-160 | 12 | Jak1 |
| Dennd1b4 | 2.95E-164 | 0.924154534 | 0.636 | 0.176 | 5.98E-160 | 12 | Dennd1b |
| Cish | 6.69E-164 | 0.603165792 | 0.366 | 0.062 | 1.35E-159 | 12 | Cish |
| Ripor23 | 1.18E-163 | 1.083622113 | 0.615 | 0.164 | 2.39E-159 | 12 | Ripor2 |
| Fam78a1 | 3.32E-163 | 0.560010423 | 0.273 | 0.036 | 6.72E-159 | 12 | Fam78a |
| Trim143 | 3.53E-162 | 0.785310574 | 0.501 | 0.114 | 7.14E-158 | 12 | Trim14 |
| Fermt33 | 5.19E-162 | 0.74363393 | 0.49 | 0.108 | 1.05E-157 | 12 | Fermt3 |
| A930015D03Rik2 | 6.93E-162 | 1.074010131 | 0.462 | 0.1 | 1.40E-157 | 12 | A930015D03Rik |
| Dennd1c2 | 1.33E-161 | 0.605202701 | 0.329 | 0.053 | 2.69E-157 | 12 | Dennd1c |
| Gem2 | 2.03E-161 | 1.135378662 | 0.46 | 0.097 | 4.11E-157 | 12 | Gem |
| Ncr12 | 2.62E-160 | 1.329119164 | 0.456 | 0.098 | 5.31E-156 | 12 | Ncr1 |
| Malt12 | 8.16E-160 | 0.978250545 | 0.561 | 0.142 | 1.65E-155 | 12 | Malt1 |
| Chn22 | 1.58E-159 | 1.411123358 | 0.639 | 0.19 | 3.21E-155 | 12 | Chn2 |
| Lyst2 | 2.57E-159 | 1.249527825 | 0.46 | 0.103 | 5.21E-155 | 12 | Lyst |
| Herc32 | 4.62E-159 | 0.704037514 | 0.37 | 0.066 | 9.35E-155 | 12 | Herc3 |
| Ubac23 | 9.53E-159 | 0.859553092 | 0.636 | 0.18 | 1.93E-154 | 12 | Ubac2 |
| Rps34 | 2.70E-158 | 0.972042748 | 0.996 | 0.891 | 5.47E-154 | 12 | Rps3 |
| Sytl32 | 5.59E-157 | 0.696924111 | 0.325 | 0.053 | 1.13E-152 | 12 | Sytl3 |
| Rps113 | 8.95E-157 | 1.001687465 | 1 | 0.906 | 1.81E-152 | 12 | Rps11 |

| | | | | | | | |
|---|---|---|---|---|---|---|---|
| Ugcg2 | 1.38E-156 | 1.012810196 | 0.604 | 0.164 | 2.79E-152 | 12 | Ugcg |
| Id22 | 1.41E-156 | 1.216889464 | 0.837 | 0.312 | 2.85E-152 | 12 | Id2 |
| Pfn14 | 1.63E-156 | 1.206817233 | 0.991 | 0.677 | 3.29E-152 | 12 | Pfn1 |
| Btbd112 | 2.34E-156 | 0.728661209 | 0.32 | 0.051 | 4.74E-152 | 12 | Btbd11 |
| AW1120104 | 2.98E-154 | 1.307388415 | 0.972 | 0.561 | 6.03E-150 | 12 | AW112010 |
| Rpl18a3 | 4.27E-154 | 0.912798783 | 0.998 | 0.93 | 8.65E-150 | 12 | Rpl18a |
| Fhl22 | 2.46E-152 | 0.560886311 | 0.271 | 0.038 | 4.99E-148 | 12 | Fhl2 |
| Ttc39b3 | 3.53E-152 | 0.880255306 | 0.518 | 0.129 | 7.14E-148 | 12 | Ttc39b |
| Atp11b2 | 5.43E-152 | 0.965603935 | 0.607 | 0.173 | 1.10E-147 | 12 | Atp11b |
| Rps134 | 5.74E-151 | 0.930047404 | 0.996 | 0.922 | 1.16E-146 | 12 | Rps13 |
| Ezr2 | 6.85E-151 | 0.59398375 | 0.535 | 0.129 | 1.39E-146 | 12 | Ezr |
| Rps104 | 8.11E-151 | 0.895911594 | 0.998 | 0.943 | 1.64E-146 | 12 | Rps10 |
| Gm203885 | 1.68E-150 | 1.08378702 | 0.91 | 0.392 | 3.40E-146 | 12 | Gm20388 |
| Arnt21 | 9.50E-150 | 0.593064275 | 0.28 | 0.041 | 1.92E-145 | 12 | Arnt2 |
| Nabp12 | 6.59E-149 | 0.716439808 | 0.391 | 0.076 | 1.33E-144 | 12 | Nabp1 |
| Akna3 | 1.27E-148 | 0.710635582 | 0.411 | 0.086 | 2.56E-144 | 12 | Akna |
| Plcl24 | 1.90E-147 | 0.865502582 | 0.465 | 0.108 | 3.85E-143 | 12 | Plcl2 |
| Samd32 | 2.86E-147 | 0.775432283 | 0.406 | 0.081 | 5.79E-143 | 12 | Samd3 |
| Oprm12 | 5.64E-147 | 0.598525208 | 0.327 | 0.055 | 1.14E-142 | 12 | Oprm1 |
| Rpl37a4 | 1.46E-146 | 0.88494038 | 1 | 0.937 | 2.95E-142 | 12 | Rpl37a |
| Satb12 | 1.82E-146 | 0.834205631 | 0.333 | 0.058 | 3.68E-142 | 12 | Satb1 |
| Akap135 | 6.56E-145 | 1.095398193 | 0.884 | 0.385 | 1.33E-140 | 12 | Akap13 |
| Rpl124 | 6.63E-145 | 1.01720921 | 0.994 | 0.839 | 1.34E-140 | 12 | Rpl12 |
| Ifitm102 | 9.67E-145 | 0.583961405 | 0.325 | 0.056 | 1.96E-140 | 12 | Ifitm10 |
| Rps183 | 2.50E-144 | 0.905947756 | 0.994 | 0.941 | 5.06E-140 | 12 | Rps18 |
| Jpt13 | 6.92E-144 | 0.783944699 | 0.692 | 0.214 | 1.40E-139 | 12 | Jpt1 |
| Gzmc1 | 8.44E-144 | 2.291243037 | 0.383 | 0.077 | 1.71E-139 | 12 | Gzmc |
| Klrb1c2 | 4.37E-143 | 0.824852989 | 0.432 | 0.091 | 8.84E-139 | 12 | Klrb1c |
| Dock52 | 7.67E-143 | 0.826812934 | 0.469 | 0.109 | 1.55E-138 | 12 | Dock5 |
| Vps37b2 | 1.02E-141 | 0.865336258 | 0.514 | 0.13 | 2.07E-137 | 12 | Vps37b |
| Anxa62 | 1.07E-141 | 0.813719394 | 0.551 | 0.15 | 2.16E-137 | 12 | Anxa6 |
| Mta33 | 3.15E-141 | 0.730264479 | 0.548 | 0.145 | 6.37E-137 | 12 | Mta3 |
| Tasp1 | 4.90E-141 | 0.70471124 | 0.348 | 0.064 | 9.93E-137 | 12 | Tasp1 |
| Rpl134 | 9.44E-141 | 0.828665442 | 1 | 0.977 | 1.91E-136 | 12 | Rpl13 |
| Fam49b3 | 1.01E-140 | 0.861255117 | 0.716 | 0.228 | 2.05E-136 | 12 | Fam49b |
| Ssbp23 | 1.32E-139 | 0.892937195 | 0.52 | 0.134 | 2.67E-135 | 12 | Ssbp2 |
| Rab19 | 4.67E-139 | 0.475372388 | 0.25 | 0.035 | 9.45E-135 | 12 | Rab19 |
| Vav13 | 2.54E-138 | 0.575018002 | 0.376 | 0.076 | 5.15E-134 | 12 | Vav1 |
| Stk262 | 9.79E-138 | 0.476911252 | 0.271 | 0.041 | 1.98E-133 | 12 | Stk26 |
| Xist6 | 1.80E-137 | 1.091205573 | 0.948 | 0.455 | 3.64E-133 | 12 | Xist |
| Rps54 | 2.67E-137 | 0.880110722 | 0.998 | 0.919 | 5.41E-133 | 12 | Rps5 |
| Gimap92 | 3.11E-137 | 0.76290053 | 0.557 | 0.152 | 6.31E-133 | 12 | Gimap9 |
| Mapkapk32 | 4.54E-137 | 0.580717264 | 0.379 | 0.077 | 9.19E-133 | 12 | Mapkapk3 |

| | | | | | | | |
|---|---|---|---|---|---|---|---|
| Gfod12 | 5.83E-137 | 0.641904147 | 0.342 | 0.064 | 1.18E-132 | 12 | Gfod1 |
| Chd32 | 6.90E-137 | 0.745285711 | 0.544 | 0.149 | 1.40E-132 | 12 | Chd3 |
| Fau3 | 6.67E-136 | 0.780912233 | 0.996 | 0.952 | 1.35E-131 | 12 | Fau |
| Znrf22 | 8.01E-136 | 0.786552468 | 0.548 | 0.152 | 1.62E-131 | 12 | Znrf2 |
| Apobec32 | 8.22E-136 | 0.84386356 | 0.671 | 0.218 | 1.66E-131 | 12 | Apobec3 |
| Fmnl13 | 2.88E-135 | 0.66427875 | 0.4 | 0.087 | 5.84E-131 | 12 | Fmnl1 |
| Gpr1712 | 3.12E-135 | 0.521161733 | 0.277 | 0.044 | 6.31E-131 | 12 | Gpr171 |
| Ncf43 | 5.46E-135 | 0.712810932 | 0.449 | 0.106 | 1.11E-130 | 12 | Ncf4 |
| Ifi2092 | 1.45E-134 | 0.607945842 | 0.305 | 0.053 | 2.94E-130 | 12 | Ifi209 |
| St6galnac33 | 4.54E-134 | 1.571449148 | 0.637 | 0.209 | 9.19E-130 | 12 | St6galnac3 |
| Cyth43 | 5.84E-134 | 0.703048143 | 0.503 | 0.127 | 1.18E-129 | 12 | Cyth4 |
| Rbl22 | 1.72E-133 | 0.785981968 | 0.495 | 0.129 | 3.49E-129 | 12 | Rbl2 |
| Rab8b3 | 2.09E-133 | 0.984700841 | 0.753 | 0.276 | 4.23E-129 | 12 | Rab8b |
| Rpl384 | 8.92E-133 | 0.868636119 | 0.994 | 0.892 | 1.81E-128 | 12 | Rpl38 |
| Rpl304 | 9.67E-133 | 0.800208517 | 0.998 | 0.937 | 1.96E-128 | 12 | Rpl30 |
| 1810011H11Rik2 | 6.26E-132 | 0.732591061 | 0.348 | 0.069 | 1.27E-127 | 12 | 1810011H11Rik |
| Gimap65 | 5.14E-131 | 1.161906704 | 0.845 | 0.357 | 1.04E-126 | 12 | Gimap6 |
| Rpl184 | 7.10E-131 | 0.850041589 | 1 | 0.899 | 1.44E-126 | 12 | Rpl18 |
| Gimap84 | 2.91E-130 | 0.826291889 | 0.563 | 0.162 | 5.89E-126 | 12 | Gimap8 |
| Rpl343 | 7.02E-130 | 0.804136222 | 0.996 | 0.916 | 1.42E-125 | 12 | Rpl34 |
| Btg24 | 9.18E-130 | 1.11758519 | 0.826 | 0.336 | 1.86E-125 | 12 | Btg2 |
| Slc38a23 | 2.10E-129 | 1.052966855 | 0.774 | 0.303 | 4.25E-125 | 12 | Slc38a2 |
| Rps74 | 3.18E-129 | 0.853962103 | 0.994 | 0.915 | 6.43E-125 | 12 | Rps7 |
| H2afy2 | 1.44E-128 | 0.876852823 | 0.813 | 0.329 | 2.92E-124 | 12 | H2afy |
| Esyt12 | 1.72E-128 | 0.720516096 | 0.493 | 0.13 | 3.49E-124 | 12 | Esyt1 |
| Pim12 | 4.41E-128 | 0.948159533 | 0.48 | 0.126 | 8.92E-124 | 12 | Pim1 |
| Srgap32 | 7.93E-128 | 0.68260277 | 0.406 | 0.091 | 1.61E-123 | 12 | Srgap3 |
| Fam105a3 | 8.04E-128 | 0.592397879 | 0.482 | 0.119 | 1.63E-123 | 12 | Fam105a |
| Tes2 | 8.26E-128 | 0.662300635 | 0.482 | 0.123 | 1.67E-123 | 12 | Tes |
| Peli13 | 2.23E-127 | 0.977588129 | 0.654 | 0.225 | 4.51E-123 | 12 | Peli1 |
| Myo1g1 | 2.73E-127 | 0.508674348 | 0.267 | 0.044 | 5.54E-123 | 12 | Myo1g |
| Eml43 | 5.54E-127 | 0.926526916 | 0.697 | 0.251 | 1.12E-122 | 12 | Eml4 |
| AC149090.13 | 5.89E-127 | 0.910458603 | 0.669 | 0.224 | 1.19E-122 | 12 | AC149090.1 |
| Gimap53 | 8.84E-127 | 0.781525244 | 0.566 | 0.164 | 1.79E-122 | 12 | Gimap5 |
| Sept114 | 9.64E-127 | 0.883073748 | 0.624 | 0.196 | 1.95E-122 | 12 | 11-Sep |
| Fyn5 | 4.31E-126 | 0.9989126 | 0.716 | 0.259 | 8.73E-122 | 12 | Fyn |
| Cyba4 | 6.30E-126 | 0.842964831 | 0.933 | 0.432 | 1.28E-121 | 12 | Cyba |
| Dgkd3 | 5.13E-125 | 0.840371092 | 0.535 | 0.155 | 1.04E-120 | 12 | Dgkd |
| Prkcb4 | 8.87E-125 | 0.731381318 | 0.544 | 0.15 | 1.79E-120 | 12 | Prkcb |
| Suco2 | 2.11E-124 | 0.77144601 | 0.52 | 0.147 | 4.27E-120 | 12 | Suco |
| Efhd23 | 4.61E-124 | 0.730960957 | 0.6 | 0.18 | 9.34E-120 | 12 | Efhd2 |
| Nfatc32 | 4.75E-124 | 0.800690749 | 0.561 | 0.168 | 9.62E-120 | 12 | Nfatc3 |
| Neurl33 | 2.71E-123 | 0.689967969 | 0.443 | 0.11 | 5.48E-119 | 12 | Neurl3 |

| | | | | | | | |
|---|---|---|---|---|---|---|---|
| Cdk63 | 9.41E-123 | 0.927538087 | 0.43 | 0.106 | 1.91E-118 | 12 | Cdk6 |
| Rps216 | 2.85E-122 | 0.754171131 | 0.996 | 0.926 | 5.77E-118 | 12 | Rps21 |
| Slc9a3r12 | 4.14E-122 | 0.5804849 | 0.43 | 0.104 | 8.39E-118 | 12 | Slc9a3r1 |
| Diaph12 | 4.21E-122 | 0.822096683 | 0.675 | 0.235 | 8.53E-118 | 12 | Diaph1 |
| Nsd34 | 1.02E-121 | 0.912380748 | 0.763 | 0.3 | 2.07E-117 | 12 | Nsd3 |
| Lbh2 | 1.36E-121 | 0.677436511 | 0.479 | 0.126 | 2.76E-117 | 12 | Lbh |
| Wipf14 | 2.94E-121 | 0.910608608 | 0.712 | 0.26 | 5.95E-117 | 12 | Wipf1 |
| Slc2a3 | 3.06E-121 | 0.570768926 | 0.273 | 0.047 | 6.19E-117 | 12 | Slc2a3 |
| Rpl394 | 3.31E-121 | 0.815050462 | 0.998 | 0.899 | 6.70E-117 | 12 | Rpl39 |
| Ptpn42 | 6.06E-121 | 0.747204083 | 0.471 | 0.125 | 1.23E-116 | 12 | Ptpn4 |
| Tmsb4x5 | 7.06E-121 | 0.762489632 | 0.998 | 0.933 | 1.43E-116 | 12 | Tmsb4x |
| Retreg13 | 2.52E-120 | 0.613940022 | 0.398 | 0.092 | 5.09E-116 | 12 | Retreg1 |
| Znrf14 | 2.56E-120 | 0.840967232 | 0.583 | 0.182 | 5.18E-116 | 12 | Znrf1 |
| Rpl194 | 4.64E-120 | 0.704894573 | 1 | 0.963 | 9.39E-116 | 12 | Rpl19 |
| Arhgap43 | 7.58E-120 | 0.507508746 | 0.293 | 0.054 | 1.54E-115 | 12 | Arhgap4 |
| Rpl324 | 1.78E-119 | 0.794268211 | 0.996 | 0.953 | 3.61E-115 | 12 | Rpl32 |
| Sulf23 | 1.92E-119 | 0.700956904 | 0.398 | 0.094 | 3.90E-115 | 12 | Sulf2 |
| Atp10a1 | 3.66E-119 | 0.516719883 | 0.262 | 0.044 | 7.41E-115 | 12 | Atp10a |
| Ppp1r183 | 1.03E-118 | 0.710082559 | 0.512 | 0.148 | 2.09E-114 | 12 | Ppp1r18 |
| Was3 | 1.64E-118 | 0.581220159 | 0.331 | 0.068 | 3.31E-114 | 12 | Was |
| Arhgap253 | 3.70E-118 | 0.651331085 | 0.419 | 0.103 | 7.49E-114 | 12 | Arhgap25 |
| Actb4 | 5.39E-118 | 0.920994749 | 0.998 | 0.84 | 1.09E-113 | 12 | Actb |
| Rasa32 | 9.00E-118 | 0.606172033 | 0.312 | 0.061 | 1.82E-113 | 12 | Rasa3 |
| Gm268272 | 1.15E-116 | 0.538321713 | 0.271 | 0.047 | 2.33E-112 | 12 | Gm26827 |
| Tmem131l3 | 1.31E-115 | 0.567043756 | 0.35 | 0.076 | 2.65E-111 | 12 | Tmem131l |
| Prkcz1 | 2.08E-115 | 0.430019113 | 0.288 | 0.053 | 4.21E-111 | 12 | Prkcz |
| Npm32 | 6.38E-115 | 0.575900785 | 0.473 | 0.128 | 1.29E-110 | 12 | Npm3 |
| Cntrl2 | 8.70E-115 | 0.646571023 | 0.449 | 0.118 | 1.76E-110 | 12 | Cntrl |
| Ptger43 | 9.92E-115 | 0.628268969 | 0.359 | 0.08 | 2.01E-110 | 12 | Ptger4 |
| B4galt53 | 1.69E-114 | 0.645813737 | 0.379 | 0.089 | 3.43E-110 | 12 | B4galt5 |
| Actr34 | 2.69E-114 | 0.823486234 | 0.869 | 0.383 | 5.45E-110 | 12 | Actr3 |
| Stk242 | 4.60E-114 | 0.790577061 | 0.581 | 0.189 | 9.31E-110 | 12 | Stk24 |
| Ifngr17 | 3.22E-113 | 1.121365565 | 0.822 | 0.386 | 6.51E-109 | 12 | Ifngr1 |
| Ccnd25 | 4.79E-113 | 0.904090938 | 0.735 | 0.278 | 9.69E-109 | 12 | Ccnd2 |
| Cdk2ap22 | 2.35E-112 | 0.621387451 | 0.596 | 0.192 | 4.77E-108 | 12 | Cdk2ap2 |
| Armc71 | 4.01E-112 | 0.438967313 | 0.26 | 0.045 | 8.12E-108 | 12 | Armc7 |
| Rpl27a4 | 8.96E-112 | 0.674456447 | 1 | 0.959 | 1.81E-107 | 12 | Rpl27a |
| Unc1192 | 4.20E-111 | 0.544088052 | 0.353 | 0.079 | 8.50E-107 | 12 | Unc119 |
| Rps203 | 1.22E-110 | 0.739355457 | 0.998 | 0.95 | 2.46E-106 | 12 | Rps20 |
| Pvt13 | 2.28E-110 | 0.588690026 | 0.34 | 0.074 | 4.61E-106 | 12 | Pvt1 |
| Fam172a5 | 4.13E-110 | 0.828170452 | 0.723 | 0.277 | 8.37E-106 | 12 | Fam172a |
| Rpl84 | 4.25E-110 | 0.714311955 | 0.996 | 0.924 | 8.60E-106 | 12 | Rpl8 |
| Cnn23 | 4.59E-110 | 0.762859145 | 0.555 | 0.176 | 9.29E-106 | 12 | Cnn2 |

| | | | | | | | |
|---|---|---|---|---|---|---|---|
| Gpr1831 | 1.31E-109 | 0.720443914 | 0.292 | 0.056 | 2.66E-105 | 12 | Gpr183 |
| Gramd1a2 | 1.44E-109 | 0.618552971 | 0.389 | 0.096 | 2.92E-105 | 12 | Gramd1a |
| Fam102a2 | 3.43E-109 | 0.542747781 | 0.391 | 0.095 | 6.94E-105 | 12 | Fam102a |
| Rcsd12 | 4.16E-109 | 0.66556076 | 0.454 | 0.124 | 8.41E-105 | 12 | Rcsd1 |
| AU0202065 | 1.06E-108 | 0.911581565 | 0.718 | 0.283 | 2.14E-104 | 12 | AU020206 |
| Mgat4a3 | 1.88E-108 | 0.664166134 | 0.398 | 0.1 | 3.80E-104 | 12 | Mgat4a |
| D16Ertd472e2 | 2.21E-108 | 0.613628183 | 0.402 | 0.101 | 4.48E-104 | 12 | D16Ertd472e |
| Vps13a2 | 1.12E-107 | 0.718793101 | 0.533 | 0.166 | 2.26E-103 | 12 | Vps13a |
| 4930523C07Rik5 | 1.72E-107 | 0.780218845 | 0.619 | 0.212 | 3.49E-103 | 12 | 4930523C07Rik |
| Rnf1662 | 7.20E-107 | 0.5757603 | 0.4 | 0.102 | 1.46E-102 | 12 | Rnf166 |
| Rpl174 | 9.56E-107 | 0.697269282 | 1 | 0.915 | 1.94E-102 | 12 | Rpl17 |
| Tagap1 | 1.15E-106 | 0.495351817 | 0.26 | 0.047 | 2.32E-102 | 12 | Tagap |
| Eya22 | 1.95E-106 | 0.517540568 | 0.393 | 0.095 | 3.94E-102 | 12 | Eya2 |
| Tle44 | 2.11E-106 | 0.802983549 | 0.617 | 0.219 | 4.27E-102 | 12 | Tle4 |
| Ppp1r12a5 | 2.62E-106 | 0.936265117 | 0.864 | 0.418 | 5.30E-102 | 12 | Ppp1r12a |
| Tecpr12 | 3.77E-106 | 0.673362524 | 0.449 | 0.125 | 7.63E-102 | 12 | Tecpr1 |
| Tmem1632 | 3.93E-106 | 0.652918204 | 0.364 | 0.083 | 7.96E-102 | 12 | Tmem163 |
| Mycbp26 | 1.07E-105 | 0.881400256 | 0.817 | 0.367 | 2.17E-101 | 12 | Mycbp2 |
| Tnfrsf92 | 1.31E-105 | 0.560397513 | 0.307 | 0.063 | 2.65E-101 | 12 | Tnfrsf9 |
| Ogt4 | 1.49E-105 | 0.751749711 | 0.589 | 0.2 | 3.01E-101 | 12 | Ogt |
| Prkacb2 | 8.21E-105 | 0.71365535 | 0.615 | 0.211 | 1.66E-100 | 12 | Prkacb |
| Rpl364 | 1.55E-104 | 0.709618166 | 0.998 | 0.902 | 3.13E-100 | 12 | Rpl36 |
| Slco3a11 | 1.88E-104 | 0.677909883 | 0.436 | 0.117 | 3.81E-100 | 12 | Slco3a1 |
| Lncpint5 | 1.89E-104 | 0.881387651 | 0.682 | 0.267 | 3.83E-100 | 12 | Lncpint |
| Tab22 | 2.31E-104 | 0.755417234 | 0.624 | 0.224 | 4.68E-100 | 12 | Tab2 |
| Vav34 | 3.27E-104 | 0.881734229 | 0.656 | 0.249 | 6.61E-100 | 12 | Vav3 |
| Zc3hav13 | 4.44E-104 | 0.69081412 | 0.587 | 0.198 | 8.99E-100 | 12 | Zc3hav1 |
| Atxn15 | 6.44E-104 | 0.972309674 | 0.716 | 0.288 | 1.30E-99 | 12 | Atxn1 |
| Mob3a2 | 7.62E-104 | 0.527612558 | 0.336 | 0.077 | 1.54E-99 | 12 | Mob3a |
| Rplp25 | 1.03E-103 | 0.676099257 | 0.998 | 0.888 | 2.09E-99 | 12 | Rplp2 |
| Arhgef18 | 1.54E-103 | 0.822622635 | 0.647 | 0.242 | 3.13E-99 | 12 | Arhgef1 |
| Rpl115 | 2.42E-103 | 0.680425031 | 1 | 0.926 | 4.91E-99 | 12 | Rpl11 |
| Rpl235 | 3.65E-103 | 0.66101738 | 1 | 0.97 | 7.39E-99 | 12 | Rpl23 |
| Ube2h2 | 7.02E-103 | 0.758171593 | 0.669 | 0.258 | 1.42E-98 | 12 | Ube2h |
| Crlf33 | 1.15E-102 | 0.748015863 | 0.495 | 0.151 | 2.32E-98 | 12 | Crlf3 |
| Stk104 | 5.97E-102 | 0.75039089 | 0.542 | 0.177 | 1.21E-97 | 12 | Stk10 |
| Inpp5d3 | 7.02E-102 | 0.634158216 | 0.598 | 0.197 | 1.42E-97 | 12 | Inpp5d |
| Prr132 | 1.17E-101 | 0.607552358 | 0.594 | 0.203 | 2.38E-97 | 12 | Prr13 |
| Rps194 | 1.65E-101 | 0.73717448 | 0.996 | 0.926 | 3.35E-97 | 12 | Rps19 |
| Plekha53 | 2.46E-101 | 0.786666755 | 0.488 | 0.15 | 4.99E-97 | 12 | Plekha5 |
| Ifi2061 | 4.10E-101 | 0.571658666 | 0.312 | 0.068 | 8.29E-97 | 12 | Ifi206 |
| Dnajc17 | 4.95E-101 | 0.902691067 | 0.74 | 0.316 | 1.00E-96 | 12 | Dnajc1 |
| Tra2b3 | 8.81E-101 | 0.799910318 | 0.708 | 0.289 | 1.78E-96 | 12 | Tra2b |

| | | | | | | | |
|---|---|---|---|---|---|---|---|
| Lrmp3 | 8.99E-101 | 0.478560673 | 0.28 | 0.057 | 1.82E-96 | 12 | Lrmp |
| Zcchc113 | 9.09E-101 | 0.777354596 | 0.677 | 0.26 | 1.84E-96 | 12 | Zcchc11 |
| Ankrd116 | 1.75E-100 | 0.916832132 | 0.806 | 0.374 | 3.54E-96 | 12 | Ankrd11 |
| Rpl13a4 | 1.88E-100 | 0.674788178 | 0.701 | 0.277 | 3.80E-96 | 12 | Rpl13a |
| Itpr23 | 2.25E-100 | 0.742718263 | 0.641 | 0.238 | 4.56E-96 | 12 | Itpr2 |
| Cbx42 | 2.96E-100 | 0.581294896 | 0.402 | 0.106 | 5.99E-96 | 12 | Cbx4 |
| Socs11 | 3.34E-100 | 0.576612629 | 0.376 | 0.093 | 6.75E-96 | 12 | Socs1 |
| Cdc42se23 | 5.63E-100 | 0.793371936 | 0.626 | 0.233 | 1.14E-95 | 12 | Cdc42se2 |
| Gzmb1 | 2.92E-99 | 1.195974993 | 0.338 | 0.078 | 5.92E-95 | 12 | Gzmb |
| Esyt25 | 3.14E-99 | 0.836071948 | 0.626 | 0.236 | 6.37E-95 | 12 | Esyt2 |
| Ahcyl21 | 4.61E-99 | 0.761266154 | 0.426 | 0.121 | 9.33E-95 | 12 | Ahcyl2 |
| Rps3a15 | 5.78E-99 | 0.693534529 | 0.998 | 0.934 | 1.17E-94 | 12 | Rps3a1 |
| Nfatc13 | 1.04E-98 | 0.655257829 | 0.406 | 0.111 | 2.10E-94 | 12 | Nfatc1 |
| Ccr22 | 1.11E-98 | 0.663868682 | 0.348 | 0.082 | 2.25E-94 | 12 | Ccr2 |
| Chd25 | 3.35E-98 | 0.771079849 | 0.647 | 0.247 | 6.79E-94 | 12 | Chd2 |
| Ust4 | 4.61E-98 | 0.674775421 | 0.338 | 0.082 | 9.34E-94 | 12 | Ust |
| Capza13 | 5.10E-98 | 0.695802324 | 0.686 | 0.268 | 1.03E-93 | 12 | Capza1 |
| Nedd95 | 2.08E-97 | 0.817382754 | 0.751 | 0.312 | 4.22E-93 | 12 | Nedd9 |
| Hmgb22 | 2.58E-97 | 0.595252221 | 0.778 | 0.321 | 5.22E-93 | 12 | Hmgb2 |
| B930036N10Rik | 2.83E-97 | 0.497805549 | 0.26 | 0.051 | 5.73E-93 | 12 | B930036N10Rik |
| Arhgef64 | 6.16E-97 | 0.59780403 | 0.432 | 0.123 | 1.25E-92 | 12 | Arhgef6 |
| Gab32 | 8.35E-97 | 0.606196102 | 0.359 | 0.09 | 1.69E-92 | 12 | Gab3 |
| Sec11c4 | 1.99E-96 | 0.621175406 | 0.746 | 0.303 | 4.03E-92 | 12 | Sec11c |
| mt-Co17 | 1.06E-95 | 0.763853714 | 1 | 0.715 | 2.14E-91 | 12 | mt-Co1 |
| Pip5k1b2 | 2.07E-95 | 0.549874979 | 0.37 | 0.095 | 4.20E-91 | 12 | Pip5k1b |
| Prkch5 | 4.51E-95 | 0.923682809 | 0.841 | 0.392 | 9.13E-91 | 12 | Prkch |
| Rps235 | 8.56E-95 | 0.600196172 | 1 | 0.952 | 1.73E-90 | 12 | Rps23 |
| Ago22 | 2.37E-94 | 0.659011585 | 0.499 | 0.162 | 4.80E-90 | 12 | Ago2 |
| Rgs23 | 2.59E-94 | 0.643683632 | 0.436 | 0.125 | 5.24E-90 | 12 | Rgs2 |
| Rfx72 | 5.14E-94 | 0.697539292 | 0.475 | 0.15 | 1.04E-89 | 12 | Rfx7 |
| Rpl95 | 1.07E-93 | 0.611449179 | 0.998 | 0.944 | 2.17E-89 | 12 | Rpl9 |
| Kif13b3 | 3.43E-93 | 0.792965456 | 0.581 | 0.21 | 6.95E-89 | 12 | Kif13b |
| Rps94 | 5.86E-93 | 0.633255653 | 1 | 0.919 | 1.19E-88 | 12 | Rps9 |
| Stap12 | 6.04E-93 | 0.276043558 | 0.426 | 0.119 | 1.22E-88 | 12 | Stap1 |
| Rps144 | 8.66E-93 | 0.571758591 | 0.998 | 0.942 | 1.75E-88 | 12 | Rps14 |
| Klrb1f2 | 2.83E-92 | 0.55527815 | 0.279 | 0.059 | 5.72E-88 | 12 | Klrb1f |
| Tnrc6b4 | 2.92E-92 | 0.773842569 | 0.796 | 0.364 | 5.90E-88 | 12 | Tnrc6b |
| Jak22 | 3.36E-92 | 0.691214765 | 0.419 | 0.122 | 6.81E-88 | 12 | Jak2 |
| Saraf5 | 2.35E-91 | 0.71705544 | 0.716 | 0.3 | 4.75E-87 | 12 | Saraf |
| H2-Q65 | 2.85E-91 | 0.650953235 | 0.639 | 0.239 | 5.78E-87 | 12 | H2-Q6 |
| Ccm22 | 2.96E-91 | 0.546991096 | 0.421 | 0.121 | 6.00E-87 | 12 | Ccm2 |
| Ldlrad43 | 6.27E-91 | 0.772629703 | 0.639 | 0.244 | 1.27E-86 | 12 | Ldlrad4 |
| Snrpg3 | 1.18E-90 | 0.591849923 | 0.75 | 0.317 | 2.40E-86 | 12 | Snrpg |

| | | | | | | | |
|---|---|---|---|---|---|---|---|
| Ccr53 | 1.52E-90 | 0.644556262 | 0.437 | 0.128 | 3.07E-86 | 12 | Ccr5 |
| H2-K16 | 2.88E-89 | 0.815829138 | 1 | 0.637 | 5.83E-85 | 12 | H2-K1 |
| Ctsd4 | 3.80E-89 | 0.771932106 | 0.714 | 0.307 | 7.70E-85 | 12 | Ctsd |
| Kansl16 | 1.01E-88 | 0.824167114 | 0.742 | 0.333 | 2.05E-84 | 12 | Kansl1 |
| Isg202 | 1.44E-88 | 0.521132601 | 0.396 | 0.111 | 2.92E-84 | 12 | Isg20 |
| Ptprj3 | 3.47E-88 | 0.567575551 | 0.473 | 0.146 | 7.02E-84 | 12 | Ptprj |
| Asxl22 | 4.42E-88 | 0.675911297 | 0.501 | 0.169 | 8.95E-84 | 12 | Asxl2 |
| Usp32 | 5.12E-88 | 0.607427439 | 0.518 | 0.177 | 1.04E-83 | 12 | Usp3 |
| Ccl43 | 7.10E-88 | 0.872083373 | 0.428 | 0.125 | 1.44E-83 | 12 | Ccl4 |
| Ssh24 | 1.51E-87 | 0.661027007 | 0.598 | 0.22 | 3.07E-83 | 12 | Ssh2 |
| 4933406I18Rik3 | 1.91E-87 | 0.614403843 | 0.303 | 0.072 | 3.86E-83 | 12 | 4933406I18Rik |
| Man1a5 | 4.05E-87 | 0.704065723 | 0.488 | 0.162 | 8.21E-83 | 12 | Man1a |
| Notch23 | 4.20E-87 | 0.537231866 | 0.312 | 0.077 | 8.51E-83 | 12 | Notch2 |
| Rps284 | 4.93E-87 | 0.674765561 | 0.998 | 0.896 | 9.98E-83 | 12 | Rps28 |
| Rpl22l14 | 4.32E-86 | 0.723055056 | 0.95 | 0.572 | 8.74E-82 | 12 | Rpl22l1 |
| Srpk11 | 5.62E-86 | 0.514725379 | 0.48 | 0.156 | 1.14E-81 | 12 | Srpk1 |
| Baz1a3 | 6.78E-86 | 0.621113973 | 0.497 | 0.167 | 1.37E-81 | 12 | Baz1a |
| Creb12 | 1.62E-85 | 0.72513392 | 0.553 | 0.202 | 3.28E-81 | 12 | Creb1 |
| Fryl3 | 2.34E-85 | 0.766797553 | 0.645 | 0.262 | 4.73E-81 | 12 | Fryl |
| Rps27a4 | 2.47E-85 | 0.562716074 | 0.998 | 0.949 | 4.99E-81 | 12 | Rps27a |
| Mapre23 | 3.55E-85 | 0.751351102 | 0.619 | 0.244 | 7.19E-81 | 12 | Mapre2 |
| Nsmaf3 | 6.04E-85 | 0.586653673 | 0.346 | 0.093 | 1.22E-80 | 12 | Nsmaf |
| Ptpn23 | 9.50E-85 | 0.555298882 | 0.441 | 0.138 | 1.92E-80 | 12 | Ptpn2 |
| Xrn22 | 9.50E-85 | 0.600833459 | 0.647 | 0.255 | 1.92E-80 | 12 | Xrn2 |
| Cbl3 | 1.18E-84 | 0.673241786 | 0.555 | 0.202 | 2.39E-80 | 12 | Cbl |
| Mtmr122 | 1.52E-84 | 0.566477347 | 0.417 | 0.126 | 3.09E-80 | 12 | Mtmr12 |
| Elmo15 | 1.89E-84 | 0.81462316 | 0.779 | 0.359 | 3.83E-80 | 12 | Elmo1 |
| Tcf204 | 2.22E-84 | 0.628214233 | 0.434 | 0.136 | 4.49E-80 | 12 | Tcf20 |
| Fosb4 | 3.42E-84 | 0.790615634 | 0.521 | 0.184 | 6.92E-80 | 12 | Fosb |
| Rps6ka13 | 3.56E-84 | 0.536850553 | 0.383 | 0.11 | 7.21E-80 | 12 | Rps6ka1 |
| Rbpj3 | 3.92E-84 | 0.673324689 | 0.576 | 0.215 | 7.94E-80 | 12 | Rbpj |
| Hvcn11 | 5.02E-84 | 0.388060085 | 0.25 | 0.053 | 1.02E-79 | 12 | Hvcn1 |
| Pycard4 | 5.79E-84 | 0.488170821 | 0.695 | 0.27 | 1.17E-79 | 12 | Pycard |
| Trip42 | 1.06E-83 | 0.56433714 | 0.391 | 0.114 | 2.15E-79 | 12 | Trip4 |
| Traf33 | 1.47E-83 | 0.580747848 | 0.37 | 0.105 | 2.98E-79 | 12 | Traf3 |
| Zfp36l23 | 4.27E-83 | 0.720847579 | 0.695 | 0.299 | 8.65E-79 | 12 | Zfp36l2 |
| Sun22 | 6.40E-83 | 0.472705477 | 0.301 | 0.074 | 1.30E-78 | 12 | Sun2 |
| Cap13 | 9.48E-83 | 0.618637444 | 0.656 | 0.265 | 1.92E-78 | 12 | Cap1 |
| Eef1a14 | 1.04E-82 | 0.52671904 | 0.998 | 0.959 | 2.11E-78 | 12 | Eef1a1 |
| Tsc22d43 | 1.69E-82 | 0.539971872 | 0.69 | 0.288 | 3.42E-78 | 12 | Tsc22d4 |
| Sms2 | 1.96E-82 | 0.454943865 | 0.456 | 0.145 | 3.97E-78 | 12 | Sms |
| Rpl274 | 2.59E-82 | 0.639469331 | 0.951 | 0.577 | 5.23E-78 | 12 | Rpl27 |
| Rbm32 | 3.69E-82 | 0.647820262 | 0.899 | 0.47 | 7.46E-78 | 12 | Rbm3 |

| | | | | | | | |
|---|---|---|---|---|---|---|---|
| Rpl375 | 4.30E-82 | 0.584183028 | 0.996 | 0.926 | 8.70E-78 | 12 | Rpl37 |
| Nfkb15 | 7.67E-82 | 0.696982348 | 0.649 | 0.265 | 1.55E-77 | 12 | Nfkb1 |
| Bcl2l112 | 1.82E-81 | 0.614190275 | 0.385 | 0.113 | 3.68E-77 | 12 | Bcl2l11 |
| Mgat53 | 3.84E-81 | 0.679088128 | 0.486 | 0.165 | 7.78E-77 | 12 | Mgat5 |
| Bach23 | 5.21E-81 | 0.770774143 | 0.464 | 0.156 | 1.06E-76 | 12 | Bach2 |
| Rpl224 | 8.73E-81 | 0.582535775 | 0.994 | 0.85 | 1.77E-76 | 12 | Rpl22 |
| Sep-94 | 1.01E-80 | 0.548035355 | 0.488 | 0.165 | 2.04E-76 | 12 | 9-Sep |
| Trmt1122 | 1.44E-80 | 0.553939259 | 0.652 | 0.268 | 2.91E-76 | 12 | Trmt112 |
| Uba525 | 2.69E-80 | 0.576888567 | 0.933 | 0.514 | 5.44E-76 | 12 | Uba52 |
| Sik34 | 5.93E-80 | 0.712630878 | 0.639 | 0.264 | 1.20E-75 | 12 | Sik3 |
| Zcchc76 | 1.31E-79 | 0.687063293 | 0.682 | 0.295 | 2.65E-75 | 12 | Zcchc7 |
| Def61 | 7.47E-79 | 0.334334002 | 0.335 | 0.089 | 1.51E-74 | 12 | Def6 |
| Ighm3 | 9.56E-79 | 0.478971237 | 0.402 | 0.122 | 1.93E-74 | 12 | Ighm |
| Rps4x4 | 1.41E-78 | 0.555950168 | 0.998 | 0.94 | 2.85E-74 | 12 | Rps4x |
| Arid1a5 | 2.58E-78 | 0.648921054 | 0.572 | 0.225 | 5.22E-74 | 12 | Arid1a |
| Rasa23 | 3.18E-78 | 0.547586572 | 0.421 | 0.134 | 6.44E-74 | 12 | Rasa2 |
| Dtnb2 | 5.45E-78 | 0.487940607 | 0.376 | 0.11 | 1.10E-73 | 12 | Dtnb |
| Itpk12 | 5.60E-78 | 0.458038005 | 0.279 | 0.068 | 1.13E-73 | 12 | Itpk1 |
| Stambpl12 | 6.13E-78 | 0.494718282 | 0.275 | 0.066 | 1.24E-73 | 12 | Stambpl1 |
| Rras22 | 6.21E-78 | 0.516997423 | 0.312 | 0.081 | 1.26E-73 | 12 | Rras2 |
| Atp2a32 | 8.62E-78 | 0.333684193 | 0.26 | 0.059 | 1.74E-73 | 12 | Atp2a3 |
| Clk14 | 8.65E-78 | 0.663150982 | 0.667 | 0.286 | 1.75E-73 | 12 | Clk1 |
| Birc32 | 8.81E-78 | 0.526740556 | 0.34 | 0.095 | 1.78E-73 | 12 | Birc3 |
| Rack15 | 9.72E-78 | 0.623768277 | 0.989 | 0.789 | 1.97E-73 | 12 | Rack1 |
| Tpp23 | 1.71E-77 | 0.599512284 | 0.574 | 0.222 | 3.46E-73 | 12 | Tpp2 |
| Fcer1g4 | 1.82E-77 | 1.091214991 | 0.632 | 0.264 | 3.68E-73 | 12 | Fcer1g |
| Rsbn1l3 | 3.69E-77 | 0.612204471 | 0.643 | 0.266 | 7.46E-73 | 12 | Rsbn1l |
| Stim14 | 7.57E-77 | 0.572781943 | 0.512 | 0.185 | 1.53E-72 | 12 | Stim1 |
| Rpl44 | 1.84E-76 | 0.633010657 | 0.97 | 0.653 | 3.73E-72 | 12 | Rpl4 |
| Vasp2 | 2.63E-76 | 0.608762239 | 0.579 | 0.232 | 5.32E-72 | 12 | Vasp |
| Plekhj12 | 2.78E-76 | 0.457823479 | 0.475 | 0.163 | 5.63E-72 | 12 | Plekhj1 |
| Rpl215 | 4.06E-76 | 0.538008562 | 0.998 | 0.922 | 8.22E-72 | 12 | Rpl21 |
| Psmb87 | 6.00E-76 | 0.679037186 | 0.953 | 0.551 | 1.21E-71 | 12 | Psmb8 |
| Zfp2925 | 6.34E-76 | 0.642492359 | 0.602 | 0.246 | 1.28E-71 | 12 | Zfp292 |
| Capzb4 | 8.37E-76 | 0.598355209 | 0.779 | 0.362 | 1.69E-71 | 12 | Capzb |
| Dennd4c3 | 1.28E-75 | 0.506952228 | 0.393 | 0.122 | 2.60E-71 | 12 | Dennd4c |
| Ttc72 | 1.34E-75 | 0.372099681 | 0.35 | 0.099 | 2.71E-71 | 12 | Ttc7 |
| Kcnq52 | 1.68E-75 | 0.72998101 | 0.273 | 0.065 | 3.40E-71 | 12 | Kcnq5 |
| Cers63 | 2.43E-75 | 0.668629371 | 0.368 | 0.112 | 4.92E-71 | 12 | Cers6 |
| Cd843 | 1.09E-74 | 0.501557311 | 0.32 | 0.087 | 2.21E-70 | 12 | Cd84 |
| Thada4 | 1.16E-74 | 0.551247448 | 0.376 | 0.116 | 2.34E-70 | 12 | Thada |
| Ptpn63 | 1.74E-74 | 0.480423878 | 0.406 | 0.126 | 3.53E-70 | 12 | Ptpn6 |
| Vmp13 | 2.70E-74 | 0.508808316 | 0.551 | 0.21 | 5.47E-70 | 12 | Vmp1 |

| | | | | | | | |
|---|---|---|---|---|---|---|---|
| Rpl65 | 3.29E-74 | 0.55611854 | 0.996 | 0.88 | 6.66E-70 | 12 | Rpl6 |
| Gas73 | 4.08E-74 | 0.859554313 | 0.479 | 0.178 | 8.25E-70 | 12 | Gas7 |
| Rpl315 | 4.12E-74 | 0.57030646 | 1 | 0.908 | 8.34E-70 | 12 | Rpl3 |
| Ncor15 | 6.07E-74 | 0.650954949 | 0.854 | 0.431 | 1.23E-69 | 12 | Ncor1 |
| Cdk175 | 6.18E-74 | 0.613113607 | 0.596 | 0.239 | 1.25E-69 | 12 | Cdk17 |
| Snrpf4 | 7.21E-74 | 0.45164923 | 0.798 | 0.372 | 1.46E-69 | 12 | Snrpf |
| Mkl14 | 7.43E-74 | 0.594091058 | 0.383 | 0.122 | 1.50E-69 | 12 | Mkl1 |
| Plekha23 | 1.00E-73 | 0.491186237 | 0.406 | 0.13 | 2.03E-69 | 12 | Plekha2 |
| Clcn31 | 1.11E-73 | 0.595789268 | 0.4 | 0.129 | 2.26E-69 | 12 | Clcn3 |
| Me22 | 1.97E-73 | 0.447421906 | 0.282 | 0.072 | 3.98E-69 | 12 | Me2 |
| Grap1 | 2.23E-73 | 0.528747605 | 0.37 | 0.111 | 4.52E-69 | 12 | Grap |
| Trp53inp11 | 2.93E-73 | 0.391307274 | 0.258 | 0.061 | 5.93E-69 | 12 | Trp53inp1 |
| mt-Atp86 | 3.51E-73 | 0.628106194 | 0.869 | 0.456 | 7.10E-69 | 12 | mt-Atp8 |
| Junb5 | 5.43E-73 | 0.725864825 | 0.793 | 0.386 | 1.10E-68 | 12 | Junb |
| mt-Nd55 | 5.51E-73 | 0.683397395 | 0.897 | 0.489 | 1.12E-68 | 12 | mt-Nd5 |
| Phf62 | 5.67E-73 | 0.548468274 | 0.406 | 0.131 | 1.15E-68 | 12 | Phf6 |
| Klre12 | 8.93E-73 | 0.624022892 | 0.351 | 0.1 | 1.81E-68 | 12 | Klre1 |
| Spcs22 | 1.75E-72 | 0.600261213 | 0.794 | 0.395 | 3.55E-68 | 12 | Spcs2 |
| Chst114 | 2.26E-72 | 0.611367531 | 0.475 | 0.171 | 4.57E-68 | 12 | Chst11 |
| Rpl10a5 | 2.59E-72 | 0.591905059 | 0.994 | 0.833 | 5.24E-68 | 12 | Rpl10a |
| mt-Co38 | 5.67E-72 | 0.639063833 | 1 | 0.722 | 1.15E-67 | 12 | mt-Co3 |
| Tnip1 | 6.61E-72 | 0.442145436 | 0.271 | 0.068 | 1.34E-67 | 12 | Tnip1 |
| Eif3e3 | 6.71E-72 | 0.528374555 | 0.815 | 0.406 | 1.36E-67 | 12 | Eif3e |
| Rngtt2 | 6.80E-72 | 0.548786941 | 0.402 | 0.131 | 1.38E-67 | 12 | Rngtt |
| Cog54 | 7.59E-72 | 0.61663819 | 0.482 | 0.178 | 1.54E-67 | 12 | Cog5 |
| Slfn83 | 1.17E-71 | 0.416059218 | 0.282 | 0.072 | 2.36E-67 | 12 | Slfn8 |
| Camk2g2 | 1.24E-71 | 0.515248605 | 0.404 | 0.131 | 2.51E-67 | 12 | Camk2g |
| Fchsd24 | 1.31E-71 | 0.557110844 | 0.45 | 0.155 | 2.65E-67 | 12 | Fchsd2 |
| Kpna46 | 1.33E-71 | 0.644473734 | 0.701 | 0.317 | 2.69E-67 | 12 | Kpna4 |
| Zbp11 | 1.74E-71 | 0.462609847 | 0.267 | 0.066 | 3.52E-67 | 12 | Zbp1 |
| Fgf132 | 3.66E-71 | 0.646402348 | 0.29 | 0.075 | 7.41E-67 | 12 | Fgf13 |
| Styk12 | 5.94E-71 | 0.672413624 | 0.335 | 0.094 | 1.20E-66 | 12 | Styk1 |
| Pacs13 | 6.82E-71 | 0.57461776 | 0.525 | 0.2 | 1.38E-66 | 12 | Pacs1 |
| Rps294 | 8.72E-71 | 0.477149615 | 0.998 | 0.971 | 1.77E-66 | 12 | Rps29 |
| Sipa1l13 | 9.45E-71 | 0.610447209 | 0.501 | 0.185 | 1.91E-66 | 12 | Sipa1l1 |
| Fam204a2 | 1.15E-70 | 0.46838365 | 0.465 | 0.165 | 2.33E-66 | 12 | Fam204a |
| Hcls13 | 1.28E-70 | 0.392886079 | 0.389 | 0.121 | 2.60E-66 | 12 | Hcls1 |
| Smad32 | 2.26E-70 | 0.496504017 | 0.325 | 0.093 | 4.58E-66 | 12 | Smad3 |
| Pla2g162 | 2.51E-70 | 0.42908925 | 0.594 | 0.237 | 5.08E-66 | 12 | Pla2g16 |
| Dnajc152 | 3.54E-70 | 0.321611931 | 0.721 | 0.315 | 7.16E-66 | 12 | Dnajc15 |
| Bin13 | 4.18E-70 | 0.404326245 | 0.477 | 0.171 | 8.46E-66 | 12 | Bin1 |
| Rab27a2 | 5.43E-70 | 0.356185416 | 0.303 | 0.081 | 1.10E-65 | 12 | Rab27a |
| Dapk23 | 7.13E-70 | 0.482595974 | 0.357 | 0.106 | 1.44E-65 | 12 | Dapk2 |

| | | | | | | | |
|---|---|---|---|---|---|---|---|
| Exoc22 | 1.23E-69 | 0.545144775 | 0.421 | 0.143 | 2.50E-65 | 12 | Exoc2 |
| Fam129a3 | 1.54E-69 | 0.557982602 | 0.421 | 0.141 | 3.11E-65 | 12 | Fam129a |
| Ambra15 | 5.79E-69 | 0.609592172 | 0.505 | 0.193 | 1.17E-64 | 12 | Ambra1 |
| Rbm256 | 1.59E-68 | 0.559900485 | 0.716 | 0.328 | 3.21E-64 | 12 | Rbm25 |
| Rps125 | 1.65E-68 | 0.520797618 | 0.994 | 0.897 | 3.34E-64 | 12 | Rps12 |
| Ppm1b2 | 1.69E-68 | 0.459177534 | 0.406 | 0.134 | 3.42E-64 | 12 | Ppm1b |
| Nckap1l3 | 2.19E-68 | 0.370318114 | 0.308 | 0.085 | 4.43E-64 | 12 | Nckap1l |
| Otulin2 | 2.21E-68 | 0.44681057 | 0.514 | 0.195 | 4.48E-64 | 12 | Otulin |
| 4932438A13Rik4 | 2.74E-68 | 0.600494532 | 0.536 | 0.214 | 5.56E-64 | 12 | 4932438A13Rik |
| Tln16 | 2.78E-68 | 0.572706497 | 0.817 | 0.398 | 5.63E-64 | 12 | Tln1 |
| Rps217 | 3.22E-68 | 0.560972129 | 0.998 | 0.919 | 6.52E-64 | 12 | Rps2 |
| Vps282 | 4.97E-68 | 0.453811403 | 0.673 | 0.295 | 1.01E-63 | 12 | Vps28 |
| Rpl35a4 | 5.66E-68 | 0.468749642 | 0.998 | 0.948 | 1.15E-63 | 12 | Rpl35a |
| Cdkn2d2 | 5.81E-68 | 0.432924752 | 0.314 | 0.089 | 1.18E-63 | 12 | Cdkn2d |
| Ccdc1072 | 7.03E-68 | 0.434949649 | 0.383 | 0.124 | 1.42E-63 | 12 | Ccdc107 |
| Mettl232 | 8.03E-68 | 0.409284956 | 0.52 | 0.198 | 1.63E-63 | 12 | Mettl23 |
| Phf21a5 | 1.06E-67 | 0.649684978 | 0.585 | 0.248 | 2.14E-63 | 12 | Phf21a |
| Ncoa26 | 1.28E-67 | 0.668255095 | 0.557 | 0.234 | 2.60E-63 | 12 | Ncoa2 |
| Arrb23 | 1.88E-67 | 0.353878592 | 0.316 | 0.089 | 3.81E-63 | 12 | Arrb2 |
| Ddx212 | 1.95E-67 | 0.46912325 | 0.533 | 0.208 | 3.94E-63 | 12 | Ddx21 |
| Tnfaip8l21 | 2.46E-67 | 0.37714668 | 0.267 | 0.069 | 4.98E-63 | 12 | Tnfaip8l2 |
| Ptpn16 | 2.82E-67 | 0.539596828 | 0.563 | 0.231 | 5.71E-63 | 12 | Ptpn1 |
| Cnot6l2 | 2.95E-67 | 0.577393263 | 0.572 | 0.239 | 5.98E-63 | 12 | Cnot6l |
| Smc42 | 2.99E-67 | 0.630599536 | 0.441 | 0.159 | 6.06E-63 | 12 | Smc4 |
| Cobll12 | 3.28E-67 | 0.463897206 | 0.299 | 0.082 | 6.64E-63 | 12 | Cobll1 |
| Irak23 | 3.40E-67 | 0.519197523 | 0.409 | 0.138 | 6.89E-63 | 12 | Irak2 |
| Zdhhc205 | 4.84E-67 | 0.557883838 | 0.559 | 0.225 | 9.79E-63 | 12 | Zdhhc20 |
| Aoah2 | 7.51E-67 | 0.466283211 | 0.271 | 0.07 | 1.52E-62 | 12 | Aoah |
| Cdc42ep34 | 1.21E-66 | 0.528447236 | 0.467 | 0.17 | 2.45E-62 | 12 | Cdc42ep3 |
| Csk3 | 1.36E-66 | 0.482141755 | 0.379 | 0.126 | 2.76E-62 | 12 | Csk |
| Galnt10 | 1.77E-66 | 0.487828016 | 0.335 | 0.101 | 3.58E-62 | 12 | Galnt10 |
| Atad22 | 5.75E-66 | 0.428538973 | 0.312 | 0.09 | 1.16E-61 | 12 | Atad2 |
| Crip15 | 6.17E-66 | 0.666575611 | 0.936 | 0.529 | 1.25E-61 | 12 | Crip1 |
| Arpc24 | 6.31E-66 | 0.567357843 | 0.946 | 0.538 | 1.28E-61 | 12 | Arpc2 |
| Gm118081 | 1.55E-65 | 0.283139207 | 0.436 | 0.149 | 3.13E-61 | 12 | Gm11808 |
| Ablim14 | 1.61E-65 | 0.544305919 | 0.807 | 0.396 | 3.25E-61 | 12 | Ablim1 |
| Ddx102 | 1.74E-65 | 0.396952131 | 0.391 | 0.128 | 3.52E-61 | 12 | Ddx10 |
| Nktr3 | 1.99E-65 | 0.599209009 | 0.591 | 0.252 | 4.02E-61 | 12 | Nktr |
| Fam173a2 | 2.10E-65 | 0.342862693 | 0.51 | 0.193 | 4.25E-61 | 12 | Fam173a |
| Pkn14 | 2.62E-65 | 0.493933219 | 0.482 | 0.182 | 5.30E-61 | 12 | Pkn1 |
| Arhgap264 | 2.65E-65 | 0.682327079 | 0.536 | 0.217 | 5.36E-61 | 12 | Arhgap26 |
| Pced1b1 | 1.04E-64 | 0.514904746 | 0.359 | 0.114 | 2.10E-60 | 12 | Pced1b |
| Eef1b25 | 1.12E-64 | 0.54742798 | 0.987 | 0.732 | 2.26E-60 | 12 | Eef1b2 |

| | | | | | | | |
|---|---|---|---|---|---|---|---|
| Arsb2 | 1.17E-64 | 0.594594564 | 0.372 | 0.123 | 2.36E-60 | 12 | Arsb |
| Zfp6542 | 1.19E-64 | 0.529589269 | 0.363 | 0.117 | 2.42E-60 | 12 | Zfp654 |
| Stk392 | 1.62E-64 | 0.346185617 | 0.434 | 0.15 | 3.29E-60 | 12 | Stk39 |
| Cln32 | 1.90E-64 | 0.387426257 | 0.256 | 0.066 | 3.84E-60 | 12 | Cln3 |
| Heca2 | 2.58E-64 | 0.491499532 | 0.402 | 0.139 | 5.23E-60 | 12 | Heca |
| Hnrnpf2 | 3.16E-64 | 0.514939463 | 0.901 | 0.489 | 6.39E-60 | 12 | Hnrnpf |
| Clic14 | 3.82E-64 | 0.572007364 | 0.959 | 0.595 | 7.73E-60 | 12 | Clic1 |
| Usp482 | 9.53E-64 | 0.419954149 | 0.379 | 0.125 | 1.93E-59 | 12 | Usp48 |
| Sp41 | 1.09E-63 | 0.464261408 | 0.314 | 0.093 | 2.20E-59 | 12 | Sp4 |
| Rps27rt2 | 1.21E-63 | 0.319594154 | 0.34 | 0.104 | 2.45E-59 | 12 | Rps27rt |
| Tbl1xr13 | 1.31E-63 | 0.561539615 | 0.432 | 0.16 | 2.66E-59 | 12 | Tbl1xr1 |
| Scfd21 | 1.88E-63 | 0.403357777 | 0.293 | 0.083 | 3.80E-59 | 12 | Scfd2 |
| Rpl294 | 2.66E-63 | 0.567198464 | 0.978 | 0.765 | 5.38E-59 | 12 | Rpl29 |
| Lnpep6 | 3.52E-63 | 0.592998097 | 0.587 | 0.255 | 7.14E-59 | 12 | Lnpep |
| Itgb16 | 4.55E-63 | 0.871863781 | 0.85 | 0.473 | 9.22E-59 | 12 | Itgb1 |
| Hectd14 | 6.85E-63 | 0.593615831 | 0.639 | 0.295 | 1.39E-58 | 12 | Hectd1 |
| Vars2 | 8.93E-63 | 0.417784758 | 0.28 | 0.079 | 1.81E-58 | 12 | Vars |
| Arid4b5 | 9.85E-63 | 0.56962616 | 0.639 | 0.287 | 1.99E-58 | 12 | Arid4b |
| Zyx3 | 1.06E-62 | 0.567607471 | 0.52 | 0.211 | 2.15E-58 | 12 | Zyx |
| Plgrkt3 | 1.20E-62 | 0.503642654 | 0.52 | 0.208 | 2.43E-58 | 12 | Plgrkt |
| Myh95 | 1.43E-62 | 0.621652934 | 0.824 | 0.431 | 2.90E-58 | 12 | Myh9 |
| Srpk21 | 2.24E-62 | 0.451170511 | 0.488 | 0.189 | 4.54E-58 | 12 | Srpk2 |
| Sh3gl12 | 2.56E-62 | 0.427393694 | 0.368 | 0.121 | 5.19E-58 | 12 | Sh3gl1 |
| Ubash3b3 | 2.76E-62 | 0.568411469 | 0.439 | 0.164 | 5.59E-58 | 12 | Ubash3b |
| Virma2 | 2.79E-62 | 0.46914375 | 0.378 | 0.128 | 5.66E-58 | 12 | Virma |
| Rin33 | 5.01E-62 | 0.541179472 | 0.46 | 0.175 | 1.01E-57 | 12 | Rin3 |
| Srrm25 | 7.94E-62 | 0.609246179 | 0.854 | 0.463 | 1.61E-57 | 12 | Srrm2 |
| Esco13 | 8.23E-62 | 0.48927603 | 0.462 | 0.175 | 1.67E-57 | 12 | Esco1 |
| Rnf1382 | 9.31E-62 | 0.394818071 | 0.389 | 0.133 | 1.88E-57 | 12 | Rnf138 |
| Smg16 | 9.71E-62 | 0.586922471 | 0.544 | 0.23 | 1.97E-57 | 12 | Smg1 |
| Sp1102 | 1.63E-61 | 0.414765978 | 0.426 | 0.153 | 3.31E-57 | 12 | Sp110 |
| Tex143 | 2.98E-61 | 0.623931467 | 0.407 | 0.146 | 6.03E-57 | 12 | Tex14 |
| Pitpnc15 | 3.47E-61 | 0.669034209 | 0.809 | 0.423 | 7.03E-57 | 12 | Pitpnc1 |
| Senp64 | 4.14E-61 | 0.519912002 | 0.645 | 0.293 | 8.38E-57 | 12 | Senp6 |
| Haus31 | 6.61E-61 | 0.343211303 | 0.262 | 0.07 | 1.34E-56 | 12 | Haus3 |
| Ranbp93 | 6.69E-61 | 0.557270748 | 0.49 | 0.196 | 1.35E-56 | 12 | Ranbp9 |
| Supt4a3 | 9.21E-61 | 0.384057264 | 0.684 | 0.316 | 1.86E-56 | 12 | Supt4a |
| Nsa24 | 9.21E-61 | 0.435919924 | 0.867 | 0.48 | 1.86E-56 | 12 | Nsa2 |
| Rnaseh2c2 | 1.07E-60 | 0.406182247 | 0.626 | 0.28 | 2.17E-56 | 12 | Rnaseh2c |
| Emg1 | 1.24E-60 | 0.400166421 | 0.538 | 0.221 | 2.50E-56 | 12 | Emg1 |
| Eif3f2 | 1.47E-60 | 0.469759908 | 0.877 | 0.476 | 2.97E-56 | 12 | Eif3f |
| Rasa14 | 2.09E-60 | 0.547757363 | 0.568 | 0.242 | 4.23E-56 | 12 | Rasa1 |
| Gm472834 | 2.18E-60 | 0.610646762 | 0.609 | 0.276 | 4.42E-56 | 12 | Gm47283 |

| | | | | | | | |
|---|---|---|---|---|---|---|---|
| Sec11a3 | 3.13E-60 | 0.478571211 | 0.619 | 0.28 | 6.33E-56 | 12 | Sec11a |
| Trim12a4 | 3.32E-60 | 0.475185518 | 0.464 | 0.177 | 6.73E-56 | 12 | Trim12a |
| Pan36 | 6.86E-60 | 0.534314463 | 0.533 | 0.224 | 1.39E-55 | 12 | Pan3 |
| Rps85 | 6.94E-60 | 0.427866374 | 0.998 | 0.964 | 1.40E-55 | 12 | Rps8 |
| Rpl154 | 1.48E-59 | 0.471711915 | 0.994 | 0.9 | 2.99E-55 | 12 | Rpl15 |
| Tpt14 | 1.53E-59 | 0.363825508 | 1 | 0.99 | 3.09E-55 | 12 | Tpt1 |
| Lta4h2 | 1.84E-59 | 0.423759716 | 0.407 | 0.145 | 3.73E-55 | 12 | Lta4h |
| Itgax3 | 2.07E-59 | 0.542205237 | 0.327 | 0.105 | 4.19E-55 | 12 | Itgax |
| Exoc47 | 2.34E-59 | 0.660315346 | 0.66 | 0.318 | 4.73E-55 | 12 | Exoc4 |
| Taok33 | 2.55E-59 | 0.411196091 | 0.436 | 0.159 | 5.15E-55 | 12 | Taok3 |
| Odc11 | 2.69E-59 | 0.382223756 | 0.477 | 0.182 | 5.45E-55 | 12 | Odc1 |
| Taf102 | 2.80E-59 | 0.435390035 | 0.757 | 0.378 | 5.66E-55 | 12 | Taf10 |
| Sqstm14 | 3.19E-59 | 0.8342291 | 0.621 | 0.3 | 6.46E-55 | 12 | Sqstm1 |
| Tpr4 | 3.46E-59 | 0.526422771 | 0.815 | 0.419 | 7.00E-55 | 12 | Tpr |
| Tbc1d13 | 3.51E-59 | 0.462406961 | 0.434 | 0.161 | 7.11E-55 | 12 | Tbc1d1 |
| Sf3b14 | 4.73E-59 | 0.482907727 | 0.772 | 0.387 | 9.57E-55 | 12 | Sf3b1 |
| Klhl63 | 6.23E-59 | 0.456094231 | 0.353 | 0.117 | 1.26E-54 | 12 | Klhl6 |
| Exosc8 | 7.05E-59 | 0.3280249 | 0.387 | 0.132 | 1.43E-54 | 12 | Exosc8 |
| Rap1gds16 | 7.44E-59 | 0.556976255 | 0.585 | 0.256 | 1.51E-54 | 12 | Rap1gds1 |
| Rbm395 | 8.74E-59 | 0.563040592 | 0.953 | 0.568 | 1.77E-54 | 12 | Rbm39 |
| Vps13b4 | 1.39E-58 | 0.559600981 | 0.514 | 0.215 | 2.82E-54 | 12 | Vps13b |
| Snrnp702 | 1.40E-58 | 0.448381098 | 0.602 | 0.267 | 2.83E-54 | 12 | Snrnp70 |
| Sppl35 | 1.70E-58 | 0.565385138 | 0.514 | 0.214 | 3.44E-54 | 12 | Sppl3 |
| Smap23 | 2.01E-58 | 0.433108782 | 0.508 | 0.207 | 4.07E-54 | 12 | Smap2 |
| Snd14 | 2.69E-58 | 0.472300559 | 0.443 | 0.169 | 5.45E-54 | 12 | Snd1 |
| Lrba3 | 3.86E-58 | 0.56332276 | 0.471 | 0.187 | 7.81E-54 | 12 | Lrba |
| Car22 | 6.95E-58 | 0.483614611 | 0.357 | 0.119 | 1.41E-53 | 12 | Car2 |
| Rps6ka35 | 7.18E-58 | 0.550894795 | 0.579 | 0.258 | 1.45E-53 | 12 | Rps6ka3 |
| Nipbl7 | 1.62E-57 | 0.510349 | 0.736 | 0.36 | 3.29E-53 | 12 | Nipbl |
| Ppp1r16b5 | 2.15E-57 | 0.565942972 | 0.49 | 0.198 | 4.35E-53 | 12 | Ppp1r16b |
| Tor1aip22 | 2.25E-57 | 0.529805592 | 0.45 | 0.178 | 4.56E-53 | 12 | Tor1aip2 |
| Bola23 | 3.27E-57 | 0.359425497 | 0.585 | 0.257 | 6.63E-53 | 12 | Bola2 |
| Clint12 | 4.15E-57 | 0.339348453 | 0.74 | 0.368 | 8.39E-53 | 12 | Clint1 |
| Smad75 | 5.21E-57 | 0.574254181 | 0.677 | 0.321 | 1.05E-52 | 12 | Smad7 |
| Usp153 | 6.66E-57 | 0.526982136 | 0.523 | 0.227 | 1.35E-52 | 12 | Usp15 |
| Maz1 | 6.73E-57 | 0.338355302 | 0.295 | 0.089 | 1.36E-52 | 12 | Maz |
| Phf35 | 9.65E-57 | 0.519933222 | 0.561 | 0.248 | 1.95E-52 | 12 | Phf3 |
| Bhlhe403 | 1.15E-56 | 0.464944142 | 0.421 | 0.154 | 2.33E-52 | 12 | Bhlhe40 |
| Abhd2 | 1.33E-56 | 0.373363438 | 0.264 | 0.075 | 2.69E-52 | 12 | Abhd2 |
| Ptbp36 | 1.35E-56 | 0.551420527 | 0.755 | 0.388 | 2.73E-52 | 12 | Ptbp3 |
| Tec3 | 2.36E-56 | 0.399649899 | 0.329 | 0.107 | 4.78E-52 | 12 | Tec |
| Herc44 | 2.90E-56 | 0.506533548 | 0.398 | 0.148 | 5.87E-52 | 12 | Herc4 |
| Afg3l21 | 3.38E-56 | 0.269355024 | 0.265 | 0.075 | 6.84E-52 | 12 | Afg3l2 |

| | | | | | | | |
|---|---|---|---|---|---|---|---|
| Ube2i2 | 3.63E-56 | 0.387513377 | 0.682 | 0.325 | 7.34E-52 | 12 | Ube2i |
| Sri2 | 4.38E-56 | 0.427277961 | 0.787 | 0.415 | 8.87E-52 | 12 | Sri |
| B4galt15 | 5.60E-56 | 0.494863478 | 0.443 | 0.173 | 1.13E-51 | 12 | B4galt1 |
| Itch5 | 6.24E-56 | 0.552710908 | 0.591 | 0.269 | 1.26E-51 | 12 | Itch |
| Rpl36a5 | 8.73E-56 | 0.476701997 | 0.981 | 0.748 | 1.77E-51 | 12 | Rpl36a |
| Ptp4a12 | 9.66E-56 | 0.413079813 | 0.387 | 0.141 | 1.96E-51 | 12 | Ptp4a1 |
| Eif3h3 | 1.44E-55 | 0.406621831 | 0.847 | 0.474 | 2.92E-51 | 12 | Eif3h |
| Dapp12 | 1.56E-55 | 0.430088326 | 0.267 | 0.079 | 3.15E-51 | 12 | Dapp1 |
| Slc44a21 | 1.70E-55 | 0.445646219 | 0.346 | 0.118 | 3.45E-51 | 12 | Slc44a2 |
| St8sia45 | 1.78E-55 | 0.577859744 | 0.572 | 0.255 | 3.60E-51 | 12 | St8sia4 |
| Ncl2 | 2.24E-55 | 0.476859491 | 0.89 | 0.51 | 4.54E-51 | 12 | Ncl |
| Pspc12 | 2.82E-55 | 0.406432964 | 0.342 | 0.115 | 5.70E-51 | 12 | Pspc1 |
| Smg64 | 2.86E-55 | 0.610103361 | 0.564 | 0.255 | 5.78E-51 | 12 | Smg6 |
| Arid1b5 | 3.01E-55 | 0.548879778 | 0.564 | 0.254 | 6.09E-51 | 12 | Arid1b |
| Mri11 | 3.73E-55 | 0.336822286 | 0.262 | 0.075 | 7.55E-51 | 12 | Mri1 |
| mt-Atp68 | 4.26E-55 | 0.568992637 | 1 | 0.75 | 8.62E-51 | 12 | mt-Atp6 |
| Zdhhc184 | 6.58E-55 | 0.422003276 | 0.439 | 0.166 | 1.33E-50 | 12 | Zdhhc18 |
| Gbp82 | 8.05E-55 | 0.445411957 | 0.378 | 0.135 | 1.63E-50 | 12 | Gbp8 |
| Tecr3 | 8.53E-55 | 0.430673921 | 0.516 | 0.22 | 1.73E-50 | 12 | Tecr |
| Slfn24 | 1.02E-54 | 0.544875908 | 0.57 | 0.256 | 2.07E-50 | 12 | Slfn2 |
| Pld32 | 1.19E-54 | 0.361434172 | 0.277 | 0.083 | 2.41E-50 | 12 | Pld3 |
| Grk23 | 1.57E-54 | 0.356377005 | 0.353 | 0.121 | 3.18E-50 | 12 | Grk2 |
| Rpl23a5 | 1.59E-54 | 0.518451371 | 0.957 | 0.675 | 3.22E-50 | 12 | Rpl23a |
| Slc2a12 | 1.80E-54 | 0.382806558 | 0.28 | 0.085 | 3.65E-50 | 12 | Slc2a1 |
| Prrc2c6 | 1.98E-54 | 0.526605754 | 0.787 | 0.415 | 4.01E-50 | 12 | Prrc2c |
| Arl6ip15 | 2.16E-54 | 0.47528169 | 0.735 | 0.364 | 4.37E-50 | 12 | Arl6ip1 |
| Emid12 | 2.95E-54 | 0.449175523 | 0.321 | 0.106 | 5.96E-50 | 12 | Emid1 |
| Rab8a2 | 3.50E-54 | 0.340676029 | 0.542 | 0.232 | 7.09E-50 | 12 | Rab8a |
| F2r1 | 4.62E-54 | 0.396940036 | 0.288 | 0.088 | 9.35E-50 | 12 | F2r |
| Smc52 | 5.38E-54 | 0.37711216 | 0.342 | 0.115 | 1.09E-49 | 12 | Smc5 |
| Rpl245 | 6.23E-54 | 0.393275205 | 0.994 | 0.906 | 1.26E-49 | 12 | Rpl24 |
| Vrk23 | 6.90E-54 | 0.443736494 | 0.35 | 0.121 | 1.40E-49 | 12 | Vrk2 |
| Tcf126 | 7.21E-54 | 0.622908989 | 0.714 | 0.371 | 1.46E-49 | 12 | Tcf12 |
| Glipr13 | 1.43E-53 | 0.324915578 | 0.299 | 0.093 | 2.90E-49 | 12 | Glipr1 |
| Phf145 | 3.16E-53 | 0.471527629 | 0.544 | 0.24 | 6.41E-49 | 12 | Phf14 |
| Sub13 | 3.55E-53 | 0.415841329 | 0.88 | 0.51 | 7.19E-49 | 12 | Sub1 |
| Rpl144 | 3.94E-53 | 0.448984336 | 0.985 | 0.799 | 7.98E-49 | 12 | Rpl14 |
| Arl6ip55 | 4.69E-53 | 0.470405462 | 0.508 | 0.217 | 9.50E-49 | 12 | Arl6ip5 |
| Cnot12 | 5.16E-53 | 0.427515502 | 0.432 | 0.169 | 1.04E-48 | 12 | Cnot1 |
| Srsf117 | 6.22E-53 | 0.434708506 | 0.71 | 0.345 | 1.26E-48 | 12 | Srsf11 |
| Snhg12 | 9.06E-53 | 0.287287427 | 0.335 | 0.111 | 1.83E-48 | 12 | Snhg12 |
| Ern12 | 9.87E-53 | 0.474453599 | 0.269 | 0.082 | 2.00E-48 | 12 | Ern1 |
| Rps265 | 1.10E-52 | 0.417342158 | 0.993 | 0.864 | 2.23E-48 | 12 | Rps26 |

| | | | | | | | |
|---|---|---|---|---|---|---|---|
| Sltm4 | 1.39E-52 | 0.482540751 | 0.578 | 0.266 | 2.81E-48 | 12 | Sltm |
| Arap24 | 1.52E-52 | 0.505428178 | 0.439 | 0.174 | 3.08E-48 | 12 | Arap2 |
| Sipa13 | 1.55E-52 | 0.466601183 | 0.449 | 0.182 | 3.13E-48 | 12 | Sipa1 |
| Zfp5122 | 2.69E-52 | 0.397170524 | 0.275 | 0.084 | 5.45E-48 | 12 | Zfp512 |
| Gm17018 | 3.11E-52 | 0.274335859 | 0.303 | 0.097 | 6.29E-48 | 12 | Gm17018 |
| Ankrd126 | 3.19E-52 | 0.510983091 | 0.634 | 0.303 | 6.45E-48 | 12 | Ankrd12 |
| Vti1a4 | 3.77E-52 | 0.521128209 | 0.551 | 0.251 | 7.64E-48 | 12 | Vti1a |
| Vps543 | 4.85E-52 | 0.498953704 | 0.449 | 0.183 | 9.81E-48 | 12 | Vps54 |
| Pbxip13 | 7.93E-52 | 0.368754522 | 0.333 | 0.114 | 1.61E-47 | 12 | Pbxip1 |
| Gls4 | 9.67E-52 | 0.477405703 | 0.548 | 0.247 | 1.96E-47 | 12 | Gls |
| Atp1b34 | 1.02E-51 | 0.41527239 | 0.553 | 0.244 | 2.07E-47 | 12 | Atp1b3 |
| Elmsan12 | 1.23E-51 | 0.377813336 | 0.31 | 0.103 | 2.50E-47 | 12 | Elmsan1 |
| Rsrc14 | 1.71E-51 | 0.458973981 | 0.561 | 0.256 | 3.46E-47 | 12 | Rsrc1 |
| Nt5c3 | 1.75E-51 | 0.317499973 | 0.559 | 0.245 | 3.54E-47 | 12 | Nt5c |
| Ets15 | 2.98E-51 | 0.422378789 | 0.798 | 0.397 | 6.04E-47 | 12 | Ets1 |
| Sesn13 | 4.06E-51 | 0.552920285 | 0.486 | 0.208 | 8.22E-47 | 12 | Sesn1 |
| Actr23 | 4.36E-51 | 0.388061218 | 0.695 | 0.338 | 8.83E-47 | 12 | Actr2 |
| Ankib13 | 6.06E-51 | 0.45898569 | 0.366 | 0.135 | 1.23E-46 | 12 | Ankib1 |
| Ralgapa22 | 7.43E-51 | 0.457662954 | 0.393 | 0.15 | 1.50E-46 | 12 | Ralgapa2 |
| Runx23 | 9.03E-51 | 0.398855253 | 0.282 | 0.089 | 1.83E-46 | 12 | Runx2 |
| Maml26 | 1.09E-50 | 0.554403745 | 0.677 | 0.338 | 2.21E-46 | 12 | Maml2 |
| Cep1921 | 1.33E-50 | 0.36863469 | 0.267 | 0.083 | 2.69E-46 | 12 | Cep192 |
| Npm14 | 1.56E-50 | 0.465554673 | 0.959 | 0.675 | 3.17E-46 | 12 | Npm1 |
| Polr2a5 | 1.68E-50 | 0.476941102 | 0.594 | 0.276 | 3.39E-46 | 12 | Polr2a |
| Dcaf52 | 1.68E-50 | 0.387739657 | 0.353 | 0.127 | 3.40E-46 | 12 | Dcaf5 |
| Dbnl2 | 2.29E-50 | 0.334094837 | 0.462 | 0.188 | 4.63E-46 | 12 | Dbnl |
| Ascc35 | 2.45E-50 | 0.496581037 | 0.475 | 0.203 | 4.97E-46 | 12 | Ascc3 |
| H2afj4 | 3.25E-50 | 0.260409614 | 0.832 | 0.447 | 6.58E-46 | 12 | H2afj |
| Sirt72 | 4.00E-50 | 0.410442054 | 0.355 | 0.131 | 8.10E-46 | 12 | Sirt7 |
| Mitd1 | 5.71E-50 | 0.356473831 | 0.396 | 0.151 | 1.16E-45 | 12 | Mitd1 |
| Cerk3 | 5.88E-50 | 0.447186825 | 0.32 | 0.112 | 1.19E-45 | 12 | Cerk |
| Galnt17 | 7.58E-50 | 0.456802777 | 0.441 | 0.181 | 1.53E-45 | 12 | Galnt1 |
| Dennd1a5 | 8.36E-50 | 0.476656631 | 0.546 | 0.249 | 1.69E-45 | 12 | Dennd1a |
| Bclaf13 | 1.06E-49 | 0.459804414 | 0.548 | 0.249 | 2.15E-45 | 12 | Bclaf1 |
| Parp43 | 1.78E-49 | 0.427526298 | 0.389 | 0.149 | 3.60E-45 | 12 | Parp4 |
| Mfng | 1.85E-49 | 0.352489155 | 0.264 | 0.081 | 3.75E-45 | 12 | Mfng |
| Rev3l3 | 2.45E-49 | 0.474152119 | 0.393 | 0.154 | 4.96E-45 | 12 | Rev3l |
| Hivep33 | 3.64E-49 | 0.48296804 | 0.404 | 0.158 | 7.38E-45 | 12 | Hivep3 |
| Chd73 | 5.68E-49 | 0.416315445 | 0.387 | 0.148 | 1.15E-44 | 12 | Chd7 |
| Rplp05 | 9.39E-49 | 0.430029088 | 1 | 0.944 | 1.90E-44 | 12 | Rplp0 |
| Dmtf13 | 9.73E-49 | 0.376315249 | 0.333 | 0.119 | 1.97E-44 | 12 | Dmtf1 |
| Atp2b15 | 9.79E-49 | 0.533756082 | 0.664 | 0.328 | 1.98E-44 | 12 | Atp2b1 |
| Nek74 | 1.01E-48 | 0.438237191 | 0.536 | 0.244 | 2.04E-44 | 12 | Nek7 |

| | | | | | | | |
|---|---|---|---|---|---|---|---|
| mt-Nd17 | 1.10E-48 | 0.499856367 | 0.968 | 0.593 | 2.23E-44 | 12 | mt-Nd1 |
| Rps65 | 1.14E-48 | 0.435035451 | 0.981 | 0.809 | 2.32E-44 | 12 | Rps6 |
| Rfx32 | 1.20E-48 | 0.442063935 | 0.307 | 0.105 | 2.43E-44 | 12 | Rfx3 |
| Rel3 | 1.22E-48 | 0.467817659 | 0.361 | 0.135 | 2.47E-44 | 12 | Rel |
| Ggnbp24 | 1.48E-48 | 0.460838115 | 0.525 | 0.238 | 3.00E-44 | 12 | Ggnbp2 |
| Setbp13 | 1.97E-48 | 0.544663461 | 0.335 | 0.119 | 3.99E-44 | 12 | Setbp1 |
| Reps12 | 2.46E-48 | 0.35608731 | 0.295 | 0.098 | 4.98E-44 | 12 | Reps1 |
| Slc28a21 | 3.21E-48 | 0.392618339 | 0.31 | 0.104 | 6.50E-44 | 12 | Slc28a2 |
| Nt5dc12 | 3.55E-48 | 0.359459582 | 0.331 | 0.117 | 7.19E-44 | 12 | Nt5dc1 |
| Vsir1 | 3.82E-48 | 0.370427328 | 0.301 | 0.101 | 7.74E-44 | 12 | Vsir |
| Arih14 | 4.68E-48 | 0.499681198 | 0.626 | 0.308 | 9.47E-44 | 12 | Arih1 |
| Fndc3a3 | 5.90E-48 | 0.525967368 | 0.574 | 0.273 | 1.19E-43 | 12 | Fndc3a |
| Abr3 | 8.54E-48 | 0.44016001 | 0.394 | 0.155 | 1.73E-43 | 12 | Abr |
| Trp532 | 9.15E-48 | 0.366880705 | 0.396 | 0.155 | 1.85E-43 | 12 | Trp53 |
| Commd72 | 1.09E-47 | 0.318678114 | 0.465 | 0.197 | 2.22E-43 | 12 | Commd7 |
| Borcs81 | 1.19E-47 | 0.267345805 | 0.307 | 0.104 | 2.40E-43 | 12 | Borcs8 |
| Rpl56 | 1.21E-47 | 0.42021121 | 0.979 | 0.706 | 2.44E-43 | 12 | Rpl5 |
| Taf152 | 1.23E-47 | 0.427285041 | 0.523 | 0.233 | 2.50E-43 | 12 | Taf15 |
| Rictor3 | 1.56E-47 | 0.413369033 | 0.391 | 0.153 | 3.16E-43 | 12 | Rictor |
| Cdyl22 | 1.66E-47 | 0.406783889 | 0.275 | 0.089 | 3.35E-43 | 12 | Cdyl2 |
| Arhgef35 | 2.03E-47 | 0.484901793 | 0.523 | 0.236 | 4.11E-43 | 12 | Arhgef3 |
| Ppp6r12 | 2.46E-47 | 0.31334588 | 0.293 | 0.098 | 4.99E-43 | 12 | Ppp6r1 |
| Fhit3 | 3.05E-47 | 0.480777472 | 0.25 | 0.077 | 6.17E-43 | 12 | Fhit |
| Cnot44 | 3.16E-47 | 0.459146023 | 0.559 | 0.264 | 6.41E-43 | 12 | Cnot4 |
| Ddx39b2 | 3.41E-47 | 0.34583765 | 0.516 | 0.229 | 6.90E-43 | 12 | Ddx39b |
| Gpcpd11 | 3.60E-47 | 0.417372028 | 0.32 | 0.114 | 7.29E-43 | 12 | Gpcpd1 |
| Gmip3 | 5.72E-47 | 0.366868485 | 0.254 | 0.08 | 1.16E-42 | 12 | Gmip |
| Rnf19b2 | 5.87E-47 | 0.334587598 | 0.26 | 0.082 | 1.19E-42 | 12 | Rnf19b |
| Rpl355 | 6.24E-47 | 0.40581247 | 0.987 | 0.826 | 1.26E-42 | 12 | Rpl35 |
| Gsap3 | 6.76E-47 | 0.414956679 | 0.26 | 0.083 | 1.37E-42 | 12 | Gsap |
| Ccnl13 | 7.45E-47 | 0.395405258 | 0.479 | 0.206 | 1.51E-42 | 12 | Ccnl1 |
| Ppp3ca6 | 7.47E-47 | 0.502403762 | 0.723 | 0.388 | 1.51E-42 | 12 | Ppp3ca |
| Crtc34 | 7.71E-47 | 0.440231684 | 0.4 | 0.161 | 1.56E-42 | 12 | Crtc3 |
| Atad2b4 | 7.78E-47 | 0.423643153 | 0.464 | 0.199 | 1.58E-42 | 12 | Atad2b |
| Arhgap103 | 8.81E-47 | 0.365077751 | 0.342 | 0.124 | 1.78E-42 | 12 | Arhgap10 |
| Cdk135 | 9.13E-47 | 0.436772059 | 0.499 | 0.223 | 1.85E-42 | 12 | Cdk13 |
| Kdm6a5 | 1.35E-46 | 0.425265536 | 0.482 | 0.21 | 2.74E-42 | 12 | Kdm6a |
| Klf64 | 1.46E-46 | 0.497318576 | 0.564 | 0.263 | 2.97E-42 | 12 | Klf6 |
| Chka3 | 1.50E-46 | 0.408727391 | 0.37 | 0.141 | 3.05E-42 | 12 | Chka |
| Ankrd176 | 1.78E-46 | 0.457762501 | 0.544 | 0.257 | 3.61E-42 | 12 | Ankrd17 |
| Uimc12 | 2.11E-46 | 0.330519415 | 0.379 | 0.146 | 4.27E-42 | 12 | Uimc1 |
| Nup1532 | 2.45E-46 | 0.391405223 | 0.318 | 0.114 | 4.96E-42 | 12 | Nup153 |
| Aprt1 | 2.60E-46 | 0.284195235 | 0.535 | 0.241 | 5.26E-42 | 12 | Aprt |

| | | | | | | | |
|---|---|---|---|---|---|---|---|
| Myo9b3 | 2.83E-46 | 0.400218008 | 0.321 | 0.117 | 5.73E-42 | 12 | Myo9b |
| Psmd141 | 3.51E-46 | 0.286824916 | 0.49 | 0.213 | 7.10E-42 | 12 | Psmd14 |
| Cnp | 4.22E-46 | 0.274540219 | 0.284 | 0.094 | 8.54E-42 | 12 | Cnp |
| Jund3 | 5.30E-46 | 0.538725396 | 0.95 | 0.641 | 1.07E-41 | 12 | Jund |
| Litaf3 | 7.18E-46 | 0.454451202 | 0.383 | 0.155 | 1.45E-41 | 12 | Litaf |
| Rae1 | 9.81E-46 | 0.250016627 | 0.293 | 0.099 | 1.99E-41 | 12 | Rae1 |
| Wac5 | 1.02E-45 | 0.452113533 | 0.632 | 0.318 | 2.06E-41 | 12 | Wac |
| Akr1b32 | 1.03E-45 | 0.368834146 | 0.437 | 0.186 | 2.08E-41 | 12 | Akr1b3 |
| Rbm56 | 1.06E-45 | 0.461970607 | 0.512 | 0.236 | 2.14E-41 | 12 | Rbm5 |
| Foxo33 | 1.13E-45 | 0.341488257 | 0.318 | 0.114 | 2.30E-41 | 12 | Foxo3 |
| Luc7l25 | 1.32E-45 | 0.457911721 | 0.738 | 0.4 | 2.67E-41 | 12 | Luc7l2 |
| Hdac42 | 1.63E-45 | 0.38115797 | 0.297 | 0.103 | 3.30E-41 | 12 | Hdac4 |
| Hnrnpll | 1.73E-45 | 0.452040215 | 0.308 | 0.111 | 3.50E-41 | 12 | Hnrnpll |
| Ccl33 | 1.82E-45 | 0.535856092 | 0.252 | 0.08 | 3.67E-41 | 12 | Ccl3 |
| Dusp113 | 1.82E-45 | 0.365427246 | 0.579 | 0.277 | 3.68E-41 | 12 | Dusp11 |
| Gna131 | 2.59E-45 | 0.363062467 | 0.409 | 0.168 | 5.24E-41 | 12 | Gna13 |
| Farp12 | 2.72E-45 | 0.316198577 | 0.305 | 0.105 | 5.50E-41 | 12 | Farp1 |
| Nr4a22 | 2.95E-45 | 0.566636137 | 0.338 | 0.128 | 5.98E-41 | 12 | Nr4a2 |
| Nin2 | 4.10E-45 | 0.38274932 | 0.355 | 0.136 | 8.30E-41 | 12 | Nin |
| Wnk16 | 4.31E-45 | 0.462418629 | 0.695 | 0.358 | 8.73E-41 | 12 | Wnk1 |
| Fam120b1 | 5.94E-45 | 0.347946297 | 0.267 | 0.087 | 1.20E-40 | 12 | Fam120b |
| Kdm5a2 | 8.11E-45 | 0.396602051 | 0.4 | 0.164 | 1.64E-40 | 12 | Kdm5a |
| Cfl14 | 8.70E-45 | 0.483996214 | 0.942 | 0.603 | 1.76E-40 | 12 | Cfl1 |
| Rbm331 | 9.30E-45 | 0.372441884 | 0.361 | 0.139 | 1.88E-40 | 12 | Rbm33 |
| Pnn5 | 1.27E-44 | 0.416910595 | 0.553 | 0.263 | 2.57E-40 | 12 | Pnn |
| Itgav4 | 1.33E-44 | 0.426435195 | 0.393 | 0.158 | 2.69E-40 | 12 | Itgav |
| Zbtb1 | 1.56E-44 | 0.293216705 | 0.277 | 0.092 | 3.16E-40 | 12 | Zbtb1 |
| Fbxw42 | 1.62E-44 | 0.313827294 | 0.316 | 0.113 | 3.28E-40 | 12 | Fbxw4 |
| Galnt73 | 2.09E-44 | 0.377984 | 0.383 | 0.152 | 4.23E-40 | 12 | Galnt7 |
| Cers4 | 2.30E-44 | 0.383814606 | 0.305 | 0.107 | 4.65E-40 | 12 | Cers4 |
| Hdac83 | 2.67E-44 | 0.474824232 | 0.346 | 0.133 | 5.41E-40 | 12 | Hdac8 |
| Krtcap23 | 3.24E-44 | 0.363924834 | 0.787 | 0.437 | 6.55E-40 | 12 | Krtcap2 |
| Celf13 | 3.52E-44 | 0.390039855 | 0.484 | 0.216 | 7.14E-40 | 12 | Celf1 |
| Rtf13 | 3.53E-44 | 0.313298666 | 0.578 | 0.275 | 7.14E-40 | 12 | Rtf1 |
| Mfsd101 | 3.63E-44 | 0.316935346 | 0.277 | 0.092 | 7.35E-40 | 12 | Mfsd10 |
| Pak24 | 3.67E-44 | 0.36924075 | 0.787 | 0.429 | 7.43E-40 | 12 | Pak2 |
| Atf7ip4 | 3.78E-44 | 0.483800652 | 0.456 | 0.204 | 7.65E-40 | 12 | Atf7ip |
| Tmem1313 | 4.33E-44 | 0.437463608 | 0.464 | 0.204 | 8.76E-40 | 12 | Tmem131 |
| Rnf1493 | 4.58E-44 | 0.350413783 | 0.29 | 0.102 | 9.27E-40 | 12 | Rnf149 |
| Lrrfip15 | 4.71E-44 | 0.446574231 | 0.703 | 0.374 | 9.54E-40 | 12 | Lrrfip1 |
| Tgfbr25 | 5.01E-44 | 0.37613507 | 0.63 | 0.306 | 1.02E-39 | 12 | Tgfbr2 |
| Senp73 | 7.78E-44 | 0.399571154 | 0.379 | 0.154 | 1.57E-39 | 12 | Senp7 |
| Eef1d3 | 7.97E-44 | 0.31708292 | 0.899 | 0.544 | 1.61E-39 | 12 | Eef1d |

| | | | | | | | |
|---|---|---|---|---|---|---|---|
| Susd66 | 8.45E-44 | 0.470302548 | 0.564 | 0.279 | 1.71E-39 | 12 | Susd6 |
| Ss182 | 8.66E-44 | 0.27737945 | 0.417 | 0.171 | 1.75E-39 | 12 | Ss18 |
| Kmt2c6 | 9.35E-44 | 0.405850075 | 0.611 | 0.302 | 1.89E-39 | 12 | Kmt2c |
| Eef1g4 | 9.44E-44 | 0.286857233 | 0.871 | 0.505 | 1.91E-39 | 12 | Eef1g |
| Smap12 | 1.08E-43 | 0.29260861 | 0.686 | 0.356 | 2.19E-39 | 12 | Smap1 |
| Rif11 | 1.14E-43 | 0.374173316 | 0.379 | 0.152 | 2.30E-39 | 12 | Rif1 |
| Med10 | 1.51E-43 | 0.308039371 | 0.447 | 0.193 | 3.06E-39 | 12 | Med10 |
| Impact1 | 1.64E-43 | 0.450508207 | 0.321 | 0.12 | 3.32E-39 | 12 | Impact |
| Twf22 | 1.68E-43 | 0.349038396 | 0.406 | 0.167 | 3.40E-39 | 12 | Twf2 |
| Pde4b2 | 2.20E-43 | 0.405209027 | 0.447 | 0.192 | 4.46E-39 | 12 | Pde4b |
| Agfg11 | 2.46E-43 | 0.352654109 | 0.394 | 0.16 | 4.99E-39 | 12 | Agfg1 |
| Khdrbs11 | 2.65E-43 | 0.323944725 | 0.551 | 0.261 | 5.37E-39 | 12 | Khdrbs1 |
| Wdr432 | 2.90E-43 | 0.322750076 | 0.372 | 0.148 | 5.87E-39 | 12 | Wdr43 |
| Zfand61 | 3.19E-43 | 0.326983365 | 0.594 | 0.295 | 6.46E-39 | 12 | Zfand6 |
| Zfp1484 | 3.60E-43 | 0.416912922 | 0.55 | 0.261 | 7.30E-39 | 12 | Zfp148 |
| Rcbtb2 | 5.59E-43 | 0.37753207 | 0.321 | 0.12 | 1.13E-38 | 12 | Rcbtb2 |
| Dis3l24 | 6.01E-43 | 0.387684239 | 0.35 | 0.136 | 1.22E-38 | 12 | Dis3l2 |
| Mgea53 | 9.02E-43 | 0.335245887 | 0.402 | 0.166 | 1.83E-38 | 12 | Mgea5 |
| Kcnq1ot13 | 1.09E-42 | 0.457713084 | 0.342 | 0.131 | 2.21E-38 | 12 | Kcnq1ot1 |
| Nudt211 | 1.10E-42 | 0.311265779 | 0.383 | 0.154 | 2.22E-38 | 12 | Nudt21 |
| mt-Co29 | 1.18E-42 | 0.493126529 | 1 | 0.724 | 2.40E-38 | 12 | mt-Co2 |
| Krit15 | 1.19E-42 | 0.450998969 | 0.417 | 0.179 | 2.41E-38 | 12 | Krit1 |
| Zc3h7a4 | 1.35E-42 | 0.363471513 | 0.389 | 0.159 | 2.73E-38 | 12 | Zc3h7a |
| Fars21 | 1.42E-42 | 0.337119429 | 0.454 | 0.2 | 2.87E-38 | 12 | Fars2 |
| Mndal5 | 1.48E-42 | 0.441961875 | 0.744 | 0.4 | 3.00E-38 | 12 | Mndal |
| Lrch35 | 1.63E-42 | 0.387647391 | 0.503 | 0.229 | 3.31E-38 | 12 | Lrch3 |
| Ppil22 | 2.42E-42 | 0.302965149 | 0.37 | 0.147 | 4.90E-38 | 12 | Ppil2 |
| Xpr12 | 2.76E-42 | 0.383409416 | 0.372 | 0.151 | 5.59E-38 | 12 | Xpr1 |
| Atxn2l | 3.16E-42 | 0.296899989 | 0.307 | 0.111 | 6.41E-38 | 12 | Atxn2l |
| Sp1004 | 3.30E-42 | 0.420329775 | 0.764 | 0.427 | 6.69E-38 | 12 | Sp100 |
| Psmb32 | 3.72E-42 | 0.301476394 | 0.757 | 0.414 | 7.53E-38 | 12 | Psmb3 |
| Zc3h132 | 4.00E-42 | 0.43984932 | 0.467 | 0.212 | 8.10E-38 | 12 | Zc3h13 |
| Rnf23 | 4.70E-42 | 0.343118757 | 0.288 | 0.103 | 9.51E-38 | 12 | Rnf2 |
| Ssbp4 | 4.89E-42 | 0.28803896 | 0.443 | 0.189 | 9.90E-38 | 12 | Ssbp4 |
| Plekhm32 | 5.83E-42 | 0.404196564 | 0.252 | 0.084 | 1.18E-37 | 12 | Plekhm3 |
| Dad14 | 6.34E-42 | 0.331916088 | 0.809 | 0.453 | 1.28E-37 | 12 | Dad1 |
| Atxn7l12 | 8.17E-42 | 0.391442507 | 0.336 | 0.131 | 1.65E-37 | 12 | Atxn7l1 |
| Osbpl84 | 9.56E-42 | 0.413832012 | 0.441 | 0.196 | 1.94E-37 | 12 | Osbpl8 |
| Pigx | 9.59E-42 | 0.276793153 | 0.279 | 0.097 | 1.94E-37 | 12 | Pigx |
| Mvb12b2 | 1.09E-41 | 0.351431254 | 0.394 | 0.164 | 2.21E-37 | 12 | Mvb12b |
| Gm40701 | 1.16E-41 | 0.333486931 | 0.279 | 0.096 | 2.34E-37 | 12 | Gm4070 |
| Ppp1ca2 | 1.26E-41 | 0.354579509 | 0.714 | 0.392 | 2.56E-37 | 12 | Ppp1ca |
| Cdkal14 | 1.31E-41 | 0.372023147 | 0.417 | 0.177 | 2.65E-37 | 12 | Cdkal1 |

| | | | | | | | |
|---|---|---|---|---|---|---|---|
| Nasp2 | 1.68E-41 | 0.35865301 | 0.402 | 0.169 | 3.40E-37 | 12 | Nasp |
| Nsun6 | 1.97E-41 | 0.380856749 | 0.275 | 0.096 | 3.99E-37 | 12 | Nsun6 |
| Tmem1643 | 2.00E-41 | 0.406071244 | 0.387 | 0.161 | 4.05E-37 | 12 | Tmem164 |
| Rpl76 | 2.12E-41 | 0.361979832 | 0.994 | 0.891 | 4.29E-37 | 12 | Rpl7 |
| Hivep26 | 2.16E-41 | 0.521241084 | 0.583 | 0.294 | 4.38E-37 | 12 | Hivep2 |
| Mlxip2 | 2.17E-41 | 0.372212236 | 0.325 | 0.125 | 4.39E-37 | 12 | Mlxip |
| Zeb14 | 2.49E-41 | 0.40987812 | 0.634 | 0.317 | 5.03E-37 | 12 | Zeb1 |
| Birc66 | 2.89E-41 | 0.453070119 | 0.639 | 0.337 | 5.84E-37 | 12 | Birc6 |
| Ifi208 | 3.00E-41 | 0.346002985 | 0.252 | 0.083 | 6.07E-37 | 12 | Ifi208 |
| Zfp4074 | 3.28E-41 | 0.413893244 | 0.381 | 0.158 | 6.64E-37 | 12 | Zfp407 |
| Rock15 | 3.79E-41 | 0.415914062 | 0.781 | 0.427 | 7.68E-37 | 12 | Rock1 |
| Larp14 | 4.79E-41 | 0.389191417 | 0.447 | 0.2 | 9.70E-37 | 12 | Larp1 |
| Usp243 | 4.87E-41 | 0.361513493 | 0.34 | 0.134 | 9.85E-37 | 12 | Usp24 |
| Kdm5c | 5.57E-41 | 0.36363099 | 0.344 | 0.135 | 1.13E-36 | 12 | Kdm5c |
| Map3k23 | 5.66E-41 | 0.36045554 | 0.368 | 0.151 | 1.15E-36 | 12 | Map3k2 |
| Morc31 | 6.25E-41 | 0.319955137 | 0.383 | 0.159 | 1.26E-36 | 12 | Morc3 |
| Ubr24 | 6.51E-41 | 0.414909739 | 0.488 | 0.226 | 1.32E-36 | 12 | Ubr2 |
| Nub11 | 6.69E-41 | 0.314361396 | 0.422 | 0.183 | 1.35E-36 | 12 | Nub1 |
| Zfp6522 | 7.68E-41 | 0.446825074 | 0.51 | 0.244 | 1.55E-36 | 12 | Zfp652 |
| Zfp3981 | 8.43E-41 | 0.298759235 | 0.267 | 0.092 | 1.71E-36 | 12 | Zfp398 |
| Slc4a71 | 8.86E-41 | 0.357784701 | 0.256 | 0.087 | 1.79E-36 | 12 | Slc4a7 |
| R3hdm1 | 9.34E-41 | 0.311582713 | 0.434 | 0.188 | 1.89E-36 | 12 | R3hdm1 |
| Mark2 | 9.48E-41 | 0.320809512 | 0.434 | 0.189 | 1.92E-36 | 12 | Mark2 |
| Cnot32 | 1.36E-40 | 0.385387 | 0.417 | 0.182 | 2.76E-36 | 12 | Cnot3 |
| Syncrip2 | 1.57E-40 | 0.332571889 | 0.469 | 0.212 | 3.19E-36 | 12 | Syncrip |
| Pdcd6ip2 | 1.72E-40 | 0.360276954 | 0.486 | 0.226 | 3.49E-36 | 12 | Pdcd6ip |
| Tex22 | 2.15E-40 | 0.317037592 | 0.256 | 0.087 | 4.36E-36 | 12 | Tex2 |
| Acin14 | 2.32E-40 | 0.414295486 | 0.607 | 0.312 | 4.70E-36 | 12 | Acin1 |
| Hspa56 | 2.52E-40 | 0.407643321 | 0.778 | 0.43 | 5.10E-36 | 12 | Hspa5 |
| Tagln24 | 3.05E-40 | 0.41388773 | 0.757 | 0.424 | 6.18E-36 | 12 | Tagln2 |
| Mdfic3 | 3.27E-40 | 0.462559635 | 0.507 | 0.242 | 6.61E-36 | 12 | Mdfic |
| Nsmce23 | 3.37E-40 | 0.355047276 | 0.45 | 0.203 | 6.82E-36 | 12 | Nsmce2 |
| Scaper4 | 3.39E-40 | 0.404173567 | 0.409 | 0.176 | 6.87E-36 | 12 | Scaper |
| Zfp3182 | 3.64E-40 | 0.369036489 | 0.271 | 0.095 | 7.36E-36 | 12 | Zfp318 |
| Supt202 | 4.33E-40 | 0.404704675 | 0.378 | 0.158 | 8.77E-36 | 12 | Supt20 |
| Gabpb2 | 4.63E-40 | 0.256479295 | 0.31 | 0.116 | 9.37E-36 | 12 | Gabpb2 |
| Tpm35 | 5.24E-40 | 0.363022372 | 0.836 | 0.474 | 1.06E-35 | 12 | Tpm3 |
| Csnk1g14 | 5.34E-40 | 0.432744277 | 0.366 | 0.155 | 1.08E-35 | 12 | Csnk1g1 |
| Med15 | 5.72E-40 | 0.332833476 | 0.335 | 0.13 | 1.16E-35 | 12 | Med15 |
| Mllt105 | 6.41E-40 | 0.43626596 | 0.542 | 0.271 | 1.30E-35 | 12 | Mllt10 |
| mt-Nd47 | 7.84E-40 | 0.447558438 | 0.989 | 0.67 | 1.59E-35 | 12 | mt-Nd4 |
| Rad501 | 8.32E-40 | 0.342317703 | 0.348 | 0.14 | 1.68E-35 | 12 | Rad50 |
| mt-Nd26 | 9.13E-40 | 0.447550346 | 0.996 | 0.684 | 1.85E-35 | 12 | mt-Nd2 |

| | | | | | | | |
|---|---|---|---|---|---|---|---|
| Stt3b3 | 9.64E-40 | 0.375186588 | 0.385 | 0.162 | 1.95E-35 | 12 | Stt3b |
| Fbxl177 | 9.93E-40 | 0.48373901 | 0.598 | 0.307 | 2.01E-35 | 12 | Fbxl17 |
| Asb31 | 1.20E-39 | 0.362976843 | 0.292 | 0.108 | 2.43E-35 | 12 | Asb3 |
| Gm372403 | 1.26E-39 | 0.351419523 | 0.282 | 0.102 | 2.55E-35 | 12 | Gm37240 |
| Xpa1 | 1.57E-39 | 0.271210423 | 0.318 | 0.121 | 3.18E-35 | 12 | Xpa |
| Fyco12 | 1.66E-39 | 0.349195212 | 0.316 | 0.121 | 3.35E-35 | 12 | Fyco1 |
| Ptpre3 | 1.91E-39 | 0.418252857 | 0.376 | 0.158 | 3.86E-35 | 12 | Ptpre |
| Prkdc1 | 2.39E-39 | 0.344914232 | 0.256 | 0.088 | 4.84E-35 | 12 | Prkdc |
| Gbf14 | 2.73E-39 | 0.418672765 | 0.407 | 0.179 | 5.53E-35 | 12 | Gbf1 |
| Jarid23 | 2.75E-39 | 0.352067592 | 0.443 | 0.2 | 5.58E-35 | 12 | Jarid2 |
| Pnisr5 | 3.03E-39 | 0.372304851 | 0.536 | 0.26 | 6.14E-35 | 12 | Pnisr |
| Nsd15 | 3.88E-39 | 0.373642407 | 0.654 | 0.345 | 7.86E-35 | 12 | Nsd1 |
| Cd1643 | 4.02E-39 | 0.284039505 | 0.465 | 0.211 | 8.13E-35 | 12 | Cd164 |
| Spata53 | 4.73E-39 | 0.347301791 | 0.314 | 0.121 | 9.58E-35 | 12 | Spata5 |
| Tmem1231 | 5.26E-39 | 0.292631927 | 0.368 | 0.151 | 1.07E-34 | 12 | Tmem123 |
| Chd64 | 7.64E-39 | 0.327031238 | 0.467 | 0.214 | 1.55E-34 | 12 | Chd6 |
| Avl92 | 7.66E-39 | 0.353057109 | 0.336 | 0.135 | 1.55E-34 | 12 | Avl9 |
| Coq10b2 | 8.30E-39 | 0.399101551 | 0.363 | 0.151 | 1.68E-34 | 12 | Coq10b |
| Efr3a3 | 8.43E-39 | 0.348557914 | 0.434 | 0.194 | 1.71E-34 | 12 | Efr3a |
| Gon4l2 | 8.57E-39 | 0.346386827 | 0.357 | 0.146 | 1.73E-34 | 12 | Gon4l |
| Adipor23 | 8.59E-39 | 0.342796724 | 0.359 | 0.148 | 1.74E-34 | 12 | Adipor2 |
| Nr2c24 | 9.66E-39 | 0.42076558 | 0.368 | 0.156 | 1.96E-34 | 12 | Nr2c2 |
| Phf20l15 | 1.34E-38 | 0.37876485 | 0.564 | 0.283 | 2.72E-34 | 12 | Phf20l1 |
| Gm268872 | 1.68E-38 | 0.404511866 | 0.271 | 0.098 | 3.41E-34 | 12 | Gm26887 |
| Lrif11 | 1.69E-38 | 0.28944677 | 0.342 | 0.136 | 3.43E-34 | 12 | Lrif1 |
| Flna2 | 1.83E-38 | 0.342170155 | 0.325 | 0.129 | 3.71E-34 | 12 | Flna |
| Dnajb1 | 1.94E-38 | 0.278729127 | 0.301 | 0.113 | 3.92E-34 | 12 | Dnajb1 |
| Bcas35 | 2.20E-38 | 0.384913266 | 0.407 | 0.179 | 4.46E-34 | 12 | Bcas3 |
| Ttc146 | 2.52E-38 | 0.339027306 | 0.462 | 0.212 | 5.10E-34 | 12 | Ttc14 |
| Foxp14 | 2.56E-38 | 0.426564649 | 0.699 | 0.385 | 5.19E-34 | 12 | Foxp1 |
| Rbm65 | 3.05E-38 | 0.396143305 | 0.458 | 0.214 | 6.18E-34 | 12 | Rbm6 |
| H2-D16 | 3.58E-38 | 0.456964978 | 0.994 | 0.664 | 7.26E-34 | 12 | H2-D1 |
| Csnk1g33 | 3.61E-38 | 0.377936524 | 0.406 | 0.18 | 7.31E-34 | 12 | Csnk1g3 |
| Psip12 | 3.78E-38 | 0.317901015 | 0.432 | 0.191 | 7.66E-34 | 12 | Psip1 |
| Lat23 | 4.30E-38 | 0.299217161 | 0.331 | 0.129 | 8.70E-34 | 12 | Lat2 |
| Kdm4c4 | 4.48E-38 | 0.353246934 | 0.387 | 0.166 | 9.07E-34 | 12 | Kdm4c |
| Lrp104 | 4.81E-38 | 0.390491126 | 0.415 | 0.185 | 9.73E-34 | 12 | Lrp10 |
| Rpa11 | 6.46E-38 | 0.251776076 | 0.264 | 0.093 | 1.31E-33 | 12 | Rpa1 |
| Appl12 | 6.68E-38 | 0.320950336 | 0.409 | 0.178 | 1.35E-33 | 12 | Appl1 |
| Zfp1821 | 6.77E-38 | 0.367183225 | 0.286 | 0.107 | 1.37E-33 | 12 | Zfp182 |
| Nop562 | 9.22E-38 | 0.303559541 | 0.327 | 0.129 | 1.87E-33 | 12 | Nop56 |
| Ifnar14 | 1.04E-37 | 0.348240323 | 0.346 | 0.142 | 2.10E-33 | 12 | Ifnar1 |
| Lbr4 | 1.04E-37 | 0.322203321 | 0.364 | 0.15 | 2.11E-33 | 12 | Lbr |

| | | | | | | | |
|---|---|---|---|---|---|---|---|
| Srgn6 | 1.09E-37 | 0.391315627 | 0.955 | 0.538 | 2.20E-33 | 12 | Srgn |
| Hspa4 | 1.42E-37 | 0.297687017 | 0.607 | 0.312 | 2.88E-33 | 12 | Hspa4 |
| Mef2a4 | 1.48E-37 | 0.569579854 | 0.725 | 0.412 | 2.99E-33 | 12 | Mef2a |
| Immp2l3 | 3.37E-37 | 0.467939901 | 0.331 | 0.133 | 6.83E-33 | 12 | Immp2l |
| Nlk2 | 3.79E-37 | 0.433235172 | 0.342 | 0.143 | 7.68E-33 | 12 | Nlk |
| Rgs34 | 5.37E-37 | 0.411887367 | 0.415 | 0.184 | 1.09E-32 | 12 | Rgs3 |
| Hnrnpa31 | 6.15E-37 | 0.288155967 | 0.768 | 0.44 | 1.24E-32 | 12 | Hnrnpa3 |
| Babam24 | 6.96E-37 | 0.400924794 | 0.488 | 0.235 | 1.41E-32 | 12 | Babam2 |
| Rcor12 | 8.75E-37 | 0.342846664 | 0.406 | 0.18 | 1.77E-32 | 12 | Rcor1 |
| Faf13 | 8.86E-37 | 0.322552701 | 0.437 | 0.198 | 1.79E-32 | 12 | Faf1 |
| Thrap33 | 1.04E-36 | 0.289496038 | 0.574 | 0.29 | 2.10E-32 | 12 | Thrap3 |
| Bcap291 | 1.18E-36 | 0.273569537 | 0.273 | 0.1 | 2.39E-32 | 12 | Bcap29 |
| Ddx46 | 1.23E-36 | 0.270020858 | 0.473 | 0.222 | 2.50E-32 | 12 | Ddx46 |
| Zfp6385 | 1.33E-36 | 0.338771223 | 0.555 | 0.278 | 2.68E-32 | 12 | Zfp638 |
| Antxr22 | 1.37E-36 | 0.359768598 | 0.25 | 0.089 | 2.78E-32 | 12 | Antxr2 |
| Casp83 | 1.93E-36 | 0.314273102 | 0.454 | 0.211 | 3.90E-32 | 12 | Casp8 |
| Atp9b3 | 2.08E-36 | 0.305849944 | 0.366 | 0.155 | 4.21E-32 | 12 | Atp9b |
| Kdm3b3 | 2.54E-36 | 0.348432592 | 0.329 | 0.134 | 5.13E-32 | 12 | Kdm3b |
| Rab3gap11 | 3.08E-36 | 0.304476479 | 0.376 | 0.161 | 6.23E-32 | 12 | Rab3gap1 |
| Nufip22 | 3.21E-36 | 0.389863342 | 0.475 | 0.229 | 6.50E-32 | 12 | Nufip2 |
| Xrn14 | 3.46E-36 | 0.337375829 | 0.436 | 0.2 | 7.01E-32 | 12 | Xrn1 |
| Dhx362 | 3.71E-36 | 0.2675663 | 0.488 | 0.232 | 7.52E-32 | 12 | Dhx36 |
| Hnrnpl2 | 4.08E-36 | 0.303712324 | 0.484 | 0.232 | 8.27E-32 | 12 | Hnrnpl |
| Mbd55 | 4.26E-36 | 0.435867636 | 0.456 | 0.217 | 8.62E-32 | 12 | Mbd5 |
| Rptor3 | 4.68E-36 | 0.35300516 | 0.329 | 0.136 | 9.48E-32 | 12 | Rptor |
| Mapk62 | 4.80E-36 | 0.258391644 | 0.273 | 0.1 | 9.71E-32 | 12 | Mapk6 |
| Plcg23 | 5.62E-36 | 0.304682832 | 0.303 | 0.118 | 1.14E-31 | 12 | Plcg2 |
| Itsn25 | 5.84E-36 | 0.396992018 | 0.544 | 0.274 | 1.18E-31 | 12 | Itsn2 |
| Ube2j12 | 6.92E-36 | 0.281364573 | 0.37 | 0.159 | 1.40E-31 | 12 | Ube2j1 |
| Snrnp481 | 8.22E-36 | 0.251313248 | 0.411 | 0.182 | 1.66E-31 | 12 | Snrnp48 |
| Scai1 | 8.87E-36 | 0.30306495 | 0.271 | 0.1 | 1.80E-31 | 12 | Scai |
| Gm269171 | 1.05E-35 | 0.307529972 | 0.301 | 0.118 | 2.12E-31 | 12 | Gm26917 |
| Btbd96 | 1.15E-35 | 0.41663792 | 0.484 | 0.234 | 2.33E-31 | 12 | Btbd9 |
| Cnot23 | 1.51E-35 | 0.299229075 | 0.449 | 0.207 | 3.05E-31 | 12 | Cnot2 |
| Scaf85 | 1.53E-35 | 0.39008426 | 0.398 | 0.182 | 3.09E-31 | 12 | Scaf8 |
| Zufsp3 | 1.73E-35 | 0.347740912 | 0.327 | 0.134 | 3.50E-31 | 12 | Zufsp |
| Fam13b4 | 2.05E-35 | 0.305662203 | 0.346 | 0.145 | 4.15E-31 | 12 | Fam13b |
| Tra2a6 | 2.57E-35 | 0.414728803 | 0.531 | 0.271 | 5.20E-31 | 12 | Tra2a |
| Rad54l22 | 2.76E-35 | 0.287482413 | 0.256 | 0.093 | 5.60E-31 | 12 | Rad54l2 |
| 5031425E22Rik3 | 2.86E-35 | 0.343709059 | 0.363 | 0.157 | 5.78E-31 | 12 | 5031425E22Rik |
| Trim30a5 | 2.89E-35 | 0.343306718 | 0.465 | 0.216 | 5.84E-31 | 12 | Trim30a |
| Rftn15 | 3.55E-35 | 0.321877372 | 0.516 | 0.248 | 7.18E-31 | 12 | Rftn1 |
| Sema6d3 | 3.59E-35 | 0.502278134 | 0.443 | 0.211 | 7.26E-31 | 12 | Sema6d |

| | | | | | | | |
|---|---|---|---|---|---|---|---|
| Cast4 | 3.84E-35 | 0.34133301 | 0.591 | 0.308 | 7.78E-31 | 12 | Cast |
| Iqgap16 | 3.96E-35 | 0.399010291 | 0.733 | 0.407 | 8.02E-31 | 12 | Iqgap1 |
| Sfpq4 | 4.48E-35 | 0.325156794 | 0.574 | 0.294 | 9.08E-31 | 12 | Sfpq |
| Zfand36 | 4.49E-35 | 0.38197526 | 0.576 | 0.298 | 9.09E-31 | 12 | Zfand3 |
| Dnajc35 | 5.97E-35 | 0.365002378 | 0.673 | 0.366 | 1.21E-30 | 12 | Dnajc3 |
| Ikbkb1 | 6.36E-35 | 0.354735949 | 0.336 | 0.142 | 1.29E-30 | 12 | Ikbkb |
| Stat5b2 | 6.73E-35 | 0.352041025 | 0.342 | 0.144 | 1.36E-30 | 12 | Stat5b |
| Helz4 | 6.87E-35 | 0.353510747 | 0.378 | 0.166 | 1.39E-30 | 12 | Helz |
| Chfr2 | 7.89E-35 | 0.320219812 | 0.292 | 0.114 | 1.60E-30 | 12 | Chfr |
| Ythdf34 | 8.51E-35 | 0.280066445 | 0.419 | 0.193 | 1.72E-30 | 12 | Ythdf3 |
| Nptn5 | 1.10E-34 | 0.396125192 | 0.495 | 0.246 | 2.22E-30 | 12 | Nptn |
| Ahnak5 | 1.10E-34 | 0.259077384 | 0.858 | 0.465 | 2.24E-30 | 12 | Ahnak |
| Tnks25 | 1.11E-34 | 0.346366503 | 0.544 | 0.276 | 2.25E-30 | 12 | Tnks2 |
| Nudcd33 | 1.19E-34 | 0.263843236 | 0.501 | 0.243 | 2.40E-30 | 12 | Nudcd3 |
| 2610035D17Rik1 | 1.20E-34 | 0.346423732 | 0.288 | 0.112 | 2.43E-30 | 12 | 2610035D17Rik |
| Ifi474 | 1.36E-34 | 0.298059257 | 0.398 | 0.176 | 2.76E-30 | 12 | Ifi47 |
| Fam208b3 | 2.11E-34 | 0.338503819 | 0.348 | 0.148 | 4.28E-30 | 12 | Fam208b |
| Pi4ka3 | 2.40E-34 | 0.311713537 | 0.308 | 0.125 | 4.85E-30 | 12 | Pi4ka |
| Aebp2 | 2.45E-34 | 0.279892522 | 0.28 | 0.107 | 4.96E-30 | 12 | Aebp2 |
| Taf1d2 | 3.12E-34 | 0.30355027 | 0.336 | 0.142 | 6.32E-30 | 12 | Taf1d |
| Ppp2r5a1 | 3.27E-34 | 0.269533356 | 0.507 | 0.247 | 6.62E-30 | 12 | Ppp2r5a |
| Fli15 | 3.47E-34 | 0.320241413 | 0.733 | 0.397 | 7.03E-30 | 12 | Fli1 |
| Dnaja13 | 3.96E-34 | 0.308278073 | 0.86 | 0.511 | 8.01E-30 | 12 | Dnaja1 |
| Riok12 | 4.09E-34 | 0.289448031 | 0.29 | 0.113 | 8.29E-30 | 12 | Riok1 |
| Twistnb | 4.29E-34 | 0.258722564 | 0.336 | 0.141 | 8.69E-30 | 12 | Twistnb |
| Xpo72 | 5.09E-34 | 0.302021324 | 0.351 | 0.151 | 1.03E-29 | 12 | Xpo7 |
| Snrnp40 | 5.95E-34 | 0.263890186 | 0.338 | 0.141 | 1.20E-29 | 12 | Snrnp40 |
| Scamp31 | 6.25E-34 | 0.279727167 | 0.273 | 0.104 | 1.27E-29 | 12 | Scamp3 |
| Ifrd12 | 6.37E-34 | 0.325376044 | 0.469 | 0.227 | 1.29E-29 | 12 | Ifrd1 |
| Ifi2036 | 7.69E-34 | 0.321195744 | 0.677 | 0.363 | 1.56E-29 | 12 | Ifi203 |
| Rsf12 | 1.29E-33 | 0.36541486 | 0.452 | 0.221 | 2.61E-29 | 12 | Rsf1 |
| Prpf40a5 | 1.37E-33 | 0.35224158 | 0.512 | 0.264 | 2.77E-29 | 12 | Prpf40a |
| Camkmt2 | 1.39E-33 | 0.415893335 | 0.323 | 0.137 | 2.81E-29 | 12 | Camkmt |
| Sf11 | 1.80E-33 | 0.269482444 | 0.404 | 0.185 | 3.64E-29 | 12 | Sf1 |
| Ccdc823 | 1.89E-33 | 0.281335729 | 0.398 | 0.178 | 3.82E-29 | 12 | Ccdc82 |
| mt-Nd4l5 | 1.91E-33 | 0.26969 | 0.821 | 0.486 | 3.86E-29 | 12 | mt-Nd4l |
| Prpf38b5 | 2.19E-33 | 0.302072084 | 0.417 | 0.193 | 4.44E-29 | 12 | Prpf38b |
| Smchd14 | 2.36E-33 | 0.402830962 | 0.51 | 0.261 | 4.79E-29 | 12 | Smchd1 |
| Neat15 | 2.41E-33 | 0.337078605 | 0.434 | 0.206 | 4.88E-29 | 12 | Neat1 |
| Nfkbiz2 | 2.63E-33 | 0.356663498 | 0.323 | 0.135 | 5.32E-29 | 12 | Nfkbiz |
| Wwp23 | 2.81E-33 | 0.298178612 | 0.305 | 0.125 | 5.70E-29 | 12 | Wwp2 |
| Hltf1 | 2.90E-33 | 0.250609699 | 0.31 | 0.126 | 5.88E-29 | 12 | Hltf |
| Elf14 | 2.97E-33 | 0.329420291 | 0.673 | 0.372 | 6.01E-29 | 12 | Elf1 |

| | | | | | | | |
|---|---|---|---|---|---|---|---|
| Cep1201 | 3.02E-33 | 0.33283254 | 0.288 | 0.115 | 6.11E-29 | 12 | Cep120 |
| Gpatch86 | 3.03E-33 | 0.352154903 | 0.507 | 0.255 | 6.13E-29 | 12 | Gpatch8 |
| Bcor1 | 3.50E-33 | 0.268075585 | 0.297 | 0.119 | 7.08E-29 | 12 | Bcor |
| Orai11 | 4.19E-33 | 0.283372744 | 0.308 | 0.126 | 8.48E-29 | 12 | Orai1 |
| Hnrnpa2b12 | 4.45E-33 | 0.311294249 | 0.843 | 0.51 | 9.01E-29 | 12 | Hnrnpa2b1 |
| Sik24 | 4.95E-33 | 0.347643394 | 0.292 | 0.118 | 1.00E-28 | 12 | Sik2 |
| Vgll44 | 5.30E-33 | 0.34869797 | 0.49 | 0.247 | 1.07E-28 | 12 | Vgll4 |
| Stag16 | 5.50E-33 | 0.396475921 | 0.647 | 0.357 | 1.11E-28 | 12 | Stag1 |
| Chd44 | 5.94E-33 | 0.299565767 | 0.604 | 0.324 | 1.20E-28 | 12 | Chd4 |
| Maf2 | 5.96E-33 | 0.589522701 | 0.284 | 0.116 | 1.21E-28 | 12 | Maf |
| Kdm2a6 | 6.12E-33 | 0.342192727 | 0.479 | 0.237 | 1.24E-28 | 12 | Kdm2a |
| Filip1l1 | 1.02E-32 | 0.263857049 | 0.346 | 0.15 | 2.06E-28 | 12 | Filip1l |
| G3bp13 | 1.06E-32 | 0.276124436 | 0.57 | 0.299 | 2.14E-28 | 12 | G3bp1 |
| Ints63 | 1.11E-32 | 0.333186159 | 0.329 | 0.14 | 2.24E-28 | 12 | Ints6 |
| Ppp1r15a1 | 1.29E-32 | 0.3064053 | 0.351 | 0.153 | 2.61E-28 | 12 | Ppp1r15a |
| Smg72 | 1.42E-32 | 0.300337194 | 0.351 | 0.154 | 2.87E-28 | 12 | Smg7 |
| Myo1e5 | 1.56E-32 | 0.519670443 | 0.51 | 0.278 | 3.16E-28 | 12 | Myo1e |
| Akt35 | 1.67E-32 | 0.359162964 | 0.501 | 0.248 | 3.39E-28 | 12 | Akt3 |
| Rlf3 | 2.04E-32 | 0.317004274 | 0.318 | 0.135 | 4.14E-28 | 12 | Rlf |
| Usp346 | 2.58E-32 | 0.34882988 | 0.619 | 0.341 | 5.22E-28 | 12 | Usp34 |
| Brd46 | 3.15E-32 | 0.379111571 | 0.508 | 0.264 | 6.38E-28 | 12 | Brd4 |
| Kmt2e4 | 3.51E-32 | 0.303430094 | 0.723 | 0.402 | 7.10E-28 | 12 | Kmt2e |
| Cyth12 | 3.86E-32 | 0.325381708 | 0.394 | 0.185 | 7.80E-28 | 12 | Cyth1 |
| Kmt2a6 | 4.17E-32 | 0.362034242 | 0.512 | 0.264 | 8.44E-28 | 12 | Kmt2a |
| Taf32 | 4.34E-32 | 0.356232849 | 0.333 | 0.145 | 8.79E-28 | 12 | Taf3 |
| Sf3b22 | 4.57E-32 | 0.262962238 | 0.645 | 0.355 | 9.25E-28 | 12 | Sf3b2 |
| Cbfb4 | 4.92E-32 | 0.332091082 | 0.346 | 0.153 | 9.95E-28 | 12 | Cbfb |
| Irf25 | 5.42E-32 | 0.311324671 | 0.621 | 0.338 | 1.10E-27 | 12 | Irf2 |
| Rpl7a5 | 5.61E-32 | 0.281108499 | 0.966 | 0.749 | 1.14E-27 | 12 | Rpl7a |
| Ube2d3 | 5.75E-32 | 0.264105386 | 0.723 | 0.414 | 1.16E-27 | 12 | Ube2d3 |
| Wdr442 | 6.06E-32 | 0.321137039 | 0.293 | 0.121 | 1.23E-27 | 12 | Wdr44 |
| Os94 | 6.12E-32 | 0.367924443 | 0.394 | 0.186 | 1.24E-27 | 12 | Os9 |
| Serbp14 | 7.42E-32 | 0.278453736 | 0.897 | 0.585 | 1.50E-27 | 12 | Serbp1 |
| Phc31 | 8.47E-32 | 0.313196185 | 0.314 | 0.133 | 1.72E-27 | 12 | Phc3 |
| Bbx4 | 8.56E-32 | 0.298932473 | 0.452 | 0.22 | 1.73E-27 | 12 | Bbx |
| Ddx54 | 9.18E-32 | 0.408055983 | 0.953 | 0.65 | 1.86E-27 | 12 | Ddx5 |
| Ctcf3 | 9.36E-32 | 0.274013511 | 0.531 | 0.273 | 1.89E-27 | 12 | Ctcf |
| Thoc25 | 9.95E-32 | 0.291311388 | 0.529 | 0.27 | 2.01E-27 | 12 | Thoc2 |
| Rlim2 | 1.22E-31 | 0.280815433 | 0.415 | 0.195 | 2.46E-27 | 12 | Rlim |
| Usp253 | 1.28E-31 | 0.256726298 | 0.544 | 0.278 | 2.60E-27 | 12 | Usp25 |
| Grb21 | 1.32E-31 | 0.279712317 | 0.536 | 0.279 | 2.67E-27 | 12 | Grb2 |
| Wdr334 | 1.33E-31 | 0.284824128 | 0.473 | 0.236 | 2.70E-27 | 12 | Wdr33 |
| Rps153 | 1.55E-31 | 0.310801109 | 0.968 | 0.745 | 3.15E-27 | 12 | Rps15 |

| | | | | | | | |
|---|---|---|---|---|---|---|---|
| Ap3b14 | 1.60E-31 | 0.307781851 | 0.439 | 0.215 | 3.24E-27 | 12 | Ap3b1 |
| B2m8 | 1.78E-31 | 0.404584921 | 0.996 | 0.686 | 3.59E-27 | 12 | B2m |
| Cdyl3 | 1.89E-31 | 0.257862479 | 0.307 | 0.127 | 3.83E-27 | 12 | Cdyl |
| Gtpbp2 | 2.06E-31 | 0.290992748 | 0.303 | 0.127 | 4.17E-27 | 12 | Gtpbp2 |
| Jmjd1c7 | 2.66E-31 | 0.335674718 | 0.572 | 0.304 | 5.39E-27 | 12 | Jmjd1c |
| Tank1 | 2.86E-31 | 0.279562173 | 0.327 | 0.141 | 5.78E-27 | 12 | Tank |
| Bbs93 | 2.87E-31 | 0.376929276 | 0.256 | 0.1 | 5.81E-27 | 12 | Bbs9 |
| Fam193a3 | 3.02E-31 | 0.295189042 | 0.387 | 0.177 | 6.12E-27 | 12 | Fam193a |
| Wdr70 | 3.30E-31 | 0.271771599 | 0.338 | 0.149 | 6.67E-27 | 12 | Wdr70 |
| Trim261 | 3.59E-31 | 0.29742629 | 0.271 | 0.108 | 7.27E-27 | 12 | Trim26 |
| Mia22 | 3.83E-31 | 0.313477974 | 0.424 | 0.205 | 7.76E-27 | 12 | Mia2 |
| Arid4a6 | 3.89E-31 | 0.270570789 | 0.497 | 0.248 | 7.88E-27 | 12 | Arid4a |
| Ccnt12 | 4.72E-31 | 0.289833173 | 0.368 | 0.165 | 9.55E-27 | 12 | Ccnt1 |
| Crebbp5 | 4.86E-31 | 0.33457466 | 0.563 | 0.301 | 9.84E-27 | 12 | Crebbp |
| Spata6 | 6.51E-31 | 0.284990086 | 0.288 | 0.118 | 1.32E-26 | 12 | Spata6 |
| Abcb72 | 6.82E-31 | 0.28733645 | 0.275 | 0.11 | 1.38E-26 | 12 | Abcb7 |
| Cul3 | 7.53E-31 | 0.262006838 | 0.54 | 0.283 | 1.52E-26 | 12 | Cul3 |
| Cdk124 | 8.46E-31 | 0.363325948 | 0.456 | 0.227 | 1.71E-26 | 12 | Cdk12 |
| Fus4 | 9.73E-31 | 0.256143433 | 0.69 | 0.384 | 1.97E-26 | 12 | Fus |
| Huwe15 | 1.05E-30 | 0.336777292 | 0.531 | 0.282 | 2.12E-26 | 12 | Huwe1 |
| Nop582 | 1.05E-30 | 0.268270666 | 0.316 | 0.136 | 2.12E-26 | 12 | Nop58 |
| Chd84 | 1.16E-30 | 0.319563603 | 0.361 | 0.166 | 2.35E-26 | 12 | Chd8 |
| 2310035C23Rik4 | 1.45E-30 | 0.347788899 | 0.316 | 0.138 | 2.94E-26 | 12 | 2310035C23Rik |
| Tbc1d55 | 1.78E-30 | 0.34748192 | 0.501 | 0.258 | 3.59E-26 | 12 | Tbc1d5 |
| Trip125 | 1.79E-30 | 0.283206045 | 0.497 | 0.253 | 3.63E-26 | 12 | Trip12 |
| Lsm14a1 | 1.84E-30 | 0.274509605 | 0.355 | 0.161 | 3.73E-26 | 12 | Lsm14a |
| Foxn35 | 2.94E-30 | 0.457613874 | 0.692 | 0.407 | 5.95E-26 | 12 | Foxn3 |
| Papd52 | 3.07E-30 | 0.264684874 | 0.284 | 0.116 | 6.21E-26 | 12 | Papd5 |
| Ubxn43 | 3.62E-30 | 0.291121466 | 0.516 | 0.269 | 7.33E-26 | 12 | Ubxn4 |
| Dnajb91 | 5.19E-30 | 0.305319766 | 0.321 | 0.141 | 1.05E-25 | 12 | Dnajb9 |
| Pdpk12 | 5.65E-30 | 0.286202405 | 0.323 | 0.141 | 1.14E-25 | 12 | Pdpk1 |
| Asxl11 | 6.31E-30 | 0.31651282 | 0.327 | 0.144 | 1.28E-25 | 12 | Asxl1 |
| Bod1l3 | 6.69E-30 | 0.290264103 | 0.516 | 0.27 | 1.35E-25 | 12 | Bod1l |
| Rfc22 | 7.28E-30 | 0.263286719 | 0.293 | 0.123 | 1.47E-25 | 12 | Rfc2 |
| Mtmr36 | 7.40E-30 | 0.339087543 | 0.409 | 0.198 | 1.50E-25 | 12 | Mtmr3 |
| Ppp2r5c1 | 7.43E-30 | 0.285885996 | 0.48 | 0.244 | 1.50E-25 | 12 | Ppp2r5c |
| Setd56 | 8.36E-30 | 0.300547504 | 0.505 | 0.261 | 1.69E-25 | 12 | Setd5 |
| Ankhd14 | 8.83E-30 | 0.333201312 | 0.473 | 0.241 | 1.79E-25 | 12 | Ankhd1 |
| Tpk13 | 1.13E-29 | 0.272231289 | 0.26 | 0.103 | 2.30E-25 | 12 | Tpk1 |
| Ncoa14 | 1.16E-29 | 0.34208536 | 0.422 | 0.207 | 2.35E-25 | 12 | Ncoa1 |
| Eif4g34 | 1.25E-29 | 0.297235177 | 0.531 | 0.283 | 2.52E-25 | 12 | Eif4g3 |
| Psen13 | 1.26E-29 | 0.275706503 | 0.25 | 0.099 | 2.54E-25 | 12 | Psen1 |
| Mcu | 1.40E-29 | 0.286955419 | 0.254 | 0.1 | 2.84E-25 | 12 | Mcu |

| | | | | | | | |
|---|---|---|---|---|---|---|---|
| Stxbp52 | 1.56E-29 | 0.311430954 | 0.299 | 0.128 | 3.16E-25 | 12 | Stxbp5 |
| Atl31 | 1.60E-29 | 0.293924496 | 0.295 | 0.126 | 3.24E-25 | 12 | Atl3 |
| Lcor4 | 1.61E-29 | 0.32644398 | 0.394 | 0.188 | 3.26E-25 | 12 | Lcor |
| Rps6ka53 | 1.74E-29 | 0.256710182 | 0.265 | 0.106 | 3.53E-25 | 12 | Rps6ka5 |
| Tmcc16 | 1.90E-29 | 0.387141922 | 0.432 | 0.215 | 3.84E-25 | 12 | Tmcc1 |
| Ier55 | 2.02E-29 | 0.318403938 | 0.581 | 0.314 | 4.10E-25 | 12 | Ier5 |
| Akap96 | 2.12E-29 | 0.289601689 | 0.465 | 0.234 | 4.29E-25 | 12 | Akap9 |
| 2810474O19Rik2 | 2.86E-29 | 0.286238925 | 0.424 | 0.208 | 5.79E-25 | 12 | 2810474O19Rik |
| Prkd33 | 3.96E-29 | 0.300930582 | 0.359 | 0.166 | 8.01E-25 | 12 | Prkd3 |
| Tnfrsf1b2 | 4.75E-29 | 0.259676846 | 0.279 | 0.115 | 9.62E-25 | 12 | Tnfrsf1b |
| Dyrk1a4 | 5.35E-29 | 0.312165196 | 0.376 | 0.177 | 1.08E-24 | 12 | Dyrk1a |
| Rai12 | 5.52E-29 | 0.255108829 | 0.265 | 0.108 | 1.12E-24 | 12 | Rai1 |
| Ddx583 | 5.85E-29 | 0.285772576 | 0.346 | 0.158 | 1.18E-24 | 12 | Ddx58 |
| Lasp12 | 9.29E-29 | 0.252094937 | 0.417 | 0.206 | 1.88E-24 | 12 | Lasp1 |
| Nrf13 | 1.01E-28 | 0.299514709 | 0.307 | 0.133 | 2.05E-24 | 12 | Nrf1 |
| Tspan142 | 1.10E-28 | 0.26368048 | 0.297 | 0.126 | 2.22E-24 | 12 | Tspan14 |
| Gatad2b3 | 1.13E-28 | 0.288546329 | 0.407 | 0.2 | 2.30E-24 | 12 | Gatad2b |
| Tet31 | 1.20E-28 | 0.306066675 | 0.297 | 0.129 | 2.42E-24 | 12 | Tet3 |
| Bptf6 | 1.24E-28 | 0.257279712 | 0.551 | 0.29 | 2.52E-24 | 12 | Bptf |
| Sf3b34 | 1.85E-28 | 0.289128387 | 0.346 | 0.158 | 3.75E-24 | 12 | Sf3b3 |
| Trim12c1 | 2.34E-28 | 0.292382716 | 0.308 | 0.136 | 4.75E-24 | 12 | Trim12c |
| Pias12 | 2.42E-28 | 0.269003589 | 0.389 | 0.186 | 4.91E-24 | 12 | Pias1 |
| Srsf73 | 2.57E-28 | 0.254657057 | 0.593 | 0.329 | 5.20E-24 | 12 | Srsf7 |
| Map2k41 | 3.37E-28 | 0.275975541 | 0.34 | 0.157 | 6.82E-24 | 12 | Map2k4 |
| Lims14 | 4.27E-28 | 0.351384561 | 0.542 | 0.297 | 8.64E-24 | 12 | Lims1 |
| Fnbp42 | 5.51E-28 | 0.255520685 | 0.293 | 0.127 | 1.12E-23 | 12 | Fnbp4 |
| Ywhaz2 | 8.81E-28 | 0.263791593 | 0.798 | 0.488 | 1.78E-23 | 12 | Ywhaz |
| Pum15 | 9.08E-28 | 0.265394161 | 0.492 | 0.256 | 1.84E-23 | 12 | Pum1 |
| Fbxo381 | 1.25E-27 | 0.272071941 | 0.273 | 0.116 | 2.52E-23 | 12 | Fbxo38 |
| Trappc93 | 1.39E-27 | 0.320327654 | 0.389 | 0.192 | 2.82E-23 | 12 | Trappc9 |
| Setd26 | 1.63E-27 | 0.271278324 | 0.394 | 0.19 | 3.30E-23 | 12 | Setd2 |
| Mapk16 | 1.97E-27 | 0.263426866 | 0.452 | 0.233 | 3.99E-23 | 12 | Mapk1 |
| Pcsk7 | 2.04E-27 | 0.255912439 | 0.269 | 0.112 | 4.13E-23 | 12 | Pcsk7 |
| Ube4a | 2.10E-27 | 0.258828512 | 0.262 | 0.108 | 4.26E-23 | 12 | Ube4a |
| Ndfip24 | 2.69E-27 | 0.253450601 | 0.467 | 0.242 | 5.44E-23 | 12 | Ndfip2 |
| Emsy1 | 2.71E-27 | 0.270023237 | 0.288 | 0.125 | 5.50E-23 | 12 | Emsy |
| Ep4004 | 3.20E-27 | 0.291455988 | 0.415 | 0.205 | 6.48E-23 | 12 | Ep400 |
| Tpst22 | 3.51E-27 | 0.262401022 | 0.452 | 0.23 | 7.10E-23 | 12 | Tpst2 |
| Malat17 | 5.07E-27 | 0.324545905 | 1 | 0.721 | 1.03E-22 | 12 | Malat1 |
| mt-Nd37 | 6.19E-27 | 0.286334419 | 0.738 | 0.442 | 1.25E-22 | 12 | mt-Nd3 |
| Gmds1 | 9.39E-27 | 0.302170064 | 0.366 | 0.176 | 1.90E-22 | 12 | Gmds |
| Rnf1695 | 1.04E-26 | 0.334041573 | 0.37 | 0.181 | 2.10E-22 | 12 | Rnf169 |
| Zzef13 | 1.06E-26 | 0.272438091 | 0.321 | 0.146 | 2.16E-22 | 12 | Zzef1 |

| | | | | | | | |
|---|---|---|---|---|---|---|---|
| Mdm42 | 1.72E-26 | 0.25170251 | 0.338 | 0.158 | 3.49E-22 | 12 | Mdm4 |
| Ppp4r3b3 | 1.77E-26 | 0.259799763 | 0.447 | 0.23 | 3.58E-22 | 12 | Ppp4r3b |
| Fam69a2 | 2.35E-26 | 0.306319849 | 0.284 | 0.125 | 4.76E-22 | 12 | Fam69a |
| Kidins2203 | 2.53E-26 | 0.299714689 | 0.331 | 0.153 | 5.13E-22 | 12 | Kidins220 |
| Selenot2 | 2.57E-26 | 0.259353278 | 0.417 | 0.209 | 5.21E-22 | 12 | Selenot |
| Abi15 | 2.69E-26 | 0.326970412 | 0.492 | 0.267 | 5.45E-22 | 12 | Abi1 |
| Tnrc6c4 | 3.39E-26 | 0.254296617 | 0.572 | 0.312 | 6.87E-22 | 12 | Tnrc6c |
| Trpm74 | 7.95E-26 | 0.259352871 | 0.402 | 0.202 | 1.61E-21 | 12 | Trpm7 |
| Naa152 | 8.75E-26 | 0.259791122 | 0.344 | 0.164 | 1.77E-21 | 12 | Naa15 |
| Kat6a2 | 1.05E-25 | 0.271618626 | 0.436 | 0.225 | 2.13E-21 | 12 | Kat6a |
| Son7 | 1.29E-25 | 0.250042322 | 0.871 | 0.521 | 2.61E-21 | 12 | Son |
| Birc22 | 1.52E-25 | 0.252118554 | 0.256 | 0.108 | 3.07E-21 | 12 | Birc2 |
| Stat13 | 1.80E-25 | 0.32759797 | 0.465 | 0.247 | 3.65E-21 | 12 | Stat1 |
| Sik13 | 2.22E-25 | 0.325679376 | 0.28 | 0.125 | 4.49E-21 | 12 | Sik1 |
| Nrip14 | 2.61E-25 | 0.37347952 | 0.449 | 0.241 | 5.29E-21 | 12 | Nrip1 |
| Cmip5 | 2.86E-25 | 0.283168 | 0.52 | 0.283 | 5.79E-21 | 12 | Cmip |
| Setx2 | 3.76E-25 | 0.250721763 | 0.353 | 0.172 | 7.61E-21 | 12 | Setx |
| Akirin1 | 4.98E-25 | 0.251979693 | 0.318 | 0.149 | 1.01E-20 | 12 | Akirin1 |
| Fbxo115 | 5.36E-25 | 0.311776662 | 0.434 | 0.23 | 1.08E-20 | 12 | Fbxo11 |
| Ccny4 | 6.92E-25 | 0.266816895 | 0.49 | 0.265 | 1.40E-20 | 12 | Ccny |
| Lcorl4 | 8.54E-25 | 0.254043295 | 0.344 | 0.165 | 1.73E-20 | 12 | Lcorl |
| Arid22 | 8.94E-25 | 0.262537853 | 0.327 | 0.155 | 1.81E-20 | 12 | Arid2 |
| Kif2a2 | 1.15E-24 | 0.250690199 | 0.359 | 0.176 | 2.33E-20 | 12 | Kif2a |
| AW5549183 | 1.69E-24 | 0.312128301 | 0.342 | 0.169 | 3.42E-20 | 12 | AW554918 |
| Bmt23 | 2.13E-24 | 0.254569283 | 0.269 | 0.119 | 4.32E-20 | 12 | Bmt2 |
| Samd9l3 | 3.65E-24 | 0.284736354 | 0.469 | 0.254 | 7.39E-20 | 12 | Samd9l |
| Msi23 | 5.43E-24 | 0.256207478 | 0.521 | 0.29 | 1.10E-19 | 12 | Msi2 |
| Zfp36l14 | 9.06E-24 | 0.31507254 | 0.71 | 0.429 | 1.83E-19 | 12 | Zfp36l1 |
| Epc12 | 1.09E-23 | 0.275902451 | 0.389 | 0.201 | 2.20E-19 | 12 | Epc1 |
| Ubn25 | 1.24E-23 | 0.293462549 | 0.447 | 0.242 | 2.52E-19 | 12 | Ubn2 |
| Rnf1305 | 1.44E-23 | 0.250390326 | 0.505 | 0.281 | 2.92E-19 | 12 | Rnf130 |
| Arfgef13 | 2.86E-23 | 0.253971035 | 0.421 | 0.225 | 5.78E-19 | 12 | Arfgef1 |
| Mbtd12 | 5.53E-23 | 0.302464749 | 0.284 | 0.133 | 1.12E-18 | 12 | Mbtd1 |
| Brd14 | 1.38E-22 | 0.261555873 | 0.437 | 0.238 | 2.80E-18 | 12 | Brd1 |
| mt-Cytb6 | 1.49E-22 | 0.284674105 | 0.998 | 0.696 | 3.03E-18 | 12 | mt-Cytb |
| Elp41 | 2.46E-22 | 0.255379822 | 0.273 | 0.126 | 4.98E-18 | 12 | Elp4 |
| Uhrf21 | 3.05E-22 | 0.269328228 | 0.331 | 0.164 | 6.18E-18 | 12 | Uhrf2 |
| Lpp5 | 5.13E-22 | 0.265930039 | 0.503 | 0.283 | 1.04E-17 | 12 | Lpp |
| Pou2f11 | 5.99E-21 | 0.271431111 | 0.265 | 0.125 | 1.21E-16 | 12 | Pou2f1 |
| Phip5 | 7.72E-21 | 0.282743471 | 0.452 | 0.254 | 1.56E-16 | 12 | Phip |
| Nfkbia4 | 2.42E-20 | 0.292754808 | 0.645 | 0.397 | 4.90E-16 | 12 | Nfkbia |
| Hip14 | 5.54E-19 | 0.312336788 | 0.548 | 0.332 | 1.12E-14 | 12 | Hip1 |
| Ski5 | 4.67E-17 | 0.261266061 | 0.374 | 0.213 | 9.45E-13 | 12 | Ski |

| | | | | | | | |
|---|---|---|---|---|---|---|---|
| Wwox5 | 5.41E-17 | 0.297645384 | 0.321 | 0.175 | 1.10E-12 | 12 | Wwox |
| Ccl53 | 0 | 2.312230883 | 0.927 | 0.143 | 0 | 13 | Ccl5 |
| Xcl13 | 0 | 2.126694102 | 0.927 | 0.121 | 0 | 13 | Xcl1 |
| Nkg73 | 0 | 2.000130004 | 0.933 | 0.118 | 0 | 13 | Nkg7 |
| Cd75 | 0 | 1.95287604 | 0.864 | 0.104 | 0 | 13 | Cd7 |
| Klrd13 | 0 | 1.728671124 | 0.897 | 0.114 | 0 | 13 | Klrd1 |
| Ltb3 | 0 | 1.719198201 | 0.918 | 0.127 | 0 | 13 | Ltb |
| Ctsw3 | 0 | 1.663536102 | 0.894 | 0.119 | 0 | 13 | Ctsw |
| Il2rb3 | 0 | 1.619457536 | 0.855 | 0.112 | 0 | 13 | Il2rb |
| Ptprcap3 | 0 | 1.49689113 | 0.894 | 0.123 | 0 | 13 | Ptprcap |
| Ms4a4b3 | 0 | 1.441154423 | 0.87 | 0.114 | 0 | 13 | Ms4a4b |
| Ncr13 | 0 | 1.392948138 | 0.764 | 0.096 | 0 | 13 | Ncr1 |
| Klre13 | 0 | 1.357351476 | 0.776 | 0.096 | 0 | 13 | Klre1 |
| Klrb1c3 | 0 | 1.259176982 | 0.758 | 0.09 | 0 | 13 | Klrb1c |
| Gzmb2 | 5.06E-305 | 2.184618029 | 0.658 | 0.076 | 1.02E-300 | 13 | Gzmb |
| Klrk13 | 1.88E-300 | 1.377746891 | 0.739 | 0.094 | 3.81E-296 | 13 | Klrk1 |
| Skap13 | 9.94E-287 | 1.495318697 | 0.852 | 0.126 | 2.01E-282 | 13 | Skap1 |
| Hcst3 | 1.23E-284 | 1.435264435 | 0.87 | 0.138 | 2.49E-280 | 13 | Hcst |
| S100a42 | 1.90E-282 | 1.732260588 | 0.809 | 0.121 | 3.85E-278 | 13 | S100a4 |
| Clnk3 | 2.86E-281 | 1.374213591 | 0.764 | 0.104 | 5.80E-277 | 13 | Clnk |
| Dusp24 | 2.72E-274 | 1.217114149 | 0.709 | 0.095 | 5.51E-270 | 13 | Dusp2 |
| Gzmc2 | 7.80E-268 | 2.266717157 | 0.615 | 0.077 | 1.58E-263 | 13 | Gzmc |
| Trbc13 | 1.57E-261 | 1.070140971 | 0.636 | 0.08 | 3.19E-257 | 13 | Trbc1 |
| Klra82 | 5.51E-256 | 1.563008201 | 0.606 | 0.074 | 1.11E-251 | 13 | Klra8 |
| Ikzf33 | 1.81E-253 | 1.184809089 | 0.761 | 0.113 | 3.66E-249 | 13 | Ikzf3 |
| Txk3 | 5.45E-253 | 1.015725749 | 0.718 | 0.1 | 1.10E-248 | 13 | Txk |
| Tnfrsf93 | 5.06E-252 | 0.883977522 | 0.545 | 0.062 | 1.02E-247 | 13 | Tnfrsf9 |
| Rac23 | 2.24E-246 | 1.491982688 | 0.909 | 0.17 | 4.53E-242 | 13 | Rac2 |
| Klra72 | 3.38E-245 | 1.353715838 | 0.573 | 0.069 | 6.85E-241 | 13 | Klra7 |
| Samd33 | 2.46E-243 | 0.936008538 | 0.621 | 0.081 | 4.98E-239 | 13 | Samd3 |
| Styk13 | 7.03E-243 | 1.180917209 | 0.664 | 0.091 | 1.42E-238 | 13 | Styk1 |
| Klra92 | 4.17E-234 | 1.342294507 | 0.576 | 0.073 | 8.45E-230 | 13 | Klra9 |
| 14-Sep | 6.39E-223 | 1.052422302 | 0.806 | 0.139 | 1.29E-218 | 13 | 1-Sep |
| Lck3 | 6.57E-220 | 0.851830945 | 0.664 | 0.097 | 1.33E-215 | 13 | Lck |
| Gimap33 | 1.99E-219 | 0.911234401 | 0.645 | 0.094 | 4.04E-215 | 13 | Gimap3 |
| Tox3 | 3.02E-218 | 1.325420654 | 0.709 | 0.115 | 6.12E-214 | 13 | Tox |
| Klra42 | 2.84E-217 | 1.495085192 | 0.564 | 0.074 | 5.76E-213 | 13 | Klra4 |
| Stat43 | 4.92E-216 | 1.044614739 | 0.664 | 0.101 | 9.96E-212 | 13 | Stat4 |
| Serpina3g3 | 1.29E-212 | 0.768632246 | 0.512 | 0.063 | 2.60E-208 | 13 | Serpina3g |
| Klri22 | 3.88E-206 | 0.93942322 | 0.533 | 0.07 | 7.85E-202 | 13 | Klri2 |
| Cd23 | 1.69E-205 | 0.857167214 | 0.57 | 0.079 | 3.43E-201 | 13 | Cd2 |
| Cst73 | 2.75E-203 | 0.812427402 | 0.527 | 0.069 | 5.56E-199 | 13 | Cst7 |
| Ptpn223 | 2.07E-198 | 1.048013487 | 0.676 | 0.113 | 4.19E-194 | 13 | Ptpn22 |

| | | | | | | | |
|---|---|---|---|---|---|---|---|
| Ptprc4 | 5.54E-194 | 1.480959206 | 0.882 | 0.191 | 1.12E-189 | 13 | Ptprc |
| Runx34 | 2.40E-191 | 1.212853479 | 0.745 | 0.143 | 4.86E-187 | 13 | Runx3 |
| AC163354.12 | 1.97E-190 | 0.993390009 | 0.461 | 0.058 | 4.00E-186 | 13 | AC163354.1 |
| Gem3 | 4.38E-190 | 1.095044148 | 0.612 | 0.098 | 8.86E-186 | 13 | Gem |
| Ipcef14 | 5.91E-187 | 1.06305569 | 0.652 | 0.112 | 1.20E-182 | 13 | Ipcef1 |
| Cd2262 | 3.02E-184 | 1.135441393 | 0.564 | 0.085 | 6.11E-180 | 13 | Cd226 |
| Inpp4b3 | 1.30E-183 | 1.567088881 | 0.842 | 0.197 | 2.62E-179 | 13 | Inpp4b |
| Gm26823 | 2.88E-183 | 1.155669875 | 0.527 | 0.076 | 5.84E-179 | 13 | Gm2682 |
| Cd2443 | 5.90E-182 | 0.797177226 | 0.57 | 0.087 | 1.19E-177 | 13 | Cd244 |
| Padi23 | 1.41E-181 | 0.774075496 | 0.582 | 0.091 | 2.86E-177 | 13 | Padi2 |
| Tbx213 | 2.20E-179 | 0.708430001 | 0.47 | 0.062 | 4.46E-175 | 13 | Tbx21 |
| Stk17b3 | 1.01E-178 | 1.071187762 | 0.767 | 0.156 | 2.04E-174 | 13 | Stk17b |
| Atp8b43 | 5.53E-178 | 1.026480641 | 0.576 | 0.092 | 1.12E-173 | 13 | Atp8b4 |
| Dock24 | 7.39E-177 | 1.289361976 | 0.824 | 0.179 | 1.50E-172 | 13 | Dock2 |
| Selplg4 | 1.13E-175 | 0.970759059 | 0.748 | 0.149 | 2.28E-171 | 13 | Selplg |
| Mir142hg4 | 1.62E-175 | 1.037689698 | 0.8 | 0.164 | 3.29E-171 | 13 | Mir142hg |
| Cd963 | 1.93E-174 | 0.649631964 | 0.476 | 0.065 | 3.90E-170 | 13 | Cd96 |
| Cd374 | 3.27E-173 | 0.70500382 | 0.633 | 0.109 | 6.62E-169 | 13 | Cd37 |
| Sulf24 | 6.13E-173 | 0.776997522 | 0.576 | 0.094 | 1.24E-168 | 13 | Sulf2 |
| Aoah3 | 1.08E-172 | 0.754420606 | 0.488 | 0.069 | 2.19E-168 | 13 | Aoah |
| Rgs15 | 1.06E-171 | 0.989787089 | 0.709 | 0.135 | 2.15E-167 | 13 | Rgs1 |
| Klrb1f3 | 1.18E-170 | 0.595208825 | 0.448 | 0.059 | 2.38E-166 | 13 | Klrb1f |
| Coro1a5 | 3.41E-167 | 1.142979137 | 0.912 | 0.212 | 6.91E-163 | 13 | Coro1a |
| Cyfip23 | 8.95E-167 | 0.720141039 | 0.545 | 0.087 | 1.81E-162 | 13 | Cyfip2 |
| Dennd4a4 | 1.06E-165 | 1.181480267 | 0.794 | 0.182 | 2.14E-161 | 13 | Dennd4a |
| Limd24 | 1.54E-165 | 0.905829831 | 0.709 | 0.143 | 3.13E-161 | 13 | Limd2 |
| Osbpl33 | 2.73E-164 | 1.006885081 | 0.621 | 0.115 | 5.52E-160 | 13 | Osbpl3 |
| Acap13 | 1.36E-163 | 0.656895209 | 0.47 | 0.067 | 2.76E-159 | 13 | Acap1 |
| Epsti14 | 3.66E-163 | 0.996002986 | 0.779 | 0.169 | 7.42E-159 | 13 | Epsti1 |
| Car23 | 1.97E-162 | 0.778620775 | 0.642 | 0.117 | 3.99E-158 | 13 | Car2 |
| Pglyrp13 | 8.34E-162 | 0.672412247 | 0.548 | 0.088 | 1.69E-157 | 13 | Pglyrp1 |
| Chsy13 | 1.76E-161 | 0.92193996 | 0.642 | 0.122 | 3.57E-157 | 13 | Chsy1 |
| Sytl33 | 2.95E-161 | 0.53912114 | 0.418 | 0.054 | 5.97E-157 | 13 | Sytl3 |
| Tbc1d10c3 | 3.76E-161 | 0.696331131 | 0.476 | 0.069 | 7.62E-157 | 13 | Tbc1d10c |
| Cd525 | 4.75E-161 | 1.516522132 | 0.991 | 0.275 | 9.62E-157 | 13 | Cd52 |
| Ccr23 | 7.81E-161 | 0.790197098 | 0.521 | 0.082 | 1.58E-156 | 13 | Ccr2 |
| Prf12 | 1.35E-160 | 0.680032177 | 0.436 | 0.06 | 2.74E-156 | 13 | Prf1 |
| Itgad1 | 4.21E-160 | 0.68830416 | 0.327 | 0.035 | 8.53E-156 | 13 | Itgad |
| Rasal33 | 1.49E-159 | 0.701085387 | 0.509 | 0.079 | 3.02E-155 | 13 | Rasal3 |
| Tmem1633 | 8.34E-157 | 0.829464291 | 0.521 | 0.083 | 1.69E-152 | 13 | Tmem163 |
| Klrb1a1 | 1.35E-156 | 0.696898219 | 0.348 | 0.04 | 2.72E-152 | 13 | Klrb1a |
| Il2rg3 | 5.46E-155 | 0.794432044 | 0.609 | 0.112 | 1.10E-150 | 13 | Il2rg |
| Cxcr62 | 9.69E-155 | 0.831123156 | 0.43 | 0.06 | 1.96E-150 | 13 | Cxcr6 |

| | | | | | | | |
|---|---|---|---|---|---|---|---|
| Itgax4 | 1.81E-154 | 0.862876738 | 0.576 | 0.103 | 3.67E-150 | 13 | Itgax |
| Mctp23 | 7.43E-154 | 0.656576928 | 0.506 | 0.08 | 1.50E-149 | 13 | Mctp2 |
| Tnfrsf183 | 8.28E-154 | 0.682630489 | 0.467 | 0.07 | 1.68E-149 | 13 | Tnfrsf18 |
| Ppp3cc3 | 1.20E-153 | 0.894669653 | 0.673 | 0.136 | 2.44E-149 | 13 | Ppp3cc |
| Fcer1g5 | 4.66E-153 | 1.46816552 | 0.97 | 0.263 | 9.44E-149 | 13 | Fcer1g |
| Unc1193 | 5.96E-153 | 0.56682919 | 0.503 | 0.08 | 1.21E-148 | 13 | Unc119 |
| Fasl3 | 4.22E-151 | 0.485113712 | 0.406 | 0.054 | 8.55E-147 | 13 | Fasl |
| Klra32 | 4.90E-151 | 0.867112173 | 0.358 | 0.043 | 9.93E-147 | 13 | Klra3 |
| Ugcg3 | 9.60E-151 | 1.097858701 | 0.724 | 0.166 | 1.94E-146 | 13 | Ugcg |
| Arhgap454 | 2.15E-150 | 0.866240489 | 0.718 | 0.153 | 4.34E-146 | 13 | Arhgap45 |
| Dock53 | 2.60E-148 | 0.860856207 | 0.588 | 0.11 | 5.26E-144 | 13 | Dock5 |
| Cd693 | 4.72E-148 | 0.748059545 | 0.558 | 0.098 | 9.56E-144 | 13 | Cd69 |
| Itk3 | 6.86E-148 | 0.560300258 | 0.479 | 0.074 | 1.39E-143 | 13 | Itk |
| Arhgdib5 | 2.04E-147 | 1.050998247 | 0.882 | 0.227 | 4.12E-143 | 13 | Arhgdib |
| Ptk2b4 | 6.25E-147 | 0.904035368 | 0.685 | 0.145 | 1.26E-142 | 13 | Ptk2b |
| Tnik4 | 1.17E-146 | 0.810056569 | 0.582 | 0.108 | 2.37E-142 | 13 | Tnik |
| Dscam3 | 2.06E-146 | 1.081370272 | 0.352 | 0.044 | 4.17E-142 | 13 | Dscam |
| Itgae2 | 3.13E-143 | 0.89342269 | 0.427 | 0.063 | 6.33E-139 | 13 | Itgae |
| Ikzf14 | 3.77E-142 | 0.912765191 | 0.697 | 0.151 | 7.63E-138 | 13 | Ikzf1 |
| Coro2a3 | 3.29E-141 | 0.867078297 | 0.588 | 0.113 | 6.67E-137 | 13 | Coro2a |
| Sema4d4 | 1.85E-139 | 0.84214463 | 0.615 | 0.125 | 3.75E-135 | 13 | Sema4d |
| Chn23 | 2.88E-139 | 1.332850333 | 0.745 | 0.193 | 5.83E-135 | 13 | Chn2 |
| Srgap33 | 6.54E-139 | 0.798379454 | 0.515 | 0.092 | 1.32E-134 | 13 | Srgap3 |
| Prkcq3 | 1.83E-138 | 0.625665589 | 0.497 | 0.084 | 3.70E-134 | 13 | Prkcq |
| Tyrobp4 | 2.00E-137 | 1.230942537 | 0.945 | 0.255 | 4.05E-133 | 13 | Tyrobp |
| Gimap73 | 2.13E-137 | 0.589321312 | 0.394 | 0.056 | 4.30E-133 | 13 | Gimap7 |
| Klrc23 | 1.58E-136 | 0.554191156 | 0.37 | 0.05 | 3.19E-132 | 13 | Klrc2 |
| Tmsb104 | 2.45E-136 | 1.665779182 | 0.994 | 0.496 | 4.97E-132 | 13 | Tmsb10 |
| S100a64 | 2.20E-135 | 1.375322061 | 0.897 | 0.265 | 4.46E-131 | 13 | S100a6 |
| B4galnt13 | 3.37E-135 | 0.607626642 | 0.515 | 0.091 | 6.82E-131 | 13 | B4galnt1 |
| Lax13 | 5.39E-135 | 0.558940034 | 0.418 | 0.063 | 1.09E-130 | 13 | Lax1 |
| Rinl3 | 4.03E-134 | 0.709345394 | 0.621 | 0.127 | 8.16E-130 | 13 | Rinl |
| Bcl23 | 1.98E-133 | 1.282665476 | 0.761 | 0.202 | 4.01E-129 | 13 | Bcl2 |
| Ccl44 | 4.44E-133 | 1.117683551 | 0.606 | 0.125 | 9.00E-129 | 13 | Ccl4 |
| Cd1602 | 6.02E-132 | 0.575187068 | 0.327 | 0.041 | 1.22E-127 | 13 | Cd160 |
| Itgal3 | 2.79E-131 | 0.615169845 | 0.479 | 0.083 | 5.64E-127 | 13 | Itgal |
| Dapk24 | 8.36E-131 | 0.714613159 | 0.552 | 0.106 | 1.69E-126 | 13 | Dapk2 |
| Card113 | 1.36E-130 | 0.584549253 | 0.467 | 0.079 | 2.76E-126 | 13 | Card11 |
| Sh2d1a2 | 3.36E-130 | 0.532715861 | 0.376 | 0.054 | 6.81E-126 | 13 | Sh2d1a |
| Pik3cd4 | 3.44E-129 | 0.735552193 | 0.539 | 0.104 | 6.97E-125 | 13 | Pik3cd |
| Trbc23 | 1.05E-128 | 0.82604532 | 0.409 | 0.063 | 2.12E-124 | 13 | Trbc2 |
| Lsp15 | 2.11E-128 | 0.673297126 | 0.63 | 0.134 | 4.27E-124 | 13 | Lsp1 |
| Jaml3 | 2.86E-128 | 0.640074517 | 0.379 | 0.055 | 5.80E-124 | 13 | Jaml |

| | | | | | | | |
|---|---|---|---|---|---|---|---|
| Ifitm103 | 1.81E-127 | 0.538153258 | 0.385 | 0.057 | 3.67E-123 | 13 | Ifitm10 |
| Sla23 | 8.17E-127 | 0.421702373 | 0.342 | 0.046 | 1.65E-122 | 13 | Sla2 |
| Sema4a3 | 2.86E-126 | 0.523804943 | 0.439 | 0.073 | 5.79E-122 | 13 | Sema4a |
| Ccr54 | 3.78E-126 | 0.755468384 | 0.606 | 0.128 | 7.66E-122 | 13 | Ccr5 |
| Dock104 | 5.59E-126 | 1.128485558 | 0.776 | 0.203 | 1.13E-121 | 13 | Dock10 |
| Acot73 | 2.12E-125 | 0.551991652 | 0.439 | 0.073 | 4.29E-121 | 13 | Acot7 |
| Arsb3 | 2.24E-125 | 0.816187838 | 0.576 | 0.123 | 4.53E-121 | 13 | Arsb |
| Rhof2 | 7.19E-125 | 0.372923301 | 0.294 | 0.035 | 1.46E-120 | 13 | Rhof |
| Zap703 | 7.52E-125 | 0.487381123 | 0.37 | 0.054 | 1.52E-120 | 13 | Zap70 |
| Cd3g2 | 8.21E-124 | 1.048361716 | 0.321 | 0.042 | 1.66E-119 | 13 | Cd3g |
| Id23 | 1.24E-123 | 1.206313511 | 0.915 | 0.316 | 2.51E-119 | 13 | Id2 |
| Lcp14 | 1.64E-123 | 0.973918431 | 0.833 | 0.232 | 3.32E-119 | 13 | Lcp1 |
| Shisa53 | 7.97E-123 | 1.181326225 | 0.915 | 0.323 | 1.61E-118 | 13 | Shisa5 |
| Lat24 | 7.32E-122 | 0.646614261 | 0.597 | 0.127 | 1.48E-117 | 13 | Lat2 |
| Cdh14 | 1.23E-121 | 0.61903372 | 0.424 | 0.071 | 2.48E-117 | 13 | Cdh1 |
| Il12rb23 | 2.47E-120 | 0.475819181 | 0.336 | 0.047 | 5.01E-116 | 13 | Il12rb2 |
| Gimap45 | 1.43E-119 | 1.02091194 | 0.833 | 0.251 | 2.90E-115 | 13 | Gimap4 |
| Arhgap154 | 1.06E-118 | 0.7951326 | 0.742 | 0.183 | 2.14E-114 | 13 | Arhgap15 |
| Gprin33 | 1.74E-118 | 0.568648377 | 0.382 | 0.06 | 3.51E-114 | 13 | Gprin3 |
| Gata33 | 2.61E-118 | 0.361941734 | 0.345 | 0.049 | 5.29E-114 | 13 | Gata3 |
| Arnt22 | 5.04E-117 | 0.496092269 | 0.315 | 0.043 | 1.02E-112 | 13 | Arnt2 |
| Laptm55 | 2.12E-116 | 0.673776159 | 0.806 | 0.205 | 4.30E-112 | 13 | Laptm5 |
| Fam189b3 | 2.61E-116 | 0.48199908 | 0.361 | 0.054 | 5.29E-112 | 13 | Fam189b |
| Ccdc88c3 | 6.34E-116 | 0.542604449 | 0.5 | 0.096 | 1.28E-111 | 13 | Ccdc88c |
| Pde7a3 | 7.77E-116 | 0.758603935 | 0.642 | 0.151 | 1.57E-111 | 13 | Pde7a |
| Vipr21 | 9.93E-116 | 0.402047917 | 0.261 | 0.03 | 2.01E-111 | 13 | Vipr2 |
| Evl4 | 1.23E-115 | 0.924399666 | 0.764 | 0.216 | 2.50E-111 | 13 | Evl |
| Efhd24 | 6.25E-114 | 0.723992928 | 0.715 | 0.182 | 1.27E-109 | 13 | Efhd2 |
| Gpr1743 | 2.97E-113 | 0.455463871 | 0.339 | 0.05 | 6.01E-109 | 13 | Gpr174 |
| Gm163412 | 3.46E-113 | 0.493472983 | 0.276 | 0.034 | 7.00E-109 | 13 | Gm16341 |
| Gpr682 | 8.93E-113 | 0.366388832 | 0.315 | 0.043 | 1.81E-108 | 13 | Gpr68 |
| Il21r4 | 2.51E-112 | 0.512819312 | 0.473 | 0.09 | 5.09E-108 | 13 | Il21r |
| Cotl15 | 2.71E-111 | 0.904446089 | 0.788 | 0.219 | 5.48E-107 | 13 | Cotl1 |
| Sep-63 | 3.14E-111 | 0.48013885 | 0.491 | 0.095 | 6.36E-107 | 13 | 6-Sep |
| Hivep34 | 4.48E-111 | 0.723354861 | 0.639 | 0.157 | 9.07E-107 | 13 | Hivep3 |
| Slc38a13 | 1.43E-110 | 0.555044874 | 0.452 | 0.085 | 2.90E-106 | 13 | Slc38a1 |
| Nup2103 | 2.61E-109 | 0.439194726 | 0.385 | 0.064 | 5.29E-105 | 13 | Nup210 |
| AW1120105 | 9.03E-109 | 1.359915219 | 0.982 | 0.565 | 1.83E-104 | 13 | AW112010 |
| Hopx3 | 1.00E-108 | 0.44650379 | 0.412 | 0.072 | 2.03E-104 | 13 | Hopx |
| Ptpn73 | 1.63E-108 | 0.491503305 | 0.352 | 0.056 | 3.30E-104 | 13 | Ptpn7 |
| Adcy75 | 5.14E-108 | 0.500137732 | 0.527 | 0.11 | 1.04E-103 | 13 | Adcy7 |
| Traf3ip33 | 2.36E-107 | 0.538126746 | 0.418 | 0.076 | 4.78E-103 | 13 | Traf3ip3 |
| Fam107b3 | 1.00E-106 | 0.546039084 | 0.542 | 0.117 | 2.02E-102 | 13 | Fam107b |

| | | | | | | | |
|---|---|---|---|---|---|---|---|
| Rhoh4 | 1.54E-106 | 0.643512528 | 0.488 | 0.1 | 3.12E-102 | 13 | Rhoh |
| Ncf44 | 3.80E-106 | 0.648678916 | 0.512 | 0.109 | 7.70E-102 | 13 | Ncf4 |
| Bin24 | 4.10E-106 | 0.634025273 | 0.555 | 0.123 | 8.30E-102 | 13 | Bin2 |
| Smyd33 | 4.21E-105 | 1.109465893 | 0.727 | 0.222 | 8.53E-101 | 13 | Smyd3 |
| Tiam22 | 5.56E-105 | 0.480923534 | 0.327 | 0.05 | 1.12E-100 | 13 | Tiam2 |
| Kcnip32 | 1.63E-104 | 0.445888724 | 0.309 | 0.045 | 3.30E-100 | 13 | Kcnip3 |
| Setbp14 | 2.34E-104 | 0.732857834 | 0.53 | 0.118 | 4.73E-100 | 13 | Setbp1 |
| Prex14 | 2.90E-104 | 0.785539376 | 0.645 | 0.169 | 5.86E-100 | 13 | Prex1 |
| Ighm4 | 3.18E-104 | 1.913399389 | 0.555 | 0.123 | 6.43E-100 | 13 | Ighm |
| Traf13 | 4.81E-104 | 0.480193987 | 0.361 | 0.06 | 9.73E-100 | 13 | Traf1 |
| Sorl14 | 5.17E-104 | 0.560991862 | 0.503 | 0.108 | 1.05E-99 | 13 | Sorl1 |
| Ptpn185 | 1.71E-103 | 0.981994716 | 0.915 | 0.31 | 3.46E-99 | 13 | Ptpn18 |
| Arhgap93 | 1.86E-103 | 0.397068385 | 0.512 | 0.108 | 3.76E-99 | 13 | Arhgap9 |
| Pde3b4 | 2.84E-103 | 0.766060572 | 0.542 | 0.124 | 5.75E-99 | 13 | Pde3b |
| Gas74 | 8.47E-103 | 0.94725527 | 0.658 | 0.178 | 1.71E-98 | 13 | Gas7 |
| Cd484 | 4.04E-102 | 0.717502701 | 0.564 | 0.131 | 8.18E-98 | 13 | Cd48 |
| Cers64 | 2.67E-101 | 0.600400353 | 0.512 | 0.112 | 5.41E-97 | 13 | Cers6 |
| Pglyrp23 | 3.74E-101 | 0.369758574 | 0.294 | 0.042 | 7.57E-97 | 13 | Pglyrp2 |
| Chl12 | 4.14E-101 | 0.509575862 | 0.324 | 0.05 | 8.38E-97 | 13 | Chl1 |
| Satb13 | 1.09E-100 | 0.455857045 | 0.361 | 0.06 | 2.21E-96 | 13 | Satb1 |
| Myo1f4 | 1.57E-100 | 0.639828502 | 0.57 | 0.134 | 3.17E-96 | 13 | Myo1f |
| Psd43 | 2.56E-100 | 0.39124127 | 0.367 | 0.062 | 5.18E-96 | 13 | Psd4 |
| Sytl23 | 5.45E-99 | 0.397221235 | 0.333 | 0.053 | 1.10E-94 | 13 | Sytl2 |
| Itgb24 | 5.88E-99 | 0.585724727 | 0.467 | 0.098 | 1.19E-94 | 13 | Itgb2 |
| Ccnd33 | 3.22E-98 | 0.785324593 | 0.764 | 0.235 | 6.52E-94 | 13 | Ccnd3 |
| Rnf432 | 5.93E-97 | 0.389970664 | 0.309 | 0.048 | 1.20E-92 | 13 | Rnf43 |
| Gramd1b3 | 9.36E-97 | 0.508038751 | 0.433 | 0.087 | 1.90E-92 | 13 | Gramd1b |
| Ssbp24 | 9.91E-97 | 0.726355347 | 0.561 | 0.136 | 2.01E-92 | 13 | Ssbp2 |
| Eomes2 | 2.21E-96 | 0.424564761 | 0.312 | 0.049 | 4.46E-92 | 13 | Eomes |
| Apbb1ip4 | 2.29E-96 | 0.575712067 | 0.655 | 0.169 | 4.64E-92 | 13 | Apbb1ip |
| Gng24 | 2.43E-96 | 0.665173127 | 0.676 | 0.182 | 4.91E-92 | 13 | Gng2 |
| Nlrc33 | 7.47E-95 | 0.420018526 | 0.333 | 0.055 | 1.51E-90 | 13 | Nlrc3 |
| Rabgap1l3 | 1.55E-92 | 1.021050457 | 0.755 | 0.252 | 3.13E-88 | 13 | Rabgap1l |
| Grap24 | 4.24E-92 | 0.49588143 | 0.473 | 0.103 | 8.59E-88 | 13 | Grap2 |
| Apol7e3 | 7.97E-92 | 0.343952503 | 0.306 | 0.048 | 1.61E-87 | 13 | Apol7e |
| Dok61 | 8.71E-92 | 0.521577884 | 0.279 | 0.042 | 1.76E-87 | 13 | Dok6 |
| Sh2d2a3 | 2.43E-91 | 0.436636137 | 0.297 | 0.047 | 4.91E-87 | 13 | Sh2d2a |
| Sept115 | 2.79E-91 | 0.752418888 | 0.682 | 0.199 | 5.64E-87 | 13 | 11-Sep |
| Hsd11b14 | 3.67E-91 | 0.440289209 | 0.433 | 0.089 | 7.44E-87 | 13 | Hsd11b1 |
| Dock115 | 9.50E-91 | 0.470557641 | 0.494 | 0.111 | 1.92E-86 | 13 | Dock11 |
| Rasgrp13 | 1.00E-90 | 0.368177308 | 0.342 | 0.06 | 2.02E-86 | 13 | Rasgrp1 |
| Pkp33 | 1.93E-90 | 0.442700417 | 0.364 | 0.067 | 3.91E-86 | 13 | Pkp3 |
| Jakmip13 | 2.84E-90 | 0.296714479 | 0.3 | 0.047 | 5.75E-86 | 13 | Jakmip1 |

| | | | | | | | |
|---|---|---|---|---|---|---|---|
| Itgb73 | 3.17E-90 | 0.39787144 | 0.339 | 0.059 | 6.42E-86 | 13 | Itgb7 |
| Mob3a3 | 6.50E-90 | 0.444487442 | 0.397 | 0.078 | 1.32E-85 | 13 | Mob3a |
| Esyt13 | 9.00E-90 | 0.606532413 | 0.536 | 0.132 | 1.82E-85 | 13 | Esyt1 |
| Ccl34 | 1.25E-89 | 0.675607003 | 0.394 | 0.08 | 2.53E-85 | 13 | Ccl3 |
| Gpsm34 | 1.54E-89 | 0.578682406 | 0.57 | 0.144 | 3.11E-85 | 13 | Gpsm3 |
| P2ry101 | 2.85E-89 | 0.3643152 | 0.258 | 0.037 | 5.77E-85 | 13 | P2ry10 |
| Gimap54 | 6.55E-89 | 0.596072423 | 0.627 | 0.167 | 1.33E-84 | 13 | Gimap5 |
| Sash33 | 7.66E-89 | 0.439664353 | 0.33 | 0.058 | 1.55E-84 | 13 | Sash3 |
| Dock84 | 9.45E-89 | 0.821067033 | 0.703 | 0.214 | 1.91E-84 | 13 | Dock8 |
| Rab373 | 1.59E-88 | 0.348255913 | 0.276 | 0.042 | 3.21E-84 | 13 | Rab37 |
| Znrf34 | 4.58E-88 | 0.532859159 | 0.464 | 0.104 | 9.27E-84 | 13 | Znrf3 |
| Vim6 | 7.59E-88 | 0.785708993 | 0.852 | 0.301 | 1.54E-83 | 13 | Vim |
| Arhgap304 | 1.13E-87 | 0.514275603 | 0.47 | 0.107 | 2.28E-83 | 13 | Arhgap30 |
| Vps37b3 | 1.57E-87 | 0.510700984 | 0.539 | 0.133 | 3.19E-83 | 13 | Vps37b |
| Prkcb5 | 1.02E-86 | 0.56628289 | 0.591 | 0.153 | 2.07E-82 | 13 | Prkcb |
| Kcnq53 | 1.12E-85 | 0.6464228 | 0.352 | 0.066 | 2.28E-81 | 13 | Kcnq5 |
| Lcp24 | 2.20E-85 | 0.472692147 | 0.427 | 0.092 | 4.45E-81 | 13 | Lcp2 |
| Fyb4 | 6.54E-85 | 0.405808852 | 0.667 | 0.18 | 1.32E-80 | 13 | Fyb |
| Ctsd5 | 1.22E-84 | 0.896808992 | 0.815 | 0.31 | 2.48E-80 | 13 | Ctsd |
| Gimap17 | 1.64E-84 | 0.777020522 | 0.848 | 0.298 | 3.32E-80 | 13 | Gimap1 |
| Rassf53 | 2.85E-84 | 0.358929703 | 0.352 | 0.066 | 5.77E-80 | 13 | Rassf5 |
| Fgl24 | 2.03E-83 | 0.641590203 | 0.448 | 0.105 | 4.12E-79 | 13 | Fgl2 |
| Akna4 | 2.48E-83 | 0.395495414 | 0.415 | 0.088 | 5.03E-79 | 13 | Akna |
| Celf25 | 2.86E-83 | 0.47614275 | 0.688 | 0.198 | 5.79E-79 | 13 | Celf2 |
| Eya23 | 1.52E-82 | 0.414827912 | 0.436 | 0.097 | 3.08E-78 | 13 | Eya2 |
| Neurl34 | 3.67E-82 | 0.520739297 | 0.473 | 0.112 | 7.43E-78 | 13 | Neurl3 |
| Cytip3 | 6.88E-82 | 0.443053978 | 0.415 | 0.089 | 1.39E-77 | 13 | Cytip |
| Spn2 | 9.39E-82 | 0.365172067 | 0.267 | 0.042 | 1.90E-77 | 13 | Spn |
| Pstpip13 | 6.16E-81 | 0.356536319 | 0.306 | 0.054 | 1.25E-76 | 13 | Pstpip1 |
| Dok22 | 7.20E-81 | 0.427345135 | 0.258 | 0.04 | 1.46E-76 | 13 | Dok2 |
| 1810011H11Rik3 | 8.42E-81 | 0.443440104 | 0.358 | 0.071 | 1.70E-76 | 13 | 1810011H11Rik |
| Armc72 | 1.07E-80 | 0.336107005 | 0.282 | 0.047 | 2.17E-76 | 13 | Armc7 |
| Camk2g3 | 1.31E-80 | 0.501673261 | 0.518 | 0.131 | 2.65E-76 | 13 | Camk2g |
| Ptpn64 | 1.36E-80 | 0.512018719 | 0.509 | 0.127 | 2.76E-76 | 13 | Ptpn6 |
| Plcl25 | 2.75E-80 | 0.517956142 | 0.467 | 0.111 | 5.56E-76 | 13 | Plcl2 |
| Spry23 | 3.32E-80 | 0.425477372 | 0.382 | 0.079 | 6.73E-76 | 13 | Spry2 |
| Rps15a4 | 3.51E-80 | 0.932048177 | 1 | 0.924 | 7.12E-76 | 13 | Rps15a |
| Gramd1a3 | 4.65E-80 | 0.474503209 | 0.43 | 0.098 | 9.41E-76 | 13 | Gramd1a |
| Oprm13 | 5.23E-80 | 0.347773146 | 0.318 | 0.058 | 1.06E-75 | 13 | Oprm1 |
| Emid13 | 1.24E-79 | 0.445819523 | 0.455 | 0.106 | 2.51E-75 | 13 | Emid1 |
| Rps6ka14 | 2.07E-79 | 0.423074557 | 0.467 | 0.111 | 4.20E-75 | 13 | Rps6ka1 |
| Ripor24 | 2.31E-79 | 0.556139599 | 0.606 | 0.168 | 4.67E-75 | 13 | Ripor2 |
| Sh3bgrl35 | 3.82E-79 | 1.039907744 | 0.982 | 0.556 | 7.74E-75 | 13 | Sh3bgrl3 |

| | | | | | | | |
|---|---|---|---|---|---|---|---|
| Fhl23 | 3.88E-79 | 0.300418474 | 0.258 | 0.04 | 7.85E-75 | 13 | Fhl2 |
| Fermt34 | 2.12E-78 | 0.45073977 | 0.467 | 0.111 | 4.30E-74 | 13 | Fermt3 |
| Cd823 | 3.88E-78 | 0.510358374 | 0.421 | 0.098 | 7.85E-74 | 13 | Cd82 |
| Lgals13 | 9.08E-78 | 0.90080496 | 0.503 | 0.132 | 1.84E-73 | 13 | Lgals1 |
| Herc33 | 2.17E-77 | 0.358457694 | 0.348 | 0.069 | 4.39E-73 | 13 | Herc3 |
| Ankrd444 | 5.35E-77 | 0.86952305 | 0.779 | 0.29 | 1.08E-72 | 13 | Ankrd44 |
| Pla2g163 | 5.47E-77 | 0.496967309 | 0.742 | 0.239 | 1.11E-72 | 13 | Pla2g16 |
| Tes3 | 5.76E-77 | 0.39296574 | 0.503 | 0.126 | 1.17E-72 | 13 | Tes |
| Mllt33 | 9.96E-77 | 0.664000676 | 0.536 | 0.148 | 2.02E-72 | 13 | Mllt3 |
| Matk1 | 1.44E-76 | 0.347791662 | 0.252 | 0.04 | 2.91E-72 | 13 | Matk |
| Vegfc | 3.37E-76 | 0.473080439 | 0.448 | 0.106 | 6.81E-72 | 13 | Vegfc |
| Aim24 | 1.17E-75 | 0.42224861 | 0.43 | 0.1 | 2.37E-71 | 13 | Aim2 |
| Rps114 | 2.02E-75 | 0.888716393 | 1 | 0.906 | 4.09E-71 | 13 | Rps11 |
| Ern13 | 2.61E-75 | 0.42017132 | 0.379 | 0.082 | 5.28E-71 | 13 | Ern1 |
| Mtmr123 | 1.12E-74 | 0.521673193 | 0.491 | 0.127 | 2.26E-70 | 13 | Mtmr12 |
| Sh2d1b13 | 1.44E-74 | 0.408655141 | 0.358 | 0.075 | 2.91E-70 | 13 | Sh2d1b1 |
| Rnf1663 | 1.48E-73 | 0.429840517 | 0.433 | 0.104 | 2.99E-69 | 13 | Rnf166 |
| Apobec33 | 1.91E-73 | 0.531984411 | 0.694 | 0.221 | 3.86E-69 | 13 | Apobec3 |
| Peli14 | 1.92E-73 | 0.761280903 | 0.679 | 0.228 | 3.89E-69 | 13 | Peli1 |
| S100a103 | 9.28E-73 | 0.935400307 | 0.961 | 0.465 | 1.88E-68 | 13 | S100a10 |
| Pycard5 | 1.34E-72 | 0.465969039 | 0.794 | 0.272 | 2.71E-68 | 13 | Pycard |
| Malt13 | 2.13E-72 | 0.541165696 | 0.527 | 0.146 | 4.32E-68 | 13 | Malt1 |
| Lrrk14 | 7.27E-72 | 0.620440561 | 0.609 | 0.187 | 1.47E-67 | 13 | Lrrk1 |
| 4933406I18Rik4 | 9.96E-72 | 0.471346157 | 0.345 | 0.073 | 2.02E-67 | 13 | 4933406I18Rik |
| Lta4h3 | 2.24E-71 | 0.535964845 | 0.524 | 0.145 | 4.54E-67 | 13 | Lta4h |
| Zyx4 | 4.09E-71 | 0.607137887 | 0.658 | 0.212 | 8.28E-67 | 13 | Zyx |
| Camkmt3 | 4.34E-71 | 0.546807099 | 0.503 | 0.136 | 8.79E-67 | 13 | Camkmt |
| Pim13 | 4.70E-71 | 0.479028498 | 0.494 | 0.129 | 9.51E-67 | 13 | Pim1 |
| Hvcn12 | 7.54E-71 | 0.277990213 | 0.291 | 0.054 | 1.53E-66 | 13 | Hvcn1 |
| Fnbp15 | 1.83E-70 | 0.628932233 | 0.679 | 0.226 | 3.70E-66 | 13 | Fnbp1 |
| Plcg24 | 2.47E-70 | 0.388052368 | 0.467 | 0.117 | 5.00E-66 | 13 | Plcg2 |
| Pfn15 | 6.66E-70 | 0.985801082 | 0.973 | 0.68 | 1.35E-65 | 13 | Pfn1 |
| Ggct2 | 7.65E-70 | 0.306950182 | 0.315 | 0.062 | 1.55E-65 | 13 | Ggct |
| Adgre54 | 9.19E-70 | 0.743371595 | 0.679 | 0.233 | 1.86E-65 | 13 | Adgre5 |
| Parp85 | 1.56E-69 | 0.586350743 | 0.524 | 0.146 | 3.15E-65 | 13 | Parp8 |
| Vasp3 | 2.08E-69 | 0.512702465 | 0.7 | 0.233 | 4.22E-65 | 13 | Vasp |
| Napsa2 | 2.14E-69 | 0.266600054 | 0.276 | 0.05 | 4.33E-65 | 13 | Napsa |
| Tespa13 | 2.75E-69 | 0.353879745 | 0.348 | 0.075 | 5.57E-65 | 13 | Tespa1 |
| Slc9a94 | 3.26E-69 | 0.524554303 | 0.53 | 0.148 | 6.60E-65 | 13 | Slc9a9 |
| Gtpbp12 | 3.49E-69 | 0.369507642 | 0.333 | 0.069 | 7.07E-65 | 13 | Gtpbp1 |
| Cd534 | 3.91E-69 | 0.339866775 | 0.455 | 0.114 | 7.92E-65 | 13 | Cd53 |
| Cyth44 | 6.99E-69 | 0.411410552 | 0.491 | 0.131 | 1.42E-64 | 13 | Cyth4 |
| Cited41 | 9.68E-69 | 0.263643487 | 0.497 | 0.131 | 1.96E-64 | 13 | Cited4 |

| | | | | | | | |
|---|---|---|---|---|---|---|---|
| Eef2k1 | 1.43E-68 | 0.344215455 | 0.352 | 0.076 | 2.90E-64 | 13 | Eef2k |
| H2-Q75 | 2.16E-68 | 0.829206737 | 0.894 | 0.393 | 4.37E-64 | 13 | H2-Q7 |
| Slc38a24 | 3.24E-68 | 0.728051537 | 0.794 | 0.307 | 6.55E-64 | 13 | Slc38a2 |
| Fam49b4 | 3.69E-68 | 0.531527963 | 0.709 | 0.233 | 7.48E-64 | 13 | Fam49b |
| Otulin3 | 5.48E-68 | 0.455637829 | 0.63 | 0.196 | 1.11E-63 | 13 | Otulin |
| Cd2473 | 5.93E-68 | 0.270126052 | 0.258 | 0.045 | 1.20E-63 | 13 | Cd247 |
| Gimap93 | 1.68E-67 | 0.418562793 | 0.548 | 0.155 | 3.40E-63 | 13 | Gimap9 |
| Plek3 | 2.26E-67 | 0.417727268 | 0.47 | 0.122 | 4.58E-63 | 13 | Plek |
| B4galt54 | 2.37E-67 | 0.379396577 | 0.388 | 0.092 | 4.80E-63 | 13 | B4galt5 |
| Mbnl14 | 2.64E-67 | 0.994613976 | 0.87 | 0.456 | 5.34E-63 | 13 | Mbnl1 |
| Chka4 | 1.25E-66 | 0.430540488 | 0.506 | 0.141 | 2.53E-62 | 13 | Chka |
| Dgkd4 | 1.50E-66 | 0.44900945 | 0.542 | 0.158 | 3.03E-62 | 13 | Dgkd |
| AC149090.14 | 1.64E-66 | 0.584626421 | 0.673 | 0.228 | 3.32E-62 | 13 | AC149090.1 |
| Npm33 | 1.66E-66 | 0.392021076 | 0.479 | 0.13 | 3.37E-62 | 13 | Npm3 |
| Gimap66 | 2.07E-66 | 0.751521835 | 0.882 | 0.361 | 4.20E-62 | 13 | Gimap6 |
| Mgat54 | 3.15E-66 | 0.56430538 | 0.558 | 0.167 | 6.37E-62 | 13 | Mgat5 |
| Rin34 | 4.25E-66 | 0.495447147 | 0.582 | 0.176 | 8.60E-62 | 13 | Rin3 |
| Fmnl14 | 4.26E-66 | 0.379670779 | 0.382 | 0.09 | 8.62E-62 | 13 | Fmnl1 |
| Slc14a13 | 4.36E-66 | 0.329805085 | 0.252 | 0.045 | 8.82E-62 | 13 | Slc14a1 |
| Stk44 | 5.66E-66 | 0.42306514 | 0.579 | 0.172 | 1.15E-61 | 13 | Stk4 |
| Btbd113 | 7.02E-66 | 0.309317915 | 0.282 | 0.054 | 1.42E-61 | 13 | Btbd11 |
| Smc43 | 1.17E-65 | 0.429871493 | 0.545 | 0.16 | 2.37E-61 | 13 | Smc4 |
| Ppm1b3 | 1.33E-65 | 0.420841088 | 0.488 | 0.135 | 2.69E-61 | 13 | Ppm1b |
| Jpt14 | 1.55E-65 | 0.439231505 | 0.67 | 0.218 | 3.14E-61 | 13 | Jpt1 |
| Slc9a3r13 | 1.64E-65 | 0.266717457 | 0.427 | 0.107 | 3.32E-61 | 13 | Slc9a3r1 |
| Gpr1713 | 1.85E-65 | 0.324060651 | 0.255 | 0.046 | 3.74E-61 | 13 | Gpr171 |
| Leprotl13 | 3.04E-65 | 0.541046067 | 0.645 | 0.214 | 6.15E-61 | 13 | Leprotl1 |
| Nr4a23 | 6.71E-65 | 0.582025405 | 0.464 | 0.128 | 1.36E-60 | 13 | Nr4a2 |
| Tex144 | 1.05E-64 | 0.494680389 | 0.512 | 0.146 | 2.12E-60 | 13 | Tex14 |
| Pip5k1b3 | 1.23E-64 | 0.339807379 | 0.397 | 0.097 | 2.48E-60 | 13 | Pip5k1b |
| Mmgt21 | 1.48E-64 | 0.255971335 | 0.309 | 0.064 | 2.99E-60 | 13 | Mmgt2 |
| Ndfip13 | 1.71E-64 | 0.707376823 | 0.785 | 0.317 | 3.46E-60 | 13 | Ndfip1 |
| Csk4 | 2.46E-64 | 0.433262725 | 0.467 | 0.127 | 4.98E-60 | 13 | Csk |
| Trip43 | 1.02E-63 | 0.426737062 | 0.439 | 0.116 | 2.06E-59 | 13 | Trip4 |
| Gfod13 | 1.21E-63 | 0.330967548 | 0.315 | 0.067 | 2.44E-59 | 13 | Gfod1 |
| Tab23 | 2.15E-63 | 0.573195946 | 0.655 | 0.227 | 4.34E-59 | 13 | Tab2 |
| Sesn34 | 3.76E-63 | 0.388213135 | 0.461 | 0.126 | 7.62E-59 | 13 | Sesn3 |
| Tec4 | 3.83E-63 | 0.368809329 | 0.421 | 0.107 | 7.75E-59 | 13 | Tec |
| Mxd12 | 1.55E-62 | 0.310197045 | 0.297 | 0.062 | 3.14E-58 | 13 | Mxd1 |
| Tnfaip34 | 1.61E-62 | 0.381639557 | 0.367 | 0.088 | 3.27E-58 | 13 | Tnfaip3 |
| N4bp2l11 | 1.69E-62 | 0.285476849 | 0.273 | 0.053 | 3.41E-58 | 13 | N4bp2l1 |
| Fxyd56 | 2.34E-62 | 0.723126585 | 0.906 | 0.407 | 4.75E-58 | 13 | Fxyd5 |
| Zfp3183 | 2.78E-62 | 0.38676268 | 0.385 | 0.095 | 5.63E-58 | 13 | Zfp318 |

| | | | | | | | |
|---|---|---|---|---|---|---|---|
| Ubac24 | 3.21E-62 | 0.492242717 | 0.579 | 0.185 | 6.49E-58 | 13 | Ubac2 |
| Kif13b4 | 5.66E-62 | 0.606208016 | 0.621 | 0.212 | 1.15E-57 | 13 | Kif13b |
| Ahdc12 | 6.78E-62 | 0.293211895 | 0.424 | 0.109 | 1.37E-57 | 13 | Ahdc1 |
| Ust5 | 7.82E-62 | 0.326786988 | 0.358 | 0.084 | 1.58E-57 | 13 | Ust |
| Sec11c5 | 9.43E-62 | 0.49483431 | 0.809 | 0.306 | 1.91E-57 | 13 | Sec11c |
| Rgs24 | 2.66E-61 | 0.381562415 | 0.464 | 0.127 | 5.38E-57 | 13 | Rgs2 |
| A930015D03Rik3 | 2.80E-61 | 0.437314439 | 0.4 | 0.104 | 5.66E-57 | 13 | A930015D03Rik |
| Gimap85 | 5.37E-61 | 0.418072554 | 0.552 | 0.166 | 1.09E-56 | 13 | Gimap8 |
| Arhgap254 | 8.46E-61 | 0.34952453 | 0.409 | 0.106 | 1.71E-56 | 13 | Arhgap25 |
| Aak14 | 1.59E-60 | 0.547387015 | 0.564 | 0.182 | 3.22E-56 | 13 | Aak1 |
| Cblb4 | 3.63E-60 | 0.540685827 | 0.536 | 0.168 | 7.35E-56 | 13 | Cblb |
| Pik3ap13 | 4.95E-60 | 0.331866542 | 0.364 | 0.088 | 1.00E-55 | 13 | Pik3ap1 |
| Magohb2 | 5.36E-60 | 0.350641965 | 0.439 | 0.119 | 1.09E-55 | 13 | Magohb |
| Actb5 | 7.29E-60 | 0.947496158 | 0.994 | 0.842 | 1.47E-55 | 13 | Actb |
| Rnf1494 | 7.66E-60 | 0.327086169 | 0.397 | 0.102 | 1.55E-55 | 13 | Rnf149 |
| Vps13a3 | 8.09E-60 | 0.471467158 | 0.545 | 0.169 | 1.64E-55 | 13 | Vps13a |
| Ptpre4 | 8.25E-60 | 0.48837735 | 0.524 | 0.158 | 1.67E-55 | 13 | Ptpre |
| Btg13 | 9.53E-60 | 0.764242408 | 0.915 | 0.456 | 1.93E-55 | 13 | Btg1 |
| Rps35 | 2.29E-59 | 0.755181139 | 0.988 | 0.892 | 4.63E-55 | 13 | Rps3 |
| Znrf23 | 2.65E-59 | 0.364724947 | 0.521 | 0.156 | 5.37E-55 | 13 | Znrf2 |
| Jak15 | 2.67E-59 | 0.774518638 | 0.8 | 0.362 | 5.40E-55 | 13 | Jak1 |
| Sh3bp11 | 5.17E-59 | 0.296811053 | 0.255 | 0.05 | 1.05E-54 | 13 | Sh3bp1 |
| Rfx73 | 5.38E-59 | 0.420784889 | 0.506 | 0.152 | 1.09E-54 | 13 | Rfx7 |
| Prkacb3 | 7.13E-59 | 0.523336257 | 0.624 | 0.215 | 1.44E-54 | 13 | Prkacb |
| Cdyl23 | 8.44E-59 | 0.35710174 | 0.364 | 0.089 | 1.71E-54 | 13 | Cdyl2 |
| Lpxn3 | 8.87E-59 | 0.29968393 | 0.37 | 0.091 | 1.80E-54 | 13 | Lpxn |
| Ptprj4 | 9.58E-59 | 0.32684009 | 0.506 | 0.148 | 1.94E-54 | 13 | Ptprj |
| Sla5 | 2.84E-58 | 0.321153708 | 0.315 | 0.072 | 5.76E-54 | 13 | Sla |
| Atp11b3 | 2.91E-58 | 0.464481929 | 0.555 | 0.178 | 5.89E-54 | 13 | Atp11b |
| H2-Q66 | 3.86E-58 | 0.525946676 | 0.676 | 0.242 | 7.81E-54 | 13 | H2-Q6 |
| Lrmp4 | 4.14E-58 | 0.358152581 | 0.279 | 0.059 | 8.38E-54 | 13 | Lrmp |
| Diaph13 | 6.51E-58 | 0.523917865 | 0.661 | 0.239 | 1.32E-53 | 13 | Diaph1 |
| Rapgef64 | 1.84E-57 | 0.665062216 | 0.73 | 0.293 | 3.72E-53 | 13 | Rapgef6 |
| Mta34 | 2.34E-57 | 0.421198812 | 0.497 | 0.149 | 4.73E-53 | 13 | Mta3 |
| Chst115 | 2.49E-57 | 0.415860619 | 0.552 | 0.173 | 5.05E-53 | 13 | Chst11 |
| Klk82 | 2.93E-57 | 0.362680601 | 0.339 | 0.081 | 5.94E-53 | 13 | Klk8 |
| Rpsa5 | 5.83E-57 | 0.800831112 | 0.997 | 0.902 | 1.18E-52 | 13 | Rpsa |
| Abr4 | 7.84E-57 | 0.516040194 | 0.503 | 0.155 | 1.59E-52 | 13 | Abr |
| Baz1a4 | 7.88E-57 | 0.492172689 | 0.536 | 0.169 | 1.59E-52 | 13 | Baz1a |
| Vars3 | 8.95E-57 | 0.314473042 | 0.336 | 0.08 | 1.81E-52 | 13 | Vars |
| Mkl15 | 1.69E-56 | 0.342451994 | 0.439 | 0.123 | 3.41E-52 | 13 | Mkl1 |
| Rras23 | 2.98E-56 | 0.311119678 | 0.342 | 0.083 | 6.04E-52 | 13 | Rras2 |
| Sp1103 | 3.54E-56 | 0.289935488 | 0.512 | 0.154 | 7.17E-52 | 13 | Sp110 |

| | | | | | | | |
|---|---|---|---|---|---|---|---|
| Btg25 | 1.73E-55 | 0.666038735 | 0.8 | 0.34 | 3.51E-51 | 13 | Btg2 |
| Ptpn43 | 1.74E-55 | 0.4434838 | 0.442 | 0.129 | 3.52E-51 | 13 | Ptpn4 |
| Sms3 | 2.13E-55 | 0.290907721 | 0.491 | 0.148 | 4.31E-51 | 13 | Sms |
| Fndc3a4 | 2.17E-55 | 0.669699597 | 0.691 | 0.274 | 4.40E-51 | 13 | Fndc3a |
| Ccnd26 | 2.34E-55 | 0.648310522 | 0.712 | 0.283 | 4.73E-51 | 13 | Ccnd2 |
| Capzb5 | 4.85E-55 | 0.571939449 | 0.842 | 0.365 | 9.82E-51 | 13 | Capzb |
| Bcl2l113 | 7.54E-55 | 0.414291349 | 0.412 | 0.115 | 1.53E-50 | 13 | Bcl2l11 |
| Ptger44 | 9.64E-55 | 0.348573876 | 0.336 | 0.083 | 1.95E-50 | 13 | Ptger4 |
| Tmsb4x6 | 1.21E-54 | 0.631251788 | 1 | 0.934 | 2.46E-50 | 13 | Tmsb4x |
| Itga44 | 1.23E-54 | 0.326572606 | 0.315 | 0.075 | 2.48E-50 | 13 | Itga4 |
| Rbpj4 | 2.29E-54 | 0.495384447 | 0.615 | 0.217 | 4.63E-50 | 13 | Rbpj |
| Samsn14 | 3.04E-54 | 0.323634824 | 0.324 | 0.079 | 6.15E-50 | 13 | Samsn1 |
| Rgs35 | 4.16E-54 | 0.430528673 | 0.558 | 0.184 | 8.42E-50 | 13 | Rgs3 |
| H2afy3 | 5.58E-54 | 0.472291027 | 0.8 | 0.334 | 1.13E-49 | 13 | H2afy |
| Cnn24 | 8.54E-54 | 0.41834199 | 0.548 | 0.179 | 1.73E-49 | 13 | Cnn2 |
| Chst121 | 1.13E-53 | 0.254559888 | 0.291 | 0.066 | 2.28E-49 | 13 | Chst12 |
| Cyba5 | 1.78E-53 | 0.510938153 | 0.924 | 0.436 | 3.61E-49 | 13 | Cyba |
| Suco3 | 1.98E-53 | 0.465498225 | 0.485 | 0.151 | 4.00E-49 | 13 | Suco |
| Metrnl3 | 1.99E-53 | 0.317396695 | 0.379 | 0.101 | 4.03E-49 | 13 | Metrnl |
| Trmt1123 | 2.47E-53 | 0.45849847 | 0.715 | 0.27 | 5.00E-49 | 13 | Trmt112 |
| Fryl4 | 2.54E-53 | 0.631322768 | 0.679 | 0.265 | 5.15E-49 | 13 | Fryl |
| Cetn22 | 2.61E-53 | 0.254760531 | 0.6 | 0.2 | 5.29E-49 | 13 | Cetn2 |
| Rps6ka54 | 3.64E-53 | 0.332324887 | 0.388 | 0.106 | 7.36E-49 | 13 | Rps6ka5 |
| Ldlrad44 | 4.35E-53 | 0.501921035 | 0.67 | 0.247 | 8.81E-49 | 13 | Ldlrad4 |
| Tmem1314 | 4.88E-53 | 0.459731044 | 0.591 | 0.205 | 9.88E-49 | 13 | Tmem131 |
| Ppm1h4 | 5.35E-53 | 0.426652484 | 0.5 | 0.159 | 1.08E-48 | 13 | Ppm1h |
| Riok13 | 1.15E-52 | 0.308870942 | 0.409 | 0.113 | 2.32E-48 | 13 | Riok1 |
| Ttc39b4 | 3.78E-52 | 0.442395681 | 0.439 | 0.133 | 7.65E-48 | 13 | Ttc39b |
| Cbx43 | 5.01E-52 | 0.329560109 | 0.394 | 0.109 | 1.01E-47 | 13 | Cbx4 |
| Tecr4 | 6.56E-52 | 0.366895026 | 0.63 | 0.221 | 1.33E-47 | 13 | Tecr |
| Trim144 | 8.47E-52 | 0.328563591 | 0.415 | 0.119 | 1.71E-47 | 13 | Trim14 |
| Slamf74 | 8.69E-52 | 0.297751223 | 0.367 | 0.097 | 1.76E-47 | 13 | Slamf7 |
| H2-K17 | 1.05E-51 | 0.759178277 | 0.973 | 0.641 | 2.13E-47 | 13 | H2-K1 |
| Dennd4c4 | 1.25E-51 | 0.363896543 | 0.424 | 0.124 | 2.53E-47 | 13 | Dennd4c |
| Ramp13 | 1.39E-51 | 0.353139519 | 0.388 | 0.108 | 2.82E-47 | 13 | Ramp1 |
| Gm472835 | 2.48E-51 | 0.58999963 | 0.694 | 0.278 | 5.02E-47 | 13 | Gm47283 |
| Junb6 | 3.20E-51 | 0.567519181 | 0.845 | 0.389 | 6.48E-47 | 13 | Junb |
| Tmem24 | 3.45E-51 | 0.420308151 | 0.461 | 0.14 | 6.99E-47 | 13 | Tmem2 |
| Grina3 | 3.77E-51 | 0.337275898 | 0.452 | 0.137 | 7.63E-47 | 13 | Grina |
| Rpl13a5 | 3.98E-51 | 0.447032352 | 0.721 | 0.281 | 8.06E-47 | 13 | Rpl13a |
| Rsbn1l4 | 6.67E-51 | 0.4860737 | 0.691 | 0.269 | 1.35E-46 | 13 | Rsbn1l |
| Thy13 | 9.09E-51 | 0.354560452 | 0.264 | 0.059 | 1.84E-46 | 13 | Thy1 |
| Prkch6 | 9.61E-51 | 0.836408357 | 0.803 | 0.397 | 1.95E-46 | 13 | Prkch |

| | | | | | | | |
|---|---|---|---|---|---|---|---|
| Chd74 | 1.13E-50 | 0.420199232 | 0.473 | 0.149 | 2.29E-46 | 13 | Chd7 |
| Ppp1r184 | 1.16E-50 | 0.383914059 | 0.479 | 0.151 | 2.35E-46 | 13 | Ppp1r18 |
| Zfp36l24 | 1.23E-50 | 0.535661651 | 0.745 | 0.302 | 2.50E-46 | 13 | Zfp36l2 |
| Tbl1xr14 | 2.21E-50 | 0.398588746 | 0.497 | 0.162 | 4.48E-46 | 13 | Tbl1xr1 |
| Cdk2ap23 | 3.43E-50 | 0.251389137 | 0.579 | 0.196 | 6.95E-46 | 13 | Cdk2ap2 |
| Sipa1l14 | 3.72E-50 | 0.439828169 | 0.545 | 0.187 | 7.54E-46 | 13 | Sipa1l1 |
| Dnajc18 | 1.15E-49 | 0.623996636 | 0.733 | 0.32 | 2.34E-45 | 13 | Dnajc1 |
| Ppp1r16b6 | 1.90E-49 | 0.420615332 | 0.576 | 0.199 | 3.84E-45 | 13 | Ppp1r16b |
| Runx14 | 1.99E-49 | 0.31402539 | 0.433 | 0.129 | 4.03E-45 | 13 | Runx1 |
| Senp74 | 2.00E-49 | 0.311320793 | 0.488 | 0.154 | 4.05E-45 | 13 | Senp7 |
| Eml44 | 2.01E-49 | 0.529275914 | 0.658 | 0.256 | 4.08E-45 | 13 | Eml4 |
| Emp35 | 2.19E-49 | 0.336342552 | 0.694 | 0.255 | 4.44E-45 | 13 | Emp3 |
| Zfp5123 | 2.32E-49 | 0.289653078 | 0.33 | 0.085 | 4.69E-45 | 13 | Zfp512 |
| Fam102a3 | 2.54E-49 | 0.347362127 | 0.361 | 0.098 | 5.14E-45 | 13 | Fam102a |
| Usp483 | 3.21E-49 | 0.287059805 | 0.427 | 0.127 | 6.51E-45 | 13 | Usp48 |
| Pag13 | 3.52E-49 | 0.39902133 | 0.406 | 0.119 | 7.14E-45 | 13 | Pag1 |
| Rps184 | 4.00E-49 | 0.656227349 | 1 | 0.941 | 8.10E-45 | 13 | Rps18 |
| Elmsan13 | 4.36E-49 | 0.284993084 | 0.376 | 0.104 | 8.82E-45 | 13 | Elmsan1 |
| Rbl23 | 4.68E-49 | 0.316610095 | 0.436 | 0.133 | 9.47E-45 | 13 | Rbl2 |
| Gab33 | 5.35E-49 | 0.333267303 | 0.345 | 0.093 | 1.08E-44 | 13 | Gab3 |
| Mlxip3 | 6.35E-49 | 0.305776913 | 0.421 | 0.125 | 1.29E-44 | 13 | Mlxip |
| Fyn6 | 8.56E-49 | 0.50170703 | 0.676 | 0.263 | 1.73E-44 | 13 | Fyn |
| Sik35 | 1.85E-48 | 0.528020308 | 0.676 | 0.267 | 3.74E-44 | 13 | Sik3 |
| Myo1e6 | 2.20E-48 | 0.625543542 | 0.673 | 0.278 | 4.45E-44 | 13 | Myo1e |
| Strbp2 | 2.23E-48 | 0.280438771 | 0.552 | 0.189 | 4.51E-44 | 13 | Strbp |
| Mgat4a4 | 2.33E-48 | 0.303131556 | 0.37 | 0.103 | 4.72E-44 | 13 | Mgat4a |
| Rps245 | 2.44E-48 | 0.63050071 | 1 | 0.972 | 4.95E-44 | 13 | Rps24 |
| Gsap4 | 2.72E-48 | 0.276900343 | 0.324 | 0.084 | 5.50E-44 | 13 | Gsap |
| Rab8b4 | 4.42E-48 | 0.526175122 | 0.688 | 0.281 | 8.94E-44 | 13 | Rab8b |
| Sema6d4 | 6.16E-48 | 0.533939692 | 0.576 | 0.211 | 1.25E-43 | 13 | Sema6d |
| Bin14 | 6.75E-48 | 0.292938678 | 0.515 | 0.173 | 1.37E-43 | 13 | Bin1 |
| Nedd96 | 7.02E-48 | 0.607597034 | 0.742 | 0.316 | 1.42E-43 | 13 | Nedd9 |
| Vwa82 | 7.91E-48 | 0.316689492 | 0.4 | 0.118 | 1.60E-43 | 13 | Vwa8 |
| Bhlhe404 | 8.68E-48 | 0.319053991 | 0.485 | 0.156 | 1.76E-43 | 13 | Bhlhe40 |
| Rps105 | 1.38E-47 | 0.617974642 | 0.994 | 0.944 | 2.79E-43 | 13 | Rps10 |
| AU0202066 | 1.39E-47 | 0.516378955 | 0.706 | 0.287 | 2.81E-43 | 13 | AU020206 |
| Ctla2a6 | 1.69E-47 | 0.453623938 | 0.882 | 0.394 | 3.42E-43 | 13 | Ctla2a |
| Chd26 | 1.92E-47 | 0.538617864 | 0.642 | 0.251 | 3.89E-43 | 13 | Chd2 |
| Litaf4 | 2.05E-47 | 0.343733975 | 0.479 | 0.155 | 4.14E-43 | 13 | Litaf |
| Dcaf53 | 2.09E-47 | 0.36226797 | 0.418 | 0.128 | 4.23E-43 | 13 | Dcaf5 |
| Nabp13 | 2.30E-47 | 0.263898673 | 0.312 | 0.08 | 4.65E-43 | 13 | Nabp1 |
| Cdk64 | 2.48E-47 | 0.369948416 | 0.379 | 0.109 | 5.03E-43 | 13 | Cdk6 |
| Vps283 | 5.75E-47 | 0.406838683 | 0.745 | 0.297 | 1.17E-42 | 13 | Vps28 |

| | | | | | | | |
|---|---|---|---|---|---|---|---|
| Ddx103 | 6.07E-47 | 0.261415786 | 0.427 | 0.129 | 1.23E-42 | 13 | Ddx10 |
| Was4 | 9.99E-47 | 0.286820975 | 0.291 | 0.071 | 2.02E-42 | 13 | Was |
| Inpp5d4 | 1.05E-46 | 0.360383313 | 0.57 | 0.201 | 2.13E-42 | 13 | Inpp5d |
| Rps164 | 1.39E-46 | 0.613180889 | 0.994 | 0.927 | 2.81E-42 | 13 | Rps16 |
| D16Ertd472e3 | 1.82E-46 | 0.297598617 | 0.367 | 0.104 | 3.69E-42 | 13 | D16Ertd472e |
| Ralgapa23 | 2.15E-46 | 0.379956026 | 0.464 | 0.152 | 4.35E-42 | 13 | Ralgapa2 |
| Cdc42ep35 | 2.29E-46 | 0.308870703 | 0.515 | 0.172 | 4.63E-42 | 13 | Cdc42ep3 |
| Hectd15 | 3.61E-46 | 0.515302728 | 0.712 | 0.297 | 7.30E-42 | 13 | Hectd1 |
| Cntrl3 | 4.85E-46 | 0.33190425 | 0.403 | 0.122 | 9.81E-42 | 13 | Cntrl |
| Kcnq1ot14 | 5.50E-46 | 0.41355919 | 0.424 | 0.132 | 1.11E-41 | 13 | Kcnq1ot1 |
| Mfsd102 | 6.44E-46 | 0.270229205 | 0.339 | 0.093 | 1.30E-41 | 13 | Mfsd10 |
| Rcc21 | 8.15E-46 | 0.28737982 | 0.379 | 0.11 | 1.65E-41 | 13 | Rcc2 |
| Ubash3b4 | 1.04E-45 | 0.331228583 | 0.497 | 0.165 | 2.11E-41 | 13 | Ubash3b |
| Nfatc14 | 1.22E-45 | 0.301967609 | 0.388 | 0.114 | 2.46E-41 | 13 | Nfatc1 |
| Vav35 | 1.26E-45 | 0.51849113 | 0.639 | 0.253 | 2.56E-41 | 13 | Vav3 |
| Iqsec12 | 1.59E-45 | 0.291714942 | 0.403 | 0.12 | 3.22E-41 | 13 | Iqsec1 |
| Dennd1b5 | 1.61E-45 | 0.402390318 | 0.521 | 0.182 | 3.25E-41 | 13 | Dennd1b |
| Rbm34 | 2.11E-45 | 0.56163519 | 0.912 | 0.473 | 4.28E-41 | 13 | Rbm3 |
| Wwp24 | 2.12E-45 | 0.310085494 | 0.412 | 0.125 | 4.29E-41 | 13 | Wwp2 |
| Ube2h3 | 2.36E-45 | 0.454175225 | 0.648 | 0.262 | 4.78E-41 | 13 | Ube2h |
| Fam204a3 | 2.47E-45 | 0.288894777 | 0.506 | 0.167 | 4.99E-41 | 13 | Fam204a |
| Mef2a5 | 2.52E-45 | 0.750835269 | 0.8 | 0.414 | 5.10E-41 | 13 | Mef2a |
| Agpat52 | 2.69E-45 | 0.363553216 | 0.439 | 0.14 | 5.45E-41 | 13 | Agpat5 |
| Cdc42se24 | 3.06E-45 | 0.388650652 | 0.627 | 0.236 | 6.20E-41 | 13 | Cdc42se2 |
| Eif3d | 4.71E-45 | 0.262978872 | 0.594 | 0.215 | 9.54E-41 | 13 | Eif3d |
| Tgfb15 | 5.32E-45 | 0.472762391 | 0.612 | 0.233 | 1.08E-40 | 13 | Tgfb1 |
| Arhgef19 | 6.99E-45 | 0.413876656 | 0.639 | 0.245 | 1.41E-40 | 13 | Arhgef1 |
| Itgb17 | 7.06E-45 | 0.672423593 | 0.876 | 0.476 | 1.43E-40 | 13 | Itgb1 |
| Ptpn24 | 8.11E-45 | 0.331852766 | 0.439 | 0.141 | 1.64E-40 | 13 | Ptpn2 |
| Ifi475 | 1.02E-44 | 0.337813073 | 0.521 | 0.176 | 2.06E-40 | 13 | Ifi47 |
| Arid1a6 | 1.13E-44 | 0.38549764 | 0.603 | 0.227 | 2.30E-40 | 13 | Arid1a |
| Rpl135 | 1.24E-44 | 0.570436853 | 0.997 | 0.977 | 2.51E-40 | 13 | Rpl13 |
| Itpr24 | 1.34E-44 | 0.41335007 | 0.618 | 0.241 | 2.71E-40 | 13 | Itpr2 |
| Rev3l4 | 1.36E-44 | 0.334400306 | 0.473 | 0.155 | 2.76E-40 | 13 | Rev3l |
| Rcor13 | 1.37E-44 | 0.341550963 | 0.518 | 0.181 | 2.77E-40 | 13 | Rcor1 |
| Creb13 | 1.87E-44 | 0.323026836 | 0.57 | 0.205 | 3.79E-40 | 13 | Creb1 |
| Rps275 | 1.92E-44 | 0.625024092 | 0.994 | 0.897 | 3.88E-40 | 13 | Rps27 |
| Rps204 | 2.33E-44 | 0.594754466 | 0.997 | 0.951 | 4.72E-40 | 13 | Rps20 |
| Prkca4 | 3.12E-44 | 0.262640709 | 0.348 | 0.099 | 6.32E-40 | 13 | Prkca |
| Rpl325 | 3.64E-44 | 0.606530792 | 0.997 | 0.953 | 7.37E-40 | 13 | Rpl32 |
| Lasp13 | 3.70E-44 | 0.283337028 | 0.567 | 0.205 | 7.49E-40 | 13 | Lasp1 |
| Tpp24 | 4.34E-44 | 0.388389281 | 0.597 | 0.225 | 8.78E-40 | 13 | Tpp2 |
| Plgrkt4 | 4.41E-44 | 0.348017232 | 0.576 | 0.21 | 8.93E-40 | 13 | Plgrkt |

| | | | | | | | |
|---|---|---|---|---|---|---|---|
| Cdk176 | 4.90E-44 | 0.476108091 | 0.624 | 0.242 | 9.92E-40 | 13 | Cdk17 |
| Elf44 | 5.45E-44 | 0.337808475 | 0.394 | 0.12 | 1.10E-39 | 13 | Elf4 |
| Cerk4 | 6.11E-44 | 0.258562036 | 0.379 | 0.113 | 1.24E-39 | 13 | Cerk |
| Dtnb3 | 9.02E-44 | 0.32180468 | 0.376 | 0.113 | 1.83E-39 | 13 | Dtnb |
| Rel4 | 1.04E-43 | 0.313380655 | 0.427 | 0.136 | 2.11E-39 | 13 | Rel |
| Parp44 | 1.19E-43 | 0.264814001 | 0.464 | 0.15 | 2.41E-39 | 13 | Parp4 |
| Sdf2l13 | 1.86E-43 | 0.323969032 | 0.491 | 0.165 | 3.77E-39 | 13 | Sdf2l1 |
| Msh31 | 1.97E-43 | 0.250644606 | 0.267 | 0.066 | 3.99E-39 | 13 | Msh3 |
| Cd1644 | 2.25E-43 | 0.333970701 | 0.576 | 0.211 | 4.55E-39 | 13 | Cd164 |
| Asxl23 | 2.86E-43 | 0.388691819 | 0.491 | 0.172 | 5.79E-39 | 13 | Asxl2 |
| Crlf23 | 2.92E-43 | 0.288962247 | 0.473 | 0.159 | 5.91E-39 | 13 | Crlf2 |
| Sep-95 | 3.45E-43 | 0.329726434 | 0.494 | 0.168 | 6.98E-39 | 13 | 9-Sep |
| Rasa15 | 4.48E-43 | 0.477578299 | 0.609 | 0.244 | 9.06E-39 | 13 | Rasa1 |
| Nol4l1 | 5.26E-43 | 0.268385907 | 0.285 | 0.073 | 1.06E-38 | 13 | Nol4l |
| Nasp3 | 8.43E-43 | 0.251411469 | 0.497 | 0.17 | 1.71E-38 | 13 | Nasp |
| Sipa14 | 9.36E-43 | 0.327337871 | 0.524 | 0.183 | 1.89E-38 | 13 | Sipa1 |
| Ddx392 | 9.47E-43 | 0.262881554 | 0.4 | 0.124 | 1.92E-38 | 13 | Ddx39 |
| Atp6v0a21 | 1.07E-42 | 0.271346504 | 0.37 | 0.11 | 2.16E-38 | 13 | Atp6v0a2 |
| Itgav5 | 1.29E-42 | 0.333068099 | 0.47 | 0.159 | 2.61E-38 | 13 | Itgav |
| Man1a6 | 1.53E-42 | 0.383508959 | 0.482 | 0.165 | 3.10E-38 | 13 | Man1a |
| Sh2d3c4 | 1.88E-42 | 0.338404403 | 0.461 | 0.155 | 3.81E-38 | 13 | Sh2d3c |
| Zdhhc206 | 2.05E-42 | 0.382192921 | 0.597 | 0.227 | 4.15E-38 | 13 | Zdhhc20 |
| Fosb5 | 2.53E-42 | 0.458935446 | 0.512 | 0.187 | 5.13E-38 | 13 | Fosb |
| Nup1533 | 3.13E-42 | 0.316822873 | 0.376 | 0.115 | 6.34E-38 | 13 | Nup153 |
| Brd47 | 3.86E-42 | 0.441531553 | 0.648 | 0.264 | 7.82E-38 | 13 | Brd4 |
| Rpl344 | 3.95E-42 | 0.54221759 | 0.994 | 0.917 | 7.99E-38 | 13 | Rpl34 |
| Mvb12b3 | 4.33E-42 | 0.342888097 | 0.473 | 0.165 | 8.78E-38 | 13 | Mvb12b |
| Cdyl4 | 4.40E-42 | 0.316279494 | 0.403 | 0.127 | 8.91E-38 | 13 | Cdyl |
| Zcchc114 | 5.94E-42 | 0.434694995 | 0.645 | 0.264 | 1.20E-37 | 13 | Zcchc11 |
| Capg3 | 7.42E-42 | 0.469690635 | 0.336 | 0.099 | 1.50E-37 | 13 | Capg |
| Slc2a13 | 1.20E-41 | 0.259176744 | 0.312 | 0.086 | 2.44E-37 | 13 | Slc2a1 |
| Rpl18a4 | 1.24E-41 | 0.579239025 | 0.994 | 0.93 | 2.51E-37 | 13 | Rpl18a |
| Cbl4 | 2.97E-41 | 0.382328328 | 0.552 | 0.205 | 6.01E-37 | 13 | Cbl |
| Prr133 | 3.82E-41 | 0.304794016 | 0.558 | 0.207 | 7.73E-37 | 13 | Prr13 |
| 2900026A02Rik4 | 4.49E-41 | 0.393330463 | 0.57 | 0.217 | 9.09E-37 | 13 | 2900026A02Rik |
| Itch6 | 4.70E-41 | 0.431613212 | 0.661 | 0.271 | 9.51E-37 | 13 | Itch |
| Atxn16 | 5.49E-41 | 0.477561017 | 0.691 | 0.292 | 1.11E-36 | 13 | Atxn1 |
| Nsd35 | 8.63E-41 | 0.487038932 | 0.7 | 0.305 | 1.75E-36 | 13 | Nsd3 |
| Cyld3 | 8.64E-41 | 0.267343756 | 0.482 | 0.168 | 1.75E-36 | 13 | Cyld |
| Mapre24 | 9.97E-41 | 0.385380853 | 0.624 | 0.247 | 2.02E-36 | 13 | Mapre2 |
| St8sia46 | 1.76E-40 | 0.420915952 | 0.636 | 0.257 | 3.55E-36 | 13 | St8sia4 |
| Kpna47 | 1.80E-40 | 0.474381022 | 0.721 | 0.32 | 3.64E-36 | 13 | Kpna4 |
| Stim15 | 1.81E-40 | 0.362927894 | 0.518 | 0.188 | 3.67E-36 | 13 | Stim1 |

| | | | | | | | |
|---|---|---|---|---|---|---|---|
| Cog55 | 2.51E-40 | 0.355031667 | 0.5 | 0.181 | 5.08E-36 | 13 | Cog5 |
| Ggnbp25 | 3.06E-40 | 0.318423183 | 0.612 | 0.239 | 6.19E-36 | 13 | Ggnbp2 |
| Esyt26 | 3.23E-40 | 0.458310325 | 0.591 | 0.24 | 6.54E-36 | 13 | Esyt2 |
| Immp2l4 | 3.77E-40 | 0.34070685 | 0.409 | 0.134 | 7.62E-36 | 13 | Immp2l |
| Sesn14 | 3.78E-40 | 0.370038557 | 0.552 | 0.21 | 7.66E-36 | 13 | Sesn1 |
| Ago23 | 4.13E-40 | 0.252824209 | 0.476 | 0.165 | 8.36E-36 | 13 | Ago2 |
| Exoc64 | 4.32E-40 | 0.270924572 | 0.482 | 0.168 | 8.74E-36 | 13 | Exoc6 |
| Tnfrsf1b3 | 4.68E-40 | 0.294951713 | 0.37 | 0.115 | 9.48E-36 | 13 | Tnfrsf1b |
| Supt4a4 | 4.73E-40 | 0.266507094 | 0.745 | 0.318 | 9.57E-36 | 13 | Supt4a |
| Ppm1g2 | 4.84E-40 | 0.272165816 | 0.455 | 0.154 | 9.80E-36 | 13 | Ppm1g |
| Arap25 | 5.10E-40 | 0.301144771 | 0.497 | 0.176 | 1.03E-35 | 13 | Arap2 |
| Stt3b4 | 5.33E-40 | 0.261534163 | 0.476 | 0.163 | 1.08E-35 | 13 | Stt3b |
| Smg73 | 5.48E-40 | 0.275929877 | 0.452 | 0.154 | 1.11E-35 | 13 | Smg7 |
| Rpl37a5 | 7.22E-40 | 0.583666683 | 1 | 0.937 | 1.46E-35 | 13 | Rpl37a |
| Gbp83 | 1.15E-39 | 0.298534494 | 0.418 | 0.136 | 2.32E-35 | 13 | Gbp8 |
| Capza14 | 1.72E-39 | 0.3683576 | 0.661 | 0.273 | 3.49E-35 | 13 | Capza1 |
| Slx4ip1 | 2.03E-39 | 0.271125908 | 0.261 | 0.067 | 4.11E-35 | 13 | Slx4ip |
| Pam5 | 2.30E-39 | 0.348233943 | 0.558 | 0.211 | 4.66E-35 | 13 | Pam |
| Bach24 | 2.47E-39 | 0.450121317 | 0.452 | 0.159 | 4.99E-35 | 13 | Bach2 |
| Mycbp27 | 2.50E-39 | 0.514398951 | 0.782 | 0.371 | 5.06E-35 | 13 | Mycbp2 |
| Socs12 | 2.91E-39 | 0.318643379 | 0.33 | 0.097 | 5.89E-35 | 13 | Socs1 |
| Snd15 | 3.12E-39 | 0.273494482 | 0.485 | 0.17 | 6.32E-35 | 13 | Snd1 |
| Tra2b4 | 4.63E-39 | 0.417949411 | 0.679 | 0.294 | 9.38E-35 | 13 | Tra2b |
| Ccm23 | 6.17E-39 | 0.266466882 | 0.391 | 0.124 | 1.25E-34 | 13 | Ccm2 |
| Csnk1g15 | 8.83E-39 | 0.260183985 | 0.452 | 0.155 | 1.79E-34 | 13 | Csnk1g1 |
| Syk3 | 1.03E-38 | 0.253612264 | 0.258 | 0.066 | 2.08E-34 | 13 | Syk |
| Rps3a16 | 1.13E-38 | 0.546090127 | 1 | 0.935 | 2.29E-34 | 13 | Rps3a1 |
| Atad23 | 1.16E-38 | 0.307044288 | 0.315 | 0.091 | 2.34E-34 | 13 | Atad2 |
| Zcchc77 | 1.58E-38 | 0.445168006 | 0.685 | 0.298 | 3.21E-34 | 13 | Zcchc7 |
| Fhit4 | 1.73E-38 | 0.2998805 | 0.282 | 0.078 | 3.49E-34 | 13 | Fhit |
| Trp533 | 1.80E-38 | 0.256544658 | 0.455 | 0.156 | 3.64E-34 | 13 | Trp53 |
| Cdk136 | 2.02E-38 | 0.326190845 | 0.573 | 0.225 | 4.10E-34 | 13 | Cdk13 |
| Nfkb16 | 2.72E-38 | 0.39171879 | 0.648 | 0.268 | 5.51E-34 | 13 | Nfkb1 |
| Smg65 | 2.85E-38 | 0.470292098 | 0.621 | 0.257 | 5.77E-34 | 13 | Smg6 |
| Ccdc1073 | 3.01E-38 | 0.268558528 | 0.388 | 0.126 | 6.10E-34 | 13 | Ccdc107 |
| Plekha54 | 3.06E-38 | 0.301619818 | 0.442 | 0.153 | 6.20E-34 | 13 | Plekha5 |
| Rps55 | 3.94E-38 | 0.575880615 | 1 | 0.919 | 7.97E-34 | 13 | Rps5 |
| Rad54l23 | 4.95E-38 | 0.279225401 | 0.318 | 0.093 | 1.00E-33 | 13 | Rad54l2 |
| Nme14 | 5.10E-38 | 0.284116728 | 0.776 | 0.352 | 1.03E-33 | 13 | Nme1 |
| Ankrd117 | 5.14E-38 | 0.596229975 | 0.736 | 0.379 | 1.04E-33 | 13 | Ankrd11 |
| Kdm5a3 | 5.28E-38 | 0.2533177 | 0.47 | 0.165 | 1.07E-33 | 13 | Kdm5a |
| Nfatc33 | 5.29E-38 | 0.314663407 | 0.482 | 0.173 | 1.07E-33 | 13 | Nfatc3 |
| Rpl22l15 | 5.70E-38 | 0.53476616 | 0.955 | 0.575 | 1.15E-33 | 13 | Rpl22l1 |

| | | | | | | | |
|---|---|---|---|---|---|---|---|
| Lgals34 | 5.80E-38 | 0.330424877 | 0.555 | 0.215 | 1.17E-33 | 13 | Lgals3 |
| Ptpn17 | 7.08E-38 | 0.315016098 | 0.591 | 0.233 | 1.43E-33 | 13 | Ptpn1 |
| Ddx213 | 7.88E-38 | 0.295189033 | 0.548 | 0.211 | 1.60E-33 | 13 | Ddx21 |
| Rftn16 | 8.68E-38 | 0.445788281 | 0.612 | 0.249 | 1.76E-33 | 13 | Rftn1 |
| Ranbp94 | 9.72E-38 | 0.296228791 | 0.527 | 0.198 | 1.97E-33 | 13 | Ranbp9 |
| Rbm257 | 1.01E-37 | 0.400333538 | 0.742 | 0.331 | 2.04E-33 | 13 | Rbm25 |
| Znrf15 | 1.15E-37 | 0.373630355 | 0.497 | 0.187 | 2.32E-33 | 13 | Znrf1 |
| Saraf6 | 1.29E-37 | 0.367642549 | 0.709 | 0.304 | 2.62E-33 | 13 | Saraf |
| Ttc7b2 | 1.36E-37 | 0.287280744 | 0.373 | 0.12 | 2.75E-33 | 13 | Ttc7b |
| Snrpg4 | 1.49E-37 | 0.320062422 | 0.736 | 0.321 | 3.01E-33 | 13 | Snrpg |
| Lyst3 | 2.14E-37 | 0.363825987 | 0.345 | 0.108 | 4.33E-33 | 13 | Lyst |
| Faf14 | 2.66E-37 | 0.262972143 | 0.533 | 0.199 | 5.39E-33 | 13 | Faf1 |
| Avl93 | 3.11E-37 | 0.288482884 | 0.406 | 0.135 | 6.31E-33 | 13 | Avl9 |
| Clcn32 | 3.52E-37 | 0.288503984 | 0.394 | 0.132 | 7.12E-33 | 13 | Clcn3 |
| Zfp6523 | 4.02E-37 | 0.308671668 | 0.603 | 0.245 | 8.14E-33 | 13 | Zfp652 |
| Tnrc6b5 | 4.23E-37 | 0.503435865 | 0.761 | 0.368 | 8.56E-33 | 13 | Tnrc6b |
| Lncpint6 | 5.97E-37 | 0.49131608 | 0.627 | 0.272 | 1.21E-32 | 13 | Lncpint |
| Vti1a5 | 7.22E-37 | 0.386704641 | 0.606 | 0.252 | 1.46E-32 | 13 | Vti1a |
| Rps75 | 8.37E-37 | 0.52823854 | 0.985 | 0.916 | 1.69E-32 | 13 | Rps7 |
| Rasa24 | 8.61E-37 | 0.303188859 | 0.409 | 0.136 | 1.74E-32 | 13 | Rasa2 |
| Rnaseh2c3 | 1.06E-36 | 0.276915462 | 0.667 | 0.283 | 2.15E-32 | 13 | Rnaseh2c |
| Kansl17 | 1.17E-36 | 0.500182301 | 0.718 | 0.337 | 2.37E-32 | 13 | Kansl1 |
| Rsf13 | 1.18E-36 | 0.337287977 | 0.558 | 0.222 | 2.38E-32 | 13 | Rsf1 |
| Rfx33 | 1.40E-36 | 0.255878633 | 0.339 | 0.106 | 2.83E-32 | 13 | Rfx3 |
| Pacs14 | 1.52E-36 | 0.321429349 | 0.527 | 0.203 | 3.08E-32 | 13 | Pacs1 |
| Lrp105 | 1.60E-36 | 0.300447868 | 0.5 | 0.186 | 3.24E-32 | 13 | Lrp10 |
| Wipf15 | 1.75E-36 | 0.384588653 | 0.636 | 0.265 | 3.55E-32 | 13 | Wipf1 |
| Sqstm15 | 2.22E-36 | 0.384272122 | 0.694 | 0.301 | 4.50E-32 | 13 | Sqstm1 |
| Rpl185 | 2.48E-36 | 0.521781551 | 0.994 | 0.9 | 5.03E-32 | 13 | Rpl18 |
| Srrm26 | 2.62E-36 | 0.503002146 | 0.861 | 0.466 | 5.29E-32 | 13 | Srrm2 |
| Kmt2a7 | 5.20E-36 | 0.399184143 | 0.618 | 0.265 | 1.05E-31 | 13 | Kmt2a |
| Smc53 | 6.99E-36 | 0.310141019 | 0.358 | 0.117 | 1.42E-31 | 13 | Smc5 |
| Rnf1306 | 7.01E-36 | 0.310547557 | 0.667 | 0.281 | 1.42E-31 | 13 | Rnf130 |
| Zc3h133 | 7.30E-36 | 0.319705339 | 0.542 | 0.214 | 1.48E-31 | 13 | Zc3h13 |
| Akap136 | 8.07E-36 | 0.574120409 | 0.761 | 0.391 | 1.63E-31 | 13 | Akap13 |
| Rngtt3 | 8.14E-36 | 0.261004396 | 0.394 | 0.134 | 1.65E-31 | 13 | Rngtt |
| Phactr25 | 8.29E-36 | 0.345931419 | 0.588 | 0.244 | 1.68E-31 | 13 | Phactr2 |
| Rps6ka36 | 9.92E-36 | 0.474100099 | 0.606 | 0.26 | 2.01E-31 | 13 | Rps6ka3 |
| Myh96 | 1.01E-35 | 0.532991852 | 0.824 | 0.434 | 2.05E-31 | 13 | Myh9 |
| Fbxl178 | 1.14E-35 | 0.518985574 | 0.664 | 0.308 | 2.31E-31 | 13 | Fbxl17 |
| Fau4 | 1.14E-35 | 0.497694421 | 0.994 | 0.953 | 2.32E-31 | 13 | Fau |
| Ntan13 | 1.32E-35 | 0.256232658 | 0.718 | 0.315 | 2.67E-31 | 13 | Ntan1 |
| Clk15 | 1.40E-35 | 0.384895518 | 0.667 | 0.289 | 2.83E-31 | 13 | Clk1 |

| | | | | | | | |
|---|---|---|---|---|---|---|---|
| Mark32 | 2.16E-35 | 0.266934821 | 0.539 | 0.21 | 4.38E-31 | 13 | Mark3 |
| Crlf34 | 2.16E-35 | 0.320014584 | 0.436 | 0.155 | 4.38E-31 | 13 | Crlf3 |
| Rps135 | 2.38E-35 | 0.55058836 | 0.997 | 0.923 | 4.83E-31 | 13 | Rps13 |
| Sec11a4 | 2.52E-35 | 0.260564627 | 0.67 | 0.282 | 5.09E-31 | 13 | Sec11a |
| Lbr5 | 3.54E-35 | 0.256708497 | 0.433 | 0.151 | 7.17E-31 | 13 | Lbr |
| Spata135 | 3.99E-35 | 0.272961319 | 0.533 | 0.204 | 8.09E-31 | 13 | Spata13 |
| Irak24 | 4.42E-35 | 0.280798635 | 0.406 | 0.14 | 8.95E-31 | 13 | Irak2 |
| Atxn103 | 4.97E-35 | 0.284107538 | 0.467 | 0.173 | 1.01E-30 | 13 | Atxn10 |
| Tcf127 | 7.30E-35 | 0.515190372 | 0.748 | 0.374 | 1.48E-30 | 13 | Tcf12 |
| N4bp2l23 | 7.96E-35 | 0.264531208 | 0.597 | 0.245 | 1.61E-30 | 13 | N4bp2l2 |
| Tln17 | 8.45E-35 | 0.457229217 | 0.8 | 0.402 | 1.71E-30 | 13 | Tln1 |
| Ran3 | 1.41E-34 | 0.317675659 | 0.827 | 0.396 | 2.86E-30 | 13 | Ran |
| Gls5 | 1.51E-34 | 0.312006235 | 0.597 | 0.249 | 3.07E-30 | 13 | Gls |
| Xist7 | 1.57E-34 | 0.536114756 | 0.867 | 0.46 | 3.18E-30 | 13 | Xist |
| Rptor4 | 1.60E-34 | 0.291872566 | 0.397 | 0.137 | 3.25E-30 | 13 | Rptor |
| Eif3f3 | 1.88E-34 | 0.494954696 | 0.888 | 0.48 | 3.81E-30 | 13 | Eif3f |
| Cnot45 | 1.91E-34 | 0.269678702 | 0.633 | 0.266 | 3.87E-30 | 13 | Cnot4 |
| Ubr25 | 2.47E-34 | 0.286436228 | 0.576 | 0.227 | 5.01E-30 | 13 | Ubr2 |
| Lman22 | 2.67E-34 | 0.329286811 | 0.527 | 0.204 | 5.41E-30 | 13 | Lman2 |
| Sppl36 | 2.73E-34 | 0.38787357 | 0.53 | 0.216 | 5.52E-30 | 13 | Sppl3 |
| Hdac84 | 2.82E-34 | 0.265158865 | 0.388 | 0.134 | 5.70E-30 | 13 | Hdac8 |
| Taf103 | 2.92E-34 | 0.258150213 | 0.821 | 0.381 | 5.92E-30 | 13 | Taf10 |
| Ppp1ca3 | 3.22E-34 | 0.388759725 | 0.8 | 0.394 | 6.52E-30 | 13 | Ppp1ca |
| Gbf15 | 4.74E-34 | 0.30026539 | 0.473 | 0.18 | 9.59E-30 | 13 | Gbf1 |
| Ube2i3 | 6.94E-34 | 0.264219547 | 0.727 | 0.327 | 1.40E-29 | 13 | Ube2i |
| Rpl385 | 7.14E-34 | 0.526237331 | 0.988 | 0.893 | 1.44E-29 | 13 | Rpl38 |
| Iqgap17 | 1.06E-33 | 0.459123103 | 0.797 | 0.409 | 2.14E-29 | 13 | Iqgap1 |
| Ablim15 | 1.24E-33 | 0.414647299 | 0.794 | 0.4 | 2.51E-29 | 13 | Ablim1 |
| Rcsd13 | 1.43E-33 | 0.252156432 | 0.376 | 0.128 | 2.89E-29 | 13 | Rcsd1 |
| Eif3e4 | 1.75E-33 | 0.299054568 | 0.839 | 0.409 | 3.54E-29 | 13 | Eif3e |
| Rpl45 | 1.82E-33 | 0.48574435 | 0.961 | 0.656 | 3.68E-29 | 13 | Rpl4 |
| Scaper5 | 1.84E-33 | 0.321255307 | 0.464 | 0.178 | 3.73E-29 | 13 | Scaper |
| Xrn23 | 1.85E-33 | 0.25112788 | 0.618 | 0.259 | 3.74E-29 | 13 | Xrn2 |
| Zfpm11 | 2.14E-33 | 0.255000607 | 0.424 | 0.15 | 4.33E-29 | 13 | Zfpm1 |
| Nsmaf4 | 2.38E-33 | 0.253741788 | 0.306 | 0.096 | 4.81E-29 | 13 | Nsmaf |
| Rpl27a5 | 2.52E-33 | 0.458611384 | 0.994 | 0.96 | 5.10E-29 | 13 | Rpl27a |
| Arid1b6 | 2.66E-33 | 0.321972124 | 0.609 | 0.256 | 5.39E-29 | 13 | Arid1b |
| Thrap34 | 3.04E-33 | 0.303590389 | 0.664 | 0.291 | 6.15E-29 | 13 | Thrap3 |
| Smg17 | 3.37E-33 | 0.358551931 | 0.561 | 0.232 | 6.82E-29 | 13 | Smg1 |
| Jarid24 | 4.10E-33 | 0.350957107 | 0.512 | 0.202 | 8.30E-29 | 13 | Jarid2 |
| Ogt5 | 5.41E-33 | 0.251077152 | 0.527 | 0.204 | 1.09E-28 | 13 | Ogt |
| Plec2 | 6.98E-33 | 0.257631042 | 0.461 | 0.172 | 1.41E-28 | 13 | Plec |
| Mndal6 | 7.74E-33 | 0.425168432 | 0.794 | 0.403 | 1.57E-28 | 13 | Mndal |

| | | | | | | | |
|---|---|---|---|---|---|---|---|
| Anxa25 | 8.67E-33 | 0.407148638 | 0.833 | 0.413 | 1.75E-28 | 13 | Anxa2 |
| Tnfsf104 | 9.07E-33 | 0.317274893 | 0.803 | 0.371 | 1.84E-28 | 13 | Tnfsf10 |
| Huwe16 | 1.02E-32 | 0.340144242 | 0.648 | 0.282 | 2.07E-28 | 13 | Huwe1 |
| Pitpnc16 | 1.15E-32 | 0.535703694 | 0.794 | 0.427 | 2.34E-28 | 13 | Pitpnc1 |
| Vps544 | 1.48E-32 | 0.287260792 | 0.476 | 0.185 | 2.99E-28 | 13 | Vps54 |
| Atad2b5 | 1.67E-32 | 0.261812925 | 0.509 | 0.201 | 3.38E-28 | 13 | Atad2b |
| Pum22 | 1.73E-32 | 0.318515214 | 0.618 | 0.267 | 3.49E-28 | 13 | Pum2 |
| Nktr4 | 1.76E-32 | 0.346246772 | 0.588 | 0.255 | 3.55E-28 | 13 | Nktr |
| Lnpep7 | 2.05E-32 | 0.358286181 | 0.603 | 0.257 | 4.15E-28 | 13 | Lnpep |
| Polr2a6 | 2.42E-32 | 0.299161901 | 0.639 | 0.279 | 4.89E-28 | 13 | Polr2a |
| Gm265101 | 2.69E-32 | 0.270372052 | 0.333 | 0.109 | 5.44E-28 | 13 | Gm26510 |
| Tle45 | 3.22E-32 | 0.374273769 | 0.536 | 0.224 | 6.51E-28 | 13 | Tle4 |
| Eif3h4 | 3.88E-32 | 0.301956849 | 0.9 | 0.477 | 7.86E-28 | 13 | Eif3h |
| Os95 | 4.39E-32 | 0.309618071 | 0.476 | 0.187 | 8.89E-28 | 13 | Os9 |
| Lims15 | 4.52E-32 | 0.324575485 | 0.664 | 0.298 | 9.16E-28 | 13 | Lims1 |
| 1700025G04Rik4 | 4.84E-32 | 0.318287311 | 0.482 | 0.187 | 9.81E-28 | 13 | 1700025G04Rik |
| Rpl395 | 5.47E-32 | 0.503292677 | 0.997 | 0.9 | 1.11E-27 | 13 | Rpl39 |
| Ppp4r3b4 | 5.51E-32 | 0.279515905 | 0.561 | 0.23 | 1.12E-27 | 13 | Ppp4r3b |
| Rps236 | 5.88E-32 | 0.473057546 | 0.994 | 0.952 | 1.19E-27 | 13 | Rps23 |
| Ncoa27 | 6.15E-32 | 0.300438002 | 0.567 | 0.236 | 1.25E-27 | 13 | Ncoa2 |
| Abhd17b4 | 6.41E-32 | 0.263161542 | 0.491 | 0.19 | 1.30E-27 | 13 | Abhd17b |
| Ankrd127 | 6.52E-32 | 0.351138197 | 0.67 | 0.305 | 1.32E-27 | 13 | Ankrd12 |
| Fli16 | 6.91E-32 | 0.520247017 | 0.761 | 0.399 | 1.40E-27 | 13 | Fli1 |
| Clic15 | 8.55E-32 | 0.506204433 | 0.958 | 0.598 | 1.73E-27 | 13 | Clic1 |
| Lrrfip16 | 8.98E-32 | 0.367171024 | 0.767 | 0.376 | 1.82E-27 | 13 | Lrrfip1 |
| Wrn3 | 1.00E-31 | 0.263081653 | 0.406 | 0.145 | 2.03E-27 | 13 | Wrn |
| Zdhhc185 | 1.01E-31 | 0.269715792 | 0.452 | 0.169 | 2.05E-27 | 13 | Zdhhc18 |
| Rpl305 | 1.03E-31 | 0.466620997 | 0.994 | 0.937 | 2.09E-27 | 13 | Rpl30 |
| Babam25 | 1.04E-31 | 0.341059362 | 0.561 | 0.236 | 2.10E-27 | 13 | Babam2 |
| Rpl125 | 1.24E-31 | 0.482178599 | 0.982 | 0.841 | 2.51E-27 | 13 | Rpl12 |
| Phf36 | 1.30E-31 | 0.311055791 | 0.585 | 0.25 | 2.64E-27 | 13 | Phf3 |
| Twf23 | 1.41E-31 | 0.284860183 | 0.452 | 0.169 | 2.85E-27 | 13 | Twf2 |
| Btbd97 | 1.46E-31 | 0.337793575 | 0.555 | 0.235 | 2.95E-27 | 13 | Btbd9 |
| Larp15 | 1.62E-31 | 0.259346172 | 0.503 | 0.201 | 3.27E-27 | 13 | Larp1 |
| Ep3004 | 1.77E-31 | 0.271467081 | 0.509 | 0.202 | 3.58E-27 | 13 | Ep300 |
| Ppp1r12a6 | 1.78E-31 | 0.531607454 | 0.785 | 0.423 | 3.61E-27 | 13 | Ppp1r12a |
| Vps13b5 | 2.17E-31 | 0.310244856 | 0.533 | 0.217 | 4.38E-27 | 13 | Vps13b |
| Rbm66 | 2.52E-31 | 0.277186097 | 0.53 | 0.215 | 5.10E-27 | 13 | Rbm6 |
| Prrc2b4 | 2.71E-31 | 0.28207568 | 0.555 | 0.227 | 5.48E-27 | 13 | Prrc2b |
| Aqp11 | 3.37E-31 | 0.409069996 | 0.291 | 0.092 | 6.82E-27 | 13 | Aqp1 |
| Actr35 | 4.11E-31 | 0.272673967 | 0.797 | 0.388 | 8.33E-27 | 13 | Actr3 |
| Ier56 | 4.16E-31 | 0.274277591 | 0.7 | 0.314 | 8.42E-27 | 13 | Ier5 |
| Acin15 | 4.70E-31 | 0.309896087 | 0.685 | 0.314 | 9.51E-27 | 13 | Acin1 |

| | | | | | | | |
|---|---|---|---|---|---|---|---|
| Arid4a7 | 5.17E-31 | 0.319436948 | 0.585 | 0.249 | 1.05E-26 | 13 | Arid4a |
| Phf21a6 | 5.24E-31 | 0.351078869 | 0.57 | 0.251 | 1.06E-26 | 13 | Phf21a |
| Zfp6445 | 5.26E-31 | 0.267072508 | 0.6 | 0.253 | 1.07E-26 | 13 | Zfp644 |
| B4galt16 | 5.59E-31 | 0.322957131 | 0.448 | 0.175 | 1.13E-26 | 13 | B4galt1 |
| Ahnak6 | 7.49E-31 | 0.454437703 | 0.879 | 0.469 | 1.52E-26 | 13 | Ahnak |
| Fnbp43 | 7.93E-31 | 0.250844641 | 0.361 | 0.127 | 1.61E-26 | 13 | Fnbp4 |
| Rreb13 | 1.07E-30 | 0.269211942 | 0.494 | 0.2 | 2.17E-26 | 13 | Rreb1 |
| Srgn7 | 1.08E-30 | 0.528252917 | 0.93 | 0.542 | 2.20E-26 | 13 | Srgn |
| Sf3b15 | 1.14E-30 | 0.31277302 | 0.788 | 0.39 | 2.31E-26 | 13 | Sf3b1 |
| Nin3 | 1.24E-30 | 0.342249502 | 0.379 | 0.137 | 2.50E-26 | 13 | Nin |
| Mllt106 | 1.38E-30 | 0.336513348 | 0.609 | 0.272 | 2.80E-26 | 13 | Mllt10 |
| Rpl275 | 1.45E-30 | 0.424380815 | 0.939 | 0.58 | 2.94E-26 | 13 | Rpl27 |
| Tmcc17 | 1.95E-30 | 0.319124478 | 0.518 | 0.216 | 3.95E-26 | 13 | Tmcc1 |
| Itsn26 | 1.98E-30 | 0.361837978 | 0.609 | 0.276 | 4.01E-26 | 13 | Itsn2 |
| Psmb104 | 2.19E-30 | 0.355656477 | 0.803 | 0.397 | 4.44E-26 | 13 | Psmb10 |
| Lcor5 | 2.43E-30 | 0.257187385 | 0.473 | 0.188 | 4.91E-26 | 13 | Lcor |
| Fam172a6 | 2.47E-30 | 0.422869533 | 0.612 | 0.282 | 5.00E-26 | 13 | Fam172a |
| Stat36 | 2.50E-30 | 0.274836348 | 0.609 | 0.264 | 5.07E-26 | 13 | Stat3 |
| Herc14 | 2.86E-30 | 0.25966997 | 0.594 | 0.255 | 5.79E-26 | 13 | Herc1 |
| Rap1gds17 | 3.54E-30 | 0.301888941 | 0.594 | 0.259 | 7.16E-26 | 13 | Rap1gds1 |
| Eef1a15 | 3.75E-30 | 0.38317833 | 1 | 0.959 | 7.60E-26 | 13 | Eef1a1 |
| Rpl85 | 6.60E-30 | 0.437578188 | 0.997 | 0.925 | 1.34E-25 | 13 | Rpl8 |
| Prpf38b6 | 6.94E-30 | 0.256319232 | 0.485 | 0.194 | 1.40E-25 | 13 | Prpf38b |
| 4932438A13Rik5 | 7.30E-30 | 0.309910948 | 0.518 | 0.217 | 1.48E-25 | 13 | 4932438A13Rik |
| Sri3 | 7.67E-30 | 0.291870084 | 0.824 | 0.417 | 1.55E-25 | 13 | Sri |
| Arhgap265 | 7.83E-30 | 0.264848615 | 0.533 | 0.22 | 1.59E-25 | 13 | Arhgap26 |
| Rpl316 | 8.11E-30 | 0.448218518 | 0.994 | 0.909 | 1.64E-25 | 13 | Rpl3 |
| Spcs23 | 1.26E-29 | 0.353648127 | 0.803 | 0.398 | 2.55E-25 | 13 | Spcs2 |
| Rapgef15 | 1.95E-29 | 0.272560952 | 0.558 | 0.239 | 3.94E-25 | 13 | Rapgef1 |
| Ncor16 | 1.95E-29 | 0.370676096 | 0.824 | 0.435 | 3.95E-25 | 13 | Ncor1 |
| H2-D17 | 2.08E-29 | 0.539447586 | 0.955 | 0.667 | 4.21E-25 | 13 | H2-D1 |
| Eef1b26 | 2.34E-29 | 0.487586654 | 0.979 | 0.734 | 4.74E-25 | 13 | Eef1b2 |
| Lrba4 | 2.69E-29 | 0.286320583 | 0.47 | 0.19 | 5.44E-25 | 13 | Lrba |
| Hip15 | 2.83E-29 | 0.396853033 | 0.682 | 0.332 | 5.73E-25 | 13 | Hip1 |
| Reep52 | 3.36E-29 | 0.270194264 | 0.87 | 0.478 | 6.80E-25 | 13 | Reep5 |
| Supt61 | 3.39E-29 | 0.252982965 | 0.418 | 0.158 | 6.87E-25 | 13 | Supt6 |
| Traf34 | 3.75E-29 | 0.323531072 | 0.315 | 0.108 | 7.59E-25 | 13 | Traf3 |
| Rpl96 | 3.79E-29 | 0.437204101 | 0.994 | 0.944 | 7.67E-25 | 13 | Rpl9 |
| Pnisr6 | 4.73E-29 | 0.269349476 | 0.594 | 0.262 | 9.58E-25 | 13 | Pnisr |
| Ncoa15 | 5.12E-29 | 0.261423108 | 0.503 | 0.208 | 1.04E-24 | 13 | Ncoa1 |
| Rpl175 | 5.43E-29 | 0.434681369 | 1 | 0.916 | 1.10E-24 | 13 | Rpl17 |
| Lpp6 | 8.82E-29 | 0.296349504 | 0.621 | 0.284 | 1.79E-24 | 13 | Lpp |
| Rps195 | 9.75E-29 | 0.453251794 | 0.988 | 0.927 | 1.97E-24 | 13 | Rps19 |

| | | | | | | | |
|---|---|---|---|---|---|---|---|
| Ccnl14 | 1.30E-28 | 0.267973617 | 0.503 | 0.208 | 2.63E-24 | 13 | Ccnl1 |
| Tpk14 | 1.43E-28 | 0.25585516 | 0.306 | 0.104 | 2.90E-24 | 13 | Tpk1 |
| Exoc48 | 1.60E-28 | 0.332313858 | 0.67 | 0.32 | 3.25E-24 | 13 | Exoc4 |
| Dnajc36 | 1.94E-28 | 0.355845358 | 0.745 | 0.368 | 3.93E-24 | 13 | Dnajc3 |
| Smad76 | 2.32E-28 | 0.260051721 | 0.697 | 0.323 | 4.70E-24 | 13 | Smad7 |
| Tmed95 | 2.77E-28 | 0.276339781 | 0.806 | 0.4 | 5.60E-24 | 13 | Tmed9 |
| Phip6 | 2.79E-28 | 0.281799424 | 0.573 | 0.254 | 5.64E-24 | 13 | Phip |
| Pan37 | 3.06E-28 | 0.299168731 | 0.527 | 0.227 | 6.19E-24 | 13 | Pan3 |
| Zfp2926 | 3.72E-28 | 0.303623289 | 0.558 | 0.249 | 7.53E-24 | 13 | Zfp292 |
| Psmb88 | 5.33E-28 | 0.425395758 | 0.939 | 0.554 | 1.08E-23 | 13 | Psmb8 |
| Akt36 | 5.46E-28 | 0.286240827 | 0.573 | 0.249 | 1.11E-23 | 13 | Akt3 |
| Bclaf14 | 5.81E-28 | 0.298722866 | 0.57 | 0.252 | 1.18E-23 | 13 | Bclaf1 |
| Nipbl8 | 7.58E-28 | 0.333054605 | 0.73 | 0.364 | 1.54E-23 | 13 | Nipbl |
| Luc7l26 | 8.70E-28 | 0.446744642 | 0.755 | 0.403 | 1.76E-23 | 13 | Luc7l2 |
| Ptbp37 | 9.52E-28 | 0.373720753 | 0.761 | 0.391 | 1.93E-23 | 13 | Ptbp3 |
| Tapbp4 | 1.20E-27 | 0.276540433 | 0.667 | 0.302 | 2.43E-23 | 13 | Tapbp |
| Eif4a13 | 1.22E-27 | 0.345665113 | 0.815 | 0.428 | 2.47E-23 | 13 | Eif4a1 |
| Gm203886 | 1.39E-27 | 0.386247537 | 0.779 | 0.398 | 2.82E-23 | 13 | Gm20388 |
| Vgll45 | 2.29E-27 | 0.300337031 | 0.561 | 0.248 | 4.64E-23 | 13 | Vgll4 |
| Rps66 | 3.06E-27 | 0.443560007 | 0.988 | 0.81 | 6.20E-23 | 13 | Rps6 |
| Bptf7 | 3.21E-27 | 0.314177422 | 0.618 | 0.292 | 6.50E-23 | 13 | Bptf |
| Ankrd177 | 3.86E-27 | 0.276135059 | 0.573 | 0.259 | 7.81E-23 | 13 | Ankrd17 |
| Dennd1a6 | 4.46E-27 | 0.273687308 | 0.567 | 0.251 | 9.03E-23 | 13 | Dennd1a |
| Tra2a7 | 6.16E-27 | 0.288104071 | 0.606 | 0.272 | 1.25E-22 | 13 | Tra2a |
| Crebbp6 | 8.22E-27 | 0.275093225 | 0.645 | 0.303 | 1.66E-22 | 13 | Crebbp |
| Rpl376 | 8.78E-27 | 0.428738982 | 1 | 0.927 | 1.78E-22 | 13 | Rpl37 |
| Sub14 | 8.92E-27 | 0.313247728 | 0.906 | 0.513 | 1.81E-22 | 13 | Sub1 |
| Rpl236 | 9.33E-27 | 0.40136339 | 0.997 | 0.971 | 1.89E-22 | 13 | Rpl23 |
| Dad15 | 1.03E-26 | 0.351790986 | 0.845 | 0.456 | 2.08E-22 | 13 | Dad1 |
| Cdk125 | 1.10E-26 | 0.290078528 | 0.53 | 0.228 | 2.22E-22 | 13 | Cdk12 |
| Birc67 | 1.15E-26 | 0.26251188 | 0.703 | 0.338 | 2.33E-22 | 13 | Birc6 |
| Rpl195 | 1.77E-26 | 0.401131135 | 1 | 0.963 | 3.58E-22 | 13 | Rpl19 |
| Rps145 | 3.05E-26 | 0.382836318 | 1 | 0.943 | 6.18E-22 | 13 | Rps14 |
| Eef1g5 | 3.31E-26 | 0.289748177 | 0.879 | 0.508 | 6.71E-22 | 13 | Eef1g |
| Susd67 | 4.56E-26 | 0.261590755 | 0.609 | 0.28 | 9.24E-22 | 13 | Susd6 |
| Itga14 | 6.37E-26 | 0.372425067 | 0.779 | 0.413 | 1.29E-21 | 13 | Itga1 |
| Rpl365 | 8.59E-26 | 0.400752524 | 0.985 | 0.903 | 1.74E-21 | 13 | Rpl36 |
| Rpl66 | 1.34E-25 | 0.404921205 | 0.994 | 0.881 | 2.71E-21 | 13 | Rpl6 |
| Arl6ip16 | 1.41E-25 | 0.271580538 | 0.724 | 0.368 | 2.85E-21 | 13 | Arl6ip1 |
| Hgsnat2 | 1.76E-25 | 0.344967938 | 0.252 | 0.083 | 3.56E-21 | 13 | Hgsnat |
| Rack16 | 1.78E-25 | 0.437146351 | 0.985 | 0.791 | 3.60E-21 | 13 | Rack1 |
| Tpr5 | 1.98E-25 | 0.338105027 | 0.785 | 0.423 | 4.02E-21 | 13 | Tpr |
| Slc3a25 | 2.13E-25 | 0.262122838 | 0.615 | 0.288 | 4.32E-21 | 13 | Slc3a2 |

| | | | | | | | |
|---|---|---|---|---|---|---|---|
| Maml27 | 2.20E-25 | 0.424154804 | 0.652 | 0.341 | 4.45E-21 | 13 | Maml2 |
| Diaph26 | 2.51E-25 | 0.336411789 | 0.727 | 0.374 | 5.08E-21 | 13 | Diaph2 |
| Ifngr18 | 2.51E-25 | 0.34183241 | 0.758 | 0.39 | 5.08E-21 | 13 | Ifngr1 |
| Tbc1d56 | 3.51E-25 | 0.250646823 | 0.567 | 0.26 | 7.10E-21 | 13 | Tbc1d5 |
| Kmt2c7 | 3.92E-25 | 0.316025371 | 0.624 | 0.304 | 7.93E-21 | 13 | Kmt2c |
| Krtcap24 | 4.71E-25 | 0.318874471 | 0.812 | 0.439 | 9.53E-21 | 13 | Krtcap2 |
| Ncl3 | 5.16E-25 | 0.425381014 | 0.873 | 0.514 | 1.04E-20 | 13 | Ncl |
| Psmb33 | 5.50E-25 | 0.25162236 | 0.806 | 0.417 | 1.11E-20 | 13 | Psmb3 |
| Rpl155 | 7.45E-25 | 0.422561807 | 0.985 | 0.901 | 1.51E-20 | 13 | Rpl15 |
| Arpc25 | 7.95E-25 | 0.378994174 | 0.903 | 0.542 | 1.61E-20 | 13 | Arpc2 |
| Rpl225 | 8.56E-25 | 0.365800304 | 0.985 | 0.851 | 1.73E-20 | 13 | Rpl22 |
| Ccdc88a6 | 9.96E-25 | 0.261828308 | 0.47 | 0.203 | 2.02E-20 | 13 | Ccdc88a |
| Uba526 | 1.02E-24 | 0.296993516 | 0.903 | 0.519 | 2.07E-20 | 13 | Uba52 |
| Kdm6a6 | 1.72E-24 | 0.28724836 | 0.479 | 0.212 | 3.48E-20 | 13 | Kdm6a |
| Peak16 | 1.73E-24 | 0.340263438 | 0.727 | 0.385 | 3.51E-20 | 13 | Peak1 |
| Foxp15 | 2.98E-24 | 0.305008018 | 0.727 | 0.387 | 6.03E-20 | 13 | Foxp1 |
| Lrrc8c4 | 4.43E-24 | 0.288267393 | 0.645 | 0.309 | 8.97E-20 | 13 | Lrrc8c |
| St6galnac34 | 4.48E-24 | 0.361485311 | 0.494 | 0.215 | 9.07E-20 | 13 | St6galnac3 |
| Rplp26 | 5.03E-24 | 0.363997584 | 1 | 0.889 | 1.02E-19 | 13 | Rplp2 |
| Jund4 | 9.04E-24 | 0.400348192 | 0.942 | 0.644 | 1.83E-19 | 13 | Jund |
| mt-Nd56 | 1.25E-23 | 0.425998588 | 0.827 | 0.494 | 2.54E-19 | 13 | mt-Nd5 |
| Rps95 | 3.32E-23 | 0.365947899 | 0.985 | 0.92 | 6.72E-19 | 13 | Rps9 |
| Sep-77 | 3.88E-23 | 0.267273903 | 0.845 | 0.459 | 7.86E-19 | 13 | 7-Sep |
| Rps4x5 | 4.67E-23 | 0.371162463 | 0.991 | 0.941 | 9.46E-19 | 13 | Rps4x |
| Zeb15 | 4.83E-23 | 0.2888714 | 0.633 | 0.32 | 9.78E-19 | 13 | Zeb1 |
| Rpl116 | 5.53E-23 | 0.366734892 | 0.994 | 0.927 | 1.12E-18 | 13 | Rpl11 |
| Rps27a5 | 5.70E-23 | 0.327191468 | 0.997 | 0.95 | 1.15E-18 | 13 | Rps27a |
| Wnk17 | 5.93E-23 | 0.254082667 | 0.709 | 0.361 | 1.20E-18 | 13 | Wnk1 |
| mt-Co18 | 6.27E-23 | 0.469221734 | 0.967 | 0.718 | 1.27E-18 | 13 | mt-Co1 |
| Arih15 | 1.38E-22 | 0.269672631 | 0.621 | 0.311 | 2.80E-18 | 13 | Arih1 |
| Hspa57 | 1.83E-22 | 0.287294559 | 0.797 | 0.432 | 3.71E-18 | 13 | Hspa5 |
| Elmo16 | 2.78E-22 | 0.258119022 | 0.7 | 0.364 | 5.63E-18 | 13 | Elmo1 |
| Prrc2c7 | 3.20E-22 | 0.316878757 | 0.755 | 0.418 | 6.48E-18 | 13 | Prrc2c |
| Foxn36 | 3.79E-22 | 0.254663641 | 0.767 | 0.408 | 7.67E-18 | 13 | Foxn3 |
| Rpl23a6 | 5.65E-22 | 0.357542583 | 0.948 | 0.677 | 1.14E-17 | 13 | Rpl23a |
| Ppp3ca7 | 9.87E-22 | 0.305790109 | 0.721 | 0.391 | 2.00E-17 | 13 | Ppp3ca |
| Klf65 | 1.13E-21 | 0.259007536 | 0.555 | 0.266 | 2.29E-17 | 13 | Klf6 |
| B2m9 | 1.20E-21 | 0.414483065 | 0.973 | 0.689 | 2.43E-17 | 13 | B2m |
| Macf16 | 1.30E-21 | 0.376351469 | 0.842 | 0.501 | 2.64E-17 | 13 | Macf1 |
| Zbtb206 | 1.44E-21 | 0.417828758 | 0.855 | 0.535 | 2.92E-17 | 13 | Zbtb20 |
| Rps285 | 2.02E-21 | 0.339423861 | 0.997 | 0.897 | 4.09E-17 | 13 | Rps28 |
| Rpl285 | 4.38E-21 | 0.342358672 | 0.994 | 0.93 | 8.86E-17 | 13 | Rpl28 |
| Rpl295 | 8.17E-21 | 0.388557614 | 0.982 | 0.767 | 1.65E-16 | 13 | Rpl29 |

| | | | | | | | |
|---|---|---|---|---|---|---|---|
| Rps218 | 1.35E-20 | 0.37874892 | 0.991 | 0.919 | 2.74E-16 | 13 | Rps2 |
| Arpc1b4 | 1.61E-20 | 0.274852965 | 0.927 | 0.589 | 3.25E-16 | 13 | Arpc1b |
| Cfl15 | 3.66E-20 | 0.35317599 | 0.948 | 0.606 | 7.42E-16 | 13 | Cfl1 |
| Stag17 | 4.16E-20 | 0.291151146 | 0.67 | 0.36 | 8.42E-16 | 13 | Stag1 |
| Srgap26 | 6.70E-20 | 0.250786765 | 0.594 | 0.302 | 1.36E-15 | 13 | Srgap2 |
| Rps86 | 1.21E-19 | 0.312854486 | 0.994 | 0.965 | 2.45E-15 | 13 | Rps8 |
| Eif5a4 | 2.52E-19 | 0.293838288 | 0.906 | 0.563 | 5.10E-15 | 13 | Eif5a |
| Rps219 | 3.95E-19 | 0.300532207 | 0.994 | 0.926 | 8.00E-15 | 13 | Rps21 |
| Myl65 | 1.85E-18 | 0.344985765 | 0.979 | 0.775 | 3.74E-14 | 13 | Myl6 |
| Npm15 | 2.32E-18 | 0.270130312 | 0.955 | 0.677 | 4.70E-14 | 13 | Npm1 |
| Rpl57 | 3.63E-18 | 0.287529462 | 0.952 | 0.708 | 7.35E-14 | 13 | Rpl5 |
| Kmt2e5 | 3.84E-18 | 0.250691438 | 0.709 | 0.405 | 7.78E-14 | 13 | Kmt2e |
| Rpl145 | 5.89E-18 | 0.305969028 | 0.988 | 0.801 | 1.19E-13 | 13 | Rpl14 |
| mt-Nd48 | 2.97E-17 | 0.376962608 | 0.939 | 0.674 | 6.02E-13 | 13 | mt-Nd4 |
| Rpl10a6 | 3.27E-17 | 0.304031985 | 0.985 | 0.835 | 6.61E-13 | 13 | Rpl10a |
| Rpl35a5 | 4.05E-17 | 0.287699132 | 0.997 | 0.948 | 8.19E-13 | 13 | Rpl35a |
| Rpl77 | 8.89E-17 | 0.300123495 | 0.985 | 0.892 | 1.80E-12 | 13 | Rpl7 |
| Rpl216 | 1.86E-16 | 0.319029902 | 0.991 | 0.923 | 3.77E-12 | 13 | Rpl21 |
| Rps126 | 8.01E-16 | 0.291217686 | 0.994 | 0.898 | 1.62E-11 | 13 | Rps12 |
| Actg13 | 2.30E-14 | 0.286193811 | 0.97 | 0.793 | 4.66E-10 | 13 | Actg1 |
| mt-Cytb7 | 3.50E-14 | 0.30486712 | 0.964 | 0.699 | 7.08E-10 | 13 | mt-Cytb |
| Hsp90b16 | 4.24E-14 | 0.272656591 | 0.87 | 0.508 | 8.59E-10 | 13 | Hsp90b1 |
| mt-Co39 | 3.44E-13 | 0.301879832 | 0.973 | 0.725 | 6.96E-09 | 13 | mt-Co3 |
| mt-Nd18 | 1.07E-11 | 0.251472963 | 0.891 | 0.597 | 2.16E-07 | 13 | mt-Nd1 |
| mt-Co210 | 1.60E-10 | 0.292621595 | 0.97 | 0.727 | 3.24E-06 | 13 | mt-Co2 |
| Cldn51 | 0 | 4.164738018 | 0.978 | 0.023 | 0 | 14 | Cldn5 |
| Mgp1 | 0 | 3.631689663 | 0.64 | 0.054 | 0 | 14 | Mgp |
| Mecom1 | 0 | 3.58503954 | 0.989 | 0.089 | 0 | 14 | Mecom |
| Stmn21 | 0 | 3.528043876 | 0.938 | 0.02 | 0 | 14 | Stmn2 |
| Clu1 | 0 | 3.127290637 | 0.887 | 0.057 | 0 | 14 | Clu |
| Fbln52 | 0 | 3.111680724 | 0.956 | 0.03 | 0 | 14 | Fbln5 |
| Pcsk5 | 0 | 3.011733567 | 0.873 | 0.018 | 0 | 14 | Pcsk5 |
| Nebl1 | 0 | 2.909267773 | 0.822 | 0.041 | 0 | 14 | Nebl |
| Col8a11 | 0 | 2.899497985 | 0.865 | 0.02 | 0 | 14 | Col8a1 |
| Sema3g1 | 0 | 2.878685826 | 0.916 | 0.024 | 0 | 14 | Sema3g |
| Vegfc1 | 0 | 2.803294104 | 0.887 | 0.102 | 0 | 14 | Vegfc |
| Gja41 | 0 | 2.762325238 | 0.884 | 0.054 | 0 | 14 | Gja4 |
| Sox171 | 0 | 2.677801742 | 0.895 | 0.081 | 0 | 14 | Sox17 |
| Edn1 | 0 | 2.486675363 | 0.745 | 0.021 | 0 | 14 | Edn1 |
| Cdh132 | 0 | 2.467482758 | 0.833 | 0.091 | 0 | 14 | Cdh13 |
| Fn11 | 0 | 2.443438178 | 0.862 | 0.025 | 0 | 14 | Fn1 |
| Alpl1 | 0 | 2.390826166 | 0.836 | 0.066 | 0 | 14 | Alpl |
| Bmp6 | 0 | 2.33536318 | 0.596 | 0.041 | 0 | 14 | Bmp6 |

| | | | | | | | |
|---|---|---|---|---|---|---|---|
| Ptprr1 | 0 | 2.143957259 | 0.796 | 0.058 | 0 | 14 | Ptprr |
| St8sia6 | 0 | 2.067121605 | 0.702 | 0.013 | 0 | 14 | St8sia6 |
| Tspan71 | 0 | 1.998148559 | 0.738 | 0.071 | 0 | 14 | Tspan7 |
| Mannr | 0 | 1.934172623 | 0.604 | 0.007 | 0 | 14 | Mannr |
| Bmx | 0 | 1.890537748 | 0.589 | 0.005 | 0 | 14 | Bmx |
| Jag21 | 0 | 1.817397836 | 0.702 | 0.077 | 0 | 14 | Jag2 |
| Gja5 | 0 | 1.69405721 | 0.545 | 0.008 | 0 | 14 | Gja5 |
| Fbn12 | 0 | 1.62438049 | 0.684 | 0.059 | 0 | 14 | Fbn1 |
| Jag1 | 0 | 1.569444306 | 0.524 | 0.034 | 0 | 14 | Jag1 |
| Tmem100 | 0 | 1.491297919 | 0.407 | 0.013 | 0 | 14 | Tmem100 |
| Hey11 | 0 | 1.449035768 | 0.549 | 0.022 | 0 | 14 | Hey1 |
| Eps8l2 | 0 | 1.41112458 | 0.596 | 0.038 | 0 | 14 | Eps8l2 |
| Efna5 | 0 | 1.287523888 | 0.335 | 0.008 | 0 | 14 | Efna5 |
| Car71 | 0 | 1.285785607 | 0.476 | 0.029 | 0 | 14 | Car7 |
| Npnt | 0 | 1.216320642 | 0.385 | 0.014 | 0 | 14 | Npnt |
| Dkk2 | 0 | 1.212662952 | 0.251 | 0.002 | 0 | 14 | Dkk2 |
| Egfl8 | 0 | 1.137200344 | 0.36 | 0.009 | 0 | 14 | Egfl8 |
| Aif1l | 0 | 0.988864942 | 0.382 | 0.005 | 0 | 14 | Aif1l |
| Kazald1 | 0 | 0.978557422 | 0.385 | 0.018 | 0 | 14 | Kazald1 |
| Dusp26 | 0 | 0.864599923 | 0.305 | 0.007 | 0 | 14 | Dusp26 |
| Prom1 | 0 | 0.84890941 | 0.273 | 0.007 | 0 | 14 | Prom1 |
| Ace2 | 2.28E-290 | 1.993101206 | 0.796 | 0.113 | 4.62E-286 | 14 | Ace |
| Mast43 | 8.78E-277 | 2.368139832 | 0.935 | 0.18 | 1.78E-272 | 14 | Mast4 |
| Tgfb22 | 1.99E-252 | 1.909837627 | 0.655 | 0.082 | 4.04E-248 | 14 | Tgfb2 |
| Atp13a32 | 6.05E-247 | 1.819283049 | 0.687 | 0.098 | 1.22E-242 | 14 | Atp13a3 |
| Lmo71 | 2.88E-245 | 1.460003329 | 0.593 | 0.068 | 5.83E-241 | 14 | Lmo7 |
| Prdm16 | 2.94E-242 | 1.339759843 | 0.487 | 0.047 | 5.95E-238 | 14 | Prdm16 |
| Lmo1 | 5.52E-242 | 1.236489442 | 0.476 | 0.045 | 1.12E-237 | 14 | Lmo1 |
| Htra11 | 3.02E-236 | 1.262490812 | 0.469 | 0.042 | 6.11E-232 | 14 | Htra1 |
| Apba11 | 7.60E-235 | 1.53119592 | 0.491 | 0.05 | 1.54E-230 | 14 | Apba1 |
| Syt7 | 4.09E-226 | 0.733826868 | 0.276 | 0.015 | 8.27E-222 | 14 | Syt7 |
| Tppp3 | 2.79E-223 | 1.020393452 | 0.309 | 0.02 | 5.64E-219 | 14 | Tppp3 |
| Eogt1 | 3.95E-222 | 1.422254138 | 0.6 | 0.077 | 8.01E-218 | 14 | Eogt |
| Efhd11 | 1.60E-220 | 0.8066242 | 0.309 | 0.02 | 3.24E-216 | 14 | Efhd1 |
| Lima13 | 1.80E-212 | 1.789842523 | 0.796 | 0.153 | 3.64E-208 | 14 | Lima1 |
| Col18a11 | 3.87E-208 | 0.941207064 | 0.302 | 0.02 | 7.84E-204 | 14 | Col18a1 |
| Fbln25 | 9.94E-201 | 2.289801083 | 0.956 | 0.267 | 2.01E-196 | 14 | Fbln2 |
| Sncaip1 | 6.15E-198 | 1.439725241 | 0.567 | 0.077 | 1.24E-193 | 14 | Sncaip |
| Tln2 | 2.59E-197 | 1.058788538 | 0.407 | 0.039 | 5.24E-193 | 14 | Tln2 |
| Msx1 | 8.40E-193 | 0.843474924 | 0.353 | 0.03 | 1.70E-188 | 14 | Msx1 |
| Ltbp44 | 1.09E-190 | 2.273693169 | 0.964 | 0.304 | 2.20E-186 | 14 | Ltbp4 |
| Heg14 | 1.21E-190 | 2.185798069 | 0.942 | 0.269 | 2.45E-186 | 14 | Heg1 |
| Tm4sf14 | 1.84E-189 | 2.556923297 | 0.989 | 0.373 | 3.72E-185 | 14 | Tm4sf1 |

| | | | | | | | |
|---|---|---|---|---|---|---|---|
| Glul1 | 2.12E-188 | 1.908865199 | 0.844 | 0.191 | 4.29E-184 | 14 | Glul |
| Crispld14 | 6.58E-187 | 1.459811164 | 0.745 | 0.138 | 1.33E-182 | 14 | Crispld1 |
| Vim7 | 7.38E-182 | 2.786270749 | 0.931 | 0.302 | 1.49E-177 | 14 | Vim |
| Hs3st1 | 6.71E-180 | 1.090882923 | 0.455 | 0.053 | 1.36E-175 | 14 | Hs3st1 |
| Eln3 | 3.44E-179 | 1.885081532 | 0.698 | 0.132 | 6.97E-175 | 14 | Eln |
| Arl153 | 4.85E-178 | 2.708814272 | 0.967 | 0.377 | 9.82E-174 | 14 | Arl15 |
| Epb41l4a1 | 2.87E-174 | 1.352932709 | 0.542 | 0.078 | 5.80E-170 | 14 | Epb41l4a |
| Fgfr3 | 5.52E-170 | 0.692903948 | 0.262 | 0.019 | 1.12E-165 | 14 | Fgfr3 |
| Atp2a33 | 4.61E-168 | 1.258062365 | 0.455 | 0.059 | 9.33E-164 | 14 | Atp2a3 |
| Fzd61 | 5.74E-166 | 1.004101423 | 0.451 | 0.056 | 1.16E-161 | 14 | Fzd6 |
| Trpv4 | 1.66E-165 | 0.832583378 | 0.316 | 0.028 | 3.37E-161 | 14 | Trpv4 |
| Msrb32 | 1.20E-163 | 1.445247401 | 0.669 | 0.128 | 2.44E-159 | 14 | Msrb3 |
| Cdk191 | 1.50E-161 | 1.674569934 | 0.68 | 0.143 | 3.03E-157 | 14 | Cdk19 |
| Afap1l2 | 3.85E-161 | 0.80499901 | 0.291 | 0.024 | 7.79E-157 | 14 | Afap1l2 |
| Pir | 2.90E-159 | 0.918490153 | 0.375 | 0.04 | 5.87E-155 | 14 | Pir |
| Tspan2 | 8.35E-159 | 0.915685932 | 0.345 | 0.035 | 1.69E-154 | 14 | Tspan2 |
| St6galnac2 | 1.38E-158 | 0.931187404 | 0.425 | 0.052 | 2.79E-154 | 14 | St6galnac2 |
| Chp2 | 1.88E-157 | 0.833370298 | 0.298 | 0.026 | 3.81E-153 | 14 | Chp2 |
| Ntn41 | 4.29E-157 | 0.846649271 | 0.331 | 0.032 | 8.69E-153 | 14 | Ntn4 |
| Cavin13 | 1.52E-156 | 1.56314506 | 0.825 | 0.209 | 3.08E-152 | 14 | Cavin1 |
| Sulf11 | 7.44E-156 | 1.586297246 | 0.407 | 0.05 | 1.51E-151 | 14 | Sulf1 |
| Sox132 | 1.30E-154 | 1.252276298 | 0.542 | 0.087 | 2.62E-150 | 14 | Sox13 |
| Ptprb4 | 5.91E-154 | 1.905688093 | 0.993 | 0.364 | 1.20E-149 | 14 | Ptprb |
| Dll41 | 5.16E-153 | 1.233812711 | 0.498 | 0.074 | 1.04E-148 | 14 | Dll4 |
| Mylk1 | 7.49E-153 | 0.79847613 | 0.396 | 0.046 | 1.52E-148 | 14 | Mylk |
| Azin11 | 1.75E-152 | 1.438480579 | 0.695 | 0.149 | 3.55E-148 | 14 | Azin1 |
| Epas14 | 5.05E-152 | 1.856880598 | 0.971 | 0.355 | 1.02E-147 | 14 | Epas1 |
| Tanc21 | 8.97E-152 | 1.593687632 | 0.716 | 0.155 | 1.82E-147 | 14 | Tanc2 |
| Efnb23 | 9.41E-152 | 1.500064031 | 0.687 | 0.14 | 1.90E-147 | 14 | Efnb2 |
| Entpd14 | 9.50E-151 | 1.870278967 | 0.876 | 0.271 | 1.92E-146 | 14 | Entpd1 |
| Palld1 | 4.04E-147 | 1.002435526 | 0.407 | 0.05 | 8.17E-143 | 14 | Palld |
| S1pr13 | 1.13E-146 | 1.770620804 | 0.873 | 0.261 | 2.28E-142 | 14 | S1pr1 |
| Chrm32 | 3.22E-146 | 1.505676212 | 0.56 | 0.096 | 6.51E-142 | 14 | Chrm3 |
| Runx1t11 | 4.28E-146 | 0.988989875 | 0.36 | 0.041 | 8.66E-142 | 14 | Runx1t1 |
| Ptprm3 | 3.26E-145 | 1.930476447 | 0.953 | 0.325 | 6.60E-141 | 14 | Ptprm |
| Syt151 | 5.32E-145 | 1.030157883 | 0.44 | 0.06 | 1.08E-140 | 14 | Syt15 |
| Ecscr3 | 8.01E-142 | 1.527132839 | 0.825 | 0.223 | 1.62E-137 | 14 | Ecscr |
| Fstl12 | 1.89E-140 | 1.19217197 | 0.564 | 0.099 | 3.82E-136 | 14 | Fstl1 |
| Sncg3 | 2.80E-140 | 1.44285096 | 0.691 | 0.151 | 5.67E-136 | 14 | Sncg |
| Pdgfd3 | 1.83E-139 | 1.968537127 | 0.858 | 0.249 | 3.70E-135 | 14 | Pdgfd |
| Arhgef281 | 1.36E-136 | 1.067072896 | 0.491 | 0.077 | 2.76E-132 | 14 | Arhgef28 |
| Icam23 | 1.99E-135 | 1.613068985 | 0.804 | 0.225 | 4.03E-131 | 14 | Icam2 |
| Ehd23 | 2.73E-134 | 1.405782146 | 0.749 | 0.189 | 5.52E-130 | 14 | Ehd2 |

| | | | | | | | |
|---|---|---|---|---|---|---|---|
| Crip16 | 1.19E-132 | 1.947804795 | 0.975 | 0.533 | 2.40E-128 | 14 | Crip1 |
| Cdc42bpa4 | 1.33E-131 | 1.502527659 | 0.72 | 0.181 | 2.69E-127 | 14 | Cdc42bpa |
| Utrn3 | 3.24E-131 | 1.833942905 | 0.945 | 0.402 | 6.56E-127 | 14 | Utrn |
| Slc6a63 | 8.95E-131 | 1.834306261 | 0.913 | 0.323 | 1.81E-126 | 14 | Slc6a6 |
| Lamb23 | 2.75E-130 | 1.288648663 | 0.593 | 0.12 | 5.56E-126 | 14 | Lamb2 |
| Alox121 | 6.26E-130 | 1.233446143 | 0.531 | 0.094 | 1.27E-125 | 14 | Alox12 |
| She3 | 7.99E-130 | 1.268921168 | 0.658 | 0.146 | 1.62E-125 | 14 | She |
| Specc11 | 7.90E-129 | 1.224025237 | 0.469 | 0.078 | 1.60E-124 | 14 | Specc1 |
| Nuak12 | 2.06E-125 | 1.200345629 | 0.571 | 0.114 | 4.17E-121 | 14 | Nuak1 |
| Klf24 | 4.70E-125 | 1.656692626 | 0.971 | 0.363 | 9.52E-121 | 14 | Klf2 |
| Pls33 | 3.69E-124 | 1.402699853 | 0.771 | 0.21 | 7.48E-120 | 14 | Pls3 |
| Plec3 | 1.78E-122 | 1.452476244 | 0.665 | 0.171 | 3.60E-118 | 14 | Plec |
| Adarb12 | 3.54E-122 | 1.056688116 | 0.556 | 0.106 | 7.17E-118 | 14 | Adarb1 |
| Prex23 | 1.96E-121 | 1.675572927 | 0.84 | 0.269 | 3.96E-117 | 14 | Prex2 |
| Ephx13 | 4.34E-120 | 1.156388814 | 0.575 | 0.118 | 8.79E-116 | 14 | Ephx1 |
| Itgb4 | 1.37E-119 | 0.686104108 | 0.291 | 0.032 | 2.77E-115 | 14 | Itgb4 |
| Limch13 | 3.51E-119 | 1.243743101 | 0.618 | 0.136 | 7.10E-115 | 14 | Limch1 |
| Ecm21 | 3.80E-119 | 1.053138142 | 0.349 | 0.047 | 7.69E-115 | 14 | Ecm2 |
| Cd2ap2 | 4.41E-119 | 1.486905658 | 0.887 | 0.322 | 8.93E-115 | 14 | Cd2ap |
| Cd364 | 1.60E-118 | 1.751017157 | 0.822 | 0.264 | 3.24E-114 | 14 | Cd36 |
| Rnf1521 | 4.08E-118 | 1.021473434 | 0.465 | 0.078 | 8.26E-114 | 14 | Rnf152 |
| Nos3 | 1.23E-117 | 1.037402176 | 0.44 | 0.072 | 2.48E-113 | 14 | Nos3 |
| Samd123 | 1.62E-117 | 1.504523254 | 0.793 | 0.225 | 3.28E-113 | 14 | Samd12 |
| Klf125 | 6.90E-117 | 1.603128739 | 0.716 | 0.204 | 1.40E-112 | 14 | Klf12 |
| Tm6sf14 | 7.34E-117 | 1.23702156 | 0.6 | 0.138 | 1.49E-112 | 14 | Tm6sf1 |
| Cav13 | 6.83E-116 | 1.540152754 | 0.884 | 0.3 | 1.38E-111 | 14 | Cav1 |
| Rasgrf23 | 5.77E-115 | 1.237987828 | 0.698 | 0.173 | 1.17E-110 | 14 | Rasgrf2 |
| Rras4 | 8.85E-115 | 1.454824513 | 0.851 | 0.294 | 1.79E-110 | 14 | Rras |
| Ssfa21 | 1.46E-113 | 1.006543723 | 0.491 | 0.094 | 2.95E-109 | 14 | Ssfa2 |
| Dock93 | 3.74E-113 | 1.342068978 | 0.727 | 0.202 | 7.56E-109 | 14 | Dock9 |
| Flnb2 | 1.03E-112 | 1.261514208 | 0.64 | 0.158 | 2.09E-108 | 14 | Flnb |
| Jcad2 | 1.34E-112 | 1.161488222 | 0.538 | 0.112 | 2.72E-108 | 14 | Jcad |
| Kalrn2 | 2.59E-112 | 1.126338395 | 0.52 | 0.103 | 5.24E-108 | 14 | Kalrn |
| Mmrn23 | 6.06E-112 | 1.368833304 | 0.778 | 0.228 | 1.23E-107 | 14 | Mmrn2 |
| Slc9a3r24 | 1.06E-111 | 1.549973241 | 0.945 | 0.366 | 2.15E-107 | 14 | Slc9a3r2 |
| Adgrf54 | 1.68E-111 | 1.481385336 | 0.945 | 0.333 | 3.40E-107 | 14 | Adgrf5 |
| Id14 | 2.39E-110 | 1.460316992 | 0.935 | 0.321 | 4.85E-106 | 14 | Id1 |
| Mgst14 | 5.03E-110 | 1.1569665 | 0.662 | 0.163 | 1.02E-105 | 14 | Mgst1 |
| Map4k34 | 2.11E-109 | 1.370072299 | 0.753 | 0.226 | 4.27E-105 | 14 | Map4k3 |
| Myo1d2 | 2.12E-108 | 1.154930689 | 0.513 | 0.104 | 4.28E-104 | 14 | Myo1d |
| Gfod14 | 3.00E-108 | 1.032822855 | 0.4 | 0.067 | 6.07E-104 | 14 | Gfod1 |
| Emp23 | 6.66E-108 | 1.395203864 | 0.76 | 0.232 | 1.35E-103 | 14 | Emp2 |
| Ldb24 | 2.53E-107 | 1.502031086 | 0.931 | 0.326 | 5.13E-103 | 14 | Ldb2 |

| | | | | | | | |
|---|---|---|---|---|---|---|---|
| S100a65 | 1.52E-106 | 1.062419657 | 0.898 | 0.267 | 3.07E-102 | 14 | S100a6 |
| Dach12 | 9.15E-106 | 1.199163145 | 0.502 | 0.101 | 1.85E-101 | 14 | Dach1 |
| Ccm2l2 | 1.25E-105 | 0.928910253 | 0.476 | 0.09 | 2.54E-101 | 14 | Ccm2l |
| Abcb1a2 | 7.63E-104 | 1.093473393 | 0.513 | 0.107 | 1.54E-99 | 14 | Abcb1a |
| Tek3 | 7.76E-104 | 1.276891811 | 0.825 | 0.256 | 1.57E-99 | 14 | Tek |
| Cdh53 | 7.76E-104 | 1.377470859 | 0.924 | 0.327 | 1.57E-99 | 14 | Cdh5 |
| Fryl5 | 3.05E-101 | 1.482461538 | 0.764 | 0.265 | 6.18E-97 | 14 | Fryl |
| Podxl4 | 4.50E-101 | 1.280140535 | 0.96 | 0.341 | 9.11E-97 | 14 | Podxl |
| Timp33 | 7.97E-101 | 1.592988052 | 0.971 | 0.381 | 1.61E-96 | 14 | Timp3 |
| C1qtnf9 | 4.77E-100 | 0.830871679 | 0.316 | 0.044 | 9.66E-96 | 14 | C1qtnf9 |
| Myo103 | 8.50E-100 | 1.473898642 | 0.705 | 0.216 | 1.72E-95 | 14 | Myo10 |
| Cyth33 | 1.24E-99 | 1.133789391 | 0.531 | 0.122 | 2.51E-95 | 14 | Cyth3 |
| Klf43 | 3.36E-99 | 1.37014391 | 0.807 | 0.266 | 6.81E-95 | 14 | Klf4 |
| Tmem441 | 4.89E-99 | 0.859785176 | 0.364 | 0.058 | 9.90E-95 | 14 | Tmem44 |
| Foxp16 | 9.33E-99 | 1.455670316 | 0.895 | 0.386 | 1.89E-94 | 14 | Foxp1 |
| Ramp24 | 1.62E-98 | 1.277250552 | 0.913 | 0.328 | 3.28E-94 | 14 | Ramp2 |
| Unc13b | 2.02E-98 | 0.745624583 | 0.4 | 0.067 | 4.09E-94 | 14 | Unc13b |
| Aplp24 | 1.69E-97 | 1.436655321 | 0.935 | 0.413 | 3.42E-93 | 14 | Aplp2 |
| Kitl3 | 2.27E-97 | 1.4334131 | 0.898 | 0.332 | 4.59E-93 | 14 | Kitl |
| Smad6 | 2.45E-97 | 0.836238652 | 0.36 | 0.058 | 4.96E-93 | 14 | Smad6 |
| Cst33 | 5.06E-97 | 1.352488684 | 0.993 | 0.6 | 1.03E-92 | 14 | Cst3 |
| Ly6c14 | 5.58E-97 | 1.393569977 | 0.993 | 0.396 | 1.13E-92 | 14 | Ly6c1 |
| Cavin24 | 8.64E-97 | 1.237924618 | 0.949 | 0.337 | 1.75E-92 | 14 | Cavin2 |
| Plcb13 | 9.28E-97 | 1.495862829 | 0.938 | 0.336 | 1.88E-92 | 14 | Plcb1 |
| Ehd1 | 5.87E-96 | 0.881132599 | 0.389 | 0.069 | 1.19E-91 | 14 | Ehd1 |
| Filip13 | 6.78E-96 | 1.20519406 | 0.68 | 0.191 | 1.37E-91 | 14 | Filip1 |
| Ly6a4 | 1.08E-95 | 1.432111159 | 0.985 | 0.456 | 2.19E-91 | 14 | Ly6a |
| Bsg5 | 1.12E-95 | 1.496727625 | 0.92 | 0.446 | 2.26E-91 | 14 | Bsg |
| Sept116 | 6.19E-95 | 1.181505014 | 0.673 | 0.2 | 1.25E-90 | 14 | 11-Sep |
| Cp1 | 1.75E-94 | 0.975413888 | 0.313 | 0.045 | 3.55E-90 | 14 | Cp |
| Uaca3 | 3.02E-94 | 1.283051277 | 0.716 | 0.223 | 6.11E-90 | 14 | Uaca |
| Amd1 | 3.40E-94 | 0.863017654 | 0.375 | 0.065 | 6.89E-90 | 14 | Amd1 |
| St6galnac35 | 1.60E-93 | 1.450408587 | 0.716 | 0.213 | 3.24E-89 | 14 | St6galnac3 |
| Ebf14 | 6.12E-93 | 1.375616088 | 0.96 | 0.371 | 1.24E-88 | 14 | Ebf1 |
| Epha43 | 7.12E-93 | 1.054866733 | 0.509 | 0.114 | 1.44E-88 | 14 | Epha4 |
| Lama33 | 7.35E-93 | 1.243252209 | 0.727 | 0.218 | 1.49E-88 | 14 | Lama3 |
| Gcnt23 | 7.65E-93 | 1.077298202 | 0.567 | 0.14 | 1.55E-88 | 14 | Gcnt2 |
| Sox184 | 1.89E-92 | 1.149930487 | 0.844 | 0.272 | 3.82E-88 | 14 | Sox18 |
| 9530026P05Rik3 | 8.59E-92 | 1.40051548 | 0.756 | 0.238 | 1.74E-87 | 14 | 9530026P05Rik |
| Gata22 | 1.33E-91 | 1.005920795 | 0.48 | 0.102 | 2.69E-87 | 14 | Gata2 |
| Slc27a3 | 1.60E-90 | 0.665337304 | 0.251 | 0.031 | 3.24E-86 | 14 | Slc27a3 |
| Gulp14 | 2.72E-90 | 1.229586272 | 0.607 | 0.163 | 5.51E-86 | 14 | Gulp1 |
| Igfbp41 | 1.12E-89 | 0.986374267 | 0.458 | 0.095 | 2.27E-85 | 14 | Igfbp4 |

| | | | | | | | |
|---|---|---|---|---|---|---|---|
| Nhsl22 | 1.72E-89 | 1.132554498 | 0.538 | 0.132 | 3.48E-85 | 14 | Nhsl2 |
| Cd933 | 2.37E-89 | 1.185658963 | 0.782 | 0.253 | 4.79E-85 | 14 | Cd93 |
| Akap123 | 4.36E-89 | 1.24744957 | 0.593 | 0.158 | 8.83E-85 | 14 | Akap12 |
| Bcam4 | 9.67E-89 | 1.192281348 | 0.767 | 0.25 | 1.96E-84 | 14 | Bcam |
| Fam13c2 | 3.52E-88 | 1.030435447 | 0.487 | 0.109 | 7.13E-84 | 14 | Fam13c |
| Lifr4 | 4.39E-88 | 1.138299163 | 0.655 | 0.189 | 8.90E-84 | 14 | Lifr |
| Clec2d4 | 8.47E-88 | 1.397752792 | 0.935 | 0.525 | 1.71E-83 | 14 | Clec2d |
| Esam3 | 1.31E-87 | 1.109064261 | 0.869 | 0.306 | 2.64E-83 | 14 | Esam |
| Adamts16 | 1.94E-86 | 1.079486591 | 0.447 | 0.095 | 3.93E-82 | 14 | Adamts1 |
| Hspa12b2 | 2.00E-86 | 0.960461263 | 0.462 | 0.1 | 4.05E-82 | 14 | Hspa12b |
| Cxcl125 | 2.08E-86 | 1.563981991 | 0.84 | 0.318 | 4.22E-82 | 14 | Cxcl12 |
| Palmd3 | 2.22E-86 | 1.133241743 | 0.72 | 0.23 | 4.49E-82 | 14 | Palmd |
| Crybg32 | 5.55E-86 | 1.117566853 | 0.535 | 0.137 | 1.12E-81 | 14 | Crybg3 |
| Pgrmc23 | 1.66E-85 | 1.051557441 | 0.538 | 0.14 | 3.35E-81 | 14 | Pgrmc2 |
| Actn43 | 3.18E-85 | 1.319777812 | 0.851 | 0.373 | 6.44E-81 | 14 | Actn4 |
| Cgnl1 | 5.92E-85 | 0.72041127 | 0.316 | 0.051 | 1.20E-80 | 14 | Cgnl1 |
| Mpzl14 | 9.55E-85 | 1.009729794 | 0.549 | 0.14 | 1.93E-80 | 14 | Mpzl1 |
| Sorbs23 | 1.42E-84 | 1.243718649 | 0.796 | 0.274 | 2.88E-80 | 14 | Sorbs2 |
| Ptprg3 | 1.65E-84 | 1.213980421 | 0.913 | 0.341 | 3.34E-80 | 14 | Ptprg |
| Sfxn3 | 2.16E-84 | 0.729556486 | 0.302 | 0.047 | 4.37E-80 | 14 | Sfxn3 |
| Adam153 | 8.58E-84 | 1.151319443 | 0.633 | 0.191 | 1.74E-79 | 14 | Adam15 |
| Col5a21 | 1.37E-83 | 0.831771322 | 0.284 | 0.041 | 2.78E-79 | 14 | Col5a2 |
| Map43 | 1.55E-83 | 1.230703603 | 0.771 | 0.301 | 3.14E-79 | 14 | Map4 |
| Tmod35 | 4.28E-83 | 1.293331152 | 0.822 | 0.359 | 8.66E-79 | 14 | Tmod3 |
| Pecam14 | 4.74E-83 | 1.223172812 | 0.982 | 0.407 | 9.60E-79 | 14 | Pecam1 |
| Mkl23 | 5.86E-83 | 1.088786203 | 0.738 | 0.252 | 1.19E-78 | 14 | Mkl2 |
| Jam23 | 8.88E-83 | 1.164180887 | 0.804 | 0.282 | 1.80E-78 | 14 | Jam2 |
| Adcy42 | 1.14E-81 | 0.901398269 | 0.465 | 0.105 | 2.30E-77 | 14 | Adcy4 |
| Ssu21 | 3.60E-81 | 0.763901485 | 0.284 | 0.043 | 7.28E-77 | 14 | Ssu2 |
| Rflnb4 | 4.76E-81 | 1.653137827 | 0.804 | 0.311 | 9.63E-77 | 14 | Rflnb |
| Ece14 | 6.82E-81 | 1.115627988 | 0.822 | 0.303 | 1.38E-76 | 14 | Ece1 |
| Tmem255b1 | 3.66E-80 | 0.731943519 | 0.375 | 0.071 | 7.42E-76 | 14 | Tmem255b |
| Ptpn142 | 4.37E-80 | 0.95497899 | 0.455 | 0.104 | 8.86E-76 | 14 | Ptpn14 |
| Ablim16 | 4.66E-80 | 1.252901392 | 0.88 | 0.4 | 9.43E-76 | 14 | Ablim1 |
| Eva1a | 5.43E-80 | 0.571340028 | 0.258 | 0.036 | 1.10E-75 | 14 | Eva1a |
| S100a164 | 7.36E-80 | 1.097377871 | 0.731 | 0.255 | 1.49E-75 | 14 | S100a16 |
| Cdk142 | 5.22E-79 | 1.138020474 | 0.527 | 0.139 | 1.06E-74 | 14 | Cdk14 |
| Gm16070 | 6.19E-79 | 0.884393808 | 0.309 | 0.052 | 1.25E-74 | 14 | Gm16070 |
| Smpdl3a3 | 6.65E-79 | 1.028547369 | 0.665 | 0.216 | 1.35E-74 | 14 | Smpdl3a |
| Mapk34 | 6.85E-79 | 1.099116046 | 0.72 | 0.263 | 1.39E-74 | 14 | Mapk3 |
| Phactr11 | 7.32E-79 | 0.899716945 | 0.349 | 0.065 | 1.48E-74 | 14 | Phactr1 |
| Ceacam13 | 2.81E-78 | 0.951871886 | 0.564 | 0.153 | 5.68E-74 | 14 | Ceacam1 |
| Efna13 | 6.53E-78 | 0.941892853 | 0.538 | 0.141 | 1.32E-73 | 14 | Efna1 |

| | | | | | | | |
|---|---|---|---|---|---|---|---|
| Itpr31 | 8.18E-78 | 0.843047624 | 0.367 | 0.074 | 1.66E-73 | 14 | Itpr3 |
| Emp12 | 1.13E-77 | 0.915456887 | 0.491 | 0.119 | 2.29E-73 | 14 | Emp1 |
| Enpp31 | 1.58E-77 | 0.722196434 | 0.295 | 0.048 | 3.19E-73 | 14 | Enpp3 |
| Tulp41 | 1.68E-77 | 1.120538216 | 0.596 | 0.185 | 3.39E-73 | 14 | Tulp4 |
| Lims23 | 2.98E-77 | 1.012734198 | 0.651 | 0.199 | 6.03E-73 | 14 | Lims2 |
| Notch13 | 2.81E-76 | 1.182554261 | 0.647 | 0.214 | 5.70E-72 | 14 | Notch1 |
| Tjp13 | 8.96E-76 | 1.162738834 | 0.72 | 0.261 | 1.81E-71 | 14 | Tjp1 |
| Fchsd25 | 1.67E-75 | 1.212114148 | 0.538 | 0.157 | 3.38E-71 | 14 | Fchsd2 |
| Slc50a14 | 2.01E-74 | 1.130614488 | 0.753 | 0.296 | 4.07E-70 | 14 | Slc50a1 |
| Itpr13 | 1.31E-73 | 1.258473575 | 0.64 | 0.219 | 2.66E-69 | 14 | Itpr1 |
| Cd92 | 1.65E-73 | 1.100958566 | 0.851 | 0.383 | 3.35E-69 | 14 | Cd9 |
| Magi13 | 1.71E-73 | 1.21482478 | 0.687 | 0.229 | 3.46E-69 | 14 | Magi1 |
| Rassf92 | 3.97E-73 | 0.935321452 | 0.491 | 0.124 | 8.03E-69 | 14 | Rassf9 |
| Bmpr23 | 2.20E-72 | 1.01291029 | 0.865 | 0.346 | 4.46E-68 | 14 | Bmpr2 |
| Fbxo71 | 2.26E-72 | 0.894987739 | 0.425 | 0.102 | 4.57E-68 | 14 | Fbxo7 |
| Hyal21 | 2.96E-72 | 0.819920022 | 0.389 | 0.085 | 5.99E-68 | 14 | Hyal2 |
| Crip24 | 7.01E-72 | 1.138446049 | 0.898 | 0.413 | 1.42E-67 | 14 | Crip2 |
| Acvrl14 | 8.20E-72 | 1.03147758 | 0.742 | 0.272 | 1.66E-67 | 14 | Acvrl1 |
| Lpar41 | 9.55E-72 | 0.739660505 | 0.309 | 0.055 | 1.93E-67 | 14 | Lpar4 |
| Ralgapa13 | 1.65E-71 | 1.157508156 | 0.749 | 0.301 | 3.33E-67 | 14 | Ralgapa1 |
| Arid5b5 | 2.56E-71 | 1.179757494 | 0.709 | 0.275 | 5.18E-67 | 14 | Arid5b |
| Atp1b35 | 2.63E-71 | 1.048428554 | 0.669 | 0.246 | 5.32E-67 | 14 | Atp1b3 |
| Cd59a | 6.89E-71 | 0.889895456 | 0.389 | 0.085 | 1.39E-66 | 14 | Cd59a |
| Atp2b43 | 9.48E-71 | 1.112674506 | 0.6 | 0.196 | 1.92E-66 | 14 | Atp2b4 |
| Mcam2 | 2.78E-70 | 0.892240213 | 0.444 | 0.108 | 5.62E-66 | 14 | Mcam |
| Kctd102 | 4.01E-70 | 0.898952223 | 0.451 | 0.115 | 8.12E-66 | 14 | Kctd10 |
| Shtn11 | 5.13E-70 | 0.678982254 | 0.338 | 0.066 | 1.04E-65 | 14 | Shtn1 |
| Ifitm35 | 5.40E-70 | 1.133375458 | 0.993 | 0.536 | 1.09E-65 | 14 | Ifitm3 |
| Fam20a | 6.84E-70 | 0.709856984 | 0.302 | 0.054 | 1.38E-65 | 14 | Fam20a |
| Notch42 | 1.07E-68 | 0.922664471 | 0.447 | 0.113 | 2.17E-64 | 14 | Notch4 |
| Id34 | 1.22E-68 | 1.027455442 | 0.96 | 0.405 | 2.47E-64 | 14 | Id3 |
| Arhgap51 | 1.41E-68 | 1.008828459 | 0.738 | 0.286 | 2.86E-64 | 14 | Arhgap5 |
| Tmem474 | 1.51E-68 | 1.031585739 | 0.535 | 0.154 | 3.07E-64 | 14 | Tmem47 |
| Sh3bp4 | 4.77E-68 | 0.690540694 | 0.324 | 0.063 | 9.65E-64 | 14 | Sh3bp4 |
| Luzp13 | 9.94E-68 | 1.074325308 | 0.811 | 0.354 | 2.01E-63 | 14 | Luzp1 |
| Ccdc88c4 | 3.08E-67 | 0.827230117 | 0.407 | 0.098 | 6.24E-63 | 14 | Ccdc88c |
| Fam171a12 | 6.20E-67 | 0.846502887 | 0.425 | 0.104 | 1.25E-62 | 14 | Fam171a1 |
| Nfia4 | 3.56E-66 | 1.040709275 | 0.749 | 0.285 | 7.21E-62 | 14 | Nfia |
| Tpst11 | 4.45E-66 | 0.672591829 | 0.284 | 0.051 | 9.00E-62 | 14 | Tpst1 |
| 4930578C19Rik2 | 1.30E-65 | 0.955283354 | 0.425 | 0.105 | 2.62E-61 | 14 | 4930578C19Rik |
| Eng3 | 5.22E-65 | 0.931699189 | 0.898 | 0.35 | 1.06E-60 | 14 | Eng |
| Dst4 | 8.24E-64 | 1.099316799 | 0.571 | 0.189 | 1.67E-59 | 14 | Dst |
| Emp36 | 1.22E-63 | 1.120470779 | 0.669 | 0.257 | 2.46E-59 | 14 | Emp3 |

| | | | | | | | |
|---|---|---|---|---|---|---|---|
| Ccnd27 | 1.39E-63 | 1.21024814 | 0.695 | 0.284 | 2.82E-59 | 14 | Ccnd2 |
| Nxpe43 | 1.75E-63 | 0.95126543 | 0.505 | 0.146 | 3.55E-59 | 14 | Nxpe4 |
| Fgd53 | 1.85E-63 | 0.933002282 | 0.662 | 0.226 | 3.76E-59 | 14 | Fgd5 |
| Egfl74 | 5.02E-63 | 0.946352745 | 0.975 | 0.389 | 1.02E-58 | 14 | Egfl7 |
| 4931406P16Rik3 | 5.72E-63 | 0.929704616 | 0.822 | 0.34 | 1.16E-58 | 14 | 4931406P16Rik |
| Cltb2 | 2.52E-62 | 0.877306084 | 0.618 | 0.217 | 5.11E-58 | 14 | Cltb |
| Pde4d2 | 2.55E-62 | 1.152403412 | 0.498 | 0.148 | 5.17E-58 | 14 | Pde4d |
| Cyyr13 | 3.36E-62 | 0.868685085 | 0.909 | 0.34 | 6.81E-58 | 14 | Cyyr1 |
| Plcl11 | 4.51E-62 | 0.798621303 | 0.276 | 0.051 | 9.13E-58 | 14 | Plcl1 |
| Fbxl73 | 1.34E-61 | 1.133576055 | 0.698 | 0.255 | 2.71E-57 | 14 | Fbxl7 |
| Rapgef23 | 3.11E-61 | 1.066808054 | 0.785 | 0.336 | 6.29E-57 | 14 | Rapgef2 |
| Rgs102 | 4.59E-61 | 0.533009591 | 0.549 | 0.159 | 9.30E-57 | 14 | Rgs10 |
| Btbd74 | 4.71E-61 | 1.062963243 | 0.673 | 0.279 | 9.53E-57 | 14 | Btbd7 |
| Pdzd21 | 6.20E-61 | 0.890451178 | 0.429 | 0.113 | 1.26E-56 | 14 | Pdzd2 |
| Igf1r3 | 1.02E-60 | 0.924228216 | 0.52 | 0.159 | 2.07E-56 | 14 | Igf1r |
| Myct13 | 1.61E-60 | 0.81873836 | 0.655 | 0.22 | 3.25E-56 | 14 | Myct1 |
| Tinagl14 | 1.85E-60 | 0.882786221 | 0.785 | 0.294 | 3.75E-56 | 14 | Tinagl1 |
| Ggta12 | 1.89E-60 | 0.832188429 | 0.476 | 0.138 | 3.82E-56 | 14 | Ggta1 |
| Lrrc3b2 | 2.01E-60 | 0.874853777 | 0.385 | 0.092 | 4.06E-56 | 14 | Lrrc3b |
| Anxa33 | 2.20E-60 | 0.910433795 | 0.8 | 0.336 | 4.45E-56 | 14 | Anxa3 |
| Rbpms3 | 2.36E-60 | 0.854036232 | 0.818 | 0.346 | 4.79E-56 | 14 | Rbpms |
| Nrarp | 2.40E-60 | 0.617556967 | 0.291 | 0.057 | 4.86E-56 | 14 | Nrarp |
| Gadd45g1 | 4.17E-60 | 0.995709408 | 0.455 | 0.129 | 8.45E-56 | 14 | Gadd45g |
| Wwc24 | 4.42E-60 | 0.907510849 | 0.585 | 0.195 | 8.95E-56 | 14 | Wwc2 |
| Tubb2a3 | 4.45E-60 | 0.867693982 | 0.524 | 0.163 | 9.00E-56 | 14 | Tubb2a |
| Ddah1 | 4.89E-60 | 0.632096182 | 0.265 | 0.048 | 9.91E-56 | 14 | Ddah1 |
| Tspo | 7.51E-60 | 1.019880408 | 0.953 | 0.645 | 1.52E-55 | 14 | Tspo |
| Sqor1 | 2.04E-59 | 0.634905389 | 0.324 | 0.07 | 4.13E-55 | 14 | Sqor |
| Pdgfa | 2.78E-59 | 0.548743682 | 0.295 | 0.058 | 5.63E-55 | 14 | Pdgfa |
| Orai12 | 3.20E-59 | 0.809156462 | 0.444 | 0.127 | 6.47E-55 | 14 | Orai1 |
| Ly6e4 | 4.80E-59 | 0.970283995 | 0.978 | 0.617 | 9.72E-55 | 14 | Ly6e |
| Cd2004 | 5.08E-59 | 0.750755386 | 0.909 | 0.346 | 1.03E-54 | 14 | Cd200 |
| Parvb2 | 8.28E-59 | 0.721828698 | 0.447 | 0.121 | 1.68E-54 | 14 | Parvb |
| Tanc13 | 8.41E-59 | 0.968008033 | 0.618 | 0.221 | 1.70E-54 | 14 | Tanc1 |
| Capg4 | 9.05E-59 | 0.690604863 | 0.393 | 0.099 | 1.83E-54 | 14 | Capg |
| Fat42 | 1.29E-58 | 0.760911495 | 0.407 | 0.104 | 2.60E-54 | 14 | Fat4 |
| Filip1l2 | 2.47E-58 | 0.952264894 | 0.487 | 0.151 | 4.99E-54 | 14 | Filip1l |
| Prdx43 | 2.64E-58 | 0.913319035 | 0.611 | 0.219 | 5.35E-54 | 14 | Prdx4 |
| Tmem1093 | 2.67E-58 | 0.813025679 | 0.484 | 0.146 | 5.40E-54 | 14 | Tmem109 |
| 8430408G22Rik | 3.17E-58 | 0.953236406 | 0.284 | 0.056 | 6.41E-54 | 14 | 8430408G22Rik |
| Pik3r3 | 3.89E-58 | 0.612066126 | 0.316 | 0.067 | 7.88E-54 | 14 | Pik3r3 |
| Cnn34 | 6.20E-58 | 0.795081333 | 0.578 | 0.191 | 1.25E-53 | 14 | Cnn3 |
| Cttnbp2 | 6.78E-58 | 0.572915886 | 0.262 | 0.048 | 1.37E-53 | 14 | Cttnbp2 |

| | | | | | | | |
|---|---|---|---|---|---|---|---|
| Camk2d5 | 3.40E-57 | 0.952634008 | 0.669 | 0.267 | 6.89E-53 | 14 | Camk2d |
| Cbfa2t32 | 3.70E-57 | 0.848946408 | 0.465 | 0.137 | 7.49E-53 | 14 | Cbfa2t3 |
| Ctsd6 | 3.86E-57 | 0.982024165 | 0.731 | 0.312 | 7.81E-53 | 14 | Ctsd |
| Marveld1 | 4.55E-57 | 0.541370148 | 0.255 | 0.047 | 9.22E-53 | 14 | Marveld1 |
| Thsd11 | 4.62E-57 | 0.693262368 | 0.364 | 0.087 | 9.34E-53 | 14 | Thsd1 |
| Enah3 | 5.22E-57 | 0.747810995 | 0.498 | 0.148 | 1.06E-52 | 14 | Enah |
| Mall | 5.44E-57 | 0.630521712 | 0.28 | 0.056 | 1.10E-52 | 14 | Mall |
| Stox23 | 1.15E-56 | 0.937368505 | 0.669 | 0.245 | 2.32E-52 | 14 | Stox2 |
| Cavin34 | 1.37E-56 | 0.872684967 | 0.76 | 0.304 | 2.78E-52 | 14 | Cavin3 |
| Malat18 | 1.71E-56 | 1.128459375 | 0.993 | 0.724 | 3.46E-52 | 14 | Malat1 |
| Atox13 | 3.35E-56 | 0.880751465 | 0.938 | 0.617 | 6.78E-52 | 14 | Atox1 |
| Pdgfb4 | 4.62E-56 | 0.89157489 | 0.742 | 0.307 | 9.35E-52 | 14 | Pdgfb |
| Trim473 | 5.20E-56 | 0.800848128 | 0.473 | 0.142 | 1.05E-51 | 14 | Trim47 |
| Itm2c3 | 8.22E-56 | 1.021269531 | 0.545 | 0.197 | 1.66E-51 | 14 | Itm2c |
| Asap22 | 9.48E-56 | 0.836509635 | 0.447 | 0.13 | 1.92E-51 | 14 | Asap2 |
| Galnt183 | 1.04E-55 | 0.753761713 | 0.465 | 0.133 | 2.10E-51 | 14 | Galnt18 |
| Fxyd57 | 2.91E-55 | 1.073430229 | 0.836 | 0.409 | 5.89E-51 | 14 | Fxyd5 |
| 9930111J21Rik24 | 3.05E-55 | 0.960384624 | 0.644 | 0.255 | 6.18E-51 | 14 | 9930111J21Rik2 |
| Zfyve93 | 3.77E-55 | 0.739129021 | 0.527 | 0.169 | 7.63E-51 | 14 | Zfyve9 |
| Tns14 | 3.85E-55 | 0.90022508 | 0.64 | 0.243 | 7.80E-51 | 14 | Tns1 |
| Lrrc321 | 4.12E-55 | 0.656428591 | 0.298 | 0.064 | 8.34E-51 | 14 | Lrrc32 |
| Ccser23 | 1.91E-54 | 0.859582507 | 0.615 | 0.229 | 3.87E-50 | 14 | Ccser2 |
| Sparcl15 | 2.10E-54 | 0.822692767 | 0.865 | 0.361 | 4.25E-50 | 14 | Sparcl1 |
| Dock63 | 2.64E-54 | 0.771014434 | 0.469 | 0.138 | 5.35E-50 | 14 | Dock6 |
| Parva1 | 4.12E-54 | 0.717006497 | 0.407 | 0.11 | 8.34E-50 | 14 | Parva |
| Nxn3 | 5.69E-54 | 0.897772827 | 0.68 | 0.266 | 1.15E-49 | 14 | Nxn |
| Cd816 | 8.60E-54 | 0.912253486 | 0.978 | 0.523 | 1.74E-49 | 14 | Cd81 |
| Afdn3 | 1.52E-53 | 0.786415637 | 0.753 | 0.312 | 3.08E-49 | 14 | Afdn |
| Lama5 | 1.76E-53 | 0.565820749 | 0.28 | 0.057 | 3.56E-49 | 14 | Lama5 |
| Ppp1r13b3 | 2.08E-53 | 0.842277896 | 0.585 | 0.214 | 4.22E-49 | 14 | Ppp1r13b |
| Gm26532 | 2.40E-53 | 0.70907768 | 0.28 | 0.059 | 4.86E-49 | 14 | Gm26532 |
| Arhgap313 | 6.03E-53 | 0.766420552 | 0.887 | 0.369 | 1.22E-48 | 14 | Arhgap31 |
| Rgs36 | 9.33E-53 | 0.94032884 | 0.527 | 0.185 | 1.89E-48 | 14 | Rgs3 |
| Slco3a12 | 1.08E-52 | 0.842309403 | 0.422 | 0.121 | 2.18E-48 | 14 | Slco3a1 |
| Pam6 | 1.32E-52 | 0.954016243 | 0.567 | 0.212 | 2.68E-48 | 14 | Pam |
| Itgb18 | 1.36E-52 | 0.944831217 | 0.876 | 0.477 | 2.74E-48 | 14 | Itgb1 |
| Smad77 | 1.85E-52 | 0.987542464 | 0.72 | 0.324 | 3.74E-48 | 14 | Smad7 |
| Alas1 | 2.35E-52 | 0.542281276 | 0.327 | 0.077 | 4.75E-48 | 14 | Alas1 |
| App4 | 2.38E-52 | 0.828276719 | 0.913 | 0.417 | 4.82E-48 | 14 | App |
| Mertk3 | 3.32E-52 | 0.725373791 | 0.444 | 0.13 | 6.73E-48 | 14 | Mertk |
| Foxn37 | 7.11E-52 | 1.031035999 | 0.818 | 0.409 | 1.44E-47 | 14 | Foxn3 |
| Cx3cl13 | 7.19E-52 | 0.70780055 | 0.385 | 0.103 | 1.46E-47 | 14 | Cx3cl1 |
| Hacd43 | 1.73E-51 | 0.766438836 | 0.404 | 0.114 | 3.50E-47 | 14 | Hacd4 |

| | | | | | | | |
|---|---|---|---|---|---|---|---|
| Pkn3 | 1.83E-51 | 0.59181197 | 0.265 | 0.054 | 3.70E-47 | 14 | Pkn3 |
| Ahnak7 | 3.69E-51 | 0.949110771 | 0.887 | 0.47 | 7.48E-47 | 14 | Ahnak |
| Prnp | 5.58E-51 | 0.442617796 | 0.313 | 0.072 | 1.13E-46 | 14 | Prnp |
| Sgms13 | 6.02E-51 | 0.732561199 | 0.433 | 0.127 | 1.22E-46 | 14 | Sgms1 |
| Tagln25 | 1.14E-50 | 0.923950664 | 0.818 | 0.427 | 2.31E-46 | 14 | Tagln2 |
| Il6st4 | 1.30E-50 | 0.808410773 | 0.611 | 0.23 | 2.64E-46 | 14 | Il6st |
| Ak31 | 1.89E-50 | 0.731785269 | 0.476 | 0.152 | 3.82E-46 | 14 | Ak3 |
| Cav23 | 2.18E-50 | 0.825446559 | 0.575 | 0.207 | 4.41E-46 | 14 | Cav2 |
| Syne23 | 2.87E-50 | 0.846614237 | 0.684 | 0.27 | 5.80E-46 | 14 | Syne2 |
| Enpp4 | 3.79E-50 | 0.638633127 | 0.327 | 0.08 | 7.67E-46 | 14 | Enpp4 |
| Rexo22 | 5.71E-50 | 0.776675422 | 0.68 | 0.295 | 1.16E-45 | 14 | Rexo2 |
| Ablim31 | 5.98E-50 | 0.62428446 | 0.313 | 0.073 | 1.21E-45 | 14 | Ablim3 |
| Tcf7l22 | 1.01E-49 | 0.727890487 | 0.425 | 0.128 | 2.04E-45 | 14 | Tcf7l2 |
| Sparc5 | 1.29E-49 | 0.849323139 | 0.902 | 0.4 | 2.60E-45 | 14 | Sparc |
| Fam198b4 | 1.51E-49 | 0.776233711 | 0.538 | 0.182 | 3.06E-45 | 14 | Fam198b |
| Rbfox24 | 1.66E-49 | 0.867628599 | 0.538 | 0.189 | 3.35E-45 | 14 | Rbfox2 |
| Laptm4a4 | 2.01E-49 | 0.813531594 | 0.898 | 0.482 | 4.06E-45 | 14 | Laptm4a |
| Tmed105 | 2.95E-49 | 0.871887821 | 0.811 | 0.434 | 5.97E-45 | 14 | Tmed10 |
| Fam129b1 | 3.63E-49 | 0.622580669 | 0.335 | 0.084 | 7.35E-45 | 14 | Fam129b |
| Ppfibp13 | 3.90E-49 | 0.784116333 | 0.524 | 0.182 | 7.89E-45 | 14 | Ppfibp1 |
| Afap1l12 | 7.20E-49 | 0.678194101 | 0.389 | 0.108 | 1.46E-44 | 14 | Afap1l1 |
| Rasip13 | 1.75E-48 | 0.738500693 | 0.56 | 0.193 | 3.54E-44 | 14 | Rasip1 |
| Pdlim71 | 1.76E-48 | 0.641287614 | 0.302 | 0.072 | 3.56E-44 | 14 | Pdlim7 |
| Clstn13 | 1.88E-48 | 0.841779199 | 0.607 | 0.235 | 3.80E-44 | 14 | Clstn1 |
| Mcfd2 | 2.39E-48 | 0.657262664 | 0.349 | 0.093 | 4.84E-44 | 14 | Mcfd2 |
| Nck14 | 5.30E-48 | 0.867900044 | 0.713 | 0.338 | 1.07E-43 | 14 | Nck1 |
| Klf33 | 8.97E-48 | 0.856347952 | 0.6 | 0.234 | 1.82E-43 | 14 | Klf3 |
| F11r5 | 1.20E-47 | 0.761263528 | 0.658 | 0.263 | 2.42E-43 | 14 | F11r |
| Cmip6 | 1.76E-47 | 0.945677175 | 0.636 | 0.284 | 3.57E-43 | 14 | Cmip |
| Ano63 | 2.95E-47 | 0.803136241 | 0.502 | 0.179 | 5.96E-43 | 14 | Ano6 |
| Fnbp1l1 | 6.54E-47 | 0.58090668 | 0.505 | 0.168 | 1.32E-42 | 14 | Fnbp1l |
| Pdlim53 | 7.53E-47 | 0.77032129 | 0.673 | 0.284 | 1.53E-42 | 14 | Pdlim5 |
| Plekhg13 | 7.95E-47 | 0.791985106 | 0.636 | 0.243 | 1.61E-42 | 14 | Plekhg1 |
| Ccny5 | 9.34E-47 | 0.934708409 | 0.622 | 0.266 | 1.89E-42 | 14 | Ccny |
| Osmr4 | 1.11E-46 | 0.773065867 | 0.436 | 0.138 | 2.25E-42 | 14 | Osmr |
| Ctnnb13 | 1.15E-46 | 0.836940217 | 0.745 | 0.362 | 2.32E-42 | 14 | Ctnnb1 |
| Ifitm25 | 1.29E-46 | 0.792894064 | 0.953 | 0.462 | 2.62E-42 | 14 | Ifitm2 |
| Atp8b13 | 1.30E-46 | 0.707811815 | 0.367 | 0.102 | 2.63E-42 | 14 | Atp8b1 |
| Ankrd29 | 3.88E-46 | 0.620870425 | 0.302 | 0.074 | 7.85E-42 | 14 | Ankrd29 |
| Hhex | 4.42E-46 | 0.717163819 | 0.305 | 0.078 | 8.95E-42 | 14 | Hhex |
| Rfk | 4.77E-46 | 0.597222359 | 0.305 | 0.076 | 9.66E-42 | 14 | Rfk |
| Swap704 | 6.89E-46 | 0.828712877 | 0.596 | 0.245 | 1.39E-41 | 14 | Swap70 |
| Aida2 | 1.47E-45 | 0.757159307 | 0.433 | 0.141 | 2.98E-41 | 14 | Aida |

| | | | | | | | |
|---|---|---|---|---|---|---|---|
| Rapgef53 | 1.52E-45 | 0.793873845 | 0.582 | 0.22 | 3.07E-41 | 14 | Rapgef5 |
| Atp1a11 | 2.85E-45 | 0.766140626 | 0.629 | 0.278 | 5.77E-41 | 14 | Atp1a1 |
| Ctnna13 | 3.71E-45 | 0.748129835 | 0.742 | 0.35 | 7.50E-41 | 14 | Ctnna1 |
| Zcchc141 | 4.05E-45 | 0.57054405 | 0.313 | 0.079 | 8.20E-41 | 14 | Zcchc14 |
| Nedd44 | 8.77E-45 | 0.694705071 | 0.847 | 0.417 | 1.77E-40 | 14 | Nedd4 |
| Dag13 | 9.56E-45 | 0.693177844 | 0.433 | 0.141 | 1.94E-40 | 14 | Dag1 |
| Msn3 | 9.93E-45 | 0.844646595 | 0.811 | 0.442 | 2.01E-40 | 14 | Msn |
| Kif26a | 1.06E-44 | 0.498585532 | 0.28 | 0.065 | 2.15E-40 | 14 | Kif26a |
| Cyb5r33 | 1.08E-44 | 0.956050058 | 0.538 | 0.215 | 2.18E-40 | 14 | Cyb5r3 |
| Ift172 | 1.61E-44 | 0.557561567 | 0.276 | 0.065 | 3.25E-40 | 14 | Ift172 |
| Mill21 | 2.29E-44 | 0.597460038 | 0.284 | 0.068 | 4.63E-40 | 14 | Mill2 |
| Tpm44 | 3.33E-44 | 0.778285264 | 0.764 | 0.378 | 6.75E-40 | 14 | Tpm4 |
| Tspan134 | 6.42E-44 | 0.871079513 | 0.644 | 0.284 | 1.30E-39 | 14 | Tspan13 |
| Clic44 | 6.96E-44 | 0.842072694 | 0.829 | 0.421 | 1.41E-39 | 14 | Clic4 |
| Slc25a32 | 4.52E-43 | 0.57285004 | 0.262 | 0.061 | 9.15E-39 | 14 | Slc25a32 |
| Apbb24 | 5.59E-43 | 0.805852552 | 0.782 | 0.365 | 1.13E-38 | 14 | Apbb2 |
| Pik3c2b1 | 7.08E-43 | 0.596494189 | 0.309 | 0.08 | 1.43E-38 | 14 | Pik3c2b |
| Nbea4 | 7.78E-43 | 0.818308052 | 0.531 | 0.195 | 1.57E-38 | 14 | Nbea |
| Socs24 | 8.80E-43 | 0.740843723 | 0.498 | 0.181 | 1.78E-38 | 14 | Socs2 |
| Ppic3 | 1.17E-42 | 0.628814323 | 0.393 | 0.119 | 2.37E-38 | 14 | Ppic |
| Usp461 | 1.23E-42 | 0.591591266 | 0.411 | 0.132 | 2.49E-38 | 14 | Usp46 |
| Dab2ip4 | 1.55E-42 | 0.68022426 | 0.629 | 0.259 | 3.13E-38 | 14 | Dab2ip |
| Igfbp34 | 1.64E-42 | 0.460972692 | 0.745 | 0.3 | 3.33E-38 | 14 | Igfbp3 |
| Lmcd12 | 4.40E-42 | 0.676746261 | 0.484 | 0.163 | 8.90E-38 | 14 | Lmcd1 |
| Hsp90b17 | 5.45E-42 | 0.853637134 | 0.865 | 0.509 | 1.10E-37 | 14 | Hsp90b1 |
| mt-Cytb8 | 7.13E-42 | 0.809826507 | 0.985 | 0.699 | 1.44E-37 | 14 | mt-Cytb |
| Csrp13 | 8.52E-42 | 0.667221417 | 0.491 | 0.181 | 1.73E-37 | 14 | Csrp1 |
| Aig13 | 1.59E-41 | 0.735226015 | 0.542 | 0.212 | 3.23E-37 | 14 | Aig1 |
| Gnaq5 | 2.47E-41 | 0.78089418 | 0.735 | 0.357 | 4.99E-37 | 14 | Gnaq |
| Sptbn14 | 2.87E-41 | 0.682083425 | 0.971 | 0.492 | 5.81E-37 | 14 | Sptbn1 |
| Myl12b3 | 2.96E-41 | 0.764896013 | 0.887 | 0.546 | 6.00E-37 | 14 | Myl12b |
| Frmd4b3 | 3.14E-41 | 0.683208392 | 0.487 | 0.176 | 6.35E-37 | 14 | Frmd4b |
| Tcf44 | 4.37E-41 | 0.698280156 | 0.949 | 0.469 | 8.85E-37 | 14 | Tcf4 |
| Cd1513 | 8.41E-41 | 0.693407538 | 0.549 | 0.216 | 1.70E-36 | 14 | Cd151 |
| Kras3 | 1.25E-40 | 0.74014075 | 0.604 | 0.268 | 2.54E-36 | 14 | Kras |
| Gng114 | 1.35E-40 | 0.575762009 | 0.833 | 0.364 | 2.74E-36 | 14 | Gng11 |
| Tsc22d14 | 1.37E-40 | 0.656747764 | 0.847 | 0.444 | 2.78E-36 | 14 | Tsc22d1 |
| H2-D18 | 1.44E-40 | 0.765035646 | 0.985 | 0.667 | 2.92E-36 | 14 | H2-D1 |
| Spag94 | 1.46E-40 | 0.809310247 | 0.716 | 0.357 | 2.95E-36 | 14 | Spag9 |
| Fam181b | 2.47E-40 | 0.346809061 | 0.251 | 0.057 | 5.01E-36 | 14 | Fam181b |
| Itm2b7 | 2.71E-40 | 0.708129093 | 0.996 | 0.724 | 5.49E-36 | 14 | Itm2b |
| Ankrd33b2 | 3.13E-40 | 0.559828236 | 0.309 | 0.082 | 6.33E-36 | 14 | Ankrd33b |
| Exoc65 | 3.30E-40 | 0.66865305 | 0.465 | 0.169 | 6.68E-36 | 14 | Exoc6 |

| | | | | | | | |
|---|---|---|---|---|---|---|---|
| Sipa1l23 | 5.17E-40 | 0.691478269 | 0.451 | 0.157 | 1.05E-35 | 14 | Sipa1l2 |
| Dnm33 | 6.98E-40 | 0.805216947 | 0.578 | 0.229 | 1.41E-35 | 14 | Dnm3 |
| Plcb43 | 7.04E-40 | 0.751961436 | 0.662 | 0.289 | 1.43E-35 | 14 | Plcb4 |
| Zfp7041 | 8.09E-40 | 0.641573108 | 0.396 | 0.129 | 1.64E-35 | 14 | Zfp704 |
| Fam189a2 | 8.11E-40 | 0.427289382 | 0.255 | 0.06 | 1.64E-35 | 14 | Fam189a2 |
| Rftn17 | 1.29E-39 | 0.824261225 | 0.582 | 0.25 | 2.60E-35 | 14 | Rftn1 |
| Bnip23 | 1.38E-39 | 0.811709511 | 0.615 | 0.282 | 2.80E-35 | 14 | Bnip2 |
| Klf73 | 1.66E-39 | 0.729010603 | 0.676 | 0.296 | 3.36E-35 | 14 | Klf7 |
| Bcl2l14 | 4.27E-39 | 0.63952854 | 0.422 | 0.146 | 8.65E-35 | 14 | Bcl2l1 |
| Sigirr | 4.39E-39 | 0.47938135 | 0.291 | 0.077 | 8.89E-35 | 14 | Sigirr |
| Lrrc584 | 4.41E-39 | 0.708181499 | 0.531 | 0.213 | 8.93E-35 | 14 | Lrrc58 |
| Pdlim1 | 4.61E-39 | 0.635787289 | 0.342 | 0.101 | 9.34E-35 | 14 | Pdlim1 |
| Hspb14 | 4.92E-39 | 0.582856717 | 0.851 | 0.378 | 9.96E-35 | 14 | Hspb1 |
| Pcp4l15 | 5.26E-39 | 0.621175012 | 0.702 | 0.311 | 1.06E-34 | 14 | Pcp4l1 |
| S100a133 | 5.67E-39 | 0.695795711 | 0.887 | 0.535 | 1.15E-34 | 14 | S100a13 |
| Smarca24 | 7.72E-39 | 0.832623767 | 0.604 | 0.279 | 1.56E-34 | 14 | Smarca2 |
| Fbxo221 | 8.08E-39 | 0.632900549 | 0.415 | 0.142 | 1.64E-34 | 14 | Fbxo22 |
| Gnai24 | 8.22E-39 | 0.763726471 | 0.927 | 0.573 | 1.66E-34 | 14 | Gnai2 |
| Wwtr14 | 8.44E-39 | 0.636204046 | 0.596 | 0.238 | 1.71E-34 | 14 | Wwtr1 |
| Fabp44 | 9.24E-39 | 0.456077061 | 0.945 | 0.432 | 1.87E-34 | 14 | Fabp4 |
| Rapgef43 | 1.22E-38 | 0.672777116 | 0.625 | 0.256 | 2.46E-34 | 14 | Rapgef4 |
| Plekhg5 | 1.30E-38 | 0.533668837 | 0.273 | 0.07 | 2.62E-34 | 14 | Plekhg5 |
| Ptk23 | 1.71E-38 | 0.692810409 | 0.629 | 0.264 | 3.46E-34 | 14 | Ptk2 |
| Map3k201 | 2.23E-38 | 0.526997008 | 0.276 | 0.072 | 4.52E-34 | 14 | Map3k20 |
| mt-Co211 | 2.30E-38 | 0.719013218 | 0.996 | 0.727 | 4.66E-34 | 14 | mt-Co2 |
| Rapgef16 | 2.33E-38 | 0.767156234 | 0.553 | 0.24 | 4.72E-34 | 14 | Rapgef1 |
| Npdc13 | 2.83E-38 | 0.598884039 | 0.396 | 0.132 | 5.73E-34 | 14 | Npdc1 |
| Sav1 | 3.06E-38 | 0.579805499 | 0.345 | 0.105 | 6.19E-34 | 14 | Sav1 |
| Tnks26 | 3.63E-38 | 0.800018611 | 0.6 | 0.279 | 7.35E-34 | 14 | Tnks2 |
| Rnf125 | 4.15E-38 | 0.534468406 | 0.28 | 0.073 | 8.40E-34 | 14 | Rnf125 |
| Epn23 | 4.49E-38 | 0.681656083 | 0.436 | 0.153 | 9.09E-34 | 14 | Epn2 |
| Etl43 | 4.66E-38 | 0.661869298 | 0.691 | 0.292 | 9.44E-34 | 14 | Etl4 |
| Arglu14 | 5.12E-38 | 0.800343928 | 0.698 | 0.353 | 1.04E-33 | 14 | Arglu1 |
| Fam126a | 7.49E-38 | 0.642921129 | 0.287 | 0.08 | 1.52E-33 | 14 | Fam126a |
| Cadps2 | 1.44E-37 | 0.479537331 | 0.269 | 0.067 | 2.92E-33 | 14 | Cadps2 |
| Nckap14 | 2.18E-37 | 0.635528313 | 0.556 | 0.225 | 4.42E-33 | 14 | Nckap1 |
| Aggf1 | 2.36E-37 | 0.658930636 | 0.375 | 0.126 | 4.77E-33 | 14 | Aggf1 |
| Ccnyl12 | 2.39E-37 | 0.669472136 | 0.353 | 0.11 | 4.84E-33 | 14 | Ccnyl1 |
| Kif1c | 2.66E-37 | 0.483393243 | 0.284 | 0.077 | 5.38E-33 | 14 | Kif1c |
| Elmo17 | 3.15E-37 | 0.795861142 | 0.735 | 0.364 | 6.38E-33 | 14 | Elmo1 |
| Capn24 | 3.89E-37 | 0.66314877 | 0.469 | 0.178 | 7.87E-33 | 14 | Capn2 |
| Card19 | 4.23E-37 | 0.598067399 | 0.498 | 0.199 | 8.57E-33 | 14 | Card19 |
| Bst26 | 4.52E-37 | 0.632183163 | 0.887 | 0.501 | 9.16E-33 | 14 | Bst2 |

| | | | | | | | |
|---|---|---|---|---|---|---|---|
| Tmem25 | 5.91E-37 | 0.678140841 | 0.407 | 0.142 | 1.20E-32 | 14 | Tmem2 |
| Pcdh13 | 5.99E-37 | 0.545996839 | 0.316 | 0.091 | 1.21E-32 | 14 | Pcdh1 |
| Nfe2l13 | 9.39E-37 | 0.711355171 | 0.607 | 0.279 | 1.90E-32 | 14 | Nfe2l1 |
| Rnf144a2 | 9.89E-37 | 0.545425851 | 0.349 | 0.106 | 2.00E-32 | 14 | Rnf144a |
| Tpgs2 | 1.07E-36 | 0.528510211 | 0.28 | 0.076 | 2.17E-32 | 14 | Tpgs2 |
| Atp2a23 | 1.35E-36 | 0.842992451 | 0.484 | 0.199 | 2.73E-32 | 14 | Atp2a2 |
| Arhgef153 | 1.99E-36 | 0.598192293 | 0.469 | 0.167 | 4.02E-32 | 14 | Arhgef15 |
| Kctd12b4 | 3.33E-36 | 0.64709901 | 0.6 | 0.248 | 6.74E-32 | 14 | Kctd12b |
| Sptan14 | 3.87E-36 | 0.70504154 | 0.702 | 0.342 | 7.83E-32 | 14 | Sptan1 |
| Ttc284 | 4.05E-36 | 0.84900294 | 0.607 | 0.284 | 8.21E-32 | 14 | Ttc28 |
| Myl12a3 | 4.28E-36 | 0.744756752 | 0.8 | 0.469 | 8.67E-32 | 14 | Myl12a |
| Lmo24 | 4.44E-36 | 0.827429905 | 0.502 | 0.207 | 9.00E-32 | 14 | Lmo2 |
| Timp22 | 7.40E-36 | 0.50135789 | 0.389 | 0.129 | 1.50E-31 | 14 | Timp2 |
| Cracr2b2 | 1.02E-35 | 0.460940589 | 0.578 | 0.237 | 2.07E-31 | 14 | Cracr2b |
| Ackr33 | 1.25E-35 | 0.563767302 | 0.436 | 0.151 | 2.52E-31 | 14 | Ackr3 |
| Serinc15 | 1.26E-35 | 0.677074491 | 0.564 | 0.248 | 2.56E-31 | 14 | Serinc1 |
| Clic54 | 1.42E-35 | 0.585998869 | 0.687 | 0.289 | 2.88E-31 | 14 | Clic5 |
| Abr5 | 1.71E-35 | 0.751636261 | 0.422 | 0.157 | 3.46E-31 | 14 | Abr |
| Tgm23 | 1.82E-35 | 0.640419175 | 0.444 | 0.161 | 3.69E-31 | 14 | Tgm2 |
| Cpd4 | 2.12E-35 | 0.664651088 | 0.669 | 0.311 | 4.28E-31 | 14 | Cpd |
| Plscr44 | 2.43E-35 | 0.601012672 | 0.491 | 0.188 | 4.91E-31 | 14 | Plscr4 |
| Lrrc8c5 | 2.89E-35 | 0.710067794 | 0.658 | 0.31 | 5.85E-31 | 14 | Lrrc8c |
| Gna114 | 3.18E-35 | 0.682830014 | 0.564 | 0.243 | 6.44E-31 | 14 | Gna11 |
| Eml13 | 4.15E-35 | 0.572684257 | 0.345 | 0.108 | 8.40E-31 | 14 | Eml1 |
| Hist1h2bc3 | 4.60E-35 | 0.625685805 | 0.571 | 0.248 | 9.30E-31 | 14 | Hist1h2bc |
| Osbpl94 | 4.66E-35 | 0.641590993 | 0.607 | 0.277 | 9.43E-31 | 14 | Osbpl9 |
| Ndrg13 | 5.60E-35 | 0.679412234 | 0.553 | 0.231 | 1.13E-30 | 14 | Ndrg1 |
| Dennd5a3 | 6.40E-35 | 0.631751678 | 0.473 | 0.186 | 1.30E-30 | 14 | Dennd5a |
| Macf17 | 7.19E-35 | 0.756867903 | 0.858 | 0.501 | 1.46E-30 | 14 | Macf1 |
| Fkbp1a3 | 8.78E-35 | 0.543816018 | 0.785 | 0.404 | 1.78E-30 | 14 | Fkbp1a |
| Galnt152 | 1.05E-34 | 0.529193446 | 0.305 | 0.089 | 2.14E-30 | 14 | Galnt15 |
| Ktn14 | 1.16E-34 | 0.690615459 | 0.636 | 0.306 | 2.35E-30 | 14 | Ktn1 |
| Ick1 | 1.51E-34 | 0.490645666 | 0.291 | 0.084 | 3.05E-30 | 14 | Ick |
| Vps13d2 | 1.63E-34 | 0.76918833 | 0.425 | 0.168 | 3.29E-30 | 14 | Vps13d |
| Mgll4 | 1.68E-34 | 0.67155911 | 0.88 | 0.447 | 3.40E-30 | 14 | Mgll |
| Capns14 | 1.85E-34 | 0.551352162 | 0.687 | 0.347 | 3.74E-30 | 14 | Capns1 |
| Elk33 | 1.96E-34 | 0.626437537 | 0.727 | 0.355 | 3.97E-30 | 14 | Elk3 |
| Shroom42 | 2.01E-34 | 0.585320797 | 0.338 | 0.104 | 4.07E-30 | 14 | Shroom4 |
| Inpp12 | 2.46E-34 | 0.590394865 | 0.349 | 0.115 | 4.98E-30 | 14 | Inpp1 |
| Plcg25 | 2.62E-34 | 0.668830384 | 0.353 | 0.119 | 5.30E-30 | 14 | Plcg2 |
| Cald14 | 2.65E-34 | 0.631821852 | 0.705 | 0.32 | 5.36E-30 | 14 | Cald1 |
| Rhoc4 | 3.17E-34 | 0.6277143 | 0.636 | 0.295 | 6.41E-30 | 14 | Rhoc |
| Efr3b2 | 3.49E-34 | 0.572270838 | 0.298 | 0.086 | 7.07E-30 | 14 | Efr3b |

| | | | | | | | |
|---|---|---|---|---|---|---|---|
| Strn34 | 3.96E-34 | 0.755312196 | 0.673 | 0.344 | 8.02E-30 | 14 | Strn3 |
| Agrn2 | 4.72E-34 | 0.523774062 | 0.287 | 0.082 | 9.55E-30 | 14 | Agrn |
| Mbnl25 | 5.02E-34 | 0.722075751 | 0.756 | 0.417 | 1.02E-29 | 14 | Mbnl2 |
| Sh3rf13 | 5.30E-34 | 0.757664471 | 0.375 | 0.13 | 1.07E-29 | 14 | Sh3rf1 |
| Tmpo3 | 5.31E-34 | 0.669301429 | 0.491 | 0.207 | 1.08E-29 | 14 | Tmpo |
| Klf133 | 5.99E-34 | 0.667720691 | 0.691 | 0.349 | 1.21E-29 | 14 | Klf13 |
| Tapbp5 | 6.22E-34 | 0.7134611 | 0.622 | 0.304 | 1.26E-29 | 14 | Tapbp |
| C130074G19Rik3 | 6.76E-34 | 0.516213291 | 0.509 | 0.198 | 1.37E-29 | 14 | C130074G19Rik |
| Erg3 | 7.11E-34 | 0.56504504 | 0.382 | 0.13 | 1.44E-29 | 14 | Erg |
| Dram23 | 7.40E-34 | 0.5916331 | 0.415 | 0.154 | 1.50E-29 | 14 | Dram2 |
| Gas63 | 7.96E-34 | 0.526944512 | 0.676 | 0.296 | 1.61E-29 | 14 | Gas6 |
| Rassf33 | 8.02E-34 | 0.598362754 | 0.549 | 0.234 | 1.62E-29 | 14 | Rassf3 |
| Tie14 | 1.11E-33 | 0.537670374 | 0.611 | 0.252 | 2.24E-29 | 14 | Tie1 |
| Anxa53 | 1.25E-33 | 0.592192713 | 0.855 | 0.492 | 2.53E-29 | 14 | Anxa5 |
| Adipor24 | 1.39E-33 | 0.644548186 | 0.404 | 0.149 | 2.81E-29 | 14 | Adipor2 |
| Dusp33 | 1.85E-33 | 0.588905546 | 0.487 | 0.191 | 3.75E-29 | 14 | Dusp3 |
| Vps361 | 1.89E-33 | 0.491473111 | 0.571 | 0.253 | 3.82E-29 | 14 | Vps36 |
| Pcmtd12 | 3.46E-33 | 0.69156479 | 0.644 | 0.326 | 7.00E-29 | 14 | Pcmtd1 |
| Akt37 | 4.08E-33 | 0.627007596 | 0.571 | 0.25 | 8.27E-29 | 14 | Akt3 |
| mt-Atp69 | 5.37E-33 | 0.676544851 | 1 | 0.753 | 1.09E-28 | 14 | mt-Atp6 |
| Tmem50a6 | 9.78E-33 | 0.628800225 | 0.705 | 0.382 | 1.98E-28 | 14 | Tmem50a |
| Slc3a26 | 1.03E-32 | 0.793633792 | 0.593 | 0.29 | 2.08E-28 | 14 | Slc3a2 |
| Slc16a21 | 1.20E-32 | 0.447537301 | 0.251 | 0.067 | 2.43E-28 | 14 | Slc16a2 |
| Sh3glb14 | 1.23E-32 | 0.672077435 | 0.833 | 0.53 | 2.50E-28 | 14 | Sh3glb1 |
| Slc12a71 | 1.54E-32 | 0.627560112 | 0.324 | 0.105 | 3.12E-28 | 14 | Slc12a7 |
| Nova22 | 1.56E-32 | 0.487194566 | 0.335 | 0.105 | 3.16E-28 | 14 | Nova2 |
| Klf66 | 1.78E-32 | 0.753495735 | 0.56 | 0.266 | 3.59E-28 | 14 | Klf6 |
| Sema7a3 | 2.40E-32 | 0.533439934 | 0.418 | 0.149 | 4.86E-28 | 14 | Sema7a |
| Cep85l3 | 2.87E-32 | 0.684248819 | 0.425 | 0.164 | 5.80E-28 | 14 | Cep85l |
| Isg153 | 2.90E-32 | 0.738238463 | 0.531 | 0.228 | 5.87E-28 | 14 | Isg15 |
| Tmem883 | 3.82E-32 | 0.615615813 | 0.52 | 0.209 | 7.73E-28 | 14 | Tmem88 |
| Vps531 | 4.88E-32 | 0.584166966 | 0.356 | 0.123 | 9.88E-28 | 14 | Vps53 |
| Flt14 | 7.67E-32 | 0.289171539 | 0.967 | 0.39 | 1.55E-27 | 14 | Flt1 |
| Esyt27 | 8.13E-32 | 0.788255065 | 0.524 | 0.242 | 1.65E-27 | 14 | Esyt2 |
| Smc1a4 | 1.05E-31 | 0.68791079 | 0.611 | 0.311 | 2.12E-27 | 14 | Smc1a |
| Anxa26 | 1.68E-31 | 0.637843442 | 0.767 | 0.415 | 3.39E-27 | 14 | Anxa2 |
| Adgre55 | 2.07E-31 | 0.598252139 | 0.545 | 0.236 | 4.19E-27 | 14 | Adgre5 |
| Gnb14 | 2.31E-31 | 0.631369696 | 0.753 | 0.426 | 4.67E-27 | 14 | Gnb1 |
| Cers41 | 2.77E-31 | 0.553978212 | 0.327 | 0.109 | 5.62E-27 | 14 | Cers4 |
| Eif3a4 | 3.57E-31 | 0.725137924 | 0.673 | 0.371 | 7.23E-27 | 14 | Eif3a |
| Hip1r1 | 3.66E-31 | 0.474330659 | 0.258 | 0.074 | 7.41E-27 | 14 | Hip1r |
| 31-Mar | 4.31E-31 | 0.64147169 | 0.295 | 0.092 | 8.73E-27 | 14 | 3-Mar |
| Diaph27 | 5.41E-31 | 0.771848625 | 0.709 | 0.375 | 1.10E-26 | 14 | Diaph2 |

| | | | | | | | |
|---|---|---|---|---|---|---|---|
| Iqgap18 | 6.02E-31 | 0.652756941 | 0.742 | 0.411 | 1.22E-26 | 14 | Iqgap1 |
| Trim242 | 6.73E-31 | 0.635829055 | 0.425 | 0.172 | 1.36E-26 | 14 | Trim24 |
| Cped14 | 3.08E-30 | 0.6980613 | 0.451 | 0.178 | 6.24E-26 | 14 | Cped1 |
| Mob2 | 3.96E-30 | 0.557581801 | 0.385 | 0.146 | 8.02E-26 | 14 | Mob2 |
| Mmp152 | 4.93E-30 | 0.512381624 | 0.313 | 0.101 | 9.99E-26 | 14 | Mmp15 |
| Tmem30a5 | 5.19E-30 | 0.66728014 | 0.585 | 0.285 | 1.05E-25 | 14 | Tmem30a |
| Plcg12 | 1.19E-29 | 0.533286918 | 0.349 | 0.122 | 2.40E-25 | 14 | Plcg1 |
| Xpr13 | 1.43E-29 | 0.649298683 | 0.389 | 0.153 | 2.90E-25 | 14 | Xpr1 |
| Sox72 | 1.49E-29 | 0.504947947 | 0.298 | 0.094 | 3.02E-25 | 14 | Sox7 |
| Abca13 | 1.85E-29 | 0.666401681 | 0.353 | 0.129 | 3.75E-25 | 14 | Abca1 |
| Arhgef22 | 2.09E-29 | 0.643993845 | 0.407 | 0.164 | 4.23E-25 | 14 | Arhgef2 |
| Wasf23 | 2.27E-29 | 0.655365286 | 0.731 | 0.424 | 4.60E-25 | 14 | Wasf2 |
| Myo1e7 | 2.85E-29 | 0.636849471 | 0.589 | 0.28 | 5.78E-25 | 14 | Myo1e |
| Rabac14 | 2.87E-29 | 0.589422353 | 0.865 | 0.563 | 5.81E-25 | 14 | Rabac1 |
| Abcc43 | 3.78E-29 | 0.643391376 | 0.364 | 0.135 | 7.65E-25 | 14 | Abcc4 |
| Zbtb207 | 4.67E-29 | 0.6246197 | 0.898 | 0.536 | 9.46E-25 | 14 | Zbtb20 |
| Son8 | 5.83E-29 | 0.594803656 | 0.876 | 0.525 | 1.18E-24 | 14 | Son |
| Plxna42 | 6.07E-29 | 0.581239986 | 0.338 | 0.118 | 1.23E-24 | 14 | Plxna4 |
| Numb3 | 8.40E-29 | 0.594346665 | 0.669 | 0.336 | 1.70E-24 | 14 | Numb |
| Amotl14 | 9.74E-29 | 0.490612854 | 0.458 | 0.185 | 1.97E-24 | 14 | Amotl1 |
| Rab124 | 1.13E-28 | 0.547637731 | 0.433 | 0.176 | 2.28E-24 | 14 | Rab12 |
| Serinc34 | 1.24E-28 | 0.59476966 | 0.833 | 0.486 | 2.51E-24 | 14 | Serinc3 |
| Plxna23 | 1.26E-28 | 0.567712697 | 0.516 | 0.218 | 2.56E-24 | 14 | Plxna2 |
| Sh2b3 | 1.72E-28 | 0.474206664 | 0.295 | 0.096 | 3.49E-24 | 14 | Sh2b3 |
| Cdkn1c3 | 1.73E-28 | 0.647305934 | 0.345 | 0.125 | 3.50E-24 | 14 | Cdkn1c |
| Itga5 | 2.15E-28 | 0.503307954 | 0.295 | 0.097 | 4.36E-24 | 14 | Itga5 |
| Plk23 | 2.74E-28 | 0.677012246 | 0.418 | 0.166 | 5.56E-24 | 14 | Plk2 |
| Abcg24 | 2.92E-28 | 0.487946608 | 0.593 | 0.269 | 5.92E-24 | 14 | Abcg2 |
| Chn24 | 3.10E-28 | 0.446111041 | 0.487 | 0.197 | 6.27E-24 | 14 | Chn2 |
| Mcc3 | 3.37E-28 | 0.494457042 | 0.622 | 0.283 | 6.82E-24 | 14 | Mcc |
| Fermt24 | 4.70E-28 | 0.412213106 | 0.731 | 0.348 | 9.51E-24 | 14 | Fermt2 |
| Cr1l3 | 4.85E-28 | 0.58523739 | 0.502 | 0.233 | 9.81E-24 | 14 | Cr1l |
| Map7d13 | 4.98E-28 | 0.573945111 | 0.407 | 0.167 | 1.01E-23 | 14 | Map7d1 |
| Kank33 | 5.02E-28 | 0.478293897 | 0.531 | 0.227 | 1.02E-23 | 14 | Kank3 |
| Ypel32 | 5.16E-28 | 0.625155221 | 0.538 | 0.266 | 1.04E-23 | 14 | Ypel3 |
| Ccrl22 | 5.26E-28 | 0.45234759 | 0.32 | 0.108 | 1.07E-23 | 14 | Ccrl2 |
| Itga15 | 6.13E-28 | 0.638636206 | 0.775 | 0.413 | 1.24E-23 | 14 | Itga1 |
| Nav14 | 6.28E-28 | 0.565513927 | 0.516 | 0.222 | 1.27E-23 | 14 | Nav1 |
| Tnrc6c5 | 6.98E-28 | 0.613898845 | 0.618 | 0.314 | 1.41E-23 | 14 | Tnrc6c |
| Pnpla2 | 7.57E-28 | 0.379659697 | 0.353 | 0.128 | 1.53E-23 | 14 | Pnpla2 |
| Atrnl13 | 7.88E-28 | 0.717034714 | 0.407 | 0.165 | 1.60E-23 | 14 | Atrnl1 |
| Rac13 | 8.31E-28 | 0.570678249 | 0.735 | 0.429 | 1.68E-23 | 14 | Rac1 |
| mt-Co19 | 8.86E-28 | 0.692301038 | 1 | 0.718 | 1.79E-23 | 14 | mt-Co1 |

| | | | | | | | |
|---|---|---|---|---|---|---|---|
| Hip16 | 1.01E-27 | 0.622407229 | 0.651 | 0.333 | 2.04E-23 | 14 | Hip1 |
| Rasgrp33 | 1.35E-27 | 0.618608745 | 0.604 | 0.281 | 2.74E-23 | 14 | Rasgrp3 |
| Maml28 | 1.38E-27 | 0.879708511 | 0.647 | 0.342 | 2.79E-23 | 14 | Maml2 |
| Golim43 | 1.38E-27 | 0.691077465 | 0.422 | 0.18 | 2.80E-23 | 14 | Golim4 |
| Nfkbia5 | 1.44E-27 | 0.559459008 | 0.72 | 0.399 | 2.92E-23 | 14 | Nfkbia |
| Raph12 | 1.48E-27 | 0.5699416 | 0.32 | 0.114 | 2.99E-23 | 14 | Raph1 |
| Rtn34 | 2.07E-27 | 0.626217443 | 0.611 | 0.323 | 4.18E-23 | 14 | Rtn3 |
| Ppp1r16b7 | 2.38E-27 | 0.615305246 | 0.462 | 0.201 | 4.81E-23 | 14 | Ppp1r16b |
| Ybx14 | 2.47E-27 | 0.551852723 | 0.949 | 0.751 | 5.00E-23 | 14 | Ybx1 |
| Map4k46 | 2.66E-27 | 0.575079586 | 0.567 | 0.276 | 5.38E-23 | 14 | Map4k4 |
| Gbp32 | 2.74E-27 | 0.506847589 | 0.331 | 0.117 | 5.54E-23 | 14 | Gbp3 |
| Hdgfl33 | 3.01E-27 | 0.520952768 | 0.316 | 0.11 | 6.09E-23 | 14 | Hdgfl3 |
| Dennd5b3 | 3.81E-27 | 0.51641963 | 0.385 | 0.146 | 7.70E-23 | 14 | Dennd5b |
| Itpkb3 | 4.29E-27 | 0.675503585 | 0.658 | 0.355 | 8.69E-23 | 14 | Itpkb |
| Oasl23 | 4.43E-27 | 0.525667188 | 0.353 | 0.131 | 8.97E-23 | 14 | Oasl2 |
| Calm14 | 5.16E-27 | 0.516003348 | 0.989 | 0.78 | 1.05E-22 | 14 | Calm1 |
| Tmem184b2 | 5.30E-27 | 0.485859388 | 0.32 | 0.113 | 1.07E-22 | 14 | Tmem184b |
| Ushbp13 | 5.31E-27 | 0.551915176 | 0.509 | 0.218 | 1.08E-22 | 14 | Ushbp1 |
| Txnip3 | 1.15E-26 | 0.554225621 | 0.873 | 0.544 | 2.32E-22 | 14 | Txnip |
| Tmem595 | 1.35E-26 | 0.559066668 | 0.793 | 0.489 | 2.73E-22 | 14 | Tmem59 |
| Trove23 | 1.60E-26 | 0.472300377 | 0.353 | 0.131 | 3.24E-22 | 14 | Trove2 |
| Ralbp12 | 1.75E-26 | 0.426011163 | 0.633 | 0.33 | 3.55E-22 | 14 | Ralbp1 |
| Dnajc102 | 2.04E-26 | 0.56247264 | 0.356 | 0.139 | 4.12E-22 | 14 | Dnajc10 |
| Fas2 | 2.16E-26 | 0.614819287 | 0.324 | 0.117 | 4.38E-22 | 14 | Fas |
| Clec14a3 | 2.97E-26 | 0.46730138 | 0.655 | 0.304 | 6.01E-22 | 14 | Clec14a |
| Dock13 | 3.17E-26 | 0.55407285 | 0.582 | 0.279 | 6.42E-22 | 14 | Dock1 |
| Dync1h15 | 3.37E-26 | 0.65818337 | 0.553 | 0.286 | 6.83E-22 | 14 | Dync1h1 |
| Map3k112 | 4.70E-26 | 0.452902688 | 0.276 | 0.091 | 9.52E-22 | 14 | Map3k11 |
| Rabgap12 | 6.18E-26 | 0.604564118 | 0.389 | 0.162 | 1.25E-21 | 14 | Rabgap1 |
| Ccdc502 | 6.90E-26 | 0.592314045 | 0.48 | 0.228 | 1.40E-21 | 14 | Ccdc50 |
| Nfib3 | 7.87E-26 | 0.380564406 | 0.916 | 0.442 | 1.59E-21 | 14 | Nfib |
| Palm1 | 8.73E-26 | 0.469962815 | 0.262 | 0.084 | 1.77E-21 | 14 | Palm |
| mt-Nd49 | 1.02E-25 | 0.621183037 | 0.978 | 0.674 | 2.07E-21 | 14 | mt-Nd4 |
| Znrf16 | 1.21E-25 | 0.551258674 | 0.433 | 0.188 | 2.45E-21 | 14 | Znrf1 |
| Pdia66 | 1.74E-25 | 0.545844829 | 0.625 | 0.336 | 3.53E-21 | 14 | Pdia6 |
| Itga63 | 1.91E-25 | 0.479023736 | 0.6 | 0.282 | 3.88E-21 | 14 | Itga6 |
| Rock23 | 2.35E-25 | 0.550951432 | 0.625 | 0.337 | 4.76E-21 | 14 | Rock2 |
| Prss235 | 2.89E-25 | 0.417462831 | 0.611 | 0.28 | 5.84E-21 | 14 | Prss23 |
| Ankrd13a1 | 3.53E-25 | 0.398453225 | 0.338 | 0.128 | 7.15E-21 | 14 | Ankrd13a |
| Rdx4 | 3.53E-25 | 0.485594675 | 0.895 | 0.55 | 7.15E-21 | 14 | Rdx |
| Limk2 | 4.09E-25 | 0.450792238 | 0.269 | 0.09 | 8.29E-21 | 14 | Limk2 |
| Lysmd23 | 4.16E-25 | 0.522230916 | 0.44 | 0.191 | 8.42E-21 | 14 | Lysmd2 |
| Grasp3 | 4.84E-25 | 0.519182492 | 0.404 | 0.166 | 9.80E-21 | 14 | Grasp |

| | | | | | | | |
|---|---|---|---|---|---|---|---|
| Nudt42 | 5.44E-25 | 0.500397366 | 0.713 | 0.409 | 1.10E-20 | 14 | Nudt4 |
| Gab23 | 5.64E-25 | 0.490935926 | 0.462 | 0.204 | 1.14E-20 | 14 | Gab2 |
| Slain21 | 6.16E-25 | 0.537699708 | 0.411 | 0.177 | 1.25E-20 | 14 | Slain2 |
| Agap12 | 6.30E-25 | 0.567411238 | 0.364 | 0.144 | 1.28E-20 | 14 | Agap1 |
| mt-Atp87 | 7.24E-25 | 0.594116588 | 0.76 | 0.462 | 1.47E-20 | 14 | mt-Atp8 |
| Stmn11 | 7.69E-25 | 0.37560068 | 0.345 | 0.13 | 1.56E-20 | 14 | Stmn1 |
| Cdk177 | 1.52E-24 | 0.613064378 | 0.505 | 0.244 | 3.08E-20 | 14 | Cdk17 |
| Rb14 | 1.93E-24 | 0.653881429 | 0.458 | 0.214 | 3.91E-20 | 14 | Rb1 |
| Pmp224 | 2.80E-24 | 0.394047891 | 0.371 | 0.143 | 5.66E-20 | 14 | Pmp22 |
| Smco44 | 2.98E-24 | 0.356063482 | 0.749 | 0.389 | 6.04E-20 | 14 | Smco4 |
| Rhoa3 | 3.24E-24 | 0.533651872 | 0.876 | 0.593 | 6.55E-20 | 14 | Rhoa |
| Mllt34 | 3.33E-24 | 0.66392568 | 0.367 | 0.151 | 6.74E-20 | 14 | Mllt3 |
| Gars | 5.06E-24 | 0.386377099 | 0.298 | 0.109 | 1.02E-19 | 14 | Gars |
| Irf71 | 6.22E-24 | 0.427029652 | 0.411 | 0.176 | 1.26E-19 | 14 | Irf7 |
| Cdc42ep36 | 7.48E-24 | 0.543511573 | 0.4 | 0.174 | 1.51E-19 | 14 | Cdc42ep3 |
| Sybu3 | 7.54E-24 | 0.520770075 | 0.313 | 0.114 | 1.53E-19 | 14 | Sybu |
| Pcm14 | 1.03E-23 | 0.565525607 | 0.505 | 0.25 | 2.09E-19 | 14 | Pcm1 |
| Ezh1 | 1.06E-23 | 0.471598881 | 0.327 | 0.127 | 2.16E-19 | 14 | Ezh1 |
| Pced1b2 | 1.16E-23 | 0.600814242 | 0.309 | 0.117 | 2.34E-19 | 14 | Pced1b |
| Tead11 | 1.54E-23 | 0.481506132 | 0.251 | 0.083 | 3.13E-19 | 14 | Tead1 |
| Dennd32 | 1.59E-23 | 0.546706415 | 0.356 | 0.146 | 3.22E-19 | 14 | Dennd3 |
| Rock16 | 1.89E-23 | 0.596314804 | 0.709 | 0.432 | 3.82E-19 | 14 | Rock1 |
| Itsn27 | 1.96E-23 | 0.641856603 | 0.524 | 0.277 | 3.97E-19 | 14 | Itsn2 |
| Arhgef125 | 2.13E-23 | 0.415180184 | 0.691 | 0.347 | 4.31E-19 | 14 | Arhgef12 |
| Tspan42 | 2.63E-23 | 0.362306295 | 0.298 | 0.106 | 5.33E-19 | 14 | Tspan4 |
| Wars3 | 2.78E-23 | 0.511778736 | 0.393 | 0.168 | 5.63E-19 | 14 | Wars |
| Klhl53 | 2.81E-23 | 0.457365327 | 0.48 | 0.224 | 5.70E-19 | 14 | Klhl5 |
| Selenof4 | 3.22E-23 | 0.459819812 | 0.793 | 0.49 | 6.52E-19 | 14 | Selenof |
| Tmco14 | 3.51E-23 | 0.573173136 | 0.564 | 0.312 | 7.11E-19 | 14 | Tmco1 |
| Plscr14 | 3.61E-23 | 0.503032666 | 0.451 | 0.204 | 7.31E-19 | 14 | Plscr1 |
| Shoc22 | 3.82E-23 | 0.544208078 | 0.465 | 0.221 | 7.74E-19 | 14 | Shoc2 |
| Atrx3 | 4.52E-23 | 0.594233553 | 0.702 | 0.419 | 9.16E-19 | 14 | Atrx |
| Tbc1d15 | 4.60E-23 | 0.493995779 | 0.382 | 0.165 | 9.31E-19 | 14 | Tbc1d1 |
| Mprip5 | 5.35E-23 | 0.490116094 | 0.571 | 0.292 | 1.08E-18 | 14 | Mprip |
| Gsk3b4 | 5.40E-23 | 0.565411005 | 0.6 | 0.33 | 1.09E-18 | 14 | Gsk3b |
| Sypl3 | 6.20E-23 | 0.548333944 | 0.491 | 0.239 | 1.25E-18 | 14 | Sypl |
| Fry3 | 8.46E-23 | 0.530243917 | 0.444 | 0.2 | 1.71E-18 | 14 | Fry |
| mt-Nd38 | 9.48E-23 | 0.551203092 | 0.735 | 0.446 | 1.92E-18 | 14 | mt-Nd3 |
| Hspa58 | 1.12E-22 | 0.589237043 | 0.72 | 0.434 | 2.26E-18 | 14 | Hspa5 |
| Cd475 | 1.13E-22 | 0.497254837 | 0.84 | 0.509 | 2.29E-18 | 14 | Cd47 |
| Baz2b5 | 1.19E-22 | 0.528558688 | 0.633 | 0.35 | 2.41E-18 | 14 | Baz2b |
| Zfp3663 | 1.22E-22 | 0.474851015 | 0.429 | 0.182 | 2.47E-18 | 14 | Zfp366 |
| Rhoj4 | 1.35E-22 | 0.492333672 | 0.52 | 0.241 | 2.74E-18 | 14 | Rhoj |

| | | | | | | | |
|---|---|---|---|---|---|---|---|
| Sema6a3 | 1.67E-22 | 0.537607263 | 0.462 | 0.209 | 3.39E-18 | 14 | Sema6a |
| Il31ra | 2.22E-22 | 0.434152913 | 0.444 | 0.208 | 4.50E-18 | 14 | Il31ra |
| Krtcap25 | 2.33E-22 | 0.446524489 | 0.727 | 0.441 | 4.72E-18 | 14 | Krtcap2 |
| Rnf2203 | 2.49E-22 | 0.533992852 | 0.502 | 0.241 | 5.03E-18 | 14 | Rnf220 |
| Rb1cc14 | 2.52E-22 | 0.588188918 | 0.469 | 0.234 | 5.11E-18 | 14 | Rb1cc1 |
| Fkbp92 | 2.79E-22 | 0.423517129 | 0.298 | 0.111 | 5.65E-18 | 14 | Fkbp9 |
| Tspan93 | 3.10E-22 | 0.458801314 | 0.545 | 0.264 | 6.28E-18 | 14 | Tspan9 |
| mt-Co310 | 3.55E-22 | 0.537814611 | 1 | 0.725 | 7.18E-18 | 14 | mt-Co3 |
| Rab11a4 | 3.61E-22 | 0.45853043 | 0.575 | 0.308 | 7.31E-18 | 14 | Rab11a |
| Dynll13 | 3.97E-22 | 0.412413308 | 0.895 | 0.604 | 8.04E-18 | 14 | Dynll1 |
| Tlk12 | 4.28E-22 | 0.463786899 | 0.458 | 0.222 | 8.67E-18 | 14 | Tlk1 |
| Tjp22 | 4.34E-22 | 0.444049045 | 0.345 | 0.14 | 8.79E-18 | 14 | Tjp2 |
| Clec1a3 | 4.66E-22 | 0.427158864 | 0.4 | 0.168 | 9.43E-18 | 14 | Clec1a |
| H2-K18 | 4.71E-22 | 0.49456412 | 0.975 | 0.642 | 9.54E-18 | 14 | H2-K1 |
| Anp32a3 | 5.01E-22 | 0.521153308 | 0.738 | 0.465 | 1.01E-17 | 14 | Anp32a |
| Lrpap13 | 5.42E-22 | 0.622180273 | 0.415 | 0.197 | 1.10E-17 | 14 | Lrpap1 |
| Bvht3 | 6.87E-22 | 0.360098548 | 0.433 | 0.186 | 1.39E-17 | 14 | Bvht |
| Bod1l4 | 7.69E-22 | 0.606455421 | 0.516 | 0.273 | 1.56E-17 | 14 | Bod1l |
| Nr3c23 | 1.02E-21 | 0.613350306 | 0.309 | 0.121 | 2.06E-17 | 14 | Nr3c2 |
| Syne13 | 1.05E-21 | 0.371395916 | 0.691 | 0.358 | 2.13E-17 | 14 | Syne1 |
| Pcnx3 | 1.21E-21 | 0.606672393 | 0.393 | 0.178 | 2.46E-17 | 14 | Pcnx |
| Ddx3x4 | 1.57E-21 | 0.50457899 | 0.571 | 0.308 | 3.17E-17 | 14 | Ddx3x |
| Herc23 | 1.77E-21 | 0.540661065 | 0.433 | 0.207 | 3.58E-17 | 14 | Herc2 |
| Phc22 | 1.85E-21 | 0.418115074 | 0.382 | 0.169 | 3.74E-17 | 14 | Phc2 |
| Spata136 | 2.01E-21 | 0.550148138 | 0.433 | 0.206 | 4.06E-17 | 14 | Spata13 |
| Pdia36 | 2.21E-21 | 0.591288685 | 0.72 | 0.443 | 4.48E-17 | 14 | Pdia3 |
| Ccdc85b4 | 2.25E-21 | 0.364261971 | 0.527 | 0.255 | 4.56E-17 | 14 | Ccdc85b |
| Ptpra5 | 2.33E-21 | 0.48212171 | 0.571 | 0.304 | 4.71E-17 | 14 | Ptpra |
| 2900026A02Rik5 | 2.84E-21 | 0.528097008 | 0.451 | 0.219 | 5.75E-17 | 14 | 2900026A02Rik |
| Card101 | 3.13E-21 | 0.360845718 | 0.251 | 0.086 | 6.34E-17 | 14 | Card10 |
| Npr11 | 3.24E-21 | 0.349299842 | 0.262 | 0.091 | 6.55E-17 | 14 | Npr1 |
| Gimap18 | 3.37E-21 | 0.480005995 | 0.578 | 0.302 | 6.83E-17 | 14 | Gimap1 |
| Ash1l4 | 3.39E-21 | 0.537571495 | 0.553 | 0.299 | 6.86E-17 | 14 | Ash1l |
| Fam219a2 | 3.74E-21 | 0.554321752 | 0.32 | 0.131 | 7.58E-17 | 14 | Fam219a |
| Klhl243 | 3.90E-21 | 0.500486559 | 0.444 | 0.215 | 7.90E-17 | 14 | Klhl24 |
| Anxa73 | 4.26E-21 | 0.455909963 | 0.404 | 0.186 | 8.62E-17 | 14 | Anxa7 |
| Pttg1ip5 | 4.43E-21 | 0.432768603 | 0.535 | 0.268 | 8.96E-17 | 14 | Pttg1ip |
| Rtn45 | 4.68E-21 | 0.452912627 | 0.611 | 0.338 | 9.48E-17 | 14 | Rtn4 |
| Trak11 | 5.07E-21 | 0.469169521 | 0.309 | 0.126 | 1.03E-16 | 14 | Trak1 |
| Nenf2 | 5.67E-21 | 0.332480783 | 0.564 | 0.295 | 1.15E-16 | 14 | Nenf |
| Gm121852 | 6.45E-21 | 0.482624058 | 0.324 | 0.133 | 1.31E-16 | 14 | Gm12185 |
| Spata61 | 6.70E-21 | 0.541208552 | 0.298 | 0.12 | 1.36E-16 | 14 | Spata6 |
| Mfsd62 | 7.35E-21 | 0.443959406 | 0.349 | 0.144 | 1.49E-16 | 14 | Mfsd6 |

| | | | | | | | |
|---|---|---|---|---|---|---|---|
| Phlpp13 | 8.24E-21 | 0.547925036 | 0.513 | 0.263 | 1.67E-16 | 14 | Phlpp1 |
| Erc12 | 8.25E-21 | 0.469342948 | 0.309 | 0.123 | 1.67E-16 | 14 | Erc1 |
| Dusp162 | 8.30E-21 | 0.496607521 | 0.298 | 0.119 | 1.68E-16 | 14 | Dusp16 |
| Prickle24 | 9.90E-21 | 0.453602805 | 0.342 | 0.139 | 2.00E-16 | 14 | Prickle2 |
| Psmd73 | 1.18E-20 | 0.427489363 | 0.56 | 0.305 | 2.40E-16 | 14 | Psmd7 |
| Tcaf12 | 1.38E-20 | 0.474265638 | 0.276 | 0.103 | 2.79E-16 | 14 | Tcaf1 |
| Cpne83 | 1.58E-20 | 0.331068804 | 0.495 | 0.228 | 3.20E-16 | 14 | Cpne8 |
| Gbp52 | 1.77E-20 | 0.413877835 | 0.32 | 0.128 | 3.59E-16 | 14 | Gbp5 |
| Senp65 | 1.87E-20 | 0.483609399 | 0.542 | 0.298 | 3.79E-16 | 14 | Senp6 |
| Hdac73 | 2.20E-20 | 0.429099521 | 0.436 | 0.202 | 4.45E-16 | 14 | Hdac7 |
| Trim30a6 | 2.24E-20 | 0.628290976 | 0.444 | 0.219 | 4.53E-16 | 14 | Trim30a |
| Ywhah4 | 2.45E-20 | 0.41816411 | 0.564 | 0.309 | 4.96E-16 | 14 | Ywhah |
| Csgalnact14 | 2.65E-20 | 0.565763449 | 0.425 | 0.195 | 5.37E-16 | 14 | Csgalnact1 |
| Pik3c2a3 | 2.82E-20 | 0.485792583 | 0.411 | 0.195 | 5.71E-16 | 14 | Pik3c2a |
| Rhob4 | 3.19E-20 | 0.45054178 | 0.52 | 0.267 | 6.46E-16 | 14 | Rhob |
| Oaz23 | 3.41E-20 | 0.345896126 | 0.498 | 0.236 | 6.91E-16 | 14 | Oaz2 |
| Add13 | 3.45E-20 | 0.458084997 | 0.433 | 0.214 | 6.98E-16 | 14 | Add1 |
| Kif5b3 | 3.47E-20 | 0.424966487 | 0.669 | 0.406 | 7.03E-16 | 14 | Kif5b |
| Sec625 | 4.09E-20 | 0.470371202 | 0.651 | 0.394 | 8.28E-16 | 14 | Sec62 |
| Ubxn44 | 4.16E-20 | 0.506310052 | 0.505 | 0.272 | 8.41E-16 | 14 | Ubxn4 |
| Atp11c3 | 4.46E-20 | 0.586270258 | 0.345 | 0.151 | 9.03E-16 | 14 | Atp11c |
| Il10rb4 | 4.82E-20 | 0.437963944 | 0.484 | 0.248 | 9.77E-16 | 14 | Il10rb |
| Zbtb7a3 | 5.15E-20 | 0.467571702 | 0.451 | 0.229 | 1.04E-15 | 14 | Zbtb7a |
| Pcdh173 | 6.02E-20 | 0.430407744 | 0.509 | 0.239 | 1.22E-15 | 14 | Pcdh17 |
| Cntln3 | 7.50E-20 | 0.46696591 | 0.353 | 0.154 | 1.52E-15 | 14 | Cntln |
| Rsu14 | 7.83E-20 | 0.415465589 | 0.495 | 0.253 | 1.59E-15 | 14 | Rsu1 |
| Adam104 | 8.93E-20 | 0.54129439 | 0.524 | 0.292 | 1.81E-15 | 14 | Adam10 |
| Frmd4a2 | 9.04E-20 | 0.401655149 | 0.342 | 0.148 | 1.83E-15 | 14 | Frmd4a |
| Usp347 | 1.02E-19 | 0.514633549 | 0.6 | 0.344 | 2.07E-15 | 14 | Usp34 |
| Cnih45 | 1.18E-19 | 0.377184138 | 0.491 | 0.253 | 2.39E-15 | 14 | Cnih4 |
| Fyn7 | 1.20E-19 | 0.432692745 | 0.524 | 0.266 | 2.42E-15 | 14 | Fyn |
| Nr3c14 | 1.29E-19 | 0.44646299 | 0.673 | 0.382 | 2.62E-15 | 14 | Nr3c1 |
| mt-Nd4l6 | 1.30E-19 | 0.522210127 | 0.767 | 0.491 | 2.64E-15 | 14 | mt-Nd4l |
| Zhx34 | 1.32E-19 | 0.515726899 | 0.429 | 0.202 | 2.66E-15 | 14 | Zhx3 |
| Arl2bp | 1.33E-19 | 0.320828287 | 0.345 | 0.15 | 2.69E-15 | 14 | Arl2bp |
| Ston12 | 1.39E-19 | 0.334420913 | 0.251 | 0.09 | 2.81E-15 | 14 | Ston1 |
| Akap97 | 1.44E-19 | 0.544400302 | 0.465 | 0.237 | 2.92E-15 | 14 | Akap9 |
| 1810037I17Rik2 | 1.46E-19 | 0.371935102 | 0.749 | 0.479 | 2.95E-15 | 14 | 1810037I17Rik |
| Myh97 | 1.52E-19 | 0.55062678 | 0.716 | 0.436 | 3.08E-15 | 14 | Myh9 |
| Phf11d2 | 1.79E-19 | 0.43656435 | 0.327 | 0.137 | 3.63E-15 | 14 | Phf11d |
| Fam13b5 | 1.86E-19 | 0.635055183 | 0.327 | 0.147 | 3.76E-15 | 14 | Fam13b |
| Gnpda21 | 1.95E-19 | 0.381164225 | 0.298 | 0.119 | 3.96E-15 | 14 | Gnpda2 |
| Ankrd128 | 2.07E-19 | 0.600424985 | 0.545 | 0.308 | 4.20E-15 | 14 | Ankrd12 |

| | | | | | | | |
|---|---|---|---|---|---|---|---|
| Glg14 | 2.17E-19 | 0.532030854 | 0.484 | 0.252 | 4.38E-15 | 14 | Glg1 |
| Klf94 | 2.36E-19 | 0.363604116 | 0.484 | 0.237 | 4.78E-15 | 14 | Klf9 |
| Grk54 | 2.45E-19 | 0.588761435 | 0.353 | 0.158 | 4.96E-15 | 14 | Grk5 |
| Plekha12 | 2.47E-19 | 0.441406102 | 0.324 | 0.138 | 5.00E-15 | 14 | Plekha1 |
| Wdr263 | 2.61E-19 | 0.467830736 | 0.422 | 0.209 | 5.29E-15 | 14 | Wdr26 |
| Fndc3b4 | 2.66E-19 | 0.463762791 | 0.48 | 0.244 | 5.38E-15 | 14 | Fndc3b |
| Camk1d1 | 3.14E-19 | 0.502259774 | 0.96 | 0.823 | 6.37E-15 | 14 | Camk1d |
| Sod14 | 3.37E-19 | 0.40263614 | 0.556 | 0.299 | 6.83E-15 | 14 | Sod1 |
| Tm9sf24 | 3.44E-19 | 0.530022879 | 0.484 | 0.255 | 6.96E-15 | 14 | Tm9sf2 |
| Ddah23 | 3.93E-19 | 0.363908474 | 0.436 | 0.205 | 7.96E-15 | 14 | Ddah2 |
| Calr5 | 4.14E-19 | 0.488520224 | 0.658 | 0.408 | 8.38E-15 | 14 | Calr |
| Lamp17 | 4.24E-19 | 0.496106433 | 0.684 | 0.417 | 8.59E-15 | 14 | Lamp1 |
| 2610035D17Rik2 | 4.46E-19 | 0.530353475 | 0.284 | 0.114 | 9.04E-15 | 14 | 2610035D17Rik |
| Calu4 | 5.39E-19 | 0.388765598 | 0.429 | 0.202 | 1.09E-14 | 14 | Calu |
| Ghr3 | 5.58E-19 | 0.513288247 | 0.345 | 0.151 | 1.13E-14 | 14 | Ghr |
| Thra3 | 5.97E-19 | 0.34847267 | 0.371 | 0.164 | 1.21E-14 | 14 | Thra |
| Lman14 | 6.32E-19 | 0.504767505 | 0.462 | 0.238 | 1.28E-14 | 14 | Lman1 |
| Git24 | 7.04E-19 | 0.50670398 | 0.458 | 0.235 | 1.43E-14 | 14 | Git2 |
| Tcf154 | 7.62E-19 | 0.263859653 | 0.535 | 0.259 | 1.54E-14 | 14 | Tcf15 |
| Grrp13 | 8.01E-19 | 0.353869107 | 0.465 | 0.219 | 1.62E-14 | 14 | Grrp1 |
| Rbm396 | 8.74E-19 | 0.445241035 | 0.898 | 0.573 | 1.77E-14 | 14 | Rbm39 |
| Golgb14 | 8.93E-19 | 0.540346917 | 0.484 | 0.262 | 1.81E-14 | 14 | Golgb1 |
| Tspan54 | 9.39E-19 | 0.490088001 | 0.415 | 0.198 | 1.90E-14 | 14 | Tspan5 |
| Ppp1r12c | 9.46E-19 | 0.358487575 | 0.262 | 0.1 | 1.91E-14 | 14 | Ppp1r12c |
| Myadm3 | 1.11E-18 | 0.440907705 | 0.291 | 0.12 | 2.24E-14 | 14 | Myadm |
| Srgn8 | 1.38E-18 | 0.396753706 | 0.92 | 0.543 | 2.79E-14 | 14 | Srgn |
| Sep-78 | 1.51E-18 | 0.389269302 | 0.749 | 0.461 | 3.06E-14 | 14 | 7-Sep |
| Usp13 | 1.85E-18 | 0.394643283 | 0.331 | 0.146 | 3.75E-14 | 14 | Usp1 |
| Fzd43 | 1.89E-18 | 0.366200985 | 0.476 | 0.229 | 3.82E-14 | 14 | Fzd4 |
| Ptpn111 | 2.26E-18 | 0.358802254 | 0.335 | 0.149 | 4.57E-14 | 14 | Ptpn11 |
| Socs52 | 2.34E-18 | 0.385146419 | 0.305 | 0.127 | 4.75E-14 | 14 | Socs5 |
| Xist8 | 2.45E-18 | 0.424312045 | 0.811 | 0.462 | 4.96E-14 | 14 | Xist |
| Ski6 | 3.17E-18 | 0.483270598 | 0.422 | 0.215 | 6.42E-14 | 14 | Ski |
| Adipor13 | 3.26E-18 | 0.447573423 | 0.484 | 0.257 | 6.59E-14 | 14 | Adipor1 |
| Slc48a1 | 3.45E-18 | 0.330170015 | 0.276 | 0.11 | 6.98E-14 | 14 | Slc48a1 |
| Csnk1e1 | 3.47E-18 | 0.393475302 | 0.313 | 0.134 | 7.03E-14 | 14 | Csnk1e |
| Rrbp15 | 3.49E-18 | 0.343875246 | 0.771 | 0.459 | 7.06E-14 | 14 | Rrbp1 |
| Tap24 | 3.62E-18 | 0.505154861 | 0.458 | 0.241 | 7.33E-14 | 14 | Tap2 |
| S100a104 | 3.80E-18 | 0.307506914 | 0.807 | 0.468 | 7.69E-14 | 14 | S100a10 |
| Stag18 | 3.80E-18 | 0.525762159 | 0.604 | 0.361 | 7.69E-14 | 14 | Stag1 |
| Slk5 | 3.95E-18 | 0.450728081 | 0.462 | 0.241 | 7.99E-14 | 14 | Slk |
| Ppme1 | 4.26E-18 | 0.342083488 | 0.291 | 0.12 | 8.63E-14 | 14 | Ppme1 |
| Pbx12 | 4.55E-18 | 0.46127969 | 0.418 | 0.198 | 9.20E-14 | 14 | Pbx1 |

| | | | | | | | |
|---|---|---|---|---|---|---|---|
| Bcl7c | 5.65E-18 | 0.319746517 | 0.469 | 0.243 | 1.14E-13 | 14 | Bcl7c |
| Pros13 | 6.59E-18 | 0.45878953 | 0.338 | 0.15 | 1.33E-13 | 14 | Pros1 |
| Gstm14 | 6.86E-18 | 0.253541638 | 0.589 | 0.322 | 1.39E-13 | 14 | Gstm1 |
| Pkd11 | 7.95E-18 | 0.419919132 | 0.251 | 0.097 | 1.61E-13 | 14 | Pkd1 |
| Bcap315 | 9.01E-18 | 0.458536257 | 0.513 | 0.285 | 1.82E-13 | 14 | Bcap31 |
| Canx6 | 9.41E-18 | 0.462734457 | 0.542 | 0.306 | 1.91E-13 | 14 | Canx |
| Spen2 | 1.02E-17 | 0.415433727 | 0.345 | 0.158 | 2.06E-13 | 14 | Spen |
| Ppib2 | 1.09E-17 | 0.379458062 | 0.825 | 0.553 | 2.21E-13 | 14 | Ppib |
| Gm424182 | 1.19E-17 | 0.405253879 | 1 | 1 | 2.42E-13 | 14 | Gm42418 |
| Atxn17 | 1.24E-17 | 0.465848306 | 0.545 | 0.295 | 2.51E-13 | 14 | Atxn1 |
| Fut83 | 1.25E-17 | 0.518685501 | 0.531 | 0.289 | 2.53E-13 | 14 | Fut8 |
| Hist1h1c | 1.32E-17 | 0.402609614 | 0.404 | 0.201 | 2.66E-13 | 14 | Hist1h1c |
| Fbxo116 | 1.35E-17 | 0.453310957 | 0.447 | 0.233 | 2.73E-13 | 14 | Fbxo11 |
| Tpm36 | 1.41E-17 | 0.43431114 | 0.738 | 0.479 | 2.86E-13 | 14 | Tpm3 |
| Ddx62 | 1.45E-17 | 0.363065578 | 0.589 | 0.341 | 2.94E-13 | 14 | Ddx6 |
| Dctn2 | 1.45E-17 | 0.350821218 | 0.487 | 0.265 | 2.94E-13 | 14 | Dctn2 |
| Mfhas12 | 1.56E-17 | 0.428584527 | 0.335 | 0.15 | 3.15E-13 | 14 | Mfhas1 |
| Ripor13 | 1.61E-17 | 0.368827934 | 0.356 | 0.16 | 3.27E-13 | 14 | Ripor1 |
| Prrg12 | 1.63E-17 | 0.469644234 | 0.269 | 0.108 | 3.29E-13 | 14 | Prrg1 |
| Setx3 | 1.88E-17 | 0.456018312 | 0.364 | 0.174 | 3.80E-13 | 14 | Setx |
| Tor1aip15 | 1.99E-17 | 0.410883458 | 0.513 | 0.281 | 4.03E-13 | 14 | Tor1aip1 |
| Xaf13 | 1.99E-17 | 0.36609423 | 0.44 | 0.221 | 4.03E-13 | 14 | Xaf1 |
| Cast5 | 2.02E-17 | 0.473750157 | 0.535 | 0.312 | 4.09E-13 | 14 | Cast |
| Irf26 | 2.26E-17 | 0.430275343 | 0.585 | 0.341 | 4.58E-13 | 14 | Irf2 |
| Phactr26 | 2.32E-17 | 0.393912319 | 0.469 | 0.246 | 4.69E-13 | 14 | Phactr2 |
| Ctnnbip13 | 2.60E-17 | 0.398945953 | 0.418 | 0.204 | 5.26E-13 | 14 | Ctnnbip1 |
| Ppp1cb1 | 2.64E-17 | 0.315655658 | 0.502 | 0.273 | 5.34E-13 | 14 | Ppp1cb |
| Arhgap104 | 2.71E-17 | 0.432749008 | 0.295 | 0.127 | 5.49E-13 | 14 | Arhgap10 |
| Rps6ka37 | 3.20E-17 | 0.508011562 | 0.476 | 0.263 | 6.48E-13 | 14 | Rps6ka3 |
| Tuba1a5 | 3.51E-17 | 0.388099683 | 0.495 | 0.256 | 7.10E-13 | 14 | Tuba1a |
| Mia33 | 3.63E-17 | 0.455039802 | 0.356 | 0.17 | 7.35E-13 | 14 | Mia3 |
| Taok14 | 3.76E-17 | 0.467539325 | 0.469 | 0.251 | 7.62E-13 | 14 | Taok1 |
| Dip2b5 | 4.19E-17 | 0.417756629 | 0.509 | 0.276 | 8.49E-13 | 14 | Dip2b |
| Nsd16 | 4.88E-17 | 0.393034401 | 0.6 | 0.349 | 9.87E-13 | 14 | Nsd1 |
| B230219D22Rik3 | 5.07E-17 | 0.399977814 | 0.589 | 0.342 | 1.03E-12 | 14 | B230219D22Rik |
| Clcn33 | 5.15E-17 | 0.450899211 | 0.302 | 0.134 | 1.04E-12 | 14 | Clcn3 |
| Nisch4 | 5.18E-17 | 0.416745758 | 0.418 | 0.211 | 1.05E-12 | 14 | Nisch |
| Evi52 | 5.29E-17 | 0.323914118 | 0.32 | 0.141 | 1.07E-12 | 14 | Evi5 |
| Rela | 5.33E-17 | 0.391156851 | 0.265 | 0.111 | 1.08E-12 | 14 | Rela |
| Hspa1a1 | 6.45E-17 | 0.335395929 | 0.258 | 0.101 | 1.31E-12 | 14 | Hspa1a |
| Lrp61 | 6.73E-17 | 0.43561759 | 0.32 | 0.144 | 1.36E-12 | 14 | Lrp6 |
| Ifi2037 | 6.89E-17 | 0.401996149 | 0.622 | 0.367 | 1.40E-12 | 14 | Ifi203 |
| Myl66 | 9.16E-17 | 0.371808308 | 0.964 | 0.775 | 1.86E-12 | 14 | Myl6 |

| | | | | | | | |
|---|---|---|---|---|---|---|---|
| Rnpepl1 | 9.76E-17 | 0.343321249 | 0.255 | 0.102 | 1.98E-12 | 14 | Rnpepl1 |
| Rspry11 | 1.13E-16 | 0.432394325 | 0.269 | 0.114 | 2.30E-12 | 14 | Rspry1 |
| Bnip3l1 | 1.15E-16 | 0.337961891 | 0.567 | 0.332 | 2.33E-12 | 14 | Bnip3l |
| Parp144 | 1.26E-16 | 0.408267563 | 0.473 | 0.252 | 2.54E-12 | 14 | Parp14 |
| mt-Nd19 | 1.31E-16 | 0.389211243 | 0.909 | 0.598 | 2.65E-12 | 14 | mt-Nd1 |
| Tnfrsf1b4 | 1.59E-16 | 0.39877522 | 0.273 | 0.117 | 3.22E-12 | 14 | Tnfrsf1b |
| Eea14 | 1.59E-16 | 0.378994702 | 0.422 | 0.218 | 3.22E-12 | 14 | Eea1 |
| Tmbim65 | 1.60E-16 | 0.432008047 | 0.665 | 0.431 | 3.24E-12 | 14 | Tmbim6 |
| Pkig1 | 2.17E-16 | 0.304478789 | 0.418 | 0.211 | 4.39E-12 | 14 | Pkig |
| Mesd2 | 2.24E-16 | 0.374479632 | 0.345 | 0.163 | 4.54E-12 | 14 | Mesd |
| Mtch12 | 2.29E-16 | 0.28420041 | 0.56 | 0.316 | 4.64E-12 | 14 | Mtch1 |
| Ddx55 | 2.35E-16 | 0.43658772 | 0.927 | 0.653 | 4.76E-12 | 14 | Ddx5 |
| Tgfbr26 | 2.35E-16 | 0.413887777 | 0.545 | 0.311 | 4.77E-12 | 14 | Tgfbr2 |
| Sri4 | 2.77E-16 | 0.320864697 | 0.673 | 0.42 | 5.60E-12 | 14 | Sri |
| Tmem245 | 2.91E-16 | 0.414145829 | 0.276 | 0.118 | 5.90E-12 | 14 | Tmem245 |
| Elovl52 | 3.14E-16 | 0.389515186 | 0.327 | 0.151 | 6.36E-12 | 14 | Elovl5 |
| Fam69a3 | 3.17E-16 | 0.461199289 | 0.287 | 0.127 | 6.42E-12 | 14 | Fam69a |
| Picalm4 | 3.35E-16 | 0.412064161 | 0.524 | 0.296 | 6.77E-12 | 14 | Picalm |
| Atp6ap12 | 3.47E-16 | 0.37056041 | 0.324 | 0.149 | 7.02E-12 | 14 | Atp6ap1 |
| Dctn11 | 3.49E-16 | 0.314502391 | 0.298 | 0.131 | 7.07E-12 | 14 | Dctn1 |
| Jup3 | 3.97E-16 | 0.286151328 | 0.498 | 0.257 | 8.03E-12 | 14 | Jup |
| Lgals94 | 4.51E-16 | 0.467803732 | 0.455 | 0.247 | 9.13E-12 | 14 | Lgals9 |
| Sppl2a4 | 5.20E-16 | 0.42371721 | 0.516 | 0.296 | 1.05E-11 | 14 | Sppl2a |
| Tmtc13 | 6.00E-16 | 0.394725533 | 0.32 | 0.142 | 1.21E-11 | 14 | Tmtc1 |
| Fcho23 | 6.53E-16 | 0.406096979 | 0.542 | 0.309 | 1.32E-11 | 14 | Fcho2 |
| Robo43 | 7.16E-16 | 0.281959159 | 0.382 | 0.176 | 1.45E-11 | 14 | Robo4 |
| Zbtb43 | 7.25E-16 | 0.407005333 | 0.276 | 0.121 | 1.47E-11 | 14 | Zbtb4 |
| Ptp4a31 | 8.12E-16 | 0.387186831 | 0.302 | 0.137 | 1.64E-11 | 14 | Ptp4a3 |
| Eif3c | 8.20E-16 | 0.368265223 | 0.618 | 0.387 | 1.66E-11 | 14 | Eif3c |
| Sel1l3 | 8.69E-16 | 0.451502983 | 0.335 | 0.162 | 1.76E-11 | 14 | Sel1l |
| Kmt2e6 | 8.90E-16 | 0.400614766 | 0.658 | 0.406 | 1.80E-11 | 14 | Kmt2e |
| Exoc6b4 | 9.27E-16 | 0.571448914 | 0.433 | 0.241 | 1.88E-11 | 14 | Exoc6b |
| Nek75 | 1.01E-15 | 0.419909261 | 0.455 | 0.248 | 2.05E-11 | 14 | Nek7 |
| Gbp23 | 1.03E-15 | 0.383982237 | 0.316 | 0.143 | 2.08E-11 | 14 | Gbp2 |
| Ubr53 | 1.12E-15 | 0.398460514 | 0.385 | 0.198 | 2.27E-11 | 14 | Ubr5 |
| Plxnd13 | 1.21E-15 | 0.382121927 | 0.375 | 0.181 | 2.45E-11 | 14 | Plxnd1 |
| Srp141 | 1.23E-15 | 0.276490093 | 0.709 | 0.461 | 2.49E-11 | 14 | Srp14 |
| Ctnnd12 | 1.41E-15 | 0.304035002 | 0.32 | 0.145 | 2.86E-11 | 14 | Ctnnd1 |
| Nptn6 | 1.45E-15 | 0.51273083 | 0.444 | 0.249 | 2.93E-11 | 14 | Nptn |
| Mfge84 | 1.54E-15 | 0.276840461 | 0.491 | 0.254 | 3.12E-11 | 14 | Mfge8 |
| Stim23 | 1.59E-15 | 0.459591506 | 0.385 | 0.199 | 3.22E-11 | 14 | Stim2 |
| Pqlc12 | 1.64E-15 | 0.327972011 | 0.291 | 0.126 | 3.32E-11 | 14 | Pqlc1 |
| mt-Nd27 | 1.72E-15 | 0.394066838 | 0.967 | 0.687 | 3.48E-11 | 14 | mt-Nd2 |

| | | | | | | | |
|---|---|---|---|---|---|---|---|
| Tpr6 | 1.80E-15 | 0.448576259 | 0.651 | 0.425 | 3.65E-11 | 14 | Tpr |
| Tubb54 | 1.80E-15 | 0.318117475 | 0.622 | 0.38 | 3.65E-11 | 14 | Tubb5 |
| Nras2 | 1.83E-15 | 0.344784889 | 0.324 | 0.15 | 3.70E-11 | 14 | Nras |
| Qser1 | 2.23E-15 | 0.3071925 | 0.255 | 0.104 | 4.51E-11 | 14 | Qser1 |
| Gjc12 | 2.27E-15 | 0.357293127 | 0.273 | 0.115 | 4.60E-11 | 14 | Gjc1 |
| Sh3bgrl5 | 2.48E-15 | 0.294908352 | 0.633 | 0.375 | 5.02E-11 | 14 | Sh3bgrl |
| Tbl1x4 | 2.50E-15 | 0.399938903 | 0.473 | 0.262 | 5.05E-11 | 14 | Tbl1x |
| Phf20l16 | 2.51E-15 | 0.468774131 | 0.491 | 0.287 | 5.08E-11 | 14 | Phf20l1 |
| Mxd45 | 2.78E-15 | 0.369722732 | 0.593 | 0.352 | 5.63E-11 | 14 | Mxd4 |
| Wdfy35 | 3.66E-15 | 0.416215547 | 0.367 | 0.182 | 7.42E-11 | 14 | Wdfy3 |
| Snx94 | 3.75E-15 | 0.318698053 | 0.367 | 0.176 | 7.59E-11 | 14 | Snx9 |
| Tpp12 | 3.82E-15 | 0.393915327 | 0.273 | 0.119 | 7.74E-11 | 14 | Tpp1 |
| Zc3h134 | 4.07E-15 | 0.473777892 | 0.4 | 0.216 | 8.24E-11 | 14 | Zc3h13 |
| Ptprj5 | 4.28E-15 | 0.43212199 | 0.324 | 0.151 | 8.66E-11 | 14 | Ptprj |
| Eif4g13 | 4.55E-15 | 0.391014475 | 0.433 | 0.236 | 9.22E-11 | 14 | Eif4g1 |
| Nfat55 | 4.62E-15 | 0.494392966 | 0.473 | 0.267 | 9.35E-11 | 14 | Nfat5 |
| Cdc42bpb3 | 4.75E-15 | 0.310422677 | 0.382 | 0.186 | 9.62E-11 | 14 | Cdc42bpb |
| Ubn26 | 4.99E-15 | 0.431560611 | 0.451 | 0.244 | 1.01E-10 | 14 | Ubn2 |
| Agpat33 | 5.22E-15 | 0.379232896 | 0.284 | 0.129 | 1.06E-10 | 14 | Agpat3 |
| Ptpn123 | 7.56E-15 | 0.438453387 | 0.404 | 0.213 | 1.53E-10 | 14 | Ptpn12 |
| Atp6v0e1 | 7.98E-15 | 0.301985658 | 0.673 | 0.425 | 1.62E-10 | 14 | Atp6v0e |
| Clock4 | 8.14E-15 | 0.414805221 | 0.389 | 0.204 | 1.65E-10 | 14 | Clock |
| Klc12 | 8.22E-15 | 0.416905976 | 0.316 | 0.149 | 1.66E-10 | 14 | Klc1 |
| Sptlc24 | 8.75E-15 | 0.458839096 | 0.44 | 0.245 | 1.77E-10 | 14 | Sptlc2 |
| Dip2c4 | 9.18E-15 | 0.388278784 | 0.433 | 0.223 | 1.86E-10 | 14 | Dip2c |
| Sp1104 | 9.52E-15 | 0.394216217 | 0.324 | 0.157 | 1.93E-10 | 14 | Sp110 |
| Cask3 | 1.10E-14 | 0.391832448 | 0.298 | 0.139 | 2.23E-10 | 14 | Cask |
| Nipbl9 | 1.25E-14 | 0.389328345 | 0.596 | 0.366 | 2.54E-10 | 14 | Nipbl |
| Grn4 | 1.38E-14 | 0.367342037 | 0.425 | 0.224 | 2.79E-10 | 14 | Grn |
| Akt1 | 1.44E-14 | 0.299433621 | 0.349 | 0.173 | 2.91E-10 | 14 | Akt1 |
| Dpp82 | 1.52E-14 | 0.405669006 | 0.36 | 0.184 | 3.07E-10 | 14 | Dpp8 |
| Cyfip14 | 1.58E-14 | 0.372147442 | 0.491 | 0.278 | 3.19E-10 | 14 | Cyfip1 |
| Zfp6446 | 1.63E-14 | 0.442481264 | 0.451 | 0.256 | 3.30E-10 | 14 | Zfp644 |
| Gls6 | 1.78E-14 | 0.427675579 | 0.447 | 0.251 | 3.60E-10 | 14 | Gls |
| Nfic2 | 1.78E-14 | 0.263223179 | 0.48 | 0.261 | 3.60E-10 | 14 | Nfic |
| Acot9 | 1.86E-14 | 0.359644867 | 0.269 | 0.119 | 3.77E-10 | 14 | Acot9 |
| Piezo13 | 2.20E-14 | 0.455246863 | 0.349 | 0.174 | 4.46E-10 | 14 | Piezo1 |
| Zmym42 | 2.23E-14 | 0.373648591 | 0.335 | 0.165 | 4.52E-10 | 14 | Zmym4 |
| Prpf38b7 | 2.31E-14 | 0.437999207 | 0.375 | 0.196 | 4.68E-10 | 14 | Prpf38b |
| Elk42 | 2.36E-14 | 0.361055953 | 0.338 | 0.167 | 4.78E-10 | 14 | Elk4 |
| Tuba1b3 | 2.44E-14 | 0.295364561 | 0.549 | 0.322 | 4.94E-10 | 14 | Tuba1b |
| Csde13 | 2.46E-14 | 0.328565004 | 0.578 | 0.36 | 4.99E-10 | 14 | Csde1 |
| Zeb16 | 2.56E-14 | 0.479887047 | 0.542 | 0.322 | 5.18E-10 | 14 | Zeb1 |

| | | | | | | | |
|---|---|---|---|---|---|---|---|
| Mum1 | 2.76E-14 | 0.337852414 | 0.258 | 0.114 | 5.59E-10 | 14 | Mum1 |
| Peak17 | 2.85E-14 | 0.404012808 | 0.625 | 0.387 | 5.76E-10 | 14 | Peak1 |
| Lats22 | 3.45E-14 | 0.340456662 | 0.273 | 0.12 | 6.99E-10 | 14 | Lats2 |
| Unc503 | 3.47E-14 | 0.317805389 | 0.4 | 0.213 | 7.02E-10 | 14 | Unc50 |
| Tmx34 | 3.70E-14 | 0.453311368 | 0.36 | 0.187 | 7.49E-10 | 14 | Tmx3 |
| Bptf8 | 3.94E-14 | 0.369477992 | 0.513 | 0.294 | 7.97E-10 | 14 | Bptf |
| P4hb3 | 5.10E-14 | 0.367349829 | 0.447 | 0.252 | 1.03E-09 | 14 | P4hb |
| Ahctf11 | 5.66E-14 | 0.426820478 | 0.269 | 0.124 | 1.15E-09 | 14 | Ahctf1 |
| Phc32 | 5.69E-14 | 0.391401722 | 0.287 | 0.135 | 1.15E-09 | 14 | Phc3 |
| Zfr5 | 5.94E-14 | 0.444697789 | 0.44 | 0.248 | 1.20E-09 | 14 | Zfr |
| Tmem1654 | 6.36E-14 | 0.4332864 | 0.349 | 0.18 | 1.29E-09 | 14 | Tmem165 |
| Ttc7b3 | 6.40E-14 | 0.332539516 | 0.269 | 0.122 | 1.30E-09 | 14 | Ttc7b |
| Psmd11 | 7.21E-14 | 0.384889423 | 0.389 | 0.21 | 1.46E-09 | 14 | Psmd1 |
| Qk4 | 7.33E-14 | 0.314687647 | 0.684 | 0.41 | 1.48E-09 | 14 | Qk |
| Emc33 | 8.19E-14 | 0.415350069 | 0.338 | 0.173 | 1.66E-09 | 14 | Emc3 |
| Marf1 | 8.57E-14 | 0.370314372 | 0.305 | 0.147 | 1.74E-09 | 14 | Marf1 |
| Spop1 | 1.10E-13 | 0.324088878 | 0.593 | 0.381 | 2.23E-09 | 14 | Spop |
| Zbtb463 | 1.17E-13 | 0.366737002 | 0.335 | 0.163 | 2.37E-09 | 14 | Zbtb46 |
| Sash14 | 1.20E-13 | 0.317430422 | 0.476 | 0.257 | 2.43E-09 | 14 | Sash1 |
| Huwe17 | 1.23E-13 | 0.451411457 | 0.476 | 0.285 | 2.49E-09 | 14 | Huwe1 |
| Reep34 | 1.24E-13 | 0.357316413 | 0.531 | 0.306 | 2.52E-09 | 14 | Reep3 |
| Pnisr7 | 1.28E-13 | 0.44567622 | 0.451 | 0.265 | 2.59E-09 | 14 | Pnisr |
| Zc2hc1a | 1.31E-13 | 0.32522707 | 0.251 | 0.11 | 2.65E-09 | 14 | Zc2hc1a |
| Pla2g164 | 1.34E-13 | 0.36510613 | 0.429 | 0.243 | 2.70E-09 | 14 | Pla2g16 |
| Ncor17 | 1.67E-13 | 0.420724337 | 0.673 | 0.438 | 3.38E-09 | 14 | Ncor1 |
| Prickle14 | 1.81E-13 | 0.326943503 | 0.313 | 0.147 | 3.66E-09 | 14 | Prickle1 |
| Cers51 | 1.83E-13 | 0.40555451 | 0.28 | 0.134 | 3.70E-09 | 14 | Cers5 |
| Gpr1462 | 1.85E-13 | 0.334302961 | 0.251 | 0.109 | 3.74E-09 | 14 | Gpr146 |
| Phf21a7 | 1.95E-13 | 0.537208465 | 0.44 | 0.253 | 3.94E-09 | 14 | Phf21a |
| Prdm22 | 2.01E-13 | 0.366535981 | 0.353 | 0.185 | 4.08E-09 | 14 | Prdm2 |
| Psma71 | 2.06E-13 | 0.250167472 | 0.796 | 0.577 | 4.16E-09 | 14 | Psma7 |
| Smurf24 | 2.07E-13 | 0.413142798 | 0.425 | 0.239 | 4.19E-09 | 14 | Smurf2 |
| Tcf128 | 2.12E-13 | 0.412130862 | 0.607 | 0.376 | 4.30E-09 | 14 | Tcf12 |
| Yes13 | 2.19E-13 | 0.34479249 | 0.32 | 0.154 | 4.43E-09 | 14 | Yes1 |
| Ryk1 | 2.38E-13 | 0.323603687 | 0.251 | 0.11 | 4.81E-09 | 14 | Ryk |
| Ext14 | 2.39E-13 | 0.479489127 | 0.331 | 0.167 | 4.84E-09 | 14 | Ext1 |
| Slc44a22 | 2.64E-13 | 0.386734639 | 0.262 | 0.122 | 5.34E-09 | 14 | Slc44a2 |
| Gphn1 | 2.76E-13 | 0.283132979 | 0.938 | 0.792 | 5.58E-09 | 14 | Gphn |
| Arhgef110 | 3.15E-13 | 0.438695553 | 0.429 | 0.249 | 6.37E-09 | 14 | Arhgef1 |
| Srsf118 | 3.22E-13 | 0.347327605 | 0.564 | 0.351 | 6.52E-09 | 14 | Srsf11 |
| Zfp6083 | 3.23E-13 | 0.406292371 | 0.295 | 0.141 | 6.55E-09 | 14 | Zfp608 |
| Vps13c2 | 3.47E-13 | 0.379360888 | 0.287 | 0.14 | 7.03E-09 | 14 | Vps13c |
| C1galt1 | 3.60E-13 | 0.268883607 | 0.291 | 0.138 | 7.30E-09 | 14 | C1galt1 |

| | | | | | | | |
|---|---|---|---|---|---|---|---|
| Snx41 | 3.73E-13 | 0.313973672 | 0.276 | 0.13 | 7.55E-09 | 14 | Snx4 |
| Ilk | 3.83E-13 | 0.354453744 | 0.331 | 0.171 | 7.75E-09 | 14 | Ilk |
| Dpm2 | 3.88E-13 | 0.361944223 | 0.291 | 0.143 | 7.85E-09 | 14 | Dpm2 |
| Nae11 | 4.01E-13 | 0.331453908 | 0.313 | 0.156 | 8.13E-09 | 14 | Nae1 |
| Nudcd34 | 4.12E-13 | 0.366330316 | 0.429 | 0.247 | 8.33E-09 | 14 | Nudcd3 |
| Grb104 | 4.20E-13 | 0.318784938 | 0.382 | 0.195 | 8.51E-09 | 14 | Grb10 |
| B4galt43 | 4.75E-13 | 0.255188634 | 0.338 | 0.162 | 9.61E-09 | 14 | B4galt4 |
| Rabep14 | 5.52E-13 | 0.34915665 | 0.411 | 0.231 | 1.12E-08 | 14 | Rabep1 |
| Sat12 | 6.12E-13 | 0.354309712 | 0.556 | 0.346 | 1.24E-08 | 14 | Sat1 |
| Rcn23 | 6.28E-13 | 0.346829763 | 0.295 | 0.143 | 1.27E-08 | 14 | Rcn2 |
| Ppp2r2d1 | 7.18E-13 | 0.299163932 | 0.305 | 0.151 | 1.45E-08 | 14 | Ppp2r2d |
| Kif1b4 | 7.75E-13 | 0.360945062 | 0.429 | 0.243 | 1.57E-08 | 14 | Kif1b |
| Specc1l2 | 7.97E-13 | 0.311053341 | 0.273 | 0.129 | 1.61E-08 | 14 | Specc1l |
| Tgoln14 | 8.27E-13 | 0.422462451 | 0.433 | 0.247 | 1.67E-08 | 14 | Tgoln1 |
| Dnaja14 | 8.34E-13 | 0.277996299 | 0.76 | 0.516 | 1.69E-08 | 14 | Dnaja1 |
| Tnfaip22 | 9.00E-13 | 0.389206999 | 0.309 | 0.152 | 1.82E-08 | 14 | Tnfaip2 |
| Jade11 | 9.32E-13 | 0.354668173 | 0.251 | 0.115 | 1.89E-08 | 14 | Jade1 |
| Tmbim12 | 1.04E-12 | 0.339631443 | 0.265 | 0.123 | 2.11E-08 | 14 | Tmbim1 |
| Rbms23 | 1.18E-12 | 0.275693295 | 0.32 | 0.16 | 2.39E-08 | 14 | Rbms2 |
| Selenow3 | 1.19E-12 | 0.284776517 | 0.851 | 0.629 | 2.40E-08 | 14 | Selenow |
| Dync1li2 | 1.28E-12 | 0.278980742 | 0.28 | 0.133 | 2.59E-08 | 14 | Dync1li2 |
| Pld12 | 1.32E-12 | 0.386702856 | 0.295 | 0.148 | 2.68E-08 | 14 | Pld1 |
| Dnajc110 | 1.37E-12 | 0.333480644 | 0.549 | 0.323 | 2.78E-08 | 14 | Dnajc1 |
| Kdm6a7 | 1.43E-12 | 0.418943259 | 0.382 | 0.214 | 2.90E-08 | 14 | Kdm6a |
| Tnrc184 | 1.46E-12 | 0.34958486 | 0.335 | 0.173 | 2.96E-08 | 14 | Tnrc18 |
| Ywhab3 | 1.48E-12 | 0.379620684 | 0.676 | 0.467 | 3.00E-08 | 14 | Ywhab |
| Diaph14 | 1.65E-12 | 0.407696472 | 0.422 | 0.243 | 3.34E-08 | 14 | Diaph1 |
| Cdc42ep12 | 1.71E-12 | 0.250891018 | 0.28 | 0.129 | 3.46E-08 | 14 | Cdc42ep1 |
| Stard93 | 1.71E-12 | 0.340274851 | 0.309 | 0.152 | 3.47E-08 | 14 | Stard9 |
| Sbno13 | 1.72E-12 | 0.340795685 | 0.378 | 0.209 | 3.48E-08 | 14 | Sbno1 |
| Rictor4 | 1.91E-12 | 0.411479382 | 0.305 | 0.157 | 3.86E-08 | 14 | Rictor |
| Zfp1485 | 1.91E-12 | 0.373722062 | 0.451 | 0.265 | 3.88E-08 | 14 | Zfp148 |
| Rap1gds18 | 2.23E-12 | 0.444673854 | 0.44 | 0.261 | 4.52E-08 | 14 | Rap1gds1 |
| Lamc13 | 2.27E-12 | 0.3254013 | 0.305 | 0.148 | 4.59E-08 | 14 | Lamc1 |
| Cdk9 | 2.29E-12 | 0.28313023 | 0.28 | 0.136 | 4.63E-08 | 14 | Cdk9 |
| Rin24 | 2.78E-12 | 0.304915742 | 0.353 | 0.182 | 5.63E-08 | 14 | Rin2 |
| Ckap51 | 2.80E-12 | 0.293196747 | 0.291 | 0.142 | 5.66E-08 | 14 | Ckap5 |
| Ppp2r2a3 | 3.32E-12 | 0.399205352 | 0.382 | 0.213 | 6.72E-08 | 14 | Ppp2r2a |
| Gpatch87 | 3.36E-12 | 0.451107528 | 0.44 | 0.258 | 6.81E-08 | 14 | Gpatch8 |
| Tm9sf35 | 3.53E-12 | 0.264675296 | 0.538 | 0.335 | 7.16E-08 | 14 | Tm9sf3 |
| Gbp43 | 3.64E-12 | 0.334717582 | 0.342 | 0.178 | 7.38E-08 | 14 | Gbp4 |
| Abi16 | 3.75E-12 | 0.317592559 | 0.458 | 0.27 | 7.59E-08 | 14 | Abi1 |
| Cds23 | 3.81E-12 | 0.330964122 | 0.342 | 0.177 | 7.70E-08 | 14 | Cds2 |

| | | | | | | | |
|---|---|---|---|---|---|---|---|
| Smchd15 | 3.98E-12 | 0.363743876 | 0.444 | 0.264 | 8.06E-08 | 14 | Smchd1 |
| Unc45b3 | 4.12E-12 | 0.337628741 | 0.291 | 0.14 | 8.34E-08 | 14 | Unc45b |
| Lgals8 | 4.31E-12 | 0.272785859 | 0.305 | 0.153 | 8.72E-08 | 14 | Lgals8 |
| Selenok4 | 4.63E-12 | 0.267148061 | 0.782 | 0.554 | 9.38E-08 | 14 | Selenok |
| Kat6a3 | 4.71E-12 | 0.340156144 | 0.404 | 0.228 | 9.54E-08 | 14 | Kat6a |
| Nrd11 | 4.71E-12 | 0.422670836 | 0.371 | 0.208 | 9.54E-08 | 14 | Nrd1 |
| Mkln12 | 4.79E-12 | 0.391164848 | 0.44 | 0.265 | 9.70E-08 | 14 | Mkln1 |
| Apc4 | 5.16E-12 | 0.394409099 | 0.422 | 0.249 | 1.04E-07 | 14 | Apc |
| Cdc37l1 | 5.70E-12 | 0.317267872 | 0.324 | 0.168 | 1.15E-07 | 14 | Cdc37l1 |
| Tmem2053 | 5.72E-12 | 0.274821036 | 0.316 | 0.159 | 1.16E-07 | 14 | Tmem205 |
| Tmed96 | 6.08E-12 | 0.350356635 | 0.6 | 0.404 | 1.23E-07 | 14 | Tmed9 |
| Stx122 | 6.10E-12 | 0.296289575 | 0.316 | 0.162 | 1.24E-07 | 14 | Stx12 |
| Lama44 | 6.32E-12 | 0.379698997 | 0.338 | 0.175 | 1.28E-07 | 14 | Lama4 |
| Tnfaip83 | 6.41E-12 | 0.342136658 | 0.349 | 0.187 | 1.30E-07 | 14 | Tnfaip8 |
| Bach25 | 6.43E-12 | 0.419693563 | 0.313 | 0.161 | 1.30E-07 | 14 | Bach2 |
| Emc4 | 6.53E-12 | 0.270408392 | 0.265 | 0.127 | 1.32E-07 | 14 | Emc4 |
| Lmbrd13 | 6.67E-12 | 0.343349762 | 0.316 | 0.163 | 1.35E-07 | 14 | Lmbrd1 |
| Flna3 | 7.51E-12 | 0.324567102 | 0.269 | 0.132 | 1.52E-07 | 14 | Flna |
| Thap32 | 8.26E-12 | 0.349843114 | 0.313 | 0.162 | 1.67E-07 | 14 | Thap3 |
| St3gal14 | 9.01E-12 | 0.338176289 | 0.338 | 0.181 | 1.82E-07 | 14 | St3gal1 |
| Lrch14 | 9.99E-12 | 0.333972277 | 0.513 | 0.312 | 2.02E-07 | 14 | Lrch1 |
| Trpm75 | 1.21E-11 | 0.427910086 | 0.36 | 0.205 | 2.44E-07 | 14 | Trpm7 |
| Dicer1 | 1.22E-11 | 0.324090522 | 0.255 | 0.123 | 2.47E-07 | 14 | Dicer1 |
| Clasp1 | 1.22E-11 | 0.343389062 | 0.269 | 0.132 | 2.47E-07 | 14 | Clasp1 |
| Lmna4 | 1.25E-11 | 0.345550777 | 0.364 | 0.198 | 2.53E-07 | 14 | Lmna |
| Cradd | 1.37E-11 | 0.287739472 | 0.251 | 0.118 | 2.77E-07 | 14 | Cradd |
| Tmed23 | 1.41E-11 | 0.259942697 | 0.575 | 0.366 | 2.86E-07 | 14 | Tmed2 |
| Wac6 | 1.43E-11 | 0.2903804 | 0.524 | 0.323 | 2.89E-07 | 14 | Wac |
| Esf1 | 1.77E-11 | 0.359849411 | 0.313 | 0.165 | 3.58E-07 | 14 | Esf1 |
| Rmnd5a | 1.84E-11 | 0.291302837 | 0.276 | 0.138 | 3.72E-07 | 14 | Rmnd5a |
| BC005561 | 2.13E-11 | 0.31684128 | 0.251 | 0.119 | 4.32E-07 | 14 | BC005561 |
| Sertad22 | 2.24E-11 | 0.375550423 | 0.316 | 0.169 | 4.54E-07 | 14 | Sertad2 |
| Zcchc63 | 2.63E-11 | 0.315440867 | 0.458 | 0.279 | 5.33E-07 | 14 | Zcchc6 |
| Atraid4 | 2.66E-11 | 0.2877928 | 0.455 | 0.271 | 5.38E-07 | 14 | Atraid |
| Med13l3 | 2.78E-11 | 0.364329471 | 0.349 | 0.194 | 5.62E-07 | 14 | Med13l |
| Vcp3 | 3.06E-11 | 0.282989837 | 0.527 | 0.344 | 6.20E-07 | 14 | Vcp |
| Chst116 | 3.10E-11 | 0.428772658 | 0.327 | 0.176 | 6.27E-07 | 14 | Chst11 |
| Larp4b2 | 3.11E-11 | 0.361956029 | 0.36 | 0.204 | 6.29E-07 | 14 | Larp4b |
| Arhgap266 | 3.61E-11 | 0.321292709 | 0.396 | 0.223 | 7.31E-07 | 14 | Arhgap26 |
| Elovl1 | 3.72E-11 | 0.289808804 | 0.251 | 0.12 | 7.53E-07 | 14 | Elovl1 |
| Zscan262 | 3.80E-11 | 0.337347476 | 0.284 | 0.145 | 7.69E-07 | 14 | Zscan26 |
| Fbxw71 | 4.25E-11 | 0.336427699 | 0.265 | 0.132 | 8.60E-07 | 14 | Fbxw7 |
| Rpn23 | 4.27E-11 | 0.344640407 | 0.371 | 0.212 | 8.65E-07 | 14 | Rpn2 |

| | | | | | | | |
|---|---|---|---|---|---|---|---|
| Ccdc473 | 4.43E-11 | 0.336983375 | 0.345 | 0.193 | 8.97E-07 | 14 | Ccdc47 |
| Tmem2083 | 4.45E-11 | 0.34157209 | 0.404 | 0.239 | 9.01E-07 | 14 | Tmem208 |
| Supt62 | 4.64E-11 | 0.306734812 | 0.305 | 0.16 | 9.40E-07 | 14 | Supt6 |
| Flii2 | 4.78E-11 | 0.337930426 | 0.298 | 0.158 | 9.68E-07 | 14 | Flii |
| Vcl2 | 4.80E-11 | 0.297653092 | 0.313 | 0.162 | 9.72E-07 | 14 | Vcl |
| Zmynd114 | 5.00E-11 | 0.312718429 | 0.473 | 0.289 | 1.01E-06 | 14 | Zmynd11 |
| Fam193a4 | 5.08E-11 | 0.364797408 | 0.324 | 0.18 | 1.03E-06 | 14 | Fam193a |
| Sbf24 | 5.23E-11 | 0.346682973 | 0.375 | 0.208 | 1.06E-06 | 14 | Sbf2 |
| Stx6 | 5.37E-11 | 0.332743104 | 0.251 | 0.122 | 1.09E-06 | 14 | Stx6 |
| Pkn24 | 5.64E-11 | 0.281649858 | 0.476 | 0.295 | 1.14E-06 | 14 | Pkn2 |
| Ttc147 | 5.84E-11 | 0.339962408 | 0.378 | 0.216 | 1.18E-06 | 14 | Ttc14 |
| Dap1 | 6.40E-11 | 0.340282364 | 0.335 | 0.187 | 1.29E-06 | 14 | Dap |
| Hnrnpc | 7.55E-11 | 0.252722642 | 0.535 | 0.349 | 1.53E-06 | 14 | Hnrnpc |
| Birc68 | 7.96E-11 | 0.309209826 | 0.545 | 0.341 | 1.61E-06 | 14 | Birc6 |
| Mat2a2 | 8.47E-11 | 0.250158325 | 0.509 | 0.309 | 1.71E-06 | 14 | Mat2a |
| Crebbp7 | 9.20E-11 | 0.32936366 | 0.491 | 0.305 | 1.86E-06 | 14 | Crebbp |
| Ifnar24 | 9.22E-11 | 0.336863065 | 0.516 | 0.325 | 1.87E-06 | 14 | Ifnar2 |
| Rsf14 | 9.23E-11 | 0.358760987 | 0.385 | 0.225 | 1.87E-06 | 14 | Rsf1 |
| Ppig4 | 9.41E-11 | 0.364339947 | 0.487 | 0.317 | 1.90E-06 | 14 | Ppig |
| Pea15a4 | 9.53E-11 | 0.255176004 | 0.418 | 0.241 | 1.93E-06 | 14 | Pea15a |
| Lap31 | 9.77E-11 | 0.281303626 | 0.273 | 0.137 | 1.98E-06 | 14 | Lap3 |
| Ephb43 | 1.03E-10 | 0.265638885 | 0.324 | 0.167 | 2.08E-06 | 14 | Ephb4 |
| Os96 | 1.07E-10 | 0.327532002 | 0.338 | 0.189 | 2.18E-06 | 14 | Os9 |
| Znfx13 | 1.08E-10 | 0.336965179 | 0.324 | 0.175 | 2.19E-06 | 14 | Znfx1 |
| Smad42 | 1.11E-10 | 0.298620892 | 0.385 | 0.223 | 2.24E-06 | 14 | Smad4 |
| Kidins2204 | 1.27E-10 | 0.395882911 | 0.295 | 0.156 | 2.56E-06 | 14 | Kidins220 |
| Mphosph82 | 1.30E-10 | 0.347579135 | 0.356 | 0.207 | 2.63E-06 | 14 | Mphosph8 |
| Fam172a7 | 1.33E-10 | 0.363825213 | 0.469 | 0.285 | 2.70E-06 | 14 | Fam172a |
| Mycbp28 | 1.33E-10 | 0.347732534 | 0.582 | 0.375 | 2.70E-06 | 14 | Mycbp2 |
| Tm9sf41 | 1.35E-10 | 0.383138599 | 0.258 | 0.132 | 2.73E-06 | 14 | Tm9sf4 |
| Lbr6 | 1.36E-10 | 0.371091286 | 0.287 | 0.153 | 2.76E-06 | 14 | Lbr |
| Xiap3 | 1.43E-10 | 0.316013687 | 0.48 | 0.3 | 2.89E-06 | 14 | Xiap |
| Thoc26 | 1.44E-10 | 0.357035982 | 0.444 | 0.274 | 2.91E-06 | 14 | Thoc2 |
| Ergic22 | 1.53E-10 | 0.334787388 | 0.324 | 0.18 | 3.10E-06 | 14 | Ergic2 |
| Ndel12 | 1.56E-10 | 0.363107061 | 0.313 | 0.175 | 3.17E-06 | 14 | Ndel1 |
| Nr2c25 | 1.58E-10 | 0.326330137 | 0.298 | 0.159 | 3.20E-06 | 14 | Nr2c2 |
| Scaf112 | 1.61E-10 | 0.348127225 | 0.425 | 0.264 | 3.25E-06 | 14 | Scaf11 |
| Tom1l22 | 1.66E-10 | 0.320461067 | 0.262 | 0.131 | 3.37E-06 | 14 | Tom1l2 |
| Gch11 | 1.68E-10 | 0.320081884 | 0.262 | 0.131 | 3.41E-06 | 14 | Gch1 |
| Ppp3ca8 | 1.76E-10 | 0.399789906 | 0.596 | 0.394 | 3.56E-06 | 14 | Ppp3ca |
| Adgrg13 | 1.77E-10 | 0.285190819 | 0.28 | 0.14 | 3.58E-06 | 14 | Adgrg1 |
| Sae1 | 1.85E-10 | 0.278201536 | 0.284 | 0.148 | 3.74E-06 | 14 | Sae1 |
| Sltm5 | 1.87E-10 | 0.329729062 | 0.433 | 0.271 | 3.78E-06 | 14 | Sltm |

| | | | | | | | |
|---|---|---|---|---|---|---|---|
| Prrc2c8 | 2.24E-10 | 0.3419875 | 0.622 | 0.421 | 4.53E-06 | 14 | Prrc2c |
| Arl6ip56 | 2.34E-10 | 0.340954468 | 0.378 | 0.222 | 4.74E-06 | 14 | Arl6ip5 |
| Tmem1281 | 2.45E-10 | 0.34196845 | 0.302 | 0.166 | 4.96E-06 | 14 | Tmem128 |
| Ifih13 | 2.52E-10 | 0.311355013 | 0.258 | 0.129 | 5.10E-06 | 14 | Ifih1 |
| Stk381 | 2.69E-10 | 0.293213036 | 0.385 | 0.227 | 5.45E-06 | 14 | Stk38 |
| Ptbp38 | 2.84E-10 | 0.349047428 | 0.593 | 0.394 | 5.74E-06 | 14 | Ptbp3 |
| Sos23 | 2.93E-10 | 0.301476364 | 0.302 | 0.162 | 5.93E-06 | 14 | Sos2 |
| Zfand37 | 3.29E-10 | 0.345110706 | 0.476 | 0.302 | 6.66E-06 | 14 | Zfand3 |
| Kmt5b3 | 3.52E-10 | 0.27508916 | 0.313 | 0.171 | 7.12E-06 | 14 | Kmt5b |
| Trim252 | 3.94E-10 | 0.261992654 | 0.287 | 0.15 | 7.97E-06 | 14 | Trim25 |
| Cflar3 | 3.98E-10 | 0.298821943 | 0.389 | 0.232 | 8.05E-06 | 14 | Cflar |
| Fam120a3 | 4.13E-10 | 0.349341507 | 0.324 | 0.182 | 8.36E-06 | 14 | Fam120a |
| Tnfsf123 | 4.39E-10 | 0.263760492 | 0.284 | 0.148 | 8.88E-06 | 14 | Tnfsf12 |
| Airn5 | 4.68E-10 | 0.308967186 | 0.418 | 0.249 | 9.47E-06 | 14 | Airn |
| Lrch36 | 4.85E-10 | 0.357669963 | 0.389 | 0.233 | 9.82E-06 | 14 | Lrch3 |
| Emc75 | 4.93E-10 | 0.291935254 | 0.436 | 0.272 | 9.97E-06 | 14 | Emc7 |
| Dab25 | 5.12E-10 | 0.275627509 | 0.371 | 0.207 | 1.04E-05 | 14 | Dab2 |
| Neat16 | 5.36E-10 | 0.42206695 | 0.356 | 0.21 | 1.08E-05 | 14 | Neat1 |
| Srgap27 | 6.27E-10 | 0.301711011 | 0.487 | 0.304 | 1.27E-05 | 14 | Srgap2 |
| Stat24 | 6.62E-10 | 0.299131943 | 0.273 | 0.141 | 1.34E-05 | 14 | Stat2 |
| Keap11 | 6.68E-10 | 0.259758457 | 0.269 | 0.139 | 1.35E-05 | 14 | Keap1 |
| Dnajb111 | 7.12E-10 | 0.32716929 | 0.364 | 0.214 | 1.44E-05 | 14 | Dnajb11 |
| Pkn15 | 8.18E-10 | 0.312306826 | 0.331 | 0.187 | 1.66E-05 | 14 | Pkn1 |
| Phldb23 | 8.35E-10 | 0.293237777 | 0.364 | 0.203 | 1.69E-05 | 14 | Phldb2 |
| Leprot2 | 8.48E-10 | 0.279240721 | 0.32 | 0.178 | 1.72E-05 | 14 | Leprot |
| Rassf14 | 9.19E-10 | 0.340882487 | 0.378 | 0.229 | 1.86E-05 | 14 | Rassf1 |
| Lpp7 | 9.27E-10 | 0.332551043 | 0.455 | 0.286 | 1.88E-05 | 14 | Lpp |
| Ext22 | 1.05E-09 | 0.336355965 | 0.269 | 0.144 | 2.12E-05 | 14 | Ext2 |
| Slc44a13 | 1.07E-09 | 0.325636221 | 0.291 | 0.157 | 2.16E-05 | 14 | Slc44a1 |
| Ap2a21 | 1.11E-09 | 0.29877158 | 0.28 | 0.15 | 2.26E-05 | 14 | Ap2a2 |
| Snrnp482 | 1.14E-09 | 0.298282324 | 0.327 | 0.186 | 2.30E-05 | 14 | Snrnp48 |
| Luc7l27 | 1.26E-09 | 0.288492249 | 0.607 | 0.405 | 2.56E-05 | 14 | Luc7l2 |
| Sos13 | 1.29E-09 | 0.320889263 | 0.324 | 0.181 | 2.61E-05 | 14 | Sos1 |
| Sec24b1 | 1.33E-09 | 0.320242311 | 0.262 | 0.139 | 2.69E-05 | 14 | Sec24b |
| Ifnar15 | 1.38E-09 | 0.392526887 | 0.265 | 0.145 | 2.79E-05 | 14 | Ifnar1 |
| Cul11 | 1.42E-09 | 0.274350403 | 0.436 | 0.277 | 2.87E-05 | 14 | Cul1 |
| Pbrm13 | 1.54E-09 | 0.332913575 | 0.447 | 0.288 | 3.13E-05 | 14 | Pbrm1 |
| Pum23 | 1.56E-09 | 0.332500144 | 0.422 | 0.27 | 3.15E-05 | 14 | Pum2 |
| Cwf19l21 | 1.61E-09 | 0.317916068 | 0.349 | 0.202 | 3.26E-05 | 14 | Cwf19l2 |
| Kmt2a8 | 1.69E-09 | 0.371436196 | 0.425 | 0.268 | 3.42E-05 | 14 | Kmt2a |
| Man1a7 | 1.70E-09 | 0.292030644 | 0.302 | 0.168 | 3.44E-05 | 14 | Man1a |
| Ccnl23 | 1.77E-09 | 0.338398702 | 0.356 | 0.215 | 3.57E-05 | 14 | Ccnl2 |
| Ankrd118 | 1.86E-09 | 0.338256699 | 0.582 | 0.381 | 3.76E-05 | 14 | Ankrd11 |

| | | | | | | | |
|---|---|---|---|---|---|---|---|
| Casp84 | 1.93E-09 | 0.255589621 | 0.364 | 0.215 | 3.90E-05 | 14 | Casp8 |
| Yif1b1 | 1.95E-09 | 0.250432002 | 0.276 | 0.149 | 3.95E-05 | 14 | Yif1b |
| Pdia42 | 1.99E-09 | 0.327595734 | 0.298 | 0.167 | 4.03E-05 | 14 | Pdia4 |
| Stat37 | 2.11E-09 | 0.294997652 | 0.429 | 0.267 | 4.27E-05 | 14 | Stat3 |
| Phactr43 | 2.39E-09 | 0.28185147 | 0.316 | 0.178 | 4.83E-05 | 14 | Phactr4 |
| Atp9b4 | 2.56E-09 | 0.276467727 | 0.287 | 0.158 | 5.19E-05 | 14 | Atp9b |
| Zdhhc207 | 2.60E-09 | 0.385963716 | 0.375 | 0.231 | 5.25E-05 | 14 | Zdhhc20 |
| 4930523C07Rik6 | 2.65E-09 | 0.287217745 | 0.371 | 0.22 | 5.36E-05 | 14 | 4930523C07Rik |
| Tcerg11 | 3.22E-09 | 0.266799058 | 0.258 | 0.137 | 6.52E-05 | 14 | Tcerg1 |
| Golga41 | 3.92E-09 | 0.274388249 | 0.287 | 0.159 | 7.93E-05 | 14 | Golga4 |
| Adam172 | 4.31E-09 | 0.283986388 | 0.28 | 0.154 | 8.72E-05 | 14 | Adam17 |
| Mbd56 | 4.69E-09 | 0.415132983 | 0.364 | 0.221 | 9.50E-05 | 14 | Mbd5 |
| Aak15 | 5.60E-09 | 0.299610327 | 0.324 | 0.185 | 0.000113439 | 14 | Aak1 |
| Kansl18 | 6.21E-09 | 0.314566581 | 0.524 | 0.34 | 0.000125764 | 14 | Kansl1 |
| Esyt14 | 7.67E-09 | 0.275022327 | 0.255 | 0.137 | 0.000155245 | 14 | Esyt1 |
| Mllt107 | 8.31E-09 | 0.346204285 | 0.429 | 0.275 | 0.000168325 | 14 | Mllt10 |
| Txndc152 | 8.40E-09 | 0.348103881 | 0.255 | 0.139 | 0.000169999 | 14 | Txndc15 |
| Chd65 | 1.07E-08 | 0.31431439 | 0.356 | 0.218 | 0.000216493 | 14 | Chd6 |
| 22-Mar | 1.11E-08 | 0.27358846 | 0.331 | 0.197 | 0.000223818 | 14 | 2-Mar |
| Magt13 | 1.20E-08 | 0.310099463 | 0.287 | 0.165 | 0.000243596 | 14 | Magt1 |
| Mtdh4 | 1.29E-08 | 0.29611837 | 0.56 | 0.387 | 0.000260737 | 14 | Mtdh |
| Dhx363 | 1.41E-08 | 0.283153983 | 0.378 | 0.236 | 0.000286257 | 14 | Dhx36 |
| Pdzd81 | 1.48E-08 | 0.272417912 | 0.255 | 0.137 | 0.000299928 | 14 | Pdzd8 |
| Nrip15 | 1.48E-08 | 0.274621046 | 0.396 | 0.244 | 0.000300553 | 14 | Nrip1 |
| Emc103 | 1.52E-08 | 0.297350705 | 0.335 | 0.201 | 0.000308746 | 14 | Emc10 |
| Fam208a2 | 1.66E-08 | 0.287389584 | 0.338 | 0.2 | 0.000336607 | 14 | Fam208a |
| Large13 | 1.68E-08 | 0.279250535 | 0.276 | 0.153 | 0.000339501 | 14 | Large1 |
| Erap11 | 1.68E-08 | 0.308418986 | 0.255 | 0.14 | 0.000340046 | 14 | Erap1 |
| Rnf19a2 | 1.71E-08 | 0.254274206 | 0.258 | 0.14 | 0.000345712 | 14 | Rnf19a |
| Hmg20a2 | 1.72E-08 | 0.343125979 | 0.258 | 0.143 | 0.000348448 | 14 | Hmg20a |
| Sec633 | 1.79E-08 | 0.29478867 | 0.345 | 0.204 | 0.000361588 | 14 | Sec63 |
| Ep3005 | 1.80E-08 | 0.296148192 | 0.338 | 0.205 | 0.000365433 | 14 | Ep300 |
| Prpf40a6 | 1.83E-08 | 0.304943712 | 0.415 | 0.268 | 0.000369565 | 14 | Prpf40a |
| Srek12 | 1.99E-08 | 0.319699396 | 0.324 | 0.194 | 0.000403395 | 14 | Srek1 |
| Dennd6a2 | 2.31E-08 | 0.298901475 | 0.313 | 0.183 | 0.000467332 | 14 | Dennd6a |
| Ssr11 | 2.84E-08 | 0.317796947 | 0.313 | 0.187 | 0.000575164 | 14 | Ssr1 |
| Foxo13 | 2.93E-08 | 0.266434912 | 0.338 | 0.2 | 0.000594156 | 14 | Foxo1 |
| Ambra16 | 3.08E-08 | 0.332364139 | 0.327 | 0.199 | 0.000624086 | 14 | Ambra1 |
| Akap137 | 3.20E-08 | 0.27505972 | 0.585 | 0.394 | 0.000647885 | 14 | Akap13 |
| Irf13 | 3.21E-08 | 0.292576222 | 0.455 | 0.299 | 0.000649353 | 14 | Irf1 |
| Tra2a8 | 3.64E-08 | 0.354501837 | 0.422 | 0.275 | 0.000737112 | 14 | Tra2a |
| Uhrf22 | 3.75E-08 | 0.357557331 | 0.284 | 0.167 | 0.000758924 | 14 | Uhrf2 |
| Crtc35 | 3.78E-08 | 0.350481452 | 0.284 | 0.165 | 0.000765419 | 14 | Crtc3 |

| | | | | | | | |
|---|---|---|---|---|---|---|---|
| Rc3h12 | 3.81E-08 | 0.340810591 | 0.335 | 0.209 | 0.000771752 | 14 | Rc3h1 |
| Fam3c2 | 3.91E-08 | 0.261594895 | 0.305 | 0.18 | 0.000791312 | 14 | Fam3c |
| Pon21 | 3.99E-08 | 0.250396504 | 0.298 | 0.172 | 0.000807943 | 14 | Pon2 |
| Dhrs74 | 4.50E-08 | 0.268309472 | 0.349 | 0.216 | 0.000911742 | 14 | Dhrs7 |
| Dstyk2 | 4.54E-08 | 0.269862862 | 0.269 | 0.15 | 0.000919557 | 14 | Dstyk |
| Clasp24 | 4.81E-08 | 0.382953895 | 0.305 | 0.183 | 0.000974018 | 14 | Clasp2 |
| Chd12 | 5.21E-08 | 0.277736512 | 0.331 | 0.202 | 0.001055612 | 14 | Chd1 |
| Dpysl24 | 5.36E-08 | 0.265024952 | 0.298 | 0.175 | 0.00108469 | 14 | Dpysl2 |
| Anapc53 | 5.43E-08 | 0.255492958 | 0.342 | 0.212 | 0.001098706 | 14 | Anapc5 |
| Brd23 | 5.63E-08 | 0.32569762 | 0.385 | 0.254 | 0.001138916 | 14 | Brd2 |
| Ssu723 | 5.72E-08 | 0.250754733 | 0.462 | 0.312 | 0.001157099 | 14 | Ssu72 |
| Mark33 | 5.74E-08 | 0.255171669 | 0.345 | 0.213 | 0.001161235 | 14 | Mark3 |
| Lgals3bp3 | 7.31E-08 | 0.310182251 | 0.258 | 0.148 | 0.001480305 | 14 | Lgals3bp |
| Cpsf61 | 7.44E-08 | 0.275645185 | 0.265 | 0.152 | 0.00150604 | 14 | Cpsf6 |
| Srpr4 | 8.31E-08 | 0.286232802 | 0.385 | 0.247 | 0.001682189 | 14 | Srpr |
| Papola2 | 9.53E-08 | 0.260957173 | 0.447 | 0.305 | 0.001929301 | 14 | Papola |
| Ranbp23 | 9.85E-08 | 0.305725207 | 0.324 | 0.199 | 0.001994334 | 14 | Ranbp2 |
| Ranbp95 | 1.29E-07 | 0.35280534 | 0.32 | 0.201 | 0.002617812 | 14 | Ranbp9 |
| Ets16 | 1.31E-07 | 0.252395779 | 0.585 | 0.404 | 0.002645745 | 14 | Ets1 |
| Phip7 | 1.52E-07 | 0.263194947 | 0.4 | 0.257 | 0.003068801 | 14 | Phip |
| Samd9l4 | 1.58E-07 | 0.281714617 | 0.396 | 0.257 | 0.003197362 | 14 | Samd9l |
| Baz1b3 | 1.82E-07 | 0.254926136 | 0.316 | 0.194 | 0.003687833 | 14 | Baz1b |
| Dnajb92 | 1.96E-07 | 0.319845107 | 0.251 | 0.144 | 0.003961709 | 14 | Dnajb9 |
| Prrc2b5 | 2.37E-07 | 0.286683209 | 0.36 | 0.23 | 0.004799969 | 14 | Prrc2b |
| Srsf41 | 2.38E-07 | 0.257840488 | 0.273 | 0.161 | 0.004812412 | 14 | Srsf4 |
| Setd57 | 2.65E-07 | 0.333274973 | 0.4 | 0.265 | 0.005362265 | 14 | Setd5 |
| Btbd98 | 2.80E-07 | 0.403748842 | 0.364 | 0.238 | 0.005673799 | 14 | Btbd9 |
| Zcchc78 | 3.11E-07 | 0.3984241 | 0.44 | 0.302 | 0.006304756 | 14 | Zcchc7 |
| Sorbs13 | 3.24E-07 | 0.255730461 | 0.302 | 0.178 | 0.006566787 | 14 | Sorbs1 |
| Rbm258 | 3.32E-07 | 0.25773517 | 0.484 | 0.335 | 0.006713846 | 14 | Rbm25 |
| Fto2 | 3.36E-07 | 0.270631453 | 0.313 | 0.192 | 0.006794345 | 14 | Fto |
| Zfp4075 | 3.63E-07 | 0.354994906 | 0.269 | 0.162 | 0.007354137 | 14 | Zfp407 |
| Ccdc88a7 | 3.68E-07 | 0.308804524 | 0.331 | 0.205 | 0.007444449 | 14 | Ccdc88a |
| Ddost2 | 3.86E-07 | 0.26723954 | 0.342 | 0.217 | 0.007805723 | 14 | Ddost |
| N4bp2l24 | 3.93E-07 | 0.274459695 | 0.378 | 0.248 | 0.007962653 | 14 | N4bp2l2 |
| Cep3506 | 4.04E-07 | 0.253243657 | 0.327 | 0.204 | 0.008178871 | 14 | Cep350 |
| Stk34 | 5.16E-07 | 0.361646911 | 0.276 | 0.167 | 0.010442535 | 14 | Stk3 |
| Ino80d2 | 5.18E-07 | 0.273344879 | 0.276 | 0.166 | 0.010487707 | 14 | Ino80d |
| Mar-71 | 6.03E-07 | 0.250701972 | 0.298 | 0.184 | 0.012209433 | 14 | 7-Mar |
| Zfp2927 | 6.10E-07 | 0.256910916 | 0.382 | 0.252 | 0.012350373 | 14 | Zfp292 |
| Fgfr22 | 6.83E-07 | 0.291011146 | 0.258 | 0.153 | 0.013833827 | 14 | Fgfr2 |
| Exoc23 | 7.31E-07 | 0.2700469 | 0.251 | 0.148 | 0.014805282 | 14 | Exoc2 |
| Ascc36 | 7.61E-07 | 0.332156711 | 0.324 | 0.208 | 0.015397125 | 14 | Ascc3 |

| | | | | | | | |
|---|---|---|---|---|---|---|---|
| Gon4l3 | 8.05E-07 | 0.30200734 | 0.251 | 0.15 | 0.016291253 | 14 | Gon4l |
| Fubp14 | 8.84E-07 | 0.279003273 | 0.4 | 0.273 | 0.017906013 | 14 | Fubp1 |
| Vti1a6 | 9.46E-07 | 0.275564309 | 0.385 | 0.256 | 0.019147033 | 14 | Vti1a |
| R3hdm11 | 9.46E-07 | 0.250017639 | 0.302 | 0.192 | 0.019159182 | 14 | R3hdm1 |
| Mon22 | 1.03E-06 | 0.310702571 | 0.255 | 0.153 | 0.020754856 | 14 | Mon2 |
| AW5549184 | 1.06E-06 | 0.329838971 | 0.276 | 0.172 | 0.021485365 | 14 | AW554918 |
| Sppl37 | 1.15E-06 | 0.273703726 | 0.335 | 0.219 | 0.023305655 | 14 | Sppl3 |
| Ap3b15 | 1.20E-06 | 0.25046795 | 0.335 | 0.218 | 0.024323279 | 14 | Ap3b1 |
| Uvrag5 | 1.25E-06 | 0.267071371 | 0.396 | 0.269 | 0.025312776 | 14 | Uvrag |
| Vps13a4 | 1.34E-06 | 0.29608269 | 0.28 | 0.173 | 0.027101449 | 14 | Vps13a |
| Fbxl33 | 1.35E-06 | 0.252794401 | 0.262 | 0.157 | 0.027251496 | 14 | Fbxl3 |
| Nfatc34 | 1.45E-06 | 0.276396496 | 0.284 | 0.176 | 0.029287 | 14 | Nfatc3 |
| Ppp6r33 | 1.46E-06 | 0.321991298 | 0.302 | 0.192 | 0.029624612 | 14 | Ppp6r3 |
| Bdp12 | 1.55E-06 | 0.268274396 | 0.273 | 0.166 | 0.031311236 | 14 | Bdp1 |
| Cdk126 | 1.69E-06 | 0.293391144 | 0.345 | 0.231 | 0.034216909 | 14 | Cdk12 |
| Scaf86 | 1.71E-06 | 0.379632275 | 0.291 | 0.186 | 0.034579595 | 14 | Scaf8 |
| Mga3 | 2.03E-06 | 0.273620989 | 0.276 | 0.172 | 0.041120833 | 14 | Mga |
| Atxn73 | 2.14E-06 | 0.294582732 | 0.269 | 0.167 | 0.043317976 | 14 | Atxn7 |
| Ankhd15 | 2.16E-06 | 0.253569458 | 0.36 | 0.245 | 0.043746867 | 14 | Ankhd1 |
| Numa13 | 2.40E-06 | 0.253359567 | 0.305 | 0.196 | 0.04855494 | 14 | Numa1 |
| Lnpep8 | 2.99E-06 | 0.259628459 | 0.385 | 0.261 | 0.060545432 | 14 | Lnpep |
| Dgkh3 | 3.55E-06 | 0.252383522 | 0.287 | 0.178 | 0.071922419 | 14 | Dgkh |
| Ubr33 | 3.70E-06 | 0.269675996 | 0.309 | 0.202 | 0.074909788 | 14 | Ubr3 |
| Scaper6 | 3.94E-06 | 0.263424643 | 0.287 | 0.18 | 0.079719799 | 14 | Scaper |
| Bclaf15 | 5.43E-06 | 0.269370618 | 0.371 | 0.255 | 0.109976107 | 14 | Bclaf1 |
| Mgat22 | 6.43E-06 | 0.257820484 | 0.251 | 0.156 | 0.130213513 | 14 | Mgat2 |
| Nemf | 7.66E-06 | 0.251909959 | 0.273 | 0.175 | 0.155150193 | 14 | Nemf |
| Man1a24 | 7.89E-06 | 0.305906837 | 0.338 | 0.228 | 0.159747286 | 14 | Man1a2 |
| Phf37 | 8.97E-06 | 0.27496846 | 0.364 | 0.254 | 0.181553243 | 14 | Phf3 |
| Fmnl34 | 9.11E-06 | 0.261571181 | 0.356 | 0.241 | 0.184396157 | 14 | Fmnl3 |
| Gigyf24 | 1.23E-05 | 0.270444682 | 0.262 | 0.169 | 0.249052379 | 14 | Gigyf2 |
| Map3k33 | 1.60E-05 | 0.25981909 | 0.276 | 0.179 | 0.324895329 | 14 | Map3k3 |
| Snrnp703 | 1.61E-05 | 0.253905941 | 0.389 | 0.274 | 0.326854372 | 14 | Snrnp70 |
| Ccar14 | 1.85E-05 | 0.288207966 | 0.305 | 0.209 | 0.374298488 | 14 | Ccar1 |
| Pan38 | 1.91E-05 | 0.303515042 | 0.335 | 0.23 | 0.38763023 | 14 | Pan3 |
| Smg18 | 2.00E-05 | 0.250282299 | 0.342 | 0.236 | 0.404877446 | 14 | Smg1 |
| Fbxl179 | 2.21E-05 | 0.329024469 | 0.429 | 0.312 | 0.447137499 | 14 | Fbxl17 |
| Ncoa16 | 4.56E-05 | 0.260606635 | 0.305 | 0.211 | 0.922545495 | 14 | Ncoa1 |
| B4galt17 | 8.32E-05 | 0.253053364 | 0.265 | 0.178 | 1 | 14 | B4galt1 |
| Dmxl14 | 0.000140367 | 0.268396127 | 0.269 | 0.186 | 1 | 14 | Dmxl1 |
| Hba-a11 | 0 | 6.42199256 | 1 | 0.12 | 0 | 15 | Hba-a1 |
| Hba-a21 | 0 | 6.393378987 | 1 | 0.103 | 0 | 15 | Hba-a2 |
| Hbb-bt1 | 0 | 6.229962574 | 0.96 | 0.073 | 0 | 15 | Hbb-bt |

| | | | | | | | |
|---|---|---|---|---|---|---|---|
| Alas2 | 0 | 2.578913692 | 0.704 | 0.019 | 0 | 15 | Alas2 |
| Snca | 0 | 2.212525344 | 0.615 | 0.019 | 0 | 15 | Snca |
| Bpgm | 4.32E-284 | 2.776451521 | 0.563 | 0.051 | 8.75E-280 | 15 | Bpgm |
| Hbb-bs1 | 3.25E-224 | 6.338532124 | 1 | 0.325 | 6.58E-220 | 15 | Hbb-bs |
| Fam220a1 | 1.72E-209 | 1.769271379 | 0.652 | 0.093 | 3.48E-205 | 15 | Fam220a |
| C1qb2 | 2.49E-175 | 1.566952478 | 0.85 | 0.15 | 5.04E-171 | 15 | C1qb |
| C1qa2 | 2.98E-160 | 1.450350339 | 0.802 | 0.143 | 6.04E-156 | 15 | C1qa |
| Aif12 | 2.98E-147 | 1.582990683 | 0.68 | 0.117 | 6.04E-143 | 15 | Aif1 |
| C1qc2 | 2.59E-146 | 1.390846353 | 0.741 | 0.13 | 5.25E-142 | 15 | C1qc |
| H2-Eb12 | 1.78E-143 | 1.511344334 | 0.862 | 0.19 | 3.61E-139 | 15 | H2-Eb1 |
| Fam46c1 | 6.84E-142 | 1.066543549 | 0.49 | 0.069 | 1.39E-137 | 15 | Fam46c |
| H2-Aa2 | 1.87E-138 | 1.418085207 | 0.838 | 0.183 | 3.79E-134 | 15 | H2-Aa |
| H2-Ab12 | 2.71E-126 | 1.396998125 | 0.83 | 0.197 | 5.49E-122 | 15 | H2-Ab1 |
| Cd742 | 1.34E-120 | 1.526549864 | 0.879 | 0.247 | 2.71E-116 | 15 | Cd74 |
| Ctss2 | 9.71E-118 | 1.317632202 | 0.741 | 0.162 | 1.97E-113 | 15 | Ctss |
| Lyz22 | 3.92E-111 | 1.193125319 | 0.579 | 0.109 | 7.93E-107 | 15 | Lyz2 |
| Ube2l61 | 7.01E-95 | 0.996670503 | 0.425 | 0.075 | 1.42E-90 | 15 | Ube2l6 |
| Mkrn11 | 4.17E-94 | 2.159637948 | 0.595 | 0.164 | 8.43E-90 | 15 | Mkrn1 |
| Ms4a72 | 8.22E-91 | 1.041302618 | 0.51 | 0.097 | 1.66E-86 | 15 | Ms4a7 |
| Slc25a37 | 2.63E-90 | 0.839799364 | 0.332 | 0.049 | 5.32E-86 | 15 | Slc25a37 |
| Ly862 | 1.39E-82 | 0.799971706 | 0.486 | 0.095 | 2.82E-78 | 15 | Ly86 |
| Isca1 | 3.81E-65 | 0.670926573 | 0.352 | 0.069 | 7.71E-61 | 15 | Isca1 |
| Tyrobp5 | 1.24E-63 | 1.071827425 | 0.806 | 0.259 | 2.51E-59 | 15 | Tyrobp |
| Cd300c22 | 8.22E-63 | 0.824652462 | 0.397 | 0.084 | 1.67E-58 | 15 | Cd300c2 |
| Isg203 | 8.80E-61 | 0.961560849 | 0.441 | 0.114 | 1.78E-56 | 15 | Isg20 |
| Fcer1g6 | 6.68E-57 | 0.803487769 | 0.818 | 0.267 | 1.35E-52 | 15 | Fcer1g |
| Bcl2a1b2 | 2.94E-56 | 0.817441517 | 0.425 | 0.105 | 5.95E-52 | 15 | Bcl2a1b |
| Lst12 | 1.51E-54 | 0.98846452 | 0.433 | 0.11 | 3.05E-50 | 15 | Lst1 |
| Wfdc172 | 2.04E-53 | 0.706544726 | 0.324 | 0.067 | 4.14E-49 | 15 | Wfdc17 |
| Rgs103 | 1.04E-51 | 0.844982692 | 0.53 | 0.159 | 2.10E-47 | 15 | Rgs10 |
| Cd682 | 1.46E-51 | 0.701981953 | 0.348 | 0.078 | 2.96E-47 | 15 | Cd68 |
| Apoe2 | 1.89E-50 | 1.355039424 | 0.87 | 0.533 | 3.83E-46 | 15 | Apoe |
| Fech | 3.53E-46 | 0.581051442 | 0.328 | 0.079 | 7.14E-42 | 15 | Fech |
| Mpeg11 | 1.44E-44 | 0.498206465 | 0.32 | 0.073 | 2.92E-40 | 15 | Mpeg1 |
| Cd526 | 2.18E-41 | 0.475829037 | 0.781 | 0.279 | 4.42E-37 | 15 | Cd52 |
| Spi12 | 9.49E-38 | 0.576996718 | 0.32 | 0.083 | 1.92E-33 | 15 | Spi1 |
| Alox5ap2 | 1.41E-37 | 0.682840729 | 0.291 | 0.073 | 2.85E-33 | 15 | Alox5ap |
| Ftl14 | 5.76E-37 | 0.993626401 | 0.996 | 0.911 | 1.17E-32 | 15 | Ftl1 |
| Cst34 | 4.18E-36 | 1.142334781 | 0.887 | 0.602 | 8.46E-32 | 15 | Cst3 |
| Basp13 | 1.21E-35 | 0.513371046 | 0.364 | 0.104 | 2.45E-31 | 15 | Basp1 |
| Hpgd2 | 3.17E-34 | 0.546225194 | 0.32 | 0.087 | 6.42E-30 | 15 | Hpgd |
| H2-DMa2 | 1.40E-33 | 0.604615688 | 0.385 | 0.119 | 2.83E-29 | 15 | H2-DMa |
| Cd633 | 1.94E-31 | 0.606835572 | 0.51 | 0.187 | 3.93E-27 | 15 | Cd63 |

| | | | | | | | |
|---|---|---|---|---|---|---|---|
| Ctsh3 | 3.71E-30 | 0.420249945 | 0.413 | 0.136 | 7.52E-26 | 15 | Ctsh |
| H2-DMb12 | 7.98E-29 | 0.366932142 | 0.312 | 0.092 | 1.62E-24 | 15 | H2-DMb1 |
| Ccndbp1 | 4.90E-27 | 0.422088738 | 0.32 | 0.105 | 9.92E-23 | 15 | Ccndbp1 |
| Scimp1 | 3.35E-26 | 0.65249533 | 0.259 | 0.076 | 6.77E-22 | 15 | Scimp |
| Cxcl162 | 2.04E-20 | 0.484239974 | 0.308 | 0.113 | 4.12E-16 | 15 | Cxcl16 |
| Lgmn2 | 1.73E-19 | 0.413886016 | 0.308 | 0.114 | 3.51E-15 | 15 | Lgmn |
| Fth13 | 2.65E-19 | 0.934508833 | 0.984 | 0.969 | 5.37E-15 | 15 | Fth1 |
| Cd722 | 1.89E-18 | 0.486977501 | 0.283 | 0.105 | 3.82E-14 | 15 | Cd72 |
| Sec61g1 | 1.48E-16 | 0.441682148 | 0.814 | 0.593 | 3.00E-12 | 15 | Sec61g |
| Ntpcr2 | 4.56E-16 | 0.434657296 | 0.267 | 0.105 | 9.24E-12 | 15 | Ntpcr |
| Ckb2 | 7.48E-16 | 0.567443409 | 0.433 | 0.216 | 1.51E-11 | 15 | Ckb |
| 23-Mar | 8.75E-16 | 0.358169581 | 0.409 | 0.197 | 1.77E-11 | 15 | 2-Mar |
| Blvrb2 | 5.32E-14 | 0.401307747 | 0.453 | 0.243 | 1.08E-09 | 15 | Blvrb |
| Tfdp2 | 4.04E-12 | 0.348101516 | 0.263 | 0.12 | 8.18E-08 | 15 | Tfdp2 |
| Bnip3l2 | 1.28E-11 | 0.485678849 | 0.518 | 0.333 | 2.59E-07 | 15 | Bnip3l |
| Epb412 | 6.87E-11 | 0.276290303 | 0.401 | 0.217 | 1.39E-06 | 15 | Epb41 |
| Npc22 | 4.38E-10 | 0.651865749 | 0.587 | 0.415 | 8.86E-06 | 15 | Npc2 |
| Cyba6 | 1.69E-08 | 0.47518567 | 0.628 | 0.441 | 0.000341611 | 15 | Cyba |
| Ctsz3 | 2.75E-08 | 0.467135628 | 0.53 | 0.361 | 0.000556255 | 15 | Ctsz |
| Creg12 | 1.15E-07 | 0.487317659 | 0.316 | 0.189 | 0.002328104 | 15 | Creg1 |
| Gpx13 | 1.83E-06 | 0.296300592 | 0.741 | 0.596 | 0.037124047 | 15 | Gpx1 |
| Ctsb3 | 5.66E-05 | 0.372583816 | 0.425 | 0.299 | 1 | 15 | Ctsb |
| Tmsb4x7 | 0.007139306 | 0.422237475 | 0.964 | 0.934 | 1 | 15 | Tmsb4x |
| Kcnq54 | 0 | 3.15602837 | 0.94 | 0.063 | 0 | 16 | Kcnq5 |
| Stab2 | 0 | 2.980308841 | 0.582 | 0.003 | 0 | 16 | Stab2 |
| Il7r2 | 0 | 2.90745803 | 0.957 | 0.034 | 0 | 16 | Il7r |
| Gata34 | 0 | 2.78771363 | 0.918 | 0.046 | 0 | 16 | Gata3 |
| Dach2 | 0 | 2.702793521 | 0.739 | 0.014 | 0 | 16 | Dach2 |
| Emb2 | 0 | 2.64769405 | 0.897 | 0.036 | 0 | 16 | Emb |
| Il1rl1 | 0 | 2.607002492 | 0.864 | 0.004 | 0 | 16 | Il1rl1 |
| Itk4 | 0 | 2.578219307 | 0.853 | 0.073 | 0 | 16 | Itk |
| Samsn15 | 0 | 2.418255876 | 0.875 | 0.076 | 0 | 16 | Samsn1 |
| Trbc24 | 0 | 2.176518223 | 0.81 | 0.062 | 0 | 16 | Trbc2 |
| Ccr24 | 0 | 2.090409251 | 0.859 | 0.082 | 0 | 16 | Ccr2 |
| Cxcr63 | 0 | 2.07463255 | 0.804 | 0.059 | 0 | 16 | Cxcr6 |
| Ptpn131 | 0 | 2.024515026 | 0.739 | 0.027 | 0 | 16 | Ptpn13 |
| Bcl11b1 | 0 | 1.954288688 | 0.772 | 0.024 | 0 | 16 | Bcl11b |
| Areg | 0 | 1.912372843 | 0.326 | 0.001 | 0 | 16 | Areg |
| 1700113H08Rik | 0 | 1.791689621 | 0.484 | 0.001 | 0 | 16 | 1700113H08Rik |
| Ltb4r1 | 0 | 1.747703537 | 0.663 | 0.007 | 0 | 16 | Ltb4r1 |
| Camk41 | 0 | 1.635185477 | 0.647 | 0.016 | 0 | 16 | Camk4 |
| Il2ra1 | 0 | 1.563365458 | 0.641 | 0.04 | 0 | 16 | Il2ra |
| Icos | 0 | 1.521654436 | 0.505 | 0.009 | 0 | 16 | Icos |

| | | | | | | | |
|---|---|---|---|---|---|---|---|
| Tspan32 | 0 | 1.438265209 | 0.609 | 0.029 | 0 | 16 | Tspan32 |
| Tcrg-C41 | 0 | 1.400296715 | 0.462 | 0.01 | 0 | 16 | Tcrg-C4 |
| Cd200r11 | 0 | 1.377078697 | 0.522 | 0.024 | 0 | 16 | Cd200r1 |
| Gm6213 | 0 | 1.320623028 | 0.457 | 0.002 | 0 | 16 | Gm6213 |
| Cntn1 | 0 | 1.289882042 | 0.418 | 0.004 | 0 | 16 | Cntn1 |
| Rnf128 | 0 | 1.25358614 | 0.44 | 0.008 | 0 | 16 | Rnf128 |
| Alox5 | 0 | 1.027070991 | 0.315 | 0.002 | 0 | 16 | Alox5 |
| Mns1 | 0 | 0.983041705 | 0.424 | 0.009 | 0 | 16 | Mns1 |
| Klrg1 | 0 | 0.971516859 | 0.315 | 0.004 | 0 | 16 | Klrg1 |
| Pcsk1 | 0 | 0.969620229 | 0.337 | 0.002 | 0 | 16 | Pcsk1 |
| Xlr4a | 0 | 0.936174957 | 0.386 | 0.006 | 0 | 16 | Xlr4a |
| Ccr8 | 0 | 0.912573653 | 0.304 | 0.002 | 0 | 16 | Ccr8 |
| Ccdc184 | 0 | 0.91246327 | 0.332 | 0.002 | 0 | 16 | Ccdc184 |
| Tnfsf11 | 0 | 0.817347935 | 0.304 | 0.003 | 0 | 16 | Tnfsf11 |
| Mboat1 | 0 | 0.780149929 | 0.315 | 0.006 | 0 | 16 | Mboat1 |
| Bmp7 | 0 | 0.755889432 | 0.31 | 0.008 | 0 | 16 | Bmp7 |
| Il17rb | 0 | 0.688924325 | 0.25 | 0.001 | 0 | 16 | Il17rb |
| Slc6a13 | 0 | 0.591272401 | 0.255 | 0.001 | 0 | 16 | Slc6a13 |
| Il9r | 0 | 0.58660032 | 0.25 | 0.001 | 0 | 16 | Il9r |
| Lpcat21 | 2.02E-305 | 1.901556803 | 0.75 | 0.063 | 4.09E-301 | 16 | Lpcat2 |
| Tespa14 | 5.34E-292 | 1.892427909 | 0.772 | 0.073 | 1.08E-287 | 16 | Tespa1 |
| Skap14 | 3.64E-291 | 2.800454762 | 1 | 0.129 | 7.36E-287 | 16 | Skap1 |
| Xlr4b | 2.22E-289 | 0.656961091 | 0.31 | 0.01 | 4.50E-285 | 16 | Xlr4b |
| Itgb74 | 2.87E-287 | 1.55346171 | 0.696 | 0.058 | 5.82E-283 | 16 | Itgb7 |
| Olfr1259 | 8.05E-285 | 0.79182291 | 0.321 | 0.011 | 1.63E-280 | 16 | Olfr1259 |
| Gpr1322 | 4.24E-267 | 1.071145495 | 0.451 | 0.025 | 8.59E-263 | 16 | Gpr132 |
| Gpr1832 | 4.53E-265 | 1.884896929 | 0.668 | 0.057 | 9.17E-261 | 16 | Gpr183 |
| Slc7a81 | 2.14E-259 | 1.357284979 | 0.533 | 0.037 | 4.33E-255 | 16 | Slc7a8 |
| Kcnn4 | 1.98E-256 | 1.026946937 | 0.484 | 0.03 | 4.00E-252 | 16 | Kcnn4 |
| Ptgir1 | 5.37E-253 | 0.685660615 | 0.321 | 0.013 | 1.09E-248 | 16 | Ptgir |
| Ltb4 | 1.50E-236 | 2.316504322 | 0.957 | 0.131 | 3.04E-232 | 16 | Ltb |
| Pard31 | 1.97E-230 | 1.437315461 | 0.549 | 0.043 | 3.99E-226 | 16 | Pard3 |
| Kit | 3.40E-230 | 0.668098394 | 0.255 | 0.009 | 6.87E-226 | 16 | Kit |
| Itga45 | 9.27E-230 | 2.085349773 | 0.69 | 0.073 | 1.88E-225 | 16 | Itga4 |
| S100a43 | 3.86E-227 | 2.385239932 | 0.913 | 0.125 | 7.82E-223 | 16 | S100a4 |
| Ramp14 | 6.48E-227 | 1.840503544 | 0.826 | 0.106 | 1.31E-222 | 16 | Ramp1 |
| Inpp4b4 | 2.26E-210 | 3.407037061 | 0.989 | 0.199 | 4.57E-206 | 16 | Inpp4b |
| Pxdc1 | 3.13E-210 | 1.148712277 | 0.37 | 0.021 | 6.34E-206 | 16 | Pxdc1 |
| Ikzf22 | 1.57E-209 | 1.398154322 | 0.56 | 0.049 | 3.17E-205 | 16 | Ikzf2 |
| Tasp11 | 2.63E-203 | 1.468303829 | 0.63 | 0.066 | 5.33E-199 | 16 | Tasp1 |
| Plxdc22 | 2.76E-199 | 1.4683904 | 0.761 | 0.09 | 5.58E-195 | 16 | Plxdc2 |
| Ptprcap4 | 8.37E-199 | 1.649535155 | 0.908 | 0.128 | 1.69E-194 | 16 | Ptprcap |
| Cd101 | 5.02E-196 | 0.782846291 | 0.277 | 0.013 | 1.02E-191 | 16 | Cd101 |

| | | | | | | | |
|---|---|---|---|---|---|---|---|
| Tnfrsf184 | 9.57E-194 | 1.442478154 | 0.636 | 0.071 | 1.94E-189 | 16 | Tnfrsf18 |
| Kbtbd11 | 4.81E-193 | 0.670779875 | 0.283 | 0.013 | 9.75E-189 | 16 | Kbtbd11 |
| Prkcq4 | 1.07E-187 | 1.567855882 | 0.69 | 0.085 | 2.17E-183 | 16 | Prkcq |
| Plcl12 | 2.27E-184 | 1.410791361 | 0.527 | 0.05 | 4.60E-180 | 16 | Plcl1 |
| Ikzf15 | 1.35E-181 | 2.047368913 | 0.88 | 0.153 | 2.73E-177 | 16 | Ikzf1 |
| Ahcyl22 | 6.97E-181 | 1.742379843 | 0.793 | 0.123 | 1.41E-176 | 16 | Ahcyl2 |
| Coro2a4 | 3.29E-180 | 1.726799792 | 0.777 | 0.115 | 6.65E-176 | 16 | Coro2a |
| Nfkbid | 5.34E-180 | 1.090208788 | 0.418 | 0.033 | 1.08E-175 | 16 | Nfkbid |
| Mbp3 | 5.68E-178 | 1.09466916 | 0.592 | 0.065 | 1.15E-173 | 16 | Mbp |
| Tmem64 | 1.68E-175 | 1.541910068 | 0.625 | 0.075 | 3.40E-171 | 16 | Tmem64 |
| Thy14 | 1.10E-173 | 1.148567979 | 0.554 | 0.058 | 2.23E-169 | 16 | Thy1 |
| Cd694 | 1.22E-173 | 2.122375173 | 0.707 | 0.099 | 2.48E-169 | 16 | Cd69 |
| Hcst4 | 2.40E-169 | 1.60612895 | 0.875 | 0.142 | 4.85E-165 | 16 | Hcst |
| Faah | 2.48E-169 | 0.736009054 | 0.353 | 0.024 | 5.03E-165 | 16 | Faah |
| St6galnac36 | 1.60E-168 | 2.517948335 | 0.957 | 0.213 | 3.24E-164 | 16 | St6galnac3 |
| Mir142hg5 | 9.10E-166 | 1.980569967 | 0.897 | 0.167 | 1.84E-161 | 16 | Mir142hg |
| Klk83 | 6.63E-165 | 1.199681558 | 0.636 | 0.08 | 1.34E-160 | 16 | Klk8 |
| Tnik5 | 2.75E-162 | 1.484729236 | 0.734 | 0.11 | 5.56E-158 | 16 | Tnik |
| Slc38a14 | 4.23E-162 | 1.362921373 | 0.641 | 0.086 | 8.57E-158 | 16 | Slc38a1 |
| Gimap34 | 2.73E-161 | 1.487332129 | 0.696 | 0.097 | 5.54E-157 | 16 | Gimap3 |
| Arhgap155 | 2.68E-160 | 1.941774279 | 0.94 | 0.185 | 5.42E-156 | 16 | Arhgap15 |
| Pde7a4 | 5.30E-158 | 1.901739715 | 0.826 | 0.153 | 1.07E-153 | 16 | Pde7a |
| Itgb3 | 1.12E-157 | 0.740117245 | 0.353 | 0.026 | 2.27E-153 | 16 | Itgb3 |
| Fam107b4 | 2.92E-156 | 1.552262149 | 0.734 | 0.118 | 5.92E-152 | 16 | Fam107b |
| Dok23 | 3.08E-152 | 0.962641692 | 0.429 | 0.04 | 6.24E-148 | 16 | Dok2 |
| Il2rg4 | 2.15E-150 | 1.449831405 | 0.723 | 0.114 | 4.35E-146 | 16 | Il2rg |
| Runx15 | 5.81E-146 | 1.508802283 | 0.755 | 0.129 | 1.18E-141 | 16 | Runx1 |
| Dock25 | 2.19E-145 | 1.680803087 | 0.929 | 0.183 | 4.43E-141 | 16 | Dock2 |
| Fosb6 | 2.88E-143 | 2.425912338 | 0.832 | 0.187 | 5.84E-139 | 16 | Fosb |
| Nabp14 | 2.96E-140 | 1.192706019 | 0.587 | 0.079 | 6.00E-136 | 16 | Nabp1 |
| Gm26802 | 1.16E-138 | 1.598084589 | 0.538 | 0.07 | 2.35E-134 | 16 | Gm26802 |
| Arhgap455 | 2.55E-134 | 1.406278173 | 0.821 | 0.156 | 5.17E-130 | 16 | Arhgap45 |
| Lcp25 | 1.93E-133 | 1.220611158 | 0.614 | 0.092 | 3.90E-129 | 16 | Lcp2 |
| Cd535 | 4.52E-133 | 1.343932399 | 0.679 | 0.114 | 9.16E-129 | 16 | Cd53 |
| Tnfaip35 | 1.14E-130 | 1.619697434 | 0.587 | 0.088 | 2.30E-126 | 16 | Tnfaip3 |
| Cnot6l3 | 2.32E-130 | 1.905508073 | 0.913 | 0.242 | 4.69E-126 | 16 | Cnot6l |
| Mgat55 | 5.27E-130 | 1.627097571 | 0.799 | 0.168 | 1.07E-125 | 16 | Mgat5 |
| Nfkbiz3 | 5.21E-129 | 1.82142383 | 0.707 | 0.135 | 1.05E-124 | 16 | Nfkbiz |
| 1810011H11Rik4 | 3.24E-126 | 1.120878622 | 0.527 | 0.072 | 6.55E-122 | 16 | 1810011H11Rik |
| Ctse1 | 5.88E-126 | 0.764242098 | 0.337 | 0.03 | 1.19E-121 | 16 | Ctse |
| Ptpn74 | 8.38E-126 | 0.935408086 | 0.467 | 0.056 | 1.70E-121 | 16 | Ptpn7 |
| Cd375 | 2.21E-124 | 1.189323219 | 0.663 | 0.112 | 4.47E-120 | 16 | Cd37 |
| Tiam1 | 3.33E-124 | 0.649862519 | 0.283 | 0.021 | 6.74E-120 | 16 | Tiam1 |

| | | | | | | | |
|---|---|---|---|---|---|---|---|
| Coro1a6 | 3.70E-124 | 1.566652134 | 0.951 | 0.216 | 7.48E-120 | 16 | Coro1a |
| Cysltr11 | 1.05E-123 | 1.100218746 | 0.484 | 0.06 | 2.13E-119 | 16 | Cysltr1 |
| Nrgn | 1.86E-121 | 0.970671996 | 0.342 | 0.031 | 3.77E-117 | 16 | Nrgn |
| Pip5k1b4 | 4.33E-121 | 1.198421415 | 0.609 | 0.097 | 8.76E-117 | 16 | Pip5k1b |
| Mdfic4 | 4.71E-121 | 2.125986437 | 0.875 | 0.243 | 9.54E-117 | 16 | Mdfic |
| St6gal11 | 2.27E-120 | 0.834454061 | 0.386 | 0.04 | 4.60E-116 | 16 | St6gal1 |
| Rac24 | 4.53E-120 | 1.386135231 | 0.859 | 0.175 | 9.18E-116 | 16 | Rac2 |
| Cd824 | 5.19E-120 | 1.210766348 | 0.603 | 0.098 | 1.05E-115 | 16 | Cd82 |
| Gmfg3 | 2.66E-118 | 1.188342111 | 0.728 | 0.137 | 5.38E-114 | 16 | Gmfg |
| Dock105 | 3.64E-118 | 1.541430538 | 0.908 | 0.206 | 7.36E-114 | 16 | Dock10 |
| Bbs94 | 8.17E-118 | 1.384004584 | 0.603 | 0.099 | 1.65E-113 | 16 | Bbs9 |
| Nfkb17 | 8.56E-118 | 1.806835799 | 0.918 | 0.269 | 1.73E-113 | 16 | Nfkb1 |
| Pim14 | 9.09E-117 | 1.472094895 | 0.679 | 0.13 | 1.84E-112 | 16 | Pim1 |
| Ptpn186 | 5.21E-115 | 1.761558017 | 0.989 | 0.313 | 1.05E-110 | 16 | Ptpn18 |
| Stat44 | 3.68E-114 | 1.176838192 | 0.636 | 0.105 | 7.45E-110 | 16 | Stat4 |
| Rora2 | 1.88E-113 | 1.519094632 | 0.826 | 0.183 | 3.80E-109 | 16 | Rora |
| Man1a8 | 2.58E-112 | 1.650414335 | 0.739 | 0.165 | 5.22E-108 | 16 | Man1a |
| Cish1 | 1.93E-109 | 0.910001963 | 0.478 | 0.066 | 3.91E-105 | 16 | Cish |
| 15-Sep | 6.22E-109 | 1.15705545 | 0.739 | 0.144 | 1.26E-104 | 16 | 1-Sep |
| Fbxo17 | 4.85E-107 | 0.543409497 | 0.261 | 0.02 | 9.81E-103 | 16 | Fbxo17 |
| Il27ra2 | 6.89E-107 | 0.715562719 | 0.37 | 0.041 | 1.40E-102 | 16 | Il27ra |
| Aff42 | 7.34E-107 | 1.671807712 | 0.848 | 0.238 | 1.49E-102 | 16 | Aff4 |
| Ap1s3 | 5.79E-106 | 0.730293467 | 0.359 | 0.038 | 1.17E-101 | 16 | Ap1s3 |
| Ptprc5 | 7.08E-106 | 1.045627227 | 0.924 | 0.195 | 1.43E-101 | 16 | Ptprc |
| Mllt35 | 3.39E-105 | 1.330130785 | 0.712 | 0.149 | 6.86E-101 | 16 | Mllt3 |
| Nck2 | 6.29E-105 | 0.905957581 | 0.457 | 0.062 | 1.27E-100 | 16 | Nck2 |
| Arhgdib6 | 6.37E-104 | 1.413978665 | 0.908 | 0.231 | 1.29E-99 | 16 | Arhgdib |
| Itpr25 | 1.23E-103 | 1.470340561 | 0.864 | 0.242 | 2.48E-99 | 16 | Itpr2 |
| Trbc14 | 2.32E-103 | 1.27520301 | 0.527 | 0.085 | 4.69E-99 | 16 | Trbc1 |
| Rps6ka55 | 8.07E-102 | 1.171361059 | 0.587 | 0.106 | 1.63E-97 | 16 | Rps6ka5 |
| Pdcd1lg2 | 1.95E-101 | 0.665055375 | 0.288 | 0.027 | 3.94E-97 | 16 | Pdcd1lg2 |
| Scml43 | 5.34E-101 | 0.791295653 | 0.359 | 0.041 | 1.08E-96 | 16 | Scml4 |
| Lgals14 | 2.22E-100 | 1.489826671 | 0.679 | 0.133 | 4.50E-96 | 16 | Lgals1 |
| Zfp362 | 3.16E-100 | 1.654262542 | 0.777 | 0.195 | 6.39E-96 | 16 | Zfp36 |
| Limd25 | 4.29E-100 | 1.235150439 | 0.69 | 0.146 | 8.69E-96 | 16 | Limd2 |
| Cd272 | 5.73E-99 | 0.808458434 | 0.342 | 0.038 | 1.16E-94 | 16 | Cd27 |
| Arg11 | 8.84E-99 | 1.038289343 | 0.652 | 0.116 | 1.79E-94 | 16 | Arg1 |
| Fam102a4 | 2.69E-98 | 1.070459752 | 0.56 | 0.098 | 5.44E-94 | 16 | Fam102a |
| Fos4 | 3.58E-97 | 2.294663949 | 0.908 | 0.315 | 7.24E-93 | 16 | Fos |
| Tox5 | 7.80E-97 | 1.045085769 | 0.652 | 0.119 | 1.58E-92 | 16 | Tox |
| Zcchc10 | 3.37E-96 | 0.865584295 | 0.5 | 0.078 | 6.82E-92 | 16 | Zcchc10 |
| Chsy3 | 6.31E-96 | 1.193080054 | 0.321 | 0.034 | 1.28E-91 | 16 | Chsy3 |
| Stk17b4 | 2.95E-95 | 1.303563263 | 0.723 | 0.16 | 5.97E-91 | 16 | Stk17b |

| | | | | | | | |
|---|---|---|---|---|---|---|---|
| Pparg | 4.36E-94 | 0.973293813 | 0.473 | 0.072 | 8.82E-90 | 16 | Pparg |
| Sh3kbp13 | 1.42E-93 | 1.132202851 | 0.62 | 0.123 | 2.87E-89 | 16 | Sh3kbp1 |
| Sorl15 | 3.59E-92 | 1.178164243 | 0.571 | 0.11 | 7.27E-88 | 16 | Sorl1 |
| Glipr22 | 4.74E-92 | 0.803122243 | 0.348 | 0.042 | 9.61E-88 | 16 | Glipr2 |
| Tnfaip84 | 6.23E-91 | 1.300621594 | 0.75 | 0.185 | 1.26E-86 | 16 | Tnfaip8 |
| Notch24 | 1.13E-90 | 0.925762537 | 0.478 | 0.079 | 2.30E-86 | 16 | Notch2 |
| Lat3 | 2.12E-90 | 0.674629354 | 0.321 | 0.036 | 4.29E-86 | 16 | Lat |
| Selplg5 | 3.95E-89 | 1.105052835 | 0.701 | 0.153 | 7.99E-85 | 16 | Selplg |
| Abcb9 | 5.79E-88 | 0.654820599 | 0.288 | 0.03 | 1.17E-83 | 16 | Abcb9 |
| Stk45 | 9.73E-88 | 1.181156958 | 0.712 | 0.173 | 1.97E-83 | 16 | Stk4 |
| Cers42 | 1.11E-87 | 0.9276747 | 0.571 | 0.108 | 2.25E-83 | 16 | Cers4 |
| Galnt121 | 2.27E-87 | 0.690349081 | 0.326 | 0.039 | 4.61E-83 | 16 | Galnt12 |
| Polg2 | 1.36E-86 | 0.895208542 | 0.429 | 0.066 | 2.76E-82 | 16 | Polg2 |
| Birc33 | 2.44E-86 | 1.032328625 | 0.527 | 0.097 | 4.95E-82 | 16 | Birc3 |
| Apbb1ip5 | 1.85E-85 | 1.066988555 | 0.728 | 0.172 | 3.75E-81 | 16 | Apbb1ip |
| Crtc36 | 2.73E-85 | 1.168506686 | 0.679 | 0.162 | 5.53E-81 | 16 | Crtc3 |
| Junb7 | 1.89E-84 | 2.036619036 | 0.951 | 0.391 | 3.82E-80 | 16 | Junb |
| Eya24 | 7.54E-84 | 0.918567975 | 0.533 | 0.099 | 1.53E-79 | 16 | Eya2 |
| Celf26 | 3.30E-83 | 1.079794962 | 0.788 | 0.2 | 6.68E-79 | 16 | Celf2 |
| Phtf2 | 6.71E-83 | 0.777336158 | 0.408 | 0.062 | 1.36E-78 | 16 | Phtf2 |
| Galnt6 | 7.56E-83 | 0.610334266 | 0.266 | 0.028 | 1.53E-78 | 16 | Galnt6 |
| Adam192 | 7.65E-83 | 1.082052055 | 0.467 | 0.079 | 1.55E-78 | 16 | Adam19 |
| Flrt21 | 1.35E-82 | 1.052924518 | 0.337 | 0.043 | 2.74E-78 | 16 | Flrt2 |
| Chn25 | 1.90E-80 | 1.034363394 | 0.777 | 0.196 | 3.85E-76 | 16 | Chn2 |
| Cd844 | 2.74E-80 | 0.91701759 | 0.489 | 0.089 | 5.55E-76 | 16 | Cd84 |
| Nrip16 | 9.18E-80 | 1.458823372 | 0.788 | 0.242 | 1.86E-75 | 16 | Nrip1 |
| Rps246 | 5.49E-78 | 1.182284923 | 1 | 0.972 | 1.11E-73 | 16 | Rps24 |
| Bzw2 | 6.40E-78 | 0.959376413 | 0.543 | 0.109 | 1.30E-73 | 16 | Bzw2 |
| Rcsd14 | 6.80E-78 | 1.05102082 | 0.582 | 0.128 | 1.38E-73 | 16 | Rcsd1 |
| P2ry102 | 9.04E-78 | 0.687563334 | 0.304 | 0.038 | 1.83E-73 | 16 | P2ry10 |
| Aak16 | 2.06E-77 | 1.202373941 | 0.685 | 0.183 | 4.17E-73 | 16 | Aak1 |
| Btg26 | 2.13E-77 | 1.639741948 | 0.908 | 0.342 | 4.31E-73 | 16 | Btg2 |
| Bach26 | 3.87E-77 | 1.382261839 | 0.636 | 0.159 | 7.83E-73 | 16 | Bach2 |
| Rassf54 | 9.45E-77 | 0.694415977 | 0.413 | 0.068 | 1.91E-72 | 16 | Rassf5 |
| Neurl35 | 1.51E-76 | 1.173644027 | 0.538 | 0.114 | 3.05E-72 | 16 | Neurl3 |
| Rinl4 | 1.97E-76 | 1.008721033 | 0.592 | 0.13 | 3.99E-72 | 16 | Rinl |
| Id24 | 3.29E-76 | 1.284904325 | 0.94 | 0.319 | 6.67E-72 | 16 | Id2 |
| Fhl24 | 4.83E-76 | 0.799571158 | 0.315 | 0.041 | 9.78E-72 | 16 | Fhl2 |
| Slco3a13 | 1.96E-73 | 0.952642952 | 0.565 | 0.121 | 3.96E-69 | 16 | Slco3a1 |
| Laptm56 | 3.01E-73 | 0.909134623 | 0.804 | 0.209 | 6.09E-69 | 16 | Laptm5 |
| Lmo4 | 3.45E-73 | 1.164138776 | 0.701 | 0.192 | 6.99E-69 | 16 | Lmo4 |
| Chd33 | 7.42E-73 | 0.963898805 | 0.63 | 0.154 | 1.50E-68 | 16 | Chd3 |
| Ankrd445 | 9.66E-73 | 1.205531575 | 0.875 | 0.292 | 1.96E-68 | 16 | Ankrd44 |

| | | | | | | | |
|---|---|---|---|---|---|---|---|
| Cd527 | 1.40E-72 | 1.034896219 | 0.989 | 0.279 | 2.84E-68 | 16 | Cd52 |
| Gm267402 | 1.96E-72 | 0.784152966 | 0.38 | 0.059 | 3.97E-68 | 16 | Gm26740 |
| Pag14 | 2.15E-72 | 0.983265524 | 0.554 | 0.12 | 4.35E-68 | 16 | Pag1 |
| Jakmip14 | 3.30E-72 | 0.672360313 | 0.337 | 0.048 | 6.68E-68 | 16 | Jakmip1 |
| Iqgap22 | 3.59E-70 | 0.871268676 | 0.511 | 0.104 | 7.27E-66 | 16 | Iqgap2 |
| Asb22 | 2.48E-69 | 0.639654522 | 0.342 | 0.051 | 5.02E-65 | 16 | Asb2 |
| Rps165 | 2.48E-69 | 1.078212797 | 1 | 0.927 | 5.02E-65 | 16 | Rps16 |
| Klf67 | 9.98E-69 | 1.497513482 | 0.788 | 0.266 | 2.02E-64 | 16 | Klf6 |
| Rapgef65 | 1.81E-68 | 1.242911451 | 0.848 | 0.295 | 3.67E-64 | 16 | Rapgef6 |
| Btg14 | 1.95E-68 | 1.529221197 | 0.973 | 0.458 | 3.95E-64 | 16 | Btg1 |
| Pard3b1 | 3.82E-68 | 1.031497635 | 0.489 | 0.1 | 7.73E-64 | 16 | Pard3b |
| Rpl37a6 | 6.69E-68 | 1.113670058 | 1 | 0.938 | 1.35E-63 | 16 | Rpl37a |
| Maf3 | 9.20E-68 | 0.924237557 | 0.538 | 0.117 | 1.86E-63 | 16 | Maf |
| Rps185 | 1.42E-67 | 1.098826941 | 1 | 0.942 | 2.87E-63 | 16 | Rps18 |
| 4930523C07Rik7 | 1.52E-67 | 1.136276629 | 0.728 | 0.218 | 3.08E-63 | 16 | 4930523C07Rik |
| Shisa54 | 2.07E-67 | 1.160618206 | 0.918 | 0.327 | 4.18E-63 | 16 | Shisa5 |
| Tmsb105 | 2.82E-67 | 1.357451967 | 0.995 | 0.499 | 5.71E-63 | 16 | Tmsb10 |
| Rps106 | 6.19E-67 | 1.04203374 | 1 | 0.944 | 1.25E-62 | 16 | Rps10 |
| Atxn18 | 3.02E-66 | 1.312222316 | 0.837 | 0.294 | 6.12E-62 | 16 | Atxn1 |
| Slc9a95 | 3.16E-66 | 0.977846763 | 0.614 | 0.15 | 6.40E-62 | 16 | Slc9a9 |
| Rpl126 | 7.01E-66 | 1.189271175 | 1 | 0.841 | 1.42E-61 | 16 | Rpl12 |
| D16Ertd472e4 | 1.10E-65 | 0.902624779 | 0.489 | 0.105 | 2.23E-61 | 16 | D16Ertd472e |
| Rpsa6 | 2.24E-64 | 1.189764182 | 1 | 0.903 | 4.54E-60 | 16 | Rpsa |
| Arhgef65 | 2.62E-64 | 0.797672596 | 0.549 | 0.127 | 5.31E-60 | 16 | Arhgef6 |
| Emp37 | 2.64E-64 | 1.090419839 | 0.793 | 0.257 | 5.35E-60 | 16 | Emp3 |
| Rps196 | 9.34E-64 | 1.040141638 | 1 | 0.927 | 1.89E-59 | 16 | Rps19 |
| Satb14 | 1.77E-63 | 0.754052548 | 0.364 | 0.062 | 3.59E-59 | 16 | Satb1 |
| Sik14 | 2.30E-63 | 1.019509354 | 0.533 | 0.125 | 4.66E-59 | 16 | Sik1 |
| Rps276 | 2.94E-63 | 1.084913909 | 1 | 0.898 | 5.96E-59 | 16 | Rps27 |
| Zcchc115 | 6.48E-63 | 1.109108017 | 0.793 | 0.266 | 1.31E-58 | 16 | Zcchc11 |
| Cd2263 | 1.04E-62 | 0.625839926 | 0.467 | 0.089 | 2.11E-58 | 16 | Cd226 |
| Fam213a | 1.05E-62 | 0.502132608 | 0.25 | 0.031 | 2.13E-58 | 16 | Fam213a |
| Gramd1b4 | 1.65E-62 | 0.786452478 | 0.44 | 0.089 | 3.33E-58 | 16 | Gramd1b |
| Gpsm35 | 1.86E-62 | 0.922557421 | 0.576 | 0.147 | 3.76E-58 | 16 | Gpsm3 |
| Pstpip14 | 2.38E-62 | 0.661187882 | 0.337 | 0.055 | 4.82E-58 | 16 | Pstpip1 |
| Fermt35 | 2.41E-62 | 0.795206666 | 0.505 | 0.113 | 4.88E-58 | 16 | Fermt3 |
| Tbc1d10c4 | 2.90E-62 | 0.658656564 | 0.397 | 0.072 | 5.87E-58 | 16 | Tbc1d10c |
| Gm446861 | 2.90E-62 | 0.564564752 | 0.359 | 0.06 | 5.88E-58 | 16 | Gm44686 |
| Rps36 | 1.06E-61 | 1.063215303 | 1 | 0.893 | 2.15E-57 | 16 | Rps3 |
| Cox10 | 1.39E-61 | 0.696075408 | 0.37 | 0.065 | 2.82E-57 | 16 | Cox10 |
| Cobll13 | 2.18E-61 | 0.855407502 | 0.424 | 0.085 | 4.42E-57 | 16 | Cobll1 |
| Mbd57 | 3.73E-61 | 1.232282766 | 0.685 | 0.219 | 7.54E-57 | 16 | Mbd5 |
| Adam121 | 1.23E-60 | 0.740607792 | 0.266 | 0.036 | 2.48E-56 | 16 | Adam12 |

| | | | | | | | |
|---|---|---|---|---|---|---|---|
| Sdc43 | 1.83E-60 | 0.630170686 | 0.386 | 0.07 | 3.71E-56 | 16 | Sdc4 |
| Gm26835 | 2.55E-60 | 0.752531142 | 0.342 | 0.058 | 5.16E-56 | 16 | Gm26835 |
| Rasgrp14 | 7.79E-60 | 0.699518516 | 0.353 | 0.062 | 1.58E-55 | 16 | Rasgrp1 |
| Rb15 | 9.21E-60 | 1.08402311 | 0.69 | 0.213 | 1.86E-55 | 16 | Rb1 |
| Bcl24 | 1.56E-59 | 0.852991952 | 0.717 | 0.206 | 3.16E-55 | 16 | Bcl2 |
| Rpl86 | 1.60E-59 | 0.982938137 | 1 | 0.925 | 3.24E-55 | 16 | Rpl8 |
| Stim16 | 1.68E-59 | 1.00783896 | 0.647 | 0.189 | 3.41E-55 | 16 | Stim1 |
| Fnbp16 | 1.89E-59 | 1.106069637 | 0.717 | 0.229 | 3.83E-55 | 16 | Fnbp1 |
| Rpl136 | 2.71E-59 | 0.945394083 | 1 | 0.977 | 5.49E-55 | 16 | Rpl13 |
| Rhoh5 | 4.40E-59 | 1.048637139 | 0.467 | 0.102 | 8.90E-55 | 16 | Rhoh |
| Cotl16 | 8.09E-59 | 0.988838916 | 0.734 | 0.223 | 1.64E-54 | 16 | Cotl1 |
| Rps136 | 9.84E-59 | 0.995825105 | 1 | 0.923 | 1.99E-54 | 16 | Rps13 |
| Arhgap305 | 1.21E-58 | 0.817564319 | 0.478 | 0.109 | 2.45E-54 | 16 | Arhgap30 |
| Usp33 | 1.78E-58 | 1.074378447 | 0.625 | 0.181 | 3.61E-54 | 16 | Usp3 |
| Dock116 | 2.43E-58 | 0.952789239 | 0.484 | 0.113 | 4.92E-54 | 16 | Dock11 |
| Jund5 | 2.93E-58 | 1.342651236 | 0.989 | 0.645 | 5.93E-54 | 16 | Jund |
| Rfx74 | 6.18E-58 | 0.961433349 | 0.571 | 0.154 | 1.25E-53 | 16 | Rfx7 |
| Rpl306 | 9.85E-58 | 0.920673759 | 1 | 0.938 | 1.99E-53 | 16 | Rpl30 |
| Zc3hav14 | 9.97E-58 | 1.146128466 | 0.658 | 0.203 | 2.02E-53 | 16 | Zc3hav1 |
| Adgre56 | 1.17E-57 | 1.358428016 | 0.712 | 0.236 | 2.37E-53 | 16 | Adgre5 |
| Rps205 | 2.71E-57 | 0.946907867 | 1 | 0.951 | 5.48E-53 | 16 | Rps20 |
| Rps15a5 | 3.94E-57 | 1.029701701 | 1 | 0.925 | 7.98E-53 | 16 | Rps15a |
| Sema4a4 | 5.19E-57 | 0.824740538 | 0.386 | 0.076 | 1.05E-52 | 16 | Sema4a |
| Tecpr13 | 9.50E-57 | 0.889125307 | 0.516 | 0.129 | 1.92E-52 | 16 | Tecpr1 |
| Large14 | 2.30E-56 | 1.594027843 | 0.538 | 0.151 | 4.66E-52 | 16 | Large1 |
| Rps56 | 2.65E-56 | 0.982175116 | 1 | 0.92 | 5.37E-52 | 16 | Rps5 |
| Stt3b5 | 1.06E-55 | 0.947269223 | 0.582 | 0.164 | 2.14E-51 | 16 | Stt3b |
| Rps76 | 1.06E-55 | 1.002103581 | 1 | 0.916 | 2.15E-51 | 16 | Rps7 |
| Hlf1 | 1.92E-55 | 0.743579881 | 0.332 | 0.057 | 3.89E-51 | 16 | Hlf |
| S100a66 | 1.97E-55 | 0.671756249 | 0.87 | 0.269 | 4.00E-51 | 16 | S100a6 |
| AI467606 | 3.27E-55 | 0.616931311 | 0.288 | 0.045 | 6.62E-51 | 16 | AI467606 |
| Rgs25 | 3.46E-55 | 0.998242762 | 0.516 | 0.129 | 7.01E-51 | 16 | Rgs2 |
| Ccnd28 | 3.83E-55 | 1.185016917 | 0.777 | 0.285 | 7.75E-51 | 16 | Ccnd2 |
| Fxyd58 | 4.41E-55 | 1.103653612 | 0.946 | 0.41 | 8.94E-51 | 16 | Fxyd5 |
| Cdc42ep37 | 1.48E-54 | 0.936995547 | 0.603 | 0.174 | 3.00E-50 | 16 | Cdc42ep3 |
| Rabgap1l4 | 1.98E-54 | 0.978307171 | 0.755 | 0.256 | 4.01E-50 | 16 | Rabgap1l |
| Isy1 | 3.23E-54 | 0.728224541 | 0.364 | 0.07 | 6.54E-50 | 16 | Isy1 |
| Elovl53 | 8.27E-54 | 1.015562378 | 0.543 | 0.15 | 1.67E-49 | 16 | Elovl5 |
| Znrf17 | 9.02E-54 | 1.04638999 | 0.62 | 0.188 | 1.83E-49 | 16 | Znrf1 |
| Fam117a | 1.20E-53 | 0.581180914 | 0.299 | 0.05 | 2.43E-49 | 16 | Fam117a |
| Atp13a33 | 1.23E-53 | 0.671548796 | 0.451 | 0.102 | 2.49E-49 | 16 | Atp13a3 |
| H2-Q76 | 1.67E-53 | 1.136869198 | 0.902 | 0.396 | 3.37E-49 | 16 | H2-Q7 |
| Rps96 | 2.28E-53 | 0.920325978 | 1 | 0.921 | 4.62E-49 | 16 | Rps9 |

| | | | | | | | |
|---|---|---|---|---|---|---|---|
| Rpl196 | 7.22E-53 | 0.830262604 | 1 | 0.963 | 1.46E-48 | 16 | Rpl19 |
| Rpl237 | 1.20E-52 | 0.839226408 | 1 | 0.971 | 2.42E-48 | 16 | Rpl23 |
| Furin | 2.59E-52 | 1.053279708 | 0.429 | 0.099 | 5.25E-48 | 16 | Furin |
| Smad33 | 3.09E-52 | 0.817275995 | 0.424 | 0.095 | 6.26E-48 | 16 | Smad3 |
| Rpl176 | 3.80E-52 | 0.892987102 | 1 | 0.916 | 7.69E-48 | 16 | Rpl17 |
| Cyld4 | 9.66E-52 | 0.79254702 | 0.587 | 0.169 | 1.96E-47 | 16 | Cyld |
| Pde7b4 | 1.62E-51 | 1.301164726 | 0.701 | 0.241 | 3.28E-47 | 16 | Pde7b |
| Socs13 | 1.69E-51 | 0.75812353 | 0.429 | 0.097 | 3.42E-47 | 16 | Socs1 |
| Ndfip14 | 1.71E-51 | 1.031355153 | 0.837 | 0.32 | 3.46E-47 | 16 | Ndfip1 |
| Rasal34 | 1.90E-51 | 0.720719727 | 0.391 | 0.083 | 3.84E-47 | 16 | Rasal3 |
| Rpl326 | 2.77E-51 | 0.89784647 | 1 | 0.953 | 5.60E-47 | 16 | Rpl32 |
| Hist1h1d | 2.78E-51 | 0.625218577 | 0.288 | 0.048 | 5.63E-47 | 16 | Hist1h1d |
| Rpl186 | 2.97E-51 | 0.884627692 | 1 | 0.9 | 6.01E-47 | 16 | Rpl18 |
| Inpp5d5 | 3.42E-51 | 0.882657586 | 0.658 | 0.203 | 6.92E-47 | 16 | Inpp5d |
| Dtnb4 | 5.59E-51 | 0.754637918 | 0.467 | 0.114 | 1.13E-46 | 16 | Dtnb |
| Rpl18a5 | 6.37E-51 | 0.873593641 | 1 | 0.931 | 1.29E-46 | 16 | Rpl18a |
| Rps3a17 | 4.77E-50 | 0.895160964 | 1 | 0.935 | 9.66E-46 | 16 | Rps3a1 |
| Gm2a2 | 8.57E-50 | 0.635878685 | 0.402 | 0.088 | 1.73E-45 | 16 | Gm2a |
| Psen21 | 9.04E-50 | 0.60142499 | 0.326 | 0.061 | 1.83E-45 | 16 | Psen2 |
| Cd442 | 9.60E-50 | 0.675897085 | 0.446 | 0.107 | 1.94E-45 | 16 | Cd44 |
| Gm203887 | 3.00E-49 | 1.022937021 | 0.886 | 0.4 | 6.07E-45 | 16 | Gm20388 |
| Rab374 | 5.17E-49 | 0.55642453 | 0.266 | 0.043 | 1.05E-44 | 16 | Rab37 |
| Nol4l2 | 5.79E-49 | 0.732710815 | 0.359 | 0.074 | 1.17E-44 | 16 | Nol4l |
| Lcp15 | 6.35E-49 | 0.876393257 | 0.734 | 0.237 | 1.29E-44 | 16 | Lcp1 |
| Csk5 | 6.44E-49 | 0.752959887 | 0.489 | 0.129 | 1.30E-44 | 16 | Csk |
| Rps115 | 7.28E-49 | 0.874805703 | 1 | 0.907 | 1.47E-44 | 16 | Rps11 |
| Cytip4 | 7.69E-49 | 0.619869152 | 0.408 | 0.091 | 1.56E-44 | 16 | Cytip |
| Cerk5 | 8.60E-49 | 0.663332255 | 0.462 | 0.114 | 1.74E-44 | 16 | Cerk |
| Gm472836 | 8.87E-49 | 1.048096085 | 0.739 | 0.28 | 1.80E-44 | 16 | Gm47283 |
| Ppm1h5 | 1.24E-48 | 0.806633425 | 0.56 | 0.16 | 2.50E-44 | 16 | Ppm1h |
| Traf3ip2 | 3.53E-48 | 0.518423735 | 0.283 | 0.048 | 7.16E-44 | 16 | Traf3ip2 |
| Leprotl14 | 4.73E-48 | 0.897955714 | 0.647 | 0.216 | 9.57E-44 | 16 | Leprotl1 |
| Ptpn91 | 5.12E-48 | 0.799545201 | 0.516 | 0.141 | 1.04E-43 | 16 | Ptpn9 |
| Hspa41 | 6.04E-48 | 1.041577277 | 0.777 | 0.315 | 1.22E-43 | 16 | Hspa4 |
| Rps2110 | 1.12E-47 | 0.821793657 | 1 | 0.927 | 2.27E-43 | 16 | Rps21 |
| Sh2d2a4 | 1.70E-47 | 0.46959036 | 0.283 | 0.049 | 3.44E-43 | 16 | Sh2d2a |
| Cysltr21 | 2.29E-47 | 0.536403533 | 0.25 | 0.039 | 4.63E-43 | 16 | Cysltr2 |
| Tnip11 | 2.42E-47 | 0.688578645 | 0.342 | 0.07 | 4.89E-43 | 16 | Tnip1 |
| Sh3bgrl36 | 4.84E-47 | 1.029388852 | 0.995 | 0.558 | 9.80E-43 | 16 | Sh3bgrl3 |
| Actr36 | 5.95E-47 | 0.92626502 | 0.891 | 0.39 | 1.20E-42 | 16 | Actr3 |
| Senp66 | 1.19E-46 | 0.950501987 | 0.75 | 0.297 | 2.40E-42 | 16 | Senp6 |
| Ern14 | 1.75E-46 | 0.737876452 | 0.375 | 0.084 | 3.53E-42 | 16 | Ern1 |
| Rpl386 | 1.97E-46 | 0.877704385 | 1 | 0.893 | 3.98E-42 | 16 | Rpl38 |

| | | | | | | | |
|---|---|---|---|---|---|---|---|
| Rpl97 | 2.27E-46 | 0.782688848 | 1 | 0.944 | 4.60E-42 | 16 | Rpl9 |
| Rpl27a6 | 2.94E-46 | 0.750182159 | 1 | 0.96 | 5.95E-42 | 16 | Rpl27a |
| Per11 | 3.88E-46 | 0.67595604 | 0.386 | 0.087 | 7.85E-42 | 16 | Per1 |
| Prkca5 | 5.34E-46 | 0.734034307 | 0.418 | 0.1 | 1.08E-41 | 16 | Prkca |
| Samhd14 | 7.36E-46 | 1.026144659 | 0.701 | 0.253 | 1.49E-41 | 16 | Samhd1 |
| Zfp36l15 | 8.06E-46 | 1.188281998 | 0.902 | 0.432 | 1.63E-41 | 16 | Zfp36l1 |
| Stk243 | 9.78E-46 | 0.850268424 | 0.592 | 0.195 | 1.98E-41 | 16 | Stk24 |
| Jak23 | 1.03E-45 | 0.754522379 | 0.473 | 0.126 | 2.09E-41 | 16 | Jak2 |
| S100a105 | 1.45E-45 | 1.0529006 | 0.967 | 0.468 | 2.93E-41 | 16 | S100a10 |
| Gpcpd12 | 1.63E-45 | 0.759468104 | 0.451 | 0.116 | 3.30E-41 | 16 | Gpcpd1 |
| Pkp34 | 1.74E-45 | 0.545185271 | 0.337 | 0.069 | 3.52E-41 | 16 | Pkp3 |
| Rpl396 | 2.03E-45 | 0.871347354 | 0.995 | 0.901 | 4.12E-41 | 16 | Rpl39 |
| Rps6ka15 | 2.78E-45 | 0.68996904 | 0.446 | 0.114 | 5.62E-41 | 16 | Rps6ka1 |
| Rpl345 | 2.96E-45 | 0.789442308 | 1 | 0.917 | 6.00E-41 | 16 | Rpl34 |
| Vars4 | 5.61E-45 | 0.653487965 | 0.364 | 0.081 | 1.14E-40 | 16 | Vars |
| Cbx44 | 5.94E-45 | 0.737903725 | 0.435 | 0.11 | 1.20E-40 | 16 | Cbx4 |
| Chd27 | 6.11E-45 | 0.894785561 | 0.701 | 0.253 | 1.24E-40 | 16 | Chd2 |
| Ptk2b5 | 8.07E-45 | 0.656606124 | 0.543 | 0.15 | 1.63E-40 | 16 | Ptk2b |
| AI5044321 | 2.67E-44 | 0.58739506 | 0.332 | 0.069 | 5.41E-40 | 16 | AI504432 |
| Jak16 | 3.54E-44 | 0.889797268 | 0.848 | 0.364 | 7.16E-40 | 16 | Jak1 |
| Ppp3ca9 | 6.34E-44 | 1.002489844 | 0.848 | 0.392 | 1.28E-39 | 16 | Ppp3ca |
| Ar4 | 7.50E-44 | 0.842742127 | 0.652 | 0.217 | 1.52E-39 | 16 | Ar |
| Ifrd13 | 1.00E-43 | 1.043776742 | 0.641 | 0.23 | 2.03E-39 | 16 | Ifrd1 |
| Ier23 | 1.12E-43 | 0.933191193 | 0.734 | 0.283 | 2.27E-39 | 16 | Ier2 |
| Cdc42se25 | 1.15E-43 | 0.8117022 | 0.679 | 0.238 | 2.33E-39 | 16 | Cdc42se2 |
| Baz1a5 | 1.28E-43 | 0.735524106 | 0.554 | 0.172 | 2.59E-39 | 16 | Baz1a |
| Tnfaip8l22 | 1.31E-43 | 0.561961113 | 0.337 | 0.071 | 2.65E-39 | 16 | Tnfaip8l2 |
| Mctp11 | 2.00E-43 | 0.861196001 | 0.342 | 0.073 | 4.06E-39 | 16 | Mctp1 |
| Nav24 | 2.01E-43 | 0.864361355 | 0.549 | 0.164 | 4.08E-39 | 16 | Nav2 |
| Gpr342 | 2.80E-43 | 0.595771342 | 0.299 | 0.059 | 5.67E-39 | 16 | Gpr34 |
| Herc45 | 1.17E-42 | 0.738345422 | 0.511 | 0.151 | 2.38E-38 | 16 | Herc4 |
| Rps27a6 | 1.57E-42 | 0.724205793 | 1 | 0.95 | 3.17E-38 | 16 | Rps27a |
| Ncoa28 | 4.94E-42 | 0.919299198 | 0.647 | 0.238 | 1.00E-37 | 16 | Ncoa2 |
| Rbl24 | 5.09E-42 | 0.700192993 | 0.478 | 0.134 | 1.03E-37 | 16 | Rbl2 |
| Gimap19 | 5.63E-42 | 0.859812697 | 0.788 | 0.302 | 1.14E-37 | 16 | Gimap1 |
| Pglyrp24 | 8.18E-42 | 0.470079994 | 0.25 | 0.044 | 1.66E-37 | 16 | Pglyrp2 |
| Pik3cd5 | 1.00E-41 | 0.623310054 | 0.424 | 0.108 | 2.03E-37 | 16 | Pik3cd |
| Rplp27 | 1.49E-41 | 0.753580672 | 1 | 0.89 | 3.03E-37 | 16 | Rplp2 |
| Dennd4a5 | 1.68E-41 | 0.650979064 | 0.609 | 0.187 | 3.41E-37 | 16 | Dennd4a |
| Spcs24 | 1.87E-41 | 0.833786796 | 0.875 | 0.4 | 3.78E-37 | 16 | Spcs2 |
| Ccm24 | 2.73E-41 | 0.65668739 | 0.457 | 0.126 | 5.52E-37 | 16 | Ccm2 |
| Srpk12 | 2.87E-41 | 0.623332755 | 0.527 | 0.161 | 5.82E-37 | 16 | Srpk1 |
| Rplp14 | 3.10E-41 | 0.700689121 | 1 | 0.983 | 6.27E-37 | 16 | Rplp1 |

| | | | | | | | |
|---|---|---|---|---|---|---|---|
| Gadd45b | 3.21E-41 | 0.839343265 | 0.413 | 0.107 | 6.50E-37 | 16 | Gadd45b |
| Rps146 | 3.34E-41 | 0.666350768 | 1 | 0.943 | 6.76E-37 | 16 | Rps14 |
| Traf35 | 5.53E-41 | 0.719912921 | 0.413 | 0.109 | 1.12E-36 | 16 | Traf3 |
| Ubash3b5 | 5.80E-41 | 0.87357184 | 0.527 | 0.167 | 1.17E-36 | 16 | Ubash3b |
| Stk105 | 9.86E-41 | 0.768537371 | 0.565 | 0.182 | 2.00E-36 | 16 | Stk10 |
| Sntb12 | 1.04E-40 | 0.692005057 | 0.31 | 0.065 | 2.11E-36 | 16 | Sntb1 |
| Pole4 | 1.22E-40 | 0.523281066 | 0.571 | 0.179 | 2.47E-36 | 16 | Pole4 |
| 4932438A13Rik6 | 1.76E-40 | 0.83340381 | 0.614 | 0.218 | 3.56E-36 | 16 | 4932438A13Rik |
| Rps4x6 | 2.34E-40 | 0.705196864 | 1 | 0.941 | 4.73E-36 | 16 | Rps4x |
| Arhgap44 | 3.13E-40 | 0.494419096 | 0.288 | 0.058 | 6.34E-36 | 16 | Arhgap4 |
| Atp10a2 | 3.64E-40 | 0.597802889 | 0.255 | 0.047 | 7.36E-36 | 16 | Atp10a |
| Slc7a11 | 5.83E-40 | 0.604652699 | 0.293 | 0.061 | 1.18E-35 | 16 | Slc7a1 |
| Rbm259 | 6.55E-40 | 0.783317717 | 0.799 | 0.333 | 1.33E-35 | 16 | Rbm25 |
| Pfn16 | 7.16E-40 | 0.908394892 | 0.995 | 0.682 | 1.45E-35 | 16 | Pfn1 |
| Tex23 | 8.74E-40 | 0.686399764 | 0.364 | 0.088 | 1.77E-35 | 16 | Tex2 |
| Clk16 | 1.32E-39 | 0.90032251 | 0.728 | 0.291 | 2.67E-35 | 16 | Clk1 |
| Arhgap255 | 1.43E-39 | 0.61888979 | 0.408 | 0.108 | 2.90E-35 | 16 | Arhgap25 |
| Fam129a4 | 1.87E-39 | 0.878797426 | 0.478 | 0.145 | 3.78E-35 | 16 | Fam129a |
| Atp2b16 | 2.12E-39 | 0.97315029 | 0.766 | 0.332 | 4.30E-35 | 16 | Atp2b1 |
| Uhrf23 | 3.48E-39 | 0.726376041 | 0.527 | 0.165 | 7.04E-35 | 16 | Uhrf2 |
| Rps286 | 3.90E-39 | 0.787466273 | 1 | 0.898 | 7.89E-35 | 16 | Rps28 |
| Ncoa17 | 5.64E-39 | 0.833855206 | 0.598 | 0.209 | 1.14E-34 | 16 | Ncoa1 |
| Apobec34 | 5.76E-39 | 0.603305767 | 0.652 | 0.225 | 1.17E-34 | 16 | Apobec3 |
| Tex145 | 6.32E-39 | 0.826609839 | 0.495 | 0.149 | 1.28E-34 | 16 | Tex14 |
| Rpl36a6 | 1.17E-38 | 0.789780824 | 0.995 | 0.752 | 2.37E-34 | 16 | Rpl36a |
| Fau5 | 1.23E-38 | 0.66794829 | 1 | 0.953 | 2.49E-34 | 16 | Fau |
| Pcgf5 | 1.38E-38 | 0.743241336 | 0.418 | 0.112 | 2.80E-34 | 16 | Pcgf5 |
| Rps67 | 1.58E-38 | 0.767792298 | 1 | 0.811 | 3.20E-34 | 16 | Rps6 |
| Lsp16 | 1.70E-38 | 0.59908507 | 0.478 | 0.139 | 3.45E-34 | 16 | Lsp1 |
| Gls7 | 1.77E-38 | 0.854094753 | 0.647 | 0.25 | 3.58E-34 | 16 | Gls |
| Fyb5 | 2.50E-38 | 0.501380214 | 0.598 | 0.183 | 5.05E-34 | 16 | Fyb |
| Acot74 | 2.54E-38 | 0.519511775 | 0.332 | 0.076 | 5.14E-34 | 16 | Acot7 |
| Grap25 | 2.71E-38 | 0.761861512 | 0.397 | 0.106 | 5.50E-34 | 16 | Grap2 |
| Lpxn4 | 3.10E-38 | 0.660675234 | 0.37 | 0.093 | 6.28E-34 | 16 | Lpxn |
| Dtnbp12 | 3.17E-38 | 0.574853766 | 0.641 | 0.227 | 6.42E-34 | 16 | Dtnbp1 |
| Rps295 | 3.33E-38 | 0.634032438 | 1 | 0.971 | 6.75E-34 | 16 | Rps29 |
| Rad502 | 3.38E-38 | 0.667943514 | 0.478 | 0.142 | 6.84E-34 | 16 | Rad50 |
| Selenot3 | 4.64E-38 | 0.799069254 | 0.592 | 0.211 | 9.40E-34 | 16 | Selenot |
| Capg5 | 6.06E-38 | 0.796047071 | 0.386 | 0.1 | 1.23E-33 | 16 | Capg |
| Cyba7 | 9.16E-38 | 0.675068359 | 0.935 | 0.439 | 1.85E-33 | 16 | Cyba |
| Sept117 | 1.03E-37 | 0.769383168 | 0.598 | 0.203 | 2.08E-33 | 16 | 11-Sep |
| Ppp1r15a2 | 1.19E-37 | 0.89661554 | 0.489 | 0.155 | 2.42E-33 | 16 | Ppp1r15a |
| Esyt15 | 3.53E-37 | 0.785298385 | 0.451 | 0.136 | 7.14E-33 | 16 | Esyt1 |

| | | | | | | | |
|---|---|---|---|---|---|---|---|
| Nlrc34 | 6.01E-37 | 0.522276645 | 0.277 | 0.058 | 1.22E-32 | 16 | Nlrc3 |
| Ppp1r12a7 | 8.46E-37 | 0.866940468 | 0.87 | 0.425 | 1.71E-32 | 16 | Ppp1r12a |
| Arid1b7 | 8.73E-37 | 0.735572208 | 0.674 | 0.258 | 1.77E-32 | 16 | Arid1b |
| Rps237 | 1.06E-36 | 0.609046971 | 1 | 0.953 | 2.14E-32 | 16 | Rps23 |
| Ccdc88c5 | 1.30E-36 | 0.658105535 | 0.38 | 0.099 | 2.63E-32 | 16 | Ccdc88c |
| Fhit5 | 1.36E-36 | 0.794470039 | 0.332 | 0.079 | 2.75E-32 | 16 | Fhit |
| Ccnd34 | 1.40E-36 | 0.74300975 | 0.647 | 0.239 | 2.83E-32 | 16 | Ccnd3 |
| Tra2b5 | 2.33E-36 | 0.772436373 | 0.712 | 0.296 | 4.72E-32 | 16 | Tra2b |
| Zfp36l25 | 2.45E-36 | 1.001100564 | 0.712 | 0.305 | 4.97E-32 | 16 | Zfp36l2 |
| Gmip4 | 2.96E-36 | 0.555152832 | 0.337 | 0.082 | 5.99E-32 | 16 | Gmip |
| Sgms14 | 4.01E-36 | 0.718169158 | 0.446 | 0.128 | 8.13E-32 | 16 | Sgms1 |
| Fam105a4 | 4.15E-36 | 0.598040716 | 0.44 | 0.125 | 8.40E-32 | 16 | Fam105a |
| Lncpint7 | 4.26E-36 | 0.902599861 | 0.674 | 0.274 | 8.62E-32 | 16 | Lncpint |
| B230307C23Rik | 4.98E-36 | 0.501720084 | 0.315 | 0.073 | 1.01E-31 | 16 | B230307C23Rik |
| Rap1gds19 | 5.19E-36 | 0.768686748 | 0.663 | 0.26 | 1.05E-31 | 16 | Rap1gds1 |
| mt-Co110 | 6.51E-36 | 0.79300173 | 1 | 0.719 | 1.32E-31 | 16 | mt-Co1 |
| Rpl366 | 9.00E-36 | 0.706466931 | 1 | 0.903 | 1.82E-31 | 16 | Rpl36 |
| Rpl217 | 1.12E-35 | 0.685356842 | 1 | 0.923 | 2.27E-31 | 16 | Rpl21 |
| Sik36 | 1.34E-35 | 0.854182968 | 0.668 | 0.269 | 2.72E-31 | 16 | Sik3 |
| Rack17 | 1.45E-35 | 0.742487401 | 1 | 0.792 | 2.93E-31 | 16 | Rack1 |
| Retreg14 | 1.58E-35 | 0.488447832 | 0.375 | 0.097 | 3.20E-31 | 16 | Retreg1 |
| Rbfa | 1.77E-35 | 0.594079798 | 0.446 | 0.132 | 3.59E-31 | 16 | Rbfa |
| Tmem1644 | 2.00E-35 | 0.707403432 | 0.5 | 0.163 | 4.05E-31 | 16 | Tmem164 |
| Lztfl1 | 2.17E-35 | 0.836972047 | 0.386 | 0.107 | 4.39E-31 | 16 | Lztfl1 |
| Trps15 | 2.24E-35 | 0.90397784 | 0.614 | 0.235 | 4.53E-31 | 16 | Trps1 |
| Tspan135 | 2.64E-35 | 0.677538367 | 0.723 | 0.285 | 5.35E-31 | 16 | Tspan13 |
| Dusp13 | 4.49E-35 | 0.833093897 | 0.56 | 0.193 | 9.08E-31 | 16 | Dusp1 |
| 4-Jun | 5.28E-35 | 1.076830187 | 0.908 | 0.514 | 1.07E-30 | 16 | Jun |
| Mbnl15 | 6.16E-35 | 0.755756507 | 0.935 | 0.458 | 1.25E-30 | 16 | Mbnl1 |
| Rps87 | 6.29E-35 | 0.599304813 | 1 | 0.965 | 1.27E-30 | 16 | Rps8 |
| Was5 | 6.48E-35 | 0.673020083 | 0.304 | 0.073 | 1.31E-30 | 16 | Was |
| Rpl117 | 7.09E-35 | 0.656812824 | 1 | 0.927 | 1.44E-30 | 16 | Rpl11 |
| Arhgef113 | 9.04E-35 | 0.740664286 | 0.647 | 0.248 | 1.83E-30 | 16 | Arhgef1 |
| Sptssa1 | 1.13E-34 | 0.638182218 | 0.728 | 0.303 | 2.30E-30 | 16 | Sptssa |
| mt-Nd57 | 1.31E-34 | 0.818219617 | 0.929 | 0.495 | 2.66E-30 | 16 | mt-Nd5 |
| Pkn16 | 2.12E-34 | 0.752196671 | 0.527 | 0.186 | 4.29E-30 | 16 | Pkn1 |
| Cyfip24 | 2.12E-34 | 0.641114755 | 0.353 | 0.091 | 4.30E-30 | 16 | Cyfip2 |
| Gm20234 | 2.75E-34 | 0.39310711 | 0.261 | 0.053 | 5.56E-30 | 16 | Gm20234 |
| Pced1b3 | 2.86E-34 | 0.613595854 | 0.413 | 0.117 | 5.80E-30 | 16 | Pced1b |
| mt-Co311 | 2.87E-34 | 0.725964915 | 1 | 0.726 | 5.80E-30 | 16 | mt-Co3 |
| Tpt15 | 3.18E-34 | 0.504669506 | 1 | 0.99 | 6.45E-30 | 16 | Tpt1 |
| Znrf35 | 3.85E-34 | 0.647881575 | 0.386 | 0.107 | 7.79E-30 | 16 | Znrf3 |
| Ifi2093 | 3.98E-34 | 0.511273527 | 0.266 | 0.057 | 8.05E-30 | 16 | Ifi209 |

| | | | | | | | |
|---|---|---|---|---|---|---|---|
| Traf14 | 4.33E-34 | 0.515207729 | 0.283 | 0.062 | 8.77E-30 | 16 | Traf1 |
| Gng25 | 4.37E-34 | 0.593738154 | 0.554 | 0.186 | 8.85E-30 | 16 | Gng2 |
| Crlf35 | 5.77E-34 | 0.664372437 | 0.484 | 0.157 | 1.17E-29 | 16 | Crlf3 |
| Rpl35a6 | 5.83E-34 | 0.597501462 | 1 | 0.948 | 1.18E-29 | 16 | Rpl35a |
| Vps37b4 | 7.56E-34 | 0.603818636 | 0.451 | 0.136 | 1.53E-29 | 16 | Vps37b |
| Abhd8 | 8.41E-34 | 0.445291774 | 0.266 | 0.056 | 1.70E-29 | 16 | Abhd8 |
| Sep-64 | 1.08E-33 | 0.555149086 | 0.37 | 0.099 | 2.19E-29 | 16 | 6-Sep |
| Ncor18 | 1.52E-33 | 0.847884621 | 0.848 | 0.437 | 3.08E-29 | 16 | Ncor1 |
| Snrpg5 | 1.56E-33 | 0.542960496 | 0.766 | 0.323 | 3.16E-29 | 16 | Snrpg |
| Trio4 | 1.69E-33 | 0.682777563 | 0.527 | 0.188 | 3.43E-29 | 16 | Trio |
| Slc38a25 | 2.74E-33 | 0.910315007 | 0.707 | 0.31 | 5.55E-29 | 16 | Slc38a2 |
| Tagln26 | 2.74E-33 | 0.784584579 | 0.853 | 0.428 | 5.56E-29 | 16 | Tagln2 |
| Rps6ka38 | 4.18E-33 | 0.743191982 | 0.652 | 0.262 | 8.47E-29 | 16 | Rps6ka3 |
| Sla6 | 4.23E-33 | 0.571069169 | 0.304 | 0.073 | 8.56E-29 | 16 | Sla |
| Nup2104 | 4.76E-33 | 0.477192154 | 0.288 | 0.067 | 9.64E-29 | 16 | Nup210 |
| Acap14 | 7.54E-33 | 0.481398397 | 0.299 | 0.071 | 1.53E-28 | 16 | Acap1 |
| H2-K19 | 7.63E-33 | 0.817607639 | 1 | 0.643 | 1.54E-28 | 16 | H2-K1 |
| Eml45 | 9.25E-33 | 0.893127847 | 0.62 | 0.258 | 1.87E-28 | 16 | Eml4 |
| Tusc32 | 1.10E-32 | 0.695956538 | 0.462 | 0.148 | 2.24E-28 | 16 | Tusc3 |
| Rpl226 | 1.13E-32 | 0.660020436 | 0.995 | 0.852 | 2.28E-28 | 16 | Rpl22 |
| Pitpnc17 | 1.22E-32 | 0.767099865 | 0.875 | 0.428 | 2.47E-28 | 16 | Pitpnc1 |
| Lcorl5 | 1.68E-32 | 0.763032649 | 0.484 | 0.166 | 3.40E-28 | 16 | Lcorl |
| Itgal4 | 1.97E-32 | 0.511335602 | 0.337 | 0.087 | 3.98E-28 | 16 | Itgal |
| Mcu1 | 2.84E-32 | 0.633919395 | 0.364 | 0.101 | 5.74E-28 | 16 | Mcu |
| Rpl22l16 | 3.61E-32 | 0.740846101 | 0.962 | 0.578 | 7.30E-28 | 16 | Rpl22l1 |
| Sash34 | 3.64E-32 | 0.477780619 | 0.266 | 0.06 | 7.37E-28 | 16 | Sash3 |
| Rnf1664 | 4.61E-32 | 0.564589083 | 0.375 | 0.107 | 9.33E-28 | 16 | Rnf166 |
| Rcn11 | 4.66E-32 | 0.486123852 | 0.386 | 0.108 | 9.43E-28 | 16 | Rcn1 |
| Orc4 | 4.94E-32 | 0.681159883 | 0.408 | 0.124 | 1.00E-27 | 16 | Orc4 |
| Rtf14 | 5.56E-32 | 0.651656541 | 0.663 | 0.279 | 1.13E-27 | 16 | Rtf1 |
| Osbpl85 | 5.61E-32 | 0.712615428 | 0.538 | 0.199 | 1.14E-27 | 16 | Osbpl8 |
| Dennd1b6 | 6.11E-32 | 0.683705891 | 0.522 | 0.184 | 1.24E-27 | 16 | Dennd1b |
| Tbc1d17 | 6.51E-32 | 0.668779096 | 0.489 | 0.165 | 1.32E-27 | 16 | Tbc1d1 |
| Hivep27 | 8.41E-32 | 0.882946131 | 0.674 | 0.298 | 1.70E-27 | 16 | Hivep2 |
| Rab8b5 | 1.01E-31 | 0.750702712 | 0.679 | 0.284 | 2.05E-27 | 16 | Rab8b |
| Frmd4b4 | 1.23E-31 | 0.662794865 | 0.516 | 0.177 | 2.49E-27 | 16 | Frmd4b |
| Ppp1cc2 | 1.45E-31 | 0.636355809 | 0.576 | 0.221 | 2.94E-27 | 16 | Ppp1cc |
| Gpr651 | 1.63E-31 | 0.402177102 | 0.342 | 0.088 | 3.31E-27 | 16 | Gpr65 |
| Slc4a72 | 1.92E-31 | 0.607349976 | 0.332 | 0.089 | 3.89E-27 | 16 | Slc4a7 |
| Prr134 | 2.08E-31 | 0.579458439 | 0.565 | 0.209 | 4.20E-27 | 16 | Prr13 |
| Vim8 | 2.11E-31 | 0.747388926 | 0.734 | 0.306 | 4.27E-27 | 16 | Vim |
| St53 | 2.20E-31 | 0.677750233 | 0.462 | 0.151 | 4.46E-27 | 16 | St5 |
| Tsc22d44 | 2.34E-31 | 0.554007922 | 0.707 | 0.294 | 4.74E-27 | 16 | Tsc22d4 |

| | | | | | | | |
|---|---|---|---|---|---|---|---|
| Psd44 | 3.02E-31 | 0.454839404 | 0.277 | 0.065 | 6.12E-27 | 16 | Psd4 |
| Tmsb4x8 | 3.14E-31 | 0.571661083 | 1 | 0.934 | 6.37E-27 | 16 | Tmsb4x |
| Ppp1r14b2 | 3.29E-31 | 0.376328813 | 0.777 | 0.326 | 6.65E-27 | 16 | Ppp1r14b |
| Galnt19 | 5.95E-31 | 0.689138546 | 0.505 | 0.184 | 1.20E-26 | 16 | Galnt1 |
| Dnaja15 | 7.49E-31 | 0.849032568 | 0.908 | 0.516 | 1.52E-26 | 16 | Dnaja1 |
| Rpl146 | 8.42E-31 | 0.629766837 | 1 | 0.802 | 1.70E-26 | 16 | Rpl14 |
| Ldha1 | 9.31E-31 | 0.643243274 | 0.837 | 0.411 | 1.88E-26 | 16 | Ldha |
| Paxbp11 | 9.36E-31 | 0.622698607 | 0.402 | 0.124 | 1.89E-26 | 16 | Paxbp1 |
| Eef1b27 | 1.02E-30 | 0.688626225 | 0.989 | 0.736 | 2.07E-26 | 16 | Eef1b2 |
| Nfkbia6 | 1.11E-30 | 0.852434366 | 0.793 | 0.4 | 2.24E-26 | 16 | Nfkbia |
| Ssbp25 | 1.20E-30 | 0.627419432 | 0.44 | 0.14 | 2.43E-26 | 16 | Ssbp2 |
| Ldlrad45 | 1.30E-30 | 0.894151523 | 0.625 | 0.25 | 2.63E-26 | 16 | Ldlrad4 |
| Card114 | 1.37E-30 | 0.626974666 | 0.315 | 0.082 | 2.78E-26 | 16 | Card11 |
| Asxl24 | 1.49E-30 | 0.632001054 | 0.5 | 0.174 | 3.01E-26 | 16 | Asxl2 |
| Ets17 | 1.96E-30 | 0.690341065 | 0.859 | 0.403 | 3.97E-26 | 16 | Ets1 |
| Capza15 | 2.89E-30 | 0.625089107 | 0.652 | 0.275 | 5.86E-26 | 16 | Capza1 |
| Snx201 | 4.72E-30 | 0.461006146 | 0.266 | 0.062 | 9.56E-26 | 16 | Snx20 |
| Taok34 | 5.53E-30 | 0.466985631 | 0.489 | 0.163 | 1.12E-25 | 16 | Taok3 |
| Herc34 | 7.96E-30 | 0.490555666 | 0.288 | 0.071 | 1.61E-25 | 16 | Herc3 |
| Tnrc6b6 | 9.49E-30 | 0.741115694 | 0.783 | 0.371 | 1.92E-25 | 16 | Tnrc6b |
| Smap13 | 1.18E-29 | 0.551028998 | 0.777 | 0.36 | 2.38E-25 | 16 | Smap1 |
| Ptpn25 | 1.25E-29 | 0.59453147 | 0.435 | 0.143 | 2.52E-25 | 16 | Ptpn2 |
| Traf3ip34 | 1.32E-29 | 0.477729212 | 0.304 | 0.079 | 2.66E-25 | 16 | Traf3ip3 |
| Dgka | 1.39E-29 | 0.480403977 | 0.261 | 0.06 | 2.82E-25 | 16 | Dgka |
| Cnn25 | 1.49E-29 | 0.615147499 | 0.505 | 0.182 | 3.01E-25 | 16 | Cnn2 |
| Ero1l | 1.56E-29 | 0.67946607 | 0.332 | 0.092 | 3.17E-25 | 16 | Ero1l |
| R3hdm12 | 1.80E-29 | 0.563366801 | 0.522 | 0.191 | 3.65E-25 | 16 | R3hdm1 |
| Atp8a13 | 1.87E-29 | 0.680642136 | 0.603 | 0.241 | 3.78E-25 | 16 | Atp8a1 |
| Rab27b | 3.08E-29 | 0.324485152 | 0.326 | 0.085 | 6.23E-25 | 16 | Rab27b |
| Hmgb23 | 3.97E-29 | 0.462309754 | 0.755 | 0.328 | 8.04E-25 | 16 | Hmgb2 |
| Xist9 | 4.40E-29 | 0.762131088 | 0.891 | 0.463 | 8.91E-25 | 16 | Xist |
| Rev3l5 | 4.60E-29 | 0.60139329 | 0.457 | 0.157 | 9.32E-25 | 16 | Rev3l |
| Serinc35 | 4.87E-29 | 0.719568687 | 0.924 | 0.486 | 9.86E-25 | 16 | Serinc3 |
| Sema4d5 | 6.09E-29 | 0.514690956 | 0.418 | 0.13 | 1.23E-24 | 16 | Sema4d |
| Ncl4 | 8.41E-29 | 0.659573694 | 0.924 | 0.516 | 1.70E-24 | 16 | Ncl |
| Rpl78 | 8.46E-29 | 0.566016166 | 0.995 | 0.893 | 1.71E-24 | 16 | Rpl7 |
| Unc1194 | 1.31E-28 | 0.497113121 | 0.31 | 0.084 | 2.65E-24 | 16 | Unc119 |
| Kalrn3 | 1.34E-28 | 0.504371309 | 0.37 | 0.106 | 2.71E-24 | 16 | Kalrn |
| Fam172a8 | 1.80E-28 | 0.87132408 | 0.636 | 0.284 | 3.64E-24 | 16 | Fam172a |
| B4galnt14 | 1.83E-28 | 0.459432894 | 0.337 | 0.095 | 3.70E-24 | 16 | B4galnt1 |
| Tmem1732 | 1.97E-28 | 0.39838128 | 0.283 | 0.071 | 3.99E-24 | 16 | Tmem173 |
| Neat17 | 2.07E-28 | 0.706884107 | 0.538 | 0.209 | 4.19E-24 | 16 | Neat1 |
| H2-Q67 | 2.61E-28 | 0.631701209 | 0.609 | 0.245 | 5.28E-24 | 16 | H2-Q6 |

| | | | | | | | |
|---|---|---|---|---|---|---|---|
| Clint13 | 3.14E-28 | 0.441468272 | 0.788 | 0.373 | 6.36E-24 | 16 | Clint1 |
| Gimap86 | 3.17E-28 | 0.520558292 | 0.489 | 0.169 | 6.42E-24 | 16 | Gimap8 |
| Sesn15 | 3.65E-28 | 0.695778048 | 0.543 | 0.212 | 7.38E-24 | 16 | Sesn1 |
| Zfp1486 | 3.75E-28 | 0.710512022 | 0.62 | 0.264 | 7.60E-24 | 16 | Zfp148 |
| Pi4ka4 | 4.14E-28 | 0.595084453 | 0.397 | 0.128 | 8.38E-24 | 16 | Pi4ka |
| Rpl356 | 5.12E-28 | 0.60378163 | 1 | 0.828 | 1.04E-23 | 16 | Rpl35 |
| Chd45 | 8.47E-28 | 0.615978887 | 0.717 | 0.327 | 1.71E-23 | 16 | Chd4 |
| Nipbl10 | 1.39E-27 | 0.762118909 | 0.734 | 0.366 | 2.81E-23 | 16 | Nipbl |
| Rpl58 | 1.39E-27 | 0.654557605 | 0.978 | 0.71 | 2.82E-23 | 16 | Rpl5 |
| Rgs16 | 1.41E-27 | 0.552035484 | 0.44 | 0.141 | 2.86E-23 | 16 | Rgs1 |
| AC149090.15 | 1.51E-27 | 0.742518816 | 0.571 | 0.231 | 3.06E-23 | 16 | AC149090.1 |
| Smyd34 | 1.58E-27 | 0.398116427 | 0.598 | 0.226 | 3.19E-23 | 16 | Smyd3 |
| mt-Atp88 | 2.24E-27 | 0.72968079 | 0.859 | 0.463 | 4.54E-23 | 16 | mt-Atp8 |
| Scfd22 | 2.89E-27 | 0.523707206 | 0.31 | 0.086 | 5.85E-23 | 16 | Scfd2 |
| Aprt2 | 3.04E-27 | 0.451670051 | 0.603 | 0.244 | 6.16E-23 | 16 | Aprt |
| Chm | 3.83E-27 | 0.420187946 | 0.38 | 0.117 | 7.76E-23 | 16 | Chm |
| Tes4 | 3.95E-27 | 0.464256509 | 0.402 | 0.129 | 7.99E-23 | 16 | Tes |
| Rpl276 | 4.06E-27 | 0.58056236 | 0.967 | 0.582 | 8.22E-23 | 16 | Rpl27 |
| Rcc22 | 4.22E-27 | 0.474356773 | 0.364 | 0.112 | 8.54E-23 | 16 | Rcc2 |
| Acvr2a3 | 5.72E-27 | 0.686889763 | 0.435 | 0.149 | 1.16E-22 | 16 | Acvr2a |
| Smc44 | 6.76E-27 | 0.60685816 | 0.457 | 0.163 | 1.37E-22 | 16 | Smc4 |
| Rpl10a7 | 7.17E-27 | 0.578381029 | 1 | 0.836 | 1.45E-22 | 16 | Rpl10a |
| Glb1 | 7.20E-27 | 0.503214715 | 0.272 | 0.07 | 1.46E-22 | 16 | Glb1 |
| Rps220 | 7.90E-27 | 0.559612193 | 1 | 0.92 | 1.60E-22 | 16 | Rps2 |
| Tbc1d9b2 | 1.06E-26 | 0.59205678 | 0.375 | 0.122 | 2.15E-22 | 16 | Tbc1d9b |
| Zfp2928 | 1.07E-26 | 0.568334047 | 0.609 | 0.251 | 2.17E-22 | 16 | Zfp292 |
| Pvt14 | 1.12E-26 | 0.485882279 | 0.293 | 0.079 | 2.26E-22 | 16 | Pvt1 |
| Rpl156 | 1.17E-26 | 0.5476522 | 1 | 0.901 | 2.36E-22 | 16 | Rpl15 |
| Esyt28 | 1.30E-26 | 0.72698187 | 0.582 | 0.242 | 2.63E-22 | 16 | Esyt2 |
| Arhgap267 | 1.61E-26 | 0.54679561 | 0.576 | 0.222 | 3.25E-22 | 16 | Arhgap26 |
| Sgf29 | 1.77E-26 | 0.420474447 | 0.299 | 0.081 | 3.59E-22 | 16 | Sgf29 |
| Jmjd1c8 | 1.78E-26 | 0.629899808 | 0.679 | 0.307 | 3.59E-22 | 16 | Jmjd1c |
| Ddx56 | 1.99E-26 | 0.730883636 | 0.978 | 0.654 | 4.03E-22 | 16 | Ddx5 |
| Kat2b3 | 2.07E-26 | 0.582741975 | 0.446 | 0.158 | 4.19E-22 | 16 | Kat2b |
| Rpl317 | 2.44E-26 | 0.534095545 | 1 | 0.91 | 4.94E-22 | 16 | Rpl3 |
| Nlrc54 | 3.06E-26 | 0.569585577 | 0.359 | 0.111 | 6.19E-22 | 16 | Nlrc5 |
| Trim145 | 3.99E-26 | 0.611082185 | 0.375 | 0.121 | 8.07E-22 | 16 | Trim14 |
| Rbpj5 | 3.99E-26 | 0.612632705 | 0.554 | 0.22 | 8.08E-22 | 16 | Rbpj |
| Egr13 | 5.05E-26 | 0.693959126 | 0.478 | 0.175 | 1.02E-21 | 16 | Egr1 |
| Ezh2 | 5.09E-26 | 0.375128064 | 0.272 | 0.07 | 1.03E-21 | 16 | Ezh2 |
| Rpl67 | 6.15E-26 | 0.536205782 | 1 | 0.881 | 1.25E-21 | 16 | Rpl6 |
| Exoc49 | 6.31E-26 | 0.676613974 | 0.685 | 0.322 | 1.28E-21 | 16 | Exoc4 |
| Fbxl1710 | 6.71E-26 | 0.762193914 | 0.668 | 0.311 | 1.36E-21 | 16 | Fbxl17 |

| | | | | | | | |
|---|---|---|---|---|---|---|---|
| Zcchc79 | 7.28E-26 | 0.706336076 | 0.663 | 0.301 | 1.47E-21 | 16 | Zcchc7 |
| Ikzf34 | 1.07E-25 | 0.403401825 | 0.397 | 0.12 | 2.18E-21 | 16 | Ikzf3 |
| Itgb25 | 1.24E-25 | 0.46288229 | 0.337 | 0.102 | 2.50E-21 | 16 | Itgb2 |
| Gm269172 | 1.57E-25 | 0.486329576 | 0.375 | 0.12 | 3.18E-21 | 16 | Gm26917 |
| mt-Atp610 | 1.65E-25 | 0.616158175 | 1 | 0.754 | 3.35E-21 | 16 | mt-Atp6 |
| Elmo18 | 1.77E-25 | 0.658707674 | 0.766 | 0.366 | 3.58E-21 | 16 | Elmo1 |
| Uba527 | 1.91E-25 | 0.512100621 | 0.946 | 0.521 | 3.87E-21 | 16 | Uba52 |
| Rpl296 | 2.16E-25 | 0.595601898 | 0.995 | 0.768 | 4.38E-21 | 16 | Rpl29 |
| Dad16 | 2.17E-25 | 0.507281876 | 0.87 | 0.458 | 4.40E-21 | 16 | Dad1 |
| Ptpn224 | 2.74E-25 | 0.490047686 | 0.38 | 0.119 | 5.56E-21 | 16 | Ptpn22 |
| Rpl13a6 | 3.06E-25 | 0.512089897 | 0.647 | 0.284 | 6.19E-21 | 16 | Rpl13a |
| Prex15 | 3.11E-25 | 0.567576593 | 0.473 | 0.174 | 6.30E-21 | 16 | Prex1 |
| Nsd36 | 3.36E-25 | 0.620572828 | 0.668 | 0.308 | 6.80E-21 | 16 | Nsd3 |
| Snrpf5 | 5.20E-25 | 0.4404554 | 0.788 | 0.379 | 1.05E-20 | 16 | Snrpf |
| Asxl12 | 6.01E-25 | 0.551868072 | 0.413 | 0.146 | 1.22E-20 | 16 | Asxl1 |
| Sun23 | 6.24E-25 | 0.415035274 | 0.283 | 0.077 | 1.26E-20 | 16 | Sun2 |
| Rnaseh2c4 | 7.06E-25 | 0.429279855 | 0.647 | 0.285 | 1.43E-20 | 16 | Rnaseh2c |
| Mapkapk33 | 7.28E-25 | 0.520333783 | 0.288 | 0.082 | 1.47E-20 | 16 | Mapkapk3 |
| Susd68 | 8.44E-25 | 0.661912927 | 0.625 | 0.282 | 1.71E-20 | 16 | Susd6 |
| Gimap94 | 8.59E-25 | 0.521003217 | 0.446 | 0.159 | 1.74E-20 | 16 | Gimap9 |
| Sec11c6 | 8.90E-25 | 0.49395104 | 0.685 | 0.31 | 1.80E-20 | 16 | Sec11c |
| Fgl25 | 1.05E-24 | 0.731005156 | 0.342 | 0.108 | 2.13E-20 | 16 | Fgl2 |
| Nr4a11 | 1.26E-24 | 0.487936631 | 0.348 | 0.106 | 2.54E-20 | 16 | Nr4a1 |
| Ralgps2 | 1.68E-24 | 0.428286735 | 0.25 | 0.063 | 3.41E-20 | 16 | Ralgps2 |
| Tcf205 | 1.71E-24 | 0.546652647 | 0.402 | 0.141 | 3.47E-20 | 16 | Tcf20 |
| mt-Nd110 | 1.87E-24 | 0.652141438 | 0.973 | 0.598 | 3.78E-20 | 16 | mt-Nd1 |
| Rsbn1l5 | 2.08E-24 | 0.528313865 | 0.63 | 0.272 | 4.22E-20 | 16 | Rsbn1l |
| Ell21 | 2.10E-24 | 0.437290785 | 0.299 | 0.085 | 4.25E-20 | 16 | Ell2 |
| Ywhaz3 | 2.17E-24 | 0.634112324 | 0.859 | 0.493 | 4.40E-20 | 16 | Ywhaz |
| Specc12 | 2.86E-24 | 0.452411672 | 0.288 | 0.081 | 5.79E-20 | 16 | Specc1 |
| Sidt2 | 3.07E-24 | 0.400828039 | 0.25 | 0.064 | 6.21E-20 | 16 | Sidt2 |
| Ankrd119 | 3.44E-24 | 0.653515446 | 0.755 | 0.381 | 6.97E-20 | 16 | Ankrd11 |
| Anxa63 | 3.71E-24 | 0.50690237 | 0.435 | 0.157 | 7.51E-20 | 16 | Anxa6 |
| Pde3b5 | 4.21E-24 | 0.631466706 | 0.38 | 0.128 | 8.52E-20 | 16 | Pde3b |
| Itpr14 | 4.79E-24 | 0.568407869 | 0.538 | 0.221 | 9.71E-20 | 16 | Itpr1 |
| Bola24 | 5.37E-24 | 0.369656355 | 0.614 | 0.262 | 1.09E-19 | 16 | Bola2 |
| Eef1g6 | 6.28E-24 | 0.475018275 | 0.908 | 0.51 | 1.27E-19 | 16 | Eef1g |
| Map2k42 | 7.27E-24 | 0.458944042 | 0.435 | 0.159 | 1.47E-19 | 16 | Map2k4 |
| Akap138 | 1.09E-23 | 0.60851871 | 0.766 | 0.394 | 2.20E-19 | 16 | Akap13 |
| Phf63 | 1.17E-23 | 0.535087614 | 0.386 | 0.135 | 2.36E-19 | 16 | Phf6 |
| Pla2g165 | 1.24E-23 | 0.416812072 | 0.576 | 0.243 | 2.52E-19 | 16 | Pla2g16 |
| Stk393 | 1.25E-23 | 0.36691619 | 0.435 | 0.154 | 2.53E-19 | 16 | Stk39 |
| Arhgap94 | 1.41E-23 | 0.322718747 | 0.353 | 0.111 | 2.86E-19 | 16 | Arhgap9 |

| | | | | | | | |
|---|---|---|---|---|---|---|---|
| Arih16 | 1.91E-23 | 0.594471339 | 0.658 | 0.313 | 3.86E-19 | 16 | Arih1 |
| Prdx62 | 2.11E-23 | 0.357880981 | 0.734 | 0.337 | 4.27E-19 | 16 | Prdx6 |
| Actb6 | 2.12E-23 | 0.621112263 | 0.995 | 0.843 | 4.30E-19 | 16 | Actb |
| Apaf1 | 2.92E-23 | 0.484264513 | 0.293 | 0.086 | 5.91E-19 | 16 | Apaf1 |
| Cnst | 3.11E-23 | 0.389915965 | 0.288 | 0.082 | 6.29E-19 | 16 | Cnst |
| Zfp6543 | 3.64E-23 | 0.600156489 | 0.353 | 0.121 | 7.36E-19 | 16 | Zfp654 |
| Esco14 | 3.82E-23 | 0.489049308 | 0.467 | 0.179 | 7.74E-19 | 16 | Esco1 |
| Ccnl15 | 4.08E-23 | 0.61361811 | 0.511 | 0.21 | 8.25E-19 | 16 | Ccnl1 |
| Trmt1124 | 4.72E-23 | 0.516881603 | 0.614 | 0.274 | 9.56E-19 | 16 | Trmt112 |
| Cdk2ap24 | 4.86E-23 | 0.25586687 | 0.516 | 0.199 | 9.84E-19 | 16 | Cdk2ap2 |
| Malt14 | 4.93E-23 | 0.785513717 | 0.402 | 0.15 | 9.98E-19 | 16 | Malt1 |
| Prrc2c9 | 5.04E-23 | 0.643813969 | 0.799 | 0.42 | 1.02E-18 | 16 | Prrc2c |
| Ppp1r185 | 5.90E-23 | 0.585379256 | 0.413 | 0.154 | 1.19E-18 | 16 | Ppp1r18 |
| Mettl233 | 6.05E-23 | 0.448124176 | 0.5 | 0.203 | 1.22E-18 | 16 | Mettl23 |
| Zeb17 | 6.11E-23 | 0.682449785 | 0.674 | 0.322 | 1.24E-18 | 16 | Zeb1 |
| Nufip23 | 8.52E-23 | 0.535215046 | 0.554 | 0.232 | 1.72E-18 | 16 | Nufip2 |
| Galnt101 | 9.15E-23 | 0.555222438 | 0.326 | 0.105 | 1.85E-18 | 16 | Galnt10 |
| Me23 | 1.23E-22 | 0.407987508 | 0.266 | 0.075 | 2.49E-18 | 16 | Me2 |
| Grb22 | 1.35E-22 | 0.490040433 | 0.625 | 0.282 | 2.73E-18 | 16 | Grb2 |
| Tbl1xr15 | 1.37E-22 | 0.535188854 | 0.435 | 0.165 | 2.78E-18 | 16 | Tbl1xr1 |
| Nt5c4 | 1.41E-22 | 0.429357644 | 0.576 | 0.25 | 2.85E-18 | 16 | Nt5c |
| Serp11 | 1.45E-22 | 0.385996852 | 0.527 | 0.217 | 2.93E-18 | 16 | Serp1 |
| Coq10b3 | 1.49E-22 | 0.497893375 | 0.418 | 0.154 | 3.02E-18 | 16 | Coq10b |
| Eif3e5 | 1.62E-22 | 0.426057312 | 0.81 | 0.412 | 3.27E-18 | 16 | Eif3e |
| Rps266 | 1.93E-22 | 0.485856603 | 0.995 | 0.865 | 3.91E-18 | 16 | Rps26 |
| Bin25 | 2.09E-22 | 0.406998585 | 0.375 | 0.128 | 4.22E-18 | 16 | Bin2 |
| Rngtt4 | 2.50E-22 | 0.515965875 | 0.38 | 0.135 | 5.07E-18 | 16 | Rngtt |
| Akna5 | 2.54E-22 | 0.446756687 | 0.299 | 0.091 | 5.15E-18 | 16 | Akna |
| Rps27rt3 | 2.57E-22 | 0.405847438 | 0.332 | 0.108 | 5.21E-18 | 16 | Rps27rt |
| Rps127 | 2.76E-22 | 0.472581003 | 1 | 0.898 | 5.59E-18 | 16 | Rps12 |
| Rps154 | 3.41E-22 | 0.554736529 | 0.995 | 0.748 | 6.91E-18 | 16 | Rps15 |
| Pak25 | 3.77E-22 | 0.518163572 | 0.799 | 0.435 | 7.63E-18 | 16 | Pak2 |
| Vav14 | 4.87E-22 | 0.368468591 | 0.277 | 0.081 | 9.87E-18 | 16 | Vav1 |
| Rptor5 | 4.98E-22 | 0.514517815 | 0.386 | 0.139 | 1.01E-17 | 16 | Rptor |
| Ankrd178 | 5.16E-22 | 0.553710472 | 0.587 | 0.261 | 1.04E-17 | 16 | Ankrd17 |
| Rbm397 | 7.13E-22 | 0.583370205 | 0.984 | 0.573 | 1.44E-17 | 16 | Rbm39 |
| Irak25 | 7.89E-22 | 0.647925496 | 0.386 | 0.142 | 1.60E-17 | 16 | Irak2 |
| Rpl46 | 8.29E-22 | 0.507454173 | 0.978 | 0.658 | 1.68E-17 | 16 | Rpl4 |
| Rplp06 | 9.10E-22 | 0.479288701 | 1 | 0.945 | 1.84E-17 | 16 | Rplp0 |
| Ptp4a13 | 1.03E-21 | 0.502076487 | 0.391 | 0.144 | 2.08E-17 | 16 | Ptp4a1 |
| Xpo73 | 1.13E-21 | 0.446202878 | 0.413 | 0.154 | 2.28E-17 | 16 | Xpo7 |
| Vps284 | 1.19E-21 | 0.39686193 | 0.652 | 0.3 | 2.40E-17 | 16 | Vps28 |
| Irf2bp21 | 1.28E-21 | 0.444929194 | 0.543 | 0.228 | 2.59E-17 | 16 | Irf2bp2 |

| | | | | | | | |
|---|---|---|---|---|---|---|---|
| Fam49b5 | 1.34E-21 | 0.397424022 | 0.576 | 0.237 | 2.71E-17 | 16 | Fam49b |
| Mum11 | 1.56E-21 | 0.415080053 | 0.342 | 0.114 | 3.15E-17 | 16 | Mum1 |
| Rai13 | 1.71E-21 | 0.442198176 | 0.332 | 0.109 | 3.47E-17 | 16 | Rai1 |
| Etv63 | 1.78E-21 | 0.49759474 | 0.625 | 0.278 | 3.61E-17 | 16 | Etv6 |
| Plp21 | 1.82E-21 | 0.358790336 | 0.321 | 0.102 | 3.68E-17 | 16 | Plp2 |
| Zfp1 | 2.07E-21 | 0.313309113 | 0.255 | 0.071 | 4.19E-17 | 16 | Zfp1 |
| Gimap67 | 2.10E-21 | 0.706346433 | 0.728 | 0.366 | 4.26E-17 | 16 | Gimap6 |
| Map2k6 | 2.38E-21 | 0.411079766 | 0.293 | 0.089 | 4.82E-17 | 16 | Map2k6 |
| Cast6 | 2.43E-21 | 0.569998161 | 0.63 | 0.312 | 4.91E-17 | 16 | Cast |
| Cebpb2 | 2.45E-21 | 0.561995963 | 0.413 | 0.157 | 4.96E-17 | 16 | Cebpb |
| Atp2a24 | 3.32E-21 | 0.582425939 | 0.484 | 0.2 | 6.73E-17 | 16 | Atp2a2 |
| Cdk137 | 3.61E-21 | 0.470605722 | 0.533 | 0.227 | 7.30E-17 | 16 | Cdk13 |
| Mycbp29 | 3.73E-21 | 0.595994593 | 0.734 | 0.374 | 7.55E-17 | 16 | Mycbp2 |
| Csf13 | 3.88E-21 | 0.489964247 | 0.337 | 0.11 | 7.85E-17 | 16 | Csf1 |
| Nckap1l4 | 4.22E-21 | 0.425891901 | 0.288 | 0.089 | 8.54E-17 | 16 | Nckap1l |
| Pan39 | 4.87E-21 | 0.547783711 | 0.522 | 0.229 | 9.86E-17 | 16 | Pan3 |
| Tshz31 | 4.94E-21 | 0.517125048 | 0.299 | 0.093 | 1.00E-16 | 16 | Tshz3 |
| Exoc3 | 4.97E-21 | 0.423284658 | 0.451 | 0.178 | 1.01E-16 | 16 | Exoc3 |
| Fmnl15 | 4.99E-21 | 0.446151779 | 0.293 | 0.093 | 1.01E-16 | 16 | Fmnl1 |
| mt-Nd410 | 5.29E-21 | 0.605333115 | 0.995 | 0.675 | 1.07E-16 | 16 | mt-Nd4 |
| Ppp1r102 | 5.53E-21 | 0.604531525 | 0.457 | 0.19 | 1.12E-16 | 16 | Ppp1r10 |
| mt-Nd28 | 5.69E-21 | 0.55934644 | 1 | 0.688 | 1.15E-16 | 16 | mt-Nd2 |
| Scaper7 | 5.70E-21 | 0.573793981 | 0.451 | 0.18 | 1.15E-16 | 16 | Scaper |
| Uhrf1bp1l | 8.37E-21 | 0.330459011 | 0.299 | 0.093 | 1.69E-16 | 16 | Uhrf1bp1l |
| Ubac25 | 1.10E-20 | 0.515375661 | 0.462 | 0.188 | 2.23E-16 | 16 | Ubac2 |
| 9130401M01Rik | 1.10E-20 | 0.321925676 | 0.266 | 0.078 | 2.24E-16 | 16 | 9130401M01Rik |
| Zbtb38 | 1.17E-20 | 0.432912538 | 0.408 | 0.154 | 2.37E-16 | 16 | Zbtb38 |
| Tspan143 | 1.20E-20 | 0.486970307 | 0.359 | 0.128 | 2.43E-16 | 16 | Tspan14 |
| Gm118082 | 1.25E-20 | 0.356581114 | 0.408 | 0.154 | 2.52E-16 | 16 | Gm11808 |
| Kansl19 | 1.36E-20 | 0.647307441 | 0.658 | 0.34 | 2.75E-16 | 16 | Kansl1 |
| Huwe18 | 1.49E-20 | 0.505013708 | 0.598 | 0.285 | 3.02E-16 | 16 | Huwe1 |
| Zhx1 | 1.64E-20 | 0.318923751 | 0.304 | 0.097 | 3.33E-16 | 16 | Zhx1 |
| Arid1a7 | 1.67E-20 | 0.492744113 | 0.527 | 0.23 | 3.39E-16 | 16 | Arid1a |
| Nsun61 | 1.74E-20 | 0.481999638 | 0.304 | 0.098 | 3.51E-16 | 16 | Nsun6 |
| Mtmr37 | 1.79E-20 | 0.494350326 | 0.484 | 0.201 | 3.63E-16 | 16 | Mtmr3 |
| Arhgef181 | 1.93E-20 | 0.394572924 | 0.25 | 0.072 | 3.90E-16 | 16 | Arhgef18 |
| Itm2c4 | 2.16E-20 | 0.465570883 | 0.484 | 0.199 | 4.37E-16 | 16 | Itm2c |
| Rbm35 | 2.51E-20 | 0.40795389 | 0.875 | 0.476 | 5.07E-16 | 16 | Rbm3 |
| Maz2 | 2.56E-20 | 0.344980055 | 0.293 | 0.093 | 5.18E-16 | 16 | Maz |
| Glipr14 | 2.61E-20 | 0.318618269 | 0.304 | 0.096 | 5.28E-16 | 16 | Glipr1 |
| Diaph15 | 2.71E-20 | 0.595641583 | 0.533 | 0.243 | 5.50E-16 | 16 | Diaph1 |
| Aim25 | 2.72E-20 | 0.472191355 | 0.31 | 0.103 | 5.51E-16 | 16 | Aim2 |
| Taf1d3 | 2.76E-20 | 0.383024433 | 0.391 | 0.145 | 5.58E-16 | 16 | Taf1d |

| | | | | | | | |
|---|---|---|---|---|---|---|---|
| Zfp6386 | 2.96E-20 | 0.584861969 | 0.592 | 0.282 | 6.00E-16 | 16 | Zfp638 |
| H2afy4 | 3.00E-20 | 0.426472991 | 0.696 | 0.337 | 6.07E-16 | 16 | H2afy |
| Osbpl34 | 3.07E-20 | 0.437148814 | 0.353 | 0.12 | 6.21E-16 | 16 | Osbpl3 |
| Cdkn2d3 | 3.19E-20 | 0.372760248 | 0.293 | 0.093 | 6.45E-16 | 16 | Cdkn2d |
| Plcl26 | 3.36E-20 | 0.541493231 | 0.332 | 0.115 | 6.80E-16 | 16 | Plcl2 |
| Nop583 | 4.34E-20 | 0.42487227 | 0.375 | 0.138 | 8.78E-16 | 16 | Nop58 |
| Ago24 | 4.36E-20 | 0.460797758 | 0.424 | 0.168 | 8.83E-16 | 16 | Ago2 |
| Thada5 | 4.84E-20 | 0.511149065 | 0.337 | 0.12 | 9.80E-16 | 16 | Thada |
| Dis3l25 | 5.02E-20 | 0.439191695 | 0.375 | 0.139 | 1.02E-15 | 16 | Dis3l2 |
| Grap3 | 7.17E-20 | 0.434398548 | 0.337 | 0.115 | 1.45E-15 | 16 | Grap |
| Gclc | 7.91E-20 | 0.6441456 | 0.31 | 0.105 | 1.60E-15 | 16 | Gclc |
| Ssh25 | 8.43E-20 | 0.452875399 | 0.533 | 0.226 | 1.71E-15 | 16 | Ssh2 |
| Cnot46 | 8.87E-20 | 0.49327324 | 0.582 | 0.269 | 1.80E-15 | 16 | Cnot4 |
| Impact2 | 1.08E-19 | 0.524833965 | 0.342 | 0.123 | 2.19E-15 | 16 | Impact |
| Ctps21 | 1.08E-19 | 0.340024219 | 0.283 | 0.089 | 2.19E-15 | 16 | Ctps2 |
| Zfp622 | 1.53E-19 | 0.336108648 | 0.315 | 0.106 | 3.10E-15 | 16 | Zfp622 |
| Snhg121 | 1.61E-19 | 0.372946735 | 0.332 | 0.115 | 3.25E-15 | 16 | Snhg12 |
| Lypla2 | 1.67E-19 | 0.347429996 | 0.293 | 0.096 | 3.39E-15 | 16 | Lypla2 |
| Setd58 | 1.93E-19 | 0.532230906 | 0.571 | 0.264 | 3.91E-15 | 16 | Setd5 |
| Adipor25 | 2.01E-19 | 0.455428687 | 0.391 | 0.15 | 4.08E-15 | 16 | Adipor2 |
| Hnrnpll1 | 2.04E-19 | 0.403308032 | 0.332 | 0.114 | 4.12E-15 | 16 | Hnrnpll |
| Nr3c15 | 2.07E-19 | 0.522591759 | 0.739 | 0.383 | 4.20E-15 | 16 | Nr3c1 |
| Slc9a3r14 | 2.18E-19 | 0.372604783 | 0.321 | 0.11 | 4.40E-15 | 16 | Slc9a3r1 |
| Psip13 | 2.32E-19 | 0.442551875 | 0.467 | 0.195 | 4.70E-15 | 16 | Psip1 |
| mt-Co212 | 2.73E-19 | 0.532176228 | 1 | 0.728 | 5.53E-15 | 16 | mt-Co2 |
| Pi4k2a | 2.75E-19 | 0.355557243 | 0.299 | 0.098 | 5.56E-15 | 16 | Pi4k2a |
| Map3k143 | 3.68E-19 | 0.411181743 | 0.31 | 0.106 | 7.44E-15 | 16 | Map3k14 |
| Rictor5 | 3.68E-19 | 0.553838988 | 0.397 | 0.157 | 7.45E-15 | 16 | Rictor |
| Wls1 | 3.77E-19 | 0.385809247 | 0.353 | 0.127 | 7.62E-15 | 16 | Wls |
| Nfatc35 | 4.22E-19 | 0.535612264 | 0.424 | 0.175 | 8.54E-15 | 16 | Nfatc3 |
| Wdr83os3 | 4.69E-19 | 0.324257658 | 0.603 | 0.283 | 9.49E-15 | 16 | Wdr83os |
| Fli17 | 5.19E-19 | 0.439958884 | 0.788 | 0.402 | 1.05E-14 | 16 | Fli1 |
| Camk2g4 | 5.94E-19 | 0.363709439 | 0.37 | 0.135 | 1.20E-14 | 16 | Camk2g |
| Lsm42 | 6.48E-19 | 0.338438098 | 0.652 | 0.322 | 1.31E-14 | 16 | Lsm4 |
| Sms4 | 8.16E-19 | 0.274661831 | 0.397 | 0.151 | 1.65E-14 | 16 | Sms |
| Gm10073 | 1.01E-18 | 0.275219068 | 0.359 | 0.133 | 2.04E-14 | 16 | Gm10073 |
| Akap98 | 1.17E-18 | 0.466538903 | 0.527 | 0.237 | 2.37E-14 | 16 | Akap9 |
| Fyco13 | 1.21E-18 | 0.416703999 | 0.342 | 0.124 | 2.45E-14 | 16 | Fyco1 |
| Kdm2b3 | 1.37E-18 | 0.344600975 | 0.277 | 0.089 | 2.78E-14 | 16 | Kdm2b |
| Akap8l1 | 1.57E-18 | 0.405372798 | 0.37 | 0.14 | 3.18E-14 | 16 | Akap8l |
| Sipa1l15 | 1.58E-18 | 0.446332226 | 0.457 | 0.19 | 3.20E-14 | 16 | Sipa1l1 |
| Rexo23 | 1.75E-18 | 0.326192557 | 0.62 | 0.297 | 3.54E-14 | 16 | Rexo2 |
| Foxn38 | 1.84E-18 | 0.441152798 | 0.793 | 0.41 | 3.72E-14 | 16 | Foxn3 |

| | | | | | | | |
|---|---|---|---|---|---|---|---|
| Pfkfb31 | 1.90E-18 | 0.377017891 | 0.272 | 0.087 | 3.84E-14 | 16 | Pfkfb3 |
| Cdkn1b2 | 2.02E-18 | 0.455900602 | 0.489 | 0.215 | 4.09E-14 | 16 | Cdkn1b |
| Gna132 | 2.12E-18 | 0.413373475 | 0.418 | 0.172 | 4.29E-14 | 16 | Gna13 |
| Mkl16 | 2.31E-18 | 0.49748202 | 0.337 | 0.126 | 4.67E-14 | 16 | Mkl1 |
| Peli15 | 2.43E-18 | 0.488741563 | 0.516 | 0.232 | 4.92E-14 | 16 | Peli1 |
| Anp32b2 | 2.45E-18 | 0.292062948 | 0.793 | 0.422 | 4.96E-14 | 16 | Anp32b |
| Eef1a16 | 2.66E-18 | 0.366442487 | 1 | 0.96 | 5.38E-14 | 16 | Eef1a1 |
| Ythdc11 | 2.82E-18 | 0.412708948 | 0.516 | 0.235 | 5.71E-14 | 16 | Ythdc1 |
| Rap1b4 | 2.85E-18 | 0.412260655 | 0.832 | 0.444 | 5.77E-14 | 16 | Rap1b |
| Nsa25 | 3.07E-18 | 0.364142326 | 0.859 | 0.486 | 6.21E-14 | 16 | Nsa2 |
| Srrm27 | 3.11E-18 | 0.508236249 | 0.826 | 0.469 | 6.29E-14 | 16 | Srrm2 |
| Celf14 | 4.09E-18 | 0.488611115 | 0.489 | 0.22 | 8.28E-14 | 16 | Celf1 |
| Galnt74 | 4.63E-18 | 0.430271302 | 0.391 | 0.156 | 9.37E-14 | 16 | Galnt7 |
| Dock85 | 5.33E-18 | 0.258583175 | 0.533 | 0.219 | 1.08E-13 | 16 | Dock8 |
| Creb14 | 5.54E-18 | 0.408725396 | 0.478 | 0.208 | 1.12E-13 | 16 | Creb1 |
| Ankib14 | 5.55E-18 | 0.39012431 | 0.364 | 0.139 | 1.12E-13 | 16 | Ankib1 |
| Srebf21 | 5.73E-18 | 0.473356517 | 0.283 | 0.097 | 1.16E-13 | 16 | Srebf2 |
| Bhlhe405 | 6.10E-18 | 0.534214683 | 0.397 | 0.159 | 1.24E-13 | 16 | Bhlhe40 |
| Ywhab4 | 6.15E-18 | 0.409330065 | 0.826 | 0.467 | 1.25E-13 | 16 | Ywhab |
| Arpc26 | 6.20E-18 | 0.488993245 | 0.908 | 0.544 | 1.26E-13 | 16 | Arpc2 |
| Nudcd35 | 6.34E-18 | 0.434140438 | 0.533 | 0.246 | 1.28E-13 | 16 | Nudcd3 |
| Srgn9 | 6.72E-18 | 0.523315915 | 0.946 | 0.544 | 1.36E-13 | 16 | Srgn |
| Ctla2a7 | 7.60E-18 | 0.401722972 | 0.804 | 0.398 | 1.54E-13 | 16 | Ctla2a |
| Dnajb12 | 8.80E-18 | 0.500921566 | 0.315 | 0.116 | 1.78E-13 | 16 | Dnajb1 |
| Ascc37 | 1.06E-17 | 0.473849061 | 0.473 | 0.208 | 2.15E-13 | 16 | Ascc3 |
| Vrk1 | 1.27E-17 | 0.351631072 | 0.266 | 0.087 | 2.57E-13 | 16 | Vrk1 |
| Tor1aip23 | 1.40E-17 | 0.531473957 | 0.424 | 0.183 | 2.84E-13 | 16 | Tor1aip2 |
| Msh2 | 1.45E-17 | 0.286991749 | 0.25 | 0.078 | 2.94E-13 | 16 | Msh2 |
| Emg11 | 1.66E-17 | 0.291415507 | 0.505 | 0.226 | 3.36E-13 | 16 | Emg1 |
| Ambra17 | 1.71E-17 | 0.501328716 | 0.451 | 0.198 | 3.47E-13 | 16 | Ambra1 |
| Crebbp8 | 1.81E-17 | 0.447627657 | 0.62 | 0.305 | 3.66E-13 | 16 | Crebbp |
| Npm16 | 1.83E-17 | 0.396006499 | 0.984 | 0.679 | 3.71E-13 | 16 | Npm1 |
| Fndc3a5 | 1.86E-17 | 0.473453055 | 0.576 | 0.277 | 3.76E-13 | 16 | Fndc3a |
| Phf21a8 | 2.08E-17 | 0.509741646 | 0.533 | 0.253 | 4.20E-13 | 16 | Phf21a |
| Ifngr22 | 2.15E-17 | 0.387142118 | 0.353 | 0.132 | 4.35E-13 | 16 | Ifngr2 |
| Sfmbt11 | 2.16E-17 | 0.362810631 | 0.261 | 0.084 | 4.37E-13 | 16 | Sfmbt1 |
| Atad2b6 | 2.35E-17 | 0.461009734 | 0.462 | 0.203 | 4.76E-13 | 16 | Atad2b |
| Tiparp2 | 2.37E-17 | 0.349176792 | 0.266 | 0.087 | 4.79E-13 | 16 | Tiparp |
| Mrpl33 | 3.03E-17 | 0.296794203 | 0.772 | 0.405 | 6.14E-13 | 16 | Mrpl33 |
| Birc69 | 3.20E-17 | 0.466622918 | 0.658 | 0.341 | 6.49E-13 | 16 | Birc6 |
| Cog56 | 3.48E-17 | 0.453339937 | 0.429 | 0.183 | 7.04E-13 | 16 | Cog5 |
| Kmt2c8 | 4.02E-17 | 0.513171398 | 0.603 | 0.306 | 8.13E-13 | 16 | Kmt2c |
| Yaf23 | 4.48E-17 | 0.427798114 | 0.484 | 0.215 | 9.06E-13 | 16 | Yaf2 |

| | | | | | | | |
|---|---|---|---|---|---|---|---|
| Cdk178 | 4.65E-17 | 0.50982025 | 0.527 | 0.245 | 9.41E-13 | 16 | Cdk17 |
| Kdm5a4 | 5.24E-17 | 0.367868907 | 0.408 | 0.168 | 1.06E-12 | 16 | Kdm5a |
| Appl13 | 5.33E-17 | 0.370280175 | 0.429 | 0.182 | 1.08E-12 | 16 | Appl1 |
| Chd75 | 6.33E-17 | 0.439270697 | 0.375 | 0.152 | 1.28E-12 | 16 | Chd7 |
| AW5549185 | 6.66E-17 | 0.563357592 | 0.402 | 0.171 | 1.35E-12 | 16 | AW554918 |
| Gch12 | 6.70E-17 | 0.536823411 | 0.342 | 0.131 | 1.36E-12 | 16 | Gch1 |
| Irf14 | 6.90E-17 | 0.452479787 | 0.62 | 0.299 | 1.40E-12 | 16 | Irf1 |
| Usp154 | 7.07E-17 | 0.443113952 | 0.495 | 0.231 | 1.43E-12 | 16 | Usp15 |
| Ppp1r9b | 7.09E-17 | 0.395964989 | 0.332 | 0.126 | 1.44E-12 | 16 | Ppp1r9b |
| mt-Nd4l7 | 7.18E-17 | 0.392435137 | 0.864 | 0.491 | 1.45E-12 | 16 | mt-Nd4l |
| Zc3h135 | 7.66E-17 | 0.460867518 | 0.473 | 0.216 | 1.55E-12 | 16 | Zc3h13 |
| Msn4 | 8.36E-17 | 0.415947156 | 0.821 | 0.443 | 1.69E-12 | 16 | Msn |
| Adcy76 | 8.44E-17 | 0.46768922 | 0.31 | 0.115 | 1.71E-12 | 16 | Adcy7 |
| Riok14 | 9.12E-17 | 0.386054902 | 0.315 | 0.116 | 1.85E-12 | 16 | Riok1 |
| Gpatch88 | 9.94E-17 | 0.511585194 | 0.533 | 0.258 | 2.01E-12 | 16 | Gpatch8 |
| Birc23 | 1.17E-16 | 0.375576263 | 0.304 | 0.11 | 2.36E-12 | 16 | Birc2 |
| Hivep12 | 1.25E-16 | 0.307711973 | 0.353 | 0.135 | 2.54E-12 | 16 | Hivep1 |
| Calm33 | 1.26E-16 | 0.337030609 | 0.554 | 0.269 | 2.54E-12 | 16 | Calm3 |
| Crem | 1.32E-16 | 0.351557708 | 0.261 | 0.087 | 2.67E-12 | 16 | Crem |
| Tcof11 | 1.40E-16 | 0.378700128 | 0.321 | 0.12 | 2.84E-12 | 16 | Tcof1 |
| Ppie | 1.45E-16 | 0.287245545 | 0.31 | 0.113 | 2.93E-12 | 16 | Ppie |
| Tgfb16 | 1.51E-16 | 0.44836289 | 0.511 | 0.236 | 3.05E-12 | 16 | Tgfb1 |
| Tecr5 | 1.57E-16 | 0.322001592 | 0.495 | 0.225 | 3.18E-12 | 16 | Tecr |
| Vti1a7 | 1.58E-16 | 0.429668412 | 0.543 | 0.255 | 3.19E-12 | 16 | Vti1a |
| Nrros1 | 1.58E-16 | 0.395972677 | 0.299 | 0.107 | 3.21E-12 | 16 | Nrros |
| Rif12 | 1.67E-16 | 0.447004129 | 0.375 | 0.155 | 3.38E-12 | 16 | Rif1 |
| Lonp2 | 1.84E-16 | 0.392735436 | 0.446 | 0.195 | 3.72E-12 | 16 | Lonp2 |
| Etv31 | 1.97E-16 | 0.383208951 | 0.283 | 0.099 | 3.98E-12 | 16 | Etv3 |
| Ier57 | 1.98E-16 | 0.488838253 | 0.625 | 0.317 | 4.00E-12 | 16 | Ier5 |
| Rpl23a7 | 2.18E-16 | 0.442439423 | 0.967 | 0.679 | 4.41E-12 | 16 | Rpl23a |
| Kidins2205 | 2.47E-16 | 0.343833342 | 0.386 | 0.156 | 5.00E-12 | 16 | Kidins220 |
| Mapk63 | 2.51E-16 | 0.424838933 | 0.288 | 0.103 | 5.08E-12 | 16 | Mapk6 |
| Elf15 | 2.89E-16 | 0.478674802 | 0.707 | 0.377 | 5.84E-12 | 16 | Elf1 |
| Rpl263 | 2.91E-16 | 0.302811782 | 1 | 0.957 | 5.90E-12 | 16 | Rpl26 |
| Fam173a3 | 3.12E-16 | 0.277966561 | 0.451 | 0.198 | 6.32E-12 | 16 | Fam173a |
| Srsf119 | 4.11E-16 | 0.389438558 | 0.668 | 0.351 | 8.31E-12 | 16 | Srsf11 |
| Tcf129 | 4.13E-16 | 0.47972247 | 0.712 | 0.376 | 8.36E-12 | 16 | Tcf12 |
| Nedd4l4 | 4.52E-16 | 0.568294674 | 0.467 | 0.218 | 9.14E-12 | 16 | Nedd4l |
| Ep4005 | 4.82E-16 | 0.423846742 | 0.457 | 0.208 | 9.76E-12 | 16 | Ep400 |
| Slc3a27 | 4.99E-16 | 0.461496186 | 0.576 | 0.291 | 1.01E-11 | 16 | Slc3a2 |
| Clic16 | 5.07E-16 | 0.402959336 | 0.967 | 0.6 | 1.03E-11 | 16 | Clic1 |
| Nmt2 | 5.21E-16 | 0.336536088 | 0.277 | 0.098 | 1.06E-11 | 16 | Nmt2 |
| Tmem176a6 | 5.81E-16 | 0.319088397 | 0.793 | 0.417 | 1.18E-11 | 16 | Tmem176a |

| | | | | | | | |
|---|---|---|---|---|---|---|---|
| Fkbp32 | 6.08E-16 | 0.252703643 | 0.734 | 0.396 | 1.23E-11 | 16 | Fkbp3 |
| Zfp3982 | 6.75E-16 | 0.332753434 | 0.272 | 0.094 | 1.37E-11 | 16 | Zfp398 |
| Dnajc111 | 7.21E-16 | 0.538468265 | 0.625 | 0.323 | 1.46E-11 | 16 | Dnajc1 |
| B4galt18 | 8.34E-16 | 0.447220375 | 0.408 | 0.177 | 1.69E-11 | 16 | B4galt1 |
| Wdr891 | 8.54E-16 | 0.276242351 | 0.293 | 0.107 | 1.73E-11 | 16 | Wdr89 |
| Capzb6 | 8.83E-16 | 0.38298081 | 0.674 | 0.369 | 1.79E-11 | 16 | Capzb |
| Bcap316 | 1.01E-15 | 0.325302998 | 0.576 | 0.285 | 2.05E-11 | 16 | Bcap31 |
| Ccar15 | 1.05E-15 | 0.442686628 | 0.457 | 0.208 | 2.13E-11 | 16 | Ccar1 |
| Rpl377 | 1.11E-15 | 0.344249481 | 1 | 0.927 | 2.26E-11 | 16 | Rpl37 |
| Tpr7 | 1.15E-15 | 0.381839584 | 0.788 | 0.425 | 2.32E-11 | 16 | Tpr |
| Polr2a7 | 1.26E-15 | 0.480470746 | 0.571 | 0.281 | 2.54E-11 | 16 | Polr2a |
| Rab21 | 1.29E-15 | 0.420964575 | 0.587 | 0.303 | 2.62E-11 | 16 | Rab21 |
| Tra2a9 | 1.37E-15 | 0.426003536 | 0.56 | 0.275 | 2.77E-11 | 16 | Tra2a |
| Hectd16 | 1.46E-15 | 0.432957737 | 0.587 | 0.301 | 2.95E-11 | 16 | Hectd1 |
| Sdhaf1 | 1.46E-15 | 0.256327742 | 0.315 | 0.117 | 2.95E-11 | 16 | Sdhaf1 |
| Ddx39b3 | 1.50E-15 | 0.264133497 | 0.505 | 0.234 | 3.04E-11 | 16 | Ddx39b |
| Ppp1r11 | 1.74E-15 | 0.337253525 | 0.592 | 0.298 | 3.52E-11 | 16 | Ppp1r11 |
| Naa153 | 1.74E-15 | 0.340140837 | 0.391 | 0.166 | 3.52E-11 | 16 | Naa15 |
| Rock17 | 1.87E-15 | 0.421126005 | 0.766 | 0.433 | 3.79E-11 | 16 | Rock1 |
| Mllt108 | 1.93E-15 | 0.452000171 | 0.549 | 0.275 | 3.90E-11 | 16 | Mllt10 |
| Sf3b16 | 1.93E-15 | 0.357585457 | 0.728 | 0.393 | 3.90E-11 | 16 | Sf3b1 |
| Lrp106 | 1.95E-15 | 0.460005594 | 0.418 | 0.189 | 3.95E-11 | 16 | Lrp10 |
| Bicra | 1.99E-15 | 0.386600565 | 0.283 | 0.103 | 4.03E-11 | 16 | Bicra |
| Sub15 | 2.06E-15 | 0.272622873 | 0.88 | 0.516 | 4.17E-11 | 16 | Sub1 |
| Wac7 | 2.13E-15 | 0.407030709 | 0.625 | 0.323 | 4.32E-11 | 16 | Wac |
| Lrba5 | 2.16E-15 | 0.386960158 | 0.429 | 0.192 | 4.36E-11 | 16 | Lrba |
| Pcna1 | 2.36E-15 | 0.383905403 | 0.315 | 0.122 | 4.78E-11 | 16 | Pcna |
| Brd15 | 2.38E-15 | 0.387435309 | 0.5 | 0.241 | 4.83E-11 | 16 | Brd1 |
| Ccdc824 | 2.63E-15 | 0.420496224 | 0.408 | 0.181 | 5.33E-11 | 16 | Ccdc82 |
| Actr24 | 2.64E-15 | 0.309143665 | 0.668 | 0.343 | 5.35E-11 | 16 | Actr2 |
| Tln18 | 2.70E-15 | 0.444474596 | 0.745 | 0.405 | 5.48E-11 | 16 | Tln1 |
| Hnrnpa2b13 | 2.90E-15 | 0.458256665 | 0.859 | 0.515 | 5.87E-11 | 16 | Hnrnpa2b1 |
| Hectd4 | 3.01E-15 | 0.342774462 | 0.293 | 0.108 | 6.08E-11 | 16 | Hectd4 |
| Sec11a5 | 3.05E-15 | 0.34253033 | 0.571 | 0.285 | 6.18E-11 | 16 | Sec11a |
| Lmna5 | 3.09E-15 | 0.510880028 | 0.44 | 0.198 | 6.25E-11 | 16 | Lmna |
| Ttc39b5 | 3.67E-15 | 0.386844312 | 0.337 | 0.136 | 7.43E-11 | 16 | Ttc39b |
| Rnf1307 | 3.97E-15 | 0.452593028 | 0.56 | 0.284 | 8.03E-11 | 16 | Rnf130 |
| Klhl64 | 4.08E-15 | 0.351472749 | 0.315 | 0.121 | 8.26E-11 | 16 | Klhl6 |
| Fam69a4 | 4.09E-15 | 0.433356379 | 0.321 | 0.127 | 8.28E-11 | 16 | Fam69a |
| Phf38 | 4.40E-15 | 0.413898993 | 0.522 | 0.253 | 8.90E-11 | 16 | Phf3 |
| Sltm6 | 4.43E-15 | 0.374377222 | 0.554 | 0.271 | 8.96E-11 | 16 | Sltm |
| Cfl16 | 4.84E-15 | 0.43660415 | 0.935 | 0.608 | 9.80E-11 | 16 | Cfl1 |
| Mapk17 | 4.93E-15 | 0.332481825 | 0.5 | 0.236 | 9.99E-11 | 16 | Mapk1 |

| | | | | | | | |
|---|---|---|---|---|---|---|---|
| Itgav6 | 5.51E-15 | 0.510874522 | 0.375 | 0.161 | 1.12E-10 | 16 | Itgav |
| Babam26 | 5.73E-15 | 0.417070636 | 0.5 | 0.239 | 1.16E-10 | 16 | Babam2 |
| Mak16 | 5.84E-15 | 0.271452273 | 0.353 | 0.145 | 1.18E-10 | 16 | Mak16 |
| Fus5 | 5.87E-15 | 0.327941005 | 0.717 | 0.389 | 1.19E-10 | 16 | Fus |
| Mia23 | 6.21E-15 | 0.386861875 | 0.446 | 0.208 | 1.26E-10 | 16 | Mia2 |
| Wdr335 | 6.90E-15 | 0.312081531 | 0.505 | 0.239 | 1.40E-10 | 16 | Wdr33 |
| Eif3f4 | 7.75E-15 | 0.303226118 | 0.832 | 0.483 | 1.57E-10 | 16 | Eif3f |
| Cpq2 | 8.07E-15 | 0.42918181 | 0.315 | 0.121 | 1.63E-10 | 16 | Cpq |
| Klf102 | 8.10E-15 | 0.520397189 | 0.375 | 0.164 | 1.64E-10 | 16 | Klf10 |
| Mpp72 | 8.32E-15 | 0.303890042 | 0.288 | 0.107 | 1.68E-10 | 16 | Mpp7 |
| Kdm6b | 8.63E-15 | 0.321260815 | 0.255 | 0.09 | 1.75E-10 | 16 | Kdm6b |
| Ubxn45 | 9.87E-15 | 0.38508267 | 0.543 | 0.272 | 2.00E-10 | 16 | Ubxn4 |
| Lbr7 | 1.02E-14 | 0.428108732 | 0.359 | 0.153 | 2.07E-10 | 16 | Lbr |
| Ssfa22 | 1.08E-14 | 0.292940495 | 0.272 | 0.097 | 2.19E-10 | 16 | Ssfa2 |
| Acin16 | 1.16E-14 | 0.402112204 | 0.598 | 0.317 | 2.35E-10 | 16 | Acin1 |
| Bcor2 | 1.18E-14 | 0.268707753 | 0.315 | 0.121 | 2.38E-10 | 16 | Bcor |
| Spata54 | 1.27E-14 | 0.417907745 | 0.315 | 0.124 | 2.57E-10 | 16 | Spata5 |
| Fam208b4 | 1.34E-14 | 0.375295922 | 0.359 | 0.151 | 2.71E-10 | 16 | Fam208b |
| Prelid1 | 1.39E-14 | 0.258428404 | 0.598 | 0.31 | 2.82E-10 | 16 | Prelid1 |
| Trip126 | 1.49E-14 | 0.433536424 | 0.511 | 0.257 | 3.02E-10 | 16 | Trip12 |
| Sp42 | 1.51E-14 | 0.349042928 | 0.266 | 0.097 | 3.06E-10 | 16 | Sp4 |
| AI3141803 | 1.56E-14 | 0.306579223 | 0.457 | 0.211 | 3.15E-10 | 16 | AI314180 |
| Spcs31 | 1.56E-14 | 0.322202424 | 0.304 | 0.119 | 3.16E-10 | 16 | Spcs3 |
| Pitpna3 | 1.62E-14 | 0.321522837 | 0.636 | 0.328 | 3.27E-10 | 16 | Pitpna |
| Casp85 | 1.71E-14 | 0.371113707 | 0.462 | 0.215 | 3.46E-10 | 16 | Casp8 |
| Camk2d6 | 1.73E-14 | 0.442699555 | 0.538 | 0.27 | 3.49E-10 | 16 | Camk2d |
| Xpo6 | 1.93E-14 | 0.29167712 | 0.25 | 0.088 | 3.91E-10 | 16 | Xpo6 |
| Ncoa33 | 2.06E-14 | 0.352242251 | 0.516 | 0.253 | 4.17E-10 | 16 | Ncoa3 |
| Ube2j13 | 2.07E-14 | 0.351962951 | 0.375 | 0.162 | 4.18E-10 | 16 | Ube2j1 |
| Ugcg4 | 2.22E-14 | 0.318145852 | 0.402 | 0.172 | 4.50E-10 | 16 | Ugcg |
| Kmt2a9 | 2.69E-14 | 0.372903285 | 0.543 | 0.267 | 5.44E-10 | 16 | Kmt2a |
| Arid4b6 | 2.99E-14 | 0.382392857 | 0.565 | 0.293 | 6.06E-10 | 16 | Arid4b |
| Fbl2 | 3.07E-14 | 0.325601253 | 0.321 | 0.131 | 6.21E-10 | 16 | Fbl |
| Rsf15 | 3.38E-14 | 0.38793006 | 0.467 | 0.225 | 6.85E-10 | 16 | Rsf1 |
| Hnrnpa01 | 3.39E-14 | 0.326278767 | 0.826 | 0.494 | 6.86E-10 | 16 | Hnrnpa0 |
| Chst117 | 3.40E-14 | 0.402715439 | 0.397 | 0.176 | 6.89E-10 | 16 | Chst11 |
| Mtdh5 | 3.50E-14 | 0.3272902 | 0.728 | 0.387 | 7.09E-10 | 16 | Mtdh |
| Akr1b33 | 3.63E-14 | 0.301908748 | 0.413 | 0.19 | 7.35E-10 | 16 | Akr1b3 |
| Erap12 | 3.83E-14 | 0.417276941 | 0.337 | 0.14 | 7.74E-10 | 16 | Erap1 |
| Herc15 | 4.17E-14 | 0.422841449 | 0.505 | 0.257 | 8.45E-10 | 16 | Herc1 |
| Taf33 | 4.49E-14 | 0.399746613 | 0.348 | 0.147 | 9.09E-10 | 16 | Taf3 |
| Krtcap26 | 4.51E-14 | 0.288453356 | 0.772 | 0.442 | 9.14E-10 | 16 | Krtcap2 |
| Snd16 | 4.56E-14 | 0.366240531 | 0.386 | 0.173 | 9.23E-10 | 16 | Snd1 |

| | | | | | | | |
|---|---|---|---|---|---|---|---|
| Nt5c31 | 4.79E-14 | 0.252335147 | 0.288 | 0.11 | 9.69E-10 | 16 | Nt5c3 |
| Ccdc1074 | 5.00E-14 | 0.413215104 | 0.31 | 0.128 | 1.01E-09 | 16 | Ccdc107 |
| Synj12 | 5.05E-14 | 0.439879069 | 0.364 | 0.16 | 1.02E-09 | 16 | Synj1 |
| Csnk1g16 | 5.06E-14 | 0.390987063 | 0.364 | 0.158 | 1.02E-09 | 16 | Csnk1g1 |
| Atxn74 | 5.42E-14 | 0.282518591 | 0.386 | 0.166 | 1.10E-09 | 16 | Atxn7 |
| Dpm1 | 5.54E-14 | 0.250736584 | 0.348 | 0.143 | 1.12E-09 | 16 | Dpm1 |
| Smpdl3a4 | 6.48E-14 | 0.349492597 | 0.467 | 0.22 | 1.31E-09 | 16 | Smpdl3a |
| Xpo1 | 8.66E-14 | 0.299153266 | 0.315 | 0.127 | 1.75E-09 | 16 | Xpo1 |
| Srsf5 | 9.64E-14 | 0.319984404 | 0.804 | 0.45 | 1.95E-09 | 16 | Srsf5 |
| Tsc22d22 | 1.03E-13 | 0.366153965 | 0.359 | 0.154 | 2.09E-09 | 16 | Tsc22d2 |
| Sp1105 | 1.03E-13 | 0.278753101 | 0.364 | 0.158 | 2.09E-09 | 16 | Sp110 |
| Ly6e5 | 1.05E-13 | 0.347723224 | 0.978 | 0.619 | 2.12E-09 | 16 | Ly6e |
| Psmb89 | 1.08E-13 | 0.362608857 | 0.918 | 0.557 | 2.19E-09 | 16 | Psmb8 |
| Map3k24 | 1.11E-13 | 0.296841222 | 0.359 | 0.154 | 2.24E-09 | 16 | Map3k2 |
| Zdhhc186 | 1.12E-13 | 0.301425382 | 0.391 | 0.171 | 2.26E-09 | 16 | Zdhhc18 |
| Arf41 | 1.15E-13 | 0.373483124 | 0.739 | 0.456 | 2.33E-09 | 16 | Arf4 |
| Ubap2l | 1.19E-13 | 0.28142205 | 0.56 | 0.287 | 2.40E-09 | 16 | Ubap2l |
| Brd48 | 1.35E-13 | 0.426319341 | 0.522 | 0.268 | 2.72E-09 | 16 | Brd4 |
| Efr3a4 | 1.47E-13 | 0.341151007 | 0.418 | 0.198 | 2.97E-09 | 16 | Efr3a |
| Fosl21 | 1.50E-13 | 0.399166565 | 0.272 | 0.102 | 3.03E-09 | 16 | Fosl2 |
| Stxbp53 | 1.55E-13 | 0.399308764 | 0.315 | 0.13 | 3.13E-09 | 16 | Stxbp5 |
| Hist1h1c1 | 1.57E-13 | 0.389025973 | 0.429 | 0.201 | 3.17E-09 | 16 | Hist1h1c |
| Vps13b6 | 1.57E-13 | 0.362262008 | 0.457 | 0.22 | 3.18E-09 | 16 | Vps13b |
| Rsbn13 | 1.67E-13 | 0.342438513 | 0.391 | 0.178 | 3.39E-09 | 16 | Rsbn1 |
| Srpk22 | 1.68E-13 | 0.278985876 | 0.424 | 0.194 | 3.41E-09 | 16 | Srpk2 |
| Arntl1 | 1.77E-13 | 0.34138835 | 0.299 | 0.119 | 3.59E-09 | 16 | Arntl |
| Cab393 | 1.83E-13 | 0.280003682 | 0.538 | 0.281 | 3.72E-09 | 16 | Cab39 |
| Ppp2r5a2 | 2.15E-13 | 0.39272364 | 0.495 | 0.251 | 4.35E-09 | 16 | Ppp2r5a |
| Rtf2 | 2.19E-13 | 0.316581097 | 0.418 | 0.198 | 4.43E-09 | 16 | Rtf2 |
| H2-D19 | 2.21E-13 | 0.425731866 | 1 | 0.669 | 4.48E-09 | 16 | H2-D1 |
| Srek13 | 2.23E-13 | 0.360653504 | 0.418 | 0.194 | 4.51E-09 | 16 | Srek1 |
| Sep-96 | 2.26E-13 | 0.349913027 | 0.386 | 0.171 | 4.58E-09 | 16 | 9-Sep |
| Snrnp704 | 2.39E-13 | 0.331921263 | 0.533 | 0.273 | 4.83E-09 | 16 | Snrnp70 |
| Sfpq5 | 2.57E-13 | 0.357156036 | 0.56 | 0.299 | 5.19E-09 | 16 | Sfpq |
| Pdia37 | 2.60E-13 | 0.362035738 | 0.772 | 0.443 | 5.26E-09 | 16 | Pdia3 |
| Pds5a2 | 2.98E-13 | 0.418465454 | 0.424 | 0.205 | 6.03E-09 | 16 | Pds5a |
| Ogt6 | 3.07E-13 | 0.370757277 | 0.435 | 0.207 | 6.21E-09 | 16 | Ogt |
| Rpl286 | 3.14E-13 | 0.304705247 | 1 | 0.93 | 6.36E-09 | 16 | Rpl28 |
| Ppp1r16b8 | 3.25E-13 | 0.435794892 | 0.435 | 0.202 | 6.57E-09 | 16 | Ppp1r16b |
| Atxn7l13 | 3.30E-13 | 0.337854879 | 0.321 | 0.134 | 6.67E-09 | 16 | Atxn7l1 |
| St3gal64 | 3.33E-13 | 0.473473515 | 0.587 | 0.318 | 6.75E-09 | 16 | St3gal6 |
| Csnk1g34 | 3.35E-13 | 0.290410342 | 0.402 | 0.183 | 6.79E-09 | 16 | Csnk1g3 |
| St8sia47 | 3.39E-13 | 0.403818604 | 0.516 | 0.26 | 6.86E-09 | 16 | St8sia4 |

| | | | | | | | |
|---|---|---|---|---|---|---|---|
| Ggnbp26 | 3.46E-13 | 0.339360545 | 0.489 | 0.242 | 7.00E-09 | 16 | Ggnbp2 |
| Cblb5 | 3.48E-13 | 0.303931592 | 0.386 | 0.171 | 7.04E-09 | 16 | Cblb |
| Ralgapa24 | 3.66E-13 | 0.340724736 | 0.353 | 0.154 | 7.40E-09 | 16 | Ralgapa2 |
| Akt38 | 4.05E-13 | 0.395825873 | 0.505 | 0.252 | 8.19E-09 | 16 | Akt3 |
| Zufsp4 | 4.22E-13 | 0.296491452 | 0.326 | 0.137 | 8.55E-09 | 16 | Zufsp |
| Zbtb11 | 4.34E-13 | 0.313012264 | 0.261 | 0.1 | 8.78E-09 | 16 | Zbtb11 |
| Rara | 4.37E-13 | 0.31744141 | 0.261 | 0.098 | 8.85E-09 | 16 | Rara |
| Ctcf4 | 4.40E-13 | 0.266234732 | 0.549 | 0.277 | 8.90E-09 | 16 | Ctcf |
| Nktr5 | 4.48E-13 | 0.440546078 | 0.505 | 0.258 | 9.06E-09 | 16 | Nktr |
| Epc13 | 4.63E-13 | 0.410478766 | 0.424 | 0.203 | 9.37E-09 | 16 | Epc1 |
| Med13l4 | 4.78E-13 | 0.295263694 | 0.424 | 0.194 | 9.68E-09 | 16 | Med13l |
| Prpf40a7 | 4.79E-13 | 0.302553858 | 0.527 | 0.267 | 9.69E-09 | 16 | Prpf40a |
| Mybbp1a | 5.03E-13 | 0.273157804 | 0.255 | 0.097 | 1.02E-08 | 16 | Mybbp1a |
| Gnl31 | 6.51E-13 | 0.278525488 | 0.283 | 0.111 | 1.32E-08 | 16 | Gnl3 |
| Srsf3 | 7.31E-13 | 0.252605127 | 0.859 | 0.519 | 1.48E-08 | 16 | Srsf3 |
| Mvb12b4 | 8.05E-13 | 0.315714322 | 0.37 | 0.168 | 1.63E-08 | 16 | Mvb12b |
| Dym | 8.47E-13 | 0.272647655 | 0.261 | 0.1 | 1.72E-08 | 16 | Dym |
| Smc64 | 8.92E-13 | 0.34413855 | 0.505 | 0.262 | 1.81E-08 | 16 | Smc6 |
| Synrg2 | 9.22E-13 | 0.302097077 | 0.332 | 0.143 | 1.87E-08 | 16 | Synrg |
| Kpna48 | 9.93E-13 | 0.434777441 | 0.587 | 0.324 | 2.01E-08 | 16 | Kpna4 |
| Dnajc37 | 1.03E-12 | 0.369638943 | 0.652 | 0.371 | 2.08E-08 | 16 | Dnajc3 |
| Son9 | 1.03E-12 | 0.352073589 | 0.891 | 0.526 | 2.09E-08 | 16 | Son |
| Arl6ip17 | 1.05E-12 | 0.316601492 | 0.663 | 0.371 | 2.12E-08 | 16 | Arl6ip1 |
| Cited21 | 1.15E-12 | 0.417671307 | 0.391 | 0.181 | 2.33E-08 | 16 | Cited2 |
| Mast44 | 1.24E-12 | 0.324639376 | 0.413 | 0.187 | 2.50E-08 | 16 | Mast4 |
| Arglu15 | 1.33E-12 | 0.250573865 | 0.652 | 0.354 | 2.70E-08 | 16 | Arglu1 |
| Avl94 | 1.43E-12 | 0.327189644 | 0.321 | 0.138 | 2.89E-08 | 16 | Avl9 |
| Mapre25 | 1.53E-12 | 0.470134007 | 0.484 | 0.251 | 3.10E-08 | 16 | Mapre2 |
| Mnat1 | 1.58E-12 | 0.273608254 | 0.315 | 0.133 | 3.19E-08 | 16 | Mnat1 |
| Sf3b35 | 1.73E-12 | 0.313045893 | 0.359 | 0.161 | 3.50E-08 | 16 | Sf3b3 |
| Exoc6b5 | 1.75E-12 | 0.398430486 | 0.473 | 0.241 | 3.54E-08 | 16 | Exoc6b |
| Rnf1696 | 1.81E-12 | 0.348975463 | 0.391 | 0.184 | 3.67E-08 | 16 | Rnf169 |
| Lpcat31 | 1.83E-12 | 0.284555307 | 0.25 | 0.094 | 3.71E-08 | 16 | Lpcat3 |
| Hcls14 | 1.88E-12 | 0.258555882 | 0.304 | 0.126 | 3.80E-08 | 16 | Hcls1 |
| Cept11 | 2.11E-12 | 0.32073808 | 0.321 | 0.138 | 4.28E-08 | 16 | Cept1 |
| Chd66 | 2.27E-12 | 0.431377923 | 0.429 | 0.218 | 4.59E-08 | 16 | Chd6 |
| Mdm43 | 2.48E-12 | 0.332668451 | 0.353 | 0.16 | 5.03E-08 | 16 | Mdm4 |
| Rbm264 | 2.92E-12 | 0.337285802 | 0.451 | 0.227 | 5.91E-08 | 16 | Rbm26 |
| Cdkal15 | 2.97E-12 | 0.32723797 | 0.386 | 0.181 | 6.01E-08 | 16 | Cdkal1 |
| Wdr433 | 2.98E-12 | 0.326311884 | 0.337 | 0.151 | 6.04E-08 | 16 | Wdr43 |
| Ap3s1 | 3.09E-12 | 0.318247301 | 0.353 | 0.159 | 6.27E-08 | 16 | Ap3s1 |
| Piezo14 | 3.15E-12 | 0.393416447 | 0.38 | 0.174 | 6.38E-08 | 16 | Piezo1 |
| Tcerg12 | 3.19E-12 | 0.339618387 | 0.315 | 0.137 | 6.45E-08 | 16 | Tcerg1 |

| | | | | | | | |
|---|---|---|---|---|---|---|---|
| Rbm67 | 3.49E-12 | 0.327516068 | 0.446 | 0.218 | 7.06E-08 | 16 | Rbm6 |
| Ptbp39 | 3.83E-12 | 0.363956717 | 0.696 | 0.394 | 7.76E-08 | 16 | Ptbp3 |
| Arhgef36 | 4.01E-12 | 0.355952441 | 0.478 | 0.241 | 8.12E-08 | 16 | Arhgef3 |
| Trp534 | 4.04E-12 | 0.316668556 | 0.348 | 0.159 | 8.17E-08 | 16 | Trp53 |
| Zfp4076 | 4.32E-12 | 0.357535302 | 0.353 | 0.162 | 8.75E-08 | 16 | Zfp407 |
| Tmem1315 | 4.35E-12 | 0.347311531 | 0.424 | 0.209 | 8.82E-08 | 16 | Tmem131 |
| Pbxip14 | 4.64E-12 | 0.351713873 | 0.283 | 0.118 | 9.40E-08 | 16 | Pbxip1 |
| Supt203 | 4.77E-12 | 0.333013336 | 0.353 | 0.162 | 9.66E-08 | 16 | Supt20 |
| Ube2h4 | 5.01E-12 | 0.366401517 | 0.505 | 0.266 | 1.01E-07 | 16 | Ube2h |
| Vps13a5 | 5.02E-12 | 0.35412956 | 0.37 | 0.173 | 1.02E-07 | 16 | Vps13a |
| Hnrnpdl1 | 5.08E-12 | 0.251586231 | 0.489 | 0.249 | 1.03E-07 | 16 | Hnrnpdl |
| Rarg3 | 5.18E-12 | 0.259106444 | 0.424 | 0.196 | 1.05E-07 | 16 | Rarg |
| Plekha55 | 5.34E-12 | 0.394454629 | 0.342 | 0.156 | 1.08E-07 | 16 | Plekha5 |
| Ptp4a21 | 5.39E-12 | 0.270216273 | 0.679 | 0.395 | 1.09E-07 | 16 | Ptp4a2 |
| Thrap35 | 5.52E-12 | 0.294183838 | 0.549 | 0.294 | 1.12E-07 | 16 | Thrap3 |
| Ngly1 | 6.17E-12 | 0.289146517 | 0.283 | 0.117 | 1.25E-07 | 16 | Ngly1 |
| Maml29 | 6.18E-12 | 0.383478381 | 0.62 | 0.343 | 1.25E-07 | 16 | Maml2 |
| Hdac85 | 6.56E-12 | 0.336079677 | 0.315 | 0.136 | 1.33E-07 | 16 | Hdac8 |
| Uri1 | 6.73E-12 | 0.253132231 | 0.391 | 0.182 | 1.36E-07 | 16 | Uri1 |
| Fyn8 | 6.92E-12 | 0.394482856 | 0.516 | 0.267 | 1.40E-07 | 16 | Fyn |
| Mbd11 | 7.02E-12 | 0.286655543 | 0.255 | 0.101 | 1.42E-07 | 16 | Mbd1 |
| Nsd17 | 7.57E-12 | 0.283249817 | 0.63 | 0.35 | 1.53E-07 | 16 | Nsd1 |
| Ints64 | 7.60E-12 | 0.299446758 | 0.326 | 0.143 | 1.54E-07 | 16 | Ints6 |
| Lfng1 | 7.64E-12 | 0.256689258 | 0.321 | 0.137 | 1.55E-07 | 16 | Lfng |
| mt-Nd39 | 8.39E-12 | 0.361270799 | 0.75 | 0.447 | 1.70E-07 | 16 | mt-Nd3 |
| Wdr701 | 8.62E-12 | 0.336095527 | 0.332 | 0.152 | 1.75E-07 | 16 | Wdr70 |
| Prkacb4 | 9.04E-12 | 0.339606629 | 0.44 | 0.219 | 1.83E-07 | 16 | Prkacb |
| Kdm3b4 | 9.22E-12 | 0.278657504 | 0.315 | 0.137 | 1.87E-07 | 16 | Kdm3b |
| Lrch37 | 9.23E-12 | 0.37486111 | 0.457 | 0.233 | 1.87E-07 | 16 | Lrch3 |
| Scaf113 | 9.52E-12 | 0.320060197 | 0.495 | 0.264 | 1.93E-07 | 16 | Scaf11 |
| Arl5b1 | 9.97E-12 | 0.309166434 | 0.283 | 0.117 | 2.02E-07 | 16 | Arl5b |
| Nr2c26 | 1.01E-11 | 0.341077552 | 0.348 | 0.159 | 2.04E-07 | 16 | Nr2c2 |
| Ddx18 | 1.25E-11 | 0.282412301 | 0.364 | 0.169 | 2.53E-07 | 16 | Ddx18 |
| Atp11b4 | 1.38E-11 | 0.318281389 | 0.38 | 0.181 | 2.79E-07 | 16 | Atp11b |
| Hist1h2bc4 | 1.38E-11 | 0.515187453 | 0.473 | 0.25 | 2.79E-07 | 16 | Hist1h2bc |
| Evl5 | 1.41E-11 | 0.296118821 | 0.451 | 0.221 | 2.86E-07 | 16 | Evl |
| Kat6a4 | 1.57E-11 | 0.273397029 | 0.457 | 0.228 | 3.18E-07 | 16 | Kat6a |
| Fryl6 | 1.65E-11 | 0.301614521 | 0.516 | 0.269 | 3.34E-07 | 16 | Fryl |
| Ppp4r3b5 | 1.70E-11 | 0.290743528 | 0.457 | 0.233 | 3.45E-07 | 16 | Ppp4r3b |
| Filip1l3 | 1.74E-11 | 0.33837201 | 0.332 | 0.153 | 3.52E-07 | 16 | Filip1l |
| Ccny6 | 2.05E-11 | 0.390177194 | 0.5 | 0.268 | 4.15E-07 | 16 | Ccny |
| Kdm4c5 | 2.07E-11 | 0.313552641 | 0.364 | 0.17 | 4.20E-07 | 16 | Kdm4c |
| Fbxw43 | 2.12E-11 | 0.278903736 | 0.277 | 0.116 | 4.29E-07 | 16 | Fbxw4 |

| | | | | | | | |
|---|---|---|---|---|---|---|---|
| Soat12 | 2.19E-11 | 0.465120592 | 0.261 | 0.108 | 4.44E-07 | 16 | Soat1 |
| Tbc1d57 | 2.25E-11 | 0.330672033 | 0.5 | 0.262 | 4.55E-07 | 16 | Tbc1d5 |
| Fcho24 | 2.45E-11 | 0.371185162 | 0.543 | 0.31 | 4.96E-07 | 16 | Fcho2 |
| Nt5dc13 | 2.49E-11 | 0.33494188 | 0.283 | 0.121 | 5.05E-07 | 16 | Nt5dc1 |
| Rftn18 | 2.61E-11 | 0.341565122 | 0.484 | 0.252 | 5.28E-07 | 16 | Rftn1 |
| Gm40702 | 2.87E-11 | 0.276067162 | 0.25 | 0.099 | 5.82E-07 | 16 | Gm4070 |
| G3bp14 | 3.04E-11 | 0.277402527 | 0.554 | 0.303 | 6.15E-07 | 16 | G3bp1 |
| Tlk13 | 3.32E-11 | 0.278956883 | 0.435 | 0.223 | 6.73E-07 | 16 | Tlk1 |
| Map2k5 | 3.48E-11 | 0.295658495 | 0.293 | 0.128 | 7.05E-07 | 16 | Map2k5 |
| Ankrd129 | 3.65E-11 | 0.387766236 | 0.56 | 0.309 | 7.38E-07 | 16 | Ankrd12 |
| Ino80d3 | 3.91E-11 | 0.264563398 | 0.359 | 0.166 | 7.92E-07 | 16 | Ino80d |
| Xpr14 | 3.92E-11 | 0.270872205 | 0.337 | 0.154 | 7.94E-07 | 16 | Xpr1 |
| Smg19 | 4.37E-11 | 0.276388711 | 0.462 | 0.235 | 8.85E-07 | 16 | Smg1 |
| Tet21 | 5.63E-11 | 0.288756285 | 0.261 | 0.109 | 1.14E-06 | 16 | Tet2 |
| Aup1 | 5.69E-11 | 0.264344415 | 0.299 | 0.133 | 1.15E-06 | 16 | Aup1 |
| Ddx421 | 5.95E-11 | 0.308859829 | 0.391 | 0.198 | 1.21E-06 | 16 | Ddx42 |
| Arid4a8 | 6.03E-11 | 0.327465685 | 0.478 | 0.252 | 1.22E-06 | 16 | Arid4a |
| Cdk11b3 | 7.56E-11 | 0.349636056 | 0.451 | 0.244 | 1.53E-06 | 16 | Cdk11b |
| Asb32 | 8.02E-11 | 0.333530654 | 0.261 | 0.111 | 1.62E-06 | 16 | Asb3 |
| Heca3 | 8.41E-11 | 0.287158167 | 0.315 | 0.144 | 1.70E-06 | 16 | Heca |
| Rlim3 | 8.43E-11 | 0.296594935 | 0.397 | 0.198 | 1.71E-06 | 16 | Rlim |
| Ppp1ca4 | 9.33E-11 | 0.259202511 | 0.668 | 0.397 | 1.89E-06 | 16 | Ppp1ca |
| Nop563 | 9.68E-11 | 0.322256896 | 0.293 | 0.132 | 1.96E-06 | 16 | Nop56 |
| Senp75 | 1.06E-10 | 0.281230165 | 0.337 | 0.157 | 2.16E-06 | 16 | Senp7 |
| Dcaf54 | 1.09E-10 | 0.372459319 | 0.288 | 0.13 | 2.20E-06 | 16 | Dcaf5 |
| Pnn6 | 1.16E-10 | 0.304407598 | 0.489 | 0.268 | 2.34E-06 | 16 | Pnn |
| Plec4 | 1.18E-10 | 0.405116863 | 0.359 | 0.175 | 2.39E-06 | 16 | Plec |
| Itsn28 | 1.19E-10 | 0.250860262 | 0.527 | 0.278 | 2.42E-06 | 16 | Itsn2 |
| Ino801 | 1.24E-10 | 0.284737834 | 0.31 | 0.14 | 2.51E-06 | 16 | Ino80 |
| Mat2a3 | 1.25E-10 | 0.360974906 | 0.549 | 0.31 | 2.54E-06 | 16 | Mat2a |
| Fam168b1 | 1.29E-10 | 0.302363216 | 0.348 | 0.167 | 2.60E-06 | 16 | Fam168b |
| Rpn24 | 1.34E-10 | 0.282210949 | 0.418 | 0.212 | 2.71E-06 | 16 | Rpn2 |
| M6pr2 | 1.34E-10 | 0.303786283 | 0.342 | 0.163 | 2.72E-06 | 16 | M6pr |
| Ube4a1 | 1.34E-10 | 0.285787189 | 0.261 | 0.11 | 2.72E-06 | 16 | Ube4a |
| Arhgap105 | 1.45E-10 | 0.32516441 | 0.288 | 0.128 | 2.94E-06 | 16 | Arhgap10 |
| Prpf4b5 | 1.48E-10 | 0.263123686 | 0.582 | 0.334 | 3.00E-06 | 16 | Prpf4b |
| Aebp21 | 1.49E-10 | 0.274100488 | 0.261 | 0.11 | 3.01E-06 | 16 | Aebp2 |
| Ikbkb2 | 1.50E-10 | 0.312515428 | 0.315 | 0.145 | 3.04E-06 | 16 | Ikbkb |
| Plgrkt5 | 1.55E-10 | 0.309736002 | 0.424 | 0.214 | 3.13E-06 | 16 | Plgrkt |
| Scamp32 | 1.62E-10 | 0.270548219 | 0.255 | 0.107 | 3.27E-06 | 16 | Scamp3 |
| Smg66 | 1.64E-10 | 0.400954606 | 0.473 | 0.261 | 3.31E-06 | 16 | Smg6 |
| Ap3b16 | 1.65E-10 | 0.291550389 | 0.424 | 0.218 | 3.35E-06 | 16 | Ap3b1 |
| Arfgef14 | 1.70E-10 | 0.278882439 | 0.435 | 0.228 | 3.44E-06 | 16 | Arfgef1 |

| | | | | | | | |
|---|---|---|---|---|---|---|---|
| Man2a14 | 1.74E-10 | 0.262205591 | 0.462 | 0.235 | 3.53E-06 | 16 | Man2a1 |
| Myo1e8 | 1.75E-10 | 0.367065348 | 0.533 | 0.282 | 3.54E-06 | 16 | Myo1e |
| Slfn25 | 1.93E-10 | 0.280049673 | 0.489 | 0.261 | 3.90E-06 | 16 | Slfn2 |
| Bod1l5 | 1.95E-10 | 0.294572259 | 0.5 | 0.274 | 3.94E-06 | 16 | Bod1l |
| Fbxw72 | 2.24E-10 | 0.308356697 | 0.293 | 0.132 | 4.53E-06 | 16 | Fbxw7 |
| Bclaf16 | 2.25E-10 | 0.360136165 | 0.467 | 0.255 | 4.56E-06 | 16 | Bclaf1 |
| Nlk3 | 2.41E-10 | 0.277015198 | 0.315 | 0.146 | 4.87E-06 | 16 | Nlk |
| Sf3b23 | 2.45E-10 | 0.28466449 | 0.614 | 0.359 | 4.95E-06 | 16 | Sf3b2 |
| Usp348 | 2.54E-10 | 0.274386391 | 0.603 | 0.345 | 5.15E-06 | 16 | Usp34 |
| Tram12 | 2.93E-10 | 0.264465757 | 0.424 | 0.22 | 5.93E-06 | 16 | Tram1 |
| Elf45 | 2.94E-10 | 0.32653809 | 0.277 | 0.123 | 5.96E-06 | 16 | Elf4 |
| Ankhd16 | 3.17E-10 | 0.304235056 | 0.457 | 0.245 | 6.42E-06 | 16 | Ankhd1 |
| Wdr264 | 3.25E-10 | 0.271447053 | 0.408 | 0.21 | 6.58E-06 | 16 | Wdr26 |
| Usp254 | 3.34E-10 | 0.266836109 | 0.511 | 0.282 | 6.76E-06 | 16 | Usp25 |
| Cnot13 | 3.77E-10 | 0.255597941 | 0.359 | 0.174 | 7.62E-06 | 16 | Cnot1 |
| Fto3 | 3.97E-10 | 0.319452492 | 0.38 | 0.192 | 8.04E-06 | 16 | Fto |
| Rbm57 | 4.14E-10 | 0.292606182 | 0.451 | 0.24 | 8.38E-06 | 16 | Rbm5 |
| Taf153 | 4.59E-10 | 0.265826836 | 0.451 | 0.238 | 9.30E-06 | 16 | Taf15 |
| Zdhhc208 | 4.61E-10 | 0.356765647 | 0.429 | 0.231 | 9.34E-06 | 16 | Zdhhc20 |
| 2610035D17Rik3 | 5.88E-10 | 0.253676122 | 0.266 | 0.115 | 1.19E-05 | 16 | 2610035D17Rik |
| Chd92 | 6.08E-10 | 0.287707809 | 0.467 | 0.251 | 1.23E-05 | 16 | Chd9 |
| Cdk127 | 6.12E-10 | 0.299748781 | 0.44 | 0.23 | 1.24E-05 | 16 | Cdk12 |
| Rprd2 | 6.70E-10 | 0.280910776 | 0.293 | 0.135 | 1.36E-05 | 16 | Rprd2 |
| Map4k47 | 7.57E-10 | 0.306979625 | 0.505 | 0.278 | 1.53E-05 | 16 | Map4k4 |
| Lcor6 | 7.61E-10 | 0.313487441 | 0.375 | 0.191 | 1.54E-05 | 16 | Lcor |
| Dennd1a7 | 8.62E-10 | 0.38898778 | 0.462 | 0.254 | 1.75E-05 | 16 | Dennd1a |
| Cep3507 | 1.02E-09 | 0.29695883 | 0.391 | 0.204 | 2.06E-05 | 16 | Cep350 |
| Arl6ip57 | 1.02E-09 | 0.328108773 | 0.418 | 0.222 | 2.06E-05 | 16 | Arl6ip5 |
| Cntrl4 | 1.08E-09 | 0.369326071 | 0.272 | 0.124 | 2.18E-05 | 16 | Cntrl |
| Cdc733 | 1.19E-09 | 0.250420062 | 0.348 | 0.171 | 2.41E-05 | 16 | Cdc73 |
| Ddx104 | 1.22E-09 | 0.304682438 | 0.288 | 0.132 | 2.47E-05 | 16 | Ddx10 |
| mt-Cytb9 | 1.22E-09 | 0.301459429 | 1 | 0.7 | 2.47E-05 | 16 | mt-Cytb |
| Clcn34 | 1.30E-09 | 0.396123676 | 0.288 | 0.134 | 2.63E-05 | 16 | Clcn3 |
| Zfand38 | 1.31E-09 | 0.291351384 | 0.527 | 0.302 | 2.66E-05 | 16 | Zfand3 |
| Bptf9 | 1.32E-09 | 0.301595099 | 0.516 | 0.294 | 2.68E-05 | 16 | Bptf |
| H131 | 1.33E-09 | 0.268512414 | 0.326 | 0.158 | 2.69E-05 | 16 | H13 |
| Eed2 | 1.36E-09 | 0.255710638 | 0.315 | 0.151 | 2.75E-05 | 16 | Eed |
| Pcm15 | 1.45E-09 | 0.252902322 | 0.462 | 0.252 | 2.93E-05 | 16 | Pcm1 |
| Mgea54 | 1.95E-09 | 0.256217305 | 0.342 | 0.17 | 3.94E-05 | 16 | Mgea5 |
| Wdr443 | 1.96E-09 | 0.267980295 | 0.272 | 0.124 | 3.98E-05 | 16 | Wdr44 |
| Orai13 | 1.99E-09 | 0.250921053 | 0.283 | 0.129 | 4.02E-05 | 16 | Orai1 |
| Kcnq1ot15 | 2.17E-09 | 0.364675302 | 0.288 | 0.135 | 4.40E-05 | 16 | Kcnq1ot1 |
| Dip2b6 | 2.97E-09 | 0.256556455 | 0.5 | 0.277 | 6.00E-05 | 16 | Dip2b |

| | | | | | | | |
|---|---|---|---|---|---|---|---|
| Abhd17b5 | 2.98E-09 | 0.264078519 | 0.375 | 0.193 | 6.04E-05 | 16 | Abhd17b |
| Scaf87 | 3.05E-09 | 0.360354471 | 0.359 | 0.186 | 6.18E-05 | 16 | Scaf8 |
| Arid23 | 3.13E-09 | 0.26738709 | 0.321 | 0.157 | 6.33E-05 | 16 | Arid2 |
| Gmds2 | 3.44E-09 | 0.276873036 | 0.353 | 0.179 | 6.97E-05 | 16 | Gmds |
| Itch7 | 3.77E-09 | 0.298430284 | 0.484 | 0.274 | 7.63E-05 | 16 | Itch |
| Lman23 | 3.79E-09 | 0.258983546 | 0.391 | 0.207 | 7.68E-05 | 16 | Lman2 |
| Tmcc18 | 4.16E-09 | 0.369586246 | 0.402 | 0.219 | 8.41E-05 | 16 | Tmcc1 |
| Phf20l17 | 4.39E-09 | 0.256889863 | 0.5 | 0.288 | 8.88E-05 | 16 | Phf20l1 |
| Rsrc15 | 4.49E-09 | 0.330536563 | 0.457 | 0.261 | 9.09E-05 | 16 | Rsrc1 |
| Cdyl5 | 4.65E-09 | 0.262017651 | 0.277 | 0.13 | 9.41E-05 | 16 | Cdyl |
| Sdccag83 | 4.90E-09 | 0.260571649 | 0.266 | 0.122 | 9.91E-05 | 16 | Sdccag8 |
| Top2b3 | 5.70E-09 | 0.251836608 | 0.495 | 0.284 | 0.000115329 | 16 | Top2b |
| Wipf16 | 6.17E-09 | 0.333256531 | 0.478 | 0.269 | 0.000124857 | 16 | Wipf1 |
| Tmem50a7 | 6.27E-09 | 0.265905304 | 0.625 | 0.384 | 0.00012687 | 16 | Tmem50a |
| Pik3ca2 | 6.75E-09 | 0.261638915 | 0.337 | 0.169 | 0.000136723 | 16 | Pik3ca |
| Btbd99 | 7.61E-09 | 0.26602003 | 0.435 | 0.238 | 0.000154083 | 16 | Btbd9 |
| Phip8 | 8.16E-09 | 0.314662645 | 0.457 | 0.257 | 0.000165276 | 16 | Phip |
| Slc10a72 | 1.11E-08 | 0.271055157 | 0.304 | 0.148 | 0.000225699 | 16 | Slc10a7 |
| Plaa | 1.28E-08 | 0.267821021 | 0.25 | 0.115 | 0.000258759 | 16 | Plaa |
| Tapbpl1 | 1.46E-08 | 0.25116066 | 0.272 | 0.129 | 0.000295929 | 16 | Tapbpl |
| Pibf11 | 1.50E-08 | 0.285525309 | 0.277 | 0.133 | 0.000303301 | 16 | Pibf1 |
| Suco4 | 1.51E-08 | 0.28175156 | 0.31 | 0.154 | 0.000306705 | 16 | Suco |
| Krit16 | 1.54E-08 | 0.261196779 | 0.353 | 0.183 | 0.000311039 | 16 | Krit1 |
| Wapl1 | 1.87E-08 | 0.265848218 | 0.435 | 0.252 | 0.000378879 | 16 | Wapl |
| Ash1l5 | 1.99E-08 | 0.250403876 | 0.516 | 0.3 | 0.000403843 | 16 | Ash1l |
| Kdm2a7 | 2.00E-08 | 0.250527233 | 0.435 | 0.241 | 0.000405725 | 16 | Kdm2a |
| 2310035C23Rik5 | 2.27E-08 | 0.286799381 | 0.288 | 0.141 | 0.000458811 | 16 | 2310035C23Rik |
| Malat19 | 2.39E-08 | 0.257763437 | 1 | 0.725 | 0.000484677 | 16 | Malat1 |
| Pds5b3 | 2.49E-08 | 0.257863317 | 0.348 | 0.181 | 0.000504261 | 16 | Pds5b |
| Ubb | 2.73E-08 | 0.334746694 | 0.995 | 0.895 | 0.000552585 | 16 | Ubb |
| Ubn27 | 3.53E-08 | 0.383916776 | 0.424 | 0.246 | 0.00071381 | 16 | Ubn2 |
| Faf15 | 3.61E-08 | 0.255086074 | 0.37 | 0.202 | 0.000731605 | 16 | Faf1 |
| Cpsf62 | 3.64E-08 | 0.267872966 | 0.299 | 0.152 | 0.00073603 | 16 | Cpsf6 |
| Rlf4 | 3.99E-08 | 0.288929751 | 0.277 | 0.138 | 0.000808119 | 16 | Rlf |
| Ehmt13 | 4.79E-08 | 0.276173461 | 0.348 | 0.186 | 0.000970438 | 16 | Ehmt1 |
| Kpna11 | 4.85E-08 | 0.261812325 | 0.255 | 0.123 | 0.000981454 | 16 | Kpna1 |
| Stat14 | 5.27E-08 | 0.437710989 | 0.429 | 0.25 | 0.001066409 | 16 | Stat1 |
| Ube3c3 | 5.32E-08 | 0.264549833 | 0.31 | 0.16 | 0.001077386 | 16 | Ube3c |
| Rnf1113 | 6.56E-08 | 0.260740648 | 0.304 | 0.157 | 0.001328062 | 16 | Rnf111 |
| Trappc94 | 7.15E-08 | 0.270004464 | 0.359 | 0.195 | 0.001447566 | 16 | Trappc9 |
| Sqstm16 | 7.21E-08 | 0.47119563 | 0.5 | 0.305 | 0.001460667 | 16 | Sqstm1 |
| Pigk1 | 8.69E-08 | 0.260167873 | 0.25 | 0.12 | 0.001759733 | 16 | Pigk |
| Zmym22 | 1.05E-07 | 0.265435883 | 0.277 | 0.14 | 0.002132228 | 16 | Zmym2 |

| | | | | | | | |
|---|---|---|---|---|---|---|---|
| Vmp14 | 1.24E-07 | 0.255044071 | 0.38 | 0.217 | 0.002502259 | 16 | Vmp1 |
| Atp9b5 | 1.29E-07 | 0.312971263 | 0.299 | 0.159 | 0.002607306 | 16 | Atp9b |
| Kdm6a8 | 1.92E-07 | 0.3235404 | 0.37 | 0.215 | 0.003888638 | 16 | Kdm6a |
| Fnip13 | 2.18E-07 | 0.260505921 | 0.283 | 0.147 | 0.004418181 | 16 | Fnip1 |
| Sppl38 | 2.19E-07 | 0.280450633 | 0.386 | 0.219 | 0.004433305 | 16 | Sppl3 |
| Ube2e22 | 2.50E-07 | 0.261620463 | 0.299 | 0.154 | 0.00506701 | 16 | Ube2e2 |
| Lnpep9 | 2.93E-07 | 0.255047278 | 0.44 | 0.261 | 0.005924883 | 16 | Lnpep |
| Elp42 | 4.78E-07 | 0.256439736 | 0.255 | 0.129 | 0.009677619 | 16 | Elp4 |
| Sf12 | 8.87E-07 | 0.263547502 | 0.332 | 0.189 | 0.017959454 | 16 | Sf1 |
| Zfc3h11 | 1.30E-06 | 0.256056318 | 0.261 | 0.137 | 0.026368782 | 16 | Zfc3h1 |
| Dnmt3a3 | 1.42E-06 | 0.467036708 | 0.277 | 0.148 | 0.028671364 | 16 | Dnmt3a |
| Ifi476 | 3.12E-06 | 0.267698038 | 0.315 | 0.18 | 0.063074755 | 16 | Ifi47 |
| Igf1r4 | 3.38E-06 | 0.273569128 | 0.293 | 0.162 | 0.068411856 | 16 | Igf1r |
| Os97 | 9.42E-06 | 0.25273261 | 0.315 | 0.19 | 0.190796903 | 16 | Os9 |
| Rgs5 | 0 | 5.898887672 | 0.988 | 0.012 | 0 | 17 | Rgs5 |
| Tagln | 0 | 4.092444837 | 0.626 | 0.011 | 0 | 17 | Tagln |
| Abcc9 | 0 | 4.066521213 | 0.819 | 0.009 | 0 | 17 | Abcc9 |
| Acta2 | 0 | 3.996626376 | 0.632 | 0.018 | 0 | 17 | Acta2 |
| Ndufa4l21 | 0 | 3.884273985 | 0.953 | 0.015 | 0 | 17 | Ndufa4l2 |
| Myl9 | 0 | 3.840197216 | 0.936 | 0.017 | 0 | 17 | Myl9 |
| Kcnj8 | 0 | 3.834907757 | 0.713 | 0.014 | 0 | 17 | Kcnj8 |
| Notch3 | 0 | 3.747323792 | 0.988 | 0.003 | 0 | 17 | Notch3 |
| Gucy1a1 | 0 | 3.688386546 | 0.982 | 0.006 | 0 | 17 | Gucy1a1 |
| Tpm2 | 0 | 3.58383982 | 0.912 | 0.024 | 0 | 17 | Tpm2 |
| Pdgfrb1 | 0 | 3.52568623 | 0.965 | 0.022 | 0 | 17 | Pdgfrb |
| Myh11 | 0 | 3.494252701 | 0.596 | 0.007 | 0 | 17 | Myh11 |
| Steap41 | 0 | 3.433287675 | 0.883 | 0.023 | 0 | 17 | Steap4 |
| Gucy1b1 | 0 | 3.283970134 | 0.936 | 0.005 | 0 | 17 | Gucy1b1 |
| Cacna1c | 0 | 3.192657456 | 0.901 | 0.011 | 0 | 17 | Cacna1c |
| Rarres21 | 0 | 3.070680443 | 0.936 | 0.032 | 0 | 17 | Rarres2 |
| Pde3a | 0 | 2.907786096 | 0.825 | 0.009 | 0 | 17 | Pde3a |
| Cox4i2 | 0 | 2.902375863 | 0.895 | 0.005 | 0 | 17 | Cox4i2 |
| Gpc61 | 0 | 2.85906919 | 0.789 | 0.027 | 0 | 17 | Gpc6 |
| Aspn | 0 | 2.802638207 | 0.865 | 0.006 | 0 | 17 | Aspn |
| Lin7a1 | 0 | 2.801733933 | 0.854 | 0.012 | 0 | 17 | Lin7a |
| Ano1 | 0 | 2.783278745 | 0.865 | 0.005 | 0 | 17 | Ano1 |
| Meg31 | 0 | 2.780386241 | 0.708 | 0.022 | 0 | 17 | Meg3 |
| Cacna2d11 | 0 | 2.678678867 | 0.836 | 0.02 | 0 | 17 | Cacna2d1 |
| Sox51 | 0 | 2.677741319 | 0.784 | 0.016 | 0 | 17 | Sox5 |
| Higd1b | 0 | 2.675974007 | 0.772 | 0.001 | 0 | 17 | Higd1b |
| Plcl13 | 0 | 2.673457565 | 0.807 | 0.048 | 0 | 17 | Plcl1 |
| Gucy1a21 | 0 | 2.619122508 | 0.819 | 0.011 | 0 | 17 | Gucy1a2 |
| Cygb1 | 0 | 2.589579401 | 0.865 | 0.03 | 0 | 17 | Cygb |

| | | | | | | | |
|---|---|---|---|---|---|---|---|
| Mylk2 | 0 | 2.586295491 | 0.784 | 0.045 | 0 | 17 | Mylk |
| Grm7 | 0 | 2.582591528 | 0.532 | 0.002 | 0 | 17 | Grm7 |
| Pcdh191 | 0 | 2.556883963 | 0.766 | 0.022 | 0 | 17 | Pcdh19 |
| Serpini1 | 0 | 2.543993029 | 0.76 | 0.02 | 0 | 17 | Serpini1 |
| Ednra1 | 0 | 2.530455565 | 0.83 | 0.012 | 0 | 17 | Ednra |
| Rasl11a | 0 | 2.522132686 | 0.76 | 0.005 | 0 | 17 | Rasl11a |
| Dmd | 0 | 2.501997432 | 0.737 | 0.056 | 0 | 17 | Dmd |
| Vtn | 0 | 2.489652365 | 0.731 | 0.006 | 0 | 17 | Vtn |
| Ebf2 | 0 | 2.480324291 | 0.772 | 0.009 | 0 | 17 | Ebf2 |
| Carmn | 0 | 2.438710874 | 0.772 | 0.004 | 0 | 17 | Carmn |
| Cpe1 | 0 | 2.394718224 | 0.825 | 0.016 | 0 | 17 | Cpe |
| Cacnb21 | 0 | 2.378431417 | 0.637 | 0.016 | 0 | 17 | Cacnb2 |
| Slc24a31 | 0 | 2.366455402 | 0.743 | 0.039 | 0 | 17 | Slc24a3 |
| Pde5a | 0 | 2.253073495 | 0.789 | 0.012 | 0 | 17 | Pde5a |
| Olfr558 | 0 | 2.182000139 | 0.696 | 0.001 | 0 | 17 | Olfr558 |
| Stac | 0 | 2.172019707 | 0.708 | 0.001 | 0 | 17 | Stac |
| Pde8b | 0 | 2.160872026 | 0.52 | 0.005 | 0 | 17 | Pde8b |
| Dgkb | 0 | 2.156668876 | 0.503 | 0.002 | 0 | 17 | Dgkb |
| Mrvi11 | 0 | 2.150963476 | 0.708 | 0.013 | 0 | 17 | Mrvi1 |
| Rgs6 | 0 | 2.148667677 | 0.661 | 0.006 | 0 | 17 | Rgs6 |
| Pde1a1 | 0 | 2.142735413 | 0.725 | 0.01 | 0 | 17 | Pde1a |
| Prrx11 | 0 | 2.129003626 | 0.784 | 0.028 | 0 | 17 | Prrx1 |
| Art3 | 0 | 2.128807996 | 0.614 | 0.024 | 0 | 17 | Art3 |
| Rgs4 | 0 | 2.108951773 | 0.655 | 0.001 | 0 | 17 | Rgs4 |
| Il34 | 0 | 2.097045483 | 0.719 | 0.008 | 0 | 17 | Il34 |
| Arhgap421 | 0 | 2.0852342 | 0.719 | 0.022 | 0 | 17 | Arhgap42 |
| Adamts121 | 0 | 2.077189977 | 0.702 | 0.036 | 0 | 17 | Adamts12 |
| B3galt11 | 0 | 2.066410465 | 0.649 | 0.018 | 0 | 17 | B3galt1 |
| Lhfp1 | 0 | 2.019662948 | 0.696 | 0.035 | 0 | 17 | Lhfp |
| Trpc6 | 0 | 2.017213497 | 0.602 | 0 | 0 | 17 | Trpc6 |
| Sgip1 | 0 | 2.016045254 | 0.655 | 0.012 | 0 | 17 | Sgip1 |
| Ppp1r14a | 0 | 1.993906462 | 0.632 | 0.008 | 0 | 17 | Ppp1r14a |
| Fap1 | 0 | 1.980743646 | 0.591 | 0.033 | 0 | 17 | Fap |
| Postn | 0 | 1.974064715 | 0.55 | 0.007 | 0 | 17 | Postn |
| Gpx81 | 0 | 1.966368751 | 0.678 | 0.047 | 0 | 17 | Gpx8 |
| Fxyd11 | 0 | 1.9619738 | 0.708 | 0.03 | 0 | 17 | Fxyd1 |
| Des | 0 | 1.951497384 | 0.626 | 0.003 | 0 | 17 | Des |
| Sema5a | 0 | 1.94050647 | 0.596 | 0.012 | 0 | 17 | Sema5a |
| Rbms31 | 0 | 1.928561249 | 0.807 | 0.049 | 0 | 17 | Rbms3 |
| Daam2 | 0 | 1.913775005 | 0.661 | 0.006 | 0 | 17 | Daam2 |
| C1s11 | 0 | 1.871398664 | 0.76 | 0.03 | 0 | 17 | C1s1 |
| Arhgef171 | 0 | 1.869263346 | 0.684 | 0.021 | 0 | 17 | Arhgef17 |
| Barx11 | 0 | 1.863808644 | 0.573 | 0.021 | 0 | 17 | Barx1 |

| | | | | | | | |
|---|---|---|---|---|---|---|---|
| Atp1b2 | 0 | 1.860161324 | 0.667 | 0.016 | 0 | 17 | Atp1b2 |
| Rem1 | 0 | 1.82055855 | 0.538 | 0.004 | 0 | 17 | Rem1 |
| Zfp804b | 0 | 1.798231685 | 0.287 | 0.001 | 0 | 17 | Zfp804b |
| Mnd1 | 0 | 1.784798132 | 0.474 | 0.019 | 0 | 17 | Mnd1 |
| Plce1 | 0 | 1.751784538 | 0.585 | 0.018 | 0 | 17 | Plce1 |
| Meox2 | 0 | 1.748973352 | 0.556 | 0.011 | 0 | 17 | Meox2 |
| Cpm | 0 | 1.727242374 | 0.497 | 0.017 | 0 | 17 | Cpm |
| Foxs1 | 0 | 1.723739188 | 0.579 | 0.007 | 0 | 17 | Foxs1 |
| Cspg41 | 0 | 1.692217203 | 0.602 | 0.011 | 0 | 17 | Cspg4 |
| Htra12 | 0 | 1.663887998 | 0.69 | 0.043 | 0 | 17 | Htra1 |
| Lgi1 | 0 | 1.661833325 | 0.439 | 0.003 | 0 | 17 | Lgi1 |
| Serping11 | 0 | 1.655471745 | 0.655 | 0.032 | 0 | 17 | Serping1 |
| Agtr1a | 0 | 1.642790865 | 0.509 | 0.006 | 0 | 17 | Agtr1a |
| Aoc31 | 0 | 1.642544183 | 0.602 | 0.013 | 0 | 17 | Aoc3 |
| Vstm41 | 0 | 1.639960337 | 0.532 | 0.012 | 0 | 17 | Vstm4 |
| Heyl | 0 | 1.638503915 | 0.561 | 0.004 | 0 | 17 | Heyl |
| Tmem1001 | 0 | 1.61336116 | 0.561 | 0.013 | 0 | 17 | Tmem100 |
| Ifitm1 | 0 | 1.611244969 | 0.398 | 0.003 | 0 | 17 | Ifitm1 |
| Serpine21 | 0 | 1.609573218 | 0.579 | 0.03 | 0 | 17 | Serpine2 |
| Egflam1 | 0 | 1.607338909 | 0.561 | 0.017 | 0 | 17 | Egflam |
| Parm1 | 0 | 1.60340128 | 0.591 | 0.005 | 0 | 17 | Parm1 |
| Rdh5 | 0 | 1.591261794 | 0.52 | 0.011 | 0 | 17 | Rdh5 |
| Nr2f11 | 0 | 1.586719387 | 0.556 | 0.023 | 0 | 17 | Nr2f1 |
| Rasl12 | 0 | 1.548710258 | 0.567 | 0.012 | 0 | 17 | Rasl12 |
| Lmod1 | 0 | 1.524940344 | 0.503 | 0.008 | 0 | 17 | Lmod1 |
| Tbx22 | 0 | 1.515554271 | 0.532 | 0.019 | 0 | 17 | Tbx2 |
| Ephx3 | 0 | 1.506766208 | 0.468 | 0.001 | 0 | 17 | Ephx3 |
| Sdc1 | 0 | 1.498470759 | 0.485 | 0.008 | 0 | 17 | Sdc1 |
| Itga7 | 0 | 1.474348096 | 0.556 | 0.001 | 0 | 17 | Itga7 |
| Rgs7bp | 0 | 1.47239471 | 0.439 | 0.008 | 0 | 17 | Rgs7bp |
| Chn1 | 0 | 1.459131714 | 0.468 | 0.002 | 0 | 17 | Chn1 |
| Synpo2 | 0 | 1.430381601 | 0.433 | 0.005 | 0 | 17 | Synpo2 |
| Rbpms21 | 0 | 1.425705947 | 0.491 | 0.006 | 0 | 17 | Rbpms2 |
| Thsd4 | 0 | 1.391300814 | 0.38 | 0.007 | 0 | 17 | Thsd4 |
| Pla2r11 | 0 | 1.341091493 | 0.48 | 0.016 | 0 | 17 | Pla2r1 |
| Hspb2 | 0 | 1.339713206 | 0.45 | 0.004 | 0 | 17 | Hspb2 |
| Kirrel1 | 0 | 1.332098309 | 0.491 | 0.019 | 0 | 17 | Kirrel |
| Casq2 | 0 | 1.330003952 | 0.363 | 0.002 | 0 | 17 | Casq2 |
| Elmod1 | 0 | 1.31801754 | 0.292 | 0.001 | 0 | 17 | Elmod1 |
| Galnt161 | 0 | 1.316139788 | 0.462 | 0.003 | 0 | 17 | Galnt16 |
| Mark1 | 0 | 1.292157264 | 0.433 | 0.007 | 0 | 17 | Mark1 |
| Naalad21 | 0 | 1.26094142 | 0.45 | 0.019 | 0 | 17 | Naalad2 |
| Slc38a111 | 0 | 1.258511244 | 0.392 | 0.001 | 0 | 17 | Slc38a11 |

| | | | | | | | |
|---|---|---|---|---|---|---|---|
| Angpt1 | 0 | 1.234020993 | 0.444 | 0.015 | 0 | 17 | Angpt1 |
| Npy1r | 0 | 1.228801883 | 0.404 | 0.001 | 0 | 17 | Npy1r |
| Nrip2 | 0 | 1.228685498 | 0.316 | 0 | 0 | 17 | Nrip2 |
| Cap2 | 0 | 1.219889718 | 0.404 | 0.002 | 0 | 17 | Cap2 |
| Ppm1e | 0 | 1.213715958 | 0.456 | 0.019 | 0 | 17 | Ppm1e |
| Tbx3os1 | 0 | 1.186922934 | 0.374 | 0.003 | 0 | 17 | Tbx3os1 |
| Klhl29 | 0 | 1.175347092 | 0.368 | 0.006 | 0 | 17 | Klhl29 |
| Pcdh181 | 0 | 1.164216569 | 0.415 | 0.011 | 0 | 17 | Pcdh18 |
| Gm13861 | 0 | 1.140716105 | 0.351 | 0.001 | 0 | 17 | Gm13861 |
| Trabd2b | 0 | 1.079298607 | 0.333 | 0.009 | 0 | 17 | Trabd2b |
| Trpc3 | 0 | 1.074308857 | 0.368 | 0.011 | 0 | 17 | Trpc3 |
| Gprc5c | 0 | 1.058303867 | 0.404 | 0.008 | 0 | 17 | Gprc5c |
| Ntm | 0 | 1.054901015 | 0.316 | 0.008 | 0 | 17 | Ntm |
| Slc2a4 | 0 | 1.052792587 | 0.398 | 0.009 | 0 | 17 | Slc2a4 |
| Dgkg | 0 | 1.050716932 | 0.386 | 0.006 | 0 | 17 | Dgkg |
| Cdh2 | 0 | 0.998086817 | 0.281 | 0.004 | 0 | 17 | Cdh2 |
| Fads3 | 0 | 0.963713065 | 0.333 | 0.009 | 0 | 17 | Fads3 |
| Ccdc3 | 0 | 0.942685778 | 0.292 | 0.001 | 0 | 17 | Ccdc3 |
| Ace21 | 0 | 0.934121495 | 0.281 | 0 | 0 | 17 | Ace2 |
| Pth1r | 0 | 0.908065834 | 0.292 | 0.006 | 0 | 17 | Pth1r |
| Ajuba | 0 | 0.851237943 | 0.31 | 0.007 | 0 | 17 | Ajuba |
| Olfml2b | 0 | 0.823210051 | 0.316 | 0.008 | 0 | 17 | Olfml2b |
| Slc4a8 | 0 | 0.794892175 | 0.251 | 0.002 | 0 | 17 | Slc4a8 |
| Jph2 | 0 | 0.729092368 | 0.281 | 0.002 | 0 | 17 | Jph2 |
| Kcnk3 | 0 | 0.701962149 | 0.263 | 0.005 | 0 | 17 | Kcnk3 |
| Angptl21 | 1.21E-299 | 1.50076835 | 0.433 | 0.019 | 2.44E-295 | 17 | Angptl2 |
| Col1a21 | 7.47E-299 | 1.002890517 | 0.614 | 0.036 | 1.51E-294 | 17 | Col1a2 |
| Nbl11 | 1.96E-298 | 1.422455169 | 0.45 | 0.021 | 3.98E-294 | 17 | Nbl1 |
| Bgn1 | 8.58E-298 | 1.378866189 | 0.649 | 0.042 | 1.74E-293 | 17 | Bgn |
| Cryab | 4.13E-295 | 1.833257365 | 0.801 | 0.067 | 8.37E-291 | 17 | Cryab |
| Gm14005 | 1.22E-294 | 1.619519554 | 0.556 | 0.033 | 2.46E-290 | 17 | Gm14005 |
| Arhgap62 | 5.13E-294 | 2.083649801 | 0.713 | 0.054 | 1.04E-289 | 17 | Arhgap6 |
| Rasgrp2 | 2.09E-291 | 1.829770267 | 0.696 | 0.053 | 4.23E-287 | 17 | Rasgrp2 |
| Tnfrsf211 | 3.78E-287 | 1.381825226 | 0.456 | 0.022 | 7.66E-283 | 17 | Tnfrsf21 |
| Gja42 | 1.70E-282 | 1.933363333 | 0.731 | 0.059 | 3.45E-278 | 17 | Gja4 |
| Ccl111 | 4.08E-282 | 2.143333417 | 0.52 | 0.029 | 8.26E-278 | 17 | Ccl11 |
| Akap6 | 4.92E-280 | 0.816954533 | 0.263 | 0.007 | 9.95E-276 | 17 | Akap6 |
| Trpc1 | 7.17E-278 | 0.998343995 | 0.281 | 0.008 | 1.45E-273 | 17 | Trpc1 |
| Nr2f21 | 7.65E-278 | 2.504615209 | 0.877 | 0.095 | 1.55E-273 | 17 | Nr2f2 |
| Adcy6 | 2.47E-274 | 1.386174135 | 0.544 | 0.034 | 5.00E-270 | 17 | Adcy6 |
| Pdgfa1 | 1.20E-268 | 1.734593851 | 0.69 | 0.056 | 2.43E-264 | 17 | Pdgfa |
| Adcyap1r11 | 4.26E-268 | 0.950283448 | 0.327 | 0.012 | 8.62E-264 | 17 | Adcyap1r1 |
| Pard32 | 9.78E-263 | 1.523880673 | 0.602 | 0.043 | 1.98E-258 | 17 | Pard3 |

| | | | | | | | |
|---|---|---|---|---|---|---|---|
| Afap1l21 | 1.70E-249 | 1.136979179 | 0.444 | 0.024 | 3.44E-245 | 17 | Afap1l2 |
| Ppfia2 | 6.96E-248 | 1.179129181 | 0.304 | 0.011 | 1.41E-243 | 17 | Ppfia2 |
| Ctnna3 | 7.71E-246 | 2.070225943 | 0.386 | 0.019 | 1.56E-241 | 17 | Ctnna3 |
| Col6a11 | 9.39E-245 | 1.391649864 | 0.503 | 0.031 | 1.90E-240 | 17 | Col6a1 |
| A830082K12Rik1 | 2.07E-244 | 1.045223106 | 0.327 | 0.013 | 4.20E-240 | 17 | A830082K12Rik |
| P3h2 | 2.63E-244 | 1.372973921 | 0.48 | 0.029 | 5.33E-240 | 17 | P3h2 |
| Adam22 | 8.26E-240 | 0.982873921 | 0.322 | 0.013 | 1.67E-235 | 17 | Adam22 |
| Sntb13 | 1.95E-235 | 1.809678955 | 0.673 | 0.062 | 3.94E-231 | 17 | Sntb1 |
| Cyp7b11 | 9.78E-232 | 1.505620156 | 0.503 | 0.034 | 1.98E-227 | 17 | Cyp7b1 |
| Bmpr1a1 | 1.12E-229 | 1.346339171 | 0.515 | 0.036 | 2.27E-225 | 17 | Bmpr1a |
| Ggt51 | 2.19E-229 | 1.168447771 | 0.439 | 0.026 | 4.43E-225 | 17 | Ggt5 |
| Ntn42 | 6.17E-227 | 1.384977901 | 0.485 | 0.032 | 1.25E-222 | 17 | Ntn4 |
| Lsamp1 | 2.66E-226 | 1.404432998 | 0.526 | 0.036 | 5.39E-222 | 17 | Lsamp |
| Ddr21 | 4.36E-224 | 1.144442167 | 0.427 | 0.025 | 8.83E-220 | 17 | Ddr2 |
| Ror11 | 8.49E-221 | 1.273279962 | 0.491 | 0.034 | 1.72E-216 | 17 | Ror1 |
| Col3a11 | 3.74E-220 | 1.241530572 | 0.526 | 0.036 | 7.57E-216 | 17 | Col3a1 |
| Myo1b2 | 7.01E-220 | 2.711315651 | 0.895 | 0.134 | 1.42E-215 | 17 | Myo1b |
| Ecm22 | 9.92E-220 | 1.562923631 | 0.567 | 0.046 | 2.01E-215 | 17 | Ecm2 |
| Etv1 | 3.53E-219 | 1.073448581 | 0.45 | 0.028 | 7.14E-215 | 17 | Etv1 |
| Pdlim3 | 6.65E-216 | 1.007850419 | 0.345 | 0.017 | 1.35E-211 | 17 | Pdlim3 |
| Jag11 | 6.72E-212 | 1.195143907 | 0.497 | 0.036 | 1.36E-207 | 17 | Jag1 |
| Zeb23 | 2.13E-210 | 2.497314135 | 0.93 | 0.14 | 4.32E-206 | 17 | Zeb2 |
| Map3k202 | 6.13E-210 | 1.935926756 | 0.667 | 0.07 | 1.24E-205 | 17 | Map3k20 |
| Cog7 | 3.44E-208 | 1.581003942 | 0.579 | 0.051 | 6.97E-204 | 17 | Cog7 |
| Klhl23 | 4.88E-207 | 0.796023176 | 0.275 | 0.011 | 9.89E-203 | 17 | Klhl23 |
| Sep-44 | 1.38E-206 | 2.63192984 | 0.942 | 0.161 | 2.79E-202 | 17 | 4-Sep |
| Adgrl3 | 2.52E-202 | 1.017528856 | 0.263 | 0.01 | 5.10E-198 | 17 | Adgrl3 |
| Cp2 | 4.77E-202 | 1.664963518 | 0.538 | 0.044 | 9.66E-198 | 17 | Cp |
| Lama21 | 3.99E-201 | 1.054602782 | 0.491 | 0.035 | 8.08E-197 | 17 | Lama2 |
| Prkg13 | 3.02E-200 | 3.260180039 | 0.982 | 0.193 | 6.12E-196 | 17 | Prkg1 |
| Arhgef251 | 1.65E-198 | 0.928955726 | 0.345 | 0.018 | 3.35E-194 | 17 | Arhgef25 |
| Dtx31 | 9.14E-198 | 1.615509096 | 0.608 | 0.06 | 1.85E-193 | 17 | Dtx3 |
| Hrct1 | 1.10E-192 | 1.070976744 | 0.368 | 0.021 | 2.22E-188 | 17 | Hrct1 |
| Filip1l4 | 1.88E-190 | 2.717202274 | 0.877 | 0.149 | 3.81E-186 | 17 | Filip1l |
| Gjc13 | 2.64E-190 | 2.135069727 | 0.789 | 0.112 | 5.35E-186 | 17 | Gjc1 |
| Olfr1033 | 1.36E-188 | 0.798072662 | 0.251 | 0.01 | 2.75E-184 | 17 | Olfr1033 |
| Sdc21 | 6.01E-188 | 1.152620703 | 0.427 | 0.029 | 1.22E-183 | 17 | Sdc2 |
| Slc12a22 | 1.45E-187 | 2.476670683 | 0.789 | 0.112 | 2.94E-183 | 17 | Slc12a2 |
| Igfbp51 | 7.15E-187 | 1.875229986 | 0.456 | 0.034 | 1.45E-182 | 17 | Igfbp5 |
| Arhgap106 | 2.69E-185 | 2.013212024 | 0.819 | 0.124 | 5.46E-181 | 17 | Arhgap10 |
| Adap22 | 3.65E-185 | 1.545497483 | 0.573 | 0.055 | 7.38E-181 | 17 | Adap2 |
| Mamdc2 | 2.72E-181 | 0.916327038 | 0.304 | 0.016 | 5.51E-177 | 17 | Mamdc2 |
| Pkia | 8.19E-181 | 0.931644237 | 0.322 | 0.017 | 1.66E-176 | 17 | Pkia |

| | | | | | | | |
|---|---|---|---|---|---|---|---|
| Twist11 | 9.01E-181 | 0.886580304 | 0.327 | 0.018 | 1.82E-176 | 17 | Twist1 |
| Lgals15 | 1.56E-178 | 2.231654163 | 0.86 | 0.132 | 3.16E-174 | 17 | Lgals1 |
| Hopx4 | 3.14E-175 | 1.744618468 | 0.632 | 0.073 | 6.35E-171 | 17 | Hopx |
| Epb41l11 | 1.07E-173 | 0.82644187 | 0.357 | 0.022 | 2.17E-169 | 17 | Epb41l1 |
| Tmem51 | 1.72E-170 | 0.879479608 | 0.333 | 0.02 | 3.47E-166 | 17 | Tmem51 |
| Ppfibp2 | 1.01E-168 | 1.044160507 | 0.415 | 0.031 | 2.05E-164 | 17 | Ppfibp2 |
| Magi21 | 6.66E-168 | 1.383489424 | 0.444 | 0.036 | 1.35E-163 | 17 | Magi2 |
| Sep-81 | 1.03E-167 | 1.188159881 | 0.485 | 0.044 | 2.08E-163 | 17 | 8-Sep |
| Ngf | 2.29E-164 | 0.995600894 | 0.48 | 0.041 | 4.65E-160 | 17 | Ngf |
| C1qtnf11 | 7.89E-162 | 0.792051875 | 0.281 | 0.015 | 1.60E-157 | 17 | C1qtnf1 |
| Phlda31 | 2.92E-160 | 1.200822901 | 0.526 | 0.052 | 5.92E-156 | 17 | Phlda3 |
| Snhg182 | 7.39E-159 | 0.997553782 | 0.45 | 0.039 | 1.50E-154 | 17 | Snhg18 |
| Cobll14 | 1.75E-157 | 1.58082124 | 0.643 | 0.083 | 3.53E-153 | 17 | Cobll1 |
| Klhdc8b | 4.95E-157 | 0.884970133 | 0.322 | 0.02 | 1.00E-152 | 17 | Klhdc8b |
| Nr1h31 | 6.62E-155 | 1.034585324 | 0.304 | 0.018 | 1.34E-150 | 17 | Nr1h3 |
| Sncg4 | 1.27E-154 | 2.520419188 | 0.825 | 0.152 | 2.58E-150 | 17 | Sncg |
| Mfge85 | 1.83E-150 | 2.604491073 | 0.971 | 0.251 | 3.70E-146 | 17 | Mfge8 |
| Rcan2 | 9.01E-150 | 1.827528884 | 0.462 | 0.044 | 1.82E-145 | 17 | Rcan2 |
| Mustn1 | 3.27E-149 | 1.725286979 | 0.363 | 0.028 | 6.62E-145 | 17 | Mustn1 |
| Runx1t12 | 3.79E-149 | 1.067478015 | 0.45 | 0.041 | 7.68E-145 | 17 | Runx1t1 |
| Vcl3 | 1.84E-148 | 1.827401298 | 0.842 | 0.159 | 3.72E-144 | 17 | Vcl |
| Gas11 | 3.12E-148 | 1.029467644 | 0.386 | 0.03 | 6.32E-144 | 17 | Gas1 |
| Gm4876 | 6.44E-148 | 1.046881753 | 0.333 | 0.023 | 1.30E-143 | 17 | Gm4876 |
| Ddit4l | 8.57E-148 | 0.855589554 | 0.404 | 0.032 | 1.73E-143 | 17 | Ddit4l |
| Ptprk3 | 2.56E-144 | 2.032136034 | 0.778 | 0.137 | 5.18E-140 | 17 | Ptprk |
| Tspan33 | 6.32E-143 | 1.558806064 | 0.684 | 0.102 | 1.28E-138 | 17 | Tspan3 |
| Lamc31 | 7.46E-141 | 1.005798339 | 0.281 | 0.017 | 1.51E-136 | 17 | Lamc3 |
| Col6a21 | 3.64E-140 | 0.95766206 | 0.363 | 0.028 | 7.36E-136 | 17 | Col6a2 |
| Sort1 | 7.46E-140 | 1.106260525 | 0.474 | 0.049 | 1.51E-135 | 17 | Sort1 |
| Lgalsl | 5.34E-139 | 1.502133529 | 0.632 | 0.089 | 1.08E-134 | 17 | Lgalsl |
| Pcolce1 | 1.24E-138 | 1.235480063 | 0.45 | 0.043 | 2.51E-134 | 17 | Pcolce |
| Atp2a25 | 2.60E-138 | 2.149742258 | 0.871 | 0.198 | 5.26E-134 | 17 | Atp2a2 |
| Phlda11 | 1.54E-137 | 1.632898806 | 0.608 | 0.084 | 3.11E-133 | 17 | Phlda1 |
| Angpt2 | 3.89E-137 | 0.756076731 | 0.263 | 0.015 | 7.87E-133 | 17 | Angpt2 |
| Wtip | 1.85E-136 | 1.841922313 | 0.602 | 0.085 | 3.74E-132 | 17 | Wtip |
| Smarcd3 | 5.72E-136 | 0.662476773 | 0.263 | 0.015 | 1.16E-131 | 17 | Smarcd3 |
| Asap23 | 5.91E-136 | 1.754904283 | 0.737 | 0.129 | 1.20E-131 | 17 | Asap2 |
| Cd634 | 1.18E-135 | 1.768354017 | 0.93 | 0.185 | 2.39E-131 | 17 | Cd63 |
| Lrp12 | 6.74E-135 | 1.058553438 | 0.503 | 0.054 | 1.37E-130 | 17 | Lrp1 |
| Ccdc141 | 1.73E-134 | 0.88031633 | 0.322 | 0.023 | 3.50E-130 | 17 | Ccdc141 |
| Slco3a14 | 5.73E-133 | 1.706745947 | 0.713 | 0.12 | 1.16E-128 | 17 | Slco3a1 |
| Limd1 | 9.50E-133 | 1.67470428 | 0.754 | 0.14 | 1.92E-128 | 17 | Limd1 |
| Ndrg21 | 6.82E-129 | 1.05079347 | 0.556 | 0.068 | 1.38E-124 | 17 | Ndrg2 |

| | | | | | | | |
|---|---|---|---|---|---|---|---|
| Col5a22 | 1.08E-126 | 1.168784732 | 0.421 | 0.041 | 2.19E-122 | 17 | Col5a2 |
| Gamt | 4.87E-124 | 1.111105668 | 0.462 | 0.052 | 9.87E-120 | 17 | Gamt |
| Ednrb2 | 8.39E-123 | 2.274359867 | 0.842 | 0.192 | 1.70E-118 | 17 | Ednrb |
| Cald15 | 3.01E-122 | 2.639681628 | 0.959 | 0.32 | 6.10E-118 | 17 | Cald1 |
| Fam110b | 1.78E-121 | 0.608896013 | 0.257 | 0.017 | 3.61E-117 | 17 | Fam110b |
| Cd2481 | 1.60E-120 | 0.738459379 | 0.263 | 0.017 | 3.24E-116 | 17 | Cd248 |
| Flna4 | 3.39E-120 | 1.724529205 | 0.69 | 0.129 | 6.87E-116 | 17 | Flna |
| Inpp4b5 | 1.71E-118 | 1.448939096 | 0.93 | 0.2 | 3.45E-114 | 17 | Inpp4b |
| Nckap5 | 9.47E-116 | 1.072564102 | 0.433 | 0.048 | 1.92E-111 | 17 | Nckap5 |
| Daam11 | 2.21E-114 | 1.484409275 | 0.632 | 0.107 | 4.46E-110 | 17 | Daam1 |
| Gm138892 | 6.10E-114 | 1.943193514 | 0.883 | 0.217 | 1.23E-109 | 17 | Gm13889 |
| Ank31 | 9.64E-112 | 1.075771931 | 0.433 | 0.049 | 1.95E-107 | 17 | Ank3 |
| Slit31 | 1.68E-111 | 1.226423152 | 0.561 | 0.08 | 3.41E-107 | 17 | Slit3 |
| Pard3b2 | 2.80E-111 | 1.443454887 | 0.608 | 0.099 | 5.67E-107 | 17 | Pard3b |
| Tpm12 | 5.86E-109 | 2.797237499 | 0.906 | 0.29 | 1.19E-104 | 17 | Tpm1 |
| Nt5dc2 | 4.21E-108 | 0.737274098 | 0.322 | 0.028 | 8.53E-104 | 17 | Nt5dc2 |
| Ift43 | 2.68E-105 | 1.286988187 | 0.667 | 0.122 | 5.42E-101 | 17 | Ift43 |
| Tbx3 | 2.72E-105 | 0.865134824 | 0.398 | 0.044 | 5.52E-101 | 17 | Tbx3 |
| Dlc14 | 1.10E-104 | 2.368798967 | 0.977 | 0.361 | 2.22E-100 | 17 | Dlc1 |
| Tmem38b | 3.49E-104 | 0.813143143 | 0.345 | 0.034 | 7.06E-100 | 17 | Tmem38b |
| Sobp1 | 1.18E-101 | 0.732369023 | 0.263 | 0.021 | 2.39E-97 | 17 | Sobp |
| Mast45 | 7.99E-100 | 1.76974753 | 0.778 | 0.185 | 1.62E-95 | 17 | Mast4 |
| Cacna1a1 | 1.50E-99 | 0.877205822 | 0.345 | 0.035 | 3.03E-95 | 17 | Cacna1a |
| Cabp1 | 2.13E-99 | 0.852582666 | 0.345 | 0.036 | 4.31E-95 | 17 | Cabp1 |
| Ccdc801 | 2.76E-99 | 0.859666167 | 0.316 | 0.03 | 5.59E-95 | 17 | Ccdc80 |
| Rbpms4 | 1.20E-96 | 1.932874623 | 0.953 | 0.348 | 2.43E-92 | 17 | Rbpms |
| Btbd31 | 1.54E-96 | 1.36936724 | 0.573 | 0.099 | 3.12E-92 | 17 | Btbd3 |
| Tbc1d18 | 2.37E-96 | 1.48393879 | 0.719 | 0.163 | 4.79E-92 | 17 | Tbc1d1 |
| Khdrbs3 | 1.02E-95 | 0.700006362 | 0.374 | 0.042 | 2.07E-91 | 17 | Khdrbs3 |
| Gnb42 | 4.09E-95 | 1.322346883 | 0.591 | 0.107 | 8.29E-91 | 17 | Gnb4 |
| Eps81 | 1.43E-94 | 1.258313207 | 0.526 | 0.085 | 2.90E-90 | 17 | Eps8 |
| Plekhh21 | 4.41E-94 | 0.760822694 | 0.269 | 0.023 | 8.94E-90 | 17 | Plekhh2 |
| Mgp2 | 2.96E-93 | 2.069954879 | 0.433 | 0.058 | 6.00E-89 | 17 | Mgp |
| Map2 | 3.40E-93 | 1.075487867 | 0.515 | 0.081 | 6.89E-89 | 17 | Map2 |
| Pmp225 | 1.06E-92 | 1.519038484 | 0.673 | 0.142 | 2.15E-88 | 17 | Pmp22 |
| Samd41 | 2.31E-92 | 1.059341356 | 0.409 | 0.053 | 4.68E-88 | 17 | Samd4 |
| Tmem94 | 3.39E-92 | 0.891812053 | 0.363 | 0.043 | 6.85E-88 | 17 | Tmem94 |
| Npas21 | 6.77E-91 | 0.759455816 | 0.281 | 0.026 | 1.37E-86 | 17 | Npas2 |
| Pawr | 1.96E-90 | 0.991959214 | 0.444 | 0.063 | 3.97E-86 | 17 | Pawr |
| Grk55 | 2.96E-90 | 1.696639571 | 0.678 | 0.156 | 6.00E-86 | 17 | Grk5 |
| Sparc6 | 6.47E-90 | 2.197659768 | 0.994 | 0.401 | 1.31E-85 | 17 | Sparc |
| Fam162a2 | 9.66E-90 | 1.690455422 | 0.854 | 0.254 | 1.96E-85 | 17 | Fam162a |
| Mprip6 | 1.19E-89 | 1.69417194 | 0.883 | 0.291 | 2.41E-85 | 17 | Mprip |

| | | | | | | | |
|---|---|---|---|---|---|---|---|
| Fry4 | 4.89E-89 | 1.657012812 | 0.766 | 0.199 | 9.90E-85 | 17 | Fry |
| Gng115 | 4.97E-88 | 1.865469195 | 0.965 | 0.365 | 1.01E-83 | 17 | Gng11 |
| Sparcl16 | 5.06E-88 | 1.80475982 | 0.977 | 0.362 | 1.02E-83 | 17 | Sparcl1 |
| Sgce1 | 6.76E-88 | 0.984178134 | 0.357 | 0.043 | 1.37E-83 | 17 | Sgce |
| Ift811 | 7.66E-88 | 0.700541302 | 0.275 | 0.026 | 1.55E-83 | 17 | Ift81 |
| Lpp8 | 3.87E-86 | 1.692793496 | 0.854 | 0.284 | 7.83E-82 | 17 | Lpp |
| Stk38l | 5.00E-86 | 0.867066083 | 0.374 | 0.048 | 1.01E-81 | 17 | Stk38l |
| Parp31 | 6.21E-86 | 0.752182873 | 0.281 | 0.028 | 1.26E-81 | 17 | Parp3 |
| Psd32 | 1.33E-85 | 1.341769423 | 0.591 | 0.116 | 2.70E-81 | 17 | Psd3 |
| Dennd2a1 | 1.71E-85 | 0.671917537 | 0.251 | 0.022 | 3.47E-81 | 17 | Dennd2a |
| Bcar3 | 2.66E-85 | 0.773126482 | 0.333 | 0.038 | 5.39E-81 | 17 | Bcar3 |
| Mcam3 | 4.95E-85 | 1.176818073 | 0.573 | 0.108 | 1.00E-80 | 17 | Mcam |
| Myof1 | 6.50E-85 | 1.093210618 | 0.409 | 0.058 | 1.32E-80 | 17 | Myof |
| Rora3 | 2.36E-84 | 1.40608603 | 0.76 | 0.184 | 4.78E-80 | 17 | Rora |
| Pid12 | 9.24E-84 | 1.152321151 | 0.45 | 0.068 | 1.87E-79 | 17 | Pid1 |
| Plekhg3 | 1.06E-83 | 0.875070601 | 0.38 | 0.051 | 2.14E-79 | 17 | Plekhg3 |
| Ebf15 | 2.45E-83 | 1.841144706 | 0.988 | 0.373 | 4.96E-79 | 17 | Ebf1 |
| Cystm12 | 4.09E-83 | 1.003475224 | 0.795 | 0.18 | 8.28E-79 | 17 | Cystm1 |
| Igfbp42 | 5.05E-83 | 0.830057456 | 0.55 | 0.096 | 1.02E-78 | 17 | Igfbp4 |
| Peli21 | 1.15E-82 | 0.955387272 | 0.433 | 0.066 | 2.34E-78 | 17 | Peli2 |
| Tinagl15 | 1.50E-82 | 1.493551488 | 0.947 | 0.295 | 3.03E-78 | 17 | Tinagl1 |
| Snx182 | 2.15E-82 | 1.028962832 | 0.503 | 0.089 | 4.35E-78 | 17 | Snx18 |
| Casc41 | 3.63E-82 | 0.829438057 | 0.304 | 0.033 | 7.35E-78 | 17 | Casc4 |
| P3h31 | 5.02E-82 | 0.730949864 | 0.263 | 0.025 | 1.02E-77 | 17 | P3h3 |
| Adcy31 | 1.59E-80 | 0.82660879 | 0.257 | 0.025 | 3.22E-76 | 17 | Adcy3 |
| Dcbld21 | 2.20E-80 | 0.84874986 | 0.327 | 0.04 | 4.46E-76 | 17 | Dcbld2 |
| S1pr31 | 3.74E-80 | 0.792297967 | 0.275 | 0.028 | 7.56E-76 | 17 | S1pr3 |
| Malat110 | 3.95E-80 | 2.024731712 | 1 | 0.725 | 8.00E-76 | 17 | Malat1 |
| Col1a11 | 1.57E-79 | 0.738565113 | 0.304 | 0.033 | 3.17E-75 | 17 | Col1a1 |
| Lbh3 | 1.79E-79 | 1.493046669 | 0.591 | 0.131 | 3.63E-75 | 17 | Lbh |
| Thy15 | 2.23E-78 | 1.050013932 | 0.398 | 0.059 | 4.52E-74 | 17 | Thy1 |
| Pten1 | 1.04E-77 | 1.689451778 | 0.86 | 0.32 | 2.11E-73 | 17 | Pten |
| Rhoj5 | 1.08E-77 | 1.431101919 | 0.819 | 0.24 | 2.18E-73 | 17 | Rhoj |
| Pdlim11 | 1.56E-77 | 0.96362469 | 0.538 | 0.101 | 3.15E-73 | 17 | Pdlim1 |
| Maml33 | 3.29E-77 | 1.502225319 | 0.749 | 0.206 | 6.67E-73 | 17 | Maml3 |
| Axl2 | 9.57E-77 | 1.008270366 | 0.462 | 0.077 | 1.94E-72 | 17 | Axl |
| Sept118 | 7.70E-76 | 1.455559764 | 0.725 | 0.202 | 1.56E-71 | 17 | 11-Sep |
| Emid14 | 8.23E-76 | 1.112910791 | 0.538 | 0.107 | 1.67E-71 | 17 | Emid1 |
| Gstm51 | 1.93E-75 | 1.1553075 | 0.409 | 0.062 | 3.91E-71 | 17 | Gstm5 |
| Dapk25 | 3.84E-75 | 1.721264207 | 0.526 | 0.109 | 7.77E-71 | 17 | Dapk2 |
| Parva2 | 1.33E-74 | 1.059090387 | 0.55 | 0.111 | 2.70E-70 | 17 | Parva |
| Fstl13 | 1.75E-74 | 1.124949982 | 0.526 | 0.102 | 3.55E-70 | 17 | Fstl1 |
| Fbn13 | 8.28E-74 | 0.947157389 | 0.409 | 0.063 | 1.68E-69 | 17 | Fbn1 |

| | | | | | | | |
|---|---|---|---|---|---|---|---|
| E130102H24Rik1 | 9.61E-74 | 0.933703528 | 0.409 | 0.065 | 1.95E-69 | 17 | E130102H24Rik |
| Cd3022 | 1.63E-73 | 0.974763404 | 0.462 | 0.08 | 3.29E-69 | 17 | Cd302 |
| C1qtnf21 | 2.90E-73 | 0.954137712 | 0.333 | 0.044 | 5.86E-69 | 17 | C1qtnf2 |
| Errfi12 | 1.47E-71 | 1.502159144 | 0.632 | 0.16 | 2.98E-67 | 17 | Errfi1 |
| Fermt25 | 4.46E-71 | 1.380675361 | 0.947 | 0.348 | 9.02E-67 | 17 | Fermt2 |
| P2ry141 | 1.81E-70 | 0.725380217 | 0.263 | 0.029 | 3.66E-66 | 17 | P2ry14 |
| Uba2 | 3.66E-70 | 1.184010415 | 0.62 | 0.153 | 7.40E-66 | 17 | Uba2 |
| Sep-79 | 1.52E-69 | 1.641373059 | 0.936 | 0.461 | 3.07E-65 | 17 | 7-Sep |
| Nhs1 | 2.56E-68 | 0.880277086 | 0.327 | 0.045 | 5.19E-64 | 17 | Nhs |
| Abi21 | 4.65E-68 | 0.942352339 | 0.474 | 0.091 | 9.41E-64 | 17 | Abi2 |
| Mapt | 6.56E-68 | 0.693749776 | 0.31 | 0.04 | 1.33E-63 | 17 | Mapt |
| Esyt16 | 7.32E-68 | 1.241189788 | 0.567 | 0.135 | 1.48E-63 | 17 | Esyt1 |
| Palld2 | 1.63E-67 | 0.908932017 | 0.357 | 0.052 | 3.30E-63 | 17 | Palld |
| Ppp1r12b1 | 5.64E-67 | 1.223933288 | 0.497 | 0.105 | 1.14E-62 | 17 | Ppp1r12b |
| Crip17 | 7.20E-67 | 2.326038115 | 0.959 | 0.535 | 1.46E-62 | 17 | Crip1 |
| Laptm4a5 | 7.31E-66 | 1.685858787 | 0.947 | 0.483 | 1.48E-61 | 17 | Laptm4a |
| Lpl2 | 8.67E-66 | 0.657112145 | 0.474 | 0.086 | 1.76E-61 | 17 | Lpl |
| Rarb1 | 9.25E-66 | 0.889260644 | 0.363 | 0.056 | 1.87E-61 | 17 | Rarb |
| Cc2d2a1 | 1.01E-65 | 0.647142845 | 0.281 | 0.035 | 2.04E-61 | 17 | Cc2d2a |
| Arhgap321 | 1.86E-65 | 0.769596392 | 0.351 | 0.053 | 3.76E-61 | 17 | Arhgap32 |
| Arhgap241 | 1.11E-64 | 0.837136841 | 0.327 | 0.047 | 2.24E-60 | 17 | Arhgap24 |
| Spry11 | 1.58E-64 | 0.972472244 | 0.415 | 0.074 | 3.20E-60 | 17 | Spry1 |
| Maged21 | 2.63E-64 | 0.796244927 | 0.409 | 0.07 | 5.33E-60 | 17 | Maged2 |
| S100a67 | 2.85E-64 | 1.03365619 | 0.895 | 0.27 | 5.78E-60 | 17 | S100a6 |
| St6gal12 | 1.21E-63 | 0.75469186 | 0.298 | 0.04 | 2.45E-59 | 17 | St6gal1 |
| Pkig2 | 4.03E-63 | 1.042311561 | 0.713 | 0.21 | 8.16E-59 | 17 | Pkig |
| Pik3r13 | 1.29E-62 | 1.373667125 | 0.754 | 0.252 | 2.60E-58 | 17 | Pik3r1 |
| Itgb19 | 2.48E-62 | 1.397536679 | 0.947 | 0.478 | 5.03E-58 | 17 | Itgb1 |
| Trpv2 | 3.62E-62 | 0.719369088 | 0.287 | 0.039 | 7.32E-58 | 17 | Trpv2 |
| Tead12 | 4.32E-62 | 0.874794731 | 0.439 | 0.083 | 8.74E-58 | 17 | Tead1 |
| Actn11 | 7.62E-62 | 1.088089057 | 0.427 | 0.083 | 1.54E-57 | 17 | Actn1 |
| Pitx1 | 3.92E-61 | 0.721243184 | 0.415 | 0.073 | 7.94E-57 | 17 | Pitx1 |
| Pip5k1b5 | 4.80E-61 | 1.092350976 | 0.468 | 0.098 | 9.71E-57 | 17 | Pip5k1b |
| Lamb24 | 7.12E-61 | 1.098316575 | 0.526 | 0.123 | 1.44E-56 | 17 | Lamb2 |
| Tspan153 | 9.92E-61 | 1.085083493 | 0.556 | 0.129 | 2.01E-56 | 17 | Tspan15 |
| Serpinh15 | 1.67E-60 | 1.206319745 | 0.801 | 0.248 | 3.38E-56 | 17 | Serpinh1 |
| St54 | 1.79E-59 | 1.084355706 | 0.591 | 0.15 | 3.62E-55 | 17 | St5 |
| Timp34 | 2.17E-59 | 1.333997349 | 0.977 | 0.384 | 4.40E-55 | 17 | Timp3 |
| Tgfb1i11 | 3.35E-59 | 0.924244068 | 0.439 | 0.087 | 6.77E-55 | 17 | Tgfb1i1 |
| Fgf1 | 5.71E-59 | 0.567808209 | 0.257 | 0.032 | 1.16E-54 | 17 | Fgf1 |
| Raph13 | 5.80E-59 | 1.283510182 | 0.491 | 0.113 | 1.17E-54 | 17 | Raph1 |
| Ecm11 | 2.12E-58 | 0.623508382 | 0.263 | 0.034 | 4.29E-54 | 17 | Ecm1 |
| Cyth34 | 2.99E-58 | 1.042406875 | 0.526 | 0.124 | 6.05E-54 | 17 | Cyth3 |

| | | | | | | | |
|---|---|---|---|---|---|---|---|
| Anxa11 | 4.41E-58 | 0.834860866 | 0.743 | 0.225 | 8.93E-54 | 17 | Anxa1 |
| Arap26 | 1.61E-57 | 1.161063914 | 0.62 | 0.177 | 3.26E-53 | 17 | Arap2 |
| Zfhx31 | 1.94E-57 | 1.280362025 | 0.632 | 0.182 | 3.92E-53 | 17 | Zfhx3 |
| Klf95 | 9.00E-57 | 1.137301954 | 0.731 | 0.236 | 1.82E-52 | 17 | Klf9 |
| Thra4 | 1.97E-56 | 0.993702123 | 0.608 | 0.163 | 3.98E-52 | 17 | Thra |
| Ppp1r12a8 | 2.35E-56 | 1.451844093 | 0.906 | 0.425 | 4.76E-52 | 17 | Ppp1r12a |
| Pbx13 | 4.89E-56 | 1.048093905 | 0.673 | 0.197 | 9.90E-52 | 17 | Pbx1 |
| Lims16 | 2.32E-55 | 1.315143952 | 0.778 | 0.3 | 4.70E-51 | 17 | Lims1 |
| Oat2 | 2.55E-55 | 0.794041226 | 0.556 | 0.135 | 5.16E-51 | 17 | Oat |
| Kalrn4 | 4.03E-55 | 1.16347615 | 0.474 | 0.105 | 8.16E-51 | 17 | Kalrn |
| Selenof5 | 1.69E-54 | 1.250546088 | 0.942 | 0.49 | 3.43E-50 | 17 | Selenof |
| Adcy9 | 3.32E-54 | 0.870371109 | 0.333 | 0.057 | 6.72E-50 | 17 | Adcy9 |
| Airn6 | 3.54E-54 | 1.373894397 | 0.743 | 0.247 | 7.18E-50 | 17 | Airn |
| Marveld11 | 3.90E-54 | 0.599399238 | 0.304 | 0.048 | 7.90E-50 | 17 | Marveld1 |
| Pdlim22 | 6.77E-54 | 0.841416723 | 0.363 | 0.068 | 1.37E-49 | 17 | Pdlim2 |
| Cep1121 | 7.00E-54 | 0.79723958 | 0.339 | 0.058 | 1.42E-49 | 17 | Cep112 |
| Specc13 | 8.45E-54 | 1.019686638 | 0.398 | 0.08 | 1.71E-49 | 17 | Specc1 |
| Epn24 | 1.95E-53 | 1.109124052 | 0.567 | 0.154 | 3.94E-49 | 17 | Epn2 |
| Tspan21 | 3.35E-53 | 0.55662268 | 0.263 | 0.037 | 6.77E-49 | 17 | Tspan2 |
| Slc16a22 | 4.24E-53 | 0.858906091 | 0.363 | 0.067 | 8.58E-49 | 17 | Slc16a2 |
| Pear13 | 5.82E-53 | 1.034105814 | 0.444 | 0.101 | 1.18E-48 | 17 | Pear1 |
| Nedd45 | 9.72E-53 | 1.24979904 | 0.906 | 0.419 | 1.97E-48 | 17 | Nedd4 |
| Ppp2r3a1 | 9.87E-53 | 0.855450619 | 0.427 | 0.091 | 2.00E-48 | 17 | Ppp2r3a |
| Ano64 | 1.67E-52 | 1.219788596 | 0.596 | 0.18 | 3.39E-48 | 17 | Ano6 |
| Tns15 | 4.31E-52 | 1.174999494 | 0.719 | 0.244 | 8.73E-48 | 17 | Tns1 |
| Wls2 | 5.41E-52 | 0.930009442 | 0.509 | 0.126 | 1.10E-47 | 17 | Wls |
| Ssbp26 | 6.60E-52 | 0.900614075 | 0.544 | 0.139 | 1.34E-47 | 17 | Ssbp2 |
| Arhgdib7 | 2.81E-51 | 1.168598535 | 0.725 | 0.233 | 5.69E-47 | 17 | Arhgdib |
| Rasal23 | 1.70E-50 | 1.12735725 | 0.848 | 0.314 | 3.45E-46 | 17 | Rasal2 |
| Amotl15 | 2.08E-50 | 1.08743779 | 0.62 | 0.185 | 4.21E-46 | 17 | Amotl1 |
| Zbtb208 | 3.06E-50 | 1.202761154 | 0.994 | 0.537 | 6.19E-46 | 17 | Zbtb20 |
| Enpep1 | 1.73E-49 | 1.122096463 | 0.392 | 0.084 | 3.49E-45 | 17 | Enpep |
| Mgst3 | 2.49E-49 | 1.112293839 | 0.532 | 0.145 | 5.04E-45 | 17 | Mgst3 |
| Dag14 | 3.25E-49 | 1.068872872 | 0.526 | 0.142 | 6.58E-45 | 17 | Dag1 |
| Kcnq55 | 5.21E-49 | 1.367462244 | 0.351 | 0.068 | 1.05E-44 | 17 | Kcnq5 |
| Itpr15 | 5.61E-49 | 1.158665925 | 0.661 | 0.221 | 1.14E-44 | 17 | Itpr1 |
| Tuba1a6 | 6.68E-49 | 1.155634153 | 0.719 | 0.255 | 1.35E-44 | 17 | Tuba1a |
| Phtf21 | 8.98E-49 | 0.7726852 | 0.333 | 0.062 | 1.82E-44 | 17 | Phtf2 |
| Tnfaip85 | 1.42E-47 | 0.923767753 | 0.614 | 0.186 | 2.88E-43 | 17 | Tnfaip8 |
| Herc16 | 1.78E-47 | 1.461561334 | 0.69 | 0.256 | 3.61E-43 | 17 | Herc1 |
| Dirc2 | 2.09E-47 | 0.692935037 | 0.298 | 0.052 | 4.22E-43 | 17 | Dirc2 |
| Frmd61 | 2.31E-47 | 0.664240387 | 0.281 | 0.046 | 4.67E-43 | 17 | Frmd6 |
| Itga16 | 3.30E-47 | 1.170828554 | 0.93 | 0.414 | 6.68E-43 | 17 | Itga1 |

| | | | | | | | |
|---|---|---|---|---|---|---|---|
| Fam20a1 | 1.22E-46 | 0.775010978 | 0.31 | 0.055 | 2.47E-42 | 17 | Fam20a |
| Dock72 | 2.05E-46 | 0.68370849 | 0.281 | 0.047 | 4.14E-42 | 17 | Dock7 |
| Nab12 | 3.26E-46 | 0.953505764 | 0.556 | 0.167 | 6.60E-42 | 17 | Nab1 |
| Ppp1r12c1 | 3.53E-46 | 0.861596816 | 0.421 | 0.1 | 7.15E-42 | 17 | Ppp1r12c |
| Rock18 | 4.30E-46 | 1.35823902 | 0.842 | 0.432 | 8.70E-42 | 17 | Rock1 |
| Ehd24 | 6.72E-46 | 1.001523642 | 0.614 | 0.192 | 1.36E-41 | 17 | Ehd2 |
| Slc8a11 | 9.13E-46 | 1.025947269 | 0.327 | 0.061 | 1.85E-41 | 17 | Slc8a1 |
| Ubr12 | 9.75E-46 | 1.063975648 | 0.538 | 0.162 | 1.97E-41 | 17 | Ubr1 |
| Gab12 | 1.61E-45 | 0.830526404 | 0.462 | 0.114 | 3.26E-41 | 17 | Gab1 |
| Kank21 | 2.34E-45 | 0.624204618 | 0.298 | 0.053 | 4.74E-41 | 17 | Kank2 |
| Prkca6 | 4.48E-45 | 0.847622998 | 0.421 | 0.1 | 9.08E-41 | 17 | Prkca |
| Slc1a52 | 7.47E-45 | 0.738531884 | 0.368 | 0.08 | 1.51E-40 | 17 | Slc1a5 |
| Frmd4a3 | 4.30E-44 | 1.029143099 | 0.509 | 0.147 | 8.71E-40 | 17 | Frmd4a |
| Plekhg2 | 1.63E-43 | 0.633822685 | 0.287 | 0.051 | 3.30E-39 | 17 | Plekhg2 |
| Ptpn92 | 2.31E-43 | 1.008882138 | 0.497 | 0.141 | 4.68E-39 | 17 | Ptpn9 |
| Nbeal14 | 4.37E-43 | 1.097710368 | 0.696 | 0.261 | 8.84E-39 | 17 | Nbeal1 |
| Crtc37 | 6.16E-43 | 1.135788157 | 0.52 | 0.164 | 1.25E-38 | 17 | Crtc3 |
| Atp2b44 | 1.13E-42 | 0.998466518 | 0.602 | 0.198 | 2.28E-38 | 17 | Atp2b4 |
| Bcas36 | 1.51E-42 | 1.014036202 | 0.567 | 0.182 | 3.05E-38 | 17 | Bcas3 |
| Myadm4 | 3.95E-42 | 0.85769273 | 0.45 | 0.119 | 8.00E-38 | 17 | Myadm |
| Msrb33 | 2.08E-41 | 0.747794112 | 0.491 | 0.132 | 4.21E-37 | 17 | Msrb3 |
| Rrad | 8.80E-41 | 0.622398694 | 0.257 | 0.045 | 1.78E-36 | 17 | Rrad |
| Stard83 | 2.79E-40 | 0.931719361 | 0.608 | 0.197 | 5.65E-36 | 17 | Stard8 |
| Psip14 | 1.00E-39 | 0.93986091 | 0.585 | 0.194 | 2.03E-35 | 17 | Psip1 |
| Zfp703 | 1.05E-39 | 0.604896582 | 0.287 | 0.055 | 2.12E-35 | 17 | Zfp703 |
| Cped15 | 1.20E-39 | 0.842360213 | 0.579 | 0.178 | 2.43E-35 | 17 | Cped1 |
| Rab11fip21 | 1.21E-39 | 0.773304455 | 0.415 | 0.106 | 2.45E-35 | 17 | Rab11fip2 |
| Inpp5a1 | 4.69E-39 | 0.746887839 | 0.468 | 0.129 | 9.49E-35 | 17 | Inpp5a |
| Ltbp12 | 8.26E-39 | 0.757101515 | 0.415 | 0.103 | 1.67E-34 | 17 | Ltbp1 |
| Gcnt24 | 1.03E-38 | 0.909268326 | 0.491 | 0.142 | 2.08E-34 | 17 | Gcnt2 |
| Gpcpd13 | 1.39E-38 | 0.944415175 | 0.427 | 0.117 | 2.81E-34 | 17 | Gpcpd1 |
| Vim9 | 2.01E-38 | 1.189533165 | 0.731 | 0.306 | 4.08E-34 | 17 | Vim |
| Ate1 | 2.96E-38 | 0.883134712 | 0.474 | 0.14 | 6.00E-34 | 17 | Ate1 |
| Csnk1e2 | 4.86E-38 | 0.840458329 | 0.462 | 0.134 | 9.84E-34 | 17 | Csnk1e |
| Pkd12 | 5.36E-38 | 0.754674723 | 0.386 | 0.096 | 1.09E-33 | 17 | Pkd1 |
| Fzd44 | 6.79E-38 | 0.857320256 | 0.667 | 0.229 | 1.38E-33 | 17 | Fzd4 |
| Pik3ca3 | 1.26E-37 | 0.994632775 | 0.52 | 0.168 | 2.55E-33 | 17 | Pik3ca |
| Emp38 | 1.80E-37 | 0.865003578 | 0.678 | 0.258 | 3.64E-33 | 17 | Emp3 |
| Col4a14 | 1.91E-37 | 1.008258962 | 0.924 | 0.391 | 3.86E-33 | 17 | Col4a1 |
| Fam69a5 | 3.82E-37 | 0.799227751 | 0.444 | 0.126 | 7.74E-33 | 17 | Fam69a |
| Tnks1bp11 | 6.36E-37 | 0.700012015 | 0.368 | 0.089 | 1.29E-32 | 17 | Tnks1bp1 |
| Nenf3 | 8.10E-37 | 0.904715492 | 0.713 | 0.295 | 1.64E-32 | 17 | Nenf |
| Ccnd29 | 9.99E-37 | 1.125792756 | 0.678 | 0.286 | 2.02E-32 | 17 | Ccnd2 |

| | | | | | | | |
|---|---|---|---|---|---|---|---|
| Ptms1 | 1.12E-36 | 0.98306504 | 0.93 | 0.583 | 2.27E-32 | 17 | Ptms |
| Rcn31 | 1.40E-36 | 0.578207877 | 0.275 | 0.053 | 2.84E-32 | 17 | Rcn3 |
| Csrp14 | 1.54E-36 | 1.037367466 | 0.538 | 0.182 | 3.12E-32 | 17 | Csrp1 |
| Il15ra | 1.97E-36 | 0.648235237 | 0.31 | 0.067 | 3.98E-32 | 17 | Il15ra |
| Actn44 | 2.22E-36 | 0.924026248 | 0.819 | 0.375 | 4.50E-32 | 17 | Actn4 |
| Gulp15 | 3.94E-36 | 0.928995514 | 0.526 | 0.166 | 7.98E-32 | 17 | Gulp1 |
| Rhob5 | 1.02E-35 | 1.052468973 | 0.684 | 0.267 | 2.06E-31 | 17 | Rhob |
| Fam129a5 | 1.22E-35 | 0.805246286 | 0.474 | 0.145 | 2.47E-31 | 17 | Fam129a |
| Gipc1 | 3.04E-35 | 0.523617122 | 0.404 | 0.106 | 6.16E-31 | 17 | Gipc1 |
| Mtus11 | 3.46E-35 | 0.838220871 | 0.503 | 0.158 | 7.01E-31 | 17 | Mtus1 |
| Cdc42bpa5 | 4.20E-35 | 0.757364758 | 0.561 | 0.184 | 8.50E-31 | 17 | Cdc42bpa |
| mt-Cytb10 | 1.04E-34 | 1.045410619 | 0.982 | 0.7 | 2.11E-30 | 17 | mt-Cytb |
| Epas15 | 1.13E-34 | 0.891104286 | 0.819 | 0.359 | 2.29E-30 | 17 | Epas1 |
| Tm4sf15 | 1.62E-34 | 0.83734207 | 0.906 | 0.377 | 3.28E-30 | 17 | Tm4sf1 |
| Armcx31 | 2.21E-34 | 0.669397057 | 0.31 | 0.071 | 4.47E-30 | 17 | Armcx3 |
| Bcam5 | 2.43E-34 | 0.830027643 | 0.678 | 0.253 | 4.93E-30 | 17 | Bcam |
| Epb41l23 | 3.56E-34 | 0.895765388 | 0.626 | 0.241 | 7.20E-30 | 17 | Epb41l2 |
| Cavin35 | 5.61E-34 | 1.043321812 | 0.719 | 0.307 | 1.14E-29 | 17 | Cavin3 |
| Ktn15 | 7.11E-34 | 0.84390292 | 0.719 | 0.307 | 1.44E-29 | 17 | Ktn1 |
| Prkar1a4 | 7.73E-34 | 0.93334288 | 0.789 | 0.403 | 1.57E-29 | 17 | Prkar1a |
| Nes3 | 1.03E-33 | 0.62978268 | 0.462 | 0.133 | 2.08E-29 | 17 | Nes |
| Tnfsf124 | 1.22E-33 | 0.903231831 | 0.468 | 0.147 | 2.47E-29 | 17 | Tnfsf12 |
| Dnajc103 | 1.58E-33 | 0.865468294 | 0.45 | 0.139 | 3.20E-29 | 17 | Dnajc10 |
| Camk2d7 | 1.90E-33 | 0.883585648 | 0.655 | 0.269 | 3.85E-29 | 17 | Camk2d |
| Glg15 | 2.10E-33 | 0.92528057 | 0.626 | 0.252 | 4.24E-29 | 17 | Glg1 |
| Suco5 | 2.32E-33 | 0.881845744 | 0.468 | 0.153 | 4.70E-29 | 17 | Suco |
| Cpne2 | 2.53E-33 | 0.765475179 | 0.298 | 0.068 | 5.13E-29 | 17 | Cpne2 |
| mt-Nd29 | 2.98E-33 | 0.974971296 | 0.994 | 0.688 | 6.02E-29 | 17 | mt-Nd2 |
| Rap2a1 | 3.48E-33 | 0.729194289 | 0.415 | 0.117 | 7.05E-29 | 17 | Rap2a |
| Lmna6 | 3.56E-33 | 0.851844456 | 0.55 | 0.198 | 7.22E-29 | 17 | Lmna |
| Mxra74 | 3.81E-33 | 0.734955567 | 0.468 | 0.141 | 7.71E-29 | 17 | Mxra7 |
| Pacsin21 | 4.42E-33 | 0.695885356 | 0.485 | 0.157 | 8.95E-29 | 17 | Pacsin2 |
| Mef2a6 | 4.76E-33 | 0.841092135 | 0.848 | 0.416 | 9.63E-29 | 17 | Mef2a |
| Ints6l | 7.15E-33 | 0.577040292 | 0.298 | 0.067 | 1.45E-28 | 17 | Ints6l |
| Mpdz2 | 7.58E-33 | 0.632835343 | 0.374 | 0.097 | 1.54E-28 | 17 | Mpdz |
| Farp13 | 8.13E-33 | 0.83191622 | 0.386 | 0.108 | 1.65E-28 | 17 | Farp1 |
| Nrarp1 | 1.72E-32 | 0.503976411 | 0.275 | 0.058 | 3.49E-28 | 17 | Nrarp |
| Unc45a | 1.78E-32 | 0.543384547 | 0.263 | 0.055 | 3.60E-28 | 17 | Unc45a |
| Ilk1 | 1.94E-32 | 0.751053338 | 0.503 | 0.17 | 3.92E-28 | 17 | Ilk |
| Rock24 | 2.83E-32 | 0.914940241 | 0.743 | 0.337 | 5.73E-28 | 17 | Rock2 |
| Asph2 | 3.63E-32 | 0.632846231 | 0.374 | 0.1 | 7.35E-28 | 17 | Asph |
| Esyt29 | 3.85E-32 | 0.906324643 | 0.608 | 0.242 | 7.80E-28 | 17 | Esyt2 |
| Atp2a34 | 5.59E-32 | 0.469479227 | 0.281 | 0.062 | 1.13E-27 | 17 | Atp2a3 |

| | | | | | | | |
|---|---|---|---|---|---|---|---|
| Soga11 | 7.68E-32 | 0.610839243 | 0.269 | 0.059 | 1.55E-27 | 17 | Soga1 |
| Atrnl14 | 8.20E-32 | 0.932423426 | 0.491 | 0.165 | 1.66E-27 | 17 | Atrnl1 |
| Cyp20a11 | 9.42E-32 | 0.64350866 | 0.333 | 0.085 | 1.91E-27 | 17 | Cyp20a1 |
| Oaz24 | 9.96E-32 | 0.688390482 | 0.632 | 0.236 | 2.02E-27 | 17 | Oaz2 |
| Zcchc241 | 1.21E-31 | 0.607295699 | 0.292 | 0.068 | 2.45E-27 | 17 | Zcchc24 |
| Mill22 | 1.42E-31 | 0.589311704 | 0.298 | 0.069 | 2.87E-27 | 17 | Mill2 |
| Ankrd50 | 1.52E-31 | 0.655670952 | 0.351 | 0.092 | 3.07E-27 | 17 | Ankrd50 |
| Cav14 | 2.32E-31 | 0.885427437 | 0.737 | 0.304 | 4.69E-27 | 17 | Cav1 |
| Efhd25 | 2.52E-31 | 0.678589761 | 0.544 | 0.187 | 5.10E-27 | 17 | Efhd2 |
| Zc2hc1a1 | 2.78E-31 | 0.629539493 | 0.392 | 0.11 | 5.64E-27 | 17 | Zc2hc1a |
| Zcchc142 | 4.08E-31 | 0.598372457 | 0.322 | 0.08 | 8.26E-27 | 17 | Zcchc14 |
| Wdfy36 | 4.96E-31 | 0.877413367 | 0.515 | 0.182 | 1.01E-26 | 17 | Wdfy3 |
| Cxcl126 | 5.10E-31 | 0.365308627 | 0.801 | 0.321 | 1.03E-26 | 17 | Cxcl12 |
| Lamc14 | 5.35E-31 | 0.678454479 | 0.468 | 0.148 | 1.08E-26 | 17 | Lamc1 |
| Hs2st11 | 6.55E-31 | 0.692274007 | 0.339 | 0.09 | 1.33E-26 | 17 | Hs2st1 |
| Tmem1316 | 7.12E-31 | 0.811530762 | 0.55 | 0.208 | 1.44E-26 | 17 | Tmem131 |
| Akap124 | 7.26E-31 | 0.801931826 | 0.491 | 0.161 | 1.47E-26 | 17 | Akap12 |
| Pdzd22 | 8.00E-31 | 0.816690594 | 0.398 | 0.115 | 1.62E-26 | 17 | Pdzd2 |
| Hic11 | 1.15E-30 | 0.617013528 | 0.292 | 0.069 | 2.33E-26 | 17 | Hic1 |
| Cd1514 | 1.81E-30 | 0.785323716 | 0.579 | 0.217 | 3.67E-26 | 17 | Cd151 |
| Lix1l1 | 2.06E-30 | 0.542886561 | 0.304 | 0.073 | 4.17E-26 | 17 | Lix1l |
| Selenom5 | 2.62E-30 | 0.84288924 | 0.901 | 0.445 | 5.31E-26 | 17 | Selenom |
| Timp23 | 2.73E-30 | 0.432026295 | 0.439 | 0.13 | 5.53E-26 | 17 | Timp2 |
| Bzw21 | 2.78E-30 | 0.631247934 | 0.386 | 0.111 | 5.62E-26 | 17 | Bzw2 |
| Parp86 | 4.17E-30 | 0.886987069 | 0.45 | 0.149 | 8.44E-26 | 17 | Parp8 |
| Cbx61 | 6.13E-30 | 0.39496066 | 0.287 | 0.065 | 1.24E-25 | 17 | Cbx6 |
| Nckap15 | 7.22E-30 | 0.755681951 | 0.591 | 0.226 | 1.46E-25 | 17 | Nckap1 |
| Stk394 | 1.03E-29 | 0.785091161 | 0.462 | 0.154 | 2.09E-25 | 17 | Stk39 |
| Pbxip15 | 1.19E-29 | 0.721960898 | 0.386 | 0.117 | 2.40E-25 | 17 | Pbxip1 |
| Ryk2 | 1.42E-29 | 0.712493382 | 0.38 | 0.11 | 2.87E-25 | 17 | Ryk |
| Ift57 | 1.46E-29 | 0.517210313 | 0.251 | 0.055 | 2.95E-25 | 17 | Ift57 |
| Plcb44 | 2.39E-29 | 0.735477535 | 0.696 | 0.291 | 4.84E-25 | 17 | Plcb4 |
| Tprgl4 | 3.62E-29 | 0.78061085 | 0.643 | 0.278 | 7.33E-25 | 17 | Tprgl |
| Lrrc585 | 4.44E-29 | 0.700761388 | 0.561 | 0.214 | 8.99E-25 | 17 | Lrrc58 |
| Utrn4 | 4.76E-29 | 1.063786927 | 0.778 | 0.406 | 9.63E-25 | 17 | Utrn |
| B230219D22Rik4 | 5.57E-29 | 0.736327044 | 0.737 | 0.342 | 1.13E-24 | 17 | B230219D22Rik |
| Rab6b | 6.19E-29 | 0.614871312 | 0.287 | 0.07 | 1.25E-24 | 17 | Rab6b |
| Marcks4 | 6.24E-29 | 0.8393217 | 0.76 | 0.341 | 1.26E-24 | 17 | Marcks |
| Tsc22d15 | 6.67E-29 | 0.751560051 | 0.865 | 0.445 | 1.35E-24 | 17 | Tsc22d1 |
| Col4a24 | 7.35E-29 | 0.789910072 | 0.836 | 0.36 | 1.49E-24 | 17 | Col4a2 |
| Eif4e3 | 9.32E-29 | 0.609148509 | 0.292 | 0.073 | 1.89E-24 | 17 | Eif4e3 |
| Tgfbr33 | 1.16E-28 | 0.747734933 | 0.509 | 0.174 | 2.34E-24 | 17 | Tgfbr3 |
| mt-Atp611 | 1.25E-28 | 0.897663947 | 0.994 | 0.754 | 2.52E-24 | 17 | mt-Atp6 |

| | | | | | | | |
|---|---|---|---|---|---|---|---|
| Slc25a42 | 1.29E-28 | 0.637746471 | 0.93 | 0.56 | 2.62E-24 | 17 | Slc25a4 |
| Bcl9l1 | 1.47E-28 | 0.560528833 | 0.292 | 0.073 | 2.98E-24 | 17 | Bcl9l |
| Tacc15 | 2.51E-28 | 0.827443831 | 0.673 | 0.305 | 5.09E-24 | 17 | Tacc1 |
| Son10 | 2.64E-28 | 0.820644863 | 0.906 | 0.526 | 5.35E-24 | 17 | Son |
| Prkacb5 | 6.43E-28 | 0.690188268 | 0.561 | 0.218 | 1.30E-23 | 17 | Prkacb |
| Itpripl23 | 7.68E-28 | 0.655597653 | 0.386 | 0.118 | 1.55E-23 | 17 | Itpripl2 |
| Rras5 | 1.10E-27 | 0.813896634 | 0.667 | 0.298 | 2.23E-23 | 17 | Rras |
| Naf1 | 1.10E-27 | 0.50204111 | 0.257 | 0.059 | 2.23E-23 | 17 | Naf1 |
| Trak12 | 1.18E-27 | 0.565516521 | 0.404 | 0.126 | 2.40E-23 | 17 | Trak1 |
| Anxa54 | 1.29E-27 | 0.84325534 | 0.883 | 0.493 | 2.61E-23 | 17 | Anxa5 |
| Eif4g35 | 1.43E-27 | 1.000113771 | 0.626 | 0.286 | 2.89E-23 | 17 | Eif4g3 |
| Thap33 | 1.96E-27 | 0.762849136 | 0.462 | 0.162 | 3.97E-23 | 17 | Thap3 |
| Pls34 | 1.98E-27 | 0.775505224 | 0.55 | 0.214 | 4.02E-23 | 17 | Pls3 |
| Mast21 | 2.53E-27 | 0.523882741 | 0.287 | 0.072 | 5.11E-23 | 17 | Mast2 |
| Ptp4a32 | 3.03E-27 | 1.035826089 | 0.404 | 0.137 | 6.13E-23 | 17 | Ptp4a3 |
| mt-Co312 | 6.29E-27 | 0.871528017 | 0.994 | 0.726 | 1.27E-22 | 17 | mt-Co3 |
| Rcsd15 | 9.29E-27 | 0.768529796 | 0.398 | 0.13 | 1.88E-22 | 17 | Rcsd1 |
| Itgb52 | 9.70E-27 | 0.602116662 | 0.444 | 0.145 | 1.96E-22 | 17 | Itgb5 |
| Aldh25 | 1.12E-26 | 0.683641412 | 0.819 | 0.396 | 2.26E-22 | 17 | Aldh2 |
| Tle11 | 1.54E-26 | 0.552137958 | 0.327 | 0.092 | 3.11E-22 | 17 | Tle1 |
| Urod | 2.14E-26 | 0.470985735 | 0.281 | 0.07 | 4.34E-22 | 17 | Urod |
| Rsrc16 | 2.16E-26 | 0.751997031 | 0.602 | 0.26 | 4.37E-22 | 17 | Rsrc1 |
| Fcgrt3 | 2.19E-26 | 0.667292258 | 0.509 | 0.185 | 4.43E-22 | 17 | Fcgrt |
| Vangl11 | 2.25E-26 | 0.505709888 | 0.269 | 0.065 | 4.55E-22 | 17 | Vangl1 |
| mt-Co213 | 2.31E-26 | 0.838852642 | 0.988 | 0.728 | 4.68E-22 | 17 | mt-Co2 |
| Id42 | 2.89E-26 | 1.096770674 | 0.392 | 0.125 | 5.85E-22 | 17 | Id4 |
| Sh3kbp14 | 3.21E-26 | 0.817429342 | 0.386 | 0.125 | 6.50E-22 | 17 | Sh3kbp1 |
| 9530026P05Rik4 | 4.24E-26 | 0.646407794 | 0.62 | 0.241 | 8.59E-22 | 17 | 9530026P05Rik |
| Sfxn11 | 4.77E-26 | 0.553540719 | 0.327 | 0.093 | 9.65E-22 | 17 | Sfxn1 |
| Syne24 | 5.11E-26 | 0.660370449 | 0.661 | 0.272 | 1.03E-21 | 17 | Syne2 |
| Ptov11 | 6.32E-26 | 0.487671235 | 0.386 | 0.118 | 1.28E-21 | 17 | Ptov1 |
| Pgrmc14 | 7.05E-26 | 0.707958822 | 0.544 | 0.218 | 1.43E-21 | 17 | Pgrmc1 |
| Akap113 | 1.07E-25 | 0.657352788 | 0.398 | 0.13 | 2.16E-21 | 17 | Akap11 |
| Cbfa2t33 | 1.20E-25 | 0.698456944 | 0.415 | 0.139 | 2.43E-21 | 17 | Cbfa2t3 |
| Pparg1 | 1.20E-25 | 0.790019805 | 0.281 | 0.074 | 2.43E-21 | 17 | Pparg |
| mt-Nd111 | 1.66E-25 | 0.800444895 | 0.936 | 0.599 | 3.36E-21 | 17 | mt-Nd1 |
| Gyg | 2.18E-25 | 0.576903751 | 0.263 | 0.066 | 4.41E-21 | 17 | Gyg |
| Top2b4 | 2.57E-25 | 0.784948553 | 0.62 | 0.283 | 5.21E-21 | 17 | Top2b |
| Zfp5211 | 2.89E-25 | 0.546246797 | 0.287 | 0.075 | 5.85E-21 | 17 | Zfp521 |
| mt-Nd411 | 3.29E-25 | 0.803538642 | 0.965 | 0.676 | 6.66E-21 | 17 | mt-Nd4 |
| Cavin14 | 4.32E-25 | 0.852177032 | 0.532 | 0.213 | 8.75E-21 | 17 | Cavin1 |
| Clstn14 | 4.65E-25 | 0.640482418 | 0.585 | 0.236 | 9.41E-21 | 17 | Clstn1 |
| Pdlim72 | 6.02E-25 | 0.525718949 | 0.281 | 0.073 | 1.22E-20 | 17 | Pdlim7 |

| | | | | | | | |
|---|---|---|---|---|---|---|---|
| Adam9 | 9.15E-25 | 0.518056099 | 0.275 | 0.072 | 1.85E-20 | 17 | Adam9 |
| Rsu15 | 1.26E-24 | 0.751359265 | 0.573 | 0.254 | 2.56E-20 | 17 | Rsu1 |
| Zfp9501 | 1.50E-24 | 0.688859788 | 0.351 | 0.112 | 3.04E-20 | 17 | Zfp950 |
| Blmh1 | 1.70E-24 | 0.626078256 | 0.433 | 0.152 | 3.45E-20 | 17 | Blmh |
| Mapre26 | 1.94E-24 | 0.846215865 | 0.567 | 0.25 | 3.94E-20 | 17 | Mapre2 |
| Arid5b6 | 2.04E-24 | 0.723992645 | 0.608 | 0.278 | 4.13E-20 | 17 | Arid5b |
| Myo1d3 | 2.61E-24 | 0.557277359 | 0.351 | 0.107 | 5.28E-20 | 17 | Myo1d |
| Syn3 | 4.04E-24 | 0.439077787 | 0.275 | 0.07 | 8.18E-20 | 17 | Syn3 |
| Mbnl26 | 6.54E-24 | 0.831750761 | 0.749 | 0.419 | 1.32E-19 | 17 | Mbnl2 |
| Msrb1 | 7.83E-24 | 0.480936331 | 0.415 | 0.142 | 1.59E-19 | 17 | Msrb1 |
| Brd31 | 7.83E-24 | 0.597437309 | 0.327 | 0.099 | 1.59E-19 | 17 | Brd3 |
| Il1r12 | 8.16E-24 | 0.563423371 | 0.357 | 0.111 | 1.65E-19 | 17 | Il1r1 |
| Tmco15 | 9.14E-24 | 0.649360757 | 0.667 | 0.312 | 1.85E-19 | 17 | Tmco1 |
| Gna115 | 9.31E-24 | 0.583401625 | 0.585 | 0.245 | 1.88E-19 | 17 | Gna11 |
| Clock5 | 1.55E-23 | 0.671415746 | 0.509 | 0.204 | 3.14E-19 | 17 | Clock |
| Pnpla81 | 2.11E-23 | 0.567181641 | 0.48 | 0.182 | 4.27E-19 | 17 | Pnpla8 |
| Actb7 | 2.11E-23 | 0.614290322 | 0.988 | 0.843 | 4.28E-19 | 17 | Actb |
| Sorbs14 | 2.19E-23 | 0.730695564 | 0.468 | 0.177 | 4.44E-19 | 17 | Sorbs1 |
| Lyst4 | 2.70E-23 | 0.645068854 | 0.345 | 0.11 | 5.46E-19 | 17 | Lyst |
| Nfia5 | 2.90E-23 | 0.768994303 | 0.632 | 0.288 | 5.87E-19 | 17 | Nfia |
| Ckb3 | 3.36E-23 | 0.590840103 | 0.544 | 0.216 | 6.79E-19 | 17 | Ckb |
| Sema6d5 | 4.09E-23 | 0.79611515 | 0.52 | 0.214 | 8.28E-19 | 17 | Sema6d |
| Rbfox25 | 4.78E-23 | 0.546266419 | 0.503 | 0.191 | 9.69E-19 | 17 | Rbfox2 |
| Rasa33 | 6.26E-23 | 0.550805455 | 0.251 | 0.066 | 1.27E-18 | 17 | Rasa3 |
| Sep-65 | 6.53E-23 | 0.629967954 | 0.322 | 0.099 | 1.32E-18 | 17 | 6-Sep |
| Rabac15 | 7.16E-23 | 0.729241639 | 0.889 | 0.564 | 1.45E-18 | 17 | Rabac1 |
| Ephx14 | 7.44E-23 | 0.555814895 | 0.374 | 0.122 | 1.51E-18 | 17 | Ephx1 |
| Lrrfip22 | 8.53E-23 | 0.616380506 | 0.439 | 0.165 | 1.73E-18 | 17 | Lrrfip2 |
| Lpcat32 | 1.13E-22 | 0.583007902 | 0.31 | 0.094 | 2.29E-18 | 17 | Lpcat3 |
| Dock106 | 1.19E-22 | 0.533031633 | 0.556 | 0.209 | 2.42E-18 | 17 | Dock10 |
| Nrp23 | 1.20E-22 | 0.651223917 | 0.485 | 0.184 | 2.43E-18 | 17 | Nrp2 |
| Myh98 | 1.27E-22 | 0.747495579 | 0.789 | 0.437 | 2.58E-18 | 17 | Myh9 |
| Sesn35 | 1.98E-22 | 0.543308893 | 0.38 | 0.129 | 4.01E-18 | 17 | Sesn3 |
| Tnrc6c6 | 2.26E-22 | 0.75710641 | 0.649 | 0.315 | 4.57E-18 | 17 | Tnrc6c |
| mt-Co111 | 2.36E-22 | 0.717259124 | 0.982 | 0.72 | 4.77E-18 | 17 | mt-Co1 |
| Cdk65 | 2.36E-22 | 0.622693538 | 0.345 | 0.112 | 4.78E-18 | 17 | Cdk6 |
| Tmem1094 | 2.48E-22 | 0.680234429 | 0.409 | 0.148 | 5.01E-18 | 17 | Tmem109 |
| Selenow4 | 2.57E-22 | 0.670954045 | 0.906 | 0.63 | 5.20E-18 | 17 | Selenow |
| Igfbp74 | 2.58E-22 | 0.505602941 | 1 | 0.466 | 5.22E-18 | 17 | Igfbp7 |
| Tcf7l23 | 2.63E-22 | 0.524713547 | 0.386 | 0.129 | 5.33E-18 | 17 | Tcf7l2 |
| Ywhaz4 | 2.92E-22 | 0.63081768 | 0.854 | 0.493 | 5.91E-18 | 17 | Ywhaz |
| Map1lc3a2 | 2.93E-22 | 0.446249313 | 0.871 | 0.481 | 5.93E-18 | 17 | Map1lc3a |
| Nbea5 | 3.23E-22 | 0.756755267 | 0.491 | 0.196 | 6.54E-18 | 17 | Nbea |

| | | | | | | | |
|---|---|---|---|---|---|---|---|
| Tnfrsf1a4 | 5.17E-22 | 0.629781603 | 0.532 | 0.222 | 1.05E-17 | 17 | Tnfrsf1a |
| Serinc16 | 6.47E-22 | 0.73506369 | 0.556 | 0.249 | 1.31E-17 | 17 | Serinc1 |
| Rmdn11 | 8.43E-22 | 0.621268272 | 0.474 | 0.186 | 1.71E-17 | 17 | Rmdn1 |
| Zbtb381 | 9.39E-22 | 0.642808739 | 0.415 | 0.154 | 1.90E-17 | 17 | Zbtb38 |
| Ifitm26 | 1.06E-21 | 0.546231033 | 0.947 | 0.464 | 2.14E-17 | 17 | Ifitm2 |
| Tmem52 | 1.35E-21 | 0.524862554 | 0.392 | 0.138 | 2.73E-17 | 17 | Tmem5 |
| Camk12 | 1.43E-21 | 0.527472261 | 0.544 | 0.227 | 2.90E-17 | 17 | Camk1 |
| Hivep28 | 1.90E-21 | 0.706598293 | 0.632 | 0.298 | 3.85E-17 | 17 | Hivep2 |
| Luzp14 | 1.95E-21 | 0.632468033 | 0.708 | 0.357 | 3.94E-17 | 17 | Luzp1 |
| Dennd1a8 | 2.25E-21 | 0.78422843 | 0.561 | 0.253 | 4.56E-17 | 17 | Dennd1a |
| Tmem1655 | 3.35E-21 | 0.620112117 | 0.45 | 0.18 | 6.78E-17 | 17 | Tmem165 |
| Tmod36 | 3.38E-21 | 0.737344157 | 0.684 | 0.362 | 6.85E-17 | 17 | Tmod3 |
| Sec24d1 | 3.40E-21 | 0.449896265 | 0.257 | 0.07 | 6.89E-17 | 17 | Sec24d |
| Pja24 | 3.55E-21 | 0.604160718 | 0.491 | 0.205 | 7.18E-17 | 17 | Pja2 |
| Lamp18 | 4.26E-21 | 0.664868321 | 0.772 | 0.417 | 8.63E-17 | 17 | Lamp1 |
| Chd93 | 4.46E-21 | 0.588040254 | 0.567 | 0.25 | 9.03E-17 | 17 | Chd9 |
| Sgms15 | 4.52E-21 | 0.538995705 | 0.374 | 0.129 | 9.14E-17 | 17 | Sgms1 |
| Snrnp705 | 5.95E-21 | 0.671677008 | 0.579 | 0.273 | 1.20E-16 | 17 | Snrnp70 |
| Dgkh4 | 6.45E-21 | 0.648475304 | 0.45 | 0.177 | 1.31E-16 | 17 | Dgkh |
| Stim17 | 6.59E-21 | 0.724876175 | 0.456 | 0.191 | 1.33E-16 | 17 | Stim1 |
| Dab2ip5 | 6.65E-21 | 0.594944777 | 0.591 | 0.261 | 1.35E-16 | 17 | Dab2ip |
| Drap11 | 6.84E-21 | 0.589615106 | 0.737 | 0.389 | 1.38E-16 | 17 | Drap1 |
| Ypel33 | 7.28E-21 | 0.5520652 | 0.579 | 0.267 | 1.47E-16 | 17 | Ypel3 |
| Myl67 | 7.55E-21 | 0.887263873 | 0.953 | 0.776 | 1.53E-16 | 17 | Myl6 |
| Cldnd11 | 7.58E-21 | 0.562235533 | 0.351 | 0.118 | 1.53E-16 | 17 | Cldnd1 |
| Cd817 | 8.14E-21 | 0.621257035 | 0.965 | 0.525 | 1.65E-16 | 17 | Cd81 |
| Smpdl3a5 | 8.55E-21 | 0.6644035 | 0.515 | 0.219 | 1.73E-16 | 17 | Smpdl3a |
| Gns2 | 1.01E-20 | 0.556124437 | 0.363 | 0.127 | 2.05E-16 | 17 | Gns |
| Ehd11 | 1.01E-20 | 0.417367927 | 0.257 | 0.072 | 2.05E-16 | 17 | Ehd1 |
| Sh3d192 | 1.09E-20 | 0.660709926 | 0.327 | 0.108 | 2.20E-16 | 17 | Sh3d19 |
| Ublcp1 | 1.10E-20 | 0.518735558 | 0.292 | 0.089 | 2.22E-16 | 17 | Ublcp1 |
| Pon22 | 1.16E-20 | 0.670697234 | 0.427 | 0.172 | 2.34E-16 | 17 | Pon2 |
| Nisch5 | 1.40E-20 | 0.613191437 | 0.503 | 0.211 | 2.83E-16 | 17 | Nisch |
| Tln19 | 1.47E-20 | 0.69326983 | 0.731 | 0.405 | 2.97E-16 | 17 | Tln1 |
| Cbx52 | 1.62E-20 | 0.58364138 | 0.462 | 0.189 | 3.28E-16 | 17 | Cbx5 |
| Ttc285 | 1.64E-20 | 0.700101735 | 0.602 | 0.285 | 3.33E-16 | 17 | Ttc28 |
| Tgfb23 | 1.76E-20 | 0.523397794 | 0.292 | 0.087 | 3.57E-16 | 17 | Tgfb2 |
| Fer2 | 1.82E-20 | 0.488513745 | 0.357 | 0.121 | 3.68E-16 | 17 | Fer |
| Trp53inp2 | 2.10E-20 | 0.33413647 | 0.269 | 0.076 | 4.25E-16 | 17 | Trp53inp2 |
| Nktr6 | 2.78E-20 | 0.689141528 | 0.561 | 0.258 | 5.62E-16 | 17 | Nktr |
| Ankmy2 | 3.74E-20 | 0.422979868 | 0.251 | 0.069 | 7.57E-16 | 17 | Ankmy2 |
| Mef2c4 | 4.89E-20 | 0.542340388 | 0.842 | 0.423 | 9.90E-16 | 17 | Mef2c |
| Ssr32 | 4.95E-20 | 0.572757836 | 0.468 | 0.196 | 1.00E-15 | 17 | Ssr3 |

| | | | | | | | |
|---|---|---|---|---|---|---|---|
| Dock64 | 5.68E-20 | 0.540746113 | 0.392 | 0.141 | 1.15E-15 | 17 | Dock6 |
| Uhrf24 | 7.83E-20 | 0.544822977 | 0.427 | 0.166 | 1.59E-15 | 17 | Uhrf2 |
| Plekha24 | 8.26E-20 | 0.677505278 | 0.363 | 0.134 | 1.67E-15 | 17 | Plekha2 |
| Cited22 | 9.18E-20 | 0.754492707 | 0.439 | 0.18 | 1.86E-15 | 17 | Cited2 |
| Ubb1 | 1.09E-19 | 0.571691324 | 0.994 | 0.895 | 2.21E-15 | 17 | Ubb |
| Exoc6b6 | 1.15E-19 | 0.679310839 | 0.532 | 0.241 | 2.33E-15 | 17 | Exoc6b |
| Calm34 | 1.44E-19 | 0.575443358 | 0.561 | 0.27 | 2.91E-15 | 17 | Calm3 |
| Nhsl23 | 1.46E-19 | 0.592150632 | 0.374 | 0.135 | 2.96E-15 | 17 | Nhsl2 |
| Smarcc22 | 2.11E-19 | 0.522954172 | 0.439 | 0.178 | 4.26E-15 | 17 | Smarcc2 |
| Srek14 | 2.29E-19 | 0.641531236 | 0.456 | 0.194 | 4.63E-15 | 17 | Srek1 |
| 4930523C07Rik8 | 2.43E-19 | 0.589108646 | 0.509 | 0.22 | 4.91E-15 | 17 | 4930523C07Rik |
| Ppp1cb2 | 2.61E-19 | 0.6226033 | 0.556 | 0.273 | 5.29E-15 | 17 | Ppp1cb |
| Xist10 | 3.07E-19 | 0.662253335 | 0.836 | 0.463 | 6.22E-15 | 17 | Xist |
| Crip25 | 3.25E-19 | 0.540746164 | 0.813 | 0.416 | 6.57E-15 | 17 | Crip2 |
| Fam13c3 | 3.85E-19 | 0.492577944 | 0.333 | 0.112 | 7.79E-15 | 17 | Fam13c |
| Arhgap1 | 4.83E-19 | 0.439380156 | 0.269 | 0.081 | 9.78E-15 | 17 | Arhgap1 |
| Coq2 | 5.09E-19 | 0.456902606 | 0.31 | 0.102 | 1.03E-14 | 17 | Coq2 |
| Selenok5 | 5.31E-19 | 0.602477431 | 0.871 | 0.554 | 1.07E-14 | 17 | Selenok |
| Strn35 | 5.97E-19 | 0.55271525 | 0.667 | 0.345 | 1.21E-14 | 17 | Strn3 |
| Ebf33 | 6.48E-19 | 0.540080162 | 0.357 | 0.126 | 1.31E-14 | 17 | Ebf3 |
| Dpysl25 | 7.27E-19 | 0.700328658 | 0.427 | 0.175 | 1.47E-14 | 17 | Dpysl2 |
| Ppp2r2a4 | 7.71E-19 | 0.669437423 | 0.48 | 0.213 | 1.56E-14 | 17 | Ppp2r2a |
| Hdgfl34 | 7.86E-19 | 0.52283727 | 0.327 | 0.111 | 1.59E-14 | 17 | Hdgfl3 |
| Cdk2ap25 | 8.80E-19 | 0.539561144 | 0.468 | 0.2 | 1.78E-14 | 17 | Cdk2ap2 |
| Nid13 | 9.20E-19 | 0.697854207 | 0.45 | 0.182 | 1.86E-14 | 17 | Nid1 |
| Actr101 | 1.01E-18 | 0.481975575 | 0.538 | 0.243 | 2.05E-14 | 17 | Actr10 |
| Hmgn12 | 1.14E-18 | 0.474436516 | 0.643 | 0.329 | 2.30E-14 | 17 | Hmgn1 |
| Gstm15 | 1.18E-18 | 0.561674599 | 0.649 | 0.323 | 2.38E-14 | 17 | Gstm1 |
| Cuedc12 | 1.32E-18 | 0.55141346 | 0.368 | 0.136 | 2.67E-14 | 17 | Cuedc1 |
| Ech11 | 1.40E-18 | 0.514363571 | 0.667 | 0.35 | 2.84E-14 | 17 | Ech1 |
| Ifitm36 | 1.47E-18 | 0.645312449 | 0.947 | 0.538 | 2.97E-14 | 17 | Ifitm3 |
| Cpne33 | 1.51E-18 | 0.55123858 | 0.485 | 0.212 | 3.06E-14 | 17 | Cpne3 |
| Mertk4 | 1.55E-18 | 0.474675213 | 0.368 | 0.132 | 3.13E-14 | 17 | Mertk |
| Cnn35 | 1.75E-18 | 0.517622771 | 0.468 | 0.194 | 3.55E-14 | 17 | Cnn3 |
| Esam4 | 1.87E-18 | 0.464786023 | 0.696 | 0.31 | 3.78E-14 | 17 | Esam |
| Ifngr23 | 2.15E-18 | 0.674896283 | 0.357 | 0.132 | 4.34E-14 | 17 | Ifngr2 |
| Spry42 | 2.37E-18 | 0.4460777 | 0.298 | 0.097 | 4.80E-14 | 17 | Spry4 |
| Eif1b1 | 2.56E-18 | 0.50770633 | 0.649 | 0.335 | 5.18E-14 | 17 | Eif1b |
| Rbms14 | 2.67E-18 | 0.523046888 | 0.871 | 0.486 | 5.40E-14 | 17 | Rbms1 |
| Klf126 | 2.95E-18 | 0.610521987 | 0.48 | 0.208 | 5.97E-14 | 17 | Klf12 |
| Hif1a1 | 3.69E-18 | 0.510362199 | 0.462 | 0.2 | 7.48E-14 | 17 | Hif1a |
| Taok35 | 3.75E-18 | 0.612731491 | 0.404 | 0.164 | 7.60E-14 | 17 | Taok3 |
| Dynll14 | 3.79E-18 | 0.635893093 | 0.871 | 0.606 | 7.68E-14 | 17 | Dynll1 |

| | | | | | | | |
|---|---|---|---|---|---|---|---|
| Atraid5 | 4.18E-18 | 0.547011805 | 0.567 | 0.271 | 8.45E-14 | 17 | Atraid |
| Itm2b8 | 4.36E-18 | 0.5254708 | 1 | 0.725 | 8.82E-14 | 17 | Itm2b |
| Cyb5r34 | 4.51E-18 | 0.598117274 | 0.491 | 0.217 | 9.12E-14 | 17 | Cyb5r3 |
| Ext15 | 5.11E-18 | 0.591335222 | 0.415 | 0.167 | 1.03E-13 | 17 | Ext1 |
| Map1b1 | 5.32E-18 | 0.469821047 | 0.31 | 0.102 | 1.08E-13 | 17 | Map1b |
| Maml210 | 6.06E-18 | 0.728907455 | 0.649 | 0.343 | 1.23E-13 | 17 | Maml2 |
| Manbal | 6.83E-18 | 0.50285041 | 0.304 | 0.105 | 1.38E-13 | 17 | Manbal |
| Luc7l28 | 7.54E-18 | 0.598013557 | 0.719 | 0.405 | 1.53E-13 | 17 | Luc7l2 |
| Tax1bp11 | 8.22E-18 | 0.498882132 | 0.743 | 0.417 | 1.66E-13 | 17 | Tax1bp1 |
| Lnpep10 | 8.94E-18 | 0.66028411 | 0.538 | 0.26 | 1.81E-13 | 17 | Lnpep |
| Spag95 | 9.40E-18 | 0.570789055 | 0.684 | 0.359 | 1.90E-13 | 17 | Spag9 |
| Nhsl1 | 9.57E-18 | 0.404698071 | 0.257 | 0.078 | 1.94E-13 | 17 | Nhsl1 |
| Wasl3 | 9.62E-18 | 0.573111745 | 0.509 | 0.237 | 1.95E-13 | 17 | Wasl |
| mt-Nd4l8 | 1.10E-17 | 0.645947994 | 0.778 | 0.492 | 2.23E-13 | 17 | mt-Nd4l |
| Jade12 | 1.14E-17 | 0.384474047 | 0.327 | 0.115 | 2.30E-13 | 17 | Jade1 |
| Arl14 | 1.66E-17 | 0.401348624 | 0.596 | 0.286 | 3.36E-13 | 17 | Arl1 |
| Ahdc13 | 1.80E-17 | 0.501285773 | 0.316 | 0.112 | 3.65E-13 | 17 | Ahdc1 |
| Zfpm21 | 1.95E-17 | 0.64406459 | 0.257 | 0.079 | 3.95E-13 | 17 | Zfpm2 |
| Hnrnpr2 | 2.08E-17 | 0.485992636 | 0.515 | 0.236 | 4.21E-13 | 17 | Hnrnpr |
| Klhdc101 | 2.32E-17 | 0.491519394 | 0.392 | 0.154 | 4.69E-13 | 17 | Klhdc10 |
| Ptk24 | 2.39E-17 | 0.518401967 | 0.579 | 0.266 | 4.85E-13 | 17 | Ptk2 |
| Ufl11 | 2.61E-17 | 0.496454607 | 0.339 | 0.127 | 5.29E-13 | 17 | Ufl1 |
| Mtdh6 | 3.01E-17 | 0.595232817 | 0.69 | 0.387 | 6.09E-13 | 17 | Mtdh |
| Gm374941 | 3.33E-17 | 0.419251652 | 0.263 | 0.084 | 6.74E-13 | 17 | Gm37494 |
| Cds24 | 3.73E-17 | 0.55236477 | 0.427 | 0.178 | 7.56E-13 | 17 | Cds2 |
| Man1a9 | 4.25E-17 | 0.604195573 | 0.404 | 0.168 | 8.61E-13 | 17 | Man1a |
| Bach1 | 4.35E-17 | 0.450676835 | 0.292 | 0.1 | 8.81E-13 | 17 | Bach1 |
| Ywhab5 | 4.59E-17 | 0.551559596 | 0.778 | 0.467 | 9.30E-13 | 17 | Ywhab |
| Vezf12 | 4.73E-17 | 0.505481081 | 0.509 | 0.234 | 9.57E-13 | 17 | Vezf1 |
| Sult1a1 | 5.06E-17 | 0.414554141 | 0.251 | 0.078 | 1.02E-12 | 17 | Sult1a1 |
| mt-Nd58 | 5.88E-17 | 0.640852489 | 0.819 | 0.496 | 1.19E-12 | 17 | mt-Nd5 |
| Aig14 | 7.42E-17 | 0.470914735 | 0.491 | 0.214 | 1.50E-12 | 17 | Aig1 |
| 1110059E24Rik | 7.73E-17 | 0.416440675 | 0.374 | 0.144 | 1.56E-12 | 17 | 1110059E24Rik |
| Siva1 | 8.41E-17 | 0.347350681 | 0.398 | 0.159 | 1.70E-12 | 17 | Siva1 |
| Ston13 | 9.36E-17 | 0.319435762 | 0.281 | 0.09 | 1.90E-12 | 17 | Ston1 |
| Ppib3 | 9.51E-17 | 0.636419786 | 0.86 | 0.554 | 1.93E-12 | 17 | Ppib |
| Fam53b | 9.95E-17 | 0.388639012 | 0.316 | 0.111 | 2.01E-12 | 17 | Fam53b |
| Rela1 | 1.05E-16 | 0.549462741 | 0.304 | 0.111 | 2.13E-12 | 17 | Rela |
| Mlf2 | 1.07E-16 | 0.506450663 | 0.444 | 0.199 | 2.16E-12 | 17 | Mlf2 |
| Nucks11 | 1.19E-16 | 0.416818448 | 0.713 | 0.39 | 2.41E-12 | 17 | Nucks1 |
| Sacm1l | 1.25E-16 | 0.436808924 | 0.415 | 0.174 | 2.53E-12 | 17 | Sacm1l |
| Cd365 | 1.31E-16 | 0.603363872 | 0.561 | 0.268 | 2.64E-12 | 17 | Cd36 |
| Pld13 | 1.45E-16 | 0.57736181 | 0.368 | 0.148 | 2.94E-12 | 17 | Pld1 |

| | | | | | | | |
|---|---|---|---|---|---|---|---|
| Camk2n13 | 1.56E-16 | 0.402173711 | 0.573 | 0.273 | 3.15E-12 | 17 | Camk2n1 |
| Baz1b4 | 1.57E-16 | 0.61317542 | 0.433 | 0.194 | 3.17E-12 | 17 | Baz1b |
| Cers43 | 1.63E-16 | 0.453334067 | 0.31 | 0.11 | 3.29E-12 | 17 | Cers4 |
| Fam76a1 | 1.74E-16 | 0.454846091 | 0.409 | 0.17 | 3.53E-12 | 17 | Fam76a |
| Rsbn1l6 | 1.77E-16 | 0.608355835 | 0.544 | 0.273 | 3.59E-12 | 17 | Rsbn1l |
| Ube2e23 | 1.80E-16 | 0.42332586 | 0.392 | 0.153 | 3.65E-12 | 17 | Ube2e2 |
| Gdi1 | 1.84E-16 | 0.418760847 | 0.287 | 0.099 | 3.72E-12 | 17 | Gdi1 |
| Ube2v2 | 1.92E-16 | 0.492387377 | 0.415 | 0.176 | 3.88E-12 | 17 | Ube2v2 |
| Sash15 | 2.00E-16 | 0.523195481 | 0.556 | 0.257 | 4.06E-12 | 17 | Sash1 |
| Krit17 | 2.17E-16 | 0.623828694 | 0.421 | 0.183 | 4.39E-12 | 17 | Krit1 |
| Fam174a1 | 2.19E-16 | 0.506555283 | 0.45 | 0.203 | 4.43E-12 | 17 | Fam174a |
| Sec634 | 2.70E-16 | 0.522266953 | 0.45 | 0.204 | 5.46E-12 | 17 | Sec63 |
| Tnrc6b7 | 2.99E-16 | 0.561885763 | 0.667 | 0.372 | 6.06E-12 | 17 | Tnrc6b |
| Lrp62 | 3.53E-16 | 0.410975188 | 0.368 | 0.145 | 7.15E-12 | 17 | Lrp6 |
| Plxnd14 | 3.54E-16 | 0.442822191 | 0.439 | 0.182 | 7.16E-12 | 17 | Plxnd1 |
| Pcdh14 | 3.58E-16 | 0.337625243 | 0.281 | 0.092 | 7.26E-12 | 17 | Pcdh1 |
| Btrc1 | 3.74E-16 | 0.5342165 | 0.304 | 0.112 | 7.58E-12 | 17 | Btrc |
| Tuba1b4 | 3.76E-16 | 0.362099961 | 0.643 | 0.323 | 7.60E-12 | 17 | Tuba1b |
| Lamp24 | 4.29E-16 | 0.581355472 | 0.591 | 0.306 | 8.69E-12 | 17 | Lamp2 |
| Mat2a4 | 4.49E-16 | 0.498173622 | 0.596 | 0.31 | 9.08E-12 | 17 | Mat2a |
| Tcaf13 | 5.48E-16 | 0.540847341 | 0.292 | 0.104 | 1.11E-11 | 17 | Tcaf1 |
| Hspb15 | 6.24E-16 | 0.377627997 | 0.778 | 0.381 | 1.26E-11 | 17 | Hspb1 |
| Epc22 | 7.04E-16 | 0.458175834 | 0.392 | 0.163 | 1.42E-11 | 17 | Epc2 |
| Fbxl1711 | 7.13E-16 | 0.639133457 | 0.585 | 0.311 | 1.44E-11 | 17 | Fbxl17 |
| Ginm14 | 7.38E-16 | 0.496676132 | 0.415 | 0.18 | 1.49E-11 | 17 | Ginm1 |
| Rab125 | 7.38E-16 | 0.431765184 | 0.421 | 0.177 | 1.49E-11 | 17 | Rab12 |
| Rtn35 | 7.44E-16 | 0.484738195 | 0.62 | 0.324 | 1.51E-11 | 17 | Rtn3 |
| Tshz25 | 8.55E-16 | 0.373736866 | 0.76 | 0.372 | 1.73E-11 | 17 | Tshz2 |
| Tmem50a8 | 8.75E-16 | 0.531466525 | 0.684 | 0.383 | 1.77E-11 | 17 | Tmem50a |
| Desi21 | 9.14E-16 | 0.430514869 | 0.404 | 0.172 | 1.85E-11 | 17 | Desi2 |
| Comt2 | 9.84E-16 | 0.450166782 | 0.439 | 0.194 | 1.99E-11 | 17 | Comt |
| Gsk3b5 | 1.02E-15 | 0.549448758 | 0.614 | 0.331 | 2.07E-11 | 17 | Gsk3b |
| Eea15 | 1.08E-15 | 0.551181345 | 0.468 | 0.219 | 2.19E-11 | 17 | Eea1 |
| Zranb2 | 1.34E-15 | 0.37651131 | 0.532 | 0.255 | 2.72E-11 | 17 | Zranb2 |
| Srpk23 | 1.34E-15 | 0.501755327 | 0.439 | 0.194 | 2.72E-11 | 17 | Srpk2 |
| Itm2c5 | 1.38E-15 | 0.535496241 | 0.439 | 0.2 | 2.79E-11 | 17 | Itm2c |
| Dad17 | 1.76E-15 | 0.459563817 | 0.772 | 0.459 | 3.57E-11 | 17 | Dad1 |
| Arhgef114 | 1.81E-15 | 0.472171899 | 0.287 | 0.101 | 3.67E-11 | 17 | Arhgef11 |
| Tm9sf25 | 1.82E-15 | 0.588247369 | 0.509 | 0.256 | 3.69E-11 | 17 | Tm9sf2 |
| Rbms24 | 2.13E-15 | 0.350089659 | 0.392 | 0.16 | 4.30E-11 | 17 | Rbms2 |
| Pnrc11 | 2.17E-15 | 0.426672752 | 0.713 | 0.411 | 4.40E-11 | 17 | Pnrc1 |
| Fkbp71 | 2.27E-15 | 0.431277876 | 0.251 | 0.082 | 4.59E-11 | 17 | Fkbp7 |
| Bloc1s1 | 2.36E-15 | 0.354633008 | 0.287 | 0.102 | 4.77E-11 | 17 | Bloc1s1 |

| | | | | | | | |
|---|---|---|---|---|---|---|---|
| Pura3 | 2.49E-15 | 0.556804601 | 0.649 | 0.366 | 5.04E-11 | 17 | Pura |
| Btbd75 | 2.75E-15 | 0.639156753 | 0.544 | 0.282 | 5.56E-11 | 17 | Btbd7 |
| Apbb25 | 2.82E-15 | 0.378671019 | 0.731 | 0.367 | 5.71E-11 | 17 | Apbb2 |
| Rhoq2 | 2.86E-15 | 0.389200687 | 0.298 | 0.109 | 5.79E-11 | 17 | Rhoq |
| As3mt | 3.28E-15 | 0.356428053 | 0.263 | 0.088 | 6.64E-11 | 17 | As3mt |
| Cnn26 | 3.73E-15 | 0.538861589 | 0.409 | 0.183 | 7.55E-11 | 17 | Cnn2 |
| mt-Nd310 | 3.80E-15 | 0.640477296 | 0.713 | 0.447 | 7.70E-11 | 17 | mt-Nd3 |
| Ctdspl1 | 4.02E-15 | 0.416501462 | 0.269 | 0.092 | 8.14E-11 | 17 | Ctdspl |
| Ppm1a | 4.37E-15 | 0.345176823 | 0.398 | 0.166 | 8.85E-11 | 17 | Ppm1a |
| Atad11 | 4.84E-15 | 0.431661265 | 0.287 | 0.104 | 9.79E-11 | 17 | Atad1 |
| D5Ertd579e | 4.91E-15 | 0.415326228 | 0.275 | 0.098 | 9.93E-11 | 17 | D5Ertd579e |
| Hmgn31 | 4.99E-15 | 0.324218192 | 0.409 | 0.171 | 1.01E-10 | 17 | Hmgn3 |
| Sec626 | 5.05E-15 | 0.530255198 | 0.678 | 0.395 | 1.02E-10 | 17 | Sec62 |
| Nr3c16 | 5.13E-15 | 0.539751257 | 0.684 | 0.384 | 1.04E-10 | 17 | Nr3c1 |
| Pbrm14 | 5.65E-15 | 0.540927123 | 0.55 | 0.288 | 1.14E-10 | 17 | Pbrm1 |
| Pkn25 | 5.89E-15 | 0.533366868 | 0.561 | 0.295 | 1.19E-10 | 17 | Pkn2 |
| Snrk4 | 6.73E-15 | 0.338733626 | 0.725 | 0.369 | 1.36E-10 | 17 | Snrk |
| Cav24 | 6.89E-15 | 0.348307069 | 0.48 | 0.21 | 1.40E-10 | 17 | Cav2 |
| C2cd21 | 7.03E-15 | 0.427434206 | 0.304 | 0.111 | 1.42E-10 | 17 | C2cd2 |
| Insr1 | 7.43E-15 | 0.344779405 | 0.339 | 0.129 | 1.50E-10 | 17 | Insr |
| Gtf2a2 | 7.52E-15 | 0.415481194 | 0.532 | 0.268 | 1.52E-10 | 17 | Gtf2a2 |
| Crim13 | 8.85E-15 | 0.326198949 | 0.725 | 0.373 | 1.79E-10 | 17 | Crim1 |
| Cisd21 | 9.76E-15 | 0.518635658 | 0.392 | 0.175 | 1.98E-10 | 17 | Cisd2 |
| Pde4d3 | 9.80E-15 | 0.822478491 | 0.357 | 0.15 | 1.98E-10 | 17 | Pde4d |
| Cfap202 | 9.97E-15 | 0.394611922 | 0.427 | 0.187 | 2.02E-10 | 17 | Cfap20 |
| Leng8 | 1.02E-14 | 0.520967116 | 0.257 | 0.09 | 2.07E-10 | 17 | Leng8 |
| Ssbp41 | 1.26E-14 | 0.400346554 | 0.433 | 0.194 | 2.56E-10 | 17 | Ssbp4 |
| Galnt184 | 1.46E-14 | 0.36418249 | 0.351 | 0.135 | 2.96E-10 | 17 | Galnt18 |
| Pkd21 | 1.47E-14 | 0.50706066 | 0.292 | 0.107 | 2.97E-10 | 17 | Pkd2 |
| Ralgapb1 | 1.47E-14 | 0.41519668 | 0.31 | 0.118 | 2.98E-10 | 17 | Ralgapb |
| Fyco14 | 1.94E-14 | 0.42551831 | 0.316 | 0.124 | 3.92E-10 | 17 | Fyco1 |
| Prdx44 | 2.08E-14 | 0.441135236 | 0.474 | 0.222 | 4.21E-10 | 17 | Prdx4 |
| Fnip22 | 2.10E-14 | 0.382693333 | 0.292 | 0.107 | 4.26E-10 | 17 | Fnip2 |
| Ahcyl11 | 2.54E-14 | 0.277135911 | 0.404 | 0.175 | 5.15E-10 | 17 | Ahcyl1 |
| Ubr26 | 3.02E-14 | 0.587953409 | 0.462 | 0.23 | 6.11E-10 | 17 | Ubr2 |
| Filip14 | 3.41E-14 | 0.508669823 | 0.433 | 0.195 | 6.90E-10 | 17 | Filip1 |
| Eva1b5 | 3.45E-14 | 0.415866585 | 0.573 | 0.286 | 6.98E-10 | 17 | Eva1b |
| Hsp90aa11 | 3.93E-14 | 0.406575031 | 0.708 | 0.426 | 7.96E-10 | 17 | Hsp90aa1 |
| Ddx1 | 3.97E-14 | 0.340105655 | 0.327 | 0.131 | 8.03E-10 | 17 | Ddx1 |
| Rcn12 | 4.01E-14 | 0.591628108 | 0.281 | 0.109 | 8.13E-10 | 17 | Rcn1 |
| Cux1 | 4.70E-14 | 0.43304298 | 0.474 | 0.231 | 9.52E-10 | 17 | Cux1 |
| Slc38a26 | 5.57E-14 | 0.427831371 | 0.591 | 0.312 | 1.13E-09 | 17 | Slc38a2 |
| Kifap31 | 5.66E-14 | 0.42445409 | 0.287 | 0.107 | 1.15E-09 | 17 | Kifap3 |

| | | | | | | | |
|---|---|---|---|---|---|---|---|
| Crebbp9 | 5.67E-14 | 0.554086093 | 0.567 | 0.306 | 1.15E-09 | 17 | Crebbp |
| Chd46 | 5.95E-14 | 0.50497587 | 0.596 | 0.328 | 1.21E-09 | 17 | Chd4 |
| Cdkn1c4 | 6.33E-14 | 0.382662226 | 0.327 | 0.127 | 1.28E-09 | 17 | Cdkn1c |
| Chd67 | 6.55E-14 | 0.569219234 | 0.444 | 0.218 | 1.33E-09 | 17 | Chd6 |
| Hnrnph32 | 6.84E-14 | 0.365520575 | 0.509 | 0.249 | 1.39E-09 | 17 | Hnrnph3 |
| Cfap36 | 7.31E-14 | 0.402201027 | 0.298 | 0.116 | 1.48E-09 | 17 | Cfap36 |
| Rheb1 | 8.15E-14 | 0.36666767 | 0.667 | 0.377 | 1.65E-09 | 17 | Rheb |
| Kmt2e7 | 8.41E-14 | 0.481494377 | 0.69 | 0.407 | 1.70E-09 | 17 | Kmt2e |
| Prpsap11 | 8.88E-14 | 0.388421646 | 0.292 | 0.111 | 1.80E-09 | 17 | Prpsap1 |
| Med13l5 | 9.23E-14 | 0.49271983 | 0.415 | 0.195 | 1.87E-09 | 17 | Med13l |
| Zdhhc172 | 9.76E-14 | 0.419459435 | 0.357 | 0.15 | 1.98E-09 | 17 | Zdhhc17 |
| Ncor19 | 1.03E-13 | 0.467099803 | 0.737 | 0.438 | 2.08E-09 | 17 | Ncor1 |
| Adprh1 | 1.20E-13 | 0.43252743 | 0.339 | 0.141 | 2.42E-09 | 17 | Adprh |
| Evi53 | 1.20E-13 | 0.453407327 | 0.339 | 0.141 | 2.44E-09 | 17 | Evi5 |
| Large15 | 1.21E-13 | 0.464091059 | 0.357 | 0.153 | 2.45E-09 | 17 | Large1 |
| Chd34 | 1.27E-13 | 0.282088809 | 0.368 | 0.156 | 2.57E-09 | 17 | Chd3 |
| Tmem2451 | 1.31E-13 | 0.573321039 | 0.298 | 0.119 | 2.66E-09 | 17 | Tmem245 |
| Ppp2r1a | 1.34E-13 | 0.502048247 | 0.374 | 0.165 | 2.71E-09 | 17 | Ppp2r1a |
| Nap1l14 | 1.42E-13 | 0.399565326 | 0.591 | 0.315 | 2.87E-09 | 17 | Nap1l1 |
| Rbm398 | 1.56E-13 | 0.54259879 | 0.889 | 0.574 | 3.16E-09 | 17 | Rbm39 |
| Tm9sf36 | 1.62E-13 | 0.519892174 | 0.596 | 0.336 | 3.29E-09 | 17 | Tm9sf3 |
| Lama45 | 1.70E-13 | 0.453584464 | 0.404 | 0.175 | 3.44E-09 | 17 | Lama4 |
| Iscu1 | 1.74E-13 | 0.290685172 | 0.515 | 0.262 | 3.52E-09 | 17 | Iscu |
| Psmd74 | 1.75E-13 | 0.402498922 | 0.573 | 0.306 | 3.53E-09 | 17 | Psmd7 |
| Tril2 | 1.80E-13 | 0.337964457 | 0.251 | 0.088 | 3.64E-09 | 17 | Tril |
| Slc29a12 | 1.80E-13 | 0.581027619 | 0.322 | 0.135 | 3.65E-09 | 17 | Slc29a1 |
| Ddhd21 | 1.82E-13 | 0.444098011 | 0.281 | 0.108 | 3.69E-09 | 17 | Ddhd2 |
| Map4k48 | 1.83E-13 | 0.455506731 | 0.538 | 0.277 | 3.70E-09 | 17 | Map4k4 |
| Vkorc11 | 1.83E-13 | 0.428142773 | 0.357 | 0.154 | 3.70E-09 | 17 | Vkorc1 |
| Zbtb44 | 1.92E-13 | 0.516208589 | 0.304 | 0.122 | 3.88E-09 | 17 | Zbtb4 |
| Srsf10 | 1.92E-13 | 0.352304713 | 0.491 | 0.245 | 3.88E-09 | 17 | Srsf10 |
| Pnisr8 | 1.96E-13 | 0.591120346 | 0.509 | 0.265 | 3.97E-09 | 17 | Pnisr |
| Fndc3b5 | 2.07E-13 | 0.461347803 | 0.503 | 0.245 | 4.20E-09 | 17 | Fndc3b |
| Dync1i23 | 2.14E-13 | 0.481934887 | 0.702 | 0.419 | 4.32E-09 | 17 | Dync1i2 |
| Kitl4 | 2.25E-13 | 0.292283645 | 0.678 | 0.336 | 4.56E-09 | 17 | Kitl |
| Gab24 | 2.38E-13 | 0.434249851 | 0.45 | 0.205 | 4.82E-09 | 17 | Gab2 |
| Ski7 | 2.41E-13 | 0.56470965 | 0.444 | 0.215 | 4.88E-09 | 17 | Ski |
| Ubap2l1 | 2.57E-13 | 0.482218426 | 0.544 | 0.288 | 5.20E-09 | 17 | Ubap2l |
| Cyhr1 | 2.62E-13 | 0.296337379 | 0.386 | 0.168 | 5.31E-09 | 17 | Cyhr1 |
| Setbp15 | 2.77E-13 | 0.505097053 | 0.31 | 0.122 | 5.60E-09 | 17 | Setbp1 |
| Rbm58 | 2.91E-13 | 0.53622189 | 0.474 | 0.24 | 5.89E-09 | 17 | Rbm5 |
| Zfp36l16 | 2.92E-13 | 0.520307304 | 0.713 | 0.433 | 5.91E-09 | 17 | Zfp36l1 |
| Rapgef24 | 3.25E-13 | 0.657332252 | 0.596 | 0.34 | 6.58E-09 | 17 | Rapgef2 |

| | | | | | | | |
|---|---|---|---|---|---|---|---|
| Ralgapa14 | 3.32E-13 | 0.537462076 | 0.561 | 0.304 | 6.73E-09 | 17 | Ralgapa1 |
| Cabin11 | 3.51E-13 | 0.367961599 | 0.269 | 0.1 | 7.11E-09 | 17 | Cabin1 |
| Nptn7 | 3.52E-13 | 0.480383868 | 0.485 | 0.25 | 7.12E-09 | 17 | Nptn |
| Prpf4b6 | 3.58E-13 | 0.479346359 | 0.591 | 0.334 | 7.25E-09 | 17 | Prpf4b |
| Supt63 | 3.76E-13 | 0.456123756 | 0.363 | 0.161 | 7.61E-09 | 17 | Supt6 |
| Igf1r5 | 3.79E-13 | 0.518436245 | 0.368 | 0.162 | 7.68E-09 | 17 | Igf1r |
| Anapc54 | 3.83E-13 | 0.440165061 | 0.444 | 0.212 | 7.76E-09 | 17 | Anapc5 |
| Mycbp210 | 4.37E-13 | 0.516497931 | 0.655 | 0.375 | 8.85E-09 | 17 | Mycbp2 |
| Morf4l12 | 4.68E-13 | 0.479299245 | 0.86 | 0.589 | 9.47E-09 | 17 | Morf4l1 |
| Stag19 | 4.71E-13 | 0.516166506 | 0.626 | 0.362 | 9.54E-09 | 17 | Stag1 |
| Snrpb1 | 5.37E-13 | 0.480709086 | 0.749 | 0.473 | 1.09E-08 | 17 | Snrpb |
| 9930021J03Rik2 | 5.51E-13 | 0.540223515 | 0.345 | 0.152 | 1.12E-08 | 17 | 9930021J03Rik |
| BC0055611 | 5.69E-13 | 0.456922267 | 0.298 | 0.119 | 1.15E-08 | 17 | BC005561 |
| Ssr21 | 5.96E-13 | 0.421676741 | 0.421 | 0.203 | 1.21E-08 | 17 | Ssr2 |
| Matr32 | 6.04E-13 | 0.388901747 | 0.509 | 0.267 | 1.22E-08 | 17 | Matr3 |
| Tra2a10 | 6.04E-13 | 0.517074228 | 0.52 | 0.275 | 1.22E-08 | 17 | Tra2a |
| Prex24 | 6.24E-13 | 0.446790924 | 0.55 | 0.274 | 1.26E-08 | 17 | Prex2 |
| Ociad1 | 6.71E-13 | 0.41044473 | 0.398 | 0.186 | 1.36E-08 | 17 | Ociad1 |
| Hbp1 | 7.16E-13 | 0.398075443 | 0.304 | 0.125 | 1.45E-08 | 17 | Hbp1 |
| Tcf7l14 | 7.81E-13 | 0.452027039 | 0.404 | 0.182 | 1.58E-08 | 17 | Tcf7l1 |
| Sh3bgrl6 | 7.85E-13 | 0.438335826 | 0.655 | 0.376 | 1.59E-08 | 17 | Sh3bgrl |
| Med31 | 7.92E-13 | 0.311047092 | 0.281 | 0.108 | 1.60E-08 | 17 | Med31 |
| Nub12 | 8.12E-13 | 0.417250348 | 0.398 | 0.187 | 1.64E-08 | 17 | Nub1 |
| Jak24 | 8.69E-13 | 0.401793504 | 0.31 | 0.128 | 1.76E-08 | 17 | Jak2 |
| Wdr1 | 8.88E-13 | 0.413068427 | 0.374 | 0.17 | 1.80E-08 | 17 | Wdr1 |
| Phlpp14 | 8.90E-13 | 0.493684584 | 0.52 | 0.264 | 1.80E-08 | 17 | Phlpp1 |
| Arhgap293 | 9.03E-13 | 0.30439547 | 0.713 | 0.366 | 1.83E-08 | 17 | Arhgap29 |
| Trpm76 | 9.44E-13 | 0.474040585 | 0.421 | 0.205 | 1.91E-08 | 17 | Trpm7 |
| Cwc271 | 9.48E-13 | 0.427417477 | 0.304 | 0.124 | 1.92E-08 | 17 | Cwc27 |
| Pcm16 | 1.02E-12 | 0.503809509 | 0.485 | 0.252 | 2.07E-08 | 17 | Pcm1 |
| Stk35 | 1.04E-12 | 0.444926839 | 0.368 | 0.167 | 2.10E-08 | 17 | Stk3 |
| Dennd4c5 | 1.04E-12 | 0.44948095 | 0.304 | 0.127 | 2.11E-08 | 17 | Dennd4c |
| Pafah1b12 | 1.07E-12 | 0.404334614 | 0.766 | 0.481 | 2.17E-08 | 17 | Pafah1b1 |
| Gkap12 | 1.10E-12 | 0.45746706 | 0.333 | 0.143 | 2.22E-08 | 17 | Gkap1 |
| Cdk143 | 1.10E-12 | 0.415918296 | 0.339 | 0.143 | 2.24E-08 | 17 | Cdk14 |
| Emd | 1.15E-12 | 0.362818847 | 0.351 | 0.153 | 2.33E-08 | 17 | Emd |
| Tmcc19 | 1.15E-12 | 0.508032711 | 0.439 | 0.219 | 2.34E-08 | 17 | Tmcc1 |
| Cask4 | 1.19E-12 | 0.466404696 | 0.327 | 0.14 | 2.40E-08 | 17 | Cask |
| Ttc148 | 1.24E-12 | 0.493379705 | 0.433 | 0.216 | 2.51E-08 | 17 | Ttc14 |
| Tnks3 | 1.24E-12 | 0.437801393 | 0.327 | 0.143 | 2.51E-08 | 17 | Tnks |
| Trim22 | 1.26E-12 | 0.320483356 | 0.404 | 0.179 | 2.55E-08 | 17 | Trim2 |
| Arhgef126 | 1.27E-12 | 0.308634028 | 0.667 | 0.349 | 2.56E-08 | 17 | Arhgef12 |
| Ash1l6 | 1.28E-12 | 0.4379312 | 0.55 | 0.3 | 2.59E-08 | 17 | Ash1l |

| | | | | | | | |
|---|---|---|---|---|---|---|---|
| Srsf1110 | 1.30E-12 | 0.534874785 | 0.614 | 0.352 | 2.63E-08 | 17 | Srsf11 |
| Klhdc2 | 1.37E-12 | 0.33374821 | 0.322 | 0.135 | 2.77E-08 | 17 | Klhdc2 |
| Peak18 | 1.37E-12 | 0.501672457 | 0.655 | 0.388 | 2.77E-08 | 17 | Peak1 |
| Sik37 | 1.39E-12 | 0.49940912 | 0.515 | 0.271 | 2.81E-08 | 17 | Sik3 |
| Kcnq1ot16 | 1.39E-12 | 0.530121647 | 0.316 | 0.135 | 2.82E-08 | 17 | Kcnq1ot1 |
| Zmynd81 | 1.40E-12 | 0.386465358 | 0.415 | 0.196 | 2.84E-08 | 17 | Zmynd8 |
| Arglu16 | 1.41E-12 | 0.438976466 | 0.608 | 0.355 | 2.86E-08 | 17 | Arglu1 |
| Rb1cc15 | 1.44E-12 | 0.566942283 | 0.456 | 0.235 | 2.92E-08 | 17 | Rb1cc1 |
| mt-Atp89 | 1.50E-12 | 0.490019359 | 0.731 | 0.464 | 3.04E-08 | 17 | mt-Atp8 |
| Tpd52l21 | 1.55E-12 | 0.318990611 | 0.38 | 0.171 | 3.15E-08 | 17 | Tpd52l2 |
| Sumo3 | 1.68E-12 | 0.288384845 | 0.421 | 0.197 | 3.39E-08 | 17 | Sumo3 |
| Senp67 | 1.89E-12 | 0.40719211 | 0.55 | 0.299 | 3.82E-08 | 17 | Senp6 |
| Gtf2h51 | 1.92E-12 | 0.276272639 | 0.596 | 0.325 | 3.88E-08 | 17 | Gtf2h5 |
| Atp6v0b2 | 1.93E-12 | 0.297664965 | 0.649 | 0.367 | 3.91E-08 | 17 | Atp6v0b |
| Ctnna14 | 1.97E-12 | 0.351968298 | 0.637 | 0.352 | 3.98E-08 | 17 | Ctnna1 |
| Macf18 | 2.01E-12 | 0.394328589 | 0.842 | 0.503 | 4.08E-08 | 17 | Macf1 |
| Pnn7 | 2.05E-12 | 0.58505996 | 0.491 | 0.268 | 4.15E-08 | 17 | Pnn |
| Fam120a4 | 2.21E-12 | 0.393626228 | 0.392 | 0.182 | 4.47E-08 | 17 | Fam120a |
| Snx17 | 2.23E-12 | 0.342788953 | 0.38 | 0.173 | 4.51E-08 | 17 | Snx17 |
| Ctdsp23 | 2.36E-12 | 0.408182219 | 0.386 | 0.179 | 4.77E-08 | 17 | Ctdsp2 |
| Sike1 | 2.43E-12 | 0.344844486 | 0.316 | 0.132 | 4.92E-08 | 17 | Sike1 |
| P4ha12 | 2.51E-12 | 0.548505313 | 0.298 | 0.124 | 5.09E-08 | 17 | P4ha1 |
| Mkln13 | 2.73E-12 | 0.461334181 | 0.491 | 0.265 | 5.52E-08 | 17 | Mkln1 |
| Emc76 | 2.78E-12 | 0.352454059 | 0.532 | 0.272 | 5.64E-08 | 17 | Emc7 |
| Dennd5a4 | 2.86E-12 | 0.451060718 | 0.398 | 0.188 | 5.79E-08 | 17 | Dennd5a |
| Crebrf2 | 3.18E-12 | 0.510979131 | 0.398 | 0.191 | 6.43E-08 | 17 | Crebrf |
| Ccdc1861 | 3.25E-12 | 0.3373568 | 0.357 | 0.157 | 6.58E-08 | 17 | Ccdc186 |
| Maf11 | 3.31E-12 | 0.34792447 | 0.292 | 0.118 | 6.69E-08 | 17 | Maf1 |
| Arid1b8 | 3.43E-12 | 0.480050573 | 0.48 | 0.259 | 6.94E-08 | 17 | Arid1b |
| Rere5 | 3.63E-12 | 0.505488726 | 0.509 | 0.272 | 7.35E-08 | 17 | Rere |
| Zfand39 | 3.74E-12 | 0.591853575 | 0.526 | 0.302 | 7.56E-08 | 17 | Zfand3 |
| Zfp5033 | 3.83E-12 | 0.479276507 | 0.339 | 0.148 | 7.76E-08 | 17 | Zfp503 |
| Kat6b2 | 4.19E-12 | 0.445167247 | 0.345 | 0.153 | 8.49E-08 | 17 | Kat6b |
| Ssb | 4.23E-12 | 0.378382798 | 0.69 | 0.414 | 8.56E-08 | 17 | Ssb |
| Pcnp1 | 4.25E-12 | 0.280952966 | 0.485 | 0.249 | 8.60E-08 | 17 | Pcnp |
| Stom2 | 4.39E-12 | 0.301041345 | 0.281 | 0.108 | 8.88E-08 | 17 | Stom |
| Zranb1 | 4.49E-12 | 0.389740177 | 0.322 | 0.14 | 9.09E-08 | 17 | Zranb1 |
| Nsfl1c | 4.55E-12 | 0.333498615 | 0.368 | 0.168 | 9.22E-08 | 17 | Nsfl1c |
| Arhgef71 | 4.56E-12 | 0.39368625 | 0.351 | 0.154 | 9.22E-08 | 17 | Arhgef7 |
| Fam107b5 | 4.56E-12 | 0.485197237 | 0.292 | 0.121 | 9.23E-08 | 17 | Fam107b |
| Celf15 | 4.95E-12 | 0.553808489 | 0.427 | 0.221 | 1.00E-07 | 17 | Celf1 |
| Sh3rf14 | 5.04E-12 | 0.478025553 | 0.31 | 0.131 | 1.02E-07 | 17 | Sh3rf1 |
| Hspa1a2 | 7.54E-12 | 0.368989875 | 0.263 | 0.101 | 1.53E-07 | 17 | Hspa1a |

| | | | | | | | |
|---|---|---|---|---|---|---|---|
| Reep35 | 7.64E-12 | 0.431295822 | 0.561 | 0.307 | 1.55E-07 | 17 | Reep3 |
| Nsd37 | 7.67E-12 | 0.525942924 | 0.556 | 0.309 | 1.55E-07 | 17 | Nsd3 |
| Gnl3l | 7.79E-12 | 0.290517529 | 0.251 | 0.096 | 1.58E-07 | 17 | Gnl3l |
| Spcs25 | 7.80E-12 | 0.361885566 | 0.667 | 0.402 | 1.58E-07 | 17 | Spcs2 |
| Hnrnpu3 | 7.86E-12 | 0.387400077 | 0.678 | 0.424 | 1.59E-07 | 17 | Hnrnpu |
| Cep85l4 | 7.95E-12 | 0.370940286 | 0.368 | 0.165 | 1.61E-07 | 17 | Cep85l |
| Sh3pxd2a3 | 9.92E-12 | 0.397730572 | 0.327 | 0.143 | 2.01E-07 | 17 | Sh3pxd2a |
| Tmem106b1 | 1.02E-11 | 0.306273752 | 0.304 | 0.128 | 2.06E-07 | 17 | Tmem106b |
| Hspa59 | 1.05E-11 | 0.467376456 | 0.731 | 0.435 | 2.12E-07 | 17 | Hspa5 |
| Clip12 | 1.06E-11 | 0.413633207 | 0.322 | 0.144 | 2.14E-07 | 17 | Clip1 |
| Cttn | 1.09E-11 | 0.282513541 | 0.251 | 0.095 | 2.20E-07 | 17 | Cttn |
| Scaf114 | 1.09E-11 | 0.461171803 | 0.497 | 0.264 | 2.21E-07 | 17 | Scaf11 |
| 4921524J17Rik | 1.15E-11 | 0.395059025 | 0.339 | 0.152 | 2.33E-07 | 17 | 4921524J17Rik |
| Rer11 | 1.22E-11 | 0.40487815 | 0.573 | 0.326 | 2.48E-07 | 17 | Rer1 |
| Tmem176b6 | 1.23E-11 | 0.494068609 | 0.737 | 0.439 | 2.49E-07 | 17 | Tmem176b |
| 1110004F10Rik2 | 1.28E-11 | 0.398184138 | 0.556 | 0.309 | 2.59E-07 | 17 | 1110004F10Rik |
| Msn5 | 1.32E-11 | 0.431559117 | 0.725 | 0.444 | 2.67E-07 | 17 | Msn |
| Vasp4 | 1.38E-11 | 0.455685794 | 0.456 | 0.238 | 2.79E-07 | 17 | Vasp |
| Phf21a9 | 1.45E-11 | 0.462496167 | 0.485 | 0.254 | 2.94E-07 | 17 | Phf21a |
| Sav11 | 1.60E-11 | 0.304898151 | 0.269 | 0.107 | 3.24E-07 | 17 | Sav1 |
| Rexo24 | 1.68E-11 | 0.291351602 | 0.544 | 0.298 | 3.40E-07 | 17 | Rexo2 |
| Hp1bp3 | 1.69E-11 | 0.342083034 | 0.526 | 0.286 | 3.43E-07 | 17 | Hp1bp3 |
| Fam104a2 | 1.70E-11 | 0.288267328 | 0.55 | 0.302 | 3.44E-07 | 17 | Fam104a |
| Cct7 | 1.76E-11 | 0.267042194 | 0.55 | 0.304 | 3.56E-07 | 17 | Cct7 |
| Spred15 | 1.77E-11 | 0.469112945 | 0.48 | 0.253 | 3.58E-07 | 17 | Spred1 |
| Pttg1ip6 | 1.80E-11 | 0.367821029 | 0.515 | 0.27 | 3.63E-07 | 17 | Pttg1ip |
| Pds5b4 | 1.87E-11 | 0.440945953 | 0.38 | 0.181 | 3.79E-07 | 17 | Pds5b |
| Ralgapa25 | 1.88E-11 | 0.486832055 | 0.333 | 0.155 | 3.80E-07 | 17 | Ralgapa2 |
| Tmx35 | 2.00E-11 | 0.479760194 | 0.386 | 0.188 | 4.05E-07 | 17 | Tmx3 |
| Ubr54 | 2.01E-11 | 0.47558644 | 0.404 | 0.199 | 4.07E-07 | 17 | Ubr5 |
| Fchsd26 | 2.04E-11 | 0.476320209 | 0.345 | 0.161 | 4.13E-07 | 17 | Fchsd2 |
| Nipbl11 | 2.17E-11 | 0.446325584 | 0.608 | 0.367 | 4.40E-07 | 17 | Nipbl |
| Spop2 | 2.24E-11 | 0.407163991 | 0.62 | 0.381 | 4.54E-07 | 17 | Spop |
| Picalm5 | 2.27E-11 | 0.480644012 | 0.52 | 0.297 | 4.60E-07 | 17 | Picalm |
| Stat61 | 2.37E-11 | 0.397836203 | 0.298 | 0.128 | 4.81E-07 | 17 | Stat6 |
| Arhgap268 | 2.39E-11 | 0.451974596 | 0.439 | 0.223 | 4.84E-07 | 17 | Arhgap26 |
| Hsd17b112 | 2.53E-11 | 0.34994379 | 0.287 | 0.119 | 5.12E-07 | 17 | Hsd17b11 |
| Zfp6447 | 2.89E-11 | 0.40887511 | 0.48 | 0.257 | 5.85E-07 | 17 | Zfp644 |
| Vps35 | 2.98E-11 | 0.308260397 | 0.415 | 0.207 | 6.04E-07 | 17 | Vps35 |
| Spcs12 | 3.07E-11 | 0.384621283 | 0.515 | 0.289 | 6.22E-07 | 17 | Spcs1 |
| Mier14 | 3.11E-11 | 0.358818885 | 0.591 | 0.332 | 6.29E-07 | 17 | Mier1 |
| Fxr1 | 3.11E-11 | 0.314943866 | 0.485 | 0.257 | 6.30E-07 | 17 | Fxr1 |
| Imp32 | 3.43E-11 | 0.306805306 | 0.561 | 0.316 | 6.95E-07 | 17 | Imp3 |

| | | | | | | | |
|---|---|---|---|---|---|---|---|
| Ankrd1110 | 3.49E-11 | 0.444083409 | 0.643 | 0.382 | 7.06E-07 | 17 | Ankrd11 |
| Arhgap52 | 3.72E-11 | 0.404958435 | 0.52 | 0.29 | 7.52E-07 | 17 | Arhgap5 |
| Pde6d | 3.86E-11 | 0.327413261 | 0.281 | 0.117 | 7.81E-07 | 17 | Pde6d |
| Ddx461 | 3.88E-11 | 0.364624618 | 0.439 | 0.226 | 7.87E-07 | 17 | Ddx46 |
| Rbm265 | 3.96E-11 | 0.484949521 | 0.433 | 0.228 | 8.02E-07 | 17 | Rbm26 |
| Galnt110 | 4.23E-11 | 0.401069746 | 0.38 | 0.185 | 8.56E-07 | 17 | Galnt1 |
| Pde4b3 | 4.64E-11 | 0.508669774 | 0.398 | 0.197 | 9.40E-07 | 17 | Pde4b |
| Srpr5 | 4.99E-11 | 0.441423453 | 0.456 | 0.247 | 1.01E-06 | 17 | Srpr |
| Map1lc3b2 | 5.00E-11 | 0.301212881 | 0.766 | 0.502 | 1.01E-06 | 17 | Map1lc3b |
| Dock117 | 5.31E-11 | 0.368716881 | 0.275 | 0.115 | 1.07E-06 | 17 | Dock11 |
| Ccnl24 | 5.67E-11 | 0.416325511 | 0.427 | 0.215 | 1.15E-06 | 17 | Ccnl2 |
| Atxn7l3b3 | 5.76E-11 | 0.27714573 | 0.69 | 0.405 | 1.17E-06 | 17 | Atxn7l3b |
| Supt204 | 5.89E-11 | 0.506782142 | 0.339 | 0.162 | 1.19E-06 | 17 | Supt20 |
| Tia11 | 5.90E-11 | 0.406518707 | 0.269 | 0.113 | 1.19E-06 | 17 | Tia1 |
| 4932438A13Rik7 | 5.94E-11 | 0.420234435 | 0.427 | 0.22 | 1.20E-06 | 17 | 4932438A13Rik |
| Rsrp11 | 6.17E-11 | 0.382798339 | 0.877 | 0.571 | 1.25E-06 | 17 | Rsrp1 |
| Tsen341 | 6.22E-11 | 0.334505758 | 0.374 | 0.178 | 1.26E-06 | 17 | Tsen34 |
| Ddx57 | 6.41E-11 | 0.441967781 | 0.918 | 0.655 | 1.30E-06 | 17 | Ddx5 |
| Rbbp63 | 6.89E-11 | 0.474111668 | 0.497 | 0.277 | 1.39E-06 | 17 | Rbbp6 |
| 4931406P16Rik4 | 6.94E-11 | 0.302460156 | 0.632 | 0.344 | 1.41E-06 | 17 | 4931406P16Rik |
| Smc1a5 | 7.20E-11 | 0.429946471 | 0.55 | 0.313 | 1.46E-06 | 17 | Smc1a |
| Rtn46 | 7.25E-11 | 0.378599053 | 0.602 | 0.34 | 1.47E-06 | 17 | Rtn4 |
| Sipa15 | 7.52E-11 | 0.477893291 | 0.38 | 0.186 | 1.52E-06 | 17 | Sipa1 |
| Smc33 | 7.75E-11 | 0.350338922 | 0.474 | 0.253 | 1.57E-06 | 17 | Smc3 |
| Tbl1x5 | 7.77E-11 | 0.419079515 | 0.485 | 0.263 | 1.57E-06 | 17 | Tbl1x |
| Kif1b5 | 8.01E-11 | 0.441538116 | 0.456 | 0.244 | 1.62E-06 | 17 | Kif1b |
| Zfp24 | 8.06E-11 | 0.258476827 | 0.275 | 0.113 | 1.63E-06 | 17 | Zfp24 |
| Setd59 | 8.16E-11 | 0.482393892 | 0.48 | 0.265 | 1.65E-06 | 17 | Setd5 |
| Trappc95 | 8.28E-11 | 0.46704182 | 0.392 | 0.195 | 1.68E-06 | 17 | Trappc9 |
| Dek | 8.60E-11 | 0.253918357 | 0.614 | 0.373 | 1.74E-06 | 17 | Dek |
| Ube3a3 | 8.82E-11 | 0.439492644 | 0.526 | 0.296 | 1.79E-06 | 17 | Ube3a |
| Gabpb21 | 1.04E-10 | 0.3608865 | 0.281 | 0.119 | 2.11E-06 | 17 | Gabpb2 |
| Pcmtd2 | 1.09E-10 | 0.297376933 | 0.257 | 0.105 | 2.21E-06 | 17 | Pcmtd2 |
| Map3k25 | 1.15E-10 | 0.344778908 | 0.333 | 0.154 | 2.32E-06 | 17 | Map3k2 |
| Aff43 | 1.21E-10 | 0.406172271 | 0.45 | 0.241 | 2.46E-06 | 17 | Aff4 |
| Capns15 | 1.32E-10 | 0.333075439 | 0.585 | 0.349 | 2.68E-06 | 17 | Capns1 |
| Rhoc5 | 1.37E-10 | 0.324119922 | 0.544 | 0.297 | 2.78E-06 | 17 | Rhoc |
| Gpx41 | 1.39E-10 | 0.315596151 | 0.901 | 0.629 | 2.82E-06 | 17 | Gpx4 |
| Ago25 | 1.49E-10 | 0.347381098 | 0.351 | 0.168 | 3.02E-06 | 17 | Ago2 |
| Rbm8a1 | 1.59E-10 | 0.261931649 | 0.544 | 0.309 | 3.21E-06 | 17 | Rbm8a |
| Dst5 | 1.74E-10 | 0.360349489 | 0.392 | 0.192 | 3.52E-06 | 17 | Dst |
| Tspan123 | 1.81E-10 | 0.250487556 | 0.474 | 0.233 | 3.67E-06 | 17 | Tspan12 |
| Igf2r3 | 1.88E-10 | 0.342937558 | 0.304 | 0.135 | 3.82E-06 | 17 | Igf2r |

| | | | | | | | |
|---|---|---|---|---|---|---|---|
| Eid12 | 1.90E-10 | 0.304871394 | 0.561 | 0.322 | 3.84E-06 | 17 | Eid1 |
| Zcchc9 | 1.91E-10 | 0.393089183 | 0.304 | 0.14 | 3.86E-06 | 17 | Zcchc9 |
| Zmym43 | 1.94E-10 | 0.418151171 | 0.345 | 0.166 | 3.92E-06 | 17 | Zmym4 |
| Ddb12 | 2.00E-10 | 0.391600662 | 0.322 | 0.149 | 4.04E-06 | 17 | Ddb1 |
| Ddx39b4 | 2.08E-10 | 0.34248107 | 0.444 | 0.234 | 4.21E-06 | 17 | Ddx39b |
| Ywhaq4 | 2.08E-10 | 0.354605561 | 0.731 | 0.489 | 4.22E-06 | 17 | Ywhaq |
| Smim12 | 2.10E-10 | 0.335423276 | 0.298 | 0.134 | 4.24E-06 | 17 | Smim12 |
| Prrc2b6 | 2.10E-10 | 0.380825009 | 0.444 | 0.23 | 4.26E-06 | 17 | Prrc2b |
| Rap1a4 | 2.13E-10 | 0.422416882 | 0.784 | 0.537 | 4.31E-06 | 17 | Rap1a |
| Bbx5 | 2.15E-10 | 0.362590749 | 0.427 | 0.224 | 4.35E-06 | 17 | Bbx |
| Tmed106 | 2.17E-10 | 0.341846935 | 0.702 | 0.437 | 4.38E-06 | 17 | Tmed10 |
| Emc34 | 2.18E-10 | 0.362577834 | 0.357 | 0.174 | 4.41E-06 | 17 | Emc3 |
| Sbf25 | 2.21E-10 | 0.345237372 | 0.421 | 0.208 | 4.48E-06 | 17 | Sbf2 |
| Dmxl15 | 2.40E-10 | 0.316410467 | 0.38 | 0.186 | 4.86E-06 | 17 | Dmxl1 |
| Asb33 | 2.41E-10 | 0.345085487 | 0.263 | 0.111 | 4.88E-06 | 17 | Asb3 |
| Ggta13 | 2.45E-10 | 0.45269452 | 0.31 | 0.141 | 4.96E-06 | 17 | Ggta1 |
| Add14 | 2.59E-10 | 0.321557843 | 0.421 | 0.215 | 5.24E-06 | 17 | Add1 |
| Eif4a21 | 2.61E-10 | 0.390073656 | 0.526 | 0.305 | 5.28E-06 | 17 | Eif4a2 |
| 1110038F14Rik | 2.62E-10 | 0.317720852 | 0.392 | 0.196 | 5.30E-06 | 17 | 1110038F14Rik |
| Ccdc88a8 | 2.70E-10 | 0.455440357 | 0.398 | 0.205 | 5.46E-06 | 17 | Ccdc88a |
| Bod1l6 | 2.81E-10 | 0.40844746 | 0.491 | 0.274 | 5.70E-06 | 17 | Bod1l |
| Dennd6a3 | 2.86E-10 | 0.310899937 | 0.374 | 0.183 | 5.79E-06 | 17 | Dennd6a |
| Erlec11 | 3.02E-10 | 0.434770077 | 0.322 | 0.152 | 6.10E-06 | 17 | Erlec1 |
| Ostc2 | 3.08E-10 | 0.390733501 | 0.474 | 0.263 | 6.24E-06 | 17 | Ostc |
| 5031425E22Rik4 | 3.08E-10 | 0.388798258 | 0.333 | 0.16 | 6.25E-06 | 17 | 5031425E22Rik |
| Mia24 | 3.22E-10 | 0.403323186 | 0.398 | 0.208 | 6.52E-06 | 17 | Mia2 |
| Zcchc710 | 3.34E-10 | 0.340204636 | 0.55 | 0.302 | 6.75E-06 | 17 | Zcchc7 |
| Zbtb7a4 | 3.41E-10 | 0.358088491 | 0.439 | 0.23 | 6.89E-06 | 17 | Zbtb7a |
| Yme1l13 | 3.41E-10 | 0.411354107 | 0.386 | 0.197 | 6.90E-06 | 17 | Yme1l1 |
| Fus6 | 3.52E-10 | 0.350134582 | 0.626 | 0.39 | 7.12E-06 | 17 | Fus |
| Snrnp483 | 3.54E-10 | 0.329896128 | 0.374 | 0.186 | 7.16E-06 | 17 | Snrnp48 |
| Bclaf17 | 3.70E-10 | 0.338941391 | 0.474 | 0.255 | 7.49E-06 | 17 | Bclaf1 |
| Tmem1403 | 3.74E-10 | 0.32956187 | 0.439 | 0.224 | 7.57E-06 | 17 | Tmem140 |
| Nae12 | 4.01E-10 | 0.324044339 | 0.333 | 0.157 | 8.12E-06 | 17 | Nae1 |
| Stk244 | 4.06E-10 | 0.445532033 | 0.38 | 0.197 | 8.21E-06 | 17 | Stk24 |
| Agpat34 | 4.10E-10 | 0.325078535 | 0.292 | 0.13 | 8.29E-06 | 17 | Agpat3 |
| Prrc2c10 | 4.15E-10 | 0.447795631 | 0.655 | 0.421 | 8.41E-06 | 17 | Prrc2c |
| Cops5 | 4.18E-10 | 0.258665268 | 0.357 | 0.17 | 8.47E-06 | 17 | Cops5 |
| Ggps13 | 4.53E-10 | 0.348327766 | 0.357 | 0.173 | 9.16E-06 | 17 | Ggps1 |
| Spata62 | 4.67E-10 | 0.372246889 | 0.275 | 0.121 | 9.46E-06 | 17 | Spata6 |
| Txndc153 | 4.82E-10 | 0.340101056 | 0.304 | 0.139 | 9.76E-06 | 17 | Txndc15 |
| Mecp22 | 4.95E-10 | 0.397382276 | 0.333 | 0.161 | 1.00E-05 | 17 | Mecp2 |
| Iah11 | 5.08E-10 | 0.278945675 | 0.304 | 0.137 | 1.03E-05 | 17 | Iah1 |

| | | | | | | | |
|---|---|---|---|---|---|---|---|
| Zfp2601 | 5.33E-10 | 0.349347838 | 0.257 | 0.109 | 1.08E-05 | 17 | Zfp260 |
| Wac8 | 5.89E-10 | 0.401873233 | 0.556 | 0.323 | 1.19E-05 | 17 | Wac |
| Frs21 | 6.01E-10 | 0.399603642 | 0.31 | 0.143 | 1.22E-05 | 17 | Frs2 |
| Yipf51 | 6.21E-10 | 0.397604606 | 0.292 | 0.133 | 1.26E-05 | 17 | Yipf5 |
| Psap3 | 6.50E-10 | 0.253647606 | 0.655 | 0.399 | 1.32E-05 | 17 | Psap |
| Csnk2a1 | 6.87E-10 | 0.400652465 | 0.38 | 0.194 | 1.39E-05 | 17 | Csnk2a1 |
| Ddx3x5 | 6.91E-10 | 0.319859264 | 0.538 | 0.309 | 1.40E-05 | 17 | Ddx3x |
| Jmjd1c9 | 6.91E-10 | 0.339697685 | 0.544 | 0.308 | 1.40E-05 | 17 | Jmjd1c |
| Appl14 | 7.11E-10 | 0.346374582 | 0.363 | 0.182 | 1.44E-05 | 17 | Appl1 |
| Fto4 | 7.12E-10 | 0.427610642 | 0.374 | 0.192 | 1.44E-05 | 17 | Fto |
| Ddx174 | 7.16E-10 | 0.375726779 | 0.491 | 0.274 | 1.45E-05 | 17 | Ddx17 |
| Nbr11 | 7.20E-10 | 0.354196409 | 0.31 | 0.141 | 1.46E-05 | 17 | Nbr1 |
| Gpatch89 | 7.58E-10 | 0.478499888 | 0.468 | 0.259 | 1.53E-05 | 17 | Gpatch8 |
| Slc23a23 | 7.97E-10 | 0.360873753 | 0.304 | 0.14 | 1.61E-05 | 17 | Slc23a2 |
| Dyrk1a5 | 8.74E-10 | 0.353637571 | 0.363 | 0.18 | 1.77E-05 | 17 | Dyrk1a |
| Kdm5b2 | 9.13E-10 | 0.383829768 | 0.287 | 0.131 | 1.85E-05 | 17 | Kdm5b |
| Mapkap1 | 9.17E-10 | 0.348176138 | 0.386 | 0.2 | 1.86E-05 | 17 | Mapkap1 |
| Ubr34 | 9.48E-10 | 0.366933052 | 0.392 | 0.202 | 1.92E-05 | 17 | Ubr3 |
| Srrm28 | 9.56E-10 | 0.401693901 | 0.719 | 0.47 | 1.94E-05 | 17 | Srrm2 |
| Mfhas13 | 9.92E-10 | 0.396145141 | 0.322 | 0.151 | 2.01E-05 | 17 | Mfhas1 |
| Txnip4 | 1.01E-09 | 0.430293495 | 0.801 | 0.546 | 2.05E-05 | 17 | Txnip |
| Tnpo13 | 1.02E-09 | 0.385143816 | 0.351 | 0.175 | 2.07E-05 | 17 | Tnpo1 |
| Nfat56 | 1.05E-09 | 0.514879601 | 0.474 | 0.268 | 2.13E-05 | 17 | Nfat5 |
| Dip2c5 | 1.07E-09 | 0.361452287 | 0.439 | 0.224 | 2.17E-05 | 17 | Dip2c |
| Ino80d4 | 1.09E-09 | 0.370820082 | 0.339 | 0.166 | 2.20E-05 | 17 | Ino80d |
| Creb3l22 | 1.11E-09 | 0.412515539 | 0.281 | 0.124 | 2.26E-05 | 17 | Creb3l2 |
| Rab11b3 | 1.12E-09 | 0.373833086 | 0.474 | 0.266 | 2.27E-05 | 17 | Rab11b |
| Unc504 | 1.27E-09 | 0.310598348 | 0.404 | 0.214 | 2.58E-05 | 17 | Unc50 |
| AW5549186 | 1.28E-09 | 0.508500609 | 0.339 | 0.172 | 2.60E-05 | 17 | AW554918 |
| Eif3a5 | 1.29E-09 | 0.416824122 | 0.591 | 0.373 | 2.61E-05 | 17 | Eif3a |
| Csnk1g17 | 1.35E-09 | 0.375082829 | 0.327 | 0.158 | 2.73E-05 | 17 | Csnk1g1 |
| Zeb18 | 1.42E-09 | 0.492410058 | 0.544 | 0.323 | 2.88E-05 | 17 | Zeb1 |
| Dctn42 | 1.47E-09 | 0.315873804 | 0.404 | 0.212 | 2.97E-05 | 17 | Dctn4 |
| Csnk1a12 | 1.48E-09 | 0.266501821 | 0.719 | 0.458 | 2.99E-05 | 17 | Csnk1a1 |
| Wbp11 | 1.49E-09 | 0.335602801 | 0.374 | 0.192 | 3.02E-05 | 17 | Wbp11 |
| Cers52 | 1.62E-09 | 0.408647027 | 0.292 | 0.135 | 3.28E-05 | 17 | Cers5 |
| Zufsp5 | 1.66E-09 | 0.415643616 | 0.292 | 0.137 | 3.36E-05 | 17 | Zufsp |
| Aaed11 | 1.67E-09 | 0.368820387 | 0.304 | 0.144 | 3.37E-05 | 17 | Aaed1 |
| Rell11 | 1.69E-09 | 0.3998443 | 0.298 | 0.138 | 3.42E-05 | 17 | Rell1 |
| Ebp1 | 1.70E-09 | 0.340276098 | 0.363 | 0.183 | 3.43E-05 | 17 | Ebp |
| Zfr6 | 1.70E-09 | 0.45649897 | 0.444 | 0.248 | 3.44E-05 | 17 | Zfr |
| Tpm45 | 2.00E-09 | 0.327939357 | 0.626 | 0.381 | 4.04E-05 | 17 | Tpm4 |
| Mtmr38 | 2.07E-09 | 0.409137781 | 0.386 | 0.201 | 4.19E-05 | 17 | Mtmr3 |

| | | | | | | | |
|---|---|---|---|---|---|---|---|
| Stx123 | 2.14E-09 | 0.356079054 | 0.333 | 0.163 | 4.34E-05 | 17 | Stx12 |
| Crk3 | 2.16E-09 | 0.330699382 | 0.38 | 0.194 | 4.36E-05 | 17 | Crk |
| Ywhag2 | 2.20E-09 | 0.362165292 | 0.363 | 0.182 | 4.45E-05 | 17 | Ywhag |
| Tmem596 | 2.33E-09 | 0.280135237 | 0.778 | 0.491 | 4.72E-05 | 17 | Tmem59 |
| Sec11a6 | 2.37E-09 | 0.4573269 | 0.485 | 0.286 | 4.80E-05 | 17 | Sec11a |
| Selenot4 | 2.61E-09 | 0.332807379 | 0.398 | 0.213 | 5.29E-05 | 17 | Selenot |
| Emc104 | 2.62E-09 | 0.327594 | 0.386 | 0.201 | 5.31E-05 | 17 | Emc10 |
| Cnbp3 | 2.65E-09 | 0.301182815 | 0.731 | 0.488 | 5.36E-05 | 17 | Cnbp |
| Cnpy21 | 2.75E-09 | 0.263113446 | 0.474 | 0.264 | 5.58E-05 | 17 | Cnpy2 |
| Ctdsp1 | 2.86E-09 | 0.294690176 | 0.275 | 0.124 | 5.79E-05 | 17 | Ctdsp1 |
| Stx81 | 2.86E-09 | 0.271425059 | 0.48 | 0.268 | 5.80E-05 | 17 | Stx8 |
| Stip1 | 2.88E-09 | 0.337302947 | 0.269 | 0.122 | 5.83E-05 | 17 | Stip1 |
| Eln4 | 2.98E-09 | 0.405115495 | 0.298 | 0.137 | 6.03E-05 | 17 | Eln |
| Iqsec13 | 3.26E-09 | 0.374654284 | 0.269 | 0.123 | 6.61E-05 | 17 | Iqsec1 |
| Amfr1 | 3.49E-09 | 0.304080731 | 0.275 | 0.125 | 7.06E-05 | 17 | Amfr |
| Abr6 | 3.58E-09 | 0.420484138 | 0.322 | 0.159 | 7.24E-05 | 17 | Abr |
| Slmap2 | 3.75E-09 | 0.276002364 | 0.38 | 0.194 | 7.59E-05 | 17 | Slmap |
| Srsf74 | 3.84E-09 | 0.337604412 | 0.556 | 0.333 | 7.77E-05 | 17 | Srsf7 |
| Thoc27 | 3.87E-09 | 0.435834179 | 0.474 | 0.275 | 7.83E-05 | 17 | Thoc2 |
| Tecr6 | 3.94E-09 | 0.377792955 | 0.409 | 0.226 | 7.98E-05 | 17 | Tecr |
| Taok15 | 3.95E-09 | 0.360722873 | 0.444 | 0.252 | 8.00E-05 | 17 | Taok1 |
| Copa2 | 4.13E-09 | 0.322694772 | 0.386 | 0.206 | 8.36E-05 | 17 | Copa |
| Cul51 | 4.51E-09 | 0.340500256 | 0.298 | 0.142 | 9.12E-05 | 17 | Cul5 |
| Trim352 | 4.61E-09 | 0.45117358 | 0.292 | 0.141 | 9.34E-05 | 17 | Trim35 |
| Dennd5b4 | 4.83E-09 | 0.380157226 | 0.316 | 0.148 | 9.77E-05 | 17 | Dennd5b |
| Ubn28 | 4.83E-09 | 0.449020729 | 0.439 | 0.245 | 9.77E-05 | 17 | Ubn2 |
| Stag23 | 4.90E-09 | 0.293938706 | 0.509 | 0.289 | 9.93E-05 | 17 | Stag2 |
| Cnot47 | 5.00E-09 | 0.372031733 | 0.462 | 0.27 | 0.000101312 | 17 | Cnot4 |
| Trip114 | 5.32E-09 | 0.326943891 | 0.415 | 0.225 | 0.000107634 | 17 | Trip11 |
| Nr3c24 | 5.43E-09 | 0.275557446 | 0.275 | 0.122 | 0.00011 | 17 | Nr3c2 |
| Tmem30a6 | 5.50E-09 | 0.36704193 | 0.491 | 0.287 | 0.000111318 | 17 | Tmem30a |
| Itgb1bp1 | 5.80E-09 | 0.285319518 | 0.257 | 0.115 | 0.000117429 | 17 | Itgb1bp1 |
| Tm9sf42 | 6.16E-09 | 0.422477187 | 0.281 | 0.133 | 0.000124784 | 17 | Tm9sf4 |
| Ppp3ca10 | 6.24E-09 | 0.464035659 | 0.632 | 0.394 | 0.000126254 | 17 | Ppp3ca |
| Dlgap4 | 6.58E-09 | 0.288994561 | 0.292 | 0.137 | 0.000133146 | 17 | Dlgap4 |
| Ccar16 | 7.02E-09 | 0.322886166 | 0.392 | 0.209 | 0.000142049 | 17 | Ccar1 |
| Zfp1487 | 7.23E-09 | 0.361947762 | 0.462 | 0.266 | 0.000146297 | 17 | Zfp148 |
| Wwox6 | 7.25E-09 | 0.437568474 | 0.345 | 0.177 | 0.000146869 | 17 | Wwox |
| Tagln27 | 7.52E-09 | 0.362111315 | 0.673 | 0.43 | 0.000152332 | 17 | Tagln2 |
| Hnrnpm | 7.68E-09 | 0.255975672 | 0.567 | 0.34 | 0.000155515 | 17 | Hnrnpm |
| Plod12 | 7.90E-09 | 0.379989391 | 0.269 | 0.122 | 0.000159994 | 17 | Plod1 |
| Ap2m1 | 8.30E-09 | 0.273107663 | 0.363 | 0.185 | 0.000168133 | 17 | Ap2m1 |
| Pum16 | 8.37E-09 | 0.407427808 | 0.45 | 0.26 | 0.000169512 | 17 | Pum1 |

| | | | | | | | |
|---|---|---|---|---|---|---|---|
| Zzz3 | 8.60E-09 | 0.29882787 | 0.31 | 0.153 | 0.000174064 | 17 | Zzz3 |
| Ccdc825 | 8.66E-09 | 0.402274181 | 0.345 | 0.182 | 0.000175331 | 17 | Ccdc82 |
| Frmd4b5 | 8.80E-09 | 0.609008209 | 0.339 | 0.178 | 0.000178265 | 17 | Frmd4b |
| Sf3b17 | 8.91E-09 | 0.285747264 | 0.626 | 0.394 | 0.000180365 | 17 | Sf3b1 |
| Lmbrd14 | 8.93E-09 | 0.477079094 | 0.322 | 0.163 | 0.000180854 | 17 | Lmbrd1 |
| Acap26 | 8.99E-09 | 0.336714688 | 0.474 | 0.27 | 0.000182092 | 17 | Acap2 |
| Atxn104 | 1.06E-08 | 0.296137032 | 0.339 | 0.176 | 0.000215027 | 17 | Atxn10 |
| Nsmce24 | 1.08E-08 | 0.4143086 | 0.38 | 0.208 | 0.000218416 | 17 | Nsmce2 |
| Psme43 | 1.16E-08 | 0.304816884 | 0.374 | 0.198 | 0.000234259 | 17 | Psme4 |
| Prrc2a2 | 1.17E-08 | 0.283590294 | 0.31 | 0.152 | 0.000236319 | 17 | Prrc2a |
| Fbxl5 | 1.23E-08 | 0.291454381 | 0.251 | 0.114 | 0.00024869 | 17 | Fbxl5 |
| Usp9x1 | 1.34E-08 | 0.395371255 | 0.421 | 0.236 | 0.000270466 | 17 | Usp9x |
| Cpsf63 | 1.35E-08 | 0.314859095 | 0.31 | 0.152 | 0.000274321 | 17 | Cpsf6 |
| Hnrnph2 | 1.36E-08 | 0.302007429 | 0.333 | 0.17 | 0.00027452 | 17 | Hnrnph2 |
| Lima14 | 1.39E-08 | 0.360390595 | 0.322 | 0.159 | 0.000280535 | 17 | Lima1 |
| Zfp911 | 1.39E-08 | 0.277413419 | 0.509 | 0.308 | 0.000282028 | 17 | Zfp91 |
| Trim443 | 1.40E-08 | 0.428246079 | 0.409 | 0.228 | 0.000282893 | 17 | Trim44 |
| Kmt2a10 | 1.45E-08 | 0.351181356 | 0.468 | 0.268 | 0.000292565 | 17 | Kmt2a |
| Pced1b4 | 1.47E-08 | 0.406785438 | 0.257 | 0.119 | 0.000297482 | 17 | Pced1b |
| Fam208a3 | 1.47E-08 | 0.318887068 | 0.38 | 0.201 | 0.000297998 | 17 | Fam208a |
| Lars | 1.48E-08 | 0.344046584 | 0.316 | 0.159 | 0.000298816 | 17 | Lars |
| Usp349 | 1.49E-08 | 0.339949413 | 0.556 | 0.345 | 0.000301529 | 17 | Usp34 |
| Pou2f12 | 1.51E-08 | 0.396501117 | 0.269 | 0.127 | 0.000305015 | 17 | Pou2f1 |
| Ube2w1 | 1.55E-08 | 0.28000797 | 0.339 | 0.172 | 0.000314236 | 17 | Ube2w |
| Atp2c14 | 1.67E-08 | 0.37987482 | 0.327 | 0.168 | 0.000338636 | 17 | Atp2c1 |
| Dctn6 | 1.74E-08 | 0.276853149 | 0.345 | 0.179 | 0.000351838 | 17 | Dctn6 |
| Atp2b17 | 1.75E-08 | 0.38964649 | 0.544 | 0.334 | 0.000354242 | 17 | Atp2b1 |
| Atf22 | 1.82E-08 | 0.308072423 | 0.322 | 0.159 | 0.000367883 | 17 | Atf2 |
| Dis3l26 | 1.89E-08 | 0.328018183 | 0.292 | 0.14 | 0.000383337 | 17 | Dis3l2 |
| Camk2g5 | 1.93E-08 | 0.40106214 | 0.281 | 0.136 | 0.00038993 | 17 | Camk2g |
| Brd81 | 2.16E-08 | 0.284094019 | 0.327 | 0.165 | 0.000436848 | 17 | Brd8 |
| Atl32 | 2.17E-08 | 0.388830883 | 0.269 | 0.129 | 0.000438853 | 17 | Atl3 |
| Chm1 | 2.23E-08 | 0.290615057 | 0.257 | 0.118 | 0.00045094 | 17 | Chm |
| Prpf39 | 2.26E-08 | 0.357681198 | 0.269 | 0.127 | 0.000457427 | 17 | Prpf39 |
| Rbm332 | 2.27E-08 | 0.329477073 | 0.292 | 0.143 | 0.000459554 | 17 | Rbm33 |
| Nf15 | 2.36E-08 | 0.409525959 | 0.398 | 0.222 | 0.000477577 | 17 | Nf1 |
| Gm472837 | 2.37E-08 | 0.445061871 | 0.48 | 0.283 | 0.000480312 | 17 | Gm47283 |
| Ddx24 | 2.43E-08 | 0.320577538 | 0.532 | 0.325 | 0.000491954 | 17 | Ddx24 |
| Arid4a9 | 2.52E-08 | 0.367822344 | 0.444 | 0.253 | 0.000511119 | 17 | Arid4a |
| Nsd18 | 2.54E-08 | 0.339531793 | 0.579 | 0.35 | 0.000513809 | 17 | Nsd1 |
| Litaf5 | 2.58E-08 | 0.316900073 | 0.316 | 0.159 | 0.000521939 | 17 | Litaf |
| Lonp21 | 2.63E-08 | 0.277534281 | 0.368 | 0.195 | 0.000532251 | 17 | Lonp2 |
| Golim44 | 2.64E-08 | 0.350729119 | 0.351 | 0.182 | 0.0005352 | 17 | Golim4 |

| | | | | | | | |
|---|---|---|---|---|---|---|---|
| Tpgs1 | 2.67E-08 | 0.261081472 | 0.257 | 0.117 | 0.000541144 | 17 | Tpgs1 |
| Ppil4 | 2.72E-08 | 0.363928101 | 0.327 | 0.168 | 0.000550054 | 17 | Ppil4 |
| Epc14 | 2.79E-08 | 0.408892087 | 0.374 | 0.204 | 0.000564758 | 17 | Epc1 |
| Hsp90b18 | 2.86E-08 | 0.424861591 | 0.731 | 0.511 | 0.000579397 | 17 | Hsp90b1 |
| Ccdc503 | 2.93E-08 | 0.30381323 | 0.409 | 0.23 | 0.000592407 | 17 | Ccdc50 |
| Mob21 | 3.13E-08 | 0.389697954 | 0.298 | 0.148 | 0.000633735 | 17 | Mob2 |
| Rab8b6 | 3.22E-08 | 0.419108944 | 0.48 | 0.285 | 0.000652792 | 17 | Rab8b |
| Cdkn1b3 | 3.25E-08 | 0.296967641 | 0.392 | 0.216 | 0.00065762 | 17 | Cdkn1b |
| Ulk23 | 3.48E-08 | 0.422878173 | 0.339 | 0.18 | 0.000705048 | 17 | Ulk2 |
| Sfpq6 | 3.75E-08 | 0.350678012 | 0.497 | 0.299 | 0.000758466 | 17 | Sfpq |
| Rmnd5a1 | 3.87E-08 | 0.33332596 | 0.281 | 0.139 | 0.00078318 | 17 | Rmnd5a |
| Sf3b24 | 4.07E-08 | 0.312141177 | 0.567 | 0.36 | 0.00082343 | 17 | Sf3b2 |
| Ubxn46 | 4.10E-08 | 0.316722358 | 0.462 | 0.273 | 0.000830244 | 17 | Ubxn4 |
| Smap14 | 4.20E-08 | 0.269168678 | 0.585 | 0.362 | 0.00085019 | 17 | Smap1 |
| Drg1 | 4.50E-08 | 0.326735472 | 0.333 | 0.177 | 0.000911422 | 17 | Drg1 |
| Ppp2r5a3 | 4.60E-08 | 0.333150411 | 0.427 | 0.252 | 0.000931483 | 17 | Ppp2r5a |
| Faf16 | 4.62E-08 | 0.321156502 | 0.368 | 0.203 | 0.000934857 | 17 | Faf1 |
| Pi4ka5 | 4.65E-08 | 0.310421742 | 0.269 | 0.129 | 0.000940713 | 17 | Pi4ka |
| Gbf16 | 4.73E-08 | 0.378154163 | 0.345 | 0.183 | 0.000956992 | 17 | Gbf1 |
| Plekha56 | 4.78E-08 | 0.294192682 | 0.31 | 0.156 | 0.00096817 | 17 | Plekha5 |
| Phc33 | 4.84E-08 | 0.298511601 | 0.281 | 0.136 | 0.000979807 | 17 | Phc3 |
| Psmd9 | 4.85E-08 | 0.292269436 | 0.269 | 0.131 | 0.000981424 | 17 | Psmd9 |
| Tom1l23 | 4.89E-08 | 0.350923449 | 0.275 | 0.132 | 0.000989751 | 17 | Tom1l2 |
| Notch14 | 4.91E-08 | 0.316947536 | 0.398 | 0.218 | 0.000993424 | 17 | Notch1 |
| Acyp21 | 4.98E-08 | 0.30943508 | 0.298 | 0.144 | 0.001007604 | 17 | Acyp2 |
| Rab101 | 5.03E-08 | 0.310183966 | 0.556 | 0.352 | 0.001018137 | 17 | Rab10 |
| Dctn21 | 5.10E-08 | 0.279267051 | 0.45 | 0.266 | 0.001032328 | 17 | Dctn2 |
| Map4k53 | 5.17E-08 | 0.285816089 | 0.275 | 0.132 | 0.001046398 | 17 | Map4k5 |
| Hnrnpk | 5.20E-08 | 0.258124907 | 0.713 | 0.496 | 0.001052393 | 17 | Hnrnpk |
| Ankrd446 | 5.45E-08 | 0.310305905 | 0.509 | 0.295 | 0.001103595 | 17 | Ankrd44 |
| Arl31 | 5.46E-08 | 0.260985786 | 0.327 | 0.171 | 0.001104691 | 17 | Arl3 |
| Brd49 | 5.48E-08 | 0.434182437 | 0.433 | 0.269 | 0.001109295 | 17 | Brd4 |
| Gpbp1l1 | 5.51E-08 | 0.264001988 | 0.339 | 0.179 | 0.001114914 | 17 | Gpbp1l1 |
| Cdc37l11 | 5.63E-08 | 0.265637308 | 0.327 | 0.169 | 0.00114052 | 17 | Cdc37l1 |
| Cbfa2t21 | 5.87E-08 | 0.343813806 | 0.251 | 0.118 | 0.001188997 | 17 | Cbfa2t2 |
| Smc65 | 5.87E-08 | 0.291026484 | 0.45 | 0.263 | 0.001189138 | 17 | Smc6 |
| Phip9 | 6.09E-08 | 0.340597558 | 0.439 | 0.257 | 0.001232899 | 17 | Phip |
| Pcp4l16 | 6.23E-08 | 0.350150017 | 0.515 | 0.314 | 0.00126164 | 17 | Pcp4l1 |
| Copb12 | 6.40E-08 | 0.307477329 | 0.368 | 0.205 | 0.001296212 | 17 | Copb1 |
| Bsg6 | 6.53E-08 | 0.270305231 | 0.684 | 0.45 | 0.001321435 | 17 | Bsg |
| Akt39 | 6.62E-08 | 0.334604224 | 0.45 | 0.252 | 0.001341084 | 17 | Akt3 |
| Ppp3cb1 | 7.14E-08 | 0.350231538 | 0.322 | 0.171 | 0.001446021 | 17 | Ppp3cb |
| Ppp2r5e3 | 7.17E-08 | 0.288398481 | 0.322 | 0.166 | 0.001451334 | 17 | Ppp2r5e |

| | | | | | | | |
|---|---|---|---|---|---|---|---|
| Caml1 | 7.23E-08 | 0.301877045 | 0.275 | 0.134 | 0.001463186 | 17 | Caml |
| Prpf6 | 8.25E-08 | 0.30276352 | 0.275 | 0.135 | 0.001669634 | 17 | Prpf6 |
| Hdac11 | 8.34E-08 | 0.326862311 | 0.322 | 0.168 | 0.001688327 | 17 | Hdac1 |
| Hook35 | 8.55E-08 | 0.360732112 | 0.485 | 0.291 | 0.001731973 | 17 | Hook3 |
| Kansl1l3 | 8.86E-08 | 0.355565916 | 0.281 | 0.14 | 0.001794609 | 17 | Kansl1l |
| Rabgap13 | 8.87E-08 | 0.453478336 | 0.31 | 0.164 | 0.001796523 | 17 | Rabgap1 |
| Sf13 | 9.31E-08 | 0.274779764 | 0.345 | 0.189 | 0.001885424 | 17 | Sf1 |
| Cebpb3 | 9.54E-08 | 0.332901175 | 0.304 | 0.158 | 0.001932053 | 17 | Cebpb |
| Tm2d1 | 9.78E-08 | 0.300030408 | 0.357 | 0.196 | 0.001979622 | 17 | Tm2d1 |
| Canx7 | 9.81E-08 | 0.289217219 | 0.497 | 0.308 | 0.001985532 | 17 | Canx |
| Mapk18 | 1.00E-07 | 0.36299372 | 0.404 | 0.237 | 0.002028717 | 17 | Mapk1 |
| Derl11 | 1.01E-07 | 0.269808668 | 0.31 | 0.158 | 0.002038791 | 17 | Derl1 |
| Golgb15 | 1.05E-07 | 0.358796569 | 0.444 | 0.264 | 0.002124928 | 17 | Golgb1 |
| Phf201 | 1.12E-07 | 0.390583911 | 0.287 | 0.145 | 0.002271444 | 17 | Phf20 |
| Bdp13 | 1.24E-07 | 0.286483332 | 0.322 | 0.167 | 0.002518771 | 17 | Bdp1 |
| Xpa2 | 1.25E-07 | 0.324274943 | 0.257 | 0.124 | 0.002536711 | 17 | Xpa |
| Mdm44 | 1.30E-07 | 0.307046561 | 0.31 | 0.161 | 0.002634569 | 17 | Mdm4 |
| Rnf2163 | 1.33E-07 | 0.279539555 | 0.339 | 0.181 | 0.002690959 | 17 | Rnf216 |
| Lrch38 | 1.37E-07 | 0.302074015 | 0.409 | 0.234 | 0.002776806 | 17 | Lrch3 |
| Zfp280d2 | 1.49E-07 | 0.357036648 | 0.339 | 0.185 | 0.003024763 | 17 | Zfp280d |
| Baz2a1 | 1.51E-07 | 0.359407385 | 0.263 | 0.133 | 0.003063281 | 17 | Baz2a |
| Dgkd5 | 1.52E-07 | 0.289450535 | 0.31 | 0.162 | 0.003074573 | 17 | Dgkd |
| Emc41 | 1.68E-07 | 0.304419937 | 0.263 | 0.128 | 0.003408632 | 17 | Emc4 |
| Elavl1 | 1.72E-07 | 0.330308722 | 0.333 | 0.18 | 0.003472974 | 17 | Elavl1 |
| Wapl2 | 1.80E-07 | 0.295455569 | 0.421 | 0.252 | 0.003651559 | 17 | Wapl |
| Papd41 | 1.81E-07 | 0.257067371 | 0.281 | 0.139 | 0.003667 | 17 | Papd4 |
| Acaa1a | 1.85E-07 | 0.260570505 | 0.322 | 0.17 | 0.003749417 | 17 | Acaa1a |
| Paip21 | 1.87E-07 | 0.25276545 | 0.602 | 0.407 | 0.003780988 | 17 | Paip2 |
| Phf39 | 1.95E-07 | 0.317313 | 0.427 | 0.254 | 0.003943807 | 17 | Phf3 |
| Sp12 | 2.02E-07 | 0.275751813 | 0.287 | 0.143 | 0.004097209 | 17 | Sp1 |
| Tnrc185 | 2.21E-07 | 0.41862042 | 0.316 | 0.174 | 0.004464864 | 17 | Tnrc18 |
| Ralb1 | 2.21E-07 | 0.258152229 | 0.304 | 0.155 | 0.004470776 | 17 | Ralb |
| Anxa64 | 2.31E-07 | 0.418896604 | 0.298 | 0.158 | 0.004672574 | 17 | Anxa6 |
| Bptf10 | 2.32E-07 | 0.255971663 | 0.491 | 0.295 | 0.004696625 | 17 | Bptf |
| Degs14 | 2.33E-07 | 0.339082174 | 0.339 | 0.186 | 0.004717945 | 17 | Degs1 |
| Ahctf12 | 2.58E-07 | 0.384239193 | 0.251 | 0.125 | 0.005229301 | 17 | Ahctf1 |
| Pam7 | 2.65E-07 | 0.329050319 | 0.38 | 0.215 | 0.005369325 | 17 | Pam |
| Smarca5 | 3.25E-07 | 0.277325808 | 0.491 | 0.303 | 0.006576815 | 17 | Smarca5 |
| Tmed97 | 3.39E-07 | 0.262843669 | 0.614 | 0.404 | 0.006857748 | 17 | Tmed9 |
| Brd92 | 3.45E-07 | 0.269891627 | 0.322 | 0.174 | 0.006991382 | 17 | Brd9 |
| Secisbp2l1 | 3.51E-07 | 0.291069368 | 0.281 | 0.144 | 0.007105379 | 17 | Secisbp2l |
| Selenos2 | 3.63E-07 | 0.281459198 | 0.404 | 0.233 | 0.007345048 | 17 | Selenos |
| Tmem176a7 | 3.82E-07 | 0.283311752 | 0.667 | 0.418 | 0.007733081 | 17 | Tmem176a |

| | | | | | | | |
|---|---|---|---|---|---|---|---|
| Stx162 | 4.09E-07 | 0.390905004 | 0.298 | 0.162 | 0.008280616 | 17 | Stx16 |
| Dazap1 | 4.10E-07 | 0.335809725 | 0.298 | 0.159 | 0.008297282 | 17 | Dazap1 |
| Eif5b3 | 4.12E-07 | 0.317154771 | 0.509 | 0.322 | 0.008338739 | 17 | Eif5b |
| Rab5b | 4.14E-07 | 0.270079265 | 0.275 | 0.14 | 0.008375723 | 17 | Rab5b |
| Map44 | 4.54E-07 | 0.269715851 | 0.491 | 0.305 | 0.009196844 | 17 | Map4 |
| Mbd58 | 4.84E-07 | 0.341459663 | 0.38 | 0.221 | 0.0097921 | 17 | Mbd5 |
| Specc1l3 | 5.01E-07 | 0.312488251 | 0.257 | 0.13 | 0.010139383 | 17 | Specc1l |
| Tmem2191 | 5.44E-07 | 0.251044543 | 0.251 | 0.124 | 0.011007044 | 17 | Tmem219 |
| Ewsr1 | 5.56E-07 | 0.272647948 | 0.363 | 0.207 | 0.011253011 | 17 | Ewsr1 |
| Kdm2a8 | 5.69E-07 | 0.305708422 | 0.404 | 0.242 | 0.011527662 | 17 | Kdm2a |
| Huwe19 | 5.73E-07 | 0.421259948 | 0.456 | 0.286 | 0.011604734 | 17 | Huwe1 |
| Anapc11 | 6.15E-07 | 0.287771629 | 0.251 | 0.125 | 0.01244686 | 17 | Anapc1 |
| Zkscan3 | 6.15E-07 | 0.360153615 | 0.251 | 0.126 | 0.012457193 | 17 | Zkscan3 |
| Orc41 | 6.47E-07 | 0.281763544 | 0.251 | 0.126 | 0.013097505 | 17 | Orc4 |
| Tnks27 | 6.56E-07 | 0.313369813 | 0.456 | 0.281 | 0.013273211 | 17 | Tnks2 |
| Trove24 | 6.58E-07 | 0.306099572 | 0.263 | 0.133 | 0.0133266 | 17 | Trove2 |
| Nrf14 | 7.00E-07 | 0.250717971 | 0.269 | 0.136 | 0.01416913 | 17 | Nrf1 |
| Hnrnpa32 | 7.02E-07 | 0.274826344 | 0.643 | 0.446 | 0.014222132 | 17 | Hnrnpa3 |
| Eif4h | 7.29E-07 | 0.313495239 | 0.468 | 0.29 | 0.014751044 | 17 | Eif4h |
| Vps13c3 | 7.47E-07 | 0.354661845 | 0.269 | 0.141 | 0.015119032 | 17 | Vps13c |
| Smg74 | 7.47E-07 | 0.325134111 | 0.292 | 0.157 | 0.015125666 | 17 | Smg7 |
| Tmed72 | 8.31E-07 | 0.262142394 | 0.287 | 0.15 | 0.016816249 | 17 | Tmed7 |
| 9930111J21Rik25 | 8.50E-07 | 0.413770272 | 0.421 | 0.258 | 0.017201093 | 17 | 9930111J21Rik2 |
| Braf3 | 9.82E-07 | 0.299876167 | 0.298 | 0.16 | 0.019871938 | 17 | Braf |
| Rprd21 | 1.01E-06 | 0.395856547 | 0.257 | 0.136 | 0.02049588 | 17 | Rprd2 |
| Dlg12 | 1.02E-06 | 0.31713076 | 0.363 | 0.213 | 0.020589239 | 17 | Dlg1 |
| Ccdc1075 | 1.07E-06 | 0.258484815 | 0.251 | 0.129 | 0.021672423 | 17 | Ccdc107 |
| Nlk4 | 1.10E-06 | 0.383225227 | 0.275 | 0.146 | 0.022178101 | 17 | Nlk |
| Nudcd36 | 1.11E-06 | 0.325606825 | 0.404 | 0.248 | 0.022464901 | 17 | Nudcd3 |
| Stt3b6 | 1.12E-06 | 0.358959013 | 0.304 | 0.166 | 0.022597233 | 17 | Stt3b |
| Itgav7 | 1.12E-06 | 0.361796697 | 0.298 | 0.162 | 0.022760318 | 17 | Itgav |
| Ankhd17 | 1.12E-06 | 0.316304563 | 0.415 | 0.245 | 0.022771463 | 17 | Ankhd1 |
| Yipf31 | 1.16E-06 | 0.250509277 | 0.281 | 0.146 | 0.023485323 | 17 | Yipf3 |
| H132 | 1.17E-06 | 0.35148566 | 0.287 | 0.158 | 0.023746531 | 17 | H13 |
| Ctdspl21 | 1.21E-06 | 0.25394577 | 0.316 | 0.173 | 0.024544841 | 17 | Ctdspl2 |
| Ino802 | 1.26E-06 | 0.284587634 | 0.269 | 0.14 | 0.025525101 | 17 | Ino80 |
| Cdc734 | 1.28E-06 | 0.338846186 | 0.304 | 0.171 | 0.025837183 | 17 | Cdc73 |
| Ncoa18 | 1.28E-06 | 0.312093041 | 0.363 | 0.211 | 0.025880695 | 17 | Ncoa1 |
| Cul12 | 1.37E-06 | 0.354534575 | 0.439 | 0.278 | 0.027774803 | 17 | Cul1 |
| Numa14 | 1.56E-06 | 0.26024358 | 0.345 | 0.196 | 0.031551798 | 17 | Numa1 |
| Irf2bp22 | 1.58E-06 | 0.291222266 | 0.392 | 0.23 | 0.031908464 | 17 | Irf2bp2 |
| Sdf23 | 1.58E-06 | 0.309583033 | 0.31 | 0.172 | 0.031931513 | 17 | Sdf2 |
| Dctn12 | 1.60E-06 | 0.298170189 | 0.251 | 0.133 | 0.032418993 | 17 | Dctn1 |

| | | | | | | | |
|---|---|---|---|---|---|---|---|
| Arnt3 | 1.62E-06 | 0.304470359 | 0.275 | 0.143 | 0.032852563 | 17 | Arnt |
| Dnajc112 | 1.70E-06 | 0.364140571 | 0.497 | 0.324 | 0.03445627 | 17 | Dnajc1 |
| Map3k34 | 1.82E-06 | 0.333960297 | 0.316 | 0.179 | 0.036832905 | 17 | Map3k3 |
| Rbm2510 | 1.84E-06 | 0.35891553 | 0.509 | 0.335 | 0.037227327 | 17 | Rbm25 |
| Arih17 | 1.95E-06 | 0.324605656 | 0.503 | 0.314 | 0.039564968 | 17 | Arih1 |
| Prpf40a8 | 2.01E-06 | 0.286884062 | 0.433 | 0.268 | 0.040610983 | 17 | Prpf40a |
| Wdr265 | 2.29E-06 | 0.282443786 | 0.357 | 0.21 | 0.046452332 | 17 | Wdr26 |
| Rlf5 | 2.34E-06 | 0.260916244 | 0.263 | 0.138 | 0.047457568 | 17 | Rlf |
| Dpm11 | 2.37E-06 | 0.270202754 | 0.269 | 0.144 | 0.047963524 | 17 | Dpm1 |
| Add32 | 2.55E-06 | 0.319787961 | 0.374 | 0.224 | 0.051570561 | 17 | Add3 |
| Rpn25 | 2.75E-06 | 0.35091091 | 0.357 | 0.213 | 0.055638381 | 17 | Rpn2 |
| Tbc1d58 | 2.97E-06 | 0.401020112 | 0.421 | 0.263 | 0.060127596 | 17 | Tbc1d5 |
| Ogt7 | 2.98E-06 | 0.44143751 | 0.339 | 0.208 | 0.060257245 | 17 | Ogt |
| Map4k35 | 3.14E-06 | 0.255968238 | 0.392 | 0.231 | 0.063632345 | 17 | Map4k3 |
| Upf3b1 | 3.18E-06 | 0.261470716 | 0.263 | 0.139 | 0.064299148 | 17 | Upf3b |
| Ep3006 | 3.45E-06 | 0.262996521 | 0.351 | 0.205 | 0.069783269 | 17 | Ep300 |
| Fbxl20 | 3.48E-06 | 0.258030076 | 0.251 | 0.13 | 0.070355623 | 17 | Fbxl20 |
| Ppig5 | 3.75E-06 | 0.253317538 | 0.48 | 0.317 | 0.075936728 | 17 | Ppig |
| Ssr12 | 3.97E-06 | 0.291135238 | 0.322 | 0.188 | 0.080401768 | 17 | Ssr1 |
| Sf3b36 | 4.36E-06 | 0.309284435 | 0.287 | 0.162 | 0.088306584 | 17 | Sf3b3 |
| AI3141804 | 4.55E-06 | 0.313460585 | 0.357 | 0.212 | 0.092147891 | 17 | AI314180 |
| Xrn15 | 4.70E-06 | 0.304643548 | 0.351 | 0.205 | 0.095092404 | 17 | Xrn1 |
| Larp16 | 4.78E-06 | 0.257894182 | 0.345 | 0.204 | 0.096846077 | 17 | Larp1 |
| Arid4b7 | 4.90E-06 | 0.274065369 | 0.456 | 0.294 | 0.099124021 | 17 | Arid4b |
| Luc7l33 | 5.14E-06 | 0.321415574 | 0.421 | 0.272 | 0.104138128 | 17 | Luc7l3 |
| Sri5 | 5.74E-06 | 0.288996288 | 0.602 | 0.422 | 0.116307079 | 17 | Sri |
| Smg67 | 5.77E-06 | 0.416381123 | 0.409 | 0.261 | 0.116746575 | 17 | Smg6 |
| Akap81 | 5.88E-06 | 0.264779951 | 0.281 | 0.157 | 0.119062847 | 17 | Akap8 |
| Akap99 | 6.02E-06 | 0.360386587 | 0.38 | 0.239 | 0.121954532 | 17 | Akap9 |
| Calr6 | 6.73E-06 | 0.303382106 | 0.602 | 0.409 | 0.136326744 | 17 | Calr |
| Ttc39b6 | 6.88E-06 | 0.334706086 | 0.251 | 0.137 | 0.139225247 | 17 | Ttc39b |
| Trip127 | 7.24E-06 | 0.261805066 | 0.409 | 0.258 | 0.146677898 | 17 | Trip12 |
| Mesd3 | 7.60E-06 | 0.255907775 | 0.292 | 0.164 | 0.153798494 | 17 | Mesd |
| Usp472 | 7.74E-06 | 0.259380303 | 0.322 | 0.189 | 0.156683064 | 17 | Usp47 |
| Nek76 | 7.76E-06 | 0.254941472 | 0.404 | 0.25 | 0.157201401 | 17 | Nek7 |
| Kmt2c9 | 7.83E-06 | 0.315984936 | 0.462 | 0.308 | 0.158461649 | 17 | Kmt2c |
| Senp76 | 7.96E-06 | 0.319540623 | 0.281 | 0.158 | 0.161141874 | 17 | Senp7 |
| Clk17 | 8.44E-06 | 0.296976696 | 0.45 | 0.293 | 0.170965479 | 17 | Clk1 |
| Zscan263 | 8.69E-06 | 0.250004943 | 0.269 | 0.146 | 0.176019617 | 17 | Zscan26 |
| Pde7b5 | 9.41E-06 | 0.554676908 | 0.374 | 0.244 | 0.190528968 | 17 | Pde7b |
| Hnrnpdl2 | 9.91E-06 | 0.250584547 | 0.398 | 0.25 | 0.200630236 | 17 | Hnrnpdl |
| Taf154 | 1.07E-05 | 0.253640549 | 0.38 | 0.239 | 0.216320024 | 17 | Taf15 |
| Sfswap1 | 1.28E-05 | 0.260794125 | 0.251 | 0.136 | 0.258453631 | 17 | Sfswap |

| | | | | | | | |
|---|---|---|---|---|---|---|---|
| Ap1s22 | 1.29E-05 | 0.379059162 | 0.251 | 0.14 | 0.261928287 | 17 | Ap1s2 |
| Sorbs24 | 1.30E-05 | 0.344057134 | 0.45 | 0.279 | 0.264092839 | 17 | Sorbs2 |
| Zmynd115 | 1.31E-05 | 0.266364306 | 0.444 | 0.291 | 0.265445471 | 17 | Zmynd11 |
| Fam172a9 | 1.43E-05 | 0.393152855 | 0.433 | 0.286 | 0.289605088 | 17 | Fam172a |
| Plec5 | 1.50E-05 | 0.268265603 | 0.304 | 0.175 | 0.3033091 | 17 | Plec |
| Hmg20a3 | 1.50E-05 | 0.274790448 | 0.257 | 0.143 | 0.304630506 | 17 | Hmg20a |
| Diaph28 | 1.58E-05 | 0.281819817 | 0.556 | 0.378 | 0.319342865 | 17 | Diaph2 |
| Rb16 | 1.59E-05 | 0.303409824 | 0.351 | 0.215 | 0.321993529 | 17 | Rb1 |
| Kdm5a5 | 1.64E-05 | 0.28759178 | 0.292 | 0.169 | 0.331889107 | 17 | Kdm5a |
| Tusc33 | 1.68E-05 | 0.299115733 | 0.269 | 0.15 | 0.339729772 | 17 | Tusc3 |
| Taf34 | 1.74E-05 | 0.263478446 | 0.263 | 0.148 | 0.352290856 | 17 | Taf3 |
| Terf11 | 1.79E-05 | 0.277282423 | 0.257 | 0.142 | 0.362862663 | 17 | Terf1 |
| Ndfip25 | 1.81E-05 | 0.252822856 | 0.392 | 0.246 | 0.366126804 | 17 | Ndfip2 |
| Tnrc6a1 | 1.83E-05 | 0.289676607 | 0.327 | 0.201 | 0.371260736 | 17 | Tnrc6a |
| Ankib15 | 1.88E-05 | 0.315879479 | 0.251 | 0.14 | 0.380389989 | 17 | Ankib1 |
| Mark34 | 1.96E-05 | 0.295984739 | 0.345 | 0.213 | 0.397582281 | 17 | Mark3 |
| Ankrd1210 | 2.21E-05 | 0.278623485 | 0.468 | 0.309 | 0.448330193 | 17 | Ankrd12 |
| Rnf1697 | 2.30E-05 | 0.31986341 | 0.31 | 0.184 | 0.464926964 | 17 | Rnf169 |
| Fam168a2 | 2.58E-05 | 0.265070691 | 0.257 | 0.144 | 0.523027669 | 17 | Fam168a |
| Raf1 | 2.61E-05 | 0.260548568 | 0.257 | 0.145 | 0.527827365 | 17 | Raf1 |
| Hectd17 | 2.87E-05 | 0.271863018 | 0.45 | 0.302 | 0.581726214 | 17 | Hectd1 |
| Ier58 | 3.99E-05 | 0.257717482 | 0.48 | 0.319 | 0.808796151 | 17 | Ier5 |
| Klf68 | 4.00E-05 | 0.28105973 | 0.409 | 0.269 | 0.8098245 | 17 | Klf6 |
| Ythdf35 | 4.02E-05 | 0.27968728 | 0.322 | 0.197 | 0.814123911 | 17 | Ythdf3 |
| Pias13 | 4.30E-05 | 0.306137844 | 0.31 | 0.19 | 0.871347097 | 17 | Pias1 |
| Herc24 | 4.97E-05 | 0.261882606 | 0.333 | 0.208 | 1 | 17 | Herc2 |
| Asxl25 | 5.09E-05 | 0.275724551 | 0.292 | 0.176 | 1 | 17 | Asxl2 |
| Birc610 | 5.31E-05 | 0.254959546 | 0.497 | 0.342 | 1 | 17 | Birc6 |
| Pcnx4 | 5.41E-05 | 0.295084141 | 0.298 | 0.18 | 1 | 17 | Pcnx |
| Ptpra6 | 6.14E-05 | 0.320513045 | 0.444 | 0.306 | 1 | 17 | Ptpra |
| Ago32 | 6.26E-05 | 0.296385034 | 0.263 | 0.156 | 1 | 17 | Ago3 |
| Nfkb18 | 7.94E-05 | 0.300781794 | 0.415 | 0.273 | 1 | 17 | Nfkb1 |
| Pan310 | 9.66E-05 | 0.25784644 | 0.357 | 0.23 | 1 | 17 | Pan3 |
| Mib12 | 0.000116351 | 0.307363366 | 0.275 | 0.167 | 1 | 17 | Mib1 |
| Eml46 | 0.00012739 | 0.339724298 | 0.386 | 0.26 | 1 | 17 | Eml4 |
| Pros14 | 0.000142816 | 0.317844791 | 0.251 | 0.151 | 1 | 17 | Pros1 |
| Trp535 | 0.000178202 | 0.296678163 | 0.263 | 0.16 | 1 | 17 | Trp53 |
| Wdr336 | 0.00018334 | 0.261314215 | 0.357 | 0.241 | 1 | 17 | Wdr33 |
| Fam193a5 | 0.000196084 | 0.263048796 | 0.287 | 0.181 | 1 | 17 | Fam193a |
| Mga4 | 0.000244989 | 0.295209158 | 0.275 | 0.173 | 1 | 17 | Mga |
| Sltm7 | 0.000299483 | 0.30742602 | 0.398 | 0.272 | 1 | 17 | Sltm |
| Tmem1645 | 0.000366139 | 0.29249043 | 0.263 | 0.165 | 1 | 17 | Tmem164 |
| Zc3h7a5 | 0.000469938 | 0.272556718 | 0.257 | 0.164 | 1 | 17 | Zc3h7a |

| | | | | | | | |
|---|---|---|---|---|---|---|---|
| Jund6 | 0.000557852 | 0.262935288 | 0.819 | 0.647 | 1 | 17 | Jund |
| Exoc410 | 0.000940816 | 0.259921004 | 0.444 | 0.324 | 1 | 17 | Exoc4 |
| Hist1h1b | 0 | 3.9440123 | 0.744 | 0.014 | 0 | 18 | Hist1h1b |
| Hist1h2ap | 0 | 3.182805956 | 0.679 | 0.046 | 0 | 18 | Hist1h2ap |
| Hist1h2ae | 0 | 2.815914497 | 0.607 | 0.026 | 0 | 18 | Hist1h2ae |
| Birc5 | 0 | 2.553495566 | 0.714 | 0.006 | 0 | 18 | Birc5 |
| Top2a | 0 | 2.413572068 | 0.613 | 0.017 | 0 | 18 | Top2a |
| Mki67 | 0 | 2.065522596 | 0.488 | 0.01 | 0 | 18 | Mki67 |
| Ube2c | 0 | 1.885630922 | 0.44 | 0.009 | 0 | 18 | Ube2c |
| Nusap1 | 0 | 1.813658606 | 0.488 | 0.01 | 0 | 18 | Nusap1 |
| Pclaf | 0 | 1.781299648 | 0.5 | 0.007 | 0 | 18 | Pclaf |
| Hist1h3c | 0 | 1.773912279 | 0.399 | 0.003 | 0 | 18 | Hist1h3c |
| Cenpf | 0 | 1.585719834 | 0.339 | 0.005 | 0 | 18 | Cenpf |
| Spc24 | 0 | 1.567421911 | 0.47 | 0.009 | 0 | 18 | Spc24 |
| Cdk1 | 0 | 1.540194481 | 0.482 | 0.014 | 0 | 18 | Cdk1 |
| Cdca8 | 0 | 1.460741345 | 0.47 | 0.009 | 0 | 18 | Cdca8 |
| Prc1 | 0 | 1.443728119 | 0.446 | 0.009 | 0 | 18 | Prc1 |
| Tpx2 | 0 | 1.377718728 | 0.435 | 0.008 | 0 | 18 | Tpx2 |
| Tk1 | 0 | 1.349862465 | 0.405 | 0.008 | 0 | 18 | Tk1 |
| Knl1 | 0 | 1.183721898 | 0.387 | 0.007 | 0 | 18 | Knl1 |
| Kif11 | 0 | 1.104814337 | 0.369 | 0.008 | 0 | 18 | Kif11 |
| Cenpe | 0 | 1.095492719 | 0.31 | 0.008 | 0 | 18 | Cenpe |
| Esco2 | 0 | 1.095204798 | 0.339 | 0.003 | 0 | 18 | Esco2 |
| Ccnb2 | 0 | 1.049871487 | 0.292 | 0.005 | 0 | 18 | Ccnb2 |
| Cdca3 | 0 | 1.014966145 | 0.357 | 0.004 | 0 | 18 | Cdca3 |
| Hist1h2ab | 0 | 0.997085156 | 0.262 | 0.003 | 0 | 18 | Hist1h2ab |
| Cdkn3 | 0 | 0.970566102 | 0.268 | 0.004 | 0 | 18 | Cdkn3 |
| Pbk | 0 | 0.961167592 | 0.286 | 0.002 | 0 | 18 | Pbk |
| Kif15 | 0 | 0.924741042 | 0.315 | 0.006 | 0 | 18 | Kif15 |
| Aurkb | 0 | 0.920399957 | 0.315 | 0.005 | 0 | 18 | Aurkb |
| Hist1h2af | 0 | 0.878008134 | 0.268 | 0.003 | 0 | 18 | Hist1h2af |
| Rad51ap1 | 0 | 0.853506375 | 0.298 | 0.008 | 0 | 18 | Rad51ap1 |
| Hmmr | 0 | 0.809204987 | 0.286 | 0.003 | 0 | 18 | Hmmr |
| Anln | 0 | 0.747675007 | 0.298 | 0.008 | 0 | 18 | Anln |
| Neil3 | 0 | 0.719880383 | 0.262 | 0.003 | 0 | 18 | Neil3 |
| Sgo1 | 0 | 0.543237799 | 0.256 | 0.003 | 0 | 18 | Sgo1 |
| Ckap2l | 1.88E-304 | 0.829584177 | 0.327 | 0.01 | 3.81E-300 | 18 | Ckap2l |
| Ndc80 | 1.28E-279 | 0.737009324 | 0.345 | 0.012 | 2.58E-275 | 18 | Ndc80 |
| Asf1b | 1.65E-273 | 0.955475201 | 0.298 | 0.009 | 3.34E-269 | 18 | Asf1b |
| Bub1b | 1.37E-268 | 0.779425129 | 0.304 | 0.01 | 2.78E-264 | 18 | Bub1b |
| Knstrn | 1.90E-266 | 0.860648753 | 0.292 | 0.009 | 3.85E-262 | 18 | Knstrn |
| Clspn | 1.44E-264 | 0.848126712 | 0.327 | 0.012 | 2.92E-260 | 18 | Clspn |
| Hist1h2bj | 3.41E-263 | 1.332649696 | 0.292 | 0.009 | 6.91E-259 | 18 | Hist1h2bj |

| | | | | | | | |
|---|---|---|---|---|---|---|---|
| Cdc20 | 4.39E-257 | 1.03696933 | 0.25 | 0.007 | 8.88E-253 | 18 | Cdc20 |
| Cenpp | 1.24E-256 | 0.953713686 | 0.357 | 0.015 | 2.51E-252 | 18 | Cenpp |
| Stmn12 | 7.62E-256 | 3.475307436 | 0.935 | 0.127 | 1.54E-251 | 18 | Stmn1 |
| Cks1b1 | 2.46E-236 | 1.659741594 | 0.554 | 0.04 | 4.98E-232 | 18 | Cks1b |
| Tacc3 | 1.13E-233 | 0.611459145 | 0.292 | 0.011 | 2.29E-229 | 18 | Tacc3 |
| Rad51 | 5.80E-229 | 0.650188068 | 0.268 | 0.009 | 1.17E-224 | 18 | Rad51 |
| Cenpm | 1.10E-228 | 0.740068283 | 0.25 | 0.008 | 2.22E-224 | 18 | Cenpm |
| Lockd | 3.09E-208 | 1.061792723 | 0.387 | 0.022 | 6.25E-204 | 18 | Lockd |
| Aif13 | 4.75E-198 | 2.367696118 | 0.875 | 0.118 | 9.62E-194 | 18 | Aif1 |
| Kif4 | 8.64E-192 | 0.587725344 | 0.274 | 0.011 | 1.75E-187 | 18 | Kif4 |
| Hist1h1a | 8.99E-192 | 0.779631317 | 0.256 | 0.01 | 1.82E-187 | 18 | Hist1h1a |
| C1qc3 | 4.16E-181 | 2.59945101 | 0.881 | 0.132 | 8.42E-177 | 18 | C1qc |
| Kif23 | 4.49E-179 | 0.933024565 | 0.315 | 0.017 | 9.08E-175 | 18 | Kif23 |
| C1qb3 | 1.00E-174 | 2.770809957 | 0.917 | 0.152 | 2.03E-170 | 18 | C1qb |
| Spi13 | 2.18E-170 | 1.719631388 | 0.679 | 0.081 | 4.42E-166 | 18 | Spi1 |
| H2afx | 1.04E-165 | 1.704363189 | 0.548 | 0.057 | 2.12E-161 | 18 | H2afx |
| Spc25 | 2.16E-164 | 0.671112748 | 0.298 | 0.016 | 4.38E-160 | 18 | Spc25 |
| Cenpa | 7.04E-155 | 1.507500049 | 0.399 | 0.031 | 1.43E-150 | 18 | Cenpa |
| C1qa3 | 1.64E-150 | 2.609635801 | 0.845 | 0.145 | 3.32E-146 | 18 | C1qa |
| Kif20b | 1.03E-149 | 0.674430216 | 0.292 | 0.017 | 2.09E-145 | 18 | Kif20b |
| Alox5ap3 | 9.57E-145 | 1.463579482 | 0.595 | 0.071 | 1.94E-140 | 18 | Alox5ap |
| Ly863 | 5.23E-144 | 1.96615926 | 0.685 | 0.095 | 1.06E-139 | 18 | Ly86 |
| Sdk11 | 1.15E-141 | 1.387353695 | 0.321 | 0.022 | 2.32E-137 | 18 | Sdk1 |
| Gatm1 | 1.06E-137 | 1.687264331 | 0.577 | 0.072 | 2.14E-133 | 18 | Gatm |
| Bcl2a1b3 | 1.21E-137 | 1.801999962 | 0.696 | 0.104 | 2.46E-133 | 18 | Bcl2a1b |
| Pak11 | 2.36E-137 | 1.047281771 | 0.435 | 0.041 | 4.79E-133 | 18 | Pak1 |
| Ube2t | 9.32E-137 | 0.596459163 | 0.25 | 0.014 | 1.89E-132 | 18 | Ube2t |
| H2-Aa3 | 1.81E-136 | 2.330606562 | 0.917 | 0.184 | 3.66E-132 | 18 | H2-Aa |
| Cd683 | 3.42E-136 | 1.527961987 | 0.601 | 0.077 | 6.93E-132 | 18 | Cd68 |
| H2-Eb13 | 3.54E-136 | 2.368632131 | 0.929 | 0.192 | 7.16E-132 | 18 | H2-Eb1 |
| Clec5a1 | 1.56E-135 | 0.884306825 | 0.369 | 0.03 | 3.16E-131 | 18 | Clec5a |
| Slamf92 | 1.69E-135 | 1.466855162 | 0.565 | 0.069 | 3.42E-131 | 18 | Slamf9 |
| Lmnb1 | 2.98E-132 | 1.36709928 | 0.476 | 0.052 | 6.04E-128 | 18 | Lmnb1 |
| Ctss3 | 5.99E-132 | 2.226532042 | 0.857 | 0.164 | 1.21E-127 | 18 | Ctss |
| Hck1 | 6.78E-132 | 1.260897894 | 0.5 | 0.055 | 1.37E-127 | 18 | Hck |
| Ncapd2 | 2.94E-131 | 0.495260907 | 0.28 | 0.018 | 5.95E-127 | 18 | Ncapd2 |
| Hist1h1d1 | 2.68E-129 | 1.686044507 | 0.446 | 0.047 | 5.43E-125 | 18 | Hist1h1d |
| H2-Ab13 | 6.07E-128 | 2.271801443 | 0.917 | 0.199 | 1.23E-123 | 18 | H2-Ab1 |
| Diaph3 | 7.24E-125 | 0.839003774 | 0.327 | 0.026 | 1.47E-120 | 18 | Diaph3 |
| Tnfsf13b1 | 1.22E-124 | 0.966542884 | 0.429 | 0.043 | 2.46E-120 | 18 | Tnfsf13b |
| Fbxo5 | 5.34E-122 | 0.750042529 | 0.25 | 0.015 | 1.08E-117 | 18 | Fbxo5 |
| Ms4a6c2 | 2.47E-121 | 1.591158015 | 0.589 | 0.082 | 5.00E-117 | 18 | Ms4a6c |
| Ctsh4 | 2.88E-121 | 1.870992459 | 0.75 | 0.134 | 5.84E-117 | 18 | Ctsh |

| | | | | | | | |
|---|---|---|---|---|---|---|---|
| Bcl2a1a1 | 3.96E-121 | 0.977171407 | 0.387 | 0.036 | 8.02E-117 | 18 | Bcl2a1a |
| Incenp | 8.75E-120 | 0.848541458 | 0.31 | 0.024 | 1.77E-115 | 18 | Incenp |
| Smc2 | 2.04E-118 | 1.34437099 | 0.506 | 0.064 | 4.13E-114 | 18 | Smc2 |
| Hist1h3e | 1.63E-116 | 1.533428578 | 0.381 | 0.037 | 3.30E-112 | 18 | Hist1h3e |
| Nlrp1b1 | 5.37E-116 | 1.083607507 | 0.393 | 0.039 | 1.09E-111 | 18 | Nlrp1b |
| Cd743 | 5.75E-115 | 2.454491178 | 0.935 | 0.249 | 1.16E-110 | 18 | Cd74 |
| Rgs104 | 1.58E-114 | 1.971094937 | 0.774 | 0.159 | 3.19E-110 | 18 | Rgs10 |
| Pld41 | 2.63E-114 | 1.116145722 | 0.482 | 0.058 | 5.32E-110 | 18 | Pld4 |
| Plbd11 | 6.47E-114 | 1.116762052 | 0.482 | 0.058 | 1.31E-109 | 18 | Plbd1 |
| Hpgd3 | 1.24E-113 | 1.619908351 | 0.583 | 0.086 | 2.51E-109 | 18 | Hpgd |
| Hmgb24 | 3.06E-110 | 2.847584815 | 0.917 | 0.327 | 6.19E-106 | 18 | Hmgb2 |
| H2-DMb13 | 2.16E-109 | 1.453659692 | 0.595 | 0.09 | 4.37E-105 | 18 | H2-DMb1 |
| Cd723 | 4.29E-109 | 1.747144744 | 0.619 | 0.104 | 8.68E-105 | 18 | Cd72 |
| Cxcl163 | 6.28E-109 | 1.606607334 | 0.649 | 0.111 | 1.27E-104 | 18 | Cxcl16 |
| Clec12a1 | 1.17E-108 | 0.947985885 | 0.345 | 0.032 | 2.37E-104 | 18 | Clec12a |
| Racgap1 | 4.80E-108 | 0.788135878 | 0.333 | 0.03 | 9.72E-104 | 18 | Racgap1 |
| Csf1r1 | 1.53E-107 | 1.440665578 | 0.53 | 0.073 | 3.09E-103 | 18 | Csf1r |
| Wfdc173 | 2.17E-107 | 1.370640783 | 0.506 | 0.067 | 4.39E-103 | 18 | Wfdc17 |
| Ms4a73 | 3.96E-107 | 1.727273564 | 0.619 | 0.098 | 8.01E-103 | 18 | Ms4a7 |
| Cks2 | 5.15E-107 | 1.437646391 | 0.5 | 0.068 | 1.04E-102 | 18 | Cks2 |
| Lyz23 | 1.02E-103 | 1.525761119 | 0.655 | 0.11 | 2.06E-99 | 18 | Lyz2 |
| Cd1801 | 2.91E-103 | 0.974334145 | 0.405 | 0.046 | 5.89E-99 | 18 | Cd180 |
| Siglece1 | 3.29E-103 | 0.670944817 | 0.292 | 0.024 | 6.66E-99 | 18 | Siglece |
| Cyp4f181 | 6.84E-102 | 0.660210664 | 0.351 | 0.035 | 1.39E-97 | 18 | Cyp4f18 |
| H2-DMa3 | 6.97E-102 | 1.694564774 | 0.649 | 0.118 | 1.41E-97 | 18 | H2-DMa |
| Cd300c23 | 6.98E-102 | 1.214757431 | 0.565 | 0.084 | 1.41E-97 | 18 | Cd300c2 |
| Basp14 | 6.19E-97 | 1.721958057 | 0.595 | 0.103 | 1.25E-92 | 18 | Basp1 |
| Ppfia41 | 3.02E-96 | 1.017745765 | 0.429 | 0.054 | 6.12E-92 | 18 | Ppfia4 |
| Scimp2 | 2.44E-95 | 1.236997916 | 0.506 | 0.075 | 4.94E-91 | 18 | Scimp |
| Lst13 | 7.22E-95 | 1.57142774 | 0.613 | 0.11 | 1.46E-90 | 18 | Lst1 |
| Pkib1 | 3.02E-94 | 0.779558705 | 0.375 | 0.042 | 6.10E-90 | 18 | Pkib |
| Batf31 | 1.04E-93 | 0.809441711 | 0.315 | 0.031 | 2.10E-89 | 18 | Batf3 |
| Ctsc2 | 5.37E-93 | 1.995384458 | 0.679 | 0.149 | 1.09E-88 | 18 | Ctsc |
| Trem22 | 2.65E-90 | 1.138723099 | 0.464 | 0.065 | 5.36E-86 | 18 | Trem2 |
| Clec4a21 | 1.00E-89 | 0.677186683 | 0.304 | 0.03 | 2.03E-85 | 18 | Clec4a2 |
| Lilra51 | 1.73E-89 | 1.06128355 | 0.405 | 0.051 | 3.50E-85 | 18 | Lilra5 |
| Cybb1 | 1.88E-89 | 1.436307652 | 0.494 | 0.074 | 3.81E-85 | 18 | Cybb |
| Lair11 | 8.57E-89 | 1.105213981 | 0.446 | 0.062 | 1.74E-84 | 18 | Lair1 |
| Csf2ra1 | 5.03E-88 | 1.06873509 | 0.476 | 0.072 | 1.02E-83 | 18 | Csf2ra |
| Fcgr11 | 9.52E-87 | 0.942558314 | 0.399 | 0.051 | 1.93E-82 | 18 | Fcgr1 |
| Trf2 | 4.56E-86 | 1.033615487 | 0.458 | 0.067 | 9.24E-82 | 18 | Trf |
| Havcr21 | 5.64E-86 | 0.681218797 | 0.304 | 0.031 | 1.14E-81 | 18 | Havcr2 |
| Cx3cr11 | 1.43E-85 | 1.701846092 | 0.506 | 0.083 | 2.89E-81 | 18 | Cx3cr1 |

| | | | | | | | |
|---|---|---|---|---|---|---|---|
| Tyms | 8.91E-85 | 1.130592923 | 0.464 | 0.07 | 1.80E-80 | 18 | Tyms |
| Lgmn3 | 1.18E-84 | 1.549998821 | 0.583 | 0.112 | 2.38E-80 | 18 | Lgmn |
| Tgfbi2 | 1.79E-84 | 0.988110969 | 0.411 | 0.055 | 3.63E-80 | 18 | Tgfbi |
| Hist2h2ac | 1.69E-83 | 1.353460102 | 0.458 | 0.071 | 3.42E-79 | 18 | Hist2h2ac |
| Adgre11 | 2.16E-82 | 0.949314112 | 0.411 | 0.056 | 4.38E-78 | 18 | Adgre1 |
| H2afz3 | 1.09E-81 | 2.297101402 | 0.958 | 0.608 | 2.22E-77 | 18 | H2afz |
| Mcm5 | 1.84E-81 | 0.693856666 | 0.262 | 0.025 | 3.73E-77 | 18 | Mcm5 |
| Rad51b | 4.25E-81 | 1.012809447 | 0.321 | 0.037 | 8.61E-77 | 18 | Rad51b |
| Cadm11 | 2.78E-80 | 1.303357808 | 0.452 | 0.071 | 5.63E-76 | 18 | Cadm1 |
| Ccl61 | 4.16E-80 | 0.631534013 | 0.304 | 0.033 | 8.43E-76 | 18 | Ccl6 |
| Mpeg12 | 1.05E-79 | 1.189712052 | 0.464 | 0.073 | 2.13E-75 | 18 | Mpeg1 |
| Lacc11 | 3.55E-79 | 0.866615712 | 0.351 | 0.044 | 7.19E-75 | 18 | Lacc1 |
| Rab7b1 | 4.29E-78 | 0.495676271 | 0.286 | 0.03 | 8.69E-74 | 18 | Rab7b |
| Tubb55 | 8.08E-76 | 2.130988335 | 0.869 | 0.38 | 1.64E-71 | 18 | Tubb5 |
| P2ry61 | 2.45E-75 | 0.755027059 | 0.369 | 0.05 | 4.96E-71 | 18 | P2ry6 |
| Pou2f21 | 5.95E-75 | 0.734755775 | 0.351 | 0.046 | 1.20E-70 | 18 | Pou2f2 |
| Fat31 | 1.24E-74 | 1.300103468 | 0.333 | 0.043 | 2.51E-70 | 18 | Fat3 |
| Slc8a12 | 1.90E-74 | 1.617260682 | 0.399 | 0.061 | 3.84E-70 | 18 | Slc8a1 |
| Phf11b1 | 1.90E-74 | 1.278142311 | 0.5 | 0.092 | 3.85E-70 | 18 | Phf11b |
| Cd861 | 2.60E-74 | 0.783406041 | 0.393 | 0.056 | 5.26E-70 | 18 | Cd86 |
| Ncf21 | 3.50E-74 | 0.81501079 | 0.375 | 0.053 | 7.08E-70 | 18 | Ncf2 |
| Gpr1833 | 1.45E-73 | 0.866124486 | 0.399 | 0.059 | 2.93E-69 | 18 | Gpr183 |
| Gm267403 | 4.50E-73 | 0.85308172 | 0.399 | 0.059 | 9.12E-69 | 18 | Gm26740 |
| Tmem1192 | 1.11E-72 | 1.007470551 | 0.393 | 0.059 | 2.24E-68 | 18 | Tmem119 |
| Ulbp1 | 1.31E-71 | 0.529629037 | 0.268 | 0.029 | 2.66E-67 | 18 | Ulbp1 |
| Rassf41 | 1.89E-70 | 0.569117796 | 0.327 | 0.042 | 3.82E-66 | 18 | Rassf4 |
| Arhgap221 | 3.54E-70 | 0.816617404 | 0.333 | 0.044 | 7.17E-66 | 18 | Arhgap22 |
| Tyrobp6 | 4.58E-70 | 1.457412328 | 0.905 | 0.26 | 9.28E-66 | 18 | Tyrobp |
| Klra21 | 8.06E-70 | 0.884980342 | 0.369 | 0.053 | 1.63E-65 | 18 | Klra2 |
| Rnf1801 | 4.98E-69 | 0.644730908 | 0.31 | 0.039 | 1.01E-64 | 18 | Rnf180 |
| Ckb4 | 7.96E-69 | 1.737818812 | 0.714 | 0.215 | 1.61E-64 | 18 | Ckb |
| Ccl121 | 9.73E-69 | 1.468283188 | 0.31 | 0.039 | 1.97E-64 | 18 | Ccl12 |
| Pla2g4a2 | 1.48E-68 | 0.720137322 | 0.321 | 0.042 | 3.00E-64 | 18 | Pla2g4a |
| Glipr15 | 4.71E-68 | 1.192153645 | 0.488 | 0.095 | 9.53E-64 | 18 | Glipr1 |
| 11-Mar | 5.60E-68 | 0.59679768 | 0.315 | 0.041 | 1.13E-63 | 18 | 1-Mar |
| Dck1 | 8.05E-68 | 0.663004116 | 0.28 | 0.033 | 1.63E-63 | 18 | Dck |
| Apobec11 | 2.61E-67 | 1.36707515 | 0.512 | 0.103 | 5.29E-63 | 18 | Apobec1 |
| Nrm | 9.24E-67 | 0.604080995 | 0.262 | 0.029 | 1.87E-62 | 18 | Nrm |
| Ms4a6d1 | 1.09E-66 | 0.687822832 | 0.292 | 0.036 | 2.21E-62 | 18 | Ms4a6d |
| Rtn11 | 4.54E-66 | 0.686467844 | 0.286 | 0.035 | 9.20E-62 | 18 | Rtn1 |
| Il2ra2 | 4.73E-66 | 0.949134989 | 0.315 | 0.043 | 9.58E-62 | 18 | Il2ra |
| Tuba1b5 | 5.40E-66 | 2.029466182 | 0.81 | 0.321 | 1.09E-61 | 18 | Tuba1b |
| Fcer1g7 | 7.41E-66 | 1.405722652 | 0.899 | 0.268 | 1.50E-61 | 18 | Fcer1g |

| | | | | | | | |
|---|---|---|---|---|---|---|---|
| Milr11 | 7.85E-66 | 0.63500132 | 0.333 | 0.046 | 1.59E-61 | 18 | Milr1 |
| Xlr1 | 5.99E-64 | 0.7804724 | 0.327 | 0.046 | 1.21E-59 | 18 | Xlr |
| Zmynd151 | 8.83E-63 | 0.905697318 | 0.345 | 0.052 | 1.79E-58 | 18 | Zmynd15 |
| Cd528 | 1.11E-62 | 1.276069003 | 0.917 | 0.28 | 2.24E-58 | 18 | Cd52 |
| Slc29a31 | 2.01E-62 | 0.904024469 | 0.399 | 0.069 | 4.08E-58 | 18 | Slc29a3 |
| Cacna1d1 | 6.05E-62 | 0.534971106 | 0.298 | 0.039 | 1.23E-57 | 18 | Cacna1d |
| Fen1 | 7.10E-62 | 0.663967092 | 0.262 | 0.032 | 1.44E-57 | 18 | Fen1 |
| Cst35 | 9.73E-61 | 2.006388482 | 0.935 | 0.602 | 1.97E-56 | 18 | Cst3 |
| Hmgn21 | 5.04E-60 | 1.523421372 | 0.673 | 0.193 | 1.02E-55 | 18 | Hmgn2 |
| Mcm7 | 7.12E-60 | 1.035830972 | 0.339 | 0.053 | 1.44E-55 | 18 | Mcm7 |
| Rrm1 | 1.03E-59 | 0.774035743 | 0.333 | 0.051 | 2.08E-55 | 18 | Rrm1 |
| Kcnk131 | 2.73E-59 | 0.354199803 | 0.25 | 0.029 | 5.53E-55 | 18 | Kcnk13 |
| Bcl2a1d1 | 2.56E-58 | 0.591638143 | 0.274 | 0.036 | 5.18E-54 | 18 | Bcl2a1d |
| Ntpcr3 | 4.86E-58 | 1.002534681 | 0.488 | 0.104 | 9.84E-54 | 18 | Ntpcr |
| Axl3 | 3.40E-57 | 0.740864743 | 0.417 | 0.077 | 6.88E-53 | 18 | Axl |
| Alcam | 7.17E-57 | 0.75248693 | 0.25 | 0.031 | 1.45E-52 | 18 | Alcam |
| Sirpa1 | 1.77E-56 | 0.618079596 | 0.327 | 0.05 | 3.58E-52 | 18 | Sirpa |
| Apoe3 | 6.71E-56 | 1.925768321 | 0.893 | 0.534 | 1.36E-51 | 18 | Apoe |
| Olfml32 | 6.99E-56 | 0.880210696 | 0.351 | 0.059 | 1.41E-51 | 18 | Olfml3 |
| Lig1 | 1.16E-55 | 0.788186753 | 0.351 | 0.059 | 2.34E-51 | 18 | Lig1 |
| Lpcat22 | 1.24E-55 | 0.862479529 | 0.375 | 0.066 | 2.50E-51 | 18 | Lpcat2 |
| Hpgds1 | 1.89E-55 | 0.787400982 | 0.351 | 0.058 | 3.83E-51 | 18 | Hpgds |
| Tmsb4x9 | 3.37E-55 | 1.509924415 | 0.988 | 0.934 | 6.82E-51 | 18 | Tmsb4x |
| Ezh21 | 1.71E-54 | 0.879045334 | 0.375 | 0.069 | 3.46E-50 | 18 | Ezh2 |
| Coro1a7 | 2.69E-54 | 1.391575162 | 0.714 | 0.219 | 5.45E-50 | 18 | Coro1a |
| Adap23 | 4.96E-54 | 0.606445333 | 0.345 | 0.057 | 1.00E-49 | 18 | Adap2 |
| Laptm57 | 1.03E-53 | 1.580972734 | 0.655 | 0.21 | 2.09E-49 | 18 | Laptm5 |
| Fcgr32 | 1.13E-53 | 0.943248201 | 0.429 | 0.087 | 2.30E-49 | 18 | Fcgr3 |
| Frmpd41 | 1.84E-53 | 1.143402083 | 0.286 | 0.041 | 3.73E-49 | 18 | Frmpd4 |
| Slc11a11 | 1.89E-53 | 0.474365447 | 0.256 | 0.034 | 3.83E-49 | 18 | Slc11a1 |
| Pid13 | 5.13E-53 | 0.874186912 | 0.375 | 0.069 | 1.04E-48 | 18 | Pid1 |
| Fth14 | 7.10E-52 | 1.482981366 | 0.994 | 0.969 | 1.44E-47 | 18 | Fth1 |
| Tbxas11 | 1.88E-51 | 0.616963405 | 0.298 | 0.046 | 3.81E-47 | 18 | Tbxas1 |
| Irf51 | 3.28E-51 | 0.704097803 | 0.327 | 0.056 | 6.64E-47 | 18 | Irf5 |
| Pf41 | 3.99E-51 | 1.019271202 | 0.28 | 0.041 | 8.07E-47 | 18 | Pf4 |
| Myo5a1 | 2.22E-50 | 0.878539448 | 0.405 | 0.084 | 4.49E-46 | 18 | Myo5a |
| Cep128 | 2.41E-50 | 0.654824611 | 0.31 | 0.051 | 4.88E-46 | 18 | Cep128 |
| Blnk1 | 1.10E-49 | 0.62242181 | 0.345 | 0.061 | 2.22E-45 | 18 | Blnk |
| Lrrc251 | 1.79E-49 | 0.39069146 | 0.262 | 0.037 | 3.63E-45 | 18 | Lrrc25 |
| Cd141 | 4.85E-49 | 0.682215099 | 0.345 | 0.062 | 9.81E-45 | 18 | Cd14 |
| Arl4c2 | 1.39E-48 | 0.777452716 | 0.369 | 0.072 | 2.82E-44 | 18 | Arl4c |
| Mb21d1 | 2.32E-48 | 0.415573931 | 0.256 | 0.037 | 4.69E-44 | 18 | Mb21d1 |
| Pmf1 | 2.59E-48 | 0.607569296 | 0.286 | 0.045 | 5.25E-44 | 18 | Pmf1 |

| | | | | | | | |
|---|---|---|---|---|---|---|---|
| Lpl3 | 3.54E-48 | 0.793161102 | 0.417 | 0.086 | 7.16E-44 | 18 | Lpl |
| Gpr652 | 4.75E-48 | 0.986210155 | 0.411 | 0.088 | 9.62E-44 | 18 | Gpr65 |
| C3ar11 | 1.11E-47 | 0.472447449 | 0.268 | 0.04 | 2.25E-43 | 18 | C3ar1 |
| Sh2d1b14 | 1.85E-47 | 0.797543391 | 0.381 | 0.077 | 3.74E-43 | 18 | Sh2d1b1 |
| Il10ra2 | 2.38E-46 | 0.692433682 | 0.393 | 0.082 | 4.82E-42 | 18 | Il10ra |
| Gtf2h21 | 2.84E-46 | 0.718091692 | 0.363 | 0.072 | 5.74E-42 | 18 | Gtf2h2 |
| Fam111a1 | 4.89E-46 | 0.836688973 | 0.411 | 0.091 | 9.90E-42 | 18 | Fam111a |
| Ncf11 | 8.55E-46 | 0.491800894 | 0.274 | 0.043 | 1.73E-41 | 18 | Ncf1 |
| Tbc1d91 | 1.24E-45 | 0.454848038 | 0.256 | 0.039 | 2.50E-41 | 18 | Tbc1d9 |
| Smc45 | 2.97E-45 | 1.380461036 | 0.53 | 0.162 | 6.01E-41 | 18 | Smc4 |
| Atad24 | 8.00E-45 | 1.080422846 | 0.399 | 0.092 | 1.62E-40 | 18 | Atad2 |
| Cotl17 | 8.06E-45 | 1.246102715 | 0.661 | 0.224 | 1.63E-40 | 18 | Cotl1 |
| Evi2a1 | 8.88E-45 | 0.530038662 | 0.274 | 0.044 | 1.80E-40 | 18 | Evi2a |
| Cyba8 | 1.27E-44 | 1.485489643 | 0.815 | 0.441 | 2.57E-40 | 18 | Cyba |
| Creb51 | 1.74E-44 | 0.682695657 | 0.327 | 0.061 | 3.52E-40 | 18 | Creb5 |
| Ciita1 | 2.07E-44 | 0.5807992 | 0.25 | 0.038 | 4.18E-40 | 18 | Ciita |
| Gm2a3 | 2.13E-44 | 0.782746579 | 0.399 | 0.088 | 4.30E-40 | 18 | Gm2a |
| Brip1 | 2.39E-44 | 0.528970843 | 0.286 | 0.048 | 4.84E-40 | 18 | Brip1 |
| Sap301 | 4.48E-44 | 0.774553656 | 0.405 | 0.09 | 9.07E-40 | 18 | Sap30 |
| Tuba1c1 | 1.91E-43 | 1.035102972 | 0.464 | 0.12 | 3.87E-39 | 18 | Tuba1c |
| Ms4a6b3 | 4.16E-43 | 0.885361951 | 0.405 | 0.092 | 8.42E-39 | 18 | Ms4a6b |
| Cfh2 | 2.33E-42 | 0.385079659 | 0.351 | 0.069 | 4.72E-38 | 18 | Cfh |
| Fgd21 | 2.53E-42 | 0.32593221 | 0.268 | 0.044 | 5.12E-38 | 18 | Fgd2 |
| Rbm471 | 3.21E-42 | 0.582759233 | 0.363 | 0.075 | 6.51E-38 | 18 | Rbm47 |
| Dut | 3.54E-42 | 1.027535787 | 0.482 | 0.129 | 7.17E-38 | 18 | Dut |
| Hcls15 | 5.07E-42 | 0.990613597 | 0.47 | 0.125 | 1.03E-37 | 18 | Hcls1 |
| Rfc4 | 8.03E-42 | 0.558082256 | 0.28 | 0.048 | 1.63E-37 | 18 | Rfc4 |
| Snx202 | 1.05E-41 | 0.499257566 | 0.321 | 0.062 | 2.13E-37 | 18 | Snx20 |
| Timp24 | 3.71E-41 | 0.758212613 | 0.494 | 0.129 | 7.52E-37 | 18 | Timp2 |
| Tep11 | 5.89E-41 | 0.652092903 | 0.304 | 0.058 | 1.19E-36 | 18 | Tep1 |
| Qpct2 | 9.01E-41 | 0.707784623 | 0.363 | 0.079 | 1.82E-36 | 18 | Qpct |
| Actb8 | 1.72E-40 | 1.058189403 | 0.982 | 0.843 | 3.48E-36 | 18 | Actb |
| Gusb1 | 2.45E-40 | 0.658425906 | 0.411 | 0.099 | 4.96E-36 | 18 | Gusb |
| Nckap1l5 | 4.04E-40 | 0.660423492 | 0.381 | 0.088 | 8.19E-36 | 18 | Nckap1l |
| Hk21 | 5.58E-40 | 0.529161296 | 0.321 | 0.064 | 1.13E-35 | 18 | Hk2 |
| 5031439G07Rik1 | 5.92E-40 | 0.387696019 | 0.369 | 0.079 | 1.20E-35 | 18 | 5031439G07Rik |
| Ran4 | 1.32E-39 | 1.278571625 | 0.804 | 0.4 | 2.67E-35 | 18 | Ran |
| Fgf133 | 1.72E-39 | 0.815245898 | 0.357 | 0.078 | 3.49E-35 | 18 | Fgf13 |
| Hirip3 | 1.81E-39 | 0.599000672 | 0.256 | 0.044 | 3.67E-35 | 18 | Hirip3 |
| Shtn12 | 1.95E-39 | 0.438178311 | 0.333 | 0.067 | 3.95E-35 | 18 | Shtn1 |
| Prim2 | 2.49E-39 | 0.583461647 | 0.262 | 0.046 | 5.03E-35 | 18 | Prim2 |
| Slc15a31 | 3.22E-39 | 0.579879084 | 0.351 | 0.075 | 6.53E-35 | 18 | Slc15a3 |
| Cyth45 | 1.44E-38 | 0.893582274 | 0.476 | 0.133 | 2.91E-34 | 18 | Cyth4 |

| | | | | | | | |
|---|---|---|---|---|---|---|---|
| Abhd121 | 1.86E-38 | 0.954983536 | 0.458 | 0.129 | 3.76E-34 | 18 | Abhd12 |
| Jpt15 | 3.00E-38 | 1.239218275 | 0.607 | 0.222 | 6.08E-34 | 18 | Jpt1 |
| Pfn17 | 3.23E-38 | 1.184661943 | 0.958 | 0.682 | 6.55E-34 | 18 | Pfn1 |
| Col14a12 | 3.51E-38 | 0.407111459 | 0.298 | 0.057 | 7.11E-34 | 18 | Col14a1 |
| Arrb24 | 1.67E-37 | 0.625998609 | 0.381 | 0.092 | 3.37E-33 | 18 | Arrb2 |
| Dctpp1 | 1.77E-37 | 0.818307565 | 0.488 | 0.139 | 3.58E-33 | 18 | Dctpp1 |
| Prkcb6 | 2.77E-37 | 1.300785194 | 0.494 | 0.156 | 5.60E-33 | 18 | Prkcb |
| Man2b11 | 1.26E-36 | 1.026925345 | 0.506 | 0.164 | 2.56E-32 | 18 | Man2b1 |
| Skap21 | 1.38E-36 | 1.037479966 | 0.518 | 0.167 | 2.80E-32 | 18 | Skap2 |
| Hist1h4d1 | 2.19E-36 | 1.870660812 | 0.619 | 0.257 | 4.44E-32 | 18 | Hist1h4d |
| Ctsz4 | 2.58E-36 | 1.322936665 | 0.726 | 0.36 | 5.22E-32 | 18 | Ctsz |
| Clic17 | 6.94E-36 | 1.217401008 | 0.875 | 0.601 | 1.41E-31 | 18 | Clic1 |
| Pvt15 | 7.21E-36 | 0.826868765 | 0.339 | 0.078 | 1.46E-31 | 18 | Pvt1 |
| Ncapd3 | 7.63E-36 | 0.579119299 | 0.333 | 0.075 | 1.55E-31 | 18 | Ncapd3 |
| Haus8 | 1.41E-35 | 0.471821303 | 0.28 | 0.054 | 2.85E-31 | 18 | Haus8 |
| Fam105a5 | 3.87E-35 | 0.795985925 | 0.44 | 0.125 | 7.84E-31 | 18 | Fam105a |
| Gmnn | 8.02E-35 | 0.826250379 | 0.351 | 0.085 | 1.62E-30 | 18 | Gmnn |
| Tns31 | 1.21E-34 | 0.423245331 | 0.327 | 0.072 | 2.44E-30 | 18 | Tns3 |
| Xylt11 | 1.43E-34 | 0.545358684 | 0.339 | 0.077 | 2.90E-30 | 18 | Xylt1 |
| Tmem371 | 1.45E-34 | 0.453085876 | 0.256 | 0.048 | 2.93E-30 | 18 | Tmem37 |
| Gmfg4 | 3.17E-34 | 0.8264812 | 0.476 | 0.139 | 6.41E-30 | 18 | Gmfg |
| Pmepa13 | 6.37E-34 | 1.238898728 | 0.613 | 0.252 | 1.29E-29 | 18 | Pmepa1 |
| Hexb2 | 8.94E-34 | 1.278744608 | 0.81 | 0.518 | 1.81E-29 | 18 | Hexb |
| Bmp2k2 | 1.21E-33 | 0.540219991 | 0.417 | 0.11 | 2.45E-29 | 18 | Bmp2k |
| Nsd21 | 1.51E-33 | 0.808331042 | 0.363 | 0.093 | 3.07E-29 | 18 | Nsd2 |
| Rab3il12 | 5.29E-33 | 0.539071513 | 0.31 | 0.069 | 1.07E-28 | 18 | Rab3il1 |
| Mcm3 | 6.46E-33 | 0.82290061 | 0.298 | 0.066 | 1.31E-28 | 18 | Mcm3 |
| Arhgap391 | 7.33E-33 | 0.498632287 | 0.28 | 0.058 | 1.48E-28 | 18 | Arhgap39 |
| Ifi27l2a3 | 2.09E-32 | 0.882015989 | 0.792 | 0.366 | 4.23E-28 | 18 | Ifi27l2a |
| Ifi2072 | 2.43E-32 | 0.658735353 | 0.399 | 0.107 | 4.92E-28 | 18 | Ifi207 |
| Ftl15 | 2.47E-32 | 1.10476697 | 0.988 | 0.911 | 5.01E-28 | 18 | Ftl1 |
| Rfc3 | 2.52E-32 | 0.644397721 | 0.262 | 0.052 | 5.11E-28 | 18 | Rfc3 |
| Fam49b6 | 3.23E-32 | 1.065987822 | 0.595 | 0.237 | 6.55E-28 | 18 | Fam49b |
| Fermt36 | 3.93E-32 | 0.775916608 | 0.411 | 0.114 | 7.96E-28 | 18 | Fermt3 |
| Hjurp1 | 4.42E-32 | 0.621141815 | 0.357 | 0.091 | 8.94E-28 | 18 | Hjurp |
| Ccdc34 | 6.82E-32 | 0.682043028 | 0.345 | 0.084 | 1.38E-27 | 18 | Ccdc34 |
| Cryl1 | 1.20E-31 | 0.400203235 | 0.28 | 0.059 | 2.42E-27 | 18 | Cryl1 |
| Itgb53 | 1.54E-31 | 0.862397984 | 0.458 | 0.145 | 3.11E-27 | 18 | Itgb5 |
| Sh3kbp15 | 2.72E-31 | 0.700819156 | 0.423 | 0.125 | 5.51E-27 | 18 | Sh3kbp1 |
| Mcub1 | 8.24E-31 | 0.446639969 | 0.298 | 0.067 | 1.67E-26 | 18 | Mcub |
| Lat25 | 8.96E-31 | 0.800308589 | 0.44 | 0.132 | 1.81E-26 | 18 | Lat2 |
| Unc93b11 | 9.74E-31 | 0.882567857 | 0.47 | 0.156 | 1.97E-26 | 18 | Unc93b1 |
| Cdkn2c | 1.12E-30 | 0.827766964 | 0.274 | 0.06 | 2.28E-26 | 18 | Cdkn2c |

| | | | | | | | |
|---|---|---|---|---|---|---|---|
| Zeb24 | 1.63E-30 | 0.956161137 | 0.452 | 0.143 | 3.30E-26 | 18 | Zeb2 |
| H2afv1 | 1.69E-30 | 1.103855299 | 0.75 | 0.379 | 3.43E-26 | 18 | H2afv |
| Tmem176b7 | 1.73E-30 | 1.180465745 | 0.786 | 0.439 | 3.50E-26 | 18 | Tmem176b |
| Apbb1ip6 | 2.69E-30 | 0.954844746 | 0.5 | 0.174 | 5.45E-26 | 18 | Apbb1ip |
| Erp292 | 2.84E-30 | 1.309004348 | 0.667 | 0.328 | 5.76E-26 | 18 | Erp29 |
| Topbp1 | 3.20E-30 | 0.531182097 | 0.363 | 0.095 | 6.48E-26 | 18 | Topbp1 |
| Rac25 | 3.72E-30 | 0.792831525 | 0.548 | 0.178 | 7.54E-26 | 18 | Rac2 |
| Npc23 | 5.30E-30 | 1.154192642 | 0.75 | 0.414 | 1.07E-25 | 18 | Npc2 |
| Vrk11 | 9.13E-30 | 0.587211171 | 0.339 | 0.086 | 1.85E-25 | 18 | Vrk1 |
| Cd845 | 1.03E-29 | 0.652991932 | 0.345 | 0.09 | 2.09E-25 | 18 | Cd84 |
| Snx242 | 1.23E-29 | 0.336148945 | 0.274 | 0.059 | 2.48E-25 | 18 | Snx24 |
| Psap4 | 1.25E-29 | 1.313759641 | 0.696 | 0.399 | 2.53E-25 | 18 | Psap |
| Rap1gap21 | 1.51E-29 | 0.413670681 | 0.333 | 0.082 | 3.05E-25 | 18 | Rap1gap2 |
| Tnfrsf11a1 | 1.70E-29 | 0.447469718 | 0.268 | 0.059 | 3.44E-25 | 18 | Tnfrsf11a |
| Vrk24 | 2.59E-29 | 0.725053425 | 0.411 | 0.124 | 5.24E-25 | 18 | Vrk2 |
| Rfwd3 | 2.72E-29 | 0.447761788 | 0.31 | 0.075 | 5.51E-25 | 18 | Rfwd3 |
| Arhgap46 | 2.77E-29 | 0.316578145 | 0.268 | 0.058 | 5.60E-25 | 18 | Arhgap4 |
| Selenoh | 3.64E-29 | 1.054753077 | 0.53 | 0.194 | 7.36E-25 | 18 | Selenoh |
| Cdk144 | 3.75E-29 | 1.065999968 | 0.429 | 0.142 | 7.60E-25 | 18 | Cdk14 |
| Dnajc9 | 4.05E-29 | 0.667867739 | 0.381 | 0.105 | 8.21E-25 | 18 | Dnajc9 |
| Alyref | 1.02E-28 | 0.9522723 | 0.56 | 0.213 | 2.07E-24 | 18 | Alyref |
| Hpf1 | 1.23E-28 | 0.867304679 | 0.524 | 0.19 | 2.48E-24 | 18 | Hpf1 |
| Pola1 | 1.57E-28 | 0.869203175 | 0.286 | 0.069 | 3.18E-24 | 18 | Pola1 |
| Ptma3 | 2.58E-28 | 0.853772024 | 0.988 | 0.94 | 5.23E-24 | 18 | Ptma |
| Specc14 | 3.20E-28 | 0.605135109 | 0.315 | 0.08 | 6.47E-24 | 18 | Specc1 |
| Ramp15 | 4.65E-28 | 0.74423579 | 0.375 | 0.11 | 9.42E-24 | 18 | Ramp1 |
| Mctp12 | 5.54E-28 | 0.516387592 | 0.304 | 0.074 | 1.12E-23 | 18 | Mctp1 |
| Apba12 | 6.26E-28 | 0.403109062 | 0.25 | 0.053 | 1.27E-23 | 18 | Apba1 |
| Lcp16 | 6.36E-28 | 1.095456733 | 0.577 | 0.238 | 1.29E-23 | 18 | Lcp1 |
| Cpq3 | 1.06E-27 | 0.709850918 | 0.399 | 0.121 | 2.14E-23 | 18 | Cpq |
| Sh3bgrl37 | 1.25E-27 | 1.148095711 | 0.851 | 0.56 | 2.54E-23 | 18 | Sh3bgrl3 |
| Ube2s | 1.37E-27 | 1.120276474 | 0.655 | 0.311 | 2.77E-23 | 18 | Ube2s |
| Ptpn65 | 2.13E-27 | 0.698173126 | 0.423 | 0.13 | 4.30E-23 | 18 | Ptpn6 |
| Ptprj6 | 2.46E-27 | 1.000227535 | 0.44 | 0.151 | 4.99E-23 | 18 | Ptprj |
| Cd485 | 2.81E-27 | 0.697058214 | 0.423 | 0.135 | 5.69E-23 | 18 | Cd48 |
| Lap32 | 3.06E-27 | 0.707835765 | 0.435 | 0.137 | 6.19E-23 | 18 | Lap3 |
| Retreg15 | 4.02E-27 | 0.687854777 | 0.345 | 0.097 | 8.14E-23 | 18 | Retreg1 |
| Tgfbr11 | 5.67E-27 | 1.620672306 | 0.494 | 0.206 | 1.15E-22 | 18 | Tgfbr1 |
| Ctsb4 | 7.75E-27 | 1.247548438 | 0.625 | 0.298 | 1.57E-22 | 18 | Ctsb |
| Arhgdib8 | 9.33E-27 | 0.788428938 | 0.613 | 0.233 | 1.89E-22 | 18 | Arhgdib |
| Dapp13 | 1.57E-26 | 0.592098796 | 0.31 | 0.081 | 3.18E-22 | 18 | Dapp1 |
| Arhgap306 | 1.72E-26 | 0.42570432 | 0.381 | 0.11 | 3.48E-22 | 18 | Arhgap30 |
| Anp32e | 2.22E-26 | 0.929525952 | 0.464 | 0.164 | 4.50E-22 | 18 | Anp32e |

| | | | | | | | |
|---|---|---|---|---|---|---|---|
| Aim26 | 2.50E-26 | 0.606682694 | 0.357 | 0.103 | 5.06E-22 | 18 | Aim2 |
| Snrpd1 | 2.80E-26 | 0.902021127 | 0.667 | 0.307 | 5.67E-22 | 18 | Snrpd1 |
| Plek4 | 2.93E-26 | 0.638671426 | 0.405 | 0.125 | 5.93E-22 | 18 | Plek |
| Tmpo4 | 2.94E-26 | 0.918251779 | 0.524 | 0.208 | 5.95E-22 | 18 | Tmpo |
| Fyb6 | 3.15E-26 | 0.975257327 | 0.5 | 0.184 | 6.37E-22 | 18 | Fyb |
| Cd3023 | 4.22E-26 | 0.588097534 | 0.315 | 0.081 | 8.54E-22 | 18 | Cd302 |
| Rgs26 | 4.39E-26 | 0.751106021 | 0.405 | 0.13 | 8.88E-22 | 18 | Rgs2 |
| Creg13 | 6.27E-26 | 0.841641396 | 0.506 | 0.188 | 1.27E-21 | 18 | Creg1 |
| Crlf24 | 1.45E-25 | 0.747596762 | 0.458 | 0.161 | 2.93E-21 | 18 | Crlf2 |
| Naip21 | 1.80E-25 | 0.384023684 | 0.25 | 0.058 | 3.65E-21 | 18 | Naip2 |
| Tmem176a8 | 2.44E-25 | 1.040374035 | 0.738 | 0.418 | 4.95E-21 | 18 | Tmem176a |
| Cd635 | 2.45E-25 | 0.653513441 | 0.53 | 0.188 | 4.96E-21 | 18 | Cd63 |
| Vav15 | 3.19E-25 | 0.487892062 | 0.304 | 0.081 | 6.46E-21 | 18 | Vav1 |
| Nap1l15 | 3.22E-25 | 1.222796905 | 0.631 | 0.315 | 6.53E-21 | 18 | Nap1l1 |
| Acer32 | 3.49E-25 | 0.732652119 | 0.423 | 0.141 | 7.06E-21 | 18 | Acer3 |
| Actr37 | 6.24E-25 | 1.126452361 | 0.685 | 0.392 | 1.26E-20 | 18 | Actr3 |
| Hist1h1e2 | 8.08E-25 | 1.640554147 | 0.679 | 0.372 | 1.64E-20 | 18 | Hist1h1e |
| Snx53 | 1.09E-24 | 0.734166889 | 0.536 | 0.211 | 2.21E-20 | 18 | Snx5 |
| Rhog1 | 1.17E-24 | 0.580478558 | 0.405 | 0.129 | 2.36E-20 | 18 | Rhog |
| Prkcd1 | 1.50E-24 | 0.370878038 | 0.321 | 0.088 | 3.04E-20 | 18 | Prkcd |
| Bin26 | 1.54E-24 | 0.591396235 | 0.399 | 0.128 | 3.12E-20 | 18 | Bin2 |
| Cmtm73 | 3.56E-24 | 0.689129371 | 0.542 | 0.213 | 7.20E-20 | 18 | Cmtm7 |
| Lsm2 | 3.79E-24 | 0.684663659 | 0.446 | 0.153 | 7.67E-20 | 18 | Lsm2 |
| Myof2 | 7.14E-24 | 0.381939581 | 0.25 | 0.059 | 1.44E-19 | 18 | Myof |
| Ifngr24 | 7.16E-24 | 0.490185993 | 0.411 | 0.132 | 1.45E-19 | 18 | Ifngr2 |
| Asph3 | 8.18E-24 | 0.518975046 | 0.339 | 0.1 | 1.66E-19 | 18 | Asph |
| Cd536 | 9.36E-24 | 0.515808087 | 0.375 | 0.117 | 1.89E-19 | 18 | Cd53 |
| Tmem2192 | 1.53E-23 | 0.517168512 | 0.387 | 0.123 | 3.10E-19 | 18 | Tmem219 |
| Tipin | 1.65E-23 | 0.544814947 | 0.315 | 0.089 | 3.35E-19 | 18 | Tipin |
| Irf82 | 1.79E-23 | 0.646049973 | 0.351 | 0.109 | 3.62E-19 | 18 | Irf8 |
| Tnf3 | 1.83E-23 | 0.742663243 | 0.274 | 0.073 | 3.71E-19 | 18 | Tnf |
| Cenpc1 | 2.10E-23 | 0.396563429 | 0.262 | 0.066 | 4.26E-19 | 18 | Cenpc1 |
| Tnfaip8l23 | 2.97E-23 | 0.588463283 | 0.274 | 0.072 | 6.02E-19 | 18 | Tnfaip8l2 |
| Bmyc1 | 3.43E-23 | 0.807839206 | 0.47 | 0.17 | 6.94E-19 | 18 | Bmyc |
| Plxdc23 | 4.42E-23 | 0.53317706 | 0.327 | 0.093 | 8.95E-19 | 18 | Plxdc2 |
| Cdkn1a1 | 5.77E-23 | 0.809600065 | 0.387 | 0.127 | 1.17E-18 | 18 | Cdkn1a |
| Commd81 | 8.14E-23 | 0.643147505 | 0.435 | 0.154 | 1.65E-18 | 18 | Commd8 |
| Rps6ka56 | 8.16E-23 | 0.537758827 | 0.351 | 0.108 | 1.65E-18 | 18 | Rps6ka5 |
| Ncf45 | 1.01E-22 | 0.61725953 | 0.363 | 0.112 | 2.05E-18 | 18 | Ncf4 |
| Tanc22 | 1.10E-22 | 1.213395663 | 0.417 | 0.16 | 2.23E-18 | 18 | Tanc2 |
| Hexa3 | 1.17E-22 | 0.889536265 | 0.53 | 0.221 | 2.36E-18 | 18 | Hexa |
| Ighm5 | 1.25E-22 | 0.640473856 | 0.381 | 0.127 | 2.53E-18 | 18 | Ighm |
| Sfmbt12 | 1.85E-22 | 0.584707273 | 0.292 | 0.084 | 3.75E-18 | 18 | Sfmbt1 |

| | | | | | | | |
|---|---|---|---|---|---|---|---|
| Myo1f5 | 2.78E-22 | 0.614810781 | 0.405 | 0.138 | 5.62E-18 | 18 | Myo1f |
| Calm35 | 3.41E-22 | 0.900831933 | 0.589 | 0.269 | 6.90E-18 | 18 | Calm3 |
| Atox14 | 6.98E-22 | 0.913537835 | 0.875 | 0.619 | 1.41E-17 | 18 | Atox1 |
| Ptpn187 | 7.70E-22 | 0.857154629 | 0.661 | 0.316 | 1.56E-17 | 18 | Ptpn18 |
| Hmgn13 | 8.53E-22 | 0.820232565 | 0.661 | 0.329 | 1.73E-17 | 18 | Hmgn1 |
| Exosc81 | 8.75E-22 | 0.46166746 | 0.405 | 0.136 | 1.77E-17 | 18 | Exosc8 |
| Acsl11 | 1.09E-21 | 0.50927043 | 0.268 | 0.073 | 2.21E-17 | 18 | Acsl1 |
| Prcp1 | 1.10E-21 | 0.412043403 | 0.304 | 0.087 | 2.22E-17 | 18 | Prcp |
| Vsir2 | 1.24E-21 | 0.411934616 | 0.339 | 0.104 | 2.51E-17 | 18 | Vsir |
| Anp32b3 | 1.41E-21 | 1.011031354 | 0.72 | 0.423 | 2.85E-17 | 18 | Anp32b |
| Hat1 | 1.82E-21 | 0.612235508 | 0.363 | 0.12 | 3.69E-17 | 18 | Hat1 |
| Mast22 | 2.02E-21 | 0.39647897 | 0.268 | 0.072 | 4.09E-17 | 18 | Mast2 |
| Cdk66 | 2.65E-21 | 0.471667531 | 0.351 | 0.112 | 5.36E-17 | 18 | Cdk6 |
| Pde3b6 | 3.37E-21 | 0.581251289 | 0.375 | 0.128 | 6.83E-17 | 18 | Pde3b |
| Mafb2 | 4.27E-21 | 0.707246104 | 0.423 | 0.155 | 8.65E-17 | 18 | Mafb |
| Syngr21 | 5.42E-21 | 0.478208118 | 0.405 | 0.142 | 1.10E-16 | 18 | Syngr2 |
| Arhgap156 | 7.10E-21 | 1.230293288 | 0.458 | 0.189 | 1.44E-16 | 18 | Arhgap15 |
| Sat13 | 8.12E-21 | 0.92173723 | 0.649 | 0.346 | 1.64E-16 | 18 | Sat1 |
| Ranbp1 | 9.97E-21 | 0.984709631 | 0.631 | 0.323 | 2.02E-16 | 18 | Ranbp1 |
| Epsti15 | 1.21E-20 | 0.636764183 | 0.464 | 0.176 | 2.46E-16 | 18 | Epsti1 |
| Hacd22 | 2.32E-20 | 0.519670626 | 0.357 | 0.117 | 4.69E-16 | 18 | Hacd2 |
| Runx16 | 2.88E-20 | 0.68227646 | 0.375 | 0.132 | 5.83E-16 | 18 | Runx1 |
| Fxyd59 | 3.79E-20 | 0.89595646 | 0.714 | 0.412 | 7.68E-16 | 18 | Fxyd5 |
| Plcg26 | 4.04E-20 | 0.560753473 | 0.351 | 0.12 | 8.17E-16 | 18 | Plcg2 |
| Gns3 | 4.21E-20 | 0.503453474 | 0.375 | 0.127 | 8.51E-16 | 18 | Gns |
| Ppia2 | 5.33E-20 | 0.64358668 | 1 | 0.938 | 1.08E-15 | 18 | Ppia |
| Gna121 | 5.33E-20 | 0.69364952 | 0.405 | 0.153 | 1.08E-15 | 18 | Gna12 |
| Sdf2l14 | 7.30E-20 | 0.870031865 | 0.429 | 0.168 | 1.48E-15 | 18 | Sdf2l1 |
| Itga91 | 2.33E-19 | 0.496068278 | 0.321 | 0.102 | 4.71E-15 | 18 | Itga9 |
| Hps31 | 2.35E-19 | 0.353156062 | 0.25 | 0.069 | 4.75E-15 | 18 | Hps3 |
| Tnfaip86 | 3.41E-19 | 0.693038121 | 0.458 | 0.187 | 6.90E-15 | 18 | Tnfaip8 |
| Ckap52 | 3.71E-19 | 0.581878218 | 0.387 | 0.142 | 7.50E-15 | 18 | Ckap5 |
| 4933406I18Rik5 | 4.43E-19 | 0.610539885 | 0.262 | 0.076 | 8.96E-15 | 18 | 4933406I18Rik |
| Lyn2 | 4.44E-19 | 1.052841945 | 0.542 | 0.273 | 8.99E-15 | 18 | Lyn |
| Rap2a2 | 4.52E-19 | 0.60113269 | 0.345 | 0.118 | 9.15E-15 | 18 | Rap2a |
| Ap1s23 | 6.76E-19 | 0.570500058 | 0.387 | 0.139 | 1.37E-14 | 18 | Ap1s2 |
| Rfc23 | 1.01E-18 | 0.465615733 | 0.357 | 0.125 | 2.04E-14 | 18 | Rfc2 |
| Slamf75 | 1.20E-18 | 0.547247771 | 0.31 | 0.1 | 2.42E-14 | 18 | Slamf7 |
| Cep295 | 1.46E-18 | 0.34774371 | 0.262 | 0.075 | 2.95E-14 | 18 | Cep295 |
| Pdcd2l1 | 1.53E-18 | 0.290783815 | 0.28 | 0.083 | 3.10E-14 | 18 | Pdcd2l |
| Zfhx32 | 2.03E-18 | 1.019667725 | 0.423 | 0.183 | 4.12E-14 | 18 | Zfhx3 |
| Tec5 | 2.26E-18 | 0.341975035 | 0.333 | 0.11 | 4.57E-14 | 18 | Tec |
| Hsp90aa12 | 2.35E-18 | 0.88286857 | 0.696 | 0.426 | 4.75E-14 | 18 | Hsp90aa1 |

| | | | | | | | |
|---|---|---|---|---|---|---|---|
| Taok36 | 2.54E-18 | 0.661938405 | 0.411 | 0.164 | 5.14E-14 | 18 | Taok3 |
| Selplg6 | 2.71E-18 | 0.49111746 | 0.417 | 0.155 | 5.48E-14 | 18 | Selplg |
| Lpxn5 | 2.95E-18 | 0.367856215 | 0.298 | 0.093 | 5.97E-14 | 18 | Lpxn |
| Hmgb11 | 5.33E-18 | 0.808310415 | 0.893 | 0.717 | 1.08E-13 | 18 | Hmgb1 |
| Glrx2 | 5.86E-18 | 0.255799247 | 0.25 | 0.071 | 1.19E-13 | 18 | Glrx |
| Gng26 | 6.01E-18 | 0.688347071 | 0.458 | 0.187 | 1.22E-13 | 18 | Gng2 |
| Capza21 | 7.15E-18 | 0.741221482 | 0.744 | 0.462 | 1.45E-13 | 18 | Capza2 |
| Fkbp22 | 7.70E-18 | 0.706310529 | 0.565 | 0.277 | 1.56E-13 | 18 | Fkbp2 |
| Cmc2 | 8.42E-18 | 0.465090186 | 0.339 | 0.117 | 1.71E-13 | 18 | Cmc2 |
| Blvrb3 | 1.35E-17 | 0.66935539 | 0.524 | 0.243 | 2.72E-13 | 18 | Blvrb |
| Fau6 | 1.40E-17 | 0.603672534 | 0.988 | 0.953 | 2.83E-13 | 18 | Fau |
| Dock107 | 1.85E-17 | 0.891276096 | 0.476 | 0.209 | 3.75E-13 | 18 | Dock10 |
| Zfp7042 | 1.92E-17 | 0.374646537 | 0.363 | 0.13 | 3.88E-13 | 18 | Zfp704 |
| Arl6ip18 | 2.24E-17 | 1.067350819 | 0.607 | 0.371 | 4.54E-13 | 18 | Arl6ip1 |
| Dek1 | 3.05E-17 | 0.827321937 | 0.637 | 0.373 | 6.17E-13 | 18 | Dek |
| Sdc31 | 3.83E-17 | 0.369208238 | 0.262 | 0.078 | 7.76E-13 | 18 | Sdc3 |
| Hist1h1c2 | 4.06E-17 | 0.999236657 | 0.452 | 0.201 | 8.22E-13 | 18 | Hist1h1c |
| Trappc1 | 5.78E-17 | 0.458777448 | 0.321 | 0.11 | 1.17E-12 | 18 | Trappc1 |
| Fam129a6 | 7.38E-17 | 0.540687568 | 0.375 | 0.146 | 1.49E-12 | 18 | Fam129a |
| Cdkn2d4 | 7.39E-17 | 0.599274823 | 0.28 | 0.093 | 1.50E-12 | 18 | Cdkn2d |
| Arpc27 | 8.22E-17 | 0.73367198 | 0.774 | 0.545 | 1.66E-12 | 18 | Arpc2 |
| Tle12 | 8.41E-17 | 0.378314749 | 0.286 | 0.092 | 1.70E-12 | 18 | Tle1 |
| Ucp24 | 8.58E-17 | 0.540142161 | 0.643 | 0.331 | 1.74E-12 | 18 | Ucp2 |
| Cfl17 | 1.01E-16 | 0.788264649 | 0.833 | 0.609 | 2.04E-12 | 18 | Cfl1 |
| Gpsm36 | 1.59E-16 | 0.562531027 | 0.381 | 0.148 | 3.23E-12 | 18 | Gpsm3 |
| Grap26 | 1.64E-16 | 0.423485404 | 0.31 | 0.106 | 3.32E-12 | 18 | Grap2 |
| Meaf6 | 2.18E-16 | 0.259681009 | 0.31 | 0.103 | 4.41E-12 | 18 | Meaf6 |
| Ifi2042 | 2.39E-16 | 0.320228843 | 0.292 | 0.095 | 4.85E-12 | 18 | Ifi204 |
| Gab25 | 2.42E-16 | 1.146893961 | 0.435 | 0.206 | 4.90E-12 | 18 | Gab2 |
| Twf24 | 3.61E-16 | 0.460923273 | 0.417 | 0.171 | 7.30E-12 | 18 | Twf2 |
| Grb23 | 4.96E-16 | 0.716636984 | 0.56 | 0.283 | 1.00E-11 | 18 | Grb2 |
| Ddx393 | 5.44E-16 | 0.592631172 | 0.339 | 0.126 | 1.10E-11 | 18 | Ddx39 |
| Gsap5 | 6.60E-16 | 0.306726757 | 0.268 | 0.086 | 1.34E-11 | 18 | Gsap |
| Prdx45 | 6.70E-16 | 0.671670501 | 0.488 | 0.222 | 1.36E-11 | 18 | Prdx4 |
| Pih1d1 | 7.59E-16 | 0.524727334 | 0.292 | 0.098 | 1.54E-11 | 18 | Pih1d1 |
| Pnp4 | 9.12E-16 | 0.627762317 | 0.536 | 0.254 | 1.85E-11 | 18 | Pnp |
| Rpsa7 | 9.67E-16 | 0.593182794 | 0.97 | 0.903 | 1.96E-11 | 18 | Rpsa |
| Cep1922 | 1.12E-15 | 0.399984413 | 0.262 | 0.086 | 2.27E-11 | 18 | Cep192 |
| Efhd26 | 1.17E-15 | 0.675024889 | 0.446 | 0.187 | 2.37E-11 | 18 | Efhd2 |
| Hspa8 | 1.18E-15 | 0.661807124 | 0.863 | 0.71 | 2.39E-11 | 18 | Hspa8 |
| Cebpzos1 | 1.57E-15 | 0.322129431 | 0.476 | 0.202 | 3.17E-11 | 18 | Cebpzos |
| Hnrnpa14 | 1.59E-15 | 0.828955092 | 0.613 | 0.325 | 3.22E-11 | 18 | Hnrnpa1 |
| Cyfip15 | 1.90E-15 | 0.695091341 | 0.548 | 0.279 | 3.85E-11 | 18 | Cyfip1 |

| | | | | | | | |
|---|---|---|---|---|---|---|---|
| Mgat4a5 | 2.25E-15 | 0.363723258 | 0.298 | 0.106 | 4.56E-11 | 18 | Mgat4a |
| Zfp7102 | 2.50E-15 | 0.45702958 | 0.387 | 0.156 | 5.05E-11 | 18 | Zfp710 |
| Fuca12 | 2.89E-15 | 0.458360776 | 0.47 | 0.208 | 5.86E-11 | 18 | Fuca1 |
| Bub3 | 3.55E-15 | 0.749188036 | 0.458 | 0.206 | 7.18E-11 | 18 | Bub3 |
| Dnajb142 | 3.81E-15 | 0.268661784 | 0.321 | 0.115 | 7.71E-11 | 18 | Dnajb14 |
| Dock86 | 5.37E-15 | 0.808442519 | 0.458 | 0.219 | 1.09E-10 | 18 | Dock8 |
| Pcna2 | 5.43E-15 | 0.332480937 | 0.327 | 0.122 | 1.10E-10 | 18 | Pcna |
| Timm50 | 5.78E-15 | 0.36005059 | 0.268 | 0.088 | 1.17E-10 | 18 | Timm50 |
| Dock26 | 9.19E-15 | 0.435963743 | 0.446 | 0.187 | 1.86E-10 | 18 | Dock2 |
| Ywhah5 | 9.20E-15 | 0.807868339 | 0.554 | 0.31 | 1.86E-10 | 18 | Ywhah |
| Slc9a96 | 9.22E-15 | 0.868112294 | 0.357 | 0.152 | 1.87E-10 | 18 | Slc9a9 |
| Fmnl16 | 9.27E-15 | 0.348544038 | 0.274 | 0.093 | 1.88E-10 | 18 | Fmnl1 |
| Inpp5d6 | 9.42E-15 | 0.859691506 | 0.423 | 0.205 | 1.91E-10 | 18 | Inpp5d |
| Frmd4b6 | 9.54E-15 | 0.705972872 | 0.405 | 0.178 | 1.93E-10 | 18 | Frmd4b |
| Celf27 | 1.00E-14 | 0.566243885 | 0.446 | 0.203 | 2.03E-10 | 18 | Celf2 |
| Nubp11 | 1.09E-14 | 0.546508214 | 0.286 | 0.101 | 2.21E-10 | 18 | Nubp1 |
| Asah13 | 1.15E-14 | 0.459645011 | 0.435 | 0.189 | 2.32E-10 | 18 | Asah1 |
| Arpc41 | 1.54E-14 | 0.647300966 | 0.56 | 0.291 | 3.11E-10 | 18 | Arpc4 |
| Pip4k2a4 | 1.55E-14 | 1.03759214 | 0.458 | 0.242 | 3.14E-10 | 18 | Pip4k2a |
| Pik3ap14 | 1.72E-14 | 0.276757742 | 0.268 | 0.091 | 3.49E-10 | 18 | Pik3ap1 |
| Dtymk | 2.29E-14 | 0.576009339 | 0.345 | 0.135 | 4.64E-10 | 18 | Dtymk |
| Rps296 | 2.54E-14 | 0.567667253 | 1 | 0.971 | 5.14E-10 | 18 | Rps29 |
| Rnpep1 | 3.23E-14 | 0.413744286 | 0.315 | 0.116 | 6.55E-10 | 18 | Rnpep |
| Eif1ad | 3.29E-14 | 0.304169696 | 0.321 | 0.118 | 6.65E-10 | 18 | Eif1ad |
| Tkt2 | 3.38E-14 | 0.428872582 | 0.524 | 0.253 | 6.84E-10 | 18 | Tkt |
| Tm6sf15 | 3.51E-14 | 0.51099276 | 0.351 | 0.142 | 7.11E-10 | 18 | Tm6sf1 |
| Atpif13 | 3.92E-14 | 0.530822311 | 0.81 | 0.561 | 7.93E-10 | 18 | Atpif1 |
| Ncaph2 | 3.99E-14 | 0.318359073 | 0.25 | 0.082 | 8.07E-10 | 18 | Ncaph2 |
| Sae11 | 4.11E-14 | 0.529827641 | 0.363 | 0.148 | 8.32E-10 | 18 | Sae1 |
| Nmt12 | 5.57E-14 | 0.551508356 | 0.56 | 0.291 | 1.13E-09 | 18 | Nmt1 |
| Rps57 | 5.68E-14 | 0.562096708 | 0.994 | 0.92 | 1.15E-09 | 18 | Rps5 |
| Derl21 | 6.64E-14 | 0.291275708 | 0.333 | 0.127 | 1.34E-09 | 18 | Derl2 |
| Rbm36 | 6.67E-14 | 0.72038842 | 0.732 | 0.478 | 1.35E-09 | 18 | Rbm3 |
| Ccr55 | 7.22E-14 | 0.345766073 | 0.345 | 0.133 | 1.46E-09 | 18 | Ccr5 |
| Soat13 | 7.35E-14 | 0.380605179 | 0.292 | 0.108 | 1.49E-09 | 18 | Soat1 |
| Cers65 | 7.60E-14 | 0.424737412 | 0.31 | 0.116 | 1.54E-09 | 18 | Cers6 |
| Runx35 | 1.55E-13 | 0.298823792 | 0.375 | 0.15 | 3.13E-09 | 18 | Runx3 |
| Mir142hg6 | 1.96E-13 | 0.493702073 | 0.399 | 0.171 | 3.97E-09 | 18 | Mir142hg |
| Rtcb1 | 2.04E-13 | 0.529126104 | 0.393 | 0.171 | 4.12E-09 | 18 | Rtcb |
| Rps206 | 2.34E-13 | 0.522924196 | 0.988 | 0.951 | 4.74E-09 | 18 | Rps20 |
| Cerk6 | 2.52E-13 | 0.25723645 | 0.31 | 0.115 | 5.09E-09 | 18 | Cerk |
| Nudt212 | 2.80E-13 | 0.504339141 | 0.375 | 0.158 | 5.66E-09 | 18 | Nudt21 |
| Epb41l24 | 2.84E-13 | 0.691370936 | 0.464 | 0.242 | 5.74E-09 | 18 | Epb41l2 |

| | | | | | | | |
|---|---|---|---|---|---|---|---|
| Cnih46 | 2.87E-13 | 0.674942515 | 0.488 | 0.254 | 5.81E-09 | 18 | Cnih4 |
| Hba-a22 | 2.87E-13 | 1.578438913 | 0.286 | 0.112 | 5.82E-09 | 18 | Hba-a2 |
| Rbpj6 | 2.90E-13 | 0.620125471 | 0.464 | 0.221 | 5.88E-09 | 18 | Rbpj |
| Scamp22 | 3.14E-13 | 0.368227598 | 0.345 | 0.142 | 6.35E-09 | 18 | Scamp2 |
| Dnmt1 | 3.45E-13 | 0.284841553 | 0.286 | 0.105 | 6.98E-09 | 18 | Dnmt1 |
| Pa2g4 | 3.74E-13 | 0.504272825 | 0.429 | 0.198 | 7.58E-09 | 18 | Pa2g4 |
| Psd33 | 4.05E-13 | 0.392019782 | 0.31 | 0.118 | 8.19E-09 | 18 | Psd3 |
| Pip5k1c1 | 4.52E-13 | 0.289929901 | 0.274 | 0.099 | 9.15E-09 | 18 | Pip5k1c |
| Pbx33 | 4.64E-13 | 0.553044585 | 0.333 | 0.14 | 9.39E-09 | 18 | Pbx3 |
| Trmt1125 | 6.84E-13 | 0.599055654 | 0.542 | 0.274 | 1.39E-08 | 18 | Trmt112 |
| Extl32 | 6.95E-13 | 0.363070771 | 0.345 | 0.14 | 1.41E-08 | 18 | Extl3 |
| Ctsa1 | 7.03E-13 | 0.441309346 | 0.363 | 0.154 | 1.42E-08 | 18 | Ctsa |
| Tmem55b1 | 8.54E-13 | 0.294818018 | 0.28 | 0.103 | 1.73E-08 | 18 | Tmem55b |
| Psmb22 | 9.16E-13 | 0.645574826 | 0.655 | 0.403 | 1.85E-08 | 18 | Psmb2 |
| Gpx14 | 9.64E-13 | 0.447783967 | 0.827 | 0.596 | 1.95E-08 | 18 | Gpx1 |
| Brk11 | 1.40E-12 | 0.650794217 | 0.685 | 0.443 | 2.83E-08 | 18 | Brk1 |
| Parp87 | 1.42E-12 | 0.463786225 | 0.351 | 0.15 | 2.88E-08 | 18 | Parp8 |
| Tubb2a4 | 1.49E-12 | 0.662865753 | 0.375 | 0.165 | 3.01E-08 | 18 | Tubb2a |
| Ppp1r212 | 1.88E-12 | 0.512448822 | 0.292 | 0.113 | 3.82E-08 | 18 | Ppp1r21 |
| Mgat56 | 2.02E-12 | 0.543696013 | 0.375 | 0.171 | 4.09E-08 | 18 | Mgat5 |
| Orc6 | 2.16E-12 | 0.476241772 | 0.274 | 0.104 | 4.38E-08 | 18 | Orc6 |
| Cenpx3 | 2.27E-12 | 0.533416689 | 0.56 | 0.301 | 4.59E-08 | 18 | Cenpx |
| Hdac92 | 2.34E-12 | 0.56893507 | 0.363 | 0.159 | 4.74E-08 | 18 | Hdac9 |
| Camk13 | 3.02E-12 | 0.555133885 | 0.47 | 0.228 | 6.12E-08 | 18 | Camk1 |
| Ccl45 | 3.62E-12 | 0.625016355 | 0.315 | 0.13 | 7.34E-08 | 18 | Ccl4 |
| Nudt5 | 3.99E-12 | 0.34402502 | 0.28 | 0.107 | 8.09E-08 | 18 | Nudt5 |
| Ccdc122 | 4.29E-12 | 0.524105026 | 0.673 | 0.415 | 8.68E-08 | 18 | Ccdc12 |
| Arhgef66 | 5.19E-12 | 0.296520688 | 0.321 | 0.129 | 1.05E-07 | 18 | Arhgef6 |
| Rbbp8 | 5.97E-12 | 0.382101381 | 0.268 | 0.102 | 1.21E-07 | 18 | Rbbp8 |
| Ppt11 | 6.13E-12 | 0.2921425 | 0.321 | 0.13 | 1.24E-07 | 18 | Ppt1 |
| Filip1l5 | 7.20E-12 | 0.524276152 | 0.351 | 0.153 | 1.46E-07 | 18 | Filip1l |
| Pitpna4 | 7.32E-12 | 0.645861233 | 0.571 | 0.329 | 1.48E-07 | 18 | Pitpna |
| Emp39 | 1.01E-11 | 0.588888048 | 0.5 | 0.26 | 2.04E-07 | 18 | Emp3 |
| Bin15 | 1.10E-11 | 0.39037282 | 0.393 | 0.176 | 2.22E-07 | 18 | Bin1 |
| Grn5 | 1.15E-11 | 0.47820525 | 0.452 | 0.225 | 2.33E-07 | 18 | Grn |
| Limd26 | 1.23E-11 | 0.274918137 | 0.351 | 0.149 | 2.49E-07 | 18 | Limd2 |
| Gdi22 | 1.37E-11 | 0.576414748 | 0.625 | 0.385 | 2.77E-07 | 18 | Gdi2 |
| Rel5 | 1.86E-11 | 0.352518118 | 0.327 | 0.139 | 3.76E-07 | 18 | Rel |
| Lsm3 | 1.89E-11 | 0.355647876 | 0.411 | 0.188 | 3.83E-07 | 18 | Lsm3 |
| Klhl65 | 2.05E-11 | 0.478342377 | 0.298 | 0.121 | 4.14E-07 | 18 | Klhl6 |
| Cep571 | 2.11E-11 | 0.327328641 | 0.28 | 0.109 | 4.27E-07 | 18 | Cep57 |
| Sdccag84 | 2.14E-11 | 0.432812825 | 0.298 | 0.122 | 4.32E-07 | 18 | Sdccag8 |
| Itgav8 | 2.22E-11 | 0.376960128 | 0.363 | 0.162 | 4.49E-07 | 18 | Itgav |

| | | | | | | | |
|---|---|---|---|---|---|---|---|
| Nav25 | 2.97E-11 | 0.519894984 | 0.363 | 0.166 | 6.02E-07 | 18 | Nav2 |
| B2m10 | 3.78E-11 | 0.59515186 | 0.905 | 0.691 | 7.65E-07 | 18 | B2m |
| Larp7 | 3.78E-11 | 0.357756979 | 0.304 | 0.122 | 7.65E-07 | 18 | Larp7 |
| Smchd16 | 4.02E-11 | 0.684916249 | 0.476 | 0.265 | 8.14E-07 | 18 | Smchd1 |
| Pag15 | 4.46E-11 | 0.264208728 | 0.298 | 0.122 | 9.03E-07 | 18 | Pag1 |
| Ppp1r186 | 4.70E-11 | 0.326105783 | 0.357 | 0.155 | 9.51E-07 | 18 | Ppp1r18 |
| Adcy77 | 4.86E-11 | 0.283101756 | 0.286 | 0.115 | 9.83E-07 | 18 | Adcy7 |
| 1700025G04Rik5 | 5.51E-11 | 0.465232555 | 0.399 | 0.189 | 1.12E-06 | 18 | 1700025G04Rik |
| Baz1a6 | 6.56E-11 | 0.404522377 | 0.375 | 0.173 | 1.33E-06 | 18 | Baz1a |
| Pycard6 | 6.68E-11 | 0.372923096 | 0.536 | 0.278 | 1.35E-06 | 18 | Pycard |
| Rnaset2a1 | 6.76E-11 | 0.383643347 | 0.28 | 0.113 | 1.37E-06 | 18 | Rnaset2a |
| Dnmt3a4 | 7.89E-11 | 0.260392193 | 0.339 | 0.148 | 1.60E-06 | 18 | Dnmt3a |
| Gltp | 8.62E-11 | 0.4242554 | 0.327 | 0.142 | 1.74E-06 | 18 | Gltp |
| Plekho14 | 8.82E-11 | 0.511690896 | 0.452 | 0.225 | 1.79E-06 | 18 | Plekho1 |
| Ola1 | 1.44E-10 | 0.304400288 | 0.423 | 0.199 | 2.92E-06 | 18 | Ola1 |
| Lsm51 | 1.54E-10 | 0.542109308 | 0.446 | 0.221 | 3.11E-06 | 18 | Lsm5 |
| Fam214a2 | 1.54E-10 | 0.293854079 | 0.298 | 0.125 | 3.12E-06 | 18 | Fam214a |
| Rps267 | 1.66E-10 | 0.592644759 | 0.958 | 0.866 | 3.36E-06 | 18 | Rps26 |
| Cbfb5 | 1.66E-10 | 0.419533683 | 0.345 | 0.156 | 3.37E-06 | 18 | Cbfb |
| Arhgap456 | 1.82E-10 | 0.285896867 | 0.363 | 0.159 | 3.69E-06 | 18 | Arhgap45 |
| Prdx13 | 1.88E-10 | 0.510201115 | 0.863 | 0.693 | 3.81E-06 | 18 | Prdx1 |
| Srsf92 | 1.89E-10 | 0.444819662 | 0.577 | 0.33 | 3.83E-06 | 18 | Srsf9 |
| Bst27 | 1.91E-10 | 0.503906845 | 0.75 | 0.504 | 3.86E-06 | 18 | Bst2 |
| Rpl35a7 | 2.10E-10 | 0.445053539 | 0.988 | 0.948 | 4.25E-06 | 18 | Rpl35a |
| Nde1 | 2.18E-10 | 0.439331503 | 0.256 | 0.102 | 4.40E-06 | 18 | Nde1 |
| Gapdh2 | 2.46E-10 | 0.438678987 | 0.881 | 0.708 | 4.99E-06 | 18 | Gapdh |
| Psmb810 | 2.55E-10 | 0.579682495 | 0.798 | 0.558 | 5.17E-06 | 18 | Psmb8 |
| Osgep | 2.76E-10 | 0.346746092 | 0.369 | 0.171 | 5.59E-06 | 18 | Osgep |
| Impdh2 | 3.31E-10 | 0.394472505 | 0.274 | 0.114 | 6.71E-06 | 18 | Impdh2 |
| Hspa1a3 | 3.39E-10 | 0.263304515 | 0.256 | 0.101 | 6.85E-06 | 18 | Hspa1a |
| Myo9b4 | 3.52E-10 | 0.311922741 | 0.286 | 0.12 | 7.13E-06 | 18 | Myo9b |
| Slfn26 | 3.67E-10 | 0.458486108 | 0.488 | 0.262 | 7.44E-06 | 18 | Slfn2 |
| Hint12 | 3.93E-10 | 0.537607034 | 0.851 | 0.696 | 7.95E-06 | 18 | Hint1 |
| Ssr43 | 4.52E-10 | 0.619701978 | 0.679 | 0.469 | 9.15E-06 | 18 | Ssr4 |
| Ddost3 | 4.61E-10 | 0.499673741 | 0.423 | 0.217 | 9.33E-06 | 18 | Ddost |
| Hsd17b4 | 4.71E-10 | 0.318365365 | 0.304 | 0.131 | 9.54E-06 | 18 | Hsd17b4 |
| Cttnbp2nl2 | 5.45E-10 | 0.268184389 | 0.28 | 0.116 | 1.10E-05 | 18 | Cttnbp2nl |
| Ilk2 | 6.15E-10 | 0.380004054 | 0.369 | 0.171 | 1.25E-05 | 18 | Ilk |
| Lsp17 | 6.22E-10 | 0.724348362 | 0.304 | 0.14 | 1.26E-05 | 18 | Lsp1 |
| Cdc423 | 6.56E-10 | 0.605935273 | 0.786 | 0.617 | 1.33E-05 | 18 | Cdc42 |
| Magt14 | 7.22E-10 | 0.365168413 | 0.351 | 0.165 | 1.46E-05 | 18 | Magt1 |
| Lhfpl22 | 7.27E-10 | 0.354427808 | 0.333 | 0.151 | 1.47E-05 | 18 | Lhfpl2 |
| Hmgxb4 | 7.52E-10 | 0.278901301 | 0.256 | 0.102 | 1.52E-05 | 18 | Hmgxb4 |

| | | | | | | | |
|---|---|---|---|---|---|---|---|
| Nsmce1 | 8.19E-10 | 0.544455818 | 0.327 | 0.15 | 1.66E-05 | 18 | Nsmce1 |
| Ext16 | 9.27E-10 | 0.43405662 | 0.351 | 0.168 | 1.88E-05 | 18 | Ext1 |
| Wdr444 | 9.78E-10 | 0.255473469 | 0.292 | 0.124 | 1.98E-05 | 18 | Wdr44 |
| Ptk2b6 | 9.82E-10 | 0.254232666 | 0.339 | 0.151 | 1.99E-05 | 18 | Ptk2b |
| Coro1b1 | 1.05E-09 | 0.323526673 | 0.292 | 0.124 | 2.13E-05 | 18 | Coro1b |
| Erh | 1.12E-09 | 0.336989228 | 0.381 | 0.184 | 2.27E-05 | 18 | Erh |
| Lcorl6 | 1.17E-09 | 0.434524083 | 0.345 | 0.167 | 2.38E-05 | 18 | Lcorl |
| Tor1a1 | 1.21E-09 | 0.297501711 | 0.262 | 0.107 | 2.44E-05 | 18 | Tor1a |
| Snrpb2 | 1.25E-09 | 0.583445935 | 0.685 | 0.473 | 2.53E-05 | 18 | Snrpb |
| Hipk21 | 1.25E-09 | 0.275434601 | 0.274 | 0.118 | 2.53E-05 | 18 | Hipk2 |
| Hnrnpa33 | 1.27E-09 | 0.50691081 | 0.685 | 0.446 | 2.57E-05 | 18 | Hnrnpa3 |
| Ccdc88a9 | 1.29E-09 | 0.538959376 | 0.399 | 0.205 | 2.61E-05 | 18 | Ccdc88a |
| Rps6ka16 | 1.34E-09 | 0.309372668 | 0.274 | 0.115 | 2.72E-05 | 18 | Rps6ka1 |
| Banf11 | 1.35E-09 | 0.514701591 | 0.464 | 0.249 | 2.73E-05 | 18 | Banf1 |
| Nhp22 | 1.38E-09 | 0.336171839 | 0.375 | 0.178 | 2.79E-05 | 18 | Nhp2 |
| H2afy5 | 1.40E-09 | 0.508357265 | 0.571 | 0.339 | 2.84E-05 | 18 | H2afy |
| Itgb26 | 1.41E-09 | 0.339763267 | 0.25 | 0.103 | 2.85E-05 | 18 | Itgb2 |
| Ppm1h6 | 1.46E-09 | 0.325763096 | 0.345 | 0.162 | 2.96E-05 | 18 | Ppm1h |
| Nxt1 | 1.49E-09 | 0.255242806 | 0.25 | 0.102 | 3.02E-05 | 18 | Nxt1 |
| Svbp | 1.58E-09 | 0.316769262 | 0.47 | 0.245 | 3.20E-05 | 18 | Svbp |
| Tgs11 | 1.62E-09 | 0.454988357 | 0.321 | 0.149 | 3.28E-05 | 18 | Tgs1 |
| Mapk142 | 1.65E-09 | 0.252787031 | 0.375 | 0.18 | 3.35E-05 | 18 | Mapk14 |
| Ebp2 | 1.68E-09 | 0.397348033 | 0.375 | 0.183 | 3.40E-05 | 18 | Ebp |
| Rab311 | 1.78E-09 | 0.352649192 | 0.25 | 0.101 | 3.60E-05 | 18 | Rab31 |
| F11r6 | 1.84E-09 | 0.475750977 | 0.482 | 0.267 | 3.72E-05 | 18 | F11r |
| Ddx39b5 | 1.93E-09 | 0.41017054 | 0.446 | 0.234 | 3.90E-05 | 18 | Ddx39b |
| Dst6 | 2.06E-09 | 0.383466091 | 0.381 | 0.192 | 4.17E-05 | 18 | Dst |
| Usp14 | 2.78E-09 | 0.264010373 | 0.321 | 0.147 | 5.64E-05 | 18 | Usp1 |
| Ormdl2 | 2.87E-09 | 0.260927675 | 0.262 | 0.11 | 5.81E-05 | 18 | Ormdl2 |
| Ggh | 2.93E-09 | 0.403868405 | 0.351 | 0.168 | 5.92E-05 | 18 | Ggh |
| Ssh26 | 3.21E-09 | 0.558849364 | 0.411 | 0.227 | 6.50E-05 | 18 | Ssh2 |
| Lage3 | 3.43E-09 | 0.289786555 | 0.375 | 0.179 | 6.94E-05 | 18 | Lage3 |
| Prex16 | 3.64E-09 | 0.30935169 | 0.363 | 0.175 | 7.36E-05 | 18 | Prex1 |
| Ahcyl23 | 3.90E-09 | 0.295856666 | 0.286 | 0.127 | 7.90E-05 | 18 | Ahcyl2 |
| Pold43 | 4.06E-09 | 0.359129011 | 0.399 | 0.201 | 8.22E-05 | 18 | Pold4 |
| Snrpg6 | 4.11E-09 | 0.486591725 | 0.554 | 0.325 | 8.31E-05 | 18 | Snrpg |
| Kpna33 | 4.78E-09 | 0.268889677 | 0.333 | 0.157 | 9.68E-05 | 18 | Kpna3 |
| Tmed33 | 4.80E-09 | 0.556289334 | 0.435 | 0.244 | 9.73E-05 | 18 | Tmed3 |
| Lipa3 | 4.92E-09 | 0.320848206 | 0.363 | 0.179 | 9.97E-05 | 18 | Lipa |
| Pkig3 | 5.62E-09 | 0.369231518 | 0.417 | 0.212 | 0.000113818 | 18 | Pkig |
| Traf36 | 6.55E-09 | 0.334837424 | 0.256 | 0.11 | 0.000132576 | 18 | Traf3 |
| Ap2s1 | 6.85E-09 | 0.299594998 | 0.47 | 0.248 | 0.00013868 | 18 | Ap2s1 |
| Cap14 | 7.04E-09 | 0.451661755 | 0.488 | 0.273 | 0.000142482 | 18 | Cap1 |

| | | | | | | | |
|---|---|---|---|---|---|---|---|
| Slbp2 | 7.15E-09 | 0.305906505 | 0.327 | 0.152 | 0.000144716 | 18 | Slbp |
| Arpc1b5 | 7.42E-09 | 0.47108673 | 0.774 | 0.592 | 0.000150175 | 18 | Arpc1b |
| Fnip14 | 7.74E-09 | 0.252673191 | 0.315 | 0.147 | 0.000156686 | 18 | Fnip1 |
| Clta3 | 8.04E-09 | 0.494214728 | 0.738 | 0.559 | 0.00016269 | 18 | Clta |
| Apobec35 | 8.26E-09 | 0.418826394 | 0.423 | 0.226 | 0.000167136 | 18 | Apobec3 |
| Fnbp17 | 8.32E-09 | 0.461545762 | 0.423 | 0.231 | 0.000168368 | 18 | Fnbp1 |
| Chd94 | 8.37E-09 | 0.483747282 | 0.446 | 0.251 | 0.000169367 | 18 | Chd9 |
| Snrpe3 | 8.50E-09 | 0.428784267 | 0.708 | 0.478 | 0.000172106 | 18 | Snrpe |
| Rps27a7 | 9.91E-09 | 0.378849326 | 0.994 | 0.95 | 0.000200648 | 18 | Rps27a |
| Lamtor11 | 1.09E-08 | 0.355743647 | 0.548 | 0.317 | 0.000220141 | 18 | Lamtor1 |
| Anapc111 | 1.10E-08 | 0.32755615 | 0.423 | 0.223 | 0.00022279 | 18 | Anapc11 |
| Agps1 | 1.16E-08 | 0.321326218 | 0.339 | 0.169 | 0.000235675 | 18 | Agps |
| Nasp4 | 1.20E-08 | 0.533219426 | 0.351 | 0.173 | 0.000243729 | 18 | Nasp |
| Rpl327 | 1.25E-08 | 0.433178238 | 0.994 | 0.953 | 0.000253538 | 18 | Rpl32 |
| Necap2 | 1.28E-08 | 0.273491733 | 0.381 | 0.191 | 0.000258281 | 18 | Necap2 |
| Plcl27 | 1.39E-08 | 0.279521224 | 0.262 | 0.115 | 0.000280601 | 18 | Plcl2 |
| Dnajc21 | 1.39E-08 | 0.289825508 | 0.321 | 0.149 | 0.000282382 | 18 | Dnajc2 |
| Grk24 | 1.49E-08 | 0.312165518 | 0.28 | 0.125 | 0.000301306 | 18 | Grk2 |
| Brd82 | 1.55E-08 | 0.301588013 | 0.345 | 0.165 | 0.000313881 | 18 | Brd8 |
| Cep1702 | 1.62E-08 | 0.322556221 | 0.286 | 0.131 | 0.000327941 | 18 | Cep170 |
| Snrpf6 | 1.63E-08 | 0.480793377 | 0.613 | 0.38 | 0.000329018 | 18 | Snrpf |
| Rdm1 | 1.64E-08 | 0.2660585 | 0.298 | 0.136 | 0.000332493 | 18 | Rdm1 |
| Ophn12 | 1.95E-08 | 0.263081279 | 0.292 | 0.133 | 0.000395165 | 18 | Ophn1 |
| Hba-a12 | 2.00E-08 | 1.6106652 | 0.268 | 0.129 | 0.000405569 | 18 | Hba-a1 |
| Ube2d2a | 2.34E-08 | 0.453230206 | 0.506 | 0.285 | 0.000474576 | 18 | Ube2d2a |
| U2af1 | 2.36E-08 | 0.460069317 | 0.518 | 0.312 | 0.000477088 | 18 | U2af1 |
| Ccdc504 | 2.54E-08 | 0.45685349 | 0.423 | 0.23 | 0.000513436 | 18 | Ccdc50 |
| Snx171 | 2.99E-08 | 0.252881362 | 0.357 | 0.174 | 0.000605891 | 18 | Snx17 |
| Rps287 | 3.24E-08 | 0.426736979 | 0.964 | 0.898 | 0.000655663 | 18 | Rps28 |
| Msrb11 | 3.46E-08 | 0.286725979 | 0.304 | 0.143 | 0.000700564 | 18 | Msrb1 |
| Atf63 | 3.57E-08 | 0.370980897 | 0.327 | 0.163 | 0.000723214 | 18 | Atf6 |
| Rad212 | 3.87E-08 | 0.348003249 | 0.375 | 0.192 | 0.000783364 | 18 | Rad21 |
| Cstb1 | 4.77E-08 | 0.427928407 | 0.44 | 0.25 | 0.000965727 | 18 | Cstb |
| Cep831 | 4.80E-08 | 0.36441526 | 0.256 | 0.114 | 0.000972204 | 18 | Cep83 |
| Rps27l1 | 4.81E-08 | 0.547308623 | 0.762 | 0.592 | 0.000973995 | 18 | Rps27l |
| Fbl3 | 4.83E-08 | 0.270660518 | 0.286 | 0.131 | 0.000976934 | 18 | Fbl |
| Sptssa2 | 5.07E-08 | 0.324645074 | 0.53 | 0.305 | 0.001026643 | 18 | Sptssa |
| Eif4a14 | 5.72E-08 | 0.475856528 | 0.625 | 0.432 | 0.001158357 | 18 | Eif4a1 |
| Lbr8 | 5.93E-08 | 0.425894303 | 0.315 | 0.154 | 0.001199808 | 18 | Lbr |
| Actg14 | 6.85E-08 | 0.419145107 | 0.905 | 0.795 | 0.001387327 | 18 | Actg1 |
| H3f3a2 | 7.49E-08 | 0.504311003 | 0.911 | 0.862 | 0.001517353 | 18 | H3f3a |
| Rps166 | 7.56E-08 | 0.372457271 | 0.982 | 0.927 | 0.001530558 | 18 | Rps16 |
| Rps247 | 7.57E-08 | 0.352359275 | 1 | 0.972 | 0.001532527 | 18 | Rps24 |

| | | | | | | | |
|---|---|---|---|---|---|---|---|
| Cisd22 | 7.78E-08 | 0.420508427 | 0.345 | 0.175 | 0.001575445 | 18 | Cisd2 |
| Psmb95 | 8.13E-08 | 0.409221201 | 0.625 | 0.407 | 0.00164538 | 18 | Psmb9 |
| Timm10b1 | 8.71E-08 | 0.507124824 | 0.446 | 0.248 | 0.001762971 | 18 | Timm10b |
| Skil2 | 1.09E-07 | 0.302229801 | 0.345 | 0.174 | 0.00221517 | 18 | Skil |
| Rps97 | 1.11E-07 | 0.360703292 | 0.97 | 0.921 | 0.002251671 | 18 | Rps9 |
| Capzb7 | 1.15E-07 | 0.540389554 | 0.571 | 0.37 | 0.002329273 | 18 | Capzb |
| Ldlrad46 | 1.21E-07 | 0.68037088 | 0.423 | 0.252 | 0.002442874 | 18 | Ldlrad4 |
| Arpp192 | 1.42E-07 | 0.501159649 | 0.643 | 0.442 | 0.002870796 | 18 | Arpp19 |
| Hnrnpab1 | 1.70E-07 | 0.466253702 | 0.56 | 0.369 | 0.003437758 | 18 | Hnrnpab |
| Papd42 | 1.83E-07 | 0.262269914 | 0.292 | 0.139 | 0.00371208 | 18 | Papd4 |
| Cebpb4 | 2.05E-07 | 0.29084841 | 0.315 | 0.158 | 0.004145664 | 18 | Cebpb |
| Fam96a | 2.25E-07 | 0.394889206 | 0.363 | 0.196 | 0.004565075 | 18 | Fam96a |
| Stk46 | 2.67E-07 | 0.309288589 | 0.345 | 0.176 | 0.005409448 | 18 | Stk4 |
| Ptms2 | 2.76E-07 | 0.532523872 | 0.762 | 0.584 | 0.005584648 | 18 | Ptms |
| Clasp25 | 2.96E-07 | 0.295118599 | 0.351 | 0.183 | 0.005999164 | 18 | Clasp2 |
| Fbxw111 | 3.41E-07 | 0.255291928 | 0.333 | 0.172 | 0.00690553 | 18 | Fbxw11 |
| Pea15a5 | 3.74E-07 | 0.359576109 | 0.435 | 0.241 | 0.0075627 | 18 | Pea15a |
| Siva11 | 3.88E-07 | 0.318584819 | 0.315 | 0.16 | 0.007859229 | 18 | Siva1 |
| Arhgdia1 | 4.02E-07 | 0.373733362 | 0.619 | 0.404 | 0.008140601 | 18 | Arhgdia |
| Dbi2 | 4.04E-07 | 0.40598443 | 0.649 | 0.433 | 0.008171693 | 18 | Dbi |
| Rps137 | 4.10E-07 | 0.274360691 | 0.988 | 0.924 | 0.008303575 | 18 | Rps13 |
| Snrpa1 | 4.21E-07 | 0.251022275 | 0.315 | 0.158 | 0.008528088 | 18 | Snrpa1 |
| Tmem2432 | 4.29E-07 | 0.331819217 | 0.321 | 0.164 | 0.008689593 | 18 | Tmem243 |
| Nudc1 | 4.68E-07 | 0.34805169 | 0.565 | 0.352 | 0.009468411 | 18 | Nudc |
| Rpl357 | 4.85E-07 | 0.420408783 | 0.917 | 0.829 | 0.009811129 | 18 | Rpl35 |
| Atp6v0b3 | 5.27E-07 | 0.400559607 | 0.554 | 0.368 | 0.010677879 | 18 | Atp6v0b |
| Rap1gds110 | 6.57E-07 | 0.491680242 | 0.429 | 0.262 | 0.013305514 | 18 | Rap1gds1 |
| Hnrnpf3 | 7.01E-07 | 0.467049044 | 0.685 | 0.497 | 0.014184621 | 18 | Hnrnpf |
| Rnaseh2c5 | 7.21E-07 | 0.444656415 | 0.476 | 0.287 | 0.014604766 | 18 | Rnaseh2c |
| Rps37 | 7.61E-07 | 0.333130484 | 0.946 | 0.893 | 0.015416509 | 18 | Rps3 |
| Nsmce25 | 8.08E-07 | 0.365555445 | 0.369 | 0.208 | 0.016356831 | 18 | Nsmce2 |
| Rnh1 | 9.93E-07 | 0.347477835 | 0.458 | 0.271 | 0.020097135 | 18 | Rnh1 |
| Ssr13 | 1.05E-06 | 0.25322132 | 0.357 | 0.187 | 0.021310344 | 18 | Ssr1 |
| Cntln4 | 1.10E-06 | 0.276387389 | 0.31 | 0.156 | 0.022226105 | 18 | Cntln |
| Galnt111 | 1.21E-06 | 0.278215153 | 0.351 | 0.185 | 0.024410318 | 18 | Galnt1 |
| Snrnp401 | 1.21E-06 | 0.38243313 | 0.286 | 0.145 | 0.024488677 | 18 | Snrnp40 |
| Lgals35 | 1.32E-06 | 0.277168517 | 0.393 | 0.219 | 0.026820634 | 18 | Lgals3 |
| G3bp15 | 1.59E-06 | 0.299596416 | 0.5 | 0.303 | 0.032279626 | 18 | G3bp1 |
| Pgp | 1.80E-06 | 0.304966967 | 0.375 | 0.21 | 0.036497866 | 18 | Pgp |
| Rps197 | 1.83E-06 | 0.413723706 | 0.976 | 0.927 | 0.037118703 | 18 | Rps19 |
| Vps13b7 | 1.98E-06 | 0.415489957 | 0.375 | 0.22 | 0.040104014 | 18 | Vps13b |
| Eif2s1 | 2.03E-06 | 0.303823263 | 0.369 | 0.204 | 0.041127272 | 18 | Eif2s1 |
| Trps16 | 2.08E-06 | 0.394824825 | 0.393 | 0.237 | 0.042138492 | 18 | Trps1 |

| | | | | | | | |
|---|---|---|---|---|---|---|---|
| Srsf31 | 2.32E-06 | 0.27373795 | 0.714 | 0.521 | 0.047070097 | 18 | Srsf3 |
| Tcerg13 | 2.39E-06 | 0.253482613 | 0.274 | 0.137 | 0.04848011 | 18 | Tcerg1 |
| Nucks12 | 2.46E-06 | 0.405284097 | 0.554 | 0.392 | 0.049782473 | 18 | Nucks1 |
| Rplp28 | 2.94E-06 | 0.375590078 | 0.946 | 0.891 | 0.059489893 | 18 | Rplp2 |
| Rps4x7 | 3.53E-06 | 0.340090598 | 0.982 | 0.941 | 0.071367641 | 18 | Rps4x |
| Mrpl521 | 3.76E-06 | 0.402882009 | 0.631 | 0.442 | 0.076063246 | 18 | Mrpl52 |
| Lsm62 | 4.88E-06 | 0.292881412 | 0.512 | 0.316 | 0.098734896 | 18 | Lsm6 |
| Ppp1ca5 | 5.58E-06 | 0.333151179 | 0.583 | 0.398 | 0.113068641 | 18 | Ppp1ca |
| Ccdc1076 | 5.77E-06 | 0.301513409 | 0.256 | 0.129 | 0.116731921 | 18 | Ccdc107 |
| Anxa55 | 5.84E-06 | 0.372740169 | 0.685 | 0.495 | 0.118173137 | 18 | Anxa5 |
| Atp6v0e2 | 6.11E-06 | 0.437481835 | 0.613 | 0.427 | 0.123694552 | 18 | Atp6v0e |
| Dennd1b7 | 6.48E-06 | 0.257372265 | 0.333 | 0.186 | 0.131290128 | 18 | Dennd1b |
| Supt16 | 7.77E-06 | 0.289235788 | 0.411 | 0.238 | 0.15726297 | 18 | Supt16 |
| Smc66 | 8.69E-06 | 0.476447802 | 0.417 | 0.263 | 0.17587692 | 18 | Smc6 |
| Rtn47 | 9.02E-06 | 0.337991902 | 0.5 | 0.34 | 0.182664259 | 18 | Rtn4 |
| Tmem1646 | 9.23E-06 | 0.329583992 | 0.304 | 0.165 | 0.186771306 | 18 | Tmem164 |
| Srsf75 | 9.37E-06 | 0.364619071 | 0.512 | 0.334 | 0.189622492 | 18 | Srsf7 |
| Rgs17 | 9.60E-06 | 0.39920212 | 0.268 | 0.142 | 0.194418265 | 18 | Rgs1 |
| Elmo19 | 1.03E-05 | 0.778103161 | 0.476 | 0.368 | 0.208029485 | 18 | Elmo1 |
| Esd3 | 1.03E-05 | 0.401433313 | 0.565 | 0.381 | 0.208368657 | 18 | Esd |
| Tbc1d59 | 1.11E-05 | 0.524583117 | 0.405 | 0.263 | 0.223995172 | 18 | Tbc1d5 |
| Stk382 | 1.15E-05 | 0.251865269 | 0.387 | 0.228 | 0.231875372 | 18 | Stk38 |
| Rpl397 | 1.21E-05 | 0.349602828 | 0.935 | 0.901 | 0.245907766 | 18 | Rpl39 |
| Akr1a11 | 1.26E-05 | 0.275882115 | 0.595 | 0.405 | 0.255691087 | 18 | Akr1a1 |
| Hprt1 | 1.31E-05 | 0.309085986 | 0.452 | 0.287 | 0.264347042 | 18 | Hprt |
| Far13 | 1.40E-05 | 0.278484056 | 0.298 | 0.166 | 0.283595843 | 18 | Far1 |
| Gm472838 | 1.65E-05 | 0.806167038 | 0.411 | 0.283 | 0.334280935 | 18 | Gm47283 |
| Rpl118 | 1.66E-05 | 0.260732497 | 0.982 | 0.927 | 0.335483789 | 18 | Rpl11 |
| Tagln28 | 1.81E-05 | 0.385869369 | 0.607 | 0.43 | 0.367223392 | 18 | Tagln2 |
| Lman24 | 1.89E-05 | 0.313280901 | 0.363 | 0.208 | 0.38317379 | 18 | Lman2 |
| Snrpc | 2.03E-05 | 0.250646836 | 0.351 | 0.201 | 0.410176602 | 18 | Snrpc |
| Suz121 | 2.06E-05 | 0.307733823 | 0.327 | 0.186 | 0.417881917 | 18 | Suz12 |
| Psip15 | 2.14E-05 | 0.40922952 | 0.333 | 0.196 | 0.432612191 | 18 | Psip1 |
| Tma74 | 2.38E-05 | 0.4335995 | 0.72 | 0.525 | 0.481447639 | 18 | Tma7 |
| Bax4 | 2.43E-05 | 0.326835443 | 0.482 | 0.315 | 0.492590979 | 18 | Bax |
| Nfkbiz4 | 2.46E-05 | 0.254786124 | 0.256 | 0.139 | 0.498719339 | 18 | Nfkbiz |
| Txn12 | 2.66E-05 | 0.522628376 | 0.815 | 0.678 | 0.537811118 | 18 | Txn1 |
| Ppib4 | 2.75E-05 | 0.387544044 | 0.696 | 0.555 | 0.556017552 | 18 | Ppib |
| Gngt24 | 2.78E-05 | 0.300397154 | 0.554 | 0.361 | 0.562649308 | 18 | Gngt2 |
| Rpn26 | 2.96E-05 | 0.334664586 | 0.363 | 0.213 | 0.598757522 | 18 | Rpn2 |
| mt-Co313 | 3.32E-05 | 0.342287873 | 0.869 | 0.727 | 0.671461558 | 18 | mt-Co3 |
| Dennd1a9 | 3.36E-05 | 0.291035891 | 0.405 | 0.254 | 0.680161899 | 18 | Dennd1a |
| Rpl37a7 | 3.55E-05 | 0.301837795 | 0.976 | 0.938 | 0.719248641 | 18 | Rpl37a |

| | | | | | | | |
|---|---|---|---|---|---|---|---|
| Baz1b5 | 3.68E-05 | 0.286014528 | 0.327 | 0.195 | 0.745757262 | 18 | Baz1b |
| Crip18 | 3.73E-05 | 0.285360152 | 0.726 | 0.537 | 0.755329631 | 18 | Crip1 |
| Tpm37 | 3.80E-05 | 0.327451552 | 0.643 | 0.481 | 0.768583928 | 18 | Tpm3 |
| Serinc36 | 4.30E-05 | 0.470422731 | 0.601 | 0.489 | 0.870011217 | 18 | Serinc3 |
| Rplp07 | 4.41E-05 | 0.272928716 | 0.994 | 0.945 | 0.893561754 | 18 | Rplp0 |
| Rpl227 | 4.60E-05 | 0.363995432 | 0.946 | 0.852 | 0.930534132 | 18 | Rpl22 |
| Cct51 | 4.61E-05 | 0.257367939 | 0.47 | 0.296 | 0.934205806 | 18 | Cct5 |
| Gtf2b | 4.93E-05 | 0.286441483 | 0.274 | 0.149 | 0.997143032 | 18 | Gtf2b |
| Cd818 | 4.99E-05 | 0.278612096 | 0.75 | 0.527 | 1 | 18 | Cd81 |
| Commd31 | 5.18E-05 | 0.348703849 | 0.417 | 0.256 | 1 | 18 | Commd3 |
| Rps128 | 5.26E-05 | 0.372872019 | 0.964 | 0.899 | 1 | 18 | Rps12 |
| P4hb4 | 5.33E-05 | 0.292341682 | 0.417 | 0.254 | 1 | 18 | P4hb |
| Serf23 | 5.45E-05 | 0.284460338 | 0.899 | 0.82 | 1 | 18 | Serf2 |
| Cdk4 | 5.64E-05 | 0.28165154 | 0.357 | 0.208 | 1 | 18 | Cdk4 |
| Ppm1g3 | 5.65E-05 | 0.392724699 | 0.28 | 0.157 | 1 | 18 | Ppm1g |
| Reep53 | 6.24E-05 | 0.274740649 | 0.649 | 0.482 | 1 | 18 | Reep5 |
| Rpl10a8 | 7.87E-05 | 0.27771624 | 0.94 | 0.836 | 1 | 18 | Rpl10a |
| Rps254 | 8.24E-05 | 0.386833594 | 0.911 | 0.831 | 1 | 18 | Rps25 |
| Pdia67 | 8.53E-05 | 0.317332067 | 0.5 | 0.339 | 1 | 18 | Pdia6 |
| Rpl218 | 9.50E-05 | 0.300857581 | 0.958 | 0.923 | 1 | 18 | Rpl21 |
| Etv64 | 9.99E-05 | 0.31900432 | 0.411 | 0.28 | 1 | 18 | Etv6 |
| Psme16 | 0.000109977 | 0.369713369 | 0.732 | 0.57 | 1 | 18 | Psme1 |
| Rps155 | 0.000116782 | 0.360477377 | 0.917 | 0.749 | 1 | 18 | Rps15 |
| Nsa26 | 0.000117737 | 0.440543887 | 0.649 | 0.488 | 1 | 18 | Nsa2 |
| Med8 | 0.000183241 | 0.308408409 | 0.286 | 0.169 | 1 | 18 | Med8 |
| Snrpd22 | 0.000193037 | 0.283154024 | 0.577 | 0.41 | 1 | 18 | Snrpd2 |
| mt-Co214 | 0.000221825 | 0.320869877 | 0.857 | 0.729 | 1 | 18 | mt-Co2 |
| Rack18 | 0.000229592 | 0.297424098 | 0.917 | 0.793 | 1 | 18 | Rack1 |
| Eid13 | 0.000241056 | 0.262257847 | 0.464 | 0.323 | 1 | 18 | Eid1 |
| Sf3b25 | 0.000264951 | 0.254068156 | 0.524 | 0.36 | 1 | 18 | Sf3b2 |
| Dpysl26 | 0.000267145 | 0.279466637 | 0.298 | 0.176 | 1 | 18 | Dpysl2 |
| Pdia38 | 0.000286099 | 0.40299632 | 0.577 | 0.445 | 1 | 18 | Pdia3 |
| Rbm8a2 | 0.000291622 | 0.302551783 | 0.476 | 0.31 | 1 | 18 | Rbm8a |
| Taf104 | 0.000302121 | 0.352236043 | 0.56 | 0.386 | 1 | 18 | Taf10 |
| Snrpd31 | 0.000317659 | 0.274383529 | 0.464 | 0.312 | 1 | 18 | Snrpd3 |
| Sarnp | 0.000357037 | 0.26547318 | 0.458 | 0.309 | 1 | 18 | Sarnp |
| Gm100765 | 0.000398079 | 0.264984587 | 0.714 | 0.547 | 1 | 18 | Gm10076 |
| Wnk18 | 0.000531358 | 0.397825579 | 0.47 | 0.365 | 1 | 18 | Wnk1 |
| Mydgf3 | 0.00055249 | 0.415773544 | 0.357 | 0.233 | 1 | 18 | Mydgf |
| mt-Co112 | 0.000570711 | 0.264340932 | 0.845 | 0.721 | 1 | 18 | mt-Co1 |
| Med281 | 0.000717182 | 0.289733749 | 0.417 | 0.281 | 1 | 18 | Med28 |
| Tmcc34 | 0.000719278 | 0.410865099 | 0.44 | 0.321 | 1 | 18 | Tmcc3 |
| Hsp90b19 | 0.00078428 | 0.366308859 | 0.631 | 0.512 | 1 | 18 | Hsp90b1 |

| | | | | | | | |
|---|---|---|---|---|---|---|---|
| Cdc371 | 0.000852647 | 0.262090938 | 0.464 | 0.323 | 1 | 18 | Cdc37 |
| Prelid11 | 0.001033725 | 0.382108005 | 0.44 | 0.312 | 1 | 18 | Prelid1 |
| Rbbp41 | 0.001129485 | 0.272899822 | 0.399 | 0.264 | 1 | 18 | Rbbp4 |
| Litaf6 | 0.001342968 | 0.262219627 | 0.262 | 0.159 | 1 | 18 | Litaf |
| AU0202067 | 0.001510771 | 0.308817249 | 0.405 | 0.293 | 1 | 18 | AU020206 |
| Tmem14c1 | 0.00155297 | 0.294981839 | 0.518 | 0.375 | 1 | 18 | Tmem14c |
| Ube2i4 | 0.001783268 | 0.30855827 | 0.47 | 0.332 | 1 | 18 | Ube2i |
| mt-Atp612 | 0.001847942 | 0.269921854 | 0.875 | 0.755 | 1 | 18 | mt-Atp6 |
| Hnrnpm1 | 0.002044088 | 0.318553122 | 0.482 | 0.341 | 1 | 18 | Hnrnpm |
| mt-Cytb11 | 0.003810062 | 0.277873608 | 0.792 | 0.702 | 1 | 18 | mt-Cytb |
| Xist11 | 0.003951197 | 0.970014424 | 0.47 | 0.466 | 1 | 18 | Xist |
| mt-Nd112 | 0.005629657 | 0.25677749 | 0.69 | 0.601 | 1 | 18 | mt-Nd1 |
| Bcap317 | 0.008422934 | 0.257576299 | 0.399 | 0.287 | 1 | 18 | Bcap31 |
| Hbb-bs2 | 0.00885143 | 1.546858821 | 0.411 | 0.331 | 1 | 18 | Hbb-bs |
| Napsa3 | 0 | 2.059340461 | 0.722 | 0.049 | 0 | 19 | Napsa |
| 12-Mar | 0 | 1.628778933 | 0.696 | 0.038 | 0 | 19 | 1-Mar |
| Gm15987 | 0 | 1.051768938 | 0.361 | 0.001 | 0 | 19 | Gm15987 |
| H2-DMb2 | 0 | 1.049035629 | 0.323 | 0.002 | 0 | 19 | H2-DMb2 |
| H2-Ob | 0 | 1.037038242 | 0.297 | 0.007 | 0 | 19 | H2-Ob |
| Btla | 0 | 0.997993722 | 0.253 | 0.003 | 0 | 19 | Btla |
| Dpp4 | 0 | 0.95780826 | 0.386 | 0.012 | 0 | 19 | Dpp4 |
| Flt3 | 0 | 0.761772239 | 0.285 | 0.001 | 0 | 19 | Flt3 |
| Wdfy41 | 5.19E-303 | 1.483047114 | 0.468 | 0.021 | 1.05E-298 | 19 | Wdfy4 |
| Cd79b | 9.23E-303 | 2.054389023 | 0.272 | 0.006 | 1.87E-298 | 19 | Cd79b |
| Alcam1 | 1.08E-274 | 1.596811671 | 0.532 | 0.029 | 2.19E-270 | 19 | Alcam |
| Cd831 | 2.27E-267 | 1.46117242 | 0.589 | 0.037 | 4.60E-263 | 19 | Cd83 |
| Pmaip11 | 1.16E-259 | 0.97741811 | 0.437 | 0.021 | 2.36E-255 | 19 | Pmaip1 |
| Pirb1 | 1.39E-248 | 0.685125643 | 0.386 | 0.017 | 2.82E-244 | 19 | Pirb |
| Rnase61 | 1.96E-241 | 1.276723618 | 0.608 | 0.044 | 3.98E-237 | 19 | Rnase6 |
| Syk4 | 5.54E-236 | 1.630759366 | 0.722 | 0.065 | 1.12E-231 | 19 | Syk |
| Pou2f22 | 3.37E-231 | 1.309103221 | 0.601 | 0.044 | 6.83E-227 | 19 | Pou2f2 |
| Kynu | 1.64E-221 | 0.641000058 | 0.259 | 0.008 | 3.32E-217 | 19 | Kynu |
| Itga46 | 1.53E-220 | 1.618499208 | 0.747 | 0.073 | 3.10E-216 | 19 | Itga4 |
| Spi14 | 6.42E-207 | 1.362149247 | 0.785 | 0.08 | 1.30E-202 | 19 | Spi1 |
| Gm267404 | 3.03E-203 | 1.472395393 | 0.646 | 0.058 | 6.13E-199 | 19 | Gm26740 |
| Gm205131 | 1.23E-202 | 0.843665778 | 0.348 | 0.017 | 2.49E-198 | 19 | Gm20513 |
| Rassf42 | 1.37E-198 | 0.989590927 | 0.538 | 0.041 | 2.77E-194 | 19 | Rassf4 |
| Pld42 | 5.54E-197 | 0.95968481 | 0.646 | 0.057 | 1.12E-192 | 19 | Pld4 |
| Arl5c | 7.37E-184 | 0.638471881 | 0.304 | 0.014 | 1.49E-179 | 19 | Arl5c |
| Plbd12 | 4.54E-182 | 1.587099747 | 0.608 | 0.057 | 9.20E-178 | 19 | Plbd1 |
| Hck2 | 6.06E-182 | 0.939610129 | 0.608 | 0.055 | 1.23E-177 | 19 | Hck |
| Nfam11 | 2.14E-177 | 0.648178557 | 0.367 | 0.021 | 4.34E-173 | 19 | Nfam1 |
| Sell1 | 2.61E-175 | 0.970216831 | 0.266 | 0.011 | 5.29E-171 | 19 | Sell |

| | | | | | | | |
|---|---|---|---|---|---|---|---|
| Cd862 | 2.93E-175 | 1.080637756 | 0.595 | 0.055 | 5.94E-171 | 19 | Cd86 |
| Ly864 | 1.50E-174 | 1.161661993 | 0.81 | 0.094 | 3.04E-170 | 19 | Ly86 |
| Map4k1 | 7.36E-173 | 0.576812621 | 0.361 | 0.021 | 1.49E-168 | 19 | Map4k1 |
| Ciita2 | 8.44E-173 | 0.927443031 | 0.475 | 0.037 | 1.71E-168 | 19 | Ciita |
| Btk1 | 1.44E-172 | 0.763102813 | 0.38 | 0.024 | 2.91E-168 | 19 | Btk |
| Gpr1834 | 7.19E-167 | 1.330119262 | 0.589 | 0.058 | 1.46E-162 | 19 | Gpr183 |
| Cybb2 | 3.49E-166 | 1.500229443 | 0.677 | 0.073 | 7.06E-162 | 19 | Cybb |
| Snx203 | 1.17E-156 | 0.949346089 | 0.582 | 0.06 | 2.37E-152 | 19 | Snx20 |
| Cd1802 | 5.37E-155 | 0.840630749 | 0.506 | 0.045 | 1.09E-150 | 19 | Cd180 |
| Lsp18 | 3.50E-153 | 2.394583543 | 0.823 | 0.137 | 7.08E-149 | 19 | Lsp1 |
| Evi2a2 | 5.81E-153 | 0.71802488 | 0.487 | 0.043 | 1.18E-148 | 19 | Evi2a |
| Cd300a | 7.19E-152 | 0.807994504 | 0.297 | 0.016 | 1.46E-147 | 19 | Cd300a |
| Ncf22 | 6.59E-149 | 1.003615003 | 0.525 | 0.052 | 1.33E-144 | 19 | Ncf2 |
| Aff32 | 5.11E-147 | 1.777197609 | 0.411 | 0.033 | 1.03E-142 | 19 | Aff3 |
| Irf52 | 2.82E-145 | 0.841937061 | 0.538 | 0.055 | 5.70E-141 | 19 | Irf5 |
| Cytip5 | 3.83E-145 | 1.308031543 | 0.684 | 0.09 | 7.75E-141 | 19 | Cytip |
| Csf2rb | 8.11E-144 | 0.625063162 | 0.304 | 0.018 | 1.64E-139 | 19 | Csf2rb |
| Dapp14 | 1.50E-141 | 1.04306087 | 0.639 | 0.079 | 3.04E-137 | 19 | Dapp1 |
| Ctsh5 | 1.14E-139 | 1.550706115 | 0.848 | 0.134 | 2.32E-135 | 19 | Ctsh |
| Ctss4 | 1.61E-138 | 1.185565413 | 0.962 | 0.163 | 3.25E-134 | 19 | Ctss |
| Ncf12 | 5.39E-138 | 0.74002049 | 0.462 | 0.042 | 1.09E-133 | 19 | Ncf1 |
| Mir142hg7 | 1.03E-136 | 1.616088525 | 0.924 | 0.168 | 2.08E-132 | 19 | Mir142hg |
| H2-Oa | 5.52E-135 | 0.659854754 | 0.272 | 0.016 | 1.12E-130 | 19 | H2-Oa |
| H2-DMa4 | 7.12E-135 | 1.46118438 | 0.772 | 0.118 | 1.44E-130 | 19 | H2-DMa |
| Dock27 | 3.64E-134 | 1.850414646 | 0.949 | 0.183 | 7.37E-130 | 19 | Dock2 |
| Myo1g2 | 7.95E-134 | 0.856898729 | 0.468 | 0.046 | 1.61E-129 | 19 | Myo1g |
| Runx17 | 9.86E-134 | 1.524771345 | 0.797 | 0.129 | 2.00E-129 | 19 | Runx1 |
| Ccl9 | 2.67E-133 | 0.698575803 | 0.291 | 0.018 | 5.42E-129 | 19 | Ccl9 |
| Batf32 | 5.37E-131 | 0.718653599 | 0.38 | 0.031 | 1.09E-126 | 19 | Batf3 |
| Alox5ap4 | 6.47E-130 | 1.385854422 | 0.589 | 0.072 | 1.31E-125 | 19 | Alox5ap |
| Il6ra1 | 1.74E-128 | 0.659619202 | 0.373 | 0.03 | 3.53E-124 | 19 | Il6ra |
| Ptpro1 | 5.01E-127 | 0.544014873 | 0.354 | 0.027 | 1.01E-122 | 19 | Ptpro |
| Tbc1d92 | 2.55E-126 | 0.644650316 | 0.418 | 0.038 | 5.17E-122 | 19 | Tbc1d9 |
| Adap1 | 8.15E-126 | 0.460509987 | 0.323 | 0.023 | 1.65E-121 | 19 | Adap1 |
| Csf2ra2 | 8.84E-125 | 0.811809371 | 0.589 | 0.072 | 1.79E-120 | 19 | Csf2ra |
| Pid14 | 5.62E-124 | 1.427946117 | 0.557 | 0.067 | 1.14E-119 | 19 | Pid1 |
| Plxnc11 | 6.85E-124 | 1.201912911 | 0.329 | 0.025 | 1.39E-119 | 19 | Plxnc1 |
| Bcl11a | 2.49E-123 | 0.852084384 | 0.272 | 0.017 | 5.05E-119 | 19 | Bcl11a |
| Ighd | 1.21E-122 | 1.574837141 | 0.31 | 0.023 | 2.45E-118 | 19 | Ighd |
| Fgd22 | 1.97E-122 | 0.529738536 | 0.443 | 0.043 | 3.99E-118 | 19 | Fgd2 |
| St6gal13 | 1.68E-121 | 1.02344267 | 0.418 | 0.04 | 3.41E-117 | 19 | St6gal1 |
| H2-DMb14 | 9.22E-121 | 1.230472594 | 0.652 | 0.09 | 1.87E-116 | 19 | H2-DMb1 |
| Fgr1 | 1.31E-120 | 0.850861618 | 0.43 | 0.042 | 2.65E-116 | 19 | Fgr |

| | | | | | | | |
|---|---|---|---|---|---|---|---|
| Notch25 | 1.53E-120 | 1.086504796 | 0.589 | 0.079 | 3.09E-116 | 19 | Notch2 |
| Slc8a13 | 3.76E-118 | 1.092677345 | 0.519 | 0.06 | 7.61E-114 | 19 | Slc8a1 |
| Plek5 | 1.55E-117 | 1.37913629 | 0.728 | 0.123 | 3.14E-113 | 19 | Plek |
| Rel6 | 2.64E-117 | 1.480840616 | 0.766 | 0.136 | 5.35E-113 | 19 | Rel |
| Ptprc6 | 2.97E-116 | 1.624887333 | 0.962 | 0.196 | 6.01E-112 | 19 | Ptprc |
| H2-Aa4 | 4.44E-116 | 1.825960915 | 0.911 | 0.185 | 8.99E-112 | 19 | H2-Aa |
| Plac81 | 7.31E-116 | 2.432772479 | 0.304 | 0.023 | 1.48E-111 | 19 | Plac8 |
| Ccl62 | 2.26E-115 | 1.359348052 | 0.367 | 0.032 | 4.58E-111 | 19 | Ccl6 |
| Coro1a8 | 6.50E-115 | 1.77069222 | 0.956 | 0.217 | 1.32E-110 | 19 | Coro1a |
| Ms4a4c1 | 3.04E-113 | 0.990439083 | 0.354 | 0.031 | 6.15E-109 | 19 | Ms4a4c |
| Bmp2k3 | 8.39E-112 | 1.424379609 | 0.677 | 0.109 | 1.70E-107 | 19 | Bmp2k |
| Tmem131l4 | 9.78E-112 | 1.204685287 | 0.57 | 0.079 | 1.98E-107 | 19 | Tmem131l |
| Grk31 | 2.09E-111 | 0.708295946 | 0.399 | 0.039 | 4.24E-107 | 19 | Grk3 |
| Arhgap457 | 7.86E-111 | 1.307839341 | 0.823 | 0.156 | 1.59E-106 | 19 | Arhgap45 |
| H2-Eb14 | 2.77E-110 | 1.750405671 | 0.911 | 0.193 | 5.60E-106 | 19 | H2-Eb1 |
| Gmfg5 | 1.16E-109 | 1.212309483 | 0.759 | 0.137 | 2.35E-105 | 19 | Gmfg |
| Arhgap157 | 3.77E-109 | 2.104584693 | 0.848 | 0.186 | 7.64E-105 | 19 | Arhgap15 |
| Samsn16 | 2.17E-108 | 1.680053279 | 0.557 | 0.079 | 4.39E-104 | 19 | Samsn1 |
| Pkib2 | 2.55E-108 | 0.769198332 | 0.411 | 0.042 | 5.15E-104 | 19 | Pkib |
| Cd744 | 3.19E-106 | 1.871024131 | 0.968 | 0.249 | 6.45E-102 | 19 | Cd74 |
| Tbc1d81 | 1.19E-105 | 0.999350993 | 0.481 | 0.059 | 2.41E-101 | 19 | Tbc1d8 |
| Aim27 | 3.92E-105 | 0.852159645 | 0.646 | 0.101 | 7.94E-101 | 19 | Aim2 |
| Cd537 | 6.88E-104 | 1.168802032 | 0.671 | 0.115 | 1.39E-99 | 19 | Cd53 |
| Mpeg13 | 1.68E-103 | 0.945474122 | 0.544 | 0.072 | 3.41E-99 | 19 | Mpeg1 |
| Slamf76 | 1.00E-102 | 0.970398624 | 0.62 | 0.097 | 2.03E-98 | 19 | Slamf7 |
| Dock118 | 1.30E-102 | 1.161385046 | 0.665 | 0.112 | 2.64E-98 | 19 | Dock11 |
| Ero1lb1 | 9.08E-102 | 0.957539306 | 0.481 | 0.062 | 1.84E-97 | 19 | Ero1lb |
| Sirpa2 | 1.41E-101 | 0.726258151 | 0.437 | 0.05 | 2.85E-97 | 19 | Sirpa |
| Clec12a2 | 1.92E-101 | 0.533908291 | 0.348 | 0.032 | 3.89E-97 | 19 | Clec12a |
| Blnk2 | 2.06E-101 | 1.014355069 | 0.481 | 0.06 | 4.16E-97 | 19 | Blnk |
| Ptprj7 | 9.97E-101 | 1.332451343 | 0.766 | 0.149 | 2.02E-96 | 19 | Ptprj |
| H2-Ab14 | 2.16E-100 | 1.731207831 | 0.88 | 0.199 | 4.37E-96 | 19 | H2-Ab1 |
| Dock108 | 3.15E-100 | 1.596150193 | 0.892 | 0.207 | 6.37E-96 | 19 | Dock10 |
| Tmem511 | 1.86E-99 | 0.417657867 | 0.272 | 0.021 | 3.76E-95 | 19 | Tmem51 |
| Il10ra3 | 4.34E-99 | 0.844588162 | 0.557 | 0.081 | 8.78E-95 | 19 | Il10ra |
| Unc93b12 | 6.39E-99 | 1.094692681 | 0.778 | 0.154 | 1.29E-94 | 19 | Unc93b1 |
| Ms4a6c3 | 7.13E-99 | 1.134570031 | 0.563 | 0.083 | 1.44E-94 | 19 | Ms4a6c |
| Fam49a2 | 7.19E-99 | 1.054525313 | 0.418 | 0.049 | 1.46E-94 | 19 | Fam49a |
| Prcp2 | 9.15E-99 | 0.759405329 | 0.576 | 0.085 | 1.85E-94 | 19 | Prcp |
| Apbb1ip7 | 3.15E-98 | 1.317507363 | 0.816 | 0.172 | 6.37E-94 | 19 | Apbb1ip |
| Nckap1l6 | 6.08E-98 | 0.787548587 | 0.576 | 0.087 | 1.23E-93 | 19 | Nckap1l |
| Gm2a4 | 1.12E-97 | 1.343333289 | 0.557 | 0.087 | 2.27E-93 | 19 | Gm2a |
| Plcg27 | 1.22E-97 | 0.999955055 | 0.671 | 0.118 | 2.47E-93 | 19 | Plcg2 |

| | | | | | | | |
|---|---|---|---|---|---|---|---|
| Arhgap307 | 4.52E-97 | 0.903969572 | 0.646 | 0.108 | 9.15E-93 | 19 | Arhgap30 |
| Gpr1323 | 1.38E-96 | 0.578947962 | 0.304 | 0.027 | 2.79E-92 | 19 | Gpr132 |
| Birc34 | 2.08E-96 | 1.073006622 | 0.601 | 0.097 | 4.21E-92 | 19 | Birc3 |
| Ptpn66 | 2.80E-96 | 1.089659588 | 0.703 | 0.129 | 5.67E-92 | 19 | Ptpn6 |
| Ifi2094 | 4.26E-96 | 0.767319342 | 0.449 | 0.056 | 8.62E-92 | 19 | Ifi209 |
| Snx301 | 8.53E-96 | 0.844115749 | 0.411 | 0.048 | 1.73E-91 | 19 | Snx30 |
| Stk17b5 | 7.71E-95 | 1.307058373 | 0.785 | 0.16 | 1.56E-90 | 19 | Stk17b |
| Ighm6 | 9.54E-95 | 1.901157265 | 0.671 | 0.125 | 1.93E-90 | 19 | Ighm |
| Laptm58 | 2.03E-94 | 1.512921984 | 0.861 | 0.209 | 4.10E-90 | 19 | Laptm5 |
| Arhgdib9 | 1.79E-93 | 1.362591363 | 0.937 | 0.231 | 3.62E-89 | 19 | Arhgdib |
| Arhgap222 | 1.04E-92 | 0.640729044 | 0.392 | 0.044 | 2.11E-88 | 19 | Arhgap22 |
| Sh3kbp16 | 1.98E-92 | 1.136307938 | 0.671 | 0.123 | 4.01E-88 | 19 | Sh3kbp1 |
| Tiam11 | 3.14E-92 | 0.433633201 | 0.266 | 0.021 | 6.35E-88 | 19 | Tiam1 |
| Pik3ap15 | 6.85E-92 | 0.85070835 | 0.563 | 0.089 | 1.39E-87 | 19 | Pik3ap1 |
| Ikzf16 | 3.02E-91 | 1.434678408 | 0.741 | 0.155 | 6.11E-87 | 19 | Ikzf1 |
| Cers66 | 1.18E-90 | 1.207071066 | 0.633 | 0.114 | 2.39E-86 | 19 | Cers6 |
| Dennd4a6 | 1.79E-90 | 1.3663073 | 0.823 | 0.186 | 3.61E-86 | 19 | Dennd4a |
| Zeb25 | 2.78E-90 | 1.617991034 | 0.709 | 0.142 | 5.63E-86 | 19 | Zeb2 |
| Cd376 | 1.41E-87 | 1.131499696 | 0.627 | 0.113 | 2.86E-83 | 19 | Cd37 |
| Cd684 | 2.78E-87 | 0.426403245 | 0.538 | 0.078 | 5.63E-83 | 19 | Cd68 |
| Prkcd2 | 3.06E-87 | 0.788796747 | 0.551 | 0.086 | 6.20E-83 | 19 | Prkcd |
| Ap1s31 | 4.72E-87 | 0.607962976 | 0.354 | 0.039 | 9.56E-83 | 19 | Ap1s3 |
| Celf28 | 6.79E-87 | 1.117952182 | 0.867 | 0.2 | 1.37E-82 | 19 | Celf2 |
| S100a44 | 1.00E-86 | 2.009539372 | 0.652 | 0.128 | 2.03E-82 | 19 | S100a4 |
| Slc38a15 | 2.45E-86 | 0.867950169 | 0.544 | 0.087 | 4.96E-82 | 19 | Slc38a1 |
| Itgb27 | 8.13E-86 | 0.782491481 | 0.589 | 0.1 | 1.65E-81 | 19 | Itgb2 |
| Prkcb7 | 2.04E-85 | 1.46436877 | 0.715 | 0.155 | 4.14E-81 | 19 | Prkcb |
| Rgs27 | 2.39E-85 | 1.080188367 | 0.665 | 0.129 | 4.85E-81 | 19 | Rgs2 |
| Irf83 | 9.94E-85 | 1.282992867 | 0.601 | 0.108 | 2.01E-80 | 19 | Irf8 |
| Nuak2 | 6.37E-84 | 0.427822357 | 0.266 | 0.023 | 1.29E-79 | 19 | Nuak2 |
| Rnf1501 | 6.54E-84 | 0.476061646 | 0.304 | 0.03 | 1.32E-79 | 19 | Rnf150 |
| Ncf46 | 1.22E-83 | 0.773133625 | 0.62 | 0.111 | 2.47E-79 | 19 | Ncf4 |
| Xylt12 | 1.54E-83 | 1.007843026 | 0.5 | 0.076 | 3.11E-79 | 19 | Xylt1 |
| Nfkbid1 | 2.26E-83 | 0.47440189 | 0.323 | 0.034 | 4.57E-79 | 19 | Nfkbid |
| Tep12 | 2.82E-83 | 0.662760307 | 0.43 | 0.058 | 5.70E-79 | 19 | Tep1 |
| Rbm472 | 1.51E-82 | 0.608853716 | 0.5 | 0.074 | 3.05E-78 | 19 | Rbm47 |
| Slc9a71 | 2.12E-81 | 0.911811177 | 0.348 | 0.041 | 4.30E-77 | 19 | Slc9a7 |
| Rasgef1b1 | 2.31E-81 | 0.693470437 | 0.329 | 0.036 | 4.68E-77 | 19 | Rasgef1b |
| Apobec12 | 5.43E-81 | 0.929432378 | 0.589 | 0.102 | 1.10E-76 | 19 | Apobec1 |
| Myo5a2 | 9.15E-81 | 0.73063664 | 0.519 | 0.083 | 1.85E-76 | 19 | Myo5a |
| Cd486 | 1.18E-80 | 0.868461098 | 0.671 | 0.133 | 2.38E-76 | 19 | Cd48 |
| Pik3r51 | 1.55E-80 | 0.432034947 | 0.297 | 0.03 | 3.14E-76 | 19 | Pik3r5 |
| Cd529 | 3.26E-80 | 1.632948273 | 0.975 | 0.28 | 6.60E-76 | 19 | Cd52 |

| | | | | | | | |
|---|---|---|---|---|---|---|---|
| Mbp4 | 3.85E-80 | 0.652171436 | 0.456 | 0.066 | 7.79E-76 | 19 | Mbp |
| Nlrp1b2 | 8.38E-80 | 0.42024236 | 0.348 | 0.039 | 1.70E-75 | 19 | Nlrp1b |
| Bcl2a1d2 | 1.00E-79 | 0.819782384 | 0.323 | 0.036 | 2.03E-75 | 19 | Bcl2a1d |
| Lyz24 | 1.58E-78 | 2.94751544 | 0.601 | 0.111 | 3.21E-74 | 19 | Lyz2 |
| Hk22 | 1.77E-78 | 0.678889074 | 0.443 | 0.063 | 3.59E-74 | 19 | Hk2 |
| Rac26 | 2.35E-78 | 1.066569808 | 0.791 | 0.176 | 4.75E-74 | 19 | Rac2 |
| Naaa | 5.11E-78 | 0.928935841 | 0.323 | 0.036 | 1.03E-73 | 19 | Naaa |
| Taok37 | 8.11E-78 | 0.988662565 | 0.728 | 0.162 | 1.64E-73 | 19 | Taok3 |
| Epsti16 | 2.26E-77 | 1.720742656 | 0.759 | 0.174 | 4.58E-73 | 19 | Epsti1 |
| Wfdc174 | 7.13E-77 | 0.908688865 | 0.456 | 0.067 | 1.44E-72 | 19 | Wfdc17 |
| Lcp17 | 7.87E-77 | 1.340123382 | 0.867 | 0.236 | 1.59E-72 | 19 | Lcp1 |
| Rassf22 | 5.13E-76 | 0.739491395 | 0.392 | 0.053 | 1.04E-71 | 19 | Rassf2 |
| Fam49b7 | 5.49E-76 | 1.31498862 | 0.854 | 0.235 | 1.11E-71 | 19 | Fam49b |
| Rasa34 | 1.26E-75 | 0.683632401 | 0.437 | 0.065 | 2.55E-71 | 19 | Rasa3 |
| Neurl36 | 2.10E-75 | 0.809522962 | 0.601 | 0.114 | 4.26E-71 | 19 | Neurl3 |
| Icosl1 | 8.86E-75 | 0.395685035 | 0.297 | 0.032 | 1.79E-70 | 19 | Icosl |
| Dock87 | 1.24E-74 | 1.361359073 | 0.816 | 0.217 | 2.51E-70 | 19 | Dock8 |
| Cyp27a11 | 1.25E-74 | 0.44944225 | 0.285 | 0.029 | 2.52E-70 | 19 | Cyp27a1 |
| Ctsc3 | 7.83E-74 | 0.950225008 | 0.69 | 0.149 | 1.58E-69 | 19 | Ctsc |
| Dusp25 | 1.19E-73 | 0.871020738 | 0.557 | 0.101 | 2.41E-69 | 19 | Dusp2 |
| Lyn3 | 1.35E-73 | 1.55473364 | 0.88 | 0.27 | 2.73E-69 | 19 | Lyn |
| Plxdc1 | 1.42E-73 | 0.318046899 | 0.253 | 0.024 | 2.87E-69 | 19 | Plxdc1 |
| Plcl28 | 3.59E-73 | 0.903908459 | 0.589 | 0.113 | 7.27E-69 | 19 | Plcl2 |
| Traf37 | 6.87E-73 | 0.864383744 | 0.57 | 0.108 | 1.39E-68 | 19 | Traf3 |
| Themis21 | 3.04E-72 | 0.361671927 | 0.259 | 0.025 | 6.15E-68 | 19 | Themis2 |
| Mctp13 | 7.91E-72 | 0.808979345 | 0.462 | 0.073 | 1.60E-67 | 19 | Mctp1 |
| Retreg16 | 1.18E-71 | 0.824707416 | 0.532 | 0.096 | 2.39E-67 | 19 | Retreg1 |
| Stap13 | 1.88E-71 | 0.659162918 | 0.614 | 0.123 | 3.81E-67 | 19 | Stap1 |
| Atf31 | 4.45E-71 | 0.83691257 | 0.5 | 0.086 | 9.01E-67 | 19 | Atf3 |
| Cd300c24 | 4.82E-71 | 0.56077562 | 0.513 | 0.085 | 9.76E-67 | 19 | Cd300c2 |
| Limd27 | 5.38E-71 | 0.916902711 | 0.677 | 0.147 | 1.09E-66 | 19 | Limd2 |
| Tnfaip8l24 | 2.10E-70 | 0.572624266 | 0.449 | 0.071 | 4.24E-66 | 19 | Tnfaip8l2 |
| Malt15 | 2.24E-70 | 1.083508577 | 0.671 | 0.148 | 4.53E-66 | 19 | Malt1 |
| Card115 | 3.14E-70 | 0.930302791 | 0.475 | 0.082 | 6.35E-66 | 19 | Card11 |
| Selplg7 | 3.19E-70 | 0.906700952 | 0.69 | 0.154 | 6.45E-66 | 19 | Selplg |
| Aif14 | 3.83E-70 | 0.533523392 | 0.633 | 0.12 | 7.76E-66 | 19 | Aif1 |
| Bri3bp | 4.07E-70 | 0.428465778 | 0.291 | 0.032 | 8.23E-66 | 19 | Bri3bp |
| Fchsd27 | 5.08E-70 | 1.675698583 | 0.671 | 0.158 | 1.03E-65 | 19 | Fchsd2 |
| Sema4d6 | 5.20E-70 | 0.894261212 | 0.62 | 0.129 | 1.05E-65 | 19 | Sema4d |
| Cx3cr12 | 5.75E-70 | 0.974553561 | 0.494 | 0.083 | 1.16E-65 | 19 | Cx3cr1 |
| Sla7 | 6.42E-70 | 0.694063323 | 0.449 | 0.073 | 1.30E-65 | 19 | Sla |
| Ccr25 | 1.14E-69 | 1.100454652 | 0.487 | 0.085 | 2.31E-65 | 19 | Ccr2 |
| Cd443 | 1.66E-69 | 1.083468602 | 0.551 | 0.107 | 3.35E-65 | 19 | Cd44 |

| | | | | | | | |
|---|---|---|---|---|---|---|---|
| Arhgef67 | 1.87E-69 | 0.79255157 | 0.614 | 0.127 | 3.79E-65 | 19 | Arhgef6 |
| Ptk2b7 | 1.90E-69 | 1.011412959 | 0.671 | 0.149 | 3.85E-65 | 19 | Ptk2b |
| Baz1a7 | 1.99E-69 | 0.910526127 | 0.722 | 0.171 | 4.02E-65 | 19 | Baz1a |
| Emb3 | 4.10E-69 | 0.680602388 | 0.329 | 0.041 | 8.30E-65 | 19 | Emb |
| Rhoh6 | 6.23E-69 | 0.881192201 | 0.538 | 0.102 | 1.26E-64 | 19 | Rhoh |
| Pip4k2a5 | 6.84E-69 | 1.313900955 | 0.804 | 0.24 | 1.39E-64 | 19 | Pip4k2a |
| Stk47 | 7.27E-69 | 0.964591383 | 0.722 | 0.174 | 1.47E-64 | 19 | Stk4 |
| Cyth46 | 8.70E-69 | 0.769728317 | 0.627 | 0.132 | 1.76E-64 | 19 | Cyth4 |
| Lst14 | 2.09E-68 | 0.77720696 | 0.576 | 0.11 | 4.23E-64 | 19 | Lst1 |
| Gsap6 | 7.00E-68 | 0.65584183 | 0.487 | 0.085 | 1.42E-63 | 19 | Gsap |
| Shtn13 | 7.52E-68 | 0.541465885 | 0.43 | 0.067 | 1.52E-63 | 19 | Shtn1 |
| A430078G23Rik1 | 9.81E-68 | 0.579391099 | 0.259 | 0.027 | 1.99E-63 | 19 | A430078G23Rik |
| Zfp363 | 1.02E-67 | 1.104463518 | 0.772 | 0.196 | 2.07E-63 | 19 | Zfp36 |
| Bcl2a1b4 | 2.39E-67 | 1.096477254 | 0.551 | 0.105 | 4.84E-63 | 19 | Bcl2a1b |
| Fam129a7 | 2.70E-67 | 0.877108655 | 0.652 | 0.144 | 5.47E-63 | 19 | Fam129a |
| AB1246111 | 2.87E-67 | 0.522556935 | 0.272 | 0.03 | 5.81E-63 | 19 | AB124611 |
| Cotl18 | 3.18E-67 | 1.106533916 | 0.81 | 0.223 | 6.44E-63 | 19 | Cotl1 |
| Phf11b2 | 7.14E-67 | 0.605377697 | 0.513 | 0.092 | 1.45E-62 | 19 | Phf11b |
| Filip1l6 | 7.76E-67 | 0.927425058 | 0.665 | 0.151 | 1.57E-62 | 19 | Filip1l |
| Rps6ka57 | 8.82E-67 | 0.760189963 | 0.551 | 0.107 | 1.79E-62 | 19 | Rps6ka5 |
| Nrros2 | 1.05E-66 | 0.700614212 | 0.551 | 0.106 | 2.13E-62 | 19 | Nrros |
| Fam105a6 | 1.33E-66 | 0.827736619 | 0.595 | 0.125 | 2.70E-62 | 19 | Fam105a |
| Skap22 | 4.23E-66 | 0.844946323 | 0.696 | 0.166 | 8.57E-62 | 19 | Skap2 |
| Tnfaip87 | 1.55E-65 | 0.814049582 | 0.747 | 0.185 | 3.15E-61 | 19 | Tnfaip8 |
| Man2b12 | 2.13E-65 | 0.75373636 | 0.696 | 0.163 | 4.31E-61 | 19 | Man2b1 |
| Inpp5d7 | 2.32E-65 | 1.124064496 | 0.753 | 0.203 | 4.69E-61 | 19 | Inpp5d |
| Fam111a2 | 3.09E-65 | 0.662019039 | 0.494 | 0.09 | 6.26E-61 | 19 | Fam111a |
| Cd695 | 4.26E-65 | 0.945780374 | 0.525 | 0.101 | 8.62E-61 | 19 | Cd69 |
| Plekhm33 | 5.55E-65 | 0.772940441 | 0.475 | 0.085 | 1.12E-60 | 19 | Plekhm3 |
| Fmnl17 | 9.56E-65 | 0.626258699 | 0.5 | 0.092 | 1.94E-60 | 19 | Fmnl1 |
| Ltb5 | 1.53E-64 | 1.237318796 | 0.62 | 0.135 | 3.10E-60 | 19 | Ltb |
| Lrrc252 | 2.78E-64 | 0.348262462 | 0.304 | 0.037 | 5.64E-60 | 19 | Lrrc25 |
| Csf1r2 | 5.79E-64 | 0.715521962 | 0.449 | 0.074 | 1.17E-59 | 19 | Csf1r |
| Klra22 | 1.19E-63 | 0.624224083 | 0.367 | 0.053 | 2.42E-59 | 19 | Klra2 |
| N4bp2l12 | 1.22E-63 | 0.611010979 | 0.367 | 0.054 | 2.47E-59 | 19 | N4bp2l1 |
| Tnfrsf11a2 | 1.23E-63 | 0.562122388 | 0.386 | 0.058 | 2.49E-59 | 19 | Tnfrsf11a |
| Tnf4 | 1.50E-63 | 0.624354585 | 0.43 | 0.072 | 3.03E-59 | 19 | Tnf |
| Pik3cd6 | 2.83E-63 | 0.72727567 | 0.538 | 0.108 | 5.72E-59 | 19 | Pik3cd |
| Sorl16 | 3.31E-63 | 0.832034087 | 0.544 | 0.11 | 6.70E-59 | 19 | Sorl1 |
| Cyfip25 | 5.47E-63 | 0.607646881 | 0.494 | 0.09 | 1.11E-58 | 19 | Cyfip2 |
| Unc1195 | 9.00E-63 | 0.571564866 | 0.468 | 0.083 | 1.82E-58 | 19 | Unc119 |
| Cyba9 | 9.42E-63 | 1.466756694 | 0.956 | 0.44 | 1.91E-58 | 19 | Cyba |
| Fermt37 | 1.35E-62 | 0.7417075 | 0.551 | 0.113 | 2.72E-58 | 19 | Fermt3 |

| | | | | | | | |
|---|---|---|---|---|---|---|---|
| Adcy78 | 1.95E-62 | 0.738745516 | 0.551 | 0.113 | 3.95E-58 | 19 | Adcy7 |
| Atp8b44 | 2.07E-62 | 0.765438621 | 0.513 | 0.096 | 4.20E-58 | 19 | Atp8b4 |
| Zeb2os1 | 3.18E-62 | 0.312607503 | 0.272 | 0.031 | 6.44E-58 | 19 | Zeb2os |
| Cd846 | 1.19E-61 | 0.810427969 | 0.475 | 0.09 | 2.41E-57 | 19 | Cd84 |
| Syngr22 | 1.95E-61 | 0.908134188 | 0.608 | 0.141 | 3.96E-57 | 19 | Syngr2 |
| Vav16 | 2.74E-61 | 0.493828332 | 0.456 | 0.08 | 5.54E-57 | 19 | Vav1 |
| Il2rg5 | 3.12E-61 | 0.626904737 | 0.57 | 0.116 | 6.31E-57 | 19 | Il2rg |
| Igsf61 | 3.90E-61 | 0.479967915 | 0.266 | 0.031 | 7.90E-57 | 19 | Igsf6 |
| Ankrd447 | 4.99E-61 | 1.171256891 | 0.886 | 0.293 | 1.01E-56 | 19 | Ankrd44 |
| Mkl17 | 6.03E-61 | 0.774636224 | 0.576 | 0.125 | 1.22E-56 | 19 | Mkl1 |
| Adgre12 | 1.19E-60 | 0.638402879 | 0.373 | 0.056 | 2.41E-56 | 19 | Adgre1 |
| Rilpl21 | 1.40E-60 | 0.633525523 | 0.424 | 0.073 | 2.84E-56 | 19 | Rilpl2 |
| Pag16 | 1.57E-60 | 0.998496867 | 0.557 | 0.121 | 3.17E-56 | 19 | Pag1 |
| Vrk25 | 1.85E-60 | 0.681467282 | 0.576 | 0.123 | 3.75E-56 | 19 | Vrk2 |
| Traf3ip35 | 2.15E-60 | 0.711620522 | 0.443 | 0.078 | 4.35E-56 | 19 | Traf3ip3 |
| Fgd31 | 2.60E-60 | 0.409960443 | 0.342 | 0.049 | 5.27E-56 | 19 | Fgd3 |
| Mob3b | 5.30E-60 | 0.50944465 | 0.361 | 0.054 | 1.07E-55 | 19 | Mob3b |
| Bcl61 | 2.53E-59 | 0.523695879 | 0.361 | 0.055 | 5.13E-55 | 19 | Bcl6 |
| Slc29a32 | 3.66E-59 | 0.36331493 | 0.418 | 0.069 | 7.40E-55 | 19 | Slc29a3 |
| Slamf93 | 4.71E-59 | 0.423645104 | 0.424 | 0.07 | 9.54E-55 | 19 | Slamf9 |
| Arhgap256 | 6.66E-59 | 0.683965274 | 0.525 | 0.107 | 1.35E-54 | 19 | Arhgap25 |
| Tec6 | 9.04E-59 | 0.744237019 | 0.525 | 0.109 | 1.83E-54 | 19 | Tec |
| Tuba1c2 | 1.44E-58 | 0.4592271 | 0.57 | 0.119 | 2.92E-54 | 19 | Tuba1c |
| Basp15 | 1.49E-58 | 0.968171144 | 0.519 | 0.104 | 3.02E-54 | 19 | Basp1 |
| Tank2 | 2.35E-58 | 0.980591256 | 0.601 | 0.142 | 4.76E-54 | 19 | Tank |
| Slc43a21 | 2.85E-58 | 0.327793961 | 0.278 | 0.035 | 5.77E-54 | 19 | Slc43a2 |
| Man1a10 | 3.19E-58 | 1.187125138 | 0.652 | 0.166 | 6.47E-54 | 19 | Man1a |
| Myo1f6 | 3.24E-58 | 0.958544451 | 0.595 | 0.137 | 6.57E-54 | 19 | Myo1f |
| Fam107b6 | 5.20E-58 | 0.768880837 | 0.557 | 0.12 | 1.05E-53 | 19 | Fam107b |
| Parp88 | 7.57E-58 | 1.010644613 | 0.614 | 0.148 | 1.53E-53 | 19 | Parp8 |
| Fnbp18 | 1.94E-57 | 1.191605128 | 0.753 | 0.229 | 3.92E-53 | 19 | Fnbp1 |
| Rassf55 | 1.98E-57 | 0.422686142 | 0.405 | 0.068 | 4.01E-53 | 19 | Rassf5 |
| Plekho21 | 2.20E-57 | 0.566928348 | 0.443 | 0.082 | 4.45E-53 | 19 | Plekho2 |
| Bcl2a1a2 | 3.30E-57 | 0.593007845 | 0.285 | 0.037 | 6.69E-53 | 19 | Bcl2a1a |
| B4galnt15 | 3.89E-57 | 0.54911398 | 0.487 | 0.095 | 7.88E-53 | 19 | B4galnt1 |
| Itgb75 | 5.41E-57 | 0.653228804 | 0.373 | 0.061 | 1.10E-52 | 19 | Itgb7 |
| Gpr137b | 8.93E-57 | 0.268462586 | 0.253 | 0.029 | 1.81E-52 | 19 | Gpr137b |
| Sp1401 | 1.38E-56 | 0.508550675 | 0.449 | 0.083 | 2.80E-52 | 19 | Sp140 |
| Arid3b1 | 3.51E-56 | 0.441054577 | 0.291 | 0.039 | 7.11E-52 | 19 | Arid3b |
| Slc15a4 | 3.70E-56 | 0.41272755 | 0.253 | 0.03 | 7.49E-52 | 19 | Slc15a4 |
| Tiparp3 | 3.80E-56 | 0.546266724 | 0.456 | 0.085 | 7.69E-52 | 19 | Tiparp |
| Soat14 | 5.03E-56 | 0.543198219 | 0.519 | 0.107 | 1.02E-51 | 19 | Soat1 |
| Uvrag6 | 9.14E-56 | 1.231368052 | 0.804 | 0.267 | 1.85E-51 | 19 | Uvrag |

| | | | | | | | |
|---|---|---|---|---|---|---|---|
| Glipr16 | 9.33E-56 | 0.581675056 | 0.481 | 0.095 | 1.89E-51 | 19 | Glipr1 |
| Fam69a6 | 1.07E-55 | 0.871692802 | 0.557 | 0.126 | 2.17E-51 | 19 | Fam69a |
| Rp21 | 1.36E-55 | 0.362773592 | 0.348 | 0.053 | 2.75E-51 | 19 | Rp2 |
| Nfkbie | 1.40E-55 | 0.445326271 | 0.285 | 0.038 | 2.84E-51 | 19 | Nfkbie |
| Traf15 | 1.45E-55 | 1.050609738 | 0.367 | 0.062 | 2.93E-51 | 19 | Traf1 |
| Prex17 | 1.56E-55 | 0.962157715 | 0.658 | 0.173 | 3.16E-51 | 19 | Prex1 |
| Lyst5 | 2.02E-55 | 0.892708943 | 0.513 | 0.109 | 4.09E-51 | 19 | Lyst |
| Bach27 | 2.35E-55 | 1.790062864 | 0.614 | 0.16 | 4.76E-51 | 19 | Bach2 |
| Crlf36 | 5.14E-55 | 0.813509664 | 0.633 | 0.156 | 1.04E-50 | 19 | Crlf3 |
| Nup2105 | 6.60E-55 | 0.388574797 | 0.392 | 0.066 | 1.34E-50 | 19 | Nup210 |
| Map3k144 | 8.17E-55 | 0.941422854 | 0.494 | 0.105 | 1.65E-50 | 19 | Map3k14 |
| Jak25 | 8.90E-55 | 0.847120127 | 0.551 | 0.126 | 1.80E-50 | 19 | Jak2 |
| Acer33 | 9.21E-55 | 0.857739096 | 0.589 | 0.14 | 1.86E-50 | 19 | Acer3 |
| Gmip5 | 1.19E-54 | 0.513800056 | 0.437 | 0.082 | 2.41E-50 | 19 | Gmip |
| Rtn12 | 1.77E-54 | 0.423362255 | 0.272 | 0.035 | 3.59E-50 | 19 | Rtn1 |
| Wipf17 | 2.27E-54 | 1.027233578 | 0.823 | 0.267 | 4.59E-50 | 19 | Wipf1 |
| Plekha25 | 3.11E-54 | 0.683828359 | 0.576 | 0.133 | 6.29E-50 | 19 | Plekha2 |
| Nfkbiz5 | 3.61E-54 | 0.762699245 | 0.582 | 0.136 | 7.32E-50 | 19 | Nfkbiz |
| Hcls16 | 4.62E-54 | 0.574912561 | 0.557 | 0.124 | 9.35E-50 | 19 | Hcls1 |
| Mthfd2 | 6.56E-54 | 0.31152166 | 0.304 | 0.043 | 1.33E-49 | 19 | Mthfd2 |
| Il21r5 | 8.06E-54 | 0.74228783 | 0.462 | 0.093 | 1.63E-49 | 19 | Il21r |
| Actr38 | 1.13E-53 | 1.09544423 | 0.937 | 0.39 | 2.28E-49 | 19 | Actr3 |
| Adgre57 | 1.95E-53 | 1.196926782 | 0.759 | 0.236 | 3.94E-49 | 19 | Adgre5 |
| Akna6 | 2.17E-53 | 0.488776682 | 0.462 | 0.091 | 4.40E-49 | 19 | Akna |
| Casp11 | 2.31E-53 | 0.380858279 | 0.361 | 0.059 | 4.67E-49 | 19 | Casp1 |
| Hist1h2ap1 | 4.19E-53 | 0.276656012 | 0.323 | 0.048 | 8.48E-49 | 19 | Hist1h2ap |
| Ikbkb3 | 4.24E-53 | 0.651763425 | 0.595 | 0.143 | 8.59E-49 | 19 | Ikbkb |
| Tmsb106 | 6.47E-53 | 1.434780537 | 0.975 | 0.5 | 1.31E-48 | 19 | Tmsb10 |
| P2ry62 | 1.17E-52 | 0.363467592 | 0.329 | 0.05 | 2.37E-48 | 19 | P2ry6 |
| Pde7a5 | 1.25E-52 | 0.877765946 | 0.62 | 0.155 | 2.53E-48 | 19 | Pde7a |
| Ttc73 | 1.29E-52 | 0.500401934 | 0.494 | 0.102 | 2.61E-48 | 19 | Ttc7 |
| Marcksl1 | 1.46E-52 | 0.667066244 | 0.266 | 0.035 | 2.96E-48 | 19 | Marcksl1 |
| Slc4a73 | 2.57E-52 | 0.687757295 | 0.443 | 0.088 | 5.21E-48 | 19 | Slc4a7 |
| Immp2l5 | 2.67E-52 | 1.013384409 | 0.57 | 0.135 | 5.40E-48 | 19 | Immp2l |
| Ezr3 | 3.06E-52 | 0.705871455 | 0.57 | 0.135 | 6.20E-48 | 19 | Ezr |
| Gpsm37 | 5.86E-52 | 0.656657595 | 0.601 | 0.147 | 1.19E-47 | 19 | Gpsm3 |
| Samhd15 | 6.11E-52 | 1.150138458 | 0.791 | 0.253 | 1.24E-47 | 19 | Samhd1 |
| Fcgr2b2 | 6.56E-52 | 0.413284367 | 0.285 | 0.04 | 1.33E-47 | 19 | Fcgr2b |
| Arrb25 | 3.04E-51 | 0.410801675 | 0.462 | 0.092 | 6.16E-47 | 19 | Arrb2 |
| Cyp4f182 | 5.13E-51 | 0.433643117 | 0.266 | 0.036 | 1.04E-46 | 19 | Cyp4f18 |
| Rap1gap22 | 5.19E-51 | 0.556317353 | 0.43 | 0.082 | 1.05E-46 | 19 | Rap1gap2 |
| Scpep11 | 6.64E-51 | 0.468712558 | 0.329 | 0.052 | 1.34E-46 | 19 | Scpep1 |
| Dusp51 | 1.46E-50 | 0.661654212 | 0.285 | 0.042 | 2.96E-46 | 19 | Dusp5 |

| | | | | | | | |
|---|---|---|---|---|---|---|---|
| Grap27 | 1.49E-50 | 0.830050715 | 0.481 | 0.105 | 3.02E-46 | 19 | Grap2 |
| Elf46 | 1.94E-50 | 0.628480553 | 0.532 | 0.121 | 3.92E-46 | 19 | Elf4 |
| Cxcl164 | 2.00E-50 | 0.893433258 | 0.506 | 0.112 | 4.04E-46 | 19 | Cxcl16 |
| Tagap2 | 2.04E-50 | 0.352932148 | 0.323 | 0.05 | 4.13E-46 | 19 | Tagap |
| Icam11 | 2.90E-50 | 0.408593764 | 0.405 | 0.075 | 5.87E-46 | 19 | Icam1 |
| Fgd42 | 4.69E-50 | 0.51276755 | 0.285 | 0.041 | 9.49E-46 | 19 | Fgd4 |
| Ahcyl24 | 7.66E-50 | 0.669782299 | 0.538 | 0.125 | 1.55E-45 | 19 | Ahcyl2 |
| Ikzf35 | 7.85E-50 | 1.262634249 | 0.513 | 0.12 | 1.59E-45 | 19 | Ikzf3 |
| Atxn7l14 | 8.13E-50 | 0.69930615 | 0.551 | 0.133 | 1.65E-45 | 19 | Atxn7l1 |
| Bin27 | 1.08E-49 | 0.641356137 | 0.538 | 0.127 | 2.19E-45 | 19 | Bin2 |
| Tgfbi3 | 1.27E-49 | 0.400900321 | 0.342 | 0.056 | 2.58E-45 | 19 | Tgfbi |
| Tbxas12 | 1.28E-49 | 0.476748891 | 0.304 | 0.046 | 2.59E-45 | 19 | Tbxas1 |
| Lat26 | 1.92E-49 | 0.644163794 | 0.551 | 0.131 | 3.88E-45 | 19 | Lat2 |
| Rnf1495 | 2.23E-49 | 0.531843126 | 0.481 | 0.103 | 4.51E-45 | 19 | Rnf149 |
| Setdb2 | 5.80E-49 | 0.473291259 | 0.335 | 0.056 | 1.17E-44 | 19 | Setdb2 |
| Tbc1d19 | 7.10E-49 | 0.723254552 | 0.627 | 0.164 | 1.44E-44 | 19 | Tbc1d1 |
| Rnf19b3 | 1.36E-48 | 0.517515194 | 0.424 | 0.084 | 2.75E-44 | 19 | Rnf19b |
| Was6 | 2.36E-48 | 0.466272081 | 0.386 | 0.072 | 4.79E-44 | 19 | Was |
| Sbno2 | 2.97E-48 | 0.351877036 | 0.367 | 0.065 | 6.01E-44 | 19 | Sbno2 |
| Trps17 | 3.93E-48 | 1.073841811 | 0.734 | 0.235 | 7.96E-44 | 19 | Trps1 |
| Ust6 | 4.41E-48 | 0.601485446 | 0.424 | 0.085 | 8.93E-44 | 19 | Ust |
| Tctex1d21 | 4.96E-48 | 0.270459516 | 0.361 | 0.063 | 1.00E-43 | 19 | Tctex1d2 |
| Net1 | 7.84E-48 | 0.669623462 | 0.266 | 0.038 | 1.59E-43 | 19 | Net1 |
| Itgal5 | 1.54E-47 | 0.690720133 | 0.424 | 0.086 | 3.12E-43 | 19 | Itgal |
| Sh3bp12 | 1.93E-47 | 0.291288848 | 0.316 | 0.051 | 3.92E-43 | 19 | Sh3bp1 |
| Ms4a6b4 | 2.25E-47 | 0.709851512 | 0.437 | 0.092 | 4.56E-43 | 19 | Ms4a6b |
| Arl4c3 | 2.82E-47 | 0.381546758 | 0.386 | 0.072 | 5.70E-43 | 19 | Arl4c |
| Jaml4 | 3.60E-47 | 0.585319033 | 0.335 | 0.058 | 7.29E-43 | 19 | Jaml |
| Mcub2 | 5.52E-47 | 0.411305405 | 0.367 | 0.066 | 1.12E-42 | 19 | Mcub |
| Adap24 | 6.73E-47 | 0.440879734 | 0.335 | 0.057 | 1.36E-42 | 19 | Adap2 |
| Snx292 | 7.20E-47 | 0.830865691 | 0.487 | 0.112 | 1.46E-42 | 19 | Snx29 |
| Dapk12 | 8.74E-47 | 0.507263935 | 0.354 | 0.063 | 1.77E-42 | 19 | Dapk1 |
| Kdm6b1 | 8.76E-47 | 0.556748099 | 0.43 | 0.089 | 1.77E-42 | 19 | Kdm6b |
| Fgl26 | 9.34E-47 | 0.475494425 | 0.487 | 0.107 | 1.89E-42 | 19 | Fgl2 |
| Vim10 | 1.12E-46 | 1.46307312 | 0.81 | 0.306 | 2.27E-42 | 19 | Vim |
| Tnfaip36 | 1.12E-46 | 0.566908713 | 0.43 | 0.089 | 2.28E-42 | 19 | Tnfaip3 |
| Pstpip15 | 1.24E-46 | 0.370115373 | 0.329 | 0.055 | 2.52E-42 | 19 | Pstpip1 |
| Tns32 | 2.01E-46 | 0.295032225 | 0.386 | 0.072 | 4.06E-42 | 19 | Tns3 |
| Ubac26 | 2.55E-46 | 0.716397011 | 0.658 | 0.187 | 5.16E-42 | 19 | Ubac2 |
| Ssh27 | 2.91E-46 | 0.934565452 | 0.709 | 0.225 | 5.90E-42 | 19 | Ssh2 |
| Sep-66 | 3.48E-46 | 0.518332811 | 0.456 | 0.098 | 7.04E-42 | 19 | 6-Sep |
| Grb24 | 7.66E-46 | 0.817388747 | 0.816 | 0.281 | 1.55E-41 | 19 | Grb2 |
| Hipk22 | 1.36E-45 | 0.396557381 | 0.513 | 0.117 | 2.75E-41 | 19 | Hipk2 |

| | | | | | | | |
|---|---|---|---|---|---|---|---|
| Lrmp5 | 1.40E-45 | 0.587331024 | 0.335 | 0.06 | 2.83E-41 | 19 | Lrmp |
| Akap139 | 1.46E-45 | 1.004496289 | 0.949 | 0.393 | 2.95E-41 | 19 | Akap13 |
| Ralgps21 | 2.26E-45 | 1.267391677 | 0.335 | 0.063 | 4.57E-41 | 19 | Ralgps2 |
| Rabgap1l5 | 2.32E-45 | 1.185891793 | 0.759 | 0.256 | 4.70E-41 | 19 | Rabgap1l |
| Trf3 | 2.46E-45 | 0.362268778 | 0.367 | 0.068 | 4.99E-41 | 19 | Trf |
| Csnk1g35 | 3.52E-45 | 0.74755944 | 0.633 | 0.182 | 7.13E-41 | 19 | Csnk1g3 |
| Btg15 | 4.00E-45 | 1.184795628 | 0.975 | 0.458 | 8.09E-41 | 19 | Btg1 |
| Il17ra | 5.42E-45 | 0.482483173 | 0.253 | 0.037 | 1.10E-40 | 19 | Il17ra |
| Arap12 | 7.98E-45 | 0.32466574 | 0.335 | 0.058 | 1.62E-40 | 19 | Arap1 |
| Rbpj7 | 8.70E-45 | 1.025595625 | 0.684 | 0.22 | 1.76E-40 | 19 | Rbpj |
| Efhd27 | 9.25E-45 | 0.802604405 | 0.639 | 0.186 | 1.87E-40 | 19 | Efhd2 |
| Sh3bgrl38 | 1.17E-44 | 1.085804155 | 0.981 | 0.559 | 2.37E-40 | 19 | Sh3bgrl3 |
| Ptprcap5 | 1.18E-44 | 0.874221792 | 0.538 | 0.131 | 2.38E-40 | 19 | Ptprcap |
| Flna5 | 1.38E-44 | 0.68980068 | 0.519 | 0.131 | 2.80E-40 | 19 | Flna |
| Pim15 | 2.45E-44 | 0.640247173 | 0.532 | 0.132 | 4.96E-40 | 19 | Pim1 |
| Rgs105 | 2.65E-44 | 0.535727261 | 0.608 | 0.16 | 5.37E-40 | 19 | Rgs10 |
| Apobec36 | 2.95E-44 | 0.646346962 | 0.734 | 0.224 | 5.98E-40 | 19 | Apobec3 |
| Ppp1r15a3 | 4.68E-44 | 0.732598702 | 0.582 | 0.154 | 9.48E-40 | 19 | Ppp1r15a |
| Actb9 | 6.81E-44 | 1.244923387 | 0.994 | 0.843 | 1.38E-39 | 19 | Actb |
| Grk25 | 6.91E-44 | 0.479392932 | 0.513 | 0.124 | 1.40E-39 | 19 | Grk2 |
| Gpr18 | 1.07E-43 | 0.439808279 | 0.253 | 0.037 | 2.17E-39 | 19 | Gpr18 |
| Coro2a5 | 1.09E-43 | 0.461893676 | 0.5 | 0.117 | 2.21E-39 | 19 | Coro2a |
| Cep1703 | 1.55E-43 | 0.516629681 | 0.532 | 0.129 | 3.13E-39 | 19 | Cep170 |
| Glipr23 | 1.67E-43 | 0.464502576 | 0.272 | 0.043 | 3.38E-39 | 19 | Glipr2 |
| Sgk32 | 1.67E-43 | 0.43585926 | 0.532 | 0.127 | 3.39E-39 | 19 | Sgk3 |
| Capg6 | 2.14E-43 | 0.669374898 | 0.443 | 0.1 | 4.34E-39 | 19 | Capg |
| Cadm12 | 2.89E-43 | 0.689710094 | 0.367 | 0.071 | 5.84E-39 | 19 | Cadm1 |
| Slc9a97 | 2.97E-43 | 0.784520849 | 0.563 | 0.151 | 6.02E-39 | 19 | Slc9a9 |
| Pak12 | 3.54E-43 | 0.283141648 | 0.272 | 0.042 | 7.17E-39 | 19 | Pak1 |
| Mob3a4 | 3.91E-43 | 0.475009221 | 0.392 | 0.081 | 7.92E-39 | 19 | Mob3a |
| Sema4a5 | 4.05E-43 | 0.354761182 | 0.386 | 0.076 | 8.20E-39 | 19 | Sema4a |
| Kif21b2 | 4.22E-43 | 0.410424803 | 0.304 | 0.052 | 8.54E-39 | 19 | Kif21b |
| Cblb6 | 4.42E-43 | 1.093834342 | 0.589 | 0.17 | 8.96E-39 | 19 | Cblb |
| Slc14a14 | 5.80E-43 | 0.375360594 | 0.285 | 0.046 | 1.17E-38 | 19 | Slc14a1 |
| Atg2a | 8.13E-43 | 0.360295198 | 0.348 | 0.065 | 1.65E-38 | 19 | Atg2a |
| Dennd1b8 | 1.26E-42 | 0.594872422 | 0.639 | 0.184 | 2.55E-38 | 19 | Dennd1b |
| Nfkb19 | 1.67E-42 | 0.914076939 | 0.759 | 0.271 | 3.39E-38 | 19 | Nfkb1 |
| Ptpn19 | 1.74E-42 | 0.799445425 | 0.715 | 0.235 | 3.53E-38 | 19 | Ptpn1 |
| Herc46 | 2.86E-42 | 0.783289617 | 0.557 | 0.151 | 5.80E-38 | 19 | Herc4 |
| Cdk145 | 3.08E-42 | 0.768238804 | 0.538 | 0.141 | 6.25E-38 | 19 | Cdk14 |
| P2ry103 | 3.12E-42 | 0.403836079 | 0.253 | 0.038 | 6.31E-38 | 19 | P2ry10 |
| Ddi21 | 3.21E-42 | 0.41009047 | 0.494 | 0.117 | 6.49E-38 | 19 | Ddi2 |
| Ramp16 | 3.49E-42 | 0.508796631 | 0.468 | 0.109 | 7.07E-38 | 19 | Ramp1 |

| | | | | | | | |
|---|---|---|---|---|---|---|---|
| Serp12 | 8.34E-42 | 0.619681752 | 0.703 | 0.216 | 1.69E-37 | 19 | Serp1 |
| Lgals36 | 9.13E-42 | 1.027191843 | 0.69 | 0.217 | 1.85E-37 | 19 | Lgals3 |
| AC149090.16 | 1.19E-41 | 0.738825293 | 0.715 | 0.231 | 2.40E-37 | 19 | AC149090.1 |
| Stat45 | 1.22E-41 | 0.62365808 | 0.462 | 0.107 | 2.47E-37 | 19 | Stat4 |
| Fosb7 | 1.24E-41 | 0.812267225 | 0.633 | 0.189 | 2.50E-37 | 19 | Fosb |
| 3110043O21Rik1 | 1.57E-41 | 0.405678453 | 0.291 | 0.049 | 3.17E-37 | 19 | 3110043O21Rik |
| Atf64 | 2.11E-41 | 0.654547413 | 0.589 | 0.161 | 4.28E-37 | 19 | Atf6 |
| Sgpl12 | 2.24E-41 | 0.369134602 | 0.323 | 0.059 | 4.53E-37 | 19 | Sgpl1 |
| Gpcpd14 | 2.26E-41 | 0.61586127 | 0.475 | 0.116 | 4.58E-37 | 19 | Gpcpd1 |
| Erp293 | 2.74E-41 | 0.814250737 | 0.823 | 0.327 | 5.55E-37 | 19 | Erp29 |
| Ripor25 | 2.80E-41 | 1.223636834 | 0.589 | 0.172 | 5.68E-37 | 19 | Ripor2 |
| Cnn27 | 2.87E-41 | 0.704375151 | 0.62 | 0.182 | 5.81E-37 | 19 | Cnn2 |
| Irak26 | 3.30E-41 | 0.667612944 | 0.538 | 0.141 | 6.68E-37 | 19 | Irak2 |
| Osbpl86 | 4.07E-41 | 0.614673751 | 0.665 | 0.198 | 8.23E-37 | 19 | Osbpl8 |
| Mbtd13 | 4.30E-41 | 0.537342458 | 0.519 | 0.134 | 8.70E-37 | 19 | Mbtd1 |
| Dclre1c1 | 5.60E-41 | 0.438007653 | 0.316 | 0.058 | 1.13E-36 | 19 | Dclre1c |
| Tbc1d141 | 5.80E-41 | 0.45804445 | 0.342 | 0.067 | 1.17E-36 | 19 | Tbc1d14 |
| Atp7a1 | 6.12E-41 | 0.434304822 | 0.386 | 0.08 | 1.24E-36 | 19 | Atp7a |
| Synj13 | 7.55E-41 | 0.633469812 | 0.582 | 0.159 | 1.53E-36 | 19 | Synj1 |
| Gng27 | 8.51E-41 | 0.634371867 | 0.633 | 0.186 | 1.72E-36 | 19 | Gng2 |
| Hvcn13 | 1.13E-40 | 0.646345768 | 0.304 | 0.056 | 2.29E-36 | 19 | Hvcn1 |
| 4930523C07Rik9 | 1.31E-40 | 1.088346526 | 0.684 | 0.219 | 2.66E-36 | 19 | 4930523C07Rik |
| Pik3cg3 | 1.42E-40 | 0.348870744 | 0.304 | 0.054 | 2.88E-36 | 19 | Pik3cg |
| Etv65 | 1.88E-40 | 1.065036342 | 0.741 | 0.277 | 3.80E-36 | 19 | Etv6 |
| Scamp23 | 2.17E-40 | 0.52880859 | 0.532 | 0.141 | 4.39E-36 | 19 | Scamp2 |
| Gm203888 | 2.45E-40 | 0.95829065 | 0.924 | 0.4 | 4.95E-36 | 19 | Gm20388 |
| Tbl1xr16 | 2.49E-40 | 0.55494307 | 0.595 | 0.164 | 5.04E-36 | 19 | Tbl1xr1 |
| Chd28 | 2.90E-40 | 0.847133686 | 0.753 | 0.253 | 5.87E-36 | 19 | Chd2 |
| Lair12 | 3.04E-40 | 0.39874506 | 0.335 | 0.063 | 6.15E-36 | 19 | Lair1 |
| Lbh4 | 3.25E-40 | 0.612817645 | 0.513 | 0.132 | 6.57E-36 | 19 | Lbh |
| Fau7 | 3.37E-40 | 0.846104814 | 1 | 0.953 | 6.82E-36 | 19 | Fau |
| Fyb7 | 3.42E-40 | 1.182663146 | 0.595 | 0.184 | 6.92E-36 | 19 | Fyb |
| Tyrobp7 | 3.62E-40 | 1.038486607 | 0.778 | 0.261 | 7.33E-36 | 19 | Tyrobp |
| Cerk7 | 3.72E-40 | 0.643629973 | 0.468 | 0.114 | 7.53E-36 | 19 | Cerk |
| Ucp25 | 3.77E-40 | 0.701328016 | 0.854 | 0.33 | 7.63E-36 | 19 | Ucp2 |
| Psd45 | 4.05E-40 | 0.37804271 | 0.335 | 0.065 | 8.20E-36 | 19 | Psd4 |
| Stk245 | 5.52E-40 | 0.635216545 | 0.652 | 0.195 | 1.12E-35 | 19 | Stk24 |
| Il161 | 5.77E-40 | 0.385526889 | 0.291 | 0.051 | 1.17E-35 | 19 | Il16 |
| H133 | 1.12E-39 | 0.493923818 | 0.57 | 0.157 | 2.27E-35 | 19 | H13 |
| Lrrk15 | 1.52E-39 | 0.829996841 | 0.62 | 0.19 | 3.07E-35 | 19 | Lrrk1 |
| Itgax5 | 1.77E-39 | 0.549834819 | 0.449 | 0.107 | 3.59E-35 | 19 | Itgax |
| Kdm7a2 | 1.84E-39 | 0.606382506 | 0.576 | 0.161 | 3.72E-35 | 19 | Kdm7a |
| Antxr23 | 1.85E-39 | 0.436218279 | 0.411 | 0.09 | 3.75E-35 | 19 | Antxr2 |

| | | | | | | | |
|---|---|---|---|---|---|---|---|
| Pot1b | 1.98E-39 | 0.577077335 | 0.367 | 0.076 | 4.02E-35 | 19 | Pot1b |
| Csk6 | 2.15E-39 | 0.447642643 | 0.506 | 0.129 | 4.35E-35 | 19 | Csk |
| Tbc1d4 | 2.52E-39 | 1.578421357 | 0.272 | 0.047 | 5.11E-35 | 19 | Tbc1d4 |
| Vrk12 | 2.93E-39 | 0.545457866 | 0.386 | 0.086 | 5.92E-35 | 19 | Vrk1 |
| Nfkbib | 3.13E-39 | 0.34387469 | 0.399 | 0.087 | 6.34E-35 | 19 | Nfkbib |
| Specc15 | 3.86E-39 | 0.562481129 | 0.373 | 0.08 | 7.82E-35 | 19 | Specc1 |
| Lgals16 | 4.67E-39 | 0.755975704 | 0.519 | 0.134 | 9.46E-35 | 19 | Lgals1 |
| Cbl5 | 5.08E-39 | 0.631791085 | 0.671 | 0.207 | 1.03E-34 | 19 | Cbl |
| Junb8 | 6.13E-39 | 0.960542246 | 0.924 | 0.392 | 1.24E-34 | 19 | Junb |
| Dgkd6 | 7.03E-39 | 0.708524528 | 0.563 | 0.16 | 1.42E-34 | 19 | Dgkd |
| Gm268873 | 7.09E-39 | 0.558543349 | 0.424 | 0.1 | 1.44E-34 | 19 | Gm26887 |
| Ifngr25 | 7.77E-39 | 0.460719855 | 0.513 | 0.131 | 1.57E-34 | 19 | Ifngr2 |
| Arhgap392 | 8.86E-39 | 0.411403061 | 0.31 | 0.058 | 1.79E-34 | 19 | Arhgap39 |
| Wwp25 | 1.28E-38 | 0.438336472 | 0.5 | 0.127 | 2.58E-34 | 19 | Wwp2 |
| 5031439G07Rik2 | 1.49E-38 | 0.387725617 | 0.373 | 0.079 | 3.02E-34 | 19 | 5031439G07Rik |
| Atp6v1b21 | 1.81E-38 | 0.331562147 | 0.462 | 0.11 | 3.67E-34 | 19 | Atp6v1b2 |
| Elmsan14 | 2.20E-38 | 0.440793947 | 0.443 | 0.106 | 4.45E-34 | 19 | Elmsan1 |
| Relb | 2.28E-38 | 1.031972668 | 0.373 | 0.084 | 4.62E-34 | 19 | Relb |
| Pde4b4 | 3.47E-38 | 0.84006959 | 0.627 | 0.195 | 7.03E-34 | 19 | Pde4b |
| Prkag2 | 6.60E-38 | 0.35900186 | 0.361 | 0.075 | 1.34E-33 | 19 | Prkag2 |
| B4galt55 | 6.61E-38 | 0.375944654 | 0.411 | 0.094 | 1.34E-33 | 19 | B4galt5 |
| Bmt24 | 7.47E-38 | 0.682207625 | 0.468 | 0.12 | 1.51E-33 | 19 | Bmt2 |
| Fcgr33 | 9.58E-38 | 0.435354432 | 0.392 | 0.087 | 1.94E-33 | 19 | Fcgr3 |
| Ly6d | 1.02E-37 | 1.738216899 | 0.272 | 0.05 | 2.07E-33 | 19 | Ly6d |
| Scimp3 | 1.07E-37 | 0.276564853 | 0.367 | 0.076 | 2.17E-33 | 19 | Scimp |
| Apba13 | 1.13E-37 | 0.449570172 | 0.291 | 0.053 | 2.30E-33 | 19 | Apba1 |
| Cdk67 | 1.25E-37 | 0.556267263 | 0.449 | 0.111 | 2.53E-33 | 19 | Cdk6 |
| Pnpla72 | 1.73E-37 | 0.409269376 | 0.323 | 0.064 | 3.50E-33 | 19 | Pnpla7 |
| Rhog2 | 1.80E-37 | 0.447704397 | 0.494 | 0.128 | 3.65E-33 | 19 | Rhog |
| Ext17 | 1.87E-37 | 0.815403766 | 0.57 | 0.166 | 3.78E-33 | 19 | Ext1 |
| Ptpn75 | 2.09E-37 | 0.25863137 | 0.31 | 0.058 | 4.24E-33 | 19 | Ptpn7 |
| Atp6ap22 | 2.23E-37 | 0.524197499 | 0.595 | 0.174 | 4.51E-33 | 19 | Atp6ap2 |
| Ccnl16 | 2.28E-37 | 0.648685351 | 0.665 | 0.209 | 4.61E-33 | 19 | Ccnl1 |
| Ccnd35 | 2.72E-37 | 0.714230903 | 0.709 | 0.239 | 5.51E-33 | 19 | Ccnd3 |
| Hdac93 | 3.13E-37 | 0.647708655 | 0.557 | 0.158 | 6.34E-33 | 19 | Hdac9 |
| Dok1 | 3.14E-37 | 0.332489791 | 0.297 | 0.056 | 6.36E-33 | 19 | Dok1 |
| Psap5 | 3.20E-37 | 1.330303849 | 0.816 | 0.398 | 6.48E-33 | 19 | Psap |
| Zdhhc141 | 4.34E-37 | 0.516688977 | 0.259 | 0.045 | 8.78E-33 | 19 | Zdhhc14 |
| Washc42 | 4.60E-37 | 0.47752681 | 0.494 | 0.128 | 9.31E-33 | 19 | Washc4 |
| Crybg1 | 5.02E-37 | 0.319974032 | 0.266 | 0.046 | 1.02E-32 | 19 | Crybg1 |
| Zc3h12a | 6.60E-37 | 0.362403936 | 0.278 | 0.05 | 1.34E-32 | 19 | Zc3h12a |
| Lactb | 6.76E-37 | 0.315613313 | 0.43 | 0.102 | 1.37E-32 | 19 | Lactb |
| Myo9b5 | 7.22E-37 | 0.460881207 | 0.475 | 0.119 | 1.46E-32 | 19 | Myo9b |

| | | | | | | | |
|---|---|---|---|---|---|---|---|
| Gadd45b1 | 7.60E-37 | 1.081599108 | 0.424 | 0.107 | 1.54E-32 | 19 | Gadd45b |
| Dennd1c3 | 8.80E-37 | 0.296709208 | 0.304 | 0.057 | 1.78E-32 | 19 | Dennd1c |
| Sap302 | 1.03E-36 | 0.297969702 | 0.399 | 0.09 | 2.09E-32 | 19 | Sap30 |
| Rapgef66 | 1.09E-36 | 0.742846433 | 0.804 | 0.296 | 2.21E-32 | 19 | Rapgef6 |
| Runx36 | 1.18E-36 | 0.693817951 | 0.532 | 0.149 | 2.38E-32 | 19 | Runx3 |
| St3gal31 | 1.38E-36 | 0.417210138 | 0.348 | 0.073 | 2.78E-32 | 19 | St3gal3 |
| Slfn27 | 1.38E-36 | 0.772192771 | 0.734 | 0.26 | 2.80E-32 | 19 | Slfn2 |
| Btg27 | 1.83E-36 | 0.940359812 | 0.823 | 0.344 | 3.70E-32 | 19 | Btg2 |
| Ehd12 | 2.35E-36 | 0.381740811 | 0.342 | 0.071 | 4.75E-32 | 19 | Ehd1 |
| Cd724 | 3.73E-36 | 0.526114502 | 0.43 | 0.105 | 7.56E-32 | 19 | Cd72 |
| Zufsp6 | 3.96E-36 | 0.484149162 | 0.506 | 0.136 | 8.02E-32 | 19 | Zufsp |
| Ddhd1 | 4.78E-36 | 0.706815196 | 0.354 | 0.079 | 9.67E-32 | 19 | Ddhd1 |
| Mcl15 | 5.03E-36 | 0.799305883 | 0.867 | 0.372 | 1.02E-31 | 19 | Mcl1 |
| Ccdc88a10 | 5.81E-36 | 1.015030709 | 0.614 | 0.204 | 1.18E-31 | 19 | Ccdc88a |
| Peli16 | 8.52E-36 | 0.753383542 | 0.665 | 0.231 | 1.72E-31 | 19 | Peli1 |
| Aftph1 | 8.82E-36 | 0.416720465 | 0.582 | 0.168 | 1.79E-31 | 19 | Aftph |
| Emp310 | 8.92E-36 | 0.827055273 | 0.715 | 0.258 | 1.81E-31 | 19 | Emp3 |
| Jpt16 | 9.20E-36 | 0.49619862 | 0.677 | 0.222 | 1.86E-31 | 19 | Jpt1 |
| Cyld5 | 9.50E-36 | 0.519264599 | 0.576 | 0.17 | 1.92E-31 | 19 | Cyld |
| Nsf | 9.55E-36 | 0.43983077 | 0.513 | 0.139 | 1.93E-31 | 19 | Nsf |
| Sulf25 | 9.59E-36 | 0.399510676 | 0.418 | 0.099 | 1.94E-31 | 19 | Sulf2 |
| Pde3b7 | 1.02E-35 | 0.794934988 | 0.468 | 0.127 | 2.07E-31 | 19 | Pde3b |
| Cyth13 | 1.18E-35 | 0.554247662 | 0.614 | 0.187 | 2.38E-31 | 19 | Cyth1 |
| Abhd122 | 1.23E-35 | 0.496756629 | 0.481 | 0.129 | 2.49E-31 | 19 | Abhd12 |
| Cask5 | 1.27E-35 | 0.671618187 | 0.506 | 0.139 | 2.58E-31 | 19 | Cask |
| Milr12 | 1.29E-35 | 0.30246377 | 0.266 | 0.047 | 2.62E-31 | 19 | Milr1 |
| Jmjd1c10 | 1.30E-35 | 0.720256179 | 0.791 | 0.306 | 2.63E-31 | 19 | Jmjd1c |
| Slc38a27 | 1.59E-35 | 0.809876101 | 0.791 | 0.31 | 3.23E-31 | 19 | Slc38a2 |
| Atg7 | 1.65E-35 | 0.326693077 | 0.329 | 0.068 | 3.33E-31 | 19 | Atg7 |
| Chka5 | 1.65E-35 | 0.667671222 | 0.506 | 0.144 | 3.34E-31 | 19 | Chka |
| Tcf206 | 1.75E-35 | 0.468540746 | 0.513 | 0.141 | 3.54E-31 | 19 | Tcf20 |
| Klrd14 | 1.89E-35 | 0.452628568 | 0.487 | 0.122 | 3.83E-31 | 19 | Klrd1 |
| Srebf22 | 1.90E-35 | 0.41083645 | 0.405 | 0.097 | 3.85E-31 | 19 | Srebf2 |
| Gm268351 | 2.09E-35 | 0.28623547 | 0.304 | 0.059 | 4.24E-31 | 19 | Gm26835 |
| Pitpna5 | 2.69E-35 | 0.73254809 | 0.791 | 0.327 | 5.44E-31 | 19 | Pitpna |
| Arid1b9 | 2.96E-35 | 0.671045844 | 0.741 | 0.258 | 6.00E-31 | 19 | Arid1b |
| Sirt73 | 3.37E-35 | 0.317118561 | 0.506 | 0.133 | 6.82E-31 | 19 | Sirt7 |
| Litaf7 | 3.42E-35 | 0.509447699 | 0.544 | 0.157 | 6.93E-31 | 19 | Litaf |
| Foxp17 | 4.73E-35 | 1.397329542 | 0.886 | 0.389 | 9.57E-31 | 19 | Foxp1 |
| Dbnl3 | 4.82E-35 | 0.436376321 | 0.62 | 0.191 | 9.76E-31 | 19 | Dbnl |
| Ldlr1 | 4.86E-35 | 0.413365151 | 0.278 | 0.052 | 9.83E-31 | 19 | Ldlr |
| Fgfr1op | 4.97E-35 | 0.303967705 | 0.272 | 0.05 | 1.01E-30 | 19 | Fgfr1op |
| Cop12 | 4.99E-35 | 0.540960262 | 0.589 | 0.179 | 1.01E-30 | 19 | Cop1 |

| | | | | | | | |
|---|---|---|---|---|---|---|---|
| Bloc1s2 | 5.97E-35 | 0.418195716 | 0.475 | 0.127 | 1.21E-30 | 19 | Bloc1s2 |
| Ube2f1 | 6.65E-35 | 0.307446683 | 0.494 | 0.131 | 1.35E-30 | 19 | Ube2f |
| Rnf2164 | 7.49E-35 | 0.47718944 | 0.601 | 0.18 | 1.52E-30 | 19 | Rnf216 |
| Cd2444 | 9.21E-35 | 0.330250024 | 0.399 | 0.092 | 1.86E-30 | 19 | Cd244 |
| Tcof12 | 9.71E-35 | 0.484813277 | 0.456 | 0.12 | 1.97E-30 | 19 | Tcof1 |
| Fos5 | 9.76E-35 | 1.045729217 | 0.797 | 0.316 | 1.98E-30 | 19 | Fos |
| Gpr653 | 1.15E-34 | 0.446862493 | 0.38 | 0.088 | 2.34E-30 | 19 | Gpr65 |
| Pbxip16 | 1.16E-34 | 0.400888 | 0.456 | 0.117 | 2.34E-30 | 19 | Pbxip1 |
| Lrrc8d2 | 1.26E-34 | 0.559126839 | 0.487 | 0.133 | 2.55E-30 | 19 | Lrrc8d |
| Plcl14 | 1.34E-34 | 0.418379444 | 0.278 | 0.052 | 2.72E-30 | 19 | Plcl1 |
| Pold44 | 1.36E-34 | 0.523647153 | 0.633 | 0.2 | 2.75E-30 | 19 | Pold4 |
| Rcsd16 | 1.39E-34 | 0.730730123 | 0.475 | 0.129 | 2.82E-30 | 19 | Rcsd1 |
| Atp6v0a11 | 1.42E-34 | 0.399317918 | 0.386 | 0.09 | 2.88E-30 | 19 | Atp6v0a1 |
| Slc35c2 | 1.45E-34 | 0.2872598 | 0.297 | 0.058 | 2.94E-30 | 19 | Slc35c2 |
| Iqgap23 | 1.57E-34 | 0.636915132 | 0.418 | 0.105 | 3.17E-30 | 19 | Iqgap2 |
| Tmcc110 | 2.28E-34 | 0.819157026 | 0.646 | 0.217 | 4.63E-30 | 19 | Tmcc1 |
| Ern15 | 2.40E-34 | 0.560040031 | 0.367 | 0.084 | 4.86E-30 | 19 | Ern1 |
| Fbl4 | 2.41E-34 | 0.318265302 | 0.487 | 0.13 | 4.89E-30 | 19 | Fbl |
| Slc15a32 | 2.45E-34 | 0.434747073 | 0.348 | 0.075 | 4.96E-30 | 19 | Slc15a3 |
| Rps6ka17 | 2.47E-34 | 0.411073712 | 0.443 | 0.114 | 4.99E-30 | 19 | Rps6ka1 |
| Sgms16 | 3.09E-34 | 0.816267379 | 0.462 | 0.128 | 6.26E-30 | 19 | Sgms1 |
| Ptpn188 | 3.21E-34 | 0.746760392 | 0.804 | 0.315 | 6.50E-30 | 19 | Ptpn18 |
| Kpna49 | 3.42E-34 | 0.839288135 | 0.797 | 0.323 | 6.93E-30 | 19 | Kpna4 |
| Snx54 | 3.46E-34 | 0.62085377 | 0.633 | 0.21 | 7.01E-30 | 19 | Snx5 |
| Zfyve26 | 3.73E-34 | 0.31333434 | 0.266 | 0.049 | 7.55E-30 | 19 | Zfyve26 |
| Atp13a34 | 3.87E-34 | 0.267220174 | 0.424 | 0.103 | 7.84E-30 | 19 | Atp13a3 |
| Asph4 | 3.96E-34 | 0.386504077 | 0.411 | 0.1 | 8.02E-30 | 19 | Asph |
| Ppp1r103 | 4.56E-34 | 0.488218012 | 0.614 | 0.189 | 9.23E-30 | 19 | Ppp1r10 |
| Plekho15 | 4.59E-34 | 0.455445606 | 0.703 | 0.224 | 9.28E-30 | 19 | Plekho1 |
| Tbc1d510 | 4.64E-34 | 0.741308818 | 0.715 | 0.261 | 9.39E-30 | 19 | Tbc1d5 |
| Susd31 | 4.95E-34 | 0.312250448 | 0.253 | 0.045 | 1.00E-29 | 19 | Susd3 |
| Rngtt5 | 6.42E-34 | 0.446564725 | 0.494 | 0.135 | 1.30E-29 | 19 | Rngtt |
| Ctsz5 | 6.80E-34 | 0.71546142 | 0.823 | 0.36 | 1.38E-29 | 19 | Ctsz |
| Ptger45 | 7.68E-34 | 0.525209964 | 0.361 | 0.085 | 1.56E-29 | 19 | Ptger4 |
| Mta35 | 8.56E-34 | 0.396439098 | 0.532 | 0.152 | 1.73E-29 | 19 | Mta3 |
| Iqgap19 | 1.03E-33 | 0.896957732 | 0.886 | 0.411 | 2.09E-29 | 19 | Iqgap1 |
| Wsb12 | 1.13E-33 | 0.503699084 | 0.582 | 0.175 | 2.30E-29 | 19 | Wsb1 |
| Zfp217 | 1.45E-33 | 0.319636795 | 0.297 | 0.06 | 2.94E-29 | 19 | Zfp217 |
| Sat14 | 1.49E-33 | 0.721737569 | 0.848 | 0.345 | 3.02E-29 | 19 | Sat1 |
| Plekhm21 | 1.50E-33 | 0.256123935 | 0.354 | 0.078 | 3.04E-29 | 19 | Plekhm2 |
| Gab26 | 1.51E-33 | 0.865882678 | 0.614 | 0.204 | 3.05E-29 | 19 | Gab2 |
| Colgalt13 | 1.65E-33 | 0.409547068 | 0.513 | 0.143 | 3.34E-29 | 19 | Colgalt1 |
| Clk18 | 1.67E-33 | 0.619758466 | 0.785 | 0.291 | 3.39E-29 | 19 | Clk1 |

| | | | | | | | |
|---|---|---|---|---|---|---|---|
| Elmo110 | 1.91E-33 | 0.800512493 | 0.861 | 0.365 | 3.86E-29 | 19 | Elmo1 |
| Xist12 | 2.75E-33 | 0.9191695 | 0.93 | 0.463 | 5.56E-29 | 19 | Xist |
| Klf69 | 2.77E-33 | 0.816853185 | 0.728 | 0.267 | 5.62E-29 | 19 | Klf6 |
| S100a68 | 3.32E-33 | 0.873234682 | 0.728 | 0.271 | 6.71E-29 | 19 | S100a6 |
| Il4ra | 3.38E-33 | 0.369867243 | 0.304 | 0.063 | 6.83E-29 | 19 | Il4ra |
| Lgmn4 | 3.41E-33 | 0.547282966 | 0.437 | 0.113 | 6.90E-29 | 19 | Lgmn |
| Zmynd152 | 3.94E-33 | 0.584526326 | 0.272 | 0.052 | 7.97E-29 | 19 | Zmynd15 |
| Ier59 | 4.24E-33 | 0.713704378 | 0.816 | 0.316 | 8.58E-29 | 19 | Ier5 |
| Lpxn6 | 4.87E-33 | 0.503639595 | 0.38 | 0.093 | 9.87E-29 | 19 | Lpxn |
| Cntrl5 | 5.29E-33 | 0.505963624 | 0.456 | 0.123 | 1.07E-28 | 19 | Cntrl |
| Eml47 | 5.32E-33 | 0.608854085 | 0.728 | 0.258 | 1.08E-28 | 19 | Eml4 |
| Usp121 | 5.39E-33 | 0.419638696 | 0.494 | 0.138 | 1.09E-28 | 19 | Usp12 |
| Jarid25 | 6.41E-33 | 0.818453361 | 0.608 | 0.203 | 1.30E-28 | 19 | Jarid2 |
| Inpp4b6 | 6.59E-33 | 0.906827145 | 0.608 | 0.203 | 1.34E-28 | 19 | Inpp4b |
| Gusb2 | 6.97E-33 | 0.320633601 | 0.405 | 0.099 | 1.41E-28 | 19 | Gusb |
| Eps82 | 7.50E-33 | 0.319160744 | 0.373 | 0.086 | 1.52E-28 | 19 | Eps8 |
| Sik15 | 7.63E-33 | 0.607034926 | 0.462 | 0.126 | 1.54E-28 | 19 | Sik1 |
| Zcchc711 | 7.98E-33 | 0.757396325 | 0.778 | 0.3 | 1.62E-28 | 19 | Zcchc7 |
| Birc24 | 8.31E-33 | 0.622859649 | 0.418 | 0.11 | 1.68E-28 | 19 | Birc2 |
| Sdccag85 | 9.30E-33 | 0.37082876 | 0.456 | 0.121 | 1.88E-28 | 19 | Sdccag8 |
| Atp11b5 | 1.06E-32 | 0.396575474 | 0.595 | 0.18 | 2.14E-28 | 19 | Atp11b |
| Ncoa19 | 1.24E-32 | 0.586469327 | 0.633 | 0.209 | 2.50E-28 | 19 | Ncoa1 |
| Lcp26 | 1.29E-32 | 0.446141406 | 0.386 | 0.094 | 2.61E-28 | 19 | Lcp2 |
| Fcer1g8 | 1.43E-32 | 0.965243919 | 0.741 | 0.269 | 2.90E-28 | 19 | Fcer1g |
| Nabp15 | 1.49E-32 | 0.392816316 | 0.354 | 0.082 | 3.03E-28 | 19 | Nabp1 |
| Mapk143 | 2.63E-32 | 0.520910837 | 0.57 | 0.178 | 5.32E-28 | 19 | Mapk14 |
| Eif2ak32 | 2.90E-32 | 0.492482516 | 0.354 | 0.084 | 5.86E-28 | 19 | Eif2ak3 |
| Cbfa2t34 | 3.40E-32 | 0.724076169 | 0.475 | 0.138 | 6.88E-28 | 19 | Cbfa2t3 |
| Cmah1 | 3.51E-32 | 0.667721446 | 0.31 | 0.067 | 7.10E-28 | 19 | Cmah |
| Slc20a1 | 4.67E-32 | 0.259511832 | 0.266 | 0.051 | 9.45E-28 | 19 | Slc20a1 |
| Hps32 | 6.45E-32 | 0.304739535 | 0.316 | 0.068 | 1.31E-27 | 19 | Hps3 |
| Myof3 | 6.57E-32 | 0.300258767 | 0.291 | 0.059 | 1.33E-27 | 19 | Myof |
| Coro1b2 | 7.84E-32 | 0.255772523 | 0.462 | 0.123 | 1.59E-27 | 19 | Coro1b |
| Ankrd1111 | 9.01E-32 | 0.784573886 | 0.873 | 0.38 | 1.82E-27 | 19 | Ankrd11 |
| Snrpb21 | 9.44E-32 | 0.29378348 | 0.475 | 0.131 | 1.91E-27 | 19 | Snrpb2 |
| Pan311 | 9.63E-32 | 0.57764822 | 0.665 | 0.228 | 1.95E-27 | 19 | Pan3 |
| Ppm1h7 | 1.10E-31 | 0.588250746 | 0.525 | 0.161 | 2.22E-27 | 19 | Ppm1h |
| Pfkfb32 | 1.63E-31 | 0.519681271 | 0.361 | 0.086 | 3.31E-27 | 19 | Pfkfb3 |
| Zfp4077 | 1.71E-31 | 0.416340851 | 0.544 | 0.161 | 3.46E-27 | 19 | Zfp407 |
| Mxd13 | 1.82E-31 | 0.645588629 | 0.297 | 0.064 | 3.68E-27 | 19 | Mxd1 |
| Me24 | 1.87E-31 | 0.258445189 | 0.335 | 0.075 | 3.79E-27 | 19 | Me2 |
| P2rx42 | 2.72E-31 | 0.290402098 | 0.304 | 0.065 | 5.52E-27 | 19 | P2rx4 |
| Rab8b7 | 2.95E-31 | 0.790493523 | 0.722 | 0.284 | 5.98E-27 | 19 | Rab8b |

| | | | | | | | |
|---|---|---|---|---|---|---|---|
| Prr135 | 3.82E-31 | 0.421056858 | 0.633 | 0.209 | 7.72E-27 | 19 | Prr13 |
| Rpsa8 | 3.91E-31 | 0.90435292 | 1 | 0.903 | 7.91E-27 | 19 | Rpsa |
| Dnajb143 | 4.04E-31 | 0.269811542 | 0.437 | 0.114 | 8.18E-27 | 19 | Dnajb14 |
| Mbnl16 | 4.28E-31 | 0.920057602 | 0.886 | 0.459 | 8.67E-27 | 19 | Mbnl1 |
| Fnip15 | 4.70E-31 | 0.512172114 | 0.494 | 0.146 | 9.52E-27 | 19 | Fnip1 |
| H2afy6 | 4.73E-31 | 0.645539434 | 0.81 | 0.337 | 9.59E-27 | 19 | H2afy |
| Rbm17 | 4.98E-31 | 0.272526204 | 0.5 | 0.142 | 1.01E-26 | 19 | Rbm17 |
| Cdc42se26 | 5.01E-31 | 0.546084967 | 0.684 | 0.239 | 1.01E-26 | 19 | Cdc42se2 |
| Mapkapk34 | 5.07E-31 | 0.325110071 | 0.348 | 0.082 | 1.03E-26 | 19 | Mapkapk3 |
| Trim146 | 5.08E-31 | 0.386507525 | 0.443 | 0.121 | 1.03E-26 | 19 | Trim14 |
| Creg14 | 5.10E-31 | 0.428556342 | 0.589 | 0.188 | 1.03E-26 | 19 | Creg1 |
| Sfmbt13 | 5.17E-31 | 0.347986905 | 0.354 | 0.084 | 1.05E-26 | 19 | Sfmbt1 |
| Hhex1 | 5.25E-31 | 0.390007091 | 0.342 | 0.079 | 1.06E-26 | 19 | Hhex |
| Fam102b2 | 6.23E-31 | 0.27778021 | 0.266 | 0.052 | 1.26E-26 | 19 | Fam102b |
| Rfx75 | 7.26E-31 | 0.524183702 | 0.513 | 0.155 | 1.47E-26 | 19 | Rfx7 |
| Srgap34 | 7.63E-31 | 0.50398762 | 0.38 | 0.096 | 1.54E-26 | 19 | Srgap3 |
| Traf52 | 7.73E-31 | 0.447024585 | 0.437 | 0.118 | 1.57E-26 | 19 | Traf5 |
| Chd76 | 1.03E-30 | 0.450453939 | 0.506 | 0.151 | 2.09E-26 | 19 | Chd7 |
| Vps13b8 | 1.09E-30 | 0.635787594 | 0.633 | 0.219 | 2.20E-26 | 19 | Vps13b |
| Ptpre5 | 1.11E-30 | 0.633233432 | 0.513 | 0.161 | 2.25E-26 | 19 | Ptpre |
| Daglb1 | 1.13E-30 | 0.314050322 | 0.259 | 0.051 | 2.28E-26 | 19 | Daglb |
| Hpgd4 | 1.19E-30 | 0.358128575 | 0.367 | 0.088 | 2.40E-26 | 19 | Hpgd |
| Rgs110 | 1.25E-30 | 0.947302704 | 0.468 | 0.141 | 2.54E-26 | 19 | Rgs1 |
| Gatm2 | 1.26E-30 | 0.363236143 | 0.329 | 0.073 | 2.56E-26 | 19 | Gatm |
| Tex146 | 1.26E-30 | 0.588926309 | 0.506 | 0.149 | 2.56E-26 | 19 | Tex14 |
| Ccm25 | 1.29E-30 | 0.517223781 | 0.449 | 0.126 | 2.60E-26 | 19 | Ccm2 |
| Limd11 | 1.45E-30 | 0.356774377 | 0.494 | 0.142 | 2.93E-26 | 19 | Limd1 |
| Znrf24 | 1.46E-30 | 0.407929274 | 0.525 | 0.159 | 2.96E-26 | 19 | Znrf2 |
| Otulin4 | 1.50E-30 | 0.414864391 | 0.608 | 0.199 | 3.05E-26 | 19 | Otulin |
| Snd17 | 1.54E-30 | 0.457206515 | 0.551 | 0.172 | 3.11E-26 | 19 | Snd1 |
| Creb52 | 1.63E-30 | 0.379063381 | 0.291 | 0.061 | 3.31E-26 | 19 | Creb5 |
| Slx4ip2 | 2.00E-30 | 0.322649153 | 0.31 | 0.068 | 4.04E-26 | 19 | Slx4ip |
| Slc38a91 | 2.68E-30 | 0.289590789 | 0.348 | 0.082 | 5.43E-26 | 19 | Slc38a9 |
| Tnfrsf1b5 | 3.03E-30 | 0.556961464 | 0.43 | 0.116 | 6.13E-26 | 19 | Tnfrsf1b |
| Stk106 | 3.07E-30 | 0.70557178 | 0.557 | 0.183 | 6.21E-26 | 19 | Stk10 |
| Mkrn12 | 3.45E-30 | 0.315052226 | 0.538 | 0.166 | 6.99E-26 | 19 | Mkrn1 |
| Mapkapk21 | 3.58E-30 | 0.410154922 | 0.481 | 0.139 | 7.25E-26 | 19 | Mapkapk2 |
| Map2k11 | 3.72E-30 | 0.273993025 | 0.437 | 0.116 | 7.53E-26 | 19 | Map2k1 |
| Mast31 | 4.05E-30 | 0.255566547 | 0.272 | 0.055 | 8.20E-26 | 19 | Mast3 |
| Cd24 | 4.61E-30 | 0.539680779 | 0.348 | 0.084 | 9.34E-26 | 19 | Cd2 |
| Arhgap47 | 4.98E-30 | 0.310148113 | 0.278 | 0.058 | 1.01E-25 | 19 | Arhgap4 |
| Paxbp12 | 5.07E-30 | 0.413340969 | 0.443 | 0.124 | 1.03E-25 | 19 | Paxbp1 |
| Tmem1733 | 5.53E-30 | 0.282770291 | 0.316 | 0.071 | 1.12E-25 | 19 | Tmem173 |

| | | | | | | | |
|---|---|---|---|---|---|---|---|
| Tmsb4x10 | 5.66E-30 | 0.975172964 | 1 | 0.934 | 1.15E-25 | 19 | Tmsb4x |
| Baz2b6 | 5.89E-30 | 0.598300368 | 0.867 | 0.35 | 1.19E-25 | 19 | Baz2b |
| Ctsa2 | 6.69E-30 | 0.463050258 | 0.5 | 0.153 | 1.35E-25 | 19 | Ctsa |
| Sash35 | 6.71E-30 | 0.322523975 | 0.285 | 0.06 | 1.36E-25 | 19 | Sash3 |
| Wnk19 | 6.86E-30 | 0.720092385 | 0.835 | 0.363 | 1.39E-25 | 19 | Wnk1 |
| Crlf25 | 7.48E-30 | 0.415500556 | 0.525 | 0.161 | 1.52E-25 | 19 | Crlf2 |
| Nap1l16 | 7.62E-30 | 0.643212918 | 0.785 | 0.314 | 1.54E-25 | 19 | Nap1l1 |
| Gna133 | 9.72E-30 | 0.386673002 | 0.551 | 0.171 | 1.97E-25 | 19 | Gna13 |
| Vmp15 | 1.01E-29 | 0.430563344 | 0.633 | 0.215 | 2.04E-25 | 19 | Vmp1 |
| Arhgef182 | 1.10E-29 | 0.496171485 | 0.31 | 0.072 | 2.24E-25 | 19 | Arhgef18 |
| Shisa55 | 1.26E-29 | 0.712092494 | 0.804 | 0.328 | 2.55E-25 | 19 | Shisa5 |
| Strbp3 | 1.49E-29 | 0.471381237 | 0.589 | 0.191 | 3.03E-25 | 19 | Strbp |
| Pepd | 1.90E-29 | 0.261007245 | 0.392 | 0.101 | 3.85E-25 | 19 | Pepd |
| Tgfbr12 | 1.91E-29 | 0.912203682 | 0.57 | 0.205 | 3.87E-25 | 19 | Tgfbr1 |
| Rras24 | 2.12E-29 | 0.363852594 | 0.348 | 0.084 | 4.30E-25 | 19 | Rras2 |
| Pfn18 | 2.37E-29 | 0.832588538 | 0.968 | 0.682 | 4.80E-25 | 19 | Pfn1 |
| Trim5 | 2.44E-29 | 0.426486982 | 0.259 | 0.053 | 4.95E-25 | 19 | Trim5 |
| Dmxl16 | 2.45E-29 | 0.597842206 | 0.563 | 0.185 | 4.95E-25 | 19 | Dmxl1 |
| Ptp4a14 | 2.45E-29 | 0.297188141 | 0.494 | 0.144 | 4.97E-25 | 19 | Ptp4a1 |
| mt-Co113 | 2.53E-29 | 0.795171652 | 1 | 0.72 | 5.12E-25 | 19 | mt-Co1 |
| Map3k82 | 2.59E-29 | 0.316364362 | 0.348 | 0.084 | 5.23E-25 | 19 | Map3k8 |
| Htt1 | 3.05E-29 | 0.342914683 | 0.392 | 0.102 | 6.17E-25 | 19 | Htt |
| Hpgds2 | 3.30E-29 | 0.354899039 | 0.278 | 0.059 | 6.68E-25 | 19 | Hpgds |
| Nubp12 | 3.80E-29 | 0.342958776 | 0.386 | 0.101 | 7.69E-25 | 19 | Nubp1 |
| Rnf24 | 4.19E-29 | 0.325610034 | 0.399 | 0.105 | 8.47E-25 | 19 | Rnf2 |
| Cdc42se1 | 4.76E-29 | 0.331835851 | 0.475 | 0.138 | 9.64E-25 | 19 | Cdc42se1 |
| Usp155 | 5.05E-29 | 0.372650578 | 0.671 | 0.23 | 1.02E-24 | 19 | Usp15 |
| Acox31 | 6.26E-29 | 0.297801109 | 0.266 | 0.055 | 1.27E-24 | 19 | Acox3 |
| Dpy19l11 | 1.01E-28 | 0.290423186 | 0.335 | 0.08 | 2.04E-24 | 19 | Dpy19l1 |
| Pvt16 | 1.02E-28 | 0.359792304 | 0.329 | 0.079 | 2.07E-24 | 19 | Pvt1 |
| Vsir3 | 1.05E-28 | 0.424833117 | 0.392 | 0.104 | 2.13E-24 | 19 | Vsir |
| Larp4b3 | 1.06E-28 | 0.460808797 | 0.601 | 0.204 | 2.14E-24 | 19 | Larp4b |
| Tes5 | 1.10E-28 | 0.311040137 | 0.456 | 0.129 | 2.23E-24 | 19 | Tes |
| Cklf | 1.10E-28 | 0.316870317 | 0.297 | 0.067 | 2.24E-24 | 19 | Cklf |
| Rps98 | 1.27E-28 | 0.690567966 | 1 | 0.921 | 2.57E-24 | 19 | Rps9 |
| Rap2a3 | 1.28E-28 | 0.36765234 | 0.424 | 0.117 | 2.60E-24 | 19 | Rap2a |
| Kcnq56 | 1.29E-28 | 0.728318261 | 0.297 | 0.068 | 2.61E-24 | 19 | Kcnq5 |
| Rara1 | 1.36E-28 | 0.420178064 | 0.373 | 0.097 | 2.75E-24 | 19 | Rara |
| Nsd38 | 1.51E-28 | 0.586016205 | 0.772 | 0.307 | 3.05E-24 | 19 | Nsd3 |
| Adam193 | 1.55E-28 | 0.351701013 | 0.335 | 0.08 | 3.14E-24 | 19 | Adam19 |
| Arid4a10 | 1.81E-28 | 0.64518634 | 0.677 | 0.251 | 3.66E-24 | 19 | Arid4a |
| Cap15 | 1.93E-28 | 0.498264886 | 0.709 | 0.271 | 3.90E-24 | 19 | Cap1 |
| Sesn36 | 2.13E-28 | 0.335983591 | 0.449 | 0.129 | 4.32E-24 | 19 | Sesn3 |

| | | | | | | | |
|---|---|---|---|---|---|---|---|
| Rps167 | 2.37E-28 | 0.644836745 | 1 | 0.927 | 4.79E-24 | 19 | Rps16 |
| Lncpint8 | 2.54E-28 | 0.590053751 | 0.722 | 0.274 | 5.14E-24 | 19 | Lncpint |
| Ccnt13 | 2.69E-28 | 0.435032542 | 0.532 | 0.167 | 5.45E-24 | 19 | Ccnt1 |
| Gramd1b5 | 2.95E-28 | 0.4941477 | 0.354 | 0.09 | 5.97E-24 | 19 | Gramd1b |
| Fam46a1 | 3.59E-28 | 0.283605457 | 0.475 | 0.14 | 7.27E-24 | 19 | Fam46a |
| Orai14 | 3.87E-28 | 0.336129324 | 0.449 | 0.128 | 7.83E-24 | 19 | Orai1 |
| Rin35 | 4.63E-28 | 0.549755146 | 0.544 | 0.179 | 9.38E-24 | 19 | Rin3 |
| Myadm5 | 4.71E-28 | 0.412899207 | 0.43 | 0.12 | 9.54E-24 | 19 | Myadm |
| Ncoa29 | 4.83E-28 | 0.489492782 | 0.658 | 0.238 | 9.78E-24 | 19 | Ncoa2 |
| Cpq4 | 5.06E-28 | 0.462895868 | 0.43 | 0.121 | 1.02E-23 | 19 | Cpq |
| Nptn8 | 5.29E-28 | 0.517963018 | 0.677 | 0.249 | 1.07E-23 | 19 | Nptn |
| Hcst5 | 6.39E-28 | 0.314287658 | 0.494 | 0.145 | 1.29E-23 | 19 | Hcst |
| Sepsecs | 7.35E-28 | 0.362384824 | 0.259 | 0.054 | 1.49E-23 | 19 | Sepsecs |
| Tor1aip24 | 8.47E-28 | 0.372533566 | 0.563 | 0.182 | 1.71E-23 | 19 | Tor1aip2 |
| mt-Nd59 | 9.23E-28 | 0.783844524 | 0.968 | 0.496 | 1.87E-23 | 19 | mt-Nd5 |
| Sdf2l15 | 9.91E-28 | 0.257377563 | 0.538 | 0.167 | 2.01E-23 | 19 | Sdf2l1 |
| Mob1a | 9.98E-28 | 0.361061855 | 0.576 | 0.191 | 2.02E-23 | 19 | Mob1a |
| Stat5b3 | 1.04E-27 | 0.334802717 | 0.487 | 0.146 | 2.10E-23 | 19 | Stat5b |
| Capzb8 | 1.19E-27 | 0.531215023 | 0.835 | 0.368 | 2.42E-23 | 19 | Capzb |
| Phf21a10 | 1.31E-27 | 0.756122004 | 0.652 | 0.253 | 2.66E-23 | 19 | Phf21a |
| Gls8 | 1.36E-27 | 0.474873708 | 0.684 | 0.251 | 2.75E-23 | 19 | Gls |
| Prpf38b8 | 1.38E-27 | 0.475446503 | 0.576 | 0.196 | 2.80E-23 | 19 | Prpf38b |
| Dusp163 | 1.64E-27 | 0.412568112 | 0.418 | 0.119 | 3.31E-23 | 19 | Dusp16 |
| Tmem1232 | 1.84E-27 | 1.389050992 | 0.475 | 0.154 | 3.72E-23 | 19 | Tmem123 |
| Calhm2 | 1.93E-27 | 0.308248937 | 0.291 | 0.066 | 3.90E-23 | 19 | Calhm2 |
| Arpc28 | 2.14E-27 | 0.794515304 | 0.937 | 0.544 | 4.33E-23 | 19 | Arpc2 |
| Nlrc55 | 2.72E-27 | 0.373572376 | 0.405 | 0.111 | 5.51E-23 | 19 | Nlrc5 |
| Wdr434 | 2.77E-27 | 0.2967212 | 0.494 | 0.15 | 5.60E-23 | 19 | Wdr43 |
| Cdyl24 | 2.77E-27 | 0.372018652 | 0.354 | 0.092 | 5.62E-23 | 19 | Cdyl2 |
| Tmem1351 | 2.93E-27 | 0.346512399 | 0.532 | 0.166 | 5.93E-23 | 19 | Tmem135 |
| Coq10b4 | 2.98E-27 | 0.381435075 | 0.494 | 0.153 | 6.04E-23 | 19 | Coq10b |
| Tbc1d22a1 | 3.01E-27 | 0.2920284 | 0.437 | 0.123 | 6.09E-23 | 19 | Tbc1d22a |
| Rps198 | 3.41E-27 | 0.737901153 | 1 | 0.927 | 6.91E-23 | 19 | Rps19 |
| Capza16 | 3.53E-27 | 0.443262386 | 0.728 | 0.275 | 7.15E-23 | 19 | Capza1 |
| Runx24 | 3.54E-27 | 0.373051357 | 0.361 | 0.092 | 7.17E-23 | 19 | Runx2 |
| Htra2 | 3.83E-27 | 0.291609724 | 0.278 | 0.062 | 7.76E-23 | 19 | Htra2 |
| Crip19 | 4.07E-27 | 1.549077553 | 0.905 | 0.536 | 8.24E-23 | 19 | Crip1 |
| Rps116 | 4.08E-27 | 0.687291934 | 0.994 | 0.907 | 8.26E-23 | 19 | Rps11 |
| Satb15 | 4.50E-27 | 0.504483087 | 0.278 | 0.063 | 9.10E-23 | 19 | Satb1 |
| Arhgef115 | 4.75E-27 | 0.524794576 | 0.671 | 0.248 | 9.61E-23 | 19 | Arhgef1 |
| Mycbp211 | 4.92E-27 | 0.755731592 | 0.823 | 0.374 | 9.95E-23 | 19 | Mycbp2 |
| Cst36 | 5.22E-27 | 1.425958919 | 0.886 | 0.603 | 1.06E-22 | 19 | Cst3 |
| Abi17 | 5.83E-27 | 0.502203291 | 0.696 | 0.269 | 1.18E-22 | 19 | Abi1 |

| | | | | | | | |
|---|---|---|---|---|---|---|---|
| Mgat4a6 | 5.93E-27 | 0.351408707 | 0.386 | 0.105 | 1.20E-22 | 19 | Mgat4a |
| Tbc1d10c5 | 6.02E-27 | 0.315422833 | 0.31 | 0.073 | 1.22E-22 | 19 | Tbc1d10c |
| Nfatc36 | 6.38E-27 | 0.652899388 | 0.525 | 0.175 | 1.29E-22 | 19 | Nfatc3 |
| Pias14 | 6.46E-27 | 0.378235688 | 0.57 | 0.188 | 1.31E-22 | 19 | Pias1 |
| Npc24 | 6.70E-27 | 0.846813508 | 0.791 | 0.414 | 1.36E-22 | 19 | Npc2 |
| Sec11c7 | 6.96E-27 | 0.352479697 | 0.785 | 0.31 | 1.41E-22 | 19 | Sec11c |
| Aff12 | 7.04E-27 | 0.505662486 | 0.494 | 0.16 | 1.43E-22 | 19 | Aff1 |
| Trim30d1 | 8.27E-27 | 0.336431773 | 0.329 | 0.082 | 1.68E-22 | 19 | Trim30d |
| Ppp3ca11 | 8.94E-27 | 0.741635027 | 0.88 | 0.393 | 1.81E-22 | 19 | Ppp3ca |
| Kdm2b4 | 1.00E-26 | 0.315423654 | 0.348 | 0.089 | 2.03E-22 | 19 | Kdm2b |
| Zfc3h12 | 1.08E-26 | 0.476456663 | 0.449 | 0.136 | 2.18E-22 | 19 | Zfc3h1 |
| Fam172a10 | 1.31E-26 | 0.548262698 | 0.722 | 0.284 | 2.66E-22 | 19 | Fam172a |
| Arhgap175 | 1.37E-26 | 0.624687144 | 0.69 | 0.27 | 2.78E-22 | 19 | Arhgap17 |
| Pgd1 | 1.37E-26 | 0.269288945 | 0.361 | 0.095 | 2.78E-22 | 19 | Pgd |
| Mia25 | 1.38E-26 | 0.456645588 | 0.589 | 0.207 | 2.79E-22 | 19 | Mia2 |
| Cdk138 | 1.43E-26 | 0.442527428 | 0.633 | 0.227 | 2.89E-22 | 19 | Cdk13 |
| Arpc42 | 1.43E-26 | 0.485565951 | 0.709 | 0.291 | 2.89E-22 | 19 | Arpc4 |
| Agps2 | 1.48E-26 | 0.315875602 | 0.525 | 0.168 | 2.99E-22 | 19 | Agps |
| Galnt112 | 1.55E-26 | 0.486082632 | 0.544 | 0.184 | 3.14E-22 | 19 | Galnt1 |
| Ifi2073 | 1.66E-26 | 0.542224993 | 0.38 | 0.107 | 3.37E-22 | 19 | Ifi207 |
| Phtf22 | 1.70E-26 | 0.305862868 | 0.278 | 0.063 | 3.44E-22 | 19 | Phtf2 |
| Ambra18 | 2.10E-26 | 0.502549067 | 0.576 | 0.198 | 4.26E-22 | 19 | Ambra1 |
| Abcg12 | 2.30E-26 | 0.33356479 | 0.43 | 0.123 | 4.66E-22 | 19 | Abcg1 |
| Supt32 | 2.59E-26 | 0.331749912 | 0.43 | 0.127 | 5.24E-22 | 19 | Supt3 |
| Ccl54 | 3.00E-26 | 1.647989945 | 0.462 | 0.152 | 6.07E-22 | 19 | Ccl5 |
| Nfatc15 | 3.11E-26 | 0.421155221 | 0.405 | 0.116 | 6.31E-22 | 19 | Nfatc1 |
| Rap1gds111 | 3.70E-26 | 0.543457309 | 0.671 | 0.261 | 7.49E-22 | 19 | Rap1gds1 |
| Fxyd510 | 3.71E-26 | 0.954986432 | 0.778 | 0.411 | 7.51E-22 | 19 | Fxyd5 |
| Plekhj13 | 3.83E-26 | 0.373452687 | 0.513 | 0.168 | 7.76E-22 | 19 | Plekhj1 |
| Zfp6084 | 4.42E-26 | 0.522604083 | 0.456 | 0.141 | 8.94E-22 | 19 | Zfp608 |
| Smap24 | 4.45E-26 | 0.451307142 | 0.595 | 0.212 | 9.01E-22 | 19 | Smap2 |
| Lcorl7 | 4.53E-26 | 0.413201524 | 0.513 | 0.166 | 9.16E-22 | 19 | Lcorl |
| Ago26 | 4.60E-26 | 0.55125955 | 0.506 | 0.167 | 9.32E-22 | 19 | Ago2 |
| Tmem1647 | 4.94E-26 | 0.411691742 | 0.506 | 0.164 | 1.00E-21 | 19 | Tmem164 |
| Cmip7 | 5.78E-26 | 0.677810647 | 0.709 | 0.285 | 1.17E-21 | 19 | Cmip |
| Smg110 | 6.15E-26 | 0.491387757 | 0.639 | 0.234 | 1.25E-21 | 19 | Smg1 |
| Rap2b1 | 6.37E-26 | 0.497878836 | 0.304 | 0.075 | 1.29E-21 | 19 | Rap2b |
| Taf1d4 | 6.57E-26 | 0.320070197 | 0.468 | 0.144 | 1.33E-21 | 19 | Taf1d |
| Ddx541 | 7.04E-26 | 0.3619359 | 0.437 | 0.13 | 1.43E-21 | 19 | Ddx54 |
| Wdr445 | 8.18E-26 | 0.410991491 | 0.418 | 0.123 | 1.66E-21 | 19 | Wdr44 |
| Neat18 | 8.39E-26 | 0.568892311 | 0.582 | 0.209 | 1.70E-21 | 19 | Neat1 |
| Foxn21 | 8.86E-26 | 0.359725493 | 0.449 | 0.136 | 1.79E-21 | 19 | Foxn2 |
| Ptpn225 | 9.10E-26 | 0.421553943 | 0.411 | 0.119 | 1.84E-21 | 19 | Ptpn22 |

| | | | | | | | |
|---|---|---|---|---|---|---|---|
| Atad2b7 | 1.01E-25 | 0.566325507 | 0.563 | 0.203 | 2.05E-21 | 19 | Atad2b |
| Xrn24 | 1.04E-25 | 0.445180407 | 0.684 | 0.261 | 2.10E-21 | 19 | Xrn2 |
| Dennd1a10 | 1.06E-25 | 0.570093195 | 0.658 | 0.253 | 2.14E-21 | 19 | Dennd1a |
| Mtss12 | 1.06E-25 | 0.646408121 | 0.361 | 0.1 | 2.15E-21 | 19 | Mtss1 |
| Supt4a5 | 1.45E-25 | 0.424846503 | 0.766 | 0.321 | 2.93E-21 | 19 | Supt4a |
| Gem4 | 1.48E-25 | 0.727477471 | 0.367 | 0.104 | 3.01E-21 | 19 | Gem |
| Polg21 | 1.50E-25 | 0.296040776 | 0.285 | 0.067 | 3.04E-21 | 19 | Polg2 |
| Fbxo117 | 1.53E-25 | 0.603302266 | 0.633 | 0.232 | 3.10E-21 | 19 | Fbxo11 |
| Etv32 | 1.56E-25 | 0.682142814 | 0.354 | 0.099 | 3.16E-21 | 19 | Etv3 |
| Trmt1126 | 1.58E-25 | 0.366126892 | 0.709 | 0.273 | 3.20E-21 | 19 | Trmt112 |
| Lrba6 | 1.76E-25 | 0.526045011 | 0.551 | 0.191 | 3.56E-21 | 19 | Lrba |
| Fuca13 | 1.82E-25 | 0.41742594 | 0.589 | 0.207 | 3.69E-21 | 19 | Fuca1 |
| Hlcs | 1.85E-25 | 0.34996623 | 0.304 | 0.075 | 3.74E-21 | 19 | Hlcs |
| Kdm6a9 | 1.88E-25 | 0.381485485 | 0.608 | 0.213 | 3.82E-21 | 19 | Kdm6a |
| Ccr56 | 1.95E-25 | 0.424659535 | 0.437 | 0.133 | 3.95E-21 | 19 | Ccr5 |
| Smchd17 | 2.07E-25 | 0.560625125 | 0.684 | 0.264 | 4.19E-21 | 19 | Smchd1 |
| Qpct3 | 2.12E-25 | 0.283145529 | 0.316 | 0.08 | 4.29E-21 | 19 | Qpct |
| Rps15a6 | 2.36E-25 | 0.655410982 | 0.994 | 0.925 | 4.78E-21 | 19 | Rps15a |
| Actr25 | 2.46E-25 | 0.508329855 | 0.785 | 0.343 | 4.97E-21 | 19 | Actr2 |
| Exoc24 | 2.68E-25 | 0.396241222 | 0.468 | 0.147 | 5.42E-21 | 19 | Exoc2 |
| Bin16 | 2.81E-25 | 0.266579297 | 0.532 | 0.175 | 5.69E-21 | 19 | Bin1 |
| Gimap35 | 2.83E-25 | 0.359771101 | 0.367 | 0.1 | 5.73E-21 | 19 | Gimap3 |
| Ythdf36 | 2.98E-25 | 0.420135623 | 0.576 | 0.195 | 6.03E-21 | 19 | Ythdf3 |
| Gns4 | 3.05E-25 | 0.357086888 | 0.424 | 0.126 | 6.17E-21 | 19 | Gns |
| Ccdc505 | 3.35E-25 | 0.368077895 | 0.627 | 0.228 | 6.78E-21 | 19 | Ccdc50 |
| Morc32 | 3.53E-25 | 0.309657877 | 0.5 | 0.162 | 7.16E-21 | 19 | Morc3 |
| Enox22 | 3.94E-25 | 0.584946706 | 0.449 | 0.143 | 7.97E-21 | 19 | Enox2 |
| Nub13 | 3.99E-25 | 0.414687583 | 0.538 | 0.186 | 8.08E-21 | 19 | Nub1 |
| Psen14 | 4.05E-25 | 0.371451651 | 0.361 | 0.1 | 8.20E-21 | 19 | Psen1 |
| Skil3 | 4.79E-25 | 0.354137347 | 0.519 | 0.173 | 9.70E-21 | 19 | Skil |
| Rftn19 | 5.10E-25 | 0.706728901 | 0.652 | 0.251 | 1.03E-20 | 19 | Rftn1 |
| Mitf2 | 5.19E-25 | 0.391349904 | 0.304 | 0.075 | 1.05E-20 | 19 | Mitf |
| Ehbp1l1 | 6.13E-25 | 0.284498525 | 0.367 | 0.1 | 1.24E-20 | 19 | Ehbp1l1 |
| Rps248 | 7.40E-25 | 0.633230773 | 1 | 0.972 | 1.50E-20 | 19 | Rps24 |
| Cdk2ap26 | 7.78E-25 | 0.308430072 | 0.57 | 0.199 | 1.58E-20 | 19 | Cdk2ap2 |
| Itpr16 | 8.26E-25 | 0.64083281 | 0.589 | 0.221 | 1.67E-20 | 19 | Itpr1 |
| Fnbp44 | 8.50E-25 | 0.314937994 | 0.43 | 0.129 | 1.72E-20 | 19 | Fnbp4 |
| Kansl110 | 8.63E-25 | 0.622005305 | 0.778 | 0.34 | 1.75E-20 | 19 | Kansl1 |
| Arsb4 | 9.56E-25 | 0.449152141 | 0.418 | 0.127 | 1.93E-20 | 19 | Arsb |
| Stx163 | 9.63E-25 | 0.331855648 | 0.5 | 0.16 | 1.95E-20 | 19 | Stx16 |
| Polr2a8 | 1.06E-24 | 0.524494186 | 0.715 | 0.281 | 2.14E-20 | 19 | Polr2a |
| Lcor7 | 1.08E-24 | 0.422568406 | 0.551 | 0.19 | 2.19E-20 | 19 | Lcor |
| Smc46 | 1.12E-24 | 0.336686658 | 0.506 | 0.163 | 2.26E-20 | 19 | Smc4 |

| | | | | | | | |
|---|---|---|---|---|---|---|---|
| Lrrfip23 | 1.23E-24 | 0.29180539 | 0.506 | 0.164 | 2.49E-20 | 19 | Lrrfip2 |
| Gm472839 | 1.40E-24 | 0.471689457 | 0.709 | 0.281 | 2.84E-20 | 19 | Gm47283 |
| Metrnl4 | 1.41E-24 | 0.382915823 | 0.367 | 0.103 | 2.86E-20 | 19 | Metrnl |
| Rbbp81 | 1.59E-24 | 0.255194717 | 0.367 | 0.101 | 3.21E-20 | 19 | Rbbp8 |
| M6pr3 | 1.66E-24 | 0.364943451 | 0.494 | 0.162 | 3.37E-20 | 19 | M6pr |
| Sub16 | 1.74E-24 | 0.681032068 | 0.924 | 0.516 | 3.52E-20 | 19 | Sub1 |
| Mapre27 | 2.13E-24 | 0.60383677 | 0.627 | 0.25 | 4.32E-20 | 19 | Mapre2 |
| Sms5 | 2.17E-24 | 0.305637222 | 0.468 | 0.15 | 4.39E-20 | 19 | Sms |
| C2cd31 | 2.41E-24 | 0.315075876 | 0.31 | 0.079 | 4.87E-20 | 19 | C2cd3 |
| Nup1534 | 2.46E-24 | 0.308101065 | 0.399 | 0.116 | 4.98E-20 | 19 | Nup153 |
| Rpl18a6 | 2.56E-24 | 0.635719555 | 1 | 0.931 | 5.19E-20 | 19 | Rpl18a |
| Arih18 | 2.77E-24 | 0.500006252 | 0.747 | 0.312 | 5.60E-20 | 19 | Arih1 |
| Arfgef15 | 2.78E-24 | 0.338744284 | 0.627 | 0.227 | 5.63E-20 | 19 | Arfgef1 |
| Stxbp3 | 3.04E-24 | 0.32873484 | 0.5 | 0.162 | 6.15E-20 | 19 | Stxbp3 |
| 16-Sep | 3.18E-24 | 0.408958516 | 0.462 | 0.146 | 6.43E-20 | 19 | 1-Sep |
| Atp6v0b4 | 3.19E-24 | 0.412801208 | 0.829 | 0.366 | 6.46E-20 | 19 | Atp6v0b |
| Tra2b6 | 3.35E-24 | 0.448696063 | 0.734 | 0.296 | 6.79E-20 | 19 | Tra2b |
| Mgat57 | 3.62E-24 | 0.637709051 | 0.487 | 0.17 | 7.33E-20 | 19 | Mgat5 |
| Spata55 | 3.81E-24 | 0.268365829 | 0.418 | 0.123 | 7.70E-20 | 19 | Spata5 |
| Cnot24 | 3.99E-24 | 0.414509544 | 0.582 | 0.21 | 8.08E-20 | 19 | Cnot2 |
| Rbm333 | 4.08E-24 | 0.304286267 | 0.456 | 0.142 | 8.26E-20 | 19 | Rbm33 |
| Ywhag3 | 5.51E-24 | 0.348285444 | 0.532 | 0.181 | 1.12E-19 | 19 | Ywhag |
| Tmem1634 | 5.56E-24 | 0.606466387 | 0.323 | 0.088 | 1.13E-19 | 19 | Tmem163 |
| Ubash3b6 | 6.18E-24 | 0.385170864 | 0.5 | 0.168 | 1.25E-19 | 19 | Ubash3b |
| Ipcef15 | 6.21E-24 | 0.454461881 | 0.392 | 0.118 | 1.26E-19 | 19 | Ipcef1 |
| Tra2a11 | 6.27E-24 | 0.495856433 | 0.69 | 0.274 | 1.27E-19 | 19 | Tra2a |
| Zcchc116 | 6.47E-24 | 0.523538733 | 0.684 | 0.267 | 1.31E-19 | 19 | Zcchc11 |
| H2-M31 | 6.77E-24 | 0.340471612 | 0.462 | 0.147 | 1.37E-19 | 19 | H2-M3 |
| Ccl35 | 7.12E-24 | 0.629945145 | 0.31 | 0.083 | 1.44E-19 | 19 | Ccl3 |
| Mtmr39 | 7.30E-24 | 0.479991019 | 0.557 | 0.2 | 1.48E-19 | 19 | Mtmr3 |
| Rptor6 | 7.92E-24 | 0.416424625 | 0.437 | 0.139 | 1.60E-19 | 19 | Rptor |
| Papd43 | 8.48E-24 | 0.414648459 | 0.43 | 0.138 | 1.72E-19 | 19 | Papd4 |
| Camk2d8 | 8.67E-24 | 0.659523538 | 0.665 | 0.27 | 1.76E-19 | 19 | Camk2d |
| Vps37b5 | 8.91E-24 | 0.377657773 | 0.437 | 0.137 | 1.80E-19 | 19 | Vps37b |
| Cnpy32 | 9.40E-24 | 0.322672853 | 0.361 | 0.102 | 1.90E-19 | 19 | Cnpy3 |
| Zc3hav15 | 9.74E-24 | 0.412293161 | 0.57 | 0.204 | 1.97E-19 | 19 | Zc3hav1 |
| Fam126a1 | 1.09E-23 | 0.301114001 | 0.31 | 0.081 | 2.21E-19 | 19 | Fam126a |
| Zfp3184 | 1.11E-23 | 0.363768038 | 0.348 | 0.098 | 2.25E-19 | 19 | Zfp318 |
| Aebp22 | 1.13E-23 | 0.391274194 | 0.373 | 0.109 | 2.28E-19 | 19 | Aebp2 |
| Sf3b18 | 1.16E-23 | 0.533652769 | 0.848 | 0.393 | 2.36E-19 | 19 | Sf3b1 |
| Ldlrad47 | 1.19E-23 | 0.690609444 | 0.614 | 0.251 | 2.41E-19 | 19 | Ldlrad4 |
| Brd410 | 1.23E-23 | 0.438866893 | 0.684 | 0.267 | 2.48E-19 | 19 | Brd4 |
| Edem11 | 1.24E-23 | 0.272016199 | 0.329 | 0.089 | 2.51E-19 | 19 | Edem1 |

| | | | | | | | |
|---|---|---|---|---|---|---|---|
| Epc15 | 1.24E-23 | 0.378366463 | 0.57 | 0.203 | 2.52E-19 | 19 | Epc1 |
| Arhgap269 | 1.31E-23 | 0.643673781 | 0.582 | 0.222 | 2.65E-19 | 19 | Arhgap26 |
| Eml61 | 1.81E-23 | 0.412382104 | 0.361 | 0.102 | 3.66E-19 | 19 | Eml6 |
| Sgpp11 | 2.06E-23 | 0.283761224 | 0.297 | 0.076 | 4.18E-19 | 19 | Sgpp1 |
| Babam27 | 2.39E-23 | 0.403209075 | 0.633 | 0.238 | 4.84E-19 | 19 | Babam2 |
| Med13l6 | 2.40E-23 | 0.561644867 | 0.544 | 0.194 | 4.87E-19 | 19 | Med13l |
| Ppt12 | 2.45E-23 | 0.570556136 | 0.418 | 0.13 | 4.95E-19 | 19 | Ppt1 |
| Ndfip15 | 2.48E-23 | 0.47591293 | 0.741 | 0.321 | 5.01E-19 | 19 | Ndfip1 |
| Smyd35 | 2.53E-23 | 0.667613135 | 0.576 | 0.227 | 5.11E-19 | 19 | Smyd3 |
| Rnf1151 | 2.68E-23 | 0.418313279 | 0.538 | 0.189 | 5.43E-19 | 19 | Rnf115 |
| Rasa25 | 2.71E-23 | 0.426119978 | 0.43 | 0.138 | 5.49E-19 | 19 | Rasa2 |
| Zbp12 | 2.75E-23 | 0.420201179 | 0.278 | 0.07 | 5.56E-19 | 19 | Zbp1 |
| Ptbp21 | 2.83E-23 | 0.264373474 | 0.424 | 0.129 | 5.73E-19 | 19 | Ptbp2 |
| Dyrk1a6 | 2.98E-23 | 0.399611421 | 0.519 | 0.179 | 6.03E-19 | 19 | Dyrk1a |
| Fam168b2 | 3.35E-23 | 0.300808601 | 0.494 | 0.166 | 6.79E-19 | 19 | Fam168b |
| Tle46 | 3.78E-23 | 0.544009036 | 0.595 | 0.225 | 7.66E-19 | 19 | Tle4 |
| Chsy14 | 3.79E-23 | 0.388998035 | 0.411 | 0.128 | 7.68E-19 | 19 | Chsy1 |
| Nsmaf5 | 4.11E-23 | 0.26854786 | 0.348 | 0.097 | 8.32E-19 | 19 | Nsmaf |
| Lpp9 | 4.14E-23 | 0.443669093 | 0.709 | 0.286 | 8.37E-19 | 19 | Lpp |
| Atp2b18 | 4.20E-23 | 0.689631669 | 0.741 | 0.333 | 8.50E-19 | 19 | Atp2b1 |
| Rps38 | 4.83E-23 | 0.564729874 | 0.987 | 0.893 | 9.78E-19 | 19 | Rps3 |
| Hmgcr | 5.62E-23 | 0.266385119 | 0.272 | 0.067 | 1.14E-18 | 19 | Hmgcr |
| Lman25 | 5.80E-23 | 0.304655675 | 0.576 | 0.206 | 1.17E-18 | 19 | Lman2 |
| Rpl37a8 | 5.89E-23 | 0.545142434 | 1 | 0.938 | 1.19E-18 | 19 | Rpl37a |
| Fam214a3 | 5.92E-23 | 0.506668282 | 0.392 | 0.124 | 1.20E-18 | 19 | Fam214a |
| Tmem1317 | 5.97E-23 | 0.463698452 | 0.563 | 0.208 | 1.21E-18 | 19 | Tmem131 |
| Otud5 | 6.83E-23 | 0.286257716 | 0.323 | 0.088 | 1.38E-18 | 19 | Otud5 |
| Cnot6l4 | 7.15E-23 | 0.317823739 | 0.652 | 0.244 | 1.45E-18 | 19 | Cnot6l |
| Atp6v0d11 | 7.47E-23 | 0.257395831 | 0.481 | 0.159 | 1.51E-18 | 19 | Atp6v0d1 |
| Ctsb5 | 8.31E-23 | 0.558273844 | 0.69 | 0.298 | 1.68E-18 | 19 | Ctsb |
| Rbm2511 | 9.51E-23 | 0.448067793 | 0.791 | 0.333 | 1.93E-18 | 19 | Rbm25 |
| Pkn17 | 9.58E-23 | 0.360120011 | 0.525 | 0.186 | 1.94E-18 | 19 | Pkn1 |
| Esyt17 | 1.16E-22 | 0.273332135 | 0.437 | 0.136 | 2.35E-18 | 19 | Esyt1 |
| AC163354.13 | 1.18E-22 | 0.263140011 | 0.259 | 0.062 | 2.40E-18 | 19 | AC163354.1 |
| Swap705 | 1.19E-22 | 0.618303632 | 0.614 | 0.246 | 2.41E-18 | 19 | Swap70 |
| Pip5k1c2 | 1.29E-22 | 0.263917029 | 0.348 | 0.098 | 2.60E-18 | 19 | Pip5k1c |
| Vps13a6 | 1.60E-22 | 0.384787583 | 0.5 | 0.173 | 3.23E-18 | 19 | Vps13a |
| Ssbp27 | 1.72E-22 | 0.359785825 | 0.437 | 0.14 | 3.48E-18 | 19 | Ssbp2 |
| Cog57 | 1.94E-22 | 0.344509808 | 0.525 | 0.183 | 3.93E-18 | 19 | Cog5 |
| Tpp25 | 2.01E-22 | 0.384540983 | 0.614 | 0.228 | 4.07E-18 | 19 | Tpp2 |
| Epb41l25 | 2.04E-22 | 0.459083248 | 0.627 | 0.241 | 4.13E-18 | 19 | Epb41l2 |
| Fam13b6 | 2.08E-22 | 0.338577548 | 0.449 | 0.147 | 4.20E-18 | 19 | Fam13b |
| Itgav9 | 2.09E-22 | 0.435549311 | 0.475 | 0.161 | 4.23E-18 | 19 | Itgav |

| | | | | | | | |
|---|---|---|---|---|---|---|---|
| Asxl26 | 2.11E-22 | 0.358276308 | 0.506 | 0.175 | 4.26E-18 | 19 | Asxl2 |
| Diaph16 | 2.37E-22 | 0.475413448 | 0.614 | 0.242 | 4.80E-18 | 19 | Diaph1 |
| Atf7ip5 | 2.38E-22 | 0.522838497 | 0.563 | 0.207 | 4.83E-18 | 19 | Atf7ip |
| Zc3h7a6 | 2.44E-22 | 0.357352368 | 0.481 | 0.162 | 4.94E-18 | 19 | Zc3h7a |
| Lnpep11 | 2.53E-22 | 0.42317928 | 0.665 | 0.259 | 5.13E-18 | 19 | Lnpep |
| Hexa4 | 2.85E-22 | 0.266662628 | 0.601 | 0.22 | 5.77E-18 | 19 | Hexa |
| 4933406I18Rik6 | 2.89E-22 | 0.31242264 | 0.291 | 0.076 | 5.85E-18 | 19 | 4933406I18Rik |
| Sp1106 | 3.02E-22 | 0.320526819 | 0.468 | 0.157 | 6.12E-18 | 19 | Sp110 |
| Zfx1 | 3.11E-22 | 0.268901199 | 0.468 | 0.151 | 6.29E-18 | 19 | Zfx |
| Arid1a8 | 3.28E-22 | 0.476628241 | 0.595 | 0.23 | 6.63E-18 | 19 | Arid1a |
| Ppp1r187 | 3.40E-22 | 0.314822283 | 0.462 | 0.154 | 6.88E-18 | 19 | Ppp1r18 |
| Sep-97 | 3.66E-22 | 0.352967438 | 0.494 | 0.171 | 7.42E-18 | 19 | 9-Sep |
| Zmynd82 | 4.22E-22 | 0.275172086 | 0.544 | 0.195 | 8.55E-18 | 19 | Zmynd8 |
| Ifrd14 | 4.69E-22 | 0.338694076 | 0.608 | 0.23 | 9.50E-18 | 19 | Ifrd1 |
| Pik3cb3 | 4.93E-22 | 0.352134399 | 0.443 | 0.146 | 9.98E-18 | 19 | Pik3cb |
| Stk383 | 4.96E-22 | 0.29438945 | 0.608 | 0.226 | 1.00E-17 | 19 | Stk38 |
| Ckb5 | 5.14E-22 | 0.703557467 | 0.551 | 0.216 | 1.04E-17 | 19 | Ckb |
| Rlim4 | 5.30E-22 | 0.427268674 | 0.544 | 0.197 | 1.07E-17 | 19 | Rlim |
| Bcl25 | 5.38E-22 | 0.548210122 | 0.538 | 0.207 | 1.09E-17 | 19 | Bcl2 |
| Pign3 | 5.86E-22 | 0.398762139 | 0.329 | 0.094 | 1.19E-17 | 19 | Pign |
| Pml | 5.91E-22 | 0.444644081 | 0.297 | 0.081 | 1.20E-17 | 19 | Pml |
| Trim30a7 | 5.93E-22 | 0.443271698 | 0.582 | 0.219 | 1.20E-17 | 19 | Trim30a |
| Vasp5 | 5.97E-22 | 0.416936448 | 0.614 | 0.237 | 1.21E-17 | 19 | Vasp |
| Vars5 | 6.18E-22 | 0.294868467 | 0.304 | 0.082 | 1.25E-17 | 19 | Vars |
| Rnf1308 | 6.27E-22 | 0.481674384 | 0.684 | 0.284 | 1.27E-17 | 19 | Rnf130 |
| Larp17 | 6.27E-22 | 0.388976166 | 0.551 | 0.203 | 1.27E-17 | 19 | Larp1 |
| Ranbp96 | 6.91E-22 | 0.427022773 | 0.544 | 0.2 | 1.40E-17 | 19 | Ranbp9 |
| Fbxw112 | 7.09E-22 | 0.292356952 | 0.5 | 0.171 | 1.44E-17 | 19 | Fbxw11 |
| Cux11 | 8.05E-22 | 0.264771595 | 0.614 | 0.23 | 1.63E-17 | 19 | Cux1 |
| Sqstm17 | 8.37E-22 | 0.526473188 | 0.709 | 0.304 | 1.69E-17 | 19 | Sqstm1 |
| Cbfb6 | 8.88E-22 | 0.29843442 | 0.468 | 0.155 | 1.80E-17 | 19 | Cbfb |
| Sun24 | 9.47E-22 | 0.334398763 | 0.291 | 0.078 | 1.92E-17 | 19 | Sun2 |
| Slc2a14 | 1.01E-21 | 0.3449006 | 0.316 | 0.088 | 2.04E-17 | 19 | Slc2a1 |
| Ap1s24 | 1.05E-21 | 0.296841491 | 0.43 | 0.139 | 2.13E-17 | 19 | Ap1s2 |
| Rps288 | 1.15E-21 | 0.562039687 | 0.987 | 0.898 | 2.33E-17 | 19 | Rps28 |
| Usp35 | 1.20E-21 | 0.294421192 | 0.519 | 0.182 | 2.44E-17 | 19 | Usp3 |
| Nipbl12 | 1.28E-21 | 0.49239917 | 0.816 | 0.366 | 2.60E-17 | 19 | Nipbl |
| Foxn39 | 1.29E-21 | 0.774046081 | 0.829 | 0.411 | 2.61E-17 | 19 | Foxn3 |
| Kpna34 | 1.33E-21 | 0.289670675 | 0.462 | 0.157 | 2.68E-17 | 19 | Kpna3 |
| Cspp11 | 1.33E-21 | 0.2943056 | 0.38 | 0.117 | 2.70E-17 | 19 | Cspp1 |
| Ralgapb2 | 1.34E-21 | 0.277735957 | 0.386 | 0.117 | 2.72E-17 | 19 | Ralgapb |
| Rps107 | 1.40E-21 | 0.558236786 | 0.994 | 0.944 | 2.83E-17 | 19 | Rps10 |
| Dock54 | 1.40E-21 | 0.376931646 | 0.373 | 0.115 | 2.83E-17 | 19 | Dock5 |

| | | | | | | | |
|---|---|---|---|---|---|---|---|
| Rasgrp15 | 1.50E-21 | 0.315357673 | 0.253 | 0.063 | 3.04E-17 | 19 | Rasgrp1 |
| Zfp7103 | 1.65E-21 | 0.556932106 | 0.443 | 0.155 | 3.34E-17 | 19 | Zfp710 |
| Tespa15 | 1.77E-21 | 0.303599819 | 0.291 | 0.077 | 3.59E-17 | 19 | Tespa1 |
| Mlxip4 | 1.79E-21 | 0.382554215 | 0.399 | 0.127 | 3.63E-17 | 19 | Mlxip |
| Foxo34 | 1.84E-21 | 0.280498241 | 0.38 | 0.117 | 3.73E-17 | 19 | Foxo3 |
| Rpl346 | 1.88E-21 | 0.543102514 | 0.994 | 0.917 | 3.80E-17 | 19 | Rpl34 |
| Zfp36l26 | 1.92E-21 | 0.575460985 | 0.684 | 0.305 | 3.89E-17 | 19 | Zfp36l2 |
| Fam193a6 | 2.18E-21 | 0.28029506 | 0.513 | 0.179 | 4.42E-17 | 19 | Fam193a |
| Tbc1d231 | 2.52E-21 | 0.260025512 | 0.392 | 0.122 | 5.10E-17 | 19 | Tbc1d23 |
| Mettl234 | 2.58E-21 | 0.258070411 | 0.551 | 0.203 | 5.22E-17 | 19 | Mettl23 |
| Dusp14 | 2.67E-21 | 0.65777049 | 0.532 | 0.193 | 5.41E-17 | 19 | Dusp1 |
| Pcgf51 | 2.74E-21 | 0.769670052 | 0.361 | 0.113 | 5.54E-17 | 19 | Pcgf5 |
| Fkbp151 | 3.01E-21 | 0.299021489 | 0.367 | 0.111 | 6.09E-17 | 19 | Fkbp15 |
| Dip2b7 | 3.03E-21 | 0.487768512 | 0.677 | 0.276 | 6.14E-17 | 19 | Dip2b |
| 2310035C23Rik6 | 3.03E-21 | 0.276698348 | 0.43 | 0.14 | 6.14E-17 | 19 | 2310035C23Rik |
| Srrm11 | 3.27E-21 | 0.287329067 | 0.81 | 0.35 | 6.61E-17 | 19 | Srrm1 |
| Rfx34 | 3.33E-21 | 0.405601361 | 0.354 | 0.108 | 6.74E-17 | 19 | Rfx3 |
| Rps277 | 3.47E-21 | 0.676474637 | 0.994 | 0.898 | 7.03E-17 | 19 | Rps27 |
| Zyx5 | 3.56E-21 | 0.460275683 | 0.57 | 0.216 | 7.20E-17 | 19 | Zyx |
| Map3k13 | 3.70E-21 | 0.367045462 | 0.538 | 0.199 | 7.49E-17 | 19 | Map3k1 |
| Hivep29 | 3.79E-21 | 0.52642028 | 0.69 | 0.298 | 7.68E-17 | 19 | Hivep2 |
| Fam60a | 4.51E-21 | 0.261622831 | 0.323 | 0.092 | 9.13E-17 | 19 | Fam60a |
| Ascc38 | 4.87E-21 | 0.42692757 | 0.551 | 0.207 | 9.86E-17 | 19 | Ascc3 |
| Rsrc17 | 5.07E-21 | 0.396636392 | 0.639 | 0.26 | 1.03E-16 | 19 | Rsrc1 |
| Klri23 | 5.32E-21 | 0.301613197 | 0.285 | 0.075 | 1.08E-16 | 19 | Klri2 |
| Aff44 | 5.71E-21 | 0.440416976 | 0.608 | 0.24 | 1.16E-16 | 19 | Aff4 |
| Exoc411 | 6.29E-21 | 0.558797861 | 0.728 | 0.323 | 1.27E-16 | 19 | Exoc4 |
| Rtn48 | 6.71E-21 | 0.485522238 | 0.759 | 0.339 | 1.36E-16 | 19 | Rtn4 |
| Setbp16 | 6.93E-21 | 0.458442877 | 0.38 | 0.122 | 1.40E-16 | 19 | Setbp1 |
| Ptpn44 | 7.53E-21 | 0.330119371 | 0.405 | 0.131 | 1.52E-16 | 19 | Ptpn4 |
| Gatad2b4 | 7.84E-21 | 0.332545735 | 0.544 | 0.202 | 1.59E-16 | 19 | Gatad2b |
| Osbpl35 | 8.43E-21 | 0.377742494 | 0.38 | 0.12 | 1.71E-16 | 19 | Osbpl3 |
| Sft2d13 | 8.44E-21 | 0.321842708 | 0.468 | 0.16 | 1.71E-16 | 19 | Sft2d1 |
| Gak1 | 8.99E-21 | 0.270095537 | 0.392 | 0.125 | 1.82E-16 | 19 | Gak |
| Nedd97 | 9.81E-21 | 0.487325017 | 0.734 | 0.319 | 1.99E-16 | 19 | Nedd9 |
| Hist1h1c3 | 9.85E-21 | 0.389326117 | 0.532 | 0.201 | 1.99E-16 | 19 | Hist1h1c |
| Itch8 | 1.09E-20 | 0.384489977 | 0.671 | 0.273 | 2.20E-16 | 19 | Itch |
| Clk41 | 1.16E-20 | 0.367015705 | 0.424 | 0.141 | 2.36E-16 | 19 | Clk4 |
| Snx2 | 1.20E-20 | 0.450880096 | 0.576 | 0.227 | 2.42E-16 | 19 | Snx2 |
| Cnot14 | 1.24E-20 | 0.275250333 | 0.494 | 0.173 | 2.50E-16 | 19 | Cnot1 |
| Tsc22d45 | 1.27E-20 | 0.272560482 | 0.703 | 0.294 | 2.57E-16 | 19 | Tsc22d4 |
| Mtpn | 1.41E-20 | 0.294761946 | 0.608 | 0.239 | 2.86E-16 | 19 | Mtpn |
| Sbno14 | 1.53E-20 | 0.260252708 | 0.563 | 0.209 | 3.09E-16 | 19 | Sbno1 |

| | | | | | | | |
|---|---|---|---|---|---|---|---|
| Rps58 | 1.53E-20 | 0.543175214 | 0.994 | 0.92 | 3.10E-16 | 19 | Rps5 |
| Prkca7 | 1.59E-20 | 0.375437774 | 0.335 | 0.101 | 3.21E-16 | 19 | Prkca |
| Stat15 | 2.03E-20 | 0.304154384 | 0.62 | 0.249 | 4.11E-16 | 19 | Stat1 |
| Acap15 | 2.11E-20 | 0.276727871 | 0.272 | 0.071 | 4.27E-16 | 19 | Acap1 |
| Rpl187 | 2.33E-20 | 0.50979011 | 0.987 | 0.9 | 4.72E-16 | 19 | Rpl18 |
| Ugcg5 | 2.34E-20 | 0.44536616 | 0.475 | 0.172 | 4.74E-16 | 19 | Ugcg |
| Gtpbp13 | 2.59E-20 | 0.256099845 | 0.272 | 0.072 | 5.25E-16 | 19 | Gtpbp1 |
| Tgfb17 | 2.74E-20 | 0.391340054 | 0.589 | 0.236 | 5.55E-16 | 19 | Tgfb1 |
| Anxa65 | 2.81E-20 | 0.266021938 | 0.456 | 0.157 | 5.68E-16 | 19 | Anxa6 |
| Irf27 | 3.10E-20 | 0.460316132 | 0.766 | 0.341 | 6.28E-16 | 19 | Irf2 |
| Nsa27 | 3.21E-20 | 0.380868918 | 0.918 | 0.486 | 6.50E-16 | 19 | Nsa2 |
| Suco6 | 3.59E-20 | 0.355872762 | 0.443 | 0.154 | 7.26E-16 | 19 | Suco |
| Btaf11 | 3.71E-20 | 0.307447805 | 0.386 | 0.126 | 7.50E-16 | 19 | Btaf1 |
| Rps207 | 3.89E-20 | 0.564022902 | 0.994 | 0.951 | 7.88E-16 | 19 | Rps20 |
| Txk4 | 4.11E-20 | 0.321633696 | 0.354 | 0.107 | 8.32E-16 | 19 | Txk |
| Wdfy22 | 4.36E-20 | 0.294478688 | 0.361 | 0.111 | 8.82E-16 | 19 | Wdfy2 |
| Nek77 | 4.53E-20 | 0.336466587 | 0.614 | 0.248 | 9.18E-16 | 19 | Nek7 |
| Uggt1 | 4.63E-20 | 0.253407274 | 0.392 | 0.126 | 9.37E-16 | 19 | Uggt1 |
| Pgap2 | 5.49E-20 | 0.260104654 | 0.449 | 0.154 | 1.11E-15 | 19 | Pgap2 |
| Padi24 | 5.63E-20 | 0.310262187 | 0.323 | 0.096 | 1.14E-15 | 19 | Padi2 |
| Gbf17 | 6.00E-20 | 0.251215798 | 0.506 | 0.182 | 1.21E-15 | 19 | Gbf1 |
| Ddx63 | 6.66E-20 | 0.4293448 | 0.759 | 0.341 | 1.35E-15 | 19 | Ddx6 |
| Ppp3cc4 | 7.14E-20 | 0.400253513 | 0.418 | 0.142 | 1.45E-15 | 19 | Ppp3cc |
| Add33 | 7.82E-20 | 0.723743823 | 0.551 | 0.223 | 1.58E-15 | 19 | Add3 |
| Nsmce26 | 7.85E-20 | 0.259986361 | 0.557 | 0.206 | 1.59E-15 | 19 | Nsmce2 |
| Far14 | 8.09E-20 | 0.346446326 | 0.462 | 0.165 | 1.64E-15 | 19 | Far1 |
| Fndc3a6 | 8.10E-20 | 0.444145193 | 0.639 | 0.277 | 1.64E-15 | 19 | Fndc3a |
| Nrf15 | 8.15E-20 | 0.311204478 | 0.411 | 0.135 | 1.65E-15 | 19 | Nrf1 |
| Mctp24 | 8.97E-20 | 0.371287374 | 0.297 | 0.084 | 1.82E-15 | 19 | Mctp2 |
| Rbm37 | 9.68E-20 | 0.486666427 | 0.873 | 0.477 | 1.96E-15 | 19 | Rbm3 |
| Ier24 | 1.06E-19 | 0.452281885 | 0.665 | 0.284 | 2.14E-15 | 19 | Ier2 |
| Fgf134 | 1.07E-19 | 0.362101999 | 0.285 | 0.079 | 2.16E-15 | 19 | Fgf13 |
| Ms4a74 | 1.09E-19 | 0.318329155 | 0.335 | 0.1 | 2.21E-15 | 19 | Ms4a7 |
| Ptbp310 | 1.15E-19 | 0.473773069 | 0.854 | 0.393 | 2.34E-15 | 19 | Ptbp3 |
| Tmem2433 | 1.17E-19 | 0.356418738 | 0.462 | 0.163 | 2.37E-15 | 19 | Tmem243 |
| Ino803 | 1.19E-19 | 0.321486999 | 0.411 | 0.14 | 2.41E-15 | 19 | Ino80 |
| Rps138 | 1.25E-19 | 0.51229412 | 0.994 | 0.924 | 2.54E-15 | 19 | Rps13 |
| Supt205 | 1.39E-19 | 0.336069032 | 0.462 | 0.161 | 2.81E-15 | 19 | Supt20 |
| Nr4a24 | 1.46E-19 | 0.381664583 | 0.392 | 0.131 | 2.96E-15 | 19 | Nr4a2 |
| Dnajb93 | 1.54E-19 | 0.375727734 | 0.418 | 0.143 | 3.12E-15 | 19 | Dnajb9 |
| Arpc51 | 1.69E-19 | 0.39884802 | 0.842 | 0.428 | 3.43E-15 | 19 | Arpc5 |
| Vav36 | 1.72E-19 | 0.442048188 | 0.614 | 0.256 | 3.49E-15 | 19 | Vav3 |
| Ankib16 | 1.81E-19 | 0.254648676 | 0.418 | 0.139 | 3.66E-15 | 19 | Ankib1 |

| | | | | | | | |
|---|---|---|---|---|---|---|---|
| Rsbn1l7 | 1.89E-19 | 0.279051988 | 0.671 | 0.272 | 3.84E-15 | 19 | Rsbn1l |
| Mllt109 | 1.90E-19 | 0.45968641 | 0.652 | 0.275 | 3.84E-15 | 19 | Mllt10 |
| Pspc13 | 2.15E-19 | 0.250983458 | 0.373 | 0.118 | 4.34E-15 | 19 | Pspc1 |
| Rreb14 | 2.21E-19 | 0.413789256 | 0.519 | 0.202 | 4.48E-15 | 19 | Rreb1 |
| Ms4a4b4 | 2.33E-19 | 0.295082189 | 0.386 | 0.123 | 4.72E-15 | 19 | Ms4a4b |
| Rsf16 | 2.51E-19 | 0.298315314 | 0.576 | 0.224 | 5.08E-15 | 19 | Rsf1 |
| Zfp6544 | 2.64E-19 | 0.332207611 | 0.373 | 0.121 | 5.35E-15 | 19 | Zfp654 |
| Parp1 | 3.09E-19 | 0.265691692 | 0.323 | 0.098 | 6.25E-15 | 19 | Parp1 |
| Tab24 | 3.40E-19 | 0.316303654 | 0.576 | 0.231 | 6.88E-15 | 19 | Tab2 |
| Txndc112 | 3.48E-19 | 0.288392541 | 0.316 | 0.095 | 7.04E-15 | 19 | Txndc11 |
| Cep3508 | 3.77E-19 | 0.333683356 | 0.532 | 0.204 | 7.63E-15 | 19 | Cep350 |
| Atl33 | 4.13E-19 | 0.4026941 | 0.38 | 0.129 | 8.37E-15 | 19 | Atl3 |
| Phip10 | 4.45E-19 | 0.403630338 | 0.614 | 0.256 | 9.02E-15 | 19 | Phip |
| Eif3f5 | 5.04E-19 | 0.419803163 | 0.886 | 0.483 | 1.02E-14 | 19 | Eif3f |
| Itgae3 | 5.07E-19 | 0.371624916 | 0.253 | 0.067 | 1.03E-14 | 19 | Itgae |
| Srsf1111 | 5.42E-19 | 0.431221229 | 0.759 | 0.351 | 1.10E-14 | 19 | Srsf11 |
| Zfand310 | 5.97E-19 | 0.360546793 | 0.709 | 0.301 | 1.21E-14 | 19 | Zfand3 |
| Scfd23 | 6.10E-19 | 0.27101497 | 0.297 | 0.086 | 1.24E-14 | 19 | Scfd2 |
| Map3k35 | 6.28E-19 | 0.31959022 | 0.487 | 0.178 | 1.27E-14 | 19 | Map3k3 |
| Cd76 | 6.32E-19 | 0.448043277 | 0.354 | 0.113 | 1.28E-14 | 19 | Cd7 |
| Abr7 | 6.54E-19 | 0.379784978 | 0.437 | 0.158 | 1.32E-14 | 19 | Abr |
| Med132 | 6.76E-19 | 0.381414467 | 0.532 | 0.205 | 1.37E-14 | 19 | Med13 |
| Tmcc35 | 7.84E-19 | 0.880584797 | 0.665 | 0.319 | 1.59E-14 | 19 | Tmcc3 |
| Bicra1 | 8.37E-19 | 0.263033208 | 0.335 | 0.103 | 1.69E-14 | 19 | Bicra |
| Nufip24 | 9.07E-19 | 0.277830262 | 0.582 | 0.232 | 1.84E-14 | 19 | Nufip2 |
| Kdm3b5 | 9.27E-19 | 0.28153382 | 0.405 | 0.136 | 1.88E-14 | 19 | Kdm3b |
| Ttc5 | 9.86E-19 | 0.258957857 | 0.354 | 0.113 | 2.00E-14 | 19 | Ttc5 |
| Usp4 | 9.87E-19 | 0.303909174 | 0.367 | 0.12 | 2.00E-14 | 19 | Usp4 |
| Adam173 | 1.00E-18 | 0.334212939 | 0.437 | 0.154 | 2.03E-14 | 19 | Adam17 |
| Kcnq1ot17 | 1.04E-18 | 0.336973744 | 0.399 | 0.134 | 2.10E-14 | 19 | Kcnq1ot1 |
| Sys1 | 1.12E-18 | 0.28334529 | 0.62 | 0.251 | 2.26E-14 | 19 | Sys1 |
| Snhg122 | 1.12E-18 | 0.261386802 | 0.354 | 0.115 | 2.27E-14 | 19 | Snhg12 |
| Exoc66 | 1.15E-18 | 0.260546569 | 0.475 | 0.171 | 2.32E-14 | 19 | Exoc6 |
| Rpl35a8 | 1.15E-18 | 0.427877852 | 1 | 0.948 | 2.34E-14 | 19 | Rpl35a |
| Xpo61 | 1.43E-18 | 0.275836033 | 0.297 | 0.088 | 2.89E-14 | 19 | Xpo6 |
| Safb2 | 1.46E-18 | 0.272007279 | 0.354 | 0.115 | 2.95E-14 | 19 | Safb2 |
| Mtmr124 | 1.46E-18 | 0.259625796 | 0.392 | 0.131 | 2.97E-14 | 19 | Mtmr12 |
| Hmgb25 | 1.58E-18 | 0.41251212 | 0.728 | 0.328 | 3.20E-14 | 19 | Hmgb2 |
| Ccl46 | 1.60E-18 | 0.696652394 | 0.373 | 0.13 | 3.25E-14 | 19 | Ccl4 |
| Btbd910 | 1.67E-18 | 0.418990902 | 0.582 | 0.237 | 3.38E-14 | 19 | Btbd9 |
| Lims17 | 1.73E-18 | 0.408605115 | 0.684 | 0.3 | 3.49E-14 | 19 | Lims1 |
| Srsf76 | 2.10E-18 | 0.283224613 | 0.759 | 0.332 | 4.25E-14 | 19 | Srsf7 |
| Kdm5a6 | 2.15E-18 | 0.276366002 | 0.462 | 0.168 | 4.36E-14 | 19 | Kdm5a |

| | | | | | | | |
|---|---|---|---|---|---|---|---|
| Rap1b5 | 2.62E-18 | 0.491434983 | 0.873 | 0.444 | 5.31E-14 | 19 | Rap1b |
| Rnf1114 | 2.71E-18 | 0.254558574 | 0.443 | 0.156 | 5.48E-14 | 19 | Rnf111 |
| Lrch15 | 2.95E-18 | 0.384039908 | 0.696 | 0.312 | 5.98E-14 | 19 | Lrch1 |
| Lmbrd15 | 3.27E-18 | 0.394430744 | 0.443 | 0.163 | 6.61E-14 | 19 | Lmbrd1 |
| Xpo11 | 3.41E-18 | 0.25192393 | 0.38 | 0.126 | 6.90E-14 | 19 | Xpo1 |
| Sfpq7 | 3.60E-18 | 0.349500832 | 0.684 | 0.298 | 7.29E-14 | 19 | Sfpq |
| Cep1923 | 3.64E-18 | 0.28692642 | 0.291 | 0.085 | 7.37E-14 | 19 | Cep192 |
| Rassf34 | 3.67E-18 | 0.42196793 | 0.576 | 0.235 | 7.43E-14 | 19 | Rassf3 |
| Ogt8 | 3.70E-18 | 0.412121123 | 0.519 | 0.206 | 7.48E-14 | 19 | Ogt |
| St8sia48 | 4.04E-18 | 0.429199236 | 0.62 | 0.259 | 8.18E-14 | 19 | St8sia4 |
| Vti1a8 | 4.25E-18 | 0.388320156 | 0.614 | 0.255 | 8.60E-14 | 19 | Vti1a |
| Akap103 | 4.26E-18 | 0.331298615 | 0.411 | 0.146 | 8.62E-14 | 19 | Akap10 |
| Rpl197 | 4.87E-18 | 0.45292652 | 1 | 0.963 | 9.87E-14 | 19 | Rpl19 |
| mt-Co314 | 5.17E-18 | 0.597499924 | 1 | 0.726 | 1.05E-13 | 19 | mt-Co3 |
| Bhlhe406 | 5.69E-18 | 0.308072088 | 0.437 | 0.159 | 1.15E-13 | 19 | Bhlhe40 |
| Rbm266 | 6.32E-18 | 0.26848696 | 0.576 | 0.227 | 1.28E-13 | 19 | Rbm26 |
| Gas75 | 6.69E-18 | 0.580053377 | 0.462 | 0.183 | 1.35E-13 | 19 | Gas7 |
| Trim12a5 | 6.74E-18 | 0.404239162 | 0.475 | 0.182 | 1.36E-13 | 19 | Trim12a |
| Nf16 | 7.00E-18 | 0.264104353 | 0.563 | 0.221 | 1.42E-13 | 19 | Nf1 |
| Zc3h136 | 7.11E-18 | 0.304955517 | 0.544 | 0.216 | 1.44E-13 | 19 | Zc3h13 |
| Rpl137 | 7.26E-18 | 0.465215678 | 1 | 0.977 | 1.47E-13 | 19 | Rpl13 |
| Heca4 | 7.53E-18 | 0.299889699 | 0.405 | 0.143 | 1.52E-13 | 19 | Heca |
| Tor1aip16 | 7.60E-18 | 0.29877614 | 0.665 | 0.281 | 1.54E-13 | 19 | Tor1aip1 |
| Clptm11 | 8.13E-18 | 0.297865533 | 0.373 | 0.125 | 1.65E-13 | 19 | Clptm1 |
| 4932438A13Rik8 | 9.40E-18 | 0.325245897 | 0.551 | 0.219 | 1.90E-13 | 19 | 4932438A13Rik |
| Fam208b5 | 1.01E-17 | 0.270748158 | 0.424 | 0.15 | 2.05E-13 | 19 | Fam208b |
| Frmd4a4 | 1.05E-17 | 0.34557538 | 0.411 | 0.148 | 2.12E-13 | 19 | Frmd4a |
| Mef2d1 | 1.09E-17 | 0.253859209 | 0.392 | 0.135 | 2.20E-13 | 19 | Mef2d |
| Sf14 | 1.11E-17 | 0.254699121 | 0.494 | 0.188 | 2.24E-13 | 19 | Sf1 |
| Sipa16 | 1.31E-17 | 0.37748459 | 0.481 | 0.186 | 2.66E-13 | 19 | Sipa1 |
| Rps297 | 1.39E-17 | 0.389820111 | 1 | 0.971 | 2.81E-13 | 19 | Rps29 |
| Gnl2 | 1.43E-17 | 0.266814052 | 0.266 | 0.076 | 2.89E-13 | 19 | Gnl2 |
| Ppp4r11 | 1.44E-17 | 0.263240817 | 0.354 | 0.117 | 2.92E-13 | 19 | Ppp4r1 |
| Phactr27 | 1.49E-17 | 0.392378829 | 0.589 | 0.246 | 3.02E-13 | 19 | Phactr2 |
| Hivep13 | 1.60E-17 | 0.40846663 | 0.38 | 0.135 | 3.24E-13 | 19 | Hivep1 |
| Kctd124 | 1.62E-17 | 0.359432868 | 0.627 | 0.258 | 3.29E-13 | 19 | Kctd12 |
| Nedd4l5 | 1.78E-17 | 0.314702523 | 0.544 | 0.217 | 3.60E-13 | 19 | Nedd4l |
| Sik38 | 1.78E-17 | 0.41813175 | 0.62 | 0.27 | 3.61E-13 | 19 | Sik3 |
| Clasp11 | 1.93E-17 | 0.272067233 | 0.38 | 0.132 | 3.91E-13 | 19 | Clasp1 |
| Tln110 | 2.02E-17 | 0.546273848 | 0.772 | 0.405 | 4.09E-13 | 19 | Tln1 |
| Ints65 | 2.06E-17 | 0.275970773 | 0.405 | 0.142 | 4.17E-13 | 19 | Ints6 |
| Arid4b8 | 2.10E-17 | 0.357092561 | 0.665 | 0.293 | 4.26E-13 | 19 | Arid4b |
| Chd95 | 2.26E-17 | 0.282595184 | 0.601 | 0.25 | 4.58E-13 | 19 | Chd9 |

| | | | | | | | |
|---|---|---|---|---|---|---|---|
| Tbcd | 2.54E-17 | 0.250285019 | 0.259 | 0.074 | 5.13E-13 | 19 | Tbcd |
| Eif4g36 | 2.75E-17 | 0.377999638 | 0.646 | 0.286 | 5.56E-13 | 19 | Eif4g3 |
| Itsn29 | 2.86E-17 | 0.465626095 | 0.627 | 0.278 | 5.79E-13 | 19 | Itsn2 |
| Sik25 | 2.93E-17 | 0.311021229 | 0.354 | 0.12 | 5.94E-13 | 19 | Sik2 |
| Tsc22d23 | 3.18E-17 | 0.297242731 | 0.424 | 0.154 | 6.44E-13 | 19 | Tsc22d2 |
| Egr14 | 3.24E-17 | 0.43726995 | 0.456 | 0.175 | 6.56E-13 | 19 | Egr1 |
| Zfp36l17 | 3.25E-17 | 0.651207197 | 0.797 | 0.433 | 6.59E-13 | 19 | Zfp36l1 |
| Xpr15 | 3.68E-17 | 0.29071738 | 0.424 | 0.154 | 7.45E-13 | 19 | Xpr1 |
| Rps27a8 | 4.00E-17 | 0.421549765 | 1 | 0.95 | 8.11E-13 | 19 | Rps27a |
| Rpl398 | 4.37E-17 | 0.435900567 | 0.994 | 0.901 | 8.85E-13 | 19 | Rpl39 |
| Zfhx33 | 4.38E-17 | 0.427923769 | 0.462 | 0.183 | 8.87E-13 | 19 | Zfhx3 |
| Gtf2a1 | 4.47E-17 | 0.288546137 | 0.323 | 0.105 | 9.05E-13 | 19 | Gtf2a1 |
| Sesn16 | 4.56E-17 | 0.469953473 | 0.506 | 0.213 | 9.23E-13 | 19 | Sesn1 |
| Taf13 | 5.01E-17 | 0.286414219 | 0.411 | 0.148 | 1.01E-12 | 19 | Taf13 |
| Ash1l7 | 5.26E-17 | 0.277383947 | 0.696 | 0.299 | 1.06E-12 | 19 | Ash1l |
| Sf3b37 | 5.56E-17 | 0.266894021 | 0.437 | 0.161 | 1.12E-12 | 19 | Sf3b3 |
| Stat38 | 5.79E-17 | 0.301347167 | 0.633 | 0.267 | 1.17E-12 | 19 | Stat3 |
| Nup981 | 6.17E-17 | 0.260719287 | 0.411 | 0.145 | 1.25E-12 | 19 | Nup98 |
| Dazap21 | 6.48E-17 | 0.270686779 | 0.671 | 0.3 | 1.31E-12 | 19 | Dazap2 |
| Trafd12 | 6.84E-17 | 0.283518987 | 0.424 | 0.154 | 1.38E-12 | 19 | Trafd1 |
| Stag110 | 7.66E-17 | 0.354413743 | 0.766 | 0.361 | 1.55E-12 | 19 | Stag1 |
| Ccdc88c6 | 7.99E-17 | 0.35411519 | 0.31 | 0.1 | 1.62E-12 | 19 | Ccdc88c |
| Sept119 | 8.16E-17 | 0.368781799 | 0.506 | 0.204 | 1.65E-12 | 19 | 11-Sep |
| Xrcc4 | 8.21E-17 | 0.271273531 | 0.361 | 0.122 | 1.66E-12 | 19 | Xrcc4 |
| Ifi2062 | 8.40E-17 | 0.297020276 | 0.253 | 0.072 | 1.70E-12 | 19 | Ifi206 |
| Rpl27a7 | 8.96E-17 | 0.409256343 | 1 | 0.96 | 1.81E-12 | 19 | Rpl27a |
| Ptp4a22 | 9.12E-17 | 0.27390755 | 0.804 | 0.394 | 1.85E-12 | 19 | Ptp4a2 |
| Prpf40a9 | 9.73E-17 | 0.264088865 | 0.62 | 0.267 | 1.97E-12 | 19 | Prpf40a |
| Hsp90aa13 | 1.02E-16 | 0.319285048 | 0.848 | 0.425 | 2.07E-12 | 19 | Hsp90aa1 |
| Zcchc64 | 1.13E-16 | 0.261829764 | 0.652 | 0.278 | 2.29E-12 | 19 | Zcchc6 |
| Clasp26 | 1.13E-16 | 0.274750076 | 0.475 | 0.183 | 2.30E-12 | 19 | Clasp2 |
| Scaf88 | 1.16E-16 | 0.258370101 | 0.481 | 0.185 | 2.36E-12 | 19 | Scaf8 |
| Rps186 | 1.40E-16 | 0.443514303 | 0.981 | 0.942 | 2.83E-12 | 19 | Rps18 |
| Scaper8 | 1.43E-16 | 0.361110025 | 0.462 | 0.18 | 2.90E-12 | 19 | Scaper |
| Clnk4 | 1.49E-16 | 0.443350096 | 0.335 | 0.112 | 3.01E-12 | 19 | Clnk |
| Stt3b7 | 1.57E-16 | 0.512723459 | 0.43 | 0.165 | 3.19E-12 | 19 | Stt3b |
| Rnf133 | 1.67E-16 | 0.255582517 | 0.468 | 0.179 | 3.37E-12 | 19 | Rnf13 |
| Tnpo14 | 1.73E-16 | 0.279226844 | 0.456 | 0.174 | 3.50E-12 | 19 | Tnpo1 |
| Rpl307 | 1.84E-16 | 0.418368916 | 1 | 0.938 | 3.73E-12 | 19 | Rpl30 |
| Ep3007 | 1.98E-16 | 0.370501885 | 0.506 | 0.204 | 4.01E-12 | 19 | Ep300 |
| Pds5a3 | 1.99E-16 | 0.273751004 | 0.519 | 0.204 | 4.02E-12 | 19 | Pds5a |
| Rpl119 | 2.10E-16 | 0.431148413 | 0.994 | 0.927 | 4.25E-12 | 19 | Rpl11 |
| Clcn35 | 2.34E-16 | 0.263238984 | 0.38 | 0.134 | 4.75E-12 | 19 | Clcn3 |

| | | | | | | | |
|---|---|---|---|---|---|---|---|
| Lmtk21 | 2.49E-16 | 0.288741444 | 0.291 | 0.091 | 5.03E-12 | 19 | Lmtk2 |
| Spag96 | 2.54E-16 | 0.427776078 | 0.734 | 0.359 | 5.15E-12 | 19 | Spag9 |
| Herpud13 | 2.68E-16 | 0.27793013 | 0.525 | 0.214 | 5.42E-12 | 19 | Herpud1 |
| Rnf2134 | 2.80E-16 | 0.388923156 | 0.582 | 0.257 | 5.67E-12 | 19 | Rnf213 |
| Psmb811 | 2.86E-16 | 0.621904617 | 0.911 | 0.558 | 5.80E-12 | 19 | Psmb8 |
| Kdm2a9 | 3.01E-16 | 0.303266516 | 0.57 | 0.241 | 6.09E-12 | 19 | Kdm2a |
| Thada6 | 3.35E-16 | 0.304338742 | 0.348 | 0.12 | 6.79E-12 | 19 | Thada |
| Esco15 | 3.46E-16 | 0.311494738 | 0.462 | 0.18 | 7.01E-12 | 19 | Esco1 |
| Hspa42 | 3.55E-16 | 0.3110238 | 0.696 | 0.316 | 7.18E-12 | 19 | Hspa4 |
| Creb15 | 3.67E-16 | 0.268406733 | 0.513 | 0.208 | 7.43E-12 | 19 | Creb1 |
| Rere6 | 3.79E-16 | 0.425163747 | 0.608 | 0.272 | 7.67E-12 | 19 | Rere |
| Tacc16 | 3.82E-16 | 0.336363341 | 0.671 | 0.305 | 7.74E-12 | 19 | Tacc1 |
| Atp6v0e3 | 4.03E-16 | 0.340497063 | 0.797 | 0.426 | 8.16E-12 | 19 | Atp6v0e |
| Ppp1ca6 | 4.09E-16 | 0.258922906 | 0.81 | 0.397 | 8.28E-12 | 19 | Ppp1ca |
| mt-Nd113 | 4.60E-16 | 0.55903398 | 0.968 | 0.599 | 9.32E-12 | 19 | mt-Nd1 |
| 9930111J21Rik26 | 5.07E-16 | 0.354914617 | 0.589 | 0.257 | 1.03E-11 | 19 | 9930111J21Rik2 |
| Tanc23 | 5.22E-16 | 0.528567986 | 0.405 | 0.16 | 1.06E-11 | 19 | Tanc2 |
| Gpatch810 | 5.92E-16 | 0.288156762 | 0.601 | 0.258 | 1.20E-11 | 19 | Gpatch8 |
| Gtdc13 | 5.93E-16 | 0.273414186 | 0.411 | 0.152 | 1.20E-11 | 19 | Gtdc1 |
| Scai2 | 6.76E-16 | 0.304810375 | 0.31 | 0.102 | 1.37E-11 | 19 | Scai |
| Calr7 | 6.95E-16 | 0.394945736 | 0.804 | 0.408 | 1.41E-11 | 19 | Calr |
| Nasp5 | 9.00E-16 | 0.280195563 | 0.449 | 0.172 | 1.82E-11 | 19 | Nasp |
| Tet32 | 9.68E-16 | 0.259145178 | 0.367 | 0.131 | 1.96E-11 | 19 | Tet3 |
| Rplp29 | 1.01E-15 | 0.406446273 | 1 | 0.89 | 2.05E-11 | 19 | Rplp2 |
| Fryl7 | 1.09E-15 | 0.312850598 | 0.614 | 0.268 | 2.21E-11 | 19 | Fryl |
| Hectd41 | 1.14E-15 | 0.270123464 | 0.323 | 0.108 | 2.31E-11 | 19 | Hectd4 |
| A930015D03Rik4 | 1.21E-15 | 0.258074779 | 0.316 | 0.107 | 2.45E-11 | 19 | A930015D03Rik |
| Tm6sf16 | 1.26E-15 | 0.275683676 | 0.386 | 0.142 | 2.55E-11 | 19 | Tm6sf1 |
| Rpl277 | 1.34E-15 | 0.438072665 | 0.924 | 0.583 | 2.71E-11 | 19 | Rpl27 |
| Cdyl6 | 1.42E-15 | 0.25871649 | 0.361 | 0.129 | 2.88E-11 | 19 | Cdyl |
| Mbnl27 | 1.44E-15 | 0.407216594 | 0.81 | 0.419 | 2.91E-11 | 19 | Mbnl2 |
| Elf16 | 1.48E-15 | 0.332060348 | 0.778 | 0.376 | 2.99E-11 | 19 | Elf1 |
| Itpr26 | 1.69E-15 | 0.373958722 | 0.557 | 0.245 | 3.43E-11 | 19 | Itpr2 |
| Cabin12 | 1.73E-15 | 0.253382764 | 0.304 | 0.1 | 3.51E-11 | 19 | Cabin1 |
| Arhgef37 | 1.89E-15 | 0.35704874 | 0.563 | 0.241 | 3.84E-11 | 19 | Arhgef3 |
| Lrrk21 | 2.10E-15 | 0.389971882 | 0.253 | 0.078 | 4.26E-11 | 19 | Lrrk2 |
| Atp8a14 | 2.17E-15 | 0.333799286 | 0.557 | 0.242 | 4.40E-11 | 19 | Atp8a1 |
| 2610035D17Rik4 | 2.32E-15 | 0.370501114 | 0.329 | 0.115 | 4.69E-11 | 19 | 2610035D17Rik |
| Tox6 | 2.42E-15 | 0.456804327 | 0.342 | 0.122 | 4.90E-11 | 19 | Tox |
| Phf64 | 2.55E-15 | 0.252689842 | 0.373 | 0.136 | 5.17E-11 | 19 | Phf6 |
| Smg68 | 2.67E-15 | 0.362378954 | 0.582 | 0.26 | 5.41E-11 | 19 | Smg6 |
| Rprd22 | 2.72E-15 | 0.26213109 | 0.373 | 0.135 | 5.51E-11 | 19 | Rprd2 |
| Ankhd18 | 2.74E-15 | 0.268673954 | 0.582 | 0.244 | 5.55E-11 | 19 | Ankhd1 |

| | | | | | | | |
|---|---|---|---|---|---|---|---|
| Tnrc6b8 | 2.78E-15 | 0.39811262 | 0.759 | 0.371 | 5.62E-11 | 19 | Tnrc6b |
| Nktr7 | 3.35E-15 | 0.301269984 | 0.582 | 0.258 | 6.79E-11 | 19 | Nktr |
| Ube2h5 | 3.38E-15 | 0.271565357 | 0.601 | 0.265 | 6.84E-11 | 19 | Ube2h |
| Ptp4a33 | 3.59E-15 | 0.368522743 | 0.373 | 0.138 | 7.28E-11 | 19 | Ptp4a3 |
| Rpl328 | 3.72E-15 | 0.404859348 | 0.994 | 0.953 | 7.53E-11 | 19 | Rpl32 |
| Dhx9 | 4.44E-15 | 0.252623685 | 0.487 | 0.199 | 8.99E-11 | 19 | Dhx9 |
| Hspa510 | 4.54E-15 | 0.429352395 | 0.829 | 0.435 | 9.19E-11 | 19 | Hspa5 |
| Klhl66 | 4.65E-15 | 0.257551781 | 0.342 | 0.121 | 9.41E-11 | 19 | Klhl6 |
| Msn6 | 5.61E-15 | 0.306941357 | 0.886 | 0.443 | 1.14E-10 | 19 | Msn |
| Prdm23 | 5.69E-15 | 0.278639832 | 0.462 | 0.185 | 1.15E-10 | 19 | Prdm2 |
| Bclaf18 | 5.78E-15 | 0.317281969 | 0.582 | 0.254 | 1.17E-10 | 19 | Bclaf1 |
| Phf146 | 6.92E-15 | 0.30533045 | 0.551 | 0.245 | 1.40E-10 | 19 | Phf14 |
| Mef2a7 | 7.42E-15 | 0.521984701 | 0.778 | 0.417 | 1.50E-10 | 19 | Mef2a |
| Frmd4b7 | 7.43E-15 | 0.341181651 | 0.443 | 0.178 | 1.50E-10 | 19 | Frmd4b |
| Arih21 | 7.55E-15 | 0.310807982 | 0.316 | 0.109 | 1.53E-10 | 19 | Arih2 |
| Dis3l27 | 8.29E-15 | 0.302356095 | 0.373 | 0.14 | 1.68E-10 | 19 | Dis3l2 |
| Fmnl24 | 8.67E-15 | 0.434453355 | 0.557 | 0.243 | 1.75E-10 | 19 | Fmnl2 |
| Maml34 | 9.59E-15 | 0.327564413 | 0.494 | 0.208 | 1.94E-10 | 19 | Maml3 |
| Il2rb4 | 9.77E-15 | 0.373709027 | 0.342 | 0.121 | 1.98E-10 | 19 | Il2rb |
| Gzmb3 | 1.09E-14 | 0.719543103 | 0.259 | 0.083 | 2.22E-10 | 19 | Gzmb |
| Rps77 | 1.20E-14 | 0.430235573 | 0.994 | 0.917 | 2.42E-10 | 19 | Rps7 |
| Cdkal16 | 1.20E-14 | 0.252276906 | 0.456 | 0.181 | 2.43E-10 | 19 | Cdkal1 |
| mt-Co215 | 1.20E-14 | 0.540719888 | 1 | 0.729 | 2.43E-10 | 19 | mt-Co2 |
| Lrp107 | 1.23E-14 | 0.273733035 | 0.468 | 0.189 | 2.49E-10 | 19 | Lrp10 |
| Kmt2c10 | 1.28E-14 | 0.311333727 | 0.665 | 0.306 | 2.58E-10 | 19 | Kmt2c |
| Srsf51 | 1.29E-14 | 0.310112787 | 0.867 | 0.45 | 2.61E-10 | 19 | Srsf5 |
| Crtc38 | 1.31E-14 | 0.250627089 | 0.424 | 0.165 | 2.65E-10 | 19 | Crtc3 |
| Srrm29 | 1.48E-14 | 0.385193402 | 0.861 | 0.469 | 3.00E-10 | 19 | Srrm2 |
| Setd27 | 1.59E-14 | 0.254580463 | 0.475 | 0.193 | 3.22E-10 | 19 | Setd2 |
| Akt310 | 1.80E-14 | 0.371936091 | 0.57 | 0.252 | 3.64E-10 | 19 | Akt3 |
| mt-Nd311 | 1.90E-14 | 0.416897792 | 0.861 | 0.446 | 3.85E-10 | 19 | mt-Nd3 |
| Tob2 | 2.04E-14 | 0.272556236 | 0.335 | 0.121 | 4.13E-10 | 19 | Tob2 |
| Cnot48 | 2.24E-14 | 0.312233539 | 0.595 | 0.269 | 4.54E-10 | 19 | Cnot4 |
| Rps3a18 | 2.28E-14 | 0.419285819 | 0.994 | 0.935 | 4.62E-10 | 19 | Rps3a1 |
| Crebbp10 | 2.40E-14 | 0.363707958 | 0.652 | 0.305 | 4.87E-10 | 19 | Crebbp |
| Skap15 | 2.51E-14 | 0.496492322 | 0.361 | 0.135 | 5.09E-10 | 19 | Skap1 |
| Atxn75 | 2.63E-14 | 0.250852489 | 0.424 | 0.166 | 5.33E-10 | 19 | Atxn7 |
| Ap3b17 | 2.96E-14 | 0.276448004 | 0.513 | 0.218 | 5.99E-10 | 19 | Ap3b1 |
| Hivep35 | 3.15E-14 | 0.307752118 | 0.405 | 0.162 | 6.38E-10 | 19 | Hivep3 |
| Pfkp1 | 3.24E-14 | 0.253542841 | 0.405 | 0.157 | 6.56E-10 | 19 | Pfkp |
| Arpc1b6 | 3.82E-14 | 0.397597756 | 0.911 | 0.592 | 7.73E-10 | 19 | Arpc1b |
| Ctsw4 | 4.19E-14 | 0.344299196 | 0.354 | 0.129 | 8.48E-10 | 19 | Ctsw |
| Luc7l29 | 4.60E-14 | 0.329604983 | 0.81 | 0.405 | 9.32E-10 | 19 | Luc7l2 |

| | | | | | | | |
|---|---|---|---|---|---|---|---|
| Rpl367 | 5.04E-14 | 0.372962347 | 0.994 | 0.903 | 1.02E-09 | 19 | Rpl36 |
| Snx95 | 5.34E-14 | 0.286915275 | 0.437 | 0.176 | 1.08E-09 | 19 | Snx9 |
| Rbm399 | 5.50E-14 | 0.466478766 | 0.956 | 0.574 | 1.11E-09 | 19 | Rbm39 |
| Cfl18 | 5.71E-14 | 0.455001841 | 0.905 | 0.609 | 1.16E-09 | 19 | Cfl1 |
| Tpr8 | 7.23E-14 | 0.304961755 | 0.842 | 0.425 | 1.46E-09 | 19 | Tpr |
| Fbxl1712 | 8.20E-14 | 0.276099702 | 0.658 | 0.311 | 1.66E-09 | 19 | Fbxl17 |
| Trim353 | 8.65E-14 | 0.260955508 | 0.367 | 0.141 | 1.75E-09 | 19 | Trim35 |
| Bcas37 | 9.84E-14 | 0.372849278 | 0.437 | 0.183 | 1.99E-09 | 19 | Bcas3 |
| Chn26 | 1.18E-13 | 0.509190029 | 0.456 | 0.199 | 2.38E-09 | 19 | Chn2 |
| Evl6 | 1.20E-13 | 0.394809788 | 0.494 | 0.221 | 2.43E-09 | 19 | Evl |
| Stim18 | 1.62E-13 | 0.296107955 | 0.456 | 0.191 | 3.27E-09 | 19 | Stim1 |
| Ppp1r12a9 | 1.69E-13 | 0.349867287 | 0.81 | 0.426 | 3.42E-09 | 19 | Ppp1r12a |
| Nt5c22 | 1.75E-13 | 0.26226221 | 0.278 | 0.096 | 3.54E-09 | 19 | Nt5c2 |
| Nr4a12 | 2.05E-13 | 0.360068334 | 0.297 | 0.107 | 4.15E-09 | 19 | Nr4a1 |
| Rpl87 | 2.23E-13 | 0.40109959 | 1 | 0.925 | 4.51E-09 | 19 | Rpl8 |
| Kmt2e8 | 2.75E-13 | 0.335398786 | 0.804 | 0.406 | 5.58E-09 | 19 | Kmt2e |
| Birc611 | 3.24E-13 | 0.269958582 | 0.696 | 0.341 | 6.56E-09 | 19 | Birc6 |
| Picalm6 | 3.55E-13 | 0.455476478 | 0.614 | 0.297 | 7.19E-09 | 19 | Picalm |
| Ppp2r5a4 | 3.56E-13 | 0.290571096 | 0.557 | 0.251 | 7.22E-09 | 19 | Ppp2r5a |
| Ankrd179 | 3.90E-13 | 0.311945877 | 0.563 | 0.261 | 7.89E-09 | 19 | Ankrd17 |
| Brd24 | 4.22E-13 | 0.254069332 | 0.557 | 0.254 | 8.55E-09 | 19 | Brd2 |
| Eif4a15 | 4.22E-13 | 0.256198602 | 0.791 | 0.431 | 8.55E-09 | 19 | Eif4a1 |
| Nsd19 | 4.60E-13 | 0.254636099 | 0.722 | 0.35 | 9.32E-09 | 19 | Nsd1 |
| Srgn10 | 5.33E-13 | 0.452587795 | 0.962 | 0.545 | 1.08E-08 | 19 | Srgn |
| Slc44a23 | 6.58E-13 | 0.272854061 | 0.323 | 0.122 | 1.33E-08 | 19 | Slc44a2 |
| Lamp25 | 7.10E-13 | 0.322649765 | 0.633 | 0.306 | 1.44E-08 | 19 | Lamp2 |
| Rpl387 | 7.67E-13 | 0.345392559 | 0.994 | 0.893 | 1.55E-08 | 19 | Rpl38 |
| Mtdh7 | 8.24E-13 | 0.279062568 | 0.753 | 0.387 | 1.67E-08 | 19 | Mtdh |
| Zeb19 | 8.38E-13 | 0.264894596 | 0.677 | 0.322 | 1.70E-08 | 19 | Zeb1 |
| Msi24 | 9.17E-13 | 0.276893263 | 0.614 | 0.293 | 1.86E-08 | 19 | Msi2 |
| Galnt75 | 1.01E-12 | 0.251203186 | 0.386 | 0.156 | 2.05E-08 | 19 | Galnt7 |
| Tpm38 | 1.04E-12 | 0.339633396 | 0.842 | 0.479 | 2.11E-08 | 19 | Tpm3 |
| Jund7 | 1.06E-12 | 0.474869351 | 0.962 | 0.646 | 2.15E-08 | 19 | Jund |
| Usp3410 | 1.11E-12 | 0.277324803 | 0.696 | 0.345 | 2.24E-08 | 19 | Usp34 |
| Nkg74 | 1.16E-12 | 0.319187764 | 0.342 | 0.128 | 2.36E-08 | 19 | Nkg7 |
| Trappc84 | 1.18E-12 | 0.256215342 | 0.405 | 0.167 | 2.38E-08 | 19 | Trappc8 |
| Jak17 | 1.18E-12 | 0.361067674 | 0.715 | 0.366 | 2.39E-08 | 19 | Jak1 |
| Brk12 | 1.24E-12 | 0.258458715 | 0.804 | 0.442 | 2.51E-08 | 19 | Brk1 |
| Styk14 | 1.46E-12 | 0.270365521 | 0.278 | 0.098 | 2.95E-08 | 19 | Styk1 |
| Psmb96 | 1.54E-12 | 0.330838879 | 0.778 | 0.406 | 3.11E-08 | 19 | Psmb9 |
| C1qc4 | 1.60E-12 | 0.373810182 | 0.348 | 0.135 | 3.25E-08 | 19 | C1qc |
| Cebpb5 | 1.70E-12 | 0.803898567 | 0.373 | 0.158 | 3.44E-08 | 19 | Cebpb |
| Abhd17b6 | 1.89E-12 | 0.347073619 | 0.443 | 0.193 | 3.82E-08 | 19 | Abhd17b |

| | | | | | | | |
|---|---|---|---|---|---|---|---|
| Rpl68 | 2.12E-12 | 0.336519103 | 0.994 | 0.882 | 4.28E-08 | 19 | Rpl6 |
| Marcks5 | 2.12E-12 | 0.288221178 | 0.69 | 0.342 | 4.28E-08 | 19 | Marcks |
| mt-Nd412 | 2.20E-12 | 0.48583289 | 0.987 | 0.676 | 4.45E-08 | 19 | mt-Nd4 |
| Fyn9 | 2.23E-12 | 0.418163557 | 0.557 | 0.267 | 4.52E-08 | 19 | Fyn |
| Map4k49 | 2.45E-12 | 0.483190889 | 0.589 | 0.277 | 4.95E-08 | 19 | Map4k4 |
| Rpl378 | 2.49E-12 | 0.312968197 | 0.994 | 0.927 | 5.04E-08 | 19 | Rpl37 |
| Chst118 | 2.50E-12 | 0.329689826 | 0.418 | 0.176 | 5.06E-08 | 19 | Chst11 |
| Clic18 | 2.58E-12 | 0.369839832 | 0.943 | 0.601 | 5.22E-08 | 19 | Clic1 |
| Rock19 | 2.86E-12 | 0.25602928 | 0.842 | 0.433 | 5.79E-08 | 19 | Rock1 |
| Bach11 | 3.18E-12 | 0.265193555 | 0.278 | 0.1 | 6.44E-08 | 19 | Bach1 |
| Rpl127 | 3.20E-12 | 0.37048478 | 1 | 0.841 | 6.47E-08 | 19 | Rpl12 |
| Ndel13 | 3.31E-12 | 0.302043025 | 0.411 | 0.175 | 6.70E-08 | 19 | Ndel1 |
| Ttc149 | 3.56E-12 | 0.251626019 | 0.487 | 0.216 | 7.21E-08 | 19 | Ttc14 |
| Rpl10a9 | 4.59E-12 | 0.32385807 | 0.987 | 0.836 | 9.29E-08 | 19 | Rpl10a |
| Msrb12 | 4.67E-12 | 0.393069406 | 0.354 | 0.142 | 9.46E-08 | 19 | Msrb1 |
| Nfe2l24 | 4.70E-12 | 0.301440057 | 0.601 | 0.302 | 9.52E-08 | 19 | Nfe2l2 |
| Acin17 | 4.79E-12 | 0.257394945 | 0.646 | 0.317 | 9.69E-08 | 19 | Acin1 |
| Xcl14 | 5.21E-12 | 0.382261937 | 0.329 | 0.131 | 1.05E-07 | 19 | Xcl1 |
| Apc5 | 5.41E-12 | 0.282739236 | 0.538 | 0.249 | 1.10E-07 | 19 | Apc |
| Asap14 | 6.77E-12 | 0.58437652 | 0.671 | 0.357 | 1.37E-07 | 19 | Asap1 |
| Brwd12 | 7.16E-12 | 0.29712705 | 0.354 | 0.142 | 1.45E-07 | 19 | Brwd1 |
| Ywhaz5 | 9.33E-12 | 0.267467183 | 0.873 | 0.493 | 1.89E-07 | 19 | Ywhaz |
| Myh99 | 9.44E-12 | 0.34596826 | 0.791 | 0.437 | 1.91E-07 | 19 | Myh9 |
| Gsr | 1.14E-11 | 0.443142956 | 0.329 | 0.13 | 2.31E-07 | 19 | Gsr |
| Senp77 | 1.18E-11 | 0.287883683 | 0.373 | 0.157 | 2.39E-07 | 19 | Senp7 |
| H3f3a3 | 1.19E-11 | 0.531042855 | 0.994 | 0.862 | 2.41E-07 | 19 | H3f3a |
| Bbx6 | 1.41E-11 | 0.253453123 | 0.487 | 0.223 | 2.86E-07 | 19 | Bbx |
| Nfkbia7 | 1.58E-11 | 0.337231907 | 0.741 | 0.4 | 3.20E-07 | 19 | Nfkbia |
| Map3k54 | 1.65E-11 | 0.278643569 | 0.437 | 0.193 | 3.34E-07 | 19 | Map3k5 |
| Ssr44 | 1.92E-11 | 0.263667307 | 0.797 | 0.468 | 3.88E-07 | 19 | Ssr4 |
| Rlf6 | 2.02E-11 | 0.255892864 | 0.342 | 0.137 | 4.09E-07 | 19 | Rlf |
| Atrnl15 | 3.59E-11 | 0.483889917 | 0.38 | 0.166 | 7.27E-07 | 19 | Atrnl1 |
| Malat111 | 3.86E-11 | 0.513284441 | 1 | 0.726 | 7.81E-07 | 19 | Malat1 |
| Prkce3 | 4.85E-11 | 0.386032274 | 0.456 | 0.215 | 9.81E-07 | 19 | Prkce |
| Cd2742 | 4.88E-11 | 0.390378675 | 0.291 | 0.115 | 9.87E-07 | 19 | Cd274 |
| Cdc424 | 4.92E-11 | 0.339093438 | 0.911 | 0.616 | 9.96E-07 | 19 | Cdc42 |
| Etnk13 | 4.98E-11 | 0.289541253 | 0.487 | 0.226 | 1.01E-06 | 19 | Etnk1 |
| Rpl358 | 5.17E-11 | 0.293948224 | 0.981 | 0.828 | 1.05E-06 | 19 | Rpl35 |
| Rpl238 | 5.78E-11 | 0.29772777 | 0.994 | 0.971 | 1.17E-06 | 19 | Rpl23 |
| Rps268 | 6.56E-11 | 0.292453785 | 0.987 | 0.866 | 1.33E-06 | 19 | Rps26 |
| mt-Atp810 | 7.33E-11 | 0.336228673 | 0.842 | 0.463 | 1.48E-06 | 19 | mt-Atp8 |
| C1qa4 | 8.53E-11 | 0.385775281 | 0.348 | 0.148 | 1.73E-06 | 19 | C1qa |
| Rack19 | 8.72E-11 | 0.319796413 | 0.975 | 0.793 | 1.77E-06 | 19 | Rack1 |

| | | | | | | | |
|---|---|---|---|---|---|---|---|
| Myo1e9 | 9.98E-11 | 0.465061078 | 0.538 | 0.282 | 2.02E-06 | 19 | Myo1e |
| Rps4x8 | 1.21E-10 | 0.28010823 | 1 | 0.941 | 2.45E-06 | 19 | Rps4x |
| H2-Q77 | 1.32E-10 | 0.293472808 | 0.734 | 0.398 | 2.68E-06 | 19 | H2-Q7 |
| Nfat57 | 1.38E-10 | 0.264485148 | 0.544 | 0.268 | 2.80E-06 | 19 | Nfat5 |
| Psme24 | 1.41E-10 | 0.43323205 | 0.753 | 0.444 | 2.85E-06 | 19 | Psme2 |
| Large16 | 1.43E-10 | 0.26095754 | 0.361 | 0.153 | 2.90E-06 | 19 | Large1 |
| Rpl98 | 1.44E-10 | 0.309454819 | 0.994 | 0.945 | 2.91E-06 | 19 | Rpl9 |
| Kif13b5 | 1.66E-10 | 0.256672944 | 0.456 | 0.216 | 3.36E-06 | 19 | Kif13b |
| Rpl177 | 2.10E-10 | 0.282767196 | 1 | 0.917 | 4.25E-06 | 19 | Rpl17 |
| Cd476 | 2.60E-10 | 0.283256461 | 0.873 | 0.51 | 5.26E-06 | 19 | Cd47 |
| Hspa81 | 2.64E-10 | 0.447176818 | 0.943 | 0.71 | 5.35E-06 | 19 | Hspa8 |
| Lamp19 | 3.40E-10 | 0.304961736 | 0.741 | 0.418 | 6.89E-06 | 19 | Lamp1 |
| Pxk2 | 4.11E-10 | 0.308153002 | 0.329 | 0.141 | 8.33E-06 | 19 | Pxk |
| Rpl228 | 4.31E-10 | 0.257756207 | 0.981 | 0.852 | 8.72E-06 | 19 | Rpl22 |
| Mef2c5 | 4.35E-10 | 0.284078993 | 0.804 | 0.424 | 8.80E-06 | 19 | Mef2c |
| Rpl23a8 | 5.47E-10 | 0.267548413 | 0.968 | 0.679 | 1.11E-05 | 19 | Rpl23a |
| Rps2111 | 5.60E-10 | 0.263275399 | 0.994 | 0.927 | 1.13E-05 | 19 | Rps21 |
| Pmepa14 | 5.85E-10 | 0.336736933 | 0.494 | 0.253 | 1.18E-05 | 19 | Pmepa1 |
| Psme17 | 1.00E-09 | 0.349493477 | 0.886 | 0.569 | 2.03E-05 | 19 | Psme1 |
| Fli18 | 1.42E-09 | 0.328179449 | 0.722 | 0.402 | 2.88E-05 | 19 | Fli1 |
| Wwox7 | 1.50E-09 | 0.271488605 | 0.38 | 0.176 | 3.03E-05 | 19 | Wwox |
| Rpl219 | 1.50E-09 | 0.275325164 | 1 | 0.923 | 3.04E-05 | 19 | Rpl21 |
| S100a106 | 2.09E-09 | 0.309174332 | 0.81 | 0.47 | 4.24E-05 | 19 | S100a10 |
| Actg15 | 2.12E-09 | 0.640101342 | 0.937 | 0.795 | 4.29E-05 | 19 | Actg1 |
| Rpl287 | 3.67E-09 | 0.27155068 | 1 | 0.93 | 7.43E-05 | 19 | Rpl28 |
| Rps255 | 6.53E-09 | 0.256702828 | 1 | 0.83 | 0.000132125 | 19 | Rps25 |
| Grap4 | 8.28E-09 | 0.315062545 | 0.272 | 0.116 | 0.000167539 | 19 | Grap |
| Tagln29 | 1.83E-08 | 0.380709098 | 0.715 | 0.43 | 0.00037082 | 19 | Tagln2 |
| H2afz4 | 3.37E-08 | 0.460887165 | 0.823 | 0.609 | 0.000681811 | 19 | H2afz |
| H2-D110 | 3.81E-08 | 0.335420004 | 1 | 0.669 | 0.000770414 | 19 | H2-D1 |
| mt-Atp613 | 4.05E-08 | 0.355033447 | 1 | 0.754 | 0.000819992 | 19 | mt-Atp6 |
| Maml211 | 1.44E-07 | 0.28866721 | 0.582 | 0.344 | 0.002913426 | 19 | Maml2 |
| Phlpp15 | 1.91E-07 | 0.263188891 | 0.5 | 0.264 | 0.003858343 | 19 | Phlpp1 |
| Tmem176a9 | 2.02E-06 | 0.293120586 | 0.658 | 0.418 | 0.040844533 | 19 | Tmem176a |
| B2m11 | 2.50E-05 | 0.275907645 | 0.956 | 0.691 | 0.505589973 | 19 | B2m |
| Thsd7a1 | 0 | 2.807763333 | 0.886 | 0.032 | 0 | 20 | Thsd7a |
| Lrg1 | 0 | 2.581135801 | 0.617 | 0.012 | 0 | 20 | Lrg1 |
| Vwf | 0 | 2.52480219 | 0.624 | 0.013 | 0 | 20 | Vwf |
| Pcdh71 | 0 | 2.390114107 | 0.617 | 0.028 | 0 | 20 | Pcdh7 |
| Selp | 0 | 2.187973797 | 0.611 | 0.007 | 0 | 20 | Selp |
| Lbp | 0 | 1.81044998 | 0.597 | 0.016 | 0 | 20 | Lbp |
| Pgm5 | 0 | 0.945759306 | 0.289 | 0.004 | 0 | 20 | Pgm5 |
| Adgrg6 | 4.72E-303 | 1.261759687 | 0.322 | 0.009 | 9.55E-299 | 20 | Adgrg6 |

| | | | | | | | |
|---|---|---|---|---|---|---|---|
| Aqp12 | 2.03E-272 | 3.256355735 | 0.886 | 0.09 | 4.11E-268 | 20 | Aqp1 |
| Slco2a11 | 6.30E-244 | 2.475097579 | 0.779 | 0.075 | 1.28E-239 | 20 | Slco2a1 |
| Myrip | 1.94E-233 | 1.175658317 | 0.416 | 0.02 | 3.93E-229 | 20 | Myrip |
| Ptn | 1.18E-227 | 1.419921519 | 0.309 | 0.011 | 2.38E-223 | 20 | Ptn |
| Vcam11 | 2.30E-222 | 1.688375564 | 0.456 | 0.026 | 4.65E-218 | 20 | Vcam1 |
| Chp21 | 4.37E-220 | 1.166931804 | 0.463 | 0.027 | 8.86E-216 | 20 | Chp2 |
| Rasa41 | 5.54E-218 | 1.967521073 | 0.651 | 0.056 | 1.12E-213 | 20 | Rasa4 |
| P3h21 | 1.15E-198 | 1.412889628 | 0.463 | 0.03 | 2.33E-194 | 20 | P3h2 |
| Mctp14 | 3.06E-194 | 2.105684811 | 0.698 | 0.072 | 6.20E-190 | 20 | Mctp1 |
| Mgp3 | 1.39E-190 | 1.646497651 | 0.644 | 0.057 | 2.82E-186 | 20 | Mgp |
| Cadm3 | 6.38E-190 | 0.878232903 | 0.322 | 0.015 | 1.29E-185 | 20 | Cadm3 |
| Cysltr12 | 4.01E-179 | 1.676132402 | 0.617 | 0.06 | 8.11E-175 | 20 | Cysltr1 |
| Myof4 | 5.46E-178 | 1.534672485 | 0.604 | 0.057 | 1.11E-173 | 20 | Myof |
| Zfp5212 | 1.05E-175 | 1.962932961 | 0.671 | 0.072 | 2.12E-171 | 20 | Zfp521 |
| Ppfia21 | 9.92E-175 | 0.878175213 | 0.275 | 0.011 | 2.01E-170 | 20 | Ppfia2 |
| Insr2 | 6.74E-173 | 2.565552989 | 0.846 | 0.126 | 1.36E-168 | 20 | Insr |
| Gm5127 | 1.24E-169 | 1.087393802 | 0.262 | 0.011 | 2.52E-165 | 20 | Gm5127 |
| Bgn2 | 1.31E-168 | 1.257457442 | 0.53 | 0.043 | 2.64E-164 | 20 | Bgn |
| Nr2f22 | 4.05E-167 | 1.83167594 | 0.765 | 0.097 | 8.19E-163 | 20 | Nr2f2 |
| Kirrel31 | 1.46E-160 | 2.421948932 | 0.624 | 0.068 | 2.95E-156 | 20 | Kirrel3 |
| Lepr | 1.87E-151 | 1.379443672 | 0.55 | 0.054 | 3.79E-147 | 20 | Lepr |
| Ctnnal1 | 8.60E-138 | 1.041129078 | 0.336 | 0.022 | 1.74E-133 | 20 | Ctnnal1 |
| Hs3st11 | 6.21E-133 | 1.141664926 | 0.523 | 0.055 | 1.26E-128 | 20 | Hs3st1 |
| C1qtnf91 | 1.29E-132 | 1.157059702 | 0.47 | 0.045 | 2.61E-128 | 20 | C1qtnf9 |
| Mecom2 | 2.74E-118 | 1.823903195 | 0.664 | 0.096 | 5.55E-114 | 20 | Mecom |
| Il6st5 | 1.04E-115 | 2.404339766 | 0.899 | 0.23 | 2.11E-111 | 20 | Il6st |
| Nrp24 | 4.48E-114 | 1.923699869 | 0.846 | 0.182 | 9.06E-110 | 20 | Nrp2 |
| Eln5 | 1.20E-105 | 1.982075729 | 0.725 | 0.135 | 2.42E-101 | 20 | Eln |
| Bmp41 | 1.23E-105 | 0.874092317 | 0.268 | 0.018 | 2.48E-101 | 20 | Bmp4 |
| Cda | 1.35E-105 | 0.694425146 | 0.275 | 0.019 | 2.73E-101 | 20 | Cda |
| Ret | 4.74E-105 | 0.758132906 | 0.262 | 0.018 | 9.59E-101 | 20 | Ret |
| Fam171a13 | 1.21E-104 | 1.601181633 | 0.638 | 0.104 | 2.45E-100 | 20 | Fam171a1 |
| Itgb41 | 5.63E-102 | 1.072339559 | 0.356 | 0.033 | 1.14E-97 | 20 | Itgb4 |
| Prkag21 | 3.13E-99 | 1.379222893 | 0.523 | 0.074 | 6.33E-95 | 20 | Prkag2 |
| Prcp3 | 3.22E-99 | 1.245960469 | 0.564 | 0.085 | 6.51E-95 | 20 | Prcp |
| AU021092 | 3.34E-97 | 0.729307446 | 0.275 | 0.021 | 6.76E-93 | 20 | AU021092 |
| Lifr5 | 5.90E-97 | 1.861040274 | 0.805 | 0.191 | 1.19E-92 | 20 | Lifr |
| Tspan72 | 4.42E-94 | 1.269969812 | 0.523 | 0.076 | 8.96E-90 | 20 | Tspan7 |
| Tln21 | 1.36E-92 | 0.877434996 | 0.383 | 0.041 | 2.74E-88 | 20 | Tln2 |
| Vegfc2 | 3.38E-92 | 1.600601433 | 0.617 | 0.108 | 6.84E-88 | 20 | Vegfc |
| Aebp11 | 8.03E-86 | 1.019691557 | 0.53 | 0.081 | 1.63E-81 | 20 | Aebp1 |
| Zfp423 | 8.95E-85 | 1.156682979 | 0.483 | 0.07 | 1.81E-80 | 20 | Zfp423 |
| Pdzd23 | 1.23E-84 | 1.602653712 | 0.611 | 0.114 | 2.49E-80 | 20 | Pdzd2 |

| | | | | | | | |
|---|---|---|---|---|---|---|---|
| Dock94 | 1.94E-83 | 1.522624172 | 0.812 | 0.204 | 3.93E-79 | 20 | Dock9 |
| Cped16 | 1.05E-82 | 1.489601907 | 0.772 | 0.178 | 2.12E-78 | 20 | Cped1 |
| Stox24 | 2.25E-80 | 1.834997242 | 0.879 | 0.245 | 4.56E-76 | 20 | Stox2 |
| Dnm34 | 2.39E-80 | 2.007397467 | 0.826 | 0.229 | 4.83E-76 | 20 | Dnm3 |
| Mras | 7.28E-80 | 0.718972323 | 0.282 | 0.026 | 1.47E-75 | 20 | Mras |
| Tgm24 | 2.40E-79 | 1.434311355 | 0.718 | 0.161 | 4.85E-75 | 20 | Tgm2 |
| Kcnb11 | 3.13E-79 | 1.12760384 | 0.503 | 0.079 | 6.34E-75 | 20 | Kcnb1 |
| Nuak13 | 4.99E-77 | 1.359755085 | 0.597 | 0.117 | 1.01E-72 | 20 | Nuak1 |
| Abcg25 | 9.33E-77 | 1.576506432 | 0.879 | 0.269 | 1.89E-72 | 20 | Abcg2 |
| Spns22 | 1.46E-76 | 0.991719403 | 0.537 | 0.092 | 2.96E-72 | 20 | Spns2 |
| Prkg14 | 1.07E-74 | 1.818596182 | 0.765 | 0.195 | 2.17E-70 | 20 | Prkg1 |
| Tll11 | 4.46E-74 | 1.421980433 | 0.443 | 0.068 | 9.03E-70 | 20 | Tll1 |
| Bace2 | 2.43E-72 | 0.867045381 | 0.416 | 0.06 | 4.93E-68 | 20 | Bace2 |
| Myzap | 2.39E-71 | 0.572380955 | 0.255 | 0.024 | 4.84E-67 | 20 | Myzap |
| Entpd15 | 4.64E-71 | 1.742878012 | 0.832 | 0.275 | 9.40E-67 | 20 | Entpd1 |
| Ptpn143 | 8.46E-71 | 1.038816894 | 0.557 | 0.106 | 1.71E-66 | 20 | Ptpn14 |
| Abcb1a3 | 1.44E-70 | 1.073575436 | 0.564 | 0.109 | 2.91E-66 | 20 | Abcb1a |
| Tshz26 | 1.25E-68 | 1.758454702 | 0.966 | 0.371 | 2.52E-64 | 20 | Tshz2 |
| Ppfibp14 | 5.25E-68 | 1.266883428 | 0.718 | 0.182 | 1.06E-63 | 20 | Ppfibp1 |
| Atp8b14 | 6.01E-68 | 1.208650037 | 0.53 | 0.102 | 1.22E-63 | 20 | Atp8b1 |
| Mylk3 | 8.23E-68 | 0.718293543 | 0.362 | 0.048 | 1.67E-63 | 20 | Mylk |
| Ralgapa26 | 3.77E-67 | 1.432030916 | 0.638 | 0.153 | 7.62E-63 | 20 | Ralgapa2 |
| Calcrl3 | 5.79E-67 | 1.596342654 | 0.846 | 0.262 | 1.17E-62 | 20 | Calcrl |
| Prss236 | 6.78E-67 | 1.599098339 | 0.872 | 0.28 | 1.37E-62 | 20 | Prss23 |
| Tspan55 | 1.01E-66 | 1.426001977 | 0.725 | 0.197 | 2.05E-62 | 20 | Tspan5 |
| Jam24 | 1.20E-66 | 1.690345097 | 0.846 | 0.285 | 2.43E-62 | 20 | Jam2 |
| Vim11 | 1.61E-66 | 1.965900048 | 0.893 | 0.305 | 3.26E-62 | 20 | Vim |
| Npr12 | 5.70E-66 | 0.899707955 | 0.503 | 0.091 | 1.15E-61 | 20 | Npr1 |
| Zfpm22 | 6.64E-66 | 1.309209865 | 0.456 | 0.078 | 1.34E-61 | 20 | Zfpm2 |
| Chrm2 | 1.26E-65 | 1.03388529 | 0.356 | 0.049 | 2.55E-61 | 20 | Chrm2 |
| Dst7 | 4.66E-65 | 1.296487086 | 0.732 | 0.19 | 9.43E-61 | 20 | Dst |
| Sema6a4 | 1.45E-64 | 1.472241133 | 0.765 | 0.208 | 2.94E-60 | 20 | Sema6a |
| Pde4d4 | 7.86E-64 | 1.401232279 | 0.631 | 0.149 | 1.59E-59 | 20 | Pde4d |
| Plec6 | 1.80E-63 | 1.286102939 | 0.664 | 0.173 | 3.65E-59 | 20 | Plec |
| Igfbp43 | 2.47E-63 | 0.942574532 | 0.517 | 0.096 | 5.00E-59 | 20 | Igfbp4 |
| Il1r13 | 2.65E-63 | 1.310968819 | 0.537 | 0.11 | 5.37E-59 | 20 | Il1r1 |
| Heg15 | 5.31E-62 | 1.565077881 | 0.826 | 0.273 | 1.08E-57 | 20 | Heg1 |
| Plekhg14 | 9.74E-62 | 1.487807208 | 0.805 | 0.244 | 1.97E-57 | 20 | Plekhg1 |
| Pam8 | 2.20E-61 | 1.420822098 | 0.725 | 0.213 | 4.45E-57 | 20 | Pam |
| Prex25 | 2.68E-61 | 1.493200802 | 0.839 | 0.272 | 5.42E-57 | 20 | Prex2 |
| Cd94 | 3.33E-61 | 1.685941763 | 0.913 | 0.385 | 6.74E-57 | 20 | Cd9 |
| Ndrg14 | 4.01E-61 | 1.316620475 | 0.779 | 0.232 | 8.11E-57 | 20 | Ndrg1 |
| Ypel2 | 4.25E-59 | 0.847151368 | 0.416 | 0.072 | 8.60E-55 | 20 | Ypel2 |

| | | | | | | | |
|---|---|---|---|---|---|---|---|
| Ldb25 | 5.73E-59 | 1.539743567 | 0.933 | 0.329 | 1.16E-54 | 20 | Ldb2 |
| Itga3 | 2.19E-58 | 0.645134692 | 0.268 | 0.032 | 4.44E-54 | 20 | Itga3 |
| Gnaq6 | 2.42E-58 | 1.507412641 | 0.906 | 0.358 | 4.90E-54 | 20 | Gnaq |
| Rhoj6 | 5.34E-58 | 1.257566783 | 0.792 | 0.241 | 1.08E-53 | 20 | Rhoj |
| Slco3a15 | 6.63E-58 | 1.117033603 | 0.55 | 0.121 | 1.34E-53 | 20 | Slco3a1 |
| Ccser11 | 1.05E-57 | 1.131985276 | 0.396 | 0.067 | 2.12E-53 | 20 | Ccser1 |
| Acer34 | 4.26E-57 | 1.171715405 | 0.584 | 0.141 | 8.62E-53 | 20 | Acer3 |
| Rora4 | 1.34E-56 | 1.551156641 | 0.678 | 0.185 | 2.72E-52 | 20 | Rora |
| Klk84 | 3.50E-56 | 0.935292329 | 0.436 | 0.082 | 7.08E-52 | 20 | Klk8 |
| Tgfbr34 | 1.05E-55 | 1.130521414 | 0.658 | 0.173 | 2.12E-51 | 20 | Tgfbr3 |
| Itpr17 | 2.89E-55 | 1.39076191 | 0.705 | 0.221 | 5.86E-51 | 20 | Itpr1 |
| Ptprb5 | 1.88E-54 | 1.475776179 | 0.953 | 0.367 | 3.80E-50 | 20 | Ptprb |
| Adcy43 | 2.18E-54 | 0.851764446 | 0.51 | 0.107 | 4.42E-50 | 20 | Adcy4 |
| Aplp25 | 5.70E-54 | 1.388846374 | 0.94 | 0.415 | 1.15E-49 | 20 | Aplp2 |
| Fcgrt4 | 1.27E-53 | 1.063755088 | 0.671 | 0.184 | 2.57E-49 | 20 | Fcgrt |
| Adgrl23 | 5.33E-53 | 1.407811062 | 0.745 | 0.236 | 1.08E-48 | 20 | Adgrl2 |
| Lmcd13 | 3.61E-52 | 1.091149215 | 0.631 | 0.164 | 7.30E-48 | 20 | Lmcd1 |
| Arhgap2610 | 9.19E-52 | 1.235176242 | 0.711 | 0.222 | 1.86E-47 | 20 | Arhgap26 |
| Ube2e24 | 1.26E-51 | 1.213313078 | 0.597 | 0.152 | 2.56E-47 | 20 | Ube2e2 |
| Flnb3 | 2.66E-51 | 1.157118141 | 0.604 | 0.161 | 5.38E-47 | 20 | Flnb |
| Mmrn24 | 5.24E-51 | 1.233364958 | 0.745 | 0.231 | 1.06E-46 | 20 | Mmrn2 |
| Net11 | 1.60E-50 | 0.766713695 | 0.275 | 0.038 | 3.23E-46 | 20 | Net1 |
| Lrrc1 | 2.51E-50 | 0.510814444 | 0.275 | 0.038 | 5.07E-46 | 20 | Lrrc1 |
| Airn7 | 5.22E-50 | 1.432106222 | 0.752 | 0.247 | 1.06E-45 | 20 | Airn |
| Thsd12 | 2.01E-49 | 0.827052706 | 0.436 | 0.088 | 4.08E-45 | 20 | Thsd1 |
| Rassf93 | 2.22E-49 | 1.062028926 | 0.53 | 0.125 | 4.49E-45 | 20 | Rassf9 |
| Eng4 | 4.27E-49 | 1.25029479 | 0.913 | 0.353 | 8.65E-45 | 20 | Eng |
| Cnn36 | 1.45E-48 | 1.048211055 | 0.664 | 0.193 | 2.93E-44 | 20 | Cnn3 |
| Ebf16 | 3.00E-48 | 1.383258772 | 0.96 | 0.374 | 6.08E-44 | 20 | Ebf1 |
| Foxo14 | 4.19E-48 | 0.996010527 | 0.671 | 0.199 | 8.49E-44 | 20 | Foxo1 |
| Ecscr4 | 5.72E-48 | 1.142714029 | 0.725 | 0.227 | 1.16E-43 | 20 | Ecscr |
| Magi14 | 7.68E-48 | 1.357450262 | 0.725 | 0.231 | 1.55E-43 | 20 | Magi1 |
| Piezo15 | 1.34E-47 | 0.956982289 | 0.631 | 0.173 | 2.72E-43 | 20 | Piezo1 |
| Tek4 | 5.88E-47 | 1.217463256 | 0.765 | 0.259 | 1.19E-42 | 20 | Tek |
| Icam12 | 7.14E-47 | 0.772977837 | 0.383 | 0.075 | 1.45E-42 | 20 | Icam1 |
| Nfat58 | 8.07E-47 | 1.245895901 | 0.752 | 0.266 | 1.63E-42 | 20 | Nfat5 |
| Ptprg4 | 9.00E-47 | 1.456050569 | 0.886 | 0.344 | 1.82E-42 | 20 | Ptprg |
| Ptgs12 | 1.01E-46 | 1.140412491 | 0.685 | 0.211 | 2.04E-42 | 20 | Ptgs1 |
| St6galnac37 | 1.38E-46 | 1.203561008 | 0.698 | 0.215 | 2.79E-42 | 20 | St6galnac3 |
| Sulf12 | 1.46E-46 | 0.725771788 | 0.322 | 0.053 | 2.95E-42 | 20 | Sulf1 |
| mt-Nd413 | 1.86E-46 | 1.156670718 | 1 | 0.676 | 3.76E-42 | 20 | mt-Nd4 |
| Actn12 | 2.63E-46 | 0.891428102 | 0.403 | 0.083 | 5.33E-42 | 20 | Actn1 |
| Eps83 | 4.03E-46 | 0.695687563 | 0.423 | 0.086 | 8.15E-42 | 20 | Eps8 |

| | | | | | | | |
|---|---|---|---|---|---|---|---|
| Cav15 | 4.54E-46 | 1.141599319 | 0.859 | 0.303 | 9.19E-42 | 20 | Cav1 |
| Ptprk4 | 5.13E-46 | 1.05611106 | 0.544 | 0.14 | 1.04E-41 | 20 | Ptprk |
| Foxp18 | 5.14E-46 | 1.380225839 | 0.859 | 0.389 | 1.04E-41 | 20 | Foxp1 |
| Arhgap314 | 1.13E-45 | 1.27499448 | 0.913 | 0.371 | 2.28E-41 | 20 | Arhgap31 |
| Pard3b3 | 1.59E-45 | 1.020232107 | 0.45 | 0.1 | 3.21E-41 | 20 | Pard3b |
| mt-Atp614 | 1.96E-45 | 1.083123928 | 1 | 0.754 | 3.96E-41 | 20 | mt-Atp6 |
| Rbp11 | 4.04E-45 | 0.985299225 | 0.45 | 0.101 | 8.18E-41 | 20 | Rbp1 |
| Cmah2 | 4.69E-45 | 1.06836516 | 0.356 | 0.067 | 9.50E-41 | 20 | Cmah |
| Hyal22 | 4.73E-45 | 0.900701361 | 0.409 | 0.086 | 9.57E-41 | 20 | Hyal2 |
| Ppp2r5a5 | 6.43E-45 | 1.181015907 | 0.698 | 0.25 | 1.30E-40 | 20 | Ppp2r5a |
| Fmo23 | 1.91E-44 | 1.236690261 | 0.792 | 0.288 | 3.87E-40 | 20 | Fmo2 |
| Nhsl24 | 3.62E-44 | 1.008697517 | 0.523 | 0.134 | 7.34E-40 | 20 | Nhsl2 |
| Nr3c25 | 4.81E-44 | 1.005928822 | 0.49 | 0.121 | 9.75E-40 | 20 | Nr3c2 |
| Fgd54 | 1.21E-43 | 0.982267455 | 0.725 | 0.228 | 2.44E-39 | 20 | Fgd5 |
| Lhx61 | 1.90E-43 | 0.660994435 | 0.342 | 0.062 | 3.85E-39 | 20 | Lhx6 |
| Emp13 | 3.41E-43 | 0.759443158 | 0.503 | 0.121 | 6.90E-39 | 20 | Emp1 |
| Adam154 | 3.57E-43 | 0.981790312 | 0.638 | 0.193 | 7.22E-39 | 20 | Adam15 |
| Myh910 | 3.72E-43 | 1.286183722 | 0.879 | 0.437 | 7.53E-39 | 20 | Myh9 |
| Fstl14 | 9.70E-43 | 0.954277937 | 0.45 | 0.103 | 1.96E-38 | 20 | Fstl1 |
| Samd124 | 9.74E-43 | 1.198700615 | 0.705 | 0.229 | 1.97E-38 | 20 | Samd12 |
| Ica13 | 1.38E-42 | 0.838815928 | 0.617 | 0.176 | 2.79E-38 | 20 | Ica1 |
| Fgl27 | 1.89E-42 | 1.58926467 | 0.443 | 0.108 | 3.83E-38 | 20 | Fgl2 |
| Pbx14 | 5.43E-42 | 1.103391462 | 0.638 | 0.198 | 1.10E-37 | 20 | Pbx1 |
| Bmpr24 | 8.56E-42 | 1.113549707 | 0.866 | 0.349 | 1.73E-37 | 20 | Bmpr2 |
| Ahnak8 | 1.16E-41 | 1.207196946 | 0.94 | 0.471 | 2.34E-37 | 20 | Ahnak |
| Prpf40b1 | 2.73E-41 | 0.668337047 | 0.376 | 0.077 | 5.53E-37 | 20 | Prpf40b |
| mt-Co114 | 3.20E-41 | 1.02803843 | 1 | 0.72 | 6.48E-37 | 20 | mt-Co1 |
| Cavin36 | 4.10E-41 | 1.090260084 | 0.805 | 0.307 | 8.30E-37 | 20 | Cavin3 |
| Ramp25 | 4.29E-41 | 1.108091903 | 0.852 | 0.331 | 8.69E-37 | 20 | Ramp2 |
| Selenop6 | 5.56E-41 | 1.436964886 | 0.832 | 0.363 | 1.13E-36 | 20 | Selenop |
| Adgrl44 | 8.00E-41 | 1.098142683 | 0.926 | 0.351 | 1.62E-36 | 20 | Adgrl4 |
| Grrp14 | 8.53E-41 | 1.085874408 | 0.664 | 0.219 | 1.73E-36 | 20 | Grrp1 |
| Rfk1 | 2.60E-40 | 0.651491922 | 0.369 | 0.077 | 5.27E-36 | 20 | Rfk |
| Tspan94 | 2.63E-40 | 1.02929065 | 0.745 | 0.265 | 5.33E-36 | 20 | Tspan9 |
| Egfl75 | 3.34E-40 | 1.128984143 | 0.953 | 0.393 | 6.76E-36 | 20 | Egfl7 |
| Elk34 | 3.37E-40 | 1.223163696 | 0.819 | 0.356 | 6.82E-36 | 20 | Elk3 |
| Msrb34 | 4.79E-40 | 0.947554154 | 0.497 | 0.132 | 9.71E-36 | 20 | Msrb3 |
| Abca14 | 6.08E-40 | 1.109617482 | 0.483 | 0.129 | 1.23E-35 | 20 | Abca1 |
| Plxna43 | 6.99E-40 | 1.086479757 | 0.463 | 0.119 | 1.42E-35 | 20 | Plxna4 |
| Flrt22 | 9.28E-40 | 0.772180889 | 0.268 | 0.044 | 1.88E-35 | 20 | Flrt2 |
| Tjp14 | 1.88E-39 | 1.083574354 | 0.725 | 0.263 | 3.80E-35 | 20 | Tjp1 |
| Cdh54 | 1.91E-39 | 0.864455168 | 0.926 | 0.33 | 3.86E-35 | 20 | Cdh5 |
| Rcn32 | 2.07E-39 | 0.599991353 | 0.302 | 0.053 | 4.19E-35 | 20 | Rcn3 |

| | | | | | | | |
|---|---|---|---|---|---|---|---|
| Fbln26 | 2.31E-39 | 1.156284953 | 0.758 | 0.272 | 4.68E-35 | 20 | Fbln2 |
| Shank33 | 3.00E-39 | 0.912738457 | 0.631 | 0.19 | 6.07E-35 | 20 | Shank3 |
| Plvap4 | 5.67E-39 | 1.198097169 | 0.913 | 0.384 | 1.15E-34 | 20 | Plvap |
| Bok | 6.67E-39 | 0.651678098 | 0.282 | 0.049 | 1.35E-34 | 20 | Bok |
| Fzd62 | 1.59E-38 | 0.593801277 | 0.315 | 0.059 | 3.22E-34 | 20 | Fzd6 |
| Gng116 | 1.63E-38 | 1.217941391 | 0.866 | 0.366 | 3.30E-34 | 20 | Gng11 |
| Cd344 | 3.15E-38 | 1.07834937 | 0.812 | 0.321 | 6.38E-34 | 20 | Cd34 |
| Cpne21 | 5.09E-38 | 0.71448655 | 0.336 | 0.068 | 1.03E-33 | 20 | Cpne2 |
| Sntb24 | 1.54E-37 | 1.104394801 | 0.705 | 0.265 | 3.12E-33 | 20 | Sntb2 |
| Cavin15 | 1.58E-37 | 0.793984754 | 0.664 | 0.213 | 3.20E-33 | 20 | Cavin1 |
| Galnt185 | 1.59E-37 | 1.204156358 | 0.49 | 0.134 | 3.21E-33 | 20 | Galnt18 |
| Cavin25 | 1.92E-37 | 1.048587202 | 0.886 | 0.341 | 3.89E-33 | 20 | Cavin2 |
| Plcb14 | 2.07E-37 | 1.031770258 | 0.893 | 0.34 | 4.18E-33 | 20 | Plcb1 |
| Ghr4 | 2.88E-37 | 0.792742341 | 0.537 | 0.151 | 5.82E-33 | 20 | Ghr |
| Rasip14 | 3.27E-37 | 0.880014006 | 0.624 | 0.195 | 6.62E-33 | 20 | Rasip1 |
| Sash16 | 7.34E-37 | 1.057583568 | 0.698 | 0.257 | 1.49E-32 | 20 | Sash1 |
| Sorbs25 | 3.77E-36 | 0.982796216 | 0.758 | 0.277 | 7.63E-32 | 20 | Sorbs2 |
| Ltbp45 | 4.20E-36 | 0.988841096 | 0.805 | 0.309 | 8.50E-32 | 20 | Ltbp4 |
| Sncaip2 | 5.96E-36 | 0.872843227 | 0.362 | 0.081 | 1.21E-31 | 20 | Sncaip |
| Rai143 | 8.54E-36 | 0.923430201 | 0.671 | 0.231 | 1.73E-31 | 20 | Rai14 |
| Esam5 | 1.01E-35 | 0.868487488 | 0.826 | 0.309 | 2.03E-31 | 20 | Esam |
| Tfpi1 | 1.06E-35 | 0.629117004 | 0.302 | 0.059 | 2.15E-31 | 20 | Tfpi |
| Arrb1 | 1.35E-35 | 0.660864471 | 0.275 | 0.051 | 2.73E-31 | 20 | Arrb1 |
| Utrn5 | 1.51E-35 | 1.014938411 | 0.879 | 0.405 | 3.06E-31 | 20 | Utrn |
| Acer21 | 1.62E-35 | 0.69366158 | 0.336 | 0.07 | 3.28E-31 | 20 | Acer2 |
| Zfp4621 | 1.85E-35 | 1.035654226 | 0.45 | 0.12 | 3.75E-31 | 20 | Zfp462 |
| Plcb45 | 2.02E-35 | 0.93320066 | 0.745 | 0.291 | 4.09E-31 | 20 | Plcb4 |
| Unc13b1 | 2.12E-35 | 0.574319807 | 0.336 | 0.069 | 4.29E-31 | 20 | Unc13b |
| Mill23 | 3.03E-35 | 0.775189781 | 0.329 | 0.069 | 6.13E-31 | 20 | Mill2 |
| Pecam15 | 3.60E-35 | 1.012250443 | 0.946 | 0.411 | 7.29E-31 | 20 | Pecam1 |
| Tie15 | 3.63E-35 | 0.742983557 | 0.745 | 0.253 | 7.35E-31 | 20 | Tie1 |
| Nbea6 | 3.72E-35 | 0.865367894 | 0.604 | 0.196 | 7.53E-31 | 20 | Nbea |
| Fli19 | 6.82E-35 | 1.20536178 | 0.859 | 0.402 | 1.38E-30 | 20 | Fli1 |
| Dock44 | 9.57E-35 | 0.95987413 | 0.779 | 0.296 | 1.94E-30 | 20 | Dock4 |
| Ptprm4 | 1.60E-34 | 0.891081063 | 0.859 | 0.329 | 3.24E-30 | 20 | Ptprm |
| Rasal24 | 2.18E-34 | 1.068786507 | 0.772 | 0.315 | 4.42E-30 | 20 | Rasal2 |
| Sptbn15 | 6.55E-34 | 0.987747075 | 0.987 | 0.494 | 1.33E-29 | 20 | Sptbn1 |
| Tbc1d41 | 9.88E-34 | 0.646473692 | 0.262 | 0.048 | 2.00E-29 | 20 | Tbc1d4 |
| Bambi | 1.21E-33 | 0.566715346 | 0.268 | 0.05 | 2.45E-29 | 20 | Bambi |
| Elmo111 | 1.64E-33 | 1.064041971 | 0.839 | 0.366 | 3.32E-29 | 20 | Elmo1 |
| Ehd43 | 1.79E-33 | 1.003602629 | 0.866 | 0.408 | 3.63E-29 | 20 | Ehd4 |
| Ctla2a8 | 2.25E-33 | 1.031105939 | 0.872 | 0.398 | 4.56E-29 | 20 | Ctla2a |
| 4931406P16Rik5 | 2.62E-33 | 0.897690885 | 0.819 | 0.343 | 5.31E-29 | 20 | 4931406P16Rik |

| | | | | | | | |
|---|---|---|---|---|---|---|---|
| Nfib4 | 4.00E-33 | 0.9612268 | 0.973 | 0.444 | 8.09E-29 | 20 | Nfib |
| Nfia6 | 5.46E-33 | 1.00713615 | 0.718 | 0.287 | 1.11E-28 | 20 | Nfia |
| Ecm23 | 6.27E-33 | 0.691451964 | 0.262 | 0.049 | 1.27E-28 | 20 | Ecm2 |
| mt-Atp811 | 1.05E-32 | 0.991785449 | 0.893 | 0.463 | 2.13E-28 | 20 | mt-Atp8 |
| Adamts93 | 1.52E-32 | 1.123427667 | 0.443 | 0.124 | 3.07E-28 | 20 | Adamts9 |
| Mical3 | 1.87E-32 | 0.618991723 | 0.322 | 0.07 | 3.79E-28 | 20 | Mical3 |
| Slfn31 | 2.02E-32 | 0.848364639 | 0.322 | 0.073 | 4.08E-28 | 20 | Slfn3 |
| Meis22 | 2.04E-32 | 1.002459498 | 0.725 | 0.282 | 4.13E-28 | 20 | Meis2 |
| Tnfrsf1a5 | 2.12E-32 | 0.953963774 | 0.611 | 0.222 | 4.29E-28 | 20 | Tnfrsf1a |
| Prkch7 | 2.37E-32 | 1.046079705 | 0.879 | 0.399 | 4.81E-28 | 20 | Prkch |
| 4930578C19Rik3 | 2.43E-32 | 0.899225386 | 0.409 | 0.106 | 4.92E-28 | 20 | 4930578C19Rik |
| Cd2005 | 3.04E-32 | 0.975762373 | 0.832 | 0.35 | 6.16E-28 | 20 | Cd200 |
| Pkp41 | 3.35E-32 | 0.813703407 | 0.725 | 0.286 | 6.78E-28 | 20 | Pkp4 |
| Timp25 | 9.09E-32 | 1.048089809 | 0.45 | 0.13 | 1.84E-27 | 20 | Timp2 |
| Lamb25 | 1.49E-31 | 0.687874043 | 0.443 | 0.124 | 3.01E-27 | 20 | Lamb2 |
| Plxnb21 | 1.60E-31 | 0.592788353 | 0.255 | 0.049 | 3.25E-27 | 20 | Plxnb2 |
| Sbf26 | 1.77E-31 | 0.879568258 | 0.597 | 0.207 | 3.57E-27 | 20 | Sbf2 |
| App5 | 1.85E-31 | 0.891065234 | 0.919 | 0.419 | 3.75E-27 | 20 | App |
| Ppic4 | 1.87E-31 | 0.622022113 | 0.443 | 0.121 | 3.79E-27 | 20 | Ppic |
| Cpd5 | 1.93E-31 | 0.92047887 | 0.752 | 0.312 | 3.91E-27 | 20 | Cpd |
| Creb3l23 | 1.93E-31 | 0.763124944 | 0.436 | 0.123 | 3.91E-27 | 20 | Creb3l2 |
| Cpne84 | 2.72E-31 | 0.853440699 | 0.644 | 0.228 | 5.52E-27 | 20 | Cpne8 |
| Btbd32 | 3.12E-31 | 0.743511749 | 0.389 | 0.101 | 6.31E-27 | 20 | Btbd3 |
| Stab11 | 3.26E-31 | 0.846008974 | 0.349 | 0.084 | 6.61E-27 | 20 | Stab1 |
| Hspa12b3 | 4.06E-31 | 0.746800565 | 0.396 | 0.102 | 8.22E-27 | 20 | Hspa12b |
| Aplnr1 | 4.66E-31 | 0.667740067 | 0.383 | 0.094 | 9.44E-27 | 20 | Aplnr |
| Limch14 | 5.08E-31 | 0.821324365 | 0.477 | 0.139 | 1.03E-26 | 20 | Limch1 |
| Erg4 | 5.39E-31 | 0.772090632 | 0.456 | 0.131 | 1.09E-26 | 20 | Erg |
| Adgrf55 | 5.55E-31 | 0.807620542 | 0.866 | 0.337 | 1.12E-26 | 20 | Adgrf5 |
| Nxpe44 | 7.48E-31 | 0.847186461 | 0.49 | 0.148 | 1.51E-26 | 20 | Nxpe4 |
| Apbb26 | 8.36E-31 | 0.965563815 | 0.832 | 0.367 | 1.69E-26 | 20 | Apbb2 |
| Klf25 | 1.01E-30 | 0.990212175 | 0.812 | 0.367 | 2.04E-26 | 20 | Klf2 |
| mt-Nd4l9 | 1.08E-30 | 0.840106012 | 0.926 | 0.491 | 2.19E-26 | 20 | mt-Nd4l |
| She4 | 1.29E-30 | 0.767933318 | 0.497 | 0.149 | 2.60E-26 | 20 | She |
| Nmt21 | 1.43E-30 | 0.785200187 | 0.369 | 0.098 | 2.90E-26 | 20 | Nmt2 |
| Sned11 | 1.46E-30 | 0.561785333 | 0.302 | 0.065 | 2.95E-26 | 20 | Sned1 |
| Shroom43 | 1.73E-30 | 0.807728806 | 0.396 | 0.105 | 3.49E-26 | 20 | Shroom4 |
| Gda | 2.06E-30 | 0.736516885 | 0.336 | 0.08 | 4.16E-26 | 20 | Gda |
| Itga51 | 2.13E-30 | 0.627839995 | 0.376 | 0.097 | 4.31E-26 | 20 | Itga5 |
| Ace3 | 3.23E-30 | 0.712995778 | 0.43 | 0.119 | 6.55E-26 | 20 | Ace |
| Ttc286 | 3.24E-30 | 0.989560894 | 0.691 | 0.285 | 6.56E-26 | 20 | Ttc28 |
| Lima15 | 3.39E-30 | 0.797366993 | 0.503 | 0.158 | 6.86E-26 | 20 | Lima1 |
| Cyyr14 | 3.87E-30 | 0.841284899 | 0.872 | 0.343 | 7.84E-26 | 20 | Cyyr1 |

| | | | | | | | |
|---|---|---|---|---|---|---|---|
| Hdac74 | 6.39E-30 | 0.81065778 | 0.57 | 0.202 | 1.29E-25 | 20 | Hdac7 |
| Tmtc23 | 9.96E-30 | 0.990854405 | 0.544 | 0.185 | 2.02E-25 | 20 | Tmtc2 |
| Lhfpl23 | 1.15E-29 | 0.802494844 | 0.483 | 0.15 | 2.33E-25 | 20 | Lhfpl2 |
| Zbtb209 | 1.27E-29 | 0.853844989 | 0.966 | 0.537 | 2.56E-25 | 20 | Zbtb20 |
| S100a165 | 1.30E-29 | 0.668224828 | 0.691 | 0.258 | 2.63E-25 | 20 | S100a16 |
| mt-Co315 | 1.36E-29 | 0.851221863 | 1 | 0.726 | 2.75E-25 | 20 | mt-Co3 |
| Lrch16 | 1.38E-29 | 1.028244052 | 0.711 | 0.312 | 2.79E-25 | 20 | Lrch1 |
| Myo1b3 | 1.49E-29 | 0.731021532 | 0.463 | 0.138 | 3.02E-25 | 20 | Myo1b |
| Nos31 | 1.77E-29 | 0.698960307 | 0.322 | 0.075 | 3.59E-25 | 20 | Nos3 |
| Tcf45 | 1.93E-29 | 0.831586958 | 0.98 | 0.472 | 3.90E-25 | 20 | Tcf4 |
| Rasgrf24 | 2.26E-29 | 0.929431902 | 0.537 | 0.177 | 4.57E-25 | 20 | Rasgrf2 |
| Tmem255b2 | 2.76E-29 | 0.680762737 | 0.315 | 0.073 | 5.59E-25 | 20 | Tmem255b |
| Ifitm37 | 3.00E-29 | 0.953802649 | 0.953 | 0.538 | 6.07E-25 | 20 | Ifitm3 |
| Mbnl28 | 3.15E-29 | 0.814673596 | 0.866 | 0.418 | 6.39E-25 | 20 | Mbnl2 |
| Fry5 | 3.40E-29 | 0.8318119 | 0.57 | 0.2 | 6.89E-25 | 20 | Fry |
| Cdc42bpa6 | 4.60E-29 | 0.799166674 | 0.55 | 0.185 | 9.31E-25 | 20 | Cdc42bpa |
| Klf44 | 4.80E-29 | 0.846622642 | 0.691 | 0.269 | 9.73E-25 | 20 | Klf4 |
| Pltp4 | 4.98E-29 | 0.839691637 | 0.752 | 0.3 | 1.01E-24 | 20 | Pltp |
| Arhgap231 | 5.86E-29 | 0.770193882 | 0.342 | 0.085 | 1.19E-24 | 20 | Arhgap23 |
| Rbpms5 | 6.67E-29 | 0.795847394 | 0.799 | 0.349 | 1.35E-24 | 20 | Rbpms |
| Gulp16 | 1.87E-28 | 0.666103314 | 0.523 | 0.166 | 3.79E-24 | 20 | Gulp1 |
| Sptlc25 | 2.77E-28 | 0.891220503 | 0.611 | 0.245 | 5.61E-24 | 20 | Sptlc2 |
| Synpo1 | 3.07E-28 | 0.677071605 | 0.376 | 0.1 | 6.21E-24 | 20 | Synpo |
| Numb4 | 3.13E-28 | 0.844142611 | 0.779 | 0.337 | 6.34E-24 | 20 | Numb |
| Ctnna15 | 3.17E-28 | 0.794900773 | 0.779 | 0.351 | 6.42E-24 | 20 | Ctnna1 |
| Piezo22 | 4.31E-28 | 0.881896387 | 0.671 | 0.253 | 8.73E-24 | 20 | Piezo2 |
| mt-Cytb12 | 5.90E-28 | 0.81213303 | 1 | 0.701 | 1.19E-23 | 20 | mt-Cytb |
| Ppp1r13b4 | 7.53E-28 | 0.914194746 | 0.577 | 0.216 | 1.52E-23 | 20 | Ppp1r13b |
| Gab27 | 7.63E-28 | 0.757930545 | 0.577 | 0.205 | 1.54E-23 | 20 | Gab2 |
| Myct14 | 9.58E-28 | 0.615188079 | 0.638 | 0.222 | 1.94E-23 | 20 | Myct1 |
| Emcn4 | 1.02E-27 | 0.766230833 | 0.94 | 0.38 | 2.07E-23 | 20 | Emcn |
| Pde7b6 | 1.22E-27 | 1.330810197 | 0.611 | 0.243 | 2.48E-23 | 20 | Pde7b |
| Etv66 | 1.60E-27 | 0.832816455 | 0.671 | 0.278 | 3.23E-23 | 20 | Etv6 |
| Abcc44 | 1.68E-27 | 0.794661501 | 0.443 | 0.136 | 3.39E-23 | 20 | Abcc4 |
| Nedd46 | 1.77E-27 | 0.818209487 | 0.846 | 0.419 | 3.59E-23 | 20 | Nedd4 |
| Slit32 | 1.80E-27 | 0.884935948 | 0.329 | 0.082 | 3.65E-23 | 20 | Slit3 |
| Cers44 | 1.88E-27 | 0.745621278 | 0.383 | 0.11 | 3.80E-23 | 20 | Cers4 |
| Rock25 | 2.00E-27 | 0.833545341 | 0.752 | 0.338 | 4.06E-23 | 20 | Rock2 |
| Wwtr15 | 2.37E-27 | 0.737347431 | 0.644 | 0.24 | 4.79E-23 | 20 | Wwtr1 |
| Tspan183 | 2.44E-27 | 0.879161813 | 0.443 | 0.138 | 4.93E-23 | 20 | Tspan18 |
| Etl44 | 2.73E-27 | 0.781393678 | 0.752 | 0.294 | 5.53E-23 | 20 | Etl4 |
| Atp11c4 | 3.16E-27 | 0.86051053 | 0.463 | 0.151 | 6.40E-23 | 20 | Atp11c |
| Ptk25 | 4.18E-27 | 0.825501239 | 0.678 | 0.266 | 8.47E-23 | 20 | Ptk2 |

| | | | | | | | |
|---|---|---|---|---|---|---|---|
| Stk36 | 5.41E-27 | 0.665387242 | 0.497 | 0.166 | 1.10E-22 | 20 | Stk3 |
| Sh3bp52 | 7.73E-27 | 0.592769369 | 0.611 | 0.216 | 1.56E-22 | 20 | Sh3bp5 |
| Macf19 | 9.17E-27 | 0.902742814 | 0.919 | 0.503 | 1.86E-22 | 20 | Macf1 |
| Phactr28 | 1.05E-26 | 0.818311364 | 0.617 | 0.246 | 2.12E-22 | 20 | Phactr2 |
| Rapgef54 | 1.49E-26 | 0.871816394 | 0.597 | 0.222 | 3.03E-22 | 20 | Rapgef5 |
| Dusp231 | 1.58E-26 | 0.491953244 | 0.403 | 0.116 | 3.19E-22 | 20 | Dusp23 |
| Cd59a1 | 1.65E-26 | 0.474275057 | 0.342 | 0.087 | 3.33E-22 | 20 | Cd59a |
| 2010111I01Rik3 | 1.84E-26 | 0.853350496 | 0.738 | 0.349 | 3.73E-22 | 20 | 2010111I01Rik |
| Diaph29 | 2.43E-26 | 0.895024304 | 0.792 | 0.377 | 4.92E-22 | 20 | Diaph2 |
| Cebpd1 | 2.71E-26 | 0.575037167 | 0.517 | 0.175 | 5.49E-22 | 20 | Cebpd |
| Tcf7l15 | 2.76E-26 | 0.816376971 | 0.523 | 0.181 | 5.60E-22 | 20 | Tcf7l1 |
| Kalrn5 | 2.97E-26 | 0.731096674 | 0.376 | 0.106 | 6.02E-22 | 20 | Kalrn |
| Ehd25 | 3.56E-26 | 0.649460707 | 0.557 | 0.193 | 7.20E-22 | 20 | Ehd2 |
| Ephb44 | 4.13E-26 | 0.716316113 | 0.497 | 0.166 | 8.36E-22 | 20 | Ephb4 |
| Ctnnb14 | 4.57E-26 | 0.771334619 | 0.765 | 0.364 | 9.26E-22 | 20 | Ctnnb1 |
| Galnt153 | 4.68E-26 | 0.648995371 | 0.342 | 0.09 | 9.48E-22 | 20 | Galnt15 |
| Mkl24 | 5.75E-26 | 0.898619621 | 0.624 | 0.255 | 1.16E-21 | 20 | Mkl2 |
| Pdlim12 | 6.08E-26 | 0.545322116 | 0.369 | 0.103 | 1.23E-21 | 20 | Pdlim1 |
| Cd934 | 7.44E-26 | 0.828790454 | 0.651 | 0.256 | 1.51E-21 | 20 | Cd93 |
| Dock14 | 8.64E-26 | 0.679155137 | 0.691 | 0.28 | 1.75E-21 | 20 | Dock1 |
| Fkbp1a4 | 1.12E-25 | 0.781330894 | 0.805 | 0.406 | 2.27E-21 | 20 | Fkbp1a |
| Socs25 | 1.45E-25 | 0.647826969 | 0.523 | 0.183 | 2.94E-21 | 20 | Socs2 |
| Wwc25 | 1.57E-25 | 0.769266139 | 0.544 | 0.197 | 3.18E-21 | 20 | Wwc2 |
| Plxna24 | 1.72E-25 | 0.699467244 | 0.597 | 0.219 | 3.49E-21 | 20 | Plxna2 |
| mt-Nd510 | 1.75E-25 | 0.796755189 | 0.906 | 0.496 | 3.54E-21 | 20 | mt-Nd5 |
| Tmem26 | 1.79E-25 | 0.664506868 | 0.443 | 0.143 | 3.62E-21 | 20 | Tmem2 |
| Gas64 | 2.09E-25 | 0.7261171 | 0.725 | 0.298 | 4.24E-21 | 20 | Gas6 |
| Amotl16 | 2.25E-25 | 0.755699983 | 0.517 | 0.186 | 4.55E-21 | 20 | Amotl1 |
| Bcr | 3.92E-25 | 0.551268581 | 0.262 | 0.059 | 7.94E-21 | 20 | Bcr |
| mt-Nd210 | 4.16E-25 | 0.777302104 | 1 | 0.689 | 8.42E-21 | 20 | mt-Nd2 |
| Tbc1d82 | 4.17E-25 | 0.494593513 | 0.262 | 0.06 | 8.45E-21 | 20 | Tbc1d8 |
| Stim19 | 4.95E-25 | 0.803413452 | 0.517 | 0.191 | 1.00E-20 | 20 | Stim1 |
| Iigp14 | 5.49E-25 | 1.081864262 | 0.591 | 0.233 | 1.11E-20 | 20 | Iigp1 |
| Plxnd15 | 6.02E-25 | 0.709609648 | 0.517 | 0.182 | 1.22E-20 | 20 | Plxnd1 |
| Mar-32 | 8.45E-25 | 0.79197369 | 0.336 | 0.093 | 1.71E-20 | 20 | 3-Mar |
| Iqgap110 | 9.23E-25 | 0.852266904 | 0.799 | 0.412 | 1.87E-20 | 20 | Iqgap1 |
| Parva3 | 1.24E-24 | 0.583845621 | 0.383 | 0.112 | 2.52E-20 | 20 | Parva |
| Gm326881 | 1.45E-24 | 0.490581123 | 0.349 | 0.094 | 2.94E-20 | 20 | Gm32688 |
| Me31 | 1.78E-24 | 0.5649591 | 0.362 | 0.1 | 3.60E-20 | 20 | Me3 |
| Clec14a4 | 1.89E-24 | 0.724469922 | 0.725 | 0.305 | 3.83E-20 | 20 | Clec14a |
| Slc12a72 | 1.90E-24 | 0.533374915 | 0.369 | 0.106 | 3.84E-20 | 20 | Slc12a7 |
| Eid14 | 1.94E-24 | 0.676193074 | 0.691 | 0.321 | 3.93E-20 | 20 | Eid1 |
| Meox13 | 2.01E-24 | 0.595396994 | 0.537 | 0.187 | 4.07E-20 | 20 | Meox1 |

| | | | | | | | |
|---|---|---|---|---|---|---|---|
| Mall1 | 2.05E-24 | 0.477385746 | 0.255 | 0.057 | 4.15E-20 | 20 | Mall |
| Csf14 | 2.28E-24 | 0.630123734 | 0.376 | 0.11 | 4.61E-20 | 20 | Csf1 |
| F11r7 | 3.04E-24 | 0.542611729 | 0.671 | 0.266 | 6.16E-20 | 20 | F11r |
| Dennd5b5 | 3.21E-24 | 0.686488112 | 0.45 | 0.147 | 6.51E-20 | 20 | Dennd5b |
| Sh3bgrl21 | 3.44E-24 | 0.569862878 | 0.295 | 0.073 | 6.96E-20 | 20 | Sh3bgrl2 |
| Tcaf14 | 3.46E-24 | 0.557566282 | 0.362 | 0.103 | 7.01E-20 | 20 | Tcaf1 |
| Rbfox26 | 3.67E-24 | 0.628447457 | 0.537 | 0.191 | 7.43E-20 | 20 | Rbfox2 |
| mt-Co216 | 3.89E-24 | 0.74485318 | 0.993 | 0.729 | 7.88E-20 | 20 | mt-Co2 |
| Hmcn13 | 4.01E-24 | 0.822940124 | 0.564 | 0.211 | 8.12E-20 | 20 | Hmcn1 |
| Stxbp11 | 4.11E-24 | 0.522865727 | 0.268 | 0.064 | 8.33E-20 | 20 | Stxbp1 |
| Pros15 | 5.20E-24 | 0.633204839 | 0.45 | 0.15 | 1.05E-19 | 20 | Pros1 |
| Klf74 | 5.74E-24 | 0.795735451 | 0.685 | 0.298 | 1.16E-19 | 20 | Klf7 |
| Slc16a91 | 6.41E-24 | 0.579395288 | 0.255 | 0.059 | 1.30E-19 | 20 | Slc16a9 |
| Mgst15 | 7.69E-24 | 0.62711952 | 0.49 | 0.167 | 1.56E-19 | 20 | Mgst1 |
| Mospd2 | 8.76E-24 | 0.581210408 | 0.362 | 0.107 | 1.77E-19 | 20 | Mospd2 |
| Dip2c6 | 9.83E-24 | 0.768678274 | 0.577 | 0.223 | 1.99E-19 | 20 | Dip2c |
| Dock65 | 1.01E-23 | 0.617839612 | 0.436 | 0.14 | 2.04E-19 | 20 | Dock6 |
| Myo1d4 | 1.07E-23 | 0.678478505 | 0.362 | 0.107 | 2.17E-19 | 20 | Myo1d |
| Hjurp2 | 1.08E-23 | 0.534591296 | 0.329 | 0.091 | 2.18E-19 | 20 | Hjurp |
| Mid21 | 1.61E-23 | 0.686472909 | 0.329 | 0.092 | 3.26E-19 | 20 | Mid2 |
| Ppp3ca12 | 1.70E-23 | 0.922241858 | 0.779 | 0.393 | 3.44E-19 | 20 | Ppp3ca |
| Ubn29 | 2.00E-23 | 0.740539397 | 0.597 | 0.245 | 4.05E-19 | 20 | Ubn2 |
| Hdgfl35 | 2.15E-23 | 0.640856835 | 0.369 | 0.111 | 4.35E-19 | 20 | Hdgfl3 |
| Slfn28 | 2.20E-23 | 0.69210085 | 0.617 | 0.261 | 4.46E-19 | 20 | Slfn2 |
| Tacc17 | 2.72E-23 | 0.786599426 | 0.671 | 0.305 | 5.50E-19 | 20 | Tacc1 |
| Tes6 | 2.79E-23 | 0.568057506 | 0.403 | 0.13 | 5.65E-19 | 20 | Tes |
| Lama46 | 3.10E-23 | 0.748314247 | 0.49 | 0.175 | 6.27E-19 | 20 | Lama4 |
| Qk5 | 3.14E-23 | 0.754862563 | 0.792 | 0.41 | 6.35E-19 | 20 | Qk |
| Cd2ap3 | 3.16E-23 | 0.549122465 | 0.732 | 0.327 | 6.40E-19 | 20 | Cd2ap |
| Spag97 | 3.38E-23 | 0.768366571 | 0.738 | 0.359 | 6.84E-19 | 20 | Spag9 |
| Hspb16 | 4.42E-23 | 0.786544043 | 0.832 | 0.381 | 8.95E-19 | 20 | Hspb1 |
| Tcn22 | 4.50E-23 | 0.677421504 | 0.497 | 0.18 | 9.10E-19 | 20 | Tcn2 |
| Stim24 | 5.80E-23 | 0.762197852 | 0.517 | 0.2 | 1.17E-18 | 20 | Stim2 |
| Igf11 | 6.06E-23 | 0.67796268 | 0.309 | 0.083 | 1.23E-18 | 20 | Igf1 |
| Zhx35 | 8.20E-23 | 0.699681162 | 0.537 | 0.202 | 1.66E-18 | 20 | Zhx3 |
| Sh3d193 | 8.84E-23 | 0.618534373 | 0.362 | 0.108 | 1.79E-18 | 20 | Sh3d19 |
| Ece15 | 9.82E-23 | 0.665920398 | 0.725 | 0.306 | 1.99E-18 | 20 | Ece1 |
| Xdh4 | 1.25E-22 | 0.579749572 | 0.523 | 0.188 | 2.53E-18 | 20 | Xdh |
| Mpdz3 | 1.48E-22 | 0.513775914 | 0.342 | 0.098 | 2.99E-18 | 20 | Mpdz |
| Rb1cc16 | 1.56E-22 | 0.757081353 | 0.564 | 0.235 | 3.17E-18 | 20 | Rb1cc1 |
| Phldb24 | 2.56E-22 | 0.687722416 | 0.537 | 0.203 | 5.17E-18 | 20 | Phldb2 |
| Nedd4l6 | 3.03E-22 | 0.782994775 | 0.55 | 0.217 | 6.14E-18 | 20 | Nedd4l |
| Tmtc14 | 3.14E-22 | 0.716564263 | 0.423 | 0.142 | 6.36E-18 | 20 | Tmtc1 |

| | | | | | | | |
|---|---|---|---|---|---|---|---|
| Tmem176a10 | 3.75E-22 | 0.743710873 | 0.819 | 0.418 | 7.58E-18 | 20 | Tmem176a |
| Ankrd33b3 | 4.18E-22 | 0.514162637 | 0.309 | 0.083 | 8.47E-18 | 20 | Ankrd33b |
| Trim474 | 5.03E-22 | 0.535822652 | 0.43 | 0.144 | 1.02E-17 | 20 | Trim47 |
| Reep36 | 6.62E-22 | 0.790324313 | 0.651 | 0.306 | 1.34E-17 | 20 | Reep3 |
| Luzp15 | 7.14E-22 | 0.669009817 | 0.752 | 0.357 | 1.45E-17 | 20 | Luzp1 |
| Adam105 | 7.59E-22 | 0.752300772 | 0.638 | 0.293 | 1.54E-17 | 20 | Adam10 |
| Fabp45 | 8.56E-22 | 0.782092988 | 0.852 | 0.436 | 1.73E-17 | 20 | Fabp4 |
| Fzd45 | 1.05E-21 | 0.61396505 | 0.591 | 0.23 | 2.12E-17 | 20 | Fzd4 |
| Taok16 | 1.05E-21 | 0.692715582 | 0.584 | 0.252 | 2.13E-17 | 20 | Taok1 |
| Kif13a3 | 1.07E-21 | 0.727728521 | 0.477 | 0.174 | 2.16E-17 | 20 | Kif13a |
| Nod1 | 1.10E-21 | 0.694003141 | 0.336 | 0.103 | 2.23E-17 | 20 | Nod1 |
| Hmgn32 | 1.12E-21 | 0.375974371 | 0.49 | 0.171 | 2.27E-17 | 20 | Hmgn3 |
| Cd1515 | 1.21E-21 | 0.653758186 | 0.55 | 0.217 | 2.45E-17 | 20 | Cd151 |
| Cds25 | 1.25E-21 | 0.569222686 | 0.483 | 0.177 | 2.53E-17 | 20 | Cds2 |
| Mafb3 | 1.80E-21 | 0.764269995 | 0.443 | 0.155 | 3.64E-17 | 20 | Mafb |
| Tnfrsf1b6 | 1.85E-21 | 0.636280427 | 0.362 | 0.117 | 3.75E-17 | 20 | Tnfrsf1b |
| Emp311 | 1.97E-21 | 0.723259593 | 0.591 | 0.259 | 4.00E-17 | 20 | Emp3 |
| Dach13 | 2.41E-21 | 0.876774227 | 0.342 | 0.104 | 4.88E-17 | 20 | Dach1 |
| Phc23 | 2.57E-21 | 0.673183592 | 0.456 | 0.17 | 5.20E-17 | 20 | Phc2 |
| Malat112 | 3.56E-21 | 0.743771035 | 1 | 0.726 | 7.21E-17 | 20 | Malat1 |
| Il13ra11 | 3.65E-21 | 0.506096224 | 0.282 | 0.076 | 7.39E-17 | 20 | Il13ra1 |
| 2900026A02Rik6 | 3.71E-21 | 0.712449465 | 0.537 | 0.22 | 7.51E-17 | 20 | 2900026A02Rik |
| Smad11 | 3.97E-21 | 0.613007798 | 0.396 | 0.133 | 8.05E-17 | 20 | Smad1 |
| Pitpnm22 | 4.31E-21 | 0.570592433 | 0.383 | 0.123 | 8.72E-17 | 20 | Pitpnm2 |
| Bcam6 | 4.33E-21 | 0.588058853 | 0.638 | 0.254 | 8.76E-17 | 20 | Bcam |
| Filip15 | 4.33E-21 | 0.648710224 | 0.523 | 0.195 | 8.76E-17 | 20 | Filip1 |
| Irak3 | 5.01E-21 | 0.630206232 | 0.262 | 0.068 | 1.01E-16 | 20 | Irak3 |
| Arntl2 | 5.91E-21 | 0.673262963 | 0.362 | 0.119 | 1.20E-16 | 20 | Arntl |
| Sbno21 | 6.32E-21 | 0.536797441 | 0.255 | 0.066 | 1.28E-16 | 20 | Sbno2 |
| Kank34 | 6.70E-21 | 0.62120621 | 0.57 | 0.228 | 1.36E-16 | 20 | Kank3 |
| Lmbr1 | 7.06E-21 | 0.48834409 | 0.255 | 0.064 | 1.43E-16 | 20 | Lmbr1 |
| Slfn55 | 7.26E-21 | 0.654119576 | 0.772 | 0.357 | 1.47E-16 | 20 | Slfn5 |
| Nos1ap | 7.49E-21 | 0.567584161 | 0.275 | 0.074 | 1.52E-16 | 20 | Nos1ap |
| Zfp36l18 | 7.90E-21 | 0.817035144 | 0.779 | 0.433 | 1.60E-16 | 20 | Zfp36l1 |
| Hip17 | 8.76E-21 | 0.698433946 | 0.691 | 0.335 | 1.77E-16 | 20 | Hip1 |
| Ifitm27 | 9.46E-21 | 0.610269873 | 0.953 | 0.464 | 1.91E-16 | 20 | Ifitm2 |
| Il31ra1 | 1.04E-20 | 0.476817193 | 0.523 | 0.209 | 2.11E-16 | 20 | Il31ra |
| Grk56 | 1.14E-20 | 0.779294249 | 0.436 | 0.158 | 2.30E-16 | 20 | Grk5 |
| Npdc14 | 1.20E-20 | 0.567457553 | 0.396 | 0.134 | 2.44E-16 | 20 | Npdc1 |
| Ptpn124 | 1.41E-20 | 0.679151493 | 0.523 | 0.213 | 2.86E-16 | 20 | Ptpn12 |
| Slc3a28 | 1.58E-20 | 0.668475137 | 0.624 | 0.291 | 3.20E-16 | 20 | Slc3a2 |
| Dhrs34 | 1.63E-20 | 0.652545661 | 0.564 | 0.232 | 3.29E-16 | 20 | Dhrs3 |
| Actn45 | 1.66E-20 | 0.63545857 | 0.758 | 0.376 | 3.35E-16 | 20 | Actn4 |

| | | | | | | | |
|---|---|---|---|---|---|---|---|
| Gm205321 | 3.19E-20 | 0.523169346 | 0.356 | 0.11 | 6.47E-16 | 20 | Gm20532 |
| Ushbp14 | 3.24E-20 | 0.551118921 | 0.564 | 0.219 | 6.56E-16 | 20 | Ushbp1 |
| Dysf3 | 3.42E-20 | 0.651233714 | 0.597 | 0.239 | 6.91E-16 | 20 | Dysf |
| Cmip8 | 3.77E-20 | 0.721723533 | 0.631 | 0.286 | 7.64E-16 | 20 | Cmip |
| Sema3f1 | 4.45E-20 | 0.46453874 | 0.322 | 0.095 | 9.00E-16 | 20 | Sema3f |
| Maml212 | 4.82E-20 | 0.801187281 | 0.705 | 0.343 | 9.76E-16 | 20 | Maml2 |
| Slc43a34 | 4.91E-20 | 0.801415439 | 0.624 | 0.27 | 9.94E-16 | 20 | Slc43a3 |
| Camk2d9 | 5.01E-20 | 0.766533605 | 0.604 | 0.27 | 1.01E-15 | 20 | Camk2d |
| Tcf1210 | 5.04E-20 | 0.775593949 | 0.738 | 0.377 | 1.02E-15 | 20 | Tcf12 |
| Adamts102 | 5.31E-20 | 0.633309458 | 0.322 | 0.101 | 1.08E-15 | 20 | Adamts10 |
| Tspan43 | 6.02E-20 | 0.506830181 | 0.342 | 0.106 | 1.22E-15 | 20 | Tspan4 |
| Fam174b1 | 6.12E-20 | 0.543770398 | 0.336 | 0.104 | 1.24E-15 | 20 | Fam174b |
| Scarf11 | 8.22E-20 | 0.422570149 | 0.268 | 0.071 | 1.66E-15 | 20 | Scarf1 |
| Syne25 | 9.04E-20 | 0.674096116 | 0.631 | 0.272 | 1.83E-15 | 20 | Syne2 |
| Notch15 | 9.36E-20 | 0.611353718 | 0.537 | 0.217 | 1.90E-15 | 20 | Notch1 |
| Golgb16 | 9.91E-20 | 0.675103691 | 0.57 | 0.263 | 2.01E-15 | 20 | Golgb1 |
| Osbpl95 | 1.20E-19 | 0.60810154 | 0.617 | 0.279 | 2.43E-15 | 20 | Osbpl9 |
| H2-T234 | 1.22E-19 | 0.772943755 | 0.819 | 0.46 | 2.46E-15 | 20 | H2-T23 |
| Pkd22 | 1.54E-19 | 0.484525774 | 0.342 | 0.107 | 3.12E-15 | 20 | Pkd2 |
| Pdlim54 | 1.55E-19 | 0.62438241 | 0.631 | 0.286 | 3.14E-15 | 20 | Pdlim5 |
| Yes14 | 1.61E-19 | 0.627758471 | 0.43 | 0.154 | 3.26E-15 | 20 | Yes1 |
| Grb105 | 1.80E-19 | 0.618402361 | 0.503 | 0.195 | 3.64E-15 | 20 | Grb10 |
| Pced1b5 | 1.82E-19 | 0.642172369 | 0.356 | 0.118 | 3.68E-15 | 20 | Pced1b |
| Uaca4 | 2.12E-19 | 0.516678445 | 0.564 | 0.226 | 4.28E-15 | 20 | Uaca |
| Sigirr1 | 2.26E-19 | 0.500855042 | 0.275 | 0.078 | 4.58E-15 | 20 | Sigirr |
| Enah4 | 2.46E-19 | 0.610821966 | 0.423 | 0.15 | 4.97E-15 | 20 | Enah |
| Sgms17 | 2.78E-19 | 0.650493238 | 0.376 | 0.129 | 5.63E-15 | 20 | Sgms1 |
| Exoc3l23 | 2.79E-19 | 0.668326162 | 0.631 | 0.271 | 5.65E-15 | 20 | Exoc3l2 |
| Prkce4 | 2.94E-19 | 0.778860155 | 0.523 | 0.214 | 5.94E-15 | 20 | Prkce |
| Rapgef44 | 3.38E-19 | 0.668569559 | 0.597 | 0.258 | 6.85E-15 | 20 | Rapgef4 |
| Pdgfd4 | 3.82E-19 | 0.553681041 | 0.611 | 0.254 | 7.72E-15 | 20 | Pdgfd |
| Ppp1r12a10 | 3.92E-19 | 0.653292398 | 0.812 | 0.426 | 7.94E-15 | 20 | Ppp1r12a |
| Flt41 | 4.05E-19 | 0.882061467 | 0.329 | 0.105 | 8.21E-15 | 20 | Flt4 |
| Flt15 | 4.24E-19 | 0.592292898 | 0.893 | 0.393 | 8.58E-15 | 20 | Flt1 |
| Pear14 | 4.47E-19 | 0.560236086 | 0.322 | 0.102 | 9.05E-15 | 20 | Pear1 |
| Ly6e6 | 4.67E-19 | 0.694081998 | 0.94 | 0.62 | 9.45E-15 | 20 | Ly6e |
| Far15 | 4.82E-19 | 0.552860674 | 0.436 | 0.165 | 9.75E-15 | 20 | Far1 |
| Rbms15 | 5.30E-19 | 0.633213523 | 0.886 | 0.486 | 1.07E-14 | 20 | Rbms1 |
| Rassf35 | 5.33E-19 | 0.513031091 | 0.564 | 0.235 | 1.08E-14 | 20 | Rassf3 |
| Galnt102 | 5.37E-19 | 0.498836823 | 0.329 | 0.105 | 1.09E-14 | 20 | Galnt10 |
| Pde8a3 | 5.41E-19 | 0.604164415 | 0.477 | 0.183 | 1.10E-14 | 20 | Pde8a |
| Mpp73 | 6.10E-19 | 0.731163927 | 0.329 | 0.107 | 1.23E-14 | 20 | Mpp7 |
| Dab2ip6 | 7.74E-19 | 0.615564042 | 0.604 | 0.262 | 1.57E-14 | 20 | Dab2ip |

| | | | | | | | |
|---|---|---|---|---|---|---|---|
| Gm268831 | 9.22E-19 | 0.517430844 | 0.282 | 0.081 | 1.87E-14 | 20 | Gm26883 |
| C130074G19Rik4 | 1.03E-18 | 0.469805234 | 0.51 | 0.2 | 2.08E-14 | 20 | C130074G19Rik |
| Gab13 | 1.11E-18 | 0.570635869 | 0.349 | 0.115 | 2.26E-14 | 20 | Gab1 |
| Pcmtd13 | 1.13E-18 | 0.508589254 | 0.671 | 0.327 | 2.30E-14 | 20 | Pcmtd1 |
| Mob22 | 1.21E-18 | 0.50309903 | 0.403 | 0.148 | 2.45E-14 | 20 | Mob2 |
| Plod31 | 1.26E-18 | 0.366657638 | 0.255 | 0.069 | 2.55E-14 | 20 | Plod3 |
| Ifi443 | 1.28E-18 | 0.736024078 | 0.409 | 0.146 | 2.60E-14 | 20 | Ifi44 |
| Laptm4a6 | 1.46E-18 | 0.622623367 | 0.852 | 0.484 | 2.96E-14 | 20 | Laptm4a |
| Ets23 | 2.37E-18 | 0.495214351 | 0.497 | 0.194 | 4.80E-14 | 20 | Ets2 |
| Ctsd7 | 2.51E-18 | 0.581660681 | 0.664 | 0.314 | 5.09E-14 | 20 | Ctsd |
| Csgalnact15 | 2.70E-18 | 0.630537353 | 0.497 | 0.196 | 5.46E-14 | 20 | Csgalnact1 |
| Cadps21 | 2.79E-18 | 0.378644586 | 0.255 | 0.068 | 5.65E-14 | 20 | Cadps2 |
| Fam69a7 | 3.10E-18 | 0.64556642 | 0.362 | 0.127 | 6.27E-14 | 20 | Fam69a |
| Igf1r6 | 3.19E-18 | 0.768938916 | 0.416 | 0.161 | 6.46E-14 | 20 | Igf1r |
| Sh3bgrl7 | 3.19E-18 | 0.513453061 | 0.738 | 0.375 | 6.47E-14 | 20 | Sh3bgrl |
| Fndc3b6 | 3.80E-18 | 0.616388992 | 0.557 | 0.245 | 7.69E-14 | 20 | Fndc3b |
| Vps13c4 | 4.41E-18 | 0.563727616 | 0.383 | 0.14 | 8.93E-14 | 20 | Vps13c |
| Ktn16 | 5.05E-18 | 0.595490699 | 0.644 | 0.308 | 1.02E-13 | 20 | Ktn1 |
| Nek78 | 5.36E-18 | 0.585894995 | 0.55 | 0.249 | 1.08E-13 | 20 | Nek7 |
| mt-Nd114 | 6.94E-18 | 0.622958066 | 0.953 | 0.599 | 1.41E-13 | 20 | mt-Nd1 |
| Lipe3 | 7.74E-18 | 0.587409238 | 0.383 | 0.138 | 1.57E-13 | 20 | Lipe |
| Wdfy37 | 8.00E-18 | 0.571718667 | 0.463 | 0.182 | 1.62E-13 | 20 | Wdfy3 |
| Large17 | 8.16E-18 | 0.611021239 | 0.409 | 0.153 | 1.65E-13 | 20 | Large1 |
| Tead13 | 8.33E-18 | 0.515522471 | 0.282 | 0.084 | 1.69E-13 | 20 | Tead1 |
| Anxa27 | 8.85E-18 | 0.63611652 | 0.772 | 0.417 | 1.79E-13 | 20 | Anxa2 |
| Cyfip16 | 9.10E-18 | 0.619970944 | 0.604 | 0.279 | 1.84E-13 | 20 | Cyfip1 |
| Atp2b45 | 9.34E-18 | 0.602439852 | 0.483 | 0.199 | 1.89E-13 | 20 | Atp2b4 |
| Arid1b10 | 9.76E-18 | 0.671439677 | 0.557 | 0.259 | 1.98E-13 | 20 | Arid1b |
| Epas16 | 1.20E-17 | 0.404656123 | 0.792 | 0.36 | 2.43E-13 | 20 | Epas1 |
| Atp8a15 | 1.31E-17 | 0.565282975 | 0.557 | 0.242 | 2.66E-13 | 20 | Atp8a1 |
| Marcks6 | 1.35E-17 | 0.574137063 | 0.725 | 0.342 | 2.72E-13 | 20 | Marcks |
| Hif1a2 | 1.49E-17 | 0.57749945 | 0.477 | 0.2 | 3.02E-13 | 20 | Hif1a |
| Leprot3 | 1.67E-17 | 0.54863607 | 0.45 | 0.178 | 3.39E-13 | 20 | Leprot |
| Lmna7 | 1.83E-17 | 0.501079871 | 0.49 | 0.198 | 3.70E-13 | 20 | Lmna |
| Gjc14 | 1.83E-17 | 0.532527934 | 0.342 | 0.115 | 3.70E-13 | 20 | Gjc1 |
| Map1b2 | 2.09E-17 | 0.483803767 | 0.322 | 0.102 | 4.24E-13 | 20 | Map1b |
| Atrx4 | 2.38E-17 | 0.601904927 | 0.779 | 0.42 | 4.82E-13 | 20 | Atrx |
| Fam208a4 | 2.64E-17 | 0.638008489 | 0.477 | 0.2 | 5.34E-13 | 20 | Fam208a |
| Ano65 | 2.72E-17 | 0.586742481 | 0.443 | 0.181 | 5.50E-13 | 20 | Ano6 |
| Rsu16 | 3.16E-17 | 0.592591819 | 0.55 | 0.254 | 6.40E-13 | 20 | Rsu1 |
| Osmr5 | 3.46E-17 | 0.509874024 | 0.389 | 0.14 | 7.01E-13 | 20 | Osmr |
| Smarca25 | 3.74E-17 | 0.604549894 | 0.604 | 0.281 | 7.58E-13 | 20 | Smarca2 |
| Atrnl16 | 4.20E-17 | 0.620838682 | 0.43 | 0.166 | 8.49E-13 | 20 | Atrnl1 |

| | | | | | | | |
|---|---|---|---|---|---|---|---|
| Arl4a | 4.25E-17 | 0.52524142 | 0.403 | 0.153 | 8.61E-13 | 20 | Arl4a |
| Gcnt25 | 4.48E-17 | 0.746457291 | 0.389 | 0.143 | 9.07E-13 | 20 | Gcnt2 |
| Ehbp11 | 4.92E-17 | 0.58127617 | 0.322 | 0.111 | 9.95E-13 | 20 | Ehbp1 |
| Tmem1648 | 5.09E-17 | 0.682126818 | 0.409 | 0.164 | 1.03E-12 | 20 | Tmem164 |
| Itga64 | 5.14E-17 | 0.489325694 | 0.631 | 0.284 | 1.04E-12 | 20 | Itga6 |
| Atp2b19 | 5.20E-17 | 0.647226804 | 0.651 | 0.333 | 1.05E-12 | 20 | Atp2b1 |
| Icam24 | 5.65E-17 | 0.37849989 | 0.564 | 0.23 | 1.14E-12 | 20 | Icam2 |
| Fmnl25 | 6.49E-17 | 0.834236515 | 0.53 | 0.243 | 1.31E-12 | 20 | Fmnl2 |
| Ralgapa15 | 7.11E-17 | 0.64319564 | 0.617 | 0.304 | 1.44E-12 | 20 | Ralgapa1 |
| Ebf34 | 7.35E-17 | 0.694857344 | 0.356 | 0.127 | 1.49E-12 | 20 | Ebf3 |
| Cav25 | 8.65E-17 | 0.459700062 | 0.517 | 0.21 | 1.75E-12 | 20 | Cav2 |
| Xiap4 | 8.69E-17 | 0.611730707 | 0.611 | 0.3 | 1.76E-12 | 20 | Xiap |
| Nav15 | 8.75E-17 | 0.489158551 | 0.537 | 0.223 | 1.77E-12 | 20 | Nav1 |
| Rab11a5 | 9.10E-17 | 0.551380742 | 0.624 | 0.309 | 1.84E-12 | 20 | Rab11a |
| Fcho25 | 9.12E-17 | 0.610854538 | 0.617 | 0.31 | 1.85E-12 | 20 | Fcho2 |
| Cbfa2t35 | 9.74E-17 | 0.609579771 | 0.376 | 0.139 | 1.97E-12 | 20 | Cbfa2t3 |
| Arhgef154 | 1.08E-16 | 0.521436509 | 0.436 | 0.169 | 2.18E-12 | 20 | Arhgef15 |
| Rtn36 | 1.19E-16 | 0.54978297 | 0.651 | 0.324 | 2.42E-12 | 20 | Rtn3 |
| Zcchc143 | 1.29E-16 | 0.390946909 | 0.268 | 0.08 | 2.61E-12 | 20 | Zcchc14 |
| Ccm2l3 | 1.47E-16 | 0.459357907 | 0.295 | 0.093 | 2.98E-12 | 20 | Ccm2l |
| Arid5b7 | 1.53E-16 | 0.598435241 | 0.591 | 0.278 | 3.09E-12 | 20 | Arid5b |
| Cdc42bpb4 | 2.15E-16 | 0.5154612 | 0.45 | 0.187 | 4.34E-12 | 20 | Cdc42bpb |
| Tm4sf16 | 2.23E-16 | 0.347784459 | 0.826 | 0.378 | 4.51E-12 | 20 | Tm4sf1 |
| Tmod37 | 2.24E-16 | 0.570153546 | 0.685 | 0.362 | 4.53E-12 | 20 | Tmod3 |
| Gpr1463 | 2.30E-16 | 0.388518414 | 0.329 | 0.11 | 4.66E-12 | 20 | Gpr146 |
| Kansl111 | 2.32E-16 | 0.686792372 | 0.658 | 0.341 | 4.71E-12 | 20 | Kansl1 |
| Fbxo118 | 2.39E-16 | 0.52637576 | 0.523 | 0.233 | 4.85E-12 | 20 | Fbxo11 |
| Pcdh174 | 2.41E-16 | 0.481056222 | 0.577 | 0.24 | 4.88E-12 | 20 | Pcdh17 |
| Arhgef72 | 2.51E-16 | 0.527176603 | 0.396 | 0.154 | 5.09E-12 | 20 | Arhgef7 |
| Rrbp16 | 2.90E-16 | 0.508095801 | 0.819 | 0.46 | 5.87E-12 | 20 | Rrbp1 |
| Ganab1 | 2.94E-16 | 0.441790061 | 0.329 | 0.116 | 5.95E-12 | 20 | Ganab |
| Msn7 | 3.04E-16 | 0.532665179 | 0.799 | 0.444 | 6.16E-12 | 20 | Msn |
| Crim14 | 3.13E-16 | 0.491160771 | 0.752 | 0.373 | 6.34E-12 | 20 | Crim1 |
| Pxn1 | 3.34E-16 | 0.540917969 | 0.409 | 0.165 | 6.76E-12 | 20 | Pxn |
| Kitl5 | 3.41E-16 | 0.477525436 | 0.718 | 0.336 | 6.91E-12 | 20 | Kitl |
| Gtf2i3 | 3.50E-16 | 0.488283531 | 0.557 | 0.251 | 7.08E-12 | 20 | Gtf2i |
| Timp35 | 3.52E-16 | 0.448771522 | 0.866 | 0.385 | 7.13E-12 | 20 | Timp3 |
| Gnai25 | 3.65E-16 | 0.617898492 | 0.893 | 0.575 | 7.39E-12 | 20 | Gnai2 |
| Ipo111 | 3.67E-16 | 0.53294427 | 0.369 | 0.137 | 7.43E-12 | 20 | Ipo11 |
| Vezt | 4.28E-16 | 0.502666513 | 0.255 | 0.078 | 8.67E-12 | 20 | Vezt |
| Cald16 | 4.36E-16 | 0.487183492 | 0.711 | 0.323 | 8.82E-12 | 20 | Cald1 |
| Plk24 | 4.54E-16 | 0.598538687 | 0.43 | 0.167 | 9.20E-12 | 20 | Plk2 |
| Ppp2r3a2 | 5.14E-16 | 0.581085899 | 0.282 | 0.092 | 1.04E-11 | 20 | Ppp2r3a |

| | | | | | | | |
|---|---|---|---|---|---|---|---|
| Ext18 | 5.21E-16 | 0.712153038 | 0.416 | 0.167 | 1.05E-11 | 20 | Ext1 |
| Nckap16 | 5.27E-16 | 0.407807796 | 0.53 | 0.227 | 1.07E-11 | 20 | Nckap1 |
| Zeb110 | 5.28E-16 | 0.697664654 | 0.638 | 0.322 | 1.07E-11 | 20 | Zeb1 |
| Robo44 | 5.55E-16 | 0.566466945 | 0.443 | 0.177 | 1.12E-11 | 20 | Robo4 |
| Tspan136 | 5.96E-16 | 0.589599089 | 0.597 | 0.286 | 1.21E-11 | 20 | Tspan13 |
| Canx8 | 6.53E-16 | 0.595219835 | 0.604 | 0.307 | 1.32E-11 | 20 | Canx |
| Plscr45 | 6.60E-16 | 0.611201941 | 0.456 | 0.19 | 1.34E-11 | 20 | Plscr4 |
| Fam13c4 | 6.92E-16 | 0.467282192 | 0.329 | 0.112 | 1.40E-11 | 20 | Fam13c |
| Gm4728310 | 7.03E-16 | 0.737755573 | 0.577 | 0.282 | 1.42E-11 | 20 | Gm47283 |
| Zfp9502 | 7.17E-16 | 0.585684399 | 0.315 | 0.112 | 1.45E-11 | 20 | Zfp950 |
| Trove25 | 7.62E-16 | 0.421150249 | 0.362 | 0.132 | 1.54E-11 | 20 | Trove2 |
| St3gal44 | 7.72E-16 | 0.898553478 | 0.523 | 0.257 | 1.56E-11 | 20 | St3gal4 |
| Gng123 | 8.41E-16 | 0.403489753 | 0.483 | 0.201 | 1.70E-11 | 20 | Gng12 |
| Zfp36l27 | 8.60E-16 | 0.612913109 | 0.604 | 0.306 | 1.74E-11 | 20 | Zfp36l2 |
| Prrc2c11 | 9.75E-16 | 0.641732237 | 0.758 | 0.421 | 1.97E-11 | 20 | Prrc2c |
| Gm138893 | 1.05E-15 | 0.54309694 | 0.51 | 0.22 | 2.12E-11 | 20 | Gm13889 |
| Glg16 | 1.30E-15 | 0.619232023 | 0.537 | 0.253 | 2.64E-11 | 20 | Glg1 |
| Anxa34 | 1.34E-15 | 0.486394806 | 0.691 | 0.339 | 2.72E-11 | 20 | Anxa3 |
| Tulp42 | 1.41E-15 | 0.508813137 | 0.443 | 0.188 | 2.85E-11 | 20 | Tulp4 |
| Fkbp93 | 1.57E-15 | 0.453880169 | 0.322 | 0.112 | 3.17E-11 | 20 | Fkbp9 |
| Phip11 | 1.58E-15 | 0.657131823 | 0.537 | 0.257 | 3.21E-11 | 20 | Phip |
| Mgat4b | 1.83E-15 | 0.351866886 | 0.282 | 0.091 | 3.70E-11 | 20 | Mgat4b |
| Sik39 | 2.10E-15 | 0.676102129 | 0.55 | 0.271 | 4.26E-11 | 20 | Sik3 |
| Kmt2e9 | 2.31E-15 | 0.569987711 | 0.745 | 0.407 | 4.68E-11 | 20 | Kmt2e |
| Mtss13 | 2.33E-15 | 0.50669828 | 0.295 | 0.1 | 4.73E-11 | 20 | Mtss1 |
| Camk1d2 | 2.59E-15 | 0.715903196 | 0.98 | 0.823 | 5.25E-11 | 20 | Camk1d |
| Ldlrad48 | 2.60E-15 | 0.396414682 | 0.564 | 0.251 | 5.26E-11 | 20 | Ldlrad4 |
| Ctsb6 | 2.86E-15 | 0.298230092 | 0.624 | 0.299 | 5.79E-11 | 20 | Ctsb |
| mt-Nd312 | 2.87E-15 | 0.607367861 | 0.758 | 0.447 | 5.81E-11 | 20 | mt-Nd3 |
| Mfge86 | 3.46E-15 | 0.46415872 | 0.564 | 0.254 | 7.00E-11 | 20 | Mfge8 |
| Prrc2b7 | 4.18E-15 | 0.465179182 | 0.51 | 0.23 | 8.46E-11 | 20 | Prrc2b |
| Fmnl35 | 4.28E-15 | 0.601422327 | 0.517 | 0.24 | 8.66E-11 | 20 | Fmnl3 |
| Nxn4 | 4.30E-15 | 0.418040551 | 0.591 | 0.269 | 8.70E-11 | 20 | Nxn |
| Stag111 | 4.49E-15 | 0.501811532 | 0.705 | 0.362 | 9.08E-11 | 20 | Stag1 |
| Ccnyl13 | 4.52E-15 | 0.507053156 | 0.315 | 0.111 | 9.15E-11 | 20 | Ccnyl1 |
| Fryl8 | 4.61E-15 | 0.520013215 | 0.564 | 0.269 | 9.33E-11 | 20 | Fryl |
| Tra2a12 | 4.65E-15 | 0.5790686 | 0.557 | 0.275 | 9.42E-11 | 20 | Tra2a |
| Itgb110 | 4.81E-15 | 0.485881705 | 0.852 | 0.479 | 9.74E-11 | 20 | Itgb1 |
| Vamp54 | 4.93E-15 | 0.460242237 | 0.758 | 0.385 | 9.98E-11 | 20 | Vamp5 |
| Apc6 | 5.22E-15 | 0.607344008 | 0.517 | 0.249 | 1.06E-10 | 20 | Apc |
| Ripply33 | 5.33E-15 | 0.525031142 | 0.383 | 0.147 | 1.08E-10 | 20 | Ripply3 |
| Fnip23 | 5.50E-15 | 0.511691869 | 0.309 | 0.107 | 1.11E-10 | 20 | Fnip2 |
| Tmem641 | 5.54E-15 | 0.367597663 | 0.255 | 0.079 | 1.12E-10 | 20 | Tmem64 |

| | | | | | | | |
|---|---|---|---|---|---|---|---|
| Card102 | 6.66E-15 | 0.415412948 | 0.268 | 0.086 | 1.35E-10 | 20 | Card10 |
| Atp1a12 | 6.82E-15 | 0.472062168 | 0.564 | 0.28 | 1.38E-10 | 20 | Atp1a1 |
| Gbp44 | 7.36E-15 | 0.635072769 | 0.423 | 0.178 | 1.49E-10 | 20 | Gbp4 |
| Gm383992 | 7.84E-15 | 0.543629605 | 0.309 | 0.108 | 1.59E-10 | 20 | Gm38399 |
| Lrp52 | 8.24E-15 | 0.43956126 | 0.282 | 0.094 | 1.67E-10 | 20 | Lrp5 |
| Kmt2a11 | 8.88E-15 | 0.591858571 | 0.544 | 0.268 | 1.80E-10 | 20 | Kmt2a |
| Arhgef101 | 9.04E-15 | 0.385429797 | 0.295 | 0.1 | 1.83E-10 | 20 | Arhgef10 |
| Phactr44 | 9.61E-15 | 0.523051881 | 0.423 | 0.179 | 1.95E-10 | 20 | Phactr4 |
| Lims24 | 1.01E-14 | 0.354416726 | 0.497 | 0.202 | 2.05E-10 | 20 | Lims2 |
| Gtdc14 | 1.11E-14 | 0.603880201 | 0.376 | 0.152 | 2.24E-10 | 20 | Gtdc1 |
| Lrrc8c6 | 1.11E-14 | 0.678493953 | 0.604 | 0.312 | 2.25E-10 | 20 | Lrrc8c |
| Wars4 | 1.16E-14 | 0.446521888 | 0.416 | 0.169 | 2.34E-10 | 20 | Wars |
| Parvb3 | 1.25E-14 | 0.417409171 | 0.342 | 0.124 | 2.52E-10 | 20 | Parvb |
| Sptan15 | 1.32E-14 | 0.454197178 | 0.678 | 0.344 | 2.67E-10 | 20 | Sptan1 |
| Myo63 | 1.42E-14 | 0.501208369 | 0.604 | 0.293 | 2.87E-10 | 20 | Myo6 |
| Afdn4 | 1.47E-14 | 0.458780848 | 0.644 | 0.315 | 2.98E-10 | 20 | Afdn |
| Nid14 | 1.50E-14 | 0.385412525 | 0.45 | 0.182 | 3.03E-10 | 20 | Nid1 |
| Gm269173 | 1.51E-14 | 0.693873917 | 0.315 | 0.121 | 3.06E-10 | 20 | Gm26917 |
| Cask6 | 1.61E-14 | 0.507399853 | 0.362 | 0.14 | 3.27E-10 | 20 | Cask |
| Mfsd63 | 1.65E-14 | 0.54631043 | 0.369 | 0.145 | 3.34E-10 | 20 | Mfsd6 |
| Arhgef127 | 1.71E-14 | 0.433342568 | 0.718 | 0.349 | 3.47E-10 | 20 | Arhgef12 |
| Nck15 | 1.80E-14 | 0.510330482 | 0.651 | 0.341 | 3.64E-10 | 20 | Nck1 |
| Crybg33 | 1.90E-14 | 0.433838617 | 0.362 | 0.14 | 3.85E-10 | 20 | Crybg3 |
| Akap910 | 1.92E-14 | 0.64368962 | 0.49 | 0.238 | 3.88E-10 | 20 | Akap9 |
| Irgm21 | 1.98E-14 | 0.552566609 | 0.336 | 0.128 | 4.00E-10 | 20 | Irgm2 |
| Gpatch811 | 2.74E-14 | 0.526291937 | 0.53 | 0.259 | 5.55E-10 | 20 | Gpatch8 |
| Bclaf19 | 2.80E-14 | 0.52026751 | 0.53 | 0.254 | 5.67E-10 | 20 | Bclaf1 |
| Ephx15 | 3.25E-14 | 0.47138437 | 0.329 | 0.123 | 6.58E-10 | 20 | Ephx1 |
| Neat19 | 3.34E-14 | 0.678569177 | 0.45 | 0.21 | 6.76E-10 | 20 | Neat1 |
| Mfng1 | 3.35E-14 | 0.417091787 | 0.255 | 0.084 | 6.78E-10 | 20 | Mfng |
| Sparc7 | 3.51E-14 | 0.306155545 | 0.859 | 0.403 | 7.11E-10 | 20 | Sparc |
| Lamc15 | 3.52E-14 | 0.468204166 | 0.376 | 0.149 | 7.13E-10 | 20 | Lamc1 |
| Atp2a26 | 3.54E-14 | 0.488072614 | 0.45 | 0.201 | 7.17E-10 | 20 | Atp2a2 |
| Pqlc13 | 3.62E-14 | 0.488327229 | 0.336 | 0.127 | 7.33E-10 | 20 | Pqlc1 |
| Tmem475 | 3.88E-14 | 0.457618446 | 0.396 | 0.157 | 7.85E-10 | 20 | Tmem47 |
| Tmem176b8 | 4.30E-14 | 0.487768399 | 0.779 | 0.439 | 8.70E-10 | 20 | Tmem176b |
| Pggt1b | 4.79E-14 | 0.373688367 | 0.289 | 0.1 | 9.71E-10 | 20 | Pggt1b |
| Exoc6b7 | 5.31E-14 | 0.548818915 | 0.51 | 0.241 | 1.08E-09 | 20 | Exoc6b |
| Smc1a6 | 5.43E-14 | 0.507195002 | 0.604 | 0.313 | 1.10E-09 | 20 | Smc1a |
| Phf21a11 | 5.50E-14 | 0.582437863 | 0.523 | 0.254 | 1.11E-09 | 20 | Phf21a |
| Lrmda7 | 5.55E-14 | 0.748623209 | 0.671 | 0.369 | 1.12E-09 | 20 | Lrmda |
| 9530026P05Rik5 | 5.95E-14 | 0.844584139 | 0.51 | 0.242 | 1.20E-09 | 20 | 9530026P05Rik |
| Id15 | 6.15E-14 | 0.254191772 | 0.705 | 0.325 | 1.24E-09 | 20 | Id1 |

| | | | | | | | |
|---|---|---|---|---|---|---|---|
| Xbp12 | 7.04E-14 | 0.427358165 | 0.443 | 0.197 | 1.43E-09 | 20 | Xbp1 |
| Mprip7 | 7.07E-14 | 0.475681339 | 0.584 | 0.294 | 1.43E-09 | 20 | Mprip |
| 4932438A13Rik9 | 7.12E-14 | 0.535058179 | 0.47 | 0.22 | 1.44E-09 | 20 | 4932438A13Rik |
| Thbd5 | 7.18E-14 | 0.424898128 | 0.564 | 0.255 | 1.45E-09 | 20 | Thbd |
| Gngt25 | 7.24E-14 | 0.428064896 | 0.698 | 0.36 | 1.47E-09 | 20 | Gngt2 |
| Wnk110 | 7.44E-14 | 0.633938324 | 0.671 | 0.364 | 1.51E-09 | 20 | Wnk1 |
| Tnfsf105 | 7.80E-14 | 0.562267739 | 0.705 | 0.375 | 1.58E-09 | 20 | Tnfsf10 |
| Smg111 | 7.93E-14 | 0.557667959 | 0.49 | 0.235 | 1.60E-09 | 20 | Smg1 |
| Ube3a4 | 8.23E-14 | 0.491530962 | 0.584 | 0.296 | 1.67E-09 | 20 | Ube3a |
| Fbxl1713 | 8.87E-14 | 0.660554434 | 0.597 | 0.312 | 1.80E-09 | 20 | Fbxl17 |
| Trio5 | 8.98E-14 | 0.579899818 | 0.423 | 0.189 | 1.82E-09 | 20 | Trio |
| Lamp26 | 9.28E-14 | 0.427564847 | 0.604 | 0.306 | 1.88E-09 | 20 | Lamp2 |
| Fnbp1l2 | 1.03E-13 | 0.277720093 | 0.416 | 0.17 | 2.09E-09 | 20 | Fnbp1l |
| P4hb5 | 1.07E-13 | 0.491988054 | 0.517 | 0.253 | 2.16E-09 | 20 | P4hb |
| Ralb2 | 1.10E-13 | 0.382817087 | 0.383 | 0.154 | 2.22E-09 | 20 | Ralb |
| Bsg7 | 1.13E-13 | 0.39426795 | 0.779 | 0.45 | 2.30E-09 | 20 | Bsg |
| Zfp6085 | 1.17E-13 | 0.526196853 | 0.356 | 0.141 | 2.37E-09 | 20 | Zfp608 |
| Nisch6 | 1.19E-13 | 0.48231189 | 0.47 | 0.212 | 2.41E-09 | 20 | Nisch |
| Rgl13 | 1.32E-13 | 0.344432646 | 0.624 | 0.29 | 2.66E-09 | 20 | Rgl1 |
| Wasf24 | 1.36E-13 | 0.455867673 | 0.752 | 0.426 | 2.75E-09 | 20 | Wasf2 |
| Taok2 | 1.53E-13 | 0.362509411 | 0.289 | 0.103 | 3.10E-09 | 20 | Taok2 |
| Scaper9 | 1.59E-13 | 0.527282507 | 0.409 | 0.18 | 3.22E-09 | 20 | Scaper |
| Strn36 | 1.63E-13 | 0.51939744 | 0.631 | 0.346 | 3.29E-09 | 20 | Strn3 |
| Xist13 | 1.63E-13 | 0.599263293 | 0.819 | 0.464 | 3.29E-09 | 20 | Xist |
| Arhgef116 | 1.78E-13 | 0.538080328 | 0.51 | 0.249 | 3.61E-09 | 20 | Arhgef1 |
| Tmpo5 | 1.79E-13 | 0.460034386 | 0.463 | 0.209 | 3.62E-09 | 20 | Tmpo |
| Ube3c4 | 2.02E-13 | 0.493345315 | 0.376 | 0.16 | 4.09E-09 | 20 | Ube3c |
| Tm6sf17 | 2.11E-13 | 0.569809951 | 0.349 | 0.142 | 4.28E-09 | 20 | Tm6sf1 |
| Ash1l8 | 2.49E-13 | 0.644431458 | 0.57 | 0.3 | 5.04E-09 | 20 | Ash1l |
| Asap24 | 2.67E-13 | 0.405414634 | 0.342 | 0.132 | 5.40E-09 | 20 | Asap2 |
| Dstyk3 | 2.67E-13 | 0.460296047 | 0.369 | 0.15 | 5.41E-09 | 20 | Dstyk |
| Nipbl13 | 2.79E-13 | 0.639682968 | 0.658 | 0.367 | 5.65E-09 | 20 | Nipbl |
| Stat39 | 3.07E-13 | 0.560614742 | 0.517 | 0.268 | 6.21E-09 | 20 | Stat3 |
| Zfyve94 | 3.10E-13 | 0.491392388 | 0.403 | 0.172 | 6.29E-09 | 20 | Zfyve9 |
| Agrn3 | 3.23E-13 | 0.327844816 | 0.255 | 0.084 | 6.55E-09 | 20 | Agrn |
| Ncoa74 | 3.29E-13 | 0.525924836 | 0.584 | 0.297 | 6.66E-09 | 20 | Ncoa7 |
| Ctnnd13 | 3.41E-13 | 0.319067289 | 0.369 | 0.146 | 6.90E-09 | 20 | Ctnnd1 |
| Emp24 | 3.47E-13 | 0.483702846 | 0.51 | 0.236 | 7.02E-09 | 20 | Emp2 |
| Tnrc6c7 | 3.75E-13 | 0.513623813 | 0.591 | 0.316 | 7.59E-09 | 20 | Tnrc6c |
| Ccnd13 | 3.79E-13 | 0.488989239 | 0.497 | 0.227 | 7.67E-09 | 20 | Ccnd1 |
| Zmiz13 | 3.82E-13 | 0.450054554 | 0.617 | 0.321 | 7.72E-09 | 20 | Zmiz1 |
| Itsn210 | 4.23E-13 | 0.579711978 | 0.537 | 0.279 | 8.56E-09 | 20 | Itsn2 |
| Scmh11 | 4.32E-13 | 0.498792783 | 0.342 | 0.139 | 8.74E-09 | 20 | Scmh1 |

| | | | | | | | |
|---|---|---|---|---|---|---|---|
| Tcf7l24 | 4.51E-13 | 0.545884995 | 0.329 | 0.13 | 9.13E-09 | 20 | Tcf7l2 |
| Exoc412 | 4.54E-13 | 0.712930286 | 0.577 | 0.324 | 9.19E-09 | 20 | Exoc4 |
| Ccser24 | 4.54E-13 | 0.442866779 | 0.497 | 0.232 | 9.20E-09 | 20 | Ccser2 |
| Vps13b9 | 4.69E-13 | 0.657842027 | 0.456 | 0.22 | 9.50E-09 | 20 | Vps13b |
| Myo1c3 | 4.92E-13 | 0.428829341 | 0.47 | 0.214 | 9.95E-09 | 20 | Myo1c |
| Crip26 | 5.08E-13 | 0.326990415 | 0.805 | 0.416 | 1.03E-08 | 20 | Crip2 |
| Sorbs15 | 5.30E-13 | 0.438078824 | 0.416 | 0.178 | 1.07E-08 | 20 | Sorbs1 |
| Anxa74 | 5.48E-13 | 0.429341346 | 0.416 | 0.187 | 1.11E-08 | 20 | Anxa7 |
| Usp255 | 5.77E-13 | 0.48452819 | 0.557 | 0.282 | 1.17E-08 | 20 | Usp25 |
| Dennd1a11 | 6.11E-13 | 0.62464672 | 0.51 | 0.254 | 1.24E-08 | 20 | Dennd1a |
| Scarb13 | 6.31E-13 | 0.47992616 | 0.57 | 0.267 | 1.28E-08 | 20 | Scarb1 |
| Smg69 | 6.35E-13 | 0.582521676 | 0.51 | 0.261 | 1.29E-08 | 20 | Smg6 |
| Gimap46 | 6.45E-13 | 0.458417184 | 0.544 | 0.257 | 1.30E-08 | 20 | Gimap4 |
| Klhl2 | 6.51E-13 | 0.412735113 | 0.295 | 0.111 | 1.32E-08 | 20 | Klhl2 |
| Sp1005 | 7.20E-13 | 0.588213657 | 0.752 | 0.433 | 1.46E-08 | 20 | Sp100 |
| Afap1l13 | 7.47E-13 | 0.396615222 | 0.302 | 0.11 | 1.51E-08 | 20 | Afap1l1 |
| Tmem2043 | 7.54E-13 | 0.325744293 | 0.483 | 0.21 | 1.53E-08 | 20 | Tmem204 |
| Etnk14 | 7.64E-13 | 0.480046476 | 0.477 | 0.226 | 1.55E-08 | 20 | Etnk1 |
| Ccdc91 | 7.73E-13 | 0.263253378 | 0.268 | 0.093 | 1.56E-08 | 20 | Ccdc91 |
| Foxn310 | 7.75E-13 | 0.631382904 | 0.705 | 0.412 | 1.57E-08 | 20 | Foxn3 |
| Pds5b5 | 7.96E-13 | 0.529175334 | 0.403 | 0.181 | 1.61E-08 | 20 | Pds5b |
| Setd510 | 8.16E-13 | 0.549009804 | 0.523 | 0.265 | 1.65E-08 | 20 | Setd5 |
| Klhl54 | 8.20E-13 | 0.516979103 | 0.477 | 0.225 | 1.66E-08 | 20 | Klhl5 |
| Msl11 | 8.44E-13 | 0.442203208 | 0.329 | 0.132 | 1.71E-08 | 20 | Msl1 |
| Arid24 | 9.78E-13 | 0.458162942 | 0.369 | 0.157 | 1.98E-08 | 20 | Arid2 |
| Wwox8 | 1.04E-12 | 0.684957682 | 0.396 | 0.176 | 2.11E-08 | 20 | Wwox |
| Tmem30a7 | 1.09E-12 | 0.577544702 | 0.544 | 0.287 | 2.21E-08 | 20 | Tmem30a |
| Smagp3 | 1.17E-12 | 0.33009599 | 0.503 | 0.232 | 2.38E-08 | 20 | Smagp |
| Kdelr12 | 1.19E-12 | 0.339180907 | 0.497 | 0.238 | 2.40E-08 | 20 | Kdelr1 |
| Prkcsh1 | 1.26E-12 | 0.365192879 | 0.255 | 0.089 | 2.55E-08 | 20 | Prkcsh |
| Gbp24 | 1.26E-12 | 0.609590794 | 0.349 | 0.144 | 2.56E-08 | 20 | Gbp2 |
| Spen3 | 1.30E-12 | 0.49686573 | 0.369 | 0.159 | 2.64E-08 | 20 | Spen |
| Add15 | 1.45E-12 | 0.391597648 | 0.456 | 0.215 | 2.94E-08 | 20 | Add1 |
| Sox185 | 1.64E-12 | 0.307103149 | 0.597 | 0.277 | 3.33E-08 | 20 | Sox18 |
| Spata63 | 1.65E-12 | 0.405024252 | 0.309 | 0.121 | 3.33E-08 | 20 | Spata6 |
| Tmem884 | 1.88E-12 | 0.439214887 | 0.463 | 0.211 | 3.80E-08 | 20 | Tmem88 |
| Kctd103 | 1.97E-12 | 0.312643938 | 0.309 | 0.117 | 3.99E-08 | 20 | Kctd10 |
| Eif4a22 | 2.05E-12 | 0.305303626 | 0.577 | 0.305 | 4.14E-08 | 20 | Eif4a2 |
| Rgs37 | 2.06E-12 | 0.534643136 | 0.409 | 0.188 | 4.16E-08 | 20 | Rgs3 |
| Crispld15 | 2.11E-12 | 0.456303196 | 0.356 | 0.144 | 4.28E-08 | 20 | Crispld1 |
| Tmem1404 | 2.17E-12 | 0.420696017 | 0.49 | 0.224 | 4.40E-08 | 20 | Tmem140 |
| Prkd13 | 2.19E-12 | 0.35921478 | 0.369 | 0.148 | 4.44E-08 | 20 | Prkd1 |
| Mesd4 | 2.21E-12 | 0.450149924 | 0.376 | 0.164 | 4.46E-08 | 20 | Mesd |

| | | | | | | | |
|---|---|---|---|---|---|---|---|
| Ctdsp24 | 2.25E-12 | 0.352828621 | 0.409 | 0.179 | 4.56E-08 | 20 | Ctdsp2 |
| Cltb3 | 2.28E-12 | 0.301243226 | 0.477 | 0.22 | 4.61E-08 | 20 | Cltb |
| Ehmt14 | 2.29E-12 | 0.449723001 | 0.409 | 0.186 | 4.63E-08 | 20 | Ehmt1 |
| Kidins2206 | 2.29E-12 | 0.486998523 | 0.362 | 0.156 | 4.64E-08 | 20 | Kidins220 |
| Sf3b38 | 2.40E-12 | 0.48175826 | 0.369 | 0.161 | 4.86E-08 | 20 | Sf3b3 |
| Agpat4 | 2.47E-12 | 0.419342038 | 0.262 | 0.094 | 5.00E-08 | 20 | Agpat4 |
| Sec24b2 | 2.59E-12 | 0.438070126 | 0.336 | 0.139 | 5.24E-08 | 20 | Sec24b |
| Ogt9 | 2.59E-12 | 0.521980779 | 0.436 | 0.207 | 5.25E-08 | 20 | Ogt |
| Arhgap176 | 2.74E-12 | 0.555642453 | 0.523 | 0.271 | 5.55E-08 | 20 | Arhgap17 |
| Jak18 | 2.75E-12 | 0.464209409 | 0.664 | 0.366 | 5.56E-08 | 20 | Jak1 |
| Parp145 | 2.75E-12 | 0.566703129 | 0.497 | 0.254 | 5.57E-08 | 20 | Parp14 |
| Nsd110 | 2.86E-12 | 0.388934389 | 0.644 | 0.35 | 5.79E-08 | 20 | Nsd1 |
| Ets18 | 3.01E-12 | 0.478809245 | 0.705 | 0.404 | 6.10E-08 | 20 | Ets1 |
| Cr1l4 | 3.03E-12 | 0.44870061 | 0.477 | 0.234 | 6.13E-08 | 20 | Cr1l |
| Pan312 | 3.23E-12 | 0.578155471 | 0.463 | 0.229 | 6.54E-08 | 20 | Pan3 |
| Ncor110 | 3.27E-12 | 0.460895219 | 0.765 | 0.438 | 6.62E-08 | 20 | Ncor1 |
| Upp11 | 3.44E-12 | 0.393357032 | 0.275 | 0.1 | 6.97E-08 | 20 | Upp1 |
| Smad43 | 3.47E-12 | 0.46449137 | 0.463 | 0.224 | 7.02E-08 | 20 | Smad4 |
| Srrm210 | 3.66E-12 | 0.439192511 | 0.805 | 0.47 | 7.40E-08 | 20 | Srrm2 |
| Gbp92 | 3.69E-12 | 0.503872563 | 0.342 | 0.142 | 7.47E-08 | 20 | Gbp9 |
| Nrip17 | 3.78E-12 | 0.501220098 | 0.49 | 0.245 | 7.66E-08 | 20 | Nrip1 |
| Pitpnc18 | 3.90E-12 | 0.598421965 | 0.738 | 0.43 | 7.89E-08 | 20 | Pitpnc1 |
| Klf96 | 3.95E-12 | 0.331930096 | 0.51 | 0.238 | 8.00E-08 | 20 | Klf9 |
| Acvrl15 | 4.11E-12 | 0.380905633 | 0.564 | 0.276 | 8.31E-08 | 20 | Acvrl1 |
| Daam12 | 4.22E-12 | 0.349910821 | 0.289 | 0.109 | 8.55E-08 | 20 | Daam1 |
| Map2k51 | 4.22E-12 | 0.407084634 | 0.315 | 0.128 | 8.55E-08 | 20 | Map2k5 |
| Cyb5r35 | 4.30E-12 | 0.548257304 | 0.45 | 0.218 | 8.71E-08 | 20 | Cyb5r3 |
| Scoc1 | 4.60E-12 | 0.286522116 | 0.383 | 0.164 | 9.31E-08 | 20 | Scoc |
| Rnf1309 | 4.78E-12 | 0.45191018 | 0.544 | 0.285 | 9.69E-08 | 20 | Rnf130 |
| Desi22 | 4.82E-12 | 0.422203076 | 0.383 | 0.172 | 9.75E-08 | 20 | Desi2 |
| Tm9sf26 | 4.87E-12 | 0.440994839 | 0.503 | 0.256 | 9.86E-08 | 20 | Tm9sf2 |
| Id35 | 5.02E-12 | 0.341500943 | 0.792 | 0.409 | 1.02E-07 | 20 | Id3 |
| Nfe2l14 | 5.18E-12 | 0.441542605 | 0.55 | 0.281 | 1.05E-07 | 20 | Nfe2l1 |
| Dram24 | 5.22E-12 | 0.340419121 | 0.369 | 0.155 | 1.06E-07 | 20 | Dram2 |
| Arhgef117 | 5.77E-12 | 0.488760379 | 0.268 | 0.101 | 1.17E-07 | 20 | Arhgef11 |
| Tpm46 | 5.95E-12 | 0.444070218 | 0.685 | 0.381 | 1.20E-07 | 20 | Tpm4 |
| Baz2b7 | 6.04E-12 | 0.52021481 | 0.617 | 0.351 | 1.22E-07 | 20 | Baz2b |
| Peak19 | 6.14E-12 | 0.700008829 | 0.658 | 0.388 | 1.24E-07 | 20 | Peak1 |
| Nfkbia8 | 6.60E-12 | 0.529855035 | 0.691 | 0.401 | 1.34E-07 | 20 | Nfkbia |
| Ahr2 | 6.99E-12 | 0.481578762 | 0.497 | 0.247 | 1.41E-07 | 20 | Ahr |
| Eif3a6 | 7.13E-12 | 0.400265186 | 0.658 | 0.373 | 1.44E-07 | 20 | Eif3a |
| Lmo25 | 7.22E-12 | 0.373778516 | 0.45 | 0.209 | 1.46E-07 | 20 | Lmo2 |
| Clcn36 | 7.50E-12 | 0.373979754 | 0.329 | 0.134 | 1.52E-07 | 20 | Clcn3 |

| | | | | | | | |
|---|---|---|---|---|---|---|---|
| Clic45 | 7.75E-12 | 0.461964083 | 0.732 | 0.424 | 1.57E-07 | 20 | Clic4 |
| Otud4 | 8.09E-12 | 0.347955313 | 0.315 | 0.126 | 1.64E-07 | 20 | Otud4 |
| Mast46 | 8.13E-12 | 0.32535964 | 0.423 | 0.187 | 1.65E-07 | 20 | Mast4 |
| Mtus12 | 8.16E-12 | 0.331899585 | 0.376 | 0.16 | 1.65E-07 | 20 | Mtus1 |
| Zmym44 | 8.27E-12 | 0.439937534 | 0.376 | 0.166 | 1.68E-07 | 20 | Zmym4 |
| Smchd18 | 8.53E-12 | 0.431939154 | 0.517 | 0.265 | 1.73E-07 | 20 | Smchd1 |
| Hdac94 | 8.55E-12 | 0.718473605 | 0.362 | 0.159 | 1.73E-07 | 20 | Hdac9 |
| Ddx175 | 8.58E-12 | 0.615096596 | 0.51 | 0.274 | 1.74E-07 | 20 | Ddx17 |
| Rps6kb12 | 8.64E-12 | 0.403656393 | 0.362 | 0.159 | 1.75E-07 | 20 | Rps6kb1 |
| Vps13d3 | 9.57E-12 | 0.526077955 | 0.376 | 0.17 | 1.94E-07 | 20 | Vps13d |
| Crebbp11 | 1.00E-11 | 0.498696299 | 0.564 | 0.306 | 2.02E-07 | 20 | Crebbp |
| Herc25 | 1.00E-11 | 0.44148134 | 0.436 | 0.208 | 2.03E-07 | 20 | Herc2 |
| Rbm267 | 1.02E-11 | 0.56430923 | 0.45 | 0.228 | 2.07E-07 | 20 | Rbm26 |
| Abi34 | 1.02E-11 | 0.402383158 | 0.497 | 0.245 | 2.07E-07 | 20 | Abi3 |
| Tm9sf37 | 1.03E-11 | 0.457188611 | 0.611 | 0.336 | 2.08E-07 | 20 | Tm9sf3 |
| Aff45 | 1.07E-11 | 0.422758555 | 0.483 | 0.241 | 2.17E-07 | 20 | Aff4 |
| Slco2b13 | 1.08E-11 | 0.406371127 | 0.409 | 0.181 | 2.19E-07 | 20 | Slco2b1 |
| Ddx3x6 | 1.12E-11 | 0.427588146 | 0.584 | 0.309 | 2.27E-07 | 20 | Ddx3x |
| Ppp1r25 | 1.22E-11 | 0.475435328 | 0.705 | 0.394 | 2.48E-07 | 20 | Ppp1r2 |
| Snx34 | 1.23E-11 | 0.411671997 | 0.738 | 0.462 | 2.50E-07 | 20 | Snx3 |
| Bvht4 | 1.25E-11 | 0.361857756 | 0.423 | 0.187 | 2.53E-07 | 20 | Bvht |
| Son11 | 1.33E-11 | 0.421586308 | 0.872 | 0.526 | 2.69E-07 | 20 | Son |
| Srek15 | 1.39E-11 | 0.534816554 | 0.403 | 0.195 | 2.82E-07 | 20 | Srek1 |
| Osbpl87 | 1.43E-11 | 0.425211062 | 0.423 | 0.2 | 2.91E-07 | 20 | Osbpl8 |
| Phf11d3 | 1.46E-11 | 0.408971731 | 0.336 | 0.138 | 2.95E-07 | 20 | Phf11d |
| Senp68 | 1.63E-11 | 0.385402687 | 0.557 | 0.299 | 3.30E-07 | 20 | Senp6 |
| Serpinb9 | 1.65E-11 | 0.330940046 | 0.336 | 0.141 | 3.35E-07 | 20 | Serpinb9 |
| Lman15 | 1.69E-11 | 0.436678518 | 0.477 | 0.239 | 3.42E-07 | 20 | Lman1 |
| Sox73 | 1.71E-11 | 0.427641854 | 0.262 | 0.096 | 3.46E-07 | 20 | Sox7 |
| Phf310 | 1.77E-11 | 0.469586236 | 0.49 | 0.254 | 3.59E-07 | 20 | Phf3 |
| Btbd76 | 1.88E-11 | 0.511991135 | 0.517 | 0.282 | 3.81E-07 | 20 | Btbd7 |
| Pls35 | 1.98E-11 | 0.344434136 | 0.47 | 0.215 | 4.01E-07 | 20 | Pls3 |
| Kctd171 | 2.16E-11 | 0.28281183 | 0.255 | 0.091 | 4.38E-07 | 20 | Kctd17 |
| Zfp6387 | 2.21E-11 | 0.507537385 | 0.517 | 0.283 | 4.48E-07 | 20 | Zfp638 |
| Rbm2512 | 2.22E-11 | 0.455484039 | 0.597 | 0.335 | 4.49E-07 | 20 | Rbm25 |
| Sncg5 | 2.24E-11 | 0.387314659 | 0.369 | 0.156 | 4.54E-07 | 20 | Sncg |
| Nucb14 | 2.25E-11 | 0.473474234 | 0.456 | 0.229 | 4.55E-07 | 20 | Nucb1 |
| Smurf25 | 2.25E-11 | 0.527730526 | 0.47 | 0.239 | 4.55E-07 | 20 | Smurf2 |
| Pkn26 | 2.33E-11 | 0.3468053 | 0.57 | 0.295 | 4.72E-07 | 20 | Pkn2 |
| D16Ertd472e5 | 2.41E-11 | 0.465758554 | 0.275 | 0.107 | 4.88E-07 | 20 | D16Ertd472e |
| Ctdspl22 | 2.69E-11 | 0.371737206 | 0.383 | 0.173 | 5.44E-07 | 20 | Ctdspl2 |
| Mcu2 | 2.71E-11 | 0.407049927 | 0.268 | 0.102 | 5.49E-07 | 20 | Mcu |
| Zfp280d3 | 2.72E-11 | 0.42212746 | 0.396 | 0.184 | 5.50E-07 | 20 | Zfp280d |

| | | | | | | | |
|---|---|---|---|---|---|---|---|
| Tns16 | 2.78E-11 | 0.352331287 | 0.51 | 0.246 | 5.62E-07 | 20 | Tns1 |
| Ddx64 | 2.84E-11 | 0.383644571 | 0.617 | 0.342 | 5.75E-07 | 20 | Ddx6 |
| Sypl4 | 2.93E-11 | 0.330311363 | 0.497 | 0.24 | 5.93E-07 | 20 | Sypl |
| Fam117b1 | 2.97E-11 | 0.499685504 | 0.369 | 0.165 | 6.00E-07 | 20 | Fam117b |
| Ubn13 | 3.09E-11 | 0.359579899 | 0.497 | 0.247 | 6.26E-07 | 20 | Ubn1 |
| Pura4 | 3.16E-11 | 0.390597413 | 0.644 | 0.367 | 6.39E-07 | 20 | Pura |
| Fbxl74 | 3.30E-11 | 0.448193325 | 0.523 | 0.258 | 6.67E-07 | 20 | Fbxl7 |
| Hook36 | 3.34E-11 | 0.44068534 | 0.537 | 0.291 | 6.76E-07 | 20 | Hook3 |
| Zc3h137 | 3.35E-11 | 0.384779458 | 0.45 | 0.217 | 6.78E-07 | 20 | Zc3h13 |
| Zbtb464 | 3.59E-11 | 0.502430648 | 0.369 | 0.163 | 7.28E-07 | 20 | Zbtb46 |
| Kdm5c1 | 3.60E-11 | 0.483533346 | 0.322 | 0.139 | 7.29E-07 | 20 | Kdm5c |
| Impad11 | 3.76E-11 | 0.429848507 | 0.262 | 0.101 | 7.62E-07 | 20 | Impad1 |
| Lpp10 | 3.78E-11 | 0.501704888 | 0.544 | 0.287 | 7.66E-07 | 20 | Lpp |
| Trim253 | 3.93E-11 | 0.420445921 | 0.349 | 0.15 | 7.96E-07 | 20 | Trim25 |
| AW5549187 | 3.95E-11 | 0.496165556 | 0.369 | 0.171 | 7.99E-07 | 20 | AW554918 |
| Ttc1410 | 3.99E-11 | 0.470633856 | 0.436 | 0.216 | 8.08E-07 | 20 | Ttc14 |
| Dph61 | 4.06E-11 | 0.481921045 | 0.315 | 0.132 | 8.22E-07 | 20 | Dph6 |
| Ubr41 | 4.27E-11 | 0.400377627 | 0.336 | 0.146 | 8.64E-07 | 20 | Ubr4 |
| Mndal7 | 4.50E-11 | 0.539440485 | 0.691 | 0.407 | 9.10E-07 | 20 | Mndal |
| Mtdh8 | 4.61E-11 | 0.492788152 | 0.651 | 0.388 | 9.34E-07 | 20 | Mtdh |
| Prkd22 | 4.82E-11 | 0.425750138 | 0.329 | 0.141 | 9.76E-07 | 20 | Prkd2 |
| Socs53 | 5.07E-11 | 0.345076368 | 0.315 | 0.128 | 1.03E-06 | 20 | Socs5 |
| Dync1h16 | 5.13E-11 | 0.47229904 | 0.537 | 0.287 | 1.04E-06 | 20 | Dync1h1 |
| Exoc67 | 5.13E-11 | 0.424284192 | 0.376 | 0.172 | 1.04E-06 | 20 | Exoc6 |
| Mfap31 | 5.16E-11 | 0.438926137 | 0.255 | 0.097 | 1.05E-06 | 20 | Mfap3 |
| Csnk1g18 | 5.25E-11 | 0.482178651 | 0.349 | 0.158 | 1.06E-06 | 20 | Csnk1g1 |
| Thra5 | 5.51E-11 | 0.358210447 | 0.376 | 0.165 | 1.12E-06 | 20 | Thra |
| Asap15 | 5.56E-11 | 0.454566993 | 0.631 | 0.357 | 1.13E-06 | 20 | Asap1 |
| Clec2d5 | 5.69E-11 | 0.509432271 | 0.805 | 0.529 | 1.15E-06 | 20 | Clec2d |
| Eif4g14 | 5.81E-11 | 0.401644753 | 0.463 | 0.237 | 1.18E-06 | 20 | Eif4g1 |
| Myo104 | 5.83E-11 | 0.349799332 | 0.463 | 0.22 | 1.18E-06 | 20 | Myo10 |
| Mrrf1 | 5.91E-11 | 0.252538036 | 0.289 | 0.112 | 1.20E-06 | 20 | Mrrf |
| Rnf2204 | 5.91E-11 | 0.380229535 | 0.503 | 0.242 | 1.20E-06 | 20 | Rnf220 |
| Ppig6 | 5.99E-11 | 0.463186211 | 0.564 | 0.317 | 1.21E-06 | 20 | Ppig |
| Tspan34 | 6.23E-11 | 0.306024922 | 0.275 | 0.106 | 1.26E-06 | 20 | Tspan3 |
| Slc31a2 | 6.27E-11 | 0.404638307 | 0.255 | 0.096 | 1.27E-06 | 20 | Slc31a2 |
| Tnfaip13 | 6.31E-11 | 0.265782998 | 0.45 | 0.204 | 1.28E-06 | 20 | Tnfaip1 |
| Tmed98 | 6.72E-11 | 0.418959918 | 0.671 | 0.404 | 1.36E-06 | 20 | Tmed9 |
| Gnb15 | 6.77E-11 | 0.341203165 | 0.725 | 0.428 | 1.37E-06 | 20 | Gnb1 |
| BC0055612 | 7.10E-11 | 0.359008236 | 0.295 | 0.12 | 1.44E-06 | 20 | BC005561 |
| Myo18a3 | 7.66E-11 | 0.343021221 | 0.517 | 0.258 | 1.55E-06 | 20 | Myo18a |
| Ablim17 | 7.94E-11 | 0.511707408 | 0.691 | 0.404 | 1.61E-06 | 20 | Ablim1 |
| Clstn15 | 8.04E-11 | 0.375292626 | 0.49 | 0.237 | 1.63E-06 | 20 | Clstn1 |

| | | | | | | | |
|---|---|---|---|---|---|---|---|
| Tanc14 | 8.07E-11 | 0.368615033 | 0.463 | 0.224 | 1.63E-06 | 20 | Tanc1 |
| Ubr35 | 8.08E-11 | 0.488769772 | 0.416 | 0.202 | 1.64E-06 | 20 | Ubr3 |
| Eva1b6 | 8.09E-11 | 0.34259504 | 0.564 | 0.286 | 1.64E-06 | 20 | Eva1b |
| Fam172a11 | 8.16E-11 | 0.609165282 | 0.53 | 0.285 | 1.65E-06 | 20 | Fam172a |
| Scaf115 | 8.32E-11 | 0.441501433 | 0.503 | 0.264 | 1.68E-06 | 20 | Scaf11 |
| Tmem184b3 | 8.55E-11 | 0.352359505 | 0.289 | 0.115 | 1.73E-06 | 20 | Tmem184b |
| Zmynd116 | 8.64E-11 | 0.39037756 | 0.544 | 0.29 | 1.75E-06 | 20 | Zmynd11 |
| Tmbim66 | 8.69E-11 | 0.394578987 | 0.711 | 0.432 | 1.76E-06 | 20 | Tmbim6 |
| Dab26 | 8.83E-11 | 0.524952745 | 0.423 | 0.207 | 1.79E-06 | 20 | Dab2 |
| Svil4 | 8.98E-11 | 0.397127667 | 0.483 | 0.244 | 1.82E-06 | 20 | Svil |
| Bod1l7 | 9.13E-11 | 0.556484671 | 0.497 | 0.274 | 1.85E-06 | 20 | Bod1l |
| Rbm334 | 9.32E-11 | 0.342202108 | 0.329 | 0.143 | 1.89E-06 | 20 | Rbm33 |
| Ptpn110 | 9.55E-11 | 0.317029568 | 0.477 | 0.237 | 1.93E-06 | 20 | Ptpn1 |
| Mark35 | 9.65E-11 | 0.486221069 | 0.423 | 0.213 | 1.95E-06 | 20 | Mark3 |
| Ythdc12 | 9.77E-11 | 0.439620588 | 0.463 | 0.236 | 1.98E-06 | 20 | Ythdc1 |
| Sfswap2 | 9.98E-11 | 0.455730929 | 0.315 | 0.136 | 2.02E-06 | 20 | Sfswap |
| Lrch39 | 1.00E-10 | 0.443681388 | 0.463 | 0.233 | 2.03E-06 | 20 | Lrch3 |
| Syncrip3 | 1.04E-10 | 0.370230199 | 0.43 | 0.216 | 2.10E-06 | 20 | Syncrip |
| Brd411 | 1.14E-10 | 0.38542192 | 0.51 | 0.268 | 2.31E-06 | 20 | Brd4 |
| Ubxn47 | 1.21E-10 | 0.402816993 | 0.517 | 0.273 | 2.45E-06 | 20 | Ubxn4 |
| Yipf52 | 1.23E-10 | 0.401010984 | 0.309 | 0.133 | 2.48E-06 | 20 | Yipf5 |
| Ubr55 | 1.23E-10 | 0.422846303 | 0.409 | 0.199 | 2.50E-06 | 20 | Ubr5 |
| Tpp13 | 1.26E-10 | 0.327546095 | 0.295 | 0.12 | 2.56E-06 | 20 | Tpp1 |
| Nrbp12 | 1.28E-10 | 0.399657014 | 0.43 | 0.212 | 2.60E-06 | 20 | Nrbp1 |
| Pnisr9 | 1.32E-10 | 0.412365968 | 0.497 | 0.265 | 2.68E-06 | 20 | Pnisr |
| Elf23 | 1.33E-10 | 0.446333231 | 0.544 | 0.301 | 2.68E-06 | 20 | Elf2 |
| Ccdc85a3 | 1.36E-10 | 0.60156245 | 0.396 | 0.182 | 2.75E-06 | 20 | Ccdc85a |
| Scn1b3 | 1.38E-10 | 0.39333921 | 0.423 | 0.196 | 2.79E-06 | 20 | Scn1b |
| Vcl4 | 1.47E-10 | 0.462151562 | 0.356 | 0.162 | 2.97E-06 | 20 | Vcl |
| Tmem1095 | 1.55E-10 | 0.397406131 | 0.336 | 0.149 | 3.13E-06 | 20 | Tmem109 |
| Usp3411 | 1.56E-10 | 0.503746481 | 0.584 | 0.345 | 3.16E-06 | 20 | Usp34 |
| Tmcc111 | 1.56E-10 | 0.537900006 | 0.436 | 0.219 | 3.17E-06 | 20 | Tmcc1 |
| Tubb56 | 1.59E-10 | 0.338086372 | 0.651 | 0.381 | 3.21E-06 | 20 | Tubb5 |
| Ppp2r5e4 | 1.63E-10 | 0.419225302 | 0.362 | 0.166 | 3.31E-06 | 20 | Ppp2r5e |
| Fus7 | 1.67E-10 | 0.337371898 | 0.685 | 0.389 | 3.38E-06 | 20 | Fus |
| Pttg1ip7 | 1.69E-10 | 0.315328101 | 0.53 | 0.27 | 3.42E-06 | 20 | Pttg1ip |
| Rnf2135 | 1.72E-10 | 0.557726563 | 0.49 | 0.258 | 3.48E-06 | 20 | Rnf213 |
| Gpatch21 | 1.73E-10 | 0.376463386 | 0.282 | 0.116 | 3.50E-06 | 20 | Gpatch2 |
| Actr1a1 | 1.76E-10 | 0.298646944 | 0.309 | 0.13 | 3.57E-06 | 20 | Actr1a |
| Arl154 | 1.79E-10 | 0.483900045 | 0.678 | 0.382 | 3.62E-06 | 20 | Arl15 |
| Slc6a64 | 1.89E-10 | 0.65647336 | 0.564 | 0.329 | 3.82E-06 | 20 | Slc6a6 |
| Erap13 | 1.96E-10 | 0.406678522 | 0.322 | 0.141 | 3.96E-06 | 20 | Erap1 |
| Vat13 | 1.99E-10 | 0.309486782 | 0.356 | 0.156 | 4.03E-06 | 20 | Vat1 |

| | | | | | | | |
|---|---|---|---|---|---|---|---|
| Rapgef25 | 2.04E-10 | 0.30292653 | 0.631 | 0.34 | 4.13E-06 | 20 | Rapgef2 |
| Xrn16 | 2.11E-10 | 0.414715218 | 0.409 | 0.204 | 4.27E-06 | 20 | Xrn1 |
| Luc7l | 2.20E-10 | 0.3675364 | 0.289 | 0.12 | 4.46E-06 | 20 | Luc7l |
| Agap13 | 2.26E-10 | 0.381518157 | 0.336 | 0.146 | 4.58E-06 | 20 | Agap1 |
| Hivep210 | 2.40E-10 | 0.534258559 | 0.544 | 0.299 | 4.86E-06 | 20 | Hivep2 |
| Chst153 | 2.45E-10 | 0.419064919 | 0.396 | 0.176 | 4.96E-06 | 20 | Chst15 |
| Ssr33 | 2.52E-10 | 0.320321912 | 0.409 | 0.197 | 5.11E-06 | 20 | Ssr3 |
| Kmt2c11 | 2.62E-10 | 0.450799614 | 0.55 | 0.307 | 5.30E-06 | 20 | Kmt2c |
| Txnip5 | 2.65E-10 | 0.30681392 | 0.852 | 0.545 | 5.36E-06 | 20 | Txnip |
| Gimap110 | 2.68E-10 | 0.421558728 | 0.57 | 0.304 | 5.43E-06 | 20 | Gimap1 |
| Man1a11 | 2.74E-10 | 0.543403903 | 0.356 | 0.168 | 5.56E-06 | 20 | Man1a |
| Add34 | 2.82E-10 | 0.334650509 | 0.456 | 0.224 | 5.71E-06 | 20 | Add3 |
| Kifap32 | 2.87E-10 | 0.331694153 | 0.268 | 0.107 | 5.82E-06 | 20 | Kifap3 |
| Hacd44 | 2.89E-10 | 0.36753067 | 0.282 | 0.116 | 5.86E-06 | 20 | Hacd4 |
| Slc29a13 | 2.98E-10 | 0.39687177 | 0.315 | 0.136 | 6.04E-06 | 20 | Slc29a1 |
| Med133 | 3.05E-10 | 0.344229418 | 0.416 | 0.206 | 6.18E-06 | 20 | Med13 |
| Dopey11 | 3.08E-10 | 0.352185627 | 0.268 | 0.108 | 6.24E-06 | 20 | Dopey1 |
| Rac14 | 3.10E-10 | 0.375839293 | 0.705 | 0.431 | 6.28E-06 | 20 | Rac1 |
| Gphn2 | 3.15E-10 | 0.397863837 | 0.973 | 0.793 | 6.37E-06 | 20 | Gphn |
| Arhgef23 | 3.21E-10 | 0.389716426 | 0.356 | 0.166 | 6.49E-06 | 20 | Arhgef2 |
| Trim354 | 3.27E-10 | 0.405797475 | 0.322 | 0.141 | 6.63E-06 | 20 | Trim35 |
| Stx124 | 3.40E-10 | 0.315783694 | 0.356 | 0.163 | 6.89E-06 | 20 | Stx12 |
| Ier31 | 3.43E-10 | 0.490605052 | 0.282 | 0.116 | 6.93E-06 | 20 | Ier3 |
| Tbl1xr17 | 3.47E-10 | 0.527719294 | 0.356 | 0.165 | 7.03E-06 | 20 | Tbl1xr1 |
| Srpk24 | 3.63E-10 | 0.378629859 | 0.403 | 0.194 | 7.36E-06 | 20 | Srpk2 |
| Zcchc712 | 3.68E-10 | 0.599174887 | 0.523 | 0.302 | 7.44E-06 | 20 | Zcchc7 |
| Ly6a5 | 3.70E-10 | 0.354331122 | 0.812 | 0.46 | 7.50E-06 | 20 | Ly6a |
| Wasl4 | 3.71E-10 | 0.31573613 | 0.463 | 0.238 | 7.52E-06 | 20 | Wasl |
| Map3k36 | 3.90E-10 | 0.418435579 | 0.376 | 0.179 | 7.90E-06 | 20 | Map3k3 |
| Ddx542 | 4.17E-10 | 0.259588217 | 0.309 | 0.131 | 8.44E-06 | 20 | Ddx54 |
| Immp2l6 | 4.18E-10 | 0.43367244 | 0.315 | 0.137 | 8.46E-06 | 20 | Immp2l |
| Colgalt23 | 4.28E-10 | 0.354246745 | 0.329 | 0.141 | 8.67E-06 | 20 | Colgalt2 |
| Mxra75 | 4.29E-10 | 0.280310074 | 0.336 | 0.143 | 8.69E-06 | 20 | Mxra7 |
| Ext23 | 4.44E-10 | 0.366172047 | 0.322 | 0.144 | 8.99E-06 | 20 | Ext2 |
| Mib13 | 4.59E-10 | 0.405821547 | 0.356 | 0.166 | 9.29E-06 | 20 | Mib1 |
| Rnf1698 | 4.90E-10 | 0.499583257 | 0.376 | 0.184 | 9.92E-06 | 20 | Rnf169 |
| Phf20l18 | 4.92E-10 | 0.436187545 | 0.523 | 0.288 | 9.97E-06 | 20 | Phf20l1 |
| Tom1l24 | 5.01E-10 | 0.390068678 | 0.302 | 0.132 | 1.01E-05 | 20 | Tom1l2 |
| Ago33 | 5.01E-10 | 0.477537365 | 0.336 | 0.156 | 1.01E-05 | 20 | Ago3 |
| Tmx36 | 5.06E-10 | 0.359225165 | 0.396 | 0.188 | 1.02E-05 | 20 | Tmx3 |
| Numa15 | 5.26E-10 | 0.36967919 | 0.403 | 0.196 | 1.06E-05 | 20 | Numa1 |
| Mga5 | 5.50E-10 | 0.448913268 | 0.362 | 0.172 | 1.11E-05 | 20 | Mga |
| Mycbp212 | 5.96E-10 | 0.453041428 | 0.631 | 0.376 | 1.21E-05 | 20 | Mycbp2 |

| | | | | | | | |
|---|---|---|---|---|---|---|---|
| Mapk1ip1l | 6.02E-10 | 0.333124179 | 0.262 | 0.106 | 1.22E-05 | 20 | Mapk1ip1l |
| Rbm68 | 6.41E-10 | 0.414779821 | 0.43 | 0.218 | 1.30E-05 | 20 | Rbm6 |
| Bnip24 | 6.44E-10 | 0.338882261 | 0.537 | 0.285 | 1.30E-05 | 20 | Bnip2 |
| Tbck | 6.80E-10 | 0.450453576 | 0.262 | 0.109 | 1.38E-05 | 20 | Tbck |
| Pum24 | 6.90E-10 | 0.329346166 | 0.503 | 0.27 | 1.40E-05 | 20 | Pum2 |
| Rabep15 | 7.31E-10 | 0.412085583 | 0.45 | 0.232 | 1.48E-05 | 20 | Rabep1 |
| Tspan124 | 7.37E-10 | 0.329148831 | 0.477 | 0.233 | 1.49E-05 | 20 | Tspan12 |
| Efna14 | 7.42E-10 | 0.285269977 | 0.336 | 0.144 | 1.50E-05 | 20 | Efna1 |
| Caskin23 | 7.52E-10 | 0.330245596 | 0.329 | 0.141 | 1.52E-05 | 20 | Caskin2 |
| Tmem2054 | 7.66E-10 | 0.265548255 | 0.356 | 0.16 | 1.55E-05 | 20 | Tmem205 |
| Dgkh5 | 7.73E-10 | 0.470217773 | 0.376 | 0.178 | 1.56E-05 | 20 | Dgkh |
| Matr33 | 7.97E-10 | 0.349711335 | 0.503 | 0.267 | 1.61E-05 | 20 | Matr3 |
| Jmjd1c11 | 8.05E-10 | 0.502555112 | 0.544 | 0.308 | 1.63E-05 | 20 | Jmjd1c |
| Dcaf81 | 8.08E-10 | 0.369370707 | 0.403 | 0.198 | 1.64E-05 | 20 | Dcaf8 |
| Trim30a8 | 8.22E-10 | 0.523208736 | 0.436 | 0.22 | 1.66E-05 | 20 | Trim30a |
| Hecw23 | 9.02E-10 | 0.403557272 | 0.497 | 0.25 | 1.83E-05 | 20 | Hecw2 |
| Cpeb41 | 9.25E-10 | 0.354453654 | 0.289 | 0.123 | 1.87E-05 | 20 | Cpeb4 |
| Shoc23 | 1.08E-09 | 0.331728772 | 0.436 | 0.222 | 2.19E-05 | 20 | Shoc2 |
| Smarcc23 | 1.13E-09 | 0.294596828 | 0.376 | 0.178 | 2.29E-05 | 20 | Smarcc2 |
| Rabgap14 | 1.18E-09 | 0.466768553 | 0.342 | 0.164 | 2.38E-05 | 20 | Rabgap1 |
| Ranbp24 | 1.20E-09 | 0.35467544 | 0.403 | 0.199 | 2.42E-05 | 20 | Ranbp2 |
| Csnk1a13 | 1.20E-09 | 0.305066307 | 0.738 | 0.458 | 2.43E-05 | 20 | Csnk1a1 |
| N4bp2l25 | 1.23E-09 | 0.388436622 | 0.47 | 0.248 | 2.50E-05 | 20 | N4bp2l2 |
| Uhrf25 | 1.28E-09 | 0.450271156 | 0.349 | 0.167 | 2.59E-05 | 20 | Uhrf2 |
| Fto5 | 1.33E-09 | 0.442769997 | 0.383 | 0.192 | 2.69E-05 | 20 | Fto |
| Nlk5 | 1.35E-09 | 0.399499554 | 0.322 | 0.146 | 2.74E-05 | 20 | Nlk |
| Cbx53 | 1.39E-09 | 0.340731649 | 0.389 | 0.189 | 2.81E-05 | 20 | Cbx5 |
| Lfng2 | 1.46E-09 | 0.384880478 | 0.309 | 0.137 | 2.95E-05 | 20 | Lfng |
| Ttc34 | 1.49E-09 | 0.464880138 | 0.463 | 0.253 | 3.03E-05 | 20 | Ttc3 |
| Yy12 | 1.50E-09 | 0.280615623 | 0.51 | 0.274 | 3.04E-05 | 20 | Yy1 |
| Mon23 | 1.51E-09 | 0.393166317 | 0.329 | 0.153 | 3.05E-05 | 20 | Mon2 |
| Tpst23 | 1.53E-09 | 0.362748784 | 0.456 | 0.233 | 3.10E-05 | 20 | Tpst2 |
| Usp473 | 1.57E-09 | 0.388431667 | 0.383 | 0.189 | 3.17E-05 | 20 | Usp47 |
| Orai15 | 1.59E-09 | 0.316192829 | 0.295 | 0.13 | 3.22E-05 | 20 | Orai1 |
| 4930402H24Rik4 | 1.63E-09 | 0.340685659 | 0.436 | 0.222 | 3.30E-05 | 20 | 4930402H24Rik |
| Ralgapb3 | 1.85E-09 | 0.368393377 | 0.275 | 0.118 | 3.75E-05 | 20 | Ralgapb |
| Akt311 | 1.86E-09 | 0.435828632 | 0.47 | 0.253 | 3.76E-05 | 20 | Akt3 |
| Stom3 | 1.86E-09 | 0.437025985 | 0.262 | 0.109 | 3.76E-05 | 20 | Stom |
| Setbp17 | 1.90E-09 | 0.441792624 | 0.289 | 0.123 | 3.86E-05 | 20 | Setbp1 |
| Capns16 | 1.93E-09 | 0.259513231 | 0.604 | 0.349 | 3.90E-05 | 20 | Capns1 |
| Uvrag7 | 1.94E-09 | 0.428477082 | 0.49 | 0.269 | 3.93E-05 | 20 | Uvrag |
| Tia12 | 1.96E-09 | 0.397346564 | 0.268 | 0.114 | 3.97E-05 | 20 | Tia1 |
| Ccnl25 | 1.98E-09 | 0.434589352 | 0.416 | 0.215 | 4.01E-05 | 20 | Ccnl2 |

| | | | | | | | |
|---|---|---|---|---|---|---|---|
| Gbp74 | 1.98E-09 | 0.470270522 | 0.497 | 0.27 | 4.01E-05 | 20 | Gbp7 |
| Spint22 | 2.07E-09 | 0.304916057 | 0.268 | 0.114 | 4.19E-05 | 20 | Spint2 |
| Znfx14 | 2.20E-09 | 0.314823381 | 0.376 | 0.176 | 4.45E-05 | 20 | Znfx1 |
| Atp9b6 | 2.25E-09 | 0.406179636 | 0.336 | 0.159 | 4.55E-05 | 20 | Atp9b |
| Srsf1112 | 2.58E-09 | 0.409268089 | 0.597 | 0.352 | 5.22E-05 | 20 | Srsf11 |
| Stard84 | 2.59E-09 | 0.349977508 | 0.416 | 0.199 | 5.24E-05 | 20 | Stard8 |
| Ppp1r9b1 | 2.76E-09 | 0.346408092 | 0.289 | 0.127 | 5.59E-05 | 20 | Ppp1r9b |
| Lrp63 | 2.83E-09 | 0.421051414 | 0.315 | 0.145 | 5.73E-05 | 20 | Lrp6 |
| Abhd17b7 | 3.00E-09 | 0.546008124 | 0.383 | 0.193 | 6.08E-05 | 20 | Abhd17b |
| Arap33 | 3.11E-09 | 0.353117412 | 0.43 | 0.219 | 6.29E-05 | 20 | Arap3 |
| Ankrd1710 | 3.11E-09 | 0.48064382 | 0.477 | 0.262 | 6.31E-05 | 20 | Ankrd17 |
| Mbd59 | 3.12E-09 | 0.560815741 | 0.416 | 0.221 | 6.32E-05 | 20 | Mbd5 |
| Prpf4b7 | 3.12E-09 | 0.379880982 | 0.57 | 0.334 | 6.32E-05 | 20 | Prpf4b |
| Dennd4c6 | 3.12E-09 | 0.40240924 | 0.282 | 0.127 | 6.33E-05 | 20 | Dennd4c |
| Bms1 | 3.15E-09 | 0.37357959 | 0.262 | 0.111 | 6.37E-05 | 20 | Bms1 |
| Pla2g166 | 3.19E-09 | 0.293312633 | 0.456 | 0.244 | 6.46E-05 | 20 | Pla2g16 |
| Zfp3664 | 3.32E-09 | 0.454116088 | 0.383 | 0.183 | 6.72E-05 | 20 | Zfp366 |
| Zscan264 | 3.37E-09 | 0.319443136 | 0.322 | 0.146 | 6.83E-05 | 20 | Zscan26 |
| Nf17 | 3.38E-09 | 0.418383231 | 0.43 | 0.222 | 6.84E-05 | 20 | Nf1 |
| Eps153 | 3.39E-09 | 0.301984184 | 0.356 | 0.17 | 6.85E-05 | 20 | Eps15 |
| Rab126 | 3.43E-09 | 0.37674281 | 0.369 | 0.178 | 6.94E-05 | 20 | Rab12 |
| Ddb13 | 3.64E-09 | 0.304687043 | 0.322 | 0.149 | 7.38E-05 | 20 | Ddb1 |
| Smc34 | 3.67E-09 | 0.275765057 | 0.477 | 0.254 | 7.42E-05 | 20 | Smc3 |
| Zzz31 | 3.75E-09 | 0.302287456 | 0.329 | 0.153 | 7.60E-05 | 20 | Zzz3 |
| mt-Nd6 | 3.76E-09 | 0.332573709 | 0.282 | 0.124 | 7.60E-05 | 20 | mt-Nd6 |
| Pum17 | 3.77E-09 | 0.390595631 | 0.477 | 0.26 | 7.64E-05 | 20 | Pum1 |
| Jak26 | 3.78E-09 | 0.32759043 | 0.289 | 0.128 | 7.65E-05 | 20 | Jak2 |
| Gigyf25 | 3.80E-09 | 0.489157459 | 0.342 | 0.169 | 7.69E-05 | 20 | Gigyf2 |
| Pea15a6 | 3.86E-09 | 0.328846104 | 0.456 | 0.241 | 7.82E-05 | 20 | Pea15a |
| Rbm59 | 3.97E-09 | 0.394086443 | 0.45 | 0.241 | 8.04E-05 | 20 | Rbm5 |
| Lamp110 | 4.07E-09 | 0.362514643 | 0.685 | 0.418 | 8.24E-05 | 20 | Lamp1 |
| Fer3 | 4.17E-09 | 0.35202369 | 0.282 | 0.122 | 8.44E-05 | 20 | Fer |
| Elk43 | 4.49E-09 | 0.354211908 | 0.349 | 0.168 | 9.09E-05 | 20 | Elk4 |
| Atxn19 | 4.49E-09 | 0.430843271 | 0.53 | 0.296 | 9.10E-05 | 20 | Atxn1 |
| Zfp6092 | 4.56E-09 | 0.388106159 | 0.295 | 0.132 | 9.24E-05 | 20 | Zfp609 |
| Cers21 | 4.61E-09 | 0.381533034 | 0.275 | 0.121 | 9.34E-05 | 20 | Cers2 |
| Armc8 | 4.64E-09 | 0.349127206 | 0.282 | 0.123 | 9.39E-05 | 20 | Armc8 |
| Map4k54 | 4.66E-09 | 0.366007479 | 0.295 | 0.132 | 9.43E-05 | 20 | Map4k5 |
| Rock110 | 4.78E-09 | 0.382030291 | 0.698 | 0.434 | 9.68E-05 | 20 | Rock1 |
| Mum12 | 4.99E-09 | 0.2835095 | 0.268 | 0.114 | 0.00010102 | 20 | Mum1 |
| Tjp23 | 5.16E-09 | 0.403776009 | 0.309 | 0.141 | 0.000104556 | 20 | Tjp2 |
| Cog58 | 5.18E-09 | 0.389325496 | 0.369 | 0.184 | 0.000104892 | 20 | Cog5 |
| Lnx21 | 5.32E-09 | 0.359518248 | 0.302 | 0.134 | 0.000107752 | 20 | Lnx2 |

| | | | | | | | |
|---|---|---|---|---|---|---|---|
| Map4k410 | 5.45E-09 | 0.422179588 | 0.503 | 0.278 | 0.00011044 | 20 | Map4k4 |
| Thoc28 | 5.47E-09 | 0.428516277 | 0.477 | 0.275 | 0.000110729 | 20 | Thoc2 |
| Adipor14 | 5.52E-09 | 0.284729695 | 0.477 | 0.258 | 0.000111839 | 20 | Adipor1 |
| Golm1 | 5.57E-09 | 0.369860058 | 0.255 | 0.108 | 0.000112733 | 20 | Golm1 |
| Frs22 | 5.69E-09 | 0.461887185 | 0.302 | 0.143 | 0.00011522 | 20 | Frs2 |
| Bcl2l15 | 5.72E-09 | 0.347821624 | 0.322 | 0.148 | 0.000115757 | 20 | Bcl2l1 |
| Rnf381 | 5.85E-09 | 0.395538964 | 0.295 | 0.134 | 0.000118523 | 20 | Rnf38 |
| Camsap21 | 5.86E-09 | 0.306639929 | 0.262 | 0.109 | 0.000118608 | 20 | Camsap2 |
| Kcnq1ot18 | 6.04E-09 | 0.454648942 | 0.295 | 0.135 | 0.000122344 | 20 | Kcnq1ot1 |
| Fam208b6 | 6.17E-09 | 0.375900258 | 0.322 | 0.151 | 0.000124852 | 20 | Fam208b |
| Pcnx5 | 6.41E-09 | 0.278506094 | 0.376 | 0.18 | 0.000129716 | 20 | Pcnx |
| Gimap55 | 6.60E-09 | 0.386337578 | 0.356 | 0.172 | 0.000133635 | 20 | Gimap5 |
| Herc17 | 6.78E-09 | 0.457522135 | 0.463 | 0.258 | 0.000137299 | 20 | Herc1 |
| Plekho16 | 7.17E-09 | 0.343871118 | 0.443 | 0.225 | 0.000145148 | 20 | Plekho1 |
| Btbd12 | 7.62E-09 | 0.271224915 | 0.45 | 0.241 | 0.000154337 | 20 | Btbd1 |
| Erc13 | 7.64E-09 | 0.403850043 | 0.282 | 0.124 | 0.000154674 | 20 | Erc1 |
| Hnrnph14 | 7.68E-09 | 0.329991948 | 0.463 | 0.251 | 0.000155536 | 20 | Hnrnph1 |
| Prpf83 | 7.69E-09 | 0.390400444 | 0.369 | 0.184 | 0.000155781 | 20 | Prpf8 |
| Derl22 | 7.75E-09 | 0.365162337 | 0.282 | 0.127 | 0.000156865 | 20 | Derl2 |
| Mcl16 | 9.45E-09 | 0.302029951 | 0.638 | 0.374 | 0.000191248 | 20 | Mcl1 |
| Ogfrl12 | 9.95E-09 | 0.267036503 | 0.302 | 0.135 | 0.00020136 | 20 | Ogfrl1 |
| Lncpint9 | 1.00E-08 | 0.513414319 | 0.483 | 0.275 | 0.000202666 | 20 | Lncpint |
| Aida3 | 1.01E-08 | 0.274726843 | 0.315 | 0.144 | 0.000204043 | 20 | Aida |
| Trpm77 | 1.02E-08 | 0.413534809 | 0.389 | 0.205 | 0.00020664 | 20 | Trpm7 |
| Btbd911 | 1.10E-08 | 0.528829466 | 0.43 | 0.238 | 0.00022299 | 20 | Btbd9 |
| Tmed107 | 1.10E-08 | 0.260005891 | 0.711 | 0.437 | 0.000223127 | 20 | Tmed10 |
| Traf53 | 1.11E-08 | 0.368960536 | 0.268 | 0.119 | 0.000224548 | 20 | Traf5 |
| Igf2r4 | 1.11E-08 | 0.369713148 | 0.295 | 0.135 | 0.000224841 | 20 | Igf2r |
| Bbx7 | 1.15E-08 | 0.365544628 | 0.423 | 0.224 | 0.000233486 | 20 | Bbx |
| Col4a3bp3 | 1.19E-08 | 0.326881102 | 0.477 | 0.259 | 0.000240004 | 20 | Col4a3bp |
| Fat43 | 1.22E-08 | 0.318154838 | 0.255 | 0.107 | 0.000246963 | 20 | Fat4 |
| Lgals81 | 1.24E-08 | 0.314361244 | 0.322 | 0.154 | 0.000250875 | 20 | Lgals8 |
| Ap3b18 | 1.25E-08 | 0.329908304 | 0.423 | 0.219 | 0.000252105 | 20 | Ap3b1 |
| Dtd1 | 1.25E-08 | 0.309753559 | 0.309 | 0.141 | 0.000253101 | 20 | Dtd1 |
| Dyrk1a7 | 1.27E-08 | 0.428715457 | 0.356 | 0.181 | 0.000257264 | 20 | Dyrk1a |
| Cdkal17 | 1.27E-08 | 0.428736259 | 0.356 | 0.181 | 0.000257584 | 20 | Cdkal1 |
| Tbl1x6 | 1.27E-08 | 0.310898341 | 0.477 | 0.263 | 0.000257646 | 20 | Tbl1x |
| Afap14 | 1.30E-08 | 0.361121226 | 0.322 | 0.151 | 0.000263985 | 20 | Afap1 |
| Irf28 | 1.32E-08 | 0.361363814 | 0.591 | 0.343 | 0.000268058 | 20 | Irf2 |
| R3hdm13 | 1.34E-08 | 0.315050551 | 0.376 | 0.192 | 0.000271323 | 20 | R3hdm1 |
| Nbeal15 | 1.35E-08 | 0.382401291 | 0.47 | 0.263 | 0.000273199 | 20 | Nbeal1 |
| Agtrap2 | 1.37E-08 | 0.399935785 | 0.282 | 0.128 | 0.000277787 | 20 | Agtrap |
| C2cd22 | 1.38E-08 | 0.316821528 | 0.262 | 0.112 | 0.000279044 | 20 | C2cd2 |

| | | | | | | | |
|---|---|---|---|---|---|---|---|
| Rere7 | 1.40E-08 | 0.431719497 | 0.483 | 0.273 | 0.000283996 | 20 | Rere |
| Lrrc586 | 1.46E-08 | 0.364873459 | 0.416 | 0.215 | 0.000295121 | 20 | Lrrc58 |
| Luc7l210 | 1.50E-08 | 0.322484447 | 0.658 | 0.406 | 0.00030299 | 20 | Luc7l2 |
| Klhl244 | 1.51E-08 | 0.34456212 | 0.409 | 0.216 | 0.000305996 | 20 | Klhl24 |
| Rbm3910 | 1.53E-08 | 0.352907371 | 0.872 | 0.575 | 0.000309768 | 20 | Rbm39 |
| Tlk21 | 1.54E-08 | 0.338605657 | 0.329 | 0.161 | 0.00031182 | 20 | Tlk2 |
| 1700025G04Rik6 | 1.54E-08 | 0.369378034 | 0.369 | 0.19 | 0.000312699 | 20 | 1700025G04Rik |
| Pdzd82 | 1.62E-08 | 0.371017131 | 0.295 | 0.137 | 0.000328668 | 20 | Pdzd8 |
| Faf17 | 1.65E-08 | 0.279813779 | 0.396 | 0.203 | 0.000334811 | 20 | Faf1 |
| Stxbp54 | 1.67E-08 | 0.391697274 | 0.282 | 0.131 | 0.000337616 | 20 | Stxbp5 |
| Adgrg14 | 1.68E-08 | 0.336435599 | 0.309 | 0.141 | 0.000341084 | 20 | Adgrg1 |
| Ddx462 | 1.69E-08 | 0.255656083 | 0.43 | 0.226 | 0.000341949 | 20 | Ddx46 |
| Tnfaip23 | 1.69E-08 | 0.287488085 | 0.329 | 0.153 | 0.000343065 | 20 | Tnfaip2 |
| Thoc11 | 1.85E-08 | 0.319710172 | 0.268 | 0.118 | 0.000374142 | 20 | Thoc1 |
| Man2a15 | 1.90E-08 | 0.380007513 | 0.443 | 0.235 | 0.000385237 | 20 | Man2a1 |
| Stk19 | 1.91E-08 | 0.251713147 | 0.268 | 0.117 | 0.000386111 | 20 | Stk19 |
| Fam168a3 | 1.93E-08 | 0.340595228 | 0.309 | 0.144 | 0.000391424 | 20 | Fam168a |
| Ccdc88a11 | 1.96E-08 | 0.386217944 | 0.396 | 0.206 | 0.00039672 | 20 | Ccdc88a |
| Tmco16 | 1.98E-08 | 0.275923194 | 0.537 | 0.313 | 0.000399993 | 20 | Tmco1 |
| Kat6a5 | 2.00E-08 | 0.328473506 | 0.43 | 0.229 | 0.000404395 | 20 | Kat6a |
| Cdc42ep38 | 2.05E-08 | 0.457069421 | 0.349 | 0.176 | 0.000415953 | 20 | Cdc42ep3 |
| Agtpbp1 | 2.19E-08 | 0.331328373 | 0.262 | 0.116 | 0.000444001 | 20 | Agtpbp1 |
| Ier5l3 | 2.30E-08 | 0.360851489 | 0.362 | 0.179 | 0.00046491 | 20 | Ier5l |
| Ep4006 | 2.31E-08 | 0.445378668 | 0.389 | 0.209 | 0.000466709 | 20 | Ep400 |
| Ppp4r12 | 2.33E-08 | 0.336602408 | 0.262 | 0.118 | 0.00047233 | 20 | Ppp4r1 |
| Tmed73 | 2.38E-08 | 0.335630361 | 0.315 | 0.15 | 0.000481152 | 20 | Tmed7 |
| Usp9x2 | 2.47E-08 | 0.372114675 | 0.43 | 0.236 | 0.000500588 | 20 | Usp9x |
| Acbd31 | 2.63E-08 | 0.277581624 | 0.396 | 0.208 | 0.000531813 | 20 | Acbd3 |
| Clasp27 | 2.68E-08 | 0.411190065 | 0.356 | 0.183 | 0.000542993 | 20 | Clasp2 |
| Zfp207 | 2.74E-08 | 0.326255063 | 0.362 | 0.184 | 0.000555563 | 20 | Zfp207 |
| Coro1c2 | 2.75E-08 | 0.426683261 | 0.295 | 0.141 | 0.000557247 | 20 | Coro1c |
| Prkd34 | 2.75E-08 | 0.367399883 | 0.336 | 0.169 | 0.000557335 | 20 | Prkd3 |
| Ncoa34 | 2.76E-08 | 0.464646285 | 0.443 | 0.253 | 0.000559301 | 20 | Ncoa3 |
| Phlpp16 | 2.85E-08 | 0.527478951 | 0.47 | 0.265 | 0.000577124 | 20 | Phlpp1 |
| Flywch11 | 2.92E-08 | 0.334114456 | 0.262 | 0.114 | 0.000591681 | 20 | Flywch1 |
| Rsf17 | 2.96E-08 | 0.392360758 | 0.416 | 0.225 | 0.000599923 | 20 | Rsf1 |
| Tor1aip17 | 2.97E-08 | 0.324576279 | 0.51 | 0.282 | 0.000601389 | 20 | Tor1aip1 |
| Serpinh16 | 3.02E-08 | 0.281696372 | 0.483 | 0.251 | 0.000612325 | 20 | Serpinh1 |
| Babam28 | 3.04E-08 | 0.448114896 | 0.43 | 0.239 | 0.000615154 | 20 | Babam2 |
| Hipk31 | 3.04E-08 | 0.317825786 | 0.315 | 0.151 | 0.000615169 | 20 | Hipk3 |
| Ier510 | 3.13E-08 | 0.419386671 | 0.537 | 0.318 | 0.00063275 | 20 | Ier5 |
| Slc44a14 | 3.17E-08 | 0.411630428 | 0.329 | 0.157 | 0.000642455 | 20 | Slc44a1 |
| Surf41 | 3.30E-08 | 0.332747892 | 0.302 | 0.145 | 0.000668864 | 20 | Surf4 |

| | | | | | | | |
|---|---|---|---|---|---|---|---|
| Pacsin22 | 3.53E-08 | 0.352281328 | 0.322 | 0.159 | 0.000714338 | 20 | Pacsin2 |
| Epb413 | 3.56E-08 | 0.43568749 | 0.396 | 0.217 | 0.000720262 | 20 | Epb41 |
| Tmem597 | 3.72E-08 | 0.31830149 | 0.758 | 0.491 | 0.000753191 | 20 | Tmem59 |
| Arih19 | 3.75E-08 | 0.316843127 | 0.544 | 0.314 | 0.00075926 | 20 | Arih1 |
| Ahctf13 | 3.88E-08 | 0.309477054 | 0.275 | 0.125 | 0.000785516 | 20 | Ahctf1 |
| Dnajc113 | 3.91E-08 | 0.426534529 | 0.557 | 0.324 | 0.000791783 | 20 | Dnajc1 |
| Serinc17 | 3.93E-08 | 0.371107779 | 0.45 | 0.25 | 0.000795697 | 20 | Serinc1 |
| Mtf21 | 4.06E-08 | 0.460102584 | 0.282 | 0.135 | 0.000821615 | 20 | Mtf2 |
| Cop13 | 4.06E-08 | 0.319718776 | 0.356 | 0.18 | 0.000821865 | 20 | Cop1 |
| Tbc1d9b3 | 4.13E-08 | 0.317059544 | 0.268 | 0.123 | 0.000837124 | 20 | Tbc1d9b |
| Selenof6 | 4.14E-08 | 0.259374431 | 0.758 | 0.492 | 0.0008385 | 20 | Selenof |
| Atxn76 | 4.21E-08 | 0.417759144 | 0.329 | 0.167 | 0.000852108 | 20 | Atxn7 |
| Pdcd6ip3 | 4.67E-08 | 0.318840161 | 0.423 | 0.231 | 0.000944758 | 20 | Pdcd6ip |
| Ripor14 | 4.72E-08 | 0.294227284 | 0.336 | 0.162 | 0.00095626 | 20 | Ripor1 |
| Wls3 | 4.75E-08 | 0.38898333 | 0.275 | 0.128 | 0.000961084 | 20 | Wls |
| Zbtb382 | 5.13E-08 | 0.334981032 | 0.315 | 0.155 | 0.001038959 | 20 | Zbtb38 |
| 5031425E22Rik5 | 5.24E-08 | 0.373325009 | 0.322 | 0.161 | 0.00106025 | 20 | 5031425E22Rik |
| Ophn13 | 5.39E-08 | 0.388857842 | 0.289 | 0.134 | 0.00109086 | 20 | Ophn1 |
| Ywhab6 | 5.49E-08 | 0.257646215 | 0.705 | 0.468 | 0.001111586 | 20 | Ywhab |
| Snx96 | 5.58E-08 | 0.353213098 | 0.349 | 0.177 | 0.001130045 | 20 | Snx9 |
| Lrrk16 | 5.65E-08 | 0.344166396 | 0.369 | 0.192 | 0.001143763 | 20 | Lrrk1 |
| Mdfic5 | 5.93E-08 | 0.471099591 | 0.43 | 0.247 | 0.00120142 | 20 | Mdfic |
| Epc16 | 5.98E-08 | 0.289457808 | 0.389 | 0.204 | 0.001211539 | 20 | Epc1 |
| Cpsf64 | 6.02E-08 | 0.392370381 | 0.309 | 0.152 | 0.001219029 | 20 | Cpsf6 |
| Ceacam14 | 6.18E-08 | 0.33135188 | 0.329 | 0.157 | 0.001251826 | 20 | Ceacam1 |
| Ddx584 | 6.20E-08 | 0.37206455 | 0.329 | 0.161 | 0.001256049 | 20 | Ddx58 |
| Nfe2l25 | 6.23E-08 | 0.263888878 | 0.53 | 0.302 | 0.001261276 | 20 | Nfe2l2 |
| Dhx151 | 6.28E-08 | 0.261318024 | 0.383 | 0.203 | 0.001270702 | 20 | Dhx15 |
| Eea16 | 6.50E-08 | 0.350866732 | 0.403 | 0.219 | 0.001315031 | 20 | Eea1 |
| Spidr | 6.64E-08 | 0.294107126 | 0.255 | 0.113 | 0.001344665 | 20 | Spidr |
| Sgk33 | 6.84E-08 | 0.433789086 | 0.275 | 0.129 | 0.00138466 | 20 | Sgk3 |
| Zfp6448 | 7.06E-08 | 0.369157284 | 0.45 | 0.257 | 0.001430012 | 20 | Zfp644 |
| Luc7l34 | 7.10E-08 | 0.375221225 | 0.477 | 0.272 | 0.001438425 | 20 | Luc7l3 |
| Dag15 | 7.20E-08 | 0.299031419 | 0.302 | 0.143 | 0.001457544 | 20 | Dag1 |
| Map7d14 | 7.34E-08 | 0.26534123 | 0.342 | 0.168 | 0.001485531 | 20 | Map7d1 |
| Zfp2929 | 8.11E-08 | 0.306378324 | 0.45 | 0.252 | 0.001641752 | 20 | Zfp292 |
| Ppp4r22 | 8.14E-08 | 0.25339293 | 0.416 | 0.225 | 0.001647473 | 20 | Ppp4r2 |
| Acap27 | 8.24E-08 | 0.406698435 | 0.463 | 0.271 | 0.001668776 | 20 | Acap2 |
| Slk6 | 8.40E-08 | 0.362598803 | 0.436 | 0.242 | 0.00170039 | 20 | Slk |
| Zfp1488 | 8.73E-08 | 0.316406899 | 0.463 | 0.266 | 0.001767195 | 20 | Zfp148 |
| Srsf42 | 9.17E-08 | 0.31757502 | 0.322 | 0.161 | 0.00185721 | 20 | Srsf4 |
| Asxl13 | 9.25E-08 | 0.414536402 | 0.302 | 0.147 | 0.001872867 | 20 | Asxl1 |
| Srgap28 | 9.45E-08 | 0.477432089 | 0.51 | 0.305 | 0.001913747 | 20 | Srgap2 |

| | | | | | | | |
|---|---|---|---|---|---|---|---|
| Tnks28 | 9.63E-08 | 0.326705469 | 0.483 | 0.281 | 0.001949775 | 20 | Tnks2 |
| Sdf45 | 9.83E-08 | 0.321421383 | 0.517 | 0.31 | 0.001989217 | 20 | Sdf4 |
| Mar-72 | 9.96E-08 | 0.352471865 | 0.349 | 0.184 | 0.0020156 | 20 | 7-Mar |
| Epc23 | 1.01E-07 | 0.395711306 | 0.322 | 0.164 | 0.002037414 | 20 | Epc2 |
| Anks13 | 1.06E-07 | 0.357865631 | 0.289 | 0.14 | 0.002136624 | 20 | Anks1 |
| Dpysl27 | 1.06E-07 | 0.430990902 | 0.342 | 0.176 | 0.00215273 | 20 | Dpysl2 |
| Huwe110 | 1.10E-07 | 0.341242319 | 0.483 | 0.286 | 0.002228462 | 20 | Huwe1 |
| Fbxw73 | 1.12E-07 | 0.282026743 | 0.282 | 0.133 | 0.002260711 | 20 | Fbxw7 |
| Baz1b6 | 1.13E-07 | 0.433352407 | 0.369 | 0.195 | 0.00228279 | 20 | Baz1b |
| Faf21 | 1.14E-07 | 0.340242951 | 0.268 | 0.127 | 0.002315817 | 20 | Faf2 |
| Anapc16 | 1.22E-07 | 0.262501887 | 0.369 | 0.194 | 0.002468999 | 20 | Anapc16 |
| Chd68 | 1.24E-07 | 0.403375767 | 0.396 | 0.218 | 0.002515956 | 20 | Chd6 |
| Fam76b1 | 1.26E-07 | 0.342142757 | 0.295 | 0.145 | 0.00255994 | 20 | Fam76b |
| Pcm17 | 1.30E-07 | 0.445557092 | 0.43 | 0.252 | 0.002635741 | 20 | Pcm1 |
| Gls9 | 1.31E-07 | 0.416724107 | 0.436 | 0.252 | 0.002658152 | 20 | Gls |
| Arhgef38 | 1.32E-07 | 0.433324421 | 0.43 | 0.242 | 0.002668454 | 20 | Arhgef3 |
| Ccdc474 | 1.34E-07 | 0.271197895 | 0.369 | 0.194 | 0.002709649 | 20 | Ccdc47 |
| Ppp2r5c2 | 1.37E-07 | 0.325011214 | 0.443 | 0.248 | 0.002766764 | 20 | Ppp2r5c |
| Xpo74 | 1.40E-07 | 0.34450475 | 0.309 | 0.155 | 0.002842647 | 20 | Xpo7 |
| Stau1 | 1.49E-07 | 0.281419314 | 0.262 | 0.122 | 0.00301499 | 20 | Stau1 |
| Mkln14 | 1.53E-07 | 0.415331186 | 0.443 | 0.266 | 0.003099058 | 20 | Mkln1 |
| Evi54 | 1.53E-07 | 0.269881896 | 0.295 | 0.142 | 0.003103328 | 20 | Evi5 |
| Fubp15 | 1.66E-07 | 0.303848516 | 0.477 | 0.274 | 0.003365374 | 20 | Fubp1 |
| Wdr337 | 1.66E-07 | 0.328899556 | 0.423 | 0.24 | 0.003369002 | 20 | Wdr33 |
| Map45 | 1.69E-07 | 0.282431682 | 0.523 | 0.305 | 0.00341156 | 20 | Map4 |
| Prickle15 | 1.70E-07 | 0.319590022 | 0.309 | 0.147 | 0.00345035 | 20 | Prickle1 |
| Parp91 | 1.74E-07 | 0.334834349 | 0.282 | 0.134 | 0.003529604 | 20 | Parp9 |
| Cdk179 | 1.76E-07 | 0.473021205 | 0.436 | 0.246 | 0.003573353 | 20 | Cdk17 |
| Cd477 | 1.77E-07 | 0.258879397 | 0.812 | 0.511 | 0.003579799 | 20 | Cd47 |
| Trip128 | 1.84E-07 | 0.437315108 | 0.436 | 0.258 | 0.003721457 | 20 | Trip12 |
| Ankhd19 | 1.85E-07 | 0.338687079 | 0.43 | 0.245 | 0.003744256 | 20 | Ankhd1 |
| Tnrc6a2 | 1.94E-07 | 0.261538329 | 0.376 | 0.2 | 0.003931474 | 20 | Tnrc6a |
| Cnot6l5 | 1.97E-07 | 0.354447642 | 0.43 | 0.246 | 0.003983852 | 20 | Cnot6l |
| Epb41l26 | 1.98E-07 | 0.319645397 | 0.436 | 0.243 | 0.004004713 | 20 | Epb41l2 |
| Cers53 | 2.09E-07 | 0.278971681 | 0.282 | 0.135 | 0.00423208 | 20 | Cers5 |
| Nup982 | 2.24E-07 | 0.288174613 | 0.295 | 0.146 | 0.004525773 | 20 | Nup98 |
| Glud11 | 2.30E-07 | 0.435706812 | 0.55 | 0.337 | 0.00464762 | 20 | Glud1 |
| Zfr7 | 2.35E-07 | 0.408561472 | 0.423 | 0.249 | 0.004762609 | 20 | Zfr |
| Cnot49 | 2.43E-07 | 0.399884516 | 0.456 | 0.27 | 0.004911153 | 20 | Cnot4 |
| Med13l7 | 2.45E-07 | 0.362168113 | 0.362 | 0.195 | 0.004956559 | 20 | Med13l |
| Stard94 | 2.45E-07 | 0.319719263 | 0.309 | 0.153 | 0.004964876 | 20 | Stard9 |
| Tnfsf125 | 2.46E-07 | 0.280718069 | 0.302 | 0.149 | 0.004980695 | 20 | Tnfsf12 |
| Fchsd28 | 2.60E-07 | 0.337369613 | 0.315 | 0.161 | 0.005269384 | 20 | Fchsd2 |

| | | | | | | | |
|---|---|---|---|---|---|---|---|
| Dicer11 | 2.64E-07 | 0.349979424 | 0.262 | 0.124 | 0.005338167 | 20 | Dicer1 |
| Sos24 | 2.67E-07 | 0.320822225 | 0.315 | 0.162 | 0.005401443 | 20 | Sos2 |
| Ppil41 | 2.68E-07 | 0.265791072 | 0.329 | 0.168 | 0.005417187 | 20 | Ppil4 |
| Tsc22d24 | 2.72E-07 | 0.357739989 | 0.309 | 0.155 | 0.005504809 | 20 | Tsc22d2 |
| Emc77 | 2.78E-07 | 0.26959286 | 0.47 | 0.273 | 0.005633891 | 20 | Emc7 |
| Bptf11 | 3.01E-07 | 0.390553367 | 0.49 | 0.295 | 0.006094707 | 20 | Bptf |
| Zufsp7 | 3.03E-07 | 0.31933087 | 0.282 | 0.137 | 0.006130771 | 20 | Zufsp |
| Hnrnpr3 | 3.09E-07 | 0.402448073 | 0.409 | 0.237 | 0.006258179 | 20 | Hnrnpr |
| Rapgef33 | 3.10E-07 | 0.342968077 | 0.336 | 0.17 | 0.00627908 | 20 | Rapgef3 |
| Ssbp42 | 3.26E-07 | 0.296779854 | 0.362 | 0.194 | 0.006602319 | 20 | Ssbp4 |
| Smc47 | 3.30E-07 | 0.352737202 | 0.315 | 0.164 | 0.006690593 | 20 | Smc4 |
| Ccnd210 | 3.36E-07 | 0.341878991 | 0.483 | 0.287 | 0.006809777 | 20 | Ccnd2 |
| Dmxl17 | 3.59E-07 | 0.300619119 | 0.349 | 0.186 | 0.007266833 | 20 | Dmxl1 |
| Trim444 | 3.66E-07 | 0.327518389 | 0.396 | 0.229 | 0.00740831 | 20 | Trim44 |
| Papd44 | 3.68E-07 | 0.285747677 | 0.282 | 0.139 | 0.007449507 | 20 | Papd4 |
| Zfp4078 | 3.76E-07 | 0.377128958 | 0.315 | 0.163 | 0.007615566 | 20 | Zfp407 |
| Phf147 | 3.77E-07 | 0.335535182 | 0.423 | 0.246 | 0.0076297 | 20 | Phf14 |
| Ggta14 | 3.78E-07 | 0.289091229 | 0.289 | 0.142 | 0.007659151 | 20 | Ggta1 |
| Prpf61 | 3.90E-07 | 0.304865676 | 0.275 | 0.135 | 0.007885832 | 20 | Prpf6 |
| Braf4 | 4.00E-07 | 0.328768686 | 0.315 | 0.16 | 0.008096418 | 20 | Braf |
| Esyt210 | 4.04E-07 | 0.39507903 | 0.423 | 0.244 | 0.008172159 | 20 | Esyt2 |
| Saraf7 | 4.14E-07 | 0.272747505 | 0.517 | 0.308 | 0.0083774 | 20 | Saraf |
| Hsd17b121 | 4.15E-07 | 0.351306426 | 0.275 | 0.136 | 0.00839766 | 20 | Hsd17b12 |
| Ulk24 | 4.21E-07 | 0.273127516 | 0.342 | 0.18 | 0.008530928 | 20 | Ulk2 |
| Enox23 | 4.48E-07 | 0.535663463 | 0.282 | 0.144 | 0.009069613 | 20 | Enox2 |
| Dhx364 | 4.65E-07 | 0.330392685 | 0.416 | 0.236 | 0.009413686 | 20 | Dhx36 |
| Zfhx34 | 4.68E-07 | 0.263642425 | 0.356 | 0.184 | 0.009482073 | 20 | Zfhx3 |
| Hdac86 | 4.71E-07 | 0.448225487 | 0.275 | 0.137 | 0.009537271 | 20 | Hdac8 |
| Ino80d5 | 4.71E-07 | 0.351288097 | 0.322 | 0.167 | 0.009541412 | 20 | Ino80d |
| Zdhhc173 | 4.74E-07 | 0.310951787 | 0.302 | 0.151 | 0.009588816 | 20 | Zdhhc17 |
| Cuedc13 | 4.79E-07 | 0.333906106 | 0.282 | 0.137 | 0.009700474 | 20 | Cuedc1 |
| Rcn24 | 4.81E-07 | 0.311197161 | 0.289 | 0.144 | 0.009732007 | 20 | Rcn2 |
| Ambra19 | 4.93E-07 | 0.389818858 | 0.356 | 0.199 | 0.009990345 | 20 | Ambra1 |
| Trip115 | 4.99E-07 | 0.285131843 | 0.403 | 0.225 | 0.010109351 | 20 | Trip11 |
| Acin18 | 5.08E-07 | 0.341697764 | 0.517 | 0.318 | 0.010276458 | 20 | Acin1 |
| Zdhhc209 | 5.09E-07 | 0.426662029 | 0.403 | 0.231 | 0.010307514 | 20 | Zdhhc20 |
| Myo9a4 | 5.12E-07 | 0.52408959 | 0.322 | 0.175 | 0.010355901 | 20 | Myo9a |
| Eif5b4 | 5.19E-07 | 0.255724724 | 0.53 | 0.322 | 0.010514977 | 20 | Eif5b |
| Pds5a4 | 5.79E-07 | 0.385355197 | 0.369 | 0.206 | 0.011729 | 20 | Pds5a |
| Ggps14 | 5.84E-07 | 0.272801156 | 0.329 | 0.173 | 0.011816344 | 20 | Ggps1 |
| Lrba7 | 5.90E-07 | 0.424435268 | 0.349 | 0.193 | 0.011938281 | 20 | Lrba |
| Pnn8 | 6.01E-07 | 0.340333349 | 0.45 | 0.268 | 0.012168907 | 20 | Pnn |
| Ptbp22 | 6.13E-07 | 0.279117594 | 0.268 | 0.13 | 0.012403663 | 20 | Ptbp2 |

| | | | | | | | |
|---|---|---|---|---|---|---|---|
| Dennd5a5 | 6.22E-07 | 0.253074113 | 0.356 | 0.188 | 0.012597582 | 20 | Dennd5a |
| Pik3c2a4 | 6.22E-07 | 0.36333931 | 0.362 | 0.197 | 0.012603069 | 20 | Pik3c2a |
| Wrn4 | 6.56E-07 | 0.324298564 | 0.289 | 0.148 | 0.01327889 | 20 | Wrn |
| Zfp4451 | 6.76E-07 | 0.337377752 | 0.262 | 0.127 | 0.013681127 | 20 | Zfp445 |
| Kat6b3 | 6.94E-07 | 0.407772472 | 0.295 | 0.153 | 0.014043286 | 20 | Kat6b |
| Triobp1 | 6.96E-07 | 0.30387188 | 0.282 | 0.138 | 0.014085886 | 20 | Triobp |
| Emc35 | 7.16E-07 | 0.310582661 | 0.329 | 0.174 | 0.014492615 | 20 | Emc3 |
| Sel1l4 | 7.70E-07 | 0.286362833 | 0.315 | 0.163 | 0.015591164 | 20 | Sel1l |
| AI3141805 | 7.76E-07 | 0.36013412 | 0.369 | 0.212 | 0.015720619 | 20 | AI314180 |
| Brd16 | 7.88E-07 | 0.273573026 | 0.423 | 0.242 | 0.015944739 | 20 | Brd1 |
| Mertk5 | 8.20E-07 | 0.265899775 | 0.275 | 0.133 | 0.016607747 | 20 | Mertk |
| Sec635 | 8.34E-07 | 0.391905 | 0.362 | 0.204 | 0.016891869 | 20 | Sec63 |
| Amfr2 | 8.42E-07 | 0.294825898 | 0.255 | 0.125 | 0.017040209 | 20 | Amfr |
| Pdia68 | 8.46E-07 | 0.280601767 | 0.55 | 0.339 | 0.017122005 | 20 | Pdia6 |
| Cyp2d224 | 8.60E-07 | 0.262624935 | 0.282 | 0.136 | 0.017410364 | 20 | Cyp2d22 |
| N4bp13 | 8.88E-07 | 0.31541226 | 0.336 | 0.177 | 0.01798413 | 20 | N4bp1 |
| Rfx76 | 8.95E-07 | 0.365380853 | 0.302 | 0.156 | 0.018117199 | 20 | Rfx7 |
| Rictor6 | 9.35E-07 | 0.334316746 | 0.302 | 0.158 | 0.018923831 | 20 | Rictor |
| Psmd15 | 9.52E-07 | 0.291366681 | 0.376 | 0.211 | 0.019264395 | 20 | Psmd1 |
| Klhdc102 | 9.77E-07 | 0.301061003 | 0.302 | 0.155 | 0.019782623 | 20 | Klhdc10 |
| Tbc1d2b2 | 9.92E-07 | 0.265824065 | 0.255 | 0.122 | 0.020087609 | 20 | Tbc1d2b |
| Tgs12 | 9.98E-07 | 0.292337979 | 0.295 | 0.149 | 0.020214658 | 20 | Tgs1 |
| Taf35 | 1.03E-06 | 0.398340203 | 0.289 | 0.148 | 0.020821788 | 20 | Taf3 |
| Cep3509 | 1.03E-06 | 0.446582036 | 0.356 | 0.205 | 0.020849543 | 20 | Cep350 |
| Kif2a3 | 1.04E-06 | 0.299319967 | 0.329 | 0.179 | 0.021085604 | 20 | Kif2a |
| Upf3b2 | 1.04E-06 | 0.299242283 | 0.275 | 0.139 | 0.021107084 | 20 | Upf3b |
| Vav37 | 1.07E-06 | 0.374301011 | 0.436 | 0.257 | 0.021739487 | 20 | Vav3 |
| Elovl11 | 1.15E-06 | 0.280851376 | 0.255 | 0.121 | 0.023230459 | 20 | Elovl1 |
| Ifnar25 | 1.15E-06 | 0.256434807 | 0.544 | 0.326 | 0.02337424 | 20 | Ifnar2 |
| Setd72 | 1.17E-06 | 0.278145162 | 0.302 | 0.155 | 0.023653727 | 20 | Setd7 |
| S100a107 | 1.24E-06 | 0.277940161 | 0.752 | 0.47 | 0.02504618 | 20 | S100a10 |
| Cdk139 | 1.24E-06 | 0.26081122 | 0.409 | 0.228 | 0.025122474 | 20 | Cdk13 |
| Gatad2b5 | 1.27E-06 | 0.320022915 | 0.356 | 0.203 | 0.02571362 | 20 | Gatad2b |
| Ap1g1 | 1.36E-06 | 0.303096374 | 0.289 | 0.152 | 0.027498807 | 20 | Ap1g1 |
| Arglu17 | 1.38E-06 | 0.267094209 | 0.557 | 0.355 | 0.028008277 | 20 | Arglu1 |
| Ppp2r2a5 | 1.43E-06 | 0.33729214 | 0.376 | 0.214 | 0.028926488 | 20 | Ppp2r2a |
| Pde4b5 | 1.46E-06 | 0.360611131 | 0.356 | 0.197 | 0.029582579 | 20 | Pde4b |
| Tecpr14 | 1.50E-06 | 0.295459813 | 0.268 | 0.132 | 0.030317975 | 20 | Tecpr1 |
| Maml35 | 1.50E-06 | 0.450335431 | 0.369 | 0.209 | 0.030349312 | 20 | Maml3 |
| Cdk128 | 1.51E-06 | 0.433067327 | 0.396 | 0.231 | 0.030572345 | 20 | Cdk12 |
| Tbc1d511 | 1.58E-06 | 0.416951162 | 0.436 | 0.263 | 0.031915486 | 20 | Tbc1d5 |
| Bdp14 | 1.67E-06 | 0.273041887 | 0.315 | 0.167 | 0.033739767 | 20 | Bdp1 |
| Emc105 | 1.74E-06 | 0.295020105 | 0.356 | 0.201 | 0.035216452 | 20 | Emc10 |

| | | | | | | | |
|---|---|---|---|---|---|---|---|
| Smarcc12 | 1.75E-06 | 0.328923327 | 0.275 | 0.143 | 0.035414703 | 20 | Smarcc1 |
| Itch9 | 1.81E-06 | 0.311847614 | 0.463 | 0.275 | 0.036567845 | 20 | Itch |
| Lbr9 | 1.81E-06 | 0.338721043 | 0.295 | 0.154 | 0.036582769 | 20 | Lbr |
| Ankrd1211 | 1.89E-06 | 0.317389877 | 0.503 | 0.309 | 0.038221957 | 20 | Ankrd12 |
| Filip1l7 | 1.91E-06 | 0.283270298 | 0.295 | 0.154 | 0.038719092 | 20 | Filip1l |
| Creb16 | 1.96E-06 | 0.254658334 | 0.369 | 0.209 | 0.03971248 | 20 | Creb1 |
| Cnot15 | 1.97E-06 | 0.316111902 | 0.322 | 0.174 | 0.03984035 | 20 | Cnot1 |
| Lnpep12 | 1.99E-06 | 0.341946716 | 0.43 | 0.261 | 0.040211597 | 20 | Lnpep |
| Rnf2165 | 2.10E-06 | 0.399569436 | 0.329 | 0.182 | 0.042430549 | 20 | Rnf216 |
| Oasl24 | 2.11E-06 | 0.514997965 | 0.262 | 0.132 | 0.042797861 | 20 | Oasl2 |
| Nrf16 | 2.19E-06 | 0.280773831 | 0.268 | 0.136 | 0.044415395 | 20 | Nrf1 |
| 9930021J03Rik3 | 2.35E-06 | 0.311485139 | 0.289 | 0.153 | 0.047537987 | 20 | 9930021J03Rik |
| Rmdn12 | 2.42E-06 | 0.260788311 | 0.342 | 0.187 | 0.049088367 | 20 | Rmdn1 |
| Atp1b36 | 2.47E-06 | 0.325586592 | 0.423 | 0.25 | 0.050086544 | 20 | Atp1b3 |
| Zfand311 | 2.69E-06 | 0.469308937 | 0.477 | 0.303 | 0.054446572 | 20 | Zfand3 |
| Mllt1010 | 2.83E-06 | 0.453399198 | 0.436 | 0.276 | 0.057368354 | 20 | Mllt10 |
| Pdia43 | 2.88E-06 | 0.309012529 | 0.309 | 0.168 | 0.058362908 | 20 | Pdia4 |
| Map3k15 | 2.97E-06 | 0.359832636 | 0.349 | 0.2 | 0.060150241 | 20 | Map3k1 |
| Gcc21 | 3.10E-06 | 0.299599384 | 0.342 | 0.193 | 0.062672327 | 20 | Gcc2 |
| Golga42 | 3.11E-06 | 0.253748157 | 0.302 | 0.159 | 0.062870477 | 20 | Golga4 |
| Stat62 | 3.12E-06 | 0.286968498 | 0.255 | 0.128 | 0.063067669 | 20 | Stat6 |
| Birc612 | 3.13E-06 | 0.254998355 | 0.55 | 0.342 | 0.063359659 | 20 | Birc6 |
| Lims18 | 3.24E-06 | 0.398698679 | 0.47 | 0.302 | 0.065561736 | 20 | Lims1 |
| Gimap87 | 3.29E-06 | 0.319720883 | 0.315 | 0.17 | 0.066578656 | 20 | Gimap8 |
| Elp43 | 3.30E-06 | 0.333538461 | 0.255 | 0.129 | 0.066842323 | 20 | Elp4 |
| Nsf1 | 3.38E-06 | 0.260345734 | 0.275 | 0.141 | 0.068336605 | 20 | Nsf |
| Tle47 | 3.38E-06 | 0.303019352 | 0.389 | 0.227 | 0.06834397 | 20 | Tle4 |
| Sppl2a5 | 3.38E-06 | 0.287856074 | 0.483 | 0.298 | 0.068385243 | 20 | Sppl2a |
| Tmem1318 | 3.41E-06 | 0.382219057 | 0.369 | 0.209 | 0.069099204 | 20 | Tmem131 |
| Sema6d6 | 3.58E-06 | 0.30967967 | 0.383 | 0.215 | 0.072512476 | 20 | Sema6d |
| Rasa26 | 3.60E-06 | 0.321751021 | 0.268 | 0.139 | 0.072794776 | 20 | Rasa2 |
| Spata137 | 3.62E-06 | 0.278544489 | 0.369 | 0.207 | 0.073210333 | 20 | Spata13 |
| Dgkz2 | 3.91E-06 | 0.349975059 | 0.255 | 0.131 | 0.079072881 | 20 | Dgkz |
| Ubr27 | 3.92E-06 | 0.280869762 | 0.396 | 0.231 | 0.079439644 | 20 | Ubr2 |
| Fnbp45 | 3.97E-06 | 0.309947925 | 0.255 | 0.13 | 0.080429964 | 20 | Fnbp4 |
| Wdr266 | 4.02E-06 | 0.306694105 | 0.362 | 0.21 | 0.081294186 | 20 | Wdr26 |
| Tpr9 | 4.07E-06 | 0.257941475 | 0.638 | 0.427 | 0.082418749 | 20 | Tpr |
| Pbx34 | 4.17E-06 | 0.400295112 | 0.268 | 0.141 | 0.084334121 | 20 | Pbx3 |
| Ccar17 | 4.21E-06 | 0.287124021 | 0.362 | 0.209 | 0.085299255 | 20 | Ccar1 |
| Rap1gds112 | 4.26E-06 | 0.283682901 | 0.436 | 0.262 | 0.086223199 | 20 | Rap1gds1 |
| Cep1704 | 4.41E-06 | 0.336042053 | 0.255 | 0.131 | 0.089304555 | 20 | Cep170 |
| Golim45 | 4.60E-06 | 0.298872991 | 0.329 | 0.182 | 0.093061837 | 20 | Golim4 |
| Nfx1 | 4.93E-06 | 0.273938647 | 0.302 | 0.163 | 0.099875536 | 20 | Nfx1 |

| | | | | | | | |
|---|---|---|---|---|---|---|---|
| Frmd4a5 | 4.95E-06 | 0.29323837 | 0.282 | 0.149 | 0.100154051 | 20 | Frmd4a |
| Agfg12 | 5.07E-06 | 0.310545901 | 0.302 | 0.165 | 0.102723568 | 20 | Agfg1 |
| Klf103 | 5.33E-06 | 0.325064144 | 0.302 | 0.165 | 0.107887856 | 20 | Klf10 |
| Chd13 | 5.47E-06 | 0.271178673 | 0.349 | 0.202 | 0.110711814 | 20 | Chd1 |
| Tbc1d151 | 5.79E-06 | 0.258621767 | 0.262 | 0.136 | 0.117125916 | 20 | Tbc1d15 |
| Eif2s3x | 5.85E-06 | 0.25273701 | 0.295 | 0.161 | 0.118387032 | 20 | Eif2s3x |
| Clk19 | 6.04E-06 | 0.300115929 | 0.463 | 0.293 | 0.122210031 | 20 | Clk1 |
| Nemf1 | 6.04E-06 | 0.253096577 | 0.315 | 0.175 | 0.122380211 | 20 | Nemf |
| Gbp53 | 6.33E-06 | 0.399831125 | 0.255 | 0.13 | 0.128139673 | 20 | Gbp5 |
| Zcchc65 | 6.36E-06 | 0.37355751 | 0.45 | 0.28 | 0.128769386 | 20 | Zcchc6 |
| Cast7 | 6.38E-06 | 0.336623405 | 0.483 | 0.313 | 0.129116088 | 20 | Cast |
| Dido1 | 6.64E-06 | 0.331512524 | 0.255 | 0.132 | 0.13436561 | 20 | Dido1 |
| Xpr16 | 6.76E-06 | 0.426406179 | 0.282 | 0.155 | 0.136936368 | 20 | Xpr1 |
| Hmg20a4 | 6.88E-06 | 0.319177527 | 0.268 | 0.144 | 0.139302787 | 20 | Hmg20a |
| Rassf15 | 7.10E-06 | 0.250843259 | 0.396 | 0.229 | 0.143842743 | 20 | Rassf1 |
| Skil4 | 7.34E-06 | 0.258380417 | 0.315 | 0.174 | 0.148695568 | 20 | Skil |
| Klf127 | 7.47E-06 | 0.282771808 | 0.362 | 0.209 | 0.151296483 | 20 | Klf12 |
| Vti1a9 | 7.59E-06 | 0.358278208 | 0.416 | 0.256 | 0.153578303 | 20 | Vti1a |
| Ppp3cb2 | 7.65E-06 | 0.253294746 | 0.309 | 0.171 | 0.154829165 | 20 | Ppp3cb |
| Phc34 | 7.69E-06 | 0.347309538 | 0.255 | 0.136 | 0.155596997 | 20 | Phc3 |
| Calu5 | 7.79E-06 | 0.267101876 | 0.356 | 0.203 | 0.157729056 | 20 | Calu |
| Wipf18 | 7.86E-06 | 0.375480273 | 0.43 | 0.269 | 0.159144465 | 20 | Wipf1 |
| Gbp84 | 7.87E-06 | 0.320252687 | 0.262 | 0.14 | 0.159302344 | 20 | Gbp8 |
| Sh3rf15 | 8.35E-06 | 0.295153813 | 0.255 | 0.132 | 0.169079031 | 20 | Sh3rf1 |
| Rif13 | 8.35E-06 | 0.275010909 | 0.289 | 0.156 | 0.169113802 | 20 | Rif1 |
| Kdm7a3 | 9.23E-06 | 0.290906496 | 0.295 | 0.163 | 0.186918539 | 20 | Kdm7a |
| Abi18 | 9.31E-06 | 0.44187225 | 0.423 | 0.271 | 0.188481136 | 20 | Abi1 |
| Rnf134 | 9.36E-06 | 0.251291981 | 0.322 | 0.18 | 0.189401856 | 20 | Rnf13 |
| Ppp1r16b9 | 9.54E-06 | 0.36193099 | 0.356 | 0.203 | 0.193107379 | 20 | Ppp1r16b |
| Appbp22 | 9.60E-06 | 0.263786637 | 0.268 | 0.141 | 0.194370677 | 20 | Appbp2 |
| Taf11 | 9.97E-06 | 0.298361941 | 0.268 | 0.144 | 0.201796452 | 20 | Taf1 |
| Scaf89 | 1.10E-05 | 0.334841086 | 0.322 | 0.186 | 0.221802348 | 20 | Scaf8 |
| Lrpap14 | 1.15E-05 | 0.250306263 | 0.342 | 0.198 | 0.23229892 | 20 | Lrpap1 |
| Car82 | 1.15E-05 | 0.289717129 | 0.342 | 0.188 | 0.233317205 | 20 | Car8 |
| Phf202 | 1.26E-05 | 0.325013662 | 0.268 | 0.146 | 0.25575746 | 20 | Phf20 |
| Chd85 | 1.33E-05 | 0.335460881 | 0.302 | 0.169 | 0.269660395 | 20 | Chd8 |
| Rprd23 | 1.34E-05 | 0.304264615 | 0.255 | 0.136 | 0.270511878 | 20 | Rprd2 |
| Supt64 | 1.42E-05 | 0.294928595 | 0.289 | 0.161 | 0.286684491 | 20 | Supt6 |
| Ar5 | 1.46E-05 | 0.260119099 | 0.376 | 0.22 | 0.295580733 | 20 | Ar |
| Eml48 | 1.48E-05 | 0.355424866 | 0.416 | 0.26 | 0.299309986 | 20 | Eml4 |
| Bcas38 | 1.51E-05 | 0.263607961 | 0.322 | 0.184 | 0.305084357 | 20 | Bcas3 |
| Mar-62 | 1.53E-05 | 0.328395352 | 0.262 | 0.144 | 0.309869128 | 20 | 6-Mar |
| Oxr13 | 1.55E-05 | 0.26310936 | 0.53 | 0.337 | 0.314281202 | 20 | Oxr1 |

| | | | | | | | |
|---|---|---|---|---|---|---|---|
| Gmds3 | 1.57E-05 | 0.351594246 | 0.315 | 0.18 | 0.31698127 | 20 | Gmds |
| Tap25 | 1.66E-05 | 0.289293765 | 0.396 | 0.242 | 0.335467833 | 20 | Tap2 |
| Erp441 | 1.67E-05 | 0.255684905 | 0.302 | 0.171 | 0.337403136 | 20 | Erp44 |
| Tapbp6 | 1.70E-05 | 0.273651885 | 0.477 | 0.306 | 0.343907342 | 20 | Tapbp |
| Psme44 | 1.82E-05 | 0.314186202 | 0.336 | 0.199 | 0.36803757 | 20 | Psme4 |
| Cntln5 | 2.01E-05 | 0.3111649 | 0.282 | 0.156 | 0.40672607 | 20 | Cntln |
| Mecp23 | 2.11E-05 | 0.279173667 | 0.289 | 0.161 | 0.427932756 | 20 | Mecp2 |
| Pias15 | 2.24E-05 | 0.301637168 | 0.322 | 0.19 | 0.453998937 | 20 | Pias1 |
| Slc10a73 | 2.41E-05 | 0.337580961 | 0.268 | 0.149 | 0.488257507 | 20 | Slc10a7 |
| Dnm23 | 2.44E-05 | 0.254715336 | 0.423 | 0.26 | 0.493230686 | 20 | Dnm2 |
| Cep85l5 | 2.62E-05 | 0.262563072 | 0.302 | 0.166 | 0.531442436 | 20 | Cep85l |
| Ccdc826 | 2.67E-05 | 0.266320251 | 0.315 | 0.182 | 0.540034134 | 20 | Ccdc82 |
| Lmbrd16 | 2.75E-05 | 0.278788502 | 0.289 | 0.164 | 0.555851207 | 20 | Lmbrd1 |
| Ankrd1112 | 2.78E-05 | 0.255086543 | 0.597 | 0.382 | 0.563093397 | 20 | Ankrd11 |
| Nsd39 | 2.91E-05 | 0.264012761 | 0.497 | 0.309 | 0.588368303 | 20 | Nsd3 |
| Vgll46 | 3.31E-05 | 0.320342716 | 0.396 | 0.251 | 0.669830027 | 20 | Vgll4 |
| 9930111J21Rik27 | 3.76E-05 | 0.304225938 | 0.416 | 0.258 | 0.76198351 | 20 | 9930111J21Rik2 |
| Rpn27 | 3.77E-05 | 0.25378408 | 0.349 | 0.213 | 0.762330436 | 20 | Rpn2 |
| Zc3hav16 | 3.80E-05 | 0.31998516 | 0.342 | 0.206 | 0.768681801 | 20 | Zc3hav1 |
| Pkn18 | 4.12E-05 | 0.345953723 | 0.315 | 0.188 | 0.834206139 | 20 | Pkn1 |
| Stt3b8 | 4.17E-05 | 0.347197769 | 0.289 | 0.166 | 0.843752772 | 20 | Stt3b |
| Naa154 | 4.38E-05 | 0.325529635 | 0.289 | 0.167 | 0.885883104 | 20 | Naa15 |
| Tial11 | 4.43E-05 | 0.259423565 | 0.275 | 0.155 | 0.896351253 | 20 | Tial1 |
| Os98 | 4.52E-05 | 0.254644206 | 0.322 | 0.19 | 0.914956006 | 20 | Os9 |
| Fam193a7 | 4.70E-05 | 0.307880124 | 0.302 | 0.181 | 0.952464115 | 20 | Fam193a |
| Trps18 | 4.90E-05 | 0.305675629 | 0.383 | 0.237 | 0.99153513 | 20 | Trps1 |
| Hsp90b110 | 5.45E-05 | 0.253814107 | 0.738 | 0.512 | 1 | 20 | Hsp90b1 |
| Brwd13 | 5.86E-05 | 0.296630663 | 0.255 | 0.143 | 1 | 20 | Brwd1 |
| Helz5 | 6.40E-05 | 0.272442245 | 0.295 | 0.17 | 1 | 20 | Helz |
| Kdm6a10 | 6.63E-05 | 0.290630813 | 0.342 | 0.215 | 1 | 20 | Kdm6a |
| Ap2a22 | 6.77E-05 | 0.282136567 | 0.268 | 0.151 | 1 | 20 | Ap2a2 |
| Senp78 | 7.51E-05 | 0.290988875 | 0.275 | 0.158 | 1 | 20 | Senp7 |
| Tnrc6b9 | 8.04E-05 | 0.278847885 | 0.537 | 0.373 | 1 | 20 | Tnrc6b |
| Rell12 | 8.07E-05 | 0.290112078 | 0.255 | 0.138 | 1 | 20 | Rell1 |
| Trappc96 | 9.24E-05 | 0.304886332 | 0.322 | 0.196 | 1 | 20 | Trappc9 |
| Atf7ip6 | 0.000114868 | 0.294308053 | 0.336 | 0.208 | 1 | 20 | Atf7ip |
| Krit18 | 0.000144859 | 0.292625451 | 0.302 | 0.184 | 1 | 20 | Krit1 |
| Cnot25 | 0.000166429 | 0.293460958 | 0.336 | 0.212 | 1 | 20 | Cnot2 |
| Kras4 | 0.000177594 | 0.310418754 | 0.403 | 0.271 | 1 | 20 | Kras |
| Fam168b3 | 0.000189953 | 0.257290236 | 0.275 | 0.168 | 1 | 20 | Fam168b |
| Smg75 | 0.000234765 | 0.257596245 | 0.268 | 0.157 | 1 | 20 | Smg7 |
| Sesn17 | 0.000247649 | 0.310420247 | 0.336 | 0.214 | 1 | 20 | Sesn1 |
| Tmem1352 | 0.000287994 | 0.257099037 | 0.275 | 0.168 | 1 | 20 | Tmem135 |

| | | | | | | | |
|---|---|---|---|---|---|---|---|
| Dlg13 | 0.000292357 | 0.594825929 | 0.322 | 0.214 | 1 | 20 | Dlg1 |
| Rasa16 | 0.000305882 | 0.256830245 | 0.383 | 0.249 | 1 | 20 | Rasa1 |
| Trim12a6 | 0.000339498 | 0.254685043 | 0.295 | 0.183 | 1 | 20 | Trim12a |
| Gm424183 | 0.000347508 | 0.278367648 | 1 | 1 | 1 | 20 | Gm42418 |
| Pbrm15 | 0.000638572 | 0.301240523 | 0.409 | 0.289 | 1 | 20 | Pbrm1 |
| 2810474O19Rik3 | 0.001089266 | 0.260909176 | 0.322 | 0.212 | 1 | 20 | 2810474O19Rik |
| Nav26 | 0.001428058 | 0.371074012 | 0.268 | 0.167 | 1 | 20 | Nav2 |
| Stat16 | 0.001756144 | 0.320223261 | 0.369 | 0.251 | 1 | 20 | Stat1 |
| Eif2ak23 | 0.003168937 | 0.286741261 | 0.255 | 0.166 | 1 | 20 | Eif2ak2 |
| Herc62 | 0.006872252 | 0.307273395 | 0.275 | 0.186 | 1 | 20 | Herc6 |
| Acta21 | 0 | 6.137687993 | 0.563 | 0.019 | 0 | 21 | Acta2 |
| Tpm21 | 0 | 5.459723592 | 0.63 | 0.027 | 0 | 21 | Tpm2 |
| Tagln1 | 0 | 5.083929468 | 0.58 | 0.012 | 0 | 21 | Tagln |
| Pcp4 | 0 | 4.360679335 | 0.471 | 0.002 | 0 | 21 | Pcp4 |
| Myh111 | 0 | 4.218909254 | 0.479 | 0.009 | 0 | 21 | Myh11 |
| Myl91 | 0 | 3.870975602 | 0.58 | 0.021 | 0 | 21 | Myl9 |
| Cnn1 | 0 | 3.406912533 | 0.454 | 0.001 | 0 | 21 | Cnn1 |
| Krt14 | 0 | 3.101740282 | 0.63 | 0.002 | 0 | 21 | Krt14 |
| Igfbp2 | 0 | 2.893379536 | 0.42 | 0.001 | 0 | 21 | Igfbp2 |
| Cck | 0 | 2.773948234 | 0.378 | 0.004 | 0 | 21 | Cck |
| Krt17 | 0 | 2.231132445 | 0.462 | 0.001 | 0 | 21 | Krt17 |
| Krt5 | 0 | 2.170384628 | 0.487 | 0.002 | 0 | 21 | Krt5 |
| Lmod11 | 0 | 1.913641253 | 0.387 | 0.01 | 0 | 21 | Lmod1 |
| Fhod3 | 0 | 1.781011853 | 0.37 | 0.002 | 0 | 21 | Fhod3 |
| Osr1 | 0 | 1.587921232 | 0.345 | 0.007 | 0 | 21 | Osr1 |
| Gm30613 | 0 | 1.537485672 | 0.303 | 0 | 0 | 21 | Gm30613 |
| Sh3bgr | 0 | 1.507562415 | 0.336 | 0.005 | 0 | 21 | Sh3bgr |
| Fbxl22 | 0 | 1.452106829 | 0.303 | 0.006 | 0 | 21 | Fbxl22 |
| Ntng1 | 0 | 1.446575683 | 0.353 | 0.003 | 0 | 21 | Ntng1 |
| Trp63 | 0 | 1.435165221 | 0.429 | 0.001 | 0 | 21 | Trp63 |
| Trim29 | 0 | 1.368377405 | 0.462 | 0.003 | 0 | 21 | Trim29 |
| Synpo21 | 0 | 1.356437315 | 0.353 | 0.006 | 0 | 21 | Synpo2 |
| Pgm51 | 0 | 1.247772447 | 0.353 | 0.004 | 0 | 21 | Pgm5 |
| Actg2 | 0 | 1.175788613 | 0.269 | 0 | 0 | 21 | Actg2 |
| Dlk2 | 0 | 1.117675508 | 0.353 | 0.008 | 0 | 21 | Dlk2 |
| Fmod | 0 | 0.998899948 | 0.294 | 0 | 0 | 21 | Fmod |
| Trnp1 | 0 | 0.985409447 | 0.261 | 0.003 | 0 | 21 | Trnp1 |
| Vsnl1 | 0 | 0.949408796 | 0.294 | 0.001 | 0 | 21 | Vsnl1 |
| Fat1 | 0 | 0.949085712 | 0.378 | 0.01 | 0 | 21 | Fat1 |
| Cdh3 | 0 | 0.937998142 | 0.311 | 0 | 0 | 21 | Cdh3 |
| Mia | 0 | 0.936902473 | 0.311 | 0.002 | 0 | 21 | Mia |
| Tfap2c | 0 | 0.888763356 | 0.303 | 0.001 | 0 | 21 | Tfap2c |
| Col17a1 | 0 | 0.846575184 | 0.311 | 0.002 | 0 | 21 | Col17a1 |

| | | | | | | | |
|---|---|---|---|---|---|---|---|
| 4631405K08Rik | 0 | 0.832778834 | 0.294 | 0 | 0 | 21 | 4631405K08Rik |
| Moxd1 | 0 | 0.811414623 | 0.294 | 0.002 | 0 | 21 | Moxd1 |
| Gm9885 | 0 | 0.771677827 | 0.252 | 0.001 | 0 | 21 | Gm9885 |
| Dgki | 0 | 0.738962177 | 0.252 | 0.002 | 0 | 21 | Dgki |
| Plch2 | 0 | 0.730186268 | 0.286 | 0.001 | 0 | 21 | Plch2 |
| Sema3d | 0 | 0.65607603 | 0.277 | 0.004 | 0 | 21 | Sema3d |
| Postn1 | 4.90E-294 | 2.057504414 | 0.361 | 0.009 | 9.92E-290 | 21 | Postn |
| Carmn1 | 7.72E-287 | 1.516963429 | 0.336 | 0.008 | 1.56E-282 | 21 | Carmn |
| Sostdc1 | 3.54E-284 | 0.954190524 | 0.261 | 0.005 | 7.17E-280 | 21 | Sostdc1 |
| Ntrk31 | 2.59E-278 | 2.225242077 | 0.403 | 0.013 | 5.25E-274 | 21 | Ntrk3 |
| Serpinb5 | 1.56E-264 | 0.694651408 | 0.261 | 0.005 | 3.15E-260 | 21 | Serpinb5 |
| Gm15867 | 4.77E-262 | 0.992099339 | 0.252 | 0.005 | 9.67E-258 | 21 | Gm15867 |
| Grhl2 | 1.95E-251 | 0.608304368 | 0.311 | 0.008 | 3.96E-247 | 21 | Grhl2 |
| Ptprf | 1.20E-242 | 0.829707569 | 0.303 | 0.008 | 2.43E-238 | 21 | Ptprf |
| Rgs61 | 7.54E-240 | 1.541427554 | 0.319 | 0.009 | 1.53E-235 | 21 | Rgs6 |
| Kcnma1 | 2.06E-239 | 1.713193927 | 0.387 | 0.013 | 4.17E-235 | 21 | Kcnma1 |
| Sox10 | 1.31E-237 | 0.770081672 | 0.261 | 0.006 | 2.66E-233 | 21 | Sox10 |
| Nectin1 | 2.16E-237 | 1.061420078 | 0.345 | 0.011 | 4.37E-233 | 21 | Nectin1 |
| Sult5a1 | 1.57E-229 | 0.953131525 | 0.303 | 0.008 | 3.17E-225 | 21 | Sult5a1 |
| Adamts17 | 1.33E-214 | 0.90810254 | 0.261 | 0.007 | 2.70E-210 | 21 | Adamts17 |
| Dsp | 3.94E-206 | 1.456377944 | 0.42 | 0.019 | 7.98E-202 | 21 | Dsp |
| Ppp1r14a1 | 1.37E-203 | 1.466188827 | 0.319 | 0.011 | 2.78E-199 | 21 | Ppp1r14a |
| Nrg2 | 3.95E-192 | 1.344428939 | 0.37 | 0.016 | 7.99E-188 | 21 | Nrg2 |
| Sfrp11 | 4.58E-192 | 1.642601225 | 0.345 | 0.014 | 9.28E-188 | 21 | Sfrp1 |
| Slc25a21 | 2.33E-185 | 1.005920939 | 0.336 | 0.013 | 4.72E-181 | 21 | Slc25a21 |
| Nrtn | 1.62E-173 | 1.582454212 | 0.437 | 0.024 | 3.28E-169 | 21 | Nrtn |
| Sema5a1 | 4.86E-173 | 1.332992903 | 0.336 | 0.014 | 9.83E-169 | 21 | Sema5a |
| Ddr1 | 2.34E-169 | 0.734738081 | 0.277 | 0.01 | 4.75E-165 | 21 | Ddr1 |
| Ptprd1 | 1.17E-166 | 1.607756751 | 0.403 | 0.022 | 2.38E-162 | 21 | Ptprd |
| Adgrl31 | 3.28E-166 | 0.825258128 | 0.286 | 0.011 | 6.64E-162 | 21 | Adgrl3 |
| Mylk4 | 2.42E-162 | 3.705509657 | 0.571 | 0.048 | 4.91E-158 | 21 | Mylk |
| Perp | 4.05E-153 | 1.679612868 | 0.504 | 0.037 | 8.20E-149 | 21 | Perp |
| Cbr2 | 5.63E-150 | 0.882939933 | 0.361 | 0.019 | 1.14E-145 | 21 | Cbr2 |
| Egfr1 | 2.99E-144 | 1.273343245 | 0.42 | 0.027 | 6.06E-140 | 21 | Egfr |
| CT025619.11 | 3.30E-134 | 0.950638038 | 0.319 | 0.017 | 6.69E-130 | 21 | CT025619.1 |
| Dapl1 | 2.93E-129 | 1.78934771 | 0.496 | 0.041 | 5.93E-125 | 21 | Dapl1 |
| Fzd7 | 9.44E-117 | 0.691011937 | 0.294 | 0.016 | 1.91E-112 | 21 | Fzd7 |
| Fgf71 | 3.37E-116 | 0.857421226 | 0.261 | 0.013 | 6.81E-112 | 21 | Fgf7 |
| Bmpr1a2 | 1.37E-115 | 1.021707765 | 0.445 | 0.037 | 2.77E-111 | 21 | Bmpr1a |
| Dcn1 | 1.55E-113 | 0.522806169 | 0.504 | 0.046 | 3.13E-109 | 21 | Dcn |
| Aldh1a11 | 3.17E-113 | 1.342177275 | 0.412 | 0.033 | 6.43E-109 | 21 | Aldh1a1 |
| Igfbp52 | 8.76E-110 | 2.043539478 | 0.42 | 0.035 | 1.77E-105 | 21 | Igfbp5 |
| Col4a51 | 2.28E-108 | 0.714857705 | 0.277 | 0.015 | 4.62E-104 | 21 | Col4a5 |

| | | | | | | | |
|---|---|---|---|---|---|---|---|
| Sox9 | 6.93E-107 | 0.973439623 | 0.378 | 0.029 | 1.40E-102 | 21 | Sox9 |
| Pax9 | 8.39E-104 | 0.853680855 | 0.353 | 0.026 | 1.70E-99 | 21 | Pax9 |
| Nrbp2 | 7.19E-100 | 0.909085338 | 0.294 | 0.019 | 1.46E-95 | 21 | Nrbp2 |
| Prelp1 | 5.53E-99 | 0.889801073 | 0.336 | 0.025 | 1.12E-94 | 21 | Prelp |
| Crlf12 | 9.22E-99 | 1.016845144 | 0.395 | 0.034 | 1.87E-94 | 21 | Crlf1 |
| Matn2 | 4.21E-97 | 0.918023774 | 0.286 | 0.018 | 8.53E-93 | 21 | Matn2 |
| Rbms32 | 2.39E-95 | 2.155405766 | 0.479 | 0.052 | 4.84E-91 | 21 | Rbms3 |
| Fst1 | 7.56E-93 | 1.357715572 | 0.336 | 0.026 | 1.53E-88 | 21 | Fst |
| Gprc5b | 9.48E-93 | 0.589317885 | 0.286 | 0.019 | 1.92E-88 | 21 | Gprc5b |
| Pard33 | 6.38E-92 | 1.339739385 | 0.437 | 0.045 | 1.29E-87 | 21 | Pard3 |
| Fxyd31 | 1.38E-90 | 2.552298095 | 0.588 | 0.087 | 2.79E-86 | 21 | Fxyd3 |
| Bnc21 | 9.15E-88 | 0.61144572 | 0.269 | 0.018 | 1.85E-83 | 21 | Bnc2 |
| Pdgfa2 | 9.92E-87 | 1.474994397 | 0.487 | 0.059 | 2.01E-82 | 21 | Pdgfa |
| Arhgef252 | 1.48E-86 | 0.891855025 | 0.277 | 0.019 | 2.99E-82 | 21 | Arhgef25 |
| Gas12 | 5.70E-86 | 1.193140606 | 0.353 | 0.031 | 1.15E-81 | 21 | Gas1 |
| Phlda32 | 5.40E-85 | 1.421259095 | 0.462 | 0.053 | 1.09E-80 | 21 | Phlda3 |
| Antxr11 | 5.82E-85 | 0.695709721 | 0.294 | 0.022 | 1.18E-80 | 21 | Antxr1 |
| Lhfp2 | 2.02E-81 | 1.527574155 | 0.378 | 0.038 | 4.09E-77 | 21 | Lhfp |
| Meis11 | 1.74E-76 | 1.287581033 | 0.361 | 0.037 | 3.52E-72 | 21 | Meis1 |
| Patj | 8.65E-76 | 0.629944982 | 0.294 | 0.025 | 1.75E-71 | 21 | Patj |
| Gpc62 | 1.22E-72 | 1.306788511 | 0.328 | 0.031 | 2.47E-68 | 21 | Gpc6 |
| Nbl12 | 1.32E-68 | 0.837005327 | 0.269 | 0.023 | 2.67E-64 | 21 | Nbl1 |
| Clmp1 | 3.98E-67 | 0.821760073 | 0.269 | 0.024 | 8.06E-63 | 21 | Clmp |
| Cp3 | 4.94E-67 | 1.082796831 | 0.387 | 0.046 | 1.00E-62 | 21 | Cp |
| Irx5 | 1.77E-66 | 0.689137867 | 0.252 | 0.02 | 3.58E-62 | 21 | Irx5 |
| Palld3 | 3.63E-66 | 1.961337645 | 0.403 | 0.053 | 7.34E-62 | 21 | Palld |
| Htra13 | 6.57E-66 | 0.935257556 | 0.387 | 0.046 | 1.33E-61 | 21 | Htra1 |
| Enpp11 | 1.55E-64 | 0.650405179 | 0.286 | 0.027 | 3.15E-60 | 21 | Enpp1 |
| Lamb31 | 4.71E-64 | 1.035225215 | 0.412 | 0.055 | 9.54E-60 | 21 | Lamb3 |
| Mdk1 | 1.04E-62 | 0.711825743 | 0.286 | 0.027 | 2.11E-58 | 21 | Mdk |
| Irx2 | 7.02E-61 | 0.726505867 | 0.294 | 0.029 | 1.42E-56 | 21 | Irx2 |
| Cryab1 | 7.59E-60 | 1.450953701 | 0.462 | 0.07 | 1.54E-55 | 21 | Cryab |
| Arhgap422 | 2.72E-59 | 0.611967978 | 0.269 | 0.026 | 5.50E-55 | 21 | Arhgap42 |
| Nebl2 | 9.39E-58 | 1.456551423 | 0.361 | 0.048 | 1.90E-53 | 21 | Nebl |
| Enho | 2.96E-57 | 0.793726101 | 0.277 | 0.029 | 5.99E-53 | 21 | Enho |
| Sdc44 | 1.71E-55 | 1.08213664 | 0.437 | 0.071 | 3.47E-51 | 21 | Sdc4 |
| Dmd1 | 2.09E-55 | 2.77152036 | 0.387 | 0.059 | 4.23E-51 | 21 | Dmd |
| Krt81 | 1.62E-53 | 1.146796301 | 0.571 | 0.111 | 3.29E-49 | 21 | Krt8 |
| Snhg183 | 2.36E-52 | 1.143337377 | 0.319 | 0.04 | 4.78E-48 | 21 | Snhg18 |
| Fbxo32 | 3.74E-51 | 0.951475689 | 0.319 | 0.041 | 7.57E-47 | 21 | Fbxo32 |
| Pitx11 | 6.06E-51 | 0.928232473 | 0.445 | 0.073 | 1.23E-46 | 21 | Pitx1 |
| Hacd1 | 6.46E-51 | 0.905074829 | 0.303 | 0.037 | 1.31E-46 | 21 | Hacd1 |
| Synm1 | 1.40E-50 | 1.413402899 | 0.303 | 0.039 | 2.84E-46 | 21 | Synm |

| | | | | | | | |
|---|---|---|---|---|---|---|---|
| Tpm13 | 4.17E-45 | 3.27155524 | 0.739 | 0.293 | 8.43E-41 | 21 | Tpm1 |
| Enc1 | 8.29E-45 | 0.567294024 | 0.277 | 0.035 | 1.68E-40 | 21 | Enc1 |
| Krt23 | 1.22E-44 | 0.827963711 | 0.429 | 0.075 | 2.46E-40 | 21 | Krt23 |
| Bgn3 | 1.34E-44 | 0.893135394 | 0.319 | 0.045 | 2.71E-40 | 21 | Bgn |
| Ank32 | 2.37E-44 | 0.929818831 | 0.336 | 0.05 | 4.80E-40 | 21 | Ank3 |
| Abi3bp1 | 2.44E-44 | 0.809365742 | 0.269 | 0.033 | 4.93E-40 | 21 | Abi3bp |
| Phgdh | 8.33E-44 | 1.053205682 | 0.328 | 0.051 | 1.69E-39 | 21 | Phgdh |
| Gpx2 | 2.73E-43 | 1.536701614 | 0.303 | 0.043 | 5.52E-39 | 21 | Gpx2 |
| Net12 | 3.56E-43 | 0.607215831 | 0.286 | 0.038 | 7.21E-39 | 21 | Net1 |
| Pawr1 | 4.46E-43 | 0.808280831 | 0.378 | 0.064 | 9.04E-39 | 21 | Pawr |
| Oat3 | 7.58E-43 | 1.512264741 | 0.546 | 0.136 | 1.54E-38 | 21 | Oat |
| Ptprs1 | 1.03E-41 | 0.617513489 | 0.277 | 0.037 | 2.09E-37 | 21 | Ptprs |
| Chrm33 | 1.30E-41 | 1.931928221 | 0.454 | 0.1 | 2.64E-37 | 21 | Chrm3 |
| Csrp15 | 1.96E-39 | 2.790701126 | 0.571 | 0.183 | 3.98E-35 | 21 | Csrp1 |
| Sntb14 | 4.97E-39 | 0.91248483 | 0.361 | 0.065 | 1.01E-34 | 21 | Sntb1 |
| Kcnq57 | 1.49E-38 | 1.736942773 | 0.361 | 0.068 | 3.02E-34 | 21 | Kcnq5 |
| Paqr6 | 3.21E-38 | 0.553932019 | 0.277 | 0.039 | 6.49E-34 | 21 | Paqr6 |
| Gstm52 | 4.27E-38 | 0.732250646 | 0.361 | 0.063 | 8.64E-34 | 21 | Gstm5 |
| Itgb42 | 4.27E-37 | 0.448953271 | 0.252 | 0.034 | 8.65E-33 | 21 | Itgb4 |
| Twsg11 | 1.77E-36 | 0.600658216 | 0.294 | 0.046 | 3.59E-32 | 21 | Twsg1 |
| Spint23 | 4.91E-36 | 1.081283259 | 0.471 | 0.113 | 9.94E-32 | 21 | Spint2 |
| Fgfr11 | 7.68E-36 | 0.64869336 | 0.303 | 0.049 | 1.55E-31 | 21 | Fgfr1 |
| Flna6 | 2.80E-35 | 2.240995681 | 0.479 | 0.132 | 5.68E-31 | 21 | Flna |
| Tns33 | 2.84E-35 | 0.874971754 | 0.37 | 0.073 | 5.74E-31 | 21 | Tns3 |
| Mtss1l | 9.02E-35 | 0.454978159 | 0.252 | 0.036 | 1.83E-30 | 21 | Mtss1l |
| Aldh7a11 | 7.42E-34 | 0.824433666 | 0.336 | 0.062 | 1.50E-29 | 21 | Aldh7a1 |
| Nexn | 1.17E-33 | 0.948045306 | 0.328 | 0.062 | 2.36E-29 | 21 | Nexn |
| Wfdc181 | 1.42E-33 | 3.026706897 | 0.319 | 0.06 | 2.87E-29 | 21 | Wfdc18 |
| Lgals17 | 1.69E-33 | 1.686960577 | 0.504 | 0.135 | 3.42E-29 | 21 | Lgals1 |
| Tgfb1i12 | 4.01E-33 | 1.416591333 | 0.387 | 0.088 | 8.12E-29 | 21 | Tgfb1i1 |
| Plxdc24 | 2.32E-31 | 0.655539471 | 0.42 | 0.093 | 4.69E-27 | 21 | Plxdc2 |
| Eps8l21 | 2.62E-31 | 0.424496143 | 0.269 | 0.043 | 5.31E-27 | 21 | Eps8l2 |
| Ank21 | 4.27E-30 | 0.632556922 | 0.261 | 0.043 | 8.65E-26 | 21 | Ank2 |
| Mrps6 | 5.48E-30 | 1.030286076 | 0.37 | 0.083 | 1.11E-25 | 21 | Mrps6 |
| Rbp12 | 1.35E-29 | 2.053595249 | 0.395 | 0.102 | 2.73E-25 | 21 | Rbp1 |
| Fam213b | 1.98E-29 | 1.075806894 | 0.353 | 0.076 | 4.01E-25 | 21 | Fam213b |
| S100a69 | 2.54E-29 | 1.973647403 | 0.681 | 0.272 | 5.14E-25 | 21 | S100a6 |
| Tead14 | 2.12E-28 | 0.95041472 | 0.361 | 0.084 | 4.29E-24 | 21 | Tead1 |
| Tspan35 | 4.06E-28 | 0.999363192 | 0.412 | 0.105 | 8.23E-24 | 21 | Tspan3 |
| Hmgn14 | 4.95E-28 | 1.273494772 | 0.723 | 0.329 | 1.00E-23 | 21 | Hmgn1 |
| Adcy91 | 2.49E-27 | 0.678411322 | 0.294 | 0.058 | 5.04E-23 | 21 | Adcy9 |
| Spats2l1 | 2.76E-27 | 0.935381592 | 0.353 | 0.081 | 5.60E-23 | 21 | Spats2l |
| Dock73 | 5.87E-27 | 0.644744884 | 0.261 | 0.048 | 1.19E-22 | 21 | Dock7 |

| | | | | | | | |
|---|---|---|---|---|---|---|---|
| Msrb13 | 1.22E-26 | 1.489355569 | 0.462 | 0.142 | 2.47E-22 | 21 | Msrb1 |
| Rbbp82 | 8.33E-26 | 0.824553598 | 0.387 | 0.101 | 1.69E-21 | 21 | Rbbp8 |
| Cd636 | 1.09E-25 | 1.024660367 | 0.571 | 0.188 | 2.22E-21 | 21 | Cd63 |
| Map3k203 | 2.46E-25 | 0.603288149 | 0.328 | 0.073 | 4.98E-21 | 21 | Map3k20 |
| Gstm16 | 3.45E-25 | 1.364566203 | 0.689 | 0.323 | 6.98E-21 | 21 | Gstm1 |
| Cdh15 | 6.79E-25 | 0.567495353 | 0.328 | 0.075 | 1.37E-20 | 21 | Cdh1 |
| Sh3d194 | 7.26E-25 | 0.968520992 | 0.395 | 0.108 | 1.47E-20 | 21 | Sh3d19 |
| Ppm1l | 1.41E-24 | 0.68601514 | 0.277 | 0.057 | 2.85E-20 | 21 | Ppm1l |
| Krt72 | 1.59E-24 | 0.813433285 | 0.563 | 0.178 | 3.21E-20 | 21 | Krt7 |
| Asb23 | 1.95E-24 | 0.764046092 | 0.261 | 0.052 | 3.94E-20 | 21 | Asb2 |
| Dstn2 | 1.04E-23 | 1.809429148 | 0.815 | 0.574 | 2.10E-19 | 21 | Dstn |
| Lmo41 | 1.52E-23 | 1.346110791 | 0.513 | 0.194 | 3.08E-19 | 21 | Lmo4 |
| Pygb | 2.07E-23 | 0.594427359 | 0.353 | 0.088 | 4.20E-19 | 21 | Pygb |
| Samd42 | 3.43E-23 | 0.981496753 | 0.261 | 0.055 | 6.93E-19 | 21 | Samd4 |
| Pdlim41 | 1.32E-22 | 0.788570181 | 0.445 | 0.134 | 2.67E-18 | 21 | Pdlim4 |
| Cbx62 | 1.48E-22 | 0.45794752 | 0.294 | 0.066 | 3.00E-18 | 21 | Cbx6 |
| Myl68 | 1.70E-22 | 1.989411593 | 0.924 | 0.777 | 3.43E-18 | 21 | Myl6 |
| Pdlim73 | 3.24E-22 | 0.951123721 | 0.303 | 0.074 | 6.56E-18 | 21 | Pdlim7 |
| Nfix1 | 4.06E-22 | 1.199980392 | 0.538 | 0.216 | 8.22E-18 | 21 | Nfix |
| Rnase42 | 5.96E-22 | 0.558305852 | 0.345 | 0.086 | 1.21E-17 | 21 | Rnase4 |
| Auts22 | 1.53E-21 | 1.553738395 | 0.454 | 0.153 | 3.09E-17 | 21 | Auts2 |
| Atp2a27 | 1.77E-21 | 2.519080435 | 0.487 | 0.201 | 3.59E-17 | 21 | Atp2a2 |
| Maged11 | 4.47E-21 | 0.737898 | 0.345 | 0.089 | 9.05E-17 | 21 | Maged1 |
| Unc45a1 | 1.25E-20 | 0.558932805 | 0.252 | 0.055 | 2.54E-16 | 21 | Unc45a |
| Urah | 1.41E-20 | 0.35566097 | 0.277 | 0.061 | 2.85E-16 | 21 | Urah |
| Ppp1r12b2 | 3.34E-20 | 0.898039618 | 0.361 | 0.107 | 6.77E-16 | 21 | Ppp1r12b |
| Kctd11 | 4.02E-20 | 0.537049389 | 0.378 | 0.105 | 8.13E-16 | 21 | Kctd1 |
| Dpysl31 | 4.56E-20 | 0.494010703 | 0.286 | 0.067 | 9.22E-16 | 21 | Dpysl3 |
| Epcam1 | 5.21E-20 | 0.652420127 | 0.311 | 0.079 | 1.05E-15 | 21 | Epcam |
| Rasa35 | 3.92E-19 | 0.579150762 | 0.269 | 0.066 | 7.94E-15 | 21 | Rasa3 |
| Shtn14 | 6.13E-19 | 0.578799153 | 0.277 | 0.068 | 1.24E-14 | 21 | Shtn1 |
| Jag22 | 6.76E-19 | 0.421147706 | 0.319 | 0.083 | 1.37E-14 | 21 | Jag2 |
| Ift431 | 1.52E-18 | 0.861230281 | 0.395 | 0.124 | 3.09E-14 | 21 | Ift43 |
| Cd59a2 | 2.90E-18 | 0.847251383 | 0.311 | 0.088 | 5.87E-14 | 21 | Cd59a |
| Wls4 | 4.99E-18 | 0.831930918 | 0.387 | 0.128 | 1.01E-13 | 21 | Wls |
| Cd95 | 5.09E-18 | 1.426255432 | 0.63 | 0.387 | 1.03E-13 | 21 | Cd9 |
| Serpinb6b | 6.85E-18 | 0.548249651 | 0.319 | 0.087 | 1.39E-13 | 21 | Serpinb6b |
| Atpif14 | 1.21E-17 | 0.896195194 | 0.798 | 0.561 | 2.44E-13 | 21 | Atpif1 |
| Pbx15 | 1.42E-17 | 1.263712575 | 0.471 | 0.199 | 2.87E-13 | 21 | Pbx1 |
| Dag16 | 2.37E-17 | 0.803706872 | 0.403 | 0.143 | 4.80E-13 | 21 | Dag1 |
| Rhou | 2.49E-17 | 0.391907098 | 0.269 | 0.066 | 5.05E-13 | 21 | Rhou |
| Nenf4 | 3.76E-17 | 1.047608901 | 0.597 | 0.296 | 7.61E-13 | 21 | Nenf |
| Eya25 | 3.95E-17 | 0.576408898 | 0.336 | 0.101 | 7.99E-13 | 21 | Eya2 |

| | | | | | | | |
|---|---|---|---|---|---|---|---|
| Zc2hc1a2 | 5.56E-17 | 0.659113298 | 0.361 | 0.111 | 1.13E-12 | 21 | Zc2hc1a |
| Map21 | 6.22E-17 | 0.521679566 | 0.303 | 0.083 | 1.26E-12 | 21 | Map2 |
| Actn13 | 1.08E-16 | 0.579619737 | 0.294 | 0.084 | 2.18E-12 | 21 | Actn1 |
| S100a113 | 1.56E-16 | 1.107523924 | 0.824 | 0.574 | 3.16E-12 | 21 | S100a11 |
| Kirrel32 | 1.63E-16 | 0.661133857 | 0.269 | 0.071 | 3.31E-12 | 21 | Kirrel3 |
| Nedd47 | 1.65E-16 | 1.488261201 | 0.63 | 0.421 | 3.35E-12 | 21 | Nedd4 |
| Snai21 | 2.10E-16 | 0.695714 | 0.311 | 0.092 | 4.26E-12 | 21 | Snai2 |
| Tgm25 | 3.95E-16 | 0.817003153 | 0.445 | 0.163 | 8.00E-12 | 21 | Tgm2 |
| Yap11 | 4.17E-16 | 0.569889681 | 0.269 | 0.072 | 8.44E-12 | 21 | Yap1 |
| Hmgn33 | 4.49E-16 | 0.860335605 | 0.454 | 0.172 | 9.08E-12 | 21 | Hmgn3 |
| Asph5 | 7.59E-16 | 0.56635497 | 0.328 | 0.101 | 1.54E-11 | 21 | Asph |
| Scd21 | 7.81E-16 | 0.519119642 | 0.286 | 0.081 | 1.58E-11 | 21 | Scd2 |
| Pik3r14 | 2.99E-15 | 1.203452871 | 0.504 | 0.255 | 6.05E-11 | 21 | Pik3r1 |
| Fam129a8 | 3.95E-15 | 1.183154289 | 0.378 | 0.147 | 8.00E-11 | 21 | Fam129a |
| Stmn13 | 5.27E-15 | 0.978008858 | 0.37 | 0.131 | 1.07E-10 | 21 | Stmn1 |
| Dst8 | 1.51E-14 | 0.99982753 | 0.445 | 0.192 | 3.07E-10 | 21 | Dst |
| Siva12 | 2.20E-14 | 0.782685975 | 0.412 | 0.16 | 4.46E-10 | 21 | Siva1 |
| Ryk3 | 3.36E-14 | 0.438541373 | 0.336 | 0.111 | 6.81E-10 | 21 | Ryk |
| Tgfbr35 | 4.23E-14 | 0.840513714 | 0.429 | 0.175 | 8.57E-10 | 21 | Tgfbr3 |
| Pfkl | 1.44E-13 | 0.537077064 | 0.277 | 0.086 | 2.92E-09 | 21 | Pfkl |
| Plcb46 | 2.88E-13 | 1.812338161 | 0.504 | 0.293 | 5.83E-09 | 21 | Plcb4 |
| Id43 | 4.36E-13 | 0.853937009 | 0.345 | 0.126 | 8.82E-09 | 21 | Id4 |
| Ctdspl3 | 4.39E-13 | 0.362364691 | 0.294 | 0.092 | 8.90E-09 | 21 | Ctdspl |
| Tcf7l25 | 5.01E-13 | 0.705793049 | 0.353 | 0.13 | 1.02E-08 | 21 | Tcf7l2 |
| Arhgap107 | 5.63E-13 | 0.71395357 | 0.353 | 0.128 | 1.14E-08 | 21 | Arhgap10 |
| Adk1 | 6.14E-13 | 0.419058332 | 0.353 | 0.125 | 1.24E-08 | 21 | Adk |
| Ndufa112 | 8.08E-13 | 0.733908735 | 0.765 | 0.525 | 1.64E-08 | 21 | Ndufa11 |
| Mgst16 | 1.06E-12 | 0.55080628 | 0.42 | 0.168 | 2.15E-08 | 21 | Mgst1 |
| Parva4 | 1.08E-12 | 0.405228777 | 0.328 | 0.113 | 2.19E-08 | 21 | Parva |
| Timp26 | 1.27E-12 | 0.562035151 | 0.353 | 0.131 | 2.56E-08 | 21 | Timp2 |
| Arl32 | 1.28E-12 | 0.620637401 | 0.42 | 0.17 | 2.59E-08 | 21 | Arl3 |
| Sfn | 1.32E-12 | 0.516914805 | 0.252 | 0.075 | 2.67E-08 | 21 | Sfn |
| Tes7 | 3.28E-12 | 0.844718406 | 0.336 | 0.131 | 6.63E-08 | 21 | Tes |
| Cebpb6 | 5.39E-12 | 0.883972036 | 0.387 | 0.158 | 1.09E-07 | 21 | Cebpb |
| Ptov12 | 5.96E-12 | 0.538150303 | 0.328 | 0.119 | 1.21E-07 | 21 | Ptov1 |
| Ptpn144 | 5.99E-12 | 0.556145352 | 0.303 | 0.107 | 1.21E-07 | 21 | Ptpn14 |
| Mid12 | 7.09E-12 | 0.657576177 | 0.261 | 0.086 | 1.44E-07 | 21 | Mid1 |
| Atp13a35 | 7.11E-12 | 0.616354422 | 0.294 | 0.104 | 1.44E-07 | 21 | Atp13a3 |
| Ephx16 | 8.21E-12 | 0.637489946 | 0.328 | 0.123 | 1.66E-07 | 21 | Ephx1 |
| Ccdc1077 | 1.16E-11 | 0.544268853 | 0.336 | 0.128 | 2.36E-07 | 21 | Ccdc107 |
| Map1lc3a3 | 1.87E-11 | 0.716963697 | 0.731 | 0.483 | 3.79E-07 | 21 | Map1lc3a |
| Ehf2 | 1.89E-11 | 0.565813877 | 0.328 | 0.117 | 3.82E-07 | 21 | Ehf |
| Kif13a4 | 1.98E-11 | 0.569716375 | 0.403 | 0.175 | 4.02E-07 | 21 | Kif13a |

| | | | | | | | |
|---|---|---|---|---|---|---|---|
| Lzts2 | 2.09E-11 | 0.358870339 | 0.277 | 0.092 | 4.23E-07 | 21 | Lzts2 |
| Cited42 | 2.29E-11 | 0.540886271 | 0.353 | 0.135 | 4.63E-07 | 21 | Cited4 |
| Ppp2r3a3 | 2.30E-11 | 0.352198828 | 0.277 | 0.092 | 4.66E-07 | 21 | Ppp2r3a |
| Msrb35 | 2.46E-11 | 0.783605344 | 0.336 | 0.133 | 4.98E-07 | 21 | Msrb3 |
| Pde4d5 | 2.99E-11 | 0.622668164 | 0.37 | 0.151 | 6.06E-07 | 21 | Pde4d |
| Naca2 | 3.03E-11 | 0.659463469 | 0.874 | 0.834 | 6.13E-07 | 21 | Naca |
| Rcc23 | 3.27E-11 | 0.479241488 | 0.311 | 0.113 | 6.62E-07 | 21 | Rcc2 |
| Atp1a13 | 3.38E-11 | 1.118729079 | 0.496 | 0.281 | 6.85E-07 | 21 | Atp1a1 |
| Vcl5 | 5.32E-11 | 0.739504134 | 0.378 | 0.163 | 1.08E-06 | 21 | Vcl |
| Hspb17 | 6.87E-11 | 0.84772892 | 0.655 | 0.382 | 1.39E-06 | 21 | Hspb1 |
| Mif2 | 1.09E-10 | 0.632418752 | 0.765 | 0.524 | 2.21E-06 | 21 | Mif |
| Rab11fip22 | 1.15E-10 | 0.575930309 | 0.294 | 0.108 | 2.33E-06 | 21 | Rab11fip2 |
| Slc25a43 | 1.20E-10 | 0.681985413 | 0.748 | 0.561 | 2.44E-06 | 21 | Slc25a4 |
| Eps84 | 1.29E-10 | 0.476055556 | 0.261 | 0.088 | 2.61E-06 | 21 | Eps8 |
| Ckb6 | 1.33E-10 | 1.704753557 | 0.403 | 0.217 | 2.69E-06 | 21 | Ckb |
| Gtf2i4 | 1.33E-10 | 0.65662002 | 0.496 | 0.252 | 2.70E-06 | 21 | Gtf2i |
| Sorl17 | 1.43E-10 | 0.450314611 | 0.303 | 0.112 | 2.90E-06 | 21 | Sorl1 |
| Maea | 1.78E-10 | 0.634642439 | 0.345 | 0.141 | 3.59E-06 | 21 | Maea |
| Csad | 1.86E-10 | 0.267628432 | 0.269 | 0.092 | 3.77E-06 | 21 | Csad |
| Gsta42 | 2.12E-10 | 0.339632655 | 0.378 | 0.151 | 4.29E-06 | 21 | Gsta4 |
| Daam13 | 2.55E-10 | 0.566932172 | 0.294 | 0.11 | 5.16E-06 | 21 | Daam1 |
| Apip | 2.64E-10 | 0.453426822 | 0.303 | 0.112 | 5.34E-06 | 21 | Apip |
| Nxn5 | 2.78E-10 | 0.79548589 | 0.504 | 0.27 | 5.63E-06 | 21 | Nxn |
| Tnrc186 | 3.15E-10 | 0.467385401 | 0.395 | 0.174 | 6.37E-06 | 21 | Tnrc18 |
| Lpp11 | 4.38E-10 | 1.09605862 | 0.479 | 0.287 | 8.86E-06 | 21 | Lpp |
| Josd2 | 5.83E-10 | 0.458279685 | 0.277 | 0.103 | 1.18E-05 | 21 | Josd2 |
| 2010111I01Rik4 | 6.00E-10 | 0.914617536 | 0.555 | 0.351 | 1.22E-05 | 21 | 2010111I01Rik |
| Rad503 | 6.76E-10 | 0.713294618 | 0.345 | 0.143 | 1.37E-05 | 21 | Rad50 |
| Ptms3 | 1.66E-09 | 0.749756474 | 0.748 | 0.585 | 3.35E-05 | 21 | Ptms |
| Rora5 | 1.75E-09 | 0.449134393 | 0.412 | 0.187 | 3.54E-05 | 21 | Rora |
| Zfp771 | 1.81E-09 | 0.360663314 | 0.336 | 0.136 | 3.66E-05 | 21 | Zfp771 |
| Cald17 | 2.14E-09 | 1.480094323 | 0.487 | 0.324 | 4.34E-05 | 21 | Cald1 |
| Gm170181 | 2.57E-09 | 0.5004574 | 0.269 | 0.101 | 5.21E-05 | 21 | Gm17018 |
| Gsto1 | 2.68E-09 | 0.472060796 | 0.261 | 0.095 | 5.43E-05 | 21 | Gsto1 |
| Svil5 | 3.06E-09 | 0.840099713 | 0.445 | 0.244 | 6.20E-05 | 21 | Svil |
| Dtymk1 | 3.12E-09 | 0.600652032 | 0.328 | 0.136 | 6.31E-05 | 21 | Dtymk |
| Frmd4b8 | 3.18E-09 | 0.605903833 | 0.387 | 0.178 | 6.44E-05 | 21 | Frmd4b |
| Fgfr23 | 3.37E-09 | 0.526221305 | 0.345 | 0.154 | 6.83E-05 | 21 | Fgfr2 |
| Rbpms6 | 3.84E-09 | 1.444610579 | 0.496 | 0.351 | 7.78E-05 | 21 | Rbpms |
| Srebf23 | 4.41E-09 | 0.434107687 | 0.261 | 0.098 | 8.93E-05 | 21 | Srebf2 |
| Nfia7 | 4.44E-09 | 0.728870307 | 0.496 | 0.289 | 8.99E-05 | 21 | Nfia |
| Scmh12 | 4.87E-09 | 0.510330912 | 0.328 | 0.139 | 9.87E-05 | 21 | Scmh1 |
| Pam9 | 5.43E-09 | 0.711705653 | 0.42 | 0.215 | 0.000109856 | 21 | Pam |

| | | | | | | | |
|---|---|---|---|---|---|---|---|
| Ctsl5 | 5.60E-09 | 1.116649907 | 0.521 | 0.348 | 0.000113338 | 21 | Ctsl |
| H3f3a4 | 5.69E-09 | 0.530483601 | 0.924 | 0.862 | 0.000115127 | 21 | H3f3a |
| Tnfaip88 | 6.62E-09 | 0.610076046 | 0.403 | 0.188 | 0.000133992 | 21 | Tnfaip8 |
| Jmy | 7.39E-09 | 0.486830608 | 0.261 | 0.099 | 0.00014962 | 21 | Jmy |
| Smarca26 | 1.04E-08 | 0.753240068 | 0.487 | 0.282 | 0.000210614 | 21 | Smarca2 |
| Smarca41 | 1.11E-08 | 0.379093379 | 0.378 | 0.167 | 0.000225703 | 21 | Smarca4 |
| S100a108 | 1.17E-08 | 0.862730721 | 0.672 | 0.471 | 0.000236271 | 21 | S100a10 |
| Cnp1 | 1.26E-08 | 0.515485073 | 0.252 | 0.097 | 0.000255598 | 21 | Cnp |
| Gstm21 | 1.91E-08 | 0.674206633 | 0.286 | 0.116 | 0.000387005 | 21 | Gstm2 |
| Acadvl | 2.26E-08 | 0.287268798 | 0.294 | 0.119 | 0.000457211 | 21 | Acadvl |
| Ext19 | 2.53E-08 | 0.656023208 | 0.353 | 0.168 | 0.000513094 | 21 | Ext1 |
| Rpl138 | 2.55E-08 | 0.621545243 | 0.983 | 0.977 | 0.000516501 | 21 | Rpl13 |
| Rexo25 | 3.16E-08 | 0.820639905 | 0.479 | 0.299 | 0.000639189 | 21 | Rexo2 |
| Slmap3 | 3.63E-08 | 0.603699094 | 0.387 | 0.194 | 0.000735176 | 21 | Slmap |
| Tsc22d16 | 4.00E-08 | 0.745921192 | 0.622 | 0.448 | 0.000809615 | 21 | Tsc22d1 |
| Bex3 | 4.18E-08 | 0.611082937 | 0.387 | 0.184 | 0.000847007 | 21 | Bex3 |
| Zfp4622 | 4.37E-08 | 0.326441683 | 0.294 | 0.121 | 0.000883863 | 21 | Zfp462 |
| Lima16 | 5.06E-08 | 0.510917911 | 0.353 | 0.16 | 0.001023823 | 21 | Lima1 |
| Itpr18 | 5.34E-08 | 1.152297696 | 0.395 | 0.223 | 0.001080137 | 21 | Itpr1 |
| Apobec37 | 5.54E-08 | 0.549246502 | 0.429 | 0.227 | 0.001120793 | 21 | Apobec3 |
| Manbal1 | 7.37E-08 | 0.4381728 | 0.261 | 0.105 | 0.0014917 | 21 | Manbal |
| Oard1 | 7.89E-08 | 0.382839084 | 0.353 | 0.159 | 0.001596876 | 21 | Oard1 |
| Gapdh3 | 8.08E-08 | 0.448035309 | 0.832 | 0.709 | 0.001635783 | 21 | Gapdh |
| Farp14 | 8.64E-08 | 0.34918439 | 0.269 | 0.109 | 0.001749586 | 21 | Farp1 |
| Cbr1 | 9.53E-08 | 0.270012415 | 0.336 | 0.146 | 0.001930198 | 21 | Cbr1 |
| Rell13 | 9.72E-08 | 0.318122487 | 0.319 | 0.138 | 0.001968418 | 21 | Rell1 |
| Lonp22 | 9.79E-08 | 0.355796301 | 0.403 | 0.195 | 0.001981569 | 21 | Lonp2 |
| Nav27 | 1.03E-07 | 0.396088416 | 0.353 | 0.166 | 0.002078875 | 21 | Nav2 |
| Pebp12 | 2.09E-07 | 0.601719755 | 0.655 | 0.474 | 0.004238005 | 21 | Pebp1 |
| Pbxip17 | 2.12E-07 | 0.427988869 | 0.269 | 0.118 | 0.004299637 | 21 | Pbxip1 |
| Ssbp28 | 2.59E-07 | 0.491152067 | 0.311 | 0.141 | 0.005245867 | 21 | Ssbp2 |
| Vkorc12 | 2.78E-07 | 0.550464489 | 0.328 | 0.154 | 0.005629114 | 21 | Vkorc1 |
| Snrpd23 | 2.97E-07 | 0.658070866 | 0.597 | 0.411 | 0.006006593 | 21 | Snrpd2 |
| Papss11 | 3.18E-07 | 0.255240749 | 0.37 | 0.176 | 0.006443442 | 21 | Papss1 |
| Rras6 | 3.22E-07 | 0.704521319 | 0.496 | 0.299 | 0.006511342 | 21 | Rras |
| Sorbs16 | 3.27E-07 | 0.766836129 | 0.345 | 0.178 | 0.006625339 | 21 | Sorbs1 |
| Phkb2 | 3.33E-07 | 0.449640662 | 0.311 | 0.146 | 0.006735032 | 21 | Phkb |
| Stim110 | 3.38E-07 | 0.528457421 | 0.378 | 0.192 | 0.006845796 | 21 | Stim1 |
| Tns17 | 3.49E-07 | 0.922827964 | 0.412 | 0.247 | 0.007059592 | 21 | Tns1 |
| Tsc22d46 | 3.57E-07 | 0.455419507 | 0.504 | 0.296 | 0.007223473 | 21 | Tsc22d4 |
| Igf1r7 | 4.91E-07 | 0.326372439 | 0.345 | 0.162 | 0.009948433 | 21 | Igf1r |
| Fam13b7 | 7.44E-07 | 0.558536262 | 0.311 | 0.148 | 0.015062418 | 21 | Fam13b |
| Cstb2 | 7.86E-07 | 0.373436582 | 0.462 | 0.251 | 0.015903321 | 21 | Cstb |

| | | | | | | | |
|---|---|---|---|---|---|---|---|
| Mocs21 | 7.87E-07 | 0.478100709 | 0.403 | 0.209 | 0.0159416 | 21 | Mocs2 |
| Itga65 | 8.70E-07 | 0.749947887 | 0.462 | 0.285 | 0.017613868 | 21 | Itga6 |
| Rpl36a7 | 8.96E-07 | 0.655363683 | 0.874 | 0.753 | 0.018149804 | 21 | Rpl36a |
| Anxa66 | 9.74E-07 | 0.454764344 | 0.328 | 0.158 | 0.01971082 | 21 | Anxa6 |
| Ube2q2 | 1.01E-06 | 0.554565602 | 0.303 | 0.143 | 0.020468116 | 21 | Ube2q2 |
| Dut1 | 1.07E-06 | 0.452504149 | 0.286 | 0.13 | 0.021721015 | 21 | Dut |
| Tanc15 | 1.20E-06 | 0.503977583 | 0.412 | 0.224 | 0.024223455 | 21 | Tanc1 |
| Rps199 | 1.29E-06 | 0.686061986 | 0.966 | 0.928 | 0.026066384 | 21 | Rps19 |
| Hist1h2bc5 | 1.31E-06 | 0.50868153 | 0.445 | 0.251 | 0.026484038 | 21 | Hist1h2bc |
| Aldh26 | 1.39E-06 | 0.551673265 | 0.58 | 0.398 | 0.02804589 | 21 | Aldh2 |
| Hspe13 | 1.39E-06 | 0.573699081 | 0.706 | 0.508 | 0.02818353 | 21 | Hspe1 |
| mt-Cytb13 | 1.41E-06 | 0.862020315 | 0.689 | 0.703 | 0.028498757 | 21 | mt-Cytb |
| Ppdpf2 | 1.56E-06 | 0.532176109 | 0.496 | 0.288 | 0.03149266 | 21 | Ppdpf |
| Adgrg15 | 1.67E-06 | 0.299281549 | 0.303 | 0.141 | 0.033768071 | 21 | Adgrg1 |
| Camta1 | 2.66E-06 | 0.335227755 | 0.336 | 0.163 | 0.053905925 | 21 | Camta1 |
| Crip110 | 3.35E-06 | 1.068562147 | 0.286 | 0.54 | 0.067871732 | 21 | Crip1 |
| Prkg15 | 3.36E-06 | 0.97351447 | 0.345 | 0.198 | 0.06810487 | 21 | Prkg1 |
| Usp484 | 3.42E-06 | 0.495246146 | 0.269 | 0.13 | 0.069203666 | 21 | Usp48 |
| Ptpre6 | 3.52E-06 | 0.482213285 | 0.319 | 0.162 | 0.071247054 | 21 | Ptpre |
| Fam162a3 | 3.83E-06 | 0.488936314 | 0.454 | 0.258 | 0.077514053 | 21 | Fam162a |
| Pink11 | 3.98E-06 | 0.538384485 | 0.277 | 0.132 | 0.080675375 | 21 | Pink1 |
| Pmp226 | 4.09E-06 | 0.536620197 | 0.294 | 0.145 | 0.082790276 | 21 | Pmp22 |
| Rpsa9 | 4.46E-06 | 0.478378286 | 0.966 | 0.903 | 0.090222256 | 21 | Rpsa |
| Tm6sf18 | 5.44E-06 | 0.667786523 | 0.269 | 0.143 | 0.110169139 | 21 | Tm6sf1 |
| AY0361183 | 5.48E-06 | 0.608642824 | 0.95 | 0.927 | 0.110932573 | 21 | AY036118 |
| Dgcr6 | 5.68E-06 | 0.41283938 | 0.277 | 0.131 | 0.115044965 | 21 | Dgcr6 |
| Pde4b6 | 5.90E-06 | 0.679717018 | 0.353 | 0.197 | 0.119524796 | 21 | Pde4b |
| Rpl7a6 | 6.43E-06 | 0.5071827 | 0.84 | 0.754 | 0.130175662 | 21 | Rpl7a |
| Chmp2b | 6.47E-06 | 0.548133776 | 0.403 | 0.223 | 0.130985862 | 21 | Chmp2b |
| Cdkn1c5 | 6.76E-06 | 0.308932889 | 0.277 | 0.127 | 0.136838037 | 21 | Cdkn1c |
| mt-Nd414 | 7.45E-06 | 0.740628418 | 0.647 | 0.678 | 0.15093089 | 21 | mt-Nd4 |
| Hsd17b102 | 7.51E-06 | 0.269852554 | 0.437 | 0.253 | 0.151989114 | 21 | Hsd17b10 |
| Neat110 | 7.94E-06 | 0.541864553 | 0.37 | 0.21 | 0.160810958 | 21 | Neat1 |
| Gclm | 8.35E-06 | 0.371944952 | 0.328 | 0.166 | 0.169017697 | 21 | Gclm |
| Prmt11 | 8.48E-06 | 0.497165309 | 0.387 | 0.217 | 0.171768187 | 21 | Prmt1 |
| Ergic32 | 8.61E-06 | 0.362948128 | 0.361 | 0.19 | 0.174221002 | 21 | Ergic3 |
| Naxd | 8.89E-06 | 0.320553394 | 0.336 | 0.169 | 0.179939396 | 21 | Naxd |
| Nbeal16 | 8.98E-06 | 0.451315024 | 0.462 | 0.264 | 0.181881688 | 21 | Nbeal1 |
| Cops51 | 9.06E-06 | 0.479530284 | 0.328 | 0.171 | 0.183376595 | 21 | Cops5 |
| Ube2v21 | 9.36E-06 | 0.476689758 | 0.345 | 0.177 | 0.189549467 | 21 | Ube2v2 |
| Camk1d3 | 1.03E-05 | 0.52663232 | 0.857 | 0.824 | 0.207652505 | 21 | Camk1d |
| Vps351 | 1.09E-05 | 0.48225681 | 0.378 | 0.207 | 0.2215527 | 21 | Vps35 |
| Cct52 | 1.21E-05 | 0.582723154 | 0.479 | 0.296 | 0.244303008 | 21 | Cct5 |

| | | | | | | | |
|---|---|---|---|---|---|---|---|
| Sparc8 | 1.25E-05 | 0.559485017 | 0.597 | 0.405 | 0.253643836 | 21 | Sparc |
| Rheb2 | 1.38E-05 | 0.519060915 | 0.563 | 0.379 | 0.278993444 | 21 | Rheb |
| Trnau1ap | 1.43E-05 | 0.575109197 | 0.328 | 0.173 | 0.288948424 | 21 | Trnau1ap |
| Lama34 | 1.49E-05 | 0.313274374 | 0.412 | 0.223 | 0.301836687 | 21 | Lama3 |
| Rpl79 | 1.53E-05 | 0.578759791 | 0.933 | 0.894 | 0.310126741 | 21 | Rpl7 |
| Dbi3 | 1.54E-05 | 0.633278743 | 0.546 | 0.434 | 0.312483333 | 21 | Dbi |
| Ppp1cb3 | 1.55E-05 | 0.447690715 | 0.454 | 0.274 | 0.313424683 | 21 | Ppp1cb |
| Rps173 | 1.71E-05 | 0.452324987 | 0.882 | 0.797 | 0.34561709 | 21 | Rps17 |
| Rps88 | 1.72E-05 | 0.619413475 | 0.95 | 0.965 | 0.348329081 | 21 | Rps8 |
| Rpl415 | 1.73E-05 | 0.399292356 | 0.966 | 0.962 | 0.350224991 | 21 | Rpl41 |
| Arhgap53 | 1.86E-05 | 0.560341231 | 0.454 | 0.291 | 0.377538437 | 21 | Arhgap5 |
| Runx18 | 1.90E-05 | 0.328064652 | 0.277 | 0.133 | 0.38375847 | 21 | Runx1 |
| Foxo35 | 1.94E-05 | 0.259991188 | 0.252 | 0.118 | 0.392527458 | 21 | Foxo3 |
| Fam3c3 | 1.95E-05 | 0.428827065 | 0.328 | 0.181 | 0.395802634 | 21 | Fam3c |
| Rsu17 | 2.02E-05 | 0.503948407 | 0.412 | 0.255 | 0.40964415 | 21 | Rsu1 |
| Meis23 | 2.03E-05 | 0.334088051 | 0.479 | 0.284 | 0.410731596 | 21 | Meis2 |
| Ccni1 | 2.11E-05 | 0.578294024 | 0.412 | 0.236 | 0.427128952 | 21 | Ccni |
| Smarcc13 | 2.16E-05 | 0.578954928 | 0.277 | 0.144 | 0.437539132 | 21 | Smarcc1 |
| Enah5 | 2.18E-05 | 0.273443498 | 0.303 | 0.151 | 0.441794648 | 21 | Enah |
| Cdc37l12 | 2.19E-05 | 0.271290703 | 0.328 | 0.169 | 0.443405849 | 21 | Cdc37l1 |
| Agpat53 | 2.23E-05 | 0.321908193 | 0.286 | 0.144 | 0.451485656 | 21 | Agpat5 |
| Npm17 | 2.25E-05 | 0.918138012 | 0.748 | 0.681 | 0.456276325 | 21 | Npm1 |
| Amotl17 | 2.26E-05 | 0.311047815 | 0.353 | 0.187 | 0.457842783 | 21 | Amotl1 |
| Zfp3261 | 2.33E-05 | 0.386333987 | 0.294 | 0.148 | 0.472119102 | 21 | Zfp326 |
| Rps78 | 2.38E-05 | 0.550665483 | 0.941 | 0.917 | 0.481001427 | 21 | Rps7 |
| Eif4a23 | 2.66E-05 | 0.407335889 | 0.496 | 0.305 | 0.538897932 | 21 | Eif4a2 |
| S100a166 | 2.94E-05 | 0.423806037 | 0.445 | 0.26 | 0.596040683 | 21 | S100a16 |
| Eif3k3 | 3.11E-05 | 0.612472384 | 0.647 | 0.503 | 0.62960473 | 21 | Eif3k |
| Cfl21 | 3.15E-05 | 0.429310809 | 0.403 | 0.231 | 0.638540072 | 21 | Cfl2 |
| Hpf11 | 3.27E-05 | 0.580362978 | 0.345 | 0.192 | 0.662274407 | 21 | Hpf1 |
| Fermt26 | 3.58E-05 | 0.589126546 | 0.513 | 0.352 | 0.725406324 | 21 | Fermt2 |
| Sox42 | 3.87E-05 | 0.497514532 | 0.361 | 0.205 | 0.783584165 | 21 | Sox4 |
| Dctn61 | 4.07E-05 | 0.423674929 | 0.336 | 0.179 | 0.824888078 | 21 | Dctn6 |
| Actg16 | 4.12E-05 | 0.383780407 | 0.84 | 0.795 | 0.834436238 | 21 | Actg1 |
| Actn46 | 4.24E-05 | 0.571197424 | 0.529 | 0.377 | 0.858658734 | 21 | Actn4 |
| Ktn17 | 4.27E-05 | 0.56560011 | 0.462 | 0.309 | 0.864746363 | 21 | Ktn1 |
| Gm424184 | 4.34E-05 | 0.421859514 | 1 | 1 | 0.87793685 | 21 | Gm42418 |
| Ehd26 | 4.44E-05 | 0.557220234 | 0.345 | 0.195 | 0.89875962 | 21 | Ehd2 |
| Nfic3 | 4.56E-05 | 0.383854898 | 0.445 | 0.263 | 0.923986735 | 21 | Nfic |
| Rps221 | 4.75E-05 | 0.594992763 | 0.95 | 0.92 | 0.96195276 | 21 | Rps2 |
| Hsp90aa14 | 4.78E-05 | 0.525225771 | 0.555 | 0.427 | 0.96758792 | 21 | Hsp90aa1 |
| 1810058I24Rik1 | 5.54E-05 | 0.54964573 | 0.479 | 0.307 | 1 | 21 | 1810058I24Rik |
| Aff46 | 6.28E-05 | 0.361538131 | 0.412 | 0.242 | 1 | 21 | Aff4 |

| | | | | | | | |
|---|---|---|---|---|---|---|---|
| Dynll15 | 6.31E-05 | 0.702340616 | 0.714 | 0.607 | 1 | 21 | Dynll1 |
| Anapc112 | 6.41E-05 | 0.591200594 | 0.378 | 0.224 | 1 | 21 | Anapc11 |
| Comt3 | 6.51E-05 | 0.329253913 | 0.345 | 0.195 | 1 | 21 | Comt |
| Rpl318 | 6.96E-05 | 0.401941547 | 0.79 | 0.685 | 1 | 21 | Rpl31 |
| Kcnq1ot19 | 7.06E-05 | 0.342384649 | 0.261 | 0.136 | 1 | 21 | Kcnq1ot1 |
| Skp1a2 | 7.32E-05 | 0.638736869 | 0.487 | 0.329 | 1 | 21 | Skp1a |
| Ndufb22 | 7.40E-05 | 0.305985183 | 0.597 | 0.406 | 1 | 21 | Ndufb2 |
| Slc3a29 | 7.69E-05 | 0.707163949 | 0.429 | 0.292 | 1 | 21 | Slc3a2 |
| Larp1b | 7.75E-05 | 0.285285457 | 0.311 | 0.17 | 1 | 21 | Larp1b |
| Rpl69 | 8.03E-05 | 0.504005834 | 0.899 | 0.882 | 1 | 21 | Rpl6 |
| Rpl157 | 8.20E-05 | 0.549817529 | 0.924 | 0.902 | 1 | 21 | Rpl15 |
| Clstn16 | 8.22E-05 | 0.51269137 | 0.387 | 0.238 | 1 | 21 | Clstn1 |
| 2810004N23Rik | 8.97E-05 | 0.284674462 | 0.261 | 0.131 | 1 | 21 | 2810004N23Rik |
| Gas65 | 9.49E-05 | 0.478063686 | 0.462 | 0.3 | 1 | 21 | Gas6 |
| Sdhc1 | 9.57E-05 | 0.260135879 | 0.387 | 0.221 | 1 | 21 | Sdhc |
| Rps278 | 9.94E-05 | 0.389343762 | 0.916 | 0.899 | 1 | 21 | Rps27 |
| Rpl2110 | 0.000101154 | 0.440425269 | 0.933 | 0.924 | 1 | 21 | Rpl21 |
| Lyar | 0.000102396 | 0.319310996 | 0.261 | 0.133 | 1 | 21 | Lyar |
| Snrpf7 | 0.000104713 | 0.465830737 | 0.538 | 0.381 | 1 | 21 | Snrpf |
| mt-Atp615 | 0.00010487 | 0.595312987 | 0.689 | 0.756 | 1 | 21 | mt-Atp6 |
| Polr2f2 | 0.000105876 | 0.484963743 | 0.504 | 0.349 | 1 | 21 | Polr2f |
| Cnpy22 | 0.000106191 | 0.483122336 | 0.412 | 0.265 | 1 | 21 | Cnpy2 |
| Ralbp13 | 0.000108077 | 0.585589606 | 0.487 | 0.332 | 1 | 21 | Ralbp1 |
| Eef1d4 | 0.000109719 | 0.462829381 | 0.672 | 0.551 | 1 | 21 | Eef1d |
| Mrps18a | 0.000111048 | 0.54818557 | 0.328 | 0.185 | 1 | 21 | Mrps18a |
| Smim8 | 0.000111936 | 0.272597712 | 0.252 | 0.124 | 1 | 21 | Smim8 |
| Hikeshi1 | 0.000111946 | 0.419691768 | 0.269 | 0.141 | 1 | 21 | Hikeshi |
| Prss237 | 0.000112771 | 0.705080914 | 0.403 | 0.283 | 1 | 21 | Prss23 |
| Acaa1a1 | 0.00011856 | 0.290327784 | 0.311 | 0.171 | 1 | 21 | Acaa1a |
| Rpl37a9 | 0.000119189 | 0.341020274 | 0.941 | 0.938 | 1 | 21 | Rpl37a |
| Minos12 | 0.000120354 | 0.271760844 | 0.681 | 0.514 | 1 | 21 | Minos1 |
| Hras1 | 0.0001309 | 0.364685663 | 0.538 | 0.362 | 1 | 21 | Hras |
| Esf11 | 0.000139142 | 0.44815824 | 0.303 | 0.166 | 1 | 21 | Esf1 |
| Rtraf1 | 0.000140636 | 0.477086967 | 0.563 | 0.423 | 1 | 21 | Rtraf |
| 2900097C17Rik1 | 0.000153259 | 0.3436009 | 0.286 | 0.153 | 1 | 21 | 2900097C17Rik |
| Thap34 | 0.000159029 | 0.29618853 | 0.303 | 0.163 | 1 | 21 | Thap3 |
| Pdha11 | 0.000161053 | 0.26086443 | 0.345 | 0.194 | 1 | 21 | Pdha1 |
| Cnbp4 | 0.000168943 | 0.451434463 | 0.647 | 0.489 | 1 | 21 | Cnbp |
| Nap1l17 | 0.00016951 | 0.41520177 | 0.479 | 0.316 | 1 | 21 | Nap1l1 |
| Ndufs72 | 0.000172819 | 0.285359995 | 0.613 | 0.459 | 1 | 21 | Ndufs7 |
| Tceal92 | 0.000181507 | 0.429140851 | 0.546 | 0.389 | 1 | 21 | Tceal9 |
| Elob2 | 0.000182073 | 0.404332263 | 0.773 | 0.697 | 1 | 21 | Elob |
| Sh3bp53 | 0.000187365 | 0.304076763 | 0.37 | 0.218 | 1 | 21 | Sh3bp5 |

| | | | | | | | |
|---|---|---|---|---|---|---|---|
| Ppia3 | 0.00019089 | 0.2951639 | 0.95 | 0.939 | 1 | 21 | Ppia |
| Rpl368 | 0.000194006 | 0.367118388 | 0.941 | 0.904 | 1 | 21 | Rpl36 |
| H3f3b2 | 0.000197832 | 0.409914065 | 0.924 | 0.907 | 1 | 21 | H3f3b |
| Suco7 | 0.000197858 | 0.315948357 | 0.286 | 0.155 | 1 | 21 | Suco |
| Sf3b51 | 0.000207195 | 0.382421272 | 0.487 | 0.312 | 1 | 21 | Sf3b5 |
| Selenow5 | 0.000221392 | 0.476447361 | 0.748 | 0.631 | 1 | 21 | Selenow |
| Dek2 | 0.000227129 | 0.453105189 | 0.529 | 0.374 | 1 | 21 | Dek |
| Vim12 | 0.00023613 | 0.777630333 | 0.445 | 0.308 | 1 | 21 | Vim |
| Cct3 | 0.0002603 | 0.518942473 | 0.387 | 0.245 | 1 | 21 | Cct3 |
| Rpl10a10 | 0.000265198 | 0.618065159 | 0.84 | 0.837 | 1 | 21 | Rpl10a |
| Rpl188 | 0.000275146 | 0.457916413 | 0.924 | 0.901 | 1 | 21 | Rpl18 |
| Eid15 | 0.000276112 | 0.455552373 | 0.471 | 0.323 | 1 | 21 | Eid1 |
| Cdk41 | 0.000280199 | 0.401231324 | 0.353 | 0.208 | 1 | 21 | Cdk4 |
| Park72 | 0.000309413 | 0.372367717 | 0.689 | 0.559 | 1 | 21 | Park7 |
| Tmem2562 | 0.000329562 | 0.448819479 | 0.513 | 0.366 | 1 | 21 | Tmem256 |
| Hcfc1r11 | 0.00034769 | 0.551405587 | 0.454 | 0.303 | 1 | 21 | Hcfc1r1 |
| Rpl147 | 0.00035384 | 0.377982802 | 0.857 | 0.803 | 1 | 21 | Rpl14 |
| Bad | 0.000358513 | 0.264071773 | 0.269 | 0.144 | 1 | 21 | Bad |
| Polr2m2 | 0.000365873 | 0.35358042 | 0.403 | 0.246 | 1 | 21 | Polr2m |
| Scand11 | 0.000366309 | 0.422906291 | 0.613 | 0.467 | 1 | 21 | Scand1 |
| Rpl22l17 | 0.000392357 | 0.515793219 | 0.706 | 0.58 | 1 | 21 | Rpl22l1 |
| Rps168 | 0.000395802 | 0.370775088 | 0.983 | 0.927 | 1 | 21 | Rps16 |
| Polr2i | 0.000397395 | 0.33599936 | 0.378 | 0.222 | 1 | 21 | Polr2i |
| Coa6 | 0.000405483 | 0.299193867 | 0.252 | 0.133 | 1 | 21 | Coa6 |
| mt-Co316 | 0.000407265 | 0.566063867 | 0.664 | 0.728 | 1 | 21 | mt-Co3 |
| Wwox9 | 0.000413403 | 0.311835189 | 0.303 | 0.177 | 1 | 21 | Wwox |
| Rps269 | 0.000430383 | 0.414924487 | 0.908 | 0.866 | 1 | 21 | Rps26 |
| Tardbp | 0.000430792 | 0.370444529 | 0.277 | 0.158 | 1 | 21 | Tardbp |
| Mrpl241 | 0.000431454 | 0.562307337 | 0.387 | 0.24 | 1 | 21 | Mrpl24 |
| Cyhr11 | 0.000436618 | 0.402552927 | 0.294 | 0.169 | 1 | 21 | Cyhr1 |
| Ugp2 | 0.000459921 | 0.264298039 | 0.277 | 0.155 | 1 | 21 | Ugp2 |
| Ddx39b6 | 0.00046333 | 0.333761351 | 0.37 | 0.235 | 1 | 21 | Ddx39b |
| Rbbp72 | 0.000467864 | 0.324267421 | 0.403 | 0.249 | 1 | 21 | Rbbp7 |
| Frmd4a6 | 0.000485445 | 0.278453963 | 0.269 | 0.149 | 1 | 21 | Frmd4a |
| Ptges32 | 0.000498596 | 0.282539009 | 0.555 | 0.388 | 1 | 21 | Ptges3 |
| Rpl379 | 0.000510131 | 0.406827238 | 0.941 | 0.928 | 1 | 21 | Rpl37 |
| Itm2c6 | 0.000513364 | 0.3792768 | 0.328 | 0.201 | 1 | 21 | Itm2c |
| Tcp1 | 0.000530317 | 0.355086949 | 0.353 | 0.216 | 1 | 21 | Tcp1 |
| Fto6 | 0.000535188 | 0.363491877 | 0.319 | 0.192 | 1 | 21 | Fto |
| Tpd52l22 | 0.000557614 | 0.278830131 | 0.303 | 0.172 | 1 | 21 | Tpd52l2 |
| Tmem176b9 | 0.000567105 | 0.59228749 | 0.563 | 0.441 | 1 | 21 | Tmem176b |
| mt-Co217 | 0.000584336 | 0.471077415 | 0.664 | 0.731 | 1 | 21 | mt-Co2 |
| Thra6 | 0.000605811 | 0.283223593 | 0.294 | 0.166 | 1 | 21 | Thra |

| | | | | | | | |
|---|---|---|---|---|---|---|---|
| Rps108 | 0.000626012 | 0.409316179 | 0.95 | 0.945 | 1 | 21 | Rps10 |
| Rpl319 | 0.000645975 | 0.485254818 | 0.924 | 0.91 | 1 | 21 | Rpl3 |
| 2310039H08Rik | 0.000667255 | 0.315195515 | 0.252 | 0.137 | 1 | 21 | 2310039H08Rik |
| Hmgb12 | 0.000699326 | 0.3472404 | 0.798 | 0.717 | 1 | 21 | Hmgb1 |
| Ndufa22 | 0.000712479 | 0.331739781 | 0.689 | 0.569 | 1 | 21 | Ndufa2 |
| Rps4x9 | 0.000713174 | 0.490257026 | 0.983 | 0.941 | 1 | 21 | Rps4x |
| Thoc72 | 0.000733395 | 0.425385886 | 0.454 | 0.3 | 1 | 21 | Thoc7 |
| Cir1 | 0.000754946 | 0.537505787 | 0.286 | 0.174 | 1 | 21 | Cir1 |
| Ano66 | 0.000767292 | 0.252569727 | 0.311 | 0.182 | 1 | 21 | Ano6 |
| Anp32b4 | 0.00076907 | 0.377912443 | 0.58 | 0.424 | 1 | 21 | Anp32b |
| Cirbp1 | 0.000801102 | 0.276943819 | 0.395 | 0.252 | 1 | 21 | Cirbp |
| Ptprk5 | 0.00085099 | 0.260948996 | 0.261 | 0.142 | 1 | 21 | Ptprk |
| Prelid12 | 0.000878163 | 0.359412729 | 0.479 | 0.312 | 1 | 21 | Prelid1 |
| Rpl246 | 0.000894524 | 0.342395923 | 0.916 | 0.908 | 1 | 21 | Rpl24 |
| Rplp08 | 0.00090511 | 0.513838199 | 0.958 | 0.945 | 1 | 21 | Rplp0 |
| mt-Nd4l10 | 0.000905607 | 0.691476195 | 0.513 | 0.494 | 1 | 21 | mt-Nd4l |
| Fryl9 | 0.000945729 | 0.27634102 | 0.412 | 0.27 | 1 | 21 | Fryl |
| Snrpd32 | 0.001057902 | 0.380655471 | 0.471 | 0.313 | 1 | 21 | Snrpd3 |
| Ckap53 | 0.001130848 | 0.25697054 | 0.252 | 0.144 | 1 | 21 | Ckap5 |
| Eif1ax1 | 0.001148875 | 0.27971069 | 0.429 | 0.277 | 1 | 21 | Eif1ax |
| Nop102 | 0.001157286 | 0.400415235 | 0.521 | 0.375 | 1 | 21 | Nop10 |
| Psma42 | 0.001239793 | 0.436342287 | 0.504 | 0.359 | 1 | 21 | Psma4 |
| Ppp1r12a11 | 0.001260408 | 0.761037573 | 0.504 | 0.428 | 1 | 21 | Ppp1r12a |
| Rps39 | 0.001271063 | 0.498824597 | 0.933 | 0.894 | 1 | 21 | Rps3 |
| Rcn25 | 0.001347308 | 0.298844515 | 0.252 | 0.144 | 1 | 21 | Rcn2 |
| Rps3a19 | 0.001354012 | 0.34488215 | 0.966 | 0.935 | 1 | 21 | Rps3a1 |
| Cct83 | 0.001420579 | 0.268994164 | 0.412 | 0.268 | 1 | 21 | Cct8 |
| Rps2112 | 0.001436623 | 0.2961002 | 0.95 | 0.927 | 1 | 21 | Rps21 |
| Rpl359 | 0.001479846 | 0.348628088 | 0.882 | 0.829 | 1 | 21 | Rpl35 |
| Atxn110 | 0.001511913 | 0.392133204 | 0.42 | 0.297 | 1 | 21 | Atxn1 |
| Btf32 | 0.001518258 | 0.345807362 | 0.773 | 0.723 | 1 | 21 | Btf3 |
| Fundc21 | 0.001585569 | 0.324192273 | 0.412 | 0.271 | 1 | 21 | Fundc2 |
| Vav38 | 0.00170848 | 0.460151303 | 0.378 | 0.258 | 1 | 21 | Vav3 |
| B230219D22Rik5 | 0.001741725 | 0.43488448 | 0.487 | 0.344 | 1 | 21 | B230219D22Rik |
| Pfn19 | 0.001770318 | 0.366167088 | 0.765 | 0.684 | 1 | 21 | Pfn1 |
| Tuba1b6 | 0.001786511 | 0.285934471 | 0.471 | 0.324 | 1 | 21 | Tuba1b |
| Ube2b2 | 0.00181462 | 0.420064236 | 0.538 | 0.397 | 1 | 21 | Ube2b |
| Chd69 | 0.00184687 | 0.253179879 | 0.345 | 0.219 | 1 | 21 | Chd6 |
| Tax1bp12 | 0.001860429 | 0.329443722 | 0.529 | 0.419 | 1 | 21 | Tax1bp1 |
| Rps208 | 0.001909006 | 0.39565063 | 0.958 | 0.951 | 1 | 21 | Rps20 |
| Zfp106 | 0.001929507 | 0.317826145 | 0.345 | 0.219 | 1 | 21 | Zfp106 |
| Mlf21 | 0.001935401 | 0.343717402 | 0.319 | 0.201 | 1 | 21 | Mlf2 |
| Anxa12 | 0.001953538 | 0.436664131 | 0.353 | 0.228 | 1 | 21 | Anxa1 |

| | | | | | | | |
|---|---|---|---|---|---|---|---|
| Tsen342 | 0.002057695 | 0.322760547 | 0.294 | 0.179 | 1 | 21 | Tsen34 |
| Rps99 | 0.002080048 | 0.373015684 | 0.941 | 0.921 | 1 | 21 | Rps9 |
| Rps59 | 0.002250684 | 0.416888336 | 0.941 | 0.92 | 1 | 21 | Rps5 |
| Pnisr10 | 0.00226441 | 0.349512175 | 0.387 | 0.266 | 1 | 21 | Pnisr |
| Arid5b8 | 0.002326707 | 0.395563109 | 0.403 | 0.28 | 1 | 21 | Arid5b |
| Nop53 | 0.002352531 | 0.255670621 | 0.37 | 0.23 | 1 | 21 | Nop53 |
| Gpx42 | 0.002403477 | 0.396181879 | 0.681 | 0.631 | 1 | 21 | Gpx4 |
| Tubb2a5 | 0.002414142 | 0.37611911 | 0.269 | 0.166 | 1 | 21 | Tubb2a |
| mt-Nd211 | 0.002577389 | 0.461189483 | 0.647 | 0.691 | 1 | 21 | mt-Nd2 |
| Uba21 | 0.002602541 | 0.515471171 | 0.261 | 0.156 | 1 | 21 | Uba2 |
| Clip13 | 0.002644893 | 0.258961628 | 0.252 | 0.145 | 1 | 21 | Clip1 |
| Map3k26 | 0.002717771 | 0.250683638 | 0.269 | 0.155 | 1 | 21 | Map3k2 |
| Tuba1a7 | 0.002816742 | 0.368243454 | 0.387 | 0.258 | 1 | 21 | Tuba1a |
| Cd1516 | 0.00282067 | 0.283871896 | 0.353 | 0.219 | 1 | 21 | Cd151 |
| Camk14 | 0.002878269 | 0.267692964 | 0.353 | 0.229 | 1 | 21 | Camk1 |
| Dctn22 | 0.002892398 | 0.363489595 | 0.395 | 0.267 | 1 | 21 | Dctn2 |
| Pkp42 | 0.002981795 | 0.288076923 | 0.42 | 0.289 | 1 | 21 | Pkp4 |
| Chmp32 | 0.003087224 | 0.268416603 | 0.479 | 0.335 | 1 | 21 | Chmp3 |
| Laptm4a7 | 0.003304694 | 0.431957184 | 0.571 | 0.486 | 1 | 21 | Laptm4a |
| Mrpl28 | 0.003364246 | 0.403128468 | 0.294 | 0.181 | 1 | 21 | Mrpl28 |
| Rpl399 | 0.003385852 | 0.394159712 | 0.933 | 0.901 | 1 | 21 | Rpl39 |
| Gm4728311 | 0.003503394 | 0.407619825 | 0.403 | 0.283 | 1 | 21 | Gm47283 |
| Prdx14 | 0.003602422 | 0.349056039 | 0.748 | 0.694 | 1 | 21 | Prdx1 |
| Ncl5 | 0.003647074 | 0.452015834 | 0.613 | 0.519 | 1 | 21 | Ncl |
| Lrpap15 | 0.003675368 | 0.415690732 | 0.311 | 0.198 | 1 | 21 | Lrpap1 |
| Ssb1 | 0.00391024 | 0.48825741 | 0.538 | 0.415 | 1 | 21 | Ssb |
| Acaa2 | 0.004060513 | 0.262673337 | 0.319 | 0.199 | 1 | 21 | Acaa2 |
| Gtf2h52 | 0.004293127 | 0.296143927 | 0.462 | 0.327 | 1 | 21 | Gtf2h5 |
| Chd47 | 0.004304782 | 0.457820861 | 0.454 | 0.33 | 1 | 21 | Chd4 |
| Rpl99 | 0.004320579 | 0.323422085 | 0.941 | 0.945 | 1 | 21 | Rpl9 |
| Cnn28 | 0.004339102 | 0.308838703 | 0.277 | 0.184 | 1 | 21 | Cnn2 |
| Lsm7 | 0.004444122 | 0.392611523 | 0.269 | 0.167 | 1 | 21 | Lsm7 |
| Atp1b12 | 0.00453184 | 0.290177755 | 0.261 | 0.163 | 1 | 21 | Atp1b1 |
| Rpl288 | 0.004566101 | 0.333950199 | 0.958 | 0.931 | 1 | 21 | Rpl28 |
| Tomm72 | 0.004665717 | 0.323971804 | 0.681 | 0.573 | 1 | 21 | Tomm7 |
| mt-Nd115 | 0.00470095 | 0.520408202 | 0.588 | 0.601 | 1 | 21 | mt-Nd1 |
| Lamtor22 | 0.004735455 | 0.43217967 | 0.597 | 0.466 | 1 | 21 | Lamtor2 |
| Gm100766 | 0.004832936 | 0.305414502 | 0.655 | 0.548 | 1 | 21 | Gm10076 |
| Hsp90ab11 | 0.004919894 | 0.2803578 | 0.832 | 0.841 | 1 | 21 | Hsp90ab1 |
| Lamp27 | 0.0050214 | 0.261894958 | 0.445 | 0.307 | 1 | 21 | Lamp2 |
| Rps156 | 0.00506533 | 0.386306919 | 0.824 | 0.75 | 1 | 21 | Rps15 |
| Fnta | 0.005165285 | 0.36252028 | 0.261 | 0.154 | 1 | 21 | Fnta |
| Ubc2 | 0.005484169 | 0.346031264 | 0.731 | 0.666 | 1 | 21 | Ubc |

| | | | | | | | |
|---|---|---|---|---|---|---|---|
| Csnk1a14 | 0.005514032 | 0.267942093 | 0.571 | 0.46 | 1 | 21 | Csnk1a1 |
| Osgep1 | 0.005634421 | 0.493928453 | 0.269 | 0.172 | 1 | 21 | Osgep |
| Sumo21 | 0.005944888 | 0.438347398 | 0.723 | 0.66 | 1 | 21 | Sumo2 |
| Erbin1 | 0.006184676 | 0.286049484 | 0.42 | 0.312 | 1 | 21 | Erbin |
| Rpl47 | 0.006210512 | 0.349593624 | 0.723 | 0.66 | 1 | 21 | Rpl4 |
| Swi51 | 0.006283608 | 0.440599763 | 0.58 | 0.443 | 1 | 21 | Swi5 |
| Uqcc22 | 0.006316885 | 0.261530871 | 0.529 | 0.406 | 1 | 21 | Uqcc2 |
| Ranbp11 | 0.006358648 | 0.366553743 | 0.454 | 0.324 | 1 | 21 | Ranbp1 |
| Ndufa72 | 0.006444729 | 0.291177787 | 0.681 | 0.576 | 1 | 21 | Ndufa7 |
| Pin1 | 0.006599689 | 0.316515204 | 0.319 | 0.207 | 1 | 21 | Pin1 |
| Timp36 | 0.006735013 | 0.295456287 | 0.521 | 0.388 | 1 | 21 | Timp3 |
| Sfr13 | 0.006747328 | 0.303860572 | 0.504 | 0.384 | 1 | 21 | Sfr1 |
| Rps68 | 0.007009343 | 0.285867824 | 0.874 | 0.813 | 1 | 21 | Rps6 |
| Nedd81 | 0.007208235 | 0.3310642 | 0.639 | 0.553 | 1 | 21 | Nedd8 |
| 4930523C07Rik10 | 0.007621751 | 0.407172408 | 0.328 | 0.221 | 1 | 21 | 4930523C07Rik |
| Knop1 | 0.007746162 | 0.275740562 | 0.277 | 0.176 | 1 | 21 | Knop1 |
| Rpl88 | 0.007833645 | 0.314197614 | 0.933 | 0.926 | 1 | 21 | Rpl8 |
| Slirp | 0.008514622 | 0.292143314 | 0.286 | 0.183 | 1 | 21 | Slirp |
| Eif5b5 | 0.008554153 | 0.422675116 | 0.437 | 0.323 | 1 | 21 | Eif5b |
| Cavin16 | 0.009059783 | 0.408537842 | 0.328 | 0.215 | 1 | 21 | Cavin1 |
| Trmt1127 | 0.009186959 | 0.272198379 | 0.403 | 0.276 | 1 | 21 | Trmt112 |
| Pacsin23 | 0.009244753 | 0.282813473 | 0.252 | 0.159 | 1 | 21 | Pacsin2 |
| Esd4 | 0.009492548 | 0.371678615 | 0.504 | 0.382 | 1 | 21 | Esd |
| Hdgf2 | 0.009828679 | 0.349603559 | 0.361 | 0.25 | 1 | 21 | Hdgf |
| Rps27l2 | 0.009863942 | 0.29019385 | 0.664 | 0.593 | 1 | 21 | Rps27l |
| Csmd1 | 0 | 5.464603289 | 0.88 | 0.006 | 0 | 22 | Csmd1 |
| Scn7a1 | 0 | 4.986506658 | 0.926 | 0.025 | 0 | 22 | Scn7a |
| Nkain2 | 0 | 4.878745252 | 0.926 | 0.008 | 0 | 22 | Nkain2 |
| Slc35f1 | 0 | 4.535489123 | 0.87 | 0.002 | 0 | 22 | Slc35f1 |
| Mpz | 0 | 4.519898801 | 0.648 | 0.004 | 0 | 22 | Mpz |
| Fxyd12 | 0 | 4.376810118 | 0.963 | 0.031 | 0 | 22 | Fxyd1 |
| Kcna1 | 0 | 4.341975912 | 0.926 | 0.001 | 0 | 22 | Kcna1 |
| Cdh191 | 0 | 4.312101343 | 0.907 | 0.013 | 0 | 22 | Cdh19 |
| Plp1 | 0 | 4.142961385 | 0.954 | 0.005 | 0 | 22 | Plp1 |
| Ank33 | 0 | 4.08809794 | 0.917 | 0.047 | 0 | 22 | Ank3 |
| Matn21 | 0 | 3.994628982 | 0.889 | 0.016 | 0 | 22 | Matn2 |
| Cadm2 | 0 | 3.873598704 | 0.852 | 0.004 | 0 | 22 | Cadm2 |
| Abca8a1 | 0 | 3.868557705 | 0.87 | 0.03 | 0 | 22 | Abca8a |
| Gpm6b1 | 0 | 3.685343925 | 0.935 | 0.03 | 0 | 22 | Gpm6b |
| Sgcd1 | 0 | 3.644390832 | 0.824 | 0.017 | 0 | 22 | Sgcd |
| Chl13 | 0 | 3.625104283 | 0.889 | 0.05 | 0 | 22 | Chl1 |
| Ptn1 | 0 | 3.620527157 | 0.833 | 0.009 | 0 | 22 | Ptn |
| Ncam1 | 0 | 3.529285887 | 0.815 | 0.003 | 0 | 22 | Ncam1 |

| | | | | | | | |
|---|---|---|---|---|---|---|---|
| Plce11 | 0 | 3.415163904 | 0.815 | 0.018 | 0 | 22 | Plce1 |
| Mal | 0 | 3.373191262 | 0.907 | 0.036 | 0 | 22 | Mal |
| Abca8b1 | 0 | 3.368825613 | 0.833 | 0.016 | 0 | 22 | Abca8b |
| C4b | 0 | 3.314870819 | 0.713 | 0.003 | 0 | 22 | C4b |
| Lgi4 | 0 | 3.282120606 | 0.843 | 0.007 | 0 | 22 | Lgi4 |
| Sostdc11 | 0 | 3.270680728 | 0.787 | 0.002 | 0 | 22 | Sostdc1 |
| Aspa | 0 | 3.216719114 | 0.824 | 0.009 | 0 | 22 | Aspa |
| Pcdh9 | 0 | 3.20349904 | 0.602 | 0.007 | 0 | 22 | Pcdh9 |
| Kcna2 | 0 | 3.197857245 | 0.861 | 0.005 | 0 | 22 | Kcna2 |
| Adam231 | 0 | 3.166070884 | 0.778 | 0.013 | 0 | 22 | Adam23 |
| Rarres22 | 0 | 3.164891394 | 0.861 | 0.035 | 0 | 22 | Rarres2 |
| Gfra3 | 0 | 3.032838282 | 0.759 | 0.001 | 0 | 22 | Gfra3 |
| Atp1a21 | 0 | 3.013467973 | 0.824 | 0.016 | 0 | 22 | Atp1a2 |
| Art31 | 0 | 3.00202251 | 0.898 | 0.024 | 0 | 22 | Art3 |
| Sfrp12 | 0 | 2.946485656 | 0.75 | 0.012 | 0 | 22 | Sfrp1 |
| Dlgap1 | 0 | 2.940422166 | 0.769 | 0.008 | 0 | 22 | Dlgap1 |
| Tenm31 | 0 | 2.821120808 | 0.75 | 0.019 | 0 | 22 | Tenm3 |
| Nrn1 | 0 | 2.811133791 | 0.685 | 0.001 | 0 | 22 | Nrn1 |
| Grik2 | 0 | 2.738682546 | 0.704 | 0.002 | 0 | 22 | Grik2 |
| Ptprz1 | 0 | 2.631882537 | 0.639 | 0.002 | 0 | 22 | Ptprz1 |
| Zfp536 | 0 | 2.628308204 | 0.75 | 0.001 | 0 | 22 | Zfp536 |
| Sox6 | 0 | 2.586046762 | 0.667 | 0.005 | 0 | 22 | Sox6 |
| Edil3 | 0 | 2.571207758 | 0.676 | 0.002 | 0 | 22 | Edil3 |
| Sox101 | 0 | 2.543834237 | 0.704 | 0.004 | 0 | 22 | Sox10 |
| Gpr37l1 | 0 | 2.536083346 | 0.722 | 0.002 | 0 | 22 | Gpr37l1 |
| Fabp7 | 0 | 2.512339568 | 0.611 | 0.001 | 0 | 22 | Fabp7 |
| Cmtm5 | 0 | 2.426552907 | 0.685 | 0.001 | 0 | 22 | Cmtm5 |
| S100b | 0 | 2.420632824 | 0.713 | 0.004 | 0 | 22 | S100b |
| Col20a1 | 0 | 2.413457178 | 0.602 | 0.01 | 0 | 22 | Col20a1 |
| Kcna6 | 0 | 2.41116649 | 0.639 | 0.001 | 0 | 22 | Kcna6 |
| Tmod2 | 0 | 2.370874533 | 0.704 | 0.025 | 0 | 22 | Tmod2 |
| Il1rapl1 | 0 | 2.359053858 | 0.583 | 0.011 | 0 | 22 | Il1rapl1 |
| Xkr4 | 0 | 2.32192304 | 0.519 | 0.005 | 0 | 22 | Xkr4 |
| Kcnh8 | 0 | 2.289825458 | 0.602 | 0 | 0 | 22 | Kcnh8 |
| Sfrp5 | 0 | 2.282493083 | 0.583 | 0 | 0 | 22 | Sfrp5 |
| Luzp2 | 0 | 2.215178129 | 0.639 | 0 | 0 | 22 | Luzp2 |
| Cdh21 | 0 | 2.188024068 | 0.583 | 0.003 | 0 | 22 | Cdh2 |
| Fign1 | 0 | 2.179861929 | 0.694 | 0.022 | 0 | 22 | Fign |
| Ntng11 | 0 | 2.170576107 | 0.472 | 0.003 | 0 | 22 | Ntng1 |
| Adgrl32 | 0 | 2.159955983 | 0.556 | 0.009 | 0 | 22 | Adgrl3 |
| Cadm4 | 0 | 2.146987462 | 0.583 | 0.005 | 0 | 22 | Cadm4 |
| L1cam | 0 | 2.146284951 | 0.63 | 0.015 | 0 | 22 | L1cam |
| Sema3b | 0 | 2.11873846 | 0.565 | 0.001 | 0 | 22 | Sema3b |

| | | | | | | | |
|---|---|---|---|---|---|---|---|
| Gjc3 | 0 | 2.114700936 | 0.5 | 0 | 0 | 22 | Gjc3 |
| Erbb3 | 0 | 2.099238081 | 0.593 | 0.007 | 0 | 22 | Erbb3 |
| Aatk | 0 | 2.051890518 | 0.602 | 0.008 | 0 | 22 | Aatk |
| Grb14 | 0 | 2.050877699 | 0.602 | 0.011 | 0 | 22 | Grb14 |
| Fndc1 | 0 | 2.04262221 | 0.463 | 0.004 | 0 | 22 | Fndc1 |
| Egfl81 | 0 | 1.989417906 | 0.574 | 0.011 | 0 | 22 | Egfl8 |
| Epb41l4b | 0 | 1.983025197 | 0.62 | 0.02 | 0 | 22 | Epb41l4b |
| Rasgef1c | 0 | 1.979986072 | 0.528 | 0.001 | 0 | 22 | Rasgef1c |
| Galnt171 | 0 | 1.963329609 | 0.509 | 0.005 | 0 | 22 | Galnt17 |
| Col16a11 | 0 | 1.940771082 | 0.556 | 0.011 | 0 | 22 | Col16a1 |
| Ttyh1 | 0 | 1.935039309 | 0.537 | 0.001 | 0 | 22 | Ttyh1 |
| Gfra2 | 0 | 1.921325449 | 0.509 | 0.002 | 0 | 22 | Gfra2 |
| Wnt6 | 0 | 1.884717637 | 0.537 | 0.002 | 0 | 22 | Wnt6 |
| Mboat2 | 0 | 1.880890912 | 0.556 | 0.006 | 0 | 22 | Mboat2 |
| Postn2 | 0 | 1.866296643 | 0.398 | 0.009 | 0 | 22 | Postn |
| Adgrg61 | 0 | 1.856931782 | 0.509 | 0.008 | 0 | 22 | Adgrg6 |
| Lrrc4c | 0 | 1.845618455 | 0.537 | 0.005 | 0 | 22 | Lrrc4c |
| Cyp2j9 | 0 | 1.839334098 | 0.519 | 0.014 | 0 | 22 | Cyp2j9 |
| Pex5l | 0 | 1.837028968 | 0.491 | 0.001 | 0 | 22 | Pex5l |
| Sorcs1 | 0 | 1.836715428 | 0.398 | 0.001 | 0 | 22 | Sorcs1 |
| Gm2115 | 0 | 1.814278315 | 0.537 | 0 | 0 | 22 | Gm2115 |
| Olfml2b1 | 0 | 1.756578255 | 0.528 | 0.008 | 0 | 22 | Olfml2b |
| Vgll3 | 0 | 1.723311948 | 0.5 | 0.007 | 0 | 22 | Vgll3 |
| Klhl291 | 0 | 1.688961344 | 0.472 | 0.006 | 0 | 22 | Klhl29 |
| Foxd3 | 0 | 1.678720437 | 0.463 | 0 | 0 | 22 | Foxd3 |
| Slitrk6 | 0 | 1.6608577 | 0.463 | 0.002 | 0 | 22 | Slitrk6 |
| Adam11 | 0 | 1.62586191 | 0.444 | 0.002 | 0 | 22 | Adam11 |
| Opcml | 0 | 1.621513499 | 0.389 | 0.007 | 0 | 22 | Opcml |
| Cnksr2 | 0 | 1.608509041 | 0.398 | 0 | 0 | 22 | Cnksr2 |
| Sema3e | 0 | 1.525883603 | 0.444 | 0.002 | 0 | 22 | Sema3e |
| Nkain4 | 0 | 1.507377789 | 0.389 | 0.001 | 0 | 22 | Nkain4 |
| Stk32a | 0 | 1.500654402 | 0.407 | 0.002 | 0 | 22 | Stk32a |
| Lgi11 | 0 | 1.487894909 | 0.37 | 0.005 | 0 | 22 | Lgi1 |
| Ncam2 | 0 | 1.480087673 | 0.296 | 0.001 | 0 | 22 | Ncam2 |
| Reln | 0 | 1.464705363 | 0.25 | 0.002 | 0 | 22 | Reln |
| Daam21 | 0 | 1.431316422 | 0.435 | 0.008 | 0 | 22 | Daam2 |
| A330049N07Rik | 0 | 1.395956864 | 0.287 | 0 | 0 | 22 | A330049N07Rik |
| Gas2l3 | 0 | 1.381665603 | 0.398 | 0.007 | 0 | 22 | Gas2l3 |
| Adamts20 | 0 | 1.342458105 | 0.361 | 0 | 0 | 22 | Adamts20 |
| Srcin1 | 0 | 1.336258765 | 0.38 | 0.001 | 0 | 22 | Srcin1 |
| Insc | 0 | 1.316943504 | 0.426 | 0.001 | 0 | 22 | Insc |
| Ephb2 | 0 | 1.314352533 | 0.37 | 0.006 | 0 | 22 | Ephb2 |
| Flrt3 | 0 | 1.311501828 | 0.343 | 0.001 | 0 | 22 | Flrt3 |

| | | | | | | | |
|---|---|---|---|---|---|---|---|
| Matn4 | 0 | 1.302920412 | 0.389 | 0.001 | 0 | 22 | Matn4 |
| Nlgn3 | 0 | 1.259016357 | 0.352 | 0.001 | 0 | 22 | Nlgn3 |
| Tenm1 | 0 | 1.255786335 | 0.278 | 0 | 0 | 22 | Tenm1 |
| Rxrg | 0 | 1.209412663 | 0.361 | 0.005 | 0 | 22 | Rxrg |
| Slc22a23 | 0 | 1.187266101 | 0.324 | 0.004 | 0 | 22 | Slc22a23 |
| Scg3 | 0 | 1.169792073 | 0.269 | 0.003 | 0 | 22 | Scg3 |
| Fam19a5 | 0 | 1.158564592 | 0.352 | 0.006 | 0 | 22 | Fam19a5 |
| Frmd3 | 0 | 1.145967686 | 0.306 | 0.001 | 0 | 22 | Frmd3 |
| Igsf11 | 0 | 1.140210409 | 0.287 | 0 | 0 | 22 | Igsf11 |
| Hmgcs2 | 0 | 1.096706297 | 0.287 | 0.001 | 0 | 22 | Hmgcs2 |
| Gal3st1 | 0 | 1.069553159 | 0.287 | 0.001 | 0 | 22 | Gal3st1 |
| Sema3d1 | 0 | 1.045984927 | 0.287 | 0.005 | 0 | 22 | Sema3d |
| Ajap1 | 0 | 0.996503811 | 0.259 | 0.002 | 0 | 22 | Ajap1 |
| Prima1 | 0 | 0.985178083 | 0.296 | 0 | 0 | 22 | Prima1 |
| Ndst3 | 0 | 0.905235287 | 0.25 | 0.002 | 0 | 22 | Ndst3 |
| C130071C03Rik | 0 | 0.892544683 | 0.259 | 0 | 0 | 22 | C130071C03Rik |
| Tmprss5 | 0 | 0.876258621 | 0.25 | 0.002 | 0 | 22 | Tmprss5 |
| Col3a12 | 1.55E-305 | 2.028357618 | 0.759 | 0.037 | 3.13E-301 | 22 | Col3a1 |
| Prnp1 | 6.55E-299 | 4.776330339 | 0.944 | 0.071 | 1.33E-294 | 22 | Prnp |
| Fez1 | 1.66E-297 | 1.355616041 | 0.389 | 0.01 | 3.35E-293 | 22 | Fez1 |
| Shpk | 5.65E-286 | 1.330975261 | 0.38 | 0.01 | 1.14E-281 | 22 | Shpk |
| Plekhb1 | 2.13E-282 | 1.418709667 | 0.398 | 0.011 | 4.31E-278 | 22 | Plekhb1 |
| Lpar11 | 5.59E-282 | 1.861160534 | 0.556 | 0.022 | 1.13E-277 | 22 | Lpar1 |
| Fads2 | 1.46E-279 | 1.002204553 | 0.296 | 0.006 | 2.96E-275 | 22 | Fads2 |
| Col18a12 | 4.89E-279 | 1.856120617 | 0.537 | 0.021 | 9.91E-275 | 22 | Col18a1 |
| Fam89a | 5.01E-279 | 1.150919214 | 0.398 | 0.011 | 1.01E-274 | 22 | Fam89a |
| Lama22 | 5.60E-276 | 1.870374442 | 0.704 | 0.035 | 1.13E-271 | 22 | Lama2 |
| Scd22 | 1.81E-259 | 3.58307169 | 0.926 | 0.078 | 3.66E-255 | 22 | Scd2 |
| Cryab2 | 2.91E-255 | 3.437645911 | 0.889 | 0.068 | 5.89E-251 | 22 | Cryab |
| Plekha4 | 1.35E-252 | 1.848970098 | 0.398 | 0.013 | 2.74E-248 | 22 | Plekha4 |
| Entpd21 | 1.17E-242 | 1.731453098 | 0.583 | 0.028 | 2.38E-238 | 22 | Entpd2 |
| Dmd2 | 8.75E-242 | 2.784411627 | 0.796 | 0.058 | 1.77E-237 | 22 | Dmd |
| Atp1b21 | 1.12E-239 | 1.412386901 | 0.472 | 0.019 | 2.27E-235 | 22 | Atp1b2 |
| Sorcs2 | 2.75E-239 | 1.033737878 | 0.296 | 0.007 | 5.57E-235 | 22 | Sorcs2 |
| Acsbg1 | 3.27E-230 | 1.192864606 | 0.315 | 0.008 | 6.62E-226 | 22 | Acsbg1 |
| Ptprf1 | 7.67E-226 | 1.032579846 | 0.306 | 0.008 | 1.55E-221 | 22 | Ptprf |
| Col1a22 | 2.64E-220 | 1.473761655 | 0.657 | 0.038 | 5.34E-216 | 22 | Col1a2 |
| Mbp5 | 3.60E-217 | 2.862794966 | 0.796 | 0.066 | 7.29E-213 | 22 | Mbp |
| Megf9 | 6.29E-215 | 2.049579503 | 0.602 | 0.036 | 1.27E-210 | 22 | Megf9 |
| Afap1l22 | 6.13E-214 | 1.727133105 | 0.509 | 0.025 | 1.24E-209 | 22 | Afap1l2 |
| Sdc45 | 4.08E-210 | 2.626517949 | 0.806 | 0.069 | 8.27E-206 | 22 | Sdc4 |
| Kirrel2 | 7.87E-209 | 1.548370025 | 0.454 | 0.02 | 1.59E-204 | 22 | Kirrel |
| Aldh1a12 | 7.96E-204 | 2.12026702 | 0.565 | 0.032 | 1.61E-199 | 22 | Aldh1a1 |

| | | | | | | | |
|---|---|---|---|---|---|---|---|
| Col5a31 | 4.76E-201 | 1.860389529 | 0.546 | 0.03 | 9.64E-197 | 22 | Col5a3 |
| Stard131 | 5.04E-201 | 2.033416385 | 0.611 | 0.04 | 1.02E-196 | 22 | Stard13 |
| Fcgr2b3 | 7.01E-196 | 2.848072777 | 0.593 | 0.039 | 1.42E-191 | 22 | Fcgr2b |
| Gpt2 | 5.65E-195 | 0.942059685 | 0.269 | 0.007 | 1.14E-190 | 22 | Gpt2 |
| Ispd | 1.49E-194 | 1.547427263 | 0.454 | 0.022 | 3.02E-190 | 22 | Ispd |
| Cd59a3 | 8.84E-192 | 2.825286381 | 0.843 | 0.085 | 1.79E-187 | 22 | Cd59a |
| Vwa12 | 6.21E-190 | 3.305303733 | 0.898 | 0.102 | 1.26E-185 | 22 | Vwa1 |
| Col14a13 | 7.16E-184 | 2.190819494 | 0.694 | 0.055 | 1.45E-179 | 22 | Col14a1 |
| Eda1 | 5.66E-183 | 1.943955602 | 0.519 | 0.03 | 1.15E-178 | 22 | Eda |
| Gsn1 | 1.01E-170 | 1.547109733 | 0.917 | 0.106 | 2.05E-166 | 22 | Gsn |
| Cadm13 | 2.33E-169 | 2.405703009 | 0.75 | 0.07 | 4.71E-165 | 22 | Cadm1 |
| Col27a11 | 1.76E-165 | 1.898459619 | 0.472 | 0.028 | 3.56E-161 | 22 | Col27a1 |
| Pdgfa3 | 2.31E-163 | 2.250273932 | 0.667 | 0.058 | 4.67E-159 | 22 | Pdgfa |
| Cnp2 | 2.33E-162 | 2.829248824 | 0.815 | 0.095 | 4.73E-158 | 22 | Cnp |
| Apod1 | 5.71E-159 | 2.053709129 | 0.537 | 0.036 | 1.16E-154 | 22 | Apod |
| Gria31 | 1.44E-156 | 1.260871776 | 0.361 | 0.017 | 2.91E-152 | 22 | Gria3 |
| Paqr61 | 1.80E-156 | 1.450805097 | 0.546 | 0.038 | 3.65E-152 | 22 | Paqr6 |
| Gatm3 | 9.33E-147 | 2.386774639 | 0.713 | 0.072 | 1.89E-142 | 22 | Gatm |
| Dock74 | 2.24E-145 | 1.875761774 | 0.565 | 0.046 | 4.54E-141 | 22 | Dock7 |
| Syt11 | 4.28E-145 | 1.609014003 | 0.463 | 0.031 | 8.66E-141 | 22 | Syt11 |
| Zeb26 | 1.09E-143 | 3.220321839 | 0.917 | 0.142 | 2.20E-139 | 22 | Zeb2 |
| Frmd5 | 9.71E-142 | 1.325893434 | 0.306 | 0.013 | 1.97E-137 | 22 | Frmd5 |
| Alcam2 | 1.74E-136 | 1.400276456 | 0.454 | 0.031 | 3.53E-132 | 22 | Alcam |
| Zfhx41 | 5.97E-134 | 1.21614772 | 0.38 | 0.022 | 1.21E-129 | 22 | Zfhx4 |
| Metrn | 5.19E-131 | 1.146527133 | 0.37 | 0.022 | 1.05E-126 | 22 | Metrn |
| Ttyh2 | 1.98E-129 | 0.967789816 | 0.296 | 0.014 | 4.01E-125 | 22 | Ttyh2 |
| Prkca8 | 4.72E-129 | 2.767350466 | 0.75 | 0.099 | 9.56E-125 | 22 | Prkca |
| Nrxn11 | 1.30E-128 | 1.635942062 | 0.324 | 0.016 | 2.63E-124 | 22 | Nrxn1 |
| Itgb43 | 3.00E-128 | 1.487428753 | 0.454 | 0.033 | 6.08E-124 | 22 | Itgb4 |
| F31 | 1.32E-126 | 1.245612793 | 0.352 | 0.02 | 2.68E-122 | 22 | F3 |
| Fads11 | 1.62E-126 | 1.672228078 | 0.565 | 0.053 | 3.29E-122 | 22 | Fads1 |
| Pmp227 | 8.24E-122 | 3.238461359 | 0.861 | 0.143 | 1.67E-117 | 22 | Pmp22 |
| Antxr12 | 1.10E-121 | 1.254182441 | 0.361 | 0.022 | 2.22E-117 | 22 | Antxr1 |
| Egflam2 | 1.21E-116 | 1.038401372 | 0.333 | 0.019 | 2.46E-112 | 22 | Egflam |
| Sntb15 | 3.45E-116 | 1.951692555 | 0.593 | 0.064 | 6.98E-112 | 22 | Sntb1 |
| Itga21 | 1.05E-115 | 1.498482757 | 0.454 | 0.037 | 2.13E-111 | 22 | Itga2 |
| Magi22 | 5.99E-114 | 1.91361503 | 0.454 | 0.037 | 1.21E-109 | 22 | Magi2 |
| Deptor | 2.74E-111 | 1.727444968 | 0.556 | 0.057 | 5.55E-107 | 22 | Deptor |
| Vcan1 | 2.12E-108 | 1.199898061 | 0.287 | 0.015 | 4.29E-104 | 22 | Vcan |
| Rftn2 | 4.71E-108 | 1.033870153 | 0.287 | 0.016 | 9.55E-104 | 22 | Rftn2 |
| Frmd4a7 | 5.76E-108 | 3.097498454 | 0.815 | 0.147 | 1.17E-103 | 22 | Frmd4a |
| Thsd7a2 | 1.44E-107 | 1.502055552 | 0.435 | 0.035 | 2.91E-103 | 22 | Thsd7a |
| Abi3bp2 | 2.84E-106 | 1.449213735 | 0.417 | 0.033 | 5.75E-102 | 22 | Abi3bp |

| | | | | | | | |
|---|---|---|---|---|---|---|---|
| Fgf2 | 1.14E-105 | 0.763412217 | 0.259 | 0.013 | 2.30E-101 | 22 | Fgf2 |
| Mtss1l1 | 4.61E-105 | 1.27210485 | 0.426 | 0.035 | 9.34E-101 | 22 | Mtss1l |
| Col5a11 | 7.79E-105 | 1.213936281 | 0.343 | 0.023 | 1.58E-100 | 22 | Col5a1 |
| Tppp31 | 2.05E-104 | 1.315870694 | 0.333 | 0.022 | 4.15E-100 | 22 | Tppp3 |
| Dst9 | 2.33E-101 | 2.881440835 | 0.88 | 0.19 | 4.73E-97 | 22 | Dst |
| Macrod2 | 2.41E-101 | 1.591138033 | 0.463 | 0.043 | 4.88E-97 | 22 | Macrod2 |
| Adamts122 | 6.63E-101 | 1.256080403 | 0.454 | 0.039 | 1.34E-96 | 22 | Adamts12 |
| Snca1 | 1.66E-99 | 1.167422561 | 0.343 | 0.024 | 3.36E-95 | 22 | Snca |
| Ppp1r9a1 | 4.05E-97 | 1.311992224 | 0.407 | 0.035 | 8.19E-93 | 22 | Ppp1r9a |
| Cyp2j6 | 9.98E-96 | 1.56028019 | 0.454 | 0.044 | 2.02E-91 | 22 | Cyp2j6 |
| Arhgef161 | 5.43E-95 | 0.751613913 | 0.25 | 0.013 | 1.10E-90 | 22 | Arhgef16 |
| Rbms33 | 4.04E-93 | 1.282207215 | 0.509 | 0.053 | 8.18E-89 | 22 | Rbms3 |
| A830082K12Rik2 | 1.80E-88 | 0.938289494 | 0.25 | 0.014 | 3.64E-84 | 22 | A830082K12Rik |
| Olfml2a1 | 6.74E-88 | 1.049801183 | 0.269 | 0.017 | 1.36E-83 | 22 | Olfml2a |
| Rora6 | 1.38E-87 | 2.773411288 | 0.833 | 0.185 | 2.79E-83 | 22 | Rora |
| Enho1 | 2.93E-87 | 0.960631584 | 0.352 | 0.029 | 5.93E-83 | 22 | Enho |
| Gulp17 | 1.82E-86 | 2.424402305 | 0.796 | 0.165 | 3.68E-82 | 22 | Gulp1 |
| Pdzd24 | 3.35E-86 | 2.030188332 | 0.685 | 0.114 | 6.77E-82 | 22 | Pdzd2 |
| Celf29 | 1.44E-85 | 2.653586009 | 0.852 | 0.201 | 2.91E-81 | 22 | Celf2 |
| Angptl22 | 1.54E-85 | 1.100198464 | 0.296 | 0.021 | 3.11E-81 | 22 | Angptl2 |
| CT025619.12 | 1.89E-85 | 0.878876626 | 0.269 | 0.017 | 3.83E-81 | 22 | CT025619.1 |
| Phlda33 | 2.50E-85 | 1.298417091 | 0.481 | 0.053 | 5.06E-81 | 22 | Phlda3 |
| Itgb81 | 1.06E-81 | 1.084694712 | 0.38 | 0.035 | 2.14E-77 | 22 | Itgb8 |
| Timp27 | 1.40E-81 | 1.584971562 | 0.731 | 0.129 | 2.84E-77 | 22 | Timp2 |
| Scd11 | 3.30E-81 | 1.142249628 | 0.315 | 0.025 | 6.69E-77 | 22 | Scd1 |
| Frmd62 | 1.31E-80 | 1.561129819 | 0.426 | 0.046 | 2.66E-76 | 22 | Frmd6 |
| Sox52 | 2.26E-79 | 0.953423068 | 0.287 | 0.021 | 4.57E-75 | 22 | Sox5 |
| Reck1 | 8.40E-79 | 1.379331225 | 0.361 | 0.034 | 1.70E-74 | 22 | Reck |
| Sdc22 | 1.03E-78 | 0.916212315 | 0.352 | 0.031 | 2.08E-74 | 22 | Sdc2 |
| Col1a12 | 1.65E-77 | 0.778707794 | 0.37 | 0.033 | 3.34E-73 | 22 | Col1a1 |
| Antxr24 | 2.31E-77 | 1.637383066 | 0.583 | 0.09 | 4.68E-73 | 22 | Antxr2 |
| Iqgap24 | 2.54E-77 | 1.862976855 | 0.63 | 0.104 | 5.14E-73 | 22 | Iqgap2 |
| Dbi4 | 2.16E-75 | 3.744139543 | 0.972 | 0.432 | 4.37E-71 | 22 | Dbi |
| Ehbp12 | 3.32E-75 | 1.907300969 | 0.63 | 0.11 | 6.73E-71 | 22 | Ehbp1 |
| Agap14 | 9.33E-74 | 2.046418552 | 0.704 | 0.144 | 1.89E-69 | 22 | Agap1 |
| Ptprs2 | 2.21E-71 | 1.15165624 | 0.361 | 0.037 | 4.47E-67 | 22 | Ptprs |
| Prdm161 | 3.13E-71 | 1.16736209 | 0.426 | 0.05 | 6.33E-67 | 22 | Prdm16 |
| Prkg16 | 2.79E-70 | 2.219661949 | 0.806 | 0.196 | 5.66E-66 | 22 | Prkg1 |
| S100a610 | 9.86E-69 | 2.118988107 | 0.944 | 0.271 | 2.00E-64 | 22 | S100a6 |
| Htra14 | 1.03E-68 | 1.170050656 | 0.407 | 0.046 | 2.08E-64 | 22 | Htra1 |
| Tln22 | 2.07E-68 | 1.211992874 | 0.38 | 0.042 | 4.19E-64 | 22 | Tln2 |
| Lbh5 | 3.27E-67 | 1.909267096 | 0.648 | 0.132 | 6.61E-63 | 22 | Lbh |
| Srebf11 | 1.29E-66 | 1.380733702 | 0.426 | 0.055 | 2.62E-62 | 22 | Srebf1 |

| | | | | | | | |
|---|---|---|---|---|---|---|---|
| Mpp74 | 1.69E-66 | 1.710554594 | 0.593 | 0.106 | 3.42E-62 | 22 | Mpp7 |
| Sgce2 | 2.31E-65 | 1.188460854 | 0.38 | 0.043 | 4.68E-61 | 22 | Sgce |
| Cd97 | 3.60E-65 | 2.832018447 | 0.926 | 0.386 | 7.29E-61 | 22 | Cd9 |
| Dag17 | 5.39E-65 | 1.787857268 | 0.667 | 0.142 | 1.09E-60 | 22 | Dag1 |
| Gas76 | 5.53E-65 | 1.986196272 | 0.75 | 0.182 | 1.12E-60 | 22 | Gas7 |
| Arhgef26 | 6.10E-65 | 0.981246999 | 0.25 | 0.019 | 1.23E-60 | 22 | Arhgef26 |
| Efemp11 | 1.34E-63 | 0.886850605 | 0.287 | 0.025 | 2.72E-59 | 22 | Efemp1 |
| Tspan36 | 2.53E-63 | 1.643257594 | 0.574 | 0.104 | 5.12E-59 | 22 | Tspan3 |
| Dpyd1 | 3.06E-63 | 1.159080919 | 0.398 | 0.048 | 6.20E-59 | 22 | Dpyd |
| Gab14 | 5.08E-63 | 1.613564658 | 0.602 | 0.114 | 1.03E-58 | 22 | Gab1 |
| Rhobtb31 | 5.40E-62 | 1.107972229 | 0.398 | 0.05 | 1.09E-57 | 22 | Rhobtb3 |
| Cpe2 | 5.74E-61 | 1.367855645 | 0.25 | 0.021 | 1.16E-56 | 22 | Cpe |
| Vim13 | 6.37E-61 | 2.35978927 | 0.88 | 0.307 | 1.29E-56 | 22 | Vim |
| Ndrg22 | 1.04E-60 | 1.163291397 | 0.481 | 0.07 | 2.11E-56 | 22 | Ndrg2 |
| Sorbs17 | 1.68E-60 | 1.839145871 | 0.731 | 0.177 | 3.40E-56 | 22 | Sorbs1 |
| Col5a23 | 2.73E-60 | 0.95674639 | 0.37 | 0.042 | 5.53E-56 | 22 | Col5a2 |
| Il1rap | 1.50E-59 | 1.142119047 | 0.361 | 0.044 | 3.04E-55 | 22 | Il1rap |
| Limch15 | 5.21E-59 | 1.810716998 | 0.648 | 0.139 | 1.05E-54 | 22 | Limch1 |
| Myl92 | 6.08E-59 | 0.559594194 | 0.259 | 0.022 | 1.23E-54 | 22 | Myl9 |
| Ptpn132 | 6.50E-59 | 0.884852729 | 0.306 | 0.031 | 1.32E-54 | 22 | Ptpn13 |
| Phgdh1 | 6.54E-59 | 1.069956257 | 0.389 | 0.05 | 1.32E-54 | 22 | Phgdh |
| Csrp16 | 7.78E-59 | 1.830451428 | 0.722 | 0.182 | 1.58E-54 | 22 | Csrp1 |
| Atp9a | 1.15E-58 | 0.923675379 | 0.315 | 0.033 | 2.34E-54 | 22 | Atp9a |
| Nid21 | 3.41E-58 | 1.349765036 | 0.481 | 0.076 | 6.91E-54 | 22 | Nid2 |
| Casc42 | 6.11E-58 | 0.927466456 | 0.315 | 0.034 | 1.24E-53 | 22 | Casc4 |
| Slc44a15 | 2.55E-57 | 1.691472953 | 0.667 | 0.156 | 5.16E-53 | 22 | Slc44a1 |
| Asap25 | 7.44E-57 | 1.819675263 | 0.611 | 0.131 | 1.51E-52 | 22 | Asap2 |
| Fbxo72 | 1.07E-56 | 1.491301306 | 0.546 | 0.104 | 2.17E-52 | 22 | Fbxo7 |
| Lrrc8b | 4.35E-56 | 0.915330548 | 0.324 | 0.037 | 8.81E-52 | 22 | Lrrc8b |
| Ppfibp21 | 6.83E-56 | 0.924658211 | 0.306 | 0.033 | 1.38E-51 | 22 | Ppfibp2 |
| Sparc9 | 3.69E-55 | 2.04031333 | 0.963 | 0.403 | 7.47E-51 | 22 | Sparc |
| Ctnnal11 | 4.38E-54 | 1.206223877 | 0.25 | 0.023 | 8.87E-50 | 22 | Ctnnal1 |
| St6gal14 | 2.29E-53 | 1.090482399 | 0.333 | 0.041 | 4.64E-49 | 22 | St6gal1 |
| Malat113 | 3.71E-53 | 2.349177317 | 0.981 | 0.726 | 7.52E-49 | 22 | Malat1 |
| Klk85 | 1.12E-52 | 1.525133206 | 0.472 | 0.082 | 2.27E-48 | 22 | Klk8 |
| Sh3d195 | 1.86E-52 | 1.472064084 | 0.546 | 0.108 | 3.77E-48 | 22 | Sh3d19 |
| P2rx71 | 3.19E-52 | 1.002390199 | 0.37 | 0.051 | 6.46E-48 | 22 | P2rx7 |
| Fasn | 3.51E-52 | 1.014027321 | 0.306 | 0.036 | 7.11E-48 | 22 | Fasn |
| Plxdc25 | 3.91E-52 | 1.449403862 | 0.528 | 0.093 | 7.91E-48 | 22 | Plxdc2 |
| Serpine22 | 4.02E-52 | 1.047098221 | 0.296 | 0.033 | 8.14E-48 | 22 | Serpine2 |
| Slc2a131 | 5.66E-52 | 0.995613262 | 0.315 | 0.037 | 1.14E-47 | 22 | Slc2a13 |
| Fgl28 | 2.58E-51 | 1.917820406 | 0.528 | 0.108 | 5.22E-47 | 22 | Fgl2 |
| Fst2 | 2.82E-51 | 0.731768226 | 0.269 | 0.027 | 5.70E-47 | 22 | Fst |

| | | | | | | | |
|---|---|---|---|---|---|---|---|
| Hbegf | 3.57E-51 | 1.198175833 | 0.417 | 0.065 | 7.23E-47 | 22 | Hbegf |
| Pard3b4 | 6.31E-51 | 1.392178913 | 0.519 | 0.101 | 1.28E-46 | 22 | Pard3b |
| Cacna2d12 | 4.73E-50 | 0.80517594 | 0.25 | 0.024 | 9.58E-46 | 22 | Cacna2d1 |
| Laptm4b | 5.63E-50 | 0.939800405 | 0.333 | 0.043 | 1.14E-45 | 22 | Laptm4b |
| Slc43a22 | 1.42E-49 | 1.004526433 | 0.296 | 0.035 | 2.88E-45 | 22 | Slc43a2 |
| Casp12 | 2.66E-49 | 0.875027463 | 0.287 | 0.033 | 5.38E-45 | 22 | Casp12 |
| Apoe4 | 3.90E-49 | 2.076425059 | 0.972 | 0.534 | 7.89E-45 | 22 | Apoe |
| Fam78b | 6.67E-48 | 1.136258598 | 0.361 | 0.052 | 1.35E-43 | 22 | Fam78b |
| Fstl15 | 2.02E-47 | 1.250693577 | 0.519 | 0.103 | 4.09E-43 | 22 | Fstl1 |
| Tspan154 | 6.26E-47 | 1.329682612 | 0.583 | 0.13 | 1.27E-42 | 22 | Tspan15 |
| Sh3pxd2a4 | 4.51E-46 | 1.728729173 | 0.583 | 0.142 | 9.13E-42 | 22 | Sh3pxd2a |
| Marcks7 | 7.02E-46 | 1.818401806 | 0.87 | 0.342 | 1.42E-41 | 22 | Marcks |
| Lpcat23 | 7.82E-46 | 1.126349286 | 0.407 | 0.067 | 1.58E-41 | 22 | Lpcat2 |
| Asrgl1 | 8.38E-46 | 1.198092095 | 0.435 | 0.078 | 1.70E-41 | 22 | Asrgl1 |
| Tanc24 | 1.08E-45 | 1.397696485 | 0.639 | 0.16 | 2.19E-41 | 22 | Tanc2 |
| Retreg17 | 1.28E-45 | 1.474209972 | 0.481 | 0.097 | 2.58E-41 | 22 | Retreg1 |
| Adk2 | 2.81E-45 | 1.06839826 | 0.574 | 0.125 | 5.70E-41 | 22 | Adk |
| Ckb7 | 3.36E-45 | 1.706132121 | 0.722 | 0.216 | 6.79E-41 | 22 | Ckb |
| S100a45 | 4.14E-45 | 1.198776526 | 0.602 | 0.129 | 8.39E-41 | 22 | S100a4 |
| Pcbp4 | 7.84E-44 | 0.944281282 | 0.25 | 0.028 | 1.59E-39 | 22 | Pcbp4 |
| Lsamp2 | 9.08E-44 | 0.706328236 | 0.306 | 0.038 | 1.84E-39 | 22 | Lsamp |
| Igf1r8 | 2.11E-43 | 1.389392921 | 0.62 | 0.161 | 4.27E-39 | 22 | Igf1r |
| Itgav10 | 3.03E-43 | 1.576734518 | 0.602 | 0.161 | 6.14E-39 | 22 | Itgav |
| Slc1a53 | 3.42E-43 | 1.186117865 | 0.426 | 0.08 | 6.92E-39 | 22 | Slc1a5 |
| Pard34 | 7.68E-43 | 0.920160576 | 0.324 | 0.046 | 1.56E-38 | 22 | Pard3 |
| Cd637 | 1.01E-42 | 1.295722848 | 0.722 | 0.188 | 2.04E-38 | 22 | Cd63 |
| Lama47 | 1.62E-42 | 1.513224811 | 0.648 | 0.175 | 3.27E-38 | 22 | Lama4 |
| Itga47 | 1.68E-42 | 1.133247881 | 0.417 | 0.076 | 3.39E-38 | 22 | Itga4 |
| Fam234a | 3.24E-42 | 0.739211703 | 0.278 | 0.035 | 6.56E-38 | 22 | Fam234a |
| Sh3pxd2b2 | 5.34E-41 | 0.957586662 | 0.296 | 0.041 | 1.08E-36 | 22 | Sh3pxd2b |
| Arhgap322 | 8.17E-41 | 1.010668402 | 0.343 | 0.054 | 1.65E-36 | 22 | Arhgap32 |
| Gstm53 | 2.90E-40 | 0.855483622 | 0.38 | 0.063 | 5.86E-36 | 22 | Gstm5 |
| Ntn43 | 1.43E-39 | 0.688684715 | 0.269 | 0.035 | 2.90E-35 | 22 | Ntn4 |
| Cep1122 | 6.28E-39 | 1.02511872 | 0.352 | 0.059 | 1.27E-34 | 22 | Cep112 |
| Park21 | 6.89E-39 | 1.175448864 | 0.398 | 0.075 | 1.39E-34 | 22 | Park2 |
| Rnd31 | 8.83E-39 | 1.170107551 | 0.463 | 0.097 | 1.79E-34 | 22 | Rnd3 |
| Col6a31 | 2.49E-38 | 0.749290415 | 0.25 | 0.03 | 5.05E-34 | 22 | Col6a3 |
| Kif21a | 5.08E-38 | 0.992753082 | 0.296 | 0.042 | 1.03E-33 | 22 | Kif21a |
| Tmtc24 | 6.64E-38 | 1.640907175 | 0.63 | 0.185 | 1.34E-33 | 22 | Tmtc2 |
| Stmn14 | 1.11E-37 | 1.363075372 | 0.537 | 0.131 | 2.25E-33 | 22 | Stmn1 |
| Plat1 | 1.52E-37 | 1.006223381 | 0.343 | 0.058 | 3.07E-33 | 22 | Plat |
| Lhfpl24 | 1.54E-37 | 1.324988983 | 0.574 | 0.15 | 3.13E-33 | 22 | Lhfpl2 |
| Ndrg15 | 3.98E-37 | 1.563455311 | 0.704 | 0.233 | 8.05E-33 | 22 | Ndrg1 |

| | | | | | | | |
|---|---|---|---|---|---|---|---|
| Plekha57 | 2.37E-36 | 1.293529896 | 0.565 | 0.156 | 4.79E-32 | 22 | Plekha5 |
| Creb53 | 4.03E-36 | 0.94089339 | 0.352 | 0.061 | 8.16E-32 | 22 | Creb5 |
| Camk2d10 | 5.87E-36 | 1.553933696 | 0.731 | 0.27 | 1.19E-31 | 22 | Camk2d |
| Sgcb1 | 1.16E-35 | 0.938988238 | 0.324 | 0.055 | 2.35E-31 | 22 | Sgcb |
| Specc16 | 1.44E-35 | 1.116546836 | 0.398 | 0.081 | 2.91E-31 | 22 | Specc1 |
| Arhgap393 | 2.31E-35 | 1.009897369 | 0.333 | 0.059 | 4.67E-31 | 22 | Arhgap39 |
| Rasa36 | 4.30E-35 | 1.071136457 | 0.352 | 0.066 | 8.71E-31 | 22 | Rasa3 |
| Prex26 | 6.05E-35 | 1.454582948 | 0.759 | 0.273 | 1.22E-30 | 22 | Prex2 |
| Nr2f23 | 6.98E-35 | 1.055644388 | 0.454 | 0.099 | 1.41E-30 | 22 | Nr2f2 |
| Srgap1 | 8.56E-35 | 1.064998537 | 0.333 | 0.059 | 1.73E-30 | 22 | Srgap1 |
| Gpx82 | 1.06E-34 | 0.920365778 | 0.306 | 0.05 | 2.14E-30 | 22 | Gpx8 |
| Timp37 | 3.41E-34 | 1.31725147 | 0.926 | 0.386 | 6.90E-30 | 22 | Timp3 |
| Qk6 | 4.10E-34 | 1.71318088 | 0.843 | 0.411 | 8.29E-30 | 22 | Qk |
| Sort11 | 4.90E-34 | 0.847783711 | 0.306 | 0.051 | 9.92E-30 | 22 | Sort1 |
| Itpk13 | 5.42E-34 | 1.077271822 | 0.361 | 0.071 | 1.10E-29 | 22 | Itpk1 |
| Selenom6 | 6.11E-34 | 1.523473863 | 0.898 | 0.446 | 1.24E-29 | 22 | Selenom |
| Synm2 | 6.36E-34 | 0.751481008 | 0.269 | 0.039 | 1.29E-29 | 22 | Synm |
| Tuba1a8 | 9.14E-34 | 1.680241845 | 0.694 | 0.257 | 1.85E-29 | 22 | Tuba1a |
| Mill24 | 1.30E-33 | 0.904948 | 0.361 | 0.069 | 2.63E-29 | 22 | Mill2 |
| Pmepa15 | 1.60E-33 | 1.551396231 | 0.685 | 0.252 | 3.25E-29 | 22 | Pmepa1 |
| Txndc16 | 5.50E-33 | 0.750340392 | 0.287 | 0.046 | 1.11E-28 | 22 | Txndc16 |
| Ank22 | 9.10E-33 | 0.915776288 | 0.278 | 0.043 | 1.84E-28 | 22 | Ank2 |
| Cnn37 | 1.51E-32 | 1.374517373 | 0.602 | 0.194 | 3.06E-28 | 22 | Cnn3 |
| Lamc16 | 3.13E-32 | 1.238531635 | 0.528 | 0.148 | 6.33E-28 | 22 | Lamc1 |
| Dock55 | 6.46E-32 | 1.104944117 | 0.463 | 0.115 | 1.31E-27 | 22 | Dock5 |
| Sat15 | 1.33E-31 | 1.666529848 | 0.759 | 0.346 | 2.70E-27 | 22 | Sat1 |
| Tnik6 | 1.93E-31 | 1.30998077 | 0.454 | 0.113 | 3.91E-27 | 22 | Tnik |
| Scarb23 | 3.75E-31 | 1.509544972 | 0.574 | 0.188 | 7.59E-27 | 22 | Scarb2 |
| Serping12 | 6.73E-31 | 0.504172029 | 0.25 | 0.036 | 1.36E-26 | 22 | Serping1 |
| Zbtb2010 | 7.52E-31 | 1.418710021 | 0.898 | 0.538 | 1.52E-26 | 22 | Zbtb20 |
| Utrn6 | 1.13E-30 | 1.461008322 | 0.843 | 0.406 | 2.30E-26 | 22 | Utrn |
| Vcl6 | 1.84E-30 | 1.325696646 | 0.537 | 0.162 | 3.73E-26 | 22 | Vcl |
| Gpcpd15 | 2.76E-30 | 1.070692443 | 0.454 | 0.117 | 5.59E-26 | 22 | Gpcpd1 |
| Hsd17b113 | 4.32E-30 | 1.146814514 | 0.454 | 0.119 | 8.74E-26 | 22 | Hsd17b11 |
| Mcam4 | 5.53E-30 | 1.031522652 | 0.444 | 0.11 | 1.12E-25 | 22 | Mcam |
| Tead15 | 1.49E-29 | 0.984481487 | 0.38 | 0.084 | 3.01E-25 | 22 | Tead1 |
| Phldb11 | 1.53E-29 | 0.811908255 | 0.333 | 0.066 | 3.11E-25 | 22 | Phldb1 |
| Lamb13 | 3.24E-29 | 1.238688563 | 0.5 | 0.137 | 6.57E-25 | 22 | Lamb1 |
| Clasp28 | 4.11E-29 | 1.299344218 | 0.556 | 0.183 | 8.33E-25 | 22 | Clasp2 |
| Rassf81 | 5.28E-29 | 0.989752355 | 0.37 | 0.082 | 1.07E-24 | 22 | Rassf8 |
| Inf2 | 1.99E-28 | 0.693457491 | 0.25 | 0.041 | 4.03E-24 | 22 | Inf2 |
| Zdhhc2 | 2.17E-28 | 0.675191096 | 0.259 | 0.042 | 4.39E-24 | 22 | Zdhhc2 |
| Map1b3 | 2.60E-28 | 1.038956385 | 0.417 | 0.102 | 5.26E-24 | 22 | Map1b |

| | | | | | | | |
|---|---|---|---|---|---|---|---|
| Agmo2 | 5.18E-28 | 0.960093399 | 0.296 | 0.056 | 1.05E-23 | 22 | Agmo |
| Lgals18 | 1.80E-27 | 1.411756465 | 0.491 | 0.135 | 3.65E-23 | 22 | Lgals1 |
| Mid1ip11 | 2.04E-27 | 0.987269774 | 0.343 | 0.075 | 4.14E-23 | 22 | Mid1ip1 |
| Snta1 | 3.57E-27 | 0.848945844 | 0.259 | 0.044 | 7.24E-23 | 22 | Snta1 |
| Lmo42 | 3.66E-27 | 1.377718555 | 0.565 | 0.194 | 7.42E-23 | 22 | Lmo4 |
| Ptprj8 | 3.77E-27 | 1.127196835 | 0.509 | 0.152 | 7.64E-23 | 22 | Ptprj |
| Tgfbr36 | 9.28E-27 | 1.323842354 | 0.537 | 0.175 | 1.88E-22 | 22 | Tgfbr3 |
| Sorbs26 | 1.34E-26 | 1.540806481 | 0.704 | 0.278 | 2.71E-22 | 22 | Sorbs2 |
| Fam129a9 | 1.39E-26 | 1.115881229 | 0.481 | 0.146 | 2.81E-22 | 22 | Fam129a |
| Samd43 | 2.16E-26 | 0.778291572 | 0.287 | 0.055 | 4.38E-22 | 22 | Samd4 |
| Ppip5k2 | 2.28E-26 | 0.935997476 | 0.324 | 0.07 | 4.61E-22 | 22 | Ppip5k2 |
| Nf18 | 3.79E-26 | 1.23918901 | 0.593 | 0.222 | 7.68E-22 | 22 | Nf1 |
| Lrp13 | 4.61E-26 | 0.759781834 | 0.296 | 0.057 | 9.34E-22 | 22 | Lrp1 |
| Fam49a3 | 5.16E-26 | 0.744021871 | 0.269 | 0.05 | 1.04E-21 | 22 | Fam49a |
| Fgfr12 | 1.01E-25 | 0.754501437 | 0.269 | 0.05 | 2.04E-21 | 22 | Fgfr1 |
| Tax1bp32 | 3.19E-25 | 1.133762125 | 0.537 | 0.176 | 6.46E-21 | 22 | Tax1bp3 |
| Pdlim42 | 5.57E-25 | 1.014765929 | 0.472 | 0.134 | 1.13E-20 | 22 | Pdlim4 |
| Arl8a | 6.29E-25 | 0.906021206 | 0.38 | 0.097 | 1.27E-20 | 22 | Arl8a |
| Snx1 | 6.43E-25 | 1.063438928 | 0.454 | 0.14 | 1.30E-20 | 22 | Snx1 |
| Negr12 | 6.64E-25 | 0.801050318 | 0.25 | 0.045 | 1.34E-20 | 22 | Negr1 |
| Ddhd11 | 2.21E-24 | 0.924926614 | 0.333 | 0.08 | 4.48E-20 | 22 | Ddhd1 |
| Map3k4 | 3.62E-24 | 1.024125134 | 0.407 | 0.114 | 7.34E-20 | 22 | Map3k4 |
| Asph6 | 9.15E-24 | 0.863375629 | 0.38 | 0.101 | 1.85E-19 | 22 | Asph |
| Cttnbp2nl3 | 4.27E-23 | 1.166790263 | 0.407 | 0.116 | 8.65E-19 | 22 | Cttnbp2nl |
| Add35 | 4.46E-22 | 1.200240541 | 0.565 | 0.224 | 9.03E-18 | 22 | Add3 |
| Plscr46 | 4.92E-22 | 1.129172319 | 0.528 | 0.19 | 9.97E-18 | 22 | Plscr4 |
| Tmem2055 | 4.98E-22 | 1.047164669 | 0.481 | 0.159 | 1.01E-17 | 22 | Tmem205 |
| Hsp90b111 | 6.63E-22 | 1.167227273 | 0.852 | 0.511 | 1.34E-17 | 22 | Hsp90b1 |
| Rgmb1 | 8.70E-22 | 0.757914928 | 0.278 | 0.061 | 1.76E-17 | 22 | Rgmb |
| Emp312 | 9.88E-22 | 1.129129574 | 0.62 | 0.26 | 2.00E-17 | 22 | Emp3 |
| Ifi27l2a4 | 2.10E-21 | 1.570371792 | 0.759 | 0.368 | 4.26E-17 | 22 | Ifi27l2a |
| Bcl26 | 2.22E-21 | 0.979152227 | 0.574 | 0.208 | 4.49E-17 | 22 | Bcl2 |
| Akap125 | 2.34E-21 | 0.930518389 | 0.491 | 0.161 | 4.73E-17 | 22 | Akap12 |
| Acaca1 | 2.36E-21 | 0.950961747 | 0.324 | 0.084 | 4.77E-17 | 22 | Acaca |
| Comt4 | 2.38E-21 | 0.997017433 | 0.519 | 0.194 | 4.82E-17 | 22 | Comt |
| Ctsl6 | 3.26E-21 | 1.094937584 | 0.75 | 0.347 | 6.60E-17 | 22 | Ctsl |
| Pmm1 | 5.95E-21 | 0.722776871 | 0.315 | 0.077 | 1.20E-16 | 22 | Pmm1 |
| Arhgef102 | 6.89E-21 | 0.913842376 | 0.361 | 0.1 | 1.40E-16 | 22 | Arhgef10 |
| Dip2b8 | 7.48E-21 | 0.996446103 | 0.62 | 0.277 | 1.52E-16 | 22 | Dip2b |
| Prdx63 | 9.12E-21 | 0.947648892 | 0.722 | 0.338 | 1.85E-16 | 22 | Prdx6 |
| Nenf5 | 1.01E-20 | 0.930251735 | 0.676 | 0.296 | 2.04E-16 | 22 | Nenf |
| Tst | 1.52E-20 | 0.63509623 | 0.333 | 0.081 | 3.07E-16 | 22 | Tst |
| Nid15 | 1.58E-20 | 0.934958852 | 0.519 | 0.183 | 3.20E-16 | 22 | Nid1 |

| | | | | | | | |
|---|---|---|---|---|---|---|---|
| Cep85l6 | 2.23E-20 | 0.93556703 | 0.481 | 0.165 | 4.51E-16 | 22 | Cep85l |
| Nfia8 | 2.37E-20 | 1.217479776 | 0.657 | 0.288 | 4.79E-16 | 22 | Nfia |
| Rcn13 | 2.74E-20 | 0.868391505 | 0.37 | 0.109 | 5.55E-16 | 22 | Rcn1 |
| Mpdz4 | 2.87E-20 | 0.935286429 | 0.352 | 0.098 | 5.80E-16 | 22 | Mpdz |
| Zfp4623 | 3.10E-20 | 0.966333101 | 0.398 | 0.121 | 6.28E-16 | 22 | Zfp462 |
| Tbc1d10a | 3.33E-20 | 0.834905004 | 0.287 | 0.069 | 6.74E-16 | 22 | Tbc1d10a |
| mt-Cytb14 | 8.50E-20 | 0.897853808 | 0.981 | 0.701 | 1.72E-15 | 22 | mt-Cytb |
| Cebpd2 | 1.39E-19 | 1.269976415 | 0.481 | 0.176 | 2.82E-15 | 22 | Cebpd |
| Tm7sf31 | 1.52E-19 | 0.769383033 | 0.287 | 0.071 | 3.07E-15 | 22 | Tm7sf3 |
| Acyp22 | 2.30E-19 | 0.731955013 | 0.444 | 0.143 | 4.66E-15 | 22 | Acyp2 |
| Palld4 | 6.78E-19 | 0.529144768 | 0.25 | 0.054 | 1.37E-14 | 22 | Palld |
| P4ha13 | 1.22E-18 | 0.855550536 | 0.389 | 0.124 | 2.48E-14 | 22 | P4ha1 |
| Zfp6388 | 1.23E-18 | 1.124836464 | 0.593 | 0.283 | 2.49E-14 | 22 | Zfp638 |
| Npdc15 | 1.29E-18 | 0.943588388 | 0.407 | 0.134 | 2.60E-14 | 22 | Npdc1 |
| Arpc1b7 | 1.39E-18 | 1.084656695 | 0.852 | 0.592 | 2.81E-14 | 22 | Arpc1b |
| Dhrs1 | 1.65E-18 | 0.735175237 | 0.343 | 0.097 | 3.35E-14 | 22 | Dhrs1 |
| mt-Nd212 | 3.03E-18 | 0.90495146 | 0.954 | 0.689 | 6.13E-14 | 22 | mt-Nd2 |
| Emp25 | 3.23E-18 | 0.952961282 | 0.583 | 0.236 | 6.54E-14 | 22 | Emp2 |
| Rarb2 | 5.67E-18 | 0.705054801 | 0.25 | 0.057 | 1.15E-13 | 22 | Rarb |
| Mmp142 | 7.82E-18 | 0.503048674 | 0.259 | 0.06 | 1.58E-13 | 22 | Mmp14 |
| Sipa1l24 | 8.70E-18 | 0.807283772 | 0.454 | 0.159 | 1.76E-13 | 22 | Sipa1l2 |
| Josd21 | 9.05E-18 | 0.75022455 | 0.343 | 0.102 | 1.83E-13 | 22 | Josd2 |
| Pik3r15 | 9.20E-18 | 1.128388112 | 0.565 | 0.254 | 1.86E-13 | 22 | Pik3r1 |
| Emp14 | 1.05E-17 | 0.840410412 | 0.389 | 0.122 | 2.13E-13 | 22 | Emp1 |
| Nedd48 | 1.12E-17 | 1.004396212 | 0.778 | 0.42 | 2.27E-13 | 22 | Nedd4 |
| Itch10 | 2.72E-17 | 1.152811325 | 0.574 | 0.275 | 5.50E-13 | 22 | Itch |
| Nrp25 | 4.41E-17 | 1.001330381 | 0.491 | 0.185 | 8.93E-13 | 22 | Nrp2 |
| Sgms18 | 9.64E-17 | 0.868071266 | 0.389 | 0.129 | 1.95E-12 | 22 | Sgms1 |
| Lamp111 | 9.96E-17 | 0.900907613 | 0.741 | 0.419 | 2.02E-12 | 22 | Lamp1 |
| Cdkn2c1 | 1.11E-16 | 0.489158538 | 0.25 | 0.061 | 2.26E-12 | 22 | Cdkn2c |
| Tns34 | 2.01E-16 | 0.715947315 | 0.278 | 0.073 | 4.07E-12 | 22 | Tns3 |
| Gsap7 | 2.25E-16 | 0.794207122 | 0.296 | 0.086 | 4.55E-12 | 22 | Gsap |
| mt-Nd4l11 | 2.29E-16 | 0.965072944 | 0.806 | 0.492 | 4.65E-12 | 22 | mt-Nd4l |
| Itm2a4 | 2.42E-16 | 0.983230531 | 0.481 | 0.181 | 4.89E-12 | 22 | Itm2a |
| Anxa56 | 2.59E-16 | 0.922354748 | 0.778 | 0.495 | 5.24E-12 | 22 | Anxa5 |
| mt-Nd415 | 2.91E-16 | 0.778179673 | 0.963 | 0.676 | 5.89E-12 | 22 | mt-Nd4 |
| Fabp52 | 3.56E-16 | 0.818562615 | 0.676 | 0.326 | 7.21E-12 | 22 | Fabp5 |
| Tox7 | 6.36E-16 | 0.584581362 | 0.389 | 0.122 | 1.29E-11 | 22 | Tox |
| Clcn37 | 6.68E-16 | 0.911787866 | 0.38 | 0.134 | 1.35E-11 | 22 | Clcn3 |
| Rasa27 | 7.28E-16 | 0.783717753 | 0.389 | 0.139 | 1.47E-11 | 22 | Rasa2 |
| Ssbp29 | 1.66E-15 | 0.899752789 | 0.398 | 0.141 | 3.36E-11 | 22 | Ssbp2 |
| Abca15 | 1.68E-15 | 0.928630922 | 0.37 | 0.13 | 3.40E-11 | 22 | Abca1 |
| Tecr7 | 1.97E-15 | 0.936863576 | 0.5 | 0.226 | 3.99E-11 | 22 | Tecr |

| | | | | | | | |
|---|---|---|---|---|---|---|---|
| Thra7 | 3.60E-15 | 0.768808389 | 0.435 | 0.165 | 7.29E-11 | 22 | Thra |
| Cald18 | 3.60E-15 | 0.722775978 | 0.694 | 0.323 | 7.29E-11 | 22 | Cald1 |
| Secisbp2l2 | 3.67E-15 | 1.299389356 | 0.389 | 0.144 | 7.43E-11 | 22 | Secisbp2l |
| Kif1b6 | 4.07E-15 | 0.894608446 | 0.528 | 0.244 | 8.24E-11 | 22 | Kif1b |
| Xist14 | 4.52E-15 | 0.979248831 | 0.787 | 0.464 | 9.15E-11 | 22 | Xist |
| Inpp5f | 5.23E-15 | 0.631215872 | 0.278 | 0.079 | 1.06E-10 | 22 | Inpp5f |
| Bhlhe407 | 5.45E-15 | 0.848121774 | 0.417 | 0.159 | 1.10E-10 | 22 | Bhlhe40 |
| Ctnna16 | 6.11E-15 | 0.802376138 | 0.676 | 0.353 | 1.24E-10 | 22 | Ctnna1 |
| Pam10 | 6.35E-15 | 0.826011588 | 0.5 | 0.215 | 1.28E-10 | 22 | Pam |
| Cpne22 | 8.15E-15 | 0.761119889 | 0.25 | 0.069 | 1.65E-10 | 22 | Cpne2 |
| Ssfa23 | 1.13E-14 | 0.852153807 | 0.306 | 0.098 | 2.28E-10 | 22 | Ssfa2 |
| Ddah24 | 1.28E-14 | 0.682166839 | 0.5 | 0.207 | 2.60E-10 | 22 | Ddah2 |
| Mical31 | 1.30E-14 | 0.753131515 | 0.259 | 0.071 | 2.63E-10 | 22 | Mical3 |
| Slc31a21 | 1.58E-14 | 0.699782858 | 0.306 | 0.096 | 3.20E-10 | 22 | Slc31a2 |
| Ick2 | 2.36E-14 | 0.599206075 | 0.287 | 0.085 | 4.78E-10 | 22 | Ick |
| Son12 | 3.09E-14 | 0.85656937 | 0.796 | 0.527 | 6.25E-10 | 22 | Son |
| Slco3a16 | 4.23E-14 | 0.82483568 | 0.352 | 0.123 | 8.57E-10 | 22 | Slco3a1 |
| Atxn23 | 4.70E-14 | 0.727508144 | 0.38 | 0.14 | 9.51E-10 | 22 | Atxn2 |
| Pdia39 | 6.15E-14 | 0.818612344 | 0.731 | 0.445 | 1.24E-09 | 22 | Pdia3 |
| Usp244 | 6.16E-14 | 0.822595126 | 0.37 | 0.138 | 1.25E-09 | 22 | Usp24 |
| mt-Atp616 | 8.80E-14 | 0.647875355 | 0.991 | 0.754 | 1.78E-09 | 22 | mt-Atp6 |
| Capg7 | 9.46E-14 | 0.610803358 | 0.315 | 0.102 | 1.92E-09 | 22 | Capg |
| Mitf3 | 1.11E-13 | 0.871452748 | 0.259 | 0.076 | 2.25E-09 | 22 | Mitf |
| Tmed108 | 1.19E-13 | 0.891778225 | 0.731 | 0.437 | 2.42E-09 | 22 | Tmed10 |
| Mar-33 | 1.42E-13 | 0.842507256 | 0.296 | 0.094 | 2.87E-09 | 22 | 3-Mar |
| Il6st6 | 1.51E-13 | 0.824891325 | 0.519 | 0.233 | 3.06E-09 | 22 | Il6st |
| Mbnl29 | 1.84E-13 | 0.827781406 | 0.704 | 0.42 | 3.73E-09 | 22 | Mbnl2 |
| Sparcl17 | 2.04E-13 | 0.819191892 | 0.731 | 0.365 | 4.13E-09 | 22 | Sparcl1 |
| Tom1l25 | 2.24E-13 | 0.948279505 | 0.352 | 0.132 | 4.53E-09 | 22 | Tom1l2 |
| Tm9sf38 | 2.38E-13 | 0.835387179 | 0.611 | 0.336 | 4.82E-09 | 22 | Tm9sf3 |
| Mmp153 | 2.68E-13 | 0.841478367 | 0.306 | 0.102 | 5.43E-09 | 22 | Mmp15 |
| Abcc52 | 2.89E-13 | 0.564443372 | 0.269 | 0.081 | 5.84E-09 | 22 | Abcc5 |
| Arpc1a3 | 3.96E-13 | 0.741832973 | 0.611 | 0.318 | 8.02E-09 | 22 | Arpc1a |
| Cyp2d225 | 4.57E-13 | 0.703383904 | 0.37 | 0.136 | 9.26E-09 | 22 | Cyp2d22 |
| Cdk5rap2 | 6.04E-13 | 0.64236184 | 0.25 | 0.075 | 1.22E-08 | 22 | Cdk5rap2 |
| Serpinh17 | 7.59E-13 | 0.7694952 | 0.556 | 0.251 | 1.54E-08 | 22 | Serpinh1 |
| Cdc42ep51 | 7.79E-13 | 0.480507325 | 0.25 | 0.072 | 1.58E-08 | 22 | Cdc42ep5 |
| Pcdh15 | 1.18E-12 | 0.616214098 | 0.287 | 0.092 | 2.39E-08 | 22 | Pcdh1 |
| Serinc18 | 1.27E-12 | 1.010891564 | 0.509 | 0.25 | 2.57E-08 | 22 | Serinc1 |
| Kiz | 1.70E-12 | 0.650354183 | 0.25 | 0.076 | 3.43E-08 | 22 | Kiz |
| Cyth35 | 1.94E-12 | 0.695786859 | 0.343 | 0.126 | 3.92E-08 | 22 | Cyth3 |
| Atp13a36 | 5.06E-12 | 0.738562222 | 0.296 | 0.104 | 1.02E-07 | 22 | Atp13a3 |
| Lmna8 | 5.65E-12 | 0.72075476 | 0.444 | 0.199 | 1.14E-07 | 22 | Lmna |

| | | | | | | | |
|---|---|---|---|---|---|---|---|
| Bsg8 | 5.84E-12 | 0.894209443 | 0.694 | 0.451 | 1.18E-07 | 22 | Bsg |
| Hipk23 | 5.95E-12 | 0.595628721 | 0.324 | 0.119 | 1.20E-07 | 22 | Hipk2 |
| Fkbp72 | 6.21E-12 | 0.639528724 | 0.259 | 0.082 | 1.26E-07 | 22 | Fkbp7 |
| Tsc22d47 | 6.47E-12 | 0.827369199 | 0.546 | 0.296 | 1.31E-07 | 22 | Tsc22d4 |
| Hspg24 | 7.10E-12 | 0.716537932 | 0.657 | 0.313 | 1.44E-07 | 22 | Hspg2 |
| Bach28 | 7.28E-12 | 0.921085782 | 0.389 | 0.162 | 1.47E-07 | 22 | Bach2 |
| Snrnp706 | 8.04E-12 | 0.772660555 | 0.519 | 0.274 | 1.63E-07 | 22 | Snrnp70 |
| Nr4a25 | 1.02E-11 | 0.728285481 | 0.343 | 0.132 | 2.06E-07 | 22 | Nr4a2 |
| Selenof7 | 1.55E-11 | 0.772295137 | 0.741 | 0.493 | 3.14E-07 | 22 | Selenof |
| Epb41l4a2 | 1.56E-11 | 0.68009457 | 0.259 | 0.083 | 3.15E-07 | 22 | Epb41l4a |
| Fam213b1 | 1.56E-11 | 0.422882575 | 0.25 | 0.076 | 3.15E-07 | 22 | Fam213b |
| Snx61 | 1.78E-11 | 0.587396428 | 0.546 | 0.276 | 3.61E-07 | 22 | Snx6 |
| Gramd1b6 | 1.82E-11 | 0.621660169 | 0.269 | 0.091 | 3.69E-07 | 22 | Gramd1b |
| Fkbp94 | 2.23E-11 | 0.717141471 | 0.306 | 0.112 | 4.51E-07 | 22 | Fkbp9 |
| Eid16 | 2.30E-11 | 0.77063593 | 0.593 | 0.322 | 4.65E-07 | 22 | Eid1 |
| Parva5 | 2.46E-11 | 0.596428653 | 0.315 | 0.113 | 4.98E-07 | 22 | Parva |
| Hspa511 | 2.68E-11 | 0.829377911 | 0.713 | 0.436 | 5.42E-07 | 22 | Hspa5 |
| Fmnl26 | 3.09E-11 | 0.822091711 | 0.5 | 0.244 | 6.25E-07 | 22 | Fmnl2 |
| Zbtb383 | 3.27E-11 | 0.826052446 | 0.361 | 0.155 | 6.63E-07 | 22 | Zbtb38 |
| Rrbp17 | 3.28E-11 | 0.693369171 | 0.75 | 0.461 | 6.63E-07 | 22 | Rrbp1 |
| 9530068E07Rik1 | 3.57E-11 | 0.617236672 | 0.278 | 0.098 | 7.23E-07 | 22 | 9530068E07Rik |
| Hfe | 3.84E-11 | 0.636581702 | 0.259 | 0.086 | 7.77E-07 | 22 | Hfe |
| mt-Co317 | 3.84E-11 | 0.579959856 | 0.981 | 0.727 | 7.78E-07 | 22 | mt-Co3 |
| Ppp2r3a4 | 4.15E-11 | 0.767420607 | 0.269 | 0.092 | 8.39E-07 | 22 | Ppp2r3a |
| Ckap54 | 4.89E-11 | 0.601253943 | 0.352 | 0.143 | 9.90E-07 | 22 | Ckap5 |
| Ahnak9 | 8.18E-11 | 0.686582386 | 0.796 | 0.473 | 1.66E-06 | 22 | Ahnak |
| Lgmn5 | 8.32E-11 | 0.348736562 | 0.324 | 0.115 | 1.68E-06 | 22 | Lgmn |
| 4930402H24Rik5 | 9.06E-11 | 0.8210262 | 0.463 | 0.223 | 1.83E-06 | 22 | 4930402H24Rik |
| Wwp1 | 1.00E-10 | 0.915205312 | 0.407 | 0.185 | 2.03E-06 | 22 | Wwp1 |
| Flna7 | 1.19E-10 | 0.85294197 | 0.324 | 0.133 | 2.40E-06 | 22 | Flna |
| Mesd5 | 1.40E-10 | 0.757537601 | 0.37 | 0.164 | 2.84E-06 | 22 | Mesd |
| Ifit32 | 1.42E-10 | 1.511949634 | 0.315 | 0.124 | 2.88E-06 | 22 | Ifit3 |
| Cdkn1a2 | 1.53E-10 | 0.962114768 | 0.324 | 0.128 | 3.11E-06 | 22 | Cdkn1a |
| Prickle25 | 1.54E-10 | 0.882062037 | 0.343 | 0.14 | 3.13E-06 | 22 | Prickle2 |
| Mvb12b5 | 1.63E-10 | 0.738919964 | 0.38 | 0.168 | 3.30E-06 | 22 | Mvb12b |
| Leng81 | 1.65E-10 | 0.661056975 | 0.259 | 0.091 | 3.33E-06 | 22 | Leng8 |
| Ryk4 | 1.72E-10 | 0.632564545 | 0.296 | 0.111 | 3.47E-06 | 22 | Ryk |
| Prkce5 | 1.89E-10 | 0.740364125 | 0.454 | 0.215 | 3.83E-06 | 22 | Prkce |
| Creb3l24 | 2.04E-10 | 0.708140387 | 0.315 | 0.124 | 4.13E-06 | 22 | Creb3l2 |
| Cdk192 | 2.25E-10 | 0.639352522 | 0.352 | 0.148 | 4.55E-06 | 22 | Cdk19 |
| Srgap29 | 3.34E-10 | 0.781445193 | 0.546 | 0.305 | 6.75E-06 | 22 | Srgap2 |
| Rock26 | 3.45E-10 | 0.881370016 | 0.565 | 0.339 | 6.98E-06 | 22 | Rock2 |
| Mocs22 | 5.43E-10 | 0.522331705 | 0.444 | 0.209 | 1.10E-05 | 22 | Mocs2 |

| | | | | | | | |
|---|---|---|---|---|---|---|---|
| Sirt2 | 5.50E-10 | 0.801239389 | 0.361 | 0.163 | 1.11E-05 | 22 | Sirt2 |
| Itgb111 | 5.56E-10 | 0.662650873 | 0.75 | 0.48 | 1.13E-05 | 22 | Itgb1 |
| Tmem53 | 6.90E-10 | 0.703441749 | 0.333 | 0.139 | 1.40E-05 | 22 | Tmem5 |
| Itfg11 | 7.23E-10 | 1.101997039 | 0.324 | 0.141 | 1.46E-05 | 22 | Itfg1 |
| Pros16 | 7.63E-10 | 0.784483579 | 0.352 | 0.151 | 1.54E-05 | 22 | Pros1 |
| Atraid6 | 8.25E-10 | 0.705899932 | 0.509 | 0.272 | 1.67E-05 | 22 | Atraid |
| Chd96 | 8.27E-10 | 0.658824824 | 0.481 | 0.251 | 1.67E-05 | 22 | Chd9 |
| Emc78 | 8.65E-10 | 0.740057536 | 0.5 | 0.273 | 1.75E-05 | 22 | Emc7 |
| Asah14 | 8.80E-10 | 0.678758919 | 0.398 | 0.189 | 1.78E-05 | 22 | Asah1 |
| Ccdc506 | 9.17E-10 | 0.614508884 | 0.454 | 0.23 | 1.86E-05 | 22 | Ccdc50 |
| S100a167 | 1.05E-09 | 0.646436762 | 0.509 | 0.259 | 2.12E-05 | 22 | S100a16 |
| Ccser25 | 1.06E-09 | 0.934790683 | 0.444 | 0.232 | 2.14E-05 | 22 | Ccser2 |
| Srpk25 | 1.06E-09 | 0.644619252 | 0.407 | 0.195 | 2.15E-05 | 22 | Srpk2 |
| Ptov13 | 1.14E-09 | 0.654861585 | 0.306 | 0.12 | 2.32E-05 | 22 | Ptov1 |
| Arl33 | 1.36E-09 | 0.447057652 | 0.389 | 0.171 | 2.76E-05 | 22 | Arl3 |
| Nfic4 | 1.37E-09 | 0.564295563 | 0.5 | 0.262 | 2.77E-05 | 22 | Nfic |
| Pten2 | 1.52E-09 | 0.684648639 | 0.546 | 0.323 | 3.08E-05 | 22 | Pten |
| Lrch310 | 1.61E-09 | 0.712021103 | 0.444 | 0.234 | 3.26E-05 | 22 | Lrch3 |
| Tgfbr27 | 3.00E-09 | 0.681056274 | 0.546 | 0.313 | 6.06E-05 | 22 | Tgfbr2 |
| mt-Atp812 | 3.45E-09 | 0.646029187 | 0.704 | 0.465 | 6.98E-05 | 22 | mt-Atp8 |
| Rhoc6 | 3.52E-09 | 0.743405227 | 0.519 | 0.298 | 7.14E-05 | 22 | Rhoc |
| Cab39l | 3.95E-09 | 0.571033943 | 0.407 | 0.199 | 8.00E-05 | 22 | Cab39l |
| Atad12 | 4.04E-09 | 0.56499567 | 0.269 | 0.104 | 8.18E-05 | 22 | Atad1 |
| Rab11fip23 | 4.67E-09 | 0.503932001 | 0.278 | 0.108 | 9.46E-05 | 22 | Rab11fip2 |
| Ndfip16 | 5.57E-09 | 0.718640712 | 0.565 | 0.323 | 0.00011283 | 22 | Ndfip1 |
| Map46 | 5.71E-09 | 0.81014189 | 0.519 | 0.305 | 0.00011553 | 22 | Map4 |
| Plin22 | 5.71E-09 | 0.65818096 | 0.259 | 0.1 | 0.000115551 | 22 | Plin2 |
| Itm2c7 | 5.72E-09 | 0.748736318 | 0.398 | 0.201 | 0.000115816 | 22 | Itm2c |
| Wls5 | 6.05E-09 | 0.625614885 | 0.306 | 0.128 | 0.000122478 | 22 | Wls |
| Trim355 | 6.38E-09 | 0.561576617 | 0.324 | 0.141 | 0.00012916 | 22 | Trim35 |
| Atrn2 | 6.73E-09 | 0.699021198 | 0.296 | 0.124 | 0.000136343 | 22 | Atrn |
| Tcaf15 | 7.18E-09 | 0.606001658 | 0.269 | 0.104 | 0.000145378 | 22 | Tcaf1 |
| Dip2c7 | 7.73E-09 | 0.657458041 | 0.444 | 0.224 | 0.000156571 | 22 | Dip2c |
| Canx9 | 7.82E-09 | 0.695224483 | 0.528 | 0.308 | 0.000158367 | 22 | Canx |
| Taok17 | 8.71E-09 | 0.62031446 | 0.472 | 0.253 | 0.000176248 | 22 | Taok1 |
| Psap6 | 8.71E-09 | 0.470214908 | 0.657 | 0.4 | 0.000176249 | 22 | Psap |
| Gtdc15 | 9.46E-09 | 0.857947345 | 0.333 | 0.153 | 0.000191484 | 22 | Gtdc1 |
| Ppp2r2a6 | 1.11E-08 | 0.690707432 | 0.417 | 0.214 | 0.000224649 | 22 | Ppp2r2a |
| Oat4 | 1.19E-08 | 0.488230044 | 0.324 | 0.138 | 0.000240558 | 22 | Oat |
| Lysmd24 | 1.30E-08 | 0.531979167 | 0.398 | 0.193 | 0.000262404 | 22 | Lysmd2 |
| Auts23 | 1.30E-08 | 0.41973316 | 0.361 | 0.154 | 0.000263917 | 22 | Auts2 |
| Sash17 | 1.40E-08 | 0.8168772 | 0.463 | 0.259 | 0.00028292 | 22 | Sash1 |
| Ddx60 | 1.44E-08 | 0.566724328 | 0.25 | 0.096 | 0.000290681 | 22 | Ddx60 |

| | | | | | | | |
|---|---|---|---|---|---|---|---|
| Anks14 | 1.67E-08 | 0.620482883 | 0.315 | 0.14 | 0.000337723 | 22 | Anks1 |
| Neat111 | 1.89E-08 | 1.142577824 | 0.389 | 0.21 | 0.000382389 | 22 | Neat1 |
| Nudt3 | 1.98E-08 | 0.500229132 | 0.296 | 0.123 | 0.000401488 | 22 | Nudt3 |
| Uaca5 | 2.94E-08 | 0.489107852 | 0.463 | 0.227 | 0.000594344 | 22 | Uaca |
| Adprh2 | 3.04E-08 | 0.568433069 | 0.315 | 0.141 | 0.000615236 | 22 | Adprh |
| Lyst6 | 3.60E-08 | 0.485620505 | 0.269 | 0.111 | 0.000729726 | 22 | Lyst |
| Pja25 | 3.84E-08 | 0.65037408 | 0.398 | 0.206 | 0.000777107 | 22 | Pja2 |
| Plec7 | 4.01E-08 | 0.889728974 | 0.352 | 0.176 | 0.000812397 | 22 | Plec |
| Glg17 | 4.46E-08 | 0.722819935 | 0.454 | 0.254 | 0.000901982 | 22 | Glg1 |
| Evi55 | 4.55E-08 | 0.748387183 | 0.315 | 0.142 | 0.000922166 | 22 | Evi5 |
| Fermt27 | 5.43E-08 | 0.465519453 | 0.63 | 0.351 | 0.001100063 | 22 | Fermt2 |
| Setbp18 | 5.70E-08 | 0.678450188 | 0.287 | 0.123 | 0.001153164 | 22 | Setbp1 |
| Thap2 | 6.35E-08 | 0.410166722 | 0.25 | 0.098 | 0.001286321 | 22 | Thap2 |
| Wdfy23 | 6.95E-08 | 0.585831537 | 0.269 | 0.112 | 0.001407122 | 22 | Wdfy2 |
| Spidr1 | 7.28E-08 | 0.508625358 | 0.269 | 0.113 | 0.001473604 | 22 | Spidr |
| Ankrd282 | 7.47E-08 | 0.630736382 | 0.296 | 0.134 | 0.001512373 | 22 | Ankrd28 |
| Mxra76 | 7.87E-08 | 0.5571124 | 0.315 | 0.143 | 0.001593577 | 22 | Mxra7 |
| Rabac16 | 8.27E-08 | 0.51058176 | 0.769 | 0.566 | 0.001673491 | 22 | Rabac1 |
| Dpysl28 | 8.80E-08 | 0.65067191 | 0.352 | 0.176 | 0.001782178 | 22 | Dpysl2 |
| Txndc121 | 8.82E-08 | 0.686420177 | 0.269 | 0.116 | 0.001785225 | 22 | Txndc12 |
| Fam102a5 | 9.58E-08 | 0.50609575 | 0.25 | 0.101 | 0.001939484 | 22 | Fam102a |
| Ddx176 | 9.80E-08 | 0.531594031 | 0.481 | 0.275 | 0.001983715 | 22 | Ddx17 |
| Fryl10 | 1.20E-07 | 0.628756924 | 0.472 | 0.27 | 0.002431525 | 22 | Fryl |
| Ptbp23 | 1.24E-07 | 0.553432308 | 0.287 | 0.13 | 0.002515819 | 22 | Ptbp2 |
| Pnisr11 | 1.33E-07 | 0.824018522 | 0.444 | 0.266 | 0.002700856 | 22 | Pnisr |
| Fkbp23 | 1.36E-07 | 0.651479607 | 0.481 | 0.278 | 0.002749057 | 22 | Fkbp2 |
| Add16 | 1.37E-07 | 0.535309354 | 0.407 | 0.216 | 0.002776242 | 22 | Add1 |
| Vapa2 | 1.41E-07 | 0.448965229 | 0.602 | 0.38 | 0.002864377 | 22 | Vapa |
| Ttc35 | 1.62E-07 | 0.550791529 | 0.454 | 0.253 | 0.003287328 | 22 | Ttc3 |
| Laptm4a8 | 1.84E-07 | 0.503045714 | 0.713 | 0.486 | 0.003730492 | 22 | Laptm4a |
| Eri31 | 2.13E-07 | 0.437798319 | 0.25 | 0.104 | 0.004311794 | 22 | Eri3 |
| Pcm18 | 2.18E-07 | 0.62694632 | 0.444 | 0.253 | 0.004411487 | 22 | Pcm1 |
| Gprasp11 | 2.38E-07 | 0.499069429 | 0.25 | 0.105 | 0.004810915 | 22 | Gprasp1 |
| Rsu18 | 2.41E-07 | 0.544084443 | 0.454 | 0.255 | 0.004877258 | 22 | Rsu1 |
| Ppp1r12b3 | 2.78E-07 | 0.662411312 | 0.25 | 0.107 | 0.00562496 | 22 | Ppp1r12b |
| mt-Nd116 | 3.13E-07 | 0.51399075 | 0.843 | 0.6 | 0.006344602 | 22 | mt-Nd1 |
| Atp6v0b5 | 3.42E-07 | 0.496419454 | 0.574 | 0.368 | 0.006933478 | 22 | Atp6v0b |
| Atp8a16 | 3.50E-07 | 0.675357856 | 0.426 | 0.243 | 0.007081954 | 22 | Atp8a1 |
| Map4k411 | 4.65E-07 | 0.635675837 | 0.472 | 0.278 | 0.009405422 | 22 | Map4k4 |
| Itga66 | 4.73E-07 | 0.419894245 | 0.519 | 0.285 | 0.009580091 | 22 | Itga6 |
| Pon23 | 4.94E-07 | 0.5565347 | 0.343 | 0.173 | 0.009997866 | 22 | Pon2 |
| Rpn28 | 4.97E-07 | 0.803025999 | 0.389 | 0.213 | 0.010055358 | 22 | Rpn2 |
| Clip14 | 5.34E-07 | 0.510426656 | 0.306 | 0.144 | 0.010803935 | 22 | Clip1 |

| | | | | | | | |
|---|---|---|---|---|---|---|---|
| Akap911 | 5.48E-07 | 0.530260084 | 0.426 | 0.239 | 0.01109285 | 22 | Akap9 |
| Cdk146 | 5.84E-07 | 0.624604246 | 0.306 | 0.143 | 0.011822485 | 22 | Cdk14 |
| Trim23 | 6.43E-07 | 0.459269822 | 0.361 | 0.18 | 0.013016648 | 22 | Trim2 |
| Cxxc53 | 6.72E-07 | 0.610501508 | 0.259 | 0.116 | 0.013605777 | 22 | Cxxc5 |
| App6 | 7.86E-07 | 0.370277182 | 0.731 | 0.421 | 0.015920278 | 22 | App |
| Itih53 | 8.09E-07 | 0.583637426 | 0.444 | 0.245 | 0.016371109 | 22 | Itih5 |
| Calr8 | 8.46E-07 | 0.542357597 | 0.602 | 0.41 | 0.017118977 | 22 | Calr |
| Dhrs75 | 8.55E-07 | 0.77479335 | 0.389 | 0.217 | 0.017315753 | 22 | Dhrs7 |
| Ifi272 | 8.67E-07 | 0.33556727 | 0.769 | 0.537 | 0.017543479 | 22 | Ifi27 |
| Mllt36 | 9.02E-07 | 0.495672605 | 0.315 | 0.153 | 0.018268467 | 22 | Mllt3 |
| Gng28 | 9.19E-07 | 0.61110458 | 0.361 | 0.188 | 0.01859789 | 22 | Gng2 |
| Iars2 | 9.47E-07 | 0.464000738 | 0.269 | 0.121 | 0.019173892 | 22 | Iars2 |
| Chd610 | 9.85E-07 | 0.706730175 | 0.389 | 0.219 | 0.019942513 | 22 | Chd6 |
| Mtdh9 | 1.01E-06 | 0.50103846 | 0.602 | 0.388 | 0.020463635 | 22 | Mtdh |
| H2-D111 | 1.05E-06 | 0.527044723 | 0.898 | 0.67 | 0.021183981 | 22 | H2-D1 |
| Arhgap54 | 1.08E-06 | 0.444656694 | 0.491 | 0.29 | 0.021795704 | 22 | Arhgap5 |
| Bcl2l16 | 1.09E-06 | 0.499851235 | 0.306 | 0.148 | 0.022168777 | 22 | Bcl2l1 |
| Dync1i24 | 1.12E-06 | 0.371716806 | 0.639 | 0.42 | 0.022614889 | 22 | Dync1i2 |
| Zfand62 | 1.13E-06 | 0.557696479 | 0.481 | 0.301 | 0.022872314 | 22 | Zfand6 |
| Cuedc14 | 1.42E-06 | 0.596645222 | 0.287 | 0.137 | 0.02874162 | 22 | Cuedc1 |
| Psmd3 | 1.43E-06 | 0.505668458 | 0.315 | 0.155 | 0.028852995 | 22 | Psmd3 |
| Rnf135 | 1.43E-06 | 0.514820053 | 0.343 | 0.18 | 0.028933968 | 22 | Rnf13 |
| Gatad2b6 | 1.43E-06 | 0.556530786 | 0.37 | 0.204 | 0.02894982 | 22 | Gatad2b |
| Top2b5 | 1.53E-06 | 0.428220521 | 0.481 | 0.285 | 0.031036163 | 22 | Top2b |
| Cfl22 | 1.56E-06 | 0.43388845 | 0.417 | 0.231 | 0.031636093 | 22 | Cfl2 |
| Acsl42 | 1.56E-06 | 0.342347035 | 0.287 | 0.134 | 0.031651487 | 22 | Acsl4 |
| Cpq5 | 1.78E-06 | 0.486817747 | 0.269 | 0.122 | 0.035951997 | 22 | Cpq |
| Cavin37 | 1.79E-06 | 0.4701286 | 0.556 | 0.309 | 0.036268949 | 22 | Cavin3 |
| Psip16 | 1.85E-06 | 0.484574426 | 0.37 | 0.196 | 0.037448629 | 22 | Psip1 |
| Senp79 | 1.87E-06 | 0.463630004 | 0.315 | 0.158 | 0.037881831 | 22 | Senp7 |
| Zfp6086 | 2.29E-06 | 0.624152673 | 0.287 | 0.142 | 0.04629531 | 22 | Zfp608 |
| Mkln15 | 2.30E-06 | 0.544542623 | 0.444 | 0.266 | 0.046608296 | 22 | Mkln1 |
| Rbm268 | 2.36E-06 | 0.599161344 | 0.389 | 0.228 | 0.047819883 | 22 | Rbm26 |
| Arl6ip19 | 2.50E-06 | 0.589621793 | 0.546 | 0.372 | 0.050671181 | 22 | Arl6ip1 |
| Ggh1 | 2.59E-06 | 0.63860649 | 0.324 | 0.168 | 0.052356874 | 22 | Ggh |
| Taf12 | 2.66E-06 | 0.597690821 | 0.287 | 0.144 | 0.053829835 | 22 | Taf1 |
| Txndc154 | 3.03E-06 | 0.460528177 | 0.287 | 0.14 | 0.061348827 | 22 | Txndc15 |
| Arntl3 | 3.14E-06 | 0.502818314 | 0.259 | 0.12 | 0.063639544 | 22 | Arntl |
| Stxbp31 | 3.62E-06 | 0.450065612 | 0.315 | 0.163 | 0.073215291 | 22 | Stxbp3 |
| Nfix2 | 3.66E-06 | 0.388045413 | 0.398 | 0.217 | 0.074045912 | 22 | Nfix |
| Myo18a4 | 3.72E-06 | 0.679239313 | 0.435 | 0.259 | 0.075318432 | 22 | Myo18a |
| Cdk68 | 3.92E-06 | 0.460973606 | 0.25 | 0.113 | 0.079403203 | 22 | Cdk6 |
| Galnt113 | 4.34E-06 | 0.47054114 | 0.343 | 0.186 | 0.087945866 | 22 | Galnt1 |

| | | | | | | | |
|---|---|---|---|---|---|---|---|
| Lamp28 | 4.77E-06 | 0.424363241 | 0.509 | 0.307 | 0.096483225 | 22 | Lamp2 |
| Zcchc713 | 4.80E-06 | 0.523235824 | 0.491 | 0.303 | 0.097244549 | 22 | Zcchc7 |
| Srek16 | 4.84E-06 | 0.455948682 | 0.352 | 0.195 | 0.097918061 | 22 | Srek1 |
| Ostc3 | 4.99E-06 | 0.622858031 | 0.435 | 0.264 | 0.100967893 | 22 | Ostc |
| Vkorc13 | 5.28E-06 | 0.609351779 | 0.296 | 0.154 | 0.106853501 | 22 | Vkorc1 |
| Appl15 | 5.31E-06 | 0.689854792 | 0.333 | 0.183 | 0.107445699 | 22 | Appl1 |
| Cd1517 | 5.33E-06 | 0.606146312 | 0.389 | 0.219 | 0.107903956 | 22 | Cd151 |
| Zfhx35 | 5.88E-06 | 0.363090703 | 0.352 | 0.184 | 0.119142421 | 22 | Zfhx3 |
| Ccdc1078 | 5.93E-06 | 0.794824244 | 0.259 | 0.129 | 0.12013755 | 22 | Ccdc107 |
| Kansl1l4 | 6.02E-06 | 0.634332427 | 0.278 | 0.14 | 0.121801119 | 22 | Kansl1l |
| Sec627 | 6.69E-06 | 0.540100227 | 0.574 | 0.396 | 0.135494819 | 22 | Sec62 |
| Clic46 | 6.91E-06 | 0.274943128 | 0.685 | 0.425 | 0.139968694 | 22 | Clic4 |
| Myo9b6 | 7.64E-06 | 0.457420986 | 0.25 | 0.121 | 0.154755266 | 22 | Myo9b |
| Fbxl34 | 7.68E-06 | 0.421406085 | 0.306 | 0.158 | 0.15547167 | 22 | Fbxl3 |
| Otud7b2 | 7.88E-06 | 0.504891122 | 0.259 | 0.126 | 0.159490698 | 22 | Otud7b |
| Selenok6 | 8.10E-06 | 0.43161116 | 0.75 | 0.556 | 0.163952855 | 22 | Selenok |
| Azin12 | 8.13E-06 | 0.275945025 | 0.306 | 0.155 | 0.164656406 | 22 | Azin1 |
| Emc36 | 8.22E-06 | 0.446652415 | 0.324 | 0.174 | 0.166429368 | 22 | Emc3 |
| Map4k36 | 8.84E-06 | 0.530854162 | 0.398 | 0.231 | 0.17904006 | 22 | Map4k3 |
| Zmynd83 | 8.88E-06 | 0.593582102 | 0.352 | 0.196 | 0.179809806 | 22 | Zmynd8 |
| M6pr4 | 9.07E-06 | 0.587657611 | 0.306 | 0.164 | 0.183650719 | 22 | M6pr |
| 1700025G04Rik7 | 1.03E-05 | 0.442555597 | 0.343 | 0.19 | 0.209432353 | 22 | 1700025G04Rik |
| mt-Co218 | 1.12E-05 | 0.408406499 | 0.981 | 0.729 | 0.226075329 | 22 | mt-Co2 |
| Pgrmc15 | 1.15E-05 | 0.408464149 | 0.389 | 0.219 | 0.233356449 | 22 | Pgrmc1 |
| Ppp3ca13 | 1.18E-05 | 0.468396688 | 0.583 | 0.395 | 0.239524074 | 22 | Ppp3ca |
| Crip27 | 1.19E-05 | 0.294369248 | 0.685 | 0.418 | 0.240515468 | 22 | Crip2 |
| Anxa67 | 1.21E-05 | 0.507503022 | 0.296 | 0.158 | 0.244765171 | 22 | Anxa6 |
| Ash1l9 | 1.22E-05 | 0.518846137 | 0.481 | 0.301 | 0.246423734 | 22 | Ash1l |
| Gpatch812 | 1.35E-05 | 0.734748483 | 0.417 | 0.26 | 0.27345684 | 22 | Gpatch8 |
| Parp146 | 1.40E-05 | 0.598737386 | 0.407 | 0.254 | 0.283436188 | 22 | Parp14 |
| Dmxl18 | 1.43E-05 | 0.526700384 | 0.333 | 0.187 | 0.290409001 | 22 | Dmxl1 |
| Adam106 | 1.54E-05 | 0.534325165 | 0.463 | 0.294 | 0.312676363 | 22 | Adam10 |
| Ociad11 | 1.57E-05 | 0.481157635 | 0.333 | 0.187 | 0.317309919 | 22 | Ociad1 |
| mt-Co115 | 1.69E-05 | 0.446792048 | 0.981 | 0.72 | 0.342805989 | 22 | mt-Co1 |
| Ift432 | 1.78E-05 | 0.38377076 | 0.259 | 0.125 | 0.359676802 | 22 | Ift43 |
| Itga17 | 1.80E-05 | 0.372380039 | 0.676 | 0.416 | 0.364598499 | 22 | Itga1 |
| Gnaq7 | 1.99E-05 | 0.54931151 | 0.537 | 0.361 | 0.403217601 | 22 | Gnaq |
| Clstn17 | 2.05E-05 | 0.44301054 | 0.407 | 0.238 | 0.415650412 | 22 | Clstn1 |
| mt-Nd511 | 2.08E-05 | 0.44086173 | 0.685 | 0.498 | 0.421714302 | 22 | mt-Nd5 |
| Hmgn15 | 2.23E-05 | 0.323143559 | 0.509 | 0.33 | 0.451448948 | 22 | Hmgn1 |
| Xaf14 | 2.26E-05 | 0.59434665 | 0.38 | 0.222 | 0.45809643 | 22 | Xaf1 |
| Ube3c5 | 2.37E-05 | 0.573995992 | 0.296 | 0.16 | 0.479168487 | 22 | Ube3c |
| Hnrnph33 | 2.68E-05 | 0.364800997 | 0.417 | 0.251 | 0.54308241 | 22 | Hnrnph3 |

| | | | | | | | |
|---|---|---|---|---|---|---|---|
| Cul13 | 2.74E-05 | 0.471534244 | 0.435 | 0.279 | 0.555396482 | 22 | Cul1 |
| Pkn27 | 2.75E-05 | 0.395602475 | 0.472 | 0.296 | 0.557490509 | 22 | Pkn2 |
| Gns5 | 2.91E-05 | 0.369949366 | 0.259 | 0.128 | 0.590112518 | 22 | Gns |
| Syne26 | 3.34E-05 | 0.499471948 | 0.454 | 0.274 | 0.675605854 | 22 | Syne2 |
| Fam212a3 | 3.34E-05 | 0.444239618 | 0.259 | 0.129 | 0.676767814 | 22 | Fam212a |
| Zeb111 | 3.53E-05 | 0.562924937 | 0.481 | 0.324 | 0.714174725 | 22 | Zeb1 |
| Psmc1 | 3.65E-05 | 0.41493316 | 0.361 | 0.208 | 0.738983232 | 22 | Psmc1 |
| Tmem50a9 | 3.80E-05 | 0.53648525 | 0.528 | 0.385 | 0.76907487 | 22 | Tmem50a |
| Srrm211 | 3.85E-05 | 0.402056521 | 0.63 | 0.471 | 0.779106222 | 22 | Srrm2 |
| Lrrc587 | 3.98E-05 | 0.548318279 | 0.361 | 0.216 | 0.80663527 | 22 | Lrrc58 |
| Slc43a35 | 4.07E-05 | 0.291670212 | 0.463 | 0.271 | 0.823399638 | 22 | Slc43a3 |
| Cdkn1b4 | 4.18E-05 | 0.76736538 | 0.361 | 0.217 | 0.846325684 | 22 | Cdkn1b |
| Pafah1b13 | 4.22E-05 | 0.371436225 | 0.639 | 0.483 | 0.854651781 | 22 | Pafah1b1 |
| Selenos3 | 4.69E-05 | 0.537010075 | 0.38 | 0.233 | 0.948807173 | 22 | Selenos |
| Mrpl331 | 4.99E-05 | 0.310762315 | 0.583 | 0.407 | 1 | 22 | Mrpl33 |
| Hes14 | 5.05E-05 | 0.408985 | 0.509 | 0.316 | 1 | 22 | Hes1 |
| Spag98 | 5.35E-05 | 0.482336899 | 0.528 | 0.361 | 1 | 22 | Spag9 |
| Hsd17b122 | 5.50E-05 | 0.403044958 | 0.259 | 0.136 | 1 | 22 | Hsd17b12 |
| Dlgap41 | 5.67E-05 | 0.418083899 | 0.259 | 0.137 | 1 | 22 | Dlgap4 |
| Foxo15 | 5.83E-05 | 0.517775214 | 0.343 | 0.201 | 1 | 22 | Foxo1 |
| Tmco17 | 5.84E-05 | 0.406536657 | 0.472 | 0.314 | 1 | 22 | Tmco1 |
| Ubc3 | 6.37E-05 | 0.354523574 | 0.815 | 0.665 | 1 | 22 | Ubc |
| Sema3c2 | 6.45E-05 | 0.326293394 | 0.306 | 0.165 | 1 | 22 | Sema3c |
| Osgep2 | 6.95E-05 | 0.336330065 | 0.306 | 0.172 | 1 | 22 | Osgep |
| mt-Nd313 | 7.03E-05 | 0.445127083 | 0.63 | 0.448 | 1 | 22 | mt-Nd3 |
| Pxn2 | 7.30E-05 | 0.461192011 | 0.296 | 0.166 | 1 | 22 | Pxn |
| Lmbrd17 | 7.31E-05 | 0.478643406 | 0.287 | 0.164 | 1 | 22 | Lmbrd1 |
| Degs15 | 7.52E-05 | 0.536836274 | 0.315 | 0.186 | 1 | 22 | Degs1 |
| Notch16 | 7.57E-05 | 0.448748265 | 0.37 | 0.219 | 1 | 22 | Notch1 |
| Mlf22 | 8.13E-05 | 0.370295522 | 0.343 | 0.201 | 1 | 22 | Mlf2 |
| Grb106 | 8.22E-05 | 0.4119396 | 0.352 | 0.196 | 1 | 22 | Grb10 |
| P4hb6 | 8.29E-05 | 0.479585225 | 0.398 | 0.254 | 1 | 22 | P4hb |
| Pkig4 | 8.59E-05 | 0.327269984 | 0.37 | 0.212 | 1 | 22 | Pkig |
| Ctdsp25 | 8.71E-05 | 0.350778168 | 0.315 | 0.18 | 1 | 22 | Ctdsp2 |
| Pik3ca4 | 8.82E-05 | 0.35870085 | 0.306 | 0.17 | 1 | 22 | Pik3ca |
| Tnrc6a3 | 9.00E-05 | 0.305502599 | 0.343 | 0.201 | 1 | 22 | Tnrc6a |
| Hmgn34 | 9.22E-05 | 0.308800085 | 0.315 | 0.172 | 1 | 22 | Hmgn3 |
| Rsrp12 | 9.59E-05 | 0.324415342 | 0.741 | 0.572 | 1 | 22 | Rsrp1 |
| Rtn49 | 9.79E-05 | 0.396541861 | 0.491 | 0.341 | 1 | 22 | Rtn4 |
| Vma21 | 9.85E-05 | 0.418796497 | 0.269 | 0.146 | 1 | 22 | Vma21 |
| Stard3nl2 | 0.000101241 | 0.379478062 | 0.278 | 0.152 | 1 | 22 | Stard3nl |
| Atp2c15 | 0.000105725 | 0.566313707 | 0.287 | 0.169 | 1 | 22 | Atp2c1 |
| Iah12 | 0.000115923 | 0.398728273 | 0.259 | 0.138 | 1 | 22 | Iah1 |

| | | | | | | | |
|---|---|---|---|---|---|---|---|
| Klf97 | 0.000119101 | 0.288919449 | 0.398 | 0.239 | 1 | 22 | Klf9 |
| Numb5 | 0.000123312 | 0.304707905 | 0.519 | 0.339 | 1 | 22 | Numb |
| Pura5 | 0.000123808 | 0.423142854 | 0.519 | 0.368 | 1 | 22 | Pura |
| Tmem476 | 0.000127506 | 0.459371059 | 0.287 | 0.158 | 1 | 22 | Tmem47 |
| Smarcc24 | 0.00012774 | 0.381917085 | 0.315 | 0.179 | 1 | 22 | Smarcc2 |
| Ddx39b7 | 0.000128445 | 0.376952887 | 0.37 | 0.235 | 1 | 22 | Ddx39b |
| Luc7l35 | 0.000139222 | 0.347962628 | 0.426 | 0.273 | 1 | 22 | Luc7l3 |
| Lrrc8d3 | 0.000139603 | 0.415201652 | 0.25 | 0.134 | 1 | 22 | Lrrc8d |
| Ddx11 | 0.000140306 | 0.311784588 | 0.25 | 0.132 | 1 | 22 | Ddx1 |
| Magt15 | 0.000141594 | 0.481286589 | 0.287 | 0.166 | 1 | 22 | Magt1 |
| Atxn7l3b4 | 0.000142202 | 0.361366823 | 0.565 | 0.406 | 1 | 22 | Atxn7l3b |
| Wwox10 | 0.00015243 | 0.584801094 | 0.306 | 0.177 | 1 | 22 | Wwox |
| Grina4 | 0.000160666 | 0.40384852 | 0.259 | 0.141 | 1 | 22 | Grina |
| Ube2e3 | 0.000172377 | 0.256012918 | 0.37 | 0.227 | 1 | 22 | Ube2e3 |
| Parp92 | 0.000174117 | 0.398338745 | 0.25 | 0.134 | 1 | 22 | Parp9 |
| Fam168a4 | 0.00017469 | 0.488706265 | 0.259 | 0.144 | 1 | 22 | Fam168a |
| Gkap13 | 0.000178174 | 0.282095881 | 0.269 | 0.144 | 1 | 22 | Gkap1 |
| Ptk26 | 0.000186155 | 0.40238086 | 0.426 | 0.268 | 1 | 22 | Ptk2 |
| Nisch7 | 0.000187409 | 0.295692488 | 0.352 | 0.213 | 1 | 22 | Nisch |
| Mbtd14 | 0.000188103 | 0.429605339 | 0.25 | 0.136 | 1 | 22 | Mbtd1 |
| Eif4h1 | 0.000192661 | 0.359233464 | 0.435 | 0.291 | 1 | 22 | Eif4h |
| Spcs26 | 0.000208898 | 0.328946035 | 0.556 | 0.403 | 1 | 22 | Spcs2 |
| Rab5b1 | 0.000214975 | 0.337727405 | 0.259 | 0.141 | 1 | 22 | Rab5b |
| Ncoa110 | 0.000223732 | 0.461297202 | 0.343 | 0.212 | 1 | 22 | Ncoa1 |
| Huwe111 | 0.000235807 | 0.384067154 | 0.426 | 0.287 | 1 | 22 | Huwe1 |
| Copa3 | 0.000243068 | 0.471902625 | 0.333 | 0.207 | 1 | 22 | Copa |
| Siva13 | 0.000245744 | 0.422693218 | 0.278 | 0.161 | 1 | 22 | Siva1 |
| Phf311 | 0.000250336 | 0.410724566 | 0.389 | 0.255 | 1 | 22 | Phf3 |
| Snw1 | 0.000253756 | 0.371206559 | 0.407 | 0.262 | 1 | 22 | Snw1 |
| Mycbp213 | 0.000259299 | 0.430379175 | 0.546 | 0.376 | 1 | 22 | Mycbp2 |
| Nbea7 | 0.00026774 | 0.410657567 | 0.333 | 0.198 | 1 | 22 | Nbea |
| Hist1h2bc6 | 0.000269536 | 0.482462933 | 0.389 | 0.251 | 1 | 22 | Hist1h2bc |
| Ifitm38 | 0.000270173 | 0.322697622 | 0.806 | 0.54 | 1 | 22 | Ifitm3 |
| Ifnar26 | 0.00027574 | 0.414712551 | 0.463 | 0.327 | 1 | 22 | Ifnar2 |
| Ahcyl12 | 0.000277008 | 0.277009572 | 0.306 | 0.176 | 1 | 22 | Ahcyl1 |
| Nipbl14 | 0.000281544 | 0.498841724 | 0.519 | 0.368 | 1 | 22 | Nipbl |
| Mysm11 | 0.000285412 | 0.361710805 | 0.296 | 0.177 | 1 | 22 | Mysm1 |
| Ppib5 | 0.000292686 | 0.327214868 | 0.694 | 0.555 | 1 | 22 | Ppib |
| Tm9sf27 | 0.00029506 | 0.480771879 | 0.389 | 0.257 | 1 | 22 | Tm9sf2 |
| Ogt10 | 0.000301687 | 0.398909383 | 0.333 | 0.208 | 1 | 22 | Ogt |
| Nptn9 | 0.000303612 | 0.392801488 | 0.389 | 0.251 | 1 | 22 | Nptn |
| Sec11a7 | 0.000305596 | 0.513574953 | 0.435 | 0.287 | 1 | 22 | Sec11a |
| Herc47 | 0.000311999 | 0.330499184 | 0.269 | 0.153 | 1 | 22 | Herc4 |

| | | | | | | | |
|---|---|---|---|---|---|---|---|
| Luc7l211 | 0.000315351 | 0.367099513 | 0.556 | 0.407 | 1 | 22 | Luc7l2 |
| Pdia69 | 0.000323018 | 0.419474608 | 0.472 | 0.339 | 1 | 22 | Pdia6 |
| Vps352 | 0.00032885 | 0.270995826 | 0.343 | 0.207 | 1 | 22 | Vps35 |
| Tns18 | 0.000362449 | 0.383027771 | 0.398 | 0.247 | 1 | 22 | Tns1 |
| Gab28 | 0.000366695 | 0.255477278 | 0.343 | 0.207 | 1 | 22 | Gab2 |
| Dync1h17 | 0.00037796 | 0.585843308 | 0.417 | 0.288 | 1 | 22 | Dync1h1 |
| Ddb14 | 0.000381899 | 0.54779647 | 0.259 | 0.15 | 1 | 22 | Ddb1 |
| Hnrnpdl3 | 0.000407389 | 0.291601387 | 0.389 | 0.25 | 1 | 22 | Hnrnpdl |
| Calu6 | 0.000411532 | 0.400067908 | 0.333 | 0.204 | 1 | 22 | Calu |
| Rere8 | 0.000428829 | 0.493662366 | 0.398 | 0.273 | 1 | 22 | Rere |
| Limd12 | 0.000459167 | 0.559245116 | 0.25 | 0.144 | 1 | 22 | Limd1 |
| Rhoj7 | 0.000478569 | 0.326350341 | 0.407 | 0.243 | 1 | 22 | Rhoj |
| Tmed74 | 0.000484583 | 0.443511473 | 0.259 | 0.151 | 1 | 22 | Tmed7 |
| Cbx54 | 0.000495825 | 0.390893237 | 0.306 | 0.19 | 1 | 22 | Cbx5 |
| Atp1a14 | 0.000542538 | 0.480823111 | 0.407 | 0.282 | 1 | 22 | Atp1a1 |
| Ahr3 | 0.00054784 | 0.318222916 | 0.389 | 0.248 | 1 | 22 | Ahr |
| Pgp1 | 0.00055445 | 0.252314397 | 0.343 | 0.211 | 1 | 22 | Pgp |
| Smc35 | 0.000585455 | 0.466035042 | 0.37 | 0.255 | 1 | 22 | Smc3 |
| Rbm510 | 0.00061606 | 0.435112992 | 0.361 | 0.241 | 1 | 22 | Rbm5 |
| Ube3a5 | 0.000640913 | 0.427512323 | 0.417 | 0.297 | 1 | 22 | Ube3a |
| Pde7b7 | 0.000671195 | 0.374497165 | 0.389 | 0.244 | 1 | 22 | Pde7b |
| Hook37 | 0.000681751 | 0.524966224 | 0.417 | 0.292 | 1 | 22 | Hook3 |
| Glo1 | 0.000700532 | 0.405209234 | 0.296 | 0.184 | 1 | 22 | Glo1 |
| Rif14 | 0.000704369 | 0.494303353 | 0.269 | 0.156 | 1 | 22 | Rif1 |
| Tmbim67 | 0.000743519 | 0.334664614 | 0.574 | 0.433 | 1 | 22 | Tmbim6 |
| Tpr10 | 0.000746713 | 0.282597884 | 0.583 | 0.427 | 1 | 22 | Tpr |
| St132 | 0.000750936 | 0.32513954 | 0.509 | 0.369 | 1 | 22 | St13 |
| Ino80d6 | 0.000753873 | 0.412159924 | 0.278 | 0.167 | 1 | 22 | Ino80d |
| Gsk3b6 | 0.0007964 | 0.324992789 | 0.463 | 0.333 | 1 | 22 | Gsk3b |
| Cdc42bpa7 | 0.000815153 | 0.447717556 | 0.306 | 0.187 | 1 | 22 | Cdc42bpa |
| Prpf4b8 | 0.000828277 | 0.401713302 | 0.454 | 0.335 | 1 | 22 | Prpf4b |
| Bcap318 | 0.000910903 | 0.392711264 | 0.417 | 0.287 | 1 | 22 | Bcap31 |
| Rbms16 | 0.000975396 | 0.393343146 | 0.648 | 0.488 | 1 | 22 | Rbms1 |
| Tnrc6c8 | 0.000997211 | 0.307796831 | 0.454 | 0.317 | 1 | 22 | Tnrc6c |
| Sbf27 | 0.001023209 | 0.57416725 | 0.324 | 0.209 | 1 | 22 | Sbf2 |
| Tmcc112 | 0.001075778 | 0.43639738 | 0.333 | 0.22 | 1 | 22 | Tmcc1 |
| Slmap4 | 0.001084124 | 0.327972694 | 0.306 | 0.195 | 1 | 22 | Slmap |
| Crlf37 | 0.001088068 | 0.45503573 | 0.259 | 0.159 | 1 | 22 | Crlf3 |
| Selenop7 | 0.001106237 | 0.269824026 | 0.528 | 0.365 | 1 | 22 | Selenop |
| Sltm8 | 0.001183612 | 0.362025011 | 0.389 | 0.273 | 1 | 22 | Sltm |
| Paics | 0.001191382 | 0.295947434 | 0.315 | 0.196 | 1 | 22 | Paics |
| Pbx16 | 0.001263077 | 0.315597328 | 0.324 | 0.2 | 1 | 22 | Pbx1 |
| Fam133b | 0.001295426 | 0.310208407 | 0.287 | 0.177 | 1 | 22 | Fam133b |

| | | | | | | | |
|---|---|---|---|---|---|---|---|
| Exoc6b8 | 0.001308131 | 0.455356386 | 0.361 | 0.243 | 1 | 22 | Exoc6b |
| Sptan16 | 0.001383388 | 0.279797549 | 0.491 | 0.346 | 1 | 22 | Sptan1 |
| Taf131 | 0.001383408 | 0.292533199 | 0.25 | 0.15 | 1 | 22 | Taf13 |
| Ktn18 | 0.001394041 | 0.338763373 | 0.444 | 0.31 | 1 | 22 | Ktn1 |
| Ddx241 | 0.001398608 | 0.383132478 | 0.454 | 0.326 | 1 | 22 | Ddx24 |
| Plekhg15 | 0.001461455 | 0.354039068 | 0.389 | 0.247 | 1 | 22 | Plekhg1 |
| Mllt1011 | 0.001462001 | 0.420169213 | 0.398 | 0.277 | 1 | 22 | Mllt10 |
| Lnpep13 | 0.001488881 | 0.405851819 | 0.38 | 0.262 | 1 | 22 | Lnpep |
| Phkb3 | 0.001492014 | 0.305344516 | 0.25 | 0.146 | 1 | 22 | Phkb |
| Myo9a5 | 0.001495869 | 0.496113663 | 0.278 | 0.175 | 1 | 22 | Myo9a |
| Setd511 | 0.001497279 | 0.473424121 | 0.38 | 0.266 | 1 | 22 | Setd5 |
| Hp1bp31 | 0.001566222 | 0.299670237 | 0.407 | 0.287 | 1 | 22 | Hp1bp3 |
| Ythdc13 | 0.001587999 | 0.3317403 | 0.361 | 0.236 | 1 | 22 | Ythdc1 |
| Ccdc475 | 0.001613974 | 0.43560115 | 0.296 | 0.194 | 1 | 22 | Ccdc47 |
| Dock15 | 0.001642623 | 0.263496379 | 0.426 | 0.282 | 1 | 22 | Dock1 |
| Eif3c1 | 0.001657796 | 0.294001734 | 0.519 | 0.389 | 1 | 22 | Eif3c |
| Klf610 | 0.001689118 | 0.453611157 | 0.389 | 0.269 | 1 | 22 | Klf6 |
| Stat17 | 0.001732194 | 0.527290415 | 0.38 | 0.251 | 1 | 22 | Stat1 |
| Mecp24 | 0.00177482 | 0.376286402 | 0.259 | 0.162 | 1 | 22 | Mecp2 |
| Ddost4 | 0.001802877 | 0.51331786 | 0.315 | 0.218 | 1 | 22 | Ddost |
| Cuedc2 | 0.00181927 | 0.517485857 | 0.315 | 0.211 | 1 | 22 | Cuedc2 |
| Golga43 | 0.001861066 | 0.348473393 | 0.259 | 0.16 | 1 | 22 | Golga4 |
| Bptf12 | 0.001868137 | 0.370068414 | 0.417 | 0.296 | 1 | 22 | Bptf |
| Ssr34 | 0.00188002 | 0.362447727 | 0.296 | 0.198 | 1 | 22 | Ssr3 |
| Nxf1 | 0.002007015 | 0.354321418 | 0.426 | 0.304 | 1 | 22 | Nxf1 |
| Spata138 | 0.002027887 | 0.629939054 | 0.306 | 0.208 | 1 | 22 | Spata13 |
| Lpcat12 | 0.002058792 | 0.331414426 | 0.269 | 0.161 | 1 | 22 | Lpcat1 |
| Spcs13 | 0.002088619 | 0.295028899 | 0.407 | 0.29 | 1 | 22 | Spcs1 |
| Dctn43 | 0.002181466 | 0.308481762 | 0.324 | 0.213 | 1 | 22 | Dctn4 |
| Scaf116 | 0.002281518 | 0.367472937 | 0.38 | 0.265 | 1 | 22 | Scaf11 |
| Prdx46 | 0.002315345 | 0.420923535 | 0.333 | 0.223 | 1 | 22 | Prdx4 |
| Cnpy23 | 0.002325814 | 0.272006738 | 0.38 | 0.265 | 1 | 22 | Cnpy2 |
| Atp6v0e4 | 0.002408141 | 0.255784624 | 0.546 | 0.427 | 1 | 22 | Atp6v0e |
| Ascc39 | 0.002465151 | 0.420097999 | 0.306 | 0.209 | 1 | 22 | Ascc3 |
| Sept710 | 0.002557643 | 0.358807242 | 0.583 | 0.464 | 1 | 22 | 7-Sep |
| Ttc1411 | 0.002588527 | 0.501332629 | 0.315 | 0.217 | 1 | 22 | Ttc14 |
| Cisd23 | 0.002679257 | 0.654922977 | 0.269 | 0.176 | 1 | 22 | Cisd2 |
| Ssr22 | 0.002702847 | 0.463857996 | 0.306 | 0.204 | 1 | 22 | Ssr2 |
| Csnk1g36 | 0.002724864 | 0.286196451 | 0.287 | 0.185 | 1 | 22 | Csnk1g3 |
| Fibp | 0.002766783 | 0.250650053 | 0.259 | 0.16 | 1 | 22 | Fibp |
| Tra2a13 | 0.002828387 | 0.289932918 | 0.398 | 0.277 | 1 | 22 | Tra2a |
| Sesn18 | 0.002932543 | 0.326932234 | 0.324 | 0.214 | 1 | 22 | Sesn1 |
| Acap28 | 0.002979819 | 0.430921349 | 0.37 | 0.271 | 1 | 22 | Acap2 |

| | | | | | | | |
|---|---|---|---|---|---|---|---|
| Kat2b4 | 0.003030796 | 0.52462836 | 0.25 | 0.16 | 1 | 22 | Kat2b |
| Dennd5a6 | 0.003165021 | 0.483546705 | 0.287 | 0.189 | 1 | 22 | Dennd5a |
| Nfat59 | 0.003289844 | 0.314687646 | 0.389 | 0.269 | 1 | 22 | Nfat5 |
| Ccnd14 | 0.003300774 | 0.304512314 | 0.343 | 0.229 | 1 | 22 | Ccnd1 |
| Ypel34 | 0.003468572 | 0.397984102 | 0.37 | 0.269 | 1 | 22 | Ypel3 |
| Kmt2e10 | 0.003502349 | 0.318567876 | 0.537 | 0.409 | 1 | 22 | Kmt2e |
| H2afv2 | 0.003524965 | 0.268078051 | 0.5 | 0.381 | 1 | 22 | H2afv |
| Fto7 | 0.003629453 | 0.459382232 | 0.287 | 0.193 | 1 | 22 | Fto |
| Dnajc7 | 0.003650343 | 0.331112412 | 0.361 | 0.26 | 1 | 22 | Dnajc7 |
| Dennd1a12 | 0.003874738 | 0.421771118 | 0.352 | 0.255 | 1 | 22 | Dennd1a |
| Adam174 | 0.003904981 | 0.367037142 | 0.25 | 0.155 | 1 | 22 | Adam17 |
| Anxa28 | 0.003975829 | 0.265652313 | 0.556 | 0.419 | 1 | 22 | Anxa2 |
| Cbfb7 | 0.003981904 | 0.271009727 | 0.25 | 0.157 | 1 | 22 | Cbfb |
| Usp9x3 | 0.004014197 | 0.340083161 | 0.333 | 0.236 | 1 | 22 | Usp9x |
| Slc38a28 | 0.00416605 | 0.310805307 | 0.435 | 0.313 | 1 | 22 | Slc38a2 |
| Lman16 | 0.004185155 | 0.321419205 | 0.343 | 0.24 | 1 | 22 | Lman1 |
| Trappc85 | 0.004192028 | 0.268770682 | 0.259 | 0.168 | 1 | 22 | Trappc8 |
| Krit19 | 0.004208765 | 0.384265535 | 0.278 | 0.184 | 1 | 22 | Krit1 |
| Trim445 | 0.004572792 | 0.359043287 | 0.333 | 0.229 | 1 | 22 | Trim44 |
| Tmed99 | 0.004618169 | 0.383004634 | 0.509 | 0.406 | 1 | 22 | Tmed9 |
| Mdm45 | 0.004799605 | 0.312857136 | 0.25 | 0.161 | 1 | 22 | Mdm4 |
| Zfp912 | 0.005005967 | 0.304815425 | 0.417 | 0.309 | 1 | 22 | Zfp91 |
| Ssr14 | 0.005017561 | 0.364278647 | 0.278 | 0.188 | 1 | 22 | Ssr1 |
| Psme45 | 0.005177631 | 0.340810335 | 0.287 | 0.199 | 1 | 22 | Psme4 |
| Tmem598 | 0.005191596 | 0.25638869 | 0.602 | 0.493 | 1 | 22 | Tmem59 |
| Cyhr12 | 0.005284182 | 0.598580601 | 0.259 | 0.169 | 1 | 22 | Cyhr1 |
| Nfx11 | 0.005321709 | 0.285554426 | 0.25 | 0.163 | 1 | 22 | Nfx1 |
| Lonp23 | 0.005625605 | 0.350112807 | 0.287 | 0.196 | 1 | 22 | Lonp2 |
| Hmgb13 | 0.005760242 | 0.293165607 | 0.852 | 0.717 | 1 | 22 | Hmgb1 |
| Naxd1 | 0.005871194 | 0.280856221 | 0.259 | 0.169 | 1 | 22 | Naxd |
| Fam172a12 | 0.006023862 | 0.32400108 | 0.389 | 0.286 | 1 | 22 | Fam172a |
| Tnrc187 | 0.006249292 | 0.335463114 | 0.269 | 0.175 | 1 | 22 | Tnrc18 |
| Ptpre7 | 0.006406626 | 0.362587106 | 0.25 | 0.163 | 1 | 22 | Ptpre |
| Eprs1 | 0.006480623 | 0.386454122 | 0.296 | 0.203 | 1 | 22 | Eprs |
| Rab127 | 0.006516465 | 0.274252321 | 0.278 | 0.179 | 1 | 22 | Rab12 |
| Sf15 | 0.006527717 | 0.272666752 | 0.278 | 0.189 | 1 | 22 | Sf1 |
| Bag53 | 0.006535704 | 0.275310832 | 0.306 | 0.203 | 1 | 22 | Bag5 |
| Dpm33 | 0.006593714 | 0.486373182 | 0.454 | 0.351 | 1 | 22 | Dpm3 |
| Vezf13 | 0.006607035 | 0.3177728 | 0.333 | 0.236 | 1 | 22 | Vezf1 |
| Tmx37 | 0.006623365 | 0.353047199 | 0.278 | 0.189 | 1 | 22 | Tmx3 |
| Mia34 | 0.006823931 | 0.373180393 | 0.259 | 0.172 | 1 | 22 | Mia3 |
| Chd48 | 0.006917151 | 0.306962454 | 0.435 | 0.33 | 1 | 22 | Chd4 |
| Ddrgk13 | 0.007311371 | 0.295850335 | 0.417 | 0.32 | 1 | 22 | Ddrgk1 |

| | | | | | | | |
|---|---|---|---|---|---|---|---|
| Atp2a28 | 0.007480856 | 0.393838228 | 0.287 | 0.202 | 1 | 22 | Atp2a2 |
| Rab102 | 0.00754243 | 0.294522009 | 0.463 | 0.352 | 1 | 22 | Rab10 |
| Cyfip17 | 0.007604767 | 0.279892419 | 0.38 | 0.28 | 1 | 22 | Cyfip1 |
| Tacc18 | 0.008005377 | 0.395864588 | 0.398 | 0.307 | 1 | 22 | Tacc1 |
| Tmem147 | 0.008048116 | 0.401343058 | 0.287 | 0.202 | 1 | 22 | Tmem147 |
| Mob41 | 0.008060573 | 0.360777525 | 0.417 | 0.308 | 1 | 22 | Mob4 |
| Bcas39 | 0.008085001 | 0.445897337 | 0.269 | 0.184 | 1 | 22 | Bcas3 |
| Tprgl5 | 0.008089679 | 0.316994561 | 0.389 | 0.281 | 1 | 22 | Tprgl |
| Isg154 | 0.009840483 | 1.037141864 | 0.324 | 0.231 | 1 | 22 | Isg15 |
| Hist1h1b1 | 0 | 3.436743908 | 0.625 | 0.017 | 0 | 23 | Hist1h1b |
| Top2a1 | 0 | 3.232573672 | 0.906 | 0.018 | 0 | 23 | Top2a |
| Pclaf1 | 0 | 2.923776489 | 0.927 | 0.007 | 0 | 23 | Pclaf |
| Mki671 | 0 | 2.840096365 | 0.906 | 0.01 | 0 | 23 | Mki67 |
| Birc51 | 0 | 2.319615365 | 0.802 | 0.008 | 0 | 23 | Birc5 |
| Cenpf1 | 0 | 2.28697116 | 0.604 | 0.005 | 0 | 23 | Cenpf |
| Nusap11 | 0 | 2.281020474 | 0.708 | 0.011 | 0 | 23 | Nusap1 |
| Ube2c1 | 0 | 2.029085608 | 0.594 | 0.01 | 0 | 23 | Ube2c |
| Prc11 | 0 | 1.919048588 | 0.698 | 0.009 | 0 | 23 | Prc1 |
| Lockd1 | 0 | 1.910652232 | 0.812 | 0.021 | 0 | 23 | Lockd |
| Spc241 | 0 | 1.810815736 | 0.885 | 0.009 | 0 | 23 | Spc24 |
| Diaph31 | 0 | 1.793906424 | 0.854 | 0.024 | 0 | 23 | Diaph3 |
| Cdca81 | 0 | 1.661349843 | 0.802 | 0.009 | 0 | 23 | Cdca8 |
| Cenpp1 | 0 | 1.628390277 | 0.75 | 0.014 | 0 | 23 | Cenpp |
| Ccnb21 | 0 | 1.610658648 | 0.625 | 0.005 | 0 | 23 | Ccnb2 |
| Rrm2 | 0 | 1.567308086 | 0.656 | 0.005 | 0 | 23 | Rrm2 |
| Tpx21 | 0 | 1.522281715 | 0.688 | 0.008 | 0 | 23 | Tpx2 |
| Kif111 | 0 | 1.518736048 | 0.688 | 0.008 | 0 | 23 | Kif11 |
| Hmmr1 | 0 | 1.484211082 | 0.562 | 0.002 | 0 | 23 | Hmmr |
| Hist1h3c1 | 0 | 1.480760729 | 0.531 | 0.004 | 0 | 23 | Hist1h3c |
| Knl11 | 0 | 1.468253698 | 0.708 | 0.007 | 0 | 23 | Knl1 |
| Clspn1 | 0 | 1.451448835 | 0.719 | 0.011 | 0 | 23 | Clspn |
| Cenpe1 | 0 | 1.424725904 | 0.615 | 0.008 | 0 | 23 | Cenpe |
| Cdk11 | 0 | 1.378827134 | 0.708 | 0.015 | 0 | 23 | Cdk1 |
| Cdca31 | 0 | 1.321690266 | 0.635 | 0.004 | 0 | 23 | Cdca3 |
| Cenpw | 0 | 1.28385779 | 0.781 | 0.017 | 0 | 23 | Cenpw |
| Asf1b1 | 0 | 1.267000054 | 0.729 | 0.008 | 0 | 23 | Asf1b |
| Hells | 0 | 1.257996573 | 0.698 | 0.023 | 0 | 23 | Hells |
| Kif231 | 0 | 1.201913183 | 0.604 | 0.016 | 0 | 23 | Kif23 |
| Kif151 | 0 | 1.190068315 | 0.583 | 0.006 | 0 | 23 | Kif15 |
| Tk11 | 0 | 1.175979675 | 0.688 | 0.008 | 0 | 23 | Tk1 |
| Fbxo51 | 0 | 1.169867361 | 0.625 | 0.014 | 0 | 23 | Fbxo5 |
| Cenpm1 | 0 | 1.140589799 | 0.729 | 0.007 | 0 | 23 | Cenpm |
| Mis18bp1 | 0 | 1.121069009 | 0.5 | 0.005 | 0 | 23 | Mis18bp1 |

| | | | | | | | |
|---|---|---|---|---|---|---|---|
| Kif20b1 | 0 | 1.117474513 | 0.646 | 0.016 | 0 | 23 | Kif20b |
| Esco21 | 0 | 1.086634416 | 0.542 | 0.003 | 0 | 23 | Esco2 |
| Ncapg2 | 0 | 1.07580155 | 0.635 | 0.018 | 0 | 23 | Ncapg2 |
| Hist1h2bj1 | 0 | 1.046920592 | 0.51 | 0.009 | 0 | 23 | Hist1h2bj |
| Kif41 | 0 | 1.031189457 | 0.656 | 0.011 | 0 | 23 | Kif4 |
| Spc251 | 0 | 1.024492331 | 0.583 | 0.016 | 0 | 23 | Spc25 |
| Neil31 | 0 | 1.016985869 | 0.49 | 0.003 | 0 | 23 | Neil3 |
| Ndc801 | 0 | 1.007182912 | 0.604 | 0.012 | 0 | 23 | Ndc80 |
| Hist1h2af1 | 0 | 0.988133042 | 0.521 | 0.002 | 0 | 23 | Hist1h2af |
| Mcm51 | 0 | 0.976959603 | 0.688 | 0.024 | 0 | 23 | Mcm5 |
| Cenpk | 0 | 0.969745757 | 0.667 | 0.009 | 0 | 23 | Cenpk |
| Ccna2 | 0 | 0.965938502 | 0.552 | 0.002 | 0 | 23 | Ccna2 |
| Ect2 | 0 | 0.953788014 | 0.531 | 0.007 | 0 | 23 | Ect2 |
| Tacc31 | 0 | 0.926509493 | 0.615 | 0.01 | 0 | 23 | Tacc3 |
| Hist1h2ab1 | 0 | 0.923778642 | 0.438 | 0.003 | 0 | 23 | Hist1h2ab |
| Cdkn31 | 0 | 0.921355914 | 0.51 | 0.004 | 0 | 23 | Cdkn3 |
| Cenph | 0 | 0.903499768 | 0.604 | 0.006 | 0 | 23 | Cenph |
| Aurkb1 | 0 | 0.863772914 | 0.51 | 0.005 | 0 | 23 | Aurkb |
| Uhrf1 | 0 | 0.862053909 | 0.573 | 0.007 | 0 | 23 | Uhrf1 |
| Ncapg | 0 | 0.860098216 | 0.552 | 0.003 | 0 | 23 | Ncapg |
| Hist1h2bm | 0 | 0.85643466 | 0.479 | 0.004 | 0 | 23 | Hist1h2bm |
| Ckap2l1 | 0 | 0.844087901 | 0.562 | 0.01 | 0 | 23 | Ckap2l |
| Kntc1 | 0 | 0.842257375 | 0.542 | 0.004 | 0 | 23 | Kntc1 |
| Bub1b1 | 0 | 0.8293389 | 0.542 | 0.01 | 0 | 23 | Bub1b |
| Brca1 | 0 | 0.825314786 | 0.5 | 0.006 | 0 | 23 | Brca1 |
| Rad511 | 0 | 0.814658435 | 0.562 | 0.009 | 0 | 23 | Rad51 |
| Cep55 | 0 | 0.810174233 | 0.521 | 0.003 | 0 | 23 | Cep55 |
| Aspm | 0 | 0.79882511 | 0.438 | 0.003 | 0 | 23 | Aspm |
| Hist1h2bb | 0 | 0.798538009 | 0.406 | 0.006 | 0 | 23 | Hist1h2bb |
| Cdca2 | 0 | 0.794931903 | 0.562 | 0.011 | 0 | 23 | Cdca2 |
| Bub1 | 0 | 0.788655975 | 0.458 | 0.002 | 0 | 23 | Bub1 |
| Cdc201 | 0 | 0.786028173 | 0.385 | 0.007 | 0 | 23 | Cdc20 |
| Mad2l1 | 0 | 0.784838662 | 0.594 | 0.015 | 0 | 23 | Mad2l1 |
| Sgo2a | 0 | 0.76675039 | 0.469 | 0.004 | 0 | 23 | Sgo2a |
| Dtl | 0 | 0.760389542 | 0.479 | 0.009 | 0 | 23 | Dtl |
| Arhgap11a | 0 | 0.760312323 | 0.5 | 0.012 | 0 | 23 | Arhgap11a |
| Dlgap5 | 0 | 0.758854735 | 0.479 | 0.006 | 0 | 23 | Dlgap5 |
| Rad51ap11 | 0 | 0.754221101 | 0.49 | 0.008 | 0 | 23 | Rad51ap1 |
| Mxd3 | 0 | 0.744782427 | 0.5 | 0.001 | 0 | 23 | Mxd3 |
| Knstrn1 | 0 | 0.738843352 | 0.448 | 0.009 | 0 | 23 | Knstrn |
| Shcbp1 | 0 | 0.738526834 | 0.438 | 0.003 | 0 | 23 | Shcbp1 |
| Ccne2 | 0 | 0.729536885 | 0.417 | 0.008 | 0 | 23 | Ccne2 |
| Ska1 | 0 | 0.721676503 | 0.479 | 0.002 | 0 | 23 | Ska1 |

| | | | | | | | |
|---|---|---|---|---|---|---|---|
| Depdc1a | 0 | 0.717916688 | 0.438 | 0.001 | 0 | 23 | Depdc1a |
| Hist1h3i | 0 | 0.708954216 | 0.438 | 0.007 | 0 | 23 | Hist1h3i |
| Kif22 | 0 | 0.700469942 | 0.438 | 0.004 | 0 | 23 | Kif22 |
| Pole | 0 | 0.697169708 | 0.49 | 0.008 | 0 | 23 | Pole |
| Sgo11 | 0 | 0.690264361 | 0.438 | 0.003 | 0 | 23 | Sgo1 |
| Ckap2 | 0 | 0.686247496 | 0.406 | 0.007 | 0 | 23 | Ckap2 |
| Plk1 | 0 | 0.677207889 | 0.406 | 0.002 | 0 | 23 | Plk1 |
| Hist1h2ai | 0 | 0.656747688 | 0.427 | 0.004 | 0 | 23 | Hist1h2ai |
| Bard1 | 0 | 0.624239713 | 0.427 | 0.008 | 0 | 23 | Bard1 |
| Pbk1 | 0 | 0.621406671 | 0.438 | 0.003 | 0 | 23 | Pbk |
| Cdc45 | 0 | 0.619795805 | 0.49 | 0.007 | 0 | 23 | Cdc45 |
| Ube2t1 | 0 | 0.605905475 | 0.49 | 0.013 | 0 | 23 | Ube2t |
| E2f8 | 0 | 0.603430439 | 0.375 | 0.004 | 0 | 23 | E2f8 |
| Melk | 0 | 0.600150928 | 0.438 | 0.003 | 0 | 23 | Melk |
| Parpbp | 0 | 0.596667493 | 0.406 | 0.004 | 0 | 23 | Parpbp |
| Ccnf | 0 | 0.593907442 | 0.396 | 0.005 | 0 | 23 | Ccnf |
| Chaf1a | 0 | 0.571891175 | 0.49 | 0.01 | 0 | 23 | Chaf1a |
| Mastl | 0 | 0.568268745 | 0.375 | 0.006 | 0 | 23 | Mastl |
| Hist1h2bn | 0 | 0.563863383 | 0.375 | 0.004 | 0 | 23 | Hist1h2bn |
| E2f7 | 0 | 0.561914697 | 0.438 | 0.008 | 0 | 23 | E2f7 |
| Zwilch | 0 | 0.554316811 | 0.396 | 0.007 | 0 | 23 | Zwilch |
| Stil | 0 | 0.543252933 | 0.5 | 0.005 | 0 | 23 | Stil |
| Ttk | 0 | 0.536883774 | 0.354 | 0.003 | 0 | 23 | Ttk |
| Kif2c | 0 | 0.526829702 | 0.365 | 0.001 | 0 | 23 | Kif2c |
| Ccnb1 | 0 | 0.524566156 | 0.271 | 0.002 | 0 | 23 | Ccnb1 |
| Ncaph | 0 | 0.519218029 | 0.396 | 0.004 | 0 | 23 | Ncaph |
| Fignl1 | 0 | 0.518255882 | 0.438 | 0.005 | 0 | 23 | Fignl1 |
| Spag5 | 0 | 0.504749262 | 0.375 | 0.003 | 0 | 23 | Spag5 |
| C330027C09Rik | 0 | 0.5028051 | 0.406 | 0.005 | 0 | 23 | C330027C09Rik |
| Hist1h2br | 0 | 0.498981105 | 0.375 | 0.004 | 0 | 23 | Hist1h2br |
| Polq | 0 | 0.494410868 | 0.375 | 0.004 | 0 | 23 | Polq |
| Rad54b | 0 | 0.485007187 | 0.323 | 0.003 | 0 | 23 | Rad54b |
| Hist1h2an | 0 | 0.474700913 | 0.302 | 0.003 | 0 | 23 | Hist1h2an |
| Kif14 | 0 | 0.468633223 | 0.281 | 0.001 | 0 | 23 | Kif14 |
| Ccne1 | 0 | 0.463541712 | 0.312 | 0.003 | 0 | 23 | Ccne1 |
| Hist1h3b | 0 | 0.4542401 | 0.312 | 0.001 | 0 | 23 | Hist1h3b |
| Pimreg | 0 | 0.453336192 | 0.302 | 0.002 | 0 | 23 | Pimreg |
| Aurka | 0 | 0.44862591 | 0.323 | 0.005 | 0 | 23 | Aurka |
| Nuf2 | 0 | 0.432735206 | 0.323 | 0.001 | 0 | 23 | Nuf2 |
| Cdca5 | 0 | 0.431869621 | 0.292 | 0.002 | 0 | 23 | Cdca5 |
| Mcm10 | 0 | 0.423276505 | 0.385 | 0.003 | 0 | 23 | Mcm10 |
| Kifc1 | 0 | 0.420136086 | 0.354 | 0.003 | 0 | 23 | Kifc1 |
| Fanci | 0 | 0.411406033 | 0.323 | 0.004 | 0 | 23 | Fanci |

| | | | | | | | |
|---|---|---|---|---|---|---|---|
| Depdc1b | 0 | 0.397803819 | 0.25 | 0.002 | 0 | 23 | Depdc1b |
| Dscc1 | 0 | 0.396614634 | 0.354 | 0.006 | 0 | 23 | Dscc1 |
| Kif20a | 0 | 0.389965738 | 0.281 | 0.002 | 0 | 23 | Kif20a |
| Nek2 | 0 | 0.370168072 | 0.25 | 0.001 | 0 | 23 | Nek2 |
| Ska3 | 0 | 0.36372472 | 0.271 | 0.002 | 0 | 23 | Ska3 |
| Spdl1 | 0 | 0.360370655 | 0.312 | 0.005 | 0 | 23 | Spdl1 |
| Aunip | 0 | 0.335432112 | 0.26 | 0.002 | 0 | 23 | Aunip |
| Exo1 | 0 | 0.328299022 | 0.281 | 0.002 | 0 | 23 | Exo1 |
| Cdc6 | 3.54E-307 | 0.337906882 | 0.271 | 0.004 | 7.16E-303 | 23 | Cdc6 |
| Cks1b2 | 2.68E-303 | 1.613259442 | 0.812 | 0.04 | 5.43E-299 | 23 | Cks1b |
| Hist1h2ak | 4.66E-302 | 0.350074206 | 0.25 | 0.003 | 9.43E-298 | 23 | Hist1h2ak |
| Incenp1 | 4.85E-299 | 1.070770387 | 0.625 | 0.024 | 9.82E-295 | 23 | Incenp |
| Efcab11 | 1.62E-298 | 0.594087308 | 0.354 | 0.007 | 3.27E-294 | 23 | Efcab11 |
| Cenpi | 7.08E-297 | 0.292560028 | 0.26 | 0.004 | 1.43E-292 | 23 | Cenpi |
| E2f1 | 1.87E-294 | 0.577338075 | 0.469 | 0.013 | 3.78E-290 | 23 | E2f1 |
| Cenpn | 7.78E-294 | 0.31508873 | 0.281 | 0.004 | 1.58E-289 | 23 | Cenpn |
| Hist1h3f | 6.55E-292 | 0.624743575 | 0.365 | 0.008 | 1.33E-287 | 23 | Hist1h3f |
| Lmnb11 | 2.36E-291 | 1.766011572 | 0.885 | 0.051 | 4.77E-287 | 23 | Lmnb1 |
| Mms22l | 7.81E-289 | 0.88550061 | 0.594 | 0.022 | 1.58E-284 | 23 | Mms22l |
| Trip13 | 2.76E-287 | 0.374796192 | 0.312 | 0.006 | 5.59E-283 | 23 | Trip13 |
| Dhfr | 9.89E-287 | 0.777100265 | 0.562 | 0.019 | 2.00E-282 | 23 | Dhfr |
| Tcf19 | 9.98E-287 | 0.648793122 | 0.469 | 0.013 | 2.02E-282 | 23 | Tcf19 |
| Ncapd21 | 1.27E-286 | 0.743195173 | 0.531 | 0.017 | 2.56E-282 | 23 | Ncapd2 |
| Rad51b1 | 6.34E-285 | 1.5202632 | 0.75 | 0.036 | 1.28E-280 | 23 | Rad51b |
| Cdc7 | 2.93E-284 | 0.522508926 | 0.417 | 0.011 | 5.93E-280 | 23 | Cdc7 |
| Anln1 | 3.85E-283 | 0.587381188 | 0.375 | 0.008 | 7.80E-279 | 23 | Anln |
| Hist1h1a1 | 2.17E-282 | 0.70789322 | 0.406 | 0.01 | 4.40E-278 | 23 | Hist1h1a |
| Hist1h2bh | 1.44E-281 | 0.361271353 | 0.26 | 0.004 | 2.92E-277 | 23 | Hist1h2bh |
| Mns11 | 4.94E-279 | 0.615115584 | 0.406 | 0.01 | 1.00E-274 | 23 | Mns1 |
| Racgap11 | 6.30E-278 | 1.166666929 | 0.677 | 0.03 | 1.28E-273 | 23 | Racgap1 |
| Psmc3ip | 3.39E-276 | 0.584626432 | 0.521 | 0.017 | 6.87E-272 | 23 | Psmc3ip |
| Rmi2 | 4.37E-271 | 0.429971075 | 0.354 | 0.008 | 8.84E-267 | 23 | Rmi2 |
| Chek1 | 5.84E-256 | 0.31064165 | 0.271 | 0.005 | 1.18E-251 | 23 | Chek1 |
| Zcwpw1 | 6.33E-256 | 0.617163257 | 0.458 | 0.014 | 1.28E-251 | 23 | Zcwpw1 |
| Hist1h2ae1 | 2.02E-254 | 2.869874322 | 0.615 | 0.028 | 4.09E-250 | 23 | Hist1h2ae |
| Hist2h2bb | 7.32E-251 | 0.980163223 | 0.479 | 0.016 | 1.48E-246 | 23 | Hist2h2bb |
| Fanca | 4.11E-246 | 0.30727879 | 0.25 | 0.004 | 8.32E-242 | 23 | Fanca |
| Cit | 4.81E-242 | 0.968067755 | 0.594 | 0.026 | 9.73E-238 | 23 | Cit |
| Nrm1 | 1.45E-235 | 0.92880704 | 0.615 | 0.029 | 2.93E-231 | 23 | Nrm |
| Hist1h2ap2 | 7.73E-235 | 3.699730868 | 0.76 | 0.047 | 1.57E-230 | 23 | Hist1h2ap |
| Cenps | 2.30E-231 | 0.515500539 | 0.469 | 0.017 | 4.66E-227 | 23 | Cenps |
| Ulbp11 | 1.69E-230 | 0.864902492 | 0.604 | 0.028 | 3.42E-226 | 23 | Ulbp1 |
| Brca2 | 3.20E-229 | 0.664081258 | 0.469 | 0.017 | 6.48E-225 | 23 | Brca2 |

| | | | | | | | |
|---|---|---|---|---|---|---|---|
| Gins2 | 6.62E-229 | 0.628647089 | 0.469 | 0.017 | 1.34E-224 | 23 | Gins2 |
| Smc21 | 2.36E-228 | 1.8973008 | 0.875 | 0.064 | 4.78E-224 | 23 | Smc2 |
| Rrm11 | 9.46E-228 | 1.365018778 | 0.781 | 0.05 | 1.91E-223 | 23 | Rrm1 |
| Ccdc18 | 2.91E-227 | 0.417703609 | 0.302 | 0.007 | 5.88E-223 | 23 | Ccdc18 |
| Plk4 | 1.34E-219 | 0.587038361 | 0.448 | 0.016 | 2.71E-215 | 23 | Plk4 |
| Hist1h3e1 | 1.40E-219 | 1.522776991 | 0.667 | 0.037 | 2.83E-215 | 23 | Hist1h3e |
| Fen11 | 1.94E-217 | 0.725921426 | 0.615 | 0.031 | 3.93E-213 | 23 | Fen1 |
| Kif24 | 2.75E-217 | 0.43442442 | 0.323 | 0.008 | 5.57E-213 | 23 | Kif24 |
| Ezh22 | 5.53E-215 | 1.607145694 | 0.885 | 0.068 | 1.12E-210 | 23 | Ezh2 |
| Cenpu | 5.39E-213 | 0.364766751 | 0.281 | 0.006 | 1.09E-208 | 23 | Cenpu |
| Prr11 | 9.71E-212 | 0.318170711 | 0.26 | 0.005 | 1.97E-207 | 23 | Prr11 |
| Mcm71 | 1.48E-210 | 1.304453467 | 0.771 | 0.052 | 3.00E-206 | 23 | Mcm7 |
| Mcm2 | 6.81E-210 | 0.506527458 | 0.5 | 0.021 | 1.38E-205 | 23 | Mcm2 |
| Hist1h4a | 6.11E-209 | 0.339892688 | 0.302 | 0.007 | 1.24E-204 | 23 | Hist1h4a |
| Lig11 | 4.62E-203 | 1.600542693 | 0.792 | 0.058 | 9.36E-199 | 23 | Lig1 |
| Hist1h3a | 4.01E-201 | 0.543574283 | 0.375 | 0.012 | 8.12E-197 | 23 | Hist1h3a |
| Spata24 | 3.52E-196 | 0.471948285 | 0.448 | 0.018 | 7.14E-192 | 23 | Spata24 |
| Prim1 | 4.30E-196 | 0.790886544 | 0.646 | 0.037 | 8.71E-192 | 23 | Prim1 |
| Pmf11 | 5.20E-196 | 0.894990711 | 0.698 | 0.044 | 1.05E-191 | 23 | Pmf1 |
| Trim59 | 8.05E-196 | 0.810325416 | 0.615 | 0.034 | 1.63E-191 | 23 | Trim59 |
| Spaca9 | 3.48E-195 | 0.622806542 | 0.479 | 0.021 | 7.05E-191 | 23 | Spaca9 |
| Chaf1b | 1.91E-186 | 0.448875081 | 0.385 | 0.014 | 3.87E-182 | 23 | Chaf1b |
| Atad5 | 4.10E-186 | 0.660418201 | 0.458 | 0.02 | 8.30E-182 | 23 | Atad5 |
| Cks21 | 2.85E-182 | 1.553617815 | 0.823 | 0.068 | 5.77E-178 | 23 | Cks2 |
| Rfc41 | 4.31E-182 | 0.892703962 | 0.698 | 0.047 | 8.72E-178 | 23 | Rfc4 |
| Hist1h1d2 | 1.69E-181 | 1.727393358 | 0.677 | 0.047 | 3.42E-177 | 23 | Hist1h1d |
| H2afx1 | 1.28E-179 | 2.051517513 | 0.74 | 0.057 | 2.59E-175 | 23 | H2afx |
| Stmn15 | 8.09E-174 | 3.914582738 | 1 | 0.129 | 1.64E-169 | 23 | Stmn1 |
| Hist1h2bg | 1.92E-173 | 0.490580667 | 0.354 | 0.013 | 3.88E-169 | 23 | Hist1h2bg |
| Poc1a | 6.54E-172 | 0.316113209 | 0.271 | 0.007 | 1.32E-167 | 23 | Poc1a |
| Kif18a | 2.02E-170 | 0.605212175 | 0.469 | 0.023 | 4.09E-166 | 23 | Kif18a |
| Cenpa1 | 2.37E-170 | 1.587307291 | 0.542 | 0.032 | 4.80E-166 | 23 | Cenpa |
| Dsn1 | 1.14E-166 | 0.271995027 | 0.281 | 0.008 | 2.30E-162 | 23 | Dsn1 |
| Pola11 | 6.41E-164 | 1.536873958 | 0.771 | 0.068 | 1.30E-159 | 23 | Pola1 |
| Serp21 | 2.47E-163 | 0.697774087 | 0.615 | 0.04 | 4.99E-159 | 23 | Serp2 |
| E2f2 | 1.97E-159 | 0.427544358 | 0.281 | 0.009 | 3.99E-155 | 23 | E2f2 |
| Brip11 | 9.24E-159 | 0.874773093 | 0.656 | 0.047 | 1.87E-154 | 23 | Brip1 |
| Mcm4 | 3.30E-158 | 0.910391062 | 0.698 | 0.054 | 6.69E-154 | 23 | Mcm4 |
| Bora | 6.01E-158 | 0.442280209 | 0.354 | 0.014 | 1.22E-153 | 23 | Bora |
| Cenpl | 2.39E-155 | 0.512764137 | 0.365 | 0.015 | 4.85E-151 | 23 | Cenpl |
| Ccdc341 | 1.40E-151 | 1.060374472 | 0.854 | 0.083 | 2.84E-147 | 23 | Ccdc34 |
| Cep1281 | 6.30E-151 | 1.040129431 | 0.656 | 0.05 | 1.27E-146 | 23 | Cep128 |
| Rbl1 | 8.91E-150 | 0.804398515 | 0.667 | 0.051 | 1.80E-145 | 23 | Rbl1 |

| | | | | | | | |
|---|---|---|---|---|---|---|---|
| Tyms1 | 1.83E-149 | 1.253841974 | 0.771 | 0.07 | 3.70E-145 | 23 | Tyms |
| Prim21 | 1.37E-147 | 0.901513226 | 0.615 | 0.045 | 2.77E-143 | 23 | Prim2 |
| Klre14 | 3.22E-147 | 1.829885931 | 0.917 | 0.102 | 6.52E-143 | 23 | Klre1 |
| Mcm31 | 4.67E-147 | 1.152881289 | 0.729 | 0.064 | 9.46E-143 | 23 | Mcm3 |
| Ncapd31 | 4.70E-145 | 1.280761556 | 0.771 | 0.074 | 9.52E-141 | 23 | Ncapd3 |
| Gmnn1 | 1.24E-143 | 1.26036545 | 0.823 | 0.084 | 2.52E-139 | 23 | Gmnn |
| Zgrf1 | 3.53E-142 | 0.445258735 | 0.312 | 0.012 | 7.14E-138 | 23 | Zgrf1 |
| Atad25 | 3.63E-141 | 1.473367584 | 0.844 | 0.092 | 7.35E-137 | 23 | Atad2 |
| Rfc5 | 2.61E-139 | 0.564256859 | 0.5 | 0.031 | 5.28E-135 | 23 | Rfc5 |
| Cd77 | 1.36E-138 | 1.973851801 | 0.948 | 0.111 | 2.76E-134 | 23 | Cd7 |
| Nrgn1 | 2.48E-138 | 0.831380201 | 0.5 | 0.032 | 5.02E-134 | 23 | Nrgn |
| Haus4 | 6.76E-138 | 0.729423222 | 0.615 | 0.047 | 1.37E-133 | 23 | Haus4 |
| Phgdh2 | 9.88E-138 | 0.836081978 | 0.625 | 0.05 | 2.00E-133 | 23 | Phgdh |
| Rad18 | 6.32E-137 | 0.580223433 | 0.49 | 0.031 | 1.28E-132 | 23 | Rad18 |
| Nup2106 | 7.13E-137 | 0.958606319 | 0.719 | 0.066 | 1.44E-132 | 23 | Nup210 |
| Dbf4 | 3.39E-136 | 0.702893067 | 0.5 | 0.033 | 6.87E-132 | 23 | Dbf4 |
| Sh2d1a3 | 8.93E-134 | 0.931863491 | 0.656 | 0.056 | 1.81E-129 | 23 | Sh2d1a |
| Ms4a4b5 | 4.06E-133 | 1.990643706 | 0.958 | 0.121 | 8.22E-129 | 23 | Ms4a4b |
| Zranb3 | 8.69E-133 | 0.414322569 | 0.323 | 0.014 | 1.76E-128 | 23 | Zranb3 |
| Hist1h2ac | 1.39E-131 | 0.503357663 | 0.406 | 0.022 | 2.82E-127 | 23 | Hist1h2ac |
| Dnajc91 | 2.36E-131 | 1.237571373 | 0.885 | 0.104 | 4.78E-127 | 23 | Dnajc9 |
| Klrk14 | 2.78E-131 | 1.607539897 | 0.875 | 0.1 | 5.64E-127 | 23 | Klrk1 |
| Clnk5 | 2.99E-131 | 1.566413534 | 0.938 | 0.11 | 6.06E-127 | 23 | Clnk |
| Rpa2 | 2.43E-130 | 0.703602179 | 0.573 | 0.044 | 4.93E-126 | 23 | Rpa2 |
| Mbnl3 | 5.17E-130 | 0.583450855 | 0.448 | 0.027 | 1.05E-125 | 23 | Mbnl3 |
| Klrd15 | 7.92E-130 | 2.011583471 | 0.948 | 0.121 | 1.60E-125 | 23 | Klrd1 |
| Haspin | 5.12E-129 | 0.298068786 | 0.26 | 0.009 | 1.04E-124 | 23 | Haspin |
| Ctsw5 | 8.41E-129 | 2.090642804 | 0.958 | 0.127 | 1.70E-124 | 23 | Ctsw |
| Gins1 | 1.38E-128 | 0.274261341 | 0.271 | 0.01 | 2.80E-124 | 23 | Gins1 |
| Wdr76 | 2.76E-128 | 0.793388625 | 0.594 | 0.048 | 5.60E-124 | 23 | Wdr76 |
| Mcm61 | 3.18E-128 | 1.196066616 | 0.76 | 0.08 | 6.43E-124 | 23 | Mcm6 |
| Psat1 | 6.68E-126 | 0.347805001 | 0.312 | 0.014 | 1.35E-121 | 23 | Psat1 |
| Txk5 | 1.03E-124 | 1.144545354 | 0.906 | 0.106 | 2.08E-120 | 23 | Txk |
| Xcl15 | 1.94E-123 | 2.541178078 | 0.938 | 0.129 | 3.93E-119 | 23 | Xcl1 |
| Il2rb5 | 6.41E-123 | 1.631780965 | 0.938 | 0.119 | 1.30E-118 | 23 | Il2rb |
| Skap16 | 2.54E-122 | 1.946496568 | 0.969 | 0.133 | 5.15E-118 | 23 | Skap1 |
| Atp8b45 | 1.12E-121 | 1.496630104 | 0.823 | 0.096 | 2.26E-117 | 23 | Atp8b4 |
| Dut2 | 2.22E-121 | 1.931144216 | 0.896 | 0.128 | 4.49E-117 | 23 | Dut |
| Lck4 | 2.52E-121 | 1.19648861 | 0.865 | 0.102 | 5.10E-117 | 23 | Lck |
| Nkg75 | 1.38E-120 | 2.033182134 | 0.948 | 0.126 | 2.80E-116 | 23 | Nkg7 |
| Ptprcap6 | 1.74E-119 | 1.671574429 | 0.958 | 0.131 | 3.52E-115 | 23 | Ptprcap |
| Blm | 2.88E-118 | 0.559957358 | 0.479 | 0.034 | 5.82E-114 | 23 | Blm |
| Ncr14 | 1.30E-117 | 1.516677331 | 0.844 | 0.103 | 2.62E-113 | 23 | Ncr1 |

| | | | | | | | |
|---|---|---|---|---|---|---|---|
| Tnfrsf185 | 2.86E-116 | 0.730275876 | 0.719 | 0.073 | 5.79E-112 | 23 | Tnfrsf18 |
| Wee1 | 2.73E-115 | 0.547573148 | 0.479 | 0.035 | 5.52E-111 | 23 | Wee1 |
| Car24 | 1.09E-114 | 1.743111869 | 0.875 | 0.121 | 2.20E-110 | 23 | Car2 |
| Pole2 | 1.44E-114 | 0.378591599 | 0.354 | 0.019 | 2.91E-110 | 23 | Pole2 |
| Tox8 | 8.70E-114 | 1.663275916 | 0.906 | 0.12 | 1.76E-109 | 23 | Tox |
| Samd34 | 1.25E-113 | 1.000990578 | 0.771 | 0.086 | 2.54E-109 | 23 | Samd3 |
| Itgal6 | 2.58E-113 | 0.948650453 | 0.76 | 0.086 | 5.23E-109 | 23 | Itgal |
| Nsd22 | 3.96E-113 | 1.236987686 | 0.771 | 0.092 | 8.01E-109 | 23 | Nsd2 |
| Smc48 | 6.22E-112 | 2.262888692 | 0.938 | 0.162 | 1.26E-107 | 23 | Smc4 |
| Cyfip26 | 1.41E-111 | 1.048100102 | 0.771 | 0.09 | 2.85E-107 | 23 | Cyfip2 |
| Cdca7l1 | 1.58E-109 | 0.612143522 | 0.521 | 0.043 | 3.20E-105 | 23 | Cdca7l |
| Sdk12 | 2.08E-109 | 0.810746582 | 0.375 | 0.023 | 4.21E-105 | 23 | Sdk1 |
| Runx37 | 5.04E-108 | 1.69773493 | 0.948 | 0.148 | 1.02E-103 | 23 | Runx3 |
| Klrb1f4 | 6.40E-108 | 0.895404802 | 0.625 | 0.062 | 1.30E-103 | 23 | Klrb1f |
| Rfc31 | 8.63E-108 | 0.668910335 | 0.573 | 0.052 | 1.75E-103 | 23 | Rfc3 |
| Eri1 | 2.95E-107 | 0.74418797 | 0.677 | 0.072 | 5.97E-103 | 23 | Eri1 |
| Pold2 | 4.41E-107 | 0.651757365 | 0.594 | 0.056 | 8.93E-103 | 23 | Pold2 |
| Tubg1 | 8.18E-106 | 0.517829195 | 0.542 | 0.046 | 1.66E-101 | 23 | Tubg1 |
| Mctp25 | 8.59E-106 | 0.974127196 | 0.729 | 0.083 | 1.74E-101 | 23 | Mctp2 |
| Acap16 | 1.01E-105 | 0.798620315 | 0.667 | 0.07 | 2.04E-101 | 23 | Acap1 |
| Ptpn76 | 7.10E-104 | 0.748568537 | 0.594 | 0.058 | 1.44E-99 | 23 | Ptpn7 |
| Rac27 | 9.48E-104 | 1.998880628 | 0.979 | 0.177 | 1.92E-99 | 23 | Rac2 |
| Arhgap19 | 1.62E-103 | 0.38677807 | 0.302 | 0.016 | 3.27E-99 | 23 | Arhgap19 |
| 17-Sep | 2.08E-103 | 1.458947011 | 0.938 | 0.145 | 4.21E-99 | 23 | 1-Sep |
| Aoah4 | 3.16E-103 | 0.741463384 | 0.677 | 0.072 | 6.40E-99 | 23 | Aoah |
| Cd25 | 5.17E-102 | 0.980184303 | 0.719 | 0.083 | 1.05E-97 | 23 | Cd2 |
| Zap704 | 7.22E-101 | 0.61189653 | 0.583 | 0.056 | 1.46E-96 | 23 | Zap70 |
| Ikzf36 | 4.48E-100 | 1.342262077 | 0.865 | 0.119 | 9.07E-96 | 23 | Ikzf3 |
| Rnaseh2b | 1.49E-99 | 0.584379553 | 0.562 | 0.053 | 3.01E-95 | 23 | Rnaseh2b |
| Osbpl36 | 2.84E-99 | 1.166055912 | 0.854 | 0.119 | 5.75E-95 | 23 | Osbpl3 |
| Fcho1 | 3.08E-99 | 0.376627981 | 0.333 | 0.02 | 6.24E-95 | 23 | Fcho1 |
| Cep57l1 | 3.18E-99 | 0.572387497 | 0.5 | 0.043 | 6.44E-95 | 23 | Cep57l1 |
| Acot75 | 5.97E-99 | 0.696637231 | 0.677 | 0.076 | 1.21E-94 | 23 | Acot7 |
| Klrc24 | 2.00E-98 | 0.782280519 | 0.552 | 0.053 | 4.05E-94 | 23 | Klrc2 |
| Itgax6 | 2.23E-98 | 1.053856404 | 0.802 | 0.107 | 4.52E-94 | 23 | Itgax |
| Ipcef16 | 8.85E-98 | 1.3336077 | 0.823 | 0.117 | 1.79E-93 | 23 | Ipcef1 |
| Pglyrp25 | 1.27E-97 | 0.572833538 | 0.5 | 0.044 | 2.58E-93 | 23 | Pglyrp2 |
| Klrb1c4 | 1.31E-97 | 1.00900668 | 0.771 | 0.096 | 2.66E-93 | 23 | Klrb1c |
| Hcst6 | 1.92E-97 | 1.321593578 | 0.938 | 0.145 | 3.88E-93 | 23 | Hcst |
| Ints7 | 2.00E-97 | 0.536674442 | 0.531 | 0.049 | 4.05E-93 | 23 | Ints7 |
| Stat46 | 2.52E-97 | 1.005084809 | 0.812 | 0.106 | 5.10E-93 | 23 | Stat4 |
| Zfand4 | 2.94E-97 | 0.292180928 | 0.25 | 0.011 | 5.96E-93 | 23 | Zfand4 |
| Rpa3 | 4.58E-97 | 0.794810958 | 0.75 | 0.094 | 9.27E-93 | 23 | Rpa3 |

| | | | | | | | |
|---|---|---|---|---|---|---|---|
| Fam78a2 | 5.41E-97 | 0.477061712 | 0.479 | 0.04 | 1.09E-92 | 23 | Fam78a |
| Nlrc35 | 5.52E-96 | 0.682463062 | 0.573 | 0.057 | 1.12E-91 | 23 | Nlrc3 |
| Cenpq | 1.25E-95 | 0.418207464 | 0.562 | 0.054 | 2.53E-91 | 23 | Cenpq |
| Styk15 | 3.79E-95 | 1.163985335 | 0.76 | 0.097 | 7.66E-91 | 23 | Styk1 |
| Hist1h4h | 3.98E-95 | 1.131282181 | 0.573 | 0.059 | 8.06E-91 | 23 | Hist1h4h |
| Cd2445 | 4.04E-95 | 0.866813147 | 0.74 | 0.091 | 8.18E-91 | 23 | Cd244 |
| Cst74 | 4.81E-95 | 0.729620219 | 0.656 | 0.073 | 9.74E-91 | 23 | Cst7 |
| Dusp27 | 5.53E-95 | 1.055197424 | 0.771 | 0.101 | 1.12E-90 | 23 | Dusp2 |
| Gpr1744 | 2.23E-94 | 0.624084345 | 0.542 | 0.052 | 4.52E-90 | 23 | Gpr174 |
| Eomes3 | 2.76E-94 | 0.715112415 | 0.531 | 0.05 | 5.59E-90 | 23 | Eomes |
| Atp10a3 | 2.79E-94 | 0.617920252 | 0.51 | 0.047 | 5.65E-90 | 23 | Atp10a |
| Ptpn226 | 6.33E-94 | 1.21418193 | 0.823 | 0.118 | 1.28E-89 | 23 | Ptpn22 |
| Dnmt11 | 7.83E-94 | 1.114904332 | 0.75 | 0.103 | 1.59E-89 | 23 | Dnmt1 |
| Zfp367 | 8.37E-94 | 0.57359911 | 0.542 | 0.052 | 1.69E-89 | 23 | Zfp367 |
| Tbc1d10c6 | 1.18E-93 | 0.751490151 | 0.646 | 0.073 | 2.39E-89 | 23 | Tbc1d10c |
| Mb21d11 | 1.78E-93 | 0.449165354 | 0.448 | 0.037 | 3.61E-89 | 23 | Mb21d1 |
| Srgap35 | 2.36E-93 | 0.927176346 | 0.74 | 0.095 | 4.77E-89 | 23 | Srgap3 |
| Hist1h2be | 4.16E-93 | 0.806006754 | 0.542 | 0.054 | 8.42E-89 | 23 | Hist1h2be |
| Klra73 | 4.28E-93 | 1.715442979 | 0.635 | 0.074 | 8.66E-89 | 23 | Klra7 |
| Cdt1 | 9.83E-93 | 0.339536259 | 0.26 | 0.013 | 1.99E-88 | 23 | Cdt1 |
| Rasgrp16 | 1.10E-92 | 0.735322994 | 0.583 | 0.062 | 2.23E-88 | 23 | Rasgrp1 |
| Gimap74 | 1.59E-92 | 0.635350258 | 0.573 | 0.058 | 3.23E-88 | 23 | Gimap7 |
| Sep-67 | 1.74E-92 | 0.857490523 | 0.75 | 0.098 | 3.52E-88 | 23 | 6-Sep |
| Nfatc2ip | 3.31E-92 | 0.400642916 | 0.396 | 0.029 | 6.71E-88 | 23 | Nfatc2ip |
| Tipin1 | 3.31E-92 | 0.928605178 | 0.698 | 0.088 | 6.71E-88 | 23 | Tipin |
| Arsb5 | 3.60E-92 | 1.335531367 | 0.833 | 0.126 | 7.28E-88 | 23 | Arsb |
| Dapk26 | 5.92E-92 | 1.04088337 | 0.792 | 0.109 | 1.20E-87 | 23 | Dapk2 |
| Tfdp1 | 2.54E-91 | 0.565655217 | 0.531 | 0.052 | 5.15E-87 | 23 | Tfdp1 |
| S100a46 | 5.49E-91 | 2.105085724 | 0.823 | 0.128 | 1.11E-86 | 23 | S100a4 |
| Cdkn2c2 | 8.92E-91 | 0.611466482 | 0.573 | 0.059 | 1.81E-86 | 23 | Cdkn2c |
| Stk17b6 | 1.10E-90 | 1.253325861 | 0.958 | 0.161 | 2.22E-86 | 23 | Stk17b |
| Cd2264 | 1.58E-90 | 1.512749108 | 0.698 | 0.09 | 3.19E-86 | 23 | Cd226 |
| Dck2 | 9.05E-90 | 0.431092198 | 0.417 | 0.033 | 1.83E-85 | 23 | Dck |
| Selplg8 | 9.18E-90 | 1.259900018 | 0.917 | 0.154 | 1.86E-85 | 23 | Selplg |
| Reep4 | 2.74E-89 | 0.392054603 | 0.365 | 0.026 | 5.54E-85 | 23 | Reep4 |
| Pold3 | 2.77E-89 | 0.69110578 | 0.698 | 0.087 | 5.61E-85 | 23 | Pold3 |
| Clec2i2 | 2.89E-89 | 0.506795669 | 0.417 | 0.034 | 5.85E-85 | 23 | Clec2i |
| Serpina3g4 | 3.59E-89 | 0.768241174 | 0.604 | 0.067 | 7.28E-85 | 23 | Serpina3g |
| Cmc21 | 5.00E-89 | 0.715321109 | 0.812 | 0.115 | 1.01E-84 | 23 | Cmc2 |
| Zfp808 | 6.24E-89 | 0.312736867 | 0.292 | 0.017 | 1.26E-84 | 23 | Zfp808 |
| Anapc15 | 8.97E-89 | 0.490750367 | 0.573 | 0.06 | 1.82E-84 | 23 | Anapc15 |
| Lax14 | 9.37E-89 | 0.697109132 | 0.594 | 0.065 | 1.90E-84 | 23 | Lax1 |
| Hist2h2ac1 | 9.61E-89 | 1.493406548 | 0.604 | 0.072 | 1.95E-84 | 23 | Hist2h2ac |

| | | | | | | | |
|---|---|---|---|---|---|---|---|
| Klra43 | 1.32E-88 | 1.780447015 | 0.646 | 0.078 | 2.68E-84 | 23 | Klra4 |
| Pkp35 | 1.78E-88 | 0.780809683 | 0.604 | 0.069 | 3.61E-84 | 23 | Pkp3 |
| Pstpip16 | 2.73E-88 | 0.601211414 | 0.542 | 0.055 | 5.52E-84 | 23 | Pstpip1 |
| Hirip31 | 4.82E-88 | 0.568016687 | 0.479 | 0.044 | 9.75E-84 | 23 | Hirip3 |
| Jaml5 | 8.41E-88 | 0.703475921 | 0.552 | 0.058 | 1.70E-83 | 23 | Jaml |
| Pglyrp14 | 9.51E-88 | 0.68864344 | 0.719 | 0.092 | 1.93E-83 | 23 | Pglyrp1 |
| Rfwd31 | 2.18E-87 | 0.831861326 | 0.625 | 0.075 | 4.41E-83 | 23 | Rfwd3 |
| Ddx43 | 3.31E-87 | 0.310388692 | 0.302 | 0.018 | 6.69E-83 | 23 | Ddx43 |
| Hmgb26 | 3.77E-86 | 3.694153717 | 1 | 0.328 | 7.63E-82 | 23 | Hmgb2 |
| Sla24 | 7.78E-86 | 0.514107727 | 0.5 | 0.048 | 1.57E-81 | 23 | Sla2 |
| Rfc24 | 1.06E-85 | 0.944036649 | 0.812 | 0.124 | 2.15E-81 | 23 | Rfc2 |
| Ttf2 | 1.12E-85 | 0.409833822 | 0.365 | 0.027 | 2.26E-81 | 23 | Ttf2 |
| Hmgn22 | 4.71E-85 | 1.779890598 | 0.948 | 0.193 | 9.54E-81 | 23 | Hmgn2 |
| Il2rg6 | 4.79E-85 | 0.844674077 | 0.802 | 0.116 | 9.69E-81 | 23 | Il2rg |
| Impa2 | 1.08E-84 | 0.400058814 | 0.594 | 0.066 | 2.19E-80 | 23 | Impa2 |
| Topbp11 | 1.12E-84 | 0.786970622 | 0.708 | 0.095 | 2.26E-80 | 23 | Topbp1 |
| Nubp13 | 1.59E-84 | 0.65674158 | 0.74 | 0.1 | 3.22E-80 | 23 | Nubp1 |
| Klra93 | 1.81E-84 | 1.487886812 | 0.625 | 0.078 | 3.66E-80 | 23 | Klra9 |
| Gem5 | 2.06E-84 | 1.024645772 | 0.74 | 0.103 | 4.16E-80 | 23 | Gem |
| Cd964 | 5.08E-84 | 0.724879109 | 0.594 | 0.068 | 1.03E-79 | 23 | Cd96 |
| Kpna2 | 6.79E-84 | 0.288604618 | 0.281 | 0.017 | 1.38E-79 | 23 | Kpna2 |
| Tnfrsf94 | 7.25E-84 | 0.743686401 | 0.583 | 0.066 | 1.47E-79 | 23 | Tnfrsf9 |
| Mturn | 1.32E-83 | 0.376251593 | 0.406 | 0.033 | 2.67E-79 | 23 | Mturn |
| Itgae4 | 4.22E-83 | 1.240545562 | 0.573 | 0.066 | 8.55E-79 | 23 | Itgae |
| Slbp3 | 4.45E-83 | 1.238292076 | 0.854 | 0.15 | 9.00E-79 | 23 | Slbp |
| Rhof3 | 5.54E-83 | 0.457634036 | 0.427 | 0.037 | 1.12E-78 | 23 | Rhof |
| Tmsb15a | 1.17E-82 | 0.447229752 | 0.25 | 0.013 | 2.38E-78 | 23 | Tmsb15a |
| Ppil1 | 2.53E-82 | 0.513219586 | 0.542 | 0.058 | 5.12E-78 | 23 | Ppil1 |
| Map4k11 | 3.64E-82 | 0.324837229 | 0.323 | 0.022 | 7.37E-78 | 23 | Map4k1 |
| Skp2 | 6.17E-82 | 0.26265152 | 0.26 | 0.015 | 1.25E-77 | 23 | Skp2 |
| Ska2 | 9.11E-82 | 0.422137142 | 0.385 | 0.031 | 1.84E-77 | 23 | Ska2 |
| Sass6 | 1.16E-81 | 0.444740919 | 0.417 | 0.036 | 2.35E-77 | 23 | Sass6 |
| Pcna3 | 2.38E-81 | 0.967610995 | 0.781 | 0.121 | 4.82E-77 | 23 | Pcna |
| Myo1g3 | 6.59E-81 | 0.433865152 | 0.479 | 0.047 | 1.34E-76 | 23 | Myo1g |
| Epsti17 | 7.78E-81 | 1.418328933 | 0.938 | 0.174 | 1.57E-76 | 23 | Epsti1 |
| Syce2 | 8.79E-81 | 0.440283564 | 0.417 | 0.037 | 1.78E-76 | 23 | Syce2 |
| Gzmc3 | 8.83E-81 | 2.380346525 | 0.625 | 0.082 | 1.79E-76 | 23 | Gzmc |
| Rangap1 | 1.20E-80 | 0.632824858 | 0.583 | 0.069 | 2.42E-76 | 23 | Rangap1 |
| Rtel1 | 1.29E-80 | 0.272500957 | 0.25 | 0.014 | 2.61E-76 | 23 | Rtel1 |
| Klri24 | 3.77E-80 | 1.055743433 | 0.604 | 0.074 | 7.62E-76 | 23 | Klri2 |
| Chsy15 | 4.15E-80 | 1.076112412 | 0.812 | 0.127 | 8.41E-76 | 23 | Chsy1 |
| Dock56 | 4.28E-80 | 1.140000695 | 0.75 | 0.114 | 8.66E-76 | 23 | Dock5 |
| Ikzf17 | 4.92E-80 | 1.321864024 | 0.885 | 0.156 | 9.97E-76 | 23 | Ikzf1 |

| | | | | | | | |
|---|---|---|---|---|---|---|---|
| Orc61 | 6.31E-80 | 0.914247366 | 0.708 | 0.102 | 1.28E-75 | 23 | Orc6 |
| Ltb6 | 4.26E-79 | 1.090461437 | 0.854 | 0.135 | 8.62E-75 | 23 | Ltb |
| Mphosph9 | 7.00E-79 | 0.540167027 | 0.49 | 0.05 | 1.42E-74 | 23 | Mphosph9 |
| Myo1f7 | 7.98E-79 | 1.060824498 | 0.844 | 0.137 | 1.62E-74 | 23 | Myo1f |
| Gm14029 | 9.72E-79 | 0.352054497 | 0.344 | 0.026 | 1.97E-74 | 23 | Gm14029 |
| Rcc1 | 1.88E-78 | 0.360320894 | 0.333 | 0.025 | 3.80E-74 | 23 | Rcc1 |
| Chn27 | 3.66E-78 | 1.551775832 | 0.958 | 0.197 | 7.42E-74 | 23 | Chn2 |
| Maz3 | 4.29E-78 | 0.643486765 | 0.677 | 0.092 | 8.68E-74 | 23 | Maz |
| Sh2d2a5 | 5.94E-78 | 0.54460061 | 0.479 | 0.049 | 1.20E-73 | 23 | Sh2d2a |
| Pold1 | 8.96E-78 | 0.33445043 | 0.344 | 0.026 | 1.81E-73 | 23 | Pold1 |
| Jpt17 | 1.15E-77 | 1.482215541 | 0.979 | 0.222 | 2.32E-73 | 23 | Jpt1 |
| Hjurp3 | 1.23E-77 | 0.925198649 | 0.656 | 0.09 | 2.49E-73 | 23 | Hjurp |
| Hvcn14 | 3.92E-77 | 0.534704247 | 0.51 | 0.056 | 7.93E-73 | 23 | Hvcn1 |
| Ptprc7 | 4.26E-77 | 1.82774849 | 0.958 | 0.198 | 8.62E-73 | 23 | Ptprc |
| Ncf47 | 4.26E-77 | 0.77446022 | 0.75 | 0.112 | 8.62E-73 | 23 | Ncf4 |
| Suv39h1 | 5.05E-77 | 0.348739841 | 0.312 | 0.022 | 1.02E-72 | 23 | Suv39h1 |
| Sema4d7 | 1.82E-76 | 0.937969369 | 0.802 | 0.13 | 3.68E-72 | 23 | Sema4d |
| Card116 | 1.86E-76 | 0.767678133 | 0.625 | 0.082 | 3.76E-72 | 23 | Card11 |
| Sema4a6 | 2.97E-76 | 0.651356528 | 0.604 | 0.076 | 6.01E-72 | 23 | Sema4a |
| Klrc11 | 4.10E-76 | 0.693681725 | 0.344 | 0.027 | 8.31E-72 | 23 | Klrc1 |
| Coro1a9 | 4.22E-76 | 1.728983815 | 0.979 | 0.219 | 8.54E-72 | 23 | Coro1a |
| Cd487 | 5.42E-76 | 0.916463568 | 0.812 | 0.134 | 1.10E-71 | 23 | Cd48 |
| Ifitm104 | 8.63E-76 | 0.535053129 | 0.531 | 0.06 | 1.75E-71 | 23 | Ifitm10 |
| Coro2a6 | 1.02E-75 | 1.089782296 | 0.74 | 0.117 | 2.06E-71 | 23 | Coro2a |
| Oprm14 | 1.03E-75 | 0.723301422 | 0.521 | 0.06 | 2.08E-71 | 23 | Oprm1 |
| Fam111a3 | 6.30E-75 | 0.68199152 | 0.656 | 0.091 | 1.28E-70 | 23 | Fam111a |
| Siva14 | 1.01E-74 | 1.121816562 | 0.844 | 0.158 | 2.05E-70 | 23 | Siva1 |
| Hist1h4c | 1.39E-74 | 0.831760445 | 0.469 | 0.05 | 2.80E-70 | 23 | Hist1h4c |
| Sorl18 | 1.39E-74 | 0.881512752 | 0.719 | 0.111 | 2.82E-70 | 23 | Sorl1 |
| Was7 | 1.64E-74 | 0.518085086 | 0.583 | 0.072 | 3.32E-70 | 23 | Was |
| Tmpo6 | 1.83E-74 | 1.515816353 | 0.938 | 0.207 | 3.70E-70 | 23 | Tmpo |
| Gprin34 | 2.55E-74 | 0.730525074 | 0.531 | 0.062 | 5.17E-70 | 23 | Gprin3 |
| 2610524H06Rik | 3.20E-74 | 0.386017953 | 0.354 | 0.029 | 6.48E-70 | 23 | 2610524H06Rik |
| Dock119 | 3.76E-74 | 0.89642692 | 0.729 | 0.113 | 7.61E-70 | 23 | Dock11 |
| Cdh16 | 3.83E-74 | 0.756101382 | 0.583 | 0.074 | 7.76E-70 | 23 | Cdh1 |
| Ccl55 | 3.88E-74 | 1.374979567 | 0.865 | 0.151 | 7.85E-70 | 23 | Ccl5 |
| Rgs111 | 7.90E-74 | 1.043565631 | 0.833 | 0.14 | 1.60E-69 | 23 | Rgs1 |
| Cntrob | 8.50E-74 | 0.283150651 | 0.271 | 0.018 | 1.72E-69 | 23 | Cntrob |
| Slc38a16 | 8.60E-74 | 0.772930304 | 0.635 | 0.088 | 1.74E-69 | 23 | Slc38a1 |
| Ccr26 | 1.00E-73 | 1.049100298 | 0.625 | 0.086 | 2.03E-69 | 23 | Ccr2 |
| Inpp4b7 | 1.27E-73 | 1.781240704 | 0.948 | 0.203 | 2.57E-69 | 23 | Inpp4b |
| Prkcq5 | 1.43E-73 | 0.702208299 | 0.646 | 0.087 | 2.91E-69 | 23 | Prkcq |
| Arhgap458 | 1.50E-73 | 1.001291223 | 0.885 | 0.158 | 3.05E-69 | 23 | Arhgap45 |

| | | | | | | | |
|---|---|---|---|---|---|---|---|
| Trbc15 | 1.53E-73 | 0.987923901 | 0.625 | 0.086 | 3.10E-69 | 23 | Trbc1 |
| Dok24 | 1.57E-73 | 0.462115866 | 0.427 | 0.041 | 3.18E-69 | 23 | Dok2 |
| Hat11 | 1.95E-73 | 0.769818088 | 0.75 | 0.119 | 3.95E-69 | 23 | Hat1 |
| Rinl5 | 5.20E-73 | 0.88761152 | 0.802 | 0.131 | 1.05E-68 | 23 | Rinl |
| Cd1603 | 7.26E-73 | 0.614379444 | 0.438 | 0.044 | 1.47E-68 | 23 | Cd160 |
| Dctpp11 | 1.74E-72 | 0.903738301 | 0.802 | 0.139 | 3.53E-68 | 23 | Dctpp1 |
| Rhoh7 | 1.99E-72 | 0.722907935 | 0.698 | 0.103 | 4.03E-68 | 23 | Rhoh |
| Wdhd1 | 2.36E-72 | 0.361380203 | 0.26 | 0.017 | 4.77E-68 | 23 | Wdhd1 |
| Sash36 | 2.52E-72 | 0.619146365 | 0.51 | 0.06 | 5.11E-68 | 23 | Sash3 |
| Dock28 | 3.15E-72 | 1.411499797 | 0.938 | 0.185 | 6.38E-68 | 23 | Dock2 |
| Chl14 | 4.56E-72 | 0.609751873 | 0.479 | 0.052 | 9.23E-68 | 23 | Chl1 |
| Tmem1635 | 4.81E-72 | 1.118775452 | 0.625 | 0.087 | 9.73E-68 | 23 | Tmem163 |
| Itgb28 | 5.37E-72 | 0.836372947 | 0.677 | 0.101 | 1.09E-67 | 23 | Itgb2 |
| Klra83 | 1.14E-71 | 1.49998797 | 0.594 | 0.079 | 2.31E-67 | 23 | Klra8 |
| Akna7 | 1.33E-71 | 0.678082812 | 0.646 | 0.091 | 2.69E-67 | 23 | Akna |
| Nelfe | 1.64E-71 | 0.547421141 | 0.635 | 0.087 | 3.32E-67 | 23 | Nelfe |
| Rasal35 | 1.75E-71 | 0.654560805 | 0.615 | 0.083 | 3.53E-67 | 23 | Rasal3 |
| Cdca4 | 2.91E-71 | 0.399064932 | 0.385 | 0.035 | 5.89E-67 | 23 | Cdca4 |
| Exosc82 | 4.22E-71 | 0.875434479 | 0.781 | 0.136 | 8.54E-67 | 23 | Exosc8 |
| Ptpn67 | 6.01E-71 | 1.087201088 | 0.76 | 0.13 | 1.22E-66 | 23 | Ptpn6 |
| Jakmip15 | 6.25E-71 | 0.535345746 | 0.458 | 0.049 | 1.27E-66 | 23 | Jakmip1 |
| C230088H06Rik | 7.42E-71 | 0.352149767 | 0.292 | 0.021 | 1.50E-66 | 23 | C230088H06Rik |
| Sytl24 | 8.96E-71 | 0.630921246 | 0.49 | 0.055 | 1.81E-66 | 23 | Sytl2 |
| Slfn9 | 1.30E-70 | 0.43924783 | 0.458 | 0.049 | 2.64E-66 | 23 | Slfn9 |
| Lsp19 | 2.75E-70 | 0.794813464 | 0.812 | 0.139 | 5.57E-66 | 23 | Lsp1 |
| Cotl19 | 3.46E-70 | 1.460481118 | 0.969 | 0.224 | 7.01E-66 | 23 | Cotl1 |
| Cdkn2d5 | 7.80E-70 | 0.947775543 | 0.625 | 0.092 | 1.58E-65 | 23 | Cdkn2d |
| Itk5 | 8.07E-70 | 0.703442393 | 0.583 | 0.077 | 1.63E-65 | 23 | Itk |
| Anp32e1 | 1.15E-69 | 1.061734156 | 0.833 | 0.164 | 2.33E-65 | 23 | Anp32e |
| Gm26824 | 2.34E-69 | 1.252778492 | 0.583 | 0.081 | 4.73E-65 | 23 | Gm2682 |
| Chd3os | 2.61E-69 | 0.307090549 | 0.26 | 0.017 | 5.28E-65 | 23 | Chd3os |
| Dennd4a7 | 2.61E-69 | 1.452097536 | 0.896 | 0.187 | 5.29E-65 | 23 | Dennd4a |
| Lta4h4 | 2.83E-69 | 0.833721865 | 0.833 | 0.148 | 5.73E-65 | 23 | Lta4h |
| Cd2474 | 3.56E-69 | 0.654124207 | 0.438 | 0.046 | 7.21E-65 | 23 | Cd247 |
| Gata35 | 4.11E-69 | 0.448465388 | 0.469 | 0.051 | 8.33E-65 | 23 | Gata3 |
| Sulf26 | 5.03E-69 | 0.812385162 | 0.656 | 0.099 | 1.02E-64 | 23 | Sulf2 |
| Cenpc11 | 9.13E-69 | 0.776897554 | 0.521 | 0.066 | 1.85E-64 | 23 | Cenpc1 |
| E2f3 | 1.91E-68 | 0.429906471 | 0.469 | 0.052 | 3.87E-64 | 23 | E2f3 |
| Sdf2l16 | 2.00E-68 | 1.023915062 | 0.854 | 0.167 | 4.05E-64 | 23 | Sdf2l1 |
| Dusp10 | 2.04E-68 | 0.387545312 | 0.375 | 0.035 | 4.12E-64 | 23 | Dusp10 |
| Arl4c4 | 2.05E-68 | 0.598763596 | 0.552 | 0.072 | 4.15E-64 | 23 | Arl4c |
| Slf1 | 2.85E-68 | 0.564966894 | 0.604 | 0.082 | 5.78E-64 | 23 | Slf1 |
| Ddx394 | 3.00E-68 | 0.79895913 | 0.75 | 0.125 | 6.08E-64 | 23 | Ddx39 |

| | | | | | | | |
|---|---|---|---|---|---|---|---|
| Ptk2b8 | 4.01E-68 | 1.018169567 | 0.823 | 0.15 | 8.12E-64 | 23 | Ptk2b |
| Padi25 | 4.45E-68 | 0.621054866 | 0.656 | 0.096 | 9.01E-64 | 23 | Padi2 |
| Mis18a | 5.60E-68 | 0.345454014 | 0.427 | 0.044 | 1.13E-63 | 23 | Mis18a |
| Hist1h4i | 6.42E-68 | 0.368517141 | 0.292 | 0.022 | 1.30E-63 | 23 | Hist1h4i |
| Lin9 | 6.65E-68 | 0.3172149 | 0.323 | 0.026 | 1.35E-63 | 23 | Lin9 |
| Tbx214 | 1.41E-67 | 0.56066433 | 0.531 | 0.066 | 2.85E-63 | 23 | Tbx21 |
| Il21r6 | 1.41E-67 | 0.608014914 | 0.646 | 0.093 | 2.86E-63 | 23 | Il21r |
| Haus5 | 2.35E-67 | 0.281338982 | 0.271 | 0.019 | 4.75E-63 | 23 | Haus5 |
| Nrarp2 | 2.53E-67 | 0.419125555 | 0.5 | 0.058 | 5.13E-63 | 23 | Nrarp |
| Pde7a6 | 2.99E-67 | 1.071628731 | 0.833 | 0.155 | 6.05E-63 | 23 | Pde7a |
| Dock109 | 3.04E-67 | 1.55878315 | 0.938 | 0.208 | 6.16E-63 | 23 | Dock10 |
| Cdk5rap21 | 3.13E-67 | 0.517057894 | 0.562 | 0.073 | 6.34E-63 | 23 | Cdk5rap2 |
| Pde3b8 | 3.86E-67 | 0.977994863 | 0.75 | 0.127 | 7.82E-63 | 23 | Pde3b |
| Iqgap25 | 4.43E-67 | 0.688974391 | 0.677 | 0.104 | 8.97E-63 | 23 | Iqgap2 |
| Plp22 | 1.17E-66 | 0.658684405 | 0.667 | 0.102 | 2.37E-62 | 23 | Plp2 |
| Itgad2 | 1.43E-66 | 0.616148272 | 0.385 | 0.038 | 2.90E-62 | 23 | Itgad |
| Samd1 | 1.62E-66 | 0.413886279 | 0.427 | 0.045 | 3.29E-62 | 23 | Samd1 |
| Cntrl6 | 3.83E-66 | 0.85901105 | 0.729 | 0.123 | 7.76E-62 | 23 | Cntrl |
| Pacsin1 | 5.56E-66 | 0.385318362 | 0.292 | 0.022 | 1.13E-61 | 23 | Pacsin1 |
| Nudt213 | 6.19E-66 | 0.863141786 | 0.823 | 0.157 | 1.25E-61 | 23 | Nudt21 |
| Nasp6 | 1.09E-65 | 0.922714741 | 0.865 | 0.171 | 2.21E-61 | 23 | Nasp |
| Ighm7 | 1.46E-65 | 0.666818158 | 0.75 | 0.126 | 2.97E-61 | 23 | Ighm |
| Cep1924 | 1.49E-65 | 0.661898119 | 0.594 | 0.085 | 3.01E-61 | 23 | Cep192 |
| Elovl62 | 2.07E-65 | 0.452550037 | 0.448 | 0.05 | 4.19E-61 | 23 | Elovl6 |
| Gfod15 | 2.19E-65 | 0.559398453 | 0.531 | 0.069 | 4.44E-61 | 23 | Gfod1 |
| Mast23 | 3.85E-65 | 0.619184897 | 0.542 | 0.072 | 7.79E-61 | 23 | Mast2 |
| Ncaph21 | 4.68E-65 | 0.556098965 | 0.583 | 0.081 | 9.48E-61 | 23 | Ncaph2 |
| Gzmb4 | 5.99E-65 | 1.74376964 | 0.573 | 0.082 | 1.21E-60 | 23 | Gzmb |
| Slc14a15 | 6.15E-65 | 0.481819524 | 0.427 | 0.046 | 1.24E-60 | 23 | Slc14a1 |
| Magohb3 | 7.26E-65 | 0.731140408 | 0.719 | 0.121 | 1.47E-60 | 23 | Magohb |
| Unc1196 | 8.92E-65 | 0.547436969 | 0.594 | 0.084 | 1.81E-60 | 23 | Unc119 |
| Hist2h2aa1 | 1.25E-64 | 0.458743653 | 0.323 | 0.028 | 2.53E-60 | 23 | Hist2h2aa1 |
| Gm268273 | 2.00E-64 | 0.546959356 | 0.448 | 0.051 | 4.04E-60 | 23 | Gm26827 |
| Tespa16 | 4.26E-64 | 0.593495853 | 0.562 | 0.076 | 8.62E-60 | 23 | Tespa1 |
| Hmgb3 | 4.59E-64 | 0.429863431 | 0.521 | 0.066 | 9.30E-60 | 23 | Hmgb3 |
| Rttn | 6.37E-64 | 0.561845178 | 0.5 | 0.063 | 1.29E-59 | 23 | Rttn |
| Katnb1 | 6.57E-64 | 0.260622877 | 0.25 | 0.017 | 1.33E-59 | 23 | Katnb1 |
| Slc16a101 | 7.69E-64 | 0.673442116 | 0.427 | 0.048 | 1.56E-59 | 23 | Slc16a10 |
| Impdh21 | 8.23E-64 | 0.504853185 | 0.708 | 0.113 | 1.67E-59 | 23 | Impdh2 |
| Mir142hg8 | 9.87E-64 | 1.013921768 | 0.875 | 0.17 | 2.00E-59 | 23 | Mir142hg |
| BC052040 | 1.09E-63 | 0.500647991 | 0.49 | 0.06 | 2.21E-59 | 23 | BC052040 |
| Gpr1714 | 1.83E-63 | 0.50453349 | 0.427 | 0.047 | 3.70E-59 | 23 | Gpr171 |
| Aim28 | 2.59E-63 | 0.673919869 | 0.656 | 0.103 | 5.24E-59 | 23 | Aim2 |

| | | | | | | | |
|---|---|---|---|---|---|---|---|
| Cd5210 | 2.65E-63 | 2.116547217 | 0.969 | 0.282 | 5.36E-59 | 23 | Cd52 |
| Arhgdib10 | 4.48E-63 | 1.355035977 | 0.969 | 0.233 | 9.06E-59 | 23 | Arhgdib |
| Kcnq58 | 5.91E-63 | 1.105766759 | 0.51 | 0.068 | 1.20E-58 | 23 | Kcnq5 |
| Rps6ka58 | 6.12E-63 | 0.895889982 | 0.656 | 0.108 | 1.24E-58 | 23 | Rps6ka5 |
| B4galnt16 | 9.42E-63 | 0.66167596 | 0.625 | 0.095 | 1.91E-58 | 23 | B4galnt1 |
| Thy16 | 1.25E-62 | 0.706161443 | 0.479 | 0.06 | 2.54E-58 | 23 | Thy1 |
| Rassf56 | 2.10E-62 | 0.500154368 | 0.521 | 0.068 | 4.25E-58 | 23 | Rassf5 |
| Mthfd21 | 2.78E-62 | 0.275031044 | 0.406 | 0.043 | 5.62E-58 | 23 | Mthfd2 |
| Ccdc88c7 | 2.87E-62 | 0.751451991 | 0.635 | 0.099 | 5.82E-58 | 23 | Ccdc88c |
| Mcph1 | 3.26E-62 | 0.329651368 | 0.406 | 0.043 | 6.60E-58 | 23 | Mcph1 |
| Fam107b7 | 3.35E-62 | 0.6050096 | 0.719 | 0.12 | 6.77E-58 | 23 | Fam107b |
| Sla8 | 3.50E-62 | 0.509059599 | 0.542 | 0.073 | 7.09E-58 | 23 | Sla |
| Emp313 | 5.18E-62 | 1.544945789 | 0.958 | 0.259 | 1.05E-57 | 23 | Emp3 |
| Ung | 5.58E-62 | 0.255939638 | 0.271 | 0.021 | 1.13E-57 | 23 | Ung |
| Prkcz2 | 6.35E-62 | 0.405943109 | 0.469 | 0.056 | 1.29E-57 | 23 | Prkcz |
| Gas77 | 1.27E-61 | 1.154097067 | 0.875 | 0.182 | 2.56E-57 | 23 | Gas7 |
| Me25 | 1.35E-61 | 0.558974257 | 0.542 | 0.075 | 2.73E-57 | 23 | Me2 |
| Ssbp210 | 1.55E-61 | 0.900609003 | 0.76 | 0.14 | 3.15E-57 | 23 | Ssbp2 |
| Mllt37 | 1.76E-61 | 0.93833627 | 0.792 | 0.151 | 3.56E-57 | 23 | Mllt3 |
| Tex9 | 2.15E-61 | 0.437026563 | 0.354 | 0.035 | 4.36E-57 | 23 | Tex9 |
| Znrf36 | 3.15E-61 | 0.751546941 | 0.656 | 0.107 | 6.38E-57 | 23 | Znrf3 |
| Limd28 | 3.55E-61 | 0.702292925 | 0.802 | 0.148 | 7.19E-57 | 23 | Limd2 |
| Selenoh1 | 4.34E-61 | 1.172206093 | 0.854 | 0.194 | 8.78E-57 | 23 | Selenoh |
| Gimap36 | 4.41E-61 | 0.647903094 | 0.646 | 0.1 | 8.92E-57 | 23 | Gimap3 |
| Arnt23 | 4.62E-61 | 0.630639251 | 0.406 | 0.045 | 9.35E-57 | 23 | Arnt2 |
| H2afv3 | 5.25E-61 | 2.112758701 | 0.99 | 0.379 | 1.06E-56 | 23 | H2afv |
| Emid15 | 5.51E-61 | 0.696351265 | 0.667 | 0.108 | 1.12E-56 | 23 | Emid1 |
| Nxt11 | 5.57E-61 | 0.612649149 | 0.635 | 0.101 | 1.13E-56 | 23 | Nxt1 |
| Vars6 | 7.54E-61 | 0.482599448 | 0.573 | 0.082 | 1.53E-56 | 23 | Vars |
| Rbbp83 | 7.83E-61 | 0.549813924 | 0.646 | 0.101 | 1.58E-56 | 23 | Rbbp8 |
| Tubb57 | 8.00E-61 | 2.439910819 | 0.99 | 0.381 | 1.62E-56 | 23 | Tubb5 |
| Ppp3cc5 | 8.13E-61 | 0.994574414 | 0.76 | 0.141 | 1.65E-56 | 23 | Ppp3cc |
| Celf210 | 1.07E-60 | 1.00452539 | 0.927 | 0.202 | 2.16E-56 | 23 | Celf2 |
| Lsm21 | 2.09E-60 | 0.689253605 | 0.802 | 0.153 | 4.24E-56 | 23 | Lsm2 |
| Prkcb8 | 3.48E-60 | 0.982824098 | 0.802 | 0.156 | 7.05E-56 | 23 | Prkcb |
| Suz122 | 4.54E-60 | 1.015712716 | 0.854 | 0.184 | 9.20E-56 | 23 | Suz12 |
| Pik3cd7 | 5.04E-60 | 0.775706441 | 0.656 | 0.108 | 1.02E-55 | 23 | Pik3cd |
| Alyref1 | 5.18E-60 | 1.105185824 | 0.906 | 0.213 | 1.05E-55 | 23 | Alyref |
| Ezr4 | 5.67E-60 | 0.624984838 | 0.75 | 0.136 | 1.15E-55 | 23 | Ezr |
| Rpa12 | 5.89E-60 | 0.559563171 | 0.615 | 0.095 | 1.19E-55 | 23 | Rpa1 |
| Esyt18 | 9.78E-60 | 0.721786438 | 0.75 | 0.135 | 1.98E-55 | 23 | Esyt1 |
| Fzr1 | 1.57E-59 | 0.3248756 | 0.406 | 0.045 | 3.17E-55 | 23 | Fzr1 |
| Haus32 | 1.61E-59 | 0.484220639 | 0.531 | 0.073 | 3.26E-55 | 23 | Haus3 |

| | | | | | | | |
|---|---|---|---|---|---|---|---|
| Gpr141 | 2.20E-59 | 0.3160575 | 0.26 | 0.02 | 4.44E-55 | 23 | Gpr141 |
| Bin28 | 2.40E-59 | 0.757111895 | 0.719 | 0.127 | 4.85E-55 | 23 | Bin2 |
| Camkmt4 | 3.00E-59 | 1.02601193 | 0.74 | 0.139 | 6.08E-55 | 23 | Camkmt |
| Bcl27 | 3.29E-59 | 1.148225735 | 0.927 | 0.207 | 6.67E-55 | 23 | Bcl2 |
| Ypel1 | 3.62E-59 | 0.256687719 | 0.26 | 0.02 | 7.34E-55 | 23 | Ypel1 |
| Lcp18 | 5.25E-59 | 1.359678303 | 0.938 | 0.238 | 1.06E-54 | 23 | Lcp1 |
| Cdc25b | 5.69E-59 | 0.296513084 | 0.25 | 0.019 | 1.15E-54 | 23 | Cdc25b |
| Ckap55 | 6.64E-59 | 0.987689433 | 0.74 | 0.142 | 1.34E-54 | 23 | Ckap5 |
| Gpr683 | 7.04E-59 | 0.428525717 | 0.406 | 0.046 | 1.42E-54 | 23 | Gpr68 |
| N4bp2l13 | 1.11E-58 | 0.427173343 | 0.448 | 0.055 | 2.25E-54 | 23 | N4bp2l1 |
| Itgb76 | 1.22E-58 | 0.537511807 | 0.479 | 0.062 | 2.48E-54 | 23 | Itgb7 |
| B4galt56 | 1.45E-58 | 0.609841213 | 0.604 | 0.094 | 2.94E-54 | 23 | B4galt5 |
| Dtymk2 | 1.47E-58 | 0.608557334 | 0.74 | 0.135 | 2.98E-54 | 23 | Dtymk |
| Ugcg6 | 2.38E-58 | 1.008261005 | 0.833 | 0.171 | 4.81E-54 | 23 | Ugcg |
| Hpf12 | 2.80E-58 | 0.795950745 | 0.875 | 0.19 | 5.67E-54 | 23 | Hpf1 |
| Apol7e4 | 3.09E-58 | 0.499488837 | 0.427 | 0.05 | 6.26E-54 | 23 | Apol7e |
| Gpr343 | 3.40E-58 | 0.35749537 | 0.469 | 0.059 | 6.89E-54 | 23 | Gpr34 |
| Rnf26 | 3.62E-58 | 0.251350734 | 0.25 | 0.019 | 7.32E-54 | 23 | Rnf26 |
| Lbr10 | 6.17E-58 | 0.881421507 | 0.771 | 0.152 | 1.25E-53 | 23 | Lbr |
| Herc35 | 8.05E-58 | 0.433963103 | 0.521 | 0.071 | 1.63E-53 | 23 | Herc3 |
| Cdkn2aipnl | 8.51E-58 | 0.390329787 | 0.521 | 0.072 | 1.72E-53 | 23 | Cdkn2aipnl |
| Cdc25a | 1.09E-57 | 0.266053641 | 0.333 | 0.032 | 2.22E-53 | 23 | Cdc25a |
| Psph1 | 1.96E-57 | 0.444906072 | 0.5 | 0.068 | 3.96E-53 | 23 | Psph |
| Cd377 | 3.40E-57 | 0.678660997 | 0.667 | 0.114 | 6.89E-53 | 23 | Cd37 |
| Fxn | 3.45E-57 | 0.351180677 | 0.469 | 0.06 | 6.98E-53 | 23 | Fxn |
| Fam105a7 | 5.37E-57 | 0.560182544 | 0.719 | 0.125 | 1.09E-52 | 23 | Fam105a |
| Apbb1ip8 | 5.79E-57 | 0.75216845 | 0.854 | 0.173 | 1.17E-52 | 23 | Apbb1ip |
| Laptm59 | 5.89E-57 | 0.897552542 | 0.958 | 0.21 | 1.19E-52 | 23 | Laptm5 |
| Rnf433 | 7.56E-57 | 0.559800465 | 0.417 | 0.05 | 1.53E-52 | 23 | Rnf43 |
| Nup62 | 8.72E-57 | 0.340640232 | 0.385 | 0.043 | 1.77E-52 | 23 | Nup62 |
| Stk263 | 1.33E-56 | 0.376712938 | 0.396 | 0.045 | 2.70E-52 | 23 | Stk26 |
| Cep572 | 1.49E-56 | 0.723105602 | 0.635 | 0.108 | 3.02E-52 | 23 | Cep57 |
| Trbc25 | 3.48E-56 | 1.064479746 | 0.479 | 0.066 | 7.04E-52 | 23 | Trbc2 |
| Fermt38 | 3.69E-56 | 0.508241003 | 0.677 | 0.114 | 7.46E-52 | 23 | Fermt3 |
| Efhd28 | 5.02E-56 | 0.9121192 | 0.865 | 0.186 | 1.02E-51 | 23 | Efhd2 |
| A430078G23Rik2 | 6.73E-56 | 0.31909653 | 0.302 | 0.028 | 1.36E-51 | 23 | A430078G23Rik |
| Traf16 | 6.83E-56 | 0.544093291 | 0.469 | 0.062 | 1.38E-51 | 23 | Traf1 |
| Psd46 | 7.21E-56 | 0.522669081 | 0.479 | 0.065 | 1.46E-51 | 23 | Psd4 |
| Adcy79 | 7.47E-56 | 0.710512264 | 0.656 | 0.114 | 1.51E-51 | 23 | Adcy7 |
| Syk5 | 8.96E-56 | 0.516249348 | 0.49 | 0.067 | 1.81E-51 | 23 | Syk |
| Haus1 | 1.27E-55 | 0.267835151 | 0.354 | 0.037 | 2.57E-51 | 23 | Haus1 |
| Arid5a | 1.30E-55 | 0.305701018 | 0.365 | 0.039 | 2.62E-51 | 23 | Arid5a |
| Fam49b8 | 2.08E-55 | 1.087515393 | 0.948 | 0.237 | 4.22E-51 | 23 | Fam49b |

| | | | | | | | |
|---|---|---|---|---|---|---|---|
| Msh21 | 2.10E-55 | 0.475220842 | 0.531 | 0.077 | 4.26E-51 | 23 | Msh2 |
| Polh | 2.40E-55 | 0.280643076 | 0.281 | 0.024 | 4.85E-51 | 23 | Polh |
| Hsd11b15 | 2.50E-55 | 0.62549907 | 0.583 | 0.092 | 5.07E-51 | 23 | Hsd11b1 |
| Rab375 | 2.97E-55 | 0.383638515 | 0.385 | 0.044 | 6.01E-51 | 23 | Rab37 |
| Fhl25 | 3.09E-55 | 0.458910103 | 0.375 | 0.042 | 6.25E-51 | 23 | Fhl2 |
| Tmsb107 | 5.90E-55 | 2.178258185 | 1 | 0.501 | 1.19E-50 | 23 | Tmsb10 |
| Ppih | 1.19E-54 | 0.339406529 | 0.438 | 0.055 | 2.41E-50 | 23 | Ppih |
| Plek6 | 2.07E-54 | 0.644738141 | 0.698 | 0.124 | 4.18E-50 | 23 | Plek |
| Ccr57 | 2.10E-54 | 0.82995038 | 0.708 | 0.132 | 4.26E-50 | 23 | Ccr5 |
| Vipr22 | 2.94E-54 | 0.341013961 | 0.323 | 0.032 | 5.96E-50 | 23 | Vipr2 |
| Dock88 | 3.56E-54 | 1.28009717 | 0.885 | 0.218 | 7.20E-50 | 23 | Dock8 |
| Grap28 | 5.00E-54 | 0.669720235 | 0.625 | 0.106 | 1.01E-49 | 23 | Grap2 |
| Tbc1d31 | 5.29E-54 | 0.302738923 | 0.281 | 0.025 | 1.07E-49 | 23 | Tbc1d31 |
| Hcls17 | 6.20E-54 | 0.535571948 | 0.698 | 0.125 | 1.25E-49 | 23 | Hcls1 |
| Cbx55 | 7.86E-54 | 0.944556524 | 0.833 | 0.188 | 1.59E-49 | 23 | Cbx5 |
| Plcl29 | 7.90E-54 | 0.704578374 | 0.646 | 0.114 | 1.60E-49 | 23 | Plcl2 |
| Arhgap308 | 8.71E-54 | 0.566894646 | 0.646 | 0.11 | 1.76E-49 | 23 | Arhgap30 |
| Mis12 | 1.03E-53 | 0.302383961 | 0.479 | 0.065 | 2.09E-49 | 23 | Mis12 |
| Nt5c3b | 1.19E-53 | 0.490027399 | 0.531 | 0.08 | 2.41E-49 | 23 | Nt5c3b |
| AB1246112 | 1.22E-53 | 0.276183249 | 0.312 | 0.03 | 2.47E-49 | 23 | AB124611 |
| Frmd8 | 1.39E-53 | 0.407646292 | 0.448 | 0.059 | 2.82E-49 | 23 | Frmd8 |
| Gm19585 | 1.79E-53 | 0.342512437 | 0.281 | 0.025 | 3.62E-49 | 23 | Gm19585 |
| Rnaseh2a | 2.28E-53 | 0.35167177 | 0.438 | 0.056 | 4.62E-49 | 23 | Rnaseh2a |
| Suco8 | 2.82E-53 | 0.697766275 | 0.771 | 0.153 | 5.71E-49 | 23 | Suco |
| Pbdc1 | 3.67E-53 | 0.456049624 | 0.667 | 0.118 | 7.43E-49 | 23 | Pbdc1 |
| Ift80 | 4.59E-53 | 0.310110782 | 0.292 | 0.027 | 9.30E-49 | 23 | Ift80 |
| H2afz5 | 5.30E-53 | 2.064759673 | 0.99 | 0.609 | 1.07E-48 | 23 | H2afz |
| Lat27 | 6.87E-53 | 0.72513331 | 0.698 | 0.132 | 1.39E-48 | 23 | Lat2 |
| Snrnp25 | 7.28E-53 | 0.278293526 | 0.51 | 0.073 | 1.47E-48 | 23 | Snrnp25 |
| Spin2c1 | 7.81E-53 | 0.327742613 | 0.344 | 0.037 | 1.58E-48 | 23 | Spin2c |
| Ran5 | 8.24E-53 | 1.536045888 | 0.99 | 0.4 | 1.67E-48 | 23 | Ran |
| Hspa14 | 9.70E-53 | 0.595460381 | 0.719 | 0.137 | 1.96E-48 | 23 | Hspa14 |
| Rif15 | 1.13E-52 | 0.850218347 | 0.75 | 0.154 | 2.28E-48 | 23 | Rif1 |
| Hmgn5 | 1.20E-52 | 0.493746444 | 0.75 | 0.145 | 2.43E-48 | 23 | Hmgn5 |
| Fmnl18 | 1.23E-52 | 0.529556596 | 0.573 | 0.092 | 2.48E-48 | 23 | Fmnl1 |
| Ttc74 | 1.24E-52 | 0.503380644 | 0.604 | 0.102 | 2.51E-48 | 23 | Ttc7 |
| Rabgap1l6 | 1.58E-52 | 1.245617251 | 0.948 | 0.257 | 3.20E-48 | 23 | Rabgap1l |
| Haus81 | 1.89E-52 | 0.293445761 | 0.427 | 0.054 | 3.83E-48 | 23 | Haus8 |
| Gpaa1 | 2.96E-52 | 0.3743665 | 0.438 | 0.057 | 5.98E-48 | 23 | Gpaa1 |
| Arhgap158 | 3.16E-52 | 1.053650711 | 0.854 | 0.188 | 6.40E-48 | 23 | Arhgap15 |
| Matk2 | 3.77E-52 | 0.412793348 | 0.365 | 0.042 | 7.63E-48 | 23 | Matk |
| Ssrp1 | 4.72E-52 | 0.742975384 | 0.812 | 0.177 | 9.56E-48 | 23 | Ssrp1 |
| Sms6 | 6.04E-52 | 0.65864491 | 0.74 | 0.15 | 1.22E-47 | 23 | Sms |

| | | | | | | | |
|---|---|---|---|---|---|---|---|
| Vav17 | 8.18E-52 | 0.430338023 | 0.531 | 0.081 | 1.66E-47 | 23 | Vav1 |
| Nup107 | 1.56E-51 | 0.296107423 | 0.333 | 0.036 | 3.16E-47 | 23 | Nup107 |
| Cysltr22 | 1.90E-51 | 0.38220739 | 0.354 | 0.04 | 3.85E-47 | 23 | Cysltr2 |
| Sh3bp13 | 2.01E-51 | 0.368642134 | 0.406 | 0.051 | 4.07E-47 | 23 | Sh3bp1 |
| Grk6 | 2.07E-51 | 0.395739065 | 0.469 | 0.065 | 4.19E-47 | 23 | Grk6 |
| Traf3ip36 | 2.37E-51 | 0.474165371 | 0.521 | 0.079 | 4.79E-47 | 23 | Traf3ip3 |
| Lpxn7 | 2.38E-51 | 0.530823911 | 0.573 | 0.093 | 4.83E-47 | 23 | Lpxn |
| Usp16 | 2.57E-51 | 0.746527041 | 0.719 | 0.146 | 5.19E-47 | 23 | Usp1 |
| Tuba1b7 | 2.64E-51 | 2.054319012 | 0.917 | 0.323 | 5.35E-47 | 23 | Tuba1b |
| Mtbp | 2.66E-51 | 0.296244989 | 0.25 | 0.021 | 5.39E-47 | 23 | Mtbp |
| Satb16 | 2.93E-51 | 0.445503483 | 0.458 | 0.063 | 5.93E-47 | 23 | Satb1 |
| Npm34 | 3.16E-51 | 0.553114211 | 0.698 | 0.133 | 6.39E-47 | 23 | Npm3 |
| Zfp8272 | 3.81E-51 | 0.408441661 | 0.458 | 0.063 | 7.72E-47 | 23 | Zfp827 |
| Tbl1xr18 | 4.78E-51 | 0.689773844 | 0.781 | 0.164 | 9.68E-47 | 23 | Tbl1xr1 |
| Smc54 | 5.46E-51 | 0.677677365 | 0.646 | 0.118 | 1.11E-46 | 23 | Smc5 |
| Fyb8 | 5.55E-51 | 0.70027549 | 0.854 | 0.184 | 1.12E-46 | 23 | Fyb |
| Rtn4rl12 | 6.35E-51 | 0.389149928 | 0.396 | 0.049 | 1.28E-46 | 23 | Rtn4rl1 |
| Lgals19 | 7.25E-51 | 1.691728814 | 0.667 | 0.135 | 1.47E-46 | 23 | Lgals1 |
| Klra33 | 7.92E-51 | 1.069042044 | 0.375 | 0.046 | 1.60E-46 | 23 | Klra3 |
| Med141 | 8.43E-51 | 0.515148516 | 0.604 | 0.103 | 1.71E-46 | 23 | Med14 |
| Cbfb8 | 8.91E-51 | 0.743786633 | 0.75 | 0.155 | 1.80E-46 | 23 | Cbfb |
| Vim14 | 9.64E-51 | 1.481190724 | 0.979 | 0.306 | 1.95E-46 | 23 | Vim |
| Mob3a5 | 1.05E-50 | 0.485488411 | 0.521 | 0.081 | 2.12E-46 | 23 | Mob3a |
| Snrnp402 | 1.23E-50 | 0.575714157 | 0.729 | 0.143 | 2.49E-46 | 23 | Snrnp40 |
| Ppat | 1.26E-50 | 0.377124163 | 0.448 | 0.061 | 2.55E-46 | 23 | Ppat |
| Pik3cg4 | 1.67E-50 | 0.411833438 | 0.417 | 0.055 | 3.39E-46 | 23 | Pik3cg |
| Cep2951 | 2.79E-50 | 0.404066596 | 0.5 | 0.074 | 5.64E-46 | 23 | Cep295 |
| Runx19 | 3.52E-50 | 0.681193138 | 0.688 | 0.131 | 7.13E-46 | 23 | Runx1 |
| Il12rb24 | 4.54E-50 | 0.380496663 | 0.396 | 0.049 | 9.19E-46 | 23 | Il12rb2 |
| Ptprj9 | 4.68E-50 | 0.648420989 | 0.75 | 0.151 | 9.48E-46 | 23 | Ptprj |
| Arhgap48 | 4.97E-50 | 0.468643458 | 0.427 | 0.058 | 1.01E-45 | 23 | Arhgap4 |
| Metrnl5 | 5.75E-50 | 0.42396308 | 0.604 | 0.103 | 1.16E-45 | 23 | Metrnl |
| Slc9a98 | 5.86E-50 | 0.775628476 | 0.74 | 0.151 | 1.19E-45 | 23 | Slc9a9 |
| Rbm38 | 6.95E-50 | 1.611669966 | 0.99 | 0.477 | 1.41E-45 | 23 | Rbm3 |
| Nfyc | 8.54E-50 | 0.388877873 | 0.625 | 0.108 | 1.73E-45 | 23 | Nfyc |
| Erg281 | 1.06E-49 | 0.614627812 | 0.635 | 0.118 | 2.15E-45 | 23 | Erg28 |
| Ttc39b7 | 1.06E-49 | 0.610187093 | 0.698 | 0.135 | 2.15E-45 | 23 | Ttc39b |
| Dek3 | 1.26E-49 | 1.630041288 | 0.969 | 0.373 | 2.55E-45 | 23 | Dek |
| Spry24 | 1.51E-49 | 0.525331904 | 0.521 | 0.082 | 3.05E-45 | 23 | Spry2 |
| Fcer1g9 | 1.57E-49 | 1.644748698 | 0.948 | 0.27 | 3.18E-45 | 23 | Fcer1g |
| Spn3 | 1.78E-49 | 0.417145757 | 0.365 | 0.044 | 3.59E-45 | 23 | Spn |
| Hdgfl21 | 2.11E-49 | 0.463237399 | 0.542 | 0.087 | 4.28E-45 | 23 | Hdgfl2 |
| Slc9a3r15 | 4.02E-49 | 0.465784541 | 0.615 | 0.109 | 8.14E-45 | 23 | Slc9a3r1 |

| | | | | | | | |
|---|---|---|---|---|---|---|---|
| Rcc24 | 4.85E-49 | 0.458285952 | 0.625 | 0.112 | 9.82E-45 | 23 | Rcc2 |
| Sept1110 | 4.87E-49 | 0.871985223 | 0.865 | 0.203 | 9.86E-45 | 23 | 11-Sep |
| Ppm1g4 | 6.44E-49 | 0.599716194 | 0.75 | 0.156 | 1.30E-44 | 23 | Ppm1g |
| Ube2s1 | 7.50E-49 | 1.699318662 | 0.906 | 0.311 | 1.52E-44 | 23 | Ube2s |
| Tnni11 | 8.11E-49 | 0.566849529 | 0.302 | 0.032 | 1.64E-44 | 23 | Tnni1 |
| Apobec38 | 1.04E-48 | 0.96294732 | 0.875 | 0.225 | 2.11E-44 | 23 | Apobec3 |
| Gramd1a4 | 1.35E-48 | 0.506552255 | 0.583 | 0.101 | 2.74E-44 | 23 | Gramd1a |
| Eif1ad1 | 1.60E-48 | 0.423578235 | 0.646 | 0.118 | 3.24E-44 | 23 | Eif1ad |
| Smarcc14 | 1.61E-48 | 0.574615634 | 0.708 | 0.142 | 3.26E-44 | 23 | Smarcc1 |
| Cxcr32 | 1.62E-48 | 0.384982975 | 0.333 | 0.038 | 3.28E-44 | 23 | Cxcr3 |
| Tiam23 | 1.69E-48 | 0.513778534 | 0.396 | 0.052 | 3.41E-44 | 23 | Tiam2 |
| Smyd36 | 3.17E-48 | 1.36025139 | 0.844 | 0.227 | 6.41E-44 | 23 | Smyd3 |
| Isy11 | 3.25E-48 | 0.332329908 | 0.479 | 0.071 | 6.57E-44 | 23 | Isy1 |
| Arhgap95 | 3.57E-48 | 0.489473514 | 0.615 | 0.111 | 7.23E-44 | 23 | Arhgap9 |
| Fnbp19 | 4.41E-48 | 0.931724564 | 0.906 | 0.23 | 8.92E-44 | 23 | Fnbp1 |
| Odf2 | 9.01E-48 | 0.480957401 | 0.521 | 0.084 | 1.82E-43 | 23 | Odf2 |
| Kcnip33 | 9.55E-48 | 0.411496551 | 0.375 | 0.047 | 1.93E-43 | 23 | Kcnip3 |
| Cytip6 | 1.18E-47 | 0.424366057 | 0.552 | 0.092 | 2.39E-43 | 23 | Cytip |
| Lsm52 | 1.41E-47 | 0.825049781 | 0.865 | 0.22 | 2.86E-43 | 23 | Lsm5 |
| Rbl25 | 1.46E-47 | 0.573907963 | 0.688 | 0.135 | 2.95E-43 | 23 | Rbl2 |
| Leprotl15 | 1.49E-47 | 0.87506051 | 0.865 | 0.217 | 3.02E-43 | 23 | Leprotl1 |
| Hist1h4d2 | 1.59E-47 | 2.104744316 | 0.854 | 0.257 | 3.23E-43 | 23 | Hist1h4d |
| Vps16 | 1.72E-47 | 0.279483282 | 0.406 | 0.054 | 3.48E-43 | 23 | Vps16 |
| A930015D03Rik5 | 1.79E-47 | 0.617211978 | 0.594 | 0.106 | 3.62E-43 | 23 | A930015D03Rik |
| Xist15 | 2.16E-47 | 1.70572959 | 0.99 | 0.464 | 4.37E-43 | 23 | Xist |
| Dtnb5 | 2.23E-47 | 0.50845431 | 0.625 | 0.114 | 4.51E-43 | 23 | Dtnb |
| Pih1d11 | 2.36E-47 | 0.364731349 | 0.573 | 0.098 | 4.78E-43 | 23 | Pih1d1 |
| Sh2d1b15 | 2.64E-47 | 0.382972482 | 0.5 | 0.078 | 5.35E-43 | 23 | Sh2d1b1 |
| Zdhhc15 | 2.66E-47 | 0.393890047 | 0.292 | 0.03 | 5.38E-43 | 23 | Zdhhc15 |
| Cd825 | 2.77E-47 | 0.535599441 | 0.573 | 0.101 | 5.60E-43 | 23 | Cd82 |
| Tnik7 | 3.19E-47 | 0.55188956 | 0.625 | 0.113 | 6.47E-43 | 23 | Tnik |
| Prkca9 | 4.09E-47 | 0.533322919 | 0.573 | 0.1 | 8.29E-43 | 23 | Prkca |
| Rad213 | 5.13E-47 | 0.937841364 | 0.792 | 0.191 | 1.04E-42 | 23 | Rad21 |
| Baz1b7 | 7.62E-47 | 0.842486643 | 0.812 | 0.193 | 1.54E-42 | 23 | Baz1b |
| Nup85 | 8.33E-47 | 0.311558853 | 0.344 | 0.041 | 1.69E-42 | 23 | Nup85 |
| Cdyl7 | 9.04E-47 | 0.536236718 | 0.667 | 0.129 | 1.83E-42 | 23 | Cdyl |
| Galnt61 | 1.11E-46 | 0.281166049 | 0.281 | 0.028 | 2.25E-42 | 23 | Galnt6 |
| Id25 | 1.15E-46 | 1.387536481 | 0.969 | 0.322 | 2.33E-42 | 23 | Id2 |
| Cdyl25 | 1.32E-46 | 0.382518851 | 0.552 | 0.091 | 2.67E-42 | 23 | Cdyl2 |
| Myo3b | 1.86E-46 | 0.269751139 | 0.26 | 0.025 | 3.77E-42 | 23 | Myo3b |
| Eya26 | 1.93E-46 | 0.462391389 | 0.573 | 0.1 | 3.90E-42 | 23 | Eya2 |
| Nup50 | 2.15E-46 | 0.375976569 | 0.542 | 0.09 | 4.34E-42 | 23 | Nup50 |
| Baz1a8 | 2.23E-46 | 0.665044306 | 0.781 | 0.172 | 4.52E-42 | 23 | Baz1a |

| | | | | | | | |
|---|---|---|---|---|---|---|---|
| Cers67 | 2.38E-46 | 0.638455797 | 0.615 | 0.115 | 4.82E-42 | 23 | Cers6 |
| Anp32b5 | 3.54E-46 | 1.368099298 | 1 | 0.423 | 7.17E-42 | 23 | Anp32b |
| Elf47 | 3.92E-46 | 0.575186135 | 0.635 | 0.122 | 7.94E-42 | 23 | Elf4 |
| Pycard7 | 4.40E-46 | 1.034255404 | 0.948 | 0.277 | 8.92E-42 | 23 | Pycard |
| Zbtb8os | 4.52E-46 | 0.469068995 | 0.75 | 0.159 | 9.16E-42 | 23 | Zbtb8os |
| Nap1l18 | 5.26E-46 | 1.221482984 | 0.917 | 0.315 | 1.06E-41 | 23 | Nap1l1 |
| Gng29 | 5.44E-46 | 0.85494093 | 0.792 | 0.187 | 1.10E-41 | 23 | Gng2 |
| Ankrd448 | 6.71E-46 | 1.286823463 | 0.948 | 0.294 | 1.36E-41 | 23 | Ankrd44 |
| Sesn37 | 7.62E-46 | 0.566659588 | 0.656 | 0.129 | 1.54E-41 | 23 | Sesn3 |
| Hivep36 | 8.57E-46 | 0.838540057 | 0.719 | 0.162 | 1.73E-41 | 23 | Hivep3 |
| Rab27a3 | 9.04E-46 | 0.323743793 | 0.521 | 0.085 | 1.83E-41 | 23 | Rab27a |
| Tcerg14 | 9.42E-46 | 0.600436497 | 0.677 | 0.136 | 1.91E-41 | 23 | Tcerg1 |
| Mbnl17 | 9.56E-46 | 1.62705719 | 0.99 | 0.459 | 1.94E-41 | 23 | Mbnl1 |
| Rbmx2 | 9.69E-46 | 0.265304635 | 0.396 | 0.053 | 1.96E-41 | 23 | Rbmx2 |
| Sun25 | 1.29E-45 | 0.566682113 | 0.479 | 0.077 | 2.62E-41 | 23 | Sun2 |
| Prex18 | 1.35E-45 | 0.885372091 | 0.76 | 0.174 | 2.73E-41 | 23 | Prex1 |
| Ubl4a | 1.50E-45 | 0.314098188 | 0.635 | 0.119 | 3.04E-41 | 23 | Ubl4a |
| Napsa4 | 1.62E-45 | 0.42793981 | 0.385 | 0.052 | 3.28E-41 | 23 | Napsa |
| Ighd1 | 1.74E-45 | 0.278824762 | 0.25 | 0.024 | 3.51E-41 | 23 | Ighd |
| Il162 | 1.79E-45 | 0.317496995 | 0.385 | 0.051 | 3.62E-41 | 23 | Il16 |
| Bid | 2.08E-45 | 0.274333348 | 0.292 | 0.031 | 4.21E-41 | 23 | Bid |
| Cd2bp2 | 2.19E-45 | 0.283542852 | 0.427 | 0.061 | 4.43E-41 | 23 | Cd2bp2 |
| Sephs1 | 2.54E-45 | 0.269694249 | 0.375 | 0.049 | 5.14E-41 | 23 | Sephs1 |
| Taf155 | 2.72E-45 | 0.850703647 | 0.885 | 0.237 | 5.51E-41 | 23 | Taf15 |
| Nsmce11 | 2.72E-45 | 0.562781509 | 0.708 | 0.149 | 5.52E-41 | 23 | Nsmce1 |
| AW1120106 | 3.02E-45 | 1.754923814 | 1 | 0.569 | 6.12E-41 | 23 | AW112010 |
| Bex31 | 3.18E-45 | 0.554730295 | 0.802 | 0.183 | 6.43E-41 | 23 | Bex3 |
| 1700017B05Rik | 4.69E-45 | 0.363375761 | 0.417 | 0.059 | 9.50E-41 | 23 | 1700017B05Rik |
| Cd696 | 5.59E-45 | 0.685400706 | 0.562 | 0.102 | 1.13E-40 | 23 | Cd69 |
| Lcp27 | 5.89E-45 | 0.538475491 | 0.542 | 0.095 | 1.19E-40 | 23 | Lcp2 |
| C2cd32 | 6.39E-45 | 0.457658149 | 0.49 | 0.079 | 1.29E-40 | 23 | C2cd3 |
| Scly | 7.17E-45 | 0.271025888 | 0.396 | 0.053 | 1.45E-40 | 23 | Scly |
| Emd1 | 7.37E-45 | 0.451977585 | 0.729 | 0.152 | 1.49E-40 | 23 | Emd |
| Cenpj | 7.72E-45 | 0.274941644 | 0.344 | 0.042 | 1.56E-40 | 23 | Cenpj |
| Evl7 | 9.12E-45 | 0.863059653 | 0.865 | 0.221 | 1.85E-40 | 23 | Evl |
| Atxn105 | 1.00E-44 | 0.631444508 | 0.771 | 0.175 | 2.03E-40 | 23 | Atxn10 |
| Eya12 | 1.08E-44 | 0.404380361 | 0.333 | 0.04 | 2.18E-40 | 23 | Eya1 |
| Rbbp42 | 1.14E-44 | 0.93355014 | 0.906 | 0.262 | 2.31E-40 | 23 | Rbbp4 |
| Paxip1 | 1.26E-44 | 0.321655948 | 0.354 | 0.045 | 2.55E-40 | 23 | Paxip1 |
| Dok62 | 1.36E-44 | 0.738247716 | 0.344 | 0.044 | 2.76E-40 | 23 | Dok6 |
| Camk2g6 | 1.37E-44 | 0.635989786 | 0.667 | 0.135 | 2.77E-40 | 23 | Camk2g |
| Plekhj14 | 1.38E-44 | 0.494317545 | 0.771 | 0.168 | 2.79E-40 | 23 | Plekhj1 |
| 4933406I18Rik7 | 1.61E-44 | 0.589316074 | 0.469 | 0.076 | 3.26E-40 | 23 | 4933406I18Rik |

| | | | | | | | |
|---|---|---|---|---|---|---|---|
| Cblb7 | 2.35E-44 | 0.644261428 | 0.771 | 0.171 | 4.77E-40 | 23 | Cblb |
| Atp2a35 | 2.43E-44 | 0.289847484 | 0.427 | 0.062 | 4.92E-40 | 23 | Atp2a3 |
| Vps37b6 | 2.85E-44 | 0.531424908 | 0.677 | 0.136 | 5.76E-40 | 23 | Vps37b |
| Litaf8 | 3.20E-44 | 0.60631192 | 0.729 | 0.157 | 6.48E-40 | 23 | Litaf |
| Cep152 | 3.68E-44 | 0.286756994 | 0.312 | 0.036 | 7.44E-40 | 23 | Cep152 |
| Sfxn12 | 3.79E-44 | 0.307095783 | 0.542 | 0.093 | 7.67E-40 | 23 | Sfxn1 |
| Rnaseh2c6 | 4.39E-44 | 0.929977912 | 0.938 | 0.286 | 8.88E-40 | 23 | Rnaseh2c |
| Dynlt1f | 4.42E-44 | 0.297035717 | 0.385 | 0.052 | 8.95E-40 | 23 | Dynlt1f |
| Sae12 | 4.57E-44 | 0.385820176 | 0.719 | 0.147 | 9.25E-40 | 23 | Sae1 |
| Med30 | 5.08E-44 | 0.493325517 | 0.76 | 0.169 | 1.03E-39 | 23 | Med30 |
| Ddb2 | 5.20E-44 | 0.28216342 | 0.292 | 0.032 | 1.05E-39 | 23 | Ddb2 |
| Fgl29 | 6.18E-44 | 0.497561093 | 0.583 | 0.108 | 1.25E-39 | 23 | Fgl2 |
| Dennd1b9 | 6.33E-44 | 0.696107583 | 0.792 | 0.184 | 1.28E-39 | 23 | Dennd1b |
| AC163354.14 | 7.22E-44 | 0.694707831 | 0.417 | 0.062 | 1.46E-39 | 23 | AC163354.1 |
| Mgat58 | 7.98E-44 | 0.813988903 | 0.74 | 0.17 | 1.61E-39 | 23 | Mgat5 |
| Sec11c8 | 8.92E-44 | 0.992500181 | 0.969 | 0.31 | 1.81E-39 | 23 | Sec11c |
| Ccdc77 | 1.09E-43 | 0.376435722 | 0.312 | 0.037 | 2.20E-39 | 23 | Ccdc77 |
| Rgs181 | 1.10E-43 | 0.342992062 | 0.292 | 0.032 | 2.23E-39 | 23 | Rgs18 |
| Mthfd1l | 1.24E-43 | 0.416019965 | 0.344 | 0.044 | 2.52E-39 | 23 | Mthfd1l |
| Slc20a11 | 1.34E-43 | 0.355285491 | 0.375 | 0.051 | 2.72E-39 | 23 | Slc20a1 |
| Rad54l24 | 1.38E-43 | 0.420042613 | 0.542 | 0.095 | 2.79E-39 | 23 | Rad54l2 |
| Cep85 | 1.59E-43 | 0.303096727 | 0.333 | 0.041 | 3.23E-39 | 23 | Cep85 |
| A430046D13Rik | 2.09E-43 | 0.296870641 | 0.25 | 0.025 | 4.24E-39 | 23 | A430046D13Rik |
| Sclt1 | 2.28E-43 | 0.463236262 | 0.542 | 0.096 | 4.61E-39 | 23 | Sclt1 |
| Haus7 | 3.22E-43 | 0.266447698 | 0.323 | 0.039 | 6.52E-39 | 23 | Haus7 |
| Snrpd11 | 3.23E-43 | 1.086144194 | 0.927 | 0.307 | 6.54E-39 | 23 | Snrpd1 |
| Tubgcp2 | 3.39E-43 | 0.329599743 | 0.438 | 0.066 | 6.86E-39 | 23 | Tubgcp2 |
| Ubac27 | 3.67E-43 | 0.767793215 | 0.781 | 0.188 | 7.43E-39 | 23 | Ubac2 |
| Rad504 | 4.47E-43 | 0.549346054 | 0.677 | 0.142 | 9.04E-39 | 23 | Rad50 |
| Gtpbp14 | 4.62E-43 | 0.347523659 | 0.458 | 0.072 | 9.35E-39 | 23 | Gtpbp1 |
| Casp8ap2 | 6.39E-43 | 0.584640191 | 0.594 | 0.116 | 1.29E-38 | 23 | Casp8ap2 |
| Psmg2 | 8.22E-43 | 0.336447391 | 0.552 | 0.098 | 1.66E-38 | 23 | Psmg2 |
| Pa2g41 | 9.17E-43 | 0.64006371 | 0.802 | 0.197 | 1.86E-38 | 23 | Pa2g4 |
| Ptpn45 | 9.26E-43 | 0.632949616 | 0.635 | 0.131 | 1.88E-38 | 23 | Ptpn4 |
| B930036N10Rik1 | 9.81E-43 | 0.388972908 | 0.385 | 0.054 | 1.99E-38 | 23 | B930036N10Rik |
| 2610020C07Rik | 9.97E-43 | 0.30026968 | 0.323 | 0.039 | 2.02E-38 | 23 | 2610020C07Rik |
| Pole41 | 1.29E-42 | 0.585259001 | 0.76 | 0.18 | 2.61E-38 | 23 | Pole4 |
| Atp11b6 | 1.45E-42 | 0.63079406 | 0.781 | 0.181 | 2.93E-38 | 23 | Atp11b |
| Raly2 | 1.47E-42 | 0.847019379 | 0.938 | 0.271 | 2.98E-38 | 23 | Raly |
| Sh3bgrl39 | 1.52E-42 | 1.500656066 | 1 | 0.56 | 3.07E-38 | 23 | Sh3bgrl3 |
| Meaf61 | 1.66E-42 | 0.362857606 | 0.562 | 0.103 | 3.37E-38 | 23 | Meaf6 |
| Nudt51 | 1.73E-42 | 0.359925942 | 0.573 | 0.106 | 3.49E-38 | 23 | Nudt5 |
| Ramp17 | 1.78E-42 | 0.615927149 | 0.573 | 0.11 | 3.61E-38 | 23 | Ramp1 |

| | | | | | | | |
|---|---|---|---|---|---|---|---|
| Cyth47 | 1.83E-42 | 0.452143541 | 0.667 | 0.134 | 3.70E-38 | 23 | Cyth4 |
| Gimap95 | 2.09E-42 | 0.632041138 | 0.719 | 0.159 | 4.24E-38 | 23 | Gimap9 |
| Dcps1 | 2.13E-42 | 0.415611536 | 0.604 | 0.117 | 4.32E-38 | 23 | Dcps |
| H2-Q78 | 2.15E-42 | 1.314975076 | 1 | 0.398 | 4.36E-38 | 23 | H2-Q7 |
| Unc13d | 2.29E-42 | 0.259231276 | 0.26 | 0.027 | 4.64E-38 | 23 | Unc13d |
| Rdm11 | 2.37E-42 | 0.601986587 | 0.646 | 0.135 | 4.80E-38 | 23 | Rdm1 |
| Kif13b6 | 2.70E-42 | 0.963819073 | 0.812 | 0.216 | 5.46E-38 | 23 | Kif13b |
| Smchd19 | 2.80E-42 | 0.851385173 | 0.917 | 0.264 | 5.68E-38 | 23 | Smchd1 |
| Smc67 | 2.99E-42 | 0.908665733 | 0.906 | 0.262 | 6.06E-38 | 23 | Smc6 |
| Rfx77 | 4.41E-42 | 0.571999667 | 0.708 | 0.155 | 8.94E-38 | 23 | Rfx7 |
| Chst122 | 4.44E-42 | 0.334256001 | 0.438 | 0.067 | 8.99E-38 | 23 | Chst12 |
| Psip17 | 4.72E-42 | 0.732234209 | 0.792 | 0.194 | 9.55E-38 | 23 | Psip1 |
| Gm140051 | 4.89E-42 | 0.374973041 | 0.302 | 0.036 | 9.90E-38 | 23 | Gm14005 |
| Lbh6 | 5.56E-42 | 0.444263246 | 0.656 | 0.132 | 1.12E-37 | 23 | Lbh |
| Pfn110 | 5.71E-42 | 1.460053642 | 1 | 0.683 | 1.16E-37 | 23 | Pfn1 |
| Agap21 | 6.04E-42 | 0.27252951 | 0.312 | 0.038 | 1.22E-37 | 23 | Agap2 |
| G2e3 | 6.60E-42 | 0.519331468 | 0.479 | 0.081 | 1.34E-37 | 23 | G2e3 |
| Rev3l6 | 6.79E-42 | 0.544894204 | 0.719 | 0.157 | 1.37E-37 | 23 | Rev3l |
| Ppm1h8 | 6.87E-42 | 0.637977186 | 0.719 | 0.161 | 1.39E-37 | 23 | Ppm1h |
| Hist1h1c4 | 7.64E-42 | 1.472692041 | 0.729 | 0.201 | 1.55E-37 | 23 | Hist1h1c |
| Strbp4 | 9.32E-42 | 0.652230991 | 0.792 | 0.191 | 1.89E-37 | 23 | Strbp |
| Odf2l | 1.01E-41 | 0.266014429 | 0.312 | 0.038 | 2.04E-37 | 23 | Odf2l |
| Ptma4 | 1.01E-41 | 1.215749371 | 1 | 0.94 | 2.04E-37 | 23 | Ptma |
| Ptpn189 | 1.04E-41 | 1.061033879 | 0.99 | 0.315 | 2.11E-37 | 23 | Ptpn18 |
| Usp37 | 1.07E-41 | 0.523876772 | 0.51 | 0.091 | 2.17E-37 | 23 | Usp37 |
| Sfmbt14 | 1.08E-41 | 0.507621375 | 0.49 | 0.084 | 2.18E-37 | 23 | Sfmbt1 |
| Arrb26 | 1.20E-41 | 0.30647076 | 0.531 | 0.093 | 2.42E-37 | 23 | Arrb2 |
| Tnf5 | 1.51E-41 | 0.508338958 | 0.448 | 0.072 | 3.07E-37 | 23 | Tnf |
| Naa40 | 2.01E-41 | 0.275495944 | 0.333 | 0.043 | 4.06E-37 | 23 | Naa40 |
| Pcnt | 2.03E-41 | 0.450552081 | 0.542 | 0.099 | 4.11E-37 | 23 | Pcnt |
| Wwp26 | 2.32E-41 | 0.456107885 | 0.635 | 0.127 | 4.70E-37 | 23 | Wwp2 |
| Nsmce27 | 2.38E-41 | 0.724202111 | 0.812 | 0.206 | 4.81E-37 | 23 | Nsmce2 |
| Ptbp1 | 2.38E-41 | 0.39429578 | 0.708 | 0.15 | 4.82E-37 | 23 | Ptbp1 |
| Bcl2l114 | 2.39E-41 | 0.551401871 | 0.594 | 0.117 | 4.83E-37 | 23 | Bcl2l11 |
| Srrt | 2.54E-41 | 0.381480854 | 0.542 | 0.098 | 5.15E-37 | 23 | Srrt |
| Bin17 | 2.83E-41 | 0.42236376 | 0.771 | 0.175 | 5.74E-37 | 23 | Bin1 |
| Gabpb22 | 2.92E-41 | 0.424379533 | 0.604 | 0.118 | 5.91E-37 | 23 | Gabpb2 |
| Ftx1 | 3.23E-41 | 0.424016434 | 0.354 | 0.048 | 6.53E-37 | 23 | Ftx |
| Cxcr64 | 3.23E-41 | 0.534832723 | 0.417 | 0.064 | 6.54E-37 | 23 | Cxcr6 |
| Zfp5124 | 6.80E-41 | 0.420750033 | 0.5 | 0.087 | 1.38E-36 | 23 | Zfp512 |
| Ppm1b4 | 9.12E-41 | 0.511831645 | 0.656 | 0.138 | 1.85E-36 | 23 | Ppm1b |
| Rapgef67 | 9.29E-41 | 0.942557172 | 0.948 | 0.297 | 1.88E-36 | 23 | Rapgef6 |
| Lsm31 | 9.72E-41 | 0.40714926 | 0.781 | 0.187 | 1.97E-36 | 23 | Lsm3 |

| | | | | | | | |
|---|---|---|---|---|---|---|---|
| Ctsd8 | 1.13E-40 | 0.963075894 | 0.979 | 0.314 | 2.29E-36 | 23 | Ctsd |
| Kmt5a | 1.18E-40 | 0.369365665 | 0.656 | 0.138 | 2.38E-36 | 23 | Kmt5a |
| Pms1 | 1.19E-40 | 0.279665831 | 0.271 | 0.03 | 2.40E-36 | 23 | Pms1 |
| Gimap47 | 1.22E-40 | 0.903634088 | 0.906 | 0.256 | 2.46E-36 | 23 | Gimap4 |
| S100a611 | 1.41E-40 | 1.234879174 | 0.906 | 0.272 | 2.84E-36 | 23 | S100a6 |
| Lin54 | 1.45E-40 | 0.286580409 | 0.354 | 0.048 | 2.93E-36 | 23 | Lin54 |
| Ust7 | 1.84E-40 | 0.738964699 | 0.479 | 0.086 | 3.73E-36 | 23 | Ust |
| Ptger46 | 2.66E-40 | 0.427205039 | 0.49 | 0.085 | 5.39E-36 | 23 | Ptger4 |
| Ddx39b8 | 2.73E-40 | 0.72044828 | 0.854 | 0.233 | 5.53E-36 | 23 | Ddx39b |
| Pla2g167 | 2.74E-40 | 0.572288807 | 0.906 | 0.243 | 5.54E-36 | 23 | Pla2g16 |
| Adgrl1 | 2.98E-40 | 0.278100757 | 0.292 | 0.034 | 6.03E-36 | 23 | Adgrl1 |
| Pds5b6 | 3.52E-40 | 0.570072645 | 0.771 | 0.18 | 7.13E-36 | 23 | Pds5b |
| Tes8 | 3.81E-40 | 0.475950668 | 0.625 | 0.13 | 7.71E-36 | 23 | Tes |
| Slc38a29 | 4.09E-40 | 1.010769862 | 0.958 | 0.311 | 8.27E-36 | 23 | Slc38a2 |
| Acd | 4.41E-40 | 0.250953126 | 0.354 | 0.048 | 8.93E-36 | 23 | Acd |
| Trdc1 | 4.57E-40 | 0.270513605 | 0.458 | 0.075 | 9.26E-36 | 23 | Trdc |
| 1810006J02Rik2 | 5.09E-40 | 0.338225444 | 0.323 | 0.042 | 1.03E-35 | 23 | 1810006J02Rik |
| Cdk2ap27 | 6.39E-40 | 0.541094959 | 0.802 | 0.199 | 1.29E-35 | 23 | Cdk2ap2 |
| Virma3 | 8.25E-40 | 0.434565503 | 0.635 | 0.131 | 1.67E-35 | 23 | Virma |
| Jazf1 | 8.29E-40 | 0.412323403 | 0.365 | 0.052 | 1.68E-35 | 23 | Jazf1 |
| A630089N07Rik | 8.51E-40 | 0.346016217 | 0.281 | 0.033 | 1.72E-35 | 23 | A630089N07Rik |
| Pms2 | 1.02E-39 | 0.284097351 | 0.312 | 0.039 | 2.07E-35 | 23 | Pms2 |
| Parvg1 | 1.03E-39 | 0.276010678 | 0.344 | 0.047 | 2.08E-35 | 23 | Parvg |
| Brip1os1 | 1.10E-39 | 0.368780083 | 0.51 | 0.091 | 2.23E-35 | 23 | Brip1os |
| Nup155 | 1.25E-39 | 0.270079516 | 0.333 | 0.044 | 2.54E-35 | 23 | Nup155 |
| Ak2 | 1.31E-39 | 0.310320479 | 0.656 | 0.137 | 2.66E-35 | 23 | Ak2 |
| Nans1 | 1.62E-39 | 0.342960674 | 0.635 | 0.134 | 3.28E-35 | 23 | Nans |
| Cmc11 | 1.67E-39 | 0.391718124 | 0.792 | 0.191 | 3.38E-35 | 23 | Cmc1 |
| Wdr702 | 1.74E-39 | 0.490449285 | 0.688 | 0.151 | 3.53E-35 | 23 | Wdr70 |
| Znrf25 | 1.80E-39 | 0.513764007 | 0.708 | 0.159 | 3.64E-35 | 23 | Znrf2 |
| Ggct3 | 1.89E-39 | 0.258069157 | 0.417 | 0.065 | 3.83E-35 | 23 | Ggct |
| Rin36 | 2.01E-39 | 0.651471309 | 0.75 | 0.18 | 4.07E-35 | 23 | Rin3 |
| Gpsm38 | 2.28E-39 | 0.579242989 | 0.667 | 0.148 | 4.62E-35 | 23 | Gpsm3 |
| Haus6 | 2.68E-39 | 0.302073254 | 0.333 | 0.045 | 5.43E-35 | 23 | Haus6 |
| Rpap21 | 2.87E-39 | 0.326918614 | 0.438 | 0.071 | 5.80E-35 | 23 | Rpap2 |
| Ripor26 | 3.15E-39 | 0.578596718 | 0.75 | 0.172 | 6.37E-35 | 23 | Ripor2 |
| Fancl | 3.71E-39 | 0.329426279 | 0.406 | 0.063 | 7.51E-35 | 23 | Fancl |
| Lin52 | 3.75E-39 | 0.284117226 | 0.375 | 0.055 | 7.60E-35 | 23 | Lin52 |
| Isg204 | 4.16E-39 | 0.414921552 | 0.583 | 0.116 | 8.42E-35 | 23 | Isg20 |
| Hmgb14 | 4.19E-39 | 1.300814805 | 1 | 0.717 | 8.48E-35 | 23 | Hmgb1 |
| Slc7a12 | 4.44E-39 | 0.412445126 | 0.396 | 0.061 | 8.99E-35 | 23 | Slc7a1 |
| Uchl5 | 4.53E-39 | 0.373468088 | 0.635 | 0.135 | 9.17E-35 | 23 | Uchl5 |
| Ppwd1 | 5.59E-39 | 0.316791511 | 0.5 | 0.088 | 1.13E-34 | 23 | Ppwd1 |

| | | | | | | | |
|---|---|---|---|---|---|---|---|
| Otulin5 | 6.28E-39 | 0.510252064 | 0.802 | 0.2 | 1.27E-34 | 23 | Otulin |
| Fbl5 | 6.71E-39 | 0.395186076 | 0.625 | 0.13 | 1.36E-34 | 23 | Fbl |
| 1700025G04Rik8 | 6.92E-39 | 0.70324641 | 0.75 | 0.189 | 1.40E-34 | 23 | 1700025G04Rik |
| Zfp8312 | 8.13E-39 | 0.288755673 | 0.281 | 0.033 | 1.65E-34 | 23 | Zfp831 |
| Snrpf8 | 8.79E-39 | 0.931762772 | 0.99 | 0.379 | 1.78E-34 | 23 | Snrpf |
| Actb10 | 9.06E-39 | 1.190516541 | 1 | 0.843 | 1.83E-34 | 23 | Actb |
| Pim16 | 9.75E-39 | 0.541544083 | 0.625 | 0.133 | 1.97E-34 | 23 | Pim1 |
| Atp6v0a22 | 9.79E-39 | 0.398642921 | 0.573 | 0.112 | 1.98E-34 | 23 | Atp6v0a2 |
| Supt161 | 1.00E-38 | 0.691460501 | 0.854 | 0.237 | 2.03E-34 | 23 | Supt16 |
| Ptpn112 | 1.05E-38 | 0.537106214 | 0.906 | 0.236 | 2.13E-34 | 23 | Ptpn1 |
| Cd538 | 1.07E-38 | 0.46797622 | 0.583 | 0.117 | 2.17E-34 | 23 | Cd53 |
| Trmt1128 | 1.22E-38 | 0.655033216 | 0.938 | 0.274 | 2.46E-34 | 23 | Trmt112 |
| Snrpc1 | 1.29E-38 | 0.44642403 | 0.812 | 0.2 | 2.62E-34 | 23 | Snrpc |
| Tra2b7 | 1.34E-38 | 0.815484889 | 0.958 | 0.296 | 2.71E-34 | 23 | Tra2b |
| Timm501 | 1.41E-38 | 0.316519233 | 0.49 | 0.088 | 2.86E-34 | 23 | Timm50 |
| Rcor14 | 1.62E-38 | 0.504344075 | 0.771 | 0.183 | 3.27E-34 | 23 | Rcor1 |
| Golt1b | 1.63E-38 | 0.292139282 | 0.365 | 0.053 | 3.30E-34 | 23 | Golt1b |
| Calm36 | 1.67E-38 | 0.758501342 | 0.885 | 0.269 | 3.38E-34 | 23 | Calm3 |
| Taf6l | 1.74E-38 | 0.354488847 | 0.708 | 0.157 | 3.51E-34 | 23 | Taf6l |
| Rngtt6 | 1.80E-38 | 0.5729184 | 0.625 | 0.135 | 3.64E-34 | 23 | Rngtt |
| Runx25 | 1.97E-38 | 0.377430495 | 0.5 | 0.092 | 3.98E-34 | 23 | Runx2 |
| Lrrk17 | 2.26E-38 | 0.696496819 | 0.76 | 0.19 | 4.58E-34 | 23 | Lrrk1 |
| Etaa1 | 2.56E-38 | 0.272148068 | 0.385 | 0.058 | 5.17E-34 | 23 | Etaa1 |
| Ormdl1 | 2.67E-38 | 0.290443567 | 0.302 | 0.038 | 5.41E-34 | 23 | Ormdl1 |
| Cep162 | 2.76E-38 | 0.270612661 | 0.458 | 0.077 | 5.59E-34 | 23 | Cep162 |
| Tuba1c3 | 2.99E-38 | 0.591434175 | 0.573 | 0.12 | 6.05E-34 | 23 | Tuba1c |
| Arrdc1 | 3.05E-38 | 0.294623853 | 0.396 | 0.061 | 6.17E-34 | 23 | Arrdc1 |
| Tmem372 | 3.16E-38 | 0.305613593 | 0.344 | 0.048 | 6.39E-34 | 23 | Tmem37 |
| Clic19 | 3.58E-38 | 1.263761627 | 1 | 0.602 | 7.25E-34 | 23 | Clic1 |
| Sp1107 | 3.87E-38 | 0.502141632 | 0.688 | 0.157 | 7.83E-34 | 23 | Sp110 |
| Abr8 | 4.19E-38 | 0.523042833 | 0.698 | 0.158 | 8.48E-34 | 23 | Abr |
| Diaph17 | 4.28E-38 | 0.700754178 | 0.875 | 0.242 | 8.66E-34 | 23 | Diaph1 |
| Ptpn26 | 4.38E-38 | 0.548574268 | 0.646 | 0.143 | 8.86E-34 | 23 | Ptpn2 |
| Shisa56 | 5.21E-38 | 0.883175246 | 0.99 | 0.329 | 1.05E-33 | 23 | Shisa5 |
| Nop584 | 5.50E-38 | 0.577472424 | 0.625 | 0.138 | 1.11E-33 | 23 | Nop58 |
| Ppp1cc3 | 5.62E-38 | 0.629153198 | 0.823 | 0.222 | 1.14E-33 | 23 | Ppp1cc |
| Helb | 6.12E-38 | 0.256138081 | 0.302 | 0.038 | 1.24E-33 | 23 | Helb |
| Orc3 | 7.19E-38 | 0.275626525 | 0.521 | 0.095 | 1.46E-33 | 23 | Orc3 |
| Glipr17 | 8.03E-38 | 0.456497862 | 0.51 | 0.096 | 1.63E-33 | 23 | Glipr1 |
| Ube2i5 | 9.43E-38 | 0.768652858 | 0.979 | 0.33 | 1.91E-33 | 23 | Ube2i |
| Gltp1 | 1.12E-37 | 0.332872373 | 0.646 | 0.141 | 2.27E-33 | 23 | Gltp |
| Nup1535 | 1.25E-37 | 0.460377396 | 0.573 | 0.116 | 2.53E-33 | 23 | Nup153 |
| Rpsa10 | 1.36E-37 | 1.258401604 | 1 | 0.903 | 2.74E-33 | 23 | Rpsa |

| | | | | | | | |
|---|---|---|---|---|---|---|---|
| Peli17 | 1.36E-37 | 0.746313121 | 0.854 | 0.232 | 2.75E-33 | 23 | Peli1 |
| Inpp5b1 | 1.42E-37 | 0.369851791 | 0.438 | 0.074 | 2.88E-33 | 23 | Inpp5b |
| Tecr8 | 1.50E-37 | 0.569681729 | 0.844 | 0.225 | 3.03E-33 | 23 | Tecr |
| Timm17b | 1.55E-37 | 0.255814967 | 0.74 | 0.168 | 3.14E-33 | 23 | Timm17b |
| Erh1 | 1.70E-37 | 0.420526995 | 0.75 | 0.183 | 3.44E-33 | 23 | Erh |
| Stk48 | 1.89E-37 | 0.62715198 | 0.719 | 0.175 | 3.82E-33 | 23 | Stk4 |
| Gm163413 | 2.06E-37 | 0.28037896 | 0.292 | 0.037 | 4.17E-33 | 23 | Gm16341 |
| Ubash3b7 | 2.10E-37 | 0.614619669 | 0.708 | 0.168 | 4.24E-33 | 23 | Ubash3b |
| Cdk193 | 2.21E-37 | 0.572631854 | 0.646 | 0.147 | 4.47E-33 | 23 | Cdk19 |
| Memo12 | 3.22E-37 | 0.428021636 | 0.521 | 0.1 | 6.52E-33 | 23 | Memo1 |
| H2afy7 | 3.81E-37 | 0.823145149 | 0.99 | 0.338 | 7.71E-33 | 23 | H2afy |
| Cdk42 | 3.92E-37 | 0.656112568 | 0.771 | 0.207 | 7.93E-33 | 23 | Cdk4 |
| Plcg28 | 5.05E-37 | 0.448109917 | 0.583 | 0.12 | 1.02E-32 | 23 | Plcg2 |
| Gab34 | 5.56E-37 | 0.48254408 | 0.5 | 0.095 | 1.13E-32 | 23 | Gab3 |
| Dazap11 | 5.68E-37 | 0.437971812 | 0.688 | 0.158 | 1.15E-32 | 23 | Dazap1 |
| Snrpg7 | 6.10E-37 | 0.867720196 | 0.938 | 0.324 | 1.23E-32 | 23 | Snrpg |
| Srpk13 | 7.32E-37 | 0.380352152 | 0.708 | 0.161 | 1.48E-32 | 23 | Srpk1 |
| Tmem131l5 | 8.31E-37 | 0.367671998 | 0.458 | 0.081 | 1.68E-32 | 23 | Tmem131l |
| Eml49 | 8.52E-37 | 0.717132726 | 0.896 | 0.259 | 1.73E-32 | 23 | Eml4 |
| Ppp1ca7 | 9.86E-37 | 1.002734971 | 0.979 | 0.397 | 2.00E-32 | 23 | Ppp1ca |
| Elmo2 | 1.01E-36 | 0.317419837 | 0.406 | 0.066 | 2.05E-32 | 23 | Elmo2 |
| Zw10 | 1.26E-36 | 0.266071861 | 0.292 | 0.037 | 2.55E-32 | 23 | Zw10 |
| Agfg2 | 1.32E-36 | 0.30930146 | 0.469 | 0.085 | 2.67E-32 | 23 | Agfg2 |
| Ppil23 | 1.33E-36 | 0.398353086 | 0.667 | 0.15 | 2.69E-32 | 23 | Ppil2 |
| Mad1l1 | 1.45E-36 | 0.393956329 | 0.396 | 0.064 | 2.93E-32 | 23 | Mad1l1 |
| Slc25a251 | 1.53E-36 | 0.302827022 | 0.406 | 0.066 | 3.09E-32 | 23 | Slc25a25 |
| Vps13a7 | 1.56E-36 | 0.599010815 | 0.708 | 0.173 | 3.15E-32 | 23 | Vps13a |
| Pde4a1 | 1.96E-36 | 0.257409668 | 0.396 | 0.063 | 3.96E-32 | 23 | Pde4a |
| Gtf2f1 | 2.01E-36 | 0.34196852 | 0.552 | 0.109 | 4.08E-32 | 23 | Gtf2f1 |
| Srbd1 | 2.08E-36 | 0.374554053 | 0.5 | 0.095 | 4.20E-32 | 23 | Srbd1 |
| Fkbp24 | 2.32E-36 | 0.695037834 | 0.896 | 0.276 | 4.70E-32 | 23 | Fkbp2 |
| Fndc3a7 | 2.55E-36 | 0.936395664 | 0.875 | 0.277 | 5.16E-32 | 23 | Fndc3a |
| Tec7 | 2.84E-36 | 0.486535217 | 0.542 | 0.11 | 5.75E-32 | 23 | Tec |
| 2700049A03Rik | 2.90E-36 | 0.261099911 | 0.333 | 0.047 | 5.88E-32 | 23 | 2700049A03Rik |
| Nde11 | 3.13E-36 | 0.392418806 | 0.521 | 0.101 | 6.35E-32 | 23 | Nde1 |
| Tyrobp8 | 3.21E-36 | 0.933740425 | 0.938 | 0.262 | 6.49E-32 | 23 | Tyrobp |
| Pbx35 | 3.42E-36 | 0.783277804 | 0.604 | 0.14 | 6.92E-32 | 23 | Pbx3 |
| Fryl11 | 3.44E-36 | 0.883917239 | 0.885 | 0.268 | 6.97E-32 | 23 | Fryl |
| Itga22 | 3.80E-36 | 0.328561529 | 0.292 | 0.038 | 7.69E-32 | 23 | Itga2 |
| Vrk26 | 3.98E-36 | 0.468483132 | 0.583 | 0.124 | 8.06E-32 | 23 | Vrk2 |
| Ranbp12 | 4.28E-36 | 0.932641608 | 0.917 | 0.323 | 8.66E-32 | 23 | Ranbp1 |
| Crlf38 | 4.36E-36 | 0.588621411 | 0.667 | 0.157 | 8.82E-32 | 23 | Crlf3 |
| Gpr181 | 6.51E-36 | 0.270142336 | 0.292 | 0.038 | 1.32E-31 | 23 | Gpr18 |

| | | | | | | | |
|---|---|---|---|---|---|---|---|
| Trim147 | 7.26E-36 | 0.496310515 | 0.573 | 0.121 | 1.47E-31 | 23 | Trim14 |
| Myg1 | 7.57E-36 | 0.32369757 | 0.448 | 0.08 | 1.53E-31 | 23 | Myg1 |
| Plcxd22 | 7.93E-36 | 0.357268096 | 0.271 | 0.034 | 1.61E-31 | 23 | Plcxd2 |
| Tango2 | 8.81E-36 | 0.258379977 | 0.396 | 0.064 | 1.78E-31 | 23 | Tango2 |
| Mbp6 | 9.34E-36 | 0.290504223 | 0.406 | 0.068 | 1.89E-31 | 23 | Mbp |
| Gins4 | 1.02E-35 | 0.321697804 | 0.448 | 0.08 | 2.06E-31 | 23 | Gins4 |
| Plgrkt6 | 1.05E-35 | 0.621002903 | 0.792 | 0.213 | 2.12E-31 | 23 | Plgrkt |
| Ilf2 | 1.07E-35 | 0.372367196 | 0.615 | 0.133 | 2.16E-31 | 23 | Ilf2 |
| Zdhhc21 | 1.12E-35 | 0.323344734 | 0.396 | 0.065 | 2.27E-31 | 23 | Zdhhc21 |
| Rps209 | 1.14E-35 | 1.064210276 | 1 | 0.951 | 2.31E-31 | 23 | Rps20 |
| Rbpj8 | 1.36E-35 | 0.709928695 | 0.802 | 0.221 | 2.75E-31 | 23 | Rbpj |
| Znrf18 | 1.50E-35 | 0.65398131 | 0.75 | 0.189 | 3.04E-31 | 23 | Znrf1 |
| Cse1l | 1.52E-35 | 0.320182199 | 0.448 | 0.08 | 3.07E-31 | 23 | Cse1l |
| Banf12 | 1.52E-35 | 0.597334711 | 0.875 | 0.248 | 3.08E-31 | 23 | Banf1 |
| Larp71 | 1.63E-35 | 0.276423512 | 0.583 | 0.121 | 3.29E-31 | 23 | Larp7 |
| Casp7 | 1.68E-35 | 0.306529336 | 0.385 | 0.062 | 3.41E-31 | 23 | Casp7 |
| Ccdc827 | 1.87E-35 | 0.518140097 | 0.729 | 0.181 | 3.79E-31 | 23 | Ccdc82 |
| Tcf207 | 1.93E-35 | 0.431542171 | 0.635 | 0.141 | 3.91E-31 | 23 | Tcf20 |
| Terf12 | 1.95E-35 | 0.421608085 | 0.635 | 0.141 | 3.95E-31 | 23 | Terf1 |
| Nckap1l7 | 2.33E-35 | 0.332956106 | 0.479 | 0.089 | 4.73E-31 | 23 | Nckap1l |
| Prr5l1 | 2.48E-35 | 0.26940599 | 0.312 | 0.043 | 5.01E-31 | 23 | Prr5l |
| Sf3a3 | 2.60E-35 | 0.314356242 | 0.625 | 0.136 | 5.26E-31 | 23 | Sf3a3 |
| Nucks13 | 3.07E-35 | 0.965652274 | 0.969 | 0.39 | 6.22E-31 | 23 | Nucks1 |
| Prkacb6 | 3.30E-35 | 0.830345992 | 0.781 | 0.218 | 6.68E-31 | 23 | Prkacb |
| Nop564 | 3.35E-35 | 0.434418227 | 0.604 | 0.132 | 6.79E-31 | 23 | Nop56 |
| Mvb12b6 | 4.43E-35 | 0.462110502 | 0.698 | 0.167 | 8.97E-31 | 23 | Mvb12b |
| Lnpk | 4.74E-35 | 0.296850008 | 0.385 | 0.062 | 9.59E-31 | 23 | Lnpk |
| Cnn29 | 4.80E-35 | 0.582040985 | 0.729 | 0.182 | 9.72E-31 | 23 | Cnn2 |
| Cdc123 | 4.81E-35 | 0.251979975 | 0.719 | 0.17 | 9.74E-31 | 23 | Cdc123 |
| Fbxl8 | 5.92E-35 | 0.261641657 | 0.333 | 0.049 | 1.20E-30 | 23 | Fbxl8 |
| Kpna35 | 6.38E-35 | 0.543822701 | 0.656 | 0.157 | 1.29E-30 | 23 | Kpna3 |
| Slx4ip3 | 7.60E-35 | 0.323751266 | 0.406 | 0.069 | 1.54E-30 | 23 | Slx4ip |
| Upf3b3 | 8.74E-35 | 0.334674552 | 0.625 | 0.138 | 1.77E-30 | 23 | Upf3b |
| Arl6ip110 | 9.45E-35 | 1.392539512 | 0.969 | 0.37 | 1.91E-30 | 23 | Arl6ip1 |
| Alms1 | 1.01E-34 | 0.308256445 | 0.26 | 0.032 | 2.05E-30 | 23 | Alms1 |
| Cops3 | 1.02E-34 | 0.28203831 | 0.594 | 0.127 | 2.06E-30 | 23 | Cops3 |
| Capg8 | 1.02E-34 | 0.751366971 | 0.49 | 0.101 | 2.07E-30 | 23 | Capg |
| Mtmr125 | 1.60E-34 | 0.549420538 | 0.583 | 0.131 | 3.23E-30 | 23 | Mtmr12 |
| Hist1h1e3 | 1.62E-34 | 2.08769594 | 0.885 | 0.372 | 3.29E-30 | 23 | Hist1h1e |
| Taf36 | 1.70E-34 | 0.508218931 | 0.635 | 0.147 | 3.44E-30 | 23 | Taf3 |
| Zcchc117 | 1.74E-34 | 0.770472127 | 0.875 | 0.267 | 3.53E-30 | 23 | Zcchc11 |
| Cep250 | 1.74E-34 | 0.305198277 | 0.458 | 0.083 | 3.53E-30 | 23 | Cep250 |
| Smc36 | 1.93E-34 | 0.7919278 | 0.823 | 0.253 | 3.91E-30 | 23 | Smc3 |

| | | | | | | | |
|---|---|---|---|---|---|---|---|
| Dnajc22 | 2.22E-34 | 0.405844384 | 0.635 | 0.149 | 4.50E-30 | 23 | Dnajc2 |
| Ccdc25 | 2.43E-34 | 0.28823698 | 0.583 | 0.125 | 4.91E-30 | 23 | Ccdc25 |
| Rfc1 | 2.79E-34 | 0.61159923 | 0.698 | 0.181 | 5.65E-30 | 23 | Rfc1 |
| Rbbp73 | 3.10E-34 | 0.491711789 | 0.865 | 0.247 | 6.27E-30 | 23 | Rbbp7 |
| Rps6ka18 | 3.72E-34 | 0.361840898 | 0.552 | 0.115 | 7.52E-30 | 23 | Rps6ka1 |
| Nsmaf6 | 3.75E-34 | 0.33541164 | 0.5 | 0.097 | 7.60E-30 | 23 | Nsmaf |
| Cnot33 | 3.97E-34 | 0.601860595 | 0.719 | 0.186 | 8.04E-30 | 23 | Cnot3 |
| Malt16 | 4.08E-34 | 0.564329159 | 0.635 | 0.15 | 8.25E-30 | 23 | Malt1 |
| Capza17 | 4.24E-34 | 0.712485964 | 0.896 | 0.275 | 8.58E-30 | 23 | Capza1 |
| Pag17 | 4.44E-34 | 0.539296024 | 0.562 | 0.122 | 8.99E-30 | 23 | Pag1 |
| Morc33 | 4.80E-34 | 0.371214866 | 0.688 | 0.162 | 9.71E-30 | 23 | Morc3 |
| Dnajc114 | 5.51E-34 | 0.875300082 | 0.948 | 0.323 | 1.12E-29 | 23 | Dnajc1 |
| Hnrnpa34 | 5.98E-34 | 0.981763049 | 0.99 | 0.445 | 1.21E-29 | 23 | Hnrnpa3 |
| Samsn17 | 6.15E-34 | 0.5156517 | 0.438 | 0.081 | 1.25E-29 | 23 | Samsn1 |
| Hdac12 | 8.53E-34 | 0.516837332 | 0.677 | 0.167 | 1.73E-29 | 23 | Hdac1 |
| Tnfaip89 | 8.68E-34 | 0.485392902 | 0.74 | 0.187 | 1.76E-29 | 23 | Tnfaip8 |
| Kin | 8.76E-34 | 0.344538155 | 0.531 | 0.109 | 1.77E-29 | 23 | Kin |
| Lyar1 | 9.27E-34 | 0.326263925 | 0.594 | 0.131 | 1.88E-29 | 23 | Lyar |
| Tcof13 | 9.57E-34 | 0.370486223 | 0.562 | 0.12 | 1.94E-29 | 23 | Tcof1 |
| Ccm26 | 1.09E-33 | 0.387832221 | 0.583 | 0.126 | 2.21E-29 | 23 | Ccm2 |
| Dars | 1.24E-33 | 0.336386517 | 0.604 | 0.134 | 2.52E-29 | 23 | Dars |
| Pnkp | 1.37E-33 | 0.256243825 | 0.427 | 0.076 | 2.78E-29 | 23 | Pnkp |
| Cenpx4 | 1.38E-33 | 0.640667164 | 0.906 | 0.3 | 2.80E-29 | 23 | Cenpx |
| Snrpa11 | 1.52E-33 | 0.407228114 | 0.656 | 0.157 | 3.08E-29 | 23 | Snrpa1 |
| Phf65 | 1.73E-33 | 0.405475132 | 0.604 | 0.135 | 3.50E-29 | 23 | Phf6 |
| AC149090.17 | 2.01E-33 | 0.643322297 | 0.812 | 0.232 | 4.07E-29 | 23 | AC149090.1 |
| Dtnbp13 | 2.10E-33 | 0.449582517 | 0.812 | 0.228 | 4.25E-29 | 23 | Dtnbp1 |
| D030056L22Rik | 2.22E-33 | 0.301767187 | 0.385 | 0.065 | 4.49E-29 | 23 | D030056L22Rik |
| Inpp5d8 | 2.35E-33 | 0.586568692 | 0.771 | 0.204 | 4.76E-29 | 23 | Inpp5d |
| Lsm63 | 2.39E-33 | 0.53877937 | 0.969 | 0.315 | 4.84E-29 | 23 | Lsm6 |
| Vrk13 | 2.74E-33 | 0.317317226 | 0.458 | 0.087 | 5.55E-29 | 23 | Vrk1 |
| Ppp1r7 | 3.26E-33 | 0.284398446 | 0.625 | 0.141 | 6.60E-29 | 23 | Ppp1r7 |
| Dap2 | 3.47E-33 | 0.834454669 | 0.688 | 0.187 | 7.02E-29 | 23 | Dap |
| Sh3kbp17 | 3.67E-33 | 0.555464655 | 0.562 | 0.125 | 7.44E-29 | 23 | Sh3kbp1 |
| Trim28 | 4.16E-33 | 0.260759382 | 0.438 | 0.08 | 8.43E-29 | 23 | Trim28 |
| Arf62 | 4.83E-33 | 0.345199934 | 0.719 | 0.178 | 9.79E-29 | 23 | Arf6 |
| Ccl47 | 5.12E-33 | 0.697006014 | 0.573 | 0.13 | 1.04E-28 | 23 | Ccl4 |
| Bcl2l131 | 5.22E-33 | 0.277067238 | 0.406 | 0.071 | 1.06E-28 | 23 | Bcl2l13 |
| Prmt12 | 5.34E-33 | 0.457196719 | 0.792 | 0.216 | 1.08E-28 | 23 | Prmt1 |
| Wrn5 | 6.37E-33 | 0.446141991 | 0.625 | 0.147 | 1.29E-28 | 23 | Wrn |
| Cdk2ap1 | 6.55E-33 | 0.269653439 | 0.583 | 0.126 | 1.33E-28 | 23 | Cdk2ap1 |
| Kif21b3 | 7.38E-33 | 0.346133647 | 0.333 | 0.052 | 1.49E-28 | 23 | Kif21b |
| Gimap111 | 7.86E-33 | 0.797227222 | 0.938 | 0.303 | 1.59E-28 | 23 | Gimap1 |

| | | | | | | | |
|---|---|---|---|---|---|---|---|
| Dda1 | 9.50E-33 | 0.265392384 | 0.542 | 0.114 | 1.92E-28 | 23 | Dda1 |
| Srsf77 | 1.11E-32 | 0.841554776 | 0.938 | 0.333 | 2.24E-28 | 23 | Srsf7 |
| Snrpe4 | 1.37E-32 | 0.924146655 | 0.99 | 0.478 | 2.78E-28 | 23 | Snrpe |
| Lsm1 | 1.39E-32 | 0.267787202 | 0.688 | 0.166 | 2.81E-28 | 23 | Lsm1 |
| Fbxo34 | 1.43E-32 | 0.273347645 | 0.375 | 0.063 | 2.90E-28 | 23 | Fbxo34 |
| Hace1 | 1.48E-32 | 0.408374781 | 0.458 | 0.089 | 3.00E-28 | 23 | Hace1 |
| Slamf77 | 1.65E-32 | 0.404587596 | 0.49 | 0.099 | 3.34E-28 | 23 | Slamf7 |
| Ptpre8 | 1.67E-32 | 0.601872702 | 0.646 | 0.161 | 3.37E-28 | 23 | Ptpre |
| Lrrcc1 | 1.67E-32 | 0.277378682 | 0.438 | 0.081 | 3.38E-28 | 23 | Lrrcc1 |
| Parp2 | 1.69E-32 | 0.282019415 | 0.479 | 0.094 | 3.43E-28 | 23 | Parp2 |
| Srsf2 | 1.84E-32 | 0.747757763 | 0.885 | 0.293 | 3.72E-28 | 23 | Srsf2 |
| Ppp4c | 2.05E-32 | 0.376727436 | 0.531 | 0.113 | 4.14E-28 | 23 | Ppp4c |
| Itga48 | 2.09E-32 | 0.651870304 | 0.406 | 0.077 | 4.23E-28 | 23 | Itga4 |
| Arhgap108 | 2.19E-32 | 0.36459951 | 0.583 | 0.127 | 4.43E-28 | 23 | Arhgap10 |
| Csk7 | 2.38E-32 | 0.412144418 | 0.573 | 0.13 | 4.83E-28 | 23 | Csk |
| Fgr2 | 2.40E-32 | 0.268350018 | 0.302 | 0.044 | 4.87E-28 | 23 | Fgr |
| Thrap36 | 2.56E-32 | 0.619247769 | 0.927 | 0.293 | 5.18E-28 | 23 | Thrap3 |
| 0610010K14Rik | 2.68E-32 | 0.272067302 | 0.521 | 0.108 | 5.43E-28 | 23 | 0610010K14Rik |
| Ms4a6b5 | 2.98E-32 | 0.390961848 | 0.469 | 0.093 | 6.04E-28 | 23 | Ms4a6b |
| Dennd1c4 | 3.21E-32 | 0.362952432 | 0.354 | 0.058 | 6.50E-28 | 23 | Dennd1c |
| Srsf1 | 3.90E-32 | 0.347807677 | 0.656 | 0.157 | 7.90E-28 | 23 | Srsf1 |
| Ctdspl23 | 4.74E-32 | 0.422231156 | 0.688 | 0.172 | 9.59E-28 | 23 | Ctdspl2 |
| P2ry104 | 5.01E-32 | 0.277843333 | 0.281 | 0.039 | 1.02E-27 | 23 | P2ry10 |
| Smc1a7 | 6.04E-32 | 0.814897623 | 0.917 | 0.312 | 1.22E-27 | 23 | Smc1a |
| Tmem1319 | 6.28E-32 | 0.623934789 | 0.76 | 0.208 | 1.27E-27 | 23 | Tmem131 |
| Lsm43 | 6.64E-32 | 0.697198013 | 0.906 | 0.322 | 1.34E-27 | 23 | Lsm4 |
| Mkl18 | 6.73E-32 | 0.445997983 | 0.562 | 0.126 | 1.36E-27 | 23 | Mkl1 |
| H2-Q68 | 6.84E-32 | 0.614250125 | 0.844 | 0.245 | 1.38E-27 | 23 | H2-Q6 |
| Utp3 | 6.89E-32 | 0.361054842 | 0.719 | 0.184 | 1.39E-27 | 23 | Utp3 |
| Mgat4a7 | 7.89E-32 | 0.319356825 | 0.51 | 0.106 | 1.60E-27 | 23 | Mgat4a |
| Snx55 | 8.20E-32 | 0.45012358 | 0.781 | 0.211 | 1.66E-27 | 23 | Snx5 |
| Chd77 | 8.70E-32 | 0.507732049 | 0.625 | 0.151 | 1.76E-27 | 23 | Chd7 |
| Psmd131 | 9.19E-32 | 0.280961679 | 0.854 | 0.235 | 1.86E-27 | 23 | Psmd13 |
| Slc25a40 | 9.41E-32 | 0.270746638 | 0.312 | 0.047 | 1.90E-27 | 23 | Slc25a40 |
| Tmem1671 | 1.02E-31 | 0.612506727 | 0.875 | 0.282 | 2.07E-27 | 23 | Tmem167 |
| Csrp17 | 1.03E-31 | 0.398745699 | 0.719 | 0.182 | 2.08E-27 | 23 | Csrp1 |
| Cnst1 | 1.11E-31 | 0.294615091 | 0.438 | 0.082 | 2.24E-27 | 23 | Cnst |
| Pcbd21 | 1.23E-31 | 0.372447673 | 0.844 | 0.248 | 2.50E-27 | 23 | Pcbd2 |
| Ccl36 | 1.24E-31 | 0.54545349 | 0.427 | 0.083 | 2.50E-27 | 23 | Ccl3 |
| Gimap56 | 1.38E-31 | 0.583600308 | 0.667 | 0.171 | 2.79E-27 | 23 | Gimap5 |
| Ipo5 | 1.41E-31 | 0.309625033 | 0.417 | 0.077 | 2.85E-27 | 23 | Ipo5 |
| Pspc14 | 1.49E-31 | 0.374638707 | 0.542 | 0.118 | 3.01E-27 | 23 | Pspc1 |
| C2cd5 | 1.49E-31 | 0.443171376 | 0.469 | 0.096 | 3.03E-27 | 23 | C2cd5 |

| | | | | | | | |
|---|---|---|---|---|---|---|---|
| Fkbp52 | 1.50E-31 | 0.41779564 | 0.573 | 0.132 | 3.04E-27 | 23 | Fkbp5 |
| Rasa37 | 1.60E-31 | 0.387312328 | 0.375 | 0.066 | 3.24E-27 | 23 | Rasa3 |
| Kpnb1 | 1.69E-31 | 0.296409758 | 0.615 | 0.141 | 3.41E-27 | 23 | Kpnb1 |
| Cnot6l6 | 1.83E-31 | 0.576680744 | 0.833 | 0.245 | 3.71E-27 | 23 | Cnot6l |
| Zfp3185 | 1.93E-31 | 0.398498463 | 0.479 | 0.098 | 3.90E-27 | 23 | Zfp318 |
| Uchl3 | 2.06E-31 | 0.280140663 | 0.646 | 0.159 | 4.16E-27 | 23 | Uchl3 |
| Trim36 | 2.12E-31 | 0.253176678 | 0.271 | 0.037 | 4.30E-27 | 23 | Trim36 |
| Itgav11 | 2.14E-31 | 0.467380337 | 0.656 | 0.161 | 4.34E-27 | 23 | Itgav |
| Hdgf3 | 2.53E-31 | 0.484644434 | 0.833 | 0.248 | 5.12E-27 | 23 | Hdgf |
| Parp89 | 3.21E-31 | 0.463237663 | 0.625 | 0.149 | 6.50E-27 | 23 | Parp8 |
| Gsap8 | 3.35E-31 | 0.268402427 | 0.448 | 0.086 | 6.77E-27 | 23 | Gsap |
| Cerk8 | 3.76E-31 | 0.352106617 | 0.531 | 0.115 | 7.61E-27 | 23 | Cerk |
| S100a109 | 3.91E-31 | 1.124876193 | 0.99 | 0.47 | 7.91E-27 | 23 | S100a10 |
| Zyx6 | 4.07E-31 | 0.587143787 | 0.771 | 0.216 | 8.24E-27 | 23 | Zyx |
| Tab25 | 4.66E-31 | 0.535980048 | 0.812 | 0.231 | 9.43E-27 | 23 | Tab2 |
| Senp1 | 5.09E-31 | 0.438515192 | 0.49 | 0.104 | 1.03E-26 | 23 | Senp1 |
| Ube2a | 6.97E-31 | 0.306968396 | 0.719 | 0.187 | 1.41E-26 | 23 | Ube2a |
| Cdc27 | 7.26E-31 | 0.424944399 | 0.49 | 0.104 | 1.47E-26 | 23 | Cdc27 |
| Myo9b7 | 7.70E-31 | 0.359468923 | 0.542 | 0.12 | 1.56E-26 | 23 | Myo9b |
| Gm170182 | 7.88E-31 | 0.333858093 | 0.479 | 0.1 | 1.60E-26 | 23 | Gm17018 |
| Pank2 | 8.26E-31 | 0.383972968 | 0.625 | 0.151 | 1.67E-26 | 23 | Pank2 |
| Lims19 | 9.45E-31 | 0.68191289 | 0.927 | 0.3 | 1.91E-26 | 23 | Lims1 |
| Tusc34 | 1.03E-30 | 0.307077922 | 0.635 | 0.149 | 2.09E-26 | 23 | Tusc3 |
| Fam189b4 | 1.04E-30 | 0.325775196 | 0.344 | 0.057 | 2.11E-26 | 23 | Fam189b |
| Ubr7 | 1.29E-30 | 0.296916148 | 0.438 | 0.085 | 2.61E-26 | 23 | Ubr7 |
| Ebp3 | 1.54E-30 | 0.429720844 | 0.698 | 0.182 | 3.13E-26 | 23 | Ebp |
| Mta36 | 1.57E-30 | 0.50668079 | 0.615 | 0.152 | 3.17E-26 | 23 | Mta3 |
| Ccnd36 | 1.62E-30 | 0.473094617 | 0.833 | 0.24 | 3.27E-26 | 23 | Ccnd3 |
| Xrn25 | 1.62E-30 | 0.46436327 | 0.885 | 0.261 | 3.27E-26 | 23 | Xrn2 |
| Ranbp97 | 1.63E-30 | 0.48998595 | 0.74 | 0.2 | 3.31E-26 | 23 | Ranbp9 |
| Glipr24 | 1.86E-30 | 0.264340399 | 0.292 | 0.043 | 3.76E-26 | 23 | Glipr2 |
| Exosc31 | 1.89E-30 | 0.298919862 | 0.604 | 0.143 | 3.82E-26 | 23 | Exosc3 |
| Mef2a8 | 2.01E-30 | 1.18141364 | 0.969 | 0.417 | 4.07E-26 | 23 | Mef2a |
| Hsp90aa15 | 2.04E-30 | 0.994477068 | 0.969 | 0.426 | 4.14E-26 | 23 | Hsp90aa1 |
| Hnrnpl3 | 2.14E-30 | 0.504685763 | 0.812 | 0.235 | 4.34E-26 | 23 | Hnrnpl |
| Bckdk | 2.23E-30 | 0.260487647 | 0.406 | 0.076 | 4.51E-26 | 23 | Bckdk |
| Ssna1 | 2.31E-30 | 0.482755744 | 0.74 | 0.209 | 4.67E-26 | 23 | Ssna1 |
| Crlf26 | 2.42E-30 | 0.411960565 | 0.646 | 0.161 | 4.89E-26 | 23 | Crlf2 |
| Rnf1496 | 2.71E-30 | 0.351811231 | 0.49 | 0.104 | 5.49E-26 | 23 | Rnf149 |
| Lmtk22 | 2.75E-30 | 0.272273877 | 0.458 | 0.091 | 5.56E-26 | 23 | Lmtk2 |
| Cklf1 | 3.10E-30 | 0.300432412 | 0.375 | 0.067 | 6.27E-26 | 23 | Cklf |
| Adss | 3.22E-30 | 0.320683877 | 0.729 | 0.194 | 6.53E-26 | 23 | Adss |
| Ubap2 | 3.23E-30 | 0.312571172 | 0.552 | 0.124 | 6.53E-26 | 23 | Ubap2 |

| | | | | | | | |
|---|---|---|---|---|---|---|---|
| Cyld6 | 3.31E-30 | 0.431462356 | 0.667 | 0.171 | 6.70E-26 | 23 | Cyld |
| Pip5k1b6 | 3.57E-30 | 0.338699959 | 0.479 | 0.1 | 7.22E-26 | 23 | Pip5k1b |
| Ppp1r35 | 3.60E-30 | 0.252170412 | 0.5 | 0.106 | 7.29E-26 | 23 | Ppp1r35 |
| Glmn1 | 3.67E-30 | 0.380558947 | 0.469 | 0.097 | 7.44E-26 | 23 | Glmn |
| Taf1d5 | 3.83E-30 | 0.270316408 | 0.615 | 0.145 | 7.76E-26 | 23 | Taf1d |
| Ppp6r13 | 4.24E-30 | 0.392863773 | 0.479 | 0.101 | 8.58E-26 | 23 | Ppp6r1 |
| Rnf168 | 4.25E-30 | 0.259205336 | 0.458 | 0.092 | 8.60E-26 | 23 | Rnf168 |
| Rab8b8 | 4.53E-30 | 0.648684147 | 0.885 | 0.284 | 9.18E-26 | 23 | Rab8b |
| Ap2s11 | 4.56E-30 | 0.461038146 | 0.823 | 0.247 | 9.24E-26 | 23 | Ap2s1 |
| Dbnl4 | 5.12E-30 | 0.404299093 | 0.719 | 0.192 | 1.04E-25 | 23 | Dbnl |
| Pds5a5 | 5.21E-30 | 0.732583033 | 0.698 | 0.205 | 1.06E-25 | 23 | Pds5a |
| Cntln6 | 5.36E-30 | 0.382935515 | 0.635 | 0.155 | 1.08E-25 | 23 | Cntln |
| Bscl2 | 6.06E-30 | 0.267621829 | 0.396 | 0.074 | 1.23E-25 | 23 | Bscl2 |
| Ppp1r188 | 6.22E-30 | 0.509409469 | 0.615 | 0.154 | 1.26E-25 | 23 | Ppp1r18 |
| Nfx12 | 6.70E-30 | 0.318043824 | 0.656 | 0.162 | 1.36E-25 | 23 | Nfx1 |
| Uimc13 | 6.74E-30 | 0.392012626 | 0.615 | 0.15 | 1.37E-25 | 23 | Uimc1 |
| Slc1a54 | 7.35E-30 | 0.337414028 | 0.417 | 0.081 | 1.49E-25 | 23 | Slc1a5 |
| Fam173a4 | 7.42E-30 | 0.440346898 | 0.708 | 0.198 | 1.50E-25 | 23 | Fam173a |
| Tmem71 | 7.80E-30 | 0.292404843 | 0.302 | 0.047 | 1.58E-25 | 23 | Tmem71 |
| Twf25 | 7.96E-30 | 0.345199207 | 0.677 | 0.171 | 1.61E-25 | 23 | Twf2 |
| Prpf40a10 | 8.22E-30 | 0.597277951 | 0.854 | 0.267 | 1.66E-25 | 23 | Prpf40a |
| Srsf6 | 8.35E-30 | 0.415506584 | 0.656 | 0.17 | 1.69E-25 | 23 | Srsf6 |
| Adgre58 | 9.57E-30 | 0.649762264 | 0.802 | 0.237 | 1.94E-25 | 23 | Adgre5 |
| Smg610 | 9.79E-30 | 0.762188727 | 0.823 | 0.26 | 1.98E-25 | 23 | Smg6 |
| Nono | 1.06E-29 | 0.51662878 | 0.771 | 0.227 | 2.15E-25 | 23 | Nono |
| Arap27 | 1.07E-29 | 0.365358333 | 0.708 | 0.178 | 2.16E-25 | 23 | Arap2 |
| Hnrnpd | 1.09E-29 | 0.723033675 | 0.906 | 0.332 | 2.21E-25 | 23 | Hnrnpd |
| Psmd142 | 1.12E-29 | 0.45286039 | 0.76 | 0.217 | 2.27E-25 | 23 | Psmd14 |
| Pnn9 | 1.26E-29 | 0.555814067 | 0.865 | 0.267 | 2.55E-25 | 23 | Pnn |
| Bub31 | 1.26E-29 | 0.573880952 | 0.719 | 0.206 | 2.56E-25 | 23 | Bub3 |
| Rgs38 | 1.34E-29 | 0.504191661 | 0.698 | 0.187 | 2.72E-25 | 23 | Rgs3 |
| Ldlrad49 | 1.38E-29 | 0.69961811 | 0.812 | 0.251 | 2.79E-25 | 23 | Ldlrad4 |
| Eefsec | 1.49E-29 | 0.282774743 | 0.354 | 0.062 | 3.03E-25 | 23 | Eefsec |
| Mfsd103 | 1.61E-29 | 0.344528017 | 0.458 | 0.095 | 3.26E-25 | 23 | Mfsd10 |
| Bcl62 | 1.75E-29 | 0.302868769 | 0.333 | 0.056 | 3.54E-25 | 23 | Bcl6 |
| Lasp14 | 1.86E-29 | 0.406522138 | 0.76 | 0.208 | 3.78E-25 | 23 | Lasp1 |
| Chka6 | 1.89E-29 | 0.400283337 | 0.594 | 0.145 | 3.82E-25 | 23 | Chka |
| Nup205 | 2.14E-29 | 0.31819629 | 0.312 | 0.051 | 4.32E-25 | 23 | Nup205 |
| Vwa83 | 2.29E-29 | 0.499431716 | 0.521 | 0.121 | 4.63E-25 | 23 | Vwa8 |
| Odc12 | 2.30E-29 | 0.37272446 | 0.698 | 0.186 | 4.66E-25 | 23 | Odc1 |
| G3bp16 | 2.57E-29 | 0.582964177 | 0.917 | 0.302 | 5.20E-25 | 23 | G3bp1 |
| Nsd310 | 2.84E-29 | 0.719618359 | 0.906 | 0.308 | 5.75E-25 | 23 | Nsd3 |
| Stip11 | 2.88E-29 | 0.326024647 | 0.531 | 0.121 | 5.84E-25 | 23 | Stip1 |

| | | | | | | | |
|---|---|---|---|---|---|---|---|
| Rnps1 | 2.96E-29 | 0.310980723 | 0.677 | 0.173 | 5.99E-25 | 23 | Rnps1 |
| Capzb9 | 3.14E-29 | 0.779105804 | 0.938 | 0.369 | 6.35E-25 | 23 | Capzb |
| Aak17 | 3.41E-29 | 0.702101762 | 0.667 | 0.185 | 6.90E-25 | 23 | Aak1 |
| Jak19 | 3.42E-29 | 0.811047093 | 0.958 | 0.366 | 6.91E-25 | 23 | Jak1 |
| Trappc101 | 3.50E-29 | 0.287042572 | 0.583 | 0.138 | 7.09E-25 | 23 | Trappc10 |
| Setx4 | 3.78E-29 | 0.307075708 | 0.688 | 0.174 | 7.64E-25 | 23 | Setx |
| Cd1645 | 3.97E-29 | 0.37602511 | 0.771 | 0.214 | 8.04E-25 | 23 | Cd164 |
| 1810011H11Rik5 | 4.14E-29 | 0.465873257 | 0.385 | 0.074 | 8.38E-25 | 23 | 1810011H11Rik |
| Exoc25 | 4.15E-29 | 0.46012221 | 0.594 | 0.147 | 8.40E-25 | 23 | Exoc2 |
| D16Ertd472e6 | 4.34E-29 | 0.405858724 | 0.49 | 0.106 | 8.79E-25 | 23 | D16Ertd472e |
| Med101 | 4.45E-29 | 0.335065961 | 0.729 | 0.196 | 9.00E-25 | 23 | Med10 |
| Ahdc14 | 5.15E-29 | 0.294221802 | 0.51 | 0.112 | 1.04E-24 | 23 | Ahdc1 |
| Bcl7c1 | 5.17E-29 | 0.368321442 | 0.812 | 0.243 | 1.05E-24 | 23 | Bcl7c |
| Hmgxb41 | 6.06E-29 | 0.267413929 | 0.479 | 0.101 | 1.23E-24 | 23 | Hmgxb4 |
| Coro1c3 | 6.22E-29 | 0.398845096 | 0.583 | 0.141 | 1.26E-24 | 23 | Coro1c |
| Brd83 | 6.44E-29 | 0.492400573 | 0.625 | 0.164 | 1.30E-24 | 23 | Brd8 |
| Rcsd17 | 6.52E-29 | 0.393811586 | 0.552 | 0.13 | 1.32E-24 | 23 | Rcsd1 |
| Adipor26 | 7.24E-29 | 0.34418133 | 0.615 | 0.15 | 1.47E-24 | 23 | Adipor2 |
| Eef2k2 | 7.70E-29 | 0.321254298 | 0.406 | 0.079 | 1.56E-24 | 23 | Eef2k |
| 4930523C07Rik11 | 7.80E-29 | 0.4595021 | 0.781 | 0.219 | 1.58E-24 | 23 | 4930523C07Rik |
| Supt33 | 8.01E-29 | 0.363149328 | 0.542 | 0.127 | 1.62E-24 | 23 | Supt3 |
| Xpo12 | 8.52E-29 | 0.387891571 | 0.542 | 0.126 | 1.72E-24 | 23 | Xpo1 |
| Rps15a7 | 9.89E-29 | 0.979503876 | 1 | 0.925 | 2.00E-24 | 23 | Rps15a |
| Magoh2 | 9.91E-29 | 0.479917604 | 0.812 | 0.257 | 2.01E-24 | 23 | Magoh |
| Ctcf5 | 1.00E-28 | 0.688620511 | 0.844 | 0.277 | 2.03E-24 | 23 | Ctcf |
| Zfp6524 | 1.13E-28 | 0.469506399 | 0.833 | 0.247 | 2.29E-24 | 23 | Zfp652 |
| Snrpd24 | 1.15E-28 | 0.666899158 | 0.979 | 0.409 | 2.33E-24 | 23 | Snrpd2 |
| Srsf32 | 1.16E-28 | 0.954448975 | 1 | 0.52 | 2.35E-24 | 23 | Srsf3 |
| Asxl14 | 1.16E-28 | 0.459314263 | 0.594 | 0.147 | 2.35E-24 | 23 | Asxl1 |
| Snrnp707 | 1.16E-28 | 0.595330928 | 0.844 | 0.273 | 2.36E-24 | 23 | Snrnp70 |
| Sf3b26 | 1.18E-28 | 0.698088509 | 0.948 | 0.359 | 2.40E-24 | 23 | Sf3b2 |
| Akr1b34 | 1.22E-28 | 0.272798486 | 0.719 | 0.189 | 2.48E-24 | 23 | Akr1b3 |
| Kdm2b5 | 1.26E-28 | 0.308182762 | 0.438 | 0.089 | 2.55E-24 | 23 | Kdm2b |
| Ilf3 | 1.51E-28 | 0.257523322 | 0.438 | 0.088 | 3.06E-24 | 23 | Ilf3 |
| Eif2s11 | 1.73E-28 | 0.392718821 | 0.719 | 0.203 | 3.50E-24 | 23 | Eif2s1 |
| Prf13 | 1.84E-28 | 0.340821365 | 0.354 | 0.064 | 3.73E-24 | 23 | Prf1 |
| Nin4 | 1.92E-28 | 0.408597679 | 0.573 | 0.139 | 3.89E-24 | 23 | Nin |
| Rpl13a7 | 2.01E-28 | 0.463050179 | 0.885 | 0.285 | 4.07E-24 | 23 | Rpl13a |
| Dgkd7 | 2.10E-28 | 0.553033747 | 0.615 | 0.161 | 4.26E-24 | 23 | Dgkd |
| Topors2 | 2.12E-28 | 0.304087873 | 0.521 | 0.117 | 4.29E-24 | 23 | Topors |
| Gramd1b7 | 2.69E-28 | 0.361135253 | 0.438 | 0.09 | 5.44E-24 | 23 | Gramd1b |
| Cnot16 | 2.75E-28 | 0.417189096 | 0.656 | 0.173 | 5.57E-24 | 23 | Cnot1 |
| Pld33 | 2.83E-28 | 0.289413838 | 0.427 | 0.086 | 5.72E-24 | 23 | Pld3 |

| | | | | | | | |
|---|---|---|---|---|---|---|---|
| Sik26 | 3.25E-28 | 0.366877717 | 0.521 | 0.12 | 6.58E-24 | 23 | Sik2 |
| Ifi2095 | 3.27E-28 | 0.263246803 | 0.333 | 0.057 | 6.61E-24 | 23 | Ifi209 |
| Cd3g3 | 3.88E-28 | 1.224104912 | 0.281 | 0.045 | 7.85E-24 | 23 | Cd3g |
| Iqsec14 | 4.16E-28 | 0.35585733 | 0.531 | 0.122 | 8.42E-24 | 23 | Iqsec1 |
| Trp536 | 4.24E-28 | 0.44157547 | 0.615 | 0.159 | 8.59E-24 | 23 | Trp53 |
| Ube2l3 | 4.27E-28 | 0.526900908 | 0.938 | 0.332 | 8.65E-24 | 23 | Ube2l3 |
| Ppia4 | 5.03E-28 | 0.872102547 | 1 | 0.938 | 1.02E-23 | 23 | Ppia |
| Rel7 | 5.46E-28 | 0.276771494 | 0.583 | 0.139 | 1.11E-23 | 23 | Rel |
| Nktr8 | 5.76E-28 | 0.569584505 | 0.823 | 0.258 | 1.17E-23 | 23 | Nktr |
| Rps117 | 5.81E-28 | 0.89904341 | 1 | 0.907 | 1.18E-23 | 23 | Rps11 |
| Srsf43 | 5.83E-28 | 0.307324426 | 0.635 | 0.16 | 1.18E-23 | 23 | Srsf4 |
| Wwox11 | 6.10E-28 | 0.462973129 | 0.656 | 0.176 | 1.24E-23 | 23 | Wwox |
| Ewsr11 | 6.47E-28 | 0.407753759 | 0.729 | 0.206 | 1.31E-23 | 23 | Ewsr1 |
| Sf3b52 | 6.64E-28 | 0.470425001 | 0.906 | 0.31 | 1.35E-23 | 23 | Sf3b5 |
| Neurl37 | 7.27E-28 | 0.330035888 | 0.51 | 0.116 | 1.47E-23 | 23 | Neurl3 |
| Tmem1734 | 7.74E-28 | 0.303569981 | 0.375 | 0.071 | 1.57E-23 | 23 | Tmem173 |
| Rnf1572 | 1.00E-27 | 0.307173828 | 0.354 | 0.065 | 2.03E-23 | 23 | Rnf157 |
| Rgs28 | 1.13E-27 | 0.32392269 | 0.552 | 0.13 | 2.29E-23 | 23 | Rgs2 |
| Dscam4 | 1.16E-27 | 0.33933263 | 0.292 | 0.047 | 2.35E-23 | 23 | Dscam |
| Pml1 | 1.20E-27 | 0.286613397 | 0.406 | 0.081 | 2.42E-23 | 23 | Pml |
| Vps285 | 1.29E-27 | 0.499535243 | 0.917 | 0.301 | 2.62E-23 | 23 | Vps28 |
| Ncald2 | 1.32E-27 | 0.415197467 | 0.458 | 0.102 | 2.67E-23 | 23 | Ncald |
| Itgb112 | 1.40E-27 | 1.191254907 | 0.969 | 0.479 | 2.84E-23 | 23 | Itgb1 |
| Nr2c27 | 1.42E-27 | 0.432898271 | 0.615 | 0.159 | 2.87E-23 | 23 | Nr2c2 |
| Hnrnpa15 | 1.45E-27 | 0.513748172 | 0.927 | 0.325 | 2.93E-23 | 23 | Hnrnpa1 |
| Cbx31 | 1.56E-27 | 0.585981717 | 0.958 | 0.377 | 3.15E-23 | 23 | Cbx3 |
| Plekha26 | 1.57E-27 | 0.41189262 | 0.552 | 0.134 | 3.19E-23 | 23 | Plekha2 |
| Grk26 | 1.59E-27 | 0.329816697 | 0.531 | 0.125 | 3.21E-23 | 23 | Grk2 |
| Lgals37 | 1.63E-27 | 0.524763886 | 0.74 | 0.218 | 3.29E-23 | 23 | Lgals3 |
| Setd28 | 2.39E-27 | 0.452262538 | 0.698 | 0.193 | 4.84E-23 | 23 | Setd2 |
| Mapre28 | 2.50E-27 | 0.59864802 | 0.792 | 0.25 | 5.06E-23 | 23 | Mapre2 |
| Prdm10 | 2.51E-27 | 0.261478437 | 0.312 | 0.053 | 5.09E-23 | 23 | Prdm10 |
| Cobll15 | 2.59E-27 | 0.355207398 | 0.417 | 0.086 | 5.25E-23 | 23 | Cobll1 |
| Sfpq8 | 2.63E-27 | 0.62219368 | 0.885 | 0.298 | 5.33E-23 | 23 | Sfpq |
| Hp1bp32 | 2.88E-27 | 0.634853439 | 0.865 | 0.285 | 5.83E-23 | 23 | Hp1bp3 |
| Uba6 | 3.18E-27 | 0.279455928 | 0.406 | 0.082 | 6.44E-23 | 23 | Uba6 |
| Soat15 | 3.38E-27 | 0.336339265 | 0.479 | 0.108 | 6.84E-23 | 23 | Soat1 |
| Rasa28 | 3.65E-27 | 0.428310075 | 0.562 | 0.139 | 7.39E-23 | 23 | Rasa2 |
| Tpk15 | 3.97E-27 | 0.389448597 | 0.469 | 0.105 | 8.04E-23 | 23 | Tpk1 |
| Pip4k2a6 | 4.10E-27 | 0.440913625 | 0.812 | 0.241 | 8.30E-23 | 23 | Pip4k2a |
| Lrmp6 | 4.71E-27 | 0.37166043 | 0.333 | 0.061 | 9.53E-23 | 23 | Lrmp |
| Actr39 | 5.13E-27 | 0.741960046 | 0.969 | 0.392 | 1.04E-22 | 23 | Actr3 |
| Atf1 | 5.59E-27 | 0.263804243 | 0.521 | 0.122 | 1.13E-22 | 23 | Atf1 |

| | | | | | | | |
|---|---|---|---|---|---|---|---|
| Hnrnpf4 | 5.60E-27 | 0.858536175 | 1 | 0.496 | 1.13E-22 | 23 | Hnrnpf |
| Tgfb18 | 5.73E-27 | 0.479498119 | 0.792 | 0.236 | 1.16E-22 | 23 | Tgfb1 |
| Tnfaip37 | 7.56E-27 | 0.346239511 | 0.427 | 0.09 | 1.53E-22 | 23 | Tnfaip3 |
| Agps3 | 8.14E-27 | 0.386758397 | 0.635 | 0.168 | 1.65E-22 | 23 | Agps |
| Usp156 | 8.14E-27 | 0.435514715 | 0.781 | 0.231 | 1.65E-22 | 23 | Usp15 |
| Srsf101 | 8.24E-27 | 0.456585401 | 0.792 | 0.244 | 1.67E-22 | 23 | Srsf10 |
| Ywhah6 | 8.26E-27 | 0.626468287 | 0.875 | 0.309 | 1.67E-22 | 23 | Ywhah |
| Wdr11 | 8.64E-27 | 0.280180106 | 0.646 | 0.169 | 1.75E-22 | 23 | Wdr1 |
| Zc3h14 | 8.70E-27 | 0.267358826 | 0.583 | 0.146 | 1.76E-22 | 23 | Zc3h14 |
| Fnip16 | 9.05E-27 | 0.344712734 | 0.583 | 0.146 | 1.83E-22 | 23 | Fnip1 |
| Phf5a | 9.74E-27 | 0.284320523 | 0.708 | 0.198 | 1.97E-22 | 23 | Phf5a |
| Arl6ip4 | 1.10E-26 | 0.250328165 | 0.677 | 0.183 | 2.23E-22 | 23 | Arl6ip4 |
| Trip44 | 1.13E-26 | 0.31549668 | 0.51 | 0.119 | 2.29E-22 | 23 | Trip4 |
| Dnajb112 | 1.14E-26 | 0.351372438 | 0.75 | 0.214 | 2.30E-22 | 23 | Dnajb11 |
| Anapc55 | 1.15E-26 | 0.350135013 | 0.74 | 0.211 | 2.33E-22 | 23 | Anapc5 |
| Efr3a5 | 1.19E-26 | 0.374111516 | 0.708 | 0.198 | 2.41E-22 | 23 | Efr3a |
| Nfatc37 | 1.23E-26 | 0.43919073 | 0.646 | 0.175 | 2.50E-22 | 23 | Nfatc3 |
| Rlf7 | 1.26E-26 | 0.308911729 | 0.562 | 0.137 | 2.55E-22 | 23 | Rlf |
| Hnrnpul1 | 1.27E-26 | 0.343004909 | 0.656 | 0.177 | 2.57E-22 | 23 | Hnrnpul1 |
| Cstf31 | 1.51E-26 | 0.316505704 | 0.531 | 0.127 | 3.05E-22 | 23 | Cstf3 |
| Kif2a4 | 1.52E-26 | 0.417012579 | 0.656 | 0.178 | 3.09E-22 | 23 | Kif2a |
| Hspa82 | 1.62E-26 | 0.827430933 | 1 | 0.71 | 3.29E-22 | 23 | Hspa8 |
| Nipa22 | 1.66E-26 | 0.413955879 | 0.531 | 0.131 | 3.37E-22 | 23 | Nipa2 |
| Rab3gap12 | 1.70E-26 | 0.359797307 | 0.625 | 0.164 | 3.45E-22 | 23 | Rab3gap1 |
| Dcaf71 | 1.73E-26 | 0.270704083 | 0.521 | 0.122 | 3.51E-22 | 23 | Dcaf7 |
| Traf2 | 1.75E-26 | 0.296645137 | 0.333 | 0.061 | 3.55E-22 | 23 | Traf2 |
| Rps310 | 1.78E-26 | 0.848522409 | 0.99 | 0.893 | 3.61E-22 | 23 | Rps3 |
| Smap25 | 1.88E-26 | 0.375283669 | 0.729 | 0.212 | 3.82E-22 | 23 | Smap2 |
| Babam1 | 2.01E-26 | 0.253854669 | 0.542 | 0.131 | 4.06E-22 | 23 | Babam1 |
| Atad2b8 | 2.06E-26 | 0.451960606 | 0.708 | 0.203 | 4.18E-22 | 23 | Atad2b |
| Ptpa | 2.07E-26 | 0.336470149 | 0.542 | 0.134 | 4.20E-22 | 23 | Ptpa |
| Prkdc2 | 2.15E-26 | 0.297839884 | 0.427 | 0.091 | 4.34E-22 | 23 | Prkdc |
| Chst119 | 2.54E-26 | 0.387775298 | 0.656 | 0.176 | 5.15E-22 | 23 | Chst11 |
| Taok38 | 2.68E-26 | 0.260350513 | 0.635 | 0.164 | 5.42E-22 | 23 | Taok3 |
| Sipa1l16 | 2.90E-26 | 0.54295329 | 0.667 | 0.19 | 5.87E-22 | 23 | Sipa1l1 |
| Pik3ap16 | 3.17E-26 | 0.273399525 | 0.427 | 0.091 | 6.42E-22 | 23 | Pik3ap1 |
| M6pr5 | 3.21E-26 | 0.385041539 | 0.615 | 0.163 | 6.50E-22 | 23 | M6pr |
| Stxbp55 | 3.52E-26 | 0.303868607 | 0.542 | 0.13 | 7.12E-22 | 23 | Stxbp5 |
| Tpr11 | 3.54E-26 | 0.762681172 | 0.979 | 0.426 | 7.16E-22 | 23 | Tpr |
| Senp710 | 3.87E-26 | 0.344801634 | 0.604 | 0.157 | 7.83E-22 | 23 | Senp7 |
| Sntb16 | 4.13E-26 | 0.42459811 | 0.344 | 0.065 | 8.37E-22 | 23 | Sntb1 |
| Sarnp1 | 4.28E-26 | 0.437748492 | 0.906 | 0.308 | 8.66E-22 | 23 | Sarnp |
| Lat4 | 4.46E-26 | 0.289377842 | 0.25 | 0.038 | 9.04E-22 | 23 | Lat |

| | | | | | | | |
|---|---|---|---|---|---|---|---|
| Hnrnpab2 | 4.48E-26 | 0.70780049 | 0.917 | 0.368 | 9.06E-22 | 23 | Hnrnpab |
| Arpc5l2 | 4.48E-26 | 0.423891784 | 0.885 | 0.305 | 9.07E-22 | 23 | Arpc5l |
| Arhgap213 | 4.71E-26 | 0.351704419 | 0.479 | 0.111 | 9.54E-22 | 23 | Arhgap21 |
| Ssr15 | 4.89E-26 | 0.343313611 | 0.677 | 0.187 | 9.90E-22 | 23 | Ssr1 |
| Tm6sf19 | 4.91E-26 | 0.361881509 | 0.562 | 0.142 | 9.93E-22 | 23 | Tm6sf1 |
| Syncrip4 | 5.08E-26 | 0.414671079 | 0.74 | 0.216 | 1.03E-21 | 23 | Syncrip |
| Nfyb | 5.23E-26 | 0.271589886 | 0.458 | 0.101 | 1.06E-21 | 23 | Nfyb |
| Faf18 | 5.55E-26 | 0.473433892 | 0.698 | 0.202 | 1.12E-21 | 23 | Faf1 |
| Usp381 | 5.66E-26 | 0.258276224 | 0.365 | 0.071 | 1.15E-21 | 23 | Usp38 |
| Eed3 | 6.20E-26 | 0.262293583 | 0.594 | 0.151 | 1.25E-21 | 23 | Eed |
| Tagln210 | 6.21E-26 | 0.858823006 | 0.958 | 0.429 | 1.26E-21 | 23 | Tagln2 |
| Nrf17 | 6.91E-26 | 0.405840994 | 0.542 | 0.136 | 1.40E-21 | 23 | Nrf1 |
| Rfx35 | 7.49E-26 | 0.413787001 | 0.469 | 0.108 | 1.52E-21 | 23 | Rfx3 |
| Herc48 | 7.62E-26 | 0.314990525 | 0.594 | 0.152 | 1.54E-21 | 23 | Herc4 |
| Fkbp8 | 7.77E-26 | 0.280633904 | 0.781 | 0.23 | 1.57E-21 | 23 | Fkbp8 |
| Naa16 | 8.73E-26 | 0.307715876 | 0.49 | 0.115 | 1.77E-21 | 23 | Naa16 |
| Smarcc25 | 8.80E-26 | 0.330287312 | 0.656 | 0.178 | 1.78E-21 | 23 | Smarcc2 |
| Mrpl571 | 9.10E-26 | 0.323353199 | 0.854 | 0.276 | 1.84E-21 | 23 | Mrpl57 |
| Fxyd511 | 9.17E-26 | 0.815283793 | 0.99 | 0.411 | 1.86E-21 | 23 | Fxyd5 |
| Nosip1 | 9.65E-26 | 0.253724147 | 0.573 | 0.144 | 1.95E-21 | 23 | Nosip |
| Luc7l36 | 1.01E-25 | 0.460168777 | 0.833 | 0.271 | 2.04E-21 | 23 | Luc7l3 |
| Rai15 | 1.21E-25 | 0.263106688 | 0.479 | 0.11 | 2.45E-21 | 23 | Rai1 |
| Stag112 | 1.29E-25 | 0.806696895 | 0.917 | 0.362 | 2.61E-21 | 23 | Stag1 |
| Cd444 | 1.37E-25 | 0.306289733 | 0.469 | 0.108 | 2.76E-21 | 23 | Cd44 |
| Coq10b5 | 1.44E-25 | 0.326699413 | 0.594 | 0.154 | 2.91E-21 | 23 | Coq10b |
| Rps510 | 1.50E-25 | 0.779423244 | 1 | 0.92 | 3.03E-21 | 23 | Rps5 |
| Fgf135 | 1.54E-25 | 0.343377371 | 0.385 | 0.079 | 3.11E-21 | 23 | Fgf13 |
| Hnrnpdl4 | 1.56E-25 | 0.428149675 | 0.792 | 0.249 | 3.16E-21 | 23 | Hnrnpdl |
| Scaf117 | 1.74E-25 | 0.47212377 | 0.812 | 0.263 | 3.53E-21 | 23 | Scaf11 |
| Pou2f13 | 1.88E-25 | 0.368382746 | 0.521 | 0.127 | 3.81E-21 | 23 | Pou2f1 |
| Tiprl2 | 2.00E-25 | 0.316525863 | 0.719 | 0.21 | 4.05E-21 | 23 | Tiprl |
| Exoc413 | 2.02E-25 | 0.767281567 | 0.885 | 0.323 | 4.09E-21 | 23 | Exoc4 |
| Atm | 2.03E-25 | 0.255514636 | 0.385 | 0.078 | 4.11E-21 | 23 | Atm |
| Dmtf14 | 2.36E-25 | 0.293008977 | 0.51 | 0.122 | 4.78E-21 | 23 | Dmtf1 |
| Rcbtb21 | 2.42E-25 | 0.286740796 | 0.51 | 0.123 | 4.91E-21 | 23 | Rcbtb2 |
| Cbx45 | 2.57E-25 | 0.27915934 | 0.479 | 0.111 | 5.20E-21 | 23 | Cbx4 |
| Ccdc1079 | 2.64E-25 | 0.306189723 | 0.521 | 0.128 | 5.34E-21 | 23 | Ccdc107 |
| Asxl27 | 2.83E-25 | 0.388666529 | 0.646 | 0.175 | 5.74E-21 | 23 | Asxl2 |
| Cdc5l | 2.85E-25 | 0.330611317 | 0.729 | 0.213 | 5.77E-21 | 23 | Cdc5l |
| Cap16 | 2.99E-25 | 0.430949759 | 0.854 | 0.272 | 6.06E-21 | 23 | Cap1 |
| Ncl6 | 3.05E-25 | 0.863664757 | 1 | 0.517 | 6.17E-21 | 23 | Ncl |
| Anxa68 | 3.18E-25 | 0.349121288 | 0.594 | 0.157 | 6.43E-21 | 23 | Anxa6 |
| Tardbp1 | 4.04E-25 | 0.320104072 | 0.594 | 0.157 | 8.17E-21 | 23 | Tardbp |

| | | | | | | | |
|---|---|---|---|---|---|---|---|
| Fam204a4 | 4.10E-25 | 0.358984898 | 0.615 | 0.17 | 8.30E-21 | 23 | Fam204a |
| Stag24 | 4.11E-25 | 0.490471035 | 0.854 | 0.288 | 8.32E-21 | 23 | Stag2 |
| Heca5 | 4.27E-25 | 0.439336491 | 0.552 | 0.143 | 8.65E-21 | 23 | Heca |
| Dnmt3a5 | 4.55E-25 | 0.294234777 | 0.573 | 0.148 | 9.21E-21 | 23 | Dnmt3a |
| Hnrnpu4 | 4.66E-25 | 0.74775647 | 0.969 | 0.423 | 9.43E-21 | 23 | Hnrnpu |
| Bola25 | 4.81E-25 | 0.384895106 | 0.802 | 0.262 | 9.74E-21 | 23 | Bola2 |
| Fuca22 | 5.16E-25 | 0.291189269 | 0.344 | 0.067 | 1.04E-20 | 23 | Fuca2 |
| Nhp23 | 5.98E-25 | 0.293115563 | 0.635 | 0.177 | 1.21E-20 | 23 | Nhp2 |
| Ss183 | 6.27E-25 | 0.325914987 | 0.635 | 0.174 | 1.27E-20 | 23 | Ss18 |
| Ehmt2 | 6.83E-25 | 0.276498783 | 0.427 | 0.094 | 1.38E-20 | 23 | Ehmt2 |
| Abcb73 | 7.53E-25 | 0.287567189 | 0.479 | 0.112 | 1.53E-20 | 23 | Abcb7 |
| Cnppd11 | 7.68E-25 | 0.26089841 | 0.427 | 0.094 | 1.56E-20 | 23 | Cnppd1 |
| Tpp26 | 7.85E-25 | 0.463791813 | 0.75 | 0.228 | 1.59E-20 | 23 | Tpp2 |
| Csnk2b2 | 7.88E-25 | 0.381485625 | 0.802 | 0.257 | 1.60E-20 | 23 | Csnk2b |
| Mphosph83 | 8.14E-25 | 0.306873634 | 0.708 | 0.207 | 1.65E-20 | 23 | Mphosph8 |
| Arhgef118 | 8.70E-25 | 0.539794325 | 0.771 | 0.249 | 1.76E-20 | 23 | Arhgef1 |
| Rpl22l18 | 8.77E-25 | 0.901739455 | 0.979 | 0.579 | 1.78E-20 | 23 | Rpl22l1 |
| Rbm2513 | 9.58E-25 | 0.620077599 | 0.917 | 0.334 | 1.94E-20 | 23 | Rbm25 |
| Actl6a | 9.66E-25 | 0.252424142 | 0.562 | 0.144 | 1.95E-20 | 23 | Actl6a |
| Emg12 | 9.77E-25 | 0.273784388 | 0.76 | 0.226 | 1.98E-20 | 23 | Emg1 |
| Atr1 | 9.99E-25 | 0.306807596 | 0.354 | 0.07 | 2.02E-20 | 23 | Atr |
| Bbs95 | 1.01E-24 | 0.32025933 | 0.448 | 0.102 | 2.05E-20 | 23 | Bbs9 |
| Retreg2 | 1.02E-24 | 0.257489664 | 0.385 | 0.08 | 2.07E-20 | 23 | Retreg2 |
| Gm4728312 | 1.05E-24 | 0.905520765 | 0.781 | 0.282 | 2.12E-20 | 23 | Gm47283 |
| Anapc113 | 1.11E-24 | 0.316355983 | 0.74 | 0.223 | 2.25E-20 | 23 | Anapc11 |
| Prr136 | 1.12E-24 | 0.329549546 | 0.719 | 0.21 | 2.27E-20 | 23 | Prr13 |
| Phf12 | 1.19E-24 | 0.295494796 | 0.542 | 0.136 | 2.41E-20 | 23 | Phf12 |
| Taf105 | 1.21E-24 | 0.566864889 | 0.958 | 0.385 | 2.45E-20 | 23 | Taf10 |
| Gm372404 | 1.27E-24 | 0.27856023 | 0.458 | 0.105 | 2.57E-20 | 23 | Gm37240 |
| Rsbn1l8 | 1.37E-24 | 0.554428421 | 0.812 | 0.273 | 2.78E-20 | 23 | Rsbn1l |
| Gm269174 | 1.39E-24 | 0.344440405 | 0.49 | 0.121 | 2.81E-20 | 23 | Gm26917 |
| Lrba8 | 1.56E-24 | 0.4125422 | 0.667 | 0.192 | 3.16E-20 | 23 | Lrba |
| Sh3gl13 | 1.65E-24 | 0.302149571 | 0.51 | 0.125 | 3.34E-20 | 23 | Sh3gl1 |
| Myh911 | 1.67E-24 | 0.782470347 | 0.99 | 0.437 | 3.37E-20 | 23 | Myh9 |
| Cog59 | 1.85E-24 | 0.324892562 | 0.667 | 0.183 | 3.74E-20 | 23 | Cog5 |
| Galnt114 | 1.93E-24 | 0.410201585 | 0.646 | 0.185 | 3.91E-20 | 23 | Galnt1 |
| Ccnl17 | 2.00E-24 | 0.361888563 | 0.719 | 0.21 | 4.04E-20 | 23 | Ccnl1 |
| Eif4a3 | 2.04E-24 | 0.317049833 | 0.719 | 0.213 | 4.14E-20 | 23 | Eif4a3 |
| Dnajc153 | 2.12E-24 | 0.290424629 | 0.917 | 0.322 | 4.30E-20 | 23 | Dnajc15 |
| Ipo9 | 2.41E-24 | 0.273411321 | 0.344 | 0.068 | 4.88E-20 | 23 | Ipo9 |
| Lman26 | 2.56E-24 | 0.404220113 | 0.698 | 0.207 | 5.18E-20 | 23 | Lman2 |
| Chordc1 | 2.56E-24 | 0.284388685 | 0.615 | 0.166 | 5.18E-20 | 23 | Chordc1 |
| Larp4b4 | 2.72E-24 | 0.367504474 | 0.698 | 0.204 | 5.51E-20 | 23 | Larp4b |

| | | | | | | | |
|---|---|---|---|---|---|---|---|
| Sipa17 | 2.77E-24 | 0.380425734 | 0.656 | 0.186 | 5.60E-20 | 23 | Sipa1 |
| Dhx91 | 2.78E-24 | 0.363290258 | 0.677 | 0.199 | 5.63E-20 | 23 | Dhx9 |
| Sytl34 | 2.80E-24 | 0.286196003 | 0.312 | 0.058 | 5.66E-20 | 23 | Sytl3 |
| Polr2a9 | 2.93E-24 | 0.455053058 | 0.865 | 0.281 | 5.93E-20 | 23 | Polr2a |
| Ccdc88a12 | 3.06E-24 | 0.572789006 | 0.677 | 0.205 | 6.19E-20 | 23 | Ccdc88a |
| Zfp6545 | 3.09E-24 | 0.424399647 | 0.49 | 0.121 | 6.25E-20 | 23 | Zfp654 |
| Akap1310 | 3.22E-24 | 0.731173011 | 0.958 | 0.394 | 6.51E-20 | 23 | Akap13 |
| Phip12 | 3.30E-24 | 0.498485346 | 0.792 | 0.256 | 6.68E-20 | 23 | Phip |
| Ddost5 | 3.41E-24 | 0.410343462 | 0.719 | 0.216 | 6.91E-20 | 23 | Ddost |
| Cbl6 | 3.94E-24 | 0.478295742 | 0.688 | 0.208 | 7.98E-20 | 23 | Cbl |
| Myef21 | 3.98E-24 | 0.302820161 | 0.375 | 0.079 | 8.06E-20 | 23 | Myef2 |
| Tmem27 | 4.05E-24 | 0.383735394 | 0.552 | 0.143 | 8.20E-20 | 23 | Tmem2 |
| Npc13 | 4.40E-24 | 0.275650373 | 0.375 | 0.078 | 8.91E-20 | 23 | Npc1 |
| Otud41 | 4.64E-24 | 0.259105537 | 0.51 | 0.126 | 9.38E-20 | 23 | Otud4 |
| Ralgapa27 | 4.79E-24 | 0.363195137 | 0.573 | 0.154 | 9.70E-20 | 23 | Ralgapa2 |
| AU0202068 | 4.97E-24 | 0.569163557 | 0.854 | 0.291 | 1.01E-19 | 23 | AU020206 |
| Ube3c6 | 5.05E-24 | 0.331232213 | 0.594 | 0.159 | 1.02E-19 | 23 | Ube3c |
| Fyn10 | 5.50E-24 | 0.640893646 | 0.792 | 0.267 | 1.11E-19 | 23 | Fyn |
| Emsy2 | 5.55E-24 | 0.290619276 | 0.51 | 0.127 | 1.12E-19 | 23 | Emsy |
| Sik310 | 6.21E-24 | 0.603760054 | 0.802 | 0.27 | 1.26E-19 | 23 | Sik3 |
| Nfkb110 | 6.55E-24 | 0.586951695 | 0.812 | 0.272 | 1.33E-19 | 23 | Nfkb1 |
| Mrpl332 | 7.09E-24 | 0.640927492 | 0.938 | 0.405 | 1.44E-19 | 23 | Mrpl33 |
| Eif4a16 | 7.36E-24 | 0.676468636 | 0.969 | 0.431 | 1.49E-19 | 23 | Eif4a1 |
| Hprt2 | 9.12E-24 | 0.427025145 | 0.833 | 0.286 | 1.85E-19 | 23 | Hprt |
| Plekhm34 | 9.77E-24 | 0.286849336 | 0.396 | 0.086 | 1.98E-19 | 23 | Plekhm3 |
| Upf12 | 1.14E-23 | 0.279926105 | 0.469 | 0.113 | 2.31E-19 | 23 | Upf1 |
| Smarca51 | 1.25E-23 | 0.502375384 | 0.854 | 0.302 | 2.53E-19 | 23 | Smarca5 |
| Cisd24 | 1.48E-23 | 0.250953883 | 0.625 | 0.175 | 2.99E-19 | 23 | Cisd2 |
| Crip111 | 1.58E-23 | 0.836969431 | 0.99 | 0.537 | 3.21E-19 | 23 | Crip1 |
| R3hdm14 | 1.59E-23 | 0.305524613 | 0.667 | 0.191 | 3.22E-19 | 23 | R3hdm1 |
| Arid1a9 | 1.60E-23 | 0.493883257 | 0.729 | 0.231 | 3.23E-19 | 23 | Arid1a |
| Arhgap257 | 1.67E-23 | 0.387598707 | 0.448 | 0.108 | 3.38E-19 | 23 | Arhgap25 |
| Mcmbp1 | 1.78E-23 | 0.288788842 | 0.531 | 0.136 | 3.61E-19 | 23 | Mcmbp |
| Arpc43 | 1.82E-23 | 0.404809908 | 0.854 | 0.291 | 3.68E-19 | 23 | Arpc4 |
| Vmp16 | 1.89E-23 | 0.370340225 | 0.708 | 0.216 | 3.82E-19 | 23 | Vmp1 |
| Usp485 | 1.91E-23 | 0.292225302 | 0.51 | 0.129 | 3.87E-19 | 23 | Usp48 |
| Luc7l212 | 1.93E-23 | 0.737391282 | 0.948 | 0.405 | 3.90E-19 | 23 | Luc7l2 |
| Ttc9c | 2.02E-23 | 0.255324143 | 0.333 | 0.066 | 4.09E-19 | 23 | Ttc9c |
| Farp15 | 2.31E-23 | 0.250268252 | 0.458 | 0.108 | 4.68E-19 | 23 | Farp1 |
| Gbf18 | 2.38E-23 | 0.307281172 | 0.646 | 0.182 | 4.81E-19 | 23 | Gbf1 |
| Ntan14 | 2.42E-23 | 0.490744909 | 0.865 | 0.318 | 4.91E-19 | 23 | Ntan1 |
| Acbd6 | 2.62E-23 | 0.261400888 | 0.75 | 0.23 | 5.30E-19 | 23 | Acbd6 |
| Pbrm16 | 2.71E-23 | 0.622363592 | 0.792 | 0.288 | 5.49E-19 | 23 | Pbrm1 |

| | | | | | | | |
|---|---|---|---|---|---|---|---|
| Ncbp1 | 2.79E-23 | 0.267492577 | 0.375 | 0.081 | 5.66E-19 | 23 | Ncbp1 |
| Gmfg6 | 2.96E-23 | 0.385142055 | 0.531 | 0.14 | 5.99E-19 | 23 | Gmfg |
| Tapt1 | 3.08E-23 | 0.253628434 | 0.333 | 0.066 | 6.23E-19 | 23 | Tapt1 |
| Ppp6r34 | 3.13E-23 | 0.307285685 | 0.667 | 0.191 | 6.34E-19 | 23 | Ppp6r3 |
| Rnaset2b1 | 3.17E-23 | 0.256838385 | 0.51 | 0.128 | 6.42E-19 | 23 | Rnaset2b |
| Plekha58 | 3.25E-23 | 0.340215016 | 0.573 | 0.156 | 6.58E-19 | 23 | Plekha5 |
| Wipf19 | 3.41E-23 | 0.434049819 | 0.823 | 0.268 | 6.90E-19 | 23 | Wipf1 |
| Rsl1d1 | 3.55E-23 | 0.255830527 | 0.688 | 0.204 | 7.18E-19 | 23 | Rsl1d1 |
| Fam208b7 | 4.04E-23 | 0.256321636 | 0.573 | 0.151 | 8.18E-19 | 23 | Fam208b |
| Zfp6449 | 4.36E-23 | 0.401464998 | 0.792 | 0.256 | 8.82E-19 | 23 | Zfp644 |
| Nlrc56 | 4.38E-23 | 0.363193219 | 0.458 | 0.112 | 8.86E-19 | 23 | Nlrc5 |
| Pdcd6ip4 | 5.23E-23 | 0.361126161 | 0.74 | 0.23 | 1.06E-18 | 23 | Pdcd6ip |
| Stt3b9 | 5.29E-23 | 0.375552806 | 0.594 | 0.165 | 1.07E-18 | 23 | Stt3b |
| Gtf2f21 | 5.50E-23 | 0.263513435 | 0.438 | 0.103 | 1.11E-18 | 23 | Gtf2f2 |
| Galnt76 | 5.53E-23 | 0.328992493 | 0.573 | 0.156 | 1.12E-18 | 23 | Galnt7 |
| Sirt74 | 5.74E-23 | 0.262197285 | 0.521 | 0.134 | 1.16E-18 | 23 | Sirt7 |
| Actg17 | 5.99E-23 | 0.832382141 | 1 | 0.795 | 1.21E-18 | 23 | Actg1 |
| Scfd24 | 6.90E-23 | 0.382757921 | 0.385 | 0.086 | 1.40E-18 | 23 | Scfd2 |
| Ankrd1711 | 7.46E-23 | 0.461206933 | 0.792 | 0.261 | 1.51E-18 | 23 | Ankrd17 |
| Casp86 | 7.53E-23 | 0.356393049 | 0.708 | 0.215 | 1.52E-18 | 23 | Casp8 |
| Atp1b37 | 7.67E-23 | 0.418960596 | 0.771 | 0.249 | 1.55E-18 | 23 | Atp1b3 |
| Dyrk1a8 | 8.17E-23 | 0.464538719 | 0.615 | 0.18 | 1.65E-18 | 23 | Dyrk1a |
| Rps249 | 8.41E-23 | 0.70940375 | 1 | 0.973 | 1.70E-18 | 23 | Rps24 |
| Sin3a | 8.54E-23 | 0.25672536 | 0.448 | 0.107 | 1.73E-18 | 23 | Sin3a |
| Vps545 | 8.61E-23 | 0.331220429 | 0.646 | 0.187 | 1.74E-18 | 23 | Vps54 |
| U2af2 | 8.70E-23 | 0.293600106 | 0.542 | 0.146 | 1.76E-18 | 23 | U2af2 |
| Supt4a6 | 8.96E-23 | 0.337879563 | 0.917 | 0.322 | 1.81E-18 | 23 | Supt4a |
| Slc9a72 | 9.82E-23 | 0.270286631 | 0.25 | 0.042 | 1.99E-18 | 23 | Slc9a7 |
| Set2 | 1.02E-22 | 0.678777465 | 0.969 | 0.474 | 2.08E-18 | 23 | Set |
| Btbd10 | 1.20E-22 | 0.262243168 | 0.396 | 0.089 | 2.43E-18 | 23 | Btbd10 |
| Sppl39 | 1.25E-22 | 0.580226751 | 0.677 | 0.219 | 2.53E-18 | 23 | Sppl3 |
| Itpk14 | 1.50E-22 | 0.284508737 | 0.344 | 0.071 | 3.03E-18 | 23 | Itpk1 |
| Hcfc1 | 1.62E-22 | 0.25284895 | 0.375 | 0.082 | 3.28E-18 | 23 | Hcfc1 |
| Gm446862 | 1.63E-22 | 0.297128708 | 0.312 | 0.061 | 3.29E-18 | 23 | Gm44686 |
| Spidr2 | 1.66E-22 | 0.314876488 | 0.458 | 0.112 | 3.36E-18 | 23 | Spidr |
| Phf148 | 1.68E-22 | 0.397449128 | 0.76 | 0.245 | 3.40E-18 | 23 | Phf14 |
| Chd14 | 1.84E-22 | 0.282077701 | 0.688 | 0.201 | 3.73E-18 | 23 | Chd1 |
| Vasp6 | 2.13E-22 | 0.394986052 | 0.74 | 0.238 | 4.32E-18 | 23 | Vasp |
| Arhgef68 | 2.22E-22 | 0.311883087 | 0.5 | 0.129 | 4.50E-18 | 23 | Arhgef6 |
| Ube2h6 | 2.30E-22 | 0.370930521 | 0.812 | 0.265 | 4.67E-18 | 23 | Ube2h |
| Jak27 | 2.33E-22 | 0.26132393 | 0.5 | 0.127 | 4.71E-18 | 23 | Jak2 |
| Rasa17 | 3.00E-22 | 0.430524853 | 0.76 | 0.247 | 6.07E-18 | 23 | Rasa1 |
| Micu22 | 3.19E-22 | 0.258291582 | 0.646 | 0.19 | 6.46E-18 | 23 | Micu2 |

| | | | | | | | |
|---|---|---|---|---|---|---|---|
| Eif4b1 | 3.27E-22 | 0.294651833 | 0.74 | 0.237 | 6.63E-18 | 23 | Eif4b |
| Reep54 | 3.37E-22 | 0.634710796 | 0.99 | 0.481 | 6.82E-18 | 23 | Reep5 |
| Zfp6389 | 3.60E-22 | 0.580580441 | 0.781 | 0.282 | 7.29E-18 | 23 | Zfp638 |
| Nme15 | 3.66E-22 | 0.586607467 | 0.885 | 0.356 | 7.42E-18 | 23 | Nme1 |
| Lnpep14 | 4.23E-22 | 0.524014708 | 0.792 | 0.26 | 8.55E-18 | 23 | Lnpep |
| Rps1910 | 5.17E-22 | 0.684463206 | 1 | 0.927 | 1.05E-17 | 23 | Rps19 |
| Asb34 | 5.78E-22 | 0.287264186 | 0.448 | 0.11 | 1.17E-17 | 23 | Asb3 |
| Polr2b | 5.88E-22 | 0.467206322 | 0.448 | 0.108 | 1.19E-17 | 23 | Polr2b |
| Vti1a10 | 5.96E-22 | 0.505356489 | 0.76 | 0.255 | 1.21E-17 | 23 | Vti1a |
| Ppp4r3a | 6.72E-22 | 0.2762076 | 0.594 | 0.17 | 1.36E-17 | 23 | Ppp4r3a |
| Kdm6a11 | 6.94E-22 | 0.356116612 | 0.698 | 0.214 | 1.41E-17 | 23 | Kdm6a |
| Ldah | 7.37E-22 | 0.272543303 | 0.385 | 0.088 | 1.49E-17 | 23 | Ldah |
| Pias16 | 8.03E-22 | 0.346748398 | 0.635 | 0.189 | 1.63E-17 | 23 | Pias1 |
| Acaca2 | 8.67E-22 | 0.251673405 | 0.375 | 0.083 | 1.75E-17 | 23 | Acaca |
| Rpn29 | 8.74E-22 | 0.334817088 | 0.688 | 0.212 | 1.77E-17 | 23 | Rpn2 |
| Cct84 | 9.18E-22 | 0.278312045 | 0.802 | 0.267 | 1.86E-17 | 23 | Cct8 |
| Taf14 | 9.49E-22 | 0.272516542 | 0.531 | 0.143 | 1.92E-17 | 23 | Taf1 |
| Psmb97 | 9.83E-22 | 0.610590173 | 0.938 | 0.406 | 1.99E-17 | 23 | Psmb9 |
| Tex147 | 1.06E-21 | 0.465253162 | 0.531 | 0.15 | 2.14E-17 | 23 | Tex14 |
| Btbd114 | 1.07E-21 | 0.294313905 | 0.292 | 0.056 | 2.16E-17 | 23 | Btbd11 |
| Ubn14 | 1.13E-21 | 0.368206879 | 0.76 | 0.246 | 2.28E-17 | 23 | Ubn1 |
| Tsc22d48 | 1.16E-21 | 0.304218188 | 0.854 | 0.294 | 2.35E-17 | 23 | Tsc22d4 |
| Vps13b10 | 1.20E-21 | 0.433274003 | 0.688 | 0.22 | 2.43E-17 | 23 | Vps13b |
| Rb17 | 1.27E-21 | 0.434439233 | 0.677 | 0.215 | 2.58E-17 | 23 | Rb1 |
| Atxn2l1 | 1.36E-21 | 0.251506587 | 0.458 | 0.114 | 2.75E-17 | 23 | Atxn2l |
| Riok15 | 1.50E-21 | 0.29423892 | 0.458 | 0.116 | 3.04E-17 | 23 | Riok1 |
| Rnf1665 | 1.70E-21 | 0.270769263 | 0.438 | 0.108 | 3.45E-17 | 23 | Rnf166 |
| Fmr11 | 1.73E-21 | 0.267429863 | 0.5 | 0.131 | 3.50E-17 | 23 | Fmr1 |
| Map2k52 | 1.74E-21 | 0.278535759 | 0.49 | 0.128 | 3.53E-17 | 23 | Map2k5 |
| Tmem2434 | 1.76E-21 | 0.334839212 | 0.573 | 0.163 | 3.57E-17 | 23 | Tmem243 |
| Stim111 | 1.87E-21 | 0.478925233 | 0.615 | 0.191 | 3.78E-17 | 23 | Stim1 |
| Ap3b19 | 1.90E-21 | 0.297108219 | 0.708 | 0.218 | 3.85E-17 | 23 | Ap3b1 |
| Gpatch813 | 1.97E-21 | 0.380663068 | 0.792 | 0.258 | 3.99E-17 | 23 | Gpatch8 |
| Sept711 | 2.04E-21 | 0.638158182 | 0.979 | 0.462 | 4.14E-17 | 23 | 7-Sep |
| Wapl3 | 2.15E-21 | 0.291915003 | 0.781 | 0.251 | 4.35E-17 | 23 | Wapl |
| Sec11a8 | 2.37E-21 | 0.406755937 | 0.812 | 0.285 | 4.80E-17 | 23 | Sec11a |
| Lsm14a2 | 2.46E-21 | 0.313826721 | 0.573 | 0.164 | 4.97E-17 | 23 | Lsm14a |
| Phb22 | 2.52E-21 | 0.288833426 | 0.844 | 0.306 | 5.10E-17 | 23 | Phb2 |
| Atxn111 | 2.65E-21 | 0.625178661 | 0.802 | 0.296 | 5.37E-17 | 23 | Atxn1 |
| Trim331 | 2.77E-21 | 0.265406248 | 0.573 | 0.163 | 5.62E-17 | 23 | Trim33 |
| Yaf24 | 2.78E-21 | 0.342933944 | 0.688 | 0.216 | 5.64E-17 | 23 | Yaf2 |
| Mxd14 | 2.93E-21 | 0.294308834 | 0.312 | 0.064 | 5.93E-17 | 23 | Mxd1 |
| Rps169 | 3.10E-21 | 0.673830152 | 1 | 0.927 | 6.28E-17 | 23 | Rps16 |

| | | | | | | | |
|---|---|---|---|---|---|---|---|
| Rpl289 | 3.10E-21 | 0.664962736 | 1 | 0.93 | 6.28E-17 | 23 | Rpl28 |
| Psen15 | 3.11E-21 | 0.271957308 | 0.417 | 0.101 | 6.29E-17 | 23 | Psen1 |
| Rtf15 | 3.52E-21 | 0.291864702 | 0.833 | 0.28 | 7.12E-17 | 23 | Rtf1 |
| Rras25 | 3.69E-21 | 0.261573066 | 0.375 | 0.085 | 7.48E-17 | 23 | Rras2 |
| Ndfip26 | 3.79E-21 | 0.331505262 | 0.76 | 0.245 | 7.67E-17 | 23 | Ndfip2 |
| Cyba10 | 3.81E-21 | 0.583300053 | 0.979 | 0.441 | 7.71E-17 | 23 | Cyba |
| Hnrnpul21 | 3.84E-21 | 0.270972624 | 0.688 | 0.212 | 7.78E-17 | 23 | Hnrnpul2 |
| Brd412 | 4.11E-21 | 0.412362841 | 0.802 | 0.268 | 8.33E-17 | 23 | Brd4 |
| Hacd23 | 4.90E-21 | 0.257608006 | 0.458 | 0.117 | 9.93E-17 | 23 | Hacd2 |
| Ing1 | 4.99E-21 | 0.287189765 | 0.521 | 0.143 | 1.01E-16 | 23 | Ing1 |
| Rsf18 | 5.13E-21 | 0.353736023 | 0.708 | 0.224 | 1.04E-16 | 23 | Rsf1 |
| Ggnbp27 | 5.52E-21 | 0.413970105 | 0.729 | 0.242 | 1.12E-16 | 23 | Ggnbp2 |
| Ppp1r16b10 | 5.71E-21 | 0.380605151 | 0.667 | 0.202 | 1.16E-16 | 23 | Ppp1r16b |
| Vav39 | 5.85E-21 | 0.5113596 | 0.75 | 0.256 | 1.18E-16 | 23 | Vav3 |
| Hint13 | 6.70E-21 | 0.675236183 | 1 | 0.696 | 1.36E-16 | 23 | Hint1 |
| Klhl67 | 6.71E-21 | 0.282482608 | 0.469 | 0.121 | 1.36E-16 | 23 | Klhl6 |
| Tmsb4x11 | 7.19E-21 | 0.708841063 | 1 | 0.934 | 1.45E-16 | 23 | Tmsb4x |
| Rpl189 | 7.33E-21 | 0.659932741 | 1 | 0.9 | 1.48E-16 | 23 | Rpl18 |
| Cdk69 | 8.28E-21 | 0.366423008 | 0.438 | 0.112 | 1.68E-16 | 23 | Cdk6 |
| Hnrnpa2b14 | 9.36E-21 | 0.722949829 | 0.979 | 0.516 | 1.90E-16 | 23 | Hnrnpa2b1 |
| Zfp2071 | 9.41E-21 | 0.333904017 | 0.604 | 0.183 | 1.91E-16 | 23 | Zfp207 |
| Esyt211 | 9.56E-21 | 0.500040124 | 0.719 | 0.243 | 1.94E-16 | 23 | Esyt2 |
| Itpr27 | 9.63E-21 | 0.380963028 | 0.74 | 0.245 | 1.95E-16 | 23 | Itpr2 |
| Hnrnpm2 | 9.64E-21 | 0.392442591 | 0.896 | 0.339 | 1.95E-16 | 23 | Hnrnpm |
| Cct22 | 9.68E-21 | 0.2556281 | 0.865 | 0.309 | 1.96E-16 | 23 | Cct2 |
| 1110008F13Rik | 9.74E-21 | 0.27137099 | 0.844 | 0.297 | 1.97E-16 | 23 | 1110008F13Rik |
| Mtf22 | 1.12E-20 | 0.266149738 | 0.5 | 0.134 | 2.26E-16 | 23 | Mtf2 |
| Ppp3cb3 | 1.12E-20 | 0.260473991 | 0.583 | 0.17 | 2.27E-16 | 23 | Ppp3cb |
| Gspt11 | 1.16E-20 | 0.254974828 | 0.74 | 0.242 | 2.35E-16 | 23 | Gspt1 |
| Mark21 | 1.17E-20 | 0.286418912 | 0.625 | 0.193 | 2.36E-16 | 23 | Mark2 |
| Mlxip5 | 1.17E-20 | 0.318815407 | 0.479 | 0.128 | 2.37E-16 | 23 | Mlxip |
| Pan313 | 1.25E-20 | 0.401759959 | 0.708 | 0.229 | 2.52E-16 | 23 | Pan3 |
| Myo1e10 | 1.67E-20 | 0.514871999 | 0.792 | 0.282 | 3.39E-16 | 23 | Myo1e |
| Agpat54 | 2.10E-20 | 0.316798224 | 0.51 | 0.143 | 4.25E-16 | 23 | Agpat5 |
| Cdc42se27 | 2.31E-20 | 0.33697577 | 0.74 | 0.24 | 4.68E-16 | 23 | Cdc42se2 |
| Tle48 | 2.52E-20 | 0.478683037 | 0.698 | 0.226 | 5.09E-16 | 23 | Tle4 |
| Acin19 | 2.54E-20 | 0.36451112 | 0.896 | 0.316 | 5.14E-16 | 23 | Acin1 |
| Atl34 | 2.73E-20 | 0.303430212 | 0.479 | 0.129 | 5.52E-16 | 23 | Atl3 |
| Mapre11 | 2.74E-20 | 0.330638815 | 0.802 | 0.285 | 5.54E-16 | 23 | Mapre1 |
| Manf2 | 3.10E-20 | 0.477592079 | 0.844 | 0.328 | 6.28E-16 | 23 | Manf |
| Fbxw74 | 3.13E-20 | 0.267898669 | 0.49 | 0.132 | 6.33E-16 | 23 | Fbxw7 |
| Krtcap27 | 3.32E-20 | 0.549095066 | 0.969 | 0.442 | 6.72E-16 | 23 | Krtcap2 |
| Khdrbs12 | 3.59E-20 | 0.26547723 | 0.802 | 0.266 | 7.28E-16 | 23 | Khdrbs1 |

| | | | | | | | |
|---|---|---|---|---|---|---|---|
| Prelid13 | 3.75E-20 | 0.420016235 | 0.823 | 0.311 | 7.59E-16 | 23 | Prelid1 |
| Nfatc16 | 4.18E-20 | 0.335339823 | 0.448 | 0.117 | 8.47E-16 | 23 | Nfatc1 |
| Gne2 | 4.31E-20 | 0.273989402 | 0.469 | 0.125 | 8.72E-16 | 23 | Gne |
| Nudc2 | 4.75E-20 | 0.367037797 | 0.896 | 0.351 | 9.61E-16 | 23 | Nudc |
| Bclaf110 | 4.87E-20 | 0.304710711 | 0.781 | 0.254 | 9.85E-16 | 23 | Bclaf1 |
| Btaf12 | 4.99E-20 | 0.308922323 | 0.469 | 0.126 | 1.01E-15 | 23 | Btaf1 |
| Ifi477 | 5.16E-20 | 0.270268834 | 0.604 | 0.179 | 1.05E-15 | 23 | Ifi47 |
| Lrp108 | 6.45E-20 | 0.290640025 | 0.625 | 0.189 | 1.31E-15 | 23 | Lrp10 |
| Ubr28 | 7.21E-20 | 0.441894348 | 0.688 | 0.23 | 1.46E-15 | 23 | Ubr2 |
| Hnrnpa02 | 7.40E-20 | 0.603487775 | 0.979 | 0.494 | 1.50E-15 | 23 | Hnrnpa0 |
| Zfp913 | 7.79E-20 | 0.285829275 | 0.854 | 0.308 | 1.58E-15 | 23 | Zfp91 |
| Fnbp46 | 8.36E-20 | 0.259065772 | 0.479 | 0.129 | 1.69E-15 | 23 | Fnbp4 |
| Hectd18 | 9.68E-20 | 0.465975084 | 0.812 | 0.301 | 1.96E-15 | 23 | Hectd1 |
| Lcor8 | 9.93E-20 | 0.38686364 | 0.604 | 0.191 | 2.01E-15 | 23 | Lcor |
| Prdx64 | 1.03E-19 | 0.276542144 | 0.885 | 0.338 | 2.09E-15 | 23 | Prdx6 |
| Ppig7 | 1.19E-19 | 0.38836839 | 0.844 | 0.316 | 2.40E-15 | 23 | Ppig |
| Cish2 | 1.19E-19 | 0.264703025 | 0.312 | 0.068 | 2.41E-15 | 23 | Cish |
| Tor1aip25 | 1.26E-19 | 0.272854938 | 0.604 | 0.183 | 2.54E-15 | 23 | Tor1aip2 |
| Pacs15 | 1.46E-19 | 0.413852882 | 0.635 | 0.205 | 2.95E-15 | 23 | Pacs1 |
| Gmip6 | 1.51E-19 | 0.2657005 | 0.354 | 0.083 | 3.05E-15 | 23 | Gmip |
| Ssr23 | 1.57E-19 | 0.276018731 | 0.646 | 0.203 | 3.18E-15 | 23 | Ssr2 |
| Ndfip17 | 1.61E-19 | 0.465202084 | 0.844 | 0.322 | 3.26E-15 | 23 | Ndfip1 |
| Sf3b19 | 1.61E-19 | 0.463096291 | 0.969 | 0.394 | 3.27E-15 | 23 | Sf3b1 |
| Ddx105 | 1.67E-19 | 0.301398186 | 0.479 | 0.132 | 3.39E-15 | 23 | Ddx10 |
| Cct53 | 1.77E-19 | 0.288057791 | 0.823 | 0.295 | 3.59E-15 | 23 | Cct5 |
| Trip116 | 1.83E-19 | 0.29072414 | 0.698 | 0.224 | 3.70E-15 | 23 | Trip11 |
| Actr26 | 1.90E-19 | 0.498385104 | 0.875 | 0.344 | 3.85E-15 | 23 | Actr2 |
| U2af11 | 2.16E-19 | 0.352983802 | 0.833 | 0.311 | 4.38E-15 | 23 | U2af1 |
| Tln111 | 2.23E-19 | 0.564703062 | 0.969 | 0.406 | 4.51E-15 | 23 | Tln1 |
| Tmem2584 | 2.32E-19 | 0.307662413 | 0.823 | 0.309 | 4.69E-15 | 23 | Tmem258 |
| Zfp36l28 | 2.47E-19 | 0.390245893 | 0.844 | 0.306 | 5.01E-15 | 23 | Zfp36l2 |
| Apaf11 | 2.50E-19 | 0.294739233 | 0.365 | 0.087 | 5.06E-15 | 23 | Apaf1 |
| Dcaf55 | 2.68E-19 | 0.359829008 | 0.469 | 0.13 | 5.42E-15 | 23 | Dcaf5 |
| Xpo75 | 2.72E-19 | 0.271337824 | 0.531 | 0.154 | 5.50E-15 | 23 | Xpo7 |
| Setd512 | 2.86E-19 | 0.365361971 | 0.771 | 0.264 | 5.78E-15 | 23 | Setd5 |
| Kansl112 | 3.13E-19 | 0.5758334 | 0.865 | 0.34 | 6.34E-15 | 23 | Kansl1 |
| Usp36 | 3.40E-19 | 0.339216316 | 0.583 | 0.183 | 6.88E-15 | 23 | Usp3 |
| Nup214 | 3.41E-19 | 0.266087883 | 0.406 | 0.104 | 6.91E-15 | 23 | Nup214 |
| Ankrd1113 | 3.77E-19 | 0.673399502 | 0.885 | 0.381 | 7.63E-15 | 23 | Ankrd11 |
| Tlk14 | 4.17E-19 | 0.252392828 | 0.688 | 0.223 | 8.43E-15 | 23 | Tlk1 |
| Ube2d2a1 | 4.17E-19 | 0.366833593 | 0.781 | 0.285 | 8.45E-15 | 23 | Ube2d2a |
| Zzef14 | 4.20E-19 | 0.286511733 | 0.521 | 0.149 | 8.51E-15 | 23 | Zzef1 |
| Rp92 | 4.25E-19 | 0.322579475 | 0.865 | 0.325 | 8.61E-15 | 23 | Rp9 |

| | | | | | | | |
|---|---|---|---|---|---|---|---|
| Npm18 | 4.67E-19 | 0.754622052 | 1 | 0.68 | 9.46E-15 | 23 | Npm1 |
| Rps3a110 | 4.78E-19 | 0.670501713 | 1 | 0.935 | 9.68E-15 | 23 | Rps3a1 |
| Dennd1a13 | 5.16E-19 | 0.420914712 | 0.729 | 0.253 | 1.04E-14 | 23 | Dennd1a |
| Tmem1233 | 5.87E-19 | 0.271764093 | 0.531 | 0.154 | 1.19E-14 | 23 | Tmem123 |
| Kcnq1ot110 | 5.94E-19 | 0.324471294 | 0.479 | 0.135 | 1.20E-14 | 23 | Kcnq1ot1 |
| B4galt19 | 5.96E-19 | 0.296388842 | 0.583 | 0.177 | 1.21E-14 | 23 | B4galt1 |
| Creb17 | 6.00E-19 | 0.296945526 | 0.646 | 0.208 | 1.22E-14 | 23 | Creb1 |
| Elavl11 | 6.34E-19 | 0.269106399 | 0.583 | 0.179 | 1.28E-14 | 23 | Elavl1 |
| Cdk1310 | 6.53E-19 | 0.358471068 | 0.688 | 0.228 | 1.32E-14 | 23 | Cdk13 |
| Zfp29210 | 7.31E-19 | 0.292048997 | 0.75 | 0.252 | 1.48E-14 | 23 | Zfp292 |
| Tbc1d512 | 7.57E-19 | 0.346365718 | 0.76 | 0.262 | 1.53E-14 | 23 | Tbc1d5 |
| Hnrnpr4 | 7.86E-19 | 0.308114375 | 0.698 | 0.236 | 1.59E-14 | 23 | Hnrnpr |
| Ern16 | 7.86E-19 | 0.26393883 | 0.354 | 0.085 | 1.59E-14 | 23 | Ern1 |
| Arid4b9 | 8.79E-19 | 0.484148716 | 0.792 | 0.293 | 1.78E-14 | 23 | Arid4b |
| Exoc68 | 9.29E-19 | 0.325437016 | 0.562 | 0.171 | 1.88E-14 | 23 | Exoc6 |
| Tnpo32 | 1.08E-18 | 0.318196452 | 0.5 | 0.146 | 2.18E-14 | 23 | Tnpo3 |
| Ranbp25 | 1.08E-18 | 0.301105922 | 0.625 | 0.199 | 2.19E-14 | 23 | Ranbp2 |
| Cct71 | 1.17E-18 | 0.311134468 | 0.812 | 0.304 | 2.37E-14 | 23 | Cct7 |
| Nr4a26 | 1.28E-18 | 0.328640695 | 0.469 | 0.131 | 2.59E-14 | 23 | Nr4a2 |
| Foxo36 | 1.29E-18 | 0.253730079 | 0.438 | 0.117 | 2.61E-14 | 23 | Foxo3 |
| Trir1 | 1.37E-18 | 0.463805139 | 0.906 | 0.418 | 2.78E-14 | 23 | Trir |
| Ep4007 | 1.57E-18 | 0.313792774 | 0.646 | 0.208 | 3.19E-14 | 23 | Ep400 |
| Fdps | 1.62E-18 | 0.260645815 | 0.312 | 0.071 | 3.29E-14 | 23 | Fdps |
| Lcorl8 | 1.78E-18 | 0.324893911 | 0.552 | 0.167 | 3.61E-14 | 23 | Lcorl |
| Pitpnc19 | 2.17E-18 | 0.68634422 | 0.938 | 0.43 | 4.39E-14 | 23 | Pitpnc1 |
| Sf3b62 | 2.25E-18 | 0.317621667 | 0.896 | 0.361 | 4.55E-14 | 23 | Sf3b6 |
| Tpm39 | 2.36E-18 | 0.629632265 | 0.969 | 0.48 | 4.78E-14 | 23 | Tpm3 |
| Lrrfip17 | 3.43E-18 | 0.444908368 | 0.938 | 0.379 | 6.94E-14 | 23 | Lrrfip1 |
| Prkch8 | 4.15E-18 | 0.651741479 | 0.927 | 0.4 | 8.40E-14 | 23 | Prkch |
| Zdhhc2010 | 4.76E-18 | 0.331215391 | 0.688 | 0.231 | 9.64E-14 | 23 | Zdhhc20 |
| Gimap88 | 4.78E-18 | 0.285369129 | 0.562 | 0.169 | 9.68E-14 | 23 | Gimap8 |
| Phf203 | 5.67E-18 | 0.302604238 | 0.49 | 0.145 | 1.15E-13 | 23 | Phf20 |
| Rps79 | 6.01E-18 | 0.646385916 | 0.99 | 0.917 | 1.22E-13 | 23 | Rps7 |
| Phf21a12 | 6.94E-18 | 0.425061441 | 0.708 | 0.253 | 1.41E-13 | 23 | Phf21a |
| Cul31 | 7.13E-18 | 0.271469328 | 0.802 | 0.287 | 1.44E-13 | 23 | Cul3 |
| Rpl320 | 7.14E-18 | 0.625393404 | 1 | 0.91 | 1.44E-13 | 23 | Rpl3 |
| Lrif12 | 7.52E-18 | 0.254553815 | 0.479 | 0.139 | 1.52E-13 | 23 | Lrif1 |
| Ncoa210 | 7.72E-18 | 0.428934779 | 0.677 | 0.239 | 1.56E-13 | 23 | Ncoa2 |
| Dbi5 | 8.26E-18 | 0.282125631 | 0.938 | 0.433 | 1.67E-13 | 23 | Dbi |
| Chd29 | 8.58E-18 | 0.516206244 | 0.698 | 0.254 | 1.74E-13 | 23 | Chd2 |
| Eif3h5 | 9.24E-18 | 0.445212924 | 0.958 | 0.481 | 1.87E-13 | 23 | Eif3h |
| Prpf4b9 | 1.02E-17 | 0.359185344 | 0.844 | 0.334 | 2.07E-13 | 23 | Prpf4b |
| Ccar18 | 1.04E-17 | 0.287864858 | 0.635 | 0.208 | 2.11E-13 | 23 | Ccar1 |

| | | | | | | | |
|---|---|---|---|---|---|---|---|
| Osbpl88 | 1.27E-17 | 0.321464747 | 0.604 | 0.2 | 2.58E-13 | 23 | Osbpl8 |
| Arpc29 | 1.38E-17 | 0.616875089 | 1 | 0.545 | 2.79E-13 | 23 | Arpc2 |
| Rps222 | 1.48E-17 | 0.719895231 | 0.99 | 0.92 | 3.00E-13 | 23 | Rps2 |
| Hnrnpc1 | 1.49E-17 | 0.315360246 | 0.896 | 0.349 | 3.01E-13 | 23 | Hnrnpc |
| Cfl19 | 1.55E-17 | 0.650280842 | 0.99 | 0.609 | 3.13E-13 | 23 | Cfl1 |
| Traf38 | 1.58E-17 | 0.265286876 | 0.406 | 0.11 | 3.21E-13 | 23 | Traf3 |
| Cdk1710 | 1.62E-17 | 0.3551343 | 0.719 | 0.245 | 3.28E-13 | 23 | Cdk17 |
| Fam172a13 | 1.65E-17 | 0.491562732 | 0.75 | 0.285 | 3.34E-13 | 23 | Fam172a |
| Rpl139 | 1.78E-17 | 0.590322304 | 1 | 0.977 | 3.60E-13 | 23 | Rpl13 |
| Ssh28 | 1.84E-17 | 0.379020388 | 0.656 | 0.227 | 3.73E-13 | 23 | Ssh2 |
| Sema6d7 | 1.89E-17 | 0.380210196 | 0.635 | 0.215 | 3.82E-13 | 23 | Sema6d |
| Sltm9 | 2.00E-17 | 0.366155672 | 0.74 | 0.271 | 4.05E-13 | 23 | Sltm |
| Ostc4 | 2.36E-17 | 0.265802957 | 0.75 | 0.263 | 4.78E-13 | 23 | Ostc |
| Rps6ka39 | 2.57E-17 | 0.351016916 | 0.74 | 0.263 | 5.20E-13 | 23 | Rps6ka3 |
| Srrm212 | 2.75E-17 | 0.559069161 | 0.99 | 0.47 | 5.58E-13 | 23 | Srrm2 |
| Rbm69 | 2.85E-17 | 0.356423987 | 0.635 | 0.218 | 5.78E-13 | 23 | Rbm6 |
| Stk246 | 2.86E-17 | 0.253722697 | 0.604 | 0.196 | 5.80E-13 | 23 | Stk24 |
| U2surp2 | 3.09E-17 | 0.296687151 | 0.74 | 0.269 | 6.25E-13 | 23 | U2surp |
| Ints66 | 3.59E-17 | 0.290100571 | 0.479 | 0.143 | 7.27E-13 | 23 | Ints6 |
| Cnot410 | 3.67E-17 | 0.290145436 | 0.771 | 0.269 | 7.43E-13 | 23 | Cnot4 |
| Ago27 | 3.83E-17 | 0.284050899 | 0.531 | 0.168 | 7.76E-13 | 23 | Ago2 |
| Clcn38 | 4.44E-17 | 0.315822335 | 0.458 | 0.134 | 9.00E-13 | 23 | Clcn3 |
| Serbp15 | 4.81E-17 | 0.56825453 | 0.99 | 0.591 | 9.73E-13 | 23 | Serbp1 |
| Sesn19 | 5.19E-17 | 0.278333628 | 0.635 | 0.213 | 1.05E-12 | 23 | Sesn1 |
| Cbfa2t22 | 5.27E-17 | 0.278698363 | 0.427 | 0.118 | 1.07E-12 | 23 | Cbfa2t2 |
| Pign4 | 5.27E-17 | 0.266057626 | 0.365 | 0.095 | 1.07E-12 | 23 | Pign |
| Tma75 | 5.77E-17 | 0.389859218 | 0.99 | 0.524 | 1.17E-12 | 23 | Tma7 |
| Psmb34 | 6.04E-17 | 0.416007289 | 0.917 | 0.42 | 1.22E-12 | 23 | Psmb3 |
| Snrpb3 | 6.57E-17 | 0.499192445 | 0.969 | 0.473 | 1.33E-12 | 23 | Snrpb |
| Arid1b11 | 6.57E-17 | 0.288471011 | 0.74 | 0.259 | 1.33E-12 | 23 | Arid1b |
| Dpm34 | 7.70E-17 | 0.353750136 | 0.865 | 0.35 | 1.56E-12 | 23 | Dpm3 |
| Eef1b28 | 8.04E-17 | 0.622712247 | 1 | 0.737 | 1.63E-12 | 23 | Eef1b2 |
| Tmcc113 | 9.44E-17 | 0.290746634 | 0.646 | 0.218 | 1.91E-12 | 23 | Tmcc1 |
| Saraf8 | 1.03E-16 | 0.455249287 | 0.781 | 0.308 | 2.09E-12 | 23 | Saraf |
| Rsrc18 | 1.04E-16 | 0.270332777 | 0.729 | 0.261 | 2.11E-12 | 23 | Rsrc1 |
| Rps187 | 1.07E-16 | 0.632345272 | 0.99 | 0.942 | 2.16E-12 | 23 | Rps18 |
| Abracl2 | 1.07E-16 | 0.255271702 | 0.927 | 0.429 | 2.17E-12 | 23 | Abracl |
| Nup983 | 1.24E-16 | 0.302876152 | 0.479 | 0.145 | 2.50E-12 | 23 | Nup98 |
| Nek79 | 1.24E-16 | 0.274442203 | 0.698 | 0.249 | 2.51E-12 | 23 | Nek7 |
| Srrm12 | 1.25E-16 | 0.322167122 | 0.896 | 0.351 | 2.54E-12 | 23 | Srrm1 |
| Ssr45 | 1.32E-16 | 0.470618187 | 0.969 | 0.468 | 2.68E-12 | 23 | Ssr4 |
| Dnajc84 | 1.34E-16 | 0.311228583 | 0.896 | 0.359 | 2.71E-12 | 23 | Dnajc8 |
| Gimap68 | 1.47E-16 | 0.39852666 | 0.938 | 0.366 | 2.97E-12 | 23 | Gimap6 |

| | | | | | | | |
|---|---|---|---|---|---|---|---|
| Sub17 | 1.50E-16 | 0.50076174 | 0.979 | 0.516 | 3.04E-12 | 23 | Sub1 |
| Btbd912 | 1.72E-16 | 0.283352472 | 0.688 | 0.238 | 3.48E-12 | 23 | Btbd9 |
| Smg112 | 1.76E-16 | 0.257234091 | 0.688 | 0.235 | 3.56E-12 | 23 | Smg1 |
| Zc3h138 | 1.78E-16 | 0.263209117 | 0.635 | 0.216 | 3.61E-12 | 23 | Zc3h13 |
| Smad2 | 1.94E-16 | 0.279691661 | 0.5 | 0.155 | 3.93E-12 | 23 | Smad2 |
| Spata56 | 1.95E-16 | 0.288716842 | 0.427 | 0.124 | 3.95E-12 | 23 | Spata5 |
| Rpl158 | 2.02E-16 | 0.575163117 | 1 | 0.901 | 4.09E-12 | 23 | Rpl15 |
| Rpl329 | 2.67E-16 | 0.602284726 | 1 | 0.953 | 5.40E-12 | 23 | Rpl32 |
| Ttc7b4 | 2.72E-16 | 0.258004772 | 0.427 | 0.123 | 5.50E-12 | 23 | Ttc7b |
| 4932438A13Rik10 | 3.12E-16 | 0.34416117 | 0.635 | 0.22 | 6.32E-12 | 23 | 4932438A13Rik |
| Atrx5 | 3.23E-16 | 0.405638119 | 0.969 | 0.42 | 6.55E-12 | 23 | Atrx |
| Psmb812 | 3.45E-16 | 0.62537161 | 0.99 | 0.558 | 6.99E-12 | 23 | Psmb8 |
| Rpl89 | 3.77E-16 | 0.536989425 | 1 | 0.925 | 7.63E-12 | 23 | Rpl8 |
| Phactr29 | 3.78E-16 | 0.42163962 | 0.667 | 0.247 | 7.65E-12 | 23 | Phactr2 |
| Eif3c2 | 4.06E-16 | 0.264584358 | 0.927 | 0.387 | 8.22E-12 | 23 | Eif3c |
| Rps109 | 4.07E-16 | 0.572331314 | 1 | 0.944 | 8.23E-12 | 23 | Rps10 |
| Fgfr1op21 | 4.36E-16 | 0.256413338 | 0.792 | 0.304 | 8.83E-12 | 23 | Fgfr1op2 |
| Phf312 | 4.84E-16 | 0.254859064 | 0.719 | 0.253 | 9.80E-12 | 23 | Phf3 |
| Gm203889 | 5.60E-16 | 0.502501674 | 0.917 | 0.401 | 1.13E-11 | 23 | Gm20388 |
| Chd49 | 6.12E-16 | 0.321761602 | 0.844 | 0.328 | 1.24E-11 | 23 | Chd4 |
| Ptbp311 | 6.45E-16 | 0.494398906 | 0.896 | 0.394 | 1.31E-11 | 23 | Ptbp3 |
| Rpl37a10 | 8.53E-16 | 0.547449077 | 1 | 0.938 | 1.73E-11 | 23 | Rpl37a |
| Spcs27 | 9.21E-16 | 0.32696019 | 0.969 | 0.401 | 1.86E-11 | 23 | Spcs2 |
| Rpl3910 | 1.04E-15 | 0.602936799 | 1 | 0.901 | 2.11E-11 | 23 | Rpl39 |
| Fau8 | 1.06E-15 | 0.479912132 | 1 | 0.953 | 2.15E-11 | 23 | Fau |
| Rps139 | 1.17E-15 | 0.541329989 | 1 | 0.924 | 2.37E-11 | 23 | Rps13 |
| Rps4x10 | 1.19E-15 | 0.532638412 | 1 | 0.941 | 2.41E-11 | 23 | Rps4x |
| Tubb4b4 | 1.22E-15 | 0.782475997 | 0.844 | 0.428 | 2.47E-11 | 23 | Tubb4b |
| Hsp90b112 | 1.42E-15 | 0.637335974 | 0.969 | 0.511 | 2.88E-11 | 23 | Hsp90b1 |
| Ankrd283 | 1.49E-15 | 0.274613516 | 0.448 | 0.134 | 3.02E-11 | 23 | Ankrd28 |
| Stk107 | 1.81E-15 | 0.30332373 | 0.552 | 0.184 | 3.67E-11 | 23 | Stk10 |
| Rpl2111 | 1.85E-15 | 0.4942856 | 1 | 0.923 | 3.75E-11 | 23 | Rpl21 |
| Mllt1012 | 2.02E-15 | 0.290001832 | 0.75 | 0.275 | 4.09E-11 | 23 | Mllt10 |
| Ppp1r12a12 | 2.09E-15 | 0.488724956 | 0.938 | 0.426 | 4.23E-11 | 23 | Ppp1r12a |
| Iws12 | 2.11E-15 | 0.251431299 | 0.49 | 0.155 | 4.27E-11 | 23 | Iws1 |
| Pdia610 | 2.24E-15 | 0.331840121 | 0.854 | 0.338 | 4.53E-11 | 23 | Pdia6 |
| Srsf1113 | 2.55E-15 | 0.383299736 | 0.854 | 0.351 | 5.16E-11 | 23 | Srsf11 |
| Pdia310 | 2.79E-15 | 0.466964637 | 0.948 | 0.444 | 5.65E-11 | 23 | Pdia3 |
| Nav28 | 2.83E-15 | 0.343526813 | 0.51 | 0.166 | 5.72E-11 | 23 | Nav2 |
| Rpl297 | 2.83E-15 | 0.566976098 | 1 | 0.769 | 5.73E-11 | 23 | Rpl29 |
| Ankrd1212 | 3.11E-15 | 0.250493935 | 0.812 | 0.309 | 6.30E-11 | 23 | Ankrd12 |
| mt-Atp813 | 3.75E-15 | 0.43828675 | 0.979 | 0.464 | 7.60E-11 | 23 | mt-Atp8 |
| Rplp09 | 3.99E-15 | 0.550173031 | 1 | 0.945 | 8.08E-11 | 23 | Rplp0 |

| | | | | | | | |
|---|---|---|---|---|---|---|---|
| Itch11 | 4.48E-15 | 0.262226128 | 0.74 | 0.274 | 9.07E-11 | 23 | Itch |
| Rpl148 | 5.05E-15 | 0.536764246 | 1 | 0.803 | 1.02E-10 | 23 | Rpl14 |
| Dis3l28 | 5.85E-15 | 0.255974378 | 0.448 | 0.14 | 1.18E-10 | 23 | Dis3l2 |
| Ankhd110 | 5.85E-15 | 0.297546006 | 0.677 | 0.244 | 1.18E-10 | 23 | Ankhd1 |
| Kdm3b6 | 6.01E-15 | 0.292532627 | 0.438 | 0.137 | 1.22E-10 | 23 | Kdm3b |
| Nsa28 | 6.73E-15 | 0.331819053 | 0.979 | 0.487 | 1.36E-10 | 23 | Nsa2 |
| Rnf13010 | 7.07E-15 | 0.298795877 | 0.74 | 0.284 | 1.43E-10 | 23 | Rnf130 |
| Eif3e6 | 8.18E-15 | 0.293817153 | 0.917 | 0.413 | 1.66E-10 | 23 | Eif3e |
| mt-Co116 | 9.44E-15 | 0.634381515 | 1 | 0.72 | 1.91E-10 | 23 | mt-Co1 |
| Eif3k4 | 1.06E-14 | 0.371587821 | 0.969 | 0.502 | 2.15E-10 | 23 | Eif3k |
| Rtn37 | 1.14E-14 | 0.298203911 | 0.812 | 0.324 | 2.31E-10 | 23 | Rtn3 |
| Btg28 | 1.29E-14 | 0.33427279 | 0.833 | 0.345 | 2.61E-10 | 23 | Btg2 |
| mt-Nd512 | 1.46E-14 | 0.519395761 | 0.979 | 0.497 | 2.95E-10 | 23 | mt-Nd5 |
| Rpl18a7 | 1.59E-14 | 0.478611343 | 1 | 0.931 | 3.22E-10 | 23 | Rpl18a |
| Akt312 | 1.62E-14 | 0.350483709 | 0.677 | 0.252 | 3.29E-10 | 23 | Akt3 |
| Btg16 | 2.17E-14 | 0.416020363 | 0.969 | 0.46 | 4.39E-10 | 23 | Btg1 |
| Eif3a7 | 2.24E-14 | 0.281278086 | 0.896 | 0.373 | 4.53E-10 | 23 | Eif3a |
| Rftn110 | 2.43E-14 | 0.331754335 | 0.667 | 0.252 | 4.91E-10 | 23 | Rftn1 |
| Psmb11 | 2.59E-14 | 0.367067447 | 0.979 | 0.512 | 5.24E-10 | 23 | Psmb1 |
| Gm100767 | 2.72E-14 | 0.476811647 | 0.958 | 0.546 | 5.51E-10 | 23 | Gm10076 |
| Fbxo119 | 3.06E-14 | 0.317802599 | 0.635 | 0.233 | 6.20E-10 | 23 | Fbxo11 |
| Rplp210 | 3.13E-14 | 0.486469007 | 1 | 0.891 | 6.34E-10 | 23 | Rplp2 |
| Birc613 | 3.22E-14 | 0.387585809 | 0.812 | 0.342 | 6.52E-10 | 23 | Birc6 |
| Arpc52 | 3.42E-14 | 0.320084597 | 0.938 | 0.429 | 6.92E-10 | 23 | Arpc5 |
| Mycbp214 | 3.64E-14 | 0.318539813 | 0.917 | 0.375 | 7.38E-10 | 23 | Mycbp2 |
| H2-K110 | 5.30E-14 | 0.685296858 | 1 | 0.644 | 1.07E-09 | 23 | H2-K1 |
| Kpna410 | 5.84E-14 | 0.272673599 | 0.812 | 0.324 | 1.18E-09 | 23 | Kpna4 |
| Rps2610 | 6.88E-14 | 0.469260762 | 1 | 0.866 | 1.39E-09 | 23 | Rps26 |
| 1810037I17Rik3 | 8.16E-14 | 0.411989341 | 0.927 | 0.48 | 1.65E-09 | 23 | 1810037I17Rik |
| Pnisr12 | 8.26E-14 | 0.265171793 | 0.698 | 0.265 | 1.67E-09 | 23 | Pnisr |
| Rps89 | 8.33E-14 | 0.469385688 | 1 | 0.965 | 1.69E-09 | 23 | Rps8 |
| Arih110 | 9.69E-14 | 0.337934468 | 0.771 | 0.313 | 1.96E-09 | 23 | Arih1 |
| Rpl48 | 9.89E-14 | 0.498228317 | 1 | 0.659 | 2.00E-09 | 23 | Rpl4 |
| Arpp193 | 1.04E-13 | 0.295193081 | 0.917 | 0.441 | 2.12E-09 | 23 | Arpp19 |
| Prrc2c12 | 1.14E-13 | 0.400342343 | 0.927 | 0.421 | 2.31E-09 | 23 | Prrc2c |
| Atf7ip7 | 1.16E-13 | 0.252938554 | 0.573 | 0.208 | 2.36E-09 | 23 | Atf7ip |
| Ncor111 | 1.18E-13 | 0.417230846 | 0.906 | 0.439 | 2.38E-09 | 23 | Ncor1 |
| Eif3f6 | 1.97E-13 | 0.433247812 | 0.948 | 0.484 | 3.99E-09 | 23 | Eif3f |
| Anxa29 | 2.43E-13 | 0.457700378 | 0.906 | 0.417 | 4.92E-09 | 23 | Anxa2 |
| Nipbl15 | 2.44E-13 | 0.303497681 | 0.875 | 0.367 | 4.94E-09 | 23 | Nipbl |
| Rpl610 | 2.56E-13 | 0.451071538 | 1 | 0.882 | 5.19E-09 | 23 | Rpl6 |
| Ablim18 | 2.64E-13 | 0.329711873 | 0.917 | 0.403 | 5.35E-09 | 23 | Ablim1 |
| S100a114 | 2.68E-13 | 0.342081956 | 0.99 | 0.574 | 5.43E-09 | 23 | S100a11 |

| | | | | | | | |
|---|---|---|---|---|---|---|---|
| Iqgap111 | 3.38E-13 | 0.391656305 | 0.906 | 0.413 | 6.85E-09 | 23 | Iqgap1 |
| Sumo22 | 4.59E-13 | 0.501117683 | 0.99 | 0.659 | 9.30E-09 | 23 | Sumo2 |
| Rpl416 | 4.63E-13 | 0.442636261 | 1 | 0.962 | 9.36E-09 | 23 | Rpl41 |
| Rock111 | 5.29E-13 | 0.382423238 | 0.927 | 0.433 | 1.07E-08 | 23 | Rock1 |
| Pcbp2 | 5.57E-13 | 0.352549827 | 0.958 | 0.488 | 1.13E-08 | 23 | Pcbp2 |
| Dad18 | 5.96E-13 | 0.322300627 | 0.969 | 0.459 | 1.21E-08 | 23 | Dad1 |
| Hspa512 | 7.01E-13 | 0.381319419 | 0.927 | 0.436 | 1.42E-08 | 23 | Hspa5 |
| Wnk111 | 7.26E-13 | 0.314475207 | 0.844 | 0.364 | 1.47E-08 | 23 | Wnk1 |
| Rpl35a9 | 9.56E-13 | 0.411252605 | 1 | 0.949 | 1.93E-08 | 23 | Rpl35a |
| Git25 | 9.68E-13 | 0.294783664 | 0.615 | 0.236 | 1.96E-08 | 23 | Git2 |
| Uba528 | 1.11E-12 | 0.285828169 | 0.979 | 0.522 | 2.25E-08 | 23 | Uba52 |
| Lrmda8 | 1.53E-12 | 0.36237259 | 0.833 | 0.369 | 3.09E-08 | 23 | Lrmda |
| mt-Co318 | 1.83E-12 | 0.545160688 | 1 | 0.727 | 3.71E-08 | 23 | mt-Co3 |
| Rpl128 | 1.90E-12 | 0.477918732 | 0.99 | 0.842 | 3.85E-08 | 23 | Rpl12 |
| Nedd98 | 2.10E-12 | 0.279996936 | 0.771 | 0.32 | 4.26E-08 | 23 | Nedd9 |
| Tcf1211 | 2.26E-12 | 0.384284411 | 0.844 | 0.377 | 4.58E-08 | 23 | Tcf12 |
| Ppp3ca14 | 2.45E-12 | 0.339176566 | 0.875 | 0.394 | 4.97E-08 | 23 | Ppp3ca |
| Elmo112 | 2.51E-12 | 0.306963595 | 0.844 | 0.367 | 5.09E-08 | 23 | Elmo1 |
| Arf51 | 2.66E-12 | 0.291861002 | 0.99 | 0.545 | 5.39E-08 | 23 | Arf5 |
| Rps129 | 2.72E-12 | 0.453508325 | 0.99 | 0.899 | 5.52E-08 | 23 | Rps12 |
| Arpc1b8 | 3.10E-12 | 0.429687496 | 0.979 | 0.592 | 6.27E-08 | 23 | Arpc1b |
| Eef1g7 | 3.26E-12 | 0.302318825 | 0.979 | 0.512 | 6.61E-08 | 23 | Eef1g |
| Rpl198 | 3.94E-12 | 0.405036204 | 1 | 0.963 | 7.98E-08 | 23 | Rpl19 |
| Rpl347 | 3.99E-12 | 0.431015683 | 0.99 | 0.918 | 8.08E-08 | 23 | Rpl34 |
| Rpl239 | 5.26E-12 | 0.425334027 | 1 | 0.971 | 1.07E-07 | 23 | Rpl23 |
| mt-Nd117 | 6.22E-12 | 0.490324892 | 1 | 0.6 | 1.26E-07 | 23 | mt-Nd1 |
| AW5549188 | 6.55E-12 | 0.301724371 | 0.469 | 0.172 | 1.33E-07 | 23 | AW554918 |
| Rps69 | 7.30E-12 | 0.460787853 | 1 | 0.812 | 1.48E-07 | 23 | Rps6 |
| Rps289 | 8.10E-12 | 0.476917213 | 1 | 0.898 | 1.64E-07 | 23 | Rps28 |
| Usp3412 | 8.18E-12 | 0.278997983 | 0.792 | 0.345 | 1.66E-07 | 23 | Usp34 |
| Rpl10a11 | 8.21E-12 | 0.45369714 | 1 | 0.836 | 1.66E-07 | 23 | Rpl10a |
| Rpl3510 | 1.01E-11 | 0.428935983 | 0.99 | 0.829 | 2.04E-07 | 23 | Rpl35 |
| Zcchc714 | 1.30E-11 | 0.284718205 | 0.708 | 0.302 | 2.63E-07 | 23 | Zcchc7 |
| Foxn311 | 1.35E-11 | 0.288702452 | 0.906 | 0.411 | 2.73E-07 | 23 | Foxn3 |
| Rpl27a8 | 1.40E-11 | 0.412243078 | 1 | 0.96 | 2.84E-07 | 23 | Rpl27a |
| mt-Nd314 | 1.52E-11 | 0.297820658 | 0.948 | 0.447 | 3.07E-07 | 23 | mt-Nd3 |
| Rps910 | 1.77E-11 | 0.399985424 | 1 | 0.921 | 3.59E-07 | 23 | Rps9 |
| Cmtm74 | 1.85E-11 | 0.277519855 | 0.542 | 0.214 | 3.74E-07 | 23 | Cmtm7 |
| Diaph210 | 2.11E-11 | 0.379404057 | 0.812 | 0.377 | 4.28E-07 | 23 | Diaph2 |
| Rps238 | 2.93E-11 | 0.403617557 | 1 | 0.953 | 5.93E-07 | 23 | Rps23 |
| Csnk1g19 | 3.05E-11 | 0.273525556 | 0.438 | 0.158 | 6.17E-07 | 23 | Csnk1g1 |
| Ywhaz6 | 3.93E-11 | 0.31849055 | 0.958 | 0.493 | 7.96E-07 | 23 | Ywhaz |
| Rpl36a8 | 4.83E-11 | 0.429310499 | 0.99 | 0.753 | 9.78E-07 | 23 | Rpl36a |

| | | | | | | | |
|---|---|---|---|---|---|---|---|
| Oaz13 | 5.89E-11 | 0.341200709 | 1 | 0.807 | 1.19E-06 | 23 | Oaz1 |
| Scmh13 | 6.56E-11 | 0.26136751 | 0.396 | 0.139 | 1.33E-06 | 23 | Scmh1 |
| Rpl1110 | 8.46E-11 | 0.393023356 | 1 | 0.928 | 1.71E-06 | 23 | Rpl11 |
| Myl69 | 1.36E-10 | 0.456479841 | 1 | 0.777 | 2.75E-06 | 23 | Myl6 |
| Rps298 | 1.36E-10 | 0.312610096 | 1 | 0.971 | 2.76E-06 | 23 | Rps29 |
| Rack110 | 2.28E-10 | 0.400932765 | 1 | 0.793 | 4.61E-06 | 23 | Rack1 |
| Rbm3911 | 3.55E-10 | 0.422536346 | 1 | 0.575 | 7.18E-06 | 23 | Rbm39 |
| Rpl23a9 | 6.04E-10 | 0.430635716 | 0.979 | 0.68 | 1.22E-05 | 23 | Rpl23a |
| Tnrc6b10 | 7.10E-10 | 0.315212459 | 0.792 | 0.372 | 1.44E-05 | 23 | Tnrc6b |
| Rpl388 | 7.53E-10 | 0.380201919 | 0.99 | 0.894 | 1.52E-05 | 23 | Rpl38 |
| Rpl3710 | 7.83E-10 | 0.37947361 | 0.99 | 0.928 | 1.58E-05 | 23 | Rpl37 |
| Srgn11 | 8.73E-10 | 0.447900767 | 1 | 0.545 | 1.77E-05 | 23 | Srgn |
| Ubb2 | 1.09E-09 | 0.510184551 | 0.99 | 0.895 | 2.21E-05 | 23 | Ubb |
| Rpl278 | 1.29E-09 | 0.263891429 | 0.99 | 0.584 | 2.61E-05 | 23 | Rpl27 |
| Calr9 | 1.49E-09 | 0.273160136 | 0.812 | 0.409 | 3.01E-05 | 23 | Calr |
| Rpl910 | 1.56E-09 | 0.340658033 | 1 | 0.945 | 3.16E-05 | 23 | Rpl9 |
| Eef1a17 | 2.03E-09 | 0.334065078 | 1 | 0.96 | 4.11E-05 | 23 | Eef1a1 |
| Rps174 | 2.16E-09 | 0.305540148 | 1 | 0.797 | 4.37E-05 | 23 | Rps17 |
| Rps27a9 | 3.29E-09 | 0.334268842 | 1 | 0.95 | 6.66E-05 | 23 | Rps27a |
| Eif5a5 | 3.73E-09 | 0.356637394 | 0.927 | 0.566 | 7.56E-05 | 23 | Eif5a |
| Rpl3110 | 4.29E-09 | 0.283045785 | 0.99 | 0.685 | 8.68E-05 | 23 | Rpl31 |
| Gapdh4 | 6.87E-09 | 0.269322013 | 0.979 | 0.708 | 0.000139081 | 23 | Gapdh |
| Rpl308 | 8.51E-09 | 0.321574618 | 0.99 | 0.938 | 0.000172344 | 23 | Rpl30 |
| Rpl229 | 1.02E-08 | 0.321233828 | 1 | 0.853 | 0.000207115 | 23 | Rpl22 |
| B2m12 | 1.05E-08 | 0.432875901 | 1 | 0.691 | 0.000211889 | 23 | B2m |
| Btf33 | 1.57E-08 | 0.294566835 | 0.99 | 0.723 | 0.00031749 | 23 | Btf3 |
| Rpl369 | 2.68E-08 | 0.312488935 | 0.99 | 0.904 | 0.000542569 | 23 | Rpl36 |
| Rhoa4 | 2.90E-08 | 0.284040418 | 1 | 0.594 | 0.000587726 | 23 | Rhoa |
| Rpl178 | 4.49E-08 | 0.273566939 | 1 | 0.917 | 0.000909541 | 23 | Rpl17 |
| Ybx15 | 1.07E-07 | 0.263802592 | 1 | 0.753 | 0.002161385 | 23 | Ybx1 |
| Rpl59 | 1.26E-07 | 0.265460606 | 1 | 0.711 | 0.0025468 | 23 | Rpl5 |
| mt-Nd416 | 5.00E-07 | 0.34908843 | 1 | 0.676 | 0.010131649 | 23 | mt-Nd4 |
| Rpl710 | 6.32E-07 | 0.258027003 | 0.99 | 0.893 | 0.012788506 | 23 | Rpl7 |
| mt-Co219 | 9.49E-07 | 0.388905569 | 1 | 0.729 | 0.019208737 | 23 | mt-Co2 |
| mt-Atp617 | 3.33E-06 | 0.374519015 | 1 | 0.755 | 0.06743579 | 23 | mt-Atp6 |
| H2-D112 | 7.71E-06 | 0.328388525 | 1 | 0.67 | 0.156156125 | 23 | H2-D1 |
| mt-Cytb15 | 6.04E-05 | 0.26140407 | 1 | 0.701 | 1 | 23 | mt-Cytb |
| C1qa5 | 2.96E-90 | 1.317352704 | 0.957 | 0.146 | 5.99E-86 | 24 | C1qa |
| C1qb4 | 2.77E-89 | 1.498785525 | 0.968 | 0.154 | 5.62E-85 | 24 | C1qb |
| C1qc5 | 2.56E-84 | 1.263847105 | 0.892 | 0.134 | 5.19E-80 | 24 | C1qc |
| H2-Eb15 | 5.82E-78 | 1.444284313 | 1 | 0.194 | 1.18E-73 | 24 | H2-Eb1 |
| H2-Aa5 | 1.61E-76 | 1.234446063 | 0.989 | 0.186 | 3.26E-72 | 24 | H2-Aa |
| H2-Ab15 | 3.21E-72 | 1.308788281 | 0.978 | 0.201 | 6.50E-68 | 24 | H2-Ab1 |

| | | | | | | | |
|---|---|---|---|---|---|---|---|
| Aif15 | 7.69E-69 | 1.094978906 | 0.763 | 0.121 | 1.56E-64 | 24 | Aif1 |
| Ctss5 | 7.43E-65 | 1.101271167 | 0.871 | 0.166 | 1.50E-60 | 24 | Ctss |
| Lyz25 | 1.66E-63 | 1.218038606 | 0.688 | 0.112 | 3.36E-59 | 24 | Lyz2 |
| Cd745 | 2.27E-63 | 1.4293034 | 0.989 | 0.251 | 4.60E-59 | 24 | Cd74 |
| Bcl2a1b5 | 1.19E-41 | 0.804126378 | 0.559 | 0.106 | 2.41E-37 | 24 | Bcl2a1b |
| H2-DMa5 | 2.15E-38 | 0.79516747 | 0.581 | 0.12 | 4.36E-34 | 24 | H2-DMa |
| Ly865 | 2.90E-37 | 0.514014774 | 0.527 | 0.097 | 5.87E-33 | 24 | Ly86 |
| Rgs106 | 4.19E-37 | 1.029231578 | 0.656 | 0.161 | 8.49E-33 | 24 | Rgs10 |
| Apoe5 | 2.50E-36 | 1.264931101 | 1 | 0.535 | 5.05E-32 | 24 | Apoe |
| Lst15 | 8.20E-36 | 0.78972501 | 0.538 | 0.112 | 1.66E-31 | 24 | Lst1 |
| Gpihbp14 | 3.82E-35 | 1.222214253 | 0.903 | 0.296 | 7.72E-31 | 24 | Gpihbp1 |
| Ramp33 | 2.95E-34 | 0.916028855 | 0.892 | 0.267 | 5.98E-30 | 24 | Ramp3 |
| Ckb8 | 3.91E-33 | 1.012530718 | 0.731 | 0.216 | 7.92E-29 | 24 | Ckb |
| Cst37 | 4.36E-33 | 1.191407704 | 1 | 0.603 | 8.84E-29 | 24 | Cst3 |
| Rbp74 | 1.22E-32 | 1.224271601 | 0.935 | 0.348 | 2.48E-28 | 24 | Rbp7 |
| Spi15 | 1.86E-32 | 0.70184016 | 0.441 | 0.084 | 3.76E-28 | 24 | Spi1 |
| Gatm4 | 7.06E-32 | 0.572636085 | 0.409 | 0.074 | 1.43E-27 | 24 | Gatm |
| Basp16 | 9.00E-32 | 0.635675888 | 0.505 | 0.105 | 1.82E-27 | 24 | Basp1 |
| Wfdc175 | 1.10E-31 | 0.673280791 | 0.387 | 0.069 | 2.22E-27 | 24 | Wfdc17 |
| Tyrobp9 | 6.36E-31 | 0.522121035 | 0.946 | 0.262 | 1.29E-26 | 24 | Tyrobp |
| Ms4a6c4 | 6.71E-31 | 0.631229229 | 0.43 | 0.084 | 1.36E-26 | 24 | Ms4a6c |
| Plcb15 | 9.71E-31 | 1.091309858 | 0.946 | 0.341 | 1.97E-26 | 24 | Plcb1 |
| Alox5ap5 | 3.53E-30 | 0.541398824 | 0.398 | 0.074 | 7.14E-26 | 24 | Alox5ap |
| Hpgd5 | 4.27E-30 | 0.62905546 | 0.441 | 0.088 | 8.64E-26 | 24 | Hpgd |
| Sox186 | 5.66E-30 | 1.011560827 | 0.849 | 0.277 | 1.15E-25 | 24 | Sox18 |
| Car42 | 7.48E-29 | 0.932336599 | 0.753 | 0.226 | 1.52E-24 | 24 | Car4 |
| Cd685 | 7.73E-29 | 0.536087658 | 0.409 | 0.08 | 1.57E-24 | 24 | Cd68 |
| Sparcl18 | 1.88E-27 | 1.060267188 | 0.957 | 0.364 | 3.80E-23 | 24 | Sparcl1 |
| Btnl93 | 3.45E-27 | 1.024667024 | 0.796 | 0.271 | 6.98E-23 | 24 | Btnl9 |
| Ms4a75 | 5.01E-27 | 0.637808412 | 0.462 | 0.1 | 1.01E-22 | 24 | Ms4a7 |
| Rgcc4 | 6.10E-27 | 1.099324009 | 0.957 | 0.405 | 1.24E-22 | 24 | Rgcc |
| Adgrl45 | 7.41E-27 | 1.090638707 | 0.957 | 0.352 | 1.50E-22 | 24 | Adgrl4 |
| Cd300c25 | 7.42E-27 | 0.539798011 | 0.419 | 0.086 | 1.50E-22 | 24 | Cd300c2 |
| Cavin26 | 7.68E-27 | 0.967537951 | 0.935 | 0.342 | 1.55E-22 | 24 | Cavin2 |
| Plpp13 | 8.43E-27 | 1.075980086 | 0.968 | 0.383 | 1.71E-22 | 24 | Plpp1 |
| Cyyr15 | 9.88E-27 | 0.923965974 | 0.957 | 0.344 | 2.00E-22 | 24 | Cyyr1 |
| Trem23 | 1.81E-26 | 0.543589656 | 0.355 | 0.067 | 3.67E-22 | 24 | Trem2 |
| Col4a25 | 1.82E-26 | 1.0834007 | 0.925 | 0.361 | 3.68E-22 | 24 | Col4a2 |
| Scarb14 | 1.88E-26 | 0.908761085 | 0.796 | 0.267 | 3.81E-22 | 24 | Scarb1 |
| Flt16 | 2.05E-26 | 1.119388257 | 1 | 0.394 | 4.15E-22 | 24 | Flt1 |
| Emcn5 | 2.75E-26 | 1.074831391 | 0.989 | 0.381 | 5.58E-22 | 24 | Emcn |
| Ly6c15 | 2.90E-26 | 1.080849269 | 1 | 0.401 | 5.87E-22 | 24 | Ly6c1 |
| Slamf94 | 3.92E-26 | 0.577941523 | 0.366 | 0.071 | 7.94E-22 | 24 | Slamf9 |

| | | | | | | | |
|---|---|---|---|---|---|---|---|
| Mpeg14 | 3.95E-26 | 0.598797273 | 0.376 | 0.074 | 8.01E-22 | 24 | Mpeg1 |
| Kdr4 | 4.24E-26 | 1.061355384 | 0.946 | 0.363 | 8.57E-22 | 24 | Kdr |
| Rgl14 | 8.69E-26 | 0.903713836 | 0.796 | 0.29 | 1.76E-21 | 24 | Rgl1 |
| Adgrf56 | 1.17E-25 | 0.909710767 | 0.935 | 0.338 | 2.37E-21 | 24 | Adgrf5 |
| Plvap5 | 1.88E-25 | 1.043632349 | 0.989 | 0.385 | 3.80E-21 | 24 | Plvap |
| Ldb26 | 3.25E-25 | 0.891059793 | 0.892 | 0.331 | 6.59E-21 | 24 | Ldb2 |
| Egfl76 | 6.56E-25 | 0.99057098 | 1 | 0.394 | 1.33E-20 | 24 | Egfl7 |
| Cd345 | 6.70E-25 | 1.001202969 | 0.86 | 0.321 | 1.36E-20 | 24 | Cd34 |
| Ptprb6 | 1.09E-24 | 0.942627432 | 0.978 | 0.369 | 2.21E-20 | 24 | Ptprb |
| Ramp26 | 1.18E-24 | 0.940384673 | 0.882 | 0.332 | 2.38E-20 | 24 | Ramp2 |
| Nostrin3 | 1.43E-24 | 0.826758585 | 0.763 | 0.259 | 2.89E-20 | 24 | Nostrin |
| Arhgap294 | 1.84E-24 | 1.032709248 | 0.892 | 0.366 | 3.72E-20 | 24 | Arhgap29 |
| Id16 | 2.20E-24 | 0.919301082 | 0.871 | 0.326 | 4.45E-20 | 24 | Id1 |
| Syne14 | 2.48E-24 | 0.872801975 | 0.903 | 0.359 | 5.01E-20 | 24 | Syne1 |
| Tmem176a11 | 2.84E-24 | 0.933422655 | 0.935 | 0.418 | 5.74E-20 | 24 | Tmem176a |
| Col13a13 | 3.93E-24 | 0.887283398 | 0.72 | 0.243 | 7.97E-20 | 24 | Col13a1 |
| Ctsh6 | 5.22E-24 | 0.694849012 | 0.527 | 0.137 | 1.06E-19 | 24 | Ctsh |
| Ppfia42 | 5.52E-24 | 0.436989678 | 0.301 | 0.055 | 1.12E-19 | 24 | Ppfia4 |
| Tmsb4x12 | 1.04E-23 | 0.803620456 | 1 | 0.934 | 2.10E-19 | 24 | Tmsb4x |
| Fabp46 | 1.12E-23 | 0.963229639 | 1 | 0.436 | 2.26E-19 | 24 | Fabp4 |
| Id36 | 1.29E-23 | 0.952644574 | 0.978 | 0.409 | 2.61E-19 | 24 | Id3 |
| Podxl5 | 1.30E-23 | 0.890932229 | 0.925 | 0.346 | 2.63E-19 | 24 | Podxl |
| Mmrn25 | 1.45E-23 | 0.825775401 | 0.71 | 0.233 | 2.93E-19 | 24 | Mmrn2 |
| Hspb18 | 1.58E-23 | 1.002451902 | 0.935 | 0.382 | 3.20E-19 | 24 | Hspb1 |
| Plpp33 | 1.63E-23 | 0.948572872 | 0.989 | 0.405 | 3.31E-19 | 24 | Plpp3 |
| Csf1r3 | 2.10E-23 | 0.50241608 | 0.366 | 0.075 | 4.24E-19 | 24 | Csf1r |
| Clic55 | 2.15E-23 | 0.761770543 | 0.828 | 0.292 | 4.35E-19 | 24 | Clic5 |
| H2-DMb15 | 2.66E-23 | 0.588848983 | 0.409 | 0.093 | 5.39E-19 | 24 | H2-DMb1 |
| Mcf2l3 | 2.76E-23 | 0.809746629 | 0.667 | 0.214 | 5.59E-19 | 24 | Mcf2l |
| Timp38 | 2.78E-23 | 0.912672507 | 0.946 | 0.386 | 5.63E-19 | 24 | Timp3 |
| Eng5 | 3.92E-23 | 0.869435784 | 0.914 | 0.355 | 7.94E-19 | 24 | Eng |
| Exoc3l24 | 4.07E-23 | 0.832479463 | 0.785 | 0.271 | 8.24E-19 | 24 | Exoc3l2 |
| Fam167b2 | 4.18E-23 | 0.818030644 | 0.656 | 0.212 | 8.47E-19 | 24 | Fam167b |
| Igfbp75 | 5.38E-23 | 1.054643943 | 1 | 0.468 | 1.09E-18 | 24 | Igfbp7 |
| Bst28 | 6.44E-23 | 1.090264372 | 0.957 | 0.504 | 1.30E-18 | 24 | Bst2 |
| Fgd55 | 8.26E-23 | 0.790873839 | 0.688 | 0.229 | 1.67E-18 | 24 | Fgd5 |
| Arhgap315 | 8.29E-23 | 0.920059443 | 0.903 | 0.373 | 1.68E-18 | 24 | Arhgap31 |
| Ctsb7 | 9.75E-23 | 0.992198596 | 0.742 | 0.299 | 1.97E-18 | 24 | Ctsb |
| Cd819 | 1.27E-22 | 0.951369129 | 1 | 0.526 | 2.58E-18 | 24 | Cd81 |
| Fcer1g10 | 1.68E-22 | 0.363151839 | 0.871 | 0.27 | 3.40E-18 | 24 | Fcer1g |
| Jam34 | 1.72E-22 | 0.862789956 | 0.763 | 0.276 | 3.47E-18 | 24 | Jam3 |
| Ushbp15 | 1.72E-22 | 0.741228887 | 0.677 | 0.219 | 3.47E-18 | 24 | Ushbp1 |
| Fmo24 | 3.02E-22 | 0.871374223 | 0.785 | 0.29 | 6.11E-18 | 24 | Fmo2 |

| | | | | | | | |
|---|---|---|---|---|---|---|---|
| Col4a15 | 3.12E-22 | 0.97977549 | 0.946 | 0.393 | 6.32E-18 | 24 | Col4a1 |
| Rsad24 | 3.36E-22 | 0.807359673 | 0.731 | 0.257 | 6.79E-18 | 24 | Rsad2 |
| Ptprm5 | 3.89E-22 | 0.89196267 | 0.849 | 0.33 | 7.87E-18 | 24 | Ptprm |
| Ece16 | 4.39E-22 | 0.804553902 | 0.806 | 0.307 | 8.89E-18 | 24 | Ece1 |
| Slc9a3r25 | 5.05E-22 | 0.808433889 | 0.925 | 0.371 | 1.02E-17 | 24 | Slc9a3r2 |
| Tmem885 | 5.57E-22 | 0.658639934 | 0.656 | 0.211 | 1.13E-17 | 24 | Tmem88 |
| Piezo23 | 5.95E-22 | 0.814957634 | 0.731 | 0.254 | 1.21E-17 | 24 | Piezo2 |
| Cdh55 | 6.90E-22 | 0.82874398 | 0.882 | 0.331 | 1.40E-17 | 24 | Cdh5 |
| Nrp13 | 9.03E-22 | 0.881690189 | 0.968 | 0.435 | 1.83E-17 | 24 | Nrp1 |
| Tcim3 | 1.10E-21 | 0.899843872 | 0.796 | 0.304 | 2.23E-17 | 24 | Tcim |
| Cd725 | 1.90E-21 | 0.549545849 | 0.419 | 0.106 | 3.84E-17 | 24 | Cd72 |
| Ebf17 | 2.18E-21 | 0.785020785 | 0.957 | 0.375 | 4.42E-17 | 24 | Ebf1 |
| Mef2c6 | 2.19E-21 | 0.889713589 | 0.946 | 0.424 | 4.43E-17 | 24 | Mef2c |
| Fbxl75 | 2.83E-21 | 0.801371052 | 0.71 | 0.258 | 5.73E-17 | 24 | Fbxl7 |
| Cd2006 | 2.92E-21 | 0.882181108 | 0.871 | 0.351 | 5.91E-17 | 24 | Cd200 |
| Aplp26 | 3.23E-21 | 0.843421944 | 0.925 | 0.417 | 6.54E-17 | 24 | Aplp2 |
| Klf75 | 3.48E-21 | 0.884226552 | 0.753 | 0.298 | 7.05E-17 | 24 | Klf7 |
| Kctd12b5 | 5.45E-21 | 0.782312339 | 0.71 | 0.25 | 1.10E-16 | 24 | Kctd12b |
| Cybb3 | 5.49E-21 | 0.307388822 | 0.355 | 0.076 | 1.11E-16 | 24 | Cybb |
| Cx3cr13 | 6.06E-21 | 0.253085243 | 0.376 | 0.085 | 1.23E-16 | 24 | Cx3cr1 |
| Pcdh175 | 6.17E-21 | 0.932682562 | 0.677 | 0.24 | 1.25E-16 | 24 | Pcdh17 |
| Pecam16 | 8.55E-21 | 0.932344142 | 0.946 | 0.412 | 1.73E-16 | 24 | Pecam1 |
| Plscr24 | 9.08E-21 | 0.809844368 | 0.849 | 0.336 | 1.84E-16 | 24 | Plscr2 |
| Ly6a6 | 9.43E-21 | 0.897534836 | 1 | 0.46 | 1.91E-16 | 24 | Ly6a |
| Hmcn14 | 9.79E-21 | 0.880589314 | 0.624 | 0.212 | 1.98E-16 | 24 | Hmcn1 |
| Lama35 | 1.01E-20 | 0.843652177 | 0.645 | 0.222 | 2.05E-16 | 24 | Lama3 |
| Cd300lg4 | 1.22E-20 | 0.676011875 | 0.817 | 0.297 | 2.47E-16 | 24 | Cd300lg |
| Meox14 | 1.33E-20 | 0.771140719 | 0.581 | 0.188 | 2.70E-16 | 24 | Meox1 |
| Tcf155 | 1.34E-20 | 0.888402698 | 0.71 | 0.261 | 2.72E-16 | 24 | Tcf15 |
| Ahr4 | 1.57E-20 | 0.739941476 | 0.677 | 0.247 | 3.18E-16 | 24 | Ahr |
| Crim15 | 2.30E-20 | 0.756588462 | 0.882 | 0.374 | 4.67E-16 | 24 | Crim1 |
| Sparc10 | 3.14E-20 | 0.790891433 | 0.968 | 0.403 | 6.37E-16 | 24 | Sparc |
| Rasgrp34 | 3.48E-20 | 0.737203904 | 0.753 | 0.283 | 7.05E-16 | 24 | Rasgrp3 |
| Kitl6 | 4.34E-20 | 0.756554189 | 0.849 | 0.337 | 8.79E-16 | 24 | Kitl |
| Nfib5 | 5.93E-20 | 0.879379852 | 0.978 | 0.445 | 1.20E-15 | 24 | Nfib |
| Itm2b9 | 6.06E-20 | 0.849067806 | 1 | 0.726 | 1.23E-15 | 24 | Itm2b |
| Thrsp4 | 6.71E-20 | 0.836951464 | 0.774 | 0.294 | 1.36E-15 | 24 | Thrsp |
| Ltbp46 | 8.86E-20 | 0.856736402 | 0.774 | 0.31 | 1.79E-15 | 24 | Ltbp4 |
| Lgmn6 | 1.03E-19 | 0.591019384 | 0.43 | 0.114 | 2.08E-15 | 24 | Lgmn |
| Gng117 | 1.53E-19 | 0.709585235 | 0.892 | 0.367 | 3.09E-15 | 24 | Gng11 |
| Rflnb5 | 1.74E-19 | 0.722987291 | 0.806 | 0.315 | 3.52E-15 | 24 | Rflnb |
| Ecscr5 | 1.76E-19 | 0.648363413 | 0.667 | 0.229 | 3.57E-15 | 24 | Ecscr |
| Chst154 | 1.81E-19 | 0.667551097 | 0.559 | 0.176 | 3.67E-15 | 24 | Chst15 |

| | | | | | | | |
|---|---|---|---|---|---|---|---|
| Rai144 | 3.02E-19 | 0.74789481 | 0.645 | 0.232 | 6.12E-15 | 24 | Rai14 |
| Clec14a5 | 3.65E-19 | 0.691539237 | 0.785 | 0.306 | 7.39E-15 | 24 | Clec14a |
| Col15a13 | 4.55E-19 | 0.607323177 | 0.688 | 0.248 | 9.21E-15 | 24 | Col15a1 |
| Sptbn16 | 4.56E-19 | 0.887292444 | 0.989 | 0.496 | 9.24E-15 | 24 | Sptbn1 |
| Esam6 | 4.68E-19 | 0.814281154 | 0.785 | 0.311 | 9.49E-15 | 24 | Esam |
| Ehd44 | 5.54E-19 | 0.747207685 | 0.892 | 0.409 | 1.12E-14 | 24 | Ehd4 |
| Selenop8 | 6.36E-19 | 0.752445235 | 0.828 | 0.364 | 1.29E-14 | 24 | Selenop |
| Tcf46 | 6.92E-19 | 0.870383058 | 0.968 | 0.473 | 1.40E-14 | 24 | Tcf4 |
| Gm120023 | 7.96E-19 | 0.734946747 | 0.57 | 0.189 | 1.61E-14 | 24 | Gm12002 |
| Npc25 | 1.12E-18 | 0.769167457 | 0.839 | 0.415 | 2.27E-14 | 24 | Npc2 |
| Igfbp35 | 1.17E-18 | 0.866947103 | 0.753 | 0.304 | 2.38E-14 | 24 | Igfbp3 |
| Cd935 | 1.31E-18 | 0.788173977 | 0.699 | 0.257 | 2.66E-14 | 24 | Cd93 |
| Tmtc25 | 1.32E-18 | 0.836006834 | 0.548 | 0.186 | 2.68E-14 | 24 | Tmtc2 |
| Calm15 | 1.51E-18 | 0.750083865 | 1 | 0.781 | 3.06E-14 | 24 | Calm1 |
| Dock45 | 2.05E-18 | 0.719284492 | 0.753 | 0.297 | 4.16E-14 | 24 | Dock4 |
| Tm4sf17 | 2.06E-18 | 0.570318463 | 0.935 | 0.378 | 4.18E-14 | 24 | Tm4sf1 |
| Etl45 | 2.43E-18 | 0.807823518 | 0.742 | 0.295 | 4.92E-14 | 24 | Etl4 |
| Psap7 | 2.58E-18 | 0.95260552 | 0.774 | 0.4 | 5.22E-14 | 24 | Psap |
| Ccdc85a4 | 3.52E-18 | 0.674796314 | 0.559 | 0.182 | 7.14E-14 | 24 | Ccdc85a |
| Cavin38 | 5.43E-18 | 0.613008927 | 0.785 | 0.308 | 1.10E-13 | 24 | Cavin3 |
| Dlc15 | 5.65E-18 | 0.731944209 | 0.86 | 0.363 | 1.14E-13 | 24 | Dlc1 |
| Plekhg16 | 5.86E-18 | 0.673126935 | 0.667 | 0.246 | 1.19E-13 | 24 | Plekhg1 |
| Tshz27 | 6.00E-18 | 0.725480213 | 0.86 | 0.373 | 1.21E-13 | 24 | Tshz2 |
| Tnfsf106 | 6.89E-18 | 0.812916115 | 0.839 | 0.375 | 1.39E-13 | 24 | Tnfsf10 |
| Ifitm39 | 9.93E-18 | 0.82391585 | 0.989 | 0.539 | 2.01E-13 | 24 | Ifitm3 |
| Rbms17 | 1.07E-17 | 0.820524426 | 0.914 | 0.487 | 2.17E-13 | 24 | Rbms1 |
| Id44 | 1.28E-17 | 0.662084284 | 0.43 | 0.126 | 2.59E-13 | 24 | Id4 |
| Cmtm83 | 1.70E-17 | 0.743665335 | 0.699 | 0.274 | 3.44E-13 | 24 | Cmtm8 |
| Myo64 | 3.22E-17 | 0.713276909 | 0.71 | 0.293 | 6.52E-13 | 24 | Myo6 |
| Pmepa16 | 3.82E-17 | 0.495415994 | 0.656 | 0.253 | 7.74E-13 | 24 | Pmepa1 |
| Cavin17 | 3.83E-17 | 0.575249836 | 0.602 | 0.214 | 7.76E-13 | 24 | Cavin1 |
| Cd638 | 4.13E-17 | 0.314082501 | 0.581 | 0.189 | 8.36E-13 | 24 | Cd63 |
| B2m13 | 5.53E-17 | 0.764964551 | 0.989 | 0.691 | 1.12E-12 | 24 | B2m |
| Ptprg5 | 5.54E-17 | 0.492497417 | 0.871 | 0.345 | 1.12E-12 | 24 | Ptprg |
| Pltp5 | 6.27E-17 | 0.791280521 | 0.731 | 0.301 | 1.27E-12 | 24 | Pltp |
| Iigp15 | 6.94E-17 | 0.677935423 | 0.624 | 0.234 | 1.41E-12 | 24 | Iigp1 |
| H2-D113 | 9.07E-17 | 0.781748311 | 1 | 0.67 | 1.84E-12 | 24 | H2-D1 |
| Trim475 | 9.15E-17 | 0.535558197 | 0.462 | 0.145 | 1.85E-12 | 24 | Trim47 |
| Klf45 | 1.01E-16 | 0.698731442 | 0.688 | 0.27 | 2.05E-12 | 24 | Klf4 |
| Hecw24 | 1.28E-16 | 0.749308226 | 0.645 | 0.25 | 2.59E-12 | 24 | Hecw2 |
| Cxcl127 | 1.37E-16 | 0.780679595 | 0.753 | 0.323 | 2.77E-12 | 24 | Cxcl12 |
| Plxna25 | 1.49E-16 | 0.7664387 | 0.581 | 0.22 | 3.01E-12 | 24 | Plxna2 |
| Nxpe21 | 1.62E-16 | 0.538490243 | 0.376 | 0.106 | 3.27E-12 | 24 | Nxpe2 |

| | | | | | | | |
|---|---|---|---|---|---|---|---|
| Gngt26 | 1.75E-16 | 0.772649221 | 0.774 | 0.361 | 3.54E-12 | 24 | Gngt2 |
| Pdgfb5 | 2.06E-16 | 0.675838169 | 0.731 | 0.31 | 4.16E-12 | 24 | Pdgfb |
| Rapgef45 | 2.10E-16 | 0.728993717 | 0.667 | 0.259 | 4.24E-12 | 24 | Rapgef4 |
| Cxcl165 | 2.16E-16 | 0.443422661 | 0.398 | 0.114 | 4.37E-12 | 24 | Cxcl16 |
| Epas17 | 2.58E-16 | 0.683341758 | 0.828 | 0.36 | 5.23E-12 | 24 | Epas1 |
| Rapgef55 | 2.82E-16 | 0.585706461 | 0.602 | 0.223 | 5.71E-12 | 24 | Rapgef5 |
| Mcam5 | 2.93E-16 | 0.570978424 | 0.387 | 0.111 | 5.92E-12 | 24 | Mcam |
| Itga18 | 3.19E-16 | 0.752933348 | 0.849 | 0.416 | 6.45E-12 | 24 | Itga1 |
| Pdgfd5 | 3.24E-16 | 0.763179599 | 0.645 | 0.254 | 6.56E-12 | 24 | Pdgfd |
| Rasal25 | 3.25E-16 | 0.725751764 | 0.753 | 0.316 | 6.59E-12 | 24 | Rasal2 |
| Tie16 | 3.38E-16 | 0.640043804 | 0.667 | 0.255 | 6.83E-12 | 24 | Tie1 |
| Bmpr25 | 4.23E-16 | 0.665688668 | 0.806 | 0.351 | 8.56E-12 | 24 | Bmpr2 |
| Fzd46 | 4.55E-16 | 0.597028927 | 0.624 | 0.23 | 9.20E-12 | 24 | Fzd4 |
| Lims25 | 4.63E-16 | 0.5958939 | 0.57 | 0.202 | 9.38E-12 | 24 | Lims2 |
| BC0285283 | 4.74E-16 | 0.754055539 | 0.602 | 0.235 | 9.61E-12 | 24 | BC028528 |
| Stard85 | 5.55E-16 | 0.61702675 | 0.559 | 0.199 | 1.12E-11 | 24 | Stard8 |
| Tinagl16 | 6.30E-16 | 0.776886952 | 0.71 | 0.298 | 1.28E-11 | 24 | Tinagl1 |
| App7 | 6.58E-16 | 0.67020571 | 0.914 | 0.421 | 1.33E-11 | 24 | App |
| Thbd6 | 6.60E-16 | 0.679214938 | 0.656 | 0.256 | 1.34E-11 | 24 | Thbd |
| Ly6e7 | 6.80E-16 | 0.717616438 | 0.989 | 0.62 | 1.38E-11 | 24 | Ly6e |
| Klf26 | 7.25E-16 | 0.634348638 | 0.828 | 0.368 | 1.47E-11 | 24 | Klf2 |
| Nfe2l15 | 8.37E-16 | 0.636367221 | 0.667 | 0.281 | 1.70E-11 | 24 | Nfe2l1 |
| Serinc37 | 8.64E-16 | 0.72982026 | 0.925 | 0.488 | 1.75E-11 | 24 | Serinc3 |
| Dgkh6 | 9.26E-16 | 0.628009007 | 0.505 | 0.178 | 1.87E-11 | 24 | Dgkh |
| Calcrl4 | 1.07E-15 | 0.712512762 | 0.656 | 0.264 | 2.17E-11 | 24 | Calcrl |
| Cd383 | 1.08E-15 | 0.625654956 | 0.602 | 0.233 | 2.20E-11 | 24 | Cd38 |
| Tmem176b10 | 1.46E-15 | 0.703376255 | 0.882 | 0.44 | 2.95E-11 | 24 | Tmem176b |
| Clic47 | 1.72E-15 | 0.678541951 | 0.86 | 0.424 | 3.47E-11 | 24 | Clic4 |
| Emp26 | 2.32E-15 | 0.605491403 | 0.613 | 0.237 | 4.69E-11 | 24 | Emp2 |
| Fth15 | 2.37E-15 | 0.511049837 | 1 | 0.969 | 4.80E-11 | 24 | Fth1 |
| Arl155 | 2.38E-15 | 0.743419773 | 0.796 | 0.382 | 4.83E-11 | 24 | Arl15 |
| Psmb813 | 2.66E-15 | 0.713468813 | 0.946 | 0.558 | 5.39E-11 | 24 | Psmb8 |
| Cav16 | 3.40E-15 | 0.656939761 | 0.731 | 0.305 | 6.89E-11 | 24 | Cav1 |
| Gas66 | 3.63E-15 | 0.584538202 | 0.731 | 0.299 | 7.35E-11 | 24 | Gas6 |
| Magi15 | 3.99E-15 | 0.566953741 | 0.613 | 0.232 | 8.07E-11 | 24 | Magi1 |
| Snai22 | 4.55E-15 | 0.473810668 | 0.333 | 0.092 | 9.21E-11 | 24 | Snai2 |
| Slc43a36 | 4.63E-15 | 0.665640802 | 0.667 | 0.271 | 9.38E-11 | 24 | Slc43a3 |
| Srgn12 | 4.68E-15 | 0.706163308 | 0.989 | 0.546 | 9.48E-11 | 24 | Srgn |
| Snrk5 | 4.84E-15 | 0.735972782 | 0.774 | 0.37 | 9.79E-11 | 24 | Snrk |
| Gbp25 | 5.41E-15 | 0.492361509 | 0.441 | 0.144 | 1.10E-10 | 24 | Gbp2 |
| S1pr14 | 5.49E-15 | 0.691398382 | 0.656 | 0.266 | 1.11E-10 | 24 | S1pr1 |
| Slfn56 | 5.74E-15 | 0.711169584 | 0.774 | 0.358 | 1.16E-10 | 24 | Slfn5 |
| Serpinh18 | 5.99E-15 | 0.532292535 | 0.645 | 0.251 | 1.21E-10 | 24 | Serpinh1 |

| | | | | | | | |
|---|---|---|---|---|---|---|---|
| Ifitm28 | 6.87E-15 | 0.675310657 | 0.978 | 0.465 | 1.39E-10 | 24 | Ifitm2 |
| Mgll5 | 7.03E-15 | 0.646279109 | 0.957 | 0.45 | 1.42E-10 | 24 | Mgll |
| Dhrs35 | 7.37E-15 | 0.649198577 | 0.591 | 0.232 | 1.49E-10 | 24 | Dhrs3 |
| Smco45 | 8.20E-15 | 0.574880804 | 0.849 | 0.391 | 1.66E-10 | 24 | Smco4 |
| Dnm35 | 8.44E-15 | 0.485461642 | 0.613 | 0.231 | 1.71E-10 | 24 | Dnm3 |
| Sptan17 | 8.68E-15 | 0.595431927 | 0.753 | 0.345 | 1.76E-10 | 24 | Sptan1 |
| Scimp4 | 9.30E-15 | 0.341565147 | 0.301 | 0.078 | 1.88E-10 | 24 | Scimp |
| Plbd13 | 1.12E-14 | 0.327509061 | 0.258 | 0.06 | 2.26E-10 | 24 | Plbd1 |
| Ripply34 | 1.17E-14 | 0.577608409 | 0.441 | 0.147 | 2.36E-10 | 24 | Ripply3 |
| Grrp15 | 1.24E-14 | 0.555522413 | 0.591 | 0.22 | 2.50E-10 | 24 | Grrp1 |
| Erp294 | 1.25E-14 | 0.627042088 | 0.71 | 0.329 | 2.54E-10 | 24 | Erp29 |
| Dip2c8 | 1.34E-14 | 0.557301517 | 0.581 | 0.224 | 2.71E-10 | 24 | Dip2c |
| Rhoc7 | 1.35E-14 | 0.612617119 | 0.677 | 0.297 | 2.74E-10 | 24 | Rhoc |
| Cyp26b14 | 1.40E-14 | 0.957809831 | 0.559 | 0.217 | 2.83E-10 | 24 | Cyp26b1 |
| Cd366 | 1.58E-14 | 0.540893698 | 0.667 | 0.269 | 3.20E-10 | 24 | Cd36 |
| Rgs122 | 1.75E-14 | 0.519676703 | 0.355 | 0.104 | 3.55E-10 | 24 | Rgs12 |
| Unc93b13 | 1.87E-14 | 0.413042661 | 0.462 | 0.157 | 3.78E-10 | 24 | Unc93b1 |
| Adam155 | 1.88E-14 | 0.579909078 | 0.527 | 0.194 | 3.80E-10 | 24 | Adam15 |
| Tspan95 | 1.93E-14 | 0.656413935 | 0.624 | 0.266 | 3.90E-10 | 24 | Tspan9 |
| Ackr34 | 1.95E-14 | 0.875768514 | 0.441 | 0.153 | 3.95E-10 | 24 | Ackr3 |
| Tjp15 | 1.96E-14 | 0.682598421 | 0.624 | 0.265 | 3.96E-10 | 24 | Tjp1 |
| Filip16 | 2.22E-14 | 0.54096026 | 0.538 | 0.196 | 4.49E-10 | 24 | Filip1 |
| Csf2ra3 | 2.29E-14 | 0.34452562 | 0.29 | 0.074 | 4.64E-10 | 24 | Csf2ra |
| Actn47 | 2.86E-14 | 0.678215535 | 0.753 | 0.377 | 5.78E-10 | 24 | Actn4 |
| 4931406P16Rik6 | 3.90E-14 | 0.709448246 | 0.742 | 0.344 | 7.89E-10 | 24 | 4931406P16Rik |
| F11r8 | 4.84E-14 | 0.491852208 | 0.656 | 0.267 | 9.80E-10 | 24 | F11r |
| Rras7 | 5.21E-14 | 0.620556598 | 0.667 | 0.299 | 1.06E-09 | 24 | Rras |
| Sema6a5 | 5.32E-14 | 0.633725649 | 0.548 | 0.21 | 1.08E-09 | 24 | Sema6a |
| Sorbs27 | 5.39E-14 | 0.594717183 | 0.677 | 0.279 | 1.09E-09 | 24 | Sorbs2 |
| Impdh12 | 5.88E-14 | 0.640893337 | 0.355 | 0.113 | 1.19E-09 | 24 | Impdh1 |
| Myct15 | 7.25E-14 | 0.585072017 | 0.581 | 0.223 | 1.47E-09 | 24 | Myct1 |
| Slc6a65 | 7.96E-14 | 0.712284957 | 0.71 | 0.329 | 1.61E-09 | 24 | Slc6a6 |
| Zfp3665 | 8.73E-14 | 0.537380397 | 0.505 | 0.183 | 1.77E-09 | 24 | Zfp366 |
| Mcc4 | 9.02E-14 | 0.784691396 | 0.656 | 0.285 | 1.83E-09 | 24 | Mcc |
| Tanc16 | 9.34E-14 | 0.562939776 | 0.57 | 0.224 | 1.89E-09 | 24 | Tanc1 |
| Itih54 | 9.74E-14 | 0.637782301 | 0.591 | 0.244 | 1.97E-09 | 24 | Itih5 |
| Tspan125 | 9.81E-14 | 0.616370412 | 0.591 | 0.233 | 1.99E-09 | 24 | Tspan12 |
| Actb11 | 1.05E-13 | 0.589099911 | 1 | 0.843 | 2.14E-09 | 24 | Actb |
| Asap16 | 1.06E-13 | 0.721802547 | 0.731 | 0.357 | 2.14E-09 | 24 | Asap1 |
| Gm138894 | 1.09E-13 | 0.548296701 | 0.581 | 0.22 | 2.21E-09 | 24 | Gm13889 |
| Grn6 | 1.17E-13 | 0.554315625 | 0.548 | 0.225 | 2.37E-09 | 24 | Grn |
| Rhbdl2 | 1.19E-13 | 0.435783854 | 0.258 | 0.065 | 2.42E-09 | 24 | Rhbdl2 |
| Bcam7 | 1.20E-13 | 0.579435861 | 0.624 | 0.255 | 2.42E-09 | 24 | Bcam |

| | | | | | | | |
|---|---|---|---|---|---|---|---|
| Prss238 | 1.28E-13 | 0.753642796 | 0.667 | 0.282 | 2.59E-09 | 24 | Prss23 |
| Eva1b7 | 1.30E-13 | 0.584684296 | 0.645 | 0.287 | 2.63E-09 | 24 | Eva1b |
| Uaca6 | 1.43E-13 | 0.614274561 | 0.57 | 0.227 | 2.89E-09 | 24 | Uaca |
| Msn8 | 1.61E-13 | 0.724126329 | 0.806 | 0.445 | 3.27E-09 | 24 | Msn |
| Lmo26 | 1.76E-13 | 0.571093446 | 0.527 | 0.209 | 3.56E-09 | 24 | Lmo2 |
| Ednrb3 | 2.12E-13 | 0.713378307 | 0.516 | 0.195 | 4.28E-09 | 24 | Ednrb |
| Ndrg16 | 2.24E-13 | 0.603811427 | 0.57 | 0.234 | 4.54E-09 | 24 | Ndrg1 |
| Fez23 | 2.30E-13 | 0.515439082 | 0.656 | 0.284 | 4.65E-09 | 24 | Fez2 |
| Arhgef155 | 2.41E-13 | 0.445359473 | 0.484 | 0.17 | 4.88E-09 | 24 | Arhgef15 |
| Rapgef26 | 2.50E-13 | 0.690869141 | 0.71 | 0.34 | 5.05E-09 | 24 | Rapgef2 |
| Meis24 | 2.56E-13 | 0.475867991 | 0.677 | 0.283 | 5.18E-09 | 24 | Meis2 |
| Irf15 | 2.58E-13 | 0.545949578 | 0.667 | 0.3 | 5.22E-09 | 24 | Irf1 |
| Ifi2038 | 2.62E-13 | 0.572248238 | 0.763 | 0.369 | 5.31E-09 | 24 | Ifi203 |
| Jup4 | 2.71E-13 | 0.588290454 | 0.613 | 0.259 | 5.49E-09 | 24 | Jup |
| Acvrl16 | 2.72E-13 | 0.650643476 | 0.645 | 0.276 | 5.51E-09 | 24 | Acvrl1 |
| Vamp55 | 2.84E-13 | 0.534612388 | 0.817 | 0.386 | 5.75E-09 | 24 | Vamp5 |
| Cd1518 | 3.38E-13 | 0.606530531 | 0.538 | 0.218 | 6.85E-09 | 24 | Cd151 |
| Csgalnact16 | 3.52E-13 | 0.635901908 | 0.505 | 0.197 | 7.12E-09 | 24 | Csgalnact1 |
| H2-T235 | 3.74E-13 | 0.765933222 | 0.806 | 0.461 | 7.58E-09 | 24 | H2-T23 |
| Stox25 | 3.78E-13 | 0.789108969 | 0.581 | 0.248 | 7.64E-09 | 24 | Stox2 |
| Ctnnb15 | 4.21E-13 | 0.563634971 | 0.753 | 0.365 | 8.53E-09 | 24 | Ctnnb1 |
| Fbln27 | 4.30E-13 | 0.580997045 | 0.645 | 0.273 | 8.70E-09 | 24 | Fbln2 |
| Gm205322 | 4.47E-13 | 0.476520763 | 0.355 | 0.11 | 9.06E-09 | 24 | Gm20532 |
| Adgrl24 | 5.83E-13 | 0.522218224 | 0.591 | 0.238 | 1.18E-08 | 24 | Adgrl2 |
| Tek5 | 6.03E-13 | 0.566755925 | 0.634 | 0.261 | 1.22E-08 | 24 | Tek |
| N4bp14 | 6.28E-13 | 0.584810277 | 0.462 | 0.177 | 1.27E-08 | 24 | N4bp1 |
| Ccnd15 | 6.38E-13 | 0.55925622 | 0.559 | 0.228 | 1.29E-08 | 24 | Ccnd1 |
| Rarg4 | 6.96E-13 | 0.562444935 | 0.505 | 0.196 | 1.41E-08 | 24 | Rarg |
| Nova23 | 7.27E-13 | 0.502857843 | 0.344 | 0.107 | 1.47E-08 | 24 | Nova2 |
| Fat44 | 7.60E-13 | 0.486873695 | 0.344 | 0.107 | 1.54E-08 | 24 | Fat4 |
| Ets24 | 7.60E-13 | 0.506112127 | 0.505 | 0.195 | 1.54E-08 | 24 | Ets2 |
| Hspg25 | 9.26E-13 | 0.58144322 | 0.71 | 0.313 | 1.88E-08 | 24 | Hspg2 |
| Hmbox13 | 1.05E-12 | 0.705982756 | 0.602 | 0.275 | 2.13E-08 | 24 | Hmbox1 |
| Ctsc4 | 1.07E-12 | 0.278973343 | 0.43 | 0.152 | 2.16E-08 | 24 | Ctsc |
| Laptm510 | 1.29E-12 | 0.613883581 | 0.538 | 0.212 | 2.62E-08 | 24 | Laptm5 |
| Cd24a2 | 1.32E-12 | 0.511094201 | 0.742 | 0.344 | 2.66E-08 | 24 | Cd24a |
| Tns19 | 1.36E-12 | 0.600829121 | 0.581 | 0.246 | 2.75E-08 | 24 | Tns1 |
| Ifit33 | 1.38E-12 | 0.453152205 | 0.376 | 0.124 | 2.80E-08 | 24 | Ifit3 |
| Cep85l7 | 1.41E-12 | 0.60690016 | 0.441 | 0.166 | 2.86E-08 | 24 | Cep85l |
| St3gal15 | 1.42E-12 | 0.532469388 | 0.473 | 0.181 | 2.87E-08 | 24 | St3gal1 |
| Ctla2a9 | 1.59E-12 | 0.60082289 | 0.806 | 0.4 | 3.21E-08 | 24 | Ctla2a |
| Tmem2044 | 1.64E-12 | 0.582975229 | 0.538 | 0.21 | 3.31E-08 | 24 | Tmem204 |
| Itga67 | 1.74E-12 | 0.724484485 | 0.634 | 0.285 | 3.53E-08 | 24 | Itga6 |

| | | | | | | | |
|---|---|---|---|---|---|---|---|
| Cald19 | 1.83E-12 | 0.486402487 | 0.731 | 0.323 | 3.70E-08 | 24 | Cald1 |
| Numb6 | 1.91E-12 | 0.646572742 | 0.699 | 0.338 | 3.87E-08 | 24 | Numb |
| Cyba11 | 1.92E-12 | 0.58368151 | 0.828 | 0.442 | 3.88E-08 | 24 | Cyba |
| Ifi444 | 1.92E-12 | 0.669874419 | 0.409 | 0.147 | 3.89E-08 | 24 | Ifi44 |
| Arpc33 | 2.11E-12 | 0.597208869 | 0.957 | 0.659 | 4.26E-08 | 24 | Arpc3 |
| Rasip15 | 2.20E-12 | 0.584555415 | 0.505 | 0.196 | 4.46E-08 | 24 | Rasip1 |
| Atox15 | 2.25E-12 | 0.561005665 | 0.946 | 0.619 | 4.56E-08 | 24 | Atox1 |
| Anxa35 | 2.35E-12 | 0.535627439 | 0.731 | 0.34 | 4.75E-08 | 24 | Anxa3 |
| Syne27 | 2.35E-12 | 0.540488247 | 0.624 | 0.273 | 4.75E-08 | 24 | Syne2 |
| Apbb27 | 2.46E-12 | 0.609315351 | 0.763 | 0.368 | 4.97E-08 | 24 | Apbb2 |
| Gnai26 | 3.11E-12 | 0.655505533 | 0.914 | 0.576 | 6.30E-08 | 24 | Gnai2 |
| Rdx5 | 3.19E-12 | 0.543459517 | 0.946 | 0.552 | 6.45E-08 | 24 | Rdx |
| Dab2ip7 | 3.19E-12 | 0.58183997 | 0.602 | 0.262 | 6.45E-08 | 24 | Dab2ip |
| Ctsz6 | 3.31E-12 | 0.583750651 | 0.72 | 0.361 | 6.70E-08 | 24 | Ctsz |
| Arhgef128 | 4.59E-12 | 0.657229332 | 0.71 | 0.35 | 9.30E-08 | 24 | Arhgef12 |
| Ptgs13 | 4.60E-12 | 0.581206239 | 0.527 | 0.213 | 9.30E-08 | 24 | Ptgs1 |
| Tuba1a9 | 4.90E-12 | 0.518118538 | 0.602 | 0.257 | 9.93E-08 | 24 | Tuba1a |
| St6galnac38 | 5.74E-12 | 0.448341605 | 0.538 | 0.217 | 1.16E-07 | 24 | St6galnac3 |
| Car83 | 5.95E-12 | 0.586608073 | 0.484 | 0.187 | 1.20E-07 | 24 | Car8 |
| Colgalt24 | 6.73E-12 | 0.542824967 | 0.398 | 0.141 | 1.36E-07 | 24 | Colgalt2 |
| Cpd6 | 6.78E-12 | 0.637207181 | 0.645 | 0.314 | 1.37E-07 | 24 | Cpd |
| Ets19 | 7.62E-12 | 0.586655011 | 0.774 | 0.405 | 1.54E-07 | 24 | Ets1 |
| Clstn18 | 7.83E-12 | 0.531906106 | 0.559 | 0.238 | 1.59E-07 | 24 | Clstn1 |
| Pea15a7 | 8.65E-12 | 0.461423979 | 0.548 | 0.242 | 1.75E-07 | 24 | Pea15a |
| Lap33 | 8.92E-12 | 0.340868982 | 0.387 | 0.138 | 1.81E-07 | 24 | Lap3 |
| Tns23 | 9.42E-12 | 0.424426499 | 0.409 | 0.146 | 1.91E-07 | 24 | Tns2 |
| Prex27 | 9.63E-12 | 0.493258433 | 0.645 | 0.274 | 1.95E-07 | 24 | Prex2 |
| Cyp2d226 | 9.75E-12 | 0.506316131 | 0.387 | 0.136 | 1.97E-07 | 24 | Cyp2d22 |
| Esm13 | 9.79E-12 | 0.944910546 | 0.323 | 0.104 | 1.98E-07 | 24 | Esm1 |
| Psmb98 | 1.19E-11 | 0.523835011 | 0.785 | 0.407 | 2.41E-07 | 24 | Psmb9 |
| Sep-45 | 1.29E-11 | 0.41582266 | 0.441 | 0.165 | 2.61E-07 | 24 | 4-Sep |
| Adcy44 | 1.41E-11 | 0.428189865 | 0.333 | 0.108 | 2.86E-07 | 24 | Adcy4 |
| Smagp4 | 1.60E-11 | 0.388318815 | 0.559 | 0.233 | 3.24E-07 | 24 | Smagp |
| H2-Q43 | 1.85E-11 | 0.595899376 | 0.645 | 0.31 | 3.74E-07 | 24 | H2-Q4 |
| Gbp75 | 2.01E-11 | 0.68456272 | 0.581 | 0.27 | 4.07E-07 | 24 | Gbp7 |
| Lysmd25 | 2.18E-11 | 0.43195603 | 0.484 | 0.193 | 4.41E-07 | 24 | Lysmd2 |
| Lamp29 | 2.22E-11 | 0.509092622 | 0.634 | 0.306 | 4.49E-07 | 24 | Lamp2 |
| Il6st7 | 2.34E-11 | 0.447317149 | 0.559 | 0.233 | 4.74E-07 | 24 | Il6st |
| Kcnq11 | 2.54E-11 | 0.358758616 | 0.312 | 0.097 | 5.15E-07 | 24 | Kcnq1 |
| Mfge87 | 2.73E-11 | 0.508181355 | 0.581 | 0.255 | 5.53E-07 | 24 | Mfge8 |
| Ahnak10 | 2.92E-11 | 0.532935531 | 0.892 | 0.473 | 5.91E-07 | 24 | Ahnak |
| Bag31 | 2.94E-11 | 0.347923357 | 0.344 | 0.115 | 5.95E-07 | 24 | Bag3 |
| Rnf2205 | 3.03E-11 | 0.538618026 | 0.548 | 0.243 | 6.13E-07 | 24 | Rnf220 |

| | | | | | | | |
|---|---|---|---|---|---|---|---|
| Nid16 | 3.06E-11 | 0.432174742 | 0.473 | 0.183 | 6.20E-07 | 24 | Nid1 |
| Plekho17 | 3.16E-11 | 0.48037831 | 0.538 | 0.226 | 6.40E-07 | 24 | Plekho1 |
| Shank34 | 3.21E-11 | 0.504752292 | 0.484 | 0.192 | 6.50E-07 | 24 | Shank3 |
| Oaz25 | 3.45E-11 | 0.608550112 | 0.548 | 0.238 | 6.99E-07 | 24 | Oaz2 |
| Itpkb4 | 3.78E-11 | 0.567577953 | 0.688 | 0.357 | 7.66E-07 | 24 | Itpkb |
| Rnf1251 | 3.80E-11 | 0.399208018 | 0.258 | 0.075 | 7.70E-07 | 24 | Rnf125 |
| Wasf25 | 4.05E-11 | 0.600449181 | 0.763 | 0.426 | 8.20E-07 | 24 | Wasf2 |
| Hexa5 | 4.17E-11 | 0.47472473 | 0.516 | 0.222 | 8.45E-07 | 24 | Hexa |
| Dab27 | 4.18E-11 | 0.428081255 | 0.505 | 0.208 | 8.46E-07 | 24 | Dab2 |
| Lipa4 | 4.38E-11 | 0.477281368 | 0.452 | 0.18 | 8.86E-07 | 24 | Lipa |
| Rhoj8 | 4.45E-11 | 0.502184554 | 0.57 | 0.243 | 9.01E-07 | 24 | Rhoj |
| Samd125 | 4.68E-11 | 0.622016016 | 0.548 | 0.231 | 9.48E-07 | 24 | Samd12 |
| Nrp26 | 4.93E-11 | 0.438922853 | 0.473 | 0.185 | 9.99E-07 | 24 | Nrp2 |
| Samhd16 | 4.95E-11 | 0.531088038 | 0.559 | 0.255 | 1.00E-06 | 24 | Samhd1 |
| Lnx22 | 5.18E-11 | 0.600213768 | 0.366 | 0.134 | 1.05E-06 | 24 | Lnx2 |
| Cd478 | 5.19E-11 | 0.584334409 | 0.86 | 0.511 | 1.05E-06 | 24 | Cd47 |
| Marcks8 | 5.50E-11 | 0.423490217 | 0.71 | 0.343 | 1.11E-06 | 24 | Marcks |
| Klf134 | 5.56E-11 | 0.540200544 | 0.699 | 0.351 | 1.13E-06 | 24 | Klf13 |
| Amotl22 | 5.59E-11 | 0.513601308 | 0.28 | 0.087 | 1.13E-06 | 24 | Amotl2 |
| Tspan137 | 5.63E-11 | 0.499176843 | 0.613 | 0.287 | 1.14E-06 | 24 | Tspan13 |
| H2-K111 | 6.01E-11 | 0.576862926 | 1 | 0.644 | 1.22E-06 | 24 | H2-K1 |
| Tcn23 | 6.42E-11 | 0.566735233 | 0.441 | 0.181 | 1.30E-06 | 24 | Tcn2 |
| Anp32a4 | 7.00E-11 | 0.47909915 | 0.839 | 0.467 | 1.42E-06 | 24 | Anp32a |
| Vat14 | 7.08E-11 | 0.471997833 | 0.409 | 0.156 | 1.43E-06 | 24 | Vat1 |
| Prkch9 | 7.19E-11 | 0.569391364 | 0.785 | 0.401 | 1.46E-06 | 24 | Prkch |
| St3gal51 | 7.28E-11 | 0.410247128 | 0.269 | 0.082 | 1.47E-06 | 24 | St3gal5 |
| Rac15 | 7.37E-11 | 0.572159983 | 0.753 | 0.432 | 1.49E-06 | 24 | Rac1 |
| Sh3glb15 | 7.53E-11 | 0.500282161 | 0.882 | 0.533 | 1.53E-06 | 24 | Sh3glb1 |
| Fut84 | 8.47E-11 | 0.502641298 | 0.613 | 0.29 | 1.71E-06 | 24 | Fut8 |
| Plxnd16 | 8.70E-11 | 0.551148584 | 0.452 | 0.183 | 1.76E-06 | 24 | Plxnd1 |
| Mkl25 | 9.65E-11 | 0.511073663 | 0.57 | 0.257 | 1.95E-06 | 24 | Mkl2 |
| Sh3bgrl8 | 1.18E-10 | 0.468092101 | 0.72 | 0.376 | 2.39E-06 | 24 | Sh3bgrl |
| Cdc42ep23 | 1.19E-10 | 0.462669189 | 0.441 | 0.174 | 2.42E-06 | 24 | Cdc42ep2 |
| Crip112 | 1.26E-10 | 0.442127414 | 0.903 | 0.537 | 2.55E-06 | 24 | Crip1 |
| Slco2b14 | 1.29E-10 | 0.527838077 | 0.452 | 0.181 | 2.62E-06 | 24 | Slco2b1 |
| Cnn38 | 1.30E-10 | 0.406782976 | 0.484 | 0.195 | 2.64E-06 | 24 | Cnn3 |
| Erg5 | 1.33E-10 | 0.465678481 | 0.366 | 0.132 | 2.69E-06 | 24 | Erg |
| Myl12a4 | 1.37E-10 | 0.48236475 | 0.817 | 0.472 | 2.77E-06 | 24 | Myl12a |
| Pls36 | 1.46E-10 | 0.457042733 | 0.516 | 0.216 | 2.95E-06 | 24 | Pls3 |
| Kank35 | 1.48E-10 | 0.508014497 | 0.538 | 0.229 | 3.00E-06 | 24 | Kank3 |
| Hexb3 | 1.60E-10 | 0.359237164 | 0.849 | 0.518 | 3.23E-06 | 24 | Hexb |
| Adam107 | 1.64E-10 | 0.602755447 | 0.591 | 0.294 | 3.32E-06 | 24 | Adam10 |
| Myo1c4 | 1.67E-10 | 0.449817115 | 0.505 | 0.215 | 3.38E-06 | 24 | Myo1c |

| | | | | | | | |
|---|---|---|---|---|---|---|---|
| Myo18a5 | 1.76E-10 | 0.447916838 | 0.57 | 0.259 | 3.57E-06 | 24 | Myo18a |
| Phldb25 | 1.77E-10 | 0.424201028 | 0.495 | 0.204 | 3.59E-06 | 24 | Phldb2 |
| Fermt28 | 1.81E-10 | 0.510229697 | 0.71 | 0.351 | 3.67E-06 | 24 | Fermt2 |
| Dysf4 | 1.91E-10 | 0.439046682 | 0.559 | 0.24 | 3.86E-06 | 24 | Dysf |
| Akt313 | 2.03E-10 | 0.637756601 | 0.538 | 0.253 | 4.12E-06 | 24 | Akt3 |
| Mfap22 | 2.04E-10 | 0.542485864 | 0.312 | 0.108 | 4.12E-06 | 24 | Mfap2 |
| Notch17 | 2.09E-10 | 0.478638647 | 0.505 | 0.218 | 4.23E-06 | 24 | Notch1 |
| Npr13 | 2.21E-10 | 0.358207312 | 0.29 | 0.093 | 4.47E-06 | 24 | Npr1 |
| Plk25 | 2.37E-10 | 0.653389366 | 0.419 | 0.168 | 4.80E-06 | 24 | Plk2 |
| Vcl7 | 2.40E-10 | 0.382734329 | 0.419 | 0.163 | 4.85E-06 | 24 | Vcl |
| Gnb43 | 2.42E-10 | 0.355491008 | 0.323 | 0.109 | 4.90E-06 | 24 | Gnb4 |
| Lrrc8c7 | 2.47E-10 | 0.513141472 | 0.645 | 0.312 | 4.99E-06 | 24 | Lrrc8c |
| Gimap69 | 2.48E-10 | 0.517253173 | 0.731 | 0.367 | 5.02E-06 | 24 | Gimap6 |
| Tspan17 | 2.55E-10 | 0.374875524 | 0.29 | 0.094 | 5.16E-06 | 24 | Tspan17 |
| Arhgap183 | 2.57E-10 | 0.617683149 | 0.656 | 0.343 | 5.19E-06 | 24 | Arhgap18 |
| Cdc42bpa8 | 2.58E-10 | 0.494565432 | 0.452 | 0.186 | 5.22E-06 | 24 | Cdc42bpa |
| Sh3bp54 | 2.82E-10 | 0.46226571 | 0.505 | 0.218 | 5.70E-06 | 24 | Sh3bp5 |
| Lmcd14 | 2.90E-10 | 0.408518288 | 0.43 | 0.166 | 5.86E-06 | 24 | Lmcd1 |
| Dll42 | 3.17E-10 | 0.429224092 | 0.258 | 0.078 | 6.42E-06 | 24 | Dll4 |
| Prrg13 | 3.30E-10 | 0.5406279 | 0.312 | 0.109 | 6.67E-06 | 24 | Prrg1 |
| 2510009E07Rik3 | 3.36E-10 | 0.436755815 | 0.344 | 0.125 | 6.80E-06 | 24 | 2510009E07Rik |
| Tap26 | 3.43E-10 | 0.441644086 | 0.527 | 0.242 | 6.95E-06 | 24 | Tap2 |
| Sema7a4 | 3.60E-10 | 0.502370825 | 0.398 | 0.151 | 7.30E-06 | 24 | Sema7a |
| Ftl16 | 3.74E-10 | 0.443177878 | 1 | 0.911 | 7.57E-06 | 24 | Ftl1 |
| C2cd23 | 3.76E-10 | 0.423900723 | 0.323 | 0.112 | 7.61E-06 | 24 | C2cd2 |
| Nxpe45 | 3.95E-10 | 0.487608382 | 0.387 | 0.149 | 7.99E-06 | 24 | Nxpe4 |
| Efnb13 | 4.07E-10 | 0.505639286 | 0.398 | 0.157 | 8.24E-06 | 24 | Efnb1 |
| Fmnl27 | 4.30E-10 | 0.586055617 | 0.527 | 0.244 | 8.71E-06 | 24 | Fmnl2 |
| Elk35 | 4.55E-10 | 0.45351742 | 0.71 | 0.358 | 9.21E-06 | 24 | Elk3 |
| Heg16 | 4.55E-10 | 0.500372893 | 0.591 | 0.275 | 9.21E-06 | 24 | Heg1 |
| Psme18 | 4.63E-10 | 0.541146609 | 0.903 | 0.57 | 9.38E-06 | 24 | Psme1 |
| Lamp112 | 4.95E-10 | 0.436474785 | 0.763 | 0.419 | 1.00E-05 | 24 | Lamp1 |
| Caskin24 | 5.14E-10 | 0.411735094 | 0.376 | 0.141 | 1.04E-05 | 24 | Caskin2 |
| Wars5 | 5.16E-10 | 0.42084231 | 0.419 | 0.169 | 1.04E-05 | 24 | Wars |
| Tbxa2r1 | 6.27E-10 | 0.353353246 | 0.28 | 0.09 | 1.27E-05 | 24 | Tbxa2r |
| Trove26 | 6.42E-10 | 0.457542607 | 0.355 | 0.133 | 1.30E-05 | 24 | Trove2 |
| Ctsl7 | 6.42E-10 | 0.324836188 | 0.71 | 0.347 | 1.30E-05 | 24 | Ctsl |
| Zhx36 | 7.30E-10 | 0.492535863 | 0.473 | 0.203 | 1.48E-05 | 24 | Zhx3 |
| Robo45 | 7.31E-10 | 0.341342786 | 0.452 | 0.178 | 1.48E-05 | 24 | Robo4 |
| Ppp1r26 | 7.61E-10 | 0.447351523 | 0.742 | 0.394 | 1.54E-05 | 24 | Ppp1r2 |
| Abi35 | 7.88E-10 | 0.570128554 | 0.516 | 0.245 | 1.60E-05 | 24 | Abi3 |
| Prkg17 | 8.51E-10 | 0.543499486 | 0.473 | 0.198 | 1.72E-05 | 24 | Prkg1 |
| Gmfg7 | 9.36E-10 | 0.371597965 | 0.366 | 0.141 | 1.89E-05 | 24 | Gmfg |

| | | | | | | | |
|---|---|---|---|---|---|---|---|
| Ctnnbip14 | 9.74E-10 | 0.365184827 | 0.484 | 0.205 | 1.97E-05 | 24 | Ctnnbip1 |
| Mprip8 | 1.01E-09 | 0.457264498 | 0.602 | 0.294 | 2.04E-05 | 24 | Mprip |
| Crip28 | 1.01E-09 | 0.414321914 | 0.817 | 0.417 | 2.04E-05 | 24 | Crip2 |
| Bvht5 | 1.14E-09 | 0.485833965 | 0.452 | 0.188 | 2.31E-05 | 24 | Bvht |
| Tnfrsf1a6 | 1.26E-09 | 0.436789534 | 0.495 | 0.223 | 2.55E-05 | 24 | Tnfrsf1a |
| Rbpms7 | 1.29E-09 | 0.397502457 | 0.699 | 0.351 | 2.62E-05 | 24 | Rbpms |
| Qk7 | 1.38E-09 | 0.551374468 | 0.731 | 0.411 | 2.79E-05 | 24 | Qk |
| Fnip24 | 1.40E-09 | 0.54845663 | 0.301 | 0.107 | 2.84E-05 | 24 | Fnip2 |
| Wwc26 | 1.49E-09 | 0.481058087 | 0.462 | 0.199 | 3.01E-05 | 24 | Wwc2 |
| Tnfaip14 | 1.54E-09 | 0.471756241 | 0.473 | 0.204 | 3.11E-05 | 24 | Tnfaip1 |
| Zdhhc174 | 1.54E-09 | 0.442783188 | 0.376 | 0.151 | 3.11E-05 | 24 | Zdhhc17 |
| Ociad12 | 1.66E-09 | 0.400730014 | 0.43 | 0.186 | 3.36E-05 | 24 | Ociad1 |
| Rapgef34 | 1.70E-09 | 0.426841751 | 0.419 | 0.17 | 3.44E-05 | 24 | Rapgef3 |
| Afap15 | 1.79E-09 | 0.379260291 | 0.387 | 0.151 | 3.62E-05 | 24 | Afap1 |
| Rasgrf25 | 2.19E-09 | 0.408623995 | 0.441 | 0.178 | 4.43E-05 | 24 | Rasgrf2 |
| Abcg26 | 2.20E-09 | 0.468204663 | 0.57 | 0.272 | 4.46E-05 | 24 | Abcg2 |
| Macf110 | 2.21E-09 | 0.617870712 | 0.839 | 0.504 | 4.47E-05 | 24 | Macf1 |
| Tpm47 | 2.32E-09 | 0.391860427 | 0.71 | 0.381 | 4.70E-05 | 24 | Tpm4 |
| Cav26 | 2.33E-09 | 0.333418282 | 0.495 | 0.21 | 4.72E-05 | 24 | Cav2 |
| Tagln211 | 2.49E-09 | 0.405113856 | 0.774 | 0.43 | 5.05E-05 | 24 | Tagln2 |
| Ntpcr4 | 2.50E-09 | 0.345392829 | 0.301 | 0.106 | 5.06E-05 | 24 | Ntpcr |
| Nfkbia9 | 2.79E-09 | 0.416726348 | 0.753 | 0.401 | 5.64E-05 | 24 | Nfkbia |
| Sod15 | 2.89E-09 | 0.395903891 | 0.591 | 0.301 | 5.86E-05 | 24 | Sod1 |
| Isg155 | 2.90E-09 | 0.403281608 | 0.505 | 0.231 | 5.87E-05 | 24 | Isg15 |
| Oxr14 | 3.06E-09 | 0.528475404 | 0.624 | 0.337 | 6.19E-05 | 24 | Oxr1 |
| Jcad3 | 3.09E-09 | 0.350045313 | 0.323 | 0.116 | 6.26E-05 | 24 | Jcad |
| Myl12b4 | 3.15E-09 | 0.401170182 | 0.882 | 0.549 | 6.39E-05 | 24 | Myl12b |
| 9530026P05Rik6 | 3.20E-09 | 0.485365134 | 0.527 | 0.243 | 6.48E-05 | 24 | 9530026P05Rik |
| Birc614 | 3.31E-09 | 0.49186183 | 0.645 | 0.342 | 6.71E-05 | 24 | Birc6 |
| Ostf15 | 3.77E-09 | 0.351784125 | 0.903 | 0.508 | 7.63E-05 | 24 | Ostf1 |
| Rab5c3 | 3.77E-09 | 0.318233389 | 0.591 | 0.292 | 7.63E-05 | 24 | Rab5c |
| Xdh5 | 4.09E-09 | 0.482684009 | 0.441 | 0.189 | 8.29E-05 | 24 | Xdh |
| Ifngr19 | 4.20E-09 | 0.476650054 | 0.731 | 0.394 | 8.51E-05 | 24 | Ifngr1 |
| Kmt2e11 | 4.21E-09 | 0.404774241 | 0.742 | 0.408 | 8.53E-05 | 24 | Kmt2e |
| Stab12 | 4.40E-09 | 0.340833256 | 0.258 | 0.085 | 8.91E-05 | 24 | Stab1 |
| Jam25 | 4.80E-09 | 0.420052668 | 0.613 | 0.287 | 9.72E-05 | 24 | Jam2 |
| Nfat510 | 5.02E-09 | 0.524721455 | 0.538 | 0.268 | 0.000101723 | 24 | Nfat5 |
| Ncoa75 | 5.15E-09 | 0.519099446 | 0.581 | 0.298 | 0.000104284 | 24 | Ncoa7 |
| Tmem1405 | 5.36E-09 | 0.665806987 | 0.473 | 0.224 | 0.000108536 | 24 | Tmem140 |
| Bsg9 | 5.65E-09 | 0.509659915 | 0.763 | 0.451 | 0.000114446 | 24 | Bsg |
| Cds26 | 5.96E-09 | 0.518657529 | 0.409 | 0.178 | 0.00012066 | 24 | Cds2 |
| Gja14 | 6.06E-09 | 0.35295285 | 0.344 | 0.129 | 0.000122705 | 24 | Gja1 |
| B230219D22Rik6 | 6.15E-09 | 0.407079578 | 0.656 | 0.344 | 0.00012448 | 24 | B230219D22Rik |

| | | | | | | | |
|---|---|---|---|---|---|---|---|
| Laptm4a9 | 6.52E-09 | 0.470475597 | 0.817 | 0.485 | 0.000132025 | 24 | Laptm4a |
| Dennd5a7 | 6.57E-09 | 0.478793483 | 0.43 | 0.188 | 0.000133019 | 24 | Dennd5a |
| Xaf15 | 6.64E-09 | 0.475090176 | 0.484 | 0.222 | 0.000134466 | 24 | Xaf1 |
| Cpne85 | 6.76E-09 | 0.448475977 | 0.505 | 0.23 | 0.000136812 | 24 | Cpne8 |
| Reep37 | 6.76E-09 | 0.565788067 | 0.591 | 0.307 | 0.000136959 | 24 | Reep3 |
| Slc48a11 | 6.96E-09 | 0.352554159 | 0.301 | 0.111 | 0.000140842 | 24 | Slc48a1 |
| Atrx6 | 7.34E-09 | 0.563354939 | 0.72 | 0.421 | 0.000148696 | 24 | Atrx |
| Gm326882 | 7.45E-09 | 0.311618255 | 0.28 | 0.095 | 0.000150844 | 24 | Gm32688 |
| Ppfibp15 | 7.48E-09 | 0.550992713 | 0.419 | 0.185 | 0.00015136 | 24 | Ppfibp1 |
| Kctd172 | 7.58E-09 | 0.32101804 | 0.269 | 0.091 | 0.000153423 | 24 | Kctd17 |
| Psme25 | 8.16E-09 | 0.422725759 | 0.774 | 0.445 | 0.000165241 | 24 | Psme2 |
| Ntf33 | 8.29E-09 | 0.421279947 | 0.43 | 0.182 | 0.00016782 | 24 | Ntf3 |
| mt-Co220 | 8.35E-09 | 0.499934701 | 1 | 0.729 | 0.000169121 | 24 | mt-Co2 |
| Zbtb465 | 8.39E-09 | 0.308471697 | 0.409 | 0.164 | 0.000169803 | 24 | Zbtb46 |
| Tubb4b5 | 8.60E-09 | 0.443428067 | 0.742 | 0.428 | 0.000174045 | 24 | Tubb4b |
| Tapbp7 | 8.67E-09 | 0.53400561 | 0.591 | 0.306 | 0.000175586 | 24 | Tapbp |
| Mapk35 | 8.68E-09 | 0.455756292 | 0.538 | 0.267 | 0.000175784 | 24 | Mapk3 |
| Tmed53 | 8.74E-09 | 0.57638042 | 0.548 | 0.29 | 0.000176945 | 24 | Tmed5 |
| 9930111J21Rik28 | 9.22E-09 | 0.459507487 | 0.527 | 0.258 | 0.000186606 | 24 | 9930111J21Rik2 |
| C130074G19Rik5 | 9.24E-09 | 0.346665961 | 0.462 | 0.201 | 0.000186992 | 24 | C130074G19Rik |
| Plscr15 | 9.32E-09 | 0.41429863 | 0.462 | 0.206 | 0.000188689 | 24 | Plscr1 |
| B4galt44 | 9.41E-09 | 0.454150246 | 0.398 | 0.164 | 0.000190431 | 24 | B4galt4 |
| Tpm310 | 9.53E-09 | 0.512542742 | 0.796 | 0.481 | 0.000192965 | 24 | Tpm3 |
| Nap1l19 | 9.87E-09 | 0.397573413 | 0.613 | 0.316 | 0.00019984 | 24 | Nap1l1 |
| Ddrgk14 | 1.02E-08 | 0.474644037 | 0.591 | 0.32 | 0.000206472 | 24 | Ddrgk1 |
| Nedd49 | 1.03E-08 | 0.384246003 | 0.774 | 0.421 | 0.000208608 | 24 | Nedd4 |
| Sipa1l25 | 1.05E-08 | 0.392837122 | 0.387 | 0.16 | 0.000212108 | 24 | Sipa1l2 |
| Arap34 | 1.12E-08 | 0.468311216 | 0.473 | 0.219 | 0.000227416 | 24 | Arap3 |
| Leprot4 | 1.14E-08 | 0.412479079 | 0.409 | 0.179 | 0.000230245 | 24 | Leprot |
| Plscr47 | 1.17E-08 | 0.461155142 | 0.43 | 0.191 | 0.000237348 | 24 | Plscr4 |
| Tmbim68 | 1.19E-08 | 0.548498368 | 0.71 | 0.433 | 0.000240692 | 24 | Tmbim6 |
| Dock16 | 1.22E-08 | 0.499956731 | 0.559 | 0.282 | 0.000247096 | 24 | Dock1 |
| Vwa3a1 | 1.24E-08 | 0.385709485 | 0.258 | 0.087 | 0.000250248 | 24 | Vwa3a |
| Nr3c17 | 1.26E-08 | 0.449211662 | 0.699 | 0.384 | 0.000254463 | 24 | Nr3c1 |
| Mafb4 | 1.40E-08 | 0.318830428 | 0.387 | 0.156 | 0.000283312 | 24 | Mafb |
| Phf11b3 | 1.46E-08 | 0.371614024 | 0.269 | 0.094 | 0.000295663 | 24 | Phf11b |
| Plcg13 | 1.53E-08 | 0.374474386 | 0.323 | 0.123 | 0.000309929 | 24 | Plcg1 |
| Proser21 | 1.58E-08 | 0.379968121 | 0.28 | 0.099 | 0.000320164 | 24 | Proser2 |
| Lipe4 | 1.66E-08 | 0.446735356 | 0.344 | 0.139 | 0.000335272 | 24 | Lipe |
| Amotl18 | 1.78E-08 | 0.346080588 | 0.441 | 0.187 | 0.000359601 | 24 | Amotl1 |
| Tspan184 | 1.83E-08 | 0.476518644 | 0.344 | 0.139 | 0.000370731 | 24 | Tspan18 |
| Nbea8 | 1.84E-08 | 0.474211329 | 0.441 | 0.198 | 0.000372456 | 24 | Nbea |
| Ctnna17 | 1.85E-08 | 0.317097173 | 0.688 | 0.353 | 0.000374139 | 24 | Ctnna1 |

| | | | | | | | |
|---|---|---|---|---|---|---|---|
| Atp2b110 | 1.85E-08 | 0.492372498 | 0.613 | 0.334 | 0.000375412 | 24 | Atp2b1 |
| Dok42 | 1.89E-08 | 0.318456421 | 0.323 | 0.12 | 0.000382376 | 24 | Dok4 |
| Limch16 | 1.94E-08 | 0.507247006 | 0.344 | 0.141 | 0.000392457 | 24 | Limch1 |
| Grb107 | 1.96E-08 | 0.453687638 | 0.441 | 0.196 | 0.000397695 | 24 | Grb10 |
| Phlpp17 | 2.02E-08 | 0.460850104 | 0.548 | 0.265 | 0.000409726 | 24 | Phlpp1 |
| Pold45 | 2.05E-08 | 0.375452584 | 0.43 | 0.202 | 0.000415103 | 24 | Pold4 |
| Atrn3 | 2.08E-08 | 0.490456859 | 0.312 | 0.124 | 0.000421177 | 24 | Atrn |
| Fkbp1a5 | 2.13E-08 | 0.432576725 | 0.71 | 0.407 | 0.000431216 | 24 | Fkbp1a |
| Myo1b4 | 2.18E-08 | 0.384277222 | 0.344 | 0.139 | 0.000441383 | 24 | Myo1b |
| 5-Jun | 2.28E-08 | 0.53078931 | 0.839 | 0.515 | 0.000462571 | 24 | Jun |
| Afdn5 | 2.36E-08 | 0.465477785 | 0.591 | 0.316 | 0.000478615 | 24 | Afdn |
| Pitpnc110 | 2.42E-08 | 0.510019175 | 0.763 | 0.43 | 0.000490663 | 24 | Pitpnc1 |
| Unc45b4 | 2.72E-08 | 0.337441459 | 0.355 | 0.141 | 0.000550438 | 24 | Unc45b |
| Utrn7 | 2.84E-08 | 0.474403054 | 0.71 | 0.407 | 0.000574634 | 24 | Utrn |
| Tsnax3 | 2.88E-08 | 0.417150613 | 0.387 | 0.167 | 0.000582365 | 24 | Tsnax |
| Sash18 | 2.89E-08 | 0.488175891 | 0.538 | 0.258 | 0.000584653 | 24 | Sash1 |
| Mecom3 | 2.90E-08 | 0.401572339 | 0.28 | 0.098 | 0.000587266 | 24 | Mecom |
| Mydgf4 | 2.98E-08 | 0.301053323 | 0.484 | 0.233 | 0.000604202 | 24 | Mydgf |
| Cfl110 | 3.21E-08 | 0.464428368 | 0.946 | 0.61 | 0.000649676 | 24 | Cfl1 |
| Zfp36l19 | 3.23E-08 | 0.493844295 | 0.742 | 0.434 | 0.000653223 | 24 | Zfp36l1 |
| Tgm26 | 3.24E-08 | 0.301500157 | 0.398 | 0.163 | 0.000655754 | 24 | Tgm2 |
| Malat114 | 3.24E-08 | 0.488868579 | 0.989 | 0.726 | 0.000656729 | 24 | Malat1 |
| Sox43 | 3.46E-08 | 0.271110193 | 0.462 | 0.205 | 0.000700897 | 24 | Sox4 |
| Nes4 | 3.52E-08 | 0.490905484 | 0.333 | 0.135 | 0.000713041 | 24 | Nes |
| Tmco18 | 3.80E-08 | 0.412444647 | 0.581 | 0.314 | 0.000769417 | 24 | Tmco1 |
| Sypl5 | 3.97E-08 | 0.417056001 | 0.495 | 0.241 | 0.000803469 | 24 | Sypl |
| Rtp42 | 4.07E-08 | 0.275638135 | 0.387 | 0.163 | 0.000823046 | 24 | Rtp4 |
| Tmed34 | 4.22E-08 | 0.35259795 | 0.495 | 0.245 | 0.00085491 | 24 | Tmed3 |
| Emp15 | 4.53E-08 | 0.260666548 | 0.323 | 0.123 | 0.000916885 | 24 | Emp1 |
| Rnf144a3 | 4.57E-08 | 0.344286637 | 0.29 | 0.108 | 0.000925382 | 24 | Rnf144a |
| Cbfa2t36 | 4.66E-08 | 0.421594141 | 0.344 | 0.14 | 0.000942518 | 24 | Cbfa2t3 |
| Tmem2591 | 4.74E-08 | 0.334042526 | 0.258 | 0.093 | 0.000959242 | 24 | Tmem259 |
| Dusp15 | 4.74E-08 | 0.420406032 | 0.43 | 0.195 | 0.000959817 | 24 | Dusp1 |
| Ralgapa16 | 5.19E-08 | 0.446891638 | 0.581 | 0.305 | 0.001050091 | 24 | Ralgapa1 |
| Myo105 | 5.21E-08 | 0.381127205 | 0.473 | 0.22 | 0.001054903 | 24 | Myo10 |
| Man2b13 | 5.30E-08 | 0.293599465 | 0.387 | 0.165 | 0.001073839 | 24 | Man2b1 |
| Klhl55 | 5.48E-08 | 0.305750995 | 0.484 | 0.226 | 0.001108645 | 24 | Klhl5 |
| Fam43a2 | 5.49E-08 | 0.32735868 | 0.355 | 0.143 | 0.001112498 | 24 | Fam43a |
| Snrpb4 | 5.74E-08 | 0.312955235 | 0.785 | 0.474 | 0.001161377 | 24 | Snrpb |
| Kdelr13 | 5.87E-08 | 0.466301971 | 0.462 | 0.239 | 0.001187574 | 24 | Kdelr1 |
| Pdia611 | 5.87E-08 | 0.51764631 | 0.602 | 0.339 | 0.00118926 | 24 | Pdia6 |
| Lats23 | 5.93E-08 | 0.338638353 | 0.312 | 0.121 | 0.001200329 | 24 | Lats2 |
| Lpcat13 | 6.06E-08 | 0.418018856 | 0.376 | 0.16 | 0.001226598 | 24 | Lpcat1 |

| | | | | | | | |
|---|---|---|---|---|---|---|---|
| Atg4a2 | 6.06E-08 | 0.507696212 | 0.323 | 0.133 | 0.001227423 | 24 | Atg4a |
| Bnip25 | 6.23E-08 | 0.450480132 | 0.548 | 0.285 | 0.001260815 | 24 | Bnip2 |
| Igbp1 | 6.26E-08 | 0.289094595 | 0.376 | 0.165 | 0.001266722 | 24 | Igbp1 |
| Lpar64 | 6.30E-08 | 0.495584081 | 0.409 | 0.185 | 0.001274798 | 24 | Lpar6 |
| Ppic5 | 6.30E-08 | 0.366883756 | 0.312 | 0.122 | 0.001274799 | 24 | Ppic |
| Fam13c5 | 6.51E-08 | 0.275496136 | 0.301 | 0.113 | 0.001317213 | 24 | Fam13c |
| Xiap5 | 6.59E-08 | 0.381546656 | 0.581 | 0.301 | 0.001333863 | 24 | Xiap |
| Phf11d4 | 6.59E-08 | 0.416305903 | 0.333 | 0.139 | 0.001334411 | 24 | Phf11d |
| Palmd4 | 6.77E-08 | 0.317709759 | 0.495 | 0.235 | 0.001371503 | 24 | Palmd |
| Clec1a4 | 6.83E-08 | 0.328381206 | 0.409 | 0.17 | 0.001382905 | 24 | Clec1a |
| Selenok7 | 6.89E-08 | 0.396877306 | 0.839 | 0.556 | 0.001395643 | 24 | Selenok |
| Stom4 | 6.92E-08 | 0.315835246 | 0.29 | 0.109 | 0.001400975 | 24 | Stom |
| Tex2611 | 6.93E-08 | 0.356674482 | 0.409 | 0.19 | 0.001403949 | 24 | Tex261 |
| Gnaq8 | 6.99E-08 | 0.374881079 | 0.656 | 0.361 | 0.00141456 | 24 | Gnaq |
| Fmnl36 | 7.44E-08 | 0.56099218 | 0.473 | 0.241 | 0.001506466 | 24 | Fmnl3 |
| St3gal65 | 7.51E-08 | 0.351877331 | 0.613 | 0.319 | 0.001521101 | 24 | St3gal6 |
| Ceacam15 | 7.73E-08 | 0.347884461 | 0.376 | 0.157 | 0.001565393 | 24 | Ceacam1 |
| Pcdh16 | 7.84E-08 | 0.329098076 | 0.258 | 0.093 | 0.001587339 | 24 | Pcdh1 |
| Ifnar27 | 8.05E-08 | 0.452577009 | 0.591 | 0.326 | 0.001630134 | 24 | Ifnar2 |
| Tbl1x7 | 8.06E-08 | 0.44564081 | 0.505 | 0.263 | 0.001631499 | 24 | Tbl1x |
| Tjp24 | 8.23E-08 | 0.294708683 | 0.344 | 0.142 | 0.001666698 | 24 | Tjp2 |
| Capn25 | 8.33E-08 | 0.497161773 | 0.387 | 0.18 | 0.001686939 | 24 | Capn2 |
| Rhob6 | 8.46E-08 | 0.376383 | 0.538 | 0.269 | 0.001712309 | 24 | Rhob |
| Tnfaip24 | 8.76E-08 | 0.304748071 | 0.366 | 0.153 | 0.001774054 | 24 | Tnfaip2 |
| Bag54 | 8.81E-08 | 0.394116868 | 0.43 | 0.203 | 0.001782764 | 24 | Bag5 |
| Lama48 | 9.06E-08 | 0.323539344 | 0.409 | 0.176 | 0.001835091 | 24 | Lama4 |
| Yes15 | 9.36E-08 | 0.402261107 | 0.366 | 0.155 | 0.001894509 | 24 | Yes1 |
| Aig15 | 9.41E-08 | 0.403859172 | 0.452 | 0.215 | 0.001905981 | 24 | Aig1 |
| Crispld16 | 9.55E-08 | 0.310712217 | 0.355 | 0.145 | 0.001932798 | 24 | Crispld1 |
| Mat2a5 | 9.61E-08 | 0.416628131 | 0.591 | 0.311 | 0.001945356 | 24 | Mat2a |
| Adgrg16 | 1.01E-07 | 0.398016495 | 0.344 | 0.141 | 0.002045577 | 24 | Adgrg1 |
| Prickle16 | 1.08E-07 | 0.379765411 | 0.355 | 0.148 | 0.002178952 | 24 | Prickle1 |
| Notch43 | 1.11E-07 | 0.366317392 | 0.301 | 0.117 | 0.002247808 | 24 | Notch4 |
| Lgals3bp4 | 1.12E-07 | 0.377339275 | 0.344 | 0.148 | 0.002270597 | 24 | Lgals3bp |
| Klf98 | 1.15E-07 | 0.348579752 | 0.495 | 0.239 | 0.002321904 | 24 | Klf9 |
| Sncg6 | 1.15E-07 | 0.439924226 | 0.366 | 0.156 | 0.002323428 | 24 | Sncg |
| Itgb54 | 1.20E-07 | 0.251063661 | 0.355 | 0.146 | 0.002423422 | 24 | Itgb5 |
| Mast47 | 1.21E-07 | 0.352671228 | 0.419 | 0.188 | 0.002446973 | 24 | Mast4 |
| Snx27 | 1.24E-07 | 0.332344208 | 0.269 | 0.103 | 0.002514931 | 24 | Snx27 |
| Smarca27 | 1.26E-07 | 0.515250629 | 0.527 | 0.282 | 0.002547835 | 24 | Smarca2 |
| Zeb112 | 1.27E-07 | 0.502795693 | 0.581 | 0.323 | 0.002572158 | 24 | Zeb1 |
| Rap1b6 | 1.31E-07 | 0.380835691 | 0.753 | 0.446 | 0.002662254 | 24 | Rap1b |
| Mndal8 | 1.33E-07 | 0.499971733 | 0.688 | 0.407 | 0.002688166 | 24 | Mndal |

| | | | | | | | |
|---|---|---|---|---|---|---|---|
| Tgfb19 | 1.33E-07 | 0.441459571 | 0.473 | 0.238 | 0.002691904 | 24 | Tgfb1 |
| Ripor15 | 1.39E-07 | 0.347256381 | 0.376 | 0.162 | 0.002817659 | 24 | Ripor1 |
| Ktn19 | 1.41E-07 | 0.431394242 | 0.57 | 0.309 | 0.00284463 | 24 | Ktn1 |
| Sgk14 | 1.41E-07 | 0.398275645 | 0.538 | 0.271 | 0.002854634 | 24 | Sgk1 |
| Rit1 | 1.42E-07 | 0.314122714 | 0.333 | 0.142 | 0.002868345 | 24 | Rit1 |
| Camk15 | 1.43E-07 | 0.27469639 | 0.473 | 0.228 | 0.002889538 | 24 | Camk1 |
| Gna116 | 1.53E-07 | 0.404624491 | 0.495 | 0.246 | 0.003101402 | 24 | Gna11 |
| Rbms25 | 1.54E-07 | 0.329997951 | 0.366 | 0.161 | 0.00312767 | 24 | Rbms2 |
| Myh912 | 1.58E-07 | 0.391209389 | 0.742 | 0.438 | 0.003194017 | 24 | Myh9 |
| Rtl8a3 | 1.66E-07 | 0.447027061 | 0.398 | 0.178 | 0.003354146 | 24 | Rtl8a |
| Pde8a4 | 1.72E-07 | 0.331474466 | 0.409 | 0.184 | 0.003483628 | 24 | Pde8a |
| Gata23 | 1.72E-07 | 0.326659143 | 0.28 | 0.106 | 0.003492035 | 24 | Gata2 |
| Tmem30a8 | 1.74E-07 | 0.396224318 | 0.548 | 0.288 | 0.003513448 | 24 | Tmem30a |
| Efna15 | 1.75E-07 | 0.369834151 | 0.344 | 0.145 | 0.003534864 | 24 | Efna1 |
| Ezh11 | 1.82E-07 | 0.328639858 | 0.312 | 0.129 | 0.003685528 | 24 | Ezh1 |
| Myadm6 | 1.87E-07 | 0.386986462 | 0.301 | 0.121 | 0.003793862 | 24 | Myadm |
| Arl6ip111 | 1.89E-07 | 0.349613239 | 0.677 | 0.372 | 0.003820627 | 24 | Arl6ip1 |
| Flywch12 | 1.90E-07 | 0.312051454 | 0.29 | 0.114 | 0.003854866 | 24 | Flywch1 |
| Calr10 | 2.04E-07 | 0.476018172 | 0.688 | 0.409 | 0.004120542 | 24 | Calr |
| Trim30a9 | 2.04E-07 | 0.402244335 | 0.452 | 0.221 | 0.004121811 | 24 | Trim30a |
| Nav16 | 2.04E-07 | 0.362168672 | 0.473 | 0.224 | 0.004124586 | 24 | Nav1 |
| Chst22 | 2.04E-07 | 0.444702608 | 0.376 | 0.167 | 0.004135982 | 24 | Chst2 |
| Prdx47 | 2.15E-07 | 0.350313173 | 0.462 | 0.223 | 0.004354389 | 24 | Prdx4 |
| Dazap22 | 2.15E-07 | 0.25053275 | 0.57 | 0.301 | 0.004359757 | 24 | Dazap2 |
| Hsp90b113 | 2.15E-07 | 0.398261911 | 0.817 | 0.512 | 0.004359884 | 24 | Hsp90b1 |
| Ace4 | 2.16E-07 | 0.326127527 | 0.301 | 0.12 | 0.004374131 | 24 | Ace |
| Selenom7 | 2.17E-07 | 0.259883379 | 0.806 | 0.447 | 0.004395556 | 24 | Selenom |
| Ipo112 | 2.17E-07 | 0.257976439 | 0.333 | 0.138 | 0.004399177 | 24 | Ipo11 |
| Ptbp312 | 2.27E-07 | 0.459154826 | 0.688 | 0.395 | 0.004589385 | 24 | Ptbp3 |
| Rnd32 | 2.39E-07 | 0.475165726 | 0.258 | 0.098 | 0.004835321 | 24 | Rnd3 |
| Abhd123 | 2.42E-07 | 0.313155558 | 0.312 | 0.13 | 0.004902833 | 24 | Abhd12 |
| 24-Mar | 2.52E-07 | 0.283336106 | 0.688 | 0.395 | 0.005106735 | 24 | 2-Mar |
| Tubb58 | 2.57E-07 | 0.502293853 | 0.645 | 0.382 | 0.005203009 | 24 | Tubb5 |
| Ctso1 | 2.59E-07 | 0.352520573 | 0.29 | 0.116 | 0.005253297 | 24 | Ctso |
| Psmb105 | 2.75E-07 | 0.411002953 | 0.667 | 0.401 | 0.005570273 | 24 | Psmb10 |
| Hspa513 | 2.87E-07 | 0.286732978 | 0.763 | 0.436 | 0.005814217 | 24 | Hspa5 |
| Lrrc3b3 | 2.88E-07 | 0.345528963 | 0.258 | 0.095 | 0.005836994 | 24 | Lrrc3b |
| Ier5l4 | 2.89E-07 | 0.510912734 | 0.387 | 0.18 | 0.005851767 | 24 | Ier5l |
| Znfx15 | 2.91E-07 | 0.319336084 | 0.387 | 0.176 | 0.005901137 | 24 | Znfx1 |
| Gab15 | 2.92E-07 | 0.359968474 | 0.29 | 0.116 | 0.005907434 | 24 | Gab1 |
| Trim254 | 3.09E-07 | 0.354617526 | 0.344 | 0.151 | 0.006250893 | 24 | Trim25 |
| Calu7 | 3.17E-07 | 0.397087669 | 0.419 | 0.204 | 0.006418447 | 24 | Calu |
| Atxn112 | 3.17E-07 | 0.358769741 | 0.57 | 0.297 | 0.00642747 | 24 | Atxn1 |

| | | | | | | | |
|---|---|---|---|---|---|---|---|
| Cyfip18 | 3.29E-07 | 0.364793031 | 0.527 | 0.28 | 0.006665727 | 24 | Cyfip1 |
| Fli110 | 3.32E-07 | 0.393954166 | 0.71 | 0.403 | 0.006728553 | 24 | Fli1 |
| Kat6b4 | 3.45E-07 | 0.382527354 | 0.344 | 0.153 | 0.006980705 | 24 | Kat6b |
| Adamts94 | 3.68E-07 | 0.414762017 | 0.301 | 0.125 | 0.007455061 | 24 | Adamts9 |
| Tubb2a6 | 3.69E-07 | 0.327375801 | 0.366 | 0.166 | 0.007472373 | 24 | Tubb2a |
| Nckap17 | 3.73E-07 | 0.27658521 | 0.473 | 0.228 | 0.007550427 | 24 | Nckap1 |
| Cx3cl14 | 3.89E-07 | 0.376121727 | 0.269 | 0.105 | 0.007873029 | 24 | Cx3cl1 |
| Parp147 | 3.89E-07 | 0.393803311 | 0.495 | 0.254 | 0.007883564 | 24 | Parp14 |
| Vps13d4 | 4.00E-07 | 0.40079674 | 0.366 | 0.17 | 0.008102813 | 24 | Vps13d |
| Inpp5a2 | 4.17E-07 | 0.381019159 | 0.312 | 0.131 | 0.008452428 | 24 | Inpp5a |
| Igtp1 | 4.26E-07 | 0.262768064 | 0.28 | 0.109 | 0.008618176 | 24 | Igtp |
| Selenow6 | 4.27E-07 | 0.30015479 | 0.925 | 0.631 | 0.00864782 | 24 | Selenow |
| Nav29 | 4.59E-07 | 0.433012113 | 0.366 | 0.167 | 0.009286032 | 24 | Nav2 |
| Foxo16 | 4.70E-07 | 0.517533997 | 0.409 | 0.201 | 0.009510294 | 24 | Foxo1 |
| Fam3c4 | 4.84E-07 | 0.443250921 | 0.376 | 0.18 | 0.009797649 | 24 | Fam3c |
| Nxn6 | 4.86E-07 | 0.298802322 | 0.527 | 0.27 | 0.009843265 | 24 | Nxn |
| Unc119b2 | 4.88E-07 | 0.336701428 | 0.333 | 0.141 | 0.009875294 | 24 | Unc119b |
| Pde7b8 | 4.90E-07 | 0.384676916 | 0.484 | 0.244 | 0.009913773 | 24 | Pde7b |
| Gimap112 | 4.98E-07 | 0.324568951 | 0.591 | 0.304 | 0.010080075 | 24 | Gimap1 |
| Kif13a5 | 5.22E-07 | 0.304756985 | 0.387 | 0.175 | 0.01055845 | 24 | Kif13a |
| Gbp33 | 5.38E-07 | 0.345239411 | 0.29 | 0.119 | 0.010883341 | 24 | Gbp3 |
| Itm2a5 | 5.50E-07 | 0.321555626 | 0.398 | 0.181 | 0.011143717 | 24 | Itm2a |
| Ccdc28b4 | 5.76E-07 | 0.374844421 | 0.516 | 0.263 | 0.011659812 | 24 | Ccdc28b |
| Ces2e2 | 5.84E-07 | 0.372362725 | 0.28 | 0.111 | 0.011829377 | 24 | Ces2e |
| Ptpra7 | 5.88E-07 | 0.420265918 | 0.548 | 0.306 | 0.011896045 | 24 | Ptpra |
| Camsap22 | 5.96E-07 | 0.380674334 | 0.269 | 0.109 | 0.012057224 | 24 | Camsap2 |
| Mfhas14 | 6.10E-07 | 0.435442983 | 0.333 | 0.151 | 0.012354277 | 24 | Mfhas1 |
| Ephb45 | 6.13E-07 | 0.353685658 | 0.376 | 0.168 | 0.012401324 | 24 | Ephb4 |
| Bbx8 | 6.15E-07 | 0.458490364 | 0.441 | 0.224 | 0.0124588 | 24 | Bbx |
| Smim7 | 6.32E-07 | 0.269195107 | 0.398 | 0.193 | 0.012788802 | 24 | Smim7 |
| She5 | 6.36E-07 | 0.37065143 | 0.344 | 0.151 | 0.012878217 | 24 | She |
| Stmn16 | 6.38E-07 | 0.435653206 | 0.312 | 0.132 | 0.012921656 | 24 | Stmn1 |
| Mdfic6 | 6.82E-07 | 0.465343652 | 0.484 | 0.247 | 0.013814656 | 24 | Mdfic |
| Lrmda9 | 7.20E-07 | 0.390766009 | 0.645 | 0.37 | 0.014584768 | 24 | Lrmda |
| mt-Nd315 | 7.72E-07 | 0.45938773 | 0.71 | 0.448 | 0.015631046 | 24 | mt-Nd3 |
| Stat18 | 7.83E-07 | 0.457011549 | 0.484 | 0.251 | 0.015848708 | 24 | Stat1 |
| Cltc4 | 8.06E-07 | 0.321195311 | 0.581 | 0.332 | 0.016311497 | 24 | Cltc |
| Rpn13 | 8.17E-07 | 0.366809677 | 0.441 | 0.223 | 0.016550453 | 24 | Rpn1 |
| mt-Nd118 | 8.35E-07 | 0.399731674 | 0.892 | 0.6 | 0.016903689 | 24 | mt-Nd1 |
| Kctd104 | 8.42E-07 | 0.266068543 | 0.29 | 0.118 | 0.017046372 | 24 | Kctd10 |
| Herc18 | 8.48E-07 | 0.401649746 | 0.484 | 0.258 | 0.017177397 | 24 | Herc1 |
| Lman17 | 9.52E-07 | 0.390903311 | 0.452 | 0.24 | 0.019276766 | 24 | Lman1 |
| St8sia49 | 9.54E-07 | 0.354185821 | 0.505 | 0.261 | 0.019312082 | 24 | St8sia4 |

| | | | | | | | |
|---|---|---|---|---|---|---|---|
| Tdrp2 | 9.78E-07 | 0.331537999 | 0.29 | 0.118 | 0.019801433 | 24 | Tdrp |
| Nhsl25 | 1.02E-06 | 0.293647613 | 0.323 | 0.136 | 0.020709934 | 24 | Nhsl2 |
| Selenof8 | 1.02E-06 | 0.378484634 | 0.763 | 0.493 | 0.020727572 | 24 | Selenof |
| Gcnt26 | 1.09E-06 | 0.34345105 | 0.333 | 0.144 | 0.022133414 | 24 | Gcnt2 |
| Kif5b4 | 1.11E-06 | 0.318726846 | 0.677 | 0.408 | 0.022377326 | 24 | Kif5b |
| Ccrl23 | 1.12E-06 | 0.37090342 | 0.269 | 0.11 | 0.022723187 | 24 | Ccrl2 |
| Pgrmc24 | 1.13E-06 | 0.293833147 | 0.323 | 0.144 | 0.022884668 | 24 | Pgrmc2 |
| Ptk27 | 1.18E-06 | 0.38433081 | 0.516 | 0.268 | 0.023939658 | 24 | Ptk2 |
| Pqlc14 | 1.19E-06 | 0.281304857 | 0.301 | 0.127 | 0.024064895 | 24 | Pqlc1 |
| Smpdl3a6 | 1.19E-06 | 0.314758069 | 0.441 | 0.221 | 0.024157475 | 24 | Smpdl3a |
| Endod12 | 1.22E-06 | 0.343838556 | 0.344 | 0.153 | 0.024602841 | 24 | Endod1 |
| Pip4k2a7 | 1.26E-06 | 0.312436546 | 0.473 | 0.243 | 0.025418271 | 24 | Pip4k2a |
| Fry6 | 1.30E-06 | 0.281490172 | 0.419 | 0.202 | 0.026248608 | 24 | Fry |
| Zfp7104 | 1.38E-06 | 0.370537473 | 0.344 | 0.156 | 0.027961802 | 24 | Zfp710 |
| Luzp16 | 1.41E-06 | 0.306702903 | 0.645 | 0.358 | 0.0285889 | 24 | Luzp1 |
| Mertk6 | 1.46E-06 | 0.456233599 | 0.301 | 0.133 | 0.029502844 | 24 | Mertk |
| Adipor15 | 1.48E-06 | 0.268097493 | 0.495 | 0.259 | 0.029972289 | 24 | Adipor1 |
| Ctsa3 | 1.50E-06 | 0.33515043 | 0.333 | 0.155 | 0.030368042 | 24 | Ctsa |
| Sec628 | 1.51E-06 | 0.40674839 | 0.645 | 0.396 | 0.03052671 | 24 | Sec62 |
| Tm9sf39 | 1.52E-06 | 0.402774367 | 0.57 | 0.337 | 0.030805642 | 24 | Tm9sf3 |
| Ifi353 | 1.55E-06 | 0.39266039 | 0.505 | 0.276 | 0.031381831 | 24 | Ifi35 |
| Elovl12 | 1.56E-06 | 0.258588537 | 0.29 | 0.121 | 0.031658217 | 24 | Elovl1 |
| Tmem1656 | 1.59E-06 | 0.305134051 | 0.376 | 0.181 | 0.032123297 | 24 | Tmem165 |
| Rabac17 | 1.61E-06 | 0.327582155 | 0.839 | 0.566 | 0.032538525 | 24 | Rabac1 |
| Pitpna6 | 1.61E-06 | 0.340991933 | 0.57 | 0.329 | 0.032667609 | 24 | Pitpna |
| Efnb24 | 1.63E-06 | 0.335981212 | 0.333 | 0.146 | 0.032913595 | 24 | Efnb2 |
| Prkar1a5 | 1.67E-06 | 0.263671065 | 0.699 | 0.404 | 0.033879413 | 24 | Prkar1a |
| Ppp1r13b5 | 1.69E-06 | 0.35188367 | 0.43 | 0.217 | 0.034249014 | 24 | Ppp1r13b |
| mt-Cytb16 | 1.81E-06 | 0.375597449 | 0.989 | 0.701 | 0.036595465 | 24 | mt-Cytb |
| Elmo113 | 1.81E-06 | 0.373945688 | 0.645 | 0.368 | 0.036638718 | 24 | Elmo1 |
| Sbf28 | 1.85E-06 | 0.386209528 | 0.419 | 0.209 | 0.037435929 | 24 | Sbf2 |
| Numa16 | 1.88E-06 | 0.31839413 | 0.398 | 0.196 | 0.0381504 | 24 | Numa1 |
| Spata139 | 1.95E-06 | 0.362916617 | 0.409 | 0.208 | 0.039459929 | 24 | Spata13 |
| Cdkn1a3 | 1.98E-06 | 0.300067362 | 0.301 | 0.128 | 0.040084589 | 24 | Cdkn1a |
| H2-Q79 | 1.99E-06 | 0.405909203 | 0.688 | 0.399 | 0.04022609 | 24 | H2-Q7 |
| Cmtm61 | 1.99E-06 | 0.375857387 | 0.28 | 0.119 | 0.040350074 | 24 | Cmtm6 |
| Rab132 | 1.99E-06 | 0.253451065 | 0.323 | 0.139 | 0.040368906 | 24 | Rab13 |
| Prdm11 | 2.02E-06 | 0.36005113 | 0.269 | 0.111 | 0.04098096 | 24 | Prdm1 |
| Hip18 | 2.03E-06 | 0.331772881 | 0.602 | 0.336 | 0.041027292 | 24 | Hip1 |
| Tox42 | 2.06E-06 | 0.372733305 | 0.376 | 0.185 | 0.041801321 | 24 | Tox4 |
| Ptprk6 | 2.08E-06 | 0.300292438 | 0.323 | 0.141 | 0.042013507 | 24 | Ptprk |
| Ier511 | 2.15E-06 | 0.414483267 | 0.559 | 0.319 | 0.043523162 | 24 | Ier5 |
| 4930523C07Rik12 | 2.16E-06 | 0.417912786 | 0.43 | 0.221 | 0.043648988 | 24 | 4930523C07Rik |

| | | | | | | | |
|---|---|---|---|---|---|---|---|
| Arpc1b9 | 2.28E-06 | 0.3262676 | 0.849 | 0.593 | 0.046138441 | 24 | Arpc1b |
| Dstyk4 | 2.29E-06 | 0.308010953 | 0.333 | 0.151 | 0.046360934 | 24 | Dstyk |
| Cped17 | 2.33E-06 | 0.313460485 | 0.387 | 0.18 | 0.047139022 | 24 | Cped1 |
| Dennd5b6 | 2.34E-06 | 0.350822135 | 0.333 | 0.148 | 0.047358546 | 24 | Dennd5b |
| Fam96b3 | 2.36E-06 | 0.3367766 | 0.452 | 0.245 | 0.047845708 | 24 | Fam96b |
| Nck16 | 2.44E-06 | 0.275462346 | 0.613 | 0.342 | 0.049316781 | 24 | Nck1 |
| Tm9sf28 | 2.47E-06 | 0.382382017 | 0.473 | 0.257 | 0.049948882 | 24 | Tm9sf2 |
| Eml14 | 2.85E-06 | 0.325981773 | 0.269 | 0.11 | 0.057704389 | 24 | Eml1 |
| Uba51 | 2.86E-06 | 0.309231949 | 0.323 | 0.151 | 0.057999531 | 24 | Uba5 |
| Wwtr16 | 2.87E-06 | 0.279748379 | 0.484 | 0.242 | 0.058185785 | 24 | Wwtr1 |
| Hnrnpf5 | 2.95E-06 | 0.373830538 | 0.753 | 0.497 | 0.059626914 | 24 | Hnrnpf |
| Slc35b12 | 2.98E-06 | 0.351077131 | 0.344 | 0.165 | 0.060292687 | 24 | Slc35b1 |
| Prkd14 | 3.00E-06 | 0.286807575 | 0.333 | 0.149 | 0.060680295 | 24 | Prkd1 |
| Fcho26 | 3.10E-06 | 0.448111522 | 0.527 | 0.311 | 0.062852867 | 24 | Fcho2 |
| Osbpl96 | 3.21E-06 | 0.406926924 | 0.495 | 0.28 | 0.065039727 | 24 | Osbpl9 |
| Fas3 | 3.23E-06 | 0.297661562 | 0.28 | 0.119 | 0.065491943 | 24 | Fas |
| Smim121 | 3.33E-06 | 0.250394549 | 0.301 | 0.134 | 0.067355107 | 24 | Smim12 |
| Ap1s25 | 3.34E-06 | 0.279400899 | 0.312 | 0.14 | 0.0676057 | 24 | Ap1s2 |
| Klf34 | 3.35E-06 | 0.27795494 | 0.462 | 0.238 | 0.067745332 | 24 | Klf3 |
| Dnajc38 | 3.37E-06 | 0.436863971 | 0.591 | 0.372 | 0.068298513 | 24 | Dnajc3 |
| Nuak14 | 3.43E-06 | 0.298274937 | 0.28 | 0.119 | 0.069489804 | 24 | Nuak1 |
| Magt16 | 3.46E-06 | 0.383581164 | 0.344 | 0.166 | 0.069967393 | 24 | Magt1 |
| Zfyve95 | 3.46E-06 | 0.258995475 | 0.366 | 0.172 | 0.069986352 | 24 | Zfyve9 |
| Grasp4 | 3.57E-06 | 0.355100625 | 0.355 | 0.168 | 0.07222255 | 24 | Grasp |
| Rock27 | 3.58E-06 | 0.441693261 | 0.559 | 0.34 | 0.072534544 | 24 | Rock2 |
| 2900026A02Rik7 | 3.59E-06 | 0.379681594 | 0.419 | 0.221 | 0.072743253 | 24 | 2900026A02Rik |
| Mxd46 | 3.62E-06 | 0.33102826 | 0.613 | 0.354 | 0.073233258 | 24 | Mxd4 |
| Clic110 | 3.72E-06 | 0.35775319 | 0.871 | 0.602 | 0.075269423 | 24 | Clic1 |
| Zbtb2011 | 3.81E-06 | 0.27537943 | 0.903 | 0.538 | 0.07706836 | 24 | Zbtb20 |
| Ccnyl14 | 3.84E-06 | 0.296555645 | 0.269 | 0.112 | 0.077787994 | 24 | Ccnyl1 |
| Lrch17 | 3.85E-06 | 0.36292532 | 0.548 | 0.313 | 0.077888956 | 24 | Lrch1 |
| Dpysl29 | 3.87E-06 | 0.436844211 | 0.355 | 0.176 | 0.078291374 | 24 | Dpysl2 |
| Tmem2344 | 4.06E-06 | 0.294569783 | 0.634 | 0.395 | 0.08222068 | 24 | Tmem234 |
| Git26 | 4.07E-06 | 0.334694104 | 0.452 | 0.237 | 0.082316702 | 24 | Git2 |
| Pole42 | 4.09E-06 | 0.287609555 | 0.366 | 0.182 | 0.082892368 | 24 | Pole4 |
| Tmem50a10 | 4.11E-06 | 0.304741112 | 0.634 | 0.384 | 0.083204916 | 24 | Tmem50a |
| Kctd125 | 4.15E-06 | 0.426053262 | 0.462 | 0.26 | 0.084012976 | 24 | Kctd12 |
| Fndc3b7 | 4.16E-06 | 0.319568792 | 0.473 | 0.246 | 0.084306184 | 24 | Fndc3b |
| Zfp6087 | 4.24E-06 | 0.404505876 | 0.312 | 0.142 | 0.085930896 | 24 | Zfp608 |
| Dennd33 | 4.33E-06 | 0.304554424 | 0.323 | 0.148 | 0.087699987 | 24 | Dennd3 |
| Fxyd512 | 4.34E-06 | 0.261816289 | 0.71 | 0.413 | 0.087850789 | 24 | Fxyd5 |
| Pttg1ip8 | 4.40E-06 | 0.434154522 | 0.484 | 0.271 | 0.089142419 | 24 | Pttg1ip |
| Cdkn1c6 | 4.44E-06 | 0.332849369 | 0.29 | 0.127 | 0.089814788 | 24 | Cdkn1c |

| | | | | | | | |
|---|---|---|---|---|---|---|---|
| Tspan155 | 4.49E-06 | 0.303380936 | 0.301 | 0.132 | 0.090836882 | 24 | Tspan15 |
| Rock112 | 4.73E-06 | 0.418505357 | 0.688 | 0.434 | 0.095761022 | 24 | Rock1 |
| Stim25 | 4.74E-06 | 0.320980232 | 0.398 | 0.201 | 0.095949748 | 24 | Stim2 |
| Samd9l5 | 4.75E-06 | 0.3040148 | 0.484 | 0.258 | 0.096128989 | 24 | Samd9l |
| Sorbs18 | 4.85E-06 | 0.362204079 | 0.366 | 0.179 | 0.098234756 | 24 | Sorbs1 |
| Sft2d14 | 5.00E-06 | 0.358230455 | 0.333 | 0.162 | 0.101222508 | 24 | Sft2d1 |
| Map47 | 5.02E-06 | 0.404045209 | 0.538 | 0.305 | 0.101609092 | 24 | Map4 |
| Hacd45 | 5.08E-06 | 0.34467896 | 0.269 | 0.117 | 0.102751666 | 24 | Hacd4 |
| Tm2d3 | 5.12E-06 | 0.287496958 | 0.301 | 0.138 | 0.10375651 | 24 | Tm2d3 |
| mt-Co319 | 5.19E-06 | 0.395111791 | 1 | 0.727 | 0.105009127 | 24 | mt-Co3 |
| Rbfox27 | 5.35E-06 | 0.422767197 | 0.376 | 0.192 | 0.108344521 | 24 | Rbfox2 |
| Sbds3 | 5.45E-06 | 0.34185118 | 0.527 | 0.312 | 0.110275085 | 24 | Sbds |
| Emc79 | 5.48E-06 | 0.418753453 | 0.473 | 0.273 | 0.111020579 | 24 | Emc7 |
| Taok18 | 5.78E-06 | 0.369594949 | 0.462 | 0.253 | 0.117041117 | 24 | Taok1 |
| mt-Nd213 | 6.14E-06 | 0.368583202 | 0.978 | 0.69 | 0.124247421 | 24 | mt-Nd2 |
| Gnas4 | 6.18E-06 | 0.275381201 | 1 | 0.863 | 0.125131333 | 24 | Gnas |
| Zfp4624 | 6.85E-06 | 0.300611561 | 0.28 | 0.122 | 0.138707315 | 24 | Zfp462 |
| Wdfy38 | 6.88E-06 | 0.372071472 | 0.366 | 0.183 | 0.139262243 | 24 | Wdfy3 |
| Rassf16 | 6.91E-06 | 0.370435121 | 0.43 | 0.23 | 0.13981537 | 24 | Rassf1 |
| Tln112 | 6.91E-06 | 0.474852663 | 0.634 | 0.407 | 0.139909694 | 24 | Tln1 |
| Nbeal17 | 7.12E-06 | 0.329492511 | 0.473 | 0.264 | 0.144077162 | 24 | Nbeal1 |
| Rtn410 | 7.18E-06 | 0.38561599 | 0.57 | 0.341 | 0.145399211 | 24 | Rtn4 |
| Fam212a4 | 7.45E-06 | 0.320459824 | 0.29 | 0.129 | 0.150834597 | 24 | Fam212a |
| Tmem1282 | 7.61E-06 | 0.350157791 | 0.333 | 0.167 | 0.153976689 | 24 | Tmem128 |
| Slc50a15 | 7.74E-06 | 0.351618041 | 0.516 | 0.301 | 0.156626636 | 24 | Slc50a1 |
| Rsbn1l9 | 7.74E-06 | 0.434126946 | 0.484 | 0.274 | 0.156780291 | 24 | Rsbn1l |
| Rmdn13 | 7.95E-06 | 0.284126408 | 0.376 | 0.187 | 0.160995497 | 24 | Rmdn1 |
| Snrpb22 | 7.99E-06 | 0.261669148 | 0.29 | 0.132 | 0.161671908 | 24 | Snrpb2 |
| Son13 | 8.29E-06 | 0.318601254 | 0.849 | 0.527 | 0.167863821 | 24 | Son |
| Adarb13 | 8.78E-06 | 0.365684274 | 0.258 | 0.111 | 0.177739623 | 24 | Adarb1 |
| Prpf40a11 | 9.16E-06 | 0.285407689 | 0.484 | 0.268 | 0.185482915 | 24 | Prpf40a |
| Klc13 | 9.43E-06 | 0.294307428 | 0.323 | 0.151 | 0.190898432 | 24 | Klc1 |
| Trp53i112 | 9.50E-06 | 0.339809845 | 0.333 | 0.162 | 0.192399395 | 24 | Trp53i11 |
| Clock6 | 9.69E-06 | 0.287663768 | 0.398 | 0.205 | 0.196275288 | 24 | Clock |
| Gulp18 | 9.79E-06 | 0.250883446 | 0.355 | 0.168 | 0.198155727 | 24 | Gulp1 |
| Tmem28 | 9.79E-06 | 0.261420545 | 0.312 | 0.144 | 0.198229636 | 24 | Tmem2 |
| Txnip6 | 9.94E-06 | 0.262659103 | 0.817 | 0.546 | 0.201226841 | 24 | Txnip |
| Il10rb5 | 1.02E-05 | 0.381925564 | 0.441 | 0.25 | 0.20570756 | 24 | Il10rb |
| Sntb25 | 1.04E-05 | 0.405578337 | 0.473 | 0.267 | 0.211408543 | 24 | Sntb2 |
| Arglu18 | 1.06E-05 | 0.361638143 | 0.581 | 0.356 | 0.214276533 | 24 | Arglu1 |
| Papola3 | 1.08E-05 | 0.252404703 | 0.527 | 0.306 | 0.219263012 | 24 | Papola |
| Kdelr21 | 1.10E-05 | 0.256931603 | 0.344 | 0.171 | 0.223061955 | 24 | Kdelr2 |
| S100a168 | 1.10E-05 | 0.255865613 | 0.495 | 0.26 | 0.223162632 | 24 | S100a16 |

| | | | | | | | |
|---|---|---|---|---|---|---|---|
| Baz2b8 | 1.11E-05 | 0.377532014 | 0.591 | 0.352 | 0.225278244 | 24 | Baz2b |
| Mcmbp2 | 1.13E-05 | 0.323758846 | 0.29 | 0.137 | 0.229501326 | 24 | Mcmbp |
| Rhoa5 | 1.15E-05 | 0.251625804 | 0.892 | 0.595 | 0.231881428 | 24 | Rhoa |
| Sgms19 | 1.17E-05 | 0.267147021 | 0.29 | 0.13 | 0.23678085 | 24 | Sgms1 |
| Txndc122 | 1.18E-05 | 0.431747637 | 0.258 | 0.116 | 0.238476772 | 24 | Txndc12 |
| Wac9 | 1.19E-05 | 0.35133606 | 0.538 | 0.324 | 0.24017626 | 24 | Wac |
| Tmed109 | 1.19E-05 | 0.271245092 | 0.699 | 0.438 | 0.241892076 | 24 | Tmed10 |
| Prdm24 | 1.21E-05 | 0.417141983 | 0.355 | 0.186 | 0.244187672 | 24 | Prdm2 |
| Afap1l14 | 1.21E-05 | 0.267729504 | 0.258 | 0.11 | 0.245242293 | 24 | Afap1l1 |
| Tmem599 | 1.23E-05 | 0.297858928 | 0.753 | 0.492 | 0.249575323 | 24 | Tmem59 |
| Nosip2 | 1.24E-05 | 0.292539903 | 0.301 | 0.145 | 0.251006401 | 24 | Nosip |
| Ggh2 | 1.27E-05 | 0.359412144 | 0.333 | 0.168 | 0.25793698 | 24 | Ggh |
| Eid17 | 1.28E-05 | 0.256215251 | 0.548 | 0.323 | 0.259561995 | 24 | Eid1 |
| Gpr1464 | 1.28E-05 | 0.262468888 | 0.258 | 0.11 | 0.259989028 | 24 | Gpr146 |
| Anxa75 | 1.32E-05 | 0.352277569 | 0.366 | 0.187 | 0.266893013 | 24 | Anxa7 |
| Stard95 | 1.33E-05 | 0.322619383 | 0.323 | 0.153 | 0.26931404 | 24 | Stard9 |
| Nucb15 | 1.36E-05 | 0.345298551 | 0.419 | 0.229 | 0.274403489 | 24 | Nucb1 |
| Asah15 | 1.42E-05 | 0.365047981 | 0.366 | 0.19 | 0.287279899 | 24 | Asah1 |
| Psmd75 | 1.42E-05 | 0.262344595 | 0.527 | 0.307 | 0.28779594 | 24 | Psmd7 |
| Hsd17b114 | 1.43E-05 | 0.255518715 | 0.269 | 0.12 | 0.289489638 | 24 | Hsd17b11 |
| Arap28 | 1.43E-05 | 0.370067005 | 0.355 | 0.179 | 0.289644847 | 24 | Arap2 |
| Rell14 | 1.45E-05 | 0.263311071 | 0.301 | 0.138 | 0.293086904 | 24 | Rell1 |
| Parvb4 | 1.48E-05 | 0.289803196 | 0.28 | 0.125 | 0.300294146 | 24 | Parvb |
| Tacc19 | 1.51E-05 | 0.271722518 | 0.527 | 0.307 | 0.305600196 | 24 | Tacc1 |
| Spred16 | 1.60E-05 | 0.260035418 | 0.473 | 0.254 | 0.324064257 | 24 | Spred1 |
| Ash1l10 | 1.61E-05 | 0.309976893 | 0.516 | 0.301 | 0.325190961 | 24 | Ash1l |
| Prickle26 | 1.76E-05 | 0.311333441 | 0.301 | 0.141 | 0.356786654 | 24 | Prickle2 |
| Srpr6 | 1.78E-05 | 0.281548693 | 0.441 | 0.247 | 0.359458319 | 24 | Srpr |
| Ywhah7 | 1.79E-05 | 0.301154331 | 0.516 | 0.311 | 0.362019882 | 24 | Ywhah |
| Tprgl6 | 1.79E-05 | 0.327612863 | 0.484 | 0.28 | 0.36338893 | 24 | Tprgl |
| Vti1a11 | 1.85E-05 | 0.320309404 | 0.462 | 0.257 | 0.374908305 | 24 | Vti1a |
| Tmed75 | 1.91E-05 | 0.263087138 | 0.312 | 0.15 | 0.385760834 | 24 | Tmed7 |
| Sema6d8 | 1.92E-05 | 0.267250556 | 0.409 | 0.216 | 0.38792282 | 24 | Sema6d |
| Flii3 | 1.96E-05 | 0.302702841 | 0.323 | 0.159 | 0.397219943 | 24 | Flii |
| Tmtc15 | 1.98E-05 | 0.354857349 | 0.301 | 0.144 | 0.400004365 | 24 | Tmtc1 |
| mt-Nd513 | 1.98E-05 | 0.385660731 | 0.774 | 0.498 | 0.400128189 | 24 | mt-Nd5 |
| Ginm15 | 2.05E-05 | 0.416416947 | 0.355 | 0.181 | 0.415319152 | 24 | Ginm1 |
| Auts24 | 2.09E-05 | 0.277000894 | 0.323 | 0.154 | 0.423895972 | 24 | Auts2 |
| Fbxw23 | 2.14E-05 | 0.251793257 | 0.43 | 0.237 | 0.433560261 | 24 | Fbxw2 |
| Arid4b10 | 2.20E-05 | 0.361384533 | 0.495 | 0.294 | 0.446090073 | 24 | Arid4b |
| Diaph211 | 2.28E-05 | 0.292313563 | 0.634 | 0.378 | 0.461259079 | 24 | Diaph2 |
| Trim243 | 2.32E-05 | 0.382902743 | 0.333 | 0.174 | 0.469817872 | 24 | Trim24 |
| Zfp5034 | 2.34E-05 | 0.271638824 | 0.312 | 0.148 | 0.472994505 | 24 | Zfp503 |

| | | | | | | | |
|---|---|---|---|---|---|---|---|
| Atrnl17 | 2.34E-05 | 0.36402134 | 0.333 | 0.167 | 0.473596995 | 24 | Atrnl1 |
| Strn37 | 2.37E-05 | 0.286039113 | 0.581 | 0.347 | 0.479814804 | 24 | Strn3 |
| Mfsd64 | 2.43E-05 | 0.445132868 | 0.301 | 0.146 | 0.492911445 | 24 | Mfsd6 |
| Rtl8b1 | 2.44E-05 | 0.282490571 | 0.258 | 0.113 | 0.493224448 | 24 | Rtl8b |
| Lrch311 | 2.49E-05 | 0.31282895 | 0.43 | 0.234 | 0.504038759 | 24 | Lrch3 |
| Hprt3 | 2.49E-05 | 0.252276438 | 0.495 | 0.287 | 0.504730786 | 24 | Hprt |
| Degs16 | 2.61E-05 | 0.304589732 | 0.355 | 0.186 | 0.528602723 | 24 | Degs1 |
| Tmed910 | 2.71E-05 | 0.336910583 | 0.634 | 0.405 | 0.549633144 | 24 | Tmed9 |
| Tsc22d33 | 2.75E-05 | 0.25491277 | 0.398 | 0.209 | 0.557033801 | 24 | Tsc22d3 |
| Srrm213 | 2.83E-05 | 0.293782873 | 0.742 | 0.471 | 0.572712204 | 24 | Srrm2 |
| Mia26 | 2.84E-05 | 0.457007149 | 0.376 | 0.209 | 0.575202477 | 24 | Mia2 |
| Fam114a13 | 2.87E-05 | 0.347174273 | 0.28 | 0.133 | 0.580401793 | 24 | Fam114a1 |
| Dock95 | 2.89E-05 | 0.34132809 | 0.398 | 0.207 | 0.584650365 | 24 | Dock9 |
| Epn25 | 2.90E-05 | 0.303591273 | 0.323 | 0.156 | 0.586554228 | 24 | Epn2 |
| Akap126 | 2.91E-05 | 0.258731068 | 0.333 | 0.162 | 0.589152489 | 24 | Akap12 |
| Saraf9 | 2.93E-05 | 0.267642805 | 0.527 | 0.309 | 0.593074313 | 24 | Saraf |
| Gatad2a5 | 3.04E-05 | 0.272843852 | 0.409 | 0.224 | 0.614910418 | 24 | Gatad2a |
| Midn3 | 3.15E-05 | 0.470925032 | 0.355 | 0.193 | 0.638525297 | 24 | Midn |
| Ehd27 | 3.31E-05 | 0.293539411 | 0.376 | 0.195 | 0.669697846 | 24 | Ehd2 |
| Stat25 | 3.33E-05 | 0.367901098 | 0.29 | 0.142 | 0.673948791 | 24 | Stat2 |
| Yy13 | 3.33E-05 | 0.36110254 | 0.462 | 0.275 | 0.67467633 | 24 | Yy1 |
| Sptlc26 | 3.55E-05 | 0.28113781 | 0.441 | 0.246 | 0.719629725 | 24 | Sptlc2 |
| Gnb16 | 3.71E-05 | 0.349848906 | 0.634 | 0.43 | 0.751539776 | 24 | Gnb1 |
| Map4k37 | 3.76E-05 | 0.311038439 | 0.419 | 0.231 | 0.761394017 | 24 | Map4k3 |
| Epha44 | 3.82E-05 | 0.360856659 | 0.258 | 0.118 | 0.77363669 | 24 | Epha4 |
| Pdia311 | 3.89E-05 | 0.256431716 | 0.699 | 0.445 | 0.788034218 | 24 | Pdia3 |
| Cdc425 | 4.09E-05 | 0.274822706 | 0.871 | 0.617 | 0.828646352 | 24 | Cdc42 |
| Serinc19 | 4.13E-05 | 0.320948511 | 0.441 | 0.251 | 0.835621704 | 24 | Serinc1 |
| Oasl25 | 4.15E-05 | 0.250007443 | 0.28 | 0.133 | 0.841150425 | 24 | Oasl2 |
| Smarca42 | 4.16E-05 | 0.258353612 | 0.323 | 0.167 | 0.842551352 | 24 | Smarca4 |
| Sdf46 | 4.17E-05 | 0.347475796 | 0.505 | 0.311 | 0.845067658 | 24 | Sdf4 |
| Ccser26 | 4.23E-05 | 0.273763236 | 0.419 | 0.233 | 0.855899524 | 24 | Ccser2 |
| Snx97 | 4.41E-05 | 0.267708414 | 0.344 | 0.177 | 0.892872956 | 24 | Snx9 |
| Airn8 | 4.56E-05 | 0.273097311 | 0.452 | 0.25 | 0.922929184 | 24 | Airn |
| Setd513 | 4.61E-05 | 0.32014119 | 0.462 | 0.265 | 0.932472628 | 24 | Setd5 |
| St3gal45 | 4.76E-05 | 0.339028336 | 0.441 | 0.258 | 0.964622352 | 24 | St3gal4 |
| Siah1a1 | 4.77E-05 | 0.293275727 | 0.269 | 0.127 | 0.965618291 | 24 | Siah1a |
| Ppp1r16b11 | 4.80E-05 | 0.275756282 | 0.387 | 0.203 | 0.972380089 | 24 | Ppp1r16b |
| Atxn77 | 4.85E-05 | 0.355309372 | 0.323 | 0.167 | 0.982932361 | 24 | Atxn7 |
| Lmbrd18 | 5.05E-05 | 0.399120076 | 0.312 | 0.164 | 1 | 24 | Lmbrd1 |
| Peak110 | 5.14E-05 | 0.27867587 | 0.634 | 0.389 | 1 | 24 | Peak1 |
| Thap35 | 5.28E-05 | 0.254744788 | 0.323 | 0.163 | 1 | 24 | Thap3 |
| Washc43 | 5.42E-05 | 0.282406382 | 0.269 | 0.13 | 1 | 24 | Washc4 |

| | | | | | | | |
|---|---|---|---|---|---|---|---|
| 1110059E24Rik1 | 5.45E-05 | 0.348387732 | 0.29 | 0.145 | 1 | 24 | 1110059E24Rik |
| Lgals95 | 5.56E-05 | 0.270268859 | 0.43 | 0.249 | 1 | 24 | Lgals9 |
| Lrrfip18 | 5.58E-05 | 0.26223579 | 0.602 | 0.38 | 1 | 24 | Lrrfip1 |
| Trim561 | 5.89E-05 | 0.265655128 | 0.269 | 0.127 | 1 | 24 | Trim56 |
| Map3k16 | 6.09E-05 | 0.324157818 | 0.366 | 0.2 | 1 | 24 | Map3k1 |
| Hdac13 | 6.15E-05 | 0.280673899 | 0.323 | 0.169 | 1 | 24 | Hdac1 |
| Srgap210 | 6.24E-05 | 0.286041842 | 0.516 | 0.306 | 1 | 24 | Srgap2 |
| Pkn28 | 6.25E-05 | 0.28317109 | 0.495 | 0.296 | 1 | 24 | Pkn2 |
| Mkln16 | 6.41E-05 | 0.294137703 | 0.462 | 0.266 | 1 | 24 | Mkln1 |
| Exoc6b9 | 6.57E-05 | 0.322446683 | 0.419 | 0.242 | 1 | 24 | Exoc6b |
| mt-Atp618 | 6.65E-05 | 0.353732823 | 1 | 0.755 | 1 | 24 | mt-Atp6 |
| Ggta15 | 6.66E-05 | 0.258293363 | 0.29 | 0.142 | 1 | 24 | Ggta1 |
| Morf4l13 | 6.79E-05 | 0.271287325 | 0.86 | 0.59 | 1 | 24 | Morf4l1 |
| Dennd6a4 | 7.09E-05 | 0.295668099 | 0.344 | 0.183 | 1 | 24 | Dennd6a |
| Gbp93 | 7.24E-05 | 0.331437216 | 0.29 | 0.143 | 1 | 24 | Gbp9 |
| S100a134 | 7.33E-05 | 0.268997248 | 0.806 | 0.538 | 1 | 24 | S100a13 |
| Tcf1212 | 7.50E-05 | 0.306776041 | 0.602 | 0.378 | 1 | 24 | Tcf12 |
| mt-Co117 | 7.59E-05 | 0.351535014 | 1 | 0.721 | 1 | 24 | mt-Co1 |
| Pja26 | 7.65E-05 | 0.256658063 | 0.376 | 0.206 | 1 | 24 | Pja2 |
| P4hb7 | 7.84E-05 | 0.255222383 | 0.441 | 0.254 | 1 | 24 | P4hb |
| Lbr11 | 8.00E-05 | 0.300139412 | 0.301 | 0.154 | 1 | 24 | Lbr |
| Snx11 | 8.02E-05 | 0.299144405 | 0.28 | 0.141 | 1 | 24 | Snx1 |
| Hook38 | 8.03E-05 | 0.344648083 | 0.484 | 0.292 | 1 | 24 | Hook3 |
| Ptpn125 | 8.31E-05 | 0.379923528 | 0.376 | 0.215 | 1 | 24 | Ptpn12 |
| H2-Q69 | 8.49E-05 | 0.4109109 | 0.419 | 0.247 | 1 | 24 | H2-Q6 |
| Map7d15 | 8.62E-05 | 0.321326018 | 0.323 | 0.169 | 1 | 24 | Map7d1 |
| Smarce1 | 8.87E-05 | 0.251882598 | 0.398 | 0.23 | 1 | 24 | Smarce1 |
| Atraid7 | 8.88E-05 | 0.254977716 | 0.462 | 0.272 | 1 | 24 | Atraid |
| Phf20l19 | 9.36E-05 | 0.404306786 | 0.462 | 0.289 | 1 | 24 | Phf20l1 |
| Rgs39 | 9.63E-05 | 0.309769418 | 0.344 | 0.189 | 1 | 24 | Rgs3 |
| Fcgrt5 | 0.00011117 | 0.412329838 | 0.344 | 0.187 | 1 | 24 | Fcgrt |
| Gcc22 | 0.000114804 | 0.419175039 | 0.344 | 0.193 | 1 | 24 | Gcc2 |
| Smad12 | 0.000116359 | 0.283614073 | 0.269 | 0.134 | 1 | 24 | Smad1 |
| Hcls18 | 0.0001226 | 0.251575614 | 0.258 | 0.127 | 1 | 24 | Hcls1 |
| Huwe112 | 0.000123248 | 0.309587367 | 0.473 | 0.287 | 1 | 24 | Huwe1 |
| Sh3rf16 | 0.000125942 | 0.293193677 | 0.269 | 0.132 | 1 | 24 | Sh3rf1 |
| Sppl2a6 | 0.000127661 | 0.285111659 | 0.495 | 0.298 | 1 | 24 | Sppl2a |
| Mgat12 | 0.000128384 | 0.339519097 | 0.312 | 0.169 | 1 | 24 | Mgat1 |
| Erc14 | 0.000130057 | 0.285609818 | 0.258 | 0.124 | 1 | 24 | Erc1 |
| Cflar4 | 0.000130626 | 0.258575713 | 0.398 | 0.233 | 1 | 24 | Cflar |
| Prkd23 | 0.000137107 | 0.288899752 | 0.28 | 0.141 | 1 | 24 | Prkd2 |
| Smchd110 | 0.000146088 | 0.293166255 | 0.441 | 0.266 | 1 | 24 | Smchd1 |
| Ssr16 | 0.000164421 | 0.285802275 | 0.333 | 0.188 | 1 | 24 | Ssr1 |

| | | | | | | | |
|---|---|---|---|---|---|---|---|
| Emc106 | 0.000166662 | 0.267585437 | 0.355 | 0.202 | 1 | 24 | Emc10 |
| Pnp5 | 0.000171186 | 0.259300107 | 0.43 | 0.256 | 1 | 24 | Pnp |
| Ints93 | 0.000174547 | 0.322201086 | 0.269 | 0.138 | 1 | 24 | Ints9 |
| Pxn3 | 0.000176277 | 0.273145056 | 0.312 | 0.166 | 1 | 24 | Pxn |
| Rc3h13 | 0.000179272 | 0.259907866 | 0.366 | 0.21 | 1 | 24 | Rc3h1 |
| Mapk19 | 0.000180518 | 0.253378235 | 0.409 | 0.237 | 1 | 24 | Mapk1 |
| Btbd77 | 0.000187743 | 0.264022065 | 0.462 | 0.283 | 1 | 24 | Btbd7 |
| Ssr24 | 0.000189158 | 0.322186002 | 0.355 | 0.204 | 1 | 24 | Ssr2 |
| Rnf13011 | 0.000189795 | 0.262832539 | 0.473 | 0.285 | 1 | 24 | Rnf130 |
| Erap14 | 0.000201163 | 0.371798705 | 0.269 | 0.141 | 1 | 24 | Erap1 |
| Txndc155 | 0.000204611 | 0.267995104 | 0.269 | 0.14 | 1 | 24 | Txndc15 |
| Lifr6 | 0.000204723 | 0.410081295 | 0.355 | 0.194 | 1 | 24 | Lifr |
| Grk57 | 0.000208569 | 0.450346586 | 0.301 | 0.16 | 1 | 24 | Grk5 |
| Sos14 | 0.000211517 | 0.294092342 | 0.333 | 0.182 | 1 | 24 | Sos1 |
| Creg15 | 0.000213425 | 0.330976047 | 0.333 | 0.19 | 1 | 24 | Creg1 |
| Hectd19 | 0.000219449 | 0.25764033 | 0.495 | 0.302 | 1 | 24 | Hectd1 |
| Ifi478 | 0.000227658 | 0.355790292 | 0.323 | 0.18 | 1 | 24 | Ifi47 |
| Spag99 | 0.000233606 | 0.256261514 | 0.559 | 0.361 | 1 | 24 | Spag9 |
| Maml36 | 0.00024636 | 0.35167814 | 0.366 | 0.21 | 1 | 24 | Maml3 |
| Mtdh10 | 0.000253593 | 0.34921109 | 0.591 | 0.388 | 1 | 24 | Mtdh |
| Slc44a16 | 0.000258964 | 0.372293155 | 0.29 | 0.158 | 1 | 24 | Slc44a1 |
| Eif4g37 | 0.000261314 | 0.251329648 | 0.473 | 0.287 | 1 | 24 | Eif4g3 |
| Elk44 | 0.000271267 | 0.324894402 | 0.312 | 0.169 | 1 | 24 | Elk4 |
| M6pr6 | 0.000275501 | 0.277183234 | 0.301 | 0.164 | 1 | 24 | M6pr |
| AW5549189 | 0.000277531 | 0.293277524 | 0.312 | 0.172 | 1 | 24 | AW554918 |
| Ncoa35 | 0.00027947 | 0.25264976 | 0.43 | 0.254 | 1 | 24 | Ncoa3 |
| Mpzl15 | 0.000288621 | 0.321574915 | 0.28 | 0.144 | 1 | 24 | Mpzl1 |
| Aph1a | 0.000289528 | 0.260692599 | 0.29 | 0.153 | 1 | 24 | Aph1a |
| Ppib6 | 0.000294476 | 0.262341824 | 0.774 | 0.555 | 1 | 24 | Ppib |
| Ccdc476 | 0.000310519 | 0.260126332 | 0.344 | 0.194 | 1 | 24 | Ccdc47 |
| Elovl54 | 0.000327022 | 0.396663101 | 0.28 | 0.152 | 1 | 24 | Elovl5 |
| Mindy21 | 0.000332099 | 0.261824869 | 0.323 | 0.182 | 1 | 24 | Mindy2 |
| Fos6 | 0.000335642 | 0.269916927 | 0.516 | 0.319 | 1 | 24 | Fos |
| Mgat23 | 0.000336803 | 0.332046353 | 0.29 | 0.157 | 1 | 24 | Mgat2 |
| Hivep211 | 0.000338163 | 0.306769215 | 0.473 | 0.3 | 1 | 24 | Hivep2 |
| Ttc287 | 0.000340784 | 0.259468962 | 0.462 | 0.287 | 1 | 24 | Ttc28 |
| mt-Nd417 | 0.000354618 | 0.265276927 | 0.989 | 0.676 | 1 | 24 | mt-Nd4 |
| Foxp19 | 0.000376189 | 0.289266986 | 0.581 | 0.391 | 1 | 24 | Foxp1 |
| Junb9 | 0.000379628 | 0.433346955 | 0.613 | 0.394 | 1 | 24 | Junb |
| Cr1l5 | 0.000382781 | 0.28570468 | 0.387 | 0.235 | 1 | 24 | Cr1l |
| Abcc45 | 0.000387116 | 0.370460547 | 0.269 | 0.138 | 1 | 24 | Abcc4 |
| Map3k55 | 0.000391653 | 0.301330787 | 0.344 | 0.194 | 1 | 24 | Map3k5 |
| Bod1l8 | 0.000415944 | 0.297684043 | 0.43 | 0.275 | 1 | 24 | Bod1l |

| | | | | | | | |
|---|---|---|---|---|---|---|---|
| Aida4 | 0.000440685 | 0.298867533 | 0.269 | 0.144 | 1 | 24 | Aida |
| Ppig8 | 0.000445868 | 0.269108854 | 0.484 | 0.318 | 1 | 24 | Ppig |
| Trafd13 | 0.000461076 | 0.397971351 | 0.28 | 0.155 | 1 | 24 | Trafd1 |
| Thoc29 | 0.00050747 | 0.268505921 | 0.441 | 0.276 | 1 | 24 | Thoc2 |
| Asap26 | 0.000537336 | 0.251855516 | 0.258 | 0.133 | 1 | 24 | Asap2 |
| Mllt1013 | 0.000572041 | 0.256439308 | 0.441 | 0.276 | 1 | 24 | Mllt10 |
| Herc63 | 0.000593914 | 0.300180219 | 0.323 | 0.186 | 1 | 24 | Herc6 |
| Plekhj15 | 0.000594059 | 0.29523762 | 0.301 | 0.17 | 1 | 24 | Plekhj1 |
| Efcab143 | 0.000606132 | 0.339177988 | 0.269 | 0.146 | 1 | 24 | Efcab14 |
| Sertad23 | 0.000680371 | 0.295727194 | 0.301 | 0.17 | 1 | 24 | Sertad2 |
| Ubn210 | 0.000710668 | 0.394525043 | 0.387 | 0.246 | 1 | 24 | Ubn2 |
| Ep4008 | 0.000728959 | 0.307369113 | 0.344 | 0.209 | 1 | 24 | Ep400 |
| Foxn312 | 0.000787099 | 0.25069976 | 0.613 | 0.413 | 1 | 24 | Foxn3 |
| Kat6a6 | 0.000801604 | 0.341080116 | 0.366 | 0.23 | 1 | 24 | Kat6a |
| Saysd1 | 0.000835451 | 0.262649735 | 0.29 | 0.165 | 1 | 24 | Saysd1 |
| Slk7 | 0.000874804 | 0.316193692 | 0.387 | 0.243 | 1 | 24 | Slk |
| Ddx585 | 0.000942921 | 0.35264433 | 0.28 | 0.162 | 1 | 24 | Ddx58 |
| Scarb24 | 0.000964558 | 0.260180628 | 0.323 | 0.189 | 1 | 24 | Scarb2 |
| Tgfbr37 | 0.001162366 | 0.252244557 | 0.312 | 0.176 | 1 | 24 | Tgfbr3 |
| Golgb17 | 0.001246096 | 0.275375952 | 0.409 | 0.264 | 1 | 24 | Golgb1 |
| Rb18 | 0.001303966 | 0.285997885 | 0.355 | 0.216 | 1 | 24 | Rb1 |
| Kmt2a12 | 0.001321623 | 0.259491255 | 0.419 | 0.269 | 1 | 24 | Kmt2a |
| Polr2a10 | 0.001324775 | 0.338028651 | 0.43 | 0.283 | 1 | 24 | Polr2a |
| Zcchc66 | 0.001339694 | 0.329093693 | 0.419 | 0.28 | 1 | 24 | Zcchc6 |
| Cyb5r36 | 0.001347577 | 0.254716758 | 0.355 | 0.219 | 1 | 24 | Cyb5r3 |
| Selenot5 | 0.001394184 | 0.261202089 | 0.344 | 0.213 | 1 | 24 | Selenot |
| Prrc2c13 | 0.001482767 | 0.26806363 | 0.602 | 0.422 | 1 | 24 | Prrc2c |
| Pcm19 | 0.001920064 | 0.275729477 | 0.387 | 0.253 | 1 | 24 | Pcm1 |
| Klhl245 | 0.001986577 | 0.304866485 | 0.344 | 0.217 | 1 | 24 | Klhl24 |
| Pdcd6ip5 | 0.002061364 | 0.291045504 | 0.355 | 0.232 | 1 | 24 | Pdcd6ip |
| Rbm2514 | 0.002112133 | 0.303497132 | 0.505 | 0.336 | 1 | 24 | Rbm25 |
| Rapgef17 | 0.002208776 | 0.325086372 | 0.376 | 0.243 | 1 | 24 | Rapgef1 |
| Cntln7 | 0.0026942 | 0.315879816 | 0.269 | 0.156 | 1 | 24 | Cntln |
| Abhd17b8 | 0.003028518 | 0.384757983 | 0.312 | 0.194 | 1 | 24 | Abhd17b |
| Atp8a17 | 0.003666209 | 0.292378702 | 0.366 | 0.244 | 1 | 24 | Atp8a1 |
| Bptf13 | 0.003813386 | 0.290298743 | 0.419 | 0.296 | 1 | 24 | Bptf |
| Mia35 | 0.004164801 | 0.275543476 | 0.28 | 0.172 | 1 | 24 | Mia3 |
| Akap912 | 0.004398447 | 0.254814847 | 0.366 | 0.239 | 1 | 24 | Akap9 |
| Swap706 | 0.004621112 | 0.25441929 | 0.376 | 0.248 | 1 | 24 | Swap70 |
| Pds5b7 | 0.004789806 | 0.253840982 | 0.29 | 0.182 | 1 | 24 | Pds5b |
| Pias17 | 0.005085908 | 0.281624135 | 0.301 | 0.19 | 1 | 24 | Pias1 |
| Nrip18 | 0.007507853 | 0.281876182 | 0.355 | 0.246 | 1 | 24 | Nrip1 |
| Rbm269 | 0.008563178 | 0.289708894 | 0.333 | 0.229 | 1 | 24 | Rbm26 |

| | | | | | | | |
|---|---|---|---|---|---|---|---|
| Alas21 | 0 | 2.276325987 | 0.795 | 0.024 | 0 | 25 | Alas2 |
| Snca2 | 1.84E-238 | 1.503726488 | 0.63 | 0.023 | 3.72E-234 | 25 | Snca |
| Hbb-bt2 | 6.24E-174 | 4.632111291 | 0.945 | 0.08 | 1.26E-169 | 25 | Hbb-bt |
| Hba-a23 | 1.22E-148 | 4.644053395 | 1 | 0.11 | 2.46E-144 | 25 | Hba-a2 |
| Hba-a13 | 2.57E-132 | 4.676725456 | 1 | 0.127 | 5.20E-128 | 25 | Hba-a1 |
| Bpgm1 | 1.60E-84 | 1.705897174 | 0.575 | 0.054 | 3.24E-80 | 25 | Bpgm |
| Fam220a2 | 6.30E-77 | 1.310542177 | 0.74 | 0.097 | 1.27E-72 | 25 | Fam220a |
| Hbb-bs3 | 2.35E-65 | 4.857791439 | 1 | 0.33 | 4.76E-61 | 25 | Hbb-bs |
| Fam46c2 | 1.72E-57 | 1.035115308 | 0.562 | 0.072 | 3.49E-53 | 25 | Fam46c |
| Slc25a371 | 3.25E-54 | 0.860982866 | 0.452 | 0.05 | 6.58E-50 | 25 | Slc25a37 |
| Ube2l62 | 2.62E-43 | 0.834969437 | 0.521 | 0.078 | 5.30E-39 | 25 | Ube2l6 |
| Mkrn13 | 2.07E-26 | 1.363854915 | 0.603 | 0.167 | 4.19E-22 | 25 | Mkrn1 |
| Rsad25 | 4.20E-26 | 1.028677271 | 0.849 | 0.257 | 8.49E-22 | 25 | Rsad2 |
| Fech1 | 5.94E-26 | 0.568008051 | 0.425 | 0.08 | 1.20E-21 | 25 | Fech |
| Clec1a5 | 4.93E-24 | 0.844150621 | 0.644 | 0.169 | 9.99E-20 | 25 | Clec1a |
| Gda1 | 1.90E-23 | 0.503907887 | 0.411 | 0.081 | 3.84E-19 | 25 | Gda |
| Kdr5 | 1.32E-22 | 1.035043151 | 1 | 0.364 | 2.68E-18 | 25 | Kdr |
| Plcb16 | 2.05E-22 | 0.815015528 | 0.986 | 0.342 | 4.14E-18 | 25 | Plcb1 |
| Stox26 | 7.44E-22 | 0.485244249 | 0.836 | 0.248 | 1.51E-17 | 25 | Stox2 |
| Hmcn15 | 8.20E-22 | 0.694608423 | 0.726 | 0.212 | 1.66E-17 | 25 | Hmcn1 |
| Kctd12b6 | 1.45E-21 | 0.812640432 | 0.808 | 0.25 | 2.94E-17 | 25 | Kctd12b |
| Isca11 | 2.35E-21 | 0.317892829 | 0.37 | 0.072 | 4.75E-17 | 25 | Isca1 |
| Btnl94 | 2.81E-21 | 0.896806879 | 0.822 | 0.271 | 5.69E-17 | 25 | Btnl9 |
| Gm120024 | 4.33E-21 | 0.587070527 | 0.685 | 0.189 | 8.77E-17 | 25 | Gm12002 |
| Rflnb6 | 6.22E-21 | 0.623627176 | 0.932 | 0.315 | 1.26E-16 | 25 | Rflnb |
| Mcc5 | 8.70E-21 | 0.679582046 | 0.863 | 0.285 | 1.76E-16 | 25 | Mcc |
| Car43 | 9.11E-21 | 0.851979641 | 0.726 | 0.226 | 1.85E-16 | 25 | Car4 |
| Rgcc5 | 1.06E-20 | 1.120874564 | 0.973 | 0.405 | 2.15E-16 | 25 | Rgcc |
| Exoc3l25 | 1.42E-20 | 0.781235903 | 0.836 | 0.271 | 2.87E-16 | 25 | Exoc3l2 |
| Cyyr16 | 3.51E-20 | 0.807588926 | 0.973 | 0.344 | 7.10E-16 | 25 | Cyyr1 |
| Myo65 | 3.65E-20 | 0.730743878 | 0.849 | 0.293 | 7.39E-16 | 25 | Myo6 |
| Tie17 | 7.31E-20 | 0.500107218 | 0.836 | 0.254 | 1.48E-15 | 25 | Tie1 |
| Ramp34 | 9.21E-20 | 0.703723317 | 0.822 | 0.268 | 1.86E-15 | 25 | Ramp3 |
| Rab133 | 9.22E-20 | 0.434357204 | 0.548 | 0.139 | 1.87E-15 | 25 | Rab13 |
| Etl46 | 1.06E-19 | 0.589076947 | 0.89 | 0.295 | 2.15E-15 | 25 | Etl4 |
| Rbp75 | 1.97E-19 | 0.713593545 | 0.945 | 0.349 | 4.00E-15 | 25 | Rbp7 |
| Rapgef46 | 2.07E-19 | 0.48662385 | 0.836 | 0.258 | 4.20E-15 | 25 | Rapgef4 |
| Gpihbp15 | 2.14E-19 | 0.832714784 | 0.863 | 0.297 | 4.34E-15 | 25 | Gpihbp1 |
| Tcim4 | 3.28E-19 | 0.764716059 | 0.89 | 0.304 | 6.65E-15 | 25 | Tcim |
| Hecw25 | 3.97E-19 | 0.642202728 | 0.781 | 0.25 | 8.03E-15 | 25 | Hecw2 |
| Cavin27 | 4.11E-19 | 0.792164557 | 0.945 | 0.342 | 8.32E-15 | 25 | Cavin2 |
| Ube2o | 4.47E-19 | 0.402797704 | 0.26 | 0.043 | 9.04E-15 | 25 | Ube2o |
| Pcdh176 | 4.69E-19 | 0.538909662 | 0.781 | 0.24 | 9.49E-15 | 25 | Pcdh17 |

| | | | | | | | |
|---|---|---|---|---|---|---|---|
| Igfbp36 | 5.67E-19 | 0.826827592 | 0.849 | 0.304 | 1.15E-14 | 25 | Igfbp3 |
| Isg205 | 7.78E-19 | 0.613161481 | 0.452 | 0.117 | 1.58E-14 | 25 | Isg20 |
| Epb414 | 9.09E-19 | 0.763178653 | 0.658 | 0.217 | 1.84E-14 | 25 | Epb41 |
| Fut41 | 9.11E-19 | 0.46080455 | 0.384 | 0.083 | 1.84E-14 | 25 | Fut4 |
| Fmo25 | 1.18E-18 | 0.726729103 | 0.836 | 0.29 | 2.39E-14 | 25 | Fmo2 |
| Thrsp5 | 1.31E-18 | 0.686472869 | 0.863 | 0.294 | 2.65E-14 | 25 | Thrsp |
| Rtp31 | 1.32E-18 | 0.419845817 | 0.384 | 0.082 | 2.67E-14 | 25 | Rtp3 |
| Clic56 | 1.70E-18 | 0.609698025 | 0.877 | 0.292 | 3.44E-14 | 25 | Clic5 |
| Tinagl17 | 1.77E-18 | 0.600012655 | 0.877 | 0.298 | 3.59E-14 | 25 | Tinagl1 |
| Adgrl46 | 1.98E-18 | 0.774374048 | 0.973 | 0.353 | 4.01E-14 | 25 | Adgrl4 |
| Plpp34 | 2.21E-18 | 0.921522505 | 1 | 0.405 | 4.47E-14 | 25 | Plpp3 |
| Ace5 | 2.28E-18 | 0.497537469 | 0.479 | 0.12 | 4.62E-14 | 25 | Ace |
| Nostrin4 | 3.84E-18 | 0.684076919 | 0.767 | 0.26 | 7.78E-14 | 25 | Nostrin |
| Tmem255b3 | 3.93E-18 | 0.391940325 | 0.356 | 0.074 | 7.95E-14 | 25 | Tmem255b |
| Scarb15 | 4.13E-18 | 0.655514025 | 0.795 | 0.268 | 8.36E-14 | 25 | Scarb1 |
| Clec14a6 | 4.20E-18 | 0.631402339 | 0.877 | 0.306 | 8.51E-14 | 25 | Clec14a |
| Chst155 | 4.52E-18 | 0.5258654 | 0.616 | 0.176 | 9.16E-14 | 25 | Chst15 |
| Syne15 | 4.55E-18 | 0.629878414 | 0.945 | 0.36 | 9.20E-14 | 25 | Syne1 |
| Ripor16 | 4.74E-18 | 0.476127773 | 0.575 | 0.161 | 9.59E-14 | 25 | Ripor1 |
| Plvap6 | 4.99E-18 | 0.878335821 | 1 | 0.385 | 1.01E-13 | 25 | Plvap |
| Crim16 | 6.24E-18 | 0.742578814 | 0.932 | 0.374 | 1.26E-13 | 25 | Crim1 |
| Plpp14 | 6.33E-18 | 0.848396066 | 0.973 | 0.384 | 1.28E-13 | 25 | Plpp1 |
| Plekhg17 | 6.50E-18 | 0.481161354 | 0.781 | 0.246 | 1.32E-13 | 25 | Plekhg1 |
| Adgrl25 | 7.28E-18 | 0.487043557 | 0.753 | 0.238 | 1.47E-13 | 25 | Adgrl2 |
| Tek6 | 7.91E-18 | 0.515948424 | 0.808 | 0.261 | 1.60E-13 | 25 | Tek |
| Plscr25 | 9.41E-18 | 0.826763999 | 0.877 | 0.336 | 1.91E-13 | 25 | Plscr2 |
| Col13a14 | 1.05E-17 | 0.542243858 | 0.753 | 0.243 | 2.13E-13 | 25 | Col13a1 |
| Sox187 | 1.32E-17 | 0.800916186 | 0.781 | 0.277 | 2.68E-13 | 25 | Sox18 |
| Ptprb7 | 1.40E-17 | 0.756778902 | 1 | 0.369 | 2.84E-13 | 25 | Ptprb |
| Prex28 | 1.62E-17 | 0.45662044 | 0.836 | 0.274 | 3.28E-13 | 25 | Prex2 |
| Hspb19 | 1.65E-17 | 0.796117794 | 0.973 | 0.382 | 3.34E-13 | 25 | Hspb1 |
| Ramp27 | 1.96E-17 | 0.58596751 | 0.932 | 0.333 | 3.97E-13 | 25 | Ramp2 |
| Eng6 | 2.24E-17 | 0.658916702 | 0.959 | 0.355 | 4.53E-13 | 25 | Eng |
| Nrp14 | 2.59E-17 | 0.852754459 | 0.986 | 0.436 | 5.25E-13 | 25 | Nrp1 |
| Ifi445 | 2.97E-17 | 0.639610371 | 0.521 | 0.147 | 6.02E-13 | 25 | Ifi44 |
| Esm14 | 3.47E-17 | 0.635424233 | 0.425 | 0.104 | 7.03E-13 | 25 | Esm1 |
| Shank35 | 4.42E-17 | 0.469822577 | 0.644 | 0.192 | 8.95E-13 | 25 | Shank3 |
| Iigp16 | 4.55E-17 | 0.802434669 | 0.699 | 0.234 | 9.21E-13 | 25 | Iigp1 |
| Ceacam16 | 5.42E-17 | 0.435408853 | 0.562 | 0.157 | 1.10E-12 | 25 | Ceacam1 |
| Tnfsf107 | 5.54E-17 | 0.740252007 | 0.918 | 0.375 | 1.12E-12 | 25 | Tnfsf10 |
| Arhgap295 | 5.57E-17 | 0.720883882 | 0.932 | 0.366 | 1.13E-12 | 25 | Arhgap29 |
| Ptprg6 | 5.69E-17 | 0.586049479 | 0.932 | 0.346 | 1.15E-12 | 25 | Ptprg |
| Flt17 | 6.77E-17 | 0.89080062 | 1 | 0.395 | 1.37E-12 | 25 | Flt1 |

| | | | | | | | |
|---|---|---|---|---|---|---|---|
| Col4a26 | 8.23E-17 | 0.816065846 | 0.918 | 0.361 | 1.67E-12 | 25 | Col4a2 |
| Timp39 | 8.30E-17 | 0.838001205 | 0.932 | 0.386 | 1.68E-12 | 25 | Timp3 |
| Jam35 | 8.30E-17 | 0.704873052 | 0.781 | 0.276 | 1.68E-12 | 25 | Jam3 |
| Cxcl128 | 1.11E-16 | 0.835964932 | 0.849 | 0.323 | 2.24E-12 | 25 | Cxcl12 |
| Fabp47 | 1.23E-16 | 0.825139491 | 1 | 0.437 | 2.49E-12 | 25 | Fabp4 |
| Dlc16 | 1.25E-16 | 0.719011792 | 0.932 | 0.363 | 2.53E-12 | 25 | Dlc1 |
| Tcf156 | 1.39E-16 | 0.82656171 | 0.726 | 0.261 | 2.81E-12 | 25 | Tcf15 |
| Thbd7 | 1.59E-16 | 0.567172952 | 0.753 | 0.256 | 3.21E-12 | 25 | Thbd |
| Cav17 | 1.69E-16 | 0.601568685 | 0.863 | 0.305 | 3.42E-12 | 25 | Cav1 |
| Tspan126 | 1.84E-16 | 0.453866117 | 0.726 | 0.233 | 3.73E-12 | 25 | Tspan12 |
| Slfn57 | 1.88E-16 | 0.685177768 | 0.918 | 0.358 | 3.81E-12 | 25 | Slfn5 |
| Meox15 | 2.29E-16 | 0.513981542 | 0.616 | 0.188 | 4.63E-12 | 25 | Meox1 |
| Prkd15 | 2.45E-16 | 0.583940112 | 0.521 | 0.149 | 4.97E-12 | 25 | Prkd1 |
| Kitl7 | 2.71E-16 | 0.563114354 | 0.904 | 0.337 | 5.49E-12 | 25 | Kitl |
| Podxl6 | 2.85E-16 | 0.70150411 | 0.918 | 0.347 | 5.78E-12 | 25 | Podxl |
| Hspg26 | 3.29E-16 | 0.494143945 | 0.89 | 0.312 | 6.65E-12 | 25 | Hspg2 |
| Tmcc2 | 3.86E-16 | 0.273528589 | 0.301 | 0.061 | 7.82E-12 | 25 | Tmcc2 |
| Sorbs28 | 3.88E-16 | 0.564912009 | 0.795 | 0.278 | 7.86E-12 | 25 | Sorbs2 |
| Pltp6 | 4.73E-16 | 0.739748095 | 0.795 | 0.301 | 9.57E-12 | 25 | Pltp |
| Mef2c7 | 5.20E-16 | 0.782666364 | 0.973 | 0.425 | 1.05E-11 | 25 | Mef2c |
| Myct16 | 6.46E-16 | 0.610842405 | 0.671 | 0.223 | 1.31E-11 | 25 | Myct1 |
| Ttll72 | 7.69E-16 | 0.459806992 | 0.411 | 0.104 | 1.56E-11 | 25 | Ttll7 |
| Emcn6 | 8.81E-16 | 0.825304815 | 0.986 | 0.382 | 1.78E-11 | 25 | Emcn |
| Gng118 | 9.46E-16 | 0.51840329 | 0.973 | 0.367 | 1.92E-11 | 25 | Gng11 |
| Adgrf57 | 1.07E-15 | 0.553448957 | 0.945 | 0.339 | 2.16E-11 | 25 | Adgrf5 |
| Cd2007 | 1.08E-15 | 0.609983928 | 0.932 | 0.351 | 2.18E-11 | 25 | Cd200 |
| Fam198b5 | 1.30E-15 | 0.471987806 | 0.603 | 0.185 | 2.63E-11 | 25 | Fam198b |
| Cd936 | 1.46E-15 | 0.484473557 | 0.767 | 0.257 | 2.96E-11 | 25 | Cd93 |
| Pecam17 | 1.61E-15 | 0.730992872 | 1 | 0.412 | 3.25E-11 | 25 | Pecam1 |
| Tmem886 | 1.89E-15 | 0.412137474 | 0.671 | 0.211 | 3.83E-11 | 25 | Tmem88 |
| Tmem2045 | 1.95E-15 | 0.592258172 | 0.644 | 0.21 | 3.95E-11 | 25 | Tmem204 |
| Igfbp76 | 2.33E-15 | 0.897111927 | 1 | 0.469 | 4.72E-11 | 25 | Igfbp7 |
| Ldb27 | 2.38E-15 | 0.516716554 | 0.877 | 0.331 | 4.83E-11 | 25 | Ldb2 |
| Tmem1406 | 2.70E-15 | 0.414042842 | 0.685 | 0.224 | 5.46E-11 | 25 | Tmem140 |
| Ccdc85a5 | 2.79E-15 | 0.643363922 | 0.575 | 0.182 | 5.66E-11 | 25 | Ccdc85a |
| Mmrn26 | 2.87E-15 | 0.613850646 | 0.685 | 0.233 | 5.81E-11 | 25 | Mmrn2 |
| Tm4sf18 | 4.13E-15 | 0.540308375 | 0.959 | 0.379 | 8.36E-11 | 25 | Tm4sf1 |
| Id37 | 4.23E-15 | 0.735502898 | 0.973 | 0.409 | 8.57E-11 | 25 | Id3 |
| Cd300lg5 | 4.28E-15 | 0.528163895 | 0.822 | 0.297 | 8.66E-11 | 25 | Cd300lg |
| Arhgap316 | 4.77E-15 | 0.610614447 | 0.932 | 0.373 | 9.65E-11 | 25 | Arhgap31 |
| Snrk6 | 4.86E-15 | 0.714420726 | 0.877 | 0.37 | 9.85E-11 | 25 | Snrk |
| Pcdh121 | 5.75E-15 | 0.270422621 | 0.397 | 0.098 | 1.16E-10 | 25 | Pcdh12 |
| Jam26 | 6.28E-15 | 0.460069582 | 0.808 | 0.287 | 1.27E-10 | 25 | Jam2 |

| | | | | | | | |
|---|---|---|---|---|---|---|---|
| Colgalt25 | 6.42E-15 | 0.396900035 | 0.493 | 0.141 | 1.30E-10 | 25 | Colgalt2 |
| Bcam8 | 6.79E-15 | 0.424583273 | 0.753 | 0.255 | 1.38E-10 | 25 | Bcam |
| Prickle17 | 7.16E-15 | 0.424834646 | 0.507 | 0.147 | 1.45E-10 | 25 | Prickle1 |
| Uaca7 | 7.30E-15 | 0.493072322 | 0.671 | 0.227 | 1.48E-10 | 25 | Uaca |
| Dysf5 | 7.36E-15 | 0.45923469 | 0.712 | 0.24 | 1.49E-10 | 25 | Dysf |
| Cdh56 | 7.46E-15 | 0.629275193 | 0.877 | 0.332 | 1.51E-10 | 25 | Cdh5 |
| Prss239 | 8.09E-15 | 0.301977992 | 0.822 | 0.282 | 1.64E-10 | 25 | Prss23 |
| Rhoj9 | 1.21E-14 | 0.484588727 | 0.712 | 0.243 | 2.45E-10 | 25 | Rhoj |
| Gbp76 | 1.72E-14 | 0.593351229 | 0.726 | 0.27 | 3.49E-10 | 25 | Gbp7 |
| Kank36 | 2.00E-14 | 0.400907014 | 0.685 | 0.229 | 4.05E-10 | 25 | Kank3 |
| Ccndbp11 | 2.21E-14 | 0.532355882 | 0.384 | 0.107 | 4.47E-10 | 25 | Ccndbp1 |
| Cmtm84 | 2.32E-14 | 0.51299137 | 0.753 | 0.274 | 4.70E-10 | 25 | Cmtm8 |
| Cd346 | 2.49E-14 | 0.420027396 | 0.877 | 0.322 | 5.05E-10 | 25 | Cd34 |
| Tspan156 | 2.59E-14 | 0.339157076 | 0.466 | 0.131 | 5.24E-10 | 25 | Tspan15 |
| B4galt45 | 3.31E-14 | 0.403117917 | 0.534 | 0.163 | 6.71E-10 | 25 | B4galt4 |
| Ly6a7 | 3.44E-14 | 0.751150508 | 0.986 | 0.461 | 6.96E-10 | 25 | Ly6a |
| Id17 | 3.44E-14 | 0.600356064 | 0.836 | 0.326 | 6.96E-10 | 25 | Id1 |
| Trib23 | 3.83E-14 | 0.479273725 | 0.575 | 0.188 | 7.75E-10 | 25 | Trib2 |
| Zbtb466 | 4.29E-14 | 0.473294273 | 0.521 | 0.164 | 8.68E-10 | 25 | Zbtb46 |
| Emp27 | 4.29E-14 | 0.363009685 | 0.699 | 0.237 | 8.68E-10 | 25 | Emp2 |
| Serpinh19 | 5.19E-14 | 0.396317744 | 0.726 | 0.251 | 1.05E-09 | 25 | Serpinh1 |
| 9530026P05Rik7 | 5.68E-14 | 0.585303744 | 0.671 | 0.243 | 1.15E-09 | 25 | 9530026P05Rik |
| Lims26 | 5.93E-14 | 0.414433757 | 0.616 | 0.203 | 1.20E-09 | 25 | Lims2 |
| Rnf144a4 | 6.13E-14 | 0.476334488 | 0.397 | 0.108 | 1.24E-09 | 25 | Rnf144a |
| Numb7 | 6.25E-14 | 0.512895189 | 0.836 | 0.338 | 1.26E-09 | 25 | Numb |
| Cavin39 | 6.87E-14 | 0.518549512 | 0.822 | 0.308 | 1.39E-09 | 25 | Cavin3 |
| Fbxl76 | 7.20E-14 | 0.299605339 | 0.74 | 0.258 | 1.46E-09 | 25 | Fbxl7 |
| Sparcl19 | 7.61E-14 | 0.68973704 | 0.89 | 0.365 | 1.54E-09 | 25 | Sparcl1 |
| Ly6c16 | 8.30E-14 | 0.649711817 | 0.986 | 0.401 | 1.68E-09 | 25 | Ly6c1 |
| Asap17 | 9.38E-14 | 0.576550488 | 0.877 | 0.357 | 1.90E-09 | 25 | Asap1 |
| Col4a16 | 1.01E-13 | 0.532713147 | 0.986 | 0.393 | 2.05E-09 | 25 | Col4a1 |
| Robo46 | 1.06E-13 | 0.411784768 | 0.562 | 0.177 | 2.14E-09 | 25 | Robo4 |
| Rapgef35 | 1.06E-13 | 0.487308662 | 0.534 | 0.17 | 2.15E-09 | 25 | Rapgef3 |
| Gcnt27 | 1.12E-13 | 0.473144879 | 0.479 | 0.144 | 2.27E-09 | 25 | Gcnt2 |
| Pdgfb6 | 1.26E-13 | 0.614755897 | 0.767 | 0.311 | 2.55E-09 | 25 | Pdgfb |
| Rasgrp35 | 1.30E-13 | 0.461012784 | 0.767 | 0.283 | 2.63E-09 | 25 | Rasgrp3 |
| Cmpk2 | 1.30E-13 | 0.44460156 | 0.274 | 0.061 | 2.63E-09 | 25 | Cmpk2 |
| Mfge88 | 1.50E-13 | 0.523952889 | 0.699 | 0.255 | 3.04E-09 | 25 | Mfge8 |
| Apbb28 | 1.53E-13 | 0.487910932 | 0.904 | 0.368 | 3.09E-09 | 25 | Apbb2 |
| Nfib6 | 1.67E-13 | 0.745807614 | 0.986 | 0.446 | 3.38E-09 | 25 | Nfib |
| Fzd47 | 1.72E-13 | 0.344294287 | 0.685 | 0.23 | 3.48E-09 | 25 | Fzd4 |
| She6 | 1.79E-13 | 0.444579989 | 0.493 | 0.15 | 3.63E-09 | 25 | She |
| Esam7 | 1.99E-13 | 0.574662934 | 0.808 | 0.311 | 4.03E-09 | 25 | Esam |

| | | | | | | | |
|---|---|---|---|---|---|---|---|
| Klf27 | 1.99E-13 | 0.491971812 | 0.877 | 0.368 | 4.03E-09 | 25 | Klf2 |
| Ltbp47 | 2.11E-13 | 0.486226741 | 0.808 | 0.311 | 4.27E-09 | 25 | Ltbp4 |
| Pdlim55 | 2.29E-13 | 0.400117823 | 0.753 | 0.287 | 4.64E-09 | 25 | Pdlim5 |
| Sncaip3 | 2.33E-13 | 0.362919185 | 0.329 | 0.082 | 4.72E-09 | 25 | Sncaip |
| BC0285284 | 2.34E-13 | 0.514391385 | 0.644 | 0.235 | 4.75E-09 | 25 | BC028528 |
| Ptprm6 | 2.54E-13 | 0.459810828 | 0.849 | 0.331 | 5.15E-09 | 25 | Ptprm |
| Cdc42ep24 | 2.60E-13 | 0.509201253 | 0.534 | 0.174 | 5.26E-09 | 25 | Cdc42ep2 |
| Itih55 | 2.68E-13 | 0.49993874 | 0.658 | 0.244 | 5.42E-09 | 25 | Itih5 |
| Slc9a3r26 | 2.94E-13 | 0.672718886 | 0.877 | 0.372 | 5.95E-09 | 25 | Slc9a3r2 |
| Zhx37 | 2.97E-13 | 0.41532831 | 0.603 | 0.203 | 6.01E-09 | 25 | Zhx3 |
| Fat45 | 3.47E-13 | 0.285522252 | 0.397 | 0.107 | 7.03E-09 | 25 | Fat4 |
| Gm205323 | 3.50E-13 | 0.433923068 | 0.397 | 0.11 | 7.08E-09 | 25 | Gm20532 |
| Gm138895 | 4.11E-13 | 0.407096542 | 0.63 | 0.22 | 8.33E-09 | 25 | Gm13889 |
| Egfl77 | 4.57E-13 | 0.613825958 | 0.986 | 0.395 | 9.26E-09 | 25 | Egfl7 |
| Cav27 | 5.08E-13 | 0.463071982 | 0.603 | 0.21 | 1.03E-08 | 25 | Cav2 |
| Dusp34 | 5.21E-13 | 0.418267031 | 0.575 | 0.193 | 1.06E-08 | 25 | Dusp3 |
| Ece17 | 5.76E-13 | 0.459928463 | 0.795 | 0.307 | 1.17E-08 | 25 | Ece1 |
| Tns110 | 6.16E-13 | 0.398628235 | 0.685 | 0.246 | 1.25E-08 | 25 | Tns1 |
| Ctnnd14 | 7.01E-13 | 0.259040649 | 0.479 | 0.146 | 1.42E-08 | 25 | Ctnnd1 |
| Tcf47 | 7.40E-13 | 0.657804667 | 1 | 0.473 | 1.50E-08 | 25 | Tcf4 |
| Rgl15 | 7.47E-13 | 0.456003798 | 0.753 | 0.29 | 1.51E-08 | 25 | Rgl1 |
| Nbeal18 | 8.42E-13 | 0.49746207 | 0.685 | 0.263 | 1.71E-08 | 25 | Nbeal1 |
| Sema3c3 | 8.44E-13 | 0.478979699 | 0.507 | 0.164 | 1.71E-08 | 25 | Sema3c |
| Rasgrf26 | 8.94E-13 | 0.42730362 | 0.548 | 0.178 | 1.81E-08 | 25 | Rasgrf2 |
| Cx3cl15 | 9.53E-13 | 0.312940814 | 0.384 | 0.105 | 1.93E-08 | 25 | Cx3cl1 |
| Tjp16 | 9.78E-13 | 0.364995947 | 0.712 | 0.265 | 1.98E-08 | 25 | Tjp1 |
| Ebf18 | 1.03E-12 | 0.380763122 | 0.973 | 0.376 | 2.08E-08 | 25 | Ebf1 |
| Bmpr26 | 1.05E-12 | 0.596267743 | 0.836 | 0.351 | 2.13E-08 | 25 | Bmpr2 |
| Rasip16 | 1.05E-12 | 0.261998962 | 0.603 | 0.196 | 2.13E-08 | 25 | Rasip1 |
| Mcf2l4 | 1.13E-12 | 0.478686296 | 0.603 | 0.215 | 2.28E-08 | 25 | Mcf2l |
| Vat15 | 1.13E-12 | 0.343583563 | 0.493 | 0.156 | 2.29E-08 | 25 | Vat1 |
| Rasal26 | 1.15E-12 | 0.480349836 | 0.795 | 0.316 | 2.33E-08 | 25 | Rasal2 |
| Tspan96 | 1.17E-12 | 0.447109626 | 0.699 | 0.266 | 2.37E-08 | 25 | Tspan9 |
| Grrp16 | 1.18E-12 | 0.449605983 | 0.616 | 0.221 | 2.39E-08 | 25 | Grrp1 |
| Dhrs36 | 1.55E-12 | 0.444691116 | 0.63 | 0.232 | 3.13E-08 | 25 | Dhrs3 |
| Rras8 | 1.69E-12 | 0.473547725 | 0.753 | 0.299 | 3.42E-08 | 25 | Rras |
| Oasl26 | 2.01E-12 | 0.304596139 | 0.438 | 0.132 | 4.07E-08 | 25 | Oasl2 |
| Sptbn17 | 2.57E-12 | 0.656491838 | 0.986 | 0.496 | 5.20E-08 | 25 | Sptbn1 |
| Pls37 | 2.68E-12 | 0.455295161 | 0.603 | 0.216 | 5.43E-08 | 25 | Pls3 |
| Dennd5b7 | 2.92E-12 | 0.301950349 | 0.479 | 0.148 | 5.92E-08 | 25 | Dennd5b |
| App8 | 3.03E-12 | 0.553383208 | 0.945 | 0.421 | 6.14E-08 | 25 | App |
| Syne28 | 3.34E-12 | 0.408587177 | 0.712 | 0.273 | 6.77E-08 | 25 | Syne2 |
| Epas18 | 3.50E-12 | 0.441649101 | 0.877 | 0.361 | 7.10E-08 | 25 | Epas1 |

| | | | | | | | |
|---|---|---|---|---|---|---|---|
| Ankrd291 | 4.01E-12 | 0.333787278 | 0.301 | 0.076 | 8.12E-08 | 25 | Ankrd29 |
| Sybu4 | 4.05E-12 | 0.379194683 | 0.397 | 0.115 | 8.20E-08 | 25 | Sybu |
| Ptgs14 | 4.14E-12 | 0.370550274 | 0.603 | 0.213 | 8.38E-08 | 25 | Ptgs1 |
| Plxnd17 | 4.34E-12 | 0.30290385 | 0.548 | 0.183 | 8.79E-08 | 25 | Plxnd1 |
| Ptk28 | 4.53E-12 | 0.392178914 | 0.712 | 0.267 | 9.18E-08 | 25 | Ptk2 |
| Gbp45 | 4.87E-12 | 0.406884362 | 0.521 | 0.179 | 9.86E-08 | 25 | Gbp4 |
| Dock46 | 5.00E-12 | 0.255473712 | 0.795 | 0.298 | 1.01E-07 | 25 | Dock4 |
| Sh3bp55 | 5.08E-12 | 0.43153545 | 0.603 | 0.218 | 1.03E-07 | 25 | Sh3bp5 |
| Lnx23 | 5.13E-12 | 0.486818317 | 0.425 | 0.134 | 1.04E-07 | 25 | Lnx2 |
| Gpr1465 | 5.32E-12 | 0.325888536 | 0.384 | 0.11 | 1.08E-07 | 25 | Gpr146 |
| Smco46 | 5.81E-12 | 0.435603244 | 0.918 | 0.391 | 1.18E-07 | 25 | Smco4 |
| Rdx6 | 6.30E-12 | 0.602251402 | 0.986 | 0.553 | 1.28E-07 | 25 | Rdx |
| Erg6 | 7.38E-12 | 0.28439965 | 0.438 | 0.132 | 1.49E-07 | 25 | Erg |
| Prdm12 | 7.98E-12 | 0.32342598 | 0.384 | 0.111 | 1.62E-07 | 25 | Prdm1 |
| Mgll6 | 8.24E-12 | 0.634231792 | 0.945 | 0.45 | 1.67E-07 | 25 | Mgll |
| Dip2c9 | 8.32E-12 | 0.341021603 | 0.63 | 0.224 | 1.69E-07 | 25 | Dip2c |
| Bnip3l3 | 8.70E-12 | 0.52760905 | 0.74 | 0.333 | 1.76E-07 | 25 | Bnip3l |
| Stard86 | 8.96E-12 | 0.36165265 | 0.575 | 0.199 | 1.81E-07 | 25 | Stard8 |
| Osmr6 | 1.00E-11 | 0.333911334 | 0.452 | 0.141 | 2.02E-07 | 25 | Osmr |
| Slc43a37 | 1.02E-11 | 0.387703797 | 0.699 | 0.271 | 2.06E-07 | 25 | Slc43a3 |
| Klf46 | 1.11E-11 | 0.5608437 | 0.671 | 0.271 | 2.24E-07 | 25 | Klf4 |
| Fbln28 | 1.17E-11 | 0.281587053 | 0.726 | 0.273 | 2.37E-07 | 25 | Fbln2 |
| Cd2743 | 1.19E-11 | 0.448458096 | 0.384 | 0.115 | 2.41E-07 | 25 | Cd274 |
| 24-Mar | 1.22E-11 | 0.455499062 | 0.534 | 0.198 | 2.47E-07 | 25 | 2-Mar |
| St6galnac39 | 1.34E-11 | 0.335617972 | 0.603 | 0.217 | 2.71E-07 | 25 | St6galnac3 |
| Piezo24 | 1.58E-11 | 0.494776766 | 0.671 | 0.254 | 3.19E-07 | 25 | Piezo2 |
| Ushbp16 | 1.66E-11 | 0.379305838 | 0.616 | 0.22 | 3.36E-07 | 25 | Ushbp1 |
| Gas67 | 1.71E-11 | 0.400022977 | 0.753 | 0.299 | 3.47E-07 | 25 | Gas6 |
| Ephb46 | 1.75E-11 | 0.337312045 | 0.507 | 0.168 | 3.55E-07 | 25 | Ephb4 |
| Sema6a6 | 1.86E-11 | 0.337989541 | 0.589 | 0.21 | 3.76E-07 | 25 | Sema6a |
| Dnm36 | 1.96E-11 | 0.383419079 | 0.63 | 0.231 | 3.97E-07 | 25 | Dnm3 |
| Nxpe46 | 1.97E-11 | 0.297169496 | 0.466 | 0.149 | 3.99E-07 | 25 | Nxpe4 |
| Arhgef129 | 2.08E-11 | 0.43702723 | 0.822 | 0.35 | 4.21E-07 | 25 | Arhgef12 |
| Ctsl8 | 2.17E-11 | 0.341358187 | 0.822 | 0.347 | 4.40E-07 | 25 | Ctsl |
| Lama36 | 2.18E-11 | 0.337742169 | 0.616 | 0.222 | 4.41E-07 | 25 | Lama3 |
| Irgm22 | 2.28E-11 | 0.314398719 | 0.411 | 0.128 | 4.61E-07 | 25 | Irgm2 |
| Klf76 | 2.30E-11 | 0.39453076 | 0.74 | 0.299 | 4.66E-07 | 25 | Klf7 |
| Ndrg17 | 2.55E-11 | 0.299830532 | 0.644 | 0.234 | 5.17E-07 | 25 | Ndrg1 |
| Itga19 | 2.61E-11 | 0.562654388 | 0.89 | 0.416 | 5.29E-07 | 25 | Itga1 |
| Gab16 | 2.64E-11 | 0.37326453 | 0.384 | 0.115 | 5.35E-07 | 25 | Gab1 |
| Ehd45 | 2.70E-11 | 0.542296346 | 0.877 | 0.41 | 5.46E-07 | 25 | Ehd4 |
| Ccnd16 | 3.35E-11 | 0.274125911 | 0.616 | 0.228 | 6.77E-07 | 25 | Ccnd1 |
| Rnf2206 | 3.43E-11 | 0.410812163 | 0.63 | 0.243 | 6.95E-07 | 25 | Rnf220 |

| | | | | | | | |
|---|---|---|---|---|---|---|---|
| Cpd7 | 3.49E-11 | 0.469881813 | 0.753 | 0.314 | 7.07E-07 | 25 | Cpd |
| Phldb26 | 3.52E-11 | 0.4585144 | 0.562 | 0.204 | 7.12E-07 | 25 | Phldb2 |
| Ecscr6 | 3.58E-11 | 0.350009701 | 0.63 | 0.229 | 7.24E-07 | 25 | Ecscr |
| Lzts21 | 3.98E-11 | 0.300791486 | 0.329 | 0.092 | 8.05E-07 | 25 | Lzts2 |
| Sptan18 | 4.17E-11 | 0.419592164 | 0.808 | 0.345 | 8.44E-07 | 25 | Sptan1 |
| Dok43 | 4.31E-11 | 0.277915479 | 0.397 | 0.12 | 8.74E-07 | 25 | Dok4 |
| Cd24a3 | 4.36E-11 | 0.489482166 | 0.781 | 0.344 | 8.84E-07 | 25 | Cd24a |
| Gngt27 | 5.26E-11 | 0.388627344 | 0.822 | 0.361 | 1.06E-06 | 25 | Gngt2 |
| Mapk36 | 5.37E-11 | 0.360239998 | 0.685 | 0.267 | 1.09E-06 | 25 | Mapk3 |
| Dock17 | 5.47E-11 | 0.324254002 | 0.712 | 0.281 | 1.11E-06 | 25 | Dock1 |
| Grb108 | 5.78E-11 | 0.40079902 | 0.548 | 0.196 | 1.17E-06 | 25 | Grb10 |
| Rtl8a4 | 5.88E-11 | 0.266973795 | 0.521 | 0.178 | 1.19E-06 | 25 | Rtl8a |
| Ints94 | 6.15E-11 | 0.361745725 | 0.425 | 0.138 | 1.24E-06 | 25 | Ints9 |
| Vamp56 | 7.03E-11 | 0.411922279 | 0.89 | 0.386 | 1.42E-06 | 25 | Vamp5 |
| Actn48 | 7.37E-11 | 0.548742965 | 0.808 | 0.377 | 1.49E-06 | 25 | Actn4 |
| Piezo16 | 7.91E-11 | 0.411026931 | 0.507 | 0.175 | 1.60E-06 | 25 | Piezo1 |
| Chst23 | 8.12E-11 | 0.332184924 | 0.493 | 0.166 | 1.64E-06 | 25 | Chst2 |
| Wwtr17 | 8.61E-11 | 0.351761115 | 0.644 | 0.241 | 1.74E-06 | 25 | Wwtr1 |
| Tmem477 | 8.65E-11 | 0.378963818 | 0.479 | 0.157 | 1.75E-06 | 25 | Tmem47 |
| Tspan44 | 8.97E-11 | 0.345474401 | 0.356 | 0.107 | 1.82E-06 | 25 | Tspan4 |
| Plxna26 | 8.97E-11 | 0.366096076 | 0.603 | 0.22 | 1.82E-06 | 25 | Plxna2 |
| Pitpnm23 | 1.06E-10 | 0.358545682 | 0.397 | 0.124 | 2.14E-06 | 25 | Pitpnm2 |
| Eva1b8 | 1.08E-10 | 0.441036451 | 0.699 | 0.287 | 2.19E-06 | 25 | Eva1b |
| Itga68 | 1.23E-10 | 0.370777595 | 0.699 | 0.285 | 2.48E-06 | 25 | Itga6 |
| 4931406P16Rik7 | 1.23E-10 | 0.497637068 | 0.767 | 0.344 | 2.49E-06 | 25 | 4931406P16Rik |
| Ackr35 | 1.24E-10 | 0.393196386 | 0.452 | 0.153 | 2.51E-06 | 25 | Ackr3 |
| Sparc11 | 1.24E-10 | 0.397688486 | 0.945 | 0.404 | 2.52E-06 | 25 | Sparc |
| Adipor16 | 1.25E-10 | 0.419630555 | 0.63 | 0.259 | 2.54E-06 | 25 | Adipor1 |
| Ces2e3 | 1.29E-10 | 0.325129244 | 0.37 | 0.111 | 2.60E-06 | 25 | Ces2e |
| Triobp2 | 1.60E-10 | 0.291075973 | 0.425 | 0.138 | 3.25E-06 | 25 | Triobp |
| Hmbox14 | 1.61E-10 | 0.334770006 | 0.685 | 0.275 | 3.26E-06 | 25 | Hmbox1 |
| Samd126 | 2.00E-10 | 0.283347644 | 0.616 | 0.231 | 4.06E-06 | 25 | Samd12 |
| Car84 | 2.08E-10 | 0.277876261 | 0.534 | 0.187 | 4.20E-06 | 25 | Car8 |
| Fgd56 | 2.52E-10 | 0.378888343 | 0.603 | 0.23 | 5.09E-06 | 25 | Fgd5 |
| Cyp26b15 | 2.54E-10 | 0.726008303 | 0.548 | 0.217 | 5.14E-06 | 25 | Cyp26b1 |
| Gata24 | 2.60E-10 | 0.255149966 | 0.356 | 0.106 | 5.27E-06 | 25 | Gata2 |
| Dab2ip8 | 2.74E-10 | 0.377574593 | 0.658 | 0.263 | 5.55E-06 | 25 | Dab2ip |
| Zfp3666 | 2.84E-10 | 0.389987456 | 0.507 | 0.184 | 5.75E-06 | 25 | Zfp366 |
| Dusp17 | 3.01E-10 | 0.56153306 | 0.507 | 0.195 | 6.09E-06 | 25 | Dusp1 |
| Cd384 | 3.07E-10 | 0.413550964 | 0.603 | 0.233 | 6.22E-06 | 25 | Cd38 |
| Ccrl24 | 3.16E-10 | 0.330571403 | 0.356 | 0.11 | 6.40E-06 | 25 | Ccrl2 |
| Fos7 | 3.26E-10 | 0.359538313 | 0.712 | 0.318 | 6.60E-06 | 25 | Fos |
| Bvht6 | 3.29E-10 | 0.324226354 | 0.521 | 0.188 | 6.65E-06 | 25 | Bvht |

| | | | | | | | |
|---|---|---|---|---|---|---|---|
| Ctnnbip15 | 3.58E-10 | 0.372498861 | 0.548 | 0.205 | 7.26E-06 | 25 | Ctnnbip1 |
| Fry7 | 3.94E-10 | 0.260295037 | 0.548 | 0.202 | 7.99E-06 | 25 | Fry |
| Lysmd26 | 4.27E-10 | 0.408958002 | 0.521 | 0.193 | 8.64E-06 | 25 | Lysmd2 |
| Nav17 | 4.27E-10 | 0.362334988 | 0.589 | 0.224 | 8.65E-06 | 25 | Nav1 |
| Gm383993 | 4.35E-10 | 0.291846831 | 0.356 | 0.109 | 8.80E-06 | 25 | Gm38399 |
| Ednrb4 | 4.40E-10 | 0.311751037 | 0.534 | 0.195 | 8.90E-06 | 25 | Ednrb |
| Msn9 | 4.54E-10 | 0.519974793 | 0.904 | 0.445 | 9.19E-06 | 25 | Msn |
| Arl156 | 4.69E-10 | 0.274498483 | 0.822 | 0.383 | 9.50E-06 | 25 | Arl15 |
| Endod13 | 4.81E-10 | 0.406221858 | 0.438 | 0.153 | 9.74E-06 | 25 | Endod1 |
| H2-Q44 | 4.94E-10 | 0.426143501 | 0.712 | 0.31 | 1.00E-05 | 25 | H2-Q4 |
| Clic48 | 5.70E-10 | 0.457603285 | 0.863 | 0.425 | 1.15E-05 | 25 | Clic4 |
| Ehd28 | 5.99E-10 | 0.295271651 | 0.534 | 0.194 | 1.21E-05 | 25 | Ehd2 |
| Ppm1f | 6.48E-10 | 0.290340618 | 0.26 | 0.068 | 1.31E-05 | 25 | Ppm1f |
| C2cd24 | 6.56E-10 | 0.351809875 | 0.356 | 0.112 | 1.33E-05 | 25 | C2cd2 |
| Cpne86 | 7.49E-10 | 0.363326932 | 0.589 | 0.23 | 1.52E-05 | 25 | Cpne8 |
| Meis25 | 7.52E-10 | 0.561953288 | 0.644 | 0.284 | 1.52E-05 | 25 | Meis2 |
| Ifi2081 | 7.99E-10 | 0.259829116 | 0.301 | 0.086 | 1.62E-05 | 25 | Ifi208 |
| Afdn6 | 8.04E-10 | 0.292695513 | 0.74 | 0.316 | 1.63E-05 | 25 | Afdn |
| Ifitm310 | 8.38E-10 | 0.576841648 | 0.986 | 0.539 | 1.70E-05 | 25 | Ifitm3 |
| Fez24 | 8.93E-10 | 0.347152118 | 0.685 | 0.284 | 1.81E-05 | 25 | Fez2 |
| Elk36 | 9.27E-10 | 0.360811858 | 0.781 | 0.358 | 1.88E-05 | 25 | Elk3 |
| Prkch10 | 9.57E-10 | 0.415163817 | 0.877 | 0.401 | 1.94E-05 | 25 | Prkch |
| Arhgef156 | 1.01E-09 | 0.281848395 | 0.479 | 0.17 | 2.05E-05 | 25 | Arhgef15 |
| Ifi354 | 1.08E-09 | 0.450119598 | 0.63 | 0.275 | 2.18E-05 | 25 | Ifi35 |
| Dgkh7 | 1.08E-09 | 0.274347382 | 0.493 | 0.178 | 2.20E-05 | 25 | Dgkh |
| Cbfa2t37 | 1.18E-09 | 0.309389999 | 0.411 | 0.14 | 2.39E-05 | 25 | Cbfa2t3 |
| Fam13c6 | 1.21E-09 | 0.322869705 | 0.356 | 0.113 | 2.46E-05 | 25 | Fam13c |
| Heg17 | 1.26E-09 | 0.272473852 | 0.671 | 0.276 | 2.55E-05 | 25 | Heg1 |
| Bcar1 | 1.27E-09 | 0.269209528 | 0.26 | 0.07 | 2.57E-05 | 25 | Bcar1 |
| Spns23 | 1.34E-09 | 0.299561246 | 0.315 | 0.094 | 2.70E-05 | 25 | Spns2 |
| Amotl19 | 1.43E-09 | 0.314592265 | 0.507 | 0.187 | 2.90E-05 | 25 | Amotl1 |
| Tnfaip15 | 1.53E-09 | 0.346110814 | 0.534 | 0.204 | 3.09E-05 | 25 | Tnfaip1 |
| Fut85 | 1.53E-09 | 0.46859865 | 0.658 | 0.29 | 3.10E-05 | 25 | Fut8 |
| Cald110 | 1.65E-09 | 0.324895267 | 0.753 | 0.324 | 3.34E-05 | 25 | Cald1 |
| Afap16 | 1.78E-09 | 0.25340644 | 0.438 | 0.151 | 3.61E-05 | 25 | Afap1 |
| Nfe2l16 | 1.83E-09 | 0.309440019 | 0.658 | 0.282 | 3.71E-05 | 25 | Nfe2l1 |
| Sypl6 | 1.84E-09 | 0.36127466 | 0.589 | 0.241 | 3.72E-05 | 25 | Sypl |
| Hlx2 | 1.88E-09 | 0.309568307 | 0.329 | 0.102 | 3.80E-05 | 25 | Hlx |
| Gbp54 | 2.00E-09 | 0.388034794 | 0.384 | 0.13 | 4.05E-05 | 25 | Gbp5 |
| Tsc22d34 | 2.03E-09 | 0.276502736 | 0.548 | 0.209 | 4.10E-05 | 25 | Tsc22d3 |
| Crip29 | 2.47E-09 | 0.384101431 | 0.904 | 0.417 | 5.01E-05 | 25 | Crip2 |
| Synpo3 | 2.76E-09 | 0.284282129 | 0.329 | 0.101 | 5.59E-05 | 25 | Synpo |
| Ginm16 | 2.82E-09 | 0.297996761 | 0.479 | 0.181 | 5.71E-05 | 25 | Ginm1 |

| | | | | | | | |
|---|---|---|---|---|---|---|---|
| Magi16 | 3.01E-09 | 0.2717981 | 0.589 | 0.233 | 6.10E-05 | 25 | Magi1 |
| Ntf34 | 3.28E-09 | 0.273871043 | 0.493 | 0.182 | 6.64E-05 | 25 | Ntf3 |
| Tanc17 | 3.74E-09 | 0.295902616 | 0.562 | 0.224 | 7.58E-05 | 25 | Tanc1 |
| Rgs123 | 3.80E-09 | 0.304898871 | 0.329 | 0.104 | 7.70E-05 | 25 | Rgs12 |
| Dock66 | 4.00E-09 | 0.298096874 | 0.411 | 0.142 | 8.10E-05 | 25 | Dock6 |
| Slc6a66 | 4.52E-09 | 0.299446263 | 0.726 | 0.329 | 9.16E-05 | 25 | Slc6a6 |
| Rapgef56 | 4.54E-09 | 0.307525866 | 0.562 | 0.223 | 9.19E-05 | 25 | Rapgef5 |
| Myo1c5 | 4.55E-09 | 0.269019456 | 0.548 | 0.215 | 9.21E-05 | 25 | Myo1c |
| S1pr15 | 5.41E-09 | 0.268486859 | 0.644 | 0.267 | 0.000109603 | 25 | S1pr1 |
| Mkl26 | 5.64E-09 | 0.324121256 | 0.616 | 0.257 | 0.000114242 | 25 | Mkl2 |
| Ncoa41 | 5.89E-09 | 0.298964771 | 0.315 | 0.102 | 0.000119327 | 25 | Ncoa4 |
| Klf135 | 6.05E-09 | 0.387570937 | 0.74 | 0.352 | 0.0001225 | 25 | Klf13 |
| Calcrl5 | 6.12E-09 | 0.298311732 | 0.63 | 0.265 | 0.000123861 | 25 | Calcrl |
| Pde8a5 | 6.65E-09 | 0.290032146 | 0.479 | 0.184 | 0.000134669 | 25 | Pde8a |
| Adarb14 | 6.80E-09 | 0.287300706 | 0.342 | 0.111 | 0.000137745 | 25 | Adarb1 |
| Klf35 | 6.98E-09 | 0.321212954 | 0.575 | 0.238 | 0.000141359 | 25 | Klf3 |
| Phlpp18 | 7.52E-09 | 0.386580173 | 0.603 | 0.265 | 0.000152153 | 25 | Phlpp1 |
| Epha45 | 8.02E-09 | 0.267286783 | 0.356 | 0.118 | 0.000162371 | 25 | Epha4 |
| Itpkb5 | 8.15E-09 | 0.337191647 | 0.753 | 0.357 | 0.000164953 | 25 | Itpkb |
| Fam43a3 | 8.21E-09 | 0.259405414 | 0.411 | 0.143 | 0.000166308 | 25 | Fam43a |
| Tspan185 | 8.68E-09 | 0.308842032 | 0.397 | 0.139 | 0.000175781 | 25 | Tspan18 |
| Stat19 | 8.69E-09 | 0.37383917 | 0.589 | 0.251 | 0.000175961 | 25 | Stat1 |
| Rhob7 | 8.74E-09 | 0.462965913 | 0.603 | 0.269 | 0.000176879 | 25 | Rhob |
| Efna16 | 9.09E-09 | 0.279742921 | 0.411 | 0.145 | 0.000184045 | 25 | Efna1 |
| Ptprk7 | 9.85E-09 | 0.298987385 | 0.397 | 0.141 | 0.00019949 | 25 | Ptprk |
| Id45 | 1.14E-08 | 0.267973265 | 0.37 | 0.126 | 0.000229982 | 25 | Id4 |
| Cdkn1a4 | 1.25E-08 | 0.252579966 | 0.37 | 0.128 | 0.000253351 | 25 | Cdkn1a |
| Rapgef27 | 1.26E-08 | 0.35014573 | 0.726 | 0.34 | 0.000254606 | 25 | Rapgef2 |
| Psmb106 | 1.26E-08 | 0.347963091 | 0.836 | 0.401 | 0.00025547 | 25 | Psmb10 |
| Nxpe22 | 1.27E-08 | 0.269587732 | 0.329 | 0.106 | 0.000257907 | 25 | Nxpe2 |
| Klhl56 | 1.48E-08 | 0.30740387 | 0.548 | 0.226 | 0.00029946 | 25 | Klhl5 |
| Gimap610 | 1.49E-08 | 0.252879848 | 0.822 | 0.367 | 0.00030217 | 25 | Gimap6 |
| Sgk15 | 1.50E-08 | 0.355398614 | 0.63 | 0.271 | 0.000302977 | 25 | Sgk1 |
| Efr3b3 | 1.52E-08 | 0.286616401 | 0.288 | 0.088 | 0.000307494 | 25 | Efr3b |
| Dusp72 | 1.54E-08 | 0.342630675 | 0.384 | 0.137 | 0.000312101 | 25 | Dusp7 |
| Csgalnact17 | 1.73E-08 | 0.268504927 | 0.507 | 0.197 | 0.000350994 | 25 | Csgalnact1 |
| Caskin25 | 1.85E-08 | 0.274083038 | 0.397 | 0.141 | 0.000374979 | 25 | Caskin2 |
| Tspan311 | 2.18E-08 | 0.266526576 | 0.356 | 0.125 | 0.000441076 | 25 | Tspan31 |
| Cyp2d227 | 2.26E-08 | 0.384586555 | 0.384 | 0.136 | 0.000457469 | 25 | Cyp2d22 |
| Tspan138 | 2.32E-08 | 0.357503481 | 0.644 | 0.287 | 0.000470049 | 25 | Tspan13 |
| Ncoa76 | 2.35E-08 | 0.34634073 | 0.658 | 0.298 | 0.000475082 | 25 | Ncoa7 |
| Nck17 | 2.35E-08 | 0.351259975 | 0.726 | 0.341 | 0.000475245 | 25 | Nck1 |
| Impdh13 | 2.40E-08 | 0.363696707 | 0.329 | 0.113 | 0.000484905 | 25 | Impdh1 |

| | | | | | | | |
|---|---|---|---|---|---|---|---|
| Rnf10 | 2.45E-08 | 0.294920733 | 0.397 | 0.15 | 0.000495228 | 25 | Rnf10 |
| Myo18a6 | 3.15E-08 | 0.262971275 | 0.616 | 0.259 | 0.000638577 | 25 | Myo18a |
| Reep38 | 3.36E-08 | 0.305369254 | 0.685 | 0.307 | 0.000680484 | 25 | Reep3 |
| Sema7a5 | 3.42E-08 | 0.336160199 | 0.411 | 0.151 | 0.000692766 | 25 | Sema7a |
| 6-Jun | 3.46E-08 | 0.36189919 | 0.918 | 0.515 | 0.000700594 | 25 | Jun |
| Ifitm29 | 3.49E-08 | 0.366958026 | 0.959 | 0.466 | 0.000706902 | 25 | Ifitm2 |
| Rnd33 | 3.53E-08 | 0.36292522 | 0.301 | 0.098 | 0.00071549 | 25 | Rnd3 |
| Bnip26 | 3.54E-08 | 0.30508144 | 0.644 | 0.285 | 0.000717034 | 25 | Bnip2 |
| Ets110 | 4.00E-08 | 0.375715664 | 0.822 | 0.405 | 0.000808848 | 25 | Ets1 |
| Tmem260 | 5.02E-08 | 0.307659639 | 0.274 | 0.088 | 0.001015842 | 25 | Tmem260 |
| Rock28 | 5.61E-08 | 0.2960751 | 0.726 | 0.339 | 0.001136214 | 25 | Rock2 |
| Rarg5 | 5.61E-08 | 0.294910735 | 0.493 | 0.197 | 0.001136447 | 25 | Rarg |
| Tnfaip25 | 5.86E-08 | 0.269468962 | 0.411 | 0.153 | 0.001185743 | 25 | Tnfaip2 |
| 9930111J21Rik29 | 6.15E-08 | 0.395424077 | 0.575 | 0.258 | 0.001245018 | 25 | 9930111J21Rik2 |
| Cntln8 | 6.80E-08 | 0.263109423 | 0.411 | 0.156 | 0.001375918 | 25 | Cntln |
| Rnf2136 | 6.81E-08 | 0.344577804 | 0.589 | 0.258 | 0.001379205 | 25 | Rnf213 |
| Oaz26 | 6.99E-08 | 0.446161823 | 0.534 | 0.238 | 0.001415902 | 25 | Oaz2 |
| Tshz28 | 7.99E-08 | 0.255639701 | 0.795 | 0.373 | 0.00161676 | 25 | Tshz2 |
| Amfr3 | 8.48E-08 | 0.392700216 | 0.342 | 0.125 | 0.001717629 | 25 | Amfr |
| Aplp27 | 8.96E-08 | 0.275873803 | 0.836 | 0.417 | 0.001813644 | 25 | Aplp2 |
| Abcg27 | 8.98E-08 | 0.251733685 | 0.616 | 0.272 | 0.001818549 | 25 | Abcg2 |
| Ppic6 | 9.33E-08 | 0.32534173 | 0.342 | 0.122 | 0.001889952 | 25 | Ppic |
| Cep85l8 | 9.52E-08 | 0.489417806 | 0.411 | 0.166 | 0.001928185 | 25 | Cep85l |
| Lipe5 | 9.72E-08 | 0.373053359 | 0.37 | 0.139 | 0.001968283 | 25 | Lipe |
| Sntb26 | 9.84E-08 | 0.374191287 | 0.603 | 0.267 | 0.001992662 | 25 | Sntb2 |
| Kif13a6 | 1.01E-07 | 0.315889556 | 0.438 | 0.175 | 0.002047274 | 25 | Kif13a |
| Oxr15 | 1.17E-07 | 0.360929287 | 0.699 | 0.337 | 0.002361157 | 25 | Oxr1 |
| Slc48a12 | 1.26E-07 | 0.280642184 | 0.315 | 0.112 | 0.002547758 | 25 | Slc48a1 |
| Mxra77 | 1.29E-07 | 0.322325444 | 0.384 | 0.143 | 0.002605724 | 25 | Mxra7 |
| Adam108 | 1.30E-07 | 0.37899396 | 0.63 | 0.294 | 0.00264183 | 25 | Adam10 |
| H2-T236 | 1.31E-07 | 0.407959844 | 0.836 | 0.461 | 0.002659513 | 25 | H2-T23 |
| Nxn7 | 1.34E-07 | 0.272471091 | 0.603 | 0.27 | 0.00271495 | 25 | Nxn |
| Gbe1 | 1.36E-07 | 0.251304964 | 0.301 | 0.103 | 0.002757927 | 25 | Gbe1 |
| Ets25 | 1.55E-07 | 0.277226481 | 0.479 | 0.195 | 0.003142021 | 25 | Ets2 |
| Fabp53 | 1.55E-07 | 0.294860345 | 0.699 | 0.326 | 0.003142757 | 25 | Fabp5 |
| Pttg1ip9 | 1.57E-07 | 0.320564038 | 0.603 | 0.27 | 0.003188371 | 25 | Pttg1ip |
| Pced1b6 | 1.67E-07 | 0.303469109 | 0.329 | 0.119 | 0.003388684 | 25 | Pced1b |
| Tnfsf126 | 1.75E-07 | 0.270680161 | 0.384 | 0.149 | 0.003533097 | 25 | Tnfsf12 |
| Anks15 | 1.78E-07 | 0.26505995 | 0.37 | 0.14 | 0.003609576 | 25 | Anks1 |
| 2900026A02Rik8 | 2.10E-07 | 0.26259633 | 0.521 | 0.221 | 0.004259649 | 25 | 2900026A02Rik |
| Gja15 | 2.11E-07 | 0.257591104 | 0.356 | 0.129 | 0.004281964 | 25 | Gja1 |
| Trim476 | 2.16E-07 | 0.379029097 | 0.37 | 0.145 | 0.004370969 | 25 | Trim47 |
| Ifi2039 | 2.16E-07 | 0.343100079 | 0.767 | 0.369 | 0.004378049 | 25 | Ifi203 |

| | | | | | | | |
|---|---|---|---|---|---|---|---|
| Nrp27 | 2.29E-07 | 0.286276165 | 0.452 | 0.185 | 0.004626288 | 25 | Nrp2 |
| Creb3l25 | 3.13E-07 | 0.278066376 | 0.342 | 0.125 | 0.006346183 | 25 | Creb3l2 |
| Ly6e8 | 3.46E-07 | 0.468638986 | 0.973 | 0.621 | 0.007001581 | 25 | Ly6e |
| Peak111 | 3.55E-07 | 0.337974841 | 0.781 | 0.389 | 0.007184349 | 25 | Peak1 |
| Sox133 | 3.67E-07 | 0.286255153 | 0.274 | 0.091 | 0.007439658 | 25 | Sox13 |
| Rb19 | 4.33E-07 | 0.448532722 | 0.479 | 0.216 | 0.00876532 | 25 | Rb1 |
| Pde7b9 | 4.37E-07 | 0.274040222 | 0.548 | 0.244 | 0.008837438 | 25 | Pde7b |
| Tom1l26 | 5.62E-07 | 0.307145336 | 0.342 | 0.132 | 0.011378809 | 25 | Tom1l2 |
| Tcf7l16 | 8.65E-07 | 0.277175302 | 0.438 | 0.182 | 0.017519598 | 25 | Tcf7l1 |
| Ifit23 | 8.79E-07 | 0.397003429 | 0.315 | 0.12 | 0.01779296 | 25 | Ifit2 |
| Mcam6 | 9.08E-07 | 0.319354057 | 0.301 | 0.111 | 0.018387023 | 25 | Mcam |
| Atf71 | 9.64E-07 | 0.317277793 | 0.288 | 0.104 | 0.019526775 | 25 | Atf7 |
| Rbms18 | 1.00E-06 | 0.344611817 | 0.877 | 0.488 | 0.020317672 | 25 | Rbms1 |
| Midn4 | 1.22E-06 | 0.319060596 | 0.438 | 0.192 | 0.024611397 | 25 | Midn |
| Gbp26 | 1.47E-06 | 0.443216824 | 0.342 | 0.145 | 0.029848299 | 25 | Gbp2 |
| Irf16 | 1.67E-06 | 0.260925407 | 0.616 | 0.3 | 0.033820732 | 25 | Irf1 |
| Ifih14 | 2.00E-06 | 0.277036126 | 0.329 | 0.13 | 0.040510099 | 25 | Ifih1 |
| Gbp94 | 2.06E-06 | 0.392290647 | 0.356 | 0.143 | 0.041624474 | 25 | Gbp9 |
| Swap707 | 2.13E-06 | 0.277190537 | 0.534 | 0.248 | 0.043091254 | 25 | Swap70 |
| Sash19 | 2.17E-06 | 0.280340815 | 0.548 | 0.259 | 0.043961812 | 25 | Sash1 |
| Tra2a14 | 2.26E-06 | 0.31832436 | 0.575 | 0.276 | 0.045832109 | 25 | Tra2a |
| Kcnq12 | 2.35E-06 | 0.256980831 | 0.274 | 0.098 | 0.04764646 | 25 | Kcnq1 |
| Sh3glb16 | 2.43E-06 | 0.295081243 | 0.918 | 0.533 | 0.049104412 | 25 | Sh3glb1 |
| Etnk15 | 2.68E-06 | 0.264123523 | 0.493 | 0.227 | 0.054311619 | 25 | Etnk1 |
| Smad78 | 2.71E-06 | 0.278203604 | 0.671 | 0.328 | 0.054940552 | 25 | Smad7 |
| Stard96 | 2.72E-06 | 0.289390376 | 0.37 | 0.153 | 0.055130755 | 25 | Stard9 |
| Flnb4 | 3.13E-06 | 0.295799047 | 0.384 | 0.163 | 0.06334892 | 25 | Flnb |
| Ahnak11 | 3.13E-06 | 0.305762346 | 0.863 | 0.473 | 0.063427932 | 25 | Ahnak |
| Nr3c26 | 3.20E-06 | 0.25415804 | 0.315 | 0.123 | 0.064732931 | 25 | Nr3c2 |
| Rhoc8 | 3.35E-06 | 0.29872231 | 0.589 | 0.298 | 0.067809278 | 25 | Rhoc |
| Gnai27 | 3.53E-06 | 0.391686654 | 0.945 | 0.576 | 0.071370375 | 25 | Gnai2 |
| Ppp1r27 | 4.14E-06 | 0.250641438 | 0.74 | 0.395 | 0.083821208 | 25 | Ppp1r2 |
| Ifit34 | 4.56E-06 | 0.413941195 | 0.315 | 0.124 | 0.092386937 | 25 | Ifit3 |
| Serinc38 | 4.70E-06 | 0.301943193 | 0.904 | 0.488 | 0.095171058 | 25 | Serinc3 |
| Mprip9 | 4.86E-06 | 0.252531768 | 0.589 | 0.295 | 0.09830388 | 25 | Mprip |
| Srgn13 | 5.27E-06 | 0.31215065 | 0.973 | 0.546 | 0.106656392 | 25 | Srgn |
| 4930402H24Rik6 | 5.43E-06 | 0.279161396 | 0.479 | 0.223 | 0.110010741 | 25 | 4930402H24Rik |
| Otud42 | 5.97E-06 | 0.253575557 | 0.315 | 0.127 | 0.120772553 | 25 | Otud4 |
| Adgrg17 | 6.95E-06 | 0.311046618 | 0.342 | 0.141 | 0.14075658 | 25 | Adgrg1 |
| Lmo27 | 8.10E-06 | 0.314986615 | 0.452 | 0.209 | 0.164028096 | 25 | Lmo2 |
| Junb10 | 1.16E-05 | 0.312513828 | 0.74 | 0.394 | 0.234463465 | 25 | Junb |
| Spred17 | 1.32E-05 | 0.272071703 | 0.521 | 0.254 | 0.266796769 | 25 | Spred1 |
| Cd8110 | 1.62E-05 | 0.298807594 | 0.959 | 0.527 | 0.327560758 | 25 | Cd81 |

| | | | | | | | |
|---|---|---|---|---|---|---|---|
| Cd479 | 1.75E-05 | 0.263584144 | 0.904 | 0.511 | 0.353767017 | 25 | Cd47 |
| Serinc110 | 1.82E-05 | 0.34448974 | 0.493 | 0.251 | 0.369420378 | 25 | Serinc1 |
| Parvb5 | 2.15E-05 | 0.26698748 | 0.301 | 0.125 | 0.435069712 | 25 | Parvb |
| Rnf167 | 2.17E-05 | 0.266493695 | 0.288 | 0.12 | 0.440238031 | 25 | Rnf167 |
| Ckap42 | 2.39E-05 | 0.325155867 | 0.26 | 0.102 | 0.483497304 | 25 | Ckap4 |
| Psmb99 | 2.40E-05 | 0.292884912 | 0.726 | 0.407 | 0.486241815 | 25 | Psmb9 |
| Mndal9 | 2.48E-05 | 0.290653864 | 0.74 | 0.407 | 0.50198787 | 25 | Mndal |
| Itm2b10 | 2.94E-05 | 0.409207056 | 0.986 | 0.726 | 0.595996937 | 25 | Itm2b |
| Man1a25 | 2.96E-05 | 0.263072292 | 0.466 | 0.228 | 0.598539231 | 25 | Man1a2 |
| Laptm4a10 | 3.75E-05 | 0.26887246 | 0.822 | 0.486 | 0.759200895 | 25 | Laptm4a |
| Exoc6b10 | 3.95E-05 | 0.277079504 | 0.493 | 0.242 | 0.799727204 | 25 | Exoc6b |
| Slc10a74 | 4.34E-05 | 0.339615716 | 0.329 | 0.149 | 0.879220042 | 25 | Slc10a7 |
| Lfng3 | 8.97E-05 | 0.293500127 | 0.301 | 0.138 | 1 | 25 | Lfng |
| Isg156 | 9.57E-05 | 0.281429947 | 0.438 | 0.231 | 1 | 25 | Isg15 |
| Bag32 | 0.000105567 | 0.278613276 | 0.274 | 0.115 | 1 | 25 | Bag3 |
| Evi56 | 0.000136348 | 0.341848651 | 0.301 | 0.142 | 1 | 25 | Evi5 |
| Egr15 | 0.000766301 | 0.27856625 | 0.342 | 0.177 | 1 | 25 | Egr1 |
| Cdk11b4 | 0.005746762 | 0.312710101 | 0.425 | 0.245 | 1 | 25 | Cdk11b |
| C31 | 0 | 5.07294757 | 1 | 0.02 | 0 | 26 | C3 |
| Pi16 | 0 | 5.001562468 | 1 | 0.003 | 0 | 26 | Pi16 |
| Igfbp61 | 0 | 4.731513301 | 1 | 0.016 | 0 | 26 | Igfbp6 |
| Mfap51 | 0 | 4.25644847 | 1 | 0.015 | 0 | 26 | Mfap5 |
| Opcml1 | 0 | 4.231190803 | 0.972 | 0.008 | 0 | 26 | Opcml |
| Dpt1 | 0 | 4.142536939 | 1 | 0.01 | 0 | 26 | Dpt |
| Fndc11 | 0 | 3.876116505 | 1 | 0.005 | 0 | 26 | Fndc1 |
| Ebf21 | 0 | 3.761954457 | 1 | 0.013 | 0 | 26 | Ebf2 |
| Smpd3 | 0 | 3.510384228 | 0.944 | 0.003 | 0 | 26 | Smpd3 |
| Cd2482 | 0 | 3.495919777 | 1 | 0.018 | 0 | 26 | Cd248 |
| Pla1a | 0 | 3.357309303 | 1 | 0.009 | 0 | 26 | Pla1a |
| Dpp41 | 0 | 3.231347362 | 1 | 0.013 | 0 | 26 | Dpp4 |
| Sdk13 | 0 | 3.140194858 | 0.972 | 0.023 | 0 | 26 | Sdk1 |
| Tnxb1 | 0 | 3.079068289 | 0.972 | 0.02 | 0 | 26 | Tnxb |
| Pcolce2 | 0 | 3.039017776 | 0.917 | 0.001 | 0 | 26 | Pcolce2 |
| Sox53 | 0 | 2.96949733 | 0.972 | 0.021 | 0 | 26 | Sox5 |
| Nova1 | 0 | 2.968243172 | 0.944 | 0.003 | 0 | 26 | Nova1 |
| Tmeff2 | 0 | 2.894684609 | 0.917 | 0.01 | 0 | 26 | Tmeff2 |
| Pcsk6 | 0 | 2.879581033 | 0.917 | 0.006 | 0 | 26 | Pcsk6 |
| Emilin21 | 0 | 2.752998856 | 0.944 | 0.011 | 0 | 26 | Emilin2 |
| Tmem1002 | 0 | 2.752416526 | 0.972 | 0.016 | 0 | 26 | Tmem100 |
| Adgrd1 | 0 | 2.738248756 | 0.972 | 0.002 | 0 | 26 | Adgrd1 |
| Loxl11 | 0 | 2.716721341 | 0.972 | 0.019 | 0 | 26 | Loxl1 |
| Efna51 | 0 | 2.631276578 | 0.833 | 0.01 | 0 | 26 | Efna5 |
| Sntg1 | 0 | 2.629331214 | 0.694 | 0.005 | 0 | 26 | Sntg1 |

| | | | | | | | |
|---|---|---|---|---|---|---|---|
| Pde3a1 | 0 | 2.515741187 | 0.806 | 0.013 | 0 | 26 | Pde3a |
| Ppp2r2b | 0 | 2.402605133 | 0.833 | 0.003 | 0 | 26 | Ppp2r2b |
| Adamts21 | 0 | 2.260837832 | 0.972 | 0.019 | 0 | 26 | Adamts2 |
| Medag1 | 0 | 2.161577805 | 0.833 | 0.012 | 0 | 26 | Medag |
| Slc4a4 | 0 | 2.155745751 | 0.833 | 0.007 | 0 | 26 | Slc4a4 |
| Tnfaip61 | 0 | 2.044465539 | 0.667 | 0.009 | 0 | 26 | Tnfaip6 |
| Sfrp2 | 0 | 1.928532965 | 0.806 | 0.004 | 0 | 26 | Sfrp2 |
| Adamts161 | 0 | 1.919757026 | 0.611 | 0 | 0 | 26 | Adamts16 |
| Scara31 | 0 | 1.913645323 | 0.972 | 0.014 | 0 | 26 | Scara3 |
| Aspn1 | 0 | 1.782455304 | 0.833 | 0.011 | 0 | 26 | Aspn |
| Dclk1 | 0 | 1.721198784 | 0.722 | 0.008 | 0 | 26 | Dclk1 |
| Dbn1 | 0 | 1.716029091 | 0.833 | 0.012 | 0 | 26 | Dbn1 |
| Vasn1 | 0 | 1.660736293 | 0.778 | 0.012 | 0 | 26 | Vasn |
| Fhl1 | 0 | 1.652443817 | 0.833 | 0.008 | 0 | 26 | Fhl1 |
| Itga111 | 0 | 1.621841131 | 0.861 | 0.004 | 0 | 26 | Itga11 |
| Ahnak21 | 0 | 1.617880841 | 0.806 | 0.008 | 0 | 26 | Ahnak2 |
| Akr1c18 | 0 | 1.60583441 | 0.611 | 0 | 0 | 26 | Akr1c18 |
| Olfml2b2 | 0 | 1.590817158 | 0.667 | 0.01 | 0 | 26 | Olfml2b |
| Nox4 | 0 | 1.557567738 | 0.528 | 0.002 | 0 | 26 | Nox4 |
| Gxylt21 | 0 | 1.529279251 | 0.75 | 0.013 | 0 | 26 | Gxylt2 |
| Aif1l1 | 0 | 1.523108475 | 0.694 | 0.009 | 0 | 26 | Aif1l |
| Rorb | 0 | 1.495559488 | 0.722 | 0.001 | 0 | 26 | Rorb |
| Mrgprg | 0 | 1.483318715 | 0.25 | 0 | 0 | 26 | Mrgprg |
| Thsd41 | 0 | 1.432202942 | 0.667 | 0.008 | 0 | 26 | Thsd4 |
| Galnt162 | 0 | 1.421848567 | 0.694 | 0.005 | 0 | 26 | Galnt16 |
| Lrrc17 | 0 | 1.413543881 | 0.806 | 0.007 | 0 | 26 | Lrrc17 |
| Pdpn1 | 0 | 1.398950973 | 0.722 | 0.011 | 0 | 26 | Pdpn |
| Ildr2 | 0 | 1.398579936 | 0.583 | 0.001 | 0 | 26 | Ildr2 |
| Dnm11 | 0 | 1.391616506 | 0.778 | 0.011 | 0 | 26 | Dnm1 |
| D630033O11Rik | 0 | 1.371307913 | 0.667 | 0.003 | 0 | 26 | D630033O11Rik |
| Osr11 | 0 | 1.334216893 | 0.639 | 0.008 | 0 | 26 | Osr1 |
| Galnt172 | 0 | 1.323268188 | 0.667 | 0.007 | 0 | 26 | Galnt17 |
| Adamtsl4 | 0 | 1.291640705 | 0.583 | 0.005 | 0 | 26 | Adamtsl4 |
| Sema3d2 | 0 | 1.263429725 | 0.611 | 0.005 | 0 | 26 | Sema3d |
| Gm46323 | 0 | 1.191764956 | 0.528 | 0 | 0 | 26 | Gm46323 |
| Pamr1 | 0 | 1.182258851 | 0.639 | 0.001 | 0 | 26 | Pamr1 |
| Lrrn4cl | 0 | 1.152538619 | 0.667 | 0.001 | 0 | 26 | Lrrn4cl |
| Chrdl1 | 0 | 1.132866686 | 0.5 | 0.001 | 0 | 26 | Chrdl1 |
| Arsi | 0 | 1.086078302 | 0.694 | 0.002 | 0 | 26 | Arsi |
| Sod3 | 0 | 1.068764363 | 0.611 | 0.007 | 0 | 26 | Sod3 |
| Vldlr | 0 | 1.050831304 | 0.639 | 0.005 | 0 | 26 | Vldlr |
| Thbs3 | 0 | 1.045820692 | 0.583 | 0.006 | 0 | 26 | Thbs3 |
| Vegfd | 0 | 1.034275668 | 0.639 | 0.005 | 0 | 26 | Vegfd |

| | | | | | | | |
|---|---|---|---|---|---|---|---|
| Adam33 | 0 | 1.033413423 | 0.5 | 0.005 | 0 | 26 | Adam33 |
| Cdh9 | 0 | 0.990415595 | 0.361 | 0.001 | 0 | 26 | Cdh9 |
| Pi15 | 0 | 0.967688229 | 0.5 | 0.002 | 0 | 26 | Pi15 |
| Duox1 | 0 | 0.960176808 | 0.556 | 0 | 0 | 26 | Duox1 |
| Dhrs9 | 0 | 0.844053029 | 0.333 | 0.001 | 0 | 26 | Dhrs9 |
| Gabra3 | 0 | 0.790293019 | 0.472 | 0.004 | 0 | 26 | Gabra3 |
| Wnt2 | 0 | 0.766724454 | 0.417 | 0.002 | 0 | 26 | Wnt2 |
| Has2 | 0 | 0.749014163 | 0.389 | 0.002 | 0 | 26 | Has2 |
| Fgf18 | 0 | 0.73993312 | 0.389 | 0.002 | 0 | 26 | Fgf18 |
| Ano3 | 0 | 0.737711204 | 0.333 | 0 | 0 | 26 | Ano3 |
| Ackr2 | 0 | 0.737613525 | 0.417 | 0.001 | 0 | 26 | Ackr2 |
| Lrfn2 | 0 | 0.723409037 | 0.361 | 0.002 | 0 | 26 | Lrfn2 |
| Prkg2 | 0 | 0.7079879 | 0.306 | 0 | 0 | 26 | Prkg2 |
| Adgrg2 | 0 | 0.7028775 | 0.389 | 0.001 | 0 | 26 | Adgrg2 |
| Gm31378 | 0 | 0.656410265 | 0.444 | 0.001 | 0 | 26 | Gm31378 |
| Adcy2 | 0 | 0.624095912 | 0.306 | 0.001 | 0 | 26 | Adcy2 |
| Gm26523 | 0 | 0.618706325 | 0.389 | 0.003 | 0 | 26 | Gm26523 |
| Duoxa1 | 0 | 0.615101746 | 0.333 | 0 | 0 | 26 | Duoxa1 |
| Folh1 | 0 | 0.591477032 | 0.306 | 0.001 | 0 | 26 | Folh1 |
| Tmem132c | 0 | 0.565250264 | 0.25 | 0.001 | 0 | 26 | Tmem132c |
| Adra1b | 0 | 0.525139396 | 0.333 | 0.002 | 0 | 26 | Adra1b |
| Nr1h4 | 0 | 0.509913053 | 0.25 | 0 | 0 | 26 | Nr1h4 |
| Sfrp4 | 0 | 0.505210787 | 0.306 | 0 | 0 | 26 | Sfrp4 |
| Bdh2 | 0 | 0.475832115 | 0.306 | 0.001 | 0 | 26 | Bdh2 |
| Plac9b | 0 | 0.331174941 | 0.25 | 0.001 | 0 | 26 | Plac9b |
| Vgll31 | 4.61E-303 | 1.290685371 | 0.611 | 0.009 | 9.33E-299 | 26 | Vgll3 |
| Adamtsl31 | 6.04E-302 | 2.352670505 | 0.806 | 0.015 | 1.22E-297 | 26 | Adamtsl3 |
| Antxr13 | 1.87E-301 | 2.691297313 | 0.944 | 0.022 | 3.78E-297 | 26 | Antxr1 |
| Gpc3 | 6.66E-299 | 1.615139331 | 0.611 | 0.009 | 1.35E-294 | 26 | Gpc3 |
| Efemp12 | 2.06E-298 | 3.410239717 | 1 | 0.025 | 4.17E-294 | 26 | Efemp1 |
| Rgs62 | 4.25E-298 | 2.427088234 | 0.639 | 0.01 | 8.61E-294 | 26 | Rgs6 |
| Wnt5a | 5.14E-297 | 0.666729574 | 0.417 | 0.004 | 1.04E-292 | 26 | Wnt5a |
| Agtr1a1 | 6.03E-296 | 0.992471494 | 0.611 | 0.009 | 1.22E-291 | 26 | Agtr1a |
| Flnc | 6.52E-294 | 1.009397663 | 0.528 | 0.007 | 1.32E-289 | 26 | Flnc |
| Esr1 | 7.00E-292 | 1.83571172 | 0.778 | 0.015 | 1.42E-287 | 26 | Esr1 |
| Col5a12 | 1.06E-290 | 2.562811491 | 0.944 | 0.023 | 2.15E-286 | 26 | Col5a1 |
| Prdm8 | 1.22E-287 | 0.612596635 | 0.278 | 0.002 | 2.47E-283 | 26 | Prdm8 |
| Ogn1 | 1.47E-287 | 3.586768108 | 1 | 0.026 | 2.97E-283 | 26 | Ogn |
| Mmp231 | 3.56E-287 | 0.949373226 | 0.583 | 0.008 | 7.21E-283 | 26 | Mmp23 |
| Epb41l31 | 2.55E-285 | 1.590759575 | 0.75 | 0.014 | 5.16E-281 | 26 | Epb41l3 |
| Cd551 | 3.71E-285 | 3.084326267 | 1 | 0.026 | 7.50E-281 | 26 | Cd55 |
| Adora2b | 1.46E-281 | 0.462978061 | 0.278 | 0.002 | 2.95E-277 | 26 | Adora2b |
| Igdcc4 | 2.12E-279 | 0.643345668 | 0.361 | 0.003 | 4.30E-275 | 26 | Igdcc4 |

| | | | | | | | |
|---|---|---|---|---|---|---|---|
| C1qtnf12 | 1.53E-278 | 1.451221407 | 0.778 | 0.016 | 3.09E-274 | 26 | C1qtnf1 |
| Adamts51 | 3.07E-278 | 3.257335534 | 0.972 | 0.025 | 6.21E-274 | 26 | Adamts5 |
| Mmp21 | 5.30E-275 | 2.425954523 | 1 | 0.026 | 1.07E-270 | 26 | Mmp2 |
| Spon21 | 1.83E-272 | 1.390273614 | 0.722 | 0.014 | 3.71E-268 | 26 | Spon2 |
| Ddr22 | 1.18E-271 | 3.161647086 | 0.972 | 0.026 | 2.38E-267 | 26 | Ddr2 |
| Arhgap201 | 2.75E-268 | 1.711026629 | 0.75 | 0.015 | 5.56E-264 | 26 | Arhgap20 |
| Abca61 | 8.42E-266 | 0.851760012 | 0.583 | 0.009 | 1.71E-261 | 26 | Abca6 |
| Akr1b8 | 1.84E-264 | 1.160147911 | 0.5 | 0.007 | 3.73E-260 | 26 | Akr1b8 |
| Cadm31 | 4.44E-262 | 1.596172508 | 0.75 | 0.015 | 9.00E-258 | 26 | Cadm3 |
| Ifi2051 | 1.14E-259 | 1.712489658 | 0.722 | 0.014 | 2.30E-255 | 26 | Ifi205 |
| Bmp8a | 2.66E-259 | 0.399099991 | 0.25 | 0.001 | 5.38E-255 | 26 | Bmp8a |
| Aff2 | 1.25E-257 | 0.779692404 | 0.361 | 0.003 | 2.53E-253 | 26 | Aff2 |
| Lgi21 | 5.52E-257 | 1.149704949 | 0.583 | 0.009 | 1.12E-252 | 26 | Lgi2 |
| Scara51 | 5.88E-256 | 2.566260349 | 0.972 | 0.027 | 1.19E-251 | 26 | Scara5 |
| 1700003F12Rik | 8.43E-256 | 0.412943235 | 0.278 | 0.002 | 1.71E-251 | 26 | 1700003F12Rik |
| Nbl13 | 3.10E-254 | 1.669033519 | 0.889 | 0.023 | 6.27E-250 | 26 | Nbl1 |
| Lmx1a | 1.12E-253 | 0.615618114 | 0.25 | 0.002 | 2.27E-249 | 26 | Lmx1a |
| Ndrg4 | 3.17E-253 | 0.33117855 | 0.25 | 0.002 | 6.42E-249 | 26 | Ndrg4 |
| Kcnn3 | 2.52E-250 | 0.988894796 | 0.472 | 0.006 | 5.10E-246 | 26 | Kcnn3 |
| Steap31 | 3.34E-249 | 1.846597788 | 0.833 | 0.02 | 6.76E-245 | 26 | Steap3 |
| Clec3b1 | 2.87E-248 | 4.456187229 | 1 | 0.031 | 5.81E-244 | 26 | Clec3b |
| Lpar12 | 5.29E-246 | 1.930650712 | 0.889 | 0.024 | 1.07E-241 | 26 | Lpar1 |
| Krt19 | 6.90E-246 | 0.808786008 | 0.333 | 0.003 | 1.40E-241 | 26 | Krt19 |
| Plce12 | 2.86E-245 | 1.830102833 | 0.833 | 0.02 | 5.79E-241 | 26 | Plce1 |
| Il331 | 6.45E-244 | 2.607956129 | 0.861 | 0.022 | 1.31E-239 | 26 | Il33 |
| Egfr2 | 1.74E-242 | 2.819515402 | 0.944 | 0.027 | 3.51E-238 | 26 | Egfr |
| Dpep11 | 7.50E-242 | 2.25582526 | 0.972 | 0.028 | 1.52E-237 | 26 | Dpep1 |
| Col6a22 | 2.48E-241 | 2.53560057 | 0.972 | 0.029 | 5.01E-237 | 26 | Col6a2 |
| Radil | 7.09E-241 | 0.732980667 | 0.444 | 0.006 | 1.44E-236 | 26 | Radil |
| Sult5a11 | 1.21E-239 | 1.041042858 | 0.556 | 0.009 | 2.44E-235 | 26 | Sult5a1 |
| Glt8d2 | 2.40E-239 | 0.869627802 | 0.444 | 0.006 | 4.85E-235 | 26 | Glt8d2 |
| Gm15675 | 5.78E-235 | 0.654469282 | 0.333 | 0.003 | 1.17E-230 | 26 | Gm15675 |
| Tubb4a | 3.94E-234 | 0.42319257 | 0.278 | 0.002 | 7.98E-230 | 26 | Tubb4a |
| Fn12 | 8.52E-234 | 4.280054355 | 1 | 0.033 | 1.73E-229 | 26 | Fn1 |
| Fam171b1 | 3.24E-233 | 0.904161818 | 0.556 | 0.009 | 6.56E-229 | 26 | Fam171b |
| Ston21 | 3.36E-232 | 1.942480426 | 0.778 | 0.019 | 6.80E-228 | 26 | Ston2 |
| Efhd12 | 4.48E-231 | 1.960923739 | 0.833 | 0.022 | 9.08E-227 | 26 | Efhd1 |
| Col1a13 | 1.01E-230 | 4.277671791 | 1 | 0.033 | 2.05E-226 | 26 | Col1a1 |
| Col6a32 | 4.51E-230 | 2.420738184 | 0.972 | 0.03 | 9.13E-226 | 26 | Col6a3 |
| Islr1 | 1.86E-229 | 2.15573169 | 0.917 | 0.027 | 3.77E-225 | 26 | Islr |
| Il1r2 | 4.93E-229 | 0.917497542 | 0.389 | 0.005 | 9.97E-225 | 26 | Il1r2 |
| C1qtnf71 | 1.14E-228 | 0.978218577 | 0.556 | 0.009 | 2.30E-224 | 26 | C1qtnf7 |
| Gpc63 | 9.79E-228 | 3.686391819 | 0.972 | 0.031 | 1.98E-223 | 26 | Gpc6 |

| | | | | | | | |
|---|---|---|---|---|---|---|---|
| Lvrn1 | 7.46E-226 | 1.297542848 | 0.556 | 0.01 | 1.51E-221 | 26 | Lvrn |
| Ttyh21 | 2.90E-225 | 1.056553959 | 0.667 | 0.014 | 5.88E-221 | 26 | Ttyh2 |
| Htra31 | 3.91E-223 | 2.197753856 | 1 | 0.033 | 7.92E-219 | 26 | Htra3 |
| Meg32 | 7.85E-222 | 2.415944161 | 0.889 | 0.026 | 1.59E-217 | 26 | Meg3 |
| Adamtsl5 | 8.81E-222 | 0.852616678 | 0.472 | 0.007 | 1.78E-217 | 26 | Adamtsl5 |
| Ccl112 | 6.46E-217 | 2.494281036 | 0.972 | 0.031 | 1.31E-212 | 26 | Ccl11 |
| Serping13 | 3.08E-215 | 3.449487147 | 1 | 0.035 | 6.24E-211 | 26 | Serping1 |
| Tubb2b | 3.59E-213 | 0.63315614 | 0.472 | 0.007 | 7.27E-209 | 26 | Tubb2b |
| Col6a12 | 3.82E-213 | 2.838175669 | 0.972 | 0.033 | 7.72E-209 | 26 | Col6a1 |
| Pdgfra1 | 1.27E-211 | 2.131137434 | 0.944 | 0.03 | 2.57E-207 | 26 | Pdgfra |
| Arhgap423 | 6.28E-210 | 1.647898244 | 0.861 | 0.026 | 1.27E-205 | 26 | Arhgap42 |
| Bnc22 | 7.38E-210 | 1.799828976 | 0.722 | 0.018 | 1.49E-205 | 26 | Bnc2 |
| Cpxm11 | 9.49E-210 | 1.680539518 | 0.833 | 0.024 | 1.92E-205 | 26 | Cpxm1 |
| Sema3e1 | 2.19E-207 | 0.673433962 | 0.333 | 0.004 | 4.44E-203 | 26 | Sema3e |
| Prrx12 | 4.53E-207 | 2.613380144 | 0.944 | 0.032 | 9.17E-203 | 26 | Prrx1 |
| Kirrel4 | 1.12E-206 | 1.333611173 | 0.778 | 0.021 | 2.27E-202 | 26 | Kirrel |
| Lum1 | 2.60E-205 | 2.174722614 | 0.944 | 0.03 | 5.26E-201 | 26 | Lum |
| CT025619.13 | 5.84E-202 | 1.527591569 | 0.694 | 0.017 | 1.18E-197 | 26 | CT025619.1 |
| Tppp32 | 1.27E-201 | 1.919201448 | 0.778 | 0.022 | 2.56E-197 | 26 | Tppp3 |
| Col3a13 | 1.19E-200 | 4.835536329 | 1 | 0.039 | 2.41E-196 | 26 | Col3a1 |
| C1s12 | 1.95E-200 | 2.160086495 | 0.972 | 0.034 | 3.95E-196 | 26 | C1s1 |
| 1700019D03Rik | 3.75E-200 | 1.090920718 | 0.556 | 0.011 | 7.59E-196 | 26 | 1700019D03Rik |
| Rarres23 | 2.78E-198 | 2.650863797 | 1 | 0.037 | 5.62E-194 | 26 | Rarres2 |
| Zfhx42 | 5.15E-197 | 1.681877801 | 0.778 | 0.022 | 1.04E-192 | 26 | Zfhx4 |
| Osr2 | 5.28E-197 | 0.503732276 | 0.306 | 0.003 | 1.07E-192 | 26 | Osr2 |
| Wnt11 | 2.07E-196 | 0.448691673 | 0.25 | 0.002 | 4.19E-192 | 26 | Wnt11 |
| Anpep1 | 3.76E-195 | 1.381109698 | 0.806 | 0.024 | 7.62E-191 | 26 | Anpep |
| Igfbp53 | 3.93E-195 | 5.146972334 | 0.944 | 0.035 | 7.96E-191 | 26 | Igfbp5 |
| Col1a23 | 4.34E-194 | 4.297886687 | 1 | 0.039 | 8.79E-190 | 26 | Col1a2 |
| Col16a12 | 3.95E-193 | 0.931732711 | 0.583 | 0.013 | 7.99E-189 | 26 | Col16a1 |
| Postn3 | 1.03E-191 | 1.748326421 | 0.528 | 0.01 | 2.08E-187 | 26 | Postn |
| Ptgis | 1.38E-188 | 1.248918884 | 0.667 | 0.017 | 2.80E-184 | 26 | Ptgis |
| Abi3bp3 | 4.34E-186 | 2.125064936 | 0.917 | 0.033 | 8.78E-182 | 26 | Abi3bp |
| Col5a32 | 3.42E-184 | 2.216329018 | 0.889 | 0.031 | 6.92E-180 | 26 | Col5a3 |
| Lama23 | 9.67E-182 | 2.292278759 | 0.972 | 0.037 | 1.96E-177 | 26 | Lama2 |
| Dact2 | 1.17E-180 | 1.900706961 | 0.75 | 0.022 | 2.37E-176 | 26 | Dact2 |
| Maob | 9.31E-178 | 0.500173146 | 0.333 | 0.004 | 1.89E-173 | 26 | Maob |
| Sema3b1 | 1.23E-177 | 0.60116158 | 0.306 | 0.004 | 2.49E-173 | 26 | Sema3b |
| Larp6 | 1.26E-177 | 0.635132839 | 0.361 | 0.005 | 2.55E-173 | 26 | Larp6 |
| Epdr1 | 7.15E-177 | 0.5697675 | 0.389 | 0.006 | 1.45E-172 | 26 | Epdr1 |
| C1ra1 | 1.98E-175 | 1.567243976 | 0.889 | 0.032 | 4.02E-171 | 26 | C1ra |
| Plac82 | 7.42E-171 | 1.507180578 | 0.75 | 0.024 | 1.50E-166 | 26 | Plac8 |
| Vcan2 | 1.05E-170 | 1.357219918 | 0.611 | 0.016 | 2.12E-166 | 26 | Vcan |

| | | | | | | | |
|---|---|---|---|---|---|---|---|
| Dact1 | 1.12E-170 | 0.778757526 | 0.389 | 0.006 | 2.27E-166 | 26 | Dact1 |
| Gm32618 | 6.31E-170 | 0.479074144 | 0.361 | 0.005 | 1.28E-165 | 26 | Gm32618 |
| Evc1 | 9.35E-170 | 0.663016894 | 0.5 | 0.01 | 1.89E-165 | 26 | Evc |
| Ecm12 | 2.58E-168 | 1.953641227 | 0.889 | 0.035 | 5.23E-164 | 26 | Ecm1 |
| Tmem178b | 3.90E-168 | 0.81443839 | 0.333 | 0.005 | 7.91E-164 | 26 | Tmem178b |
| Ndn1 | 2.03E-167 | 1.000556321 | 0.667 | 0.019 | 4.12E-163 | 26 | Ndn |
| Col5a24 | 4.93E-165 | 2.460544129 | 0.972 | 0.042 | 9.99E-161 | 26 | Col5a2 |
| Angptl23 | 1.07E-164 | 1.257807991 | 0.694 | 0.021 | 2.17E-160 | 26 | Angptl2 |
| Thbs21 | 2.37E-162 | 1.153957172 | 0.556 | 0.014 | 4.79E-158 | 26 | Thbs2 |
| Lgi12 | 1.51E-161 | 0.723500205 | 0.361 | 0.006 | 3.05E-157 | 26 | Lgi1 |
| Fxyd13 | 4.00E-160 | 1.741543802 | 0.861 | 0.034 | 8.11E-156 | 26 | Fxyd1 |
| Aldh1a3 | 6.05E-160 | 1.179740771 | 0.583 | 0.015 | 1.23E-155 | 26 | Aldh1a3 |
| Aldh1a13 | 2.01E-159 | 1.794973121 | 0.861 | 0.034 | 4.07E-155 | 26 | Aldh1a1 |
| Pygl | 4.80E-159 | 1.231292301 | 0.667 | 0.02 | 9.72E-155 | 26 | Pygl |
| Sbsn | 2.34E-156 | 1.931680068 | 0.583 | 0.016 | 4.74E-152 | 26 | Sbsn |
| Ntrk21 | 1.70E-155 | 1.314814145 | 0.639 | 0.019 | 3.44E-151 | 26 | Ntrk2 |
| Dcn2 | 1.97E-154 | 3.582237054 | 1 | 0.047 | 3.99E-150 | 26 | Dcn |
| Bmp11 | 2.34E-154 | 1.719949752 | 0.972 | 0.045 | 4.74E-150 | 26 | Bmp1 |
| Gpr1 | 2.08E-151 | 0.452892879 | 0.25 | 0.003 | 4.22E-147 | 26 | Gpr1 |
| Rab38 | 2.51E-151 | 0.563790136 | 0.25 | 0.003 | 5.07E-147 | 26 | Rab38 |
| Tmem45a | 4.10E-151 | 0.554516784 | 0.306 | 0.004 | 8.29E-147 | 26 | Tmem45a |
| Eya4 | 1.43E-150 | 1.109368391 | 0.417 | 0.008 | 2.89E-146 | 26 | Eya4 |
| Gfpt21 | 7.64E-150 | 2.613550739 | 0.917 | 0.043 | 1.55E-145 | 26 | Gfpt2 |
| Il1rl2 | 6.28E-148 | 1.057516353 | 0.556 | 0.015 | 1.27E-143 | 26 | Il1rl2 |
| Irs1 | 9.61E-148 | 1.152418708 | 0.5 | 0.012 | 1.94E-143 | 26 | Irs1 |
| Oaf1 | 2.84E-147 | 1.481452493 | 0.722 | 0.026 | 5.75E-143 | 26 | Oaf |
| Serpinf11 | 2.77E-146 | 2.400226964 | 1 | 0.05 | 5.61E-142 | 26 | Serpinf1 |
| Naaladl21 | 4.05E-146 | 2.747133501 | 0.889 | 0.041 | 8.19E-142 | 26 | Naaladl2 |
| Magi23 | 4.14E-146 | 2.698096842 | 0.861 | 0.037 | 8.39E-142 | 26 | Magi2 |
| Gan | 2.75E-145 | 2.668528825 | 0.806 | 0.034 | 5.56E-141 | 26 | Gan |
| Gpx31 | 3.17E-143 | 1.256770739 | 0.611 | 0.019 | 6.42E-139 | 26 | Gpx3 |
| Dcbld22 | 1.02E-142 | 1.398025239 | 0.889 | 0.04 | 2.06E-138 | 26 | Dcbld2 |
| Sdc23 | 8.50E-142 | 1.880762999 | 0.778 | 0.031 | 1.72E-137 | 26 | Sdc2 |
| Twist12 | 7.93E-140 | 1.282481393 | 0.611 | 0.019 | 1.61E-135 | 26 | Twist1 |
| Loxl21 | 2.56E-133 | 1.121767763 | 0.639 | 0.022 | 5.18E-129 | 26 | Loxl2 |
| Cdon1 | 4.98E-133 | 1.454397211 | 0.611 | 0.021 | 1.01E-128 | 26 | Cdon |
| Pcdh72 | 2.34E-132 | 1.808944386 | 0.75 | 0.031 | 4.75E-128 | 26 | Pcdh7 |
| Plxna1 | 3.62E-132 | 0.904013742 | 0.556 | 0.017 | 7.33E-128 | 26 | Plxna1 |
| Mtcl1 | 9.61E-131 | 0.490928934 | 0.389 | 0.008 | 1.95E-126 | 26 | Mtcl1 |
| Lrp14 | 4.85E-130 | 2.202323908 | 1 | 0.056 | 9.82E-126 | 26 | Lrp1 |
| Loxl3 | 1.04E-129 | 0.664295657 | 0.417 | 0.01 | 2.11E-125 | 26 | Loxl3 |
| Myh101 | 1.77E-129 | 1.272247416 | 0.694 | 0.027 | 3.58E-125 | 26 | Myh10 |
| Col14a14 | 5.16E-129 | 2.91583803 | 0.972 | 0.057 | 1.05E-124 | 26 | Col14a1 |

| | | | | | | | |
|---|---|---|---|---|---|---|---|
| Adamts123 | 1.62E-128 | 1.296052589 | 0.861 | 0.04 | 3.29E-124 | 26 | Adamts12 |
| Tmem1581 | 4.26E-125 | 1.501397358 | 0.667 | 0.026 | 8.62E-121 | 26 | Tmem158 |
| Creb54 | 8.05E-125 | 3.771261249 | 0.972 | 0.061 | 1.63E-120 | 26 | Creb5 |
| Pcdh91 | 1.70E-124 | 2.319045146 | 0.389 | 0.009 | 3.44E-120 | 26 | Pcdh9 |
| Flrt23 | 3.75E-124 | 2.206060401 | 0.861 | 0.044 | 7.58E-120 | 26 | Flrt2 |
| Gprc5b1 | 1.15E-123 | 0.775660999 | 0.583 | 0.02 | 2.33E-119 | 26 | Gprc5b |
| Omd | 3.32E-123 | 0.58633312 | 0.306 | 0.005 | 6.72E-119 | 26 | Omd |
| Dock75 | 9.69E-123 | 1.678727314 | 0.889 | 0.048 | 1.96E-118 | 26 | Dock7 |
| Entpd22 | 1.27E-122 | 1.199509316 | 0.722 | 0.03 | 2.57E-118 | 26 | Entpd2 |
| Adamtsl11 | 7.13E-122 | 2.315917152 | 0.889 | 0.048 | 1.44E-117 | 26 | Adamtsl1 |
| Mmp143 | 1.25E-121 | 2.257884458 | 0.972 | 0.059 | 2.52E-117 | 26 | Mmp14 |
| Fgfr13 | 1.28E-121 | 2.158144891 | 0.889 | 0.049 | 2.59E-117 | 26 | Fgfr1 |
| Sgce3 | 2.04E-121 | 1.414939025 | 0.861 | 0.044 | 4.13E-117 | 26 | Sgce |
| Fap2 | 2.13E-121 | 1.659478521 | 0.778 | 0.036 | 4.31E-117 | 26 | Fap |
| Fbn14 | 6.36E-121 | 4.44009607 | 0.972 | 0.064 | 1.29E-116 | 26 | Fbn1 |
| Fgf21 | 9.13E-121 | 0.883266575 | 0.472 | 0.013 | 1.85E-116 | 26 | Fgf2 |
| Rbms34 | 1.67E-119 | 2.248242607 | 0.944 | 0.053 | 3.38E-115 | 26 | Rbms3 |
| Il181 | 4.83E-117 | 0.979768723 | 0.611 | 0.023 | 9.78E-113 | 26 | Il18 |
| Cmah3 | 1.07E-116 | 2.419968541 | 1 | 0.067 | 2.17E-112 | 26 | Cmah |
| Gli31 | 3.03E-116 | 1.324321265 | 0.611 | 0.023 | 6.14E-112 | 26 | Gli3 |
| Fst3 | 3.93E-116 | 1.164250346 | 0.667 | 0.027 | 7.95E-112 | 26 | Fst |
| Zbtb16 | 4.41E-116 | 1.459967702 | 0.833 | 0.044 | 8.93E-112 | 26 | Zbtb16 |
| Sh3pxd2b3 | 5.10E-116 | 1.562397166 | 0.806 | 0.041 | 1.03E-111 | 26 | Sh3pxd2b |
| Pard35 | 9.81E-116 | 1.497096576 | 0.861 | 0.046 | 1.99E-111 | 26 | Pard3 |
| Rab30 | 1.33E-114 | 1.074076222 | 0.444 | 0.012 | 2.70E-110 | 26 | Rab30 |
| Clip4 | 2.56E-114 | 1.045201042 | 0.667 | 0.027 | 5.18E-110 | 26 | Clip4 |
| Klhl131 | 4.36E-112 | 1.34068818 | 0.778 | 0.039 | 8.83E-108 | 26 | Klhl13 |
| Lgi41 | 2.22E-110 | 0.611774513 | 0.389 | 0.01 | 4.49E-106 | 26 | Lgi4 |
| Srgap11 | 7.00E-110 | 1.785313015 | 0.944 | 0.059 | 1.42E-105 | 26 | Srgap1 |
| Ctsk1 | 2.16E-108 | 1.012558555 | 0.556 | 0.02 | 4.38E-104 | 26 | Ctsk |
| Ugdh1 | 5.47E-108 | 2.07466246 | 0.889 | 0.055 | 1.11E-103 | 26 | Ugdh |
| Robo11 | 5.64E-107 | 1.798517585 | 0.611 | 0.025 | 1.14E-102 | 26 | Robo1 |
| Acss31 | 2.58E-106 | 1.049804916 | 0.472 | 0.015 | 5.23E-102 | 26 | Acss3 |
| Scd12 | 7.54E-106 | 0.878665826 | 0.611 | 0.025 | 1.53E-101 | 26 | Scd1 |
| Nav31 | 1.15E-105 | 3.637742081 | 0.972 | 0.072 | 2.32E-101 | 26 | Nav3 |
| Dennd2a2 | 2.84E-105 | 0.910582896 | 0.583 | 0.023 | 5.75E-101 | 26 | Dennd2a |
| Fgf72 | 3.43E-105 | 0.657976332 | 0.444 | 0.013 | 6.95E-101 | 26 | Fgf7 |
| Fkbp141 | 4.28E-104 | 0.747296943 | 0.5 | 0.017 | 8.66E-100 | 26 | Fkbp14 |
| Clmp2 | 4.56E-103 | 1.151551038 | 0.583 | 0.024 | 9.24E-99 | 26 | Clmp |
| Ccdc802 | 1.06E-102 | 1.298351009 | 0.667 | 0.031 | 2.15E-98 | 26 | Ccdc80 |
| Efemp21 | 1.97E-102 | 0.868304907 | 0.528 | 0.02 | 3.98E-98 | 26 | Efemp2 |
| Axl4 | 4.27E-102 | 2.627740101 | 1 | 0.078 | 8.64E-98 | 26 | Axl |
| Xpnpep2 | 5.80E-101 | 0.400819924 | 0.25 | 0.004 | 1.17E-96 | 26 | Xpnpep2 |

| | | | | | | | |
|---|---|---|---|---|---|---|---|
| Megf91 | 1.37E-100 | 1.421428327 | 0.722 | 0.037 | 2.77E-96 | 26 | Megf9 |
| Cpt1c | 1.63E-100 | 0.405581371 | 0.306 | 0.007 | 3.29E-96 | 26 | Cpt1c |
| Dlg4 | 1.31E-99 | 0.546879216 | 0.417 | 0.012 | 2.65E-95 | 26 | Dlg4 |
| Pcdh192 | 1.87E-99 | 1.060078104 | 0.611 | 0.027 | 3.79E-95 | 26 | Pcdh19 |
| Ptges1 | 2.32E-99 | 0.708343056 | 0.5 | 0.018 | 4.70E-95 | 26 | Ptges |
| Glis31 | 3.33E-99 | 0.868899011 | 0.472 | 0.016 | 6.75E-95 | 26 | Glis3 |
| Aebp12 | 1.04E-98 | 3.017497122 | 1 | 0.082 | 2.10E-94 | 26 | Aebp1 |
| Fbln53 | 5.52E-98 | 1.167516937 | 0.75 | 0.04 | 1.12E-93 | 26 | Fbln5 |
| Il4ra1 | 2.42E-96 | 1.953350739 | 0.889 | 0.063 | 4.89E-92 | 26 | Il4ra |
| Sulf13 | 3.43E-95 | 1.668420741 | 0.833 | 0.053 | 6.94E-91 | 26 | Sulf1 |
| Plat2 | 2.50E-94 | 1.677083532 | 0.861 | 0.058 | 5.05E-90 | 26 | Plat |
| Zfp385a1 | 4.54E-94 | 1.123184408 | 0.556 | 0.024 | 9.19E-90 | 26 | Zfp385a |
| Ntn44 | 3.66E-93 | 1.454180028 | 0.667 | 0.035 | 7.42E-89 | 26 | Ntn4 |
| Slit33 | 1.08E-92 | 3.151579683 | 0.972 | 0.082 | 2.18E-88 | 26 | Slit3 |
| Htra15 | 6.21E-92 | 1.697889589 | 0.778 | 0.046 | 1.26E-87 | 26 | Htra1 |
| Scn7a2 | 6.23E-92 | 1.011700675 | 0.611 | 0.029 | 1.26E-87 | 26 | Scn7a |
| Gata6 | 7.41E-92 | 0.501946543 | 0.306 | 0.007 | 1.50E-87 | 26 | Gata6 |
| Rgma1 | 8.18E-92 | 0.792440367 | 0.472 | 0.017 | 1.66E-87 | 26 | Rgma |
| Sestd1 | 1.63E-91 | 1.230353182 | 0.75 | 0.044 | 3.29E-87 | 26 | Sestd1 |
| Zfpm23 | 1.14E-90 | 2.95882817 | 0.944 | 0.079 | 2.30E-86 | 26 | Zfpm2 |
| Hpse | 2.63E-90 | 0.554623804 | 0.333 | 0.009 | 5.33E-86 | 26 | Hpse |
| Pla2r12 | 1.45E-88 | 1.034846858 | 0.472 | 0.018 | 2.94E-84 | 26 | Pla2r1 |
| Antxr25 | 3.85E-88 | 2.507339128 | 1 | 0.091 | 7.79E-84 | 26 | Antxr2 |
| Rnase43 | 2.04E-87 | 2.83469796 | 0.972 | 0.086 | 4.14E-83 | 26 | Rnase4 |
| Mrc21 | 1.39E-86 | 0.870168067 | 0.583 | 0.028 | 2.81E-82 | 26 | Mrc2 |
| Ccdc8 | 1.67E-86 | 0.437570489 | 0.278 | 0.006 | 3.39E-82 | 26 | Ccdc8 |
| Bmp2 | 2.19E-86 | 0.398170819 | 0.278 | 0.006 | 4.44E-82 | 26 | Bmp2 |
| Pdgfc | 1.15E-85 | 0.319515682 | 0.278 | 0.006 | 2.32E-81 | 26 | Pdgfc |
| Fez11 | 1.12E-84 | 0.584666123 | 0.361 | 0.011 | 2.27E-80 | 26 | Fez1 |
| Olfr10331 | 1.93E-84 | 0.936196988 | 0.361 | 0.011 | 3.91E-80 | 26 | Olfr1033 |
| Dapk13 | 2.37E-84 | 1.734047139 | 0.861 | 0.064 | 4.79E-80 | 26 | Dapk1 |
| Fstl16 | 8.65E-84 | 3.573524285 | 1 | 0.103 | 1.75E-79 | 26 | Fstl1 |
| Ptpn133 | 1.50E-83 | 1.019704821 | 0.611 | 0.032 | 3.05E-79 | 26 | Ptpn13 |
| Fgf101 | 2.51E-83 | 1.184508975 | 0.611 | 0.031 | 5.09E-79 | 26 | Fgf10 |
| Plxdc26 | 5.66E-83 | 2.929515891 | 1 | 0.094 | 1.14E-78 | 26 | Plxdc2 |
| Vit | 3.22E-82 | 0.818454432 | 0.306 | 0.008 | 6.53E-78 | 26 | Vit |
| Arntl21 | 4.07E-82 | 0.455752272 | 0.278 | 0.007 | 8.23E-78 | 26 | Arntl2 |
| Dpysl32 | 5.19E-81 | 1.344448481 | 0.889 | 0.067 | 1.05E-76 | 26 | Dpysl3 |
| Mxra81 | 9.87E-81 | 1.178077049 | 0.694 | 0.042 | 2.00E-76 | 26 | Mxra8 |
| Atf51 | 1.65E-79 | 1.298618717 | 0.722 | 0.047 | 3.34E-75 | 26 | Atf5 |
| Pdgfrb2 | 4.89E-78 | 0.751182851 | 0.556 | 0.028 | 9.91E-74 | 26 | Pdgfrb |
| Abca8a2 | 8.53E-78 | 1.064607358 | 0.611 | 0.033 | 1.73E-73 | 26 | Abca8a |
| Abca5 | 1.02E-77 | 0.679990741 | 0.389 | 0.014 | 2.07E-73 | 26 | Abca5 |

| | | | | | | | |
|---|---|---|---|---|---|---|---|
| Adamts151 | 1.35E-77 | 0.588063206 | 0.361 | 0.012 | 2.73E-73 | 26 | Adamts15 |
| Zfp651 | 9.07E-76 | 0.489536227 | 0.361 | 0.012 | 1.84E-71 | 26 | Zfp651 |
| Gsn2 | 1.88E-75 | 4.145592975 | 1 | 0.108 | 3.81E-71 | 26 | Gsn |
| Rcn33 | 3.00E-74 | 1.343927448 | 0.75 | 0.054 | 6.07E-70 | 26 | Rcn3 |
| Igfbp44 | 4.84E-74 | 2.447743457 | 0.972 | 0.098 | 9.81E-70 | 26 | Igfbp4 |
| Nhsl11 | 4.88E-74 | 1.694407951 | 0.889 | 0.078 | 9.88E-70 | 26 | Nhsl1 |
| Twsg12 | 4.93E-74 | 1.131126175 | 0.694 | 0.046 | 9.97E-70 | 26 | Twsg1 |
| Snx331 | 8.42E-74 | 0.668329563 | 0.444 | 0.019 | 1.70E-69 | 26 | Snx33 |
| Igsf101 | 2.60E-73 | 0.642051543 | 0.472 | 0.021 | 5.27E-69 | 26 | Igsf10 |
| Spsb1 | 4.17E-73 | 1.063666654 | 0.583 | 0.034 | 8.45E-69 | 26 | Spsb1 |
| Zfp5213 | 4.73E-73 | 1.720115675 | 0.861 | 0.075 | 9.58E-69 | 26 | Zfp521 |
| Clip3 | 9.05E-73 | 0.395805188 | 0.25 | 0.006 | 1.83E-68 | 26 | Clip3 |
| Ubtd21 | 1.05E-72 | 1.030775977 | 0.639 | 0.04 | 2.13E-68 | 26 | Ubtd2 |
| Ntn1 | 1.38E-72 | 0.497613551 | 0.361 | 0.013 | 2.79E-68 | 26 | Ntn1 |
| Lhfp3 | 3.06E-72 | 1.128548232 | 0.639 | 0.039 | 6.19E-68 | 26 | Lhfp |
| Creb3l11 | 3.49E-72 | 1.178496394 | 0.611 | 0.037 | 7.06E-68 | 26 | Creb3l1 |
| Dsel1 | 1.11E-71 | 1.148510054 | 0.667 | 0.044 | 2.26E-67 | 26 | Dsel |
| Evc21 | 1.11E-71 | 0.397042135 | 0.333 | 0.011 | 2.26E-67 | 26 | Evc2 |
| Scrn1 | 1.36E-71 | 0.321020268 | 0.25 | 0.006 | 2.76E-67 | 26 | Scrn1 |
| Lpl4 | 6.27E-71 | 1.788007912 | 0.944 | 0.087 | 1.27E-66 | 26 | Lpl |
| Bicc12 | 7.00E-71 | 2.087963575 | 0.972 | 0.1 | 1.42E-66 | 26 | Bicc1 |
| Negr13 | 7.28E-71 | 1.573929986 | 0.667 | 0.045 | 1.47E-66 | 26 | Negr1 |
| Lrrc49 | 8.74E-71 | 0.839428774 | 0.611 | 0.037 | 1.77E-66 | 26 | Lrrc49 |
| Kdelr31 | 1.07E-70 | 0.679287767 | 0.417 | 0.018 | 2.17E-66 | 26 | Kdelr3 |
| Kcnt21 | 1.33E-70 | 1.01610605 | 0.389 | 0.016 | 2.70E-66 | 26 | Kcnt2 |
| Cachd11 | 1.53E-70 | 1.041456521 | 0.611 | 0.038 | 3.09E-66 | 26 | Cachd1 |
| Uchl1 | 2.73E-70 | 0.385848251 | 0.25 | 0.006 | 5.53E-66 | 26 | Uchl1 |
| Daglb2 | 6.33E-70 | 2.037732425 | 0.694 | 0.051 | 1.28E-65 | 26 | Daglb |
| Mmp191 | 4.52E-69 | 0.537967796 | 0.389 | 0.016 | 9.15E-65 | 26 | Mmp19 |
| Timp28 | 8.62E-69 | 4.198207006 | 1 | 0.13 | 1.74E-64 | 26 | Timp2 |
| Sirpa3 | 8.34E-66 | 1.072611699 | 0.694 | 0.051 | 1.69E-61 | 26 | Sirpa |
| Arhgap281 | 2.29E-65 | 0.780983585 | 0.528 | 0.03 | 4.64E-61 | 26 | Arhgap28 |
| Olfml11 | 3.44E-65 | 0.601544387 | 0.278 | 0.009 | 6.96E-61 | 26 | Olfml1 |
| Adam221 | 3.43E-64 | 0.578226419 | 0.361 | 0.015 | 6.94E-60 | 26 | Adam22 |
| Frmd63 | 1.44E-63 | 1.382112305 | 0.639 | 0.046 | 2.91E-59 | 26 | Frmd6 |
| Cacna2d13 | 1.47E-63 | 0.74509509 | 0.472 | 0.025 | 2.97E-59 | 26 | Cacna2d1 |
| Myof5 | 1.75E-63 | 1.36388292 | 0.722 | 0.06 | 3.54E-59 | 26 | Myof |
| Prkca10 | 2.50E-63 | 1.947299989 | 0.917 | 0.101 | 5.07E-59 | 26 | Prkca |
| Glce | 1.97E-62 | 1.440244464 | 0.806 | 0.073 | 3.99E-58 | 26 | Glce |
| Reck2 | 2.11E-62 | 0.745412127 | 0.556 | 0.035 | 4.28E-58 | 26 | Reck |
| Prelp2 | 1.17E-61 | 0.887699093 | 0.472 | 0.026 | 2.36E-57 | 26 | Prelp |
| Dzip1 | 1.53E-61 | 0.510915801 | 0.306 | 0.011 | 3.09E-57 | 26 | Dzip1 |
| Psd34 | 2.65E-61 | 2.371825405 | 0.944 | 0.118 | 5.37E-57 | 26 | Psd3 |

| | | | | | | | |
|---|---|---|---|---|---|---|---|
| Glipr25 | 5.39E-61 | 0.997292981 | 0.611 | 0.043 | 1.09E-56 | 26 | Glipr2 |
| Nipal2 | 6.40E-61 | 0.581065821 | 0.306 | 0.011 | 1.30E-56 | 26 | Nipal2 |
| Chst1 | 5.70E-60 | 0.599156426 | 0.278 | 0.009 | 1.15E-55 | 26 | Chst1 |
| Mustn11 | 9.39E-60 | 0.839011256 | 0.5 | 0.029 | 1.90E-55 | 26 | Mustn1 |
| Txndc51 | 1.61E-59 | 1.236844953 | 0.833 | 0.082 | 3.27E-55 | 26 | Txndc5 |
| Epha31 | 2.27E-59 | 0.704942682 | 0.389 | 0.018 | 4.60E-55 | 26 | Epha3 |
| Maf4 | 2.39E-59 | 1.89745354 | 0.972 | 0.119 | 4.84E-55 | 26 | Maf |
| B3gnt91 | 3.11E-59 | 0.527308206 | 0.306 | 0.011 | 6.30E-55 | 26 | B3gnt9 |
| Gsc1 | 3.76E-59 | 0.508969752 | 0.25 | 0.008 | 7.62E-55 | 26 | Gsc |
| Arhgap63 | 7.44E-59 | 1.785565927 | 0.694 | 0.058 | 1.51E-54 | 26 | Arhgap6 |
| Bmp71 | 9.05E-59 | 0.509569434 | 0.278 | 0.01 | 1.83E-54 | 26 | Bmp7 |
| Naaa1 | 1.29E-58 | 0.82217742 | 0.556 | 0.038 | 2.61E-54 | 26 | Naaa |
| Col6a61 | 3.01E-58 | 0.787840426 | 0.389 | 0.019 | 6.09E-54 | 26 | Col6a6 |
| Rab7b2 | 7.30E-58 | 0.867592392 | 0.5 | 0.031 | 1.48E-53 | 26 | Rab7b |
| Pard3b5 | 1.12E-57 | 1.854161988 | 0.889 | 0.101 | 2.26E-53 | 26 | Pard3b |
| Thy17 | 1.41E-57 | 1.499111323 | 0.694 | 0.061 | 2.86E-53 | 26 | Thy1 |
| Basp17 | 4.06E-57 | 1.762303411 | 0.944 | 0.105 | 8.22E-53 | 26 | Basp1 |
| Tmeff1 | 1.37E-56 | 0.373598988 | 0.25 | 0.008 | 2.77E-52 | 26 | Tmeff1 |
| Ntrk32 | 1.37E-56 | 0.800064822 | 0.333 | 0.014 | 2.78E-52 | 26 | Ntrk3 |
| Zc3h12b | 1.48E-56 | 0.821240191 | 0.417 | 0.022 | 3.00E-52 | 26 | Zc3h12b |
| Gpm6b2 | 1.71E-56 | 0.60993657 | 0.528 | 0.033 | 3.47E-52 | 26 | Gpm6b |
| Bicd1 | 2.76E-56 | 0.65347272 | 0.333 | 0.014 | 5.58E-52 | 26 | Bicd1 |
| Mid13 | 2.94E-56 | 1.736257523 | 0.806 | 0.085 | 5.95E-52 | 26 | Mid1 |
| Pid15 | 3.50E-56 | 1.552927942 | 0.75 | 0.07 | 7.08E-52 | 26 | Pid1 |
| Kdelc21 | 3.83E-56 | 0.904148551 | 0.667 | 0.054 | 7.76E-52 | 26 | Kdelc2 |
| Casc43 | 4.17E-56 | 0.775314351 | 0.528 | 0.034 | 8.44E-52 | 26 | Casc4 |
| Fzd21 | 6.16E-56 | 0.424198854 | 0.278 | 0.01 | 1.25E-51 | 26 | Fzd2 |
| Bmpr1a3 | 6.21E-56 | 0.91837536 | 0.556 | 0.039 | 1.26E-51 | 26 | Bmpr1a |
| Nnmt1 | 1.11E-55 | 0.447699158 | 0.306 | 0.012 | 2.25E-51 | 26 | Nnmt |
| Eda2 | 1.96E-55 | 1.034850038 | 0.5 | 0.032 | 3.96E-51 | 26 | Eda |
| Pcolce3 | 1.99E-55 | 1.022111803 | 0.611 | 0.045 | 4.04E-51 | 26 | Pcolce |
| Notch26 | 7.05E-55 | 1.508448834 | 0.778 | 0.081 | 1.43E-50 | 26 | Notch2 |
| Runx1t13 | 9.66E-55 | 0.991972411 | 0.583 | 0.043 | 1.96E-50 | 26 | Runx1t1 |
| Kank1 | 4.95E-54 | 0.648208107 | 0.389 | 0.02 | 1.00E-49 | 26 | Kank1 |
| Sh3d196 | 5.06E-54 | 1.454039269 | 0.917 | 0.108 | 1.02E-49 | 26 | Sh3d19 |
| Ace6 | 5.26E-54 | 1.868277636 | 0.944 | 0.12 | 1.06E-49 | 26 | Ace |
| Slc24a32 | 6.15E-54 | 0.935269713 | 0.583 | 0.044 | 1.25E-49 | 26 | Slc24a3 |
| Rhbdf11 | 7.90E-54 | 0.541848699 | 0.417 | 0.023 | 1.60E-49 | 26 | Rhbdf1 |
| Cd3024 | 8.33E-54 | 1.192828826 | 0.806 | 0.082 | 1.69E-49 | 26 | Cd302 |
| Sspn1 | 2.12E-53 | 0.846161754 | 0.472 | 0.03 | 4.30E-49 | 26 | Sspn |
| Ext110 | 4.69E-53 | 3.39030216 | 1 | 0.168 | 9.49E-49 | 26 | Ext1 |
| Gstt11 | 5.09E-53 | 0.685038823 | 0.528 | 0.036 | 1.03E-48 | 26 | Gstt1 |
| Tgfb24 | 6.53E-53 | 1.310006919 | 0.833 | 0.088 | 1.32E-48 | 26 | Tgfb2 |

| | | | | | | | |
|---|---|---|---|---|---|---|---|
| Lrrc8b1 | 6.83E-53 | 0.882850404 | 0.528 | 0.038 | 1.38E-48 | 26 | Lrrc8b |
| Pde1a2 | 7.48E-53 | 0.570054042 | 0.333 | 0.015 | 1.51E-48 | 26 | Pde1a |
| P3h32 | 1.46E-52 | 0.630067583 | 0.444 | 0.026 | 2.96E-48 | 26 | P3h3 |
| Ackr36 | 2.11E-52 | 2.800820183 | 0.972 | 0.153 | 4.26E-48 | 26 | Ackr3 |
| Ltbp13 | 3.27E-52 | 1.816592901 | 0.861 | 0.105 | 6.61E-48 | 26 | Ltbp1 |
| St3gal21 | 5.65E-52 | 0.872190743 | 0.667 | 0.058 | 1.14E-47 | 26 | St3gal2 |
| Nid17 | 7.18E-52 | 3.421200853 | 1 | 0.183 | 1.45E-47 | 26 | Nid1 |
| Il13ra12 | 1.15E-51 | 1.188469751 | 0.75 | 0.076 | 2.33E-47 | 26 | Il13ra1 |
| Rcan21 | 1.25E-51 | 1.361807299 | 0.583 | 0.046 | 2.54E-47 | 26 | Rcan2 |
| Meis12 | 1.46E-51 | 0.83711123 | 0.528 | 0.038 | 2.95E-47 | 26 | Meis1 |
| Snx21 | 1.64E-51 | 0.673572441 | 0.583 | 0.045 | 3.32E-47 | 26 | Snx21 |
| Sec24d2 | 2.47E-51 | 1.204584966 | 0.722 | 0.071 | 5.01E-47 | 26 | Sec24d |
| Pcsk51 | 3.07E-50 | 0.74234855 | 0.444 | 0.027 | 6.21E-46 | 26 | Pcsk5 |
| Limch17 | 8.19E-50 | 2.139880839 | 0.972 | 0.14 | 1.66E-45 | 26 | Limch1 |
| Qpct4 | 8.78E-50 | 1.306484614 | 0.75 | 0.08 | 1.78E-45 | 26 | Qpct |
| Fam102b3 | 1.05E-49 | 0.986746693 | 0.611 | 0.053 | 2.12E-45 | 26 | Fam102b |
| Ism1 | 1.53E-49 | 0.980175977 | 0.333 | 0.016 | 3.11E-45 | 26 | Ism1 |
| Gm140052 | 1.79E-49 | 1.022246086 | 0.5 | 0.036 | 3.63E-45 | 26 | Gm14005 |
| Sema3c4 | 2.44E-48 | 2.996192564 | 0.972 | 0.164 | 4.95E-44 | 26 | Sema3c |
| Itsn11 | 2.82E-48 | 1.183273794 | 0.722 | 0.076 | 5.72E-44 | 26 | Itsn1 |
| Pcdh182 | 3.41E-48 | 0.420800373 | 0.306 | 0.014 | 6.90E-44 | 26 | Pcdh18 |
| Lbp1 | 3.43E-48 | 0.732842553 | 0.361 | 0.019 | 6.94E-44 | 26 | Lbp |
| Nectin3 | 3.76E-48 | 1.165550665 | 0.639 | 0.059 | 7.62E-44 | 26 | Nectin3 |
| Coro2b | 4.64E-48 | 0.919165285 | 0.556 | 0.044 | 9.40E-44 | 26 | Coro2b |
| Abca8b2 | 8.96E-48 | 0.80317817 | 0.361 | 0.019 | 1.81E-43 | 26 | Abca8b |
| Zcchc242 | 9.40E-48 | 0.968664916 | 0.694 | 0.069 | 1.90E-43 | 26 | Zcchc24 |
| Tubb61 | 1.37E-47 | 1.104588326 | 0.583 | 0.051 | 2.77E-43 | 26 | Tubb6 |
| Zfp9 | 1.52E-47 | 0.516730865 | 0.361 | 0.019 | 3.09E-43 | 26 | Zfp9 |
| Man1a12 | 2.31E-47 | 2.697834859 | 0.944 | 0.168 | 4.67E-43 | 26 | Man1a |
| Prdm51 | 2.58E-47 | 0.708751229 | 0.417 | 0.026 | 5.22E-43 | 26 | Prdm5 |
| Slco3a17 | 2.61E-47 | 1.646038291 | 0.917 | 0.123 | 5.28E-43 | 26 | Slco3a1 |
| Lamc17 | 3.72E-47 | 2.078389869 | 0.944 | 0.149 | 7.53E-43 | 26 | Lamc1 |
| Scd23 | 4.78E-47 | 1.05674701 | 0.75 | 0.081 | 9.67E-43 | 26 | Scd2 |
| Ust8 | 8.14E-47 | 1.482149726 | 0.75 | 0.087 | 1.65E-42 | 26 | Ust |
| Ms4a4d1 | 9.10E-47 | 0.618674655 | 0.417 | 0.026 | 1.84E-42 | 26 | Ms4a4d |
| Zeb27 | 1.04E-46 | 2.246078121 | 0.944 | 0.144 | 2.11E-42 | 26 | Zeb2 |
| Lamb14 | 4.92E-46 | 1.626505494 | 0.944 | 0.137 | 9.96E-42 | 26 | Lamb1 |
| Sgcd2 | 5.92E-46 | 0.52314614 | 0.361 | 0.02 | 1.20E-41 | 26 | Sgcd |
| Mturn1 | 7.40E-46 | 0.693234365 | 0.472 | 0.034 | 1.50E-41 | 26 | Mturn |
| Celf211 | 7.58E-46 | 3.212783051 | 1 | 0.203 | 1.53E-41 | 26 | Celf2 |
| Zhx21 | 1.01E-45 | 1.024088929 | 0.75 | 0.082 | 2.04E-41 | 26 | Zhx2 |
| Ehbp13 | 1.81E-45 | 1.733046129 | 0.833 | 0.111 | 3.66E-41 | 26 | Ehbp1 |
| C1qtnf6 | 2.77E-45 | 0.651734268 | 0.417 | 0.027 | 5.61E-41 | 26 | C1qtnf6 |

| | | | | | | | |
|---|---|---|---|---|---|---|---|
| Lgals110 | 1.21E-44 | 2.124116807 | 0.917 | 0.136 | 2.45E-40 | 26 | Lgals1 |
| Procr | 4.19E-44 | 0.686834927 | 0.389 | 0.024 | 8.49E-40 | 26 | Procr |
| Arhgef282 | 4.45E-44 | 1.517586053 | 0.722 | 0.081 | 9.02E-40 | 26 | Arhgef28 |
| Mitf4 | 6.10E-44 | 1.309653491 | 0.694 | 0.076 | 1.23E-39 | 26 | Mitf |
| Ppp1r9a2 | 2.09E-43 | 0.72374004 | 0.472 | 0.036 | 4.23E-39 | 26 | Ppp1r9a |
| Snx183 | 3.33E-43 | 1.212761533 | 0.75 | 0.091 | 6.75E-39 | 26 | Snx18 |
| Naalad22 | 5.16E-43 | 0.6255878 | 0.361 | 0.021 | 1.05E-38 | 26 | Naalad2 |
| Scpep12 | 5.44E-43 | 0.737262161 | 0.583 | 0.054 | 1.10E-38 | 26 | Scpep1 |
| Nckap51 | 8.55E-43 | 1.070861157 | 0.556 | 0.05 | 1.73E-38 | 26 | Nckap5 |
| Clcn51 | 4.99E-42 | 1.028707631 | 0.556 | 0.052 | 1.01E-37 | 26 | Clcn5 |
| Vtn1 | 9.16E-42 | 0.44255171 | 0.25 | 0.011 | 1.86E-37 | 26 | Vtn |
| Hsd11b16 | 1.16E-41 | 1.362828823 | 0.75 | 0.093 | 2.35E-37 | 26 | Hsd11b1 |
| Edn11 | 1.23E-41 | 0.481664261 | 0.417 | 0.029 | 2.49E-37 | 26 | Edn1 |
| Wdr35 | 1.29E-41 | 0.325515664 | 0.25 | 0.011 | 2.62E-37 | 26 | Wdr35 |
| Atrnl18 | 2.62E-41 | 2.044594665 | 0.944 | 0.166 | 5.30E-37 | 26 | Atrnl1 |
| Prkab2 | 2.92E-41 | 0.361560029 | 0.306 | 0.016 | 5.90E-37 | 26 | Prkab2 |
| Lama49 | 3.75E-41 | 1.782806798 | 1 | 0.176 | 7.60E-37 | 26 | Lama4 |
| Kctd9 | 5.09E-41 | 0.842654309 | 0.472 | 0.038 | 1.03E-36 | 26 | Kctd9 |
| Casp121 | 5.75E-41 | 0.618141739 | 0.444 | 0.033 | 1.16E-36 | 26 | Casp12 |
| Snx243 | 1.33E-40 | 0.629581184 | 0.611 | 0.06 | 2.70E-36 | 26 | Snx24 |
| Pknox21 | 3.35E-40 | 0.425909723 | 0.278 | 0.014 | 6.78E-36 | 26 | Pknox2 |
| Jag12 | 9.86E-40 | 0.792240532 | 0.472 | 0.039 | 2.00E-35 | 26 | Jag1 |
| Plau2 | 1.08E-39 | 0.928706425 | 0.611 | 0.062 | 2.19E-35 | 26 | Plau |
| Pla2g4a3 | 2.51E-39 | 0.722584415 | 0.5 | 0.044 | 5.08E-35 | 26 | Pla2g4a |
| Gas78 | 3.05E-39 | 2.210984857 | 0.944 | 0.184 | 6.18E-35 | 26 | Gas7 |
| Arhgef253 | 4.39E-39 | 0.503834286 | 0.333 | 0.02 | 8.89E-35 | 26 | Arhgef25 |
| Lima17 | 4.72E-39 | 1.772874505 | 0.917 | 0.159 | 9.57E-35 | 26 | Lima1 |
| Nov | 5.17E-39 | 0.800018263 | 0.361 | 0.024 | 1.05E-34 | 26 | Nov |
| Lsp110 | 6.01E-39 | 1.710554408 | 0.889 | 0.14 | 1.22E-34 | 26 | Lsp1 |
| Samd44 | 9.17E-39 | 1.132046205 | 0.556 | 0.055 | 1.86E-34 | 26 | Samd4 |
| Mcu3 | 2.68E-38 | 1.307813268 | 0.75 | 0.102 | 5.43E-34 | 26 | Mcu |
| Il6ra2 | 3.01E-38 | 0.65928575 | 0.417 | 0.032 | 6.10E-34 | 26 | Il6ra |
| Cep1123 | 5.91E-38 | 0.783936347 | 0.583 | 0.059 | 1.20E-33 | 26 | Cep112 |
| Eln6 | 6.96E-38 | 1.973405547 | 0.833 | 0.137 | 1.41E-33 | 26 | Eln |
| Gpr1531 | 4.18E-37 | 0.310146874 | 0.278 | 0.015 | 8.47E-33 | 26 | Gpr153 |
| Colec121 | 5.56E-37 | 0.834110719 | 0.528 | 0.05 | 1.13E-32 | 26 | Colec12 |
| Smim13 | 6.04E-37 | 0.429236417 | 0.417 | 0.032 | 1.22E-32 | 26 | Smim1 |
| Rhoq3 | 6.46E-37 | 1.059693157 | 0.778 | 0.109 | 1.31E-32 | 26 | Rhoq |
| Snhg184 | 6.60E-37 | 0.62164963 | 0.472 | 0.041 | 1.34E-32 | 26 | Snhg18 |
| Hspb8 | 2.15E-36 | 0.432905263 | 0.333 | 0.022 | 4.34E-32 | 26 | Hspb8 |
| Srpx1 | 6.21E-36 | 0.749822461 | 0.361 | 0.025 | 1.26E-31 | 26 | Srpx |
| Wls6 | 8.07E-36 | 1.150957711 | 0.833 | 0.128 | 1.63E-31 | 26 | Wls |
| Chpf1 | 8.50E-36 | 0.496714062 | 0.389 | 0.029 | 1.72E-31 | 26 | Chpf |

| | | | | | | | |
|---|---|---|---|---|---|---|---|
| Cacnb4 | 9.49E-36 | 0.44110352 | 0.306 | 0.019 | 1.92E-31 | 26 | Cacnb4 |
| Mpdz5 | 1.23E-35 | 1.171673325 | 0.722 | 0.099 | 2.48E-31 | 26 | Mpdz |
| Cped18 | 3.75E-35 | 1.591171437 | 0.972 | 0.18 | 7.58E-31 | 26 | Cped1 |
| Tpst12 | 9.14E-35 | 0.702365398 | 0.528 | 0.053 | 1.85E-30 | 26 | Tpst1 |
| Gk51 | 9.46E-35 | 0.539866494 | 0.389 | 0.03 | 1.91E-30 | 26 | Gk5 |
| 1700029J07Rik | 1.70E-34 | 0.36754722 | 0.25 | 0.013 | 3.44E-30 | 26 | 1700029J07Rik |
| Wnt10b | 1.81E-34 | 0.493089923 | 0.25 | 0.013 | 3.66E-30 | 26 | Wnt10b |
| Tmem43 | 2.24E-34 | 0.772229694 | 0.444 | 0.04 | 4.53E-30 | 26 | Tmem43 |
| Rora7 | 3.65E-34 | 1.934234564 | 0.944 | 0.187 | 7.40E-30 | 26 | Rora |
| Tgfbr38 | 4.22E-34 | 1.555334145 | 0.917 | 0.175 | 8.54E-30 | 26 | Tgfbr3 |
| Metrnl6 | 5.46E-34 | 1.230623805 | 0.722 | 0.104 | 1.11E-29 | 26 | Metrnl |
| Hic12 | 6.15E-34 | 0.637754569 | 0.611 | 0.07 | 1.24E-29 | 26 | Hic1 |
| Morc4 | 6.84E-34 | 0.255437571 | 0.25 | 0.013 | 1.38E-29 | 26 | Morc4 |
| Pkd23 | 7.26E-34 | 1.07968303 | 0.75 | 0.108 | 1.47E-29 | 26 | Pkd2 |
| Ptgfrn1 | 1.08E-33 | 0.514279814 | 0.361 | 0.027 | 2.19E-29 | 26 | Ptgfrn |
| Pacs2 | 1.09E-33 | 0.673912314 | 0.556 | 0.061 | 2.20E-29 | 26 | Pacs2 |
| Ddhd12 | 1.49E-33 | 0.859214613 | 0.639 | 0.08 | 3.02E-29 | 26 | Ddhd1 |
| Slc1a55 | 2.63E-33 | 0.836361738 | 0.639 | 0.081 | 5.33E-29 | 26 | Slc1a5 |
| Fsd1l | 2.93E-33 | 0.598916746 | 0.389 | 0.031 | 5.93E-29 | 26 | Fsd1l |
| Sytl25 | 2.98E-33 | 0.818066614 | 0.528 | 0.057 | 6.02E-29 | 26 | Sytl2 |
| Prnp2 | 3.97E-33 | 0.762345433 | 0.611 | 0.074 | 8.04E-29 | 26 | Prnp |
| Wdpcp | 4.13E-33 | 0.682419357 | 0.444 | 0.041 | 8.37E-29 | 26 | Wdpcp |
| Tgfbr28 | 4.60E-33 | 2.578175472 | 1 | 0.313 | 9.31E-29 | 26 | Tgfbr2 |
| Kcnk5 | 4.81E-33 | 0.61568427 | 0.361 | 0.028 | 9.74E-29 | 26 | Kcnk5 |
| Uap1 | 5.83E-33 | 0.855449351 | 0.778 | 0.115 | 1.18E-28 | 26 | Uap1 |
| Cdk147 | 6.81E-33 | 1.412427395 | 0.833 | 0.143 | 1.38E-28 | 26 | Cdk14 |
| Mbp7 | 7.21E-33 | 0.743805025 | 0.583 | 0.068 | 1.46E-28 | 26 | Mbp |
| Gnpnat1 | 7.62E-33 | 0.716917456 | 0.583 | 0.068 | 1.54E-28 | 26 | Gnpnat1 |
| Neo11 | 8.09E-33 | 1.110291676 | 0.556 | 0.065 | 1.64E-28 | 26 | Neo1 |
| Cdr2l | 9.50E-33 | 0.587287038 | 0.361 | 0.028 | 1.92E-28 | 26 | Cdr2l |
| Fam129a10 | 1.24E-32 | 1.302511824 | 0.833 | 0.147 | 2.50E-28 | 26 | Fam129a |
| Serpinh110 | 1.28E-32 | 2.151830949 | 0.972 | 0.251 | 2.59E-28 | 26 | Serpinh1 |
| Anxa13 | 2.00E-32 | 2.033099322 | 0.972 | 0.228 | 4.05E-28 | 26 | Anxa1 |
| Gm162631 | 2.54E-32 | 0.551536935 | 0.25 | 0.014 | 5.14E-28 | 26 | Gm16263 |
| Rarb3 | 3.32E-32 | 0.856161154 | 0.528 | 0.057 | 6.72E-28 | 26 | Rarb |
| Plekha41 | 3.65E-32 | 0.293109007 | 0.25 | 0.014 | 7.40E-28 | 26 | Plekha4 |
| Gas13 | 3.96E-32 | 0.636521997 | 0.389 | 0.032 | 8.01E-28 | 26 | Gas1 |
| Dst10 | 4.15E-32 | 1.508771986 | 0.944 | 0.192 | 8.39E-28 | 26 | Dst |
| Marcks9 | 6.95E-32 | 2.82075432 | 1 | 0.343 | 1.41E-27 | 26 | Marcks |
| Cacna1a2 | 1.33E-31 | 0.521830834 | 0.417 | 0.037 | 2.70E-27 | 26 | Cacna1a |
| Nfix3 | 1.37E-31 | 1.569687487 | 0.972 | 0.217 | 2.76E-27 | 26 | Nfix |
| Arap13 | 1.46E-31 | 0.703926251 | 0.528 | 0.059 | 2.95E-27 | 26 | Arap1 |
| Brdt | 1.64E-31 | 0.54400812 | 0.361 | 0.029 | 3.31E-27 | 26 | Brdt |

| | | | | | | | |
|---|---|---|---|---|---|---|---|
| Abca92 | 2.08E-31 | 0.819148632 | 0.444 | 0.043 | 4.20E-27 | 26 | Abca9 |
| Plod21 | 2.34E-31 | 0.53403286 | 0.5 | 0.052 | 4.75E-27 | 26 | Plod2 |
| Wwox12 | 2.92E-31 | 2.013272669 | 0.861 | 0.177 | 5.91E-27 | 26 | Wwox |
| Lurap1l | 4.20E-31 | 0.742215042 | 0.5 | 0.053 | 8.50E-27 | 26 | Lurap1l |
| Csgalnact21 | 4.99E-31 | 0.786141571 | 0.528 | 0.06 | 1.01E-26 | 26 | Csgalnact2 |
| Tram2 | 6.33E-31 | 0.549778726 | 0.472 | 0.048 | 1.28E-26 | 26 | Tram2 |
| Csf15 | 6.98E-31 | 0.853214264 | 0.75 | 0.111 | 1.41E-26 | 26 | Csf1 |
| Kif21a1 | 8.90E-31 | 0.611687584 | 0.444 | 0.043 | 1.80E-26 | 26 | Kif21a |
| Calu8 | 1.07E-30 | 1.372305172 | 0.944 | 0.203 | 2.17E-26 | 26 | Calu |
| Pde10a2 | 1.08E-30 | 0.92803675 | 0.639 | 0.085 | 2.18E-26 | 26 | Pde10a |
| Mgst17 | 1.18E-30 | 1.457911192 | 0.889 | 0.168 | 2.38E-26 | 26 | Mgst1 |
| Rab11fip5 | 1.23E-30 | 0.498052475 | 0.361 | 0.029 | 2.49E-26 | 26 | Rab11fip5 |
| Cpq6 | 2.16E-30 | 1.190705985 | 0.75 | 0.122 | 4.37E-26 | 26 | Cpq |
| Ube2e25 | 2.32E-30 | 1.625725597 | 0.833 | 0.154 | 4.69E-26 | 26 | Ube2e2 |
| Tcf7l26 | 3.47E-30 | 1.177695178 | 0.778 | 0.13 | 7.02E-26 | 26 | Tcf7l2 |
| Gda2 | 3.66E-30 | 0.992314334 | 0.611 | 0.081 | 7.40E-26 | 26 | Gda |
| St3gal16 | 4.18E-30 | 1.393012694 | 0.889 | 0.181 | 8.47E-26 | 26 | St3gal1 |
| Sobp2 | 5.53E-30 | 0.435193967 | 0.306 | 0.022 | 1.12E-25 | 26 | Sobp |
| Vegfa | 6.54E-30 | 0.534150863 | 0.444 | 0.044 | 1.32E-25 | 26 | Vegfa |
| Ifi2043 | 1.05E-29 | 1.263231374 | 0.639 | 0.095 | 2.12E-25 | 26 | Ifi204 |
| Fam126a2 | 1.11E-29 | 0.806747166 | 0.611 | 0.082 | 2.25E-25 | 26 | Fam126a |
| Selenon1 | 1.15E-29 | 0.628625075 | 0.5 | 0.056 | 2.34E-25 | 26 | Selenon |
| Pxdn1 | 1.35E-29 | 0.860394544 | 0.583 | 0.074 | 2.74E-25 | 26 | Pxdn |
| Npr14 | 3.16E-29 | 0.739644459 | 0.667 | 0.092 | 6.40E-25 | 26 | Npr1 |
| Arhgap109 | 4.57E-29 | 0.907909364 | 0.778 | 0.128 | 9.25E-25 | 26 | Arhgap10 |
| Fkbp95 | 5.04E-29 | 0.946413434 | 0.722 | 0.112 | 1.02E-24 | 26 | Fkbp9 |
| S100a612 | 5.37E-29 | 2.167419199 | 1 | 0.273 | 1.09E-24 | 26 | S100a6 |
| Aox11 | 5.38E-29 | 0.441784742 | 0.278 | 0.019 | 1.09E-24 | 26 | Aox1 |
| Add36 | 6.70E-29 | 1.640187037 | 0.944 | 0.224 | 1.36E-24 | 26 | Add3 |
| Grb109 | 7.92E-29 | 1.539315852 | 0.917 | 0.196 | 1.60E-24 | 26 | Grb10 |
| Pde8a6 | 7.93E-29 | 1.417980532 | 0.889 | 0.184 | 1.61E-24 | 26 | Pde8a |
| Ar6 | 1.13E-28 | 1.594643435 | 0.944 | 0.22 | 2.30E-24 | 26 | Ar |
| Ccser12 | 1.26E-28 | 1.516974982 | 0.528 | 0.068 | 2.56E-24 | 26 | Ccser1 |
| Itih56 | 1.36E-28 | 1.763982827 | 0.972 | 0.244 | 2.75E-24 | 26 | Itih5 |
| Adgra21 | 1.52E-28 | 0.36747005 | 0.278 | 0.019 | 3.09E-24 | 26 | Adgra2 |
| Smurf1 | 1.76E-28 | 0.998240802 | 0.583 | 0.08 | 3.55E-24 | 26 | Smurf1 |
| Micu1 | 1.90E-28 | 0.870424379 | 0.667 | 0.1 | 3.85E-24 | 26 | Micu1 |
| Gpx83 | 2.18E-28 | 0.490129039 | 0.472 | 0.051 | 4.41E-24 | 26 | Gpx8 |
| Ispd1 | 2.49E-28 | 0.559324457 | 0.306 | 0.023 | 5.05E-24 | 26 | Ispd |
| Aldh7a12 | 2.82E-28 | 0.509579049 | 0.528 | 0.063 | 5.72E-24 | 26 | Aldh7a1 |
| Ppp1r12b4 | 3.23E-28 | 0.949882908 | 0.694 | 0.107 | 6.54E-24 | 26 | Ppp1r12b |
| Col18a13 | 3.36E-28 | 0.393472651 | 0.306 | 0.023 | 6.81E-24 | 26 | Col18a1 |
| Anxa36 | 3.47E-28 | 2.255728057 | 1 | 0.34 | 7.03E-24 | 26 | Anxa3 |

| | | | | | | | |
|---|---|---|---|---|---|---|---|
| Irak31 | 3.84E-28 | 0.547036846 | 0.556 | 0.068 | 7.77E-24 | 26 | Irak3 |
| Dock1110 | 4.15E-28 | 1.217800149 | 0.694 | 0.115 | 8.41E-24 | 26 | Dock11 |
| Fbln11 | 5.16E-28 | 0.497903294 | 0.25 | 0.016 | 1.05E-23 | 26 | Fbln1 |
| Bace11 | 5.24E-28 | 0.497512192 | 0.417 | 0.041 | 1.06E-23 | 26 | Bace1 |
| Ror12 | 5.66E-28 | 0.552476334 | 0.389 | 0.037 | 1.15E-23 | 26 | Ror1 |
| Sema3a1 | 7.77E-28 | 0.548426881 | 0.278 | 0.02 | 1.57E-23 | 26 | Sema3a |
| Agpat41 | 8.01E-28 | 1.015306798 | 0.639 | 0.094 | 1.62E-23 | 26 | Agpat4 |
| Vim15 | 8.64E-28 | 1.996139615 | 1 | 0.308 | 1.75E-23 | 26 | Vim |
| Pmp228 | 9.53E-28 | 1.153842824 | 0.806 | 0.145 | 1.93E-23 | 26 | Pmp22 |
| Ppip5k21 | 1.46E-27 | 0.537185728 | 0.556 | 0.071 | 2.95E-23 | 26 | Ppip5k2 |
| Cd639 | 1.91E-27 | 1.021203279 | 0.972 | 0.189 | 3.86E-23 | 26 | Cd63 |
| Cd347 | 1.95E-27 | 2.138703648 | 0.972 | 0.323 | 3.95E-23 | 26 | Cd34 |
| Fbxw81 | 2.03E-27 | 0.778830492 | 0.583 | 0.081 | 4.11E-23 | 26 | Fbxw8 |
| Ergic11 | 2.42E-27 | 0.741788559 | 0.583 | 0.081 | 4.90E-23 | 26 | Ergic1 |
| Kif2a5 | 3.51E-27 | 1.13824305 | 0.889 | 0.179 | 7.11E-23 | 26 | Kif2a |
| Fig4 | 4.17E-27 | 0.58505324 | 0.444 | 0.049 | 8.44E-23 | 26 | Fig4 |
| Kremen1 | 5.15E-27 | 0.47076354 | 0.444 | 0.048 | 1.04E-22 | 26 | Kremen1 |
| Serinc5 | 5.75E-27 | 0.476551292 | 0.389 | 0.038 | 1.16E-22 | 26 | Serinc5 |
| Gucy1a22 | 6.29E-27 | 0.554240182 | 0.25 | 0.017 | 1.27E-22 | 26 | Gucy1a2 |
| P3h11 | 7.18E-27 | 0.324989247 | 0.278 | 0.02 | 1.45E-22 | 26 | P3h1 |
| Sfxn13 | 9.98E-27 | 0.613455659 | 0.639 | 0.094 | 2.02E-22 | 26 | Sfxn1 |
| Plekhh22 | 1.10E-26 | 0.531257736 | 0.306 | 0.025 | 2.23E-22 | 26 | Plekhh2 |
| Ank23 | 1.22E-26 | 0.696064985 | 0.417 | 0.044 | 2.46E-22 | 26 | Ank2 |
| Triobp3 | 1.40E-26 | 0.920629529 | 0.778 | 0.138 | 2.84E-22 | 26 | Triobp |
| Ang1 | 1.46E-26 | 0.572962885 | 0.389 | 0.039 | 2.96E-22 | 26 | Ang |
| Smarcd31 | 1.68E-26 | 0.38823308 | 0.25 | 0.017 | 3.41E-22 | 26 | Smarcd3 |
| Ptov14 | 1.79E-26 | 0.47984107 | 0.75 | 0.119 | 3.62E-22 | 26 | Ptov1 |
| Trio6 | 1.95E-26 | 1.290913334 | 0.861 | 0.189 | 3.95E-22 | 26 | Trio |
| Abhd14a | 2.43E-26 | 0.339961856 | 0.389 | 0.038 | 4.92E-22 | 26 | Abhd14a |
| Arl4d | 2.60E-26 | 0.718085556 | 0.556 | 0.075 | 5.26E-22 | 26 | Arl4d |
| Galm | 2.89E-26 | 0.318058159 | 0.417 | 0.043 | 5.85E-22 | 26 | Galm |
| Airn9 | 3.24E-26 | 1.886596633 | 0.944 | 0.25 | 6.55E-22 | 26 | Airn |
| Rcn14 | 3.36E-26 | 0.923318103 | 0.667 | 0.109 | 6.81E-22 | 26 | Rcn1 |
| Nmt22 | 4.65E-26 | 0.782799028 | 0.639 | 0.099 | 9.42E-22 | 26 | Nmt2 |
| Nynrin | 5.15E-26 | 0.380112972 | 0.333 | 0.029 | 1.04E-21 | 26 | Nynrin |
| Pdzd25 | 6.24E-26 | 1.078293131 | 0.694 | 0.116 | 1.26E-21 | 26 | Pdzd2 |
| Maged12 | 6.85E-26 | 0.708236891 | 0.611 | 0.089 | 1.39E-21 | 26 | Maged1 |
| Naglu | 7.57E-26 | 0.428753081 | 0.389 | 0.039 | 1.53E-21 | 26 | Naglu |
| Lpgat11 | 8.25E-26 | 0.849615815 | 0.583 | 0.087 | 1.67E-21 | 26 | Lpgat1 |
| Dlgap42 | 9.74E-26 | 0.699017017 | 0.778 | 0.137 | 1.97E-21 | 26 | Dlgap4 |
| Hsf2 | 1.04E-25 | 0.603810361 | 0.361 | 0.035 | 2.11E-21 | 26 | Hsf2 |
| Asap31 | 1.23E-25 | 0.384642295 | 0.278 | 0.021 | 2.49E-21 | 26 | Asap3 |
| Ehd13 | 1.42E-25 | 0.776640542 | 0.528 | 0.072 | 2.88E-21 | 26 | Ehd1 |

| | | | | | | | |
|---|---|---|---|---|---|---|---|
| Dmd3 | 2.22E-25 | 1.222115988 | 0.472 | 0.06 | 4.50E-21 | 26 | Dmd |
| Zeb113 | 2.76E-25 | 2.193996206 | 0.972 | 0.323 | 5.59E-21 | 26 | Zeb1 |
| Sep-82 | 3.44E-25 | 0.644381573 | 0.417 | 0.047 | 6.97E-21 | 26 | 8-Sep |
| Daam14 | 3.45E-25 | 0.80718828 | 0.667 | 0.11 | 6.98E-21 | 26 | Daam1 |
| Nfia9 | 3.54E-25 | 1.668538828 | 1 | 0.289 | 7.17E-21 | 26 | Nfia |
| Thra8 | 6.34E-25 | 0.941522738 | 0.833 | 0.165 | 1.28E-20 | 26 | Thra |
| Mex3b | 7.04E-25 | 0.304198057 | 0.25 | 0.018 | 1.43E-20 | 26 | Mex3b |
| Gpx7 | 7.54E-25 | 0.489666846 | 0.361 | 0.035 | 1.53E-20 | 26 | Gpx7 |
| Rtn13 | 8.76E-25 | 0.692258471 | 0.361 | 0.036 | 1.77E-20 | 26 | Rtn1 |
| Ptprj10 | 9.56E-25 | 1.270257358 | 0.778 | 0.152 | 1.94E-20 | 26 | Ptprj |
| Fer4 | 1.08E-24 | 0.971395162 | 0.694 | 0.122 | 2.18E-20 | 26 | Fer |
| Flnb5 | 1.09E-24 | 1.134765398 | 0.806 | 0.163 | 2.20E-20 | 26 | Flnb |
| Tspan56 | 1.24E-24 | 1.317201225 | 0.861 | 0.2 | 2.51E-20 | 26 | Tspan5 |
| Palm2 | 1.25E-24 | 0.808704708 | 0.583 | 0.085 | 2.52E-20 | 26 | Palm |
| Galnt115 | 1.27E-24 | 1.19132755 | 0.833 | 0.185 | 2.56E-20 | 26 | Galnt1 |
| Kifap33 | 1.36E-24 | 0.696488312 | 0.667 | 0.108 | 2.75E-20 | 26 | Kifap3 |
| Arfgap3 | 1.40E-24 | 0.481902397 | 0.583 | 0.084 | 2.83E-20 | 26 | Arfgap3 |
| Cc2d2a2 | 1.74E-24 | 0.45696378 | 0.361 | 0.036 | 3.53E-20 | 26 | Cc2d2a |
| Ryk5 | 1.95E-24 | 0.806108416 | 0.667 | 0.111 | 3.96E-20 | 26 | Ryk |
| Arhgap242 | 2.17E-24 | 0.991061769 | 0.417 | 0.048 | 4.39E-20 | 26 | Arhgap24 |
| Gramd31 | 2.31E-24 | 0.512856595 | 0.361 | 0.036 | 4.68E-20 | 26 | Gramd3 |
| Epb41l27 | 2.32E-24 | 1.350460827 | 0.944 | 0.243 | 4.70E-20 | 26 | Epb41l2 |
| B3galt12 | 2.62E-24 | 0.623931722 | 0.278 | 0.022 | 5.31E-20 | 26 | B3galt1 |
| Mast48 | 2.63E-24 | 1.76743563 | 0.833 | 0.188 | 5.32E-20 | 26 | Mast4 |
| Ckap43 | 2.64E-24 | 0.761362948 | 0.639 | 0.102 | 5.34E-20 | 26 | Ckap4 |
| Chd97 | 2.95E-24 | 1.4007099 | 0.944 | 0.251 | 5.98E-20 | 26 | Chd9 |
| Fgd43 | 3.00E-24 | 0.592306388 | 0.389 | 0.042 | 6.08E-20 | 26 | Fgd4 |
| Akt21 | 3.43E-24 | 0.429540831 | 0.5 | 0.065 | 6.94E-20 | 26 | Akt2 |
| Cblb8 | 3.43E-24 | 0.979480635 | 0.833 | 0.172 | 6.95E-20 | 26 | Cblb |
| Nr2f12 | 3.55E-24 | 0.392368985 | 0.306 | 0.026 | 7.20E-20 | 26 | Nr2f1 |
| Pbx17 | 4.10E-24 | 1.651940289 | 0.833 | 0.2 | 8.29E-20 | 26 | Pbx1 |
| Vwa5a2 | 4.24E-24 | 0.753392212 | 0.611 | 0.097 | 8.59E-20 | 26 | Vwa5a |
| Tcf7l17 | 4.57E-24 | 0.997780104 | 0.889 | 0.182 | 9.25E-20 | 26 | Tcf7l1 |
| Ap2a23 | 4.67E-24 | 0.859000908 | 0.778 | 0.151 | 9.45E-20 | 26 | Ap2a2 |
| Phldb12 | 5.36E-24 | 0.741526722 | 0.5 | 0.067 | 1.08E-19 | 26 | Phldb1 |
| Nptn10 | 6.15E-24 | 1.322608969 | 0.944 | 0.25 | 1.24E-19 | 26 | Nptn |
| Akap127 | 7.08E-24 | 0.924941686 | 0.833 | 0.162 | 1.43E-19 | 26 | Akap12 |
| Lpcat33 | 7.99E-24 | 0.617760861 | 0.611 | 0.095 | 1.62E-19 | 26 | Lpcat3 |
| Sqor2 | 9.13E-24 | 0.50181581 | 0.528 | 0.072 | 1.85E-19 | 26 | Sqor |
| Slc38a101 | 9.40E-24 | 0.749865732 | 0.722 | 0.13 | 1.90E-19 | 26 | Slc38a10 |
| Hebp2 | 9.63E-24 | 0.275197797 | 0.25 | 0.019 | 1.95E-19 | 26 | Hebp2 |
| Vat16 | 9.88E-24 | 0.879115983 | 0.806 | 0.156 | 2.00E-19 | 26 | Vat1 |
| Abcc53 | 1.17E-23 | 0.685685751 | 0.556 | 0.081 | 2.37E-19 | 26 | Abcc5 |

| | | | | | | | |
|---|---|---|---|---|---|---|---|
| Snx293 | 1.32E-23 | 1.040258912 | 0.639 | 0.113 | 2.68E-19 | 26 | Snx29 |
| Npepps1 | 1.39E-23 | 0.852738131 | 0.694 | 0.127 | 2.81E-19 | 26 | Npepps |
| 9530068E07Rik2 | 1.74E-23 | 0.694518447 | 0.611 | 0.098 | 3.52E-19 | 26 | 9530068E07Rik |
| Usp2 | 2.01E-23 | 0.385905981 | 0.278 | 0.023 | 4.07E-19 | 26 | Usp2 |
| Cnrip11 | 2.10E-23 | 0.637114327 | 0.417 | 0.049 | 4.25E-19 | 26 | Cnrip1 |
| Pkia1 | 2.70E-23 | 0.581314027 | 0.25 | 0.019 | 5.48E-19 | 26 | Pkia |
| Cavin18 | 2.71E-23 | 1.159745901 | 0.917 | 0.215 | 5.48E-19 | 26 | Cavin1 |
| Zfp4625 | 2.71E-23 | 1.105704291 | 0.667 | 0.121 | 5.48E-19 | 26 | Zfp462 |
| Xdh6 | 2.72E-23 | 1.114440156 | 0.861 | 0.189 | 5.52E-19 | 26 | Xdh |
| Arhgap323 | 3.03E-23 | 0.52415942 | 0.444 | 0.054 | 6.13E-19 | 26 | Arhgap32 |
| Gaa1 | 3.11E-23 | 0.82416061 | 0.472 | 0.063 | 6.30E-19 | 26 | Gaa |
| Dennd4c7 | 3.46E-23 | 0.900042844 | 0.694 | 0.127 | 7.00E-19 | 26 | Dennd4c |
| Cep1705 | 3.97E-23 | 0.909908318 | 0.694 | 0.131 | 8.04E-19 | 26 | Cep170 |
| Golim46 | 4.29E-23 | 1.020711546 | 0.833 | 0.182 | 8.68E-19 | 26 | Golim4 |
| Hdac75 | 4.42E-23 | 1.146335587 | 0.889 | 0.204 | 8.96E-19 | 26 | Hdac7 |
| Phactr12 | 5.34E-23 | 0.593170623 | 0.5 | 0.068 | 1.08E-18 | 26 | Phactr1 |
| Tmod21 | 5.46E-23 | 0.463188438 | 0.306 | 0.028 | 1.10E-18 | 26 | Tmod2 |
| Tead16 | 1.04E-22 | 0.794053555 | 0.556 | 0.085 | 2.10E-18 | 26 | Tead1 |
| Sparc12 | 1.04E-22 | 1.989225626 | 1 | 0.405 | 2.11E-18 | 26 | Sparc |
| Ebf35 | 1.09E-22 | 0.945648074 | 0.694 | 0.127 | 2.20E-18 | 26 | Ebf3 |
| Disp1 | 1.18E-22 | 0.608626578 | 0.472 | 0.063 | 2.39E-18 | 26 | Disp1 |
| G6pdx | 1.18E-22 | 0.415203368 | 0.417 | 0.05 | 2.39E-18 | 26 | G6pdx |
| Fam20a2 | 1.29E-22 | 0.686352802 | 0.444 | 0.057 | 2.62E-18 | 26 | Fam20a |
| Prkag22 | 2.28E-22 | 0.527002743 | 0.528 | 0.076 | 4.61E-18 | 26 | Prkag2 |
| Rock29 | 2.28E-22 | 1.490551845 | 1 | 0.339 | 4.62E-18 | 26 | Rock2 |
| Man2a16 | 2.88E-22 | 1.399059154 | 0.889 | 0.236 | 5.83E-18 | 26 | Man2a1 |
| Lpp12 | 3.47E-22 | 1.47658583 | 0.917 | 0.288 | 7.03E-18 | 26 | Lpp |
| B3galnt1 | 3.59E-22 | 0.476942305 | 0.306 | 0.029 | 7.27E-18 | 26 | B3galnt1 |
| Errfi13 | 4.07E-22 | 1.772046776 | 0.722 | 0.162 | 8.23E-18 | 26 | Errfi1 |
| Adcy92 | 4.15E-22 | 0.658200043 | 0.444 | 0.059 | 8.40E-18 | 26 | Adcy9 |
| Slc7a6 | 5.84E-22 | 0.376761088 | 0.306 | 0.029 | 1.18E-17 | 26 | Slc7a6 |
| Ap3m2 | 5.90E-22 | 0.466757609 | 0.361 | 0.04 | 1.19E-17 | 26 | Ap3m2 |
| Asph7 | 7.14E-22 | 0.602314638 | 0.611 | 0.101 | 1.45E-17 | 26 | Asph |
| Zbtb2012 | 7.43E-22 | 2.065137852 | 0.972 | 0.539 | 1.50E-17 | 26 | Zbtb20 |
| Tuba1a10 | 9.07E-22 | 1.301170322 | 0.944 | 0.258 | 1.84E-17 | 26 | Tuba1a |
| Eri32 | 1.08E-21 | 0.597644145 | 0.611 | 0.104 | 2.19E-17 | 26 | Eri3 |
| Ebf19 | 1.11E-21 | 2.167652068 | 1 | 0.377 | 2.24E-17 | 26 | Ebf1 |
| P4ha14 | 1.14E-21 | 0.851009421 | 0.667 | 0.125 | 2.32E-17 | 26 | P4ha1 |
| Wdr60 | 1.39E-21 | 0.36286952 | 0.278 | 0.025 | 2.82E-17 | 26 | Wdr60 |
| Sesn38 | 1.47E-21 | 0.747657507 | 0.694 | 0.13 | 2.97E-17 | 26 | Sesn3 |
| Fyn11 | 1.81E-21 | 1.337284875 | 0.917 | 0.268 | 3.67E-17 | 26 | Fyn |
| Csrp21 | 2.30E-21 | 1.010718883 | 0.861 | 0.201 | 4.65E-17 | 26 | Csrp2 |
| Limk1 | 2.60E-21 | 0.302110631 | 0.306 | 0.03 | 5.25E-17 | 26 | Limk1 |

| | | | | | | | |
|---|---|---|---|---|---|---|---|
| Rbfox28 | 3.01E-21 | 0.906820783 | 0.861 | 0.192 | 6.10E-17 | 26 | Rbfox2 |
| Cbx63 | 3.16E-21 | 0.565979927 | 0.472 | 0.066 | 6.40E-17 | 26 | Cbx6 |
| Bmp61 | 3.22E-21 | 0.658301096 | 0.389 | 0.047 | 6.53E-17 | 26 | Bmp6 |
| Zdhhc1 | 3.34E-21 | 0.432018184 | 0.333 | 0.035 | 6.77E-17 | 26 | Zdhhc1 |
| Cmtm32 | 3.41E-21 | 0.731760954 | 0.611 | 0.107 | 6.90E-17 | 26 | Cmtm3 |
| Etfbkmt | 4.17E-21 | 0.430211526 | 0.306 | 0.03 | 8.45E-17 | 26 | Etfbkmt |
| Ocrl1 | 4.62E-21 | 0.438144392 | 0.444 | 0.059 | 9.36E-17 | 26 | Ocrl |
| Nbas1 | 5.21E-21 | 0.614770679 | 0.556 | 0.09 | 1.06E-16 | 26 | Nbas |
| Arhgap214 | 5.59E-21 | 0.886754608 | 0.611 | 0.112 | 1.13E-16 | 26 | Arhgap21 |
| Clasp12 | 6.37E-21 | 0.666745675 | 0.694 | 0.133 | 1.29E-16 | 26 | Clasp1 |
| Emp314 | 6.70E-21 | 1.220292997 | 0.917 | 0.261 | 1.36E-16 | 26 | Emp3 |
| Kazn | 7.44E-21 | 0.488077244 | 0.333 | 0.036 | 1.51E-16 | 26 | Kazn |
| Gpr137b1 | 8.90E-21 | 0.326764131 | 0.306 | 0.03 | 1.80E-16 | 26 | Gpr137b |
| Nckap18 | 9.89E-21 | 1.200151826 | 0.889 | 0.228 | 2.00E-16 | 26 | Nckap1 |
| Nav18 | 1.11E-20 | 1.28738844 | 0.861 | 0.224 | 2.24E-16 | 26 | Nav1 |
| Dag18 | 1.17E-20 | 0.858903074 | 0.722 | 0.144 | 2.37E-16 | 26 | Dag1 |
| Map1b4 | 1.18E-20 | 0.626571578 | 0.611 | 0.103 | 2.39E-16 | 26 | Map1b |
| Wdfy24 | 1.51E-20 | 0.869904203 | 0.611 | 0.112 | 3.06E-16 | 26 | Wdfy2 |
| Prss2310 | 1.51E-20 | 1.547230941 | 0.944 | 0.282 | 3.06E-16 | 26 | Prss23 |
| Slc30a4 | 1.77E-20 | 0.33384365 | 0.333 | 0.036 | 3.59E-16 | 26 | Slc30a4 |
| Kank22 | 1.85E-20 | 0.442369907 | 0.417 | 0.054 | 3.74E-16 | 26 | Kank2 |
| Tshz32 | 2.17E-20 | 0.772964164 | 0.556 | 0.094 | 4.40E-16 | 26 | Tshz3 |
| Adcy32 | 2.30E-20 | 0.398976877 | 0.278 | 0.026 | 4.66E-16 | 26 | Adcy3 |
| Gng124 | 2.39E-20 | 1.154498403 | 0.833 | 0.202 | 4.84E-16 | 26 | Gng12 |
| Dnajc251 | 2.60E-20 | 0.44793022 | 0.417 | 0.055 | 5.26E-16 | 26 | Dnajc25 |
| Slc39a11 | 2.60E-20 | 0.440936902 | 0.333 | 0.037 | 5.26E-16 | 26 | Slc39a11 |
| Adamts142 | 2.60E-20 | 0.340885492 | 0.333 | 0.036 | 5.27E-16 | 26 | Adamts14 |
| B4galt110 | 2.77E-20 | 0.851071213 | 0.806 | 0.178 | 5.62E-16 | 26 | B4galt1 |
| Stxbp62 | 2.93E-20 | 0.645430375 | 0.861 | 0.189 | 5.92E-16 | 26 | Stxbp6 |
| Sgpp12 | 3.68E-20 | 0.497650027 | 0.5 | 0.076 | 7.45E-16 | 26 | Sgpp1 |
| Rhou1 | 3.79E-20 | 0.303505742 | 0.472 | 0.067 | 7.68E-16 | 26 | Rhou |
| Pgm3 | 3.84E-20 | 0.404331302 | 0.333 | 0.037 | 7.77E-16 | 26 | Pgm3 |
| Chst123 | 3.84E-20 | 0.460852195 | 0.472 | 0.068 | 7.78E-16 | 26 | Chst12 |
| Cers68 | 3.89E-20 | 0.67422957 | 0.639 | 0.117 | 7.87E-16 | 26 | Cers6 |
| Slc44a17 | 3.99E-20 | 1.071033524 | 0.722 | 0.157 | 8.07E-16 | 26 | Slc44a1 |
| Igf2r5 | 4.29E-20 | 0.695396608 | 0.694 | 0.135 | 8.68E-16 | 26 | Igf2r |
| Sntb27 | 4.83E-20 | 1.295878173 | 0.889 | 0.267 | 9.77E-16 | 26 | Sntb2 |
| Fam171a14 | 5.20E-20 | 0.644554297 | 0.611 | 0.107 | 1.05E-15 | 26 | Fam171a1 |
| Clip15 | 5.52E-20 | 0.959648796 | 0.694 | 0.144 | 1.12E-15 | 26 | Clip1 |
| Heg18 | 6.06E-20 | 1.273100344 | 0.944 | 0.276 | 1.23E-15 | 26 | Heg1 |
| Ugp21 | 7.14E-20 | 0.751160354 | 0.722 | 0.155 | 1.45E-15 | 26 | Ugp2 |
| Aco1 | 7.26E-20 | 0.351251131 | 0.417 | 0.055 | 1.47E-15 | 26 | Aco1 |
| Lamb26 | 7.45E-20 | 0.74160711 | 0.667 | 0.125 | 1.51E-15 | 26 | Lamb2 |

| | | | | | | | |
|---|---|---|---|---|---|---|---|
| Gga2 | 7.82E-20 | 0.401396513 | 0.278 | 0.027 | 1.58E-15 | 26 | Gga2 |
| Agap15 | 8.06E-20 | 1.011294192 | 0.694 | 0.146 | 1.63E-15 | 26 | Agap1 |
| Kcnq1ot111 | 8.55E-20 | 1.690545568 | 0.639 | 0.135 | 1.73E-15 | 26 | Kcnq1ot1 |
| Unk | 8.56E-20 | 0.345573058 | 0.361 | 0.043 | 1.73E-15 | 26 | Unk |
| Samd4b | 1.05E-19 | 0.721352677 | 0.583 | 0.104 | 2.13E-15 | 26 | Samd4b |
| Zyx7 | 1.12E-19 | 1.058762536 | 0.889 | 0.217 | 2.27E-15 | 26 | Zyx |
| Rab341 | 1.22E-19 | 0.308387658 | 0.25 | 0.022 | 2.47E-15 | 26 | Rab34 |
| Il17rc | 1.23E-19 | 0.40377266 | 0.278 | 0.027 | 2.48E-15 | 26 | Il17rc |
| Efcab144 | 1.27E-19 | 0.64495536 | 0.722 | 0.146 | 2.56E-15 | 26 | Efcab14 |
| Ankrd46 | 1.32E-19 | 0.409340896 | 0.389 | 0.049 | 2.67E-15 | 26 | Ankrd46 |
| Zfand312 | 1.33E-19 | 1.276376741 | 0.944 | 0.303 | 2.69E-15 | 26 | Zfand3 |
| Calml4 | 1.42E-19 | 0.301034796 | 0.25 | 0.022 | 2.87E-15 | 26 | Calml4 |
| Nucb22 | 1.70E-19 | 0.656498833 | 0.639 | 0.122 | 3.44E-15 | 26 | Nucb2 |
| Pam11 | 1.73E-19 | 1.010819089 | 0.861 | 0.215 | 3.51E-15 | 26 | Pam |
| Tbc1d121 | 1.87E-19 | 0.930649473 | 0.583 | 0.111 | 3.78E-15 | 26 | Tbc1d12 |
| Nenf6 | 2.13E-19 | 1.060959225 | 0.972 | 0.297 | 4.31E-15 | 26 | Nenf |
| Magi32 | 2.19E-19 | 0.75035828 | 0.583 | 0.106 | 4.43E-15 | 26 | Magi3 |
| Rhobtb32 | 2.24E-19 | 0.650631512 | 0.389 | 0.051 | 4.54E-15 | 26 | Rhobtb3 |
| Park22 | 2.29E-19 | 0.97786712 | 0.472 | 0.076 | 4.65E-15 | 26 | Park2 |
| Phlda12 | 2.32E-19 | 0.511640642 | 0.528 | 0.087 | 4.71E-15 | 26 | Phlda1 |
| Adam91 | 2.47E-19 | 0.680029196 | 0.472 | 0.073 | 5.00E-15 | 26 | Adam9 |
| Dip2c10 | 2.71E-19 | 0.947607612 | 0.889 | 0.224 | 5.48E-15 | 26 | Dip2c |
| Arsk | 2.80E-19 | 0.418886671 | 0.361 | 0.044 | 5.68E-15 | 26 | Arsk |
| Prkg18 | 3.07E-19 | 1.437463439 | 0.778 | 0.198 | 6.21E-15 | 26 | Prkg1 |
| Setbp19 | 3.07E-19 | 1.171660522 | 0.611 | 0.123 | 6.23E-15 | 26 | Setbp1 |
| Npas22 | 3.13E-19 | 0.348317868 | 0.278 | 0.028 | 6.33E-15 | 26 | Npas2 |
| Tshz11 | 3.52E-19 | 0.836414509 | 0.611 | 0.119 | 7.13E-15 | 26 | Tshz1 |
| Fcgrt6 | 4.96E-19 | 0.852927494 | 0.806 | 0.187 | 1.00E-14 | 26 | Fcgrt |
| Gns6 | 5.30E-19 | 0.742483736 | 0.639 | 0.128 | 1.07E-14 | 26 | Gns |
| Krt80 | 5.49E-19 | 0.251871054 | 0.25 | 0.023 | 1.11E-14 | 26 | Krt80 |
| Wipi11 | 5.61E-19 | 0.413947151 | 0.417 | 0.057 | 1.13E-14 | 26 | Wipi1 |
| Nisch8 | 5.87E-19 | 0.941036999 | 0.833 | 0.213 | 1.19E-14 | 26 | Nisch |
| Pigk2 | 8.39E-19 | 0.581154797 | 0.639 | 0.12 | 1.70E-14 | 26 | Pigk |
| Igf1r9 | 1.03E-18 | 0.999141382 | 0.722 | 0.162 | 2.09E-14 | 26 | Igf1r |
| Abl12 | 1.06E-18 | 0.822383902 | 0.611 | 0.124 | 2.14E-14 | 26 | Abl1 |
| Cask7 | 1.26E-18 | 0.942469161 | 0.667 | 0.14 | 2.56E-14 | 26 | Cask |
| Fam172a14 | 1.28E-18 | 1.268856438 | 0.917 | 0.286 | 2.59E-14 | 26 | Fam172a |
| Gstm17 | 1.36E-18 | 1.203052039 | 0.972 | 0.324 | 2.76E-14 | 26 | Gstm1 |
| P2rx43 | 1.39E-18 | 0.449510988 | 0.444 | 0.066 | 2.81E-14 | 26 | P2rx4 |
| Smurf26 | 1.46E-18 | 1.076190988 | 0.861 | 0.24 | 2.95E-14 | 26 | Smurf2 |
| Sgsm2 | 1.49E-18 | 0.449214655 | 0.278 | 0.029 | 3.01E-14 | 26 | Sgsm2 |
| Plin3 | 1.54E-18 | 0.408770246 | 0.583 | 0.104 | 3.13E-14 | 26 | Plin3 |
| Parp32 | 1.73E-18 | 0.488661749 | 0.278 | 0.029 | 3.50E-14 | 26 | Parp3 |

| | | | | | | | |
|---|---|---|---|---|---|---|---|
| Dse2 | 1.75E-18 | 0.645987459 | 0.417 | 0.061 | 3.55E-14 | 26 | Dse |
| Rras26 | 1.89E-18 | 0.730805766 | 0.5 | 0.086 | 3.82E-14 | 26 | Rras2 |
| Il11ra11 | 1.99E-18 | 0.759609051 | 0.556 | 0.099 | 4.02E-14 | 26 | Il11ra1 |
| Plekhf1 | 1.99E-18 | 0.313059864 | 0.25 | 0.024 | 4.03E-14 | 26 | Plekhf1 |
| Ggt52 | 2.10E-18 | 0.31214644 | 0.278 | 0.028 | 4.26E-14 | 26 | Ggt5 |
| Car85 | 2.14E-18 | 0.971489113 | 0.806 | 0.188 | 4.34E-14 | 26 | Car8 |
| Sbf29 | 2.79E-18 | 0.922911612 | 0.833 | 0.209 | 5.64E-14 | 26 | Sbf2 |
| Tdrd31 | 3.21E-18 | 0.609597611 | 0.528 | 0.092 | 6.49E-14 | 26 | Tdrd3 |
| Ctsl9 | 3.40E-18 | 1.365208874 | 0.972 | 0.348 | 6.89E-14 | 26 | Ctsl |
| Pgd2 | 3.77E-18 | 0.357327748 | 0.556 | 0.096 | 7.64E-14 | 26 | Pgd |
| Lamp210 | 4.34E-18 | 1.148841176 | 0.972 | 0.307 | 8.78E-14 | 26 | Lamp2 |
| Prrc11 | 4.57E-18 | 0.629740164 | 0.528 | 0.094 | 9.26E-14 | 26 | Prrc1 |
| St71 | 4.65E-18 | 0.53848673 | 0.444 | 0.068 | 9.42E-14 | 26 | St7 |
| Ttc39c | 6.52E-18 | 0.525526339 | 0.417 | 0.061 | 1.32E-13 | 26 | Ttc39c |
| Tlr42 | 6.52E-18 | 0.572042763 | 0.389 | 0.055 | 1.32E-13 | 26 | Tlr4 |
| Ctdsp26 | 8.99E-18 | 0.662808794 | 0.778 | 0.179 | 1.82E-13 | 26 | Ctdsp2 |
| Mtmr310 | 9.68E-18 | 0.859972803 | 0.806 | 0.202 | 1.96E-13 | 26 | Mtmr3 |
| Scmh14 | 1.16E-17 | 0.892469857 | 0.639 | 0.139 | 2.34E-13 | 26 | Scmh1 |
| Ces2g1 | 1.21E-17 | 0.398136521 | 0.333 | 0.041 | 2.45E-13 | 26 | Ces2g |
| Zc3h6 | 1.22E-17 | 0.445935987 | 0.417 | 0.061 | 2.46E-13 | 26 | Zc3h6 |
| Nedd410 | 1.33E-17 | 1.402359483 | 0.972 | 0.421 | 2.70E-13 | 26 | Nedd4 |
| Setd73 | 1.41E-17 | 0.719785811 | 0.694 | 0.156 | 2.85E-13 | 26 | Setd7 |
| Stambpl13 | 1.43E-17 | 0.538933744 | 0.444 | 0.07 | 2.89E-13 | 26 | Stambpl1 |
| Fam114a14 | 1.47E-17 | 0.747509661 | 0.639 | 0.133 | 2.98E-13 | 26 | Fam114a1 |
| P4hb8 | 1.48E-17 | 0.923786178 | 0.889 | 0.254 | 3.00E-13 | 26 | P4hb |
| Ift74 | 1.49E-17 | 0.291112765 | 0.333 | 0.041 | 3.01E-13 | 26 | Ift74 |
| Atg71 | 1.58E-17 | 0.421786512 | 0.444 | 0.069 | 3.20E-13 | 26 | Atg7 |
| Ppm1a1 | 1.81E-17 | 0.523016517 | 0.75 | 0.167 | 3.67E-13 | 26 | Ppm1a |
| Hspa514 | 1.84E-17 | 1.536175233 | 1 | 0.437 | 3.72E-13 | 26 | Hspa5 |
| Tnks1bp12 | 1.95E-17 | 0.433853308 | 0.528 | 0.091 | 3.96E-13 | 26 | Tnks1bp1 |
| Bcl63 | 2.04E-17 | 0.541602096 | 0.389 | 0.057 | 4.13E-13 | 26 | Bcl6 |
| Kctd111 | 2.28E-17 | 0.282895527 | 0.25 | 0.025 | 4.62E-13 | 26 | Kctd11 |
| Stk37 | 2.41E-17 | 0.971541963 | 0.694 | 0.167 | 4.87E-13 | 26 | Stk3 |
| Trim356 | 2.49E-17 | 0.666782894 | 0.667 | 0.141 | 5.04E-13 | 26 | Trim35 |
| Cd1646 | 2.65E-17 | 0.746227581 | 0.833 | 0.215 | 5.36E-13 | 26 | Cd164 |
| Ttc36 | 3.07E-17 | 0.945837416 | 0.889 | 0.253 | 6.21E-13 | 26 | Ttc3 |
| Crtc39 | 3.09E-17 | 0.695436765 | 0.722 | 0.166 | 6.26E-13 | 26 | Crtc3 |
| Slc25a241 | 3.20E-17 | 0.40862123 | 0.5 | 0.084 | 6.47E-13 | 26 | Slc25a24 |
| Klf47 | 3.51E-17 | 1.252069821 | 0.889 | 0.271 | 7.11E-13 | 26 | Klf4 |
| Myo9a6 | 3.64E-17 | 0.933048178 | 0.722 | 0.175 | 7.36E-13 | 26 | Myo9a |
| Sash110 | 3.72E-17 | 1.192476962 | 0.833 | 0.259 | 7.54E-13 | 26 | Sash1 |
| Lrch312 | 3.88E-17 | 1.116371435 | 0.833 | 0.234 | 7.86E-13 | 26 | Lrch3 |
| Cttn1 | 4.34E-17 | 0.538808397 | 0.528 | 0.095 | 8.79E-13 | 26 | Cttn |

| | | | | | | | |
|---|---|---|---|---|---|---|---|
| Creb3l26 | 4.47E-17 | 0.728572967 | 0.611 | 0.124 | 9.04E-13 | 26 | Creb3l2 |
| Kdelr22 | 4.52E-17 | 0.880064317 | 0.694 | 0.171 | 9.15E-13 | 26 | Kdelr2 |
| Tmco3 | 4.74E-17 | 0.384630932 | 0.361 | 0.049 | 9.60E-13 | 26 | Tmco3 |
| Ddah25 | 6.52E-17 | 0.789536081 | 0.806 | 0.207 | 1.32E-12 | 26 | Ddah2 |
| Aebp23 | 7.07E-17 | 0.41604859 | 0.583 | 0.111 | 1.43E-12 | 26 | Aebp2 |
| Rcan1 | 8.33E-17 | 0.288069802 | 0.361 | 0.049 | 1.69E-12 | 26 | Rcan1 |
| Ppic7 | 9.21E-17 | 0.629815996 | 0.611 | 0.122 | 1.87E-12 | 26 | Ppic |
| Tcaf16 | 9.22E-17 | 0.562233733 | 0.556 | 0.104 | 1.87E-12 | 26 | Tcaf1 |
| Tanc25 | 9.67E-17 | 0.551130628 | 0.722 | 0.161 | 1.96E-12 | 26 | Tanc2 |
| Npdc16 | 1.04E-16 | 0.640098368 | 0.639 | 0.134 | 2.11E-12 | 26 | Npdc1 |
| Tshz29 | 1.05E-16 | 1.349350484 | 0.972 | 0.374 | 2.12E-12 | 26 | Tshz2 |
| E130307A14Rik | 1.12E-16 | 0.502817672 | 0.417 | 0.065 | 2.28E-12 | 26 | E130307A14Rik |
| Umps | 1.17E-16 | 0.391155375 | 0.306 | 0.038 | 2.36E-12 | 26 | Umps |
| Impad12 | 1.19E-16 | 0.401877505 | 0.556 | 0.102 | 2.41E-12 | 26 | Impad1 |
| Ctsf | 1.22E-16 | 0.283102919 | 0.306 | 0.037 | 2.46E-12 | 26 | Ctsf |
| Iqsec15 | 1.23E-16 | 0.611352617 | 0.611 | 0.123 | 2.48E-12 | 26 | Iqsec1 |
| Osbpl5 | 1.35E-16 | 0.265293769 | 0.25 | 0.026 | 2.74E-12 | 26 | Osbpl5 |
| Ralgps1 | 1.38E-16 | 0.31846757 | 0.25 | 0.026 | 2.78E-12 | 26 | Ralgps1 |
| Plpp35 | 1.45E-16 | 1.559526833 | 1 | 0.406 | 2.93E-12 | 26 | Plpp3 |
| Glis21 | 1.46E-16 | 0.417342137 | 0.361 | 0.05 | 2.95E-12 | 26 | Glis2 |
| Cyth36 | 1.48E-16 | 0.684431955 | 0.611 | 0.126 | 3.00E-12 | 26 | Cyth3 |
| Armcx32 | 1.50E-16 | 0.403335013 | 0.444 | 0.072 | 3.04E-12 | 26 | Armcx3 |
| Cxxc54 | 1.55E-16 | 0.447546609 | 0.583 | 0.115 | 3.14E-12 | 26 | Cxxc5 |
| Spry25 | 1.56E-16 | 0.600620532 | 0.472 | 0.083 | 3.15E-12 | 26 | Spry2 |
| C1galt11 | 1.57E-16 | 0.58132898 | 0.639 | 0.139 | 3.17E-12 | 26 | C1galt1 |
| Ncoa111 | 1.57E-16 | 0.875434683 | 0.806 | 0.211 | 3.17E-12 | 26 | Ncoa1 |
| Il17ra1 | 1.66E-16 | 0.405553516 | 0.306 | 0.038 | 3.37E-12 | 26 | Il17ra |
| Lmbr11 | 1.95E-16 | 0.487479625 | 0.417 | 0.065 | 3.94E-12 | 26 | Lmbr1 |
| Mospd21 | 2.31E-16 | 0.64611573 | 0.556 | 0.107 | 4.68E-12 | 26 | Mospd2 |
| Esyt212 | 2.38E-16 | 0.88263454 | 0.861 | 0.244 | 4.82E-12 | 26 | Esyt2 |
| Rhoj10 | 2.51E-16 | 0.927948718 | 0.889 | 0.243 | 5.08E-12 | 26 | Rhoj |
| Phf21a13 | 2.59E-16 | 1.079209226 | 0.833 | 0.254 | 5.24E-12 | 26 | Phf21a |
| Ifi2074 | 2.68E-16 | 0.652801733 | 0.556 | 0.108 | 5.43E-12 | 26 | Ifi207 |
| Farp16 | 2.73E-16 | 0.805407324 | 0.556 | 0.109 | 5.53E-12 | 26 | Farp1 |
| Abca16 | 2.81E-16 | 0.831851601 | 0.611 | 0.131 | 5.69E-12 | 26 | Abca1 |
| Plekho22 | 2.94E-16 | 0.536329131 | 0.472 | 0.083 | 5.95E-12 | 26 | Plekho2 |
| Copa4 | 3.14E-16 | 0.867293448 | 0.778 | 0.206 | 6.36E-12 | 26 | Copa |
| Alcam3 | 3.15E-16 | 0.455554784 | 0.278 | 0.032 | 6.37E-12 | 26 | Alcam |
| Dtx2 | 3.34E-16 | 0.378898125 | 0.278 | 0.032 | 6.76E-12 | 26 | Dtx2 |
| Alms11 | 3.39E-16 | 0.399212387 | 0.278 | 0.033 | 6.86E-12 | 26 | Alms1 |
| Sec24a1 | 3.69E-16 | 0.697444204 | 0.472 | 0.087 | 7.48E-12 | 26 | Sec24a |
| Afap1l23 | 4.06E-16 | 0.47851147 | 0.25 | 0.027 | 8.23E-12 | 26 | Afap1l2 |
| Fndc3b8 | 4.25E-16 | 1.120918685 | 0.833 | 0.246 | 8.61E-12 | 26 | Fndc3b |

| | | | | | | | |
|---|---|---|---|---|---|---|---|
| Vkorc14 | 4.36E-16 | 0.720885442 | 0.667 | 0.154 | 8.82E-12 | 26 | Vkorc1 |
| Itga23 | 4.54E-16 | 0.363350149 | 0.306 | 0.038 | 9.20E-12 | 26 | Itga2 |
| Zdhhc3 | 4.95E-16 | 0.473815045 | 0.5 | 0.092 | 1.00E-11 | 26 | Zdhhc3 |
| Spag910 | 5.11E-16 | 1.039112237 | 0.972 | 0.36 | 1.03E-11 | 26 | Spag9 |
| Ssbp31 | 5.40E-16 | 0.587711703 | 0.722 | 0.171 | 1.09E-11 | 26 | Ssbp3 |
| Sdccag86 | 6.43E-16 | 0.645120712 | 0.583 | 0.123 | 1.30E-11 | 26 | Sdccag8 |
| Selenos4 | 6.43E-16 | 0.900747019 | 0.806 | 0.233 | 1.30E-11 | 26 | Selenos |
| Stat5b4 | 6.46E-16 | 0.839019446 | 0.639 | 0.148 | 1.31E-11 | 26 | Stat5b |
| Rab11fip24 | 6.76E-16 | 0.451586668 | 0.556 | 0.108 | 1.37E-11 | 26 | Rab11fip2 |
| Ibtk | 6.83E-16 | 0.376495182 | 0.417 | 0.067 | 1.38E-11 | 26 | Ibtk |
| Bcr1 | 8.93E-16 | 0.52793429 | 0.389 | 0.06 | 1.81E-11 | 26 | Bcr |
| Appl2 | 9.34E-16 | 0.305982423 | 0.417 | 0.067 | 1.89E-11 | 26 | Appl2 |
| Ltbp31 | 9.39E-16 | 0.295587162 | 0.25 | 0.027 | 1.90E-11 | 26 | Ltbp3 |
| 2210408I21Rik1 | 9.60E-16 | 0.670854099 | 0.389 | 0.061 | 1.94E-11 | 26 | 2210408I21Rik |
| Hif1a3 | 9.68E-16 | 0.708059629 | 0.778 | 0.201 | 1.96E-11 | 26 | Hif1a |
| Mill25 | 1.02E-15 | 0.752865863 | 0.417 | 0.07 | 2.06E-11 | 26 | Mill2 |
| Itgb31 | 1.02E-15 | 0.465040847 | 0.25 | 0.028 | 2.07E-11 | 26 | Itgb3 |
| Erlec12 | 1.03E-15 | 0.695601655 | 0.667 | 0.152 | 2.08E-11 | 26 | Erlec1 |
| Kif13b7 | 1.05E-15 | 0.962673596 | 0.806 | 0.217 | 2.12E-11 | 26 | Kif13b |
| Nin5 | 1.07E-15 | 0.822171028 | 0.611 | 0.14 | 2.17E-11 | 26 | Nin |
| Plpp5 | 1.16E-15 | 0.500328486 | 0.333 | 0.047 | 2.35E-11 | 26 | Plpp5 |
| Ahnak12 | 1.20E-15 | 1.41173705 | 1 | 0.474 | 2.43E-11 | 26 | Ahnak |
| Rnf2171 | 1.21E-15 | 0.354588508 | 0.306 | 0.039 | 2.46E-11 | 26 | Rnf217 |
| Acaca3 | 1.33E-15 | 0.447913407 | 0.472 | 0.084 | 2.69E-11 | 26 | Acaca |
| Filip1l8 | 1.38E-15 | 0.79672326 | 0.639 | 0.154 | 2.79E-11 | 26 | Filip1l |
| Plscr48 | 1.40E-15 | 0.842621797 | 0.75 | 0.191 | 2.83E-11 | 26 | Plscr4 |
| Sec31a1 | 1.40E-15 | 0.925490277 | 0.667 | 0.173 | 2.83E-11 | 26 | Sec31a |
| Asap27 | 1.40E-15 | 0.444483342 | 0.639 | 0.133 | 2.84E-11 | 26 | Asap2 |
| Armcx21 | 1.44E-15 | 0.265218144 | 0.25 | 0.027 | 2.92E-11 | 26 | Armcx2 |
| Soga12 | 1.51E-15 | 0.354190552 | 0.389 | 0.06 | 3.07E-11 | 26 | Soga1 |
| Atp8b15 | 1.60E-15 | 0.654017885 | 0.528 | 0.104 | 3.25E-11 | 26 | Atp8b1 |
| Slc38a6 | 1.67E-15 | 0.369083315 | 0.361 | 0.053 | 3.38E-11 | 26 | Slc38a6 |
| Dennd1a14 | 1.68E-15 | 0.986144629 | 0.833 | 0.255 | 3.40E-11 | 26 | Dennd1a |
| Eif2ak33 | 1.69E-15 | 0.469026942 | 0.472 | 0.085 | 3.42E-11 | 26 | Eif2ak3 |
| Arhgap223 | 1.74E-15 | 0.325642517 | 0.333 | 0.046 | 3.53E-11 | 26 | Arhgap22 |
| Exoc6b11 | 1.88E-15 | 1.008736326 | 0.833 | 0.242 | 3.81E-11 | 26 | Exoc6b |
| Il15ra1 | 1.91E-15 | 0.427427477 | 0.417 | 0.068 | 3.86E-11 | 26 | Il15ra |
| Smad34 | 1.95E-15 | 0.653230745 | 0.5 | 0.097 | 3.95E-11 | 26 | Smad3 |
| Ksr1 | 1.99E-15 | 0.413824629 | 0.444 | 0.076 | 4.03E-11 | 26 | Ksr1 |
| Erlin1 | 2.22E-15 | 0.354620652 | 0.306 | 0.04 | 4.50E-11 | 26 | Erlin1 |
| Itgav12 | 2.29E-15 | 0.572656235 | 0.694 | 0.162 | 4.64E-11 | 26 | Itgav |
| Ssh29 | 2.30E-15 | 1.029615051 | 0.778 | 0.227 | 4.65E-11 | 26 | Ssh2 |
| Thrb | 2.35E-15 | 0.426171409 | 0.306 | 0.04 | 4.76E-11 | 26 | Thrb |

| | | | | | | | |
|---|---|---|---|---|---|---|---|
| Tfdp11 | 2.39E-15 | 0.286024164 | 0.361 | 0.053 | 4.84E-11 | 26 | Tfdp1 |
| Peli22 | 2.47E-15 | 0.382876197 | 0.417 | 0.068 | 5.00E-11 | 26 | Peli2 |
| Zfhx36 | 2.67E-15 | 0.863289875 | 0.722 | 0.184 | 5.41E-11 | 26 | Zfhx3 |
| Flna8 | 2.68E-15 | 0.793895367 | 0.583 | 0.133 | 5.43E-11 | 26 | Flna |
| Rassf23 | 3.00E-15 | 0.441500225 | 0.361 | 0.055 | 6.08E-11 | 26 | Rassf2 |
| Rgmb2 | 3.26E-15 | 0.452927288 | 0.389 | 0.062 | 6.59E-11 | 26 | Rgmb |
| Abhd21 | 3.49E-15 | 0.519231025 | 0.444 | 0.079 | 7.07E-11 | 26 | Abhd2 |
| P4ha2 | 3.53E-15 | 0.391782687 | 0.25 | 0.028 | 7.14E-11 | 26 | P4ha2 |
| Lmnb12 | 4.31E-15 | 0.344178418 | 0.361 | 0.054 | 8.72E-11 | 26 | Lmnb1 |
| Add17 | 4.35E-15 | 0.758999604 | 0.778 | 0.216 | 8.80E-11 | 26 | Add1 |
| Amotl23 | 4.76E-15 | 0.580191965 | 0.472 | 0.087 | 9.64E-11 | 26 | Amotl2 |
| Klf128 | 4.82E-15 | 0.860538018 | 0.778 | 0.209 | 9.75E-11 | 26 | Klf12 |
| Ppp1r14b3 | 5.60E-15 | 1.031646515 | 0.944 | 0.328 | 1.13E-10 | 26 | Ppp1r14b |
| Usp40 | 5.86E-15 | 0.445046623 | 0.389 | 0.063 | 1.19E-10 | 26 | Usp40 |
| Wipf110 | 6.20E-15 | 0.984403453 | 0.833 | 0.27 | 1.26E-10 | 26 | Wipf1 |
| Fbxw17 | 6.39E-15 | 0.363396103 | 0.25 | 0.029 | 1.29E-10 | 26 | Fbxw17 |
| Mast24 | 6.46E-15 | 0.564453125 | 0.417 | 0.073 | 1.31E-10 | 26 | Mast2 |
| Stxbp4 | 6.76E-15 | 0.535639186 | 0.25 | 0.029 | 1.37E-10 | 26 | Stxbp4 |
| Morf4l21 | 6.81E-15 | 0.57383439 | 0.806 | 0.215 | 1.38E-10 | 26 | Morf4l2 |
| Rusc2 | 7.52E-15 | 0.362353662 | 0.361 | 0.054 | 1.52E-10 | 26 | Rusc2 |
| Rab3il13 | 7.55E-15 | 0.405960029 | 0.417 | 0.07 | 1.53E-10 | 26 | Rab3il1 |
| Zdhhc51 | 8.27E-15 | 0.370544832 | 0.472 | 0.086 | 1.67E-10 | 26 | Zdhhc5 |
| Calcrl6 | 8.28E-15 | 1.001937155 | 0.889 | 0.265 | 1.68E-10 | 26 | Calcrl |
| Focad | 8.41E-15 | 0.298629576 | 0.278 | 0.035 | 1.70E-10 | 26 | Focad |
| Alg9 | 8.99E-15 | 0.389536227 | 0.278 | 0.035 | 1.82E-10 | 26 | Alg9 |
| Lrp64 | 9.36E-15 | 0.808867626 | 0.611 | 0.146 | 1.89E-10 | 26 | Lrp6 |
| Mkl19 | 1.01E-14 | 0.585321707 | 0.583 | 0.127 | 2.05E-10 | 26 | Mkl1 |
| Mia36 | 1.03E-14 | 0.86568441 | 0.667 | 0.171 | 2.09E-10 | 26 | Mia3 |
| Pi4kb | 1.04E-14 | 0.442080584 | 0.417 | 0.072 | 2.11E-10 | 26 | Pi4kb |
| Dnajc181 | 1.07E-14 | 0.315820829 | 0.361 | 0.055 | 2.16E-10 | 26 | Dnajc18 |
| Evi5l | 1.07E-14 | 0.346369853 | 0.25 | 0.029 | 2.17E-10 | 26 | Evi5l |
| Pdia44 | 1.12E-14 | 0.788332305 | 0.667 | 0.168 | 2.26E-10 | 26 | Pdia4 |
| Mapk9 | 1.19E-14 | 0.366221852 | 0.528 | 0.105 | 2.40E-10 | 26 | Mapk9 |
| Ttc288 | 1.27E-14 | 1.065447102 | 0.889 | 0.286 | 2.57E-10 | 26 | Ttc28 |
| Plekha59 | 1.34E-14 | 0.882534026 | 0.639 | 0.157 | 2.71E-10 | 26 | Plekha5 |
| Osbpl6 | 1.34E-14 | 0.252790796 | 0.25 | 0.029 | 2.71E-10 | 26 | Osbpl6 |
| Tek7 | 1.45E-14 | 1.009204617 | 0.861 | 0.262 | 2.94E-10 | 26 | Tek |
| Slc35a5 | 1.52E-14 | 0.389907787 | 0.278 | 0.036 | 3.07E-10 | 26 | Slc35a5 |
| Idh11 | 1.57E-14 | 0.609295451 | 0.389 | 0.065 | 3.17E-10 | 26 | Idh1 |
| Apbb1ip9 | 1.60E-14 | 0.554761831 | 0.722 | 0.175 | 3.24E-10 | 26 | Apbb1ip |
| Atic | 1.77E-14 | 0.384045536 | 0.361 | 0.057 | 3.57E-10 | 26 | Atic |
| Tmem651 | 1.81E-14 | 0.40686489 | 0.5 | 0.097 | 3.66E-10 | 26 | Tmem65 |
| Slc35f5 | 1.87E-14 | 0.371797704 | 0.306 | 0.042 | 3.79E-10 | 26 | Slc35f5 |

| | | | | | | | |
|---|---|---|---|---|---|---|---|
| Tbc1d191 | 1.89E-14 | 0.586991098 | 0.306 | 0.043 | 3.83E-10 | 26 | Tbc1d19 |
| Phlpp19 | 1.93E-14 | 1.076764872 | 0.833 | 0.265 | 3.91E-10 | 26 | Phlpp1 |
| Fam168a5 | 2.06E-14 | 0.738810495 | 0.611 | 0.144 | 4.18E-10 | 26 | Fam168a |
| Rpn210 | 2.06E-14 | 0.987275041 | 0.722 | 0.213 | 4.18E-10 | 26 | Rpn2 |
| Brwd14 | 2.12E-14 | 0.609068346 | 0.611 | 0.143 | 4.30E-10 | 26 | Brwd1 |
| Arnt4 | 2.19E-14 | 0.766431587 | 0.611 | 0.143 | 4.43E-10 | 26 | Arnt |
| Atl35 | 2.29E-14 | 0.533263004 | 0.583 | 0.13 | 4.63E-10 | 26 | Atl3 |
| Ino80c | 2.33E-14 | 0.351410226 | 0.278 | 0.036 | 4.72E-10 | 26 | Ino80c |
| Usp22 | 2.40E-14 | 0.26774437 | 0.417 | 0.071 | 4.86E-10 | 26 | Usp22 |
| Lnpep15 | 2.44E-14 | 0.882334069 | 0.833 | 0.261 | 4.95E-10 | 26 | Lnpep |
| Spats2 | 2.47E-14 | 0.425953376 | 0.389 | 0.065 | 5.01E-10 | 26 | Spats2 |
| Atp2a29 | 2.48E-14 | 0.724432104 | 0.75 | 0.202 | 5.01E-10 | 26 | Atp2a2 |
| Plxnb22 | 2.53E-14 | 0.401986347 | 0.333 | 0.05 | 5.11E-10 | 26 | Plxnb2 |
| Uggt11 | 2.71E-14 | 0.469842716 | 0.583 | 0.127 | 5.48E-10 | 26 | Uggt1 |
| Rnf170 | 2.74E-14 | 0.293372548 | 0.306 | 0.042 | 5.54E-10 | 26 | Rnf170 |
| Tmem63a1 | 2.90E-14 | 0.435582285 | 0.417 | 0.073 | 5.87E-10 | 26 | Tmem63a |
| Btbd115 | 3.02E-14 | 0.369823911 | 0.361 | 0.057 | 6.11E-10 | 26 | Btbd11 |
| Cpsf7 | 3.07E-14 | 0.361677448 | 0.472 | 0.089 | 6.21E-10 | 26 | Cpsf7 |
| Canx10 | 3.08E-14 | 0.955733946 | 0.889 | 0.308 | 6.23E-10 | 26 | Canx |
| Taok39 | 3.11E-14 | 0.730496731 | 0.667 | 0.165 | 6.30E-10 | 26 | Taok3 |
| Dram1 | 3.15E-14 | 0.313166011 | 0.306 | 0.042 | 6.37E-10 | 26 | Dram1 |
| Gfpt11 | 3.25E-14 | 0.41166339 | 0.583 | 0.131 | 6.58E-10 | 26 | Gfpt1 |
| Atp7a2 | 3.27E-14 | 0.443714529 | 0.444 | 0.082 | 6.62E-10 | 26 | Atp7a |
| Parva6 | 3.45E-14 | 0.721454001 | 0.528 | 0.113 | 6.99E-10 | 26 | Parva |
| Cnih11 | 3.50E-14 | 0.419067256 | 0.583 | 0.126 | 7.09E-10 | 26 | Cnih1 |
| Sned12 | 3.63E-14 | 0.511142832 | 0.389 | 0.066 | 7.35E-10 | 26 | Sned1 |
| Slc39a6 | 3.74E-14 | 0.273848051 | 0.278 | 0.036 | 7.57E-10 | 26 | Slc39a6 |
| Tbl1x8 | 4.45E-14 | 0.788248776 | 0.861 | 0.264 | 9.01E-10 | 26 | Tbl1x |
| Tns111 | 4.48E-14 | 0.674338932 | 0.861 | 0.246 | 9.08E-10 | 26 | Tns1 |
| 2510009E07Rik4 | 5.02E-14 | 0.455149171 | 0.583 | 0.126 | 1.02E-09 | 26 | 2510009E07Rik |
| Rps6kc1 | 5.13E-14 | 0.383155666 | 0.306 | 0.044 | 1.04E-09 | 26 | Rps6kc1 |
| Abhd5 | 5.18E-14 | 0.275504703 | 0.417 | 0.072 | 1.05E-09 | 26 | Abhd5 |
| Ifi2111 | 5.28E-14 | 0.339073588 | 0.333 | 0.05 | 1.07E-09 | 26 | Ifi211 |
| Lyst7 | 5.52E-14 | 0.478458853 | 0.528 | 0.111 | 1.12E-09 | 26 | Lyst |
| Rere9 | 5.91E-14 | 0.940814017 | 0.833 | 0.273 | 1.20E-09 | 26 | Rere |
| Itgb55 | 6.05E-14 | 0.479261062 | 0.639 | 0.146 | 1.22E-09 | 26 | Itgb5 |
| Klhl221 | 6.85E-14 | 0.375994278 | 0.306 | 0.044 | 1.39E-09 | 26 | Klhl22 |
| Pds5b8 | 6.87E-14 | 0.662842425 | 0.694 | 0.181 | 1.39E-09 | 26 | Pds5b |
| Kat6b5 | 7.20E-14 | 0.569217725 | 0.639 | 0.153 | 1.46E-09 | 26 | Kat6b |
| Rad51b2 | 7.40E-14 | 0.766465667 | 0.278 | 0.039 | 1.50E-09 | 26 | Rad51b |
| Carm1 | 7.48E-14 | 0.304985106 | 0.333 | 0.051 | 1.51E-09 | 26 | Carm1 |
| St3gal52 | 7.68E-14 | 0.446007825 | 0.444 | 0.082 | 1.55E-09 | 26 | St3gal5 |
| Prkci | 7.75E-14 | 0.276735406 | 0.278 | 0.037 | 1.57E-09 | 26 | Prkci |

| | | | | | | | |
|---|---|---|---|---|---|---|---|
| Nars2 | 7.78E-14 | 0.396008029 | 0.333 | 0.051 | 1.57E-09 | 26 | Nars2 |
| Gulp19 | 8.12E-14 | 0.772291482 | 0.667 | 0.168 | 1.64E-09 | 26 | Gulp1 |
| Sec636 | 8.86E-14 | 0.668851456 | 0.75 | 0.205 | 1.79E-09 | 26 | Sec63 |
| Zadh21 | 9.36E-14 | 0.489976709 | 0.5 | 0.104 | 1.89E-09 | 26 | Zadh2 |
| Zfp9503 | 1.08E-13 | 0.503458835 | 0.528 | 0.113 | 2.19E-09 | 26 | Zfp950 |
| Vangl12 | 1.09E-13 | 0.411183006 | 0.389 | 0.066 | 2.22E-09 | 26 | Vangl1 |
| Zfp7043 | 1.10E-13 | 0.490497765 | 0.583 | 0.131 | 2.23E-09 | 26 | Zfp704 |
| Sar1a1 | 1.26E-13 | 0.667323869 | 0.722 | 0.2 | 2.56E-09 | 26 | Sar1a |
| Smo1 | 1.31E-13 | 0.365980237 | 0.306 | 0.045 | 2.65E-09 | 26 | Smo |
| Tmem39a1 | 1.34E-13 | 0.300469502 | 0.333 | 0.051 | 2.71E-09 | 26 | Tmem39a |
| Calr11 | 1.39E-13 | 1.047559147 | 0.972 | 0.41 | 2.81E-09 | 26 | Calr |
| Gigyf26 | 1.41E-13 | 0.570182017 | 0.667 | 0.169 | 2.86E-09 | 26 | Gigyf2 |
| Zfp74 | 1.45E-13 | 0.401961887 | 0.333 | 0.052 | 2.93E-09 | 26 | Zfp74 |
| Arhgap121 | 1.47E-13 | 0.46413869 | 0.556 | 0.124 | 2.97E-09 | 26 | Arhgap12 |
| Crtap1 | 1.62E-13 | 0.6117867 | 0.417 | 0.078 | 3.29E-09 | 26 | Crtap |
| Ptprs3 | 1.73E-13 | 0.356980697 | 0.278 | 0.038 | 3.49E-09 | 26 | Ptprs |
| Rspo31 | 1.77E-13 | 0.552646275 | 0.306 | 0.045 | 3.59E-09 | 26 | Rspo3 |
| Acly | 1.88E-13 | 0.299162734 | 0.611 | 0.136 | 3.80E-09 | 26 | Acly |
| Fhit6 | 2.04E-13 | 0.982940685 | 0.417 | 0.081 | 4.12E-09 | 26 | Fhit |
| Tmem2631 | 2.13E-13 | 0.346688636 | 0.444 | 0.084 | 4.31E-09 | 26 | Tmem263 |
| Armcx1 | 2.25E-13 | 0.312424059 | 0.278 | 0.038 | 4.55E-09 | 26 | Armcx1 |
| Itga92 | 2.25E-13 | 0.471711419 | 0.5 | 0.103 | 4.56E-09 | 26 | Itga9 |
| Klf36 | 2.30E-13 | 0.824421609 | 0.806 | 0.238 | 4.65E-09 | 26 | Klf3 |
| Ammecr1 | 2.31E-13 | 0.428627543 | 0.361 | 0.061 | 4.68E-09 | 26 | Ammecr1 |
| Mbtps11 | 2.39E-13 | 0.43603733 | 0.5 | 0.104 | 4.85E-09 | 26 | Mbtps1 |
| Surf42 | 2.42E-13 | 0.718604632 | 0.583 | 0.145 | 4.90E-09 | 26 | Surf4 |
| Wdr45b | 2.53E-13 | 0.444828791 | 0.611 | 0.145 | 5.12E-09 | 26 | Wdr45b |
| Xist16 | 2.64E-13 | 1.259731994 | 0.972 | 0.465 | 5.34E-09 | 26 | Xist |
| Ttc37 | 2.64E-13 | 0.437742857 | 0.444 | 0.086 | 5.35E-09 | 26 | Ttc37 |
| Ubap1 | 2.73E-13 | 0.328010115 | 0.417 | 0.076 | 5.52E-09 | 26 | Ubap1 |
| Snx42 | 2.78E-13 | 0.615366713 | 0.556 | 0.131 | 5.63E-09 | 26 | Snx4 |
| Ambra110 | 2.94E-13 | 0.559174778 | 0.75 | 0.199 | 5.96E-09 | 26 | Ambra1 |
| Dnajc104 | 2.95E-13 | 0.62604766 | 0.583 | 0.141 | 5.98E-09 | 26 | Dnajc10 |
| Stxbp56 | 3.16E-13 | 0.660742993 | 0.556 | 0.131 | 6.39E-09 | 26 | Stxbp5 |
| Apba14 | 3.50E-13 | 0.631584447 | 0.333 | 0.054 | 7.09E-09 | 26 | Apba1 |
| Agl | 3.67E-13 | 0.27732033 | 0.333 | 0.053 | 7.43E-09 | 26 | Agl |
| Pvt17 | 3.67E-13 | 0.591258977 | 0.417 | 0.08 | 7.44E-09 | 26 | Pvt1 |
| Gpatch814 | 3.73E-13 | 1.005581029 | 0.778 | 0.26 | 7.55E-09 | 26 | Gpatch8 |
| Kdm6a12 | 3.75E-13 | 0.76597205 | 0.75 | 0.215 | 7.58E-09 | 26 | Kdm6a |
| Taok19 | 4.09E-13 | 0.546179549 | 0.861 | 0.253 | 8.27E-09 | 26 | Taok1 |
| Ophn14 | 4.34E-13 | 0.509657387 | 0.583 | 0.134 | 8.80E-09 | 26 | Ophn1 |
| Serpinb6a2 | 4.35E-13 | 0.924586265 | 0.972 | 0.41 | 8.81E-09 | 26 | Serpinb6a |
| Rab3gap13 | 4.37E-13 | 0.485198907 | 0.667 | 0.165 | 8.85E-09 | 26 | Rab3gap1 |

| | | | | | | | |
|---|---|---|---|---|---|---|---|
| Tmem127 | 4.56E-13 | 0.518875117 | 0.417 | 0.08 | 9.24E-09 | 26 | Tmem127 |
| Cul9 | 5.14E-13 | 0.429464357 | 0.306 | 0.047 | 1.04E-08 | 26 | Cul9 |
| Fam46a2 | 5.30E-13 | 0.59425394 | 0.583 | 0.142 | 1.07E-08 | 26 | Fam46a |
| Syde1 | 5.70E-13 | 0.291266618 | 0.278 | 0.039 | 1.15E-08 | 26 | Syde1 |
| Pola12 | 6.50E-13 | 0.487462332 | 0.389 | 0.07 | 1.31E-08 | 26 | Pola1 |
| Dap3 | 6.57E-13 | 0.504615461 | 0.694 | 0.188 | 1.33E-08 | 26 | Dap |
| Cdkn1c7 | 6.72E-13 | 1.435332687 | 0.528 | 0.127 | 1.36E-08 | 26 | Cdkn1c |
| Pcyt1a1 | 6.83E-13 | 0.45325071 | 0.444 | 0.088 | 1.38E-08 | 26 | Pcyt1a |
| Anks16 | 7.05E-13 | 0.595299903 | 0.583 | 0.14 | 1.43E-08 | 26 | Anks1 |
| Renbp | 8.08E-13 | 0.251198272 | 0.278 | 0.039 | 1.64E-08 | 26 | Renbp |
| Smarca28 | 8.12E-13 | 0.99310575 | 0.806 | 0.282 | 1.64E-08 | 26 | Smarca2 |
| Tmem214 | 8.64E-13 | 0.335705166 | 0.361 | 0.062 | 1.75E-08 | 26 | Tmem214 |
| Nucb16 | 9.86E-13 | 0.876834018 | 0.75 | 0.229 | 2.00E-08 | 26 | Nucb1 |
| Abca3 | 1.03E-12 | 0.373642189 | 0.389 | 0.071 | 2.08E-08 | 26 | Abca3 |
| Klc14 | 1.04E-12 | 0.514778403 | 0.611 | 0.151 | 2.10E-08 | 26 | Klc1 |
| Heatr5a1 | 1.08E-12 | 0.486152393 | 0.528 | 0.119 | 2.19E-08 | 26 | Heatr5a |
| Rpn14 | 1.09E-12 | 0.836071551 | 0.722 | 0.223 | 2.20E-08 | 26 | Rpn1 |
| Slc9a11 | 1.14E-12 | 0.481804862 | 0.389 | 0.073 | 2.31E-08 | 26 | Slc9a1 |
| Wnk112 | 1.18E-12 | 0.869987608 | 0.972 | 0.365 | 2.39E-08 | 26 | Wnk1 |
| Fam13c7 | 1.21E-12 | 0.381973215 | 0.528 | 0.113 | 2.44E-08 | 26 | Fam13c |
| Csnk1g110 | 1.22E-12 | 0.657370046 | 0.611 | 0.159 | 2.47E-08 | 26 | Csnk1g1 |
| Asb6 | 1.23E-12 | 0.250416098 | 0.25 | 0.033 | 2.49E-08 | 26 | Asb6 |
| Spg20 | 1.27E-12 | 0.326616693 | 0.361 | 0.063 | 2.56E-08 | 26 | Spg20 |
| Eprs2 | 1.28E-12 | 0.635228619 | 0.722 | 0.203 | 2.60E-08 | 26 | Eprs |
| Pan314 | 1.34E-12 | 0.666499915 | 0.778 | 0.23 | 2.72E-08 | 26 | Pan3 |
| Slc39a12 | 1.42E-12 | 0.512471712 | 0.389 | 0.073 | 2.87E-08 | 26 | Slc39a1 |
| Ncoa61 | 1.44E-12 | 0.638440651 | 0.556 | 0.135 | 2.92E-08 | 26 | Ncoa6 |
| Cfh3 | 1.55E-12 | 0.271451702 | 0.389 | 0.07 | 3.14E-08 | 26 | Cfh |
| Iqsec21 | 1.66E-12 | 0.525164239 | 0.472 | 0.101 | 3.37E-08 | 26 | Iqsec2 |
| Cdc42bpa9 | 1.68E-12 | 0.648286241 | 0.694 | 0.186 | 3.39E-08 | 26 | Cdc42bpa |
| Sipa1l3 | 1.70E-12 | 0.408170885 | 0.417 | 0.081 | 3.44E-08 | 26 | Sipa1l3 |
| Zcchc715 | 1.74E-12 | 0.954531648 | 0.861 | 0.303 | 3.52E-08 | 26 | Zcchc7 |
| Camk2d11 | 1.74E-12 | 0.896764066 | 0.806 | 0.271 | 3.53E-08 | 26 | Camk2d |
| Ghr5 | 1.76E-12 | 0.963897812 | 0.583 | 0.153 | 3.57E-08 | 26 | Ghr |
| Pdgfd6 | 1.80E-12 | 0.942635398 | 0.806 | 0.255 | 3.65E-08 | 26 | Pdgfd |
| Rrbp18 | 1.89E-12 | 1.224929878 | 0.944 | 0.461 | 3.83E-08 | 26 | Rrbp1 |
| Rbpj9 | 1.90E-12 | 0.863339385 | 0.722 | 0.222 | 3.84E-08 | 26 | Rbpj |
| Fkbp73 | 1.90E-12 | 0.548315283 | 0.417 | 0.083 | 3.84E-08 | 26 | Fkbp7 |
| Oat5 | 1.92E-12 | 0.273888098 | 0.583 | 0.138 | 3.88E-08 | 26 | Oat |
| Eif4g15 | 1.95E-12 | 0.751609359 | 0.778 | 0.237 | 3.95E-08 | 26 | Eif4g1 |
| Ccdc171 | 1.97E-12 | 0.407721972 | 0.25 | 0.034 | 3.99E-08 | 26 | Ccdc171 |
| Nadk | 2.00E-12 | 0.334363099 | 0.583 | 0.139 | 4.04E-08 | 26 | Nadk |
| Atp8b2 | 2.08E-12 | 0.305510247 | 0.278 | 0.041 | 4.21E-08 | 26 | Atp8b2 |

| | | | | | | | |
|---|---|---|---|---|---|---|---|
| Fbxo341 | 2.18E-12 | 0.339778074 | 0.361 | 0.064 | 4.41E-08 | 26 | Fbxo34 |
| Zfp7031 | 2.18E-12 | 0.333913002 | 0.333 | 0.056 | 4.42E-08 | 26 | Zfp703 |
| Man1a26 | 2.23E-12 | 0.500847227 | 0.806 | 0.228 | 4.51E-08 | 26 | Man1a2 |
| Tgfb1i13 | 2.32E-12 | 0.416879547 | 0.444 | 0.089 | 4.70E-08 | 26 | Tgfb1i1 |
| Htt2 | 2.34E-12 | 0.532578961 | 0.472 | 0.104 | 4.74E-08 | 26 | Htt |
| Elp44 | 2.40E-12 | 0.470853634 | 0.556 | 0.129 | 4.85E-08 | 26 | Elp4 |
| Tmem106b2 | 2.45E-12 | 0.410252739 | 0.556 | 0.128 | 4.96E-08 | 26 | Tmem106b |
| Slc8a14 | 2.75E-12 | 0.296783615 | 0.361 | 0.063 | 5.57E-08 | 26 | Slc8a1 |
| Zfp2602 | 2.87E-12 | 0.399075293 | 0.5 | 0.11 | 5.81E-08 | 26 | Zfp260 |
| Fto8 | 2.90E-12 | 0.712744387 | 0.667 | 0.192 | 5.87E-08 | 26 | Fto |
| Cp4 | 2.97E-12 | 0.270994248 | 0.306 | 0.047 | 6.01E-08 | 26 | Cp |
| Tbc1d32 | 3.05E-12 | 0.380244505 | 0.306 | 0.048 | 6.18E-08 | 26 | Tbc1d32 |
| Tango21 | 3.14E-12 | 0.363239276 | 0.361 | 0.065 | 6.35E-08 | 26 | Tango2 |
| Cebpd3 | 3.25E-12 | 0.510985701 | 0.667 | 0.177 | 6.57E-08 | 26 | Cebpd |
| mt-Nd119 | 3.30E-12 | 1.155860464 | 1 | 0.601 | 6.67E-08 | 26 | mt-Nd1 |
| Magt17 | 3.50E-12 | 0.476012157 | 0.639 | 0.166 | 7.08E-08 | 26 | Magt1 |
| Zfp322a | 3.87E-12 | 0.313814689 | 0.417 | 0.082 | 7.83E-08 | 26 | Zfp322a |
| Ctsb8 | 3.97E-12 | 0.601215903 | 0.889 | 0.3 | 8.05E-08 | 26 | Ctsb |
| Srek17 | 3.99E-12 | 0.780753432 | 0.667 | 0.195 | 8.08E-08 | 26 | Srek1 |
| Tbc1d8b | 4.07E-12 | 0.408389124 | 0.333 | 0.057 | 8.25E-08 | 26 | Tbc1d8b |
| Atp9a1 | 4.19E-12 | 0.294406245 | 0.25 | 0.034 | 8.48E-08 | 26 | Atp9a |
| Ftx2 | 4.25E-12 | 0.360838222 | 0.306 | 0.049 | 8.61E-08 | 26 | Ftx |
| Scyl1 | 4.32E-12 | 0.409435134 | 0.389 | 0.074 | 8.75E-08 | 26 | Scyl1 |
| Smarcc26 | 4.33E-12 | 0.513577453 | 0.667 | 0.179 | 8.76E-08 | 26 | Smarcc2 |
| Lgals3bp5 | 4.44E-12 | 0.681435717 | 0.583 | 0.148 | 8.99E-08 | 26 | Lgals3bp |
| Gm268832 | 4.45E-12 | 0.502543432 | 0.417 | 0.082 | 9.00E-08 | 26 | Gm26883 |
| Cald111 | 4.69E-12 | 0.750531034 | 0.944 | 0.324 | 9.50E-08 | 26 | Cald1 |
| Pros17 | 4.70E-12 | 0.494000122 | 0.611 | 0.152 | 9.52E-08 | 26 | Pros1 |
| Dync2h11 | 4.78E-12 | 0.422535694 | 0.306 | 0.05 | 9.68E-08 | 26 | Dync2h1 |
| Tom1l27 | 5.24E-12 | 0.448679884 | 0.556 | 0.132 | 1.06E-07 | 26 | Tom1l2 |
| Hgsnat3 | 5.38E-12 | 0.420011811 | 0.417 | 0.084 | 1.09E-07 | 26 | Hgsnat |
| Ppa1 | 5.96E-12 | 0.421882237 | 0.528 | 0.123 | 1.21E-07 | 26 | Ppa1 |
| Stard97 | 6.10E-12 | 0.484681186 | 0.611 | 0.153 | 1.24E-07 | 26 | Stard9 |
| Ssr17 | 6.35E-12 | 0.592668826 | 0.667 | 0.188 | 1.29E-07 | 26 | Ssr1 |
| Pacsin24 | 6.48E-12 | 0.432703768 | 0.611 | 0.159 | 1.31E-07 | 26 | Pacsin2 |
| Ncor21 | 6.63E-12 | 0.378621378 | 0.528 | 0.123 | 1.34E-07 | 26 | Ncor2 |
| Hook39 | 6.68E-12 | 0.741095052 | 0.833 | 0.292 | 1.35E-07 | 26 | Hook3 |
| Ckb9 | 7.19E-12 | 0.530904045 | 0.75 | 0.218 | 1.46E-07 | 26 | Ckb |
| Adgra31 | 7.26E-12 | 0.307993923 | 0.278 | 0.042 | 1.47E-07 | 26 | Adgra3 |
| Calhm21 | 7.86E-12 | 0.389199976 | 0.361 | 0.067 | 1.59E-07 | 26 | Calhm2 |
| Ano67 | 8.09E-12 | 0.816398563 | 0.639 | 0.182 | 1.64E-07 | 26 | Ano6 |
| Bet1l1 | 8.37E-12 | 0.389102335 | 0.528 | 0.124 | 1.70E-07 | 26 | Bet1l |
| Rassf82 | 8.56E-12 | 0.367657761 | 0.417 | 0.083 | 1.73E-07 | 26 | Rassf8 |

| | | | | | | | |
|---|---|---|---|---|---|---|---|
| Sel1l5 | 8.67E-12 | 0.415952113 | 0.639 | 0.163 | 1.76E-07 | 26 | Sel1l |
| Ogt11 | 8.73E-12 | 0.751253499 | 0.694 | 0.208 | 1.77E-07 | 26 | Ogt |
| Abcc1 | 8.80E-12 | 0.323783325 | 0.333 | 0.058 | 1.78E-07 | 26 | Abcc1 |
| 2900097C17Rik2 | 8.95E-12 | 0.636038914 | 0.583 | 0.153 | 1.81E-07 | 26 | 2900097C17Rik |
| Neat112 | 9.02E-12 | 1.070510983 | 0.667 | 0.211 | 1.83E-07 | 26 | Neat1 |
| Nfic5 | 9.82E-12 | 0.525066279 | 0.833 | 0.263 | 1.99E-07 | 26 | Nfic |
| Ybx32 | 9.95E-12 | 0.649471118 | 0.917 | 0.34 | 2.02E-07 | 26 | Ybx3 |
| Ahi1 | 1.03E-11 | 0.375812765 | 0.278 | 0.043 | 2.09E-07 | 26 | Ahi1 |
| Kif1b7 | 1.03E-11 | 0.839228716 | 0.75 | 0.245 | 2.09E-07 | 26 | Kif1b |
| Zfp6088 | 1.03E-11 | 0.796961703 | 0.556 | 0.142 | 2.09E-07 | 26 | Zfp608 |
| Dcun1d3 | 1.04E-11 | 0.339949707 | 0.306 | 0.05 | 2.12E-07 | 26 | Dcun1d3 |
| Purg1 | 1.07E-11 | 0.346065037 | 0.278 | 0.043 | 2.16E-07 | 26 | Purg |
| Lrba9 | 1.08E-11 | 0.782707302 | 0.667 | 0.193 | 2.19E-07 | 26 | Lrba |
| Ssh1 | 1.09E-11 | 0.49857736 | 0.417 | 0.087 | 2.21E-07 | 26 | Ssh1 |
| Cast8 | 1.11E-11 | 0.855803182 | 0.833 | 0.313 | 2.25E-07 | 26 | Cast |
| Slc37a3 | 1.19E-11 | 0.260272511 | 0.278 | 0.042 | 2.41E-07 | 26 | Slc37a3 |
| Hsp90b114 | 1.26E-11 | 1.032894345 | 1 | 0.512 | 2.56E-07 | 26 | Hsp90b1 |
| 2410089E03Rik1 | 1.30E-11 | 0.458137073 | 0.444 | 0.096 | 2.64E-07 | 26 | 2410089E03Rik |
| Sfxn31 | 1.31E-11 | 0.274330358 | 0.306 | 0.05 | 2.65E-07 | 26 | Sfxn3 |
| Acap29 | 1.33E-11 | 0.68977725 | 0.833 | 0.271 | 2.70E-07 | 26 | Acap2 |
| 1110051M20Rik1 | 1.38E-11 | 0.427359514 | 0.306 | 0.052 | 2.80E-07 | 26 | 1110051M20Rik |
| 8030462N17Rik | 1.43E-11 | 0.36754591 | 0.361 | 0.068 | 2.89E-07 | 26 | 8030462N17Rik |
| Rc3h21 | 1.45E-11 | 0.515663732 | 0.5 | 0.118 | 2.94E-07 | 26 | Rc3h2 |
| Cd99l2 | 1.49E-11 | 0.332634168 | 0.333 | 0.059 | 3.02E-07 | 26 | Cd99l2 |
| Srgap211 | 1.57E-11 | 0.780285595 | 0.861 | 0.306 | 3.17E-07 | 26 | Srgap2 |
| Lmbrd19 | 1.70E-11 | 0.380079718 | 0.639 | 0.164 | 3.45E-07 | 26 | Lmbrd1 |
| 2310035C23Rik7 | 1.72E-11 | 0.530126739 | 0.556 | 0.142 | 3.49E-07 | 26 | 2310035C23Rik |
| Atp9b7 | 1.73E-11 | 0.725750502 | 0.583 | 0.159 | 3.50E-07 | 26 | Atp9b |
| Plbd21 | 1.74E-11 | 0.350224124 | 0.389 | 0.076 | 3.53E-07 | 26 | Plbd2 |
| Ints8 | 1.75E-11 | 0.424929185 | 0.417 | 0.087 | 3.54E-07 | 26 | Ints8 |
| Fasn1 | 1.78E-11 | 0.332102247 | 0.25 | 0.037 | 3.60E-07 | 26 | Fasn |
| Pir1 | 1.82E-11 | 0.458012516 | 0.278 | 0.044 | 3.68E-07 | 26 | Pir |
| Ncoa211 | 1.90E-11 | 0.768090158 | 0.722 | 0.24 | 3.85E-07 | 26 | Ncoa2 |
| Usp245 | 1.94E-11 | 0.75758564 | 0.528 | 0.138 | 3.92E-07 | 26 | Usp24 |
| Slc35d1 | 1.95E-11 | 0.260740531 | 0.25 | 0.036 | 3.95E-07 | 26 | Slc35d1 |
| Asap18 | 1.99E-11 | 0.854641256 | 0.944 | 0.358 | 4.02E-07 | 26 | Asap1 |
| Elovl63 | 2.10E-11 | 0.302635924 | 0.306 | 0.051 | 4.26E-07 | 26 | Elovl6 |
| Stt3b10 | 2.15E-11 | 0.569243087 | 0.611 | 0.166 | 4.35E-07 | 26 | Stt3b |
| Ski8 | 2.29E-11 | 0.595376747 | 0.722 | 0.216 | 4.64E-07 | 26 | Ski |
| Leo1 | 2.31E-11 | 0.285904129 | 0.361 | 0.068 | 4.67E-07 | 26 | Leo1 |
| Ttc7b5 | 2.45E-11 | 0.597892991 | 0.5 | 0.123 | 4.96E-07 | 26 | Ttc7b |
| Dnmbp | 2.47E-11 | 0.318344452 | 0.389 | 0.076 | 4.99E-07 | 26 | Dnmbp |
| Tmem106a1 | 2.48E-11 | 0.283844932 | 0.306 | 0.051 | 5.01E-07 | 26 | Tmem106a |

| | | | | | | | |
|---|---|---|---|---|---|---|---|
| Ino804 | 2.57E-11 | 0.470886531 | 0.556 | 0.141 | 5.19E-07 | 26 | Ino80 |
| Dmxl19 | 2.77E-11 | 0.502584581 | 0.667 | 0.187 | 5.60E-07 | 26 | Dmxl1 |
| Slc33a1 | 2.78E-11 | 0.309876646 | 0.306 | 0.052 | 5.62E-07 | 26 | Slc33a1 |
| Xpnpep3 | 2.85E-11 | 0.302764413 | 0.278 | 0.044 | 5.76E-07 | 26 | Xpnpep3 |
| Mxra78 | 2.90E-11 | 0.399239743 | 0.583 | 0.143 | 5.87E-07 | 26 | Mxra7 |
| Pmm11 | 2.94E-11 | 0.274079364 | 0.389 | 0.077 | 5.95E-07 | 26 | Pmm1 |
| Ubfd1 | 3.10E-11 | 0.292951714 | 0.361 | 0.069 | 6.28E-07 | 26 | Ubfd1 |
| Ankib17 | 3.11E-11 | 0.461625102 | 0.556 | 0.14 | 6.30E-07 | 26 | Ankib1 |
| Il6st8 | 3.18E-11 | 0.433591574 | 0.806 | 0.234 | 6.43E-07 | 26 | Il6st |
| Cavin310 | 3.26E-11 | 0.629499165 | 0.917 | 0.309 | 6.60E-07 | 26 | Cavin3 |
| Hcfc2 | 3.42E-11 | 0.323792286 | 0.278 | 0.045 | 6.92E-07 | 26 | Hcfc2 |
| Slc30a7 | 3.47E-11 | 0.354810705 | 0.361 | 0.07 | 7.02E-07 | 26 | Slc30a7 |
| Arhgap351 | 3.53E-11 | 0.491335779 | 0.417 | 0.089 | 7.14E-07 | 26 | Arhgap35 |
| Tmcc114 | 3.55E-11 | 0.780213464 | 0.694 | 0.219 | 7.18E-07 | 26 | Tmcc1 |
| Gbf19 | 3.73E-11 | 0.62263387 | 0.639 | 0.183 | 7.55E-07 | 26 | Gbf1 |
| Diaph18 | 3.80E-11 | 0.802369633 | 0.722 | 0.244 | 7.69E-07 | 26 | Diaph1 |
| Rbm610 | 3.85E-11 | 0.577183163 | 0.722 | 0.219 | 7.79E-07 | 26 | Rbm6 |
| Trip129 | 3.87E-11 | 0.649165156 | 0.778 | 0.258 | 7.84E-07 | 26 | Trip12 |
| Aida5 | 4.01E-11 | 0.532519233 | 0.556 | 0.144 | 8.12E-07 | 26 | Aida |
| Tfpi2 | 4.02E-11 | 0.31739803 | 0.333 | 0.06 | 8.15E-07 | 26 | Tfpi |
| Smarcad1 | 4.20E-11 | 0.288623708 | 0.5 | 0.116 | 8.50E-07 | 26 | Smarcad1 |
| Ylpm12 | 4.30E-11 | 0.478382427 | 0.556 | 0.141 | 8.71E-07 | 26 | Ylpm1 |
| Pdia612 | 4.32E-11 | 0.762738324 | 0.917 | 0.339 | 8.74E-07 | 26 | Pdia6 |
| Nr6a1 | 4.42E-11 | 0.365322297 | 0.306 | 0.053 | 8.94E-07 | 26 | Nr6a1 |
| Glb11 | 4.42E-11 | 0.417777451 | 0.361 | 0.071 | 8.94E-07 | 26 | Glb1 |
| Ube4a2 | 4.54E-11 | 0.478654425 | 0.472 | 0.111 | 9.20E-07 | 26 | Ube4a |
| Ifi27l2a5 | 4.67E-11 | 0.461895041 | 0.944 | 0.368 | 9.46E-07 | 26 | Ifi27l2a |
| Cndp2 | 4.98E-11 | 0.39435841 | 0.361 | 0.071 | 1.01E-06 | 26 | Cndp2 |
| Alg11 | 5.28E-11 | 0.271991316 | 0.25 | 0.037 | 1.07E-06 | 26 | Alg11 |
| Aak18 | 5.52E-11 | 0.876645834 | 0.611 | 0.186 | 1.12E-06 | 26 | Aak1 |
| Pstk | 5.58E-11 | 0.266723633 | 0.389 | 0.078 | 1.13E-06 | 26 | Pstk |
| Exoc26 | 5.62E-11 | 0.480029748 | 0.583 | 0.149 | 1.14E-06 | 26 | Exoc2 |
| Inpp5f1 | 5.79E-11 | 0.360901743 | 0.389 | 0.079 | 1.17E-06 | 26 | Inpp5f |
| Arl157 | 6.24E-11 | 0.714171028 | 0.944 | 0.383 | 1.26E-06 | 26 | Arl15 |
| Stk384 | 6.62E-11 | 0.438190044 | 0.75 | 0.228 | 1.34E-06 | 26 | Stk38 |
| Golt1b1 | 6.99E-11 | 0.380151536 | 0.306 | 0.054 | 1.41E-06 | 26 | Golt1b |
| Ccdc507 | 7.00E-11 | 0.625753345 | 0.722 | 0.23 | 1.42E-06 | 26 | Ccdc50 |
| Ganab2 | 7.07E-11 | 0.30900134 | 0.5 | 0.117 | 1.43E-06 | 26 | Ganab |
| Agfg13 | 7.12E-11 | 0.659529875 | 0.583 | 0.165 | 1.44E-06 | 26 | Agfg1 |
| Setd514 | 7.88E-11 | 0.605663328 | 0.806 | 0.265 | 1.59E-06 | 26 | Setd5 |
| Dock47 | 8.21E-11 | 0.872487541 | 0.833 | 0.298 | 1.66E-06 | 26 | Dock4 |
| Lims27 | 8.70E-11 | 0.490417688 | 0.722 | 0.203 | 1.76E-06 | 26 | Lims2 |
| Aff47 | 9.09E-11 | 0.510647756 | 0.778 | 0.242 | 1.84E-06 | 26 | Aff4 |

| | | | | | | | |
|---|---|---|---|---|---|---|---|
| Dtnbp14 | 9.09E-11 | 0.437599527 | 0.75 | 0.229 | 1.84E-06 | 26 | Dtnbp1 |
| Usp9x4 | 9.35E-11 | 0.649214726 | 0.722 | 0.236 | 1.89E-06 | 26 | Usp9x |
| Nfe2l26 | 9.40E-11 | 0.671340428 | 0.833 | 0.303 | 1.90E-06 | 26 | Nfe2l2 |
| Spidr3 | 9.45E-11 | 0.475124729 | 0.472 | 0.113 | 1.91E-06 | 26 | Spidr |
| Trip101 | 9.58E-11 | 0.297465138 | 0.444 | 0.097 | 1.94E-06 | 26 | Trip10 |
| Kdm5b3 | 9.69E-11 | 0.412803359 | 0.528 | 0.132 | 1.96E-06 | 26 | Kdm5b |
| Zfp282 | 9.84E-11 | 0.326655279 | 0.306 | 0.053 | 1.99E-06 | 26 | Zfp282 |
| Atp13a1 | 9.92E-11 | 0.285126529 | 0.306 | 0.054 | 2.01E-06 | 26 | Atp13a1 |
| Sav12 | 1.03E-10 | 0.274586275 | 0.472 | 0.107 | 2.08E-06 | 26 | Sav1 |
| Adk3 | 1.04E-10 | 0.526964236 | 0.5 | 0.126 | 2.10E-06 | 26 | Adk |
| Ssbp211 | 1.08E-10 | 0.388144062 | 0.556 | 0.142 | 2.19E-06 | 26 | Ssbp2 |
| Uba61 | 1.11E-10 | 0.418370913 | 0.389 | 0.083 | 2.25E-06 | 26 | Uba6 |
| Tram13 | 1.12E-10 | 0.642763467 | 0.694 | 0.221 | 2.28E-06 | 26 | Tram1 |
| Dhx57 | 1.14E-10 | 0.429601958 | 0.389 | 0.082 | 2.30E-06 | 26 | Dhx57 |
| Sil11 | 1.15E-10 | 0.40396696 | 0.389 | 0.083 | 2.32E-06 | 26 | Sil1 |
| Vps13b11 | 1.17E-10 | 0.761679475 | 0.667 | 0.221 | 2.38E-06 | 26 | Vps13b |
| Slc2a132 | 1.21E-10 | 0.285778201 | 0.25 | 0.038 | 2.45E-06 | 26 | Slc2a13 |
| Fnip17 | 1.22E-10 | 0.473030432 | 0.556 | 0.148 | 2.48E-06 | 26 | Fnip1 |
| Trim30d2 | 1.30E-10 | 0.546003686 | 0.389 | 0.083 | 2.64E-06 | 26 | Trim30d |
| Rsrc19 | 1.31E-10 | 0.668866272 | 0.778 | 0.262 | 2.64E-06 | 26 | Rsrc1 |
| Atp6v1b22 | 1.33E-10 | 0.36323565 | 0.472 | 0.112 | 2.69E-06 | 26 | Atp6v1b2 |
| Dlc17 | 1.39E-10 | 0.813283583 | 0.972 | 0.364 | 2.80E-06 | 26 | Dlc1 |
| Ssr35 | 1.51E-10 | 0.630680572 | 0.639 | 0.197 | 3.06E-06 | 26 | Ssr3 |
| Ap3s11 | 1.55E-10 | 0.473347288 | 0.583 | 0.16 | 3.14E-06 | 26 | Ap3s1 |
| Abi19 | 1.58E-10 | 0.622824368 | 0.778 | 0.271 | 3.19E-06 | 26 | Abi1 |
| Ric8b | 1.60E-10 | 0.362900201 | 0.361 | 0.073 | 3.23E-06 | 26 | Ric8b |
| Hivep212 | 1.67E-10 | 0.890877696 | 0.806 | 0.3 | 3.38E-06 | 26 | Hivep2 |
| Rps6ka310 | 1.68E-10 | 0.726517357 | 0.778 | 0.264 | 3.40E-06 | 26 | Rps6ka3 |
| Glg18 | 1.72E-10 | 0.659590417 | 0.75 | 0.254 | 3.48E-06 | 26 | Glg1 |
| Usp331 | 1.73E-10 | 0.395379874 | 0.417 | 0.093 | 3.50E-06 | 26 | Usp33 |
| Ndst1 | 1.75E-10 | 0.284937058 | 0.333 | 0.063 | 3.54E-06 | 26 | Ndst1 |
| Twf1 | 1.75E-10 | 0.324038464 | 0.5 | 0.122 | 3.55E-06 | 26 | Twf1 |
| Ckap56 | 1.81E-10 | 0.377509319 | 0.556 | 0.143 | 3.66E-06 | 26 | Ckap5 |
| Btrc2 | 1.84E-10 | 0.424492625 | 0.472 | 0.112 | 3.72E-06 | 26 | Btrc |
| Golgb18 | 1.87E-10 | 0.60700262 | 0.778 | 0.264 | 3.79E-06 | 26 | Golgb1 |
| Parp810 | 1.92E-10 | 0.560299743 | 0.556 | 0.151 | 3.89E-06 | 26 | Parp8 |
| Sec61a11 | 1.94E-10 | 0.315913001 | 0.611 | 0.17 | 3.93E-06 | 26 | Sec61a1 |
| Fam193a8 | 1.98E-10 | 0.612911243 | 0.611 | 0.181 | 4.00E-06 | 26 | Fam193a |
| Bms11 | 2.03E-10 | 0.323041363 | 0.472 | 0.112 | 4.11E-06 | 26 | Bms1 |
| Kat2b5 | 2.10E-10 | 0.471566965 | 0.583 | 0.16 | 4.26E-06 | 26 | Kat2b |
| mt-Cytb17 | 2.21E-10 | 0.953026261 | 1 | 0.702 | 4.47E-06 | 26 | mt-Cytb |
| Synrg3 | 2.27E-10 | 0.335303056 | 0.556 | 0.144 | 4.59E-06 | 26 | Synrg |
| Stt3a1 | 2.31E-10 | 0.354581721 | 0.528 | 0.134 | 4.67E-06 | 26 | Stt3a |

| | | | | | | | |
|---|---|---|---|---|---|---|---|
| Ehd29 | 2.31E-10 | 0.400199351 | 0.694 | 0.195 | 4.68E-06 | 26 | Ehd2 |
| mt-Nd418 | 2.33E-10 | 1.018634656 | 1 | 0.677 | 4.72E-06 | 26 | mt-Nd4 |
| Copb21 | 2.40E-10 | 0.452210059 | 0.694 | 0.212 | 4.86E-06 | 26 | Copb2 |
| Manf3 | 2.41E-10 | 0.859493456 | 0.806 | 0.33 | 4.87E-06 | 26 | Manf |
| Apex2 | 2.45E-10 | 0.342265559 | 0.306 | 0.056 | 4.95E-06 | 26 | Apex2 |
| Plcb47 | 2.53E-10 | 0.527092828 | 0.861 | 0.293 | 5.11E-06 | 26 | Plcb4 |
| St3gal32 | 2.67E-10 | 0.383778497 | 0.361 | 0.075 | 5.40E-06 | 26 | St3gal3 |
| Slc12a41 | 2.91E-10 | 0.337938059 | 0.306 | 0.056 | 5.89E-06 | 26 | Slc12a4 |
| Cerk9 | 3.06E-10 | 0.468150877 | 0.472 | 0.116 | 6.20E-06 | 26 | Cerk |
| Plec8 | 3.17E-10 | 0.531862411 | 0.611 | 0.176 | 6.41E-06 | 26 | Plec |
| Fmo5 | 3.26E-10 | 0.353334421 | 0.278 | 0.048 | 6.59E-06 | 26 | Fmo5 |
| Mgat24 | 3.27E-10 | 0.564474059 | 0.556 | 0.157 | 6.62E-06 | 26 | Mgat2 |
| Zfp4231 | 3.55E-10 | 0.301738855 | 0.361 | 0.072 | 7.19E-06 | 26 | Zfp423 |
| Rnf19a3 | 3.67E-10 | 0.523622133 | 0.528 | 0.141 | 7.44E-06 | 26 | Rnf19a |
| Flot21 | 3.71E-10 | 0.329559353 | 0.389 | 0.084 | 7.51E-06 | 26 | Flot2 |
| Atxn24 | 3.80E-10 | 0.498582133 | 0.528 | 0.141 | 7.70E-06 | 26 | Atxn2 |
| Spast1 | 3.85E-10 | 0.318600456 | 0.472 | 0.114 | 7.79E-06 | 26 | Spast |
| Uba1 | 3.88E-10 | 0.549809338 | 0.583 | 0.175 | 7.86E-06 | 26 | Uba1 |
| Yipf32 | 3.89E-10 | 0.329150162 | 0.556 | 0.146 | 7.87E-06 | 26 | Yipf3 |
| Dhrs76 | 3.93E-10 | 0.527733906 | 0.694 | 0.217 | 7.97E-06 | 26 | Dhrs7 |
| Invs1 | 3.94E-10 | 0.415617656 | 0.361 | 0.074 | 7.97E-06 | 26 | Invs |
| Srsf61 | 3.95E-10 | 0.331061065 | 0.611 | 0.171 | 7.99E-06 | 26 | Srsf6 |
| Bag33 | 4.03E-10 | 0.427305178 | 0.472 | 0.115 | 8.17E-06 | 26 | Bag3 |
| Tspan37 | 4.05E-10 | 0.448719016 | 0.444 | 0.106 | 8.19E-06 | 26 | Tspan3 |
| Ddx177 | 4.11E-10 | 0.707523186 | 0.75 | 0.275 | 8.31E-06 | 26 | Ddx17 |
| Trpm78 | 4.13E-10 | 0.546086314 | 0.667 | 0.206 | 8.36E-06 | 26 | Trpm7 |
| Ccdc61 | 4.15E-10 | 0.421330511 | 0.389 | 0.087 | 8.40E-06 | 26 | Ccdc6 |
| Med134 | 4.26E-10 | 0.490003915 | 0.667 | 0.207 | 8.63E-06 | 26 | Med13 |
| mt-Co118 | 4.27E-10 | 0.891340817 | 1 | 0.721 | 8.64E-06 | 26 | mt-Co1 |
| Hsd17b123 | 4.32E-10 | 0.373516895 | 0.528 | 0.136 | 8.75E-06 | 26 | Hsd17b12 |
| Cbr31 | 4.37E-10 | 0.345579507 | 0.361 | 0.075 | 8.85E-06 | 26 | Cbr3 |
| Lars1 | 4.44E-10 | 0.320533261 | 0.583 | 0.16 | 8.98E-06 | 26 | Lars |
| Rapgef18 | 4.48E-10 | 0.630894017 | 0.722 | 0.242 | 9.06E-06 | 26 | Rapgef1 |
| Lonp24 | 4.56E-10 | 0.37048855 | 0.667 | 0.196 | 9.24E-06 | 26 | Lonp2 |
| Dopey12 | 4.62E-10 | 0.550132665 | 0.444 | 0.109 | 9.35E-06 | 26 | Dopey1 |
| Ash1l11 | 4.65E-10 | 0.625129824 | 0.833 | 0.301 | 9.41E-06 | 26 | Ash1l |
| Rabgap15 | 4.79E-10 | 0.464742912 | 0.583 | 0.165 | 9.69E-06 | 26 | Rabgap1 |
| Slmap5 | 4.91E-10 | 0.586315873 | 0.639 | 0.195 | 9.93E-06 | 26 | Slmap |
| Iah13 | 4.99E-10 | 0.297805629 | 0.528 | 0.138 | 1.01E-05 | 26 | Iah1 |
| Acvr12 | 5.48E-10 | 0.354318787 | 0.333 | 0.066 | 1.11E-05 | 26 | Acvr1 |
| Urgcp | 5.56E-10 | 0.294755871 | 0.361 | 0.075 | 1.12E-05 | 26 | Urgcp |
| Emp16 | 5.63E-10 | 0.427023069 | 0.5 | 0.123 | 1.14E-05 | 26 | Emp1 |
| Dock18 | 5.70E-10 | 0.451161733 | 0.861 | 0.282 | 1.15E-05 | 26 | Dock1 |

| | | | | | | | |
|---|---|---|---|---|---|---|---|
| Wars21 | 5.82E-10 | 0.400731304 | 0.306 | 0.058 | 1.18E-05 | 26 | Wars2 |
| Zmym45 | 5.92E-10 | 0.49176333 | 0.583 | 0.167 | 1.20E-05 | 26 | Zmym4 |
| Cyp20a12 | 5.96E-10 | 0.398967585 | 0.389 | 0.086 | 1.21E-05 | 26 | Cyp20a1 |
| D5Ertd579e1 | 6.25E-10 | 0.547583588 | 0.417 | 0.099 | 1.26E-05 | 26 | D5Ertd579e |
| Stx18 | 6.25E-10 | 0.546910579 | 0.444 | 0.113 | 1.27E-05 | 26 | Stx18 |
| S100a169 | 6.62E-10 | 0.721055852 | 0.75 | 0.26 | 1.34E-05 | 26 | S100a16 |
| Vps13a8 | 6.96E-10 | 0.456434381 | 0.611 | 0.174 | 1.41E-05 | 26 | Vps13a |
| Sgms110 | 7.01E-10 | 0.492337082 | 0.5 | 0.13 | 1.42E-05 | 26 | Sgms1 |
| Ptdss1 | 7.11E-10 | 0.276829848 | 0.333 | 0.066 | 1.44E-05 | 26 | Ptdss1 |
| Smarcc15 | 7.25E-10 | 0.406927029 | 0.528 | 0.144 | 1.47E-05 | 26 | Smarcc1 |
| Zfp532 | 7.29E-10 | 0.328203057 | 0.306 | 0.057 | 1.48E-05 | 26 | Zfp532 |
| Pigs | 7.45E-10 | 0.35022566 | 0.278 | 0.05 | 1.51E-05 | 26 | Pigs |
| Rtn4rl13 | 7.46E-10 | 0.378275651 | 0.278 | 0.05 | 1.51E-05 | 26 | Rtn4rl1 |
| Zfyve16 | 7.55E-10 | 0.532545663 | 0.361 | 0.079 | 1.53E-05 | 26 | Zfyve16 |
| Atxn106 | 7.62E-10 | 0.407123644 | 0.611 | 0.177 | 1.54E-05 | 26 | Atxn10 |
| Grk58 | 8.13E-10 | 0.59915645 | 0.556 | 0.159 | 1.65E-05 | 26 | Grk5 |
| Rap2a4 | 8.18E-10 | 0.476103925 | 0.472 | 0.119 | 1.66E-05 | 26 | Rap2a |
| Large18 | 8.24E-10 | 0.818182385 | 0.528 | 0.154 | 1.67E-05 | 26 | Large1 |
| Gabpb23 | 8.64E-10 | 0.404513827 | 0.472 | 0.12 | 1.75E-05 | 26 | Gabpb2 |
| Exoc414 | 8.81E-10 | 0.735992998 | 0.889 | 0.324 | 1.78E-05 | 26 | Exoc4 |
| Hdac43 | 9.13E-10 | 0.385564474 | 0.444 | 0.107 | 1.85E-05 | 26 | Hdac4 |
| Pdia312 | 9.31E-10 | 0.776052329 | 0.944 | 0.445 | 1.88E-05 | 26 | Pdia3 |
| Ubn211 | 9.53E-10 | 0.598372281 | 0.75 | 0.246 | 1.93E-05 | 26 | Ubn2 |
| Thada7 | 9.56E-10 | 0.484357446 | 0.472 | 0.121 | 1.94E-05 | 26 | Thada |
| Cpne87 | 9.83E-10 | 0.621415146 | 0.722 | 0.23 | 1.99E-05 | 26 | Cpne8 |
| Hdac87 | 1.01E-09 | 0.591800578 | 0.5 | 0.137 | 2.05E-05 | 26 | Hdac8 |
| Elmsan15 | 1.02E-09 | 0.34403328 | 0.444 | 0.107 | 2.06E-05 | 26 | Elmsan1 |
| Bcl7c2 | 1.03E-09 | 0.342140271 | 0.75 | 0.245 | 2.09E-05 | 26 | Bcl7c |
| Dcaf63 | 1.04E-09 | 0.267716238 | 0.528 | 0.135 | 2.11E-05 | 26 | Dcaf6 |
| Cyb561 | 1.04E-09 | 0.264711443 | 0.278 | 0.049 | 2.11E-05 | 26 | Cyb561 |
| Eps154 | 1.05E-09 | 0.531569634 | 0.583 | 0.171 | 2.13E-05 | 26 | Eps15 |
| Svil6 | 1.05E-09 | 0.652988756 | 0.722 | 0.245 | 2.13E-05 | 26 | Svil |
| Lztfl11 | 1.07E-09 | 0.378378233 | 0.444 | 0.109 | 2.17E-05 | 26 | Lztfl1 |
| Appl16 | 1.08E-09 | 0.450961637 | 0.611 | 0.183 | 2.19E-05 | 26 | Appl1 |
| Lrrc8d4 | 1.12E-09 | 0.542121289 | 0.5 | 0.134 | 2.27E-05 | 26 | Lrrc8d |
| Mbd510 | 1.12E-09 | 0.511276603 | 0.694 | 0.222 | 2.27E-05 | 26 | Mbd5 |
| Gorasp21 | 1.14E-09 | 0.26442899 | 0.5 | 0.127 | 2.31E-05 | 26 | Gorasp2 |
| Capns17 | 1.22E-09 | 0.553298826 | 0.889 | 0.35 | 2.46E-05 | 26 | Capns1 |
| Psap8 | 1.24E-09 | 0.627133916 | 0.917 | 0.4 | 2.52E-05 | 26 | Psap |
| Psen16 | 1.25E-09 | 0.458725005 | 0.417 | 0.102 | 2.52E-05 | 26 | Psen1 |
| Simc1 | 1.25E-09 | 0.33407083 | 0.25 | 0.042 | 2.53E-05 | 26 | Simc1 |
| Mon24 | 1.28E-09 | 0.396083276 | 0.556 | 0.154 | 2.58E-05 | 26 | Mon2 |
| Asb7 | 1.28E-09 | 0.320249621 | 0.25 | 0.042 | 2.59E-05 | 26 | Asb7 |

| | | | | | | | |
|---|---|---|---|---|---|---|---|
| Trak2 | 1.30E-09 | 0.370761134 | 0.389 | 0.088 | 2.63E-05 | 26 | Trak2 |
| Golga44 | 1.34E-09 | 0.457216208 | 0.556 | 0.16 | 2.72E-05 | 26 | Golga4 |
| Cant1 | 1.37E-09 | 0.289122726 | 0.306 | 0.059 | 2.77E-05 | 26 | Cant1 |
| Eps85 | 1.40E-09 | 0.429984453 | 0.389 | 0.088 | 2.84E-05 | 26 | Eps8 |
| Fam120a5 | 1.49E-09 | 0.437593483 | 0.611 | 0.183 | 3.01E-05 | 26 | Fam120a |
| mt-Nd214 | 1.51E-09 | 1.014899709 | 1 | 0.69 | 3.06E-05 | 26 | mt-Nd2 |
| Pcca2 | 1.58E-09 | 0.389316616 | 0.472 | 0.123 | 3.20E-05 | 26 | Pcca |
| Ptpn93 | 1.63E-09 | 0.464542531 | 0.528 | 0.143 | 3.29E-05 | 26 | Ptpn9 |
| mt-Co320 | 1.65E-09 | 0.848490364 | 1 | 0.728 | 3.34E-05 | 26 | mt-Co3 |
| mt-Nd316 | 1.65E-09 | 0.77722335 | 0.972 | 0.448 | 3.35E-05 | 26 | mt-Nd3 |
| Clcn41 | 1.66E-09 | 0.41126225 | 0.472 | 0.122 | 3.35E-05 | 26 | Clcn4 |
| Pias2 | 1.71E-09 | 0.505216588 | 0.444 | 0.115 | 3.46E-05 | 26 | Pias2 |
| Jpt18 | 1.71E-09 | 0.49318843 | 0.694 | 0.224 | 3.47E-05 | 26 | Jpt1 |
| Ddost6 | 1.71E-09 | 0.44526389 | 0.694 | 0.218 | 3.47E-05 | 26 | Ddost |
| Tmem184b4 | 1.75E-09 | 0.256294688 | 0.472 | 0.115 | 3.54E-05 | 26 | Tmem184b |
| Mark36 | 1.79E-09 | 0.599167425 | 0.667 | 0.214 | 3.62E-05 | 26 | Mark3 |
| Tmed35 | 1.80E-09 | 0.648362131 | 0.694 | 0.245 | 3.63E-05 | 26 | Tmed3 |
| Mettl11 | 1.80E-09 | 0.270664302 | 0.306 | 0.06 | 3.65E-05 | 26 | Mettl1 |
| Rab6b1 | 1.84E-09 | 0.406977909 | 0.333 | 0.071 | 3.73E-05 | 26 | Rab6b |
| Med151 | 1.87E-09 | 0.436836853 | 0.5 | 0.134 | 3.78E-05 | 26 | Med15 |
| Tbc1d232 | 1.89E-09 | 0.444933938 | 0.472 | 0.124 | 3.82E-05 | 26 | Tbc1d23 |
| Gorab | 1.98E-09 | 0.277374805 | 0.25 | 0.043 | 4.00E-05 | 26 | Gorab |
| Amotl110 | 2.10E-09 | 0.406648964 | 0.639 | 0.187 | 4.25E-05 | 26 | Amotl1 |
| S100a1010 | 2.11E-09 | 0.789158218 | 0.972 | 0.471 | 4.27E-05 | 26 | S100a10 |
| Strn38 | 2.12E-09 | 0.648158541 | 0.861 | 0.347 | 4.29E-05 | 26 | Strn3 |
| Acot91 | 2.13E-09 | 0.34769763 | 0.472 | 0.121 | 4.30E-05 | 26 | Acot9 |
| Tor1b | 2.30E-09 | 0.268494622 | 0.361 | 0.078 | 4.65E-05 | 26 | Tor1b |
| Prrc2b8 | 2.50E-09 | 0.541396785 | 0.694 | 0.231 | 5.07E-05 | 26 | Prrc2b |
| Dip2a | 2.54E-09 | 0.28024333 | 0.306 | 0.06 | 5.14E-05 | 26 | Dip2a |
| Fam120b2 | 2.55E-09 | 0.412771766 | 0.389 | 0.091 | 5.16E-05 | 26 | Fam120b |
| Rcn26 | 2.55E-09 | 0.395921167 | 0.528 | 0.144 | 5.16E-05 | 26 | Rcn2 |
| Sec23a1 | 2.56E-09 | 0.4018255 | 0.333 | 0.071 | 5.18E-05 | 26 | Sec23a |
| Cbfb9 | 2.57E-09 | 0.379441734 | 0.556 | 0.157 | 5.21E-05 | 26 | Cbfb |
| Cdc42ep25 | 2.61E-09 | 0.37601594 | 0.611 | 0.174 | 5.29E-05 | 26 | Cdc42ep2 |
| Mfap23 | 2.62E-09 | 0.393417479 | 0.444 | 0.108 | 5.31E-05 | 26 | Mfap2 |
| Tmem1649 | 2.67E-09 | 0.361703396 | 0.583 | 0.165 | 5.41E-05 | 26 | Tmem164 |
| Sp33 | 2.75E-09 | 0.538291482 | 0.611 | 0.192 | 5.57E-05 | 26 | Sp3 |
| Afap17 | 2.82E-09 | 0.578130743 | 0.528 | 0.152 | 5.71E-05 | 26 | Afap1 |
| Rictor7 | 2.93E-09 | 0.427807621 | 0.556 | 0.158 | 5.93E-05 | 26 | Rictor |
| Bcas310 | 2.97E-09 | 0.496978976 | 0.611 | 0.184 | 6.01E-05 | 26 | Bcas3 |
| Rps6kb13 | 3.09E-09 | 0.527594879 | 0.556 | 0.159 | 6.26E-05 | 26 | Rps6kb1 |
| Rap2c | 3.17E-09 | 0.394649219 | 0.361 | 0.082 | 6.42E-05 | 26 | Rap2c |
| Washc44 | 3.27E-09 | 0.548663802 | 0.472 | 0.13 | 6.61E-05 | 26 | Washc4 |

| | | | | | | | |
|---|---|---|---|---|---|---|---|
| Olfm11 | 3.28E-09 | 0.288835167 | 0.25 | 0.044 | 6.64E-05 | 26 | Olfm1 |
| Zbp13 | 3.30E-09 | 0.274676912 | 0.333 | 0.071 | 6.69E-05 | 26 | Zbp1 |
| Arntl4 | 3.37E-09 | 0.278387898 | 0.472 | 0.12 | 6.83E-05 | 26 | Arntl |
| Pofut1 | 3.39E-09 | 0.430791747 | 0.278 | 0.053 | 6.85E-05 | 26 | Pofut1 |
| Snx14 | 3.39E-09 | 0.258869986 | 0.528 | 0.141 | 6.87E-05 | 26 | Snx1 |
| Fip1l12 | 3.41E-09 | 0.479190536 | 0.639 | 0.201 | 6.91E-05 | 26 | Fip1l1 |
| Stam1 | 3.47E-09 | 0.275416183 | 0.333 | 0.07 | 7.02E-05 | 26 | Stam |
| Ift271 | 3.54E-09 | 0.279680957 | 0.389 | 0.091 | 7.16E-05 | 26 | Ift27 |
| Sh3gl14 | 3.62E-09 | 0.444467098 | 0.472 | 0.126 | 7.33E-05 | 26 | Sh3gl1 |
| Bmp2k4 | 3.66E-09 | 0.334717049 | 0.444 | 0.112 | 7.42E-05 | 26 | Bmp2k |
| Il31ra2 | 3.67E-09 | 0.455988151 | 0.639 | 0.21 | 7.43E-05 | 26 | Il31ra |
| Gm265612 | 3.79E-09 | 0.29176334 | 0.861 | 0.311 | 7.67E-05 | 26 | Gm26561 |
| Dpysl210 | 3.92E-09 | 0.516857831 | 0.583 | 0.176 | 7.94E-05 | 26 | Dpysl2 |
| Slc7a7 | 4.04E-09 | 0.276133985 | 0.25 | 0.044 | 8.18E-05 | 26 | Slc7a7 |
| Copg11 | 4.05E-09 | 0.25645579 | 0.417 | 0.099 | 8.20E-05 | 26 | Copg1 |
| Ppp6r21 | 4.06E-09 | 0.455065417 | 0.333 | 0.073 | 8.22E-05 | 26 | Ppp6r2 |
| Styx | 4.07E-09 | 0.323367506 | 0.25 | 0.044 | 8.23E-05 | 26 | Styx |
| Zcchc144 | 4.73E-09 | 0.369913684 | 0.361 | 0.081 | 9.58E-05 | 26 | Zcchc14 |
| Ttc1412 | 5.00E-09 | 0.527822041 | 0.667 | 0.217 | 0.000101193 | 26 | Ttc14 |
| Rptor7 | 5.18E-09 | 0.442476943 | 0.5 | 0.14 | 0.000104871 | 26 | Rptor |
| Comt5 | 5.19E-09 | 0.471291668 | 0.611 | 0.195 | 0.000105169 | 26 | Comt |
| Clip21 | 5.24E-09 | 0.298055101 | 0.278 | 0.053 | 0.000105989 | 26 | Clip2 |
| Adam175 | 5.28E-09 | 0.616983661 | 0.528 | 0.155 | 0.000106844 | 26 | Adam17 |
| Zbtb45 | 5.29E-09 | 0.317311255 | 0.472 | 0.122 | 0.000107046 | 26 | Zbtb4 |
| Ostc5 | 5.35E-09 | 0.575253644 | 0.722 | 0.264 | 0.000108315 | 26 | Ostc |
| Pik3ca5 | 5.53E-09 | 0.348526338 | 0.583 | 0.17 | 0.000111984 | 26 | Pik3ca |
| Stim26 | 5.54E-09 | 0.463393399 | 0.639 | 0.201 | 0.000112243 | 26 | Stim2 |
| Tmem87a1 | 5.55E-09 | 0.328288426 | 0.444 | 0.113 | 0.000112272 | 26 | Tmem87a |
| Tm7sf32 | 5.56E-09 | 0.309478854 | 0.333 | 0.072 | 0.000112518 | 26 | Tm7sf3 |
| Cand1 | 5.68E-09 | 0.416408672 | 0.361 | 0.084 | 0.000115 | 26 | Cand1 |
| Ptpra8 | 5.93E-09 | 0.616522185 | 0.806 | 0.306 | 0.000120032 | 26 | Ptpra |
| Ulk25 | 6.13E-09 | 0.474558276 | 0.583 | 0.18 | 0.000124047 | 26 | Ulk2 |
| Tm9sf310 | 6.21E-09 | 0.534399852 | 0.861 | 0.337 | 0.000125742 | 26 | Tm9sf3 |
| Memo13 | 6.29E-09 | 0.252697076 | 0.417 | 0.101 | 0.000127444 | 26 | Memo1 |
| Rela2 | 6.32E-09 | 0.263384232 | 0.444 | 0.112 | 0.000127878 | 26 | Rela |
| Timp310 | 6.61E-09 | 0.76790436 | 0.944 | 0.387 | 0.000133788 | 26 | Timp3 |
| Prrc2a3 | 6.64E-09 | 0.387721103 | 0.528 | 0.152 | 0.000134519 | 26 | Prrc2a |
| Prpf84 | 6.86E-09 | 0.355721045 | 0.611 | 0.184 | 0.000138925 | 26 | Prpf8 |
| Rb1cc17 | 7.02E-09 | 0.383244224 | 0.722 | 0.236 | 0.000142054 | 26 | Rb1cc1 |
| Snx98 | 7.28E-09 | 0.477474445 | 0.583 | 0.177 | 0.000147363 | 26 | Snx9 |
| Kdm4c6 | 7.34E-09 | 0.533600454 | 0.556 | 0.171 | 0.000148519 | 26 | Kdm4c |
| Blvrb4 | 7.61E-09 | 0.356853479 | 0.722 | 0.245 | 0.000154163 | 26 | Blvrb |
| Pum18 | 7.87E-09 | 0.271183653 | 0.806 | 0.26 | 0.000159319 | 26 | Pum1 |

| | | | | | | | |
|---|---|---|---|---|---|---|---|
| Ppp2r3a5 | 7.98E-09 | 0.336073968 | 0.389 | 0.093 | 0.000161655 | 26 | Ppp2r3a |
| Lrrc59 | 8.08E-09 | 0.423924541 | 0.444 | 0.118 | 0.000163487 | 26 | Lrrc59 |
| Glud12 | 8.17E-09 | 0.404102927 | 0.889 | 0.338 | 0.000165397 | 26 | Glud1 |
| Tbc1d513 | 8.41E-09 | 0.617401243 | 0.722 | 0.263 | 0.000170196 | 26 | Tbc1d5 |
| 8-Mar | 8.64E-09 | 0.282475158 | 0.389 | 0.093 | 0.000174902 | 26 | 8-Mar |
| Wdr446 | 8.66E-09 | 0.302539264 | 0.472 | 0.124 | 0.000175267 | 26 | Wdr44 |
| Tpcn11 | 8.76E-09 | 0.283528732 | 0.306 | 0.064 | 0.000177446 | 26 | Tpcn1 |
| Fam20b | 8.81E-09 | 0.313756102 | 0.306 | 0.064 | 0.000178409 | 26 | Fam20b |
| Laptm4a11 | 8.86E-09 | 0.816242686 | 0.944 | 0.486 | 0.000179464 | 26 | Laptm4a |
| Micu3 | 9.02E-09 | 0.297482591 | 0.333 | 0.073 | 0.000182579 | 26 | Micu3 |
| Cdk5rap22 | 9.02E-09 | 0.388007295 | 0.333 | 0.075 | 0.00018267 | 26 | Cdk5rap2 |
| Atp11a | 9.20E-09 | 0.350296555 | 0.278 | 0.055 | 0.000186169 | 26 | Atp11a |
| B3galnt2 | 9.33E-09 | 0.404681237 | 0.278 | 0.055 | 0.000188815 | 26 | B3galnt2 |
| Desi23 | 9.62E-09 | 0.290162397 | 0.583 | 0.173 | 0.000194817 | 26 | Desi2 |
| Srpr7 | 9.65E-09 | 0.428933073 | 0.722 | 0.247 | 0.000195425 | 26 | Srpr |
| Dnajb94 | 1.00E-08 | 0.459970103 | 0.5 | 0.144 | 0.000203098 | 26 | Dnajb9 |
| Mpp5 | 1.05E-08 | 0.430529493 | 0.361 | 0.084 | 0.00021205 | 26 | Mpp5 |
| Sae13 | 1.10E-08 | 0.274013515 | 0.528 | 0.149 | 0.000222535 | 26 | Sae1 |
| Cntln9 | 1.14E-08 | 0.490564554 | 0.528 | 0.156 | 0.000229947 | 26 | Cntln |
| Xbp13 | 1.16E-08 | 0.301528232 | 0.639 | 0.198 | 0.000235433 | 26 | Xbp1 |
| Fam222b | 1.17E-08 | 0.332455334 | 0.389 | 0.094 | 0.000236575 | 26 | Fam222b |
| Man1b1 | 1.17E-08 | 0.255741795 | 0.306 | 0.064 | 0.000236717 | 26 | Man1b1 |
| Mfsd14b | 1.19E-08 | 0.464172242 | 0.361 | 0.087 | 0.000240411 | 26 | Mfsd14b |
| Ythdf2 | 1.24E-08 | 0.268821812 | 0.444 | 0.116 | 0.000251727 | 26 | Ythdf2 |
| Tns24 | 1.27E-08 | 0.324721441 | 0.528 | 0.146 | 0.000256493 | 26 | Tns2 |
| Grn7 | 1.28E-08 | 0.484230394 | 0.667 | 0.226 | 0.00025915 | 26 | Grn |
| Psmd143 | 1.29E-08 | 0.343100771 | 0.667 | 0.219 | 0.000261257 | 26 | Psmd14 |
| Bscl21 | 1.32E-08 | 0.298660081 | 0.333 | 0.075 | 0.000266564 | 26 | Bscl2 |
| Lncpint10 | 1.34E-08 | 0.563859155 | 0.75 | 0.276 | 0.000270709 | 26 | Lncpint |
| Zfp4511 | 1.34E-08 | 0.289588209 | 0.417 | 0.105 | 0.000271304 | 26 | Zfp451 |
| Ctif | 1.37E-08 | 0.422518563 | 0.333 | 0.075 | 0.000278286 | 26 | Ctif |
| Orc31 | 1.42E-08 | 0.385100716 | 0.389 | 0.097 | 0.000286699 | 26 | Orc3 |
| Tubb2a7 | 1.44E-08 | 0.413912032 | 0.556 | 0.166 | 0.000291018 | 26 | Tubb2a |
| Akap7 | 1.52E-08 | 0.326474987 | 0.528 | 0.153 | 0.000307556 | 26 | Akap7 |
| Slc43a38 | 1.53E-08 | 0.420091374 | 0.778 | 0.272 | 0.000310347 | 26 | Slc43a3 |
| Capn26 | 1.60E-08 | 0.426140415 | 0.583 | 0.181 | 0.000322948 | 26 | Capn2 |
| Mapk1ip1l1 | 1.60E-08 | 0.314471172 | 0.417 | 0.107 | 0.00032368 | 26 | Mapk1ip1l |
| Wdfy39 | 1.60E-08 | 0.529833866 | 0.583 | 0.184 | 0.000324315 | 26 | Wdfy3 |
| Med27 | 1.65E-08 | 0.28731616 | 0.333 | 0.074 | 0.000334142 | 26 | Med27 |
| Crebbp12 | 1.67E-08 | 0.552747534 | 0.806 | 0.307 | 0.000338439 | 26 | Crebbp |
| Pitpnm24 | 1.67E-08 | 0.280215447 | 0.472 | 0.124 | 0.000339014 | 26 | Pitpnm2 |
| Pdlim74 | 1.68E-08 | 0.295427035 | 0.333 | 0.075 | 0.000340967 | 26 | Pdlim7 |
| Senp11 | 1.72E-08 | 0.275759201 | 0.417 | 0.105 | 0.000348037 | 26 | Senp1 |

| | | | | | | | |
|---|---|---|---|---|---|---|---|
| Ppp3cb4 | 1.73E-08 | 0.506520072 | 0.556 | 0.171 | 0.000350848 | 26 | Ppp3cb |
| Bin18 | 1.84E-08 | 0.316956966 | 0.583 | 0.177 | 0.000373439 | 26 | Bin1 |
| Mrpl3 | 1.87E-08 | 0.269558991 | 0.389 | 0.096 | 0.000377758 | 26 | Mrpl3 |
| Ankrd1712 | 1.89E-08 | 0.713294603 | 0.694 | 0.263 | 0.000383593 | 26 | Ankrd17 |
| 1600010M07Rik | 2.00E-08 | 0.293500429 | 0.25 | 0.047 | 0.000404293 | 26 | 1600010M07Rik |
| Ubr13 | 2.17E-08 | 0.363358427 | 0.556 | 0.164 | 0.00043957 | 26 | Ubr1 |
| Sart3 | 2.23E-08 | 0.262942463 | 0.278 | 0.056 | 0.000451063 | 26 | Sart3 |
| Enpep2 | 2.27E-08 | 0.367913059 | 0.361 | 0.085 | 0.000459587 | 26 | Enpep |
| Sik311 | 2.30E-08 | 0.916881853 | 0.667 | 0.272 | 0.000464674 | 26 | Sik3 |
| Cul52 | 2.30E-08 | 0.3385339 | 0.5 | 0.143 | 0.000465572 | 26 | Cul5 |
| Frmd4b9 | 2.36E-08 | 0.631795801 | 0.556 | 0.179 | 0.000477861 | 26 | Frmd4b |
| 1700025G04Rik9 | 2.38E-08 | 0.539140321 | 0.583 | 0.19 | 0.000481037 | 26 | 1700025G04Rik |
| Strn | 2.39E-08 | 0.269726556 | 0.417 | 0.106 | 0.000483882 | 26 | Strn |
| Srebf12 | 2.44E-08 | 0.295954099 | 0.278 | 0.057 | 0.000494325 | 26 | Srebf1 |
| Stk395 | 2.49E-08 | 0.311997357 | 0.528 | 0.156 | 0.000504064 | 26 | Stk39 |
| Ptbp24 | 2.54E-08 | 0.38481984 | 0.472 | 0.131 | 0.000513327 | 26 | Ptbp2 |
| Mib14 | 2.55E-08 | 0.51618537 | 0.528 | 0.167 | 0.00051704 | 26 | Mib1 |
| Pkd13 | 2.57E-08 | 0.368644901 | 0.389 | 0.098 | 0.000519324 | 26 | Pkd1 |
| Rubcn | 2.60E-08 | 0.281576815 | 0.278 | 0.056 | 0.000526324 | 26 | Rubcn |
| AC165953.2 | 2.63E-08 | 0.308763916 | 0.278 | 0.057 | 0.000532413 | 26 | AC165953.2 |
| Jak110 | 2.66E-08 | 0.47547325 | 0.944 | 0.367 | 0.000538143 | 26 | Jak1 |
| Tbcel | 2.66E-08 | 0.312001738 | 0.306 | 0.066 | 0.000539 | 26 | Tbcel |
| AC162302.1 | 2.66E-08 | 0.336228014 | 0.306 | 0.066 | 0.000539315 | 26 | AC162302.1 |
| Snx62 | 2.83E-08 | 0.490250522 | 0.722 | 0.277 | 0.000573104 | 26 | Snx6 |
| Arid25 | 2.86E-08 | 0.398898671 | 0.528 | 0.158 | 0.000579404 | 26 | Arid2 |
| Colgalt14 | 2.91E-08 | 0.387513768 | 0.5 | 0.145 | 0.000588162 | 26 | Colgalt1 |
| Cbx56 | 2.92E-08 | 0.311284636 | 0.611 | 0.19 | 0.000590692 | 26 | Cbx5 |
| Rnf136 | 2.93E-08 | 0.382303587 | 0.583 | 0.18 | 0.000593237 | 26 | Rnf13 |
| Stag113 | 2.99E-08 | 0.550236838 | 0.889 | 0.363 | 0.000604943 | 26 | Stag1 |
| Ube2w2 | 3.01E-08 | 0.427526947 | 0.556 | 0.173 | 0.000609655 | 26 | Ube2w |
| Rnf103 | 3.04E-08 | 0.374015481 | 0.333 | 0.078 | 0.000616309 | 26 | Rnf103 |
| Hnrnph15 | 3.07E-08 | 0.286509043 | 0.75 | 0.252 | 0.00062247 | 26 | Hnrnph1 |
| Snx271 | 3.12E-08 | 0.566462177 | 0.389 | 0.103 | 0.000631584 | 26 | Snx27 |
| Serinc111 | 3.21E-08 | 0.504712015 | 0.694 | 0.251 | 0.000649553 | 26 | Serinc1 |
| Zwint | 3.25E-08 | 0.325250489 | 0.361 | 0.088 | 0.000658599 | 26 | Zwint |
| Arfgef16 | 3.27E-08 | 0.304825797 | 0.694 | 0.229 | 0.000662434 | 26 | Arfgef1 |
| Mtch13 | 3.28E-08 | 0.266544688 | 0.861 | 0.318 | 0.000663078 | 26 | Mtch1 |
| Slc31a22 | 3.38E-08 | 0.263662501 | 0.389 | 0.097 | 0.000684336 | 26 | Slc31a2 |
| Hectd42 | 3.42E-08 | 0.363900669 | 0.417 | 0.109 | 0.000691807 | 26 | Hectd4 |
| Poglut1 | 3.42E-08 | 0.256955533 | 0.278 | 0.057 | 0.000692492 | 26 | Poglut1 |
| Zfp4079 | 3.44E-08 | 0.570417215 | 0.528 | 0.163 | 0.000695747 | 26 | Zfp407 |
| Rabep16 | 3.44E-08 | 0.498165768 | 0.667 | 0.232 | 0.000697221 | 26 | Rabep1 |
| Gmeb1 | 3.47E-08 | 0.345849128 | 0.306 | 0.068 | 0.000702382 | 26 | Gmeb1 |

| | | | | | | | |
|---|---|---|---|---|---|---|---|
| Fam208a5 | 3.47E-08 | 0.432736367 | 0.611 | 0.201 | 0.000702942 | 26 | Fam208a |
| St55 | 3.48E-08 | 0.270876728 | 0.528 | 0.153 | 0.000703891 | 26 | St5 |
| Dcaf10 | 3.56E-08 | 0.36583905 | 0.361 | 0.089 | 0.000720149 | 26 | Dcaf10 |
| Scfd25 | 3.56E-08 | 0.3173573 | 0.361 | 0.087 | 0.000720636 | 26 | Scfd2 |
| Klhl91 | 3.59E-08 | 0.260173371 | 0.389 | 0.097 | 0.000727073 | 26 | Klhl9 |
| Yme1l14 | 3.60E-08 | 0.394084932 | 0.611 | 0.197 | 0.000729754 | 26 | Yme1l1 |
| Ubr36 | 3.68E-08 | 0.458759436 | 0.611 | 0.203 | 0.000745532 | 26 | Ubr3 |
| Npc14 | 3.70E-08 | 0.401127688 | 0.333 | 0.079 | 0.000749748 | 26 | Npc1 |
| Tnpo15 | 3.76E-08 | 0.415784188 | 0.556 | 0.175 | 0.000761563 | 26 | Tnpo1 |
| Maml213 | 3.82E-08 | 0.876058425 | 0.778 | 0.345 | 0.000773323 | 26 | Maml2 |
| Pik3r16 | 3.88E-08 | 0.472813366 | 0.694 | 0.255 | 0.000786368 | 26 | Pik3r1 |
| Tnrc6a4 | 3.93E-08 | 0.38103669 | 0.611 | 0.201 | 0.000795269 | 26 | Tnrc6a |
| Usp18 | 4.01E-08 | 0.474535311 | 0.333 | 0.081 | 0.000812801 | 26 | Usp18 |
| Nf19 | 4.18E-08 | 0.519698711 | 0.639 | 0.223 | 0.000846668 | 26 | Nf1 |
| Atf23 | 4.18E-08 | 0.365771534 | 0.528 | 0.16 | 0.00084711 | 26 | Atf2 |
| Prkce6 | 4.20E-08 | 0.380058308 | 0.667 | 0.216 | 0.000849322 | 26 | Prkce |
| E130102H24Rik2 | 4.28E-08 | 0.284656088 | 0.306 | 0.067 | 0.000865665 | 26 | E130102H24Rik |
| Rtn38 | 4.32E-08 | 0.577287558 | 0.778 | 0.326 | 0.000873897 | 26 | Rtn3 |
| 9130011E15Rik | 4.33E-08 | 0.30391687 | 0.361 | 0.088 | 0.000876532 | 26 | 9130011E15Rik |
| Os99 | 4.33E-08 | 0.420243803 | 0.583 | 0.19 | 0.000876901 | 26 | Os9 |
| Dlg14 | 4.41E-08 | 0.417462097 | 0.639 | 0.214 | 0.000893483 | 26 | Dlg1 |
| Ubr56 | 4.43E-08 | 0.493452255 | 0.583 | 0.2 | 0.000897556 | 26 | Ubr5 |
| Slfn58 | 4.70E-08 | 0.401136775 | 0.944 | 0.359 | 0.000951942 | 26 | Slfn5 |
| Marf11 | 4.79E-08 | 0.395422961 | 0.5 | 0.148 | 0.000970114 | 26 | Marf1 |
| Fkbp53 | 4.80E-08 | 0.271719121 | 0.472 | 0.133 | 0.00097182 | 26 | Fkbp5 |
| Ube3c7 | 4.82E-08 | 0.397676761 | 0.528 | 0.16 | 0.00097644 | 26 | Ube3c |
| Dennd5a8 | 5.07E-08 | 0.445416293 | 0.583 | 0.189 | 0.001027436 | 26 | Dennd5a |
| Utrn8 | 5.14E-08 | 0.697249711 | 0.889 | 0.408 | 0.001040828 | 26 | Utrn |
| Nktr9 | 5.34E-08 | 0.711162686 | 0.667 | 0.259 | 0.001080227 | 26 | Nktr |
| mt-Nd514 | 5.48E-08 | 0.796822841 | 0.944 | 0.498 | 0.001109623 | 26 | mt-Nd5 |
| Rspry12 | 6.27E-08 | 0.400368222 | 0.417 | 0.115 | 0.001270309 | 26 | Rspry1 |
| Reep39 | 6.28E-08 | 0.412717924 | 0.833 | 0.308 | 0.001271413 | 26 | Reep3 |
| Babam29 | 6.31E-08 | 0.622442523 | 0.639 | 0.24 | 0.001277946 | 26 | Babam2 |
| Tmem13110 | 6.56E-08 | 0.389409944 | 0.639 | 0.21 | 0.00132749 | 26 | Tmem131 |
| Uso11 | 6.66E-08 | 0.384229923 | 0.472 | 0.141 | 0.001347386 | 26 | Uso1 |
| Hsd17b115 | 6.76E-08 | 0.302902107 | 0.444 | 0.12 | 0.00136889 | 26 | Hsd17b11 |
| Camk1d4 | 6.80E-08 | 0.705287804 | 1 | 0.824 | 0.00137748 | 26 | Camk1d |
| Anxa4 | 7.19E-08 | 0.315900644 | 0.389 | 0.103 | 0.00145596 | 26 | Anxa4 |
| Mgea55 | 7.20E-08 | 0.286478243 | 0.556 | 0.171 | 0.001457147 | 26 | Mgea5 |
| Mdn1 | 7.25E-08 | 0.336104406 | 0.306 | 0.07 | 0.001467678 | 26 | Mdn1 |
| St7l1 | 7.43E-08 | 0.334172014 | 0.361 | 0.091 | 0.001503386 | 26 | St7l |
| Pip4k2b | 7.71E-08 | 0.3444658 | 0.306 | 0.07 | 0.001561114 | 26 | Pip4k2b |
| Ecm24 | 8.24E-08 | 0.307559843 | 0.25 | 0.05 | 0.001667308 | 26 | Ecm2 |

| | | | | | | | |
|---|---|---|---|---|---|---|---|
| Sf3b39 | 8.52E-08 | 0.377916524 | 0.528 | 0.162 | 0.001725825 | 26 | Sf3b3 |
| Otud7b3 | 8.68E-08 | 0.366323921 | 0.444 | 0.126 | 0.00175651 | 26 | Otud7b |
| Dync1h18 | 8.80E-08 | 0.498371885 | 0.75 | 0.288 | 0.001781533 | 26 | Dync1h1 |
| Ints71 | 8.98E-08 | 0.258845932 | 0.25 | 0.05 | 0.001817661 | 26 | Ints7 |
| Eml15 | 9.12E-08 | 0.309362514 | 0.417 | 0.11 | 0.001847316 | 26 | Eml1 |
| Ahcyl13 | 9.21E-08 | 0.30647374 | 0.556 | 0.176 | 0.001864539 | 26 | Ahcyl1 |
| Sympk | 9.44E-08 | 0.257376324 | 0.333 | 0.08 | 0.001911478 | 26 | Sympk |
| Mga6 | 9.58E-08 | 0.519623959 | 0.528 | 0.173 | 0.001939957 | 26 | Mga |
| Copb13 | 9.66E-08 | 0.481665475 | 0.583 | 0.206 | 0.001956 | 26 | Copb1 |
| Rbms19 | 9.94E-08 | 0.642068765 | 0.972 | 0.488 | 0.002012945 | 26 | Rbms1 |
| Syncrip5 | 1.02E-07 | 0.368904948 | 0.639 | 0.217 | 0.002066529 | 26 | Syncrip |
| Cpne23 | 1.03E-07 | 0.255237161 | 0.306 | 0.069 | 0.002089658 | 26 | Cpne2 |
| Fbxw44 | 1.04E-07 | 0.393978771 | 0.417 | 0.117 | 0.002110291 | 26 | Fbxw4 |
| Agk | 1.04E-07 | 0.369193167 | 0.333 | 0.082 | 0.002110567 | 26 | Agk |
| Pdcd6ip6 | 1.06E-07 | 0.403771684 | 0.667 | 0.232 | 0.002136023 | 26 | Pdcd6ip |
| Kif3b | 1.07E-07 | 0.26036273 | 0.278 | 0.06 | 0.002157834 | 26 | Kif3b |
| Wdr111 | 1.11E-07 | 0.317414297 | 0.278 | 0.061 | 0.002247791 | 26 | Wdr11 |
| Fam111a4 | 1.13E-07 | 0.358742793 | 0.361 | 0.093 | 0.002293789 | 26 | Fam111a |
| Itfg12 | 1.21E-07 | 0.391082391 | 0.472 | 0.141 | 0.002446815 | 26 | Itfg1 |
| Lamp113 | 1.22E-07 | 0.646392157 | 0.861 | 0.419 | 0.002462016 | 26 | Lamp1 |
| Hdgfl36 | 1.23E-07 | 0.328065747 | 0.417 | 0.112 | 0.002499767 | 26 | Hdgfl3 |
| Trim8 | 1.25E-07 | 0.348398541 | 0.389 | 0.105 | 0.002535752 | 26 | Trim8 |
| Atrn4 | 1.28E-07 | 0.292848673 | 0.444 | 0.124 | 0.002600436 | 26 | Atrn |
| Fbxl1714 | 1.29E-07 | 0.559793393 | 0.778 | 0.313 | 0.002614954 | 26 | Fbxl17 |
| Raph14 | 1.31E-07 | 0.327267612 | 0.417 | 0.116 | 0.002656766 | 26 | Raph1 |
| Zfyve96 | 1.32E-07 | 0.296073741 | 0.556 | 0.172 | 0.002679145 | 26 | Zfyve9 |
| Leng82 | 1.34E-07 | 0.255741163 | 0.361 | 0.091 | 0.00270635 | 26 | Leng8 |
| Nsd311 | 1.34E-07 | 0.470006411 | 0.806 | 0.31 | 0.002721675 | 26 | Nsd3 |
| Rufy32 | 1.35E-07 | 0.405982565 | 0.333 | 0.084 | 0.002728915 | 26 | Rufy3 |
| Derl12 | 1.38E-07 | 0.434310757 | 0.5 | 0.159 | 0.002797089 | 26 | Derl1 |
| Snx12 | 1.40E-07 | 0.422539211 | 0.389 | 0.109 | 0.002824327 | 26 | Snx12 |
| Aig16 | 1.44E-07 | 0.487622446 | 0.611 | 0.215 | 0.002915008 | 26 | Aig1 |
| Tm9sf29 | 1.44E-07 | 0.447276463 | 0.694 | 0.257 | 0.002920258 | 26 | Tm9sf2 |
| H134 | 1.45E-07 | 0.398704989 | 0.5 | 0.159 | 0.002935892 | 26 | H13 |
| Arhgef39 | 1.46E-07 | 0.627439186 | 0.639 | 0.242 | 0.00295669 | 26 | Arhgef3 |
| Phkb4 | 1.47E-07 | 0.470841773 | 0.472 | 0.146 | 0.002966663 | 26 | Phkb |
| Arcn11 | 1.50E-07 | 0.309520186 | 0.583 | 0.197 | 0.003042834 | 26 | Arcn1 |
| mt-Nd4l12 | 1.53E-07 | 0.77240472 | 0.944 | 0.493 | 0.003091823 | 26 | mt-Nd4l |
| Nr3c18 | 1.55E-07 | 0.596139148 | 0.917 | 0.385 | 0.003142353 | 26 | Nr3c1 |
| Ttc172 | 1.56E-07 | 0.386153183 | 0.417 | 0.118 | 0.003157903 | 26 | Ttc17 |
| Ints67 | 1.58E-07 | 0.406172279 | 0.472 | 0.143 | 0.003197619 | 26 | Ints6 |
| Slc38a92 | 1.62E-07 | 0.364802834 | 0.333 | 0.083 | 0.003287124 | 26 | Slc38a9 |
| Rnf121 | 1.66E-07 | 0.273399911 | 0.278 | 0.061 | 0.003359429 | 26 | Rnf121 |

| | | | | | | | |
|---|---|---|---|---|---|---|---|
| Kdm5c2 | 1.66E-07 | 0.299585085 | 0.472 | 0.14 | 0.003361422 | 26 | Kdm5c |
| Ctso2 | 1.71E-07 | 0.336621798 | 0.417 | 0.116 | 0.003456951 | 26 | Ctso |
| Sec23ip | 1.72E-07 | 0.293662737 | 0.278 | 0.062 | 0.003488739 | 26 | Sec23ip |
| Ngly11 | 1.74E-07 | 0.298677313 | 0.417 | 0.118 | 0.003514609 | 26 | Ngly1 |
| Usf3 | 1.76E-07 | 0.279528379 | 0.361 | 0.093 | 0.003572866 | 26 | Usf3 |
| Tex2612 | 1.80E-07 | 0.306916412 | 0.583 | 0.19 | 0.003640847 | 26 | Tex261 |
| Ccdc174 | 1.80E-07 | 0.265426411 | 0.472 | 0.141 | 0.003653362 | 26 | Ccdc174 |
| Nbea9 | 1.85E-07 | 0.314836663 | 0.611 | 0.198 | 0.003737205 | 26 | Nbea |
| Tmsb108 | 1.88E-07 | 0.364131494 | 0.972 | 0.502 | 0.003815411 | 26 | Tmsb10 |
| Ppp1r9b2 | 1.90E-07 | 0.334929025 | 0.444 | 0.127 | 0.003847509 | 26 | Ppp1r9b |
| Cuedc15 | 1.92E-07 | 0.254398255 | 0.472 | 0.137 | 0.003895378 | 26 | Cuedc1 |
| Adam194 | 1.98E-07 | 0.310207463 | 0.333 | 0.082 | 0.004009917 | 26 | Adam19 |
| Gmds4 | 2.00E-07 | 0.644690878 | 0.528 | 0.18 | 0.004046141 | 26 | Gmds |
| Akap1311 | 2.05E-07 | 0.59617632 | 0.889 | 0.396 | 0.00415713 | 26 | Akap13 |
| Creld2 | 2.18E-07 | 0.311654491 | 0.306 | 0.073 | 0.004417142 | 26 | Creld2 |
| Usp486 | 2.20E-07 | 0.329895801 | 0.444 | 0.131 | 0.004450145 | 26 | Usp48 |
| Prkd35 | 2.25E-07 | 0.358405172 | 0.528 | 0.169 | 0.004551787 | 26 | Prkd3 |
| Cwf19l22 | 2.35E-07 | 0.317983158 | 0.611 | 0.203 | 0.004760923 | 26 | Cwf19l2 |
| Rftn111 | 2.37E-07 | 0.678312602 | 0.667 | 0.253 | 0.004795251 | 26 | Rftn1 |
| Cstf32 | 2.41E-07 | 0.27885033 | 0.444 | 0.128 | 0.004870067 | 26 | Cstf3 |
| Eea17 | 2.42E-07 | 0.46985119 | 0.611 | 0.22 | 0.004893693 | 26 | Eea1 |
| Ccl25 | 2.42E-07 | 0.272730022 | 0.25 | 0.053 | 0.004907327 | 26 | Ccl25 |
| Fbxl77 | 2.44E-07 | 0.619819365 | 0.694 | 0.259 | 0.004932298 | 26 | Fbxl7 |
| Cyld7 | 2.48E-07 | 0.507418164 | 0.5 | 0.172 | 0.005018266 | 26 | Cyld |
| Ap3s2 | 2.51E-07 | 0.317797012 | 0.278 | 0.064 | 0.005076706 | 26 | Ap3s2 |
| Gprasp12 | 2.64E-07 | 0.266483087 | 0.389 | 0.105 | 0.005335169 | 26 | Gprasp1 |
| Pogz | 2.65E-07 | 0.309516845 | 0.306 | 0.073 | 0.005357758 | 26 | Pogz |
| Tbc1d9b4 | 2.66E-07 | 0.425104653 | 0.417 | 0.123 | 0.005379092 | 26 | Tbc1d9b |
| Chic21 | 2.69E-07 | 0.437646209 | 0.667 | 0.254 | 0.005447933 | 26 | Chic2 |
| Zswim8 | 2.78E-07 | 0.266022894 | 0.361 | 0.094 | 0.00561911 | 26 | Zswim8 |
| Tmem1657 | 2.81E-07 | 0.55653974 | 0.528 | 0.181 | 0.005696642 | 26 | Tmem165 |
| mt-Atp814 | 2.88E-07 | 0.634303476 | 0.972 | 0.465 | 0.005837069 | 26 | mt-Atp8 |
| Clstn19 | 2.93E-07 | 0.32117326 | 0.694 | 0.238 | 0.005931522 | 26 | Clstn1 |
| Zbtb441 | 2.93E-07 | 0.309008535 | 0.333 | 0.085 | 0.005934226 | 26 | Zbtb44 |
| Jmjd1c12 | 2.97E-07 | 0.414281216 | 0.806 | 0.309 | 0.006016018 | 26 | Jmjd1c |
| Prpf40a12 | 3.00E-07 | 0.340971051 | 0.722 | 0.269 | 0.00606757 | 26 | Prpf40a |
| Ext24 | 3.02E-07 | 0.343398156 | 0.472 | 0.145 | 0.006122641 | 26 | Ext2 |
| Mbtd15 | 3.02E-07 | 0.448518826 | 0.444 | 0.136 | 0.006123261 | 26 | Mbtd1 |
| Zfp63810 | 3.04E-07 | 0.685685423 | 0.694 | 0.284 | 0.006160414 | 26 | Zfp638 |
| Plekhg31 | 3.10E-07 | 0.285142438 | 0.25 | 0.053 | 0.006282935 | 26 | Plekhg3 |
| Ezh12 | 3.11E-07 | 0.264345831 | 0.444 | 0.129 | 0.006301713 | 26 | Ezh1 |
| Fuca14 | 3.15E-07 | 0.469549165 | 0.583 | 0.209 | 0.006370364 | 26 | Fuca1 |
| Strap | 3.27E-07 | 0.292128961 | 0.528 | 0.167 | 0.006614522 | 26 | Strap |

| | | | | | | | |
|---|---|---|---|---|---|---|---|
| Wasl5 | 3.27E-07 | 0.318194775 | 0.667 | 0.239 | 0.006618132 | 26 | Wasl |
| Mpp75 | 3.34E-07 | 0.277394382 | 0.389 | 0.108 | 0.006756627 | 26 | Mpp7 |
| Chd611 | 3.48E-07 | 0.324705021 | 0.639 | 0.219 | 0.007053066 | 26 | Chd6 |
| Luc7l1 | 3.50E-07 | 0.307798223 | 0.417 | 0.12 | 0.00708598 | 26 | Luc7l |
| Tial12 | 3.56E-07 | 0.262452915 | 0.5 | 0.155 | 0.007214852 | 26 | Tial1 |
| Uqcc23 | 3.56E-07 | 0.35442677 | 0.861 | 0.406 | 0.007217039 | 26 | Uqcc2 |
| Usp474 | 3.66E-07 | 0.376938736 | 0.556 | 0.189 | 0.007399934 | 26 | Usp47 |
| Cd4710 | 3.68E-07 | 0.696570788 | 0.944 | 0.512 | 0.007459537 | 26 | Cd47 |
| Brd413 | 3.88E-07 | 0.337240345 | 0.75 | 0.269 | 0.007865387 | 26 | Brd4 |
| AI3141806 | 3.95E-07 | 0.59635634 | 0.583 | 0.212 | 0.008002481 | 26 | AI314180 |
| Cers22 | 3.99E-07 | 0.340659894 | 0.417 | 0.122 | 0.008081437 | 26 | Cers2 |
| Meis26 | 4.05E-07 | 0.322165361 | 0.75 | 0.284 | 0.00820004 | 26 | Meis2 |
| Sclt11 | 4.08E-07 | 0.270778558 | 0.361 | 0.097 | 0.008250833 | 26 | Sclt1 |
| Med13l8 | 4.19E-07 | 0.487921519 | 0.556 | 0.196 | 0.008479318 | 26 | Med13l |
| Hyou1 | 4.25E-07 | 0.39930786 | 0.25 | 0.055 | 0.008609994 | 26 | Hyou1 |
| Kmt2c12 | 4.25E-07 | 0.584729808 | 0.75 | 0.308 | 0.008613318 | 26 | Kmt2c |
| Yipf53 | 4.28E-07 | 0.36939529 | 0.444 | 0.134 | 0.008675077 | 26 | Yipf5 |
| Pdlim56 | 4.50E-07 | 0.274436112 | 0.778 | 0.288 | 0.00910435 | 26 | Pdlim5 |
| Ell | 4.55E-07 | 0.329505774 | 0.25 | 0.055 | 0.009220297 | 26 | Ell |
| Larp4b5 | 4.62E-07 | 0.469061043 | 0.583 | 0.206 | 0.009358239 | 26 | Larp4b |
| Slc39a71 | 4.64E-07 | 0.254583451 | 0.361 | 0.095 | 0.009392663 | 26 | Slc39a7 |
| Zhx11 | 4.72E-07 | 0.275859775 | 0.361 | 0.098 | 0.009549934 | 26 | Zhx1 |
| Snd18 | 4.91E-07 | 0.628035773 | 0.5 | 0.174 | 0.009934828 | 26 | Snd1 |
| Kdelr14 | 4.92E-07 | 0.415375792 | 0.639 | 0.239 | 0.00996491 | 26 | Kdelr1 |
| Ktn110 | 4.94E-07 | 0.315143729 | 0.806 | 0.309 | 0.009999737 | 26 | Ktn1 |
| Gab29 | 4.99E-07 | 0.557930403 | 0.556 | 0.207 | 0.010104877 | 26 | Gab2 |
| Lman18 | 5.05E-07 | 0.45756998 | 0.639 | 0.24 | 0.010232797 | 26 | Lman1 |
| Ube4b1 | 5.13E-07 | 0.286196149 | 0.389 | 0.11 | 0.010384903 | 26 | Ube4b |
| Ifngr26 | 5.41E-07 | 0.30109476 | 0.444 | 0.133 | 0.010944075 | 26 | Ifngr2 |
| Rev3l7 | 5.42E-07 | 0.630459122 | 0.472 | 0.159 | 0.010975042 | 26 | Rev3l |
| Ccdc1862 | 5.49E-07 | 0.263637168 | 0.5 | 0.158 | 0.011107181 | 26 | Ccdc186 |
| Amfr4 | 5.74E-07 | 0.376536393 | 0.417 | 0.125 | 0.011620458 | 26 | Amfr |
| Psme46 | 5.74E-07 | 0.273882787 | 0.583 | 0.199 | 0.011624915 | 26 | Psme4 |
| Lgals96 | 5.83E-07 | 0.321259459 | 0.694 | 0.249 | 0.011796536 | 26 | Lgals9 |
| Foxo17 | 5.83E-07 | 0.405646002 | 0.583 | 0.201 | 0.01180822 | 26 | Foxo1 |
| Pex14 | 5.86E-07 | 0.347289683 | 0.361 | 0.101 | 0.0118621 | 26 | Pex14 |
| Ago34 | 5.97E-07 | 0.515288511 | 0.472 | 0.156 | 0.012087472 | 26 | Ago3 |
| Ablim19 | 6.02E-07 | 0.472433262 | 0.889 | 0.405 | 0.012179961 | 26 | Ablim1 |
| Zbtb24 | 6.10E-07 | 0.250713078 | 0.25 | 0.055 | 0.012344645 | 26 | Zbtb24 |
| Ppfibp16 | 6.39E-07 | 0.320953167 | 0.556 | 0.185 | 0.01293865 | 26 | Ppfibp1 |
| Flii4 | 6.41E-07 | 0.294994844 | 0.5 | 0.16 | 0.012978884 | 26 | Flii |
| Stxbp12 | 6.45E-07 | 0.276877596 | 0.278 | 0.065 | 0.013066088 | 26 | Stxbp1 |
| Rab1a2 | 6.46E-07 | 0.275616003 | 0.75 | 0.275 | 0.013075259 | 26 | Rab1a |

| | | | | | | | |
|---|---|---|---|---|---|---|---|
| Stau11 | 6.49E-07 | 0.303481296 | 0.417 | 0.122 | 0.01314202 | 26 | Stau1 |
| Anapc4 | 6.53E-07 | 0.276066858 | 0.389 | 0.111 | 0.013220158 | 26 | Anapc4 |
| Acbd32 | 6.73E-07 | 0.365077851 | 0.583 | 0.208 | 0.01363363 | 26 | Acbd3 |
| Lipa5 | 6.81E-07 | 0.262438279 | 0.556 | 0.18 | 0.013790489 | 26 | Lipa |
| Ap3b110 | 7.33E-07 | 0.36799534 | 0.611 | 0.219 | 0.014846265 | 26 | Ap3b1 |
| Luc7l213 | 7.54E-07 | 0.582264108 | 0.861 | 0.407 | 0.015269491 | 26 | Luc7l2 |
| Rsu19 | 7.82E-07 | 0.28450043 | 0.694 | 0.256 | 0.015822454 | 26 | Rsu1 |
| Ptpn113 | 7.97E-07 | 0.285871128 | 0.472 | 0.15 | 0.016146092 | 26 | Ptpn11 |
| Rprd24 | 8.22E-07 | 0.286111723 | 0.444 | 0.136 | 0.016639545 | 26 | Rprd2 |
| Dcaf72 | 8.25E-07 | 0.300774752 | 0.417 | 0.124 | 0.016693734 | 26 | Dcaf7 |
| Dmtf15 | 8.36E-07 | 0.270919417 | 0.417 | 0.123 | 0.016923555 | 26 | Dmtf1 |
| Celf16 | 8.51E-07 | 0.378527486 | 0.611 | 0.222 | 0.017223065 | 26 | Celf1 |
| Agmo3 | 8.77E-07 | 0.360914077 | 0.25 | 0.057 | 0.017756965 | 26 | Agmo |
| Eef21 | 9.07E-07 | 0.502698162 | 1 | 0.546 | 0.018355101 | 26 | Eef2 |
| Krit110 | 9.38E-07 | 0.438822964 | 0.528 | 0.184 | 0.018994642 | 26 | Krit1 |
| Arhgef119 | 9.52E-07 | 0.348719219 | 0.361 | 0.102 | 0.01926442 | 26 | Arhgef11 |
| Rab128 | 9.52E-07 | 0.350947392 | 0.528 | 0.179 | 0.019266921 | 26 | Rab12 |
| Fam120c | 9.64E-07 | 0.291863746 | 0.25 | 0.057 | 0.019516315 | 26 | Fam120c |
| Tsc22d25 | 9.77E-07 | 0.404997752 | 0.472 | 0.155 | 0.019777183 | 26 | Tsc22d2 |
| Npc26 | 9.92E-07 | 0.269805045 | 0.917 | 0.416 | 0.020083374 | 26 | Npc2 |
| Nrf18 | 1.01E-06 | 0.286962771 | 0.444 | 0.137 | 0.020473922 | 26 | Nrf1 |
| Dpp3 | 1.02E-06 | 0.256841892 | 0.25 | 0.057 | 0.020622028 | 26 | Dpp3 |
| 9930021J03Rik4 | 1.03E-06 | 0.363576651 | 0.472 | 0.153 | 0.020860708 | 26 | 9930021J03Rik |
| Axin1 | 1.03E-06 | 0.259097194 | 0.333 | 0.089 | 0.020902956 | 26 | Axin1 |
| Tnks4 | 1.04E-06 | 0.473131098 | 0.444 | 0.144 | 0.021070573 | 26 | Tnks |
| Mlec3 | 1.07E-06 | 0.341047236 | 0.694 | 0.262 | 0.021738526 | 26 | Mlec |
| Dnajc39 | 1.09E-06 | 0.521155338 | 0.833 | 0.372 | 0.022088706 | 26 | Dnajc3 |
| St133 | 1.11E-06 | 0.266869335 | 0.889 | 0.368 | 0.022499812 | 26 | St13 |
| Sos25 | 1.13E-06 | 0.303057753 | 0.5 | 0.163 | 0.022802767 | 26 | Sos2 |
| Atp2b111 | 1.14E-06 | 0.523442013 | 0.778 | 0.335 | 0.02299333 | 26 | Atp2b1 |
| Acin110 | 1.14E-06 | 0.332336584 | 0.806 | 0.318 | 0.023076231 | 26 | Acin1 |
| Ctdnep1 | 1.14E-06 | 0.253721663 | 0.417 | 0.126 | 0.023080064 | 26 | Ctdnep1 |
| Tcf208 | 1.17E-06 | 0.398873235 | 0.444 | 0.143 | 0.023596139 | 26 | Tcf20 |
| Nufip25 | 1.18E-06 | 0.296087585 | 0.639 | 0.234 | 0.02397855 | 26 | Nufip2 |
| Sfswap3 | 1.20E-06 | 0.287573245 | 0.444 | 0.136 | 0.024310517 | 26 | Sfswap |
| Erp442 | 1.21E-06 | 0.386740898 | 0.5 | 0.171 | 0.024434723 | 26 | Erp44 |
| Aldh27 | 1.22E-06 | 0.361939655 | 0.917 | 0.399 | 0.024611299 | 26 | Aldh2 |
| Zfp4452 | 1.24E-06 | 0.33699331 | 0.417 | 0.128 | 0.025019511 | 26 | Zfp445 |
| Phip13 | 1.26E-06 | 0.41227675 | 0.667 | 0.258 | 0.025509461 | 26 | Phip |
| Ggnbp28 | 1.26E-06 | 0.276701763 | 0.667 | 0.244 | 0.025556372 | 26 | Ggnbp2 |
| Fbxw113 | 1.27E-06 | 0.422722364 | 0.5 | 0.173 | 0.025807857 | 26 | Fbxw11 |
| Cyb5a3 | 1.28E-06 | 0.351824029 | 1 | 0.501 | 0.025843677 | 26 | Cyb5a |
| Clasp29 | 1.28E-06 | 0.378779263 | 0.528 | 0.184 | 0.025914968 | 26 | Clasp2 |

| | | | | | | | |
|---|---|---|---|---|---|---|---|
| Kpnb11 | 1.29E-06 | 0.327210693 | 0.444 | 0.143 | 0.026179318 | 26 | Kpnb1 |
| Tnfsf127 | 1.33E-06 | 0.338400813 | 0.472 | 0.149 | 0.026832739 | 26 | Tnfsf12 |
| Foxp110 | 1.34E-06 | 0.328792783 | 0.917 | 0.391 | 0.027047929 | 26 | Foxp1 |
| mt-Co221 | 1.34E-06 | 0.695504874 | 1 | 0.73 | 0.027115947 | 26 | mt-Co2 |
| Kansl113 | 1.34E-06 | 0.469970618 | 0.806 | 0.342 | 0.027217979 | 26 | Kansl1 |
| Ap1g11 | 1.39E-06 | 0.333288975 | 0.472 | 0.152 | 0.028185345 | 26 | Ap1g1 |
| Elovl55 | 1.43E-06 | 0.321825795 | 0.472 | 0.152 | 0.028949164 | 26 | Elovl5 |
| Rabl61 | 1.46E-06 | 0.279274763 | 0.389 | 0.116 | 0.029592311 | 26 | Rabl6 |
| mt-Atp619 | 1.46E-06 | 0.677878982 | 1 | 0.755 | 0.029634629 | 26 | mt-Atp6 |
| Hnrnpul11 | 1.49E-06 | 0.413070841 | 0.5 | 0.179 | 0.030187771 | 26 | Hnrnpul1 |
| Pou2f14 | 1.51E-06 | 0.302761621 | 0.417 | 0.128 | 0.030494453 | 26 | Pou2f1 |
| Ppp2r2a7 | 1.51E-06 | 0.385189138 | 0.583 | 0.215 | 0.030598312 | 26 | Ppp2r2a |
| Depdc5 | 1.54E-06 | 0.310953664 | 0.306 | 0.08 | 0.031217492 | 26 | Depdc5 |
| Rgl16 | 1.58E-06 | 0.441856727 | 0.722 | 0.291 | 0.03202678 | 26 | Rgl1 |
| Cdip11 | 1.62E-06 | 0.267639435 | 0.278 | 0.069 | 0.032893693 | 26 | Cdip1 |
| Emsy3 | 1.69E-06 | 0.32423767 | 0.417 | 0.128 | 0.034315218 | 26 | Emsy |
| Nf2 | 1.70E-06 | 0.396013078 | 0.333 | 0.095 | 0.034334063 | 26 | Nf2 |
| Aven | 1.71E-06 | 0.331058845 | 0.278 | 0.07 | 0.034555945 | 26 | Aven |
| Ywhag4 | 1.72E-06 | 0.336131575 | 0.528 | 0.183 | 0.034722768 | 26 | Ywhag |
| Sh3kbp18 | 1.74E-06 | 0.303182546 | 0.417 | 0.126 | 0.035171553 | 26 | Sh3kbp1 |
| Nfkb111 | 1.74E-06 | 0.512084803 | 0.667 | 0.273 | 0.035201771 | 26 | Nfkb1 |
| Gsk3b7 | 1.77E-06 | 0.392508271 | 0.778 | 0.333 | 0.0358416 | 26 | Gsk3b |
| Pmepa17 | 1.81E-06 | 0.332867629 | 0.639 | 0.254 | 0.036546871 | 26 | Pmepa1 |
| Nsf2 | 1.81E-06 | 0.277566596 | 0.444 | 0.141 | 0.036707233 | 26 | Nsf |
| Sptan19 | 1.85E-06 | 0.316265892 | 0.833 | 0.345 | 0.037435848 | 26 | Sptan1 |
| Clptm1l1 | 1.87E-06 | 0.304570782 | 0.361 | 0.107 | 0.037767905 | 26 | Clptm1l |
| Ndel14 | 1.88E-06 | 0.30148158 | 0.528 | 0.176 | 0.038065278 | 26 | Ndel1 |
| Casc3 | 1.99E-06 | 0.266461762 | 0.389 | 0.117 | 0.040216327 | 26 | Casc3 |
| Creb18 | 2.05E-06 | 0.424650535 | 0.583 | 0.209 | 0.041426453 | 26 | Creb1 |
| Prickle27 | 2.08E-06 | 0.383689438 | 0.444 | 0.141 | 0.042024497 | 26 | Prickle2 |
| Hspg27 | 2.10E-06 | 0.43444684 | 0.778 | 0.314 | 0.042468015 | 26 | Hspg2 |
| Zmym23 | 2.12E-06 | 0.281224663 | 0.444 | 0.141 | 0.042912513 | 26 | Zmym2 |
| Aga1 | 2.21E-06 | 0.251945766 | 0.306 | 0.081 | 0.044686622 | 26 | Aga |
| Rnf2166 | 2.25E-06 | 0.287915109 | 0.528 | 0.182 | 0.045477986 | 26 | Rnf216 |
| Hecw26 | 2.27E-06 | 0.478230429 | 0.639 | 0.251 | 0.046054086 | 26 | Hecw2 |
| Crip113 | 2.29E-06 | 0.442925061 | 0.972 | 0.538 | 0.046399393 | 26 | Crip1 |
| Pxn4 | 2.35E-06 | 0.295775624 | 0.5 | 0.166 | 0.047520109 | 26 | Pxn |
| Ccnt21 | 2.35E-06 | 0.254326673 | 0.389 | 0.117 | 0.047530825 | 26 | Ccnt2 |
| Nudcd37 | 2.39E-06 | 0.294415444 | 0.667 | 0.248 | 0.048421068 | 26 | Nudcd3 |
| Copz1 | 2.49E-06 | 0.357129516 | 0.389 | 0.122 | 0.050379199 | 26 | Copz1 |
| Cnot26 | 2.51E-06 | 0.296361459 | 0.583 | 0.212 | 0.050882367 | 26 | Cnot2 |
| Brip1os2 | 2.51E-06 | 0.255556158 | 0.333 | 0.092 | 0.050897136 | 26 | Brip1os |
| Zc3h151 | 2.60E-06 | 0.319978966 | 0.694 | 0.293 | 0.052555931 | 26 | Zc3h15 |

| | | | | | | | |
|---|---|---|---|---|---|---|---|
| Numa17 | 2.62E-06 | 0.330145544 | 0.556 | 0.197 | 0.053079692 | 26 | Numa1 |
| Atad2b9 | 2.63E-06 | 0.281295711 | 0.583 | 0.205 | 0.053196256 | 26 | Atad2b |
| Acyp23 | 2.65E-06 | 0.293757396 | 0.444 | 0.144 | 0.053649752 | 26 | Acyp2 |
| Ago28 | 2.67E-06 | 0.255231301 | 0.5 | 0.169 | 0.054155015 | 26 | Ago2 |
| Hmg20a5 | 2.74E-06 | 0.317805674 | 0.444 | 0.144 | 0.055533719 | 26 | Hmg20a |
| Trf4 | 2.75E-06 | 0.271608391 | 0.278 | 0.07 | 0.055612887 | 26 | Trf |
| Gm269175 | 2.76E-06 | 0.326637247 | 0.389 | 0.122 | 0.055906859 | 26 | Gm26917 |
| Wbp1l1 | 2.79E-06 | 0.420788697 | 0.333 | 0.097 | 0.056465074 | 26 | Wbp1l |
| Chd410 | 2.79E-06 | 0.408991699 | 0.75 | 0.33 | 0.056501414 | 26 | Chd4 |
| Cdk7 | 2.81E-06 | 0.365795194 | 0.333 | 0.096 | 0.056975011 | 26 | Cdk7 |
| 4932438A13Rik11 | 2.82E-06 | 0.287538962 | 0.611 | 0.221 | 0.057076368 | 26 | 4932438A13Rik |
| Dnajc131 | 2.85E-06 | 0.344784585 | 0.417 | 0.133 | 0.057681971 | 26 | Dnajc13 |
| Trip117 | 2.88E-06 | 0.324417457 | 0.611 | 0.225 | 0.05840724 | 26 | Trip11 |
| Braf5 | 2.92E-06 | 0.414428202 | 0.472 | 0.161 | 0.059176991 | 26 | Braf |
| Dcaf56 | 2.96E-06 | 0.283732106 | 0.417 | 0.131 | 0.059871345 | 26 | Dcaf5 |
| Ddx422 | 3.00E-06 | 0.289617747 | 0.556 | 0.199 | 0.06081965 | 26 | Ddx42 |
| Pml2 | 3.13E-06 | 0.276939504 | 0.306 | 0.082 | 0.063451986 | 26 | Pml |
| Fbxo382 | 3.15E-06 | 0.361861991 | 0.389 | 0.119 | 0.063827472 | 26 | Fbxo38 |
| Pxk3 | 3.17E-06 | 0.262799821 | 0.444 | 0.142 | 0.064245724 | 26 | Pxk |
| Gapvd13 | 3.21E-06 | 0.279255003 | 0.556 | 0.195 | 0.064939127 | 26 | Gapvd1 |
| Spcs32 | 3.27E-06 | 0.31586117 | 0.389 | 0.12 | 0.066229909 | 26 | Spcs3 |
| AU040320 | 3.29E-06 | 0.264920482 | 0.333 | 0.093 | 0.06651239 | 26 | AU040320 |
| Itm2c8 | 3.35E-06 | 0.287277428 | 0.556 | 0.201 | 0.067830376 | 26 | Itm2c |
| Rbpms8 | 3.37E-06 | 0.368695581 | 0.833 | 0.351 | 0.068231722 | 26 | Rbpms |
| Pigt | 3.41E-06 | 0.314339831 | 0.306 | 0.084 | 0.068990387 | 26 | Pigt |
| Mycbp215 | 3.42E-06 | 0.480515055 | 0.889 | 0.376 | 0.06926703 | 26 | Mycbp2 |
| Ctdspl4 | 3.59E-06 | 0.278318148 | 0.333 | 0.093 | 0.07268149 | 26 | Ctdspl |
| Sltm10 | 3.62E-06 | 0.281964833 | 0.694 | 0.273 | 0.073279917 | 26 | Sltm |
| Nrd12 | 3.67E-06 | 0.348104693 | 0.556 | 0.209 | 0.0742277 | 26 | Nrd1 |
| Cbfa2t23 | 3.67E-06 | 0.277814326 | 0.389 | 0.119 | 0.074309061 | 26 | Cbfa2t2 |
| Xpo4 | 3.80E-06 | 0.281966855 | 0.306 | 0.084 | 0.076964012 | 26 | Xpo4 |
| Tnpo33 | 4.01E-06 | 0.292963617 | 0.444 | 0.147 | 0.081191934 | 26 | Tnpo3 |
| Tbck1 | 4.04E-06 | 0.398684803 | 0.361 | 0.109 | 0.081837064 | 26 | Tbck |
| Mcc6 | 4.04E-06 | 0.33809808 | 0.722 | 0.286 | 0.081856513 | 26 | Mcc |
| Abl23 | 4.26E-06 | 0.300465933 | 0.389 | 0.121 | 0.086317137 | 26 | Abl2 |
| Dido11 | 4.30E-06 | 0.287724471 | 0.417 | 0.133 | 0.087102457 | 26 | Dido1 |
| Ccdc88a13 | 4.31E-06 | 0.611552055 | 0.528 | 0.206 | 0.087330598 | 26 | Ccdc88a |
| Tnrc188 | 4.41E-06 | 0.326941928 | 0.5 | 0.174 | 0.089253699 | 26 | Tnrc18 |
| Stag25 | 4.46E-06 | 0.263260973 | 0.722 | 0.29 | 0.09021492 | 26 | Stag2 |
| Papola4 | 4.53E-06 | 0.360037818 | 0.722 | 0.306 | 0.091709728 | 26 | Papola |
| Hdlbp3 | 4.54E-06 | 0.448001366 | 0.667 | 0.299 | 0.091972088 | 26 | Hdlbp |
| Txndc113 | 4.67E-06 | 0.267030834 | 0.333 | 0.096 | 0.09464731 | 26 | Txndc11 |
| Ankrd1213 | 4.75E-06 | 0.490507873 | 0.722 | 0.31 | 0.09619695 | 26 | Ankrd12 |

| | | | | | | | |
|---|---|---|---|---|---|---|---|
| Rreb15 | 4.84E-06 | 0.328330727 | 0.556 | 0.204 | 0.09804744 | 26 | Rreb1 |
| Irf3 | 5.02E-06 | 0.265872391 | 0.389 | 0.121 | 0.101670402 | 26 | Irf3 |
| Myadm7 | 5.11E-06 | 0.31241557 | 0.389 | 0.121 | 0.103473568 | 26 | Myadm |
| Pdpk13 | 5.11E-06 | 0.310756576 | 0.444 | 0.145 | 0.103488777 | 26 | Pdpk1 |
| Picalm7 | 5.19E-06 | 0.364400994 | 0.722 | 0.298 | 0.105020864 | 26 | Picalm |
| Ube2g1 | 5.32E-06 | 0.260950656 | 0.528 | 0.195 | 0.107687187 | 26 | Ube2g1 |
| Fbxl12 | 5.36E-06 | 0.345249523 | 0.306 | 0.086 | 0.108436651 | 26 | Fbxl12 |
| Smg611 | 5.39E-06 | 0.344290325 | 0.667 | 0.262 | 0.109189264 | 26 | Smg6 |
| Wrn6 | 5.49E-06 | 0.259677203 | 0.444 | 0.148 | 0.111101351 | 26 | Wrn |
| Ly6a8 | 5.57E-06 | 0.410964505 | 1 | 0.462 | 0.112764573 | 26 | Ly6a |
| Map3k27 | 5.66E-06 | 0.38716744 | 0.444 | 0.155 | 0.114645724 | 26 | Map3k2 |
| AW55491810 | 5.79E-06 | 0.281547149 | 0.5 | 0.172 | 0.117190755 | 26 | AW554918 |
| Apbb29 | 5.91E-06 | 0.337844679 | 0.889 | 0.369 | 0.119708552 | 26 | Apbb2 |
| Smad51 | 5.95E-06 | 0.34562886 | 0.333 | 0.099 | 0.120405375 | 26 | Smad5 |
| Rfx36 | 6.06E-06 | 0.259651823 | 0.361 | 0.109 | 0.122686205 | 26 | Rfx3 |
| Kdm3b7 | 6.07E-06 | 0.291752965 | 0.417 | 0.138 | 0.12288329 | 26 | Kdm3b |
| Mecp25 | 6.16E-06 | 0.300456568 | 0.472 | 0.162 | 0.124715483 | 26 | Mecp2 |
| Zfx2 | 6.26E-06 | 0.341293826 | 0.444 | 0.153 | 0.126765473 | 26 | Zfx |
| Ubxn48 | 6.26E-06 | 0.363334709 | 0.667 | 0.274 | 0.126800068 | 26 | Ubxn4 |
| Cltc5 | 6.40E-06 | 0.370162248 | 0.75 | 0.333 | 0.12958978 | 26 | Cltc |
| Fndc3a8 | 6.42E-06 | 0.369304668 | 0.667 | 0.279 | 0.12991911 | 26 | Fndc3a |
| Klf611 | 6.46E-06 | 0.433405355 | 0.667 | 0.269 | 0.130822546 | 26 | Klf6 |
| Mllt1014 | 6.58E-06 | 0.359020383 | 0.667 | 0.277 | 0.133238633 | 26 | Mllt10 |
| Thbd8 | 6.83E-06 | 0.329558911 | 0.667 | 0.257 | 0.138279807 | 26 | Thbd |
| Stim112 | 6.96E-06 | 0.331574057 | 0.528 | 0.192 | 0.140998828 | 26 | Stim1 |
| S100a115 | 7.18E-06 | 0.361579584 | 1 | 0.575 | 0.145435232 | 26 | S100a11 |
| Arih111 | 7.42E-06 | 0.338395393 | 0.75 | 0.315 | 0.150312431 | 26 | Arih1 |
| Togaram11 | 7.56E-06 | 0.417245345 | 0.306 | 0.091 | 0.153128644 | 26 | Togaram1 |
| Rad23b | 7.61E-06 | 0.28165559 | 0.5 | 0.182 | 0.1540851 | 26 | Rad23b |
| Cln5 | 8.02E-06 | 0.265167272 | 0.25 | 0.064 | 0.162288124 | 26 | Cln5 |
| Gatad2b7 | 8.04E-06 | 0.260891791 | 0.556 | 0.204 | 0.162693398 | 26 | Gatad2b |
| Bclaf111 | 8.22E-06 | 0.3591079 | 0.639 | 0.256 | 0.166337543 | 26 | Bclaf1 |
| Tex2641 | 8.27E-06 | 0.299463942 | 0.333 | 0.1 | 0.167530521 | 26 | Tex264 |
| Pias18 | 8.60E-06 | 0.262443288 | 0.528 | 0.19 | 0.174062776 | 26 | Pias1 |
| Ube2r21 | 8.72E-06 | 0.286923093 | 0.694 | 0.29 | 0.176574836 | 26 | Ube2r2 |
| Apc7 | 8.78E-06 | 0.329511543 | 0.639 | 0.25 | 0.177807722 | 26 | Apc |
| Cers54 | 8.79E-06 | 0.253231876 | 0.417 | 0.135 | 0.177922256 | 26 | Cers5 |
| Lman27 | 9.07E-06 | 0.271727305 | 0.556 | 0.208 | 0.183724401 | 26 | Lman2 |
| Pja27 | 9.23E-06 | 0.278812993 | 0.556 | 0.206 | 0.186946319 | 26 | Pja2 |
| Fyco15 | 9.41E-06 | 0.276093317 | 0.389 | 0.125 | 0.190472755 | 26 | Fyco1 |
| Stk247 | 9.50E-06 | 0.276880794 | 0.528 | 0.197 | 0.192312325 | 26 | Stk24 |
| Gpr1071 | 9.64E-06 | 0.296680526 | 0.306 | 0.088 | 0.19518694 | 26 | Gpr107 |
| Faf22 | 9.79E-06 | 0.30365887 | 0.389 | 0.127 | 0.198271361 | 26 | Faf2 |

| | | | | | | | |
|---|---|---|---|---|---|---|---|
| Zfp131 | 1.02E-05 | 0.349863867 | 0.389 | 0.13 | 0.206223534 | 26 | Zfp131 |
| Xrn17 | 1.04E-05 | 0.417132353 | 0.528 | 0.205 | 0.210707258 | 26 | Xrn1 |
| Nipbl16 | 1.05E-05 | 0.329663588 | 0.833 | 0.368 | 0.211929922 | 26 | Nipbl |
| Nrip19 | 1.06E-05 | 0.335477495 | 0.611 | 0.246 | 0.215181702 | 26 | Nrip1 |
| Fam210a | 1.09E-05 | 0.272860007 | 0.25 | 0.065 | 0.22133806 | 26 | Fam210a |
| Gab17 | 1.11E-05 | 0.434361515 | 0.361 | 0.116 | 0.224816958 | 26 | Gab1 |
| Far16 | 1.12E-05 | 0.260571911 | 0.472 | 0.166 | 0.2267796 | 26 | Far1 |
| Gnb17 | 1.12E-05 | 0.298221059 | 0.917 | 0.43 | 0.227706629 | 26 | Gnb1 |
| Kidins2207 | 1.14E-05 | 0.425918846 | 0.444 | 0.157 | 0.229905476 | 26 | Kidins220 |
| Capg9 | 1.19E-05 | 0.274961747 | 0.333 | 0.102 | 0.240036162 | 26 | Capg |
| Zfp1489 | 1.19E-05 | 0.255136891 | 0.667 | 0.267 | 0.240450037 | 26 | Zfp148 |
| Mtdh11 | 1.19E-05 | 0.288014549 | 0.889 | 0.388 | 0.241340151 | 26 | Mtdh |
| Camkmt5 | 1.20E-05 | 0.308770822 | 0.417 | 0.141 | 0.242036162 | 26 | Camkmt |
| Atp6ap13 | 1.21E-05 | 0.270591104 | 0.444 | 0.151 | 0.245234644 | 26 | Atp6ap1 |
| Arhgef1210 | 1.21E-05 | 0.272423097 | 0.833 | 0.35 | 0.245967708 | 26 | Arhgef12 |
| Hdac21 | 1.25E-05 | 0.343010523 | 0.333 | 0.103 | 0.253182526 | 26 | Hdac2 |
| Nsd111 | 1.28E-05 | 0.390207124 | 0.778 | 0.351 | 0.260140481 | 26 | Nsd1 |
| Phf204 | 1.30E-05 | 0.353899076 | 0.417 | 0.146 | 0.262720535 | 26 | Phf20 |
| Vps13d5 | 1.34E-05 | 0.328421288 | 0.472 | 0.171 | 0.270696151 | 26 | Vps13d |
| Zranb11 | 1.34E-05 | 0.265155128 | 0.417 | 0.141 | 0.271596143 | 26 | Zranb1 |
| Pcm110 | 1.36E-05 | 0.424547428 | 0.611 | 0.253 | 0.274372697 | 26 | Pcm1 |
| Pbxip18 | 1.42E-05 | 0.343265479 | 0.361 | 0.119 | 0.287441084 | 26 | Pbxip1 |
| Scaf118 | 1.43E-05 | 0.433733377 | 0.611 | 0.265 | 0.28992983 | 26 | Scaf11 |
| Cop14 | 1.46E-05 | 0.289646208 | 0.5 | 0.181 | 0.295263675 | 26 | Cop1 |
| Emc710 | 1.47E-05 | 0.270893159 | 0.667 | 0.274 | 0.297081009 | 26 | Emc7 |
| Fermt29 | 1.49E-05 | 0.320717047 | 0.833 | 0.352 | 0.300738305 | 26 | Fermt2 |
| Lmtk23 | 1.51E-05 | 0.38950492 | 0.306 | 0.092 | 0.306388105 | 26 | Lmtk2 |
| Itsn211 | 1.55E-05 | 0.347400225 | 0.694 | 0.28 | 0.312942598 | 26 | Itsn2 |
| Atp2c16 | 1.63E-05 | 0.292037069 | 0.472 | 0.169 | 0.32978785 | 26 | Atp2c1 |
| Ehbp1l11 | 1.64E-05 | 0.264038844 | 0.333 | 0.102 | 0.331935787 | 26 | Ehbp1l1 |
| Ifih15 | 1.64E-05 | 0.308505135 | 0.389 | 0.13 | 0.332847963 | 26 | Ifih1 |
| Nudt13 | 1.78E-05 | 0.252076264 | 0.25 | 0.066 | 0.360323016 | 26 | Nudt13 |
| Zfp2661 | 1.88E-05 | 0.301942346 | 0.333 | 0.106 | 0.381006144 | 26 | Zfp266 |
| Dnajc115 | 1.90E-05 | 0.447429818 | 0.75 | 0.325 | 0.383998005 | 26 | Dnajc1 |
| Agtrap3 | 1.91E-05 | 0.258284712 | 0.389 | 0.129 | 0.387002962 | 26 | Agtrap |
| Bicd2 | 1.97E-05 | 0.319265578 | 0.25 | 0.066 | 0.397992458 | 26 | Bicd2 |
| Chd86 | 2.01E-05 | 0.272624822 | 0.472 | 0.17 | 0.406419746 | 26 | Chd8 |
| Rfx78 | 2.03E-05 | 0.268123659 | 0.444 | 0.157 | 0.410317712 | 26 | Rfx7 |
| Cbl7 | 2.06E-05 | 0.278692483 | 0.556 | 0.209 | 0.417410625 | 26 | Cbl |
| Cdk1311 | 2.11E-05 | 0.312940173 | 0.583 | 0.229 | 0.426925083 | 26 | Cdk13 |
| Sun11 | 2.18E-05 | 0.271206904 | 0.333 | 0.103 | 0.442305882 | 26 | Sun1 |
| Gpatch2l | 2.25E-05 | 0.257053977 | 0.25 | 0.067 | 0.455817577 | 26 | Gpatch2l |
| Smad13 | 2.29E-05 | 0.331536598 | 0.389 | 0.134 | 0.463596323 | 26 | Smad1 |

| | | | | | | | |
|---|---|---|---|---|---|---|---|
| Nbeal19 | 2.32E-05 | 0.304060825 | 0.639 | 0.264 | 0.469154564 | 26 | Nbeal1 |
| Phf149 | 2.35E-05 | 0.364629815 | 0.583 | 0.246 | 0.476446335 | 26 | Phf14 |
| Ppp3ca15 | 2.49E-05 | 0.504030015 | 0.833 | 0.395 | 0.505016891 | 26 | Ppp3ca |
| Ipo91 | 2.75E-05 | 0.255624079 | 0.25 | 0.069 | 0.556694324 | 26 | Ipo9 |
| Trpc4ap | 2.86E-05 | 0.260401806 | 0.472 | 0.177 | 0.580013181 | 26 | Trpc4ap |
| Clock7 | 2.87E-05 | 0.286892472 | 0.528 | 0.205 | 0.580574868 | 26 | Clock |
| Ube2k | 2.88E-05 | 0.32331867 | 0.667 | 0.305 | 0.584074148 | 26 | Ube2k |
| Atxn7l15 | 2.91E-05 | 0.335141615 | 0.389 | 0.135 | 0.588875417 | 26 | Atxn7l1 |
| Ppp4r13 | 2.93E-05 | 0.272650879 | 0.361 | 0.118 | 0.592512726 | 26 | Ppp4r1 |
| Ppig9 | 2.93E-05 | 0.267138195 | 0.722 | 0.318 | 0.592890505 | 26 | Ppig |
| Ube2d1 | 3.11E-05 | 0.258665402 | 0.361 | 0.118 | 0.628962735 | 26 | Ube2d1 |
| Osbpl11 | 3.28E-05 | 0.295942596 | 0.278 | 0.082 | 0.66404037 | 26 | Osbpl11 |
| C2cd51 | 3.30E-05 | 0.340488668 | 0.306 | 0.097 | 0.668174247 | 26 | C2cd5 |
| Bfar1 | 3.42E-05 | 0.291446615 | 0.361 | 0.12 | 0.692045598 | 26 | Bfar |
| Eya3 | 3.44E-05 | 0.27438996 | 0.306 | 0.095 | 0.696262687 | 26 | Eya3 |
| Mapkap11 | 3.44E-05 | 0.273288591 | 0.5 | 0.2 | 0.696620104 | 26 | Mapkap1 |
| Sec629 | 3.47E-05 | 0.332835852 | 0.833 | 0.396 | 0.701602628 | 26 | Sec62 |
| Ncoa77 | 3.62E-05 | 0.258103608 | 0.694 | 0.298 | 0.733705337 | 26 | Ncoa7 |
| Huwe113 | 3.82E-05 | 0.318471777 | 0.667 | 0.287 | 0.773980109 | 26 | Huwe1 |
| Ppp2r5e5 | 3.83E-05 | 0.292082104 | 0.444 | 0.167 | 0.775997245 | 26 | Ppp2r5e |
| Maml37 | 3.87E-05 | 0.411398946 | 0.528 | 0.21 | 0.782510964 | 26 | Maml3 |
| Baz2b9 | 3.92E-05 | 0.396230843 | 0.778 | 0.353 | 0.793050232 | 26 | Baz2b |
| Sfpq9 | 3.98E-05 | 0.359502292 | 0.667 | 0.3 | 0.804831801 | 26 | Sfpq |
| Cdk129 | 3.99E-05 | 0.332340448 | 0.556 | 0.232 | 0.808084153 | 26 | Cdk12 |
| Mbnl210 | 4.11E-05 | 0.267523023 | 0.889 | 0.421 | 0.831153932 | 26 | Mbnl2 |
| Setd29 | 4.14E-05 | 0.272725734 | 0.5 | 0.194 | 0.838531695 | 26 | Setd2 |
| Eif4g38 | 4.26E-05 | 0.443060147 | 0.639 | 0.288 | 0.862551969 | 26 | Eif4g3 |
| Trappc97 | 4.31E-05 | 0.29121414 | 0.5 | 0.196 | 0.872112562 | 26 | Trappc9 |
| Xpo13 | 4.41E-05 | 0.410358925 | 0.361 | 0.128 | 0.891961447 | 26 | Xpo1 |
| Gm424185 | 4.50E-05 | 0.702595463 | 1 | 1 | 0.91188026 | 26 | Gm42418 |
| Zfr8 | 4.60E-05 | 0.365630444 | 0.583 | 0.249 | 0.931370753 | 26 | Zfr |
| Egr16 | 4.66E-05 | 0.532070551 | 0.444 | 0.177 | 0.943437641 | 26 | Egr1 |
| Por1 | 4.70E-05 | 0.259983462 | 0.333 | 0.109 | 0.951524174 | 26 | Por |
| Rbm511 | 4.84E-05 | 0.260363291 | 0.583 | 0.242 | 0.980049804 | 26 | Rbm5 |
| Zmynd84 | 5.06E-05 | 0.25927143 | 0.5 | 0.197 | 1 | 26 | Zmynd8 |
| Phc35 | 5.18E-05 | 0.289946536 | 0.389 | 0.136 | 1 | 26 | Phc3 |
| Efr3a6 | 5.22E-05 | 0.264821999 | 0.5 | 0.199 | 1 | 26 | Efr3a |
| Zfp1822 | 5.22E-05 | 0.288260708 | 0.333 | 0.111 | 1 | 26 | Zfp182 |
| Ggh3 | 5.40E-05 | 0.338033661 | 0.444 | 0.169 | 1 | 26 | Ggh |
| Cog510 | 5.51E-05 | 0.351407471 | 0.472 | 0.185 | 1 | 26 | Cog5 |
| Ppp1r213 | 5.68E-05 | 0.35415752 | 0.333 | 0.114 | 1 | 26 | Ppp1r21 |
| Asb35 | 6.12E-05 | 0.276800355 | 0.333 | 0.111 | 1 | 26 | Asb3 |
| Kdm2a10 | 6.28E-05 | 0.28907816 | 0.583 | 0.242 | 1 | 26 | Kdm2a |

| | | | | | | | |
|---|---|---|---|---|---|---|---|
| Kdm5a7 | 6.33E-05 | 0.295077677 | 0.444 | 0.169 | 1 | 26 | Kdm5a |
| Birc615 | 6.65E-05 | 0.25673039 | 0.778 | 0.343 | 1 | 26 | Birc6 |
| Sptbn18 | 6.78E-05 | 0.267280573 | 1 | 0.497 | 1 | 26 | Sptbn1 |
| Snhg20 | 7.21E-05 | 0.265593061 | 0.25 | 0.073 | 1 | 26 | Snhg20 |
| Anapc12 | 7.37E-05 | 0.254405801 | 0.361 | 0.125 | 1 | 26 | Anapc1 |
| Itgb113 | 7.72E-05 | 0.276007722 | 0.972 | 0.481 | 1 | 26 | Itgb1 |
| Pi4ka6 | 7.84E-05 | 0.297615668 | 0.361 | 0.129 | 1 | 26 | Pi4ka |
| Tacc110 | 7.92E-05 | 0.269908275 | 0.694 | 0.307 | 1 | 26 | Tacc1 |
| Eif3a8 | 8.07E-05 | 0.288622611 | 0.778 | 0.374 | 1 | 26 | Eif3a |
| Cnot411 | 8.16E-05 | 0.307119392 | 0.611 | 0.271 | 1 | 26 | Cnot4 |
| Anxa210 | 8.45E-05 | 0.341640755 | 0.861 | 0.419 | 1 | 26 | Anxa2 |
| Rnf1115 | 8.51E-05 | 0.290041075 | 0.417 | 0.158 | 1 | 26 | Rnf111 |
| Tpr12 | 8.67E-05 | 0.294548856 | 0.861 | 0.427 | 1 | 26 | Tpr |
| Hnrnpll2 | 9.27E-05 | 0.285309146 | 0.333 | 0.115 | 1 | 26 | Hnrnpll |
| Anxa69 | 0.00011105 | 0.255263207 | 0.417 | 0.158 | 1 | 26 | Anxa6 |
| Mkln17 | 0.000116253 | 0.289068922 | 0.611 | 0.266 | 1 | 26 | Mkln1 |
| Ppib7 | 0.000123269 | 0.5197567 | 0.917 | 0.555 | 1 | 26 | Ppib |
| Stx164 | 0.000125083 | 0.290124492 | 0.417 | 0.162 | 1 | 26 | Stx16 |
| Agtpbp11 | 0.000130604 | 0.296087108 | 0.333 | 0.117 | 1 | 26 | Agtpbp1 |
| Epn26 | 0.000131019 | 0.281095688 | 0.417 | 0.156 | 1 | 26 | Epn2 |
| Ccm27 | 0.000131062 | 0.260698951 | 0.361 | 0.128 | 1 | 26 | Ccm2 |
| Gm4728313 | 0.000132403 | 0.259813564 | 0.639 | 0.283 | 1 | 26 | Gm47283 |
| Sdf2l17 | 0.000132611 | 0.416593749 | 0.417 | 0.169 | 1 | 26 | Sdf2l1 |
| Arf42 | 0.000134276 | 0.274740561 | 0.861 | 0.457 | 1 | 26 | Arf4 |
| Zmynd117 | 0.000135632 | 0.298622347 | 0.639 | 0.291 | 1 | 26 | Zmynd11 |
| Eml62 | 0.000137465 | 0.521527199 | 0.306 | 0.104 | 1 | 26 | Eml6 |
| Sp13 | 0.000140041 | 0.283727256 | 0.389 | 0.144 | 1 | 26 | Sp1 |
| Nfat511 | 0.000140438 | 0.278259786 | 0.611 | 0.269 | 1 | 26 | Nfat5 |
| Kif13a7 | 0.000143158 | 0.307312648 | 0.444 | 0.175 | 1 | 26 | Kif13a |
| Mdm46 | 0.000145934 | 0.260288908 | 0.417 | 0.161 | 1 | 26 | Mdm4 |
| Zc3h7a7 | 0.000147027 | 0.290033373 | 0.417 | 0.164 | 1 | 26 | Zc3h7a |
| Tmed1010 | 0.000149686 | 0.294333954 | 0.833 | 0.438 | 1 | 26 | Tmed10 |
| Aatf1 | 0.000149852 | 0.405357481 | 0.306 | 0.107 | 1 | 26 | Aatf |
| Cep35010 | 0.000167715 | 0.459323729 | 0.472 | 0.205 | 1 | 26 | Cep350 |
| Pbrm17 | 0.000178503 | 0.360625051 | 0.611 | 0.29 | 1 | 26 | Pbrm1 |
| Peak112 | 0.000179341 | 0.30491607 | 0.778 | 0.389 | 1 | 26 | Peak1 |
| Hacd24 | 0.000185999 | 0.259332042 | 0.333 | 0.118 | 1 | 26 | Hacd2 |
| Atrx7 | 0.000204059 | 0.261880254 | 0.861 | 0.422 | 1 | 26 | Atrx |
| Slc10a75 | 0.000206809 | 0.327726072 | 0.389 | 0.149 | 1 | 26 | Slc10a7 |
| Tor1aip26 | 0.000217316 | 0.319949979 | 0.444 | 0.184 | 1 | 26 | Tor1aip2 |
| Dnajb4 | 0.000224075 | 0.287800978 | 0.278 | 0.089 | 1 | 26 | Dnajb4 |
| Zfp277 | 0.00022519 | 0.322902294 | 0.306 | 0.107 | 1 | 26 | Zfp277 |
| Tra2a15 | 0.000236507 | 0.348547056 | 0.611 | 0.277 | 1 | 26 | Tra2a |

| | | | | | | | |
|---|---|---|---|---|---|---|---|
| Btbd913 | 0.00025537 | 0.261249558 | 0.556 | 0.239 | 1 | 26 | Btbd9 |
| Nr2c28 | 0.000294285 | 0.415466319 | 0.389 | 0.161 | 1 | 26 | Nr2c2 |
| Fnip25 | 0.000316572 | 0.26016008 | 0.306 | 0.108 | 1 | 26 | Fnip2 |
| Tapbp8 | 0.000325515 | 0.283534018 | 0.639 | 0.307 | 1 | 26 | Tapbp |
| Fbxo1110 | 0.000333854 | 0.267415377 | 0.528 | 0.235 | 1 | 26 | Fbxo11 |
| Crkl | 0.000371695 | 0.302139592 | 0.25 | 0.083 | 1 | 26 | Crkl |
| Herc26 | 0.000372025 | 0.307662217 | 0.472 | 0.209 | 1 | 26 | Herc2 |
| Gcn1l1 | 0.000381223 | 0.254064101 | 0.278 | 0.096 | 1 | 26 | Gcn1l1 |
| Tcf1213 | 0.000408294 | 0.428081964 | 0.75 | 0.378 | 1 | 26 | Tcf12 |
| Pkn29 | 0.000494812 | 0.322898293 | 0.583 | 0.297 | 1 | 26 | Pkn2 |
| Grina5 | 0.000499484 | 0.28827598 | 0.361 | 0.141 | 1 | 26 | Grina |
| Pip5k1a | 0.000514224 | 0.327460225 | 0.278 | 0.098 | 1 | 26 | Pip5k1a |
| Gphn3 | 0.000514249 | 0.311237922 | 1 | 0.794 | 1 | 26 | Gphn |
| Kmt2a13 | 0.000518068 | 0.323557984 | 0.556 | 0.269 | 1 | 26 | Kmt2a |
| Igf12 | 0.000533685 | 0.305719699 | 0.25 | 0.084 | 1 | 26 | Igf1 |
| Ascc310 | 0.000542952 | 0.300032203 | 0.472 | 0.209 | 1 | 26 | Ascc3 |
| Gmcl1 | 0.000614676 | 0.280419569 | 0.25 | 0.085 | 1 | 26 | Gmcl1 |
| Rbm2610 | 0.000653886 | 0.321244041 | 0.5 | 0.229 | 1 | 26 | Rbm26 |
| Itpr28 | 0.000675667 | 0.4513692 | 0.528 | 0.246 | 1 | 26 | Itpr2 |
| Scai3 | 0.000679293 | 0.409586219 | 0.278 | 0.103 | 1 | 26 | Scai |
| Tmem241 | 0.000681512 | 0.315235123 | 0.25 | 0.085 | 1 | 26 | Tmem241 |
| Ccar19 | 0.000709841 | 0.275976214 | 0.472 | 0.21 | 1 | 26 | Ccar1 |
| Tbl1xr19 | 0.000716722 | 0.269414669 | 0.389 | 0.166 | 1 | 26 | Tbl1xr1 |
| Zufsp8 | 0.000803295 | 0.362156622 | 0.333 | 0.138 | 1 | 26 | Zufsp |
| Sesn110 | 0.000905146 | 0.276719315 | 0.472 | 0.214 | 1 | 26 | Sesn1 |
| Mllt38 | 0.000906057 | 0.359696446 | 0.361 | 0.153 | 1 | 26 | Mllt3 |
| Auts25 | 0.00113729 | 0.282454462 | 0.361 | 0.154 | 1 | 26 | Auts2 |
| Wtap | 0.001485359 | 0.275273398 | 0.389 | 0.164 | 1 | 26 | Wtap |
| Golph3l | 0.001523435 | 0.271767615 | 0.25 | 0.09 | 1 | 26 | Golph3l |
| Malat115 | 0.001752154 | 0.430991616 | 1 | 0.727 | 1 | 26 | Malat1 |
| Map3k37 | 0.00194566 | 0.29809241 | 0.389 | 0.18 | 1 | 26 | Map3k3 |
| Ifrd15 | 0.002113371 | 0.368802793 | 0.472 | 0.233 | 1 | 26 | Ifrd1 |
| Son14 | 0.002258912 | 0.2876434 | 0.889 | 0.528 | 1 | 26 | Son |
| Ranbp98 | 0.002326796 | 0.25210062 | 0.444 | 0.202 | 1 | 26 | Ranbp9 |
| Enox24 | 0.002368516 | 0.452548572 | 0.333 | 0.145 | 1 | 26 | Enox2 |
| Arid1b12 | 0.002447422 | 0.255523472 | 0.528 | 0.261 | 1 | 26 | Arid1b |
| Map3k56 | 0.002773758 | 0.252201481 | 0.417 | 0.194 | 1 | 26 | Map3k5 |
| Rpl49 | 0.00341237 | 0.293767117 | 0.917 | 0.66 | 1 | 26 | Rpl4 |
| Ppt13 | 0.003886695 | 0.272144249 | 0.306 | 0.132 | 1 | 26 | Ppt1 |
| Zfp36l110 | 0.0038918 | 0.394405447 | 0.694 | 0.435 | 1 | 26 | Zfp36l1 |
| Cmip9 | 0.005931845 | 0.297713259 | 0.528 | 0.288 | 1 | 26 | Cmip |

**Appendix Table 2 Positively differentially expressed genes in the salivary gland relative to 11 other organs**

The FindAllMarkers function in Seurat was used to identify positively differentially expressed genes per organ using assay = “RNA”, min.pct = 0.25, and logfc.threshold = 0.25.

| | p_val | avg_log2FC | pct.1 | pct.2 | p_val_adj | cluster | gene |
|---|---|---|---|---|---|---|---|
| Mucl2 | 0 | 6.19306326 | 1 | 0 | 0 | Salivary_gland | Mucl2 |
| Cmss1 | 0 | 5.81259825 | 1 | 0.01 | 0 | Salivary_gland | Cmss1 |
| Gm42418 | 0 | 4.86164099 | 1 | 0.78 | 0 | Salivary_gland | Gm42418 |
| Cdk8 | 0 | 4.52567577 | 0.99 | 0.03 | 0 | Salivary_gland | Cdk8 |
| Camk1d | 0 | 3.6731806 | 0.95 | 0 | 0 | Salivary_gland | Camk1d |
| Lars2 | 0 | 3.56667033 | 0.97 | 0.04 | 0 | Salivary_gland | Lars2 |
| Cavin2 | 0 | 3.5283923 | 0.93 | 0 | 0 | Salivary_gland | Cavin2 |
| Scgb1b27 | 0 | 3.39643998 | 0.87 | 0 | 0 | Salivary_gland | Scgb1b27 |
| Klk1 | 0 | 3.35670504 | 0.81 | 0 | 0 | Salivary_gland | Klk1 |
| Gphn | 0 | 3.23970631 | 0.95 | 0.04 | 0 | Salivary_gland | Gphn |
| Plcb1 | 0 | 3.1291299 | 0.96 | 0.21 | 0 | Salivary_gland | Plcb1 |
| Rflnb | 0 | 3.06296814 | 0.85 | 0 | 0 | Salivary_gland | Rflnb |
| Prol1 | 0 | 2.95822057 | 0.83 | 0 | 0 | Salivary_gland | Prol1 |
| Tcim | 0 | 2.92360362 | 0.83 | 0 | 0 | Salivary_gland | Tcim |
| Rbp7 | 0 | 2.85723967 | 0.91 | 0.15 | 0 | Salivary_gland | Rbp7 |
| Selenow | 0 | 2.76433387 | 0.89 | 0 | 0 | Salivary_gland | Selenow |
| Rack1 | 0 | 2.72257681 | 0.86 | 0 | 0 | Salivary_gland | Rack1 |
| Fxyd2 | 0 | 2.6702393 | 0.67 | 0 | 0 | Salivary_gland | Fxyd2 |
| Fbxl7 | 0 | 2.66272345 | 0.73 | 0 | 0 | Salivary_gland | Fbxl7 |
| Itga1 | 0 | 2.61580243 | 0.91 | 0.13 | 0 | Salivary_gland | Itga1 |
| Ptprm | 0 | 2.58391148 | 0.92 | 0.15 | 0 | Salivary_gland | Ptprm |
| Arl15 | 0 | 2.57760405 | 0.81 | 0.08 | 0 | Salivary_gland | Arl15 |
| Lrmda | 0 | 2.56229646 | 0.71 | 0 | 0 | Salivary_gland | Lrmda |
| Selenom | 0 | 2.51643604 | 0.83 | 0 | 0 | Salivary_gland | Selenom |
| Cyp26b1 | 0 | 2.48228964 | 0.6 | 0.12 | 0 | Salivary_gland | Cyp26b1 |
| Scgb2b27 | 0 | 2.48186836 | 0.71 | 0 | 0 | Salivary_gland | Scgb2b27 |
| Ptprg | 0 | 2.45063504 | 0.93 | 0.23 | 0 | Salivary_gland | Ptprg |
| Dock4 | 0 | 2.41338114 | 0.77 | 0.01 | 0 | Salivary_gland | Dock4 |
| Elob | 0 | 2.4123233 | 0.83 | 0 | 0 | Salivary_gland | Elob |
| Selenok | 0 | 2.40401183 | 0.83 | 0 | 0 | Salivary_gland | Selenok |
| Cavin3 | 0 | 2.40157128 | 0.79 | 0 | 0 | Salivary_gland | Cavin3 |
| Syne1 | 0 | 2.37082409 | 0.91 | 0.17 | 0 | Salivary_gland | Syne1 |
| Xist | 0 | 2.36496961 | 0.77 | 0 | 0 | Salivary_gland | Xist |
| Piezo2 | 0 | 2.34190925 | 0.76 | 0.04 | 0 | Salivary_gland | Piezo2 |
| 9530026P05Rik | 0 | 2.32400659 | 0.67 | 0.02 | 0 | Salivary_gland | 9530026P05Rik |
| Sem1 | 0 | 2.29674015 | 0.8 | 0 | 0 | Salivary_gland | Sem1 |
| Ldb2 | 0 | 2.26853815 | 0.9 | 0.21 | 0 | Salivary_gland | Ldb2 |
| Selenop | 0 | 2.251651 | 0.75 | 0 | 0 | Salivary_gland | Selenop |
| Selenof | 0 | 2.23370026 | 0.8 | 0 | 0 | Salivary_gland | Selenof |
| Rasal2 | 0 | 2.15069251 | 0.85 | 0.15 | 0 | Salivary_gland | Rasal2 |

| | | | | | | | |
|---|---|---|---|---|---|---|---|
| Samd12 | 0 | 2.14043466 | 0.7 | 0.02 | 0 | Salivary_gland | Samd12 |
| Flt1 | 0 | 2.13767142 | 1 | 0.68 | 0 | Salivary_gland | Flt1 |
| Btnl9 | 0 | 2.13495018 | 0.81 | 0.12 | 0 | Salivary_gland | Btnl9 |
| Nfib | 0 | 2.09168691 | 0.99 | 0.53 | 0 | Salivary_gland | Nfib |
| Etl4 | 0 | 2.07615554 | 0.83 | 0.14 | 0 | Salivary_gland | Etl4 |
| Afdn | 0 | 2.07076214 | 0.71 | 0 | 0 | Salivary_gland | Afdn |
| Mcc | 0 | 2.05332349 | 0.81 | 0.13 | 0 | Salivary_gland | Mcc |
| Arhgap29 | 0 | 2.02608393 | 0.92 | 0.28 | 0 | Salivary_gland | Arhgap29 |
| Slfn5 | 0 | 1.99966528 | 0.87 | 0.19 | 0 | Salivary_gland | Slfn5 |
| Lama3 | 0 | 1.98341644 | 0.69 | 0.02 | 0 | Salivary_gland | Lama3 |
| Crim1 | 0 | 1.98243552 | 0.9 | 0.25 | 0 | Salivary_gland | Crim1 |
| Ebf1 | 0 | 1.97696027 | 0.98 | 0.37 | 0 | Salivary_gland | Ebf1 |
| Prkg1 | 0 | 1.96750434 | 0.51 | 0 | 0 | Salivary_gland | Prkg1 |
| Pde7b | 0 | 1.96596569 | 0.61 | 0.01 | 0 | Salivary_gland | Pde7b |
| Mef2c | 0 | 1.93875048 | 0.96 | 0.41 | 0 | Salivary_gland | Mef2c |
| Stox2 | 0 | 1.88770881 | 0.72 | 0.1 | 0 | Salivary_gland | Stox2 |
| Fmo2 | 0 | 1.88275695 | 0.8 | 0.13 | 0 | Salivary_gland | Fmo2 |
| Tnfsf10 | 0 | 1.85402376 | 0.86 | 0.18 | 0 | Salivary_gland | Tnfsf10 |
| Diaph2 | 0 | 1.84673372 | 0.69 | 0.06 | 0 | Salivary_gland | Diaph2 |
| Kctd12b | 0 | 1.84413964 | 0.72 | 0.09 | 0 | Salivary_gland | Kctd12b |
| Adgrl4 | 0 | 1.82311077 | 0.98 | 0.49 | 0 | Salivary_gland | Adgrl4 |
| Rapgef2 | 0 | 1.79420722 | 0.77 | 0.12 | 0 | Salivary_gland | Rapgef2 |
| Plpp3 | 0 | 1.79149768 | 0.99 | 0.6 | 0 | Salivary_gland | Plpp3 |
| Cyyr1 | 0 | 1.78625431 | 0.96 | 0.48 | 0 | Salivary_gland | Cyyr1 |
| Zfp366 | 0 | 1.75914737 | 0.59 | 0 | 0 | Salivary_gland | Zfp366 |
| Sptbn1 | 0 | 1.75522657 | 1 | 0.74 | 0 | Salivary_gland | Sptbn1 |
| Gm20388 | 0 | 1.74948648 | 0.62 | 0 | 0 | Salivary_gland | Gm20388 |
| Hecw2 | 0 | 1.74002296 | 0.73 | 0.1 | 0 | Salivary_gland | Hecw2 |
| Col4a1 | 0 | 1.73790758 | 0.97 | 0.52 | 0 | Salivary_gland | Col4a1 |
| Pdgfd | 0 | 1.71984387 | 0.72 | 0.15 | 0 | Salivary_gland | Pdgfd |
| Magi1 | 0 | 1.69980467 | 0.65 | 0.06 | 0 | Salivary_gland | Magi1 |
| Fut8 | 0 | 1.6865196 | 0.66 | 0.08 | 0 | Salivary_gland | Fut8 |
| Airn | 0 | 1.68336434 | 0.57 | 0.03 | 0 | Salivary_gland | Airn |
| Rdx | 0 | 1.68314618 | 0.98 | 0.52 | 0 | Salivary_gland | Rdx |
| Fbln2 | 0 | 1.67635836 | 0.75 | 0.17 | 0 | Salivary_gland | Fbln2 |
| Col13a1 | 0 | 1.67380473 | 0.73 | 0.11 | 0 | Salivary_gland | Col13a1 |
| Phlpp1 | 0 | 1.66963442 | 0.65 | 0.06 | 0 | Salivary_gland | Phlpp1 |
| Cavin1 | 0 | 1.66217428 | 0.6 | 0 | 0 | Salivary_gland | Cavin1 |
| Numb | 0 | 1.64722277 | 0.78 | 0.14 | 0 | Salivary_gland | Numb |
| Rapgef5 | 0 | 1.62529529 | 0.61 | 0.04 | 0 | Salivary_gland | Rapgef5 |
| Arhgef12 | 0 | 1.60027752 | 0.81 | 0.19 | 0 | Salivary_gland | Arhgef12 |

| | | | | | | | |
|---|---|---|---|---|---|---|---|
| Dysf | 0 | 1.59661564 | 0.74 | 0.12 | 0 | Salivary_gland | Dysf |
| Dlc1 | 0 | 1.58599278 | 0.91 | 0.38 | 0 | Salivary_gland | Dlc1 |
| Oxr1 | 0 | 1.584234 | 0.72 | 0.14 | 0 | Salivary_gland | Oxr1 |
| Utrn | 0 | 1.58234492 | 0.84 | 0.25 | 0 | Salivary_gland | Utrn |
| Scgb2b26 | 0 | 1.57527586 | 0.45 | 0 | 0 | Salivary_gland | Scgb2b26 |
| Dip2c | 0 | 1.57441889 | 0.56 | 0.02 | 0 | Salivary_gland | Dip2c |
| Kdr | 0 | 1.56986153 | 1 | 0.59 | 0 | Salivary_gland | Kdr |
| Rnf220 | 0 | 1.5603017 | 0.62 | 0.11 | 0 | Salivary_gland | Rnf220 |
| Macf1 | 0 | 1.5563108 | 0.9 | 0.36 | 0 | Salivary_gland | Macf1 |
| Arhgap18 | 0 | 1.55097488 | 0.67 | 0.09 | 0 | Salivary_gland | Arhgap18 |
| Jam3 | 0 | 1.54933523 | 0.8 | 0.19 | 0 | Salivary_gland | Jam3 |
| Kitl | 0 | 1.54494497 | 0.88 | 0.34 | 0 | Salivary_gland | Kitl |
| Rgcc | 0 | 1.54136379 | 0.95 | 0.4 | 0 | Salivary_gland | Rgcc |
| Hmbox1 | 0 | 1.54016268 | 0.69 | 0.11 | 0 | Salivary_gland | Hmbox1 |
| Col4a2 | 0 | 1.53272675 | 0.94 | 0.42 | 0 | Salivary_gland | Col4a2 |
| Maml2 | 0 | 1.53172723 | 0.61 | 0.09 | 0 | Salivary_gland | Maml2 |
| Dock1 | 0 | 1.51840682 | 0.69 | 0.1 | 0 | Salivary_gland | Dock1 |
| Apbb2 | 0 | 1.51612646 | 0.88 | 0.32 | 0 | Salivary_gland | Apbb2 |
| Trir | 0 | 1.50655344 | 0.58 | 0 | 0 | Salivary_gland | Trir |
| Plscr2 | 0 | 1.50422868 | 0.88 | 0.23 | 0 | Salivary_gland | Plscr2 |
| Prkch | 0 | 1.50261111 | 0.83 | 0.26 | 0 | Salivary_gland | Prkch |
| Elmo1 | 0 | 1.49978425 | 0.68 | 0.12 | 0 | Salivary_gland | Elmo1 |
| Clic5 | 0 | 1.48111609 | 0.85 | 0.26 | 0 | Salivary_gland | Clic5 |
| Itih5 | 0 | 1.47343635 | 0.56 | 0.06 | 0 | Salivary_gland | Itih5 |
| Tceal9 | 0 | 1.45924153 | 0.53 | 0 | 0 | Salivary_gland | Tceal9 |
| Tcf4 | 0 | 1.45897427 | 0.99 | 0.65 | 0 | Salivary_gland | Tcf4 |
| Tmtc2 | 0 | 1.45482411 | 0.52 | 0.06 | 0 | Salivary_gland | Tmtc2 |
| Syne2 | 0 | 1.45450755 | 0.73 | 0.16 | 0 | Salivary_gland | Syne2 |
| Exoc3l2 | 0 | 1.45299045 | 0.81 | 0.16 | 0 | Salivary_gland | Exoc3l2 |
| Dnm3 | 0 | 1.44778468 | 0.69 | 0.15 | 0 | Salivary_gland | Dnm3 |
| Rai14 | 0 | 1.44776993 | 0.67 | 0.09 | 0 | Salivary_gland | Rai14 |
| Ptk2 | 0 | 1.43682663 | 0.72 | 0.14 | 0 | Salivary_gland | Ptk2 |
| Emcn | 0 | 1.43286802 | 0.99 | 0.64 | 0 | Salivary_gland | Emcn |
| Rtl8a | 0 | 1.42937941 | 0.49 | 0 | 0 | Salivary_gland | Rtl8a |
| Gm15564 | 0 | 1.42094054 | 0.47 | 0 | 0 | Salivary_gland | Gm15564 |
| Klf12 | 0 | 1.41697374 | 0.46 | 0 | 0 | Salivary_gland | Klf12 |
| Myo6 | 0 | 1.41499326 | 0.73 | 0.16 | 0 | Salivary_gland | Myo6 |
| Btbd7 | 0 | 1.41192035 | 0.56 | 0.03 | 0 | Salivary_gland | Btbd7 |
| Plpp1 | 0 | 1.40193738 | 0.97 | 0.58 | 0 | Salivary_gland | Plpp1 |
| Fmnl2 | 0 | 1.39449651 | 0.55 | 0.05 | 0 | Salivary_gland | Fmnl2 |
| Zhx3 | 0 | 1.39407587 | 0.55 | 0.05 | 0 | Salivary_gland | Zhx3 |

| | | | | | | | |
|---|---|---|---|---|---|---|---|
| Atp5o.1 | 0 | 1.39345719 | 0.51 | 0 | 0 | Salivary_gland | Atp5o.1 |
| Fbxl17 | 0 | 1.39339283 | 0.53 | 0.03 | 0 | Salivary_gland | Fbxl17 |
| Plekhg1 | 0 | 1.38693029 | 0.73 | 0.16 | 0 | Salivary_gland | Plekhg1 |
| Igfbp3 | 0 | 1.38139528 | 0.83 | 0.22 | 0 | Salivary_gland | Igfbp3 |
| Shank3 | 0 | 1.37798434 | 0.61 | 0.07 | 0 | Salivary_gland | Shank3 |
| Ralgapa1 | 0 | 1.36816431 | 0.64 | 0.1 | 0 | Salivary_gland | Ralgapa1 |
| Plxna2 | 0 | 1.36633185 | 0.66 | 0.12 | 0 | Salivary_gland | Plxna2 |
| Thrsp | 0 | 1.35471841 | 0.84 | 0.24 | 0 | Salivary_gland | Thrsp |
| mt-Nd4l | 0 | 1.35248325 | 0.75 | 0.18 | 0 | Salivary_gland | mt-Nd4l |
| Rgl1 | 0 | 1.35058693 | 0.75 | 0.19 | 0 | Salivary_gland | Rgl1 |
| Ppp1r12a | 0 | 1.34735053 | 0.74 | 0.2 | 0 | Salivary_gland | Ppp1r12a |
| Foxn3 | 0 | 1.33921587 | 0.71 | 0.17 | 0 | Salivary_gland | Foxn3 |
| Dgkh | 0 | 1.33617274 | 0.48 | 0 | 0 | Salivary_gland | Dgkh |
| Rasgrf2 | 0 | 1.32697936 | 0.55 | 0.04 | 0 | Salivary_gland | Rasgrf2 |
| Gnaq | 0 | 1.32419282 | 0.72 | 0.18 | 0 | Salivary_gland | Gnaq |
| Hist1h1e | 0 | 1.32336745 | 0.48 | 0.01 | 0 | Salivary_gland | Hist1h1e |
| Ahnak | 0 | 1.32216597 | 0.9 | 0.42 | 0 | Salivary_gland | Ahnak |
| Rtraf | 0 | 1.32010895 | 0.51 | 0 | 0 | Salivary_gland | Rtraf |
| Nbea | 0 | 1.31565347 | 0.54 | 0.05 | 0 | Salivary_gland | Nbea |
| Col15a1 | 0 | 1.3152429 | 0.68 | 0.12 | 0 | Salivary_gland | Col15a1 |
| Car6 | 0 | 1.31207725 | 0.36 | 0 | 0 | Salivary_gland | Car6 |
| Csgalnact1 | 0 | 1.29721514 | 0.56 | 0.08 | 0 | Salivary_gland | Csgalnact1 |
| Akt3 | 0 | 1.2844984 | 0.53 | 0.05 | 0 | Salivary_gland | Akt3 |
| Podxl | 0 | 1.28294215 | 0.95 | 0.48 | 0 | Salivary_gland | Podxl |
| Hexb | 0 | 1.28103405 | 0.66 | 0.13 | 0 | Salivary_gland | Hexb |
| Iigp1 | 0 | 1.27600348 | 0.66 | 0.26 | 0 | Salivary_gland | Iigp1 |
| Rock1 | 0 | 1.27415135 | 0.75 | 0.22 | 0 | Salivary_gland | Rock1 |
| Adgrl2 | 0 | 1.26876224 | 0.72 | 0.19 | 0 | Salivary_gland | Adgrl2 |
| Snu13 | 0 | 1.26508399 | 0.49 | 0 | 0 | Salivary_gland | Snu13 |
| Nedd9 | 0 | 1.264299 | 0.64 | 0.13 | 0 | Salivary_gland | Nedd9 |
| Bmpr2 | 0 | 1.2572528 | 0.85 | 0.32 | 0 | Salivary_gland | Bmpr2 |
| Cd24a | 0 | 1.24941165 | 0.77 | 0.21 | 0 | Salivary_gland | Cd24a |
| Spred1 | 0 | 1.24855908 | 0.57 | 0.08 | 0 | Salivary_gland | Spred1 |
| Lrch1 | 0 | 1.2481393 | 0.6 | 0.1 | 0 | Salivary_gland | Lrch1 |
| Tcf12 | 0 | 1.24733648 | 0.66 | 0.15 | 0 | Salivary_gland | Tcf12 |
| Rock2 | 0 | 1.24624215 | 0.69 | 0.16 | 0 | Salivary_gland | Rock2 |
| mt-Atp8 | 0 | 1.24602913 | 0.72 | 0.18 | 0 | Salivary_gland | mt-Atp8 |
| Entpd1 | 0 | 1.24494773 | 0.63 | 0.14 | 0 | Salivary_gland | Entpd1 |
| Gm12002 | 0 | 1.24452631 | 0.59 | 0.1 | 0 | Salivary_gland | Gm12002 |
| Pitpnc1 | 0 | 1.24348235 | 0.83 | 0.3 | 0 | Salivary_gland | Pitpnc1 |
| Ramp3 | 0 | 1.24311911 | 0.78 | 0.19 | 0 | Salivary_gland | Ramp3 |

| | | | | | | | |
|---|---|---|---|---|---|---|---|
| Sorbs1 | 0 | 1.23809239 | 0.46 | 0.02 | 0 | Salivary_gland | Sorbs1 |
| Tshz2 | 0 | 1.2373759 | 0.87 | 0.35 | 0 | Salivary_gland | Tshz2 |
| Hnrnpa0 | 0 | 1.23680861 | 0.7 | 0.14 | 0 | Salivary_gland | Hnrnpa0 |
| Peak1 | 0 | 1.23513457 | 0.74 | 0.21 | 0 | Salivary_gland | Peak1 |
| Exoc4 | 0 | 1.23481342 | 0.57 | 0.09 | 0 | Salivary_gland | Exoc4 |
| Asap1 | 0 | 1.2338248 | 0.78 | 0.24 | 0 | Salivary_gland | Asap1 |
| Atxn1 | 0 | 1.23239056 | 0.58 | 0.1 | 0 | Salivary_gland | Atxn1 |
| Pcdh17 | 0 | 1.23082807 | 0.73 | 0.22 | 0 | Salivary_gland | Pcdh17 |
| 9930111J21Rik2 | 0 | 1.23035874 | 0.59 | 0.1 | 0 | Salivary_gland | 9930111J21Rik2 |
| Stag1 | 0 | 1.22004722 | 0.63 | 0.12 | 0 | Salivary_gland | Stag1 |
| Prkd1 | 0 | 1.21601359 | 0.45 | 0.03 | 0 | Salivary_gland | Prkd1 |
| Babam2 | 0 | 1.21583649 | 0.41 | 0 | 0 | Salivary_gland | Babam2 |
| Nsd3 | 0 | 1.21405998 | 0.44 | 0 | 0 | Salivary_gland | Nsd3 |
| Baz2b | 0 | 1.21224314 | 0.66 | 0.15 | 0 | Salivary_gland | Baz2b |
| Gm47283 | 0 | 1.20521483 | 0.41 | 0 | 0 | Salivary_gland | Gm47283 |
| Fli1 | 0 | 1.2038747 | 0.8 | 0.3 | 0 | Salivary_gland | Fli1 |
| Ripor1 | 0 | 1.1997159 | 0.45 | 0 | 0 | Salivary_gland | Ripor1 |
| Ncoa7 | 0 | 1.19861235 | 0.65 | 0.13 | 0 | Salivary_gland | Ncoa7 |
| Etv6 | 0 | 1.19009159 | 0.49 | 0.05 | 0 | Salivary_gland | Etv6 |
| Prkce | 0 | 1.18728448 | 0.54 | 0.08 | 0 | Salivary_gland | Prkce |
| Ccdc85a | 0 | 1.18705499 | 0.6 | 0.13 | 0 | Salivary_gland | Ccdc85a |
| Clec2d | 0 | 1.18515622 | 0.84 | 0.36 | 0 | Salivary_gland | Clec2d |
| Rbms1 | 0 | 1.17862539 | 0.94 | 0.53 | 0 | Salivary_gland | Rbms1 |
| Id4 | 0 | 1.17246696 | 0.37 | 0.02 | 0 | Salivary_gland | Id4 |
| Snrk | 0 | 1.17217055 | 0.9 | 0.44 | 0 | Salivary_gland | Snrk |
| Maml3 | 0 | 1.16601506 | 0.41 | 0.02 | 0 | Salivary_gland | Maml3 |
| Pdgfb | 0 | 1.16595937 | 0.71 | 0.2 | 0 | Salivary_gland | Pdgfb |
| Tcf7l1 | 0 | 1.16532973 | 0.5 | 0.06 | 0 | Salivary_gland | Tcf7l1 |
| Ahr | 0 | 1.16419813 | 0.65 | 0.16 | 0 | Salivary_gland | Ahr |
| Rasgrp3 | 0 | 1.16137979 | 0.8 | 0.28 | 0 | Salivary_gland | Rasgrp3 |
| Hivep2 | 0 | 1.15859454 | 0.56 | 0.1 | 0 | Salivary_gland | Hivep2 |
| Gm26561 | 0 | 1.15826916 | 0.4 | 0 | 0 | Salivary_gland | Gm26561 |
| Hip1 | 0 | 1.15803852 | 0.71 | 0.2 | 0 | Salivary_gland | Hip1 |
| Zeb1 | 0 | 1.15511229 | 0.66 | 0.18 | 0 | Salivary_gland | Zeb1 |
| Ntf3 | 0 | 1.15299416 | 0.54 | 0.08 | 0 | Salivary_gland | Ntf3 |
| Erbin | 0 | 1.15213963 | 0.42 | 0 | 0 | Salivary_gland | Erbin |
| Chst2 | 0 | 1.1518101 | 0.5 | 0.05 | 0 | Salivary_gland | Chst2 |
| Lncpint | 0 | 1.14907678 | 0.46 | 0.03 | 0 | Salivary_gland | Lncpint |
| Gm20532 | 0 | 1.14809895 | 0.37 | 0 | 0 | Salivary_gland | Gm20532 |
| Ar | 0 | 1.14717081 | 0.45 | 0.04 | 0 | Salivary_gland | Ar |
| Cpd | 0 | 1.14693328 | 0.74 | 0.21 | 0 | Salivary_gland | Cpd |

| | | | | | | | |
|---|---|---|---|---|---|---|---|
| Filip1 | 0 | 1.1452206 | 0.57 | 0.1 | 0 | Salivary_gland | Filip1 |
| Klk1b26 | 0 | 1.14513795 | 0.27 | 0 | 0 | Salivary_gland | Klk1b26 |
| Prrc2c | 0 | 1.13370414 | 0.74 | 0.23 | 0 | Salivary_gland | Prrc2c |
| Pip | 0 | 1.12876832 | 0.31 | 0 | 0 | Salivary_gland | Pip |
| Rsad2 | 0 | 1.12706138 | 0.72 | 0.21 | 0 | Salivary_gland | Rsad2 |
| Hmcn1 | 0 | 1.12414366 | 0.64 | 0.18 | 0 | Salivary_gland | Hmcn1 |
| Gulp1 | 0 | 1.12410955 | 0.43 | 0.04 | 0 | Salivary_gland | Gulp1 |
| Gab2 | 0 | 1.12013208 | 0.38 | 0.02 | 0 | Salivary_gland | Gab2 |
| Rapgef4 | 0 | 1.11837359 | 0.77 | 0.21 | 0 | Salivary_gland | Rapgef4 |
| Nr3c1 | 0 | 1.1142655 | 0.77 | 0.26 | 0 | Salivary_gland | Nr3c1 |
| Map4k3 | 0 | 1.11326496 | 0.54 | 0.1 | 0 | Salivary_gland | Map4k3 |
| Qk | 0 | 1.1132499 | 0.84 | 0.36 | 0 | Salivary_gland | Qk |
| Arhgap26 | 0 | 1.10635278 | 0.46 | 0.04 | 0 | Salivary_gland | Arhgap26 |
| Sorbs2 | 0 | 1.10632973 | 0.76 | 0.23 | 0 | Salivary_gland | Sorbs2 |
| Cast | 0 | 1.10598097 | 0.53 | 0.08 | 0 | Salivary_gland | Cast |
| Cep85l | 0 | 1.10596131 | 0.45 | 0.04 | 0 | Salivary_gland | Cep85l |
| Col4a3bp | 0 | 1.10552236 | 0.53 | 0.07 | 0 | Salivary_gland | Col4a3bp |
| Eloc | 0 | 1.10052764 | 0.42 | 0 | 0 | Salivary_gland | Eloc |
| Rora | 0 | 1.09954078 | 0.41 | 0.08 | 0 | Salivary_gland | Rora |
| Ppp4r3b | 0 | 1.0974371 | 0.4 | 0 | 0 | Salivary_gland | Ppp4r3b |
| Kansl1 | 0 | 1.08905742 | 0.56 | 0.1 | 0 | Salivary_gland | Kansl1 |
| Myh9 | 0 | 1.0853688 | 0.76 | 0.29 | 0 | Salivary_gland | Myh9 |
| Lnpep | 0 | 1.08267017 | 0.41 | 0.01 | 0 | Salivary_gland | Lnpep |
| Pdlim5 | 0 | 1.07621875 | 0.62 | 0.16 | 0 | Salivary_gland | Pdlim5 |
| Mlec | 0 | 1.07495705 | 0.54 | 0.09 | 0 | Salivary_gland | Mlec |
| AU020206 | 0 | 1.06895118 | 0.51 | 0.08 | 0 | Salivary_gland | AU020206 |
| Rab21 | 0 | 1.06763887 | 0.46 | 0.03 | 0 | Salivary_gland | Rab21 |
| Zbtb20 | 0 | 1.06413471 | 0.9 | 0.51 | 0 | Salivary_gland | Zbtb20 |
| St8sia4 | 0 | 1.06271441 | 0.53 | 0.1 | 0 | Salivary_gland | St8sia4 |
| Tbc1d5 | 0 | 1.06191554 | 0.4 | 0.02 | 0 | Salivary_gland | Tbc1d5 |
| Svil | 0 | 1.05579427 | 0.51 | 0.08 | 0 | Salivary_gland | Svil |
| Fndc3b | 0 | 1.05162479 | 0.53 | 0.1 | 0 | Salivary_gland | Fndc3b |
| Ier5l | 0 | 1.05044165 | 0.47 | 0.05 | 0 | Salivary_gland | Ier5l |
| Tnrc6c | 0 | 1.05035838 | 0.57 | 0.11 | 0 | Salivary_gland | Tnrc6c |
| Hdac9 | 0 | 1.04929022 | 0.29 | 0 | 0 | Salivary_gland | Hdac9 |
| Nav2 | 0 | 1.04855229 | 0.41 | 0.04 | 0 | Salivary_gland | Nav2 |
| Fzd4 | 0 | 1.04718146 | 0.69 | 0.2 | 0 | Salivary_gland | Fzd4 |
| Colgalt2 | 0 | 1.04691513 | 0.42 | 0.04 | 0 | Salivary_gland | Colgalt2 |
| Nxn | 0 | 1.04628798 | 0.63 | 0.2 | 0 | Salivary_gland | Nxn |
| Tpt1 | 0 | 1.04241645 | 1 | 0.93 | 0 | Salivary_gland | Tpt1 |
| Nedd4l | 0 | 1.04238136 | 0.41 | 0.04 | 0 | Salivary_gland | Nedd4l |

| | | | | | | | |
|---|---|---|---|---|---|---|---|
| Sik3 | 0 | 1.04186717 | 0.45 | 0.05 | 0 | Salivary_gland | Sik3 |
| H2-T23 | 0 | 1.04024169 | 0.84 | 0.37 | 0 | Salivary_gland | H2-T23 |
| Ttc28 | 0 | 1.03760148 | 0.62 | 0.18 | 0 | Salivary_gland | Ttc28 |
| Vti1a | 0 | 1.03572373 | 0.4 | 0.03 | 0 | Salivary_gland | Vti1a |
| Nsd1 | 0 | 1.0274014 | 0.57 | 0.11 | 0 | Salivary_gland | Nsd1 |
| Vps28 | 0 | 1.02345415 | 0.38 | 0 | 0 | Salivary_gland | Vps28 |
| Chrm3 | 0 | 1.02240807 | 0.28 | 0.02 | 0 | Salivary_gland | Chrm3 |
| Cald1 | 0 | 1.02182626 | 0.83 | 0.32 | 0 | Salivary_gland | Cald1 |
| Sos1 | 0 | 1.01926045 | 0.42 | 0.04 | 0 | Salivary_gland | Sos1 |
| Nrp1 | 0 | 1.01866192 | 0.98 | 0.63 | 0 | Salivary_gland | Nrp1 |
| Mgll | 0 | 1.01649377 | 0.96 | 0.56 | 0 | Salivary_gland | Mgll |
| Gm13889 | 0 | 1.01641465 | 0.63 | 0.18 | 0 | Salivary_gland | Gm13889 |
| Nostrin | 0 | 1.01524273 | 0.74 | 0.27 | 0 | Salivary_gland | Nostrin |
| Mkl2 | 0 | 1.01492634 | 0.65 | 0.19 | 0 | Salivary_gland | Mkl2 |
| Nbdy | 0 | 1.01273036 | 0.38 | 0 | 0 | Salivary_gland | Nbdy |
| Rex1bd | 0 | 1.01047066 | 0.39 | 0 | 0 | Salivary_gland | Rex1bd |
| Elf1 | 0 | 1.00979863 | 0.69 | 0.21 | 0 | Salivary_gland | Elf1 |
| Nedd4 | 0 | 1.00975609 | 0.81 | 0.34 | 0 | Salivary_gland | Nedd4 |
| Jcad | 0 | 1.00905886 | 0.37 | 0 | 0 | Salivary_gland | Jcad |
| Sppl2a | 0 | 1.00881262 | 0.6 | 0.14 | 0 | Salivary_gland | Sppl2a |
| Msi2 | 0 | 1.00810085 | 0.56 | 0.13 | 0 | Salivary_gland | Msi2 |
| 4930578C19Rik | 0 | 1.00691099 | 0.35 | 0.01 | 0 | Salivary_gland | 4930578C19Rik |
| Selenot | 0 | 1.00236245 | 0.37 | 0 | 0 | Salivary_gland | Selenot |
| Trps1 | 0 | 1.00213534 | 0.43 | 0.07 | 0 | Salivary_gland | Trps1 |
| Vav3 | 0 | 0.99798415 | 0.43 | 0.06 | 0 | Salivary_gland | Vav3 |
| Tnrc6b | 0 | 0.99788896 | 0.61 | 0.15 | 0 | Salivary_gland | Tnrc6b |
| Ednrb | 0 | 0.99622669 | 0.6 | 0.13 | 0 | Salivary_gland | Ednrb |
| Nop53 | 0 | 0.99447544 | 0.37 | 0 | 0 | Salivary_gland | Nop53 |
| 4931406P16Rik | 0 | 0.99072048 | 0.85 | 0.37 | 0 | Salivary_gland | 4931406P16Rik |
| Myo1e | 0 | 0.98957717 | 0.48 | 0.07 | 0 | Salivary_gland | Myo1e |
| Arih1 | 0 | 0.98746448 | 0.53 | 0.1 | 0 | Salivary_gland | Arih1 |
| Nipbl | 0 | 0.98701061 | 0.61 | 0.18 | 0 | Salivary_gland | Nipbl |
| Sntb2 | 0 | 0.98623822 | 0.61 | 0.16 | 0 | Salivary_gland | Sntb2 |
| Prickle2 | 0 | 0.98596322 | 0.4 | 0.04 | 0 | Salivary_gland | Prickle2 |
| Tmtc1 | 0 | 0.98490242 | 0.43 | 0.05 | 0 | Salivary_gland | Tmtc1 |
| Pecam1 | 0 | 0.98424269 | 1 | 0.75 | 0 | Salivary_gland | Pecam1 |
| Srgap2 | 0 | 0.98309057 | 0.56 | 0.13 | 0 | Salivary_gland | Srgap2 |
| Nav1 | 0 | 0.98223797 | 0.63 | 0.18 | 0 | Salivary_gland | Nav1 |
| Birc6 | 0 | 0.98017643 | 0.61 | 0.15 | 0 | Salivary_gland | Birc6 |
| Usp34 | 0 | 0.97973585 | 0.59 | 0.14 | 0 | Salivary_gland | Usp34 |
| Mbd5 | 0 | 0.9772964 | 0.38 | 0.02 | 0 | Salivary_gland | Mbd5 |

| | | | | | | | |
|---|---|---|---|---|---|---|---|
| Plcb4 | 0 | 0.97709952 | 0.64 | 0.2 | 0 | Salivary_gland | Plcb4 |
| Polr2a | 0 | 0.9751839 | 0.49 | 0.1 | 0 | Salivary_gland | Polr2a |
| Kmt2c | 0 | 0.97457743 | 0.54 | 0.11 | 0 | Salivary_gland | Kmt2c |
| Arhgap31 | 0 | 0.97326631 | 0.95 | 0.62 | 0 | Salivary_gland | Arhgap31 |
| Mbd2 | 0 | 0.97225829 | 0.56 | 0.1 | 0 | Salivary_gland | Mbd2 |
| Eif4ebp2 | 0 | 0.96763759 | 0.47 | 0.06 | 0 | Salivary_gland | Eif4ebp2 |
| Dip2b | 0 | 0.96692709 | 0.51 | 0.08 | 0 | Salivary_gland | Dip2b |
| Hist1h4d | 0 | 0.96148663 | 0.36 | 0 | 0 | Salivary_gland | Hist1h4d |
| Fcho2 | 0 | 0.96065471 | 0.56 | 0.13 | 0 | Salivary_gland | Fcho2 |
| St6galnac3 | 0 | 0.96034374 | 0.6 | 0.19 | 0 | Salivary_gland | St6galnac3 |
| Kif13a | 0 | 0.96014933 | 0.45 | 0.06 | 0 | Salivary_gland | Kif13a |
| Pde8a | 0 | 0.95856356 | 0.49 | 0.08 | 0 | Salivary_gland | Pde8a |
| Fam172a | 0 | 0.95835789 | 0.42 | 0.06 | 0 | Salivary_gland | Fam172a |
| Mcf2l | 0 | 0.95702463 | 0.63 | 0.17 | 0 | Salivary_gland | Mcf2l |
| Susd6 | 0 | 0.95629005 | 0.51 | 0.09 | 0 | Salivary_gland | Susd6 |
| Fgd5 | 0 | 0.94851493 | 0.69 | 0.23 | 0 | Salivary_gland | Fgd5 |
| Pde4d | 0 | 0.9451499 | 0.35 | 0.03 | 0 | Salivary_gland | Pde4d |
| Gbp7 | 0 | 0.9447835 | 0.64 | 0.24 | 0 | Salivary_gland | Gbp7 |
| Grb10 | 0 | 0.9441561 | 0.55 | 0.13 | 0 | Salivary_gland | Grb10 |
| Zfyve9 | 0 | 0.94012088 | 0.43 | 0.05 | 0 | Salivary_gland | Zfyve9 |
| Atrx | 0 | 0.9376953 | 0.73 | 0.29 | 0 | Salivary_gland | Atrx |
| Enah | 0 | 0.93690294 | 0.43 | 0.05 | 0 | Salivary_gland | Enah |
| Atrnl1 | 0 | 0.9361841 | 0.38 | 0.03 | 0 | Salivary_gland | Atrnl1 |
| Rnf213 | 0 | 0.93461524 | 0.5 | 0.12 | 0 | Salivary_gland | Rnf213 |
| Gm12185 | 0 | 0.92934605 | 0.33 | 0 | 0 | Salivary_gland | Gm12185 |
| Mbnl1 | 0 | 0.92478133 | 0.72 | 0.28 | 0 | Salivary_gland | Mbnl1 |
| Trim2 | 0 | 0.92425066 | 0.43 | 0.06 | 0 | Salivary_gland | Trim2 |
| Adamts9 | 0 | 0.92381385 | 0.37 | 0.06 | 0 | Salivary_gland | Adamts9 |
| Ankrd11 | 0 | 0.92168531 | 0.61 | 0.17 | 0 | Salivary_gland | Ankrd11 |
| Il31ra | 0 | 0.91877865 | 0.33 | 0 | 0 | Salivary_gland | Il31ra |
| Sybu | 0 | 0.91877156 | 0.39 | 0.05 | 0 | Salivary_gland | Sybu |
| Mindy2 | 0 | 0.91752452 | 0.33 | 0 | 0 | Salivary_gland | Mindy2 |
| Man2a1 | 0 | 0.91665319 | 0.52 | 0.13 | 0 | Salivary_gland | Man2a1 |
| Mndal | 0 | 0.91500407 | 0.76 | 0.29 | 0 | Salivary_gland | Mndal |
| Atp8a1 | 0 | 0.90989501 | 0.46 | 0.09 | 0 | Salivary_gland | Atp8a1 |
| Ripor2 | 0 | 0.9094816 | 0.29 | 0 | 0 | Salivary_gland | Ripor2 |
| Large1 | 0 | 0.90918045 | 0.3 | 0 | 0 | Salivary_gland | Large1 |
| Hook3 | 0 | 0.9090662 | 0.57 | 0.14 | 0 | Salivary_gland | Hook3 |
| Kmt2a | 0 | 0.90814097 | 0.5 | 0.11 | 0 | Salivary_gland | Kmt2a |
| Rb1 | 0 | 0.90796877 | 0.39 | 0.05 | 0 | Salivary_gland | Rb1 |
| Zcchc6 | 0 | 0.9060918 | 0.51 | 0.11 | 0 | Salivary_gland | Zcchc6 |

| | | | | | | | |
|---|---|---|---|---|---|---|---|
| Dab2ip | 0 | 0.90591174 | 0.66 | 0.2 | 0 | Salivary_gland | Dab2ip |
| Clic4 | 0 | 0.90446035 | 0.89 | 0.52 | 0 | Salivary_gland | Clic4 |
| Mesd | 0 | 0.9043578 | 0.33 | 0 | 0 | Salivary_gland | Mesd |
| Nr3c2 | 0 | 0.90405583 | 0.31 | 0.01 | 0 | Salivary_gland | Nr3c2 |
| Brd1 | 0 | 0.90088042 | 0.41 | 0.04 | 0 | Salivary_gland | Brd1 |
| Myo18a | 0 | 0.90039313 | 0.62 | 0.17 | 0 | Salivary_gland | Myo18a |
| Cdc42bpa | 0 | 0.89970607 | 0.5 | 0.11 | 0 | Salivary_gland | Cdc42bpa |
| Spata13 | 0 | 0.8974755 | 0.43 | 0.06 | 0 | Salivary_gland | Spata13 |
| Cebpd | 0 | 0.89689659 | 0.3 | 0 | 0 | Salivary_gland | Cebpd |
| Usp25 | 0 | 0.89492579 | 0.51 | 0.11 | 0 | Salivary_gland | Usp25 |
| Clstn1 | 0 | 0.89455344 | 0.63 | 0.18 | 0 | Salivary_gland | Clstn1 |
| Wwox | 0 | 0.89446372 | 0.28 | 0.03 | 0 | Salivary_gland | Wwox |
| Cenpx | 0 | 0.89428624 | 0.35 | 0 | 0 | Salivary_gland | Cenpx |
| Sptan1 | 0 | 0.89350878 | 0.79 | 0.35 | 0 | Salivary_gland | Sptan1 |
| Ankrd28 | 0 | 0.89350373 | 0.32 | 0 | 0 | Salivary_gland | Ankrd28 |
| Gm32688 | 0 | 0.88970857 | 0.3 | 0 | 0 | Salivary_gland | Gm32688 |
| Cops9 | 0 | 0.88629893 | 0.33 | 0 | 0 | Salivary_gland | Cops9 |
| Pak2 | 0 | 0.88276985 | 0.67 | 0.22 | 0 | Salivary_gland | Pak2 |
| Lpp | 0 | 0.88191145 | 0.46 | 0.1 | 0 | Salivary_gland | Lpp |
| Cpne8 | 0 | 0.88155341 | 0.61 | 0.18 | 0 | Salivary_gland | Cpne8 |
| Rtl8b | 0 | 0.88131373 | 0.32 | 0 | 0 | Salivary_gland | Rtl8b |
| Vwa3a | 0 | 0.8807587 | 0.29 | 0.01 | 0 | Salivary_gland | Vwa3a |
| Chst15 | 0 | 0.880099 | 0.56 | 0.14 | 0 | Salivary_gland | Chst15 |
| Lrrc8c | 0 | 0.8768669 | 0.7 | 0.27 | 0 | Salivary_gland | Lrrc8c |
| Dnajc1 | 0 | 0.87647149 | 0.53 | 0.14 | 0 | Salivary_gland | Dnajc1 |
| Itga6 | 0 | 0.87645926 | 0.73 | 0.3 | 0 | Salivary_gland | Itga6 |
| 2010111I01Rik | 0 | 0.87258959 | 0.65 | 0.2 | 0 | Salivary_gland | 2010111I01Rik |
| Stard8 | 0 | 0.87230316 | 0.58 | 0.16 | 0 | Salivary_gland | Stard8 |
| Itsn2 | 0 | 0.87226557 | 0.52 | 0.12 | 0 | Salivary_gland | Itsn2 |
| Smurf2 | 0 | 0.87223333 | 0.47 | 0.09 | 0 | Salivary_gland | Smurf2 |
| Samd9l | 0 | 0.87142513 | 0.49 | 0.1 | 0 | Salivary_gland | Samd9l |
| Purb | 0 | 0.87081159 | 0.59 | 0.17 | 0 | Salivary_gland | Purb |
| Exoc6b | 0 | 0.86641336 | 0.46 | 0.09 | 0 | Salivary_gland | Exoc6b |
| mt-Co2 | 0 | 0.86537962 | 1 | 0.93 | 0 | Salivary_gland | mt-Co2 |
| mt-Nd5 | 0 | 0.8624535 | 0.78 | 0.33 | 0 | Salivary_gland | mt-Nd5 |
| Kif13b | 0 | 0.86117186 | 0.3 | 0 | 0 | Salivary_gland | Kif13b |
| Ablim1 | 0 | 0.85859761 | 0.68 | 0.26 | 0 | Salivary_gland | Ablim1 |
| Slco2b1 | 0 | 0.85622985 | 0.44 | 0.09 | 0 | Salivary_gland | Slco2b1 |
| Cep350 | 0 | 0.84966322 | 0.37 | 0.04 | 0 | Salivary_gland | Cep350 |
| Uvrag | 0 | 0.84886799 | 0.49 | 0.1 | 0 | Salivary_gland | Uvrag |
| Rbpms | 0 | 0.84882478 | 0.76 | 0.32 | 0 | Salivary_gland | Rbpms |

| | | | | | | | |
|---|---|---|---|---|---|---|---|
| Sema6d | 0 | 0.84854998 | 0.45 | 0.1 | 0 | Salivary_gland | Sema6d |
| Phldb2 | 0 | 0.84799519 | 0.58 | 0.17 | 0 | Salivary_gland | Phldb2 |
| Ldlrad4 | 0 | 0.84607159 | 0.32 | 0.03 | 0 | Salivary_gland | Ldlrad4 |
| Myo10 | 0 | 0.84552038 | 0.6 | 0.19 | 0 | Salivary_gland | Myo10 |
| Tpr | 0 | 0.84528085 | 0.67 | 0.25 | 0 | Salivary_gland | Tpr |
| R3hdm2 | 0 | 0.84478615 | 0.46 | 0.08 | 0 | Salivary_gland | R3hdm2 |
| Apc | 0 | 0.84407879 | 0.48 | 0.1 | 0 | Salivary_gland | Apc |
| Cped1 | 0 | 0.84289062 | 0.46 | 0.1 | 0 | Salivary_gland | Cped1 |
| Ccdc82 | 0 | 0.84094806 | 0.34 | 0.02 | 0 | Salivary_gland | Ccdc82 |
| Vps13b | 0 | 0.84010835 | 0.35 | 0.03 | 0 | Salivary_gland | Vps13b |
| Pafah1b1 | 0 | 0.8385153 | 0.73 | 0.29 | 0 | Salivary_gland | Pafah1b1 |
| Pip4k2a | 0 | 0.83653103 | 0.38 | 0.05 | 0 | Salivary_gland | Pip4k2a |
| AW554918 | 0 | 0.83584545 | 0.32 | 0.02 | 0 | Salivary_gland | AW554918 |
| Timp3 | 0 | 0.8357441 | 0.97 | 0.68 | 0 | Salivary_gland | Timp3 |
| Tmcc3 | 0 | 0.83429765 | 0.64 | 0.21 | 0 | Salivary_gland | Tmcc3 |
| Egln1 | 0 | 0.83383645 | 0.43 | 0.07 | 0 | Salivary_gland | Egln1 |
| Rabgap1l | 0 | 0.83159898 | 0.35 | 0.04 | 0 | Salivary_gland | Rabgap1l |
| Ywhaz | 0 | 0.83146174 | 0.76 | 0.32 | 0 | Salivary_gland | Ywhaz |
| Smg6 | 0 | 0.83115289 | 0.43 | 0.07 | 0 | Salivary_gland | Smg6 |
| Sash1 | 0 | 0.830171 | 0.65 | 0.21 | 0 | Salivary_gland | Sash1 |
| Ivns1abp | 0 | 0.82875041 | 0.58 | 0.19 | 0 | Salivary_gland | Ivns1abp |
| Pld1 | 0 | 0.82853775 | 0.34 | 0.03 | 0 | Salivary_gland | Pld1 |
| Kctd12 | 0 | 0.82767241 | 0.59 | 0.18 | 0 | Salivary_gland | Kctd12 |
| Arid1b | 0 | 0.82663767 | 0.41 | 0.06 | 0 | Salivary_gland | Arid1b |
| Mia2 | 0 | 0.82649105 | 0.3 | 0 | 0 | Salivary_gland | Mia2 |
| Kmt5b | 0 | 0.82511019 | 0.3 | 0 | 0 | Salivary_gland | Kmt5b |
| Nf1 | 0 | 0.82456781 | 0.41 | 0.07 | 0 | Salivary_gland | Nf1 |
| Rapgef6 | 0 | 0.82399097 | 0.43 | 0.07 | 0 | Salivary_gland | Rapgef6 |
| Selenos | 0 | 0.81969331 | 0.28 | 0 | 0 | Salivary_gland | Selenos |
| Ebf3 | 0 | 0.81820629 | 0.35 | 0.03 | 0 | Salivary_gland | Ebf3 |
| Trappc9 | 0 | 0.81788299 | 0.36 | 0.03 | 0 | Salivary_gland | Trappc9 |
| Rpl30 | 0 | 0.8169952 | 0.98 | 0.76 | 0 | Salivary_gland | Rpl30 |
| Gtdc1 | 0 | 0.81672896 | 0.31 | 0.03 | 0 | Salivary_gland | Gtdc1 |
| Arid4b | 0 | 0.81390273 | 0.5 | 0.13 | 0 | Salivary_gland | Arid4b |
| Mef2a | 0 | 0.81268343 | 0.7 | 0.28 | 0 | Salivary_gland | Mef2a |
| Map3k5 | 0 | 0.81156754 | 0.34 | 0.02 | 0 | Salivary_gland | Map3k5 |
| Crebbp | 0 | 0.80915023 | 0.53 | 0.14 | 0 | Salivary_gland | Crebbp |
| Rnf169 | 0 | 0.80913756 | 0.32 | 0.03 | 0 | Salivary_gland | Rnf169 |
| Psmb8 | 0 | 0.80766887 | 0.89 | 0.5 | 0 | Salivary_gland | Psmb8 |
| Ints9 | 0 | 0.80762726 | 0.35 | 0.04 | 0 | Salivary_gland | Ints9 |
| Kcnq1 | 0 | 0.80761497 | 0.33 | 0.04 | 0 | Salivary_gland | Kcnq1 |

| | | | | | | | |
|---|---|---|---|---|---|---|---|
| Uaca | 0 | 0.80479264 | 0.63 | 0.2 | 0 | Salivary_gland | Uaca |
| Adgrf5 | 0 | 0.80458615 | 0.94 | 0.62 | 0 | Salivary_gland | Adgrf5 |
| Sipa1l1 | 0 | 0.80416384 | 0.34 | 0.03 | 0 | Salivary_gland | Sipa1l1 |
| Cop1 | 0 | 0.80367533 | 0.3 | 0 | 0 | Salivary_gland | Cop1 |
| Sipa1l2 | 0 | 0.80354873 | 0.43 | 0.08 | 0 | Salivary_gland | Sipa1l2 |
| Grk5 | 0 | 0.80217039 | 0.36 | 0.05 | 0 | Salivary_gland | Grk5 |
| Tspan9 | 0 | 0.80151577 | 0.69 | 0.26 | 0 | Salivary_gland | Tspan9 |
| Iqgap1 | 0 | 0.79938315 | 0.73 | 0.31 | 0 | Salivary_gland | Iqgap1 |
| Acap2 | 0 | 0.79929477 | 0.5 | 0.12 | 0 | Salivary_gland | Acap2 |
| Acvr2a | 0 | 0.79889519 | 0.34 | 0.03 | 0 | Salivary_gland | Acvr2a |
| Bach2 | 0 | 0.79771434 | 0.28 | 0.02 | 0 | Salivary_gland | Bach2 |
| Utp11 | 0 | 0.79754695 | 0.29 | 0 | 0 | Salivary_gland | Utp11 |
| Wdfy3 | 0 | 0.7974561 | 0.42 | 0.07 | 0 | Salivary_gland | Wdfy3 |
| Ybx3 | 0 | 0.79644056 | 0.5 | 0.12 | 0 | Salivary_gland | Ybx3 |
| Ptbp3 | 0 | 0.79526909 | 0.7 | 0.28 | 0 | Salivary_gland | Ptbp3 |
| Smco4 | 0 | 0.79508379 | 0.86 | 0.44 | 0 | Salivary_gland | Smco4 |
| Zfp462 | 0 | 0.79469591 | 0.34 | 0.04 | 0 | Salivary_gland | Zfp462 |
| Map4k4 | 0 | 0.79244189 | 0.56 | 0.16 | 0 | Salivary_gland | Map4k4 |
| Prex2 | 0 | 0.79189375 | 0.77 | 0.36 | 0 | Salivary_gland | Prex2 |
| Ankhd1 | 0 | 0.7915369 | 0.44 | 0.09 | 0 | Salivary_gland | Ankhd1 |
| Hspg2 | 0 | 0.79123352 | 0.84 | 0.36 | 0 | Salivary_gland | Hspg2 |
| Nbeal1 | 0 | 0.79114977 | 0.57 | 0.18 | 0 | Salivary_gland | Nbeal1 |
| Ndufaf8 | 0 | 0.79068918 | 0.29 | 0 | 0 | Salivary_gland | Ndufaf8 |
| Mcrip1 | 0 | 0.79062167 | 0.3 | 0 | 0 | Salivary_gland | Mcrip1 |
| Sh3bgrl | 0 | 0.78746875 | 0.73 | 0.32 | 0 | Salivary_gland | Sh3bgrl |
| Lrrc3b | 0 | 0.7863655 | 0.31 | 0.05 | 0 | Salivary_gland | Lrrc3b |
| Ncoa2 | 0 | 0.78555197 | 0.38 | 0.06 | 0 | Salivary_gland | Ncoa2 |
| Dennd1a | 0 | 0.7855113 | 0.37 | 0.06 | 0 | Salivary_gland | Dennd1a |
| Zcchc7 | 0 | 0.78445612 | 0.47 | 0.12 | 0 | Salivary_gland | Zcchc7 |
| Afap1 | 0 | 0.78432174 | 0.4 | 0.07 | 0 | Salivary_gland | Afap1 |
| Scarb1 | 0 | 0.7836618 | 0.78 | 0.38 | 0 | Salivary_gland | Scarb1 |
| Rab8b | 0 | 0.78161971 | 0.4 | 0.06 | 0 | Salivary_gland | Rab8b |
| Pkp4 | 0 | 0.78155028 | 0.54 | 0.15 | 0 | Salivary_gland | Pkp4 |
| Fut4 | 0 | 0.78117754 | 0.27 | 0.01 | 0 | Salivary_gland | Fut4 |
| Btbd9 | 0 | 0.78115505 | 0.4 | 0.08 | 0 | Salivary_gland | Btbd9 |
| Hdgfl3 | 0 | 0.78013348 | 0.28 | 0 | 0 | Salivary_gland | Hdgfl3 |
| Ddx6 | 0 | 0.77994653 | 0.57 | 0.17 | 0 | Salivary_gland | Ddx6 |
| Piezo1 | 0 | 0.77803971 | 0.47 | 0.1 | 0 | Salivary_gland | Piezo1 |
| Lrch3 | 0 | 0.77745934 | 0.43 | 0.08 | 0 | Salivary_gland | Lrch3 |
| Ppp1r13b | 0 | 0.77709522 | 0.52 | 0.13 | 0 | Salivary_gland | Ppp1r13b |
| Arhgap5 | 0 | 0.77667109 | 0.5 | 0.14 | 0 | Salivary_gland | Arhgap5 |

| | | | | | | | |
|---|---|---|---|---|---|---|---|
| Zfand3 | 0 | 0.7746164 | 0.5 | 0.13 | 0 | Salivary_gland | Zfand3 |
| Tnks2 | 0 | 0.77442417 | 0.5 | 0.12 | 0 | Salivary_gland | Tnks2 |
| Slk | 0 | 0.77423579 | 0.46 | 0.11 | 0 | Salivary_gland | Slk |
| Rbfox2 | 0 | 0.77367992 | 0.5 | 0.13 | 0 | Salivary_gland | Rbfox2 |
| Mycbp2 | 0 | 0.7720825 | 0.57 | 0.2 | 0 | Salivary_gland | Mycbp2 |
| Zfp638 | 0 | 0.77132399 | 0.46 | 0.1 | 0 | Salivary_gland | Zfp638 |
| Cul3 | 0 | 0.77019359 | 0.43 | 0.08 | 0 | Salivary_gland | Cul3 |
| Zfp608 | 0 | 0.76098179 | 0.35 | 0.05 | 0 | Salivary_gland | Zfp608 |
| Myo9a | 0 | 0.76050158 | 0.34 | 0.04 | 0 | Salivary_gland | Myo9a |
| Trim44 | 0 | 0.75802013 | 0.41 | 0.07 | 0 | Salivary_gland | Trim44 |
| Smad4 | 0 | 0.75713853 | 0.41 | 0.08 | 0 | Salivary_gland | Smad4 |
| Ephb1 | 0 | 0.75554809 | 0.29 | 0.02 | 0 | Salivary_gland | Ephb1 |
| AC149090.1 | 0 | 0.75519451 | 0.27 | 0 | 0 | Salivary_gland | AC149090.1 |
| Herc6 | 0 | 0.75290808 | 0.39 | 0.09 | 0 | Salivary_gland | Herc6 |
| Ptprk | 0 | 0.7514762 | 0.39 | 0.08 | 0 | Salivary_gland | Ptprk |
| Tmem47 | 0 | 0.74938659 | 0.46 | 0.12 | 0 | Salivary_gland | Tmem47 |
| Spag9 | 0 | 0.7465482 | 0.67 | 0.26 | 0 | Salivary_gland | Spag9 |
| AY036118 | 0 | 0.74564697 | 0.96 | 0.73 | 0 | Salivary_gland | AY036118 |
| Endod1 | 0 | 0.74516707 | 0.44 | 0.09 | 0 | Salivary_gland | Endod1 |
| Kat6a | 0 | 0.74370869 | 0.41 | 0.08 | 0 | Salivary_gland | Kat6a |
| Pacs1 | 0 | 0.74355683 | 0.33 | 0.04 | 0 | Salivary_gland | Pacs1 |
| Herc1 | 0 | 0.74249175 | 0.49 | 0.13 | 0 | Salivary_gland | Herc1 |
| Pcnx | 0 | 0.74175762 | 0.42 | 0.08 | 0 | Salivary_gland | Pcnx |
| Plxna4 | 0 | 0.74042207 | 0.29 | 0.03 | 0 | Salivary_gland | Plxna4 |
| Map3k3 | 0 | 0.73987846 | 0.34 | 0.04 | 0 | Salivary_gland | Map3k3 |
| Rere | 0 | 0.73955706 | 0.44 | 0.1 | 0 | Salivary_gland | Rere |
| Gbp5 | 0 | 0.73927626 | 0.34 | 0.06 | 0 | Salivary_gland | Gbp5 |
| Zdhhc17 | 0 | 0.7390226 | 0.35 | 0.05 | 0 | Salivary_gland | Zdhhc17 |
| Ccnyl1 | 0 | 0.73881902 | 0.33 | 0.03 | 0 | Salivary_gland | Ccnyl1 |
| Chd9 | 0 | 0.73834774 | 0.35 | 0.05 | 0 | Salivary_gland | Chd9 |
| Dpp8 | 0 | 0.73809649 | 0.36 | 0.05 | 0 | Salivary_gland | Dpp8 |
| Rtf2 | 0 | 0.73807658 | 0.28 | 0 | 0 | Salivary_gland | Rtf2 |
| Ptgs1 | 0 | 0.73622195 | 0.56 | 0.16 | 0 | Salivary_gland | Ptgs1 |
| Gm38399 | 0 | 0.7358784 | 0.32 | 0.04 | 0 | Salivary_gland | Gm38399 |
| Kpna4 | 0 | 0.73448689 | 0.52 | 0.15 | 0 | Salivary_gland | Kpna4 |
| Rmdn1 | 0 | 0.73420214 | 0.33 | 0.04 | 0 | Salivary_gland | Rmdn1 |
| Zcchc11 | 0 | 0.73216736 | 0.41 | 0.09 | 0 | Salivary_gland | Zcchc11 |
| Lrba | 0 | 0.73135456 | 0.3 | 0.02 | 0 | Salivary_gland | Lrba |
| Mosmo | 0 | 0.73125498 | 0.27 | 0 | 0 | Salivary_gland | Mosmo |
| Gcnt2 | 0 | 0.7311677 | 0.43 | 0.11 | 0 | Salivary_gland | Gcnt2 |
| Ranbp2 | 0 | 0.73043043 | 0.36 | 0.04 | 0 | Salivary_gland | Ranbp2 |

| | | | | | | | |
|---|---|---|---|---|---|---|---|
| Nfia | 0 | 0.73033216 | 0.62 | 0.23 | 0 | Salivary_gland | Nfia |
| Ifit2 | 0 | 0.72991085 | 0.3 | 0.06 | 0 | Salivary_gland | Ifit2 |
| Cog5 | 0 | 0.7299032 | 0.28 | 0.01 | 0 | Salivary_gland | Cog5 |
| Pcm1 | 0 | 0.72985969 | 0.48 | 0.12 | 0 | Salivary_gland | Pcm1 |
| Atg4a | 0 | 0.7295778 | 0.32 | 0.06 | 0 | Salivary_gland | Atg4a |
| Nxpe2 | 0 | 0.72788431 | 0.34 | 0.05 | 0 | Salivary_gland | Nxpe2 |
| Ubr3 | 0 | 0.72769445 | 0.37 | 0.06 | 0 | Salivary_gland | Ubr3 |
| Eml4 | 0 | 0.72709279 | 0.37 | 0.06 | 0 | Salivary_gland | Eml4 |
| Fam198b | 0 | 0.72580928 | 0.5 | 0.16 | 0 | Salivary_gland | Fam198b |
| Ddx50 | 0 | 0.72488596 | 0.5 | 0.13 | 0 | Salivary_gland | Ddx50 |
| Wwtr1 | 0 | 0.72432965 | 0.64 | 0.24 | 0 | Salivary_gland | Wwtr1 |
| Dock9 | 0 | 0.72421543 | 0.55 | 0.18 | 0 | Salivary_gland | Dock9 |
| Map4 | 0 | 0.72410079 | 0.62 | 0.22 | 0 | Salivary_gland | Map4 |
| Ccdc28b | 0 | 0.72361577 | 0.56 | 0.19 | 0 | Salivary_gland | Ccdc28b |
| Fmc1 | 0 | 0.72105571 | 0.25 | 0 | 0 | Salivary_gland | Fmc1 |
| St3gal4 | 0 | 0.72048593 | 0.56 | 0.19 | 0 | Salivary_gland | St3gal4 |
| Fgfr2 | 0 | 0.72030706 | 0.27 | 0.01 | 0 | Salivary_gland | Fgfr2 |
| AW112010 | 0 | 0.71916706 | 0.93 | 0.49 | 0 | Salivary_gland | AW112010 |
| Fmnl3 | 0 | 0.7180163 | 0.55 | 0.16 | 0 | Salivary_gland | Fmnl3 |
| Adam10 | 0 | 0.71756002 | 0.59 | 0.2 | 0 | Salivary_gland | Adam10 |
| Srgn | 0 | 0.71726415 | 0.97 | 0.69 | 0 | Salivary_gland | Srgn |
| Zfp91 | 0 | 0.7167238 | 0.46 | 0.12 | 0 | Salivary_gland | Zfp91 |
| Wasf2 | 0 | 0.71584488 | 0.8 | 0.4 | 0 | Salivary_gland | Wasf2 |
| Cmtm8 | 0 | 0.71469525 | 0.74 | 0.33 | 0 | Salivary_gland | Cmtm8 |
| Cdk13 | 0 | 0.71412081 | 0.37 | 0.06 | 0 | Salivary_gland | Cdk13 |
| Sp100 | 0 | 0.7129535 | 0.73 | 0.33 | 0 | Salivary_gland | Sp100 |
| Sh3rf1 | 0 | 0.7127515 | 0.33 | 0.05 | 0 | Salivary_gland | Sh3rf1 |
| Ica1 | 0 | 0.71122448 | 0.44 | 0.12 | 0 | Salivary_gland | Ica1 |
| Fry | 0 | 0.71099296 | 0.51 | 0.15 | 0 | Salivary_gland | Fry |
| Smc1a | 0 | 0.71065951 | 0.57 | 0.2 | 0 | Salivary_gland | Smc1a |
| Arid4a | 0 | 0.71037083 | 0.44 | 0.12 | 0 | Salivary_gland | Arid4a |
| Top2b | 0 | 0.7098637 | 0.5 | 0.14 | 0 | Salivary_gland | Top2b |
| Cntln | 0 | 0.70955656 | 0.34 | 0.06 | 0 | Salivary_gland | Cntln |
| Mllt10 | 0 | 0.70771838 | 0.48 | 0.13 | 0 | Salivary_gland | Mllt10 |
| Ash1l | 0 | 0.70587058 | 0.56 | 0.2 | 0 | Salivary_gland | Ash1l |
| Wdr33 | 0 | 0.70410242 | 0.37 | 0.06 | 0 | Salivary_gland | Wdr33 |
| Wwc2 | 0 | 0.70408983 | 0.5 | 0.14 | 0 | Salivary_gland | Wwc2 |
| Zfp609 | 0 | 0.70354023 | 0.31 | 0.03 | 0 | Salivary_gland | Zfp609 |
| Bod1l | 0 | 0.703288 | 0.48 | 0.16 | 0 | Salivary_gland | Bod1l |
| Gbp8 | 0 | 0.7021099 | 0.26 | 0 | 0 | Salivary_gland | Gbp8 |
| Mfsd6 | 0 | 0.70021942 | 0.41 | 0.08 | 0 | Salivary_gland | Mfsd6 |

| | | | | | | | |
|---|---|---|---|---|---|---|---|
| Zmynd11 | 0 | 0.69899652 | 0.54 | 0.17 | 0 | Salivary_gland | Zmynd11 |
| Kdm6a | 0 | 0.69864613 | 0.33 | 0.04 | 0 | Salivary_gland | Kdm6a |
| Rin2 | 0 | 0.69821156 | 0.4 | 0.09 | 0 | Salivary_gland | Rin2 |
| Bptf | 0 | 0.69728331 | 0.53 | 0.18 | 0 | Salivary_gland | Bptf |
| Mrpl58 | 0 | 0.69726194 | 0.27 | 0 | 0 | Salivary_gland | Mrpl58 |
| Chn2 | 0 | 0.69718241 | 0.29 | 0.05 | 0 | Salivary_gland | Chn2 |
| Epb41l4a | 0 | 0.69715302 | 0.25 | 0.02 | 0 | Salivary_gland | Epb41l4a |
| 2900026A02Rik | 0 | 0.69708799 | 0.5 | 0.14 | 0 | Salivary_gland | 2900026A02Rik |
| Kalrn | 0 | 0.69615333 | 0.3 | 0.04 | 0 | Salivary_gland | Kalrn |
| Selenoh | 0 | 0.69586164 | 0.26 | 0 | 0 | Salivary_gland | Selenoh |
| Zfp710 | 0 | 0.69550504 | 0.32 | 0.04 | 0 | Salivary_gland | Zfp710 |
| Crybg3 | 0 | 0.69531465 | 0.35 | 0.06 | 0 | Salivary_gland | Crybg3 |
| Stim1 | 0 | 0.69450366 | 0.3 | 0.04 | 0 | Salivary_gland | Stim1 |
| Trip11 | 0 | 0.69197456 | 0.38 | 0.08 | 0 | Salivary_gland | Trip11 |
| Cd2ap | 0 | 0.69151981 | 0.6 | 0.23 | 0 | Salivary_gland | Cd2ap |
| Lnx2 | 0 | 0.69007058 | 0.38 | 0.07 | 0 | Salivary_gland | Lnx2 |
| Gab1 | 0 | 0.69006759 | 0.32 | 0.05 | 0 | Salivary_gland | Gab1 |
| Phf20l1 | 0 | 0.6898666 | 0.53 | 0.17 | 0 | Salivary_gland | Phf20l1 |
| Znfx1 | 0 | 0.68917879 | 0.42 | 0.1 | 0 | Salivary_gland | Znfx1 |
| Ascc3 | 0 | 0.68767313 | 0.32 | 0.04 | 0 | Salivary_gland | Ascc3 |
| Nfe2l1 | 0 | 0.68719765 | 0.63 | 0.23 | 0 | Salivary_gland | Nfe2l1 |
| Nhsl2 | 0 | 0.686162 | 0.41 | 0.09 | 0 | Salivary_gland | Nhsl2 |
| Rap1gds1 | 0 | 0.68580616 | 0.4 | 0.1 | 0 | Salivary_gland | Rap1gds1 |
| Cdk17 | 0 | 0.68528467 | 0.48 | 0.14 | 0 | Salivary_gland | Cdk17 |
| Epb41 | 0 | 0.68440819 | 0.36 | 0.06 | 0 | Salivary_gland | Epb41 |
| Rassf9 | 0 | 0.68434183 | 0.39 | 0.12 | 0 | Salivary_gland | Rassf9 |
| Kmt2e | 0 | 0.68413943 | 0.71 | 0.33 | 0 | Salivary_gland | Kmt2e |
| Klhl24 | 0 | 0.68385508 | 0.45 | 0.11 | 0 | Salivary_gland | Klhl24 |
| Ppp4r3a | 0 | 0.68371694 | 0.25 | 0 | 0 | Salivary_gland | Ppp4r3a |
| Setx | 0 | 0.68364958 | 0.33 | 0.05 | 0 | Salivary_gland | Setx |
| Aig1 | 0 | 0.68283091 | 0.48 | 0.14 | 0 | Salivary_gland | Aig1 |
| Elf2 | 0 | 0.6827096 | 0.55 | 0.19 | 0 | Salivary_gland | Elf2 |
| Zfp407 | 0 | 0.68146794 | 0.27 | 0.02 | 0 | Salivary_gland | Zfp407 |
| Epha4 | 0 | 0.6797936 | 0.36 | 0.07 | 0 | Salivary_gland | Epha4 |
| Tjp1 | 0 | 0.67941001 | 0.65 | 0.27 | 0 | Salivary_gland | Tjp1 |
| Fbxo11 | 0 | 0.676906 | 0.39 | 0.09 | 0 | Salivary_gland | Fbxo11 |
| Ankrd12 | 0 | 0.67574329 | 0.54 | 0.21 | 0 | Salivary_gland | Ankrd12 |
| Sbf2 | 0 | 0.67419083 | 0.48 | 0.14 | 0 | Salivary_gland | Sbf2 |
| Kctd17 | 0 | 0.67417959 | 0.26 | 0 | 0 | Salivary_gland | Kctd17 |
| 2900097C17Rik | 0 | 0.67413609 | 0.25 | 0 | 0 | Salivary_gland | 2900097C17Rik |
| Rc3h1 | 0 | 0.67372229 | 0.39 | 0.09 | 0 | Salivary_gland | Rc3h1 |

| | | | | | | | |
|---|---|---|---|---|---|---|---|
| Ifi203 | 0 | 0.67248224 | 0.72 | 0.29 | 0 | Salivary_gland | Ifi203 |
| Larp1b | 0 | 0.67247407 | 0.28 | 0.03 | 0 | Salivary_gland | Larp1b |
| Ncor1 | 0 | 0.67244389 | 0.7 | 0.31 | 0 | Salivary_gland | Ncor1 |
| Pik3c2a | 0 | 0.67216704 | 0.42 | 0.1 | 0 | Salivary_gland | Pik3c2a |
| Med13 | 0 | 0.67196249 | 0.35 | 0.06 | 0 | Salivary_gland | Med13 |
| Atp11c | 0 | 0.67097825 | 0.34 | 0.06 | 0 | Salivary_gland | Atp11c |
| Hspa4 | 0 | 0.67045513 | 0.48 | 0.13 | 0 | Salivary_gland | Hspa4 |
| Ets1 | 0 | 0.6703708 | 0.83 | 0.43 | 0 | Salivary_gland | Ets1 |
| Stk10 | 0 | 0.67036393 | 0.32 | 0.04 | 0 | Salivary_gland | Stk10 |
| Frs2 | 0 | 0.67003858 | 0.25 | 0.01 | 0 | Salivary_gland | Frs2 |
| Phf21a | 0 | 0.66999217 | 0.38 | 0.09 | 0 | Salivary_gland | Phf21a |
| Snx9 | 0 | 0.66923314 | 0.37 | 0.07 | 0 | Salivary_gland | Snx9 |
| Smad7 | 0 | 0.66829343 | 0.66 | 0.27 | 0 | Salivary_gland | Smad7 |
| Trib2 | 0 | 0.66693695 | 0.51 | 0.15 | 0 | Salivary_gland | Trib2 |
| Fam208a | 0 | 0.66620983 | 0.39 | 0.09 | 0 | Salivary_gland | Fam208a |
| Ambra1 | 0 | 0.66591067 | 0.29 | 0.03 | 0 | Salivary_gland | Ambra1 |
| Sema6a | 0 | 0.66559406 | 0.63 | 0.25 | 0 | Salivary_gland | Sema6a |
| Parp14 | 0 | 0.66390405 | 0.52 | 0.18 | 0 | Salivary_gland | Parp14 |
| Atad2b | 0 | 0.66335367 | 0.3 | 0.03 | 0 | Salivary_gland | Atad2b |
| Ncoa1 | 0 | 0.66282398 | 0.3 | 0.04 | 0 | Salivary_gland | Ncoa1 |
| Lcor | 0 | 0.66266486 | 0.28 | 0.03 | 0 | Salivary_gland | Lcor |
| Nxpe4 | 0 | 0.66230554 | 0.45 | 0.13 | 0 | Salivary_gland | Nxpe4 |
| Scaper | 0 | 0.66140831 | 0.28 | 0.04 | 0 | Salivary_gland | Scaper |
| Tnks | 0 | 0.66030287 | 0.25 | 0.01 | 0 | Salivary_gland | Tnks |
| Braf | 0 | 0.65985445 | 0.31 | 0.05 | 0 | Salivary_gland | Braf |
| Vps13c | 0 | 0.65910581 | 0.27 | 0.02 | 0 | Salivary_gland | Vps13c |
| Fam114a1 | 0 | 0.65886071 | 0.32 | 0.05 | 0 | Salivary_gland | Fam114a1 |
| Prdm1 | 0 | 0.65858016 | 0.31 | 0.05 | 0 | Salivary_gland | Prdm1 |
| Arhgap17 | 0 | 0.65797402 | 0.48 | 0.13 | 0 | Salivary_gland | Arhgap17 |
| Ext1 | 0 | 0.65678425 | 0.29 | 0.05 | 0 | Salivary_gland | Ext1 |
| Zbtb46 | 0 | 0.65649175 | 0.46 | 0.14 | 0 | Salivary_gland | Zbtb46 |
| Zfpm1 | 0 | 0.65619229 | 0.35 | 0.06 | 0 | Salivary_gland | Zfpm1 |
| Msrb3 | 0 | 0.65596122 | 0.37 | 0.08 | 0 | Salivary_gland | Msrb3 |
| Taok1 | 0 | 0.65577043 | 0.49 | 0.15 | 0 | Salivary_gland | Taok1 |
| Fat4 | 0 | 0.65534963 | 0.33 | 0.06 | 0 | Salivary_gland | Fat4 |
| Itpr1 | 0 | 0.65478788 | 0.42 | 0.11 | 0 | Salivary_gland | Itpr1 |
| Eif4g3 | 0 | 0.65450595 | 0.47 | 0.14 | 0 | Salivary_gland | Eif4g3 |
| Me3 | 0 | 0.65408219 | 0.25 | 0.03 | 0 | Salivary_gland | Me3 |
| Phactr4 | 0 | 0.65337887 | 0.39 | 0.08 | 0 | Salivary_gland | Phactr4 |
| Chd2 | 0 | 0.65331732 | 0.36 | 0.08 | 0 | Salivary_gland | Chd2 |
| Usp15 | 0 | 0.65281636 | 0.32 | 0.05 | 0 | Salivary_gland | Usp15 |

| | | | | | | | |
|---|---|---|---|---|---|---|---|
| Aaed1 | 0 | 0.65252928 | 0.27 | 0.02 | 0 | Salivary_gland | Aaed1 |
| Smchd1 | 0 | 0.65144552 | 0.46 | 0.13 | 0 | Salivary_gland | Smchd1 |
| Ggps1 | 0 | 0.65096821 | 0.32 | 0.04 | 0 | Salivary_gland | Ggps1 |
| Glg1 | 0 | 0.65053749 | 0.48 | 0.15 | 0 | Salivary_gland | Glg1 |
| Zfp292 | 0 | 0.6504398 | 0.43 | 0.11 | 0 | Salivary_gland | Zfp292 |
| 4930402H24Rik | 0 | 0.65015474 | 0.45 | 0.13 | 0 | Salivary_gland | 4930402H24Rik |
| Calm1 | 0 | 0.64970464 | 1 | 0.9 | 0 | Salivary_gland | Calm1 |
| Hnrnpd | 0 | 0.64714584 | 0.49 | 0.15 | 0 | Salivary_gland | Hnrnpd |
| Zfp148 | 0 | 0.64542014 | 0.44 | 0.12 | 0 | Salivary_gland | Zfp148 |
| Oxct1 | 0 | 0.64540232 | 0.37 | 0.09 | 0 | Salivary_gland | Oxct1 |
| Psmb9 | 0 | 0.64498993 | 0.72 | 0.33 | 0 | Salivary_gland | Psmb9 |
| Gpatch8 | 0 | 0.64486295 | 0.45 | 0.12 | 0 | Salivary_gland | Gpatch8 |
| Chst11 | 0 | 0.64423304 | 0.31 | 0.05 | 0 | Salivary_gland | Chst11 |
| Jarid2 | 0 | 0.64184822 | 0.33 | 0.06 | 0 | Salivary_gland | Jarid2 |
| Dennd3 | 0 | 0.64108675 | 0.38 | 0.07 | 0 | Salivary_gland | Dennd3 |
| Psme4 | 0 | 0.64086524 | 0.32 | 0.05 | 0 | Salivary_gland | Psme4 |
| Luzp1 | 0 | 0.64073913 | 0.75 | 0.35 | 0 | Salivary_gland | Luzp1 |
| Klhdc10 | 0 | 0.63988977 | 0.27 | 0.03 | 0 | Salivary_gland | Klhdc10 |
| Jmjd1c | 0 | 0.63959514 | 0.52 | 0.18 | 0 | Salivary_gland | Jmjd1c |
| Swap70 | 0 | 0.63950353 | 0.51 | 0.16 | 0 | Salivary_gland | Swap70 |
| Eea1 | 0 | 0.63944516 | 0.41 | 0.11 | 0 | Salivary_gland | Eea1 |
| Zmym4 | 0 | 0.63923425 | 0.3 | 0.04 | 0 | Salivary_gland | Zmym4 |
| Myo1d | 0 | 0.63852755 | 0.29 | 0.04 | 0 | Salivary_gland | Myo1d |
| Atp8b1 | 0 | 0.63769886 | 0.29 | 0.04 | 0 | Salivary_gland | Atp8b1 |
| Rsrc1 | 0 | 0.63764019 | 0.36 | 0.08 | 0 | Salivary_gland | Rsrc1 |
| Stag2 | 0 | 0.63749488 | 0.45 | 0.12 | 0 | Salivary_gland | Stag2 |
| Csnk2a2 | 0 | 0.63667914 | 0.27 | 0.02 | 0 | Salivary_gland | Csnk2a2 |
| Stard9 | 0 | 0.63505316 | 0.37 | 0.08 | 0 | Salivary_gland | Stard9 |
| Ep300 | 0 | 0.63472428 | 0.36 | 0.06 | 0 | Salivary_gland | Ep300 |
| Rps24 | 0 | 0.63426577 | 0.99 | 0.88 | 0 | Salivary_gland | Rps24 |
| Wipf1 | 0 | 0.63145386 | 0.38 | 0.09 | 0 | Salivary_gland | Wipf1 |
| Smarca5 | 0 | 0.6311699 | 0.46 | 0.14 | 0 | Salivary_gland | Smarca5 |
| Strn3 | 0 | 0.63080134 | 0.6 | 0.24 | 0 | Salivary_gland | Strn3 |
| Slc10a7 | 0 | 0.6304034 | 0.29 | 0.04 | 0 | Salivary_gland | Slc10a7 |
| Rsf1 | 0 | 0.63023651 | 0.39 | 0.09 | 0 | Salivary_gland | Rsf1 |
| Lcorl | 0 | 0.63021928 | 0.26 | 0.03 | 0 | Salivary_gland | Lcorl |
| Rapgef1 | 0 | 0.63019483 | 0.46 | 0.14 | 0 | Salivary_gland | Rapgef1 |
| Xrn1 | 0 | 0.6299562 | 0.35 | 0.07 | 0 | Salivary_gland | Xrn1 |
| Tbl1x | 0 | 0.6281133 | 0.49 | 0.16 | 0 | Salivary_gland | Tbl1x |
| Itpkb | 0 | 0.62769202 | 0.77 | 0.38 | 0 | Salivary_gland | Itpkb |
| Ube2e2 | 0 | 0.6260058 | 0.31 | 0.08 | 0 | Salivary_gland | Ube2e2 |

| | | | | | | | |
|---|---|---|---|---|---|---|---|
| Smg1 | 0 | 0.62447002 | 0.38 | 0.08 | 0 | Salivary_gland | Smg1 |
| Ktn1 | 0 | 0.62336408 | 0.62 | 0.26 | 0 | Salivary_gland | Ktn1 |
| mt-Nd2 | 0 | 0.62222237 | 0.98 | 0.82 | 0 | Salivary_gland | mt-Nd2 |
| Xdh | 0 | 0.62168748 | 0.54 | 0.18 | 0 | Salivary_gland | Xdh |
| Scaf8 | 0 | 0.62059595 | 0.3 | 0.04 | 0 | Salivary_gland | Scaf8 |
| Irgm2 | 0 | 0.62031339 | 0.31 | 0.08 | 0 | Salivary_gland | Irgm2 |
| Ece1 | 0 | 0.62029508 | 0.82 | 0.43 | 0 | Salivary_gland | Ece1 |
| Dstyk | 0 | 0.61937815 | 0.36 | 0.07 | 0 | Salivary_gland | Dstyk |
| Ubr2 | 0 | 0.61919117 | 0.38 | 0.08 | 0 | Salivary_gland | Ubr2 |
| Ptpra | 0 | 0.61913718 | 0.6 | 0.23 | 0 | Salivary_gland | Ptpra |
| Mfhas1 | 0 | 0.61851132 | 0.39 | 0.09 | 0 | Salivary_gland | Mfhas1 |
| Tmem131 | 0 | 0.61761769 | 0.33 | 0.06 | 0 | Salivary_gland | Tmem131 |
| Man1a2 | 0 | 0.61758748 | 0.4 | 0.1 | 0 | Salivary_gland | Man1a2 |
| Rps6ka3 | 0 | 0.61592958 | 0.45 | 0.13 | 0 | Salivary_gland | Rps6ka3 |
| Trio | 0 | 0.61539388 | 0.34 | 0.07 | 0 | Salivary_gland | Trio |
| Kif1b | 0 | 0.61364748 | 0.46 | 0.14 | 0 | Salivary_gland | Kif1b |
| Osbpl8 | 0 | 0.61146959 | 0.3 | 0.05 | 0 | Salivary_gland | Osbpl8 |
| Ifi44 | 0 | 0.61125025 | 0.44 | 0.12 | 0 | Salivary_gland | Ifi44 |
| Sltm | 0 | 0.6106775 | 0.44 | 0.13 | 0 | Salivary_gland | Sltm |
| Dach1 | 0 | 0.60974872 | 0.27 | 0.06 | 0 | Salivary_gland | Dach1 |
| Pten | 0 | 0.6083583 | 0.48 | 0.15 | 0 | Salivary_gland | Pten |
| Kcnb1 | 0 | 0.60820303 | 0.27 | 0.05 | 0 | Salivary_gland | Kcnb1 |
| Shroom4 | 0 | 0.6073494 | 0.35 | 0.08 | 0 | Salivary_gland | Shroom4 |
| Mertk | 0 | 0.60722663 | 0.36 | 0.08 | 0 | Salivary_gland | Mertk |
| Tlk1 | 0 | 0.60720138 | 0.35 | 0.07 | 0 | Salivary_gland | Tlk1 |
| Kdm4c | 0 | 0.60683807 | 0.29 | 0.04 | 0 | Salivary_gland | Kdm4c |
| 4932438A13Rik | 0 | 0.60664498 | 0.37 | 0.08 | 0 | Salivary_gland | 4932438A13Rik |
| Cd274 | 0 | 0.60508376 | 0.27 | 0.04 | 0 | Salivary_gland | Cd274 |
| Mprip | 0 | 0.60460819 | 0.62 | 0.25 | 0 | Salivary_gland | Mprip |
| Ces2e | 0 | 0.60439367 | 0.36 | 0.07 | 0 | Salivary_gland | Ces2e |
| Mdfic | 0 | 0.60025854 | 0.5 | 0.17 | 0 | Salivary_gland | Mdfic |
| Dync1h1 | 0 | 0.59998062 | 0.57 | 0.21 | 0 | Salivary_gland | Dync1h1 |
| Ap3b1 | 0 | 0.59802893 | 0.35 | 0.07 | 0 | Salivary_gland | Ap3b1 |
| Ccny | 0 | 0.59784886 | 0.47 | 0.15 | 0 | Salivary_gland | Ccny |
| Kat6b | 0 | 0.59782242 | 0.3 | 0.05 | 0 | Salivary_gland | Kat6b |
| Abcc4 | 0 | 0.59738866 | 0.37 | 0.08 | 0 | Salivary_gland | Abcc4 |
| Snai2 | 0 | 0.59602658 | 0.29 | 0.04 | 0 | Salivary_gland | Snai2 |
| Slc12a2 | 0 | 0.59592041 | 0.26 | 0.03 | 0 | Salivary_gland | Slc12a2 |
| Prss23 | 0 | 0.59590145 | 0.75 | 0.28 | 0 | Salivary_gland | Prss23 |
| Tspan5 | 0 | 0.59523283 | 0.44 | 0.13 | 0 | Salivary_gland | Tspan5 |
| Spred2 | 0 | 0.59515481 | 0.37 | 0.08 | 0 | Salivary_gland | Spred2 |

| | | | | | | | |
|---|---|---|---|---|---|---|---|
| Arhgef3 | 0 | 0.59425461 | 0.52 | 0.18 | 0 | Salivary_gland | Arhgef3 |
| Cd300lg | 0 | 0.59422575 | 0.86 | 0.39 | 0 | Salivary_gland | Cd300lg |
| Kcmf1 | 0 | 0.59405531 | 0.4 | 0.1 | 0 | Salivary_gland | Kcmf1 |
| Meox1 | 0 | 0.59380163 | 0.57 | 0.19 | 0 | Salivary_gland | Meox1 |
| Tmem164 | 0 | 0.59370908 | 0.25 | 0.03 | 0 | Salivary_gland | Tmem164 |
| Rbm26 | 0 | 0.59288556 | 0.37 | 0.09 | 0 | Salivary_gland | Rbm26 |
| Prrg1 | 0 | 0.59211194 | 0.31 | 0.06 | 0 | Salivary_gland | Prrg1 |
| Dph6 | 0 | 0.59009626 | 0.27 | 0.04 | 0 | Salivary_gland | Dph6 |
| Mtmr3 | 0 | 0.58970433 | 0.32 | 0.06 | 0 | Salivary_gland | Mtmr3 |
| Tcf15 | 0 | 0.58889546 | 0.74 | 0.26 | 0 | Salivary_gland | Tcf15 |
| Ubr5 | 0 | 0.58802214 | 0.35 | 0.07 | 0 | Salivary_gland | Ubr5 |
| Prdm2 | 0 | 0.58697979 | 0.35 | 0.07 | 0 | Salivary_gland | Prdm2 |
| Nufip2 | 0 | 0.58678744 | 0.38 | 0.09 | 0 | Salivary_gland | Nufip2 |
| Nsa2 | 0 | 0.5860564 | 0.55 | 0.21 | 0 | Salivary_gland | Nsa2 |
| Socs5 | 0 | 0.5855574 | 0.3 | 0.06 | 0 | Salivary_gland | Socs5 |
| Ppp2r5c | 0 | 0.58516276 | 0.42 | 0.11 | 0 | Salivary_gland | Ppp2r5c |
| Sap30l | 0 | 0.58318772 | 0.33 | 0.06 | 0 | Salivary_gland | Sap30l |
| Fer | 0 | 0.58162042 | 0.27 | 0.04 | 0 | Salivary_gland | Fer |
| N4bp1 | 0 | 0.58109071 | 0.39 | 0.1 | 0 | Salivary_gland | N4bp1 |
| Rb1cc1 | 0 | 0.58099631 | 0.42 | 0.13 | 0 | Salivary_gland | Rb1cc1 |
| Wac | 0 | 0.58052747 | 0.55 | 0.2 | 0 | Salivary_gland | Wac |
| Spen | 0 | 0.58034352 | 0.33 | 0.06 | 0 | Salivary_gland | Spen |
| Ccdc85b | 0 | 0.579585 | 0.59 | 0.23 | 0 | Salivary_gland | Ccdc85b |
| Stk3 | 0 | 0.57906422 | 0.32 | 0.07 | 0 | Salivary_gland | Stk3 |
| Dcbld1 | 0 | 0.57897291 | 0.29 | 0.06 | 0 | Salivary_gland | Dcbld1 |
| Ndfip1 | 0 | 0.57856694 | 0.43 | 0.14 | 0 | Salivary_gland | Ndfip1 |
| Pds5b | 0 | 0.57837402 | 0.32 | 0.07 | 0 | Salivary_gland | Pds5b |
| Fnip2 | 0 | 0.5779374 | 0.26 | 0.03 | 0 | Salivary_gland | Fnip2 |
| Slfn2 | 0 | 0.57785156 | 0.46 | 0.15 | 0 | Salivary_gland | Slfn2 |
| Arfgef1 | 0 | 0.57776911 | 0.37 | 0.09 | 0 | Salivary_gland | Arfgef1 |
| Lama4 | 0 | 0.57662966 | 0.49 | 0.16 | 0 | Salivary_gland | Lama4 |
| Suz12 | 0 | 0.57539617 | 0.32 | 0.06 | 0 | Salivary_gland | Suz12 |
| Ctcf | 0 | 0.57418445 | 0.48 | 0.16 | 0 | Salivary_gland | Ctcf |
| Mier1 | 0 | 0.57327803 | 0.56 | 0.22 | 0 | Salivary_gland | Mier1 |
| Ulk2 | 0 | 0.57276166 | 0.32 | 0.06 | 0 | Salivary_gland | Ulk2 |
| Bbx | 0 | 0.57257565 | 0.4 | 0.12 | 0 | Salivary_gland | Bbx |
| Tle4 | 0 | 0.5713999 | 0.34 | 0.08 | 0 | Salivary_gland | Tle4 |
| Crispld1 | 0 | 0.57123008 | 0.42 | 0.12 | 0 | Salivary_gland | Crispld1 |
| Sppl3 | 0 | 0.57104843 | 0.37 | 0.09 | 0 | Salivary_gland | Sppl3 |
| Yy1 | 0 | 0.57078552 | 0.45 | 0.15 | 0 | Salivary_gland | Yy1 |
| Dmxl1 | 0 | 0.56984848 | 0.29 | 0.05 | 0 | Salivary_gland | Dmxl1 |

| | | | | | | | |
|---|---|---|---|---|---|---|---|
| Chd4 | 0 | 0.56891828 | 0.5 | 0.18 | 0 | Salivary_gland | Chd4 |
| Dhx36 | 0 | 0.56868028 | 0.38 | 0.09 | 0 | Salivary_gland | Dhx36 |
| Ranbp9 | 0 | 0.56823902 | 0.28 | 0.04 | 0 | Salivary_gland | Ranbp9 |
| Cmip | 0 | 0.56590574 | 0.52 | 0.18 | 0 | Salivary_gland | Cmip |
| Acyp2 | 0 | 0.5643894 | 0.3 | 0.07 | 0 | Salivary_gland | Acyp2 |
| Eepd1 | 0 | 0.56399696 | 0.34 | 0.08 | 0 | Salivary_gland | Eepd1 |
| Adarb1 | 0 | 0.56279585 | 0.34 | 0.08 | 0 | Salivary_gland | Adarb1 |
| Cyfip1 | 0 | 0.56248556 | 0.51 | 0.19 | 0 | Salivary_gland | Cyfip1 |
| C130074G19Rik | 0 | 0.56216874 | 0.52 | 0.18 | 0 | Salivary_gland | C130074G19Rik |
| Cnot4 | 0 | 0.56147109 | 0.44 | 0.14 | 0 | Salivary_gland | Cnot4 |
| Huwe1 | 0 | 0.56132735 | 0.48 | 0.15 | 0 | Salivary_gland | Huwe1 |
| Clasp2 | 0 | 0.56093935 | 0.3 | 0.05 | 0 | Salivary_gland | Clasp2 |
| Wrn | 0 | 0.560823 | 0.27 | 0.03 | 0 | Salivary_gland | Wrn |
| Slc23a2 | 0 | 0.56043067 | 0.29 | 0.05 | 0 | Salivary_gland | Slc23a2 |
| Fryl | 0 | 0.55951518 | 0.45 | 0.15 | 0 | Salivary_gland | Fryl |
| Prpf4b | 0 | 0.55896823 | 0.54 | 0.2 | 0 | Salivary_gland | Prpf4b |
| Herc2 | 0 | 0.55887688 | 0.41 | 0.11 | 0 | Salivary_gland | Herc2 |
| Nfat5 | 0 | 0.55799833 | 0.52 | 0.19 | 0 | Salivary_gland | Nfat5 |
| Rictor | 0 | 0.55765481 | 0.27 | 0.04 | 0 | Salivary_gland | Rictor |
| Tnrc18 | 0 | 0.55681924 | 0.34 | 0.07 | 0 | Salivary_gland | Tnrc18 |
| Trpm7 | 0 | 0.55656385 | 0.36 | 0.08 | 0 | Salivary_gland | Trpm7 |
| Diaph1 | 0 | 0.55653426 | 0.36 | 0.08 | 0 | Salivary_gland | Diaph1 |
| Phf14 | 0 | 0.55605281 | 0.38 | 0.1 | 0 | Salivary_gland | Phf14 |
| Lamb1 | 0 | 0.55583195 | 0.35 | 0.08 | 0 | Salivary_gland | Lamb1 |
| Nid1 | 0 | 0.5555692 | 0.5 | 0.17 | 0 | Salivary_gland | Nid1 |
| Faf1 | 0 | 0.55488616 | 0.33 | 0.08 | 0 | Salivary_gland | Faf1 |
| Ncoa3 | 0 | 0.55457196 | 0.5 | 0.16 | 0 | Salivary_gland | Ncoa3 |
| Ago3 | 0 | 0.55431568 | 0.3 | 0.06 | 0 | Salivary_gland | Ago3 |
| Tlk2 | 0 | 0.55388445 | 0.29 | 0.06 | 0 | Salivary_gland | Tlk2 |
| Cdk12 | 0 | 0.55291005 | 0.42 | 0.13 | 0 | Salivary_gland | Cdk12 |
| Vps13d | 0 | 0.55191732 | 0.31 | 0.06 | 0 | Salivary_gland | Vps13d |
| AI314180 | 0 | 0.55130977 | 0.33 | 0.07 | 0 | Salivary_gland | AI314180 |
| Hectd1 | 0 | 0.55064834 | 0.47 | 0.15 | 0 | Salivary_gland | Hectd1 |
| Gmds | 0 | 0.55050909 | 0.26 | 0.05 | 0 | Salivary_gland | Gmds |
| Etnk1 | 0 | 0.54898185 | 0.47 | 0.15 | 0 | Salivary_gland | Etnk1 |
| Lpcat1 | 0 | 0.54828457 | 0.45 | 0.13 | 0 | Salivary_gland | Lpcat1 |
| Kif2a | 0 | 0.54444522 | 0.29 | 0.05 | 0 | Salivary_gland | Kif2a |
| Fam193a | 0 | 0.54406229 | 0.31 | 0.06 | 0 | Salivary_gland | Fam193a |
| Clint1 | 0 | 0.54359647 | 0.35 | 0.09 | 0 | Salivary_gland | Clint1 |
| Rftn1 | 0 | 0.54330144 | 0.51 | 0.17 | 0 | Salivary_gland | Rftn1 |
| Il1r1 | 0 | 0.54202664 | 0.28 | 0.05 | 0 | Salivary_gland | Il1r1 |

| | | | | | | | |
|---|---|---|---|---|---|---|---|
| Cdc42bpb | 0 | 0.54136152 | 0.48 | 0.15 | 0 | Salivary_gland | Cdc42bpb |
| Trip12 | 0 | 0.54071445 | 0.42 | 0.12 | 0 | Salivary_gland | Trip12 |
| Epc1 | 0 | 0.54066322 | 0.34 | 0.08 | 0 | Salivary_gland | Epc1 |
| Snx13 | 0 | 0.54020775 | 0.31 | 0.06 | 0 | Salivary_gland | Snx13 |
| Epc2 | 0 | 0.53954872 | 0.28 | 0.05 | 0 | Salivary_gland | Epc2 |
| Eml1 | 0 | 0.53867382 | 0.31 | 0.08 | 0 | Salivary_gland | Eml1 |
| Nckap1 | 0 | 0.53562057 | 0.51 | 0.19 | 0 | Salivary_gland | Nckap1 |
| Vps54 | 0 | 0.53557161 | 0.26 | 0.04 | 0 | Salivary_gland | Vps54 |
| Map1b | 0 | 0.53371998 | 0.3 | 0.08 | 0 | Salivary_gland | Map1b |
| Larp4 | 0 | 0.53240671 | 0.3 | 0.06 | 0 | Salivary_gland | Larp4 |
| Aff4 | 0 | 0.53146454 | 0.4 | 0.12 | 0 | Salivary_gland | Aff4 |
| Gls | 0 | 0.5303222 | 0.41 | 0.12 | 0 | Salivary_gland | Gls |
| Sh3pxd2a | 0 | 0.52996744 | 0.29 | 0.06 | 0 | Salivary_gland | Sh3pxd2a |
| Ubn2 | 0 | 0.52888065 | 0.44 | 0.15 | 0 | Salivary_gland | Ubn2 |
| Fndc3a | 0 | 0.52814617 | 0.36 | 0.1 | 0 | Salivary_gland | Fndc3a |
| Ep400 | 0 | 0.52755327 | 0.38 | 0.1 | 0 | Salivary_gland | Ep400 |
| Kdm1a | 0 | 0.52664274 | 0.35 | 0.09 | 0 | Salivary_gland | Kdm1a |
| Esco1 | 0 | 0.5262063 | 0.28 | 0.05 | 0 | Salivary_gland | Esco1 |
| Dctn4 | 0 | 0.52577003 | 0.36 | 0.09 | 0 | Salivary_gland | Dctn4 |
| Wapl | 0 | 0.52483266 | 0.4 | 0.12 | 0 | Salivary_gland | Wapl |
| Rabep1 | 0 | 0.52465196 | 0.36 | 0.1 | 0 | Salivary_gland | Rabep1 |
| Enpep | 0 | 0.5240232 | 0.27 | 0.04 | 0 | Salivary_gland | Enpep |
| Prrc2b | 0 | 0.52288544 | 0.42 | 0.14 | 0 | Salivary_gland | Prrc2b |
| Dlg1 | 0 | 0.52154014 | 0.31 | 0.07 | 0 | Salivary_gland | Dlg1 |
| Cul1 | 0 | 0.5214266 | 0.45 | 0.15 | 0 | Salivary_gland | Cul1 |
| Hmg20a | 0 | 0.51955643 | 0.26 | 0.04 | 0 | Salivary_gland | Hmg20a |
| Cask | 0 | 0.51879177 | 0.26 | 0.05 | 0 | Salivary_gland | Cask |
| Agap1 | 0 | 0.51741363 | 0.33 | 0.09 | 0 | Salivary_gland | Agap1 |
| Stat2 | 0 | 0.51696696 | 0.31 | 0.07 | 0 | Salivary_gland | Stat2 |
| Acbd3 | 0 | 0.51497419 | 0.34 | 0.09 | 0 | Salivary_gland | Acbd3 |
| Ehmt1 | 0 | 0.51477902 | 0.34 | 0.08 | 0 | Salivary_gland | Ehmt1 |
| Crebrf | 0 | 0.51388167 | 0.34 | 0.09 | 0 | Salivary_gland | Crebrf |
| Rcor1 | 0 | 0.51363292 | 0.26 | 0.04 | 0 | Salivary_gland | Rcor1 |
| Sesn1 | 0 | 0.51285615 | 0.3 | 0.07 | 0 | Salivary_gland | Sesn1 |
| Trim24 | 0 | 0.51242243 | 0.35 | 0.09 | 0 | Salivary_gland | Trim24 |
| Kat2b | 0 | 0.51234808 | 0.29 | 0.06 | 0 | Salivary_gland | Kat2b |
| Trappc8 | 0 | 0.51217985 | 0.27 | 0.05 | 0 | Salivary_gland | Trappc8 |
| Cap1 | 0 | 0.50921392 | 0.35 | 0.09 | 0 | Salivary_gland | Cap1 |
| Usp47 | 0 | 0.50902923 | 0.31 | 0.06 | 0 | Salivary_gland | Usp47 |
| Erap1 | 0 | 0.50836811 | 0.28 | 0.06 | 0 | Salivary_gland | Erap1 |
| Ppp6r3 | 0 | 0.50828456 | 0.35 | 0.09 | 0 | Salivary_gland | Ppp6r3 |

| | | | | | | | |
|---|---|---|---|---|---|---|---|
| Nfatc3 | 0 | 0.5071002 | 0.26 | 0.04 | 0 | Salivary_gland | Nfatc3 |
| Ankrd44 | 0 | 0.50686681 | 0.4 | 0.12 | 0 | Salivary_gland | Ankrd44 |
| Itch | 0 | 0.50633187 | 0.41 | 0.13 | 0 | Salivary_gland | Itch |
| Pmepa1 | 0 | 0.50632606 | 0.3 | 0.07 | 0 | Salivary_gland | Pmepa1 |
| 2510009E07Rik | 0 | 0.50615822 | 0.31 | 0.07 | 0 | Salivary_gland | 2510009E07Rik |
| Cwf19l2 | 0 | 0.50560035 | 0.33 | 0.08 | 0 | Salivary_gland | Cwf19l2 |
| Spry4 | 0 | 0.50546905 | 0.28 | 0.05 | 0 | Salivary_gland | Spry4 |
| Dusp7 | 0 | 0.50535377 | 0.36 | 0.09 | 0 | Salivary_gland | Dusp7 |
| Ttll7 | 0 | 0.50482483 | 0.33 | 0.09 | 0 | Salivary_gland | Ttll7 |
| St3gal1 | 0 | 0.5043905 | 0.42 | 0.14 | 0 | Salivary_gland | St3gal1 |
| Rasa1 | 0 | 0.50312271 | 0.41 | 0.13 | 0 | Salivary_gland | Rasa1 |
| Supt16 | 0 | 0.50270235 | 0.36 | 0.1 | 0 | Salivary_gland | Supt16 |
| Erc1 | 0 | 0.50196926 | 0.32 | 0.08 | 0 | Salivary_gland | Erc1 |
| Gbp9 | 0 | 0.50169368 | 0.38 | 0.11 | 0 | Salivary_gland | Gbp9 |
| Rif1 | 0 | 0.49988604 | 0.25 | 0.04 | 0 | Salivary_gland | Rif1 |
| Lonp2 | 0 | 0.49983227 | 0.32 | 0.07 | 0 | Salivary_gland | Lonp2 |
| Igf2r | 0 | 0.49980388 | 0.29 | 0.06 | 0 | Salivary_gland | Igf2r |
| Larp1 | 0 | 0.49956453 | 0.3 | 0.06 | 0 | Salivary_gland | Larp1 |
| Cdkal1 | 0 | 0.49877889 | 0.29 | 0.06 | 0 | Salivary_gland | Cdkal1 |
| Dcaf6 | 0 | 0.49805131 | 0.27 | 0.05 | 0 | Salivary_gland | Dcaf6 |
| Ubr1 | 0 | 0.49764484 | 0.3 | 0.07 | 0 | Salivary_gland | Ubr1 |
| Asxl1 | 0 | 0.49709807 | 0.26 | 0.04 | 0 | Salivary_gland | Asxl1 |
| Ccdc186 | 0 | 0.49692633 | 0.31 | 0.08 | 0 | Salivary_gland | Ccdc186 |
| Dyrk1a | 0 | 0.4966201 | 0.28 | 0.05 | 0 | Salivary_gland | Dyrk1a |
| Pitpnm2 | 0 | 0.49525851 | 0.37 | 0.11 | 0 | Salivary_gland | Pitpnm2 |
| Rnf19a | 0 | 0.49358999 | 0.33 | 0.08 | 0 | Salivary_gland | Rnf19a |
| Slc44a1 | 0 | 0.49354327 | 0.39 | 0.12 | 0 | Salivary_gland | Slc44a1 |
| Hivep1 | 0 | 0.49329244 | 0.26 | 0.05 | 0 | Salivary_gland | Hivep1 |
| Zdhhc18 | 0 | 0.4931526 | 0.37 | 0.1 | 0 | Salivary_gland | Zdhhc18 |
| Osmr | 0 | 0.49306707 | 0.41 | 0.13 | 0 | Salivary_gland | Osmr |
| Mark3 | 0 | 0.4928636 | 0.38 | 0.11 | 0 | Salivary_gland | Mark3 |
| Zfp652 | 0 | 0.49210327 | 0.4 | 0.13 | 0 | Salivary_gland | Zfp652 |
| Rreb1 | 0 | 0.49088812 | 0.27 | 0.06 | 0 | Salivary_gland | Rreb1 |
| Med13l | 0 | 0.49017298 | 0.35 | 0.1 | 0 | Salivary_gland | Med13l |
| Mpdz | 0 | 0.48990075 | 0.28 | 0.06 | 0 | Salivary_gland | Mpdz |
| Rnf216 | 0 | 0.48432482 | 0.32 | 0.08 | 0 | Salivary_gland | Rnf216 |
| Kdm2a | 0 | 0.48287354 | 0.41 | 0.13 | 0 | Salivary_gland | Kdm2a |
| Akap11 | 0 | 0.48263536 | 0.27 | 0.05 | 0 | Salivary_gland | Akap11 |
| Efr3b | 0 | 0.48109061 | 0.29 | 0.07 | 0 | Salivary_gland | Efr3b |
| Mid2 | 0 | 0.48072044 | 0.27 | 0.06 | 0 | Salivary_gland | Mid2 |
| Lrrc8d | 0 | 0.48031371 | 0.26 | 0.05 | 0 | Salivary_gland | Lrrc8d |

| | | | | | | | |
|---|---|---|---|---|---|---|---|
| Bdp1 | 0 | 0.47895522 | 0.29 | 0.07 | 0 | Salivary_gland | Bdp1 |
| Dhx9 | 0 | 0.47816061 | 0.31 | 0.07 | 0 | Salivary_gland | Dhx9 |
| Unc119b | 0 | 0.4776283 | 0.35 | 0.09 | 0 | Salivary_gland | Unc119b |
| Plvap | 0 | 0.47590171 | 0.99 | 0.56 | 0 | Salivary_gland | Plvap |
| Rtp3 | 0 | 0.47405205 | 0.28 | 0.06 | 0 | Salivary_gland | Rtp3 |
| 9930021J03Rik | 0 | 0.47232553 | 0.26 | 0.06 | 0 | Salivary_gland | 9930021J03Rik |
| Map4k5 | 0 | 0.47171625 | 0.3 | 0.07 | 0 | Salivary_gland | Map4k5 |
| Dennd6a | 0 | 0.47110462 | 0.37 | 0.11 | 0 | Salivary_gland | Dennd6a |
| Ltbp1 | 0 | 0.46874664 | 0.28 | 0.05 | 0 | Salivary_gland | Ltbp1 |
| Mib1 | 0 | 0.46862629 | 0.33 | 0.09 | 0 | Salivary_gland | Mib1 |
| Cx3cl1 | 0 | 0.46817891 | 0.31 | 0.08 | 0 | Salivary_gland | Cx3cl1 |
| Setd7 | 0 | 0.46766653 | 0.28 | 0.06 | 0 | Salivary_gland | Setd7 |
| Chd6 | 0 | 0.46577049 | 0.35 | 0.1 | 0 | Salivary_gland | Chd6 |
| Sos2 | 0 | 0.46520195 | 0.29 | 0.07 | 0 | Salivary_gland | Sos2 |
| Iws1 | 0 | 0.46295866 | 0.29 | 0.06 | 0 | Salivary_gland | Iws1 |
| Nuak1 | 0 | 0.46256518 | 0.34 | 0.09 | 0 | Salivary_gland | Nuak1 |
| Srek1 | 0 | 0.46235342 | 0.32 | 0.09 | 0 | Salivary_gland | Srek1 |
| Arl5a | 0 | 0.46153268 | 0.3 | 0.07 | 0 | Salivary_gland | Arl5a |
| Acbd5 | 0 | 0.46147834 | 0.31 | 0.08 | 0 | Salivary_gland | Acbd5 |
| 2700081O15Rik | 0 | 0.46133804 | 0.25 | 0.05 | 0 | Salivary_gland | 2700081O15Rik |
| Uri1 | 0 | 0.46053446 | 0.31 | 0.07 | 0 | Salivary_gland | Uri1 |
| Rnf38 | 0 | 0.45992621 | 0.26 | 0.05 | 0 | Salivary_gland | Rnf38 |
| Taf15 | 0 | 0.45856665 | 0.35 | 0.1 | 0 | Salivary_gland | Taf15 |
| Cdkn1b | 0 | 0.45839273 | 0.36 | 0.1 | 0 | Salivary_gland | Cdkn1b |
| Cdc40 | 0 | 0.45752415 | 0.29 | 0.07 | 0 | Salivary_gland | Cdc40 |
| C2cd2 | 0 | 0.45665214 | 0.32 | 0.08 | 0 | Salivary_gland | C2cd2 |
| Gbf1 | 0 | 0.45580699 | 0.28 | 0.07 | 0 | Salivary_gland | Gbf1 |
| Atxn7 | 0 | 0.45498937 | 0.29 | 0.07 | 0 | Salivary_gland | Atxn7 |
| Kdm7a | 0 | 0.45391375 | 0.29 | 0.07 | 0 | Salivary_gland | Kdm7a |
| Dnajc21 | 0 | 0.45132383 | 0.25 | 0.05 | 0 | Salivary_gland | Dnajc21 |
| Mon2 | 0 | 0.45107697 | 0.26 | 0.05 | 0 | Salivary_gland | Mon2 |
| Tom1l2 | 0 | 0.44967082 | 0.27 | 0.06 | 0 | Salivary_gland | Tom1l2 |
| Wwp1 | 0 | 0.44864336 | 0.25 | 0.06 | 0 | Salivary_gland | Wwp1 |
| Creb1 | 0 | 0.44833302 | 0.33 | 0.09 | 0 | Salivary_gland | Creb1 |
| Fam117b | 0 | 0.44792279 | 0.38 | 0.1 | 0 | Salivary_gland | Fam117b |
| Arid2 | 0 | 0.44557708 | 0.29 | 0.08 | 0 | Salivary_gland | Arid2 |
| Zfp280d | 0 | 0.44471603 | 0.32 | 0.09 | 0 | Salivary_gland | Zfp280d |
| Mysm1 | 0 | 0.44428085 | 0.31 | 0.08 | 0 | Salivary_gland | Mysm1 |
| Ctdspl2 | 0 | 0.44007048 | 0.3 | 0.08 | 0 | Salivary_gland | Ctdspl2 |
| Supt6 | 0 | 0.43518962 | 0.29 | 0.07 | 0 | Salivary_gland | Supt6 |
| Smg7 | 0 | 0.43472716 | 0.27 | 0.06 | 0 | Salivary_gland | Smg7 |

| | | | | | | | |
|---|---|---|---|---|---|---|---|
| Ythdf3 | 0 | 0.43318424 | 0.29 | 0.07 | 0 | Salivary_gland | Ythdf3 |
| Nr2c2 | 0 | 0.43197463 | 0.26 | 0.06 | 0 | Salivary_gland | Nr2c2 |
| Taf1 | 0 | 0.43046605 | 0.26 | 0.06 | 0 | Salivary_gland | Taf1 |
| Rab3gap1 | 0 | 0.4298187 | 0.26 | 0.06 | 0 | Salivary_gland | Rab3gap1 |
| Pdzd8 | 0 | 0.42850568 | 0.28 | 0.07 | 0 | Salivary_gland | Pdzd8 |
| Ube3c | 0 | 0.4180548 | 0.27 | 0.07 | 0 | Salivary_gland | Ube3c |
| Meis2 | 0 | 0.41748482 | 0.67 | 0.29 | 0 | Salivary_gland | Meis2 |
| Galnt15 | 0 | 0.41422667 | 0.28 | 0.07 | 0 | Salivary_gland | Galnt15 |
| Fam76b | 0 | 0.41268174 | 0.25 | 0.06 | 0 | Salivary_gland | Fam76b |
| Ice1 | 0 | 0.41110657 | 0.27 | 0.06 | 0 | Salivary_gland | Ice1 |
| Camsap2 | 0 | 0.40399964 | 0.25 | 0.06 | 0 | Salivary_gland | Camsap2 |
| Parp4 | 0 | 0.40395778 | 0.25 | 0.06 | 0 | Salivary_gland | Parp4 |
| Snrnp200 | 0 | 0.39066498 | 0.3 | 0.08 | 0 | Salivary_gland | Snrnp200 |
| Cdc14a | 0 | 0.38571593 | 0.25 | 0.05 | 0 | Salivary_gland | Cdc14a |
| Emp2 | 0 | 0.38134449 | 0.64 | 0.17 | 0 | Salivary_gland | Emp2 |
| Fabp4 | 0 | 0.33211989 | 0.99 | 0.58 | 0 | Salivary_gland | Fabp4 |
| Numa1 | 1.91E-307 | 0.44185055 | 0.38 | 0.12 | 3.95E-303 | Salivary_gland | Numa1 |
| Nlk | 3.05E-307 | 0.43098066 | 0.26 | 0.06 | 6.32E-303 | Salivary_gland | Nlk |
| Mcl1 | 9.02E-307 | 0.60781109 | 0.62 | 0.26 | 1.87E-302 | Salivary_gland | Mcl1 |
| Ifih1 | 2.70E-306 | 0.43994592 | 0.29 | 0.08 | 5.58E-302 | Salivary_gland | Ifih1 |
| Ccar1 | 9.55E-306 | 0.47946982 | 0.35 | 0.1 | 1.98E-301 | Salivary_gland | Ccar1 |
| Nrp2 | 1.66E-305 | 0.48768824 | 0.53 | 0.2 | 3.44E-301 | Salivary_gland | Nrp2 |
| Samhd1 | 2.64E-305 | 0.5271649 | 0.5 | 0.18 | 5.46E-301 | Salivary_gland | Samhd1 |
| Rps10 | 4.84E-305 | 0.6044863 | 0.98 | 0.8 | 1.00E-300 | Salivary_gland | Rps10 |
| Cdc37l1 | 8.32E-305 | 0.3924301 | 0.26 | 0.06 | 1.72E-300 | Salivary_gland | Cdc37l1 |
| Cdc73 | 4.17E-304 | 0.4110483 | 0.28 | 0.07 | 8.64E-300 | Salivary_gland | Cdc73 |
| Phf3 | 7.95E-304 | 0.47711897 | 0.41 | 0.13 | 1.64E-299 | Salivary_gland | Phf3 |
| Kidins220 | 7.52E-303 | 0.43374214 | 0.28 | 0.07 | 1.56E-298 | Salivary_gland | Kidins220 |
| Shoc2 | 8.88E-303 | 0.44034907 | 0.37 | 0.11 | 1.84E-298 | Salivary_gland | Shoc2 |
| Ankrd17 | 1.25E-302 | 0.49588677 | 0.41 | 0.14 | 2.58E-298 | Salivary_gland | Ankrd17 |
| Rab12 | 1.77E-302 | 0.47149986 | 0.41 | 0.14 | 3.66E-298 | Salivary_gland | Rab12 |
| Rbm6 | 1.94E-302 | 0.46242656 | 0.38 | 0.12 | 4.01E-298 | Salivary_gland | Rbm6 |
| Pbrm1 | 5.47E-302 | 0.57264943 | 0.52 | 0.2 | 1.13E-297 | Salivary_gland | Pbrm1 |
| Cbx5 | 1.27E-300 | 0.45378164 | 0.37 | 0.12 | 2.63E-296 | Salivary_gland | Cbx5 |
| Cenpb | 2.16E-300 | 0.43841985 | 0.39 | 0.13 | 4.47E-296 | Salivary_gland | Cenpb |
| Eif2ak2 | 5.39E-300 | 0.46089037 | 0.29 | 0.08 | 1.12E-295 | Salivary_gland | Eif2ak2 |
| Igf1r | 5.78E-300 | 0.45447404 | 0.39 | 0.12 | 1.20E-295 | Salivary_gland | Igf1r |
| Map3k1 | 1.44E-298 | 0.42009654 | 0.36 | 0.11 | 2.98E-294 | Salivary_gland | Map3k1 |
| Tnpo1 | 9.42E-298 | 0.42015429 | 0.29 | 0.08 | 1.95E-293 | Salivary_gland | Tnpo1 |
| Sncg | 2.21E-297 | 0.46322348 | 0.45 | 0.16 | 4.57E-293 | Salivary_gland | Sncg |
| Trim25 | 7.36E-296 | 0.42241912 | 0.35 | 0.1 | 1.52E-291 | Salivary_gland | Trim25 |

| | | | | | | | |
|---|---|---|---|---|---|---|---|
| Prpf8 | 2.87E-295 | 0.43573908 | 0.36 | 0.11 | 5.94E-291 | Salivary_gland | Prpf8 |
| Setd5 | 3.68E-295 | 0.51310256 | 0.45 | 0.16 | 7.62E-291 | Salivary_gland | Setd5 |
| Ppfibp1 | 5.70E-293 | 0.47803864 | 0.5 | 0.18 | 1.18E-288 | Salivary_gland | Ppfibp1 |
| Sec24b | 1.37E-292 | 0.39484396 | 0.26 | 0.06 | 2.84E-288 | Salivary_gland | Sec24b |
| Nrip1 | 3.26E-291 | 0.62017972 | 0.45 | 0.16 | 6.74E-287 | Salivary_gland | Nrip1 |
| Tln1 | 8.74E-291 | 0.56528189 | 0.65 | 0.29 | 1.81E-286 | Salivary_gland | Tln1 |
| Sp1 | 1.96E-289 | 0.36967568 | 0.29 | 0.08 | 4.06E-285 | Salivary_gland | Sp1 |
| Acin1 | 4.91E-289 | 0.48170843 | 0.52 | 0.2 | 1.02E-284 | Salivary_gland | Acin1 |
| Ghr | 5.17E-289 | 0.53931234 | 0.37 | 0.12 | 1.07E-284 | Salivary_gland | Ghr |
| Pkn1 | 1.53E-288 | 0.3994737 | 0.32 | 0.09 | 3.16E-284 | Salivary_gland | Pkn1 |
| Usp9x | 2.22E-288 | 0.43758622 | 0.36 | 0.11 | 4.60E-284 | Salivary_gland | Usp9x |
| Otud7b | 3.03E-287 | 0.43120838 | 0.28 | 0.07 | 6.28E-283 | Salivary_gland | Otud7b |
| Nek7 | 5.28E-286 | 0.43468802 | 0.4 | 0.13 | 1.09E-281 | Salivary_gland | Nek7 |
| Arap2 | 8.77E-286 | 0.44670028 | 0.31 | 0.09 | 1.81E-281 | Salivary_gland | Arap2 |
| Prickle1 | 8.50E-285 | 0.51018337 | 0.4 | 0.13 | 1.76E-280 | Salivary_gland | Prickle1 |
| Pja2 | 4.30E-284 | 0.48347504 | 0.41 | 0.14 | 8.90E-280 | Salivary_gland | Pja2 |
| Smc3 | 7.49E-284 | 0.50355083 | 0.41 | 0.14 | 1.55E-279 | Salivary_gland | Smc3 |
| Fam168a | 7.41E-283 | 0.3925754 | 0.26 | 0.07 | 1.53E-278 | Salivary_gland | Fam168a |
| Evl | 1.79E-282 | 0.38647053 | 0.26 | 0.07 | 3.71E-278 | Salivary_gland | Evl |
| Cyp2d22 | 5.00E-282 | 0.43921903 | 0.39 | 0.13 | 1.03E-277 | Salivary_gland | Cyp2d22 |
| Arap3 | 5.92E-282 | 0.48589761 | 0.58 | 0.24 | 1.22E-277 | Salivary_gland | Arap3 |
| Gjc1 | 7.04E-282 | 0.43793208 | 0.33 | 0.1 | 1.46E-277 | Salivary_gland | Gjc1 |
| Sec63 | 1.00E-281 | 0.46865921 | 0.39 | 0.13 | 2.08E-277 | Salivary_gland | Sec63 |
| Rbm27 | 3.96E-281 | 0.42539073 | 0.32 | 0.09 | 8.19E-277 | Salivary_gland | Rbm27 |
| Tab2 | 4.88E-281 | 0.42090388 | 0.29 | 0.08 | 1.01E-276 | Salivary_gland | Tab2 |
| Lrrfip1 | 5.97E-281 | 0.57214222 | 0.61 | 0.27 | 1.24E-276 | Salivary_gland | Lrrfip1 |
| Micu2 | 1.63E-280 | 0.36771276 | 0.25 | 0.06 | 3.37E-276 | Salivary_gland | Micu2 |
| Exoc6 | 2.32E-280 | 0.41317412 | 0.27 | 0.07 | 4.80E-276 | Salivary_gland | Exoc6 |
| Epb41l2 | 2.56E-278 | 0.4441226 | 0.42 | 0.15 | 5.31E-274 | Salivary_gland | Epb41l2 |
| Setd2 | 1.98E-277 | 0.41230243 | 0.32 | 0.09 | 4.10E-273 | Salivary_gland | Setd2 |
| Plcg1 | 2.58E-277 | 0.43478019 | 0.38 | 0.13 | 5.34E-273 | Salivary_gland | Plcg1 |
| Zfp503 | 4.35E-277 | 0.50158547 | 0.4 | 0.14 | 9.01E-273 | Salivary_gland | Zfp503 |
| Tpp2 | 1.01E-276 | 0.41686591 | 0.3 | 0.09 | 2.10E-272 | Salivary_gland | Tpp2 |
| Ceacam1 | 4.17E-276 | 0.47344837 | 0.49 | 0.18 | 8.62E-272 | Salivary_gland | Ceacam1 |
| Nasp | 2.93E-274 | 0.40624482 | 0.27 | 0.07 | 6.07E-270 | Salivary_gland | Nasp |
| Tgfbr3 | 1.17E-273 | 0.4839389 | 0.4 | 0.14 | 2.42E-269 | Salivary_gland | Tgfbr3 |
| Fam13c | 1.96E-273 | 0.43884142 | 0.32 | 0.1 | 4.05E-269 | Salivary_gland | Fam13c |
| Zfr | 4.58E-273 | 0.4599256 | 0.42 | 0.15 | 9.48E-269 | Salivary_gland | Zfr |
| Txnrd1 | 8.75E-273 | 0.41241475 | 0.29 | 0.08 | 1.81E-268 | Salivary_gland | Txnrd1 |
| Ubn1 | 1.22E-272 | 0.45323904 | 0.45 | 0.16 | 2.52E-268 | Salivary_gland | Ubn1 |
| Trove2 | 1.83E-272 | 0.45396945 | 0.34 | 0.11 | 3.78E-268 | Salivary_gland | Trove2 |

| | | | | | | | |
|---|---|---|---|---|---|---|---|
| Ube2h | 2.60E-272 | 0.43151394 | 0.37 | 0.12 | 5.37E-268 | Salivary_gland | Ube2h |
| Klhl5 | 5.00E-272 | 0.44257811 | 0.55 | 0.22 | 1.03E-267 | Salivary_gland | Klhl5 |
| Dennd5b | 1.72E-271 | 0.48543229 | 0.45 | 0.17 | 3.56E-267 | Salivary_gland | Dennd5b |
| Fbxw11 | 2.47E-271 | 0.36690835 | 0.27 | 0.07 | 5.10E-267 | Salivary_gland | Fbxw11 |
| Esyt2 | 2.68E-271 | 0.42541151 | 0.4 | 0.14 | 5.55E-267 | Salivary_gland | Esyt2 |
| Clock | 4.16E-271 | 0.4409098 | 0.39 | 0.13 | 8.61E-267 | Salivary_gland | Clock |
| Slmap | 6.04E-271 | 0.41622698 | 0.33 | 0.1 | 1.25E-266 | Salivary_gland | Slmap |
| Thrap3 | 3.06E-270 | 0.46097013 | 0.46 | 0.17 | 6.33E-266 | Salivary_gland | Thrap3 |
| Senp6 | 1.39E-269 | 0.49615014 | 0.47 | 0.18 | 2.88E-265 | Salivary_gland | Senp6 |
| Scaf11 | 1.32E-268 | 0.45173929 | 0.42 | 0.15 | 2.73E-264 | Salivary_gland | Scaf11 |
| Ttc3 | 1.63E-268 | 0.49085821 | 0.44 | 0.16 | 3.37E-264 | Salivary_gland | Ttc3 |
| Jam2 | 2.46E-268 | 0.53932755 | 0.82 | 0.43 | 5.09E-264 | Salivary_gland | Jam2 |
| Sparcl1 | 6.78E-268 | 0.43868894 | 0.93 | 0.47 | 1.40E-263 | Salivary_gland | Sparcl1 |
| Itpr2 | 8.01E-268 | 0.46519641 | 0.35 | 0.11 | 1.66E-263 | Salivary_gland | Itpr2 |
| Pum1 | 1.01E-267 | 0.46891909 | 0.43 | 0.16 | 2.09E-263 | Salivary_gland | Pum1 |
| Pik3ca | 1.41E-267 | 0.38648326 | 0.3 | 0.09 | 2.92E-263 | Salivary_gland | Pik3ca |
| Mga | 1.44E-267 | 0.40133317 | 0.33 | 0.1 | 2.98E-263 | Salivary_gland | Mga |
| Ddx42 | 1.50E-265 | 0.38874062 | 0.32 | 0.1 | 3.10E-261 | Salivary_gland | Ddx42 |
| Rps20 | 1.71E-264 | 0.55172355 | 0.99 | 0.85 | 3.55E-260 | Salivary_gland | Rps20 |
| Ddx46 | 7.20E-264 | 0.39285876 | 0.35 | 0.12 | 1.49E-259 | Salivary_gland | Ddx46 |
| Pds5a | 7.21E-264 | 0.41188497 | 0.34 | 0.11 | 1.49E-259 | Salivary_gland | Pds5a |
| Aftph | 1.25E-263 | 0.37588525 | 0.26 | 0.07 | 2.58E-259 | Salivary_gland | Aftph |
| Myo1b | 1.90E-263 | 0.38511403 | 0.34 | 0.11 | 3.93E-259 | Salivary_gland | Myo1b |
| Xiap | 2.05E-263 | 0.51294001 | 0.61 | 0.27 | 4.25E-259 | Salivary_gland | Xiap |
| Atp2b1 | 3.52E-263 | 0.51960536 | 0.58 | 0.25 | 7.28E-259 | Salivary_gland | Atp2b1 |
| Zc3h13 | 2.03E-262 | 0.46117997 | 0.37 | 0.12 | 4.20E-258 | Salivary_gland | Zc3h13 |
| Pan3 | 2.23E-262 | 0.4244711 | 0.36 | 0.12 | 4.61E-258 | Salivary_gland | Pan3 |
| Atf2 | 3.44E-260 | 0.3712607 | 0.3 | 0.09 | 7.11E-256 | Salivary_gland | Atf2 |
| Pias1 | 7.81E-260 | 0.38107216 | 0.3 | 0.09 | 1.62E-255 | Salivary_gland | Pias1 |
| Hacd4 | 1.52E-259 | 0.40194483 | 0.27 | 0.07 | 3.15E-255 | Salivary_gland | Hacd4 |
| Ppp4r2 | 2.12E-259 | 0.42383975 | 0.36 | 0.12 | 4.38E-255 | Salivary_gland | Ppp4r2 |
| Plekho1 | 7.73E-259 | 0.40757331 | 0.53 | 0.21 | 1.60E-254 | Salivary_gland | Plekho1 |
| Irf2 | 5.60E-258 | 0.52050323 | 0.58 | 0.25 | 1.16E-253 | Salivary_gland | Irf2 |
| Mbnl2 | 6.06E-258 | 0.55282186 | 0.73 | 0.36 | 1.25E-253 | Salivary_gland | Mbnl2 |
| Nfx1 | 8.81E-258 | 0.33829449 | 0.25 | 0.07 | 1.82E-253 | Salivary_gland | Nfx1 |
| Dnajc3 | 1.30E-257 | 0.49495711 | 0.64 | 0.29 | 2.69E-253 | Salivary_gland | Dnajc3 |
| Rrbp1 | 1.34E-257 | 0.56266568 | 0.8 | 0.44 | 2.77E-253 | Salivary_gland | Rrbp1 |
| Zfp664 | 2.29E-257 | 0.40238085 | 0.34 | 0.11 | 4.74E-253 | Salivary_gland | Zfp664 |
| Nucks1 | 9.90E-256 | 0.47389012 | 0.54 | 0.22 | 2.05E-251 | Salivary_gland | Nucks1 |
| Akap12 | 3.56E-255 | 0.48996588 | 0.44 | 0.17 | 7.37E-251 | Salivary_gland | Akap12 |
| Gatad2a | 1.11E-253 | 0.41295067 | 0.41 | 0.15 | 2.29E-249 | Salivary_gland | Gatad2a |

| | | | | | | | |
|---|---|---|---|---|---|---|---|
| Vezf1 | 1.32E-253 | 0.44619348 | 0.45 | 0.17 | 2.73E-249 | Salivary_gland | Vezf1 |
| Zfand5 | 1.50E-253 | 0.59728999 | 0.69 | 0.34 | 3.11E-249 | Salivary_gland | Zfand5 |
| Pitpnb | 3.94E-253 | 0.38528925 | 0.38 | 0.13 | 8.15E-249 | Salivary_gland | Pitpnb |
| Gtf2i | 5.09E-253 | 0.45427818 | 0.48 | 0.19 | 1.05E-248 | Salivary_gland | Gtf2i |
| Atxn2 | 4.34E-252 | 0.35071789 | 0.26 | 0.07 | 8.98E-248 | Salivary_gland | Atxn2 |
| Rapgef3 | 4.90E-252 | 0.44958972 | 0.54 | 0.22 | 1.01E-247 | Salivary_gland | Rapgef3 |
| Gcc2 | 2.87E-251 | 0.4519985 | 0.34 | 0.11 | 5.94E-247 | Salivary_gland | Gcc2 |
| Tmod3 | 4.51E-251 | 0.50499764 | 0.66 | 0.31 | 9.32E-247 | Salivary_gland | Tmod3 |
| Gbp4 | 2.90E-250 | 0.53834905 | 0.44 | 0.17 | 6.00E-246 | Salivary_gland | Gbp4 |
| Lhfpl2 | 4.38E-250 | 0.37730063 | 0.32 | 0.1 | 9.06E-246 | Salivary_gland | Lhfpl2 |
| Insr | 1.23E-249 | 0.68864918 | 0.3 | 0.09 | 2.55E-245 | Salivary_gland | Insr |
| Slc6a6 | 3.29E-249 | 0.57233948 | 0.76 | 0.4 | 6.81E-245 | Salivary_gland | Slc6a6 |
| Rarg | 5.98E-249 | 0.44331331 | 0.52 | 0.21 | 1.24E-244 | Salivary_gland | Rarg |
| Thoc2 | 1.12E-248 | 0.49053451 | 0.47 | 0.18 | 2.32E-244 | Salivary_gland | Thoc2 |
| Zzz3 | 1.16E-248 | 0.3508828 | 0.26 | 0.07 | 2.41E-244 | Salivary_gland | Zzz3 |
| Nab1 | 6.95E-248 | 0.35385394 | 0.31 | 0.1 | 1.44E-243 | Salivary_gland | Nab1 |
| Helz2 | 1.27E-247 | 0.32322427 | 0.26 | 0.07 | 2.62E-243 | Salivary_gland | Helz2 |
| Prdx4 | 2.89E-247 | 0.46346592 | 0.51 | 0.21 | 5.98E-243 | Salivary_gland | Prdx4 |
| Sema7a | 1.06E-246 | 0.42179692 | 0.48 | 0.19 | 2.19E-242 | Salivary_gland | Sema7a |
| Atrn | 1.72E-246 | 0.35291357 | 0.26 | 0.07 | 3.56E-242 | Salivary_gland | Atrn |
| Baz1b | 3.27E-246 | 0.37934143 | 0.35 | 0.12 | 6.77E-242 | Salivary_gland | Baz1b |
| Usp7 | 1.74E-245 | 0.35982383 | 0.28 | 0.08 | 3.60E-241 | Salivary_gland | Usp7 |
| Irf2bp2 | 2.48E-245 | 0.38745205 | 0.37 | 0.13 | 5.14E-241 | Salivary_gland | Irf2bp2 |
| Zfp106 | 3.80E-245 | 0.37341564 | 0.27 | 0.08 | 7.86E-241 | Salivary_gland | Zfp106 |
| Secisbp2l | 5.15E-245 | 0.35399478 | 0.29 | 0.09 | 1.07E-240 | Salivary_gland | Secisbp2l |
| Smap1 | 2.05E-244 | 0.4162537 | 0.52 | 0.21 | 4.23E-240 | Salivary_gland | Smap1 |
| Car8 | 2.30E-244 | 0.37525391 | 0.54 | 0.21 | 4.76E-240 | Salivary_gland | Car8 |
| B230219D22Rik | 4.39E-244 | 0.47057079 | 0.64 | 0.3 | 9.09E-240 | Salivary_gland | B230219D22Rik |
| Mia3 | 6.86E-244 | 0.36821852 | 0.3 | 0.09 | 1.42E-239 | Salivary_gland | Mia3 |
| Cnot2 | 8.23E-244 | 0.40263369 | 0.33 | 0.11 | 1.70E-239 | Salivary_gland | Cnot2 |
| Proser2 | 1.87E-243 | 0.35606613 | 0.3 | 0.09 | 3.87E-239 | Salivary_gland | Proser2 |
| Anks1 | 1.93E-243 | 0.35118023 | 0.28 | 0.08 | 3.99E-239 | Salivary_gland | Anks1 |
| Eef1b2 | 3.72E-243 | 0.5890246 | 0.8 | 0.47 | 7.70E-239 | Salivary_gland | Eef1b2 |
| Mapre2 | 3.55E-242 | 0.40522186 | 0.36 | 0.12 | 7.34E-238 | Salivary_gland | Mapre2 |
| Ccdc88a | 9.27E-242 | 0.48032897 | 0.33 | 0.11 | 1.92E-237 | Salivary_gland | Ccdc88a |
| Tmx3 | 1.46E-241 | 0.40253243 | 0.38 | 0.13 | 3.03E-237 | Salivary_gland | Tmx3 |
| Helz | 1.83E-241 | 0.38429522 | 0.31 | 0.1 | 3.79E-237 | Salivary_gland | Helz |
| Rabgap1 | 1.84E-241 | 0.35339558 | 0.28 | 0.08 | 3.81E-237 | Salivary_gland | Rabgap1 |
| Slc12a7 | 4.46E-241 | 0.30524338 | 0.29 | 0.09 | 9.22E-237 | Salivary_gland | Slc12a7 |
| Tcaf1 | 5.03E-241 | 0.3704203 | 0.27 | 0.08 | 1.04E-236 | Salivary_gland | Tcaf1 |
| H2-K1 | 1.55E-239 | 0.47891716 | 0.99 | 0.84 | 3.20E-235 | Salivary_gland | H2-K1 |

| | | | | | | | |
|---|---|---|---|---|---|---|---|
| Chd1 | 1.59E-238 | 0.4133889 | 0.36 | 0.12 | 3.29E-234 | Salivary_gland | Chd1 |
| Ptpn14 | 6.26E-238 | 0.35564713 | 0.33 | 0.11 | 1.30E-233 | Salivary_gland | Ptpn14 |
| Cab39 | 1.10E-237 | 0.39938751 | 0.4 | 0.15 | 2.28E-233 | Salivary_gland | Cab39 |
| Nova2 | 2.08E-237 | 0.39835127 | 0.35 | 0.12 | 4.30E-233 | Salivary_gland | Nova2 |
| Synj1 | 2.69E-237 | 0.3406024 | 0.26 | 0.07 | 5.58E-233 | Salivary_gland | Synj1 |
| Ube2m | 3.31E-237 | 0.39321426 | 0.33 | 0.11 | 6.85E-233 | Salivary_gland | Ube2m |
| Brd4 | 4.07E-237 | 0.40588056 | 0.46 | 0.18 | 8.42E-233 | Salivary_gland | Brd4 |
| Zzef1 | 2.20E-236 | 0.36054945 | 0.26 | 0.08 | 4.55E-232 | Salivary_gland | Zzef1 |
| Ppp3ca | 1.54E-235 | 0.55693838 | 0.64 | 0.32 | 3.19E-231 | Salivary_gland | Ppp3ca |
| Pde4b | 1.07E-234 | 0.40832291 | 0.36 | 0.13 | 2.22E-230 | Salivary_gland | Pde4b |
| Ppp1r10 | 4.09E-234 | 0.39537524 | 0.33 | 0.11 | 8.47E-230 | Salivary_gland | Ppp1r10 |
| Tor1aip1 | 5.31E-234 | 0.46952669 | 0.5 | 0.21 | 1.10E-229 | Salivary_gland | Tor1aip1 |
| Stat1 | 8.23E-234 | 0.52830884 | 0.45 | 0.18 | 1.70E-229 | Salivary_gland | Stat1 |
| Hdlbp | 9.65E-234 | 0.40429017 | 0.42 | 0.16 | 2.00E-229 | Salivary_gland | Hdlbp |
| She | 3.40E-233 | 0.43436078 | 0.48 | 0.2 | 7.04E-229 | Salivary_gland | She |
| Tanc1 | 8.20E-233 | 0.49804602 | 0.56 | 0.25 | 1.70E-228 | Salivary_gland | Tanc1 |
| Efnb2 | 1.63E-232 | 0.43159811 | 0.44 | 0.17 | 3.38E-228 | Salivary_gland | Efnb2 |
| Xrn2 | 1.90E-232 | 0.41580988 | 0.35 | 0.12 | 3.93E-228 | Salivary_gland | Xrn2 |
| Cnot6l | 2.51E-232 | 0.37952012 | 0.39 | 0.14 | 5.19E-228 | Salivary_gland | Cnot6l |
| Akap13 | 4.10E-232 | 0.4335172 | 0.61 | 0.28 | 8.48E-228 | Salivary_gland | Akap13 |
| Mgst1 | 1.61E-230 | 0.39596597 | 0.38 | 0.14 | 3.34E-226 | Salivary_gland | Mgst1 |
| Fez2 | 3.80E-230 | 0.47467489 | 0.68 | 0.33 | 7.86E-226 | Salivary_gland | Fez2 |
| Stx2 | 6.09E-230 | 0.33827416 | 0.3 | 0.1 | 1.26E-225 | Salivary_gland | Stx2 |
| Fam219a | 2.49E-229 | 0.40851043 | 0.35 | 0.12 | 5.16E-225 | Salivary_gland | Fam219a |
| Prpf40a | 1.61E-228 | 0.44214206 | 0.42 | 0.16 | 3.33E-224 | Salivary_gland | Prpf40a |
| Abi1 | 8.80E-228 | 0.41473361 | 0.41 | 0.16 | 1.82E-223 | Salivary_gland | Abi1 |
| Cbl | 2.44E-227 | 0.37491478 | 0.29 | 0.09 | 5.05E-223 | Salivary_gland | Cbl |
| Eif3a | 1.46E-226 | 0.52239102 | 0.61 | 0.29 | 3.01E-222 | Salivary_gland | Eif3a |
| Tmed5 | 9.48E-226 | 0.49435374 | 0.6 | 0.28 | 1.96E-221 | Salivary_gland | Tmed5 |
| Gspt1 | 5.90E-225 | 0.34182974 | 0.34 | 0.12 | 1.22E-220 | Salivary_gland | Gspt1 |
| Zmiz1 | 6.82E-225 | 0.46736515 | 0.6 | 0.28 | 1.41E-220 | Salivary_gland | Zmiz1 |
| Dock6 | 1.58E-224 | 0.377027 | 0.42 | 0.16 | 3.28E-220 | Salivary_gland | Dock6 |
| Cggbp1 | 3.27E-224 | 0.41241079 | 0.45 | 0.18 | 6.78E-220 | Salivary_gland | Cggbp1 |
| H2-Q7 | 1.05E-223 | 0.44555226 | 0.73 | 0.36 | 2.18E-219 | Salivary_gland | H2-Q7 |
| Pkn2 | 2.22E-223 | 0.42054716 | 0.52 | 0.22 | 4.59E-219 | Salivary_gland | Pkn2 |
| Fto | 3.24E-223 | 0.37511842 | 0.34 | 0.12 | 6.70E-219 | Salivary_gland | Fto |
| Clec14a | 7.23E-223 | 0.35261027 | 0.85 | 0.48 | 1.50E-218 | Salivary_gland | Clec14a |
| Appl1 | 1.37E-222 | 0.35754807 | 0.3 | 0.1 | 2.84E-218 | Salivary_gland | Appl1 |
| Palmd | 2.93E-222 | 0.43520533 | 0.59 | 0.27 | 6.05E-218 | Salivary_gland | Palmd |
| Notch1 | 3.59E-222 | 0.43122817 | 0.57 | 0.26 | 7.42E-218 | Salivary_gland | Notch1 |
| Stk38 | 7.12E-222 | 0.3463283 | 0.34 | 0.12 | 1.47E-217 | Salivary_gland | Stk38 |

| | | | | | | | |
|---|---|---|---|---|---|---|---|
| Exoc3 | 2.62E-220 | 0.33350188 | 0.26 | 0.08 | 5.41E-216 | Salivary_gland | Exoc3 |
| Anapc1 | 1.15E-219 | 0.3381783 | 0.25 | 0.08 | 2.38E-215 | Salivary_gland | Anapc1 |
| Psme1 | 1.22E-218 | 0.53218615 | 0.84 | 0.51 | 2.52E-214 | Salivary_gland | Psme1 |
| Irf1 | 5.43E-217 | 0.36499872 | 0.61 | 0.28 | 1.12E-212 | Salivary_gland | Irf1 |
| Eid1 | 1.02E-216 | 0.45224821 | 0.48 | 0.2 | 2.11E-212 | Salivary_gland | Eid1 |
| Scn1b | 5.30E-216 | 0.3044048 | 0.4 | 0.15 | 1.10E-211 | Salivary_gland | Scn1b |
| 6-Mar | 1.32E-213 | 0.29961694 | 0.26 | 0.08 | 2.72E-209 | Salivary_gland | 6-Mar |
| Pik3r1 | 2.70E-213 | 0.31103404 | 0.31 | 0.1 | 5.60E-209 | Salivary_gland | Pik3r1 |
| Arnt | 4.72E-213 | 0.32994946 | 0.26 | 0.08 | 9.78E-209 | Salivary_gland | Arnt |
| Slc43a3 | 9.19E-213 | 0.47382172 | 0.76 | 0.4 | 1.90E-208 | Salivary_gland | Slc43a3 |
| Smarca2 | 1.20E-212 | 0.44434168 | 0.55 | 0.25 | 2.49E-208 | Salivary_gland | Smarca2 |
| Nsmce2 | 1.80E-212 | 0.36345846 | 0.26 | 0.08 | 3.73E-208 | Salivary_gland | Nsmce2 |
| Pfkp | 2.35E-212 | 0.35110615 | 0.3 | 0.1 | 4.87E-208 | Salivary_gland | Pfkp |
| Cnot1 | 5.83E-212 | 0.31793034 | 0.26 | 0.08 | 1.21E-207 | Salivary_gland | Cnot1 |
| Vcl | 5.90E-212 | 0.39989823 | 0.42 | 0.16 | 1.22E-207 | Salivary_gland | Vcl |
| 1700020I14Rik | 2.28E-211 | 0.37839544 | 0.55 | 0.25 | 4.72E-207 | Salivary_gland | 1700020I14Rik |
| Ncl | 4.19E-211 | 0.4868348 | 0.7 | 0.37 | 8.66E-207 | Salivary_gland | Ncl |
| Tacc1 | 3.70E-210 | 0.38087712 | 0.51 | 0.22 | 7.66E-206 | Salivary_gland | Tacc1 |
| Rassf3 | 3.63E-209 | 0.33695396 | 0.53 | 0.23 | 7.50E-205 | Salivary_gland | Rassf3 |
| Heg1 | 8.82E-209 | 0.51564457 | 0.72 | 0.39 | 1.83E-204 | Salivary_gland | Heg1 |
| Tmcc1 | 9.43E-209 | 0.42903983 | 0.34 | 0.12 | 1.95E-204 | Salivary_gland | Tmcc1 |
| Larp4b | 2.37E-208 | 0.36370974 | 0.32 | 0.11 | 4.91E-204 | Salivary_gland | Larp4b |
| Mapkap1 | 3.19E-208 | 0.34447479 | 0.29 | 0.1 | 6.60E-204 | Salivary_gland | Mapkap1 |
| Syap1 | 5.52E-208 | 0.30511122 | 0.27 | 0.09 | 1.14E-203 | Salivary_gland | Syap1 |
| Gna12 | 4.03E-207 | 0.28359203 | 0.25 | 0.08 | 8.33E-203 | Salivary_gland | Gna12 |
| Dennd5a | 6.12E-207 | 0.36719577 | 0.42 | 0.17 | 1.27E-202 | Salivary_gland | Dennd5a |
| Ehd4 | 7.02E-207 | 0.37039024 | 0.9 | 0.51 | 1.45E-202 | Salivary_gland | Ehd4 |
| U2surp | 4.64E-206 | 0.38177946 | 0.43 | 0.18 | 9.60E-202 | Salivary_gland | U2surp |
| Lyn | 1.81E-205 | 0.38496179 | 0.48 | 0.2 | 3.74E-201 | Salivary_gland | Lyn |
| Phip | 2.88E-205 | 0.40602562 | 0.44 | 0.18 | 5.96E-201 | Salivary_gland | Phip |
| Prkd3 | 1.55E-204 | 0.32598142 | 0.32 | 0.11 | 3.20E-200 | Salivary_gland | Prkd3 |
| Lman1 | 2.60E-204 | 0.37999659 | 0.49 | 0.21 | 5.39E-200 | Salivary_gland | Lman1 |
| Tie1 | 3.38E-204 | 0.32465357 | 1 | 0.56 | 7.00E-200 | Salivary_gland | Tie1 |
| Tax1bp1 | 8.12E-203 | 0.42295118 | 0.62 | 0.3 | 1.68E-198 | Salivary_gland | Tax1bp1 |
| Spns2 | 3.22E-202 | 0.321563 | 0.29 | 0.1 | 6.65E-198 | Salivary_gland | Spns2 |
| Lsm12 | 3.51E-202 | 0.30766063 | 0.26 | 0.08 | 7.26E-198 | Salivary_gland | Lsm12 |
| Hipk3 | 5.33E-202 | 0.31608009 | 0.3 | 0.1 | 1.10E-197 | Salivary_gland | Hipk3 |
| Chd8 | 1.85E-200 | 0.31993669 | 0.3 | 0.1 | 3.84E-196 | Salivary_gland | Chd8 |
| Ptp4a2 | 5.99E-200 | 0.41798048 | 0.54 | 0.25 | 1.24E-195 | Salivary_gland | Ptp4a2 |
| Cnot6 | 8.24E-200 | 0.32916524 | 0.33 | 0.12 | 1.70E-195 | Salivary_gland | Cnot6 |
| Mmp15 | 5.99E-198 | 0.30121361 | 0.33 | 0.12 | 1.24E-193 | Salivary_gland | Mmp15 |

| | | | | | | | |
|---|---|---|---|---|---|---|---|
| Arhgef7 | 8.61E-198 | 0.34045734 | 0.35 | 0.13 | 1.78E-193 | Salivary_gland | Arhgef7 |
| Trim56 | 1.24E-197 | 0.31230218 | 0.29 | 0.1 | 2.56E-193 | Salivary_gland | Trim56 |
| Lipa | 3.72E-197 | 0.37036273 | 0.38 | 0.15 | 7.69E-193 | Salivary_gland | Lipa |
| Phc2 | 5.82E-197 | 0.3000376 | 0.31 | 0.11 | 1.20E-192 | Salivary_gland | Phc2 |
| Tjp2 | 1.09E-196 | 0.34550493 | 0.31 | 0.11 | 2.25E-192 | Salivary_gland | Tjp2 |
| Ube2q1 | 1.50E-196 | 0.32090874 | 0.31 | 0.11 | 3.09E-192 | Salivary_gland | Ube2q1 |
| Cdk11b | 2.33E-196 | 0.35300132 | 0.44 | 0.18 | 4.82E-192 | Salivary_gland | Cdk11b |
| St5 | 3.33E-196 | 0.34313026 | 0.31 | 0.11 | 6.88E-192 | Salivary_gland | St5 |
| Exoc5 | 4.07E-196 | 0.34359992 | 0.36 | 0.14 | 8.43E-192 | Salivary_gland | Exoc5 |
| Fermt2 | 5.60E-196 | 0.44611482 | 0.83 | 0.48 | 1.16E-191 | Salivary_gland | Fermt2 |
| Lipe | 8.17E-196 | 0.30255478 | 0.41 | 0.16 | 1.69E-191 | Salivary_gland | Lipe |
| Notch4 | 6.99E-195 | 0.27286815 | 0.37 | 0.14 | 1.45E-190 | Salivary_gland | Notch4 |
| Zranb2 | 1.11E-193 | 0.33952749 | 0.37 | 0.14 | 2.30E-189 | Salivary_gland | Zranb2 |
| Terf1 | 1.12E-193 | 0.30637207 | 0.27 | 0.09 | 2.32E-189 | Salivary_gland | Terf1 |
| Sec14l1 | 1.41E-193 | 0.37208946 | 0.54 | 0.25 | 2.92E-189 | Salivary_gland | Sec14l1 |
| Epn2 | 4.65E-193 | 0.35458206 | 0.43 | 0.18 | 9.62E-189 | Salivary_gland | Epn2 |
| Ppp2r5e | 5.79E-193 | 0.32270731 | 0.28 | 0.1 | 1.20E-188 | Salivary_gland | Ppp2r5e |
| Trim30a | 6.15E-193 | 0.39678666 | 0.41 | 0.17 | 1.27E-188 | Salivary_gland | Trim30a |
| Lmcd1 | 2.43E-192 | 0.35968127 | 0.52 | 0.23 | 5.02E-188 | Salivary_gland | Lmcd1 |
| Foxo1 | 3.11E-191 | 0.35274332 | 0.49 | 0.22 | 6.43E-187 | Salivary_gland | Foxo1 |
| Pnpla8 | 4.54E-191 | 0.29496955 | 0.3 | 0.1 | 9.40E-187 | Salivary_gland | Pnpla8 |
| Smndc1 | 7.67E-191 | 0.3005929 | 0.3 | 0.11 | 1.59E-186 | Salivary_gland | Smndc1 |
| Ap2b1 | 1.31E-190 | 0.33624091 | 0.42 | 0.17 | 2.70E-186 | Salivary_gland | Ap2b1 |
| Gimap8 | 6.64E-190 | 0.29753763 | 0.33 | 0.12 | 1.37E-185 | Salivary_gland | Gimap8 |
| Capza1 | 7.67E-190 | 0.31659167 | 0.38 | 0.15 | 1.59E-185 | Salivary_gland | Capza1 |
| Nudt4 | 3.42E-189 | 0.42950342 | 0.42 | 0.17 | 7.07E-185 | Salivary_gland | Nudt4 |
| Abcg2 | 3.44E-189 | 0.27508379 | 0.7 | 0.35 | 7.12E-185 | Salivary_gland | Abcg2 |
| Itgb1 | 1.03E-188 | 0.44078533 | 0.83 | 0.51 | 2.14E-184 | Salivary_gland | Itgb1 |
| Rlim | 1.61E-188 | 0.37239737 | 0.34 | 0.13 | 3.32E-184 | Salivary_gland | Rlim |
| Grrp1 | 1.47E-187 | 0.35853428 | 0.66 | 0.33 | 3.05E-183 | Salivary_gland | Grrp1 |
| Cyth1 | 7.84E-187 | 0.2561644 | 0.32 | 0.11 | 1.62E-182 | Salivary_gland | Cyth1 |
| Prkar2a | 1.15E-186 | 0.30587706 | 0.28 | 0.09 | 2.37E-182 | Salivary_gland | Prkar2a |
| Stim2 | 1.51E-186 | 0.3552139 | 0.43 | 0.18 | 3.12E-182 | Salivary_gland | Stim2 |
| Sp3 | 1.63E-186 | 0.30330238 | 0.33 | 0.12 | 3.38E-182 | Salivary_gland | Sp3 |
| Parp9 | 1.72E-186 | 0.32429609 | 0.26 | 0.09 | 3.56E-182 | Salivary_gland | Parp9 |
| Mdm4 | 7.54E-186 | 0.27435261 | 0.26 | 0.08 | 1.56E-181 | Salivary_gland | Mdm4 |
| Rhoj | 1.81E-185 | 0.43368739 | 0.64 | 0.33 | 3.75E-181 | Salivary_gland | Rhoj |
| Atp2b4 | 5.69E-185 | 0.3134143 | 0.46 | 0.2 | 1.18E-180 | Salivary_gland | Atp2b4 |
| Ddx58 | 2.55E-184 | 0.30846474 | 0.31 | 0.11 | 5.29E-180 | Salivary_gland | Ddx58 |
| Tdrp | 3.57E-184 | 0.32383413 | 0.36 | 0.14 | 7.39E-180 | Salivary_gland | Tdrp |
| Brwd1 | 4.28E-183 | 0.31358588 | 0.25 | 0.08 | 8.86E-179 | Salivary_gland | Brwd1 |

| | | | | | | | |
|---|---|---|---|---|---|---|---|
| Hif1a | 1.77E-182 | 0.32727293 | 0.37 | 0.15 | 3.66E-178 | Salivary_gland | Hif1a |
| Prkacb | 2.37E-182 | 0.31385573 | 0.31 | 0.12 | 4.91E-178 | Salivary_gland | Prkacb |
| Tox4 | 4.56E-182 | 0.30131277 | 0.31 | 0.12 | 9.44E-178 | Salivary_gland | Tox4 |
| Ogdh | 5.72E-182 | 0.34123847 | 0.37 | 0.15 | 1.18E-177 | Salivary_gland | Ogdh |
| Smarcc2 | 6.64E-182 | 0.29137402 | 0.29 | 0.11 | 1.37E-177 | Salivary_gland | Smarcc2 |
| Zdhhc20 | 2.01E-181 | 0.31646399 | 0.42 | 0.18 | 4.16E-177 | Salivary_gland | Zdhhc20 |
| Pola2 | 2.37E-181 | 0.26643731 | 0.25 | 0.08 | 4.90E-177 | Salivary_gland | Pola2 |
| Elk4 | 8.29E-181 | 0.28466851 | 0.34 | 0.13 | 1.72E-176 | Salivary_gland | Elk4 |
| Plscr4 | 8.57E-181 | 0.33538552 | 0.49 | 0.22 | 1.77E-176 | Salivary_gland | Plscr4 |
| Ppil4 | 3.43E-180 | 0.29567834 | 0.29 | 0.1 | 7.09E-176 | Salivary_gland | Ppil4 |
| Abhd17b | 1.42E-179 | 0.34550958 | 0.36 | 0.14 | 2.93E-175 | Salivary_gland | Abhd17b |
| Fxr1 | 1.67E-178 | 0.31023317 | 0.38 | 0.15 | 3.46E-174 | Salivary_gland | Fxr1 |
| Btbd3 | 4.81E-178 | 0.29075307 | 0.26 | 0.09 | 9.95E-174 | Salivary_gland | Btbd3 |
| Ube3a | 1.08E-177 | 0.32734545 | 0.47 | 0.21 | 2.23E-173 | Salivary_gland | Ube3a |
| Glud1 | 9.74E-177 | 0.34679054 | 0.55 | 0.26 | 2.01E-172 | Salivary_gland | Glud1 |
| Rsbn1 | 2.33E-176 | 0.29525278 | 0.3 | 0.11 | 4.82E-172 | Salivary_gland | Rsbn1 |
| Stat3 | 4.61E-176 | 0.34672308 | 0.5 | 0.23 | 9.55E-172 | Salivary_gland | Stat3 |
| Ccdc50 | 5.14E-176 | 0.32112168 | 0.31 | 0.12 | 1.06E-171 | Salivary_gland | Ccdc50 |
| Hint3 | 8.79E-176 | 0.27082118 | 0.27 | 0.09 | 1.82E-171 | Salivary_gland | Hint3 |
| App | 8.94E-176 | 0.4236641 | 0.96 | 0.73 | 1.85E-171 | Salivary_gland | App |
| Reep3 | 1.02E-175 | 0.41306661 | 0.67 | 0.36 | 2.11E-171 | Salivary_gland | Reep3 |
| Nfkb1 | 3.61E-175 | 0.31024618 | 0.39 | 0.16 | 7.48E-171 | Salivary_gland | Nfkb1 |
| Ddx3x | 7.17E-175 | 0.36196257 | 0.58 | 0.29 | 1.48E-170 | Salivary_gland | Ddx3x |
| Tspan12 | 4.80E-174 | 0.37731567 | 0.65 | 0.33 | 9.93E-170 | Salivary_gland | Tspan12 |
| Fkbp3 | 1.66E-173 | 0.39543076 | 0.58 | 0.29 | 3.43E-169 | Salivary_gland | Fkbp3 |
| Fam120a | 2.48E-173 | 0.25436654 | 0.32 | 0.12 | 5.13E-169 | Salivary_gland | Fam120a |
| Ptprb | 3.34E-173 | 0.40614562 | 0.98 | 0.88 | 6.92E-169 | Salivary_gland | Ptprb |
| Rpsa | 6.56E-173 | 0.48472721 | 0.93 | 0.7 | 1.36E-168 | Salivary_gland | Rpsa |
| Ltbp4 | 6.91E-173 | 0.2928597 | 0.83 | 0.48 | 1.43E-168 | Salivary_gland | Ltbp4 |
| Ppig | 1.80E-172 | 0.40700659 | 0.51 | 0.24 | 3.72E-168 | Salivary_gland | Ppig |
| Cltc | 1.85E-172 | 0.33033562 | 0.58 | 0.28 | 3.83E-168 | Salivary_gland | Cltc |
| Adam17 | 4.91E-172 | 0.27263045 | 0.27 | 0.1 | 1.02E-167 | Salivary_gland | Adam17 |
| Usp1 | 5.30E-172 | 0.29584007 | 0.27 | 0.1 | 1.10E-167 | Salivary_gland | Usp1 |
| Mtdh | 8.08E-172 | 0.43273169 | 0.66 | 0.36 | 1.67E-167 | Salivary_gland | Mtdh |
| Marcks | 3.00E-170 | 0.36389564 | 0.66 | 0.35 | 6.20E-166 | Salivary_gland | Marcks |
| Sgk3 | 3.65E-170 | 0.27652277 | 0.26 | 0.09 | 7.56E-166 | Salivary_gland | Sgk3 |
| Pxk | 3.85E-170 | 0.30425741 | 0.3 | 0.11 | 7.96E-166 | Salivary_gland | Pxk |
| Vamp5 | 6.28E-170 | 0.3734416 | 0.87 | 0.52 | 1.30E-165 | Salivary_gland | Vamp5 |
| Dst | 7.78E-170 | 0.33168165 | 0.4 | 0.17 | 1.61E-165 | Salivary_gland | Dst |
| Rab13 | 1.48E-169 | 0.28215159 | 0.38 | 0.16 | 3.05E-165 | Salivary_gland | Rab13 |
| Calcrl | 2.04E-169 | 0.26823861 | 0.72 | 0.38 | 4.23E-165 | Salivary_gland | Calcrl |

| | | | | | | | |
|---|---|---|---|---|---|---|---|
| Eprs | 1.84E-168 | 0.30284244 | 0.31 | 0.12 | 3.81E-164 | Salivary_gland | Eprs |
| Akap9 | 2.27E-168 | 0.38586288 | 0.4 | 0.18 | 4.70E-164 | Salivary_gland | Akap9 |
| Hnrnpu | 7.96E-168 | 0.36747085 | 0.67 | 0.36 | 1.65E-163 | Salivary_gland | Hnrnpu |
| Txlna | 1.95E-167 | 0.27298951 | 0.27 | 0.1 | 4.04E-163 | Salivary_gland | Txlna |
| Cdc5l | 2.05E-167 | 0.32295552 | 0.33 | 0.13 | 4.24E-163 | Salivary_gland | Cdc5l |
| Ptpn12 | 3.35E-167 | 0.32540952 | 0.44 | 0.2 | 6.94E-163 | Salivary_gland | Ptpn12 |
| Cd47 | 3.91E-167 | 0.37143151 | 0.92 | 0.62 | 8.08E-163 | Salivary_gland | Cd47 |
| Add3 | 4.44E-167 | 0.32606979 | 0.39 | 0.16 | 9.19E-163 | Salivary_gland | Add3 |
| Eif3c | 5.38E-167 | 0.37490024 | 0.55 | 0.27 | 1.11E-162 | Salivary_gland | Eif3c |
| Ubr4 | 5.73E-167 | 0.25766934 | 0.28 | 0.1 | 1.19E-162 | Salivary_gland | Ubr4 |
| Ppp3cb | 1.14E-166 | 0.28308485 | 0.26 | 0.09 | 2.35E-162 | Salivary_gland | Ppp3cb |
| Fyn | 1.71E-166 | 0.33385349 | 0.49 | 0.23 | 3.54E-162 | Salivary_gland | Fyn |
| Mphosph8 | 2.37E-166 | 0.33012469 | 0.36 | 0.15 | 4.91E-162 | Salivary_gland | Mphosph8 |
| Far1 | 3.25E-166 | 0.25155374 | 0.28 | 0.1 | 6.73E-162 | Salivary_gland | Far1 |
| Golim4 | 1.15E-165 | 0.27649694 | 0.35 | 0.14 | 2.38E-161 | Salivary_gland | Golim4 |
| Appbp2 | 2.96E-165 | 0.29269847 | 0.27 | 0.1 | 6.12E-161 | Salivary_gland | Appbp2 |
| Top1 | 5.08E-165 | 0.40799556 | 0.66 | 0.36 | 1.05E-160 | Salivary_gland | Top1 |
| Yes1 | 1.68E-164 | 0.2841385 | 0.45 | 0.2 | 3.48E-160 | Salivary_gland | Yes1 |
| Arid5b | 2.25E-164 | 0.37707673 | 0.52 | 0.25 | 4.66E-160 | Salivary_gland | Arid5b |
| Actn4 | 5.17E-164 | 0.38621262 | 0.82 | 0.5 | 1.07E-159 | Salivary_gland | Actn4 |
| Cdv3 | 7.09E-164 | 0.30128391 | 0.46 | 0.21 | 1.47E-159 | Salivary_gland | Cdv3 |
| Sf3b3 | 8.24E-164 | 0.25620529 | 0.26 | 0.09 | 1.71E-159 | Salivary_gland | Sf3b3 |
| Stx8 | 4.92E-163 | 0.28508937 | 0.34 | 0.14 | 1.02E-158 | Salivary_gland | Stx8 |
| Zmynd8 | 5.54E-163 | 0.30284445 | 0.33 | 0.13 | 1.15E-158 | Salivary_gland | Zmynd8 |
| Synpo | 7.42E-163 | 0.25233075 | 0.29 | 0.11 | 1.53E-158 | Salivary_gland | Synpo |
| Ssbp3 | 8.86E-163 | 0.28293898 | 0.28 | 0.11 | 1.83E-158 | Salivary_gland | Ssbp3 |
| Vps4b | 1.77E-162 | 0.29372797 | 0.37 | 0.15 | 3.66E-158 | Salivary_gland | Vps4b |
| Dnm1l | 2.12E-162 | 0.26929462 | 0.26 | 0.09 | 4.39E-158 | Salivary_gland | Dnm1l |
| Rpl12 | 3.34E-162 | 0.45250858 | 0.88 | 0.6 | 6.90E-158 | Salivary_gland | Rpl12 |
| Mtf2 | 3.76E-162 | 0.26415234 | 0.26 | 0.09 | 7.77E-158 | Salivary_gland | Mtf2 |
| Gigyf2 | 3.03E-161 | 0.26595592 | 0.28 | 0.11 | 6.27E-157 | Salivary_gland | Gigyf2 |
| Camk2n1 | 1.16E-160 | 0.32228074 | 0.32 | 0.13 | 2.41E-156 | Salivary_gland | Camk2n1 |
| B4galt4 | 3.33E-160 | 0.29635715 | 0.49 | 0.23 | 6.89E-156 | Salivary_gland | B4galt4 |
| Pum2 | 4.56E-160 | 0.29433349 | 0.45 | 0.2 | 9.43E-156 | Salivary_gland | Pum2 |
| Ppm1a | 1.61E-159 | 0.28307462 | 0.29 | 0.11 | 3.34E-155 | Salivary_gland | Ppm1a |
| Amotl1 | 5.56E-159 | 0.32831653 | 0.51 | 0.24 | 1.15E-154 | Salivary_gland | Amotl1 |
| Atf7ip | 3.17E-158 | 0.26589713 | 0.33 | 0.13 | 6.57E-154 | Salivary_gland | Atf7ip |
| Sacm1l | 8.73E-158 | 0.27186662 | 0.25 | 0.09 | 1.81E-153 | Salivary_gland | Sacm1l |
| Sptlc2 | 2.26E-157 | 0.32237108 | 0.5 | 0.24 | 4.67E-153 | Salivary_gland | Sptlc2 |
| Auts2 | 2.68E-157 | 0.52797491 | 0.3 | 0.12 | 5.54E-153 | Salivary_gland | Auts2 |
| Akap8 | 5.00E-157 | 0.25642896 | 0.27 | 0.1 | 1.04E-152 | Salivary_gland | Akap8 |

| | | | | | | | |
|---|---|---|---|---|---|---|---|
| Syncrip | 7.88E-157 | 0.27587516 | 0.34 | 0.14 | 1.63E-152 | Salivary_gland | Syncrip |
| Scfd1 | 1.06E-155 | 0.2742568 | 0.25 | 0.09 | 2.20E-151 | Salivary_gland | Scfd1 |
| Scand1 | 2.00E-155 | 0.30544589 | 0.58 | 0.29 | 4.13E-151 | Salivary_gland | Scand1 |
| Ppp2r5a | 2.03E-155 | 0.27818779 | 0.44 | 0.2 | 4.20E-151 | Salivary_gland | Ppp2r5a |
| 2810474O19Rik | 1.65E-154 | 0.32075423 | 0.34 | 0.14 | 3.41E-150 | Salivary_gland | 2810474O19Rik |
| Inpp5a | 1.93E-154 | 0.26852463 | 0.34 | 0.14 | 3.98E-150 | Salivary_gland | Inpp5a |
| Flywch1 | 2.96E-154 | 0.265449 | 0.28 | 0.11 | 6.12E-150 | Salivary_gland | Flywch1 |
| Ptpn11 | 4.15E-154 | 0.26116936 | 0.26 | 0.09 | 8.58E-150 | Salivary_gland | Ptpn11 |
| Tek | 8.87E-154 | 0.36194228 | 0.78 | 0.46 | 1.84E-149 | Salivary_gland | Tek |
| Osbpl9 | 8.93E-154 | 0.31180573 | 0.55 | 0.27 | 1.85E-149 | Salivary_gland | Osbpl9 |
| Pgrmc2 | 1.65E-153 | 0.25178182 | 0.28 | 0.11 | 3.42E-149 | Salivary_gland | Pgrmc2 |
| Gapvd1 | 7.75E-153 | 0.27092769 | 0.33 | 0.13 | 1.60E-148 | Salivary_gland | Gapvd1 |
| Zfp644 | 1.25E-152 | 0.28700429 | 0.42 | 0.19 | 2.59E-148 | Salivary_gland | Zfp644 |
| Wars | 1.37E-152 | 0.29452602 | 0.42 | 0.19 | 2.83E-148 | Salivary_gland | Wars |
| Ano6 | 4.09E-152 | 0.2884502 | 0.38 | 0.17 | 8.46E-148 | Salivary_gland | Ano6 |
| Sh3bp5 | 2.01E-151 | 0.33012004 | 0.57 | 0.29 | 4.16E-147 | Salivary_gland | Sh3bp5 |
| Csde1 | 3.34E-151 | 0.29651555 | 0.59 | 0.31 | 6.92E-147 | Salivary_gland | Csde1 |
| Phactr2 | 4.45E-151 | 0.27267696 | 0.44 | 0.2 | 9.21E-147 | Salivary_gland | Phactr2 |
| Ifrd1 | 2.06E-150 | 0.26910805 | 0.35 | 0.15 | 4.25E-146 | Salivary_gland | Ifrd1 |
| Mylip | 2.18E-150 | 0.25756734 | 0.31 | 0.12 | 4.52E-146 | Salivary_gland | Mylip |
| Cux1 | 6.58E-150 | 0.25733215 | 0.36 | 0.15 | 1.36E-145 | Salivary_gland | Cux1 |
| Smc6 | 1.32E-149 | 0.31010709 | 0.42 | 0.19 | 2.74E-145 | Salivary_gland | Smc6 |
| Irf2bpl | 2.51E-149 | 0.29418999 | 0.52 | 0.26 | 5.19E-145 | Salivary_gland | Irf2bpl |
| Ubxn2a | 2.52E-149 | 0.27166926 | 0.3 | 0.12 | 5.21E-145 | Salivary_gland | Ubxn2a |
| Nrd1 | 1.60E-148 | 0.26411499 | 0.33 | 0.14 | 3.32E-144 | Salivary_gland | Nrd1 |
| Ppp2r2d | 2.91E-148 | 0.25376416 | 0.28 | 0.11 | 6.02E-144 | Salivary_gland | Ppp2r2d |
| Snrnp48 | 3.98E-148 | 0.2609854 | 0.3 | 0.12 | 8.24E-144 | Salivary_gland | Snrnp48 |
| Cebpz | 4.15E-148 | 0.28665829 | 0.3 | 0.12 | 8.60E-144 | Salivary_gland | Cebpz |
| Psmc6 | 1.01E-146 | 0.28340066 | 0.47 | 0.22 | 2.09E-142 | Salivary_gland | Psmc6 |
| Ccnt1 | 1.11E-145 | 0.25789661 | 0.31 | 0.13 | 2.29E-141 | Salivary_gland | Ccnt1 |
| Ino80d | 6.22E-145 | 0.26761816 | 0.32 | 0.13 | 1.29E-140 | Salivary_gland | Ino80d |
| Swt1 | 1.17E-144 | 0.28047145 | 0.3 | 0.12 | 2.43E-140 | Salivary_gland | Swt1 |
| Nck1 | 2.36E-144 | 0.30346158 | 0.69 | 0.38 | 4.88E-140 | Salivary_gland | Nck1 |
| Bzw1 | 2.50E-144 | 0.27225424 | 0.57 | 0.29 | 5.18E-140 | Salivary_gland | Bzw1 |
| Taf10 | 9.50E-144 | 0.27752653 | 0.46 | 0.22 | 1.96E-139 | Salivary_gland | Taf10 |
| Tgfbr2 | 2.34E-143 | 0.25275545 | 0.52 | 0.26 | 4.84E-139 | Salivary_gland | Tgfbr2 |
| Pdcd4 | 4.14E-143 | 0.28200448 | 0.32 | 0.13 | 8.56E-139 | Salivary_gland | Pdcd4 |
| Srrm1 | 6.47E-143 | 0.30750215 | 0.52 | 0.26 | 1.34E-138 | Salivary_gland | Srrm1 |
| Riok3 | 8.95E-143 | 0.25615999 | 0.32 | 0.13 | 1.85E-138 | Salivary_gland | Riok3 |
| Ndrg1 | 1.78E-142 | 0.27662018 | 0.66 | 0.36 | 3.67E-138 | Salivary_gland | Ndrg1 |
| mt-Nd3 | 3.67E-142 | 0.28505832 | 0.72 | 0.41 | 7.59E-138 | Salivary_gland | mt-Nd3 |

| | | | | | | | |
|---|---|---|---|---|---|---|---|
| Cdh5 | 1.15E-141 | 0.28643426 | 1 | 0.72 | 2.38E-137 | Salivary_gland | Cdh5 |
| Ifngr1 | 1.37E-141 | 0.28972511 | 0.7 | 0.38 | 2.83E-137 | Salivary_gland | Ifngr1 |
| Hras | 1.75E-141 | 0.26760299 | 0.52 | 0.26 | 3.63E-137 | Salivary_gland | Hras |
| Rfc1 | 4.46E-141 | 0.26967129 | 0.29 | 0.11 | 9.23E-137 | Salivary_gland | Rfc1 |
| Tcf7l2 | 8.57E-141 | 0.25925469 | 0.26 | 0.1 | 1.77E-136 | Salivary_gland | Tcf7l2 |
| Fam76a | 2.50E-140 | 0.25747164 | 0.35 | 0.15 | 5.17E-136 | Salivary_gland | Fam76a |
| Camk2d | 4.64E-140 | 0.29695212 | 0.47 | 0.23 | 9.61E-136 | Salivary_gland | Camk2d |
| Bclaf1 | 1.11E-139 | 0.2728186 | 0.41 | 0.19 | 2.30E-135 | Salivary_gland | Bclaf1 |
| Csnk1a1 | 1.78E-139 | 0.34139259 | 0.73 | 0.42 | 3.68E-135 | Salivary_gland | Csnk1a1 |
| Sox13 | 5.15E-139 | 0.26245601 | 0.27 | 0.11 | 1.07E-134 | Salivary_gland | Sox13 |
| Picalm | 8.00E-139 | 0.26166316 | 0.5 | 0.25 | 1.65E-134 | Salivary_gland | Picalm |
| Hsp90b1 | 1.34E-138 | 0.30707992 | 0.85 | 0.55 | 2.78E-134 | Salivary_gland | Hsp90b1 |
| Flnb | 1.76E-138 | 0.26896313 | 0.43 | 0.2 | 3.64E-134 | Salivary_gland | Flnb |
| Pnp | 8.27E-138 | 0.3086576 | 0.43 | 0.2 | 1.71E-133 | Salivary_gland | Pnp |
| Zc3h15 | 9.50E-138 | 0.25033634 | 0.4 | 0.18 | 1.97E-133 | Salivary_gland | Zc3h15 |
| Dusp11 | 1.42E-137 | 0.26739113 | 0.43 | 0.2 | 2.94E-133 | Salivary_gland | Dusp11 |
| Nkap | 6.82E-137 | 0.25655505 | 0.26 | 0.1 | 1.41E-132 | Salivary_gland | Nkap |
| Taf3 | 1.29E-136 | 0.2506881 | 0.26 | 0.1 | 2.67E-132 | Salivary_gland | Taf3 |
| Eif4g2 | 8.42E-136 | 0.35711527 | 0.8 | 0.51 | 1.74E-131 | Salivary_gland | Eif4g2 |
| Pura | 7.35E-135 | 0.29385286 | 0.6 | 0.32 | 1.52E-130 | Salivary_gland | Pura |
| Rsbn1l | 1.51E-134 | 0.27088781 | 0.42 | 0.2 | 3.13E-130 | Salivary_gland | Rsbn1l |
| Grb2 | 6.25E-134 | 0.27931706 | 0.38 | 0.18 | 1.29E-129 | Salivary_gland | Grb2 |
| Zfp36l1 | 6.46E-133 | 0.38068357 | 0.79 | 0.52 | 1.34E-128 | Salivary_gland | Zfp36l1 |
| Eif5 | 8.72E-133 | 0.28935287 | 0.61 | 0.33 | 1.80E-128 | Salivary_gland | Eif5 |
| Sox18 | 5.83E-132 | 0.28509492 | 0.8 | 0.49 | 1.21E-127 | Salivary_gland | Sox18 |
| Creb3l2 | 1.42E-130 | 0.25941554 | 0.32 | 0.14 | 2.94E-126 | Salivary_gland | Creb3l2 |
| Btbd1 | 2.17E-130 | 0.27356159 | 0.36 | 0.16 | 4.49E-126 | Salivary_gland | Btbd1 |
| Cxcl12 | 3.71E-130 | 0.36432002 | 0.8 | 0.48 | 7.69E-126 | Salivary_gland | Cxcl12 |
| Dusp23 | 3.79E-130 | 0.25365403 | 0.25 | 0.1 | 7.84E-126 | Salivary_gland | Dusp23 |
| Pcmtd1 | 2.55E-128 | 0.25043223 | 0.54 | 0.28 | 5.28E-124 | Salivary_gland | Pcmtd1 |
| Ccnd1 | 6.65E-128 | 0.35716157 | 0.52 | 0.28 | 1.38E-123 | Salivary_gland | Ccnd1 |
| Rbbp6 | 4.53E-127 | 0.2933863 | 0.47 | 0.24 | 9.38E-123 | Salivary_gland | Rbbp6 |
| Chchd3 | 5.09E-127 | 0.27186441 | 0.41 | 0.19 | 1.05E-122 | Salivary_gland | Chchd3 |
| Gng12 | 3.72E-126 | 0.27015842 | 0.42 | 0.2 | 7.69E-122 | Salivary_gland | Gng12 |
| Rap1a | 1.81E-125 | 0.33002344 | 0.83 | 0.54 | 3.74E-121 | Salivary_gland | Rap1a |
| Odc1 | 2.62E-123 | 0.37962587 | 0.26 | 0.11 | 5.42E-119 | Salivary_gland | Odc1 |
| Yaf2 | 7.09E-122 | 0.26718854 | 0.39 | 0.18 | 1.47E-117 | Salivary_gland | Yaf2 |
| Rras | 8.10E-120 | 0.33333688 | 0.69 | 0.4 | 1.68E-115 | Salivary_gland | Rras |
| Rab10 | 1.05E-117 | 0.25984291 | 0.55 | 0.3 | 2.18E-113 | Salivary_gland | Rab10 |
| Psmb10 | 4.62E-115 | 0.28143996 | 0.71 | 0.42 | 9.55E-111 | Salivary_gland | Psmb10 |
| 2-Mar | 6.11E-115 | 0.27974941 | 0.65 | 0.37 | 1.27E-110 | Salivary_gland | 2-Mar |

| | | | | | | | |
|---|---|---|---|---|---|---|---|
| Msn | 1.53E-111 | 0.30692049 | 0.84 | 0.57 | 3.16E-107 | Salivary_gland | Msn |
| Rps21 | 5.92E-111 | 0.30598167 | 0.98 | 0.85 | 1.22E-106 | Salivary_gland | Rps21 |
| Klf7 | 1.85E-110 | 0.30433143 | 0.76 | 0.48 | 3.83E-106 | Salivary_gland | Klf7 |
| Elk3 | 1.49E-109 | 0.28548812 | 0.82 | 0.54 | 3.09E-105 | Salivary_gland | Elk3 |
| Cox5a | 5.97E-107 | 0.31978278 | 0.81 | 0.52 | 1.23E-102 | Salivary_gland | Cox5a |
| Ctnna1 | 5.97E-107 | 0.25077584 | 0.73 | 0.44 | 1.23E-102 | Salivary_gland | Ctnna1 |
| Igtp | 2.84E-101 | 0.28663329 | 0.28 | 0.13 | 5.87E-97 | Salivary_gland | Igtp |
| Rab2a | 3.82E-100 | 0.27820542 | 0.81 | 0.53 | 7.91E-96 | Salivary_gland | Rab2a |
| Atxn7l3b | 7.25E-97 | 0.25017059 | 0.65 | 0.38 | 1.50E-92 | Salivary_gland | Atxn7l3b |
| Rpl28 | 7.73E-97 | 0.28880661 | 0.97 | 0.84 | 1.60E-92 | Salivary_gland | Rpl28 |
| Sh3glb1 | 1.32E-89 | 0.26528245 | 0.83 | 0.57 | 2.73E-85 | Salivary_gland | Sh3glb1 |
| Txn1 | 2.00E-86 | 0.26547251 | 0.88 | 0.64 | 4.13E-82 | Salivary_gland | Txn1 |
| Rpl23 | 2.49E-86 | 0.25357828 | 0.99 | 0.9 | 5.16E-82 | Salivary_gland | Rpl23 |
| Mecom | 3.16E-07 | 0.32951804 | 0.26 | 0.22 | 0.00654101 | Salivary_gland | Mecom |
| Itm2a | 0 | 5.35859669 | 0.99 | 0.13 | 0 | brain | Itm2a |
| Bsg | 0 | 4.45888513 | 1 | 0.72 | 0 | brain | Bsg |
| Slc2a1 | 0 | 4.14289451 | 0.95 | 0.07 | 0 | brain | Slc2a1 |
| Pglyrp1 | 0 | 3.81536864 | 0.89 | 0.04 | 0 | brain | Pglyrp1 |
| Slco1a4 | 0 | 3.62410292 | 0.93 | 0.06 | 0 | brain | Slco1a4 |
| Slco1c1 | 0 | 3.6218162 | 0.86 | 0 | 0 | brain | Slco1c1 |
| Spock2 | 0 | 3.60737015 | 0.91 | 0.04 | 0 | brain | Spock2 |
| Hspa1a | 0 | 3.5716609 | 0.93 | 0.26 | 0 | brain | Hspa1a |
| Foxq1 | 0 | 2.85347142 | 0.72 | 0.01 | 0 | brain | Foxq1 |
| Gkn3 | 0 | 2.72635959 | 0.41 | 0.01 | 0 | brain | Gkn3 |
| Vwa1 | 0 | 2.70409257 | 0.93 | 0.31 | 0 | brain | Vwa1 |
| Ptn | 0 | 2.63941288 | 0.88 | 0.08 | 0 | brain | Ptn |
| Pltp | 0 | 2.63788787 | 0.99 | 0.45 | 0 | brain | Pltp |
| Glul | 0 | 2.55177143 | 0.9 | 0.35 | 0 | brain | Glul |
| Sgms1 | 0 | 2.54357138 | 0.88 | 0.21 | 0 | brain | Sgms1 |
| Wfdc1 | 0 | 2.47382296 | 0.8 | 0.07 | 0 | brain | Wfdc1 |
| Lsr | 0 | 2.43720148 | 0.79 | 0.07 | 0 | brain | Lsr |
| Slc22a8 | 0 | 2.43576459 | 0.58 | 0 | 0 | brain | Slc22a8 |
| Mt1 | 0 | 2.42360429 | 0.68 | 0.09 | 0 | brain | Mt1 |
| Hbb-bs | 0 | 2.28230137 | 0.59 | 0.11 | 0 | brain | Hbb-bs |
| Gsta4 | 0 | 2.27416608 | 0.6 | 0.04 | 0 | brain | Gsta4 |
| Tsc22d1 | 0 | 2.26597117 | 0.99 | 0.64 | 0 | brain | Tsc22d1 |
| Cldn5 | 0 | 2.20253185 | 1 | 0.61 | 0 | brain | Cldn5 |
| Degs2 | 0 | 2.15673493 | 0.6 | 0.02 | 0 | brain | Degs2 |
| Prom1 | 0 | 2.15021544 | 0.6 | 0.03 | 0 | brain | Prom1 |
| Ctsh | 0 | 2.11857769 | 0.74 | 0.09 | 0 | brain | Ctsh |
| Slc7a5 | 0 | 2.09026724 | 0.67 | 0.08 | 0 | brain | Slc7a5 |

| | | | | | | | |
|---|---|---|---|---|---|---|---|
| Nebl | 0 | 2.06983409 | 0.58 | 0.02 | 0 | brain | Nebl |
| Gm694 | 0 | 2.04332127 | 0.61 | 0.05 | 0 | brain | Gm694 |
| Isyna1 | 0 | 2.02394331 | 0.69 | 0.11 | 0 | brain | Isyna1 |
| Slc7a1 | 0 | 2.01075563 | 0.46 | 0.04 | 0 | brain | Slc7a1 |
| Ocln | 0 | 1.97132307 | 0.57 | 0.04 | 0 | brain | Ocln |
| Pcp4l1 | 0 | 1.96367694 | 0.9 | 0.31 | 0 | brain | Pcp4l1 |
| Arl4a | 0 | 1.91802239 | 0.75 | 0.24 | 0 | brain | Arl4a |
| Hspb1 | 0 | 1.84289335 | 0.99 | 0.7 | 0 | brain | Hspb1 |
| Mt2 | 0 | 1.80551424 | 0.41 | 0.03 | 0 | brain | Mt2 |
| Lef1 | 0 | 1.80481191 | 0.51 | 0.03 | 0 | brain | Lef1 |
| Mpzl1 | 0 | 1.79420295 | 0.8 | 0.25 | 0 | brain | Mpzl1 |
| Fn1 | 0 | 1.79082277 | 0.56 | 0.06 | 0 | brain | Fn1 |
| Slc38a5 | 0 | 1.77293457 | 0.33 | 0 | 0 | brain | Slc38a5 |
| Foxf2 | 0 | 1.76252906 | 0.48 | 0 | 0 | brain | Foxf2 |
| Paqr5 | 0 | 1.76206506 | 0.5 | 0.03 | 0 | brain | Paqr5 |
| Slc38a3 | 0 | 1.76179512 | 0.43 | 0 | 0 | brain | Slc38a3 |
| Pllp | 0 | 1.7537671 | 0.65 | 0.1 | 0 | brain | Pllp |
| Igf1r1 | 0 | 1.65646651 | 0.54 | 0.1 | 0 | brain | Igf1r |
| Abcb1a | 0 | 1.62795904 | 0.72 | 0.18 | 0 | brain | Abcb1a |
| Abcg21 | 0 | 1.62683335 | 0.81 | 0.33 | 0 | brain | Abcg2 |
| Slc40a1 | 0 | 1.62574135 | 0.46 | 0.04 | 0 | brain | Slc40a1 |
| Alas1 | 0 | 1.61194361 | 0.55 | 0.11 | 0 | brain | Alas1 |
| Ccdc141 | 0 | 1.61001077 | 0.4 | 0.01 | 0 | brain | Ccdc141 |
| Abhd2 | 0 | 1.60147693 | 0.48 | 0.12 | 0 | brain | Abhd2 |
| Itih51 | 0 | 1.60089104 | 0.46 | 0.06 | 0 | brain | Itih5 |
| Mfsd2a | 0 | 1.60035782 | 0.49 | 0.05 | 0 | brain | Mfsd2a |
| Tpd52 | 0 | 1.59522142 | 0.56 | 0.1 | 0 | brain | Tpd52 |
| Gpcpd1 | 0 | 1.59327787 | 0.61 | 0.11 | 0 | brain | Gpcpd1 |
| Slc1a1 | 0 | 1.58012118 | 0.39 | 0.02 | 0 | brain | Slc1a1 |
| Tspan51 | 0 | 1.57841384 | 0.55 | 0.11 | 0 | brain | Tspan5 |
| Slc6a61 | 0 | 1.57012792 | 0.87 | 0.38 | 0 | brain | Slc6a6 |
| Slc39a10 | 0 | 1.50405896 | 0.49 | 0.09 | 0 | brain | Slc39a10 |
| Apold1 | 0 | 1.49365814 | 0.58 | 0.21 | 0 | brain | Apold1 |
| Ablim11 | 0 | 1.49334794 | 0.71 | 0.24 | 0 | brain | Ablim1 |
| Slc16a4 | 0 | 1.49330468 | 0.42 | 0.03 | 0 | brain | Slc16a4 |
| Dusp1 | 0 | 1.44796659 | 0.91 | 0.64 | 0 | brain | Dusp1 |
| Elovl7 | 0 | 1.40657176 | 0.37 | 0.01 | 0 | brain | Elovl7 |
| 9430020K01Rik | 0 | 1.40411221 | 0.59 | 0.17 | 0 | brain | 9430020K01Rik |
| Plat | 0 | 1.38281304 | 0.62 | 0.15 | 0 | brain | Plat |
| Tdrp1 | 0 | 1.3634905 | 0.5 | 0.12 | 0 | brain | Tdrp |
| Edn1 | 0 | 1.33370248 | 0.49 | 0.17 | 0 | brain | Edn1 |

| | | | | | | | |
|---|---|---|---|---|---|---|---|
| Nkd1 | 0 | 1.32364376 | 0.53 | 0.13 | 0 | brain | Nkd1 |
| Arl2bp | 0 | 1.30376726 | 0.69 | 0.26 | 0 | brain | Arl2bp |
| Alpl | 0 | 1.2937717 | 0.39 | 0.08 | 0 | brain | Alpl |
| Gstm7 | 0 | 1.29208307 | 0.32 | 0.02 | 0 | brain | Gstm7 |
| Rad54b | 0 | 1.26611952 | 0.3 | 0 | 0 | brain | Rad54b |
| Nr4a1 | 0 | 1.26037465 | 0.42 | 0.12 | 0 | brain | Nr4a1 |
| Mal | 0 | 1.24770378 | 0.53 | 0.1 | 0 | brain | Mal |
| Maoa | 0 | 1.24045001 | 0.51 | 0.15 | 0 | brain | Maoa |
| Jund | 0 | 1.21863742 | 0.92 | 0.71 | 0 | brain | Jund |
| Gpd2 | 0 | 1.20850333 | 0.43 | 0.09 | 0 | brain | Gpd2 |
| Fxyd5 | 0 | 1.20303577 | 0.8 | 0.45 | 0 | brain | Fxyd5 |
| Zic3 | 0 | 1.19844098 | 0.28 | 0 | 0 | brain | Zic3 |
| Car4 | 0 | 1.19732754 | 0.69 | 0.36 | 0 | brain | Car4 |
| Rasd1 | 0 | 1.19503878 | 0.29 | 0.04 | 0 | brain | Rasd1 |
| Ly6c1 | 0 | 1.19459297 | 1 | 0.84 | 0 | brain | Ly6c1 |
| Osbpl1a | 0 | 1.18653445 | 0.34 | 0.03 | 0 | brain | Osbpl1a |
| St3gal6 | 0 | 1.18227269 | 0.71 | 0.35 | 0 | brain | St3gal6 |
| Ddc | 0 | 1.16444995 | 0.26 | 0 | 0 | brain | Ddc |
| Prnp | 0 | 1.15736893 | 0.53 | 0.17 | 0 | brain | Prnp |
| Eps8l2 | 0 | 1.13027119 | 0.3 | 0.04 | 0 | brain | Eps8l2 |
| Polk | 0 | 1.12689704 | 0.29 | 0.05 | 0 | brain | Polk |
| Grb14 | 0 | 1.12580104 | 0.3 | 0.04 | 0 | brain | Grb14 |
| Aqp11 | 0 | 1.12518633 | 0.32 | 0.04 | 0 | brain | Aqp11 |
| Notum | 0 | 1.11235692 | 0.26 | 0 | 0 | brain | Notum |
| Fam63a | 0 | 1.10220622 | 0.44 | 0.13 | 0 | brain | Fam63a |
| Slc31a1 | 0 | 1.10088821 | 0.34 | 0.08 | 0 | brain | Slc31a1 |
| Ndrg11 | 0 | 1.08883448 | 0.7 | 0.35 | 0 | brain | Ndrg1 |
| Lmo2 | 0 | 1.08666706 | 0.71 | 0.37 | 0 | brain | Lmo2 |
| Ifnar1 | 0 | 1.08326535 | 0.47 | 0.15 | 0 | brain | Ifnar1 |
| Ldlrad3 | 0 | 1.07936046 | 0.34 | 0.06 | 0 | brain | Ldlrad3 |
| Tnfrsf19 | 0 | 1.07861626 | 0.28 | 0.02 | 0 | brain | Tnfrsf19 |
| Tpd52l1 | 0 | 1.07409726 | 0.29 | 0.02 | 0 | brain | Tpd52l1 |
| Car2 | 0 | 1.06574716 | 0.37 | 0.08 | 0 | brain | Car2 |
| Slc16a2 | 0 | 1.06082733 | 0.39 | 0.1 | 0 | brain | Slc16a2 |
| Il10rb | 0 | 1.05848147 | 0.72 | 0.37 | 0 | brain | Il10rb |
| Klf2 | 0 | 1.05682265 | 0.98 | 0.87 | 0 | brain | Klf2 |
| Parvb | 0 | 1.05331081 | 0.59 | 0.26 | 0 | brain | Parvb |
| Gatm | 0 | 1.05327093 | 0.51 | 0.15 | 0 | brain | Gatm |
| Tgfb2 | 0 | 1.05284264 | 0.32 | 0.07 | 0 | brain | Tgfb2 |
| St6galnac2 | 0 | 1.04922851 | 0.43 | 0.14 | 0 | brain | St6galnac2 |
| Hspa1b | 0 | 1.046429 | 0.34 | 0.08 | 0 | brain | Hspa1b |

| | | | | | | | |
|---|---|---|---|---|---|---|---|
| Stmn1 | 0 | 1.04334814 | 0.56 | 0.2 | 0 | brain | Stmn1 |
| Ptprr | 0 | 1.03759469 | 0.45 | 0.15 | 0 | brain | Ptprr |
| Flt11 | 0 | 1.03343518 | 0.96 | 0.68 | 0 | brain | Flt1 |
| Cdkn1a | 0 | 1.03210618 | 0.73 | 0.34 | 0 | brain | Cdkn1a |
| Klf6 | 0 | 1.03199567 | 0.71 | 0.42 | 0 | brain | Klf6 |
| Rhob | 0 | 1.01731723 | 0.8 | 0.53 | 0 | brain | Rhob |
| Afap1l2 | 0 | 1.01479557 | 0.32 | 0.05 | 0 | brain | Afap1l2 |
| Hrct1 | 0 | 1.0053061 | 0.32 | 0.08 | 0 | brain | Hrct1 |
| Cystm1 | 0 | 0.99950808 | 0.32 | 0.05 | 0 | brain | Cystm1 |
| Eogt | 0 | 0.99747861 | 0.43 | 0.14 | 0 | brain | Eogt |
| Tmem252 | 0 | 0.9831402 | 0.37 | 0.11 | 0 | brain | Tmem252 |
| Jun | 0 | 0.98066697 | 0.91 | 0.71 | 0 | brain | Jun |
| Sema3c | 0 | 0.98042615 | 0.57 | 0.15 | 0 | brain | Sema3c |
| Nfkbia | 0 | 0.97875292 | 0.86 | 0.61 | 0 | brain | Nfkbia |
| Slco2b11 | 0 | 0.95225191 | 0.34 | 0.09 | 0 | brain | Slco2b1 |
| Fam13c1 | 0 | 0.94553353 | 0.34 | 0.09 | 0 | brain | Fam13c |
| Tob1 | 0 | 0.94514776 | 0.43 | 0.16 | 0 | brain | Tob1 |
| Usp53 | 0 | 0.94378582 | 0.25 | 0.03 | 0 | brain | Usp53 |
| Csrp2 | 0 | 0.93659692 | 0.48 | 0.2 | 0 | brain | Csrp2 |
| Fry1 | 0 | 0.930962 | 0.42 | 0.15 | 0 | brain | Fry |
| Golim41 | 0 | 0.9280575 | 0.39 | 0.13 | 0 | brain | Golim4 |
| Sox17 | 0 | 0.9280092 | 0.74 | 0.36 | 0 | brain | Sox17 |
| Rgs12 | 0 | 0.9264604 | 0.4 | 0.13 | 0 | brain | Rgs12 |
| Slc3a2 | 0 | 0.92136677 | 0.81 | 0.52 | 0 | brain | Slc3a2 |
| Cd320 | 0 | 0.90765147 | 0.31 | 0.08 | 0 | brain | Cd320 |
| Plpp6 | 0 | 0.90357139 | 0.31 | 0.08 | 0 | brain | Plpp6 |
| Pir | 0 | 0.90231225 | 0.3 | 0.08 | 0 | brain | Pir |
| Esyt21 | 0 | 0.86689919 | 0.38 | 0.13 | 0 | brain | Esyt2 |
| Tmem123 | 0 | 0.86187468 | 0.55 | 0.26 | 0 | brain | Tmem123 |
| Cd34 | 0 | 0.85363817 | 0.86 | 0.53 | 0 | brain | Cd34 |
| Pkig | 0 | 0.84654168 | 0.51 | 0.24 | 0 | brain | Pkig |
| Hbegf | 0 | 0.84425133 | 0.31 | 0.1 | 0 | brain | Hbegf |
| Tuba1a | 0 | 0.84249912 | 0.86 | 0.63 | 0 | brain | Tuba1a |
| Gch1 | 0 | 0.83150997 | 0.37 | 0.13 | 0 | brain | Gch1 |
| Slc52a3 | 0 | 0.82779308 | 0.27 | 0.06 | 0 | brain | Slc52a3 |
| Hprt | 0 | 0.81972959 | 0.64 | 0.37 | 0 | brain | Hprt |
| Sema7a1 | 0 | 0.80146986 | 0.45 | 0.18 | 0 | brain | Sema7a |
| Fzd6 | 0 | 0.79425666 | 0.31 | 0.1 | 0 | brain | Fzd6 |
| Myl12b | 0 | 0.77450395 | 0.86 | 0.65 | 0 | brain | Myl12b |
| Pomp | 0 | 0.7686384 | 0.79 | 0.55 | 0 | brain | Pomp |
| Atox1 | 0 | 0.75599126 | 0.96 | 0.78 | 0 | brain | Atox1 |

| | | | | | | | |
|---|---|---|---|---|---|---|---|
| Fcgrt | 0 | 0.75524726 | 0.76 | 0.45 | 0 | brain | Fcgrt |
| Slc39a8 | 0 | 0.72883755 | 0.27 | 0.08 | 0 | brain | Slc39a8 |
| Tspan13 | 0 | 0.70749823 | 0.93 | 0.61 | 0 | brain | Tspan13 |
| Cxcl121 | 0 | 0.70100449 | 0.82 | 0.47 | 0 | brain | Cxcl12 |
| Efr3b1 | 0 | 0.69626919 | 0.25 | 0.07 | 0 | brain | Efr3b |
| Snx3 | 0 | 0.69613474 | 0.87 | 0.65 | 0 | brain | Snx3 |
| Ly6a | 0 | 0.66572771 | 0.99 | 0.87 | 0 | brain | Ly6a |
| Hspa8 | 0 | 0.63308981 | 0.98 | 0.9 | 0 | brain | Hspa8 |
| Rpl38 | 0 | 0.6187847 | 0.99 | 0.92 | 0 | brain | Rpl38 |
| Acvrl1 | 0 | 0.50081423 | 0.8 | 0.47 | 0 | brain | Acvrl1 |
| Ap3s1 | 1.24E-306 | 0.82455541 | 0.44 | 0.2 | 2.56E-302 | brain | Ap3s1 |
| Nampt | 8.30E-295 | 0.91900514 | 0.35 | 0.14 | 1.72E-290 | brain | Nampt |
| Pls3 | 3.73E-294 | 0.83185181 | 0.6 | 0.34 | 7.72E-290 | brain | Pls3 |
| Arf6 | 3.77E-291 | 0.773872 | 0.52 | 0.26 | 7.79E-287 | brain | Arf6 |
| Tsc22d3 | 4.51E-291 | 0.92865108 | 0.62 | 0.37 | 9.32E-287 | brain | Tsc22d3 |
| 1110034G24Rik | 6.73E-290 | 0.76652575 | 0.27 | 0.09 | 1.39E-285 | brain | 1110034G24Rik |
| 1810037I17Rik | 3.56E-286 | 0.76354633 | 0.77 | 0.55 | 7.37E-282 | brain | 1810037I17Rik |
| Gpr4 | 4.28E-285 | 0.7385813 | 0.37 | 0.15 | 8.85E-281 | brain | Gpr4 |
| Anxa3 | 9.40E-284 | 0.69255289 | 0.77 | 0.51 | 1.95E-279 | brain | Anxa3 |
| Faim | 3.44E-282 | 0.7580705 | 0.31 | 0.11 | 7.12E-278 | brain | Faim |
| Esam | 5.54E-281 | 0.59347984 | 0.93 | 0.72 | 1.15E-276 | brain | Esam |
| Ndufa7 | 1.22E-278 | 0.70538618 | 0.82 | 0.6 | 2.53E-274 | brain | Ndufa7 |
| Bnip3l | 1.25E-278 | 0.72745295 | 0.73 | 0.49 | 2.58E-274 | brain | Bnip3l |
| Ppp1cb | 2.75E-277 | 0.78203621 | 0.57 | 0.32 | 5.69E-273 | brain | Ppp1cb |
| Aspscr1 | 3.74E-276 | 0.72690182 | 0.3 | 0.1 | 7.75E-272 | brain | Aspscr1 |
| Max | 8.76E-275 | 0.79592344 | 0.41 | 0.18 | 1.81E-270 | brain | Max |
| Grap | 4.47E-272 | 0.75821719 | 0.57 | 0.3 | 9.25E-268 | brain | Grap |
| Fkbp4 | 7.52E-269 | 0.75039453 | 0.59 | 0.34 | 1.56E-264 | brain | Fkbp4 |
| Amd1 | 9.21E-269 | 0.83122039 | 0.31 | 0.11 | 1.91E-264 | brain | Amd1 |
| Klf4 | 1.24E-264 | 0.71755561 | 0.82 | 0.57 | 2.57E-260 | brain | Klf4 |
| Ldha | 3.71E-262 | 0.75125716 | 0.8 | 0.56 | 7.67E-258 | brain | Ldha |
| Btg2 | 1.06E-261 | 0.74528325 | 0.73 | 0.47 | 2.18E-257 | brain | Btg2 |
| Capg | 1.21E-260 | 0.65559107 | 0.28 | 0.1 | 2.50E-256 | brain | Capg |
| Ywhaq | 2.01E-259 | 0.64033473 | 0.83 | 0.62 | 4.16E-255 | brain | Ywhaq |
| Pik3ip1 | 2.13E-259 | 0.7178202 | 0.31 | 0.12 | 4.41E-255 | brain | Pik3ip1 |
| Cgnl1 | 1.04E-258 | 0.69277035 | 0.32 | 0.12 | 2.16E-254 | brain | Cgnl1 |
| Gltp | 1.70E-256 | 0.71610387 | 0.49 | 0.25 | 3.52E-252 | brain | Gltp |
| Sparcl11 | 2.11E-255 | 0.52498444 | 0.83 | 0.47 | 4.36E-251 | brain | Sparcl1 |
| Smox | 3.62E-250 | 0.74065232 | 0.27 | 0.09 | 7.50E-246 | brain | Smox |
| Pon2 | 1.88E-247 | 0.71896622 | 0.54 | 0.3 | 3.88E-243 | brain | Pon2 |
| Pop5 | 2.21E-247 | 0.7542305 | 0.47 | 0.24 | 4.56E-243 | brain | Pop5 |

| | | | | | | | |
|---|---|---|---|---|---|---|---|
| Atf3 | 2.92E-247 | 0.68095495 | 0.4 | 0.17 | 6.04E-243 | brain | Atf3 |
| Gnas | 8.64E-247 | 0.43177447 | 0.97 | 0.9 | 1.79E-242 | brain | Gnas |
| Sh3bp4 | 2.91E-245 | 0.61807801 | 0.25 | 0.09 | 6.03E-241 | brain | Sh3bp4 |
| Nos3 | 8.92E-245 | 0.7042199 | 0.4 | 0.19 | 1.85E-240 | brain | Nos3 |
| Fam129b | 5.07E-243 | 0.72006209 | 0.38 | 0.17 | 1.05E-238 | brain | Fam129b |
| Frmd4b | 5.17E-242 | 0.69084322 | 0.38 | 0.17 | 1.07E-237 | brain | Frmd4b |
| Cyr61 | 1.47E-237 | 1.12752906 | 0.48 | 0.25 | 3.05E-233 | brain | Cyr61 |
| Pcmtd11 | 7.59E-234 | 0.77866521 | 0.5 | 0.28 | 1.57E-229 | brain | Pcmtd1 |
| Gadd45g | 9.93E-227 | 1.07659859 | 0.51 | 0.27 | 2.06E-222 | brain | Gadd45g |
| Pkm | 1.18E-225 | 0.66933099 | 0.73 | 0.51 | 2.43E-221 | brain | Pkm |
| Ak3 | 6.49E-225 | 0.68668636 | 0.32 | 0.13 | 1.34E-220 | brain | Ak3 |
| Ucp2 | 3.53E-222 | 0.68016263 | 0.47 | 0.24 | 7.31E-218 | brain | Ucp2 |
| Actg1 | 1.27E-220 | 0.47880015 | 0.97 | 0.88 | 2.63E-216 | brain | Actg1 |
| App1 | 1.83E-220 | 0.56640981 | 0.89 | 0.73 | 3.79E-216 | brain | App |
| Htra1 | 1.99E-216 | 0.5193011 | 0.26 | 0.09 | 4.12E-212 | brain | Htra1 |
| Palm | 1.51E-215 | 0.65348402 | 0.44 | 0.22 | 3.12E-211 | brain | Palm |
| Slc30a1 | 4.21E-214 | 0.65161232 | 0.29 | 0.11 | 8.71E-210 | brain | Slc30a1 |
| Bcl2l1 | 1.12E-212 | 0.65840899 | 0.42 | 0.21 | 2.32E-208 | brain | Bcl2l1 |
| Palmd1 | 2.48E-210 | 0.69169616 | 0.5 | 0.27 | 5.14E-206 | brain | Palmd |
| Slc16a1 | 7.53E-210 | 0.87106245 | 0.28 | 0.11 | 1.56E-205 | brain | Slc16a1 |
| Mecom1 | 1.93E-200 | 0.49865504 | 0.43 | 0.2 | 3.99E-196 | brain | Mecom |
| Crem | 6.39E-198 | 0.63489099 | 0.3 | 0.13 | 1.32E-193 | brain | Crem |
| Ier2 | 1.74E-197 | 0.42263084 | 0.79 | 0.55 | 3.60E-193 | brain | Ier2 |
| Fut81 | 2.65E-197 | 0.58412543 | 0.27 | 0.11 | 5.47E-193 | brain | Fut8 |
| Fos | 3.61E-196 | 0.49518639 | 0.8 | 0.55 | 7.48E-192 | brain | Fos |
| Slc9a3r2 | 2.10E-191 | 0.44538006 | 0.94 | 0.81 | 4.34E-187 | brain | Slc9a3r2 |
| Hsp90aa1 | 5.38E-189 | 0.73223105 | 0.65 | 0.45 | 1.11E-184 | brain | Hsp90aa1 |
| Eno1 | 3.06E-188 | 0.65202506 | 0.53 | 0.32 | 6.32E-184 | brain | Eno1 |
| Sqstm1 | 2.01E-186 | 0.62847268 | 0.56 | 0.36 | 4.16E-182 | brain | Sqstm1 |
| Afap1l1 | 3.79E-185 | 0.56969303 | 0.57 | 0.34 | 7.84E-181 | brain | Afap1l1 |
| Ywhae | 1.07E-183 | 0.5163453 | 0.82 | 0.64 | 2.21E-179 | brain | Ywhae |
| Pde2a | 2.83E-183 | 0.50317946 | 0.48 | 0.25 | 5.85E-179 | brain | Pde2a |
| Hist1h2bc | 2.50E-179 | 0.54246502 | 0.52 | 0.3 | 5.17E-175 | brain | Hist1h2bc |
| Nes | 2.38E-178 | 0.58919174 | 0.35 | 0.17 | 4.93E-174 | brain | Nes |
| Stard3nl | 7.72E-177 | 0.61943973 | 0.35 | 0.17 | 1.60E-172 | brain | Stard3nl |
| Uqcc2 | 8.63E-177 | 0.56818274 | 0.61 | 0.4 | 1.79E-172 | brain | Uqcc2 |
| Nostrin1 | 1.10E-176 | 0.6679663 | 0.49 | 0.28 | 2.28E-172 | brain | Nostrin |
| Kctd10 | 1.58E-176 | 0.60651436 | 0.53 | 0.32 | 3.27E-172 | brain | Kctd10 |
| Cltb | 1.05E-174 | 0.57619105 | 0.62 | 0.41 | 2.16E-170 | brain | Cltb |
| Ap1s2 | 4.55E-172 | 0.5929726 | 0.34 | 0.16 | 9.42E-168 | brain | Ap1s2 |
| Cdkn1c | 5.71E-172 | 0.6365748 | 0.42 | 0.22 | 1.18E-167 | brain | Cdkn1c |

| | | | | | | | |
|---|---|---|---|---|---|---|---|
| Adgrl41 | 1.43E-171 | 0.42105916 | 0.75 | 0.5 | 2.95E-167 | brain | Adgrl4 |
| Map1lc3a | 3.23E-170 | 0.52922637 | 0.64 | 0.43 | 6.69E-166 | brain | Map1lc3a |
| Calm11 | 4.78E-170 | 0.39286269 | 0.97 | 0.9 | 9.89E-166 | brain | Calm1 |
| Utrn1 | 6.38E-170 | 0.54714777 | 0.48 | 0.27 | 1.32E-165 | brain | Utrn |
| Gbp3 | 1.02E-164 | 0.58289245 | 0.3 | 0.14 | 2.10E-160 | brain | Gbp3 |
| Dynll1 | 5.25E-164 | 0.40223347 | 0.94 | 0.84 | 1.09E-159 | brain | Dynll1 |
| Cnbp | 5.76E-164 | 0.48425879 | 0.83 | 0.67 | 1.19E-159 | brain | Cnbp |
| Hspa12b | 5.06E-163 | 0.58518474 | 0.44 | 0.25 | 1.05E-158 | brain | Hspa12b |
| Myl6 | 3.48E-161 | 0.32904104 | 0.97 | 0.9 | 7.20E-157 | brain | Myl6 |
| B4galt41 | 1.48E-159 | 0.56886647 | 0.42 | 0.23 | 3.06E-155 | brain | B4galt4 |
| Crip2 | 5.96E-158 | 0.36195245 | 0.95 | 0.85 | 1.23E-153 | brain | Crip2 |
| Ifnar2 | 1.10E-157 | 0.57486752 | 0.52 | 0.33 | 2.27E-153 | brain | Ifnar2 |
| Gata2 | 1.10E-156 | 0.52480048 | 0.5 | 0.3 | 2.28E-152 | brain | Gata2 |
| Ly6e | 1.52E-156 | 0.29124621 | 0.99 | 0.94 | 3.14E-152 | brain | Ly6e |
| Akr1a1 | 1.20E-155 | 0.48922384 | 0.78 | 0.6 | 2.49E-151 | brain | Akr1a1 |
| Mgp | 3.41E-155 | 0.60062208 | 0.37 | 0.18 | 7.06E-151 | brain | Mgp |
| C130074G19Rik1 | 4.04E-155 | 0.57230279 | 0.36 | 0.19 | 8.37E-151 | brain | C130074G19Rik |
| Arl6ip5 | 3.60E-154 | 0.48567438 | 0.7 | 0.5 | 7.45E-150 | brain | Arl6ip5 |
| Mff | 1.71E-151 | 0.58569424 | 0.45 | 0.27 | 3.53E-147 | brain | Mff |
| H3f3b | 1.79E-151 | 0.32441898 | 0.96 | 0.88 | 3.70E-147 | brain | H3f3b |
| Sorbs21 | 4.15E-151 | 0.43845376 | 0.46 | 0.25 | 8.59E-147 | brain | Sorbs2 |
| Osbpl91 | 9.23E-150 | 0.59555449 | 0.46 | 0.27 | 1.91E-145 | brain | Osbpl9 |
| Tbx3 | 1.68E-149 | 0.41682933 | 0.3 | 0.14 | 3.47E-145 | brain | Tbx3 |
| Mtus1 | 1.89E-149 | 0.56035093 | 0.42 | 0.24 | 3.92E-145 | brain | Mtus1 |
| Ubc | 1.10E-148 | 0.45246633 | 0.92 | 0.83 | 2.27E-144 | brain | Ubc |
| Gpx1 | 1.68E-147 | 0.55915167 | 0.72 | 0.54 | 3.47E-143 | brain | Gpx1 |
| Tpm1 | 7.69E-142 | 0.49358712 | 0.53 | 0.34 | 1.59E-137 | brain | Tpm1 |
| Gclm | 1.81E-141 | 0.5453316 | 0.32 | 0.17 | 3.75E-137 | brain | Gclm |
| Id3 | 1.10E-137 | 0.37970096 | 0.96 | 0.84 | 2.28E-133 | brain | Id3 |
| Kank3 | 2.19E-137 | 0.48772286 | 0.73 | 0.55 | 4.53E-133 | brain | Kank3 |
| Mospd1 | 7.38E-136 | 0.53149962 | 0.26 | 0.12 | 1.53E-131 | brain | Mospd1 |
| Cd59a | 4.74E-134 | 0.32673722 | 0.4 | 0.21 | 9.80E-130 | brain | Cd59a |
| Rab18 | 8.36E-134 | 0.48552146 | 0.58 | 0.4 | 1.73E-129 | brain | Rab18 |
| Grrp11 | 2.00E-133 | 0.48571599 | 0.53 | 0.33 | 4.13E-129 | brain | Grrp1 |
| Map4k2 | 9.19E-132 | 0.46748791 | 0.27 | 0.13 | 1.90E-127 | brain | Map4k2 |
| BC028528 | 1.80E-131 | 0.30620657 | 0.66 | 0.43 | 3.73E-127 | brain | BC028528 |
| Junb | 7.94E-130 | 0.42207225 | 0.81 | 0.61 | 1.64E-125 | brain | Junb |
| Ubl3 | 1.15E-129 | 0.52904999 | 0.49 | 0.31 | 2.38E-125 | brain | Ubl3 |
| Bhlhe40 | 8.85E-129 | 0.58379395 | 0.32 | 0.17 | 1.83E-124 | brain | Bhlhe40 |
| Adamts1 | 4.05E-128 | 0.635528 | 0.39 | 0.22 | 8.39E-124 | brain | Adamts1 |
| Myl12a | 5.75E-127 | 0.35472694 | 0.93 | 0.83 | 1.19E-122 | brain | Myl12a |

| | | | | | | | |
|---|---|---|---|---|---|---|---|
| Hnrnpf | 1.21E-124 | 0.39594543 | 0.79 | 0.64 | 2.50E-120 | brain | Hnrnpf |
| 8430408G22Rik | 3.81E-123 | 0.66580572 | 0.41 | 0.23 | 7.88E-119 | brain | 8430408G22Rik |
| Smpdl3a | 4.75E-123 | 0.49498349 | 0.51 | 0.33 | 9.84E-119 | brain | Smpdl3a |
| Id1 | 1.20E-122 | 0.40698948 | 0.83 | 0.64 | 2.48E-118 | brain | Id1 |
| Eif3f | 4.11E-121 | 0.43105909 | 0.77 | 0.6 | 8.51E-117 | brain | Eif3f |
| Rapgef41 | 5.01E-121 | 0.36756834 | 0.42 | 0.24 | 1.04E-116 | brain | Rapgef4 |
| Lmo7 | 5.45E-121 | 0.50326798 | 0.28 | 0.14 | 1.13E-116 | brain | Lmo7 |
| Hsp90ab1 | 8.59E-121 | 0.30405751 | 0.97 | 0.91 | 1.78E-116 | brain | Hsp90ab1 |
| Gng5 | 1.94E-119 | 0.37758013 | 0.88 | 0.75 | 4.02E-115 | brain | Gng5 |
| Cxx1a | 6.30E-119 | 0.3631402 | 0.67 | 0.44 | 1.30E-114 | brain | Cxx1a |
| Stom | 7.54E-119 | 0.48128842 | 0.3 | 0.16 | 1.56E-114 | brain | Stom |
| Tprgl | 3.97E-117 | 0.41278507 | 0.7 | 0.53 | 8.22E-113 | brain | Tprgl |
| Thsd1 | 1.88E-114 | 0.48860188 | 0.3 | 0.16 | 3.89E-110 | brain | Thsd1 |
| Rtn3 | 2.22E-114 | 0.45090458 | 0.55 | 0.38 | 4.58E-110 | brain | Rtn3 |
| Uaca1 | 4.95E-114 | 0.45331822 | 0.37 | 0.21 | 1.02E-109 | brain | Uaca |
| Eif4a2 | 4.75E-112 | 0.41549937 | 0.73 | 0.57 | 9.83E-108 | brain | Eif4a2 |
| Ctnnb1 | 3.67E-111 | 0.40840021 | 0.69 | 0.51 | 7.60E-107 | brain | Ctnnb1 |
| Nxpe41 | 1.56E-110 | 0.44656187 | 0.28 | 0.14 | 3.23E-106 | brain | Nxpe4 |
| Cct7 | 3.31E-110 | 0.44717309 | 0.53 | 0.37 | 6.85E-106 | brain | Cct7 |
| Gm10076 | 5.39E-109 | 0.3307201 | 0.65 | 0.47 | 1.11E-104 | brain | Gm10076 |
| S1pr1 | 6.90E-106 | 0.3874253 | 0.63 | 0.45 | 1.43E-101 | brain | S1pr1 |
| Sigirr | 5.88E-105 | 0.44401373 | 0.3 | 0.17 | 1.22E-100 | brain | Sigirr |
| Gabarapl2 | 5.62E-104 | 0.39438904 | 0.69 | 0.54 | 1.16E-99 | brain | Gabarapl2 |
| Dap | 1.81E-102 | 0.48678424 | 0.26 | 0.14 | 3.75E-98 | brain | Dap |
| Bnip2 | 2.07E-102 | 0.46893924 | 0.54 | 0.38 | 4.27E-98 | brain | Bnip2 |
| Hey1 | 4.56E-102 | 0.41910262 | 0.27 | 0.14 | 9.44E-98 | brain | Hey1 |
| St6galnac31 | 2.06E-100 | 0.41522245 | 0.35 | 0.21 | 4.27E-96 | brain | St6galnac3 |
| Tmbim6 | 6.18E-100 | 0.33331382 | 0.83 | 0.69 | 1.28E-95 | brain | Tmbim6 |
| Plk2 | 1.15E-99 | 0.6154304 | 0.57 | 0.42 | 2.38E-95 | brain | Plk2 |
| Ifit1 | 1.78E-98 | 0.48362427 | 0.26 | 0.14 | 3.69E-94 | brain | Ifit1 |
| Taldo1 | 2.17E-98 | 0.42307456 | 0.55 | 0.4 | 4.49E-94 | brain | Taldo1 |
| Eif4h | 2.83E-96 | 0.41116426 | 0.55 | 0.4 | 5.85E-92 | brain | Eif4h |
| Tuba1b | 1.40E-95 | 0.3431836 | 0.82 | 0.67 | 2.90E-91 | brain | Tuba1b |
| Spr | 6.48E-95 | 0.44133493 | 0.42 | 0.28 | 1.34E-90 | brain | Spr |
| Srp14 | 2.31E-94 | 0.38616432 | 0.65 | 0.5 | 4.78E-90 | brain | Srp14 |
| Capns1 | 3.50E-91 | 0.35431958 | 0.77 | 0.63 | 7.23E-87 | brain | Capns1 |
| Trim47 | 4.05E-91 | 0.35088651 | 0.49 | 0.32 | 8.39E-87 | brain | Trim47 |
| Zfand6 | 1.74E-90 | 0.40023259 | 0.47 | 0.32 | 3.59E-86 | brain | Zfand6 |
| Tmem184b | 2.83E-89 | 0.41579276 | 0.35 | 0.21 | 5.85E-85 | brain | Tmem184b |
| Zfp36 | 1.84E-88 | 0.25488438 | 0.58 | 0.41 | 3.81E-84 | brain | Zfp36 |
| Irf7 | 6.90E-88 | 0.3728411 | 0.33 | 0.19 | 1.43E-83 | brain | Irf7 |

| | | | | | | | |
|---|---|---|---|---|---|---|---|
| Kif5b | 1.14E-87 | 0.41084202 | 0.58 | 0.43 | 2.35E-83 | brain | Kif5b |
| Gde1 | 2.42E-86 | 0.44151864 | 0.29 | 0.17 | 5.02E-82 | brain | Gde1 |
| Prkar1a | 3.76E-86 | 0.36623564 | 0.7 | 0.55 | 7.79E-82 | brain | Prkar1a |
| Sri | 1.17E-85 | 0.37548628 | 0.63 | 0.49 | 2.42E-81 | brain | Sri |
| Gpi1 | 1.18E-84 | 0.41443628 | 0.51 | 0.37 | 2.44E-80 | brain | Gpi1 |
| Cacybp | 3.67E-84 | 0.41316011 | 0.42 | 0.28 | 7.60E-80 | brain | Cacybp |
| Uap1 | 1.05E-82 | 0.40986896 | 0.26 | 0.15 | 2.18E-78 | brain | Uap1 |
| Epas1 | 3.40E-81 | 0.30223865 | 0.79 | 0.64 | 7.04E-77 | brain | Epas1 |
| Gnai2 | 6.00E-80 | 0.2747038 | 0.91 | 0.8 | 1.24E-75 | brain | Gnai2 |
| Cxx1b | 1.98E-79 | 0.33072872 | 0.46 | 0.3 | 4.10E-75 | brain | Cxx1b |
| Psma6 | 3.69E-79 | 0.40916562 | 0.5 | 0.36 | 7.63E-75 | brain | Psma6 |
| Atp2c1 | 1.14E-78 | 0.41189161 | 0.29 | 0.17 | 2.35E-74 | brain | Atp2c1 |
| Isg15 | 7.36E-78 | 0.40625874 | 0.44 | 0.3 | 1.52E-73 | brain | Isg15 |
| Pttg1ip | 1.62E-77 | 0.36259563 | 0.55 | 0.41 | 3.34E-73 | brain | Pttg1ip |
| Cd2ap1 | 1.72E-77 | 0.41606297 | 0.37 | 0.24 | 3.57E-73 | brain | Cd2ap |
| Rasgrp2 | 6.47E-77 | 0.33033066 | 0.32 | 0.19 | 1.34E-72 | brain | Rasgrp2 |
| Anxa7 | 3.34E-76 | 0.38388118 | 0.42 | 0.29 | 6.92E-72 | brain | Anxa7 |
| Peli1 | 6.12E-76 | 0.41572197 | 0.27 | 0.16 | 1.27E-71 | brain | Peli1 |
| Gabarapl1 | 1.19E-75 | 0.35957765 | 0.42 | 0.28 | 2.47E-71 | brain | Gabarapl1 |
| Capn2 | 1.35E-75 | 0.38462289 | 0.45 | 0.32 | 2.79E-71 | brain | Capn2 |
| Mif4gd | 1.43E-74 | 0.38928355 | 0.25 | 0.15 | 2.96E-70 | brain | Mif4gd |
| Dnajb1 | 4.49E-74 | 0.47292179 | 0.31 | 0.19 | 9.29E-70 | brain | Dnajb1 |
| Cdc42 | 6.19E-73 | 0.28984111 | 0.84 | 0.72 | 1.28E-68 | brain | Cdc42 |
| Ctsa | 6.32E-73 | 0.37349404 | 0.32 | 0.21 | 1.31E-68 | brain | Ctsa |
| Pfkl | 7.21E-73 | 0.39465198 | 0.26 | 0.15 | 1.49E-68 | brain | Pfkl |
| Dnajb9 | 2.29E-72 | 0.4101448 | 0.3 | 0.18 | 4.73E-68 | brain | Dnajb9 |
| Impdh1 | 3.64E-71 | 0.29134725 | 0.38 | 0.25 | 7.53E-67 | brain | Impdh1 |
| Rnaset2b | 3.90E-71 | 0.3116253 | 0.52 | 0.37 | 8.06E-67 | brain | Rnaset2b |
| Vamp4 | 1.22E-70 | 0.38964444 | 0.26 | 0.15 | 2.52E-66 | brain | Vamp4 |
| Syf2 | 1.37E-70 | 0.35396699 | 0.52 | 0.4 | 2.84E-66 | brain | Syf2 |
| Tpm4 | 1.77E-70 | 0.27560247 | 0.8 | 0.67 | 3.66E-66 | brain | Tpm4 |
| Ecm1 | 4.71E-70 | 0.352102 | 0.3 | 0.18 | 9.74E-66 | brain | Ecm1 |
| Hnrnpa2b1 | 5.05E-68 | 0.30074345 | 0.77 | 0.65 | 1.05E-63 | brain | Hnrnpa2b1 |
| Dnaja1 | 6.52E-68 | 0.3018666 | 0.78 | 0.65 | 1.35E-63 | brain | Dnaja1 |
| Tmed10 | 7.09E-68 | 0.30880164 | 0.71 | 0.57 | 1.47E-63 | brain | Tmed10 |
| Grasp | 1.72E-67 | 0.39177976 | 0.38 | 0.26 | 3.56E-63 | brain | Grasp |
| Med28 | 3.56E-67 | 0.33346073 | 0.54 | 0.42 | 7.37E-63 | brain | Med28 |
| Puf60 | 6.95E-67 | 0.35047482 | 0.42 | 0.3 | 1.44E-62 | brain | Puf60 |
| Stx12 | 2.45E-66 | 0.34023287 | 0.41 | 0.29 | 5.08E-62 | brain | Stx12 |
| Ssbp4 | 7.89E-66 | 0.36047955 | 0.4 | 0.28 | 1.63E-61 | brain | Ssbp4 |
| Rrp1 | 1.02E-65 | 0.37559543 | 0.41 | 0.29 | 2.11E-61 | brain | Rrp1 |

| | | | | | | | |
|---|---|---|---|---|---|---|---|
| H2afv | 8.97E-65 | 0.3331251 | 0.53 | 0.41 | 1.86E-60 | brain | H2afv |
| Sec11c | 2.45E-64 | 0.3698441 | 0.26 | 0.16 | 5.06E-60 | brain | Sec11c |
| Aes | 3.15E-64 | 0.31773977 | 0.63 | 0.5 | 6.52E-60 | brain | Aes |
| Adipor1 | 4.11E-64 | 0.27146063 | 0.59 | 0.45 | 8.50E-60 | brain | Adipor1 |
| Maea | 7.20E-63 | 0.37784966 | 0.28 | 0.18 | 1.49E-58 | brain | Maea |
| Txn2 | 6.03E-62 | 0.36262495 | 0.4 | 0.29 | 1.25E-57 | brain | Txn2 |
| Ypel3 | 3.45E-61 | 0.29210649 | 0.56 | 0.42 | 7.14E-57 | brain | Ypel3 |
| Tcf3 | 5.07E-61 | 0.34769437 | 0.28 | 0.18 | 1.05E-56 | brain | Tcf3 |
| Fam63b | 7.99E-61 | 0.30837365 | 0.25 | 0.15 | 1.65E-56 | brain | Fam63b |
| Vamp3 | 1.35E-60 | 0.32373118 | 0.43 | 0.31 | 2.79E-56 | brain | Vamp3 |
| Pnpla2 | 3.30E-60 | 0.33477369 | 0.3 | 0.19 | 6.83E-56 | brain | Pnpla2 |
| Pink1 | 1.20E-59 | 0.34396617 | 0.36 | 0.24 | 2.49E-55 | brain | Pink1 |
| Zfp36l11 | 1.64E-59 | 0.32286422 | 0.66 | 0.53 | 3.39E-55 | brain | Zfp36l1 |
| Polr2m | 2.33E-59 | 0.34414851 | 0.4 | 0.28 | 4.83E-55 | brain | Polr2m |
| Fnta | 6.70E-59 | 0.34455113 | 0.31 | 0.21 | 1.39E-54 | brain | Fnta |
| Sypl | 1.59E-58 | 0.28576745 | 0.64 | 0.52 | 3.29E-54 | brain | Sypl |
| Pitpna | 1.67E-58 | 0.33440865 | 0.47 | 0.36 | 3.46E-54 | brain | Pitpna |
| Higd2a | 3.92E-58 | 0.31120867 | 0.54 | 0.42 | 8.12E-54 | brain | Higd2a |
| Tm6sf1 | 1.24E-57 | 0.29647421 | 0.29 | 0.18 | 2.56E-53 | brain | Tm6sf1 |
| Ube2v2 | 2.35E-57 | 0.35471864 | 0.26 | 0.17 | 4.86E-53 | brain | Ube2v2 |
| Gdi1 | 9.15E-57 | 0.32301187 | 0.3 | 0.2 | 1.89E-52 | brain | Gdi1 |
| Agfg1 | 1.12E-56 | 0.3557755 | 0.25 | 0.16 | 2.31E-52 | brain | Agfg1 |
| Ctbp2 | 1.24E-56 | 0.35076146 | 0.27 | 0.17 | 2.56E-52 | brain | Ctbp2 |
| Lsm14a | 6.25E-56 | 0.30497755 | 0.31 | 0.21 | 1.29E-51 | brain | Lsm14a |
| Irf9 | 1.36E-55 | 0.353239 | 0.27 | 0.17 | 2.81E-51 | brain | Irf9 |
| Skp1a | 1.96E-55 | 0.31476456 | 0.45 | 0.34 | 4.06E-51 | brain | Skp1a |
| Higd1a | 2.76E-55 | 0.32641616 | 0.42 | 0.31 | 5.71E-51 | brain | Higd1a |
| Mkl21 | 1.28E-54 | 0.32975836 | 0.32 | 0.21 | 2.64E-50 | brain | Mkl2 |
| Chmp5 | 2.99E-54 | 0.29892718 | 0.53 | 0.42 | 6.18E-50 | brain | Chmp5 |
| Ywhah | 1.47E-53 | 0.25259079 | 0.67 | 0.55 | 3.03E-49 | brain | Ywhah |
| Mlf2 | 2.89E-53 | 0.30359259 | 0.48 | 0.37 | 5.99E-49 | brain | Mlf2 |
| 4833439L19Rik | 4.72E-53 | 0.33638243 | 0.27 | 0.17 | 9.77E-49 | brain | 4833439L19Rik |
| Dnm2 | 6.10E-53 | 0.3165145 | 0.37 | 0.26 | 1.26E-48 | brain | Dnm2 |
| Rheb | 1.04E-52 | 0.27761701 | 0.58 | 0.47 | 2.15E-48 | brain | Rheb |
| Sec62 | 1.68E-52 | 0.27832783 | 0.7 | 0.59 | 3.48E-48 | brain | Sec62 |
| Ostf1 | 1.45E-51 | 0.28674556 | 0.68 | 0.56 | 3.00E-47 | brain | Ostf1 |
| Rexo2 | 7.54E-50 | 0.32466489 | 0.42 | 0.32 | 1.56E-45 | brain | Rexo2 |
| Ccnd3 | 1.72E-49 | 0.27036606 | 0.42 | 0.3 | 3.56E-45 | brain | Ccnd3 |
| Ctbp1 | 1.76E-49 | 0.28669483 | 0.45 | 0.34 | 3.65E-45 | brain | Ctbp1 |
| Copz2 | 2.42E-49 | 0.30085513 | 0.28 | 0.18 | 5.01E-45 | brain | Copz2 |
| Tmem9b | 3.09E-49 | 0.29061623 | 0.38 | 0.28 | 6.39E-45 | brain | Tmem9b |

| | | | | | | | |
|---|---|---|---|---|---|---|---|
| Hmgn1 | 4.24E-49 | 0.28416005 | 0.45 | 0.34 | 8.78E-45 | brain | Hmgn1 |
| Znrf1 | 2.18E-48 | 0.28065851 | 0.35 | 0.25 | 4.50E-44 | brain | Znrf1 |
| Bbip1 | 3.15E-48 | 0.30548409 | 0.25 | 0.16 | 6.52E-44 | brain | Bbip1 |
| Map7d1 | 3.18E-48 | 0.25291781 | 0.47 | 0.36 | 6.58E-44 | brain | Map7d1 |
| N4bp3 | 4.75E-48 | 0.26405754 | 0.3 | 0.2 | 9.83E-44 | brain | N4bp3 |
| Smim14 | 5.90E-48 | 0.27030958 | 0.52 | 0.41 | 1.22E-43 | brain | Smim14 |
| M6pr | 1.91E-47 | 0.32304682 | 0.34 | 0.25 | 3.94E-43 | brain | M6pr |
| Cct5 | 5.26E-47 | 0.31282489 | 0.44 | 0.34 | 1.09E-42 | brain | Cct5 |
| Ikbkb | 1.71E-46 | 0.26398812 | 0.31 | 0.21 | 3.54E-42 | brain | Ikbkb |
| Stra13 | 7.25E-46 | 0.2909095 | 0.32 | 0.22 | 1.50E-41 | brain | Stra13 |
| Ddit4 | 1.08E-45 | 0.31359517 | 0.27 | 0.18 | 2.24E-41 | brain | Ddit4 |
| Hadh | 1.94E-45 | 0.30668717 | 0.26 | 0.17 | 4.02E-41 | brain | Hadh |
| Cyb5b | 2.25E-45 | 0.29163192 | 0.25 | 0.16 | 4.66E-41 | brain | Cyb5b |
| Rbbp7 | 2.12E-44 | 0.3078216 | 0.32 | 0.23 | 4.38E-40 | brain | Rbbp7 |
| Psma1 | 2.68E-44 | 0.25566861 | 0.48 | 0.38 | 5.54E-40 | brain | Psma1 |
| Arhgef2 | 1.40E-43 | 0.28811496 | 0.3 | 0.21 | 2.90E-39 | brain | Arhgef2 |
| Hes1 | 1.72E-43 | 0.41159345 | 0.6 | 0.49 | 3.55E-39 | brain | Hes1 |
| Tpi1 | 1.77E-43 | 0.29705446 | 0.37 | 0.28 | 3.66E-39 | brain | Tpi1 |
| Ifi27 | 2.92E-43 | 0.27648046 | 0.62 | 0.52 | 6.04E-39 | brain | Ifi27 |
| Mea1 | 7.46E-43 | 0.28341857 | 0.26 | 0.18 | 1.54E-38 | brain | Mea1 |
| Fam174a | 7.54E-43 | 0.28314448 | 0.37 | 0.27 | 1.56E-38 | brain | Fam174a |
| Psmd2 | 1.22E-42 | 0.28859853 | 0.31 | 0.22 | 2.53E-38 | brain | Psmd2 |
| Wbp2 | 1.42E-42 | 0.29610371 | 0.32 | 0.23 | 2.93E-38 | brain | Wbp2 |
| Smim10l1 | 7.39E-42 | 0.3027124 | 0.42 | 0.32 | 1.53E-37 | brain | Smim10l1 |
| Mat2b | 1.68E-41 | 0.28164936 | 0.37 | 0.27 | 3.47E-37 | brain | Mat2b |
| Mbd3 | 9.18E-41 | 0.28871224 | 0.3 | 0.22 | 1.90E-36 | brain | Mbd3 |
| Thra | 1.49E-40 | 0.27729975 | 0.34 | 0.24 | 3.09E-36 | brain | Thra |
| Vps36 | 1.85E-40 | 0.29322163 | 0.32 | 0.23 | 3.82E-36 | brain | Vps36 |
| Eif4e | 2.27E-40 | 0.26703511 | 0.3 | 0.21 | 4.70E-36 | brain | Eif4e |
| Tmem33 | 1.26E-39 | 0.26194371 | 0.3 | 0.21 | 2.60E-35 | brain | Tmem33 |
| Ube2g1 | 3.66E-39 | 0.30091284 | 0.27 | 0.19 | 7.58E-35 | brain | Ube2g1 |
| Akap121 | 1.47E-38 | 0.30010958 | 0.26 | 0.18 | 3.03E-34 | brain | Akap12 |
| Pqlc1 | 2.01E-38 | 0.26953845 | 0.32 | 0.23 | 4.16E-34 | brain | Pqlc1 |
| Card19 | 5.45E-38 | 0.25387795 | 0.36 | 0.27 | 1.13E-33 | brain | Card19 |
| Cldnd1 | 1.89E-37 | 0.28348085 | 0.27 | 0.19 | 3.92E-33 | brain | Cldnd1 |
| Chmp3 | 2.10E-37 | 0.27398198 | 0.39 | 0.3 | 4.34E-33 | brain | Chmp3 |
| Hmcn11 | 3.13E-37 | 0.28598453 | 0.29 | 0.21 | 6.48E-33 | brain | Hmcn1 |
| Rnaseh2c | 4.34E-37 | 0.30466596 | 0.31 | 0.23 | 8.98E-33 | brain | Rnaseh2c |
| 2610001J05Rik | 1.51E-36 | 0.26412428 | 0.37 | 0.28 | 3.13E-32 | brain | 2610001J05Rik |
| Lmbrd1 | 1.58E-36 | 0.29315175 | 0.28 | 0.2 | 3.27E-32 | brain | Lmbrd1 |
| Psmd7 | 1.18E-35 | 0.25629382 | 0.43 | 0.34 | 2.44E-31 | brain | Psmd7 |

| | | | | | | | |
|---|---|---|---|---|---|---|---|
| Ccdc85b1 | 1.30E-35 | 0.26051771 | 0.34 | 0.25 | 2.69E-31 | brain | Ccdc85b |
| Sae1 | 1.80E-35 | 0.25205898 | 0.25 | 0.18 | 3.71E-31 | brain | Sae1 |
| Cers2 | 3.74E-35 | 0.25733847 | 0.34 | 0.25 | 7.74E-31 | brain | Cers2 |
| Tspan3 | 6.70E-34 | 0.25955167 | 0.25 | 0.18 | 1.39E-29 | brain | Tspan3 |
| Mien1 | 1.33E-33 | 0.2702881 | 0.3 | 0.22 | 2.76E-29 | brain | Mien1 |
| Nap1l4 | 2.14E-33 | 0.26315278 | 0.31 | 0.23 | 4.44E-29 | brain | Nap1l4 |
| Chmp2b | 5.62E-33 | 0.26232175 | 0.32 | 0.24 | 1.16E-28 | brain | Chmp2b |
| Eci2 | 3.11E-31 | 0.25327128 | 0.25 | 0.18 | 6.44E-27 | brain | Eci2 |
| Dusp111 | 7.83E-31 | 0.25837503 | 0.28 | 0.21 | 1.62E-26 | brain | Dusp11 |
| Rgs5 | 0 | 1.92578243 | 0.29 | 0.06 | 0 | heart | Rgs5 |
| Hbb-bs1 | 0 | 1.87075695 | 0.49 | 0.11 | 0 | heart | Hbb-bs |
| Fabp41 | 0 | 1.84123538 | 1 | 0.56 | 0 | heart | Fabp4 |
| Cpe | 0 | 1.83012378 | 0.28 | 0.05 | 0 | heart | Cpe |
| Cd36 | 0 | 1.8206595 | 0.9 | 0.49 | 0 | heart | Cd36 |
| C1qtnf9 | 0 | 1.64080174 | 0.6 | 0.15 | 0 | heart | C1qtnf9 |
| Aqp7 | 0 | 1.62028962 | 0.47 | 0.09 | 0 | heart | Aqp7 |
| Mgp1 | 0 | 1.48980268 | 0.46 | 0.17 | 0 | heart | Mgp |
| Cdkn1c1 | 0 | 1.45707119 | 0.5 | 0.2 | 0 | heart | Cdkn1c |
| Fabp5 | 0 | 1.45163222 | 0.76 | 0.41 | 0 | heart | Fabp5 |
| Lpl | 0 | 1.40931388 | 0.48 | 0.13 | 0 | heart | Lpl |
| Tcf151 | 0 | 1.36873321 | 0.66 | 0.25 | 0 | heart | Tcf15 |
| Meox2 | 0 | 1.35226309 | 0.33 | 0.06 | 0 | heart | Meox2 |
| Rbp71 | 0 | 1.21695723 | 0.48 | 0.17 | 0 | heart | Rbp7 |
| Slc28a2 | 0 | 1.1267322 | 0.27 | 0.08 | 0 | heart | Slc28a2 |
| Timp4 | 0 | 1.00216337 | 0.37 | 0.09 | 0 | heart | Timp4 |
| Cav1 | 0 | 0.93956592 | 0.86 | 0.64 | 0 | heart | Cav1 |
| Cd81 | 0 | 0.90544637 | 0.93 | 0.81 | 0 | heart | Cd81 |
| Ptrf | 0 | 0.90010571 | 0.7 | 0.46 | 0 | heart | Ptrf |
| Sepw1 | 0 | 0.82757307 | 0.83 | 0.7 | 0 | heart | Sepw1 |
| Tmsb4x | 0 | 0.79238598 | 1 | 0.99 | 0 | heart | Tmsb4x |
| Gng11 | 0 | 0.77136598 | 0.92 | 0.84 | 0 | heart | Gng11 |
| Gpihbp1 | 0 | 0.76584317 | 0.84 | 0.54 | 0 | heart | Gpihbp1 |
| Crip1 | 0 | 0.76005585 | 0.88 | 0.69 | 0 | heart | Crip1 |
| Sdpr | 0 | 0.72030442 | 0.87 | 0.69 | 0 | heart | Sdpr |
| Rpl18a | 0 | 0.57996082 | 0.97 | 0.95 | 0 | heart | Rpl18a |
| Hspb11 | 3.85E-291 | 0.67951566 | 0.87 | 0.71 | 7.98E-287 | heart | Hspb1 |
| Ablim3 | 1.35E-289 | 1.038751 | 0.26 | 0.09 | 2.79E-285 | heart | Ablim3 |
| Rps29 | 1.77E-281 | 0.48616812 | 0.99 | 0.98 | 3.67E-277 | heart | Rps29 |
| AW1120101 | 2.59E-279 | 0.97305081 | 0.7 | 0.5 | 5.35E-275 | heart | AW112010 |
| Rgcc1 | 4.72E-277 | 0.77353193 | 0.67 | 0.41 | 9.76E-273 | heart | Rgcc |
| Lims2 | 5.05E-260 | 0.91844273 | 0.65 | 0.48 | 1.04E-255 | heart | Lims2 |

| | | | | | | | |
|---|---|---|---|---|---|---|---|
| Rps5 | 4.22E-259 | 0.54152744 | 0.94 | 0.91 | 8.74E-255 | heart | Rps5 |
| Gm100761 | 4.19E-252 | 0.94015959 | 0.63 | 0.46 | 8.67E-248 | heart | Gm10076 |
| Rps14 | 3.63E-250 | 0.45536055 | 0.97 | 0.96 | 7.51E-246 | heart | Rps14 |
| Aqp1 | 1.06E-249 | 0.60119528 | 0.76 | 0.46 | 2.20E-245 | heart | Aqp1 |
| Rpl35 | 1.62E-249 | 0.58358993 | 0.9 | 0.87 | 3.34E-245 | heart | Rpl35 |
| Sparcl12 | 2.74E-247 | 0.76721476 | 0.71 | 0.48 | 5.67E-243 | heart | Sparcl1 |
| Ehd2 | 3.19E-247 | 0.86764245 | 0.63 | 0.45 | 6.60E-243 | heart | Ehd2 |
| Ctnnbip1 | 3.34E-247 | 1.01576877 | 0.46 | 0.27 | 6.90E-243 | heart | Ctnnbip1 |
| Hspa81 | 3.31E-243 | 0.55517292 | 0.93 | 0.91 | 6.85E-239 | heart | Hspa8 |
| 8430408G22Rik1 | 7.57E-239 | 1.1911419 | 0.43 | 0.23 | 1.57E-234 | heart | 8430408G22Rik |
| Car81 | 7.14E-237 | 0.82789548 | 0.42 | 0.21 | 1.48E-232 | heart | Car8 |
| Tspan131 | 3.47E-236 | 0.60588125 | 0.81 | 0.62 | 7.17E-232 | heart | Tspan13 |
| Hist1h2bc1 | 2.22E-234 | 1.04933299 | 0.49 | 0.3 | 4.59E-230 | heart | Hist1h2bc |
| Unc45b | 1.65E-223 | 1.04859607 | 0.48 | 0.3 | 3.42E-219 | heart | Unc45b |
| Mgll1 | 1.98E-221 | 0.88268552 | 0.7 | 0.57 | 4.09E-217 | heart | Mgll |
| Ehd1 | 7.13E-221 | 0.78635076 | 0.46 | 0.26 | 1.48E-216 | heart | Ehd1 |
| Fscn1 | 2.42E-212 | 0.98639949 | 0.38 | 0.2 | 5.01E-208 | heart | Fscn1 |
| Ptma | 3.56E-199 | 0.45600093 | 0.96 | 0.95 | 7.37E-195 | heart | Ptma |
| Rps15 | 5.66E-199 | 0.4843848 | 0.93 | 0.91 | 1.17E-194 | heart | Rps15 |
| Ybx1 | 1.15E-198 | 0.52844104 | 0.89 | 0.84 | 2.39E-194 | heart | Ybx1 |
| Rps27 | 1.93E-195 | 0.40296151 | 0.99 | 0.98 | 4.00E-191 | heart | Rps27 |
| Fam101b | 4.32E-193 | 0.90162904 | 0.44 | 0.25 | 8.94E-189 | heart | Fam101b |
| Cyr611 | 4.64E-192 | 1.09555418 | 0.43 | 0.25 | 9.61E-188 | heart | Cyr61 |
| Aplnr | 5.10E-192 | 0.96032856 | 0.36 | 0.19 | 1.06E-187 | heart | Aplnr |
| Rpl13 | 5.48E-191 | 0.44302825 | 0.97 | 0.95 | 1.13E-186 | heart | Rpl13 |
| Klk8 | 1.67E-190 | 0.86573984 | 0.26 | 0.11 | 3.46E-186 | heart | Klk8 |
| Rpl41 | 7.27E-186 | 0.39759192 | 0.99 | 0.96 | 1.50E-181 | heart | Rpl41 |
| Cav2 | 4.09E-185 | 0.62107005 | 0.77 | 0.67 | 8.46E-181 | heart | Cav2 |
| Rpl32 | 1.72E-181 | 0.43523958 | 0.94 | 0.93 | 3.57E-177 | heart | Rpl32 |
| Fos1 | 1.04E-180 | 1.21903033 | 0.7 | 0.55 | 2.14E-176 | heart | Fos |
| Myl12a1 | 3.78E-178 | 0.48709145 | 0.88 | 0.84 | 7.82E-174 | heart | Myl12a |
| Tpm41 | 4.51E-163 | 0.73696756 | 0.73 | 0.67 | 9.33E-159 | heart | Tpm4 |
| Rplp0 | 1.35E-161 | 0.4494627 | 0.89 | 0.86 | 2.79E-157 | heart | Rplp0 |
| Serpinh1 | 1.24E-160 | 0.67002764 | 0.74 | 0.68 | 2.57E-156 | heart | Serpinh1 |
| Rps27rt | 6.57E-157 | 0.45397271 | 0.92 | 0.83 | 1.36E-152 | heart | Rps27rt |
| Rpl23a-ps3 | 4.14E-156 | 0.61463688 | 0.74 | 0.64 | 8.56E-152 | heart | Rpl23a-ps3 |
| Rps19 | 1.89E-155 | 0.37577574 | 0.97 | 0.96 | 3.92E-151 | heart | Rps19 |
| Fxyd6 | 1.68E-151 | 0.55150662 | 0.31 | 0.15 | 3.48E-147 | heart | Fxyd6 |
| Myl61 | 3.67E-151 | 0.45608413 | 0.92 | 0.9 | 7.59E-147 | heart | Myl6 |
| Rpl23a | 5.65E-147 | 0.43166008 | 0.93 | 0.91 | 1.17E-142 | heart | Rpl23a |
| Eef2 | 1.51E-142 | 0.49364717 | 0.82 | 0.79 | 3.13E-138 | heart | Eef2 |

| | | | | | | | |
|---|---|---|---|---|---|---|---|
| Klf21 | 2.05E-137 | 0.6166456 | 0.88 | 0.88 | 4.23E-133 | heart | Klf2 |
| Id11 | 2.38E-136 | 0.79083357 | 0.73 | 0.65 | 4.92E-132 | heart | Id1 |
| Hspa1a1 | 2.52E-135 | 0.50600627 | 0.47 | 0.3 | 5.21E-131 | heart | Hspa1a |
| Gm10073 | 6.13E-134 | 0.74323267 | 0.53 | 0.4 | 1.27E-129 | heart | Gm10073 |
| Hsp90ab11 | 1.36E-133 | 0.42729071 | 0.91 | 0.91 | 2.81E-129 | heart | Hsp90ab1 |
| Egr1 | 8.89E-132 | 0.66923643 | 0.48 | 0.33 | 1.84E-127 | heart | Egr1 |
| Cst3 | 2.74E-129 | 0.28293077 | 0.9 | 0.88 | 5.66E-125 | heart | Cst3 |
| Klf61 | 1.08E-121 | 0.8750618 | 0.54 | 0.43 | 2.23E-117 | heart | Klf6 |
| Cd300lg1 | 3.66E-120 | 0.61920498 | 0.56 | 0.41 | 7.58E-116 | heart | Cd300lg |
| Rpl37a | 5.98E-120 | 0.32884964 | 0.97 | 0.97 | 1.24E-115 | heart | Rpl37a |
| Dusp12 | 6.19E-118 | 0.60335263 | 0.74 | 0.66 | 1.28E-113 | heart | Dusp1 |
| Rps28 | 4.18E-117 | 0.36930008 | 0.93 | 0.93 | 8.64E-113 | heart | Rps28 |
| Rps6 | 1.08E-115 | 0.38944388 | 0.92 | 0.9 | 2.24E-111 | heart | Rps6 |
| Mllt4 | 5.31E-115 | 0.70806587 | 0.39 | 0.26 | 1.10E-110 | heart | Mllt4 |
| Rplp1 | 2.75E-114 | 0.37275503 | 0.94 | 0.93 | 5.68E-110 | heart | Rplp1 |
| Tm4sf1 | 8.30E-113 | 0.7162009 | 0.86 | 0.86 | 1.72E-108 | heart | Tm4sf1 |
| Prr13 | 8.70E-111 | 0.68884838 | 0.47 | 0.34 | 1.80E-106 | heart | Prr13 |
| Gm9843 | 3.31E-109 | 0.53366479 | 0.7 | 0.61 | 6.85E-105 | heart | Gm9843 |
| Rps7 | 1.02E-108 | 0.42936484 | 0.87 | 0.86 | 2.11E-104 | heart | Rps7 |
| Rps4x | 2.41E-106 | 0.37337143 | 0.92 | 0.9 | 4.99E-102 | heart | Rps4x |
| Gm8730 | 2.66E-106 | 0.63886558 | 0.53 | 0.41 | 5.50E-102 | heart | Gm8730 |
| S100a11 | 5.91E-104 | 0.50976398 | 0.67 | 0.59 | 1.22E-99 | heart | S100a11 |
| Myadm | 1.78E-103 | 0.6819704 | 0.36 | 0.24 | 3.69E-99 | heart | Myadm |
| Cdh13 | 1.61E-102 | 0.51190438 | 0.28 | 0.16 | 3.34E-98 | heart | Cdh13 |
| Btg21 | 4.29E-101 | 0.79811389 | 0.58 | 0.48 | 8.87E-97 | heart | Btg2 |
| Rpl37 | 9.63E-101 | 0.32995152 | 0.96 | 0.96 | 1.99E-96 | heart | Rpl37 |
| Rpl13a | 2.02E-100 | 0.3359785 | 0.95 | 0.89 | 4.18E-96 | heart | Rpl13a |
| Rps26 | 1.10E-99 | 0.44877289 | 0.83 | 0.81 | 2.28E-95 | heart | Rps26 |
| 4-Sep | 3.57E-99 | 0.63896262 | 0.45 | 0.32 | 7.38E-95 | heart | 4-Sep |
| Rpl21 | 7.40E-99 | 0.35327561 | 0.92 | 0.92 | 1.53E-94 | heart | Rpl21 |
| Fau | 7.46E-99 | 0.35600687 | 0.94 | 0.93 | 1.54E-94 | heart | Fau |
| Atp5j | 1.18E-94 | 0.43526311 | 0.79 | 0.78 | 2.44E-90 | heart | Atp5j |
| Rpl4 | 5.37E-94 | 0.40981123 | 0.82 | 0.81 | 1.11E-89 | heart | Rpl4 |
| Aplp2 | 1.83E-91 | 0.54825374 | 0.7 | 0.67 | 3.79E-87 | heart | Aplp2 |
| Rps23 | 1.11E-90 | 0.31240318 | 0.96 | 0.95 | 2.29E-86 | heart | Rps23 |
| Ptp4a3 | 8.72E-88 | 0.61918906 | 0.49 | 0.39 | 1.80E-83 | heart | Ptp4a3 |
| Apoe | 1.49E-86 | 0.53180976 | 0.68 | 0.52 | 3.08E-82 | heart | Apoe |
| Zfp361 | 5.05E-86 | 0.76175759 | 0.51 | 0.41 | 1.04E-81 | heart | Zfp36 |
| Gm2000 | 1.14E-82 | 0.65627038 | 0.35 | 0.24 | 2.36E-78 | heart | Gm2000 |
| Eef1a1 | 1.32E-82 | 0.27581032 | 0.95 | 0.95 | 2.72E-78 | heart | Eef1a1 |
| Gm11808 | 2.80E-82 | 0.51850036 | 0.65 | 0.58 | 5.80E-78 | heart | Gm11808 |

| | | | | | | | |
|---|---|---|---|---|---|---|---|
| Rpl8 | 2.09E-81 | 0.30903994 | 0.91 | 0.9 | 4.33E-77 | heart | Rpl8 |
| Eif1 | 2.06E-80 | 0.26193435 | 0.93 | 0.93 | 4.26E-76 | heart | Eif1 |
| Rplp2 | 3.10E-80 | 0.31751859 | 0.91 | 0.91 | 6.42E-76 | heart | Rplp2 |
| Pdzd2 | 2.18E-75 | 0.58320426 | 0.29 | 0.19 | 4.51E-71 | heart | Pdzd2 |
| Rps25 | 1.69E-74 | 0.35321771 | 0.89 | 0.89 | 3.50E-70 | heart | Rps25 |
| Rpl14 | 2.12E-74 | 0.33421028 | 0.89 | 0.89 | 4.38E-70 | heart | Rpl14 |
| Gm9493 | 1.56E-73 | 0.54488362 | 0.58 | 0.51 | 3.22E-69 | heart | Gm9493 |
| Rpl35a | 2.03E-73 | 0.29120799 | 0.95 | 0.94 | 4.20E-69 | heart | Rpl35a |
| Ltbp41 | 2.14E-73 | 0.50120285 | 0.59 | 0.49 | 4.42E-69 | heart | Ltbp4 |
| Gltscr2 | 8.24E-71 | 0.52046407 | 0.54 | 0.45 | 1.70E-66 | heart | Gltscr2 |
| Cdc42ep3 | 9.50E-71 | 0.56079798 | 0.3 | 0.21 | 1.96E-66 | heart | Cdc42ep3 |
| Palm1 | 1.46E-70 | 0.54389989 | 0.33 | 0.23 | 3.02E-66 | heart | Palm |
| Rps18 | 1.24E-68 | 0.2743442 | 0.93 | 0.93 | 2.56E-64 | heart | Rps18 |
| Rpl36 | 5.09E-67 | 0.3176401 | 0.9 | 0.91 | 1.05E-62 | heart | Rpl36 |
| S100a16 | 9.13E-67 | 0.39237057 | 0.72 | 0.7 | 1.89E-62 | heart | S100a16 |
| Uba52 | 2.14E-64 | 0.41319963 | 0.75 | 0.72 | 4.43E-60 | heart | Uba52 |
| Fam43a | 6.03E-64 | 0.58854441 | 0.39 | 0.3 | 1.25E-59 | heart | Fam43a |
| Rps26-ps1 | 2.21E-62 | 0.56346179 | 0.34 | 0.24 | 4.57E-58 | heart | Rps26-ps1 |
| Ldha1 | 5.06E-62 | 0.43583942 | 0.63 | 0.58 | 1.05E-57 | heart | Ldha |
| Rps241 | 3.88E-61 | 0.32806355 | 0.9 | 0.89 | 8.04E-57 | heart | Rps24 |
| Tmem88 | 1.26E-60 | 0.57063392 | 0.5 | 0.44 | 2.61E-56 | heart | Tmem88 |
| Pea15a | 1.16E-59 | 0.5705177 | 0.54 | 0.48 | 2.40E-55 | heart | Pea15a |
| Rps17 | 4.80E-58 | 0.28594138 | 0.89 | 0.89 | 9.93E-54 | heart | Rps17 |
| 4930523C07Rik | 2.60E-57 | 0.60234185 | 0.32 | 0.24 | 5.37E-53 | heart | 4930523C07Rik |
| H3f3b1 | 7.48E-57 | 0.33571422 | 0.86 | 0.89 | 1.55E-52 | heart | H3f3b |
| Rpl9 | 4.09E-56 | 0.27702125 | 0.92 | 0.91 | 8.47E-52 | heart | Rpl9 |
| Rps12-ps3 | 1.99E-55 | 0.45038245 | 0.63 | 0.57 | 4.11E-51 | heart | Rps12-ps3 |
| Rps16 | 4.18E-55 | 0.26229686 | 0.93 | 0.93 | 8.64E-51 | heart | Rps16 |
| Rpl9-ps6 | 7.13E-55 | 0.49106461 | 0.51 | 0.44 | 1.47E-50 | heart | Rpl9-ps6 |
| Rps2 | 1.34E-54 | 0.30484375 | 0.85 | 0.85 | 2.77E-50 | heart | Rps2 |
| Gas6 | 2.31E-53 | 0.45211362 | 0.55 | 0.49 | 4.79E-49 | heart | Gas6 |
| Cltb1 | 3.65E-53 | 0.53806863 | 0.48 | 0.42 | 7.56E-49 | heart | Cltb |
| Wdr89 | 6.19E-53 | 0.35411712 | 0.76 | 0.72 | 1.28E-48 | heart | Wdr89 |
| 1810011O10Rik | 9.15E-53 | 0.38486739 | 0.55 | 0.46 | 1.89E-48 | heart | 1810011O10Rik |
| Tsnax | 1.01E-52 | 0.61964452 | 0.26 | 0.18 | 2.10E-48 | heart | Tsnax |
| Rps9 | 8.47E-52 | 0.27910847 | 0.88 | 0.89 | 1.75E-47 | heart | Rps9 |
| Rpl26 | 8.74E-52 | 0.26519377 | 0.92 | 0.93 | 1.81E-47 | heart | Rpl26 |
| Rpl17 | 3.16E-50 | 0.26119811 | 0.89 | 0.9 | 6.54E-46 | heart | Rpl17 |
| Nudt41 | 4.06E-50 | 0.55316998 | 0.26 | 0.18 | 8.40E-46 | heart | Nudt4 |
| Rpl11 | 2.33E-49 | 0.26760335 | 0.89 | 0.9 | 4.82E-45 | heart | Rpl11 |
| Jam21 | 2.22E-48 | 0.44305753 | 0.52 | 0.45 | 4.59E-44 | heart | Jam2 |

| | | | | | | | |
|---|---|---|---|---|---|---|---|
| Sult1a1 | 3.31E-48 | 0.44961045 | 0.25 | 0.17 | 6.85E-44 | heart | Sult1a1 |
| Ahnak1 | 1.28E-47 | 0.44103224 | 0.52 | 0.45 | 2.65E-43 | heart | Ahnak |
| Adipor11 | 1.62E-47 | 0.50398902 | 0.5 | 0.46 | 3.36E-43 | heart | Adipor1 |
| Apold11 | 2.03E-46 | 0.4155173 | 0.32 | 0.24 | 4.20E-42 | heart | Apold1 |
| Rpl13-ps3 | 4.26E-46 | 0.48706683 | 0.43 | 0.36 | 8.82E-42 | heart | Rpl13-ps3 |
| Selm | 1.28E-45 | 0.42149187 | 0.5 | 0.43 | 2.64E-41 | heart | Selm |
| Zyx | 2.28E-45 | 0.48308966 | 0.26 | 0.18 | 4.71E-41 | heart | Zyx |
| Rftn11 | 1.41E-43 | 0.50900628 | 0.26 | 0.19 | 2.91E-39 | heart | Rftn1 |
| Junb1 | 1.49E-42 | 0.51812701 | 0.67 | 0.63 | 3.09E-38 | heart | Junb |
| Rpl31 | 1.38E-39 | 0.26589261 | 0.88 | 0.88 | 2.86E-35 | heart | Rpl31 |
| Ccdc85b2 | 1.68E-39 | 0.52361023 | 0.32 | 0.25 | 3.48E-35 | heart | Ccdc85b |
| Dnaja11 | 8.35E-39 | 0.42577389 | 0.66 | 0.66 | 1.73E-34 | heart | Dnaja1 |
| Rpl29 | 9.80E-39 | 0.28581921 | 0.82 | 0.83 | 2.03E-34 | heart | Rpl29 |
| Klf41 | 2.22E-38 | 0.49825083 | 0.61 | 0.59 | 4.60E-34 | heart | Klf4 |
| Trp53i11 | 4.14E-38 | 0.55115183 | 0.3 | 0.24 | 8.57E-34 | heart | Trp53i11 |
| Rpl10-ps3 | 1.32E-37 | 0.48564539 | 0.3 | 0.23 | 2.73E-33 | heart | Rpl10-ps3 |
| Cox7a2l | 1.98E-37 | 0.4259616 | 0.61 | 0.61 | 4.09E-33 | heart | Cox7a2l |
| Sgk1 | 7.57E-37 | 0.45520374 | 0.58 | 0.54 | 1.57E-32 | heart | Sgk1 |
| Ier21 | 1.78E-36 | 0.26905666 | 0.62 | 0.56 | 3.68E-32 | heart | Ier2 |
| Ifngr11 | 2.06E-35 | 0.44793988 | 0.45 | 0.4 | 4.26E-31 | heart | Ifngr1 |
| Rassf31 | 3.93E-35 | 0.66359029 | 0.31 | 0.25 | 8.13E-31 | heart | Rassf3 |
| Palmd2 | 5.03E-35 | 0.47998966 | 0.35 | 0.28 | 1.04E-30 | heart | Palmd |
| Ndel1 | 8.87E-35 | 0.42538359 | 0.32 | 0.26 | 1.84E-30 | heart | Ndel1 |
| Tmem1231 | 5.22E-34 | 0.5029943 | 0.34 | 0.28 | 1.08E-29 | heart | Tmem123 |
| Dusp3 | 9.08E-34 | 0.50252624 | 0.53 | 0.51 | 1.88E-29 | heart | Dusp3 |
| Socs3 | 1.23E-33 | 0.31131693 | 0.36 | 0.28 | 2.55E-29 | heart | Socs3 |
| Rpl6l | 4.65E-33 | 0.39477952 | 0.52 | 0.47 | 9.63E-29 | heart | Rpl6l |
| Pim3 | 7.54E-33 | 0.43736558 | 0.41 | 0.35 | 1.56E-28 | heart | Pim3 |
| Rasgrp21 | 1.15E-32 | 0.44818307 | 0.26 | 0.19 | 2.38E-28 | heart | Rasgrp2 |
| Gstm1 | 5.20E-32 | 0.33001933 | 0.58 | 0.53 | 1.08E-27 | heart | Gstm1 |
| Plod1 | 8.98E-31 | 0.44813989 | 0.27 | 0.21 | 1.86E-26 | heart | Plod1 |
| Tubb2a | 7.91E-30 | 0.33754647 | 0.39 | 0.32 | 1.64E-25 | heart | Tubb2a |
| Rpl39 | 9.04E-29 | 0.25894011 | 0.86 | 0.89 | 1.87E-24 | heart | Rpl39 |
| Rps12 | 2.62E-28 | 0.27037136 | 0.81 | 0.83 | 5.41E-24 | heart | Rps12 |
| Anxa2 | 6.73E-25 | 0.3095991 | 0.64 | 0.65 | 1.39E-20 | heart | Anxa2 |
| Ppp1r2 | 3.11E-24 | 0.28544798 | 0.63 | 0.63 | 6.44E-20 | heart | Ppp1r2 |
| Adamts11 | 9.72E-23 | 0.42884319 | 0.28 | 0.23 | 2.01E-18 | heart | Adamts1 |
| Pfdn5 | 1.23E-21 | 0.28550769 | 0.72 | 0.76 | 2.55E-17 | heart | Pfdn5 |
| Cyb5r3 | 1.49E-21 | 0.39117588 | 0.5 | 0.48 | 3.08E-17 | heart | Cyb5r3 |
| Ccnd11 | 7.96E-21 | 0.50259124 | 0.33 | 0.29 | 1.65E-16 | heart | Ccnd1 |
| Rhoc | 1.02E-20 | 0.40972128 | 0.53 | 0.54 | 2.11E-16 | heart | Rhoc |

| | | | | | | | |
|---|---|---|---|---|---|---|---|
| Ywhah1 | 1.57E-20 | 0.35390748 | 0.56 | 0.56 | 3.25E-16 | heart | Ywhah |
| Smco41 | 3.95E-20 | 0.40427952 | 0.49 | 0.47 | 8.17E-16 | heart | Smco4 |
| Trim471 | 1.24E-19 | 0.3492445 | 0.37 | 0.33 | 2.57E-15 | heart | Trim47 |
| Calm3 | 2.07E-19 | 0.45867577 | 0.37 | 0.34 | 4.29E-15 | heart | Calm3 |
| Arpc1b | 4.94E-19 | 0.29486645 | 0.63 | 0.65 | 1.02E-14 | heart | Arpc1b |
| Hnrnpf1 | 1.49E-18 | 0.31934893 | 0.63 | 0.65 | 3.07E-14 | heart | Hnrnpf |
| Aldoa | 5.66E-18 | 0.25871369 | 0.65 | 0.67 | 1.17E-13 | heart | Aldoa |
| Ctnnb11 | 5.86E-18 | 0.35325429 | 0.54 | 0.53 | 1.21E-13 | heart | Ctnnb1 |
| Rpl36-ps3 | 6.75E-18 | 0.33415841 | 0.41 | 0.37 | 1.40E-13 | heart | Rpl36-ps3 |
| Cdkn1a1 | 1.10E-16 | 0.27934059 | 0.42 | 0.37 | 2.28E-12 | heart | Cdkn1a |
| Hsp90aa11 | 3.17E-16 | 0.34341715 | 0.48 | 0.47 | 6.57E-12 | heart | Hsp90aa1 |
| Ets11 | 1.55E-15 | 0.37890201 | 0.47 | 0.45 | 3.20E-11 | heart | Ets1 |
| Kctd101 | 1.88E-14 | 0.45924879 | 0.36 | 0.33 | 3.89E-10 | heart | Kctd10 |
| Plk21 | 5.92E-14 | 0.39123274 | 0.45 | 0.43 | 1.23E-09 | heart | Plk2 |
| Tnfaip1 | 7.56E-14 | 0.39737593 | 0.38 | 0.36 | 1.56E-09 | heart | Tnfaip1 |
| Hspa12b1 | 1.11E-13 | 0.37231936 | 0.29 | 0.26 | 2.29E-09 | heart | Hspa12b |
| Fkbp41 | 3.31E-13 | 0.32895584 | 0.38 | 0.36 | 6.85E-09 | heart | Fkbp4 |
| Btg1 | 6.37E-13 | 0.29284491 | 0.52 | 0.52 | 1.32E-08 | heart | Btg1 |
| Tprgl1 | 2.53E-12 | 0.31654661 | 0.53 | 0.54 | 5.24E-08 | heart | Tprgl |
| Triobp | 4.06E-12 | 0.38443493 | 0.31 | 0.29 | 8.41E-08 | heart | Triobp |
| Arpc2 | 4.19E-12 | 0.25124055 | 0.63 | 0.65 | 8.67E-08 | heart | Arpc2 |
| Tax1bp3 | 7.88E-12 | 0.37683326 | 0.37 | 0.36 | 1.63E-07 | heart | Tax1bp3 |
| Socs2 | 8.67E-12 | 0.35732418 | 0.29 | 0.26 | 1.79E-07 | heart | Socs2 |
| Dhrs3 | 1.14E-11 | 0.27155992 | 0.39 | 0.37 | 2.36E-07 | heart | Dhrs3 |
| Itgb11 | 2.77E-11 | 0.31344735 | 0.53 | 0.54 | 5.72E-07 | heart | Itgb1 |
| Gm10036 | 6.63E-11 | 0.31377271 | 0.36 | 0.34 | 1.37E-06 | heart | Gm10036 |
| Tmod31 | 1.06E-10 | 0.37007076 | 0.35 | 0.33 | 2.19E-06 | heart | Tmod3 |
| Ypel31 | 2.93E-10 | 0.30354297 | 0.44 | 0.43 | 6.06E-06 | heart | Ypel3 |
| Ushbp1 | 3.55E-10 | 0.33693532 | 0.44 | 0.44 | 7.35E-06 | heart | Ushbp1 |
| Wnk1 | 7.91E-10 | 0.34512141 | 0.35 | 0.34 | 1.64E-05 | heart | Wnk1 |
| Ilk | 1.64E-09 | 0.29981485 | 0.34 | 0.32 | 3.39E-05 | heart | Ilk |
| Nck11 | 1.76E-09 | 0.37650115 | 0.4 | 0.4 | 3.65E-05 | heart | Nck1 |
| Hspd1 | 6.60E-08 | 0.35766547 | 0.25 | 0.23 | 0.00136628 | heart | Hspd1 |
| Smpdl3a1 | 1.35E-07 | 0.33031143 | 0.36 | 0.35 | 0.00279331 | heart | Smpdl3a |
| Lrrc8a | 2.75E-07 | 0.30796123 | 0.31 | 0.3 | 0.00569063 | heart | Lrrc8a |
| Sri1 | 5.47E-07 | 0.29083953 | 0.48 | 0.5 | 0.01132619 | heart | Sri |
| Arhgef15 | 6.81E-07 | 0.29681678 | 0.37 | 0.37 | 0.01408966 | heart | Arhgef15 |
| Gdi2 | 8.69E-07 | 0.27908943 | 0.49 | 0.51 | 0.0179872 | heart | Gdi2 |
| Pls31 | 1.14E-06 | 0.29382096 | 0.37 | 0.36 | 0.02350781 | heart | Pls3 |
| Pabpc1 | 1.76E-06 | 0.2503875 | 0.48 | 0.5 | 0.03640992 | heart | Pabpc1 |
| Zfp36l2 | 2.21E-06 | 0.28044831 | 0.29 | 0.28 | 0.04573355 | heart | Zfp36l2 |

| | | | | | | | |
|---|---|---|---|---|---|---|---|
| Anp32a | 2.84E-06 | 0.28922621 | 0.62 | 0.66 | 0.05866248 | heart | Anp32a |
| Vasp | 1.05E-05 | 0.29098341 | 0.34 | 0.34 | 0.21710333 | heart | Vasp |
| Wwtr11 | 1.27E-05 | 0.30634525 | 0.28 | 0.27 | 0.26262245 | heart | Wwtr1 |
| Capn21 | 1.58E-05 | 0.30132092 | 0.34 | 0.33 | 0.32676358 | heart | Capn2 |
| Rpl27-ps3 | 2.26E-05 | 0.25857742 | 0.34 | 0.33 | 0.46724195 | heart | Rpl27-ps3 |
| H1f0 | 2.82E-05 | 0.30588177 | 0.39 | 0.4 | 0.58278919 | heart | H1f0 |
| Fez21 | 2.83E-05 | 0.33244466 | 0.35 | 0.35 | 0.58567421 | heart | Fez2 |
| Rab6a | 0.00049309 | 0.29371406 | 0.35 | 0.36 | 1 | heart | Rab6a |
| Fosb | 0.0007635 | 0.30538343 | 0.3 | 0.29 | 1 | heart | Fosb |
| Polr2m1 | 0.00094086 | 0.28611591 | 0.29 | 0.29 | 1 | heart | Polr2m |
| Med281 | 0.00411609 | 0.28296227 | 0.41 | 0.43 | 1 | heart | Med28 |
| Ahsa1 | 0.00509531 | 0.25078522 | 0.3 | 0.31 | 1 | heart | Ahsa1 |
| Plvap1 | 0 | 2.10831001 | 0.95 | 0.56 | 0 | intestine_colon | Plvap |
| Nkx2-3 | 0 | 2.00408839 | 0.6 | 0.08 | 0 | intestine_colon | Nkx2-3 |
| Col15a11 | 0 | 1.40273448 | 0.55 | 0.13 | 0 | intestine_colon | Col15a1 |
| Tppp3 | 0 | 1.3848812 | 0.44 | 0.17 | 0 | intestine_colon | Tppp3 |
| Lmna | 0 | 1.28022549 | 0.71 | 0.29 | 0 | intestine_colon | Lmna |
| Crip11 | 0 | 1.2620356 | 0.96 | 0.69 | 0 | intestine_colon | Crip1 |
| Esm1 | 0 | 1.25576095 | 0.48 | 0.11 | 0 | intestine_colon | Esm1 |
| Prss231 | 0 | 1.23477391 | 0.65 | 0.28 | 0 | intestine_colon | Prss23 |
| Cxcl122 | 0 | 1.21343954 | 0.79 | 0.47 | 0 | intestine_colon | Cxcl12 |
| Ptprr1 | 0 | 1.1980744 | 0.55 | 0.15 | 0 | intestine_colon | Ptprr |
| Ncoa71 | 0 | 1.18612508 | 0.48 | 0.14 | 0 | intestine_colon | Ncoa7 |
| Igfbp7 | 0 | 1.1710652 | 0.98 | 0.87 | 0 | intestine_colon | Igfbp7 |
| Cxx1a1 | 0 | 1.16914018 | 0.79 | 0.44 | 0 | intestine_colon | Cxx1a |
| Sox181 | 0 | 1.16384628 | 0.8 | 0.49 | 0 | intestine_colon | Sox18 |
| Fam101b1 | 0 | 1.14941755 | 0.64 | 0.24 | 0 | intestine_colon | Fam101b |
| Ssu2 | 0 | 1.14519343 | 0.36 | 0.05 | 0 | intestine_colon | Ssu2 |
| Ace | 0 | 1.13796254 | 0.62 | 0.2 | 0 | intestine_colon | Ace |
| Tmem255b | 0 | 1.08483179 | 0.43 | 0.09 | 0 | intestine_colon | Tmem255b |
| Lamb11 | 0 | 1.07979839 | 0.39 | 0.07 | 0 | intestine_colon | Lamb1 |
| S100a6 | 0 | 1.05105646 | 0.67 | 0.28 | 0 | intestine_colon | S100a6 |
| Rasa4 | 0 | 1.02503967 | 0.36 | 0.09 | 0 | intestine_colon | Rasa4 |
| Col13a11 | 0 | 1.00587531 | 0.41 | 0.13 | 0 | intestine_colon | Col13a1 |
| Thrsp1 | 0 | 1.00096966 | 0.6 | 0.26 | 0 | intestine_colon | Thrsp |
| Sncg1 | 0 | 0.97189238 | 0.47 | 0.15 | 0 | intestine_colon | Sncg |
| Cd24a1 | 0 | 0.96361666 | 0.59 | 0.22 | 0 | intestine_colon | Cd24a |
| Jam31 | 0 | 0.94243552 | 0.52 | 0.21 | 0 | intestine_colon | Jam3 |
| S100a111 | 0 | 0.93744283 | 0.86 | 0.57 | 0 | intestine_colon | S100a11 |
| Cxx1b1 | 0 | 0.920311 | 0.63 | 0.29 | 0 | intestine_colon | Cxx1b |
| Mgll2 | 0 | 0.88809652 | 0.84 | 0.56 | 0 | intestine_colon | Mgll |

| | | | | | | | |
|---|---|---|---|---|---|---|---|
| Podxl1 | 0 | 0.88120961 | 0.8 | 0.49 | 0 | intestine_colon | Podxl |
| Mall | 0 | 0.87458265 | 0.42 | 0.12 | 0 | intestine_colon | Mall |
| Ccl21a | 0 | 0.85679048 | 0.62 | 0.22 | 0 | intestine_colon | Ccl21a |
| Ramp31 | 0 | 0.85113638 | 0.51 | 0.2 | 0 | intestine_colon | Ramp3 |
| Alox12 | 0 | 0.84853651 | 0.32 | 0.08 | 0 | intestine_colon | Alox12 |
| Synpo1 | 0 | 0.81028825 | 0.35 | 0.1 | 0 | intestine_colon | Synpo |
| Bcr | 0 | 0.80341591 | 0.33 | 0.08 | 0 | intestine_colon | Bcr |
| Gas61 | 0 | 0.80252833 | 0.77 | 0.47 | 0 | intestine_colon | Gas6 |
| Slfn21 | 0 | 0.80231708 | 0.41 | 0.15 | 0 | intestine_colon | Slfn2 |
| Sparcl13 | 0 | 0.79385796 | 0.81 | 0.48 | 0 | intestine_colon | Sparcl1 |
| Cd811 | 0 | 0.78036507 | 0.93 | 0.82 | 0 | intestine_colon | Cd81 |
| Ebf11 | 0 | 0.77660758 | 0.74 | 0.38 | 0 | intestine_colon | Ebf1 |
| Ly6a1 | 0 | 0.76943393 | 0.95 | 0.87 | 0 | intestine_colon | Ly6a |
| Ly6c11 | 0 | 0.76811554 | 0.96 | 0.84 | 0 | intestine_colon | Ly6c1 |
| Pdlim4 | 0 | 0.7373323 | 0.47 | 0.15 | 0 | intestine_colon | Pdlim4 |
| Endod11 | 0 | 0.73618522 | 0.32 | 0.1 | 0 | intestine_colon | Endod1 |
| Proser21 | 0 | 0.73453313 | 0.32 | 0.09 | 0 | intestine_colon | Proser2 |
| Capg1 | 0 | 0.73376324 | 0.32 | 0.1 | 0 | intestine_colon | Capg |
| Rbp72 | 0 | 0.7226806 | 0.5 | 0.18 | 0 | intestine_colon | Rbp7 |
| Hoxa7 | 0 | 0.67466996 | 0.31 | 0.07 | 0 | intestine_colon | Hoxa7 |
| Itm2b | 0 | 0.60332372 | 0.99 | 0.96 | 0 | intestine_colon | Itm2b |
| Mfge8 | 9.47E-307 | 0.83952942 | 0.72 | 0.43 | 1.96E-302 | intestine_colon | Mfge8 |
| Hlx | 1.27E-295 | 0.82141734 | 0.43 | 0.17 | 2.63E-291 | intestine_colon | Hlx |
| Vwa11 | 7.93E-289 | 0.6951282 | 0.65 | 0.34 | 1.64E-284 | intestine_colon | Vwa1 |
| Pqlc11 | 1.05E-287 | 0.82558307 | 0.48 | 0.21 | 2.18E-283 | intestine_colon | Pqlc1 |
| Pcp4l11 | 2.90E-286 | 0.52664554 | 0.69 | 0.34 | 6.01E-282 | intestine_colon | Pcp4l1 |
| Cst31 | 3.20E-284 | 0.84622721 | 0.96 | 0.88 | 6.61E-280 | intestine_colon | Cst3 |
| Ehd41 | 5.10E-284 | 0.91686009 | 0.78 | 0.51 | 1.05E-279 | intestine_colon | Ehd4 |
| Igfbp31 | 5.46E-282 | 0.70592191 | 0.54 | 0.24 | 1.13E-277 | intestine_colon | Igfbp3 |
| Tgfbr21 | 2.10E-276 | 0.75177762 | 0.54 | 0.26 | 4.34E-272 | intestine_colon | Tgfbr2 |
| Calm12 | 6.43E-274 | 0.55627382 | 0.96 | 0.9 | 1.33E-269 | intestine_colon | Calm1 |
| S100a10 | 1.27E-265 | 0.72495701 | 0.87 | 0.65 | 2.63E-261 | intestine_colon | S100a10 |
| Plpp31 | 1.90E-261 | 0.85654709 | 0.85 | 0.61 | 3.93E-257 | intestine_colon | Plpp3 |
| Cyb561 | 2.07E-259 | 0.67180287 | 0.33 | 0.12 | 4.29E-255 | intestine_colon | Cyb561 |
| Trim472 | 2.66E-256 | 0.74370391 | 0.6 | 0.31 | 5.50E-252 | intestine_colon | Trim47 |
| Prkcdbp | 4.77E-254 | 0.61242141 | 0.8 | 0.49 | 9.87E-250 | intestine_colon | Prkcdbp |
| Col4a11 | 1.47E-250 | 0.75315005 | 0.79 | 0.53 | 3.04E-246 | intestine_colon | Col4a1 |
| Pecam11 | 5.29E-242 | 0.59704175 | 0.91 | 0.76 | 1.10E-237 | intestine_colon | Pecam1 |
| Anxa21 | 9.42E-241 | 0.66432264 | 0.83 | 0.63 | 1.95E-236 | intestine_colon | Anxa2 |
| Ctsz | 4.17E-239 | 0.72427766 | 0.62 | 0.35 | 8.63E-235 | intestine_colon | Ctsz |
| Col4a21 | 1.05E-238 | 0.78501433 | 0.71 | 0.44 | 2.17E-234 | intestine_colon | Col4a2 |

| | | | | | | | |
|---|---|---|---|---|---|---|---|
| Serpinh11 | 2.07E-235 | 0.68713072 | 0.84 | 0.67 | 4.29E-231 | intestine_colon | Serpinh1 |
| Magix | 2.11E-226 | 0.57707022 | 0.25 | 0.08 | 4.36E-222 | intestine_colon | Magix |
| Fxyd51 | 1.55E-225 | 0.70212493 | 0.73 | 0.46 | 3.21E-221 | intestine_colon | Fxyd5 |
| Meox11 | 4.87E-223 | 0.59059628 | 0.45 | 0.2 | 1.01E-218 | intestine_colon | Meox1 |
| Upp1 | 5.75E-222 | 0.62284934 | 0.35 | 0.14 | 1.19E-217 | intestine_colon | Upp1 |
| Cd9 | 1.03E-219 | 0.77018891 | 0.84 | 0.61 | 2.14E-215 | intestine_colon | Cd9 |
| Plcb11 | 4.47E-213 | 0.26420897 | 0.52 | 0.24 | 9.25E-209 | intestine_colon | Plcb1 |
| Mef2c1 | 1.18E-212 | 0.76404902 | 0.69 | 0.43 | 2.43E-208 | intestine_colon | Mef2c |
| Aplnr1 | 3.25E-210 | 0.9594165 | 0.42 | 0.2 | 6.73E-206 | intestine_colon | Aplnr |
| Mmrn2 | 2.78E-208 | 0.66316105 | 0.75 | 0.5 | 5.76E-204 | intestine_colon | Mmrn2 |
| Tril | 7.17E-206 | 0.535912 | 0.26 | 0.09 | 1.48E-201 | intestine_colon | Tril |
| Tagln2 | 2.15E-205 | 0.55286508 | 0.89 | 0.73 | 4.45E-201 | intestine_colon | Tagln2 |
| Arl6ip51 | 2.04E-203 | 0.71067704 | 0.72 | 0.5 | 4.22E-199 | intestine_colon | Arl6ip5 |
| Osmr1 | 7.73E-200 | 0.62706044 | 0.34 | 0.14 | 1.60E-195 | intestine_colon | Osmr |
| Ldha2 | 1.73E-197 | 0.62128165 | 0.77 | 0.57 | 3.59E-193 | intestine_colon | Ldha |
| Cd341 | 4.57E-197 | 0.65665658 | 0.77 | 0.54 | 9.45E-193 | intestine_colon | Cd34 |
| Lamc1 | 5.14E-195 | 0.5803972 | 0.34 | 0.14 | 1.06E-190 | intestine_colon | Lamc1 |
| Tnfaip2 | 3.28E-190 | 0.35712964 | 0.36 | 0.15 | 6.78E-186 | intestine_colon | Tnfaip2 |
| Hspg21 | 9.69E-190 | 0.71377362 | 0.64 | 0.38 | 2.01E-185 | intestine_colon | Hspg2 |
| Orai1 | 1.10E-185 | 0.6093843 | 0.45 | 0.22 | 2.27E-181 | intestine_colon | Orai1 |
| Acvrl11 | 5.24E-184 | 0.39043441 | 0.75 | 0.47 | 1.08E-179 | intestine_colon | Acvrl1 |
| Thbd | 2.13E-182 | 0.64486388 | 0.74 | 0.5 | 4.41E-178 | intestine_colon | Thbd |
| Anxa1 | 9.27E-181 | 0.51985896 | 0.33 | 0.13 | 1.92E-176 | intestine_colon | Anxa1 |
| Ptrf1 | 9.79E-181 | 0.45526474 | 0.75 | 0.47 | 2.03E-176 | intestine_colon | Ptrf |
| Gramd1a | 1.84E-179 | 0.48036296 | 0.28 | 0.1 | 3.81E-175 | intestine_colon | Gramd1a |
| Lysmd2 | 2.57E-177 | 0.57238707 | 0.47 | 0.24 | 5.32E-173 | intestine_colon | Lysmd2 |
| Slc43a31 | 2.71E-174 | 0.51282657 | 0.68 | 0.41 | 5.61E-170 | intestine_colon | Slc43a3 |
| Bhlhe401 | 1.33E-173 | 0.4915319 | 0.38 | 0.17 | 2.75E-169 | intestine_colon | Bhlhe40 |
| Mcam | 7.86E-173 | 0.57368383 | 0.39 | 0.19 | 1.63E-168 | intestine_colon | Mcam |
| Anxa6 | 3.59E-170 | 0.4924354 | 0.39 | 0.18 | 7.43E-166 | intestine_colon | Anxa6 |
| Adgrg1 | 1.98E-168 | 0.57495159 | 0.47 | 0.24 | 4.09E-164 | intestine_colon | Adgrg1 |
| Txnip | 3.18E-167 | 0.52707592 | 0.83 | 0.64 | 6.57E-163 | intestine_colon | Txnip |
| Fam43a1 | 1.39E-164 | 0.51453299 | 0.53 | 0.29 | 2.88E-160 | intestine_colon | Fam43a |
| Gapdh | 5.00E-164 | 0.54391746 | 0.8 | 0.62 | 1.04E-159 | intestine_colon | Gapdh |
| Tspan2 | 5.39E-164 | 0.47586325 | 0.27 | 0.11 | 1.12E-159 | intestine_colon | Tspan2 |
| Tubb4b | 7.55E-163 | 0.6263212 | 0.71 | 0.49 | 1.56E-158 | intestine_colon | Tubb4b |
| Ivns1abp1 | 8.65E-163 | 0.60148015 | 0.41 | 0.2 | 1.79E-158 | intestine_colon | Ivns1abp |
| St6galnac21 | 6.72E-161 | 0.5270413 | 0.33 | 0.15 | 1.39E-156 | intestine_colon | St6galnac2 |
| Kctd12b1 | 1.24E-158 | 0.3943635 | 0.3 | 0.12 | 2.56E-154 | intestine_colon | Kctd12b |
| Flt12 | 4.50E-158 | 0.41623065 | 0.84 | 0.69 | 9.31E-154 | intestine_colon | Flt1 |
| Vat1 | 4.92E-158 | 0.5778655 | 0.57 | 0.34 | 1.02E-153 | intestine_colon | Vat1 |

| | | | | | | | |
|---|---|---|---|---|---|---|---|
| Scn1b1 | 3.09E-154 | 0.41950994 | 0.34 | 0.15 | 6.39E-150 | intestine_colon | Scn1b |
| AW1120102 | 4.25E-154 | 0.51168872 | 0.74 | 0.5 | 8.80E-150 | intestine_colon | AW112010 |
| Scarb11 | 5.06E-152 | 0.66117253 | 0.6 | 0.39 | 1.05E-147 | intestine_colon | Scarb1 |
| Lrrc8c1 | 2.63E-151 | 0.53951119 | 0.5 | 0.28 | 5.44E-147 | intestine_colon | Lrrc8c |
| Rdx1 | 3.01E-150 | 0.51451477 | 0.76 | 0.54 | 6.23E-146 | intestine_colon | Rdx |
| Plscr21 | 2.94E-148 | 0.34046636 | 0.5 | 0.26 | 6.08E-144 | intestine_colon | Plscr2 |
| Socs21 | 3.50E-146 | 0.53299797 | 0.45 | 0.24 | 7.25E-142 | intestine_colon | Socs2 |
| Bcam | 7.63E-146 | 0.59060557 | 0.73 | 0.55 | 1.58E-141 | intestine_colon | Bcam |
| Fam129b1 | 5.16E-144 | 0.47287926 | 0.36 | 0.17 | 1.07E-139 | intestine_colon | Fam129b |
| Clu | 1.82E-140 | 0.34424162 | 0.61 | 0.35 | 3.77E-136 | intestine_colon | Clu |
| Tmem176b | 1.40E-137 | 0.51884004 | 0.7 | 0.49 | 2.90E-133 | intestine_colon | Tmem176b |
| Adrb2 | 3.21E-137 | 0.40246858 | 0.28 | 0.12 | 6.63E-133 | intestine_colon | Adrb2 |
| Eef1a11 | 2.39E-136 | 0.31309928 | 0.98 | 0.95 | 4.94E-132 | intestine_colon | Eef1a1 |
| Tmem176a | 6.81E-132 | 0.48980087 | 0.65 | 0.41 | 1.41E-127 | intestine_colon | Tmem176a |
| Vim | 2.36E-130 | 0.6045163 | 0.84 | 0.65 | 4.89E-126 | intestine_colon | Vim |
| Cd200 | 7.22E-130 | 0.40030164 | 0.85 | 0.65 | 1.49E-125 | intestine_colon | Cd200 |
| Cav11 | 1.57E-129 | 0.26411428 | 0.88 | 0.65 | 3.25E-125 | intestine_colon | Cav1 |
| Lpcat11 | 5.60E-129 | 0.42751253 | 0.3 | 0.14 | 1.16E-124 | intestine_colon | Lpcat1 |
| Sparc | 8.61E-129 | 0.32681837 | 0.89 | 0.86 | 1.78E-124 | intestine_colon | Sparc |
| Emp3 | 1.75E-127 | 0.49579616 | 0.45 | 0.25 | 3.61E-123 | intestine_colon | Emp3 |
| Ybx11 | 3.38E-127 | 0.38077845 | 0.92 | 0.84 | 6.99E-123 | intestine_colon | Ybx1 |
| Nrp11 | 2.02E-124 | 0.58840964 | 0.79 | 0.65 | 4.19E-120 | intestine_colon | Nrp1 |
| Kdr1 | 1.52E-123 | 0.40198599 | 0.81 | 0.61 | 3.15E-119 | intestine_colon | Kdr |
| Oaz2 | 1.93E-123 | 0.54115116 | 0.75 | 0.58 | 3.98E-119 | intestine_colon | Oaz2 |
| Slco2a1 | 3.42E-123 | 0.62438949 | 0.5 | 0.3 | 7.07E-119 | intestine_colon | Slco2a1 |
| Hmgn3 | 3.15E-122 | 0.41243509 | 0.29 | 0.14 | 6.53E-118 | intestine_colon | Hmgn3 |
| Ahnak2 | 1.52E-121 | 0.4590832 | 0.66 | 0.44 | 3.15E-117 | intestine_colon | Ahnak |
| Grrp12 | 2.30E-121 | 0.55312358 | 0.53 | 0.33 | 4.76E-117 | intestine_colon | Grrp1 |
| Ifitm3 | 1.52E-120 | 0.31830367 | 0.98 | 0.96 | 3.15E-116 | intestine_colon | Ifitm3 |
| Emp1 | 6.66E-120 | 0.27499445 | 0.41 | 0.21 | 1.38E-115 | intestine_colon | Emp1 |
| Cd151 | 8.18E-118 | 0.43532265 | 0.74 | 0.55 | 1.69E-113 | intestine_colon | Cd151 |
| Selm1 | 5.64E-115 | 0.34539373 | 0.65 | 0.41 | 1.17E-110 | intestine_colon | Selm |
| Sntb21 | 1.25E-114 | 0.41153549 | 0.34 | 0.18 | 2.59E-110 | intestine_colon | Sntb2 |
| Mcf2l1 | 5.08E-114 | 0.40893337 | 0.37 | 0.19 | 1.05E-109 | intestine_colon | Mcf2l |
| Ly6e1 | 1.69E-113 | 0.27666897 | 0.98 | 0.94 | 3.49E-109 | intestine_colon | Ly6e |
| Ifngr12 | 7.49E-113 | 0.44365634 | 0.59 | 0.39 | 1.55E-108 | intestine_colon | Ifngr1 |
| Lama41 | 2.47E-112 | 0.41591024 | 0.33 | 0.17 | 5.10E-108 | intestine_colon | Lama4 |
| Ltbp42 | 4.22E-112 | 0.4330677 | 0.71 | 0.48 | 8.72E-108 | intestine_colon | Ltbp4 |
| B4galt42 | 8.56E-112 | 0.43837588 | 0.42 | 0.23 | 1.77E-107 | intestine_colon | B4galt4 |
| Fez22 | 3.48E-111 | 0.45586002 | 0.54 | 0.34 | 7.20E-107 | intestine_colon | Fez2 |
| Myadm1 | 1.14E-110 | 0.36712142 | 0.43 | 0.24 | 2.37E-106 | intestine_colon | Myadm |

| | | | | | | | |
|---|---|---|---|---|---|---|---|
| Atox11 | 2.89E-110 | 0.38555664 | 0.89 | 0.79 | 5.98E-106 | intestine_colon | Atox1 |
| Lipe1 | 1.03E-109 | 0.42547437 | 0.33 | 0.17 | 2.14E-105 | intestine_colon | Lipe |
| Wars1 | 7.26E-107 | 0.48043466 | 0.36 | 0.19 | 1.50E-102 | intestine_colon | Wars |
| Ddah1 | 1.36E-105 | 0.38539893 | 0.25 | 0.12 | 2.82E-101 | intestine_colon | Ddah1 |
| Irf2bpl1 | 4.70E-105 | 0.42954582 | 0.45 | 0.26 | 9.73E-101 | intestine_colon | Irf2bpl |
| Lrrfip11 | 7.36E-105 | 0.42074269 | 0.47 | 0.28 | 1.52E-100 | intestine_colon | Lrrfip1 |
| Lmcd11 | 9.55E-105 | 0.38234658 | 0.43 | 0.24 | 1.98E-100 | intestine_colon | Lmcd1 |
| Mgst3 | 6.61E-103 | 0.43630286 | 0.39 | 0.22 | 1.37E-98 | intestine_colon | Mgst3 |
| Actn41 | 2.72E-100 | 0.42749482 | 0.7 | 0.51 | 5.63E-96 | intestine_colon | Actn4 |
| Pdlim1 | 2.12E-98 | 0.36731031 | 0.5 | 0.3 | 4.39E-94 | intestine_colon | Pdlim1 |
| Tmem881 | 3.46E-98 | 0.42595689 | 0.62 | 0.43 | 7.16E-94 | intestine_colon | Tmem88 |
| Crip21 | 1.51E-97 | 0.28863214 | 0.93 | 0.85 | 3.13E-93 | intestine_colon | Crip2 |
| Eef21 | 1.66E-96 | 0.32857571 | 0.9 | 0.79 | 3.44E-92 | intestine_colon | Eef2 |
| Fstl1 | 1.72E-96 | 0.38239353 | 0.3 | 0.16 | 3.55E-92 | intestine_colon | Fstl1 |
| Ifi35 | 3.70E-95 | 0.45551761 | 0.54 | 0.36 | 7.66E-91 | intestine_colon | Ifi35 |
| App2 | 9.18E-95 | 0.36121539 | 0.85 | 0.74 | 1.90E-90 | intestine_colon | App |
| Tap2 | 3.49E-93 | 0.40974884 | 0.48 | 0.3 | 7.22E-89 | intestine_colon | Tap2 |
| Fmo21 | 1.93E-92 | 0.43446768 | 0.31 | 0.16 | 3.98E-88 | intestine_colon | Fmo2 |
| Robo4 | 1.72E-91 | 0.42974541 | 0.51 | 0.33 | 3.55E-87 | intestine_colon | Robo4 |
| Aplp21 | 6.94E-91 | 0.39642332 | 0.81 | 0.66 | 1.44E-86 | intestine_colon | Aplp2 |
| Nrarp | 1.85E-90 | 0.40713876 | 0.28 | 0.14 | 3.84E-86 | intestine_colon | Nrarp |
| Csrp1 | 1.67E-88 | 0.36076634 | 0.6 | 0.41 | 3.46E-84 | intestine_colon | Csrp1 |
| Efna1 | 6.42E-88 | 0.35063464 | 0.55 | 0.35 | 1.33E-83 | intestine_colon | Efna1 |
| Iqgap11 | 1.13E-86 | 0.4184843 | 0.51 | 0.33 | 2.33E-82 | intestine_colon | Iqgap1 |
| Rpn2 | 3.09E-86 | 0.36504483 | 0.55 | 0.36 | 6.40E-82 | intestine_colon | Rpn2 |
| Arhgap291 | 2.03E-85 | 0.25337861 | 0.5 | 0.31 | 4.21E-81 | intestine_colon | Arhgap29 |
| Nfib1 | 1.06E-84 | 0.29263857 | 0.73 | 0.54 | 2.19E-80 | intestine_colon | Nfib |
| Esam1 | 3.93E-84 | 0.35180297 | 0.83 | 0.73 | 8.13E-80 | intestine_colon | Esam |
| Klf10 | 5.17E-84 | 0.3393751 | 0.39 | 0.23 | 1.07E-79 | intestine_colon | Klf10 |
| Spop | 1.10E-83 | 0.37427737 | 0.52 | 0.34 | 2.27E-79 | intestine_colon | Spop |
| Eno11 | 1.06E-81 | 0.35519003 | 0.5 | 0.32 | 2.19E-77 | intestine_colon | Eno1 |
| Rpl27 | 3.52E-81 | 0.27488897 | 0.91 | 0.82 | 7.29E-77 | intestine_colon | Rpl27 |
| Aprt | 4.46E-80 | 0.33383498 | 0.3 | 0.16 | 9.23E-76 | intestine_colon | Aprt |
| Entpd11 | 6.65E-80 | 0.36819605 | 0.3 | 0.17 | 1.38E-75 | intestine_colon | Entpd1 |
| Msn1 | 6.11E-79 | 0.35349291 | 0.74 | 0.58 | 1.26E-74 | intestine_colon | Msn |
| Stom1 | 1.99E-77 | 0.33537891 | 0.29 | 0.16 | 4.11E-73 | intestine_colon | Stom |
| Dnajc8 | 2.42E-77 | 0.37509125 | 0.58 | 0.41 | 5.02E-73 | intestine_colon | Dnajc8 |
| Azin1 | 4.43E-77 | 0.50395267 | 0.28 | 0.15 | 9.18E-73 | intestine_colon | Azin1 |
| Cds2 | 8.37E-77 | 0.35411954 | 0.38 | 0.23 | 1.73E-72 | intestine_colon | Cds2 |
| Fkbp31 | 5.06E-76 | 0.38779155 | 0.46 | 0.3 | 1.05E-71 | intestine_colon | Fkbp3 |
| Pkig1 | 5.57E-75 | 0.35479554 | 0.41 | 0.25 | 1.15E-70 | intestine_colon | Pkig |

| | | | | | | | |
|---|---|---|---|---|---|---|---|
| Rftn12 | 1.52E-74 | 0.26048897 | 0.33 | 0.19 | 3.14E-70 | intestine_colon | Rftn1 |
| Slc48a1 | 1.72E-74 | 0.33150077 | 0.37 | 0.22 | 3.57E-70 | intestine_colon | Slc48a1 |
| Gja4 | 3.15E-74 | 0.47471523 | 0.39 | 0.24 | 6.51E-70 | intestine_colon | Gja4 |
| Cdipt | 5.92E-74 | 0.37342884 | 0.39 | 0.24 | 1.23E-69 | intestine_colon | Cdipt |
| Camk1 | 6.19E-73 | 0.37379722 | 0.48 | 0.32 | 1.28E-68 | intestine_colon | Camk1 |
| Hsp90ab12 | 5.04E-72 | 0.25071673 | 0.96 | 0.91 | 1.04E-67 | intestine_colon | Hsp90ab1 |
| Arpc1b1 | 5.31E-71 | 0.33014925 | 0.77 | 0.64 | 1.10E-66 | intestine_colon | Arpc1b |
| Hmox2 | 2.53E-70 | 0.33070646 | 0.33 | 0.19 | 5.24E-66 | intestine_colon | Hmox2 |
| Mtx2 | 4.39E-70 | 0.2970682 | 0.28 | 0.16 | 9.08E-66 | intestine_colon | Mtx2 |
| Slc25a5 | 2.51E-69 | 0.34496904 | 0.68 | 0.53 | 5.19E-65 | intestine_colon | Slc25a5 |
| Hexa | 1.25E-68 | 0.31204831 | 0.38 | 0.23 | 2.58E-64 | intestine_colon | Hexa |
| Adam15 | 6.53E-67 | 0.34790045 | 0.6 | 0.43 | 1.35E-62 | intestine_colon | Adam15 |
| Ctnnb12 | 3.24E-66 | 0.30192846 | 0.69 | 0.51 | 6.70E-62 | intestine_colon | Ctnnb1 |
| Pttg1 | 1.10E-65 | 0.35568779 | 0.52 | 0.36 | 2.28E-61 | intestine_colon | Pttg1 |
| Map1lc3b | 6.68E-65 | 0.33054699 | 0.75 | 0.62 | 1.38E-60 | intestine_colon | Map1lc3b |
| Tinagl1 | 1.24E-64 | 0.26860535 | 0.8 | 0.64 | 2.57E-60 | intestine_colon | Tinagl1 |
| Ano61 | 3.87E-64 | 0.31971278 | 0.3 | 0.17 | 8.01E-60 | intestine_colon | Ano6 |
| Map1lc3a1 | 3.93E-64 | 0.32411912 | 0.6 | 0.43 | 8.13E-60 | intestine_colon | Map1lc3a |
| Cyb5r31 | 1.42E-62 | 0.34842808 | 0.62 | 0.47 | 2.94E-58 | intestine_colon | Cyb5r3 |
| St3gal61 | 3.52E-62 | 0.27340744 | 0.55 | 0.37 | 7.29E-58 | intestine_colon | St3gal6 |
| Aldoa1 | 4.49E-61 | 0.28149555 | 0.78 | 0.65 | 9.29E-57 | intestine_colon | Aldoa |
| Krtcap2 | 5.05E-61 | 0.32355735 | 0.68 | 0.52 | 1.05E-56 | intestine_colon | Krtcap2 |
| Slc9a3r21 | 5.64E-61 | 0.25903292 | 0.9 | 0.81 | 1.17E-56 | intestine_colon | Slc9a3r2 |
| Nid11 | 9.74E-60 | 0.28322343 | 0.31 | 0.19 | 2.01E-55 | intestine_colon | Nid1 |
| Gnb2l1 | 2.12E-59 | 0.2710261 | 0.75 | 0.56 | 4.39E-55 | intestine_colon | Gnb2l1 |
| Mob4 | 1.59E-58 | 0.28344244 | 0.42 | 0.27 | 3.30E-54 | intestine_colon | Mob4 |
| Capns11 | 5.28E-58 | 0.28632913 | 0.76 | 0.63 | 1.09E-53 | intestine_colon | Capns1 |
| Degs1 | 8.98E-58 | 0.28101073 | 0.54 | 0.39 | 1.86E-53 | intestine_colon | Degs1 |
| Ppp1r12a1 | 3.14E-56 | 0.25304851 | 0.36 | 0.23 | 6.49E-52 | intestine_colon | Ppp1r12a |
| Fam174b | 4.24E-56 | 0.32052603 | 0.27 | 0.16 | 8.78E-52 | intestine_colon | Fam174b |
| Hspa82 | 5.45E-56 | 0.27995709 | 0.95 | 0.91 | 1.13E-51 | intestine_colon | Hspa8 |
| Smpdl3a2 | 9.87E-56 | 0.27458191 | 0.49 | 0.34 | 2.04E-51 | intestine_colon | Smpdl3a |
| Myct1 | 1.26E-55 | 0.2542059 | 0.66 | 0.49 | 2.60E-51 | intestine_colon | Myct1 |
| Rrbp11 | 1.54E-55 | 0.30969756 | 0.61 | 0.45 | 3.19E-51 | intestine_colon | Rrbp1 |
| Tm2d3 | 2.74E-55 | 0.27980734 | 0.3 | 0.18 | 5.67E-51 | intestine_colon | Tm2d3 |
| Capn22 | 5.50E-55 | 0.25004752 | 0.47 | 0.32 | 1.14E-50 | intestine_colon | Capn2 |
| Zfp36l21 | 5.97E-55 | 0.30218167 | 0.41 | 0.27 | 1.23E-50 | intestine_colon | Zfp36l2 |
| Atp5b | 9.97E-55 | 0.27060653 | 0.81 | 0.7 | 2.06E-50 | intestine_colon | Atp5b |
| G3bp1 | 3.66E-54 | 0.29725933 | 0.5 | 0.35 | 7.57E-50 | intestine_colon | G3bp1 |
| Rnaset2a | 1.68E-53 | 0.27130701 | 0.49 | 0.33 | 3.49E-49 | intestine_colon | Rnaset2a |
| Ndrg12 | 2.87E-53 | 0.26886936 | 0.53 | 0.36 | 5.95E-49 | intestine_colon | Ndrg1 |

| | | | | | | | |
|---|---|---|---|---|---|---|---|
| Tpi11 | 5.55E-52 | 0.27550265 | 0.41 | 0.27 | 1.15E-47 | intestine_colon | Tpi1 |
| Ccdc28b1 | 3.42E-51 | 0.30338442 | 0.33 | 0.21 | 7.07E-47 | intestine_colon | Ccdc28b |
| Tnfaip11 | 3.46E-51 | 0.25728653 | 0.5 | 0.35 | 7.16E-47 | intestine_colon | Tnfaip1 |
| Ppp2r1a | 5.61E-50 | 0.25354676 | 0.38 | 0.25 | 1.16E-45 | intestine_colon | Ppp2r1a |
| Fbxo22 | 1.29E-49 | 0.25570437 | 0.25 | 0.15 | 2.68E-45 | intestine_colon | Fbxo22 |
| Taldo11 | 1.51E-49 | 0.26987193 | 0.55 | 0.4 | 3.13E-45 | intestine_colon | Taldo1 |
| Tsc22d31 | 2.12E-49 | 0.27960824 | 0.52 | 0.38 | 4.39E-45 | intestine_colon | Tsc22d3 |
| Cpne3 | 5.71E-49 | 0.26734867 | 0.3 | 0.19 | 1.18E-44 | intestine_colon | Cpne3 |
| Atp2b41 | 1.43E-48 | 0.25989094 | 0.32 | 0.21 | 2.95E-44 | intestine_colon | Atp2b4 |
| Stat31 | 2.50E-47 | 0.26128177 | 0.36 | 0.24 | 5.17E-43 | intestine_colon | Stat3 |
| Txn11 | 1.62E-46 | 0.26772746 | 0.76 | 0.65 | 3.36E-42 | intestine_colon | Txn1 |
| Gbp2 | 1.09E-44 | 0.28533963 | 0.34 | 0.22 | 2.25E-40 | intestine_colon | Gbp2 |
| Nfe2l2 | 1.37E-44 | 0.25903838 | 0.45 | 0.31 | 2.83E-40 | intestine_colon | Nfe2l2 |
| Itga61 | 2.90E-44 | 0.28238207 | 0.45 | 0.32 | 6.00E-40 | intestine_colon | Itga6 |
| Tmed101 | 4.18E-43 | 0.26165678 | 0.7 | 0.58 | 8.65E-39 | intestine_colon | Tmed10 |
| Slc25a3 | 2.86E-42 | 0.27228132 | 0.65 | 0.52 | 5.91E-38 | intestine_colon | Slc25a3 |
| Lpar6 | 4.48E-40 | 0.25055898 | 0.37 | 0.25 | 9.27E-36 | intestine_colon | Lpar6 |
| Nudt42 | 7.48E-40 | 0.27523628 | 0.28 | 0.18 | 1.55E-35 | intestine_colon | Nudt4 |
| Mast4 | 9.46E-28 | 0.25394485 | 0.39 | 0.29 | 1.96E-23 | intestine_colon | Mast4 |
| Fth1 | 0.00179919 | 0.2637733 | 0.94 | 0.91 | 1 | intestine_colon | Fth1 |
| AY0361181 | 0 | 3.02799603 | 0.99 | 0.74 | 0 | intestine_small | AY036118 |
| Scgb3a1 | 0 | 3.01379891 | 0.42 | 0.04 | 0 | intestine_small | Scgb3a1 |
| Mmrn1 | 0 | 2.836298 | 0.46 | 0.06 | 0 | intestine_small | Mmrn1 |
| Rnd1 | 0 | 2.32193388 | 0.42 | 0.11 | 0 | intestine_small | Rnd1 |
| Reln | 0 | 2.28459558 | 0.35 | 0.04 | 0 | intestine_small | Reln |
| Atf31 | 0 | 2.28126133 | 0.56 | 0.18 | 0 | intestine_small | Atf3 |
| Thbs1 | 0 | 2.20689083 | 0.37 | 0.03 | 0 | intestine_small | Thbs1 |
| Bcr1 | 0 | 2.15253873 | 0.44 | 0.09 | 0 | intestine_small | Bcr |
| Fgl2 | 0 | 1.79934364 | 0.32 | 0.06 | 0 | intestine_small | Fgl2 |
| Cyp26b11 | 0 | 1.6148465 | 0.54 | 0.15 | 0 | intestine_small | Cyp26b1 |
| Gm424181 | 0 | 1.07955705 | 1 | 0.79 | 0 | intestine_small | Gm42418 |
| Ccl21a1 | 0 | 1.04137862 | 0.81 | 0.23 | 0 | intestine_small | Ccl21a |
| Apold12 | 2.68E-281 | 2.04002985 | 0.59 | 0.23 | 5.54E-277 | intestine_small | Apold1 |
| Nkx2-31 | 2.27E-269 | 1.79367174 | 0.4 | 0.11 | 4.70E-265 | intestine_small | Nkx2-3 |
| Egr11 | 9.47E-263 | 2.13407792 | 0.68 | 0.33 | 1.96E-258 | intestine_small | Egr1 |
| Zfp362 | 2.28E-260 | 2.15155885 | 0.72 | 0.41 | 4.72E-256 | intestine_small | Zfp36 |
| 8430408G22Rik2 | 3.15E-244 | 2.0048343 | 0.58 | 0.24 | 6.52E-240 | intestine_small | 8430408G22Rik |
| Rps27rt1 | 3.01E-218 | 0.87196535 | 0.93 | 0.84 | 6.22E-214 | intestine_small | Rps27rt |
| Nr4a11 | 3.99E-169 | 1.63472866 | 0.38 | 0.14 | 8.26E-165 | intestine_small | Nr4a1 |
| Rasa41 | 1.99E-148 | 1.37506973 | 0.31 | 0.1 | 4.12E-144 | intestine_small | Rasa4 |
| Nfkbiz | 6.69E-142 | 1.49825911 | 0.31 | 0.11 | 1.38E-137 | intestine_small | Nfkbiz |

| | | | | | | | |
|---|---|---|---|---|---|---|---|
| Gm100762 | 1.80E-141 | 1.10776528 | 0.67 | 0.48 | 3.73E-137 | intestine_small | Gm10076 |
| Hspb12 | 4.68E-141 | 1.05974553 | 0.9 | 0.72 | 9.68E-137 | intestine_small | Hspb1 |
| Bhlhe402 | 6.60E-135 | 1.42669084 | 0.4 | 0.18 | 1.37E-130 | intestine_small | Bhlhe40 |
| Tgfbr22 | 1.54E-134 | 1.35304039 | 0.51 | 0.27 | 3.18E-130 | intestine_small | Tgfbr2 |
| Gm42726 | 1.06E-132 | 1.07460513 | 0.37 | 0.15 | 2.19E-128 | intestine_small | Gm42726 |
| Fos2 | 9.75E-130 | 1.42355179 | 0.78 | 0.56 | 2.02E-125 | intestine_small | Fos |
| Gm100731 | 2.33E-127 | 1.01912846 | 0.63 | 0.41 | 4.82E-123 | intestine_small | Gm10073 |
| Prss232 | 5.07E-126 | 1.02371219 | 0.56 | 0.31 | 1.05E-121 | intestine_small | Prss23 |
| H2-K11 | 1.02E-120 | 0.75412736 | 0.91 | 0.84 | 2.12E-116 | intestine_small | H2-K1 |
| Gm10263 | 7.92E-112 | 0.92027527 | 0.28 | 0.1 | 1.64E-107 | intestine_small | Gm10263 |
| Gramd1a1 | 6.32E-111 | 1.11416822 | 0.3 | 0.11 | 1.31E-106 | intestine_small | Gramd1a |
| Lyve1 | 1.83E-104 | 1.67605798 | 0.33 | 0.14 | 3.79E-100 | intestine_small | Lyve1 |
| Gm20001 | 3.52E-104 | 1.0027319 | 0.46 | 0.25 | 7.29E-100 | intestine_small | Gm2000 |
| Irf11 | 6.70E-102 | 1.50124524 | 0.49 | 0.3 | 1.39E-97 | intestine_small | Irf1 |
| Gm98431 | 1.06E-99 | 0.7800167 | 0.76 | 0.62 | 2.20E-95 | intestine_small | Gm9843 |
| Junb2 | 9.77E-99 | 0.93520992 | 0.81 | 0.63 | 2.02E-94 | intestine_small | Junb |
| Hsp90ab13 | 1.18E-98 | 0.59314701 | 0.92 | 0.91 | 2.44E-94 | intestine_small | Hsp90ab1 |
| Ier22 | 1.06E-97 | 1.1149639 | 0.73 | 0.56 | 2.19E-93 | intestine_small | Ier2 |
| H2-Q4 | 1.56E-96 | 1.03681195 | 0.61 | 0.44 | 3.22E-92 | intestine_small | H2-Q4 |
| Rpl23a-ps31 | 3.06E-96 | 0.75688969 | 0.77 | 0.65 | 6.32E-92 | intestine_small | Rpl23a-ps3 |
| Nrp21 | 2.57E-95 | 1.41056024 | 0.41 | 0.22 | 5.32E-91 | intestine_small | Nrp2 |
| Tppp31 | 1.10E-94 | 1.35880789 | 0.38 | 0.18 | 2.28E-90 | intestine_small | Tppp3 |
| Timp31 | 2.04E-93 | 1.01875951 | 0.79 | 0.7 | 4.22E-89 | intestine_small | Timp3 |
| Gadd45g1 | 5.21E-92 | 1.34776118 | 0.49 | 0.29 | 1.08E-87 | intestine_small | Gadd45g |
| Flt4 | 4.71E-84 | 1.33364232 | 0.39 | 0.21 | 9.75E-80 | intestine_small | Flt4 |
| Ubc1 | 3.04E-82 | 0.72521245 | 0.87 | 0.84 | 6.29E-78 | intestine_small | Ubc |
| Rpl36-ps31 | 3.61E-82 | 0.89781988 | 0.54 | 0.37 | 7.46E-78 | intestine_small | Rpl36-ps3 |
| Plvap2 | 1.15E-81 | 0.83168366 | 0.79 | 0.58 | 2.37E-77 | intestine_small | Plvap |
| Cyr612 | 1.85E-79 | 1.35815305 | 0.45 | 0.27 | 3.84E-75 | intestine_small | Cyr61 |
| Rpl13-ps31 | 2.72E-78 | 0.91472825 | 0.53 | 0.36 | 5.63E-74 | intestine_small | Rpl13-ps3 |
| Wdr891 | 3.40E-78 | 0.61520702 | 0.8 | 0.72 | 7.03E-74 | intestine_small | Wdr89 |
| Ncoa72 | 2.53E-77 | 1.13206015 | 0.33 | 0.16 | 5.24E-73 | intestine_small | Ncoa7 |
| Igfbp4 | 3.54E-77 | 0.91602252 | 0.58 | 0.38 | 7.32E-73 | intestine_small | Igfbp4 |
| Mall1 | 3.74E-76 | 0.93283542 | 0.29 | 0.13 | 7.74E-72 | intestine_small | Mall |
| Gm118081 | 1.05E-75 | 0.68002841 | 0.71 | 0.58 | 2.18E-71 | intestine_small | Gm11808 |
| Mast41 | 3.45E-75 | 1.24621042 | 0.46 | 0.3 | 7.15E-71 | intestine_small | Mast4 |
| Lcn2 | 5.11E-75 | 0.86853865 | 0.25 | 0.1 | 1.06E-70 | intestine_small | Lcn2 |
| Tnfaip21 | 3.15E-73 | 0.94314518 | 0.32 | 0.16 | 6.51E-69 | intestine_small | Tnfaip2 |
| Nfkbia1 | 1.34E-72 | 1.27928928 | 0.71 | 0.63 | 2.76E-68 | intestine_small | Nfkbia |
| Socs31 | 3.00E-71 | 1.10120388 | 0.48 | 0.28 | 6.20E-67 | intestine_small | Socs3 |
| Col15a12 | 5.93E-71 | 1.03529802 | 0.32 | 0.16 | 1.23E-66 | intestine_small | Col15a1 |

| | | | | | | | |
|---|---|---|---|---|---|---|---|
| Camk2n11 | 1.41E-69 | 1.17171622 | 0.28 | 0.14 | 2.92E-65 | intestine_small | Camk2n1 |
| Ier3 | 2.07E-68 | 1.0130376 | 0.56 | 0.38 | 4.28E-64 | intestine_small | Ier3 |
| Ltbp43 | 1.47E-67 | 0.91884259 | 0.65 | 0.5 | 3.04E-63 | intestine_small | Ltbp4 |
| Plk22 | 4.12E-65 | 0.99065252 | 0.58 | 0.43 | 8.53E-61 | intestine_small | Plk2 |
| Osmr2 | 1.31E-64 | 0.97788317 | 0.29 | 0.15 | 2.71E-60 | intestine_small | Osmr |
| Lama42 | 3.17E-64 | 0.97107391 | 0.33 | 0.18 | 6.55E-60 | intestine_small | Lama4 |
| Ifi27l2a | 1.68E-63 | 0.96752592 | 0.26 | 0.12 | 3.48E-59 | intestine_small | Ifi27l2a |
| Sox182 | 2.98E-63 | 0.84717872 | 0.63 | 0.51 | 6.16E-59 | intestine_small | Sox18 |
| Rpl10-ps31 | 8.99E-63 | 0.81155433 | 0.39 | 0.23 | 1.86E-58 | intestine_small | Rpl10-ps3 |
| Gm9844 | 1.50E-62 | 0.85048907 | 0.27 | 0.13 | 3.10E-58 | intestine_small | Gm9844 |
| Hspa83 | 1.78E-60 | 0.40017527 | 0.92 | 0.91 | 3.68E-56 | intestine_small | Hspa8 |
| Uba521 | 2.41E-57 | 0.53804278 | 0.77 | 0.72 | 4.98E-53 | intestine_small | Uba52 |
| Rps26-ps11 | 4.87E-57 | 0.77293114 | 0.4 | 0.25 | 1.01E-52 | intestine_small | Rps26-ps1 |
| Cd91 | 1.52E-55 | 0.49506307 | 0.77 | 0.62 | 3.14E-51 | intestine_small | Cd9 |
| Rpl351 | 4.51E-55 | 0.41069956 | 0.85 | 0.87 | 9.32E-51 | intestine_small | Rpl35 |
| Rpl27-ps31 | 4.04E-54 | 0.77788658 | 0.47 | 0.33 | 8.36E-50 | intestine_small | Rpl27-ps3 |
| Btg22 | 2.49E-53 | 0.93349619 | 0.6 | 0.49 | 5.15E-49 | intestine_small | Btg2 |
| Rpl9-ps61 | 7.46E-53 | 0.70763147 | 0.56 | 0.44 | 1.54E-48 | intestine_small | Rpl9-ps6 |
| H2-Q71 | 3.66E-52 | 0.82552601 | 0.53 | 0.39 | 7.56E-48 | intestine_small | H2-Q7 |
| Pim31 | 5.88E-52 | 1.05019372 | 0.48 | 0.36 | 1.22E-47 | intestine_small | Pim3 |
| H2-D1 | 4.52E-51 | 0.32093318 | 0.96 | 0.96 | 9.34E-47 | intestine_small | H2-D1 |
| Gm87301 | 5.49E-51 | 0.6892507 | 0.55 | 0.42 | 1.14E-46 | intestine_small | Gm8730 |
| Scn1b2 | 1.04E-50 | 0.90948037 | 0.29 | 0.16 | 2.15E-46 | intestine_small | Scn1b |
| Grrp13 | 1.23E-50 | 0.95286864 | 0.47 | 0.34 | 2.54E-46 | intestine_small | Grrp1 |
| Ace1 | 7.35E-48 | 0.76127759 | 0.38 | 0.23 | 1.52E-43 | intestine_small | Ace |
| Mmrn21 | 2.16E-47 | 0.68198054 | 0.62 | 0.52 | 4.47E-43 | intestine_small | Mmrn2 |
| Gm94931 | 3.59E-46 | 0.63637861 | 0.62 | 0.52 | 7.42E-42 | intestine_small | Gm9493 |
| Cav12 | 8.17E-45 | 0.3704365 | 0.8 | 0.66 | 1.69E-40 | intestine_small | Cav1 |
| Tagln21 | 1.65E-42 | 0.49753971 | 0.78 | 0.74 | 3.42E-38 | intestine_small | Tagln2 |
| Gm8186 | 5.02E-42 | 0.74628804 | 0.3 | 0.18 | 1.04E-37 | intestine_small | Gm8186 |
| Txnip1 | 1.15E-40 | 0.66775468 | 0.71 | 0.66 | 2.38E-36 | intestine_small | Txnip |
| Neat1 | 1.43E-40 | 0.74810811 | 0.55 | 0.45 | 2.95E-36 | intestine_small | Neat1 |
| Rps12-ps31 | 1.74E-39 | 0.50828665 | 0.67 | 0.58 | 3.60E-35 | intestine_small | Rps12-ps3 |
| Psme2b | 3.28E-39 | 0.65925249 | 0.31 | 0.19 | 6.79E-35 | intestine_small | Psme2b |
| Gm6133 | 2.02E-36 | 0.64832569 | 0.39 | 0.28 | 4.17E-32 | intestine_small | Gm6133 |
| Tsc22d11 | 1.67E-34 | 0.45847806 | 0.74 | 0.67 | 3.45E-30 | intestine_small | Tsc22d1 |
| Esm11 | 2.42E-34 | 0.51302395 | 0.25 | 0.14 | 5.00E-30 | intestine_small | Esm1 |
| Igfbp32 | 1.07E-31 | 0.32343594 | 0.4 | 0.26 | 2.21E-27 | intestine_small | Igfbp3 |
| Ece11 | 1.46E-31 | 0.64608601 | 0.54 | 0.46 | 3.02E-27 | intestine_small | Ece1 |
| Dusp13 | 4.33E-31 | 0.54084201 | 0.74 | 0.66 | 8.95E-27 | intestine_small | Dusp1 |
| Col4a12 | 6.07E-31 | 0.56153583 | 0.64 | 0.55 | 1.26E-26 | intestine_small | Col4a1 |

| | | | | | | | |
|---|---|---|---|---|---|---|---|
| Aqp12 | 1.17E-29 | 0.34520247 | 0.67 | 0.49 | 2.42E-25 | intestine_small | Aqp1 |
| S100a112 | 4.46E-29 | 0.46155253 | 0.65 | 0.59 | 9.24E-25 | intestine_small | S100a11 |
| Hspa5 | 4.83E-29 | 0.66374059 | 0.62 | 0.57 | 1.00E-24 | intestine_small | Hspa5 |
| Tmsb10 | 5.64E-29 | 0.42154119 | 0.78 | 0.75 | 1.17E-24 | intestine_small | Tmsb10 |
| Serinc3 | 1.65E-28 | 0.33685799 | 0.84 | 0.85 | 3.41E-24 | intestine_small | Serinc3 |
| Tspan18 | 3.69E-27 | 0.83525118 | 0.34 | 0.24 | 7.63E-23 | intestine_small | Tspan18 |
| Nfib2 | 9.20E-27 | 0.48084669 | 0.63 | 0.56 | 1.90E-22 | intestine_small | Nfib |
| H2-Q6 | 9.34E-27 | 0.6708466 | 0.33 | 0.24 | 1.93E-22 | intestine_small | H2-Q6 |
| Mfge81 | 2.23E-26 | 0.58942745 | 0.52 | 0.45 | 4.60E-22 | intestine_small | Mfge8 |
| Rpl6l1 | 5.87E-26 | 0.53647801 | 0.55 | 0.48 | 1.21E-21 | intestine_small | Rpl6l |
| Gm100361 | 6.25E-26 | 0.56268404 | 0.43 | 0.34 | 1.29E-21 | intestine_small | Gm10036 |
| Hist1h1c | 1.75E-25 | 0.80120816 | 0.25 | 0.17 | 3.61E-21 | intestine_small | Hist1h1c |
| Aplp22 | 2.20E-25 | 0.57270361 | 0.69 | 0.67 | 4.55E-21 | intestine_small | Aplp2 |
| Ackr3 | 9.26E-25 | 0.59498822 | 0.4 | 0.3 | 1.92E-20 | intestine_small | Ackr3 |
| Stab1 | 1.23E-24 | 0.8522378 | 0.37 | 0.28 | 2.54E-20 | intestine_small | Stab1 |
| Cxcl123 | 1.29E-24 | 0.45571353 | 0.61 | 0.5 | 2.68E-20 | intestine_small | Cxcl12 |
| Actn42 | 2.27E-24 | 0.56065051 | 0.58 | 0.52 | 4.70E-20 | intestine_small | Actn4 |
| Podxl2 | 4.80E-24 | 0.5988652 | 0.59 | 0.51 | 9.93E-20 | intestine_small | Podxl |
| Kdr2 | 3.42E-23 | 0.44028416 | 0.7 | 0.62 | 7.08E-19 | intestine_small | Kdr |
| Fam43a2 | 8.63E-23 | 0.67808782 | 0.39 | 0.31 | 1.79E-18 | intestine_small | Fam43a |
| Ptrf2 | 2.89E-22 | 0.47398714 | 0.58 | 0.48 | 5.97E-18 | intestine_small | Ptrf |
| Cd2001 | 4.30E-21 | 0.41099047 | 0.7 | 0.67 | 8.90E-17 | intestine_small | Cd200 |
| Wars2 | 6.44E-21 | 0.69376534 | 0.28 | 0.2 | 1.33E-16 | intestine_small | Wars |
| Ahnak3 | 7.37E-21 | 0.50343421 | 0.53 | 0.45 | 1.53E-16 | intestine_small | Ahnak |
| S1pr11 | 9.05E-21 | 0.54543149 | 0.53 | 0.46 | 1.87E-16 | intestine_small | S1pr1 |
| Robo41 | 9.55E-21 | 0.5858625 | 0.41 | 0.34 | 1.98E-16 | intestine_small | Robo4 |
| Tgm2 | 5.40E-20 | 0.65502093 | 0.46 | 0.4 | 1.12E-15 | intestine_small | Tgm2 |
| Iigp11 | 6.99E-19 | 0.7246042 | 0.36 | 0.28 | 1.45E-14 | intestine_small | Iigp1 |
| Myadm2 | 1.04E-18 | 0.61649512 | 0.33 | 0.25 | 2.14E-14 | intestine_small | Myadm |
| Maf | 2.73E-18 | 1.11249173 | 0.27 | 0.2 | 5.65E-14 | intestine_small | Maf |
| Cd24a2 | 2.93E-18 | 0.43719813 | 0.34 | 0.25 | 6.06E-14 | intestine_small | Cd24a |
| Rps18-ps3 | 9.48E-18 | 0.46050463 | 0.25 | 0.18 | 1.96E-13 | intestine_small | Rps18-ps3 |
| H3f3b2 | 9.52E-18 | 0.58128347 | 0.85 | 0.89 | 1.97E-13 | intestine_small | H3f3b |
| Ddit41 | 4.11E-17 | 0.64922601 | 0.26 | 0.19 | 8.50E-13 | intestine_small | Ddit4 |
| mt-Atp81 | 4.40E-17 | 0.53505845 | 0.29 | 0.21 | 9.11E-13 | intestine_small | mt-Atp8 |
| Tcf41 | 1.31E-16 | 0.34096771 | 0.71 | 0.67 | 2.72E-12 | intestine_small | Tcf4 |
| Sparcl14 | 2.60E-16 | 0.29482216 | 0.6 | 0.5 | 5.38E-12 | intestine_small | Sparcl1 |
| Comt | 2.90E-16 | 0.6711463 | 0.33 | 0.27 | 6.00E-12 | intestine_small | Comt |
| Sdpr1 | 2.99E-16 | 0.41658942 | 0.77 | 0.71 | 6.19E-12 | intestine_small | Sdpr |
| Dock91 | 5.45E-16 | 0.71728383 | 0.27 | 0.2 | 1.13E-11 | intestine_small | Dock9 |
| Ccnl1 | 3.73E-15 | 0.67854107 | 0.31 | 0.25 | 7.72E-11 | intestine_small | Ccnl1 |

| | | | | | | | |
|---|---|---|---|---|---|---|---|
| Ebf12 | 5.07E-15 | 0.32604495 | 0.5 | 0.41 | 1.05E-10 | intestine_small | Ebf1 |
| Slc43a32 | 6.55E-15 | 0.37922125 | 0.5 | 0.43 | 1.36E-10 | intestine_small | Slc43a3 |
| Tmem882 | 9.65E-15 | 0.46882172 | 0.49 | 0.45 | 2.00E-10 | intestine_small | Tmem88 |
| Nfat51 | 1.40E-14 | 0.63032021 | 0.28 | 0.21 | 2.90E-10 | intestine_small | Nfat5 |
| Gja1 | 1.46E-14 | 0.5186569 | 0.31 | 0.24 | 3.02E-10 | intestine_small | Gja1 |
| Gm26917 | 2.01E-13 | 0.39477546 | 0.36 | 0.29 | 4.16E-09 | intestine_small | Gm26917 |
| Ppfibp11 | 2.01E-13 | 0.63359009 | 0.26 | 0.2 | 4.17E-09 | intestine_small | Ppfibp1 |
| Lmna1 | 3.76E-13 | 0.55464848 | 0.38 | 0.32 | 7.77E-09 | intestine_small | Lmna |
| Ptpn18 | 4.30E-13 | 0.54412567 | 0.26 | 0.2 | 8.91E-09 | intestine_small | Ptpn18 |
| Pde2a1 | 4.70E-13 | 0.42789548 | 0.34 | 0.27 | 9.72E-09 | intestine_small | Pde2a |
| Notch11 | 5.12E-13 | 0.61592963 | 0.33 | 0.28 | 1.06E-08 | intestine_small | Notch1 |
| Gm8797 | 6.81E-13 | 0.46067072 | 0.28 | 0.22 | 1.41E-08 | intestine_small | Gm8797 |
| Ccnl2 | 1.63E-12 | 0.5345088 | 0.32 | 0.27 | 3.37E-08 | intestine_small | Ccnl2 |
| Tns1 | 2.99E-12 | 0.59501674 | 0.43 | 0.39 | 6.20E-08 | intestine_small | Tns1 |
| Ppp1r15a | 3.60E-12 | 0.67206595 | 0.33 | 0.28 | 7.45E-08 | intestine_small | Ppp1r15a |
| Ndrg13 | 3.66E-12 | 0.55023065 | 0.41 | 0.38 | 7.58E-08 | intestine_small | Ndrg1 |
| Stip1 | 4.74E-12 | 0.48298011 | 0.28 | 0.23 | 9.81E-08 | intestine_small | Stip1 |
| Prex21 | 2.09E-11 | 0.56000939 | 0.43 | 0.39 | 4.32E-07 | intestine_small | Prex2 |
| Zfp36l12 | 2.84E-11 | 0.53221683 | 0.55 | 0.54 | 5.87E-07 | intestine_small | Zfp36l1 |
| Orai11 | 7.38E-11 | 0.50931261 | 0.29 | 0.24 | 1.53E-06 | intestine_small | Orai1 |
| Mcf2l2 | 1.76E-10 | 0.5247629 | 0.26 | 0.2 | 3.65E-06 | intestine_small | Mcf2l |
| Pdlim11 | 2.37E-10 | 0.39998018 | 0.37 | 0.31 | 4.90E-06 | intestine_small | Pdlim1 |
| Midn | 8.61E-10 | 0.57455549 | 0.27 | 0.22 | 1.78E-05 | intestine_small | Midn |
| Vwf | 1.00E-09 | 0.27426285 | 0.28 | 0.21 | 2.08E-05 | intestine_small | Vwf |
| Lims1 | 2.36E-09 | 0.67920331 | 0.3 | 0.26 | 4.88E-05 | intestine_small | Lims1 |
| Zfp36l22 | 2.95E-09 | 0.53363435 | 0.32 | 0.28 | 6.11E-05 | intestine_small | Zfp36l2 |
| Gbp21 | 5.15E-09 | 0.50352879 | 0.27 | 0.23 | 0.00010661 | intestine_small | Gbp2 |
| Klf62 | 5.38E-09 | 0.41089339 | 0.48 | 0.44 | 0.00011137 | intestine_small | Klf6 |
| Lrrc8a1 | 5.80E-09 | 0.55014887 | 0.33 | 0.3 | 0.00012008 | intestine_small | Lrrc8a |
| Cyb5r32 | 6.37E-09 | 0.43278604 | 0.49 | 0.48 | 0.00013188 | intestine_small | Cyb5r3 |
| 4931406P16Rik1 | 7.09E-09 | 0.4832663 | 0.43 | 0.41 | 0.0001467 | intestine_small | 4931406P16Rik |
| Lrrc58 | 1.16E-08 | 0.44430915 | 0.39 | 0.36 | 0.00024027 | intestine_small | Lrrc58 |
| Cst32 | 1.40E-08 | 0.27073566 | 0.86 | 0.89 | 0.00028983 | intestine_small | Cst3 |
| Nr2f2 | 1.54E-08 | 0.61983424 | 0.27 | 0.22 | 0.00031898 | intestine_small | Nr2f2 |
| Lrrc8c2 | 5.25E-08 | 0.5299663 | 0.32 | 0.3 | 0.00108606 | intestine_small | Lrrc8c |
| Mecom2 | 6.38E-08 | 0.49843929 | 0.26 | 0.22 | 0.00132004 | intestine_small | Mecom |
| Col4a22 | 7.43E-08 | 0.36985518 | 0.48 | 0.46 | 0.00153827 | intestine_small | Col4a2 |
| Elf11 | 7.98E-08 | 0.48183573 | 0.28 | 0.24 | 0.00165163 | intestine_small | Elf1 |
| Irf2bpl2 | 8.74E-08 | 0.51694322 | 0.31 | 0.28 | 0.00180852 | intestine_small | Irf2bpl |
| Tap21 | 9.53E-08 | 0.45655124 | 0.34 | 0.31 | 0.00197264 | intestine_small | Tap2 |
| Stat32 | 1.80E-07 | 0.45611119 | 0.28 | 0.25 | 0.0037349 | intestine_small | Stat3 |

| | | | | | | | |
|---|---|---|---|---|---|---|---|
| Arhgap292 | 2.56E-07 | 0.40572235 | 0.36 | 0.32 | 0.00529046 | intestine_small | Arhgap29 |
| Thrsp2 | 2.67E-07 | 0.38335278 | 0.32 | 0.28 | 0.00552584 | intestine_small | Thrsp |
| Hdac7 | 4.63E-07 | 0.552674 | 0.33 | 0.31 | 0.00958518 | intestine_small | Hdac7 |
| Myh91 | 4.91E-07 | 0.48532142 | 0.34 | 0.32 | 0.01016922 | intestine_small | Myh9 |
| Emp11 | 5.42E-07 | 0.34736813 | 0.27 | 0.23 | 0.01120601 | intestine_small | Emp1 |
| Tshz21 | 6.84E-07 | 0.52140929 | 0.41 | 0.39 | 0.01414663 | intestine_small | Tshz2 |
| Rapgef42 | 8.06E-07 | 0.49424136 | 0.29 | 0.25 | 0.0166746 | intestine_small | Rapgef4 |
| Pabpc11 | 8.17E-07 | 0.36802099 | 0.5 | 0.49 | 0.01690224 | intestine_small | Pabpc1 |
| Akap131 | 8.52E-07 | 0.47100816 | 0.33 | 0.3 | 0.01763863 | intestine_small | Akap13 |
| Hsp90aa12 | 9.97E-07 | 0.3801366 | 0.47 | 0.47 | 0.02062337 | intestine_small | Hsp90aa1 |
| Hspg22 | 1.01E-06 | 0.37567271 | 0.44 | 0.4 | 0.02088545 | intestine_small | Hspg2 |
| Nme2 | 1.08E-06 | 0.30828843 | 0.29 | 0.25 | 0.02240636 | intestine_small | Nme2 |
| 1810011O10Rik1 | 1.55E-06 | 0.30844692 | 0.5 | 0.47 | 0.03198832 | intestine_small | 1810011O10Rik |
| Nr3c11 | 2.30E-06 | 0.48990969 | 0.32 | 0.29 | 0.04760424 | intestine_small | Nr3c1 |
| AW1120103 | 2.90E-06 | 0.2878401 | 0.54 | 0.52 | 0.05990474 | intestine_small | AW112010 |
| Tns2 | 3.48E-06 | 0.47692317 | 0.25 | 0.22 | 0.0719684 | intestine_small | Tns2 |
| Son | 4.89E-06 | 0.26079694 | 0.64 | 0.67 | 0.10117103 | intestine_small | Son |
| Prpf4b1 | 5.33E-06 | 0.39724981 | 0.26 | 0.22 | 0.11029888 | intestine_small | Prpf4b |
| Iqgap12 | 5.36E-06 | 0.4830965 | 0.36 | 0.34 | 0.11096185 | intestine_small | Iqgap1 |
| Nfe2l21 | 6.53E-06 | 0.41656425 | 0.34 | 0.32 | 0.13501908 | intestine_small | Nfe2l2 |
| Dab2 | 7.24E-06 | 0.4838905 | 0.3 | 0.27 | 0.14988611 | intestine_small | Dab2 |
| Sox171 | 7.36E-06 | 0.41809008 | 0.42 | 0.39 | 0.15237288 | intestine_small | Sox17 |
| Scarb12 | 8.39E-06 | 0.5341722 | 0.41 | 0.41 | 0.17350756 | intestine_small | Scarb1 |
| Fkbp42 | 1.02E-05 | 0.37938845 | 0.37 | 0.36 | 0.21110316 | intestine_small | Fkbp4 |
| Tra2b | 1.07E-05 | 0.40159766 | 0.43 | 0.43 | 0.22191452 | intestine_small | Tra2b |
| Fgd51 | 1.14E-05 | 0.48789079 | 0.29 | 0.26 | 0.23655872 | intestine_small | Fgd5 |
| Ptms | 1.16E-05 | 0.29172074 | 0.57 | 0.59 | 0.23979913 | intestine_small | Ptms |
| Mllt41 | 1.20E-05 | 0.31784891 | 0.3 | 0.27 | 0.24741736 | intestine_small | Mllt4 |
| Ucp21 | 1.42E-05 | 0.48928485 | 0.28 | 0.26 | 0.2935775 | intestine_small | Ucp2 |
| Ifi351 | 1.51E-05 | 0.36293376 | 0.38 | 0.37 | 0.31198835 | intestine_small | Ifi35 |
| Efna11 | 1.54E-05 | 0.27395117 | 0.39 | 0.37 | 0.31814401 | intestine_small | Efna1 |
| Socs22 | 1.70E-05 | 0.40369431 | 0.28 | 0.26 | 0.35247106 | intestine_small | Socs2 |
| Acly | 1.79E-05 | 0.34425466 | 0.29 | 0.26 | 0.37033012 | intestine_small | Acly |
| Mat2a | 1.79E-05 | 0.36248179 | 0.38 | 0.37 | 0.3707706 | intestine_small | Mat2a |
| Lrrfip12 | 2.89E-05 | 0.42708579 | 0.31 | 0.29 | 0.59853131 | intestine_small | Lrrfip1 |
| Psmb81 | 4.24E-05 | 0.30503608 | 0.51 | 0.53 | 0.87637308 | intestine_small | Psmb8 |
| Mcl11 | 5.31E-05 | 0.48241909 | 0.3 | 0.29 | 1 | intestine_small | Mcl1 |
| Atf4 | 9.48E-05 | 0.43817759 | 0.32 | 0.31 | 1 | intestine_small | Atf4 |
| Clk1 | 0.00018617 | 0.46275774 | 0.33 | 0.32 | 1 | intestine_small | Clk1 |
| Xbp1 | 0.00021138 | 0.33162315 | 0.38 | 0.37 | 1 | intestine_small | Xbp1 |
| Mbnl11 | 0.00025766 | 0.41176143 | 0.32 | 0.31 | 1 | intestine_small | Mbnl1 |

| | | | | | | | |
|---|---|---|---|---|---|---|---|
| Ier5 | 0.00027766 | 0.33822928 | 0.48 | 0.5 | 1 | intestine_small | Ier5 |
| Sema6a1 | 0.0003009 | 0.39078026 | 0.3 | 0.28 | 1 | intestine_small | Sema6a |
| Lgals9 | 0.0005345 | 0.31207805 | 0.4 | 0.4 | 1 | intestine_small | Lgals9 |
| St13 | 0.00054772 | 0.27203612 | 0.43 | 0.44 | 1 | intestine_small | St13 |
| Csde11 | 0.00077018 | 0.38666266 | 0.33 | 0.33 | 1 | intestine_small | Csde1 |
| Utrn2 | 0.00088971 | 0.31888775 | 0.31 | 0.29 | 1 | intestine_small | Utrn |
| Klf101 | 0.00100899 | 0.41258959 | 0.25 | 0.24 | 1 | intestine_small | Klf10 |
| Zbtb201 | 0.00115779 | 0.29040805 | 0.52 | 0.54 | 1 | intestine_small | Zbtb20 |
| Sat1 | 0.00182261 | 0.54205018 | 0.4 | 0.42 | 1 | intestine_small | Sat1 |
| Hspe1 | 0.00191621 | 0.378455 | 0.36 | 0.37 | 1 | intestine_small | Hspe1 |
| Hnrnpl | 0.0023854 | 0.34630232 | 0.25 | 0.24 | 1 | intestine_small | Hnrnpl |
| Fosb1 | 0.00240816 | 0.9056644 | 0.3 | 0.29 | 1 | intestine_small | Fosb |
| Msn2 | 0.00275306 | 0.31100251 | 0.55 | 0.59 | 1 | intestine_small | Msn |
| Hnrnph1 | 0.00284939 | 0.43291555 | 0.29 | 0.29 | 1 | intestine_small | Hnrnph1 |
| H2-T231 | 0.00325487 | 0.28212397 | 0.4 | 0.41 | 1 | intestine_small | H2-T23 |
| Wnk11 | 0.0037197 | 0.35027401 | 0.34 | 0.34 | 1 | intestine_small | Wnk1 |
| Arap31 | 0.0048385 | 0.38727512 | 0.27 | 0.26 | 1 | intestine_small | Arap3 |
| Btg11 | 0.00538388 | 0.26942164 | 0.49 | 0.52 | 1 | intestine_small | Btg1 |
| Actg11 | 0.00579411 | 0.35794541 | 0.85 | 0.89 | 1 | intestine_small | Actg1 |
| Tmem176a1 | 0.00715946 | 0.28016746 | 0.43 | 0.43 | 1 | intestine_small | Tmem176a |
| Sdcbp | 0.00783639 | 0.29641016 | 0.47 | 0.5 | 1 | intestine_small | Sdcbp |
| Itpkb1 | 0.00802124 | 0.36001159 | 0.4 | 0.41 | 1 | intestine_small | Itpkb |
| Eif2s2 | 0.00805308 | 0.32998206 | 0.45 | 0.48 | 1 | intestine_small | Eif2s2 |
| Hes11 | 0.0099268 | 0.44550728 | 0.47 | 0.5 | 1 | intestine_small | Hes1 |
| Igfbp5 | 0 | 4.41610501 | 0.78 | 0.05 | 0 | kidney | Igfbp5 |
| Pbx1 | 0 | 3.81941681 | 0.89 | 0.15 | 0 | kidney | Pbx1 |
| Meis21 | 0 | 3.15317743 | 0.9 | 0.25 | 0 | kidney | Meis2 |
| Esm12 | 0 | 2.82369266 | 0.51 | 0.1 | 0 | kidney | Esm1 |
| Rsad21 | 0 | 2.53480512 | 0.69 | 0.2 | 0 | kidney | Rsad2 |
| Igfbp33 | 0 | 2.35801075 | 0.58 | 0.23 | 0 | kidney | Igfbp3 |
| Ptpru | 0 | 2.22486937 | 0.38 | 0.04 | 0 | kidney | Ptpru |
| Plscr22 | 0 | 2.19603105 | 0.67 | 0.23 | 0 | kidney | Plscr2 |
| Tnfaip22 | 0 | 2.06990979 | 0.5 | 0.13 | 0 | kidney | Tnfaip2 |
| Dkk2 | 0 | 1.92539541 | 0.26 | 0 | 0 | kidney | Dkk2 |
| Exoc3l21 | 0 | 1.89634114 | 0.51 | 0.18 | 0 | kidney | Exoc3l2 |
| Malat1 | 0 | 1.84891199 | 1 | 0.97 | 0 | kidney | Malat1 |
| Emcn1 | 0 | 1.84178206 | 0.92 | 0.64 | 0 | kidney | Emcn |
| Ifi441 | 0 | 1.82045968 | 0.4 | 0.11 | 0 | kidney | Ifi44 |
| Irx3 | 0 | 1.70252812 | 0.31 | 0.06 | 0 | kidney | Irx3 |
| Gm269171 | 0 | 1.59314753 | 0.48 | 0.27 | 0 | kidney | Gm26917 |
| Gm427261 | 0 | 1.44424738 | 0.35 | 0.13 | 0 | kidney | Gm42726 |

| | | | | | | | |
|---|---|---|---|---|---|---|---|
| Ramp32 | 0 | 1.43534513 | 0.43 | 0.21 | 0 | kidney | Ramp3 |
| Rapgef43 | 0 | 1.41940025 | 0.45 | 0.23 | 0 | kidney | Rapgef4 |
| 1810011O10Rik2 | 0 | 1.41464685 | 0.71 | 0.45 | 0 | kidney | 1810011O10Rik |
| Fam117b1 | 0 | 1.41219724 | 0.3 | 0.1 | 0 | kidney | Fam117b |
| Plac8 | 0 | 1.37026314 | 0.26 | 0.06 | 0 | kidney | Plac8 |
| Ehd3 | 0 | 1.28762412 | 0.42 | 0.14 | 0 | kidney | Ehd3 |
| Plpp11 | 0 | 1.12571571 | 0.8 | 0.58 | 0 | kidney | Plpp1 |
| Plvap3 | 0 | 1.03532313 | 0.89 | 0.56 | 0 | kidney | Plvap |
| Timp32 | 0 | 1.03197049 | 0.8 | 0.69 | 0 | kidney | Timp3 |
| S100a61 | 6.66E-284 | 0.96851829 | 0.54 | 0.29 | 1.38E-279 | kidney | S100a6 |
| Cd24a3 | 3.04E-282 | 1.21578516 | 0.45 | 0.23 | 6.29E-278 | kidney | Cd24a |
| Ifi2031 | 8.62E-264 | 1.31042255 | 0.49 | 0.31 | 1.78E-259 | kidney | Ifi203 |
| Jup | 8.53E-253 | 1.29133356 | 0.55 | 0.39 | 1.77E-248 | kidney | Jup |
| Lmo21 | 4.18E-246 | 1.24213931 | 0.54 | 0.39 | 8.65E-242 | kidney | Lmo2 |
| B2m | 1.28E-240 | 0.43828648 | 0.99 | 0.98 | 2.65E-236 | kidney | B2m |
| Fam167b | 7.33E-234 | 0.92204143 | 0.48 | 0.27 | 1.52E-229 | kidney | Fam167b |
| 9530082P21Rik | 2.01E-227 | 1.21681896 | 0.27 | 0.12 | 4.16E-223 | kidney | 9530082P21Rik |
| Ly6c12 | 3.35E-216 | 0.49505243 | 0.95 | 0.84 | 6.93E-212 | kidney | Ly6c1 |
| Nrp12 | 6.72E-216 | 0.76349082 | 0.78 | 0.64 | 1.39E-211 | kidney | Nrp1 |
| AU021092 | 1.42E-212 | 0.99598773 | 0.51 | 0.32 | 2.94E-208 | kidney | AU021092 |
| Gimap4 | 2.44E-211 | 1.10301328 | 0.56 | 0.44 | 5.05E-207 | kidney | Gimap4 |
| Limch1 | 5.47E-211 | 1.23944045 | 0.38 | 0.22 | 1.13E-206 | kidney | Limch1 |
| Selm2 | 2.51E-204 | 1.00040833 | 0.55 | 0.42 | 5.20E-200 | kidney | Selm |
| Ebf13 | 9.24E-193 | 0.89463898 | 0.57 | 0.39 | 1.91E-188 | kidney | Ebf1 |
| Myo1c | 3.74E-192 | 1.13226203 | 0.47 | 0.34 | 7.73E-188 | kidney | Myo1c |
| Irf71 | 4.54E-192 | 1.27465452 | 0.34 | 0.19 | 9.39E-188 | kidney | Irf7 |
| Rsrp1 | 8.60E-178 | 0.83207651 | 0.67 | 0.64 | 1.78E-173 | kidney | Rsrp1 |
| Kdr3 | 1.81E-177 | 0.8627832 | 0.7 | 0.61 | 3.74E-173 | kidney | Kdr |
| Rgcc2 | 4.28E-173 | 0.78366017 | 0.61 | 0.42 | 8.85E-169 | kidney | Rgcc |
| Ly6e2 | 6.44E-164 | 0.49452771 | 0.94 | 0.95 | 1.33E-159 | kidney | Ly6e |
| Il10rb1 | 1.37E-160 | 1.01202653 | 0.5 | 0.39 | 2.83E-156 | kidney | Il10rb |
| Arhgap311 | 5.47E-157 | 0.75099812 | 0.69 | 0.64 | 1.13E-152 | kidney | Arhgap31 |
| Gimap6 | 2.04E-153 | 0.77206515 | 0.67 | 0.66 | 4.23E-149 | kidney | Gimap6 |
| Gnas1 | 5.94E-153 | 0.44102056 | 0.91 | 0.91 | 1.23E-148 | kidney | Gnas |
| Sncg2 | 1.46E-147 | 1.02892575 | 0.3 | 0.16 | 3.03E-143 | kidney | Sncg |
| Flt13 | 1.94E-145 | 0.45198286 | 0.8 | 0.69 | 4.02E-141 | kidney | Flt1 |
| Cd812 | 7.06E-128 | 0.36629836 | 0.88 | 0.82 | 1.46E-123 | kidney | Cd81 |
| Sgk11 | 1.61E-122 | 0.80055735 | 0.6 | 0.54 | 3.32E-118 | kidney | Sgk1 |
| Podxl3 | 7.92E-120 | 0.80123096 | 0.58 | 0.5 | 1.64E-115 | kidney | Podxl |
| Zbtb202 | 4.43E-119 | 0.89255984 | 0.57 | 0.54 | 9.16E-115 | kidney | Zbtb20 |
| S100a101 | 7.02E-114 | 0.57509754 | 0.7 | 0.67 | 1.45E-109 | kidney | S100a10 |

| | | | | | | | |
|---|---|---|---|---|---|---|---|
| Cald11 | 2.35E-113 | 0.85519772 | 0.45 | 0.35 | 4.87E-109 | kidney | Cald1 |
| Plat1 | 1.95E-112 | 1.56560487 | 0.3 | 0.18 | 4.04E-108 | kidney | Plat |
| Hspb13 | 1.28E-110 | 0.26113429 | 0.83 | 0.72 | 2.64E-106 | kidney | Hspb1 |
| Arglu1 | 2.40E-110 | 1.01762812 | 0.41 | 0.32 | 4.96E-106 | kidney | Arglu1 |
| Neat11 | 2.10E-109 | 0.92346275 | 0.52 | 0.45 | 4.34E-105 | kidney | Neat1 |
| F2r | 4.55E-109 | 1.03115811 | 0.31 | 0.2 | 9.41E-105 | kidney | F2r |
| Ehd42 | 4.45E-105 | 0.53042376 | 0.64 | 0.52 | 9.20E-101 | kidney | Ehd4 |
| Tmem204 | 6.49E-103 | 0.83953892 | 0.48 | 0.42 | 1.34E-98 | kidney | Tmem204 |
| Gimap5 | 2.59E-101 | 0.91416653 | 0.41 | 0.33 | 5.35E-97 | kidney | Gimap5 |
| Plpp32 | 2.63E-100 | 0.4483231 | 0.71 | 0.62 | 5.45E-96 | kidney | Plpp3 |
| Ifitm31 | 6.33E-100 | 0.35387346 | 0.95 | 0.96 | 1.31E-95 | kidney | Ifitm3 |
| Rbm39 | 1.62E-97 | 0.54859074 | 0.74 | 0.77 | 3.35E-93 | kidney | Rbm39 |
| Vamp51 | 1.83E-87 | 0.66145138 | 0.58 | 0.54 | 3.78E-83 | kidney | Vamp5 |
| Ly6a2 | 9.90E-86 | 0.33034609 | 0.91 | 0.88 | 2.05E-81 | kidney | Ly6a |
| Ptprb1 | 5.54E-83 | 0.28852989 | 0.89 | 0.89 | 1.15E-78 | kidney | Ptprb |
| Mef2c2 | 2.81E-82 | 0.62145249 | 0.51 | 0.45 | 5.82E-78 | kidney | Mef2c |
| Slc9a3r22 | 1.46E-81 | 0.40047286 | 0.81 | 0.82 | 3.02E-77 | kidney | Slc9a3r2 |
| Btnl91 | 9.74E-81 | 0.83534335 | 0.26 | 0.16 | 2.01E-76 | kidney | Btnl9 |
| Hspg23 | 4.63E-77 | 0.78077749 | 0.46 | 0.39 | 9.58E-73 | kidney | Hspg2 |
| Gas62 | 9.01E-76 | 0.72942399 | 0.52 | 0.49 | 1.86E-71 | kidney | Gas6 |
| Laptm4a | 1.88E-75 | 0.37303188 | 0.81 | 0.88 | 3.90E-71 | kidney | Laptm4a |
| Txn12 | 1.48E-71 | 0.54835112 | 0.63 | 0.66 | 3.07E-67 | kidney | Txn1 |
| Rhob1 | 2.14E-70 | 0.5390431 | 0.59 | 0.55 | 4.43E-66 | kidney | Rhob |
| Cd2002 | 9.03E-69 | 0.44702222 | 0.7 | 0.67 | 1.87E-64 | kidney | Cd200 |
| Tie11 | 2.54E-67 | 0.67272757 | 0.57 | 0.6 | 5.26E-63 | kidney | Tie1 |
| Oasl2 | 1.99E-66 | 0.9775801 | 0.25 | 0.17 | 4.11E-62 | kidney | Oasl2 |
| 2010111I01Rik1 | 3.49E-65 | 0.88520591 | 0.3 | 0.23 | 7.23E-61 | kidney | 2010111I01Rik |
| Cd300lg2 | 1.01E-64 | 0.54735578 | 0.5 | 0.42 | 2.09E-60 | kidney | Cd300lg |
| Bst2 | 1.27E-64 | 0.50335276 | 0.7 | 0.74 | 2.63E-60 | kidney | Bst2 |
| Prkcdbp1 | 4.54E-63 | 0.46184798 | 0.58 | 0.51 | 9.39E-59 | kidney | Prkcdbp |
| Meox12 | 5.35E-62 | 0.80798277 | 0.29 | 0.21 | 1.11E-57 | kidney | Meox1 |
| Atox12 | 1.22E-61 | 0.36919427 | 0.75 | 0.8 | 2.52E-57 | kidney | Atox1 |
| Efnb21 | 4.75E-57 | 0.89606545 | 0.26 | 0.18 | 9.82E-53 | kidney | Efnb2 |
| Eng | 7.43E-57 | 0.43471739 | 0.82 | 0.86 | 1.54E-52 | kidney | Eng |
| Tek1 | 1.03E-56 | 0.73615536 | 0.49 | 0.49 | 2.13E-52 | kidney | Tek |
| Plscr1 | 2.58E-56 | 0.81715816 | 0.3 | 0.23 | 5.33E-52 | kidney | Plscr1 |
| Btg12 | 2.73E-56 | 0.68314698 | 0.52 | 0.52 | 5.66E-52 | kidney | Btg1 |
| Pde2a2 | 6.72E-56 | 0.71577652 | 0.34 | 0.26 | 1.39E-51 | kidney | Pde2a |
| Mndal1 | 5.26E-52 | 0.73693703 | 0.38 | 0.32 | 1.09E-47 | kidney | Mndal |
| Srgn1 | 1.03E-51 | 0.3822684 | 0.7 | 0.71 | 2.13E-47 | kidney | Srgn |
| Txnip2 | 1.32E-48 | 0.4894938 | 0.64 | 0.66 | 2.73E-44 | kidney | Txnip |

| | | | | | | | |
|---|---|---|---|---|---|---|---|
| S100a113 | 2.07E-48 | 0.42027421 | 0.61 | 0.6 | 4.27E-44 | kidney | S100a11 |
| Tm4sf11 | 2.07E-45 | 0.30608279 | 0.83 | 0.86 | 4.28E-41 | kidney | Tm4sf1 |
| Nisch | 2.68E-43 | 0.7660652 | 0.4 | 0.38 | 5.55E-39 | kidney | Nisch |
| Lamb2 | 4.78E-42 | 0.85561921 | 0.26 | 0.21 | 9.88E-38 | kidney | Lamb2 |
| Selk | 2.61E-41 | 0.37978963 | 0.65 | 0.68 | 5.41E-37 | kidney | Selk |
| Nfat52 | 7.63E-41 | 0.81479781 | 0.26 | 0.21 | 1.58E-36 | kidney | Nfat5 |
| Gimap1 | 3.88E-39 | 0.72425808 | 0.45 | 0.46 | 8.03E-35 | kidney | Gimap1 |
| Tcf42 | 2.72E-35 | 0.36140899 | 0.65 | 0.68 | 5.64E-31 | kidney | Tcf4 |
| Apbb21 | 4.62E-35 | 0.64624109 | 0.38 | 0.35 | 9.55E-31 | kidney | Apbb2 |
| Gng51 | 2.11E-33 | 0.32904052 | 0.7 | 0.77 | 4.36E-29 | kidney | Gng5 |
| Son1 | 9.35E-33 | 0.45196306 | 0.6 | 0.68 | 1.93E-28 | kidney | Son |
| S100a1 | 2.21E-32 | 0.61997129 | 0.4 | 0.38 | 4.56E-28 | kidney | S100a1 |
| Pnisr | 2.59E-32 | 0.75878695 | 0.34 | 0.32 | 5.35E-28 | kidney | Pnisr |
| Ndel11 | 2.06E-31 | 0.71558461 | 0.3 | 0.26 | 4.26E-27 | kidney | Ndel1 |
| Rdx2 | 4.13E-30 | 0.44241493 | 0.54 | 0.56 | 8.55E-26 | kidney | Rdx |
| Ttc14 | 1.32E-27 | 0.73077486 | 0.26 | 0.23 | 2.73E-23 | kidney | Ttc14 |
| Mat2a1 | 1.84E-27 | 0.68898065 | 0.37 | 0.37 | 3.80E-23 | kidney | Mat2a |
| Efna12 | 9.08E-27 | 0.61437269 | 0.38 | 0.37 | 1.88E-22 | kidney | Efna1 |
| Itpkb2 | 2.90E-26 | 0.67139312 | 0.41 | 0.41 | 6.00E-22 | kidney | Itpkb |
| Uqcc21 | 4.70E-26 | 0.61658906 | 0.41 | 0.42 | 9.72E-22 | kidney | Uqcc2 |
| Adgrl42 | 7.66E-25 | 0.36010792 | 0.53 | 0.53 | 1.59E-20 | kidney | Adgrl4 |
| Sp1001 | 3.35E-24 | 0.65966441 | 0.36 | 0.36 | 6.94E-20 | kidney | Sp100 |
| H1f01 | 3.50E-24 | 0.68974885 | 0.39 | 0.4 | 7.23E-20 | kidney | H1f0 |
| Isg151 | 4.51E-22 | 0.74095405 | 0.33 | 0.31 | 9.34E-18 | kidney | Isg15 |
| Slc6a62 | 1.11E-20 | 0.30951025 | 0.46 | 0.42 | 2.30E-16 | kidney | Slc6a6 |
| Rbms11 | 3.65E-20 | 0.45124354 | 0.52 | 0.57 | 7.55E-16 | kidney | Rbms1 |
| Heg11 | 7.35E-20 | 0.59447379 | 0.4 | 0.41 | 1.52E-15 | kidney | Heg1 |
| Cmtm81 | 7.75E-20 | 0.5376048 | 0.37 | 0.36 | 1.60E-15 | kidney | Cmtm8 |
| Fus | 1.33E-18 | 0.4956581 | 0.49 | 0.55 | 2.75E-14 | kidney | Fus |
| Atrx1 | 2.53E-18 | 0.62830229 | 0.33 | 0.32 | 5.23E-14 | kidney | Atrx |
| Cd38 | 2.54E-15 | 0.52456437 | 0.33 | 0.32 | 5.26E-11 | kidney | Cd38 |
| Krit1 | 2.73E-15 | 0.68414989 | 0.26 | 0.24 | 5.65E-11 | kidney | Krit1 |
| Dlc11 | 5.25E-15 | 0.53037172 | 0.41 | 0.42 | 1.09E-10 | kidney | Dlc1 |
| Xaf1 | 5.83E-15 | 0.59809902 | 0.31 | 0.3 | 1.21E-10 | kidney | Xaf1 |
| Lims21 | 6.83E-15 | 0.31583948 | 0.48 | 0.5 | 1.41E-10 | kidney | Lims2 |
| Gngt2 | 9.22E-15 | 0.30052448 | 0.54 | 0.58 | 1.91E-10 | kidney | Gngt2 |
| Eva1b | 2.58E-14 | 0.50395747 | 0.42 | 0.47 | 5.33E-10 | kidney | Eva1b |
| Tspan15 | 1.05E-13 | 0.55966013 | 0.26 | 0.24 | 2.18E-09 | kidney | Tspan15 |
| Tshz22 | 5.45E-13 | 0.53847723 | 0.39 | 0.39 | 1.13E-08 | kidney | Tshz2 |
| Tgfbr23 | 7.51E-13 | 0.57979328 | 0.29 | 0.28 | 1.55E-08 | kidney | Tgfbr2 |
| Hes12 | 1.64E-12 | 0.48765692 | 0.47 | 0.5 | 3.39E-08 | kidney | Hes1 |

| | | | | | | | |
|---|---|---|---|---|---|---|---|
| Id2 | 2.42E-12 | 0.45972757 | 0.34 | 0.33 | 5.00E-08 | kidney | Id2 |
| Akap132 | 8.70E-12 | 0.57301499 | 0.3 | 0.3 | 1.80E-07 | kidney | Akap13 |
| Arhgef1 | 1.79E-10 | 0.62268755 | 0.33 | 0.35 | 3.70E-06 | kidney | Arhgef1 |
| Cbfa2t3 | 1.02E-09 | 0.64022992 | 0.27 | 0.27 | 2.11E-05 | kidney | Cbfa2t3 |
| Luzp11 | 4.38E-09 | 0.53911598 | 0.35 | 0.38 | 9.07E-05 | kidney | Luzp1 |
| Rasgrp31 | 5.90E-09 | 0.48957773 | 0.31 | 0.32 | 0.00012202 | kidney | Rasgrp3 |
| Cox17 | 1.65E-08 | 0.38752276 | 0.43 | 0.48 | 0.00034074 | kidney | Cox17 |
| Tmbim1 | 3.48E-08 | 0.54136759 | 0.28 | 0.28 | 0.00072045 | kidney | Tmbim1 |
| Peak11 | 4.37E-08 | 0.55732994 | 0.25 | 0.25 | 0.00090432 | kidney | Peak1 |
| Asap11 | 6.03E-08 | 0.5128768 | 0.28 | 0.28 | 0.00124689 | kidney | Asap1 |
| Xiap1 | 1.20E-07 | 0.60660797 | 0.28 | 0.3 | 0.00249331 | kidney | Xiap |
| Marcks1 | 2.49E-07 | 0.55753211 | 0.35 | 0.38 | 0.00514601 | kidney | Marcks |
| Rbm25 | 5.06E-07 | 0.56523237 | 0.33 | 0.36 | 0.01047721 | kidney | Rbm25 |
| Notch12 | 6.95E-07 | 0.29253287 | 0.21 | 0.29 | 0.014384 | kidney | Notch1 |
| PISD | 1.32E-06 | 0.50026346 | 0.25 | 0.25 | 0.02724197 | kidney | PISD |
| Irf21 | 1.34E-06 | 0.26057082 | 0.21 | 0.28 | 0.02768179 | kidney | Irf2 |
| Klf3 | 1.57E-06 | 0.51147286 | 0.34 | 0.37 | 0.03257068 | kidney | Klf3 |
| Rpn1 | 2.57E-06 | 0.33549336 | 0.21 | 0.28 | 0.05307522 | kidney | Rpn1 |
| Ifnar21 | 2.60E-06 | 0.52476099 | 0.32 | 0.35 | 0.05373367 | kidney | Ifnar2 |
| Elk31 | 2.86E-06 | 0.39145305 | 0.48 | 0.56 | 0.05917696 | kidney | Elk3 |
| Wnk12 | 5.79E-06 | 0.26757093 | 0.26 | 0.35 | 0.11982385 | kidney | Wnk1 |
| Clec1a | 1.03E-05 | 0.3552217 | 0.29 | 0.29 | 0.21342785 | kidney | Clec1a |
| Arap32 | 1.07E-05 | 0.31816111 | 0.2 | 0.27 | 0.22134564 | kidney | Arap3 |
| Vgll4 | 1.10E-05 | 0.51419152 | 0.3 | 0.33 | 0.2269129 | kidney | Vgll4 |
| Hnrnph11 | 1.11E-05 | 0.31237262 | 0.22 | 0.29 | 0.22919856 | kidney | Hnrnph1 |
| Ephb4 | 1.85E-05 | 0.32312768 | 0.24 | 0.31 | 0.38342955 | kidney | Ephb4 |
| Rtp4 | 2.42E-05 | 0.45589966 | 0.27 | 0.28 | 0.50120409 | kidney | Rtp4 |
| Sat11 | 2.67E-05 | 0.26522909 | 0.33 | 0.43 | 0.5527328 | kidney | Sat1 |
| Pura1 | 3.98E-05 | 0.32890121 | 0.26 | 0.35 | 0.82336666 | kidney | Pura |
| Pcdh171 | 5.10E-05 | 0.62810991 | 0.25 | 0.26 | 1 | kidney | Pcdh17 |
| Git2 | 6.56E-05 | 0.32034686 | 0.21 | 0.28 | 1 | kidney | Git2 |
| Prex22 | 6.78E-05 | 0.44598298 | 0.36 | 0.4 | 1 | kidney | Prex2 |
| Sec14l11 | 6.87E-05 | 0.30514645 | 0.21 | 0.28 | 1 | kidney | Sec14l1 |
| Csnk2a1 | 9.98E-05 | 0.25348077 | 0.21 | 0.28 | 1 | kidney | Csnk2a1 |
| Cebpg | 0.00010993 | 0.27693708 | 0.21 | 0.28 | 1 | kidney | Cebpg |
| Dync1i2 | 0.00011884 | 0.53670669 | 0.38 | 0.44 | 1 | kidney | Dync1i2 |
| Tln11 | 0.00012305 | 0.28061658 | 0.25 | 0.32 | 1 | kidney | Tln1 |
| Fubp1 | 0.00014157 | 0.53720504 | 0.26 | 0.28 | 1 | kidney | Fubp1 |
| B230219D22Rik1 | 0.00021443 | 0.25684931 | 0.25 | 0.33 | 1 | kidney | B230219D22Rik |
| Sertad2 | 0.0003333 | 0.30526671 | 0.2 | 0.27 | 1 | kidney | Sertad2 |
| Fam43a3 | 0.00046417 | 0.40608941 | 0.29 | 0.31 | 1 | kidney | Fam43a |

| | | | | | | | |
|---|---|---|---|---|---|---|---|
| Clec2d1 | 0.00049208 | 0.43373165 | 0.36 | 0.4 | 1 | kidney | Clec2d |
| Wbp5 | 0.00050229 | 0.38170071 | 0.37 | 0.42 | 1 | kidney | Wbp5 |
| Ncor11 | 0.00079446 | 0.480905 | 0.31 | 0.34 | 1 | kidney | Ncor1 |
| Fgd52 | 0.00092231 | 0.33084558 | 0.21 | 0.27 | 1 | kidney | Fgd5 |
| Slc50a1 | 0.00171014 | 0.26196885 | 0.31 | 0.4 | 1 | kidney | Slc50a1 |
| Npr1 | 0.00219967 | 0.27693129 | 0.24 | 0.3 | 1 | kidney | Npr1 |
| Cpd1 | 0.00246344 | 0.35697847 | 0.2 | 0.25 | 1 | kidney | Cpd |
| Ppp1r11 | 0.00361101 | 0.2614473 | 0.29 | 0.38 | 1 | kidney | Ppp1r11 |
| Zfp207 | 0.00433251 | 0.32190539 | 0.21 | 0.26 | 1 | kidney | Zfp207 |
| Smad71 | 0.00590566 | 0.36043952 | 0.24 | 0.31 | 1 | kidney | Smad7 |
| Mrpl33 | 0.00594293 | 0.25266541 | 0.32 | 0.42 | 1 | kidney | Mrpl33 |
| Ccdc12 | 0.00713703 | 0.2630505 | 0.32 | 0.42 | 1 | kidney | Ccdc12 |
| Mgat1 | 0.00863242 | 0.33451559 | 0.21 | 0.26 | 1 | kidney | Mgat1 |
| Brd2 | 0.00954903 | 0.32611145 | 0.2 | 0.25 | 1 | kidney | Brd2 |
| Dnase1l3 | 0 | 4.83095891 | 0.88 | 0 | 0 | liver | Dnase1l3 |
| Clec4g | 0 | 4.67595181 | 0.97 | 0.07 | 0 | liver | Clec4g |
| Rspo3 | 0 | 3.58055458 | 0.39 | 0.02 | 0 | liver | Rspo3 |
| Ehd31 | 0 | 3.47348461 | 0.91 | 0.08 | 0 | liver | Ehd3 |
| Fcgr2b | 0 | 3.36727506 | 0.76 | 0.06 | 0 | liver | Fcgr2b |
| Apoe1 | 0 | 3.12936289 | 0.99 | 0.49 | 0 | liver | Apoe |
| Gm12840 | 0 | 3.07846298 | 0.55 | 0.04 | 0 | liver | Gm12840 |
| Oit3 | 0 | 2.8506738 | 0.67 | 0.01 | 0 | liver | Oit3 |
| Bmp2 | 0 | 2.73175087 | 0.69 | 0.06 | 0 | liver | Bmp2 |
| Wnt2 | 0 | 2.7142094 | 0.59 | 0.04 | 0 | liver | Wnt2 |
| Gpr182 | 0 | 2.70970981 | 0.86 | 0.2 | 0 | liver | Gpr182 |
| Mrc1 | 0 | 2.67689036 | 0.63 | 0.02 | 0 | liver | Mrc1 |
| Sepp1 | 0 | 2.6466532 | 1 | 0.74 | 0 | liver | Sepp1 |
| Lgmn | 0 | 2.48773952 | 0.74 | 0.14 | 0 | liver | Lgmn |
| Gatm1 | 0 | 2.45270865 | 0.66 | 0.12 | 0 | liver | Gatm |
| Serpina3g | 0 | 2.44139352 | 0.52 | 0.01 | 0 | liver | Serpina3g |
| Maf1 | 0 | 2.3946961 | 0.79 | 0.14 | 0 | liver | Maf |
| Lgals1 | 0 | 2.35709849 | 0.67 | 0.17 | 0 | liver | Lgals1 |
| Cd55 | 0 | 2.29540229 | 0.76 | 0.11 | 0 | liver | Cd55 |
| Bgn | 0 | 2.16335008 | 0.73 | 0.14 | 0 | liver | Bgn |
| Cd59a1 | 0 | 2.12303841 | 0.74 | 0.17 | 0 | liver | Cd59a |
| Fam167b1 | 0 | 2.11986859 | 0.79 | 0.23 | 0 | liver | Fam167b |
| Il6st | 0 | 2.08441307 | 0.8 | 0.29 | 0 | liver | Il6st |
| Slc29a1 | 0 | 2.07168292 | 0.8 | 0.3 | 0 | liver | Slc29a1 |
| Mt11 | 0 | 2.04022598 | 0.59 | 0.1 | 0 | liver | Mt1 |
| Adam23 | 0 | 2.01935631 | 0.4 | 0 | 0 | liver | Adam23 |
| F8 | 0 | 2.00407866 | 0.56 | 0.04 | 0 | liver | F8 |

| | | | | | | | |
|---|---|---|---|---|---|---|---|
| Tpbgl | 0 | 1.99124443 | 0.5 | 0.01 | 0 | liver | Tpbgl |
| Ctsl | 0 | 1.98814136 | 0.86 | 0.53 | 0 | liver | Ctsl |
| Igfbp41 | 0 | 1.96389637 | 0.89 | 0.33 | 0 | liver | Igfbp4 |
| Tmem106a | 0 | 1.94256417 | 0.51 | 0.05 | 0 | liver | Tmem106a |
| Clec1b | 0 | 1.9231632 | 0.41 | 0 | 0 | liver | Clec1b |
| Plxnc1 | 0 | 1.85712803 | 0.43 | 0.01 | 0 | liver | Plxnc1 |
| Lamp2 | 0 | 1.84196262 | 0.8 | 0.38 | 0 | liver | Lamp2 |
| Dpp4 | 0 | 1.83797019 | 0.5 | 0.04 | 0 | liver | Dpp4 |
| Fcgrt1 | 0 | 1.78959794 | 0.84 | 0.43 | 0 | liver | Fcgrt |
| Msr1 | 0 | 1.68891269 | 0.3 | 0 | 0 | liver | Msr1 |
| Marcksl1 | 0 | 1.68468728 | 0.71 | 0.21 | 0 | liver | Marcksl1 |
| Fcer1g | 0 | 1.68461384 | 0.36 | 0.05 | 0 | liver | Fcer1g |
| Gpihbp11 | 0 | 1.66137071 | 0.94 | 0.53 | 0 | liver | Gpihbp1 |
| Degs11 | 0 | 1.65580259 | 0.78 | 0.36 | 0 | liver | Degs1 |
| Stab11 | 0 | 1.64909696 | 0.7 | 0.24 | 0 | liver | Stab1 |
| Lifr | 0 | 1.61173301 | 0.67 | 0.24 | 0 | liver | Lifr |
| Calm2 | 0 | 1.61096803 | 0.94 | 0.74 | 0 | liver | Calm2 |
| Spint2 | 0 | 1.61077352 | 0.52 | 0.07 | 0 | liver | Spint2 |
| Cotl1 | 0 | 1.57104305 | 0.59 | 0.13 | 0 | liver | Cotl1 |
| Tcn2 | 0 | 1.56260188 | 0.8 | 0.46 | 0 | liver | Tcn2 |
| Cxcl16 | 0 | 1.54984804 | 0.61 | 0.15 | 0 | liver | Cxcl16 |
| Ctsb | 0 | 1.54679126 | 0.86 | 0.51 | 0 | liver | Ctsb |
| Cd381 | 0 | 1.54556778 | 0.73 | 0.27 | 0 | liver | Cd38 |
| Ftl1 | 0 | 1.51549025 | 0.99 | 0.92 | 0 | liver | Ftl1 |
| Ccdc80 | 0 | 1.50531857 | 0.44 | 0.03 | 0 | liver | Ccdc80 |
| Stab2 | 0 | 1.49554109 | 0.64 | 0.11 | 0 | liver | Stab2 |
| Cd302 | 0 | 1.49026167 | 0.47 | 0.07 | 0 | liver | Cd302 |
| Tinagl11 | 0 | 1.45556583 | 0.91 | 0.63 | 0 | liver | Tinagl1 |
| Aqp13 | 0 | 1.4542168 | 0.96 | 0.44 | 0 | liver | Aqp1 |
| Car82 | 0 | 1.43240953 | 0.61 | 0.2 | 0 | liver | Car8 |
| Dab21 | 0 | 1.43117826 | 0.62 | 0.24 | 0 | liver | Dab2 |
| Emilin1 | 0 | 1.41686759 | 0.47 | 0.09 | 0 | liver | Emilin1 |
| Hmha1 | 0 | 1.41117419 | 0.34 | 0 | 0 | liver | Hmha1 |
| Slc43a33 | 0 | 1.41063006 | 0.75 | 0.39 | 0 | liver | Slc43a3 |
| Nr2f21 | 0 | 1.40943197 | 0.61 | 0.18 | 0 | liver | Nr2f2 |
| Gja41 | 0 | 1.3958675 | 0.6 | 0.21 | 0 | liver | Gja4 |
| Rcn1 | 0 | 1.39094356 | 0.52 | 0.16 | 0 | liver | Rcn1 |
| Ptgs11 | 0 | 1.38395588 | 0.51 | 0.16 | 0 | liver | Ptgs1 |
| Dhrs31 | 0 | 1.37281554 | 0.68 | 0.33 | 0 | liver | Dhrs3 |
| Bace2 | 0 | 1.36808572 | 0.54 | 0.14 | 0 | liver | Bace2 |
| Tmem2 | 0 | 1.35469726 | 0.52 | 0.14 | 0 | liver | Tmem2 |

| | | | | | | | |
|---|---|---|---|---|---|---|---|
| Manf | 0 | 1.3448869 | 0.74 | 0.44 | 0 | liver | Manf |
| Sh3bp51 | 0 | 1.33583324 | 0.65 | 0.27 | 0 | liver | Sh3bp5 |
| Plpp12 | 0 | 1.33568804 | 0.87 | 0.58 | 0 | liver | Plpp1 |
| Aass | 0 | 1.31911872 | 0.26 | 0 | 0 | liver | Aass |
| Ifitm2 | 0 | 1.31705435 | 0.99 | 0.92 | 0 | liver | Ifitm2 |
| Lhfpl21 | 0 | 1.31440257 | 0.42 | 0.08 | 0 | liver | Lhfpl2 |
| Gata4 | 0 | 1.30812926 | 0.31 | 0 | 0 | liver | Gata4 |
| Clta | 0 | 1.30645048 | 0.83 | 0.55 | 0 | liver | Clta |
| Arhgdib | 0 | 1.3062311 | 0.52 | 0.15 | 0 | liver | Arhgdib |
| Hspa51 | 0 | 1.29441857 | 0.8 | 0.55 | 0 | liver | Hspa5 |
| F2r1 | 0 | 1.27959684 | 0.55 | 0.17 | 0 | liver | F2r |
| Sdf2l1 | 0 | 1.27214304 | 0.39 | 0.1 | 0 | liver | Sdf2l1 |
| Jak1 | 0 | 1.25413518 | 0.64 | 0.31 | 0 | liver | Jak1 |
| Man2a11 | 0 | 1.25119807 | 0.44 | 0.12 | 0 | liver | Man2a1 |
| Vcam1 | 0 | 1.24383889 | 0.4 | 0.13 | 0 | liver | Vcam1 |
| Snx2 | 0 | 1.22356477 | 0.52 | 0.2 | 0 | liver | Snx2 |
| Hn1 | 0 | 1.20328617 | 0.49 | 0.18 | 0 | liver | Hn1 |
| Id21 | 0 | 1.20198695 | 0.57 | 0.31 | 0 | liver | Id2 |
| Adgrg3 | 0 | 1.18456556 | 0.34 | 0.03 | 0 | liver | Adgrg3 |
| Snx5 | 0 | 1.18378954 | 0.47 | 0.17 | 0 | liver | Snx5 |
| Lpar61 | 0 | 1.15449751 | 0.53 | 0.23 | 0 | liver | Lpar6 |
| Scarb2 | 0 | 1.15417869 | 0.48 | 0.18 | 0 | liver | Scarb2 |
| Dapk2 | 0 | 1.14673762 | 0.38 | 0.08 | 0 | liver | Dapk2 |
| Kit | 0 | 1.1460336 | 0.32 | 0.06 | 0 | liver | Kit |
| Lgals3bp | 0 | 1.14286733 | 0.49 | 0.19 | 0 | liver | Lgals3bp |
| Ly96 | 0 | 1.14034391 | 0.45 | 0.14 | 0 | liver | Ly96 |
| Gm10605 | 0 | 1.13464827 | 0.26 | 0.01 | 0 | liver | Gm10605 |
| Psen2 | 0 | 1.11422278 | 0.37 | 0.09 | 0 | liver | Psen2 |
| Agpat5 | 0 | 1.10850287 | 0.32 | 0.08 | 0 | liver | Agpat5 |
| Csrp21 | 0 | 1.09550853 | 0.49 | 0.2 | 0 | liver | Csrp2 |
| Golga7 | 0 | 1.09496498 | 0.54 | 0.25 | 0 | liver | Golga7 |
| Sgpp1 | 0 | 1.09433387 | 0.34 | 0.08 | 0 | liver | Sgpp1 |
| Cyp4b1 | 0 | 1.09407231 | 0.7 | 0.26 | 0 | liver | Cyp4b1 |
| Ap1b1 | 0 | 1.09091136 | 0.3 | 0.04 | 0 | liver | Ap1b1 |
| Pde2a3 | 0 | 1.09025372 | 0.59 | 0.24 | 0 | liver | Pde2a |
| Calr | 0 | 1.08027216 | 0.82 | 0.6 | 0 | liver | Calr |
| Hyal2 | 0 | 1.06939673 | 0.6 | 0.31 | 0 | liver | Hyal2 |
| Hsp90b11 | 0 | 1.06838377 | 0.78 | 0.55 | 0 | liver | Hsp90b1 |
| 2200002D01Rik | 0 | 1.06571242 | 0.29 | 0.06 | 0 | liver | 2200002D01Rik |
| Ap2s1 | 0 | 1.06491557 | 0.66 | 0.36 | 0 | liver | Ap2s1 |
| Cdh131 | 0 | 1.06096032 | 0.39 | 0.14 | 0 | liver | Cdh13 |

| | | | | | | | |
|---|---|---|---|---|---|---|---|
| Npl | 0 | 1.05297407 | 0.27 | 0.03 | 0 | liver | Npl |
| Akr1b8 | 0 | 1.04713249 | 0.31 | 0.06 | 0 | liver | Akr1b8 |
| Slc39a81 | 0 | 1.01888459 | 0.33 | 0.07 | 0 | liver | Slc39a8 |
| Bst21 | 0 | 1.01683512 | 0.92 | 0.72 | 0 | liver | Bst2 |
| Pdia3 | 0 | 0.99595355 | 0.8 | 0.59 | 0 | liver | Pdia3 |
| Cd471 | 0 | 0.98781666 | 0.84 | 0.62 | 0 | liver | Cd47 |
| Man2b1 | 0 | 0.98715695 | 0.34 | 0.1 | 0 | liver | Man2b1 |
| Ctsd | 0 | 0.98076631 | 0.77 | 0.53 | 0 | liver | Ctsd |
| Cyp26b12 | 0 | 0.9725694 | 0.42 | 0.13 | 0 | liver | Cyp26b1 |
| Pros1 | 0 | 0.95212801 | 0.48 | 0.24 | 0 | liver | Pros1 |
| Ccnd2 | 0 | 0.94600471 | 0.64 | 0.35 | 0 | liver | Ccnd2 |
| Col13a12 | 0 | 0.94051848 | 0.38 | 0.13 | 0 | liver | Col13a1 |
| Fam124a | 0 | 0.93685375 | 0.27 | 0.05 | 0 | liver | Fam124a |
| Npr11 | 0 | 0.93675221 | 0.55 | 0.26 | 0 | liver | Npr1 |
| Nqo2 | 0 | 0.93030696 | 0.27 | 0.07 | 0 | liver | Nqo2 |
| Hfe | 0 | 0.92755089 | 0.4 | 0.14 | 0 | liver | Hfe |
| Arrb2 | 0 | 0.92684949 | 0.25 | 0.05 | 0 | liver | Arrb2 |
| Tmem64 | 0 | 0.91511319 | 0.28 | 0.06 | 0 | liver | Tmem64 |
| Cd300lg3 | 0 | 0.909879 | 0.74 | 0.39 | 0 | liver | Cd300lg |
| Slc26a10 | 0 | 0.908485 | 0.36 | 0.11 | 0 | liver | Slc26a10 |
| Cyb5a | 0 | 0.90845305 | 0.83 | 0.62 | 0 | liver | Cyb5a |
| Prdx1 | 0 | 0.88481495 | 0.92 | 0.79 | 0 | liver | Prdx1 |
| H2afz | 0 | 0.86696686 | 0.78 | 0.59 | 0 | liver | H2afz |
| Calml4 | 0 | 0.86486124 | 0.32 | 0.08 | 0 | liver | Calml4 |
| Atpif1 | 0 | 0.8595729 | 0.71 | 0.48 | 0 | liver | Atpif1 |
| Itm2b1 | 0 | 0.85802813 | 0.99 | 0.96 | 0 | liver | Itm2b |
| Lbp | 0 | 0.85264481 | 0.33 | 0.08 | 0 | liver | Lbp |
| Neurl3 | 0 | 0.8309804 | 0.25 | 0.05 | 0 | liver | Neurl3 |
| Serf2 | 0 | 0.82949857 | 0.94 | 0.84 | 0 | liver | Serf2 |
| Man1a | 0 | 0.82400226 | 0.31 | 0.09 | 0 | liver | Man1a |
| Zeb2 | 0 | 0.81954633 | 0.3 | 0.08 | 0 | liver | Zeb2 |
| Plpp33 | 0 | 0.81670652 | 0.86 | 0.61 | 0 | liver | Plpp3 |
| Ldhb | 0 | 0.80024302 | 0.25 | 0.06 | 0 | liver | Ldhb |
| St3gal5 | 0 | 0.79342015 | 0.26 | 0.08 | 0 | liver | St3gal5 |
| Psap | 0 | 0.78963889 | 0.86 | 0.72 | 0 | liver | Psap |
| Fam174b1 | 0 | 0.78836393 | 0.37 | 0.14 | 0 | liver | Fam174b |
| Vamp52 | 0 | 0.77799682 | 0.77 | 0.52 | 0 | liver | Vamp5 |
| B2m1 | 0 | 0.77640921 | 1 | 0.98 | 0 | liver | B2m |
| Mfsd5 | 0 | 0.76589338 | 0.29 | 0.09 | 0 | liver | Mfsd5 |
| Lamp1 | 0 | 0.73040653 | 0.86 | 0.68 | 0 | liver | Lamp1 |
| Tspan7 | 0 | 0.72371807 | 0.85 | 0.48 | 0 | liver | Tspan7 |

| | | | | | | | |
|---|---|---|---|---|---|---|---|
| Prelp | 0 | 0.70492608 | 0.27 | 0.06 | 0 | liver | Prelp |
| Srgn2 | 0 | 0.68986159 | 0.86 | 0.69 | 0 | liver | Srgn |
| Igfbp71 | 0 | 0.59961956 | 1 | 0.87 | 0 | liver | Igfbp7 |
| Tfpi | 0 | 0.5901038 | 0.53 | 0.2 | 0 | liver | Tfpi |
| Ptprb2 | 0 | 0.58249642 | 0.94 | 0.89 | 0 | liver | Ptprb |
| Cfh | 0 | 0.56168321 | 0.28 | 0.06 | 0 | liver | Cfh |
| H2-D11 | 0 | 0.50379894 | 0.97 | 0.95 | 0 | liver | H2-D1 |
| Hspa1a2 | 0 | 0.39067789 | 0.58 | 0.29 | 0 | liver | Hspa1a |
| Meis22 | 0 | 0.3821244 | 0.64 | 0.28 | 0 | liver | Meis2 |
| Wbp51 | 3.39E-301 | 0.83156267 | 0.64 | 0.39 | 7.01E-297 | liver | Wbp5 |
| Rfk | 3.34E-299 | 0.85187718 | 0.42 | 0.19 | 6.92E-295 | liver | Rfk |
| St3gal41 | 9.64E-296 | 0.82729076 | 0.43 | 0.2 | 1.99E-291 | liver | St3gal4 |
| Smap2 | 4.57E-292 | 0.78490529 | 0.31 | 0.11 | 9.46E-288 | liver | Smap2 |
| Flt41 | 7.00E-290 | 0.78177934 | 0.43 | 0.2 | 1.45E-285 | liver | Flt4 |
| Kdelr2 | 2.40E-287 | 0.92111638 | 0.5 | 0.27 | 4.97E-283 | liver | Kdelr2 |
| Ralgapa2 | 1.19E-285 | 0.7670007 | 0.27 | 0.09 | 2.47E-281 | liver | Ralgapa2 |
| Cmtm82 | 2.96E-285 | 0.83496161 | 0.59 | 0.34 | 6.13E-281 | liver | Cmtm8 |
| Mcfd2 | 4.20E-282 | 0.83930255 | 0.5 | 0.26 | 8.68E-278 | liver | Mcfd2 |
| Blvrb | 1.39E-280 | 0.84104333 | 0.39 | 0.17 | 2.88E-276 | liver | Blvrb |
| Rcn3 | 7.16E-278 | 0.70499813 | 0.36 | 0.15 | 1.48E-273 | liver | Rcn3 |
| Hexa1 | 2.27E-277 | 0.83313701 | 0.45 | 0.22 | 4.71E-273 | liver | Hexa |
| Gimap41 | 1.39E-270 | 0.77188633 | 0.67 | 0.43 | 2.87E-266 | liver | Gimap4 |
| S100a12 | 7.00E-263 | 0.75753532 | 0.6 | 0.35 | 1.45E-258 | liver | S100a1 |
| Camk11 | 9.22E-252 | 0.80641043 | 0.54 | 0.31 | 1.91E-247 | liver | Camk1 |
| PISD1 | 4.01E-243 | 0.72174589 | 0.46 | 0.23 | 8.29E-239 | liver | PISD |
| Upp11 | 6.28E-242 | 0.67489409 | 0.33 | 0.14 | 1.30E-237 | liver | Upp1 |
| Pdlim12 | 2.17E-241 | 0.64223504 | 0.54 | 0.29 | 4.48E-237 | liver | Pdlim1 |
| Gpx4 | 6.15E-235 | 0.57497489 | 0.87 | 0.76 | 1.27E-230 | liver | Gpx4 |
| Dnajc31 | 3.48E-233 | 0.79417125 | 0.51 | 0.3 | 7.20E-229 | liver | Dnajc3 |
| Mylip1 | 1.28E-231 | 0.80446612 | 0.29 | 0.12 | 2.66E-227 | liver | Mylip |
| Pdia6 | 6.64E-229 | 0.81260758 | 0.65 | 0.45 | 1.37E-224 | liver | Pdia6 |
| Clec14a1 | 7.86E-225 | 0.51918908 | 0.73 | 0.48 | 1.63E-220 | liver | Clec14a |
| Cpne2 | 2.74E-223 | 0.74942256 | 0.33 | 0.14 | 5.67E-219 | liver | Cpne2 |
| Nucb1 | 5.96E-220 | 0.76457817 | 0.53 | 0.33 | 1.23E-215 | liver | Nucb1 |
| Fyn1 | 1.27E-219 | 0.70807385 | 0.43 | 0.23 | 2.62E-215 | liver | Fyn |
| Nt5c3 | 1.99E-218 | 0.74782213 | 0.29 | 0.12 | 4.11E-214 | liver | Nt5c3 |
| Fkbp1a | 4.20E-218 | 0.4563085 | 0.92 | 0.82 | 8.70E-214 | liver | Fkbp1a |
| Tnfrsf1a | 9.19E-215 | 0.71203173 | 0.62 | 0.42 | 1.90E-210 | liver | Tnfrsf1a |
| Jam22 | 5.62E-213 | 0.79990507 | 0.63 | 0.44 | 1.16E-208 | liver | Jam2 |
| Ap2a2 | 1.11E-210 | 0.74478501 | 0.42 | 0.22 | 2.30E-206 | liver | Ap2a2 |
| Eng1 | 5.22E-210 | 0.47549115 | 0.9 | 0.85 | 1.08E-205 | liver | Eng |

| | | | | | | | |
|---|---|---|---|---|---|---|---|
| Tpp1 | 3.62E-203 | 0.70588504 | 0.37 | 0.18 | 7.49E-199 | liver | Tpp1 |
| Rnase4 | 9.22E-200 | 0.60073136 | 0.28 | 0.11 | 1.91E-195 | liver | Rnase4 |
| Pitpna1 | 1.35E-199 | 0.72158227 | 0.55 | 0.35 | 2.80E-195 | liver | Pitpna |
| Id31 | 2.11E-194 | 0.49904517 | 0.92 | 0.85 | 4.36E-190 | liver | Id3 |
| Gm138891 | 1.14E-191 | 0.75846898 | 0.38 | 0.19 | 2.36E-187 | liver | Gm13889 |
| Mxra8 | 1.47E-190 | 0.60073474 | 0.29 | 0.13 | 3.04E-186 | liver | Mxra8 |
| Hdac1 | 1.45E-189 | 0.74641711 | 0.41 | 0.23 | 3.00E-185 | liver | Hdac1 |
| Mtch1 | 2.50E-182 | 0.67014383 | 0.59 | 0.4 | 5.17E-178 | liver | Mtch1 |
| Fam3c | 3.14E-181 | 0.72279603 | 0.39 | 0.21 | 6.49E-177 | liver | Fam3c |
| Atp1b3 | 6.75E-177 | 0.64061279 | 0.55 | 0.35 | 1.40E-172 | liver | Atp1b3 |
| Wls | 3.28E-175 | 0.59941128 | 0.3 | 0.14 | 6.79E-171 | liver | Wls |
| Efnb1 | 1.55E-170 | 0.91489138 | 0.41 | 0.23 | 3.21E-166 | liver | Efnb1 |
| Clic1 | 3.49E-167 | 0.49156951 | 0.83 | 0.71 | 7.23E-163 | liver | Clic1 |
| H2-K12 | 1.88E-166 | 0.4274834 | 0.91 | 0.84 | 3.88E-162 | liver | H2-K1 |
| Pnp1 | 2.14E-160 | 0.62821347 | 0.37 | 0.2 | 4.42E-156 | liver | Pnp |
| Nrp22 | 6.45E-160 | 0.4784945 | 0.39 | 0.21 | 1.33E-155 | liver | Nrp2 |
| Plxnd1 | 1.06E-159 | 0.66482922 | 0.46 | 0.28 | 2.19E-155 | liver | Plxnd1 |
| Ecm11 | 3.84E-159 | 0.56194928 | 0.34 | 0.17 | 7.94E-155 | liver | Ecm1 |
| Rnf130 | 2.31E-156 | 0.66026321 | 0.44 | 0.27 | 4.78E-152 | liver | Rnf130 |
| Atp5h | 2.02E-152 | 0.50117391 | 0.81 | 0.71 | 4.19E-148 | liver | Atp5h |
| Epb41l21 | 1.33E-151 | 0.63413881 | 0.3 | 0.15 | 2.75E-147 | liver | Epb41l2 |
| Cmtm3 | 5.20E-151 | 0.61017382 | 0.39 | 0.22 | 1.08E-146 | liver | Cmtm3 |
| Rbfa | 1.55E-146 | 0.57615151 | 0.28 | 0.13 | 3.20E-142 | liver | Rbfa |
| Raph1 | 6.82E-146 | 0.57763237 | 0.29 | 0.14 | 1.41E-141 | liver | Raph1 |
| Ckap4 | 3.87E-145 | 0.5561374 | 0.26 | 0.12 | 8.00E-141 | liver | Ckap4 |
| Canx | 3.46E-144 | 0.63189729 | 0.54 | 0.37 | 7.15E-140 | liver | Canx |
| Arl6ip1 | 6.30E-143 | 0.60048754 | 0.61 | 0.45 | 1.30E-138 | liver | Arl6ip1 |
| Serpinb6a | 2.50E-142 | 0.43911916 | 0.63 | 0.44 | 5.18E-138 | liver | Serpinb6a |
| Eif4ebp1 | 1.56E-141 | 0.57029355 | 0.33 | 0.17 | 3.22E-137 | liver | Eif4ebp1 |
| Atp6ap1 | 2.12E-139 | 0.58464952 | 0.48 | 0.31 | 4.38E-135 | liver | Atp6ap1 |
| Grina | 1.91E-138 | 0.55170627 | 0.36 | 0.2 | 3.95E-134 | liver | Grina |
| Creld2 | 1.53E-137 | 0.63203272 | 0.29 | 0.15 | 3.17E-133 | liver | Creld2 |
| H2-T232 | 9.88E-137 | 0.55970304 | 0.57 | 0.39 | 2.04E-132 | liver | H2-T23 |
| Tpt11 | 1.27E-136 | 0.33876942 | 0.96 | 0.93 | 2.63E-132 | liver | Tpt1 |
| Txndc12 | 2.58E-136 | 0.55134824 | 0.34 | 0.19 | 5.34E-132 | liver | Txndc12 |
| Snap23 | 4.80E-136 | 0.59752144 | 0.37 | 0.22 | 9.94E-132 | liver | Snap23 |
| Acer2 | 1.42E-135 | 0.48138926 | 0.37 | 0.2 | 2.94E-131 | liver | Acer2 |
| Rpl10 | 6.40E-132 | 0.38016036 | 0.94 | 0.88 | 1.32E-127 | liver | Rpl10 |
| Dnajb11 | 4.77E-131 | 0.6262505 | 0.44 | 0.29 | 9.87E-127 | liver | Dnajb11 |
| Cdc42ep2 | 1.32E-130 | 0.55294977 | 0.42 | 0.25 | 2.72E-126 | liver | Cdc42ep2 |
| Smagp | 8.06E-129 | 0.57504187 | 0.62 | 0.46 | 1.67E-124 | liver | Smagp |

| | | | | | | | |
|---|---|---|---|---|---|---|---|
| Utp11l | 1.49E-126 | 0.5284457 | 0.35 | 0.2 | 3.08E-122 | liver | Utp11l |
| Mgat11 | 5.05E-126 | 0.57806548 | 0.39 | 0.24 | 1.05E-121 | liver | Mgat1 |
| Rbp1 | 7.01E-126 | 0.64815906 | 0.27 | 0.14 | 1.45E-121 | liver | Rbp1 |
| Atp6v0e | 5.35E-125 | 0.51793897 | 0.65 | 0.51 | 1.11E-120 | liver | Atp6v0e |
| Kdr4 | 5.55E-123 | 0.39619499 | 0.77 | 0.61 | 1.15E-118 | liver | Kdr |
| Aldh2 | 3.14E-122 | 0.451248 | 0.73 | 0.58 | 6.50E-118 | liver | Aldh2 |
| Serinc31 | 3.73E-121 | 0.41751745 | 0.9 | 0.84 | 7.72E-117 | liver | Serinc3 |
| Mpp1 | 6.35E-120 | 0.51401423 | 0.25 | 0.13 | 1.31E-115 | liver | Mpp1 |
| Ggh | 1.95E-119 | 0.52440043 | 0.33 | 0.19 | 4.04E-115 | liver | Ggh |
| Tgfb1 | 3.23E-119 | 0.53896302 | 0.58 | 0.43 | 6.69E-115 | liver | Tgfb1 |
| Smad6 | 1.27E-115 | 0.33266135 | 0.34 | 0.18 | 2.63E-111 | liver | Smad6 |
| Txnip3 | 9.20E-115 | 0.3950083 | 0.78 | 0.65 | 1.90E-110 | liver | Txnip |
| S100a102 | 3.19E-114 | 0.42724722 | 0.78 | 0.66 | 6.61E-110 | liver | S100a10 |
| Vsir | 1.33E-112 | 0.56468349 | 0.35 | 0.21 | 2.75E-108 | liver | Vsir |
| Dlc12 | 3.67E-112 | 0.43469488 | 0.58 | 0.4 | 7.59E-108 | liver | Dlc1 |
| Cltc1 | 3.92E-112 | 0.56760831 | 0.44 | 0.29 | 8.11E-108 | liver | Cltc |
| Gimap61 | 5.03E-112 | 0.42051752 | 0.77 | 0.65 | 1.04E-107 | liver | Gimap6 |
| Arhgap23 | 2.25E-111 | 0.52180631 | 0.25 | 0.13 | 4.67E-107 | liver | Arhgap23 |
| Tmed102 | 2.07E-109 | 0.4920844 | 0.7 | 0.57 | 4.29E-105 | liver | Tmed10 |
| Fkbp2 | 6.85E-109 | 0.53192753 | 0.4 | 0.25 | 1.42E-104 | liver | Fkbp2 |
| Rps61 | 1.11E-108 | 0.36536158 | 0.94 | 0.89 | 2.30E-104 | liver | Rps6 |
| Ccdc53 | 8.26E-108 | 0.49106599 | 0.26 | 0.13 | 1.71E-103 | liver | Ccdc53 |
| Lrrc32 | 9.97E-108 | 0.47271027 | 0.29 | 0.16 | 2.06E-103 | liver | Lrrc32 |
| Tmem140 | 1.05E-107 | 0.49118633 | 0.53 | 0.37 | 2.18E-103 | liver | Tmem140 |
| Anxa5 | 9.35E-104 | 0.40853409 | 0.75 | 0.65 | 1.93E-99 | liver | Anxa5 |
| Camk2n12 | 1.71E-102 | 0.42897144 | 0.25 | 0.13 | 3.53E-98 | liver | Camk2n1 |
| Rrbp12 | 2.42E-102 | 0.50177309 | 0.6 | 0.45 | 5.00E-98 | liver | Rrbp1 |
| Nrp13 | 2.93E-102 | 0.26672175 | 0.83 | 0.64 | 6.06E-98 | liver | Nrp1 |
| Elk32 | 2.96E-102 | 0.42985758 | 0.67 | 0.54 | 6.13E-98 | liver | Elk3 |
| Ostc | 7.41E-102 | 0.54336035 | 0.43 | 0.29 | 1.53E-97 | liver | Ostc |
| Mesdc1 | 1.79E-101 | 0.45962293 | 0.37 | 0.23 | 3.70E-97 | liver | Mesdc1 |
| Ormdl2 | 2.45E-101 | 0.49228052 | 0.27 | 0.14 | 5.07E-97 | liver | Ormdl2 |
| Asah1 | 5.30E-101 | 0.49315973 | 0.4 | 0.25 | 1.10E-96 | liver | Asah1 |
| Ppia | 3.16E-100 | 0.2529819 | 0.97 | 0.96 | 6.54E-96 | liver | Ppia |
| Rab5c | 1.77E-99 | 0.52322143 | 0.57 | 0.43 | 3.65E-95 | liver | Rab5c |
| Slco3a1 | 3.59E-98 | 0.51597035 | 0.26 | 0.14 | 7.44E-94 | liver | Slco3a1 |
| Cdk4 | 3.40E-97 | 0.51100779 | 0.42 | 0.28 | 7.04E-93 | liver | Cdk4 |
| Nfe2l22 | 3.05E-96 | 0.48117029 | 0.45 | 0.31 | 6.31E-92 | liver | Nfe2l2 |
| Ralb | 3.43E-96 | 0.51817344 | 0.43 | 0.29 | 7.10E-92 | liver | Ralb |
| Rps3 | 2.86E-93 | 0.30655524 | 0.94 | 0.9 | 5.91E-89 | liver | Rps3 |
| Sema6a2 | 2.47E-92 | 0.50168929 | 0.41 | 0.27 | 5.11E-88 | liver | Sema6a |

| | | | | | | | |
|---|---|---|---|---|---|---|---|
| Fundc2 | 1.68E-91 | 0.48845774 | 0.42 | 0.28 | 3.48E-87 | liver | Fundc2 |
| Pdia4 | 7.30E-90 | 0.5009516 | 0.32 | 0.2 | 1.51E-85 | liver | Pdia4 |
| Ctso | 2.91E-88 | 0.48521803 | 0.3 | 0.18 | 6.02E-84 | liver | Ctso |
| Akr1b3 | 4.47E-87 | 0.47072998 | 0.33 | 0.21 | 9.26E-83 | liver | Akr1b3 |
| Rtp41 | 2.00E-86 | 0.48453024 | 0.4 | 0.27 | 4.13E-82 | liver | Rtp4 |
| Rpn11 | 3.35E-86 | 0.48191743 | 0.39 | 0.26 | 6.93E-82 | liver | Rpn1 |
| Ephb41 | 3.75E-86 | 0.4228042 | 0.43 | 0.29 | 7.77E-82 | liver | Ephb4 |
| Sptlc21 | 1.15E-85 | 0.49992735 | 0.37 | 0.24 | 2.38E-81 | liver | Sptlc2 |
| Atp6v1d | 2.48E-82 | 0.49179113 | 0.31 | 0.19 | 5.12E-78 | liver | Atp6v1d |
| Pcdh12 | 5.81E-82 | 0.41566946 | 0.26 | 0.15 | 1.20E-77 | liver | Pcdh12 |
| Plin2 | 1.16E-80 | 0.48085968 | 0.31 | 0.19 | 2.40E-76 | liver | Plin2 |
| Add31 | 3.14E-80 | 0.46784954 | 0.28 | 0.17 | 6.49E-76 | liver | Add3 |
| Chst151 | 5.43E-80 | 0.48142343 | 0.27 | 0.16 | 1.12E-75 | liver | Chst15 |
| Tmem30a | 6.21E-80 | 0.44788489 | 0.48 | 0.35 | 1.29E-75 | liver | Tmem30a |
| Eef1g | 1.26E-78 | 0.41817903 | 0.73 | 0.63 | 2.60E-74 | liver | Eef1g |
| Pvrl2 | 1.89E-78 | 0.3157815 | 0.28 | 0.16 | 3.90E-74 | liver | Pvrl2 |
| Fis1 | 2.00E-78 | 0.43619705 | 0.6 | 0.48 | 4.13E-74 | liver | Fis1 |
| Tsc22d4 | 8.55E-77 | 0.44161027 | 0.26 | 0.16 | 1.77E-72 | liver | Tsc22d4 |
| Fkbp7 | 4.32E-76 | 0.4104075 | 0.29 | 0.17 | 8.94E-72 | liver | Fkbp7 |
| Tmem9b1 | 7.96E-76 | 0.44234535 | 0.4 | 0.27 | 1.65E-71 | liver | Tmem9b |
| Tpst2 | 8.98E-76 | 0.40144018 | 0.58 | 0.46 | 1.86E-71 | liver | Tpst2 |
| Cstb | 1.07E-75 | 0.35657013 | 0.48 | 0.34 | 2.22E-71 | liver | Cstb |
| Mgst11 | 4.53E-75 | 0.5046292 | 0.25 | 0.15 | 9.37E-71 | liver | Mgst1 |
| Dhrs7 | 1.81E-74 | 0.44126703 | 0.37 | 0.25 | 3.74E-70 | liver | Dhrs7 |
| Cd164 | 3.27E-74 | 0.45307579 | 0.32 | 0.21 | 6.76E-70 | liver | Cd164 |
| Sash11 | 1.25E-73 | 0.43235284 | 0.35 | 0.23 | 2.59E-69 | liver | Sash1 |
| Mndal2 | 3.16E-73 | 0.37013874 | 0.45 | 0.31 | 6.55E-69 | liver | Mndal |
| Stard81 | 5.86E-73 | 0.40606283 | 0.29 | 0.18 | 1.21E-68 | liver | Stard8 |
| Akr1a11 | 6.14E-73 | 0.35363864 | 0.72 | 0.61 | 1.27E-68 | liver | Akr1a1 |
| Snrk1 | 6.76E-73 | 0.37593791 | 0.6 | 0.46 | 1.40E-68 | liver | Snrk |
| Sft2d1 | 3.84E-71 | 0.41675637 | 0.39 | 0.27 | 7.94E-67 | liver | Sft2d1 |
| Gstm2 | 6.28E-71 | 0.31700177 | 0.29 | 0.18 | 1.30E-66 | liver | Gstm2 |
| Npc2 | 1.05E-70 | 0.40095937 | 0.61 | 0.49 | 2.17E-66 | liver | Npc2 |
| Slc31a2 | 1.56E-70 | 0.45265619 | 0.33 | 0.22 | 3.22E-66 | liver | Slc31a2 |
| Cdc421 | 2.42E-70 | 0.34720392 | 0.79 | 0.72 | 5.00E-66 | liver | Cdc42 |
| Scarf1 | 2.73E-70 | 0.38305165 | 0.27 | 0.17 | 5.66E-66 | liver | Scarf1 |
| Tgm21 | 2.56E-69 | 0.39384482 | 0.51 | 0.39 | 5.30E-65 | liver | Tgm2 |
| Prr131 | 3.74E-68 | 0.36244851 | 0.47 | 0.35 | 7.73E-64 | liver | Prr13 |
| Tmed9 | 1.33E-67 | 0.37589476 | 0.62 | 0.51 | 2.74E-63 | liver | Tmed9 |
| Icam1 | 3.74E-67 | 0.54176181 | 0.29 | 0.18 | 7.73E-63 | liver | Icam1 |
| Tmem205 | 3.72E-66 | 0.42479915 | 0.33 | 0.22 | 7.71E-62 | liver | Tmem205 |

| | | | | | | | |
|---|---|---|---|---|---|---|---|
| Glmp | 4.67E-66 | 0.40856392 | 0.44 | 0.32 | 9.66E-62 | liver | Glmp |
| Sgk12 | 6.24E-66 | 0.29173536 | 0.65 | 0.53 | 1.29E-61 | liver | Sgk1 |
| Yif1b | 1.20E-65 | 0.41682962 | 0.36 | 0.25 | 2.49E-61 | liver | Yif1b |
| Bax | 6.50E-65 | 0.4176455 | 0.54 | 0.42 | 1.34E-60 | liver | Bax |
| Atp6v1g1 | 1.15E-63 | 0.3790215 | 0.67 | 0.57 | 2.37E-59 | liver | Atp6v1g1 |
| Tbxa2r | 1.46E-63 | 0.37169746 | 0.3 | 0.19 | 3.02E-59 | liver | Tbxa2r |
| Spcs2 | 6.33E-63 | 0.40938769 | 0.57 | 0.46 | 1.31E-58 | liver | Spcs2 |
| Atp6v1f | 1.77E-62 | 0.37844271 | 0.59 | 0.49 | 3.67E-58 | liver | Atp6v1f |
| Atp6ap2 | 1.92E-62 | 0.37855446 | 0.32 | 0.21 | 3.97E-58 | liver | Atp6ap2 |
| Hmox21 | 2.75E-62 | 0.45683796 | 0.29 | 0.2 | 5.69E-58 | liver | Hmox2 |
| Grn | 4.44E-62 | 0.36750441 | 0.52 | 0.4 | 9.19E-58 | liver | Grn |
| Ran | 4.76E-62 | 0.40414147 | 0.55 | 0.45 | 9.84E-58 | liver | Ran |
| Atp6v0b | 5.35E-62 | 0.3962385 | 0.57 | 0.46 | 1.11E-57 | liver | Atp6v0b |
| Tspan4 | 6.60E-61 | 0.38390854 | 0.42 | 0.3 | 1.37E-56 | liver | Tspan4 |
| Mmrn22 | 1.09E-60 | 0.33619615 | 0.63 | 0.51 | 2.25E-56 | liver | Mmrn2 |
| Mydgf | 6.62E-60 | 0.41836815 | 0.4 | 0.29 | 1.37E-55 | liver | Mydgf |
| Gipc1 | 6.92E-60 | 0.38526904 | 0.25 | 0.16 | 1.43E-55 | liver | Gipc1 |
| Spag91 | 8.27E-60 | 0.38727294 | 0.39 | 0.28 | 1.71E-55 | liver | Spag9 |
| Herpud1 | 1.05E-59 | 0.47274071 | 0.38 | 0.28 | 2.17E-55 | liver | Herpud1 |
| 2700094K13Rik | 2.64E-59 | 0.39809015 | 0.28 | 0.18 | 5.46E-55 | liver | 2700094K13Rik |
| Cnn2 | 1.08E-57 | 0.35714753 | 0.4 | 0.29 | 2.24E-53 | liver | Cnn2 |
| Nid12 | 2.29E-57 | 0.42018794 | 0.28 | 0.19 | 4.75E-53 | liver | Nid1 |
| Tmem59 | 4.19E-57 | 0.29978528 | 0.78 | 0.72 | 8.68E-53 | liver | Tmem59 |
| Tuba1b1 | 4.36E-57 | 0.30990417 | 0.76 | 0.68 | 9.03E-53 | liver | Tuba1b |
| 11-Sep | 1.31E-56 | 0.32333768 | 0.25 | 0.16 | 2.71E-52 | liver | 11-Sep |
| Rnaset2b1 | 1.43E-56 | 0.33162048 | 0.49 | 0.37 | 2.96E-52 | liver | Rnaset2b |
| Rps121 | 1.11E-55 | 0.26479483 | 0.89 | 0.82 | 2.30E-51 | liver | Rps12 |
| Derl1 | 1.36E-55 | 0.38783884 | 0.31 | 0.21 | 2.82E-51 | liver | Derl1 |
| Shisa5 | 2.06E-55 | 0.29600348 | 0.67 | 0.55 | 4.26E-51 | liver | Shisa5 |
| Lrrc581 | 5.11E-54 | 0.40933413 | 0.45 | 0.35 | 1.06E-49 | liver | Lrrc58 |
| Vps29 | 1.03E-53 | 0.37352364 | 0.46 | 0.36 | 2.13E-49 | liver | Vps29 |
| Znrf11 | 3.62E-52 | 0.34577752 | 0.35 | 0.25 | 7.49E-48 | liver | Znrf1 |
| Bcap31 | 4.07E-52 | 0.34548147 | 0.47 | 0.37 | 8.42E-48 | liver | Bcap31 |
| Marcks2 | 4.38E-52 | 0.25950029 | 0.48 | 0.36 | 9.06E-48 | liver | Marcks |
| Tex264 | 6.47E-52 | 0.37054282 | 0.3 | 0.2 | 1.34E-47 | liver | Tex264 |
| Nptn | 2.76E-51 | 0.40947696 | 0.34 | 0.25 | 5.71E-47 | liver | Nptn |
| Casp8 | 4.19E-51 | 0.36963255 | 0.31 | 0.22 | 8.66E-47 | liver | Casp8 |
| Fuca1 | 1.92E-50 | 0.3599961 | 0.31 | 0.22 | 3.96E-46 | liver | Fuca1 |
| Mesdc2 | 2.30E-49 | 0.33127896 | 0.27 | 0.18 | 4.76E-45 | liver | Mesdc2 |
| Sumo3 | 2.46E-49 | 0.35985544 | 0.42 | 0.32 | 5.08E-45 | liver | Sumo3 |
| Emc7 | 2.00E-48 | 0.34769991 | 0.46 | 0.36 | 4.14E-44 | liver | Emc7 |

| | | | | | | | |
|---|---|---|---|---|---|---|---|
| Cuta | 6.13E-48 | 0.32722872 | 0.46 | 0.36 | 1.27E-43 | liver | Cuta |
| 15-Sep | 6.51E-48 | 0.25793236 | 0.77 | 0.66 | 1.35E-43 | liver | 15-Sep |
| Ncoa31 | 9.47E-48 | 0.4130284 | 0.26 | 0.17 | 1.96E-43 | liver | Ncoa3 |
| Vimp | 9.62E-48 | 0.36793915 | 0.25 | 0.17 | 1.99E-43 | liver | Vimp |
| Tmem256 | 2.16E-47 | 0.33548572 | 0.6 | 0.51 | 4.47E-43 | liver | Tmem256 |
| Cept1 | 3.93E-47 | 0.35129113 | 0.29 | 0.2 | 8.14E-43 | liver | Cept1 |
| Tmed3 | 6.19E-47 | 0.36008828 | 0.39 | 0.29 | 1.28E-42 | liver | Tmed3 |
| Npdc1 | 8.44E-47 | 0.34499386 | 0.44 | 0.34 | 1.75E-42 | liver | Npdc1 |
| Ssr4 | 6.91E-46 | 0.35889042 | 0.49 | 0.39 | 1.43E-41 | liver | Ssr4 |
| Cnn3 | 1.26E-45 | 0.29551118 | 0.61 | 0.52 | 2.62E-41 | liver | Cnn3 |
| Pkm1 | 1.80E-45 | 0.28903471 | 0.61 | 0.52 | 3.72E-41 | liver | Pkm |
| Elf12 | 1.83E-45 | 0.29855491 | 0.33 | 0.23 | 3.79E-41 | liver | Elf1 |
| Ifi271 | 2.15E-45 | 0.29482184 | 0.62 | 0.52 | 4.45E-41 | liver | Ifi27 |
| Tgoln1 | 2.53E-45 | 0.33187486 | 0.37 | 0.28 | 5.24E-41 | liver | Tgoln1 |
| Rps12-ps32 | 3.07E-45 | 0.25887505 | 0.69 | 0.57 | 6.35E-41 | liver | Rps12-ps3 |
| Eif2s21 | 3.73E-45 | 0.32854951 | 0.56 | 0.47 | 7.72E-41 | liver | Eif2s2 |
| Trappc5 | 4.95E-45 | 0.35523794 | 0.26 | 0.18 | 1.02E-40 | liver | Trappc5 |
| Sp1002 | 6.75E-45 | 0.284971 | 0.45 | 0.35 | 1.40E-40 | liver | Sp100 |
| Sod1 | 1.92E-43 | 0.32742467 | 0.52 | 0.43 | 3.98E-39 | liver | Sod1 |
| Cnpy2 | 3.32E-43 | 0.34671404 | 0.36 | 0.27 | 6.87E-39 | liver | Cnpy2 |
| G3bp2 | 3.39E-43 | 0.34030798 | 0.36 | 0.27 | 7.02E-39 | liver | G3bp2 |
| Smc61 | 5.83E-43 | 0.34255461 | 0.28 | 0.2 | 1.21E-38 | liver | Smc6 |
| Ap2m1 | 1.81E-42 | 0.32138109 | 0.56 | 0.47 | 3.75E-38 | liver | Ap2m1 |
| Adipor12 | 4.44E-42 | 0.31001771 | 0.54 | 0.45 | 9.18E-38 | liver | Adipor1 |
| Arpc3 | 8.90E-42 | 0.26898933 | 0.76 | 0.7 | 1.84E-37 | liver | Arpc3 |
| Eva1b1 | 1.56E-41 | 0.28532733 | 0.55 | 0.45 | 3.22E-37 | liver | Eva1b |
| Xiap2 | 2.91E-41 | 0.30976316 | 0.38 | 0.29 | 6.01E-37 | liver | Xiap |
| Plekhg11 | 5.91E-41 | 0.2810386 | 0.28 | 0.19 | 1.22E-36 | liver | Plekhg1 |
| Ddah2 | 1.16E-40 | 0.267424 | 0.5 | 0.4 | 2.39E-36 | liver | Ddah2 |
| Xbp11 | 1.18E-40 | 0.27122416 | 0.46 | 0.36 | 2.44E-36 | liver | Xbp1 |
| Prdx41 | 1.64E-40 | 0.38191597 | 0.3 | 0.22 | 3.40E-36 | liver | Prdx4 |
| Tbcb | 2.13E-40 | 0.32771384 | 0.43 | 0.34 | 4.41E-36 | liver | Tbcb |
| Adgrl21 | 1.98E-39 | 0.25368454 | 0.31 | 0.22 | 4.09E-35 | liver | Adgrl2 |
| Prcp | 3.30E-39 | 0.26416174 | 0.29 | 0.2 | 6.82E-35 | liver | Prcp |
| Saraf | 5.29E-39 | 0.32297143 | 0.55 | 0.47 | 1.10E-34 | liver | Saraf |
| Amotl11 | 8.30E-39 | 0.34589885 | 0.34 | 0.25 | 1.72E-34 | liver | Amotl1 |
| Slbp | 1.16E-38 | 0.33372929 | 0.26 | 0.18 | 2.41E-34 | liver | Slbp |
| Cldnd11 | 1.28E-38 | 0.31571692 | 0.27 | 0.19 | 2.65E-34 | liver | Cldnd1 |
| Tnfaip12 | 1.88E-38 | 0.29326194 | 0.44 | 0.35 | 3.89E-34 | liver | Tnfaip1 |
| Amotl2 | 1.91E-38 | 0.25109198 | 0.27 | 0.18 | 3.95E-34 | liver | Amotl2 |
| Gabarap | 6.02E-38 | 0.26864517 | 0.73 | 0.67 | 1.25E-33 | liver | Gabarap |

| | | | | | | | |
|---|---|---|---|---|---|---|---|
| Ndufa8 | 1.36E-37 | 0.37418501 | 0.58 | 0.52 | 2.81E-33 | liver | Ndufa8 |
| Tceal8 | 2.41E-37 | 0.31662244 | 0.3 | 0.22 | 4.98E-33 | liver | Tceal8 |
| Prkar1a1 | 4.00E-37 | 0.27499544 | 0.64 | 0.56 | 8.27E-33 | liver | Prkar1a |
| Hnrnpa3 | 4.05E-37 | 0.32749096 | 0.56 | 0.49 | 8.37E-33 | liver | Hnrnpa3 |
| Pcbp1 | 4.87E-37 | 0.29245664 | 0.45 | 0.37 | 1.01E-32 | liver | Pcbp1 |
| Ddost | 1.89E-36 | 0.28857179 | 0.37 | 0.28 | 3.92E-32 | liver | Ddost |
| 1110038F14Rik | 6.86E-36 | 0.32572986 | 0.34 | 0.26 | 1.42E-31 | liver | 1110038F14Rik |
| Lman11 | 1.29E-35 | 0.30512439 | 0.3 | 0.22 | 2.68E-31 | liver | Lman1 |
| Cnih1 | 1.92E-35 | 0.31049484 | 0.33 | 0.25 | 3.97E-31 | liver | Cnih1 |
| Gas63 | 2.84E-35 | 0.28392103 | 0.57 | 0.49 | 5.87E-31 | liver | Gas6 |
| Psenen | 6.61E-34 | 0.25746476 | 0.51 | 0.42 | 1.37E-29 | liver | Psenen |
| Slc39a1 | 9.13E-34 | 0.3385618 | 0.26 | 0.19 | 1.89E-29 | liver | Slc39a1 |
| Pet100 | 2.62E-33 | 0.31954049 | 0.35 | 0.27 | 5.42E-29 | liver | Pet100 |
| Hspe11 | 7.06E-33 | 0.31343957 | 0.44 | 0.36 | 1.46E-28 | liver | Hspe1 |
| Krtcap21 | 1.19E-32 | 0.29155429 | 0.6 | 0.53 | 2.46E-28 | liver | Krtcap2 |
| Sptssa | 3.20E-32 | 0.27998342 | 0.52 | 0.44 | 6.62E-28 | liver | Sptssa |
| Tubb5 | 9.64E-32 | 0.25605093 | 0.71 | 0.65 | 1.99E-27 | liver | Tubb5 |
| Cdc42ep31 | 2.57E-31 | 0.33107511 | 0.28 | 0.21 | 5.32E-27 | liver | Cdc42ep3 |
| Tacc11 | 3.48E-31 | 0.31243731 | 0.31 | 0.23 | 7.20E-27 | liver | Tacc1 |
| Tmbim4 | 5.33E-31 | 0.28056638 | 0.44 | 0.36 | 1.10E-26 | liver | Tmbim4 |
| Lamtor1 | 1.06E-30 | 0.34270888 | 0.34 | 0.27 | 2.19E-26 | liver | Lamtor1 |
| Gdi21 | 1.10E-30 | 0.2802381 | 0.57 | 0.5 | 2.27E-26 | liver | Gdi2 |
| Imp3 | 3.44E-30 | 0.26861952 | 0.37 | 0.29 | 7.12E-26 | liver | Imp3 |
| Nenf | 5.20E-30 | 0.2623798 | 0.49 | 0.41 | 1.08E-25 | liver | Nenf |
| Ndufa5 | 1.36E-29 | 0.2787285 | 0.51 | 0.43 | 2.82E-25 | liver | Ndufa5 |
| Ltbr | 5.63E-29 | 0.28309449 | 0.26 | 0.19 | 1.16E-24 | liver | Ltbr |
| Arf4 | 5.81E-29 | 0.25617181 | 0.51 | 0.44 | 1.20E-24 | liver | Arf4 |
| Stip11 | 7.98E-29 | 0.2589977 | 0.3 | 0.22 | 1.65E-24 | liver | Stip1 |
| Sec11a | 8.45E-29 | 0.267721 | 0.36 | 0.29 | 1.75E-24 | liver | Sec11a |
| Psmb2 | 8.54E-29 | 0.26163452 | 0.55 | 0.48 | 1.77E-24 | liver | Psmb2 |
| 2010107E04Rik | 1.10E-28 | 0.27048132 | 0.55 | 0.47 | 2.27E-24 | liver | 2010107E04Rik |
| Nav11 | 1.34E-28 | 0.27276981 | 0.27 | 0.21 | 2.76E-24 | liver | Nav1 |
| Tm9sf2 | 5.54E-28 | 0.27814328 | 0.33 | 0.26 | 1.15E-23 | liver | Tm9sf2 |
| Gm16286 | 1.01E-27 | 0.26828705 | 0.29 | 0.22 | 2.09E-23 | liver | Gm16286 |
| Myeov2 | 1.67E-27 | 0.28444682 | 0.48 | 0.39 | 3.45E-23 | liver | Myeov2 |
| Dnajb91 | 1.24E-26 | 0.29155246 | 0.25 | 0.19 | 2.56E-22 | liver | Dnajb9 |
| Psma4 | 1.56E-26 | 0.27896408 | 0.43 | 0.36 | 3.22E-22 | liver | Psma4 |
| Flnb1 | 2.86E-26 | 0.27845383 | 0.27 | 0.21 | 5.92E-22 | liver | Flnb |
| Erp44 | 8.05E-26 | 0.2696398 | 0.28 | 0.21 | 1.66E-21 | liver | Erp44 |
| Tmed7 | 8.66E-26 | 0.25046634 | 0.29 | 0.22 | 1.79E-21 | liver | Tmed7 |
| Sirt2 | 4.94E-25 | 0.26272378 | 0.32 | 0.26 | 1.02E-20 | liver | Sirt2 |

| | | | | | | | |
|---|---|---|---|---|---|---|---|
| Snrpb2 | 1.12E-23 | 0.255198 | 0.29 | 0.22 | 2.31E-19 | liver | Snrpb2 |
| Capza2 | 1.52E-23 | 0.26047731 | 0.54 | 0.48 | 3.15E-19 | liver | Capza2 |
| Strn4 | 2.08E-23 | 0.28735715 | 0.25 | 0.19 | 4.30E-19 | liver | Strn4 |
| Tm9sf3 | 7.22E-23 | 0.28556168 | 0.4 | 0.33 | 1.49E-18 | liver | Tm9sf3 |
| Trappc3 | 6.46E-22 | 0.26196268 | 0.29 | 0.23 | 1.34E-17 | liver | Trappc3 |
| Erp29 | 2.37E-19 | 0.25226483 | 0.36 | 0.31 | 4.91E-15 | liver | Erp29 |
| Cfdp1 | 1.00E-16 | 0.26599187 | 0.36 | 0.31 | 2.07E-12 | liver | Cfdp1 |
| Sftpc | 0 | 4.97033829 | 0.78 | 0 | 0 | lung | Sftpc |
| Tmem100 | 0 | 4.76202736 | 0.94 | 0.04 | 0 | lung | Tmem100 |
| Hpgd | 0 | 3.5445675 | 0.9 | 0.12 | 0 | lung | Hpgd |
| Foxf1 | 0 | 3.49354251 | 0.87 | 0.09 | 0 | lung | Foxf1 |
| Sema3c1 | 0 | 2.73421249 | 0.71 | 0.11 | 0 | lung | Sema3c |
| Gm12216 | 0 | 2.72964305 | 0.72 | 0.2 | 0 | lung | Gm12216 |
| Hilpda | 0 | 2.63494802 | 0.67 | 0.08 | 0 | lung | Hilpda |
| Cyp4b11 | 0 | 2.56963423 | 0.84 | 0.22 | 0 | lung | Cyp4b1 |
| Emp21 | 0 | 2.53699281 | 0.55 | 0.16 | 0 | lung | Emp2 |
| Mfap2 | 0 | 2.43513952 | 0.66 | 0.05 | 0 | lung | Mfap2 |
| Ier31 | 0 | 2.34095537 | 0.83 | 0.32 | 0 | lung | Ier3 |
| Ace2 | 0 | 2.30486565 | 0.77 | 0.15 | 0 | lung | Ace |
| Tspan71 | 0 | 2.26332453 | 0.95 | 0.45 | 0 | lung | Tspan7 |
| Socs32 | 0 | 2.24599291 | 0.73 | 0.22 | 0 | lung | Socs3 |
| Acvrl12 | 0 | 2.19750573 | 0.9 | 0.43 | 0 | lung | Acvrl1 |
| Pmp22 | 0 | 2.18090828 | 0.61 | 0.21 | 0 | lung | Pmp22 |
| Myzap | 0 | 2.15652792 | 0.64 | 0.07 | 0 | lung | Myzap |
| Fibin | 0 | 2.13187429 | 0.32 | 0 | 0 | lung | Fibin |
| Tmcc2 | 0 | 2.10657482 | 0.57 | 0.08 | 0 | lung | Tmcc2 |
| BC0285281 | 0 | 2.05926496 | 0.88 | 0.39 | 0 | lung | BC028528 |
| Cd74 | 0 | 2.05046744 | 0.36 | 0.04 | 0 | lung | Cd74 |
| Calcrl1 | 0 | 2.03047791 | 0.84 | 0.34 | 0 | lung | Calcrl |
| Fmo1 | 0 | 2.01570425 | 0.71 | 0.18 | 0 | lung | Fmo1 |
| Gpx3 | 0 | 2.00971811 | 0.31 | 0.02 | 0 | lung | Gpx3 |
| Rasgef1a | 0 | 1.96334817 | 0.5 | 0 | 0 | lung | Rasgef1a |
| Scn7a | 0 | 1.94845753 | 0.46 | 0 | 0 | lung | Scn7a |
| Nkd11 | 0 | 1.94527723 | 0.63 | 0.1 | 0 | lung | Nkd1 |
| Tspan8 | 0 | 1.92595108 | 0.55 | 0.04 | 0 | lung | Tspan8 |
| Clec14a2 | 0 | 1.91589528 | 0.87 | 0.45 | 0 | lung | Clec14a |
| Prx | 0 | 1.90837713 | 0.43 | 0 | 0 | lung | Prx |
| Adgre5 | 0 | 1.87068603 | 0.71 | 0.2 | 0 | lung | Adgre5 |
| Ramp2 | 0 | 1.86299767 | 0.97 | 0.82 | 0 | lung | Ramp2 |
| Smad61 | 0 | 1.82945586 | 0.6 | 0.14 | 0 | lung | Smad6 |
| H2-Ab1 | 0 | 1.77981684 | 0.41 | 0.04 | 0 | lung | H2-Ab1 |

| | | | | | | | |
|---|---|---|---|---|---|---|---|
| Car41 | 0 | 1.75792197 | 0.69 | 0.34 | 0 | lung | Car4 |
| Emp12 | 0 | 1.74270193 | 0.61 | 0.17 | 0 | lung | Emp1 |
| Nsg1 | 0 | 1.70232301 | 0.54 | 0.08 | 0 | lung | Nsg1 |
| Thbd1 | 0 | 1.68917754 | 0.86 | 0.47 | 0 | lung | Thbd |
| Casc4 | 0 | 1.68095068 | 0.41 | 0.01 | 0 | lung | Casc4 |
| Clec1a1 | 0 | 1.67616436 | 0.69 | 0.23 | 0 | lung | Clec1a |
| Stmn11 | 0 | 1.67032537 | 0.64 | 0.17 | 0 | lung | Stmn1 |
| Ier23 | 0 | 1.66868194 | 0.85 | 0.53 | 0 | lung | Ier2 |
| Stmn2 | 0 | 1.61188814 | 0.66 | 0.16 | 0 | lung | Stmn2 |
| Pllp1 | 0 | 1.60075587 | 0.54 | 0.09 | 0 | lung | Pllp |
| H2-Aa | 0 | 1.58039591 | 0.34 | 0.02 | 0 | lung | H2-Aa |
| Icam2 | 0 | 1.56810308 | 0.89 | 0.63 | 0 | lung | Icam2 |
| Aard | 0 | 1.5636947 | 0.37 | 0 | 0 | lung | Aard |
| Junb3 | 0 | 1.54034915 | 0.87 | 0.6 | 0 | lung | Junb |
| Cyth3 | 0 | 1.49271292 | 0.57 | 0.17 | 0 | lung | Cyth3 |
| Lyve11 | 0 | 1.47754254 | 0.56 | 0.09 | 0 | lung | Lyve1 |
| Phlda3 | 0 | 1.46247768 | 0.46 | 0.13 | 0 | lung | Phlda3 |
| Tbx31 | 0 | 1.4621683 | 0.48 | 0.1 | 0 | lung | Tbx3 |
| Adrb1 | 0 | 1.45546595 | 0.39 | 0.03 | 0 | lung | Adrb1 |
| Mapt | 0 | 1.45389452 | 0.39 | 0.02 | 0 | lung | Mapt |
| Ankrd33b | 0 | 1.42047853 | 0.48 | 0.1 | 0 | lung | Ankrd33b |
| Sox11 | 0 | 1.41863439 | 0.31 | 0 | 0 | lung | Sox11 |
| Lhfp | 0 | 1.41668586 | 0.39 | 0.03 | 0 | lung | Lhfp |
| Acer21 | 0 | 1.40204554 | 0.53 | 0.17 | 0 | lung | Acer2 |
| Cdkn1a2 | 0 | 1.38916741 | 0.7 | 0.32 | 0 | lung | Cdkn1a |
| Impdh11 | 0 | 1.37708984 | 0.58 | 0.21 | 0 | lung | Impdh1 |
| Ehd43 | 0 | 1.37425217 | 0.88 | 0.48 | 0 | lung | Ehd4 |
| Tspan132 | 0 | 1.36659101 | 0.91 | 0.6 | 0 | lung | Tspan13 |
| 1-Jun | 0 | 1.3623037 | 0.93 | 0.7 | 0 | lung | Jun |
| Fosb2 | 0 | 1.34449501 | 0.59 | 0.25 | 0 | lung | Fosb |
| Ctla2a | 0 | 1.32566444 | 0.91 | 0.61 | 0 | lung | Ctla2a |
| Adrb21 | 0 | 1.31047199 | 0.42 | 0.09 | 0 | lung | Adrb2 |
| Ripply3 | 0 | 1.26220138 | 0.51 | 0.15 | 0 | lung | Ripply3 |
| Csrnp1 | 0 | 1.252027 | 0.36 | 0.06 | 0 | lung | Csrnp1 |
| Nhlrc2 | 0 | 1.23420567 | 0.34 | 0.04 | 0 | lung | Nhlrc2 |
| Galnt18 | 0 | 1.21745137 | 0.55 | 0.21 | 0 | lung | Galnt18 |
| Foxp1 | 0 | 1.21643295 | 0.72 | 0.45 | 0 | lung | Foxp1 |
| Cish | 0 | 1.20333366 | 0.35 | 0.04 | 0 | lung | Cish |
| Pcdh1 | 0 | 1.20099567 | 0.49 | 0.17 | 0 | lung | Pcdh1 |
| Gltp1 | 0 | 1.19919223 | 0.57 | 0.23 | 0 | lung | Gltp |
| Egr12 | 0 | 1.19762285 | 0.64 | 0.3 | 0 | lung | Egr1 |

| | | | | | | | |
|---|---|---|---|---|---|---|---|
| Hoxa5 | 0 | 1.19305166 | 0.36 | 0.07 | 0 | lung | Hoxa5 |
| Fendrr | 0 | 1.18764006 | 0.31 | 0.01 | 0 | lung | Fendrr |
| Hs3st1 | 0 | 1.18691614 | 0.44 | 0.13 | 0 | lung | Hs3st1 |
| Casz1 | 0 | 1.17755115 | 0.37 | 0.06 | 0 | lung | Casz1 |
| Afap1l11 | 0 | 1.17253392 | 0.65 | 0.32 | 0 | lung | Afap1l1 |
| Pde4b1 | 0 | 1.17202443 | 0.4 | 0.1 | 0 | lung | Pde4b |
| Jund1 | 0 | 1.1606653 | 0.88 | 0.7 | 0 | lung | Jund |
| Pltp1 | 0 | 1.15882904 | 0.89 | 0.44 | 0 | lung | Pltp |
| Klhl51 | 0 | 1.15576106 | 0.52 | 0.2 | 0 | lung | Klhl5 |
| Ecscr | 0 | 1.14873179 | 0.83 | 0.61 | 0 | lung | Ecscr |
| Ncald | 0 | 1.14470382 | 0.29 | 0.02 | 0 | lung | Ncald |
| Kit1 | 0 | 1.13643192 | 0.33 | 0.05 | 0 | lung | Kit |
| Slco2a11 | 0 | 1.12874951 | 0.67 | 0.26 | 0 | lung | Slco2a1 |
| Lamp11 | 0 | 1.12762073 | 0.88 | 0.67 | 0 | lung | Lamp1 |
| Gata21 | 0 | 1.1201587 | 0.6 | 0.28 | 0 | lung | Gata2 |
| Cldn51 | 0 | 1.11806254 | 0.97 | 0.59 | 0 | lung | Cldn5 |
| Cav21 | 0 | 1.11441809 | 0.91 | 0.65 | 0 | lung | Cav2 |
| Cadm1 | 0 | 1.10692079 | 0.4 | 0.12 | 0 | lung | Cadm1 |
| Plaur | 0 | 1.10191134 | 0.36 | 0.08 | 0 | lung | Plaur |
| Epas11 | 0 | 1.10091564 | 0.87 | 0.63 | 0 | lung | Epas1 |
| Ghr1 | 0 | 1.09615503 | 0.4 | 0.1 | 0 | lung | Ghr |
| Klf42 | 0 | 1.09309243 | 0.82 | 0.55 | 0 | lung | Klf4 |
| Sep-41 | 0 | 1.09257015 | 0.55 | 0.31 | 0 | lung | 4-Sep |
| Lysmd21 | 0 | 1.07887631 | 0.51 | 0.21 | 0 | lung | Lysmd2 |
| Tubb2a1 | 0 | 1.07835487 | 0.62 | 0.29 | 0 | lung | Tubb2a |
| Plscr11 | 0 | 1.07374304 | 0.53 | 0.19 | 0 | lung | Plscr1 |
| Tmem2521 | 0 | 1.0711398 | 0.31 | 0.11 | 0 | lung | Tmem252 |
| Tuba1a1 | 0 | 1.06750513 | 0.85 | 0.62 | 0 | lung | Tuba1a |
| Aldh21 | 0 | 1.06697408 | 0.82 | 0.56 | 0 | lung | Aldh2 |
| Icam11 | 0 | 1.0605177 | 0.44 | 0.16 | 0 | lung | Icam1 |
| Meis1 | 0 | 1.05052965 | 0.29 | 0.02 | 0 | lung | Meis1 |
| Gstm11 | 0 | 1.04599871 | 0.79 | 0.5 | 0 | lung | Gstm1 |
| Scn3b | 0 | 1.04326855 | 0.25 | 0.01 | 0 | lung | Scn3b |
| Hmcn12 | 0 | 1.02205911 | 0.42 | 0.18 | 0 | lung | Hmcn1 |
| Fkbp1a1 | 0 | 1.02151765 | 0.95 | 0.81 | 0 | lung | Fkbp1a |
| Cd92 | 0 | 1.01743375 | 0.91 | 0.59 | 0 | lung | Cd9 |
| Timp2 | 0 | 1.01169949 | 0.6 | 0.23 | 0 | lung | Timp2 |
| Cd361 | 0 | 1.00703764 | 0.92 | 0.48 | 0 | lung | Cd36 |
| Tspan181 | 0 | 0.99277552 | 0.51 | 0.21 | 0 | lung | Tspan18 |
| Enho | 0 | 0.97908255 | 0.28 | 0.06 | 0 | lung | Enho |
| Dpysl2 | 0 | 0.97599852 | 0.57 | 0.33 | 0 | lung | Dpysl2 |

| | | | | | | | |
|---|---|---|---|---|---|---|---|
| Atp13a2 | 0 | 0.97281723 | 0.3 | 0.05 | 0 | lung | Atp13a2 |
| Gja42 | 0 | 0.96544371 | 0.5 | 0.21 | 0 | lung | Gja4 |
| Plscr3 | 0 | 0.96370154 | 0.49 | 0.21 | 0 | lung | Plscr3 |
| Rgs121 | 0 | 0.9566275 | 0.36 | 0.12 | 0 | lung | Rgs12 |
| Ddah21 | 0 | 0.95598604 | 0.64 | 0.38 | 0 | lung | Ddah2 |
| Gm138892 | 0 | 0.94716206 | 0.41 | 0.18 | 0 | lung | Gm13889 |
| Wfdc11 | 0 | 0.93948072 | 0.38 | 0.1 | 0 | lung | Wfdc1 |
| Itm2b2 | 0 | 0.93160403 | 0.99 | 0.96 | 0 | lung | Itm2b |
| Bmpr21 | 0 | 0.92431421 | 0.6 | 0.32 | 0 | lung | Bmpr2 |
| Xbp12 | 0 | 0.92277321 | 0.59 | 0.33 | 0 | lung | Xbp1 |
| Il4ra | 0 | 0.92203163 | 0.35 | 0.09 | 0 | lung | Il4ra |
| Ppp1r15a1 | 0 | 0.91947847 | 0.51 | 0.25 | 0 | lung | Ppp1r15a |
| Cd24a4 | 0 | 0.90868601 | 0.53 | 0.21 | 0 | lung | Cd24a |
| Cdk14 | 0 | 0.90853279 | 0.26 | 0.03 | 0 | lung | Cdk14 |
| Ehd11 | 0 | 0.89508652 | 0.54 | 0.24 | 0 | lung | Ehd1 |
| Elovl1 | 0 | 0.88879994 | 0.53 | 0.27 | 0 | lung | Elovl1 |
| S1pr12 | 0 | 0.88852773 | 0.7 | 0.43 | 0 | lung | S1pr1 |
| Pdlim2 | 0 | 0.88586404 | 0.34 | 0.09 | 0 | lung | Pdlim2 |
| Qsox1 | 0 | 0.8857781 | 0.36 | 0.13 | 0 | lung | Qsox1 |
| Ddah11 | 0 | 0.88113629 | 0.33 | 0.1 | 0 | lung | Ddah1 |
| Crip22 | 0 | 0.87548872 | 0.96 | 0.84 | 0 | lung | Crip2 |
| Agpat4 | 0 | 0.87145497 | 0.42 | 0.17 | 0 | lung | Agpat4 |
| Ankrd441 | 0 | 0.87036525 | 0.36 | 0.11 | 0 | lung | Ankrd44 |
| Fas | 0 | 0.86972342 | 0.49 | 0.23 | 0 | lung | Fas |
| Cxx1a2 | 0 | 0.86467531 | 0.7 | 0.43 | 0 | lung | Cxx1a |
| Btg23 | 0 | 0.86085568 | 0.7 | 0.46 | 0 | lung | Btg2 |
| Tcn21 | 0 | 0.85839257 | 0.75 | 0.45 | 0 | lung | Tcn2 |
| Vat11 | 0 | 0.85745138 | 0.56 | 0.33 | 0 | lung | Vat1 |
| Car21 | 0 | 0.84811777 | 0.31 | 0.08 | 0 | lung | Car2 |
| Carhsp1 | 0 | 0.8461918 | 0.49 | 0.22 | 0 | lung | Carhsp1 |
| Atf32 | 0 | 0.84468825 | 0.44 | 0.15 | 0 | lung | Atf3 |
| Adk | 0 | 0.83697574 | 0.27 | 0.06 | 0 | lung | Adk |
| Pxdc1 | 0 | 0.8359624 | 0.37 | 0.15 | 0 | lung | Pxdc1 |
| Arl2bp1 | 0 | 0.82946762 | 0.53 | 0.27 | 0 | lung | Arl2bp |
| Hyal21 | 0 | 0.81672722 | 0.57 | 0.31 | 0 | lung | Hyal2 |
| Cav13 | 0 | 0.81670431 | 0.92 | 0.63 | 0 | lung | Cav1 |
| Pitpnc11 | 0 | 0.81452204 | 0.57 | 0.31 | 0 | lung | Pitpnc1 |
| Efnb11 | 0 | 0.81431945 | 0.49 | 0.21 | 0 | lung | Efnb1 |
| Nostrin2 | 0 | 0.81074338 | 0.52 | 0.27 | 0 | lung | Nostrin |
| Tppp32 | 0 | 0.80756206 | 0.43 | 0.15 | 0 | lung | Tppp3 |
| Cd472 | 0 | 0.80145232 | 0.8 | 0.62 | 0 | lung | Cd47 |

| | | | | | | | |
|---|---|---|---|---|---|---|---|
| Shisa51 | 0 | 0.79453033 | 0.74 | 0.54 | 0 | lung | Shisa5 |
| Rnf144a | 0 | 0.79121091 | 0.36 | 0.15 | 0 | lung | Rnf144a |
| Rap1a1 | 0 | 0.78665211 | 0.72 | 0.53 | 0 | lung | Rap1a |
| S1pr4 | 0 | 0.7833396 | 0.3 | 0.1 | 0 | lung | S1pr4 |
| Aph1a | 0 | 0.77916961 | 0.41 | 0.19 | 0 | lung | Aph1a |
| Ifitm10 | 0 | 0.7788004 | 0.26 | 0.06 | 0 | lung | Ifitm10 |
| Ccrl2 | 0 | 0.77503706 | 0.35 | 0.14 | 0 | lung | Ccrl2 |
| Grap1 | 0 | 0.77466924 | 0.54 | 0.3 | 0 | lung | Grap |
| Cxx1b2 | 0 | 0.77374599 | 0.54 | 0.28 | 0 | lung | Cxx1b |
| Egfl7 | 0 | 0.77243518 | 0.97 | 0.91 | 0 | lung | Egfl7 |
| Mapk3 | 0 | 0.76970491 | 0.72 | 0.51 | 0 | lung | Mapk3 |
| Pard6g | 0 | 0.76333776 | 0.35 | 0.06 | 0 | lung | Pard6g |
| Anxa31 | 0 | 0.74317386 | 0.71 | 0.5 | 0 | lung | Anxa3 |
| H2-D12 | 0 | 0.73749713 | 0.98 | 0.95 | 0 | lung | H2-D1 |
| Klf22 | 0 | 0.73061168 | 0.96 | 0.87 | 0 | lung | Klf2 |
| Stxbp6 | 0 | 0.7281494 | 0.26 | 0.06 | 0 | lung | Stxbp6 |
| Ly6a3 | 0 | 0.71404869 | 0.97 | 0.86 | 0 | lung | Ly6a |
| Lurap1l | 0 | 0.71398027 | 0.26 | 0.07 | 0 | lung | Lurap1l |
| Tmem176a2 | 0 | 0.71304563 | 0.7 | 0.39 | 0 | lung | Tmem176a |
| Sdpr2 | 0 | 0.70674722 | 0.91 | 0.68 | 0 | lung | Sdpr |
| Ybx12 | 0 | 0.70338673 | 0.94 | 0.83 | 0 | lung | Ybx1 |
| Bin1 | 0 | 0.70103349 | 0.37 | 0.14 | 0 | lung | Bin1 |
| Cdh51 | 0 | 0.68890028 | 0.86 | 0.72 | 0 | lung | Cdh5 |
| Arl4d | 0 | 0.68461385 | 0.31 | 0.12 | 0 | lung | Arl4d |
| Tmbim61 | 0 | 0.68251169 | 0.83 | 0.69 | 0 | lung | Tmbim6 |
| P2rx4 | 0 | 0.68070835 | 0.25 | 0.08 | 0 | lung | P2rx4 |
| Arap21 | 0 | 0.67954297 | 0.27 | 0.08 | 0 | lung | Arap2 |
| Slc3a21 | 0 | 0.6642824 | 0.76 | 0.51 | 0 | lung | Slc3a2 |
| Ttyh3 | 0 | 0.65706327 | 0.27 | 0.08 | 0 | lung | Ttyh3 |
| Ly6e3 | 0 | 0.64895456 | 0.98 | 0.94 | 0 | lung | Ly6e |
| Npr3 | 0 | 0.6485229 | 0.25 | 0.05 | 0 | lung | Npr3 |
| Fos3 | 0 | 0.63956085 | 0.79 | 0.53 | 0 | lung | Fos |
| Esam2 | 0 | 0.61294273 | 0.88 | 0.72 | 0 | lung | Esam |
| Vim1 | 0 | 0.61117195 | 0.87 | 0.63 | 0 | lung | Vim |
| Slc43a34 | 0 | 0.60560399 | 0.68 | 0.39 | 0 | lung | Slc43a3 |
| Dusp14 | 0 | 0.60187121 | 0.84 | 0.64 | 0 | lung | Dusp1 |
| Zfp363 | 0 | 0.59038623 | 0.64 | 0.39 | 0 | lung | Zfp36 |
| Gmfg | 0 | 0.56375193 | 0.27 | 0.09 | 0 | lung | Gmfg |
| Ubb | 0 | 0.47089553 | 0.99 | 0.96 | 0 | lung | Ubb |
| Eif11 | 0 | 0.47000589 | 0.97 | 0.92 | 0 | lung | Eif1 |
| Snrpb | 8.53E-307 | 0.70117987 | 0.76 | 0.6 | 1.77E-302 | lung | Snrpb |

| | | | | | | | |
|---|---|---|---|---|---|---|---|
| Mid1ip1 | 1.00E-302 | 0.7198944 | 0.31 | 0.12 | 2.07E-298 | lung | Mid1ip1 |
| S100a161 | 2.34E-301 | 0.62021871 | 0.83 | 0.68 | 4.85E-297 | lung | S100a16 |
| Lxn | 1.54E-295 | 0.79795889 | 0.41 | 0.2 | 3.18E-291 | lung | Lxn |
| Tmem243 | 6.29E-293 | 0.68640845 | 0.37 | 0.16 | 1.30E-288 | lung | Tmem243 |
| Gpr146 | 6.21E-291 | 0.73395672 | 0.4 | 0.19 | 1.29E-286 | lung | Gpr146 |
| Cd93 | 9.55E-291 | 0.85129621 | 0.68 | 0.53 | 1.98E-286 | lung | Cd93 |
| Trim8 | 4.35E-288 | 0.69796261 | 0.38 | 0.17 | 9.00E-284 | lung | Trim8 |
| Map1lc3a2 | 1.84E-282 | 0.81208314 | 0.61 | 0.42 | 3.81E-278 | lung | Map1lc3a |
| Cdc42ep1 | 7.33E-282 | 0.69600227 | 0.48 | 0.26 | 1.52E-277 | lung | Cdc42ep1 |
| Smagp1 | 3.74E-278 | 0.73910061 | 0.63 | 0.45 | 7.75E-274 | lung | Smagp |
| Gpihbp12 | 1.03E-275 | 0.3785752 | 0.84 | 0.53 | 2.14E-271 | lung | Gpihbp1 |
| Ifngr2 | 3.30E-274 | 0.66601857 | 0.35 | 0.15 | 6.83E-270 | lung | Ifngr2 |
| Atf41 | 1.40E-271 | 0.71003275 | 0.49 | 0.28 | 2.91E-267 | lung | Atf4 |
| Clic41 | 1.78E-269 | 0.62696551 | 0.71 | 0.52 | 3.69E-265 | lung | Clic4 |
| Hn11 | 3.67E-265 | 0.5991349 | 0.4 | 0.19 | 7.60E-261 | lung | Hn1 |
| Edn11 | 1.35E-264 | 0.67617481 | 0.38 | 0.17 | 2.79E-260 | lung | Edn1 |
| Rcsd1 | 2.64E-263 | 0.60552505 | 0.36 | 0.16 | 5.46E-259 | lung | Rcsd1 |
| Sertad1 | 1.40E-261 | 0.73026555 | 0.39 | 0.19 | 2.90E-257 | lung | Sertad1 |
| Hey11 | 2.58E-260 | 0.71460018 | 0.31 | 0.13 | 5.33E-256 | lung | Hey1 |
| Mfng | 5.93E-256 | 0.64539844 | 0.36 | 0.17 | 1.23E-251 | lung | Mfng |
| N4bp31 | 1.21E-255 | 0.63071183 | 0.38 | 0.18 | 2.50E-251 | lung | N4bp3 |
| Tmcc31 | 5.87E-254 | 0.64922908 | 0.42 | 0.22 | 1.22E-249 | lung | Tmcc3 |
| Dynll11 | 9.14E-253 | 0.45463761 | 0.91 | 0.84 | 1.89E-248 | lung | Dynll1 |
| Tgfb1i1 | 1.30E-251 | 0.69897415 | 0.38 | 0.19 | 2.70E-247 | lung | Tgfb1i1 |
| Gcc21 | 1.41E-246 | 0.61437813 | 0.27 | 0.11 | 2.91E-242 | lung | Gcc2 |
| Stx3 | 2.79E-246 | 0.58221758 | 0.27 | 0.11 | 5.78E-242 | lung | Stx3 |
| Myl12a2 | 6.35E-245 | 0.44400128 | 0.91 | 0.83 | 1.31E-240 | lung | Myl12a |
| Tmem176b1 | 4.16E-243 | 0.43954065 | 0.72 | 0.47 | 8.62E-239 | lung | Tmem176b |
| Myl12b1 | 2.35E-237 | 0.54782342 | 0.8 | 0.65 | 4.86E-233 | lung | Myl12b |
| Map7d11 | 2.74E-236 | 0.65612957 | 0.54 | 0.34 | 5.67E-232 | lung | Map7d1 |
| Sgce | 2.15E-235 | 0.5194265 | 0.25 | 0.1 | 4.45E-231 | lung | Sgce |
| Jup1 | 3.55E-233 | 0.50772766 | 0.6 | 0.38 | 7.35E-229 | lung | Jup |
| Zfp5031 | 4.02E-233 | 0.67317297 | 0.3 | 0.13 | 8.32E-229 | lung | Zfp503 |
| Prickle11 | 2.28E-231 | 0.60095752 | 0.3 | 0.13 | 4.72E-227 | lung | Prickle1 |
| Tmem21 | 2.92E-229 | 0.62163595 | 0.33 | 0.16 | 6.04E-225 | lung | Tmem2 |
| Myct11 | 5.47E-228 | 0.6069298 | 0.67 | 0.48 | 1.13E-223 | lung | Myct1 |
| Dpf2 | 6.62E-223 | 0.57191914 | 0.31 | 0.14 | 1.37E-218 | lung | Dpf2 |
| Lfng | 2.58E-219 | 0.59903928 | 0.33 | 0.15 | 5.33E-215 | lung | Lfng |
| Ppp1ca | 6.02E-219 | 0.59512481 | 0.69 | 0.54 | 1.25E-214 | lung | Ppp1ca |
| Sox7 | 2.23E-218 | 0.62290994 | 0.36 | 0.18 | 4.61E-214 | lung | Sox7 |
| Plvap4 | 3.87E-218 | 0.30236783 | 0.84 | 0.55 | 8.00E-214 | lung | Plvap |

| | | | | | | | |
|---|---|---|---|---|---|---|---|
| Pdlim13 | 8.04E-217 | 0.52114106 | 0.51 | 0.29 | 1.66E-212 | lung | Pdlim1 |
| Guk1 | 5.02E-215 | 0.71142592 | 0.42 | 0.24 | 1.04E-210 | lung | Guk1 |
| H2-Q72 | 3.54E-213 | 0.59318005 | 0.58 | 0.36 | 7.33E-209 | lung | H2-Q7 |
| Hspa84 | 5.88E-211 | 0.34538151 | 0.96 | 0.9 | 1.22E-206 | lung | Hspa8 |
| Sec621 | 8.77E-211 | 0.58353519 | 0.73 | 0.58 | 1.81E-206 | lung | Sec62 |
| Vimp1 | 9.01E-211 | 0.56109304 | 0.32 | 0.15 | 1.86E-206 | lung | Vimp |
| Plk23 | 5.96E-208 | 0.72489285 | 0.6 | 0.41 | 1.23E-203 | lung | Plk2 |
| Dusp6 | 3.55E-206 | 0.72530937 | 0.42 | 0.24 | 7.35E-202 | lung | Dusp6 |
| Tspan91 | 4.95E-206 | 0.61276934 | 0.46 | 0.27 | 1.02E-201 | lung | Tspan9 |
| Tmem1401 | 1.23E-203 | 0.67116738 | 0.54 | 0.36 | 2.55E-199 | lung | Tmem140 |
| Clic11 | 2.98E-203 | 0.47807311 | 0.82 | 0.71 | 6.16E-199 | lung | Clic1 |
| Ppt2 | 3.04E-203 | 0.54641562 | 0.3 | 0.14 | 6.29E-199 | lung | Ppt2 |
| Tmem109 | 1.65E-201 | 0.61044699 | 0.55 | 0.36 | 3.40E-197 | lung | Tmem109 |
| Adgrf51 | 8.30E-200 | 0.48615727 | 0.76 | 0.63 | 1.72E-195 | lung | Adgrf5 |
| Itpripl2 | 2.05E-197 | 0.54939399 | 0.26 | 0.11 | 4.25E-193 | lung | Itpripl2 |
| Tmco1 | 3.52E-197 | 0.62533591 | 0.61 | 0.45 | 7.29E-193 | lung | Tmco1 |
| Skap2 | 1.49E-196 | 0.61366846 | 0.32 | 0.16 | 3.08E-192 | lung | Skap2 |
| Slc9a3r23 | 4.62E-195 | 0.38901602 | 0.89 | 0.81 | 9.55E-191 | lung | Slc9a3r2 |
| Arf1 | 1.22E-193 | 0.50767592 | 0.76 | 0.63 | 2.52E-189 | lung | Arf1 |
| Rac1 | 2.84E-191 | 0.47815733 | 0.78 | 0.69 | 5.87E-187 | lung | Rac1 |
| Mylk | 9.15E-190 | 0.45395006 | 0.26 | 0.11 | 1.89E-185 | lung | Mylk |
| Adcy4 | 1.80E-189 | 0.57854199 | 0.39 | 0.22 | 3.73E-185 | lung | Adcy4 |
| Hlx1 | 6.81E-189 | 0.52520099 | 0.34 | 0.17 | 1.41E-184 | lung | Hlx |
| Gna13 | 1.80E-188 | 0.49675999 | 0.26 | 0.11 | 3.72E-184 | lung | Gna13 |
| Ubald2 | 8.21E-188 | 0.56885477 | 0.58 | 0.4 | 1.70E-183 | lung | Ubald2 |
| Ifitm32 | 6.23E-185 | 0.39534534 | 0.97 | 0.96 | 1.29E-180 | lung | Ifitm3 |
| Hpcal1 | 7.92E-182 | 0.58103336 | 0.52 | 0.35 | 1.64E-177 | lung | Hpcal1 |
| Ecm12 | 1.22E-180 | 0.56225767 | 0.33 | 0.17 | 2.53E-176 | lung | Ecm1 |
| Arhgef31 | 1.95E-180 | 0.53689748 | 0.35 | 0.19 | 4.04E-176 | lung | Arhgef3 |
| Ctsd1 | 5.00E-180 | 0.46714096 | 0.7 | 0.53 | 1.03E-175 | lung | Ctsd |
| Ccnl11 | 5.06E-180 | 0.61851509 | 0.4 | 0.23 | 1.05E-175 | lung | Ccnl1 |
| Hmgn2 | 4.31E-179 | 0.50556015 | 0.29 | 0.14 | 8.93E-175 | lung | Hmgn2 |
| Pon21 | 5.60E-178 | 0.56707669 | 0.47 | 0.3 | 1.16E-173 | lung | Pon2 |
| Tra2b1 | 1.41E-176 | 0.63277928 | 0.57 | 0.41 | 2.91E-172 | lung | Tra2b |
| Ppp1r16b | 2.44E-176 | 0.66234525 | 0.44 | 0.27 | 5.05E-172 | lung | Ppp1r16b |
| Cracr2b | 2.89E-174 | 0.5534856 | 0.56 | 0.38 | 5.98E-170 | lung | Cracr2b |
| Klf71 | 6.14E-173 | 0.55478852 | 0.63 | 0.48 | 1.27E-168 | lung | Klf7 |
| Stard3nl1 | 2.23E-171 | 0.50756329 | 0.32 | 0.17 | 4.62E-167 | lung | Stard3nl |
| Pecam12 | 8.39E-170 | 0.38123057 | 0.86 | 0.76 | 1.74E-165 | lung | Pecam1 |
| Ccnd31 | 1.40E-169 | 0.55289334 | 0.46 | 0.29 | 2.89E-165 | lung | Ccnd3 |
| Rhob2 | 3.21E-169 | 0.50606582 | 0.69 | 0.54 | 6.65E-165 | lung | Rhob |

| | | | | | | | |
|---|---|---|---|---|---|---|---|
| Tor3a | 2.83E-168 | 0.5490479 | 0.28 | 0.14 | 5.85E-164 | lung | Tor3a |
| Cnn31 | 1.70E-167 | 0.54312285 | 0.65 | 0.51 | 3.52E-163 | lung | Cnn3 |
| Rasip1 | 1.97E-167 | 0.53671116 | 0.66 | 0.51 | 4.08E-163 | lung | Rasip1 |
| Cyb5611 | 4.93E-166 | 0.50536634 | 0.25 | 0.12 | 1.02E-161 | lung | Cyb561 |
| Mcam1 | 7.58E-165 | 0.560975 | 0.34 | 0.18 | 1.57E-160 | lung | Mcam |
| Ifi47 | 3.30E-162 | 0.64191178 | 0.33 | 0.18 | 6.83E-158 | lung | Ifi47 |
| Homer3 | 4.97E-162 | 0.47831834 | 0.26 | 0.12 | 1.03E-157 | lung | Homer3 |
| Atxn10 | 1.76E-161 | 0.49772073 | 0.25 | 0.12 | 3.64E-157 | lung | Atxn10 |
| Tecpr1 | 6.23E-161 | 0.48850169 | 0.36 | 0.2 | 1.29E-156 | lung | Tecpr1 |
| Gnai21 | 1.18E-160 | 0.37872772 | 0.86 | 0.81 | 2.45E-156 | lung | Gnai2 |
| Dnajc81 | 6.04E-158 | 0.57959336 | 0.56 | 0.4 | 1.25E-153 | lung | Dnajc8 |
| Dram2 | 8.70E-157 | 0.52549327 | 0.46 | 0.29 | 1.80E-152 | lung | Dram2 |
| Syngr2 | 8.03E-154 | 0.51683541 | 0.39 | 0.22 | 1.66E-149 | lung | Syngr2 |
| Adamts12 | 1.23E-153 | 0.54545319 | 0.37 | 0.21 | 2.54E-149 | lung | Adamts1 |
| Ict1 | 1.42E-152 | 0.50388823 | 0.34 | 0.18 | 2.93E-148 | lung | Ict1 |
| Trib21 | 1.96E-152 | 0.50229166 | 0.3 | 0.16 | 4.05E-148 | lung | Trib2 |
| Mxra7 | 9.19E-152 | 0.53268979 | 0.41 | 0.25 | 1.90E-147 | lung | Mxra7 |
| Rassf1 | 2.35E-151 | 0.53843438 | 0.45 | 0.29 | 4.87E-147 | lung | Rassf1 |
| Dapk3 | 2.99E-151 | 0.46711098 | 0.27 | 0.13 | 6.19E-147 | lung | Dapk3 |
| Ldb21 | 2.83E-150 | 0.3388824 | 0.42 | 0.24 | 5.86E-146 | lung | Ldb2 |
| Klf9 | 1.33E-149 | 0.58621057 | 0.52 | 0.36 | 2.76E-145 | lung | Klf9 |
| Ubl7 | 7.04E-146 | 0.45400649 | 0.27 | 0.13 | 1.46E-141 | lung | Ubl7 |
| St3gal11 | 2.09E-144 | 0.44972989 | 0.28 | 0.14 | 4.32E-140 | lung | St3gal1 |
| Eng2 | 7.33E-144 | 0.31982225 | 0.9 | 0.84 | 1.52E-139 | lung | Eng |
| Add32 | 2.07E-143 | 0.442706 | 0.3 | 0.16 | 4.28E-139 | lung | Add3 |
| Inpp5k | 7.97E-143 | 0.4101473 | 0.27 | 0.13 | 1.65E-138 | lung | Inpp5k |
| Dgkz | 2.30E-142 | 0.47090666 | 0.3 | 0.16 | 4.76E-138 | lung | Dgkz |
| Saraf1 | 7.95E-142 | 0.49214528 | 0.6 | 0.46 | 1.64E-137 | lung | Saraf |
| Bicd2 | 1.14E-141 | 0.46036264 | 0.3 | 0.16 | 2.35E-137 | lung | Bicd2 |
| Tmbim11 | 8.06E-141 | 0.48350985 | 0.42 | 0.26 | 1.67E-136 | lung | Tmbim1 |
| Dad1 | 1.39E-140 | 0.46466266 | 0.71 | 0.59 | 2.87E-136 | lung | Dad1 |
| Rnf187 | 1.05E-137 | 0.49675664 | 0.53 | 0.37 | 2.18E-133 | lung | Rnf187 |
| Tapbp | 4.06E-137 | 0.50312495 | 0.62 | 0.47 | 8.40E-133 | lung | Tapbp |
| Kdm6b | 1.66E-136 | 0.54759836 | 0.27 | 0.14 | 3.44E-132 | lung | Kdm6b |
| Irf12 | 9.59E-133 | 0.5052074 | 0.44 | 0.29 | 1.98E-128 | lung | Irf1 |
| Rela | 1.32E-132 | 0.44601044 | 0.33 | 0.18 | 2.74E-128 | lung | Rela |
| Pnpla21 | 5.08E-131 | 0.4575963 | 0.33 | 0.19 | 1.05E-126 | lung | Pnpla2 |
| St3gal62 | 1.29E-130 | 0.44757817 | 0.52 | 0.36 | 2.67E-126 | lung | St3gal6 |
| Derl2 | 4.10E-130 | 0.4693199 | 0.29 | 0.16 | 8.48E-126 | lung | Derl2 |
| Srsf5 | 1.67E-129 | 0.5237587 | 0.64 | 0.52 | 3.46E-125 | lung | Srsf5 |
| Smad72 | 7.66E-129 | 0.48044431 | 0.43 | 0.28 | 1.59E-124 | lung | Smad7 |

| | | | | | | | |
|---|---|---|---|---|---|---|---|
| Ednrb1 | 1.28E-128 | 0.65705649 | 0.27 | 0.15 | 2.65E-124 | lung | Ednrb |
| Ucp22 | 3.05E-128 | 0.43200371 | 0.4 | 0.24 | 6.32E-124 | lung | Ucp2 |
| Tapbpl | 4.26E-128 | 0.43441304 | 0.26 | 0.14 | 8.81E-124 | lung | Tapbpl |
| Slc48a11 | 1.21E-127 | 0.51838085 | 0.36 | 0.22 | 2.50E-123 | lung | Slc48a1 |
| Selk1 | 1.91E-127 | 0.36103544 | 0.78 | 0.66 | 3.95E-123 | lung | Selk |
| Hist3h2a | 4.40E-126 | 0.44672076 | 0.33 | 0.19 | 9.11E-122 | lung | Hist3h2a |
| Mgst31 | 5.03E-126 | 0.53675667 | 0.35 | 0.21 | 1.04E-121 | lung | Mgst3 |
| Plekho11 | 1.37E-125 | 0.4850755 | 0.36 | 0.22 | 2.83E-121 | lung | Plekho1 |
| Elovl5 | 5.83E-125 | 0.50265352 | 0.44 | 0.29 | 1.21E-120 | lung | Elovl5 |
| Ddx5 | 1.79E-124 | 0.3421187 | 0.89 | 0.84 | 3.71E-120 | lung | Ddx5 |
| Efnb22 | 2.67E-123 | 0.45739733 | 0.31 | 0.17 | 5.53E-119 | lung | Efnb2 |
| Card191 | 3.03E-123 | 0.48584716 | 0.4 | 0.26 | 6.26E-119 | lung | Card19 |
| Tspan121 | 5.34E-123 | 0.49296222 | 0.48 | 0.34 | 1.11E-118 | lung | Tspan12 |
| Ifngr13 | 2.34E-122 | 0.49024775 | 0.53 | 0.38 | 4.85E-118 | lung | Ifngr1 |
| Dnajb12 | 3.55E-122 | 0.45316202 | 0.32 | 0.19 | 7.35E-118 | lung | Dnajb1 |
| Akr1b31 | 2.91E-121 | 0.42721584 | 0.34 | 0.2 | 6.01E-117 | lung | Akr1b3 |
| Anp32e | 3.80E-121 | 0.47540038 | 0.26 | 0.14 | 7.87E-117 | lung | Anp32e |
| Rab1a | 4.40E-121 | 0.47543968 | 0.56 | 0.42 | 9.10E-117 | lung | Rab1a |
| Zcrb1 | 1.38E-118 | 0.52914356 | 0.43 | 0.28 | 2.85E-114 | lung | Zcrb1 |
| Trim35 | 4.17E-117 | 0.47679221 | 0.27 | 0.15 | 8.62E-113 | lung | Trim35 |
| Manf1 | 4.58E-117 | 0.37080622 | 0.6 | 0.45 | 9.47E-113 | lung | Manf |
| Rbm391 | 5.71E-117 | 0.33569143 | 0.82 | 0.76 | 1.18E-112 | lung | Rbm39 |
| Csrp11 | 1.65E-116 | 0.48564525 | 0.55 | 0.4 | 3.41E-112 | lung | Csrp1 |
| Snx21 | 1.80E-116 | 0.42592211 | 0.25 | 0.13 | 3.73E-112 | lung | Snx21 |
| Rtn4 | 3.53E-116 | 0.45620024 | 0.56 | 0.42 | 7.31E-112 | lung | Rtn4 |
| Atp2b11 | 1.11E-115 | 0.47844619 | 0.39 | 0.25 | 2.30E-111 | lung | Atp2b1 |
| Vamp8 | 5.84E-115 | 0.47793068 | 0.54 | 0.39 | 1.21E-110 | lung | Vamp8 |
| Klf63 | 9.08E-115 | 0.37253123 | 0.58 | 0.42 | 1.88E-110 | lung | Klf6 |
| Zfp706 | 1.39E-114 | 0.47450573 | 0.52 | 0.39 | 2.88E-110 | lung | Zfp706 |
| Anxa61 | 1.56E-114 | 0.44658076 | 0.31 | 0.18 | 3.22E-110 | lung | Anxa6 |
| Ubc2 | 2.12E-114 | 0.30182747 | 0.89 | 0.83 | 4.39E-110 | lung | Ubc |
| Agfg11 | 9.50E-112 | 0.41458904 | 0.27 | 0.15 | 1.97E-107 | lung | Agfg1 |
| Mtus11 | 9.88E-111 | 0.410965 | 0.37 | 0.23 | 2.04E-106 | lung | Mtus1 |
| Ube2d3 | 6.87E-110 | 0.3591542 | 0.72 | 0.62 | 1.42E-105 | lung | Ube2d3 |
| H2-Q61 | 9.22E-110 | 0.53412763 | 0.35 | 0.22 | 1.91E-105 | lung | H2-Q6 |
| Vcl1 | 1.13E-109 | 0.42010612 | 0.29 | 0.17 | 2.33E-105 | lung | Vcl |
| Ypel32 | 1.15E-108 | 0.4189665 | 0.55 | 0.41 | 2.39E-104 | lung | Ypel3 |
| Raph11 | 2.80E-108 | 0.3297073 | 0.26 | 0.14 | 5.79E-104 | lung | Raph1 |
| Top11 | 3.13E-106 | 0.45332637 | 0.5 | 0.36 | 6.47E-102 | lung | Top1 |
| Abi3 | 3.90E-106 | 0.44541111 | 0.42 | 0.29 | 8.06E-102 | lung | Abi3 |
| Tnfrsf1a1 | 5.20E-106 | 0.40150124 | 0.56 | 0.42 | 1.08E-101 | lung | Tnfrsf1a |

| | | | | | | | |
|---|---|---|---|---|---|---|---|
| H3f3b3 | 1.07E-105 | 0.27362177 | 0.91 | 0.88 | 2.22E-101 | lung | H3f3b |
| Mesdc11 | 1.36E-105 | 0.42182126 | 0.36 | 0.22 | 2.82E-101 | lung | Mesdc1 |
| Tspo | 3.17E-105 | 0.38407303 | 0.7 | 0.59 | 6.56E-101 | lung | Tspo |
| Pelo | 2.52E-104 | 0.3874206 | 0.25 | 0.14 | 5.22E-100 | lung | Pelo |
| Rhoa | 2.57E-104 | 0.32661308 | 0.84 | 0.77 | 5.32E-100 | lung | Rhoa |
| Tmed4 | 5.10E-103 | 0.43119773 | 0.35 | 0.22 | 1.06E-98 | lung | Tmed4 |
| Pdia61 | 1.30E-102 | 0.42997178 | 0.59 | 0.45 | 2.70E-98 | lung | Pdia6 |
| Tubb51 | 6.08E-102 | 0.38328785 | 0.74 | 0.64 | 1.26E-97 | lung | Tubb5 |
| Ctbp21 | 3.06E-101 | 0.40958555 | 0.29 | 0.17 | 6.33E-97 | lung | Ctbp2 |
| Atp1a1 | 2.24E-98 | 0.52631282 | 0.47 | 0.35 | 4.63E-94 | lung | Atp1a1 |
| Dhrs71 | 4.49E-97 | 0.4439361 | 0.38 | 0.25 | 9.28E-93 | lung | Dhrs7 |
| Ppdpf | 1.54E-96 | 0.42479831 | 0.44 | 0.31 | 3.18E-92 | lung | Ppdpf |
| Tmem591 | 1.68E-96 | 0.32662712 | 0.8 | 0.71 | 3.48E-92 | lung | Tmem59 |
| Spcs1 | 1.41E-95 | 0.39403517 | 0.57 | 0.45 | 2.92E-91 | lung | Spcs1 |
| Sgta | 1.33E-94 | 0.38508333 | 0.31 | 0.19 | 2.76E-90 | lung | Sgta |
| Tmem50a | 1.74E-94 | 0.36912716 | 0.68 | 0.58 | 3.60E-90 | lung | Tmem50a |
| Gnb2 | 3.42E-94 | 0.32595058 | 0.78 | 0.71 | 7.07E-90 | lung | Gnb2 |
| Cyb5a1 | 3.61E-94 | 0.32666782 | 0.73 | 0.62 | 7.46E-90 | lung | Cyb5a |
| Vgll41 | 5.02E-94 | 0.46780414 | 0.44 | 0.31 | 1.04E-89 | lung | Vgll4 |
| Msn3 | 1.16E-93 | 0.37810882 | 0.69 | 0.57 | 2.40E-89 | lung | Msn |
| Rhoc1 | 1.92E-93 | 0.36715343 | 0.63 | 0.52 | 3.97E-89 | lung | Rhoc |
| Rasgrp22 | 9.84E-93 | 0.34548172 | 0.31 | 0.19 | 2.04E-88 | lung | Rasgrp2 |
| Tmem9b2 | 1.87E-92 | 0.40649271 | 0.4 | 0.27 | 3.88E-88 | lung | Tmem9b |
| Fermt21 | 5.09E-92 | 0.40837442 | 0.6 | 0.49 | 1.05E-87 | lung | Fermt2 |
| Ppp2ca | 3.74E-90 | 0.3954202 | 0.44 | 0.31 | 7.74E-86 | lung | Ppp2ca |
| Lyn1 | 2.54E-89 | 0.39707037 | 0.32 | 0.21 | 5.25E-85 | lung | Lyn |
| Ctnna11 | 6.58E-89 | 0.36934095 | 0.58 | 0.45 | 1.36E-84 | lung | Ctnna1 |
| Stx7 | 8.52E-89 | 0.44287085 | 0.39 | 0.26 | 1.76E-84 | lung | Stx7 |
| Cd1511 | 8.04E-88 | 0.36925422 | 0.65 | 0.55 | 1.66E-83 | lung | Cd151 |
| Il2rg | 1.00E-87 | 0.28635926 | 0.29 | 0.17 | 2.07E-83 | lung | Il2rg |
| Eif4a1 | 1.44E-87 | 0.37991294 | 0.69 | 0.57 | 2.98E-83 | lung | Eif4a1 |
| Aup1 | 2.70E-87 | 0.36565452 | 0.3 | 0.18 | 5.59E-83 | lung | Aup1 |
| H2-K13 | 2.12E-86 | 0.25304156 | 0.91 | 0.84 | 4.39E-82 | lung | H2-K1 |
| Arhgdia | 1.96E-85 | 0.34491429 | 0.68 | 0.6 | 4.06E-81 | lung | Arhgdia |
| Tmem258 | 5.30E-84 | 0.39785428 | 0.48 | 0.36 | 1.10E-79 | lung | Tmem258 |
| Cmtm31 | 1.74E-83 | 0.36634355 | 0.34 | 0.22 | 3.60E-79 | lung | Cmtm3 |
| Cers21 | 9.19E-83 | 0.37541585 | 0.37 | 0.25 | 1.90E-78 | lung | Cers2 |
| Sypl1 | 1.22E-81 | 0.34308772 | 0.63 | 0.51 | 2.53E-77 | lung | Sypl |
| Serpinh12 | 3.34E-81 | 0.25003667 | 0.78 | 0.67 | 6.92E-77 | lung | Serpinh1 |
| Vps361 | 5.05E-81 | 0.37909828 | 0.34 | 0.23 | 1.04E-76 | lung | Vps36 |
| Tmbim41 | 4.42E-80 | 0.37445294 | 0.47 | 0.35 | 9.15E-76 | lung | Tmbim4 |

| | | | | | | | |
|---|---|---|---|---|---|---|---|
| Stx5a | 6.06E-80 | 0.36765286 | 0.34 | 0.22 | 1.25E-75 | lung | Stx5a |
| Ahr1 | 1.02E-79 | 0.35833659 | 0.29 | 0.18 | 2.10E-75 | lung | Ahr |
| Pcbp11 | 1.18E-79 | 0.37320664 | 0.48 | 0.36 | 2.44E-75 | lung | Pcbp1 |
| Gipc11 | 2.85E-79 | 0.35827579 | 0.26 | 0.15 | 5.89E-75 | lung | Gipc1 |
| Srpr | 6.51E-79 | 0.39755039 | 0.35 | 0.24 | 1.35E-74 | lung | Srpr |
| Sdhb | 6.52E-79 | 0.38253598 | 0.48 | 0.37 | 1.35E-74 | lung | Sdhb |
| Parvb1 | 3.04E-78 | 0.34883135 | 0.4 | 0.28 | 6.28E-74 | lung | Parvb |
| Sep151 | 1.10E-77 | 0.28984669 | 0.77 | 0.65 | 2.28E-73 | lung | 15-Sep |
| Pttg1ip1 | 3.50E-77 | 0.36928236 | 0.52 | 0.4 | 7.24E-73 | lung | Pttg1ip |
| Abhd16a | 5.02E-77 | 0.38319029 | 0.33 | 0.22 | 1.04E-72 | lung | Abhd16a |
| Ssr2 | 5.22E-77 | 0.38530537 | 0.48 | 0.36 | 1.08E-72 | lung | Ssr2 |
| Gadd45g2 | 1.16E-76 | 0.29115274 | 0.4 | 0.28 | 2.41E-72 | lung | Gadd45g |
| Sipa1 | 2.17E-76 | 0.3268366 | 0.39 | 0.27 | 4.50E-72 | lung | Sipa1 |
| Tax1bp11 | 3.18E-76 | 0.48172623 | 0.43 | 0.31 | 6.59E-72 | lung | Tax1bp1 |
| Arid5b1 | 7.56E-76 | 0.33282593 | 0.37 | 0.25 | 1.57E-71 | lung | Arid5b |
| Atp6v0b1 | 1.50E-75 | 0.3274566 | 0.56 | 0.46 | 3.11E-71 | lung | Atp6v0b |
| BC031181 | 4.81E-75 | 0.38564828 | 0.46 | 0.35 | 9.96E-71 | lung | BC031181 |
| Spcs21 | 9.22E-75 | 0.34032136 | 0.57 | 0.46 | 1.91E-70 | lung | Spcs2 |
| Ptp4a31 | 1.14E-74 | 0.64913646 | 0.48 | 0.39 | 2.35E-70 | lung | Ptp4a3 |
| Tmed91 | 2.23E-74 | 0.34071365 | 0.61 | 0.51 | 4.62E-70 | lung | Tmed9 |
| 1110008F13Rik | 3.72E-74 | 0.38271261 | 0.39 | 0.28 | 7.69E-70 | lung | 1110008F13Rik |
| Srsf6 | 6.69E-74 | 0.34522124 | 0.33 | 0.22 | 1.38E-69 | lung | Srsf6 |
| Orai12 | 2.73E-73 | 0.35162759 | 0.34 | 0.23 | 5.65E-69 | lung | Orai1 |
| Sat12 | 6.39E-71 | 0.32283427 | 0.52 | 0.4 | 1.32E-66 | lung | Sat1 |
| Npdc11 | 3.26E-70 | 0.3564971 | 0.45 | 0.34 | 6.76E-66 | lung | Npdc1 |
| Ctdsp1 | 3.61E-69 | 0.31539202 | 0.35 | 0.24 | 7.47E-65 | lung | Ctdsp1 |
| Hnrnpf2 | 2.75E-68 | 0.28395452 | 0.72 | 0.64 | 5.70E-64 | lung | Hnrnpf |
| Atp6v0d1 | 2.81E-68 | 0.34240813 | 0.33 | 0.22 | 5.81E-64 | lung | Atp6v0d1 |
| Ltbr1 | 3.04E-68 | 0.31597162 | 0.29 | 0.19 | 6.29E-64 | lung | Ltbr |
| Npc21 | 6.32E-68 | 0.42226255 | 0.59 | 0.49 | 1.31E-63 | lung | Npc2 |
| Map1lc3b1 | 2.95E-67 | 0.29204899 | 0.7 | 0.62 | 6.09E-63 | lung | Map1lc3b |
| Ociad1 | 3.01E-66 | 0.3395693 | 0.45 | 0.34 | 6.23E-62 | lung | Ociad1 |
| Sema6a3 | 1.82E-65 | 0.26743367 | 0.38 | 0.27 | 3.78E-61 | lung | Sema6a |
| Nhp2l1 | 2.40E-65 | 0.3383755 | 0.5 | 0.38 | 4.98E-61 | lung | Nhp2l1 |
| Clk11 | 3.32E-65 | 0.35122428 | 0.41 | 0.3 | 6.87E-61 | lung | Clk1 |
| Csnk1a11 | 3.40E-65 | 0.31493029 | 0.54 | 0.43 | 7.04E-61 | lung | Csnk1a1 |
| Tpst21 | 4.70E-65 | 0.33491524 | 0.56 | 0.46 | 9.73E-61 | lung | Tpst2 |
| Yipf4 | 5.05E-65 | 0.35489378 | 0.44 | 0.33 | 1.05E-60 | lung | Yipf4 |
| Bcam1 | 1.00E-64 | 0.39253819 | 0.63 | 0.56 | 2.08E-60 | lung | Bcam |
| Fuca11 | 1.10E-64 | 0.3280645 | 0.32 | 0.21 | 2.28E-60 | lung | Fuca1 |
| Mrps18a | 2.13E-64 | 0.35711491 | 0.32 | 0.22 | 4.40E-60 | lung | Mrps18a |

| | | | | | | | |
|---|---|---|---|---|---|---|---|
| Tspan17 | 3.69E-64 | 0.31953044 | 0.26 | 0.16 | 7.64E-60 | lung | Tspan17 |
| Pdia31 | 4.60E-64 | 0.27420175 | 0.7 | 0.6 | 9.51E-60 | lung | Pdia3 |
| Rtn31 | 5.74E-64 | 0.3364187 | 0.49 | 0.38 | 1.19E-59 | lung | Rtn3 |
| Sbds | 7.74E-64 | 0.35353097 | 0.46 | 0.35 | 1.60E-59 | lung | Sbds |
| Emc3 | 8.46E-64 | 0.33556804 | 0.39 | 0.28 | 1.75E-59 | lung | Emc3 |
| Cnbp1 | 2.57E-63 | 0.26333535 | 0.75 | 0.67 | 5.32E-59 | lung | Cnbp |
| Tagln22 | 2.86E-63 | 0.25601259 | 0.8 | 0.73 | 5.92E-59 | lung | Tagln2 |
| Hibadh | 2.96E-63 | 0.34144533 | 0.25 | 0.16 | 6.12E-59 | lung | Hibadh |
| Nsmce1 | 1.07E-62 | 0.31752612 | 0.29 | 0.19 | 2.21E-58 | lung | Nsmce1 |
| Wars3 | 5.33E-62 | 0.2834581 | 0.29 | 0.19 | 1.10E-57 | lung | Wars |
| Vapa | 9.78E-62 | 0.35496906 | 0.5 | 0.39 | 2.02E-57 | lung | Vapa |
| Rab5a | 1.56E-61 | 0.34532748 | 0.37 | 0.26 | 3.24E-57 | lung | Rab5a |
| Rap1b | 6.59E-60 | 0.32100115 | 0.6 | 0.52 | 1.36E-55 | lung | Rap1b |
| 1110038F14Rik1 | 8.32E-60 | 0.3401725 | 0.36 | 0.26 | 1.72E-55 | lung | 1110038F14Rik |
| Sox172 | 1.54E-59 | 0.25613919 | 0.51 | 0.38 | 3.19E-55 | lung | Sox17 |
| Ddx39b | 2.61E-58 | 0.32263656 | 0.39 | 0.28 | 5.39E-54 | lung | Ddx39b |
| Tspan31 | 8.04E-58 | 0.29779035 | 0.31 | 0.21 | 1.66E-53 | lung | Tspan31 |
| Tmem2041 | 1.23E-57 | 0.32854449 | 0.5 | 0.41 | 2.55E-53 | lung | Tmem204 |
| Mocs2 | 1.36E-57 | 0.31400092 | 0.3 | 0.2 | 2.81E-53 | lung | Mocs2 |
| Tmed71 | 3.38E-57 | 0.30398215 | 0.31 | 0.22 | 7.00E-53 | lung | Tmed7 |
| Emc4 | 3.40E-57 | 0.31652945 | 0.33 | 0.24 | 7.03E-53 | lung | Emc4 |
| Ubl31 | 8.13E-57 | 0.31765013 | 0.42 | 0.32 | 1.68E-52 | lung | Ubl3 |
| Eif4h1 | 3.04E-56 | 0.33466143 | 0.49 | 0.4 | 6.29E-52 | lung | Eif4h |
| Crlf2 | 3.26E-56 | 0.28165217 | 0.27 | 0.18 | 6.75E-52 | lung | Crlf2 |
| Rhog | 5.65E-56 | 0.2945495 | 0.28 | 0.19 | 1.17E-51 | lung | Rhog |
| Tmx1 | 6.38E-56 | 0.30565082 | 0.28 | 0.19 | 1.32E-51 | lung | Tmx1 |
| Serinc1 | 2.40E-55 | 0.32122102 | 0.48 | 0.39 | 4.97E-51 | lung | Serinc1 |
| Tex2641 | 3.11E-55 | 0.29313973 | 0.29 | 0.2 | 6.43E-51 | lung | Tex264 |
| Cbfa2t31 | 9.61E-55 | 0.25057294 | 0.36 | 0.26 | 1.99E-50 | lung | Cbfa2t3 |
| Akr1a12 | 1.83E-54 | 0.27312897 | 0.7 | 0.61 | 3.79E-50 | lung | Akr1a1 |
| Ube2s | 2.98E-54 | 0.31044937 | 0.45 | 0.35 | 6.17E-50 | lung | Ube2s |
| Git21 | 8.51E-54 | 0.27103318 | 0.36 | 0.26 | 1.76E-49 | lung | Git2 |
| Bag5 | 5.23E-53 | 0.2947638 | 0.33 | 0.24 | 1.08E-48 | lung | Bag5 |
| Polr2c | 8.70E-53 | 0.3140076 | 0.39 | 0.29 | 1.80E-48 | lung | Polr2c |
| Cnih11 | 1.49E-52 | 0.31578757 | 0.34 | 0.24 | 3.08E-48 | lung | Cnih1 |
| Alyref | 1.54E-52 | 0.27871431 | 0.25 | 0.17 | 3.20E-48 | lung | Alyref |
| Klf31 | 1.96E-52 | 0.30143315 | 0.45 | 0.35 | 4.05E-48 | lung | Klf3 |
| Vdac2 | 5.62E-52 | 0.30727382 | 0.5 | 0.41 | 1.16E-47 | lung | Vdac2 |
| Ppp4c | 6.29E-52 | 0.27297126 | 0.36 | 0.26 | 1.30E-47 | lung | Ppp4c |
| Psmb101 | 1.36E-51 | 0.28083251 | 0.53 | 0.43 | 2.82E-47 | lung | Psmb10 |
| Grina1 | 1.64E-51 | 0.31563296 | 0.29 | 0.2 | 3.39E-47 | lung | Grina |

| | | | | | | | |
|---|---|---|---|---|---|---|---|
| Mrpl4 | 2.91E-51 | 0.27623379 | 0.34 | 0.25 | 6.01E-47 | lung | Mrpl4 |
| Ggh1 | 3.69E-51 | 0.29003133 | 0.28 | 0.19 | 7.64E-47 | lung | Ggh |
| Cops4 | 8.45E-51 | 0.32114608 | 0.26 | 0.17 | 1.75E-46 | lung | Cops4 |
| Arpc21 | 1.71E-50 | 0.2541686 | 0.7 | 0.64 | 3.54E-46 | lung | Arpc2 |
| Rab7 | 1.78E-50 | 0.31041728 | 0.54 | 0.45 | 3.68E-46 | lung | Rab7 |
| Smarca21 | 1.23E-49 | 0.33083724 | 0.35 | 0.26 | 2.55E-45 | lung | Smarca2 |
| Scamp3 | 1.38E-49 | 0.26532625 | 0.26 | 0.17 | 2.85E-45 | lung | Scamp3 |
| Tmem165 | 1.62E-49 | 0.29868422 | 0.36 | 0.27 | 3.36E-45 | lung | Tmem165 |
| Tmem331 | 2.12E-49 | 0.25318406 | 0.3 | 0.21 | 4.39E-45 | lung | Tmem33 |
| Glrx5 | 3.04E-49 | 0.31872697 | 0.3 | 0.22 | 6.28E-45 | lung | Glrx5 |
| Mcl12 | 1.04E-48 | 0.29515715 | 0.37 | 0.28 | 2.15E-44 | lung | Mcl1 |
| Actn1 | 3.42E-48 | 0.2538443 | 0.25 | 0.17 | 7.08E-44 | lung | Actn1 |
| Ier51 | 4.22E-48 | 0.40340144 | 0.56 | 0.49 | 8.73E-44 | lung | Ier5 |
| Rhoj1 | 6.37E-48 | 0.27917702 | 0.44 | 0.34 | 1.32E-43 | lung | Rhoj |
| Prdx6 | 1.96E-47 | 0.51603982 | 0.33 | 0.25 | 4.05E-43 | lung | Prdx6 |
| Dazap2 | 3.00E-47 | 0.29817541 | 0.5 | 0.42 | 6.21E-43 | lung | Dazap2 |
| Adrm1 | 7.58E-47 | 0.2525279 | 0.27 | 0.19 | 1.57E-42 | lung | Adrm1 |
| Dnajb92 | 4.46E-46 | 0.31122927 | 0.26 | 0.18 | 9.22E-42 | lung | Dnajb9 |
| Traf7 | 1.00E-45 | 0.28836316 | 0.25 | 0.17 | 2.07E-41 | lung | Traf7 |
| Tmem2561 | 1.85E-45 | 0.26151943 | 0.59 | 0.51 | 3.83E-41 | lung | Tmem256 |
| Ube2a | 3.99E-45 | 0.26728235 | 0.29 | 0.2 | 8.26E-41 | lung | Ube2a |
| Ptov1 | 7.97E-45 | 0.270454 | 0.32 | 0.23 | 1.65E-40 | lung | Ptov1 |
| Nol7 | 1.14E-44 | 0.28475137 | 0.42 | 0.32 | 2.37E-40 | lung | Nol7 |
| Puf601 | 6.21E-44 | 0.26075765 | 0.39 | 0.29 | 1.29E-39 | lung | Puf60 |
| Arf61 | 8.08E-44 | 0.25223818 | 0.37 | 0.28 | 1.67E-39 | lung | Arf6 |
| Sdf2 | 1.22E-43 | 0.27387785 | 0.29 | 0.21 | 2.53E-39 | lung | Sdf2 |
| Tmem184b1 | 1.42E-43 | 0.29382293 | 0.3 | 0.21 | 2.94E-39 | lung | Tmem184b |
| Mbnl21 | 1.61E-43 | 0.27448103 | 0.47 | 0.38 | 3.32E-39 | lung | Mbnl2 |
| Anp32a1 | 1.62E-43 | 0.25925936 | 0.71 | 0.64 | 3.36E-39 | lung | Anp32a |
| Rbpms1 | 3.56E-43 | 0.25705483 | 0.44 | 0.34 | 7.36E-39 | lung | Rbpms |
| Leprot | 3.69E-42 | 0.29021302 | 0.43 | 0.35 | 7.64E-38 | lung | Leprot |
| Iscu | 9.36E-42 | 0.29740929 | 0.45 | 0.36 | 1.94E-37 | lung | Iscu |
| Thra1 | 9.96E-42 | 0.25990497 | 0.33 | 0.24 | 2.06E-37 | lung | Thra |
| Tek2 | 1.12E-41 | 0.31269262 | 0.55 | 0.48 | 2.32E-37 | lung | Tek |
| Vdac3 | 2.48E-41 | 0.30570169 | 0.48 | 0.4 | 5.13E-37 | lung | Vdac3 |
| Cops8 | 2.09E-40 | 0.25989806 | 0.29 | 0.21 | 4.33E-36 | lung | Cops8 |
| Klf13 | 3.32E-40 | 0.26466886 | 0.54 | 0.47 | 6.86E-36 | lung | Klf13 |
| Pcmt1 | 3.63E-40 | 0.26726536 | 0.33 | 0.25 | 7.51E-36 | lung | Pcmt1 |
| Chmp4b | 4.55E-40 | 0.26965069 | 0.41 | 0.33 | 9.41E-36 | lung | Chmp4b |
| Naa38 | 5.96E-40 | 0.25599897 | 0.37 | 0.28 | 1.23E-35 | lung | Naa38 |
| Bri3 | 5.88E-39 | 0.26746834 | 0.56 | 0.49 | 1.22E-34 | lung | Bri3 |

| | | | | | | | |
|---|---|---|---|---|---|---|---|
| Sf3b5 | 4.76E-38 | 0.25533082 | 0.36 | 0.28 | 9.85E-34 | lung | Sf3b5 |
| Nras | 5.47E-38 | 0.25330827 | 0.35 | 0.27 | 1.13E-33 | lung | Nras |
| Sec11a1 | 1.47E-37 | 0.27488073 | 0.37 | 0.29 | 3.04E-33 | lung | Sec11a |
| Trappc4 | 1.63E-37 | 0.28896161 | 0.34 | 0.26 | 3.37E-33 | lung | Trappc4 |
| Ndfip2 | 1.29E-36 | 0.27126736 | 0.34 | 0.26 | 2.66E-32 | lung | Ndfip2 |
| Ddost1 | 1.55E-35 | 0.25222454 | 0.36 | 0.28 | 3.21E-31 | lung | Ddost |
| Abracl | 2.04E-33 | 0.28134659 | 0.27 | 0.2 | 4.22E-29 | lung | Abracl |
| Paip2 | 1.51E-30 | 0.25132402 | 0.52 | 0.45 | 3.12E-26 | lung | Paip2 |
| Vwf1 | 1.89E-19 | 0.29150209 | 0.26 | 0.21 | 3.90E-15 | lung | Vwf |
| Timp41 | 0 | 2.88094456 | 0.78 | 0.1 | 0 | muscle_edl | Timp4 |
| C1qtnf91 | 0 | 2.34561718 | 0.83 | 0.18 | 0 | muscle_edl | C1qtnf9 |
| Tcf152 | 0 | 2.28032042 | 0.88 | 0.28 | 0 | muscle_edl | Tcf15 |
| Aqp14 | 0 | 2.17791892 | 0.94 | 0.48 | 0 | muscle_edl | Aqp1 |
| Aqp71 | 0 | 2.11048955 | 0.68 | 0.12 | 0 | muscle_edl | Aqp7 |
| S100a62 | 0 | 1.89447678 | 0.81 | 0.3 | 0 | muscle_edl | S100a6 |
| Fabp42 | 0 | 1.85040824 | 1 | 0.6 | 0 | muscle_edl | Fabp4 |
| Lpl1 | 0 | 1.73600732 | 0.61 | 0.16 | 0 | muscle_edl | Lpl |
| Cav14 | 0 | 1.62833685 | 0.96 | 0.65 | 0 | muscle_edl | Cav1 |
| Meox21 | 0 | 1.50389775 | 0.42 | 0.08 | 0 | muscle_edl | Meox2 |
| Tmsb4x1 | 0 | 1.30305643 | 1 | 0.99 | 0 | muscle_edl | Tmsb4x |
| Gng111 | 0 | 1.27296946 | 0.97 | 0.85 | 0 | muscle_edl | Gng11 |
| Ptrf3 | 7.90E-308 | 1.3879999 | 0.85 | 0.48 | 1.64E-303 | muscle_edl | Ptrf |
| Gpihbp13 | 1.04E-307 | 1.30550372 | 0.93 | 0.56 | 2.15E-303 | muscle_edl | Gpihbp1 |
| Cd362 | 9.85E-307 | 1.2285336 | 0.92 | 0.52 | 2.04E-302 | muscle_edl | Cd36 |
| Sparcl15 | 1.89E-304 | 1.32761711 | 0.89 | 0.49 | 3.91E-300 | muscle_edl | Sparcl1 |
| Tspan133 | 7.39E-297 | 1.2585477 | 0.92 | 0.63 | 1.53E-292 | muscle_edl | Tspan13 |
| Rgs51 | 1.99E-295 | 1.50219908 | 0.37 | 0.08 | 4.12E-291 | muscle_edl | Rgs5 |
| Gm98432 | 3.11E-290 | 1.34113169 | 0.87 | 0.61 | 6.43E-286 | muscle_edl | Gm9843 |
| Cav22 | 2.93E-283 | 1.30730498 | 0.89 | 0.67 | 6.06E-279 | muscle_edl | Cav2 |
| Cst33 | 3.67E-277 | 0.99265215 | 0.97 | 0.88 | 7.60E-273 | muscle_edl | Cst3 |
| Fam101b2 | 4.10E-270 | 1.43604301 | 0.65 | 0.26 | 8.48E-266 | muscle_edl | Fam101b |
| Ptprb3 | 9.39E-267 | 0.84560497 | 0.97 | 0.89 | 1.94E-262 | muscle_edl | Ptprb |
| Mllt42 | 1.61E-259 | 1.56476028 | 0.62 | 0.26 | 3.34E-255 | muscle_edl | Mllt4 |
| Sult1a11 | 4.68E-258 | 1.53036073 | 0.5 | 0.17 | 9.68E-254 | muscle_edl | Sult1a1 |
| Uba522 | 1.53E-257 | 1.14320775 | 0.9 | 0.72 | 3.17E-253 | muscle_edl | Uba52 |
| Rpl36-ps32 | 1.66E-255 | 1.46803666 | 0.7 | 0.36 | 3.44E-251 | muscle_edl | Rpl36-ps3 |
| Rpl13-ps32 | 3.96E-255 | 1.47211576 | 0.69 | 0.36 | 8.18E-251 | muscle_edl | Rpl13-ps3 |
| Rgcc3 | 1.20E-253 | 1.25687784 | 0.83 | 0.43 | 2.49E-249 | muscle_edl | Rgcc |
| Rps27rt2 | 8.76E-243 | 0.92948269 | 0.96 | 0.84 | 1.81E-238 | muscle_edl | Rps27rt |
| Lims22 | 1.61E-234 | 1.27935651 | 0.8 | 0.49 | 3.34E-230 | muscle_edl | Lims2 |
| Ctnnbip11 | 1.28E-230 | 1.34390152 | 0.62 | 0.28 | 2.65E-226 | muscle_edl | Ctnnbip1 |

| | | | | | | | |
|---|---|---|---|---|---|---|---|
| Ybx13 | 2.73E-228 | 0.95865234 | 0.95 | 0.84 | 5.65E-224 | muscle_edl | Ybx1 |
| Rpl9-ps62 | 4.32E-218 | 1.31528899 | 0.73 | 0.44 | 8.95E-214 | muscle_edl | Rpl9-ps6 |
| Gm118082 | 3.19E-215 | 1.20117099 | 0.82 | 0.58 | 6.60E-211 | muscle_edl | Gm11808 |
| Rpl27-ps32 | 1.09E-210 | 1.38163794 | 0.63 | 0.32 | 2.26E-206 | muscle_edl | Rpl27-ps3 |
| Rpl23a-ps32 | 1.92E-206 | 1.07696048 | 0.85 | 0.65 | 3.98E-202 | muscle_edl | Rpl23a-ps3 |
| Cxcl124 | 4.20E-202 | 1.03691348 | 0.85 | 0.49 | 8.70E-198 | muscle_edl | Cxcl12 |
| Cdkn1c2 | 1.06E-197 | 1.38381722 | 0.55 | 0.23 | 2.20E-193 | muscle_edl | Cdkn1c |
| AW1120104 | 3.16E-197 | 1.22212911 | 0.81 | 0.51 | 6.53E-193 | muscle_edl | AW112010 |
| Sepw11 | 2.39E-191 | 0.97262836 | 0.89 | 0.71 | 4.94E-187 | muscle_edl | Sepw1 |
| Slc28a21 | 2.86E-178 | 1.10305595 | 0.32 | 0.09 | 5.91E-174 | muscle_edl | Slc28a2 |
| Crip12 | 1.56E-169 | 0.94113594 | 0.92 | 0.7 | 3.24E-165 | muscle_edl | Crip1 |
| Gm87302 | 1.76E-168 | 1.12500247 | 0.68 | 0.42 | 3.63E-164 | muscle_edl | Gm8730 |
| Gm61331 | 1.51E-166 | 1.19918407 | 0.55 | 0.27 | 3.13E-162 | muscle_edl | Gm6133 |
| Ehd21 | 2.51E-163 | 1.15675696 | 0.71 | 0.46 | 5.20E-159 | muscle_edl | Ehd2 |
| Wdr892 | 4.63E-161 | 0.87784294 | 0.87 | 0.72 | 9.58E-157 | muscle_edl | Wdr89 |
| Cdh132 | 3.92E-156 | 1.07422436 | 0.42 | 0.16 | 8.10E-152 | muscle_edl | Cdh13 |
| Gm100732 | 2.16E-154 | 1.12289824 | 0.67 | 0.41 | 4.47E-150 | muscle_edl | Gm10073 |
| Ly6c13 | 7.29E-154 | 0.66999678 | 0.99 | 0.85 | 1.51E-149 | muscle_edl | Ly6c1 |
| Gm94932 | 6.02E-152 | 1.02838869 | 0.74 | 0.51 | 1.25E-147 | muscle_edl | Gm9493 |
| Gm100362 | 1.08E-149 | 1.16527294 | 0.59 | 0.33 | 2.23E-145 | muscle_edl | Gm10036 |
| Hrct11 | 1.94E-147 | 1.12019903 | 0.3 | 0.09 | 4.01E-143 | muscle_edl | Hrct1 |
| Fabp51 | 3.57E-147 | 0.97647648 | 0.75 | 0.44 | 7.39E-143 | muscle_edl | Fabp5 |
| Lmo1 | 1.69E-145 | 0.96552977 | 0.3 | 0.09 | 3.50E-141 | muscle_edl | Lmo1 |
| Prr132 | 1.42E-142 | 1.18744154 | 0.6 | 0.35 | 2.94E-138 | muscle_edl | Prr13 |
| Tubb2a2 | 1.66E-142 | 1.17360401 | 0.59 | 0.32 | 3.43E-138 | muscle_edl | Tubb2a |
| Gm120021 | 5.52E-141 | 1.10204121 | 0.36 | 0.13 | 1.14E-136 | muscle_edl | Gm12002 |
| Ppp1r21 | 4.60E-138 | 0.96747741 | 0.79 | 0.63 | 9.51E-134 | muscle_edl | Ppp1r2 |
| Ablim31 | 7.25E-135 | 0.97362483 | 0.31 | 0.1 | 1.50E-130 | muscle_edl | Ablim3 |
| Cd2003 | 4.93E-134 | 0.89474412 | 0.85 | 0.66 | 1.02E-129 | muscle_edl | Cd200 |
| Ercc1 | 2.10E-133 | 1.10349768 | 0.37 | 0.15 | 4.34E-129 | muscle_edl | Ercc1 |
| Rps26-ps12 | 2.36E-130 | 1.08811005 | 0.5 | 0.25 | 4.89E-126 | muscle_edl | Rps26-ps1 |
| Ehd12 | 3.64E-124 | 1.03699921 | 0.53 | 0.27 | 7.53E-120 | muscle_edl | Ehd1 |
| Gm87971 | 7.38E-124 | 1.06784208 | 0.45 | 0.21 | 1.53E-119 | muscle_edl | Gm8797 |
| Cd300lg4 | 1.23E-117 | 0.88895395 | 0.69 | 0.42 | 2.55E-113 | muscle_edl | Cd300lg |
| Sdpr3 | 2.19E-112 | 0.66471148 | 0.9 | 0.71 | 4.54E-108 | muscle_edl | Sdpr |
| Rps291 | 2.11E-107 | 0.40684955 | 0.99 | 0.98 | 4.37E-103 | muscle_edl | Rps29 |
| Rpl6l2 | 4.20E-107 | 0.89048932 | 0.68 | 0.47 | 8.69E-103 | muscle_edl | Rpl6l |
| Rpl411 | 6.35E-106 | 0.42232086 | 0.99 | 0.97 | 1.31E-101 | muscle_edl | Rpl41 |
| Atp5j1 | 9.15E-106 | 0.68513413 | 0.86 | 0.78 | 1.89E-101 | muscle_edl | Atp5j |
| Rps12-ps33 | 2.99E-101 | 0.82414422 | 0.75 | 0.57 | 6.18E-97 | muscle_edl | Rps12-ps3 |
| Sep-42 | 2.83E-97 | 0.92616522 | 0.56 | 0.33 | 5.85E-93 | muscle_edl | 4-Sep |

| | | | | | | | |
|---|---|---|---|---|---|---|---|
| Rps18-ps31 | 5.68E-97 | 0.95928781 | 0.37 | 0.17 | 1.17E-92 | muscle_edl | Rps18-ps3 |
| Sparc1 | 8.10E-97 | 0.56114915 | 0.94 | 0.86 | 1.68E-92 | muscle_edl | Sparc |
| Myadm3 | 1.06E-96 | 1.01639152 | 0.46 | 0.25 | 2.20E-92 | muscle_edl | Myadm |
| Hspb14 | 7.56E-96 | 0.47723255 | 0.91 | 0.72 | 1.56E-91 | muscle_edl | Hspb1 |
| Mustn1 | 9.82E-89 | 0.83538634 | 0.26 | 0.1 | 2.03E-84 | muscle_edl | Mustn1 |
| Ptp4a32 | 1.01E-86 | 0.83814051 | 0.6 | 0.4 | 2.10E-82 | muscle_edl | Ptp4a3 |
| Jam23 | 7.85E-86 | 0.84067473 | 0.65 | 0.45 | 1.62E-81 | muscle_edl | Jam2 |
| Chek2 | 2.12E-83 | 0.88053085 | 0.32 | 0.15 | 4.40E-79 | muscle_edl | Chek2 |
| Car83 | 4.00E-83 | 0.85342462 | 0.44 | 0.23 | 8.27E-79 | muscle_edl | Car8 |
| Fbln21 | 9.56E-82 | 0.86909537 | 0.41 | 0.2 | 1.98E-77 | muscle_edl | Fbln2 |
| Myl12a3 | 9.65E-81 | 0.4974764 | 0.9 | 0.84 | 2.00E-76 | muscle_edl | Myl12a |
| Smco42 | 3.94E-80 | 0.92900447 | 0.63 | 0.47 | 8.16E-76 | muscle_edl | Smco4 |
| Oat | 7.43E-79 | 0.92155762 | 0.34 | 0.17 | 1.54E-74 | muscle_edl | Oat |
| Cdh52 | 1.76E-78 | 0.66615721 | 0.83 | 0.73 | 3.63E-74 | muscle_edl | Cdh5 |
| Ifitm21 | 3.14E-77 | 0.38811486 | 0.96 | 0.93 | 6.50E-73 | muscle_edl | Ifitm2 |
| Tspan6 | 2.55E-76 | 0.8825659 | 0.29 | 0.13 | 5.27E-72 | muscle_edl | Tspan6 |
| Cd813 | 9.23E-74 | 0.49311928 | 0.92 | 0.82 | 1.91E-69 | muscle_edl | Cd81 |
| Psme2b1 | 2.09E-73 | 0.9151836 | 0.35 | 0.18 | 4.32E-69 | muscle_edl | Psme2b |
| Gm427262 | 1.47E-69 | 0.76992185 | 0.32 | 0.15 | 3.04E-65 | muscle_edl | Gm42726 |
| Pdlim14 | 3.64E-69 | 0.8114831 | 0.5 | 0.31 | 7.54E-65 | muscle_edl | Pdlim1 |
| Ppic | 1.26E-64 | 0.84430875 | 0.52 | 0.36 | 2.62E-60 | muscle_edl | Ppic |
| Gm20002 | 2.23E-64 | 0.77186892 | 0.42 | 0.25 | 4.62E-60 | muscle_edl | Gm2000 |
| Tspan72 | 1.03E-62 | 0.3332753 | 0.75 | 0.51 | 2.13E-58 | muscle_edl | Tspan7 |
| Serinc32 | 3.36E-61 | 0.50201332 | 0.9 | 0.84 | 6.95E-57 | muscle_edl | Serinc3 |
| Hmgb1 | 7.18E-61 | 0.45211179 | 0.87 | 0.83 | 1.49E-56 | muscle_edl | Hmgb1 |
| Rpl352 | 9.32E-61 | 0.4058166 | 0.91 | 0.87 | 1.93E-56 | muscle_edl | Rpl35 |
| Gm98441 | 3.26E-58 | 0.73251349 | 0.27 | 0.13 | 6.75E-54 | muscle_edl | Gm9844 |
| Nme21 | 3.34E-58 | 0.8348535 | 0.4 | 0.25 | 6.92E-54 | muscle_edl | Nme2 |
| Tmsb101 | 7.77E-58 | 0.42792354 | 0.85 | 0.74 | 1.61E-53 | muscle_edl | Tmsb10 |
| Ccdc85a1 | 6.76E-57 | 0.73574861 | 0.3 | 0.15 | 1.40E-52 | muscle_edl | Ccdc85a |
| Calm13 | 7.63E-57 | 0.3808781 | 0.94 | 0.9 | 1.58E-52 | muscle_edl | Calm1 |
| mt-Nd1 | 7.79E-57 | 0.39727946 | 0.97 | 0.95 | 1.61E-52 | muscle_edl | mt-Nd1 |
| Actb | 2.73E-55 | 0.68542114 | 0.96 | 0.95 | 5.64E-51 | muscle_edl | Actb |
| Id12 | 5.69E-55 | 0.67952119 | 0.76 | 0.65 | 1.18E-50 | muscle_edl | Id1 |
| Aplnr2 | 9.36E-54 | 0.66174956 | 0.37 | 0.21 | 1.94E-49 | muscle_edl | Aplnr |
| Gstm12 | 1.20E-52 | 0.70644725 | 0.66 | 0.53 | 2.49E-48 | muscle_edl | Gstm1 |
| S100a114 | 7.92E-52 | 0.5870553 | 0.71 | 0.59 | 1.64E-47 | muscle_edl | S100a11 |
| Fau1 | 1.57E-51 | 0.35275982 | 0.95 | 0.93 | 3.24E-47 | muscle_edl | Fau |
| Cyth11 | 4.21E-51 | 0.73178346 | 0.25 | 0.13 | 8.72E-47 | muscle_edl | Cyth1 |
| Tmem883 | 2.56E-50 | 0.72327305 | 0.58 | 0.44 | 5.29E-46 | muscle_edl | Tmem88 |
| Slc9a3r24 | 7.32E-50 | 0.45503278 | 0.88 | 0.82 | 1.51E-45 | muscle_edl | Slc9a3r2 |

| | | | | | | | |
|---|---|---|---|---|---|---|---|
| 1810011O10Rik3 | 1.03E-49 | 0.60336744 | 0.63 | 0.47 | 2.14E-45 | muscle_edl | 1810011O10Rik |
| Eif12 | 1.72E-49 | 0.34757072 | 0.93 | 0.93 | 3.55E-45 | muscle_edl | Eif1 |
| Myct12 | 2.62E-49 | 0.65340182 | 0.62 | 0.5 | 5.41E-45 | muscle_edl | Myct1 |
| Rpl10-ps32 | 1.48E-47 | 0.71225337 | 0.37 | 0.23 | 3.05E-43 | muscle_edl | Rpl10-ps3 |
| Tsfm | 1.72E-47 | 0.73942237 | 0.26 | 0.13 | 3.55E-43 | muscle_edl | Tsfm |
| Marcks3 | 1.70E-46 | 0.6685771 | 0.52 | 0.37 | 3.51E-42 | muscle_edl | Marcks |
| Fscn11 | 1.91E-45 | 0.62931025 | 0.36 | 0.22 | 3.96E-41 | muscle_edl | Fscn1 |
| Sub1 | 8.69E-45 | 0.69381009 | 0.64 | 0.58 | 1.80E-40 | muscle_edl | Sub1 |
| Jup2 | 2.28E-44 | 0.63471614 | 0.54 | 0.4 | 4.71E-40 | muscle_edl | Jup |
| Tubb4b1 | 6.19E-43 | 0.67205437 | 0.61 | 0.51 | 1.28E-38 | muscle_edl | Tubb4b |
| Nrarp1 | 1.38E-41 | 0.71373376 | 0.27 | 0.15 | 2.86E-37 | muscle_edl | Nrarp |
| S100a103 | 5.40E-41 | 0.46331451 | 0.76 | 0.67 | 1.12E-36 | muscle_edl | S100a10 |
| Rpl361 | 1.56E-40 | 0.31353778 | 0.92 | 0.91 | 3.24E-36 | muscle_edl | Rpl36 |
| Dhrs32 | 4.93E-40 | 0.62413672 | 0.5 | 0.36 | 1.02E-35 | muscle_edl | Dhrs3 |
| Sypl2 | 1.26E-39 | 0.59798173 | 0.62 | 0.53 | 2.61E-35 | muscle_edl | Sypl |
| Dusp15 | 1.49E-39 | 0.47660501 | 0.75 | 0.66 | 3.08E-35 | muscle_edl | Dusp1 |
| Hspa12b2 | 1.74E-39 | 0.69385162 | 0.39 | 0.26 | 3.61E-35 | muscle_edl | Hspa12b |
| Palmd3 | 2.35E-39 | 0.67079678 | 0.42 | 0.29 | 4.86E-35 | muscle_edl | Palmd |
| Thrsp3 | 2.51E-39 | 0.59396216 | 0.43 | 0.28 | 5.19E-35 | muscle_edl | Thrsp |
| Esam3 | 5.64E-39 | 0.42257889 | 0.82 | 0.74 | 1.17E-34 | muscle_edl | Esam |
| Mecom3 | 7.08E-39 | 0.65908649 | 0.35 | 0.22 | 1.47E-34 | muscle_edl | Mecom |
| Anxa32 | 9.02E-39 | 0.557429 | 0.63 | 0.53 | 1.87E-34 | muscle_edl | Anxa3 |
| Ahnak4 | 1.00E-38 | 0.57649728 | 0.58 | 0.45 | 2.08E-34 | muscle_edl | Ahnak |
| Pim32 | 3.47E-38 | 0.63645385 | 0.48 | 0.36 | 7.18E-34 | muscle_edl | Pim3 |
| Tpr1 | 5.31E-38 | 0.81487514 | 0.4 | 0.28 | 1.10E-33 | muscle_edl | Tpr |
| Col4a23 | 8.62E-38 | 0.56740454 | 0.58 | 0.45 | 1.78E-33 | muscle_edl | Col4a2 |
| Anp32a2 | 1.68E-37 | 0.48975997 | 0.71 | 0.65 | 3.48E-33 | muscle_edl | Anp32a |
| Sertad11 | 3.48E-37 | 0.69273917 | 0.34 | 0.22 | 7.21E-33 | muscle_edl | Sertad1 |
| Col4a13 | 2.06E-36 | 0.47527025 | 0.66 | 0.55 | 4.26E-32 | muscle_edl | Col4a1 |
| H3f3a | 3.28E-35 | 0.35586676 | 0.84 | 0.83 | 6.79E-31 | muscle_edl | H3f3a |
| Uqcrb | 8.37E-35 | 0.43211885 | 0.75 | 0.72 | 1.73E-30 | muscle_edl | Uqcrb |
| Crip23 | 1.37E-34 | 0.30427898 | 0.9 | 0.86 | 2.84E-30 | muscle_edl | Crip2 |
| Gm81861 | 2.06E-34 | 0.57461173 | 0.3 | 0.18 | 4.27E-30 | muscle_edl | Gm8186 |
| Ubc3 | 4.52E-33 | 0.3869143 | 0.85 | 0.84 | 9.36E-29 | muscle_edl | Ubc |
| Rbm3 | 1.20E-32 | 0.57813465 | 0.52 | 0.42 | 2.48E-28 | muscle_edl | Rbm3 |
| Prnp1 | 1.73E-32 | 0.63665128 | 0.31 | 0.2 | 3.59E-28 | muscle_edl | Prnp |
| Rftn13 | 1.98E-32 | 0.61948871 | 0.31 | 0.19 | 4.09E-28 | muscle_edl | Rftn1 |
| Bloc1s1 | 4.49E-31 | 0.64031075 | 0.43 | 0.32 | 9.29E-27 | muscle_edl | Bloc1s1 |
| Shfm1 | 5.37E-31 | 0.42249204 | 0.75 | 0.68 | 1.11E-26 | muscle_edl | Shfm1 |
| Dynll12 | 5.52E-31 | 0.34808222 | 0.86 | 0.85 | 1.14E-26 | muscle_edl | Dynll1 |
| Plcb41 | 1.53E-30 | 0.62821169 | 0.34 | 0.23 | 3.17E-26 | muscle_edl | Plcb4 |

| | | | | | | | |
|---|---|---|---|---|---|---|---|
| Id32 | 2.76E-30 | 0.39112331 | 0.9 | 0.85 | 5.72E-26 | muscle_edl | Id3 |
| Ifitm33 | 3.39E-30 | 0.30387065 | 0.97 | 0.96 | 7.01E-26 | muscle_edl | Ifitm3 |
| Rpl271 | 4.86E-30 | 0.30637048 | 0.85 | 0.82 | 1.01E-25 | muscle_edl | Rpl27 |
| Cfl1 | 7.49E-30 | 0.32733235 | 0.85 | 0.83 | 1.55E-25 | muscle_edl | Cfl1 |
| Grcc10 | 1.96E-29 | 0.62308709 | 0.5 | 0.4 | 4.06E-25 | muscle_edl | Grcc10 |
| Pdzd21 | 3.66E-29 | 0.58371861 | 0.3 | 0.19 | 7.57E-25 | muscle_edl | Pdzd2 |
| Unc45b1 | 4.80E-29 | 0.55886997 | 0.43 | 0.32 | 9.93E-25 | muscle_edl | Unc45b |
| Lmcd12 | 8.63E-29 | 0.6041966 | 0.36 | 0.25 | 1.79E-24 | muscle_edl | Lmcd1 |
| Dusp31 | 1.01E-28 | 0.56474405 | 0.58 | 0.51 | 2.10E-24 | muscle_edl | Dusp3 |
| Ctsl1 | 7.79E-28 | 0.37141245 | 0.63 | 0.56 | 1.61E-23 | muscle_edl | Ctsl |
| Klf64 | 1.25E-27 | 0.58949201 | 0.53 | 0.44 | 2.60E-23 | muscle_edl | Klf6 |
| Tprgl2 | 3.32E-27 | 0.50184766 | 0.61 | 0.54 | 6.88E-23 | muscle_edl | Tprgl |
| Tpm42 | 9.66E-27 | 0.41954198 | 0.71 | 0.68 | 2.00E-22 | muscle_edl | Tpm4 |
| Cyb5r33 | 1.03E-26 | 0.48668988 | 0.56 | 0.48 | 2.13E-22 | muscle_edl | Cyb5r3 |
| Myl62 | 3.83E-26 | 0.26742641 | 0.92 | 0.9 | 7.92E-22 | muscle_edl | Myl6 |
| Pls32 | 5.50E-26 | 0.56779385 | 0.46 | 0.36 | 1.14E-21 | muscle_edl | Pls3 |
| Pabpc12 | 5.79E-26 | 0.59183677 | 0.55 | 0.49 | 1.20E-21 | muscle_edl | Pabpc1 |
| Klf23 | 2.52E-25 | 0.39048816 | 0.91 | 0.88 | 5.20E-21 | muscle_edl | Klf2 |
| Gm100763 | 5.62E-25 | 0.44252779 | 0.56 | 0.48 | 1.16E-20 | muscle_edl | Gm10076 |
| S100a13 | 8.27E-25 | 0.37890552 | 0.75 | 0.73 | 1.71E-20 | muscle_edl | S100a13 |
| Rps251 | 1.14E-24 | 0.25833631 | 0.9 | 0.89 | 2.37E-20 | muscle_edl | Rps25 |
| Efna13 | 1.59E-24 | 0.54036763 | 0.46 | 0.37 | 3.30E-20 | muscle_edl | Efna1 |
| Ier24 | 2.12E-24 | 0.37065094 | 0.65 | 0.57 | 4.39E-20 | muscle_edl | Ier2 |
| Egr13 | 7.85E-24 | 0.42531521 | 0.45 | 0.34 | 1.62E-19 | muscle_edl | Egr1 |
| Mgll3 | 5.71E-23 | 0.28920274 | 0.69 | 0.58 | 1.18E-18 | muscle_edl | Mgll |
| S100a162 | 2.00E-22 | 0.3457656 | 0.74 | 0.7 | 4.14E-18 | muscle_edl | S100a16 |
| Lbr | 2.87E-22 | 0.55574578 | 0.29 | 0.2 | 5.93E-18 | muscle_edl | Lbr |
| Cdc42ep32 | 5.76E-22 | 0.52028182 | 0.31 | 0.21 | 1.19E-17 | muscle_edl | Cdc42ep3 |
| Atox13 | 1.06E-21 | 0.28673272 | 0.82 | 0.79 | 2.20E-17 | muscle_edl | Atox1 |
| Adgrg11 | 2.45E-21 | 0.56305366 | 0.34 | 0.26 | 5.06E-17 | muscle_edl | Adgrg1 |
| Tshz23 | 5.42E-21 | 0.41954047 | 0.49 | 0.39 | 1.12E-16 | muscle_edl | Tshz2 |
| Nck12 | 8.84E-21 | 0.58082204 | 0.46 | 0.4 | 1.83E-16 | muscle_edl | Nck1 |
| Dok4 | 9.61E-21 | 0.5586056 | 0.27 | 0.19 | 1.99E-16 | muscle_edl | Dok4 |
| Prkcdbp2 | 3.56E-20 | 0.27816959 | 0.63 | 0.51 | 7.37E-16 | muscle_edl | Prkcdbp |
| Ctnnb13 | 3.88E-20 | 0.45107642 | 0.59 | 0.53 | 8.02E-16 | muscle_edl | Ctnnb1 |
| Tppp33 | 4.12E-20 | 0.33880678 | 0.28 | 0.19 | 8.52E-16 | muscle_edl | Tppp3 |
| Rassf32 | 8.89E-20 | 0.53152716 | 0.33 | 0.25 | 1.84E-15 | muscle_edl | Rassf3 |
| Lrrc8a2 | 2.15E-19 | 0.54320488 | 0.37 | 0.29 | 4.45E-15 | muscle_edl | Lrrc8a |
| Cdkn1a3 | 2.29E-19 | 0.46817707 | 0.47 | 0.37 | 4.73E-15 | muscle_edl | Cdkn1a |
| Bcas3 | 5.45E-19 | 0.47376366 | 0.26 | 0.18 | 1.13E-14 | muscle_edl | Bcas3 |
| Bcam2 | 1.79E-18 | 0.41109037 | 0.62 | 0.56 | 3.71E-14 | muscle_edl | Bcam |

| | | | | | | | |
|---|---|---|---|---|---|---|---|
| Hnrnpa31 | 5.10E-18 | 0.43274183 | 0.54 | 0.49 | 1.05E-13 | muscle_edl | Hnrnpa3 |
| Ptms1 | 3.61E-17 | 0.39892565 | 0.62 | 0.59 | 7.47E-13 | muscle_edl | Ptms |
| Tsnax1 | 8.07E-17 | 0.51681536 | 0.26 | 0.18 | 1.67E-12 | muscle_edl | Tsnax |
| Scarb13 | 2.92E-16 | 0.4368894 | 0.47 | 0.4 | 6.04E-12 | muscle_edl | Scarb1 |
| Tmem1232 | 4.18E-16 | 0.46623338 | 0.36 | 0.29 | 8.66E-12 | muscle_edl | Tmem123 |
| Kitl1 | 8.22E-16 | 0.39115447 | 0.46 | 0.37 | 1.70E-11 | muscle_edl | Kitl |
| Zfp364 | 8.60E-16 | 0.44141964 | 0.49 | 0.42 | 1.78E-11 | muscle_edl | Zfp36 |
| Pxdc11 | 9.19E-16 | 0.40304335 | 0.25 | 0.17 | 1.90E-11 | muscle_edl | Pxdc1 |
| Cox6a1 | 1.80E-15 | 0.25824526 | 0.78 | 0.77 | 3.72E-11 | muscle_edl | Cox6a1 |
| Ywhaq1 | 2.07E-15 | 0.31828098 | 0.66 | 0.64 | 4.27E-11 | muscle_edl | Ywhaq |
| Wasf21 | 2.53E-15 | 0.47475501 | 0.48 | 0.43 | 5.23E-11 | muscle_edl | Wasf2 |
| F11r | 2.77E-15 | 0.38771442 | 0.58 | 0.54 | 5.72E-11 | muscle_edl | F11r |
| Sumo2 | 2.93E-15 | 0.25252914 | 0.74 | 0.73 | 6.06E-11 | muscle_edl | Sumo2 |
| Ndel12 | 4.06E-15 | 0.46984316 | 0.33 | 0.26 | 8.40E-11 | muscle_edl | Ndel1 |
| Tcf43 | 4.88E-15 | 0.25789201 | 0.72 | 0.67 | 1.01E-10 | muscle_edl | Tcf4 |
| Ywhah2 | 9.16E-15 | 0.39129334 | 0.58 | 0.56 | 1.90E-10 | muscle_edl | Ywhah |
| Rasgrp23 | 9.41E-15 | 0.44247661 | 0.27 | 0.2 | 1.95E-10 | muscle_edl | Rasgrp2 |
| 4930523C07Rik1 | 1.33E-14 | 0.46287053 | 0.32 | 0.25 | 2.75E-10 | muscle_edl | 4930523C07Rik |
| Gltscr21 | 1.40E-14 | 0.35415041 | 0.52 | 0.46 | 2.91E-10 | muscle_edl | Gltscr2 |
| Fli11 | 1.44E-14 | 0.4665999 | 0.4 | 0.34 | 2.98E-10 | muscle_edl | Fli1 |
| Adipor13 | 1.61E-14 | 0.43369543 | 0.49 | 0.46 | 3.34E-10 | muscle_edl | Adipor1 |
| Uaca2 | 2.39E-14 | 0.46139906 | 0.3 | 0.23 | 4.95E-10 | muscle_edl | Uaca |
| Capza21 | 3.02E-14 | 0.41246361 | 0.51 | 0.48 | 6.26E-10 | muscle_edl | Capza2 |
| Cltb2 | 3.56E-14 | 0.38921863 | 0.48 | 0.43 | 7.36E-10 | muscle_edl | Cltb |
| Sgk13 | 4.11E-14 | 0.34787325 | 0.59 | 0.54 | 8.50E-10 | muscle_edl | Sgk1 |
| Ets12 | 5.26E-14 | 0.39670999 | 0.51 | 0.45 | 1.09E-09 | muscle_edl | Ets1 |
| Itga62 | 1.09E-13 | 0.45872565 | 0.39 | 0.33 | 2.26E-09 | muscle_edl | Itga6 |
| Aplp23 | 1.53E-13 | 0.27594819 | 0.7 | 0.67 | 3.17E-09 | muscle_edl | Aplp2 |
| Myl12b2 | 6.66E-13 | 0.25712359 | 0.69 | 0.67 | 1.38E-08 | muscle_edl | Myl12b |
| Psap1 | 1.45E-12 | 0.27263295 | 0.75 | 0.73 | 2.99E-08 | muscle_edl | Psap |
| Gimap62 | 1.51E-12 | 0.28351396 | 0.67 | 0.66 | 3.12E-08 | muscle_edl | Gimap6 |
| Dpysl21 | 1.52E-12 | 0.38538647 | 0.42 | 0.36 | 3.14E-08 | muscle_edl | Dpysl2 |
| Btg13 | 2.93E-12 | 0.41434898 | 0.54 | 0.52 | 6.05E-08 | muscle_edl | Btg1 |
| Mif | 6.35E-12 | 0.45332011 | 0.37 | 0.32 | 1.31E-07 | muscle_edl | Mif |
| Klf131 | 1.27E-11 | 0.35577115 | 0.52 | 0.48 | 2.62E-07 | muscle_edl | Klf13 |
| Ushbp11 | 1.33E-11 | 0.4296423 | 0.47 | 0.44 | 2.75E-07 | muscle_edl | Ushbp1 |
| Slc25a51 | 1.42E-11 | 0.31352833 | 0.55 | 0.54 | 2.94E-07 | muscle_edl | Slc25a5 |
| Anxa22 | 3.21E-11 | 0.29866789 | 0.65 | 0.65 | 6.64E-07 | muscle_edl | Anxa2 |
| Ndufa3 | 3.85E-11 | 0.29164928 | 0.63 | 0.61 | 7.96E-07 | muscle_edl | Ndufa3 |
| Csrp12 | 5.98E-11 | 0.39667464 | 0.46 | 0.42 | 1.24E-06 | muscle_edl | Csrp1 |
| Myo101 | 9.00E-11 | 0.52726157 | 0.27 | 0.22 | 1.86E-06 | muscle_edl | Myo10 |

| | | | | | | | |
|---|---|---|---|---|---|---|---|
| Cd931 | 9.39E-11 | 0.32713479 | 0.58 | 0.55 | 1.94E-06 | muscle_edl | Cd93 |
| Chmp2a | 2.90E-10 | 0.3544472 | 0.53 | 0.52 | 6.01E-06 | muscle_edl | Chmp2a |
| Cxx1b3 | 3.40E-10 | 0.33594385 | 0.37 | 0.31 | 7.04E-06 | muscle_edl | Cxx1b |
| Itpkb3 | 4.02E-10 | 0.42230004 | 0.44 | 0.41 | 8.32E-06 | muscle_edl | Itpkb |
| Snrpg | 4.06E-10 | 0.3244692 | 0.59 | 0.58 | 8.39E-06 | muscle_edl | Snrpg |
| Fkbp32 | 1.48E-09 | 0.42110349 | 0.36 | 0.31 | 3.06E-05 | muscle_edl | Fkbp3 |
| Gja11 | 1.90E-09 | 0.51734526 | 0.29 | 0.24 | 3.94E-05 | muscle_edl | Gja1 |
| Xdh1 | 2.10E-09 | 0.45517236 | 0.25 | 0.2 | 4.34E-05 | muscle_edl | Xdh |
| N4bp32 | 2.24E-09 | 0.36407407 | 0.26 | 0.2 | 4.64E-05 | muscle_edl | N4bp3 |
| Spop1 | 2.28E-09 | 0.4275583 | 0.4 | 0.36 | 4.71E-05 | muscle_edl | Spop |
| Atf42 | 2.31E-09 | 0.42578173 | 0.35 | 0.31 | 4.77E-05 | muscle_edl | Atf4 |
| Vasp1 | 2.66E-09 | 0.43305131 | 0.37 | 0.33 | 5.50E-05 | muscle_edl | Vasp |
| Kank31 | 2.77E-09 | 0.34576641 | 0.58 | 0.57 | 5.73E-05 | muscle_edl | Kank3 |
| Eif4a11 | 2.78E-09 | 0.27826321 | 0.6 | 0.59 | 5.76E-05 | muscle_edl | Eif4a1 |
| Higd2a1 | 3.16E-09 | 0.42046553 | 0.45 | 0.43 | 6.55E-05 | muscle_edl | Higd2a |
| Hmgb2 | 3.53E-09 | 0.43262378 | 0.25 | 0.21 | 7.30E-05 | muscle_edl | Hmgb2 |
| Btg24 | 3.84E-09 | 0.26772118 | 0.54 | 0.49 | 7.95E-05 | muscle_edl | Btg2 |
| Tspan122 | 4.02E-09 | 0.34503046 | 0.4 | 0.35 | 8.33E-05 | muscle_edl | Tspan12 |
| Ablim12 | 6.03E-09 | 0.296559 | 0.34 | 0.28 | 0.00012475 | muscle_edl | Ablim1 |
| Eno12 | 1.10E-08 | 0.35802188 | 0.38 | 0.34 | 0.00022856 | muscle_edl | Eno1 |
| S1pr13 | 1.74E-08 | 0.30474765 | 0.5 | 0.47 | 0.00035996 | muscle_edl | S1pr1 |
| Gnb1 | 3.89E-08 | 0.2974378 | 0.62 | 0.62 | 0.00080499 | muscle_edl | Gnb1 |
| Eif6 | 4.14E-08 | 0.33414041 | 0.41 | 0.38 | 0.00085737 | muscle_edl | Eif6 |
| Mndal3 | 6.76E-08 | 0.3428077 | 0.37 | 0.32 | 0.00139785 | muscle_edl | Mndal |
| Ypel33 | 2.50E-07 | 0.35091754 | 0.45 | 0.43 | 0.00516459 | muscle_edl | Ypel3 |
| Mast42 | 3.21E-07 | 0.36926018 | 0.34 | 0.3 | 0.00663671 | muscle_edl | Mast4 |
| Fam43a4 | 4.30E-07 | 0.29844733 | 0.35 | 0.31 | 0.00890191 | muscle_edl | Fam43a |
| Hes13 | 4.79E-07 | 0.40434149 | 0.51 | 0.5 | 0.00990822 | muscle_edl | Hes1 |
| 9430020K01Rik1 | 5.28E-07 | 0.30232033 | 0.25 | 0.21 | 0.01093552 | muscle_edl | 9430020K01Rik |
| Dram21 | 5.42E-07 | 0.38674431 | 0.34 | 0.31 | 0.01121741 | muscle_edl | Dram2 |
| Adgrf52 | 6.91E-07 | 0.29519606 | 0.64 | 0.64 | 0.01430468 | muscle_edl | Adgrf5 |
| Ifngr14 | 6.93E-07 | 0.30886243 | 0.43 | 0.4 | 0.01433409 | muscle_edl | Ifngr1 |
| Psme2 | 7.07E-07 | 0.34519384 | 0.51 | 0.51 | 0.01463948 | muscle_edl | Psme2 |
| Srsf2 | 8.83E-07 | 0.32900899 | 0.49 | 0.49 | 0.01826439 | muscle_edl | Srsf2 |
| Sri2 | 8.98E-07 | 0.28748273 | 0.51 | 0.5 | 0.01859141 | muscle_edl | Sri |
| Atp5k | 1.04E-06 | 0.32132307 | 0.55 | 0.55 | 0.02157452 | muscle_edl | Atp5k |
| Gemin7 | 1.05E-06 | 0.3378718 | 0.35 | 0.32 | 0.02180488 | muscle_edl | Gemin7 |
| Ostf11 | 1.11E-06 | 0.26552212 | 0.57 | 0.57 | 0.02296956 | muscle_edl | Ostf1 |
| Tmod32 | 1.85E-06 | 0.35420831 | 0.36 | 0.33 | 0.03825642 | muscle_edl | Tmod3 |
| Bag51 | 1.97E-06 | 0.34289127 | 0.28 | 0.25 | 0.04073509 | muscle_edl | Bag5 |
| Cd382 | 2.50E-06 | 0.25610511 | 0.36 | 0.32 | 0.05166299 | muscle_edl | Cd38 |

| | | | | | | | |
|---|---|---|---|---|---|---|---|
| Tmem1402 | 2.53E-06 | 0.32339166 | 0.41 | 0.39 | 0.05225278 | muscle_edl | Tmem140 |
| Ikbkb1 | 4.80E-06 | 0.38585063 | 0.26 | 0.22 | 0.09932999 | muscle_edl | Ikbkb |
| Bvht | 5.56E-06 | 0.30645814 | 0.35 | 0.32 | 0.11511884 | muscle_edl | Bvht |
| Hypk | 6.06E-06 | 0.35838015 | 0.32 | 0.29 | 0.12546497 | muscle_edl | Hypk |
| Anxa11 | 7.74E-06 | 0.31839799 | 0.27 | 0.24 | 0.16006495 | muscle_edl | Anxa11 |
| Hspg24 | 8.57E-06 | 0.28056919 | 0.44 | 0.4 | 0.17741749 | muscle_edl | Hspg2 |
| Klf102 | 1.20E-05 | 0.38431499 | 0.27 | 0.24 | 0.24793479 | muscle_edl | Klf10 |
| Pgrmc1 | 1.27E-05 | 0.34594003 | 0.38 | 0.37 | 0.26361411 | muscle_edl | Pgrmc1 |
| Xaf11 | 1.35E-05 | 0.32139133 | 0.33 | 0.3 | 0.27851528 | muscle_edl | Xaf1 |
| Vamp81 | 1.43E-05 | 0.28235096 | 0.43 | 0.41 | 0.29488561 | muscle_edl | Vamp8 |
| Arf5 | 1.70E-05 | 0.2599748 | 0.58 | 0.6 | 0.35130758 | muscle_edl | Arf5 |
| Krtcap22 | 2.02E-05 | 0.26344913 | 0.53 | 0.54 | 0.41757215 | muscle_edl | Krtcap2 |
| Usp50 | 2.48E-05 | 0.33277485 | 0.27 | 0.24 | 0.51240573 | muscle_edl | Usp50 |
| Tax1bp31 | 3.09E-05 | 0.28486852 | 0.38 | 0.36 | 0.63962599 | muscle_edl | Tax1bp3 |
| Srsf3 | 3.14E-05 | 0.29158904 | 0.52 | 0.53 | 0.64942813 | muscle_edl | Srsf3 |
| Lamtor5 | 4.88E-05 | 0.30880606 | 0.41 | 0.41 | 1 | muscle_edl | Lamtor5 |
| Wwtr12 | 6.46E-05 | 0.28603575 | 0.3 | 0.27 | 1 | muscle_edl | Wwtr1 |
| Myo1c1 | 8.10E-05 | 0.27392212 | 0.37 | 0.35 | 1 | muscle_edl | Myo1c |
| Hcfc1r1 | 8.39E-05 | 0.32528557 | 0.34 | 0.33 | 1 | muscle_edl | Hcfc1r1 |
| 1110038F14Rik2 | 9.57E-05 | 0.34130189 | 0.29 | 0.27 | 1 | muscle_edl | 1110038F14Rik |
| Ctbp11 | 0.00010611 | 0.30891947 | 0.36 | 0.35 | 1 | muscle_edl | Ctbp1 |
| Fus1 | 0.00011146 | 0.26121923 | 0.53 | 0.54 | 1 | muscle_edl | Fus |
| Slc50a11 | 0.00012392 | 0.35087828 | 0.4 | 0.39 | 1 | muscle_edl | Slc50a1 |
| Tmem2562 | 0.00015344 | 0.28692342 | 0.51 | 0.52 | 1 | muscle_edl | Tmem256 |
| Mbnl22 | 0.00019626 | 0.27416713 | 0.39 | 0.39 | 1 | muscle_edl | Mbnl2 |
| Wnk13 | 0.0002207 | 0.35218397 | 0.35 | 0.34 | 1 | muscle_edl | Wnk1 |
| Pink11 | 0.00024645 | 0.31170451 | 0.27 | 0.25 | 1 | muscle_edl | Pink1 |
| Ndufc2 | 0.00035588 | 0.25514039 | 0.52 | 0.54 | 1 | muscle_edl | Ndufc2 |
| Timm10b | 0.00038808 | 0.31831613 | 0.33 | 0.32 | 1 | muscle_edl | Timm10b |
| Cracr2b1 | 0.0003909 | 0.26317293 | 0.41 | 0.4 | 1 | muscle_edl | Cracr2b |
| Psmc3 | 0.00040243 | 0.35425619 | 0.38 | 0.38 | 1 | muscle_edl | Psmc3 |
| Fos4 | 0.00049238 | 0.28551551 | 0.57 | 0.57 | 1 | muscle_edl | Fos |
| Vamp31 | 0.00063745 | 0.32295599 | 0.33 | 0.32 | 1 | muscle_edl | Vamp3 |
| Rarg1 | 0.00080894 | 0.30688724 | 0.25 | 0.23 | 1 | muscle_edl | Rarg |
| Ilk1 | 0.00103639 | 0.31111024 | 0.33 | 0.32 | 1 | muscle_edl | Ilk |
| Ppp1cb1 | 0.00108726 | 0.25354773 | 0.35 | 0.34 | 1 | muscle_edl | Ppp1cb |
| U2af1 | 0.00132763 | 0.29252094 | 0.37 | 0.37 | 1 | muscle_edl | U2af1 |
| Dpm3 | 0.0014725 | 0.26582568 | 0.44 | 0.45 | 1 | muscle_edl | Dpm3 |
| Nfia1 | 0.00147423 | 0.26916902 | 0.27 | 0.26 | 1 | muscle_edl | Nfia |
| Pdcl3 | 0.00167372 | 0.27981276 | 0.3 | 0.29 | 1 | muscle_edl | Pdcl3 |
| Kif5b1 | 0.00169446 | 0.25350558 | 0.44 | 0.45 | 1 | muscle_edl | Kif5b |

| | | | | | | | |
|---|---|---|---|---|---|---|---|
| Picalm1 | 0.00189394 | 0.2908471 | 0.28 | 0.26 | 1 | muscle_edl | Picalm |
| Pdcd10 | 0.00274045 | 0.26437153 | 0.33 | 0.33 | 1 | muscle_edl | Pdcd10 |
| Zfand61 | 0.00343428 | 0.28802386 | 0.34 | 0.34 | 1 | muscle_edl | Zfand6 |
| Tspan151 | 0.0037609 | 0.28712403 | 0.26 | 0.24 | 1 | muscle_edl | Tspan15 |
| Snapc5 | 0.00419247 | 0.30967551 | 0.28 | 0.27 | 1 | muscle_edl | Snapc5 |
| Hras1 | 0.00441635 | 0.29900514 | 0.29 | 0.28 | 1 | muscle_edl | Hras |
| Fez23 | 0.00455927 | 0.26937742 | 0.35 | 0.35 | 1 | muscle_edl | Fez2 |
| 2-Mar | 0.00609059 | 0.3137189 | 0.32 | 0.31 | 1 | muscle_edl | 2-Mar |
| Chmp51 | 0.00643633 | 0.25921014 | 0.41 | 0.43 | 1 | muscle_edl | Chmp5 |
| 2-Sep | 0.0082994 | 0.26843433 | 0.31 | 0.31 | 1 | muscle_edl | 2-Sep |
| Timp42 | 0 | 2.60762732 | 0.74 | 0.08 | 0 | muscle_sol | Timp4 |
| Lpl2 | 0 | 2.28865923 | 0.72 | 0.13 | 0 | muscle_sol | Lpl |
| Tcf153 | 0 | 2.19643186 | 0.89 | 0.25 | 0 | muscle_sol | Tcf15 |
| C1qtnf92 | 0 | 2.15411225 | 0.84 | 0.16 | 0 | muscle_sol | C1qtnf9 |
| Aqp72 | 0 | 2.12226972 | 0.75 | 0.09 | 0 | muscle_sol | Aqp7 |
| Fabp43 | 0 | 2.11540694 | 1 | 0.58 | 0 | muscle_sol | Fabp4 |
| S100a63 | 0 | 1.92985262 | 0.88 | 0.27 | 0 | muscle_sol | S100a6 |
| Rgs52 | 0 | 1.81620513 | 0.38 | 0.07 | 0 | muscle_sol | Rgs5 |
| Ptrf4 | 0 | 1.77262628 | 0.92 | 0.46 | 0 | muscle_sol | Ptrf |
| Aqp15 | 0 | 1.74134659 | 0.9 | 0.46 | 0 | muscle_sol | Aqp1 |
| Cxcl125 | 0 | 1.70882322 | 0.92 | 0.47 | 0 | muscle_sol | Cxcl12 |
| Cd363 | 0 | 1.43187396 | 0.96 | 0.51 | 0 | muscle_sol | Cd36 |
| Cav15 | 0 | 1.43004568 | 0.96 | 0.64 | 0 | muscle_sol | Cav1 |
| Ctnnbip12 | 0 | 1.42597966 | 0.69 | 0.26 | 0 | muscle_sol | Ctnnbip1 |
| Ehd13 | 0 | 1.41137129 | 0.71 | 0.25 | 0 | muscle_sol | Ehd1 |
| Lims23 | 0 | 1.40077343 | 0.85 | 0.47 | 0 | muscle_sol | Lims2 |
| Fam101b3 | 0 | 1.39676254 | 0.7 | 0.24 | 0 | muscle_sol | Fam101b |
| Rgcc4 | 0 | 1.38526729 | 0.87 | 0.41 | 0 | muscle_sol | Rgcc |
| Sparcl16 | 0 | 1.36616152 | 0.92 | 0.47 | 0 | muscle_sol | Sparcl1 |
| Fabp52 | 0 | 1.28411227 | 0.86 | 0.42 | 0 | muscle_sol | Fabp5 |
| Crip13 | 0 | 1.26099635 | 0.95 | 0.69 | 0 | muscle_sol | Crip1 |
| Mllt43 | 0 | 1.24767727 | 0.64 | 0.25 | 0 | muscle_sol | Mllt4 |
| Myadm4 | 0 | 1.22595061 | 0.59 | 0.23 | 0 | muscle_sol | Myadm |
| Meox22 | 0 | 1.21959923 | 0.44 | 0.07 | 0 | muscle_sol | Meox2 |
| Cdkn1c3 | 0 | 1.21594139 | 0.58 | 0.21 | 0 | muscle_sol | Cdkn1c |
| Tubb2a3 | 0 | 1.18369513 | 0.66 | 0.31 | 0 | muscle_sol | Tubb2a |
| Ehd22 | 0 | 1.15837378 | 0.79 | 0.45 | 0 | muscle_sol | Ehd2 |
| Slc26a101 | 0 | 1.14980859 | 0.41 | 0.12 | 0 | muscle_sol | Slc26a10 |
| Fbln22 | 0 | 1.14428941 | 0.55 | 0.18 | 0 | muscle_sol | Fbln2 |
| Gpihbp14 | 0 | 1.1239641 | 0.93 | 0.55 | 0 | muscle_sol | Gpihbp1 |
| Cdh133 | 0 | 1.06113013 | 0.48 | 0.15 | 0 | muscle_sol | Cdh13 |

| | | | | | | | |
|---|---|---|---|---|---|---|---|
| Car84 | 0 | 1.04970584 | 0.55 | 0.22 | 0 | muscle_sol | Car8 |
| Gng112 | 0 | 1.04646819 | 0.95 | 0.84 | 0 | muscle_sol | Gng11 |
| Mgll4 | 0 | 1.02650352 | 0.86 | 0.57 | 0 | muscle_sol | Mgll |
| Slc28a22 | 0 | 1.0212869 | 0.38 | 0.08 | 0 | muscle_sol | Slc28a2 |
| Cd300lg5 | 0 | 1.01444741 | 0.78 | 0.4 | 0 | muscle_sol | Cd300lg |
| Rftn14 | 0 | 0.99113089 | 0.47 | 0.18 | 0 | muscle_sol | Rftn1 |
| Lmo11 | 0 | 0.98749221 | 0.35 | 0.08 | 0 | muscle_sol | Lmo1 |
| Ablim32 | 0 | 0.97680256 | 0.37 | 0.09 | 0 | muscle_sol | Ablim3 |
| Ybx14 | 0 | 0.9589341 | 0.95 | 0.84 | 0 | muscle_sol | Ybx1 |
| Cav23 | 0 | 0.93938448 | 0.9 | 0.66 | 0 | muscle_sol | Cav2 |
| Fam89a | 0 | 0.93242987 | 0.27 | 0.03 | 0 | muscle_sol | Fam89a |
| 1810011O10Rik4 | 0 | 0.92223898 | 0.78 | 0.45 | 0 | muscle_sol | 1810011O10Rik |
| Tmsb4x2 | 0 | 0.88612148 | 1 | 0.99 | 0 | muscle_sol | Tmsb4x |
| Tspan134 | 0 | 0.88516977 | 0.91 | 0.62 | 0 | muscle_sol | Tspan13 |
| Cst34 | 0 | 0.81003255 | 0.96 | 0.88 | 0 | muscle_sol | Cst3 |
| Slc9a3r25 | 0 | 0.78907775 | 0.94 | 0.81 | 0 | muscle_sol | Slc9a3r2 |
| Kcna5 | 0 | 0.745458 | 0.26 | 0.06 | 0 | muscle_sol | Kcna5 |
| Pdlim15 | 5.84E-299 | 1.00605678 | 0.61 | 0.29 | 1.21E-294 | muscle_sol | Pdlim1 |
| Sparc2 | 3.08E-296 | 0.65086085 | 0.97 | 0.86 | 6.37E-292 | muscle_sol | Sparc |
| Hrct12 | 3.81E-292 | 0.8202557 | 0.31 | 0.09 | 7.87E-288 | muscle_sol | Hrct1 |
| Oat1 | 4.59E-275 | 0.89091625 | 0.41 | 0.16 | 9.49E-271 | muscle_sol | Oat |
| Sdpr4 | 1.36E-272 | 0.73795345 | 0.92 | 0.7 | 2.82E-268 | muscle_sol | Sdpr |
| Cd814 | 3.83E-265 | 0.63216325 | 0.95 | 0.82 | 7.93E-261 | muscle_sol | Cd81 |
| Jam24 | 5.15E-263 | 0.83684388 | 0.74 | 0.44 | 1.07E-258 | muscle_sol | Jam2 |
| Ercc11 | 3.58E-261 | 0.86889715 | 0.37 | 0.14 | 7.42E-257 | muscle_sol | Ercc1 |
| Cd2004 | 2.08E-260 | 0.81132663 | 0.88 | 0.65 | 4.31E-256 | muscle_sol | Cd200 |
| Tppp34 | 3.87E-240 | 0.82790183 | 0.43 | 0.17 | 8.01E-236 | muscle_sol | Tppp3 |
| Hspa12b3 | 3.94E-238 | 0.84826868 | 0.52 | 0.25 | 8.16E-234 | muscle_sol | Hspa12b |
| Sepw12 | 3.90E-236 | 0.68069999 | 0.9 | 0.7 | 8.06E-232 | muscle_sol | Sepw1 |
| Tubb4b2 | 1.80E-229 | 0.86176762 | 0.74 | 0.49 | 3.72E-225 | muscle_sol | Tubb4b |
| Cdkn1a4 | 5.11E-227 | 0.88444575 | 0.64 | 0.35 | 1.06E-222 | muscle_sol | Cdkn1a |
| Gm120022 | 2.87E-226 | 0.74345319 | 0.34 | 0.12 | 5.94E-222 | muscle_sol | Gm12002 |
| Ly6c14 | 1.08E-225 | 0.52408178 | 0.98 | 0.84 | 2.23E-221 | muscle_sol | Ly6c1 |
| Unc45b2 | 5.38E-221 | 0.82264497 | 0.57 | 0.3 | 1.11E-216 | muscle_sol | Unc45b |
| Klf43 | 4.43E-219 | 0.82946083 | 0.8 | 0.57 | 9.16E-215 | muscle_sol | Klf4 |
| Cyth12 | 1.99E-218 | 0.79542373 | 0.32 | 0.12 | 4.12E-214 | muscle_sol | Cyth1 |
| 2610305D13Rik | 5.84E-217 | 0.67747373 | 0.27 | 0.09 | 1.21E-212 | muscle_sol | 2610305D13Rik |
| Sgk14 | 4.30E-213 | 0.86947122 | 0.76 | 0.53 | 8.90E-209 | muscle_sol | Sgk1 |
| Id13 | 6.69E-213 | 0.80609953 | 0.85 | 0.64 | 1.38E-208 | muscle_sol | Id1 |
| Myl12a4 | 1.29E-212 | 0.54688636 | 0.93 | 0.84 | 2.67E-208 | muscle_sol | Myl12a |
| Ppp1r22 | 8.81E-212 | 0.72856099 | 0.81 | 0.62 | 1.82E-207 | muscle_sol | Ppp1r2 |

| | | | | | | | |
|---|---|---|---|---|---|---|---|
| Stmn21 | 3.21E-203 | 0.94872605 | 0.46 | 0.21 | 6.65E-199 | muscle_sol | Stmn2 |
| Mustn11 | 1.41E-202 | 0.69362155 | 0.27 | 0.09 | 2.91E-198 | muscle_sol | Mustn1 |
| Ptma1 | 1.57E-201 | 0.43467371 | 0.98 | 0.95 | 3.24E-197 | muscle_sol | Ptma |
| Ptp4a33 | 2.27E-199 | 0.71204858 | 0.66 | 0.38 | 4.70E-195 | muscle_sol | Ptp4a3 |
| Slc6a63 | 8.87E-195 | 0.75267628 | 0.69 | 0.41 | 1.84E-190 | muscle_sol | Slc6a6 |
| Lmna2 | 9.71E-195 | 0.81364467 | 0.56 | 0.3 | 2.01E-190 | muscle_sol | Lmna |
| Smco43 | 1.08E-194 | 0.80630877 | 0.69 | 0.46 | 2.23E-190 | muscle_sol | Smco4 |
| Cdh53 | 1.92E-193 | 0.64118201 | 0.88 | 0.73 | 3.97E-189 | muscle_sol | Cdh5 |
| Rassf33 | 5.01E-188 | 0.7805327 | 0.47 | 0.24 | 1.04E-183 | muscle_sol | Rassf3 |
| Pim33 | 5.68E-187 | 0.86866643 | 0.58 | 0.34 | 1.17E-182 | muscle_sol | Pim3 |
| Pxdc12 | 1.94E-183 | 0.67105234 | 0.38 | 0.16 | 4.01E-179 | muscle_sol | Pxdc1 |
| Adgrg12 | 6.60E-181 | 0.75954555 | 0.48 | 0.24 | 1.37E-176 | muscle_sol | Adgrg1 |
| AW1120105 | 7.08E-177 | 0.60766936 | 0.78 | 0.5 | 1.46E-172 | muscle_sol | AW112010 |
| Dok41 | 1.62E-175 | 0.72952758 | 0.39 | 0.18 | 3.34E-171 | muscle_sol | Dok4 |
| Tuba1a2 | 1.46E-174 | 0.67022719 | 0.83 | 0.63 | 3.02E-170 | muscle_sol | Tuba1a |
| Ahnak5 | 5.75E-173 | 0.70408132 | 0.69 | 0.44 | 1.19E-168 | muscle_sol | Ahnak |
| 8430408G22Rik3 | 2.77E-167 | 0.75644673 | 0.47 | 0.23 | 5.73E-163 | muscle_sol | 8430408G22Rik |
| S100a104 | 1.16E-163 | 0.58837822 | 0.85 | 0.66 | 2.39E-159 | muscle_sol | S100a10 |
| Sult1a12 | 3.13E-161 | 0.64307076 | 0.38 | 0.17 | 6.48E-157 | muscle_sol | Sult1a1 |
| Ckb | 6.08E-161 | 0.81924645 | 0.34 | 0.15 | 1.26E-156 | muscle_sol | Ckb |
| Gm98433 | 1.18E-160 | 0.57680965 | 0.82 | 0.61 | 2.45E-156 | muscle_sol | Gm9843 |
| Cyb5r34 | 3.86E-155 | 0.64010472 | 0.68 | 0.47 | 7.98E-151 | muscle_sol | Cyb5r3 |
| Fbxo221 | 2.80E-150 | 0.65166409 | 0.33 | 0.15 | 5.80E-146 | muscle_sol | Fbxo22 |
| Anxa33 | 3.33E-149 | 0.60160117 | 0.74 | 0.51 | 6.88E-145 | muscle_sol | Anxa3 |
| Aplnr3 | 6.95E-149 | 0.69617436 | 0.41 | 0.2 | 1.44E-144 | muscle_sol | Aplnr |
| Myct13 | 1.56E-148 | 0.64734016 | 0.7 | 0.49 | 3.23E-144 | muscle_sol | Myct1 |
| Prr133 | 9.52E-148 | 0.6487754 | 0.57 | 0.34 | 1.97E-143 | muscle_sol | Prr13 |
| Col4a24 | 2.13E-147 | 0.6205701 | 0.68 | 0.44 | 4.41E-143 | muscle_sol | Col4a2 |
| Palmd4 | 4.09E-146 | 0.66526882 | 0.5 | 0.28 | 8.46E-142 | muscle_sol | Palmd |
| S100a115 | 5.38E-144 | 0.53875544 | 0.79 | 0.58 | 1.11E-139 | muscle_sol | S100a11 |
| Ctnnb14 | 6.27E-144 | 0.61197744 | 0.72 | 0.51 | 1.30E-139 | muscle_sol | Ctnnb1 |
| Tmem255b1 | 8.25E-144 | 0.5622269 | 0.27 | 0.1 | 1.71E-139 | muscle_sol | Tmem255b |
| Tspan61 | 1.15E-143 | 0.59410476 | 0.3 | 0.13 | 2.38E-139 | muscle_sol | Tspan6 |
| Rbm31 | 1.22E-143 | 0.62024773 | 0.63 | 0.41 | 2.53E-139 | muscle_sol | Rbm3 |
| Ccdc85a2 | 5.45E-141 | 0.62017065 | 0.33 | 0.15 | 1.13E-136 | muscle_sol | Ccdc85a |
| Pdzd22 | 8.14E-140 | 0.66276112 | 0.38 | 0.18 | 1.68E-135 | muscle_sol | Pdzd2 |
| Sep-43 | 1.72E-138 | 0.60518855 | 0.55 | 0.32 | 3.56E-134 | muscle_sol | 4-Sep |
| Tmem1233 | 2.29E-138 | 0.61733049 | 0.49 | 0.27 | 4.75E-134 | muscle_sol | Tmem123 |
| Cltb3 | 3.11E-138 | 0.59300903 | 0.63 | 0.42 | 6.43E-134 | muscle_sol | Cltb |
| Anxa23 | 1.05E-135 | 0.57205415 | 0.8 | 0.63 | 2.17E-131 | muscle_sol | Anxa2 |
| Pls33 | 8.76E-134 | 0.68177592 | 0.56 | 0.35 | 1.81E-129 | muscle_sol | Pls3 |

| | | | | | | | |
|---|---|---|---|---|---|---|---|
| Calm14 | 5.55E-133 | 0.38690603 | 0.95 | 0.9 | 1.15E-128 | muscle_sol | Calm1 |
| Jup3 | 1.08E-131 | 0.56305349 | 0.61 | 0.39 | 2.23E-127 | muscle_sol | Jup |
| Gm94933 | 4.02E-130 | 0.56011111 | 0.72 | 0.51 | 8.33E-126 | muscle_sol | Gm9493 |
| Gstm13 | 3.87E-129 | 0.5973801 | 0.72 | 0.52 | 8.00E-125 | muscle_sol | Gstm1 |
| Ptprr2 | 4.24E-129 | 0.56478926 | 0.35 | 0.17 | 8.77E-125 | muscle_sol | Ptprr |
| Gnas2 | 2.87E-128 | 0.36399658 | 0.94 | 0.9 | 5.94E-124 | muscle_sol | Gnas |
| Tsfm1 | 1.52E-127 | 0.61337483 | 0.28 | 0.13 | 3.15E-123 | muscle_sol | Tsfm |
| Socs33 | 3.24E-126 | 0.56010089 | 0.49 | 0.28 | 6.69E-122 | muscle_sol | Socs3 |
| Ppic1 | 4.55E-125 | 0.61205254 | 0.56 | 0.36 | 9.42E-121 | muscle_sol | Ppic |
| Chek21 | 2.35E-124 | 0.52332912 | 0.31 | 0.14 | 4.86E-120 | muscle_sol | Chek2 |
| Fam198b1 | 1.43E-122 | 0.59367645 | 0.35 | 0.17 | 2.97E-118 | muscle_sol | Fam198b |
| Sertad12 | 3.50E-121 | 0.68358022 | 0.39 | 0.21 | 7.24E-117 | muscle_sol | Sertad1 |
| Tmem884 | 3.86E-121 | 0.5971438 | 0.63 | 0.43 | 8.00E-117 | muscle_sol | Tmem88 |
| Tmod33 | 9.73E-121 | 0.6358223 | 0.51 | 0.32 | 2.01E-116 | muscle_sol | Tmod3 |
| Hspg25 | 5.09E-120 | 0.53830984 | 0.62 | 0.38 | 1.05E-115 | muscle_sol | Hspg2 |
| Uaca3 | 2.96E-116 | 0.56231072 | 0.41 | 0.22 | 6.12E-112 | muscle_sol | Uaca |
| Nck13 | 1.41E-115 | 0.65880469 | 0.58 | 0.39 | 2.92E-111 | muscle_sol | Nck1 |
| Atp5j2 | 3.09E-113 | 0.43473134 | 0.88 | 0.78 | 6.39E-109 | muscle_sol | Atp5j |
| Kras | 6.98E-112 | 0.60561683 | 0.42 | 0.24 | 1.44E-107 | muscle_sol | Kras |
| Sox173 | 3.53E-111 | 0.65744124 | 0.59 | 0.38 | 7.31E-107 | muscle_sol | Sox17 |
| Klf91 | 1.48E-110 | 0.59287905 | 0.56 | 0.37 | 3.06E-106 | muscle_sol | Klf9 |
| Arhgef151 | 3.11E-110 | 0.56843362 | 0.55 | 0.36 | 6.44E-106 | muscle_sol | Arhgef15 |
| Xdh2 | 2.73E-109 | 0.63929132 | 0.36 | 0.19 | 5.64E-105 | muscle_sol | Xdh |
| Itpkb4 | 7.52E-109 | 0.56838903 | 0.59 | 0.4 | 1.56E-104 | muscle_sol | Itpkb |
| Lzts2 | 9.21E-109 | 0.56035517 | 0.36 | 0.19 | 1.91E-104 | muscle_sol | Lzts2 |
| Gadd45g3 | 7.36E-107 | 0.65297843 | 0.47 | 0.28 | 1.52E-102 | muscle_sol | Gadd45g |
| Lipe2 | 1.39E-106 | 0.55930473 | 0.33 | 0.17 | 2.88E-102 | muscle_sol | Lipe |
| Col4a14 | 3.18E-106 | 0.46724199 | 0.73 | 0.54 | 6.58E-102 | muscle_sol | Col4a1 |
| Fscn12 | 1.40E-105 | 0.47733526 | 0.39 | 0.21 | 2.89E-101 | muscle_sol | Fscn1 |
| Sdc3 | 2.21E-104 | 0.5014248 | 0.29 | 0.13 | 4.57E-100 | muscle_sol | Sdc3 |
| Ier25 | 3.84E-103 | 0.46802713 | 0.74 | 0.56 | 7.94E-99 | muscle_sol | Ier2 |
| Fos5 | 3.88E-103 | 0.75842951 | 0.71 | 0.56 | 8.04E-99 | muscle_sol | Fos |
| Palm2 | 5.10E-103 | 0.51031356 | 0.41 | 0.23 | 1.06E-98 | muscle_sol | Palm |
| Rps12-ps34 | 8.81E-103 | 0.48183188 | 0.75 | 0.57 | 1.82E-98 | muscle_sol | Rps12-ps3 |
| Adipor14 | 2.24E-102 | 0.54446089 | 0.63 | 0.45 | 4.63E-98 | muscle_sol | Adipor1 |
| Klk81 | 1.78E-101 | 0.54941506 | 0.26 | 0.12 | 3.69E-97 | muscle_sol | Klk8 |
| Rpl23a-ps33 | 9.27E-101 | 0.42330746 | 0.81 | 0.64 | 1.92E-96 | muscle_sol | Rpl23a-ps3 |
| 4930523C07Rik2 | 1.73E-100 | 0.55047951 | 0.41 | 0.24 | 3.58E-96 | muscle_sol | 4930523C07Rik |
| Ndel13 | 4.16E-100 | 0.52000766 | 0.43 | 0.25 | 8.61E-96 | muscle_sol | Ndel1 |
| Btg25 | 6.82E-100 | 0.51827169 | 0.67 | 0.48 | 1.41E-95 | muscle_sol | Btg2 |
| Ywhaq2 | 8.07E-100 | 0.47907455 | 0.77 | 0.63 | 1.67E-95 | muscle_sol | Ywhaq |

| | | | | | | | |
|---|---|---|---|---|---|---|---|
| Id33 | 1.17E-97 | 0.44005852 | 0.92 | 0.85 | 2.42E-93 | muscle_sol | Id3 |
| Wdr893 | 1.48E-97 | 0.39763139 | 0.87 | 0.71 | 3.06E-93 | muscle_sol | Wdr89 |
| Arpc22 | 1.44E-96 | 0.46012954 | 0.78 | 0.64 | 2.99E-92 | muscle_sol | Arpc2 |
| Tagln23 | 1.08E-95 | 0.46123073 | 0.84 | 0.73 | 2.23E-91 | muscle_sol | Tagln2 |
| Dhrs33 | 1.02E-94 | 0.4738507 | 0.55 | 0.36 | 2.11E-90 | muscle_sol | Dhrs3 |
| Klf65 | 1.64E-93 | 0.5865025 | 0.61 | 0.43 | 3.40E-89 | muscle_sol | Klf6 |
| Mmp151 | 4.98E-92 | 0.48348103 | 0.26 | 0.12 | 1.03E-87 | muscle_sol | Mmp15 |
| H3f3a1 | 6.11E-92 | 0.37866695 | 0.9 | 0.83 | 1.27E-87 | muscle_sol | H3f3a |
| Sox71 | 4.15E-90 | 0.55472566 | 0.35 | 0.19 | 8.60E-86 | muscle_sol | Sox7 |
| Ywhah3 | 8.52E-90 | 0.4870938 | 0.7 | 0.55 | 1.76E-85 | muscle_sol | Ywhah |
| Sptbn11 | 2.53E-89 | 0.32977424 | 0.87 | 0.75 | 5.24E-85 | muscle_sol | Sptbn1 |
| Anp32a3 | 4.58E-89 | 0.43887137 | 0.77 | 0.64 | 9.48E-85 | muscle_sol | Anp32a |
| Tprgl3 | 5.93E-89 | 0.47148797 | 0.69 | 0.53 | 1.23E-84 | muscle_sol | Tprgl |
| Ets13 | 2.27E-88 | 0.49771765 | 0.63 | 0.44 | 4.70E-84 | muscle_sol | Ets1 |
| Klf24 | 7.00E-88 | 0.4457608 | 0.93 | 0.88 | 1.45E-83 | muscle_sol | Klf2 |
| Picalm2 | 2.06E-87 | 0.53506254 | 0.41 | 0.25 | 4.26E-83 | muscle_sol | Picalm |
| Crip24 | 9.35E-86 | 0.30112127 | 0.93 | 0.85 | 1.93E-81 | muscle_sol | Crip2 |
| Kitl2 | 6.23E-85 | 0.45708817 | 0.55 | 0.36 | 1.29E-80 | muscle_sol | Kitl |
| Apold13 | 1.76E-84 | 0.43863338 | 0.4 | 0.24 | 3.64E-80 | muscle_sol | Apold1 |
| N4bp33 | 2.61E-84 | 0.48480262 | 0.35 | 0.19 | 5.41E-80 | muscle_sol | N4bp3 |
| Vasp2 | 6.35E-84 | 0.5158065 | 0.49 | 0.33 | 1.31E-79 | muscle_sol | Vasp |
| Lmcd13 | 1.11E-82 | 0.50054468 | 0.4 | 0.24 | 2.30E-78 | muscle_sol | Lmcd1 |
| Fam43a5 | 3.42E-82 | 0.4652559 | 0.47 | 0.3 | 7.07E-78 | muscle_sol | Fam43a |
| Nrarp2 | 7.01E-82 | 0.43470879 | 0.28 | 0.15 | 1.45E-77 | muscle_sol | Nrarp |
| Adam151 | 6.61E-81 | 0.47930502 | 0.6 | 0.43 | 1.37E-76 | muscle_sol | Adam15 |
| Nes1 | 7.66E-81 | 0.53309577 | 0.31 | 0.17 | 1.59E-76 | muscle_sol | Nes |
| Gltscr22 | 2.05E-80 | 0.48547951 | 0.62 | 0.45 | 4.24E-76 | muscle_sol | Gltscr2 |
| Gna11 | 3.19E-80 | 0.49793891 | 0.5 | 0.34 | 6.60E-76 | muscle_sol | Gna11 |
| Tpm43 | 3.98E-80 | 0.40141556 | 0.79 | 0.67 | 8.23E-76 | muscle_sol | Tpm4 |
| Plcb42 | 1.65E-79 | 0.48556211 | 0.37 | 0.22 | 3.41E-75 | muscle_sol | Plcb4 |
| Tinagl12 | 2.59E-79 | 0.39754542 | 0.79 | 0.65 | 5.35E-75 | muscle_sol | Tinagl1 |
| Mecom4 | 1.12E-77 | 0.42546924 | 0.37 | 0.21 | 2.32E-73 | muscle_sol | Mecom |
| Rasgrp24 | 3.88E-77 | 0.44892435 | 0.34 | 0.19 | 8.03E-73 | muscle_sol | Rasgrp2 |
| Cxx1a3 | 8.73E-77 | 0.43553205 | 0.63 | 0.45 | 1.81E-72 | muscle_sol | Cxx1a |
| Litaf | 9.12E-77 | 0.41810557 | 0.3 | 0.16 | 1.89E-72 | muscle_sol | Litaf |
| Bvht1 | 6.59E-76 | 0.51101585 | 0.47 | 0.31 | 1.36E-71 | muscle_sol | Bvht |
| Dusp32 | 2.97E-75 | 0.5062232 | 0.65 | 0.5 | 6.14E-71 | muscle_sol | Dusp3 |
| Ltbp44 | 3.46E-75 | 0.33858095 | 0.69 | 0.49 | 7.15E-71 | muscle_sol | Ltbp4 |
| Cyr613 | 6.35E-75 | 0.269592 | 0.43 | 0.26 | 1.31E-70 | muscle_sol | Cyr61 |
| Nxpe42 | 8.83E-75 | 0.46711928 | 0.28 | 0.15 | 1.83E-70 | muscle_sol | Nxpe4 |
| Ubc4 | 7.41E-74 | 0.35639319 | 0.9 | 0.83 | 1.53E-69 | muscle_sol | Ubc |

| | | | | | | | |
|---|---|---|---|---|---|---|---|
| Clic51 | 7.84E-74 | 0.42146529 | 0.45 | 0.29 | 1.62E-69 | muscle_sol | Clic5 |
| Egr14 | 2.28E-73 | 0.48897786 | 0.5 | 0.33 | 4.72E-69 | muscle_sol | Egr1 |
| Cdc42ep33 | 7.96E-73 | 0.44522266 | 0.35 | 0.21 | 1.65E-68 | muscle_sol | Cdc42ep3 |
| Zfp365 | 5.82E-72 | 0.57696648 | 0.56 | 0.41 | 1.20E-67 | muscle_sol | Zfp36 |
| Dusp16 | 2.38E-71 | 0.32960565 | 0.8 | 0.66 | 4.92E-67 | muscle_sol | Dusp1 |
| Tuba1b2 | 2.44E-71 | 0.39394298 | 0.8 | 0.68 | 5.05E-67 | muscle_sol | Tuba1b |
| Fam177a | 1.18E-70 | 0.45168662 | 0.35 | 0.2 | 2.43E-66 | muscle_sol | Fam177a |
| Cd1512 | 6.14E-70 | 0.41399751 | 0.7 | 0.55 | 1.27E-65 | muscle_sol | Cd151 |
| Gm87303 | 3.19E-69 | 0.39042014 | 0.58 | 0.41 | 6.60E-65 | muscle_sol | Gm8730 |
| Spop2 | 1.16E-68 | 0.48073849 | 0.5 | 0.35 | 2.40E-64 | muscle_sol | Spop |
| Mall2 | 1.92E-68 | 0.3801956 | 0.25 | 0.13 | 3.97E-64 | muscle_sol | Mall |
| Zfp6521 | 6.17E-67 | 0.51662259 | 0.25 | 0.14 | 1.28E-62 | muscle_sol | Zfp652 |
| Uba523 | 2.10E-66 | 0.32588745 | 0.83 | 0.71 | 4.34E-62 | muscle_sol | Uba52 |
| Rassf91 | 3.12E-66 | 0.40678603 | 0.25 | 0.14 | 6.45E-62 | muscle_sol | Rassf9 |
| Tm4sf12 | 7.43E-66 | 0.3298741 | 0.91 | 0.85 | 1.54E-61 | muscle_sol | Tm4sf1 |
| Zfp36l23 | 7.94E-66 | 0.51352589 | 0.41 | 0.27 | 1.64E-61 | muscle_sol | Zfp36l2 |
| Junb4 | 1.64E-65 | 0.47144512 | 0.74 | 0.63 | 3.39E-61 | muscle_sol | Junb |
| Atf43 | 3.64E-65 | 0.49533531 | 0.44 | 0.3 | 7.53E-61 | muscle_sol | Atf4 |
| Wwtr13 | 4.47E-65 | 0.45699338 | 0.41 | 0.26 | 9.25E-61 | muscle_sol | Wwtr1 |
| Mcf2l3 | 5.40E-65 | 0.43090402 | 0.33 | 0.19 | 1.12E-60 | muscle_sol | Mcf2l |
| Hbb-bs2 | 6.20E-65 | 1.0943691 | 0.28 | 0.15 | 1.28E-60 | muscle_sol | Hbb-bs |
| Fmo11 | 6.61E-65 | 0.29756344 | 0.39 | 0.24 | 1.37E-60 | muscle_sol | Fmo1 |
| Psap2 | 9.18E-65 | 0.33616476 | 0.83 | 0.73 | 1.90E-60 | muscle_sol | Psap |
| Rpl13-ps33 | 1.06E-64 | 0.36095284 | 0.52 | 0.36 | 2.20E-60 | muscle_sol | Rpl13-ps3 |
| Ldha3 | 1.23E-64 | 0.35729101 | 0.72 | 0.57 | 2.55E-60 | muscle_sol | Ldha |
| St6galnac4 | 1.08E-63 | 0.48416562 | 0.25 | 0.14 | 2.23E-59 | muscle_sol | St6galnac4 |
| Cxx1b4 | 1.32E-63 | 0.41066881 | 0.46 | 0.3 | 2.73E-59 | muscle_sol | Cxx1b |
| Polr2e | 2.70E-63 | 0.4568367 | 0.48 | 0.34 | 5.58E-59 | muscle_sol | Polr2e |
| Marcks4 | 4.13E-63 | 0.40600579 | 0.52 | 0.36 | 8.55E-59 | muscle_sol | Marcks |
| Adgrf53 | 7.79E-63 | 0.4142892 | 0.76 | 0.64 | 1.61E-58 | muscle_sol | Adgrf5 |
| Actg12 | 1.44E-62 | 0.30244103 | 0.94 | 0.88 | 2.98E-58 | muscle_sol | Actg1 |
| Akap2 | 3.38E-62 | 0.40151723 | 0.28 | 0.16 | 6.99E-58 | muscle_sol | Akap2 |
| Tsnax2 | 3.52E-62 | 0.47225339 | 0.3 | 0.18 | 7.29E-58 | muscle_sol | Tsnax |
| Rapgef11 | 3.70E-62 | 0.48599559 | 0.27 | 0.15 | 7.66E-58 | muscle_sol | Rapgef1 |
| Bag52 | 5.46E-62 | 0.4376672 | 0.38 | 0.24 | 1.13E-57 | muscle_sol | Bag5 |
| Rassf11 | 1.47E-61 | 0.41558647 | 0.44 | 0.3 | 3.03E-57 | muscle_sol | Rassf1 |
| Thrsp4 | 1.90E-61 | 0.3627924 | 0.43 | 0.27 | 3.92E-57 | muscle_sol | Thrsp |
| Bcam3 | 2.93E-61 | 0.38108033 | 0.7 | 0.56 | 6.07E-57 | muscle_sol | Bcam |
| Pkp41 | 2.04E-60 | 0.46864914 | 0.29 | 0.17 | 4.21E-56 | muscle_sol | Pkp4 |
| Bcl2l11 | 4.06E-60 | 0.44932403 | 0.35 | 0.22 | 8.40E-56 | muscle_sol | Bcl2l1 |
| Klf132 | 4.97E-60 | 0.42042678 | 0.61 | 0.47 | 1.03E-55 | muscle_sol | Klf13 |

| | | | | | | | |
|---|---|---|---|---|---|---|---|
| Lmo71 | 6.68E-60 | 0.37994162 | 0.26 | 0.15 | 1.38E-55 | muscle_sol | Lmo7 |
| Fez24 | 1.85E-59 | 0.41955112 | 0.48 | 0.34 | 3.82E-55 | muscle_sol | Fez2 |
| Eef22 | 3.59E-59 | 0.29314425 | 0.88 | 0.79 | 7.43E-55 | muscle_sol | Eef2 |
| Acot9 | 6.37E-59 | 0.45212972 | 0.29 | 0.17 | 1.32E-54 | muscle_sol | Acot9 |
| Ggta1 | 5.67E-58 | 0.39352482 | 0.3 | 0.18 | 1.17E-53 | muscle_sol | Ggta1 |
| Kctd102 | 8.37E-58 | 0.39076824 | 0.47 | 0.33 | 1.73E-53 | muscle_sol | Kctd10 |
| Lrrc8c3 | 1.37E-57 | 0.41742611 | 0.43 | 0.29 | 2.83E-53 | muscle_sol | Lrrc8c |
| Trim473 | 3.72E-57 | 0.36864652 | 0.48 | 0.33 | 7.70E-53 | muscle_sol | Trim47 |
| Itga63 | 5.77E-57 | 0.44229801 | 0.46 | 0.32 | 1.19E-52 | muscle_sol | Itga6 |
| Hnrnpf3 | 1.65E-56 | 0.33745725 | 0.76 | 0.64 | 3.41E-52 | muscle_sol | Hnrnpf |
| Gm61332 | 1.24E-55 | 0.3811189 | 0.41 | 0.27 | 2.57E-51 | muscle_sol | Gm6133 |
| Myo102 | 1.50E-55 | 0.44040805 | 0.33 | 0.21 | 3.11E-51 | muscle_sol | Myo10 |
| Ppp1cb2 | 3.72E-55 | 0.37117316 | 0.47 | 0.33 | 7.70E-51 | muscle_sol | Ppp1cb |
| Ceacam11 | 5.88E-54 | 0.39369463 | 0.32 | 0.2 | 1.22E-49 | muscle_sol | Ceacam1 |
| Dst1 | 6.98E-54 | 0.37815361 | 0.29 | 0.18 | 1.44E-49 | muscle_sol | Dst |
| Jund2 | 9.12E-54 | 0.35450391 | 0.8 | 0.72 | 1.89E-49 | muscle_sol | Jund |
| Cnn21 | 1.67E-53 | 0.3850233 | 0.43 | 0.29 | 3.47E-49 | muscle_sol | Cnn2 |
| Higd2a2 | 3.89E-53 | 0.3918514 | 0.55 | 0.42 | 8.04E-49 | muscle_sol | Higd2a |
| Tcf44 | 8.12E-53 | 0.26943341 | 0.79 | 0.67 | 1.68E-48 | muscle_sol | Tcf4 |
| Tpr2 | 8.67E-53 | 0.44829905 | 0.4 | 0.27 | 1.79E-48 | muscle_sol | Tpr |
| Wasf22 | 1.02E-52 | 0.40887149 | 0.55 | 0.42 | 2.11E-48 | muscle_sol | Wasf2 |
| Cdc42ep11 | 3.94E-52 | 0.39623527 | 0.41 | 0.28 | 8.14E-48 | muscle_sol | Cdc42ep1 |
| Podxl4 | 1.64E-51 | 0.25917738 | 0.67 | 0.5 | 3.40E-47 | muscle_sol | Podxl |
| Cd932 | 1.89E-51 | 0.34912581 | 0.67 | 0.54 | 3.91E-47 | muscle_sol | Cd93 |
| Lbr1 | 3.02E-51 | 0.42267761 | 0.31 | 0.19 | 6.25E-47 | muscle_sol | Lbr |
| Fkbp33 | 3.43E-51 | 0.48397355 | 0.43 | 0.31 | 7.11E-47 | muscle_sol | Fkbp3 |
| Cfl11 | 7.50E-51 | 0.26567551 | 0.89 | 0.83 | 1.55E-46 | muscle_sol | Cfl1 |
| Ifngr15 | 8.64E-51 | 0.3695006 | 0.53 | 0.39 | 1.79E-46 | muscle_sol | Ifngr1 |
| Csrp13 | 2.88E-50 | 0.38946093 | 0.55 | 0.41 | 5.96E-46 | muscle_sol | Csrp1 |
| 21-Mar | 3.58E-50 | 0.36591936 | 0.43 | 0.31 | 7.41E-46 | muscle_sol | 2-Mar |
| F11r1 | 5.42E-50 | 0.36155164 | 0.65 | 0.53 | 1.12E-45 | muscle_sol | F11r |
| Oaz1 | 1.43E-49 | 0.26853887 | 0.86 | 0.79 | 2.96E-45 | muscle_sol | Oaz1 |
| Hmgb21 | 2.45E-49 | 0.37898308 | 0.32 | 0.2 | 5.08E-45 | muscle_sol | Hmgb2 |
| Elovl51 | 3.21E-49 | 0.37435011 | 0.43 | 0.3 | 6.65E-45 | muscle_sol | Elovl5 |
| Ablim13 | 3.79E-49 | 0.29145708 | 0.42 | 0.28 | 7.85E-45 | muscle_sol | Ablim1 |
| Cald12 | 6.23E-49 | 0.32192055 | 0.49 | 0.35 | 1.29E-44 | muscle_sol | Cald1 |
| Capn23 | 6.47E-49 | 0.38656912 | 0.45 | 0.32 | 1.34E-44 | muscle_sol | Capn2 |
| Ier52 | 9.88E-49 | 0.43743275 | 0.6 | 0.49 | 2.04E-44 | muscle_sol | Ier5 |
| Plod11 | 1.13E-48 | 0.38908905 | 0.33 | 0.21 | 2.35E-44 | muscle_sol | Plod1 |
| Aplp24 | 1.91E-48 | 0.28333811 | 0.78 | 0.67 | 3.95E-44 | muscle_sol | Aplp2 |
| Tax1bp32 | 6.72E-48 | 0.35401598 | 0.48 | 0.36 | 1.39E-43 | muscle_sol | Tax1bp3 |

| | | | | | | | |
|---|---|---|---|---|---|---|---|
| Vamp32 | 1.05E-47 | 0.38777433 | 0.44 | 0.32 | 2.18E-43 | muscle_sol | Vamp3 |
| Ddx3x1 | 1.85E-47 | 0.42295574 | 0.42 | 0.3 | 3.84E-43 | muscle_sol | Ddx3x |
| Itgb12 | 2.09E-47 | 0.35310877 | 0.65 | 0.53 | 4.32E-43 | muscle_sol | Itgb1 |
| Fmo22 | 1.16E-46 | 0.32645159 | 0.28 | 0.17 | 2.40E-42 | muscle_sol | Fmo2 |
| Btg14 | 2.09E-46 | 0.41742111 | 0.62 | 0.51 | 4.32E-42 | muscle_sol | Btg1 |
| Sox183 | 2.55E-46 | 0.35681297 | 0.63 | 0.51 | 5.28E-42 | muscle_sol | Sox18 |
| Slc25a52 | 4.10E-46 | 0.35450756 | 0.65 | 0.53 | 8.49E-42 | muscle_sol | Slc25a5 |
| Rhoa1 | 7.24E-46 | 0.28682008 | 0.85 | 0.77 | 1.50E-41 | muscle_sol | Rhoa |
| Gm118083 | 6.14E-45 | 0.27833458 | 0.71 | 0.58 | 1.27E-40 | muscle_sol | Gm11808 |
| Ptms2 | 8.05E-45 | 0.31784434 | 0.7 | 0.58 | 1.67E-40 | muscle_sol | Ptms |
| Arid5b2 | 9.56E-45 | 0.33699765 | 0.38 | 0.26 | 1.98E-40 | muscle_sol | Arid5b |
| 9430020K01Rik2 | 1.07E-44 | 0.33094134 | 0.31 | 0.2 | 2.22E-40 | muscle_sol | 9430020K01Rik |
| Eif4a12 | 1.08E-44 | 0.30016673 | 0.7 | 0.58 | 2.24E-40 | muscle_sol | Eif4a1 |
| Rpl9-ps63 | 1.85E-44 | 0.30770611 | 0.58 | 0.44 | 3.83E-40 | muscle_sol | Rpl9-ps6 |
| Smim141 | 2.01E-44 | 0.36804592 | 0.53 | 0.41 | 4.16E-40 | muscle_sol | Smim14 |
| Wnk14 | 3.37E-44 | 0.38226945 | 0.45 | 0.33 | 6.97E-40 | muscle_sol | Wnk1 |
| Ddit42 | 3.79E-44 | 0.38719791 | 0.29 | 0.18 | 7.85E-40 | muscle_sol | Ddit4 |
| Sh3glb11 | 8.34E-44 | 0.32526456 | 0.7 | 0.59 | 1.73E-39 | muscle_sol | Sh3glb1 |
| Anxa51 | 1.36E-43 | 0.30332653 | 0.75 | 0.65 | 2.81E-39 | muscle_sol | Anxa5 |
| Klf32 | 2.23E-43 | 0.3707144 | 0.48 | 0.36 | 4.61E-39 | muscle_sol | Klf3 |
| Ctnna12 | 4.89E-43 | 0.34464922 | 0.58 | 0.46 | 1.01E-38 | muscle_sol | Ctnna1 |
| Sypl3 | 7.92E-43 | 0.32461534 | 0.63 | 0.52 | 1.64E-38 | muscle_sol | Sypl |
| Cmpk1 | 8.38E-43 | 0.3869782 | 0.4 | 0.29 | 1.73E-38 | muscle_sol | Cmpk1 |
| Tnfaip13 | 1.16E-42 | 0.34366944 | 0.47 | 0.35 | 2.41E-38 | muscle_sol | Tnfaip1 |
| Gpr41 | 1.60E-42 | 0.28761222 | 0.26 | 0.16 | 3.30E-38 | muscle_sol | Gpr4 |
| Lasp1 | 4.74E-42 | 0.35877118 | 0.36 | 0.26 | 9.81E-38 | muscle_sol | Lasp1 |
| Hdac2 | 1.49E-41 | 0.34852404 | 0.27 | 0.17 | 3.08E-37 | muscle_sol | Hdac2 |
| Serpinh13 | 2.30E-41 | 0.25404927 | 0.78 | 0.68 | 4.75E-37 | muscle_sol | Serpinh1 |
| Ushbp12 | 3.54E-41 | 0.33623633 | 0.55 | 0.44 | 7.33E-37 | muscle_sol | Ushbp1 |
| Rpl6l3 | 1.09E-40 | 0.29116295 | 0.6 | 0.47 | 2.25E-36 | muscle_sol | Rpl6l |
| Psme2b2 | 3.06E-40 | 0.30745201 | 0.28 | 0.18 | 6.32E-36 | muscle_sol | Psme2b |
| Atf33 | 6.34E-40 | 0.53343196 | 0.28 | 0.18 | 1.31E-35 | muscle_sol | Atf3 |
| Rpl36-ps33 | 2.36E-39 | 0.25460449 | 0.5 | 0.36 | 4.89E-35 | muscle_sol | Rpl36-ps3 |
| Gm100363 | 4.65E-39 | 0.29142614 | 0.46 | 0.33 | 9.62E-35 | muscle_sol | Gm10036 |
| Lama43 | 8.16E-39 | 0.2710586 | 0.27 | 0.17 | 1.69E-34 | muscle_sol | Lama4 |
| Sptan11 | 1.07E-38 | 0.33123524 | 0.49 | 0.37 | 2.21E-34 | muscle_sol | Sptan1 |
| Slc38a2 | 3.02E-38 | 0.39190954 | 0.44 | 0.34 | 6.24E-34 | muscle_sol | Slc38a2 |
| Dab2ip1 | 3.08E-38 | 0.37930861 | 0.33 | 0.23 | 6.38E-34 | muscle_sol | Dab2ip |
| Arhgdia1 | 6.61E-38 | 0.28806568 | 0.7 | 0.6 | 1.37E-33 | muscle_sol | Arhgdia |
| Ilk2 | 1.31E-37 | 0.32804934 | 0.43 | 0.31 | 2.70E-33 | muscle_sol | Ilk |
| Capza22 | 1.60E-37 | 0.31025271 | 0.59 | 0.48 | 3.31E-33 | muscle_sol | Capza2 |

| | | | | | | | |
|---|---|---|---|---|---|---|---|
| Arpc1b2 | 2.79E-37 | 0.28423503 | 0.73 | 0.64 | 5.77E-33 | muscle_sol | Arpc1b |
| Zfos1 | 3.20E-37 | 0.3409055 | 0.35 | 0.24 | 6.62E-33 | muscle_sol | Zfos1 |
| Srsf51 | 8.10E-37 | 0.31956668 | 0.63 | 0.53 | 1.68E-32 | muscle_sol | Srsf5 |
| Tubb52 | 1.77E-36 | 0.26746108 | 0.74 | 0.65 | 3.67E-32 | muscle_sol | Tubb5 |
| Neat12 | 1.95E-36 | 0.28775095 | 0.56 | 0.45 | 4.04E-32 | muscle_sol | Neat1 |
| Tns11 | 2.16E-36 | 0.31883206 | 0.5 | 0.38 | 4.47E-32 | muscle_sol | Tns1 |
| Tmem1403 | 2.26E-36 | 0.31704664 | 0.49 | 0.38 | 4.67E-32 | muscle_sol | Tmem140 |
| Tspan123 | 3.14E-36 | 0.27977175 | 0.47 | 0.35 | 6.49E-32 | muscle_sol | Tspan12 |
| Rarg2 | 3.29E-36 | 0.30969839 | 0.33 | 0.23 | 6.80E-32 | muscle_sol | Rarg |
| Myo18a1 | 3.76E-36 | 0.29796328 | 0.3 | 0.2 | 7.79E-32 | muscle_sol | Myo18a |
| Efnb12 | 9.20E-36 | 0.25179288 | 0.35 | 0.24 | 1.90E-31 | muscle_sol | Efnb1 |
| Txnip4 | 9.94E-36 | 0.27329491 | 0.76 | 0.65 | 2.06E-31 | muscle_sol | Txnip |
| B4galt43 | 1.63E-35 | 0.36402589 | 0.34 | 0.24 | 3.37E-31 | muscle_sol | B4galt4 |
| Lrrc8a3 | 2.00E-35 | 0.36036087 | 0.39 | 0.29 | 4.13E-31 | muscle_sol | Lrrc8a |
| Pabpc13 | 3.19E-35 | 0.34838684 | 0.58 | 0.49 | 6.60E-31 | muscle_sol | Pabpc1 |
| Rpl27-ps33 | 5.28E-35 | 0.26130163 | 0.45 | 0.33 | 1.09E-30 | muscle_sol | Rpl27-ps3 |
| Ube2d31 | 5.67E-35 | 0.29539813 | 0.72 | 0.63 | 1.17E-30 | muscle_sol | Ube2d3 |
| S1pr14 | 3.03E-34 | 0.28145984 | 0.59 | 0.46 | 6.27E-30 | muscle_sol | S1pr1 |
| Arf62 | 5.34E-34 | 0.30074512 | 0.39 | 0.28 | 1.10E-29 | muscle_sol | Arf6 |
| Srsf31 | 1.89E-33 | 0.31167531 | 0.62 | 0.53 | 3.91E-29 | muscle_sol | Srsf3 |
| Anp32b | 5.07E-33 | 0.33600065 | 0.42 | 0.32 | 1.05E-28 | muscle_sol | Anp32b |
| Pdgfb1 | 8.49E-33 | 0.28716821 | 0.33 | 0.23 | 1.76E-28 | muscle_sol | Pdgfb |
| Gdi22 | 1.34E-32 | 0.27494169 | 0.6 | 0.5 | 2.78E-28 | muscle_sol | Gdi2 |
| Kif5b2 | 3.44E-32 | 0.29572557 | 0.54 | 0.44 | 7.11E-28 | muscle_sol | Kif5b |
| Gm122161 | 5.23E-32 | 0.32144865 | 0.36 | 0.26 | 1.08E-27 | muscle_sol | Gm12216 |
| Srsf21 | 9.36E-32 | 0.28521315 | 0.58 | 0.48 | 1.94E-27 | muscle_sol | Srsf2 |
| Mob2 | 1.17E-31 | 0.29381456 | 0.31 | 0.22 | 2.41E-27 | muscle_sol | Mob2 |
| Zfp36l13 | 3.28E-31 | 0.29747484 | 0.63 | 0.53 | 6.80E-27 | muscle_sol | Zfp36l1 |
| Gm87972 | 4.97E-31 | 0.30155629 | 0.31 | 0.22 | 1.03E-26 | muscle_sol | Gm8797 |
| Actb1 | 1.51E-29 | 0.28149122 | 0.96 | 0.95 | 3.12E-25 | muscle_sol | Actb |
| Pink12 | 4.53E-29 | 0.2976936 | 0.34 | 0.25 | 9.36E-25 | muscle_sol | Pink1 |
| Bnip21 | 6.26E-29 | 0.27703735 | 0.49 | 0.39 | 1.30E-24 | muscle_sol | Bnip2 |
| Pgrmc11 | 1.23E-28 | 0.26276752 | 0.46 | 0.36 | 2.54E-24 | muscle_sol | Pgrmc1 |
| Ndufa81 | 5.93E-28 | 0.28960206 | 0.61 | 0.52 | 1.23E-23 | muscle_sol | Ndufa8 |
| Mast43 | 3.12E-26 | 0.27653509 | 0.39 | 0.3 | 6.45E-22 | muscle_sol | Mast4 |
| Kank32 | 3.12E-26 | 0.25419547 | 0.65 | 0.56 | 6.46E-22 | muscle_sol | Kank3 |
| Bloc1s11 | 4.77E-26 | 0.28570578 | 0.41 | 0.32 | 9.86E-22 | muscle_sol | Bloc1s1 |
| Cfap20 | 8.00E-26 | 0.29036556 | 0.28 | 0.2 | 1.66E-21 | muscle_sol | Cfap20 |
| Irf2bpl3 | 3.65E-25 | 0.29301537 | 0.36 | 0.27 | 7.54E-21 | muscle_sol | Irf2bpl |
| Nfe2l11 | 3.66E-25 | 0.28106122 | 0.34 | 0.26 | 7.57E-21 | muscle_sol | Nfe2l1 |
| Tob11 | 3.85E-25 | 0.31802126 | 0.25 | 0.18 | 7.96E-21 | muscle_sol | Tob1 |

| | | | | | | | |
|---|---|---|---|---|---|---|---|
| Mprip1 | 4.86E-25 | 0.29228227 | 0.36 | 0.27 | 1.00E-20 | muscle_sol | Mprip |
| Socs23 | 1.38E-24 | 0.25340377 | 0.34 | 0.25 | 2.85E-20 | muscle_sol | Socs2 |
| Copz21 | 2.05E-24 | 0.25440959 | 0.26 | 0.19 | 4.25E-20 | muscle_sol | Copz2 |
| Cebpg1 | 3.74E-24 | 0.2807553 | 0.35 | 0.27 | 7.74E-20 | muscle_sol | Cebpg |
| Trp53i111 | 5.01E-24 | 0.30338035 | 0.32 | 0.24 | 1.04E-19 | muscle_sol | Trp53i11 |
| Klf103 | 5.66E-24 | 0.27128991 | 0.32 | 0.23 | 1.17E-19 | muscle_sol | Klf10 |
| Nptn1 | 6.34E-22 | 0.28029031 | 0.33 | 0.25 | 1.31E-17 | muscle_sol | Nptn |
| Irf13 | 1.48E-21 | 0.2733675 | 0.38 | 0.3 | 3.06E-17 | muscle_sol | Irf1 |
| Hnrnpu1 | 7.60E-21 | 0.26777444 | 0.46 | 0.38 | 1.57E-16 | muscle_sol | Hnrnpu |
| Isg152 | 4.62E-16 | 0.30749989 | 0.38 | 0.31 | 9.56E-12 | muscle_sol | Isg15 |
| Ltc4s | 0 | 3.78898096 | 0.91 | 0.04 | 0 | spleen | Ltc4s |
| Cxcl10 | 0 | 3.65847611 | 0.81 | 0.03 | 0 | spleen | Cxcl10 |
| Star | 0 | 3.54222716 | 0.85 | 0 | 0 | spleen | Star |
| Ankrd2 | 0 | 3.44610258 | 0.87 | 0 | 0 | spleen | Ankrd2 |
| Stab21 | 0 | 3.25846466 | 0.93 | 0.08 | 0 | spleen | Stab2 |
| Tfpi1 | 0 | 3.22661657 | 0.9 | 0.16 | 0 | spleen | Tfpi |
| Mafb | 0 | 3.20076834 | 0.86 | 0.06 | 0 | spleen | Mafb |
| Serpinb6b | 0 | 2.85371248 | 0.83 | 0.07 | 0 | spleen | Serpinb6b |
| Serpinb1a | 0 | 2.74607071 | 0.79 | 0.1 | 0 | spleen | Serpinb1a |
| Dstn | 0 | 2.55802134 | 0.98 | 0.57 | 0 | spleen | Dstn |
| Efhd1 | 0 | 2.49898513 | 0.79 | 0.06 | 0 | spleen | Efhd1 |
| Tmem35 | 0 | 2.46314105 | 0.7 | 0.03 | 0 | spleen | Tmem35 |
| Sema3a | 0 | 2.44191586 | 0.67 | 0.01 | 0 | spleen | Sema3a |
| Adamts5 | 0 | 2.42468789 | 0.63 | 0.01 | 0 | spleen | Adamts5 |
| Ackr31 | 0 | 2.31042058 | 0.85 | 0.24 | 0 | spleen | Ackr3 |
| Sema3d | 0 | 2.30853393 | 0.69 | 0.03 | 0 | spleen | Sema3d |
| Pcolce | 0 | 2.28691126 | 0.74 | 0.09 | 0 | spleen | Pcolce |
| Lrg1 | 0 | 2.27917935 | 0.8 | 0.14 | 0 | spleen | Lrg1 |
| Pdlim41 | 0 | 2.26036519 | 0.78 | 0.11 | 0 | spleen | Pdlim4 |
| Tgfbi | 0 | 2.21079051 | 0.72 | 0.07 | 0 | spleen | Tgfbi |
| Sema4b | 0 | 2.11264888 | 0.58 | 0.02 | 0 | spleen | Sema4b |
| Tmem215 | 0 | 2.10261774 | 0.57 | 0 | 0 | spleen | Tmem215 |
| Clu1 | 0 | 2.08226762 | 0.95 | 0.31 | 0 | spleen | Clu |
| Irf8 | 0 | 2.0580177 | 0.63 | 0.04 | 0 | spleen | Irf8 |
| Nupr1 | 0 | 2.05047228 | 0.62 | 0.04 | 0 | spleen | Nupr1 |
| Gm37350 | 0 | 2.020194 | 0.47 | 0.03 | 0 | spleen | Gm37350 |
| Serpinb6a1 | 0 | 1.99231348 | 0.91 | 0.41 | 0 | spleen | Serpinb6a |
| AU0210921 | 0 | 1.96093096 | 0.87 | 0.28 | 0 | spleen | AU021092 |
| Cd551 | 0 | 1.94117568 | 0.77 | 0.11 | 0 | spleen | Cd55 |
| Tmem176a3 | 0 | 1.88564744 | 0.9 | 0.38 | 0 | spleen | Tmem176a |
| Slc16a11 | 0 | 1.88006764 | 0.63 | 0.07 | 0 | spleen | Slc16a1 |

| | | | | | | | |
|---|---|---|---|---|---|---|---|
| Tst | 0 | 1.8635362 | 0.57 | 0.04 | 0 | spleen | Tst |
| Tmem176b2 | 0 | 1.86024856 | 0.93 | 0.46 | 0 | spleen | Tmem176b |
| Rcn31 | 0 | 1.85929562 | 0.69 | 0.11 | 0 | spleen | Rcn3 |
| Dhcr24 | 0 | 1.8403645 | 0.56 | 0.03 | 0 | spleen | Dhcr24 |
| Akr1b81 | 0 | 1.78313121 | 0.58 | 0.04 | 0 | spleen | Akr1b8 |
| Timp21 | 0 | 1.77271364 | 0.83 | 0.22 | 0 | spleen | Timp2 |
| Sh3bgrl3 | 0 | 1.77104964 | 0.95 | 0.67 | 0 | spleen | Sh3bgrl3 |
| Crispld11 | 0 | 1.75879023 | 0.61 | 0.09 | 0 | spleen | Crispld1 |
| Lrrn1 | 0 | 1.75565613 | 0.5 | 0 | 0 | spleen | Lrrn1 |
| Dbi | 0 | 1.75049022 | 0.88 | 0.44 | 0 | spleen | Dbi |
| St18 | 0 | 1.71481386 | 0.42 | 0 | 0 | spleen | St18 |
| Bin11 | 0 | 1.7145496 | 0.66 | 0.12 | 0 | spleen | Bin1 |
| Marcksl11 | 0 | 1.70857578 | 0.77 | 0.2 | 0 | spleen | Marcksl1 |
| C1qtnf7 | 0 | 1.68960305 | 0.49 | 0.01 | 0 | spleen | C1qtnf7 |
| Prkcdbp3 | 0 | 1.6821382 | 0.93 | 0.47 | 0 | spleen | Prkcdbp |
| Bok | 0 | 1.66766366 | 0.62 | 0.14 | 0 | spleen | Bok |
| Tpm2 | 0 | 1.66603034 | 0.46 | 0.02 | 0 | spleen | Tpm2 |
| Slco2a12 | 0 | 1.64274142 | 0.78 | 0.27 | 0 | spleen | Slco2a1 |
| Sorbs22 | 0 | 1.60914119 | 0.75 | 0.22 | 0 | spleen | Sorbs2 |
| Cd63 | 0 | 1.58176608 | 0.68 | 0.13 | 0 | spleen | Cd63 |
| Cstb1 | 0 | 1.57891786 | 0.77 | 0.31 | 0 | spleen | Cstb |
| Cyp21a1 | 0 | 1.51427707 | 0.35 | 0 | 0 | spleen | Cyp21a1 |
| Ipo11 | 0 | 1.50031535 | 0.69 | 0.2 | 0 | spleen | Ipo11 |
| Il2rg1 | 0 | 1.49169262 | 0.62 | 0.14 | 0 | spleen | Il2rg |
| Ninj1 | 0 | 1.48629603 | 0.69 | 0.24 | 0 | spleen | Ninj1 |
| Cotl11 | 0 | 1.48432592 | 0.65 | 0.13 | 0 | spleen | Cotl1 |
| Cxcl161 | 0 | 1.46618745 | 0.65 | 0.14 | 0 | spleen | Cxcl16 |
| Unc119 | 0 | 1.46398941 | 0.41 | 0.02 | 0 | spleen | Unc119 |
| Met | 0 | 1.45508578 | 0.41 | 0.01 | 0 | spleen | Met |
| Timp1 | 0 | 1.43503204 | 0.4 | 0.01 | 0 | spleen | Timp1 |
| Nr5a1 | 0 | 1.4264369 | 0.4 | 0.01 | 0 | spleen | Nr5a1 |
| Hmgcs1 | 0 | 1.3919529 | 0.47 | 0.08 | 0 | spleen | Hmgcs1 |
| Maf2 | 0 | 1.38015902 | 0.77 | 0.14 | 0 | spleen | Maf |
| Jag1 | 0 | 1.37461947 | 0.47 | 0.09 | 0 | spleen | Jag1 |
| Rgs2 | 0 | 1.37446162 | 0.39 | 0.04 | 0 | spleen | Rgs2 |
| Pvrl21 | 0 | 1.36832657 | 0.59 | 0.13 | 0 | spleen | Pvrl2 |
| Nkx2-32 | 0 | 1.34864538 | 0.59 | 0.07 | 0 | spleen | Nkx2-3 |
| Serpinb9 | 0 | 1.33978975 | 0.59 | 0.17 | 0 | spleen | Serpinb9 |
| Itm2c | 0 | 1.33965204 | 0.82 | 0.4 | 0 | spleen | Itm2c |
| Acly1 | 0 | 1.33771325 | 0.66 | 0.21 | 0 | spleen | Acly |
| Emp31 | 0 | 1.33700881 | 0.69 | 0.22 | 0 | spleen | Emp3 |

| | | | | | | | |
|---|---|---|---|---|---|---|---|
| Gpr1821 | 0 | 1.33451554 | 0.79 | 0.21 | 0 | spleen | Gpr182 |
| Cdk2ap2 | 0 | 1.33441098 | 0.62 | 0.19 | 0 | spleen | Cdk2ap2 |
| Fcer1g1 | 0 | 1.32774157 | 0.36 | 0.05 | 0 | spleen | Fcer1g |
| Ptpn181 | 0 | 1.32414518 | 0.59 | 0.16 | 0 | spleen | Ptpn18 |
| Asrgl1 | 0 | 1.32368101 | 0.41 | 0.04 | 0 | spleen | Asrgl1 |
| Plxnb1 | 0 | 1.32016206 | 0.35 | 0 | 0 | spleen | Plxnb1 |
| Vsir1 | 0 | 1.3108041 | 0.62 | 0.18 | 0 | spleen | Vsir |
| Grhpr | 0 | 1.29714117 | 0.43 | 0.05 | 0 | spleen | Grhpr |
| C1qtnf6 | 0 | 1.28724877 | 0.39 | 0.03 | 0 | spleen | C1qtnf6 |
| Gngt21 | 0 | 1.27892023 | 0.9 | 0.54 | 0 | spleen | Gngt2 |
| Mcfd21 | 0 | 1.27791903 | 0.66 | 0.25 | 0 | spleen | Mcfd2 |
| Anxa12 | 0 | 1.26395587 | 0.49 | 0.11 | 0 | spleen | Anxa1 |
| Igfbp72 | 0 | 1.25553416 | 1 | 0.87 | 0 | spleen | Igfbp7 |
| Fam213b | 0 | 1.25437505 | 0.37 | 0.03 | 0 | spleen | Fam213b |
| Bmp21 | 0 | 1.24364811 | 0.57 | 0.08 | 0 | spleen | Bmp2 |
| H2-Q73 | 0 | 1.23377507 | 0.77 | 0.35 | 0 | spleen | H2-Q7 |
| Nr2f22 | 0 | 1.23168944 | 0.63 | 0.18 | 0 | spleen | Nr2f2 |
| Gata6 | 0 | 1.23093454 | 0.38 | 0.03 | 0 | spleen | Gata6 |
| Bace21 | 0 | 1.22939649 | 0.55 | 0.15 | 0 | spleen | Bace2 |
| Synm | 0 | 1.19571522 | 0.42 | 0.07 | 0 | spleen | Synm |
| Plekho2 | 0 | 1.19394801 | 0.43 | 0.07 | 0 | spleen | Plekho2 |
| Fabp53 | 0 | 1.1925139 | 0.88 | 0.41 | 0 | spleen | Fabp5 |
| Atp2b42 | 0 | 1.18563815 | 0.56 | 0.18 | 0 | spleen | Atp2b4 |
| Phlda1 | 0 | 1.17758434 | 0.46 | 0.11 | 0 | spleen | Phlda1 |
| Fxyd61 | 0 | 1.17118077 | 0.57 | 0.13 | 0 | spleen | Fxyd6 |
| Fam167b2 | 0 | 1.15515315 | 0.75 | 0.24 | 0 | spleen | Fam167b |
| Dll1 | 0 | 1.15335183 | 0.42 | 0.08 | 0 | spleen | Dll1 |
| Ttyh2 | 0 | 1.15082551 | 0.33 | 0.03 | 0 | spleen | Ttyh2 |
| Cyba | 0 | 1.13990126 | 0.79 | 0.45 | 0 | spleen | Cyba |
| Extl1 | 0 | 1.13705745 | 0.29 | 0 | 0 | spleen | Extl1 |
| Rtn4rl1 | 0 | 1.13146244 | 0.3 | 0.01 | 0 | spleen | Rtn4rl1 |
| Arhgap231 | 0 | 1.09939961 | 0.45 | 0.11 | 0 | spleen | Arhgap23 |
| Trp53i112 | 0 | 1.09814597 | 0.61 | 0.21 | 0 | spleen | Trp53i11 |
| Arfgap3 | 0 | 1.09497764 | 0.41 | 0.09 | 0 | spleen | Arfgap3 |
| Ube2e21 | 0 | 1.09469778 | 0.38 | 0.07 | 0 | spleen | Ube2e2 |
| Sigmar1 | 0 | 1.09309374 | 0.48 | 0.14 | 0 | spleen | Sigmar1 |
| Camk4 | 0 | 1.08744328 | 0.29 | 0 | 0 | spleen | Camk4 |
| Mmp16 | 0 | 1.08235862 | 0.29 | 0 | 0 | spleen | Mmp16 |
| Flrt2 | 0 | 1.0818582 | 0.33 | 0.03 | 0 | spleen | Flrt2 |
| Calml41 | 0 | 1.07161843 | 0.4 | 0.08 | 0 | spleen | Calml4 |
| Tgfbr31 | 0 | 1.04721746 | 0.45 | 0.12 | 0 | spleen | Tgfbr3 |

| | | | | | | | |
|---|---|---|---|---|---|---|---|
| Tbxa2r1 | 0 | 1.03554914 | 0.54 | 0.16 | 0 | spleen | Tbxa2r |
| Gja12 | 0 | 1.03393684 | 0.54 | 0.21 | 0 | spleen | Gja1 |
| Akr1b10 | 0 | 1.0252515 | 0.33 | 0.04 | 0 | spleen | Akr1b10 |
| Tmed31 | 0 | 1.02430121 | 0.6 | 0.27 | 0 | spleen | Tmed3 |
| Msmo1 | 0 | 1.02273624 | 0.36 | 0.08 | 0 | spleen | Msmo1 |
| Ppp1r14b | 0 | 1.01616069 | 0.36 | 0.06 | 0 | spleen | Ppp1r14b |
| Ccdc8 | 0 | 1.00568237 | 0.3 | 0.02 | 0 | spleen | Ccdc8 |
| Tpm11 | 0 | 0.99930688 | 0.68 | 0.32 | 0 | spleen | Tpm1 |
| Ralgapa21 | 0 | 0.99587121 | 0.38 | 0.08 | 0 | spleen | Ralgapa2 |
| Vamp53 | 0 | 0.99519378 | 0.86 | 0.51 | 0 | spleen | Vamp5 |
| Prox1 | 0 | 0.9883259 | 0.33 | 0.03 | 0 | spleen | Prox1 |
| Cox6c | 0 | 0.98601172 | 0.95 | 0.76 | 0 | spleen | Cox6c |
| Gimap51 | 0 | 0.97144044 | 0.65 | 0.3 | 0 | spleen | Gimap5 |
| Nfic | 0 | 0.97121529 | 0.64 | 0.32 | 0 | spleen | Nfic |
| Fam174b2 | 0 | 0.97105212 | 0.46 | 0.13 | 0 | spleen | Fam174b |
| Tnip1 | 0 | 0.95815376 | 0.36 | 0.08 | 0 | spleen | Tnip1 |
| Zmiz11 | 0 | 0.95100083 | 0.6 | 0.27 | 0 | spleen | Zmiz1 |
| B2m2 | 0 | 0.94957853 | 1 | 0.98 | 0 | spleen | B2m |
| Mrpl52 | 0 | 0.94781742 | 0.74 | 0.4 | 0 | spleen | Mrpl52 |
| mt-Co1 | 0 | 0.94712609 | 0.99 | 0.97 | 0 | spleen | mt-Co1 |
| Rps31 | 0 | 0.94644013 | 0.98 | 0.89 | 0 | spleen | Rps3 |
| Cited2 | 0 | 0.9416622 | 0.36 | 0.11 | 0 | spleen | Cited2 |
| Fam181b | 0 | 0.93742406 | 0.41 | 0.11 | 0 | spleen | Fam181b |
| Sumo1 | 0 | 0.93563602 | 0.77 | 0.47 | 0 | spleen | Sumo1 |
| Hint1 | 0 | 0.93530202 | 0.9 | 0.65 | 0 | spleen | Hint1 |
| Proser22 | 0 | 0.93389707 | 0.36 | 0.08 | 0 | spleen | Proser2 |
| Cmtm7 | 0 | 0.92662303 | 0.4 | 0.11 | 0 | spleen | Cmtm7 |
| Maged2 | 0 | 0.91860205 | 0.36 | 0.08 | 0 | spleen | Maged2 |
| Ndn | 0 | 0.91756275 | 0.32 | 0.05 | 0 | spleen | Ndn |
| Cdc42ep5 | 0 | 0.90795048 | 0.34 | 0.08 | 0 | spleen | Cdc42ep5 |
| Pfn1 | 0 | 0.90105047 | 0.96 | 0.82 | 0 | spleen | Pfn1 |
| Ccdc184 | 0 | 0.89206978 | 0.27 | 0.03 | 0 | spleen | Ccdc184 |
| Tcn22 | 0 | 0.8918613 | 0.83 | 0.46 | 0 | spleen | Tcn2 |
| Tspan62 | 0 | 0.88735939 | 0.4 | 0.11 | 0 | spleen | Tspan6 |
| Minos1 | 0 | 0.87995818 | 0.81 | 0.53 | 0 | spleen | Minos1 |
| Rab131 | 0 | 0.87359506 | 0.43 | 0.15 | 0 | spleen | Rab13 |
| Rab14 | 0 | 0.86949593 | 0.77 | 0.49 | 0 | spleen | Rab14 |
| 2700094K13Rik1 | 0 | 0.86331863 | 0.46 | 0.16 | 0 | spleen | 2700094K13Rik |
| P2ry6 | 0 | 0.84926577 | 0.26 | 0.03 | 0 | spleen | P2ry6 |
| Zfhx3 | 0 | 0.84915891 | 0.41 | 0.12 | 0 | spleen | Zfhx3 |
| Rnaset2b2 | 0 | 0.84773716 | 0.66 | 0.35 | 0 | spleen | Rnaset2b |

| | | | | | | | |
|---|---|---|---|---|---|---|---|
| Ndufc1 | 0 | 0.84316492 | 0.72 | 0.41 | 0 | spleen | Ndufc1 |
| Rps3a1 | 0 | 0.84131932 | 0.97 | 0.86 | 0 | spleen | Rps3a1 |
| H2-Q62 | 0 | 0.8410298 | 0.52 | 0.21 | 0 | spleen | H2-Q6 |
| Tpst22 | 0 | 0.83796026 | 0.74 | 0.44 | 0 | spleen | Tpst2 |
| mt-Cytb | 0 | 0.83476343 | 0.99 | 0.96 | 0 | spleen | mt-Cytb |
| Gm2a | 0 | 0.83459929 | 0.35 | 0.11 | 0 | spleen | Gm2a |
| mt-Nd4 | 0 | 0.82751233 | 0.98 | 0.9 | 0 | spleen | mt-Nd4 |
| Gypc | 0 | 0.82745643 | 0.31 | 0.06 | 0 | spleen | Gypc |
| Fstl11 | 0 | 0.82325798 | 0.43 | 0.14 | 0 | spleen | Fstl1 |
| Homer31 | 0 | 0.82220703 | 0.38 | 0.12 | 0 | spleen | Homer3 |
| Flna | 0 | 0.81796063 | 0.39 | 0.13 | 0 | spleen | Flna |
| Gpx11 | 0 | 0.81032998 | 0.8 | 0.53 | 0 | spleen | Gpx1 |
| Cped11 | 0 | 0.80852752 | 0.36 | 0.1 | 0 | spleen | Cped1 |
| Nos1ap | 0 | 0.80628199 | 0.29 | 0.06 | 0 | spleen | Nos1ap |
| Cib1 | 0 | 0.80570518 | 0.43 | 0.17 | 0 | spleen | Cib1 |
| Rnase41 | 0 | 0.80482012 | 0.39 | 0.1 | 0 | spleen | Rnase4 |
| Npl1 | 0 | 0.80233544 | 0.27 | 0.03 | 0 | spleen | Npl |
| Fdps | 0 | 0.800708 | 0.33 | 0.11 | 0 | spleen | Fdps |
| Phpt1 | 0 | 0.79830191 | 0.41 | 0.15 | 0 | spleen | Phpt1 |
| Zfp521 | 0 | 0.79183169 | 0.27 | 0.05 | 0 | spleen | Zfp521 |
| Glo1 | 0 | 0.79057736 | 0.51 | 0.24 | 0 | spleen | Glo1 |
| Fkbp5 | 0 | 0.78980468 | 0.26 | 0.04 | 0 | spleen | Fkbp5 |
| Nme22 | 0 | 0.78971629 | 0.52 | 0.23 | 0 | spleen | Nme2 |
| Lrrfip13 | 0 | 0.7876953 | 0.55 | 0.27 | 0 | spleen | Lrrfip1 |
| mt-Co3 | 0 | 0.77270004 | 0.99 | 0.98 | 0 | spleen | mt-Co3 |
| Sec61b | 0 | 0.77123992 | 0.72 | 0.45 | 0 | spleen | Sec61b |
| Sema3f | 0 | 0.76849715 | 0.48 | 0.2 | 0 | spleen | Sema3f |
| H2afz1 | 0 | 0.76704224 | 0.82 | 0.59 | 0 | spleen | H2afz |
| H2-T22 | 0 | 0.76580683 | 0.62 | 0.33 | 0 | spleen | H2-T22 |
| Cadm11 | 0 | 0.7625047 | 0.41 | 0.13 | 0 | spleen | Cadm1 |
| Pdlim21 | 0 | 0.76244796 | 0.34 | 0.1 | 0 | spleen | Pdlim2 |
| Slf1 | 0 | 0.76223186 | 0.25 | 0.05 | 0 | spleen | Slf1 |
| 1110001J03Rik | 0 | 0.75952878 | 0.46 | 0.19 | 0 | spleen | 1110001J03Rik |
| Ndufb4 | 0 | 0.74793047 | 0.76 | 0.5 | 0 | spleen | Ndufb4 |
| Mxra81 | 0 | 0.7469066 | 0.37 | 0.12 | 0 | spleen | Mxra8 |
| Phlda31 | 0 | 0.74336392 | 0.45 | 0.14 | 0 | spleen | Phlda3 |
| Aes1 | 0 | 0.74332436 | 0.75 | 0.49 | 0 | spleen | Aes |
| PISD2 | 0 | 0.74288552 | 0.52 | 0.23 | 0 | spleen | PISD |
| Dpysl3 | 0 | 0.7415589 | 0.3 | 0.08 | 0 | spleen | Dpysl3 |
| Rpl101 | 0 | 0.73847491 | 0.97 | 0.88 | 0 | spleen | Rpl10 |
| Nrip11 | 0 | 0.73751271 | 0.4 | 0.16 | 0 | spleen | Nrip1 |

| | | | | | | | |
|---|---|---|---|---|---|---|---|
| Mmp14 | 0 | 0.73449891 | 0.26 | 0.05 | 0 | spleen | Mmp14 |
| Dtymk | 0 | 0.73345367 | 0.33 | 0.11 | 0 | spleen | Dtymk |
| Ctnnd1 | 0 | 0.73156561 | 0.48 | 0.22 | 0 | spleen | Ctnnd1 |
| H2-K14 | 0 | 0.73074859 | 0.97 | 0.83 | 0 | spleen | H2-K1 |
| Rpl3 | 0 | 0.72808083 | 0.96 | 0.84 | 0 | spleen | Rpl3 |
| Birc2 | 0 | 0.72680481 | 0.28 | 0.09 | 0 | spleen | Birc2 |
| Cnp | 0 | 0.72347586 | 0.34 | 0.11 | 0 | spleen | Cnp |
| Jak11 | 0 | 0.72242169 | 0.6 | 0.31 | 0 | spleen | Jak1 |
| Vcam11 | 0 | 0.71776154 | 0.48 | 0.12 | 0 | spleen | Vcam1 |
| Rpl34 | 0 | 0.71703068 | 0.98 | 0.9 | 0 | spleen | Rpl34 |
| Rab3il1 | 0 | 0.71309961 | 0.27 | 0.06 | 0 | spleen | Rab3il1 |
| Clec4g1 | 0 | 0.71037572 | 0.61 | 0.11 | 0 | spleen | Clec4g |
| Nudt19 | 0 | 0.71021581 | 0.26 | 0.06 | 0 | spleen | Nudt19 |
| Peak12 | 0 | 0.7059397 | 0.51 | 0.22 | 0 | spleen | Peak1 |
| Stard4 | 0 | 0.70423127 | 0.32 | 0.11 | 0 | spleen | Stard4 |
| Marveld1 | 0 | 0.69987685 | 0.28 | 0.07 | 0 | spleen | Marveld1 |
| Tsc22d12 | 0 | 0.69807269 | 0.93 | 0.64 | 0 | spleen | Tsc22d1 |
| Ccdc107 | 0 | 0.69172381 | 0.32 | 0.1 | 0 | spleen | Ccdc107 |
| Fcgrt2 | 0 | 0.68459808 | 0.78 | 0.44 | 0 | spleen | Fcgrt |
| Fnbp1 | 0 | 0.68045043 | 0.26 | 0.07 | 0 | spleen | Fnbp1 |
| Scn1b3 | 0 | 0.6803598 | 0.4 | 0.14 | 0 | spleen | Scn1b |
| Parp1 | 0 | 0.67494158 | 0.28 | 0.08 | 0 | spleen | Parp1 |
| Cnrip1 | 0 | 0.67009855 | 0.27 | 0.07 | 0 | spleen | Cnrip1 |
| Rps15a | 0 | 0.64792246 | 0.97 | 0.88 | 0 | spleen | Rps15a |
| Chchd2 | 0 | 0.64524585 | 0.95 | 0.83 | 0 | spleen | Chchd2 |
| Cox7c | 0 | 0.64326222 | 0.9 | 0.73 | 0 | spleen | Cox7c |
| Rps27rt3 | 0 | 0.63952458 | 0.98 | 0.83 | 0 | spleen | Rps27rt |
| Clca3a1 | 0 | 0.63747509 | 0.26 | 0.05 | 0 | spleen | Clca3a1 |
| Rpl111 | 0 | 0.63621051 | 0.97 | 0.89 | 0 | spleen | Rpl11 |
| Rps91 | 0 | 0.61353659 | 0.97 | 0.88 | 0 | spleen | Rps9 |
| Gatm2 | 0 | 0.61012526 | 0.5 | 0.14 | 0 | spleen | Gatm |
| Rps13 | 0 | 0.61004688 | 0.96 | 0.84 | 0 | spleen | Rps13 |
| Rps11 | 0 | 0.608437 | 0.98 | 0.91 | 0 | spleen | Rps11 |
| Rpl6 | 0 | 0.60223764 | 0.97 | 0.89 | 0 | spleen | Rpl6 |
| Cox8a | 0 | 0.59843089 | 0.91 | 0.74 | 0 | spleen | Cox8a |
| Rpl371 | 0 | 0.59803044 | 0.99 | 0.95 | 0 | spleen | Rpl37 |
| Ttyh31 | 0 | 0.59702349 | 0.29 | 0.09 | 0 | spleen | Ttyh3 |
| Stab12 | 0 | 0.58831672 | 0.57 | 0.25 | 0 | spleen | Stab1 |
| Rps191 | 0 | 0.58199901 | 0.99 | 0.95 | 0 | spleen | Rps19 |
| Gmpr | 0 | 0.57735615 | 0.25 | 0.07 | 0 | spleen | Gmpr |
| Rps271 | 0 | 0.57448507 | 0.99 | 0.98 | 0 | spleen | Rps27 |

| | | | | | | | |
|---|---|---|---|---|---|---|---|
| Sgpp11 | 0 | 0.5597858 | 0.28 | 0.08 | 0 | spleen | Sgpp1 |
| Rpl35a1 | 0 | 0.54239821 | 0.99 | 0.94 | 0 | spleen | Rpl35a |
| Rpl13a1 | 0 | 0.53377011 | 0.98 | 0.89 | 0 | spleen | Rpl13a |
| Rpl24 | 0 | 0.53315599 | 0.96 | 0.87 | 0 | spleen | Rpl24 |
| Rpl211 | 0 | 0.50668788 | 0.97 | 0.91 | 0 | spleen | Rpl21 |
| Rpl23a1 | 0 | 0.48913851 | 0.98 | 0.9 | 0 | spleen | Rpl23a |
| Rpl91 | 0 | 0.48813314 | 0.97 | 0.9 | 0 | spleen | Rpl9 |
| Rps8 | 0 | 0.48372529 | 0.98 | 0.93 | 0 | spleen | Rps8 |
| mt-Co21 | 0 | 0.48061271 | 0.98 | 0.93 | 0 | spleen | mt-Co2 |
| Rps27a | 0 | 0.46618545 | 0.98 | 0.91 | 0 | spleen | Rps27a |
| Rps231 | 0 | 0.44454903 | 0.99 | 0.94 | 0 | spleen | Rps23 |
| Plvap5 | 0 | 0.44049271 | 0.97 | 0.55 | 0 | spleen | Plvap |
| Wnt21 | 0 | 0.38162414 | 0.31 | 0.07 | 0 | spleen | Wnt2 |
| Rasa42 | 2.30E-305 | 0.50907457 | 0.29 | 0.09 | 4.75E-301 | spleen | Rasa4 |
| Vsig10 | 1.98E-300 | 0.58588922 | 0.26 | 0.08 | 4.09E-296 | spleen | Vsig10 |
| Tnfaip8l1 | 9.71E-300 | 0.6265219 | 0.26 | 0.08 | 2.01E-295 | spleen | Tnfaip8l1 |
| Atpif11 | 4.09E-294 | 0.67427522 | 0.75 | 0.48 | 8.46E-290 | spleen | Atpif1 |
| mt-Nd11 | 2.34E-292 | 0.40498529 | 0.98 | 0.95 | 4.85E-288 | spleen | mt-Nd1 |
| Rps161 | 1.09E-288 | 0.4230576 | 0.98 | 0.92 | 2.25E-284 | spleen | Rps16 |
| Hmgn11 | 2.15E-288 | 0.71810016 | 0.6 | 0.33 | 4.45E-284 | spleen | Hmgn1 |
| Ctsb1 | 3.22E-285 | 0.68716842 | 0.78 | 0.52 | 6.67E-281 | spleen | Ctsb |
| Mpc2 | 2.16E-283 | 0.73388051 | 0.52 | 0.27 | 4.47E-279 | spleen | Mpc2 |
| Rps141 | 6.86E-283 | 0.39896112 | 0.99 | 0.95 | 1.42E-278 | spleen | Rps14 |
| mt-Atp6 | 2.18E-282 | 0.34482564 | 0.99 | 0.99 | 4.51E-278 | spleen | mt-Atp6 |
| Reep31 | 5.94E-281 | 0.71322809 | 0.63 | 0.36 | 1.23E-276 | spleen | Reep3 |
| Rpl171 | 6.78E-280 | 0.45702206 | 0.97 | 0.89 | 1.40E-275 | spleen | Rpl17 |
| Zbtb461 | 1.55E-279 | 0.68329627 | 0.36 | 0.14 | 3.21E-275 | spleen | Zbtb46 |
| Idh1 | 1.19E-278 | 0.60228648 | 0.28 | 0.09 | 2.46E-274 | spleen | Idh1 |
| Serp1 | 3.06E-278 | 0.69344897 | 0.42 | 0.19 | 6.34E-274 | spleen | Serp1 |
| Rps71 | 1.61E-273 | 0.47959355 | 0.96 | 0.85 | 3.33E-269 | spleen | Rps7 |
| Prdx3 | 1.93E-273 | 0.70931108 | 0.37 | 0.15 | 4.00E-269 | spleen | Prdx3 |
| Arsa | 5.00E-273 | 0.6037388 | 0.25 | 0.08 | 1.03E-268 | spleen | Arsa |
| Rpl5 | 1.02E-270 | 0.61160383 | 0.85 | 0.65 | 2.11E-266 | spleen | Rpl5 |
| Rpl37a1 | 1.05E-268 | 0.36824974 | 0.99 | 0.97 | 2.18E-264 | spleen | Rpl37a |
| Ndufa4 | 1.44E-268 | 0.56124696 | 0.89 | 0.73 | 2.99E-264 | spleen | Ndufa4 |
| Cox4i1 | 2.92E-267 | 0.4819903 | 0.94 | 0.83 | 6.04E-263 | spleen | Cox4i1 |
| Abca3 | 9.31E-266 | 0.60724766 | 0.3 | 0.11 | 1.93E-261 | spleen | Abca3 |
| Ccdc88a1 | 1.38E-261 | 0.59931975 | 0.3 | 0.11 | 2.85E-257 | spleen | Ccdc88a |
| Cox5b | 1.77E-261 | 0.6106286 | 0.83 | 0.62 | 3.66E-257 | spleen | Cox5b |
| Prcp1 | 2.56E-261 | 0.63696716 | 0.42 | 0.19 | 5.30E-257 | spleen | Prcp |
| Mrpl38 | 2.74E-260 | 0.61470095 | 0.34 | 0.13 | 5.68E-256 | spleen | Mrpl38 |

| | | | | | | | |
|---|---|---|---|---|---|---|---|
| Rps51 | 8.91E-260 | 0.41723737 | 0.97 | 0.91 | 1.84E-255 | spleen | Rps5 |
| Rpl10a | 2.85E-259 | 0.54950992 | 0.9 | 0.73 | 5.89E-255 | spleen | Rpl10a |
| Cox20 | 4.04E-258 | 0.64137405 | 0.43 | 0.19 | 8.35E-254 | spleen | Cox20 |
| Rfk1 | 2.54E-257 | 0.62508048 | 0.42 | 0.19 | 5.27E-253 | spleen | Rfk |
| Ifitm22 | 1.39E-256 | 0.3697097 | 0.98 | 0.93 | 2.88E-252 | spleen | Ifitm2 |
| Rpl36a | 1.69E-256 | 0.49254222 | 0.94 | 0.8 | 3.51E-252 | spleen | Rpl36a |
| Tecr | 7.52E-256 | 0.69945318 | 0.53 | 0.28 | 1.56E-251 | spleen | Tecr |
| Mcee | 3.93E-255 | 0.64743895 | 0.37 | 0.15 | 8.12E-251 | spleen | Mcee |
| Polr3gl | 4.98E-253 | 0.62950752 | 0.39 | 0.17 | 1.03E-248 | spleen | Polr3gl |
| Ndufb5 | 7.55E-253 | 0.66909155 | 0.65 | 0.4 | 1.56E-248 | spleen | Ndufb5 |
| Clta1 | 3.19E-252 | 0.56137389 | 0.79 | 0.56 | 6.60E-248 | spleen | Clta |
| Sdpr5 | 1.58E-246 | 0.46327184 | 0.93 | 0.69 | 3.26E-242 | spleen | Sdpr |
| Frmd4b1 | 6.50E-245 | 0.62029266 | 0.39 | 0.17 | 1.35E-240 | spleen | Frmd4b |
| Rps4x1 | 8.70E-245 | 0.43121466 | 0.97 | 0.89 | 1.80E-240 | spleen | Rps4x |
| Eln | 3.28E-242 | 0.48964604 | 0.42 | 0.19 | 6.79E-238 | spleen | Eln |
| Edf1 | 1.60E-241 | 0.60510793 | 0.81 | 0.61 | 3.31E-237 | spleen | Edf1 |
| Yes11 | 8.37E-241 | 0.63139298 | 0.42 | 0.19 | 1.73E-236 | spleen | Yes1 |
| Atp5j21 | 4.17E-238 | 0.5294697 | 0.87 | 0.7 | 8.63E-234 | spleen | Atp5j2 |
| Zeb21 | 5.02E-238 | 0.48972511 | 0.26 | 0.09 | 1.04E-233 | spleen | Zeb2 |
| Rps281 | 1.35E-237 | 0.38409629 | 0.99 | 0.92 | 2.79E-233 | spleen | Rps28 |
| Slbp1 | 1.86E-237 | 0.63983138 | 0.39 | 0.17 | 3.84E-233 | spleen | Slbp |
| Relb | 2.36E-237 | 0.57433365 | 0.28 | 0.1 | 4.88E-233 | spleen | Relb |
| Snx51 | 1.62E-236 | 0.58856167 | 0.4 | 0.18 | 3.35E-232 | spleen | Snx5 |
| Gnb2l11 | 2.42E-236 | 0.5566789 | 0.8 | 0.55 | 5.00E-232 | spleen | Gnb2l1 |
| Rpl391 | 1.21E-235 | 0.42468513 | 0.96 | 0.87 | 2.50E-231 | spleen | Rpl39 |
| Rps181 | 9.95E-234 | 0.37201086 | 0.98 | 0.92 | 2.06E-229 | spleen | Rps18 |
| Psen21 | 1.28E-233 | 0.50915009 | 0.28 | 0.1 | 2.65E-229 | spleen | Psen2 |
| Pgam1 | 5.72E-231 | 0.66786594 | 0.53 | 0.29 | 1.18E-226 | spleen | Pgam1 |
| Romo1 | 9.25E-231 | 0.60454409 | 0.67 | 0.43 | 1.91E-226 | spleen | Romo1 |
| 2010107E04Rik1 | 1.18E-230 | 0.58199201 | 0.7 | 0.45 | 2.44E-226 | spleen | 2010107E04Rik |
| Dnajc15 | 5.76E-229 | 0.56250922 | 0.3 | 0.11 | 1.19E-224 | spleen | Dnajc15 |
| Gbas | 9.46E-227 | 0.53866935 | 0.26 | 0.09 | 1.96E-222 | spleen | Gbas |
| Abca1 | 3.23E-226 | 0.53285873 | 0.25 | 0.09 | 6.68E-222 | spleen | Abca1 |
| Nop10 | 1.02E-225 | 0.60758633 | 0.69 | 0.45 | 2.10E-221 | spleen | Nop10 |
| Anapc11 | 2.51E-225 | 0.6173463 | 0.64 | 0.4 | 5.19E-221 | spleen | Anapc11 |
| 6430548M08Rik | 8.40E-223 | 0.50354732 | 0.26 | 0.1 | 1.74E-218 | spleen | 6430548M08Rik |
| Cyb5a2 | 1.34E-221 | 0.52682456 | 0.83 | 0.62 | 2.78E-217 | spleen | Cyb5a |
| Rps171 | 3.03E-221 | 0.40979443 | 0.96 | 0.88 | 6.27E-217 | spleen | Rps17 |
| Atp5l | 1.11E-219 | 0.50549765 | 0.88 | 0.72 | 2.30E-215 | spleen | Atp5l |
| Pcdh121 | 7.82E-218 | 0.54700801 | 0.33 | 0.14 | 1.62E-213 | spleen | Pcdh12 |
| Ttc281 | 3.16E-217 | 0.59920301 | 0.4 | 0.19 | 6.53E-213 | spleen | Ttc28 |

| | | | | | | | |
|---|---|---|---|---|---|---|---|
| Rpl311 | 1.23E-216 | 0.3981985 | 0.96 | 0.87 | 2.54E-212 | spleen | Rpl31 |
| Lmo4 | 1.50E-215 | 0.61265686 | 0.35 | 0.16 | 3.11E-211 | spleen | Lmo4 |
| Rpl7 | 1.98E-215 | 0.43356543 | 0.95 | 0.85 | 4.09E-211 | spleen | Rpl7 |
| Cox6a11 | 3.09E-215 | 0.49037536 | 0.9 | 0.76 | 6.40E-211 | spleen | Cox6a1 |
| Ndufa6 | 9.26E-210 | 0.55766804 | 0.79 | 0.58 | 1.92E-205 | spleen | Ndufa6 |
| Rpl22 | 2.02E-209 | 0.49695637 | 0.88 | 0.72 | 4.18E-205 | spleen | Rpl22 |
| Pcdh172 | 4.29E-209 | 0.56546145 | 0.47 | 0.24 | 8.87E-205 | spleen | Pcdh17 |
| Rpl18 | 7.43E-208 | 0.42110313 | 0.95 | 0.84 | 1.54E-203 | spleen | Rpl18 |
| Cmtm32 | 1.69E-207 | 0.5428004 | 0.44 | 0.22 | 3.50E-203 | spleen | Cmtm3 |
| Fundc21 | 4.75E-207 | 0.5966424 | 0.5 | 0.27 | 9.83E-203 | spleen | Fundc2 |
| Med10 | 8.89E-207 | 0.62721465 | 0.43 | 0.22 | 1.84E-202 | spleen | Med10 |
| Slc39a11 | 1.15E-204 | 0.84257803 | 0.37 | 0.17 | 2.39E-200 | spleen | Slc39a1 |
| Rpl15 | 7.44E-204 | 0.44109995 | 0.93 | 0.82 | 1.54E-199 | spleen | Rpl15 |
| Trmt112 | 1.16E-202 | 0.56215595 | 0.48 | 0.26 | 2.40E-198 | spleen | Trmt112 |
| Jmjd1c1 | 9.99E-201 | 0.5733572 | 0.38 | 0.18 | 2.07E-196 | spleen | Jmjd1c |
| Ndufb6 | 6.01E-200 | 0.57972352 | 0.55 | 0.32 | 1.24E-195 | spleen | Ndufb6 |
| Brk1 | 5.30E-199 | 0.55996397 | 0.7 | 0.48 | 1.10E-194 | spleen | Brk1 |
| Rps27l | 8.90E-199 | 0.5305528 | 0.78 | 0.58 | 1.84E-194 | spleen | Rps27l |
| Prdx2 | 1.28E-197 | 0.58093304 | 0.68 | 0.46 | 2.66E-193 | spleen | Prdx2 |
| Hoxa51 | 3.05E-197 | 0.42527528 | 0.25 | 0.09 | 6.30E-193 | spleen | Hoxa5 |
| Rpl27a | 9.49E-196 | 0.36378203 | 0.97 | 0.91 | 1.96E-191 | spleen | Rpl27a |
| Cnpy21 | 3.91E-195 | 0.5771589 | 0.48 | 0.26 | 8.10E-191 | spleen | Cnpy2 |
| Rps101 | 1.45E-193 | 0.43672822 | 0.92 | 0.8 | 3.01E-189 | spleen | Rps10 |
| Cox6b1 | 3.29E-193 | 0.4645146 | 0.88 | 0.73 | 6.80E-189 | spleen | Cox6b1 |
| Atp5g2 | 7.30E-193 | 0.53337176 | 0.77 | 0.56 | 1.51E-188 | spleen | Atp5g2 |
| Shfm11 | 6.97E-191 | 0.45716441 | 0.87 | 0.66 | 1.44E-186 | spleen | Shfm1 |
| Bloc1s12 | 9.33E-191 | 0.5042653 | 0.53 | 0.3 | 1.93E-186 | spleen | Bloc1s1 |
| Rpl36al | 2.19E-190 | 0.50921154 | 0.77 | 0.56 | 4.53E-186 | spleen | Rpl36al |
| Mtfr1l | 3.01E-189 | 0.50469144 | 0.33 | 0.15 | 6.22E-185 | spleen | Mtfr1l |
| Gm269172 | 3.77E-182 | 0.36531722 | 0.51 | 0.27 | 7.81E-178 | spleen | Gm26917 |
| Odc11 | 5.32E-182 | 0.51604282 | 0.26 | 0.1 | 1.10E-177 | spleen | Odc1 |
| Eef1g1 | 5.38E-181 | 0.4841379 | 0.82 | 0.62 | 1.11E-176 | spleen | Eef1g |
| Agfg12 | 1.52E-179 | 0.51839873 | 0.33 | 0.15 | 3.15E-175 | spleen | Agfg1 |
| mt-Nd21 | 1.65E-179 | 0.41200475 | 0.93 | 0.82 | 3.41E-175 | spleen | mt-Nd2 |
| Ly961 | 2.12E-179 | 0.48584016 | 0.34 | 0.16 | 4.39E-175 | spleen | Ly96 |
| Pigp | 2.27E-179 | 0.53078724 | 0.37 | 0.18 | 4.70E-175 | spleen | Pigp |
| Rps22 | 1.17E-178 | 0.39094798 | 0.94 | 0.84 | 2.43E-174 | spleen | Rps2 |
| Pald1 | 2.24E-178 | 0.45345615 | 0.26 | 0.1 | 4.63E-174 | spleen | Pald1 |
| Rpl131 | 1.10E-177 | 0.31532548 | 0.98 | 0.95 | 2.28E-173 | spleen | Rpl13 |
| Rps261 | 1.75E-177 | 0.41381254 | 0.91 | 0.8 | 3.63E-173 | spleen | Rps26 |
| Fkbp1a2 | 2.02E-177 | 0.37651235 | 0.94 | 0.82 | 4.17E-173 | spleen | Fkbp1a |

| | | | | | | | |
|---|---|---|---|---|---|---|---|
| Wdr894 | 3.22E-177 | 0.41037526 | 0.9 | 0.71 | 6.66E-173 | spleen | Wdr89 |
| Abhd17b1 | 2.17E-176 | 0.50864517 | 0.32 | 0.14 | 4.50E-172 | spleen | Abhd17b |
| Arrb1 | 9.19E-176 | 0.45322547 | 0.28 | 0.12 | 1.90E-171 | spleen | Arrb1 |
| Rab11a | 9.60E-176 | 0.50103319 | 0.82 | 0.64 | 1.99E-171 | spleen | Rab11a |
| Smarca22 | 1.80E-174 | 0.5451283 | 0.46 | 0.25 | 3.72E-170 | spleen | Smarca2 |
| Mgat4b | 2.87E-174 | 0.48210324 | 0.3 | 0.13 | 5.94E-170 | spleen | Mgat4b |
| Pigyl | 3.62E-173 | 0.49704529 | 0.33 | 0.15 | 7.48E-169 | spleen | Pigyl |
| mt-Nd51 | 9.78E-173 | 0.52446735 | 0.57 | 0.34 | 2.02E-168 | spleen | mt-Nd5 |
| Rpl22l1 | 1.50E-172 | 0.51017844 | 0.76 | 0.55 | 3.10E-168 | spleen | Rpl22l1 |
| Ube2e1 | 3.44E-172 | 0.55418056 | 0.41 | 0.22 | 7.12E-168 | spleen | Ube2e1 |
| Rpl231 | 1.43E-171 | 0.33950669 | 0.97 | 0.9 | 2.96E-167 | spleen | Rpl23 |
| Rcn11 | 2.03E-171 | 0.42306138 | 0.36 | 0.17 | 4.21E-167 | spleen | Rcn1 |
| Rps62 | 9.44E-170 | 0.34006919 | 0.96 | 0.89 | 1.95E-165 | spleen | Rps6 |
| Cox14 | 8.53E-169 | 0.55803968 | 0.54 | 0.33 | 1.77E-164 | spleen | Cox14 |
| Anapc13 | 5.14E-168 | 0.50347809 | 0.47 | 0.26 | 1.06E-163 | spleen | Anapc13 |
| Rpl261 | 1.43E-167 | 0.32372755 | 0.97 | 0.92 | 2.96E-163 | spleen | Rpl26 |
| Grcc101 | 1.60E-167 | 0.4887238 | 0.62 | 0.39 | 3.31E-163 | spleen | Grcc10 |
| Naca | 9.68E-166 | 0.44597382 | 0.88 | 0.74 | 2.00E-161 | spleen | Naca |
| Rpl141 | 1.36E-165 | 0.35614613 | 0.95 | 0.89 | 2.82E-161 | spleen | Rpl14 |
| Cox171 | 6.14E-164 | 0.47862598 | 0.68 | 0.46 | 1.27E-159 | spleen | Cox17 |
| Rpl7a | 1.18E-162 | 0.43698505 | 0.87 | 0.7 | 2.43E-158 | spleen | Rpl7a |
| Uqcrh | 1.72E-162 | 0.45152485 | 0.83 | 0.64 | 3.56E-158 | spleen | Uqcrh |
| Fam173a | 6.20E-160 | 0.49266626 | 0.29 | 0.13 | 1.28E-155 | spleen | Fam173a |
| Serf21 | 4.17E-157 | 0.34612861 | 0.93 | 0.84 | 8.63E-153 | spleen | Serf2 |
| Rpl19 | 2.30E-156 | 0.31055464 | 0.97 | 0.91 | 4.75E-152 | spleen | Rpl19 |
| 0610007P14Rik | 3.38E-156 | 0.41953946 | 0.26 | 0.11 | 6.99E-152 | spleen | 0610007P14Rik |
| Zfyve21 | 1.02E-155 | 0.43765622 | 0.26 | 0.11 | 2.11E-151 | spleen | Zfyve21 |
| Tln12 | 1.18E-155 | 0.50685746 | 0.5 | 0.3 | 2.45E-151 | spleen | Tln1 |
| Snrpf | 1.65E-155 | 0.50233363 | 0.54 | 0.34 | 3.42E-151 | spleen | Snrpf |
| Tmem14c | 8.07E-155 | 0.51516473 | 0.59 | 0.39 | 1.67E-150 | spleen | Tmem14c |
| Rpl412 | 1.61E-154 | 0.25088814 | 0.99 | 0.96 | 3.33E-150 | spleen | Rpl41 |
| Uqcr11 | 6.55E-153 | 0.42598217 | 0.82 | 0.65 | 1.36E-148 | spleen | Uqcr11 |
| Tomm7 | 1.60E-151 | 0.41815146 | 0.79 | 0.61 | 3.31E-147 | spleen | Tomm7 |
| Myeov21 | 2.22E-151 | 0.43249382 | 0.6 | 0.38 | 4.59E-147 | spleen | Myeov2 |
| Zfp36l14 | 3.81E-151 | 0.45016888 | 0.72 | 0.52 | 7.89E-147 | spleen | Zfp36l1 |
| Mrps36 | 2.57E-148 | 0.47045611 | 0.36 | 0.19 | 5.31E-144 | spleen | Mrps36 |
| Sec61g | 8.24E-148 | 0.43368161 | 0.79 | 0.61 | 1.70E-143 | spleen | Sec61g |
| Mrps12 | 1.80E-146 | 0.47850126 | 0.38 | 0.2 | 3.72E-142 | spleen | Mrps12 |
| Mycbp21 | 4.65E-146 | 0.48551051 | 0.39 | 0.21 | 9.62E-142 | spleen | Mycbp2 |
| Pfdn51 | 5.78E-146 | 0.4001669 | 0.88 | 0.74 | 1.20E-141 | spleen | Pfdn5 |
| Kcnq1ot1 | 2.92E-145 | 0.50979983 | 0.32 | 0.16 | 6.05E-141 | spleen | Kcnq1ot1 |

| | | | | | | | |
|---|---|---|---|---|---|---|---|
| Fam171a1 | 3.79E-144 | 0.43323288 | 0.33 | 0.17 | 7.84E-140 | spleen | Fam171a1 |
| Prnp2 | 4.16E-144 | 0.62698893 | 0.35 | 0.18 | 8.62E-140 | spleen | Prnp |
| Pgap2 | 4.25E-144 | 0.44649008 | 0.27 | 0.12 | 8.80E-140 | spleen | Pgap2 |
| Park7 | 5.34E-144 | 0.45615111 | 0.7 | 0.5 | 1.10E-139 | spleen | Park7 |
| Arpc5l | 5.45E-142 | 0.49241763 | 0.41 | 0.23 | 1.13E-137 | spleen | Arpc5l |
| Pomp1 | 2.34E-141 | 0.43546797 | 0.75 | 0.55 | 4.83E-137 | spleen | Pomp |
| Ndufb9 | 2.81E-141 | 0.45898884 | 0.68 | 0.49 | 5.82E-137 | spleen | Ndufb9 |
| Rps201 | 3.00E-141 | 0.32612466 | 0.95 | 0.86 | 6.21E-137 | spleen | Rps20 |
| Tpd52l2 | 6.10E-141 | 0.48615056 | 0.41 | 0.23 | 1.26E-136 | spleen | Tpd52l2 |
| Ndufb2 | 3.29E-140 | 0.44535 | 0.55 | 0.35 | 6.80E-136 | spleen | Ndufb2 |
| Wsb2 | 5.84E-140 | 0.42536583 | 0.27 | 0.13 | 1.21E-135 | spleen | Wsb2 |
| Rps151 | 1.51E-138 | 0.29936458 | 0.97 | 0.9 | 3.13E-134 | spleen | Rps15 |
| Atp5g1 | 2.10E-137 | 0.44305573 | 0.64 | 0.44 | 4.35E-133 | spleen | Atp5g1 |
| Rpl281 | 7.40E-137 | 0.34206079 | 0.94 | 0.84 | 1.53E-132 | spleen | Rpl28 |
| Rnaset2a1 | 3.61E-136 | 0.47435683 | 0.52 | 0.33 | 7.48E-132 | spleen | Rnaset2a |
| Ccdc531 | 6.80E-136 | 0.40110429 | 0.28 | 0.13 | 1.41E-131 | spleen | Ccdc53 |
| Rpl301 | 1.63E-135 | 0.35494332 | 0.91 | 0.76 | 3.38E-131 | spleen | Rpl30 |
| mt-Nd31 | 1.79E-134 | 0.4348302 | 0.61 | 0.41 | 3.70E-130 | spleen | mt-Nd3 |
| Scand11 | 2.63E-134 | 0.45004952 | 0.49 | 0.3 | 5.43E-130 | spleen | Scand1 |
| Golgb1 | 3.41E-134 | 0.46155845 | 0.41 | 0.23 | 7.06E-130 | spleen | Golgb1 |
| Hmcn13 | 5.12E-134 | 0.31007993 | 0.38 | 0.2 | 1.06E-129 | spleen | Hmcn1 |
| Ogt | 6.05E-134 | 0.46654299 | 0.43 | 0.24 | 1.25E-129 | spleen | Ogt |
| Nudt43 | 2.64E-133 | 0.41167892 | 0.34 | 0.18 | 5.45E-129 | spleen | Nudt4 |
| Fam162a | 1.16E-132 | 0.45763834 | 0.37 | 0.2 | 2.40E-128 | spleen | Fam162a |
| Slirp | 6.96E-132 | 0.44158495 | 0.48 | 0.29 | 1.44E-127 | spleen | Slirp |
| Nenf1 | 9.71E-131 | 0.44969095 | 0.6 | 0.4 | 2.01E-126 | spleen | Nenf |
| Rpl272 | 1.76E-130 | 0.32842153 | 0.93 | 0.81 | 3.64E-126 | spleen | Rpl27 |
| Aldh22 | 1.34E-128 | 0.38085692 | 0.77 | 0.58 | 2.77E-124 | spleen | Aldh2 |
| Ifngr21 | 1.62E-128 | 0.42378764 | 0.32 | 0.16 | 3.36E-124 | spleen | Ifngr2 |
| Eid11 | 6.84E-128 | 0.46442379 | 0.37 | 0.21 | 1.41E-123 | spleen | Eid1 |
| Cd1641 | 4.59E-126 | 0.42739022 | 0.37 | 0.2 | 9.50E-122 | spleen | Cd164 |
| Fkbp21 | 6.61E-126 | 0.47069478 | 0.42 | 0.25 | 1.37E-121 | spleen | Fkbp2 |
| Capzb | 9.02E-126 | 0.43364334 | 0.72 | 0.54 | 1.87E-121 | spleen | Capzb |
| Mrps16 | 3.56E-125 | 0.43866199 | 0.39 | 0.22 | 7.36E-121 | spleen | Mrps16 |
| Sod11 | 1.23E-124 | 0.41868921 | 0.61 | 0.42 | 2.54E-120 | spleen | Sod1 |
| Actb2 | 4.10E-124 | 0.56425674 | 0.98 | 0.95 | 8.48E-120 | spleen | Actb |
| Aurkaip1 | 5.63E-124 | 0.44193077 | 0.5 | 0.32 | 1.16E-119 | spleen | Aurkaip1 |
| Slc38a21 | 6.28E-124 | 0.43440492 | 0.51 | 0.32 | 1.30E-119 | spleen | Slc38a2 |
| Ppp2r5a1 | 9.44E-123 | 0.42414438 | 0.36 | 0.2 | 1.95E-118 | spleen | Ppp2r5a |
| Rnf13 | 2.14E-122 | 0.41822779 | 0.36 | 0.2 | 4.43E-118 | spleen | Rnf13 |
| Atp5k1 | 2.88E-122 | 0.38843543 | 0.72 | 0.54 | 5.97E-118 | spleen | Atp5k |

| | | | | | | | |
|---|---|---|---|---|---|---|---|
| Lamtor2 | 2.97E-121 | 0.43119829 | 0.53 | 0.35 | 6.14E-117 | spleen | Lamtor2 |
| Dynll2 | 3.82E-121 | 0.41954514 | 0.35 | 0.19 | 7.89E-117 | spleen | Dynll2 |
| Ndufa31 | 4.32E-121 | 0.36063774 | 0.78 | 0.59 | 8.94E-117 | spleen | Ndufa3 |
| Ccnt2 | 1.37E-120 | 0.39241133 | 0.32 | 0.16 | 2.84E-116 | spleen | Ccnt2 |
| Ifi272 | 2.77E-120 | 0.41066006 | 0.7 | 0.51 | 5.74E-116 | spleen | Ifi27 |
| Eef1d | 3.27E-120 | 0.43427891 | 0.65 | 0.46 | 6.78E-116 | spleen | Eef1d |
| Rpl362 | 7.96E-120 | 0.27321753 | 0.96 | 0.9 | 1.65E-115 | spleen | Rpl36 |
| Atp5e | 3.10E-119 | 0.35318817 | 0.83 | 0.68 | 6.42E-115 | spleen | Atp5e |
| Tbca | 2.76E-118 | 0.39827745 | 0.68 | 0.49 | 5.71E-114 | spleen | Tbca |
| Ndufb8 | 6.61E-118 | 0.41978631 | 0.64 | 0.46 | 1.37E-113 | spleen | Ndufb8 |
| Atraid | 2.94E-117 | 0.44591459 | 0.57 | 0.39 | 6.09E-113 | spleen | Atraid |
| Bola3 | 9.05E-117 | 0.41076047 | 0.37 | 0.21 | 1.87E-112 | spleen | Bola3 |
| Rps211 | 1.12E-116 | 0.28526576 | 0.95 | 0.85 | 2.33E-112 | spleen | Rps21 |
| Jam32 | 1.93E-116 | 0.41158471 | 0.39 | 0.21 | 3.99E-112 | spleen | Jam3 |
| Churc1 | 8.96E-116 | 0.40335942 | 0.44 | 0.26 | 1.85E-111 | spleen | Churc1 |
| Rpl42 | 1.98E-115 | 0.31282553 | 0.93 | 0.8 | 4.10E-111 | spleen | Rpl4 |
| Gpx41 | 4.83E-115 | 0.34423412 | 0.89 | 0.76 | 1.00E-110 | spleen | Gpx4 |
| Swi5 | 7.45E-115 | 0.40865045 | 0.64 | 0.46 | 1.54E-110 | spleen | Swi5 |
| Uqcrq | 1.94E-114 | 0.36673366 | 0.72 | 0.55 | 4.01E-110 | spleen | Uqcrq |
| Ngfrap1 | 6.10E-114 | 0.36101397 | 0.27 | 0.13 | 1.26E-109 | spleen | Ngfrap1 |
| Ttc31 | 1.98E-113 | 0.40821274 | 0.31 | 0.17 | 4.09E-109 | spleen | Ttc3 |
| Echs1 | 3.58E-113 | 0.41428589 | 0.31 | 0.16 | 7.41E-109 | spleen | Echs1 |
| Sox4 | 4.22E-113 | 0.36102252 | 0.45 | 0.27 | 8.74E-109 | spleen | Sox4 |
| Prkaca | 5.31E-113 | 0.3782799 | 0.28 | 0.14 | 1.10E-108 | spleen | Prkaca |
| Hif1a1 | 5.70E-113 | 0.38313237 | 0.29 | 0.15 | 1.18E-108 | spleen | Hif1a |
| Mmrn23 | 6.58E-113 | 0.37598697 | 0.69 | 0.5 | 1.36E-108 | spleen | Mmrn2 |
| Fam195b | 8.52E-112 | 0.40389193 | 0.45 | 0.27 | 1.76E-107 | spleen | Fam195b |
| Tomm5 | 6.39E-111 | 0.39279906 | 0.41 | 0.24 | 1.32E-106 | spleen | Tomm5 |
| Pdrg1 | 1.16E-110 | 0.38015868 | 0.4 | 0.23 | 2.41E-106 | spleen | Pdrg1 |
| Rpl121 | 1.90E-110 | 0.36341432 | 0.79 | 0.61 | 3.93E-106 | spleen | Rpl12 |
| Eif4a21 | 3.84E-110 | 0.37730784 | 0.75 | 0.56 | 7.95E-106 | spleen | Eif4a2 |
| Rpl81 | 1.69E-109 | 0.27352332 | 0.96 | 0.9 | 3.50E-105 | spleen | Rpl8 |
| Smim11 | 2.61E-109 | 0.4052288 | 0.5 | 0.32 | 5.41E-105 | spleen | Smim11 |
| Pet1001 | 2.85E-109 | 0.38877668 | 0.43 | 0.27 | 5.91E-105 | spleen | Pet100 |
| Swap701 | 6.47E-109 | 0.40586301 | 0.32 | 0.17 | 1.34E-104 | spleen | Swap70 |
| Snrpg1 | 1.59E-108 | 0.33776488 | 0.75 | 0.56 | 3.30E-104 | spleen | Snrpg |
| Hmgb11 | 2.74E-108 | 0.3039566 | 0.91 | 0.82 | 5.66E-104 | spleen | Hmgb1 |
| Cux11 | 3.02E-108 | 0.40517273 | 0.29 | 0.15 | 6.24E-104 | spleen | Cux1 |
| Lsm5 | 7.34E-108 | 0.3864926 | 0.41 | 0.25 | 1.52E-103 | spleen | Lsm5 |
| Psmb91 | 2.85E-107 | 0.41278106 | 0.52 | 0.34 | 5.89E-103 | spleen | Psmb9 |
| Ndufb3 | 3.45E-107 | 0.4078929 | 0.54 | 0.36 | 7.13E-103 | spleen | Ndufb3 |

| | | | | | | | |
|---|---|---|---|---|---|---|---|
| Msrb1 | 1.27E-106 | 0.39463438 | 0.26 | 0.13 | 2.63E-102 | spleen | Msrb1 |
| Snrpd2 | 1.75E-106 | 0.37673568 | 0.58 | 0.4 | 3.62E-102 | spleen | Snrpd2 |
| Srp141 | 3.64E-106 | 0.38594414 | 0.67 | 0.5 | 7.53E-102 | spleen | Srp14 |
| Amotl12 | 3.87E-106 | 0.38547693 | 0.41 | 0.25 | 8.01E-102 | spleen | Amotl1 |
| Cpne21 | 6.48E-106 | 0.31900062 | 0.29 | 0.15 | 1.34E-101 | spleen | Cpne2 |
| 1810058I24Rik | 8.46E-106 | 0.46901303 | 0.42 | 0.26 | 1.75E-101 | spleen | 1810058I24Rik |
| Maoa1 | 1.13E-105 | 0.31804927 | 0.31 | 0.17 | 2.33E-101 | spleen | Maoa |
| 2410015M20Rik | 3.80E-104 | 0.3865181 | 0.52 | 0.35 | 7.87E-100 | spleen | 2410015M20Rik |
| Ost4 | 2.54E-103 | 0.35628887 | 0.7 | 0.52 | 5.26E-99 | spleen | Ost4 |
| Zeb11 | 2.32E-102 | 0.33530459 | 0.35 | 0.2 | 4.80E-98 | spleen | Zeb1 |
| Uba524 | 8.19E-102 | 0.26146842 | 0.87 | 0.71 | 1.69E-97 | spleen | Uba52 |
| Snrpe | 9.24E-102 | 0.3809954 | 0.54 | 0.37 | 1.91E-97 | spleen | Snrpe |
| Degs12 | 6.59E-101 | 0.30995364 | 0.57 | 0.38 | 1.36E-96 | spleen | Degs1 |
| Nagk | 7.42E-101 | 0.34612354 | 0.28 | 0.15 | 1.54E-96 | spleen | Nagk |
| Skap21 | 1.12E-100 | 0.34979559 | 0.31 | 0.17 | 2.32E-96 | spleen | Skap2 |
| Rps122 | 2.58E-100 | 0.2846461 | 0.92 | 0.82 | 5.34E-96 | spleen | Rps12 |
| Cnn22 | 4.64E-100 | 0.38485389 | 0.45 | 0.28 | 9.61E-96 | spleen | Cnn2 |
| Tia1 | 1.07E-99 | 0.35579351 | 0.28 | 0.15 | 2.21E-95 | spleen | Tia1 |
| Tns12 | 2.43E-99 | 0.34379047 | 0.55 | 0.37 | 5.04E-95 | spleen | Tns1 |
| Reep5 | 6.82E-99 | 0.38479076 | 0.59 | 0.42 | 1.41E-94 | spleen | Reep5 |
| Tmem147 | 1.38E-98 | 0.3799314 | 0.42 | 0.27 | 2.87E-94 | spleen | Tmem147 |
| Cgnl11 | 2.82E-98 | 0.34243695 | 0.25 | 0.13 | 5.83E-94 | spleen | Cgnl1 |
| Psmb7 | 5.16E-98 | 0.44577668 | 0.48 | 0.32 | 1.07E-93 | spleen | Psmb7 |
| Mrps21 | 1.22E-97 | 0.36404664 | 0.57 | 0.4 | 2.53E-93 | spleen | Mrps21 |
| Mrpl54 | 1.33E-97 | 0.35636983 | 0.33 | 0.19 | 2.76E-93 | spleen | Mrpl54 |
| Pura2 | 3.74E-97 | 0.39087416 | 0.49 | 0.32 | 7.74E-93 | spleen | Pura |
| Hspe12 | 3.84E-97 | 0.35361919 | 0.52 | 0.35 | 7.95E-93 | spleen | Hspe1 |
| Serinc33 | 1.04E-96 | 0.27647045 | 0.93 | 0.84 | 2.15E-92 | spleen | Serinc3 |
| Agpat41 | 5.03E-96 | 0.36075916 | 0.34 | 0.19 | 1.04E-91 | spleen | Agpat4 |
| Arhgdib1 | 7.41E-96 | 0.31847283 | 0.32 | 0.17 | 1.53E-91 | spleen | Arhgdib |
| Fam53b | 1.34E-95 | 0.32436118 | 0.31 | 0.17 | 2.77E-91 | spleen | Fam53b |
| Abi31 | 4.12E-95 | 0.34167435 | 0.46 | 0.29 | 8.52E-91 | spleen | Abi3 |
| St3gal42 | 3.74E-94 | 0.37369312 | 0.35 | 0.21 | 7.74E-90 | spleen | St3gal4 |
| 1110008F13Rik1 | 4.00E-94 | 0.43719939 | 0.43 | 0.28 | 8.29E-90 | spleen | 1110008F13Rik |
| Eif3e | 7.97E-94 | 0.39822115 | 0.59 | 0.43 | 1.65E-89 | spleen | Eif3e |
| Ift43 | 3.45E-92 | 0.34217772 | 0.25 | 0.13 | 7.13E-88 | spleen | Ift43 |
| Dab22 | 7.59E-92 | 0.263538 | 0.43 | 0.26 | 1.57E-87 | spleen | Dab2 |
| Glrx51 | 3.85E-91 | 0.37121876 | 0.35 | 0.21 | 7.96E-87 | spleen | Glrx5 |
| 2700060E02Rik | 2.37E-90 | 0.34318601 | 0.57 | 0.39 | 4.91E-86 | spleen | 2700060E02Rik |
| Imp31 | 3.13E-90 | 0.38448074 | 0.43 | 0.28 | 6.47E-86 | spleen | Imp3 |
| Adprh | 3.47E-90 | 0.33217082 | 0.32 | 0.18 | 7.17E-86 | spleen | Adprh |

| | | | | | | | |
|---|---|---|---|---|---|---|---|
| Myh92 | 6.71E-90 | 0.34967579 | 0.47 | 0.3 | 1.39E-85 | spleen | Myh9 |
| Cox7b | 7.60E-89 | 0.35685183 | 0.59 | 0.42 | 1.57E-84 | spleen | Cox7b |
| Pdcd41 | 1.67E-88 | 0.35046343 | 0.25 | 0.13 | 3.46E-84 | spleen | Pdcd4 |
| Naa381 | 4.54E-88 | 0.34313667 | 0.43 | 0.28 | 9.39E-84 | spleen | Naa38 |
| Sdhaf4 | 7.01E-88 | 0.37401765 | 0.27 | 0.15 | 1.45E-83 | spleen | Sdhaf4 |
| Jtb | 1.47E-87 | 0.34821262 | 0.34 | 0.21 | 3.04E-83 | spleen | Jtb |
| Ndufa11 | 6.58E-87 | 0.34636334 | 0.58 | 0.42 | 1.36E-82 | spleen | Ndufa11 |
| Arl3 | 2.91E-86 | 0.3300009 | 0.31 | 0.18 | 6.01E-82 | spleen | Arl3 |
| Pam | 5.25E-86 | 0.35130288 | 0.38 | 0.24 | 1.09E-81 | spleen | Pam |
| Tgfb11 | 1.47E-85 | 0.32214149 | 0.6 | 0.43 | 3.05E-81 | spleen | Tgfb1 |
| Ndufa1 | 1.84E-84 | 0.31657164 | 0.6 | 0.44 | 3.81E-80 | spleen | Ndufa1 |
| Btf3 | 2.78E-84 | 0.3087817 | 0.84 | 0.7 | 5.75E-80 | spleen | Btf3 |
| Lamtor4 | 8.86E-84 | 0.35101255 | 0.38 | 0.24 | 1.83E-79 | spleen | Lamtor4 |
| 9-Sep | 1.64E-83 | 0.31053165 | 0.28 | 0.15 | 3.40E-79 | spleen | 9-Sep |
| 1500011K16Rik | 1.93E-83 | 0.32490623 | 0.31 | 0.18 | 3.99E-79 | spleen | 1500011K16Rik |
| Ets2 | 2.22E-83 | 0.33471642 | 0.5 | 0.34 | 4.58E-79 | spleen | Ets2 |
| Mrpl17 | 2.34E-83 | 0.35794731 | 0.46 | 0.31 | 4.83E-79 | spleen | Mrpl17 |
| 1810043H04Rik | 2.50E-83 | 0.30145355 | 0.29 | 0.17 | 5.17E-79 | spleen | 1810043H04Rik |
| Ndufs6 | 3.51E-83 | 0.33213975 | 0.56 | 0.4 | 7.27E-79 | spleen | Ndufs6 |
| Cdv31 | 6.95E-83 | 0.35272503 | 0.35 | 0.22 | 1.44E-78 | spleen | Cdv3 |
| Ndufb11 | 7.19E-83 | 0.34408059 | 0.66 | 0.5 | 1.49E-78 | spleen | Ndufb11 |
| Gm100364 | 2.70E-82 | 0.25525243 | 0.49 | 0.32 | 5.59E-78 | spleen | Gm10036 |
| Arhgef11 | 9.93E-82 | 0.34568309 | 0.49 | 0.34 | 2.06E-77 | spleen | Arhgef1 |
| Pcbd2 | 2.39E-81 | 0.29876316 | 0.31 | 0.18 | 4.95E-77 | spleen | Pcbd2 |
| Apoa1bp | 2.78E-81 | 0.30237975 | 0.36 | 0.22 | 5.76E-77 | spleen | Apoa1bp |
| Usmg5 | 3.89E-81 | 0.29956978 | 0.69 | 0.53 | 8.06E-77 | spleen | Usmg5 |
| Tmem208 | 4.87E-81 | 0.35602837 | 0.39 | 0.25 | 1.01E-76 | spleen | Tmem208 |
| Arpp19 | 1.29E-80 | 0.3034141 | 0.69 | 0.54 | 2.67E-76 | spleen | Arpp19 |
| Zfand51 | 2.91E-80 | 0.33250478 | 0.51 | 0.35 | 6.03E-76 | spleen | Zfand5 |
| Sf3b51 | 3.85E-80 | 0.35658997 | 0.42 | 0.28 | 7.96E-76 | spleen | Sf3b5 |
| Nono | 2.27E-79 | 0.33969146 | 0.44 | 0.3 | 4.69E-75 | spleen | Nono |
| Carhsp11 | 3.23E-79 | 0.31114032 | 0.39 | 0.25 | 6.69E-75 | spleen | Carhsp1 |
| Ostc1 | 5.51E-79 | 0.36742768 | 0.44 | 0.29 | 1.14E-74 | spleen | Ostc |
| Morf4l2 | 1.29E-78 | 0.34331821 | 0.49 | 0.34 | 2.67E-74 | spleen | Morf4l2 |
| Zfos11 | 1.66E-78 | 0.29998099 | 0.38 | 0.24 | 3.44E-74 | spleen | Zfos1 |
| Lage3 | 1.93E-78 | 0.32366786 | 0.28 | 0.16 | 3.99E-74 | spleen | Lage3 |
| Bola2 | 4.21E-78 | 0.31387674 | 0.4 | 0.26 | 8.72E-74 | spleen | Bola2 |
| Mrpl14 | 4.23E-78 | 0.31668055 | 0.4 | 0.25 | 8.75E-74 | spleen | Mrpl14 |
| Uqcrfs1 | 9.13E-78 | 0.34143222 | 0.43 | 0.29 | 1.89E-73 | spleen | Uqcrfs1 |
| Atp5d | 9.20E-78 | 0.3185434 | 0.68 | 0.52 | 1.90E-73 | spleen | Atp5d |
| Erh | 1.78E-77 | 0.30585126 | 0.55 | 0.38 | 3.68E-73 | spleen | Erh |

| | | | | | | | |
|---|---|---|---|---|---|---|---|
| Cisd2 | 2.81E-77 | 0.31215864 | 0.33 | 0.2 | 5.81E-73 | spleen | Cisd2 |
| C1qbp | 3.64E-77 | 0.33790515 | 0.29 | 0.17 | 7.53E-73 | spleen | C1qbp |
| Ubl5 | 4.87E-77 | 0.28524883 | 0.74 | 0.59 | 1.01E-72 | spleen | Ubl5 |
| Stx4a | 1.14E-76 | 0.30919477 | 0.36 | 0.22 | 2.35E-72 | spleen | Stx4a |
| Lsm4 | 3.95E-76 | 0.29479041 | 0.37 | 0.23 | 8.18E-72 | spleen | Lsm4 |
| Dpm31 | 3.61E-75 | 0.3101392 | 0.59 | 0.44 | 7.47E-71 | spleen | Dpm3 |
| Snrpd1 | 4.91E-75 | 0.30085131 | 0.38 | 0.24 | 1.02E-70 | spleen | Snrpd1 |
| Npr12 | 5.37E-75 | 0.27112363 | 0.43 | 0.28 | 1.11E-70 | spleen | Npr1 |
| Ndufs8 | 6.66E-75 | 0.31233257 | 0.48 | 0.33 | 1.38E-70 | spleen | Ndufs8 |
| 2810428I15Rik | 1.68E-74 | 0.28957145 | 0.41 | 0.26 | 3.49E-70 | spleen | 2810428I15Rik |
| Slc31a21 | 2.25E-74 | 0.32715246 | 0.35 | 0.22 | 4.65E-70 | spleen | Slc31a2 |
| Actn11 | 4.06E-74 | 0.30267955 | 0.29 | 0.17 | 8.41E-70 | spleen | Actn1 |
| Etfb | 6.51E-74 | 0.32566252 | 0.43 | 0.28 | 1.35E-69 | spleen | Etfb |
| Asap12 | 6.72E-74 | 0.27557747 | 0.42 | 0.27 | 1.39E-69 | spleen | Asap1 |
| Tsen34 | 9.03E-74 | 0.3295697 | 0.46 | 0.31 | 1.87E-69 | spleen | Tsen34 |
| Sub11 | 1.16E-73 | 0.27552627 | 0.72 | 0.57 | 2.39E-69 | spleen | Sub1 |
| Hsd17b10 | 1.36E-73 | 0.33506718 | 0.29 | 0.18 | 2.82E-69 | spleen | Hsd17b10 |
| Tspan152 | 2.02E-73 | 0.32417992 | 0.37 | 0.23 | 4.18E-69 | spleen | Tspan15 |
| Hnrnpa1 | 2.92E-73 | 0.35294888 | 0.4 | 0.26 | 6.04E-69 | spleen | Hnrnpa1 |
| Ppp2r4 | 3.18E-73 | 0.29496321 | 0.27 | 0.15 | 6.58E-69 | spleen | Ppp2r4 |
| Hsbp1 | 8.03E-73 | 0.29401837 | 0.62 | 0.47 | 1.66E-68 | spleen | Hsbp1 |
| Ndufa12 | 9.62E-73 | 0.29842936 | 0.54 | 0.38 | 1.99E-68 | spleen | Ndufa12 |
| Ift20 | 3.11E-72 | 0.29557389 | 0.43 | 0.29 | 6.43E-68 | spleen | Ift20 |
| Toporsos | 4.42E-72 | 0.28249592 | 0.28 | 0.16 | 9.16E-68 | spleen | Toporsos |
| Atp5g3 | 4.92E-72 | 0.3257639 | 0.5 | 0.35 | 1.02E-67 | spleen | Atp5g3 |
| Cisd1 | 5.79E-72 | 0.32071935 | 0.28 | 0.17 | 1.20E-67 | spleen | Cisd1 |
| Ptms3 | 6.10E-72 | 0.25334838 | 0.73 | 0.58 | 1.26E-67 | spleen | Ptms |
| Rnf7 | 6.90E-72 | 0.29749149 | 0.59 | 0.44 | 1.43E-67 | spleen | Rnf7 |
| Ndufa2 | 7.38E-72 | 0.2615381 | 0.75 | 0.6 | 1.53E-67 | spleen | Ndufa2 |
| Ndufc21 | 1.04E-71 | 0.29563253 | 0.67 | 0.52 | 2.15E-67 | spleen | Ndufc2 |
| Vkorc1 | 1.28E-71 | 0.30400559 | 0.34 | 0.21 | 2.66E-67 | spleen | Vkorc1 |
| Cirbp | 2.06E-71 | 0.35758752 | 0.34 | 0.22 | 4.26E-67 | spleen | Cirbp |
| Trim351 | 3.47E-71 | 0.30541381 | 0.27 | 0.16 | 7.18E-67 | spleen | Trim35 |
| Prrc2a | 5.17E-71 | 0.2859148 | 0.28 | 0.16 | 1.07E-66 | spleen | Prrc2a |
| Uqcr10 | 6.05E-70 | 0.28671535 | 0.71 | 0.57 | 1.25E-65 | spleen | Uqcr10 |
| Sf3b6 | 1.13E-69 | 0.30007338 | 0.45 | 0.31 | 2.33E-65 | spleen | Sf3b6 |
| Ei24 | 1.67E-69 | 0.29592404 | 0.36 | 0.23 | 3.46E-65 | spleen | Ei24 |
| Mafg | 2.87E-69 | 0.27888009 | 0.27 | 0.15 | 5.95E-65 | spleen | Mafg |
| Tceb1 | 5.59E-69 | 0.2707396 | 0.47 | 0.32 | 1.16E-64 | spleen | Tceb1 |
| Commd6 | 2.10E-68 | 0.29385932 | 0.28 | 0.17 | 4.34E-64 | spleen | Commd6 |
| Sntb22 | 2.24E-68 | 0.31754573 | 0.29 | 0.18 | 4.63E-64 | spleen | Sntb2 |

| | | | | | | | |
|---|---|---|---|---|---|---|---|
| Pam16 | 3.05E-68 | 0.25742403 | 0.27 | 0.16 | 6.31E-64 | spleen | Pam16 |
| Wdr83os | 5.77E-68 | 0.32754619 | 0.39 | 0.26 | 1.19E-63 | spleen | Wdr83os |
| Mrfap1 | 1.03E-67 | 0.27610123 | 0.71 | 0.56 | 2.13E-63 | spleen | Mrfap1 |
| Tspan41 | 1.09E-67 | 0.32546331 | 0.44 | 0.3 | 2.25E-63 | spleen | Tspan4 |
| 1110059E24Rik | 2.42E-67 | 0.29872944 | 0.26 | 0.15 | 5.01E-63 | spleen | 1110059E24Rik |
| Pdcd5 | 3.14E-67 | 0.27343334 | 0.5 | 0.35 | 6.49E-63 | spleen | Pdcd5 |
| Mrpl36 | 5.75E-67 | 0.29107059 | 0.32 | 0.2 | 1.19E-62 | spleen | Mrpl36 |
| 0610012G03Rik | 2.39E-66 | 0.29200483 | 0.46 | 0.32 | 4.94E-62 | spleen | 0610012G03Rik |
| Nfkb11 | 3.22E-66 | 0.29060649 | 0.28 | 0.17 | 6.66E-62 | spleen | Nfkb1 |
| Ubn21 | 3.72E-66 | 0.30504784 | 0.27 | 0.16 | 7.69E-62 | spleen | Ubn2 |
| Tomm20 | 3.05E-65 | 0.30307071 | 0.56 | 0.41 | 6.30E-61 | spleen | Tomm20 |
| Eif3f1 | 6.76E-65 | 0.26143628 | 0.76 | 0.6 | 1.40E-60 | spleen | Eif3f |
| Ndufs3 | 8.79E-65 | 0.30807022 | 0.41 | 0.28 | 1.82E-60 | spleen | Ndufs3 |
| Coa3 | 1.22E-64 | 0.26426886 | 0.5 | 0.35 | 2.52E-60 | spleen | Coa3 |
| Cox7a2l1 | 1.75E-64 | 0.25973144 | 0.75 | 0.6 | 3.62E-60 | spleen | Cox7a2l |
| R3hdm1 | 4.54E-64 | 0.29123895 | 0.26 | 0.15 | 9.39E-60 | spleen | R3hdm1 |
| Tceal81 | 6.24E-64 | 0.30469789 | 0.33 | 0.21 | 1.29E-59 | spleen | Tceal8 |
| mt-Nd4l1 | 6.81E-64 | 0.26103442 | 0.33 | 0.2 | 1.41E-59 | spleen | mt-Nd4l |
| Phldb21 | 8.92E-64 | 0.27883139 | 0.3 | 0.19 | 1.85E-59 | spleen | Phldb2 |
| Tnfsf12 | 1.06E-63 | 0.28128993 | 0.33 | 0.21 | 2.19E-59 | spleen | Tnfsf12 |
| Prdx5 | 1.09E-63 | 0.2773877 | 0.42 | 0.28 | 2.25E-59 | spleen | Prdx5 |
| Mien11 | 1.78E-63 | 0.29781579 | 0.34 | 0.22 | 3.68E-59 | spleen | Mien1 |
| Set | 5.07E-63 | 0.27956242 | 0.49 | 0.36 | 1.05E-58 | spleen | Set |
| H13 | 2.39E-62 | 0.29328034 | 0.35 | 0.23 | 4.94E-58 | spleen | H13 |
| Tspan171 | 2.72E-62 | 0.28669626 | 0.27 | 0.16 | 5.63E-58 | spleen | Tspan17 |
| Cuta1 | 1.94E-61 | 0.2773561 | 0.5 | 0.36 | 4.01E-57 | spleen | Cuta |
| Gatad2b | 2.89E-61 | 0.26497691 | 0.3 | 0.19 | 5.98E-57 | spleen | Gatad2b |
| Cnbp2 | 4.14E-61 | 0.27222236 | 0.8 | 0.67 | 8.57E-57 | spleen | Cnbp |
| Mrps17 | 8.38E-61 | 0.27401292 | 0.31 | 0.2 | 1.73E-56 | spleen | Mrps17 |
| Chchd7 | 1.96E-60 | 0.276123 | 0.27 | 0.16 | 4.07E-56 | spleen | Chchd7 |
| Fkbp71 | 2.39E-60 | 0.27440458 | 0.28 | 0.18 | 4.96E-56 | spleen | Fkbp7 |
| Dad11 | 3.79E-60 | 0.26729537 | 0.73 | 0.59 | 7.84E-56 | spleen | Dad1 |
| Ndufab1 | 1.03E-59 | 0.26361405 | 0.48 | 0.35 | 2.13E-55 | spleen | Ndufab1 |
| Psmd4 | 1.43E-59 | 0.2897806 | 0.48 | 0.35 | 2.95E-55 | spleen | Psmd4 |
| Lsm6 | 8.02E-59 | 0.27530519 | 0.35 | 0.23 | 1.66E-54 | spleen | Lsm6 |
| Btf3l4 | 1.52E-58 | 0.26848097 | 0.26 | 0.16 | 3.15E-54 | spleen | Btf3l4 |
| Smdt1 | 2.02E-58 | 0.25693799 | 0.69 | 0.55 | 4.17E-54 | spleen | Smdt1 |
| Tm2d2 | 2.05E-58 | 0.3026614 | 0.4 | 0.28 | 4.23E-54 | spleen | Tm2d2 |
| Ndufa51 | 2.09E-58 | 0.26339981 | 0.56 | 0.43 | 4.33E-54 | spleen | Ndufa5 |
| Nxf1 | 6.02E-58 | 0.28320928 | 0.43 | 0.3 | 1.25E-53 | spleen | Nxf1 |
| Gadd45gip1 | 7.30E-58 | 0.27038022 | 0.3 | 0.19 | 1.51E-53 | spleen | Gadd45gip1 |

| | | | | | | | |
|---|---|---|---|---|---|---|---|
| Tnrc6b1 | 7.71E-58 | 0.26293956 | 0.28 | 0.18 | 1.59E-53 | spleen | Tnrc6b |
| Nfu1 | 1.81E-57 | 0.29637729 | 0.28 | 0.18 | 3.74E-53 | spleen | Nfu1 |
| 1110004F10Rik | 2.08E-57 | 0.28787074 | 0.46 | 0.33 | 4.31E-53 | spleen | 1110004F10Rik |
| Psmb21 | 2.98E-57 | 0.29684563 | 0.61 | 0.48 | 6.16E-53 | spleen | Psmb2 |
| Mrpl24 | 5.27E-57 | 0.27019773 | 0.33 | 0.22 | 1.09E-52 | spleen | Mrpl24 |
| Rpp21 | 6.86E-57 | 0.25103882 | 0.3 | 0.19 | 1.42E-52 | spleen | Rpp21 |
| Slc25a31 | 2.92E-55 | 0.28092715 | 0.65 | 0.52 | 6.05E-51 | spleen | Slc25a3 |
| Npm1 | 1.09E-53 | 0.28540396 | 0.63 | 0.49 | 2.25E-49 | spleen | Npm1 |
| Dnajc19 | 5.54E-53 | 0.26613653 | 0.32 | 0.21 | 1.15E-48 | spleen | Dnajc19 |
| Cdc26 | 7.25E-53 | 0.26633576 | 0.35 | 0.24 | 1.50E-48 | spleen | Cdc26 |
| Anxa52 | 1.42E-52 | 0.28302785 | 0.77 | 0.64 | 2.94E-48 | spleen | Anxa5 |
| Mrps34 | 5.81E-52 | 0.27276486 | 0.26 | 0.16 | 1.20E-47 | spleen | Mrps34 |
| Ube3a1 | 6.36E-52 | 0.2602222 | 0.33 | 0.22 | 1.32E-47 | spleen | Ube3a |
| Eif3k | 7.60E-51 | 0.25116235 | 0.65 | 0.51 | 1.57E-46 | spleen | Eif3k |
| Mrps14 | 4.60E-50 | 0.2509038 | 0.46 | 0.34 | 9.52E-46 | spleen | Mrps14 |
| Ube2r2 | 1.74E-49 | 0.26289274 | 0.39 | 0.28 | 3.59E-45 | spleen | Ube2r2 |
| Psma5 | 4.19E-48 | 0.25626723 | 0.45 | 0.33 | 8.66E-44 | spleen | Psma5 |
| Stat11 | 5.70E-45 | 0.26950883 | 0.29 | 0.19 | 1.18E-40 | spleen | Stat1 |
| Ccl21a2 | 0 | 4.32903207 | 0.97 | 0.17 | 0 | testis | Ccl21a |
| Lcn21 | 0 | 3.09468207 | 0.81 | 0.03 | 0 | testis | Lcn2 |
| Ptn1 | 0 | 2.63240351 | 0.72 | 0.09 | 0 | testis | Ptn |
| Slc7a51 | 0 | 2.5230348 | 0.7 | 0.08 | 0 | testis | Slc7a5 |
| Mmrn11 | 0 | 2.09504456 | 0.35 | 0.04 | 0 | testis | Mmrn1 |
| Mfsd2a1 | 0 | 2.01392175 | 0.51 | 0.05 | 0 | testis | Mfsd2a |
| Wt1 | 0 | 2.00440051 | 0.54 | 0.02 | 0 | testis | Wt1 |
| Htra11 | 0 | 1.96970572 | 0.54 | 0.06 | 0 | testis | Htra1 |
| Mal1 | 0 | 1.86017459 | 0.56 | 0.1 | 0 | testis | Mal |
| Mgp2 | 0 | 1.85371801 | 0.59 | 0.16 | 0 | testis | Mgp |
| Ptgis | 0 | 1.85331485 | 0.48 | 0.02 | 0 | testis | Ptgis |
| Gpcpd11 | 0 | 1.79863141 | 0.57 | 0.11 | 0 | testis | Gpcpd1 |
| Tmsb102 | 0 | 1.78760663 | 0.95 | 0.73 | 0 | testis | Tmsb10 |
| Vwf2 | 0 | 1.7729829 | 0.68 | 0.17 | 0 | testis | Vwf |
| Clu2 | 0 | 1.72396394 | 0.81 | 0.32 | 0 | testis | Clu |
| Fgl21 | 0 | 1.68773357 | 0.27 | 0.04 | 0 | testis | Fgl2 |
| Abcb1a1 | 0 | 1.67504055 | 0.64 | 0.19 | 0 | testis | Abcb1a |
| Pcp4l12 | 0 | 1.55291917 | 0.77 | 0.32 | 0 | testis | Pcp4l1 |
| Prss233 | 0 | 1.52597281 | 0.75 | 0.27 | 0 | testis | Prss23 |
| Reln1 | 0 | 1.46371668 | 0.26 | 0.03 | 0 | testis | Reln |
| Sparc3 | 0 | 1.46245421 | 0.92 | 0.86 | 0 | testis | Sparc |
| Gstm21 | 0 | 1.44254039 | 0.52 | 0.15 | 0 | testis | Gstm2 |
| Fxyd62 | 0 | 1.44136895 | 0.39 | 0.15 | 0 | testis | Fxyd6 |

| | | | | | | | |
|---|---|---|---|---|---|---|---|
| Aoc3 | 0 | 1.37762313 | 0.37 | 0 | 0 | testis | Aoc3 |
| Rbm32 | 0 | 1.35637794 | 0.79 | 0.39 | 0 | testis | Rbm3 |
| Lsr1 | 0 | 1.35470136 | 0.55 | 0.09 | 0 | testis | Lsr |
| Abcg22 | 0 | 1.34720658 | 0.73 | 0.34 | 0 | testis | Abcg2 |
| Cldn52 | 0 | 1.33618171 | 0.99 | 0.61 | 0 | testis | Cldn5 |
| Slc3a22 | 0 | 1.2886492 | 0.81 | 0.52 | 0 | testis | Slc3a2 |
| Hmgb3 | 0 | 1.27439618 | 0.39 | 0.04 | 0 | testis | Hmgb3 |
| 21-Mar | 0 | 1.25713685 | 0.69 | 0.36 | 0 | testis | 2-Mar |
| Cthrc1 | 0 | 1.24608157 | 0.36 | 0.03 | 0 | testis | Cthrc1 |
| Slco1a41 | 0 | 1.2389013 | 0.56 | 0.1 | 0 | testis | Slco1a4 |
| Vim2 | 0 | 1.22977116 | 0.88 | 0.64 | 0 | testis | Vim |
| Fth11 | 0 | 1.22023191 | 0.99 | 0.91 | 0 | testis | Fth1 |
| Atp1b1 | 0 | 1.21663247 | 0.3 | 0.01 | 0 | testis | Atp1b1 |
| Col18a1 | 0 | 1.20217296 | 0.33 | 0.04 | 0 | testis | Col18a1 |
| C130074G19Rik2 | 0 | 1.15002344 | 0.52 | 0.17 | 0 | testis | C130074G19Rik |
| Tgfbi1 | 0 | 1.14586725 | 0.43 | 0.1 | 0 | testis | Tgfbi |
| Hoxd9 | 0 | 1.13633549 | 0.33 | 0.02 | 0 | testis | Hoxd9 |
| Igfbp42 | 0 | 1.13476737 | 0.77 | 0.35 | 0 | testis | Igfbp4 |
| Slc39a82 | 0 | 1.11477058 | 0.36 | 0.07 | 0 | testis | Slc39a8 |
| Ndn1 | 0 | 1.04665964 | 0.32 | 0.05 | 0 | testis | Ndn |
| Lmo12 | 0 | 1.03160745 | 0.37 | 0.07 | 0 | testis | Lmo1 |
| Cadm12 | 0 | 1.00965879 | 0.44 | 0.13 | 0 | testis | Cadm1 |
| Timp22 | 0 | 0.97866764 | 0.64 | 0.24 | 0 | testis | Timp2 |
| Olfml2a | 0 | 0.95436781 | 0.26 | 0.02 | 0 | testis | Olfml2a |
| Isyna11 | 0 | 0.94397177 | 0.47 | 0.13 | 0 | testis | Isyna1 |
| Itm2c1 | 0 | 0.93395715 | 0.71 | 0.41 | 0 | testis | Itm2c |
| Rbp11 | 0 | 0.92964078 | 0.4 | 0.12 | 0 | testis | Rbp1 |
| Rplp01 | 0 | 0.92365146 | 0.94 | 0.86 | 0 | testis | Rplp0 |
| Rpl353 | 0 | 0.91695588 | 0.95 | 0.86 | 0 | testis | Rpl35 |
| Pdia5 | 0 | 0.90236958 | 0.36 | 0.09 | 0 | testis | Pdia5 |
| mt-Co11 | 0 | 0.89610464 | 0.98 | 0.97 | 0 | testis | mt-Co1 |
| Fam198b2 | 0 | 0.87810466 | 0.43 | 0.16 | 0 | testis | Fam198b |
| Cd2005 | 0 | 0.87495812 | 0.84 | 0.65 | 0 | testis | Cd200 |
| Rps282 | 0 | 0.85611765 | 0.98 | 0.92 | 0 | testis | Rps28 |
| Tgm22 | 0 | 0.85023328 | 0.68 | 0.37 | 0 | testis | Tgm2 |
| Fbln5 | 0 | 0.84643382 | 0.31 | 0.09 | 0 | testis | Fbln5 |
| Rab11fip5 | 0 | 0.8435915 | 0.33 | 0.07 | 0 | testis | Rab11fip5 |
| Syt15 | 0 | 0.8418736 | 0.29 | 0.06 | 0 | testis | Syt15 |
| Kazald1 | 0 | 0.83398394 | 0.26 | 0.04 | 0 | testis | Kazald1 |
| Slc16a41 | 0 | 0.82618022 | 0.29 | 0.04 | 0 | testis | Slc16a4 |
| Gm98442 | 0 | 0.82175146 | 0.35 | 0.11 | 0 | testis | Gm9844 |

| | | | | | | | |
|---|---|---|---|---|---|---|---|
| Pkig2 | 0 | 0.81875051 | 0.52 | 0.24 | 0 | testis | Pkig |
| Nos1ap1 | 0 | 0.80244536 | 0.29 | 0.06 | 0 | testis | Nos1ap |
| Hoxd3os1 | 0 | 0.78155623 | 0.28 | 0.06 | 0 | testis | Hoxd3os1 |
| Efhd11 | 0 | 0.76565865 | 0.41 | 0.11 | 0 | testis | Efhd1 |
| Rpl36a1 | 0 | 0.76357177 | 0.92 | 0.8 | 0 | testis | Rpl36a |
| Rps212 | 0 | 0.74942166 | 0.94 | 0.85 | 0 | testis | Rps21 |
| Gm2a1 | 0 | 0.74286105 | 0.34 | 0.11 | 0 | testis | Gm2a |
| Rps111 | 0 | 0.74033149 | 0.98 | 0.91 | 0 | testis | Rps11 |
| Sorbs23 | 0 | 0.72774635 | 0.53 | 0.24 | 0 | testis | Sorbs2 |
| Rpl321 | 0 | 0.72080646 | 0.97 | 0.92 | 0 | testis | Rpl32 |
| 1110008P14Rik | 0 | 0.71922442 | 0.34 | 0.11 | 0 | testis | 1110008P14Rik |
| Paqr51 | 0 | 0.71036648 | 0.27 | 0.05 | 0 | testis | Paqr5 |
| mt-Nd12 | 0 | 0.70108633 | 0.97 | 0.95 | 0 | testis | mt-Nd1 |
| Rpl122 | 0 | 0.69333763 | 0.82 | 0.6 | 0 | testis | Rpl12 |
| Rps182 | 0 | 0.69066098 | 0.97 | 0.92 | 0 | testis | Rps18 |
| Cd3201 | 0 | 0.68654529 | 0.29 | 0.08 | 0 | testis | Cd320 |
| Rpl273 | 0 | 0.68074877 | 0.92 | 0.81 | 0 | testis | Rpl27 |
| Rpl312 | 0 | 0.67824186 | 0.95 | 0.87 | 0 | testis | Rpl31 |
| Rpl232 | 0 | 0.67713317 | 0.96 | 0.9 | 0 | testis | Rpl23 |
| Rps202 | 0 | 0.66992617 | 0.95 | 0.86 | 0 | testis | Rps20 |
| Rpl13a2 | 0 | 0.65871219 | 0.98 | 0.89 | 0 | testis | Rpl13a |
| Rpl392 | 0 | 0.64080201 | 0.95 | 0.88 | 0 | testis | Rpl39 |
| Rpl172 | 0 | 0.63228786 | 0.96 | 0.89 | 0 | testis | Rpl17 |
| Rps192 | 0 | 0.62941323 | 0.98 | 0.96 | 0 | testis | Rps19 |
| Sepp11 | 0 | 0.62401132 | 0.97 | 0.75 | 0 | testis | Sepp1 |
| Rpl23a2 | 0 | 0.62348165 | 0.97 | 0.91 | 0 | testis | Rpl23a |
| Rpl37a2 | 0 | 0.61940906 | 0.99 | 0.97 | 0 | testis | Rpl37a |
| Rpl61 | 0 | 0.60681515 | 0.96 | 0.89 | 0 | testis | Rpl6 |
| AY0361182 | 0 | 0.60598525 | 0.95 | 0.73 | 0 | testis | AY036118 |
| Rpl291 | 0 | 0.60363484 | 0.91 | 0.82 | 0 | testis | Rpl29 |
| Rpl241 | 0 | 0.60016098 | 0.95 | 0.87 | 0 | testis | Rpl24 |
| Cox4i11 | 0 | 0.57815735 | 0.92 | 0.84 | 0 | testis | Cox4i1 |
| Rps172 | 0 | 0.5768637 | 0.95 | 0.88 | 0 | testis | Rps17 |
| Rpl142 | 0 | 0.5542404 | 0.95 | 0.89 | 0 | testis | Rpl14 |
| Rps63 | 0 | 0.5502202 | 0.95 | 0.89 | 0 | testis | Rps6 |
| Rpl112 | 0 | 0.54567167 | 0.95 | 0.89 | 0 | testis | Rpl11 |
| Rpl381 | 0 | 0.54220673 | 0.96 | 0.92 | 0 | testis | Rpl38 |
| Rpl363 | 0 | 0.53674683 | 0.95 | 0.9 | 0 | testis | Rpl36 |
| Rps15a1 | 0 | 0.52941476 | 0.95 | 0.89 | 0 | testis | Rps15a |
| Rps292 | 0 | 0.52006389 | 0.99 | 0.98 | 0 | testis | Rps29 |
| Rpl262 | 0 | 0.51059697 | 0.97 | 0.92 | 0 | testis | Rpl26 |

| | | | | | | | |
|---|---|---|---|---|---|---|---|
| Rplp21 | 0 | 0.50817394 | 0.95 | 0.9 | 0 | testis | Rplp2 |
| Rps27a1 | 0 | 0.50797205 | 0.96 | 0.91 | 0 | testis | Rps27a |
| Rpl413 | 0 | 0.50402118 | 0.98 | 0.97 | 0 | testis | Rpl41 |
| Rps272 | 0 | 0.48817246 | 0.99 | 0.98 | 0 | testis | Rps27 |
| Rps142 | 0 | 0.48358908 | 0.98 | 0.95 | 0 | testis | Rps14 |
| Rpl372 | 0 | 0.46066238 | 0.98 | 0.95 | 0 | testis | Rpl37 |
| Fcgr2b1 | 0 | 0.43423909 | 0.4 | 0.1 | 0 | testis | Fcgr2b |
| mt-Co31 | 2.38E-300 | 0.52934177 | 0.98 | 0.98 | 4.93E-296 | testis | mt-Co3 |
| Rpl33 | 3.45E-299 | 0.58049047 | 0.92 | 0.84 | 7.14E-295 | testis | Rpl3 |
| Rpl132 | 1.64E-297 | 0.45400556 | 0.98 | 0.95 | 3.39E-293 | testis | Rpl13 |
| Rps232 | 5.93E-297 | 0.44565318 | 0.97 | 0.95 | 1.23E-292 | testis | Rps23 |
| Rpl212 | 9.63E-295 | 0.46283506 | 0.96 | 0.91 | 1.99E-290 | testis | Rpl21 |
| Tm6sf11 | 7.60E-293 | 0.66383938 | 0.41 | 0.17 | 1.57E-288 | testis | Tm6sf1 |
| Rps92 | 5.76E-290 | 0.50758741 | 0.94 | 0.88 | 1.19E-285 | testis | Rps9 |
| Rpl10a1 | 4.59E-284 | 0.60903278 | 0.87 | 0.74 | 9.49E-280 | testis | Rpl10a |
| Lbp1 | 6.15E-283 | 0.9004354 | 0.27 | 0.09 | 1.27E-278 | testis | Lbp |
| Hmgn31 | 2.48E-279 | 0.62685374 | 0.35 | 0.13 | 5.13E-275 | testis | Hmgn3 |
| Rpl18a1 | 2.02E-278 | 0.44063066 | 0.97 | 0.95 | 4.17E-274 | testis | Rpl18a |
| Cgnl12 | 1.18E-275 | 0.61764052 | 0.33 | 0.12 | 2.44E-271 | testis | Cgnl1 |
| Rps81 | 2.70E-274 | 0.46656154 | 0.96 | 0.93 | 5.58E-270 | testis | Rps8 |
| Spock21 | 5.10E-273 | 0.41319047 | 0.31 | 0.1 | 1.05E-268 | testis | Spock2 |
| Rps162 | 1.28E-272 | 0.44221736 | 0.96 | 0.93 | 2.65E-268 | testis | Rps16 |
| Rplp11 | 1.87E-270 | 0.45027095 | 0.96 | 0.92 | 3.88E-266 | testis | Rplp1 |
| Rps4x2 | 4.59E-268 | 0.4896186 | 0.94 | 0.89 | 9.50E-264 | testis | Rps4x |
| Prkcdbp4 | 2.20E-265 | 0.58979759 | 0.79 | 0.48 | 4.55E-261 | testis | Prkcdbp |
| Gm87304 | 6.08E-265 | 0.68722512 | 0.66 | 0.4 | 1.26E-260 | testis | Gm8730 |
| Atp1b31 | 2.14E-258 | 0.6960385 | 0.61 | 0.34 | 4.43E-254 | testis | Atp1b3 |
| Rpl181 | 2.47E-254 | 0.49531868 | 0.91 | 0.84 | 5.11E-250 | testis | Rpl18 |
| Smtn | 2.32E-253 | 0.63782499 | 0.36 | 0.15 | 4.79E-249 | testis | Smtn |
| Rps252 | 4.83E-251 | 0.44884106 | 0.94 | 0.89 | 1.00E-246 | testis | Rps25 |
| Stt3b | 6.85E-247 | 0.70801843 | 0.32 | 0.12 | 1.42E-242 | testis | Stt3b |
| Fau2 | 4.89E-240 | 0.39634228 | 0.96 | 0.93 | 1.01E-235 | testis | Fau |
| Pam1 | 4.60E-239 | 0.94584981 | 0.46 | 0.23 | 9.51E-235 | testis | Pam |
| Rpl71 | 1.45E-238 | 0.45413259 | 0.93 | 0.85 | 3.00E-234 | testis | Rpl7 |
| Dnm31 | 1.29E-237 | 0.57796897 | 0.39 | 0.17 | 2.67E-233 | testis | Dnm3 |
| Mgst12 | 8.37E-237 | 0.62881888 | 0.34 | 0.14 | 1.73E-232 | testis | Mgst1 |
| Cd94 | 1.04E-235 | 0.58464617 | 0.87 | 0.6 | 2.15E-231 | testis | Cd9 |
| Rpl102 | 1.19E-233 | 0.46254906 | 0.94 | 0.88 | 2.47E-229 | testis | Rpl10 |
| Selm3 | 3.95E-232 | 0.58380833 | 0.69 | 0.41 | 8.18E-228 | testis | Selm |
| Ddit43 | 1.23E-230 | 0.60237648 | 0.39 | 0.17 | 2.54E-226 | testis | Ddit4 |
| Gm100764 | 6.89E-227 | 0.6155134 | 0.69 | 0.46 | 1.43E-222 | testis | Gm10076 |

| | | | | | | | |
|---|---|---|---|---|---|---|---|
| Rps123 | 1.07E-225 | 0.48858727 | 0.91 | 0.82 | 2.22E-221 | testis | Rps12 |
| Cirbp1 | 4.54E-222 | 0.63657065 | 0.44 | 0.21 | 9.40E-218 | testis | Cirbp |
| Maoa2 | 6.03E-220 | 0.58455395 | 0.37 | 0.16 | 1.25E-215 | testis | Maoa |
| Nmt2 | 2.50E-219 | 0.54765625 | 0.27 | 0.1 | 5.18E-215 | testis | Nmt2 |
| Rpl35a2 | 2.04E-218 | 0.38249655 | 0.97 | 0.94 | 4.22E-214 | testis | Rpl35a |
| Rps242 | 6.22E-218 | 0.39732389 | 0.95 | 0.89 | 1.29E-213 | testis | Rps24 |
| Rpl341 | 3.66E-217 | 0.41054549 | 0.94 | 0.9 | 7.58E-213 | testis | Rpl34 |
| Tmsb4x3 | 4.17E-211 | 0.28445923 | 1 | 0.99 | 8.63E-207 | testis | Tmsb4x |
| Rps210 | 5.43E-207 | 0.44183314 | 0.91 | 0.84 | 1.12E-202 | testis | Rps2 |
| Rps72 | 1.60E-206 | 0.44714066 | 0.92 | 0.86 | 3.31E-202 | testis | Rps7 |
| Fzd61 | 3.25E-204 | 0.51559743 | 0.27 | 0.1 | 6.73E-200 | testis | Fzd6 |
| Hhex | 4.56E-204 | 0.64574266 | 0.39 | 0.18 | 9.44E-200 | testis | Hhex |
| Ssbp41 | 9.22E-203 | 0.58813336 | 0.49 | 0.27 | 1.91E-198 | testis | Ssbp4 |
| Rps32 | 1.16E-200 | 0.38211752 | 0.95 | 0.9 | 2.40E-196 | testis | Rps3 |
| Rpl191 | 1.51E-197 | 0.36796146 | 0.95 | 0.91 | 3.13E-193 | testis | Rpl19 |
| Rpl302 | 1.50E-193 | 0.44734006 | 0.88 | 0.77 | 3.11E-189 | testis | Rpl30 |
| Eogt1 | 3.27E-193 | 0.50834562 | 0.34 | 0.15 | 6.77E-189 | testis | Eogt |
| Lgals11 | 1.27E-191 | 0.28744728 | 0.43 | 0.2 | 2.63E-187 | testis | Lgals1 |
| Mfge82 | 2.00E-191 | 0.74045928 | 0.65 | 0.43 | 4.14E-187 | testis | Mfge8 |
| Prdx51 | 2.16E-190 | 0.59721689 | 0.5 | 0.28 | 4.47E-186 | testis | Prdx5 |
| Prdx31 | 3.18E-187 | 0.53336022 | 0.35 | 0.16 | 6.58E-183 | testis | Prdx3 |
| Pdcd42 | 1.06E-184 | 0.73559047 | 0.3 | 0.13 | 2.20E-180 | testis | Pdcd4 |
| Igfbp73 | 1.46E-184 | 0.57571217 | 0.98 | 0.87 | 3.01E-180 | testis | Igfbp7 |
| mt-Atp61 | 2.70E-181 | 0.38439223 | 0.98 | 0.99 | 5.58E-177 | testis | mt-Atp6 |
| Eef23 | 8.03E-181 | 0.47061181 | 0.88 | 0.79 | 1.66E-176 | testis | Eef2 |
| Rps102 | 1.54E-180 | 0.43605556 | 0.9 | 0.8 | 3.18E-176 | testis | Rps10 |
| Rps131 | 1.64E-178 | 0.4166009 | 0.91 | 0.84 | 3.39E-174 | testis | Rps13 |
| Rps3a11 | 2.08E-178 | 0.39640212 | 0.93 | 0.86 | 4.31E-174 | testis | Rps3a1 |
| Emp32 | 4.22E-175 | 0.57984415 | 0.46 | 0.24 | 8.73E-171 | testis | Emp3 |
| Rpl43 | 2.67E-174 | 0.42437572 | 0.89 | 0.81 | 5.52E-170 | testis | Rpl4 |
| Sema7a2 | 3.47E-173 | 0.53099235 | 0.39 | 0.19 | 7.18E-169 | testis | Sema7a |
| Eno13 | 4.36E-173 | 0.6000147 | 0.53 | 0.32 | 9.03E-169 | testis | Eno1 |
| AU0210922 | 1.14E-172 | 0.47013762 | 0.56 | 0.32 | 2.36E-168 | testis | AU021092 |
| Cox6c1 | 1.49E-172 | 0.43768005 | 0.87 | 0.77 | 3.09E-168 | testis | Cox6c |
| Rpl27a1 | 2.87E-171 | 0.34690757 | 0.95 | 0.91 | 5.94E-167 | testis | Rpl27a |
| Hjurp | 2.05E-168 | 0.69697744 | 0.25 | 0.1 | 4.25E-164 | testis | Hjurp |
| Rps262 | 1.68E-164 | 0.410789 | 0.9 | 0.8 | 3.47E-160 | testis | Rps26 |
| Abca31 | 2.37E-164 | 0.511661 | 0.27 | 0.11 | 4.91E-160 | testis | Abca3 |
| Smarca23 | 2.82E-162 | 0.56780077 | 0.45 | 0.25 | 5.84E-158 | testis | Smarca2 |
| Tpi12 | 1.61E-159 | 0.53692561 | 0.47 | 0.27 | 3.33E-155 | testis | Tpi1 |
| Bgn1 | 6.70E-158 | 0.31516741 | 0.39 | 0.18 | 1.39E-153 | testis | Bgn |

| | | | | | | | |
|---|---|---|---|---|---|---|---|
| Tob12 | 6.62E-157 | 0.51229594 | 0.34 | 0.17 | 1.37E-152 | testis | Tob1 |
| Tshz24 | 3.17E-156 | 0.4410222 | 0.62 | 0.37 | 6.56E-152 | testis | Tshz2 |
| Rpl23a-ps34 | 1.26E-152 | 0.4148153 | 0.81 | 0.64 | 2.60E-148 | testis | Rpl23a-ps3 |
| Uba525 | 2.21E-152 | 0.41077378 | 0.84 | 0.71 | 4.57E-148 | testis | Uba52 |
| Arfgap31 | 3.05E-152 | 0.46959744 | 0.25 | 0.11 | 6.30E-148 | testis | Arfgap3 |
| Oaz21 | 2.54E-151 | 0.55174481 | 0.73 | 0.58 | 5.26E-147 | testis | Oaz2 |
| Cd2ap2 | 9.35E-151 | 0.53767334 | 0.43 | 0.24 | 1.94E-146 | testis | Cd2ap |
| mt-Cytb1 | 3.93E-150 | 0.37511069 | 0.96 | 0.96 | 8.12E-146 | testis | mt-Cytb |
| Pgp | 9.73E-149 | 0.51188179 | 0.32 | 0.16 | 2.01E-144 | testis | Pgp |
| Il11ra1 | 1.95E-148 | 0.57736295 | 0.27 | 0.12 | 4.04E-144 | testis | Il11ra1 |
| Rps12-ps35 | 8.02E-148 | 0.43232141 | 0.76 | 0.56 | 1.66E-143 | testis | Rps12-ps3 |
| Nenf2 | 2.19E-147 | 0.5183533 | 0.61 | 0.4 | 4.53E-143 | testis | Nenf |
| Gm20003 | 1.15E-146 | 0.41420057 | 0.44 | 0.24 | 2.38E-142 | testis | Gm2000 |
| Gm100733 | 2.04E-146 | 0.41561357 | 0.61 | 0.39 | 4.22E-142 | testis | Gm10073 |
| Tspan42 | 2.22E-146 | 0.58723421 | 0.5 | 0.3 | 4.59E-142 | testis | Tspan4 |
| Abhd17a | 2.19E-145 | 0.51346715 | 0.53 | 0.33 | 4.53E-141 | testis | Abhd17a |
| Rpl221 | 4.37E-145 | 0.40336437 | 0.85 | 0.73 | 9.04E-141 | testis | Rpl22 |
| Eif3f2 | 5.06E-145 | 0.47321399 | 0.75 | 0.61 | 1.05E-140 | testis | Eif3f |
| Serinc34 | 4.83E-143 | 0.46240126 | 0.93 | 0.84 | 9.98E-139 | testis | Serinc3 |
| Eef1a12 | 3.07E-141 | 0.3163146 | 0.97 | 0.95 | 6.36E-137 | testis | Eef1a1 |
| Tmem2522 | 4.74E-138 | 0.30233167 | 0.28 | 0.12 | 9.81E-134 | testis | Tmem252 |
| Sh3glb2 | 5.20E-138 | 0.45722225 | 0.35 | 0.18 | 1.08E-133 | testis | Sh3glb2 |
| Ehd44 | 1.16E-137 | 0.45163411 | 0.72 | 0.52 | 2.40E-133 | testis | Ehd4 |
| Irak1 | 5.18E-137 | 0.42059815 | 0.28 | 0.13 | 1.07E-132 | testis | Irak1 |
| Rps52 | 1.26E-134 | 0.30117079 | 0.94 | 0.91 | 2.60E-130 | testis | Rps5 |
| Trappc6a | 4.61E-128 | 0.41436375 | 0.28 | 0.14 | 9.54E-124 | testis | Trappc6a |
| Sult1a13 | 3.19E-127 | 0.53970148 | 0.33 | 0.17 | 6.59E-123 | testis | Sult1a1 |
| Ndufa71 | 7.01E-126 | 0.45692877 | 0.75 | 0.61 | 1.45E-121 | testis | Ndufa7 |
| Aldoa2 | 1.68E-125 | 0.43575995 | 0.8 | 0.65 | 3.48E-121 | testis | Aldoa |
| Mif1 | 1.13E-124 | 0.44430781 | 0.49 | 0.3 | 2.33E-120 | testis | Mif |
| Pfdn52 | 1.87E-124 | 0.37518545 | 0.85 | 0.74 | 3.88E-120 | testis | Pfdn5 |
| Bambi | 1.33E-122 | 0.46562092 | 0.3 | 0.15 | 2.75E-118 | testis | Bambi |
| Ppib | 1.44E-121 | 0.43276542 | 0.79 | 0.64 | 2.98E-117 | testis | Ppib |
| Kctd121 | 4.02E-121 | 0.48120216 | 0.35 | 0.19 | 8.32E-117 | testis | Kctd12 |
| Pkm2 | 8.79E-121 | 0.55532503 | 0.67 | 0.51 | 1.82E-116 | testis | Pkm |
| Rnh1 | 1.20E-120 | 0.46643348 | 0.39 | 0.22 | 2.48E-116 | testis | Rnh1 |
| Pebp1 | 1.24E-119 | 0.52669214 | 0.61 | 0.44 | 2.57E-115 | testis | Pebp1 |
| Rpl282 | 6.42E-117 | 0.31967458 | 0.91 | 0.84 | 1.33E-112 | testis | Rpl28 |
| Rps152 | 2.16E-116 | 0.25871904 | 0.94 | 0.91 | 4.48E-112 | testis | Rps15 |
| Ctsz1 | 1.25E-115 | 0.46896143 | 0.53 | 0.35 | 2.58E-111 | testis | Ctsz |
| Ptms4 | 1.03E-114 | 0.4163061 | 0.74 | 0.58 | 2.12E-110 | testis | Ptms |

| | | | | | | | |
|---|---|---|---|---|---|---|---|
| Adam152 | 1.18E-113 | 0.58133252 | 0.59 | 0.43 | 2.45E-109 | testis | Adam15 |
| Ndufc22 | 5.83E-112 | 0.47583321 | 0.69 | 0.52 | 1.21E-107 | testis | Ndufc2 |
| Rps27rt4 | 5.74E-111 | 0.28291455 | 0.94 | 0.83 | 1.19E-106 | testis | Rps27rt |
| Pvrl22 | 4.57E-110 | 0.46459394 | 0.31 | 0.16 | 9.45E-106 | testis | Pvrl2 |
| Rpl151 | 6.49E-110 | 0.33684669 | 0.89 | 0.83 | 1.34E-105 | testis | Rpl15 |
| Ankrd13a | 1.91E-109 | 0.38554234 | 0.29 | 0.14 | 3.96E-105 | testis | Ankrd13a |
| Hif1a2 | 5.73E-109 | 0.38856521 | 0.29 | 0.15 | 1.19E-104 | testis | Hif1a |
| Rpl6l4 | 4.78E-107 | 0.38346015 | 0.65 | 0.46 | 9.89E-103 | testis | Rpl6l |
| Uqcrc1 | 8.35E-107 | 0.45193942 | 0.54 | 0.37 | 1.73E-102 | testis | Uqcrc1 |
| S100a163 | 5.33E-106 | 0.33968669 | 0.82 | 0.69 | 1.10E-101 | testis | S100a16 |
| Fam181b1 | 8.59E-105 | 0.38547536 | 0.26 | 0.13 | 1.78E-100 | testis | Fam181b |
| Spg21 | 3.15E-104 | 0.34719239 | 0.26 | 0.13 | 6.53E-100 | testis | Spg21 |
| Polr2l | 3.40E-104 | 0.35547371 | 0.35 | 0.19 | 7.03E-100 | testis | Polr2l |
| Crip14 | 5.77E-102 | 0.29770201 | 0.87 | 0.69 | 1.19E-97 | testis | Crip1 |
| Fryl1 | 7.49E-102 | 0.37141926 | 0.3 | 0.16 | 1.55E-97 | testis | Fryl |
| Uqcrh1 | 1.06E-101 | 0.37320787 | 0.77 | 0.65 | 2.19E-97 | testis | Uqcrh |
| Reep51 | 1.10E-101 | 0.46485678 | 0.59 | 0.42 | 2.28E-97 | testis | Reep5 |
| Gnb2l12 | 3.61E-101 | 0.36778145 | 0.73 | 0.55 | 7.47E-97 | testis | Gnb2l1 |
| Sin3b | 8.44E-100 | 0.39821036 | 0.38 | 0.22 | 1.75E-95 | testis | Sin3b |
| Hint11 | 2.21E-99 | 0.37490572 | 0.8 | 0.66 | 4.57E-95 | testis | Hint1 |
| Rpl82 | 3.35E-99 | 0.25846265 | 0.93 | 0.9 | 6.93E-95 | testis | Rpl8 |
| mt-Co22 | 8.62E-99 | 0.29822699 | 0.95 | 0.93 | 1.78E-94 | testis | mt-Co2 |
| Rpl51 | 1.88E-98 | 0.35715947 | 0.78 | 0.66 | 3.90E-94 | testis | Rpl5 |
| Ltbp45 | 7.13E-98 | 0.53063146 | 0.67 | 0.48 | 1.48E-93 | testis | Ltbp4 |
| Iqgap13 | 3.11E-97 | 0.42458455 | 0.5 | 0.33 | 6.43E-93 | testis | Iqgap1 |
| Glul1 | 4.43E-97 | 0.65498766 | 0.55 | 0.39 | 9.17E-93 | testis | Glul |
| Atp5e1 | 7.33E-97 | 0.34695155 | 0.8 | 0.68 | 1.52E-92 | testis | Atp5e |
| Eci21 | 1.37E-96 | 0.35540291 | 0.31 | 0.17 | 2.82E-92 | testis | Eci2 |
| Ecscr1 | 1.56E-95 | 0.35541192 | 0.8 | 0.62 | 3.22E-91 | testis | Ecscr |
| Cox5b1 | 1.78E-95 | 0.3579242 | 0.76 | 0.63 | 3.68E-91 | testis | Cox5b |
| Pink13 | 1.57E-94 | 0.36860392 | 0.4 | 0.24 | 3.24E-90 | testis | Pink1 |
| Palm3 | 3.61E-94 | 0.32718758 | 0.39 | 0.23 | 7.46E-90 | testis | Palm |
| Flna1 | 2.22E-92 | 0.4024852 | 0.27 | 0.14 | 4.59E-88 | testis | Flna |
| Socs24 | 3.23E-92 | 0.35876797 | 0.41 | 0.25 | 6.67E-88 | testis | Socs2 |
| Atp5l1 | 7.32E-91 | 0.33171985 | 0.82 | 0.72 | 1.51E-86 | testis | Atp5l |
| Pqlc12 | 2.36E-90 | 0.37424541 | 0.37 | 0.22 | 4.88E-86 | testis | Pqlc1 |
| Gm81862 | 1.70E-89 | 0.26965237 | 0.31 | 0.17 | 3.52E-85 | testis | Gm8186 |
| Anxa111 | 1.75E-89 | 0.36893589 | 0.38 | 0.22 | 3.62E-85 | testis | Anxa11 |
| Actn43 | 2.91E-89 | 0.41046961 | 0.68 | 0.51 | 6.02E-85 | testis | Actn4 |
| Tgfbr24 | 6.45E-89 | 0.50953464 | 0.42 | 0.26 | 1.33E-84 | testis | Tgfbr2 |
| Card192 | 7.76E-89 | 0.36234269 | 0.42 | 0.26 | 1.61E-84 | testis | Card19 |

| | | | | | | | |
|---|---|---|---|---|---|---|---|
| Orai13 | 3.21E-87 | 0.37511728 | 0.38 | 0.23 | 6.63E-83 | testis | Orai1 |
| Wdr895 | 1.11E-86 | 0.26908504 | 0.85 | 0.71 | 2.30E-82 | testis | Wdr89 |
| Plec | 2.01E-86 | 0.39772587 | 0.4 | 0.25 | 4.16E-82 | testis | Plec |
| Rpl27-ps34 | 2.43E-85 | 0.29551606 | 0.49 | 0.32 | 5.04E-81 | testis | Rpl27-ps3 |
| Ndufv3 | 8.05E-84 | 0.35523927 | 0.56 | 0.4 | 1.67E-79 | testis | Ndufv3 |
| Uqcc3 | 1.08E-82 | 0.37597306 | 0.46 | 0.31 | 2.24E-78 | testis | Uqcc3 |
| Sox184 | 1.94E-82 | 0.28628314 | 0.67 | 0.5 | 4.02E-78 | testis | Sox18 |
| Capns12 | 4.43E-82 | 0.33276916 | 0.77 | 0.63 | 9.17E-78 | testis | Capns1 |
| Ctnnb15 | 1.49E-81 | 0.30360178 | 0.68 | 0.51 | 3.08E-77 | testis | Ctnnb1 |
| Mrpl541 | 1.81E-81 | 0.35242759 | 0.33 | 0.19 | 3.74E-77 | testis | Mrpl54 |
| Zyx1 | 2.76E-81 | 0.44967111 | 0.31 | 0.18 | 5.70E-77 | testis | Zyx |
| Mospd2 | 1.19E-80 | 0.32061641 | 0.28 | 0.15 | 2.47E-76 | testis | Mospd2 |
| Ssr41 | 2.98E-80 | 0.36749883 | 0.55 | 0.39 | 6.17E-76 | testis | Ssr4 |
| Eif4a13 | 1.58E-79 | 0.3653179 | 0.71 | 0.58 | 3.26E-75 | testis | Eif4a1 |
| Mpzl11 | 2.43E-79 | 0.28917125 | 0.45 | 0.29 | 5.03E-75 | testis | Mpzl1 |
| Gapdh1 | 2.82E-79 | 0.38984249 | 0.74 | 0.62 | 5.83E-75 | testis | Gapdh |
| Atpif12 | 2.08E-78 | 0.31083642 | 0.66 | 0.49 | 4.31E-74 | testis | Atpif1 |
| Phpt11 | 2.55E-78 | 0.28580087 | 0.29 | 0.16 | 5.27E-74 | testis | Phpt1 |
| H2afj | 1.51E-77 | 0.34977619 | 0.44 | 0.29 | 3.12E-73 | testis | H2afj |
| Gltscr23 | 3.85E-77 | 0.30847535 | 0.62 | 0.45 | 7.97E-73 | testis | Gltscr2 |
| Rpsa1 | 1.26E-76 | 0.30110443 | 0.8 | 0.71 | 2.62E-72 | testis | Rpsa |
| Snrpg2 | 1.83E-75 | 0.32572642 | 0.7 | 0.57 | 3.79E-71 | testis | Snrpg |
| Eif3e1 | 2.66E-75 | 0.33538906 | 0.58 | 0.43 | 5.50E-71 | testis | Eif3e |
| Sptssa1 | 1.20E-74 | 0.37433632 | 0.59 | 0.43 | 2.49E-70 | testis | Sptssa |
| 2410015M20Rik1 | 3.76E-74 | 0.34085399 | 0.5 | 0.35 | 7.78E-70 | testis | 2410015M20Rik |
| Ndufa14 | 1.82E-73 | 0.32589579 | 0.59 | 0.44 | 3.76E-69 | testis | Ndufa1 |
| Eif3i | 1.10E-72 | 0.33750273 | 0.44 | 0.29 | 2.28E-68 | testis | Eif3i |
| Gm94934 | 2.30E-72 | 0.2606928 | 0.67 | 0.51 | 4.75E-68 | testis | Gm9493 |
| Rpl36al1 | 3.04E-71 | 0.29859263 | 0.7 | 0.56 | 6.29E-67 | testis | Rpl36al |
| Ndufa61 | 4.08E-71 | 0.32589245 | 0.71 | 0.58 | 8.43E-67 | testis | Ndufa6 |
| Cox7a2l2 | 4.56E-71 | 0.31460777 | 0.71 | 0.6 | 9.43E-67 | testis | Cox7a2l |
| Traf71 | 9.55E-71 | 0.30642275 | 0.29 | 0.17 | 1.98E-66 | testis | Traf7 |
| Eif3k1 | 1.21E-70 | 0.33213655 | 0.65 | 0.52 | 2.50E-66 | testis | Eif3k |
| Use1 | 1.56E-70 | 0.31575969 | 0.46 | 0.31 | 3.23E-66 | testis | Use1 |
| mt-Nd52 | 2.05E-69 | 0.33515202 | 0.5 | 0.35 | 4.24E-65 | testis | mt-Nd5 |
| Cd631 | 3.26E-69 | 0.65803086 | 0.28 | 0.17 | 6.75E-65 | testis | Cd63 |
| Mrpl521 | 8.24E-69 | 0.31468026 | 0.58 | 0.42 | 1.71E-64 | testis | Mrpl52 |
| Rab11a1 | 1.18E-68 | 0.30329203 | 0.77 | 0.65 | 2.44E-64 | testis | Rab11a |
| Clstn11 | 3.08E-68 | 0.34613648 | 0.33 | 0.2 | 6.38E-64 | testis | Clstn1 |
| Hmgn12 | 8.07E-68 | 0.30870779 | 0.49 | 0.34 | 1.67E-63 | testis | Hmgn1 |
| Ndufa21 | 3.83E-66 | 0.30852342 | 0.72 | 0.6 | 7.92E-62 | testis | Ndufa2 |

| | | | | | | | |
|---|---|---|---|---|---|---|---|
| Atp5g31 | 9.54E-66 | 0.30900278 | 0.5 | 0.35 | 1.97E-61 | testis | Atp5g3 |
| Arl4a1 | 3.72E-65 | 0.35257334 | 0.41 | 0.27 | 7.70E-61 | testis | Arl4a |
| Lxn1 | 3.75E-65 | 0.32507643 | 0.35 | 0.22 | 7.75E-61 | testis | Lxn |
| Atp5o | 3.85E-65 | 0.27596371 | 0.59 | 0.43 | 7.96E-61 | testis | Atp5o |
| Pfkl1 | 4.75E-64 | 0.28610213 | 0.26 | 0.15 | 9.83E-60 | testis | Pfkl |
| Hnrnpa11 | 1.08E-63 | 0.29557459 | 0.4 | 0.26 | 2.24E-59 | testis | Hnrnpa1 |
| Ndufb91 | 1.53E-63 | 0.31294892 | 0.63 | 0.49 | 3.17E-59 | testis | Ndufb9 |
| Sox174 | 5.63E-63 | 0.25016027 | 0.54 | 0.38 | 1.17E-58 | testis | Sox17 |
| Timm10b1 | 6.22E-63 | 0.32401122 | 0.44 | 0.31 | 1.29E-58 | testis | Timm10b |
| Ogt1 | 1.06E-62 | 0.32853494 | 0.38 | 0.25 | 2.20E-58 | testis | Ogt |
| Rnf71 | 1.68E-62 | 0.31977969 | 0.58 | 0.44 | 3.47E-58 | testis | Rnf7 |
| Atp2b43 | 2.37E-62 | 0.26792403 | 0.33 | 0.2 | 4.90E-58 | testis | Atp2b4 |
| Cdk2ap21 | 1.60E-61 | 0.29142961 | 0.34 | 0.22 | 3.32E-57 | testis | Cdk2ap2 |
| Ppic2 | 1.84E-61 | 0.32209318 | 0.5 | 0.36 | 3.82E-57 | testis | Ppic |
| Cd342 | 1.87E-61 | 0.43653948 | 0.66 | 0.55 | 3.87E-57 | testis | Cd34 |
| Eif3h | 4.24E-61 | 0.26280141 | 0.64 | 0.5 | 8.76E-57 | testis | Eif3h |
| Gnb4 | 5.05E-60 | 0.28846181 | 0.27 | 0.16 | 1.04E-55 | testis | Gnb4 |
| Myl63 | 4.75E-59 | 0.26022709 | 0.93 | 0.9 | 9.82E-55 | testis | Myl6 |
| Ndufs81 | 4.78E-59 | 0.30144613 | 0.47 | 0.34 | 9.89E-55 | testis | Ndufs8 |
| Fuca12 | 1.73E-58 | 0.30193754 | 0.34 | 0.22 | 3.59E-54 | testis | Fuca1 |
| Ptov11 | 2.73E-58 | 0.28107641 | 0.35 | 0.23 | 5.65E-54 | testis | Ptov1 |
| Ndufa13 | 1.78E-57 | 0.26329552 | 0.69 | 0.56 | 3.68E-53 | testis | Ndufa13 |
| Gpi11 | 5.08E-57 | 0.31347963 | 0.5 | 0.37 | 1.05E-52 | testis | Gpi1 |
| Hprt1 | 2.06E-56 | 0.28328265 | 0.52 | 0.38 | 4.27E-52 | testis | Hprt |
| Snrpd21 | 2.72E-56 | 0.26801277 | 0.55 | 0.41 | 5.63E-52 | testis | Snrpd2 |
| Lamb21 | 6.15E-56 | 0.25237679 | 0.32 | 0.2 | 1.27E-51 | testis | Lamb2 |
| Apobec3 | 1.73E-55 | 0.33824576 | 0.3 | 0.19 | 3.59E-51 | testis | Apobec3 |
| Cope | 1.11E-53 | 0.26814542 | 0.54 | 0.39 | 2.31E-49 | testis | Cope |
| Ndufa111 | 1.82E-53 | 0.28589786 | 0.56 | 0.42 | 3.76E-49 | testis | Ndufa11 |
| Ndufa121 | 2.06E-53 | 0.26378143 | 0.52 | 0.39 | 4.27E-49 | testis | Ndufa12 |
| Aprt1 | 3.55E-53 | 0.27431732 | 0.27 | 0.17 | 7.35E-49 | testis | Aprt |
| Ncl1 | 4.39E-53 | 0.28359903 | 0.52 | 0.38 | 9.09E-49 | testis | Ncl |
| Eif3m | 1.43E-52 | 0.28495165 | 0.43 | 0.31 | 2.96E-48 | testis | Eif3m |
| Slc6a64 | 1.49E-51 | 0.58419638 | 0.51 | 0.42 | 3.09E-47 | testis | Slc6a6 |
| Pigyl1 | 1.95E-51 | 0.25540455 | 0.26 | 0.16 | 4.03E-47 | testis | Pigyl |
| Smad1 | 7.66E-51 | 0.44360184 | 0.28 | 0.18 | 1.59E-46 | testis | Smad1 |
| Mrpl20 | 8.12E-51 | 0.26358914 | 0.43 | 0.31 | 1.68E-46 | testis | Mrpl20 |
| Pgls | 1.09E-50 | 0.25780085 | 0.39 | 0.27 | 2.26E-46 | testis | Pgls |
| Tulp4 | 1.60E-50 | 0.26959345 | 0.27 | 0.17 | 3.32E-46 | testis | Tulp4 |
| Erh1 | 1.96E-50 | 0.2546816 | 0.52 | 0.39 | 4.06E-46 | testis | Erh |
| Rpl22l11 | 4.91E-49 | 0.26109282 | 0.68 | 0.55 | 1.02E-44 | testis | Rpl22l1 |

| | | | | | | | |
|---|---|---|---|---|---|---|---|
| Lama44 | 1.84E-47 | 0.2770024 | 0.27 | 0.17 | 3.82E-43 | testis | Lama4 |
| Rps27l1 | 2.48E-47 | 0.26704131 | 0.69 | 0.59 | 5.13E-43 | testis | Rps27l |
| Eln1 | 2.55E-47 | 0.59299537 | 0.3 | 0.2 | 5.28E-43 | testis | Eln |
| Ddrgk1 | 1.78E-46 | 0.25911142 | 0.48 | 0.35 | 3.68E-42 | testis | Ddrgk1 |
| Tomm201 | 1.93E-46 | 0.2523074 | 0.54 | 0.42 | 3.98E-42 | testis | Tomm20 |
| Cracr2b2 | 2.29E-46 | 0.25982045 | 0.52 | 0.39 | 4.73E-42 | testis | Cracr2b |
| Kank33 | 9.15E-46 | 0.25788377 | 0.68 | 0.56 | 1.89E-41 | testis | Kank3 |
| Lims11 | 1.94E-45 | 0.27766891 | 0.36 | 0.25 | 4.00E-41 | testis | Lims1 |
| Ndufv1 | 8.77E-44 | 0.25294904 | 0.33 | 0.22 | 1.82E-39 | testis | Ndufv1 |
| Grn1 | 2.09E-43 | 0.27013297 | 0.53 | 0.4 | 4.33E-39 | testis | Grn |
| Phb2 | 3.12E-43 | 0.26294696 | 0.42 | 0.3 | 6.46E-39 | testis | Phb2 |
| Flt42 | 2.63E-41 | 0.53002574 | 0.3 | 0.21 | 5.45E-37 | testis | Flt4 |
| Lrg11 | 4.33E-41 | 0.55519595 | 0.28 | 0.19 | 8.95E-37 | testis | Lrg1 |
| Comt1 | 2.42E-36 | 0.27022778 | 0.37 | 0.27 | 5.02E-32 | testis | Comt |
| Scarb14 | 5.02E-36 | 0.32211013 | 0.5 | 0.4 | 1.04E-31 | testis | Scarb1 |
| Lmcd14 | 1.79E-33 | 0.27850001 | 0.34 | 0.25 | 3.70E-29 | testis | Lmcd1 |
| Slc25a4 | 1.85E-33 | 0.26640888 | 0.64 | 0.53 | 3.83E-29 | testis | Slc25a4 |
| Ptpn182 | 3.28E-33 | 0.25251709 | 0.28 | 0.2 | 6.79E-29 | testis | Ptpn18 |
| Ppfibp12 | 4.71E-33 | 0.25126718 | 0.28 | 0.2 | 9.76E-29 | testis | Ppfibp1 |
| Nrp23 | 5.79E-11 | 0.25798018 | 0.27 | 0.22 | 1.20E-06 | testis | Nrp2 |

References:

Kalucka J, de Rooij LPMH, Goveia J, Rohlenova K, Dumas SJ, Meta E, Conchinha NV, Taverna F, Teuwen L-A, Veys K, et al. 2020. Single-Cell Transcriptome Atlas of Murine Endothelial Cells. Cell. 180(4):764-779.e20. doi:10.1016/j.cell.2020.01.015.
